# Adaptive immunity: from CRISPR to CRIHSP?

**DOI:** 10.1101/2023.09.25.559221

**Authors:** Zhiyuan Niu, Yanfeng Wang, Bingqian Xu, Xinru Jin, Linwei Ning, Yuekai Hao, Yangchun Yan, Mengjun Wang, Wuling Zhu, Lingtong Zhi, Changjiang Guo

## Abstract

Clustered regularly interspaced short palindromic repeats (CRISPR) confer adaptive immunity in prokaryotes against viral infection and plasmid transformation. Here, we identified abundant clustered regularly interspaced homologous stem-loop pairs (CRIHSP) within human adaptive immunity-related genes, such as antigen receptor germline genes. By examining genomic stability under activation-induced cytidine deaminase (AID) overexpression, we found that CRIHSP structures are preferred targets for AID. Through collecting and analyzing cancer cell genomic mutation databases, we discovered that these mutation sites tend to cluster toward stem-loop structures. Importantly, the frequency of stem-loop sequences within immunoglobulin heavy chain variable (*IGHV*) gene segments was over 3-fold higher compared to that within other regions. We concurrently observed stem-loop structures frequently flanking RSS motifs. Conservative estimates indicate a 74.07% probability of immunoglobulin heavy chain diversity (*IGHD*) gene segment enclosure within stem-loop. These RSS-proximal stem-loops may assemble into homologous stem-loop pair (HSP)-like intermediates and subsequent CRIHSP arrays. Our results implicate CRIHSP architectures as critical enhancer of antigen receptor gene recombination and diversification. Further molecular dissection of these processes could potentially elucidate mechanisms underlying cancer cell genomic instability, chromatin folding and dynamic regulation, gene expression control, etc.

## Introduction

Clustered Regularly Interspaced Short Palindromic Repeats (CRISPR) is an adaptive immune system found in the majority of archaea and about half of the bacteria which stores pieces of infecting viral DNA as spacers in genomic CRISPR arrays to reuse them for specific virus destruction upon a second wave of infection. The short palindromic repeats in the CRISPR locus can bind with tracrRNA and form gRNA, an important component of the immune effector of archaea and bacteria, after being cleaved by RNaseL. This can eliminate their main threat: the invading rapidly expanding DNA or RNA. Higher organisms, such as humans and other mammals, are exposed to more complex and diverse foreign pathogens, and the mere excision of DNA or RNA within cells by the CRISPR/Cas system no longer guarantees their safety. So these higher organisms evolved multi-functional and diverse antigen receptors – antibodies or BCRs and TCRs. Variable-(Diversity)-Joining (V(D)J) fragments rearrangement in their germline gene is the process by which antigen receptor diversity is attained, then enables the recognition of a nearly limitless array of potential antigens. In the evolutionary “arms race” from prokaryotes to eukaryotes, traces of Short Palindromic Repeats (SPR) have been conserved in the antigen receptor germline genes, for example, the CAC(A/T)GTG short palindrome sequences are preserved adjacent to the recombination sites of V(D)J gene segments^1^. In a sense, the antigen receptor germline genes of higher organisms also have CRISPR sequence features and are evolved higher-level acquired immune-related genes. However, unlike CRISPR, which exerts adaptive immunity by transcribing into RNA, the antigen receptor germline genes need to undergo rearrangement and eventually form antibodies or BCRs and TCRs to mediate humoral and cellular immunity, respectively.

SPR (CAC(A/T)GTG) are considered to play an important role in the antigen receptor germline gene recombination process, SPR constitute inverted repeats at both ends of the sequences to be joined. These palindromes can be written as a hypothetical stem structure that draws variable and J regions together, providing a possible molecular basis for the DNA joining event^1^. Later studies found that each rearranging gene segment is flanked by a recombination signal sequence (RSS). RSS not only includes the SPR heptamer (CAC(A/T)GTG) but also a nonamer (consensus ACAAAAACC), separated by a 12 or 23-bp spacer of conserved length but not conserved sequence. For recombination, Max, E. E. *et al.* proposed the hypothetical stem structure asymmetrically formed between RSSs partners. In these stem-loop structures that draw V- and J-regions together, the high-efficiency cleavage and rejoining occur only between a 12/23 RSS pair, this is referred to as the “12/23” rule^2–4^. However, the hypothetical stem structure formed between RSS sites cannot explain why V(D)J gene segments combine in a supposedly stochastic fashion. Because the VJ sequences at the periphery have the highest probability of being recombined and retained by RSS sequences. Also, the hypothetical stem-loop structure formed between inverted repeats located next to germ-line V- and J-region genes on the same strand, presumably many thousands of bases away, how are such long-distance stem-loop structures formed? Also, this model cannot explain the ordered rearrangement process, such as D_H_-J_H_ rearrangements preceding V_H_-DJ_H_ recombination in the *IGH* locus, and the *IGK* locus rearranges before the *IGL* locus^5^. Gene arrangement also occurs between joining (J) and constant (C) genes, but we did not find any RSS sites between them. Moreover, not every pair of 12- and 23-RSS is compatible; the factors that govern the “beyond 12/23” restriction are not completely understood^6^. Other mysteries include allelic exclusion, isotype exclusion, and so on. Therefore, we must explore other unknown details in the antigen receptor germline gene recombination process.

In this study, we found a large number of Clustered Regularly Interspaced Homologous Stem-loop Pairs (CRIHSP) sequences in the antigen receptor germline genes, such as antibody or B cell receptor (BCR) and T cell receptor (TCR) germline genes. The CRIHSP sequences we found in eukaryotic cells are longer and more diverse than the CRISPR sequences in prokaryotic genomes. We found that these CRIHSP sequences are highly related to AID-mediated gene mutation. RSS flanking stem-loops also form CRIHSP structures and might be an important mechanism in V(D)J recombination, class switch recombination (CSR), and somatic hypermutation (SHM) of the antigen receptor.

## Results

### The discovery of CRIHSP in antigen receptor germline genes and its structural features

As mentioned above, although the hypothetical stem-loop structure based on RSS is not a perfect model to explain the rearrangement of antigen receptor germline genes^1, 4^, it gives us an important hint that other hypothetical stem-loop structures hiding in the gene sequence may help us explore the secrets therein. Therefore, we analyzed the distribution of theoretical stem-loop structures on the whole chromosome and found that the stem-loop structures were evenly distributed on chromosome 14 (Fig S1A), and the number of stem-loop structures on antigen receptor germline genes, such as Immunoglobulin Heavy Chain Gene (*IGH*), Kappa Chain Gene (*IGK*), Lambda Chain Gene (*IGL*), and T Cell Receptor Alpha Chain Gene (*TCRA*), Beta Chain Gene (*TCRB*), Gamma Chain Gene (*TCRG*), Delta Chain Gene (*TCRD*), is significantly lower than the other regions (Fig S1A and S1B). Thus, this result does not give us more hints.

Rao *et al*. reported that the CCCTC-binding factor (CTCF) binds to the consensus DNA sequence 5’-CCACNAGGTGGCAG-3’, which contains a stem-loop structure sequence, on chromatin to facilitate chromatin looping and folding^7^, also, CTCF has been implicated in *IGH* locus contraction and recombination^8, 9^. Then we came up with a reasonable conjecture, that is, sequence-similar stem-loop structures may produce a “convergent effect” under the action of auxiliary factors, then mediate the antigen receptor germline genes locus contraction and engage in gene rearrangement.

To comprehensively analyze genome-wide sequence-similar stem-loop structures, we developed a Python program. The reference human genome assembly GRCh38 was examined to identify all stem-loops with stem size ≥7 bp and overall length about 60 bp. Stem-loop pairs exhibiting >89% similarity in their first 10 bp were designated as homologous stem-loop pairs (HSPs). The 5′ positions of these HSP stem-loops were plotted on graphs using horizontal and vertical coordinates, with each HSP represented by a dot. For sequences with an odd number of total HSPs, the unpaired stem-loop was plotted using its 5′ position for both axes. Following this criterion, we utilized Plotly, a data visualization tool in Python, to plot the locations of these HSPs on the graph. As a result, we found that there are a large number of Clustered Regularly Interspaced Homologous Stem-loop Pairs (CRIHSP) in the Ig germline gene V(D)JC regions. As shown in Figure 1, we analyzed the *IGH* (Figure 1A), *IGK* (Figure 1B), and *IGL* (Figure 1C) germline genes. The red arrows in the figure indicate the CRIHSP structures, which are arranged in a seemingly "parallel" or "vertical" fashion (the visual non-verticality is an artifact of different scaling ratios between the horizontal and vertical axes) with respect to the diagonal line representing the entire gene region. We also randomly analyzed the non-Ig gene region on chromosome 14 and found that the overall number of HSPs was not significantly different from the Ig germline gene region, but there were no CRIHSP structures (Figure 1D). In addition, we found that the stem-loop structures were uniformly distributed across all the gene regions studied above, as evidenced by the red barcodes in Figure 1 which depict their distribution.

**Figure 1.**
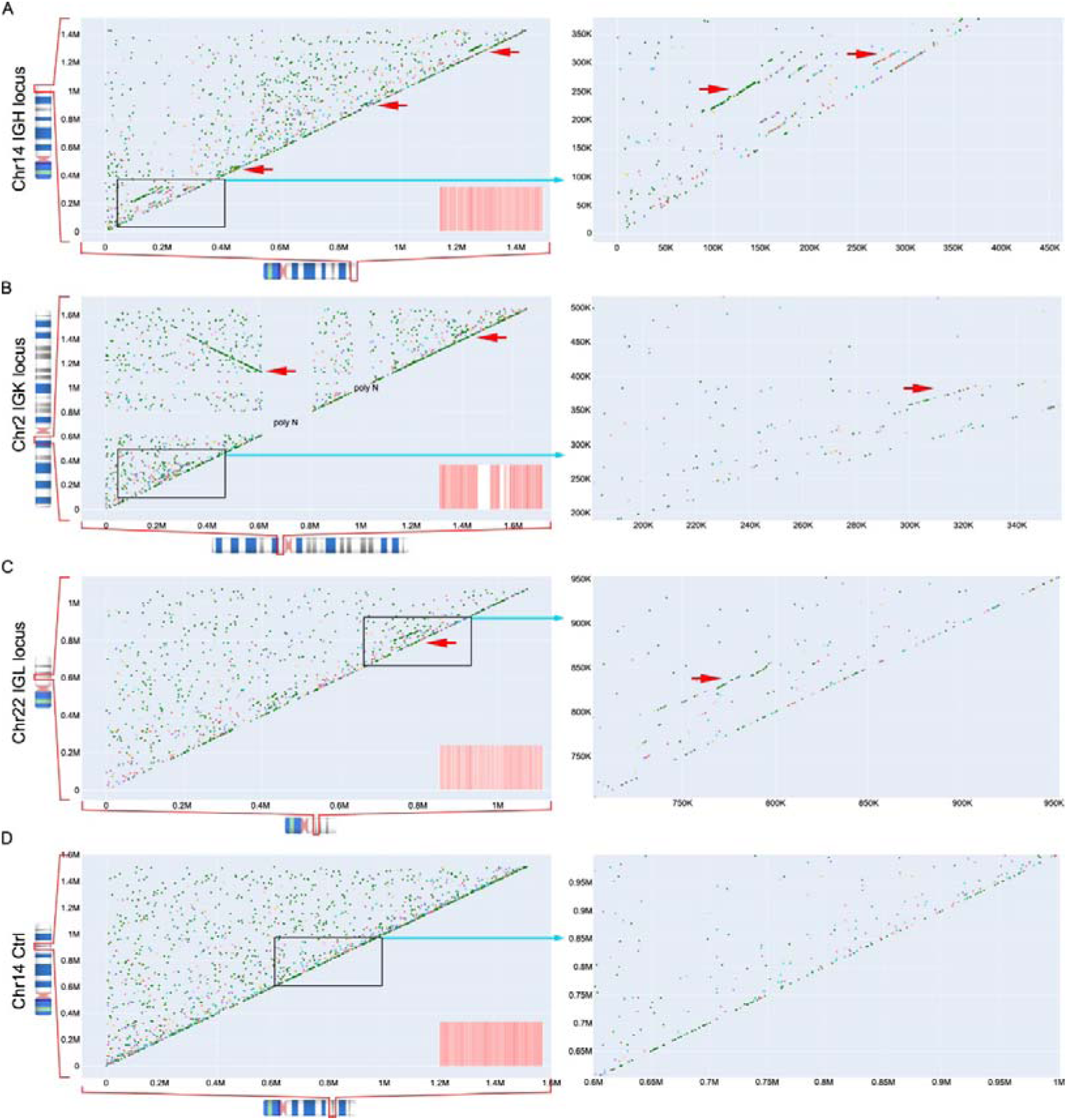
CRIHSP structures in antibody germline genes. The distribution of CRIHSP in the *IGH* heavy chain germline gene located on chromosome 14 (A), the *IGK* (B) and *IGL* (C) light chain germline genes located on chromosomes 2 and 22, respectively, and a random control region on chromosome 14 (D). The red arrows indicate the identified CRIHSP structures. The right panels show enlarged views depicting parts of the left figures, while the red barcodes on the left represent the overall stem-loops distribution across each corresponding gene region.

In addition to antibody germline genes, T cell receptor (TCR) germline genes can also undergo rearrangement. Thus, we analyzed CRIHSP in the TRA (Figure S2A), TRB (Figure S2B), TRG (Figure S2C), and TRD (Figure S2D) TCR genes. We identified abundant CRIHSP structures in the TRB and TRG germline regions. Notably, TCR genes are known to rarely undergo somatic hypermutation (SHM). Therefore, further studies are needed to elucidate the roles of these highly structured patterns in gene rearrangement and mutation, as well as the underlying molecular mechanisms.

After expanding the analysis of HSP distribution to the chromosome level, we discovered new CRIHSP structures formed in certain antibody germline gene regions and more distal DNA regions (Figure 2A). Concurrently, abundant CRIHSP were present in the centromeric region (Figure 2B), while no such structures occurred in other chromosomal areas. We also detected CRIHSP structures flanking the antibody germline genes. Intriguingly, some closely situated regions contained both “parallel” and “vertical” (as mentioned above, the visual non-verticality is an artifact of disproportionate horizontal and vertical axis scaling) arrangements (Figures 2C-E).

**Figure 2.**
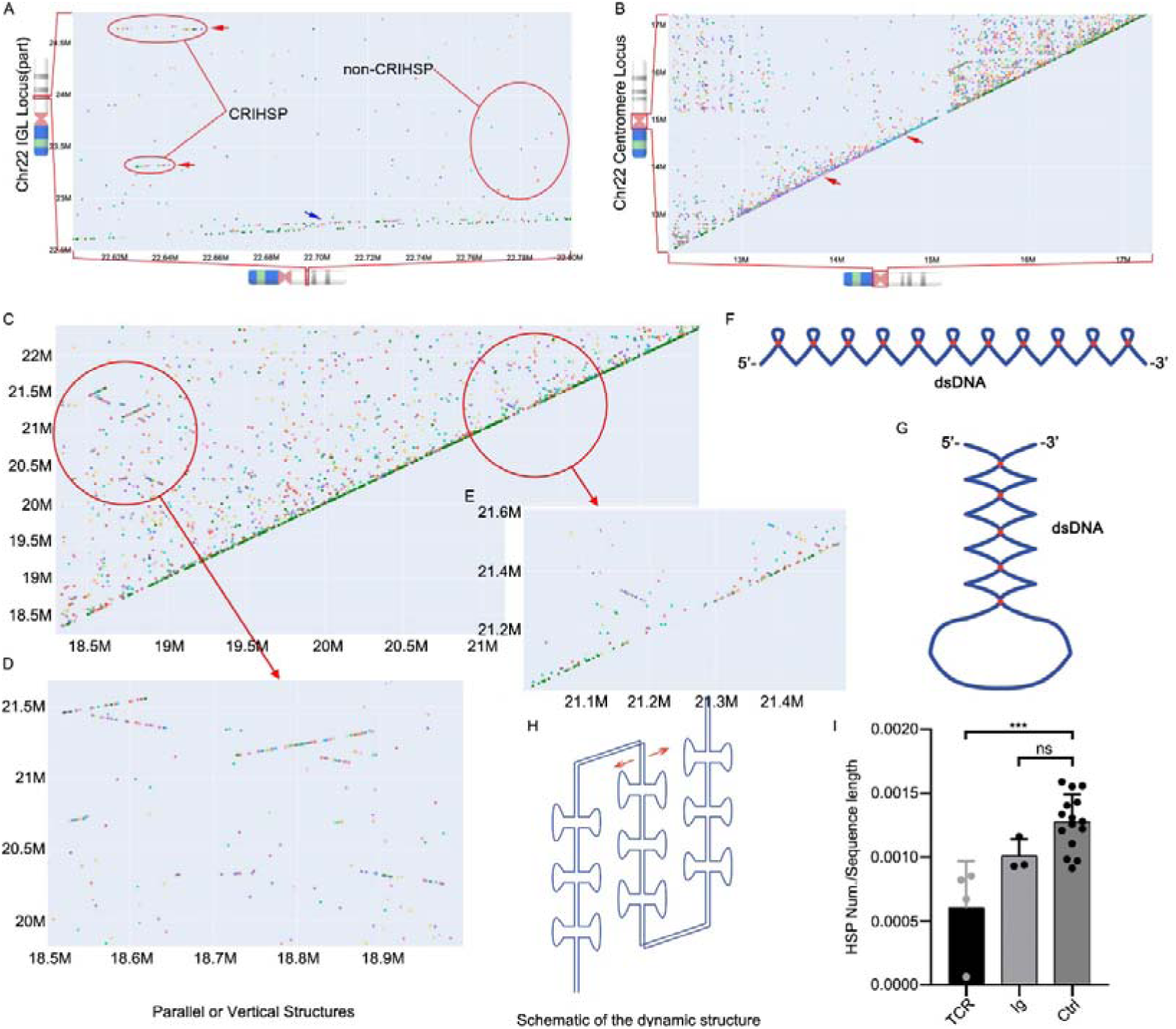
Putative static and dynamic DNA conformations mediated by CRIHSP. (A) CRIHSP formed between the *IGL* locus (21.9-23.0 M) and flanking *non-IGL* areas, denoted by red arrows, while the blue arrow marks CRIHSP within *IGL*. A non-CRIHSP region was framed in red. (B) CRIHSP arrangement in the centromeric region of chromosome 22. (C) CRIHSP distribution adjoining the 5’ end of *IGL*. (D-E) Enlarged parts of Figure C. (F-G) Schematics of chromatin folding patterns for “parallel” (F) or “vertical” (G) CRIHSP structures. Red dots symbolize HSPs. (H) Possible dynamic chromatin architecture elicited by CRIHSP. (I) HSP quantities in poly-CRIHSP (TCR, Ig) and non-CRIHSP (Ctrl) loci (>1 Mbp). (***P < 0.001, ns, not significant, one way ANOVA for multiple comparisons).

Schematic diagrams illustrating how the “parallel” (Figure 2F) and “vertical” (Figure 2G) arrangements could affect DNA folding under the “convergent effect”. Given their physical proximity, we postulate these patterns may precipitate dynamic chromatin architectures, as depicted in Figure 2H. Comprehensively mapping genome structure and dynamics would substantially improve our knowledge of cell biology. We quantified and analyzed HSP numbers in poly-CRIHSP-containing regions within *IGH*, *IGK*, *IGL,* and *TCRA/B/G/D* germline genes, alongside randomly selected non-CRIHSP areas across chromosomes. Intriguingly, poly-CRIHSP regions exhibited fewer HSPs compared to non-CRIHSP regions (Figure 2I). Moreover, random shuffling of *IGH* and *TRB* germline sequences maintained overall stem-loop totals, whereas CRIHSP configurations disappeared (Figure S3). This implicates CRIHSP as a non-random and likely biologically vital structure.

**Figure 3.**
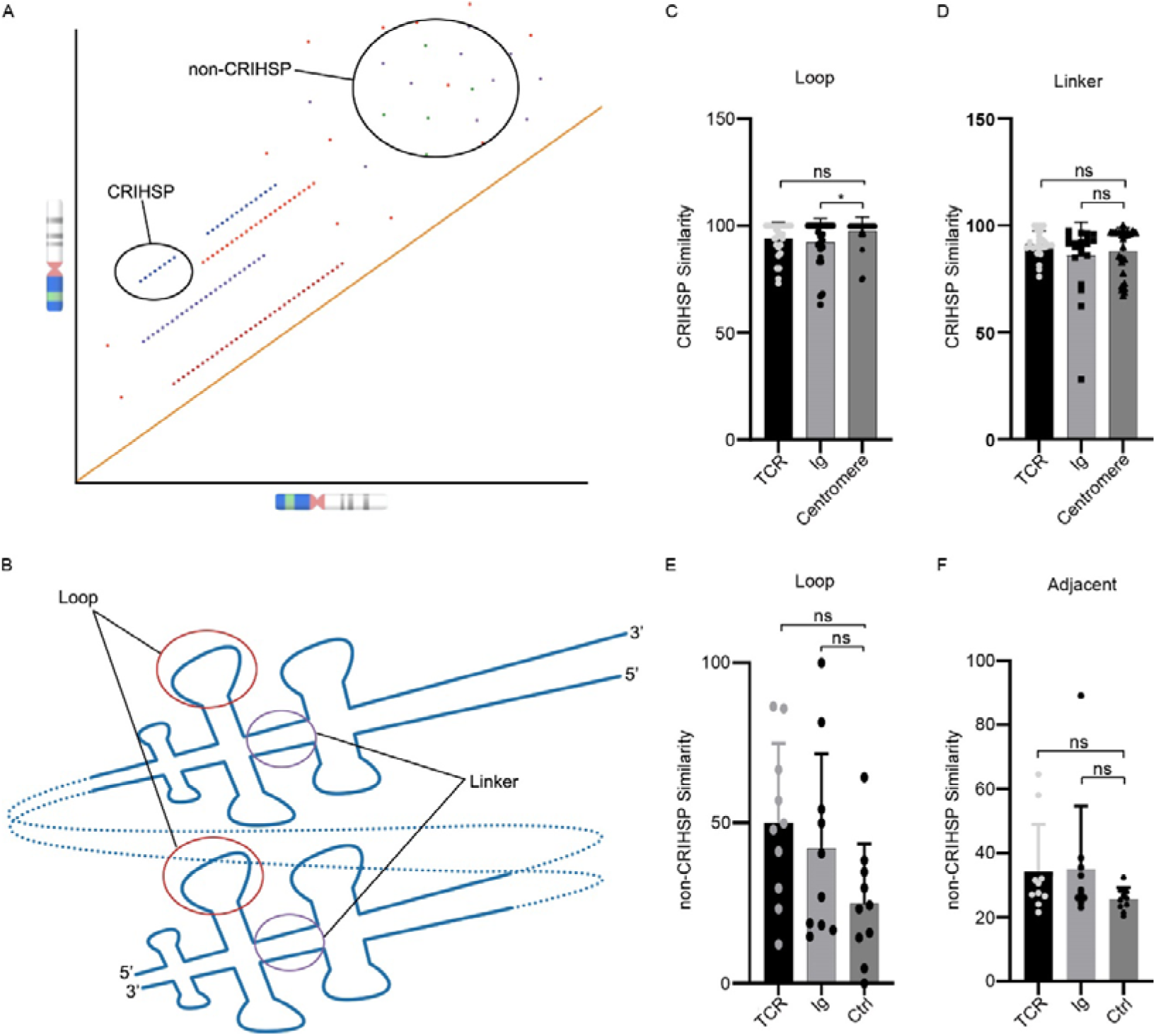
Sequence similarity of HSPs within CRIHSP versus non-CRIHSP loci. (A) Schematic of CRIHSP and non-CRIHSP regions. (B) Tandem HSP array within a CRIHSP locus. (C, D) Sequence similarity analysis of Loop (C) and Linker (D) regions for CRIHSP-associated HSPs. (E, F) Similarity analysis of Loop (E) and adjacent (F) segments for non-CRIHSP HSPs. The adjacent regions in (F) denote the 200 bp sequences flanking the 3’ end of randomly chosen HSPs. Ctrl regions denote randomly selected non-CRIHSP and non-antigen receptor gene loci. (*P < 0.05, ns, not significant, one way ANOVA for multiple comparisons).

We compared the sequence similarity of HSPs situated within CRIHSP and non-CRIHSP loci, as illustrated in Figure 3A. CRIHSP regions were selected from Ig, TCR germline sequences, and centromeric DNA. For CRIHSP-associated HSPs, we focused on pairs arranged as depicted in Figure 3B. These corresponding HSPs exhibited homology not only in the Stem but also in the Loop (Figure 3C) and inter-HSP linker regions (Figure 3D). In contrast, HSP sequence similarity in non-CRIHSP areas was markedly reduced in the Loop (Figure 3E) and flanking segments (Figure 3F). Interestingly, Loop similarity was significantly higher for centromeric versus Ig region HSPs (Figure 3C).

### CRIHSP are involved in the diversification of Ig gene loci

Ig gene locus diversification has been a topic of intensive study. AID is a central factor in generating antibody repertoires, which enhances antibody affinity through somatic hypermutation (SHM) and alters the effector function of antibodies by causing DNA breaks that trigger class switch recombination (CSR)^10, 11^. The expression of AID alone is sufficient to reproduce the salient features of SHM in both B cells and other mammalian cells^12, 13^. The mechanism by which AID preferentially mutates Ig genes is still incompletely understood. This issue has implications for immunity as well as for the oncogenic mutations introduced by AID outside the Ig loci^14^. A popular view is that AID preferentially targeted to WRCH^15^, RGYW/WRCY^16^, or WGCW^17^ (W = A/T, R = A/G, H = A/C/T, Y = C/T) motifs, however, this view can not explain why most of the genome is spared from AID activity. Also, there is no evidence of a specific AID targeting factor^10^.

Research carried out over the past two decades has provided a fundamental understanding of AID by uncovering its biochemical and structural properties. AID has been suggested to act on ssDNA by deaminating deoxycytidines (dC) to deoxyuridine (dU) within immunoglobulin genes, leading to SHM^16^. Researchers also visualized the ssDNA scanning motion of AID and suggested that the enzyme was able to move bi-directionally while remaining bound to the same ssDNA^18^ (Figure 4A). Qiao *et al.* demonstrated a bifurcated substrate-binding surface of AID, which includes two branched single-stranded DNA binding grooves, so AID has a higher affinity for forked DNA (Figure 4B)^19^. Recent studies show that AID deaminates ssDNA more robustly in the context of structured substrates, such as G-quadruplex (G4)^19^, for G4 has more branched structures, and AID easily recognizes ssDNA overhangs on these structures (Figure 4C). G4 DNA additionally promotes AID oligomerization, which would facilitate double-strand breaks (DSB) formation by creating mutation clusters at the S regions^19^, and this may also help to recruit CSR cofactors^20^. AID typically displays greater enzymatic efficiency on single-stranded substrates with the following order: G4 > branched substrate > linear substrate. Based on these facts, we propose a speculative model that HSP share the same structure of G4 DNA leads to ssDNA exposure, and favors AID jumping to ssDNA, and the two ssDNAs “extruded” from dsDNA, making them perfect substrates for AID deaminase (Figure 4D). However, our model posits that not all CRIHSP-enriched areas necessarily constitute hotspots. As depicted in Figure 4E, loops with substantial sequence differences can readily form single-stranded motifs (left), enabling accessible AID binding and ensuing mutations. In contrast, excessively homologous stem-loops may anneal complementary loop structures without single-strand exposure (right), impeding mutagenesis. Moreover, our data revealed higher centromeric versus *IGH* HSP homology (Figure 3C), consistent with greater centromeric sequence stability as predicted by the speculative model.

**Figure 4.**
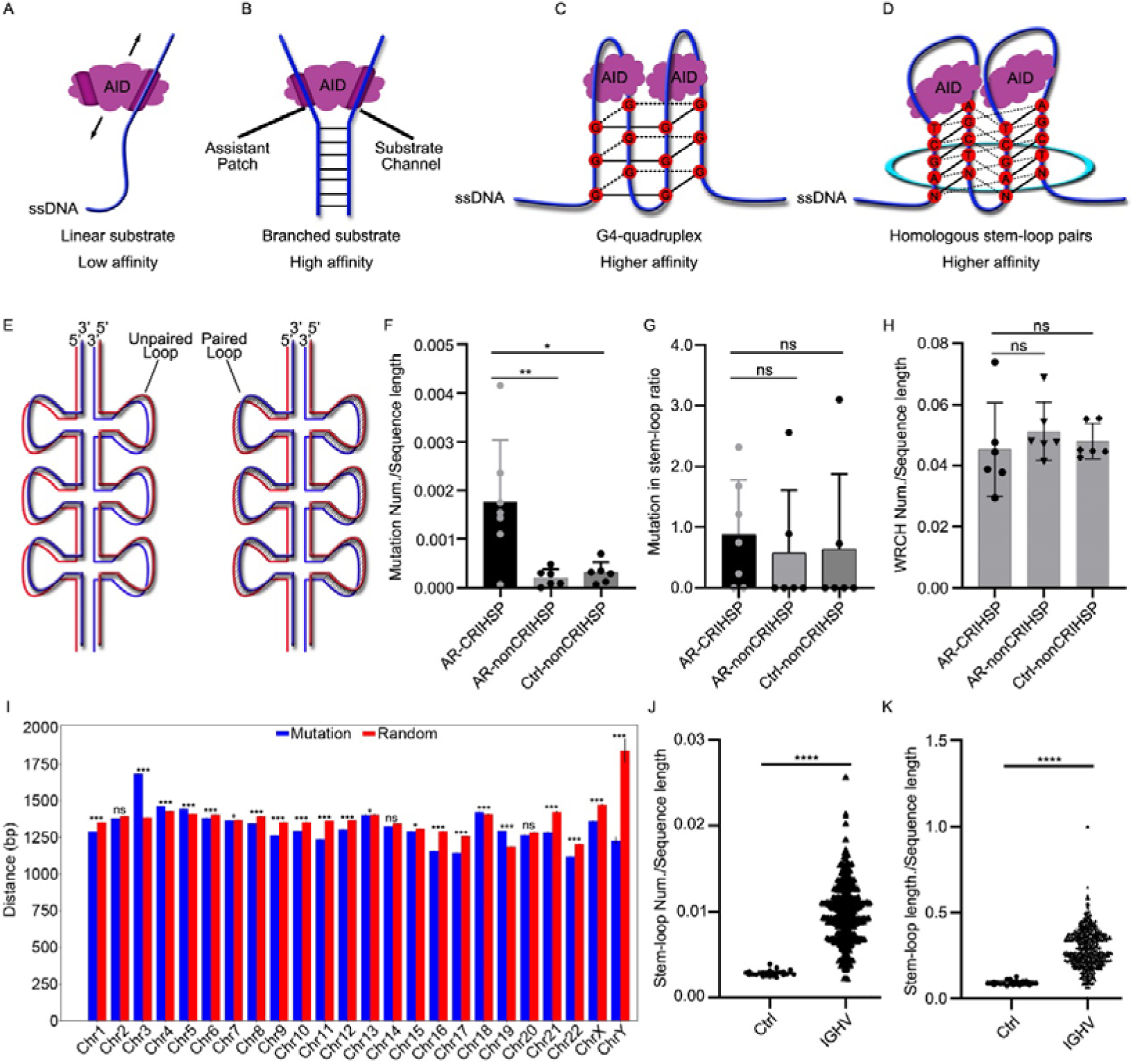
Association between stem-loop structures and mutation hotspots. (A-C) The efficiency of AID-mediated deamination on single-stranded substrates including linear DNA (A), forked double-stranded DNA (B), and G4 exposed single-strands (C). (D) Schematic of AID targeting HSP substrates. (E) Model of HSPs with non-complementary (left) or complementary (right) loop sequences. (F) Analysis of mutation frequency in CRIHSP and non-CRIHSP regions of IGH versus non-immunoglobulin non-CRIHSP loci under AID. (G) Examination of mutation clustering proximal to stem-loops. (H) Assessment of WRCH motif enrichment within CRIHSP regions. (I) Comparison of mean minimum distance from mutations or random sites to adjacent stem-loops using Wilcoxon rank sum test. (J-K) Quantification of stem-loop numbers (J) and length (K) in IGH “V_gene_segments” versus other genomic regions. (*P < 0.05, ** < 0.01, *** < 0.001, **** < 0.0001, ns, not significant, one way ANOVA for multiple comparisons).

To investigate the potential involvement of germline CRIHSP in shaping AID activity and locus preference, we examined mutations in three types of target regions, the CRIHSP loci and non-CRIHSP loci within *IGH*, and randomly selected non-CRIHSP non-immunoglobulin loci–under AID-deficient or -overexpressing conditions. Additionally, it is important that all selected sites contain approximately the same number of stem-loop structures. Our results demonstrate that the CRIHSP region is a hotspot for mutations, while the non-CRIHSP regions within or out of the *IGH* locus are not hotspots for mutations (Figure 4F). We calculated the ratio of mutations localized to stem-loop structures over total mutations and the ratio of stem-loop length to total sequence length. Comparing these ratios allowed us to determine any preferential accumulation of mutations within stem-loop motifs. Results demonstrated that although CRIHSP loci were mutation hotspots, mutations showed no overt preference for stem-loop sequences (Figure 4G). We speculate that this is due to the complex folding of DNA that exposes the single strands of the non-stem-loop structure regions, essentially, the mutations are still caused by the stem-loop structures and tend to cluster towards the stem-loops. Our results also demonstrate intrinsic instability within CRIHSP gene regions. Both G4 (Figure S4A) and CRIHSP structures (Figures S4B and S4C) are susceptible to mutations and indels. While G4 motifs have known links with nuclear DNA instability in cancer^21^, it remains uncertain whether CRIHSP induces mutations via shared mechanisms and thereby contributes to cancer genomic instability. Moreover, we excluded spontaneous mutation sites (mutations generated without overexpression of AID) from analysis in Figures 4F and 4G.

Our results demonstrate that CRIHSP regions accumulate more mutations, while WRCH constitutes a preferred AID mutational motif. We hypothesized that this overlap may derive from WRCH enrichment within CRIHSP. However, analysis of WRCH frequencies in CRIHSP, non-CRIHSP structures in *IGH,* and random non-CRIHSP structures in non-immunoglobulin loci indicated no preferential WRCH incorporation within these structures (Figure 4H). Notably, the palindromic WRCH variants AGCT and TGCA represent optimal AID hotspots^17, 22^, and are overrepresented within stem-loop sequences. Given the multitude of reported AID-biased motifs though, precisely correlating these motifs and CRIHSP configurations remains challenging.

If genomic regions adopt CRIHSP configurations and expose single-stranded motifs susceptible to mutations under physiological conditions, we can validate this hypothesis by examining correlations between mutation site data and stem-loop positioning on a genome-wide scale. To validate this, we obtained genomic mutations by downloading Genome Screen Mutants (release v98) from the COSMIC Cell Line Project. After excluding centromeric and gap regions, we calculated the minimum distance from each mutation site to neighboring stem-loops across the remaining hg38 intervals. In parallel, equally sized sets of random chromosomal coordinates were generated and their minimal distance to adjacent stem-loops was computed. Averaged minimum distances were then compared between the two groups. Large-scale analysis revealed only 5 of 24 chromosomes lacked the posited association between mutations and proximity to stem-loops. The remaining 19 chromosomes exhibited mutation sites closer than expected to stem-loops by chance, supporting the hypothesis (Figure 4I). This phenomenon also has important significance for revealing the intrinsic laws of tumor genome instability.

Although *IGH* germline regions exhibited fewer stem-loops (Figure S1B) and HSPs (Figure 2I) compared to other loci. We found stem-loop numbers and length within *IGH* “V_gene_segments” surpassed average genome levels (in non-special structure regions) by 3.4- and 3.2-fold, respectively (Figure 4J and 4K). Notably, abundant CRIHSP structures flanking *IGL* locus, facilitated stem-loops within *IGL* remaining proximal to CRIHSP after rearrangement (Figure 2C), potentially promoting SHM. Due to the large amount of data produced, we have not yet fully analyzed the adjacent regions of *IGH* and *IGK*. This fact also confirms the clear association between stem-loop structures and hypervariable gene regions, but the underlying mechanisms still need further exploration.

### HSP-mediated contraction might be the potential mechanism that underpins antigen receptor germline gene rearrangement

Another strong evidence supporting HSP involvement in V(D)J recombination is the frequent co-localization of RSSs with stem-loops, put it another way, the presence of RSS motifs results in adjacent stem-loop structures capable of forming HSP-like configurations with other co-localized RSSs and stem-loops. The diverse antigen receptor repertoire of lymphocytes is generated by V(D)J recombination. V(D)J gene segments are flanked by RSSs that function as recognition sites for the V(D)J recombinase consisting of recombination activating gene 1 (RAG1) and RAG2 proteins. After pairing two compatible RSSs, the RAG1-RAG2 complex introduces double-strand DNA breaks, followed by processing and ligation of the DNA ends by repair factors of the nonhomologous end-joining machinery^23, 24^.

Beyond RSS, we identified stem-loop structures as additional conserved elements at immunoglobulin recombination junctions, with positional conservation exceeding that of RSS sites. For example, we analyzed 664 “V_gene_segment”, 61 “D_gene_segment”, 128 “J_gene_segment”, and 44 “C_gene_segment” on *IGH*, *IGL*, *IGK*, *TCRA*, *TCRB*, *TCRG*, *TCRD* germline genes. We searched for the presence of stem-loop structures at the positions adjacent to heptamers and nonamers, and then calculated the proportion of stem-loop structures at each position (Figure 5A, Supplementary Data 1-5). Take the *IGH* gene as an example, most of the “V_gene_segment” (*IGHV*) either nonamer or heptamer were bordered by a stem-loop sequence. To be more specific, for all RSSs at the 3’ end of the *IGHV* genes, 55.95% heptamers and 49.21% nonamers were flanked by a stem-loop sequence. Surprisingly, the probability of “D_gene_segment” being located in the stem-loop structure is 74.07%, most D fragments containing stem-loops are flanked by two RSSs. According to the average length of the D segment (∼20 bp), under natural conditions, the probability of “D_gene_segment” being located in a stem-loop structure is only about 4% (In our research, the average length of the stem-loop structure is ∼33 bp, and there is a stem-loop every 350 bp, as shown in Figure S1B). According to our research, the Diversity (D) segment is only diverse in sequence, but very conservative in structure. D segment itself being part of a stem-loop structure might be the important structural basis for the D segment to be retained in the V(D)J rearrangement process. Importantly, our analysis was limited to stem-loops within 5 bp to RSS heptamers and nonamers; relaxing proximity criteria further increases stem-loop adjacency frequencies. Likewise, lowering stem-loop structural constraints results in near-universal encapsulation of “D_gene_segment” within stem-loops.

**Figure 5.**
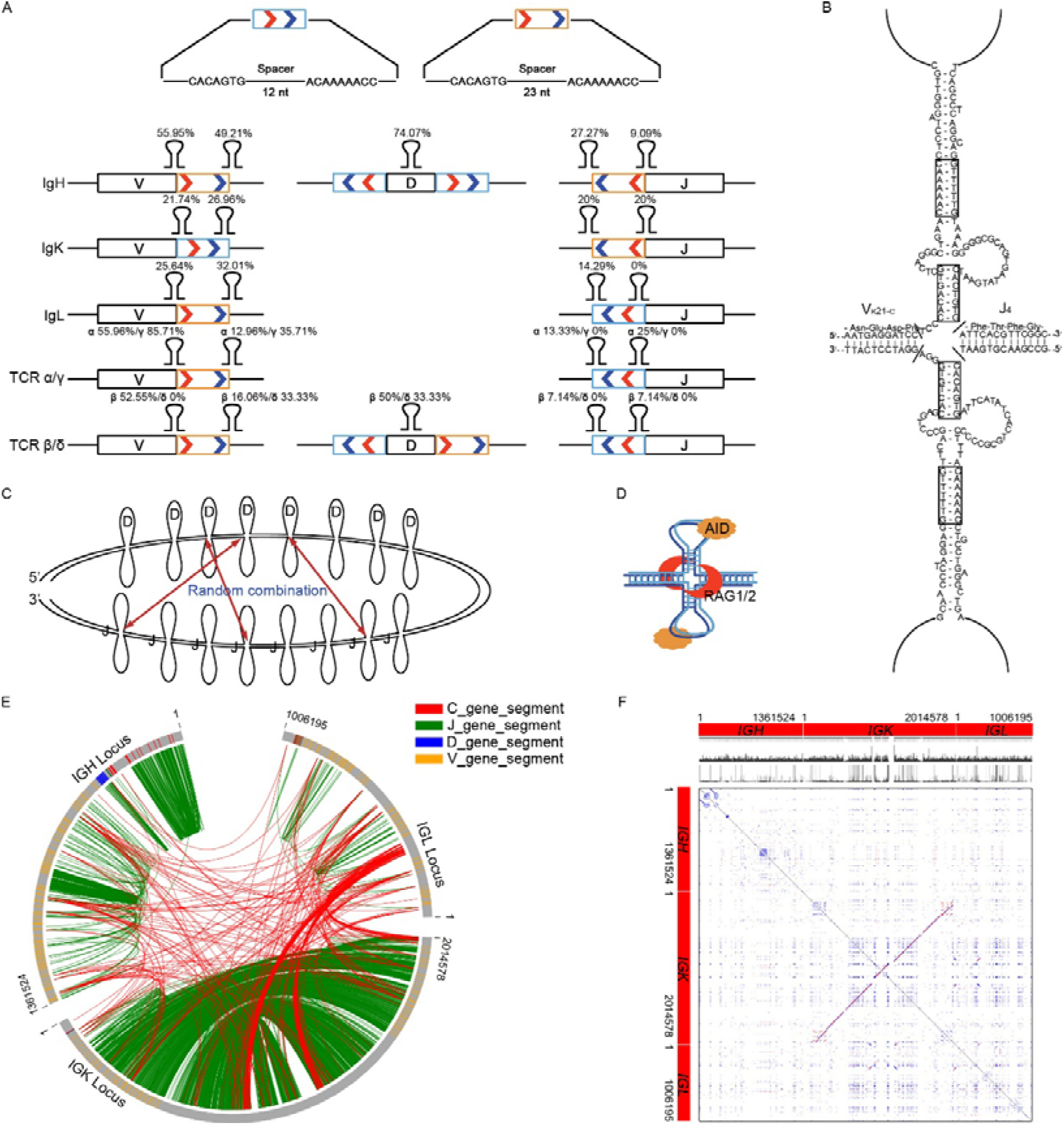
HSPs are widely discovered in antigen receptor gene loci. (A) We marked the stem-loop structures adjacent to heptamers and nonamers and calculated the proportions of heptamers or nonamers with adjacent stem-loop structures. (B) Diagram of hypothetical stem-loop structure formed between inverted repeats of two RSSs, that mediated the locus contraction between V and J segment, proposed by Tonegawa, S. *et al*. in 1979. (C) We proposed the hypothetical stem-loop structure formed by D and J segments, and HSP mediated the locus contraction between D and J segments. (D) Schematic diagram of the interaction mode of HSP with AID and RAG1/2. (E) A circular diagram of the human *IGH*, *IGK*, and *IGL* locus indicates the potential intra- and inter-gene connections of HSPs. Positions of the V (yellow), D (blue), J (dark green), and C (red) gene segments were marked. The intragenic connections were shown by dark green tracks, the intergenic connections were shown by red tracks. (F) We connected the *IGH*, *IGK*, and *IGL* genes, and then analyzed the distribution of HSPs and CRIHSP.

Additionally, despite detecting RSS elements flanking most V(D)J segments, this relies on the 63 heptamer and 90 nonamer motifs documented in current publications^3, 4, 25–27^, we provided these heptamer and nonamer sequences in Supplementary Data 6. The heterogeneity of RSS elements, comprising distinct heptamers, nonamers, and intervening spacers, probably precludes their identification as conserved sequences. However, the stem-loop structures adjacent to RSSs are relatively conservative structures, and warrant continued investigation of their biological significance.

Figure 5B illustrates a theoretical stem-loop structure formed between two RSS sites^4^, as we mentioned in the Introduction part, this is not a convincing antigen receptor rearrangement model. However, RSS proximity facilitates many stem-loops formation of HSP-like conformations with other co-localized RSSs and stem-loops (Figure 5A). HSPs might be producing a convergent effect on each other according to our proposed “convergent effect”. Although speculative, we establish a new paradigm for depicting the antigen receptor locus contraction and recombination (Figure 5C). In this model, stem-loops and RSSs work together to facilitate the rearrangement process, V(D)J segments are able to aggregate between the same type or different types of segments through HSPs, without the need for long-distance unwinding as shown in Figure 5B. V(D)J can recombine in a random way that is more consistent with thermodynamics. This model also leads to a pleasing mechanistic simplification and unification in the recombination events. Further, it was shown that RAGs could cleave symmetric bubbles, hairpins, heterologous loops, and potential G4 structures under physiological conditions^28^. These facts provide the molecular basis for our G4-like HSP to recruit RAG1/2 and AID (Figure 5D), and then mediate the antigen receptor germline gene rearrangement.

If the *IGK* rearrangement fails, RSS located 3’ end to the *IGKC* genes (KDE sequence) will mediate *IGK* locus deletion, either via another RSS located immediately 3’ end to the *IGKV* genes by 12/23 joining rule or via recombining element (RE) located between *IGKJ*-*IGKC* locus (KDE recombination)^29, 30^. RE is composed of an isolated conserved heptamer (CACAGTG) motif without an obvious nonamer sequence, however, it can form a perfect stem-loop structure with the heptamer in the KDE sequence (CACTGTG), this phenomenon also suggests the existence of a rearrangement mechanism beyond RSS 12/23 joining rule.

Early studies in human and mouse showed that during B-cell differentiation, the immunoglobulin light chain gene rearrangements occur in an orderly fashion starting with the kappa chain and proceeding to the lambda chain^31, 32^. If the *IGK* rearrangement fails, KDE recombination can delete part of the *IGK* cluster, this process correlates with *IGL* recombination and expression, and KDE mutation made fewer B cells expressing lambda chain^33^. V(D)J recombination at *IGH* and *IGK* loci also takes place sequentially during successive stages in B cell development, researchers identified an interchromosomal association between 3′ *IGK* enhancer and *IGH* locus, that marks the transition between *IGH* and *IGK* recombination^34^. Based on the above facts, we speculate that, besides intragenic interaction, antigen receptor germline genes have HSP-mediated intergenic interactions. So we use circular diagrams to show HSPs, HSPs were marked by lines. We found that there is a strong connection between *IGK* and *IGL* locus, as shown by red lines (Figure 5E). However, whether these interconnected sites are associated with the sequential rearrangement of *IGK* and *IGL* requires further investigation. We also connected the *IGH*, *IGK*, and *IGL* genes, and then analyzed the distribution of HSP and CRIHSP by dots (Figure 5F), we can see similar intra- and inter-genic HSP distributions as in Figure 5E. Indeed, recent observations have indicated that the α-globin locus is similarly organized into a rosette-like structure. Determining the spectrum of conformations adopted by the chromatin fiber of antigen receptor loci in developing lymphocytes is now critical, because it is the ensemble of trajectories that determines whether V(D)J elements will find each other with the appropriate probabilities^35^.

Large-scale contraction of the immunoglobulin locus is important for rearrangements and even allelic exclusion^36^, the robust D-to-J_H_ but little V_H_-to-DJ_H_ rearrangements, also, presumably due to locus contraction^37, 38^. Some researchers have also proposed the concept of homologous pairing, which usually occurs at the *IGH* and *IGK* loci^39^. Homologous pairing is also a form of locus contraction, but its exact structure is still unclear, and our proposed HSP model can well explain how homologous pairing occurs. Now we know that these long-range interactions are mediated by the ‘looping’ of individual *IGH* subdomains by many factors, which juxtaposes distal V_H_ genes next to proximal D_H_ segments to facilitate V_H_-DJ_H_ rearrangements. For example, CTCF orchestrates long-range cohesin-driven V(D)J recombinational scanning^40^. Other regulators that promote *IGH* locus contraction in pro-B cells include Pax5, E2A, YY1, and cohesin by binding to multiple sites in the V_H_ gene cluster, such as Pax5-activated intergenic repeat (PAIR) elements ^9, 41^. Factors that interact with HSPs and maintain these bundles or loops are currently unknown, and a detailed analysis is required.

### HSP as a single factor to predict chromosome conformation

The preceding models and experimental designs presume HSPs elicit a “convergent effect”. This means the more HSPs formed by two loci on a chromosome, the greater the possibility of spatial contacts between these two loci. Based on this premise, we simulate the ensemble of chromosome conformation based on HSPs and comprehensively map the chromatin interactions at a genome-wide scale. According to the algorithm that the more HSPs in a region, the stronger the interaction, after computing the distribution of the HSPs group and visualization plotting, results demonstrate that the genome is partitioned into numerous domains, large squares of enhanced contact frequency tiling the diagonal of the contact matrices, with loci in the same compartment showing more frequent interaction. Intriguingly, a subset of these squares showed pronounced concordance with the conformation predicted by the high-throughput chromosome conformation capture technique (Hi-C)^42, 43^ (Figure 6). Notably, HSPs accurately predicted the centromere topology of all chromosomes, while these repetitive regions are unmappable by Hi-C techniques. Collectively, these computational and experimental observations provide indirect evidence supporting the hypothesized HSP “convergent effect”.

**Figure 6.**
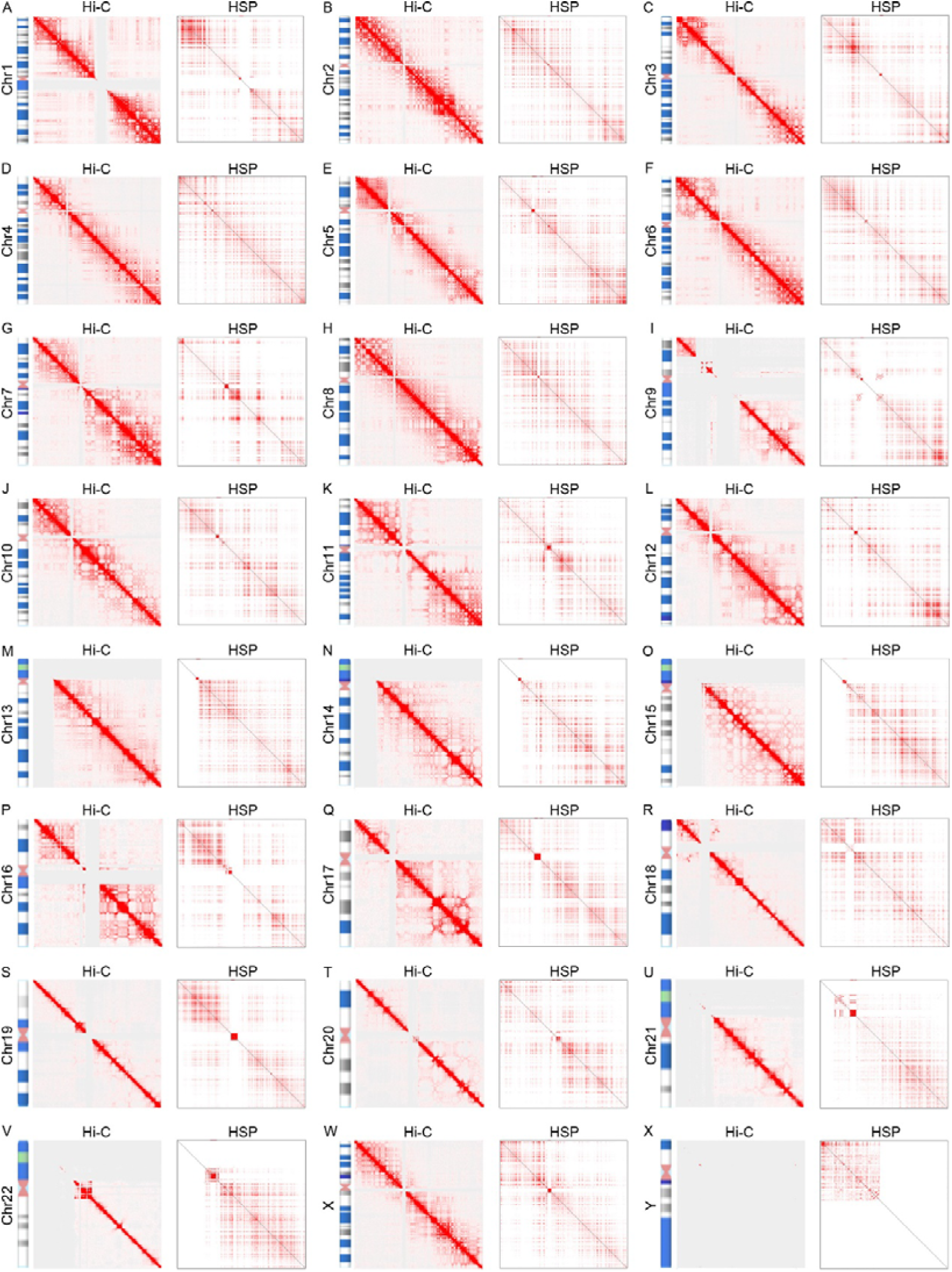
Comparison of chromosome conformation predicted by Hi-C and HSPs. (A-X) Contact matrices of chromosomes from Hi-C (left) and HSPs (right). The clustered red pixels in both figures indicate regions with stronger interactions.

We already know that, in addition to the canonical double helix, DNA can fold into various other inter- and intramolecular secondary structures under physiologically relevant conditions, which include G4 as well as Z-DNA, cruciforms, and triplexes. bioinformatics demonstrates that DNA sequences capable of forming these structures are conserved throughout evolution, and a growing body of work suggests that the resolution of DNA secondary structures is critical for genome integrity, stability, and cellular processes, such as transcription^44^. In future work, we still need to confirm the existence of these structures by using biophysical techniques (for example, circular dichroism, cryo-electron microscopy, *et al*.). Moving forward, the variable “convergent effects” of distinct HSPs could be inferred from empirical findings to refine algorithm parameters and achieve enhanced chromatin mapping resolution. And in the future, simulations of the ensemble of chromosome conformations based on the HSPs may not only accurately reproduce experimental contact probabilities, but also provide a picture of chromosome dynamics and topology.

## Discussion

Despite decades of work and intense effort from numerous labs, we are still left with an incomplete picture of how antigen receptor loci are regulated. Beyond the DGYW/WRCH motifs as predictors of Ig hypermutation hotspots^45^, several mechanisms that can provide ssDNA as AID substrates have been suggested. For example, DNA–RNA R loops or G4 DNA formed during transcription can stall RNAPII and are preferred AID substrates^46^. Transcription-induced DNA supercoiling also provides ssDNA substrate, and AID gains access to the gene body by traveling with transcription elongation^47^. Cis elements and genome topology act together with trans-acting factors, which biochemically copurify with AID, involved in transcription and RNA processing to determine AID activity at specific Ig regions^14, 48, 49^. Other loci sharing genomic and transcriptional features with the Ig are collaterally targeted during SHM and CSR. However, these views are currently controversial, as such structures are rare, especially over diverse Ig V exons, also, their presence at other AID off-targets, is unlikely and uncertain. In addition, the dissociation of AID from Ig loci to prevent mutation of constant regions must be somehow regulated – a step that has received little attention. Genomic targeting of AID appears to be multilayered, with inbuilt redundancy, but robust enough to ensure that most of the genome is spared from AID activity. However, there is no perfect hypothesis that can explain productive and precise AID targeting. Our results suggest that CRIHSP are the cis-acting targeting elements that render the target gene a suitable platform for AID-mediated mutation, and thus define a mechanistic basis for locus-specific targeting of SHM in the genome. Combining the results of ourselves and other researchers, we conclude, AID targeting must be integrated into a comprehensive picture that in conjunction with chromatin architecture, transcription quality, cis- and trans-acting factors, and AID regulation all contribute to defining whether a locus is mutated or not, and to what extent.

The essence of antigen receptor germline gene recombination is gene contraction, breakage, and joining. Interestingly, in these phenomena, we usually find traces of stem-loop structures. For example, the previously mentioned chromatin looping by CTCF and its stem-loop containing binding site^7^. In 2010, Xu *et al.* found that IgH switch (S) regions contain 5′-AGCT-3′ repeats in their core. 14-3-3 proteins specifically bind 5′-AGCT-3′ repeats and are recruited together with AID to the S regions involved in CSR events^50^, Multiple 5′-AGCT-3′ sequences can form stem-loops and then HSP structures. Other well-known phenomenon that can undergo gene breakage and insertion, such as CRISPR/Cas mediated genome expansion in Archaea and Bacteria, also have a large number of sequences that can form stem-loop structures by themselves or with each other^51^. However, the mechanism by which the stem-loop structure plays a role in this is unknown. Our results show that RSSs and RSS flanked stem-loops widely present in antigen receptor germline genes recombination sites might form HSP-like structures, and facilitate rearrangement, but we only speculate on how they work, whether HSP-like structures can form under physiological conditions, and whether there are trans-acting factors that assist these structures to form will be interesting challenges for future work.

The discovery of CRISPR structure in prokaryotes has sparked a wave of gene editing research, and CRIHSP structure in eukaryotes is also very likely to be a new key that can help us understand the secrets of how eukaryotes fine-tune their genes. In our proposed model, CRIHSP can also influence gene expression over large genomic distances. For example, CRIHSP structures bring distant enhancer elements into the proximity of promoter regions by compaction and looping. The single strand exposed by the CRIHSP structure might be the recognition sites of various trans-acting factors, and greatly reduce the energy consumption of unwinding double-stranded DNA during the initiation process of transcription, replication, gene recombination, or even heterochromatin interactions and homologous pairing of sister chromatids. Our results also clearly demonstrate the strong correlation between stem-loop structures and mutation sites in the cancer cell genome, further study of these molecular mechanisms may also help us uncover the potential mechanisms of genomic instability in cancer cells and find corresponding solutions.

## Materials and Methods

### Reagents

Puromycin (P8230), Trytone (T8490), Agar Powder (A8190), Yeast Extract Powder (01-012), DNA purification kit (D1300), NaCl (S8210), Trypsin 0.25% (T1360), PBS (D1040) were from Solarbio, Beijing, China. Stbl3 Chemically Competent Cells (KTSM110L) were from AlpaLife, Shenzhen, China. Polybrene (40804ES76) and Hieff Canace® High-Fidelity DNA Polymerase (10148ES10) were from Yeasen Biotech, Shanghai, China. Tissue gDNA Isolation Kit (BW-GD2211) was from BIOMIGA, Zhe Jiang, China. EcoR L (1040S), Xba L (1093A), T4 ligase (2011A) were form Takara, Da Lian, China. SanPrep Column DNA Gel Extraction Kit (B518131), and Pfu DNA Polymerase were purchased from Sangon Biotech, Shang Hai, China. E.Z.N.A.® Plasmid Midi Kit I (D6926-03) was purchased from Omega Bio-Tek, Guangzhou, China. Green Taq Mix (P131-01) was from Vazyme Biotech Co., Ltd, Nanjing, China. Calcium Phosphate Cell Transfection Kit (C0508) was from Beyotime, Haimen, PR China.

### Cell culture

Dulbecco’s modified Eagle’s medium (DMEM), and Fetal bovine serum (FBS) were from Gibco, Thermo Fisher, Shanghai, China. The HEK293T (GDC0187) cell line was obtained from the China Center for Type Culture Collection (CCTCC, Wuhan, China) and cultured according to their instructions. All media were supplemented with 1% penicillin-streptomycin (C0222, Beyotime, Haimen, China) and 10% FBS. All experiments were performed with mycoplasma-free cells.

### HEK239T infection or transfection and gene locus sequencing

LaG-2/G4 sequence and the lentivirus packaging, infection, and sequencing methods were described by us previously^52^. Transfection of HEK293T was performed by using Calcium Phosphate Cell Transfection Kit according to the manufacturer’s instructions. 1×10^6^ HEK293T cells were seeded in 6-well plates. At one day post-seeding, cells were transfected with 4 µg plasmids expressing AID. After 12 hours, replace the cell culture medium with fresh medium. 2 days later, genomic DNA was isolated from HEK293T cells by using the Tissue gDNA Isolation Kit according to the manufacturer’s instructions. The PCR primers we used to amplify the CRIHSP, non-CRIHSP and Ctrl sites in Figures 4F, 4G and 4H were provided in Supplementary Data 7. The reaction conditions consisted of initial denaturation at 98°C for 5 min; 35Lcycles of 98°C for 10 s, 68°C for 50 s; and a final extension step for 5 min at 72°C. We performed sequencing at Sangon Biotech (Shanghai) Co., Ltd.

### Databases

For most of our analysis, the latest version of the human reference genome assembly GRCh38.p13 was used, for Figures 5E and 5F, version T2T-CHM13v2.0 was used. The NCBI-provided human genome annotation file (GFF3 format) was retrieved and parsed using gffutils, a Python package for processing GFF and GTF files, to extract “V_gene_segment”, “D_gene_segment”, “J_gene_segment” and “C_gene_segment” features. Genome Screen Mutants (release v98) was downloaded from the Cell line project, a sub-database of the COSMIC database (https://cancer.sanger.ac.uk/cosmic/download), to obtain all genomic mutation information. Mutation profiles of over 1,000 cell lines used in cancer research were collected and analyzed in our study.

### Experimental repeats and data analysis

All assays had been performed at least thrice to ensure the repeatability of the experiments. The representative pictures and statistical analysis were shown in the final figures. Data for most experiments were analyzed with the Python applets we wrote. In Figure 4I-K, Figure 5, Figure 6, and Figure S1, the criteria for defining stem-loop structure sequences are: minimum size of a stem is 8, maximum size of a stem is 29, minimum size of a loop is 3, maximum size of a loop is 20, the number of mismatches allowed in the stem is 1. Maximum free energy threshold: -5.0 kcal/mol in Figure 4I. Python applet we used for analyzing Figure 1, Figure 2, Figure S2, and Figure S3 was provided as Supplementary Data 8, in these figures, the criteria for defining stem-loop structure sequences are: minimum size of a stem is 7, the number of mismatches allowed in the stem is 1, the maximum length of the stem-loop being about 60 bp. We use the random.shuffle () Python function to shuffle *IGH* and *TRB* gene sequence in Figure S3.

## Authors’ contributions

ZYN discovered the CRIHSP structures and initiated this program, CJG wrote most of the Python applets and analyzed data. ZYN, YFW, BQX, XRJ, LWN, YKH, YCY and LTZ carried out the experiments and analyzed data. ZYN, WLZ, CJG, and LTZ acquired funding. ZYN and WLZ supervised the study. All authors were involved in writing the paper and had final approval of the submitted and published versions.

## Acknowledgments

This work was supported by the National Natural Science Foundation of China (Grant 81902916), Natural Science Foundation of Henan Province (Grant 222300420513 and 232300420060), National Key R&D Program of China (2019YFA0906000), Major Science and Technology Projects in Xinxiang (21ZD007), Henan Program for Science and Technology Development (232102311042) and National Natural Science Foundation of China (Grant 81903187 and 82003261). We thank members of Zhu’s laboratory for helpful discussions, technical assistance, and critical reading of the manuscript.

---

**Figure S1.**
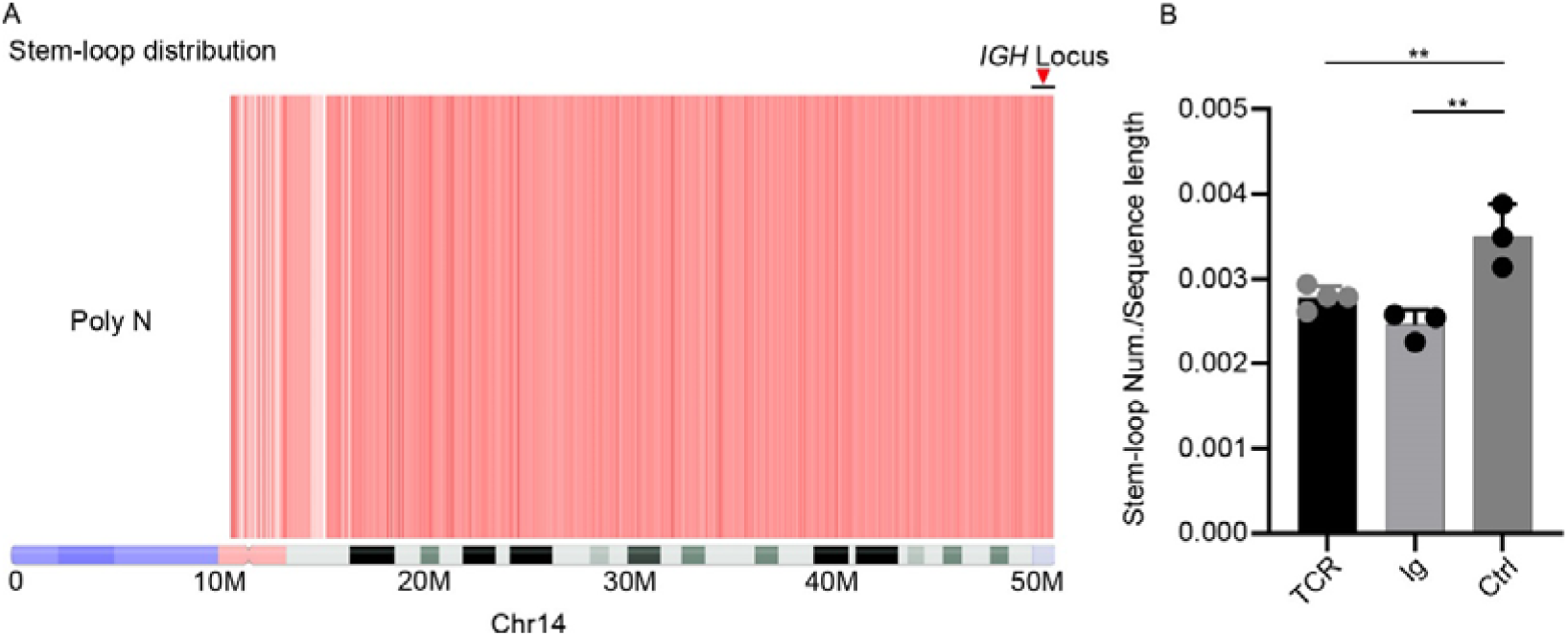
The distribution of stem-loop sequences on chromosomes. (A) The distribution of stem-loop sequences on chromosome 14. The location of the *IGH* germline gene was marked with red triangles and black lines. (B) Comparing the density of stem-loop sequences in the *TCR*, immunoglobulin (Ig) region with other control regions (Ctrl). The average length of the sequences selected for the Ctrl region is greater than 13 Mbp. (**P < 0.01, ns, not significant, one way ANOVA for multiple comparisons).

**Figure S2.**
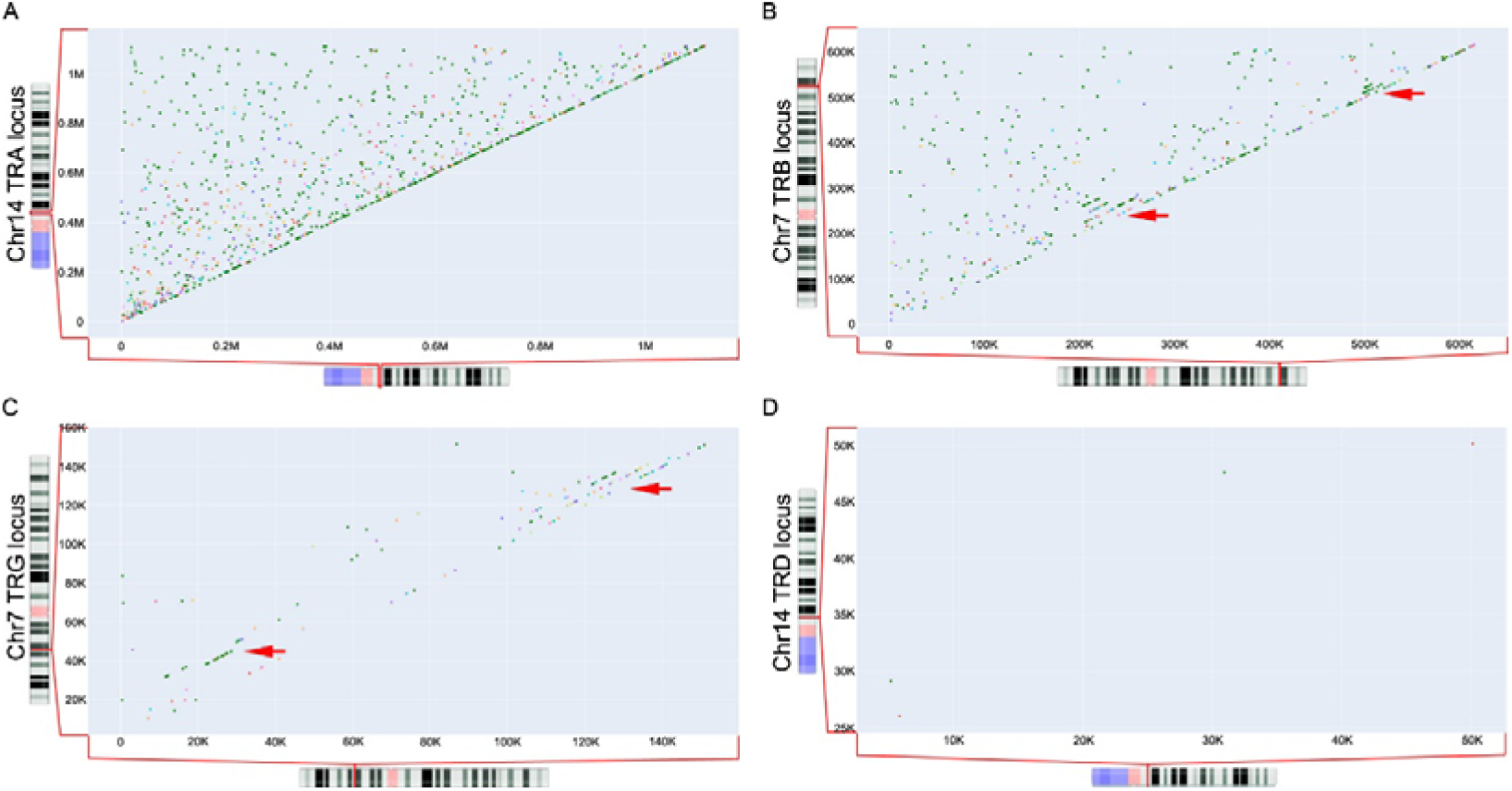
CRIHSP structures in TCR germline genes. Distribution of CRIHSP in the TCR α chain germline gene on chromosome 14 (A), the TCR β chain (B) and TCR γ chain (C) germline genes on chromosome 7, and the TCR δ chain germline gene on chromosome 14 (D), as indicated by the red arrows.

**Figure S3.**
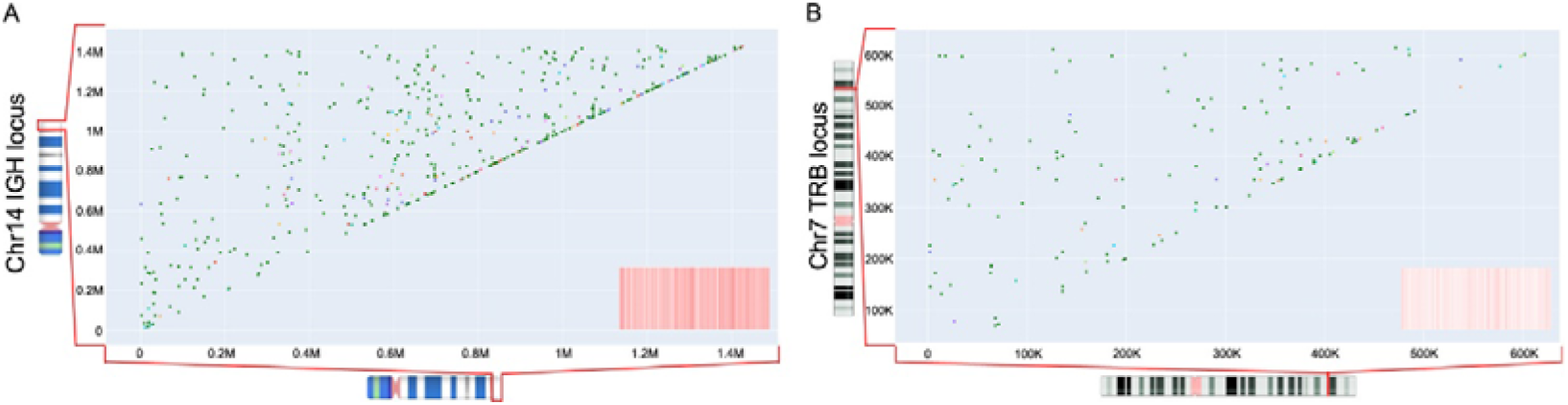
CRIHSP distribution in shuffled *IGH* and *TRB* germline sequences. (A,B) The *IGH* (A) and *TRB* (B) germline sequences were randomly shuffled prior to the analysis of CRIHSP distribution.

**Figure S4:**
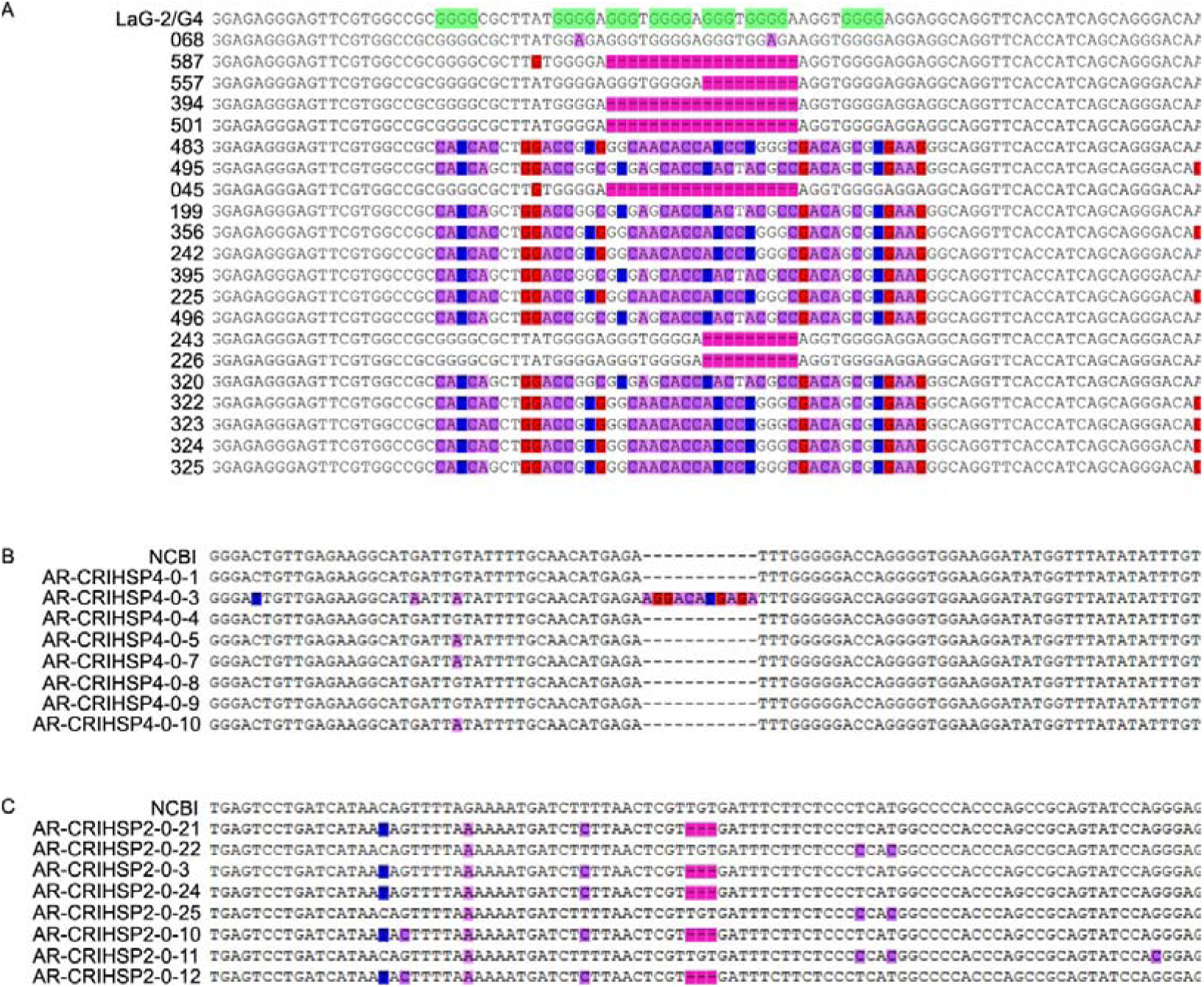
Sequences of LaG-2/G4 and two CRIHSP sites on *IGH* germline gene locus and their spontaneously mutated sequences. We integrated the LaG-2/G4 sequence into the genome of 293T cells by lentiviral transduction, then LaG-2/G4 sequences or CRIHSP sequences in the genome of 293T cells were amplified by Hieff Canace High-Fidelity DNA Polymerase, and subcloned to pCDH vectors, then transformed into Stbl3 Competent Cell, and LaG-2/G4 loci or CRIHSP sites were sequenced. The top sequence is the original sequence, bases marked with colored backgrounds are the spontaneously mutated bases, and “-” marks the position where the base deletion mutation occurred.

