## Supplementary Data 1 for "Adaptive immunity: from CRISPR to CRIHSP?"

id-IGHV3-9[V_gene_segment]

ctct**aagctgagcagcagctt**tatttgaaatacacagaaactgaaaacaatccaaatattaatcagcaccttcataagtaaacaaattgtggaaaagtcatttattgggatagtacccactactacaatcagtaaatgttggatacgatcaacagcatggttcaattcacaagta**catatgatgagtatagtgagccaaagcacatacatatg**attccgttttataaactgtacaaagtgaatactcattggaaattacataacaaagatcactaactga**cttctccatagtaagagaag**cgaaggtataggaggagaaattgcgagag**acaaagagaaaattgagaggcgaattcatttgttttctctgtgt**atggtcattaggtcaatgtttgtcaaatggtga**acatattgtgtgaagattatatgt**cagtat**tacctcattaaagttattataaataaaagtctaatgtggta**gaaaaagatgaagagagaaataaaaataatacaagaaaagtcatgaactcctgaatgaattaaccctta**gtttttctctattacttacaaaaac**accaagatacagccaaataatatcacgatatcattataagaagagtgttttgtaaacctcactgggaatttctagctctttcctagagttaattttgggaacagttggatccaattgtgagaaacgcaggccggacactgagactggctcttatgag**atgtgagctcttgtctatgtcacat**ggtccctccatacttggggg**tttacattcacatctgtaaa**tgaaggaaacattgactctcaaagaacatatttcatgtgcatgtaaaagtatgaatgctaatgagaattaattacttatgaagtataatcacccacatccactcttggacacagcccactctgaggcatccgttacagaactcattatatagtaggagacatgcaaatagggtcctccctctgctgatgaaaaccagcccagccctgACCCTGCAGCT**CTGGGAAAGGAGCCCCAGCCCTGAGATTCCCAG**GTGTTTCCATTCGGTGATCAGCACTGAACACAGAACTCACCATGGAGTTTGGACTGAGCTGGGTTTTCCTTGTTGCTATTTTAAAAGGTGATTCATGGATAAATAGAGATGTTGAGTGTGAGTGAACATGAGTGAGAGAAACAGTGGATATGTGTGGCAGTGTCTGACCAGGGTGTCTCTGTGTTTGCAGGTGTCCAGTGTGAAGTGCAGCTGGTGGAGTCTGGGGGAGTCGTGGTACAGCCTGGGGGGTCCCTGAGACTCTCCTGTGCAGCCTCTGGATTCACCTTTGATGATTATGCCATGCACTGGGTCCGTCAAGCTCCGGGGAAGGGTCTGGAGTGGGTCTCTCTTATTAGTTGGGATGGTGGTAGCACCTACTATGCAGACTCTGTGAAGGGTCGATTCACCATCTCCAGAGACAACAGCAAAAACTCCCTGTATCTGCAAATGAACAGTCTGAGAGCTGAGGACACCGCCTTGTATT**ACTGTGCAAAAGATACACAGT**GAGGAGAAGTCAGTGAGagcccagacaaaaacctcc**ctgcaggaagacaggaggggcctgggctgcag**aggccgctcaagacaca**ctgagcatagggttaactctgggacaagttgctcag**gaaggttaagagctggtttcctttcagagtcttcacaatttctccatctaacagtttcctcaggaaccctgtctagatctgtgatctggatctgctgaaactgcctgtgtcaccttcctcacctgtgattttgggggagctgattgtggacactccagtgtgtgggatttcttgatgacagcaattgtgtcttctgtctaggcatgtctagggctggccatcaggaag**ggcaggctggaatttttggaaagaggcgcacctgcc**atccaccaggaaattttgttgtcttttgttc**tgctagaattaaatcagacacaccaaggttaactagca**ctatcttcctagctcgagaaacttgatagcaagcttgaataacacttgtatgaaacc**atcagagcaacacctagaatagtgtctgat**ttaataattgagactatggtctagccaaggagacacataaaacgtgatttccaaggttggttcatattttatatttatcaatagaatctggctggtattataaacagctttgccaaaatatgtgtgttagtgttggtcttcggagaagcaaactctgtgactggattagcttcg**tgtacagtttccttcgtgtaca**gtttccctgtccactcttctgagaggaaatgcagaggtggtggaaggggtctggggtagctgaatgcatgtgagggaagtcggtccctgagtgaaggagaaagggaggaggactgggtggaactttcctaaacttctgtgttgttctatgaaggtccagcaaggtcactgaatcagagtcgtgtcacagtttcccatcggggacccagtgactcccagcagtg**gctctgctcagatcagcgcagagc**ttgttcatctcctaagagtggagcacaggacgt

Uppercase: IGHV3-9

Lowercase: Flanking sequence[1000bp]

Red & Bold & Underline: Stem-loop [18]

Blue: Heptamer[35]

Green: Nonamer [6]

id-IGHV2-70[V_gene_segment]

gaattcagccaagacagagccccatttaggagcaggtgcatcaaggaaccagccagagaacctctgactacaatctagggttgtcttcttttgcttcacagcatccagtcctaaggagaccacccactatatcatc**tgtaactcagaactttcttgttaca**aaggaaaacacaaatgaatataactctgcagttaacttcttgaacctggattgatgagtttccacggccttatgagcacagagaccacatccagtgcaggtgcagagca**aggctgcaacagtcagcct**gctgctccctcaggagtcttccaa**ttcccttgtccaaagcatccgctcattcatgggctgagaaaagggaa**gtcattcatgcagtctactctctccacagaactgacggggcacaaggacaacatcgatttttcatggaacatgcctctaggaatgcagc**tgtgtgcacacacactgctaagcacaca**cttctttacataattacttgtaactgtattttcttatttattctctccaatttttttacacaaattcatcacttttccccata**atcaaagaggattttgat**cagaatgcttgtggggagccccttgccccccagatgcccaccatcattattcttgaagggaggaggaacggcagctctcttgatttctactctaatcctctaggactaaaaccagaaggttgcatgtccggtgcgggagcatcgaagaagatcctgtctgtagaagcaggagcgtcaagacttgactgagagccatggtgctgaaatgagatagattccctgatggagagcacacgtggacccccacacctgagggctcactgctcctcaccacaga**tgcactcccctactgagtcctgagacctgagtgca**ccccatagagtagggctcagatg**aggggatgcaaatctccaccagctccaccctcccct**gggttcaaaagacgaggacagggcctcgctcagtgaatcctgctctccaccATGGACATACTTTGTTCCACGCTCCTGCTACTGACTGTCCCGTCCTGTGAGTGCTGTGGTCAGGTAGTACTTCAGAAGCAAAAAATCTATTCTCTCCTTTGTGGGCTTCATCTTCTTATGTCTTCTCCACAGGGGTCTTATCCCAGGTCACCTTGAGGGAGTCTGGTCCTGCGCTGGTGAAACCCACACAGACCCTCACACTGACCTGCACCTTCTCTGGGTTCTCACTCAGCACTAGTGGAATGTGTGTGAGCTGGATCCGTCAGCCC**CCAGGGAAGGCCCTGG**AGTGGCTTGCACTCATTGATTGGGATGATGATAAATACTACAGCACATCTCTGAAGACCAGGCTCACCATCTCC**AAGGACACCTCCAAAAACCAGGTGGTCCTT**ACAATGACCAACATGGACC**CTGTGGACACAGCCACGTATTACTGTGCACGGATACcacag**agacacagcccagggcgcctcctgtacaagaacccaggctgcttctcagtggtgctccctccccacctctgcagaacaggatagtgtggctgagatgccatttcctgccagggcctgcgtttcccatccccatctgactcagagccttgttttcctccctcttctttactaataaatggcatgtcccctgttagtggttcatgcaagcagaagctgtatcctgtttgacaaagattcagcatgaaaggtcctgctacctaa**aaaaaaatagacagatgagacttaattaacctaaataattttttt**cacaacaacagagtgaatacgcaatttacagaatgacagaaaacttttgcacactttgcctgtgacagggaactaatatgaagaatttgcaaggaactcaaacaactctacaacaacaacagcaacaagaaccaaataactccgtt**aaaatgagcaaaggacatgagtagacatttt**caaaagaacacatagaaatggataataaatatataaacaatactcaacatcactaaccatcagggaaatgcaaattaaaaccacaataagatatcatcttccaccagtcacaatgactattactaaaaactcaaataatatcagatgttgctgaggatgggaaatgaaggcaactcttagacattgttgatgaggatgtagatgagtacaacctctgtggaaaatggtatggagatttcccagaaaactagaaatagaactgtcatttggtccagcaatcccactactgggtaactacccaaagga**aaataaactattattt**caaaaagatacccaccttctatgcttaccataaaactactctcaatagcacatatgtcaaactgagtgtctgccaaccgatgattttataaaagaatatagcatgtatgcacaattcaatactagtcagccacaataaggaatgaaactgtgtcttttgcagcaagatgcatagaagtgggggacaatat

Uppercase: IGHV2-70

Lowercase: Flanking sequence[1000bp]

Red & Bold & Underline: Stem-loop [13]

Blue: Heptamer[41]

Green: Nonamer [4]

id-TRAV32[V_gene_segment]

attttctttgactatgagtttatctttcctgccagatccaaaatgactaataaaatgagagatacagtcagtct**agactcgaaccaggaagagtct**ctaaaatccaaatgagtaactctaatggtctctctgaagactctgcactgttgcttctttaaagataaacattaaaatagaaacccgcatctctaataaaatatttgtctttc**ccagactttcttgagagtgagtctgg**agtgaccattttgtggtaacagatcatcatttggggttattatttaagcaccagttccctcatatttgtgctgtcagtcacttcctcaaagcttcccattcagtatgttttaacttaagactcagaaggacacattgatgataagaagtttctcagtgggtctgaagcttagctccaacactgcttgtcttagaatagagactttatcttactaggacacc**aaagaagactgttactacatggcatgaatatctgcttcttcttt**atttaaaaaaaaaaagtgcagaaggcatcaatggataacattctctatgttttcatttgagtctagtagttttctgttcttactttgtactgtcaggctttttttccccttatatctaccttaaaccctataaagaactac**tacagatcagaaactttaaatgcaagatctgta**acttaatcaccagaaatataaaccagaaatatgactttctactaatttcctacttgtgaagtaagatgttctaagggataggcagacatgccagctctccttctaaagacagaatgtgtgttatacacgtggccttggcaattagaggtctgtacacaggatggcagcacttcattttcacatgcagagaaatggaatgttcaactctgttgggcagagcatagcttgtctcctattggcttatagccaggttaagaaagagcaagaatacactgagaatagaatatgttctccttcccagatagtatagatcattgatgattctaATGGCAAGAAGAATGGAAAAGTCCCTGGGAGCTTTATTCAAATTCAGCTGAAGCGTAAGTGCAGGGCTTCTGGAGGGGTCCATCACAAGTCTGGGGAAGGAGGGGGATTCATTTTAGTTCTCCCCTG**GTTTTCTGCTGATACTCTGCAAAGAAAAC**GTACATTC**GGAAAATGACCCAGTAACCCATTTTCC**TTGTTTCAACGGTGATGACTGGCCAAGAAAAGGATGTGATACAGAGTTATTCAAATCTAAATGTCTAGGAGAGAGAAATGGCCGTTATTAATGACAGTTATACAGATGGAGCTTTGAATTATTTCTGTTGGTACAAGAAGAAAACGGGGAAGGCCCTAATATCTTAATGGAGATTCATTCAAATGTGGATAGAA**AACAGGACAGAAGGCTCACTGTACTGTT**GAATAAAAATGCTAAACAT**GTCTCCCTGCATATTACAGCCACCCAACCAGGAGAC**TCATTCCTGTAC**TTCTGTGCAGTGAGAACACAgaa**ctcttcaggcacctgcaacctgtactcaaacctgcaactgggagtccagtca**cattctttgtctttgaacgggttttgggttagaatg**gtttaccataatgtgcttgtttctaaattgaaatgaatgcagaaatctaataattgtgattacaaaatcattacgttt**aaagtaactcaaaatgagctggtatacttt**gtactaactccaatgtattcgagaggagaagtagtgttgttttaaatagatatgtaaataataaagat**aacctcattggaatatctgaggttgtttcattgagatcctacacttccataataaaacaa**ggacacctaaaactcttctgggtggcaatctgctggccttcagaaaaaaaaagaagattcatttttgaaaaa**attttaaaatttttaaatcatcatattttaaaat**catcacacactcattatttattataagtattttcattccaggtgtgtctgcatatgccctaattcatagcaacttaacatatcaagtatatttatcatataacattt**atgtccattctcctctggacat**ccaccttcatagtacgggtattcagtcataaaagttacaattcaaatagccaccagaagaattcctgcaaagatttttgtcttgcaactgttctacattcagaaaatatttaccaactgcctactctgtgccagacattgttcttggtgttgaggattgagcaggatgaacagaaagacatgacaaaatttaattgtattctgcttctctcctccaatggatgtctgctcaccagaaattaactggggaaacattttgatgacagtgtatatccagggatcaatgacaccacctaggagttttaggattttgcccaaattcaccaattccttcatacctcataggtctgtatgtacagtagcaacagtgtgatctttctcttttttccaatg**ttccttcaagaaaggaa**gataatagttattttagtcaaactgttca

Uppercase: TRAV32

Lowercase: Flanking sequence[1000bp]

Red & Bold & Underline: Stem-loop [16]

Blue: Heptamer[23]

Green: Nonamer [4]

id-IGLV2-11[V_gene_segment]

tcagtgttgcagggttgagagaggtcagctttaggttggatgtgctcagatgaggaaaagcaagaaagcaggaaacacgcggct**ctgtgacctgacacataaatcacag**taaaatggaaataaatctttcaataaatgaaacattcaaaataaacgtcactctggtgcacacagaaaggcctattagcaaactttccctttgtcctgctggctcatatccccatggatcacctgtgtgcagagtggacactgtagtccccat**ttctcaggctgagaa**ctagaggctgggaaggacaagtcacctgcttgtgagtgggaggggcaggattgggagcagtggactcgggctcaggactgcggccctggtcggtgctctgtgttccctccacaggatccct**cccacatgtgcccgtgtggatgtggg**tgcactgcctggggtggctga**tgtgctccagggattcgatggtgcagatgcttcagcaca**aagcttgggggtggggatgggatgaggctgca**gctggggagaccagggaccaatgccacccagc**agaggtttccacggggtgggcagccccatttcctggcagtagagacagcctggaggtgggtgccaagcaga**gggcactgcagggtgtgccc**aggcctggagggtcacaaagacacaggcaccacacagcagagtcactgaggaccagaagctca**ctcaggtaagggatttcagcagaactttctctcctgag**caaatcaggtacaaaaaaattaagatcctgccatcagaaaagacaaacaagaactttttatctcctcagctgagccccactgtccagggaagcagaagtctctgagcccagcccaggtgagggtggggtgaggagaggagctc**agggtggagatttgcatggaggccccaccctcctctgaggcagagg**ggataagacagggctgggggcaggccc**agtgctggggtctcaggaggcagcact**ctcgggacgtctccaccATGGCCTGGGCTCTGCTCCTCCTCAGCCTCCTCACTCAGGGCACAGGTGACACCTCCAGGGAAAGGGTCACAGGGGTCTCTGGGCTGATCCTTGGTCTCCTGCTCCTCAGGCTCACCTGGGCCCAGCACTGACTCACTAGAGTGTGTTTCTCCCTCTTTCCAGGATCCTGGGCTCAGTCTGCCCTGACTCAGCCTCGCTCAGTGTCCGGGTCTCCTGGACAGTCAGTCACCATCTCCTGCACTGGAACCAGCA**GTGATGTTGGTGGTTATAACTATGTCTCCTGGTACCAACAGCAC**CCAGGCAAAGCCCCCAAACTCATGATTTATGATGTCAGTAAGCGGCCCTCAGGGGTCCCTGATCGCTTCTCTGGCTCCAAGTCTGGCAACACGGCCTCCCTGACCATCTCTGGGCTCCAGGCTGAGGATGAGGCTGATTATTACTGCTGCTCATATGCAGGCAGCTACACTTTCcacagtggtcca**agttcctggggaactgagaccaaaacccaccctgggctctcag**gctcccttttgctctgaagatgcttcctcaccctgtgcaaggggcttcctgcagcacggccttgagaattcttctctctcagctcctctcctttcccaccatgaaatccaacag**gaaacctgccctgtggtttc**tcatcccggacagggacagcttcctggtgcttgtgtgctgtg**gtccctgaatgtgcagctcttcccagctcttcaaatgcagggacagtgacaagcagctgcctgattggtgcagtcactg**cttttttttca**gagatgtcttcaccctacatgcatcaccatctc**ccacactgtgggtagaattttagccaaatatg**gctgggcgcagtggctcacacctgtaatcccagc**actttgggaggccaaggctggcagatcatgaggtcaggagatcgagaccatcctggctaatacggtgaaaccccatctctattaaaaatacaaaaaattaaccgggcacggtggcgggcgcctgtagtcccagctacttgggaggctgaggcaggagaatggcatgaacccgggaggcggagcttgcagggagccaagattcgtcactgcactccagcctgggtgacagagcaagactccatctcaaaaaaaaaaaaaaaaaaaagaattttagcaactacattctaatagttatcgccacaactttgatcttaaataacagtgcagtgaacatccctatgcagcctcctttgagttcctgtgtgaatatgaccataggatacatttctaaaactgaaattgcgggtcagaaagatgtgtgtttgtaattttcacccattgttgtcagattcccctccagaggggttctaccaactt**acaatgccaaccataagtatttgttcagagtacattgt**tgaaagtgtaa**atttttgccaacctaccagactgaaaaat**gttatcttgttacagttttattttt

Uppercase: IGLV2-11

Lowercase: Flanking sequence[1000bp]

Red & Bold & Underline: Stem-loop [20]

Blue: Heptamer[53]

Green: Nonamer [1]

id-IGHVIII-67-3-2[V_gene_segment]

cctgtaccccataaatatatacgtgtactatttacccacaaagattcaaaattaaagtgtttatgtgtctaatattctgtaactatattacattaaacctaagaacacactggcttttacacactactgtcatccacatagacattcatagccttc**tttctctgcttgtccataatcccacactccaaaatgagaaa**acctctcctaagatatgacattaatttacttagtgcacaatttcaaaatacacaaatcgtctttaggattgttaaactgacccttgttgaaaatatgtttatcaactagaatacagtgcttatgtggagcttatttatgttttacaatcgcaacatcaaattctttccaagtctcttgtgtccaatatttatgccccaccttcttcagtgtaattagttcaaaccttctgattgttgttagatatttttatacagctgtctttttttgtgtttcatgacctgctaaatactttcgaagttgtatgcatttaggttaacactttgtctcaaaaaatatgagttttcagaaattctttatgccatgtgtccatcattgaatatcataaaaatagtttcattggccg**ggcgtgttggctcacgcc**tgtaatcccagcactttgggaggccgaggtgggcggatcacgaggtcaggagatcgagaccatcctggctaacacaatgaaaccccatctctactaaaaatacaaaaattagccgggagtggtggtgggcacctgtagtcccaactactcgggaggctgaggcaggagaa**tggcgcgaacccgggaggcagagcttgctatgagcagagatcgcgcca**ctgcactccagcctgggtgacagagcaagactccctctcaaaaaaaaaaaatagtttcattgggattaaaaa**gtgtgtgtttcacctaatccacacac**cttctccccaccttcctgggaaacctcaaatgtttactggatctatatttttatctttggcaa**aaCAGCCTGGCTGTT**CTCTTAGCCTCTTTTGCAAAGCATCAAGATTCACCTTCACTGACTACAGCATAAATTGAGCCCAGATGGCTGGGGGACAGAGGCTGGAGTGGGTGGTAACAGTGATTGATTCAAGTGGAAGTTCTCAGTGATATTCTGCATCAGCATAATGAAGATTCACAATTCCCAGGGACACCAATTACCAGCACAGTCTCCCTTAAAATAATCTACTTGGAAGCTGAGGG**GGCTCTCACAGGGGTAGGCAGTGTATTACTGTGAGAGAc**acagcgagggaca**tttctgtgagtccagacagaaa**cctccctgcagggagacaagagaggactttgtgataaatggtgcttaggacaccagggggcactcaggacagcagagggtgctcagcagagcagggtgcgctcaggacatggtgtgtgttgtagggggcattgctcaggatcaccagggggctctcagaaccaccaagggtcactcaggaaaccagggggcgctctggacaccagggggagctcagaaccaccaggtggtactcaggacaccagggggcgctcatgaccagcagg**gcatgcttgggactctatgaggcatgc**aggagaagcaggccgtgctcagggcaacagggggctcacagaaccacctgggggtgctcagaaccactaggggtcactcaggacagcagcaggcattcgggacaccatggggccctcaggacctgcagggggcgctcagaacaccaggggaaggtcagaaagacctgggggcacttggaaccaccagggggcactcaggacagcagggggcgctcaggaccactagggggcgctcaggaccagcagggggtgctcaggacaccatggggcatgcaggagcag**caggctgttctcagggtaacagggggctctcagaacaacctg**g**gggtgctcagaaccaccagggggcaccc**aggacatcatgcggccctcagaacctgcagggggcgctcaggacaccaggggaagctcagaatg**acctgggggcactcagaaccaccaggt**ggcactcaggacagcagggggcgctcagtaccacggggtgcactcaggacagcagggggcgctcaagacacagtgggatgctcaggaccagcaagaggcactcaggaccaccagggggcactcgggggagcaggaggcactcaggacaccagaagaggatcagaaccactcgggggcactcagaaccaccagggggcgctcaggacactgggggcattcagaaccacc

Uppercase: IGHVIII-67-3-2

Lowercase: Flanking sequence[1000bp]

Red & Bold & Underline: Stem-loop [11]

Blue: Heptamer[32]

Green: Nonamer [4]

id-IGKV1-5-3[V_gene_segment]

atgtccagtttcatctggatccaactgatttctccatgtacatagacaattgcttgataagagattgagtatgtttttcctaaaggtgttaacagggaggctggtgtctgggtcaggatgatgtccccatgcactgataaaaagtataagaagaaagtgtcattgatggtgcatggcagggacatgctccgtgcagtggccaccctcactaagacagatgaactttgggaaataatacccaatggcagaaaagaaggtagactatgaaggtacccaaaacaagaataaggtgcacctcattta**gtctctgggtattaaagagac**ctgcagttcttgatagtggtggatctgtgagtgctgcatgcatggagacaacacggtatcatctttgtatatctgtaataaattgcttgatctaatactagtaagaacaaaggcataacaccattacctaatacttacaaatatatagcatcatgccgatacattttatttttaattttttttagaaaggaacaatgttaaactcacagaaatgttgcaggtatagcacaattacccccttccctacccggaatcttatgag**agtcttttgaagact**tgagaatcctaccatctaacattttactatgtgtttcctacaaacaagaatattctcctaaataatcctgatacaccaatgaaatacattactctatcggctcctgaggaatatttaaaattctcaaaaaaatacctaaa**aattgtttctcataataaaatagtccccagtagaaacacatt**ctctgcagacaaatttgtgctaccctggtcttacctgggacacctggggacact**gagctggtgctgagttactgagatgagccagctc**tgcagctgtgcccagcctgccccatcccctgctcatttgcatgttcccagagcacaacctcctgccctgaagccttattaataggctggtcacactttgtgcaggagtcagacccagtcaggacacagcATGGACATGAGGGTCCCCGCTCAGCT**CCTGGGGCTCCTGCTGCTCTGGCTCCCAGG**TAAGGAAGGAGAACACTAGGAATTTACTCAGCCCAGTGTGCTCAGTACTGCCTGGTTATTCAGGGAAGTCTTCCTATAATATGATCAATAGTATGAATATTTGTGTTTCTATTTCCAATCTCAGGTGCCAAATGTGACATCCAGATGACCCA**GTCTCCTTCCACCCTGTCTGCATCTGTAGGAGAC**AGAGTCACCATCACTTGCCGGGCCAGTCAGAGTATTAGTAGCTGGTTGG**CCTGGTATCAGCAGAAACCAGG**GAAAGCCCCTAAGCTCCTGATCTATAAGGCGTCTAGTTTAGAAAGTGGG**GTCCCATCAAGGTTCAGCGGCAGTGGATCTGGGAC**AGAATTCACTCTCAC**CATCAGCAGCCTGCAGCCTGATG**ATTTTGCAACTTATTACTGCCAACAGTATAATAGTTATTCTcccacagtgttacacacccgaacataaacccccagggaagcagatgtgtgaggcagggctgccc**cagctgctcctcctgatgcctctatcagctgagagtggtcctcagatgcagccacactct**gatggtgttggtagagggggacatggagtcacctctgcaccctaattcttttctctttctcagccccaactgcacagatctagcaatgcctctcctgatttaataaagacagagatcatgacac**ctgaagagtctagtttatggcttcag**ctggactttatataacagagaagaggccactatagatattcaaagcagg**aattgtcttaatacagacaatt**agagtctaaactactgaagtctaaata**aaatgtagagatgaatctctaaattt**aatgttttacgtgcaaagaaatatttgccaaatggggcatacaggaaaactcagtggtcttcaatatgttggaagaacaaagagaaggttagagttttacgaaaaagggaacatgttacctatggctctttgagaaagttcatgggcactaggaagggttgggagctggcaagctcagactgggaagcagtggtggacaaagtgaatcctacagttatatcaagttatctcagaagttgtggataaatttgatttcaggttacaataagccaaaggcagtgaagcttgcagagaattttgtt**actgaaatgccagggattcagt**gtagatcctgctgctcaccacacagaaaaacaatcactaagacaagtattgccaaggaacaggctttaatcaggtgctgcagccga**ggagatgggacaccattctcaaatgtatctccctgaactactaaaatgaggggtatatatatcaggga**agaaatgtgggaaaacaggaattagggaggggtaaggaagatgatttggtcaacaggaagcaggaggtcagttaggcaatcacaatgggtgaagagtctgatgtctcactgtcccgattcagtg

Uppercase: IGKV1-5-3

Lowercase: Flanking sequence[1000bp]

Red & Bold & Underline: Stem-loop [17]

Blue: Heptamer[24]

Green: Nonamer [4]

id-IGKV1D-27[V_gene_segment]

atcaagtgccctaaacccagtcattaagaactgttt**tctcttaggagctggaagaga**ttggtgatttggaaatgtgcaggtataagaaacagagtagtcacagggatagagggtgacaacttggtttagaggacacctcagcttctgaaggggaatggcttggataagagaaataaaaggcataaataaaattcagggagcacagggaaata**tctagcatgagactgtaggatggcatacagagctaga**atatagctgagaactttcagaagtaaagggagaaaatttatcatgttggctggcccagctgaaagaggtaggaaagaacattcagatatggaggataacaattatgtgtctggagatgggagattagctatgccagatagccagtggcaggacccttccttgctgtggcattattttcaagtattagggtttttttaagttttttatttcttttttaatgagcaaatctatctatctatctatctatctatctatctatctatctatctatctatcattcatgattataccttaatgcatccattgttggtagcactgacaatttacagcactggtgtttccagggaattggaccaaaaaggaagtctctctgaccttaatagtactcatctgtatcaaatgcag**gaaacttctaaaatttcttgagtttc**tagagatgtttttccctagcagaccttatcataaatagaaagctagcaagagaagcatgtcatgaaacatgaagagagcaaaagaacactccacacataggacagtaggctgattctttcctgtagc**ctgcagggagaaacacatgctctgcag**actttggacacctgggaggcactgggcctgtgcagtgttattgagataagtcatctttgcagctgtgcagatttgcatgtcccacagagcaacgcctactgccctgaacatttatcaataggctggtgacatcctgtgcagaagtctctctcagtccggacacagcATGGACATGAGGGCCCCCGCTCAGCT**CCTGGGACTCCTGCTGCTCTGGCTCCCAGG**TAAGGAGGGAAACAACAAAAATTTTATTCAGCCAGTGTAGCCATTAATGCCTGGCACTTCAGGAAATTCTTCTTA**GAACATTACTAATCATGTGGATATGTGTTTTTATGTTC**CTAATATCAGATACCAGATGTGACATCCAGATGACCCAGTCTCCATCCTCC**CTGTCTGCATCTGTAGGAGACAG**AGTCACCATCACTTGCCGGGCGAGTCAGGGCATTAGCAATTATTTAG**CCTGGTATCAGCAGAAACCAGG**GAAAGTTCCTAAGCT**CCTGATCTATGCTGCATCCGCTTTGCAATCAGG**GG**GTCCCATCTCGGTTCAGTGGCAGTGGATCTGGGAC**AGATTTCACTCTCACCATCAGCAGCCTGCAGCCTGAAGATGTTGCAACTTATTACTGTCAAAA**GTATAACAGTGCCCCTcccactgtgatac**aagcctgaacataaaccatggagggaagtagatgtgtgagtctgggctgccccagctgctcctcctggtgccgccgtctgctgacagcagttctcagatgcagccaaggtttgaagctcactgggaagttttggtagaaggggtcagggaggcacatttacattctatctctctttatcc**tcagctccatcagctga**tatgcaagtatctctcctgattattattaataaaggacaaagaaaattaaacctaggaggtctagattgcagcaaaagtcagacttatacagaaaaggagaaggtactctacatatttttaaagaatttttttttgatacaggggattagagtctaaattatgacctttcaaagatcagacataagt**acatacacatataagtatgt**tcagccaacaatatatcagataaacgtattgcacctatatgtatactgtaaacatttatgctccgatcccaccttcttccaacactacccagatcaagatccacactcttcccagcatctcaaatatctccccaggttacttccctgtcaattccgtgttccatctatcatcatcctaggcttccgctgtattttatttcaatatttattaatttttcctatacttaagctttatataatttcaatcatacagtatttactctttgacgttttgcttattttga**tcaaaatatttttga**gtttcatctgtgttgtttctctggcagtaat**ttgtgcctttttattgaggcacaa**ttttcc**atgaatatgccaaaatatattcat**ttttcaactaatgattt**gtttttggtttaagaatattactggtaaaaac**cttaagaacattcttgtgcttgtcttttgtgaacttatgcactcatttctcttgggaatattcctaagaatggaagtgttgagttataaagttagcatagattttgtcttagtaaacattgacaatagtttactgaaatactgaacatat

Uppercase: IGKV1D-27

Lowercase: Flanking sequence[1000bp]

Red & Bold & Underline: Stem-loop [17]

Blue: Heptamer[21]

Green: Nonamer [5]

id-IGHV5-10-1[V_gene_segment]

tgatgctgacagtca**ttgcatagcatgcaa**attacctaggtcatagcctgtgttgtgtgtttatgttgatggtaatggtcattgcttagcatgcaccttacccaggtcatagcctgtgctgtgataaagagaagtcattttgcttctgcaccccaggtatacaattcctgtgggtaatctggcaaatatttt**agaaactgtgtagtttct**gttgccttggcttt**tcatttggatcttttagatgttaaggaaatga**gagtggttcacaggaaaatggggcaaatacaggggacagtaaaaagccccctccgagcaatgtggatatcataaaataaaaaaaataagaataaaagaactaaacagatt**gtgctggggtggaattaaaacagcac**taccaggagttgggtgcagcaatggggatcccttcaggtccatcagcaatacagag**acctaaacaaatatttcctatttaggt**tgaattggcagagtgtaaaatctagaagatgaaatggccaggaaacatttgtccttgattttgcatgggtgaaggtctggcaggttaatcaaattgagggttgttaaa**ggcctggggtggtcctgccctttctgggtgcccaggcc**atagtcacattcatgggaaattgccatggaataaagagtagttttttgtggtagttcttgttatcaggcataagtgcatttgaattttctgttcacggcctttcctggctctatttctcatttttttaacagacatgattttatctagatttgtcaactttcacagggtcacagagaagggtggaaggagggatgccctggtttgggtcttgcaggaccacagacaagagtg**gggaggggcaagaagaggttgtaaggaattattgagacctattctgcccctccc**aggagg**ctcagttcagccttttctctgcaactgag**gttctgggttataaaccctgtagactcttcccttcaggtcagggcggcaactatgcaaATGC**AAGTGGGGGCCTCTCCACTT**AAACCCAGGGCTCCCCTCCACAGTGAGTCTCCTTCACCACCCAGCTGGGATCTCAGGGCTTCCTTTTCTGTCCTCCTCC**AGGATGGGGTCAACCGCCATCCT**CGCCCTCCTCCTGGCTGTTCTCCAAGGTCAGTCCTGCCGAGGGCTTGAGGTCACAGAGGAGAACGGGTGGAAAGCA**ACTCCTGATTCAAATTTTGTGTCTTCCATACAGGAGT**CTGTGCCGAAGTGCAGCTGGTGCAGTCCGGAGCAGAGGTGAAAAAGCCCGGGGAGTCTCTGAGGATCTCCTGTAAGGGTTCTGGATACAGCTTTA**CCAGCTACTGGATCAGCTGG**GTGCGCCAGATGCCCGGGAAAGGCCTGGAGTGGATGGGGAGGATTGATCCTAGTGACTCTTATACCAACTACAGCCCGTCCTTCCAAGGCCACGTCACCATCTCAGCTGACAAGTCCATCAG**CACTGCCTACCTGCAGTG**GAGCAGCCTGAAGGCCTCGGACACCGCCATGTATT**ACTGTGCGAGACAcacagt**gagagaaaccagcactgagcccgtctaaaaccctccacagcacaggtgcagaggaagctgccagagacacactccccaggggcctctctattcatccagggaggaaacactggctgtgtcctgaggagtgagaaccagagaacaacgtgggagggttcccagtccctaaggcaactagatggggcacctgctcctgggaggtggattctctgagggggctcttgttctacaaggttgttcatggtgtatattacatggttaaaataa**aaaagctgtctaatatgcacctcttaagcatgacagtctttt**aattagtgaaaagtctacacagttaatcattgctggcatgctttcctccctcctatcttctcttgctccctacatctattattttctacttaatttttcaaaatcatttgatccctttgtgaactattactaggttctgttgtttttgttgttgttttcctttcaataatatgtactgaataattcatctttgtgccaattcataagttcttgggtataataaatacttctctcataaaaattagataaa**ttaaaataaataaattttaa**aaaacatacaacctatcaaaactgaaccattaagaaataaaaaatctgggctggttgtggtgggtcatgcctgtaatcccagcagttttggaggctgaagcaggtagatcaggagttctatgtgaatctgtggtcaggagttcaagatcagcctggccgacatggtgacaccaagtctctactaaaaatataaaaatta**gctgggtgtggttgtgttcacctgtagtcccagc**cactcgggatgctgaggcaggagaatggcttgaacctgagaggcagaggt**tgcagtaagctgaaattgagccactgca**ctccagactgggaaacagagtgagactccatctcaacaacaacaacaaaaaaatctgaacagaccaatgagtaaagagattgagtcagtgatttttcaaacatc

Uppercase: IGHV5-10-1

Lowercase: Flanking sequence[1000bp]

Red & Bold & Underline: Stem-loop [18]

Blue: Heptamer[44]

Green: Nonamer [6]

id-TRBV26OR9-2[V_gene_segment]

aaaggaaaaccaaaaccaagccaaattgagtactctaacagttacagcaccttactattgcactt**attctttcagaccctaacccaaagccaggactgaaataat**ctaaccaggacctccctagggtaggattcaaacccatgaagctacacaaaaataggactctaatctacactcctagaccaaagataatttaaggtgtaagcacatattaccaaggtactgactccaaaagacatcttatccagaagcaatttctttcctcaaaagagaaagtagcactcaaagggcctgaaatgtcatgacagagaataaggacctcacagctaagcctctggataaactggtctaggtagcaccactcaggg**tcagtgaagaaactttagttcccactga**ggggctacagtgccttggcaggcattggcctcacattgggcacctatatttgttatcaagcaaatgggtttgcacacagaagccaataactatggcaccagcttttgagaaaagaaaaggctttgttgaaaaaccagtcagcaaaaagaaaggagtgagct**caaatctgtttccctgatttg**gggtctagggcaggttttaagggatcagagagcaagggaaaggatttaggaatgttgatttggtagggtctgattggagggcttcaaatatgactatctatggtaagatatgttgaggcacatttta**gcccaggagcttctgggc**caatggacccctcacta**ctgaaagaggtccggctttcag**gttccagtcatgtcttggtcttcttgtttctaaggggaggaatctttggttctgggtattgttagaagtcaaagcttttactgttgcacatgccttg**gctacatgatttgcagttatgtagc**tgttggcagtgtgagagacaagcgttgtttctctggccaggaagtgaatgaaaagtgctggtatcagagactaaattaatca**cagcaccaagcaacatcctgcttttggtgctg**ccATGAGCAACAGGCTTCTCTGCTGTGTGATCATTTGTCTCCTAAGAGCAGGTGAGTCCTGGGCACAGGTGAGGAATCCCTATTACTGAATTCACAGGATCTAACTATAAAATGTTTCTTCATAAAAGTCACATTAGTCTCCTCTTTCCCAGGCTCTGTCTTTATTTGATCACCCATTTTTCCCAACAGGCCTCAAGGATGCTGTAGTTACACAAT**TCCCAAGACACAGAATCATTGGGACAGGAAAGGAATTCATTCTACTGTGTCCC**CAGAATATGAATCATGTTGCAATGTACTGGTATCGACAGAACCCAGGACTTGGACTGAAGCTAGTCTATTATTCACCTGGCACTGGGAGCATTGAAAAAGGAGATGTCTCTGAGAGGTATCATGTTTCTTGAAATACTATAGCATCTTTTCTCCTGACCCTGAAGTCTGCTAGCACCAACCAGACATGTGTGTATCTCTGCGCCAGCAGTTCATCcacagcactgcatagctgccacatcctctccacataaaaaaaggtgcataccaaa**gaggaaaagcctgccctcaaaattcctc**accgcaaataagagaagttacctcacaggtattgacaaaaagagccaaactctgtaaaatatttgaagagatttattc**tgagccaaacatgagtgagcatggccagaggcagaggctca**agggcctaagaacgtgtgtccaaggtggtcagcctacagcttggttttatacctttttggagacatcagacatcaatcaatacatgtaagctgtacattgtttcaatttggaaaggtgggacaactcaaagagggggcttccgggtcataggtggattcaaagatttttcagactggcagttggttgaaagagttaagttatcgtctaaagatttgaaatcaatataaagaagtatctgggttgtggggaccaaggttcttattatgcaga**tgaagcctccagggagcaggcttca**gagagaatagattgtatttcttaatcaaacttttaaaaggtgccagactc**ataattctcttctggatcaggggaaaaacctgaaaaggaaagagaattatctacagaatgtagat**ttccctcatgatagacagctttgtagggccgtttaaaaatatgtcggaaaatacattttggggtaaaatattgatttttttcagggcctgctaggagtcatgttggtgtcttattgctacaaagggtctgttttgtcactcataaggtttccattgtaatgttaatgctggttggttgtgcctgaattctaaagtgaagagggtataatgaggcatgtctgaccccacttcccactaagaactgaactggtttttcatgttaactttggaatgctcttggccaagagaaggggtccacacagttggttgtggggcttaggattttatttttggttcacacaaaagtataccaatgtcctcccctaggtttgatatgccctgcttccc

Uppercase: TRBV26OR9-2

Lowercase: Flanking sequence[1000bp]

Red & Bold & Underline: Stem-loop [14]

Blue: Heptamer[22]

Green: Nonamer [5]

id-TRBV23OR9-2[V_gene_segment]

ttttaagtggtcattttgttggtatgttatcaagtt**attgtctatatattcaaagacaat**cttgtaatgtggaggaaacaacctcactgaacaattaaataagtaaggagaaatctatggaaagatatacattaaactgttaacattacttacctgaaatgaggatgagatggaggaagaaaaaagtaaaaaaataaataaatcaccaaaaatgc**actatttaaaataaatagt**tacagaaagaaaagcaaaaaagggaggttatctaaaagaaacactagtaaatgaacaagct**aaaatacaccaaaaagagctaaaaggtgaagtatttt**tcaaagcagtggctagtttcagaatcataaaaatagcaattacaatagag**atctatattatactaaaagataattagat**aattaatagccagaagaaatgtctagtcgtgagttagaaactgatgctcaaattaatggcaaaggacatttttgaaacaactgtaaaactctgcataaaaactaaatagcagataagatgaaactgcattgatgagaaaggatgatgatcattaactgaaaaaaaaagcttatcaaacagaatttcaagaatattgtttataataaaaaat**aatgtgtctatatatctacacatt**tatatatggggagagaaggatctagaaagactctcactaatgtgttaacagtgataattgctttctggcagtggtatatttttattt**tgttattttattttctaactttctaaaatatca**gtgc**tttgcttatattagcaaaaatagacatacaatatttt**aaatgaatacaaaataacttagttatatttactgagttaggaataaaagctttttttcccgataccatgaagtgtttttgctcagaaacatgaagcagtaacaagatgattatgaatctgtggagcaatcacgtcacaagaaggtgcagacg**gggatgaaaaagcctcatccc**tttgcgacgtcaatgcgaccACGGGCACCAGGCTTCTCGGCTGGGTAGCTCTTTGTCTCCTGGCAGCAGGTGAGCTCTGAGCACAGAATAAAAACCCTTGTCCTGGGCTGGCCAACCCCCAGATCCAAGCTTTGCTGTTGTTGTTAAGGTCTTTCCTGGACTCTGTCTACAGCATCTGTTTCTTTCTCTTACAGGCTGTTTTCATGCCAAAGTCACACAGACTCCAGGATATTTGGTCAAAGGAAAAGGAAGGAAAACAAAGATGTATTGTACCCCCAAAAACGGACATACTTTTGTTTGTTGGTATCAGCAGAATCAGAATAAAGAGTTTATGTTTTTGATTTCCTTTCAGAATGAACAAGTTCTTCAAGAAATG**GAGATGCACAAGAAGCGATTCTCATCTC**AATGCCCCAAGAACCCACCCTGCAGCCTGGCAATCCTGTCCTCGGAACCGGGAGACACCGCACTGTATCTCTGTGCCACCAGTCCGTCcacagcactgaaatgtcagttcctgttagcacacaaacttgccacagacccagctcaggaagcaggtgatgtattaggttggaagggagtaacagaaaataactggagccagcttaagccacagtggaatttaacacaaggataaagggtgtttagcagaagtcaagaggtgatggacagacagttctgaaaaggacagacagttctgaaaaggcaccactagggccaggctctagaatgactatctctccagctctgatttttattcagttttgcagctgcatgcttttctctccctgtacaccatccctttttctccaggcctcgtggtgagaatgagatggtcactaaagattacaaattcagttgccataatttccctacagtgacaaataccaccccagctctttctaagcctgagttcaaaatcccaggagaaaaaaacagatccgtcatgatccagccatgaaccttctcatagagcactcagcagtaggcatgtatgtatgtatttaaaatcacataaacgttacttttccactcagctcaagcaagggaaatgctctgtttttcagtatgtacattcggtctgttggttct**gtatgatagagagtactcatac**ccatttcttcagaaatacatacccag**attttctctggtttccggccagaattgaaaat**tattgtacaaataccttcatagacttgtatgattgctttataggattatataccaagaacaagttgtgattgctgagacatgtaattctatgcatattttatctatctaaatgccaccacaggttcttcaggatcaccgtatacatctgcaatcccacataaaagtgaatagggtgttttcctacatcctaatacctcctttaatccttatccagtgatct**agattgttatcagtatagcgatgtaaagaggttccaatct**ccctctctctccatcccctcagccctgctcacccaacaccagaatatcatcagtgttta

Uppercase: TRBV23OR9-2

Lowercase: Flanking sequence[1000bp]

Red & Bold & Underline: Stem-loop [13]

Blue: Heptamer[18]

Green: Nonamer [5]

id-IGHVIII-67-2-2[V_gene_segment]

gttgtcatgttcctttattctcttctggcaaatcacagtttgtgtttctgaagttttctagtttaatattttatagataaaaatt**tatcaagtagattgata**gtcctatttaggacacttatatctttacttatttg**aacactgcttgtcaatcacaatgagtgtt**gatttctccgtcttctatacatgttattatattttctg**ttccaaatccattcattaagtatttttaaatgtattttggaa**agttgatcc**tattactttatgtaata**tttctcttgagtcccaaaggtcttctttttatgattattttgt**cttaaattaacatagctatttaag**ttttcttttggtttatattttcatgatatatatttttccatctttattttttct**atttctgaatatgtaaagaaat**tttcttggagataaata**atagttgggtgtttttttttagtcaactat**g**aagaattattttttaatgatgttactcttttgtccaattattctt**gttttaatttgtattttt**acaataaagtttattgt**ttctattaacttattatttaatcattctgtgactaacatt**ttatgggttaccataa**ggataacaataaaaagtttctataa**ttgaattctaattcaa**aaatttaattcccattcatatatactacag**agatattataactctatatatct**aattcctctttcctatcccatgttttttattttctgtttgtatctatacatgctat**aaaatgtaatattggcatttt**tataattgctttccacacttacataatgaaca**aaatacaaatgcaaaaactgtattt**caacctcatttttccattgttgaccatcttctttgtgcatattttcaggtgttggatctatgtcacatggctattcatctagagactctgctaaaatgtctacggaaagattcatgtgctggaaataggttttctcaggttttattggtccgacagaatatttatttgtctttGATTT**ATTGTCTCCAGAGACAAT**GTCAAGAATATGCTATATCTGCAAATGGGCGATCTGTAAACCAAGAACACATCAGTATATCACTGTGCAAGAGGAGcacatgagggaaagccagtgtgtgcccagacataaacctccagggacacatgggcgaattagctgcagggggagcttgggacccactgatgagagtcgaccccagagggaggtgcacagaaaggttaagaactgctttcctgccagtatatttttaaaaaaacaaaacttccatctgacagtttctccagagaacttccctaattttagcattctgtgcctaccaatgccatcactaaataggtttttaaaaaatattgta**atatgaggacatattctcatat**ggacaaaacgcaaattgacatttacagaaatgaaaagtcctcaaccgtggtcgctagaattggagacctgtagcagctcagtgggacctgttgagtcttctccaatcacactcagaacagagaccttagtgcatctccctgactagaatacc**ctttaggtatcgagataacagcctaaag**atggtaaatcatgagagtctcagatgtccagatgacagacatgggtctatgggaatag**tactgtaactgacagta**actaattttccataccgccagatgtcctattcaggagcagctacagcagtcatgcctag**gtgtgaagatcacac**actgacctcacccatgctgtctctggccacttcatcacaaccaatgcttaatattggacgtggatctgccagtccccggggaatgggttgaatggataaaatgcatggctag**tgctgctggaaacccattcctactgtggcaaatggcagca**tctcttttaaaaggctaaataatattctattctgtatacataccacattgccattatcctttttgtttgtttgtttgtttgtttgtttgagatggagattcactcactctgtcacccaggctggagtgcagtggcgtgatttcaggtcactgtaatccctgccttctgggctcaaacaattctcctgcttcagcctccggagtatctgggattacaggcatgcaccaccactcccg

Uppercase: IGHVIII-67-2-2

Lowercase: Flanking sequence[1000bp]

Red & Bold & Underline: Stem-loop [20]

Blue: Heptamer[18]

Green: Nonamer [3]

id-TRBV10-3-2[V_gene_segment]

aatccaggggattgctagtgcaagtcccggaatccaaaggtaggagaatctagaattctgatatccaagggcagtagaagacacatgtctcatctctgagagagaatgcaaattcaccctttctgtgcctttttgtttc**atccaggcccccagctgactggat**attgccatacacattgagagtagatctttcccacacagttcatcaatttacaagccagtctcctccagaaataccctcacaga**cataccaagaagtgttgctgtaccagctatccaggtatg**cc**ttaatctagtcagattaa**cacctaaaattaaccatcaaactaccagcgggaacacacagaggtctgagctctgatgaggaggctcctgtgaggtgaactaatgtgacagagcataaaacaatatacaaattgtattgat**cctgaccttaagactttttgaaaccttcccagatgtcagg**tggctcttttaacagaggccagcactgtaggctgacatccagttatccagagtttgagaaaagataaaatctagtaaaaggtacttagaggcaaatttgaggagatgagttataaaggacaccagactgccacgagaatgtcccttagactacataaatattagaaacaattaaaacagc**atacaaatgaggaattgtat**cctaatgggaataaaatatgcagagaggaatgcattttctttctaataacctgcaaagagcatatgcacttatgtctttaaaaatagtctacaaattattcccattctgagctgttaaattgtatatatgtcc**tataaaatattgttttatataccataaattacatatatgtatata**tttgatttatttttttggagaatgtaaggggtgtggcaggttaactttccagctcaactatttgtcaggggcactgatgtcatcgagtcactgagaacctaagttctatttcc**ccaggcagggctgggagagatgagatcctggcctgg**acctgaaATGGGCACAAGGTTGTTCTTCTATGTGGCCCTTTGTCT**CCTGTGGACAGGTGAGGGCTGGTCACAGG**TGGGCTTCCTTCCCTAGAAT**TCCCAAGGCCTCAATACAAGTCTTTTTCTTGGGA**TT**ACAACATCAGGGTCTGTTGT**TTTCTATTACAGGACACATGGATGCTGGAATCACCCAGAGCCCAAGACACAAGGTCACAGAGACAGGAACACCAGTGACTCTGAGATGTCATCAGACTGAGAACCACCGCTATATGTACTGGTATCGACAAGACCCGGGGCATGGGCTGAGGCTGATCCATTACTCATATGGTGTTAAAGATACTGACAAAGGAGAAGTCTCAGATGGCTATAGTGTCTCTAGATCAAAGACAGAGGATTTCCTCCTCACTCTGGAGTCCGCTACCAGCTCCCAGACATCTGTGTACTTCTGTGCCATCAGTGAGTCcacagtgctgcatggctgcctcctctctgcacgtaaacagcagttagaaagactgaggttgctctgtgtctatccccacccttggaag**tccaggcctccatagaagtcagagggccctggccagcctgga**agccatagagcaggggccttatgaccctcagtgctgacgtccattcctaccccagtctcagaccaactggaggtcaccccaacacacttagcttgccaagtctctc**ttctgcagctctcttttgctgcttgcagaa**aggaaaaggcatcattagttgaggttagcgatgattctcttaaccccaaagcctggactcccttctctcccctgtgggcttcagtgacttcttcatctgcccctttccccacgctccactatccttccactcatagtcatgaaccttcactcctgacctctgttcctggctgcctttcttctatgggccaatagcctcatggggccctctatcatttgctcttcacctacatctccatcatcattttgcacaagtctcccctggctctctctgctccagccatgctcctttatgttcagttatttcctcatatgtgttatgctgtttcccaccctggggacgtgtgtttttacaaattatttcttctacactgaacatatgttcctccctcttattcctaagtctctacttcatttcttacatgaaagtatatttcaggtacatttactcttaactataaacagaaaaacttaaat**ttcactaaaagaaattaaggaaaatatttttatcctttgagtgaa**gacttttttagtggaaacacaaaagaaca**catgccataaaaaataatactggatgggcatg**gtggctcacacctttaattccagcactttgggaggcctaggcgggtggatcacttga**gcccaggagttcaagaccagcctgggc**aacatggcaaaacctcgtctctactaaaaacacataaattagccgggggtagtggcgtttgcctgtaa

Uppercase: TRBV10-3-2

Lowercase: Flanking sequence[1000bp]

Red & Bold & Underline: Stem-loop [16]

Blue: Heptamer[33]

Green: Nonamer [3]

id-TRBV11-2-2[V_gene_segment]

ggtggaatatctctgagtgcacctcactgaacacatatatcctcattcccacctccctggacgtgcatctccttgtgttattgtcttatgccaactctgaggagcgccatcccctaactc**cttttatttcttattactcaggcaaggaataaaag**tgcgtttgtga**ctaatgttgccccattag**aaacaaaaaagagataatgaccactactgttcaactgtaatttacactagaggtatagctataaggttatttttagatttcagcaagtctctttgattttgcatttcttacaatatatactgatattttcagggataagagaattccccacctatcttcgataaaaatgagaacttactttcattgcactggtgacccaaaactacaggcagaac**catgatgaaggcttggtacacaatgtatcatg**aatgat**aacagtaaagaggaaactgttcatcggccttcactaaaaatgaac**ccttccaactctgtgcccaggttattgttttatccct**ttagcttacagcgcgtcagctaa**ttcaacaagcttctaggataaacttgtgtcaattaccccagcttctaagtgcataggacaaggaagatagagaacctagtaaaagagaaacacccaaagagaaggataattgctgataagacaaagaggtcactaactgatatctc**tgtttcgagttgccttcaactcgaaaca**tccagcagaggatgggcctagagatggagtaggagatccagtccccaagccctagagatgtgagtgaggacacttggctgaacttacacagtctgtgaatacggtaaagcaacttcgtgctctcttttagcttataataattggtttctaacaatgtaggcatttgtggaggcaatgatgtcactgtgggaactgccatgagaggacagggatgtccctcctc**ctttgctgttgctcacagtgaccctgattgggcaaag**ctcccatccttccctgaccctgccATGGGCACCAGGCTCCTCTGCTAGGCGGCCCTCTGTCTCCTGGGAGCAGGTGAGTCCTAGGAACACCATGATCACATTGGATCTCTCTGATTATTTCAATCATTTCCTTCTGTTTTCAAA**TTCTGTCTTTTTCCTTCTCAGAACTCACAGAA**GCTGGAGTTGCCCAGTCTCCCGGATATAAGATTATAGAGAAAAGGCAGAGTGTGGCTTTTTGGTGCAATCCTATATCTGGCCATGCTACCCTTTACTGGTACCAGCAGATCCTGGGACAGGGCCCAAAGC**TTCTGATTCAGTTTCAGAA**TAACGGTGTAGTGGATGATTCACAGTTGCCTAAGGATCGATTTTCTGCAGAGAGGCTCAAAGGAGTAGACTCCACT**CTCAAGATCCAGCCTGCAAAGCTTGAG**GACTCGGCCGTGTA**TCTCTGTGCCAGCAGCTTAGAcacagtgtagcagaga**cacttccctcctgtgcagaaaaccagaaaaccgcaggactctctcctctctactcagctcacagcagcct**ttccttattcctcatcctcccaaggaa**gaagtgagttttcagatatagctaggactcatatagtgggaggaaataaactttttcttttcttttctttttttctttttgtttttgag**atggagtcttgctccat**tgcccaggctg**gagtgcagtggtgcgatctgggctcactgcagtc**tctgcctcctgagttcaagtgattgtcctgcctcagcctccccag**tagctgggatgacaggcatgtgccaccatgcccagcta**atttctgtatttttagtagagatgggattttgatatgttggccaggctgatttcgaactcctgacctcaagtgattcgcccttctcggcatctcaaagcgctggaattacaagcttgagccactgtgcctggctggagataaactatttcttaaaacatgaaggctgcagttgttatttgaaaatatgtctaaggaatctgaaacacctatcggtgagaattcaagaaaccaaaactgaaagtgacttatcacagacttagtcttctggggtactaatagttgttc**ttgtatttaaaaaatgcatacaatattttatatcaattaaaaaata**cgtaaatttgaaaaataaaaacatgcagtcaataaacatgtgtgataaactttaataatacagataa**tgattgtatgtgagtaacaatca**gcaggttttctctctgcagttgtcagcatatgaataatttgaatcctatccttggtgacaccctggctctggagcctcctctagacctctcattggatgtattatgcaaccaaatctc**aacaggtactatgtgtttggaaatctttttcccctgtt**tttgcggaaatatgctaataagagtaatgttaaagatgatgatgttgataatttcaactatttcattcagcaccc

Uppercase: TRBV11-2-2

Lowercase: Flanking sequence[1000bp]

Red & Bold & Underline: Stem-loop [20]

Blue: Heptamer[37]

Green: Nonamer [4]

id-IGKV1-27[V_gene_segment]

atcaagtgccctaaacccagtcactaagaactgttt**tctcttaggagctggaagaga**ttggtgatttggaaatatgcaggtataagaaacagagtagtcacagggatagagggtgacaacttggtttagaggacacctcagcttctgaaggggaatggcttggataagagaaataaaaggcataaataaaattcagggagcacagggaaata**tctagcatgagactgtaggatggcatacagagctaga**atatagctgagaactttcagaagtaaagggagaaaatttatcatgttggctggcccagctgaaagaggtaggaaagaacattcagatatggaggataacaattatgtgtctggagatgggagattagctatgccagatagccagtggcaggacccttccttgctgtggcactattttcaagtattaggttttttttaagttttttatttcttttttaatgagcaaatctatctatctatctatctatctatctatctatctatctatctatctatcattcatgattataccttaatgcatccattgttggtagcactgacaatttacagcactggtggttccagggaattggaccaaaaaggaagtctctctgaccttaatagtactcatctgtatcaaatgcag**gaaacttctaaaatgtcttgagtttc**tagagatgtttttccctagcagaccttgtcataaatagaaagctagcaagagaagcatgtcatgaaacatgaagagagcaaaagaacactccacatat**aggacagtaggctgattctgtcct**gtagc**ctgcagggagaaacacatgctctgcag**actttggacacctgggaggcactgggcctgtgcagtgttattgagataagacatctttgcagctgtgcagatttgcatgtcccacagagcaacgcctactgccctgaacatttatcaataggctggtgacatcctgtgcagaagtctctctcagtcaggacacagcATGGACATGAGGGTCCCTGCTCAGCT**CCTGGGACTCCTGCTGCTCTGGCTCCCAGG**TAAGGAGGGAAACAACAAAAATTTTATTCAGCCAGTGTAGCCACTAATGCCTGGCACTTCAGGAAATTCTTCTTA**GAACATTACTAATCATGTGGATATGTGTTTTTATGTTC**CTAATATCAGATACCAGATGTGACATCCAGATGACCCAGTCTCCATCCTCC**CTGTCTGCATCTGTAGGAGACAG**AGTCACCATCACTTGCCGGGCGAGTCAGGGCATTAGCAATTATTTAG**CCTGGTATCAGCAGAAACCAGG**GAAAGTTCCTAAGCT**CCTGATCTATGCTGCATCCACTTTGCAATCAGG**G**GTCCCATCTCGGTTCAGTGGCAGTGGATCTGGGAC**AGATTTCACTCTCACCATCAGCAGCCTGCAGCCTGAAGATGTTGCAACTTATTACTGTCAAAA**GTATAACAGTGCCCCTcccactgtgatac**aagcccgaacataaaccatggagggaagtagatgtgtgagtctgggctgccccagctgctcctcctggtgccgccctctgctgacagcagttctcagatgcagccaaggtttgaagctcactgggaagttttggtagaaggggtcagggaggcacatttacattctatctctctttatcc**tcagctccatcagctga**tatgcaagtatctctcctgattattattaataaaggacaaagaaaattaaacctaggaggtctagattgcagcaaaagtcagacttatacagaaaaggagaaggtactctacatatttttaaagaattttttttgatacaggggattagagtctaaattatgacctttcaaagatcagacataagt**acatacacatataagtatgt**tcagccaacaacatatcagataa**acatattgcacctatatgt**atactgtaaacatttatgctccgatcccaccttcttccaacactacccagatcaagatccacactcttcccagcatctcaaatatctccccagtttacttccctgtcaattccgtgttccatctatcatcatcctaggcttccactgtattttatatcaatatttattaatttttcctatacttaagctttatataatttcaatcatacagtatttactctttgacgtcttgcttattttgg**tcaaaatatttttga**gcttcatctgcgttgtttctctggcagtaatttgtgcctttttattgaggaacaattttcc**atgaatatgccaaaatatattcat**ttttcaactaattattt**gtttttggtttaagaatattactggtaaaaac**cttaagaacattcttgtgcttgtcttttgtgaacttatgcactcatttctcttgggaatattcctaagaatggaagtgttgagttataaagttagcatagaatttgtcttagtaaacattgacaatagtttactgaaatgctgaacatgtt

Uppercase: IGKV1-27

Lowercase: Flanking sequence[1000bp]

Red & Bold & Underline: Stem-loop [18]

Blue: Heptamer[21]

Green: Nonamer [5]

id-IGKV6-21-3[V_gene_segment]

tgtttattttgggggcatttcaggatcttgctcaagaactgtagagatttttttctgtgactcttttttggtgcttgcatggaggtttacagagtttcctcatctaatatagattatctagcaccaggcaatgtgctggatctcatggctgaagtgacagaggcatttgcatta**aaactcaaacttactacagaatattttctttctcagagttt**attcataaaagacagccttccaagttagctgataaa**tgggttggtatagtaaaccca**agtgcaaaatgcattgtcaacactctaggatggcttaaccagtaatgtgcttcattgctagtggttggaagtacaaggtgcaattatttttccttactttggaggggataagccagcatgactcatactccttttataaacacttgacatcttctctaatgtgacaagcccttgatgttttggggcgtgcatcccaccctctagagcacatgtgttttcacaagaaattcagagttcttacaatgtccagctcatcacgtctaattaccatgatgt**catcaatatagtgttgatgctttgtggaacgttcacaaagc**tttttcagcctacattgtgacagagagcaggagagttaacatagccctgggacgagactgaggatgtgagctgttattcaccccagataactgcagactctcccagagatggcgatggactctgccttcactctgcagctgtgccctggggtcttggtcaagccctgccagagcctcagcggagctcgtctgcaggtgccagcagagggcgcttcacacccctcatggaaggggccgggagggcgctctcctggcaacagtgatttctgtttatttaaaccagcaggacatccccataatttgcatgtatcgttcctcctatatgtga**agaggccctgcctct**cggtatcttaaaagaggttctttctctgggatgtggcatgagcaaaactgacaagtcaaggcaggaagATGTTGCCATCACAACTCATTGGGTTTCTGCTGCTCTGGGTTCCAGGTG**AGAATATTTCCACAAACCTAGGCGGAGATATTCT**TTCAATCTGTAATTTCTTTCATTGGGGACTCTGCAATAGGTGATTTTTGGCTTGA**TTTTAAAATCCTAATTTTAAAA**ATGTAATGCATATTCTTTCTTCATGTCTAGCAAGATTAAAGGTGATTTTCATACACAGATATTTATGTTGTACTGATGTTTGCTGTATATTTTCAGCCTCCAGGGGTGAAATTGTGCTGACTCAGTCTC**CAGACTTTCAGTCTGTGACTCCAAAGGAGAAAGTCAC**CATCACCTGCCGGGCCAGTCAGAGCATTGGTAGTAGCTTACACTGGTACCAGCAGAAACCAGATCAGTCTCCAAAGCTCCTCATCAAGTATGCTTCCCAGTCCTTCTCAGGGGTCCCCTCGAGGTTCAGTGGCAGTGGATCTGGGACAGATTTCACCCTCACCATCAATAGCCTGGAAGCTGAAGATGCTGCAACGTATTACTGTCATCAGAGTAGTAGTTTACCTcacactgtgttacaacccagaacaaaaacta**gttcagcctggctgaac**agagaaactgggtgataccctagaatacttctgattgttacaggtgctttggaggcaatgagttaaccaatacaatgaagtctggctcacc**cagcagagaggaaactagagtcactgctg**catactttcatctttttaaaaatgaattattttatttatttcaatagtttttgggggtataggtggtttttagttacatggataagttctttagtggtgatgtctgagattttggtggacctgttacctgagcagtgcatactgtgcccaatatgttgtcttctagccttcacctccccttctatccttcctccccagtccccaaagtccattatatcattcttacgcctttgcatcctcatagcttagctcccacttatagat**gaaaacatataggttttc**cattcctgagttacttcatttagaataatagcctccagcttcatccatgttgctgcaaagttcattattttgttctgttctgttttatggctgagaagtattt**cgtggtgtatatacaccacg**ttttctttatccactcgttgcttgattggcacttatggtggttccatatttttgaaatggagaaatgtgctggaat**aaacatgcatgtgcatgttt**ctttttcctataataactttt**tttttctttgggtagataagaaaaa**taagtactgggattgctggactgaatggtatttctacttttagttctttaaggaatctccatactgtttttcatagtggttgtactag**tttacattcccaccagctgtgtaaa**agtgttccctcttcaccacatccacgccaatatctattattttttgacattttaattatggccattcttgcatgagtaaggtggtatttcaaggctatggttaccaaaacagcatggttctagtataaaaataggcacatagatcaatggaacacaatagagaacacagaaa

Uppercase: IGKV6-21-3

Lowercase: Flanking sequence[1000bp]

Red & Bold & Underline: Stem-loop [16]

Blue: Heptamer[26]

Green: Nonamer [8]

id-IGHV3-16-2[V_gene_segment]

gtgaggtgacatggggtatatgtgaggtgacccccctgacgtgtg**aggtgacctgagcacatgtgaggtcacct**aagg**cacatctgagttgacctgggggatgtg**tgaggtgacctagggaaatgtatgagctgacaaagggcatgtgtgcagtgacctgggtgcatgtgaggtgacctggtgcatgtgttagtttacctgggggatgtgtgaggtgaacaggggcaggtgtgaagtgcctgaggaatgcatgaggtgacctggtgcatgtgtgtggtgacctagcagatttacgaggtgacctgcgggatgtgtgaggtgaatagggacttgtgtgaggtggccacaggaatgtgtgaggtgacctgggggatgtgtaaggtgacattggggatgtgtgaggtgactcagggcacatgtgaggtgacctgggggatgtg**caaggtcaccttgggtgagtgtgaggtgaccttg**gggatgtgcaagg**tcacactgtgtgagtgtga**ggtcatctgggatatttcaaacttcgtaaggaaagcctgcaacagtatatggcactaaaagcttcacactccaacaccgttaatgcgcacatcctcaaggaactctaagtgtttccaggttagcttaaatgccatggagcgacacctgc**cccaggcacactcatatatgcatcctggg**tctccctatcaacccctatcctcaaattttcagtttatgtttgctccatg**atatcaactctgatat**gttgaaggtttttttttttcttttatttgtagttgttcaggtttgctatttcactctcattactctgggctcagtcctctcctcaggtgtcccacttcagagcttgctatataataggagacatgcaaataggaccctcc**cttttctgatgaaaag**cagcgcagccctgaccctgcagccctgggagagaagccccagccctgggattctcaggtgtttctattgggtcaacagcaataaa**caaattaccATGGAATTTG**GGCTGAGCTGGGTTTTTCTTGCTGGTATTTTAAAAGGTGATTCATGGAGAACTAAGGATATTGAGTGAGTGGACATGAGTGAG**AGAAACAGTGGATATGTGTGGCAGTTTCT**GACCAGGGTGTCTCTGTGTTTGCAGGTGTCCAGTGTGAGGTGCAGCTGGTGGAGTCTGGGGGAGGCTTGGTACAGCCTGGGGGGTCCCTGAGACTCTCCTGTGCAGCCTCTGGATTCACCTTCAGTAACAGTGACATGAACTGGGCCCGCAAGG**CTCCAGGAAAGGGGCTGGAG**TGGGTATCGGGTGTTAGTTGGAATGGCAGTAGGACGCACTATGTGGACTCCGTGAAGCGCCGATTCATCATCTCCAGAGACAATTCCAGGAACTCCCTGTATCTGCAAAAGAACAGACGGAGAGCCGAGGACATGGCTGTGTATTACTGTGTGAGAAAtcctgtgaggggacacaagtgcgagcccagacacaa**acctcctgcaggaacactgggcgaaatcagctgcagggggt**gctcagggcccactcatcagagtc**aaccccagagcaggtgcacatggaggctggggtt**tgtttc**ctgtcaggatttgggacttcctctgcttctgacagtttccctagggaaact**ctttaattttagatttctgtgcccaccaatgtcatctctacatatatttttttaaatcattataatacgaggactcattctcacatgcacaatatgtatattgccacctacgggaatgaaaac**tcctcaaccatggtcaccagcatcagagttgtgagga**agctcaggggtgcctgg**tgagtcttctccagtcagactca**ggacagtaacctcaatgggattccctgactagaatggcttttagggatttcgattacagccaagagagaggctggaccaggttcag**tgtcatgtaggacctcacagattttatgtatgaca**tttctcctgacaataaagtatgcaaatgagtatcagcactgatctggtgcttcctgactttcatttttagtggttctcgcttttccatttgattttcctgctttctggataaaaggatgttgtccctgtggtctaaatcctggggctcaagccctttccctggagcttaggtggggctcaggctgtgactcct**gcagccattgggagaggctgc**tgagactttcttctctctcattgttcaggatactccactctgttttctggag**acgcgtctgagaacgcgt**gtggccattagtaatgagggcacaagcttctttggtcaaaatggggtgaggatgtggaattgatcctgtgctgtgtaaactgtcacagagtcaccttcttcaccagtagtgttagaagagcttgtaaaatttgtcggaatccaactggagtcccctgtgttcaaaccctgacaaatggagctggggaaggccatggatggaag

Uppercase: IGHV3-16-2

Lowercase: Flanking sequence[1000bp]

Red & Bold & Underline: Stem-loop [19]

Blue: Heptamer[36]

Green: Nonamer [4]

id-IGHV7-4-1[V_gene_segment]

aaaggaactgacaagagattaggtgttttcccaaagtccctgggagttcctggactcactatgagtgtggactgatccagtgcttccggagct**ccagggaaggggctccctgg**tggtttcatagaatccttgcttggggtgtttctgcagagttcactggctttcctacggccaatttactattgtaagagatggtgtcagcagtacatggtgtcacgtgtcactaaaggagcattccgagccaggacacagccacttcatacttgggggaaaatgctctggg**agcccagacaggagcctctctgcagtgcaagggctgggct**gcaggggacgctcaaagcccacccagtacaagtttcag**ccccagagcaggtgcacaggaggctgggg**agggattcctctcgagatctgtgtcattctttaaaaaatctaaaatacatatttgacaactgaacattctataaatatcctattcaattgcgagcatttatcaaacttgatgttgtaatgagaaccacttttacactggagatttctaacccttctagatatcttaatagtatgcagctggaggttaaggaaactacttttcttttctataaataagtgaaacttttggagaaacacactcatcccccaaa**taatacattcatgtatta**aagcctagaaatgcttatattacctctgagccatttgagtgtgggttcccagtgaagtcctgttctagggaaaattgttagccaatgggag**aagctctgtcaacatagagctt**agggatatggcagggctcacatggcctctaaggggattacagcttgaaccttgagcatcctccgtgtgtcatctctctgctctttctcatgcaatatcaggtatgaaataggatcactcatgaatatgcaaataactgaggtaaatatagctatctttgggccctgagagcatcacccaacaaccacacccctcctaagaagaagcccctagaccacagctccacaccATGGACTGGACCTGGAGGATCCTCTTCTTGGTGGCAGCAGCAACAGGTAAGGGGCTCCCCAGTCACTGGGCTGAGGGAGAAACCAGCACAGTCAAGTGAGACTTCATGCACTCCCATCTCCTCTCCACAGGTGCCCACTCCCAGGTGCAGCTGGTGCAATCTGGGTCTGAGTTGAAGAAGCCTGGGGCCTC**AGTGAAGGTTTCCTGCAAGGCTTCTGGATACACCTTCACT**AGCTATGCTATGAATTGGGTGCGACAGGCCCCTGGACAAGGGCTTGAGTGGATGGGATGGATCAACACCAACACTGGGAACCCAACGTATGCCCAGGGCTTCACAGGACGGTTTGTCTTCTCCTTGGACACCTCTGTCAGCACGGCATATCTGCAGATCTGCAGCCTAAAGGCTGAGGACACTGCCGTGTATT**ACTGTGCGAGAGAcacagt**gtggaaa**cccacatcctgagagtgtcaaaaaccctgtggg**aggaggcagctgtgctgagctgaggcagtgacagggacaacgtggctgcaccctttgtagggttataaacatacatatatatgcacccaacagtggagtacccagatatatcaagcaaacattattagagctaaacagagagatagacccaatacagtaaaagctggagactttaacaccccaccctcagcattggacagatcttccagacagaaaattaacaatgaaatcttagactaaatgtacatgacagaccaaatggacataatggatattgacagaacacttcatccaatggctgtagagtgcacagtcttttcctaaac**atatggatcagtctcaaggatagaccatat**agtagaccagaaaattagtcattttttaatt**gagaagatatcaagtaccttctc**tgatcataatggaagaaaactaaattaataagaggaatcttggaaaactatgcaaacacatggaaattaaacaatatgctccctaattaaacaatgtgctcccaaattgaaccacatgctccaaaatgggttaataaagacataaaaaattttaaaatttattgaaagaaatgataaagaagacaaagcatatgaaaacctatgaaatacagtgaaagcagtattaagaggaaagtttatagctataagcact**tactttaaaaaagta**gacaaccataaaataaacatcctaggaatgtcttttaaaaaatagaaaagcaagagcaaatcaaacccaaacacagtaaaaaaaaaaaaaggaaagatcaaaacgttaatacataaaattgaaataaagaaaacaatacaaaagatggatcaaatgaaaagctgttttttaaaaagataaaagacatctacaaacctttaaccagactaagaaaaaaagaaagaagaccaaagtaaatgaaatcagagagaaaaatggaaatattacaaccaatgg

Uppercase: IGHV7-4-1

Lowercase: Flanking sequence[1000bp]

Red & Bold & Underline: Stem-loop [11]

Blue: Heptamer[42]

Green: Nonamer [1]

id-TRBV18-2[V_gene_segment]

aaaagggaggaaggaagatctctcaaacacaggcatcccatgcatgactacagagaatgatacagataggtatgcagaataacgggacctgaggtcagagggggaagtctttttagagtttgttctttgaa**tcacttatattgaaaattcaagtggaataagtga**aggtgacaaaagtgaattacaaatcttaaagttccctgaaaatgcttgaagccatcccttagctgaggtgtgggagtgaaaaagcaaactgagacaatgtgaaatcattttctatctcgaattttccctccagcaatgttggtgtccacttccctgtctcttaatgggtgcatctatatatcacctcttgtcaatacctctactcagcaacagccagaaaatagaaagattatataagccaataaaaggaaaataaaggaacaaaaaataagaatagtggagagatcatctacttgggaggctgaggtggggaggattgcttga**gcccaggaatttgggaccagcctgggc**aaacagcaagaccccatttatgaaaaaaatgaataacattatctgggtatggtggcttgt**agtcctagctacttaggaggctgagatgaaggact**cacttgactgatcctaggagattg**aggctgtaatgagatatgattgcactgctgcactccagcct**gggcagcggagctagatccagtcttgaaaaaaaaagaaaaaaaaggtgaagagggaaaaaaaagaacgagtaatagaggtgcttacttcacaacactaacctcacaa**acacacactaagctaggttgtgtgtataggtcagaacaaaagatctaccatttaactgacatata**agattggttctcgaaaccataggacccactggaaatggt**gacctcacaggaagatgcatcttgtaggaggcagctgtgaggtc**tggttccccgacgtgctgcagcaagtgcctttgccctgcctgtgggctccctccatggcca**actctgctATGGACACCAGAGT**ACTCTGCTGTGCGGTCATCTGTCTTCTGGGGGCAGGTGAGTTTTCAGCTAAAGGATCACCATCACTGGGCTTTGTTTTGGGCTCTACAGCAGATTCTCAAATATACCCTGGACCTCTGTGTTGACCAAATATTTTGTGAAAGGTATTGACATTTATATGAAAACAAATAATTGCATTATGGAGTAAAAAGCAAAATTCATGTGAGTTTTTAAATGAACGCAGGAGACTTCAAACAGGCTCCTTCACTGTAAGACATTCATCCTTCTGCCATTACAACTCTGTCTCATGCAGTCTCTTCTCCTCTGAGGTCTCTCAAATGCCGGCGTCATGCAGAACCCAAGACACCTGGTCAGGAGGAGGGGACAGGAGGCAAGACTGAGATGCAGCCCAATGAAAGGACACAGTCATGTTTACTGGTATCGGCAGCTCCCAGAGGAAGGTCTGAAATTCATGGTTTATCTCCAGAAAGAAAATATCATAGATGAGTCAGGAATGCCAAAGGAACGATTTTCTGCTGAATTTCCCAAAGAGGGCCCCAGCATCCTGAGGATCCAGCAGGTAGTGCGAGGAGATTCGGCAGCTTATTTCTGTGCCAGCTCACCACCcacattgatgcagagccacatcctctcagtccacaaacatcctccagacctgccttggaaacagcggtgggccaggaagggaaacgcgtt**acctgtacagtgaacaggt**cagctctacggtattctgcaagttgagggtggaaataggggaggattgtaggaatagcatgaaacatcccaggctacgctcagaaactaagactctgtcatccaaagaacataaagatttttaagaactagacataatttttcagtttcaacattgacttaatttgtcagtttcaaaacacttag**cagtgtaactctgttttttactccacactg**tatgtcaagttctatatttcatgctttacatcaataattttatttaatcaccaagacacacaagc**aaaggaagtgctttcttccttt**tcaccaagag**aaattatggtcctaggagttaattt**ttagaaagaaagagacagagagatgggggaagaaaggggaggggaggggacaggaggagaagggaagggaggggaagggaggggaggggaggggaggggagggaagggagaaggaaacctagtaggtggcaaaaagcatttggctcaaagggttatgatttttatctcacatcatgctgcttagcattcttttaggcataatccttgttctgttccttgctgagtacccgtgcaggtttttacggatgctaggttcaagaattcctgctataaaattatagaacatctgagagatgatggaaatagagttgga**taaatttgtctagaattta**gggaaaaggtctggcccacagatattgttcattagaaatgtgtcacctcaactttttgtcttaatgtttgtttttattctcatcactcaacagatataggccacctacagaaaaagatttgttataatttgc**tttgatgaaacaatttcataaaa**tcaaaaggcagacaaaagcaaagtcatttcaaatatatgtgcagactaagtccagcctcaccagctt

Uppercase: TRBV18-2

Lowercase: Flanking sequence[1000bp]

Red & Bold & Underline: Stem-loop [14]

Blue: Heptamer[22]

Green: Nonamer [3]

id-TRBV5-4[V_gene_segment]

gggaggagcagactagcaaaagcctagactcctgaaaacagagcattaagggcaattctggtgcagtgtcagggggaaatgagggacaagctatcggaaattgaagcaaataccatccttgttataaatagcaaaaaccttggcaaaattgtgtcctgttctaggactttatggaatataaaattatgagccattcactaggatatcaactgaaagaaatatctaagcggcaaagcattcaggctactgtgtgactactttcgggcaccaggaaatttcacccagcaagaagggagccatgggaatagattttgcaaaccagcacagatggtaaccctacctccctctgctgtcctgtctcccataagccaaaccctgtgctgtgagctcttaaaatcctagaattatttctggggaa**tctatactcttaataatttataga**caacccaggtctgctttggatctgatcagacggactaaatcttggggactctgcaccactggccactgaagaaaggggctggg**aatgttgatgggacaggaaaatataataagaaacatt**ggtgtgaatctagtatcagaaagatgatgtgaggacaacaaggaagagctggaat**gtgagggtttaatcacaggctcctcac**cctccgctgatgggtaggtgtgtta**gctccagcatggagc**accaccgcactaggtgggggaagggtgagagggcaatggggcagcctgtgagctggggcaacgt**aggcagagaaggaactgtgtcaccacagaaacttctgcct**tcacccatcccttcagctctgcaggacaggtagagactccaggatcatccactgagcactggacataaggaaggctgcatggggaggacacaggacagtgacatcacaggatacccctcctattaggaaaatcaaggcccagaattcactcggctcttccccaggaggaccaagccctgaatcaggtgcagtgctgcctgccccactgtgccATGGGCCCTGGGCTCCTCTGCTGGGCGCTGCTTTGTCTCCTGGGAGCAGGTGA**GTCCTGGGCACAGGAC**AGCAGCCCCATTCTCAGCTTTCCCACCCAT**GTCCTCCACTTTACCTTGGGGAGGAC**CTCCAGGCTTTCTGCTGTGCTCATCCTCCATCTGCTTTTCCCACAGGCTCAGTGGAGACTGGAGTCACCCAAAGTCCCACACACCTGATCAAAACGAGAGGACAGCAAGTGACT**CTGAGATGCTCTTCTCAG**TCTGGGCACAACACTGTGTCCTGGTACCAACAGGCCCTGGGTCAGGGGCCCCAGTTTATCTTTCAGTATTATAGGGAGGAA**GAGAATGGCAGAGGAAACTTCCCTCCTAGATTCTC**AGGTCTCCAGTTCCCTAATTATAGCTCTGAGCTGAATGTGAACGCCTTGGAGCTGGACGACTCGGCCCTGTATCT**CTGTGCCAGCAGCTTGGcacag**cccggcagagtctctgacattctatacataaacttcctgccttagctttgacttgagagc**tgcaggccccacccaggtttcactcctgca**agggaagcttttagttgtatggaaggcatgtgttgtgtcctactgagcacagagctctcccaaacagagcccaggtttcctatgccctgagtgtgccggcttctgtgcagcatcttcttgcagctt**gtcacttcctgggtaacttcagtagaagagtgac**tgcggagccccagatgtgtgctagattctttgaatttgttatatagcttaaagtctttcaacaaccttgcacatcaatcattcttattcttccttgacagatggaagtctcagggacattcagtcattttccccagtgtctcctggcttgtaaggaccagaagtaggaaacaaagtagtgcatccattttccacctacccccctgctccatcatcaccttctgtgtcctggtcagtaagtcagagccctcaaact**gccatctagtgaccagcagggcctcactagaggc**ttacatgtcttcaggggtcattactctggcctcttcattagaaactttgaggaatgttaaatttaacactttttaaaaatcatttatatgcattccctttgtctatccccagtctgcagcctacactttcatttttatggtgttttacacaagtttaatttaatgcaacaataactaaaaacaa**ttttgtttgtgtgtagacaaaa**actgctttctcatataaatgtctaatgatatttttctcaatttattctactaaaatttaagtttggattttcaccaataagaataaatgaacctgaaatatatctcatgtgagaaataagatagagacctaggaattggagggaagcacatatgttcagtgaagaagcaatcgt**ctgtcaaaaagagctctaaagtggacacctcgtgctgacag**tggagagataattggaaggcaaattatcatgaacc

Uppercase: TRBV5-4

Lowercase: Flanking sequence[1000bp]

Red & Bold & Underline: Stem-loop [15]

Blue: Heptamer[34]

Green: Nonamer [6]

id-TRBV6-7[V_gene_segment]

agagggacctgggggcagggtgttggctgggtctaggagaagatgatgagctcagaaaatagacaaggagaaaccccaggggaaagcttgtaaaatcctggaggaatcacaagatatattatttgagctcctttttatttattactaaattaatggc**aggaatcccatataaagggatattcct**tatgaaacatgacatctaaagtacaaacag**aggattcttccttaccctaaatcct**gccctctctccctgagtctcttacgcatcaggccacctttg**tgcccataagtcatgggca**acgca**gtgtggtcaccttcatccacac**ctagaggacagtcagcaaagtgaggtggttctgcctgctgcggtctcaccccagacatggaaagcaagagccctgggtgg**agctgaaggtgctcagct**gggtttatcaggagtctcatctgtcagtggattgacaagaaacagagcaaaacgactcctccaatgttgatgagcctgcccctg**ggttttggaaacctgataacagagaaaacc**aatatagacaaaggattttaaacaggattatggtcaattaagcaaattagaaaaagatacttgaaagaatatttgggacacaggagtcaaaaacaccaggaagacatgaggagtgtccctaagactctagactacagcactgtgtagacaactaacaccag**aatattattatattatcgaagtaataaaatt**tacagtgaattacaaactgtttccaaaaggtcatgaaaatcttgtagacttgtttcaattcagacaaatgtgttcttctcattctcagctgttcactggtgcatttatcttgaatttgaccatctggggaatgggcgtggcctctcctgacaggaaggctctgggggccaggc**agggagaatgatgtctcagaatgactccct**tgagagtccagttcccctttcatcaatgcacagacccagaagacccctccgtcctgcagcccctgccATGAGCCTCGGGCTCCTGTGCTGTGTGGCCTTTTCTCTCCTGTGGGCAGGTGGGTCCT**GGGCAGGGCCCCTTGTGTGGATATCAAGGCCCAGCCC**CTTTCCACTGGAGCTGTAGCATCAGCTTTGTCCTTCCCTGCAGGTCCAATGAATGCTGGTGTCACTCAGACCCCAAAATTCCAC**GTCCTGAAGACAGGAC**AGAGCATGACTCTGCTGTGTGCCCAGGATATGAACCATGAATACATGTATCGGTATCGACAAGACCCAGGCAAGGGGCTGAGGCTGATTTACTACTCAGTTGCTGCTGCTCTCACTGACAAAGGAGAAGTTCCCAATGGCTACAATGT**CTCCAGATCAAACACAGAGGATTTCCCCCTCAAGCTGGAG**TCAGCTGCTCCCTCTCAGACTTCTGTTTACTTCTGTGCCAGCAGTTACTCcacagcgctgcaaggctgtctcctctctgcacataaaggcaagggaaggtgctgccctcctcccccacccaagactcaaggatgc**cctgtgcagagatctctgcaccagg**aaccttggaacccagagtggccccaagtggcccggacagtatgagcctcgctctgtgccacgtgcctctgcaggcatctcagccaggcctggactggtcccaggtcctcagatgtctcccttgttgctctctctggtctgtcctccgaggtctcctttcgggttgtggccagggcttccccagctcctacttttctactcatcatcctgagtccgaggcccccagggtggaagaggatttgtatttcagatccatctagactcctgtctcttcctggtgaccatgttgcttcctctctctagggttttcccagcccccaccctcacgtagtctctcctgtggcccacctttcccatctgggcagtcaccctccaaggccttgctgggtctctcctcccctcacttccccg**cccctttctactgcagctataagggg**agcc**tctcttctgtgcctccttccttcccatcaaagaga**tttcaaaggccattccctctgccctaggctagaggctaccttttctactggccagcgcctacctgtgcttcagatctcagcataatctgcccctcctgcaggaagcactcccccgcctcccagctgagatcaactttccactcaaaagctctcatagaatcatgtgtctgttggtaacctttagcacagttgggctttctcaggtacttgtctgataatttttgtactccacccattttttttccctatttttccttttttttttga**gacagagtctcgctctgtc**accaggctggaatacagtggcactacctcggctcactgcaacctccgactctccgactctctggttcaagcgattctcctgcctcagcctcccaagtagctgggattacaggcacacaccaccacaccc

Uppercase: TRBV6-7

Lowercase: Flanking sequence[1000bp]

Red & Bold & Underline: Stem-loop [15]

Blue: Heptamer[34]

Green: Nonamer [3]

id-TRBV9[V_gene_segment]

tgcactgtctaaa**atttagtaataatgaacctaaat**acaaaatatgaaaacatatgggtgtaattaactgggattagctgtggaatatagggtttccaggctgggtttccttgatagcatcacactgtttaattgtatggcctttataattctggctggtctctaccatcttaacatattatttgcaattatgtaacaattgttt**attttgtaatcaaaat**gttggttcatgtgtcctttgactccattggcta**agggtagatttgaaaatcaaccct**aaattgactcttctgggaaagcatggacggggtgtgtaagagggtgggggtggcaggagtgggggcatctctctttctgatttggacaatttgagcatatcacgtcagagggtatgaatccatgagactgtagactgttggccacttggaggcgctgtggctccacagtgctgcagagaactcagcggaggggtgggagagtggccaagagaggaggactgaaggg**gatgccctggctgcctgctggcatc**agtagggtggatggaattccctgagtggggcctggccatgtagagttaagcacagactatcattgccttgaaagagttggggtgccagaggcctcagtttgcatcaaggagggagtgtctctgctgctgcacgggcagtggagggtgaaggggctggggtggcaagggctggggaatcagtgtcagggcagggacagggcgggcagaggaaagtaaggtgtgatgtgggtatgttggtactgtgttcagtcctcctcacctttagcttctggcctgtgcaagacaggaatagtacatggagtagcagaacctgaggcctgccaaagagtacagtgggctggagtggcctgcaggggaaagt**atgtcagtgtggtgacat**cacaggtcaatgctcctatcaggaagaagggagcttaggaacttcagaatgcttactacagagacaccagccccaagctaggagatcctgccATGGGCTTCAGGCTCCTCTGCTGTGTGGCCTTTTGTCTCCTGGGAGCAGGTGAGTCCTGGGCACAGCTTGAAAGTCTCCGATCTT**CATTTCTTGTCCCTGAAATG**CATGTGGGCCAACGATGGCTTCAGCAGGAGGCTTTCTTCTGTGCCTTATGGTTAACTTTTGTCTTCTGACACACAGGCCCAGTGGATTCTGGAGTCACACAAACCCCAAAGCACCTGATCACAGCAACTGGACAGCGAGTGACGCTGAGATGCTCCCCTAGGTCTGGAGACCT**CTCTGTGTACTGGTACCAACAGAG**CCTGGACCAGGGCCTCCAGTTCCTCATTCACTATTATAATGGAGAAGAGAGAGCAAAAGGAAACATTCTTGAACGATTCTCCGCACAACAGTTCCCTGACTTGCACTCTGAACTAAACCTGAGCTCTCTGGAGCTGGGGGACTCAGCTTTGTATTT**CTGTGCCAGCAGCGTAGcacag**ccctgcatgagcatcagccttctgtgcaataaca**ttcctgccccactcaggaa**gtgacggtgaggggagggctgccagccagaggggctcaggccctg**gagagtggacaggcctttcccaggcactctc**tggtgcttctgtatgtgatctggtttactgctcactgaagctgtgtaagcgaaatattattttacctgtttatatatctgagggacagcagatgtcctggggctcatatttagtagctagctgaagtctattgtttcaacacctctgcactgtccctcctcactccggagatctgtccttttacctggtgtcccctttttgtgtccccaagagcaggatgctaaggtcatatatgaaactgtgatcatgata**acttgttccctgacaagt**ccaatggacacatgatttccaggcaaaacatatcca**tctattctttattgtgatttaaaaaaaaaagaaagaaaaga**aaggtgcagttttctgctctgtagccctcctatgggcaaggccaggtcatctcttttatggagactgtatgaggagggagc**tttcctaggttcacagcaaatagcaggaaa**cgttttttatggggcgtgtagtcttcctgtggcggatgcaatctgccatcagtgaaaggacatgaaaaattgttattcattagttaatattttttattcttttttttttttgagatagagtttcactcttgttgcccaggctggag**tgcagtggtgctatcttggctcactgca**acctccacctcttgggttcaagcaattctcctgccttagcctcccgggtggcctccaccacacctggccagttttgtgtttttagtagaggtggggtttctctttggtggccaggctggtctcaaactcccaatctcaggtgatcacctgccttggcttcccaacgtgctgggattatagagccattgcactaggccaattttttattctttatcaataaatcttttggggcat

Uppercase: TRBV9

Lowercase: Flanking sequence[1000bp]

Red & Bold & Underline: Stem-loop [14]

Blue: Heptamer[38]

Green: Nonamer [5]

id-IGKV1D-42[V_gene_segment]

gataaagaaaatgtggtacatatgcaacatggaatgcttaatatacagctatgaaaaatgagatcatgtcttttgcaacactgtggatggtgctggaggccattatcctaagcaaactaacacaaagattcaagtttatgctaatcaaggctcctatagtagaagaaaaaaaattgttttaaagtaatcagatttagcattgctagtcagacttttatgctgatggaccgtgggaaataattcacaatgacagaaaagggggtagaatatgggggtccccaaacagagcataaaatgaatcccattaaaactcaagcattaaagagacttatagctctggacaatgcaggaactgtggatgtcatatgctttaaggaatcacaaaatcatcctctttgtcaatctgcag**taaacctcttcagcttagactggtaataacattggttta**gggccattac**aaaatgcttttgagaacatttt**acttgctcatgactaag**tgttcttttttacttaaaaaaagatca**atttcatgcttacaaaaatgtagtatgtatgtcacaaagtatcgctcccaactggaacaatttcacagtgtgttgaagacctgatagccccactctctaagactttattaagcactctctacaaacaaggatattctcctacttattcccaatacagcattgaaatatattattccatctagttctcacaaccacttcgagttttgccaataaatgctcaaaaatgtacttggtaacaaaatatcctttaggaagaaacattctctgcaggcaaatctaggtgccctggtctgacctgggacactggggacactgcccctgtgctgagttactgagatgagccagccacgcagctgtatccagcctgccccaccccctgccgttttgcttgttcccagagcaccaccccctgccctaaagacttcttaataggctggtcacacctgtgcaggagtcagtcccagtcaggacacagcATGGACATGAGGGTCCCCGCTCAGCT**CCTGGGGCTCCTGCTGCTCTGGCTCCCAGG**TAAGGAAGGAGAACACTAGGAATTTTCTTAGCCCACTGTGCTCTGGCACTTCTGGGAAGTTCTCTTAT**ACCATGATTCATGGT**GTGGATATTTGTTTTTATGTTTCCAATCTCAGGTGTCAGATTTGACATCCAGATGACCCAGTCTCCATCTTTC**CTGTCTGCATCTGTAGGAGACAG**AGTCAGTATCATTTGCTGGGCAAGTGAGGGCATTAGCAGTAATTTAG**CCTGGTATCTGCAGAAACCAGG**GAAATCCCCTAAGCTCTTCCTCTATGA**TGCAAAAGATTTGCA**CCCTGGGGTCTCATCGAGGTTCAGTGGCAGGGGATCTGGGACGGATTTCACTCTCACCATCATCAGC**CTGAAGCCTGAAGATTTTGCAGCTTATTACTGTAAACAGGACTTCAG**TTACCCTcccacagggttacaagcctgaacataagccccttagtgaaacaaatgtatgaagttgggttgcacagaggctccacctggtgcctc**catctgctgaaagcatttctcagatg**tatacaagtttttaagcttttggcacaagggatcagggagtctcccctgttttctaactctctttcttcatcctcagtcccaggagcacagacatgacaacgtctccctgatttagtaatgatagcaattatgatacctgaggaatctgtgttgttgcatcagtcagggttcatacatcaaaatagaaaccactctacagatttcaagaaggaattgttttaaaacagggaattagagtcaaacatccatacctgggggtgtggtaaa**cagagaagctgatactagaaatacgggactctctggtggtccaccaga**agcctgaacgcatctgaccctgaatgtatgggctgcttaaacatgtggtcctctgccctgtctaggaagtagagatgttagggtgctgattctcttagctgcctgtagcacgtcactcggtgattctccagtcctcacctcagtccatgtgtctacctgcaggtgtcaatgaatattgaatcatcctcctctgactcttacgtatgacatgagggcccattattgagcaactctacaagaaaccatagtgggaaaagggtttttgtgaaatgtgtttccagaaatgatggtgataatgaggaatggagagctgacaaccaattaaggtcaatatttc**cccaaatctcagaattttccagatttggg**atgacccaagaatatccttgcatgtgcttccacgtcttattagc**aagtttgggattattgcaagaaaactt**cctgtgtcataaag**taaacagaaaaaggagaaaattgttta**gaatacctagcaattctttgacaaaaaaatgaaacactttcttgataaaacaaccaagacacctaattgttcacaaagtatattttttccttaa

Uppercase: IGKV1D-42

Lowercase: Flanking sequence[1000bp]

Red & Bold & Underline: Stem-loop [15]

Blue: Heptamer[28]

Green: Nonamer [6]

id-TRAV8-5[V_gene_segment]

aacaggaacaaatgaatcaagaacccaaggagagaactgggcttatgctttcaaatcaaaggaaagaagtattctgactttgtcaagggtcccattagccaca**gagaaagcagtgactttctgcattctc**gtgttttcagagttttatttttttccccatgcactatctacatttattattatttgtttttatttttaaatttcaacttttatttctcgcttggtgtggttgtatttccttcagccccacaggtgtctgtagttttattttctttacccctagaggcaaagccttcccatcatgagttacgccttgtcattagttgcatggaggcactactctccttatgtgaaatcttttcctctatggctgcacaagcatcatccgagggaaatatctctaccatcctccatcagcactgtggcaacactgaaccatccaaaactc**agtccagggcaattggtctgaacccaaactctcatggactcatatctaaattctgatatg**cttttcaataacgattcctcttgatgtattactccatctttacatgaaatcttattgctaatggactctaggactaaaacaccatcatagatttggtcaggaagaaacacgtcttgtggaagttttgatacatctcagaaaacaggaagaacacactaatgaacagtctgtgctaaa**ctctggttctgcacttggcctccagag**agcgatgctgcacacactggagcttttgtttctgttgaagatcagtccactgctcagtttcttcttcctgcagctggttgagttctttccagacaaagacaagtgacaagaatcagagtttt**aaaagcaaccagattcatctctgcagctttt**gtagttttatataaacaaagagcttgtccagcaaaccttccccacacttcttgttcttggtctcttcagccgctttccttcctgttctctggagatcttgcagaaaagagcctgcagtgtttccattgctcagccATGCTCCTGGTGCTCATCCCACTGCTGGGGATACATTTTGTCCTGAGTGAGTAAAATTTCTTTAT**CATCTCTAGTTCCACAGGTTCTGACTAGAAATG**CTTGCTTTTTATACTGAGTCTGCACTGCTTTCACTGATAGTACATTACTTTTTCAGGAACTGTCAGAGCCCAGT**CAGTGACCCAGCCTGACATCCGCATCACTG**TCTCTGAAGGAGCCTCACTGGAGTTGAGATGTAACTATTCCTATGGGGCGATGTTGTGGGAAGTCAGGGACCCCAAACGGAGGGACCGGCTGAAGCCATGGCAGAAGAATGTGGATTGTGAAGATTTCATGGACATTTATTAGTTCCCC**AAATTAATACTTTTATAATTT**CTTATGCCTC**TCTTTACTGCAATCTCTAAACATAAATTGTAAAGA**TTTCATGGACACTTATCACTTCCCCAATCAATACCCCTGTGATTTCCTATGCCTGTCTTTACTTTAATCTCTTAATCCTGTCAG**CTGAGGAGGATGTATGTCACCTCAG**GACCCTGTG**ATAATTGTGTTAACTGCACAAATTAT**AGAGCATGTGTGTTCAAACAATATGAAATCTGGGCACCTTGAAAAAAGAACAGGATAACAGCAATGTTCAGGGAATAAGAGAGATAACCTTAAACTCTGACTGCCAGTGAGCCAGGCGGAACAG**AGCCATATTTCTCTTCTTTCAAAAGCAAATGGGAGAAATATGGCT**GAATTCTTTTTCTCAGCAAGGAACATCCCTGGGGAAGAGAATACGTGCCTGGGGGTGGGT**CTATAGACGGCCCCCTTGGGTGTGGCCATCTTCTATAG**TCAAAACTGTAGGGGTGAAATAAATCCCAGTCTCCCATAGCACTCCCAGGC**TTATTAGGAAGAGGAAATTCCCACCTAATAA**ATTTTGGTCAGACTGGCTGCTCTCAAAACCCTGTCTCC**TGATAAGATGTTATCA**ATGACAATGGTGCCCGAAACTTCATTAGCAATTTTAATTTCGCCCCAGTCCTGTGGTCCTGTGATCTCGCCCTGCCTCCATTTGCCTTGTGATATTCTATTACCTTGTGAAGTACATGATCTCTGTGACCCACACCTATTCATACACTCCCTCCCCTTTTGAAAATCAGTAATAAAAACTTGCTGGTTTTGCGGCTTGTGGGGCATCACGGAACCTACCGACATGTGATTTCTCCCCCGGACTCCCAGCTTTAAAATTTCTCTCTTTTGTACTCTGTCCCTTTATTTCTCAAGCCAGCCAAAACTTAGGGAAAATAGAAAA**GAACCTATGTGACTATCGGGGCAGGTTC**CCCAATACGGCAACACCTTCTCTCTTCTGGCATGTCCAGTCCCCCGGCCAAGGCCTCCAGCTGCTCCTAAAGTACTTTTCAGGAGACACTCTGGTTCAAGGCATTGAAGGCT**TTGAAGCTGAATTTAAGAGGAGTCAATCTTCCTTCAA**CCTGAGGAGACCCTCTGTGCATTGGAGTGATGCTGCTGAGTACTTCTGTG**CTGCAGGTACcacagtgcctgggtctgcag**gggagctgaacacaaactgcctatgctagagaagttttcagagactcaatgtatcttcctgtggcattttcaaaga**tctcttatttttatgatggtgagtgaggcaaagaga**gctgagcacatccagccatttctatataattctgcaattcctgctatgtcccatgtcagataaactttgttttagaagctttttgttaattgaagaaactgcataataggaaatagtattctagaatgatggaggagacatacttcttatcacctaacaatcaggttc**tcaaggacctttccttga**ctgtgactgcattaagtggttgactatgggaattttgatatggtctaggagaaccttg**gttttcatttatcatttaatctctcaactaacccagaaaac**aaaacataagtttctcatgttcttaccttgcaagttgcatccttgagctggtcattcacaggatattttcttagaatttatttttctggctctaactgcgcaattcagatttcaggatattttg**tctcatacctctatttaggaggaattatgaga**aattgttagctaacagtgaaagtttcattagaatcacgaaggacttagaaaattgtcacagtgatgtgagttctggcaactaggaatagcacacaa**ggagaatgaagatttagaattcatcagtgtttctcc**aaagactttttcagttataaaaatagcttctattctattggtatattaattatgcaaatctacattgtaagtttgtctatggaattccattcatctcttctagctactgacacttttacttgttgctttcactctgttgttacatggtttcactattgaaatattaaagattgattctcttttttgagatagagttttactctttcacctaggctggaa**tgcagtggcatgatcatagctcactgca**gccttgacccatgcccccaggctcaagcaattctcccacctcagtcag

Uppercase: TRAV8-5

Lowercase: Flanking sequence[1000bp]

Red & Bold & Underline: Stem-loop [24]

Blue: Heptamer[42]

Green: Nonamer [4]

id-IGHVIII-13-1[V_gene_segment]

catgagatgcaggcaacaaagaatgaagtaaccttccccatttgcacataagatgtttccattcacaagagacctttcttttatcaggttcatgtgcaaccaagtttccctgctgacaagcatttaagccagatgattcgcaacttcctttg**actgagaaagatttccctcaaacttcagctcagt**gcaggcacacaccatctctgaa**tgggcatttaccatcagacaatgccca**cacctgtccccacatggacctttccctcagacaaacaaacacatcctcaggttgactcttccctcagacaagcacccctgtctt**catgtgaactcttccctcagataagcacacatg**tccccacattgactttttcctcagacaag**cacatatagctgacaacgaacagttatgtg**gcaaaatgagctcaggatagtggtaattatggactccagctctgatagtttgtagaaattgtcatttttaaaattctgactgaagactttcctttattgtagaagacagtcctttacagctctaattgcacagcctacaggcaggagtccatttcctctgggcaaggtttatttttatttgtttactgtacttatttgttgataaatattgatactacaaagatagcctatagggtccacatacgagaaaaaaagagtaatgggcagatcaaccctgcaaatccagtcccaggagtctttgaccctgccctccctggaatccagagacagagatggaagaggcctgctgagcagtgcact**catgtccccagggagaaagacatg**gaaatgaggcccctcctctgcaaatgaaaagtagctcatcccctgttcctgtagatcctgg**tgaggagccatcccacatctgtgcccttcctca**gtgtccacaccatggggtctgtgcggatctgggctgctcttgtcatcactctcaatgtttaggttccccgtggatcaggccctgctgtggctgttgcctctgtgtttgcagAAGTCCCCTGTGAAGTTAACTAATGGAGTCAGACAGAGAAATACTACAGACCAGGAATTCTGCCTTTTCTGCA**AAGCCTCTGGATTCACTTTCACTGAAAACAGCATAAGCTTGATCCAGCAGGCTT**CATGACAGGGGTGGGTGTGGGTAATAACAATAATTCAAATAGAAGTTCTCAGTGGGACTCTCCTTGAGTAAAAAGATGATTAACAATCCTCAAATACACTCAGTTCAGGAGATTCTCTTTTAAGATGATTAACCTGAGAGCTCAGGAAAAGTCCGTGTATTACTTTGAGGG**Acacagtgaggggacatctgtgt**gaactcagacaccaacctccctgcagggagacaggaggggactg**cctggtagatgcttctcagaaccaccagg**gggtgctcaggacatcagggggcactcagaaccatcaggggatgctcaggacaccagagggcactcaggacactggggggcggggggtcactcagaaccaccagggggcgctcaggacactgggggggggggggtcactcagaaccaccagggggcgctcaggacactatgaggggg**ggtggtcactcagaaccacc**agggggcactccggacacggtggtgaggggtagctcaggatagcaggggtgctcagaaccaccagggggcgctcaggacactgagggggggtcactcagaaccaccaggggacactcgagacaccagggagccctcaggacactagggggagctcagaaacaccaaagggcaatcaagacaccagggggatctcagaaccaccaggaggtgctcaggacacca**ggggtctctgaaccactagggggagattaggacccc**agggggctcagaaccactagggggttttcaggagaccagggggcgctcaggacaccagggggcgatagggaca**ccaggtggttcagaaccactagagtgtgctgagaaccaccagg**ggtgctcagaaccccagggggctgctaaggacaatagggagtgctgagaaccaccagtgggcgctcaggacaccagggggatctcagaatcaccaggctgtgctcaggacaccggggg**gtgctcaggacctccaggagcac**acacgacacaaaggatagctcaggacctacagggggcgatcagaacgccagggtgcgttgaggacaacaaggggctcacaggacacaacgatgtgcttagtaaaccgggggacactcacaaccaccagggggtactcaggacaccagggggtgcccaggaaaccagggagcactgaggacaccaccgctccc

Uppercase: IGHVIII-13-1

Lowercase: Flanking sequence[1000bp]

Red & Bold & Underline: Stem-loop [13]

Blue: Heptamer[42]

Green: Nonamer [2]

id-IGHVIII-16-1-2[V_gene_segment]

aacccagataattcttt**tctttccttggttgagaaaga**ttttccccaaacttcagctcag**ttcaggcacacactgtccctgaa**tgggcat**ttaccctcagatgggtaa**cacacctgtcaacatggggactcttctgtcagacaaacacacctttactcacgtggattcttccctcagaccaacacacctgtccccacatggactctttcctcagaccaccacat**atgtccttacatttactctttcctctagtaatgacat**ttcctcaagtggactcttgtct**cagacaaacaaacatgtctg**catgtgaactcttcactctgataagtacaaatatttccacagtgactgtttcctgacacaagcacatatatccaatgttgaactgttttatggcaaaatgatctcaagataataattataaaatccctcctgacaaggtgtagatctgcatttttttattgtaccttaactttgcct**tattgtcaagaacaata**gtttgcagctctatatgcaccgattagagattggtgtccattttctctggaaatgtatttttatattcttactggatgtatttttttgataatgtttgctactgtgaagatacctgaacagggtccacactagaaaataaaaaagactaatcggcagattaaccctgtgcatccagacccacgagtcctttgatcctgcccccctgaaatggagacacagaggacagatgagcaatgctgagcggtgcacccaagaccacaaaaagaaagacaaggaaatgtgtcccctcccctcctcatgaaaggcagctcatcccctgttccttcaggccctggtgaggagccaa**cccatgtctgttcccttcctcggtgtccacaccatggg**gtctgccctgatttgggcttcccttctcatcaccctcaatattagtgtcccttgtgaatcaggtccagctgcggctgctccacatggggccgttcttccatttcctctgtgtttgcagAAGTCCTGTGTGAAATTTACTGATAGAGTCAGGGGGGAAAAATTGTACAGCCCAGTGGTTCACTGAGACTCTCATGCAAAGCCTCTGATTTCACCTTTACTGGCTACAGCATGAGCTTGGTCCAGCAGGCTTCATGACAGGGATTGGTGTGGGTGGAAACAGTGAGTGATCAAGTGGGAGTTCTCAGAGTTACTCTCCATGAGTACAAATAAATTAACAGTCCCAAGTGACACCTTTTCATGTGCAGTCTACCTTAAAGGGACCAAACTGAAAGTCAAGGACAAGGCCTTGTAATACTGTGAGAGAcacaggagagggaatatctgcgtgagcccagacagaaaaatctctgcaggaagacaggagggagctgcatggtagatgctcctcagaaccaccagggcaccttggggacaacctgggggcactcagaaccaccagggtgtgcttaggaccacgggacactcaggatattagggggtgctcaggatcacgagggggcgctcatgacaccaggggacactcagaatcacaagggggcgctcaggacacgagggtt**tgcctgggaccaccagcaggca**ctcaggtcaccagggggtgctcaggaccaccagggggcgctcaggacaccagggggcgctcagaaccaccagggggcgctcaggacacctgggggcgctcagaagcatcaggggtgctcaggatatcagagtgccctcagtaccacgagggggcgctcatgacaccaggggcactc**agaaccaccagggggcgctcagtacaggggggttct**cttaggaggcagctccaaatcagaagcctgagaggctgtggttttcttttaaaccttggtgattcccgacctggtcaagcaaaagtcttccccaggatctctcaccatttcttccttgtaagtccatgattacttttacctacaaaacattaacttagaacaggaatttaattcaacttttaatgctgcatattttccaagtaatactagcaatgatctctcaggacaatttttacaataggttatttata**tattttcttgattaaaaata**a**tactattattaaaatagta**aagttataaaaatcacacctgtaatcccagcactttgggaggctaaggcaggcagatcacttgagctcaggacttttagaacagcctggcagcacagtgaggtcttttctatatatatatataaaataataataataataataaatacctgggtatggttccacctgtggtagcagctacttgaaaggctgaggcaggaggatcccttaagcctgggaggtt

Uppercase: IGHVIII-16-1-2

Lowercase: Flanking sequence[1000bp]

Red & Bold & Underline: Stem-loop [11]

Blue: Heptamer[35]

Green: Nonamer [5]

id-IGHV3-75-2[V_gene_segment]

atacacacaacaaactaaaatatctaggtatttatgctgagtaaaataagccaaaaaaataagtatatatgctatattattccatttctataaattctggaaaatgaaaactaacctacaataatttaatgaagaacagttgttgcctgaggaaatggtagaagaagggaaagggataggaggaaatgtggaacaggagaaaaaattgaggcgaattcacttgtccactttc**ttgataatggtgatagttatgtcatgtttatcaa**tg**tatactttaagtata**tgaagtttattatctctcaattagaccttataaag**tttattacaagcaaacaaataaa**aaattagacaaaaagaactgatagaaaatg**ggtaacatatgttaaatgccagttacctgaaaatttatctgcgtgaaccttagttctctccataattttaggt**gaacgctggagtgtaacagagtcacaatattttcattaaggcagtaagttctgcaagccacactagacatgtcccgctctgtcctggagttagttcagggaacacagt**gatgaggaacacgggctcagataccggaactcactcatc**tcagacatgagcacttagacacatacataggcactctgtgtgtgggtttactttcacatccgtacatggagaaacattgagtctcacagacataatttacacacatatttaaaatataataaggtgtttga**tgctgtttatcacagca**caatttcctcgtaaaagtgcaaatcttctaaataccataagttaccaaactcctgcaggacaccatcatcctctctcggcaatgccttctcctcacgtgtcccaccctagagttttttatgtagttggaaatatgcaaaggaccctccttctttggtgaaaaccagccctgccctcaccctacagttctgggatgggagacccagccctggaatttcaaggttttt**ccacatggtgattagcactgaacagagaactcaccATGGGG**TTTGTGCTGACTTGGGATTTACTTGTTACTATTTTAAAAGTAGATTCAGGGTGAACTAGAG**ATTTTGAGTGTGAGTGGATAACAGTTAAGGAAACACTGAAAAT**ATGTGCCAGTTTCTGACCAGGATGCCCCTGTGTTTTTGGGTGTCCAGTGTGAGGCGCAGTTGTTGGAGCCTGGGGGAGACTTGGTACAGCCTGGCGCGGTGGGGGTGGGGGTCCCTGAGACTGTCCTGTGCAGCCTCTGGATTTGCCTTCAGTAGCCACTGGATGCACTGGGCCTGCCAGG**CTCCAGGGAACAGACTGGAG**TGGGTCTCATGTATTAGTACTGATGGGAGTAGCATACGCTATGCACA**TTCTGGGAAGTTCTGATTCCCCATCTCCAGAA**ACAATGCCAAGAACACACTCTTATCTGGAAATGAACAGTTTGAGAGCTGAGTACACAGCCACGTATTACTGTGCTGGAGAcgcagtgaagggaagtcaggttgagcccagacacaaacctctctgcagaaacacatgaagtaaatcagctacgggggcgctcaggacccactgatcagattcaacctcagaggcagatgcagatacagggtaggagctggttttctcatagtgtttgtaacttccctctccaactgacagtttcccagagaacctcactaatttagagttctgtgcctaccaaaaaaatctccaaatatacaccttttaaaaggtagtaatataaaggcctagtctcacatgcacaaaatgcagatcaccacttccaggcatgagaagttctcaaccacggtcaccaagatcagagt**cctgaggaagctcagg**ggtgccttgtgggtcttctccaatcacattcgggacaggaaactaag**tttaattccctgattaaa**aaaggctataattccctgatttaaaacgtttcattccctgattaaaaaa**ggctggaatttttctgattccagcc**aagaaagaggctgggccagagtcagtgtcacgtacaagctcacaggtttcacgtctgccccttctc**ctgatatgagactatgcaaagcagtatcag**cactcatctgctgcactttttgctgtcaatccattttctttcttttcagctgttgtcactttttcgt**tttcttttctgctccctgaaaaagaaa**gatgtggtccatgcggtctaaattccaaggctcaagctcttccgctggagctcaggtggggctcaggctgtggctcctgcagccgcatgggagaggctgatggggctttcttctt**tccccagtgcttgggaccctccaccgtgtttcatgggga**ctcatgtggaaatttgagtggcccaggggaactgaagggcataagcttgtttggtgagagtgaaacgtgactgtggaattgattctttgttgtgcaaaccagaagaaa**ttttgaattttgtcagaatcaaaa**cggagtcacttttgttaaaact

Uppercase: IGHV3-75-2

Lowercase: Flanking sequence[1000bp]

Red & Bold & Underline: Stem-loop [18]

Blue: Heptamer[35]

Green: Nonamer [4]

id-IGHV3-73-2[V_gene_segment]

tctctctaaagggcgtgaccagatttgcagtagaatgagaatagtttcaaccctaaagttcatctaaaatgttagtggttt**tcatggtaagtaattaatagaaacctgacagctccatga**cagcaatatagaaatcaatagtccactcattttaacagagagaattagaaaacctataatttaagaataaccagctgggcactgtggctctgcctgcagttgaatttacttgggagtttgaggtgagaggatcacttgagcccaggatttcaagcgcagcctggaaacatagctagacttaatttcttgaaaaaaatcagtgctattttggggtcagaacaatcctcaaaaatggtccccggaaagaagcccaaatttaattgcaccagattgttgaacgatttatgctacaaaacagtgtcacatgtcagaaatgagcacagtgtaacatctacatggtttgttaagagacacaaatggtcaagtagaacaatcaggtaattag**gctgtcgaaggcaacactgccaaatgacagc**atttccgtggaaactgcgtgtacatccaggactgcac**ctgtgaacgatgacatcatacccttcacag**tgtcgaggaaagagacatcactcaaacagacaagccaagggacttcagaaaatataaggggaaatacagtgtgcaaatatgtaaaaaatgcaataagatgattactcctaaatgaatatcaagacacaatcacataatatgaaattaaattttcctgaatgataggattactaccaatcaccccccaggacaccctcatctactctgtgcacagccttctcgtcaggcgtcccagcccagaccttgctatgtagcagaagacatgcaaataagacccccctttttgctgatgaaaagcagcccagccctgaccctgcagct**ctgggagaggagctccagccttgggattcccag**ct**gtctccactcggtgatcggcactgaatacaggagac**tcaccATGGAGTTTGGGCTGAGCTGGGTTTTCCTTGTTGCTATTTTAAAAGGTGATTCATGGGGAACTAGAGATACTGAGTGTGAGTGGACATGAGTGAGAGAAACAGTGGACGTGTGTGGCACTTTCTGACCAGGGTGTCTCTGTGTTTGCAGGTGTCCAGTGTGAGGTGCAGCTGGTGGAGTCCGGGGGAGGCTTGGTCCAGCCTGGGGGGTCCCTGAAACTCTCCTGTGCAGCCTCTGGGTTCACCTTCAGTGGCTCTGCTATGCACTGGGTCCGCCAGGCTTCCGGGAAAGGGCTGGAGTGGGTTGGCCGTATTAGAAGCAAAGCTAACAGTTACGCGACAGCATATGCTGCGTC**GGTGAAAGGCAGGTTCACC**ATCTCCAGAGATGATTCAAAGAACACGGCGTATCTGCAAATGAACAGCCTGAAAACCGAGGACACGGCCGTGTATTACTGTACTAGACAcacagtgag**gggaggtcagtgtgagcccggacacaaacctccc**tgcaggggcgcgcggggctaccagggggcgctcgggactcactgagggcgggacaggtcccaggaacaggtgcagcggtaggttttct**ttctcctcagctggagaa**gtcaggtttgtgttttcagaactctggagtcttacaggttgctacattttcatacagttattagtatgtatttattatcattggtattta**agttttaataattttaaaccttttatgtagtgttattttttaaaact**gtttactttcatttgcagttattcttccagagtttcattaacatctatt**gctatgagcaactacatagc**tatgagagcataa**atttacacctgtagacgtaggtctaaat**gccacaaacctgtgcataaatgtatagtgacttatatttaacattataataagataatttttaaaatatatcctaaacgatcaaacttactgatgaacta**aatataaattatcagagtaatgcataattgatttcaataattttatatt**gtttatattaattaatatctatttctttactgaaacataatatattggtcatttcaaaatagctacgatacatttcaaatggccttgatgctaataatgaaaagattttgaggtgattaatatgc**taattagttatatttaatta**ttccatattgtattaatatatcaaaacattgctttgtacctcataaatagatacagtcgtaatttgtcaattttcaagacagttatttaatccatcctgggtcataatctttttttcccctgtccatggctgattggattgtcctgataaacccatccatac**gcctgcctcccgcaggcttctgagacgggctccatgaaggccagaagc**aggcaccctgtgagagtccacacgacctggagcctccctctccttggattaggccatctcctcgggatcgcagggctcttcattatcctcacccccttgt

Uppercase: IGHV3-73-2

Lowercase: Flanking sequence[1000bp]

Red & Bold & Underline: Stem-loop [15]

Blue: Heptamer[32]

Green: Nonamer [6]

id-IGHV3-23-2[V_gene_segment]

gctttgtcgtcc**aggctggagctccgtggtgagatcatagctctctgcagcct**cagaccctgggtctccagcaatcctcctccctctgccctatgtagctgggactacaggggtgcaccaccatgctcggcttcagtgcgtattaaaccgcacacttcaattatgcagtatttattatatagcaataatgcctcaataagggtatt**acaaataagtggataatttgt**taaagattgatggaaagatagataccaacatgagaaatgtatgacactc**aagaaaataaaactgtaggaaacttgcttttctt**tatatttgttaggtaatcaccacagtgtgtacacatcacaccatgttcccattacagagaaa**aggttctgcgaacct**cacgagctgtgacccctgtgtgctgggcttggttcagggagaagtcaggtccagtggtgagaagcacaggcccagatg**cccaggctcactctgaccaaatgtgagcactggg**gacattgtaaaacccacctgtgcttttgctgataatttttcatctttaacatggaaataatattgatactatataccatggtttctctgcgtatgtaaaaataaaagatgattggtgctaactttaaaaatatg**cagtttatgtagatctatggtacctcaataaaactg**ttttaaaa**taaaaattacaaaattataagattttta**ggttttaaggtttaagtttatcacaaaacaaactgacaatag**gaaagcacaatttcccaatgctttc**aatatcacagatctccccgaggacattctgacatgctctgagccccactatctccaaaggcctctcaccccagagcttactatatagtaggagatatgcaaatagagccctccgtctgctgatgaaaaccagcccagccctgaccctgcagctctgagagaggagcccagccctgggattttcaggtgttttcatt**tggtgatcaggactgaacagagagaactcaccA**TGGAGTTTGGGCTGAGCTGGCTTTTTCTTGTGGCTATTTTAAAAGGTAATTCATGGAGAAATAGAAAAATTGAGTGTGAATGGATAAGAGTGAG**AGAAACAGTGGATACGTGTGGCAGTTTCT**GACCAGGGTTTCTTTTTGTTTGCAGGTGTCCAGTGTGAGGTGCAGCTGGTGGAGTCTGGGGGAGGCTTGGTACAGCCTGGGGGGTCCCTGAGACTCTCCTGTGCAGCCTCTGGATTCACCTTTAGCAGCTATGCCATGAGCTGGGTCCGCCAGG**CTCCAGGGAAGGGGCTGGAG**TGGGTCTCAGCTATTAGTGGTAGTGGTGGTAGCACATACTACGCAGACTCCGTGAAGGGCCGGTTCACCATCTCCAGAGACAATTCCAAGAACACGCTGTATCTGCAAATGAACAGCCTGAGAGCCGAGGACACGGCCGTATATT**ACTGTGCGAAAGAcacagt**gaggggaagtcattgtgagcccagacacaaacctccctgcaggaacgatgggggggaa**atcagcggcagggggcgctcaggacccgctgat**cagagtcatccccagaggcaggtgcagatggaggctgtttc**ctgtcagggtgtgggacttcatcttcttctgacag**tttctctagtgaacctctctaacctcagaattctgtgcttactaatgtcatctctacgtatttt**ttaaaagatcattttaa**tatgagcacctattctcacacgcaccaaatgcagattgacgcttacagagatgaaaagttctcaaccatg**gtcaccaggatgagagtcctgaggaaactcaggggtgcctggtgac**tcttccgcaatcagtcctgggacagaaacctcagggacagcctttatgagttttgatcacatctaacagagaggatgggccagggccagagtcatgta**gaacctcacaggttc**cacgtctgacccttctcctgacactaaagccaatcagcatcagcactgacctggtgctccttttgctcccaatccatgttctttctttggagagtttcttctcccttttttatttgcttttc**ctgtttcctgaaaaagaaacag**atggtccccttggtccacattccagggctcaaggcattttcttggagctcaggtggggctcaggct**gtggctcctgcagccac**gt**gggagaggctgatgggacttccttctctccc**cattttctcaggaccctccactgtgttgtgtgtaggctcatctgggaatgcagttgaccattaatactgaaggggacaaactcatttgatcaaaatgggatgtggatgtggagttaaccctgttccgtgcacagtgtcagagtcgtctgcttcagaagtagtgttagaaagagctggtgaaatttatcagaatcaagatggagccacttgtgttaaagccctaatgaatgaagctggggaaggccatgaaggagggt

Uppercase: IGHV3-23-2

Lowercase: Flanking sequence[1000bp]

Red & Bold & Underline: Stem-loop [20]

Blue: Heptamer[42]

Green: Nonamer [6]

id-IGHV6-1[V_gene_segment]

tgaggagtggtggcacccccgttcccac**cccacagatgcaacgcctgtggg**tgacgcatgtgagtactgaggagtagagggtagaactgtaggc**cccgagaaccacagaaactcggg**tgttacactctggggccatgtaaggagaa**agtgtcactggacagaaacaggcccctcctagacact**gtgtgcgccatagtcacctgtcattagctctcactcttgcagattcatgattgaggtggttaaaaaaaaaaaagctcctactcacccatcca**accccatcctggggt**gtttccaccacccttggggtttgggatgagctgcccttgcccactgtgctctgtggacctccctttagaagctcacagctccctgcactcggctccatcctgccccaccacacagaagcaaaacccctctcctttccactgcaggcttttcctggaccagaatgctgacctgctgcccttcactcccgaagtggtgggactgcctggggtggtgtgggtgttgagccttcttactctagggacctggcacctggccccaggggcacagggatggtgcatctgcctagggatgcctcctcatgccagggggtgggggttagtaccatcggccctcaggatttgttgcatgaatgagtgaatgggtgaataaatgaaggggatctgatctatgaataagggtatatagactttggttgatgtaggacgccaaatgctggaatt**tcagagtcatcacacccaggggccctgcctctga**gctcctctttgcatccaatctgctgaagaacatggctctagggaaacccagttgtagacctgag**ggccccggctcttcaatgagccatctccgtcccggggcc**ttatatcagcaagtgacgcacacaggcaaatgccagggtgtggtttcctgtttaaatgtagcctcccccgctgcagaactgcagagcctgctgaattctggctgaccagggcagtcaccagagctccagacaATGTCTGTCTCCTTCCTCATCTTCCTGCCCGTGCTGGGCCTCCCATGGGGTCAGTGTCAGGGAGATGCCGTATTC**ACAGCAGCATTCACAGACTGAGGGGTGTTTCACTTTGCTGT**TTCCTTTTGTCTCCAGGT**GTCCTGTCACAGGTACAGCTGCAGCAGTCAGGTCCAGGAC**TGGTGAAGCCCTCGCAGACCCTCTCACTCACCTGTGCCATCTCCGGGGACAGTGTCTCTAGCAACAGTGCTGCTTGGAACTGGATCAGGCAGTCCCCATCGAGAGGCCTTGAGTGGCTGGGAAGGACATACTACAGGTCCAAGTGGTATAATGATTATGCAGTATCTGTGAAAAGTCGAATAACCATCAACCCAGACACATCCAAGAACCAGTTCTCCCTGCAGCTGAACTCTGTGACTCCCGAGGACACGGCTGTGTATT**ACTGTGCAAGAGAcacagt**gaggggaagtcagtgtgagcccagacacaaacctccctgcagggatgctcaggaccccagaaggcacccagcactaccagcgcagggcccagaccaggagcaggtgtggagttaagcaaaaatggaacttcttgctgtgtcttaaa**ctgttgttgttttttttttttttttggctcagcaacag**agatcatagaaaaccctttttcatatttttgaaatctgttcttagtctaatggagattctctaatatgtgacaatgtttttctcttgctgtttttggaattctttgtctttgacttttgacaacttgacttttgacagtgtgcctcaaagaagttctattttgggttctgtgaacctcctggatctgggaagttttcagctatgatttcattaaacgtgttttctacaccatttccctactcttttggaatacccataatgcaaatatttgttcacttaattgtgtcccataaatgctggggattttcttcattcctttttactctttttttctttttattcatctgcctgaattattt**caaaagatctgtcttcaacttcagaaactcttttg**cttggcctagtctaatcttgaaggtctcaattgtacttttaatttcattcattgaattcttcaactctggaatttctgttggttcttttttatgatacttatctctttg**ttgaattcctcattcaa**atgataaattgttttcctgatttcactgaattttctatctgtacactattgtatctccctgagtttcttagagattatccttttgaattatttttctgacattctgtatatttccttatgattggggtctgctactggagaatgactgttgtctttttcaggtgtcgtgtttcctggccttttcatgtttt**atgtgttcctacgttgatttctacacat**ctggcggaccagtcatcccttgcaatttaatggagtaggttttgcaggaaaagacttcctagtacagacgg

Uppercase: IGHV6-1

Lowercase: Flanking sequence[1000bp]

Red & Bold & Underline: Stem-loop [13]

Blue: Heptamer[34]

Green: Nonamer [5]

id-TRBV11-3[V_gene_segment]

gacatgtgtcccctaggttttaggaacttggatttctttaggctgagttgtactattccctttctctgaacacacatgtcattggttagttcattcatattgattctcaggacttctccaatccctcaagccttcccactctaccactcagataaggagaagtgtgtcttaactaatgttgcat**aattttttaaaaattgctggatactaacactgcaatt**caattgtaaaatatgatagaggtgtaagtgttttagatcccagaagtcctgctcaattttgcatttattgttagatattcaccctgatattttcaggaattagagtgttcttcatccatttgggttaagatatcagaacttactttcaattcattggtgaccagagactacaggcagaatcatagtgaaggcttgatacataatacataatgaagattgataacagcaaaaagggaactagtcattggccttcagataaaatgaaacctcccaactttgtgctcagggaattattttacccctttaggttagagtatgtcagctagttcaatgcagt**tctaggataaagctgtgtcaattaccctaga**ggactgggagcatggaggacctggtagaaaagaaatacccaaagagaaggaaaattacacataaaatcaagaggtcactaattgatatatctattttgagttgcctttgagacttccacgtagaa**acatctagcacaggataggcctagagatgt**agtgacagatctaatccc**ccaagccctgaatatgtgagtgaggacgcttgg**atgaacttccagtctatgaatacattaaacaaacttttagctctgttttagtttacaacaactgttttctaagaacttaagcatttgtggagacaatgatgtcactgtaggaacttctctgtaaggacagcaacatcccacttcctctgctcctgctcacagtgaccctgatctggcaaagcttccatcctgccctgaccctgccATGGGT**ACCAGGCTCCTCTGCTGGGTGGCCTTCTGTCTCCTGGT**GGAAGGTGAGTCCTAGGAACACCATGATCCTCAC**ATAATCTCCCAGTGATTAT**TCTAGTCATTTC**CTTCTATTTTAAAATTCTTTTTCCCCCACAGAACTCATAGAAG**CTGGAGTGGTTCAGTCTCCCAGATATAAGATTATAGAGAAAAAACAGCC**TGTGGCTTTTTGGTGCAATCCTATTTCTGGCCACA**ATACCCTTTACTGGTACCTGCAGAACTTGGGACAGGGCCCGGAGCTTCTGATTCGATATGAGAATGAGGAAGCAGTAGACGATTCACAGTTGCCTAAGGATCGATTTTCTGCAGAGAGGCTCAAAGGAGTAGACTCCACTCTCAAGATCCAGCCTGCAGAGCTTGGGGACTCGGCCGTGTA**TCTCTGTGCCAGCAGCTTAGAcacagtgtagcagaga**cacttccctcctgtgcagaaaaccgcaggactctctcctctctactcagctcacagcagc**ctttccttattcctcatcctcccaggaaag**aagtgagttttcagatatagctaggattcatatagtgggaggaaatgaactatttcttacaacatgagcgctataattgttgcttgaaaatgtgtctcagggatttgaaacac**ttcttggggacaactcaagaa**acctaaagtgacttatcagagattgagtcctcaggggtagtgatggtttatctcccataaaagtatgtgatcatgaaacatgtacaataaacttaataatacagatagtgattataccaagcagcaaacagcaggctccccttatgtagttgtcagcatgtgaatgacttgttacccttagtgacaccccggctgtctagcaaagtctc**aacacacagtatgtgtt**tggaaatctttttccttctgttttatttttggtaaagtatgctaataatgatgttgttaatgatagtgattataatttcaggtatgttattcagccctttttccagttcaaagaaatgcacacaaatggataccccggcaaccaggtgcaaatcttagctccacatgaattctacttctcctgaactctctaactcttctatgtaatcatcccagcccaagaatatatctagacctggaccagtcagttcagaggacatttctacgtgtctgccttgtatattaa**aagttccttagtggattgaaatattccttgtaagaactt**ctgaagcaatcaattgccctctctgataatatacactctgtgtgtctagctccatattcattactctacttctgcctaaactctct**aatactagaaatctttctgaagtatt**ggttagaattacttaaaaagcagcatactggaaattgtctgaaagcactgtcactgtccatttgcccctcctgacttccctcccctccttatccaatc

Uppercase: TRBV11-3

Lowercase: Flanking sequence[1000bp]

Red & Bold & Underline: Stem-loop [15]

Blue: Heptamer[38]

Green: Nonamer [1]

id-IGHVIII-11-1[V_gene_segment]

aagaagatccatgctacattattcattttcttgttcaaatctccacagctttattaggtgctgggacctcatttagtttggatcctgcatccttacagcaaagctcatccttttgtttttgaacacttccctgtttcctgatattacgatagattctaagcttgtttctttatcacctttttcatacatagaattagccatttatataaagattgcttgtttctg**attttaaagagtagtgttaaaat**aaaatattgtgataatgggtatgtgtgttgttaatgtggtataagtacttctgggacctctcaaccttctgttctagcaaatgtgcatgtt**tatatgaatcatgtcatgtccatata**tccaatgttgaactatgttgcgaaaagtgatctcaa**gataatggtaattatc**aaccccctccctaacaaggtgtagatctgcatttttttttcattgtaactcaactttgcctcatggtcaggaacagtggtttccagctctaaatgtactgattacagcgagatgtccgttttccctggaaatgtatttttaagttcttactggatgtatttgttgataatgttttctactatgaagatacctgaacagtgtccgcactagagagtaagaaagagtaatgggcagattaaccctgtgcatccagacccagga**atcctctgatcctgccctccctgaaacggagacacagaggat**ggatgagcaatgctgagcggtgcacccaagaccacaaaaagaaagacatggaaatatgtcccctcccctcctcaagaaaggcagctcatcccctgttccctcaggcccgggcgaggagccaccccatgtctcttcccttcc**tcagtgtccacactgtggggtctgcactga**tctgggcttcccttctcatcaccctcgtattagtggcccttgtgaatcaggtccagctgtggctgctgctcatggggctgctctcagtctgctctctctgtgtttgcagAAGTCCTGTGTGAAGTTTACTGATGGAGTCAGAGGGAGAAAGATTGTACAGCCCAGCGGTTCACTCAGCCACTC**CTGCAAAGACTCTGGATCACCTTCACTGATTGCAG**CATAAGCT**TGGTCCAGCAGGCTCCAGGACCA**GAGTTGGCGTGGGCAGCAACAGGGAGAAATTCAAGAGGAAGTTCTTAcatgcacccttacgtgcacggtctcactgagatctttacttcctttatcacgtttgttctgtaaatcacaacgaatggtgcattcttcatctattatacacttgttaagtct**ttttttggcatcttttaaaaaa**ctggtaactttatcctgtgtaatatccctgttaagtcctaaaagtcttttttgatgtctattttttcttaactttacacagctactatagatttattttggttcatatttttgtaatccatgttttctcatctttagtttttacatttgtgaatacgtacagtaattttctaatacatagcttc**tagttggggcttgctttttaaaatcaacta**taataagctctatt**tttaaactaatattatttttctgttatattgtttaaa**ttagcatttcacaatgatgtttatttctctactaacatataatctaattcacttttataaatattatattggttaccataagttt**tacaataagaattgta**tataattagattctataagaattgtatataattggattctaatgcagatactgtgatggccttgatatgaagaacagagacagtgtaactgtgtgctttgaattcctccttctcactgctctttcttgtttattcccagtttgcacttacatatactataaaacataatatataaattttttattcctttcatagttatataatatagcaattacaaagataaaaactgcacttcatcttgattttcccattttgtacagtcttaatttcttcgtatagatttatgttttgaatgtatatcacacggctactagcagagaaagtttgttaaaatgtgcactgaagcatgaatgtgctgacaataaattgtctcaagatctttttgtaatgacagaattttatttgcctttcacttttaatgaaaatgt**aatgcacgtagaattccagtttgcatt**tcacttgtaatttattttcttgtgtttattattttgttcatggag

Uppercase: IGHVIII-11-1

Lowercase: Flanking sequence[1000bp]

Red & Bold & Underline: Stem-loop [12]

Blue: Heptamer[40]

Green: Nonamer [4]

id-TRDV3[V_gene_segment]

gaaatgaaggaccttctagaaatactcctaggaa**cactcctgcattttggggagtg**agttagtcagtgttctgtttttcacagcagtgcctcccaactgttgtcatattgatggcttacatggaaaactgtatttgtagtgctcctgactgggagg**ggctggctcaggggctccagcc**accccaggtacatgggcactctgaacactatctcagcaagcttataacccacttatggcagttcacctgccgggaagctctgttcaggattttccagctggagccaagtgtgtgctgcagtgggcccagtgagagatggaaagtcaagctggtg**aagtgcaagaagcactt**aggatcaggacactccctcaagcatct**gaaatcaaagatttc**tggaattcaggtctagga**cagattttagaaagaatctg**gtccaaccccctttcct**tcctctttacacagagga**gtaagggagttgtccaaggtactctttcacttattccatttattgaatagaacaaagtgtgtggagccgtgctagatgctatgagacagtgtcactcaaagtacgacatgtgattaatggactagtacaccagtctgtgaccagaggatacagaaatggagagtgaactacttttgtagcagttggatatccaagaacgccatcattagtcttgcagttttctttcctccctcacttcctctctctctcccttcctttgtttttcttttcttttcttttttaaaagt**aatccatttttcattgtcatatgtgttagttcaagatggatt**ggggccaaaatcataatagcaatagcagtttccttggctgagcgcatttgcctctagaagtcacagctgaatcagtcacatgaccgtcagcgctgccagtctcctcttgttttcctcccgtcttctttcattagaaccgtcagatagtctctgagatcttaggaaccctgttttgttgtatatcctctgccctgatatgaagaaaagATGATTCTTACTGTGGGCTTTAGCTTTTTGTTTTTCTGTAAGTAGTTTCATTTGTTTACATTCCACTTGCTCTATAACCCCATTTCTCCTTCTCTGACTCTTGAAGGCTTTTTCTCTTGCACTCTTGGTTTTTGGTTATTTGCTGCCTTTGGCTTTATTTCTGTGTCCTTCTCTTGCCTCTGCTCATTGAGTTCTTCTGGGCTAGGCATGTGCTGTACTCACTGTCTGGGGGGTGCAGAATTCACTATTTCCTCATTGTCTTTTTCCCAGACAGGGGCACGCTGTGTGACAAAGTAACCCAGAGTTCCCCGGACCAGACGGTGGCGAGTGGCAGTGAGGTGGTACTGCTCTGCACTTACGACACTGTATATTCAAATCCAGATTTATTCTGGTACCGGATAAGGCCAGATTATTCCTTTCAGTTTGTCTTTTATGGGGATAACAGCAGATCAGAAGGTGCAGATTTTACTCAAGGACGGT**TTTCTGTGAAACACATTCTGACCCAGAAA**GCCTTTCACTTGGTGATCTCTCC**AGTAAGGACTGAAGACAGTGCCACTTACT**ACTGTGCCTTTAGcactatgatgcag**gtgcccaggaagtcataacacaaactcctggggcacagctcagcagagctg**cctcttagggcaggtcatgtctgggacttggcatccttctcttagccattttgggttccagcctggc**ctttagaagtgacctaaag**tgtcttctgagaaataaatggcctggtcagat**ccaggggagcaccctgg**cttcagagaagcagagtatctccccacttctgactcacggggcccttgccacctgaatgactcctggtaggcaaaggagaactgcatgtttacagagcgctgtcttcgt**gcccctggtgccctggtcgtcccgatacagaaaggggc**ttgacaagcctgtg**ctgcattttctgtgagtgtttcaggatgcag**gttcaggtgcaggtaaagggcccatctaaggcgaggccgtcagccgtgtctttattttagtcaccaccaccatttatccattattatgaattgatctgttttcattgcagaattaaattagggccagatgtggtggctcatgccaataattgcaacactttgggaggctgaggcgggcagatcacttgaggccaggagttcaagaccagcttggccaacatggcgaagacccgtctctactaaaaatacaagaat**tagccgggcgtggtggctcgcgcccgtagtcccggcta**ctcgggaggctgaggcacaagaatcgcttgaacccaggaggcagaggt**tgcagtgagccgaggtggtgccactgca**ttccagcctgggt**gacagagcgagattctgtc**tcaaaaaaaaaaaaaaaa**ttaagttagacacaaagttatttacttaa**aagacttttcaccaatttactttttttttcccgcatctttttatatccagccaccttctccttgatgttctttgacacctccattttacttattttatttggagaacattttcttttttttcactgcagagcactgaggcactcctggaaagttgtgactt

Uppercase: TRDV3

Lowercase: Flanking sequence[1000bp]

Red & Bold & Underline: Stem-loop [19]

Blue: Heptamer[33]

Green: Nonamer [5]

id-IGLV3-7[V_gene_segment]

atcacaag**gtcaggagattgagaccatcctgac**acggtgaaaccccgtctctactaaaaatacaaaaaattagccgggcatgatggcaggtgcctttagtcccagctacttgggaggctgaggcaggagaatggcgtgaacccgggaggcagcgat**tgcagtgagccgagatcgcgtcactgca**ctccagcctgggtgacagagcgagactccgtctcaaaaaaaaaaaaaaaaaaaacagataaataaaaaataaatctgagagatggctgagcaatttcactgagtttcatcatggaattctggcctcccaaacaccagcatggcccctgttcctacaatctcctgcttcaagatcaacccaaccctgctacccagaccagatgctagatggtcaagacagggctgtgagagctaaagaatcttcagggtccacgatgaatttacattggagcttccttttccacggatattcacatcaatagagaagatgactgagttcttgcctacctctgtgaatacctgagacatgaactgagatgtgtgtgtgcatatggtgcgggcagcac**ttcctcactcaatactagggagagtgtgaggaa**actggatacagacctggagatagggcacatttttacactctctattaagccctgtccttctgtgctcaccaaggagtcacccaacaaggccacaacagg**aagagaagcaggtttctatgaagagacttctctt**tttgtgatgagggcccaggaagtccactgtctagctatgaac**tcagaggcagaactctga**tgcatctccaccatgacctgaatccctctcctctactcacccttcctgctgtcggcacaggttctgccttc**cagggctcagccccataagactagggatcagcctggccctg**accttcagctcagcacagggagtgacgcagtgtgtggggctctgagaatgagaccctcatcttcagactcacctctcctgtcctatcttgtag**GCTCAGTGGCCTCCTCTGAGC**GGACTCAGTTGCCTGCAGCGTCTGTGGCTTTGGGACAGAGGGCCAGGATCACCTACCAGGGAGACAGCATAGAATACTTTTATGCAAACTTTTG**TACCAGCAGAAGCTAGGACAGGTCCCTGTGCTGGTA**ATCTATGGTGACAGCAACTGGCACTCAGTGATTCCTGAACAAC**TCTCTGACTCCATATCAGAGA**ACATGGCCACCCTGATAATCAATGGGCCCCAGGCTGGAAACAAGGCTATTACTGTCAATCATGAGACAGCACTGATACTCA**TCTcacagtgacacaggcagacaggaaagtgaga**cacaaaccccttcccacctatgtcaccctccccctccaacctcaggaggacagtggtcagagctataagcaggtctggcccagttcactcagatttgagaccccc**aggctgcccttccctccagcct**tccaggcaggctcagcagagtgtaaattaagagtggatacacaatgaattattagaattgtatttcagtttttcaagaagtagaggaatgattgaacatgttaagtggaagtatagaataccaaaaaaggactaattgagcttgtactgctaataattacagcatctgagacattaaaaacacactggatgtgattaacaacagatgatgcaacacaaaagattagcaaactcaataatgtagcactagaaaatattcaaaatgcagcacataaagaaaaaataatgatgaaaatgaagagagtgttacagtatatttatagaaaaaatgaagagtgagttataggacaacttcaaacagcctgaggttg**gctgggaggggtggctcacacctgtaatcccagc**actttgggaggtctcggagggagaatccttttgagactagaagctcaacaccagactagacaacataacgagacttcatctctaca**aattttttttaagaaaatt**aactaggcatcatagcctgcagctgtagtcctacctacttaggagcctgagacaggaagattgcttgagcccagaaattcaagatg**gcagtgagtgaggtacgatcatgccactgc**actctaacctagga**cacagagcaagttcctgggtctcactctgtg**gttc**aggctggagtgccatggtgcaattacagctcactgcagcct**tgacttcccaggcccaagagatcc**tcccagctaagcctcccaaagtttctggga**cta**caggcatgcatcacatgcctg**gttaattctttgtattttttgtagagatggggttctccctatgttgccc

Uppercase: IGLV3-7

Lowercase: Flanking sequence[1000bp]

Red & Bold & Underline: Stem-loop [18]

Blue: Heptamer[43]

Green: Nonamer [5]

id-IGLV2-28[V_gene_segment]

atccattaatattta**ttttaatatggaacattaaaa**gagttttgtaataggcaatgggaaaaggtaaaaaagataaaagc**tttttgagaaaaaaaacaaaaa**caaacttatagaatgtctatgtgtcttcagtgttgcagggttgagagagatcagctttatggtg**ccccatggatgtgctgagatgggg**aaaatcaagaaagaaggaagtatgtaccactgtgactcaacaaataaatcaaagtaaaatg**gacttaaatctttcaataagtc**attcacaataaatatcactctggtgcacacagaaatgcctgttagcaaactttcccttccccctggccatgtctccatggatcacctgtgtgcagagtggctccggtcagtgctctgtgttccctc**cacagggctccccacaacatgtgccctgtg**tggatgtgggt**gcatcgcctggggtggctgatgtgctccagggattcgatgc**tgcagatgcctcagcaccaagggtgggggtgggaatgggacaaggctgcagctggggagaccaaggatcaaccacacacagcaggggtttccacggggtgagcagcctcatttcctgacagtagagcgagcctaaaggtaggtg**ccaggcagtggtatctgtggggtggcccaggcctgg**agggtcacagagacacaggcaccacacagcagagacactgagggccaggagctcactcaggtgagggacttcagcagatctttctctcctggtacaaagagatcaagttcctaccatcagaatagacaaacaaggactttaatctcctcagctgagccccactgtccagggaagaagtctctgagcccaggcccaggtgagagtgggatgaggagaggagccaggacacagattttcatggatgccccgccctcctctgaggcagagcggataagacagggctgggggcatacccagtgcttgagtctcaggaggcagcgctctcgggacgtctccaccATGGCCTGGGCTCTGCTCCTCCTCACCCTCCTCACTCAGGGCACAGGTGATGCCTCCAGGAAAGGGGCCTCAGGGACCTCTGGGCTGATTCTTAGTCTCCTGCTCCTCAGGCTCACCTGGGCCCAGCACTGACTCACTAGAGTGTGTTTTTTCCTTTTTCCGGGCTCTCGGGCTCAGTCTGCCCTGACTCAGCCTCACTCAGTACCCGGGTCTCCTGGACAGTCGGTCACCATCTCCTGCACTGGAACCAGCAGTGACGTTGGGCATTATAACTATGTCTCCTGGTGCCAGCAGCACCCAGGCACATGATTTATGAAGTCAGTTATAGGCTCTCTGGAGTCTCTGATCACTTCTCTGGCTCCCAGCTCTGGCAACATGGCCTCCATGACCATCTCTGGGCTCCAGGCTGAGGACGAGGCTGATTATTACTGCAGTTCATATACAAGCAGCAACATTTTCcacagtggacataagattgattctcaggctccaagtctggccagtgagcttctttgagactccctgggatcccagcagtgacactg**atcactattgctgtcccacacatcccaagtgat**gaggaagtaagaccaaaaccctcctgagattcctgacttgtgtcctgacaccagggctgttgggattcctgtctttccttcaagattgttcaaataagcagtgacagtcacttccatgtgagatatgagaagaagaataattccatatgaaccaaatccacgaagcctacactgaagcctgccaagttctctgcagtccagcgcgatgaagggagcattttcgtgggaacttttgggaatatgaggggagccaggctcagctatgggttccagagacaaagctcctggatgtttctccccggccttggtccatcacctgctacctctcctcactctttccacagacactgccccaggctaagcccccacaggaccctaagcctattcctgaccctgagctcagcccaggtgatgct**tcaggagatgggatcctga**gaatgagccccaacatctctttaaccta**aggtgtggggtcatgagacacct**gttagccataagacacctgtccccca**cctgctcaggagacagtggagcagg**caacagtgcagagagaaccagcaggagccaggccaggccctgcatccaaagcaggccccagtccccacagattctccagctct**agcctgtggtacagcccccactttctgcaggct**gaaatgggcttctcaggatgggcaggccatgatgggggatggcag**tgtgtccacaggacaca**gccctgctgcctgcctgaccctcatgccccctccttccccaccctggttaatctccaccagaaagggc**ccaggttcagaactcaggacacctgg**atcagctcctggacttgcccccactgctctctgttctcctgagccacttagtctcttttggcctcagtgtta

Uppercase: IGLV2-28

Lowercase: Flanking sequence[1000bp]

Red & Bold & Underline: Stem-loop [14]

Blue: Heptamer[42]

Green: Nonamer [4]

id-IGKV3-31-3[V_gene_segment]

ttttagcctgatacaaag**agaagccagggaaagcttct**gagctcctgatctacgatgcatccaatttgcaaacctgggtcccattgcagttatgtggcattggatccaggacagatttgattctcaccattagcatcctccagtct**gaagttgctgcaacttc**ttattattggtcaacagtataaaagtgaccctctcacagtgttacaaacccaataagctccccaag**gaagcagatatgtgagggtgggctgccccagctgcttc**tcctgtttcctccatct**gctgagagtgtttctcagactcagc**cacactctgaaggtcactgagcagttttcatagaagggg**taagggaagtgtctctacaccctta**gtttcttttatcctcctcagccccagcagcagacatggcaatgcctctcctaatttcataga**agagtcattacctatgaggactct**gggttacagcactggtccaggttcatacaacaaaagagaagctattgtaggtaacccaaatagaaagtttttcctaatatgggaaatccatgtctaaattacaacttttcaaagaccagaggatataatggttaggaaaccagaaaggcaaaaggagtgctgggaaatctacttcaagaagcaatggtggccgcgcacagtggttcaggcctgtaatcccagcactttgggaggccgaggcaggcggatcacctgagatcaggagttcaagaccagcctggccaacatggcgaaaccccgtctctactaaaaatacaaaaattagccgggcatagtggtgggtgcctgtaatcctagctactcagaaggctgaggcaggagaatcatttgaaccaggcaggtggtctctgcagtttcgaagcccagcgccatct**gtggctgttatgcatgtctctcccagccac**cctgctgtccagagcccatatcaatccgtgggggtaggtctgtgaaaga**gcagctcagttaggacccagagggaaccATGGAAGCTGC**AGCTCAGCTGCTTTGATTCTTGTTACTCTGGCTCCCAGGTGCAGGAAACACCGGATGGTTCTGCATGTCAGTGAAACTTT**CTCAACCTTGTTGAG**TCCTGTTACCTGGCACACCTGCTGGGAAGGCACAATGATTAAAGCTCAATGTAGATCAATGGTCCTGGATGCACTGGGAAGACAATAGGTATGATGTAGTGTACGTGTGTGACATTTCTGTTTTTATTCCAATTTCAGATACCACCAGAGAACTTGTGATGACAGTCTCCAGCCTCCCTGTCTTTATCTCTAGGGGAAAAAGCCACCCTCACCTACAGAGCCAAGTAGACTATTAGCAGCTCCTTAGCAGGGTATCAGTGGAAACCTGGACAGGCTCTCAGGCTCCTCATCCATGGTGCA**TCCACCAGGACCACCAATGTCCCAGCCTGGTGGA**GTGGCAGTGGGTTCGGGGAAAACTTCAGTCTCATTATCAGCAGGCTGGAGCATGAAGATTTTGCACTTTAA**CACTGTTATCAGCATAGTGGTGGGTATtccacagtg**attccac**aggaaaccaaacctccacaagacagctggtgttttttcct**caagccttctgtttacttatgggaagctactatggtggctgct**tagttattgagagaaaacaatggagacttcaataaata**g**aagtttataccacattcatgcattataaaactt**aatattgtgagtaacataattctcccttaa**ttaattcaatacagtgacaattaa**aatcccagaggacttttgtgaaattgaaatgatgtttcatatatatatagtatatatatatatatatatatatatgtactcacataccacgcactatatata**tatatatatacacacacactatatatggtgtatgtatatatata**gtgatatgtatgtgtataatatatatatagagagtg**tgtgtgtatatatatagatacatacatatcacaca**tgcatttaaaaaggacaaaaatagccattggaatcttgcataagaataacaggtcgg**gctgggcgtggtggatcactcctgtaatcccagc**aatttgggaggctgaggtgggcaggtcacctga**ggtcaggaggttgagactagcctgacc**aatatggagaaaccccatctctactaaaaatacaaaattagtcaggtgtggtggcacatgcctgtaatcccagctactcaggaggctgaggcaggagaatcatttgaacccgggtggtggaggttgcagtgagctgagatcgcaccattgcactccagcctggggaacagagccagaatctatctcaaaaaaacaaaaacaaaaaacaacaacaaaacaaaacaaaaaaagaataacagggcaggcatggtgtctcacgcctataatcccagtactttgagaggctgaggcaagtgtatcgcttgaggtcaggagttagaccaccctggccaacatggtgaaaccttgactctactaaaaaaatacaaaaaattagccaggcatggttgcacattcctgtagtccca

Uppercase: IGKV3-31-3

Lowercase: Flanking sequence[1000bp]

Red & Bold & Underline: Stem-loop [19]

Blue: Heptamer[36]

Green: Nonamer [1]

id-IGLV3-10[V_gene_segment]

tagaggagatgctctccagagggaagggggttctcacctgattcaggccagagctctgaccacaaggctgtccttggctcagtgacaagagtctgctgcattgcagagaccagggaccacccaccatccagagctcacac**tctgtgcaaaggctgtgtccctgggatatttgcagaga**ccaccaggcctaaagttgaatttcccatcaatatttca**tgtaaacttgtaaaaagcaccagaaaaacacactggactgagaaccccgatgcttttctggggccttttacaagtttaca**caaaatactcataggaaaaaataaataaataatggaagagctgaggatggcaatggctgatctatgtggggatgggggatggcctaaaaggagcggcaccatggctgaaatgagagcagacgtccagatgatgccacaagggtgt**gtggttgcgtaaatgaaccac**ctactcctactgaaataattaacaagtatcctgacgacacagtacgttatttagatgaaatgaccaggatgggactaagtaatgctgggggacccagagaggagggggaatatggtgggttcttggtccacctg**ctgctgagtcaggtttcatcaatgatctctcagcag**ggccaaactcagccccaa**gccaggtttctagaaggcccgcagccctgctgagcctggc**ccaggaca**gaccacaggtcagagaggtttgggacagtggtc**atatggccagtccccaggaaggtgactgtgatcacaggcctcagaggtcgaggctaattttctgaccaaagacccaactccttcttcagtcacctgtgctgcccccacagctagtttgagtgacatctttcaatgaggagtcccagaggatagaaatttgcatgaacaccaagcacttactgccccatagggcctgggaaggaataagagaggcctggg**gagcccagctgtgctgtgggctc**aggaggcagagctctgggaatctcaccATGGCCTGGACCCCTCTCCTGCTCCCCCTCCTCACTTTCTGCACAGGTGCTTCTCCCAGGCCCTGCCCCAGGCTCAGTGCCCATAGACCCCAAGTTGGCCCTGCCCTGAACCCTGTGCAAAGCCCAGACACAGTCTTAGGGTAGGACCCCTGGGAATGGGCTCTTGATCTTCAAGCCCCCTCTCCTGTTTTCCTTGCAGTCTCTGAGGCCTCCTATGAG**CTGACACAGCCACCCTCGGTGTCAG**TGTCCCCAGGACAAACGGCCAGGATCACCTGCTCTGGAGATGCATTGCCAAAAAAATATGCTTATTGGT**ACCAGCAGAAGTCAGGCCAGGCCCCTGTGCTGGT**CATCTATGAGGACAGCAAACGACCCTCCGGGA**TCCCTGAGAGATTCTCTGGCTCCAGCTCAGGGA**CAATGG**CCACCTTGACTATCAGTGGGGCCCAGGTGG**AGGATGAAGCTGACTACTACTGTTACTCAACAGACAGCAGTGGTAATCATAGcacagtgacactggcagatggggaagtgagacacaaaccccttcttcatctattttaccctctccctccagccccaggaccgctgtggaccaacccataagcaggtctggcagaattcacctggttaggagatccttaagctgcttttccc**tccactcctctaggcagactttgcaggaatgagtcaggagtgga**tttgggcctgg**cagggccagactgccctg**tgtttccttgcccacggtttgagttgtgaatggagagtatgagtggaaagacagacacagcgggacatccatgcaggcatgaacagcccagagcgacaggggaaactttcagctctcagttcaatgcagtcattgttttatctcttggaagaagcatttctccagtgcaactaagactctagactcacagagcccgaggttgtgggggaggacattaaattttgctcatggatcactaaggacgtttttgagtggagccactacaatttattcattgattcctggactcatggatcctgactgtgggagaaatagtactgcata**ttgggcactgctgtacagcgcatgccccatcctcaaagtcattgcccaa**ccttacatgtttttgccatcctgctgccacagtaactgagtcttcaagcagacactgagccattcatataagccagatgccacattggtgaagatggtaagacacttgaatgccatgaggatgggcccatggttgctgtttatttgctgtgatatgagttccttggtcaaagccccctgtggaataccatcatggtgggtgaagcattctgtagtccagagataaaaggtttggcagaagcactgcagtcaaagaaagcaaatccctgtctatactaagtgtgtattctggcaagagcaaagagctatccctgccatgaaggaacagctttaatgaaattaatatcaaccaggcatcctatagggatcttcctgttggtcttcactgttggcagatggtcact

Uppercase: IGLV3-10

Lowercase: Flanking sequence[1000bp]

Red & Bold & Underline: Stem-loop [14]

Blue: Heptamer[57]

Green: Nonamer [3]

id-IGHVIV-44-1[V_gene_segment]

ataatggagggaatctcttggtcccaggaagatgacaatttaatatgtctcaggcctgtggagctctctttaggagctttgtgtgtggcttctttgtatactctggctccaggaactccagtgagggaagagattcgtcacctcctgtgcgaagggggctgaagcgaggaagacaagtggcctggcttccacagaacccataagctaaaacactccgagttcccttgactgcaagaaatgcagctgtcatcacggtttcaagtgccagagtctttaatgattatgtttcaactcatgagagtgccaggaaaattccaccagtatgaactactcaaatatgtcattttgtaacattacttcattgcttgctctcatatcctccatgtttc**tgaaaaagcaaagtttccatgctttatca**tctgcaccaatgaggccatatttaagctatattttcatatgaacgtctcttgaagattttaaaatagaacaaatgttgaacaatatgcttaaaagaaaaaaaataaagaaagcagtctatgttgactcatgtgacattgataatgtatattagattatattggttgcttatttagttggtcatgggagaagagcc**ttctttccagccaaagaa**tactaagaactccctcttctaatgacgaatttcagctga**tcttgctgagaatatttagcaaga**agacagaaacagagactcctaaagggagctccagagtcctgagatacatttaagttttcattttcctggaacccagagattccccactgttctacaagacaagaccccattctatgaccctgtctga**ggagagctcaggggcagtgagctcccc**ctcacattaccccttcaccctatgcccttggggctctgtccatgcaaataagtcccgggttcaagaacgaatagaggcagggctgtgagtgttatgctgtgtgctggctctgagctggggtctctagtagcactctccacatccatagatacATGGCCCCTCCGCTACTCCTTCTCTGCCTGCTGGCTGCCCTCTGTGGTGAGTTACTCATGATTTCATACGTGGGG**AGACACGGGATGAGGCAAATGTGTCTGTGACTCACAGA**TGTTCTCTATTCTCAGGGGTTCACAGTGA**GGACCACCTTGTGCAATGGGAGGAAGAAGTAGTGGTCC**CCTTGGTCATGCTCAGCCTCACCTATGCCGCCTATGCACATTCAATTTCTGAACATTCTGTTTCCTGGATCCACCATCTCCCATCAAAAGGTCTTCAGTGTGTTGGTGTGATATGGGTTAAGGGAAACACTAAGCCCCAACCTTCAGGGCAGAGCTAGCATCTCCAGAAACACATAGTAAAAAACAAGAAAACTTACAGCTGAGAAGTGTGATGGCTGGGGAGGCAGGCGTGTATTACTGTGCTCAAGGcactgtgactcgaatccagagtgaactcagacacaaacctgccctgcaggggttcttgggaccacaaggggaaggatcaggtcaccagggtgtacttaggaaccactgaactgggtcaggcacaggaggtgggggtcagggctcctccccagggaagggcttttaatctccatgccttgtaaacctctgacaagccaggcagacacagacatatttttaaactatggaaacagatgtac**ctttcattttggaggaaaaagatcataagtgtgaaag**caaaacttaccaagggccaggtactggaataactgtaatctacataatcactctaagaattctaagaatttacaagtgcaatcatcctcatttcataaactcaccttaaataagagctttctaactgatcccatatattctgattttgtttatttcttcaacactttattatcttcaaggcatttcactgatgtcaattttagaagaatcatatgcaaacagcctctcgtggtggttgaaagc**ccacagcctcgccagtcctgtgg**atatttctggatgttctcagagtctccattgtctgtagagaagaattgtcccaatagttatttcacttttgtcttccacaaagttggaaagatggggatcactcaggggaatcgagatcaacaaacagaaagcctgaacccaccatatacaagataggatgtatttttttaggaatatattaattcccagaccctcaggctcacataccccatgtggaggggaacacacatccactccttcactcaagcc**aagctggccatgcagctcagctt**ctgcaggctcaggactgcaagctctccagtgtgggagtggagctaatgggag**gtgaggccaagtctctgccctcac**agagtctcatgatgaagatctgatgaaatatttccacagta**acatttctttggatatataaatgt**agaatatattttgaaataatgaatgatgtatcttctgatggatt

Uppercase: IGHVIV-44-1

Lowercase: Flanking sequence[1000bp]

Red & Bold & Underline: Stem-loop [12]

Blue: Heptamer[35]

Green: Nonamer [3]

id-IGHV3-21-2[V_gene_segment]

aaatatctaagtatttataccgagtaaaatatgccaaacaaataagaatatatactatgtcattt**catttttataaatgctgatgaataaaaaatg**catctgaagtaaaataatgaggatcagtagttgctgggggaaatggtagaagggaaggggagaggaggaggaatacagcagaataagaggaagtgttgagaagaattcacttgtccactttcttgataatgacgacagttacatcacgtttattaattgtacattttaaa**tatgtgaagtttattatctttcaattaagccacata**aaatgtattacaagcaaatggaaatttagacaagcacggagtaatagaa**agatacaaacatatgttaaatgtcagaagtatctgaaaattagtgtgtctggaccctagttctctccatattttcag**gtgagtgctggtggcaacatcacacatgctcttactacagaaagtgggttctgaaaaccacaccaggcacgtccagctttgtcctggagttggtttagggggatgtcacagccagtgacgaggagcacagggccagataccagcgttcacttatcccagacatgaactcctggatgcatacagagccccctccacttgtgggtttacttccac**ttctgtaaaaggagaaaatattgactcctacagaa**cataatttacacaaatatttacagatgaaatagggtgatcagggcaaagtgtttatcacagcacaatttcataagacagcatattttccaaataccatcgtcagcaaacatctgcagggcaccgtcttattatctgggtacagcctattcctccagcgtcccaccctagagcttgttatatagtaggagatatgcaaatagggacctccctctactgatgaaaaccaaccgaaccctgaccctgcagctctgagagaggagccttagccctggattccaaggcctatccact**tggtgatcagcactgagcaccgaggattcaccA**TGGAACTGGGGCTCCGCTGGG**TTTTCCTTGTTGCTATTTTAGAAGGTGAATCATGGAAAA**GTAGAGAGATTTAGTGTGTGTGGATATGAGTGAG**AGAAACGGTGGATGTGTGTGACAGTTTCT**GACCAATGTCTCTCTGTTTGCAGGTGTCCAGTGTGAGGTGCAGCTGGTGGAGTCTGGGGGAGGCCTGGTCAAGCCTGGGGGGTCCCTGAGACTCTCCTGTGCAGCCTCTGGATTCACCTTCAGTAGCTATAGCATGAACTGGGTCCGCCAGG**CTCCAGGGAAGGGGCTGGAG**TGGGTCTCATCCATTAGTAGTA**GTAGTAGTTACATATACTAC**GCAGACTCAGTGAAGGGCCGATTCACCATCTCCAGAGACAACGCCAAGAACTCACTGTATCTGCAAATGA**ACAGCCTGAGAGCCGAGGACACGGCTGT**GTATT**ACTGTGCGAGAGAcacagt**gaggggaagtcagtgtgagcccagacacaaacctccctgcaggggtccccaggaccaccagggggcgcccgggacactgtgcacggggctgtctccagggcaggtgcaggtgctgctgagggctggcttcctgtcatggcctgcggcggcctcattgtcaaatttccccagggaacttctccagatttacaattctgtactg**acatttcatgtctctaaatgt**aaaacttttt**tgtcctttttgtttctttgtttttgtaacaggaggaca**caccctcacctccacagaa**accacagtgtcactttgggggcagatgatccttctgtggt**cagcaggatgaaagttccgaggaatctcaggggaacccgaagagtgtttgccagttagactcagggcagcgacctccacgggaatctctgattagaacaggctttgagttctgataggagccaagagagacgctcacacagggtcagggtccttaaagcctgatggttttcacagctatcccccctggtcttgtaaaactgtgtacatatgactcagactgattcacttgaccccctttctgctaatccattttccttctctgcatacttgattctcacaattccctttcttctcttcccc**gaaaactgaggatgtgttttc**tgtagtctaaattccagggct**caggtctgcaggacctg**ggtaggctacggggactttctcactcaccattgtctggactctcctgttgtcgtctgtgaacagagatgtttggaagatgaagtggacattagtcatgaagggaataatactagttttctccaatgggatactgatgtagagctgatcttgtgcttctcacactgtcacagagtttggactctcacccgtgactttgagg**agagctgaggatggacactccattgtgctgtgagctct**gggtaacaatagtagggcctggctaggcagcctaaggtcaatactgctggccttcaggaaagacag

Uppercase: IGHV3-21-2

Lowercase: Flanking sequence[1000bp]

Red & Bold & Underline: Stem-loop [18]

Blue: Heptamer[44]

Green: Nonamer [7]

id-IGHV3-13-2[V_gene_segment]

attctcagtatt**tctattgagaaaaataagccaaataga**tgagtacatactatattatttcattcttataaattctagagaataaaaactagtctaaagaaatatgaaaacatcagtacttttataaagaaatggtagaagaaaaggagaaaaacaaaaatatatctataaaagagcaagaggaattctg**aggtgagttgacttgtcacct**tcttgaaaatagagactttctttatcaaagtttactattgtacaggttaaatatgtgaattttatcatctgtcaattaaaactca**taaaatttattacaagtaaacaagtgaaatttta**gacaaaaaagggatgataagaaggaacaaataaatacattaaatgt**cagatacaccaaaaatgtatctg**cctgacgcctagttgtctctgtatttttaggtaaatgcagcaaaatcacacaggttgtcgtggcaggaagtggattctgcaaaccacactaggccgtttatctctgtcctgtagttggttcagagcaactgaggccagctgtgaggcgcataggcccaggtactaggactcactcatgccagatataagccctgagacacgtacatagcccctccatgtgtgggttcacttttacatctgtacatgaagaaaccactgactcctaaataacatcatctatacacataggtaaaaaaattaaaaatatgatagttgttaaatgtttatcgcagaacaatttcaaaataaggcagcattttcccaaatacaatcattgtcatcaaaatcccccaggacgctctcatctactctgcgccctgccttcacctcagatgtcccaccccagagcttgctatatagtaacagacatgcaaatagttgactccctctcctgatgaaaaccagcccagccctgaccctgcagctctgggagtggagccccagc**cttgggattcccaag**tgtt**tgtattcagtgatcaggactgaacaca**caggactcaccATGGAGTTGGGGCTGAGCTGGGTTTTCCTTGTTGCTATATTAGAAGGTG**ATTCATGGAGAACTAGAGATATTGAGTGTGAATGGGCATGAAT**GAGAGAAACAGTGGGTATGTGTGGCAATTTCTGACTTTTGTGTCTCTGTGTTTGCAGGTGTCCAGTGTGAGGTGCAGCTGGTGGAGTCTGGGGGAGGCTTGGTACAGCCTGGGGGGTCCCTGAGACTCTCCTGTGCAGCCTCTGGATTCACCTTCA**GTAGCTACGACATGCACTGGGTCCGCCAAGCTAC**AGGAAAAGGTCTGGAG**TGGGTCTCAGCTATTGGTACTGCTGGTGACCCA**TACTATCCAGGCTCCGTGAAGGGCCGATTCACCATCTCCAGAGAAAATGCCAAGAACTCCTTGTATCTTCAAATGA**ACAGCCTGAGAGCCGGGGACACGGCTGT**GTATT**ACTGTGCAAGAGAcacagt**gaggggaagtcagtatgagcccagacacaaacct**ccctgcagaatgcctgggggaaatcagctgcaggg**ggcgcacaggacccactgatcagagtcatccccagaggcaggtgcagatggaggctggtttcctgtcaggatgtgggacttcatcttt**ttagagtttctctagggaacctctctaa**gttcagaattctgtgcttaccaatgccatccctacatattt**ttaaaatgattattttaa**tatgaaaacctattctcttatgcacgaaacacagactgatgcttacagagatgaaaagccctcaaccattgt**caccaggatcagagtattgaggaaactcaggggtacctggtg**ggtcttctccactcagactcaggacagaaacctcagtgagtttccctgactaggacggtctttaggaattgtgatcgcagccaatagagagtctgggcaaggatcagtgtcatgtagaacttcacaggtttcacttctgacccttctc**ctgacactaaagtatacaacttagtgtcag**cactgatctggggccccttttgctcttagcccattctatttctttttatttgttgttgttgttcttgcttttccttgtagtgttcctgctccctgtaaagtggggatgtggctcttgctgccaaagctccaggtctcaagcccattccctgcagctcaggtggggctcaggct**gtggctcctgcagccac**gtggaagaggctg**atgggactttcctctctcccat**tgctcagcaccctccagtgtgtcacgtggagactcacctgcgaatggaagtggccaacagtagtgaaggggatgagcttgtgtggacaaaatgggatgtggatgtgaaatttatc**ctgtgctgtgcaaagtagcacag**agtgagtcaccttcctcaccagtagtgttagaaagagggtgtgaaagttgtcagaatcaaaatagatccacttgtgttaaaaccctgacaaatgg

Uppercase: IGHV3-13-2

Lowercase: Flanking sequence[1000bp]

Red & Bold & Underline: Stem-loop [19]

Blue: Heptamer[38]

Green: Nonamer [7]

id-TRBV13[V_gene_segment]

atgtgtttatttctccagcttagatgattgtttcactgatagcatgtcactgtcagctgcctccaggaattatctttggttaaag**ggaatgctttgcccaggtctcaccccattcc**tatggcagctggcatttaatgactggctgatatctcccttcctacaccattatttgatga**aatttaaaatttaaatt**aaaatgaaaaatttttctttcttgtttaaaa**agacatttgctttatgtct**ccctcaccaacctgtgcaaaatttgagtttgaggcaggtaaacctgaaatcagcaagagctgtgctgtgtcctgagagatcctgagggtctaattt**aggaagcacccagtctccctagtcagggtccttagagcttcct**catagtgactacacactggccacttgggggcactgtggatccactgagtgggtcactgatagcg**ctgtctgagagagagagggttaagggagacag**tcttgatttcgtctctgtatcagggatattaccagatgaaccagaggtactgctggcacaggaagagttaatcatagtct**tatcagatcaactgata**tgcaacatgtatttgtcatgacaggaagccatgaactgaagccttgtgtggcctctctagcaacctaggtttgatggggagtgtgatgagggtttcagcagggcagcctggataacccaagccaggtggagataaaggag**catctgcctcagatg**aaaatctcagaagctgtgcggatggctgtctgcccggaagcctgccccctctcgctcagcacagct**agcttcccctgctctgcaggaagct**ggacaggatgggggaaagcctgagttagctgagcta**gtcctggagtacagggtgcctatgggagcctacaggac**gatgacatcagaacagtgacatcacagtaaaaacctccaaaaacggtgaggaggagcaaaag**ccctgctttctcaccccaggagaccagcaacctgagcaggg**agATGCTTAGTCCTGACCTGCCTGACTCT**GCCTGGAACACCAGGC**TCCTCTGCCGTGTCATGCTTTGTCTCCTGGGAGCAGGTGAGTCCTGAG**TACAGGTGGGACATCCCTGTA**TCCACAGTGTTCAATTGTTGCTGAAGTGTCAAACTCTCCCG**AGCTGAGTCTTCAGCT**TCTGTCTCCTTCCTCCACAGGTTCAGTGGCTGCTGGAGTCATCCAGTCCCCAAGACATCTGATCAAAGAAAAGAGGGAAACAGCCACTCTGAAATGCTATCCTATCCC**TAGACACGACACTGTCTA**CTGGTACCAGCA**GGGTCCAGGTCAGGACCC**CCAGTTCCTCATTTCGTTTTATGAAAAGATGCAGAGCGATAAAGGAAGCATCCCTGATCGATTCTCAGCTCAA**CAGTTCAGTGACTATCATTCTGAACTG**AACATGAGCTCCTTGGAGCTGGGGGACTCAGCCCTGTACTT**CTGTGCCAGCAGCTTAGGcacag**ccctggagaattactggctttctgtacccaaac**cctcctatctcacttgaggatgtaatagggagaaggagg**tg**ggggctgccacacaactttagccaagcccc**agagatgcttctattcttttctaacattttcccctgccctgct**gagctcagtgagagctc**ctgcacttgtgggctccagacccactggaagttctcacatcttagccagtactttttaattcctagcaagtggcgggagcttctactctgtgcc**aacataggggtatatgtt**ttagtgtgtttcctgttgccatacag**aatacctgagactgggtatt**ttata**aagaaatgaaatttgtttctt**acagctctggaggctgggaagtcc**aaggtcaaggacccacatctagtgaggacctt**cttgctggtga**ggactctgcagagtcc**ccaggtggcatagggcattacatatgagggggctcatgaaagatggtcaaactggcttttataacagacccaacctcataactaactaattccttctataaccca**ttaatccatgaatggattaa**cctttatgagggcagagtccttaagatccagtcaccttccaaaggtcctacctctcaacgctgctgcattgggaaccaagttttcaatatataaattatttaggaacacagtcaaatcatagcagtacacattgcgggttttttttttttttcttttt**tgagacagggtctca**ctctgtcacccaggctgaagtgcagtggtgctgtgatcatggctcattgcaaccttgagcgcgtgggttcaagcaatcctcctgcctcagcggcctgagtagctatgactaca**gggacacatcactacaaggtgtgtccc**cagttcctcagttcctacaaggtgatgtgtccccttgcctggctaattttttttaatttttatttttgtagagatgaggccttgtcatgttgtccagattagtctcaaactcctcaaatgatcctc

Uppercase: TRBV13

Lowercase: Flanking sequence[1000bp]

Red & Bold & Underline: Stem-loop [28]

Blue: Heptamer[43]

Green: Nonamer [3]

id-IGKV2-19-2[V_gene_segment]

atcctcttgtaagaagtccccactccacccaggaagtctagctggcctcacctctcactatcaatgactgaaatgatcattagtagcagtggctttggttaaatgtataccagtagttacatacagggtagaaaaatggtctgaagaggtatattcagcatcaaatgaaaaaattacaccttgtgacccctttctctccctgtatgatcccttaagattgtgctgaaacctcagagcatcaggtcagcctcatattgctccttcctaaatcttggtccctgctcaacctgagcctatgtttggccatccacgggcatccatgtatacaactttgaaggcaaaactccctgcttggagtggctcattagatgaccagggtccagatagcattgccaaggaggctcacagatggcttcatactgagtctctcccctcagcctcccagcatcaccagccctgaaggaatgtccctgttatccgtggcacccagagcac**agtcagccctgccctgact**tcactgcccaattcacagcctcatcgcagattccactgcaaaatagacagtgtgagatctcaaagacaggctgatttgtagtcctgaaaagcaacaacccatcgttccctcgatctctttggtggaatttcaaagttcgtcataagctgaatttctgtacaaacatactctcttaacaatggaaactatttgcaatatgggaaacataaaacatgggaaaagtttaaatctctcaagagcatcttctagtcagattcatattcagcttccctgattctgaactttctgtcttgcggtcagcttcttttggtccatttgtttcaccctttccagctcagatcttctctccctggctacccttagtctcaggttgtccttacgtttgataggcttct**accttatggatgctgtaaggt**tatcctctcatttgctggattcttgtgggaaaaaaaatccctccaagttcacatcctgagacttcagactCCACTCTTCCTTGGAGAGCCATCC**TCCATTTCCTGCAGGTCTGGTCAGAGCCCCTTTCATAGAAATGGA**AACACTTACTTGAATTGGTTTCTGCAGAAGCCAGACCAACCTCTAAGGTGCCTGATTTATGAAGTTTCTAACAGGAAGTCTGGGGTCTCGGGCAGGTTCAGCAGCAGTGGTTCAGGGACAGATTTCATATTGAAAATCAGCAGGGTAGAGGCTGAGGACGTTGGGGTTTATTACTGCCTGCAAGGTACACAAGTGCCTcccacagtggtacaaccctgaatacaaacctccctgcttggagtggcccagctgctcaaata**tgttgtttatctggggagtggcacaaca**gaatctctgctgtataagatgaagatgttgggccgaggcgggcagatcatgaggtcaaaagtttgagaccagcccgaccaacatggtgaaaccccatctctactaaaaatacaaaaattagctggatgtggtggtgcacgcctgtaatcctagttactcaggaggctgaggcaggagaatagcttgaacccgggagatggagcc**tgcagtgagccaagatcacaccactgca**ctccagcctggatgacagaaggagactccatctcaaaaaaaaaaaaaaaaaaaaaaaaaatgaagatgttggtgagctcagggtaaaaggttgcagctgaatgacctgtcccatgggggactcagcagtacatcaggtaaaacccattcatggtcctgtcagctcccacagccttggcatggcataagccaaaggaaaacagagataattcaagtgccttcagagtaagcagacaggactgagggagagcgg**agggaaatctcacactaatcttccct**gccttgcctacattcgacaatgagacttcaaatagcttaatagccagacaagtaacacagattcgtgtcaacacgtgttgaatatctcttgaagtttaggtcttttgtgtatatttttaagagggtagtatttggtagtatttagaaactggtatttttcaacttttccaatttccttcttctcctttttactagttctcttcccattgcaccacataacaaatagtgaaaagagcattctacacaagc**tgtcctcagggagagctggctgaggaca**atcatgaaaaagcttgaatttgcacctccaaatagacttttgtgacgtcatggaagacagaagatcctgatgttaaaactctttcattcatttcaattacttcttgctaataaaaaaggataa

Uppercase: IGKV2-19-2

Lowercase: Flanking sequence[1000bp]

Red & Bold & Underline: Stem-loop [7]

Blue: Heptamer[23]

Green: Nonamer [6]

id-IGHV5-10-1-2[V_gene_segment]

tgatgctgacagtca**ttgcatagcatgcaa**attacctaggtcatagcctgtgttgtgtgtttatgttgatggtaatggtcattgcttagcatgcaccttacccaggtcatagcctgtgctgtgataaagagaagtcattttgcttctgcaccccaggtatacaattcctgtgggtaatctggcaaatatttt**agaaactgtgtagtttct**gttgccttggcttt**tcatttggatcttttagatgttaaggaaatga**gagtggttcacaggaaaatggggcaaatacaggggacagtaaaaagccccctccgagcaatgtggatatcataaaataaaaaaaataagaataaaagaactaaacagatt**gtgctggggtggaattaaaacagcac**taccaggagttgggtgcagcaatggggatcccttcaggtccatcagcaatacagag**acctaaacaaatatttcctatttaggt**tgaattggcagagtgtaaaatctagaagatgaaatggccaggaaacatttgtccttgattttgcatgggtgaaggtctggcaggttaatcaaattgagggttgttaaa**ggcctggggtggtcctgccctttctgggtgcccaggcc**atagtcacattcatgggaaattgccatggaataaagagtagttttttgtggtagttcttgttatcaggcataagtgcatttgaattttctgttcacggcctttcctggctctatttctcatttttttaacagacatgattttatctagatttgtcaactttcacagggtcacagagaagggtggaaggagggatgccctggtttgggtcttgcaggaccacagacaagagtg**gggaggggcaagaagaggttgtaaggaattattgagacctattctgcccctccc**aggagg**ctcagttcagccttttctctgcaactgag**gttctgggttataaaccctgtagactcttcccttcaggtcagggcggcaactatgcaaATGC**AAGTGGGGGCCTCTCCACTT**AAACCCAGGGCTCCCCTCCACAGTGAGTCTCCTTCACCACCCAGCTGGGATCTCAGGGCTTCCTTTTCTGTCCTCCTCC**AGGATGGGGTCAACCGCCATCCT**CGCCCTCCTCCTGGCTGTTCTCCAAGGTCAGTCCTGCCGAGGGCTTGAGGTCACAGAGGAGAACGGGTGGAAAGCA**ACTCCTGATTCAAATTTTGTGTCTTCCATACAGGAGT**CTGTGCCGAAGTGCAGCTGGTGCAGTCCGGAGCAGAGGTGAAAAAGCCCGGGGAGTCTCTGAGGATCTCCTGTAAGGGTTCTGGATACAGCTTTA**CCAGCTACTGGATCAGCTGG**GTGCGCCAGATGCCCGGGAAAGGCCTGGAGTGGATGGGGAGGATTGATCCTAGTGACTCTTATACCAACTACAGCCCGTCCTTCCAAGGCCACGTCACCATCTCAGCTGACAAGTCCATCAG**CACTGCCTACCTGCAGTG**GAGCAGCCTGAAGGCCTCGGACACCGCCATGTATT**ACTGTGCGAGACAcacagt**gagagaaaccagcactgagcccgtctaaaaccctccacagcacaggtgcagaggaagctgccagagacacactccccaggggcctctctattcatccagggaggaaacactggctgtgtcctgaggagtgagaaccagagaacaacgtgggagggttcccagtccctaaggcaactagatggggcacctgctcctgggaggtggattctctgagggggctcttgttctacaaggttgttcatggtgtatattacatggttaaaataa**aaaagctgtctaatatgcacctcttaagcatgacagtctttt**aattagtgaaaagtctacacagttaatcattgctggcatgctttcctccctcctatcttctcttgctccctacatctattattttctacttaatttttcaaaatcatttgatccctttgtgaactattactaggttctgttgtttttgttgttgttttcctttcaataatatgtactgaataattcatctttgtgccaattcataagttcttgggtataataaatacttctctcataaaaattagataaa**ttaaaataaataaattttaa**aaaacatacaacctatcaaaactgaaccattaagaaataaaaaatctgggctggttgtggtgggtcatgcctgtaatcccagcagttttggaggctgaagcaggtagatcaggagttctatgtgaatctgtggtcaggagttcaagatcagcctggccgacatggtgacaccaagtctctactaaaaatataaaaatta**gctgggtgtggttgtgttcacctgtagtcccagc**cactcgggatgctgaggcaggagaatggcttgaacctgagaggcagaggt**tgcagtaagctgaaattgagccactgca**ctccagactgggaaacagagtgagactccatctcaacaacaacaacaaaaaaatctgaacagaccaatgagtaaagagattgagtcagtgatttttcaaacatc

Uppercase: IGHV5-10-1-2

Lowercase: Flanking sequence[1000bp]

Red & Bold & Underline: Stem-loop [18]

Blue: Heptamer[44]

Green: Nonamer [6]

id-IGKV1-16[V_gene_segment]

tctggaac**caactgatttctccatgtacctgggcagttg**cttgataaaagattgagtgcctctttccaaaagatg**ttaacagggatcttggtgtctgggtcaggatgatgtccctgataa**aaagtaaaagaagaaagtgtcacct**atggtgcatggcagggacatgctccat**gcagtgatcaccctcactaagagagatgaatgttgggaaataatacttaatggcagaaaagaaggtagactatgaaggtgcccaaaacaggaataaggtgcaaaacaggaataaggtgcagcccattta**gtctctggatattaaagagac**ctagagctcttgataatggtggatctgtaattgctgcatgcattgaggaaacacggtatcaacttagtgtatctgaagtaaattgc**ttgatcttacagtggtaagaacaa**tggcataacaccattacctaacatttacaaatatatgtagcatcatgtgaataaattttatttttaatttttttttagaaaggagcaatgttaaactcacagaaatgttgcaagtatatgataagtaccaccttccctaacagaaatcacatgagagtcttttgaagacctgaaaa**ttgtagggtctaaaattttactatgtgtttcctacaa**aaaagaatattctcctaaataatccccatacaccaa**tgaaatacattgctccatcgactcccaaggaatatttca**aattgtcaaaaagaaaccataaaa**tgtttcccataacaaaatagtccccagtagaaaca**cattctctgcagacaaatttgagctaccttgatcttacctgggacacgtggggacact**gagctggtgctgagttactgagatgcgccagctc**tgcagctgtgcccagcctgccccatcccctgctcatttgcatgttcccagagcacaacctcctgccctgaagccttattaataggctggtcagactttgtgcaggaatcagacccagtcaggacacagcATGGACATGAGAGTCCTCGCTCAGCT**CCTGGGGCTCCTGCTGCTCTGTTTCCCAGG**TAAGGATGGAGAACACT**AGCAGTTTACTCAGCCCAGGGTGCTCAGTACTGCT**TTACTATTCAGGGAAATTCTCTTACAACATGATTAATTGTGTGGACATTTGTTTTTATGTTTCCAATCTCAGGTGCCAGATGTGACATCCAGATGACCCAGTCTCCATCCTCA**CTGTCTGCATCTGTAGGAGACAG**AGTCACCATCACTTGTCGGGCGAGTCAGGGCATTAGCAATTATTTAG**CCTGGTTTCAGCAGAAACCAGGGAAAGCCCCTAAGTCCCTG**ATCTATGCTGCATCCAGTTTGCAAAGTGGG**GTCCCATCAAAGTTCAGCGGCAGTGGATCTGGGAC**AGATTTCACTCTCACCATCAGCAGCCTGCAGCCTGAAGATTTTGCAACTTATTACTGCCAACAGTATAATAGTTACCCTcccacagtgttacacacccaaacataaacccccagggaagcagatgtgtgag**gctgggctgccccagc**agcttctc**ctgatgcctccatcag**ctgagagtgttcctcagatgcagccacgctctgatggtgttggtagatggggacatgaagtcacctctgcaccctaattcttttctctttctcagcaccaactgcacagacatagcaatgcttctccttatttaataaaaacagagatcatgacacctgaggagtctagttta**tggcttcagttggaaatcataatgcagaagaagcca**ctataga**tattctaagcaggaatagtcttaatacagagaattagaata**aactgctgaagtctaaata**aaatgtagagatgaatctctaaattt**aatgttttatttgaagagaaatatttgccaaatgaggcatacaggaaaactcagtggtcttcagtacgttggaagaacaaagaaaaagtcagtgttccatgaaaaagggaaatattacctattgcactttgagaaagttcattggcactaggaagggttgggagctggcaagctcagactggtaaacagtggtggacaaagtgaatcctagagttatatcaagttatctcagaaattgtggataaatttgatttcaggttacaataagccaaagcactgaaggttgcagagaattttcttactgaaataccagggacttggtgtagatcctgcagctcaccacacagaaagccaatctcgaagacaacaagtattgccaaggaacaggctttaatcaggtgctgcagccgaggagatgggacgccattctcaattgtatctctctgacaaaattaattaggggtttatataggagggaagaaatgttgaaaacaggaattagggaggggtaaggaagataatttggtcaagaagaagcaggaggtcagttatgcaatc**acagtgggtgaaggttctgatgtctcactgt**cccaattcagtgatatg

Uppercase: IGKV1-16

Lowercase: Flanking sequence[1000bp]

Red & Bold & Underline: Stem-loop [21]

Blue: Heptamer[25]

Green: Nonamer [3]

id-IGHV7-56[V_gene_segment]

tcccc**aaacccctgaagagtgtttgggttt**accgtgtgtccagtgattcagtgcctctagagctccaggaaggggctccccggtgatgcctgagatcttttcttcaggtctccctgcagagttctctgggtttcctaagggcaattcactatttcaaaagatggtgtgagaagcacatgctgtcattaaaggagaattctgagccacggcacagccactttatactgggctagagacactggtatgaatatactctgtaagtttagacagaaagcctcgtgcatggtaggggctgggctgcagggggtgctcaggataaacgcagcacagtct**cccgccccagagcaggtgcacaggaggctggggaggg**gttcctctcagggcctgggacttcctttgaaaatatctaaaataagtatttcacaagggctgctgtt**gtttgtataaatatcctattcaattgtgagcatttatcaaac**tggatg**ttgtaatgacaaccacttttacaa**tggggatttcaaactcccctagatatcttaatagtaagcagctggaggtcaagaagagatcctttcttttaaataagtgcaat**ttttggagaaacatactaattcccaaaa**taacgcattcacatattaaggtctagaaatggctcaagttgtccctggtgcattcgaatgtgggttcaaagtgaggtgcgtgtcctgagg**gagcttgttctccagtggaggaagctctgtcaacacagag**ttcagggatgtgtaggggtcgcatcgcctctaacaggattacggcttgaaccctcagcatgtacaattgtgtcgtccatctgtcatgtatttgctctatctcatcctggctcaggaattgggc**tattcaatagcatccttcgtgaata**tgcaaatcactaaggttaatacagatatctctgtgccgtgagagcatcacccaacaaccacacccctccttgggagaatcccctagatcacagctcctcaccATGGACTGGACCTGGAGCATCCTCTTCTTGGTGGCAGCAGCAACAGGTAA**GGGACTCCCCAGTCCC**AGGGCTGAGGGAGAAACCAGGCCAGTCATGTGAGACTTCACCCACTGCTGTCTCCTCTCCACAGGTGCCCACTCCCGAGTGCAGCTGGTGCAGTCTGGGCCTGAGGTGAAGCAGCCTGGGGCCTCGGCGAAGGTCTCCTGCAAGGTGTCTGGTTAAACTGTCATCACCTATGGTATGAATTGGATACGACAGACCCCAGGACAGGGGCTTGAGTGGATGGGATGGATCATCCTACCCTGGTGAACCCAATGTATGCCCACAGATTCACACACGGTTTGTCTTCTCCATGGACACCTCTGTCAGCACGGCGGATCTGCAGACTAGCTGCCTAAAGACTGAGGATGCAGCCATTTATTACTGTGTGAGGTAcaccgtgtggaaacccacatcccgagagttttagaaaccctgaggaaggaggcagctgtgatgagctgaggcagtggtgcagcatgtctct**aaacttccattttatctaagttt**gcattgagttccgctttaatattagccaggaatgtgggatagacgggtgctcctaagaggtccttaatttgcccattttgatgggttttccagaagacgtgagaagccactttgttaacaaagcatcccaaagccatgccctgctccagaaacacgtgtacccatttcctggtctttggttaactgacaagctctcatcagcgcacctgggctaatttctcatcaggtagaaaaatgtgttgtaaagcaaggctaacgttgtgatagcaattcctgctcaataaccttcagcatcgttgttgttgtgttctatcaactaattacttgacttcaaggttctcattgggagtgtcttataaatttaagggatatatagaagttcccctaattaaaataaaacaattgtgagcacaacctcagtgttcaaccatgtctccacccttcccaccattcaccccaaagaaatgttcacctctcctggaagtcgggttcattttcaaattagttattttttattttactatatcaagattattgtatgtgactactgtagcagaaagtgaattaggggaacttgaagtaaccaacgaaagataaattcagaactaattaaacaagatgtcagaacgtgattggctctagtcttttaaaattcagcaggttatgtaaccaggctttaaatttacacatcttcctgttaccttcacggcacagtcaactcccattatgtaagaaatggcaactgcattcccaagcgtcatccaaaattgtaaaaatagactgggtgaggtgag**gagttgattgtttaaattccgctctgaagaagcagcatcaactc**aacaaaccaccgcttttccctcagtgactagagctatgtcgcaggcca

Uppercase: IGHV7-56

Lowercase: Flanking sequence[1000bp]

Red & Bold & Underline: Stem-loop [11]

Blue: Heptamer[31]

Green: Nonamer [3]

id-TRGVB[V_gene_segment]

ttctgaagtccaaggtgagttcccacagactgccctgtcggtccctccagcaatttcaatcaagtttcctcttgaggtgcaccctctttataagtgaaattggatttctctgagttataatctgcattgtgacatcccaattaattataaacc**aaatactttactgtattt**caaacagaaatgactatagctttttaaatgaggtatgaataatgctagtgtaaaatttcagtttgccttttttaaaattgtaaaatactgtcc**atgttttatactcctgagtcctgcatccccaaaacat**taacagagggatcaatttttacgtgaggtaaagaaagttatatgcttatatatctggaaaaatattgtacagtctgaaaatcaaggagagaacagctaagtttgaagcagaattagttcaactccagagctgctcccatccctttccacagacacattcttgtcaggtaatgcaccctacaccccgtcagtggtcagtgtaatgctcctgggaactgctcagctcttcagtaggccatgaagtccccagagggcacctgacccaccacatttttctctctcaccaagtcatgccatgctgtctccccactgactggggccagccagcaaccagggtccttctcctggggatatggctcagcaga**cagcaggatgtccctgctg**agaatattcatatttttctccctctgcagcggtgagttgctcctcaattatttcaacatagatgagagatgaaagtgattctttttttcttttattttaagacagtctcgttctttcacccaggctggagtgcagtggtgtgatctcagctcacttcaacctctgcctcctgggttcaagcaattctctcttgtgcttcagcctccaaagtagctaagattacaggcatgcgccactgtgcctagcttatttttttatttttagtagaggcagggttttgccatgttggtcaggctggtctcgatctcctggcctcATGTGATCTGCCCTCCTTGGCCTCCCAAAGTACTGTGATTACAGTCATGAGCCACCACACCTGGCCAG**ATGAAAGTGTTTCAT**TATCATTTCCTCTTTCTATGTCCCAGGTTCATGGGGATTTCTCTTTTTTGTTTGCAGTTGAACTTGGTAAGTTAAATTTAAAGCAATA**AAAAATGTCAACTACATTTTT**GTCAACAGAGCAACAGATAAAAGTGTCTAGGTATCTTGTGTGGTGTCCACTGAAGACTTTGTAAATATAGTTATACACTGGTACCAGCAGAAACTGAATCTAAATTATGAGCCATCTGACAAATATCACTTCAATAACAAGCCCCAGTTCGAGTTTGCTTAGGTGAGAAAAACAAGAAACTTGAGGCAAGAACAAATTTTCAAATGTCTACTTCAGTCTTTACCATAAACTTCATAGGAAAGGAAGATGAGGCCATTTACTACT**GCACTGCTTAGGACCcacagcatcagtgc**cacactgtcccacacaacaacctct**gttgggtctctgcccaac**cacatccttccca**tgggagcaaactctatggactcctagctgggctccca**ccctcagccttgccttctctgcggcacttctggatctttctgggaagccatcttgggtta**atgctttaagcaggaagcat**gcctgatcatgacgttcccagat**attcattgaacactacttcctaatgaat**taaggtttggcctg**ggaatgagtggacacaggctaaggtcccattcc**tgtcagcatccagataattctattaactgcaaacggcttaggccaagggagagaatcgttgtgtataggaatcccatatataattatttcaacacctctgcaacagaactattatttgcccat**tttaataaagaaattaaa**gcttacaagttcaaaatgactggtcgaagttgagaagtgcatgttttttgttttttaaaactggatcatgaggccaccttgaatatcta**aaagtcacagttgtacaacagactctttgaggtgacttt**ggggtggagcaagtgtcttttttctgtagagaagaagatggttcct**cacttcacactaaggggtgtgacagtg**tgatatcatcagaaaagagccccaggcatcggact**gttaaatgagccatactttgcatttattaactcctttaac**ctctggcaacctctgagg**tctgttctgttatgaacaga**agtgcattttactcaggtaagtgagatctatttgtcctagccactcatccaggtcacac**agctggttcatggcagtgtaggattggacccagct**agtctggctccagagtc**tgtactcctcagtcacagaagggtaaagagaagtaca**ttgttcctactcccatggagaagacaatactgacaacttctgtaaaacatttactggcacaattattattttgaactgaatcaggtgaccagaatttgagcaattaggaatc

Uppercase: TRGVB

Lowercase: Flanking sequence[1000bp]

Red & Bold & Underline: Stem-loop [18]

Blue: Heptamer[20]

Green: Nonamer [3]

id-TRBV6-7-2[V_gene_segment]

agagggacctgggggcagggtgttggctgggtctaggagaagatgatgagctcagaaaatagacaaggagaaaccccaggggaaagcttgtaaaatcctggaggaatcacaagatatattatttgagctcctttttatttattactaaattaatggc**aggaatcccatataaagggatattcct**tatgaaacatgacatctaaagtacaaacag**aggattcttccttaccctaaatcct**gccctctctccctgagtctctcacgcatcaggccacctttg**tgcccataagtcatgggca**acgca**gtgtggtcaccttcatccacac**ctagaggacagtcagcaaagtgaggtggttctgcctgctgcggtctcaccccagacatggaaagcaagagccctgggtgg**agctgaaggtgctcagct**gggtttatcaggagtctcatctgtcagtggattgacaagaaacagagcaaaacgactcctccaatgttgatgagcctgcccctg**ggttttggaaacctgataacagagaaaacc**aatatagacaaaggattttaaacaggattatggtcaattaagcaaattagaaaaagatacttgaaagaatatttgggacacaggagtcaaaaacaccaggaagacatgaggagtgtccctaagactctagactacagcactgtgtagacaataaacaccag**aatattattatattatcgaagtaataaaatt**tacagtgaattacaaactgtttccaaaaggtcatgaaaatcttgtagacttgtttcaattcagacaaatgtgttcttctcattctcagctgttcactggtgcatttatcttgaatttgaccatctggggaatgggcgtggcctctcctgacaggaaggctctgggggccaggc**agggagaatgatgtctcagaatgactccct**tgagagtccagttcccctttcatcaatgcacagacccagaagacccctccgtcctgcagcccctgccATGAGCCTCGGGCTCCTGTGCTGTGTGGCCTTTTCTCTCCTGTGGGCAGGTGGGTCCT**GGGCAGGGCCCCTTGTGTGGATATCAAGGCCCAGCCC**CTTTCCACTGGAGCTGTAGCATCAGCTTTGTCCTTCCCTGCAGGTCCAATGAATGCTGGTGTCACTCAGACCCCAAAATTCCAC**GTCCTGAAGACAGGAC**AGAGCATGACTCTGCTGTGTGCCCAGGATATGAACCATGAATACATGTATCGGTATCGACAAGACCCAGGCAAGGGGCTGAGGCTTATTTACTACTCAGTTGCTGCTGCTCTCACTGACAAAGGAGAAGTTCCCAATGGCTACAATGT**CTCCAGATCAAACACAGAGGATTTCCCCCTCAAGCTGGAG**TCAGCTGCTCCCTCTCAGACTTCTGTTTACTTCTGTGCCAGCAGTTACTCcacagcgctgcaaggctgtctcctctctgcacataaaggcaagggaaggtgctgccctcctcccccacccaagactcaaggatgc**cctgggcagagatctctgcaccagg**aaccttggaacgcagagtggccccaagtggcccggacagtatgagcctcgctctgtgccacgtgcctctgcaggcatctcagccaggcctggactggtcccaggtcctcagatgtctcccttgttgctctctctggtctgtcctccgaggtctcctttcgggttgtggccagggcttccccagctcctacttttctactcatc**atcctgagtccgaggcccccagggtggaacaggat**ttgtatttcagatccatctagactcctgtctcttcctggtgaccatgttgcttcctctctctagggttttcccagcccccaccctcacgtagtctctcctgtggcccacctttcccatctgggcagtcaccctccaaggccttgctgggtctctcctcccctcacttccccg**cccctttctactgcagctataagggg**agcc**tctcttctgtgcctccttccttcccatcaaagaga**tttcaaaggccattccctctgccctaggctagaggctaccttttctactggccagcgcctacctgtgcttcagatctcagcataatctgcccctcctgcaggaagcactcccccgcctcccagctgagatcaactttccactcaaaagctctcatagaatcatatgtctgttggtaacctttagcacagttgggctttctcaggtacttgtctgatactttttgtactccacccattttttttccctatttttccttttttttttga**gacagagtctcgctctgtc**accaggctggaatacagtggcactacctcagctcactgcaacctccgactctccgactctctggttcaagcgattctcctgcctcagcctcccaagtagctgggattacaggcacacaccaccacaccc

Uppercase: TRBV6-7-2

Lowercase: Flanking sequence[1000bp]

Red & Bold & Underline: Stem-loop [16]

Blue: Heptamer[31]

Green: Nonamer [3]

id-IGHVIII-13-1-2[V_gene_segment]

catgagatgcaggcaacaaagaatgaagtaaccttccccatttgcacataagatgtttccattcacaagagacctttcttttatcaggttcatgtgcaaccaagtttccctgctgacaagcatttaagccagatgattcgcaacttcctttg**actgagaaagatttccctcaaacttcagctcagt**gcaggcacacaccatctctgaa**tgggcatttaccatcagacaatgccca**cacctgtccccacatggacctttccctcagacaaacaaacacatcctcaggttgactcttccctcagacaagcacccctgtctt**catgtgaactcttccctcagataagcacacatg**tccccacattgactttttcctcagacaag**cacatatagctgacaacgaacagttatgtg**gcaaaatgagctcaggatagtggtaattatggactccagctctgatagtttgtagaaattgtcatttttaaaattctgactgaagactttcctttattgtagaagacagtcctttacagctctaattgcacagcctacaggcaggagtccatttcctctgggcaaggtttatttttatttgtttactgtacttatttgttgataaatattgatactacaaagatagcctatagggtccacatacgagaaaaaaagagtaatgggcagatcaaccctgcaaatccagtcccaggagtctttgaccctgccctccctggaatccagagacagagatggaagaggcctgctgagcagtgcact**catgtccccagggagaaagacatg**gaaatgaggcccctcctctgcaaatgaaaagtagctcatcccctgttcctgtagatcctgg**tgaggagccatcccacatctgtgcccttcctca**gtgtccacaccatggggtctgtgcggatctgggctgctcttgtcatcactctcaatgtttaggttccccgtggatcaggccctgctgtggctgttgcctctgtgtttgcagAAGTCCCCTGTGAAGTTAACTAATGGAGTCAGACAGAGAAATACTACAGACCAGGAATTCTGCCTTTTCTGCA**AAGCCTCTGGATTCACTTTCACTGAAAACAGCATAAGCTTGATCCAGCAGGCTT**CATGACAGGGGTGGGTGTGGGTAATAACAATAATTCAAATAGAAGTTCTCAGTGGGACTCTCCTTGAGTAAAAAGATGATTAACAATCCTCAAATACACTCAGTTCAGGAGATTCTCTTTTAAGATGATTAACCTGAGAGCTCAGGAAAAGTCCGTGTATTACTTTGAGGG**Acacagtgaggggacatctgtgt**gaactcagacaccaacctccctgcagggagacaggaggggactg**cctggtagatgcttctcagaaccaccagg**gggtgctcaggacatcagggggcactcagaaccatcaggggatgctcaggacaccagagggcactcaggacactggggggcggggggtcactcagaaccaccagggggcgctcaggacactgggggggggggggtcactcagaaccaccagggggcgctcaggacactatgaggggg**ggtggtcactcagaaccacc**agggggcactccggacacggtggtgaggggtagctcaggatagcaggggtgctcagaaccaccagggggcgctcaggacactgagggggggtcactcagaaccaccaggggacactcgagacaccagggagccctcaggacactagggggagctcagaaacaccaaagggcaatcaagacaccagggggatctcagaaccaccaggaggtgctcaggacacca**ggggtctctgaaccactagggggagattaggacccc**agggggctcagaaccactagggggttttcaggagaccagggggcgctcaggacaccagggggcgatagggaca**ccaggtggttcagaaccactagagtgtgctgagaaccaccagg**ggtgctcagaaccccagggggctgctaaggacaatagggagtgctgagaaccaccagtgggcgctcaggacaccagggggatctcagaatcaccaggctgtgctcaggacaccggggg**gtgctcaggacctccaggagcac**acacgacacaaaggatagctcaggacctacagggggcgatcagaacgccagggtgcgttgaggacaacaaggggctcacaggacacaacgatgtgcttagtaaaccgggggacactcacaaccaccagggggtactcaggacaccagggggtgcccaggaaaccagggagcactgaggacaccaccgctccc

Uppercase: IGHVIII-13-1-2

Lowercase: Flanking sequence[1000bp]

Red & Bold & Underline: Stem-loop [13]

Blue: Heptamer[42]

Green: Nonamer [2]

id-TRBVA-2[V_gene_segment]

gtactcttatgttccttgaagcattattcacatagccaaaatacagaattaacctaggtttccaccaagggatgactagatgaagaatatatgatataaattctgtatatgt**atatataaatatatatat**tatttatac**gtatatgtgtatacatacacacatac**acacacacacacaatggaataatactcaaccctacagaaaaaggaaatcatataatttgcaacaacatgtttggacctggagacattatgctaagtgaaataattcaggcacagaaagacaaataatgcatggtctcacttattcatggaatccaacaaagtcaaacccacagaagcagagagtagaagggtaattgccaagcactgggatggggtttcaggaagggaggcataggaaatagggaaatgttggtcaaaaggtacatgttttaatcagacagaaagaagttctagagacctaatatgcagcatgattactatagttaataatatatgtactcttgaaaattgctaagaatgtagattttaaatgttcctaccaaaaaaaagtcaaaacaaaataagtatgtgagattatgcatatgttagcttgatttaattattccacaa**tgtatacatatattaaaatgtcacaatatcacattgtataca**ataaatatatataatttctatttgtcaattttaaataaataataaagaaatgagatagctttcaactaagc**ttatatagcagtataatataa**aaatttttgtagtaa**caaaattatagtaattttg**gcttatattgaaatgtgcattaaacaagtgtttgcataaaatataattcaaattactattcatagcccacagccatgaaaacatgtgtgccttactcttggtacagaaacaaatagcttttactcctaaggaaaaaagaaaattcctgccttttgatgagaattgtttgagttttatacttattttatggggaactttgctgacctcttcattccacCATAGCTGGTTATTCCTCCCTTGCATGTACTTT**TGTTTCCTGGGGGCAGTGAGTCTTGGAAACA**TTTGGGAAATTCTTATTCTGAGTTTGCCTGCCCACAGCCCAGGTTCCTTCCTGGAGTCTGCAGCATCAGGCTCTCTCCTCGGCTCTCTTTCTGTCTTCTGTCCCC**TCAGCTCATAAACCCCAGCTCGTAGAAGCTGA**AGCCACCTAGACTCTAAGACACCTGATTGCAGAGACAGGAAAGGAGTTCTCAAGATAAGTGCCAAGATTTCATACTGGTTTTCACAAGAATCAGGGTTGGAATTAGAGCCGACTCATTATTCAGTTAACATTGACATTGTTGATTAGAGACATATCCCTATTGAAAATATTTCCTGGCAAAAAATAGAAGTTCTCTTTGGCT**CTGAAATCTGCAACTCCCTTTCAG**GTGTC**CCTGTGTCCTTGTACCGTCACTCcacagcactgcacagg**catgtgctcacctcacaaaatggcagtctcaaagggaggagtgcccacccacaagaggctccacccta**ttctgagaaagaacttctttcagaggaggagagaatagaactttccagaa**ccttttacatgagatttttttttttgggtag**gaagtctgacctgagcagcaaccttgcatcagacttc**aaaaggcaatatttcctatacctccttgtgctctactagtgagtacagaagtcgagaaaatgttcctaagtctatggccagttc**aaatagtctcattttctattt**ctgagattaaattatatgtattaagatccttactctgccctttcaggtttattgctcctcagacaagatatcccctcttcaggtcccaaacacattcattgcaatgcctttcctcatccttgtcccagtcctggctctctagcttcttgagacaataacccaccctaaggtatgtt**gtgacatgcatcttctgtgtcac**tgggatccctaagtggtacaagcattcactcaattgctttagctagagtccccagaatatcttcacctcttctgtcacattcttcttccacattggtcc**ataaacacatctttaacatttctgtttat**taaaacctctcaaattcaaccactattttctatcccaacgaatactgccttagctcaggcaaccactgtcttcgatctgaattacttctttgacctgaattacttcaactcctaccaaatatccacctacctgatgtcttactctcttttctaactcagttcattctctgcagctgaaaacacttcctttccattgactttagggttatgaccagtcttctttatatctaaatgttagcctataa**gttttttcgtaaaaaaaaaaac**aaacttccaaactacaaattcttaaaaccctccatgatttggcccttgcatacctatccagtcttgtgaaattcacctactctttacccactttttgca

Uppercase: TRBVA-2

Lowercase: Flanking sequence[1000bp]

Red & Bold & Underline: Stem-loop [15]

Blue: Heptamer[29]

Green: Nonamer [7]

id-IGHVII-43-1-2[V_gene_segment]

accaaaagtgtgtaagtggtccgttatctctatagccttgcctgcatctgttaatttctgatctttcagtgatagcaattctgacatatgtgagaaagtatatcatttggttttgattcgcacttatctgatgactgaatatgctgagcatttttataggttagttggcc**acttctatgtttgtatttgagaagt**ttctgtgcatgtcctttctccatttttaatggggttatttggtttatgctggtgattaagttccctatgaattaccaccttacacacactaagataatcagcattcaaaaaagaaaaatgaatagaaaaacataaccacttgtaggtatagacatactgaatttggaatggtcatggcttgcatgttggtattgaaatgggacagcaactttggaaaatggtcttcacagaaaacctcataaatgtagctagtgtcaccactcacaga**agtagatggagaatacaactcaagtgtccatcgact**gccagagggatga**agacgccgcggtgcgtct**gtaacccacagggatga**agacgccgcggtgcgtct**gtaac**ccacagggatgaagacgctgtggtgctttaaaaaggagtgaagcac**tgacataggctgcaactcggatgggccttgtaaacattgggtgagtgaactgagggagacacggaagtctacatcctgaattgtcctattgacatgaagtatgcagaggaggaaaatccctagagacagatcacagggggtgagtggaggggaaatggagagtgacgcttaatggagctgaggttttcttttctgtgatgaaaatgttcccaaaccaaatactggtgat**atttgcacaacattgcaaat**gtgtaaagtatcactgacccgtacacttaatcgtggc**taaaattgtaaatttta**tgtcatgtgtatctcaacacaatagaaatgagcaccacattttgtttagtgactcaacaacaataacatttaggtttaaatTCTGGATTCCCCAACAGAACCAGTGCTTCCTGCTGGAGCTGGATCCATCAGCCCCCAGGGAAGGGACT**GGAGTGGGTCAGGTGCACAGGTCATGAAGGGAGCACAAATTCTAACCCACTCC**TCAAGAGTCCAGTCACCACCTCCAGATCTATGTCCAAAAACAGCTCTTCGT**ATGGCTGAGTGACATTAGCAACAAGCACACAGCCAT**GTttgtttttttgttttgttttggtgtgtgtgtgtgtgtttttgatagagtcttgtgtcacccaggctggagtgcagtggggcaatcaatcttggctcactacgagctccacctcctgggttcaagtgattctccagcctcagcc**tccaagtagcttgga**ctgcaggcactcaccaccacacctggctaatttttgtatttttagtagagacagggtttcactatgttggccaggctggtcttgaactcctgaccttctgatccacccgactcggcctccaaaagt**gctgggattacaggtataagccactgtgcccagc**cacaaacaagtatttttaagcaaaagacacagtgaagagacttcagtatgag**cccacacacaaaccttcctgtggg**agtttacagaacagcag**tgggtgctgaggacagaagccagcaccca**ggaaccagcagggaaacccagggggcatttggcacc**gcctgaaggctcaggaccattgtggggctcagtggtcaggc**aggctcaaggttcagcc**tcagagcaggtgtagcaggcggggaaaggctctga**agacggagtttagtgtcaccttctcatttccactacta**aacaccctccagcacatctattctaatgtgtatgggtgtt**catgtgtttagagaatattatttatgttatgaatctatagccatcttgtgggtgcatcaagttaaccacttcaacctatgtggaccctgttcattaggaataagtccctgtatttgaggacctcacaaattaataattatgtagaatcactttctttttcagtctcctttccttcctctttctttctttctcactcacacactgacaaacacacggggtgccataacattaattacctgatgtattaaagaatattgattcatttgcaacttgaccagtttagctgttgtt**catgttgttgtaagatcaggaacatg**tttctcagctgtgtactcctctaagctgagcagcag

Uppercase: IGHVII-43-1-2

Lowercase: Flanking sequence[1000bp]

Red & Bold & Underline: Stem-loop [18]

Blue: Heptamer[36]

Green: Nonamer [5]

id-IGHV3-35-2[V_gene_segment]

gaggtgacatggggaatgcgtgaggtgacctggggcatgtgtgaggtgacaaagggcatgtgtgcagtgacctgggtgcatgtgaggtgacctggtgcatgtgttaggttacctgggggatgtgtgaggtgaacaggggcaggtgtgaagtgcctgaggaatgtgtgaggtgaactggtgcatgtatgtggtgacctgggggatttacgaggtgacctgcgggatgtgtgaggtgaccttgggtgagcgtgaggtgacctgacgtatgtatgaggtgaccagggtgtgtctgaggtgatgtggggtatatgtgcggtgacctgggaaatgtttgaggtgacctggtggatgtgtgaggtgacctgcaggatgtgtaaggtgacactggggatgtgtgaggtgactcagggcacatgtgaggtgaaattggggatgcgtgaggtgattcagggcatatgtg**aggtgaccttggggatgtgtgaggtcacct**tgggtgagtgtgtggtcatctgggatatttcccaacttcgtaaggaaagcctgcaacaacacatg**gcattagaagcttcacactctagcaccattaatgc**gtacat**cctcaaggaactctaaatgttttgagg**ttagcttaaatgccatggagcggcaccttc**cccaggcacactcatatatgcatcctggg**tctccctgccaagctatatctcctcaaatttacagtttatgtttgctctgtgacatcaactctgatatgttcaagtgtgtttttttttctttatttgtagttgttcaggcttgttgtttcactctcattactctgggctcagtcctctcctcaggtgtcccacttcagagctcgctgtgaaataggagacatgcaaatagggccccccctttcctgataaaaagcagcccagtcctgaccctgcagccctgggagagaagcaccagccctgggattctcaggtgtttccactttgtcatcagcaacaaa**caaattaccATGGAATTTG**GGCTGAGCTGGGTTTTTCTTGCTGCTATTTTAAAAGGTGATTCATGAAGAACTAAGGATATTGAGTGAGTGGACATGAGTGAG**AGAAACAGTGGATTTGTGTGGCAGTTTCT**GACCAGGGTGTCTCTGTGTTTGCAGGTGTCCAGTGTGAGGTGCAGCTGGTGGAGTCTGGGGGAGGCTTGGTACAGCCTGGGGGATCCCTGAGACTCTCCTGTGCAGCCTCTGGATTCACCTTCAGTAACAGTGACATGAACTGGGTCCATCAGG**CTCCAGGAAAGGGGCTGGAG**TGGGTATCGGGTGTTAGTTGGAATGGCAGTAGGACGCACTATGCAGACTCTGTGAAGGGCCGATTCATCATCTCCAGAGACAATTCCAGGAACACCCTGTATCTGCAAACGAATAGCCTGAGGGCCGAGGACACGGCTGTGTATTACTGTGTGAGAAAcactgtgagaggtcggaagtgtgagcccagacacaaacct**cctgcaggaacgttgggggaaatcagctgcagg**gggcgctcaggacccactcatcagagtc**aaccccagagcaggtgcacatggaggctggggtt**tgtttc**ctgtcaggatttgggacttcctctgcttctgacagtttctctagggaaact**ctttaattttagatttctgtgcccaccaatgtcatctctacatttttttaatcattgtatatgaggactcgttctcacatgcacaatatgtatattgccacctatgggaatgaaagg**tcctcaaccatggtcaccagcatcagagtcgtgagga**agctcaggggtgcctgg**tgagtcttctccagtcagactca**ggacagtaacct**caaggggattcccttg**tgagaactcacacattttcatgagaacagcaccaggagtcagttctaaaccattcatgaaagacccactccatga**cccagtcacctcccaccaggtcaccccttcacaactggg**gattataatacaacatga**gatttggggcaggacacaaatc**caaaccatatcagatacacattgt**gaaatacgcatggtggtcaggtaatttgtatttc**tatcacctcacagcgctaccattttattattttttttaatttactgcatgagtgagtgtctgatgagaacacctaagatctaccctttcagcaacaatcatttttacaatacagtattaactataggaccattgctgtacgttagatctccagaactcatccaacctgcacaactgaaactctgtacaatttaacacacatcacccaatttccctcacc**tcccaggtcctggga**cccactattctactctctgctttcaagagcttgaatattttagatcccacatgtaaatgagatc**atgcagcatttgtctttctgcat**ctggcttattcaactcagcatcatgtcctccaggcccatccgtgttgttgcaa

Uppercase: IGHV3-35-2

Lowercase: Flanking sequence[1000bp]

Red & Bold & Underline: Stem-loop [19]

Blue: Heptamer[31]

Green: Nonamer [5]

id-IGLV4-69[V_gene_segment]

actgatgtaaaaatcattgttttatgtactcagttttcctttgagtttaccctcttttctctactttttaaaatatttatttctaatttggggagctactaactgagatta**tttttcatctcactgaaaaa**cagttttagaatttcctgtagagcagttctgctggtggcaaattccatcgggttttgtctgaaaagtagccattttctcctattttttctgttta**ttatatagaaagataacttatataa**agtaaaattcacaggtcttaattatacagtttgatggtttttacaaatgcagatgctcatgtagccaacagtccgatccattctcacaaca**tctccattactgcagaatggaga**cattctgttccagtcaatgtaaattatcccattacacctaccaaatagaacgtgtatgagagacacctttctcctgaggacttttgcaa**agtggggtggatcatgtgtcccgctcccact**gaaaagggctaaatggaaaactaaagtctgaaataaaataggaggctgccctgacgaggggtcccactttgcccttggacagagaacaggccgtggtcaaggccctggtccgggcagaagcctctgtcaggacccactggcatctggtcacagacacgatggacctgggcctaggcagaagggggtgctgttggtctgct**gctgagggctctgtgggtttctcagc**tgggaaaccaaacacttgaacttggtctccacgcagggttcactggggccagcag**ctgggctctctctgcacccttggagagcctcag**gccaggcccagcccaggtaacccc**tcccagaaatgtcaccccaccactggga**ctgacactcaggcacacggagtgatttggttggg**cagaggaagaggagcacatttgcatgaagggcccctctctctcctctg**ggactacagggtgggtaagaaatacctgcaactgtcagcctcagcagagctctggggagtctgcaccATGGCTTGGACCCCACTCCTCTTCCTCACCCTCCTCCTCCACTGCAC**AGGTCAGGATGGCCCTCAGCACCCTGACCT**CCAGCTCACTGATACCACCTCCCAAACTTATGCCAGGAATGTCCTTCCCTCTTTTCTTGACTCCAGCCGGTAATGGGTGTCTGTGTTTTCAGGGTCTCTCTCCCAGCTTGTGCTGACTCAATCGCCCTCTGCCTCTGCCTCCCTGGGAGCCTCGGTCAAGCTCACCTGCACTCTGAGCAGTGGGCACAGCAGCTACGCCATCGCATGGCATCAGCAGCAGCCAGAGAAGGGCCCTCGGTACTTGATGAAGCTTAACAGTGATGGCAGCCACAGCAAGGGGGACGGGATCCCTGATCGCTTCTCAGGCTCCAGCTCTGGGGCTGAGCGCTACCTCACCATCTC**CAGCCTCCAGTCTGAGGATGAGGCTG**ACTATTACTGTCAGACCTGGGGCACTGGCATTCAcacagtgacacaggcagatgaggaagtgggacagaaacctcagcctgctcagggtcttgttatatgacaagttttaaattttcaaatatgttttatatgtacacactccacttggaagcaccgtctggtttaaaatgtttctactctcaatttctcattca**agttttctgaagtctcaggaaaact**aaaaacagaaacaaaaatatcatgatgactatgaaatattacctaattttccaaatatgctgaattcagtgcctgtggagctaac**cattttctcttgacagaagaaaaatg**agccttagaaattgtttccaaa**tcctgaaagagtcatcattccatgtactcagga**cagatttagagagtgctgaccctgccctttgaaagatg**ttatttgcctcttagtgcccaaataa**ggaaggataattaggaatcagaagcctacctagggctcccctctctcctctcaaaattgcacttggtgtcatattgccctagaatttatgcctggtgctggcaaatatttttaaatttattttttaatcttttccttttctgcaatcaa**atttcttgaggggctgagaagaagaaat**agg**cccagagagaagcaaggcagtctggg**tgcaggctctagacagcccagggagaagaaagagggagca**gctccctgtgtcctggagggagc**cctgtccccattgggcttcctgctctgatgtcccctagt**ggaggctcctcagggagtccagcctcc**cagaactgccctataaacctcacagtgagctaagcaagtatatccaggaaggagggtctgtggagatgtgagactaaggtgtgttatggcctaaacactgcccatctccatccttcttggggatgacaaagtgaacagtggaggagacaggatacatgggtgcatctggacctcaagcagtaggctgttctgtggtctgtccctaaatcttgtgctggagaccccatacgagagccactccctgaccct

Uppercase: IGLV4-69

Lowercase: Flanking sequence[1000bp]

Red & Bold & Underline: Stem-loop [18]

Blue: Heptamer[37]

Green: Nonamer [3]

id-IGHV5-78-2[V_gene_segment]

gtgataaagagaagtccttttgcttctgcaccccaggtacaactcctgtgagcaatctggcaaatattttagaaactgtgtagtttgtattgccttgtcttt**tcatttgaatcttttagatgataaggaaatga**gagtggttcacaagataatggggcagacacaggggatagttaagggcctcctccaagcaatgtggatatcataaaatgaatgttaaaaaatgaaagaactaaacaagaaggtgctggggttgacacaaaggacttaaaacagaactaccaggagttgggtgcagcaatggcaatcccttcagttccatcagcaattcagagacctaaacaaatattccctatttatgttgaattggcagactgtagaaatgtagaagacaaaatgaccagaaaaaatgtgtgatttggcatgggtgaaggtctggcaagttaatcaagttgaggactgttgaagccctagggtggtcctgcctttgctgggtgcccagaccataatggcactcatgggaaattgccagcaagtagaaactaggtgttttttttttttttgtagttcttattatcaggcatatgtgcatttgaattttgtcttcatagcatttcctggttctatttctcattttttaaaacacacgtgatttcatctagatttatcaccttcacagggtcacagagaagggtaggaagaagggaggccctggtttggg**tcttgtaggaccatggacaaga**gtgcggagggacaacagggggttgtaagacattattaagaccttattct**gcctctcccaagaggc**tcagttcagccgtttttctgcagttgcggttctgggttataaaccttgtagactcttcccttcaggccagggcggcaactatgcaaattc**aagtgggggcctccccactt**aaacccagggctctcctccacagtgagtctccttcaccacccagctgggatctcagtgctttcttttctgtcctcctcc**aggATGGGGTCAACCGTCATCCT**TTCCCTCGTCCTGGCTGTTCTCCAAGGTCAGTC**CTGCCCAGGGTTTGAGGTCACAGAGAAGAATGGGCAG**AAGGGAGCCCCTGATGCAAATTTTGTGTCTCCCACACAGGTGTCTTTGCCGAGGTGCAGCTGTTGCAGTCTGCAGCAGAGGTGAAAAGACCCGGGGAGTCTCTGAGGATCTCCTGTAAGACTTCTGGATACAGCTTTACCAGCTACTGGATCCACTGGGTGCGCC**AGATGCCCGGGAAAGAACTGGAGTGGATGGGGAGCATCT**ATCCTGGGAACTCTGATACCAGATACAGCCCATCCTTCCAAGGCCACGTCACCATCTCAGCCGACA**GCTCCAGCAGCACCGCCTACCTGCAGTGGAGC**AGCCTGAAGGCCTCGGACGCCGCCATGTATTATTGTGTGAGAGggaccatctaaaaccttccgcggtgcaggtgcagagtgagctgccagacacac**cctccccaggggcctctctattcatccggggagg**aaacactggctgtttgtgtcctcaggagcaaaaaccagagaacaacatgggagcgttcctaacccctaaggc**aactggatgggagacctgacccatccagtt**ctctgagggggctcttgtgtgttctacaaggttgttcatggtgtatattacatggttaacatcaaaaggctgcctaataggcacctcttcaatataatagtcttttaattagtgaaaattttacacagttcatcattgcttgcttgccttcctcccttctgtccgctcttactccctccttcttttattttctacttacttttccaaaatcatttaacccctttttgtaccattaataagttatcttgtttttgttgttgttttccttttaacaatatgcactgaataattca**tctttgtgccaattcgtaaatattttgatataacaaaga**cttctgtcataaatagatttcctgtgagtaatcttgcaaatattttagaacctgtttggttaagaataaa**ttaaaataaatagacaaatttttaa**agacatacaacctatcaaaactg**aaccaccaagaaataaaaaatctgggctgggcgtggtt**gctcatatgtgtattctcagcactttgggagaccgatgcgggtggatcacctcaggtcaggagttcaagatcagcctggccaacatggcaaaaccccgtctctactaaaaatacacagattagctgggtgtggtggtgttcacccgcagtcccatagtcccggctactcgggaggctgaggtaggagaatgacttgaacctgggaggcagaggttgcagaaagccaaaatcgagccactgcactccagactgagcaacagagtgagactccaattaaaaaaaatctgaacagaccaatgagtaaagagatcgagtcagtgatt

Uppercase: IGHV5-78-2

Lowercase: Flanking sequence[1000bp]

Red & Bold & Underline: Stem-loop [13]

Blue: Heptamer[32]

Green: Nonamer [7]

id-IGHV3-35[V_gene_segment]

gaggtgacatggggaatgcgtgaggtgacctggggcatgtgtgaggtgacaaagggcatgtgtgcagtgacctgggtgcatgtgaggtgacctggtgcatgtgttaggttacctgggggatgtgtgaggtgaacaggggcaggtgtgaagtgcctgaggaatgtgtgaggtgaactggtgcatgtatgtggtgacctgggggatttacgaggtgacctgcgggatgtgtgaggtgaccttgggtgagcgtgaggtgacctgacgtatgtatgaggtgaccagggtgtgtctgaggtgatgtggggtatatgtgcggtgacctgggaaatgtttgaggtgacctggtggatgtgtgaggtgacctgcaggatgtgtaaggtgacactggggatgtgtgaggtgactcagggcacatgtgaggtgaaattggggatgcgtgaggtgattcagggcatatgtg**aggtgaccttggggatgtgtgaggtcacct**tgggtgagtgtgtggtcatctgggatatttcccaacttcgtaaggaaagcctgcaacaacacatg**gcattagaagcttcacactctagcaccattaatgc**gtacat**cctcaaggaactctaaatgttttgagg**ttagcttaaatgccatggagcggcaccttc**cccaggcacactcatatatgcatcctggg**tctccctgccaagctatatctcctcaaatttacagtttatgtttgctctgtgacatcaactctgatatgttcaagtgtgtttttttttctttatttgtagttgttcaggcttgttgtttcactctcattactctgggctcagtcctctcctcaggtgtcccacttcagagctcgctgtgaaataggagacatgcaaatagggccccccctttcctgataaaaagcagcccagtcctgaccctgcagccctgggagagaagcaccagccctgggattctcaggtgtttccactttgtcatcagcaacaaa**caaattaccATGGAATTTG**GGCTGAGCTGGGTTTTTCTTGCTGCTATTTTAAAAGGTGATTCATGAAGAACTAAGGATATTGAGTGAGTGGACATGAGTGAG**AGAAACAGTGGATTTGTGTGGCAGTTTCT**GACCAGGGTGTCTCTGTGTTTGCAGGTGTCCAGTGTGAGGTGCAGCTGGTGGAGTCTGGGGGAGGCTTGGTACAGCCTGGGGGATCCCTGAGACTCTCCTGTGCAGCCTCTGGATTCACCTTCAGTAACAGTGACATGAACTGGGTCCATCAGG**CTCCAGGAAAGGGGCTGGAG**TGGGTATCGGGTGTTAGTTGGAATGGCAGTAGGACGCACTATGCAGACTCTGTGAAGGGCCGATTCATCATCTCCAGAGACAATTCCAGGAACACCCTGTATCTGCAAACGAATAGCCTGAGGGCCGAGGACACGGCTGTGTATTACTGTGTGAGAAAcactgtgagaggtcggaagtgtgagcccagacacaaacct**cctgcaggaacgttgggggaaatcagctgcagg**gggcgctcaggacccactcatcagagtc**aaccccagagcaggtgcacatggaggctggggtt**tgtttc**ctgtcaggatttgggacttcctctgcttctgacagtttctctagggaaact**ctttaattttagatttctgtgcccaccaatgtcatctctacatttttttaatcattgtatatgaggactcgttctcacatgcacaatatgtatattgccacctatgggaatgaaagg**tcctcaaccatggtcaccagcatcagagtcgtgagga**agctcaggggtgcctgg**tgagtcttctccagtcagactca**ggacagtaacct**caaggggattcccttg**tgagaactcacacattttcatgagaacagcaccaggagtcagttctaaaccattcatgaaagacccactccatga**cccagtcacctcccaccaggtcaccccttcacaactggg**gattataatacaacatga**gatttggggcaggacacaaatc**caaaccatatcagatacacattgt**gaaatacgcatggtggtcaggtaatttgtatttc**tatcacctcacagcgctaccattttattattttttttaatttactgcatgagtgagtgtctgatgagaacacctaagatctaccctttcagcaacaatcatttttacaatacagtattaactataggaccattgctgtacgttagatctccagaactcatccaacctgcacaactgaaactctgtacaatttaacacacatcacccaatttccctcacc**tcccaggtcctggga**cccactattctactctctgctttcaagagcttgaatattttagatcccacatgtaaatgagatc**atgcagcatttgtctttctgcat**ctggcttattcaactcagcatcatgtcctccaggcccatccgtgttgttgcaa

Uppercase: IGHV3-35

Lowercase: Flanking sequence[1000bp]

Red & Bold & Underline: Stem-loop [19]

Blue: Heptamer[31]

Green: Nonamer [5]

id-IGKV2-18-3[V_gene_segment]

gggcaacagagcaagactccatctcaaaaaaaaaaagaaaaacaaaaatctatattccatagttccaccttgacagttaatata**cacaaacttcattcagttacactcttaatatttgtg**ccctttactctggggattttgtcaactaaaatgaaactaatttaaaaacctagatgctaaaataaatatgaaaataataaataagaataactggccaggtgtggtggcacatgcctgtaatcccagcactttgggaggtggaggcaggaggatcacttgaagtcaagagttcgagaccggcctggccaacatatgaaaccccatctctactaaaaatacaaaaattaaccaggcatggtggtgcacgccagtaatcccagctagctcgggaggctgaggcaggaaaatcgctttaaccttggaggcggaggttgcagtgagccgagatcctgacactgtactccagtctgggt**gacagagtaagtgaaactctgtc**tcaaaatatatatatatatatttatacacaaaat**aatatatatatttatatatatttatatatatcaaaaatatatatatt**t**atacatatatacatatgtgtatatatatgtat**gtgtgtgtatatatataataaaat**taaataagaataaccaaatattta**tggaaactcagcctattaaagacaagtgtaaaatgtgtctgggaaagtgaaaacaatgtagaaatctcttagaggaaatagaagaacatgattaaatatgtccctttttccactcaggcagcacaaactttattaggttccctctaaacacagaggacacattatgtcactgtgctcacactagatgtccccatactctccttggctctttccaccccactgcacccacc**aggggatttgcatactgtcccct**agggaggaccttccattgtgagtctgagataaaagctcagctgtaaccttgccttgactgatcagaactcctcagctcaccttctcacaATAAGGGTCCCTGCTCAGCTCCTGGGGCTGCTAATGCTTTGGGTTCCTGGTAAGGACAGAAGGGAGATAAGGGAGGAGAATGGAGTGTGAGGGTGAGCTCTGGGGGTCCCACTGCCTGTCCACATGCACATCTTGACCTGCAAGAAAAGGTGTATA**AAGTTTAGAACTGCAAGAGTCAGAAAGAGAAGATTCCTCACTCTTATAGTTCTAAACTT**CATATCCTAGAAGGACAAAGGACTTGTGCTCTCATGAAGAGTGTCACACAGGAAGAGGATAATAGTGTAGGCGACTTCTTGAGTCTCTTTTGTGCCTTGTGAATG**TTGGTTCTTTTTATGCCTGGATGTTTAGGGGTATGAACCAA**AGTCACACAAAAAAATCACTTAGCATAAAGTAGATAACAGAAAGAATATCATTTCAATGGTTCCCAATATTTGTACATAACTTTGCACTTCTCTCACTATTTCAGGATCCA**GTGGAGATATTGTGATGACCCAGACTCCAC**CCTCCCTGCCCGTCAACCCTGGAGAGCCGGCCTCCATCTCT**TGCAGGTCTAGTCAGAGCCTCCTGCA**TAGTAATGGATATACCTATTTGCATTGGTACCTGCAGAAGCCAGGGCAGTCTCCACAGCTCCTGATTTATAGGGTTTCCAATCATCTTTCTGGGGTCCCAGACAGGTTTAGTGGCAGTGGGTCAGGTAG**TGATTTCACACTGAAAATCA**GCTGGGTGGAGGCTGAGGATGTTGGGGTTTATTACTGCATGCAAGCTACACAGTTTCCTaacacagtggtacaacc**ctgaacagaaacctcccttcttgctgtggttcag**ctgcccaaatgtgttgtttatctggaaagcagacactgtctattatcttgggagagtaaagaggaagatgatggagaattcaggagaatacattacagctgagggctcttgaccatcaatatctccgttacatctcaggtaccacaatttagtccccatcagt**tgcaagagtcttggcctggcagaacttgca**ggagaatggagggaggtcaagtgccctctgaacaaccagtcatgttgcaagggaaagctgaatcaaagctcatcgtaatcctctctgccttgcctacgttcattcactaactaaa**ttcattcagcctgacagccacctaaatgaa**acagattagtggcaata**caaaactaatacgttcttggttttg**atttggtttagcagttactagtgtacatgtaccttgacaagatttggtgatactaaaacagtttctctcccatcttctaccttccactttacacttttcttaacatatatctctctctgggacagcagg**aagcacagcattttatcctatttttttttcagggagtgctt**gcgtgtatttacattctagactttgtaactgctagatatattttggtaagtttatgcaaaaa**caaagacctagtgacaaaaatctttg**aacaatcttattcacagtatgttgaaatgaaaagaaaattctatgaaatccaaagctgtgttttaaaaataaatatccaagtttcaaaagacaaaagaaaaaacgactaaaaatgtatgccatagtcaaaaatgtatgccatattcaatggaatctggatt**aatataaaataatatataagcttatattttgccccttgaaaaata**ataatctatagactttgggcagagggcagtaaatgagaatgtgagggcagctgttaactgtgttagtggtggtgtgttggagttgatatatgaacccaagtggaa

Uppercase: IGKV2-18-3

Lowercase: Flanking sequence[1000bp]

Red & Bold & Underline: Stem-loop [19]

Blue: Heptamer[30]

Green: Nonamer [12]

id-IGHV3-37-2[V_gene_segment]

aactaaattcacaagtacatataagtaaaataagccaaaaaaaaaaaactacatcccgtatgatga**cacttttatacattctacagaatagaaactaaagtg**atatgaaggttgggagttgaatggagaaaaggtgagcgaaaagagggaggacggaggagttgcaagagaacaag**agaaaatgttgagggtaattgatctgttttct**atcttaataatggttatggttacatcaataatttctaaattgtacattttgtgtgttaattataattttatattattcaagctattacaaataaatacctgtataaatttcttagggacggtttaagggagatggataaatcacataaaaagtcagatatttctgaatgtacaatgaataacccttagctctatctttattttttggtaattctaaaatgcagcagatttttttaatgtttttattacaagaagggtttcttcaaaccacaccaggtgtgtgtaactctgggctaaagttggttaagggagcagtgggatccggtggagaggagcacaggcctcgatcctggagcccattcttgtcacatgtgagctcctggacacatctcatagctccctcatgcttctgcg**tttacattctcatctgtaaa**tggaggaccaatgggcatctgcctctaacaatgtattttcatgtgttaagatgatgtatgttaagtgtttactatagcacaatcttttaataagaaattacatttttttccaaatactatcattgtcttcaagccccaccacccaccagaaaacttaatatgctctgtgccctgccctgtcctcagacatccagcccaagagacttttatatagtaggagatatgcaaatagttccctccctctgctgatgatgaaaaccagcccagccgtgaccgtgcagctctgggagacgagctcaagtgccaggattc**ccaagtgttttcacttggtgatcagaacttaacacagaggaatcaccA**TGTTGTTTGGGCTGAGCTGGGCTTTCCTTTTTACTATTTTAAGAGGTGATACATGAAGAACTACAGATATTGTTCGTGAGTGGATATTAG**AGAAACAGTGGATATGTGTGGCAGTTTCT**GACCAGGATGTCTCTGTGTTTGCAGGTGTGCAGTATGAGGTG**CAGCTGGTAGAGTCTGGGGGAGACTTGGTACAGCTG**TGGTGGGGTTGGGGGTGTCCTGAGACTCTCATGTGCAGCCTGCGGATTCATCTTGAGAAGCAATGACATGAACTGGGCCCACCGGGCTTCAAGAAAGGGGCTGGAATGGGTCTCATACATTAGTGCTAGTGGTGGTAGCCTATACTATGCAGACACTGAAGGGTAGATTCACCATCTCTAGAGACAATGGCAAGAACATGCTGTACTTGCAAATGAACAGTCTGAGAGATGAGGACTCGGCTGTGTGAGAGAcatggtgaggggaaatcagtatgagcccagccagaaacct**ccctgcaggaaccctggggtggggggaaatcagctgcaggg**ggcactcaggacccactgatcagaatcaacccca**gaaggaggtgcacatggagattaggggctggattccttc**caggatgtgggacttcaacttcttctgacagtttccccagggaaccgctctaagtttagaattctgtgtcatctctatgtat**ttaaaaaagatcattttaatgtgaggaactaactattctcacat**gcacaaaatgcagattcacactcacagagatgaaaagtcctcaactgtggtcactaggatcagagt**cctgaggaatctcagg**cgt**tcctggtgagtcttctccaatcagacccagga**cagacacctcagtgagattccatgactagaacaggctttatggattttaaaaaaaatagactgagcc**agggtcagtgtcatgtagaacctcacaggtttcacatctgaccct**tctcctgacactaaagtgtgcaaatcagcatcaatactgatctggtgctgcttttgctcccaatctgtgttctttcttgagttgtccc**tttttcatttgcttttcctgctccctgaaaaa**agaaagatgttgtctctgtggtctaaattccagggctcaagcccttttcttggagctcaggtgggtctcaggtt**gtggctcctgcagccac**gctggggaggctg**atgggactttcttctctcccat**tgctcagggcc**ctccactatattgtgtggag**actcatctcagaatagatgtggccagtgggagctgaaggggacaagcttgtttggtcaaaatgagatacaaatatggaattcaccctgtgctgtacaaactgtcacagagtc**actttctgaaacaatagtgttagaaagaacttgagaaagt**tgtcagaatcaaaattgggtcacttgtgttaaaactctgctgaatgaagctgggaaaggccg

Uppercase: IGHV3-37-2

Lowercase: Flanking sequence[1000bp]

Red & Bold & Underline: Stem-loop [19]

Blue: Heptamer[30]

Green: Nonamer [6]

id-IGHVIII-5-2-2[V_gene_segment]

ctgaactcgtggagatagagagtgaaaggatggtcaccagaggctgggaaggtctatggtagatggggattattggggatgctcaatggatgaaaaaagtggttagaaataa**caaataatacatagtatttg**ataacacaacagactaagtataatccttaataacataattatagattgtaaaataactaaaagtgtataaatggaatgttggtaactcaaagaattaatacttgaaaggatgtaggaaccattcat**cataatgtgagtattatg**cagttcatgctatatcaaagtcaggtaccccataaatatacacatataccatttacccacaaaaattcaaaattaaagtatttatgtatctc**atcttctgtatctatattacattaaacctaagaagat**actggtttcttcacactaccaagatccacatagacatccgtagccttctttctttgcctgtccataatctgacattgaaaaatgaggaaacctttcctatcatatgacattaatttacttagtg**cacaatttcaaaatacacaaattgtg**ttagaattattaaactgacccttgttgaaaatatgtttatcgactagaataccgtgcttatatgcagcttatttatattttacaatcacaacatcaaattatttccaaagatacttgtttctgatattcatgcccccctttctgcagtgaaattagtacaaacctgtttatcattgttagatatttttatacagctttcattttattctgtgtttcatgaccagctaaatacttttgaagttgtatgcatttaggttaacactttgtctcaaaaaattatgagttttcacaaattatttatgccatgtgtttatcattgaa**caccatgaaaatagtttcattgtg**atta**aaaagtgtgtgtttcacctaatccacacaccttttccccagattcctggaaaa**ccacaaaggtttactggatcta**tatttttgtctttggcaaaatAGCCTGGCTATT**CTCTTAGCTTCTTCTGCAAAGCATCAAGATTCACCTTCACTGACTGCAGCATTTATTGGGCCCTG**ATGGCTGGAGGAGAGGGGCTGGAGTGGGTGGTAGCCAT**GAGTGACCCAAGTGGAAGTTCTCAGTGGTGCTCTGCTCCAGCACAAAGAAGATTCACAGTTCCTGGGGACAACACTTAACATCACAATCTCCCTTAAAATTATCTACTGGAAAGCTGAGGAGTAGGCAGTGTATT**ACTGTGAGAGAcacagt**gagggacatatgggtgagtctagacgcaaacttccctgcaggaagacaagaggggactttgtggtaaatggtgctcagaaccactaggggacactcagaaccactaggagacactcaggatagcagaaggtgctcaggatgccgcagggtgctcaaaacactaggggtgctcaagaccagcagggggcactcaggacaccaggagacacttaggacatcagggggcgctcaggacaccagggggcgctcagcacaagcaggggccacagaatgacaagggtgcattccagaccaccaggaaacactcaggaccaacaagagattctcagaaaccaccaggtgtaacaggaccacctaaaggcccacaagacaaccaggagacactca**tggccactgaagaacattgaggacaccaggggttcagggcca**acagggagtgctcggggccaccatggggctttcagtaaccacgagggaatcaatcactcaggacaccacggttttcttaggaggcagctccacatcaggtgcctggggaaggt**gacgtttgcttttagacctagctgattcctgagctggtcaagcaaaagtc**ttcccacgatctctcactatttcttccttgtaacccatggtttctttcacatacaaatcattaacttaggataggaattcaattcaacttttaaccttgcatatttttagaataataatagcaatagtaattcaatgtgttttgataataagaaattgtgtatgcctaattcaaactctggttccacgcatgggtttttgtttcttgtgctgtgccagtcacactacagtcaatgcttttctaataataactcaacaatgtcacggtgttaagtttcctttcattttgtctgccattt**gtggagatgaaacaccactttcacgggtcagttcctccac**tttggttgggttctggtcttctgctcttcacactgtttctccaccttcccttctgtcaaatcct

Uppercase: IGHVIII-5-2-2

Lowercase: Flanking sequence[1000bp]

Red & Bold & Underline: Stem-loop [14]

Blue: Heptamer[31]

Green: Nonamer [6]

id-IGHV3-53[V_gene_segment]

tctcaaagaattgtgctgagtgaaagaagctaag**gaattaagagtaaattttatataattc**attgtagaaatattagaagatgccactaccataaattaaaatgaagaagacttaaatttttctgagaaaatggtgttgggaatgatgcggatgtgatttaagtttcagaggaataagaaaaagatttagggattaatttaattattcaaaagttgattgaagtgccgagtgaatggctgcaaacatagctctacattt**ttcaaatcattccctataaatttgaa**ttaatt**atttatttttatacttgaataaagcaataacaaagaaataaat**gaatatttttgctaaaatggagcaataaaaagactgatattga**cagaagaaatatgactgacttctg**aaaatacacacacatgagccgtggttctctctacatatttagataaattacagaaagttgtcataactgatggggaatcctgcagacttcactaggcatagtccacactgcc**ctggagttgtctcaggggagctgcctcctccag**tggttagagcacaggcccaggtaataggactcatttttttagatgtgtaattttagacacac**tgcacaactgctgtgttctctgtgca**aattatctcctgtaaaatgtaacattgaaacctgccttaaatatattgtgtaaatatgtaaaaataaaatcagattgtgagagctaaatgctaatcaaggcgcaatcacgt**aatatacaattatatt**ttcctgaatgatggaattaataccaatctcccccaggacactt**catctgcacggagcccggcctctcctcagatgtcccaccccagagcttgctatatagtcggggacat**gcaaatagggccctccctctgctgatgaaaaccagcccagctgaccctgcagctctgggagaggagcccagcactgggattccgaggtgtttccattcggtgatcagcactgaacacagaggactcaccATGGAGTTTTGGCTGAGCTGGGTTTTCCTTGTTGCTATTTCAAAAGGTGATTCATGGAGAACTAGAGATATCGAGTGTGAGTGAACACGAGTGAG**AGAAACAGTGGATATGTGTGGCAGTTTCT**AACCAATGTCTCTGTGTTTGCAGGTG**TCCAGTGTGAGGTGCAGCTGGTGGAGACTGGA**GGAGGCTTGATCCAGCCTGGGGGGTCCCTGAGACTCTCCTGTGCAGCCTCTGGGTTCACCGTCAGTAGCAACTACATGAGCTGGGTCCGCCAGG**CTCCAGGGAAGGGGCTGGAG**TGGGTCTCAGTTATTTATAGCGGTGGTAGCACATACTACGCAGACTCCGTGAAGGGCCGATTCACCATCTCCAGAGACAATTCCAAGAACACGCTGTATCTTCAAATGAACAGCCTGAGAGCCGAGGACACGGCCGTGTATT**ACTGTGCGAGAGAcacagt**gag**gggaggccattgtgcgcccagacacaaacctccctgcaggaacgctggggaaatcagcggcaggg**ggcgctcaggagccactgatcagag**tcagccccggaggcaggtgcagatggaggctga**tttcctgtcaggatgtgggactctgtcttcttctgac**ggttccccagggaacc**tctctaagtttagcattctgtgcctatgaacgtcttctctaagtatttgaaagagattattttaatatgaagagcagttctcactcgcacaaaatgtggattgatgcttactgggatgaaaagtcctcaaacatggtcaccacgatcagagtctgagtgagctcagggcttcctgctgagtctcctcctatcagaccaaggacagggacctcagtgaggttccccgtctagaacagtctttatggatactgattgtgggcggcaagccacccaggtgccgacgcaagagaccgaggacacgagctgttccagtacaataaaatataaaacaagaatagttataccagatatagatcttagat**atgattatatatgaatatcattaatcat**tagttggtagcaattactctttattccaatattataataatcctcactctacaatcataacctaggaaaagccaggccatacagagataggagctgaggggacatagtgagaagtgaccagaagacaagagtgc**gagccttctgttatgcctggacagggcgaccagagggctc**cttggtctagcagtaatgccagcatctgggaagacgcctgttgccaagcggaccatggtctagtggtagactcagtgtcaaggaaaaacacctgctacttagcagacca**ggaaagggagtctccctttcc**ccggggagtttagagaagactctgct**cctccacctcctgtggagg**gcctgatatcagtcagacccgcccgcacttatccggaggcctaacagtctccctgtgatgctgtgcttcagtggcc

Uppercase: IGHV3-53

Lowercase: Flanking sequence[1000bp]

Red & Bold & Underline: Stem-loop [21]

Blue: Heptamer[40]

Green: Nonamer [6]

id-IGKV2-10[V_gene_segment]

aaatcattggctaaattttcaccattacaaatggaggtgagaagacagtggaatgatatttttaaatactgagccacaaattctcagccacaaattctatatccagaaaaacaatcgtttaaatataaaagcaaggtagaaatattccttcatacacaaagattaattcattaatagcatatatgccttgtaagagattcaagagaaaattcttcagaataacagagaatgacagcagacagcaactcaaattgacaagaaggaaccaaggcctcaaaaaatgaaaaaagttagaaagaagctacaaaaaatgtttactttacctgatataaagattaagtccaggactaaaacaattatatgtcagaatatcatttacattgattgtgttttgagcagcagtttttgactgg**gaaggtgcataatgaaagcagcgagagaattggaaaccttc**tgcgtttctatgtg**gatcatgatatgaaatgatc**tggatcttgacattcatttgcacccttaagagttgcaccctttgcactgtgtgttttaccttagataaaataaaataaaataaaataaaataaaataaaataaaataaacaaacctagactcttttaaataaacctaaacaaagaaataagaatcatcaaatatttatgaaagatcaagcttattaaaaagaaacataaaacatgcccgagaaagtcctaaagaactagaagtctagtagaggaaacagcaggaaaatctaaatacgtcccttctgccactcaggcaacacagattttacctctattgatttccttaaaa**acataaaatatatttatgt**cattttgctcacaagagacgcccccaccttctccttggctctttccaccccactgcacccacc**aggggatttgcatactgtcccct**agggaggactttccttgtgagtctgagacgaaagctcactctaaccttgccttgactgatcaggactcctcagttcaccttctcacaGTGAGGCTCCCTGCTCAGCTCCTGGGGCTGCTAATGCTCTGGGTCCCTGGTAAGGGCAGAAGGGAAATGAGGGAGGATGATGGGG**TGGGAGGGTGAACTCTGTGGACCCCGCCACCTCCCA**TGTGTGTCCTGTCCTCGTGTTAGATGTGTCTTGTCCTCCAGGATGGGGCATGTGATGTCTAGATCTGTGAGAGTGAGGAAGATTCCAGAAGGAGCAAGGATATGTAAATATTGGTTCTTTTTAAATCTGTATGTTTTGGGAGGATTAATCAAAATCACACACACAAAAATAATTGAGCAAAACATAAATAACAGACAGAAAATGATTAAAATGACTCACAATGTTTGCACATAACCTTGCACTTCTCTCTCATTATTTCATGATCCAATGGAGATGCTGTGATGACCCAACCTCCA**CTCTCCCTGCCCATCACCCCTGGAGAG**CCAGCCTCCATCTCCTGCAGGTCTAGTCACAGCCCCCTTCACAGTAATGGATACATCTATTTCAACTGGTACCTACAGAAGACAGGCCAGCCTCCTTGGCTCCCAATCTATTTGGTTTCCAATCACGACCCTGGAGTCCCAGACAGGTTCAGTGGCAGTGGGTTGGGGACA**GATTTCATGCTGAAATC**AGGAGGATGGATGCTGAGGATGTTGGGGTTTATTGCTGCCAGCAAAGTACACATTATCCTCCcacaatggtatagtcttgaacacaaacccccccacttgctgtggcctagctgcccagatgtgctgtttctgtggagagcaggcactgtggattctcttagatgcctaaag**accaagatgttggagaactcagaggacttggt**gcag**ctgagggctcatgaccataaatttcttggctacacctcag**gcaccacattttaaggtcccgtcag**ctgcagcagcctttgcatgacagagtctgcag**tatggaggaggtccacgtgccctctgagcaatgagacac**aagagaagagaaggctcaatgagagctcattctaatcctctcttcctt**ccctatgttcattcatcaactaaattcattctgcataacaggcacctaattgagactgattactggcaacacaaaactaacacattcttttgatttggtttagcagtt**accagagtacatgtacattgataagatctggt**gatattaaatcagtttccctcccctcctccacctcccccctccactttttgtaaggccgacccttgaccagaacagcagcaagcactgtattttatcctacttttttcagggagaagccaaataaaactcttcacagtctagattttttaattactagatatgttgtggcaaatgta**tgcaaaaataaagacctagttgctaaagtgtttgca**cagtctcactcacagcatattgaagtacaaaagtag**ctttctgaactccaaagctgggtttcaaaaataaggacaaagttcgaaag**aataaaggaacatatgtatgttatttaaacagaatttgaatgaatacaagataatatataagctaactttttaccccatgaaaaattacaatgtgaagattttgagcagagggcagcaagtgagagactgg**tgggttgtgtttgagatatgaaccca**gatggaag**ttctcacgtatttcctgggaagaatcacagagttgagaa**gcgcctgcctggctttcaaggcagtaa

Uppercase: IGKV2-10

Lowercase: Flanking sequence[1000bp]

Red & Bold & Underline: Stem-loop [16]

Blue: Heptamer[35]

Green: Nonamer [8]

id-IGKV1-32[V_gene_segment]

aacccagtcattaagaactgctt**tctcttaaaaactggaagaga**ttggtaatatagagatatacagtttatggaacagagtagattcaagagt**agatgtgagcactgggtctagataacatct**cagtttccggaagatattagcttaagaatgacgtaaggcacacatattaatccaaggagctcaggaaacatctagcataggaatgcatgatggcacacagaggtagagagtagctgacacttt**tcagaagtggagggagaagatttctga**tgttggcctagatgaaagaagtaggaaagaaaatgtagaaatacaggaagatgatcacatgtctgagataagagatttgccatgtcgtttacccagtgta**caaggattacccaatccttg**caaagcattacactactttcacatattttgtgttaacattttcctttttc**ttatttttaataaataa**atctgtccttctatccatccatctatgtatccatccatctttcttatacc**tcatcattgcatctctggctgatagtgctgatga**tttacaacatcaagggctctagaggatcagagcaaagaagaagtctctctggcctcaataatattcatcagtaccaaacgcgggacacttctaaaatttattgcagattc**tggagatgtgcttctcca**acaagccaggtcttagacgggaagctagcaagaa**aagtaggtcatgacctacttctgcagagcagaa**aagcaatctccttgcaggacagtagacagattctgttctgtggcccccagagagaa**acttgttctctgctaacaagt**gtatg**caggcctgatcaatcttaggacccctgggggacactaggcctg**tgcaatattatgcaggtaaagtcattcttgcatctg**ttgaaattttcattttcaa**aaaaacacagccaacttcctgaagacttgtacacaggctggtcataccacttgcaagagtcagtcccagtcaagacacagcATGGACATGAGGGTCCCTGCTCAGCTCCTGGGGCTCCTGCTGCTCTGGCTCTCAGGCAAGGAAGGAGAACACTAGGAATTTACTG**AGCCAGTGTGGTCAGTACTACCTGGCT**ATTCAGGAAAGTCCTCTCATAACATGGTTAATAGTGTTAATAGTTTTGTTTCCCATCTCAGGTGCCAGATATGACATGCAGATGACCCAGTCTCCATCCTCGCTGTC**TTCCTGTCTAGGTGAGAGTCACCATCACATGCCAGGAA**AG**TCAGGCAGGGCATTAGCCATGTTTTAGCCTGA**TACAAAG**AGAAGCCAGGGAAAGCTTCT**GAGCTCCTGATCTACGATGCATCCAATTTGCAAACCTGGGTCCCATTGCAGTTATGTGGCATTGGATCCAGGACAGATTTGATTCTCACCATTAGCATCCTCCAGTCT**GAAGTTGCTGCAACTTC**TTATTATTGGTCAACAGTATAAAAGTGACCCTctcacagtgttacaaacccaataagctccccaag**gaagcagatatgtgagggtgggctgccccagctgcttc**tcctgtttcctccatct**gctgagagtgtttctcagactcagc**cacactctgaaggtcactgagcagttttcatagaagggg**taagggaagtgtctctacaccctta**gtttcttttatcctcctcagccccagcagcagacatggcaatgcctctcctaatttcataga**agagtcattacctatgaggactct**gggttacagcactggtccaggttcatacaacaaaagagaagctattgtaggtaacccaaatagaaagtttttcctaatatgggaaatccatgtctaaattacaacttttcaaagaccagaggatataatggttaggaaaccagaaaggcaaaaggagtgctgggaaatctacttcaagaagcaatggtggccgcgcacagtggttcaggcctgtaatcccagcactttgggaggccgaggcaggcggatcacctgagatcaggagttcaagaccagcctggccaacatggcgaaaccccgtctctactaaaaatacaaaaattagccgggcatagtggtgggtgcctgtaatcctagctactcagaaggctgaggcaggagaatcatttgaaccaggcaggtggtctctgcagtttcgaagcccagcgccatct**gtggctgttatgcatgtctctcccagccac**cctgctgtccagagcccatatcaatccgtgggggtaggtctgtgaaaga**gcagctcagttaggacccagagggaaccatggaagctgc**agctcagctgctttgattcttgttactctggctcccaggtgcaggaaacaccggatggttctgcatgtcagtgaaacttt**ctcaaccttgttgag**tcctgttacctggcacacctgctgggaaggcacaatgattaaagctcaatgtagatcaatggtcctggatgcactgggaagacaatagg

Uppercase: IGKV1-32

Lowercase: Flanking sequence[1000bp]

Red & Bold & Underline: Stem-loop [24]

Blue: Heptamer[26]

Green: Nonamer [3]

id-TRBV20-1[V_gene_segment]

cctagaaactattttcactgtccagtacttagtatgacctccattacaat**agaataaatttaattattct**cacaacctcaacagtgattaacacgg**aaacatacaagctaaattgagaaatattattagaatgttt**agaattaagtccctttatatatgagggaaattgtc**atgagatacacatgaggtaactatctcatttagaaaatga**tgaagtggcttcaacatgattatgaagaaacattggacttccagccga**ggcactggcctgtgctttatcaatatttatggtagtgcc**aacctgtgggtctagtgattaacaaccatgagttgtcttggtgttattactttgaggatatgaaacagaagcagcaaacagactccaatgccattcattacctaaacctacactcc**tttccttataggaaa**ctcacttaagtcattacaatgctttatgtctcagtgtatctttaataaatttggaagttgaattgggaagtctttaggccactttaggcaatgtatgggttttttgttttcttttgatatggggtctcactctgtcacccaggctggagtgcagtggcatgatttcagctcattgcaacctccgcctcctgggttcaagcaattctctccctcagcctcctgagtagctgggattacaggtgtccaccaccacgcctggctaatttttgtgtttttagtagagatgggttttcaccatgttggccagattggtcttgaactcctaaccttgtgatccacccgcctcggactcccaaagttctggaat**tacaggcgtgagccacctcacccgacctgta**tggttttttgacaggcagagatggacgtaacatcagtcatgggcaaagattaccaccagggggcagactagggcatccttgggattctgtgatcagtcatccctcctcgctggtgaatggaggcagtggtcacaactctccccagagaaggtggtgtgaggccatcacggaagATGCTGCTGCTTCTGCTGCTTCTGGGG**CCAGGTATAAGCCTCCTTCTACCTGG**GAGCTTGGGTGGGCATGTGCGTGTGTTGGCATGGTCAAGTGGTGGCCAGCAGGGTTGCAATGTGGATTGTTTATGCTCATCGAAGGGAGAGGGAGAGGCCCTGCTCTCTAGAGGTGTAAATGGTAAGGTGAAAGCCGCCGGTCAGAGGAGATGGGGGATTATGGCCCTAGGGAGATGACGGGAAGATTGCACAAAACAAACAGGACTCTCCAG**GAGCTGGGAGCACAGGGAGGGAGTGAGGCTCAGCTC**TGCCTGGCGTCCTGTCTGACTCGGCTCCCACTGGGCTCTCCTCTCTCTCTGGCTTCTGTCTCAGCAGGCTCCGGGCTTGGTGCTGTCGTCTCTCAACATCCGAGCAGGGTTATCTGTAAGAGTGGAACCTCTGTGAAGATCGAGTGCCGTTCCCTGGACTTTCAGGCCACAACTATGTTTTGGTATCGTCAGTTCCCGAAACAGAGTCTCATGCTGATGGCAACTTCCAATGAGGGCTCCAAGGCCACATACGAGCAAGGCGTCGAGAA**GGACAAGTTTCTCATCAACCATGCAAGCCTGACCTTGTCC**ACTCTGACAGTGACCAGTGCCCATCCTGAAGACAGCAGCTTCTACATCTGCAGTGCTAGAGAcacagcgccaggaggggatcagacaccgcggcaagaacccctgcagctgccctccgccccagcgggccccctgagtgctgagaggggaagcgtggagaatggaaaaccacagctttcctgactgagacatctgggagtgtgtgtgcagggggtggggcgggggggagggggagaaaaagaaaagaagcactgtgagtgtggagtatggaggaggtgtagtgtttgagacccacgaaatggcaatagagttgggcctaaagtggtcggcggacatggaaagttccctg**aaaattataaaacctggaatttt**gccctgactctgccacattagtcatgtattcttgaatag**atctacacataatcctagaggtagat**atgaagacagttgtataatcactgtgtatttcctataataaacttcagcgattggctttcttgtcatcatgctgggctctcaagacaaagataaccaagaggcagccaagtagagagttatgcatgaatacgttaccttaggggaggatagcaacagcattgaatagcattcactccaaatttgtgttctgtcctccttagaccaggccaccaaaataaaatcctatc**acatttcctgtaaatgt**tgaacgccatcatgttctccatggtgtttcggttgatgaatgaacagagtaagactcggtacaaagggaggggagcaccctctatgcctaggtacacaccatacggaatgtatcgctagggaaggtgctggtgagctggctcctgggaaaggtgtcaaggggaagagggggcagaggtacctagaaggaagacctc**agacaacatggcttgtct**acagagttggaaagttctgccagtagttagaagacagtaagtggaacagtgaaaatcctttcctaggagaaaccaaacatttacattgaccacaaatctgtatgaaccaacagaaaatgagcaaaacatatgaccatacatttaatagaataaacatatatgacta

Uppercase: TRBV20-1

Lowercase: Flanking sequence[1000bp]

Red & Bold & Underline: Stem-loop [14]

Blue: Heptamer[40]

Green: Nonamer [4]

id-TRBV25-1[V_gene_segment]

agagatataatagtaaaaacccaaagcctatgtcagcatcatgtgaacaaaaattttagtaaataccaactgtagatcttagtgaactagggcaaaggaagtggacaacactgggagaaattctgggattaa**tctttgtatctcacaatcaaaaccaaaga**cttttctgttaagaagaaatgggggaaagggcggagggagaatggcacttggtttagacattgtt**tactgcagcagcacccaaaccatttgtagcagta**ttaccaaagcatgtattggattatattttctgttctgtcatgattgatagtgcaattaaaggcagatggactaaaatcatcctaggcaaaggctaagttgataaagctgataaagcctgaacaccaagtcagtgaaaaatgagactgagaaa**gttattatcacacaataataac**accattctgaggactcagtgaattccagttactgctccaagctctttacatgtgtctatataaatagattgccccgtttaatgctcacaagttctcctgctccttgtctgatcaagccacgctgttcactggttccctct**tctgcatgttcttctctcttgggtttgcaga**gttcattcctgttcatccttcagaacccaactcaaatgccacttcctttcccaacctttttaattaggtaaaatcttgctattacaggcactca**aagctatgtagtttttatgttcatagcttacaaaacagttgcaatggaatgttttgtaag**aatattaaagacttgagttaagataagcaaagcatcaagcctactttggatgagggaagtcaacaaaaatattactcatttgatgggttaaaaggatatagatgaaactgtttcaacaggatggctgtgggacccaggaagtttgcaggggctg**tgtcatcacacacacacaatgaca**ttatcaagagcccatcctgcttccccactactgggagacatcctctctagccccaactgtgccATGACTATCAGGCTCCTCTGCTACGTGGGCTTTTATTTTCTGGGGGCAGGTAAGTCATAGACACAGTTCATTCCAAAATCTAGAAGGAATAATTCCTCTGTGTTGGGTTTGTGCCTGGCTCAGCATCAAAGTCCATCGTGAACTCTGTTACCAATTTTTGTCTCTTCCCATAGGCCTCATGGAAGCTGACATCTACCAGACCCCAAGATACCTTGTTATAGGGACAGGAAAGAAGATCACTCTGGAATGTTCTCAAACCATGGGCCATGACAAAATGTACTGGTATCAACAAGATCC**AGGAATGGAACTACACCTCATCCACTATTCCT**ATGGAGTTAATTCCACAGAGAAGGGAGATCTTTCCTCTGAGTCAA**CAGTCTCCAGAATAAGGACGGAGCATTTTCCCCTGACCCTGGAGTCTG**CCAGGCCCTCACATACCTCTCAGTACCTCTGTGCCAGCAGTGAATAcacagtgctacatggataccgacactccgcacagaaagggtcgcctctaatgtgaggacatcttgcctccagaaacctcatcttaaactacagaaacccctacaaatcttcccagactcctcagccttgagggaccagtgtgctttaagtaacagtctggactaaagactatctctggctcagtctctaacaagctgagaaatgacccagccccacagtcatcctagctgactttttccttgcatctagttgttgttgttcaattgagcttgca**caaattttatccttcaagataaatgcgataatatttg**acagttaccttatggc**tagctgaagaaagtctttgtcagcta**agaaacaggaaaaggatcctttgactttctacacaggctgtctttatctctggttcatggtttcttcctcatgttaaacttgttccttcatacacagccacacaaagtccctgccctttcctgaccccagtcctgcacctgacggcttcagttcctcaatctatgactttgaacttagaataaaattttattt**tcacttgaaataagtga**ttaatgattattttcactaatatccctctaaataaatgtaattttagctatatctgcatgaggcctaaggaaaagaaaaggctccaactttggcaaagaatagtctttaacaatgtccctggattcagccctaaatctctttggagactacccttagcattggaagactatcccacggctaggtgattcttctgtcttggatatgctga**atctcagggaagtgttctctatccacatgagat**cagacctgggaccctgaatagatttatttttcacagaagtatagactgctgtggggccctgggtgctgagctcagatgtgaaat**ctctgttttgaggttaccaaaaagtaaagtcacagag**tctctgtcctggttttgcctccttaaaaagtagctttaatatgttagaaaagaatctagaaaacatcatacata

Uppercase: TRBV25-1

Lowercase: Flanking sequence[1000bp]

Red & Bold & Underline: Stem-loop [14]

Blue: Heptamer[40]

Green: Nonamer [8]

id-IGLV8OR8-1[V_gene_segment]

gttctgagaaaggcatagtgtctcgtatctacca**ttatgtgccctacagaatagttctcacataa**aaatcctgtgtgccaccaatttatcccttctctccgacactcattttaagaaagaaaactctgctgatggaggaactgctgagggat**tcacggcatcaggtgttttaagctgctccgtga**tgctcacactaacttgttcacacacagtgccatttctatttccggttatgtgatgcgtgttttcctagtttgctatttttgatgttgg**gataaatcccataatataatggtaaatttatc**atctttctcctcaaaatctgcaattctgctggaggtcctccttaccattgatggcttcttagtgtgtgagagagttcacccccagcctggcagtgctgctccgtgcctaggtcc**cctcagccctgggccccacactgagg**ttactgaagacaacgcctgggacagagtcagaccctgctttgggacgtggttag**agagcccaccccagacacggctct**gtggattcccagtgtgccctctgctggaggaaggaggccc**tgctactcactcccaaagaaagtagca**atcatttcactggggagggaagaaaatgctcccaaggatggggttaggtttgccataaacaagattatgtggc**aggaaggagaaaacacctatctccctcct**tcagcccttgag**ctttccagagaaggaaag**ttggattgaggtcaccaagggcttggaagagctgatttggggagaggcatcaacactcattcctcagtgtctcagccctgtgtggtcgtgggtctgtgtgaggtctcagcatctgcctctctcaaggtctctcttagggga**cagcagggtgaaaactgctg**caggaaggg**aggggaagctgatttgcataaataactttttcccct**gttgggtctgtagagccatgaaaaggccatgaggacaacaaaccccagcggagaagcctggcccacgtgactGTCCCCACCATGGCCTGCATGATGCTTCTCCTTGGGCTCCTTGCTTATGGGTCAGGT**CAGGGAAAGGGACTCTGCATCCCTGGAGGGACCACACAAAGCAGGATCCAGG**TTACCCTTATCCCCATGACCATAACTCTGTCTCTCTTGTTGGTTTTAGGGGTGGATTCTCAGTCTGTGGTGAC**CCAGGAGCCATCACTCTCAGGGTCTCCTGG**AGGGACGGTCACACTCACCTGTGCC**CTGAGCTCTGGCTCAG**TCTCTACCAGTCACTACCCCAGGTGGTACCAGCAGACCCCAGGCCAGGCTCCACACATGCTCATCTGCAGCCCAAACACCTGCCCTTCTGGGGTCCCTGGTCGCTTCTCTGGCTCCATCCTTGGGAACAAAGCTGCCCTCACCATCACGGGGACT**CAGTAGATGATGACTCTGATCATTACTG**TGTGCTGTA**CATGGGTAGTGGCAATTCCACAGTGATTTAAACCCATG**AGGAGGTGCAACTAAAACCTCTTTACATACTCAGAAAGATTCAGCCCTTAGAAGCAAGAGAGAAGTTGAGAGGGTGGGAATGTCAACACCATGAGCTGGGAACCT**CCTTGCCCCAGTGTCCTGTGCAAGG**GTTCACTGCCAGGTGCCTTATCTAGTCTCTCCCAAGGTGTGTCCATCTCCCCAG**TGTGACGCTTCCATGAATATGTAAACTCACGGTGAGTCACA**GAGACGCCTGTGCTCCAGGAAGACTCTTCTTATTCAAAGAGAATCAACTGACATATGCTCATTCATTAATTTGTCCATGGAGACCAATTTAGTATTTCATTTATTACTGAAGTAAGGGCCTTGCCAAGTGCCTTGTTAAAAATGCAATTCTCCAACCTAGACCTACTGAATCAGAAATTTGTGTTGGTGCCTAGTAATTTGTATT**TAACAAGATGTACATGAGATTTTTGTTA**GCATTAATATTTGGGAAGAGCCTACTTTTATTATTATGAAATTATTCCTGGGGAATCTCTGTTTAAAATGATCTCAGGTAATATTCCATGACCCCCCTCTCCTAAATTCCCATTCAAAACCTCCAGAATGCTCAGAAACAGACTTAAAATCTCAAGCCATGGACAAATTTATAACTTTATTTCTGCAGACAGAGATGGGAGGAATTTCCAAAAACATACGTTCTTAGGTGCCAGGTGTGGTGACTCATGCCTATAATAACTAATACTTTGGGAGGCTGAGGCAC**GAGGATTGCTTGAGGCCAAGAGTTTGAGACCATCCTC**TTCCACA**TAGTGAGATTCCATCACTA**CAAAAAAAAAAAAAAAAAAGACATAAGCCCCTTTAGGAATTTCCAAGACCATAAGTTTTCTTAGGGTACATTTAGTAGCTCATATGAATATTTCTGTTATTAGAGACCATCTCCTGTAGGATTGGCTCCAGAATCTCTTCTCCCTCTCACT**GAGCTGACCTACCTGCAGCTC**TGGTTCTTAGGATCTTTGTAAGACATAGGGATCAGCCTTCCCCATCCTCTACATCTTATCACAGATTATCCCAGC**GGATCACAGTTCCACTGAGATCC**ACTCAGAACATATATATACTTGTCCTGAACAGTGTCTCTTAACATGACGCTGGCAAGATGAGGAGGACTGAGCGGTGACAAAGACTTGGAGCCACCTGGGGATGACCTGTGACTGCACGGCTGAGCTTTGCTGAGCCGAC**CTCAGGCTCCCTCCCTGCTCCCCTCCCCTGAG**TCTGT**CTCCTGGCCACCCTAGATCCAGGAG**CTTCCTCTACTTGCCTG**GGTTCTGGGGTCACCTCAGAGAACC**TCTCAGTGAGATAAAGGAGGTGCTGTGGGGGATCTAACACACCTGAGGGCAAATTTCAAGACTGTCCTG**CACAGGAAGCCCTGTG**AAAAGAAACCAACCAGCCCTACCTGTGTCTGGCACCTGAGTTTTTAGTCCTGACTCCCTGGTCTCATGGCTGTGTTGAGATTGCAAATGTGCGGTTTATTATCCAGGTCAGGAGAAATCCATGACAGTTTATGAGGACAAAGAAAACAACTACCCCGTGTACTTATCACACAATTAAGACACTGGCCTACTTTGGGAGGCCGAGGTGGGTGGATCACCTGAGGTCA**GGAGTTCAAGACCAGCTTGGCCAACATGATGAAACTCC**ATCTCTACTAAAAATACAAAAAATTAGTCAGGCACTGTGGCGGGCTCCTGTAATCCCAGCTACTTGGGAAGCTGAGGCACGAGAATCGCTTGAACCCGGGAGGCAGA**GGTTGCAGTGAGCCGAGATTGTGCCACTGCACCC**CAGCCTGGGCAACAAGAGTGAAAGTCTGTCTCAAAAAAAAAAAAAAAAAAAAAAAAGACACTGGCCTTTGATTTGATTTGTCAGTGCCCTGGAAGAATCCTGGAAGGAGAGGGTCAAGGTTACTTAGAAGGTGTTGGAGCTCAAA**CATGAGGATCAGTGGAGAAGACACCTCATG**GTGACCCAGAGAGATAAAGGTGGGATCAGTTGAGAGGGTGGGAGATGGGGATTTCTTAAGGTCATTTTGTTGGCTTCTTTCTCATTGCAAAAGATTCAAATTGAAATACACACATGATTTTCCTAATAAATGCTTATGAGGCTTCC**CATTAAGTTTAATTATCTTTCTTGCCTTAATG**AAAATAACCTAGAAA**GTAAGAAATGAAATATACTTCTGTAATTTCTTAC**AGATTGTGTATT**TTTAGACTAGAGGAAATCATTACTTCTAAA**GGAAAACTAC**TAGGATTGTTTGCATTTTAATCTGGGATCCTA**CATGATGGCTACAGATACAGCTATTGTGCAAGATTTTAACCTGGAATTTCTGCCAATTCAATAACAGAGGCAATAAAAATGGAT**ATGAATATTATAAAGACAGAATTCAT**TTATTTTAAAAGTTTTGAATGCTATGAATATCTACCAAGAATAGAATACATGGTACTG**AAAATTTATTAGAATGAATTTT**AGAATTGCCAAATTTGCAACTAATGAAATAAAATGTATTATAAAGAGTTCAAAATCC**TTATTTTTTGGTCTCAGTAGACAAAAATAA**ACATTGAATTTATCTATTAGAAAATAACTAGT**AAATTTAGTAAATTATATAAAAGAAAATTT**GAACAAATGGTACA**ACAGGCCAGGTGTGGTGGCTCAGGCCTGT**GAATCCCAGCACTTTGGGAGGCCTAGGCGGGCAGAT**CTCCTGAGGTCAGGAG**TTTGAGACCAGCCTGTCCAACCTGGTGAAACCCCATCTCTATTAAAAATACAAAAATTAGCCGGGTGTGATGGTGAGCACCTGTAATCCCAGCTACTTGGGAGGCTGAGGCAGGAGAATCGCTTGAACCTGGGAGGCACACTT**TGCAGTGAGCCGAGATTATGCCACTGCA**CTCCAGCCTGGGCAACACAGCAAGATTCTGTCTCAAAAAAAAAAATTGGTAAAAC**ATTTTCTTTTATCTGAGGACATCAAATAAAAAAAAT**TCAGGCCAGATGCAGTGGCTCACACCTGTAATCCCAACTACTTTACGAGGTCAA**GGTGGTAGGATCTCTTGATCCCAGGAGTTCAGTACCACC**CAGGGCAACATAGTGAGATCTAATCTCCAGTGAAAATTATTAAATTTAAATGTAACTAAAATTA**TTTAATTAAAAATAATTAAA**ATTAGCCAGGCAT**GCTGGGCTGTGCCTGTAGTCCCAGC**TACTAAGGAGGCTGAGATTGGAGGATTGCTC**AAGCCTGGAAGGTCGAGGCTT**CC**TGCTGTGAGCTGAGATCCTGCCACTGCACCACAGCA**TGGGCAACATGGTGCTCAAAAAACAAAACAAAACAAAAATTCAGCAAGTAGATTGGTGATGTAAATGCATTGTTATTATTTTTATCATTGTTTGCTTGTGTAGGTTT**TCCTCTGAGAAGGTGGACGGAGGGTGTTAGCAGAGGA**GATAGCGCAGCCCCCTTAGATATCAGAAACAGGGATCTATGGTAAATGGAAAATGCCAGTCACTTGTCGCCATG**CACAGGCGGCTATACCCTCCCTGTG**TCAGGCTCTGTGCCAGCAGTGATGGGCCCCATGTGCCCCCTTCAGGGTCCTGCATGCCATCTGTGCTACCCTCCAAACTCCCAAGGGCTGGGCATTGGTGCCTGTGTTGCTATGACGACTCAGCACTGTGTCCTCCCTGAGAGGGGCAGAGTCTGTAGAGGGTTCTAGTATCTGCTTCCTGTCTGATCCTCCCATGGGACAAACTTTGCATCTCACCCACCAGCCTCCTGTGTCTAAAGAATAAATGTCAGACTGCAGCTGTCTTCTCAGGTGGAACAGAGATCACC**AGGAGGAGAACACACTTGTATAAACATCTCCTCCTCTTTCAGAGCAACAAAGATAAAAAGAGG**CCTGAACCAGCAGCCTTCATTACAGGCAAGAGTACTG**TGTCCACTGTGGCCTGGACA**ACTGTCTTCTTCATGCTCCACCATTATTGCACAGGTGAGGACAGAATTTGGTTGTCCAGGGTGAGCTCCTTAGCTTGTCAGCCCCTCTGGCTCAGACCCCCAAGGGACTTCACCCTGGCCCCGCCCCAGCCCCCATGTGAGTCTGCGTCTGCAGTTCTTTCTCCCAGCCTGTGCTGACTCAGCGCCCTCCCTCACTGCATTGCCTGGAGCATCAGCCACACTCACCTGCACCCTG**AGCAGTGGGGCTTCAGTGTTGGTGACTGCAATGTACTGCT**TCCAGCAGAGGCCAGGGAGTCCTCCCAGGTTTCTCTTGAGATACTCCTCAAACTCTAGTAAGCTACAGGGCTCTGGGGTCCCATCTGCTTCTCTGGATCCAAAGATGCCTCAGCCAATGCAGGGATTCTGCACATCCTTGGGCTGCAGCCTGAGGACAAGGCTTACTATTATTGTTGTAGCAGCTTGCAGGCTTACACAGTGCCCCAGACCCAT**GGGGAAGTGAGGCGCTAAcctactctcttttcccc**ttgcctggcgc**agtcaccccgctagtggttctgaccaagtccagctcagggggtgact**tctg**ttgtcagatctttaaatagcaactgattttcccaaatggatctgacaa**tgtggtaccattttcttcagtttaacttttaaatcagttcacttct**catttcttttcactctgttcatggttattcaggaagaaatg**ttttgtttaaagctaaaaatctctcagtggaga**ctttggaggaggcccaaag**aggctggttcaggccatgcccagctcagggtccaggagtgatatggagatgcttctgctcagagggcgtcagtctgggccctggatacagcacccgcatgcaggagagagctctattctctatattagcgatgaaccatcactcattttaaaataatcttcctggaggaacagaaaaccatggggtgaggctgctctaagagtcacagaaataaacttggctggagggcagagaggggagccagacaccagtggacccagtcattagtgctgctcatg**accttgagcccctgaccatgtgagattgccaaggt**gtgggataaatctaagacattttaggtgggaaaataagacaagcatcctcatttcttcatgactcagtgaaggcaccagccttggctctgtgggtttcctggaagatgctggaaatgggggagtatg**tcatgttgcctagcaggtgttgggactcaaacatga**agatcagtgataaaaataaattaacagagccaaaataaataaaattaaatgtttataataagttaagaataatataatacctttatttaaagacagggtaagtcataagggtggaaacagtggatttacctcaaggccttttttttttttttttttgcaatggagttttgctctgtcgctcaggctggagtgcagcggtgtgatcttggctcaccacaacctccacctactgggttcaagtgattctcctgcctcagc

Uppercase: IGLV8OR8-1

Lowercase: Flanking sequence[1000bp]

Red & Bold & Underline: Stem-loop [60]

Blue: Heptamer[114]

Green: Nonamer [9]

id-TRAV40[V_gene_segment]

gagaaatatttccatttctgagaaaccagataaaattatgccaggactgaggaagaaaaaaagagaaaaaaaaaaagagcagtcaggaaaaaaaaagcagtcaaaaaaaaaaaaaaagaa**caaagttaagtagactttg**caaaatgacttcaattgaattacaccatcattatcaaataaatcttcaagtccagataa**aaaaggaataagtgtaccctcagcactaccctttt**ctagaagagagaaccaaaaatatcttga**ttcactaccgttttcagtgaa**gaatattgcaaatgtgaaagatggtaataacaggagcaatcaaacaatagacagtcctgagaggttactgctcttttactgtaaagctaaactttgaaatgtgaatgctaagtctttgttaacac**aattataataataatt**acattttag**atagaatttcatatttattcttgcattgttgaagttaattctat**tttttgatattcagattcagctttagcttcccccagatattgactttaacatctcaccattcggcatttggtcacaatgtgtgttcagtggaccatgttttaaggaaataattagacgtatgaagagggtttatgtaagcatagatttatcacaat**gttgtttgtaaaaactgaaaatattgaaacaac**tgtctaaaatactgacatgttgtgatgtctaagttctagtggaaa**ttattctcccttagaataa**gtgttgtagcgataacagacttaggagtgtggtgagtcccctttgcacaaaaagaccacctcacacccaacatgactcttaagaatgaggacagtagtgaggacacactgcagaacactgagaggcagatgacgtcttataacctgttcattaacttgccacttgggggtgctattcaccagtgaaatcataatttatgcatgaattctgagtacaactgtgaatcctcacttcaacagtgatgccctctgct**aggccagagacactaacaATGAACTCCTCTCTGGACT**TTCTAATTCTGATCTTAATGTTTGGTAAGTACATTTTATTAAAATTTTTTG**TATTATTTCAAAATAATA**AATACTTCTCCAAAAGATACAACAACTCTGGCCAAACCCATAAAATCAGTACTTTTTTCCCCCTACAGGAGGAACCAGCAGCAATTCAGTCAAGC**AGACGGGCCAAATAACCGTCT**CGGAGGGAGCATCTGTGACTATGAACTGCACATACACATCCAC**GGGGTACCCTACCCTTTTCTGGTATGTGGAATACCCC**AGCAAACCTCTGCAGCTTCTTCAGAGAGAGACAATGGAAAACAGCAAAAACTTCGGAGGCGGAAATATTAAAGACAAAAACTCCCCCATTGTGAAATATTCAGTCCAGGTATCAGACTCAGCCGTGTACTACTGTCTTCTGGGAGA**cactgtgttaaaagcacagtg**ggagctatacaaaaacctcaaaggctcagaggaagtatgtagtgaggctggaaaacccaggttgtagagccctgttctctctttcacagacagtcctgttaagacattattagaagataaggaaaatcaaaatcagatataactgcctctaattccagggggcacaggggtttctttgacagaaaatggggt**cagggaagagggtctcagatgatatccctg**ctctcaattaggtcagttctctcttgtccttcaggctttcttgcatgatgtgtaggaattagaatatgatgacttggatgtcagtcctggctctactacttattagctatgcaaatttggccaagttaattgatgtctcagagctttagtttttttcacttgtgaagtagaaataaaaataatcacacaaatttagcatggagataaagtggaaaaaaggtaggttaggaaggggcagattagtactcaaagcatttctggaacttgcaagtcccaccgtagcagttctgctctctcacagcaatctctctgcctagtgctgaccatcctaacagggcttggcatgcccactgctttgtcacaccacaccacaccccacgccacgcccagccccaggccattatgatcaacatatcatcttttctctcacaggctcctactgacactacacacaaaaattaactcaaagtggatcatagacctaaatgtaaaaactaaaattgtaaaacccttagaagaaaacatagcagtaaattttcatgatcttgggttaagcactgatttcttagatgtgacaccaaaagcacaagcaataaaatagaacattgataagttggacttcaacaaaattaaacttgtttatgcttcaaaggacaccatcaagaaaatgaaatgataacccacagaatgggagaaaacatttgcaaatcatatgtactatgtacccgatgataagggacttgtatttagaatatat

Uppercase: TRAV40

Lowercase: Flanking sequence[1000bp]

Red & Bold & Underline: Stem-loop [13]

Blue: Heptamer[28]

Green: Nonamer [5]

id-TRAV26-1[V_gene_segment]

tctcttattaagtgtgttatggcttcaaaagttctggggcccttacactgatggactaagtcttttcagagcacctcccttaacactgctttctccaagtttcacttccctaatcatctcttacctcccctattccttacacctttttcttcctttcctttacctattcttaccaccgtcaccccaaacttctttgcttttcctggtctcttttctttccgcctcagtgtccttggtgcacacagtaggcataagcacgggcatagaaattaggaaatgaacatgaaggccatgagtttaaggaggatgagagccagcaccatttatgtcattag**tcactatatcaagaacagatatattga**gaaatggctacttagacaacagtcaacacatgattccagggacctctatagaaaacccgagtttggggagacttagctttcagtgtcaatgttatcaatttttattttccccctgagcacatcacttcctacttttgaacttcacttccgttttctagaatgtagatgggaatttccaactccttacaacacagagctcactataggcttctgggatttaatgacactttccacaagctccatggagcagaggaaactttatttggtacattgaggaagaaatgatggaagaaaaggtctggatagatacatccttatgatcagttttcctcattctgtgccagcctcttagactttaaacctcataaatgatttcagtagacaccccctgagt**tgaggatgtctggttgatatcctca**cc**gtcttctcaggaacatgactaaacctggggaggaagac**atagaaaagcaactgctcta**tcattggctgacaagatcctgacaacaacacaatga**gagctgcactccttgcaagtctttactagtagctgttgcttctgttctctggaaattctttaaacctagaatcagacacaaaaactgaactctgggtccacaat**cctcatttgtccttgaagtATGAGG**CTGGTGGCAAGAGTAACTGTGTTTCTGACCTTTGGTAAGTGCTCTGTGTCTACCTAGGGAGGGCTTCTGAGGACAAGGGACCCCGAAGGGTGGTTTGCCCTC**TATGCTCCCTGTGCTTTATCCAGCATA**CTAGATATGACTGAGAACTGTTCTGTGCATTTGTCTAGGAAGTACATACATCTACATACCAGGCTAGATCATAACCATAATCCCAGAAGAGCCCTTGGCAGTATTCAGTTTTAAGAAATCTATGGGTTTACTAAATCAAGCTCCCATGGAATACAGCCTAGAGGGTCTCTGATTATTGGCATGTATCTTTGGTCTTTATAATTAACAGCAATGATATTTTTTGTGTTCTTGCAGGACATAAAAGTTTTGGAGTGTTAAGGGAGTGTTGTATCCTTGATCCATAGAATTGCATTGGTTTGAGAGTCAGGAGCTAATCTGATTTTCTTGACCCCTTAGGAACTATAATTGATGCTAAGACCACCCAGCCCACCTCCATGGATTGCGCTGAAGGAAGAGCTGCAAACCTGCCTTGTAATCACTCTACCATCAGTGGAAATGAGTATGTGTATTGGTATCGACAGATTCACTCCCAGGGGCCACAGTATATCATTCATGGTCTAAAAAACAATGAAACCAATGAAATGGCCTCTCTGATCATCACAGAAGACAGAAAGTCCAGCACCTTGATCCTGCCCCACGCTACGCTGAGAGACACTGCTGTGTACTATTGCATCGTCAGAGTCGcacactgggacagatggggctgcacctgtgcaatatctccctggtggcaagtgaggaggagggtagcattcacctagagcaaaatgtcgataggagtcaaaaagtaacaagaaaagaggaaaaggaagggaaggagagaaggaggaaaataggtaaagtaaataagggagatacgatgaagggtaaaaagataagaataagaactttattgttgatgcggctgagaaaaatgataaagaggctgtgaatcataatttgaacatagcaataaa**atgaagagggttaagaattttcaagactcttcttcat**gtcaaaaaaaaa**tcatttcagtttgttagtaaaaatga**atggataaggaaaatcaaaataattctgcttgtcccaaaggcttttttcgtctggatccaggggcaagtccaagctttgaaatgcatgctgttatagttgttccaattac**cagaaacaagttgatgtttctg**c**aagaaaaatctgaaatgttctttctt**actagtgc**agtaacatcagttact**agagcatgcatatttgtgatgaaggtt**tttctttttggaaagaaa**ca**aaaatcaaatgaatattaatttgttttt**atttttcttgaagtcttgaatctttaaatagtcttcatcacgttttatttcctaagcagcacacaaatccatcttaccataaaagactcagataaagcaaaagtgaaagacctagctgcatttcctcattcttccctatctcacaacactcttt**tcaaagaacactagctttga**cagtttaaggtataccatgatagccagtttaatttcagttatcta**tgtttgcatataactttatgcaatca**cacacaaac**ttacatacatacacacagttttgtcatgtaa**attggatttattagattttttaacttgcc**atttatttttcatttaataaat**tgtcctagagatctttctatatcaatagagaagttcactttattctttttggtgcttat

Uppercase: TRAV26-1

Lowercase: Flanking sequence[1000bp]

Red & Bold & Underline: Stem-loop [17]

Blue: Heptamer[22]

Green: Nonamer [8]

id-IGHV4-30-2[V_gene_segment]

ttctggtctccaacaatactcctgcatctgccatctaagtagctggaactaccgttgtgtgccaggaggtttggcttggcttgagtacttattaaaccacacactttttgcaatatttaatgtatagtaataatgtctcaataagactact**acaaatcaatgaatgaataatttgt**tcagtacag**attcatggaaaaatagacactaacatgatgaat**gtctg**acatttatgaaaatacaactgcatgaaatgt**gcttctctttacattcattaggtaaacacaatagtgcatacacatcacaccgtgctttcattacaggaagaaagatctgaaaatgtcactgggg**tgaaccacattgtgctgggcttggttca**gggagcagtcaggcccagtgttgtgaccttcaaccacagaatccttaaaaaataataaaagaggctcccccaaagtccccatcagttcccggactcgctatgtttctggatcgtatcagtgcatccggagctccctggtggctttagtgattccttgcttgccatgctgaggtctccctgtagattatgttgggttttctgaggccgttttgctatttaagacccactccctggcacagagcgattc**cctctaaacctgatagagg**ttctgaactaaaaataacattaagtgaatcctggtgtgtctgaactcaagtgattgttacattaagctgctgttgcaatctgtttcctcacctgggaaaagaggagccaggacatagtgagttgaggccccaggaagataactgaattctcagagggcacagccagcatcctcctccc**agggagagtctaaaagactggggcctccct**catcccttttcacttctccatacagaggcaccacccccatgcaaatctcacttaggcacccac**aggaaaccaccacacatttcct**taaattcag**ggtccagctcacatgggaaatactttctgagagtcctggacc**tcctgtgc**aagaacATGAAACACCTGTGGTTCTT**CCTCCTGCTGGTGGCAGCTCCCAGATGTGAGTGTCTCAAGGCTGCAGACATGGAGATATGGGAGGTGCCTC**TGAGCCCAGGGCTCA**CTGTGGGTCTCTCTGTTCACAGGG**GTCCTGTCCCAGCTGCAGCTGCAGGAGTCCGGCTCAGGAC**TGGTGAAGCCTTCACAGACCCTGTCCCTCACCTGCGCTGTCTCTGGTG**GCTCCATCAGCAGTGGTGGTTACTCCTGGAGC**TGGATCCGGCAGCCACCAGGGAAGGGCCTGGAGTGGATTGGGTACATCTATCATAGTGGGAGCACCTACTACAACCCGTCCCTCAAGAGTCGAGTCACCATATCAGTAGACAGGTCCAAGAACCAGTTCTCCCTGAAGCTGAGCTCTGTGACCGCCGCGGACACGGCCGTGTATTACTGTGCCAGAGAcacaatgaggggaggtgagtgtgagcccagacacaaa**cctccctgcagggagg**cggagggggcgggcgcaggtgctgctcaggaccagcagggggcgcgcggggcccacagagcatgaggccgggtcaggagcaggtgcag**ggagggcggggcttcctcatcagctcagtgctctccctcc**tcgccagcacctcagctgtccccaggactcctctttctttattatctgtggttctg**cttcctcacatccttgtggtaggaaaggaaggaggaag**gcaaattttcctcttagagtcaaagtgtcactaattact**aggaactttcctacaagttcct**gaatgtccc**atttttccttcttaattaaaaaaaat**atatattctaatacttctcaccatctcttgatttgtgtcat**cagttgaattgtgctgtctttgaaattcaaatg**ctga**aaccttaaatccaattgatctatattggaattttaaggatggaattaaggtt**aaatgtgatcataagtctgagattctaatgcaatagatctgttgtctttataagaagtggaagagtcaccagagacctctcacttttcccgtgcacgcagagaagaggccatgtggagacatagtggactagaaggtgcaagccaagaagaagccgcaccaagaaccaaccctgccagcacctggaccttggacattcagacttcagaatggtgagaaaatcaatgtttgttgtttaagccacccactcctgtt**gtcttcttatgaagac**ccagacagactaataccacataactctgttagctcctgg**agggagaagcagctccct**gaggctgggcacatctctcagatatccatatgaagtaggcagaaatagtagttctcatataaaaatgtgtc**atggccctgttggccat**ttttttttgacggagtctcgctctgtggccaggctggagtgcagtggtgcaatctcagctgactacaaaccccgtctcctgggt

Uppercase: IGHV4-30-2

Lowercase: Flanking sequence[1000bp]

Red & Bold & Underline: Stem-loop [22]

Blue: Heptamer[42]

Green: Nonamer [4]

id-IGKV1D-22[V_gene_segment]

ggacaagaacttcactaacaattgagtcactgaagacttatggccctgtgtgacgcaccacataaccgtgagtttgcagtggttgcaggtcagggacagattttatgcttaagatcagtagggtggaggct**gaggatcttggctattacaactgccaccacactctacaatatcctc**ccacaatggttcagcaccaaacaaaagcctcctgcttggattgtcccagctgcccaaattagttccttcactgaggagtagacagggtatattctctaaatctatgtaacaggaagatgttggtgaactcaggggattagtatgaagctacacctcaggcatcacacataagatcacttcagcagtcgcagccttagcatgggcagaacctacagaagatgcaagtgccctctgagccaggagacaggaggaaggaggaagggaaaggtgacttagctcatctcaatcctctctcctttgcatacatttgtcaaccag**atgtattcagcctaccagtcacacaactgaggctgatacat**gacaacatagcactggtatattcttggtattgtttggcttagcagttactagtatatatttaat**gggagaatatttggtggtgttaacacattgcttatctccc**ttaccccagttgtactttacacttgttctcggcacacattctcctccaggactggagcattcacagggttttatgttactgttcttatgggagtaaaaagaaaaacgattcacattcttgctactgagctaggctgggatgtcctgggccaagctgaaaatgtgaaaaataagagtatgaatatttattaagttttatctggatctaagatacttatccatgaaccagtcctgcagctgtgcccagcctgctccattccctgctgatttgcatgttcccagagcacaaccccctgttctgaagacttcttaataggctggtcacaccctgtgcaggagtcagtctcagtcaggacacagcATGGACATGAGGGTCCCCACTCAGCTCCAGGGGCTCCTGCTGCTCCGGCTCCCAGGTAAGGATGGAGAACACTAGGAATTTACTCAGCCAATGTGCTCAGTACAGCCTGGCCTTTCAGGGAAATCATCTT**ACAAATAGTTGTGTGGATTATTTGT**TTTTATGTCCCAG**GAGTCAGATGTGATTTCCAGATGACTC**AGTCTCCATCCTCCCTGACTGCATCTGTAGGAGAGAGAGTCACCATCACTTGCTGGGCGAGTCAGGGCATTTGCAATTATTTAAGCTAGTATCAGTAGAAACTAGAGAATCCTCCTAAGCTCCTGATCTATGCTGCATCCAGTTTGCAATCTGGGGTCCCGTCACGGTTCAGTGGCAGTAGGTCTGGGACACATTTCACACATTCTCACCATCAGGAGCCTGCAACCTGAAGATGTTATAACTTATTACTGTCTATAGACTTACAGCAGCCATcctagagtgttacaggtcataaaataaaccc**ccagggaagcagaagtatgactcatggctgccccaggtgcttccactgg**tgcctccatctg**ctgagagtgtttctcag**gtgcagccaagatttaaaggtttttgtaggaatggtcagaagtctcatctgcattctaattctttttcttcctgcttagccccagcagcacagacatgacactatctctcctgatttaataaaggatagcatttacgatacctgaagaatctgtgttattgcatccatctgggtcatagattaaaagagaaaccactctacagattgccagaaggcattgttttaatacagggaattagagttgaatatacaaaactgggagtgtggtagttagggaagctgacactagaaacacgggagt**ctctggaggtctgccagaagccagag**ttcatcagccgctaaaggcatgggctatctaaccatatagtcttctttgtctaggaagtccgtatgcgaagatgctgatgctatcagttgttgcagcacctcaccaggtgattctccagtccttatctcagtgaacatgtttgcctaccggtgtcaaagaatattgaatcgccttcttcttaccttcaaatatgatgagaggtcttctctttgagtaactctacaa**gaaaccatagagggtttaatgggtttc**aggaaaggtgcttttagaaatcatggtgaatatgaggaattacagccaagtgggataagtatttcccaaaatctcagaattttccaggtatggggtggcttcag**aatacatttggatgttcttacatgtatt**attagaaagtttggtattattgcaagaaaattttattaagtcgtaaagtaaaagaaaaaaatgacaac**attgcttgaaatacatagcaat**cctttgacaaatgaaaaaaaaattgacaaaacaaacaagaacacctataggtgcatgtagcatactttttccttaatataagagcactt

Uppercase: IGKV1D-22

Lowercase: Flanking sequence[1000bp]

Red & Bold & Underline: Stem-loop [11]

Blue: Heptamer[34]

Green: Nonamer [5]

id-TRBV25OR9-2[V_gene_segment]

tagtagagatataatcataaaaacccaaaccccatgtcagcatcgtgtgaacaaaaattttagtaaataccaactgtaaatctcaatgaactagggcaaaggaagtgg**acaaagctgggagaaattctgggattaatctttgt**atttcacaatcaaaaccaaaggcttttctgttaagaagaaatgggggaaagggcggagggaaaatgacact**tggtttagacattgtttaccgcagcagcacccaaacca**tttttagcagtattaccaaagcatgtattggattgtattttctgttctgtcatgattgatggtgaaattaaaaggcagatggactaaaatcatcctaggcaaaggctaagttgattaagctgataaaggctgaacaccaagtcaatgaaaaatgagactgagaaa**gttattatcacacaataataac**accatattttctgaggactcagtgaatcccagttactgctccaagctctttacatgtgtctatatagattgccccatttaatgctaacaagttctcttgctccttgtctgatcaagccacgctgttcactggttctctct**tctgcatgttcttctctcttgggtttgcagagttcattcctgttcatccttcagaactc**aactcaaatgccacttcctttcccaatctttttaattaggtaaaatcccgctattataggcattca**aagctatgtagtttttatgctcatagcttataaaacagttgcaatggaatattttataag**aatattaaagacttgagttaagcaaagcatgaagcctactttggatgagggaagtgaacaaaaatattagtcatttgatgggttaaaaggatatagatgaaactgtttcaacaggatggctgtgggacccaggaagtttgcaggggctgtgtcatcacacacacacaatgactttatcaagagcccatcctgcctccccactactgggagacatcctctgtagccccaactgtgccATGACTATCAGGTTCCTCCGCCAAGTGGCCTTTTATTTTCTGGAGGCAGGTAAGTAATAGGCACAGTTCATTCCAAAATCTAGAAGGAACACTTCCTCTGTGTTGGGTTTGTGCATGGCTCAGCCTCAA**AGTTCATCATGAACT**CTGTTGCCAGTTTTTGTCTCTTCCCATAGGCCTTAAGGAAGCTGAAATCTACCAGACCCCAAGACACCGTGTTATAGGGGCAGGAAAGAAGATCACTCTGGAATGTTCTCAAACCATGGGCCATGACAAAATGTACTGGTCAAGATCC**AGGAATGGAATTACACCTCATCCACCATTCCT**ATGGGTTAATTCCACAGAGAAGGGAGATCTTTGCTCTGAGTCAA**CAGTCTCCAGAATAAGGATAGAGCGTTTTCCCCTGACCCTGGAGTCTG**CCAGCCCCTCACATACCTCTCAGTACCTCTGTGCCAGCAGTGAATAcacagtgctacatagataccgacactctgcacagaaagggtcgcctc**taaggtgaggacatcttgccttcagaaacctta**tcttaaactacagaaacccctgcaaatcttcccagactcctcagccttgagggaccagtgtgctttaagtaacagtctggactaaagactatctctggctcagtctctaacaagctgagaaatgacccagccccacagtcatcctggctgacttcttccttgcatctagttcttgctgttcaattgagcttgcacaaat**tttatccttcaagataaa**tgtgatgatatttgacagttgccttatggc**tagctgaagaaagtctttgtcagcta**agaaacaggaaaaggatcctttaactttctgcacaggctttctttatctctggttcatggttccttcctcatgttaaactcgttccttcatacacagccacacaaagtccttgccctttcctgaccccagtcctgcacctgatgacttcagttcctctctatgactttgaacttggaataaatttttattt**tcacttgaaataagtga**ttaatgattattttcactaatctccctctaaataaatgtaatttgagctaaatctgcatgaggcctaaggaagagaaaaggctccaactttggaaaagaatagtcttcaacaatgtccctggattcagccctaaatctctttggagactacccttaccagtgggagactatcctacagatagatgattcttctctcttggatatgctga**atctcagggaagtgttctctatccacatgagat**cagacctgggaccttgaatagattgatttatttttcacaaaagcacagactactgtggggccctgggtgctgagctcagatgtgaaacttctgttttgaggttaccaaaaagtaaagtcacagagtctctgtcctggttttgcctccttaaaaagtagctttaatatgttagaaaagaatctagaaaacataataca

Uppercase: TRBV25OR9-2

Lowercase: Flanking sequence[1000bp]

Red & Bold & Underline: Stem-loop [15]

Blue: Heptamer[40]

Green: Nonamer [11]

id-IGHV1-68[V_gene_segment]

ggaagcagagtt**tgcagtgagctgagattgtgccactgca**ctccagcttgggcaacaaagcgagattccatctccaaaaaaaaaattcagatttcagtgttaagtaaagttgcctacattgtgtgagtgacagggcagtggtggatccgagagtgtggatctgagagtgagtcagaaatcagcatgtaaagatgaggatctatgca**catgaactgaaagtatgtaaacagttcatg**aaattctaataaatccagtaggaaataaaacccaaacttatccaaaacacaaattcccttgaaattattatgggagcat**gagttcataaagaactc**ctaactcctgtttcaacttctgaatcctagtgtccatgacataagaaaatcatctccaattatgcatcacagggcaaatctgtaaactaagagtttttctgttgacga**tcctggggaatcaggacaccagga**aggtgctggagaaactgtctcaggagcgccccagggatctcagaggaacgtgctggccactcacgtgggacatcagcgtcacttgctcagagtcatcagtgagctg**tgctggtgtctgacgggtccagca**tagggccaaggcacctgctctgtgtcatggaccgagatggtccccagaatgatccaagtggtctctgtgctaatctaatgtagtttcacagtgagaggccgttctgagggggcttcttcttcagtgaaaggacct**ctgtccacaaatattcctaaatggacag**gggcatgcatttcctcaagcaggattagggcttg**gaccatcagcatctcactcttgcaaggctgatgtgtc**atttgtcttccctttcttatcatggatcaggctttgagctatgaaatgccctgtctcatgaatatgtaaatacctgagatccactgaggtaaatatggtctgtgccctgagagcttcacccaacaatcacatcccgcctctagagaatcccccgagagcatggctcctcaccATGGACAGGACCTAGAGCTAATCTTCCTGGTGGCAGCAGCTACAGGTAAGGTGCTCCCAAGTCCCAGTGATGAGAAGGGGATTGAGTACAGTCAAGGAGGCTTTCATCCACTCCTGTGTCCCACCCTCCAATGGGTGTCTGCTCCCAGGTGCAGCTGGGGCAGT**CTGAGGCTGAGGTAAAGAAGCCTGGGGCCTCAGTGAAGGTCTCCTGCAAGGCTTCCGGATACACCTTCACT**TGCTGCTCCTTGCACTGGTTGCAACAGGCCCCTGGACAAGGGCTTGAAAGGATGAGATGGATCACACTTTACAATGGTAACACCAACTATGCAAAGAAGTTCCAGGGCAGAGTCACCATTAC**CAGGGACATGTCCCTG**AGGACAGCCTACATAGAGCTGAGCAGCCTGAGATCTGAGGACTCGGCT**GTGTATTACTGGGCAAGATACAC**GGTGCGAGAACCCACA**TCCTGAGAGAGTCAGGA**ACCccagggaggaggcagctgtgctggcatggaggagatgacaaagattattagattgaagactttcttagaaaatagcattaagt**catttacgaaaaggaacaatataaatg**tgtatttg**agaaattgtaattatttgagagatttct**tatgcaacatttattctgtaagca**aattctagggattggagaattaatcaaattaat**aaagctgacat**agaaattcctctgaaggtatctttgtaaacatcaatttct**gaatcagtgttgt**aaatgttttggaacacagacacaagatcacattt**taactctacttttgtctctataa**aaaatgccaaaaagagtctcatttt**gaccatgtacctctttgaattcccaccatcaatgaatgattgttcttgggtttccacattatattaccatttatcattatgagaattgtgtgtttta**accattataataggtgagtaatggt**atctaatttttgcttaaatgcacatgcccctaat**aaaattcatatttaacaattttcatataatttt**tggtgagatgcctctcctgatatttggttcattttcaatagcactgttttcttttgattagttgtaagtttacttgcatattgattataaaag**ttaattaacaaattaa**aagaattcatttaacaaatatgtgacttggaagt**tgcaggggtttgtcctgca**caccctgacacaatgacgaatgaataaagtgcactgacacaaagatattctgctttgccagctcgactgagcatctgggccacttagtcacagccaccgcctcgatcagtcagggagacttacatttattcagtaaagattaattgacaaaggccgtgagtaaacaccactagaaggtaattgacattgtggacctcttgagtggaaagcaattaagcacccgcggtagatcaaaggttagtcttaggaacacatgagtaaacaagctagttagataagctcctcacattcctttgtttc

Uppercase: IGHV1-68

Lowercase: Flanking sequence[1000bp]

Red & Bold & Underline: Stem-loop [23]

Blue: Heptamer[45]

Green: Nonamer [4]

id-IGHV4-61-2[V_gene_segment]

agggtatgttcatgcaattcaatggaggagaaagtc**tttcagagacagatggatctgaaatgataaatatgtgggtaaggactctgggcttgagtatcatt**gtccagccatgtttcacaagtg**tgtcctgtcagggaaggaca**agagttccttgtgttctcagagggaaggggtcacagagttcctctctggttcccaggaaagataatcgcactaatct**tcatgatcttcatga**gactatcctccagtgctgacctgttatagagtttttgtctgaagttctcactgcaatccccaatctacatattttcaatcagaagtgtttagaggcca**ggacatatcttcacggtcacacattgagaaggatgtagatatgtcc**cactaccttctcctgagatctcagacagaatcccagatttcaaaaggacacagaaggacagctctcaggtgcttttaaaaaatgacccacttccagggacagggagcttccctataaccatggtggatgttctgaactacaataaacattggatggatccaggattgtttgaagtcactgtcattattacattcagctgctgtttcaatgtgtctgaagtagtaaatgacaatttagatgacaatttatatgaatcttcaagggtagaacaatattgaccatattccaaaatctgtccttgatccatgatcacactcatctcccagaccaggtccttcagcacgtctctttacctgaaagaagaggactctgggcttggagaggggagaccccaagaagacaactgagttctcaaagggcacagccagcatcctactcccagggcgagcccaaaagactggggcctccctcctcctttttcacctctccgtacaaaggcaccacccacatgcaaatccttacttaagcacccac**aggaaaccaccacacatttcct**taaattcaggttccagctcacatgggaaatactttctgagagtcctggacctcctgtgc**aagaacATGAAACACCTGTGGTTCTT**CCTCCTCCTGGTGGCAGCTCCCAGATGTGAGTGTCTCA**GGGATCCAGACATGGGGGTATGGGAGGTGCCTCTGATCCC**AGGGCTCACTGTGGGTCTCTCTGTTCACAGGG**GTCCTGTCCCAGGTGCAGCTGCAGGAGTCGGGCCCAGGAC**TGGTGAAGCCTTCGGAGACCCTGTCCCTCACCTGCACTGTCTCTGGTGGCTCCGTCAGCAGTGGTAGTTACTACTGGAGCTGGATCCGGCAGCCCCCAGGGAAGGGACTGGAGTGGATTGGGTATATCTATTACAGTGGGAGCACCAACTACAACCCCTCCCTCAAGAGTCGAGTCACCATATCAGTAGACACGTCCAAGAACCAGTTCTCCCTGAAGCTGAGCTCTGTGACCGCTGCGGACACGGCCGTGTATT**ACTGTGCGAGAGAcacagt**gag**gggaggtgagtgtgagcccagacacaaacctccc**tgcatggacgcggaggggaccggcgcaggtgctgctcaggaccagcaggtggcgcgcggggcccccagagcatgaggccgggtcag**gagcaggtgcagggagggcggggcttcctcatctgctc**agtggtctccgtcctcgccagcacctcgctgtcaccagggctcctctttctttattatctgtggttctg**cttcctcacattcttgtgccaggaaagaaacgaggaag**acaaattttcgtctatagttgaagcttcaccaattact**aggaacttgcctacaagttcct**gcatgacccattataacttatcgattaaaaaatatatattctaatgcttctcaccatctcttgatttgtatcatcaac**tgaattgtaccctctttgaaattca**tatgatgaaacctt**aaattcaatggatctatattggaattt**taatgaaataattaaggttaaatgtggtcataattgtaagaccctaatgcaatagacgtgttgtctttataagaagaggaagagacaccagagacctctcacttttcacgtgcaggcagagaagaggccatgtggagacatag**tgcactagaaggtggcccagtgca**agccaggaagaagccgcgccaagaaccagccctgccagcacactaatcttcaacattcagactgcagaattttaagaaaatcaatatttgttgtttaagccacccactcctgttgtcttcttatgaagatcca**gacagactaataccacataactctgttagtgctgtc**ccctggatggagaattagcctcctgaggctgggcacatctctcagatttccacataaaacaggtaaaaaatagtagttctgatataaaaacttgtcatgtccctgttggccaatttctgggcaaggtcttttaaataagccaagtttgcggggaaatggagaccatatgtttgtgggactctaacc

Uppercase: IGHV4-61-2

Lowercase: Flanking sequence[1000bp]

Red & Bold & Underline: Stem-loop [18]

Blue: Heptamer[40]

Green: Nonamer [4]

id-IGHV3-15[V_gene_segment]

tgatgttcaggaaaataaagc**aaatgaataagaatatgtactatgttattccattttaataaactctgataaattaaa**ttgaatctacagcaatataaagaagaacagaatt**taccatttggggaaatggta**gaagaagggaagaggaaaggaggaggaatatagaagaatgagagggaaattttgtgaattttctggttcaccttggtaactaggatggttacatcaggtttatcaattgtacactttaaatatgtga**agtttattatcagtaaact**g**aaatttataaaattt**attaccagcaaacaaatgaaaacttgcacaagaaataagtgacataaagatagaaaaaatattaaatttcagaaa**cacctaataatttatcttcgtgaaccctagttctcaccatatttttaggtg**aatgctagaatgcagcaaaattacacatgctctcaatacagaaagtgggtttcacaaaccacactaggcatgctcagctctgtcctggagttgggttagggagtaatatagggccagtggatgaggagcacaggcctagatactggggctcactaacctcaggtatgagctcttagatacatacaaagcccctccacgcatgggtttacttccccatctgtaaattgagaaaccattgacccctaaaaatatga**tttacacaaatatgtaaa**aatgtaagagagtgattagtgcaaagtgtttatcacagcacaatttcataacaagacagcaagttttccaaacagcatcattgtcattagattcct**gcagggcatcattaccttatctgggccctgc**cctctgctcaggcatcccaccccagagcttgctatatagtaggtgacatgcaaatagggccctccctctcctgatgaaaaccagcccagtcctgaccctgcagctctgggagaggagccccagc**cttgggattcccaag**tgttttca**ttcagtgatcaggactgaa**cacagaggactcaccATGGAGTTTGGGCTGAGCTGGATTTTCCTTGCTGCTATTTTAAAAGGTGATTTATGGAGAACTAGAGAGATTAAGTGTGAGTGGACGTGAGTGAG**AGAAACAGTGGATATGTGTGGCAGTTTCT**GATCTTAGTGTCTCTGTGTTTGCAGGTGTCCAGTGTGAGGTGCAGCTGGTGGAGTCTGGGGG**AGGCTTGGTAAAGCCT**GGGGGGTCCCTTAGACTCTCCTGTGCAGCCTCTGGATTCACTTTCAGTAAC**GCCTGGATGAGCTGGGTCCGCCAGGCTCCAGGGAAGGGGCTGGAG**TGGGTTGGCCGTATTAAAAGCAAAACTGATGGTGGGACAACAGACTACGCTGCACCCGTGAAAGGCAGATTCACCATCTCAAGAGATGATTCAAAAAACACGCTGTATCTGCAAATGAACAGCCTGAAAACCGAGGACACAGCCGTGTATTACTGTACCACAGAcacagtgag**gggaggtcagtgtgagcccggacacaaacctccc**tgcaggggcgcgcggggccaccagggggcgctcgtgacccactgagggcgggacaggtcccaggagcaggtgccgggagaggtttcctttctcctcagctggaaaagtcaggtttatcttcgcaggactctggagtcttctaggctgtgatattttgttacttatatttattatgaattttatca**ttaatacttaaattttagtaattattaa**cattctacatattattatatttttaagtatatactttcaagaaataaacattcctaattgcat**aataattattgtaacataattatt**gta**acatataattatatcctaaatgcataataattattgt**aacaatcacattctcattttgtatttgaattttcatatttattttactaagatttta**attttaatataaaat**atttttctacaattgtctgttttc**catttttgtgagataaaattagcatatacaaaaatg**cacagattctccaggtacagttagagggtttcaggcaaatgtgcacatacttgtctacagcagctaagtgagggtgaggaacaggtgaggtccatctccccacaaagtgtcctcttagtgcttccagttagttctcacataaggatttttttttatttcaactttataatttagatacagagggttcatgtgtggatttgttacatgggattactgagtgatgctgaggtttggaatacagatttcttcaccccctccctccatggtctagcagtccacagtgtctgttgctcccatatttatgtctatgtgtgctcaatgctgaggtcccacttacaagaatatgtggtgttcagttttattttcctgaattaatttgtttaggattaagaactccagctccatttgttttgctgcaaaggacatgatttcattcttttttatggctgtgtagtattatatattttacatgtaacacattttat

Uppercase: IGHV3-15

Lowercase: Flanking sequence[1000bp]

Red & Bold & Underline: Stem-loop [20]

Blue: Heptamer[33]

Green: Nonamer [7]

id-IGHVII-74-1-2[V_gene_segment]

tttaactgcctaagaatttataccgtgaagtaattgcttcatttagtgttaatattaatgcccttttgacaatctttgaagctctttttttctgatatttcttccaacataaagaatgggagctttttcaccatataaagaatttttaggaagatatgatggaaagggaaactcaatgtattgttcagtgtttctgcctaacacatcaatctcttaaaatattttgctgtgtctgcatgtgataaagcagatgctaacactggagataaaagaaaaaaatgcacagaccttccacattccaaatcataaagtggtactggatgtgtttctgatgaagtgaaccatgggtcataagtgctagtgcaagtagctttggaaaatggagttcatagc**ttcttaaagttttgctagttttaatttaagaa**tgttatttactatttctttttagtacaagctttccagaag**attcttgatggtaatggaagtgtcgcatttaagaattacaaatgactcacagagttactttgtaatt**ttttctgtagggtgtacatgtcagtcact**tgacataaaagctcatatgtca**cagatagaaaaatg**aaaattgagccaatttt**aataactgcagtggaaatagtttttggcaatgagtgcttcaagacaaccacaaaataaatcggcaaaatattaaaatacagcaggaatctgatctcagtttctctgccttataagaacaaggctgtcccacatcaaaattattattcaagctctaggaaaaaatacacttttggattctggaagctaaaagctctctccttagatagtgctaaagtacttgtgaaatgcagagttgtgttcatgaaaaagatgactttattgtgatttttttttttttttttttgcctgttgacaactcaccaatctgtctcagatgtaaagccctgctctatgggttgaatgtggttgcacatgactggcagagactgattattcatatctccagGTGCCCTGTCTTTGGAGCAGCTACAGGAGTCAGGCCCAGACTAGAAGAGCCCACACTGACCTTCTGCATCACTTATGCTCTCTCTGGCCACTACATCACAAACAGTGCTTATGACTGGGCCTGAATCTG**CCAGCCCAGGAAAGGGCTGG**AATGGATGGGGTGCATTGGTagtgatggtgggggtcctactagcctgtggcaaatggaagcatctcttttttatcagactgaataatattgtagtgttttcttataccacatttacttcatccctttgtgcattaacacttaggttgtttttatatattggctgccataaatagagctgcaattaatataagagcacaggtgtctttacaagttcatgattttatttcctttccctgtatttccagagaagcgtttgctgagtcatattgtatttccatttttgatttatttatgagctgctat**actgtttttcataatggtgcaactaaaaaatggtgtagtcatcatgaaaaacagt**tttagtttttgacataatc**caaacacaatgtttg**aatctggttatctataggagtttctagtaaaattggtgaatgtaaattaaaaagcagtaagaaaaatctgcaattcagaaatagtgtacccagatggtctttctctccgaaatgaaa**gagaaataaactatttctc**aggtgagataaactaagatattttcttcacacttgaccagtc**atacaagaaatattcaagggaatttttgtat**aaaagccagcatatgaaaaccatcagcttggaaaaaaacacacaaaacctatatttcaatggtagaaacaattaaaaaaggagaaaagaaacaaacattattactacaaaacaacatcaaatcataaaaccaaacagaaagagaataaacaaagatagagatatatacatatatacaaacaatcataaaacaataaagaaaataatttgagtaaatacttataaataataaccttatgtaataaaattcctcaattaaaaatacagt**ataaaaattgattttat**taaagtcccaactctatgctgcttacaagagactcaaatcatctgagaatctacacacagactgaaagtgagtgggtggaaaaaagatatgctacaaaaatagaaaccaacagcacaaatagctctacttatat

Uppercase: IGHVII-74-1-2

Lowercase: Flanking sequence[1000bp]

Red & Bold & Underline: Stem-loop [11]

Blue: Heptamer[10]

Green: Nonamer [5]

id-TRBV19[V_gene_segment]

aatgtaagtctacactccaaacagaaataaatacactcaaggaagcagctgaggatataagtgaccataaccagaccaggactttaaaaactcagactacaaatggccaaaatgggaagacatcagaaaaaaacacaaattgattgaacccaggagggacatgaaagaaaaagacaaagttatcccagtcatgaaaaataaaagatgaccaagggagaataaacccaaatgaaaatttagtcaggacattgaacaaaggcaaagaaacaagtagcagaatgagaaggacatgagtgaaaccagccagttagaaggcggtcaaagaaggcgaggtcttggtatgattgcatgaggggcctggaacagcagcactgggctctgtgcagcaggacggctaagcagcaggttgtcacag**tggccacagcactactggcca**gaccccaggagtcacagtttccctgagagagctgagggcataaacaatgataagttctccaggggatggacttcttaaatactctgagttcatcattacacattctatgcatgtagcaaaataccacaggcacccacaaataggtacacatattatgtatcaatgtttttttaaaaaaaggaaaaataaatgaattccacattcttgtataatcagagtatctacagcgctcctggagtaacccaaaccttatggttgctggaaagataaaaaaccaat**gggccaattggggtgaaactctggccc**ggaaacgtggggtgccagccccctcagcacagatgcaat**gcagagttatgggaggtgcgaatgactctgc**tctctgtcctgtctcctcatctgcaaaattaggaagcctgtcttgattatctccaggaacctcccacctcttcattccagcctctgacaaactctgcacattaggccaggagaagcccccgagccaagtctcttttctcattctcttccaacaagtgcttggagctccaagaaggccccctttgcactATGAGCAACCAGGTGCTCTGCTGTGTGGTCCTTTGTCTCCTGGGAGCAAGTGAGTCCTGGGTTCAGGGAGAAAATTCCTATCTGGAGTGCTGCAAGTTCCAAATCTAAGGCCTCCCCGAGGGACAGCAGCATCAGGCGCCTCCCTGGGCTGTACTCAGGCATGCCTCTCTTTCTTTTCCAGACACCGTGGATGGTGGAATCACTCAGTCCCCAAAGTAC**CTGTTCAGAAAGGAAGGACAGAATGTGACCCTGAGTTGTGAACAG**AATTTGAACCACGATGCCATGTACTGGTACCGACAGGACCCAGGGCAAGGGCTGAGATTGATCTACTACTCACAGATAGTAAATGACTTTCAGAAAGGAGATATAGCTGAAGGGTACAGCGTCTCTCGGGAGAAGAAGGAATCCTTTCCTCTCACTGTGACATCGGCCCAAAAGAACCCGACAGCTTTCTATCTCTGTGCCAGTAGTATAGAcacagtgaagcacggatgtcgcctctctgtgcataaatgtgcccagtcctgcttccccgaccaggtggcagggctcctctgcactctatgatggcaggaaacgccactcagccactaagcaggtttaaaaaactgtatataactaaaactctatatataaactatg**catataaaactaaactatatatg**tatacatttatatataac**taaaactttatatgagactaaaatgatagtttta**tatagttttttaaagtatagtttt**atatatatataactatatatatatat**catttttt**aaactatagttttgtagttt**tatttatttatttactttttggagatggagtctcactctgttgcccaggctgcag**tgcagtggcatgatctcagctcactgca**atctctgcctcctgggttcaagcaattttcctgcctcagcctcccaagtagctgggatta**caggtgcataccaccacacctg**gctaatttttgtattttagtagaggcagggtttcaccatgttggccaggatggtctcgaactcctgacctcaggtgatccacccacctcggcctcccaaagtgctgggatta**caggcatgagtcaccgtgcctg**gctgttttatagtttcatatagttattttt**aaaaactatataaaagttttt**gtat**agccacttagtggct**gagtgaagtttcctg**tcatatatgatatagatatagatatgtttatatatga**tatagatacagatatgtta**tatatatgttatatgttgtatataagatatata**catagagagagagtcagagagacacagagaagaaggaaaagttaatcttacaacccccagcttttt**tctattttaccgaatgctagatgaataga**tggatgaatgaatagaaatgaatgaatgaatagaaaattttccct**tctttcaggaagataaaaaagcctgtggaaaga**gagatgtagaagacaaaccctgtaatcacaatttc

Uppercase: TRBV19

Lowercase: Flanking sequence[1000bp]

Red & Bold & Underline: Stem-loop [17]

Blue: Heptamer[32]

Green: Nonamer [2]

id-TRAV12-1[V_gene_segment]

aaggcaaagagaaagaccaaattc**ataaattttggagatttat**tcctagatataaacttaaaacaagaaaagaaataagaggaattagacaaatgaacagacatataaaagtctagatagtgaaggtaggtccatctcttttgcaacagcccaaacaaatactgttt**tgacatgacatgcacatgtcattggtgtaagacagaatgac**gaattccaagctccagtccttggctttgttaaaaaatgttggccaggctgtgagctcaaatcatatttctttatctagattttaggtcattct**attttgaaagaattcttttgaattataggaaatttctttaaaaat**tggagataacagaatgctatggggatgtatggcctgacatggagaagagctacgtaatgtgaaatgctccttaaattgcaagaatctcactcaaagttttggaatagaatgcttcatgactgattgttgtgtgcaggaaagtggctattagaaataattcctcttactcatacaagagttgccatttgatatcaaacacatgcatgcataaacatcccccacagacattgacttggagcatttgcttctcaaatttttaatttaactaatatgtcagatttttacattt**acctgttcacataatacaggt**ctacatcttgagtacgtctttgattactatcatgattccacaagatggcagtgtgttcttagggaccctgaggaaagctctcaagtgctcgtgtgtgtgtgtgtctgtgtgtgtgtgtgtgtgcttgagagagagagaaggagagaaagagagagaatgggaagggcgataagaggaggggttagtgatataccaggggttgtgaaaataacctcttttttctaattggtaggacagattctttttacgattcctaaagtgg**aagaaataaagtatctctgctatgttcatttctt**tttggattgaaaattttaatcctcagtgaaccagggcagaaaagaatgATGATATCCTTGAGAGTTTTACTGGTGATCCTGTGGCTTCAGTTAAGCTGTGAGTTGGGCATCTCTATAGGAACATAATGT**ACAATATTTTGAGATATTGT**CTATCCTAGGCTGGGGGCTAGAAGCCTGAATTTTTTCTCTAACTAAGAGAGTAGAAAGCCTGTAAAAACATGATTTGAATTCTGGGGAGTAAGCATTTACCTCCCAATTTGTCTTCTCTGTCTGACCACTGTCTTTTTCACAGGGGTTTGGAGCCAACGGAAGGAGGTGGAGCAGGATCCTGGACCCTTCAATGTTCCAGAGGGAGC**CACTGTCGCTTTCAACTGTACTTACAGCAACAGTG**CTTCTCAG**TCTTTCTTCTGGTACAGACAGGATTGCAGGAAAGA**ACCTAAGTTGCTGATGTCCGTATACTCCAGTGGTAATGAAGATGGAAGGTTTACAGCACAGCTCAATAGAGCCAGCCAGTATATTTCCCTGCTCATCAGAGACTCCAAGCTCAGTGATTCAGCCACCTACCT**CTGTGTGGTGAACAcacag**tgctccccagacacctgcagtctgtacccaaacctgctgggccccaggaatgcctgatgtagagcttagactgcagggcagtaaagtcctctttgctctctagattcaagtggaaattaagagtactagtatggaggactgattgattagggaagt**attattataattctgaaaataat**tgagaccctgaagaaaaatgaaagatacagtcttggtctggtt**aaaatattttccagtattctctttctactttagttttaaaatatttt**ttatatttcttctggagagatt**aaaaaaataaattttttt**gagatcattgacaaaccagaaggtttacatattcatttcatacatacatcacaatctatgccataagcatatccatcatctctaaaagtttacttcctccctccttttttaaaaaatttaggtatgat**tgatatacagcaaggtgcctatatca**cataatgtatacattttaatgctcactctagttttatgccacatctatgctttattagatcggttttatgattccttttatttatatcaagaaaacattttcatacaataacagatattctgtctctgtctgtaggacatcttattctttccccatttagttacccaagaagtttgatagacgttgctttaggaatgcaacagaaaaacacaaatgcaaaataaaaaagatcagcaaactactgtgaaaatcctgtaagtcataaattcttttagttatagaaccctcaccccatctccagccttagctgcagctccactcctgagttatataagggcacatttgatgagtagatggcagaactagagggagagggtgagaacacgttaaaaaaaagcctagatatttgcagagattgaagattagagagggtctatattgccattttggggtaaagtctgaattgaactcaaaaccttgtggaagagtatattgatttctatgtaactgagtgtggcttggagatggtagg

Uppercase: TRAV12-1

Lowercase: Flanking sequence[1000bp]

Red & Bold & Underline: Stem-loop [14]

Blue: Heptamer[39]

Green: Nonamer [7]

id-IGKV2D-19[V_gene_segment]

atcctcttgtaagaagtccccactccacccaggaagtccagctggcctcacctctcactatcaatgactgaaatgatcattagtagcagtggctttggttaaatgtataccagtagttacatacagggtagaaaaatggtctgaagaggtatattcagcatcaaatgaaaaaattacaccttgtgacccctttctctccctgtatgatcccttaagattgtgctgaaacctc**ggagcatcaggtcagcctcatattgctcc**ttcctaaatcttggtccctgctcaacctgagcctatgtttggccatccacgggcatccatgtatacaactttgaaggcaaaactccctgcttggagtggctcattcgatgaccagggtccagatagcattgccaaggaggctcacagatggcttcatactgagtctctcccctcagcctcccagcatcaccggccctgaaggaatgtccctgttatccgtggcacccagagcac**agtcagccctgccctgact**tcactgcccaattcacagcctcatcgcagattccactgcaaaatagacagtgtcagatctcaaagacaggctgatttgtagtcctgaaaagcaacaacccatcgttccctcgatctctttggtggaatttcaaagttcgtcataagctgaatttctgtacaaacatattctcttaacaatggaaactatttgcaatatgagaaacataaaacatgggaaaagtttaaatctctcaagagcatcttctagtcagattcatattcagcttccctgattctgaacttcctgtcttgcggtcagcttcttttggtccatttgtttcaccctttccagctcagatcttctctccctggctacccttagtctcaggttgtccttacgtttgataggcttct**accttatggatgctgtaaggt**tatcctctcatttgctggattcttgtgggaaaaaaaatccctccaagttcacatcctgagacttcacactCCACTCTTCCTTGGAGAGCCATCC**TCCATTTCCTGCAGGTCTGGTCAGAGCCCCTTTCATAGAAATGGA**AACACTTACTTGAATTGGTTTCTGCAGAAGCCAGGCCAACCTCTAAAGTGCCTGATTTATGAAGTTTCTAACAGGAAGTCTGGGGTCTCAGGCAGGTTCAGCAGCAGTGGTTCAGGGACAGATTTCATATTGAAAATCAGCAGGGTAGAGGCTGAGGACGTTGGGGTTTATTACTGCCTGCAAGGTACACAAGTGCCTcccacagtggtacaaccctgaatacaaacctccctgtttggagtggcccagctgctgaaata**tgttgtttatctggggagtggcacaaca**gaatctctgctgtataagatgaagatgttgggccgaggtgggcagatcatgaggtcaaaagtttgagaccagcccgaccaacatggtgaaaccccatctctactaaaaatacaaaaattagctggacgtggtggtgcacgcctgtaatcctagttactcaggaggctgaggcaggagaatagcttgaacccgggagatggagcc**tgcagtgagccaagatcacaccactgca**ctacagcctggatgacagagggagactccatctcaaaaaaaaaaaaaaaaaaaaagatgaagatgttggcgagctcagggta**acaggttgcagctgaatgacctgt**cccatgggggactccgcagtacatcaggtaaaacccattcatggtcctgtcagctcccacagccttggcatggcataagccaaaggaaaacagagataattcaagtgccttcagagtaagcagacaggactg**agggagagtggagggaaatctcacactaatcctccct**gccttgcctacattcgacaatgagacttcaaatagcttaatagccagacaagtaacacagattcgt**gtcaacatgtgttgactatctcttgaagtttaggtcttttgtgtatatttttaagaggatagt**atttggtagtatttagaaactggtatttttcaacttttccaatttccttcttctcctttttactagtt**ctctttccattgcaacacataacaaatagtgaaaagag**cattctacacaagc**tgtcctcagggagagctggctgaggaca**atcattaaaaagcttgaatttgcacctccaaatagacttttgtgacgtcatggaagacagaagatcctgatgttaaaactctttcattcatttcaattacttcttgctaataaaaaaggataatat

Uppercase: IGKV2D-19

Lowercase: Flanking sequence[1000bp]

Red & Bold & Underline: Stem-loop [12]

Blue: Heptamer[21]

Green: Nonamer [6]

id-IGHVII-60-1-2[V_gene_segment]

ggcggggcttcctcatctgctcagtggtctccgtcctcgccagcacctcgctgtcaccagggctcctctttctttattatctgtggttctg**cttcctcacattcttgtgccaggaaagaaacgaggaag**acaaattttcgtctatagttgaagcttcaccaattact**aggaacttgcctacaagttcct**gcatgacccattataacttatcgattaaaaaatatatattctaatgcttctcaccatctcttgatttgtatcatcaac**tgaattgtaccctctttgaaattca**tatgatgaaacctt**aaattcaatggatctatattggaattt**taatgaaataattaaggttaaatgtggtcataattgtaagaccctaatgcaatagacgtgttgtctttataagaagaggaagagacaccagagacctctcacttttcacgtgcaggcagagaagaggccatgtggagacatag**tgcactagaaggtggcccagtgca**agccaggaagaagccgcgccaagaaccagccctgccagcacactaatcttcaacattcagactgcagaattttaagaaaatcaatatttgttgtttaagccacccactcctgttgtcttcttatgaagatcca**gacagactaataccacataactctgttagtgctgtc**ccctggatggagaattagcctcctgaggctgggcacatctctcagatttccacataaaacaggtaaaaaatagtagttctgatataaaaacttgtcatgtccctgttggccaatttctgggcaaggtcttttaaataagccaagtttgcggggaaatggagaccatatgtttgtgggactctaaccgtggaatctactgcattgctctagcagaatcaaaaattgaacaccagtgagctctttattctcattaaaatgtgtgtctttggaaatgttaacattctgtccagaatctgtgctcacaaacaactgtgtttcttagtgcactcttGGCCTGGTGGAGCCCTCGCAGACCCTCTCCCTCACCTGTGCTGTCTCTGGATTTTCCATCACAACCAGTGTTTCCTGCTGGAGCTGGATCCACGA**GTCCACATGGGAAGGACTGGAGTGGAC**CAGGCGCACACGTCATGAAGGGAGCAAAAATTCCCACCCACTCCTTATGAATCCAGTCACCATCTCCAAATTCGGGTCCA**AAAAACACTTGTTTTT**ACAGTGGAGCTA**TGTGAGCAACAAGCTCACA**GCCATGTTTTAAAGAAGAGAcagagtgaggggaccacggtgcgagctcacacccaaaccttcctggaggggtgcacaggacagcaggagtgccgatgatggaaggggg**tggtctggattccagggcacacgcaagacca**ttgtagaggcactcaggtcacccgggggctctcaggaaccatcgaggaaaatcaggacaccagaagagactcggtacagcagggggctcaggaccattgtggggactcagaaacagcaggttctaggctcagcctcagggcaggtgcagatggggtgaaaaggggctggatgaggggttttgtgtcaccatcatatttcaccactagacaccctccactacatgtgttctactgcatatgagt**gtatgattagaaaatcatac**ttctataaatatataaccatacgaaggtgtgtcaagttttcctcttcaacttctatcggccttgttcgtaaggactaattccctgtaattacttgaggacctcataaattgtggtcagttttgtaggtttcctctctttccttgtctccctttccccttattctctctcacacaaatggacataatccccactccacacgcaaagaaatctataactttcattacctgatgtattgaataaaattgattaatgtgcagctttcccatctttgttgttgttcatgctgctgtaagtataagaaccatgagatt**ctcagctgtgtacttctccaagctgag**tggcagctttgcttatcatactgagatggccaacaatccaaatgtcaatcatcagcttaactggtaaacaaattgtggaaaagtcattcattggcatactactgactgtgaccacaatatactaatgttggatacgctcaacagtatggttaaattc**tcatatggtgagtaaaatgagccaaacaaataaaagtacacacatatga**taccacttttatacattctcccatccaagtactaaccaggcccgaccctgcttagcttccgagatcagacgag

Uppercase: IGHVII-60-1-2

Lowercase: Flanking sequence[1000bp]

Red & Bold & Underline: Stem-loop [13]

Blue: Heptamer[28]

Green: Nonamer [3]

id-IGHV1-24-2[V_gene_segment]

gtctggaccgtatcagtggatccggagctc**cagggaaggggctccctg**atggcttttatgattccttgcttgccgtgctgaggtctccctgtagattatgttgggttttctaaggcctattttctattgtaagaggtggtgggggaaacaatttctgtcactgaaagagcattctgagccagggcacagccccttcctactgagcttgagattctgggaggaaaaagttctctgacaggaggaaa**aagttcccagacaggaactt**ccctgcagggcaggag**ctgagcggcagggggcgctcag**ggagcacccagcacagga**cccagccctggggcaggtgcacaggaggctggg**gcagggttttctctcaggaattgaatgttc**cttatttcaaagcaataatgacctaacatttaaataag**aatttagcaagtactgatgtgcctgtaagtattttattgtatacgcagcctatacctaacccaataatttaatgcaaaccggatttaaaagg**agaaatgtctaggtctttcaaatgtatttct**agttaggaattgaggggtggttttattaattcaatgggtgttactgtccggagacacactcatcccagaagttagatgtgcagagggcaa**ggctcaggaaaagttcaggtagacagtgagcc**atatgaacagggaatcacattgaggaccatgtcctgtgagattctgggtttctgtgagaggagttctgtcttcatggacttctgggcatgtcagaggacaaatattaaacaaagttcagggcagggagctcagcatcccactgtggcgtggtccacatgtcacctatcttcttc**ctcagggtggcgtggcctgag**ctatgaaaaacctgcctcatgaatatgcaaatgcactgctgtctaccgaggtaaatacagatctttccttgcccagagaccatcacacaacagccacatccctcccctacagaagcccccagagcgcagcacctcaccATGGACTGCACCTGG**AGGATCCTCTTCTTGGTGGCAGCAGCTACAGGCAAGAGAATCCT**TAGTTCCAGGGCT**GATGAGGGGACTGGGTCCAGTTAAGTGGTGTCTCATC**CACTCCTCTGTCCTCTCCACAGGCACCCACGCCCAGGTCCAGCTGGTACAGTCTGGGG**CTGAGGTGAAGAAGCCTGGGGCCTCAGTGAAGGTCTCCTGCAAGGTTTCCGGATACACCCTCACTGA**ATTATCCATGCACTGGGTGCGACAGGCTCCTGGAAAAGGGCTTGAGTGGATGGGAGGTTTTGATCCTGAAGATGGTGAAACAATCTACGCACAGAAGTTCCAGGGCAGAGTCACCATGACCGAGGACACATCTACAGACACAGCCTACATGGAGCTGAGCAGCCTGAGATCTGAGGACACGGCCGTGTATT**ACTGTGCAACAGAcacagt**gtgaaaacccacatcctgagagcgtcagaaaccctgaggaatgaggcagctgtgctgaggctgaggagatgacag**gatttatgaagtttaaatctgtttagaaaatgggttgagtaattgagtaaaacag**caatggaaagatgtgtacactgtcattatgtaggaaatagtattttcaactgtcaccctataagtaacactcacagagtgggaaaggcagccatcaatcaagctgatgcaaacattcccatgggggctttgtggggacatacat**tttataattggatctataaa**tcattagaaacaggattgcttgatctcatggtaagactaaatat**aatttccaagaagtggccgaaatt**tcttccaaaatgtctttgccct**tccttttacactgaatttattttaaagga**gttttaggatcacaacaagtttgagtagaagaaacagagttcccatgcactcctgccccccgacgctcagccttctctgctatcaacaccctgcaccagtgtcataaatcggttacagaggatgaatctccacagacacattggttgtttccttttctggtggtccctggtataacaagcctcaactat**cttgaagcaccccaggttcaagtggattcttcaag**tgggttactgggaatgatgccaggtaga**aggaaagtgggtgggaccattcctctttgcactctttcct**caaggatccataaaatgtacatttacatggagctcctctttctcctggttttatatgcctcttccccaatggtaaggtttcccagtgtttacagcgggggtccccagtccccgtgctgtggatccgttaaggtgtgcagtgcgttaggaaccgggccacacagctggagatgagtgggggcgagccagcatcgctgcctgagctctgtctcctgtcggatcagctgtgcattagattctcctacaagccgaattcactttttttatatatatactttaattctagggta

Uppercase: IGHV1-24-2

Lowercase: Flanking sequence[1000bp]

Red & Bold & Underline: Stem-loop [20]

Blue: Heptamer[53]

Green: Nonamer [5]

id-TRGV9[V_gene_segment]

ggtagttatgtcatacagggaactcggaagaagatatggtttgggggaagggaatgagttcactctggctatgagcagactttgtttgagatcccatgtcagacgtacttgaaaatgtccaagagttagctggatatgggtacagacagctttcgatttaactgaggtccgacttaagatttttttttactttatgatggtgcaaaagtgacacgcatttggtagaaaacataatttgacctttgctgtttccccaggacagtgatataaggttatgatactctcttgcgatgctgggcagtggcagtgagctgcagctcccagtgggtaaattaagcatggaaatctggaacacagcagcttttaagggatgcactggggagagcagtttgtaaaaataccttgggacggattaaatgtcttggcttagagaaagaacaacagacaataggaaactgtcattgaggagaagccgggattcaggagatgtacatt**attcaggtagctgagggaggggacaggaacctgaat**cttcaacagaagcaaacaaactgggatggagggcccagcctcatgaaggtgaccagaaggacccggtgctgcaggctgtgtgggtagctgagcagagctaagcggcttgacggaccaacatctctccagctggttgaagacaagctctcagaagacaatgctgcatgtca**cagccccagcaaccaacaacaccagcctgacaacttgctggggtg**gccgccttgtggtctgaggtggccgtctaaactatgt**ggtctgatctcaggctgcagacc**ttgcaggactgtcttcacacagactggaagtgctaacaggtggtgaggacaccgctttacaacgatgcagggggc**cccatgtcaccctcacccatggg**aagtttgacttggtggactcagccaagccacagaggtctaacgcttctctgcggtgatttcaggctgccctggcagaaagcacagtgcctgcagacATGCTGTCACTGCTCCACACATCAACGCTGGCAGTCC**TTGGGGCTCGTAAGTAGTTTTGCTCCCCCAA**TCACTTTGGACTGATATTCTCTATTGTTATATATTTTTATAAATTCCAAATTCTTGGTTTATTTACTCCCTCCCATTTTTTTCCCCAGTGTGTGTATATGGTGCAGGTCACCTAGAGCAACCTCAAAT**TTCCAGTACTAAAACGCTGTCAAAAACAGCCCGCCTGGAA**TGTGTGGTGTCTGGAATAACAATTTCTGCAACATCTGTATATTGGTATCGAGAGAGACCTGGTGAA**GTCATACAGTTCCTGGTGTCCATTTCATATGACGGCACTGTCAGAAAGGAATCCGGCATTCCGTCA**GGCAAATT**TGAGGTGGATAGGATACCTGAAACGTCTACATCCACTCTCA**CCATTCACAATGTAGAGAAACAGGACATAGCTACCTACTA**CTGTGCCTTGTGGGAGGTGcacag**cagcagacagtttgagccatcccat**tcaataaatgtttattga**gtctttgtttataattacgaattgggaagccacagttaccaccagtgtgcttgtaaacagtttttaagataaacattc**atgtggtgactctacttggagttgccaaattccacat**attttct**caaagcagaagctttg**cataagtctatacaagtgaccagcctcctctttttcagggttgtactctctcaccaacagcttttctaggtcacccatgcttatttttctggttcaacctctcctactcat**tatatattttttttctggtataatatata**cacatttttagtgtgattcttttgtactagtcaatatgcattacataagttttctctgaccttcctatttctctcgatta**aaaaatggatatgacattaaatggtattcattttt**ctggaggttaacatgtaacaaacgcagcatgatctaaccacgagcagggcagactgcaatgacaaatctatcacatgcatgccttgagtggtgatgttt**gattttgtacaaaatc**cgtaaacacttccactaatcgaatctaagtcaaagtccactttggattggggtttgagaaaaaattcatttgctgcaacctctgcctcccgggttcaagcgattcttctacctcagtttcccaagtagctgagactacaggtgcgtgccaccacgcccatctaatttttttgtattttta**atagagacggggtttcgccatgttggccaggatggtctctat**ctcctcctgacctcgtgatctgcccacctcagccttccaaagtgctgggattacaggcgtgagccaccg**tgcctggtcatccaaccatgagcttttgatagactactctgagccaggca**ttatgcacagcagaaattacaaggcccatacgctctgcaaaggcccgaggttgcttctgttgaaccacattctgctactgggctgccattcaccactaaaatg

Uppercase: TRGV9

Lowercase: Flanking sequence[1000bp]

Red & Bold & Underline: Stem-loop [18]

Blue: Heptamer[26]

Green: Nonamer [3]

id-LOC112268316[V_gene_segment]

aataaacaatacaacataaaaggtttaattctcggtatttctactgagaaaaataagccaaatagataagagtacatactatattgtttcattcttataaattctagagaataaaaactagtctaaagaaatatgaaaacatcagtacttttataaagaaatggtagaagaaaagaggagaaaaacaaaattatgtctgtaaaagagcaagaggaattctg**aggtgagttgacttgtcacct**tctt**gaaaatagagattttc**tttatcaaagtttactattatgcaggttaaatatgtga**attttatcatctgtcaattaaaactcataaaat**gtattacaagtaaacaagtgaaattttagacaaaaaagggatgataagaaggaacaaatgaatacattaaatat**cagatacaccaaaaatttatctg**cctgatgcctagatgtttccgtatttttaggtaaatgcagcaaaatcacacaggttctcgtggcaggaagtggattctgcaaaccacactaggcccatttagctctgtccaatagttggttaagagagcaattgaggccagctgtgaggagcataggcccaggtactaggactcactcatgccagatataagcccttag**acacatacatagcccctccatgtgt**gggttcacttttacatctgtaaacgaagaaaccactgagtgctaaataacatcatttatacacataggtaaaaataattaaaaatatgatagttgttaaatgtttatcgcacaacaatttcacattaagacagcattttcccaaacacaatcattgtcatcaaaatcccccaggacgctctcatctactctgggccctgccctctcctcaggagtcccaccccatagcttgctatatagtaggtgacatgcaaatagagccctccctctcctgatgaaaaccagcccagccctgaccctgcagct**ctgggagtggagccccagccttgggattcccag**gtgtttccA**TTCAGTGATCAGGACTGAA**CACACAGGAATCACCATGGAGTTTGTGCTGAGCTGGGTTTTCCTTGTTGCTATATTAAAAGGTG**ATTCATGGAGAACTAGAGATATTGAGTGTGAATGGGCATGAAT**GAGAGAAACAGTGGGTATGTGATGTGTGGCAATTTCTGACCTTTGTGTCTCTCTGTTTGCAGGTGTCCAGTGTGAGGTTCAGCTGGTGCAGTCTGGGGGAGGCTTGGTACATCCTGGGGGGTCCCTGAGACTCTCCTGTGCAGGCTCTGGATTCACCTTCAGTAGCTATGCTATGCACTGGGTTCGCCAGG**CTCCAGGAAAAGGTCTGGAG**TGGGTATCAGCTATTGGTACTGGTGGTGGCACATACTATGCAGACTCCGTGAAGGGCCGATTCACCATCTCCAGAGACAATGCCAAGAACTCCTTGTATCTTCAAATGA**ACAGCCTGAGAGCCGAGGACATGGCTGT**GTATT**ACTGTGCAAGAGACACAGT**GAGGGGAAGTCAGTGTGAGCCCAGACACAAACCTCTCTGCAGAATGCTTGGGGGAAATCAGCTGCGGGGGGCACACAGGACCCACTGATCAGAGTCATCGCCAGAGGCAAGTTGCAGATGGAGGCTGGTTTCCTGTCAGGATGTGGGACTTCATCTTT**TTAGAGTTTCTCTAGGGAATCTCTCTAA**GTTCAGAATTCTGTGCTTACCAATGTCATCTCTACATATTT**TTAAAATGATTATTTTAA**TATGAAAACCTATTCTCCTATGCACAAAACACAGATTGATGCTTACAGAGATGAAAAGCCCTCAACCATTG**TCACCAGGATCAGAGTATTGAGGAAACTCAGGGATACCTGGTGAGTCTTCTCCAGTCAGACTCA**GGACAGAAACCTCAGTGAGATTCCCTGACTAGGACGGTCTTTAGGAATTGTGATCACAGCCAATAGAGTCTGGGCC**AGGGTCAGTGTCGTGTAGAACCTCACAGTTTTCATTCCTGACCCT**TCTC**CTGACACTAAAGTATGCAACTTAGTATCAG**CACTGATCTGGGGCCCCTTTTGCTCTTAGCCCACTCTATTTCTTTTTATTTGTTGTTGTTGTTCTTGCTCTTCCTTGTGCTGTTCGTGCTTCCTGTAAAGTGGGGATGTGGTTCTTGCTGCCAAAGCTCGAGGTCTCAAGCCCATTCC**CTGCAGCTGAGGTGGGGCTCAGGCTGTGGCTCCTGCAG**CCATG**TGGGAGAGGCTGATAGGACTTTTCTCTCTCCCA**TTGCTCAGCACCCTCCAGTGTGTCATGTGGAGACTCACCTGGGAATGCAAGTGGCCAACAGTAGTGAAGAGGATGAGCTTGTGTGGTCAAAATGGGATGTGGATGTGAAATTTATCCTGTGCTGTGCAAAGTACCACAGAGTGAGTCACCTTCCTCACCAGTAGTGTTAGAAAGAGGGTGTGAAAGTTGTCAGAATCAAAATAGATCCACTTGTGTTAAAACCCTGACAAAAGGAACTAGGAATGACCATGAAGGAGGTTTCCCATGCACATACTCCTGATAACAAGAACGACCATGAATGGATTCTGCTTAACCACAACCTTTGATAGAAGCCACCATGACCTTATAAAaatcacatctacaaggacatcttcccagcaaatcactgtttaaccctatattgatgccaaccttggtattgactctacaagcaaggaaaatactctcaaaacaatttatgtaacccacctcattttcactaataaacctatggattgacatcctggagtcactgctgcatttgttgtta**attgtaattagccccttttacaat**gtttgtgactgtttttctccgatgtctcttggaaaataaataatttacaagttgatggcagtaaagaagctatttaggacatttttaacatcctgttgaatatttctgcatagcacatcgatcccctaaaatacttcgctgtattggcatgtgataaatcagagtataatgctgaaggtaaaatggaaaatacatgggcttttgatgaatcaagtcatagggtgatattgtctttgcccttgaggaagcagaccatgggctgttaagttctagtgggagtacctttggcaaaaggatttcatgagtttctgaatgttatgctactttc**aatttaagaatgcaacttgtcatttatttttacttaaatt**tttccagaagatatttggcagtaaggacagggtagcattcgtgtgatactgatgacttagagaattattttgtaatttctcctgtaaggtatgcacattgctcactcgatacagaaggtcaaatgtcacaggtgggaaaatagg**aataaagcaaattttatt**aaatgtcatgactgtagtttttggcaaggaagtgcttcatgtcaacctgaaaacagacagacaacaataaaacatattcaaacccacagggagtcagacctatgtccttctctcgtataagtacaaggccttgccacatccaaactatcctttaggctccagggtataaaatgcttttggactgtggaagctaacagctctcccctcaggcagggctaaggtatctggggaatgcagagttgtgttcacgaagaagatggcattatg

Uppercase: LOC112268316

Lowercase: Flanking sequence[1000bp]

Red & Bold & Underline: Stem-loop [22]

Blue: Heptamer[60]

Green: Nonamer [8]

id-IGLVV-66[V_gene_segment]

aattctatttttttgatct**gaaaaaatggtattttttc**tcttttttatttttaaaacttatcttgaagcataatatgtaa**agtgaaatttgcagatcttaattgtaaatttcagt**gtgctttttatgaacacatatactcatgtgccccatgcgtactcacaacatttccattgctgcagaattaaaacattttgttctagtcaatgaaattattccattacagaataga**tatgtgtaccacata**gaatgtgtacctgagacacacttcccctggggactccttataaagtgcggtgaattgtgtgtctt**ttttcccattgaggaggccagtgtggaaaa**ctaaagactgagatgatgtagaaaactgccctgagaggga**acctcactctgccctgaggt**agggagccagcactggtcaaggcccc**ggtggaggatggcaactctgttgggtccacc**tggcctctggtcacaggcacgatggacctgggcctgggcagcagggggcgctgtggggctggcacttagagtctgtggggctcttagctgggaaatgatac**ctggccctggcctctgcctagggctcactggggccag**cagctgggtcctctctgggccctttgggagcctcagtccaggtacatcccaggcctactct**tcccagaaatgacccagccactggga**ctgcacacgggcccagggaggggcacgcttgggcaggggaagaggagcacatttgcatgaaaggcccatctctct**cctctaaggctagagg**gtgaacacgagagacctgccgcgtggcttcctc**tgcagagatccggggagtctgca**ccatggccttgaccccactcctcctcctcat**tgtcctccttcactgcacagtccaggaca**gcacaaagcatcctgatcccccagctcactgacaccacctcccaactcatgc**cagaactgtccttccctgttctg**ctttattcagtgtataatggctgtctgtgttttcggGATCCCTCTGGCAGCCCGTGCCGACTCAGTCGACCTCTGCATCTGCCTCCCTGGGAGCCCTGGCCAAGCTCGCCCACACTCAGCAGTCA**GCATAGCAGCTATGC**CATTGAATGCTTTCGGCAGCAGCCAGGGAAGGCC**CATCAGTACCTGATG**CAGCTTAAGAGTGATGGAAGCCACACCAAGGGGGCTGGGATTC**CCAGTCACTAGTTCTCAGTCTCCAGGACTGG**AGCTGACCACTATAGTGTCATTTCTACAATCCCGTCTGAGGATGGAGCTGACTAT**ATCTGTGGTACAGAT**TGTAGCATTGGTGTGccgtttgggtaaagcacagataaatggggaaatgagacaaaaactg**tttttctactctgctaccaaggttgaaaaa**tggctctcagaaccagtgtctgctgacctgcatactcaaatatgttatgtgtgtgtgtgtgtgaatgcgtgtgtgtgtgtactttaattttacgagtagagagccagatattttctgattcctgcttcaagttcccttggaaacagctttttaggccaggcctctagggagtgcactctgatctgtaactattgcgcaaatattacaacagagcctgtctcctcctgctggtctggtgtattgtggctatataggggggtgcaaagaaattctccttttcatattcagacactgggactcagctgcatacctcagtcctgtcttattga**gatcatttaaatgatc**aatcttagaagaaa**acttctgacttaaataaattttagaagt**ctatcaacatgctagaaagcatattccagataaataaaaatagaataaagatttatagagattgagatgaaaagcttt**aaaccctttgggggttt**ctgaaggtattctcaggtgttcagtgcctgggatttgggagtaagggataaggaggagaacttatttccaacacactgaagatgacaacagataagcagaaagttaccaaagaaggaggaatagaaatagccatgaatatcatcctggtaaccctgatctgg**aatgaaggatttcatt**tatt**tacttatttatttatttattttcaaaataagta**catctcatggagtccttggcaaaag**ctgaaggaagtattgcttcag**gaaatatctgtccagtaattaggaaactacta**atgtgattgagcagagacttcattaacttcacat**gcaaagaatacagacttgcagaaccctcctggaaaaccatgaaacaatacatcatcaacaataacaactatagcaaaaacagcaacaacatcaaccaaaacgctttacaaaaaag

Uppercase: IGLVV-66

Lowercase: Flanking sequence[1000bp]

Red & Bold & Underline: Stem-loop [24]

Blue: Heptamer[22]

Green: Nonamer [3]

id-IGHVIII-82[V_gene_segment]

ttggagggtaacagcactgggttgttagggatgtatgttggcattcatggtagtcattggtgctgttctccaaatattttcagttcattttttatgaatgcattctgactcttccatcccacctacttaaattttctaatggccacatgagtttttttttgtttttttttttttgtttttttttttttttgccaatggaggtgagaagaaataacatgtgactttttcaggagaaatctccaagaaacagagttctattctgcatgcttttttctcttttctatagcaatggggatcttattgatggtccctccttccgtctggattc**ctgtgttaggatgacacag**cacagagctacctctcacctgacccgtga**tgagatgttttgtggcaacatgatctca**agataatgataattataaaaccctttac**tgacaaaatgtagatctccattttttca**ttgtaatttaactttgccttactttaagaacagtagtttgcagctccaaatgcactgattaaagacagacgtccattttctctggaaatgtatttttatgttcttactggatgtatttgctgttaatgtttgctactatgaagatacctgaacagtgtccacattagagaataagagaaagtaatgggcagattaaccctgtgcatctagacccgggaatcctttgaccctaccctccctgaaatggagacacagaggatggatgagaaaaactgagcagtgcacccatgatcacaaaaagaatgacatggaaatgtgcctcttcccatctcatgaaacgcagctcatcctctgttctttcaggcccttggtgaagagccaccccatgtctgtcccttcc**tcagtgtccacaccatgaggtctgcactga**tctgggcttcccttctcatcactctcagtattagtgtgcattgtgaatcaggtccagctgtggctgctctgcatggggctgcagtcagtacgtttcctctgtgttcatgtAAGTCCTGTTGAAGCTTACTGATGGAGTCAGAGGGGGAAACATTGTACAGCCCAGCGGTTCTCACAGACTCTCCTGCAAAGCCTCTGATTTCACTTGTACTGGCTACAGCATGAGCTTGGTCCAGCAGGCTTCATGACAGGGATTTGTGTGGGTGGCAACAGTGAGTTATCAGGGTTACTCTCCATGAGTACAAGTAAATTAACAGTCCCAAGCAACACC**CTTTCAAGTGCAGTCTACCTTAAAATGACCAATGTGAAAG**CCAAGGACAAGACCTTGTATTACTGTGAGTGAcataggagcaggaacatctgcgtgagcccagacacaaaatcctctgcagggagacaggagggaat**cgcatggtagatgctgattggaactaccatggg**tcgctcagaactaccaggaggtactcagaaccactagggggcgctcaggacaccaggggacgctcaggacaaccagggggtgttcaggacaccagggggtgctccgagccaccagggggcactcaggacaccaggggacgctcagaaccattacttcc**ttgtaaatccatgatttctttacaaatgttacttccaaaacatt**aacttagaaccgggaatttttttttttttgacaaagtctccctcttgtcgcacaggctggagtgcagtggcatgatcttggctcatggcaacctccaactcccgggttcaagcaattctcctgcctcatacttctgagtagctgccaccacgtgcctgccaccatgcctggctaatgtttgtacttttagtagagacggggtttcaccatgttgaccaggctagtctcgaattcatgacctcaggtgatctgccccgctaggcttcccaaagt**gctgggattacaggcgtgagccaccatgcccagc**ctatttgaagttttaatgctgcgtgtt**ttctgagtaacgctagcaatgatctgtcagaa**caat**ttttaaaataggttatttatatattttattaaaatactattattaaaa**gagtaaaattgtaaaaattgcacctgcaatctcagcactttgggaggttgaggcaggcagatcacttgagctcaggagttttagaacagcctggcaacacaatgaaatcctttctatacaaaaaaaatagctgggtgtgatgctgcctgtggtcccagatacttggaaggctgaggtggggggatcccttaatcctgggaggttg**aggctgtagtgagatgtgattgtaccactgcactccagcct**ggttgacagagtgagaccctgtcttaaaaaatgtgt

Uppercase: IGHVIII-82

Lowercase: Flanking sequence[1000bp]

Red & Bold & Underline: Stem-loop [12]

Blue: Heptamer[36]

Green: Nonamer [6]

id-IGHV7-27-2[V_gene_segment]

aggtgtt**ctcccaaagcccctgggag**ctcctggactcattgtgag**tgtggactgatccattgcttccggagctccaca**gaaggggctccctggtggtttcatagaatccttgcttggggtctttctgcagagttc**actggctttcctacggccagt**ttagtattgtaagagatggtgtcagcagcacatggtatcacatgtcactaaatgagcattctgagccaggacacagcatcttcatactggggggagggaaatgctttggg**agcccagacaggagcctctctgcagtgcaagggctgggct**gcagggggcgctcaaggcccacccatcacaggctccac**ccccagagcaggtgcaccggaggctgggg**agggattcctctcaagatctgtgtcattcttctaaaaaaaatctaaaatgagtatttgacaaagactgctgaacattctataaatatcctattcaattgtgagcatttatcaaatttgatgttgtaatgagaaccacttttatact**agatatttctaaccctgctagatatct**taatagtatgcagctggaggttaagcaagcttttcttttatataaataagtgcaactttggagaaacacactcatcccccgaataagacattcatgta**ttaaagcctagaaatactttaa**attacctctgagccattcaagtgtgtgttcacagtgaagtcctgttctagggaaaattg**ttctccagggggagaa**gctttgtcaacatagagcttagggatgtatcaggaggcacatgacct**ctaaggggattatagcttgaacccttag**catcctcctgttggtaatacatgtgtcactctccaatctttctcatgcagtatcaggtatgaaataggatcgctcatgaatatgcaaataactgaggtaaatatagatatctttgtgccttgagagcatcatccaacaaccacaactctcctcagaagaagcccctagaccacagcacctcaacATGTACTGGACCTGGAGGATCCTCTTCTTGGTGGCAGCAGCAACAGGTAAGGGACCTCCCAGTCACCGGGCTGAGAGAGAAACCAGGCCAGTCAAGTGAGACTTCACGCACTCCTGTCTCCTCTCCACAGGTGTCCACTCACAGGTGCAGCTGGTGCAGTCTGGGCCTGAGGTGA**AGAAGCCTGGAGCCTCATTGAAGGTTTCCTGCAAGGCTTCT**GGATACACCTTCACAAGCTATGCTATCAGCTGGGTATGACAGGCCCATGGACAAGGGCTTGAGGAAATGGGATGGATCAACACCAACACTGGGAACCTAACGTATGCCCAGGGCTTCACAGGACGGTTTGTCTTCTCCATGGACACCTCCGTCAGCATGGCATATCTTCATATCAGCAGCCTAAAGGCTGAGGACACGtgcaagaggcacagtgtggaaacccacatcctgagagaaccagaaatcctgagggaggaggcagctgtgctgagctgaggcagtgacagggacaacgtggctgcaccctttgttgacatagggttatacacatacatatatatgcacccaacactggagcacccaggtatatcaagcaaatacaattagagataaacagagagatagacccaataccataatagctggagactttaacaccccactttcagcattggacagatcttccagacagaaaattaacaatgaaatcttagactaaatctgcatgacagaccaagtggacataatggatattgatggaacacttcatccaatggctgcagagtacacaatcttttcctaagcatatggatcattctcagggatagaccacatagtagaccagaaaatttgtcatttttaattaaaaagatatcatgtaccttctctgatagtaatgaaagaaaactaaaataataagaggaattttgaaaacacatgga**aattaaacagtatcctccttaatt**aaacaatgtgcttccaaactgaactatatgctctaaaatgggttaataaagaaacaaacaagaaaattttacaatttattgaaagaaatgataatgaaaacacagcatatgaacacctatgaaataca**gtgaacatagtactaagaggaaagttcac**agctataagcacc**tatgttaaaaaagtagacaattataacata**aacatcttaggaatatctctcaaaaaaaactagaaaagcaagagcaaatcaaatccaaacatagtaaaataaaaaagttaagatcagaacgttaataaataaaattgaaataaat**aaaacaataccaaagatcgatcaaatgaaaagttgtttt**ttaaaaagataaaagacatctataaacctttaaccagaccaagaaaaaaaacaaagaagaccaaagtaaataaaatcagaaagaaaactggaaatatt

Uppercase: IGHV7-27-2

Lowercase: Flanking sequence[1000bp]

Red & Bold & Underline: Stem-loop [14]

Blue: Heptamer[27]

Green: Nonamer [2]

id-IGHV2-5[V_gene_segment]

aaaattggagacatgtagaagctcactgggacctgttgagtcttctccaatcacactcagaaca**gagaccttagtgggtctc**ccttagagtagtcattaaggatggtgataacagcctaaagatggtagaccatggtcagtgtcatatagaatatggggaccctcacaagttttttgtctgacccttctcctgacactaaattatgcaaattaataacactgatctggtgcttcttttgattctaatttattttatttttagttgtcgttctca**cttttcctttggattttcctgctccctggaaaag**gtaaatgtggtctccgtgacctcaattc**aagggctgaagccctt**tccctgtagctcagctggggctcaggctgtggctactgcagccatgtggaagaggctgaagggactttcttcactctccttgctcaggaccatccactgtattgtgtataggcttctctggaaatgcaagtgg**ccatttgtagtgaaagaaatatgtttgtctggttaaaatgg**gaggtgg**atgtagagttaattggctgctacat**aaactgtccttctccaccagtgcttttaggatgagattgtgaaatttgtaagaatcaaaatggagtcacatatgttaaaaccctgacaaatggattcaggaagtgtagggagaattcttacacacatatccctgacaa**caagaactatcataaaatagttcttg**caaaaagaccaacatgacctcataatcatgacttctgcaaagacttctactcag**aatctacttgcccagccttagatt**aatgccatctgaattacactgatcatgttactatcactgctcctcaccacagatgcaacaccctcctgagtcctgaaacctgactccatcccatagagtagggca**cagatgaggggaatgcaaatctccaccagctccaccctcctctg**ggttgaaaaagccga**gcacaggtcccagctcagtgactcctgtgc**cccaccATGGACACACTTTGCTCCACGCTCCTGCTG**CTGACCATCCCTTCATGTGAGTGCTGTGGTCAG**GGACTCCTTCACGGGTGAAACATCAGTTTTCTTGTTTGTGGGCTTCATCTTCTTATGCTTTCTCCACAGGGGTCTTGTCCCAGATCACCTTGAAGGAGTCTGGTCCTACGCTGGTGAAACCCACACAGACCCTCACGCTGACCTGCACCTTCTCTGGGTTCTCACTCAGCACTAGTGGAGTGGGTGT**GGGCTGGATCCGTCAGCCC**CCAGGAAAGGCCCTGGAGTGGCTTGCACTCATTTATTGGGATGATGATAAGCGCTACAGCCCATCTCTGAAGAGCAGGCTCACCATCACC**AAGGACACCTCCAAAAACCAGGTGGTCCTT**ACAATGACCAACATGGACCCTGTGGACACAGCCACATATTACTGTGCACACAGACcacaaagacacagcccagggcacctcctgtacaaaaacccaggctgcttctcattggtgctccctccccacctctgcagaaca**ggaaagtgcagctgagatacgttttcctgccagggcctgcatttcc**catccccattagactcagagccctgtcttcctccttcttctttaataataaatggcatgactcctgttaatagttcatagaagcagaagctgagtcctgtttgtcaaacattcag**catgaaatgttcatg**ttacctgggccagatgcatcactggtatgt**ggccgccagtttaatgggctcatactactccaagcggcc**gaatacaaaacagcagaacatatgcctttattaaactgcaggaaatgacactggagaaaaatgtggggataagggagcaggaggaattgtgaaaattcagataatagct**tttctcgtagagagaaa**ggtgcgtta**ataaaataattttat**aaaacaataaaacagatggaatgtgtccccatcacggagtgtctcacagtaacagtgcagaaataaatttctaaaaatgtcgtagggtaggtgtgatagacaaggcttatttccacacagggagcccataaatacccatgtaattaaaaatccagctcttagaaagggagtgcctcaacttgacatcagaaacccacgtcagtgagtctgaggagcaatgtggaa**aagaaatggaaaattcaacaatacctttctt**atgaaaatcctagctcac**gtttcaacatgaaac**ataactgtctacaattttaggaaaaccatcccatggcaacaagtcactacactcaaaaggtactgagctaacagagttccagcagtgagcaggtcagtgctgaaatccccggagaatcaaggctctcaggtacgttgtacaaaacccagaactgagccacacaatctatggtcaagtgatctttgacaaagtcaacaaaaatatgcacggggaagaggccacgctcttcatgc

Uppercase: IGHV2-5

Lowercase: Flanking sequence[1000bp]

Red & Bold & Underline: Stem-loop [19]

Blue: Heptamer[39]

Green: Nonamer [6]

id-IGKV2D-24[V_gene_segment]

aaactgaatgcaatatgggatccagaaattttatcctaaaacattaaaagtaaaaatggtaaatacatgtatcagtggaaagctcagtgaaattcaaatgtagattgtaacttc**gttaataatagtggattaac**cattaatgttaaagctatttgaagtactaga**aaaatcagtttaaaatgatttt**atattcagcaaaactatccttaaagaaaagaaaagaagccgtgactagcatatatgtcctataagaaactcaggaagaaatccttcagaattcagaatcacagtaaatgacaatgaacagta**atttaaatccatgaaattaaat**gaaagcttcataaatatacttacctcaactcatatgttgttgatgttcacgaaaactgaatctttgtgatagatatcagagttgcagttcccttggtaggttagaggcagaagctattgactagaaagg**tgaatgaaggcagcatgtggagaatttcaaatcattca**tatttgtatctgggtagtgaatgtgagtactttatttggttgagcagtgaacatgtttgcactttactcagggcacaatttattttgatttataaaattaacagcaaaccaagaccctttcaacacacatgaagaaaaaaataagaagcaccaaatatttacagaaactcagccgtattaaagagaagtgtaacaagcactgggaaaatactaggaagtaaaaaaattgacagtaaacacagtaaacatagaaatatatcctgtcccaatcaggctgcatagattgttatttctgccagttttttctcaagcatacaaaatatgttgttcataggaaaggcccccatacccctgcacatatcatgttatttctataccactgcacccaccaggggatttgcatattgtcccccagggaggaccttcccttg**caagtctgagataaaagctcagcaccaaccttgacttg**actaattaggactcctcaggtcaccttctcacaATGAGGCTCCTTGCTCAGCTTCTGGGGCTGCTAATGCTCTGGGTCCCTGGTGAGGACAGAAGAGAGATGAGGGAGGAGAATGGGG**TGGGAGGGTGAACTCTGGGGGCCCCATTGCCTCCCA**TGTGTGTTCTGTCCTCATGTTAGATGTGTACGTCTTGTACTCCAGGATGGGGCTTGTAACTTTTATATCTGCGTGAGTAAGGCATGTGAGGTTTAGATCTGTAAGAATGAGGAAGATTCCAGAAGGAACAAAGACCAGTGCTCCGGTGAAGACTCTAACAGAGAAAGAGGGAATGGTAGAGGAAACTTCTAGCAC**TCAAAGCACTCTGCTGTGCTTTGAAAATATGTTTTTATTTT**GAAATTATATATTACTAGGGTCTG**AATCAAATTATAAAAATTGATT**TAGCCTGAAAT**AAATAACAGAAGAAAAATTATTT**TAAAATTGTGCTTAAAGTTTCTACATAACCTTGCACTTCTCTCTCATTATTTCAGGATCCAGTGGGGATATTGTGATGACCCAGACTCCACTCTCCTCGCCTGTCACCCTTGGACAGCCGGCCTCCATCTCCTTCAGGTCTAGTCAAAGCCTCGTACACAGTGATGGAAACACCTACTTGAGTTGGCTTCAGCAGAGGCCAGGCCAGCCTCCAAGACTCCTAATTTATAAGGTTTCTAACCGGTTCTCTGGGGTCCCAGACAGA**TTCAGTGGCAGTGGGGCAGGGACAGATTTCACACTGAA**AATCAGCAGGGTGGAAGCTGAGGATGTCGGGGTTTATTACTGCACGCAAGCTACACAATTTCCTcacacagtggtacagccctgaacaaaaacctcccgctggagtggcccag**ctgctcaagtgtgttgtttctctggggagcag**ttgaacagaatctctatctgtatgagataaacatgttggagaactcagggcaacaggttgcat**ctgagggttctgtcccatgggtgcctcag**ttgtacgtcaggcaaaacctgttcacagccctgtcagctgcaacagccttggcatggcataagccataggaaaccagaggtgatcccagtgcctgcacaggtaatagactgccctgagggagagcttaagaaaatcctattccaatcttccctgccttgcctgcattgggaaataagacttaaagaggtaaataaccagacaagtaacccagatttgttgcaacacttgaatatatcttgaggtttagcaatttaaagtctatatttaggaggataatatgtggtaatatcccaaaattgaacttttcaactttcctaacttcttatttttctctttcaccaccta**tcttcccaccacatattgatggtggaaaga**gccttccgcacaagctgtcatcatgaggagctggatgagggcaattagtgaaaatcttggatttcagcctcagaatggacttttgtaaattggtgagagatag**aaaatatgaatgctaaaattatttt**attcgcttcaattgtgtcttgctgacagaaaaggatagtttttgaaatttcagaagttg**agtttcataaacagaaact**taaactagaagacataggttatagaatttacctcatagaacactgaaataacacagaatgatgtgcgatttctttccccaaaatgtaagagt**ttgaagacagtgggccgacttcaa**gaatgggagaattaatggaagatagtggaggtcaactatggcccaataacctgctctttgacttacattaggtacagttgtggatgacagtgactgttgggggttggtgatataaactcagaaaggagc

Uppercase: IGKV2D-24

Lowercase: Flanking sequence[1000bp]

Red & Bold & Underline: Stem-loop [17]

Blue: Heptamer[32]

Green: Nonamer [9]

id-IGLVV-58[V_gene_segment]

ctcagctaccagagaatgcagtgaaccatccaatccaaataaattatcctgttaaacattgaccagtaggtctcctttgtgcggatctgaggaatcaagcagggagaaagaacctgactttgtgagctgc**ctgagaaatgagttcatggctgaagtgtcccttttctcag**ctaacatgttttatttcatgcacagggtgtatacagagacaatcttcccctgaggactccataaaagatggggagaat**tgtgtgtccagttcccattgaggagacacaca**cagaaaactaaaggctgaggtgaggtaggaggctg**ccctgaggacgggccttgccctgcccttggtcaggg**agtgagcagtgttcaagaccctggtggggacaggaggctctgtcaggaccccggcatctggtcagagacacggtggacctggacttg**agcagcaggggatgctgtggccctgctgct**gagccctctgtggggctgtcagctgggaaaccagaaccctgtgtggtctgtgctcagagatcactgggccaagtagctggatcctctgtg**ggctcctggggagcc**tcaggccaggctggatcctctgtg**ggctcctggggagcc**tcaggccaggcccaagttaggcccacccctcccagaaatgacacagccattgccgctgcatacagccccagggaggggtacggttggg**cagaagagagacaagtttgcatgaagggcccctctctctcttctg**ggactaccggatggataagagggacctgcatcatgtcagcctcagcagagctc**tggggtgtttgcaccatggcctggacccca**ctcttcctcctcctcaccctcctcctccacca**ctgcagaggtcaggatggcccttagcaccctgatcctgcag**ctcactcacactgcctcacaaactcataccaggaatgtccttccctcttatccttacttcaggccataatgggcgtctatgtcttcagggtccctctccCAGCCTGTGCTGACTCAGTCACCCACTGCATCTGCCTCCCTGTGGGAGCCTCGGTCAAGCTTACCT**CCACTCTGAGCAGCAAATACAGCACCTACATTATTGAGTGG**CATCAGCAGTAGCCAGGGAAGGTCCCTTGGTACTTGATGTATGTTACCAGTGATGGAAGCCACAAAAATG**GGATCCCCAGTTACAGTTCTCAGGATCC**AGCTATGGGGCTGATCGGTAGGTCACCATCTCCAACATCCAGTTTGAGGATGAAGCTGATTGTATCTGTGGTGCAGATCATAGCATTGGTGTGACATATGGGTaaggcaaagtgaccccagtgaatgaggaagcaggacaaaaactgttttctctgctccactatgaaggctgccacgtggccctgag**aaacagtgcctgttt**tccttactactcaagaaagatagatagatagataggcagatag**ctagctagatagatagctagctagacagataggcagatagctagctagctagctagctag**acagatagataggcaga**tagctagctagatgctagatagacagatacctagatagcta**gatagattttttgctttgggt**tccttaggaacaagtttgctaggccaagctcctaagga**gtacacaagtctgtattaatgtggccaatattaaggcagatcctgtcaccccctgcatgtcgatacattgtca**ccacatgggaactgcaaggacatttttccctgtgg**atttaaacactggaactctgctgcatggcacacccctgccacatcaagtttcattttaacaatcacccataagatataacttcttaagaaa**attttgaattctaacaaaat**gctagaaatcacactgttgatacacagcgatggaaataatg**aaaatttatagacagtgaaataaaaagatttaaaatttt**tggtggtttataaaggttttctcagttgttcagtgcgtgggttttgggattaggagaaaatgaggagaaactgcctccaccacattgaacatgacaactgatttgcaataatctaacaaagaagaaggatataaataatcattaatgtcatc**ctgacagccctgatcctaaatgtcag**aattttatttcttaac**ttttttctccaaaaaaa**aaatgtagcttcagaaaaacttctgga**atggtggtatgaggccatcagaaaacctgattcaccat**acaagcaatgagaacagcagtaaaaatgatataaatctacttttcagaactctgtacattaccaaaagcttacaaaattctgggaatcataaaaaatagtaataa

Uppercase: IGLVV-58

Lowercase: Flanking sequence[1000bp]

Red & Bold & Underline: Stem-loop [21]

Blue: Heptamer[38]

Green: Nonamer [2]

id-IGHV3-66-2[V_gene_segment]

ctcaaagaattgtgctgagtgaaagaagctaag**gaattaagagtaaattttatataattc**attgtagaaatattagaagatgccactaccataaattaaaatgaagaagacttaaatttttctgagaaaatggtgttgggaatgatgcggatgtgatttaagtttcagaggaataaggaaaaagatttagggattaatttaattattcaaaagttgattgaagtgccgagtgaatggctgcaaacatagctctacatttttcaaatcattccctataaatctgaattaatt**atttatttattatacttgaataaagcaataacgaagaaataaat**gaatatttttgctaaaatggagcaataaaaagactgatattga**cagaagaaatatgactgacttctg**aaaacacacatgaaccatggttctctctgcatatttaggtgaattacagaaagttgtcataacagatggggaatcctgcagacttcactaggcatggtccacgctgccctggagttgtctcaggggagctgcctcctccggtgattagagcacaggcccagataataggattacatttttttagatgtgtaaacttagacgcactgcacagctgctgtattctctatgtaaattatcttctgtaaaatacaacattaaaggctgcattaaatatattgtgtaaatatgtaaaaataaaatcagattatgagagctaaatgttaatcaaggcacaatcacat**aatataaaattatatt**ttcctgaatgatggaattactaccaatctcccccaggagactt**catctgcactgggcccggcctctcctcagatgtcccatcacagagcttgctatataatgggggacat**gcaaatagggccctccctctgctgatgaaaaccagcccagccctgaccctgcagctctgggagaggagcccagcactgggattccgaggtgtttcca**ttcagtgatctgcactgaa**cacagaggactcgccATGGAGTTTGGGCTGAGCTGGGTTTTCCTTGTTGCTATTTTAAAAGGTGATTCATGGAGAACTAGAGATATTGAGTGTGAGTGAACACGAGTGAG**AGAAACAGTGGATATGTGTGGCAGTTTCT**AACCAATGTCTCTGTGTTTGCAGGTGTCCAGTGTGAGGTGCAGCTGGTGGAGTCTGGAGGAGGCTTGATCCAGCCTGGGGGGTCCCTGAGACTCTCCTGTGCAGCCTCTGGGTTCACCGTCAGTAGCAACTACATGAGCTGGGTCCGCCAGG**CTCCAGGGAAGGGGCTGGAG**TGGGTCTCAGTTATTTATAGCTGTGGTAGCACATACTACGCAGACTCCGTGAAGGGCCGATTCACCATCTCCAGAGACAATTCCAAGAACACGCTGTATCTTCAAATGA**ACAGCCTGAGAGCTGAGGACACGGCTGT**GTATT**ACTGTGCGAGAGAcacagt**gag**gggaggccattgtgcgcccagacacaaacctccctgcaggaacgctgggggaatcagcggcaggg**ggcgctcaggagccactgatcagagtcagccccggaggcaggtgcaggtggaggctgtttcctgtcaggatgtgggacttcatcttcttccaacagtttctctaatgaacctctctaattttagaattctgtggttcctaatgtcatctctacatattttcaaaagatcattttaa**catgaggacataacctctcatg**caccaaatgcacattgatgcttacaaagatgaaaagttctcaaccattgtgaccaggatcgcagtcctgaggaagctcacgggtgtctgatgagtctcctccattcaggc**ccaggacagaaacctcagagggattcctgg**accagaacggcagggattctgatcacagccaatagagaggctgggcc**agggtcagtgtcctgtagaagctcacagggttcaagtctgaccct**tctcctgaccctaaagccaatgagcat**cagcactgatctggtgctg**cttttgctcccaatacatgttctttctttggagtgtttgttctcccttttttatttgct**tttctttcttcctgaaaaagaaa**cacatggtctctgtgatctacactccagggctcaaggcattttcttagaactcaggcaaggctcaggcttggctactccagccacgtg**ggagaggctgacgggatttccttctctctcc**atattctcagggccctcctctgtgttgtgtgtagactcatctgggaatgcaattggctgttagtagtgaaggggatgaactcatttgatcaaaatgggatgtggatgtggaattaaccctgttctatgcacactgtcagagtcatcttcttcagaagtagtgttagaaagagcttgtgaaatttatcggcaacaaaatggatccccttgtgttaaaaccctaatgaatgaagctggggaaggccatgaa

Uppercase: IGHV3-66-2

Lowercase: Flanking sequence[1000bp]

Red & Bold & Underline: Stem-loop [19]

Blue: Heptamer[42]

Green: Nonamer [10]

id-IGHVIII-22-2[V_gene_segment]

ctaacttttagaggttttattatgagatcttgaattttttaaatgttattttctatgtctggtgagaagttaa**ctgatgcttttcctttattctcttaatttaccacatcag**atttattgatttgaagatattgattcatcagcgcatctcaaaaataatttggagcttgtcatggtgtacaatccttttaaagtcttatagaacttagttgactatgaattgggagtagcttgtttacatatactaagaacatttgtcagaagtttggttttcttcttttgccttgtctattactggtaatacagtgattggagcctcttagaaagagattagaaggtggggacctcacggatt**ttttaaaaagttagagaacggttgtcatcacttttttaaa**cgctgagagtcatttgttatctaatatataacccatcatgaggaatatctctagtgtgcttgggaagcatgcatattacgctgctgttggttggaatgttccgtagatgtctgttagggcagtttgttcgagagtgcttaagtccacgggcttcctaagaatttttctgtggatgctctgtacattattgaaggcgaggttttgaagtctcctaatttttttatagttttctatttctccttttatacatcttaaagtttcataaatgtagttatatttctagttttt**caaatatgtaaaagagaaataaaaatatttg**cttagtaattttaatga**gaaaatcacataacaagaaattatattttc**caaatgctgccatcaacactaaactcctccaggagtctcacatctgccctgggctctgctct**ctcctcaggcatcccacctcagcatttgctgcagaggaggggacgtgaaaatagggccctcc**ctttcctgatggaaaccatcgcagccctgagcctacagctctgggagaagagccccagtccagcattaccaggagttttgatttgacaatcagctctgaacacacatggctcaccATAGAGTTAGGGCTGAGCTGGGTTTTCTTGTTGCTATTTTAAAAGGCAAATAATGGAGAACTAGAGATATTAAGCATGAGTGGATGTGAGTGAAAATACAGTGATTATGTGTGGCAGGTTCGGACCAAAATGTCCCTATGTTTGTAGGTGTCTAGTGTGAGGTGCAGGTGGTGGAGTGTAGGGGAGAGacaagaggcagaaaagtgtgcaggaggccgggtgaggctgtagacactgtcagcccactatgccaa**tcccaccacgagtgctggagaaggtggga**gtctgatgaagcttactaacaaccctgtggtccaagctaattccatcaaggccattggttcttcccagagcatagctgtccatcgggaatctcccatgtgcccagcagcagccacgcgttagcaacttcactgtgcacagtcattgtttgggaggagctccaggatgtgtctctttgtcaaaaactgggtgatgg**tgtcagagagctgtgccgggtgcctggtccacgtgctccctgaca**ttggaggtataggaggtgcgttctcagggtggaaacaccttttatggaaatgtacataataaatcacgattttc**cttggttctcttagacgacatggagaagcaag**catgcaccctgcaaacaattgtaatttcctgctttaccccaagttcatgatttcttgtaaactttgatttctgtccaagtacagagatactgattagagtgagtttggttctttccacacactaaccctca**cctcccccagatagagcacattgtcctcactctgagtctgagggagg**agctgttcctgtacaactcagggactgcagagacccccgtgtgcagctttgcagagtcagatctttcctcatgtgaggcgacctccactgcccgtgattgctgctcaggtctaattgtggattcgggattatgacaccctttgggctatcacaggtcaatcttattctgaaaatcaccgtaattata**gacagaggtaataatccagtacccattcttctgtc**cgcagtttgtc**tttcttatttggttgaaagtttgaagaaa**aaatgtgaccatatattcatctaattctaacctctgaatctgctatttgctctcggggacctacaaattgcaaaacagtgagttcttcattttcatcaaaatgtgcaaatttgtgaatt

Uppercase: IGHVIII-22-2

Lowercase: Flanking sequence[1000bp]

Red & Bold & Underline: Stem-loop [12]

Blue: Heptamer[27]

Green: Nonamer [4]

id-IGHVII-30-1-2[V_gene_segment]

ccagcaagaccattggtgcctcccagagcacagttgtccatcagggatgtccaatgcttcccagcagcagccatgcctcagtgtctccactgtgcacagccactgt**ctgggaggagctcccag**gatgggtgtctttggcacacaccaggtggcgggtgttaagagtgcagtgcagcagctggctgcctggtctgttgggcctg**atgctggagagatgggaggtgcattcttggggccagcat**gctgtttgtgaatttttatataaaaactctgattttacttcattttc**gcagatgacatagataactaaaaacagaatctgc**aaagaaattgtaattttcaactttaccccaaattcattgtttcttaattctgtgcaagatccagacatattattgccttcctcatgataatttattctatgtaaaaccgaaatcattttttctcctactctttgtttctgttcaagtacagagatcttgagcaaagtaagttgggttctttccacacactaaccctcaccttccccagagaatgagcagagattgtcctcagtctgagtctaagggaggagctgtt**tctgcacaactcagagcctgcaga**gacccccaggtgcagcttcagtgagtcagacatttctccatgtgggcgacctccagtgcctgtgactgctgctcagg**cctaattgttggttaagcattagg**acactcttta**ggtgatcacatctcaagcctattctgaaaatcacc**atgatcagagatagttcaatggccattctcctaacataagtttctctttat**tacttggttccaagta**tggagaaaaatgtgaccctatatttgtctgaatccaacatcagcatcaactgcattatgctaggaga**ctcactaattgagcacaagtgag**ctcgttattctcatagaaatgtaagtatttggaaatttcagtgtgttctccagaacctgtggctgccaacaactgtatttctcggtgcactcttGGCCTGGTGAAGCCTTCACAGAACCTCTCCCTCACCTGTGCCATCTCTGCATTCTCCATCACAACCAGTGTTTCCTTCTGGAGCTGCATCCATCAGCGCCCCCATGGGAGGTACTGGAGTGGATCGGGTGCATAGGTCATGGAGGGAGCACAAATTACTCCCCTCTTCTCAAGAGTCCAGTCACCATCTCCAGATCCATGTCCAAAAAG**TAGTTCTTCTTACAGCTGAACTATGTGAGGAACAAACACATAG**CCATGTATTTTAGAGCAAAAGA**cacagtgaggaaaccacagtgtgaactcatacccaagcctccctgtg**ggggtgcataggacagccagggttactcaggacaccaggctccctcaggacaccaaggggcactcaagaccattgtagaggcatgcaggtagctgggggctctcaggaaccatgggggaaaatcaggacaccaaagggtgcttggtacagcggggggctcaggattattgtggcgattcagaaagagcaggttcaaggctcaacctcagggcatgtgcagctggtgtgaaaaggagctggatgaggcgttttgtgtcaccatcatgtttcaccaccagacaccctccactacctcta**ttctaatgcatgtgtttgtatgattagaa**aatgatattgatataaatatataaccatagctaggtgtgtcaagttgtcctctccatcttatatcagccttgtctgtaaggactaattccc**cataattacttgagaacctcataaattgtggtcaattatg**taggattcctctctttttctgccttccttcctccctcattctctctctctcacacagaaacttatatacagccaccccacaacacacataaatcta**taacttttattacctgatgtattgataagtaatctaaagtta**cataaaaatcagtagttcactgtcaatattgtaggagaagggaaggtctaggaaggaggaattacag**aacaagaggatattttgagggtaattgacttgtt**atctatgtggataatgataatgtctatgaccatatttgtaaaattgaacacttcgtgtggagactattatttttttatttaaccccatgccattattttgccgtattctag**catttctctaaaaatacagaaatg**ttcagcactcattcatgtgtatattcaggagtttctgacttttcatgtattttatttatctctgtctaattgctttgataccaaattttacctagtataattagtactacctttagcactga

Uppercase: IGHVII-30-1-2

Lowercase: Flanking sequence[1000bp]

Red & Bold & Underline: Stem-loop [16]

Blue: Heptamer[28]

Green: Nonamer [4]

id-IGKV1D-43[V_gene_segment]

gtgtccagtttcatctggatccaactgatttctccatgtacatgggaaattgcttgat**aagagattgactgtctctt**tcctaaaaatggtaacaagcaggctggtgtctgggtcatgatgatgtccccagtcactgataaaatgtaaaagaggaaagtgtcattgatggtgcatggcagggacatgctctgtgcagtggccgacctcactaagagagataaactttaggaaaagatactcaatgacagaaaagaaggtagactatgaaggtgcccaaaacaagaatacggtgcagcgcatttagtctctggtattaaaaagacctgtagctcttgataatggtggatgtgtgagtgctgcatgcattgaggaaacacggtatcatctctgtgtatctgtagtaaattgcttgatcttatagtggtaagaacaatggcataataccattacctaatacttacaaatatatatagcatcatgtcaataaattttatttttaatttttttttagaaagaaacaatgttaaactcacagtaatgttgcaaatgtagcacaaagtacccccttgcctaaccggaatcatgtgagagtcctttgaagacctgaga**atagtaccgtttaacattttactat**gtatttcctacaaacaagaatattctcctaaataatccccatacaccaatgaaacacattactccttcgactcctgaggaatatttcaaattgtcaaaaaatacctaaaaaatacttctcataacaaaatagtccaaagtagaacacatcactgcagacaaatttgtgctaccctggtcttttctgggacacctggggacactga**gctggtgctgagttactgagatgaaccagc**cctgcagctgtgcccagcctgccttgccccctgctaatttgcatgttcccagagcacatcctcctaccctgaagacttattaatgcgctggtcacacttcatgcaggagtcagacccagtcaggacacagcATGGACATGAGGGTGCCCGCTCAGCG**CCTGGGGCTCCTGCTGCTCTGGTTCCCAGG**TAAGGAAGGAGAACCCTAGCAGTTTACTCAGCCCAGTGTGTTCCGTACAGCCTGGCTCTTGAGGGAAGTTCTCTTACAACATGATTAATTGTATGGACATTTGTGTTTATATTTCCAATCTCAGGTGCCAGATGTGCCATCCGGATGACCCAGTCTCCATTCTCC**CTGTCTGCATCTGTAGGAGACAG**AGTCACCATCACTTGCTGGGCCAGTCAGGGCATTAGCAGTTATTTAGCCTGGTATCAGCAAAAACCAGCAAAAGCCCCTAAGCTCTTCATCTATTATGCATCCAGTTTGCAAAGTGGG**GTCCCATCAAGGTTCAGCGGCAGTGGATCTGGGAC**GGATTACACTCTCACCATCAGCAGCCTGCAGCCTGAAGATTTTGCAACTTATTACTGTCAACAGTATTATAGTACCCCTcccacagtgttacacacccaaacaaaaacccccagggaaagagatgtgtgag**gctgggctgccccagc**tgctcctcctgatgtctccatcggcta**agagtgttcctcagatgcagccacactct**gatggtgttggtagagg**gggatgtgaagtcacctctgcatccc**aatttctttttctttcttagcaccaggtgcacagacataacagttcctctcctgatttaaaaaaggcagggatcatgacacctgaggagtctagttta**tggcttcacttggaattcaagtaacagagaagaagcca**ctatagatattctaagcaggaattgtcttgatacagagaaatagagtataaaccatggaagtct**aaataaaaatatagagatgaatctcaaatttcatgttttattt**gctaagaaatatttgctaaatggggcatacagaaaaactcaatggtcttcaatatgttgaaagaacaaagagaagcttagagtttcatgaaaaaaggaaaatgttacctattgctctttgagaaagtttattggctccagaaagttttgggagctggcaagctcagaatggtaagcactgatggacaaa**ctgaatcctagaattatgttaaattattcag**aagttggaggtgaatttgatttcaggtt**acaagaggccaaagcagtgaagcttgcagagaatcttgt**tactgaagtgcgagggatttggtgtagatcctgctgctcaccacatataaaaccaatcagtaagacaacaagtattgccaagaaacaggctttaatcaggtgctgcagctgaggaaatgggaccccattctcaaatgtatc**tccctgacaaactaaaattagggggttatatagcaggga**agaaatgtgggcaaacaggaattagagaggggtaatgaagttaattttgtcaacagggagcagaaggt**cagtgaggcaatcacagtgtgtgaagggtctgacgtctcactg**tcctgattcagt

Uppercase: IGKV1D-43

Lowercase: Flanking sequence[1000bp]

Red & Bold & Underline: Stem-loop [15]

Blue: Heptamer[25]

Green: Nonamer [3]

id-TRBV7-1-2[V_gene_segment]

atggaggaggctggggcatcaatggggatggcacaggggactctgacttgcaggaaagacaatgagctcaccttttggtgctttgtgttggcggagctgttgacacatcctagagaacatgcccagcagacagaggagcggctgtgggatgaggagataaactcagagatgcagcgtgaggcctccgggtccagacagcatgagagcccaaagcgatgatacatgcattgatgttgttaaaaaggatttttttttttttttga**gacagattctcgctctgtc**gcccaggctggagtacagtggcgcaatctcggctcactgcaagctccgcctcccgggttcaagccattctcctgcctcagcctcccaagtagcttggactagaggcgcccgccacaacgcctggctcattttttgtatttttagaagagacagggtttcaccatattagccaggatggtctcgatctcctgacctcatgatccacctgcctccgc**ctcccaaagtgctgggctcacgcctgtaatcccagcactttgggag**accaaggtgggcggatcacttgaggtcaggagtttgagaccagccggggcaatgtggtgaaaccccgtctctactaaaaatacaaaaattagcctggtatggtggtgtgagtctgtaatcccagctactcaggaggctgaggcaggagaatcacttgaacttgggaggcagagttt**gcagtgagccgagatcacactgc**tgca**ctccattctgggcaatggag**caaggctctgtctcaaagaacaaacaagcaaaaagcaaggaactcataaatattt**aaaggagtcatttaagtaggtcctaaaataaatccttt**gttcctgtcattgcgtggattgagagaggatgt**gatgtcactatgggaccttctgtgtgggaacaaggacatc**cctcctcctctgctcctactcacagtgactctgatctggtaaagctcccatcctgccctgaccctgccATGGGCACAAGGCTCCTCTGCTGGGCAGCCATATGTCTCCTGGGGGCAGGTGATTCCTCAGATGCCAAGCAGTCTCCTGTGTGTGTGTGTGTGTGTGTGTGTATGTGTGTGAGATGTGTGTGTGAGAGAGAGAGAGAGAGAGCTGACTATAGTTGTTTTTCTCATTCAGTTCCCAATTTCTGTCTCCACAGATCACACAGGTGCTGGAGTCTCCCAGTCCCTGAGACACAAGGTAGCAAAGAAGGGAAAGGATGTAGCTCTCAGATATGATCCAATTTCAGGTCATAATGCCCTTTATTGGTACCGACAGAGCCTGGGGCAGGGCCTGGAGTTTCCAATTTACTTCCAAGGCAAGGATGCAGCAGACAAATCGGGGCTTCCCCGTGATCGGTTCTCTGCACAGAGGTCTGAGGGATCCATCTCCACTCTGAAGTTCCAGCGC**ACACAGCAGGGGGACTTGGCTGTGT**ATCTCTGTGCCAGCAGCTCAGCcacagcactactgctccagtgtcagcttggttccctaggaaatggggtttctagaacctgaatgctgacaaataagagttgtatatgtgtataccatgcaacctgcgtttaaaaatgtgtgtacatagtgcaatgactaaatctagctagttaacatatgcagtacctcaatccttatcctttccagtggtgagaatacttaaaatgtactct**gttagcattttcccagaatacaatccactgttgctaac**tctagtcatttgttgtataatagatctttgtaacatcttcgtcttatctgagtgagattttgtat**ctgttgaccaacatcttcccaacag**tccatctctactccagcctttgct**aattactgctctagtctttgaatttgaagtaatt**tccctgaggtctttagctcaacaagatgggggaattttctct**ataatgatgcttacattat**ttttgctttgtttttgtttcgttattttatgtcttgcaatatagtgtttattgta**aatataaataagtatatt**gtaataaaaatccttcacctctctttgggtaaagctacagttcacatattctaaatttcatgctgtgaccaacgctgacataattatggatcatg**ggtttctgggagtcctcagagacagccccataaacc**aagtttaagatagaatattccagacccagc**caggacagaagtgcatggtcctg**catagctccttggaaaattatagttccccaaattcagttattggatattgtggtgccgcagacaccaaacgtctttctctagcaaatgatggtccttggcagggattgttctgaacccattacaattttgccatcatcaaa**tcactccttgctgttaccttgtgtctcctgggagtga**ggacgccccgggcacagatggaaattccctgaccttcggatgctatttcaaggacttcctaagaccttgtgtccatctttttccacctttatccacatgactcct

Uppercase: TRBV7-1-2

Lowercase: Flanking sequence[1000bp]

Red & Bold & Underline: Stem-loop [15]

Blue: Heptamer[38]

Green: Nonamer [4]

id-TRBV2[V_gene_segment]

tttctttctttctttctttctttgtttctttgtttctttctttccttctttcttttttg**agacagagtctcactgtct**ggagtgcagtggcgcgatcttggctcacttccacctccatcttccagattcaagcgattctcctgcctcagcctcccgggtagctggaactacaggcgcccagcaccacgactggctaatttttgtatttttagtagagacagggtttcaccatgttggccaggatggtctcaatctcttaacttcttgatttgcccccctcgggctcccaaagtgctgggattacaggcgtgagccaccgcgcccggcaggtccttttc**tttttctttctttcttttttttttgaaaaa**ctaaagaatacccaatcccc**tcctcaggtggctctatcactgcctcaccctgagga**cagtatttccccatggggtttggatcactgttctctagttccctctccctggggcatgtgcagagtttcaacttcttgttggctgctctcttccactcaggatttaaaacattcacacttgattctttttccttctatctctgtaatcctaataccaagttttcttggcattcattatcagccgtggaatctgttg**tacctgaattgttactcacagcaggta**ataagagaacaagacctcctggaatttctgagccagactttgtactcagggttaccttgctgacccccatattttccagcattgctcttattaattaatggccctcagtcctgcacacaccatcagatcccag**gggctgagcacagccc**tgcattgtcaggttctgcaacaacacaaaggctccctcctctctcctatagagggcgctccagggatggtgggtgttgccagagacaccagtaattctgccagaccttgcctgtggggccatgggagctcaaaatgcccct**cctttcctccacaggaccagatgcctgagctaggaaagg**cctcattcctgctgtgatcctgccATGGATACCTGGCTCGTATGCTGGGCAATTTTTAGTCTCTTG**AAAGCAGGTCGATGCTTAGACTCTAGGAAATTCTTGCTTT**GAACTTACCTAAGACAATTCTAAACCATTCTCTTAATCTTCTTCTTTTCTCACAGGACTCACAGA**ACCTGAAGTCACCCAGACTCCCAGCCATCAGGT**CACAC**AGATGGGACAGGAAGTGATCTTGCGCTGTGTCCCCATCT**CTAATCACTTATACTTCTATTGGTACAGACAAATCTTGGGG**CAGAAAGTCGAGTTTCTG**GTTTCCTTTTATAATAATGAAATCTCAGAGAAGTCTGAAATATTCGATGATCAATTCTCAGTTGAAAGGCCTGATGGATCAAATTTCACTCTGAAGATCCGGTCCACAAAGCTGGAGGACTCAGCCATGTACTTCTGTGCCAGCAGTGAAGCcacagccttgcaaagacaactccagcctgtgcaaaatccctcaca**gagctgcctccctcccagccgccagctc**cc**acttcctgcctaagaaaaggaagt**ctctggttgggtttgttcttgcag**ctgtctccaaatagacag**atatctgaaattaaattaccaaaggtatcttgggcagaaatgaccagaca**ctgtaggatttgaagaatgtttacagatactacag**agtgtgagtgtgatggtgggctctgaatatgagagaattgtacacaaccaagtaacttatatcttctcctcattcccatttggccatctttattagaattgttt**tgttttagttgcttaggaaaaca**tattttaacttctagggtggcgagatgattcctgagtgggagttcaggtatttccatgagaatatagagggtgctcacagaaatcttcagaagaaggggcttggtacaaagaactcaattatcaatga**ctcttgctcagcatgtggtcaagag**cagaactatctgatgaccctgtgaacctggtgtgatgcatttgggctaatgatgctagctcacaaaataacaagttaagagagtagaaaacatggagtctaatatttcacaccatttcaaggtcaacagagttctagaccagcatgagaatcgccagaataaagcaaggcagtgcaggaacagaattcagcagcaaaagtcaagagacacctagaggcagaggcaggaatgaatttgtcactttcaacttaacctgaagcagccagattaacttcttgagtttcatttaagaaattattcctcttt**taaacatacctattgaagtctaaattttaggtatttta**tatatttcagtt**ctggaatttccatttaaatcttgctacaatttccag**ttgtttgctaaaattgcatattttgtcatttactttgttgactatattaacatctgtcattttaaagccactttttgataactttaacatctgaatctgttt

Uppercase: TRBV2

Lowercase: Flanking sequence[1000bp]

Red & Bold & Underline: Stem-loop [18]

Blue: Heptamer[30]

Green: Nonamer [0]

id-IGHV2-70D-2[V_gene_segment]

gaactcagccaagacagagccccatttaggagcaggtgcatcaaggaaccagccagagaacctctgactacaatc**tagggttgtcttcttttgcttcacagcatccaacccta**aggagaccacccactatatcatc**tgtaactcagaactttcttgttaca**aaggaaaacacaaatgaatataactctgcagttaacttcttgaacctggattgatgagtttccatggcctcatgagcacagagaccacatccagtgcaggtgcagagca**aggctgcaacagtcagcct**gctgctccctcaggagtcttccaa**ttcccttgtccaaagcatccgctcattcatgggctgagaaaagggaa**gtcattcatgcagtctactctctccacagaactgacggggcacaaggacaacatcgatttttcatggaacatgcctctaggaatgcagc**tgtgtgcacacacactgctaagcacaca**cttctttacataattacttgtaactgtattttcttatttattctctccaatttttttacacaaattcatcacttttccccata**atcaaagaggattttgat**cagaatgcttgtggggagccccttgcctgccagatgcccaacatcactactcttgaagggaggaggaacggcagctctcttgatttctactctaatcctctaggactaaaaccagaaggttgcatgtccagtgcgggagcatcgaagaagatcctgtctgtagaagcaggagcgtcaagacttgactgagagccatggtgctgaaatgagatagattccctgatggagagcacacgtggacccccacacctgagggctcactgctcctcaccacaga**tgcactcccctactgagtcctgagacctgagtgca**ccccatagagtagggctcagatg**aggggatgcaaatctccaccagctccaccctcccct**gggttcaaaagacgaggacagggcctcgctcagtgaatcctgctctccaccATGGACATACTTTGTTCCACGCTCCTGCTACTGACTGTCCCGTCCTGTGAGTGCTGTGGTCAGGTAGTACTTCAGAAGCAAAAAATCTATTCTCTCCTTTGTGGGCTTCATCTTCTTATGTCTTCTCCACAGGGGTCTTATCCCAGGTCACCTTGAAGGAGTCTGGTCCTGCGCTGGTGAAACCCACACAGACCCTCACACTGACCTGCACCTTCTCTGGGTTCTCACTCAGCACTAGTGGAATGCGTGTGAGCTGGATCCGTCAGCCC**CCAGGGAAGGCCCTGG**AGTGGCTTGCACGCATTGATTGGGATGATGATAAATTCTACAGCACATCTCTGAAGACCAGGCTCACCATCTCC**AAGGACACCTCCAAAAACCAGGTGGTCCTT**ACAATGACCAACATGGACC**CTGTGGACACAGCCACGTATTACTGTGCACGGATACcacag**agacacagcccagggcgcctcctgtacaagaacccaggctgcttctcagtggtgctccctccccacctctgcagaacaggatagtgtggctgagatgccatttcctgccagggcctgcgtttcccatccccatctgactcagagccttgttttcctccctcttctttactaataaatggcatgtcccctgttagtggttcgtgcaagcagaagctgtatcctgtttgacaaagattcagcatgaaaggttcctgttacctaa**aaaaaaatagacagatgagacttaattaacctaaataattttttt**cacaacaacagagtgaatacacaatttgcagaatgacagaaaacttttgcacactttgtctgtgacagggaactaatatgaagaatttgcaaggaactcaaacaactctacaacaacaacagcaacgagaaccaaataaccccatt**aaaatgagcaaagaacatgagtagacatttt**caaaagaacacatagaaatggataataaatatataaacaatgctcaacatcactaaccatcagggaaatgcaaattaaaaccacaataagatatcatcttccac**cagtcacaatgactg**ttactaaaaactcaaataatatcagatgttgctgaggatgggaaataaaggcaactcttagacattgttgatgaggatgtagacgagtacaacctctgtggaaaatggtatggagatttcccagaaaactagaaatagaactgccatttggtccagcaatcccactactgggtaactacccaaagga**aaataaactattattt**caaaaagatacccaccttctatgcttaccataaaactactctcaatagcacatatgtcaaactgagtgtctgccaacaaatgattttataaaagaatatagcacgtatgcacaattcaatactagtcagtcacaataaggaatgaaactgtgtcttttgcagcaagatgcatagaagtgggggacaata

Uppercase: IGHV2-70D-2

Lowercase: Flanking sequence[1000bp]

Red & Bold & Underline: Stem-loop [15]

Blue: Heptamer[39]

Green: Nonamer [3]

id-IGKV1-39[V_gene_segment]

ctgaacactcacaaacattatggtgggaaaagcagatcagagcattaggcaagttgcattaccttggccttcttcctttggagacaattgatgtggggttctagattgacccagagtttcaagtttatcctgattcagg**cttcaacagctggaggaagaaacagagatgttttttgaag**taaacagatctagcattactaatcaacccttcatactgatgacctatgggaaataatacccaagggcagaaa**aatgggcagaataaggggagccccaaaccaagacgaagctgctgcccatt**gagaccctgggtattacagagacctatagctctggataatggaagatctatgagtggcacaggcgctgaggaatcacagcatcattatcgtgcatctgcagggaattgc**ttgtaaatatactggtaattacaa**atgtttaaggtcactacaaatactttggagtgtattaaatatgcttctgataaagactgtttttc**tcacatgaaacaatgggaaccatgtga**caatcacagaggtgttgttactatagca**aaagggattgttactctccacatcccttt**aagtaacttgaaggcctgatagacccaccctctaagacttcattagacattccctacgaatggttatactctcctgtatactcccaatacaactctaaaatatattattccatatagtccttaggtttgtattaaagtttgacttttttccttcaaaatatctcttgtcacaacagcggctctagagagaaatacattccctccaggcaaatctatgctgcgctggtctgacctgggaccctggggacattgcccctgtgctgagttactaagatgagccagccctgcagctgtgctcagcctgccccatgccctgctgattgatttgcatgttccagagca**cagccccctgccctgaagacttttttatgggctg**gtcgcaccctgtgcaggagtcagtctcagtcaggacacagcATGGACATGAGGGTCCCCGCTCAGCTCCTGGGGCTCCTGCTACTCTGGCTCCGAGGTAAGGATGGAGAACACTAGGAATTTACTCAGCCAGTGTGCTCAGTACTGACTG**GAACTTCAGGGAAGTTC**TCTGATAACATGATTAATAGTAAGAATATTTGTTTTTATGTTTCCAATCTCAGGTGCCAGATGTGACATCCAGATGACCCAGTCTCCATCCTCC**CTGTCTGCATCTGTAGGAGACAG**AGTCACCATC**ACTTGCCGGGCAAGT**CAGAGCATTAGCAGCTATTTAAATTGGTATCAGCAGAAACCAGGGAAAGCCCCTAAGCTCCTGATCTATGCTGCATCCAGTTTGCAAAGTGGG**GTCCCATCAAGGTTCAGTGGCAGTGGATCTGGGAC**AGATTTCACTCTCACCATCAGCAGTCTGCAACCTGAAGATTTTGCAACTTAC**TACTGTCAACAGAGTTACAGTA**CCCCTcccacagtgttacaagtcataacataaacctccaaggaagcagatgtgtgaggacgagccaccccag**atgctcctcctggtgcctccatctgctgagagcat**ttctcaaactcagtcaggttttg**aaagtcattgggagacttt**tgtagaggggaccagg**gaggctcctctgaactctaagcctcttttgcccctatccccaggagaaaaga**tgtgacaatgcctgtcctgattgaataaggaagagatacaagtccacctgaggagt**ctgtgttatgggataattggaatttacacag**caaaagagaagctattctcggtatttcaaggagaaattgttcaagttgaataaattagagtctaaactagtctttttgaagcctacggtatgttattcgtgaagcagccactagagacaggggattct**caggtgctcctgcagaagtcagagtgcacctg**cccctggtggtatgtgctgagtaccgtgtgatgatcctcagacctgtctgggaagccgagggctggg**gtgctgatgctctcagctgcctgcagcac**gtctccaggtgattctccagtccacaaacaattccacatgttttacttca**gatgtcagagtacatgaatccaccactctgacttc**ccaatctcatgggagtgcctctcattaagcaactctaaagaaaccatagagagaaaaggagttttggaaaatgtgcgtccagaagtgatagtagcgatggggaattgacagctgacaggtcagtaaggttgctctttccacaaggctcaaaactttgccagttacagattgtcccagaatatactcgaatg**tgctatcacgtcttacgagcaactctgggttcatagca**agaaaacttcattaagtcatagatgaaacagaaaaatcaggaaactgtatgaaatacattataaggctgtgtgtggtagctcatgcctgtaattcccagcactttgagaggctaaggcaggagaattgctt

Uppercase: IGKV1-39

Lowercase: Flanking sequence[1000bp]

Red & Bold & Underline: Stem-loop [20]

Blue: Heptamer[26]

Green: Nonamer [7]

id-TRAV13-1[V_gene_segment]

agttacatatgcttacattttttgaatctgcagcctgatgttttctgagaatttcttcttgttttttcaacactttgcagaaagcaagccacatagccataggaaaagacttcttaaaacatgccagatttctccttgcaaaccagcccagtctatgggtcaaactattcctattgttggagtattaataactcatggaacttgtccccctcctcctttatcattgta**ctttaattctttaccttttattaaag**ccatctttactgaagaaatattcaacattttgattgtcggtaccgtcaattctctgggaatagcaaggtcatggcccaggcttgagaagcagaacctgagataatacgcactcgaaatattgagtggtcctctctctcttgctctgatctttggcttacatattgccta**aagaaagagttttctt**cataacaaacgagtcatgatagtagacttaacctaaaaggaaggcattattttttcttgtttatgagagcgaattatggtattttcttaatttcaaacataaactcagcccatcattgaataaatatcaacagcatctacttctt**tataatttatgaataatgactcagtagtgtgcataattata**gagttagatctctacattcttactgtgcatttaatctctgattgaacacccagtctgtttgactcattgctgcttttgtgcacctgagtgcctcatagggttagagaaaaacattcaaccagcaagttcagtagttagggatgtggccacaagatggcagtgctcctctgctgaggcagaatacagggttcactttagtgtgtcctgaatgaataggtttatggtagcagcagagcctttttcttattggttggctacacagtgtgagaaacccctatggctgccagaggagagaagaga**caacctgatgatagaagtaactcttataactggaggttg**caggtcaatgactgatcttaattgggaagaacaaggATGACATCCATTCGAGCTGTATTTATATTCCTGTGGCTGCAGCTGGACTGTGAGTTGAGGGTATTTGGGTAACAGAGTATATTAGTGAATATTCTTCAGCAGTTCAA**AGAGGAGACTTGCTTTATTCAGGTTTCCTCT**CGTTACTATGAAAAATAATTCTAGAGAGTAATGACCTAACAATCCCTCTTCTTTCTCTGATTTTTTCTTTCTGCATAGTGGTGAATGGAGAGAATGTGGAGCAGCATCCTTCAACCCTGAGTGTCCAGGAGGGAGACAGCGCTGTTATCAAGTGTACTTATTCAGACAGTGCCTCAAACTACTTCCCTTGGT**ATAAGCAAGAACTTGGAAAAGGACCTCAGCTTAT**TATAGAC**ATTCGTTCAAATGTGGGCGAAAAGAAAGACCAACGAAT**TGCTGTTACATTGAACAAGACAGCCAAACATTTCTCCCTGCACAT**CACAGAGACCCAACCTGAAGACTCGGCTGTCTACTTCTGTG**CAGCAAGTAcacattgcttcccaggcacctgctacccgtacacaaacctgagactggagctgaagctgcaccccctttcctttgtcatagatcgtcaattatagcatttgtcatattgtttgtttgcaagttgaaactaacatattgacatttaaagcatataataaaaaggttaggttcaatgtttttgtatttttagtagagactgggtttcacatgctcagctaattttttctatttttagtagagatggggtttcaccatgttggctaggatggtcttgatctcctgacctcgtgatcctcccgcctcggcccccgaaagt**gctgggattacaggcctaagccactgcgcccagc**ctaatttttgtatttttagtagagacgggatttcactatgttagccaggctggtctcaaactcctgacctcaagtgatccacccgcctcggcctcccaaaatgctgggattacaggcgcgagccaccgcaaccggccctatcttacacattctattagtagattacaacaggactttggaccaatggttaatttacagacagcttgctaaaaatctccctgcctcattttc**atgttttgactgaattaaacat**tactaccacaatttcttt**taaaagagttatatctttta**gtgttgtagataaatcagtggaaataaagaaggtgcataaataattaattccctatttctcccacctgtggt**caagagtaggctaatttcaagatctcttg**tattcttcctaaggatggatctaatcagcagttggaatgggagaacaattcctcctggctgagttttggctgttatggcttgttcttatagggaaagtagttagattaaattagggttaa**aacagctggacaagaaaggtcagctgtt**ccttgtctcttccggctctgagggctgcctgcatttcttggcttgtggctccatcactctgatctctgcctctatggttgcactgcttctcttcttctgtcatctccctc

Uppercase: TRAV13-1

Lowercase: Flanking sequence[1000bp]

Red & Bold & Underline: Stem-loop [13]

Blue: Heptamer[22]

Green: Nonamer [3]

id-IGHVIV-44-1-2[V_gene_segment]

ataatggagggaatctcttggtcccaggaagatgacaatttaatatgtctcaggcctgtggagctctctttaggagctttgtgtgtggcttctttgtatactctggctccaggaactccagtgagggaagagattcgtcacctcctgtgcgaagggggctgaagcgaggaagacaagtggcctggcttccacagaacccataagctaaaacactccgagttcccttgactgcaagaaatgcagctgtcatcacggtttcaagtgccagagtctttaatgattatgtttcaactcatgagagtgccaggaaaattccaccagtatgaactactcaaatatgtcattttgtaacattacttcattgcttgctctcatatcctccatgtttc**tgaaaaagcaaagtttccatgctttatca**tctgcaccaatgaggccatatttaagctatattttcatatgaacgtctcttgaagattttaaaatagaacaaatgttgaacaatatgcttaaaagaaaaaaaataaagaaagcagtctatgttgactcatgtgacattgataatgtatattagattatattggttgcttatttagttggtcatgggagaagagcc**ttctttccagccaaagaa**tactaagaactccctcttctaatgacgaatttcagctga**tcttgctgagaatatttagcaaga**agacagaaacagagactcctaaagggagctccagagtcctgagatacatttaagttttcattttcctggaacccagagattccccactgttctacaagacaagaccccattctatgaccctgtctga**ggagagctcaggggcagtgagctcccc**ctcacattaccccttcaccctatgcccttggggctctgtccatgcaaataagtcccgggttcaagaacgaatagaggcagggctgtgagtgttatgctgtgtgctggctctgagctggggtctctagtagcactctccacatccatagatacATGGCCCCTCCGCTACTCCTTCTCTGCCTGCTGGCTGCCCTCTGTGGTGAGTTACTCATGATTTCATACGTGGGG**AGACACGGGATGAGGCAAATGTGTCTGTGACTCACAGA**TGTTCTCTATTCTCAGGGGTTCACAGTGA**GGACCACCTTGTGCAATGGGAGGAAGAAGTAGTGGTCC**CCTTGGTCATGCTCAGCCTCACCTATGCCGCCTATGCACATTCAATTTCTGAACATTCTGTTTCCTGGATCCACCATCTCCCATCAAAAGGTCTTCAGTGTGTTGGTGTGATATGGGTTAAGGGAAACACTAAGCCCCAACCTTCAGGGCAGAGCTAGCATCTCCAGAAACACATAGTAAAAAACAAGAAAACTTACAGCTGAGAAGTGTGATGGCTGGGGAGGCAGGCGTGTATTACTGTGCTCAAGGcactgtgactcgaatccagagtgaactcagacacaaacctgccctgcaggggttcttgggaccacaaggggaaggatcaggtcaccagggtgtacttaggaaccactgaactgggtcaggcacaggaggtgggggtcagggctcctccccagggaagggcttttaatctccatgccttgtaaacctctgacaagccaggcagacacagacatatttttaaactatggaaacagatgtac**ctttcattttggaggaaaaagatcataagtgtgaaag**caaaacttaccaagggccaggtactggaataactgtaatctacataatcactctaagaattctaagaatttacaagtgcaatcatcctcatttcataaactcaccttaaataagagctttctaactgatcccatatattctgattttgtttatttcttcaacactttattatcttcaaggcatttcactgatgtcaattttagaagaatcatatgcaaacagcctctcgtggtggttgaaagc**ccacagcctcgccagtcctgtgg**atatttctggatgttctcagagtctccattgtctgtagagaagaattgtcccaatagttatttcacttttgtcttccacaaagttggaaagatggggatcactcaggggaatcgagatcaacaaacagaaagcctgaacccaccatatacaagataggatgtatttttttaggaatatattaattcccagaccctcaggctcacataccccatgtggaggggaacacacatccactccttcactcaagcc**aagctggccatgcagctcagctt**ctgcaggctcaggactgcaagctctccagtgtgggagtggagctaatgggag**gtgaggccaagtctctgccctcac**agagtctcatgatgaagatctgatgaaatatttccacagta**acatttctttggatatataaatgt**agaatatattttgaaataatgaatgatgtatcttctgatggatt

Uppercase: IGHVIV-44-1-2

Lowercase: Flanking sequence[1000bp]

Red & Bold & Underline: Stem-loop [12]

Blue: Heptamer[35]

Green: Nonamer [3]

id-IGKV2D-23[V_gene_segment]

atgcagtcaaaattttaatgattacatactcacaatcacttctgtggggcctggagatactgcacatacgactgttagcaagacactcactgggacgctgcgttgtgtgatggccccacatacaaacctcaaggaggctcagcctctcaatgcagcaggagcagctggggtacccaggccacacgtccataccaggtgggctcagttagagatggctggagagccttccag**gaagaggccatgaggtttcagtcacaaacactggctcctcttc**tgtgtaaacaggggctagagccctccaggacaattcctagagcctctccctttctctccaattagtgcgctgacaccctacagactctccaggaagtggttgtcatgtcctccctgcaacagccactaaagttccctactgctgtcatgaatgcagggacacttagtcacatcactgggaggcgaccctagtgtatcctgacctcacctgctgccactgatgactttcagggcacctctttctccctttgct**gagtgactctcactc**tcaccaaccatcaggagaatggaaagctgcctgcaatgcatgatgttggctgttgagcaaatcaaagctcacaggagtctcaaacatgtacaccacataataatattttctgataatactatttggacttttcttcctt**tcaattctggaagtaattga**gaatattttttgaactcttagaaacacttagtatatatgtgtagtaggtagtaactagttttgtctactggtttattttgtttgcttgtttcaggccatgatgcggcatgttaaaatactgaagacaaagatacattttagaattaagcatactgtacattggctctttccacaccactgcaaccaccaggggatgtgcatattgtcccttaggaatgaccttcccttgtgagtctgggagtaaagctcagctgtaaccttgcctt**aactgatcaggactcctcagtt**caccttctcacaGTGAGGTTCCCTGCTCAGCTCCTGGGGCTGCTAATGCTTTGGGTTCCTGGTAAGGACAGAGGAGATGAGGGAGGAGAATGGGGTGGGA**GGGTGAGCTCTGGGGGCCCCACTGTCACCCATGTGTGTTCCGTCCACATG**TTAGATGCACGTGTCTTGTGCTCCAGG**ATAAAATGTATGGTGGCACTTTTAT**ATGTGAAAGAGTGAGGAAGATTCCAGAAAAAGCAAAGACCTGTGCTCTGGTGCAGATTCTGACATAGAAAGAGGAGGGTAGCATAAGTGACTTCCATAGGGCAACTTGGGCCTTCAAAATGTCTGTTTTTTTTTTTAATTG**AATTTTTTTGGTGCATGAATCAAAATT**ACACACACACTCACACACACACACACACACACACACGCCGCAA**TACAATTATTTAGCATTAAATAATTGTA**GAGAAATTATGATAATGTCTCATGATTTACATAACATTGTACTTCTTTTTTATATTACTTTAGGATCCTGTGGGAATATTGTGATGACCCAGACTCCACTCTCTCTGCCCGTCACCA**ATGGAGAGCCGGCCTCCAT**CTCCTGCAGGTCTAGTCAGAACCTTTTACATGGTAATGGATACACCTATTTGTATTAGTTCCTGCAGAAGCCAGGCCACTCTCCACAGCTCCTGATCTGTAGGACTTCCAATCAGTTTTCTGCCTTCCCACACAGGTTCTCCCCAATGGGAGGAGAGAGTAGACCAGTCATCCCCAGATATATCACAGGACTAGTTTCAACCTTTGGAAGCTGGTCTATATCCTATGGTTAAATAGGCATTTGTGATACGACCTGAAATACATTTGGACAAGAACTTCACTAACAATTGAGTCACTGAAGACTTATGGCCCTGTGTGACGCACCACATAACCGTGAGTTTGCAGTGGTTGCAGGTCAGGGACAGATTTTATGCTTAAGATCAGTAGGGTGGAGGCT**GAGGATCTTGGCTATTACAACTGCCACCACACTCTACAATATCCTc**ccacaatggttcagcaccaaacaaaagcctcctgcttggattgtcccagctgcccaaattagttccttcactgaggagtagacagggtatattctctaaatctatgtaacaggaagatgttggtgaactcaggggattagtatgaagctacacctcaggcatcacacataagatcacttcagcagtcgcagccttagcatgggcagaacctacagaagatgcaagtgccctctgagccaggagacaggaggaaggaggaagggaaaggtgacttagctcatctcaatcctctctcctttgcatacatttgtcaaccag**atgtattcagcctaccagtcacacaactgaggctgatacat**gacaacatagcactggtatattcttggtattgtttggcttagcagttactagtatatatttaat**gggagaatatttggtggtgttaacacattgcttatctccc**ttaccccagttgtactttacacttgttctcggcacacattctcctccaggactggagcattcacagggttttatgttactgttcttatgggagtaaaaagaaaaacgattcacattcttgctactgagctaggctgggatgtcctgggccaagctgaaaatgtgaaaaataagagtatgaatatttattaagttttatctggatctaagatacttatccatgaaccagtcctgcagctgtgcccagcctgctccattccctgctgatttgcatgttcccagagcacaaccccctgttctgaagacttcttaataggctggtcacaccctgtgcaggagtcagtctcagtcaggacacagcatggacatgagggtccccactcagctccaggggctcctgctgctccggctcccaggtaaggatggagaacactaggaatttactcagccaatgtgctcagtacagcctggcctttcagggaaatcatctt**acaaatagttgtgtggattatttgt**ttttatgtcccaggagtcaga

Uppercase: IGKV2D-23

Lowercase: Flanking sequence[1000bp]

Red & Bold & Underline: Stem-loop [14]

Blue: Heptamer[47]

Green: Nonamer [5]

id-IGKV1-35-3[V_gene_segment]

ggagctgcacctgagacactaggaacatgctggtagggagcacagcttggagcattacccaggggatatgatctttgtcttcctactttggacacatttgattataagagtctggattgatacagggtttcaagtttatcctaatcaaggctccta**tagttcaatgaagaaacagagaatcttgttgaacta**at**cagatctggcatcactagccagccctttatgcgatgatctg**taagaaatattacacagtgacagaaaaggtggtagaatattctc**cccaaggcaagaataaggtgctatccatttaatccttggg**tattaaagtgaccgatagctctggaaaatggagggtctgtgcgtggtacatgcatgagaaatcatggtattatctttgtgaaactgcagtcaattccttgtccttataccagtaacaacaatggcttaaagttatgacaaact**atttttgaatattgttaaaaat**gcttctgataaagtcttattttcattttaaaaaagacaattcctcactgacag**aaatattgttaacacaacacaaaatattt**ttacactcaccaaatacatttgccagtaacttgaagacccaagagctctactttctaagattttattatgcattccctgcaaacaaggatagtctcctacatactgccaatacaactataaaatgcattactcaatgtagtccttaggttcgtttaaaa**ttttggaaattgtcccaaaa**atacgttttgtaatagtatggccaccaaagagaaatgtattatctgcaggtaagttgatgctaccctgctctgatctgggaccctggggacactgcccctgtgctgagttactgagatgagccagccttgcagctgtgcccagccttccccatcctctgctgatttgcatgtcccagagcacagctcccttcctgaagacttattaattggctggtcacaccccgtgcaggagtcagtttttgtctggacacagcATGGACATGAAGACCCTCATTCAGCTCCCGGGACTTCTGCCATCCTGGTTTCCAGTTAAGAAAAGAAATTACTAGCCTCTTACTCAGCCAGTGTGCTAGGTACTGCCTGGCCATTCAGGGAAGTCTTCTTATAACATGATTAATTGTGTGGATATTTGTTTTTCTGTTTGCAATTTCAGGGACCAGAAGTGACATCCAGTCACCCAGTCTCCATCC**ACACTGTCTGCATCTGTAAGGAGACAGTGT**CACCATCACTTGCCGGGCGAGTCAGAGCATTGGTAGTAATTTAGACTGGTATCAGCAGAAACCAGGGA**AGGCCCCTAAGGGCCT**GATCTATGCTGCATCCAGTTTGCAATCTGGGGCTCCTTCGCGGTTCGGTGGCAGTGGATCTGGGACAGATTTTACTCTCACCATCAGAATCCTGCAGCCTAAAGATGTTGCAACTTATTACTGTCAACAGTATAAAAATTACCCTatcacagtgttacaaaccataacaaaccccccccaggaaagcagacatgtgacgctg**ggctgccccacctgctcttctttgtgcagcc**atctggtgacaacacttctcagactcagcctgagttttgagggttattgggagattacggaagagggg**gccatgaaggtgtctctccaccctaagtctctttcctctcatggc**aatgtctcttcgaatttagtaaagaaaggagattagtctacctgaggagtctgtgttatggggtaattgaaattcttacagtaaaatggaagccacacattccaagcaggaattgttttaactgagtgaattagaatctaaactatagcctttgaaagcctggcatatgttattcatgaagcagatactagagatggggttctctgatgcaaaaatataaatatttattgatctg**tttttaaagatactaagataaaacttaaaaa**tgttgtcactgtatcaaatatattgcatcatatatctataaaaacagtaaaatttgtgtcatttatgaaaaatgcacatatttgagtgtatataacatacatattaatccatata**tatttatttgataacttgttagctataaata**ctaag**tacacatattcatgtgta**gataagattatttaaatatgt**ttatatatatatatataa**gcaacacatgcatacgtatgaaattattgtataaattaaatctgttaagataattccagttactcccatgaaaaaaactgatgtccaggactatatatgttacctgctctggctacagggatttcctttgtaatctttatttctaacactcaccaaatgggtatgtgcacctgaaa**atatcttttagtggaggggtaaaggatat**gaagaaaggta**attttgcctaatgttaaaattaggcaaaat**gttaaaatttcacagtgatgcctgctgggcagagttctaaattgggtgctggaccaaagccacacccctgccatcgtca

Uppercase: IGKV1-35-3

Lowercase: Flanking sequence[1000bp]

Red & Bold & Underline: Stem-loop [16]

Blue: Heptamer[31]

Green: Nonamer [6]

id-TRBV7-9[V_gene_segment]

atatgaaagagggtgcccatgaagttttaagagttaagttaggaatgaaaatgagaaattaattgctcataatgcagactgttaggaaccccaaggattaattcaactccctattttttaattgtactactcccatcacaatgtcaagggtaaaacagttctgactttttctaaaggctatggaacatttattgatttcagaggtgacttagaaacacaggaggtgtcgtgataagtagtagagacaaaaaagaaaaagtattgagattaatgaattcatatggggaattgtttaactgttctttttttaacctttagaatgagttgaagatcccaacttagcttggtgcctgctggaggcaacgtttcttcccttgcccctctgagctgt**gaattgaaactctttcgggaaaaagacgcggcaattc**tctccacttcccaaacacctcccactcctacccagaccgtggatgggcaggaaatgcaggaacagagccagaaacaggagatctccaaggaaggttga**cagtcagcactgggatcgtctgtgtaaagtgctgctg**aagcagccaggtggcatgtccagccgacaatgcgaaaggaaaaagtgagaagacttcccgaaggcggagggtggaatgcgggcag**cagccccctggagggctg**agtggggaaaacaaaatggacctca**cagaagctgtgtgtgtggaaacccacttctg**acttatcacttgt**catgaattctatgcttcatg**gtgttacaccgtttattgtttctgatgagtgacagtaattattttctttcttgctggtacataataaagtggtgcacatcagagttgctgccatcttagacttaactcatcagtatcaggtgatcctgaggctcagtgatgtcactgtgggaactgctctgtggcgacaaggacgtccctcatcctctgctcctgctcacagtgaccctgatctggtaaagctcccatcctgccctgaccctgccATGGGCACCAGCCTCCTCTGCTGGATGGCCCTGTGTCTCCTGGGGGCAGGTGAGTCCTCAGAACACCAAGCAATCTCATTGTGTCTGTGTATGTCTGTGTGTGTGTGCGTGTGTGTGTGTGTGTGTGTGATGACTACAATTGTTTTCCTCCTGTTCCCAACTT**GTATCTCCACAGATCACGCAGATAC**TGGAGTCTCCCAGAACCCCAGACACAA**GATCACAAAGAGGGGACAGAATGTAACTTTCAGGTGTGATC**CAATTTCTGAACACAACCGCCTTTATTGGTACCGACAGACCCTGGGGCAGGGCCCAGAGTTTCTGACTTACTTCCAGAATGAAGCTCAACTAGAAAAATCAAGGCTGCTCAGTGATCGGTTCTCTGCAGAGAGGCCTAAGGGATCTT**TCTCCACCTTGGAGA**TCCAGCGCACAGAGCAGGGGGACTCGGCCATGTATCT**CTGTGCCAGCAGCTTAGCcacagcatggcacag**tcgcctccttcctgctcacaaaccctcaggcacttacttctccttc**cagctctcagaagccctgaacaaaggagctg**ccctgctctttcctcagcaaggagaatgaatgcatttggaactgcaggtgttc**ttctgatactaggaggtcagaa**aataacctctgaaatacaggaacagggaatactgggtagtaataattttgacttatggatttctgggattccttatatatagttcaaatttccataattaggatataacagag**cttagtctcatggatagtgactaag**taaatattcttttatagaactatgaagtttcagcacatttatattaaactactgttaccacatgtcaccaactcagacctataatctaccaaggagtggggaagccaaacgcaaacactgctaaggccatttacagctaccacccttggaag**aaaattggaatctttagtaagaatttccatcaaaatcaatttt**ttagaaataactattacccacaataatgcacattaactcatgttcatacattgctttttttccttctgaaaatcttgtagtca**ttctatgtttaatttccttcttctccagtctaacttaaaaaaatagaa**atacaaattgagattgtttcataagataaaccccacctgggaaggtgttaaaaggacacaattcaaagaaaacctgaataaaattatatcaaccacgtagga**ttctttgaggtttccaaagaa**gatgtaccagtttacattcccacgagaggtatatgagttttggatggaatttttttttttctctctctctctctgtag**attagtttttccctctactaat**agtttctcttgacaaacagaagttcttaagtgacaatcattccatgtttctcccatttcagttagtgttttgtattttggtgtccactttaggaaatccttgcttacttcaaaatgatcatgttttcttacatttccttctaaaat

Uppercase: TRBV7-9

Lowercase: Flanking sequence[1000bp]

Red & Bold & Underline: Stem-loop [16]

Blue: Heptamer[39]

Green: Nonamer [4]

id-IGHV3-43-2[V_gene_segment]

gtggaaaagtcgtttattgggatagtacccactactacaatcagtaaatgttggatacgatcaacagcatggttcaattcacaagta**catatgatgagtatagtgagccaaagcacatacatatg**attccgttttataaactgtacaaagtgaatactcattggaaattacataacaaagatcactaactga**cttctccatagtaagagaag**cgaaggtataggagggagaaattgtgagaga**caaagagaaaattgagaggcgaattgatttgttttctctgtg**aatggtcattaggtcaatgtttgtcaaatggtga**acatattgtgtgaagattatatgt**ctgtattacttcattaaagctattataaataaaagtctaatgtggtagaaaaagatgaagagagaaataaaaataatacaagaaaagtcatgaactcctgaatgaattaaccctta**gtttttctctattacttataaaaac**accaagatacagccaaataatatcacgatatcattataagaagagtgttttgtaaacctcactgggaatttatagctctttcctagagttaattttgggaacagttggatccaattgtgagaaatgcaggctggacactgagactggctcttatgag**atgtgagctcttgtctatgtcacat**ggtccttccatacttggggg**tttacattcacatctgtaaa**tgaaggaaacattgactctcaaagaacatatttcatgtgcatgtaaaagtatgaatgctagtgagaattaattacttatgaagtataatcacccacatccactcttggacacagcccactctgaggcatctgttacagaactcattatatagtaggagacatgcaaatagggtcctccctctgctgatgaaaaccagcccagccctgaccctgcagct**ctgggagaggagccccagccctgagattcccag**gtgtttccattcggtgatcagcactgaacacagagaacgcaccATGGAGTTTGGACTGAGCTGGGTTTTCCTTGTTGCTATTTTAAAAGGTGATTCATGGATAAATAGAGATGTTGAGTGTGAGTGAACATGAGTGAGAGAAACAGTGGATATGTGTGGCAGTGTCTGACCAGGGTGTCTCTGTGTTTGCAGGTGTCCAGTGTGAAGTGCAGCTGGTGGAGTCTGGGGGAGTCGTGGTACAGCCTGGGGGGTCCCTGAGACTCTCCTGTGCAGCCTCTGGATTCACCTTTGATGATTATACCATGCACTGGGTCCGTCAAGCTCCGGGGAAGGGTCTGGAGTGGGTCTCTCTTATTAGTTGGGATGGTGGTAGCACATACTATGCAGACTCTGTGAAGGGCCGATTCACCATCTCCAGAGACAACAGCAAAAACTCCCTGTATCTGCAAATGAACAGTCTGAGAACTGAGGACACCGCCTTGTATT**ACTGTGCAAAAGATAcacagt**gaggggaagtcagcgagagcccagacaaaaaccttg**ctgcaggaagacaggaggggcctgggctgcag**aggccactcaagacaca**ctgagcatagggttaactctgggacaagttgctcag**gaaggttaagagctggtttcctttcagagtcttcacaatttctccatctaacagtttccccaggaaccctgtctagatctgtgatctggatctgctgaaactgcctgtgtcaccttcctcacctgtgactttgggggagctgattgtggacactccagtgtgtgggatttcttggtgacagcaattgtgtcttctgtctaggcatgtctagggctggccatcaggaag**ggcaggctggaatttttggaaagaggcgcacctgcc**atccaccaggaaattttgttgtcttttgttc**tgctagaattaaatcagacacaccaaggttaactagca**ctatcttcctagcttgagaaact**tgatggcaggcttgaataacacctgtatgaagccatca**gagcaacaactagattaatgtc**tgctagaattaaatcaggcacaccaaggttaactagca**ctatcttcctagctcgagaaacttgatagcaggcttgaataacacttgtatgaaacc**atcagagcaacacctagaatagtgtctgat**ttaataattgagactatggtctagccaaggagacac**ataaaacatgatttccatggttggttcatattttat**atttattaatagaatctggctggtattataaacagctttgccaaaatatatgtgttagtgttggtcttcggagaagcaaactctgtgactggattagcttcg**tgtacagtttccttcgtgtaca**gtttccctgtccactcttctgagaggaaatgcagaggtggtggaaggggtctggggtagctgaatgcatgtgagggaagtcggtccctgagtgaaggagaaagggaggaggactgggtggaactttcctaaacttc

Uppercase: IGHV3-43-2

Lowercase: Flanking sequence[1000bp]

Red & Bold & Underline: Stem-loop [18]

Blue: Heptamer[30]

Green: Nonamer [3]

id-IGKV1-37[V_gene_segment]

aaaccaaaaataagatgcagcttatataatctctgggtattgaggtggcttacaggtttaaataatggtgaatctgtgagtgataccggccttgaggacttgtggtttcatttttatgcatctgtagtgaattgcgtgaccatacagtcatgtaagaatggcttaaggtcattacaaaatactttcaaatatatttagcattatcttgtaaaattgtctcttttttttaagaaggaac**catttcactctcacagaaatg**ttataagtatagaacaaagtaccctctcctttgcagagtatattcaagacttgatgttcccaccctctaaaactttactgagtttcctacaaataaaaatattcccctggctggacgtggtggctcatgcctgtaatcccagcactttgggaggctgaggcgagggggtcacctgaggttaggagtttgagaccagcttggccaacatggtgaaaccccatttctactactaaaaatacaaaattagcctggcatggtggtgcgtacctgtagtcccacctaccctggaggctgaggcaggagaaatcgatggtacctggcaggtggaggt**tgcagtgaggtgagatcgtgccactgca**ctccagcctaggt**gacagaacgagactctgtc**tcaaaaaaaaaaaaccttaattaatccccatatacaaatagaaaacattacttcatcgagtactcaagagtatttcaaatat**taacaattattaaaaaaaaaagttctttgtta**caaaacggccccaagaggaaacatgttcttacagacaaatctgtgctaccctcctctgacttgggacactggggacacga**gggactggctcagagatgagccagcccc**gcagctgtgcccagcctgccccaacccctgctgatttgcatgttcccagagcacagccccctgccctgaagtcttcttaacaggctggtcacaccccgtgcagcagtcagtcccagtcaggacacagcATGGACATGAGGGTCCCCGCTCAGCT**CCTGGGGCTCCTACTGCTCTGGGTCCCAGG**TAAGGAAGGAGAACACTAGGAATTTACTCAGCTGGTGCGCTCAGTATAGCCTGGCTCTTCAGGGAGGTCTTCTTATAACATGATGGATTGAATGGATGTTTGTTTTTATATTTCCAATCTCAGGTGCCAGATGTGACATCCAGTTGACCCAGTCTCCATCCTCC**CTGTCTGCATCTGTAGGAGACAG**AGTCACCATCACTTGCCGGGTGAGTCAGGGCATTAGCAGTTATTTAAATTGGTATCGGCAGAAACCAGGGAAAGTTCCTAAGCTCCTGATCTATAGTGCATCCAATTTGCAATCTGGA**GTCCCATCTCGGTTCAGTGGCAGTGGATCTGGGAC**AGATTTCACTCTCACTATCAGCAGCCTGCAGCCTGAAGATGTTGCAACTTATTACGGTCAACGGACTTACAATGCCCCTcccacagtgttacaaggcataacataaaccccccaaggaagcagatgta**tggggctggcctgcccca**gatactcctcctactgcctccagctgctcagagggtttctcatattccagtcaagctttgaaagtcactgaaaagttttgatggaaggggccatgaagactctgaaactgcctttgcaaaaa**tcataactgagaaaattatga**cggtggaagatatt**acacagccagctctgtgt**gaattactctttcactactgctattgccgtgtcttgataaatcgactctgtctaagcagcaggcaaggtgaaccccttgggcgattacaactcctctataccttaactgtttttccttctcatcccctgcagcacaaacatggcaatactgttcctgatttcattaagaaaagagatgattacacctgagtcttggctatggtgtgagttggaatcaataccacaaaggaaaacctactcttggaattccaagtctatttttttttttctaaatacacgcaatgagagtctaaactac**agcctctgggaggct**tgggagcatggtactcaaagatgcaaataattagaaaacgat**tctctggatctccccaagaatccagaga**gcatctggtgcagaagctataattgccaatcatgtggtcctcagacatgtctgggaagcccatgttcggggctgttgatgctctgcctggaggaccttcctgattttctatggcatctgcttacaattgaccagccccctattcaaagcttgccgtgttgcatctcaactcttgtccgtgggaccccaagtctctagcc**acatccagttcttcacagaagctggatgt**atgtgccagggcaaggagaactgggtgggaggtgagt**ctgtgggctccacag**cttcacgtgtttatgaggtaggagttattaagaaattattttaggcagatagagaggaaaaaagggatccttgggagacttttcacagctgttgt

Uppercase: IGKV1-37

Lowercase: Flanking sequence[1000bp]

Red & Bold & Underline: Stem-loop [15]

Blue: Heptamer[29]

Green: Nonamer [4]

id-IGKV2D-28[V_gene_segment]

gtggatccagttcgtgctcttcttacatagagaatttccactcaaattgtgaactcatggctgtggctgcaccacttacaggcccaggggacactcagatctcacttggtagttgacgaaaaactggaagtcctgatagga**cctcattccacatgatgagg**aagactgtgggaagagcttttgttaacaccattcag**aaaaacatagtgcaaagttagttttt**gttcattctatataattatcctagaaaagtcttcccttaataaatccttcaggttaatctagcatatgtgagatgattatgatggaatatataccagattgaacaattggtcaccaggaattttatattcactgcct**gaggaataaattgtttcccactttcctc**ttacctgcactgggctcttgaatctaaatatagagacccacattattttccctatgaggcccttggacagagcgctcttatggggctcactcaccaggtgccaggggagggtagattccaacacttgctatgaacattcttgaacagttatcctggaaaccgcagataccagaccac**tcttgaactggctcaaga**ctatgttttatttgtaggctgtctgctcatcagtgtttgt**aggaaagggtaacgtttcct**tttttagattagctgggagggagccaaga**agaatggcattcatccatattcattct**agacatatctcta**cattgttagggttgttatgctttcctagagttgcatatcctacacaatg**ggacctacccaagattcaaactgtcacagtcagatccttcctcccattttatatcacattgctcacaggagagacatatcccctgcccgcctgccccattgactctttccacaccactgcatgcaccaggggatttgcatattgtcccacagggaggaccttcccttgtgagtctgagataaaagctcagctgtaactgtgccttgactgatcaggactcctcagttcaccttctcacaATGAGGCTCCCTGCTCAGCTCCTGGGGCTGCTAATGCTCTGGGTCTCTGGTAAGAAAAGAAGGAGATGAGGAAGGAGAATAGGGTGGGAGGGTGAGCTCTAGGGCTCCACAGCATCCCATGATCCCATGTTTAG**TCTAACCCTGTGTTAGA**GGAGTATAATCTGTGCTGTAGAAAAGGGAACTTGATATTTTGCTCTGTGAATAATTAGAAGCCTCATAAGAAATATGACGTCTGGTGCTCCGATTAAGATTTTCAAAATATAAAGGTCTCTTATACTTTACAAAAATTGAATTCATTTTAGAATGTGTATTTTTATGGCATAAATCAGT**ATTTTTTAAAATTAAGTTTAAATAAAT**GACATAAGATAAATTATGAA**AATTGCTCATTAGGTTTGTACATAACTTTGCAATT**CATTATTTCAGGATCCAGTGGGGATATTGTGATGACTCAGTCTCCA**CTCTCCCTGCCCGTCACCCCTGGAGAG**CCGGCCTCCATCTCC**TGCAGGTCTAGTCAGAGCCTCCTGCA**TAGTAATGGATACAACTATTTGGATTGGTACCTGCAGAAGCCAGGGCAGTCTCCACAGCTCCTGATCTATTTGGGTTCTAATCGGGCCTCCGGGGTCCCTGACAGG**TTCAGTGGCAGTGGATCAGGCACAGATTTTACACTGAA**AATCAGCAGAGTGGAGGCTGAGGATGTTGGGGTTTATTACTGCATGCAAGCTCTACAAACTCCtcccacagtggta**caacccctaacagaaacctcctcctggggttg**cccagttgctcacatgt**gctgcttgtctggagagcagc**tcagcagggtctctgagtctgcagaagaggaggctgttg**gagacctcagggcagaggttgctgctgaggactc**tggctcatgatagcctcagctgtacttcagtcccacatgttaaggccccattaggtgaaaaataaa**tgattccaaaaactgagatgaaataccaaggagaatca**gagtacaattaaggctgttacaaagaagcctcaaaatatggtggactaaatgtgacatggtttctgtgtctgttgcctgacagtgcagaggcaggtgggtggctttggtggtgtcggtggctcttctccaagaggtcactcagatgggcaggagacacgaccaccctcagacacagccttcctccttcctcagagtcacttgcccccatgcccatcctcgacagcatgaagtggaacgactagagagaaagctgt**tttcttctaaagaccaaagaaa**tctagagtttcccatcagggataaatgttcttttactttaagcacgcattggagaaattttccatttagtggatctgctgataaacccactttgttttgtttgtttgttttgttgtgttgtgttttcagatggagtctctctctgtcacccaggctagagc**gcagtgacatgatctcggctcactgc**aaactgcctcctgggttcaagagattctcatgcctcagcctccggagcagctgggattacaggcacccaccaccattccaggctaatttttgtatttttagtagagacagggtttcaccatgttggccagggtggtgtcgaactcctgatctgaagcgatccacctgccttagccttccaaagtgctgggattataggcgtgagccactgcacctggcagttatttgctcttatatgataatgtaaaattattttggaaaataattttttgagtat

Uppercase: IGKV2D-28

Lowercase: Flanking sequence[1000bp]

Red & Bold & Underline: Stem-loop [19]

Blue: Heptamer[43]

Green: Nonamer [8]

id-TRAV37[V_gene_segment]

caagaaatatagctgagaaaatgtgggagggta**ataggtaggcacctat**gctccacaggatgagacctggatgcaaagaac**agatgagagctttcttccctatcctctcctcatct**tgagtccctagtcattgctgcaagtacaggaaaccataaaacagagtttcagaaaactaatatgtaaaacatgagaaattaccagtgtgcacaggtgcccagagct**aaatataaagcacttggatattt**gcaaaagcactagcatgctcagagttagccagttactagactg**ctttctcacacctgaaaacatcctaaaagaaaggatactacattttcctccttt**gtctttttaacgagaatttggaaaaggtctaactcttttaacttatttaaagctgctgagattgagacatataatacagctctggtagaaaaataatcaattaagggaaataatcagatcttctctcaacctattcctacccccctcagaatgtactacagtgccatagaggcaaaaagcctgcataattctttaaaggtgaggggagtgagggagtttcaggtgt**aaaggattttcagacagtcataactggagaaactttccttt**taacactgtttgctttctttcttttttgtttgctcctcacctctctgtatgttccgtctacc**tttcatttcaacataacgagttactttatattaagggggaaatgaaaaaacaattttttt**gagttcagaccacctggg**ggcagtgcttcctttgacagaggcactgcc**agaaactagaacaaggggaagggaacagcaggaggagggcattcaatgatgtgtgtgatgtaacagtgaggttttgattatccgtaaaaacaatgaagattccccaaaggctacccaaccaacagaaccagaagactctagaacagtgaactaggcccagtgaacacggccaaatctaggtctccaaactgaatccacagaggctccaggaaaaggaaagaATGGAAACTCCACTGAGCACTCTGCTGCTGCTCCTCTGTGTGCAGCTGACCTGTGAGTCAGGCGAGTCATGAGGAGGGGAGGAATCAAGTAGGGGCGGCCAGGGCATTGAAATTTAGAAAGATCCCGGGGACAATCTTGGGTCTAGTCTCAGGAATGTAAAACAAATAATGAAACTGATGAACAAACACACTATATAT**ATGAAAATGTTTCAT**TCCTCCTAAGCAGGGTCAAATGGACAACTGCCAGTGGAACAGAATGCT**CCTTCCCTGAAAGTCAAGGAAGG**TGACAGCGTCACACTGAACTGCAGTTACAGAGACAGCCCTTCAGATTTCTTCAGTGGTTCAGGCAGGATCCTGAGGAAGGCCTCATTTCCCTGATACAAATGCTATCAA**CTGTGAGAGAGAAGATCAGTGGAAGATTCACAG**CCAGGCTTAAAAAAGGAGACCAGCACATTTCCCTGCACATACAGGATTCCCAGCTCCATGACTCAACCACATTCTTCTGCGCAGCAAGCACACAGTGCCCACAGTCACCTGCACCCGGTACCTAAAGCTTGCTGAGGGGCCTGGGCACACCTCCTTTTATAAGGGCCCTGGGGCACTGACTATAACTCTGCTGCATACAAAGGGAAATATGAGGGACTGTTTTATAACATAATATCAAACATTTC**ATTTGAAGACCATGTGAGGAAGGATAGTGTTTTCAAAT**ACCATCTAAAGTACTTTGTTTCTCAGTGTGAGGCCCT**TCCTTTGTGAAAACTAGCCCAAAAACTAGAGGCTAAATATCAAAAAGGA**GAGAGCACTAAAGAA**CAAAGAAAGAACAGGGTAAAATTCTTGTCTTTG**AGCT**TCTTTGTAGACTTTTACAAAGA**TATAGAAAATACAAAATCTTGAAAGCCATGACCATCAAATTCAGTAGTGAAAGGAAAAATATTTTGACATGCCAAAAAGATTGGCTTAGAAGATCAAGTACCAAAATCTGAGGGAAAATTGTACAAAGCATTTA**AAGGAAAAGAGAACACAACATGGTAGAAAAGCTGTCTTCTCCTT**AAAAGGGATAAAATGCCAACCAGCTAGCTAGAAAGGAAGTCAATGATGTGAGGAGAGGTCATAAACAGTGAGTCCAGCAT**CCTGCTCTTTTCTAGGGTCAGAGCTCTGAGCAGG**TAGGGTGGGAGGAGGGT**GAGTCACATTTAGCAATGACTC**TGAACTGAAGG**AACAGTGAATTTTGGCAAAAACTGTT**TTGATCATTGTGTCTTATATCAATACTCACAAGAAAACGCCCATTACAGAGGAAGCATTCAACAACCATGTGGACAAGACGGTCGGTCCAAAGCATGTCAGCCAGCCAGAGAATGTCTGTGCTTGTCAATTTCTTGAACAATAGACCCATGAATAGATTAGCTGAGGTGACAGATTGAGTTTCTTTATaagcccagaagcatgggttttctcccaccaagacagattcagctacttctgctgctgtcacaggacttttccttagttcaaagacagggttcttttcccatgggcacgaaaattcaggctcgcaaacaatttgaatggtgagtaagacacggttttattgggtgaagaggaagaaaagggggaaa**cagggactcttacaaggccagagtccctg**ctagagcacttcccaccccgcagttcgaatcccaggttccacacaggaaga**ggaggagcctggctcctcc**ccactgcaaaggatgccaacttctgcggctccaccccagtgcacaggctgcttggaatttctctagggaccccctcccacctggctgtctcactaccaaatgtccaccctgtcagcagtaaatacaacactgagccttcgatgtggaatcatccttcaaaggaaccagccacacactagttggcaagttgattacattggaccacttccaacctggag**gtattaggaattttttttcttcttaatac**atattctgggtgtgaatctgtggtctcctccaaagtgccacacatagttgcatgaactgatggctcagagtgtctcat**tgatcattctgatca**tcaaaggatttcacat**cccttgcatcaacagacccactttatatcaaggg**cgtgtgacagtagtcatgtcatatagcatatagcttcatcatacagcattagccatgaaccactagcctgatagaactttggaatgaccacttaagggcccagctaaagcaccagcttagggacagcaccctgaggggctgaaaagctattctttgagatagtgttgtgtccctgttacataaaatacatggacctgggtagcaagggatggaagtaggactatcaccccatcaaccaccactttcagtaacccacttttgtgcttcctgttcttataactataggttctgctggactagggattctgtttccctttgagggggaa

Uppercase: TRAV37

Lowercase: Flanking sequence[1000bp]

Red & Bold & Underline: Stem-loop [25]

Blue: Heptamer[51]

Green: Nonamer [5]

id-IGKV2-38-2[V_gene_segment]

cttctctgtcctttctgattatcgttagttttgttttgttttgttttgttttgtttttt**gagacagagtctctctctgtctc**cc**aggttgcagtgcagtggcgcgatctcggctcactgcaagct**ccgccccctgggttcacgctattctcctgcctcagcctcccaagtagctgggactacaggcgcctgccaccacgctcggctattttttttgtatttttaatagagacaaggtttcatcgtgttagccagaatggtctcgatttcctgacctcgtgatctacccg**ctttggcctcccaaag**tcctgggattacaggtgtgagccaccgcgcccagccatgatcattgttagttttaagtctgttgtatctgaaataagaatagcaacccctgctcttttttgctttccatttgtttgatagatctttctttctccctttactttgagcctatgtgtaatcattagaaaccacctgaaagaacaatgaccagtgccttgaataatatcttttaataaagggtcgcttgtacctag**taaatactaagcttattta**tttatttatttatttatttatttatttatttattttgttgcccagtttcgctcttgttgcccaggatggagcgcaatggtgcgatctcggctcaccaaaacctccgcctcccggattcaagtgattctcctgcctcagcctcccaagtagctgggattacaggcataagcccccaacccccaggtgattttatgtatttttagtagagatggggtttttccatgttggtcaggctgaactcccgacctcgggtgatc**cgcctgccgcggaccaagaagtgctgggattacaggcg**tgagccaccacaaccggcctaagctcatttttaaaaacgt**attttttatgatgtgaataaaaaat**cttttaaaagataa**atgataaatccatatcat**aagggaaatcgtgatacttgctcataatatttgtacataactttgtacttcATCTCC**TGCAGGTCTAGTCAGACCCTCCTGCA**TAGCAATGGAAACACCTATTTGTATTGGTAACTATAGAAGCCAGGCGAGTCTCCACAGCTCCTGATCTATGGGGTTTCCAATGGGGCCTCTGGAGTCCCAGACAGGTTCAATAGCAGTGGGTCAGGCACATATTTTAAACTCAAAATTAGCAGGGTGGAGGCTGAGGATATTCGACTTTATTAATACA**TGCAAGCTACATAATATCCTtgca**cagtggtaacgccttgaacacaaacctccttgctggggtgtcccaactgcccacat**gagctgctgtctgggagcagctc**tgcagggtct**ctgagtctgcggaagaggaggctgttggggaactcag**ggaaagggtttgctgctgaagactctggctcatgagagcct**cagctgtgcctcagtcacatatgttaaggtcccatcagctg**ccatggcctggggaagacatcatcaggaggagaattcttgggggccatgagccctgggaacaaccagctatgatga**agggagagtgaatgaaagctcatccctccct**atcttgcctacatttttcaaggacatttac**ttgtcaaaacaaccagaccgtggatgcagattagtgacaacacaagtggaatacgtgttg**gaaatgaccagcttggggtttattttatacaagatgatacttggtcctattgagaaatcatttttttcatttcattctcccaaccttgtatcttttttagcactcgtacactaaactg**ccctctccagtgtcatgctggagaagg**tattctacacaagctgtcctcaggggagtgtggcaaaaaataattattaaaaa**tgtttaggtttgactattcatagacttggtaaaca**catggcaaataaaagatctcaatgccaaaatacttcaattctcctcaattaccttcatcccagtgaaagttcttccagttacaaaagtgagctttgaaaataaacagaatgaactcacagatggacaacatagactcactaaatatgtcacatgaataatgcagaattacataatattttttcttcttaccccag**ttttaagaatttaaaa**agcatttaacctaagtactgttttagaggtgacaaatttagtggtgtgtccccatgagaaaatatttc**aaacaataccaaaattccttgttt**acatattacatcatcgttcctagaaaatatatctaaatcctctattttaggtata

Uppercase: IGKV2-38-2

Lowercase: Flanking sequence[1000bp]

Red & Bold & Underline: Stem-loop [19]

Blue: Heptamer[14]

Green: Nonamer [4]

id-TRBV22OR9-2[V_gene_segment]

tatacatatgtaacaaacctgcacgttgtgcacatgtaccctaaaacttaaagtataataataataataataaagaaatcatctaggatcctatttttatttactaaatctattgggaagaacctatgttatacagtgaatttacactcatacaaatttctactcgtgttggaaaatgatactttagtcagctggtacaaccctatgaccataaatgtctgatttcacttgatatagtattgacttcctaacttcctgtgaattagaatatcaggtccagaacgtaataatagc**aaacacaaatatagtgtttcttatgtactaagaa**ctgttctaaatgctttatttatgtaatgttcatcacaggtttgggggtagttacttattatctcaatt**ttacaggtaaagtcagattgaagagaggctctgtaa**cttttacaaggttacgtagatgctaaatggcacagacaggactaaagcccatgccgtgtcccaagtccctgctctaatccattgtcacaaggggaggggcacttcagacgctttgctattcctaaaattacaggagtagcgcctaccccat**ctctactgtattaattatttatagtaaatacagagag**tgtcgagaattggttctttaaaaaaaaacaaccacaaaaaactaaacttccaaaccccttcttgtgttgagttgagcacaggtgcatggcactctgtgtatgactctgaccaggaaatgagggattggctcttctgaccatgcaggcaggcacttgtccatt**tagaagggatataaaaagcttcta**taagcagaagaggaggtgcatgtgacttgatctctctaattcttagtctcaccatctgcaaaatcagctgtgacaatcagtcatcatgtagcaatgacaaattctttatattaagccctggggaggggctagatctcagcctctc**tctctctcagagcctgtgtctgtaacttcagagaga**cttcatc**gctgccATGGGCAGC**TGTGTCCTCTGCTATGTGACCCTGTGTCTCCTGGGAGCAGGTGAGCTCAGCACACATCCAGTTGGTCAATTCCTATCCTGGGCTTGCCAGGCCCCTCCCCGGCCAGCACCTCCCTTCTGGGCATGATCTCTAGCCTTTGTCTCTTGGACCCTTGGATGCTGACATCTATCAGACGCCAT**TCCAGCTCACTGGGGCTGGA**TGGGATGTGACCCTGGAGTAGAAACAATTTGAGACACAATGACATGTACTGGTACTGGTACTGGCAGGACCCAAAGCAAAATCTGAGACTGATCTATTACTCAAGGGTTGAAAAGGATATTCAGAGA**GGAGACCTAACTGAAGGCTACGGTGTCTCC**CGAGAG**GAGAAGGGGCTGTTTCTTCTC**ATGGTGAAGCTGGCCCACACCAGCCAAACAGCTCTGTACTTCTGTCCTGGGAGTGCACcacagtggaa**gcacagccattgtctctctgtgc**ggaaatgtgtcctagccctgtagtccccaccacatcctctagtttaattttttcatttttaatattttcttgagattttcctatgtcctgttacttttgcaaacac**aaaggatacaaaactgaaaagaaaatagatccttt**atccaaggagctaagagtctagcaagaaagtaattataatagaacggaaaatatcatatataatactacaatcagaagtcatttgggagcttcagggaggacatgcacaattctgtttgattgaatgagaaaaagcaaagaggaagtcccatt**agaacctcaccagatagacctggttct**gagtagagcattccaggcaaagagaacaacttttgcaaaggcatattaaagagtttgacactttggggaacagtttacaaagagaaagatgatgatcaatgtatatgaatcaaaagaaatggaatatatctgaggaagtagaaaggagtttttgaaaaagttgagtttttggttccatgaatataataagtatgaaagtcctttgtaggctgttgctcaaaagagagagctgaa**gcttgttatcagcaaacaagcaaatgatacaatttttcaattttcctcatataactttgatcattt**acaaagggcaca**atttcaaacaatatgaaat**cacgttagcagtagcagttcccatttgggataatagggttcaactgtctatggaacaatcagataaaaactagaagtacgcaactgcatagacaagatgtgaaattgagatgccataaagctggactagaggttgagattctgcagattacagcatgtcaataattgaactgtggggacagaaaattccacccaaatatagatttagatcacagagataaaatgcatgggaattaagactgtag**tattctggttccttatggagaata**aagaagcctgagagtgaggagtgattagaacatgggatgtaaaaaca

Uppercase: TRBV22OR9-2

Lowercase: Flanking sequence[1000bp]

Red & Bold & Underline: Stem-loop [17]

Blue: Heptamer[32]

Green: Nonamer [2]

id-IGHV4-61[V_gene_segment]

agggtatgttcatgcaattcaatggaggagaaagtc**tttcagagacagatggatctgaaatgataaatatgtgggtaaggactctgggcttgagtatcatt**gtccagccatgtttcacaagtg**tgtcctgtcagggaaggaca**agagttccttgtgttctcagagggaaggggtcacagagttcctctctggttcccaggaaagataatcgcactaatct**tcatgatcttcatga**gactatcctccagtgctgacctgttatagagtttttgtctgaagttctcactgcaatccccaatctacatattttcaatcagaagtgtttagaggcca**ggacatatcttcacggtcacacattgagaaggatgtagatatgtcc**cactaccttctcctgagatctcagacagaatcccagatttcaaaaggacacagaaggacagctctcaggtgcttttaaaaaatgacccacttccagggacagggagcttccctataaccatggtggatgttctgaactacaataaacattggatggatccaggattgtttgaagtcactgtcattattacattcagctgctgtttcaatgtgtctgaagtagtaaatgacaatttagatgacaatttatatgaatcttcaagggtagaacaatattgaccatattccaaaatctgtccttgatccatgatcacactcatctcccagaccaggtccttcagcacgtctctttacctgaaagaagaggactctgggcttggagaggggagaccccaagaagacaactgagttctcaaagggcacagccagcatcctactcccagggcgagcccaaaagactggggcctccctcctcctttttcacctctccgtacaaaggcaccacccacatgcaaatccttacttaagcacccac**aggaaaccaccacacatttcct**taaattcaggttccagctcacatgggaaatactttctgagagtcctggacctcctgtgc**aagaacATGAAACACCTGTGGTTCTT**CCTCCTCCTGGTGGCAGCTCCCAGATGTGAGTGTCTCA**GGGATCCAGACATGGGGGTATGGGAGGTGCCTCTGATCCC**AGGGCTCACTGTGGGTCTCTCTGTTCACAGGG**GTCCTGTCCCAGGTGCAGCTGCAGGAGTCGGGCCCAGGAC**TGGTGAAGCCTTCGGAGACCCTGTCCCTCACCTGCACTGTCTCTGGTGGCTCCGTCAGCAGTGGTAGTTACTACTGGAGCTGGATCCGGCAGCCCCCAGGGAAGGGACTGGAGTGGATTGGGTATATCTATTACAGTGGGAGCACCAACTACAACCCCTCCCTCAAGAGTCGAGTCACCATATCAGTAGACACGTCCAAGAACCAGTTCTCCCTGAAGCTGAGCTCTGTGACCGCTGCGGACACGGCCGTGTATT**ACTGTGCGAGAGAcacagt**gag**gggaggtgagtgtgagcccagacacaaacctccc**tgcatggacgcggaggggaccggcgcaggtgctgctcaggaccagcaggtggcgcgcggggcccccagagcatgaggccgggtcag**gagcaggtgcagggagggcggggcttcctcatctgctc**agtggtctccgtcctcgccagcacctcgctgtcaccagggctcctctttctttattatctgtggttctg**cttcctcacattcttgtgccaggaaagaaacgaggaag**acaaattttcgtctatagttgaagcttcaccaattact**aggaacttgcctacaagttcct**gcatgacccattataacttatcgattaaaaaatatatattctaatgcttctcaccatctcttgatttgtatcatcaac**tgaattgtaccctctttgaaattca**tatgatgaaacctt**aaattcaatggatctatattggaattt**taatgaaataattaaggttaaatgtggtcataattgtaagaccctaatgcaatagacgtgttgtctttataagaagaggaagagacaccagagacctctcacttttcacgtgcaggcagagaagaggccatgtggagacatag**tgcactagaaggtggcccagtgca**agccaggaagaagccgcgccaagaaccagccctgccagcacactaatcttcaacattcagactgcagaattttaagaaaatcaatatttgttgtttaagccacccactcctgttgtcttcttatgaagatcca**gacagactaataccacataactctgttagtgctgtc**ccctggatggagaattagcctcctgaggctgggcacatctctcagatttccacataaaacaggtaaaaaatagtagttctgatataaaaacttgtcatgtccctgttggccaatttctgggcaaggtcttttaaataagccaagtttgcggggaaatggagaccatatgtttgtgggactctaacc

Uppercase: IGHV4-61

Lowercase: Flanking sequence[1000bp]

Red & Bold & Underline: Stem-loop [18]

Blue: Heptamer[40]

Green: Nonamer [4]

id-TRBV19-2[V_gene_segment]

aatgtaagtctacactccaaacagaaataaatacactcaaggaagcagctgaggatataagtgaccataaccagaccaggactttaaaaactcagactacaaatggccaaaatgggaagacatcagaaaaaaacacaaattgattgaacccaggagggacatgaaagaaaaagacaaagttatcccagtcatgcaaaataaaagatgaccaagggagaataaacccaaatgaaaatttagtcaggacattgaacaaaggcaaagaaacaagtagcagaatgagaaggacatgagtgaaaccagccagttagaaggcggtcaaagaaggcgaggtcttggtatgattgcatgaggggcctggaacagcagcactgggctctgtgcagcaggacggctaagcagcaggttgtcacag**tggccacagcactactggcca**gaccccaggagtcacagtttccctgagagagctgagggcataaacaatgataagttctccaggggatggacttcttaaatactctgagttcatcattacacattctatgcatgtagcaaaataccacaggcacccacaaataggtacacatattatgtatcaatgtttttttaaaaaaaggaaaaataaatgaattccacattcttgtataatcagagtatctacagcgctcctggagtaacccaaaccttatggttgctggaaagataaaaaaccaat**gggccaattggggtgaaactctggccc**ggaaacgtggggtgccagccccctcagcacagatgcaat**gcagagttatgggaggtgcgaatgactctgc**tctctgtcctgtctcctcatctgcaaaattaggaagcctgtcttgattatctccaggaacctcccacctcttcattccagcctctgacaaactctgcacattaggccaggagaagcccccgagccaagtctcttttctcattctcttccaacaagtgcttggagctccaagaaggccccctttgcactATGAGCAACCAGGTGCTCTGCTGTGTGGTCCTTTGTTTCCTGGGAGCAAGTGAGTCCTGGGTTCAGGGAGAAAATTCCTATCTGGAGTGCTGCAAGTTCCAAATCTAAGGCCTCCCCGAGGGACAGCAGCATCAGGCGCCTCCCTGGGCTGTACTCAGGCATGCCTCTCTTTCTTTTCCAGACACCGTGGATGGTGGAATCACTCAGTCCCCAAAGTAC**CTGTTCAGAAAGGAAGGACAGAATGTGACCCTGAGTTGTGAACAG**AATTTGAACCACGATGCCATGTACTGGTACCGACAGGACCCAGGGCAAGGGCTGAGATTGATCTACTACTCACAGATAGTAAATGACTTTCAGAAAGGAGATATAGCTGAAGGGTACAGCGTCTCTCGGGAGAAGAAGGAATCCTTTCCTCTCACTGTGACATCGGCCCAAAAGAACCCGACAGCTTTCTATCTCTGTGCCAGTAGTATAGAcacagtgaagcacggatgtcgcctctctgtgcataaatgtgcccagtcctgcttccccgaccaggtggcagggctcctctgcactctatgatggcaggaaacgccactcagccactaagcaggtttaaaaaactgtatataactaaaactctatatataaactatg**catataaaactaaactatatatg**tatacatttatatataac**taaaactttatatgagactaaaatgatagtttta**tatagttttttaaagtatagtttt**atatatatataactatatatatatat**catttttt**aaactatagttttgtagttt**tatttatttatttactttttggagatggagtctcactctgttgcccaggctgcag**tgcagtggcatgatctcagctcactgca**atctctgcctcctgggttcaagcaattttcctgcctcagcctcccaagtagctgggatta**caggtgcataccaccacacctg**gctaatttttgtattttagtagaggcagggtttcaccatgttggccaggatggtctcgaactcctgacctcaggtgatccacccacctcggcctcccaaagtgctgggatta**caggcatgagtcaccgtgcctg**gctgttttatagtttcatatagttattttt**aaaaactatataaaagttttt**gtat**agccacttagtggct**gagtgaagtttcctg**tcatatatgatatagatatagatatgtttatatatga**tatagatacagatatgtta**tatatatgttatatgttgtatataagatatata**catatagagagagtcagagagacacagagaagaaggaaaagttaatcttacaacccccagcttttt**tctattttaccgaatgctagatgaataga**tggatgaatgaatagaaatgaatgaatgaatagaaaattttccct**tctttcaggaagataaaaaagcctgtggaaaga**gagatgtagaagacaaaccctgtaatcacaatttc

Uppercase: TRBV19-2

Lowercase: Flanking sequence[1000bp]

Red & Bold & Underline: Stem-loop [17]

Blue: Heptamer[32]

Green: Nonamer [2]

id-IGHVII-43-1[V_gene_segment]

accaaaagtgtgtaagtggtccgttatctctatagccttgcctgcatctgttaatttctgatctttcagtgatagcaattctgacatatgtgagaaagtatatcatttggttttgattcgcacttatctgatgactgaatatgctgagcatttttataggttagttggcc**acttctatgtttgtatttgagaagt**ttctgtgcatgtcctttctccatttttaatggggttatttggtttatgctggtgattaagttccctatgaattaccaccttacacacactaagataatcagcattcaaaaaagaaaaatgaatagaaaaacataaccacttgtaggtatagacatactgaatttggaatggtcatggcttgcatgttggtattgaaatgggacagcaactttggaaaatggtcttcacagaaaacctcataaatgtagctagtgtcaccactcacaga**agtagatggagaatacaactcaagtgtccatcgact**gccagagggatga**agacgccgcggtgcgtct**gtaacccacagggatga**agacgccgcggtgcgtct**gtaac**ccacagggatgaagacgctgtggtgctttaaaaaggagtgaagcac**tgacataggctgcaactcggatgggccttgtaaacattgggtgagtgaactgagggagacacggaagtctacatcctgaattgtcctattgacatgaagtatgcagaggaggaaaatccctagagacagatcacagggggtgagtggaggggaaatggagagtgacgcttaatggagctgaggttttcttttctgtgatgaaaatgttcccaaaccaaatactggtgat**atttgcacaacattgcaaat**gtgtaaagtatcactgacccgtacacttaatcgtggc**taaaattgtaaatttta**tgtcatgtgtatctcaacacaatagaaatgagcaccacattttgtttagtgactcaacaacaataacatttaggtttaaatTCTGGATTCCCCAACAGAACCAGTGCTTCCTGCTGGAGCTGGATCCATCAGCCCCCAGGGAAGGGACT**GGAGTGGGTCAGGTGCACAGGTCATGAAGGGAGCACAAATTCTAACCCACTCC**TCAAGAGTCCAGTCACCACCTCCAGATCTATGTCCAAAAACAGCTCTTCGT**ATGGCTGAGTGACATTAGCAACAAGCACACAGCCAT**GTttgtttttttgttttgttttggtgtgtgtgtgtgtgtttttgatagagtcttgtgtcacccaggctggagtgcagtggggcaatcaatcttggctcactacgagctccacctcctgggttcaagtgattctccagcctcagcc**tccaagtagcttgga**ctgcaggcactcaccaccacacctggctaatttttgtatttttagtagagacagggtttcactatgttggccaggctggtcttgaactcctgaccttctgatccacccgactcggcctccaaaagt**gctgggattacaggtataagccactgtgcccagc**cacaaacaagtatttttaagcaaaagacacagtgaagagacttcagtatgag**cccacacacaaaccttcctgtggg**agtttacagaacagcag**tgggtgctgaggacagaagccagcaccca**ggaaccagcagggaaacccagggggcatttggcacc**gcctgaaggctcaggaccattgtggggctcagtggtcaggc**aggctcaaggttcagcc**tcagagcaggtgtagcaggcggggaaaggctctga**agacggagtttagtgtcaccttctcatttccactacta**aacaccctccagcacatctattctaatgtgtatgggtgtt**catgtgtttagagaatattatttatgttatgaatctatagccatcttgtgggtgcatcaagttaaccacttcaacctatgtggaccctgttcattaggaataagtccctgtatttgaggacctcacaaattaataattatgtagaatcactttctttttcagtctcctttccttcctctttctttctttctcactcacacactgacaaacacacggggtgccataacattaattacctgatgtattaaagaatattgattcatttgcaacttgaccagtttagctgttgtt**catgttgttgtaagatcaggaacatg**tttctcagctgtgtactcctctaagctgagcagcag

Uppercase: IGHVII-43-1

Lowercase: Flanking sequence[1000bp]

Red & Bold & Underline: Stem-loop [18]

Blue: Heptamer[36]

Green: Nonamer [5]

id-TRAV17[V_gene_segment]

ttatgggagccctggtaaactaatac**agcctactggttaggct**tgatgtgggcacatggcttgggctcc**aacaattagatgaattgtt**ccctaactttcactcaggagttacttttgcaaaaaaacaaaaacaaaaaacaaa**caaaaaatgagtaagagtttcctttttg**agagtggtggtaattgcaacaaggtggacatgagagtttacag**gctacaaggcagtggaattgtagc**tgcatacattgtcctgcgtgaacagtcttgcataaactacgggc**agtatctggagctcaggaaaggagccagtgggatact**cagaggagtgcttctccagagtgattttctgtactgctgctgacttcatagcttttgagttcagttctctgaccc**tactggagattctgaggtaaatccagta**ttcattaatttattccttttctacttaatttaggcagagtccatttctgttgcccatcagtaaaaaccttgacttgtatataatctcattccagggaaaattttttaaaaggaagctcgtaggtacatgatcagaggtcatagtaactacagggaatttaacagtgcttctcagaggtagtctgaaaatattgagcaccaactgtctttcgtagttgagttcttacctga**attgaaatgtatctgttcctattctaagttagaattagatttcaat**aaaggcagagactggaatagctttgtaatttcttctgagtcagctctgaacacacgtcctgatggcagcaggatgatgttacgatcaca**agagggcagtgcctccctct**cctcagactaacttggctttaaatcagcctatctgcattgaaaggaaaggactgagcttgcctgtgactggctagggaggaacctgagactaggggacagaaagactagggattcacccagtaaagagagctcatctgtgactgaggagccttgctccatttcagg**tcttctgtgatttcaataaggaagaaga**ATGGAAACTCTCCTGGGAGTGTCTTTGGTGATTCTATGGCTTCAACTGGCTAGTGAGTTGGGGAT**TTTGGTGATGGGAATGGCAAGACCAAA**TTTTAGAGCCCACAGGGCTGAGGGGTGAAGTGTTCTTATTAATTTGGAAATGTAATTTGAAAATGTAATGACTGACTCCTGGTCCCGGACCTGTGTTTCTGCCTAGGGGTGAACAGTCAACAGGGAGAAGAGGATCCTCAGGCCTTGAGCATCCAGGAGGGTGAAAATGCCACCATGAACTGCAGTTACAAAACTAGTATAAACAATTTACAGTGGTATAGAC**AAAATTCAGGTAGAGGCCTTGTCCACCTAATTTT**AATACGTTCAAATGAAAGAGAGAAACACAG**TGGAAGATTAAGAGTCACGCTTGACACTTCCA**AGAAAAGCAGTTCCTTGTTGATCACGGCTTCCCGGGCAGCAGACACTGCTTCTTACTT**CTGTGCTACGGACGcacag**tgttccccaggaacctgcagcctctacgcaaaccctgccaaagcagcttcttagaagccctaatagtgggtagaattagtggttatgtctttcagtcaag**aagagtctacaaacagctggcaaagtaaaaggggaactctt**tcaaattcgtgattttttaaaaagcatcttggactaggggaaaaggcatgaaatttggaacaggaaatggaaat**gccaggctcagttctgctaatcctggctacatagggacattgtcagtgtagc**agccttcaatttcctcacttgaaataaaggatatggatcaatgagcccaaaatattacttaa**tccctaattttattaggga**aaaaaatcactgggcacttgtggaagagga**ggaaggctgcacagttttagaaggtctcatttgtcctatgtgcagccctcc**caagacatcagtgggtctttaaggtgcatgaggtggctcttaggttcc**tctaattcttgtataagaagcagtagattcattaagcattaga**attggaagaggtcctaggactctcaaccaagcttttcattttagagctgaggagtaaaaccattgcataataagacagacaattttctgaatctttggagtgatagtgacaaatatctggatctgatttcaccatttattagttgtatg**attcccaagcttcaatgtctctattaggcaaaatgggaat**ttataagaaagtaacaacttttccccatccaaaggagagataggtgtttgacactacca**ggtcctaaaacgctgtgttttgaaagcaggcatttttagtacc**aaatacataacagtttt**tcagctatttattcttgctcctactgagaaataaatactga**ctatagttttcttgacacagtttagatactactgaaaatgtgtttgtggagaaaattagagcatttgtatttaggttcatgctttggtgagaaaagtgggggagggcagggaaaggttgtgtcaga

Uppercase: TRAV17

Lowercase: Flanking sequence[1000bp]

Red & Bold & Underline: Stem-loop [22]

Blue: Heptamer[15]

Green: Nonamer [3]

id-IGKV3D-34[V_gene_segment]

tcagataataaatatttaggtttggtgaggcaagatgcaaaattgaatatattatgtgagtacttagagaagagagaaaacaaatttctgcaatgtctattaatgaactcagatggaatgtaacccaaagagcttctcagcctgtccagtgaactgtcacttaccaccaaatgctca**caattttaaggaacacaaggaaaccccaaattg**aattattgtttagatcagaaacatgagcagaggtttccaaaattattaaagtgagaaaggcctgagtctgattgaatggatgtgttgtgcaggatggggtggcatagaaagtagggcatcacccaaggaattgttcctggcctgctggacagcattgtcgtctgtttgcagggaatggaattacacacactgaagggtctgagcccggggcctgctcatctgtgcagccgtgatgatcatcacacccacagtgctgggcctaattgcgtgttgccctttaatagtttttacaaagaatgaggaggcttcagggctcataggaaagatggctaagctgcaaaggaaaagtgtgtgtggaagatgaacttctgatgaactagttttcagccaggtacattcatgcaccaatggagatgactc**ggttctttggagaaataaggagaggaaaagaacc**acaggcttctattttattcccaacaggtttgtactaaataccatctggctgggtctaggtgactagtgagacgggaagaaacaggggctatagtctgtgaccacagggaagaagttctacca**tcaggctgagccaatggacttttctggcctga**ccacctgggcaggggctgctgag**tgcagagaggaggaggcagggggtctctgca**gctggaagcccagcacccactccaactactttgcatatccttccagccactctgctgtccagagcccatatcaatgcctgggtcag**agctctcgggaagagct**gctcggttaggacccagagggaaccATGGAAGTTCCAGCTCAGCTCCTCTCCTTTCTGGTACTCTGGCTCCCAGGTGAGGGGAACACGGGATGATTCTGCATGCCAGTGAAACTCTGTCAATGTTATTGGGTCCTGTGACCTGCTCAGCAAGAACAATAATTAATATTCAATGTACATCAATGA**TCCCAGCTGTACTGGGA**AGACAGTGGATTTG**ATCTAGATTGCACAATTGACTTTCCTGTTTTATTCCAATCTCAGAT**ACCACTGGAGAAGTTGTGCTGACATGGTCCCCAATCACCCTGTGTTTGTCCCCAGGGGAAGGAGCCACCCTCACCTGAAAGGCCAGTCAGATCATTGACAACTATTC**AGCCTGGGACCAGTAGAAACCTGTTCTGGTTCCCAGGCT**CCTTATTTATAGGGCATCTACCAAGGCTACCAGCATCCCAGCCCGGTTCAGTGGTGGTGGGCCTGAGGCAGACTTTACCCCAACCATCAACAGCCTAGACCCTGAAGATGTCACAATTTTATTACCCTCATCAGTACAGCAGTGGGTGTCCcacagtaattcaacat**gaaacaaaaactttcacaaaaccattgattttttttttc**taaaacc**agcagctttatgggctgcagctatgatggctgct**cagttttagcaactgtgcctctatttggaaattttgaaattctgaaaagtaactcgttgtcaagatggtgactccaatg**tctgtatttttctcagacttatttacaga**gttaggttttgaatcaattttactgcaaggccttcagacagact**cacctgcatggatgcctttaaagacacaggtg**ccgctgggcacagtggctcactctttttttttccaatagagtagaacaaacatagcttttcatttaataaattttcacttaatggcagcattttcaggaacaagtaaaggccaccgagtgaaggttaggtgcacgtgagtaaataagacaattttgggtcccaattcagtaggggaaagagccatgaaggaagaataaccacacaataaaagtgcgtagaataatttctgtagttgacaaattgtgtaaaaagcaggaatggtctgtggtatctggaggctgaggccatatatttgtgctgatcagggaaggccccagggtcatactgaagattcatctgagtggtgaggaggcagccaggtgaccacaaaaacagcaaagaccacgggatcattatgggaagggcagggaccagtgtgtactgaaaaagtcagcaagtgctaaact**aatcagatgccttctctgatt**tttattcccgta**aagataaaacttactgtaatatctt**acatcatacaccttggtctgggtcactcccatggcatagaatgttccctggttcagaagttggaagcccaggtctttcctttgttgacagtttgcaggctccccagagcccttccctgcc**actcaagcccttatcactgggtcagtctctgtctttgagt**tgggatcaccataggcaggggcagcactagacactggacatggggctctggccct

Uppercase: IGKV3D-34

Lowercase: Flanking sequence[1000bp]

Red & Bold & Underline: Stem-loop [15]

Blue: Heptamer[23]

Green: Nonamer [7]

id-LOC102725101[V_gene_segment]

aataaacaatacaacataaaaggtttaattctcggtatttctactgagaaaaataagccaaatagataagagtacatactatattgtttcattcttataaattctagagaataaaaactagtctaaagaaatatgaaaacatcagtacttttataaagaaatggtagaagaaaagaggagaaaaacaaaattatgtctgtaaaagagcaagaggaattctg**aggtgagttgacttgtcacct**tctt**gaaaatagagattttc**tttatcaaagtttactgttatgcaggttaaatatgtga**attttatcatctgtcaattaaaactcataaaat**gtattacaagtaaacaagtgaaattttagacaaaaaagggatgataagaaggaacaaatgaatacattaaatat**cagatacaccaaaaatttatctg**cctgatgcctagatgtttccgtatttttaggtaaatgcagcaaaatcacacaggttctcgtggcaggaagtggattctgcaaaccacactaggcccatttagctctgtccaatagttggttaagagagcaattgaggccagctgtgaggagcataggcccaggtactaggactcactcatgccagatataagcccttag**acacatacatagcccctccatgtgt**gggttcacttttacatctgtaaacgaagaaaccactgagtgctaaataacatcatttatacacataggtaaaaataattaaaaatatgatagttgttaaatgtttatcgcacaacaatttcacattaagacagcattttcccaaacacaatcattgtcatcaaaatcccccaggacgctctcatctactctgggccctgccctctcctcaggagtcccaccccatagcttgctatatagtaggtgacatgcaaatagagccctccctctcctgatgaaaaccagcccagccctgaccctgcagct**ctgggagtggagccccagccttgggattcccag**gtgtttccA**TTCAGTGATCAGGACTGAA**CACACAGGAATCACCATGGAGTTTGTGCTGAGCTGGGTTTTCCTTGTTGCTATATTAAAAGGTG**ATTCATGGAGAACTAGAGATATTGAGTGTGAATGGGCATGAAT**GAGAGAAACAGTGGGTATGTGATGTGTGGCAATTTCTGACCTTTGTGTCTCTCTGTTTGCAGGTGTCCAGTGTGAGGTTCAGCTGGTGCAGTCTGGGGGAGGCTTGGTAC**AGCCTGGGGGGTCCCTGAGACTCTCCTGTGCAGGCT**CTGGATTCACCTTCAGTAGCTATGCTATGCACTGGGTTCGCCAGG**CTCCAGGAAAAGGTCTGGAG**TGGGTATCAGCTATTGGTACTGGTGGTGGCACATACTATGCAGACTCCGTGAAGGGCCGATTCACCATCTCCAGAGACAATGCCAAGAACTCCTTGTATCTTCAAATGA**ACAGCCTGAGAGCCGAGGACATGGCTGT**GTATT**ACTGTGCAAGAGACACAGT**GAGGGGAAGTCAGTGTGAGCCCAGACACAAACCTCTCTGCAGAATGCTTGGGGGAAATCAGCTGCGGGGGGCACACAGGACCCACTGATCAGAGTCATCCCCAGAGGCAAGTTGCAGATGGAGGCTGGTTTCCTGTCAGGATGTGGGACTTCATCTTT**TTAGAGTTTCTCTAGGGAATCTCTCTAA**GTTCAGAATTCTGTGCTTACCAATGTCATCTCTACATATTT**TTAAAATGATTATTTTAA**TATGAAAACCTATTCTCCTATGCACAAAACACAGATTGATGCTTACAGAGATGAAAAGCCCTCAACCATTG**TCACCAGGATCAGAGTATTGAGGAAACTCAGGGATACCTGGTGAGTCTTCTCCAGTCAGACTCA**GGACAGAAACCTCAGTGAGATTCCCTGACTAGGACGGTCTTTAGGAATTGTGATCACAGCCAATAGAGTCTGGGCC**AGGGTCAGTGTCGTGTAGAACCTCACAGTTTTCATTCCTGACCCT**TCTC**CTGACACTAAAGTATGCAACTTAGTATCAG**CACTGATCTGGGGCCCCTTTTGCTCTTAGCCCACTCTATTTCTTTTTATTTGTTGTTGTTGTTCTTGCTCTTCCTTGTGCTGTTCGTGCTTCCTGTAAAGTGGGGATGTGGTTCTTGCTGCCAAAGCTCGAGGTCTCAAGCCCATTCC**CTGCAGCTGAGGTGGGGCTCAGGCTGTGGCTCCTGCAG**CCATG**TGGGAGAGGCTGATAGGACTTTTCTCTCTCCCA**TTGCTCAGCACCCTCCAGTGTGTCATGTGGAGACTCACCTGGGAATGCAAGTGGCCAACAGTAGTGAAGAGGATGAGCTTGTGTGGTCAAAATGGGATGTGGATGTGAAATTTATCCTGTGCTGTGCAAAGTACCACAGAGTGAGTCACCTTCCTCACCAGTAGTGTTAGAAAGAGGGTGTGAAAGTTGTCAGAATCAAAATAGATCCACTTGTGTTAAAACCCTGACAAAAGGAACTAGGAATGACCATGAAGGAGGTTTCCCATGCACATACTCCTGATAACAAGAACGACCATGAATGGATTCTGCTTAACCACAACCTTTGATAGAAGCCACCATGACCTTATAAAaatcacatctacaaggacatcttcccagcaaatcactgtttaaccctatattgatgccaaccttggtattgactctacaagcaaggaaaatactctcaaaacaatttatgtaacccacctcattttcactaataaacctatggattgacatcctggagtcactgctgcatttgttgtta**attgtaattagccccttttacaat**gtttgtgactgtttttctccgatgtctcttggaaaataaataatttacaagttgatggcagtaaagaagctatttaggacatttttaacatcctgttgaatatttctgcatagcacatcgatcccctaaaatacttcgctgtattggcatgtgataaatcagagtataatgctgaaggtaaaatggaaaatacatgggcttttgatgaatcaagtcatagggtgatattgtctttgcccttgaggaagcagaccatgggctgttaagttctagtgggagtacctttggcaaaaggatttcatgagtttctgaatgttatgctactttc**aatttaagaatgcaacttgtcatttatttttacttaaatt**tttccagaagatatttggcagtaaggacagggtagcattcgtgtgatactgatgacttagagaattattttgtaatttctcctgtaaggtatgcacattgctcactcgatacagaaggtcaaatgtcacaggtgggaaaatagg**aataaagcaaattttatt**aaatgtcatgactgtagtttttggcaaggaagtgcttcatgtcaacctgaaaacagacagacaacaataaaacatattcaaacccacagggagtcagacctatgtccttctctcgtataagtacaaggccttgccacatccaaactatcctttaagctccagggtataaaatgcttttggactgtggaagctaacagctctcccctcaggcagggctaaggtatctggggaatgcagagttgtgttcacgaagaagatggcattatg

Uppercase: LOC102725101

Lowercase: Flanking sequence[1000bp]

Red & Bold & Underline: Stem-loop [23]

Blue: Heptamer[60]

Green: Nonamer [8]

id-IGHV3OR16-17[V_gene_segment]

ctgaaatacaatgaacaattgaaatcaaggcatgcctcag**tcacataagaatgtga**ggacattatcaaataattgtgctgagtgtaggaagctaaa**gaattcacagtaaatctcctgtgatttc**atttgtataaattgtagaaaatgcaactattctaaattaacatggagaagatctgaatttttctgagaaaagtgtggtgagagtaacaagatggtgaaataaaattacagagaagtgagagaaa**aaattagaggttaattt**aattgctaatagcatgattgaagtgctgattcaaaggctgtacacatataccaacattttc**caaattgtacactataaatttgaattcactatggattgaatttttgaataaagcagtaacaaaaa**atgagtatattggctgaggaagaggaagaaagagatgaatattgacacttgaataatcacggact**cctgaaaatacacacatgtgaacactgattgcatatttcagg**taaacactagaaaaagcaaagtcacataaagttgttatgacag**gtgggacatcctgaaaacctcactaggcatgtcccac**atcgcc**ctggagctgtctcaggggagcagtctcctccag**tgtttagaggcacagacacaggtaatagggctaactctggccag**atgtgtgatattggacacat**tgcacaactgctctgttatgtatgtaattcatcttctctacaaatgtaacattgacacttgcactgaatatattctgcaaatgtgtaaacattaaataaga**tgatgactgctaattgatcatca**aggcacaatcacataatctgaagttatattttcctgagagataggattacctccaatgttttctgggatgctctcatcttctctgggcactgccctctcctcagctgtcccaccacagagcttgctatatagtaggagacatgcaaatagggccctccgtctgctgataaaaaccagcccagccctgaccctgcaAC**TCTGGGAGAAGAGCCCCAGCCCCAGA**ATTCCCAGGAATTTCCATC**TGGTGATCAGCACTGAACACAGAGGACTCACCA**TGGAGTTTGGGCTGAACTGGGTTTTCCTTGTTGCTATTATAAAAGGTGATTTATGGCGAACTAGAGACACTGAGAGGACGTGAGTGAGATAAGCAGTGAATATATGTGGCAGTTTCTGACCAGGTTGTCTCTGTGTTTGCAGGTGCCCAGTGTGAGGTGCAGCTGGTGGAGTCTGGGGGAGGCTTGGTACAGCCTGGGGGGTCCCTGAGACTGTCCTGTCCAGACTCTGGATTCACCTTCAGTAACCACTACATGAGCTGGGTCCGCCAGG**CTCCAGGGAAGGGACTGGAG**TGGATTTCATACATTAGTGGTGATAGTGGTTACACAAACTACGCAGACTCTGTGAAGGGCCGATTCACCATCTCCAGGGACAACGCCAATAACTCACCGTATCTGCAAATGAACAGCTTGAGAGCTGAGGACACGGCTGTGTATTA**CTGTGTGAAACACACAG**CGAGGGGAAGTCAGTGAgagcccaggcacaaacctccctgaaagggtcccagagaccaccagggggcgccaggacactgtgcacggggctgtctccagggcaggtgcaggtgctgctgagggctggctttctgtcatgg**cctggggcagcctcatcgtcaaatttccccagg**gaacttctccagatttacaattctatatta**acatttgatgtctctaaatgt**aaaacgtttttttttg**ttcctttgtttttgtaacaagaggaaacaccctcacctccacagaagccagggtgt**cactttgggggcagaaataatcctttcatggtcaggatgagagt**cctgaggaatctcagg**gaaacctggagagtgttttccaattagactcagag**cagagacctccatgggaatctctg**attagaacaggccttgagctctgatgggagccaagagagaggctcacccagggtcagggtcctt**aaaacctgatggtttt**cacagctgtcccctctcaacttgtaaaactgtgcccatctgactcagactgattcagctgaccctctttctgctgatccattttccatctctgtagacttgattctcacagttccctttcttcttctcttccct**gaaaacagaagatgtgttttc**tgtagtcaaaattccagggcttgggtctgcaggacctgggtaggctgaggggactttctcactcaccattgtctggacactcctgttgtcttctgtgcatggaggcatttggaaaatgtagtggacattagccatgaagagaataatactagttttctccaatgggatattgatgtagagctgatcttgtgcttctcacactatctgagtttggactctcacctgtgactttgaga**agagctgaggatgggcactccattgtgctgtgagctct**gggtaacaataattgtagaatctggctaggcagtttaaggtcaatactactggccttcgggaaagacaggctggaattcctgggaagatctgca

Uppercase: IGHV3OR16-17

Lowercase: Flanking sequence[1000bp]

Red & Bold & Underline: Stem-loop [24]

Blue: Heptamer[41]

Green: Nonamer [9]

id-IGHV7-40-2[V_gene_segment]

taagaaagacagcatgctactagtagaaaaaaatagcccagaaagaaagccaaatatctaaaaccaact**gttgtttgacaaaactgacaaaaatatacactggagaaacaac**cctctattcaataagtggtgccgggaaaattaggtggcttatg**tggaagaatagaacgagacttcca**tatcaccgtagacacaaatta**actgaatatggattcagt**gtttaaa**tttataaactaaaactataaa**aatacttgaataaaatctaaaaagagtcctctggacattggtctaagcaaacaatatacgactaagacttcaaaagcaaatgcaataaaaacagaagtagacaaacagga**tttagttgaactaaa**ggtcttctgcacagcaaaagaaataaccaacatggtgaccagacaatctgcaaatggaaaaaatatctggaatctattcatcttgcaaagggctaatatatagaatctacaagtaactcaaataagtcaactaaaaattacaaataacttcattaaagaatagacataaacagacatttatcaaaagaatacacagaagtggccaacacaaataagaaaatatactcagcatcaccaatcatctgataaatgtaaattagaaacaacatgatacggcatcttccaccagtcagaatg**gctgttattacaaataaaaacagc**aggttttgcagacaaacataggaaaaataatgatttatatatgcttggtgagaatgtaaattagtacaacctccatgaaaaacaacatagaaatttctcaaagaactaaaattagaattaccatttcttccagtaagctgtcccagtaggcatgttcctcccaaacttttatatcagagaatgttgcctgcactcatatgtttatttcaacaccattttcaatagaaaagtcaaataatctaagtgtcaatcagtgg**atgattagataaaatatgatatatgtaaatcat**ggaatactatgcagccagTATGGTATGAATTCAGTGTGACCAGCCCCTGGACAAGGGCTTGAGTGGATGGGATGGATCATCACCTACACTGGGAACCCAACATATACCAACGGCTTCACAGGACGGTTTCTATTCTCCATGGACACCTCTGTCAGCATGGCGTATCTGCAGATCAGCAGCCTAAAGGCTGAGGACACGGCCGTGTATGACTGTATGAGAGAcacagtgtggaaacccacatcccgagggagtcagaaaccccgagggaggaggcagctgtgctgagctgaggcagtggtgcagcagtttctttaacttccatatgatctcattttgcatcatcttctacttttatattagctaagaacttggggtagacaggtgctcctaagagatccttaacttgcccattttgatgggttttccagaagacgtgagaagccactttgttagcaaagcatcccaaatccatgccctgttctagatacatgtgagcccatttcctggtctttgcttaactgacaagctctcatcagtgcacctgggctaatttcacatcaggtagaggaacgcgttataaaggaaagctaatgttgtaatagcaattcctgctt**aaaaaccttcagcttcattgttttt**gtgtaatccatcagcaaattatgttagttcaaggttctcaatgggag**tttctaataaatagaaa**ggatgtatagagcttcccctaattaaaatgaaacaattgtgaacacaacctcggtattcagccatgtctccacccttcacacccttcgccacaaaggaattttcacctctcctggaagctgggttcattttcaaattagttatttttttcaatgtaatatctcaagattatcgtatatgactattttagcagaaagtgaattatgggaacttgaactaaacaactgaaaacaaattcacaactaattaaacaagatgccagaatgtgattggctccaggctttgtaattcagcagtt**catgtacccagactggaaatttacatg**tcttcttgttaccttcacagca**cagtcaactcccattatgtaagaaatggtgactg**cattcccaagggttatgcatagatatgaaaatagactgggtaaggtgag**gagttgattgtttaaattcccctctgaagaagcagcatcaactc**aacaaaccacctctcttcactctgtgactagagctatgtcacagg

Uppercase: IGHV7-40-2

Lowercase: Flanking sequence[1000bp]

Red & Bold & Underline: Stem-loop [12]

Blue: Heptamer[24]

Green: Nonamer [4]

id-TRBV5-8[V_gene_segment]

gaactgggaggagcagactagcaaaagcctagactcctgaaaacagagcattaagggcaattctggtgaaggctcagggggaaatgagga**acaaggtatcggaaattgaagtaaatgccatccttgt**ggtaagtagcaaaatccttggcaaaactgtgttctgttctaggaatttattgaatataaaaattatgagtcattcgctaggatatctgctgaaagaaatatctaagtggctaagcattcaggctaacgtgtgactactttcaggcaccaggaaatttaaccaagcaagaagggagccaagggaatagattttgcaaaccagcac**agatggtgaccctacctccctctgctgtcccatct**ccaataagaca**aattctgtgctgttagctgttaaagccctagaatt**atttctgaggaa**tctgtactcttcataatttacaga**caacccagctctgctttggatctgatcagatggactaaatcttggggactctgcaccactggccactgaggaaagggaagag**aatgttgcctgggacaggaaaatatagtaagaaacatt**ggtgtgaatctagcatcagaaagatgatgtgaggacagcaaggaagagc**aggaacctgaagtttaatcaggctcct**cgccctctgctgatgggcatgtgtgtga**gctccagcatggagc**accgcagcactaggtcagaggagagtaagagggcgatggggcagcctgtgagctggggcag**tgtaggcagaggagtaactgtatcatcacagaagcttctgccttca**accatccctccagctctgcaggacaggtagagagtccagggtctgtggggcactagacctaaggaaggcttcctg**gggaggacacaggacagccacatcacaggatacccctccc**atcaggaaaatgaaggcccagaactcactcggc**tcttccccaggaaga**ccaagccctgaatcaggtgcagtgcc**gcctggcccactgtgccATGGGACCCAGGC**TCCTCTTCTGGGCACTGCTTTGTCTCCTCGGAACAGGTGA**GTCCTGGGCACAGGAC**AGCAGCCCCATTCTCAGCTTTCCCACCCCAGT**GTCCTCCACTTTACCTTGGGGAGGAC**CTCTGGGCTGTCTCCTGAGCTCAT**CCTCCATCTGCTTTTCCTGCAGGCCCAGTGGAGG**CTGGAGTCACACAAAGTCCCACACACCTGATCAAAACGAGAGGACAGCAAGCGACTCTGAGATGCTCTCCTATCTCTGGGCAC**ACCAGTGTGTACTGGT**ACCAAC**AGGCCCTGGGTCTGGGCCT**CCAGTTCCTCCTTTGGTATGACGAGGGTGAAGAGAGAAACAGAGGAAACTTCCCTCCTAGATTTTCAGGTCGCCAGTTCCCTAATTATAGCTCTGAGCTGAATGTGAACGCCTTGGAGCTGGAGGACTCGGCCCTGTATCT**CTGTGCCAGCAGCTTGGcacag**cccag**cagaatcactgacattctg**tatataaacttcctgccttagctttgacttgagagctgcaggccccactcagtcttcactccttcaagggaagcttttagttgtttggaaggcatgtcttgtgtcct**actgagggcacagctctcccaaccaacagagcccaggtttcctgtgccctgagt**gtgcccacttctcc**gctgcatcttcttgcagc**tt**gtcacttcctgggtaacttcagtagaagagtgac**tgctgagccccagatgtgtgctagattctttgtatttgttatgggcttaaagccttgcaacaatcttgtacatcaagcattcttattcttccttgacagatgggatgctcagggacattgagtcattttccccgttgtctcctggcttgtaaggatcagaagtaggaaacaaagtagtccatccattttccacctacccccctgctccatcatcaccttctgcatcctggtcagagtaagtcagaaccctcacactgccctctagtgaccagcagggccgcagcagaggctcagatgccttcaggggtcattactatggcc**tcctcattagcaactttgagga**atgttaaatttcacactttctaaaaatcatttatatgcattccctgtgtctttccccagtctgcagcttatattttcatttaatggtgt**tttttgatgcacaagagtttaatttaatgcaacaaaaa**ctaaaaacaattttgtttgtgtg**tagacaaaaaccgctttcccatataagtgtcta**aatatatttttctca**atttattctaggaaaatttaagtttggattttcatcactaagaataaat**gtatctgaaatatatctcatgtaaaaaataaggtggaaacctaggaatgggagcgaaggacatatgttcagtgaagatgaaatagtgtgtcaaaaagagctggaaagtagacacccggagctgatggtggagaggtaggtggagggcaaata

Uppercase: TRBV5-8

Lowercase: Flanking sequence[1000bp]

Red & Bold & Underline: Stem-loop [25]

Blue: Heptamer[22]

Green: Nonamer [5]

id-IGKV1-12-3[V_gene_segment]

ctgtccagtttcatctggatccaagtgatttctccatgtacataggc**aactgcttgataagagattcagtgtctctttcctaaagcagtt**aacagggaggctgttgcctgggtcaggatgatttccccaatcactgataaaaagtaaaagaggaaagtgtcattg**atggtgcatggcagggacatgctccat**gcagtggtcaccc**tcagtaagagagatgaactttgggaaataatactga**atggcagaaaagaaggtagactatgaaggtgcccaaaacaagaataaggtgcaacccatttagtgtctgagtactatagagacctgtcgctcttgataattgtggatctgtgactgctgcatgcatcgagaaaacacggtatcatctttgtgtatctatagtacatagtttgatctcatactggtaagaacaatggcataacaccattacctgatacttacaaatgtatgtaccatcatgtcaataaattttatttttaatttttttacataggaacaatgttaaactcacagaaatgttgcatgtatacgacaaatacccccttccctagccggaatcatataag**agtcttttgaagact**tgagaattgtaccgtctaacattttactatgtgtttcctacaaacaacaatattctcctaaataatccccatacaccaa**tgaaatacattagtccatcaactcctgaggaatatttca**aatcgtcaaaaagaaaccaaaaaatgttactcacaacaaaatagttcccagtagaaacacattctctgcagacaaatttgtgctaccctggtcttacctgggacacctggggacactgagctggtgctgagttactgagatgagccagccctgcagctgcgcccagcctgccccatcccctgctcatttgcatgttcccagagcacagtctcctgacctgaagacttattaacaggctgatcacaccctgtgcaggagtcagacccagtcaggacacagcATGGACATGAGGGTCCCCGCTCAGCT**CCTGGGGCTCCTGCTGCTCTGGTTCCCAGG**TAAGAAAGGAGAACACTAGGATTATACTCGGTCAGTGTGCTGAGTACTGCTTTACTATTCAGGGAACTTCTCTTACAGCATGATTAATTGTGTGGACATTTGTTTTTATGTTTCCAATCTCAGGTTC**CAGATGCGACATCCAGATGACCCAGTCTCCATCTTCCGTGTCTGCATCTG**TAGGAGACAGAGTCACCATCACTTGTCGGGCGAGTCAGGGTATTAGCAGCTGGTTAG**CCTGGTATCAGCAGAAACCAGG**GAAAGCCCCTAAGCTCCTGATCTATGCTGCATCCAGTTTGCAAAGTGGG**GTCCCATCAAGGTTCAGCGGCAGTGGATCTGGGAC**AGATTTCACTCTCACCATCAGCAGCCTGC**AGCCTGAAGATTTTGCAACTTACTATTGTCAACAGGCT**AACAGTTTCCCTcccacagtgttaccaacccgaacataaacccccagggaagcagatgtgtga**agctgggctgccccagct**gctcctcctgatgcctccattggctg**agagtgttgctcagatgcagccacactct**gatggtgttggtagaggggtacgtgaaatcgcctctgcaccctaattcttttctctttctcagccccaactgcacagacatagcaatgcatctcctgatttgataaatacagagatcatgacacttgaggagtctagttta**tggcttcagcttgaattcatataacacagaagaagcca**ctatagatattctaagcaggaattgtcttaatacagagaattaaagtaaactactaaagtcaaaata**aaatgtagagatgaatctctaaattt**aagttttatttgcaaagaaatatttgccaggtggggcatacaggaaaactcagtggtcttcaagatattggaagaacgaagagaaaattag**cattttatgaaaaagggaaaatg**ttacctgtggctctttgagaaagttcattggcactaggaagggttgggagctggcaagctcagactggtaagcagtggtggacaaaatgaatcctagaattatatcaagttatctcagaagttgtggacaaatttgatttcaggttacaataagccaaagcagtgaaggttgcagagaattttgtt**actgaaatgccagggattcagt**gaagactctgcagcttaccccacagaaagccaatcactaagacaacaagcattgccaaggaacaggctttaatcaggtgctgcagccta**ggagatgtgataccattctcaaatgtatctccctgaccaactgaaattaggagtttatatagcaggga**agaaatgtgggaaaacaggaattagggaggggtaaggaagataatttggtcaacaggaagcagga**ggtcagtgaggcaatcataatgggtgaaggttctgatgtctcactgtcc**cgattcagtga

Uppercase: IGKV1-12-3

Lowercase: Flanking sequence[1000bp]

Red & Bold & Underline: Stem-loop [19]

Blue: Heptamer[25]

Green: Nonamer [5]

id-IGKV1D-37[V_gene_segment]

aaaccaaaaataagatgcagcttatataatctctgggtattaaggtggcttacaggtttaaataatggtgaatctgtgagtgataccggccttgaggacttgtggtttcatttttatgcatctgtagtgaattgcgtgaccatacagtcatgtaagaatggcttaaggtcattacaaaatactttcaaatatatttagcattatcttgtaaaattgtctcttttttttaagaaggaac**catttcactctcacagaaatg**ttataagtatggaacaaagtatcctctcctttgcagagtatattcaagacttgatgttcccaccctctaaaactttactgagtttcctacaaataaaaatattcccctggctggacgtggtggctcatgcctgtaatcccagcactttgggaggctgaggcgagggggtcatctgaggttaggagttcgagaccagcttggccaacatggtgaaaccccatttctactactaaaaatacaaaattagcctggcatggtggtgcgcacctgtagtcccacctaccctggaggctgaggcaggagaaatcgatggtacctggcaggtggaggt**tgcagtgaggtgagatcgtgccactgca**ctccagcctaggt**gacagaacgagactctgtc**tcaaaaaaaaaaaaccttaattaatccccatatacaaatagaaaacattacttcatcgagtactcaagagtatttcaaatat**taacaattattaaaaaaaaaagttctttgtta**caaaacggccccaagaggaaacatgttcttacagacaaatctgtgctaccctcctctgacttgggacactggggacacga**gggactggctcagagatgagccagcccc**gcagctgtgcccagcctgccccaacccctgctgatttgcatgttcccagagcacagccccctgccctgaagtcttcttaacaggctggtcacaccccgtgcagcagtcagtcccagtcaggacacagcATGGACATGAGGGTCCCCGCTCAGCT**CCTGGGGCTCCTACTGCTCTGGGTCCCAGG**TAAGGAAGGAGAACACTAGGAATTTACTCAGCTGGTGCGCTCAGTATAGCCTGGCTCTTCAGGGAGGTCTTCTTATAACATGATGGATTGAATGGATGTTTGTTTTTATATTTCCAATCTCAGGTGCCAGATGTGACATCCAGTTGACCCAGTCTCCATCCTCC**CTGTCTGCATCTGTAGGAGACAG**AGTCACCATCACTTGCCGGGTGAGTCAGGGCATTAGCAGTTATTTAAATTGGTATCGGCAGAAACCAGGGAAAGTTCCTAAGCTCCTGATCTATAGTGCATCCAATTTGCAATCTGGA**GTCCCATCTCGGTTCAGTGGCAGTGGATCTGGGAC**AGATTTCACTCTCACTATCAGCAGCCTGCAGCCTGAAGATGTTGCAACTTATTACGGTCAACGGACTTACAATGCCCCTcccacagtgttacaaggcataacataaaccccccaaggaagcagatgta**tggggctggcctgcccca**gatactcctcctactgcctccagctgctcagagcgtttctcatattccagtcaagctttgaaagtcactgaaaagttttgatggaaggggccatgaagactctgaaactgcctttgcaaaaa**tcataactgagaaaattatga**cggtggaagatatt**acacagccagctctgtgt**gaattactctttcactactgctattgccgtgtcttgataaatcgactctgtctaagcagcaggcaaggtgaaccccttgggcgattacaactcctctataccttaactgtttttccttctcatcccctgcagcacaaacatggtaatactgttcctgatttcattaagaaaagagacgattacacctgagtcttggctatggtgtgagttggaatcaataccacaaaggaaaacctactcttggaattccaagtctatttttttttttctaaatacacgcaatgagagtctaaactac**agcctctgggaggct**tgggagcatggtactcaaagatgcaaataattagaaaacgat**tctctggatctccccaagaatccagaga**gcatctggtgcagaaggtataattgccaatcatgtggtcctcagacatgtctgggaagcccatgttcggggctgttgatgcgctgcctggaggaccttcctgattttctatggcatctgcttacaattgaccagccccctattcaaagcttgccgtgttgcatctcaactcttgtccgtgggaccccaagtctctagcc**acatccaattcttcacagaagctggatgt**atgtgccagggcaaggagaactgggtgggaggtgagt**ctgtgggctccacag**cttcacgtgtttatgaggtaggagttattaagatattattttaggcagatagagaggaaaaaagggatccttgggagacttttcacagctgttgt

Uppercase: IGKV1D-37

Lowercase: Flanking sequence[1000bp]

Red & Bold & Underline: Stem-loop [15]

Blue: Heptamer[29]

Green: Nonamer [4]

id-IGHV3-54[V_gene_segment]

tgtctacattctcaaataactctgctgagtgaaagtagctgaagaattaagagtgcaattcataaacttctaattgtataaactgcaaaaggtccaactattctaaagtaacagagcagatttgaaatttgtgagaaacgggtgttgaaa**gtaattggctggtgagatgaaattac**agagaagtgacagaaagatttaggggttaacttaattgtacacaacctgattaaagtt**tgcacacatacgttaccattttccaaattgtgca**gtgtagatttg**aattaattattaatt**gtacttaaaaaaagcagtaacaaa**taaacacatgaatatgttta**ctgaggaggaacaaaaaatagatgggtatgaacactggaaacatctcagactcttgaaagtacacaggcttgaacactggttctctccgtatacttccggtaaacggctgaatacactaaaagaaaacagagatgtcctggcaggggtggaatcctgcagacctcactaggtgtgtcccacactgccctggagttgtctcaggggagcagtctcctctagtggtcagaggcacaggctgagataatggggttaactctgtccagctgtgtgactttgaatgcattgtataaacactctgttctgtatgtaatttatcttccttaaaatgcaaca**ttgacacttacattaaatgtattctacaaatatgtcaa**aaagaagatgatgactgctaaatgattatcaaggcacaatcacataatataatgatattttcctgagtgataagatgactaccaatctcg**ggggcactttgtctgctctgagccctgcccc**tcctcaggattcccatcccagagcttgctatacagtaggagacatgcaaataggtttctccctctgctgatgaccagtcctgaccccatagctctgggagagaagcgccag**ccctgggattcccaggg**gtttccatttggtgatcaggactaaagacagaggacccaccATGGAGCTTGGGCTGAGCTGGGTTTTCACTGTTGCTGTTTTAAAAGGTGAACTAGAGAGATTGAGTGTGAATGGATACACTTGAGAGAAACAGTGGATATGTCTGGAACTTTCTGACCA**GGACACCTACAAAGTTTGCAGGTGTCC**AGTGTGAGGTACAGCTGGTGGAGTCTGAAGAAAACCAAAGACAACTTGACAACTTGGGGGATCCCTGAGACTCTCCTGTGCAGACTCTGGATTAACC**TTCAGTAGCTACTGAA**TGAGCTCAGATTCCCAGG**CTCCAGGGAAGGGGCTGGAG**TGAGTAGTAGATATATAGTACGATAGAAGTCAGATATGTTATGCAC**AATCTGTGAAGAGCAGATT**CACCATCTCCAAAGAAAATGCCAAGAACTCACTCTGTTTGCAAATGAACAGTCTGAGAGCAGAGGGCACAGCTGTGTGTTACTGTATGTGAGGCAccaggtaagaagacatcagtgtgaacacagacacagaatttcctgaaataagggaggagtctgggctaaaagggcactcaggacccacagaaaacaggggaagctctagggcaggtgcagatggtcatcatgggctgctttccttgagggtct**gaggcttcctctgcatctaacagtttccctgggagcctc**tctacatttatgcttctgtggccacccctgaggtctctggacattctcatttgttgcaaaggcagatgtaagtattggaggcataaaaatgcacaggaggccagggagtctgtagacattgttaccccagaaggtcaatctcaccactagtgctggaggagggtgggagtttgatgaagctgccctaagtatcctgtggtctaagctaagtccaacgaggccatttgtgcctccctgagcacagttgtccatcagagatgtcccatgtgtcccagcagcagccatgtctcagtgtcttcactgtgcacagccattgt**ctgggaggagctcccag**gatgggtgtctttggcacacaccaggtggcgggtgttagagtgcggtgcagcagctggctgcctgttctattgggctccctga**tgctggagagatgggaggtgcattctcaggtccagca**ccctgtttgtgaatttttatataaaaccatgattttacttcattttctcagatgacatagataattaggaacagaacctgcaaagaaattgtaattttcaactttaccccaaatttattgtttcttaattctgtgtaagatccagacatattattgccttcctca**tgagaaattgttctatttaaaatgaaattagttttttctca**cattctttgtttctgttcaagtacagagatctttattaaagtaagttgggt**tctttccacacactaaccctcacctcccccagagaaaga**gcagagattttcctcactctgagtctaagggaggagctgttcctgcacgactcagagc

Uppercase: IGHV3-54

Lowercase: Flanking sequence[1000bp]

Red & Bold & Underline: Stem-loop [16]

Blue: Heptamer[41]

Green: Nonamer [3]

id-IGLVIV-66-1[V_gene_segment]

tgtaccatttctcccagtacagtctgtcaggcttcccttggctagga**aaggggaatcccccgacccctt**gtgcttcccgggtgaggtgacaccccaccatgcttcagctctttgggttgcacccactg**tccaactagtcccaatgagatgaaccaggtacctcagttgga**aat**gcagaaatcacctgtcttctgc**gtcgatctcactgggagctgcagaccagagctgttcctgtttggccatcttggaagagaccttccctgtgtattccgagccttctgcattctgccatcactgcagcctcctcatgttggatggacctgatatttttgcttttctagtaattcagaaatcctcagtggaaaggctgtttattaatatgagttatctccagacagggcaaaggtgtggtctatggatgctgtggattagcaatgaccaggctctcaggtgactccactcatgtgattttcacagaaagactgctgtctggccagtgagcagcacaggg**cactgtgagtcacactcagtgtctgcacagtg**aaaacccaaaacttgccaccccccccaaactccccagggctgggtagatggagtctgcactcactgacacagatttagcagctagatatggtcccagcacccccaccttcttccaagtcctgtgtgccagacaccccgacaacagctcccagatactgataaatccatcgggagagttgggggtaggcagtgggagcaacacacttgcatccataacccctcctttgtgaggccactgaaggggataagagaggcctggagaagccattccagctctggacctcaggagggctgtgtccactgtgacctgggcacctattctcctgtgatattctatcattt**ctcaggtagggacaggtctcacgtcctggggaagccccctagcctgag**ttcttgccctgtgtttcactctcaccctttgtcctgattcctcacccacgatcatctttgcttgcagGTTCCCTCTCTCAGCCTTTGCTTATTCAGCTGCCCTCCCTCTCTGCACATCTGAGAAATGCTATAAGACTTCCCTGTACCCCAAGTAGTGGCTGCATTGTTGGAGGCTGTCACATATTCTTGTACCAGCAGAAGTCACGGAGCCTTCCCC**AGTATCTCCTGAGATACT**ACTCAGACTCAAGTAAGCACCAGGACTCTGGGGTCCCCAGTCACTTCTGTGAATCCAAAGATCCCTCGGGC**AATGTGCAGGGATTCTGCACATT**TCTGAGCAGCCTGAGATCAAGTCCGACTATTACTATTTTACATATCACAGCA**ACAGTGGCACTTTcactgt**gctccagacttacggggaagtgagattagaacctcc**cctgcattctctctgccttgtgcagg**caacaatacactgtctgggaccgagtgtggctcatcagtagcagctttgttgtttgtatttttcattcttcagtacagggtgggacctttctactggcctccattaatagtga**gagtcaaacaagacagtctgtgggtgactc**caaagt**ctaactcagggtttccttatgttagttag**cttggactactcttggctcctcc**ccagagatctcaacttctctgg**aattctcc**aggtgccacaagcactgatttcagtcttggccttgcacct**cccctgagcagcttctgactctcctctcattacatctccatttatggcctgtcacccactcaatgtgttaaaaggtaaagcattttgcaaagtacttatcctgaactttagccccaaatccacccagcttccctagcccaggccagctctgcttcttatgaggctgatgagaggta**caagggctgcgacccttg**ctgagtccattcctgaaaatgcccccaataaagcctgattagtgccaatgccacaggacattgttctgtttcctttgcaggtggtgcccagt**gaggaaagttcttcctc**ctgaatatgaatattttggagcatgaaacagctgagctgatctggggaggggaggcaaccccttgtgatggtggtcagattgcttggttgggattgtcacagacca**tgtgataatgggattatattgcatatcaca**gatgccctggaccctcaagtgtgctggggaactctggttaaagaatagaagtgatccctttgttcatattcactatgctggatgcccttttgagatcc**ctgcaggacgctaaccctctgactgcag**cagccttgccctttgatctgaagcttttatcatacattgtcctgctcaggggatttgggtggggcgttactcctatatatcatcacaagaaaggga

Uppercase: IGLVIV-66-1

Lowercase: Flanking sequence[1000bp]

Red & Bold & Underline: Stem-loop [17]

Blue: Heptamer[53]

Green: Nonamer [5]

id-LOC124905475[V_gene_segment]

atgtatttgttttcccccatgtcgcatgtgagacctgtcagtcaggaccagggttccttgtgcactcagaggtgagggctcacagagttcctctctggtttccaggaaaggtaactgcagtaatcttggtgatgagaatatcctccagtgctggcctattatagagtttacatatgaaattgtcactgcaattcacaatctactctttcacacagaagtgtacagaggtcaggccacatcctcagggtcacacattgagaatgatgaagatatgtcccacgagtctctcctaaggtctcagaaagaattccaggactcaaaaggtctcagagggcagctcccagtgccttagttaaaatggtggctcaggcctgtaatcccagcactttgggaggccaaggtgggtggatcacctgatgtcgggacgttgagactagcttggccaacatggtgaaaccttatctctactaaaaatataaaaat**tagatgggagtggttgtgcgtgtctgtactcccagcta**cttgggaggctgaggcaggagaatcacttgaacccaggaggtggaggttgcggtaagctgagatcgggccactgcactctagcctgcgcaaaggagcaaaagttcatctaaaaaatttattttaatttaaa**cacttttaaaaagtg**gcccactccctagaacagagagattccctctaaacatgatggatgtcctgaactataaattacattaagtgaatcctggtgtgtctgaactcacatgattattacgttaagctgctgttccaatctacttcctcacctgggaaaagaggagccaggacatggctagttgaggccccaggaa**gagaactgagttctc**aaagggcaaagcaagcatcctcatcccagggtgagcctaaaagactggggcctccctcatcccttttcacctctttatacaaaggcactacctacatgcaaatcctcgcttaggcacccac**aggaaaccaccacacaTTTCCT**TAAATTCAGGGTCCAGCTCACATGGGAAATACTTTCTGAGACTCATGGACCTCCTGCAC**AAGAACATGAAACACCTGTGGTTCTT**TCTCCTCCTGGTGGCAGCTCCCAGATGTGAGTATCTCA**GGGATCCAGACATGAAGATAGGGGAGGCTGCCTCTGATCCC**AGGGCTCACTGTGGGTCTCTCTGTTCACAGGG**GTCCTGTCCCAGGTGCAGCTGCAGGAGTCGGGCCCAGGAC**TGGTGAAGCCTTCGGAGACCCTGTCCCTCACCTGCGTTGTCTCTGGTG**GCTCCATCAGCAGTAGTAACTGGTGGAGC**TGGGTC**CGCCAGCCCCCAGGGAAGGGGCTGGAG**TGGA**TTGGGGAAATCTATCATAGTGGGAGCCCCAA**CTACAACCCATCCCTCAAGAGTCGAGTCACCATATCAGTAGACAAGTCCAAGAACCAGTTCTCCCTGAAGCTGAGCTCTGTGACCGCC**GCTGGGAGGTGAGTGTGAGCCCAGACACAAACCTCCCTGC**CAGGAGGCGGAGGGCGCGGACGCAGGTGCTGCTCAGGACCAGCAGGGGGCGCGGGGCCCACAGAGCATGAGGCCGGGTCAGGAGCAGGGGCAG**GGAGGGCGGGGCTTCCTCATCTGCTCAGTGGTCTCCCTCC**TCGCCAGCACCTCAGCTGTCCCCAGGGCTCCTCTTTCTTTATTGTTTGTGGTTCTGCTTCCTCACATCCTCACTGCAGGCAAGAAAGAAGGGAGAAATTTCTCTGTTTACCATTAAGGTTTTAACAGTCACTGGCACCTTTTTTTCAACAAATTCCCACAAGGCCC**ATTTTACCTTCTTGACAAATAAAAT**ATT**TCAAGGCTTCTCACTATCCCTTGA**TTGACGTCATTGGCTGCATCGTGTCCTTCTCTTAAATTCATATGCTGAAACCCC**AATCCCAGTGGCTCTGTATTTGGAGTGAGGGATT**TTAAGAGGGCGAATAAGTTAACTGTGGTCATAAATGTGGGACCCTAATCCAGTAGGACTCTTGTCCTTATGGGAAGTGGAAGACATCAGAGACCTCTCTCT**CCACATGCACACAGAGGAGAGGCCATGTGG**GGATGCACTGCATGAGAAGGTGGCCCTTTGCAAGCCTGGAAGAGGCCTCTCAAAAAACAAAATACTATCCCTTTCTGCACCCAGATCTTGGACATCAAGACTCCAGAACAATAAGAAAATTAATATTTGTTATTTGAAACAACCTAG**TCTGTGGTATCTTCTGATGGAAAGCCAAGCGGATTCACAGA**TACAATTGTGTTAGCTCTGTCTCCTGGAGGGAGAAACAGCCCACTGAGGCTGGACACATCTCTCA**GATTATTTTTATAAAGAAAAATAGATC**CAAGATGAAAACCCCACCACATTTCTGCTGAGTCATTCACTTAGCAGGCGTCTCTGAGAT**CAGCCCTGGGGGCTG**TGTTCTAGGTCATGTCTCTCACTTTCCACTATGAGAATCCATGTAGATAATGAGAACCAGCCAGATGGAGTCCTGATCCCGGCCCACCTTCTGCTGCCCCAAGTATCCCAAAGAAAACCACTCCAGCATCACCCCTGTGTCTTCTACTCTCTAAAATCCCACTAAAGCTGACCCTAAACCAGGAGCTATTTGGGGCTGGACTCTTGTTCTCTTCTTCTTAGTCTTCCATAAGGTCCAGGTGCCGAGTAAAGCCTCAGTAAAGATTGAGCTTCAGTTCCCCTAATCCGCAGCATTCCCTTACATCCTTGTCATGCATCTGTCCTCTGTGGGCGCCCAGGGCAGGTTACACAGGAATTGCCTGAAGC**TGGCCTCAGCTGATGTGCTGAGACCA**CGGGTCATGCACACGTATGATTCCAGGTCATGCGGGCTCTACTG**CAGGACAGACCTGTGTCCTG**TGGGGCACAGCCACCTGCTGACCCCGGGCCTCCTCTAATAACTCACACACCTGGTGTTTGTGTATGTCCAGATGGCCCCATGAGCACAGCACACTAGGGTGGCCAACCTCACAGGTGGTGCCGTCAGATGGTGGACAGAAAATGGAATGGACAGGCTGGTCACTCCCATCATCAGTGATTTCCCAGTG**AATTTTAAATAAATTAAACAAAATT**AACAACAAAGGGGAAGGATATGTA**CCTGTATCTGGGAGGTATACAGG**GTCCTAGAACCGTGCTGGGAGTGGGTGTGAAAGGCCCTTTTGGGTGGAAAGCCCTCCGGCCTGAGTGAAAGCTGAAGGAATGATCACTGTGCAGGTGGAGGCTGATAGGCAGCTGGCAGGGTACATTTCCTGAAGCTGTTCCTGGGTGCATGATGGTTGGATGGCCCGTGGTGATGAGGGCTGGACTCACATAGGAACAAGGAAATTATCCCATGGGAATGGCAGAAAAGAAAACGGAGATGGACACCCCTGGAGACAGCTTATTAGAATTGGTGGAGGAAAAATAGAAATTTATTCCAATGACCCCAGCATTGTCAGCTAAATATGAGTTGCATTAATAGACAGAATTATGAAAGGAAGGGAGAAGGAGTTTATGGAGGGAAAAAACAGAAGAGAAATTAAAATTAAAAACTATGCAAATCTATCAATTGGCAGATAACAGAGTGGAGCTGATGCAGCTACTTCCTTGTT**CAGGAAACCTGCCAGAGACACTTGGCATTCCTG**TAAGGCCGGCCTAGTGGTGATGAACTTGTCGACTTTTTACTTTTTTGGGAAACTTCTCAAACATCCTGCATTACTGAAGAAAAGCTTTAATCCCTGTAAGATATTTATTTGGAAGTTTTTGGCTTTTTTTTTTCAGCAACTTGAATAGAATCATGCACATTTTGATATTTGGGTGAACAACAGGCCACATACATGACAAAGGTCTCATTAGATTATAATACTGTATTTTCACTGCATTTTTTCCATGCTTAGATATGTTCATCTGTATAAACATCTACTATGGTGATTCAGCTGCCCACAGTATTTAGTACACTTACATGACATACAGATTTGTTACCTAGGAGCACTATGCTATCCCACACAGCCTGGGTGTAGCAGGGTATTTCATGTAGGTTTGTGTAAACACACTCTATGACCCTTGCACAATGACAAATTGCCTGAGGACACATTGCTCAGACGTATCCCTACTGTTTTTCATTTTTTTTTAGATGGAGTCTCACTCTGTCACCCAGGCTGGAG**TGCAGTGGCACAATCTTGGCTCACTGCA**ACCTCCACCTCCGAGGCTCAAGCGATTCTCCTG**CCTCAGCCTTCTGAGG**AGATGGGATTACAGGCATGTACCACCACGCCCAGCTAATTCTTTTTTTTCTATTTTTAGGAGAGATGGGGTTTCACCATGTTGCCCTGGCTGGTCTTGAACTCGTGACCTAGGTGATCCACACT**CTTTGGCCTCCCAAAG**TGCTGGGATTACAAGCATGAGCCACTGCGCCTGGCCAATGTATCCCTACTGTTAAGCCACAAATCATTGTATCTCATCTCTCTCCCTTTTTTGTCCCTGTTAATACTTTCATGTTGTTCGTGCATTCTTCCCCTAGCTCAGTGAACATATTTATAAGGTTTACTTTGAATTTTCTGCTAGGTAATCAATATATCTCCTTTTAATTTGTGTCAGTTTCTAGTGCTTTTTTTTTTGTTAAACAATTTGGGCCACTTTTTTCTGTTTCGTGATTATTATTATTATTTTTTTTTACTTTCTCTGTTGCTTTCTCCTCATTAGAAGGAAAAGCTACCATAATCACTCTTCACAGAATATGCTCAGGGAAA**ATGCCATCACCTATCAACCCAGGTTGTCATTGTGGCAT**CTCTTAAAACTTCATAATGGTCGAAATTTTTTTTGGTTGTTAGTGGCACTCAGACCTATAGAGGATTTTAAGTTTTGCTCCTACTCTAAATTAGATGGCATTGAAATTCATCCTGCAGGCAGTCTCTGATAAAGTGGAGGAATTGGACAAAGTTTTGTTATGCATGTGGTTCATAAAAATAATATGAGTTTAAGGTTTTCTCAACCAAGTGCTACATACTGATACTTTATAATCACTTCTTTGAATCTGTCTTGAGTTTATCCTTCGAAAATTTAGATCAGAATATAAACC**ATGTTTGAGGCTTCTTGTTTGGTGTAAACAT**CTCCTACAGGTCATTGTGTGAAAAGGCACAGAGTCTATCTGAGGAGGAGAACTCAGGGAAGCCTCACAACAACAGGGGAGACAAAACCGAGGTCACTAAAAAAATATGAAGCTTCTGGCTCCTGCAGCCTCCAGACATGCCCTGCACCCTCTCATTGCATACAAACGAATTCCTTTCTCTTTCCTGTGTGATATTTGCCAAGATTCCTCTCCTCCACTTTGACACTTTGGTATATCCATGTGTGTCCATT**CTGGTTCAGCCAGGCACACAGAACCAG**GAAGGCATCATGGCTTTTCTAGAGGGAGCTGACCTGGGTAATCAACGGTTGGAGTAGAAAGTGTCTTTGTCCATGTTTCACAGCCCTGAGGGCCATGGGAAGAAGGTCAGGAAGAGAGGATGACCAGACGGCCGGGGAAACCAGTATGAACAAAGCCACTTTCTTTCCTGGTGAAGAGGGGCATTTGGAAGCCTGGTGTCCCTGTAGGGGATGAACCACACTCACCTACCCACTGTTACTTGCCAACGA**GGTGACCCGCAGAGCAGCAGCTCCTGAGTCACC**GACAGTGGCGCCTTCCTCCTCCAAACCCTCCA**AGATCTGTCCTTCCAGGAGTGTGAACCCCAGCAGATCT**GCATGTTTTCAATGGGTCCACTTAGCTAGTGTCTGTCCTCAGCCCCATCCCTGGGATCCTTCCCTGGTTCTGTGTACAG**TGTCACAATAGTGGTGACA**TTGACAGCTGGTGCCTGCACTTTCCAGGCCAAGAAATTTGAAAGAAGCAAGTTACAGGACCAGCTTGGATTTGGGGCAAACAGACACATTTCACTCTGTGGGGAAGTGCTGGGA**AGCAGCCTTGGGTCCTCCTCAGCTCACGTGTGGCTGCT**GCTTCTCATCCCAGGTGGGACACATGAGCCCTCCCCATGCCCAGGATGGAGCTGTTTCAATCACAGCAGGATGGGAATGTGTCTTCTCTATGGGGGCTTTTGAAGCACAGCTCTCCTGTTCCTGAGATTTGAGATTTATAAAGACAAGTCCCACAGAGTGGACCTGCACAAATAAGACAGTACAGGATCCCCAGGAGAGACATCCCATATGAGAAGGGACAGGCTGGTCCCATCACTGACACTCGTCCTCAGAAACCCTGACCAGCTACTGACCTTCCCAGGTCCC**CTGTTCCTGGAACAG**TTCCTGTGTCTGCTCCTGAAAAGAT**GCCCATCAAGAGTCCCAAGGGTTCTGCCCCCTTCCTTGATGTGC**CTCTGGGAGTGGGGCTGGCTGGGCTGTGGGT**CTGCTGTGACCCTCTCCACAGGTCAGCAG**TTCAGTGCAGGCTGCACCAACTCGGTGGGGCTGGGCTGTGGCTGAccctcctgggggagcctggactcttcttaggggtatctgcagga**tcccaggttctggga**gaacagggaagccacagtgtgtgcagggccccatctccagtaaagtgtttgctgttgttctgcttgacaagggcatatccccaggcctagaggcactatgggcacacagccaatgcgctagacacagggagtaagacacagttccctctcatt**actgtgaggactctagacacagt**gtgtgtgtgtgtgtgtgtgtgtgtgtgtgtgtgtgtgtgtgtgtgtccattactgtggggactctaaccacccgtgtgtgtgtgtgtgtgtgtgtgtgtgtgtgtattaggagtttcgcagtgggctctgagacccatgattttgtaggtttagaaattatctgatgagatgttcatccaaaggaaccgacaagagattaggtgtt**ctcccaaagcccctgggag**ctcctggactcattgtgagtgtggactgatccagtgcttccggagctccagagaaggggctccctggtggtttcatagaatccttgtttggggtgtttctgcagagttcactggctttcctacaaccaatttactattgcaagcgatggtgtcagcaggacatggtgtcacgtgtcactaaagaagcattctgagccaggacacagccacttcatacagggaggaaaatgctctggg**agcccagacaggagcctctctgcagtgcaagggctgggct**gcagggggcgctcaaggcccacccagcaaagg**ccccagccccagagcaggtgcacaggaggctgggg**agggattcctctcaagatctgtgtcattcttctaaaaaaatctaaaataagtatttgacaaagactgctgaagattccataaatatcctattcaattgcaagaatttatcaatttacactggagatttctaaccctgctacataccttaatagtaagcatctggagatcaattaagcttttattttatataaataa

Uppercase: LOC124905475

Lowercase: Flanking sequence[1000bp]

Red & Bold & Underline: Stem-loop [42]

Blue: Heptamer[130]

Green: Nonamer [10]

id-IGHV3-63[V_gene_segment]

tgttagagaccagttctttacactatggcatgtaataaattccaaatggcagtacgtcattgtttacttagcttttgtgcttata**tttttcagaggaaaaa**a**aatactgtaaattgtaaatacccaatacctaacagtatt**gtatgcaa**atctgtgactgttggcagtgtcatctctgagaaacagat**aaagtttatttactatatataaaaaaagagtttggaaggtggactcctcaccaatttttgaaagagttagagaagcattagcattaattct**ttaaatgttaagactaattttatgatgtaacatataa**cctatcatggagaatgttcaatgggtgcttgagaaggatgtgtattacatggctcttggttggaaggttctgtaaatgtctttcaggtaaatttgttcaatactgttgttcaagttcagaggcttattaagaattttctttctggatttgctatacattattctaagtgaggtattaaggttttcccttatttt**tatatagttttctatttctctctatata**tatatcttaaagtttgcttaatgcattta**tatttgtatttgtactgtgtaaaaagtaaaataaaata**ataattgcttagtgagtttcatggcacaatcacgttatgaataatcatattttcccaaatgctgtcattgccactaactcctccagatacatggtatattatcca**attctgtgagttggaattatttcgttgctttgcagaat**ctttttttgttaatctagttccacttgttcaattatgctcctccgggagtctcacatctgctctgggcactgccttcttctcaggcatcccacactggagcttgcta**tagaggaggaggcatgaaaacagggccctccctcta**ctggtgaaaacaag**cccagacctggccctgcagctctggg**agaagagccacagcactgggattcccaggggtttccatgtggtaatcagggctgaacacag**agctcactATGGGGTTTGAGCT**AACCAGAATTTTTCTTGTTGCTATTTTAAAAGGTGACTCATAGAGAAATAGAGTGAGTGAGAGTGAGTGGATATAAGTGAGAAAAACAGTAGATGTGTTTGGCAGTTTCTGACCAGGACGTTTGTGTATTTTCAGGTGTTCAGTGTGAGGTGGAGCTGATAGAGTCCATAGAGGGCCTGAGACAACTTGGGAAGTTCCTGAGACTCTCCTGTGTAGCCTCTGGATTCACC**TTCAGTAGCTACTGAA**TGAGCTGGGTCAATGAGACTCTAGGGAAGGGGCTGGAGGGAGTAATAGATGTAAAATATGATGGAAGTCAGATATACCATGCAGACTCTGTGAAGGGCAGATTCACCATCTCCAAAGACAATGCTAAGAACTCACCGTATCTGCAAACGAACAGTCTGAGAGCTGAGGACATGACCATGCATGGCTGTACATAAGGTTCCAAGTG**AGGaaacatcggtgtgagtccagacacaaaatttcct**gcagaaagaagaaaggattctgggccgaaggggacactcagcactcacaaaacaggtggagccccagggcaggtacaga**aaagcagtcaagggctgctgtccttcaggatctgtgcttt**cctctgcatgtagcagttcccctcgaatcctctgcacttttatgtttctgtgcccaccatgaggtccctggattacaaaactttaatttaaaagaggaaacattcttatatgtcccaaaaacaaaagtaagtattagaggcacaaaagtgcacaggcggccaggtgaggctgtagacactgccaccccgc**aatgccagtctcacaactagcactggagagtagtgggcatt**cagtggagcttcctacctatcctgtggtccgagctaagtccagcaaggccattggtgccttccagagcacagtagtccatcaggaatctcccatgtgtcccagcagcagccttgcctcagtatctccactgtgcacagccattgt**ctgggaggagctcccag**gatgagtgtctttggcacacaccaggtggcgggtgttagagtgcagtacagcagctggctgcctggtctattgggctccctgatgttggagggatttgaggtgcattctcaggcccagcaccctctttgtgaatttttatataaaaaccctgattttacttcattttctcagatgacatagataactaaaaacagaatctgcaaagaaattgtaattttcaactttaccccaaattcattgtttcttaattctgtgcaagatccaaacatatttttgccttccac**atgagaaattgttctgtttaaactgaaatcattttttctcat**attctttctttttgcccaagtacagagatcttgttgaaaataagttgggt**tctttccacacactaaccctcacctcccctagagaaaga**gcagagattgtcctcactctgagtctaggggaggagct

Uppercase: IGHV3-63

Lowercase: Flanking sequence[1000bp]

Red & Bold & Underline: Stem-loop [17]

Blue: Heptamer[31]

Green: Nonamer [4]

id-IGHV3-29-2[V_gene_segment]

taagtttgtttttagacgtatgcaatacattttgaggtgaaacctggtggaa**ttttttctaataaattagaaaaaaataaatcatttaattgactattttattcaggttgatttgttt**aatatttgctaaaggccagttctttaagctatgacacataataaatcccaaatggcagtacctcattgtttacttagcttttgtacttatatttctcagaggaagaaccactactgtaaattgtaaatagc**caatacataattgtattg**tatgcaaatctgtgactgtttacagtgtcatctctgagaaacataaagtttatttactatatatatataaagagtttggaaggtggactcctcaccaatttttgaaagagttacagaagggctggcattacttctttaaatgttaaggttcatttt**atgatgtgacatataacccatcat**ggagaatgttcaatgtgttcttgagaaggatgtgtattacgtgactcttggttggaaggttctgtaaatatcattcagataaatttgttcaatagtgttgttcaagttcagaggcacattaggaattttctttccggatttgctatacattatggtcatgaggtattaaggtcttgtgttatttttgtattgttttctatttctttatatctcttacagtttgcttaatgcagttatatttgtatttgtacacatgtgtaaaaacaaaaatcataattgctaaatgagttttatggcacagtcacattataagtaatatttttccaaatgctaccattgccactaaactcctcctggagtctgacgtctgctttgggcactgccttcttctcaggcatcccacactggagcttgctatagaggaggagggagggcccctctcctggtgaaaacaag**cccagacctgaccctgcagctctggg**agaagagccacag**ccctgggattcccaggg**gtttccatttggtaatcaggactgaacacagagaactcactATGGGGTGTGAATTAAGCTGAATTTTTCTTGTTGGTATTTTAAAAGGTA**ACTCATAGGGAACTAGAGTGAGTGAGAGTGAGTGGATATGAGT**GAG**AGAAACAGTGGATATGTTTGCCAGTTTCT**GACCAGGATTTGTTTGTATTTTCAGGTGTTCAGTGTGAGGTGGAGCTGATAGAGCCCACAGAGGACCTGAGACAACCTGGGAAGTTCCTGAGACTCTCCTGTGTAGCCTCTAGATTCGCCTTCAGTAGCTTCTGAATGAGCCCAGTTCACCAGTCTGCAGGCAAGGGGCTGGAGTGAGTAATAGATATAAAAGATGATGGAAGTCAGATACACCATGCAGACTCTGTGAAGGGCAGATTCTCCATCTCCAAAGACAATGCTAAGAAC**TCTCTGTATCTGCAAATGAACAGTCAGAGA**ACTGAGGACATGGCTGTGTATGGCTGTACATAAGGTTccaagtgaggaaacatcggtgtgagt**ccagacacaaaatttcctgcaaaaagaagaaaggagtctgg**gccaaaggggacactcagcactcacaaaacaggt**gcagccccaaggcaggtgcagatggagggagggtaagggctgc**tttccttcaggatctgtggtttcctctgcttctaatatttcccctctgagcctctgtacatttatattttgtgcccaccatgaggtcgctggattagaaaactaatttgaaagaggaaatattctcatatgtcctaaaaacagatgtaagtattggaggcataaaaatgcataggagccgggtgtgtctgtagacactgccacccccaatgccagacccacaactagtgctggagaagggtgggagtttgatggagcttccctgatgaccccgtggtccaagctaagtccagcaaggccattggtgcctcgctgagcacagttgtccatcagggatctcccatgtgtcccagcagcagccatgcctcaatctctccactgtgcacagccattgt**ctgggaggagctcccag**gatgggtgtctttggcacacacagttgatgggtgttagagtgcagtgcagcagctggctgcctggtctattgggctcctggatattggagggatttgaggtgcattctcagggccagcacgctttgtcaatttttatatgaaaaccatgtgatttaattcattttctcagatgacatagataattaataacacaatctgc**aaacaattataattttcaactttaccccaagttcattgttt**ctgaattctgtgcaggatccagacatggtactgcccttctcat**gagaaactgttcgatctaaactgaaaccagttgtttctc**gtatactttggttctccccacatgcagagatcttgattagagcaagtttggtactttccacgcactcaccctcacctccccagataaagagcagaagttctccttagactgagtctgaggga

Uppercase: IGHV3-29-2

Lowercase: Flanking sequence[1000bp]

Red & Bold & Underline: Stem-loop [14]

Blue: Heptamer[36]

Green: Nonamer [3]

id-TRAV20[V_gene_segment]

aaagaacagatgcatagatgcctctataatgagaactgtgatactgaaggacagatgaggaggaaagacttacttttgctgtgtattatttatattgtgtaaaatatttataatttgcatggaataatctccctcgccacacacacacacacacacacacacacacacacacacacacacagtgactttc**aggtctctctgaggtccagaaatggagacct**cttttttctctctgttatgctttttcaaaacttccttccttctaatatctaccaattt**tgtgaccctgaattttggtcaca**gttggtctcaatcctgtagtcacacagaaaattatttgaagacatggatttcttaatttatgttccaaat**tctctgcacattgcatcagaga**ggtttctagacacatccacaagctatttgatcccctcaactccatcttttattttcagagctctagtctttttaaggtttcacctgctgaggttttgaggttttaaggttttaagg**tttcacctgctgaggttcttgtgcattctagtgaaa**aa**tatattttagtgagaaatata**attacagctaccagtcggattcaactcctttgtaacttctggttgcagcttagctttaaaaaccattcaaaaaacaatctagagaattaagagtccaagtaaaataaggcttttctcattgccactttaaacctcatatgcactatgcccacc**aaatatgaagaatattt**ttctttgtaggtggcataaagtgggtaccgtgggacagtactgtggccacgaggt**ggcagtggttcaatttactctcattctacaactgcc**tt**taatttcaaaagcatgaaaatta**ctagaggacttgcatttgattggttgagcagccaacagaaaggctctttcctgcattacagaatatgaactaagatacagaagtggcgcctctgagaaaagaaggttggaattatcgtaatttgtttctaggctgagataccagcATGGAGAAAATGTTGGAGTGTGCATTCATAGTCTTGTGGCTTCAGCTTGGCTGTAAGTTGGAGGTTAAGAAATGAGAACCATTCGTGCAA**AGCTGAGGAAGAGAGATGAAGATTTAATCTCAGCT**CATTCAGTTTTCCTGTTGGAATTTTT**CAAAGCTGCTAAACTAAAAGCTTTG**TATAATTTCTGATTTTGTTTTCCTGAACAGGGTTGAGTGGAGAAGACCAGGTGACGCAGAGTCCCGAGGCCC**TGAGACTCCAGGAGGGAGAGAGTAGCAGTCTCA**ACTGCAGTTACACAGTCAGCGGTTTAAGAGGGCTGTTCTGGTATAGGCAAGATCCTGGGAAAGGCC**CTGAATTCCTCTTCACCCTGTATTCAG**CTGGGGAAGAAAAGGAGAAAGAAAGGCTAAAAGCCACATTAACAAAGAAGGAAAGCTTTCTGCACATCACAGCCCCTAAACCTGAAGACTCAGCCACTTATCT**CTGTGCTGTGCAGGcacag**cgttccccaggcacctgcaacttgtatcaaaaccctgcagctgaggatctgaaatgatg**gcagaggtatctctgc**tgttcttcctcttgaaggagtatttatttaatgccc**aggacctttttccaaatggtcct**ttggagaatgaagtcatgagacaaagccagagggc**agggactactgttcattctgtccaggcatcctagacagtccct**tgtgcatgttcatgggtcagtctactcttaataacatcccagaacttccctcacataaattcagaagggccagatgtaatatgaagagtcctgg**tttccagttgcggactacttggaaa**aagactgcacaggtg**catacactccctgccgaaaaaaaaaaaatgtgtgtatg**tgtctgtgttgtctcacaataatctggtcacgttaatataagtaaatctttttcattttaatatagtcttcttgaaacccacatctccaccagccacatctctatatttctcatcccttcatcatcaaattacttgaaagaattgtctggactcattctcttaatttccattcttcagtctcattcaaacctggcccctgttagcattactccaagggaagttctcactttaacaccagcggttttggtattgataaggctatcagatatttttcagtactcatcttgtctaagatttgcagcagtctttcctcccttctc**attaaaataaatgttactttaat**caaaataatactcacatatatttacaaaactcaacagcactgtctcacaccccagtcatgtctctccattctcaggtgatggtactcacatttatctgaacatttaaagatgctgtttcttcgctttcttctctttctttttgaaattgaggtacaacacataataaggtgaataaaactttagcttacagctcaatagaattttacacttctttgtatctgggtaacaccatccagacctacatattgaaattttccattactcca

Uppercase: TRAV20

Lowercase: Flanking sequence[1000bp]

Red & Bold & Underline: Stem-loop [19]

Blue: Heptamer[31]

Green: Nonamer [2]

id-IGKV1-8-3[V_gene_segment]

caaatccatgtccagtttcatctggatccagctgatttctccatgtaagttggcaattgcttgataagggattgtctctttcctaaagatg**ttaacagggaggatggtgtctgggtcaggatgatgtccctgataa**aacgtaaaagaagaaagtggcattgttgg**tgcatgacagggacatgctccatgca**gtggtcaccctcaataagagagaagaactttggaaagtaatactcaatgacagaaaagaaggtagacaatgaaggtgcccaaaacaagaataaggtgaagtgaattta**gtctctgggtattaaagagac**ctgtagctcttgataatggtggatgtgtgagtgctgcatgcattgaggaaactcggtatcatctctctgtatctgtagtaaattgcttgatcttatagtggtaagaacaatggcataacaccattacctaatacttacaaatatgtatagcatcatgtcaataaattttat**ttttaatatttttagaaaggaacaatgttaaaa**ctcacagaaatgttgaagtataggacaaagtacctccttccctaacccgaatcatatgagagtcttttgaagtcctgagaatcatactgtttaacattttactatgtatttcctacaaacaagatattccaaataatccccatgtgccaa**tgaaatacattactccgtcgactcctgaggaatatttca**aattttcaaaaaaatacataaaaaa**tgtttctcataacaaactactctccagtagaaaca**cattcaccacagacaaatttgtgctaccctggtctttcctgggacacctggggatactgagctggtgctgagttactgagatgagccagccctgcagctgtgcccagtcagccccatcccctgctcatttgcatgttcc**cagggcacaacctcctgccctg**aagacttattaataggctggacacacttcatgcaggagtca**gaccctgtcaggacacagcatagacATGAGGGTC**CCCGCTCAGCT**CCTGGGGCTCCTGCTGCTCTGGCTCCCAGG**TAAGGAAGGAGAACACTAGGAATTTACTCAGCCCAGTGTGCTTGGTACAGCCTGGCCCTTCAGGGAAGTTCTCTTACAACATGATTAATTGTATGGACATTTGTTTTTATGTTTCCAATCTCAGGTGCCAGATGTGCCATCCGGATGACCCA**GTCTCCATCCTCATTCTCTGCATCTACAGGAGAC**AGAGTCACCATCACTTGTCGGGCGAGTCAGGGTATTAGCAGTTATTTAG**CCTGGTATCAGCAAAAACCAGG**GAAAGCCCCTAAGCTCCTGATCTATGCTGCAT**CCACTTTGCAAAGTGG**G**GTCCCATCAAGGTTCAGCGGCAGTGGATCTGGGAC**AGATTTCACTCTCACCATCAGCTGCCTGCAGTCTGAAGATTTTGCAACTTATTACTGTCAACAGTATTATAGTTACCCTcccacagtgttacacacccaaacaaaaacccccagggaagcaaatgtgtgag**gctgggctgccccagc**tgctcctcctgatgcctccatcgcctg**agagtgttcctcagatgcaaccacactct**gatggtgttggtagagggggacctgaaatcacctctgcaacccaattcctttttctttgtaagccccagctgcacagacataacaatgcctctcctgatttaataaaggcagagatcttgacacctaaggagtctagtttagggctttggttggaattcaaataacagagaagaaaccactatagatattctaagcaggaattgtcttaatacagaaaattagagtctaaactactgaagtctaaata**aaatgtagagcggaatctctaaattt**catgttttatttgcaaagaaatgtttgccaaattggacacacaggaaaactcagtggtcttcaatatattggaagtacgaagagaaggttagtgtttt**atgaaaaagggaaaatattaccttttgcat**tttcagaaagttcactggcactagtaagggttgggagctggcaagctcagactggtaagcagtggtggacaa**agtgaattaggattcact**aattatatcaattatatcaagttgtctcagaagttgcagttaaatgtgaattcaggttacaagaagccaaagcagtgaaggttgcagagaattttcttactaaaatgccaaggattcattgtacaccctgctgctcaccacgcagaaagccaatctctaaaacaagtattgccaagaaacaggctttaatcaggtactgcagccga**ggagatgggacaccatttgcgaatgtatctccctgaccaactaaaattaagactttatataacaggga**caaaatgtgggaaaacaggaatttgggaggggtaaggaagataatttggtcaacaggaagcaagaggtcagttatgcaatcataatgggtgaaggttctgatgtctcagt

Uppercase: IGKV1-8-3

Lowercase: Flanking sequence[1000bp]

Red & Bold & Underline: Stem-loop [20]

Blue: Heptamer[15]

Green: Nonamer [2]

id-IGLV3-12[V_gene_segment]

atacttgctccttaatcttttgttgacttaagtttttatccattgatatattttcccagcaagtgaagacaacttagttataataaacattcacctccagggtcttggagtttgcagccccctctcattctctcacaaagccaacatttccttcacctcctgatgtgtccctggccctgacagcaccctggggatactgaggcacagcacctatgacctggcattagaactcacagccaagtagaaacttatcacatcccatggatcttccaaaactctgtgtgcacttggcaaccaagaattgcattctcctctagcactgaggggctgtgccctggggtggggcctgtactggtgcaaggcttgtgctgtcccctgtagagtgggaagtaaaaggtccctgcattgggttctggattgtgtcttcaaaatgaatttaggcctataccaaacatctcatgtctga**cataaaatatttatg**accaggacatttctagagaagcagaagaaagttaaccacct**ccctcctgagccaggatggaatggaggaggg**gactgtggaccccagataattcccccatcaccactgtgactctggtaacctcttgaacagggccaaccactatcccataggaaggactttatatcccctagaaaatacagag**aaactcagctctgagttt**ttccatgaccaactcagcccaggggcaaaagtgggcacaatctgggtaaagatgtgaa**ctcagaccatgggaccagtgggtggaggagaatcccatgggctgagggggtgggcaagcagtggtcacccc**ttttctctctctctctgtgtcccttgggactgagcccttctctggaaaccacaaagctcct**ccagcagcaacccctgactctgctgg**tttgcatcatgggctgttctctccagcaaggggataagagaggtctgggaggaacctgcctagcctgggcctcaggaagcagcatcagcagtgcctcagccATGGCCTGGACCCCTCTCCTCCTCAGCCTCCTCGCTCACTGCACAGGTGCTCTGCCCAGGGTATCACCAACCTGCCCATCCCCAGGGCTCTGGGTCCAGTGTGGCCATGACTATGAGCTCAGGAGGGC**CCTGCCTGTGGTGGGCAGG**ATGCTCATG**ACCCTGCTGCAGGGT**GA**GGGACTGGCGGAGCTGAAGTCCCCTCAAACTCTGCTCAGAGGCTTGTGAGAGCCTGAGGGG**CTGCACCTGCCAGGAGAGAGTACTGGGTTTTCAGTTCAAAGGCTCCATGCAGAGGGAAAGTCCATGGGCCACTGGGGCTAGGGCTGATTGCAGGGGATACCCTGAGGGTTCACAGACTCTCTGAAGCTTTTCCAGGACAGCAGGGCAGGGGATTTCATACGGATCTTTTACCTAAAAGCCATCCTCTCCTTTTTTTTTTTTTTTAATCTTTGCAGGCTCTGCGACCTCCTATGAGCTGACTCAGCCACACTCAGTGTCAG**TGGCCACAGCACAGATGGCCA**GGATCACCTGTGGGGGAAACAACATTGGAAGTAAAGCTGTGCACTGGT**ACCAGCAAAAGCCAGGCCAGGACCCTGTGCTGGT**CATCTATAGCGATAGCAACCGGCC**CTCAGGGATCCCTGAG**CGATTCTCTGGCTCCAACCCAGGGAACACCGCCACCCTAACCATCAGCAGGATCGAGGCTGGGGATGAGGCTG**ACTATTACTGTCAGGTGTGGGACAGTAGTAGT**GATCATCCcacggtgacacaggcagatggggaagtgagacaaaaacac**cctcccagcctcggtcaccctcttgctccagccccgggagg**cctatggatagagccatgagtgaatctggtccagttcacctggatctgagcctcccaggccgcccttccctccagccccctccaggagtctctatagagcatacatcaggcatacacatggcctggaaggaccagaatcatctaatgacttggggctgtcgtgtgagttagagaatgagggcctgggt**ggaaagacagacagaagcaacctctgtcctctgtcttacc**cctggatggtaatgggg**tggggatgaggccagtccttagaccagctatctggctcagtcccca**gaaccaaccctgctgaggtcctggcccccaggctgtgtggca**gtctgtgattcccaacagac**caaaccagag**gaggggacactgtgaagtctgcccaggtcccctc**ttcaatgtgacccacctagca**ctgctgagaagcccagcag**ctcagagctgtgcccacaatg**ggaagcgctgttggttgcagaaagcttcc**tcaagtttatgtcctttctcagatggtt**ttggtttaatcaaccaa**gatctcaagtccctgccttaatgtaagatgccactgaatgaagggcctcccagctccagagctccctgtgtggacggacacctgaggcctcagtggcaaccccattatgagtcagggtctccttctgcccactgttgcctccctcactctcttccaaatcttctgtgcatggaaatcattgcagtcaccttccagggaaccccatctaagatggccagctgtccccaacatgggctgtcagcgacccgaggagtccactatcca**ctgaaaattctggtttctgtcctaaattttcag**agtacatgttcaaatggccaatctgatggttccttgaatttttatggaatgaaaagggagcctgacatgctctaggttgagagatagttg

Uppercase: IGLV3-12

Lowercase: Flanking sequence[1000bp]

Red & Bold & Underline: Stem-loop [23]

Blue: Heptamer[48]

Green: Nonamer [6]

id-TRBV15[V_gene_segment]

aaaagaaataatatccttaggaaggccaccatggaattcattctatctgaacaatccttacataatatcatttcttataatatgtcctcaacatcgtgctctgcatatatttctccaccatctctaaaaatcctcttagcccaggcctcctcaaactttaatgtgcatgtaaattatgaccttgttaaaatgcagattctaatccaatcagtcagggttggagctaaggttccctgtattttgaacaagagtccgtccaagtgatgccaaagctgccagtccatggaccacaatgagcagccaagctctagatg**acacctacaataattacaggtgt**aaggcccctcttctcaagtatcattttgcacagttcctggactaggtc**agatacagaccctacttaccacatgtatct**ctgtattccaagtattacgtaatcatcagttttcctaaatctaggcaccaatattttaacatatattttcatggtataa**atagaaattaattttctat**agtaaatacactaatac**ctaatcacattcatgcatctttaatgtgagtag**caaaaagccaaggcactatcttaatggttattgagtgagactttagcaaatactatctttatggagaaacctctgggctgctttttaggatgtgagaaaagtgaggtaaagtgaggagaactggatgaacacgggaca**gagacagggacaggggcaaatatggggacacctgtctc**aaggaagcagcaaatgatatagaaaataaatatctgttccatccctgttccagacaagcccatgtacctgccaagtaggaagctgtgtatcacattgcaacaaggaatgaccccggccctggtaaagtcaacagcaacagtcatcacaggccaatctgcctatcagggactggagactctctaaactcccacctctcaacc**caggaatcagagcctgagacagacagatgcttcattcctg**tatggggtggtattcctgccATGGGTCCTGGGCTTCTCCACTGGATGGCCCTTTGTCTCCTTGGAACAGGTGAGTACCGGGCAGAAAGGAAATCTTTGAGCAAAGCTATCTTGTCCTCAGTCTGCACCTTTCATTCACAGCAGTAACACTGTCCTCCTTAACTCTGACTCCAAATTTGTCTTCTTTCTCTACAGGTCATGGGGATGCCATGGTCATCCAGAACCCAAGATACCAGGTTACCCAGTTTGGAAAGCCAGTGACCCTGAGTTGTTCTCAGACTTTGAACCATAACGTCATGTACTGGTAC**CAGCAGAAGTCAAGTCAGGCCCCAAAGCTGCTG**TTCCACTACTATGACAAAGATTTTAACAATGAAGCAGACACCCCTGATAACTTCCAATCCAGGAGGCCGAACACTTCTTTCTGCTTTCTTGACATCCGCTCACCAGGCCTGGGGGACGCAGCCATGTAC**CTGTGTGCCACCAGCAGAGAcacagagctgcagtgcttcctgctctctgt**tcataaacctcattgtttcccagatccaggtgctttctctaggacttctccctcaccacctcttacaacaataggaagtgggttggtggctgtcaatatctgtagacagaagttgagcacaaaccaataaaaaccattgacagtt**atgccaagagtggaaaagtggttacacatgcctggcat**tcagttggtggtattttcccaggatcacatttagaacccatgctgtccttttcagaagacaaataagaactttttctttctttttttcattgttttaa**tctaagtcagaggcttaga**aatataagaggtgatatgtgaagaattatggacacccaatgtctataaagtccagtctctggcattcgatgaactcttataggatttaatcctcctggagaatccacatgtttttagcctagacctggagcaactacctcattaatgggagaatgaaagcacaagtatcccaagataaaaa**tttctcagaggtaaaaccacattggagagaaa**aaaagagaagaccatacattaatc**tgctcatatctggagttggaggtactattgagatgagca**aatggaaaccataaatacataaagaaatgatattgaaaatagagtacatttggagatgaagatcatggagccatctgtatagaagagagtaataatgtcattaaggtgcataagagtgccagcaaatgtcaataagaaaaactacaggagggaagaagatgggagatcctgctccccttaacaaagataaataaaatgctgagtgagaaatattgcaacctcataaattgtcacatggaagtatgtat**ttataaattaccttataa**ttaacatccctcaatgagaacgatattctgagaaaaaaaagcagaaaatatagttatatggctatctaaaaatgtaaaattttgcacattaaattaagataatctaggaaaaatatgacaagacctat

Uppercase: TRBV15

Lowercase: Flanking sequence[1000bp]

Red & Bold & Underline: Stem-loop [14]

Blue: Heptamer[24]

Green: Nonamer [2]

id-IGHV4-4[V_gene_segment]

cagaggtgagggctcacagaattcctctctggtttccaggaaaggtaactgcagtaatcttggtaatgagaatatcctccagtgctgacctattatagagtttacatatgaaattttcactgcaattcacaatctactctttcacacagaagtgtacagaggtcaggacacatcctcagggtcacacattgagaatgatgaagatatgtcccacgagtttttcctaaggtctcagaaagaattccaggactcaaaaggtctcagagggcagctcccagtgccttagtt**aaaatggtggctcaggcctgtaatcccagcatttt**gggaggctaaggcaagtggatcacctgaggtcgggacattgagactagcttggccaacatggtgaaaccttatctctactaaaaatataaaaattagacgggggtggttgtgcgtgcctgtactcccagctactcgggaggctgaggcaggagaatcacttgaacccaggaggcggaggttgcggtgagccgagatcgggacactgcactctagcctgggcaaaggagcaaaagttcatctaa**aaaatttattttaatttaaaaatttt**aaaaaaatggcccactccctagaacagagagattccctctaaacatgatggaggtcccgaactatacattaagtgaatcctggtgtgtctgaactcacatgattattacgttaagctgctgttccaatctacttcctcacctgggaaaagaggagccagggcatggctagttgaggccccaggaa**gagaactgagttctc**aaagggcaaagcaagcatcctcatcccagggcgagcctaaaagactggggcctccctcatcccttttcacctctttatacaaaggcaccacctacatgcaaatcctcacttaggcacccac**aggaaaccaccacacatttcct**taaattcagggtccagctcacatgggaaatactttctgagagtcatggacctcctgcac**aagaacATGAAACACCTGTGGTTCTT**CCTCCTCCTGGTGGCAGCTCCCAGATGTGAGTGTCTCAAGGCTGCAGACATGGGGGTATGGGAGGTGCCTCTGATCCCAGGGCTCACTGTGGGTCTCTCTGTTCACAGGG**GTCCTGTCTCAGGTGCAGCTGCAGGAGTCGGGCCCAGGAC**TGGTGAAGCCTTCGGGGACCCTGTCCCTCACCTGCGCTGTCTCTGGTGGCTCCATCAGCAGTAGTAACTGGTGGAGTTGGGTC**CGCCAGCCCCCAGGGAAGGGGCTGGAG**TGGATTGGGGAAATCTATCATAGTGGGAGCACCAACTACAACCCGTCCCTCAAGAGTCGAGTCACCATATCAGTAGACAAGTCCAAGAACCAGTTCTCCCTGAAGCTGAGCTCTGTGACCGCCGCGGACACGGCCGTGTATT**ACTGTGCGAGAGAcacagt**gaggggaggtgagtgtgagcccagacacaaa**cctccctgcagggagg**cggaggggaccggcgcaggtgctgctcagagccagcagggggcgcgcggggcccacagagcaagaggccgggtctggagcaggtgcag**ggagggcggggcttcctcatcagctcagtgctctccctcc**tcgccaggacctcagctgtccccaggcctcctctttctttattatctgtggttctgcttcctcacatcctcgctgcaggcaagaaagaagggagaaatttctctgtttaccattaaggttttaacagtcactggcacctttttttcaacaaattcccacaaggcccattttacctt**cttgaaaaaatatttcaag**gcttcccaccatctcttgattggcgtcatcggctgcatcgtgtcctgctcttaaattcatgtgatgaaacccc**aatcccagtggctctgtatttggagagagggatt**ttaagaaggcaaataagttaactgtggtcataaatgtgggaccctaatccagtaggactcttgtccttatgggaagtggaagacatcaga**ggcctctctctccacatgcacacagaggagaggcc**atgtggggacgcgctgcatgagaaggtggccctttgcaagcctggaagagggctctcaaaaaactaaccctttctgcacccagatcttggacgtcaagactccagaacaataagaaaattaatatttgttatttgaaactacctag**tctgtggtatcttctgatggaaagccaagcagattcacaga**tacaattgtgttagctccgtctcctggagggagaagcagcccaccgaggctgacacatctctca**gattatttttataaagaaaaatagatc**caagatgaaaaccccaccacatttctgctgagtcattcacttagcaggcttctctgagat**cagccctggggggctg**tgttctaggtcatgtctctcactttccactatgagaat

Uppercase: IGHV4-4

Lowercase: Flanking sequence[1000bp]

Red & Bold & Underline: Stem-loop [16]

Blue: Heptamer[38]

Green: Nonamer [3]

id-TRBV18[V_gene_segment]

aaagggaggaaggaagatctctcaaacacaggcatcccatgcatgactacagagaatgatacagataggtatgcagaataacgggacctgaggtcagagggggaagtctttttagagtttgttctttgaa**tcacttatattgaaaattcaagtggaataagtga**aggtgacaaaagtgaattacaaatcttaaagttccctgaaaatgcttgaagccatcccttagctgaggtgtgggagtgaaaaagcaaactgagacaatgtgaaatcattttctatctcgaattttccctccagcaatgttggtgtccacttccctgtctcttaatgggtgcatctatatatcacctcttgtcaatacctctactcagcaacagccagaaaatagaaagattatataagccaataaaaggaaaataaaggaacaaaaaataagaatagtggagagatcatctacttgggaggctgaggtggggaggattgcttga**gcccaggaatttgggaccagcctgggc**aaacagcaagaccccatttatgaaaaaaatgaataacattatctgggtatggtggcttgt**agtcctagctacttaggaggctgagatgaaggact**cacttgactgatcctaggagattg**aggctgtaatgagatatgattgcactgctgcactccagcct**gggcagcggagctagatccagtcttgaaaaaaaaaaaaaaaaaaaggtgaagagggaaaaaaaagaacgagtaatagaggtgcttacttcacaacactaacctcacaa**acacacactaagctaggttgtgtgtataggtcagaacaaaagatctaccatttaactgacatata**agattggttctcgaaaccataggacccactggaaatggt**gacctcacaggaagatgcatcttgtaggaggcagctgtgaggtc**tggttccccgacgtgctgcagcaagtgcctttgccctgcctgtgggctccctccatggccaactctgctATGGACACCAGACTACTCTGCTGTGCGGTCATCTGTCTTCTGGGGGCAGGTGAGTTTTCAGCTAAAGGATCACCATCACTGGGCTTTGTTTTGGGCTCTACAGCAGATTCTCAAATATACCCTGGACCTCTGTGTTGACCAAATATTTTGTGAAAGGTATTGACATTTATATGAAAACAAATAATTGCATTATGGAGTAAAAAGCAAAATTCATGTGAGTTTTTAAATGAACGCAGGAGACTTCAAACAGGCTCCTTCACTGTAAGACATTCATCCTTCTGCCATTACAACTCTGTCTCATGCAGTCTCTTCTCCTCTGAGGTCTCTCAAATGCCGGCGTCATGCAGAACCCAAGACACCTGGTCAGGAGGAGGGGACAGGAGGCAAGACTGAGATGCAGCCCAATGAAAGGACACAGTCATGTTTACTGGTATCGGCAGCTCCCAGAGGAAGGTCTGAAATTCATGGTTTATCTCCAGAAAGAAAATATCATAGATGAGTCAGGAATGCCAAAGGAACGATTTTCTGCTGAATTTCCCAAAGAGGGCCCCAGCATCCTGAGGATCCAGCAGGTAGTGCGAGGAGATTCGGCAGCTTATTTCTGTGCCAGCTCACCACCcacattgatgcagagccacatcctctcagtccacaaacatcctccagacctgccttggaaacagcggtgggccaggaagggaaacgcgtt**acctgtacagtgaacaggt**cagctctacggtattctgcaagttgagggtggaaataggggaggattgtaggaatagcatgaaacatcccaggctacgctcagaaactaagactctgtcatccaaagaacataaagatttttaagaactagacataatttttcagtttcaacattgacttaatttgtcagtttcaaaacacttag**cagtgtaactctgttttttactccacactg**tatgtcaagttctatatttcatgctttacatcaataattttatttaatcaccaagacacacaagc**aaaggaagtgctttcttccttt**tcaccaagag**aaattatggtcctaggagttaattt**ttagaaagaaagagacagagagatgggggaagaaaggggaggggaggggacaggaggagaagggaagggaggggaagggaggggaggggaggggaggggagggaagggagaaggaaacctagtaggtggcaaaaagcatttggctcaaagggttatgatttttatctcacatcatgctgcttagcattcttttaggcataatccttgttctgttccttgctgagtacccgtgcaggtttttacggatgctaggttcaagaattcctgctataaaattatagaacatctgagagatgatggaaatagagttgga**taaatttgtctagaattta**gggaaaaggtctggcccacagatattgttcattagaaatgtgtcacctcaactttttgtcttaatgtttgtttttattctcatcactcaacagatataggccacctacagaaaaagatttgttataatttgc**tttgatgaaacaatttcataaaa**tcaaaaggcagacaaaagcaaagtcatttcaaatatatgtgcagactaagtccagcctcaccagctt

Uppercase: TRBV18

Lowercase: Flanking sequence[1000bp]

Red & Bold & Underline: Stem-loop [13]

Blue: Heptamer[22]

Green: Nonamer [3]

id-TRBV7-6[V_gene_segment]

cagctgagtacccaggatatcacacatcccagcatcttctaacaagggaaaattcttcacatgccagccttcttcctgcaccacattgactaacc**ctggaaacttcgcctttccag**aaaaaagctgtcttctcttggcccagtctagcccagtaggtttaattcatgatttatgtgaccttttaa**tttactcagaggagaacatatgatttcagagtaaa**gaatat**tcattttactttgtataagttggacctaaacagtaaactga**gtacttctcttccatcagtgtcttaaattagacaacggtaaaacaaatctttgatgcaagaactttaggtcctggtgtttttcatatggaagatcccagctatccagtccagggcatggaatagatgagcgttaataagtcactctatgcctgagatggttttctattggcatcttttctgagttacagttaacaaaggtgggaaatgatcccagaagaaaggcacgagggcagttcttgagggaatccatgtgatagga**cagtcattgggcacgtgtgactg**ggggaatggaggaggctggagcatgaatggggatggcactggggaccctgacttgcaggaaatgcaatgagctcaccactttgtgccctatgttaggggcggtgttggtgcatcctagagtaaatgtccagcagacaaaggagcaagaagcggctgtgggatgacaagataaactcagagatacagcatgagacctccggg**tccagacagctctgga**gcccaaggcgatgagccatgcattgatgttgttaaaaaggagctgataaatatttaaagcagcacccaactgtgttctaatagaaatgctgtgatcctgaggtcctggggattgagagaggaagt**gatgtcactgtgggaactgccctgtggagacaaggacatc**cctcatcctctgctgctgctcacagtgacactgatctggtaaagccctcatcctgtcctgaccctgccATGGGCACCAGTCTCCTATGCTGGGTGGTCCTGGGTTTCCTAGGGACAGGTGAGTCCTCAGAACACAAAGTAGTTTCATTTTTTTTTCTGTGTGTAGGCGTGTGGGCGTGTGTGTGTGTGTGTGTGTGTGTGTGTGTGTGTGTGTGTGTGCTGACGACAAATGTTTTCCTTATTCTGTTGCCAAATTCTATTTCCACAGATCACACAGGTGCTGGAGTCTCCCAGTCTCCCAGGTACAAAGTCACAAAGAGGGGACAGGATGTAGCTCTCAGGTGTGATCCAATTTCGGGTCATGTATCCCTTTATTGGTACCGACA**GGCCCTGGGGCAGGGCC**CAGAGTTTCTGACTTACTTCAATTATGAAGCCCAACAAGACAAATCAGGGCTGCCCAATGATCGGTTCTCTGCAGAGAGGCCTGAGGGATCCATCTCCACTCTGACGATCCAGC**GCACAGAGCAGCGGGACTCGGCCATGTATCGCTGTGC**CAGCAGCTTAGCcacagtgtggcatagtcgcctccttcctgttcacaaacctcatccttctctctccttgcacctcctagagacccttaacagaggcctctctttgctcctcacttttgatgggaaagaagtagatttggacatcggctgtcctttgggtagaaagagaccacagattcattcctgaaacacagtgactgcaaatgtaggtggtgaaaacaatcacgtcccact**gccctctaggagggc**tcggagccagctcactgctcgaaacagggagtgggtgtcttagccttggccttcagggcagacatgcatcttctataggtcttgggggctgctgtg**ctgcccacatacatgaggttgtcatgggcag**gaaacatgcctctttttcatatgttggggcatctggaaggtct**gaggctacatccccagtaacatctttcttctgaagcctc**ttctatccctgtcaccttggaagtttctgcaacaaaata**tcaaacctctcttcctgtttga**agtaaaggtctgtgcaacttttgtggtccttacgtgataaaagcaatcatgataacaataagacttcatttcttctgtctactttaagccaagagtatccttcattttgtttccatttgccattgctgctgttctgatacacacaagcatgcattcaccactgctgccggttcaccttgattcgctcagcaaatctgattattagactctcgagtgttttcattgctgtc**aaactcatttgatttgaaataagttt**cctgaggccttgaactcaagaggtgttttacttataatattgaatctattcctttttaattttcacat**tatatattgttatatagtacttgttataaatagaagtac**aatgattatattgcagtagaatcttccacctatctgtgggtgatgctgcagtttgtatccatgaaagtgaatgcactggtcagagatcatgtgatcatggatcatgggtttctgagagtcctcagggaca

Uppercase: TRBV7-6

Lowercase: Flanking sequence[1000bp]

Red & Bold & Underline: Stem-loop [15]

Blue: Heptamer[39]

Green: Nonamer [6]

id-IGHVII-74-1[V_gene_segment]

tttaactgcctaagaatttataccgtgaagtaattgcttcatttagtgttaatattaatgcccttttgacaatctttgaagctctttttttctgatatttcttccaacataaagaatgggagctttttcaccatataaagaatttttaggaagatatgatggaaagggaaactcaatgtattgttcagtgtttctgcctaacacatcaatctcttaaaatattttgctgtgtctgcatgtgataaagcagatgctaacactggagataaaagaaaaaaatgcacagaccttccacattccaaatcataaagtggtactggatgtgtttctgatgaagtgaaccatgggtcataagtgctagtgcaagtagctttggaaaatggagttcatagc**ttcttaaagttttgctagttttaatttaagaa**tgttatttactatttctttttagtacaagctttccagaag**attcttgatggtaatggaagtgtcgcatttaagaattacaaatgactcacagagttactttgtaatt**ttttctgtagggtgtacatgtcagtcact**tgacataaaagctcatatgtca**cagatagaaaaatg**aaaattgagccaatttt**aataactgcagtggaaatagtttttggcaatgagtgcttcaagacaaccacaaaataaatcggcaaaatattaaaatacagcaggaatctgatctcagtttctctgccttataagaacaaggctgtcccacatcaaaattattattcaagctctaggaaaaaatacacttttggattctggaagctaaaagctctctccttagatagtgctaaagtacttgtgaaatgcagagttgtgttcatgaaaaagatgactttattgtgatttttttttttttttttttgcctgttgacaactcaccaatctgtctcagatgtaaagccctgctctatgggttgaatgtggttgcacatgactggcagagactgattattcatatctccagGTGCCCTGTCTTTGGAGCAGCTACAGGAGTCAGGCCCAGACTAGAAGAGCCCACACTGACCTTCTGCATCACTTATGCTCTCTCTGGCCACTACATCACAAACAGTGCTTATGACTGGGCCTGAATCTG**CCAGCCCAGGAAAGGGCTGG**AATGGATGGGGTGCATTGGTagtgatggtgggggtcctactagcctgtggcaaatggaagcatctcttttttatcagactgaataatattgtagtgttttcttataccacatttacttcatccctttgtgcattaacacttaggttgtttttatatattggctgccataaatagagctgcaattaatataagagcacaggtgtctttacaagttcatgattttatttcctttccctgtatttccagagaagcgtttgctgagtcatattgtatttccatttttgatttatttatgagctgctat**actgtttttcataatggtgcaactaaaaaatggtgtagtcatcatgaaaaacagt**tttagtttttgacataatc**caaacacaatgtttg**aatctggttatctataggagtttctagtaaaattggtgaatgtaaattaaaaagcagtaagaaaaatctgcaattcagaaatagtgtacccagatggtctttctctccgaaatgaaa**gagaaataaactatttctc**aggtgagataaactaagatattttcttcacacttgaccagtc**atacaagaaatattcaagggaatttttgtat**aaaagccagcatatgaaaaccatcagcttggaaaaaaacacacaaaacctatatttcaatggtagaaacaattaaaaaaggagaaaagaaacaaacattattactacaaaacaacatcaaatcataaaaccaaacagaaagagaataaacaaagatagagatatatacatatatacaaacaatcataaaacaataaagaaaataatttgagtaaatacttataaataataaccttatgtaataaaattcctcaattaaaaatacagt**ataaaaattgattttat**taaagtcccaactctatgctgcttacaagagactcaaatcatctgagaatctacacacagactgaaagtgagtgggtggaaaaaagatatgctacaaaaatagaaaccaacagcacaaatagctctacttatat

Uppercase: IGHVII-74-1

Lowercase: Flanking sequence[1000bp]

Red & Bold & Underline: Stem-loop [11]

Blue: Heptamer[10]

Green: Nonamer [5]

id-TRAV38-1[V_gene_segment]

tttagtgctgatgagaacttctgcaatagttatttcagagcttcttgatatagc**ataaacaaggtggttttgtttat**ttgtttgtttgcgtttgttttttgagatggagtttcactcttgtcgcccaggctggagtgtggtggcacaaactcagctcactgcaacctttgcctctcaggttcaagcgattctctgacctcagcctcct**aagtagctgggattacaggcatgtactaccatgcccagctaatt**tttgtatttttaatagagat**ggagtttaaccatgttgcccgggatggtctcgaactcc**tgacctcaagtgatctgcccacctcggcctcccaaagtgctgggattacaggtgtgagccaccatgcctggccaacaagattatttttttaatccatgtaaaatgtgaacacaattacacactacatattctgcttttacttatctctgaatttagtcgatgctcttttcagccacaagcatatatcttt**tttctcattctttattttgagaaa**ttataaaaatgtaaaaaaatt**gaatataaataggcatattc**ttatcactcaaaattaagtgtccttatttcataagatttgctttgta**attttttttaataagaagaaaaat**aaaatcacagtttgccatttgttgt**gaatggagtcagaggcttctgctcaccattc**tggatcacgctctcctctccaagattaggtcgtattaaaaacaaatttgccagctccattttctctttatttttcttctaggctaactaaaagcactccagaaacattttttattttcatttgaaattataaccagaggcctat**tttctgagagaccaaaaagtattcagaaa**tgtttccctccttgaaacctagtcacccacccacctatcatagcatttcctgccctgaaggagaattctcaccaagcacagaggagaacccatcagagcaggagacttttcactctgcaggggagcgctgtcagcATGACACGAGTTAGCTTGCTGTGGGCAGTCGTGGTCTCCACCTGTCTTGGTAAGGAAGATGTGGTTCTTTCTTTATAAAGGCATCTAATGGGGCTGTGAAGCAGGAGGGATTAACAGAACAGCAACTGCATTGCCTCTTAGAGCCCAGCTACCTGG**AGGCAGAAATATGCTGCCT**GAGGCTAGAATCACAAAGAAAGGTGTGTGGGAATGGGGAATGGGAATCAGCT**TGTGAGTCTGTTTCTCACTCACA**CGTGATATCCCAGTGAAGTCCTTTCTCTCCTATTTTCCACAGAATCCGGCATGGCCCAGACAGTCACT**CAGTCTCAACCAGAGATGTCTGTGCAGGAGGCAGAGACTG**TGACCCTGAGTTGCACATATGACACCAGTGAGAATAATTATTATTTGTTCTGGTACAAGCAGCCTCCCAGCAGGCAGATGATTCTCGTTATTCGCCAAGAAGCTTATAAGCAACAGAATGCAACGGAGAATCGTTTCTCTGTGAACTTCCAGAAAGCAGCCAAATCCTTCAGTCTCAAGATCTCAGACTCACAGCTGGGGGACACTGCGATGTATTTC**TGTGCTTTCATGAAGCAca**caatgagatgagcagcagggagaggcttacagaaacctcagacctcagcatctgtgcaaaggtcacagggtgagagggaagtggtagggtaataggtatagaaaatcattgacttctcttggaaatgaaagcttctccta**tttcttctcctgtaagaaa**ctcagagagatggctgtctgcaagcttcaatccttaaagttacataaagataatgacgtgtgcagtgcattccagttttacaaggtgccttcactaaggtacttattacaggtcagatttcctaagc**attttctctgtccagaaaat**ttttataatttgcctgggcattcttacctccaggaattttaaaccttttccatttgttaattgttgcttgtccttttatttgtccatctggtccaaaactgcatctcctccgaataaggtacccattttagaacacaacaggtgtgacaacggtggcataaccacgaaggggacttgctgggcccaccagactcaagagatgac**ctaggtccactgcagtagaacctag**aagaatgatgaaaaggctttaaaatgctccgctgaagggccagctcgtgggcaacacctcagtggttggaatgctatgcttcagaaaatgataatattcgttaaaccactggctgattcatggtgtcgtggcaaaagtttagaaaccaagaaatagacatagaatagagtcttctcctatcactccctataagtcattttcagaattcgtgtttccctttcccacaaccctaggttctgctagattagaggtcctagttcccaaagaaggatgctcccttcggggattcagtaacctgcagctatggctgccacttttaggtcactctgaa**ctctctgtgccaatagcccagcaggcaaagagag**gagctattagaccagaaggggtcattgaccctgactgtcatggagcaccaggttgttgtgacacaaggagggtgaggaggaatgtggctataatgtaa

Uppercase: TRAV38-1

Lowercase: Flanking sequence[1000bp]

Red & Bold & Underline: Stem-loop [16]

Blue: Heptamer[36]

Green: Nonamer [5]

id-IGHV3-65-2[V_gene_segment]

gaattacatatgttaagtaaaataagccaa**acaaataagaatatatactgcattgatttgt**ataaattttaataaactaaatgactcagccgcaatgtatagctcaggagttgccagagaaatggtagaagaatgaaagggaaaaggaggaagaatgtagaagaacaaaaggaaatgttgagaattctcttgtccacc**ttgataacaatgatggttacattatatttatcaa**ttgtacaatttaaatatgtgaa**agtttattatctgtaaact**aaaactctcaaaatttattacaagcaaacaaatggaaacttagacaaagaaggagtgatagaaagatagaaaaaatgtatattaaatttcagaaataccta**agaacttatctgcctgaaccctagttct**caccatatttttaggtgaatgctagaatgcagcaaaatcacacgtgttctcactacagaaagtgggttccacaaccacactaggcacgcccagctctgtcctggagttggctcagggagtaattggggccagt**gatgaggagcacaggctcaggtactggggcttactcatc**cagacgtgagctcttagacacatacttagccc**tttctccatgcgtggttgacttccacatctgtacatggagaaa**caattgactccgacaaaacataatttgcacaaatatgtaaaaataaaataggatgatgaattcaaaaagtttatcacagcataattttataataagacagcatattttctgagtaccatcgttgtcaccaaattcttgcagggcacagtcatttta**tctgggtactgccttctcgtcaggcttcccaccccaga**gcttgctatatagtaggagacatgcaaatagggccatctctctgctgaagaaaaccagcccagccctgactctgcagctctgggagaggggccccagc**cctgggatgcccaggtgtttccacttggtgatcagcactgaacacc**gactaccaaccatATGAAGTCTGGGCTGAGCCGGTTTTTCCTTGTTGCTATTTTAAAAGATAATTAATGGTGAACTAGAGATACTTAGTGTGAGGGGACATGAGTGAG**AGAAACAGTGGATAGTGTGGCAGTTTCT**GATCACGATGTCTCTGGAGGTGTCCAGTGTGAGGTGCAGCTGGTGGAGTCTGGGGGAGGCTTGGTACAGCCTGGGGGGTCCCTGAGACTCTCCTGTGCAGCCTCTGGATTCACCTTCTGTTAACTACTGCATGCAGTGGATCTGCCGGG**CTCCAGGAAAGGGGCTGGAG**TGGGTAGGTTTCACTAAAAACAAAACTAATCGTGGAACAACAGAATACGCCGCGTCTGTGAAAGGCAGATTCACCATCTCAA**GCGATGATTCCAAAAGCATCGC**CTATCTGCAAATGAACAGCCTGAAAACCGAGGACACGGCCGTGTATTACTATACCAGAGACACAGTGAG**GGGAGGTCAGTGTGAGCCCAGACACAAACCtccc**tgcaggggcgcgcggggccaccgggggcggtcaaaacccactgaggacaggacaggtcccaggagcagatgctggggtagttttcctttctcatcagctggagaagtcaggtttgtatttgcaggactctgcagcattctaggctgtgacattttatcacttgtatttattatgactttattatcgttagtattta**aattttagtaattattaaaatt**atatat**aagtacattttaaatatgtactt**tgaagatataaatattcctaattatttgcactgattcttccagagttttattaacatttgttgacatcagcaactacatagctatagggacataaatttataaacatagaaatatgtatcggcaagacatggtggtctgtgcctgtaaccc**cagcttgggaagctg**aggcagaagtactgcttgaacccagaaggcggaggttgcaatgagccgtgatcgcgccactgcactccagccgggcaactgagactccatctcaaaaaactaaataaataaaaagaaagaaagaaaaaaagatgtacaaa**tacacagacctatgcatatatgtgta**ggcattcatattaaacattaagataaaataattctaaaaacatgtcctgaagaatcaaaattaatgatgaactaaatata**aattattagagtaattcataatt**gattttggttatttttaattgtttacatgttgtgattttacacaaaatattagtcatttatcc**tgataaaacaggtcaaaatgttatatca**agagtttatttactttgtatta**ctaataatataaaatgttccacatatttttag**ccatgttttacttatctgtgaaatgtgtt**tattaatttttcattttaata**tgtcacctttatctttctaatttcctggatttaattccactacagctgaatgcattttaataattattgtaatcacatttacattttttatttgaattttcatatatattttattagga

Uppercase: IGHV3-65-2

Lowercase: Flanking sequence[1000bp]

Red & Bold & Underline: Stem-loop [21]

Blue: Heptamer[29]

Green: Nonamer [4]

id-TRBV29OR9-2[V_gene_segment]

aggctggag**tgcagtggcgcaatcttggctcactgca**agctccgcctcccgggttcaccccattctcctgcctcagcctcccaagtagctgggactacaggcgcccgccactacgcccggctaattttttgtatttttagtagagacggggtttcaccgttttagccgggatggtctcgatctcctgacctcgtgatccgcccgcctcggcctcccaaagtgctgggattacaggcgtgagccaccgcgcccggcccattgtggtcaagttttacacaggcatgatttgtttcgtggcctagactgagttgaattagaatagaagaaaccatattacaagtattcaggaattaagcaaagacaaaaaaat**agttatgactgtgataact**gaaacaaaggaaaacatataagagacaatgcatttaatttgcaaattgacaaataaggtaggtgaggaaaaaaagaggcttatggaa**ttcttgagcctaacgtgcaagaa**gaacaaggatgccgtttgttgggagaggggaattgaggaataagagcccatttattggtgtgtgtatgtgttgggggaggaatgagttcagtttgggacaaccatgaagacccagcagacagttgaaatactgttatggagttcccttatattctaacctattacctttagttctttgaagtcagttctgcaccccactttttgctctacacccagacagtagcacagactggtgttcagtaatggcagatggaaaagtaatgggagaacggtggtaagtggccggtgcatgctgcaaggcttgtgcatgaagactcatcctgagtgaagagccacacagttccaccagggggaagactggtttttgcatatcctgcagaaactcaaccacctattagaggagagaaacagagcgggaaagaactacgctgtaggaaggaaagatgaacttgagtttcacttcttagtgccttttctcaggggagaggccatcacttgaagATGCTGTCTCCTGCTCCTTCTCCGGGGATTAGGTATGAGCCTTGACTTGTGAGGAGTCTGGGGTTAATGTTAATGCTGGGTA**GAGAGGTGGAAAGACTTCCTCTC**CGGAGAGTCTGAAGCCACCAGACAGAAGATGAAGGGGATATGTTAGATTAAAGGACCATAACAGAGTAATTGTCGGCACAATAA**AAAGGAGGCAGTGCCCTGGGGTCCTTT**GGGAGCTGAGGTGGTATTGGAGAGATAAAGCAGAGACTTTCCCGTCTGACCCCAAGCTTGTCGGCCCTCATGAATATCTCTCTGGAACTAGGCTCTGTGTTCAGTGCTGTCATCTCTCAAAAGCCAAGCAGGGATATCTGTCAACGTGGAACCTCCATGATGATCCAGTGTCAAGTCGACAGCCAAGTCACCATGATGTTCTGGTACTGTCAGCAACCTGGACAGAGCGTGACA**CTGATTGCAACTGCAAATCAG**GGCTCTGAGGCCACATATGAGAGCAGATTTGTCATTGACAAGTTTCCCATCAGCCGCCCAAACCTAACATTCTCAACTCTGACTGTGAGCAACAGGAGACCTGAAGACAGCAGCATATACCTCTGCAGCGTTGAAGAcacagtgcagggcacagatcaaagatctaagcaagaacctcagctcccttctacccagctc**ccctcacatgaacctgaggg**ccctgtcaaggtgggacagaagaggaaaccacagctcttgggtagacacagctgcttctgtgcttgtaggtgggcatgggtatgaatggatagactggggatgaggagagcat**ctgcaggactagctgcag**tgttctatccacagtcaggaagagtggctcctggaaagctgagggtaaa**atttcttctagaagggtgctcagaaat**acccaataaaaacaatcagatacagtttccttcaagtggcagggtcagtaat**agcctttaactgttttatctcaaggct**ctatttagctggttttctatcataatatattttggaagaactttagtcatcttgaaacaaagctggtattacattaatgacatctggttgttctccgtggataggaattataaagccctctagatgcctcagtatgacaatatgacacatgcatcccagaccaatataaaatctgcaaagttcaggaaccttctatactggtgagttttggagtggtccagaggtttggggcatgttgaaatatctcctctaatgtgagaagtaagctgctgcaccttagatcatctaccacagagaaagagatgcaatgcttagtctttggactttgaattaaactttggatatactgcatttgattgtgctactctattccgtttactatgtaatctgtaaagctggggccagggaaagagagg**gttctgtggcagaac**aagcccatgataaaagcttcctg**ctgcttgggccatagactaagcag**acccagttgctcctgaagagcctatggtagatacagatgctctgtggagctcctgacattccccagtaagagaatcacaaggcagtgtggaacaggttcactgtgccctgatttgcacttcagaccactgcatcttgggctacacttttgttgct

Uppercase: TRBV29OR9-2

Lowercase: Flanking sequence[1000bp]

Red & Bold & Underline: Stem-loop [12]

Blue: Heptamer[32]

Green: Nonamer [4]

id-TRAV26-2[V_gene_segment]

tcaaatggcatcatttgatttccctctccaaacctcttcccatttcttcctttctccttcaatgtatgccactcctaggcatcatgaagatgagaaagcatatgttaaagaggagttcaagtggcatagaaataggggtcattcaagccattggctcctgtctcaaaaccagaggcacaggc**aatgtctatctagaagacacttaggacatt**attccaggagccagtgtgggagacctgagtaccaggaaacttggcttctaacctgagttctaaatttgttctttcctttcacacttagcacagcaatattatacaattatgaaattacttttgttatatatgttgcaaataccttctttcattgtgtcatttgattttcaattttgctta**tggtaattttttttacca**aaaagaagtttctaactgttgaaattaaattatatgctactgcttcttctaagacttactcgccctgaagatgtggatagaaattcccttctttcccttacagcacagggctcacatagacttcttggaataaagggctgttcttaccagctcaatgggttgtcaaactaccaattaatggtgggttttgtccttaccataataagggatgggtaatcccatttacttctttagatgtagcctacttg**agtttcagccttgaaaatgatcttaatggaaact**ttctgaattaaagctagat**ttaaagaaaatattcttcccctctttttaaagttttatttttttaa**ttaacaaataataatcgtattttatcttct**tttaaaatgtatgtttaaa**cctgggggtgggctagacctaaccaggaagctgctctatctttagctgacgatttctggggagcactcaaattgaaacctgcctgatgtgggatgtgctgtggctgctgctttgttgcttgggacctcctctgacctagga**tcagacacagagtctga**gttctggggcctggaacctcaatgtgcacttgaacaATGAAGTTGGTGACAAGCATTACTGTACTCCTATCTTTGGGTAAGTGTTCTGGGCCTAAGAAAGCATAG**TTTTGTGGCTAATGACAAAA**GAGAGTTATCTGCTGTCTCCATGTTTTCTACCATGATTTACCAATTGCGGGAAAAAATTTGTTTCTATGTGTTTCTTGAGGGTGTCTGAGCGTCAGTGTGAGGGGTTGGTCATATCCACTACTCAGAGAAGATTCTCCAGGTCTC**ACAGCTCTCAAGCTGT**AAGGAGTTTTAGGTTCATTGAGCTCAAGGGTCATGGAAGATGGGAAATTTATTTTTGGCCTTGTGGCTCTGCATAGCTCAGGA**ACAAATAAATACCAGTAATATTTATCTTGT**CCTGTGCAACACACATAAAAACACTCAAATATGGAGTAGGGCAGGGAGGACAGTTCTGGAGCAGTGGTGAACCTGACCTTCAGATGCACCAAGAGCTGATGATCCAGAGCTAATCTGCTGTTTTTCAAC**CCCATAGGTATTATGGG**TGATGCTAAGACCACACAGCCAAATTCAATGGAGAGTAACGAAGAAGAGCCTGTTCACTTGCCTTGTAACCACTCCAC**AATCAGTGGAACTGATT**ACATACATTGGTATCGACAGCTTCCCTCCCAGGGTCCAGAGTACGTGATTCATGGTCTTACAAGCAATGTGAACAACAGAATGGCCTCTCTGGCAATCGCTGAAGACAGAAAGTCCAGTACCTTGATCCTGCACCGTGCTACCTTGAGAGATGCTGCTGTGTACTACTGCATCCTGAGAGACcacagtgggacagatggggctgcagctgtgcaatatctccctggtgatgaaagggaaggcatctaacgaggccactgcacaagaaggagcagaagtttaatagaggaagaagaaaatttaaagaagaaaagggaaaagaagtaaagaggaggtggtaaggaaatcaatgagaggatgagaggattaaaggaatgaagaaactgagaaagaatagctaaga**agtcctgaagaaaggact**gagaagcttggcgaagagatgaagaatcacaattggaattctgcagtcagg**aagaaaggaaataagaattttcccggacgctttctt**ttaatcaactcttgctttccatgtagacctcactataaa**agatgagaaccatcatct**ggtagtcccaaagcatttcttcatctagc**cccaggcaagtgcctggg**ctttgaatattatatcacattagtccttccaacctctcaataattg**aaatgaatttcagaaattgattcattt**gcccttacga**aaaatttgaaatttt**atttgttactcatccatagcacatactcttgggatatagatgatttctggaagggagcaagaaagtggtggatatgtgaatattg**atttcattttttaaatctatgttgaaat**atctctattattaaacaaaaatacaca**aatcttttttcattgtaaaagatt**caaactattaagtatgcgaagtaaaaccttagaggcccaaactgactgtctccatctccatcccattttctcttcaattcctacaataacccattttgacagtttggagagtgccctgatgatcatactccttttc**atttatgtggatactcacataaat**atctacccaattttattacat**aagttggaacataccatgtgtatttgtctgcaactt**gcttt**tatttttatttaaaaata**gttttttgaggtgtgttcataccgagatacatgtgtatgcatgtgtgtataaactggccttattctttt

Uppercase: TRAV26-2

Lowercase: Flanking sequence[1000bp]

Red & Bold & Underline: Stem-loop [22]

Blue: Heptamer[37]

Green: Nonamer [5]

id-IGLV7-43[V_gene_segment]

tatacagtcaaataattttaaacaaggttgtc**aagacttttcagtggaaacagtacagtctt**tcacaaatatagacgttctgggaaataatactgaatggcttttgctgttgttgtttttgaatggtctcaa**atgttctctcaaagatgctacagaacat**agcttattgattttgtaagttaaaaggctgagtaagactgtgagagactccctgctcatcatcattcttcttccacttccagtgactactcttatattctgtgccataagccaacagctactgaccccaggatcttttaatccttctgtaaaactgtcattaaa**aaaaaattgtctaaagtaaatttttt**aaaaaattctcacttaaaccagctgtttatccaatataaggctcaacacaaatagttacagaaagccagtaggtcattaaaacaattaatttagtcattcaaataaaaatgaacaccatgtaacactaatatcaaattatagtcacaggggcagaatttcagcaacagcaagagaaagtagcacaaataggaatgagcaaagaatagacaattaatgagtgtgaattggcctggaaaatattgtggcaatgtccaaagataaatccacagttatttgtgaaacaagtgaaagaccacgcattctcagtccaggaaagccaccagggggtggtctgggtctc**ttggactgtaagaagctcaagaggtccaa**acttcatgaaccctgcacaggtgctcattgactcatgtaaatgttagagcagctgcttcctcccacaggacaaatccacagcacacagcctccctgggctggcccctcacagtgtagctgctctcaggcctgtggacccaggtggtgataaaggcagtaaagatttgcataaagcagcacacagcacaccccctccatggagagagctcaataggagataaagagccatcagaatccagccccagctctggcgccaggggtcccttccaatatcagcaccATGGCCTGGACTC**CTCTCTTTCTGTTCCTCCTCACTTGCTGCCCAGGTTAAGAGAG**ATTTCAAATACCAGCCTTTGGAGGGATCCTTCTGTCTGCCCTTCTAATTTCTAACATGTGTCTGTTTTTTGTTTCAGGGTCCAATTCTCAGACTGTGGTGACTCAGGAGCCCTCACTGACTG**TGTCCCCAGGAGGGACA**GTCACTCTCACCTGTGCTTCCAG**CACTGGAGCAGTCACCAGTG**GTTACTATCCAAACTGGTTCCAGCAGAAACCTGGACAAGCACCCAGGGCACTGATTTATAGTACAAGCAACAAACACTCCTGGACCCCTGCCCGGTTCTCAGGCTCCCTCCTTGGGGGCAAAGCTGC**CCTGACACTGTCAGG**TGTG**CAGCCTGAGGACGAGGCTGAGTATTACTGCCTGCTCTACTATGGTGGTGCTCAGc**acagtgacagactcataagaggaaccaagacataaacctccctcggcccttgtgatgtggagattgtgtgatcatacacaccagctctcaagacagcctacatgtggaccagcc**atagaaaggggaaggaaagggtctgaattgatttctat**ccctccttgtgccctgaagtggaggaaatgtgagagtgatttgcagtaattgaatgagacaaagcaaaagttatttgttttatatgaaaaaaaaaaaccagaaacagcaggatcagatctaaaggctgagtctaaatgcatttcctccagacagaagcttcttcaaacgatgggctttctgagctaagagca**aagaaaataaactctccatgggtatattattaaagtttattttatt**gagttactttcaaagcaatccatgactattatataaagtcagaaagtattaaaaatcaccaagttctctgctaagctaccttatcccatgcaatcaaaataagtacttttcttcatttggatgcattttttatttctgtttttaatatttccacaatggtgattaaacctggtgctcattcctctggtttgatccacctttatgctcttgtcatttaaagttaatgtaagtagtgttccatttat**ttaaaaggataaaatattttaa**tagttgatcattgcattttgaactgagtaccagtgaagtattctagcttttaggtaatttcaattgtttgctagtcaatgatgttatacgttgttgtcattaaaccattcctggtcatttctgtggattttgtgaggtcagagaaaatgtctgtgcttttcaatggaaagccttgggtacattattcctgacttcccaatgcacaaaatagagaaacacaaatttcctgctcatgttagttagagcataagagattcaggtacgaggaacaatttataccccaaactacatacagtgtcttcaggcatgtattagaagtgtacttctcaaactctctgatac

Uppercase: IGLV7-43

Lowercase: Flanking sequence[1000bp]

Red & Bold & Underline: Stem-loop [13]

Blue: Heptamer[36]

Green: Nonamer [9]

id-IGLV3-1[V_gene_segment]

cagcaaagttttctttcagcatctaagc**tgtacctgaccatgaccacatactgggggtaca**ta**ggcacagcacctgtgcc**ctaccctaggagctcacagccaaggccaggaacttacagcatctcctgagtctttcaacactccgtgtgcacatgacaagggtgaagtttgattgtggaaagcaccactcagaagcaatggcaggtccctgcatgtgtgccagccttacggtgtcacctgtagagtggggtcatgagggtcactgcactgggttgaaaagtgccc**tccagagggggagctagaaccacacctaacttctggattttgccacaaaat**atttagggacaggacacccctggagtcctcaattacccaagttattctgagccagtattcaacagaggaagtaccttagatctcagaataatccctcagtcgccattgtaagtcagtccc**tggccatctccacgcaggacaaggaatggcca**catgggcaggacatcatactacctggaaaacgcacaaagaattcctctcagagttctgc**atggccagatcagctcaggagtgaggccat**aacacaacctacagtgacgatgtcaacccagatgatgggaccagaaggagaatgagaattctgtgtgctgagggtgggtctttaggggccccctctctctctgtcccttggggctgagcccttctctggaaaccacacagctcctcctg**cagcagcccctgactgctg**atttgcatcacg**ggccgctctttccagcaaggggataagagaggcc**tggaagaacctgcccagcctgggcctcaggaagcagcatcggaggtgcctcagccatggcatggatccctctcttcctcggcgtccttgcttactgcacaggtgctgcccctagggtcctagccactggtccagtcccagggctctgggtccagcct**ggccctgactctgagctcagcagggcc**cccgcctgtggtgggcaggatgctcATGACCCTGCTGCAGGTGGATGGGCTCGGCGGGGCTGAAATCCCCCCACACAGTGCTCATGTGCTCACACTGCCTTAGGGCTCTTTCATC**CCTGGATCTGTGTCCAGG**CCAGGCACGTGGGAAGATTTACTTGGAGTTCAGCTCCTCAGTTTCAAGCCTTTTCTCTCCCGTTTTCTCTCCTGTAGGATCCGTGGCCTCCTATGAGCTGACTCAGCCACCCTCAGTGTCCGTGTCCCCAGGACAGACAGCCAGCATCACCTGCTCTGGAGATAAATTGGGGGATAAATATGCTTGCTGGTATCAGCAGAAGCCAGGCCAGTCCCCTGTGCTGGTCATCTATCAAGATAGCAAGCGGCC**CTCAGGGATCCCTGAG**CGATTCTCTGGCTCCAACTCTGGGAACACAGCCACTCTGACCATCAGCGGGACCCAGGCTATGGATGAGGCTGACTATTACTGTCAGGCGTGGGACAGCAGCACTGCAcacagtgacacaggcagatgcggaagtgagacagaaaccagccacctcggcctggctcacaagacccttcc**ctctctcctgccctgtcacactgagcaggagggag**c**cttccatgtggaatggaag**tttccagtcctatccctgcccttatgttcctgagagacgggagcaagttcctgcccacctctag**gctcagcttatcccagaataaactgagc**tagtcattttgatgatcaaatgccagctcccaaaagaccccagaaaccctgatatctaagtagcaccgactctattagtatcaagggagactagccctagggtggaatcattttagtgtctcagaaggcacagggcaatggaaagtgtttatgaggtttcaggatatgcacgtgagcagttaaaggcaggtcttacaaggaaggaacctactagaattggggccca**tctgtgacatcatagcacagcctggtggacacaga**gaagggaaggtcctgaatcaagtcttgatcagtaaatatttattggataagtgagcaatttacataggtgagaactgtgtgctctcttgagcagaacacttacctggataattggtt**ttcaggaattccctgaa**gcaatgagtgacattctttattgttttcaccctcatccacctgggaaagagtatcctgga**accagcagttaacattgacacagctggt**ctcggtcctcagcacaaacattcattgcaggctgaaaagtgacaacggaagagaaaggagtttattaaatccctagacacaaacaaatccataagcagagatgagagatgcgggctcagctggcccagtcccacagg**ggtcattcctcttgtgatggaaatgacc**acatgagggt**cccccaagcggtgttggggg**gcagtcatggggaactggcctcccagggctacctgctgcttgggctgggcagaggttagagggatggaagtctggtccagtccttcccagcagcatctccaggc

Uppercase: IGLV3-1

Lowercase: Flanking sequence[1000bp]

Red & Bold & Underline: Stem-loop [19]

Blue: Heptamer[47]

Green: Nonamer [5]

id-IGHV6-1-2[V_gene_segment]

tgaggagtggtggcacccccgttcccac**cccacagatgcaacgcctgtggg**tgacgcatgtgagtactgaggagtagagggtagaactgtaggc**cccgagaaccacagaaactcggg**tgttacactctggggccatgtaaggagaa**agtgtcactggacagaaacaggcccctcctagacact**gtgtgcgccatagtcacctgtcattagctctcactcttgcagattcatgattgaggtggttaaaaaaaaaaaagctcctactcacccatcca**accccatcctggggt**gtttccaccacccttggggtttgggatgagctgcccttgcccactgtgctctgtggacctccctttagaagctcacagctccctgcactcggctccatcctgccccaccacacagaagcaaaacccctctcctttccactgcaggcttttcctggaccagaatgctgacctgctgcccttcactcccgaagtggtgggactgcctggggtggtgtgggtgttgagccttcttactctagggacctggcacctggccccaggggcacagggatggtgcatctgcctagggatgcctcctcatgccagggggtgggggttagtaccatcggccctcaggatttgttgcatgaatgagtgaatgggtgaataaatgaaggggatctgatctatgaataagggtatatagactttggttgatgtaggacgccaaatgctggaatt**tcagagtcatcacacccaggggccctgcctctga**gctcctctttgcatccaatctgctgaagaacatggctctagggaaacccagttgtagacctgag**ggccccggctcttcaatgagccatctccgtcccggggcc**ttatatcagcaagtgacgcacacaggcaaatgccagggtgtggtttcctgtttaaatgtagcctcccccgctgcagaactgcagagcctgctgaattctggctgaccagggcagtcaccagagctccagacaATGTCTGTCTCCTTCCTCATCTTCCTGCCCGTGCTGGGCCTCCCATGGGGTCAGTGTCAGGGAGATGCCGTATTC**ACAGCAGCATTCACAGACTGAGGGGTGTTTCACTTTGCTGT**TTCCTTTTGTCTCCAGGT**GTCCTGTCACAGGTACAGCTGCAGCAGTCAGGTCCAGGAC**TGGTGAAGCCCTCGCAGACCCTCTCACTCACCTGTGCCATCTCCGGGGACAGTGTCTCTAGCAACAGTGCTGCTTGGAACTGGATCAGGCAGTCCCCATCGAGAGGCCTTGAGTGGCTGGGAAGGACATACTACAGGTCCAAGTGGTATAATGATTATGCAGTATCTGTGAAAAGTCGAATAACCATCAACCCAGACACATCCAAGAACCAGTTCTCCCTGCAGCTGAACTCTGTGACTCCCGAGGACACGGCTGTGTATT**ACTGTGCAAGAGAcacagt**gaggggaagtcagtgtgagcccagacacaaacctccctgcagggatgctcaggaccccagaaggcacccagcactaccagcgcagggcccagaccaggagcaggtgtggagttaagcaaaaatggaacttcttgctgtgtcttaaa**ctgttgttgttttttttttttttttggctcagcaacag**agatcatagaaaaccctttttcatatttttgaaatctgttcttagtctaatggagattctctaatatgtgacaatgtttttctcttgctgtttttggaattctttgtctttgacttttgacaacttgacttttgacagtgtgcctcaaagaagttctattttgggttctgtgaacctcctggatctgggaagttttcagctatgatttcattaaacgtgttttctacaccatttccctactcttttggaatacccataatgcaaatatttgttcacttaattgtgtcccataaatgctggggattttcttcattcctttttactctttttttctttttattcatctgcctgaattattt**caaaagatctgtcttcaacttcagaaactcttttg**cttggcctagtctaatcttgaaggtctcaattgtacttttaatttcattcattgaattcttcaactctggaatttctgttggttcttttttatgatacttatctctttg**ttgaattcctcattcaa**atgataaattgttttcctgatttcactgaattttctatctgtacactattgtatctccctgagtttcttagagattatccttttgaattatttttctgacattctgtatatttccttatgattggggtctgctactggagaatgactgttgtctttttcaggtgtcgtgtttcctggccttttcatgtttt**atgtgttcctacgttgatttctacacat**ctggcggaccagtcatcccttgcaatttaatggagtaggttttgcaggaaaagacttcctagtacagacgg

Uppercase: IGHV6-1-2

Lowercase: Flanking sequence[1000bp]

Red & Bold & Underline: Stem-loop [13]

Blue: Heptamer[34]

Green: Nonamer [5]

id-TRBV20OR9-2[V_gene_segment]

ctagaaactagtttcactgtccagtacttagtatgacctccattacaa**gagaataaatgtaactaattattctc**acaacctcaacagtgattaacacag**aaacatacaagctaaatcgagaaatgttattagaatgttt**agaattaagtccatctatatgtgagggaaattgtc**atgagatactcacgaggtaactatctcatttagaaaatga**cgaagtggcttcaacatgattatgaagaaacattggacttccagccga**ggcactggcctgggcattatcaatatttatggtagtgcc**aacctgtaggtctagtgattaacaaccatgagttgtcttggtgtaattactttgaggatatgaaacagaagcagcagactccaatgccattcattacccaaacatacactcctttccttacaagaaactcgcttaagtcattgcaatgctctacgtctcagtgtatctttaataaatttggaagttgaattgggaaatctttaggccactttagacactgtatgggttttttgttttcttt**tgagacggagtctca**ttctgtcgcccaggctggag**tgcagtggcacgatctcagctcactgca**acctctgcctcccgggttcaagcaattctctgcctcagcctcccgaggagccgggattataggcgcccgccaccacgcctggctaatttttgtgtttttagtagagacggggtttcaccatgttggccaggttggtcttgaattcctaaccttgtgatctacccgcctcggcctcccaaagtgctggaat**tacaggcgtgagccacctcgccgacctgta**tggttctttgacaggcagagagatggatgtaacacgggtcatgagcaaagattaccaccagggggcagactagagcatccttgggattctgtgatcagtcatccctcctcgctggtgaatggagacagtggtcacaactctccccagagaaggtggtgtggggccatcacggaagATGCTGCTGCTTCTGCTGCTTCTGGGG**CCAGGTATGAGCCTCCTTCTACCTGG**GAGCTTGGGTGGGCATGTGCGTGTGTTGCCATGGTCAAGCGGTGGCCAGGCAATGTGGATTGTTTATGCTCATCGAAGGGAGAGGGAGCGGCCCT**GCTCTCTAGAGGTGTAAAGGGTAAGGTGAGAGC**CGCCAGTCAGAGGAGATAGGGGATTATAGCCCTAGGGAGATGACAGGAAGATTGCACAAAACAAACAGGATTCTCCAGGAGCTGGGAGCACGGGGAGGGAGGGAGGCTGAGCTCGGCCTGGCATCCTGTCTGACTCGGCTCCCACTGGGCTCTCCTCTCTCTGGCTTCTGTTTCAGCAGGCTCCGGGCTTAGTGCTGTCGTCTCTCAACATCCGAGCAGGGTTATCTGTAAGAGTGGAACCTCTGTGAACATCGAGTGCCGTTCCCTGGACTTTCAGGCCACAACTATGTTTTGGTATCGTCAGCTCCGGAAACAGAG**CCTCATGCTGATGGCAACTTCCAATGAGG**GCTCCGAGGTCACATACGAGCAAGGCGTCAAGAAGGACAAGTTTCCCATCAACCATCCAAACCTGACCTTCTCCGCTCTGACAGTGACCAGTGCCCATCCTGAAGACAGCAGCTTCTACATCTGCAGTGCTAGAGAcacagcgccaggaggggatcagagaccgcggcaagaacccctgcagctgctcctccaccccagcaggcccactgagtgctgagaagggaggcgtggagaatggaaaaccacagctttcctgactgagacatctgggagtgtgtgtgcagggggtggggcaggggatgggtagggggagaaaaagaaaagaagcactgtgagtgtggagtatggaggaggtgtagtgtttgagacccacgaaatggcaatagagttgggcctaaagaggtaagtgaacatggaaagttccctg**aaaattataaaacctagaatttt**gccctgactctgccacattagtcatgtgttcttgaatagatctacatataatcctagaggtacatatgaagacagttgtataaccactgtgtacttcctataataaacttcagcaattggctctctttcttgtcatcatgctgggctctcaagacaaagataaccaagaggcagccaagtagagagttatgcatgaacacattaccttaggggaggatggcaacagcattgaatagcatccactccaaatttgtgttctgtcctccttggaccaggaccacgaaaataaaatcctatc**acatttcctgtaaatgt**tgaatgccatcatgttctccacggtgtttcagttgatgaatgaacagggtaagactcggtacaaagggaggggagcacctctatg**cctaggtacacaccatacggaacgtatccctagg**gaaggtgctggtgacctggctcctgggaaaggtgtcaagggaaagagggggcagaggtacctagaaggaagacctc**agacaacatggcttgtct**aaagagttggaaagttctgccagtagctagaagacagtaagtggaacagtgaaaatcctttcctaggagaaaccaaacatttacattgaccacaaatc**tgtatgagccaatagaaaatgagcaaaacatatgatcataca**tttaatagaagaaacat

Uppercase: TRBV20OR9-2

Lowercase: Flanking sequence[1000bp]

Red & Bold & Underline: Stem-loop [16]

Blue: Heptamer[39]

Green: Nonamer [6]

id-TRBV12-3-2[V_gene_segment]

ccagtaacacaagctagtctgcaaaccaatgcccttttactttgggaagtgcttttgactgcaggggactcagaagcatgcctctg**tgccaacagcaaaaatgtgcccttctcttttgttggca**agtaacttaaccaaccaacccaaaaaaaagatctttc**tctcagctttccataatctctgaga**cgaagtaggtttggagaagtggggttacaggggaaaaagccaggtgttaatgatgaaaaaacattgaacttttctaggggtagtaat**aagatttaattcaagactagaacattttagctgcaaatctt**caagaataag**acaatattatcccctttctgttttattgt**gggactaga**gaatgtgagagaggttacattc**catgggctttgggaatttaatatggttcaaggataaacacacccaggtttttcactgcagagaagagcttcaaatataatcagttttcaggtcatcagctcagctcttgtatccctaacaatgcagttgacatgcgtcttctcagatgtctaactcctaactcactgagggatactttaagtacatataaaggactagaagcaccaagctaccagtgagaggaagaggagagtttgcagagaagctggcttgaaataagacaatgagttcatctttaaatacttgccatttgaggtgcagatggatatagttggcaggctcctatgtaaggcatgttatggagaagctaccgtgaattgataatatcaaaacaaatatccagggagcctctgcaagtgtgcatctctatttcacaccaattatagttgagttaattcctgcctgatt**catctcccagagatg**cagcctcctcttaaagaagttgggggtggtggcccat**tcagtgatgtcactga**cagatgcattctgtggggataaaatgtcacaaaattcatttctttgctcatgctcacagagggcctggtctagaatattccacatctgctctcactctgccATGGACTCCTGGACCTTCTGCTGTGTGTCCCTTTGCATCCTGGTAGCGAGTGAGTCTTCAGAATATTTGCC**ATCATCAGGCTGGGCTTCTGCATGGATGAT**CTCATATATTTTCCTTATTCTGACGCCCAATTCTGTCTTCTTTCATAGAGCATACAGATGCTGGAGTTATCCA**GTCACCCCGCCATGAGGTGAC**AGAGATGGGACAAGAAGTGACTCTGAGATGTAAACCAATTTCAGGCCACAACTCCCTTTTCTGGTACAGACAGACCATGATGCGGGGACTGGAGTTGCTCATTTACTTTAACAACAACGTTCCGATAGATGATTCAGGGATGCCCGAGGATCGATTCTCAGCTAAGATGCCTAATGCATCATTCTCCACTCTGAAGATCCAGCCCTCAGAACCCAGGGACTCA**GCTGTGTACTTCTGTGCCAGCAGTTTAGCcacagc**gctgcagaatcacccctttcctgtgcagaaaacccggtgtttccccttctccttctacctcc**cagcagtcctgggcaaagtctctgctg**ttcctccctccctatgagaaaaaagtggtttgggggtgtgaaaaagacagaaaatgagaagggat**caacataggaaaccttatgttg**gtttgaggattacaaa**atgggttttgaggattccttaaaaattgtctctgctcaaaacacat**aggagtaagataaaccttggctactgacactggagatttccctgccctcctgcatttgccatcccatgagaatggtgggggctcttgagaagggctgcattttctgaactgtgaggccctcttcattctctcctaactctaagctgcaaacagaaatttccctcacacgttttctagattgta**aaagaaagttcttcttt**actatgattgtggacgttcctttataatgccaatttcaactttacattacttcaggatttttcactactcctaaagagtgtctcaaatgtggctagagcaagcaggttagtacactagatgtaagctacctggcctggaatctaaggatccatttgtctctgttctgcgtaagatgagccgggtgctggccaaaggctgtgcacactcacagagcactgatgacgcctcctggtaaggacccacactggggtatctaaaagcagacaggcatgtccagtcttctgttgccctgtttcctttctgattatatgtccttaacacacaaatttacattttccttcttatttatatgagaagtttctatacaatacctgcaatccattctgagtggttataatttctgtgtgatattcatatttacatgctgattccttctaaatacctatcatggtatcattgacaactgaggcaaaagacccctatattttgagtgcccaaggccattgaggttttttggagctctgccataagcccaattccactgtgtcattttcc

Uppercase: TRBV12-3-2

Lowercase: Flanking sequence[1000bp]

Red & Bold & Underline: Stem-loop [14]

Blue: Heptamer[34]

Green: Nonamer [4]

id-TRAV34[V_gene_segment]

attcagtgaaaccctagagaact**ttcttgttctttctgtcatgtgaggacaccacaagaa**gactgccatctatgaaccaggaagcaggccctcaccagacatgagatcttccagcccctcgatcttagacttcccagccttcagaaccatgagaaataaatttccattgttttagtcacacagtctatggtat**tttgttatggcagcccaagctaacaaa**cacaatctcaaaataaatatgctgaaagaaaccggacg**acagaaagcagatacggtatgattctgt**ttacataaaactctagaa**aatctatagtaagagaaggcagactagagatt**gcctggggttgaataggcattgggatgatatattggaaaggtatacagaaaggattaacaaggaacacaaggaatagagacacaaggaaactttggggagtggtgcatatgttctttatcttgattgtgatgatgatttcacatttgtatgaccatgtcaa**aaattatcatattggaccttgtaaacatatgtaattt**gttatatgcaaagtgtacctcagaaaagtagtttttaagaaggctcaaaacaaaagtag**acagagataaatattgaaaaataatcactctgt**actat**agagaacagaattttttactttctct**taattatggagggtaaactttctcaagcatgcaagggtaacatatg**ttatgagattatattttctttatctcataa**gtggcatattttaatgcaataaatagatctatgtcatatataaattaactccagaaaacctgtccattattcagcatttctcagtctggccaaagagtggcagtgttaaaaaagtatttgagtaagattgtacagtgctgaacaaacaggaatccaggtatggctt**ctgattggtgcaatctcctgcaccaatgag**caaagtaacttctgctggggaagctcattcagtaaaatctgatttaactgtgttttctaaatagctaagggATGGAGACTGTTCTGCAAGTACTCCTAGGGATATTGGGGTTCCAAGCAGCCTGTGAGTATGTGGTTAGTCTGGGAGAATTGCGAGATGGTGACAGCATCCTGAGGGCAAGAGTCGAGTCATCTTCTGTTCTGGGAAGGCCCGTTTGAGAGCCAGATATGTGTTGGGAAAAAAAATGACAATACTTTGTTTTAATGTGACAATTCTGCTGCCTAGGGGAAGTGGGGAGGGGGAGAAAAACATCATCCCTACCATGACCAT**GAAAATCTTATGAGGGTCTTCTGTATTTTC**TTTACATTGTCTTTTTGAACAGGGGTCAGTAGCCAAGAACTGGAGCAGAGTCCTCAGTCCTTGATCGTCCAAGAGGGAAAGAATCTCACCATAAACTGCACGTCATCAAAGACGTTATATG**GCTTATACTGGTATAAGC**AAAAGTATGGTGAAGGTCTTATCTTCTTGATGATGCTACAGAAAGGTGGGGAAGAGAAAAGTCATGAAAAGATAACTGCCAAGTTGGATGAGAAAAAGCAGCAAAGTTC**CCTGCATATCACAGCCTCCCAGCCCAGCCATGCAGG**CATCTACCTCTGTGGAGCAGACAcacagcgatcttcaggcctctat**cagctgtctccaaacctgcagctg**g**gccacatatgctcttctgacatggggctcctgagatgtggc**tgggacctttgcc**aagacatgaagtctcagaacgttgcaattgtctt**attggaaaaatggggaggagagtaggaatgatacaagggggagctatatcactaatcccaaatttaatttgactttattgtctggtagttatgttactttttagtagatggaacatc**aaattcgaatgagaatagcatgaattt**cagtgatttatgcaagaactatgcacgaatattatgtccctgttgtgccaaggaacttctttccctttgctatttacccaccttgttaataa**aaagaaattgtgcattgtttgtcagattcttt**gtcacagattaacattgtaccaagagtgttagaatatccatactgaggagtcatgggcaatagt**tttttctaagtttgtagtatagatatagaaaaa**gacaaatgtttagacagataaataggtgatagatagattataggaagctagattattagatagatgataggtagatagatagataataggtagattagatagagtgagctagagagaggggaatatattaataaatatctaatgcagtcttagactggcctccacttctcttctcattgtgctttcattcatattctagtctcagtattcttctggccactccaaattatcagcttaaagaattgtttcccagttgtctcagta**tgaagtggagaagatgttaccttcatcctccactccacatca**gaatagtgctggcctacagtgggtgtttacaagtattaaactgatactgaataactgataagatcataggttagagagaaataat**tcacatgtttcttaaaagctatgtga**tgttgaccaacaactttaa**ctttacaagtctttattttccctctgtaaag**atgatacaatcatattattttattcatatgtt

Uppercase: TRAV34

Lowercase: Flanking sequence[1000bp]

Red & Bold & Underline: Stem-loop [21]

Blue: Heptamer[25]

Green: Nonamer [2]

id-TRBV7-8[V_gene_segment]

aggaagaaaaaaatggatgtttcctatgtattaga**atatgcatatttttcataatttctcattcttttcttgcatat**g**atttatgtttcaaaaagataaat**aggaagaatgttatgaggtgatatttatgagatgttagctgagaatttttagacgtgtttcaggacctaagttcagagtttccagaatcaaaatgaatctcaaataagacagtcaagtcatagggaaattgtagaacatgagggaaaaagagaagattttgaaagcagtcagaaaggaaagagaaatacgtcccatagaatagatggagaacaacagtaagtcaagatagtagttaaaaataaccattagctcaattatcattctccttcatttaaggtgagattgattaaaaaaacttttagaagtggaatatgaaaaatgactgctgtgagtttgctaccaaggtctctttactgaattacagttaagaaaggtgggaaatgatcccagaagaaaggcatgagggcagttcgtgagggaacccatgtg**atgggacagtcccat**tggacacgtgtgactgtgggaatggaggaggctggggcatcagtgaggatggcagaggggaccctgaattgcaggatagacaatgagctcatgccttggtgccttgtgttgggggtgctgttagtgcatcgacaggacatgcccagcagacagaggagtggctgtaggatgagaaggtgaactcagagatgtagtgtgaggcccacgggtccagaca**gcatgggagcccaaaaaatgaaccatgc**attgatagtggtaaaaagctgatacatatttaaagcagcacccaagtgtgttctaatagaaatgctgtgaccctgaggtcctggggattgagagaggaagtgatgtcactgtgggaactgccctgtggagacaaggacggcccttatcctctgcttctgttcacagtgacactgatctggtaaagcccccatcctggcctgaccctgccATGGGCACCAGGCTCCTCTGCTGGGTGGTCCTGGGTTTCCTAGGGACAGGTGAGTCCTCAGAACACCAAGTAGTTTCATTTTTTCTGTTTGTAGGTGTGTGTGTGTGAGAGAGTGGTTGTGTGTGTGTGTGTGTATGATGACTACAAATATTTTCCTTATTCTGTTGCCAAATTCTATTTCCACAGATCACACAGGTGCTGGAGTCTCCCAGTCCCCTAGGTACAAAGTCGCAAAGAGAGGACAGGATGTAGCTCTCAGGTGTGATCCAATTTCGGGTCATGTATCCCTTTTTTGGTACCAACAGGCCCTGGGGCAGGGGCCAGAGTTTCTGACTTATTTCCAGAATGAAGCTCAACTAGACAAATCGGGGCTGCCCAGTGATCGCTTCTTTGCAGAAAGGCCTGAGGGATCCGTCTCCACTCTGAAGATCCAGCGCACACAGCAGGAGGACTCCGCCGTGTATCT**CTGTGCCAGCAGCTTAGCcacagcatggcacag**ttgcctccttcctgttcacaaacctcatccttctctctccttgcagctcctagagaccctaaacagaggcctctctttgctcctcacttttcatggg**aaagaattacatctggacttcagctgttcttt**gggtagaaagagaccacagattcattcctgaaacacagtgactgaaaatgtaggtggtgaaaacaatcatgggagtc**cttggagccagctcactgctccaag**caaggagtgggtgtcttagtcttggccttcagggcagacatgcatcttctataggtcttggaggctgctgtg**ctgcccacatccatgaggttgtcatgggcaggaaacacgctcttctcctgc**atatgttggggcatctggaaggtctgaggttacatcccaaggaacatctttcttctgaagcctcttctatccgtgtcacatt**agaggttttctgcaacaaaatatcgaacctct**cttcctgtttgaagtaaaggtctttgcaacttttgaaatccttacttgataaatacagtcaggataacaataagacttcatttcttctgcctactttaagccaggtgtatccttcattttatttccatttgctattgctgctgtcctgatagacagaagtatgcattcaccaccactgccagttcaccttgattctctcaggaaatctgatttctagactctgagggttttcattgctgtccaactcatttgatttgaaataatttttctgaggccgttaagaagtgttatatt**tataatattgattctattgctgtttatttttcatattatatattgttatatagtacttgttatagatagaagtac**aatgattatattgcaatagaatcttccacctgtctgtgggtgccgctgcagtttgtatctatgaaagcgaatgcactggtcagagctgatgtgattatgaatcatgggtttctgagagtcctcggcgacagaccacttctcccagtct

Uppercase: TRBV7-8

Lowercase: Flanking sequence[1000bp]

Red & Bold & Underline: Stem-loop [13]

Blue: Heptamer[31]

Green: Nonamer [7]

id-TRBV25-1-2[V_gene_segment]

agagatataatagtaaaaacccaaagcctatgtcagcatcatgtgaacaaaaattttagtaaataccaactgtagatcttagtgaactagggcaaaggaagtggacaacactgggagaaattctgggattaa**tctttgtatctcacaatcaaaaccaaaga**cttttctgttaagaagaaatgggggaaagggcggagggagaatggcacttggtttagacattgtt**tactgcagcagcacccaaaccatttgtagcagta**ttaccaaagcatgtattggattatattttctgttctgtcatgattgatagtgcaattaaaggcagatggactaaaatcatcctaggcaaaggctaagttgataaagctgataaagcctgaacaccaagtcagtgaaaaatgagactgagaaa**gttattatcacacaataataac**accattctgaggactcagtgaattccagttactgctccaagctctttacatgtgtctatataaatagattgccccgtttaatgctcacaagttctcctgctccttgtctgatcaagccacgctgttcactggttccctct**tctgcatgttcttctctcttgggtttgcaga**gttcattcctgttcatccttcagaacccaactcaaatgccacttcctttcccaacctttttaattaggtaaaatcttgctattacaggcactca**aagctatgtagtttttatgttcatagcttacaaaacagttgcaatggaatgttttgtaag**aatattaaagacttgagttaagataagcaaagcatcaagcctactttggatgagggaagtcaacaaaaatattactcatttgatgggttaaaaggatatagatgaaactgtttcaacaggatgtctgtgggacccaggaagtttgcaggggctg**tgtcatcacacacacacaatgaca**ttatcaagagcccatcctgcttccccactactgggagacatcctctctagccccaactgtgccATGACTATCAGGCTCCTCTGCTACATGGGCTTTTATTTTCTGGGGGCAGGTAAGTCATAGACACAGTTCATTCCAAAATCTAGAAGGAATAATTCCTCTTTGTTGGGTTTGTGCCTGGCTCAGCATCAAAGTCCATCGTGAACTCTGTTACCAATTTTTGTCTCTTCCCATAGGCCTCATGGAAGCTGACATCTACCAGACCCCAAGATACCTTGTTATAGGGACAGGAAAGAAGATCACTCTGGAATGTTCTCAAACCATGGGCCATGACAAAATGTACTGGTATCAACAAGATCC**AGGAATGGAACTACACCTCATCCACTATTCCT**ATGGAGTTAATTCCACAGAGAAGGGAGATCTTTCCTCTGAGTCAA**CAGTCTCCAGAATAAGGACGGAGCATTTTCCCCTGACCCTGGAGTCTG**CCAGGCCCTCACATACCTCTCAGTACCTCTGTGCCAGCAGTGAATAcacagtgctacatggataccgacactccgcacagaaagggtcgcctctaatgtgaggacatcttgcctccagaaacctcatcttaaactacagaaacccctacaaatcttcccagactcctcagccttgagggaccagtgtgctttaagtaacagtctggactaaagactatctctggctcagtctctaagaagctgagaaatgacccagccccacagtcatcctagctgactttttccttgcatctagttgttgttgttcaattgagcttgca**caaattttatccttcaagataaatgcgataatatttg**acagttaccttatggc**tagctgaagaaagtctttgtcagcta**agaaacaggaaaaggatcctttgactttctacacaggctgtctttatctctggttcatggtttcttcctcatgttaaacttgttccttcatacacagccacacaaagtccctgccctttcctgaccccagtcctgcacctgacggcttcagttcctcaatctatgactttgaacttagaataaaattttattt**tcacttgaaataagtga**ttaatgattattttcactaatatccctctaaataaatgtaattttagctatatctgcatgaggcctaaggaaaagaaaaggctccaactttggcaaagaatagtctttaacaatgtccctggattcagccctaaatctctttggagactacccttagcattggaagactatcccacggctaggtgattcttctgtcttggatatgctga**atctcagggaagtgttctctatccacatgagat**cagacctgggaccctgaatagatttatttttcacagaagtatagactgctgtggagccctgggtgctgagctcagatgtgaaat**ctctgttttgaggttaccaaaaagtaaagtcacagag**tctctgtcctggttttgcctccttaaaaagtagctttaatatgttagaaaagaatctagaaaacatcatacata

Uppercase: TRBV25-1-2

Lowercase: Flanking sequence[1000bp]

Red & Bold & Underline: Stem-loop [14]

Blue: Heptamer[37]

Green: Nonamer [7]

id-IGKV2-40[V_gene_segment]

nnnnnnnnnnnnnnnnnnnnnnnnnnnnnnnnnnnnnnnnnnnnnnnnnnnnnnnnnnnnnnnnnnnnnnnnnnnnnnnnnnnnnnnnnnnnnnnnnnnnnnnnnnnnnnnnnnnnnnnnnnnnnnnnnnnnnnnnnnnnnnnnnnnnnnnnnnnnnnnnnnnnnnnnnnnnnnnnnnnnnnnnnnnnnnnnnnnnnnnnnnnnnnnnnnnnnnnnnnnnnnnnnnnnnnnnnnnnnnnnnnnnnnnnnnnnnnnnnnnnnnnnnnnnnnnnnnnnnnnnnnnnnnnnnnnnnnnnnnnnnnnnnnnnnnnnnnnnnnnnnnnnnnnnnnnnnnnnnnnnnnnnnnnnnnnnnnnnnnnnnnnnnnnnnnnnnnnnnnnnnnnnnnnnnnnnnnnnnnnnnnnnnnnnnnnnnnnnnnnnnnnnnnnnnnnnnnnnnnnnnnnnnnnnnnnnnnnnnnnnnnnnnnnnnnnnnnnnnnnnnnnnnnnnnnnnnnnnnnnnnnnnnnnnnnnnnnnnnnnnnnnnnnnnnnnnnnnnnnnnnnnnnnnnnnnnnnnnnnnnnnnnnnnnnnnnnnnnnnnnnnnnnnnnnnnnnnnnnnnnnnnnnnnnnnnnnnnnnnnnnnnnnnnnnnnnnnnnnnnnnnnnnnnnnnnnnnnnnnnnnnnnnnnnnnnnnnnnnnnnnnnnnnnnnnnnnnnnnnnnnnnnnnnnnnnnnnnnnnnnnnnnnnnnnnnnnnnnnnnnnnnnnnnnnngaattcaggtttgtcttatgaataatcaggattcacctccagggaacgatgaccagtgctctgattaagaacttgaaaaaaaagagttcccttgtggctaataaataatgggtctattttagaaagtctacttttca**tgatataaatcaaaactttaaaaatgtaactgtaaatttatatca**caagagaa**attatgaaagttgctcataat**gtatctatataaacttgcacttctctgttattatttcagGATCCAGTGAGGATATTGTGATGACCCAGACTCCA**CTCTCCCTGCCCGTCACCCCTGGAGAG**CCGGCCTCCATCTCCTGCAGGTCTAGTCAGAGCCTCTTGGATAGTGATGATGGAAACACCTATTTGGACTGGTACCTGCAGAAGCCAGGGCAGTCTCCACAGCTCCTGATCTATACGCTTTCCTATCGGGCCTCTGGAGTCCCAGACAGGT**TCAGTGGCAGTGGGTCAGGCACTGATTTCACACTGAAAATCAG**CAGGGTGGAGGCTGAGGATGTTGGAGTTTATTACTGCATGCAACGTATAGAGTTTCCTt**ccacagtggtacagccctgaacagaaacctccctgctgtgg**tg**ccccagctgctcacatgcactgcttgtctgggg**agcaggtcagcagcgtctctgagtctgcaaaagaggaggctgttggagaatacagggcagggtttgcttctgaggactctgcctgggactacaggtgcatgccactaaacatggctaatttttctatttttttgtagagtcggtgcttcaccatgttgcccagcctgttgtcaaaatcatgggctcaagccacccacctgacttggcctcccaacgtgctggcagt**acagtgtgagccactgtggcaggtcagcacccctgtttatgttcctgtcacctgccacag**ccttgactctcataaccaacaggaaaatgaggaggttctagggccctgtgagtaaaaaactgggatgataggga**aaggagaatggaatctcatctgaatcctcctt**ccttgcctacatttgtttaaatttattgagcaaaagg**gccagactactgattatttctggcaaaacatgttgagtacattttagggtttaacagttttg**ggtacctttcaaa**gaaaatatttggttatatgtaaaattggtattttc**ccactttttaaattcctacttctcctgtttgccattcttctccactccatgagacagtagagacaggattattcactaattctcctctggcggagctggctgaggacaatcagtaagatcttggttgtgagtgtcaaatagattttgtaatttcatagcagacgcaagttcctaatactaaaactctttgattacaattacctcttgctgatagaaaaagggagttcttgaaattttgaaagttggttttaaaaataaaatgcatacactggaagatgcagtatactaaagatgta**gtattttccagggatcactgagaaaatac**aggatgaggtaaacagttttattttccaaagttcagaatttgagattgggcagactgca

Uppercase: IGKV2-40

Lowercase: Flanking sequence[1000bp]

Red & Bold & Underline: Stem-loop [14]

Blue: Heptamer[17]

Green: Nonamer [3]

id-TRAV28[V_gene_segment]

agcatccctcatctctgcctactagatgccagtaagattctcacctgtggcaaacacaaatgt**ctccagacactgtcaaatgtccctggag**gcaaatagcccctggttgaaaaccaatggtctatatgtctacatattgaatgttgaattgatagagcaggaaaaacttttagaagg**tatatatatgtgtatatatata**tttagtatatatatgt**atatatatttattatatat**gtatatatttat**tatatatgtgtatatata**tttat**tatatatgtgtatatata**tttat**tatatatgtgtatatata**tttattatata**tgtgtatatatatttttattacttatatattcatatacacaaaatgaggatgttaacatttt**tctactattataacacatttttattcctatggtggtcatacgtcctaagttgcta**ttatttcatttggctttaataaattaaataa**gataatttcaatgtaaaaacgggaattgta**atatatatatataaacatatgtattttatatataaatat**atataactttttatacaattcattcacaatgcaaaaaatttttcagtctgatcacgttccagatttcaccttggtcaatatcttttatctctgactcctcaaagacatatagactgtttctctattctttctctaagctactgtgttcctgttcttccccaaaatgattcttaaaccctaaaaagatgacccctccattaccccccacccacacacaaacacacacaaggactcacgcatacgcaccacagtgtgcagaaggtgaactaaaccacaggtaggt**ggagttttccactaggaagctgaggggcaactcc**cgggagctgctgccgctgaatctgtctcatgaaatttctcagccatgggaacaaaattagttccaaattcaagatatgatacttaccaatatttaaagagaatagatatcaaggtgtgcccctgtagcaaaaaaagtaaagaatcATGAAGGCATTAATAGGAATCTTG**CTGGGCTTCCTGTGGATACAGATTTGCTGTGAGTTAAGGACAGCCCAG**AGGAAACCTGGAGGAGTATCACAACTGAATTTGGAAAGGAGCAATGGAGGAACGGGAGAGATGGAGAAAGAGAAAGAACGCAACTCTCAATTAAGCTGTGTCCTCTATCATAAGGA**AGAAAACTACAGAGAAATTTTCT**ACACTTGGAAACAGCTACAGTGATTTTGTGATGACTCTCATCTGCATCTACTTTTCTCTTTCTGAACGGGGGTAAAAGCGCAAATGAAAGTGGAGCAGAGTCCTCACGTCCTGATCCTCCAAGAGGGAAGAAATTCATTCCTGGTGTGCAGTTGTTCTATTTACATGATCCGTGTGCAGTGGTTTCATCAAAAGCCTGGAGGACCCCTCATGTCCTTATTTAACATTAATTCAGGAATACAGCAAAAAAGAAGACTAAAATCCGCAGTCAAAGCTGAGGAACTTTATGGCCACCTATACATCAGATTCCCAGCCTGAGGACTCAGCTATTTACTTCTGTGCTGTGGGGAcactgtgct**cttcagacacctgcagcctatacatgaaaccatagctgaag**gcctaacccatccccgagagtggcagtaggtcccgatgtgattagcattgcattcccactgcctacatctccttgcatgtcctgattcatgtggaaagtatttctaaatgagttgtaacttgacactatgg**tgcaggtgcataaatggttcccctgca**gatatctgtgggaatggtattactaggcaaaatctattgattgatcaactctttcatatattatcatat**ctctctggataagaatgagcagagtcctcaatcctccatatccaagag**caagaaagcaccagtctcacctgtacttccttttccacatcttttttatggcttactttagtacagacagaatcctaggaaattttccatgatattttttctaacagacttaagtgaagatgaaaaggaagagggaaaatagatggcctctactaccatgaagga**tggtcagagctttctgcatacttgacca**gagactcagccacttacctctatgctgtggagacacagtgcttcccaggcacctgaagcctgaaatcagacaagtacctgcaattctaagactcaacaataagttagagatgt**ccatgtgttttgctgtaacatgg**ttttgagaagcatacacttgat**aaatcaaagtgattt**catttccaaaaaagattctgatgaatacttactctagtaactatctctgtgctaaacagaacaattcctaacatcctaatccctaaatcaggga**atcaatgagtcagaaattctagataaaggatccctcaaactcaatgat**aaggctttctgtcaaaaagggtatttgcaccccgaatgctacttattcttatcttacattatctatgtataaaggaaaatcttgagaaagtcaaaccaaaatgtgtcaaaaaagacatgctaagctctctaatccttcattatttttttttccaacttctttttttt

Uppercase: TRAV28

Lowercase: Flanking sequence[1000bp]

Red & Bold & Underline: Stem-loop [20]

Blue: Heptamer[29]

Green: Nonamer [1]

id-IGHV3-22-2[V_gene_segment]

ttgctttgtcatcc**aggctggagctcagtggtgagatcacagctctctgcagcct**cagaccctgggtctccaatcctcctccctctgccctatgtagctgggactacagttgtgcaccaccatgctcggcttc**agtgtgtattcaactgcacact**ttaattatgcaatatttattatata**gcaataatgcctcaataagggtattgc**aaataagtgaatatataatttgttagataaagactgatggaaagatagacacgaacatgagagatgtatgacacttaagaaaataaaactgtgggaaacatgctgttctttacaattgttaggtaatcacaaaagagcatacacatcacaccatgttccca**ttacagagaaatggttctgtaa**acctcactaggtgcaacccctgtgtgctgggcttggttcagggagaagtcaggtccagtggtgagaagcacagg**cccagatgcccaggtcactctgaccaaatgtgagctctggg**gacactgtacaacccatctgtgcttctgctggtaatttctcatttgtaacatggaaataacattgatactacataccatggtttctctgcatgtgtaaaaataaaagattattggtgtgaacttcaaaaatatgcagtttatgtagatctattgtacctcaataaaacttttaaaataaaaattacaaaattataagatttataggttttaagggtttaccacagaacaaacttacaataagtaaccacaattcccaaatgctatcaatatcacaaatctcccccaggacgct**ctcacatgctgtgag**ccccactctctcctcaggcctctcaccccagagcttactatatagtaggagacatgcaaatagagccctccctctgctgatgaaaaccagcccagccctgaccctgcagctctgggagaggagccccagccctgggattttcagatgttttcatt**tggtgatcaggactgaacacagaggactcaccA**TGGAGTCATGGCTGAGCTGGGTTTTTCTTGCCGCTATTTTAAAAGGTAATTCATTGAGAACTATTGAAATTGAGTGTGAGCGGATAAGAGTGAG**AGAAACAGTGGATACGTGTGGCAGTTTCT**GACCAGGGTTTCTTTTTGTTTGCAGGTGTCCAGTGTGAGGTGCAGCTGGTGGAGTCTGGGGGAGGCTTGGTACAGCCTGGGGGGTCCCTGAGACTCTCCTGTGCAGCCTCTGGATTCACCTTCAGTTACTACTACATGAGCGGGGTCCGCCAGGCTCCCGGGAAGGGGCTGGAATGGGTAGGTTT**CATTAGAAACAAAGCTAATG**GTGGGACAACAGAATAGACCACGTC**TGTGAAAGGCAGATTCACA**ATCTCAAGAGATGATTCCAAAAGCATCACCTATCTGCAAATGAAGAGCCTGAAAACCGAGGACACGGCCGTGTATTACTGTTCCAGAGAcacagtgag**gggaggtcagtgtgagcccggacacaaacctccc**tgcagggccgcgcggggccaccagggggcgctagggacccactgagggcgggacaggagcaggtgccgggagaggtttcctttctcctcagctggaaaagtcaggtttgtgtttgccggactctggagctttctaggctgtgacgttttattacttgtgtttattatgaatttattatcgttagta**tttaaattttagtaatttaaa**aattatatatatatgtttacactttaatgaaataagcattcctatttgcaccgattcttccagagttttattaacatttgttgacgtcagcagctacgaagctatagggacataaatttataaccatagaaagatgtataggccaggcgcggtggctcacgcatgtaatcccagcacgttggaagccataggtgggctgatcacttgaggcca**ggagttcgagaccagcctgggcaaaatagtgaaactcc**atctctactaaaaatgcaaaaatcagctgtgcgtattggcatgtgcctttaatcccagctactcggtaggctgaagcaggagaatcacttgaacacggaaggcagagat**tgcagtgagctaaatggcgccactgca**ctccagcctgggtgacagagcaagattcgatctcaaaaaaacaacaaaaaacaaaaacaaaacaaagaaagaaaaaatatataaatacaaag**acatatgcatatatgt**gtaggaattcataataaacattacaataaagtaattctaaaaatatgtcctaaagaatcaaacttaatgatgagctaaatgtaaattatttgagtatttca**caattgattttggttatttttaattgtttacatcaattg**atttatatttgttc**atttaactactgttaaataatggttatttcaaatgtggttgtcacaataaataataagaatttgaa**ttgataaatgataattatcttaatcatttcttaat

Uppercase: IGHV3-22-2

Lowercase: Flanking sequence[1000bp]

Red & Bold & Underline: Stem-loop [19]

Blue: Heptamer[37]

Green: Nonamer [8]

id-TRBV6-1[V_gene_segment]

tcaccatgtttgtcagactggtcttgaactcctgacctcatgatccgcccacctcagccttccaatgtgttgggatta**caggcgtgatccactgcgcctg**gccttttttttttttttttttaattgttactgaattagtggtgagaatcctgaccaagggaatattccctatgaaatatgacacttcaaataaaatgagagcgctttttcttatcctggagcctgcctcttctctcctctccccacatccctcacatgtcaagacacctttg**tgcccacagatcatgggcagcgttgggctgcc**actatcatccacacagagagggcagtcagcagcgtgaggtggttc**tgcctgctgcggtctcaccccaggca**cagaaagcaagagccctgggtgg**agctgaaggtgctcagct**gggcttgtcaggagtcccatctgtcggtgaattgaaaagaaacagagcaaaatgactcctccaatggtgatgagcctgcccctgggatttggaaacttggtaacagagaaaaccaatatagacaaaggattttaaacaggattatggtcaattaagcaaattagaaaaggatacttgaagggggatttgggacacaggagtcaaaaacaccgggaagacatgagaagtttccctaagag**tctaaattagagaaatatttaga**taactgacaccagtgtatgaatgaggaattatatcatcacagggataagagttccattgagttacaaactgcttccaaaaaggttaagaaaaactcgtaaggctgtgtcaattcagacaaaggcattcttccc**attcaaacggttcaccggtgcatgaatcttgaat**ttgaccatctggggaaggggcgtggcctctcctgacaggaaggctctggggcccaggcagggagaatgaagtctcagaatgacccccttgagagtactgttcccctatcaccgatgcacagacccagaagacccctccatcctgtagcacctgccATGAGCATCGGGCTCCTGTGCTGTGTGGCCTTTTCTCTCCTGTGGGCAAGTAAGTC**CTGGGCAGGGCCCCATGTGTGGATTTCAAGGCCCAG**CCTGTTTCCATTGTGG**CTGCAGCATCAGCTTTGTTCTTCTCTGCAG**GTCCAGTGAATGCTGGTGTCACTCAGACCCCAAAATTCCAG**GTCCTGAAGACAGGAC**AGAGCATGACACTGCAGTGTGCCCAGGATATGAACCATAACTCCATGTACTGGTATCGACAAGACCCAGGCATGGGACTGAGGCTGATTTATTACTCAGCTTCTGAGGGTACCACTGACAAAGGAGAAGTCCCCAATGGCTACAATGT**CTCCAGATTAAACAAACGGGAGTTCTCGCTCAGGCTGGAG**TCGGCTGCTCCCTCCCAGACATCTGTGTACTTCTGTGCCAGCAGTGAAGCcacagcgctgcatggccgtctcctctctgcacataaaggc**agggaggctctgccctcctccct**caccccagactcagtgatgc**cctgggcagagttctctgcaccagg**aaacttgaaacccca**tcatcatgggtctgaggcccccaggatga**aacaggatttgtatttcagatccatctagactctcgtctctccctggggaccatgttgcttcttctctctagggttcc**ccccagcccccaccctcatgttgtctctccctgggg**accatgttgcttcttctctctagggtttcccccagcccccaccctcatgttgtctctcctgtggcccacctttcccatctgggcagtcaccctccaaggccttgctgggtctctc**ctcccctcacttccccacacctctccacagcagccatgaggggag**cccctct**tctgtgcctccttccttcccatcacagagacttcaaagtccattttctctg**ccctgggctggaggcttcctttctgcagtggccaactcctacctg**tgcttcacatctcagcatgatcagccctcctgtgggaagca**tgcccttgcctcccagctgagatcaatttttctgttat**aagcactgatggtaacatgtgctt**gttagtcaacagagttgagcacagttggagttttccagttacttgtctggatatttttctgctcatgtcgattttaaactccataaaggggaaggctgtgcttgttacatttaccaacagctgttcctggaatgaagggagacctgtttcacagataatggatgagttaatgaatgacaggctgattgagtgatgagcgggtggatgaaccaataacaggaacactccaggtccgctgtaccctggcagaaatctagagttaactgtgcctgagatgctctgctatgagctcttcaaagggcact**cgcttgttgaacaagcg**ttggaccattcattgtgcaagccagtttgtcagttcatctaggagt

Uppercase: TRBV6-1

Lowercase: Flanking sequence[1000bp]

Red & Bold & Underline: Stem-loop [21]

Blue: Heptamer[42]

Green: Nonamer [2]

id-IGHVIII-38-1[V_gene_segment]

aaaatttatagagcttcttcattgtggatg**aatatttgcttcccttctccaaatttggaaatatt**ttttgct**attattttgaataat**ttatctcttctctctctctctctctctctctctctctctctctctctctctccctctctctctctctctctctctctctctctctctctctctctccccctctctcccttcactaccccttttgtcttgtctttgtataactcctcaaatgcgtagttgagtgtttagtgtctttcctggcccataagtcttgaactcttttttatttctttcacttaaatttttcttctctgtctctactatttcaaatgatctttctttgagtttactgtttcttctattacttgatctagtttggtctaaaatcttgttaaaatttttggctgtattcttcaggatcaatatttttgtttagtcatc**ttttaaatgttttatctttattaaaa**ttctcacattgttcttgcatgctgccactagctcattgaacatgtttctaacatttactttaaagtttctgtcaggtaatccatgttcttccttttcattgatgtcagtttatggaccttttggttgttcttctgtttggaccatgttcccctgcttcttcattttccttgactgtctttg**ttgttatctgttctttataacaa**aagccatcatatacttcttcaaagactaggctcatagagatcaccctcaccaatcagcccagacagtaattgtgggtctctcaaaacttgataggattcaaacttctgtctgttcttagtggcccctaggttttagaggatgatagattgtgccaggactcacaggtaagggaggtagaaactatgtactcatgaccatggggtctgtgctgatctgagcttctcttgtcatcactctcaatattcaacttccc**cgtggatcaggctcagctgtggctgctccacg**tggggctgttctcagtctgttgcctctgtgtgtgcagAAGTCCTCTGTGAAGTTAACTAGTGGAGTCAGACAGGAAAATACTACAGACCAAGAATTCTCAGAC**TGTTCTGCAAAGCCCCTGGATTCACTGAAAAAAGAACA**AGTTTGGTCCAGCAGGATTCATGGCAAGTGTTGGTGAGGGAGATAACAGTAATTCAAGTGGAAGTTCTCAATGGGACTCGCCTTCAGTACAAAGAAGATT**AACAGTCCTCAGAGACACTGTT**CAGAAGATTCTCTTTTAAGATAATAAAACTGAGAGCCCAAGACAAGTCTGTGTATT**ACTGTGAGGGAcacagt**gtggggacatctatgtgagcacagacacaaa**agtccctgcagggagacaggaggggact**gcatgttagttgctgctcagaaccaccagggggcattcaggacaccagtaggcactcagaaccaccaaggagagctcagaatcaccagggggcactcaggaaactggcaggtgctcagaaccaccaaggggcgctcaagatcccagcgggagctcataaccaccagggggcgctcaggacactagggggctcagaaccactagggg**gcgcccaggaccaccaggtctctcaagacaggagggcgc**actcaggacactagggggcgctcagaaccatcagggggcaatcaggacaccagagggacttcagaaccaccagggggcgctgagaaccaccagggggtgctcaggacatgaccggccctcaggaccaccaggggatgttcagaacaccaggaagcactcaggacaccagagagcgatcagaacaccagggggcactcagatcctccagggggcactcaggacaacagg**gagctctcaggacaccagggtgagctc**agtaaaccagtgttcgctcagaaccacc**agggagcactgaggacaccaccgctccct**taggaggcagctccacatcagg**tccctggatccgggcaggga**gggcggttccttttggatcttgccactaaccttttgggagttttcctgcttcctttgtggtttc**aagaatcatttgcagattctt**ctcatgtacaaagctctgctttcttggattatgtaatgtttttggctttggatgctaccagaattacgttgtactgtgagagga**ttcattcgtggtgtgtgcaatagtgaatgaa**agctccaatgttaggggtggctttgaaagctacgttaggggtggctgagggcagttagcaga**aaatgatcatcactatagaaggctactcattt**ctttgcacatttgcataaacaattgtagtttatgccctaaaa

Uppercase: IGHVIII-38-1

Lowercase: Flanking sequence[1000bp]

Red & Bold & Underline: Stem-loop [16]

Blue: Heptamer[22]

Green: Nonamer [6]

id-IGHV3-11-2[V_gene_segment]

ctgagtgaaggaagct**aaagaattcacagtaaatctcctgtgatttcttt**tgtataaattgtagaaaatgcaactattctaaattaacatggagaagatctgaatttttctgagaaaagtgtggtgaggtaacaagatggtgaaataaaattacagagaagtgagagaaaaaaatttggaggttaatttaattgctaatagcacgattgaagtgctgattcaaaggctgcacacatataccaacattttccatatcgtacactataaatttg**aattcactattgattgaatttttgaataaagcagtaacaaaaat**gagtatattggctgaggaagagcaagaaagagatgaatattgacacttgaataatcacggact**cctgaaaatacacacatgtgaacactgattgcatatttcagg**taaatactagaaaaagcaaagtcacataaagtcgttatgacag**gtgggataccctgcaaatcacactaggcatgtcccac**actgcc**ctggagctgtctcaggggagcagtctcctccag**tgtttagaggcacaggcacggataatagggctgacgc**tgtccagatgtgtgatattggaca**cattgcacaactgctctgttatgtatgtaattcatcttctctaaaaatgtaacattgacacttgcactgaatatattctgcaaatatgtaaacattaaataaga**tgatgactgctaattgatcatca**agtcacaatcacataatctgaagttatattttcctgagagataggattacctccagtgttttccgggaccctctcatctgctctgggcactgccctctcctccagcgtcccactagagcttgctatatagtaggagacatgcaaatagggccctccctctgctgataaaaaccagccgag**cccagaccctgcagctctggg**agaagagccccagccccagaattcccaggagtttccattcggtgatcagcactgaacacagaggactcaccATGGAGTTTGGGCTGAGCTGGGTTTTCCTTGTTGCTATTATAAAAGGTGATTTATGGAGAACTAGAGACATTGAGTGGACGTGAGTGAGATAAGCAGTGAATATATGTGGCAGTTTCTGACCAGGTTGTCTCTGTGTTTGCAGGTGTCCAGTGTCAGGTGCAGCTGGTGGAGTCTGGGGG**AGGCTTGGTCAAGCCT**GGAGGGTCCCTGAGACTCTCCTGTGCAGCCTCTGGATTCACCTTCAGTGACTACTACATGAGCTGGATCCGCCAGG**CTCCAGGGAAGGGGCTGGAG**TGGGTTTCATACATTAGTAGTAGTA**GTAGTTACACAAACTAC**GCAGACTCTGTGAAGGGCCGATTCACCATCTCCAGAGACAACGCCAAGAACTCACTGTATCTGCAAATGA**ACAGCCTGAGAGCCGAGGACACGGCTGT**GTATT**ACTGTGCGAGAGAcacagt**gag**gggaggtcagtgtgagcccagacacaaacctccc**tgcaggggtccccaggaccaccagggggcgcccgggacactgtgcacggggctgtctccagggcaggtgcaggtgctgctgagggctggcttcctgtcatgg**cctggggctgcctcattgtcaaatttccccagg**aacttctccagatttacaattctgtactaacatttgatgtctctaaatgcaatactttttttgtcctttttgtttctttgtttttttgcaacaggagtacatatcctcagctccacagaagccagggtgtcactttgggggcagaaataatcctttcatggttaccaggataagagtcctgaggaatcccagggaaacctggagagtgttttccagttagactcaggg**cagagacctccatgggaatctctg**attagaacaggccttgag**ctctgacgggagccaagagagaggctcgcccagggtcagag**tcctt**aaaacctgatggtttt**cacagctatcccccctcgtcttgtaaactcagactgattcagttgaccctctttctgctaatccatttccttctctgtaggtttgattctcacagttcgctttcttcttctcttccct**gaaaacagacgatgtgttttc**tgtagtcaaaatcccagggct**caggtctgcaggacctg**ggtaggctacggggactttctcactcaccattgtccggacactcttgtcttctgtgcatggaggcatttggaaaatgaagtggacattagccatgaagggaataatactagttttctccaataggatattgatgtagagctgatcttgtgcttctcacactgtctcagagtttggactctcacctgtgactttgaga**agagctggggatgggcactccattgtgctgtgagctct**gggtaacgataattgtagaatctgcctaggcagtctaaggtcaatactactcgtcatcaggaaagacagctggaattc

Uppercase: IGHV3-11-2

Lowercase: Flanking sequence[1000bp]

Red & Bold & Underline: Stem-loop [22]

Blue: Heptamer[35]

Green: Nonamer [8]

id-IGLV10-67[V_gene_segment]

aagctagttgtcaaacacaaggggaatttataacctgc**agttagctgatggaatgctaact**agttggaag**taataaaataaaagtattatta**tgagtttcaaaggcaataaatatataaataaaattaaaaagccaagagctaaag**aaataaattattttaaaatatgaacatgcactataattttttt**taagtttataccatgagctatttgaaagttagactggaagcacattatatacacttatgtaagagcctctgaaaaagatggaaacacatacgaaggttaaaagataatttaaaacaatagagagttatttcagtagctgaagatatgagtgccaataaaatgaatgcgattcaaccctgtgagcattgctttcataaaaaaattaaaacagaaaatgaatattaatagatgacttaagtgaaatcacaatgtatgtggtgatacagctgaaagacaaacgttttgttagtaagaagggcatcgctaagcctgatgagtgctttgtttagaattcccagctaccctggattcctctccaacctcatgaacagcttcagaaaagaggatcaggacaaccgatgct**cctctcattgaaatgcccggatgtggcagagagg**caggaaatattttgggtctgaggacagtg**aggatcatcactgagatgatcct**gcccaaatgaaaaccactcagggcctccaagggagttcagccatcagctcactggactaggtttgggatgcaacactgacctgccccgacccagagtcata**atgagccaggccacctccacccactgggtcagctcat**ctcctctttatcttttctccttctgacctgtgagtacccagatcagggtggacaagtaagaga**cacatagcaaggtcttatgtg**aatacagtgactc**taccttacgacgtttttgggagatacaaaggta**cttagaggagctcag**tttccaaagcagtgctccaggaagctggaaa**tctgcactATGCTCTGGGCTCCTCTCCTCCATCTCCACCCTCACTATTCAGGTGAGAAAGAACCTTATGTCGGGGGATCCACACCCAGCTCTGCTCCATACCCCCTCACTCTCTGGCTCACTGGGTTGTAAACCTGCTCTCATGGCCCTCTCTGTCC**CCCTGCAGTGTCTGTGGCCCAGGCAGGG**CAGACTCAGCAGCTCTCTGTATCCAAGGGCTTGGGATAGACAACCACACTCACCTGCACTGGG**GACAGCAACAATGTTGGCAAAGATGGAGCTGTC**TGGCCGCAGCAGCACCAGGGCCATGCCCCCAAACACCTGATCAACAGGAAAAACAGCCAACTCT**CAGGGATCTCAGAGAGATTCCCTG**GCTCCCGGTTAGGAACATGGCATCTCTGACCATCTCTGGCCTCCAGACCAAGGACAAGCCTGCCTATTACTGCTCAGCCTGGGACAGCAGCCTCAGTGCTCAcacagcgcctc**aggggaagtgagacgaaaactcaggagctcccct**agcttcactcggtatgcgggggcgtcatagagcactgtttaaact**aaaccaaaaatgacaagggctggttt**ccacgtgacaaagtgtgtgctattcaaaactattaaaaataatatcataagttaaaataactaaatctgcagtattaatgaatatttataggaaacattgttgtgaatatatttgtggctagatacttacatgaacatacaca**tgcattaatcagtagtaaaaatgaatgca**agggtaaact**aaggaaatatttattttcccagattttcctt**ctcttttatttggagatgctggactaaatgctcagaggttccggtggtctgcctttgagtgtgtgttcagcagggtcacctcaagcttatcccaccttcttatgacctgggcccatgcctgtcctctcattagaatcccctttccctctgtcacagagctctctttattccttctacagacccagacttgttcctcacacttagttgaggcttcctgggccccacgtccctgtttgcccccaacacc**cacacaagacaatgatcccatgtgtg**tgctcttgttgtgccctttccttatcctctgacatacctcaggcagtatctggaaagaacaaatgaagtgggagcgttagcttgtagttgaaaatgatttttgatattttgg**ggaaaatatctggtttaaagatagttttcc**atcgagataaaaatgcatgtgtgacagcaccacatttctaatggcggaccaaagccccccacatctttgccccccacatggctccagggacattgacttgttaatatacaacccaggagagagcactgtgtcctatgcaactgcaatgcagaggcaggtttctgcccgctcctccaatgaagtgagtagcaaccccaaccccatcttcacccatcatttagatgacccccatacaaccacaacatagaaacacatt

Uppercase: IGLV10-67

Lowercase: Flanking sequence[1000bp]

Red & Bold & Underline: Stem-loop [18]

Blue: Heptamer[36]

Green: Nonamer [2]

id-TRAV11[V_gene_segment]

acaggcattgaaatgctctatctatcacttactaaattgcattactctggacaaaattacataacttcagagcacctacgcgccatcaatggtaattacacgtacatcacatggtcataagg**attgaattaagttaagtgaatttaattaaat**gattgaattgcattggtagaaaataatttttcaactaatggtagttattcttggacataaatgataaaacaaaggtaatttaaaataagacaataagaaactttaagtatagattaaggtttgcctttccactcttcttctgttgattatattcttttactacagttatggccgagagtcaccctttatttcacattggccttattccattcaattatttgaactgacagtattggctttcttgtttgtttgtttgtttgtttgtttttatctcatagg**ttttctgcacagaaaa**tgaagctttcttgcttccggtgt**aatttaatgtaaatt**ctatgatgaatccacatgagctttatggagcaggactccgatataacaccattatactgacctacagtgccaataactagatccatacccag**gagctcatttaacagagctc**tgcattggaaaggag**agaaagtaacctttct**aggtttgggggagaaatccttttt**agttattcaatttataact**tgtacatacatggtagatgcccaggaagtattacagaagaaaaaaggaaagagaaaaatctaagggacaaatgctagatgcagctgactgagaatcagcacctggttctgccagggtccttttccatgctacctcttcag**agatcaggaggagctgtgtgatct**gtgcaccggggagtaaatcttgtttctggctggttggggagactttcagcttcctgcaacaaagaactatgaatgttaactcagagaaaccagagatttggcttgatagtgtttgagggctcccagaccccagccagagacctcactgagtctaagtgataaaaACGGAGAAGCCCTTGGGAGTTTCATTCTTGATTTCCTCCTGGCAGCTGTGCTGTGAGTTGCTGGGCTCTGGAAATATC**ATAAAAAAGAGCAGATTGTAGTTTCTTCTTTAT**AAGTCTGGAAGTTATAGAAGGGTGGTTAGGGTCTCAGTGCTCACCTGTGGGGGCTGTGAAAAGAGAGTATATCTTCCAAATGAATGAATTTCCGTGAATGAATGTCCTCTGTCTGTGGTTCTCTGTTTAAACAGGGGTGAATAGACTACATACACTGGAGCAGAGTCCTTCAT**TCCTGAATATTCAGGA**GGGAATGCATGCCGTTCTTAATTGTACTTATCAGGAGAGAACACTCTTCAATTTCCACTGGTTCCGGCAGGATCCGGGGAGAAGACTTGTGTCTTTGACCTTAATTCAATCAAGCCAGAAGGAGCAGGGAGACAAATATTTTAAAGAACTGCTTGGAAAAGAAAAATTTTATAGTGTTTGGAATATCGCAGCCTCTCATCTGGGAGATTCAGCCACCTACTTCTGTGCTTTGcagtagtgtctccccagcacctgcagcctgtaccata**acctgcagccgggacccttgacacaggctagccttgcaggt**gggagtgaagatttttttttttttttgtatagagggaactttctttatgacccgcatctccaaaatagaagtgaaggataatgtatctttcttatgtaatgaaactg**aatttatattaaattaaatttccttttgttattatttaaatttcatt**ccttgatgcagtttgtacagattctgagcctgtttgcccacagattccttccactgctttactctttaagaggaaagacccctttgtggtgcaggctctgagtccatgaagcaagttttctccaggctataccctcatcaggaacattttacccatttgatcatatcttcatgatgtttttgtaccatctataccattatttactt**aatttaatattaaatt**gatgtttatgttaaatagactttaaaaataact**gtaaaattgtcttcattttac**cagataccatctgtgaaaccatgtatatgata**taaataatatattta**aagtgctgc**ttatatttttcccaatataa**attaaaatagcatcatatatgtaacaattgaaaaggctgtctgtatactgcctaataggacctcttgtatcaatcacactttaagaaatactgatccagcccatacccc**tcatttcataaatga**aagaattgagatccacaagtatggta**tcatttactcaagcacgacttacagtaaaatga**aagctataaaaagagtctaggctttatgatcccttgtcagttgcttcttccaccaatgatgcaaagcaggatctaaagtttgaagaaaagagttgcgtgttttacattta**aattaaattaaaagttttatttaatt**gtgtatgttcaacc**taggtaaattagaagatacgactaccta**tcaaagtagagcattctcataacttcatcgcctagaaataacaagtatttatttt

Uppercase: TRAV11

Lowercase: Flanking sequence[1000bp]

Red & Bold & Underline: Stem-loop [20]

Blue: Heptamer[19]

Green: Nonamer [3]

id-IGKV3-34[V_gene_segment]

tcagataataaatatttaggtttggtgaggcaagatgcaaaattgaatatattatgtgagtacttagagaagagagaaaacaaatttctgcaatgtctattaatgaactcagatggaatgtaacccaaagagcttctcagcctgtccagtgaactgtcacttaccaccaaatgctca**caattttaaggaacacaaggaaaccccaaattg**aattattgtttagatcagaaacatgagcagaggtttccaaaattattaaagtgagaaaggcctgagtctgattgaatggatgtgttgtgcaggatggggtggcatagaaagtagggcatcacccaaggaattgttcctggcctgctggacagcattgtcgtctgtttgcagggaatggaattacacacactgaagggtctgagcccggggcctgctcatctgtgcagccgtgatgatcatcacacccacagtgctgggcctaattgcgtgttgccctttaatagtttttacaaagaatgaggaggcttcagggctcataggaaagatggctaagctgcaaaggaaaagtgtgtgtggaagatgaacttctgatgaactagttttcagccaggtacattcatgcaccaatggagatgactc**ggttctttggagaaataaggagaggaaaagaacc**acaggcttctattttattcccaacaggtttgtactaaataccatctggctgggtctaggtgactagtgagacgggaagaaacaggggctatagtctgtgaccacagggaagaagttctacca**tcaggctgagccaatggacttttctggcctga**ccacctgggcaggggctgctgag**tgcagagaggaggaggcagggggtctctgca**gctggaagcccagcacccactccaactactttgcatatccttccagccactctgctgtccagagcccatatcaatgcctgggtcag**agctctcgggaagagct**gctcggttaggacccagagggaaccATGGAAGTTCCAGCTCAGCTCCTCTCCTTTCTGGTACTCTGGCTCCCAGGTGAGGGGAACACGGGATGATTCTGCATGCCAATGAAACTCTGTCAATGTTATTGGGTCCTGTGACCTGCTCAGCAAGAACAATAATTAATATTCAATGTACATCAATGA**TCCCAGCTGTACTGGGA**AGACAGTGGATTTG**ATCTAGATTGCACAATTGACTTTCCTGTTTTATTCCAATCTCAGAT**ACCACTGGAGAAGTTGTGCTGACATGGTCCCCAATCACCCTGTGTTTGTCCCCAGGGGAAGGAGCCACCCTCACCTGAAAGGCCAGTCAGATCATTGACAACTATTC**AGCCTGGGACCAGTAGAAACCTGTTCTGGTTCCCAGGCT**CCTTATTTATAGGGCATCTACCAAGGCTACCAGCATCCCAGCCCGGTTCAGTGGTGGTGGGCCTGAGGCAGACTTTACCCCAACCATCAACAGCCTAGACCCTGAAGATGTCACAATTTTATTACCCTCATCAGTACAGCAGTGGGTGTCCcacagtaattcaacat**gaaacaaaaactttcacaaaaccattgattttttttttc**taaaacc**agcagctttatgggctgcagctatgatggctgct**cagttttagcaactgtgcctctatttggaaattttgaaattctgaaaagtaactcgttgtcaagatggtgactccaatg**tctgtatttttctcagacttatttacaga**gttaggttttgaatcaattttactgcaaggccttcagacagact**cacctgcatggatgcctttaaagacacaggtg**ccgctgggcacagtggctcactctttttttttccaatagagtagaacaaacatagcttttcatttaataaattttcacttaatggcagcattttcacgaacaagtaaaggccaccgagtgaaggttaggtgcacgtgagtaaataagacaattttgggtcccaattcagtaggggaaagagccatgaaggaagaataaccacacaataaaagtgcgtagaataatttctgtagttgacaaattgtgtaaaaagcaggaatggtctgtggtatctggaggctgaggccatatatttgtgctgatcagggaaggccccagggtcatactgaagattcatctgagtggtgaggaggcagccaggtgaccacaaaaacagcaaagaccacgggatcattatgggaagggcagggaccagtgtgtactgaaaaagtcagcaagtgctaaact**aatcagatgccttctctgatt**tttattcccgta**aagataaaacttactgtaatatctt**acatcatacaccttggtctgggtcactcccatggcatagaatgttccctggttcagaagttggaagcccaggtctttcctttgttgacagtttgcaggctccccagagcccttccctgcc**actcaagcccttatcactgggtcagtctctgtctttgagt**tgggatcaccataggcaggggcagcactagacactggacatggggctctggccct

Uppercase: IGKV3-34

Lowercase: Flanking sequence[1000bp]

Red & Bold & Underline: Stem-loop [15]

Blue: Heptamer[22]

Green: Nonamer [7]

id-IGKV3-20[V_gene_segment]

tagtagtctcagaacagggagagacaga**ctgagtccctccactaactcag**cccaacacctgtggatgagtaggggatgcctgagatcctggggtggatgggaggtggggcactgatctgtcaa**tctgctttttcttcaaggatcaggcagcaga**gacccagaagcttcatgtctttgtaaggctcttccaagccaaca**cagataatttgacaaaacactgtatctg**catcccagacctcaccctgtcacagactgaagtggttgtttcatcctactaaggg**taaactataccagctatacagaataaaaagactggatagttta**gaggatcacccaagaaatagttctttgcctgcatggacaaaaccatcttctgtctttagggaaatggtatcactaccctgaggatttggagcccaggtctcacttatctgtgcagttgtgaaagtcctcacacccacagtgctgaggttaattgaatgttactcttttaatttctgcaaagaatgagacagcttctggaccctcaggaaagatcactaacaagtaaatacaagtatatcc**ggaagataaagttgtaatagactcttcc**tttcaacctgatccatcatgcatttagggagctgactgggcacaagttggagcagaaagagaaaaatgaaaccacagccttctattttgtttctaacagacttgtaccaaacattctgtggctcaatctaggtgatggtgagacaagaggacacaggggttaaattctgtggccgcaggggagaagttctaccctcagactgagccaacggccttttctggcctgatcacctgggcatgg**gctgctgagagcagaaaggggaggcagattgtctctgcagc**tgcaagcccagcacccgccccagctgctttgcatgtccctcccagccgccctgcag**tccagagcccatatcaatgcctgggtcagagctctgga**gaagagctgctcagttaggacccagagggaaccATGGAAACCCCAGCGCAGCTTCTCTTCCTCCTGCTACTCTGGCTCCCAGGTGAGGGGAACATGGGAT**GGTTTTGCATGTCAGTGAAAACC**CTCTCAAGTCCTGTTACCTGGCAACTCTGCTCAGTCAATACAATAATTAAAGCTCAATATAAAGCAATAATTCTGGCTCTTCTGGGAAGACAATGGGTTTGATTTAGATTACATGGGTGACTTTTCTGTTTTATTTCCAATCTCAGATACCACCGGAGAAATTGTGTTGACGCAGTCTCCAGGCACCCTGTCTTTGTCTCCAGGGGAAAGAGCCACCCTCTCCTGCAGGGCCAGTCAGAGTGTTAGCAGCAGCTACTT**AGCCTGGTACCAGCAGAAACCTGGCCAGGCT**CCCAGGCTCCTCATCTATGGTGCATCCAGCAGGGCCACTGGCA**TCCCAGACAGGTTCAGTGGCAGTGGGTCTGGGA**CAGACTTCACTCTCACCATCAGCAGACTGGAGCCTGAAGATTTTGCAGTGTATTACTGTCAGCAGTATGGTAGCTCACCTcccacagtgattcagcttg**aaacaaaaacctctgcaagaccttcattgttt**actagattataccagctgcttcctttacagatagctgctgcaatgacaactcaattttagcatctcttctctgcttgggcattttggggatcttaaaaaagtaatcccttgatatatttttgactctgattcctgcatttttcctcagaccaagatggacagccaggtttaagcacagtttcacagtaatgg**ccactggatcagatttacatcagtgg**atgtcagtaaaggtcccaaccagagccataaggcaacaacaatagcaacaaataatcaaaattggaaaagaa**gaattaaagctgtcataattc**actgatgaaggattgtgtgcagataaaattcaaatttgtctacagagaaactactaaaattgacatgagaaatagaaaatcattagattcaagatcaatttattaattcatagattcaaaaatcaatttcatttctgcataataaaaaatgctaaaaattaacattataaaacaaacacaccatttacaaaaacatcaaagtatcaattatttaaaaaaaatagactaaatacactgacgtctccagaata**ttattttgaaaaataa**aagaaaacctaagtaaat**agaaattcagttcaaagactgaatgtct**cagtactataaaaatgtcaattcttcaaagattaaaata**ttgattacatataagaaaaatcaa**aatcctaaagtatactccaatttaaattaagaagctaatctaaatattatatgggaatgtcaaggatgtagaatagccacagtgaacctgaagaaacaccaaaatgagaac**ttccagtgcctgaatacctggaa**tatagtgtgggtggcagtatggtgatggtgagatcagaagtttaaaaatttgcaaac**gtgcttatttttggaaataatcac**tacgcagatgtagccaaaccctcttcaactgtgccaccagaatctca

Uppercase: IGKV3-20

Lowercase: Flanking sequence[1000bp]

Red & Bold & Underline: Stem-loop [18]

Blue: Heptamer[19]

Green: Nonamer [7]

id-IGKV6D-21[V_gene_segment]

ttgtttattttgggggcatttcaggatcttgctcaagaactgtagagatttttttctgtgactcttttttggtgcttgcatggaggtttacagagtttcctcatctaatatagattatctagcaccaggcaatgtgctggatctcatggctgaagtgacagaggcatttgcatta**aaactcaaacttactacagaatattttctttctcagagttt**attcataaaagacagccttccaagttagctgataaatgggatggtatagtaaacccaagtgcaaaatgcattgtcaacactctaggatggcttaaccagtaatgtgcttcattgctagtggttggaagtacaaggtgcaattatttttccttactttggaggggataagccagcatgactcataccccttttataaacacttgacatcttctctaatgtgacaagcccttgatgttttggggcgtgcatcccaccctctagagcacatgtgttttcacaagaaattcagagttcttacaatgtccagctcatcacgtctaattaccatgatgt**catcaatatagtgttgatgctttgtggaacgttcacaaagc**tttttcagcctacattgtgacagagagcaggagagttaacatagccctgggacgagactgaggatgtgagctgttattcaccccagataactgcagactctcccagagatggcgatggactctgccttcactctgcagctgtgccctggggtctggtcaagccctgccagagcctcagcggagctcgtctgcaggtgccagcagagggcgcttcacacccctcatggaaggggccgggagggcgctctcctggcaacagtgatttctgtttatttaaaccagcaggacatccccataatttgcatgtatcgttcctcctatatgtga**agaggccctgcctct**cggtatcttaaaagaggttctttctctgggatgtggcatgagcaaaactgacaagtcaaggcaggaagATGTCGCCATCACAACTCATTGGGTTTCTGCTGCTCTGGGTTCCAGGTG**AGAATATTTCCACAAACCTAGGCGGAGATATTCT**TTCAATCTGTAATTTCTTTCATTGGGGACTCTGCAATAGGTGATTTTTGGCTTGA**TTTTAAAATCCTAATTTTAAAA**ATGTAATGCATATTCTTTCTTCATGTCTAGCAAGATTAAAGGTGATTTTCATACACAGATATTTATGTTGTACTGATGTTTGCTGTATATTTTCAGCCTCCAGGGGTGAAATTGTGCTGACTCAGTCTC**CAGACTTTCAGTCTGTGACTCCAAAGGAGAAAGTCAC**CATCACCTGCCGGGCCAGTCAGAGCATTGGTAGTAGCTTACACTGGTACCAGCAGAAACCAGATCAGTCTCCAAAGCTCCTCATCAAGTATGCTTCCCAGTCCATCTCAGGGGTCCCCTCGAGGTTCAGTGGCAGTGGATCTGGGACAGATTTCACCCTCACCATCAATAGCCTGGAAGCTGAAGATGCTGCAGCGTATTACTGTCATCAGAGTAGTAGTTTACCTcacactgtgttacaacccagaacaaaaacta**gttcagcctggctgaac**ggagaaactgggtgataccctagaatacttctgattgttgcaggtgctttgggggcaatgagttaaccaatacaatgaagtctggctcacc**cagcagagaggaaactagagtcactgctg**catactttcatctttttaaaaatgatttatttcaatagtttttgggggtataggtggtttttatttacatggataagttctttagtggtgatgtctgagattttggtggacctgttacttgagcagtgcatactgtgcccaatatgttgtcttctagccttcacctccccttctatccttcctccccagtccccaaagtccattatatcattcttacgcctttgcatcctcatagcttagctcccacttacagat**gaaaacatataggttttc**cattcctgagttacttcatttagaataatagcctccagcttcatccatgttgctgcaaaggtcattattttgttctgttctgttttatggctgagaagtatttc**gtggtgtatatacaccac**attttctttatccacccgttgcttgattggcacttatggtggttccatatttttgaaatggagaaatgtgctggact**aaacatgcatgtgcatgttt**ctttttcctatactaactttt**tttttctttgggtagataagaaaaa**taagtactggaattgctgaactgaatggtatttctacttttagttctttaaggaatctccatactgtttttcatagtggttgtattag**tttacattcccaccagctgtgtaaa**agtgttccctcttcaccacatccatgccaatatctattattttttgacattttaattatggccattcttgcatgagtaaggtggtatttcaaggctatggttaccaaaacagcatggttctagtataaaaataggcacatagatcaatggaacacaatagagaacacagaaataaacccaa

Uppercase: IGKV6D-21

Lowercase: Flanking sequence[1000bp]

Red & Bold & Underline: Stem-loop [15]

Blue: Heptamer[26]

Green: Nonamer [9]

id-TRBV10-2-2[V_gene_segment]

cttctgattctgtgtcctctctcactggagcatgattgtgggaattacccatccagg**ttgcatcaagatatgcaatagataatctattg**tatctgaaataa**caatagcagtgctattg**actggatgtaagtttggtgccaaagcacgtttacacctcctccaatcttctcccatttagtgaatacaagatcttctgaat**ggaaatagaacatttcc**ctgtgaccctagagactccagatagttgtagttctgggtgaacatttcatccacattcttaatgagcctcctcttttcttcataggaagtacagtcacttatgggacttaggtggtgagtgtttgtggcttcacctgaagcaaagggaaaaatcatcctcatgagaacacaccgcagcctgagcttggatggtgaggctcccatcaggtgcaatgaggtgctacagtgtgaaacaagac**ttcaaatgttaaagaccttgaatccacaactttgaa**acctttctggatgtcagttgactcttttaaccaagggtatctctgtaggctgaaatacaaatatccagaggttaaacaaagtggaaaat**aattcaacttggagttgaatt**ttgtacagagaa**gtcataaaagtcaccaggactgctatgacaatgctctttagactacagaaatattcataaatgttaaaacagcatt**caaatgaagaatttagcataatgggagtaaaatacgcatagaagaacaa**attcccctattaaaatcagggaat**gagattacacattttaagatttctttcaaaacagatctacaaatttttcccaatctaagcttctcaagtatgtctataataaatgatttgttcctgtggggaagagaacagatgtggcaggttagctctgaggttcaagcatttagcaggga**cagtgacatcatcaagtcactg**agagctcaacttcaatttgcccacag**cagggctgggagacacaagatcctgccctg**gagctgaaATGGGCACCAGGCTCTTCTTCTATGTGGCCCTTTGTCTGCTGTGGGCAGGTGAGGGCTGGTCACAGGAGGGCCTCCTTCCCTGGAAT**TCCCAAGGACTCAATGCAAGCTTTTCTGTTGGGA**TGACAGCATCAGTGTCTGTTGTTCTCTATTACAGGACACAGGGATGCTGGAATCACCCAGAGCCCAAGATACAAGATCACAGAGACAGGAAGGCA**GGTGACCTTGATGTGTCACCAGACTTGGAGCCACAGCTATATGTTCTGGT**ATCGACAAGACCTGGGACATGG**GCTGAGGCTGATCTATTACTCAGC**AGCTGCTGATATTACAGATAAAGGAGAAGTCCCCGATGGCTATGT**TGTCTCCAGATCCAAGACAGAGAATTTCCCCCTCACTCTGGAGTCA**GCTACCCGCTCCCAGACATCTGTGTATTTCTGCGCCAGCAGTGAGTCcacagtgctgcacagctgcctcctctctgcacggaaacggcagttagaaaaactgaggttgcctgtgcacccaagtctgggccccaccctgggacgtctcag**cccccataggagtcacagagtcctgcccaggttggggg**ggtgtct**tagagcaggggccatatgaccctcagtgctgtggctcta**ttcctacccctgtctcagaccatctggagtttgccccaacacactaaccttgctgggtctttc**ttctgcagctctctttttctgcttgcagaa**tggaagagatattttgttgatattcctaattcccttcataatttcaaagccaacatattcttctctcccttatgggatcagtgacttcttcgtgtaaaccttaccgtatcctctactacccttcaactcacagacacaaatcttcccccatgtggtctctgctcctggctgcctttcttct**aagggcaatagccctacaaggtcatctatcatttgccctt**cacctgcttctccatcatcattttttttttttttttttttttttgagacggagtcttgctctgttgcccaggctggagtgcag**tggcgcgatctcggctcactgcaagcccgcctcccgggttcgcgcca**ttctcctgcctcagcctcccaagtagctgggactacaggtgcctgccacggcgcccggctaattttttgtatttttagtagagatggggtttcacggtgttagccaggatggtctcgatttcctgacctcgttattcacccgccttggcctcccaaagt**gctgggattacaggcgtgagccactgcgcccagc**ctccatcatcattttgcattaagtttcccctggctttctgctttagccacactcccctacattcagttacagcttcgtaggtgtagtgttgttttccaccctggggattgtttatgaataatagattatttctacttcactgagcatatgtgcctccaacttattcccaggcctcta

Uppercase: TRBV10-2-2

Lowercase: Flanking sequence[1000bp]

Red & Bold & Underline: Stem-loop [22]

Blue: Heptamer[42]

Green: Nonamer [2]

id-IGLV2-23[V_gene_segment]

agtgttgcagggttcagagaggtcagttttatggttcactaaggaggggctaagatggaggaaaccaagaaagccaaaaacacacatctctgccactcaaaacataaatcaaagtaaaatggaaaaaatctttcaataaatcattcccaataaatatcactctggtgcacata**gaaaggcctgttagcaaactttccctttc**ccctgctggctcatatccccatggatcacctgtgtgcagagtggactctgtggtccccat**ttctcaggctgagaa**ccagagactgggaaggacaagtcac**ctgctcatgagtgggaagggcaggattgggagcag**ctggcttgggttcaggactgtggccctggtcggtgctctgtgtccctccagagggtccct**cccacatgtgccccgtgtggatgtggg**tgcactgcctggggtggctga**tgtgctccagggattcgatggtgcagatgcttcagcaca**aaacgtggaggtggggatgggatgaggctgca**gctggggagaccagagaccagcgccacccagc**agaggtttccacggggtgggcagccccatttcctgacagtagagacaggctggaggtgggtgccaagcagagggcgctgtggggtatgcccaggcctagagagtcacagagacacaggcactacacagcagaggcactgagggccaggagctca**ctcaggtgagagacttcagcagatccttctctcctgag**caaatcaggtacaaaaaaaattaagttcctgccatcagcatgcacaaacaagaacattcatctcttcagctgagccctgctgtccagggaagcaaaagtctctgagcccaggcccacgtgagggtggggtgaggagaggagcccaggatgctgattttcatggag**gccccgccctcctctgaggcaaaggggataagacagggctggggc**agggccagtgctggggtcacaagaggcagcgctctcgggacgtctccaccATGGCCTGGGCTCTGCTGCTCCTCACTCTCCTCACTCAGGACACAGGTGACGCCTCCAGGGAAGGGGTCTTGGGGACCTCTGGGCTGATCCTTGGTCTCCTGCTCCTCAGGCTCACCGGGGCCCAGCACTGACTCACTGGCATGTGTTTCTCCCTCTTTCCAGGGTCCTGGGCCCAGTCTGCCCTGACTCAGCCTGCCTCCGTGTCTGGGTCTCCTGGACAGTCGATCACCATCTCCTGCACTGGAACCAGCA**GTGATGTTGGGAGTTATAACCTTGTCTCCTGGTACCAACAGCAC**CCAGGCAAAGCCCCCAAACTCATGATTTA**TGAGGGCAGTAAGCGGCCCTCA**GGGGTTTCTAATCGCTTCTCTGGCTCCAAGTCTGGCAACACGGCCTCCCTGACAATCTCTGGGCTCCAGGCTGAGGACGAGGCTGATTATTACTGCTGCTCATATGCAGGTAGTAGCACTTTCcacagtggtccaagttcatggggaactgagaccaaaacctgcccagggc**cttcagacttcctccttgctctgaag**atgcttcctcacccggtgcaagaggcttgctgcagcgcggccttgagaattcttctctctcagctccttccctttccaccatgaattccaaca**ggaaacctgccctgtggtttcc**catccaggacagggacagcttcctgatgcttgtgtgctgtg**gtccctgaatgtgcaactcttcccagctcttcaaatgcagggacagtgacaaggagctgcctgattggtgcagtcactg**cttttttca**gggatgtcttcaccctacatgtatcatcatccc**ctacactgtggg**tagaattttagcaactacattcta**atggttatcgccacaactttgatcttagaaataacagtgcagtgaacatccctatgcaggctcctttgagttcctgtgtgaatacgaccataggattcatttctaaaagtgaaattgcgggtcagaaagatgtgtgtttgtgattttcacccaatgttgtcagattcccctccagagggattgtaccaattttcaatgccagctggaagtacttgttcagagtacattgttgaatgtggaaatttttgccaacgtatc**agagtgaaaaatggtaccttgttacagttttatttttcagtct**cccgttctgcttaaggttgggccttttcacaaatagttcagtactcttcatttttcatgatttggtaaactttataaaaaatatccagataagaagaattttagtctttgtggccagacacggtggctcatgcctgtaatcccagcattttggaagactaagacgggtgaatcgcttgat**cccaggagtttgggattagcctggg**ccacacagtgaaccctcatctcaattaaattaaagaaaa**taaaaataattttta**aaaaa**atcctttgtggtccaaggat**caactttggggatattacctatggact

Uppercase: IGLV2-23

Lowercase: Flanking sequence[1000bp]

Red & Bold & Underline: Stem-loop [20]

Blue: Heptamer[48]

Green: Nonamer [8]

id-TRBV6-5[V_gene_segment]

acaacttattagcatcccgttgg**ttcaaaaaatcataagaatttgaa**gaacatgacaatgttatactgactgtggcgtcctaataagtgtgggactctgtgcaccaagatatgttatttga**attcctttttattgattactaaattaatggcaggaat**cccacacaaagggatactccctatgaaacatgacaactcaaattaaaaaagagcactcttccttactctaaagtctgccctctctccccgagtccctcacgcataaggccacatttgtgcccctaggtcatgggcagcgctggttggccaccatcgtccacatgtaga**aggcagtcagcaacgtgaagtggttctgcct**actgcggtctcaccccagacatggaaagcaagagccctgggtgg**agctgaaggtgctcagct**gggt**ttgtcagaagtctcatctgtcagtgagttgacaa**gaaacagagcaaaacgactcctccaatgttgacgagcctgcccctgggatttggaaacttcataacagaaaaaaccaatatagacaaaggattttaaacaggattaaggtcaattaagcaaattagaaaaggacacttgaaggagtatttgggacaggggaatcaaaagcaccaggaagacatgaggagtttccctaagactctagactacagcactagcactatgtagataactaacactaaaatattaattatatcattgaaaaaataaaatttacagtgaattacaaactgtttccaaaaggtcatgaaaatcttgtagacttgtttcaattcagacagatgtgttcttctcattctcagctgttcactggtgcatttattttggatttgaccatctggggaatgggtgtggcctctcctggcc**tctccctccctggggcccaggcagggaga**atgtctcagaatgacttccttgagagtcctgctcccctttcatcaatgcacagatacagaagacccctccgtcatgcagcatctgccATGAGCATCGGCCTCCTGTGCTGTGCAGCCTTGTCTCTCCTGTGGGCAGGTGGGTC**CTGGGCAGAGCCCCTTGTGTGAATTTCAAGGCCCAG**CGCCTTCCCATTGGGG**CTGCAGCATCAGCTTTGTTCTCCTTCTCTGCAG**GTCCAGTGAATGCTGGTGTCACTCAGACCCCAAAATTCCAG**GTCCTGAAGACAGGAC**AGAGCATGACACTGCAGTGTGC**CCAGGATATGAACCATGAATACATGTCCTGG**TATCGACAAGACCCAGGCATGGGGCTGAGGCTGATTCATTACTCAGTTGGTGCTGGTATCACTGACCAAGGAGAAGTCCCCAATGGCTACAATGTCTCCAGATCAACCACAGAGGATTTCCCGCTCAGGCTGCTGTCGGCTGCTCCCTCCCAGACATCTGTGTACTTCTGTGCCAGCAGTTACTCcacagcgctacaaggccgtctcctctctgcacataaaggca**gggaggttctgccctcctccc**ccacccaagactcagggatgc**cctgggcagagatctctgcgccagg**aaccttggaacccacagtggccccaagtggcctggacagtatgagccttgctctgtgccaggtgcctctgcaggcatctcagccaggcctggactggtcccaggtcctcagatgtctcctttgttgctttctctggtctatccttggagctttccttttggggtggggccagggctttcccagctcctacttttctactca**ttatcctgagtccgaggcccccaggataa**aacaggatttgcatttcagatccatctagactcccgtctctccctggtgaccctgttgcttcctctctctaatgtttcccccagcccccaccctcatgttttctctcctgtggcccacctttcccatctgggcagtcaccctccaaggcgttgctgggtctctcctccccccacctcctcg**cccctctctactgcagccatgagggg**agcccctgt**tctgtgcctccttccttcccatcacaga**gacttcaaaggccatttcctctgcccttgcatggagccttcctttctgcagtggccagctcctacctgtccttcatatctcagcatgatcaacccctcttgtgggaagcgctccctcccctcccatctgaaatcaactttcctctcataagctgtcacagattcatgtgtctgttagtaacctttagcacagcttggctttctcagatacttgtctggttatttttgtactccacccaatttttaagcccatgagagctaaggctgtgcctgctgcatttaccaacattggtgtctggcatgtagggagacccatttcacaaagaatggatgagtgaatgacaggctgagtgagtgatgagtgggtggataaaccaataataaaaacacactacatctactgtaacctggcagagatc

Uppercase: TRBV6-5

Lowercase: Flanking sequence[1000bp]

Red & Bold & Underline: Stem-loop [15]

Blue: Heptamer[36]

Green: Nonamer [3]

id-IGHVII-40-1[V_gene_segment]

gctactaaatagtaggtgttattcattctttctaactctgtattttgtacccattaaccatatccacctgttctctggcccccactagcctcccagactccagttaccatccttctaccatccatgttcctgagtttaattctgttgatttttagagcacacacataagtaagaacctgcaatgctttttttttcagtgccttgtctttttcacttaacatagtgacctccagttcca**tccatgttgttgcaaatgactggatctcatgga**atacctaattttattcggttctttcatatgacgtagagaagcaagcatgcagtctacaaacaactgtaattttcacatttaccccaaatgaattattacttatttctgtgcagggtccaggcacatagtagcctttctcttgagggactgttaaatttagagtgagatcacttgtttctcatattatttgattcctgtcca**tgtgcagagatcttaagtaatctgcttctttctgcaca**ctaactctgaccttctcgacaaaaagagtagagattttcatgtttcctgagtctggg**gtaggagttagtcctac**acaactcagagcctgcagagacctcccagtgcagttttatagacacaggtgctttctcatggggggtgagacctccacttcctgtgattgctgctcagatctaactgtgatttcagatttaggacaccattaggctctcaaacca**tttctattctaaaaatcagcctcatcatagaaaatggaaataattcaatatccattt**ccctgaccgtatttcatccttgtttttaatttggtgcaagtttattacatatttgcctgaatctaacttcagaatcaactgaattggtctagaaatctcaaaaattgaacagaagtggattttattatttttaccaaaatgtgtgtaggtagggtttccaatgtatcgtccagaacttgtgcatgctaacaactgtttttctcagtcaattatcAGCCTGGTGAAGCCCTTGCAAACCCCCTCACTCACCTGTGCTGCCTCTGGATTCTCTGTCACAAT**CAGTGCTTCCTGtaggcactg**gatggaaagcacaggagtgggtcaggtgcatacg**tgatgagtggaggatgaattccagcccacttatca**tgaattcagacaagcccacatgttcccacatgcactatatctt**cagacaagcagatgtgtctg**cacatgaattcttctttcagacaaacacgtctgtcccctcatggactcttgccaca**cacaagcacacgtggcttcttgtg**gactgtactttcagacaagcacacattgtttcatgtggacatcttcctcccacaaatgcattttcccctgagtagactcttctcccacaagcccat**atgtccgcacatggacat**ttctcacagacaaacacccatttccccatgtagtcccttccctcagacaagcacacgtattcccaggtagactcttcccttagacaagcacagatgttcc**catatgaacccttccatatg**acaagca**cacatgttcccatgtg**aactcccgtataacaagcacacatatttcctcacgggctcatcactcagataggtctgcatgtccacatgtggactctactctcagacaagcccacaggtcctcacatggactcttctctcaaacacccgttctccattgtagactcttcccttagacaagcacgcatt**ttcccatgtgaactcttctgcatgggaa**gcacacatattcccacagggactcttctctgaaacacccattctccattgtagactcttcccttagacaagcacgcattttcccatgtgaactcttccgcatggaaagcacacatattcccacagggattcttcacta**agacaagcatgattgtct**ccactgtgacccttccctcagagaagtaca**cttgtctacatatggactcttctctcagacaag**caaacatgtccc**cacgtggactctttcttctgacaagcacacgtg**tccccgtgttgttataatcattttctcacctctgtctcacaaagttccctgtcctccaaga

Uppercase: IGHVII-40-1

Lowercase: Flanking sequence[1000bp]

Red & Bold & Underline: Stem-loop [16]

Blue: Heptamer[23]

Green: Nonamer [2]

id-TRBV21-1[V_gene_segment]

ttataagtgtgtcaaaagctactatttgtaaagaccgagttgtatgattccttacctgagatggaagggaggggaagaagcaaactgacaagttaaaaaagtaaatagagtgtaaattagttcaaccattgtggaagacagtgtggag**atttctcaaggatctagaaccagaaat**accatttg**acccagcaatcccattactgggt**atatacccaaaggattataaatcattctactataaagagacatgcacatacatgtttattgcaccactgttcacaatagcaaagacttggaaccaaccaaaatgcccatcaatgatagactggataaagaaaatgtgacacat**atattccgtggaatat**tatgcagccataaaaaaggatgagttc**atgtcctttgcagggacat**agatgaagttggaaaccatcattctcagcaatctatcacaagaacagaaaaccaaacaccgca**tgttctccctcataagtgggagctgaacaatgagaaca**catggacacagggaggggaacatcacacatggggcctgtcgggggatggagggctaggggagggatag**cattaggagaaatacctaatg**tagatgatgggttgataggtgcagcaaaccaccatggcatgtgtatacttatgtaacaaaagtgcacattctgcacatgtaccccagaacttaaagtataataa**ttttttaagtaaataaatgtatgagtgattttacttaaaaaa**aatagcaaaacagtgagtaacagaaaaaaaaaaaactaaaagcaggctgaaacttacactatttttaacagctaaaacaaagtctgccctcctacattctttgaaattgaaatga**ctctgtcccatgtacagag**catctacaaaggctatggaaagtgatcacgtcacaaggaggttgctgacgggggctaggagagtgggtgaagaagcctta**cagaaaaagctaccactacgatttctttctgagccaaccATGTGCCTCAGA**CTTCTCTGCTGTGTGGCCATTTCTTTCTGGGGAGCCAGGTAAGGCCCTGTTCT**GAACTGGTTGAATTTCAGTTC**CAAGACTCTCTCCGGTGGCTGCAGTATCAAGCTCCCTCCTGCGCTTTTTCTACAGCTGCTCTCTCCTTCCTCCACAGGCTCCACGGACACCAAGGTCACCCAGAGACCTAGACTTCTGGCCAAAGCAAGTGAACAGAAAGCAAAGATGGATTGTGTTCCTATAAAAGCACATAGTTATGTTTACTGGTATCGTAAGAAGCTGGAAGAAGAGCTCAAGTTTTTGGTTTACTTTCAGAACGAAGAACTTATTCAGAAAGCAGAAATAATCAATGAGCGATTTTTAGCCCAATG**CTCCAAAAACTCATCCTGTACCTTGGAG**ATCCAGTCCACGGAGTCAGGGGACACA**GCACTGTATTTCTGTGCCAGCAGCAAAGCcacagtgc**cgaatgttagcccttcttagaacacaaactcattatggacccagctcaggaaataagtgtgtagcaggttggtaggcactacgtaacagaaacccaacttgaaagacaataaatcagaaggaaaaacttgtaaataacagctctaagtgagcaaaaaacaaagg**ggggttatttaaaacaactgatggggacaaaacccc**agggaaggggccagaaggcataaaaagcttggattcctcagtgttc**agcatattaggtaagatatgct**aggttaccctctctctctctctacatgagctttatggatttctaacaatctttatttaaagaaggggattctgtgcttttcacttggtaaagttgtgggtgcacaaataagaatgacaagcggatcaactgtgtctttcaggtagaccctgatcttctgacccagcacatttccttgaactagagacagactctaaagggctttttgact**gcttctcacagctctccctgccatagaagc**ctagtgggaaccattagcaattcctgccctatggaatgttacaatcacaaatttttatttttatttatttatttatatttattttaatttttttttgagacggagtctcgctctgtcatccaggctggagtgcag**tggcgtgatctcggctcactacaagttccgcctcctgggttcacgcca**ttctcctgcctcagcctcccgag**tagctgggactacaggcgcccgccaccacgcccagcta**attttttgtatttcttttagtggag**acggtgtttcaccgt**gttagccaggatggtctcgatctcctgacctcgtaatccgcctgcctcggcc**tcccagagtgctggga**ttacaggcatgagccaccacgcctggccaacaattacgaacatttttaaagcaactctattgagacataagatatatatggaatatgtatgcaataaacaaatccaactgctgcaataaaacacacccgcat

Uppercase: TRBV21-1

Lowercase: Flanking sequence[1000bp]

Red & Bold & Underline: Stem-loop [20]

Blue: Heptamer[31]

Green: Nonamer [7]

id-IGHV3-11[V_gene_segment]

ctgagtgaaggaagct**aaagaattcacagtaaatctcctgtgatttcttt**tgtataaattgtagaaaatgcaactattctaaattaacatggagaagatctgaatttttctgagaaaagtgtggtgaggtaacaagatggtgaaataaaattacagagaagtgagagaaaaaaatttggaggttaatttaattgctaatagcacgattgaagtgctgattcaaaggctgcacacatataccaacattttccatatcgtacactataaatttg**aattcactattgattgaatttttgaataaagcagtaacaaaaat**gagtatattggctgaggaagagcaagaaagagatgaatattgacacttgaataatcacggact**cctgaaaatacacacatgtgaacactgattgcatatttcagg**taaatactagaaaaagcaaagtcacataaagtcgttatgacag**gtgggataccctgcaaatcacactaggcatgtcccac**actgcc**ctggagctgtctcaggggagcagtctcctccag**tgtttagaggcacaggcacggataatagggctgacgc**tgtccagatgtgtgatattggaca**cattgcacaactgctctgttatgtatgtaattcatcttctctaaaaatgtaacattgacacttgcactgaatatattctgcaaatatgtaaacattaaataaga**tgatgactgctaattgatcatca**agtcacaatcacataatctgaagttatattttcctgagagataggattacctccagtgttttccgggaccctctcatctgctctgggcactgccctctcctccagcgtcccactagagcttgctatatagtaggagacatgcaaatagggccctccctctgctgataaaaaccagccgag**cccagaccctgcagctctggg**agaagagccccagccccagaattcccaggagtttccattcggtgatcagcactgaacacagaggactcaccATGGAGTTTGGGCTGAGCTGGGTTTTCCTTGTTGCTATTATAAAAGGTGATTTATGGAGAACTAGAGACATTGAGTGGACGTGAGTGAGATAAGCAGTGAATATATGTGGCAGTTTCTGACCAGGTTGTCTCTGTGTTTGCAGGTGTCCAGTGTCAGGTGCAGCTGGTGGAGTCTGGGGG**AGGCTTGGTCAAGCCT**GGAGGGTCCCTGAGACTCTCCTGTGCAGCCTCTGGATTCACCTTCAGTGACTACTACATGAGCTGGATCCGCCAGG**CTCCAGGGAAGGGGCTGGAG**TGGGTTTCATACATTAGTAGTAGTA**GTAGTTACACAAACTAC**GCAGACTCTGTGAAGGGCCGATTCACCATCTCCAGAGACAACGCCAAGAACTCACTGTATCTGCAAATGA**ACAGCCTGAGAGCCGAGGACACGGCTGT**GTATT**ACTGTGCGAGAGAcacagt**gag**gggaggtcagtgtgagcccagacacaaacctccc**tgcaggggtccccaggaccaccagggggcgcccgggacactgtgcacggggctgtctccagggcaggtgcaggtgctgctgagggctggcttcctgtcatgg**cctggggctgcctcattgtcaaatttccccagg**aacttctccagatttacaattctgtactaacatttgatgtctctaaatgcaatactttttttgtcctttttgtttctttgtttttttgcaacaggagtacatatcctcagctccacagaagccagggtgtcactttgggggcagaaataatcctttcatggttaccaggataagagtcctgaggaatcccagggaaacctggagagtgttttccagttagactcaggg**cagagacctccatgggaatctctg**attagaacaggccttgag**ctctgacgggagccaagagagaggctcgcccagggtcagag**tcctt**aaaacctgatggtttt**cacagctatcccccctcgtcttgtaaactcagactgattcagttgaccctctttctgctaatccatttccttctctgtaggtttgattctcacagttcgctttcttcttctcttccct**gaaaacagacgatgtgttttc**tgtagtcaaaatcccagggct**caggtctgcaggacctg**ggtaggctacggggactttctcactcaccattgtccggacactcttgtcttctgtgcatggaggcatttggaaaatgaagtggacattagccatgaagggaataatactagttttctccaataggatattgatgtagagctgatcttgtgcttctcacactgtctcagagtttggactctcacctgtgactttgaga**agagctggggatgggcactccattgtgctgtgagctct**gggtaacgataattgtagaatctgcctaggcagtctaaggtcaatactactcgtcatcaggaaagacagctggaattc

Uppercase: IGHV3-11

Lowercase: Flanking sequence[1000bp]

Red & Bold & Underline: Stem-loop [22]

Blue: Heptamer[35]

Green: Nonamer [8]

id-IGHV3-60-2[V_gene_segment]

aaaaatatctaagtgtttatgctgagtaaaatacgccagacaaataag**aatatgtaccatatt**actccatttatgcaat**ttcttttaagtgaaaaagaa**tctaaagcaatatccagaagatcagtagttacctggaaaaagggtagaccaaaggaaggggaaaggaggaaacttacagaagaacaagagaaaatgttgaggggagttcacttgtccagcttggaa**atgatgggttacatcat**gttgatcaattgcacactttaaatatgtgaagtctattatctgccaattaacactcgcaaaatttattgcaagcagacaaatgaaaaattagacaaagaaaggatggtataaagatagaaaatatatattaaatgtcagaaatgtctgagaatttaactcctgaccctagttccgtccttatttttaggtgaatggtagcgtgcaccaaaatcacacacattctcagtacaggaagtgggttccacaaagcacacgaggtatgtccaattcttaccaagatttggttcagggagtaacagt**gatgaggaatcacaggcccagataccggggctcactcatc**tcagacatgacctcgtggacacacacttagcccctc**ctccatgtgtaggttgacttccacatatgtaaatggag**aaaccattgactcctacagaacataatttacagaaatatacaaaagataaaatagtgcaaatacttatcacaacaaaatttcctaataagacagt**gtattttccaaatac**cgtaattgtcacccaactcctgtggggccgtgtcatttta**tctggggtctgccgtctcctcaggattcccaccccaga**gctctctatgtagtaggagacaagcaaatagggccctccctctgctgctgaaaatcagccaaatcctgaccctgcagct**ctgggagaggagcccccgccccgggattcccag**ctgtctccacttggtcatgaacactgaacacagaagacacaccATGGAGTCTGGGCTGAGCTGGATTTTCCTTGTTGCAGTTTTAAAAGGTGATTTATGGAGAATGAGACACACTGAGTGTGACTGGACATAAGTGAG**AGAAACAGTGGATTTGTGTGGCAGTTTCT**GACCAGGGTGTCTCCGTGTTTGCAGGTGTCCAGTGTGAGGTGCAGCTGGTGGAGTCTGGGGGAGGCTTAGTAAAGACTGGAGGGGTCTCTGAGACTCTCCTGTGCAGCCTCTGGATTCACCTTCAGTAGCTCTGCTATGCACTGGGTCCACCAGGCTCCAGGAAAGGGTTTGGAGTGGGTCTCAGTTATTAGTACAAGTGGTGATACCGTACTCTACACAGACTCTGTGAAGGGCTGATTCACCATCTCT**AGAGACAATGCCCAGAATTCACTGTCTCT**GCAAATGAACAGCCTGAGAGCCGAGGG**CACAGTTGTGTACTACTGTG**TGAAAGAcgcagtgagaagtcagtgtgagcccagacacaaacctcctgcagggtacctgggacaatcagggaaagcctgggacactgtatactgggctgtc**cccaggggcaagtccaggtgatataagcctggg**tttcctgtcatgatctagggttgcattgttagcaaattaccccagggaccatctctagatttccaattctgtaataacatttgatgtcgtctctgactgcacaatgtcccctcaactttgtatcttttttttttttgtaacaggaggacacatcctcaccctgcagaagcctgagtgtcactttgggggcagaaatgacctgcct**tgatcacattgatca**ctgtcctgaggaaaataccccacaggggaccccgatgactccagcaaaggctctgcctcaaaaccattgaagagtccttcctttcattagaattgaccacagcacctgggcttcagcacaagccataccacagacgtcacaaagcagcagcttga**cacctgatccaggtg**cattttctc**acctttagaagctgagagaggggtgtattctcaaggt**caacacactctttgtg**ggtttttacacagaaaacc**tgcttttactttat**ttcatttgaagataaatgaacaaatgtgcatctataaatgattgtaattttcacatttattc**caaatccaatatttctgaattctgcataatgtcctggcacaaggttgtg**ttttctataggtatttttcaacagaccaagaaaa**tgcattttccaatttccctgtttttcctccatgtgcagagaccttgagtagagcacgtctgactgtgtttgtctgttttcacgctgctgatgaagacataccagagacgaggaaacttacaatagaaagagttttattggacttacag**ttccacgtggctggaaaggtctcacaatcatggtggaa**ggtgaaaggcacatctgacatggcggcagacaagagaagagagcttgtgctgggtct

Uppercase: IGHV3-60-2

Lowercase: Flanking sequence[1000bp]

Red & Bold & Underline: Stem-loop [20]

Blue: Heptamer[46]

Green: Nonamer [5]

id-IGHV3-66[V_gene_segment]

ctcaaagaattgtgctgagtgaaagaagctaag**gaattaagagtaaattttatataattc**attgtagaaatattagaagatgccactaccataaattaaaatgaagaagacttaaatttttctgagaaaatggtgttgggaatgatgcggatgtgatttaagtttcagaggaataaggaaaaagatttagggattaatttaattattcaaaagttgattgaagtgccgagtgaatggctgcaaacatagctctacatttttcaaatcattccctataaatctgaattaatt**atttatttattatacttgaataaagcaataacgaagaaataaat**gaatatttttgctaaaatggagcaataaaaagactgatattga**cagaagaaatatgactgacttctg**aaaacacacatgaaccatggttctctctgcatatttaggtgaattacagaaagttgtcataacagatggggaatcctgcagacttcactaggcatggtccacgctgccctggagttgtctcaggggagctgcctcctccggtgattagagcacaggcccagataataggattacatttttttagatgtgtaaacttagacgcactgcacagctgctgtattctctatgtaaattatcttctgtaaaatacaacattaaaggctgcattaaatatattgtgtaaatatgtaaaaataaaatcagattatgagagctaaatgttaatcaaggcacaatcacat**aatataaaattatatt**ttcctgaatgatggaattactaccaatctcccccaggagactt**catctgcactgggcccggcctctcctcagatgtcccatcacagagcttgctatataatgggggacat**gcaaatagggccctccctctgctgatgaaaaccagcccagccctgaccctgcagctctgggagaggagcccagcactgggattccgaggtgtttcca**ttcagtgatctgcactgaa**cacagaggactcgccATGGAGTTTGGGCTGAGCTGGGTTTTCCTTGTTGCTATTTTAAAAGGTGATTCATGGAGAACTAGAGATATTGAGTGTGAGTGAACACGAGTGAG**AGAAACAGTGGATATGTGTGGCAGTTTCT**AACCAATGTCTCTGTGTTTGCAGGTGTCCAGTGTGAGGTGCAGCTGGTGGAGTCTGGAGGAGGCTTGATCCAGCCTGGGGGGTCCCTGAGACTCTCCTGTGCAGCCTCTGGGTTCACCGTCAGTAGCAACTACATGAGCTGGGTCCGCCAGG**CTCCAGGGAAGGGGCTGGAG**TGGGTCTCAGTTATTTATAGCTGTGGTAGCACATACTACGCAGACTCCGTGAAGGGCCGATTCACCATCTCCAGAGACAATTCCAAGAACACGCTGTATCTTCAAATGA**ACAGCCTGAGAGCTGAGGACACGGCTGT**GTATT**ACTGTGCGAGAGAcacagt**gag**gggaggccattgtgcgcccagacacaaacctccctgcaggaacgctgggggaatcagcggcaggg**ggcgctcaggagccactgatcagagtcagccccggaggcaggtgcaggtggaggctgtttcctgtcaggatgtgggacttcatcttcttccaacagtttctctaatgaacctctctaattttagaattctgtggttcctaatgtcatctctacatattttcaaaagatcattttaa**catgaggacataacctctcatg**caccaaatgcacattgatgcttacaaagatgaaaagttctcaaccattgtgaccaggatcgcagtcctgaggaagctcacgggtgtctgatgagtctcctccattcaggc**ccaggacagaaacctcagagggattcctgg**accagaacggcagggattctgatcacagccaatagagaggctgggcc**agggtcagtgtcctgtagaagctcacagggttcaagtctgaccct**tctcctgaccctaaagccaatgagcat**cagcactgatctggtgctg**cttttgctcccaatacatgttctttctttggagtgtttgttctcccttttttatttgct**tttctttcttcctgaaaaagaaa**cacatggtctctgtgatctacactccagggctcaaggcattttcttagaactcaggcaaggctcaggcttggctactccagccacgtg**ggagaggctgacgggatttccttctctctcc**atattctcagggccctcctctgtgttgtgtgtagactcatctgggaatgcaattggctgttagtagtgaaggggatgaactcatttgatcaaaatgggatgtggatgtggaattaaccctgttctatgcacactgtcagagtcatcttcttcagaagtagtgttagaaagagcttgtgaaatttatcggcaacaaaatggatccccttgtgttaaaaccctaatgaatgaagctggggaaggccatgaa

Uppercase: IGHV3-66

Lowercase: Flanking sequence[1000bp]

Red & Bold & Underline: Stem-loop [19]

Blue: Heptamer[42]

Green: Nonamer [10]

id-IGKV2D-38[V_gene_segment]

atgctcttctcagtcctttctgattatcgttagttttgttttgttttgttttgtttttt**gagacagagtctctctctgtctc**cc**aggttgcagtgcagtggcgcgatctcggctcactgcaagct**ccgccccccgggttcacgctattctcctgcctcagcctcccaagtagctgggactacaggcgcccgccaccacgctcggctattttttttgtatttttaatagagacaaggtttcatcgtgttagccagaatggtctcgatttcctgacctcgtgatctacccg**ctttggcctcccaaag**tcctgggattacaggtgtgagccaccgcgcccagccatgatcattgttagttttaagtctgttgtatctgaaataagaatagcaacccctgctcttttttgccttccatttgtttgatagatctttctttctccctttactttgagcctatgtgtaatcattagaaaccacctgaaagaacaatgaccagtgccttgaataatatcttttaataaagggtctcttgtacctag**taaatactaagcttattta**tttatttatttatttatttatttatttatttattttgttgcccagtttcgctcttgttgcccaggatggagcgcaatggtgcgatctcggctcaccaaaacctccgcctcccagattcaagtgattctcctgcctcagcctcccaagtagctgggattacaggcataagcccccaacccccaggtgattttatgtatttttagtagagatggggtttttccatgttggtcagtctgaactcccgacctcgggtgatc**cgcctgccgcggaccaagaagtgctgggattacaggcg**tgagccaccacaaccggcctaagctcatttttaaaaacgt**attttttatgatgtgaataaaaaat**cttttaaaagataa**atgataaatccatatcat**aagggaaatcgtgatacttgctcataatatttgtacataactttgtacttcATCTCC**TGCAGGTCTAGTCAGACCCTCCTGCA**TAGCAATGGAAACACCTATTTGTATTGGTAACTATAGAAGCCAGGCGAGTCTCCACAGCTCCTGATCTATGGGGTTTCCAATGGGGCCTCTGGAGTCCCAGACAGGTTCAATAGCAGTGGGTCAGGCACATATTTTAAACTCAAAATTAGCAGGGTGGAGGCTGAGGATATTCGACTTTATTAATACA**TGCAAGCTACATAATATCCTtgca**cagtggtaacgccttgaacacaaacctccttgctggggtgtcccaactgcccacat**gagctgctgtctgggagcagctc**tgcagggtct**ctgagtctgcggaagaggaggctgttggggaactcag**ggaaagggtttgctgctgaagactctggctcatgagagcct**cagctgtgcctcagtcacatatgttaaggtcccatcagctg**ccatggcctggggaagacatcatcaggaggagaattcttgggggccatgagccttgggaacaaccagctatgatga**agggagagtgaatgaaagctcatccctccct**atcttgcctacatttttcaaggacatttac**ttgtcaaaacaaccagaccgtggatgcagattagtgacaacacaagtggaatacgtgttg**gaaatgaccagcttggggtttattttatacaagatgatacttggtcctattgagaaatcatttttttcatttcattctcccaaccttgtatcttttttagcactcgtacactaaactg**ccctctccagtgtcatgctggagaagg**tattctacacaagctgtcctcaggggagtgtggcaaaaaataattagtaaaaa**tgtttaggtttgactattcatagacttggtaaaca**catggcaaataaaagatctcaatgccaaaatacttcaattctcctcaattaccttcatcccagtgaaagttcttccagttacaaaagtgagctttgaaaataaacagaatgaactcacagatggacaacatagactcactaaatatgtcacatgaataatgcagaattacataatattttttcttcttaccccag**ttttaagaatttaaaa**agcatttaacctaagtactgttttagaggtgacaaatttagtggtgtgtccccatgagaaaatatttcaaacattaccaaaattccttgtttacatattacatcatcgttcctagaaaatatatctaaatcctctattttaggtata

Uppercase: IGKV2D-38

Lowercase: Flanking sequence[1000bp]

Red & Bold & Underline: Stem-loop [18]

Blue: Heptamer[14]

Green: Nonamer [4]

id-IGHV1-2-2[V_gene_segment]

ctccctttgtttccagtgatccagtgcatctagacctccaggaagtggactccctggtgattttagcgattcttctcttggagccaccctgaagaggacattgggtttccaaaggcccattcactatttcaagaggtggtgtcatcagctcatgttgtcactgaaggagcattctgagccaggtcacagtcacttcctagtgagct**acagaggctgagagaaaaatgctctgt**gagacccaatgggaa**gctccctgcagtgcaaggtctgggtggcagggagc**gctagggcctcgcccagcacaggctgcagccctggagcaggtgcaagggaggctg**gggaggggttcctccc**agggtctgatgtcttccttttctcggacaaacatgctttaataagttaaacaagactttagt**aaagactattgatgtgtcttt**gtgtctttcagtatacagttctatttgtaggatttatctaacctaacaagtcaatgagaatcacatgtaaaaggagaaatttctaggattttcagatatcttaataggtaggagatggagaaaagggatggttttattaattcagtgcttgccaatcttaacagagacagtagtaagacatgcagaaagcaaagcccagaaaagtatgaaggtgtcaaagtgccatttaagtatgggttcacttggaggaccatgttctgcgggaacttgttttcagcagacaatctattttagcagagttctgggcatacaaggggacacacatcattaaacaaggat**tgggacagggacttcagcgtccca**ctgttgcatggcccataaattatgtgtgttctctttctcatcttggatcaagtctagagctatgaaatagtatccctcatgaatatgcaaataacctgag**atttactgaagtaaat**acagatctgtcctgtgccctgagagcatcacccagcaaccacatctgtcctctagagaatcccctgagagctccgttcctcaccATGGACTGGACCTGGA**GGATCCTCTTCTTGGTGGCAGCAGCCACAGGTAAGAGGCTCC**CTAGTCCC**AGTGATGAGAAAGAGATTGAGTCCAGTCCAGGGAGATCTCATCCACT**TCTGTGTTCTCTCCACAGGAGCCCACTCCCAGGTGCAGCTGGTGCAGTCTGGGGCTGAGGTGA**AGAAGCCTGGGGCCTCAGTGAAGGTCTCCTGCAAGGCTTCT**GGATACACCTTCACCGGCTACTATATGCACTGGGTGCGACAGGCCCCTGGACAAGGGCTTGAGTGGATGGGATGGATCAACCCTAACAGTGGTGGCACAAACTATGCACAGAAGTTTCAGGGCTGGGTCACCATGACCAGGGACACGTCCATCAGCA**CAGCCTACATGGAGCTGAGCAGGCTG**AGATCTGACGACACGGCCGTGTATT**ACTGTGCGAGAGAcacagt**gtgaaaacccacatcctgagggtgtcagaaa**ccccagggaggaggcagctgtgctgggg**ctgagaaatgaaagggattact**atttttaatgttgtttacagtatgtcattaataaat**tgaaaaaaagtaacaatagaagtatatactctaattatatgggaactttgttttttcagttttttcatttttttttttttttttggtttgttt**gtgacagagtctcactctgccac**ccaggctggagtgtaacggcacaatctcagctcactacaacctccacctcccaggttcaagcaattctcctgcctcggcctccagagtagttgggattacaggcacccgccaccatgccc**ggtgaatttttgtatttttagtagagacggggtttcacc**atgttagctaggctggtctcaaactgctgatctcaggtgatctaccctcctcagcctcccaaagtcctgggatta**caggcgtgagccactgcgcctg**gccc**aattatatgggaattgtttatataatt**atcaccctataagcaaaattcatggaggaggaaaagctctactgaagaaagctgataccggcattcccatgaaagtatctgtgtagaagtaagtattaaaatcagttga**ataggcaaggcatggtggctcacgcctat**aatcccagcactttgggagaccgaggcaggtggatcacaaggtaaggagttcaagatcagcctgcccaagatggtgaaaccccttctctactaaaaatacaaagaat**tagctgggcgtggtggtgggtgcctgtaatcgcagcta**ttcgggaggctgaggcagagaattgcttgaacctgggaggtgaaggt**tgcagtgagccgagatcacgccactgca**ctccagcctgggcgacagagtgaaactccatctcaaaacaaaacaaaacaaaacaaaacaaaaaaacagttgaataaagtaccttagagtcatctgttcaattaacatgtttaact

Uppercase: IGHV1-2-2

Lowercase: Flanking sequence[1000bp]

Red & Bold & Underline: Stem-loop [20]

Blue: Heptamer[43]

Green: Nonamer [4]

id-TRAV9-1[V_gene_segment]

tatcttatcccgtggtaaaaata**actattggtcaatagt**attggtggatggacatatgtagggggtatataatcaaggctaaatttccattaaaaggttagtatcacaacactttgcctcatgcacataggccagtcacattatctggga**atcatttcttctcctacaggagaagaaagtgat**gccagataattttgtagggatt**taatgactttagttactatattttttcatta**taagagt**tatatattctttactttttaatcaatatata**cacaagatataatttccttttttctttgaaaagaattt**attaattggaaattacaattaat**ttatttctgtaattgatgccttaaatgtgaagcaatatgatatttcaagtagtaacaattcctcaaccaaagaccattcatttattcatgcatttgtgtatttaagaagtgtttcctaaccacctgctgagtgtttgacattgtactaggtatgggaatacagcagtgaaaaaaagcttaagtctctctcatctcaaataatctactaatgtttaaacatgtgctactgacattcctctggttttatc**ctaaaaaagtgctaatacggtattttag**tggatcatgggttgggcaaagatggctaaataagaaaaggtaaaatgaattattctcagaaatgccatggtatgaaaagaacttgcaagtcagacagtgtgctgttaaaaaaaacagagttataaaaaagaccaacta**ttctttagatatacaaagaa**ctatcagacagctaataggtgtcttagaaaaaaaaaaaaagccagaagtgaagatac**aggattgacttttgaaagaacagtcaacaatcctttgagtttaggatcttgcccacaaagg**ggtgcatcttcctaacacattcacatttcctgctcacgctcat**ttcttcagtgtgaccttaacttttcctcacactaagaa**gaca**agacccaagggcaccagagggtct**aaaaATGAATTCTTCTCCAGGACCAGCGATTGCACTATTCTTAATGTTTGGTAAGTAAAATGGCACTTAAAACTTAGAATCTTTTGTTCCATTATGTTGATGTAATCTGTAACCACAGTTTTCTCTAACGGGTACAGCTAAGATCACGTCTAAGGTAACTTGGTATAATTTGTTTATTTTTTTTTCAGGGGGAATCAATGGAGATTCAGTGGTCCAGACAGAAGGCCAAGTGC**TCCCCTCTGAAGGGGA**TTCCCTGATTGTGAACTGCTCCTATGAAACCACACAGTACCCTTCCCTTTTTTGGTATGTCCAATATCCTGGAGAAGGTCCACAGCTCCACCTGAAAGCCATGAAGGCCAATGACAAGGGAAGGAACAAAGGTTTTGAAGCCATGTACCGTAAAGAAACCACT**TCTTTCCACTTGGAGAAAGA**CTCAGTTCAAGAGTCAGACTCCG**CTGTGTACTTCTGTGCTCTGAGTGAcacag**tgacagggactgcaggggaagctgagcacaaactctg**agcagcacgaggggcctggctgct**gagtgtaagccactgtgatcccctctggttagggaccaggaactactctactattgttgcccaccacccaccctaggcactcc**tagtgtgaaagccatccaaataaactctaaaacacta**tagaataactaaaagtaaaaatacctcacaccagatgctttaacaatgagagtgaaaacagaatgtcaccatagttcctcaaccttgtttgaaatttgtgattttaaaagtataaaaccttgagataaatgaaatggtctagctgctcatccctgtataaaatgtagcctcctgaccttgtctcaggccatccaaggaaatagtatgcttacagacacagctgtgacagatgaggttgcaaatgctt**aatgcatacctgagtggggtatacatt**cagctggggcaagatggggtg**atttcttaggaaagcttctgcccaagaaat**aataggttcttaaagaaaaaaagaattataaatggaagaagtcaaa**ttttccttgtttgcagatgatatgattttatatttggaaaa**acatagagactccacaaaaaaaccttagaactaaaaattcagtaaagttgcaaaatacagaatcaacctataaaaatcagtagcatttctatatgtcaacaataaacaacccgaaaaagaaataaaaaagtaatcgcatttatgatagccatgaataaaattaaataactagggattaacttaaccgaagaagtgaagatctctataatgaaaagtataaaacattgatgaaagaaatcgaagaaaacaccaaaaaaaatactccgtgttcatggattgtaagaattaatattgataaaatgtccatactaccccaaacagtctacagattcagtacaatccctaaaaaaataccaatgacattcttcacagaaatagaaaaaaaatcctaaaatttatatg

Uppercase: TRAV9-1

Lowercase: Flanking sequence[1000bp]

Red & Bold & Underline: Stem-loop [19]

Blue: Heptamer[32]

Green: Nonamer [1]

id-TRAV24[V_gene_segment]

tagatcttccttttctgaacatttcatataaatggaatcactcaatatgtaatcttttgtggctggattctttttcttaaaatagtatttttgaggctcatctgtgttgtagcatatattatgttttattttatttatttcatttttgcttatttattttattttatttattatttatttatttttgttgccgaatatcattcacatcttgtatgaacatactgcattttgtttattcattcaccagctgatggacatttgggttgttctcatttttttagccttaataaagaatatccacaaattagacttccattatttttagaacatatctaacatgtaaaacattcatggtgcatcacaaatgtaggtggcatatcagattccaattataaagtctagagctagacagatatatgttctaattcttaatatgccacttaggtaattgaccttgagtgaattatttaatctccttgagctttgggttattttttgtttgtaaattagaaacagtaatgtctctattgcatgac**tttcttattagatagatgtacataaagaaa**aaggtaatttagatgtgtgcacatttataatttgaggggga**ttcaaaagttgttttttaatgtttgaa**gattcaaagaaaaatgcttttcctgccacaaatgccatcccacctgtaactccctctaaatttgaataatagcagtccacagttaggcagcgcacttctgaaagggtgtcaattctcgttgtgttccacactcatgggggatttcctcaaagtttatgaagttggcctgaagaaagcagttacttcgactcctctttgtgaaccacaggaaactgaacatagggcagagcgtttctaggacatttatgggagctggtctctgattggctaggcaaccagatgcagaatcctgttttctgaagcagaatattt**ccacagattttggctgcaaaacgtttttctgctgtgg**gtacgtgagcaggaaacATGGAGAAGAATCCTTTGGCAGCCCCATTACTAATCCTCTGGTTTCATCTTGA**CTGTAAGTCCAGGGGTTACAG**GAGACATTCTTAGTCTGGGGTGATGATGAGGAAGGAGGAGGATGCAGTCAGACACTCTAGGTTTTATTTGGGGATGTGTGAAGAAAATGCCAACTCTGTAGAGAGACCAATTTCTATGTCTGACTCTTTTTTAAAAAAACAGGCGTGAGCAGCATA**CTGAACGTGGAACAAAGTCCTCAGTCACTGCATGTTCAG**GAGGGAGACAGCACCAATTTCACCTGCAGCTTCCCTTCCAGCAATTTTTATGCCTTACACTGGTACAGATGGGAAACTGCAAAAAGCCCCGAGGCCTTGTTTGTAATGACTTTAAATGGGGATGAAAAGAAGAAAGGACGAATAAGTGCCACTCTTAATACCAAGGAGGGTTACAGCTATTTGTACATCAAAGGATCCCAGCCTGAAGACTCAGCCACATACCTCT**GTGCCTTTAcacagtgctgttcaggcac**ctgcagcccatacgcaaacctgtgtctggtgttgcactgttaccagcattgacaaagaaccatgagtaggatggaaaagacaagttcgttgaattacagttt**tttttcaagaggattgtgcaaacatagaaaaa**taagcatgctagtgatcataactaactaccacaagagcaaattaagtaaacatattccttatgacagatagatgtcagagtgctaatgaagcataacacatgacctggtaactgggccgttactaaacaatactaaccccagctcccaacagctcattaaccac**atcatggagtcagacccattgcctgctcttttggcatgat**tctgaaagaacccaatagttactcaaatctatctaaaacaatct**gggtaaacaaaggcgatatttaccc**cctcattgttgctatgtcctgaaaacgaattcttctattatactttatcacagcacaaaaggagcatgtcggcatgcccctccctcacacacctggcctggaaatatgcatgtttaataccagaacaggtgatgctgcctcctttaatccattggggattttggttttgctttagttcaacttttccc**ccacatgaacatttgtatttttatttcttctcacatgtgg**ctattttcatttttaaatatcactgataatatcataaaggagcactgatgaaggaccttcactattttacgtggtctataccccagcttgtactgtgattgtgatattggatgactaactgaactctattatcaaagtctcttcatttgtaatttccaggtatattaagtaggctgag**tgtttggtgctctcctccttttgcagatttccaaaca**acccaccagtttaaggtcccacttgtactctagaggcatctaattctaagcgttcttgggggctgtta**agatgaagtaagttgcttctgggcctcctcatct**aaaagctagaatttcaactttctctaaac

Uppercase: TRAV24

Lowercase: Flanking sequence[1000bp]

Red & Bold & Underline: Stem-loop [12]

Blue: Heptamer[21]

Green: Nonamer [2]

id-IGHV4-80[V_gene_segment]

ataaccttcagcttcattgtttttgtgtaatctatcaacaaattatgttagttcaaggttctcaatgggagtttctaataaatagaagggatgtatagaagttcccctaattaaaacaattgtgaacacaatcttggtattcagctgtgtctccacccttcttaccattcaccacaaagtaattctcacttctggaagctgggttcattttcaaattagttttttttttcattttaatatctcaaggttattgtatgtgactattttagcagaaat**tgaattttgggaaattgaactaaccaactgaaaataaattca**caactaaataacaagatgccagaatataattggctccaggctttgttaattcagcagtttatctacctaggcattatatttacacatcttcatgctatttttacagtcaactc**ccattatgtaagaaatggtcatacaccatttcttacatatgg**taagatatctacgatggatattcagaactacgataaacattggatgcatccaggtttgtctgaactcactgagattattacactcagctgctgtttcaatgtgtctgaaggtataaatgacaattcgaatgacttgggtgtgtggatggttttacatgagtctttaagtgtagaacaatattgaccctgttccaaaatctgtccttgatccatgatcacactcatctcccagaacagctccttcaacacatttcctatctggatgaagaggactctgtgcttggtgaggggaggccacaggta**gagaactgagttctc**agagggcacagccagcatcctcctcccagggtgaacccgaaagactgcggcctccctcatcccttttcatctctccatacaaaggcaccacccacatgcaaatcctcacttaagcgcccac**aggaaaccaccacacatttcct**taaattcggcgttctgcttacacgggaaatgctttctgagagtcatggacctcgtgtgaa**aagaacATGAAAAACCTGTGGTTCTT**CCTCCCCCTGGTGGCAGCTCCCAGAAGTGAGTGTCTCCAG**GATGCAGATATGAAGATATGGGAGGCTGCATCTGAGCCCAGGGCTCA**CAGTGGGTCTCTGTGTTCACAGGG**GTCCTGTCCCAGGTGCAGCTGCAGGAGTGGGGCCCAGGAC**TGGTGAAGCCTTCGGAGACCCTGTCCCTCACATGCGCTGTCTCTGGTGACTCCATCAGCAGTGGTAACTGGTGGAGCTGGGTCCGCCAGTCCCCAGGGAAGGGGCTGGAATGGATTGGATACATCTATTATAGTGGGAGGAGCTACTACACCCCGTCCCTCAGGAGTTGAGTCACCATGTCAATAGAAACGTCCAAGAACCAGTTT**TCCCTGAAGCTGAGCTCTGTGACCGCAggga**ggcggagggggcgggcgcaggtgccgctcaggaccagcagggggcgcgcggggcccacagagcatgaggccgggtcaggagcaggtgcag**ggagggcggggcttcctcatctgctcagggctctccctcc**tcgccagcacctcagctgtctccagggctcctctttctttcttatctgtggttctgcttcctcacatccttgtggcaggcaagaaaggaggacgacaggcctg**gtgcggtggtgtatgcctgtaatccccgcac**tttgggaggccgaggcgggtggatcacctgacgtctggagttcgagaccagcctgaccaacatggagaaaccccgtctctactaaaagtacaaaattagccaggcgtggtggcgcatgcctgtaatcccagctactccggaggctgaggcaggagaacggcttgaacctggggggcggaggttgctgtgagccgagatcgtgccattgcactccagcctgggcaacaagagcaaaactccgtctaaaaaaaaataaaaggaagaagacaaattttcctcttacagtcgaagtatcaacaatact**aggaactttcctacgagttcct**gaatggccc**cttttttccttcttgattaaaaaaaag**ttctaatgcttctcaccatctctttatttgtgtcgtcaat**tgaattgtgccctctttgaaattcatatactgaaatcttgaatccagtggatctatattggaattttaagtatat**aattaaggttaaatgtggtcatacgtgtgagaccctaaagcaatagacatgttgtctttataagaagaggaagagacaccagagacctctcacttttcacgtgcacatagaaaggaggccatgtggacacacagtggactagaaggtggccctgggcaagccaggaagaagccgca**ccaagaaccaccctgccagcaccgtgatcttgg**acattcagactgcagaattgtaagaaagtcagtatttgttgtttat

Uppercase: IGHV4-80

Lowercase: Flanking sequence[1000bp]

Red & Bold & Underline: Stem-loop [16]

Blue: Heptamer[29]

Green: Nonamer [4]

id-TRBV17-2[V_gene_segment]

ctgaccatgccctaagtc**ctgagcattaatttttgctcag**tttcatttacatcattttcaactagtttctcattttctctctattctttttgttttcaatgcagctcccagaaactaaatctatccatttaatattctttttcccaggaaaattcttctcattctatttaaagtgtaatattctagtttaatgggaaagttttttgttgttgttgtttagccttttaaaaatctctactgaaaa**aattttggatatgaaaatt**agccaggcatggtggcgggagcctgtaatcccagctactcaggaggctgaggcaggagaatcgcttgaaacttggaggcggaggt**tgcagtgagctgagactgca**ccactgtgctccagcctgggca**atggagagagactccat**ctcaaaaaaaaaaaaaaaagaaaagaaaagaaaaaattgtgaaatgtggt**ctgagacaacttttctcag**tgtcacctgaggcatgggttaaacaggtatattctaggacaggttcag**aatttcagagtcgaaatt**tcttgttggtggactttaggaatacgtatcattgctcctgagtatttctgttagtgctgaagttctgccccaaggacttcgt**aagattaataacatcgcaaatctt**ctgtagtgctttaaaattttgttcttcatttttttattctcactttcttctctttccttttccacttcctcttatgttttccttaatcttcaactttcctaagcacctgcaagtgggattggagccttgtttaacatcgtccatgtagcaaaataaggatgaggccaaatatttgaaccaaggatccccatctcctatggaaggtgccctgaggttgtgggtgttgctgggga**catgatgtcatggccagatcctacatcatg**cggccaagggaacccagaactttcactgctctttgctactgcacatcagaacccatcgctgggagtgtcttgcactgcctgacctcaccATGGATATCTGGCTCCTCTGCTGGGTGACCCTGTGTCTCTTGGCGGCAGGTGGGTCCAGGTATACTT**AAACATTTGCATAAAGATGTTT**TTGGCTGGGCGTGGTGGCTCACAGCCGTAATCCCACCTTTTTGGGAGGTTGAGGTGAGTAGATCACCAGAGGTCAAGAGTTCGAGACCAGCCTGGTCAACGTGGTGAAACCCCTTCT**CTACCAAAAAATACAAAAATTAGCCAGGCGTGGTAG**TGTGCTCCTGTAGTCCCAGCTACTTGGGAGGCTGAGGTGGGAGGATCACTTGAATCTGGGAGGTAG**AGGCTGCAGTGAGCAGAGATCACGACATTTCACTCCAGCCT**GGGCAACACAGAGAGACCCTATCTCAAAAAAAAAAGATGTTTTCTTTGGGCTTCCCTTCACCTTCTATGGCTTCCGTCTTCTTCCACAGGACACTCGGAGCCTGGAGTCAGCCAGACCCCCAGACACAAGGTCACCAACATGGGACAGGAGGTGATTCTGAGGTGCGATCCATCTTCTGGTCACATGTTTGTTCACTGGTACCGACAGAATCTGAGGCAAGAAATGAAGTTGCTGATTTCCTTCCAGTACCAAAACATTGCAGTTGATTCAGGGATGCCCAAGGAACGATTCACAGCTGAAAGACCTAACGGAACGTCTTCCACGCTGAAGATCCATCCCGCAGAGCCGAGG**GACTCAGCCGTGTATCTCTACAGTAGCGGTGGcactgcatggctgagtc**agttccctccagggtgcaaaccctctggctgctcttctcccagttgaactccaagaaaacatttgaaaaagcctcttccttatcttcctaccccagaagaaagaagcgagttgattgttgtggctgcagctgctaccgggagagtac**aagaccatgaattaaggtctt**aaatggtcatgatgggcacactggacaatgggctcctgaaactacccaaatacaaaatgagacattc**tgtggatcaggaggaatccaca**tgtttagaaggaagggccccagaccaatttccaagttcagagaccagctatctgaggttgatattacttaccaaaacacaaaaacattcctactgattttatctcaaacacagttctcctgcaaactcttcccccaccaacgtaccccagcagaggcaatgacatgtacatttatggaacagcctgttgactactgcagatccatctcttcaataagaccaactttccctggaggtcagtgactaaaccaaaccgcaaggcagacagcaacacgaccctctta**ggggatggatgccagagaaacactgactaaaatcccc**actcggggactgtcctagcacattcagttcctcacacagctgcccaggggagcgcgacgttttattctgttacttcaaacactcatctgttcctctctctctcttgcaggccctagactcacagacac**tggaattagagtcagaaaaattgctttctaatccca**aaactggttttgtttactccccttcagctgtgaaatcttggcaaatgctcttacttctctgggactcagtgtcctcctcttcaaaa**ttagagacttgctccacttagtctctaa**gatctgata**caaatatgattatttg**acttgctatttatttatttatacaaatattaga**atatttatgtatacatttaacataaatat**atttatatttaataaaaacataaaatttataacattttata

Uppercase: TRBV17-2

Lowercase: Flanking sequence[1000bp]

Red & Bold & Underline: Stem-loop [19]

Blue: Heptamer[36]

Green: Nonamer [5]

id-TRBV10-1[V_gene_segment]

tgggcctcctccctgtctccacttctgattctgtgtcctctctcactggagcatgattgtgggaattacccatccaggttgcatcaagatg**tgcaatagatagcctattgtatctgaaataactattgcagtgctatggactacatgtgagttttgcaccatagcac**gtttacaaatccatcaaccttc**tcccatttggtgaatataagatctgctggatggga**atagaggattttcctgtgaccctagagactccaggtaggtgtagttcttggtgaacatttcatccatattcctaatgagcctcctcttttcttcacaggaagtacagtcactcaggggacttagatggggca**gtgtttgtggcttcacctgaagcaaagggagaaaacac**ccccatgagaatacacagcagcctgagcttggatggtgacattcccatcaggtgtaatgaggtgctacagcatgtaacaagtc**ttcaagcggtcatgaccttgaa**cctggaacttggaacttggagacatttctggatgtcatttaaccgagggtatccc**tgtatgctggcataca**agtgtccagaggttaaacgaagaggaaaat**aattcaacttggagttgaatt**ttgtgcaggaa**gtcataaaagtcaccaggactgctatgac**aatgttctttagcctacagaaatattcataaaagtgaaaatagtattcaaataaagaatttctcatgatggaaataaaatatacatagaagaacaaattcccctattaaaatcagg**gaaagagcttaggagtttattagatttctttc**aaaacagacctaa**aaattattcaactctgttataaataattt**gttc**ctgcggggaagagaaggggcatggcaggttagctccgcag**ttcaagcattcagcaggaa**cagtgacatcatcaagtcactg**agagcccaacttcagtctgcccacag**cagggctgggagacacaagatcctgccctg**gagctgaaATGGGCACGAGGCTCTTCTTCTATGTGGCCCTTTGTCTGCTGTGGGCAGGTGAGGGCTGGTCACAGGAGGGCCTCCTTCCCTGGAAT**TCCCAAGGCCTCAGTGCAGGCTTTTCTGTTGGGA**TGACAGCA**TCAGCATCTGTTGTTCTCTATTACAGGACACAGGGATGCTGA**AATCACCCAGAGCCCAAGACACAAGATCACAGAGACAGGAAGGCA**GGTGACCTTGGCGTGTCACCAGACTTGGAACCACAACAATATGTTCTGGT**ATCGACAAGACCTGGGACATGGGCTGAGGCTGATCCATTACTCATATGGTGTTCAC**GACACTAACAAAGGAGAAGTCTCAGATGGCTACAGTGTC**TCTAGATCAAACACAGAGGACCTCCCCCTCACTCTGGAGTCTGCTGCCTCCTCCCAGACATCTGTATATTTCTGCGCCAGCAGTGAGTCcacagtgctgcacagctgcctcctctctgcacataaagggcagttagaatgactgaggttg**cctgtgctcccaagtcccagccttcacagg**agtcggagagccctggctagcctgggggccatagagcaggggccatataaaccttgatgttgaggtccattcctaccccagtctcagaccaactggaggtcaccccaacacccttagcttctgcagttccctcttt**ccttttgtaaaatggaaaagg**catctttagtcgagattcctaatttcgttgaaaatttgaaacccaccatattcttctctaccctaagggctgggtgacttttccacataccctttcccctaccctctactgcctttcaactcccaatcataaatctttgcccatgtggtctctgctcctgggg**gccctttattctaagggc**aatagacccacgaggtcctatatcattagccatttacgtacctcttcatcatcattttgcaccatgttttccctggctctctgctccagccacactcccttacatgcagttatttcttcatatgtgtctgttgttttccaccctggggattgtttatataaattctttcttctgcactgagcatacataccacccacttagtcctacatctctacttcatttcttaaatta**aagtgtatttcaggcacactt**gctcttaattgtaaaaagcaaaaaagaaagtttactagaaaaatttaagaaaagcattttatgcattcagttgagggatgacatttttagaggagacaaaaagccacaagccaaaagaacaacattcataacgctagttgcattaaattgtttcaagccaaaacacaccataaaaaataaagttaagctacagcctgggaaaggcatgaggaatgcatacaaatgatgaaaaatcagtgtgacagagaac**attcagtgtaacagagaacattcagaacatcactgaat**acctgtagctcctccatggtttctggtctcc

Uppercase: TRBV10-1

Lowercase: Flanking sequence[1000bp]

Red & Bold & Underline: Stem-loop [23]

Blue: Heptamer[43]

Green: Nonamer [5]

id-IGLV3-27[V_gene_segment]

atttgggatcctgatcccttcatggggcatccccagcatccttaaggagacactct**tcagagggacagggcttctcacctgcctcagctctga**ccatggggctgtccacac**tcagtgacaggacactga**ccagggatctgactggagtgaccaggaacctggctggggcaaccagggaccacccaccctctggagctcacatgctgtgcagagaccatgaccctgggaccc**cttcagagaccatggggcctgaag**ttagattttcagtccagtgtggttttctcctgctttttgcaagtttacataaaacattgatgagaaaaacatacataaataacagacgagctgagggtgccaattgctaatctcttgtggcaatgaggggtgatgtagcaggagcagcaccatggctgaaatgagaccagagg**ccctgttgactccacaggg**gtgtgtggttgcttaaatgacccaactattcccactgaataatcagcaagtgtcccaatgacacagcacagcatttagatgaaatgaacaagaggggactgagtctg**agggaccagggggaggaagatcatgggtccct**gggccacctgc**tgctgagtcaggtgtcagca**agatctctccgcagagcccagctcagcccatggccaacctcctggaaggtccacagctct**gctgagcctggcccaggacagaccccaggtcagc**gaggtttgggacagtggtcatgagattaattcccagaaaggtgactgtgatcacagggcttgaaggctgcagctgattttctaatctgagaccctgcttcttcctcctgttgcctgtgctgccctcacagctggtttgagtgacatctctccaggaggagtcgcagaggatgtaaatttgcataaacgccaaacactgactgcccccaaa**agctggagagggaataagagaggcctggggagcccagct**gtgctgtaggctcaggaggcagagctctgaatgtctcaccATGGCCTGGATCCCTCTCCTGCTCCCCCTCCTCATTCTCTGCACAGGTGCTGCCCCT**AGGCTCAGTCTCCACAGACCCCAAGTTGAGCCT**GACCTGAATCCTGAGCAAAGCCCTGCCACTGCCTCTGGGGGGGATTCCTGGCAATGCGTCCTTTGTCCTCAAGCCCCCTCTCCTGTCTTTTCTTGCAGTCTCTGTGGCCTCCTATGAG**CTGACACAGCCATCCTCAGTGTCAG**TGTCTCCGGGACAGACAGCCAGGATCACCTGCTCAGGAGATGTACTGGCAAAAAAATATGCTCGGTGGTT**CCAGCAGAAGCCAGGCCAGGCCCCTGTGCTGG**TGATTTATAAAGACAGTGAGCGGCCCTCAGGGA**TCCCTGAGCGATTCTCCGGCTCCAGCTCAGGGA**CCACAGTCACCTTGACCATCAGCGGGGCCCAGGTTGAGGATGAGGCTGACTATTACTGTTACTCTGCGGCTGACAACAATCTcacagtgacacagg**cagatggagaaatgagacacaaaccccttgccatctg**tgtcaccttct**tcctccagccccaggagga**ctgtggacaaagccacgaacaggtctggcccagtacacccagatcagagactctga**ggcctccttccttccagatctctacggaggcc**ctacagaaggtggatcaataataaattcggggttggcaggtccagtaggtcctggtgttttctgggctggagtgtgacttggggaaaaatgacttgaatggaagtacagtcagtgggagacatctattgagtggtgatggtcccagtatttccttgtgtggtgactg**aacctagggcagtgccactactctgtggcccaaatgccctgggtt**attcctctgagttccccgtgtattattgagatgctccagcccccactctcatggtatcctgatctgaagaatcatggccagtgcccattgcagtgtctccatctaggtgccattttcaggactcatgcatccagcatccaaacaag**gctgtgtgggagaccgtgacttgtcacagc**tgttccccactggaaaatgccactgagctgcagagaactgctctcccagtgatgttcccccttcacagtgtggttcaggtgcaatgcctgcctggtccctag**attttaatgccatgttttcatttaaaat**aaccacgaagtgtctcttgctgagcgctccggtgtaacacattcccagtcatgcctctcac**aattttctttctgacaaaatt**tcctgaaaccctattgcaatgaatt**ctccctctcagagtaggattgcagggag**ttcacaaattaacaaacaaaccaaatactgtctagaagaaagaggatcaaagaggagagagctgcacagggagcgcactctggagtctgcagatattcatgagcaccgatgagcatgtgcatgggaggaaaccttctgtgatagagaaaataaccattcaaagacatttgaggaaaca

Uppercase: IGLV3-27

Lowercase: Flanking sequence[1000bp]

Red & Bold & Underline: Stem-loop [20]

Blue: Heptamer[54]

Green: Nonamer [4]

id-IGHV3-41-2[V_gene_segment]

gctttgtcatcc**aggctggagctccgtggtgacatcatagctctctgcagcct**cagaccctgggtctccagcaatcctcctccctctgccctatctagctgggactacagctgtgcaccaccatgctcggcttcagtgcttattcaaccagacactttaattatgcagtatttattacgtg**gcaataatgcctcaataagggtattgc**aaataagtggatggataatttgttcaattaagattgatggaaatatagacacttacatgacaaatgtatgacaatcaagaaaataaaactgtaggaaacatactttactttttgttaggtaatcgcaacagtgcatacacatcacaccatgttctcattacagagaaaaggttctgcaaacctcacttggtgtgacctcctgtatgctgggcttggttcagggagaagtcaggaccag**tggttagaagcacgggcccagataacca**gactccctctgaccaaatgtgagtgctggggacattgtacaatccatctgtgttttgctgatattttttcatctgtaacatggaaataacattgatactacataccatggtttctgtgcatatggaaaaataaaagatgataggtgcaaactttaaatacac**acagtttatgtagatcaattgtacctcaataaaactgt**tttaaaataaaaattac**aaaactataaattttataggttt**taagggtttatcatagaacaaacttacaataagaaaccagaatttccaaatgctatcaatatcacaaatctcccccaggacactctcacatgctctgagccccactctctcctcaggcgtcccatcccagagcttg**gcatgtattaggagacatgc**aaatagggccctccctctgcttatgaaaaccagcccagccctgaccctgcagctgtgggagaggagccccagccctgggattttcaggtgctttcattttgtgatcaggactgaacacagaggattcaccATGGAGTCATGGCTGAGCTGGGTTTTTCTTGCCGCTATTTTAAAAGGTAATTCATTGAGAACTATTGAAATTGAGTGTGAGTGGATAAGAGTGAGATAAACAGTGGATTTGTGTGGAAGTTTCTGACCAGGGTTTCTTTTTGTTTGCAGGTGTCCAGTGTGAGGTGCAGCTGGTGGAGTCTGGGGGAGGCTTGGTCCAGCCTGGGGGGTC**CCTGAGACTCTCCTGTGCAGCCTCAGG**ATTCTCCTTTAGTAGCTATGGCATGAGCTGGGTCCGCCAGGCTCCAGGGAAGGGGCTGGACTGAGTGGCACATATCTGGAATGATGGAAGTCAGAAATACTATGCAGACTC**TGTGAAGGGCCGATTCACA**ATCTCCAGAGACAATTCTAAGAGCATGCTCTATCTGCAAATGGACAGTCTGAAAGCTAAGGACACGGCCATGTATTACTGTACCAGACAcagtgagaggaagtccgtgtgagcccagacacaaacctccctgcag**gggcacgcggggccaccagagggtgccc**aggatcccctgaagacagggacagcccaaaggcaggtgcagatggatgtcaagagggtctgtggcttcgtctacatctaactgtttcctggtgagcctctgtatatttatttttctgtgcccactaatgaggttgctaagtttaaacacttttttttttgagatgaagtctcgtgcttgtgccccaggctggagtgcaatggcatgatctcagctcactgcaacatccgcctcatgaattcaagcgattttcctgcctcagcctcccaagtagctgggattccagacaactgccaccacgcccggctaatttttgtatttttagtagagacagggtttcaccatgttggccaggctagtctcgaa**ctcctgacctcaggag**accctcccaccttggcctcccaaagtgctgggattacaggcatgagccactgtgcctagccaaaaaagacttcattattataggaggaaaaattctcatctgcaagtgtcactgatggaagcagaaaaattgcccaggaagtcaggtgaggctgtagacactgtcaacccaggatgcaaatctcaccacaagtgcaggagaaagggcagggggttgatggatcatcctaactaacgtatggtttaagctgagcccagcaagaccctcagtgcctccctaaccacagttgcccttcagggataacccatgtgctcaacagcagccacaccttagtatctccactgtgcagtcattgtctaggacaagctcccaggatgtggggctttggcacaaaccaggtcatggatgtcagagaggggcagctgggtgtctggtccataggctccctg**atgttggagggttgagaggtgcattctctgggccaacat**atactctagggatatttacttcagaacacatacttttatttgttggtttatggttaa

Uppercase: IGHV3-41-2

Lowercase: Flanking sequence[1000bp]

Red & Bold & Underline: Stem-loop [11]

Blue: Heptamer[39]

Green: Nonamer [11]

id-IGHV3-36-2[V_gene_segment]

tgtatttctattttgggtagagtttttatggtgaaaccttgaatttttttaatgttattttctgtctgagaagttaatgggaccttcttcgtattta**atttggcataccaaat**gtattgatttgaggattttgaatcatctgtgcat**ctcaaaaataaatttgag**ttggtcatggtgtatgatccttctaaagtcctgtggaacttagttgactattatttggtgtgtggagggggctcatttatgtctactaggaatattggtctgtagtttacttttttgtgaccttgtctattactgaaaatacggtaatttgggccttatagaaagactggaaggttggcatctcaccaatttttgaaaaagttagagaatggttggcattacttctttaaatgttgagagtcctttttttacttaacatgtgacctatcatggagaatatttagtctttgcttcggaagcatacatattacactgctcttggttggaaggttctgtagatgtctattaggtcaatttgttcaatagtgttgttcaagtccactgtcttc**ttaagaattctcttaatgatgttctatccatcatt**gaaagtgaggtgctaaagtcttctcctaatttttattattttctatttatctctccatatctcttaaggtttgcttaatgcagttat**atttatttttgtacaaatatgtaaaaaataaat**gaaataataattgctaagtaattttaatggcaaaatcacataataagaaattatattttcccaaatgctgtcattgacactaagctcctccaggaatctcacatct**gctctggacacggctctcctggtgtcctaccccagagc**ttgctatagaggaggagacatgcaaatagggccctcactgtcctgatgaaaaccagccttgcctgcgtctacgggagaagagccccagtccagaagtaccaggggtttccatttggtggtcaggtctctgaacacagaggactcactATGGAGTTTGGGCTGAGCTGGGTTTTCCTTGTTGCTAATGTAAAAGGTGACTCATGGAGAACTAGAGATATTGAGTGTGAGTGGACACAAGTGAGAGAAACAGTGGATATGTGTGGCAGGTTCTGACCAGGGTGTCTGTGTGTGTTTGCAGGTGTCCAGTGTGAGGTGCAGCTGGTGGAGTCTTTGGGAGGCTTGTTATAGCCTGGGGGTCCCTGAGACTTTCTTTTGCAGCCTCTGGATTCACCTTT**AGTACCTTTATTAGGTACT**GGATGAGCTGGGTCCATCAGGCTCCTGGGAAAGGGCTGGAGTAGGTCTCATTTATGAGTTGTTGTGTAGGTAGCACAAGCTATGCAGACTCTGTGAAGGGTCGATTCACCCTCTCCAGAGATGATGCCAAGAAATCACTGTATCTGCAAATGAACAGCGTCAGAGCCGAGGATAGGTCTGTGTATTACTGTGGTGGcattgtgtgcatcccttgtttaggtacatgcagagatgctgct**ttggtgtgttcaggggctcctgttttggggacaccaa**ttttggagtttgcagtatccttgagtccagtacgttcatggtggcattgggtctggggtgtgcagtggcaatgagtccaggatacatgcagctgatgccgta**tcttgtgaggacaaga**acaactacaaaacgat**cctattaataaatagg**caaatgacttgaatacacatttttcaaaaaatacatacaaatgacatatagatatataaaatgttaaatatgagtaataatttttaaaattcaaatgtaatctacaataaattatcaccccatacctgttagaatagctatttttaaagaagatgaaagataaccagtattggggcagaagtggagtaaagggagcctatatatattgttactgaa**aatgtcaattagtacagacatt**atgga**gaatactatggaggttcctcaaaaaataaaaatagaactaccatagtttc**caacaagcccactactggctatgtatccacaggaagtgcaatcagtatgtcaaagagacatctgtgcccccacgttcattgcaggaatatttatgatagtaatgatgtgggaacatcctaaattaccataaatggatgaattactaaagaaaatgtagtatgtacccaatggagtaatgttcatcctaaaataagaagggaatcctgcatttgcaacagcaggcatgactctagaggaccatatgtgaagttccacaggcaggcgcagaagacaagtaattcgcgatttcactgtagtttgtttaaatggtagactcacagaagtagagtggagtgttggttaccaggggctggaggggtggactggaaaaggcgagattttggtcaaagggtgaaatgttccagttagacagaaggagaaagttacagtgttctaatgcacaagatggaaaatatggctaataatgcattatatat

Uppercase: IGHV3-36-2

Lowercase: Flanking sequence[1000bp]

Red & Bold & Underline: Stem-loop [12]

Blue: Heptamer[32]

Green: Nonamer [2]

id-IGHV3-50[V_gene_segment]

aggggtgctaagttgacaagaggtgggcttttgatgcttgatactaggtgtcaacttgattggattaaaggatgcctacatggctgggaaagtattgtttcttggtgtgt**ctctgggggttttgccagag**gaggctgacatttgagtcaatggactggaaatttgagtttgtcatgttgcatgatccttttaaagtcttatagaacttagttgactattattttggtgtagttcatttgtataaatatttctttgaagtttaattttcttcctttgtgtttgtctgtcactggcaacacggtaattgtagccttttagaaatagcttagaaggttggcgactcatcattt**tttaaaaaagttagagaacagttggcatgacttttttaaa**ttctgagagccattttttatctaacatataggctatcatgaagaatatttatagtgtgcttgagaagcatgcgtattacactgcgcttggttggaatgttctgtagatgtctattaggtcagtttgttcaatagtgttgtttaagtccatgggcttcctaagaatttttttatggatgttctatacattattgagggtgaggtgttgaagtctcctcctttcttattgttttctatttctcacttcatatctcttaaactcttgttaatgcggtta**tatttctatttttataactatgtaaaatagaaataaaataataattgcttagtaatttta**a**tgggaaaatcacataataagaaattatattttccca**aatgctgccatcaccactaaactcctccaggagtctcacatctgctctgggctctgctctctcctaaggggtcccacatcagagtttgctacagagg**aggagacaagaaaatagggccctccctctcct**gatgaaaaccagccctgccctgaccctgcaactctgggagaagatctctagtccagaattaccaggagtttggatttgatgatcagctctgtacaaacatggctcaccATAGAGTTAGGGCTGAGCTGGGTTTCCCTTGT**CATTATTTTAAAAGGCGAATAATG**GAGAACTTGAGATATGGAGTGTGAGTGGATATGAGTGAAAAAACAGTGATTCTGTGTGGCAGGTTCTGACTCAGATGTCTCTGTGCTTGTAGGTGTCTAGTGTGGGGTGCAGATGGTGGAGTCTTGGGGAGAGTT**GGCACAAGCTGAATGTGCC**TGAGACTCTGCCGTGCATCCTCTGAATCCACCTTCTGTAGCTACTAGATCAGCTGAATCTGCCAGGCTCCAGGAAAGGGGCTGCAGTGAGTAGTAGATATAATGTACGATGGAAGTCAGACATAATATGCAGACTCTGTGAAGGTCAGATTCACCATCTCCAAAGACAATGCCAAGCACAGGTTGTATCTGCAAATGAACAGTCTGAGAGCTGAGAATATGGCTCTGTATTATTGAGTCAAAGGTAccaaatgaagggacatcagtgtgaacccagacacaaaatttc**ctgcagggaggagggaggaggctgggctgcag**tgggcactcagcacacacaaaaggcagggacagttccaggggcaggtacaggtgcaggtgaaggcaaaggtctactttccttccagatctgtggattcctctgcatccaacagttcgcctgggcctctgtctttatggatctgcgcctaccactgatgtctctgggttagtaaagtttgctactataggaggaaacattgtcatttgtcagaaaggcgaataatggagaacaaaagatattgagtgtgagtggatatgagtgaaaaaacagtgattatgtgtggcaggttctgaccaaaatgtctctgtgtttgtaagtgtctagtgtgaggtgcaggtggtggagtgtaggggagaggcaagaggcagaaaagcgcgcaggaggccgggtgaggctgtagacattgtcagctcactatgccaatctcacaacagtgctggagaaggtgggagtc**tgatggagcttcccaacaaccctgtggtctaagctaagtccatca**aggccgctggttcctcctggaacacagctgtccatcaggaatcccccatgtgcccagcagcagccacgcgttagcatcttcactgtgcacagtcattgtttgggaggagctccaggatgggtctctttgtcaaaaaccgggtgatggt**gtcagagagctgtgccgggtgcctggtccaggggctccctgac**gttggaggtataggaggtgcgttctcagggtgggaacaccttttatggaaatgtacagaagaaaacatgattttccttggtcctctcagatgacatggagaa**gcaagcgcgcagcttgc**aaacaattgtaatgtccttgtaatttcctactttatcccaagttcattgattcttgtaaaccttggtttctgtccaagtacagagatattgattagagtgagtttggttctttcca

Uppercase: IGHV3-50

Lowercase: Flanking sequence[1000bp]

Red & Bold & Underline: Stem-loop [12]

Blue: Heptamer[33]

Green: Nonamer [7]

id-IGLV5-45[V_gene_segment]

agctcactgcaagctccacctcccgggttcccgggttcacgccattctcctgcatcagcctcccgag**tagctgggactacaggcgcctgccaccacgcccagcta**atatgttgtatttttagtagagacggggtttcaccgcgttagccaggatggtcttgatctactgacctcgtgatccgcccgccttggcttccc**aaaggaattgctgcctttaacatgtcattgatccttt**atttttcctggaaatttttcattattcaccatctttttgaacaaccc**cagctgcccttcaacccagctcctcagctg**accccttttccatga**tgcccactctgatgacatatgtgggca**aatagatttttctaaaatgccctgttggggtgctgacacctataaaagttatgtatgtaca**ggtctgcacatcccacagacc**aagactacttctgggagtgagaaatcgtagtaataatgcatagcatgatagaagcag**tgtctgggaatcagaccatggatatgtgaatacagaca**gtgactaaag**agctgcagagaccagctgggttctgaagtctctgagct**ttgggaaagtgtgtttcctgggaccagcgtcagatgttgtggggctgccctgtggttccgacacagc**tgctcagtaatgagca**ttcactcagagaaatccaggccttgaggctgggccctgtgctcactgat**ggacactcggcagctgtgtcc**cctctggcctggcagagttccctgagcaggg**acctgtgtatggtctcaacaggt**gcttcctcccagggccttgcaaaaagaaaagcaaaccttcctcct**gctcagtctgcagagagagctgagc**tgcatc**cccaggtctcaaggaggggcccggcagtccctggg**tggaaacacatttgcatgagcaacccctcctctgcaa**cagtgggaagaaaaggaggcctggggcagcccagtcccactg**cgggggtaagaggttgtgtccaccATGGCCTGGACTCCTCTCCTCCTCCTGTTCCTCTCTCACTGCACAGGTAGGAATAGACTTCAGAGACCAGGGTCAGCCACCCAGCCTGATTCTGACTCTTCTGGCAAAGATCCCTGAAAAACTTTACCCTGGTTTCTGCCTTAGCACCCATTAATGTCTGTGTTTCCAGGTTCCCTCTCGCAGGCTGTGCTGACTCAGCCGTCTTCCCTCTCTGCATCTCCTGGAGCATCAGCCAGTCTCAC**CTGCACCTTGTGCAG**TGGCATCAATGTTGGTACCTA**CAGGATATACTGGTACCAGCAGAAGCCAGGGAGTCCTCCCCAGTATCTCCTG**AGGTACAAATCAGACTCAGATAAGCAGCAGGGCTCTGGAGTCCCCAGCCGCTTCTCTGGATCCAAAGATGCTTCGGCCAATGCAGGGATTTTA**CTCATCTCTGGGCTCCAGTCTGAGGATGAG**GCTGACTATTACTGTATGATTTGGCACAGCAGCGCTTCTcacagtgacacacacagatggggaagtgggacaaaaacctcaccctgctctgggtcttgctctgtaccaat**ttttaaattttaaaa**taactggcctaggcacaaactatatttggaagtactttctagttgtaaaaggctcttctctcaattttctatccatttttctgaagtctcagtaaaacaaaacaaaacaaaacaaaaccacctgatgaccctgaagtgttggctggttg**tccacatgtgctaacaccagtgcctgtgga**gctggacgtttttttcttgaaagaagcaaaatgaggctttcctagctcacatcatggtcagagcctggggacaacatagcaggtgaacagtgccctgcagcatgga**ctctgctcttccagcgtacgggcagag**acctccccttccatccccagtagtctgttctcagggctgtctgaggctcactgtgtcccaggaacaaggcactcaaatatgaaatgctgcctgaggact**caggtcagggaattctggatatagtgctgacctg**gctcagtcctgggtacctgcatttcagtggcaatgtcaggagatggcgttagatattagggtaactgcgggcccgatgccagtaaacctctatgtaggacatgacagaatcaaggggaggc**tccaccatggctgagacaagccagaggctgctccttggccatggggga**agtagatcccatggatgtgactcctgtagctcatcccactcaggaccacggggactg**ctgagggtgggtttcctgatccctcag**ggcctgactgtgttctcttctcctctcttacaccattggagcttct**cactgactcaggtctagtggtgagtctgtg**gaatatctggatacagaaactgcccaatatttgcttcaagtgactcctccctgggaaact**cttgaggaggcccccacctcaag**gtcatctttttc**tccatggggagcagcagaaacattggccattcccatgga**aatggc

Uppercase: IGLV5-45

Lowercase: Flanking sequence[1000bp]

Red & Bold & Underline: Stem-loop [25]

Blue: Heptamer[38]

Green: Nonamer [7]

id-TRBV5-5-2[V_gene_segment]

aattgtgtcctgttctaggactttctggaataaaaatattatgagcta**ttagctaggatatctactgaaggaaatatctaagcagctaa**gcattcaggctattgtggactactttcaggcaccaggaaatttaacccagcaagaaaggagccaagggaatagattttgcaaaccagcacagatggtgacactacctccctctgctgtcctgtctccaataagacaaactcgctctggagttagagaaaagaggagccagttaacataatacactggggtcagcttctgaaggcagggtggatctgaagggaataacataaactgtccagagcactacgtataggatgaccttgagggtgctgagtcccaagctggctaggtctgtgctatgagctgagaaggccctagaattatttctggggaatctgaactcttgataatttacagacaacacaggtctgctttgattctgatcagatggactaaatcttggggactctgcaccgctggcca**ctcaggaaatgggttgggaatgttgcctgag**acaggaaaatatgataagaaacatggtgtgaatctagcatcagaaagatgatgtgaggacagcaaggaagaactgcaacttgaggtttaatcacaggctcct**catcctcccctgatgggcaggtgtgtgagctccaggatg**gagtaccacagcactaggtgggagaagggtgatagggtggtggggctgcctgtgagctggggcagtgtaggtagaggagcaactgtatcaccacagaagcttctgccttcacacattcctccagctctgcaggacaggttgagtccagggtccgtagtgcgctagacttaaggaaggctgcatgaggaggacacaggacagtgacatcacaggatacccctcccatcaggaaaatcaaggctcagaactcactgggttcttccccaggaggaccaagccctgaatcaggtgcagtgctgcctgccccactgtgccATGGGCCCTGGGCTCCTCTGCTGGGTGCTGCTTTGTCTCCTGGTAGCAGGTGA**GTCCTGGGCACAGGAC**AGCAGCCCCATTCTCAGCTTTTCCACCCGTGTCCTCCACTTTACCATGGAGAAGACCTCCAGGCTGTCTCCTGAGCTCATCCTCCATCTGCTTTTCCCACAGGCCCAGTGGACGCTGGAGTCACCCAAAGTCCCACACACCTGATCAAAACGAGAGGACAGCAAGTGACTCTGAGATGCTCTCCTATCTCTGGGCACAAGAGTGTGTCCTGGTACCAACAGGTCCTGGGTCAGGGGCCCCAGTTTA**TCTTTCAGTATTATGAGAAAGA**AGAGAGAGGAAGAGGAAACTTCCCTGATCGATTC**TCAGCTCGCCAGTTCCCTAACTATAGCTCTGAGCTGA**ATGTGAACGCCTTGTTGCTGGGGGACTCGGCCCTGTATCT**CTGTGCCAGCAGCTTGGcacag**cc**cggcagaatcactgacattctgtatataaacttcctgccg**taactttgacttgagagctgcaggccccacccaggtttcactccttcaagggaagcttttagttgcttggaaggcatgtcttgtgtcctactgagggcagacctttcccaaccaatagagcccaggtttcctgtgccctgagtgtgcctgcttctgt**gctgcatcttcttgcagc**ttgtcccttcctgggtaacttcagtataagagtgactgctgagcgcttggtgtgtgctagatttttgtatatgttacatatcttaaagcttttccacaaccttgcaaatcaaacattctcatccttcctttacagatgggaggctcagggacattgagtcattttccccggtgtctcctggcttgtaaggatcagaagtgggaaacaaactagtccatccattttccacctacccccctgtttcatcaccaccttctgtatcctgttcagtaaaccagagccctcacactgccctctagtgaccagcagggccgcagcagaggctcagatgtcttcagggggttattactatggcctcctaattagcaattttgaggaatgttaaatttcacactttttaaaaaatcatttatatgcattccctttatctttcccatgtctgtagcttacattttcatttttatggtgttttttgatgtacaagagtttaatttaatgcc**acaaaaatgaaaaacaattttgt**ttg**tgtgtagacaaaaacttctttctcatataaatgtctaaaca**tatttttcttaatttattctagtaaaatttaagtttggcttttctccattaagaagaaatgtacctgaaatatatctcacgtcagaaataacatagagacctaggaatgggagggaagcacatatgttcagtgaagaagcaacagcctgtcaaaaagatcccaaaatggacaccctgcgctgacggtggagaggtagctggagggtaaatg

Uppercase: TRBV5-5-2

Lowercase: Flanking sequence[1000bp]

Red & Bold & Underline: Stem-loop [11]

Blue: Heptamer[20]

Green: Nonamer [3]

id-IGHVIII-26-1-2[V_gene_segment]

aagc**aaataacccacaggattattttctttccttgattgagaaaga**ttttccccaaacttcagctcagttcaggcatatactgtccctaaatgggcatttaccctcagatgagtccacacacctgtcaacacgtggagtcttctgtcagacaaacacacctttactg**atgtggattcttccctcagacaaaaacacatgtccccacatggacactttcctcagactaccatgt**atgtccttacatttacgctttcctcagaaaacagacttttcctcatgtggact**cttgtctcagacaag**caaatatgtctccatgtgaactcttcactcagataagtacacatatgtccacatttactgtttccttacacaggcacatatattcaatgttgaattgttttgtggcaaaatgatctcaagataatggtaattataaatcccctccctaataaagtgtagatctgtattt**tttttgttgtaacctaactttgccttattgtcaaaaa**cagtaatttgcag**ctctaaatgcaccagttagag**acaggtgtccattttctctggaaatgtatttttattttcttactggacatatttgttgataatgttgctactatgaagatacctgaacagtatccacattaaagaataaaaaagagtaatgggcagaataaccctgtgcatccagacccagaaatcctttgaccctgccctcccgaaatgtagacacagaggatggatgagcagtgctgagcagtgtacccatgaccacaaacatgaagacatggaaatgtgtccccactcctcctcatgaaaggcagctcatgccctcttccttcaggccctggtatggagcc**accccatgtctctgcccttccttagtgtccacaccatggggt**ctgcactgatctggattcccttctcatccccctcaatattagtgtgccttgtgaatcaggtccagctgtggctgttctcagtctgtttcctctgtattcacagAAGTCCTGTGTGATGTTTACTGATGGAGTCAGAAGGAGAAAAATTTAACAGCCCAGCAGTTCACTGAGACTGTCCTGCAAAGCCTCTGGTTTCGGGTTTACTGGGTGCAGCATGAGCTTGGTCCAGCACACTTCACAACAGGAATTGGTGTGGGTAGCAACAGTGAGAGATCATATGGGAGTTCTAAGGGTTACTCTCCATGAGTACAGATAAATCAACATTCCCAAGTGACACC**CTTTCAAGTGCAGTCTACCTTACAAGGACCAACCTGAAAG**CCAAGGGCAAGGCCGTATATTACAGTGAGGGAcacaggggagggaagatc**tgtctgagcccagaca**caaaaatctctgcagagagacaggagggaactgcgtggtagatgctgctcagaaccaccatggggtggccaggaaacaaggggtgcttaggacctctagggagcgctcaggacgccaggggggtgctcagaaccactaggggacgctcaggacactaggtggcgctcaagacaacactactgcactcatgtcaccacggggtactcagaaccaccagggggcgctcagaaccaccagggggcgctcagaaccaccaggtggcgctcagcaccaccagggggcgctcaggacaccagggggcgctcagaaacactcggggacgctcaggacaagagggggtactcagaaccaacagggggcgctcaggaccaccaatgggtgctcagaaccacaagggggcgctcaggacataggtggtgctaaagacaacaattctgggctcaggtcaccagggggcgctc**agaaccaccaggggacgctcaggacatgcaggttct**cttaggaggcagctgcacatcaggagcttcagaggctgtg**atttcctttcagaccctggaaat**tcctgacctggtcaagcaaaagtcttccccaggatgtctgatttttccttttaattccatgatttcttttacctacaaaacactaacttagaacagggatgtaatgcaactttt**attgctgcgtattttctgagtaatactagcaat**gatctctcgggac**aattattaaaataggttatttatatattttaattatt**aaaaataa**tactattattaaaatagta**aaattttttaaatcacacttgtaatcccagcacttggggagactgaggcagacagattacttgagctcaggagttttagaacagcctggcaacatggtgagatgctttctatacataaaagaagaaa**tagctgagattggtgcaaccagcta**cttaaaaggctgaggagggaggattcctta

Uppercase: IGHVIII-26-1-2

Lowercase: Flanking sequence[1000bp]

Red & Bold & Underline: Stem-loop [16]

Blue: Heptamer[34]

Green: Nonamer [5]

id-IGHV2-70D[V_gene_segment]

gaactcagccaagacagagccccatttaggagcaggtgcatcaaggaaccagccagagaacctctgactacaatc**tagggttgtcttcttttgcttcacagcatccaacccta**aggagaccacccactatatcatc**tgtaactcagaactttcttgttaca**aaggaaaacacaaatgaatataactctgcagttaacttcttgaacctggattgatgagtttccatggcctcatgagcacagagaccacatccagtgcaggtgcagagca**aggctgcaacagtcagcct**gctgctccctcaggagtcttccaa**ttcccttgtccaaagcatccgctcattcatgggctgagaaaagggaa**gtcattcatgcagtctactctctccacagaactgacggggcacaaggacaacatcgatttttcatggaacatgcctctaggaatgcagc**tgtgtgcacacacactgctaagcacaca**cttctttacataattacttgtaactgtattttcttatttattctctccaatttttttacacaaattcatcacttttccccata**atcaaagaggattttgat**cagaatgcttgtggggagccccttgcctgccagatgcccaacatcactactcttgaagggaggaggaacggcagctctcttgatttctactctaatcctctaggactaaaaccagaaggttgcatgtccagtgcgggagcatcgaagaagatcctgtctgtagaagcaggagcgtcaagacttgactgagagccatggtgctgaaatgagatagattccctgatggagagcacacgtggacccccacacctgagggctcactgctcctcaccacaga**tgcactcccctactgagtcctgagacctgagtgca**ccccatagagtagggctcagatg**aggggatgcaaatctccaccagctccaccctcccct**gggttcaaaagacgaggacagggcctcgctcagtgaatcctgctctccaccATGGACATACTTTGTTCCACGCTCCTGCTACTGACTGTCCCGTCCTGTGAGTGCTGTGGTCAGGTAGTACTTCAGAAGCAAAAAATCTATTCTCTCCTTTGTGGGCTTCATCTTCTTATGTCTTCTCCACAGGGGTCTTATCCCAGGTCACCTTGAAGGAGTCTGGTCCTGCGCTGGTGAAACCCACACAGACCCTCACACTGACCTGCACCTTCTCTGGGTTCTCACTCAGCACTAGTGGAATGCGTGTGAGCTGGATCCGTCAGCCC**CCAGGGAAGGCCCTGG**AGTGGCTTGCACGCATTGATTGGGATGATGATAAATTCTACAGCACATCTCTGAAGACCAGGCTCACCATCTCC**AAGGACACCTCCAAAAACCAGGTGGTCCTT**ACAATGACCAACATGGACC**CTGTGGACACAGCCACGTATTACTGTGCACGGATACcacag**agacacagcccagggcgcctcctgtacaagaacccaggctgcttctcagtggtgctccctccccacctctgcagaacaggatagtgtggctgagatgccatttcctgccagggcctgcgtttcccatccccatctgactcagagccttgttttcctccctcttctttactaataaatggcatgtcccctgttagtggttcgtgcaagcagaagctgtatcctgtttgacaaagattcagcatgaaaggttcctgttacctaa**aaaaaaatagacagatgagacttaattaacctaaataattttttt**cacaacaacagagtgaatacacaatttgcagaatgacagaaaacttttgcacactttgtctgtgacagggaactaatatgaagaatttgcaaggaactcaaacaactctacaacaacaacagcaacgagaaccaaataaccccatt**aaaatgagcaaagaacatgagtagacatttt**caaaagaacacatagaaatggataataaatatataaacaatgctcaacatcactaaccatcagggaaatgcaaattaaaaccacaataagatatcatcttccac**cagtcacaatgactg**ttactaaaaactcaaataatatcagatgttgctgaggatgggaaataaaggcaactcttagacattgttgatgaggatgtagacgagtacaacctctgtggaaaatggtatggagatttcccagaaaactagaaatagaactgccatttggtccagcaatcccactactgggtaactacccaaagga**aaataaactattattt**caaaaagatacccaccttctatgcttaccataaaactactctcaatagcacatatgtcaaactgagtgtctgccaacaaatgattttataaaagaatatagcacgtatgcacaattcaatactagtcagtcacaataaggaatgaaactgtgtcttttgcagcaagatgcatagaagtgggggacaata

Uppercase: IGHV2-70D

Lowercase: Flanking sequence[1000bp]

Red & Bold & Underline: Stem-loop [15]

Blue: Heptamer[39]

Green: Nonamer [3]

id-TRBV3-1-2[V_gene_segment]

aaaggggactgtggtcctcaacattagataattcacaaaacgcgaaggcattttctgtcgtcacattgccatcaggtgggtagggccca**aggatgccgccaagcatcct**ccctacacaggacagctccccacaacaaacaatcacctgatccaacatggcagtagtgctgaaattgagaaaccccgcaataagcttcggttgtaaactttgtaccaagaaggagaattctctgcagacaactcaaggggcaatacc**gtcaactacttttgttttctctgactgccgtgttgttgac**cttcatcgccattctttcccagcacatggtgtctgcatattttccagcagcctcctgtttcctcataaatctgccccttggacagataagaacatcacagggagcagggctgcagttcactcccaataataacgcctcagcagggtttgctctgcagtagcatcacagtcatgtggagtaggtggctatggctggt**attcaggtgaaatctcaggccaatctctcctgaat**caaggggaccactgatcattaatgagaccttcaagtgagcaaagattcaagagagcaaattaatgtcctgcgctattgtgaagggacgcaaaggatccagagaggaaaccagctgactctgagagaagggcctgaaaaggagaggaagggaggaggtggtcagatccccgggaaggagcagggagggagggggctgctggc**ccaggaatgactgtcagagacacatcctgg**aggtcacaattccaccctctcaattattcccagaaattccaaaaactcctttaaaggagcaagtgcagggcatgaaagggaggagccatgctagaggagaccctgggaatgggggaatgattacaggctgtgacctcactggcgcagcacctctcagcg**gcagtggaaaccacagcctagtcctctcaccactgcagaccagaatcctgccctgggccttgcctggtctgc**ctcactctgccATGGGCTGCAGGCTCCTCTGCTGTGTGGTCTTCTGCCTCCTCCAAGCAGGTGAGTCCCGGGCCCAGGTGACATGATCCTATTGGAGTCCCTAAGCCTTTTCACCATGACAACAACAGCAGGCCGTCTCCTAGGATTTGCCTGAATTCTGCTTCTTTCCTTTGCAGGTCCCTTGGACACAGCT**GTTTCCCAGACTCCAAAATACCTGGTCACACAGATGGGAAAC**GACAAGTCCATTAAATGTGAACAAAATCTGGGCCATGATACTATGTATTGGTATAAACAGGACTCTAAGAAATTTCTGAAGATAATGTTTAGCTACAATAATAAGGAGCTCATTATAAATGAAACAGTTCCAAATCGCTTCTCACCTAAATCTCCAGACA**AAGCTCACTTAAATCTTCACATCAATTCCCTGGAGCTT**GGTGACTCT**GCTGTGTATTTCTGTGCCAGCAGCCAAGAcacagc**cctgcaaagtcactgcatccctgtgcacaaacctcccggctcagccaggaagc**tgcagggcagcgtgtgcacctgca**cccagggctccagt**ctccattccctgatggcctctgatggag**tttcagtctgtagtacagccagctagtgcacccagtggagaagtcctccatcttatgcacacaaaagtctcacagaattgtttatcagctagagcagggcttcacaacaaagccactaaagtattagggctccacaaacaaaaaagagcctgtcctatgcactgatgatcttaactgatgggggaagctctttcttttctttctgtgtggcacatattggtacgtattttatataacattatgaaaaaaacacgattgctgaaaatggttgatgagcacaagcataaaggagggggaaggaaatatttccaagatcaagagacaattctgatgatcagtgagtactttattcaacagaaaaatatttggagaggcatgaaattccatggcaataggtaaactggaaagtgttgaggagatgccggtggaggtagccttgtgcgttgtgggct**ccttctcttcaagaagg**gtgaccagcaagtgtctctcccaattctgatgaata**tcatcttgaactgtcctcaacagcaggcaatagccagatga**ctggggatattgggagacctgaaaggactttggacacaaatctctaggcagtggtgagccattggtcctactccaagtgaaacaacatgatgaaactgatattttggtagaggaacactggtagtagtatggatcagaaaggagatagatttttgggtggcatgaatgattaggagac**ttctaaatatcctacatgaaagattagaa**agggttgatgtgaagcattgtggacaatataggcttgaggctctgcattatgttatttcttggaaatgtattgggttttattctgggagggttgaatgtaggacttgctagggtctatttca

Uppercase: TRBV3-1-2

Lowercase: Flanking sequence[1000bp]

Red & Bold & Underline: Stem-loop [14]

Blue: Heptamer[48]

Green: Nonamer [6]

id-IGKV1D-16[V_gene_segment]

atccctgtccagtttcatctggaac**caactgatttctccatgtacatgggcagttg**cttgataaaagattgagtgcctctttccaaaagatg**ttaacagggatcttggtgtctgggtcaggatgatgtccctgataa**aaagtaaaagaagaaagtgtcacctatggtgcgtggcagggacatgctccatgcgatgatcaccctcactaagagagatgaatgttgggaaataatacttaatggcagaaaagaaggtagactatgaaggtgcccaaaacaggaataaggtgcagcccattta**gtctctgggtattaaagagac**ctagagctcttgataatggtggatctgtaattgctgcatgcattgaggaaacacggtatcaacttagtgtatctgaagtaaattgc**ttgatcttacagtggtaagaacaa**tggcataacaccattacctaacatttacaaatatatgtagcatcatgtgaataaattttatttttaattttttttttagaaaggaacaatgttaaactcacagaaatgttgcaagtatatgataagtaccaccttccctaacagaaatcacatgagagtcttttgaagacctgaaaa**ttgtagggtctaaaattttactatgtgtttcctacaa**aaaagaatattctcctaaataatccccatacaccaa**tgaaatacattgctccatcgactcccaaggaatatttca**aattgtcaaaaagaaaccataaaa**tgtttcccataacaaaatagtccccagtagaaaca**cattctctgcagacaaatttgagctaccttgatcttacctgggacaggtggggacact**gagctggtgctgagttactcagatgcgccagctc**tgcagctgtgcccagcctgccccatcccctgctcatttgcatgttcccagagcacaacctcctgccctgaagccttattaataggctggtcagactttgtgcaggaatcagacccagtcaggacacagcATGGACATGAGGGTCCTCGCTCAGCT**CCTGGGGCTCCTGCTGCTCTGTTTCCCAGG**TAAGGATGGAGAACACT**AGCAGTTTACTCAGCCCAGGGTGCTCAGTACTGCT**TTACTATTCAGGGAAATTCTCTTACAACATGATTAATTGTGTGGACATTTGTTTTTATGTTTCCAATCTCAGGTGCCAGATGTGACATCCAGATGACCCAGTCTCCATCCTCA**CTGTCTGCATCTGTAGGAGACAG**AGTCACCATCACTTGTCGGGCGAGTCAGGGTATTAGCAG**CTGGTTAGCCTGGTATCAGCAGAAACCAG**AGAAAGCCCCTAAGTCCCTGATCTATGCTGCATCCAGTTTGCAAAGTGGG**GTCCCATCAAGGTTCAGCGGCAGTGGATCTGGGAC**AGATTTCACTCTCACCATCAGCAGCCTGCAGCCTGAAGATTTTGCAACTTATTACTGCCAACAGTATAATAGTTACCCTcccacagtgttacacacccaaacataaacccc**cagggaagcagatgtgtgaggctgggctgccccagctgcttctcctgatgcctccatcag**ctg**agagtgttcctcagatgcagccacactct**gatggtgtttgtagatggggacatgaagtcacctctgcaccctaattcttttctctttctcagcaccaacttcacagacatagcaatgcttctccttatttaataaaaacagagatcatgacacctgaggagtctagttta**tggcttcagttggaaatcataatgcagaagaagcca**ctataga**tattctaagcaggaatagtcttaatacagagaattagaata**aactactgaagtctaaata**aaatgtagagatgaatctctaaattt**aatgttttatttgaagagaaatatttgccaaatgaggcatacaggaaaattcagtggtcttcagtatgttggaagaacaaagaaaaagtcagtgttccatgaaaaagggaaatattacctat**tgaactttgagaaagttca**ttggcactaggaagggttgggagctggcaagctcagactggtaaacagtggtggacaaagtgaatcctagagttatatcaagttatctcagaaattgtggataaatttgatttcaggttacaataagccaaagcactgaaggttgcagagaattttcttactgaaataccagggacttggtgtagatcctgcagctcaccacacagaaagccaatctcgaagacaacaagtattgccaaggaacaggctttaatcaggtgctgcagccgaggagatgggacgccattctcaattgtatctctctgacaaaataaattaggggtttatataggagggaagaaatgttgaaaacaggaattagggaggggtaaggaagataatttggtcaagaagaagcaggaggtcagttatgcaatcacagcgggtgaaggttctgttgtctcactgtcccgattcagtgatatg

Uppercase: IGKV1D-16

Lowercase: Flanking sequence[1000bp]

Red & Bold & Underline: Stem-loop [20]

Blue: Heptamer[24]

Green: Nonamer [3]

id-IGKV1D-32[V_gene_segment]

aacccagtcattaagaactgctt**tctcttaaaaactggaagaga**ttggtaatatagagatatacagtttatggaacagagtagattcaagagt**agatgtgagcactgggtctagataacatct**cagtttccggaagatattagcttaagaatgacgtaaggcacacatattaatccaaggagctcaggaaacatctagcataggaatgcatgatggcacacagaggtagagagtagctgacacttt**tcagaagtggagggagaagatttctga**tgttggcctagatgtaagaagtaggaaagaaaatgtagaaatacaggaagatgatcacatgtctgagataagagatttgccatgtcgtttacccagtgta**caaggattacccaatccttg**caaagcattacactactttcacatattttgtgttaacattttcctttttc**ttatttttaataaataa**atctgtccttctatccatccatctatgtatccatccatctttcttatacc**tcatcattgcatctctggctgatagtgctgatga**tttacaacatcaagggctctagaggatcagagcaaagaagaagtctctgtggcctcaataatattcatcagtaccaaacgcaggacacttctaaaatttattgcagattc**tggagatgtgcttctcca**acaagccaggtcttagacgggaagctagcaagaa**aagtaggtcatgacctacttctgcagagcagaa**aagcaatctccttgcaggacagtagacagattctgttctgtggcccccagagagaa**acttgttctctgctaacaagt**gtatg**caggcctgatcaatcttaggacccctgggggacactaggcctg**tgcaatattatgcagataaagtcattcttgcatctg**ttgaaattttcattttcaa**aaaaacacagccaacttcctgaagacttgtacacaggctggtcataccacttgcaagagtcagtcccagtcaagacacagcATGGACATGAGGGTCCCTGCTCAGCTCCTGGGGCTCCTGCTGCTCTGACTCTCAGGCAAGGAAGGAGAACACTAGGAATTTACTG**AGCCAGTGTGGTCAGTACTACCTGGCT**ATTCAGGAAAGTCCTCTCATAACATGGTTAATAGTGTTAATAGTTTTGTTTCCCATCTCAGGTGCCAGATATGACATGCAGATGACCCAGTCTCCATCCTCGCTGTC**TTCCTGTCTAGGTGAGAGTCACCATCACATGCCAGGAA**AGTCAGGTAGGGCATTAGCCATGTTTTAGCCTGATACAAAG**AGAAGCCAGGGAAAGCTTCT**GAGCTCCTGATCTACGATGCATCCAATTTGCAAACCTGGGTCCCATTGCAGTTATGTGGCATTGGATCCAGGACAGATTTGATTCTCACCATTAGCATCCTCCAGTCT**GAAGTTGCTGCAACTTC**TTATTATTGGTCAACAGTATAAAAGTGACCCTctcacagtgttacaaacccaataagctccccaag**gaagcagatatgtgagggtgggctgccccagctgcttc**tcctgtttcctccatct**gctgagagtgtttctcagactcagc**cacactctgaaggtcactgagcagttttcatagaagcgg**taagggaagtgtctctacaccctta**gtttcttttatcctcctcagccccagcagcagacatggcaatgcctctcctaatttcataga**agagtcattacctatgaggactct**gggttacagcactggtccaggttcatacaacaaaagagaagctattgtaggtaacccaaatagaaagtttttcctaatatgggaaatccatgtctaaattacaacttttcaaagaccagaggatataatggttaggaaaccagaaaggcaaaaggagtgctgggaaatctacttcaagaagcaatggtggccgcgcacggtggttcaggcctgtaatcccagcactttgggaggccgaggcaggcggatcacctgagatcaggagttcaagaccagcctggccaacatggtgaaaccccgtctctactaaaaatacaaaaattagccgggcatagtggtgggtgcctgtaatcctagctactcagaaggctgaggcaggagaatcatttgaaccaggcaggtggtctctgcagtttcgaagcccagcgccatct**gtggctgttatgcatgtctctcccagccac**cctgctgtccagagcccatatcaatccatgggggtaggtctgtgaaaga**gcagctcagttaggacccagagggaaccatggaagctgc**agctcagctgctctgattcttgttactctggctcccaggtgcaggaaacatcggatggttctgcatgtcagtgaaacttt**ctcaaccttgttgag**tcctgttacctggcacacctgctgggaaggcacaatgattaaagctcaatgtagatcaatggtcctggatgcactgggaagacaatagg

Uppercase: IGKV1D-32

Lowercase: Flanking sequence[1000bp]

Red & Bold & Underline: Stem-loop [23]

Blue: Heptamer[29]

Green: Nonamer [3]

id-TRAV8-7[V_gene_segment]

gcaaagagagtggccaaaaccagctaaaaccaagatggagatgaaagtgacctctgattggccccactgctc**attatacactaattataat**acatttgcatgctaagtgaaactcccaccaccaccatgacagtttacaaatgccatgacaatccccagaagttaccttatatggcttaaaagggg**aggaaccctcagtttagggttcct**tttccaggtaattcatgaatttatttattatttagaatataatcaaaaaatagccatatatatagctagccaacaatccacaagggctgtaactgctgctgctattactctgtcta**tggagcaaccactttcctgtactctgtagctcca**atacacttgccttcactttgctctgtcagctcactcatgaattc**tttcctgtgtgaagccaggaaa**cctgctgggctggttttggggatcacctacctcactatcacttactatgacttggacaaattgaatcttttcttccatgctttagtttccccgtttataaaatgacgatg**ataatagtatttacttcataggattaaataatattat**ttactttataagattaaaatagcttcctcata**agcagacccctgaaaacactgtcatcagattattctgct**aagaaattcaaaccaaggaaatggagcccaaacccaaaagactcagtgtgtttccacatttgacctccagggggcaatgtctcacagaccggaaatgtctctgctaagaagagttctgtccttcctgagcggctttatctcacagctcactgaggtgcatattctgagtaggatcaacaccagagtggttgttgtcaagagagcagctcttactagtggtctctaggccttacctaagcagggaactcttgaatcccagattttcctcattatagggactcaccatcctctggcacattccttcccactcaggagatcttctagaatagagctctcagcttcctcactgcctagccATGCTCTTAGTGGTCATTCTGCTGCTTGGAATGTTCTTCACACTGAGTAAGTATAATAATATTGTTTCCCTGCTCCCGTCTCCATACAAGGAACATTGTTCCAATCGGAATCTAAATAGAGACTCTGTCTCATTACAGGTAGATCT**GTTCTGGTCTCACTAACAGAGTGTTGATGTTGCAGGAGGAACCAGAAC**CCAGTCGGTGACCCAGCTTGATGGCCACATCACTGTCTCTGAAGAAGCCCCTCT**GGAACTGAAGTGCAACTATTCCTATAGTGGAGTTCC**TTCTCTCTTCTGGTATGTCCAATACTCTAGCCAAAGCCTCCAGCTTCTCCTCAAAGACCTAACAAAGGCCACCCAGGTTAAAGGCATCA**GAGGTTTTGAGGCTGAATTTAAGAAGAGCGAAACCTC**CTTCTACCTGAGGAAACCATCAACCCATGTGAGTGATGCTGCTGAGTACTTCTGTGCTGTGGGTGACAGGAGgactgtgcctgggactgcaggaggagctgaacacaaac**ttcctgagacactgaggttttcaggaa**ctcaagggcacagcctgacctatttgtagcaaggtctctcatttgatgaaagtgagaaaggtggaagaacagagccatgaggagtgcttgactccagagaagaaacagagacttcatcacgttccttcttttactactgggt**cacatcagacaagatgtg**gaagagacagacttctatgactgttctccttctgcttggaaataccacactgaagagggtcaagtttttttttattatactttaagttttagggtacatgtgcaagattttaaattagtgtgaccagtactggactcctgatcctctccttctcactgccaac**atgctgcttctattaccttttgtttttcaatcgacagcat**ctcattctttcagaatctgcccatagcctcagagttatttttgactcttattt**tttctttcactattcacattgaatccatcaagaaa**ttcagttggccctaactaaaagatcgacccatagt**ttgaccactgctatctctttggtcaa**gccaccctcatctcaatgaattaccgtagtagccttccaaatggtttccctgcttgtcctcctacagaacattcttaatacagcagccaatgagatctttcaaatatagataggtcactcatctactccaatccttgcaatagcttcccatttctcttagagta**acatcaaagtacttacagtgacctagaggaatgcatgatgt**atctctcgttac**atttttaaaacaccaacagaaaaat**aactttggcaaccctgaattctataccagtaaaaatatcacaaagaaattatagtaaagac**attctcgaaaggaaaactgagagaattcgttgtgagcagacctgctataaacgaa**gtcctaaaggaagttattcaggctgaagggaaaggataccagaggaaaatttgcaacttcaaata

Uppercase: TRAV8-7

Lowercase: Flanking sequence[1000bp]

Red & Bold & Underline: Stem-loop [18]

Blue: Heptamer[39]

Green: Nonamer [3]

id-IGHV7-27[V_gene_segment]

aggtgtt**ctcccaaagcccctgggag**ctcctggactcattgtgag**tgtggactgatccattgcttccggagctccaca**gaaggggctccctggtggtttcatagaatccttgcttggggtctttctgcagagttc**actggctttcctacggccagt**ttagtattgtaagagatggtgtcagcagcacatggtatcacatgtcactaaatgagcattctgagccaggacacagcatcttcatactggggggagggaaatgctttggg**agcccagacaggagcctctctgcagtgcaagggctgggct**gcagggggcgctcaaggcccacccatcacaggctccac**ccccagagcaggtgcaccggaggctgggg**agggattcctctcaagatctgtgtcattcttctaaaaaaaatctaaaatgagtatttgacaaagactgctgaacattctataaatatcctattcaattgtgagcatttatcaaatttgatgttgtaatgagaaccacttttatact**agatatttctaaccctgctagatatct**taatagtatgcagctggaggttaagcaagcttttcttttatataaataagtgcaactttggagaaacacactcatcccccgaataagacattcatgta**ttaaagcctagaaatactttaa**attacctctgagccattcaagtgtgtgttcacagtgaagtcctgttctagggaaaattg**ttctccagggggagaa**gctttgtcaacatagagcttagggatgtatcaggaggcacatgacct**ctaaggggattatagcttgaacccttag**catcctcctgttggtaatacatgtgtcactctccaatctttctcatgcagtatcaggtatgaaataggatcgctcatgaatatgcaaataactgaggtaaatatagatatctttgtgccttgagagcatcatccaacaaccacaactctcctcagaagaagcccctagaccacagcacctcaacATGTACTGGACCTGGAGGATCCTCTTCTTGGTGGCAGCAGCAACAGGTAAGGGACCTCCCAGTCACCGGGCTGAGAGAGAAACCAGGCCAGTCAAGTGAGACTTCACGCACTCCTGTCTCCTCTCCACAGGTGTCCACTCACAGGTGCAGCTGGTGCAGTCTGGGCCTGAGGTGA**AGAAGCCTGGAGCCTCATTGAAGGTTTCCTGCAAGGCTTCT**GGATACACCTTCACAAGCTATGCTATCAGCTGGGTATGACAGGCCCATGGACAAGGGCTTGAGGAAATGGGATGGATCAACACCAACACTGGGAACCTAACGTATGCCCAGGGCTTCACAGGACGGTTTGTCTTCTCCATGGACACCTCCGTCAGCATGGCATATCTTCATATCAGCAGCCTAAAGGCTGAGGACACGtgcaagaggcacagtgtggaaacccacatcctgagagaaccagaaatcctgagggaggaggcagctgtgctgagctgaggcagtgacagggacaacgtggctgcaccctttgttgacatagggttatacacatacatatatatgcacccaacactggagcacccaggtatatcaagcaaatacaattagagataaacagagagatagacccaataccataatagctggagactttaacaccccactttcagcattggacagatcttccagacagaaaattaacaatgaaatcttagactaaatctgcatgacagaccaagtggacataatggatattgatggaacacttcatccaatggctgcagagtacacaatcttttcctaagcatatggatcattctcagggatagaccacatagtagaccagaaaatttgtcatttttaattaaaaagatatcatgtaccttctctgatagtaatgaaagaaaactaaaataataagaggaattttgaaaacacatgga**aattaaacagtatcctccttaatt**aaacaatgtgcttccaaactgaactatatgctctaaaatgggttaataaagaaacaaacaagaaaattttacaatttattgaaagaaatgataatgaaaacacagcatatgaacacctatgaaataca**gtgaacatagtactaagaggaaagttcac**agctataagcacc**tatgttaaaaaagtagacaattataacata**aacatcttaggaatatctctcaaaaaaaactagaaaagcaagagcaaatcaaatccaaacatagtaaaataaaaaagttaagatcagaacgttaataaataaaattgaaataaat**aaaacaataccaaagatcgatcaaatgaaaagttgtttt**ttaaaaagataaaagacatctataaacctttaaccagaccaagaaaaaaaacaaagaagaccaaagtaaataaaatcagaaagaaaactggaaatatt

Uppercase: IGHV7-27

Lowercase: Flanking sequence[1000bp]

Red & Bold & Underline: Stem-loop [14]

Blue: Heptamer[27]

Green: Nonamer [2]

id-TRBV21-1-2[V_gene_segment]

ttataagtgtgtcaaaagctactatttgtaaagaccgagttgtatgattccttacctgagatggaagggaggggaagaagcaaactgacaagttaaaaaagtaaatagagtgtaaattagttcaaccattgtggaagacagtgtggag**atttctcaaggatctagaaccagaaat**accatttg**acccagcaatcccattactgggt**atatacccaaaggattataaatcattctactataaagagacatgcacatacatgtttattgcaccactgttcacaatagcaaagacttggaaccaaccaaaatgcccatcaatgatagactggataaagaaaatgtgacacat**atattccgtggaatat**tatgcagccataaaaaaggatgagttc**atgtcctttgcagggacat**agatgaagttggaaaccatcattctcagcaatctatcacaagaacagaaaaccaaacaccgca**tgttctccctcataagtgggagctgaacaatgagaaca**catggacacagggaggggaacatcacacatggggcctgtcgggggatggagggctaggggagggatag**cattaggagaaatacctaatg**tagatgatgggttgataggtgcagcaaaccaccatggcatgtgtatacttatgtaacaaaagtgcacattctgcacatgtaccccagaacttaaagtataataa**ttttttaagtaaataaatgtatgagtgattttacttaaaaaa**aatagcaaaacagtgagtaacagaaaaaaaaaaaactaaaagcaggctgaaacttacactatttttaacagctaaaacaaagtctgccctcctacattctttgaaattgaaatga**ctctgtcccatgtacagag**catctacaaaggctatggaaagtgatcacgtcacaaggaggttgctgacgggggctaggagagtgggtgaagaagcctta**cagaaaaagctaccactacgatttctttctgagccaaccATGTGCCTCAGA**CTTCTCTGCTGTGTGGCCATTTCTTTCTGGGGAGCCAGGTAAGGCCCTGTTCT**GAACTGGTTGAATTTCAGTTC**CAAGACTCTCTCCGGTGGCTGCAGTATCAAGCTCCCTCCTGCGCTTTTTCTACAGCTGCTCTCTCCTTCCTCCACAGGCTCCACGGACACCAAGGTCACCCAGAGACCTAGACTTCTGGTCAAAGCAAGTGAACAGAAAGCAAAGATGGATTGTGTTCCTATAAAAGCACATAGTTATGTTTACTGGTATCGTAAGAAGCTGGAAGAAGAGCTCAAGTTTTTGGTTTA**CTTTCAGAATGAAGAACTTATTCAGAAAG**CAGAAATAATCAATGAGCGATTTTTAGCCCAATG**CTCCAAAAACTCATCCTGTACCTTGGAG**ATCCAGTCCACGGAGTCAGGGGACACA**GCACTGTATTTCTGTGCCAGCAGCAAAGCcacagtgc**cgaatgttagcccttcttagaacacaaactcattatggacccagctcaggaaataagtgtgtagcaggttggtaggcactacgtaacagaaacccaacttgaaagacaataaatcagaaggaaaaacttgtaaataacagctctaagtgagcaaaaaacaaagg**ggggttatttaaaacaactgatggggacaaaacccc**agggaaggggccagaaggcataaaaagcttggattcctcagtgttc**agcatattaggtaagatatgct**aggttaccctctctctctctctacatgagctttatggatttctaacaatctttatttaaagaaggggattctgtgcttttcacttggtaaagttgtgggtgcacaaataagaatgacaagcggatcaactgtgtctttcaggtagaccctgatcttctgacccagcacatttccttgaactagagacagactctaaagggctttttgact**gcttctcacagctctccctgccatagaagc**ctagtgggaaccattagcaattcctgccctatggaatgttacaatcacaaatttttatttttatttatttatttatatttattttaatttttttttgagacggagtctcgctctgtcatccaggctggagtgcag**tggcgtgatctcggctcactacaagttccgcctcctgggttcacgcca**ttctcctgcctcagcctcccgag**tagctgggactacaggcgcccgccaccacgcccagcta**attttttgtatttcttttagtggag**acggtgtttcaccgt**gttagccaggatggtctcgatctcctgacctcgtaatccgcctgcctcggcc**tcccagagtgctggga**ttacaggcatgagccaccacgcctggccaacaattacgaacatttttaaagcaactctattgagacataagatatatatggaatatgtatgcaataaacaaatccaactgctgcaataaaacacacccgcat

Uppercase: TRBV21-1-2

Lowercase: Flanking sequence[1000bp]

Red & Bold & Underline: Stem-loop [21]

Blue: Heptamer[31]

Green: Nonamer [7]

id-TRAV41[V_gene_segment]

ttaaagctctagttacactgtctaactgcggtttctgtacctctagttcaaattaatgagcgtagtatcttcccagatccaataagctattgccagaaaaaggaggttatttggtacaagcatgacctctatgtcatctctttagaaaggaatcccaaaagtggaaaaggactacatgctcaaaataattcaccaatgtgataaaacaactgaaactatcatagataacaataaaacaagtggaaaccagatgagtacagtaatactctgacagggcaacaatatttgaatcacatttgaagagacgtgaacaaactgtagcaagtctaaaaataagaaagataaaactaataaagtagtctgaaataataccatgtgcagaaagaaacaaggatgtcacataaagaaaccagagatatttataaggacacctttacaaaaaagaacagatctggccttgagttttcaaaagatataatgtgaaatg**cagctactctacttgatctatcatagctg**gctgaagtctaacaatcaacaagcttctgttctcccacactaccctcacctttcttctaggactcctagtaggaggatgctaagtttggttctctatctgtattgtgggtgagaggggctgtttgtttgttttagagacagggtcttgctctgttgcccaggctagagtggagtaacatgatcatatctcactgtatcctcaaactcctgggctcaagtatttgacctgcataataaatgtctacgcctca**tgccactaggtggca**atgtgggtgttatactgaaaagatcacagatggttcactttgcaagtaaaactgtaaatgttcttaagtgtgcatttctgctgcttctgatgggctgaaaatcccctttgatttctaaagtaaatgtagagacgttttaaaaataaaggactcctttgtccaagatatattccgaaatcctccaacagagacctgtgtgagcttctgctgcagtaataATGGTGAAGATCCGGCAATTTTTGTTGGCTATTTTGTGGCTTCAGCTAAGCTGTAAGTTGGGGAGTTTCAGGACATGAGATATTTTTCAAAATTCTGGGGAGGGGAGAGA**AGGGACTCCAAGTGGCTAATCTGGAGGCCCT**CCTGAAAATGTGTAAAGAAATGTGTATA**AATATTGCAATAATTGTAATATT**CTGTATCTGATGATGTCTTTGAGAACAGGTGTAAGTGCCGCCAAAAATGAAGTGGAGCAGAGTCCTCAGAACCTGACTGCCCAGGAAGGAGAATTTATCACAATCAACTGCAGTTACTCGGTAGGAATAAGTGCCTTACACTGGCTGCAA**CAGCATCCAGGAGGAGGCATTGTTTCCTTGTTTATGCTG**AGCTCAGGGAAGAAGAAGCATGGAAGATTAATTGCCACAATAAACATACAGGAAAAGCACAGCTCCCTGCACATCACAGCCTCCCATCCCAGAGACTCTGCCGTCTACATCTGTGCTGTCAGAcacagtgctccccaggcacctggagcccgtacctaaactctaaagttgaggcatcatttcttactcctgtctttcagacttgtctgtctctatccttggtcagatgatgtaaaatgtttaagtttttaaaatataatattctccccttctagaccatctctcttctctacaacatcatcaccaattatgttaaccagacagtatgt**gatttttatcttgaaagtaaaatatacagggtataaaaatc**ccgattctgctacttattaactggatgatcttgacaaagttacttaacttctctgtgcttcagttttccctcagctataaaaaggaaacagtgttagccatggtacgtaaccccaattgtggggattacatgaggtaatatacacaaagggcttgacacatttcctggcacgtagtgttacataaggggtatcaattattagtattagttttagatgtaccatgagtttgctaatattcactatggattaataccacagatcattggagaagtatgtggaatataaataataagcaccttataataattttagtttttagattctttcctaatgatgtccagcacttggcctttttaatagtgttaagaaaaacagggagtagctgtagaaaacgtaatagaaggatgactctatggtttggtttgtggtgtggaaatgaaagactggtaagagatgtttctgaggaagctcagctgcttctttggaaaaaaggacaggttct**aagttttcatgtggagatctctaatggaaactt**gcatttgtggaaattcatgaagcaaaatatcaaacccagtgtatctaaatcaacggttctgtatatcccccaaggattctttacattactattcccaatcccagtggaattcttatatcaaaacaagttttatgattataattgttcttattactcaacaacatcaatcacatatattaaattacagctacattcttttttaaaatcaaaggctttt

Uppercase: TRAV41

Lowercase: Flanking sequence[1000bp]

Red & Bold & Underline: Stem-loop [7]

Blue: Heptamer[27]

Green: Nonamer [4]

id-IGLV7-46[V_gene_segment]

gggcagagttcaggagaattccaagcacaagcatccatttgtcctctcccagtggagtaatagacaacgctgcattctgccggcaacaatgtgtgattctatgtaggaagcattgccaaccagcaaagctcacccaggcttggtgtccagaaaaattttgattggtggggtgggtgagagtggggttgttatgtagaccttgatgaccatcctcatggctcacttgagtttccactgaattgacaataaaacttcaatcttcaacctttggtcattttc**ttttaaaaaattttctaaagaaaattattttaaaa**agtctcat**ataaacgagctgtttat**ccaatatgaggctcaacgcaaatagtttcaaaaagtcagtaggtcattaaaacaat**tatttagtcattcaaactaaata**tttagtcattcaaa**cgttgttatctggtgttcagatagcaacg**aacaccatgtagcactaaaatcgcattatagtcacaggggcaggatttcagcaacagcaagagaaagtaatacaaacaggaatgagcaaagaatagacaagtaatgggtatgaattggcctgggaaatattgtggcaatgtccaaag**acaaatcaatagttatttgt**taaagaagtgaaagacaacacattctcagtccaggaaagccaccagggggtggtctgggt**ctcttggattgtaagaagctcaagaggttcaaacttaatgaacc**ctgcacagatgctcactgactcatggaaatgttagagcagctgcttcctcccacaggacaaatccacagcacacagccaccctgggctggcccc**ccacagtgtagctgctctcagggctgtgg**acccaggtggtgataaaggcagtaaagatttgcataaagcagcacacagcacaccccctccgtgcggagagctcaataggagataaagagccatcagaatccagccccagctctggcaccaggggtcccttccaatatcagcaccATGGCCTGGACTC**CTCTCTTTCTGTTCCTCCTCACTTGCTGCCCAGGTTAAGAGAG**ATTTCAAATACCAGCCTTTGGAGGGATCCCTTTTTCTCCCTTTCTAATTCCTAATATATGTCTGTTTTTTTTGTTTCAGGGTCCAATTCCCAGGCTGTGGTGACTCAGGAGCCCTCACTGACTG**TGTCCCCAGGAGGGACA**GTCACTCTCACCTGTG**GCTCCAGCACTGGAGC**TGTC**ACCAGTGGTCATTATCCCTACTGGT**TCCAGCAGAAGCCTGGCCAAGCCC**CCAGGACACTGATTTATGATACAAGCAACAAACACTCCTGG**ACACCTGCCCGGTTCTCAGGCTCCCTCCTTGGGGGCAAAGCTGCCCTGACCCTTTTGGGTGCG**CAGCCTGAGGATGAGGCTG**AGTATTACTGCTTGCTCTCCTATAGTGGTGCTCGGcacagtgacagacccatgagaggaaccaagacataaacctccctcggccct**tgtgatgtggagatcaca**tgatcagacatgccagatcccaagatagcctacatgtggaccagccatagaaaggggaaggaaagggtttgaattga**ttcctatttctacttaagtcttgatgtgtaggaa**atgtaagaatgatttgggagtaactggatcaaaaaaaaaaaaggaaaagttaattgttttctattctaaaaggaacactaacagcaggatcacatataaaggctgggtcaaaatgcatttcctc**cagaaagaagcatcttcagtgtataggctttctg**agctaagagc**aaagaaactttttttgcatattactgaagtttgttt**tattgagttactttcaaaccaatacctc**cagaaagaagcatcttcagtgtataggctttctg**agctaagagc**aaagagaagaaactcttt**ttttgtatatgactaaaatttgctttattgatttactttcaaaccaatacatgcctattatataaaataaaaatattaaaaaccaccaagttcactgcaaagctgtctcatcagaaaca**atccaatttcttcatttggat**gcattctttcatttttgttctcaatatttccacaatgggggtcataccttgtgctttttctctctggttttacacatctttatgcctcctcccacaccctgtgcctctct**cctgactctcagggtcctgggcagtcagg**gctcacttggtagcctgcacagtctgtctgtgacaaaaccagtcaccatctcctgcactggaatcagcagcaataaggttcttggaattgtgtcctggtaccaacaatgacaaggaagtgcctccaaacttctgatgtatagaagcaaatatctgcctgcagggacccaggatg**gatgctcgggctaccagtttggcaaggaggccttcccaagcatc**tctgggctccaggctgaggacaacgctga

Uppercase: IGLV7-46

Lowercase: Flanking sequence[1000bp]

Red & Bold & Underline: Stem-loop [23]

Blue: Heptamer[42]

Green: Nonamer [10]

id-TRAV3[V_gene_segment]

ttgctgaatacgtacttcatttgaagtctgttttaagtaggctctgcatgtggaaagtacctgttttgcaaccccttctcagagtttcatttttttgttttatatgtactgtttaatatgacaaatgctccaaaaatatttataacacctattcgtacttttctcactctaaaaatgtttaaagagaagattcaagtttctttttgccctaaagatcattatgggatgggctcctaataggcccaacgtggagaaaattctgagttatttttggtcaaagaggtttacacagaagccagctctgaattacttctaac**gagaagagcaaggctcctgttttcgcctgtcatttatcttctc**ttgccttactcttttctcctattttgctttttcttatctctcactttaaatttttctctccctcttgatgcaaagctatattttcctttctttttctgtgcattttcccatcctatttcccttctttctgtctctacattatcttaagagagcacaaatcatttacagcttgatccactcagcc**tgaatcttttccactttgattca**caaacttctatttcttctcctttgaacccctaatttgctccctatgtccctcatggagcgtgtaaccatccaaaggttggcccagggataaggccaggaagc**accagctgcttgacagtaaagtcgctggt**cctacgtgtgggctctagggggcgatgcagtgtaaggtctgtcctgggcataggcggagcggcagtctt**gctcctcacagagctttgaggagc**tggatcaaaattgtgctccacagagagaagataccggtgtcgggaagcaccagtgccctgaggaagggccatttccaaaag**ccctgtgctgacacaggg**ttgctggttcctcttcaag**agcccactctctggggtggggccatatctccagcagaggtgggct**ggaaaggacccccccaatcccgcccgccgtgagcttagctggagccATGGCCTCTGCACCCATCTCGATGCTTGCGATGCTCTTCACATTGAGTGAGTAATATCCCCTTCCCATTTTCCACTCGCATGATCCACTTA**CATGCTCTCCAGCAGCTTCACTGATGCAGCATG**ATTAATTTACAGGTG**GGCTGAGAGCTCAGTCAGTGGCTCAGCC**GGAAGATCAGGTCAACGTTGCTGAAGGGAATCCTCTGACTGTGAAATGCACCTATTCAGTCTCTGGAAACCCTTATCTTTTTTGGTATGTTCAATACCCCAACCGAGGCCTCCAGTTCCTTCTGAAATACATCACAGGGGATAACCTGGTTAAAGGCAGCTATGGCTTTGAAGCTGAATTTAACAAGAGCCAAACCTCCTTCCACCTGAAGAAACCATCTGCCCTTGTGAGCGACTCCGCTTTGTACTTCTGTGCTGTGAGAGACAcacactgataggggctgcagggggagcagaacacaaactcttgagtctggtaaagcccattttcttgaagtctttgttccttcacatgagaacggtgtgcttccaggatatgtcacttattatgttaaactacaatttaaaaccatagaatttgctcattcagcaagtgtttattgagtcctgatgtgccaggtactgggcaaagcaccggtcataaaaaggtgaaagaagtgtaaaacaaatagaaaaaaacataggggaaaattaccatgacattggtctgggcaagaagttccttccttccttccttccttccttccttccttccttccttccttccttcctccctccttccctccctgcctctctctctttctttctttc**tgaaaaggggtttattttca**aca**gacatttctttttataccgaggtgaaatgtc**atcatcataataattttgcaatgttacataccctgttgagaccattaaaaaattaggttcatggtatacacgtccaagtttgttacaaaagcatattgtgtgatgctgtggtttgggcatctattgatcctgccacccgaagagtgaacatagtacccaggagaaagcttttcatctcttgtgagcaaggaat**ttttggatatgaccccaaaa**gcacaggcaacaaaagcaaaaatagacaaatggggctatattaaactaaaaagcttctgcacagcaaaagaaacaataaatagagtgaagggaaagcctatggaatgggaggaaatatttgcaaactctacatctgataaggggttaatattcaaa**atgtttaagaagctcacataccaaaaacat**ccaaataacctgattaaaaaataggccaaggaactgaatagtcgtttctcaaatgaaggcataaaaatggccaagaggtatataaaaaatgctcaacatca**ttaattatcaggaacgtgcaattaa**gaccaccattagatatcacctcacaacctattaggtt

Uppercase: TRAV3

Lowercase: Flanking sequence[1000bp]

Red & Bold & Underline: Stem-loop [13]

Blue: Heptamer[21]

Green: Nonamer [4]

id-IGHV4-39[V_gene_segment]

caattcaatggaggagaaa**atctttcagagacagatggatctgaaatgat**aaatatgtgggtaaggactctgggcttgagtatcattgtccagccatgttt**cacaagtgtgtcctgtcagggaaggatcagagttccttgtg**ctgtcagagggaaggggtcacagagttcctctctggttcccaggaaaggtaatcgcactaatct**tcatgatcttcatga**gactatcctccagtgctgacctgttacggagtttttgtctgaagttctcactgcaatccccaatctacatattttcaatcagaagtgtttagaggccaggacacatcttcaaggtcacacatt**gagaaggatggagatatgtcccactaccttctc**ctacgatctcagacagaatcccaaatttcaaaaggacacagaaggacagctctcaggtgcttttaaaaaatgacccacttc**cagggacagtgagcttccctg**taaccatggtggatgttctgaact**acaataaacattggatggatccagtattgt**ttgaagtcactgtcattattacattcagctgttgtttcaatgtgtctgaaagggtaaatgactatttagatggcctgggtgtgtggttggttttatatgaatctttaagggttgaacagtactgaccctattccaaaatctgtccttgatccaggatcacactcatctctcagaccagctccttcagcacatctctttacctggaagaagaggactctgggcttggagaggggaggccccaagaa**gagaactgagttctc**aaagggcacagccagcattctcctcccagggtgagctcaaaagactggcgcctctctcatcccttttcactgctccgtacaaacgcaccacccccatgcaaatcctcacttaggcgcccacaggaagccaccaca**catttccttaaattcaggtccaactcataagggaaatg**ctttctgagagtcatggatctcatgtgcaagaaaATGAAGCACCTGTGGTTCTTCCTCCTGCTGGTGGCGGCTCCCAGATGTGAGTGTTTCTAGGATGCAGACATGGAGATATGGGAGGCTGCCTCTGATCCCAGGGCTCACTGTGGGTTTTTCTGTTCACAGGG**GTCCTGTCCCAGCTGCAGCTGCAGGAGTCGGGCCCAGGAC**TGGTGAAGCCTTCGGAGACCCTGTCCCTCACCTGCACTGTCTCTGGTGGCTCCATCAG**CAGTAGTAGTTACTACTGGGGCTGGATCCGCCAGCCCC**CAGGGAAGGGGCTGGAGTGGATTGGGAGTATCTATTATAGTGGGAGCACCTACTACAACCCGTCCCTCAAGAGTCGAGTCACCATATCCGTAGACACGTCCAAGAACCAGTTCTCCCTGAAGCTGAGCTCTGTGACCGCCGCAGACACGGCTGTGTATT**ACTGTGCGAGACAcacagt**gaggggaggtgagtgtgagcccagacaaaaa**cctccctgcagggagg**ctgagggggcgggcgcaggtgcagctcagggccagcagggggcgtgcggagctcacggaatacaaggccgggtcaggagcaggtgcagggtgagcggggcttgctcatcttctcagagatcatcatctccctcctcgccagcacctcagctttccgtagaggtcctctttctttattgtgtgtggttctacttcctcacatccttgtgc**caggaaagaaaggagtaaggcaaattttcctg**ttacaattgaagtttcaccaattactaagaactttcctgcaagtacctgcacagcccattataccttatttatatatgtatatattctaatgcttctcaccatctcttgatttgtgtcatcaatttaattgtgccctttttgaaattcatatgctgaaactttaaatccaatggatctatatcggaattttaatggtataattaatgttaaatgtggtcataaatgagaccctaatgcaatagagctgttgtctttataagaagaggaagagacaccagatacctctca**cttctcacatgcactcagagaagaggccacgtggagacatagtgcactagaaggtgggcctct**gcaagccaggaagaagccgcaccaagaaccaaccctgccagcacc**ttgatcttctacattcagactgcagaattgtaagaaaatcaa**tatttgttgtttaagccacccactcctttt**gtcttcttacgaagac**ccagacaggctaataccacacaactctgttagctccatctcctggagggagaagcagccccctgaagctgggcacatcgctcagattttcacatgaattaggcaaaaacagtagctctcatataaaaactgtcacgtccctgttgggacaaggtcttttaaacaagccctggggctttgtcacaaatgttgcattttatcctttattaggacttaattaat

Uppercase: IGHV4-39

Lowercase: Flanking sequence[1000bp]

Red & Bold & Underline: Stem-loop [18]

Blue: Heptamer[41]

Green: Nonamer [7]

id-IGKV1D-13[V_gene_segment]

ttaaatgaattttgtttgccttttctcttattaatct**tccttttgtcagtttattttcagcaaacatttagaggacaaatgga**gttttcttcctttctcccaatataagcaagttcccttaaaattcaggcggcttacaaagcagcaagaaggtttgtgcacgggctatggcactgtgattt**ggctcccctactcaggcatcagtaaaattttgtggagcc**ctaggctacagcccactgatgctaatatagttggatccacttcccctgctactgagctaggctgggacagttttgg**gcacattagatatgtgc**gatat**aatgattgcaaatcatt**tc**cagtttcatctgtatcaaactg**ctttctccatgtacataggcatcatctctgtggatctgtagtaaattgcttgatcttatagtggtaagaacaatggcataacaccattaccgaatac**tgacatgtatatatagcatcatgtca**ataaattttatttttgatttttttagaaa**ggaacaatgttaaactcacagaaatgttcc**aagtataggacaaagtaaccccttccctaaccgggatcatatgagagtcttttggagacctgataatcataccgtctaacattttattatatatttcctacaaacaagaatattctcctaaataatccccatacaccaa**tgaaatacattactccatcaactcctgaggaatatttca**aatcgtcaaaaaaaaactaaaaaatgtctctcataataaaatagttcccagtagaaacacattctctggagacaaatttgtgctaccctggtcttacctgggacacctggggacactgaactggtgctgagttactgagatgagccagccctgcagctgtgcccagcctgccccatcctctgctcatttgcatattcccagaacacaacctcctgccctgaagacttcttaataggctggtcacacttcttgcaggagtcagacccactcaggacacagcATGGACATGAGGGTCCCCGCTCAGCT**CCTGGGGCTTCTGCTGCTCTGGCTCCCAGG**TAAGGAAGGAGAACACTAGCAGTT**TACTCAGCCCAGGGGGCTCAGTA**CAGCCTGGCTATTCAGGGAAATTCTCTTACTACATGATTAATTGTGTGGACCATTTGTGTTTATGCTTCCAATCTCAGGTGCCAGATGTGCCATCCAGTTGACCCAGTCTCCATCCTCC**CTGTCTGCATCTGTAGGAGACAG**AGTCACCATC**ACTTGCCGGGCAAGT**CAGGGCATTAGCAGTGCTTTAG**CCTGGTATCAGCAGAAACCAGG**GAAAGCTCCTAAGCTCCTGATCTATGATGCCTCCAGTTTGGAAAGTGGG**GTCCCATCAAGGTTCAGCGGCAGTGGATCTGGGAC**AGATTTCACTCTCACCATCAGCAGCCTGCAGCCTGAAGATTTTGCAACTTATTACTGTCAACAGTTTAATAGTTACCCTcacacagtgttacaaacccgaacataaacccccagg**gaagcagatgtgtgagactgggctgccccagctgcttc**tccttatgcctccat**tggctgagagtgttcctcagatgcagcca**cactctgatggtgttggtagaggaggatatgagatcacctctgcatcccaatttctttttcttttctcagccccagctgcacagacataacaatgcctctgctgatttaataaagatagagatcatgacacctgaagagtctagttt**atggctttggttagaattcatataacagagaagaagccat**tatagatatcctaagcaggaatagtcttaatagatagaattggagtctaaagtattgaagtct**aaataaaatgtacagataaatttaatgttttattt**gttaagaaatttttgccaaatggggcatacagg**aaaactcaatggtcttcaatatgttggaagagcaaagagtttt**ataaaaagggaaattattacctattgttctttgagaaattttgttggctgtagtaagggttgggagctggcaagctcagactggtaagcagtggtggtcaaactgaatcctagaattatattaagttatctcagaagttgtgggtaaatt**tgctttcaggttacaataagccaaagca**gtgaagcttgcagagaattttgt**tactgaaatgccagggattcagta**tagatcctgcggctcaccacacagaaagccaatcactaa**gacaacaagtgttgtc**aaagaacaggctttaatcaggtgctgcagccgaggagacgggacaccattctcaaatgtatc**tccctgacaaaataaattaggggtttatatagcaggga**agaaatgtggaaaacaggaattagagaggggtaaggaagataatttagtcaacaggaagctggaggtcagttaggcaatcat**aatgggtgaagggtctgatgtctcactgtccccatt**cagtgatatataactttcagctc

Uppercase: IGKV1D-13

Lowercase: Flanking sequence[1000bp]

Red & Bold & Underline: Stem-loop [24]

Blue: Heptamer[18]

Green: Nonamer [4]

id-IGLV2-8[V_gene_segment]

tgtcgaagggttgagagaggtcagctttaggttgtcctgaggatgtgctcagatgggggaaagtaagaaagccggaaacgtgcatct**ctgtgacctgacacataaatcacag**taaaatggaagtaaatctttcaataaa**cctttcaagataacatcactctggtgcacatagaaagg**cctgttagcaaactttcccttttccctgctggctcatatccccatggatcacctgtatgcagaacggactctgtggtctccatttatcaggctgagaaccagaggctgggaaggacaagtcac**ctgctcatgagtgggaggggcaggattgggagcag**cggactcaggctcaggactgtggccctggttggtgctccttgttcccttcacagggtcc**cacccacatgtgcccgtgtggatgtgggtg**cactgcctggggtggctgatgtgctccagggatttgatggtgcag**atgcttcaggacaaagcat**gggagtggggatgggacgaggctgccgctgggcagaccaggga**ccaacgccacccagcagaggcttccagtggggttgg**gcagccccatttcctgagaatagagagaac**ctggcaatgagtgccag**aaagaggttacttcgaggtgtgcccaggcctggagggtcacagagacacaggcaccacacagcagagacactgagggccaagagctca**ctcaggtgaggggcttcagcagatttttctctcctgag**caaatcacgtgcaaagaaatcaacttcctgtcatcagaatagacaatcagggactttagtctccttggctgagccccgctgtcaagggaagcagaagtctctaagcccaggcccaagtgagggtggggtgagaagaggagctcaggatgcagatttgcatggaggtcccgcccttctctgaggcagaggggataagacagggctgggggcaggcccagtgctggggtctcaggaggcagcgctctcaggacgtcaccaccATGGCCTGGGCTCTGCTCCTCCTCACCCTCCTCACTCAGGGCACAGGTGATGCCTCCAGGGAAGGGGCCACAGGGACCTCTGGGCTGATCCTTGGTCTCCTGCTCCTCAGGCTCACCTGGGCCCAGCACTGACTCACT**AGACTGTGTTTCTCCCTTTCCAGGGTCCTGGGCCCAGTCT**GCCCTGACTCAGCCTCCCTCCGCGTCCGGGTCTCCTGGACAGTCAGTCACCATCTCCTGCACTGGAACCAGCAGTGAC**GTTGGTGGTTATAACTATGTCTCCTGGTACCAAC**AGCACCCAGGCAAAGCCCCCAAACTCATGATTTATGAGGTCAGTAAGCGGCCCTCAGGGGTCCCTGATCGCTTCTCTGGCTCCAAGTCTGGCAACACGGCCTCCCTGACCGTCTCTGGGCTCCAGGCTGAGGATGAGGCTGATTATTACTGCAGCTCATATGCAGGCAGCAACAATTTCcacagtgttttaagtcaatgaggaagtaagatcaaaacctgccctgggctctcaggcccccttcttgct**ctgcagatgcttcctcacgctgtataagggtttcctgcag**gatggccttgacaattctcctctctcagctcctc**tcctttcccaccatgaggtctaaaagga**aacctgctcgtgatttctcgttcaggactgtggcaacttccttttgcttgtgtgctctggtcccttaacgtgcaactattcctagctcttcaatgcagggacgt**agggacaaggagtttactgcttggtgcagtccct**cctgttttcaggaacatcctcattctaaatgcatccccca**tctgtcacactatgcagatcaatctggacagaagccatcagggggatggcttc**cagtttccaggaattgcatcttgttccactctgtgtccaccacgctctaatgaagatggccctcctag**cttaaagtgaccactttaag**aagacttgaagatgtttttgagggattaagaacaaagaggatgctgctgtttttctttttattgctcctgctctttgccgaaattcctcaggttg**ctgagctggggagattttgagtgacaggctcag**tgctctcgcagaattctcctcccctcacatcgctgaggccctgtcctggaaactcctcacaagtggacggtcttcccataggatgggggattccaaaatggctccacaggaaagagttaacctgagtccacctcccttcctcatggacatcgagcatttctaattttcatggctgtcaatacttttgtacctggaatccctaataatctaatggtgagaattgatttagaacacattcggacattagctgggtccaa**tatttaaattttctgagccagttgttaaata**cagctattatcatatat**agtttaggctccttaaact**tcgattatacagattatatttaaaacaaagtaactagtttactatatatatttacaaa

Uppercase: IGLV2-8

Lowercase: Flanking sequence[1000bp]

Red & Bold & Underline: Stem-loop [19]

Blue: Heptamer[42]

Green: Nonamer [2]

id-IGLV3-29[V_gene_segment]

ccctttgggccagcagttcagggatca**agaaatcctagacctgaggagtaatttct**gaatctacacctagaccaaggtggtgtctgagcctccagcagtcagtgaagagcaaccagcagtggtcaacagagagtcacatggcaaagccgcctggagctgggctgga**cctgcatgtcagggagatgctgacatgcagg**gagggtcagtgaccatgacctaacactggggg**gaaaaatgtccttctttttc**tagagtatcatcgttaacatt**gatgtacaaactcttttgccccatgtctcttacatc**tggctcttttagaatttgcttatttcacagcaacaagttcaagtcttatactacattgcatcccagcccacccaattatgctaaaactctggatctgtcaaacactttcagagcagcagatttgtacctggccctaaacgcatagtggggacactgaggcacagctcctagggccttc**ccctgaagctccaccaagggcaaaaacttatacaagcttcatgg**atattaaaaacctccatgttcacttagcaagggagaaattttattcaagttctgcagctcaggagctgtgctcaagagcaagacccagaacttctcaaagcctgg**ttttggcacctgtagaatggagcccaaaa**acgcagacagacttacctcggagctctatagggccaggacagcccaggagtaaagccagtttgagacacaatttggggtgaagatgtcggcccagaatagaaccaggagaggaagaagaaacccgtgtgctga**gagtgggtctgctgggaccactc**ctctccttcctgggccctttggggctgagcccttctctaaaaaccacagagctactccag**cagcagcccctgactctgctg**atttgcatcatggccgct**ttatccagcaaggggataa**gagaggcctgggaggaacctgctcagcctgggtcttgggaagcagcatcgggggtgcctcagccATGGCCTGGACCCCTTTCCTCCTCGGCCTCCTCCCTCACTGCACAGGTGCTGCCCCCAGGGTCTCACTCACTTGCCCAGCC**CCAGGCCTCTGGGTCCAGCCTGGCCCTAACACTGAGCTCAGTAGGGCC**CTACCTGTGGTGGGCAGGATGCTCATGAA**CCTGCCGCAGGGCAGG**GGGCTG**GTGGGGCAGAAATCCCCCCAC**ACTGTGCTCCTGTTCTCTTG**GTGCCCTAAGGGCAC**TTTTATTCCTGGGGCTCAGGTCAGTTCAGGCAAGTGGCAAGATATTCTTTGGTTAATCCCCAGAGTTTCAGGCCTTTTCGCTCCTCTTTTCTCTCCTTGCAGGCTCTGTGGCCTCATATGTG**CTGACACAGCCACCCTCAGTGTCAG**TGGCCCCAGGACAGACACCCAGGATCACCTGTAGGGGAAATAATATTGGAATTAAACCTGCTTAATGGTACCAGCAGAAACCAGGCCAGGCCCTTGTGCAGGTCATCTATGGGGATAGCAGATGGCCTTCAGGGATCTCTAAGTGATTCTCCAGCTCCAACTCGGGGAAC**ATGGTCACCCTGACCAT**CAGTGGAGCCCAGGCTGGGGACGAGGCTTTCCTCTCAGGTGTGGGGCAGTGGCACTGCAcacagtgacagaggcagacaaggaagtaaga**cacagaccccttccccatctgtg**ctgctgtcgtcctccagcccggcaacactgtggacaaagccatgagcatgcatgacccagttcacctggatctgagacccccagtctttacttccttcagccctccaggcaggctgtgcagagggggcatccagaatggatttagggctgaggtgaccaggatgtattggggttttgtggactccactgtgattcagggaaattagatcttccaagcagaccaaggtcaatatcctcagttatctctggatttccataggtggtgataggccagtccttggccgaaatgtcctgaatcagtccccagaactgcccagctgaggctcctgtcattctctacagaaaaactagtggagtggacactgt**gggatcctcccagatccc**ctcttatgagctcacccacacagcactcctgggagcttctggcagctcagagctgttcggtgcagaaatctg**cctcccccaggtttatgtctcttctcagggagg**tttggtttcatgactgcctgtggccaaggtctcaagactcttcctcagtataacaggacctgaaaatcctccaagatccatagctccttgtagggctgactgaagcctcaatatgtgtcagggcctccttctgcctggtgttgcctccatcactttcttaatcatctgtgtatgatattctccatctcagagtctgtttccagggaacccaaactaagatcaccagccatccccaacacgggccatgagggagctgaagaggccactgtacactagaaagcatcagtttactcttaaatt**accactgcgagtctgcaaatgcccagcgagacctgagtggt**ttttt**ttttcatttgtgtgaaatgaaaa**gggagactgacatgccagatgctctgggttctgagggagagttggagtcagatcctcctgcaggaatccaggggcagcaggagcagcc

Uppercase: IGLV3-29

Lowercase: Flanking sequence[1000bp]

Red & Bold & Underline: Stem-loop [21]

Blue: Heptamer[39]

Green: Nonamer [7]

id-IGHV4-39-2[V_gene_segment]

caattcaatggaggagaaa**atctttcagagacagatggatctgaaatgat**aaatatgtgggtaaggactctgggcttgagtatcattgtccagccatgttt**cacaagtgtgtcctgtcagggaaggatcagagttccttgtg**ctgtcagagggaaggggtcacagagttcctctctggttcccaggaaaggtaatcgcactaatct**tcatgatcttcatga**gactatcctccagtgctgacctgttacggagtttttgtctgaagttctcactgcaatccccaatctacatattttcaatcagaagtgtttagaggccaggacacatcttcaaggtcacacatt**gagaaggatggagatatgtcccactaccttctc**ctacgatctcagacagaatcccaaatttcaaaaggacacagaaggacagctctcaggtgcttttaaaaaatgacccacttc**cagggacagtgagcttccctg**taaccatggtggatgttctgaact**acaataaacattggatggatccagtattgt**ttgaagtcactgtcattattacattcagctgttgtttcaatgtgtctgaaagggtaaatgactatttagatggcctgggtgtgtggttggttttatatgaatctttaagggttgaacagtactgaccctattccaaaatctgtccttgatccaggatcacactcatctctcagaccagctccttcagcacatctctttacctggaagaagaggactctgggcttggagaggggaggccccaagaa**gagaactgagttctc**aaagggcacagccagcattctcctcccagggtgagctcaaaagactggcgcctctctcatcccttttcactgctccgtacaaacgcaccacccccatgcaaatcctcacttaggcgcccacaggaagccaccaca**catttccttaaattcaggtccaactcataagggaaatg**ctttctgagagtcatggatctcatgtgcaagaaaATGAAGCACCTGTGGTTCTTCCTCCTGCTGGTGGCGGCTCCCAGATGTGAGTGTTTCTAGGATGCAGACATGGAGATATGGGAGGCTGCCTCTGATCCCAGGGCTCACTGTGGGTTTTTCTGTTCACAGGG**GTCCTGTCCCAGCTGCAGCTGCAGGAGTCGGGCCCAGGAC**TGGTGAAGCCTTCGGAGACCCTGTCCCTCACCTGCACTGTCTCTGGTGGCTCCATCAG**CAGTAGTAGTTACTACTGGGGCTGGATCCGCCAGCCCC**CAGGGAAGGGGCTGGAGTGGATTGGGAGTATCTATTATAGTGGGAGCACCTACTACAACCCGTCCCTCAAGAGTCGAGTCACCATATCCGTAGACACGTCCAAGAACCAGTTCTCCCTGAAGCTGAGCTCTGTGACCGCCGCAGACACGGCTGTGTATT**ACTGTGCGAGACAcacagt**gaggggaggtgagtgtgagcccagacaaaaa**cctccctgcagggagg**ctgagggggcgggcgcaggtgcagctcagggccagcagggggcgtgcggagctcacggaatacaaggccgggtcaggagcaggtgcagggtgagcggggcttgctcatcttctcagagatcatcatctccctcctcgccagcacctcagctttccgtagaggtcctctttctttattgtgtgtggttctacttcctcacatccttgtgc**caggaaagaaaggagtaaggcaaattttcctg**ttacaattgaagtttcaccaattactaagaactttcctgcaagtacctgcacagcccattataccttatttatatatgtatatattctaatgcttctcaccatctcttgatttgtgtcatcaatttaattgtgccctttttgaaattcatatgctgaaactttaaatccaatggatctatatcggaattttaatggtataattaatgttaaatgtggtcataaatgagaccctaatgcaatagagctgttgtctttataagaagaggaagagacaccagatacctctca**cttctcacatgcactcagagaagaggccacgtggagacatagtgcactagaaggtgggcctct**gcaagccaggaagaagccgcaccaagaaccaaccctgccagcacc**ttgatcttctacattcagactgcagaattgtaagaaaatcaa**tatttgttgtttaagccacccactcctttt**gtcttcttacgaagac**ccagacaggctaataccacacaactctgttagctccatctcctggagggagaagcagccccctgaagctgggcacatcgctcagattttcacatgaattaggcaaaaacagtagctctcatataaaaactgtcacgtccctgttgggacaaggtcttttaaacaagccctggggctttgtcacaaatgttgcattttatcctttattaggacttaattaat

Uppercase: IGHV4-39-2

Lowercase: Flanking sequence[1000bp]

Red & Bold & Underline: Stem-loop [18]

Blue: Heptamer[41]

Green: Nonamer [7]

id-IGHV7-56-2[V_gene_segment]

tcccc**aaacccctgaagagtgtttgggttt**accgtgtgtccagtgattcagtgcctctagagctccaggaaggggctccccggtgatgcctgagatcttttcttcaggtctccctgcagagttctctgggtttcctaagggcaattcactatttcaaaagatggtgtgagaagcacatgctgtcattaaaggagaattctgagccacggcacagccactttatactgggctagagacactggtatgaatatactctgtaagtttagacagaaagcctcgtgcatggtaggggctgggctgcagggggtgctcaggataaacgcagcacagtct**cccgccccagagcaggtgcacaggaggctggggaggg**gttcctctcagggcctgggacttcctttgaaaatatctaaaataagtatttcacaagggctgctgtt**gtttgtataaatatcctattcaattgtgagcatttatcaaac**tggatg**ttgtaatgacaaccacttttacaa**tggggatttcaaactcccctagatatcttaatagtaagcagctggaggtcaagaagagatcctttcttttaaataagtgcaat**ttttggagaaacatactaattcccaaaa**taacgcattcacatattaaggtctagaaatggctcaagttgtccctggtgcattcgaatgtgggttcaaagtgaggtgcgtgtcctgagg**gagcttgttctccagtggaggaagctctgtcaacacagag**ttcagggatgtgtaggggtcgcatcgcctctaacaggattacggcttgaaccctcagcatgtacaattgtgtcgtccatctgtcatgtatttgctctatctcatcctggctcaggaattgggc**tattcaatagcatccttcgtgaata**tgcaaatcactaaggttaatacagatatctctgtgccgtgagagcatcacccaacaaccacacccctccttgggagaatcccctagatcacagctcctcaccATGGACTGGACCTGGAGCATCCTCTTCTTGGTGGCAGCAGCAACAGGTAA**GGGACTCCCCAGTCCC**AGGGCTGAGGGAGAAACCAGGCCAGTCATGTGAGACTTCACCCACTGCTGTCTCCTCTCCACAGGTGCCCACTCCCGAGTGCAGCTGGTGCAGTCTGGGCCTGAGGTGAAGCAGCCTGGGGCCTCGGCGAAGGTCTCCTGCAAGGTGTCTGGTTAAACTGTCATCACCTATGGTATGAATTGGATACGACAGACCCCAGGACAGGGGCTTGAGTGGATGGGATGGATCATCCTACCCTGGTGAACCCAATGTATGCCCACAGATTCACACACGGTTTGTCTTCTCCATGGACACCTCTGTCAGCACGGCGGATCTGCAGACTAGCTGCCTAAAGACTGAGGATGCAGCCATTTATTACTGTGTGAGGTAcaccgtgtggaaacccacatcccgagagttttagaaaccctgaggaaggaggcagctgtgatgagctgaggcagtggtgcagcatgtctct**aaacttccattttatctaagttt**gcattgagttccgctttaatattagccaggaatgtgggatagacgggtgctcctaagaggtccttaatttgcccattttgatgggttttccagaagacgtgagaagccactttgttaacaaagcatcccaaagccatgccctgctccagaaacacgtgtacccatttcctggtctttggttaactgacaagctctcatcagcgcacctgggctaatttctcatcaggtagaaaaatgtgttgtaaagcaaggctaacgttgtgatagcaattcctgctcaataaccttcagcatcgttgttgttgtgttctatcaactaattacttgacttcaaggttctcattgggagtgtcttataaatttaagggatatatagaagttcccctaattaaaataaaacaattgtgagcacaacctcagtgttcaaccatgtctccacccttcccaccattcaccccaaagaaatgttcacctctcctggaagtcgggttcattttcaaattagttattttttattttactatatcaagattattgtatgtgactactgtagcagaaagtgaattaggggaacttgaagtaaccaacgaaagataaattcagaactaattaaacaagatgtcagaacgtgattggctctagtcttttaaaattcagcaggttatgtaaccaggctttaaatttacacatcttcctgttaccttcacggcacagtcaactcccattatgtaagaaatggcaactgcattcccaagcgtcatccaaaattgtaaaaatagactgggtgaggtgag**gagttgattgtttaaattccgctctgaagaagcagcatcaactc**aacaaaccaccgcttttccctcagtgactagagctatgtcgcaggcca

Uppercase: IGHV7-56-2

Lowercase: Flanking sequence[1000bp]

Red & Bold & Underline: Stem-loop [11]

Blue: Heptamer[31]

Green: Nonamer [3]

id-TRBV4-3[V_gene_segment]

tttcctcctggcccagcacatctccaagttaaaaaaatctctaaattctcaacccatagtttaatcatggtaactcccttctacccccaacatagatttttatgtaactgatttgtaactacaaccctaaccctacaatcacaattgtcattacaaaatactaacaaacatgtgctattaccaatagatggttaaaatcacttagggattatgaatcatgtcattcttgtatttatttgtacttattctttgtgtttatgtattatg**tcagcttcaagctga**tttccctttcccttgctccccacaccttttgttgttccttttcagcatcctgtgtatgttagctctaagcccagccaagccccatgccctccacttcagggacagagggtggctcatccctttcgtgctccctcagctccctccccagtgcctgg**tggctggttggcatcacagcca**tgcccttgaataatgggcgctggtccacactgct**tctgtggtcatgcagtcatcctccctccacaga**gctgggccaaa**aagggctccactgccctt**aggcattgagggaatggacattctgagtggca**ctgtcttgtgtgatcccacaggacag**aaggagattccatcaatgggagtagtctgttgctgagag**ggatcctgaaagatagggggacagagacacactgggaggatcc**acataaagaggaagcagctgagaacaaaggagcccctgccagaggaatatcct**tggggagtaaaaagctgactctgccctttctcccca**gcccattcatcccagcccagacaattcaaatctaccttctatcaggacctagaaaggatgtaaaacggctgggtataaatatcccctgggtctggggaaactgtcaggagcagtgacatcacaggaaaaaccaccaaccaaggccaaggagaccagagcccagcacctcacccagaggaccccagtcagaggccccatctcagacccgaggctagcATGGGCTGCAGGCTGCTCTGCTGTGCGGTTCTCTGTCTCCTGGGAGCGGGTGAGTTGGGTTCACATCAGCTGTCCTTGAATTCCAAGCTTTTTCCTTGTGATTTCAGCAACAAGCCTCCTCCTGGGCTCTGCCTGAATTTTGTCCCTTTCCCCCCACAGTCCCCATGGAAACGGGAGTTACGCAGACACCAAGACACCTGGTCATGGGAATGACAAATAAGAAGTCTTTGAAATGTGAACAACATCTGGGTCATAACGCTATGTATTGGTACAAGCAAAGTGCTAAGAAGCCACTGGAGCTCATGTTTGTCTACAGTCTTGAAGAACGGGTTGAAAACAACAGTGTGCCAAGTCGCTTCTCACCTGAATGCCCCAACAGCTCTCACTTATTCCTTCACCTACACACCCTGCAGCCAGAAGACTCGGCCCTGTAT**CTCTGCGCCAGCAGCCAAGAcacagccttgcagag**tcaccgct**ttcctgtgcagaaaccttaggggccctccaggaa**gctgtgggggccaccaaagccttcggtgaacatttcctgcaagagccccgacagaagattcagaacatcatagcacctgctcattcatccatgtggcaacgttacatcctatgctcatatttagagtgtgggttttctttagcgtgtgactctgccctgttaactgatttgaaaaatgataaacacattcagatctagtctt**tttaaaatccctttttaaa**ttgtaaaataaagcaagcatgctataaggggattgtttggtttaagatgttataagaccctctcaacaaccacctgcgtcactacagagtcataccctgcactggaccacag**agtgtcctgtgaggagccaagttcttctgacact**ttatg**cctggtggtctccgtgcctctctttctcagtaaccagg**ggtccattctgtcataggttgctgacacccaaggtcaagcagcactgggtttgggggaatcatagaggcaacttccaggcaagggtcccagggcgggagccagctggagagaacagtagacaactgagcttttggggaa**tgggtttgagaaatatctgtcaaccca**attctattattctatcttccatcttgtctcttctctattgttcttatttaaactccaagctagagcgtaaattccagccctgatcaaagctccatatccctcacagacaccttctctgaaccccgcaggcag**caaatgtgcctctgtccactcccatttg**tgtgggcggtaagccacccaagtcctgaggcaagagaccgagggcacgagctgttcc**agtataataaagaaaatatataaaataagaatagttatact**agtaatagattatagatatgatggta**tacaaatattatgaataatcagtttgta**gcattactcttcattccaatattataataatccttgctctacaattataacctag

Uppercase: TRBV4-3

Lowercase: Flanking sequence[1000bp]

Red & Bold & Underline: Stem-loop [16]

Blue: Heptamer[35]

Green: Nonamer [4]

id-TRAV31[V_gene_segment]

cagatccctctgaatagattaatgctctggagtggagagaatgaata**agttctcactctgttagttcccaagagaact**ggttgttgaaaagagcctcgtatgtccccctctctattgcttcctctctcactttgtgatctctgcaaacaacagctcccctttg**ccttccaccatgagtggaagg**agccggaggccct**caccagaagcagatgctggtg**ccattcttcttgtatactctacaaaatcatgagccaaata**aatttattttctttataaatt**acccagtctcagtattcctttatagtaac**ataaaatggactacgatgatgggaaataatattttat**caatggaagtgatc**ttctattataactgagccaagagaaataaaattgaatagaa**tggaagttttcttttgtcttaacaaagttaccaaaatcacacattgggaggataagcaga**agtctaagataatcagaaaacagagtagact**gcaaaagaattgtttgaaatcatcaattaagatggaaaggaggcagggaaaacatcatagtttctagtatcaccctgaaattttgtgtgtaacaatatatgtatgtctatctacatccaaggatacaaaaccacatctg**tattttaaagtaaaatatattttcttcactcaaaaaaatatat**aggtatctttttaatctcagtctaaatctgggaataaa**attttgtggccccaaaat**gtctggagaataactaagcaaactttgctagggaaagagctgtcttaaacacatctagtggctcaatttagtctgtgcaagtatgagcaggatatgatttaacaacttgtccctgcccctgccttccccactgtaccactttctttcctattctgagacagatgcattgctcacctccaataattctaggtggagctttatac**aaaactgaataaagtaggtctcaggctctgagcaggcagtttt**ttttcctagaaataga**ggtgccaacATGACTGTTGGCAGC**ATATTACGGGCACTCATGGCCTCTGCCTTCCTTGGTAAGGACAGTAGCTGATAGGCTGAGGACAGTCTAGGACTGGGAGAAGACAGTGGGGAGGGAAGAGGGGATGAACAACTGGTCATATGTGACAAACACTCAGGTGTAATAAGAAGGAAGAGGAAAGGAACCAGATTCTGACTCCTGCCTTTACTGAAGGTTTAACCCCTGCCCCAAAAGTTCTTGGAAGCTCTCTTACAGGTCAGCTCAGGGGACCAATCTTGTTATTTCTTCTCATTTACGGGATCCAGCATGTCACAGAGGGTCATTCAATCCCAACCAGCAATATCTACGCAGGAGGGTGAGACCGTGAAACTGGACTGTGC**ATACAAAACTAATATTGTAT**ATTACATATTGTATTGGTACAAAAGGTCTCCCAATGGGAAGATTATTTTCCTCATTTATCAGCAAACAGATGCAGAAACCAATGCGACACAGGGTCAATATTCTGTGAGCTTCCAGAAAGCAACTAAAACTATTCAGCTTATCATATCATCATCACAGCCAGAAGACCTGCAACATATTTTTGTTGTCTCAAAGAGCCcactgtgaagaacatgttagaagagccttacaaaaagatcggaactcaa**cctgaggcaattgcctattcccacattctcagg**aaaaactcacaaaccttacccaggcatttgttagcagctggtggttagggtgcataagataagacccagaaaggaagtcattgagttcattccaggaagcacaccctcacaactgccttagaaaccacacaggaccacatgtaaaagccatactgctcggatgacatcacatctctatttcttcccatggagtatgccg**tttaaaaaagcaacatgtgaaacatggaattttaaa**gcttaattgattgtacaggactttatatctgctcaaccaaatctcttgaggaaaatagtatggaagacactagggaaaaaaaacacgcatacaaaataaagatttttatggcaggatccctaaacactggccagagagatgcaatcttggtgagactatagactggagtacttttggaccatgtagaacagcagcctattctggaccagatgcatttttctatctccttggctacatttcactctaatcctgccaactccagcttggcttaccatgctttctgattctcatgttccttgcagctgggtctgctagttgggtcctccctgcagtcacatgcctatcacaagatttggttaaccacaacagagaaaggagatatgtggggagacactg**ctacagatgctaggatctgtagttaatcttgtagccacctaacta**attcatg**gaaactccagaaatgagactgcagtttc**taaatcccgtatttctttattcagtccccaaacccaattgtttcggtcaagcacataagactagtcttc**tgactctgatgagtca**tttttcttccctacagattctcctaaaaatccctcctatgaatcctctgctaaggatgctaagtgaaccctatcacatgtggctgcaggcctgacacagtgcctaagccttaaccttcctct

Uppercase: TRAV31

Lowercase: Flanking sequence[1000bp]

Red & Bold & Underline: Stem-loop [19]

Blue: Heptamer[26]

Green: Nonamer [9]

id-IGHVII-1-1[V_gene_segment]

ttacac**ttaagaaatttcttaa**tagttttttcctccttaaggtgaactacagtcagataacatacttatcaattgtctctagctcttgtcagaaaaacatatagatgtgtgtgtgcgtgtgtcttggcctttccaatgatgaattaagatgtgcattgagaaggcattcactttatttgacgttaaggaagtaccaagaagacgctctccacagaccctgggaaagccagcagctgcaccccgaggctgtgccaggcagggaacaaggaggcagcaccacctgctg**ggcagggaaaatgtcctcccagtccctgcc**gcttctctgcagaggcacaaagagctgccccttctcctgggccttctcctgggctgatgagattgctccccgatatgccaaatcagggttgtgcatctgaggctctgtctagactctcagctccttcctactcctgcaaagtgaagaaaacaatgcca**aggggtcctggaggcgtctctacccct**g**gagagttttgactctc**ttcaatagtctccactaccctgccctcactccatgtcctccgtttctccctaaagcggtgcccagtctgattgcactgtggcagggataacgaggggccaggacatcaggggagagaagtttctacctgagtcacagcagcggctgccctgcagactcctgaagacacaagaca**catttccatcccagagacccagcgaaatg**caacctcaaactagagacagccagttattttttcttgttctgtcctggagaggccactgagaaagtcgagccccttgttgaggaaaacatgagatctctgtgtgtcgtcctctgcctgatggctgtacctccatgtgagtgtctcagagatttcagaacgggggctgtgggctgtggtgtccgcttgtgactcatctctttgcttcttgtccctgagtgtc**ctgcatcagatgcag**ctactggagtcatgccc**agggctggtgaggtcctcacagacctCTGGGCCTGGACCCAGCAGCCCT**CTGGGAAGGCGCTGGGGCACCTCAGCTCCAGGGGCAGCACACACTTCA**GCCCAGCCTTTCTGGGC**CAACTCTCCATCTGTAGAGACACATCCAAGGCCCAGTTAT**CCCTGCAGCTGAGCTCCGTGATGGCCAAGGGCAGGG**CCGCACATTCCCGTGGGAGACAGAATGGGGACCTCAGCGTGAGCCCAGACACAAACCTCCCTGCAGGGAAGCACAAGACCACCAGGCGGCGCTCCAGACCACACAGCGGCCCCAGAAGCAGGTTTTAGGGGGCGGGGCAGACGTGTCCGCGTTGAGTCAGGTCATTGGTTTTACTTTCCCTGAGCAAACGGCCTCTGCCAAGGACTCACTGCACCTCTCACCTTCACAGTTGTTTTTTTTTTTTTTTTTTAATCACCCTGTAGAGTTTTGCTAGCTAATTTAGATATTGAggagtgcttcatacttccttgggcctctgcttgcagaaacatagcaattgtaaggaggcacgtgggaaag**ccccggctcggtgacccgggg**gatgctgctgtggccctggcaagagggcgtcgggccgcagtaacaaaggtgcagacggctctcagcctgcgcccgcggagtacaacacataagggctgtaacctaacgaaaaaagaatcgcagtgcaactgtcctgcatttgagtttgtgatcagttttgccctttgtctttaacaggttctaacataaaattttgaatgttggttcaagccc**tgtgggtaaaatgcacttacccaca**ttccttaaacaaatagaacactgaggtggaaatgttt**tgaaaaagtagttttca**gacatttggaaacaagcatcacaggatcataacccctgagaaaagaaaaacaaatgaacgaatcctgctattgcctgaaagcagctgccaggacacacggaaaggcttagtgagctgagcggacagagagcagagttcaag**gcagcagcagcccgaggggaggagcaccggggagcaggctgctgtgc**agccaggatgggccggggtggggcggggggagaacagctggagacttgccgcagggagggggatccctcaggtttggggctgagaactgacttatgcctgacttatgcctgcatgaaaagaaactactcgatatcagggggaaatcaccagaaacctgtggacccaaaactacacagagcct**acacaaggaaagcattgtttgtgt**tctcccagccagggtggaaagaccttgagatatgtaaagcttcaagcaatcttccgaagtaatctcgtgagtagtggtgccacattaattcaggactaaaggctgctctgaactgaacctaagaaatgcttcaagtgtagcctggagcccgggtgcagtggctcacacctgtaatcccagcactgtgggaggccgaggcaggcggatcacttgaggtcaggac

Uppercase: IGHVII-1-1

Lowercase: Flanking sequence[1000bp]

Red & Bold & Underline: Stem-loop [14]

Blue: Heptamer[39]

Green: Nonamer [5]

id-IGKV1-6[V_gene_segment]

cgttcatcatggtacagttaacccttgaataacttgggtttgaaatttgcagatacacttatattttgttctgtgtgtgtcacctgtaagcaaatgcttaataagtagtaaatatattttctctattttatggtttccttaataacatttttttctttagc**atactttattggaataatagagtatataatatatataacattaaaaaacatgtta**atcaactatttatttt**attgctaaggcttccaatcaaaagcaat**ctattagtacttaagttagggatgcaaaaattctacacaggtgcagcccatttagtctctgggtattaaaaagacctgtagctcttgataatggtggggctgttagtgccacatgcattgaagaagcacggcatcatctttgtgtatctgtagtaa**attgtttgatcttatactggtaagaacaat**ggcataacaccattacctaatacttacaaatatatatagcatcatgtcaataaattttatttttaattttttagaaaggaacaatgttaaactcacagaaatgtttcaagtatagcacaattacccccttccctaaatggaatcttataacagtcttttgaagacctgagaatcgtaccatctaatatgtactgtgtttcctacaaacaagaatattctccaaaataatccccatacaccaatgaaatatattaccccattgactctcgaggaatattgtcaaaaaatatctttaaaa**tgtttctcataacaaaatagtccccagtagaaaca**cattctctgcagacaaatttgtgctaccctggtcttacctgggacacctggggacactgagccggtgctgagttactgagatgagccagccctgcagctgtgcccagcctgccccttcccctgctcatttgcatgttcccagagcacagtctcctgacctgaagacttattaacaggctgatcacaccctgtgcaggagtcagacccactcaggacacagcATGGACATGAGGGTCCCCGCTCAGCT**CCTGGGGCTCCTGCTGCTCTGGCTCCCAGG**TAAGGATGGAGAACACTGGCAGTTTACTCAGCCCAGGGTGCTCAGCACAGCCTGGCTATTCAGGGAAATTCTCTTACTACATGATTAATTGTGTGGACCATTTGTTTTTGTGTTTCCAATCTCAGGTGCCAGATGTGCCATCCAGATGACCCA**GTCTCCATCCTCCCTGTCTGCATCTGTTGGAGAC**AGAGTCACCATC**ACTTGCCGGGCAAGT**CAGGGCATTAGAAATGATTTAGGCTGGTATCAGCAGAAACCAGGGAAAGCCCCTAAGCTCCTGATCTATGCTGCATCCAGTTTACAAAGTGGGGTCCCATCAAGGTTCAGCGGCAGTGGATCTGGCACAGATTTCACTCTCACCATCAGCAGCCTGCAGCCTGAAGATTTTGCAACTTATTACTGTCTACAAGATTACAATTACCCTcccacagtgttacacacccgaacagaaacccccagggaagcagatgtgtgagactgggctgccccagctgctcctcctgatgcctccat**tggctgagattgttcctcagatgcagcca**cactctgatggtgttggtagagggggacacgaagtcacctctgcaccctaattcttttctctttctcagccccaactgcacagttctagcaatgcctctcttgatttaataaagacagagatcatgacacctgaagagtctagttta**tggcttcagttggaattcgtataacagagaagaagccactatagatatttgaagcagatatagt**cttaatacagagaattagagtaaactactgaagtctaaata**aaatgtagagatgaatctctaaattt**aatgttttatttgcaaagaaatatttgccaaatggggcatacaggaaaactcagtggtcttcaatatgttggaagaacaaagagaaatttag**cattttatgaaaaagggaaaatg**ttacctatggctcttttgagaaagctcattggcactaggaagggttgggagctggcaagctcagactggtaagcagtggtggacaaagtgaatcctagaattatatcaagttatctcagaa**gttttggataaatttgatttcaggttacaataaaccaaaac**ggtgaagcttgcagagaattttgttactgaaatgccagggattcggtgtagatcctgcagctcgtcacacagaaagccaatcactaagacaagtattttcaaggaacaggctttaatcaggtgctgcagctgaggagatgggatgccattctcaaatgtatctctctgaccaactgaaattaggggtttatataacaaggaagaaatgtggaaaaacaggagttagggagggttaagaaagataattgggtcaacaggaagcaggaggtcagttaggcaatcataatgggtgaagggtctgatgtctcactgtcccgattcagtgat

Uppercase: IGKV1-6

Lowercase: Flanking sequence[1000bp]

Red & Bold & Underline: Stem-loop [15]

Blue: Heptamer[27]

Green: Nonamer [6]

id-TRBV8-2-2[V_gene_segment]

attatcatcctaatcttttgctaatctttatgtttacggtaacattgattgttattttgggggacacg**tatttcatctttggtaatgaaatgaaata**actatgcacttcaattcctgtatgttctgcctacttaagagtaaatccaattttcttccattcctcccattctatccca**ctccctttatatgatagttttaagggag**aaaatattaattagactctaactgcagctatgtgataagagttatttagaatgagggtgggatattaagctttgagtaaagcaatagtcagggttagaactagtgttggaatcagggtttagggaagttgctcataaaacctgcaggatgacacttctagaatagtctctcactcactctgcagacacttcccagcatcccttgggc**cattgcaaaagcaatg**atgaaacttcacgtgttggccacaagatggcactgtggtccactgggatctaagggactctggggagagcctgggagagcagcctaggaagggaagggttaagaaaaattagggctggcatccaatattatgcagatgttgcagcagttttcagtcattgctaggctacctacagctatgcaagatgcaggaaatacctctaatctttaatgagatccacagttgaagaattttggcctggcagccttggtgtcaagcgcaggtgcaagcagaggagcaactgcctcaaaggaacgtgggaaagctaggggcgggctgtgccctaagctcagtatgtctctgctgcaccccatcttccctgcagacctcgccgggcaacagccttcatgagaggaggtggatcctgcctgagaccagtcagaaggccctggaccagctggactccgtaggcatggggggtgacaccacaggcccaccagactattccctagggagcagcctccaagccccagctcacagtcc**tgaggatcactgcacttagttgggccagcccctcctca**ctctgaccctaccATGAACCCCAAACTCTTCTGTGTGACCCTTTGTCTCCTGGGAGCAGATAAGTTCTGGAAACAGCTGCA**AAACCCTGTCATGGGTTT**GCAATGTCTGGGCTACAAGCCTTTTTTGTCCTTTCCTCTGCAGAGTCCCCTTATTTCTTAAGTCTTGTCTTCACTTTCTGTCTGCCTCT**CCAGCAGGCTCTATTGATGCTGG**GATCACCCAGATGCCAAGATAT**CACATTGTACAGAAGAAAGAGATGATCCTGGAATGTG**CTCAGGTTAGGAACAGTGTTCTGATATCGACAGGACCCAAGACGGGGGCTGAAGCTTATCCACTATTCAGGCAGTGGTCACAGCAGGACCAAAGTTGATGTCACAGAGGGGTACTGTGTTTCTTGAAACAAGCTTGAGCATTTCCCCAATCCTGGCATCCACCAGCACCAGCCAGACCTATCTGTACCACTGTGGCAGCACATCcgcagccctgcacagccag**ctgccctctgcacaaaaagggcag**tcacaggctggaggtgggcactccttatggaagcccgtgtctcaaccagaagaaaaagctgccctttctgaagctcttcccagacttcccagcaatgagatctcattctatgtgtgctcctgcagcataaaacttagagtgagctgggtctgtcactgtagtgcggatgtgtaaatgcaaacaacgtaatgttat**tttgctcaatttggaatattctggtaattgtgcaaa**ggaatggtagcttctaacaagaaattcatatttcac**tttcttttcaggagagaagaaa**aaagcaga**aagcagctgggaagctttagcctgcttgagtgatctgggattctcaag**tagcatacataacatcaccttggctttggttcctttagataccaagcatgtgatctgtggaggacaagaaaatccatttttttttgttataaagttataaacacacaagagaacaatccagttagggagaatgttaataaatacaacaaactctgccattagacctgaa**gataatcagctagtagaattatc**aaagagattgtaagactcttactgagtacctgtttgaggtcctctgaggatagatgaaggaaggaacttaattcttaccatcactgtggcttggattagaatccatgagttccttaacttgtttttcttgtaaa**atattataaaataatat**agaacacacattttattttcctgtagcctgtagtcct**ctttagcacctaaagggagaagcgcatatcccggtgcaaaag**ggagctggttgcttgcttggcaaaaaaatgaattcaccaattggccaatctgctgaaagtcaaaatgaaaggctaatgtatgagactgatgacagttcatggttcttggtataacttcaacagcgagtgaacatttttttctttgtctatagctttctttcagccatcgatcctcagttgtgtcttagcc

Uppercase: TRBV8-2-2

Lowercase: Flanking sequence[1000bp]

Red & Bold & Underline: Stem-loop [15]

Blue: Heptamer[24]

Green: Nonamer [0]

id-IGLV1-50[V_gene_segment]

gaatgttcttggtcttctcaatgaataacg**tctgaaaggttcaga**gtatatgtttcaaaaaggtaagtgcataactttcaaaaaaatgtagtattaacatgtcaacaattttaattggtatatatatttctgaaaaaaaaagcagtatgatggtaaaagaaatgtaaaaaagcctttgtaaaacccagtgcttacagtccaccaggatagcacctgtga**agcaaatttcctttgct**gaagatacaccgccttttagaataatagcaagatagtttgaatttgtcttcttctctcacagcaatctaaaaattaacttgatcccatttatatgacatatctagaaatattcctgtacatgtggacattcccatcgcacacctcccatccaggctgtcctaaagtggcatctcctcttcttctgctcatgttctccccctctcctcacctctcctgctgacacgtggctcctccctgccttcctctccagtagccccagcagatcaggcccaaccagtgaacaagctcctggacttcttggatttctgaggacttgccctgccagggtgtctttctcaatcacaatctctccttcccgggagtccagttctcctagatcctgttatttatctaatgttatacatctgtatctacttggtggacttccctacatcttcacccgaaccccaggaatcttc**aggtcatgcccatatgacct**tagttctcctggagtcccttcttccttcttcccctgtgatcccaccaagtggagatctcttctcagaccctggagtgcagcagggacttctctggtctctgccccagggattaagggggtggaattgaggactcctgtgcccttcctatgcctgtgttttcatcagggtcagctgcacaaggtagggattcccactctcagagtatgaagaggggcagggagagatttggggaagctctgcttcagctgtgggcacaggaggcagcactcaggacaatctccagcATGGCCTGGTCTTCTCTCCTCCTCACTCTCCTCGCTCACTGCACAGGTGACTGGATGCAGATCGAGGGGAGGGTCCCTGGGAAGCCTATG**GATTCTTGCTTTCTCCTCTTGTCTCTAGAAGCAGAATC**ATGATGCCTGTGTCTCTCCCACTTCCAGGGTCCTGGGCCCAGTCTGTGCTGACGCAGCCGCCCTCAGTGTCTGGGGC**CCCAGGGCAGAGGGTCACCATCTCCTGCACTGGG**AGCAGCTC**CAACATTGGGGCGGGTTATGTTG**TACATTGGTACCAGCAGCTTCCAGGAACAGCCCCCAAACTCCTCATCTATGGTAACAGCAATCGGCCCTCAGGGGTCCCTGACCAATTCTCTGGCTCCAAGTCTGGCA**CCTCAGCCTCCCTGGCCATCACTGGACTCCAGTCTGAGGATGAGGCTGATTATTACTGCAAAGCATGGGATAACAGCCT**GAATGCTCAcacagtgctccaggccc**agggggaggtgagagaagaaccccct**t**cctcctctaccaggagg**gtgagcgctcagctgctactgctcaggcgtggcctgtggcttctgctgctgcagcttcccccatggtccaggggcatccagggccctgc**ctcagagtagaggcttctctttttttctgttctgag**aatcatgcacagcagctccttcctaggaacagggcctccagggaccagaacatccatctctttgtaccttgggacacagggtctctgcagtgaagtcctgggctgaggtg**tcaggtccaactgtcatcacaaacctga**tgcagaacctgtgtctccatcaacctcatttcctgatttctggaagttccgactgttggctttctgtccctgctcctcagggtgaatcgcttggcttctttcctccccagcctactctttttaataaaaccctttccaaggaggaataatgtccatacgagaaaacacc**tatatttataatata**ccatgacacccattttggcatgtttctggatccttgaaagcatcaccaccaccgagctatgcaagctgtgaccaaccccagcagctcctcatgaccctttgtcatctccctttctaccccaatgccctcccgggaagccacagatcctcctctgtcattagtgtttggtttccattttctggaaggtatcagtgttcacaaacattctcttgccatttgttgaggcttctttcactccgcacactttt**tgagattctccctgttgttgctgaatgacatctca**tcccataaagggctcatcagtacctttgagggcccctagccattttgcctttcatgcccattcctgcatttgtatatacac**atatagaatccagggtagcctatat**tcc**ttttaatctgaattttaaaa**aatcaatacattttctggggcccctaacagcgtcgagtcctgtaggcatttgatcttctctctccagtgatgctcagccc

Uppercase: IGLV1-50

Lowercase: Flanking sequence[1000bp]

Red & Bold & Underline: Stem-loop [16]

Blue: Heptamer[26]

Green: Nonamer [6]

id-IGHVIII-44[V_gene_segment]

ataaacgcactaaattcacgagtacatataagtaaaataagccaaaaaaaaaaaactacatcctgtatgatgacacttttatacattctacagaatagaaactgaagtgacatgaaggtcgggagttgaatggagaaaaggtgagagaaaagagggaggaaggcgttgcaagagaataag**agaaaatgttgagggtaattgatttgttttct**atcttaataatggtgatggttacatcaataatttccaaattgtacattttgtgtgttaatt**agaattttatattattcaagctattacaaataaatacctgtataaatttct**tagggacggtgtaagggagatggataaatcacataaaaagtcagatatttctgaatgtacaatgaataacccttagctctatctctattttttggtaattctaaaatgcagcagttttttttaatgtttttattacaagaagggtttcttcaaaccacaccaggtatgcgtaactctgggataaagttggctaa**gggagctgtgggatccggtggagaggagcacaggtccc**ga**tcctggagcccattcttgtcacatgtgagctccagga**cacatctcatagctccctcatgcttctggg**tttacattcacatctgtaaa**tggaggaccaatgggcatctgcctctaacaatgtattttcgtgtgttaagataatgtatgctaaatgtttaccacagcacaatcttttactaagaaattacattttttccaaatattatcattgtcttcaagccccaccacccactagaaaacttcatctgctctgtgccttgccctgtcctcagacatcctgcccaagagacttctatatagtaggagacatgcaaatagttccctccctctgctgatgaaaaccagcccagccttgaccctgcagct**ctgggagaggagctcaagtgccaggattcccag**gtgttttcact**tggtgatcagaacttaacacagaggactcaccA**TGTTGTTTGGGCTGAGCTGGGCTTTCCTTGTTACTATTTTAAGAGGTGATTCATGAAGAACTACAGATATTGTTTGTGAGTGGATATTAGAGAAACAGTGGATATGTGTGGCAGTTGCTGACCAGGATTTCTCTGTGTTTGCAGGTGTGCAGTATGAGGTGCAGCTGGTAGAGTCTtttttttttttttttttcactttttagcgaacatccatgggttacaaaataatgggttggcttttcttccaacactttacagacaccatcaattttcctcttgcttataaggttttaaccagaagaatgctgtcatcatctttcctgttcttttagaaggaatgccccctcaactcatctccacttgtctgcatgtatttctatttgtctttggttttcagcagttttaataagattcacctaaatgtgtgtgtgtgtgtcgaggggtgttatgctattgttctgtgttctctgagatgcatggattcaccgtttactctgtctccatttttgtgaaaacaattagaaaaaaagtcagtgtgagcctagaaacaagcctccctgaagtgggcactggaccacctgggggc**gctcaggacccactgagc**acaagagccagccccagggcaggtgcagaggggtgttaaggtctggtttcctgtcagccctgtggcttcctctccataaaacagtttcctttgtggcacatctctggattccttatcctttttttcctgtgaagtctgaagaagaaacatttgtcgtaacaa**gagaaaaactttctc**atatgcaccaaaggcagagtcacctacagtcatttactcctgtttctgaatgtcaataaggtgtcaatgcttctgaagttaatcagctaaatctataaaaggtgcggtgtttaactcagcatggcagcccagctcaacagaactccaagggccagtgagcaagcaggcaggataaagtgcatgctgggcattggggcagagggagttagcatccagtgcaagagaagaaagcccccgtggtggtcattgtcaggactccaagcccacagttccaattgtaggtgatactgggcaaaggaagagagaccccaccaatggtt**agtgtggatttcgagtctgatggttccacact**cacactccaggtgaatatgaaaacatttattaactctattttt

Uppercase: IGHVIII-44

Lowercase: Flanking sequence[1000bp]

Red & Bold & Underline: Stem-loop [10]

Blue: Heptamer[22]

Green: Nonamer [3]

id-IGHV3-30-2-2[V_gene_segment]

aataactgtgctgagtgaaagtagcta**acaaattaacagtgcacttcataaacttctatttgt**ataaactgcagaaggtacagctattctaaagcaacagagaagattacatattggtgggaactgggttttgaaagtaatggggtggtgagatgaaattacagagaagtgacacaaagatttaggtgtgaattgtacacaaattaattgaagctctgattaaagggttgcacacatgttaccattttccaaattatgcagtatatatttg**aattaattattaatt**gtatttaaataaagcagtaacaaagaaacaaatgaatacatttattgaggtggagccaaaactaaatgggaatgaacactcgaaacatctctgactcttga**aaatacacaagcgtgaacactggttctctctatgtattt**caggtcaatagcagaatacactaaaagaaaacagagatgtcctggcaggggtggaatcctgcagacctcactaggcatgtctcacactgccctggagttgtctcaggggagcagtctcctctagtggtcagaggcacaggccgagataatggggctaactctgtccagctgtgtgaccttgaatgcattgtataaacactctgttctctgtgtgatttatcttccttaaaatgtgaca**ttgacacttgcattaaatgtattctacaaatatgtcaa**aaagatgatgactgctaa**atgattatcaaggcacaatcat**ataatataatgatattttcctgagtgataagatgactaccaatctcccc**cagggcactttgtctgctctgagccctg**cacctcctcaggattcccatcccagagcttgctatacagtaggagacatgcaaataggtttctccctctgctgatgaaaaccagcccagtcctgaccgcacagctctgggagagaagcgccag**ccctgggattcccaggg**gtttctatttggtgatcaggactaaagacagaggacccaccATGGAGCTTGGGCTGAGCTGGGTTTTCACTGTTGCTGTTTTAAAAGGTGAACTAGAGAGATTGAGCGTGAGTGGATACCCTTGAGAGAAATGGTGGATTATGTCTGGGAGTTTCTGACCAGGATGTCTACGAGTTTGCAGGTGTCTAGTGAGAGGTACAGCTCGTGGAGTCCGGAGAGGACCCAAGACAACCTGGGGGATCCCTGAGACTCTCCTGTGCAGACTCTGGATTAA**CCTTCAGTAGCTACTGAAGG**AACTCGGTTTCCCAGG**CTCCAGGGAAGGGGCTGGAG**TGAGTAGTAGATATACAGTGTGATGGAAGTCAGATATGTTATGCATAA**TCTTTGAAGAGCAAATTCACCATCTCCAAAGA**AAATGCCAAGAACTCACTGTATTTGCTAATGAACAGTCTGAGAGCAGCGGGCACAGCTGTGTGTTACTGTATGTGAGGCAccaggtaagaagacatcagtgtgaacacagacacagtttcctgcaatgataagggaggaggctgggctaaaaggggcactcaggacccactgaaaacgggcagctctagggcaggtacagatggttatcatgggctgctttccttcggggtctgtggcttcctctgcatctaccagttcccctcggagcctctggacatttatgtttctgtggccacccctgacatctctggattagaaaagtttattataggaagaggaaacattttcatttgtcccaaagcagatgtaagtaatggaagcaaaaaatgcacaggaggccaggtggggctgtagatactgtcaccccagaatgtcaatctcaccactagtactggagaggcatgggagtttgatggagcttccctaacaactctgtcttccaagataagtccagcaagaccattggtgcctcccagagcacagttgtccatcagggatgtccaatgcttcccagcagcagccatgcctcagtgtctccactgtgcacagccactgt**ctgggaggagctcccag**gatgggtgtctttggcacacaccaggtggcgggtgttaagagtgcagtgcagcagctggctgcctggtctgttgggcctg**atgctggagagatgggaggtgcattcttggggccagcat**gctgtttgtgaatttttatataaaaactctgattttacttcattttc**gcagatgacatagataactaaaaacagaatctgc**aaagaaattgtaattttcaactttaccccaaattcattgtttcttaattctgtgcaagatccagacatattattgccttcctcatgataatttattctatgtaaaaccgaaatcattttttctcctactctttgtttctgttcaagtacagagatcttgagcaaagtaagttgggttctttccacacactaaccctcaccttccccagagaatgagcagagattgtcctcagtctgagtctaaggg

Uppercase: IGHV3-30-2-2

Lowercase: Flanking sequence[1000bp]

Red & Bold & Underline: Stem-loop [13]

Blue: Heptamer[35]

Green: Nonamer [3]

id-IGHVII-33-1[V_gene_segment]

aggaagaccattggtgcctcccagagcacagttgtccatcagggatgtccaatgcgtcccagcagcagccatgcctcagtgtctccactgtgcacagccactgt**ctgggaggagctcccag**gatgggtgtctttggcacacaccaggtggcgggtgttagagtgcagtgcagcagctggctgcctgttctattgggctccctga**tgctggagagatgggaggtgcattcgcagggccagca**ccctgtttgtgaatttttatataaaaaccctgattttacttcatt**tttgcagatgacatagataactaaaaacagaatctgcaaa**gaaattgtaattttcaactttaccccaaattcattgtttcttaattctgtgcaagatccagacatattattgccttcctcatgataatttattctatgtaaaaccgaaatcattttttctcctactctttgtttctgttcaagtacagagatcttgagcaaagtaagttgggttctttccacacactaaccctcaccttccccagagaatgagcagagattgtcctcagtctgagtctaagggaggagctgtt**tctgcacaactcagagcctgcaga**gacccccaggtgcagcttcagtgagtcagacatttctccatgtgggcgacctccagtgcctgtgactgctgctcaggcctaattgttggttaagtattacgacacccttta**ggtgatcacatctcaagcctattctgaaaatcacc**atgaacagagatagttcaatggccattctcctgacataagtttctctttat**tacttggttccaagta**tggagaaaaatgtgaccctatatttgtctgaatccaacatcagcatccacggcattacgctaggaga**ctcactaattgagcacaagtgag**ctcgttattctcatagaaatgtaagtatttggaaatttcagcgtgttctccagaacctgtggctgccaacaactgtatttctcggtgcactcttGGCCTGGTGAAGCCTTCACAGAACCTCTCCCTCACCTGTGCCATCTCTGCATTCTCCATCACAACCAGTGTTTCCTTCTGGAGCTGCATCCATCAGCGCCCCCATGGGAGGTACTGGAGTGGATCGGGTGCATAGGTCATGGAGGGAGCACAGATTACTCCCCTCTTCTCAAGAGTCCAGTCACCATCTCCAGATCCATGTCCAAAAAGTACTTCTTCTTACAGGTGAA**CTATGTGAGCAACAAACACATAG**CCATGTATTTTAGAGCAAAAGA**cacagtgaggaaaccacagtgtgaactcatacccaagcctccctgtg**ggggtgcataggacagccagggttactcaggacaccaggctccctcaggacaccaaggggcactcaagaccattgtagaggcatgcaggtagctgggggctctcaggaaccatgggggaaaatcaggacaccaaagggtgcttggtacagcagggggctcaggattattgtggggattcagaaagagcaggttcaaggctcaacctcagggcatgtgcagctggtgtgaaaaggggctggatgaggcgttttgtgtcaccatcatgtttcaccaccagacaccctccactacctcta**ttctaatgcatgtgtttgtatgattagaa**aatgatattgatataaatatataaccatagctaggtgtgtcaagttgtcctctccatcttatatcagccttgtctgtaaggactaattccc**cataattacttgagaacctcataaattgtggtcaattatg**taggattcctctctttttctgccttccttcctccctccttctctctctcacacagaaacttatatacagccaccccacaacacacataaatcta**taacttttattacctgatgtattgataagtaatctaaagtta**cataaaaatcagtagttcactgtcaatattgtaggagaagggaaggtctaggaaggaggaattacag**aacaagaggatattttgagggtaattgacttgtt**atctatgtggataatgataatgtctatgaccatatttgtaaaattgaacacttcgtgtggagactattatttttttatttaaccccatgccattattttgccgtattctag**catttctctaaaaatacagaaatg**ttcagcactcattcatgtgtatattcaggagtttctgacttttcatgtattttatttatctctgtctaattgctttgataccaaattttacctagtataattagtactacctttagcactgatt

Uppercase: IGHVII-33-1

Lowercase: Flanking sequence[1000bp]

Red & Bold & Underline: Stem-loop [14]

Blue: Heptamer[29]

Green: Nonamer [5]

id-TRAV19[V_gene_segment]

tgagc**atggaatgttcttccat**ttgtttgtatcctctttgatttcattgagcagtggtttgtagttctccttgaagaggtccttcacgtcccttgtaagttgggttcctaggtattttattctctttgaagctg**ttgtgaatgggagttcgctcacaa**tttggctctctgtctgttattggtgtataagaatgcttgtgatttttgcacactgattttgtatcctgagactttgctgaaattgcctatcagcttaaggagattttgggctgagacgatggggttttctagatgtacaatcatgtcatctgcaaacagggacaatttgacttgcgtaataagagctatctatgacaaacccacagccagtatcatactgaatgggcaaaaactagaagcattccctttgaaaactggcacaagacagggataccctctctcaccacccctattcaacatagtgttggaagttctggccacggcaatcaggcaggagaaggaaataaaccacgact**ggcaaagtagtaacaaagattgagactaaaagttttgcc**ttcctgctagaaagtgaaaactccgtatggttagtttgtaccttgattctatctggagaaaaagacatatgtaagaccaaaaggtgcataactgtatcatcccatgtattcactatagctcataataaattgtaagtctcacattttccaccaaagatgaattaactaaccgtgattcaattttacc**actgtatatttacagt**ccagtaagttatattgagacaaaattggtacttaggacagtctagcatcttccatctctacaggaagtatgtgacatgacacagtcaaactcatcatatatctgttaactatacttcctgtaaaaggcagaagcctcacacagcccagtaactttgctagtacctcttgagtgcaaggtggagaattaagatctggatttgagacggagcacggaacatttcactcaggggaagagctatgaacATGC**TGACTGCCAGCCTGTTGAGGGCAGTCA**TAGCCTCCATCTGTGTTGGTAAGCGTGACCACTAGC**ACAGGACTTAGGCTCCTGT**TAGTTCAGAAGGATGTGGAGAGAGGCAGATCTTTTTAGTCTAATGAGGCTTAGAAAGAAGGGGTAAAGGGCAATGAAATCAGAGCACTAGGTCTGAAGCCACTTAATCTATTTTATCCCCAGAGTTACAAATAAGGGATCAGTCTATTTTTTCCTCTTTTTCACAGTATCCAGCATGGCTCAGAAGGTAACTCAAGCGCAGACTGAAATTTCTGTGGTGGAGAAGGAGGATGTGACCTTGGACTGTGTGTATGAAACCCGTGATACTACTTATTACTTATTCTGGTACAAGCAACCACCAAGTGGAGAATTGGTTTTCCTTATTCGTCGGAACTCTTTTGATGAGCAAAATGAAATAAGTGGTCGGTATTCTT**GGAACTTCCAGAAATCCACCAGTTCC**TTCAACTTCACCATCACAGCCTCACAAGTCGTGG**ACTCAGCAGTATACTTCTGTGCTCTGAGT**GAGG**Ccacagtgagatgggtgcctgtgggagccctacaaaaacctcaacaagaggcagggctcc**tggggagagact**ctgtcacagacaggaagaagcaaggagggtctgtgtcag**cacaggtggtttggcaaggaaacctgagttcaattcatgtacaatgtctagatataacatttat**tagaagcatgtgtatcttctattctttagaataga**ttaagacagaatgtatagcacatatttttatatcacatatagaattatgcaacaaaaagtgcaataacaaaatatattaatcttgtgttggttttattgagggtgggaaactatccaattatcttcagacca**aatggatgattccatt**cataagctgttaagtttttagaatcacttatacatttatgcattaattactatgtagaatgctttaatccaagagtaatagaaatgcctttcatcgatttttcttaagaatggtatcagtactcaatgaggtaaaaaaattaaatttaaacttctttccataatcatgccagcaacctcttgtatatcagaatagatgcgaaatgtttctcaaatttcctttctctggctcctctgtttttgtgacatgattatgatattttaaaagta**tctaaatattttgctgtcatttaga**ggctgcctactaagagagatgtctaaggtggaaatttgtcctaacggattgaatatcttgctttcaccctttcccctttgag**ttctggccagtttgaagctgagaagtggttagggccagaa**ttacagctgccatggcaaccagaacccacatggtgtgtccagtgtctacttcctgtctaaacacatatttc**tccttggatgttacaacaggtgctcaggaacaagga**tttttatttcttttgctccacacaagtttttagaaac**ctgagtttttaacattaaaagaaaacagacatcacattaaaaaccag**atttctagcttacctataaacatgggaagagctggg

Uppercase: TRAV19

Lowercase: Flanking sequence[1000bp]

Red & Bold & Underline: Stem-loop [18]

Blue: Heptamer[29]

Green: Nonamer [6]

id-LOC112268314[V_gene_segment]

gtcccaggtcctctctggctgttaccttcctccctccctattgtaaggacccttgtgattatgatggtctcacccagataat**tcaggatactctcctga**cccccaaattctcaaccatgtctgccaagttatttttgacatattcataagtaatgatcatagattccagatattaggacaatgatgtctttagtgggtgtattatttattccacaaacaactctcatcatccacacaatggtcttcccctaaggtagaataaaaatatcacaaggcagatttatgaggcgatcga**cctagaaaaaacctgagactctagg**actgtctgatgtgtggatgtcaaatcctgggagattctgagtctctgctctatgtggactctatgttgtgtagccatttgtggaaggcttctg**tgattttgtgacctagagaaaatgaatctctgctaaaatca**aatctaagaaagattggcaa**agggaatttaaagatttcctaaatttttggaatttccct**aggcattaaaacaggagaagtggcaataattcaaaccaatgatgccctccaagaatgagaatttttccaatgcattaggttgggtcccctcagtgagaaggatgccaaagattcgcatgcaggcagtatatttacaaagtgcgggaaacaagcaagtgagcaagggaggggaggagggaaagggaaagtgaaaggtgt**ctcagaaggagccacctctgag**gatgacgagagctcaagcctacatagaaacacaggaaaaatgcctctgttattccacctgagaggtg**agggagctgggggatgtatacacctccct**tgtcatcactgattgacagccgtcctaggggttgctaattccaggcc**atgaggtctgcctcat**ttgcagcctgagctgcttccccaggttcagatagagcagtgaaggggagaaagggccatagagagtcggctgaagtataatgactagaatccccaaggcgtagtaacaATGACTGCAAAAATTATGCACAAAGAAGAAGCGCATTTGA**ATCCAGAGATGTATCTCTCTGAATCTGGAT**ATATGGATCCTGGCAGCCTGTTCAGTAGCCATTTCCCAGAAATCCAGTCCTCTGGAAAAGCAGCAGGAGGTTTGTGCACAGGCTGC**ACTACCTTGGTCTGGCCACTGGTAGT**CGTGCA**TGAGAACTACTCCCTGGAGTGTTTCTCA**GTCCACTGACACTGACGTAAT**TGGCTCCACTTCCCCTGCTGTTGAGCCA**GGCTGACACACCCTGGACAAAGACATCTGTGTGAAGTATTGAGGTTCAAATCAGTGCTTAAGATATGTTTGGATGCAAAATACTTTTTCATCTACATGGGCAGTGTCTTGGCAGAAGATGGAGATTCTCTCTAAATGGATGTGAGACAGGGTGGCTGGCATCTGGGTCAGGATGTTGCCCTGG**TGCATGGCAAGGACATGCA**TT**GGGCAGCAACTGCCC**TCGCTAAGGAGAGAGGTTCACTGAC**CTGGCTTTTCCCCCCTCACCTGCTCTCCAGAAAGCCAG**ACTCTAGGGCAAATGCTCCTGAGACCCCAAGAACAGGCTGGTGGGGAGCGCAGCTCAGAGCATTACTCAGGGGATGTGGCCTTTGTCATCCTACTTTGAAACAATTGACTATTT**GAGCCTAGATTGATAGAGGGCTTCAAGTTGATTTTAATCCAGGCTC**CTATAGTCAGCGAGTGAAACAGAGATTTTAGTTGAAATAATGAGACCTGGTATTACTAGTCAGC**TCTCCATGCTGGAGA**ACCATAAGAAATTATACCAAAGGCAGGAAAGGGGATAGAATATGGGGATCATCACACCAAGAATAAGGTGCAGCCCATTTAGCCCC**TGGGTCTTAAAGAGACCCATAGCTCTGGATAATGGCAGATCTATG**CGTGACACAGTTATCATCTTTGTGCATCTTCAGAGA**ATTGTTTTTCCTTTTACTCCTAGGAACAAT**GTCTTAAGTTTGTTAG**TAAATTCTATTGAATTTA**TTAAAGATTCTTCTGATAAA**TTCTTTTTATATTCATTTCAAAAAAGAA**GAAATTTCACACTGACAGAGACATTGTTATTATAGCACTAAATACTTTTACACTCTTCAAATTCCTTTGAGGCTAACTGAAATTTCTGACAGCCCCACATTCTACAACTTTATTGTAAATTTTCTGCCAAAAGTGATGCTTCCCTATACACTCCTAATACAAGTA**TAAATATATTATTTA**ATCTAGGCTTAGGTTGATTTAAAATTTTGAAAATTCACTCCAAAAATATGTTCTGTAACCGTATGGCCACCAATGAGAAATGTATTCTTTCAAGGTAAATCTGTGCTGCCCTGGTCTGACCTGGGACTCTGGGGATACTGCGCCCCTGTGCTGAGTTACTGAAATGAGCCAGCCCCGCAGCTGTGCTCAGCCTGCCCCATCCCCTGCTGATTTGCCTGTTCCTAAAGCACAACCCCCTGCCCTGAAGACTTTTTATAGGCTGGTCACACCCGGTGCAGGAGTCAGTCCCAGTCAGGACACAGCACAGATGTGAGGGCCCCCACTCAGCTCCTGGGGCTCCTGGTGCTCTGGCTGCCAGGTAAGGAAGGAGAACACTAGGATTATACTTGGTCAGTGTGCTCAGTACTGTCTGGAACTTCAGGGAAGTCCTCTGATAACATGATTAATTGCAAGAATATTTGTTTTTATGTTTCCAACTTCAGGTGCCAGATGTGACATCCAGATGACCCAGTCTCCATCCTCC**CTGTCTGCATCTGTAGGAGACAG**AGTCACCATCACTTGCCGGGCGAGTCAGGGCATTAGCAATAATTTAAATTGGTATCAGCAGAAACCAGGGAAAACTCCTAAGCTCCTGATCTATGCTGCATCCAGTCTGCAAAGTGGGATTCCCTCTCGGTTCAGTGACAGTGGATCTGGGACAGATTACACTCTCACCATCAGCAGCCTGCAGCCTGAAGATTTTGCAACTTAT**TACTGTCAACAGAGTGACAGTA**CCCCTCCCACAGTGTTACAAGTCATAACATAAaccccaaggaagcagatgcatgaggctgggctgccccaatgctccttctggtgcctctgtctgc**tgagaaaagttctca**aactcagtcaggtttggaaagtcatagggagattttcct**agaggaggccagggaggttcctct**gaaccctaagcctctttcatcctcatccccagcagaaaagacgtgacaatgactgtcctgactgaataaagaagagagataagtccagctgaggagtctgtgttatgggataatcggaatttgtacagcaaaagagaagctattctcagtatttcaaggagaaa**ttattcaagttgaataa**attagagtctaaaccacagtctttctgaagcctatggagtgttattcatgaagcaggtactagacacaggggattctcaggtgttacttcagaag**ccagggttcacctgcccctgg**tggtatgtgctgaacaccgtgtgatgatcctcagtcctg**tctgggaagcccaga**gctgggg**gtgctgatgctctcagctgcctgcagcac**atctccaggtgattctccagtcc**acacctaacagcatgtgctttacttcaggtgt**cagtgtacatgaatccaccactctgacttcccaatctcatgacagtaattagttgtaacttattgtgacctcgtggagcaactctaaagaaaccatagagagaaaaggagttttggaaaatgtgctcccagaagtgatagtagtgatggggaattgacagctgacggggaagtaaggtgactctttccacaaggctcaac**attttgccagttacgaattgttgcaaaat**acatttgaatgtgctttcaagtattaacagtttggggtcatagctgaaaaactttattaagtcacagataaaatgggaaaatcaggaa**attgtatgaaatatacaat**aacactatgtgtgatggctcagg**tctgtaatcctgtgatagttaatacaga**ttgtcaacttgattacattgaaggatgta

Uppercase: LOC112268314

Lowercase: Flanking sequence[1000bp]

Red & Bold & Underline: Stem-loop [34]

Blue: Heptamer[38]

Green: Nonamer [6]

id-TRBV23-1-2[V_gene_segment]

tgtttttttaagtgatcattctgttggtattttatcaagttattatctatatattcaaagtccattttgtaatgtggagaaaacaacctcactgaacaattaaattagtaaggagaaatctatggaaagatatacattaagctgttaacattacttacctgaaatgaggatgagatggaggaagagaaaagtaagcgaaaaaaaaaatcatcaaaaaatgcacaatttaaaataaatagttacagagagaaaaagcaaaaaagggaggttatgtaaaagaagcactagtaaatgaacaagct**aaaatacaccagaaagagctagaaggtgaagtatttt**tcaaagcagtcactag**tttcagaatcataaagatagcaattacaacagagatctgaaa**tatataaaagataattagataactaatagccagaaggaatgtccagtcttgagttagaaactgatgctcaaattaatg**gcaaaggacgtttttgaaacaactgtgaaactttgc**ataaaaactaaatagtagataagatgaaaatgcatcgatgagaaaggatgatcattaactgaaaaaaagcttatcaaacagaatttcaagaatattgtttactaaaaaaagtgtctatatatctacacatttatatatggggcagagaaggatct**agaaagactcactaaggtgttaacagtgataattgctttct**ggcagtggta**tcatatttttattttgttattttattttctaactttctaaaatatga**gtgtattgcttatattagc**aaaaatagacataacatattttt**aaatgaatacaaaataactgagttatatttcctgagttaggataaaagctttttttccctgatactatgaag**tgtttttgctcaggaacatgaagcaataaca**tgatgattataagtcagtggagggatcacgtcacaagaaggtgcagaaag**gggatgaaaaagcctcatccc**tttgcaacgtcaatgcgatcATGGGCACCAGGCTCCTCGGCTGTGCAGCCCTGTGTCTCCTGGCAGCAGATGCGCCCTGAGCACAAAAGAAAAATCCTTGTCCTGGGCTTGCCAACCAGCAGATCCAAGCTTTGTTTTGTTTTGTTTTGTTTTGTTTTGTTTTGTTTGTTAAGCTCTTTCCTGGACTCTTTCTACAGCATCTGTTTCTTTCTCTTACAGACTCTTTTCATGCCAAAGTCACACAGACTCCAGGACATTTGGTCAAAGGAAAAGGACAGAAAACAAAGATGGATTGTACCCCCGAAAAAGGACATACTTTTGTTTATTGGTATCAACAGAATCAGAATAAAGAGTTTATGCTTTTGATTTCCTTTCAGAATGAACAAGTTCTTCAAGAAACG**GAGATGCACAAGAAGCGATTCTCATCTC**AATGCCCCAAGAACGCACCCTGCAGCCTGGCAATCCTGTCCTCAGAACCGGGAGACACGGCACTGTATCTCTGCGCCAGCAGTCAATCcacagcactgaaatgtcagttcctcttagcacacaaacttgtcacagacccagctcaggaagcaggtgatgtattaggctggaagggagtaacagaaaataactggagccagcttaagccacagtgtaatttaacacaaggataaagggtgtttagcagaagtcaagaggtgatagaagttctgaaaaggcaccactagggctgggctctagaatggctatctctccagcgctgatttttattcaattttggctgctgaatgcttttctctccctgtac**accatctctttttctccaggcctcgtggtgagaatgagatggt**cactaaagattacaaattcagttgccatcatttccctacactgataaataccaggctggctctttctaagcctcaattcaaaatcccaagagaaaaaaacagatcagtcatgatccagccattaaccttctcacagagcactcagcaggaggcatgtat**gtatatatttaaaatcacataaatattac**ttttccactcagctcaagcaagggaagtgctccgtttttctgtatgtacatttggtcttgttgcttctacatgatagagagtactcatacccatttctt**cagaaatacatagccagattttctccagttttctg**cc**agaattgaaaattct**tgtacaaataccttcatagacttgtatgattgctttataggattatatacccagagcaagtcatgattgctgagacatggaattctatgcatattttatctatctaaataccaccacgggttcttcaggatcactgtatacatctgcaatcccacataaaagtgaatagggtgttttcctacatcctaatatctcctttaatccttatccagtgatctagattgttctcagtctagcaatgt**aaagagattccaatctctgtctctcttt**ctctccaccccccacccctgcaaccctctcacctactgccagaatatcatcagtgtttaccataaagat

Uppercase: TRBV23-1-2

Lowercase: Flanking sequence[1000bp]

Red & Bold & Underline: Stem-loop [14]

Blue: Heptamer[24]

Green: Nonamer [4]

id-TRBV7-3-2[V_gene_segment]

aatatagaatatagaatatattatatag**aatatataaatatatatatt**tccca**tttttcttatagataattgaagtttcagaaaaa**gaaagtagagggaataagtgattgtgatgagacattagc**tgagaattttccagaggtgtttcaggacctaaattctcaatttcaagaattcaaatgaatatgaaat**aagataaatgtttatattataatgagattgtggaacataagtgacaaatagatgattctaaaagctgtcaaaaagaaaagataaatctctcctagtggaatagactgaaaaaatcaatgaaagaaagaggatcagaaataaccatttagtcggaattataatccccattcattaaacactgagatggaggaaaatatatttacaag**tggaaaaattaaaaaaaaagactatgagttttcca**ataaggtatcttttctgaattatagtcaacaaaggtgggaagtggtcccagaagaaaggcatgaggtgcagcttgtgagggaacccatgtgttgggacagccccgttgggcacgtgtgactgggggatggaggaggctggggcatcaatggggatggcataggggaccctgacttgcagaaaagacaatgagctcatcccttggtgccttgtgttgggggcgctgttggcacatcctagagaacatgcccagcagacagaggagcgcctgtgtggtgaggagataaactcagaaatgcagcatgaggcctgtagctccagacagctctagagcacaaagtgaagaccgatgcattgatgttgtttaaaaggagctaataaatatctaaagcagtcatccaagtgtgttctaatataaatcctgtgttcctgaggttgtggggattgagagaggaagtgatgtcactgtgggtactgttctgtgtcaggacaaggacgtccctcctcctctgctcctgctcacagtgaccctgatctggtaaagctcccatcctgccctgactctgtcATGGGCACCAGGCTCCTCTGCTGGGCAGCCCTGTGCCTCCTGGGGGCAGGTGAGTCCT**CAGACACCAACCAGTCTCATTCTGTGTGTCTG**TATGTGTGTGTGTGTGTGT**GATGAGTACAACTGTTTTCCTCATC**CTGTTCTCAACTTGTGTCTCCGCAGATCACACAGGTGCTGGAGTCTCCCAGACCCCCAGTAACAAGGTCACAGAGAAGGGAAAATATGTAGAGCTCAGGTGTGATCCAATTTCAGGTCATA**CTGCCCTTTACTGGTACCGACAAAGCCTGGGGCAG**GGCCCAGAGTTTCTAATTTACTTCCAAGGCACGGGTGCGGCAGATGA**CTCAGGGCTGCCCAACGATCGGTTCTTTGCAGTCAGGCCTGAG**GGATCCGTCTCTACTCTGAAGATCCAGC**GCACAGAGCGGGGGGACTCAGCCGTGTATCTCTGTGC**CAGCAGCTTAACcacagcatgacacaatcgcctccttcctgctcataaacctcctcctctctctccttgcttccttatgatactattttgcaccaggggatcctcatctcacaccactccactgcctcttccagaatcattgcttgcatgtccgctgctcagatcagcagatgtgcgttgtatatgttaatcattttctgtggaaggctcgttgctggtcttcat**tatccatccttattttccacattggata**ctgc**tcccttcaagattacttgttgcctacaaggga**tcctactttttacat**tcaaattttactttttattaatacgtatttga**tgccatcttgaaggcaatctcattatcatcctaatcttttgctaatctttatgtttacggtaacattgattgttattttgggggacacg**tatttcatctttggtaatgaaatgaaata**actatgcacttcaattcctgtatgttctgcctacttaagagtaaatccaattttcttccattcctcccattctatccca**ctccctttatatgatagttttaagggag**aaaatattaattagactctaactgcagctatgtgataagagttatttagaatgagggtgggatattaagctttgagtaaagcaatagtcagggttagaactagtgttggaatcagggtttagggaagttgctcataaaacctgcaggatgacacttctagaatagtctctcactcactctgcagacacttcccagcatcccttgggc**cattgcaaaagcaatg**atgaaacttcacgtgttggccacaagatggcactgtggtccactgggatctaagggactctggggagagcctgggagagcagcctaggaagggaagggttaagaaaaattagggctggcatccaatattatgcagatgttgcagcagttttcagtcattgctaggctacctacagctatgcaagatgcaggaaatacctctaatctttaatgagatccacagttgaag

Uppercase: TRBV7-3-2

Lowercase: Flanking sequence[1000bp]

Red & Bold & Underline: Stem-loop [16]

Blue: Heptamer[28]

Green: Nonamer [4]

id-IGKV1-22-3[V_gene_segment]

ggacaagaacttcactaacaattgagtcactgaagacttacggccctgtgtgacgcaccacataactgtgagtttgcagtggttgcaggtcagggacagattttatgcttaagatcagtagggtggaggct**gaggatcttggctattacaactgccaccacactctacaatatcctc**ccacaatggttcagcaccaaacaaaagcctcctgcttggattgtcccagctgcccaaattagttccttcactgaggagtagacagggtatattctctaaatctatgtaacaggaagatgttggtgaactcaggggattagtatgaagctacacctcaggcatcacacataagatcacttcagcagtcgcagccttagcatgggcagaacctacagaagatgcaagtgccctctgagccaggagacaggaggaaggaggaagggaaaggtgacttagctcatctcaatcctctctcctttgcatacatttgtcaaccag**atgtattcagcctaccagtcacacaactgaggctgatacat**gacaacatagcactggtatattcttggtattgtttggcttagcagttactagtatatatttaat**gggagaatatttggtggtgttaacacattgcttatctccc**tttccccagttgtactttacacttgttcttggcacacattctcctccaggactggagcattcacagggttttatgttactgttcttatgggagtaaaaagaaaaacgattcacattcttgctactgagctaggctgggatgccctgggccaagctgaaaatgtgaaaaataagagtatgaatatttattaagttttatctggatctaagatacttatccatgaaccagtcctgcagctgtgcccagcctgctccattccctgctgatttgcatgttcccagagcacaaccccctgttctgaagacttcttaataggctggtcacaccctgtgcaggagtcagtctcagtcaggacacagcATGGACATGAGGGTCCCCACTCAGCTCCAGGGGCTCCTGCTGCTCTGGCTCCCAGGTACGGATGGAGAACACTAGGAATTTACTCAGCCAATGTGCTCAGTACAGCCTGGCCTTTCAGGGAAATCATCTTATAAATAGTTGTGTGGATTATTTGTTTTTATGTCCCAG**GAGTCAGATGTAATTTCCAGATGACTC**AGTCTCCATCCTCCCTGACTGCATCTGTAGGAGAGAGAGTCACCATCACTTGCTGGGCGAGTCAGGGCATTTGCAATTATTTAAGCTAGTATCAGTAGAAACTAGAGAATCCTCCTAAGCTCCTGATCTATGCTGCATCCAGTTTGCAATCTGGGGTCCCGTCACGGTTCAGTGGCAGTAGGTCTGGGACACATTTCACACATTCTCACCATCAGGAGCCTGCAACCTGAAGATGTTATAACTTAT**TGCTGTCTATAGACTTACAGCA**GCCATcctagagtgttacaggtcataaaataaaccc**ccagggaagcagaagtatgactcatggctgccccaggtgcttccactgg**tgcctccatctg**ctgagagtgtttctcag**gtgcagccaagatttaaaggtttttgtaggaatggccagaagtctcatctgcattctaactctttttcttcctgcttagccccagcagcacagacatgacactatctctcctgatttaataagggatagcatttacgatacctgaagaatctgtgttattgcatccatctgggtcatagattaaaagagaaaccactctacagattgccagaaggcattgttttaatacagggaattagagttgaatatacaaaactgggagtgtggtagttagggaagctgacactagaaacacgggagt**ctctggaggtctgccagaagccagag**ttcatcagccgctaaaggcatgggctatctaaccatatagtcttctttgtctaggaagtccgtatgcgaagatgctgatgctatcagttgttgcagcacctcaccaggtgattctccagtccttatctcagtgaacatgtttgcctaccggtgtcaaagaatattgaatcgccttcttcttaccttcaaatatgatgagaggtcttctctttgagtaactctacaagaaaccatagagggtttaatgtgtttcaggaaaggtgcttttagaaatcatggtgaatatgaggaattacagccaag**tgagataagtatttcccaaaatctca**gaattttccaggtatggggtggcttcag**aatacatttggatgttcttacatgtatt**attagaaagtttggtattattgcaagaaaattttattaagtcgtaaagtaaaagaaaaaaatgacaac**attgcttgaaatacatagcaat**cctttgacaaatgaaaaaaaaattgacaaaacaaacaagaacacctataggtgcacgtagcatactttttccttaatataagagcactt

Uppercase: IGKV1-22-3

Lowercase: Flanking sequence[1000bp]

Red & Bold & Underline: Stem-loop [11]

Blue: Heptamer[35]

Green: Nonamer [5]

id-IGKV1-32-3[V_gene_segment]

aacccagtcattaagaactgctt**tctcttaaaaactggaagaga**ttggtaatatagagatatacagtttatggaacagagtagattcaagagt**agatgtgagcactgggtctagataacatct**cagtttccggaagatattagcttaagaatgacgtaaggcacacatattaatccaaggagctcaggaaacatctagcataggaatgcatgatggcacacagaggtagagagtagctgacacttt**tcagaagtggagggagaagatttctga**tgttggcctagatgaaagaagtaggaaagaaaatgtagaaatacaggaagatgatcacatgtctgagataagagatttgccatgtcgtttacccagtgta**caaggattacccaatccttg**caaagcattacactactttcacatattttgtgttaacattttcctttttc**ttatttttaataaataa**atctgtccttctatccatccatctatgtatccatccatctttcttatacc**tcatcattgcatctctggctgatagtgctgatga**tttacaacatcaagggctctagaggatcagagcaaagaagaagtctctctggcctcaataatattcatcagtaccaaacgcgggacacttctaaaatttattgcagattc**tggagatgtgcttctcca**acaagccaggtcttagacgggaagctagcaagaa**aagtaggtcatgacctacttctgcagagcagaa**aagcaatctccttgcaggacagtagacagattctgttctgtggcccccagagagaa**acttgttctctgctaacaagt**gtatg**caggcctgatcaatcttaggacccctgggggacactaggcctg**tgcaatattatgcaggtaaagtcattcttgcatctg**ttgaaattttcattttcaa**aaaaacacagccaacttcctgaagacttgtacacaggctggtcataccacttgcaagagtcagtcccagtcaagacacagcATGGACATGAGGGTCCCTGCTCAGCTCCTGGGGCTCCTGCTGCTCTGGCTCTCAGGCAAGGAAGGAGAACACTAGGAATTTACTG**AGCCAGTGTGGTCAGTACTACCTGGCT**ATTCAGGAAAGTCCTCTCATAACATGGTTAATAGTGTTAATAGTTTTGTTTCCCATCTCAGGTGCCAGATATGACATGCAGATGACCCAGTCTCCATCCTCGCTGTC**TTCCTGTCTAGGTGAGAGTCACCATCACATGCCAGGAA**AG**TCAGGCAGGGCATTAGCCATGTTTTAGCCTGA**TACAAAG**AGAAGCCAGGGAAAGCTTCT**GAGCTCCTGATCTACGATGCATCCAATTTGCAAACCTGGGTCCCATTGCAGTTATGTGGCATTGGATCCAGGACAGATTTGATTCTCACCATTAGCATCCTCCAGTCT**GAAGTTGCTGCAACTTC**TTATTATTGGTCAACAGTATAAAAGTGACCCTctcacagtgttacaaacccaataagctccccaag**gaagcagatatgtgagggtgggctgccccagctgcttc**tcctgtttcctccatct**gctgagagtgtttctcagactcagc**cacactctgaaggtcactgagcagttttcatagaagggg**taagggaagtgtctctacaccctta**gtttcttttatcctcctcagccccagcagcagacatggcaatgcctctcctaatttcataga**agagtcattacctatgaggactct**gggttacagcactggtccaggttcatacaacaaaagagaagctattgtaggtaacccaaatagaaagtttttcctaatatgggaaatccatgtctaaattacaacttttcaaagaccagaggatataatggttaggaaaccagaaaggcaaaaggagtgctgggaaatctacttcaagaagcaatggtggccgcgcacagtggttcaggcctgtaatcccagcactttgggaggccgaggcaggcggatcacctgagatcaggagttcaagaccagcctggccaacatggcgaaaccccgtctctactaaaaatacaaaaattagccgggcatagtggtgggtgcctgtaatcctagctactcagaaggctgaggcaggagaatcatttgaaccaggcaggtggtctctgcagtttcgaagcccagcgccatct**gtggctgttatgcatgtctctcccagccac**cctgctgtccagagcccatatcaatccgtgggggtaggtctgtgaaaga**gcagctcagttaggacccagagggaaccatggaagctgc**agctcagctgctttgattcttgttactctggctcccaggtgcaggaaacaccggatggttctgcatgtcagtgaaacttt**ctcaaccttgttgag**tcctgttacctggcacacctgctgggaaggcacaatgattaaagctcaatgtagatcaatggtcctggatgcactgggaagacaatagg

Uppercase: IGKV1-32-3

Lowercase: Flanking sequence[1000bp]

Red & Bold & Underline: Stem-loop [24]

Blue: Heptamer[26]

Green: Nonamer [3]

id-IGLVIV-53[V_gene_segment]

atcctccctggctccacacatgtccagccttcccccttatcaacgggtcccatcagaaatgtacatgtcctgtaattgatgaaccacattgacaggtcattatcactacaagtcagtattttccttaaggaatttttctcgctga**tgcacatcctatggctatgggtttgtgca**a**atatataatgacatatat**ccaccattacagtatcatgcagtgtatttcccttccctacaaatcttctggctttactgattcatctctctctctcccctaacctttggaagtcgatgacttttctacttttttcctttgtttgactgtaatacacttgaaattatttccccctttttggactgtaatatacttggaatcacgtatcatgtcatttgcaggtcagctccttttacttactaataagcatttaagtttcttccatgtcttttcgtggtttgatagtgtgttgtccataaacatgtagaatatattaatataaagtactttggagctgatcacatcactggc**cattcaggtttttcaattactgtttgttgaatg**acagtgaccagtcagatgacaggaagtcttagacc**ccctgagaacacagaagatggactgaggatgtcaggg**tcaccgaggactgcactgggtgctctcatggag**ggctcagtaggggtgtcctctagttccagtgagcc**tggactgtgatgggcactcagagaagggtcctctctagcctagcagagcccctgctcacaggcctaccagctgtcccagcagctttccctcccaggtccttccgtggggcagacagagtctcactgcctgttagcccttcctacccacaagaccctcagcaaaagatccttattcagtgctcagggagaggtggggcagatagc**agctggggccacacctgcatgaccagct**cctcctcgctgaggcctacagcaggtaataaactcaatccatcccactggaccaaagggagctgtgtctaccATGGCCTGGACCCCTCTCGTGCTTGTCCTCACTCCCTGCACATGTAGGGAAAAGGCCTAGAGACCTGGGGCAGTTCCCAGCTTGTGTCAGCCCCTCTGGCTCAGACTTGCCAACAGCTTAACCCTGACACCTGCTCCAACAATGTAAATGTTTGCATTTCCAGGTTCCCTCTTCCAG**GCTGGGCTGACCCAGGTGTCCTCACTCCCAGC**ATCTTCATGCAGATCAAACACTCACCTGCACTCAGAGCATATCGGCTTTCATATCGGTGGCAAAAATACCTTCTTTTCCCTACAGAATCCAGGGAGCCCTCCAACTATGTTCTGAGATTATGCTCAGGCTAAAATGAGTACCAGGCC**TCTGGGGTCCCCAGA**CAATT**CTCTGGGTTGAGAGGCTCCTCCAGAG**T**CTCAAGTTATTTGGTCGTCTCTGGCCTTCACCTTGAG**GATGGAGCAGATCATCTCTCTCAGATGGGCTGACAGGGCATGGCTTAcactgtgctctaggccaatgggaaaatcccctctgcttgtgctgcctgggctcccactaggcccct**gctgtttgtgacaacagc**cagcactggtggtgacgcttcagccatgtatgccctacggtgagagtctgcagcccttccacgcaactatcact**tgcagaatgtctgca**tcattgctaatagcacattctgagcctcctcacgtttctgtgggcaacaatttccttcccaagacgtgcacatgatacaggagat**ttatttaccaaataa**gtttattaggaatccaattataattagaagatattcatcagtgatcgggggaaatacctggtataattaaaatagaatttgatgaaaatataaataaagaactaaattatgaaggtgtgggcagagagtctggacaccctaagggagatggaaaac**cattcatccttgccaggaatgaatg**tctctgggatgcttctctaacaggaggaacctggagtgagaccaggaacctggagaactagcagctgggtgactggaaaggaaaccatctggggtggggttgggtctgcaaggcaggaggtgtctgggcagagaaagcagggctggggcggagggtcctgcctcatcccattctcctgctcactccttcctagatccctgtcagggcagtaaccatgagcaaggcttgaggctacttcttat**aaactgtggccctgggcacattccttagctcacctgagcctcagttt**cttgatgagaaaaaatcgggacaatagcaaccaacctatctcagagggctgctgtgagaccctgagccactgtttctcaaactgcccctgagatca**cctacaacagaatggctgtagg**gtgaagtttgagaaccgcagtataggtgaggtggtgcgtggggcaggccctc**ccacactgtaaagtgcagtgtgg**tgcggtaaaagctgatattcccgcaggcatcctcagagtgggcccaattagactcttgagcact

Uppercase: IGLVIV-53

Lowercase: Flanking sequence[1000bp]

Red & Bold & Underline: Stem-loop [17]

Blue: Heptamer[20]

Green: Nonamer [1]

id-IGLV1-62[V_gene_segment]

agaagttga**ccacagcatctgaatagctattaccccaagtctgtgg**aaattagagaacaaaattttaccagaaatgccccccagag**cttctccccataacctcaccacccttagagaag**agagctcctgtt**ctcctcagtgctgcagggtccacagaggag**gaagagctctggccagcagggaggagccacagaagaaagga**cctggtgtctcccgtgaccagg**gtagtgagtcatgggaaccagtcactgctctggacttgacctgacctaa**gaggacttggtatagagtcaccacttcagtcctc**accaaggaacatccaaaaagccctggacagattcacccttaaggagaccagtaaagcatgaactgccctgagttaatgagaaggtcacaaaaagggtctgtctggcctggtgctgagcctgggggcagtgat**cagagcagggggaatgcacagctggctctg**gggacatctggtcatcatgtcacaggagagtgtgtctggatgtggacacctggggacatgaagggctcattaccaag**tgatctaggggagatca**gagctgcgatcagctaactgcctgcatggtgccctctgctcagtgatcacagtggtgagctcctgatgccccggaacacacagtttcacccttgcctgcatccctgatatccacgagaggggcctgacccaaggccagcgtgtcatcagaaggctggggggctctaatttgcatggatggaccc**tcattctcagagaatga**agaggggaagggagagatttggggaagctctgtctcagctgtgggcacagaaggaaggcaggactcaggacaatctccagcatggtctggtttcctctcctcctcactctccttgcacactgcacaggtgactagatataagg**ccatggctaaagccctaggaagcccatgg**gccctgctttctccttttgtctctcaacccagaatcaccatgactgtttctctcccacttccagGATCCTGGGCTCAGTCTGTGCTGACTCAGCCACCCTCAGTGTCTTGGGCCACAAGGCAGAGGCTCACTGTCTCCTGCACTGGAAGCAGCT**CCAACACTGGGACTGGCTATAACGTAAACTGTTGG**CAGTAGCTCCCAAGAACTGACCCCAAACTCCTCAGGCATGGTGATA**AGAATTGGGCCTCCTGGGTATCTGACCAATTCT**CTGGTTCCAAGTCTGGCAGCTTGGCCTCCCTGGGCACCACTGGGCTCTGGGCTGAGGACAAGACTGATTATCACTGCCAGTCCCGTGACATCTGCTGAGTGCTTGtacagtgctccaggcttgcaggggagtgagacaagaaccccctt**cctcctttcccaggagg**gtgagtgcccag**cagctactgcacaggcctggcctgtggcttctgcagttgctg**tttccccagtgggcctggg**agcatccagggccccacctgggagtagatgct**tctcctcccctccgtcctcagagtcacccacagcagt**ccctttctaggaaaaggg**cctccagaaaacagaatgtcctgt**accctgcaacttgggacacagggt**ctctgctctaaagtccaggg**gctggagtgacaaatccagc**agtgaaatc**tcagatctacatctgatggggaacctgcatctccatca**tccttttgctatttccaggagttttcaaagcagtgtcttcctctcactggtcctcagtgtgttttcctgttctttttgtttcagcctattttttttttttttttgagatggagtctcgctctgtcgcccaggctggagc**gcagtggcgcgatcttggctcactgc**aaactccgcctcccgggttcacgccattctcctgcctcagcctcccgagtagctgggactacaggcgcccgccaccatgcccggctaattttttgtatttttagtagggatggggtttcaccattcacaggattgtctcgatctcctgaccttaagtgatccgcccacctcggcctcccaaagtgctgggattacaggcgtgagccaccatgccgggtctgtttcagcctattttttatgtgtctttttattgaggaataatta**acaaaataaacccctgatatttaaagtgcccaattttgt**gtgttttaagatatttatacacccttggat**aatcaccacaatgaatcagtgatt**ccatcagaagaaatcggtgactggttctggcagaatccagggaaccctatctaatatctcgtgaacttc**tacttggactcaagta**agaaacagggctctgaggctcccagccacatcttggg

Uppercase: IGLV1-62

Lowercase: Flanking sequence[1000bp]

Red & Bold & Underline: Stem-loop [23]

Blue: Heptamer[47]

Green: Nonamer [4]

id-IGKV2-40-2[V_gene_segment]

tatttcttatttgttctgtatcattaatctagaagattcttcccttaagacagagtcctctgggccaggcacaatggctcacacctgtaatcccactaccttaggggccaaggcaggcagat**ctcctgaggtcaggag**ttcaagacaagcccagacaacttggtgaatgaaaccccatctctactaaaaacaaaaacagaaacaacaaaaaagaaagagccctctggttaaccttgtatgtgtgagacgattatgatgagatagatcccagattgaacaactggtcaccaggaatttt**aaatttgctgctggaggcacaaaattt**tgtctctctttcctttttcttacactgggctcttggctctaaatgtagagg**ctcacatcattctccctgtgag**gcgcttggacagagagctcttatgctgttcactcaccaggtgccaaggcagagtagattctaatatttgagttgaacattcttgaacagttatcctggaaacagtagataccagacagcccttgaactggctccaggccgctttttatttgcaggctctcagttcagcagtgcttgtggggatgggcctgtttcatactctagattga**ctgggagggaatcaagccagatggcattcacctcccag**agatgtatcctagacacacatttccacattgtcagggttctggtgctttcttacagtcatgccctacacagtgtgtccctacaaaaggtccgaactttcaccttcagatccttcttcccttgattgtgggcaaacttggctgaatctagttctgttttattccaaaggacaatttatatcacattgttcacagaagagacattccccctgccccgtcaaccttttccacaccactgcacccaccaggtgatttgcatattgtcccctagggtggaccttcccttgtgagtctgagataaaagctcagctctatccttgccttgactgatcaggactcctcagttcaccttctcaccATGAGGCTCCCTGCTCAGCTCCTGGGGCTGCTAATGCTCTGGGTCCCTGGTAAGGACAGAAAGAGATGAGGGAGGACAACTGGGTGGGAGGTGAGCT**CTGTGGGCTCCACAG**CT**TCACATGTTTATTCCAATAATGTGA**TAGAGGCACATGGTCTATGCTCCAGGGAATGGAATTCAGGTTTGTCTTATGAATAATCAGGATTCACCTCCAGGGAACGATGACCAGTGCTCTGATTAAGAACTTGAAAAAAAAGAGTTCCCTTGTGGCTAATAAATAATGGGTCTATTTTAGAAAGTCTACTTTTCA**TGATATAAATCAAAACTTTAAAAATGTAACTGTAAATTTATATCA**CAAGAGAA**ATTATGAAAGTTGCTCATAAT**GTATCTATATAAACTTGCACTTCTCTGTTATTATTTCAGGATCCAGTGAGGATATTGTGATGACCCAGACTCCA**CTCTCCCTGCCCGTCACCCCTGGAGAG**CCGGCCTCCATCTCCTGCAGGTCTAGTCAGAGCCTCTTGGATAGTGATGATGGAAACACCTATTTGGACTGGTACCTGCAGAAGCCAGGGCAGTCTCCACAGCTCCTGATCTATACGCTTTCCTATCGGGCCTCTGGAGTCCCAGACAGGT**TCAGTGGCAGTGGGTCAGGCACTGATTTCACACTGAAAATCAG**CAGGGTGGAGGCTGAGGATGTTGGAGTTTATTACTGCATGCAACGTATAGAGTTTCCTt**ccacagtggtacagccctgaacagaaacctccctgctgtgg**tg**ccccagctgctcacatgcactgcttgtctgggg**agcaggtcagcagcgtctctgagtctgcaaaagaggaggctgttggagaatacagggcagggtttgcttctgaggactctgcctgggactacaggtgcatgccactaaacatggctaatttttctatttttttgtagagtcggtgcttcaccatgttgcccagcctgttgtcaaaatcatgggctcaagccacccacctgacttggcctcccaacgtgctggcagt**acagtgtgagccactgtggcaggtcagcacccctgtttatgttcctgtcacctgccacag**ccttgactctcataaccaacaggaaaatgaggaggttctagggccctgtgagtaaaaaactgggatgataggga**aaggagaatggaatctcatctgaatcctcctt**ccttgcctacatttgtttaaatttattgagcaaaagg**gccagactactgattatttctggcaaaacatgttgagtacattttagggtttaacagttttg**ggtacctttcaaa**gaaaatatttggttatatgtaaaattggtattttc**ccactttttaaattcctacttctcctgtttgccattcttctccactccatgagacagtagagacaggattattcactaattctcctctggcggagctggctgaggacaatcagtaagatcttggttgtgagtgtcaaatagattttgtaatttcatagcagacgcaagttcctaatactaaaactctttgattacaattacctcttgctgatagaaaaagggagttcttgaaattttgaaagttggttttaaaaataaaatgcatacactggaagatgcagtatactaaagatgta**gtattttccagggatcactgagaaaatac**aggatgaggtaaacagttttattttccaaagttcagaatttgagattgggcagactgca

Uppercase: IGKV2-40-2

Lowercase: Flanking sequence[1000bp]

Red & Bold & Underline: Stem-loop [20]

Blue: Heptamer[38]

Green: Nonamer [5]

id-IGHV3-38-2[V_gene_segment]

cagaagttaaactgtgc**tttcttatagctagacatcttatactctaaagaaa**ata**aaatcaatgacatgattt**ggctctgtgtccccaccccaatctcatcttggattgt**aataattcccacatgtcaagtggaaaaaggtgggaataatt**gaat**catggaggctgttttctccatg**ctgttctcctgttattctcacaagatctgatggttgtataaggggcctccctcttcccttggcacttctctcttgcagccatgtgaagaagcacgtgtttgctaccccttccatcatgattgtaaatttctt**gtggctttctcagccac**gcagaactgtaagacaattaaaccttttttctttataatttacccagcctcaggtatgtccttatagcagcatgaaaacggacaaatgtaagcaacttaattatatacaaaaactcttttatacctgcctactgtgaaaggaaaataagtcttgagaccgcaaaatcaccaagctaaatggaagagtcaaactggtgttacaggagatagaa**agaaattattttggtagatagttaggatgacagaatttct**ggcaaaaacatttcttctaacaaaaatcagcttggcaattacttattttctattcacacagagttcaaagaaatcacttctaacaaaaag**cagcctgaaagatcaggctg**taaaacatataagcaactctgccacagagagggtgtttctgggtgtaatcaccaaacttcacatacatacactgggccctagtaaaaacagtgggccttaatgagcacattcctttcct**tttctgggggcactataacatcagaaa**gctgg**aaacttgcacaagttt**aggatgcctgcacttgcaaggaggtacctgggacctagcatggaaaatcctcttcccctttttagcact**ctgggagtggagccccagccttgggattcccag**gtgtttccc**ttcagtgatcaggactgaa**cacacacaactcatcATGCAGTTTGTGCTGAGCTGGGTTTTCCTTGTTGGTATTTTAAAAGGTGATTCATGGAGAACTACAGATGTTGAGTGTGAGTGGACATGAGTGAGCAAAACAGTGGGTTTGTGTGGCAGTTTCTGACCTTGGTGTCTCTGTGTTTGCAGGTGTCCAGTGTGAGGTGCAGCTGGTGGAGTCTGGGGGAGGCTTGGTACAGCCTAGGGGGTCCCTGAGACTCTCCTGTGCAGCCTCTGGATTCACCGTCAGTAGCAATGAGATGAGCTGGATCCGCCAGG**CTCCAGGGAAGGGGCTGGAG**TGGGTCTCATCCATTAGTGGTGGTAGCACATACTACGCAGACTCCAGGAAGGGCAGATTCACCATCTCCAGAGACAATTCCAAGAACACGCTGTATCTTCAAATGAACAACCTGAGAGCTGAGGGCACGGCC**GTGTATTACTGTGCCAGATAtacac**agaggggaagtcattgtgcgcccagacacaaacct**ccctgcaggaacgctggggggaaatcagcggcaggg**ggcgctcaggagccactgatcagagtcagtcccggaggcaggtgcagatggaggctgatttcctgtcaggatgtgggacttcatcttcttccaacagtttctctaatgaacctccctaattttagaattctgtggttcctaatgtcatctcta**catattttcaaaagatcattttaatatg**aggacataacctctcatgtaccaaatgcacattgatgcttacaaagatgaaaagttctcaaccattgtcaccaggatcgcagtcctgaggaagctcacgggtgtctgat**gagtctcctccattcaggcccaggacagaaacctcagggagactc**ctggactagaacggcagggattctgatcacagccaatagagagcctgggcc**agggtcagtgtcctgtagaagctcacagggttcacatctgaccct**tctcctgaccctaaagccaatccgcatcagcactgatctggtgctccttttgctcccaatacatgttctttctttggagtgtttgttctcccttttttatttgct**tttctttcttcctgaaaaagaaa**cacatggtctctatgctctacactccagggctcaaggcattttcttagaactcaggcaaggctcaggcttggctactccagcccacgt**gggagaggctgatgggatttccttctctccc**catattctcagggccctcctctgtgttgtgtgtagactcatctgggaatgcaattggccgttaatagtgaaggggataaactcatttgatcaaaatgggatgtggatgtggaattaaccctgttctatgcacactgtcagagtcatcttcttcagaagtagtgttagaaagagcttgtgaactttatcagaatcaaaatggatccacttgtgttaaaaccctaatgaatgaagctggggaaggcca

Uppercase: IGHV3-38-2

Lowercase: Flanking sequence[1000bp]

Red & Bold & Underline: Stem-loop [19]

Blue: Heptamer[36]

Green: Nonamer [14]

id-TRGV3[V_gene_segment]

ttcttacagaggtggtgataatc**tgtgaatgccttactgaattttcaca**ataatattactgtacttgatttgtccccttattttcatctgtgaagccaggtgcactgcagatattagtctgctttctgagggtagggatgacaaataaaatatacaataccgggggaattggaagttcagataaataacaagtttttagtttaagtatgttccaaatattgcatgacatggcacccagttctactttaaaaacaaacaagcaaacaaacaaattcacactcctactttaatgaacaaaaatcttctaataaggaggtcacacacacaggattacaggcacaacccccacacacaaacacacacacacatttaccttagagcagagtccaggacacaccaacacacatgatcctcagtccttttaatttaaagcacgttcaacccttctcagagtggccttgagtggttgcctccatgagtctgcagatgcgaaa**aggaagttggttccgggccctaatcagaggcttcct**gtcaggacaaggttctggcatcagcattacctgctgcaactctgaa**gggcagggtgggagcctgctatgggactgccc**cctccaaggtgtgagcctgttaattccagggtcagcttacccaacatctgcagtccctcagct**ggaggcctgagctcttaatgcccctgcctcc**cttctgccctgggcctgctctcagctgccacaagagggcgccgaagaaagggcct**gaggagatgcaggagctcctc**tgcagg**tcctcccaggcaccaccccttctgggagga**aggcttgggagtcc**cctaagacacatcttcagtcacttctctctgcctgtgtctcagg**aaaccagctcctcctactgtcttctgtgctagggatcacttccttgttgagtggggcctgagtttt**aagaggatcttctgctcctctt**catctggtccgttt**ccttccaaggcccccgagaggaagg**cATGCGGTGGGCCCTACTGGTGCTTCTAGCTTTCCTGTCTCCTGGTGAGTACGCTGCCTACAGAGAGGCTCACAGGTTGGGTTTTGTTTTGTTTTCTTCTTGAAAGGGGTGCC**ATACAAAGGAATACCTCATTGTAT**TTTGTGTTGTTCCCATTGCAGCCAGTCAGAAAT**CTTCCAACTTGGAAG**GGAGAACGAAGTCAGTCACCAGGCAGACTGGGTCATCTGCTGAAATCACTTGCGATCTTACTGTAACAAATACCTTCTACATCCACTGGTACCTACACCAGGAGGGGAAGGCCCCACAGCGTCTTCTGTACTATGACGTCTCCACCGCAAGGGATGTGTTGGAATCAGGACTCAGTCCAGGAAAGTATTATACTCATACACCCAGGAGGTGGAGCTGGATATTGAGACTGCAAAATCTAATTGAAAATGATTCTGGGGTCTATT**ACTGTGCCACCTGGGACAGGcacagt**gattcagacctgtcctacaccacactgaaaatctgc**cttgtggctgcctctggtacacaag**atagagccgccccctctcatttcctgccaccaaatttcctgtattctgaacaagagaaagacagcttaactcctgatctccctcctaatatcacactgtcctggcagcagctgcatcctgttccccacccctcccccacaactttcctgaagatcaagctgccatct**ccaggcctcagctaagcagcctggctgagagcaaggttctctcagc**tctcctaggacatgggggaggcccactcactctgcttcctatgacacacaggtacaactagggtccagctgtgaagcgagatattttggacagcaaatgggaagggtatttatactgaatcatgtat**ccactcatggtccagagatcacagctgagagtgg**tgct**tattccttatgtctataacaacataagcaata**ctataatgaccactaaaacactaggcccaggtatgcagttgaacatggtctccctctgccctcccttttattctctctctcttttcttacttctttgcttgtttttatttttctctccctccctaactccctccctccatccatcctttcctcccttcctcccttgtttccttccatccattctccttccttgttttccttccttccttccttgcttttctctctctttctttcctttttgggtacataatcagaatactttttccacaaaatcatttgatataatttttctctatatgttcatgccaacattcatacagctagtttaattgttcagttctctttttcacattttggaattttgttgtattactttgttttgcttcatatc**tgttttgtatatataaaaca**tttacatgaaact**gaagccaaaattataaaagatgatgcatttgaagattttagcttc**cattgtagatttctcgttcctcttttctccctgacactataggtaacaatattt

Uppercase: TRGV3

Lowercase: Flanking sequence[1000bp]

Red & Bold & Underline: Stem-loop [19]

Blue: Heptamer[19]

Green: Nonamer [5]

id-IGLV3-21[V_gene_segment]

acattttcccagcaacagaagacaacttagttataataaacacttacttcctgggtcttgga**gtttgcagccccctctcattttctcatgaagcaaac**atttccttcacctcctgatctgtccctggtcctgacagcaccctggtgatactgaggcacagcacccatgacctgacactagaactcgcagccaaggtagaaacttaccgcatcccatggatcttccaa**aattctatgtgcccttggcaaccaagaatt**gcatt**ctcctctagcacagaggag**ctgtgccctggaatggggcctgtacctgtccaaggcttgtgccgtcccctgtgggagatgagaagcgtccctgcattgggctcttg**gggacccgtcttggacatgagtgagaatgaagagggtccc**tgcattgggctctggcatgtgactttaaatggatttaggcctgtac**cagacatctcatgtctg**acataaaatatttacaatcaggacattactagagaagcagaaaaaagctaaccacctcccttctgagccaggatggaatgaagg**aggggactgtggaccccagataattcccct**gtcaccactgtgactctaacaacctcttaaatcacggccaacatctatcccataggaaggtctttatatcccctagaaaatacagaggaagtcagctctgagcttttccacgaccaa**cccagccaaggagcaaggctggg**cacaacctgggtaaagatgtgagcccagaccatgggaccagtgggtgaaggaaaatcgcatgggctgagggggtgggtaagcaggggc**cagccctcctctctctgtttcctttggggctg**agtccttctctggaaaccacagatctcctccag**cagcagcctctgactctgctg**atttgcatcatgggccg**ctctctccagcaaggggataagagag**gcctgggaggaacctgctcagtctgggcctaaggaagcagcactggtggtgcctcagccATGGCCTGGACCGTTCTCCTCCTCGGCCTCCTCTCTCACTGCACAGGTGATCCCCCCAGGGTCTCACCAACCTGCCCAGCCCAAGGGTTCTGGGTCCAGCGTGTCCTTGATTCTGAGCTCAGGAGGGCCCTTCCTGTGGTGGGCAGGATGCTCATG**ACCCTGCTGCAGGGT**GGGAGGCTGGTGGGGCTGAACTCCCCCCAAACTGTGCTCAAAGGCTTGTGAGAGCCTGAGGGACTGCACCTGCCAGGAGAGAGTAGTGAGTTTTCAGTTCAAAGTCTCCATACAACAGGAAAGTCATGGGCCACTGGGGCTGGGGCTGATTGCAGGGGATACCCTGAGGGTTCACAGACTCTCTGGAGCTTGTCTGGGACAGCAGGGCAAGGGATTTCATAAGAAGCATCTTTCACCTGCAAGCCAACCTCTCTCTTATTTATTTATTTATTTATTTATTTATTTATTTATTTATTTTTATCTTTGCAGGCTCTGTGACCTCCTATGTGCTGACTCAGCCACCCTCGGTGTCAGTGGCCCCAGGACAGACGGCCAGGATTACCTGTGGGGGAAACAACATTGGAAGTAAAAGTGTGCACTGGT**ACCAGCAGAAGCCAGGCCAGGCCCCTGTGCTGGT**CGTCTATGATGATAGCGACCGGCC**CTCAGGGATCCCTGAG**CGATTCTCTGGCTCCAACTCTGGGAACACGGCC**ACCCTGACCATCAGCAGGGT**CGAAGCCGGGGATGAGGCCGACTATTACTGTCAGGT**GTGGGATAGTAGTAGTGATCATCCcac**ggtgacacaggcagatgaggaagtgagacaaaaacaccctcccagcctcggtcaccctcttgctccagccccgggaagcctgttgataaagccatgagtgaatctggcccagttcacctggatctgagcctttcaggttgcccttccctccagccccctccaggagtctctacagaagatacatcaggcataaatatggcctggaagggccagaatcatctggtgacttggggctgttgtgtgagttagagaatgaaggcttgggtggaaagaca**gacagaggcaacctctgtc**ca**ctgtcctacccctggatggtcatatggtggggacag**ggcaagtccttagaccaactgtctggatcaggccc**cagaactactgcccagttctg**ctgaggtcctggcccc**caggctgtgtggcagcctg**tgattcccaacagagcaaaccagaggaatggacactgtgaagtctgcccagatcccctcctcaatgtgacccacctggca**ctgctgagaagcccagcag**ctcagagctgtgccctcactgggaagtgctgttggttgcagaaagcttcctcaagtttgtgtcccttttcagaggggttcggtttaatcaaccaagatctcaaatccttgcctcaatttaagatgccactgaatgaagggcctcccagctccagagctccctgtgtggatacctgaggcctcaatgtcaactccatcacgagtcagggtctccttctgccccgtgttgcctcccccactcccttctgaatcttctgtgcatggacatctctattgcagagttagcttccagagaaccccatctaagatggccagctgtccccaacatgggtcatcagggacctgagtaggccactataaactgaaa**actctggtttctgtccaaatttgcagagt**aaatgttgaaatgcccaatctgatggttccttgaatttttatggaatgaaaagggagcctgacatgccaggtgctctggg

Uppercase: IGLV3-21

Lowercase: Flanking sequence[1000bp]

Red & Bold & Underline: Stem-loop [21]

Blue: Heptamer[42]

Green: Nonamer [5]

id-IGHV1-45-2[V_gene_segment]

atcctgtgtgagagcagcacttgagtgatttcaataacaacttcccaggagaatcagctgaaaactacttgtcccattttccatacagatataacccc**tctattttcctgaagaaataga**aagagctgaatgctgaatacactgaatgtctgctggttttgcaagtttgtgactatatcactttctaatttctgacctgtgcagaccactgtacagacttttctcactggtgggaccagccttccagatgtcaaatataact**gagcttcttcatataaaagtcaacacaagctc**ctcatggt**ttcagtgctcactgaa**tggagttggaaataaaacccaca**attatccataacacaaattccttggaagttattttgggataat**gagttcataacctgtagaccaagagtccaagagtgtttctattgaaga**gcctgggggatcaagacaccaggc**aggtgatgcagacactgtctaaagagtgcccagcggctctcagagggacctactggatactcacgtgggacatcagcaatcactttctcagagtcaccagtgagctgtgctggttcctgaagggtccaggatagggccaaggcacctgctctgtgtcgtggagagtgattgttccagaaat**catagaggtggtctctatg**cttataaaatctatgttcacagtgagaagtctgttctgagagggcttattcttcagtgaaaggacctctgcccacaaatgttcataaatggagcagggcattcatttc**ctcaagcaggatcagggcttgag**t**catcagcatctcactcttgcaaggctgatg**tgtcgtttgtcttccctttcttatcatcgaccaggctttgagctatgaaatgccctgtctcatcaatatgcaaataacctgagatcgactgaggtaaatatggatatgtctgtgccctgagagcatcacccaacaaccacatccctcctctagagaatcccctgaaagcacagctcctcaccATGGACTGGACCTGGAGAATCCTCTTCTTGGTGGCAGCAGTCACAGGTAAGGGGCTCCCAAGTCCCAGTGATG**AGGAGGGGATTGAGTCCAGTCAAGGTGGCTTTTATCCACTCCT**GTGTCCCCTCCACAGATGCCTACT**CCCAGATGCAGCTGGTGCAGTCTGGG**G**CTGAGGTGAAGAAGACTGGGTCCTCAGTGAAGGTTTCCTGCAAGGCTTCCGGATACACCTTCAC**CTACCGCTACCTGCACTGGGTGCGACAGGCCCCCGGACAAGCGCTTGAGTGGATGGGATGGATCACACCTTTCAATGGTAACACCAACTACGCACAGAAATTCCAGGACAGAGTCACCATTACCAGGGACAGGTCTATGAGCACAGCCTACATGGAGCTGAGCAGCCTGAGATCTGAGGACACAGCCATGTATT**ACTGTGCAAGATAcacagt**gtgaaaacccacatcctgagaccgtcagaaaccccaaggaggaggcagcttcactgaatgaggaggttacagggcttacgatgtttaaagttgttcagaaaataggctaagcaattgaggaatatgagtaa**tagaaatatgtatgcactctatacaggaaatatttcta**ataattgtcaccctatatgcaaaattcgcagagaggtaaaagcagaaatcagtcaagctgatgcaaagttccccacgtaggctttgtgcagatgtaagttctaaaatcagatagataaataatttggagcaagattgcttgataacatggctaatgctgaatatgattcctaaaaactggccaaaatatattccaatttgtctctgccacctctcttacataaaatgta**ttaaaaagtagttttaa**gaccacagcaaaattgaacagaaggtgcagagatttcctatgtgcccctgcttcacacatgcacagccttccccactgtcaccatcctgccccagagtcatcaa**taagttacaatggatgaactta**catggatggattggttctttcctcttccggcggtcccttggcataccaagtctaaactatcttgaagcacaacaggttcttcgagtgggttcctgggaatgaagccagttagaggaaaagtgggtggggctattccta**ttaggagtcttttttagaagactcataa**aatgtatatgttcctatagattctgtgactcctgacttagtatcccttcccagacggtaagcttcctaaatgtttagaggcagatccatatctatggaaagaaagcaagttctagtgaatcccataaggaatgtcctttaatgagaagtggagaccttggtcatgaggcacatcatgtatgattttctataattcctttagattcactgtaagcttttgagggtg**tttctggatgaggccctttgtacagaaa**atgaaaactcaggcatgagttccaggcacaacca

Uppercase: IGHV1-45-2

Lowercase: Flanking sequence[1000bp]

Red & Bold & Underline: Stem-loop [18]

Blue: Heptamer[42]

Green: Nonamer [5]

id-IGHV8-51-1-2[V_gene_segment]

agtcaccatctccagatccacgtccaagaaccaaattttcttttagctgagttctgtgaccaacaatgccacaaccttgtattactgtgagaggaacagaagagatgtcagtgtgatcccagacacaaacttccctggagaggggcccaggaccaccaaagagcactcaggctccatgaaaacagggcccaagctggagaacgggtttcctgtcaccctcacgtttcaccattagatactctacactatgcctatgctgaggtgtgtgtttaatataattagaaaatggtatttatatacgtgtataatcatatctagtgtaagaatttgtagtttccaccttatgtgaccattaaagacaacatctctcacgttattcaaggaccttat**aaagtataatacttt**tacataagatttatctccttttttgtttctccttctgtctatattctctctgtctctctcttcctccctctttctctctctctctccatctctctctctctgtctctctctcacacacaaatacacacacatgcaatcgttgacagtatataggctaattaagaatatattcccatatgatttaaaaataacagttttacaaatatttgcc**tttaatttttcaaagaaaattaaa**tccaaatgccagtgattttttatgtgactttt**aaaatttaaattgttaatttt**tgttttcatcaatgcttgtatgtgtgcttttgtattctttcagattatttggtgcct**tcacagaattaaacatgtgaaactctccacttgaatttctgtgactcaggactggtgagcaaaaagttagaagtcac**tctcctgatccttccctacagctgcagattcctgaaggtaatgacctcacagtagaccaaggccctcaaaggtgaccttgagttccagcacatcttgggaggccaagcggatgtttgtgaaagaaatagtggctatgtgacagtttctaaccagaatatctctgtgttttcagGTATCCAGGGTGAGGCCCAGCTTACAGAGTCTGGGGGAGACTTGGTACACTTAGAGGGGCCCCT**GAGGCTCTCCTGTGCAGCCTC**TTGGTTCACCTTCAGTATCTATGAGATTCACTGGGTTTGCCAGGCCTCAGGGAAGGGGCTGGAATGGGTTGCAGTTATATGGCGTGGTGAAAGTCATCAATACAATGCAGACTATGTTAGGGGCAGACTCACCACTTCCAGAGACAACACCAAGTACATGCTGTACATGCAAATGATCAGCCTGAGAACCCAGAACATGGCAGCATTTAACTGTGCAGGAAAcatcgtgatgggaagtccacgtgggctcagagacagactgccatgcaggacacagggggtggcttggctgaagggggcactcagcacccacagaagacagg**agcagcccagggcaggggcaggtggagttcaagggctgct**ttcctgtcaggttctgtggcttcttctgcatcaaatgcgttctcctgggagcctctctatatttatggttctctgcctaccactgaggcctctggattggaaaagattactactagaaaaaaaaattctcatatacccctgaagaatacatcattaattgaggcagaagatgtcacaggaggccagggaggctgtggaaactgtgcgacgtggatgcacatctgacaacaaagacataaaaaatcagggaacactgataaaacctcctaattatgccatggccctctctaagcctagtaaagccattgatgccttcctgggccaaacccacccatcagggatcactcctgtgtcccagcagcagacatgccttagaatctccactgtatgtaatcactgt**ctgggaggagctcccag**ggcatgtgtctttgcattaaccaggtgatgggtgtcagagaacagcagctggatgcctggtctatgggttccctgatgttgaagaaatgggaggtgcagtctcagagccaatacactgtttatgcatttttatatagaaaccatgat**ttttcttgcttctctcagatgacacagaggatcaagaaaa**cagtctgtaaacaattgtaatgtccccatttcttctgaataaatgcattatttcttaattctgagtgggggtttgtcagaccatgcccttctcctaagcaattgttcaatccaggctg**aaaccacatggggatcacatactttggttt**ggttcaggtgcagaggtcttcagtagagcaaatttggtccttcccacacagtaagcctcaccttcctcagagaaagagctgaaattgatccaactttgtctgaggtg

Uppercase: IGHV8-51-1-2

Lowercase: Flanking sequence[1000bp]

Red & Bold & Underline: Stem-loop [10]

Blue: Heptamer[35]

Green: Nonamer [4]

id-TRBV11-3-2[V_gene_segment]

gacatgtgtcccctaggttttaggaacttggatttctttaggctgagttgtactattccctttctctgaacacacatgtcattggttagttcattcatattgattctcagggcttctccaatccctcaagccttcccactctaccactcagataaggagaagtgtgtcttaactaatgttgcat**aattttttaaaaattgctggatactaacactgcaatt**caattgtaaaatatgatagaggtgtaagtgttttagatcccagaagtcctgctcaattttgcatttattgttagatattcaccctgatattttcaggaattagagtgttcttcacccatttgggttaagatatcagaacttactttcaattcattggtgaccagagactacaggcagaatcatagtgaaggcttgatacataatacataatgaagattgataacagcaaaaagggaactagtcattggccttcagataaaatgaaacctcccaactttgtgctcagggaattattttacccctttaggttagagtatgtcagctagttcaatgcagt**tctaggataaagctgtgtcaattaccctaga**ggactgggagcatggaggacctggtagaaaagaaatacccaaagagaaggaaaattacacataaaatcaagaggtcactaattgatatatctattttgagttgcctttgagacttccacgtagaa**acatctagcacaggataggcctagagatgt**agtgacagatctaattcc**ccaagccctgaatatgtgagtgaggacgcttgg**atgaacttccagtctatgaatacattaaacaaacttttagctctgttttagtttacaacaactgttttctaagaacttaagcatttgtggagacaatgatgtcactgtaggaacttctctgtaaggacagcaacatcccacttcctctgctcctgctcacagtgaccctgatctggcaaagcttccatcctgccctgaccctgccATGGGT**ACCAGGCTCCTCTGCTGGGTGGCCTTCTGTCTCCTGGT**GGAAGGTGAGTCCTAGGAACACCATGATCCTCAC**ATAATCTCCCAGTGATTAT**TCTAGTCATTTC**CTTCTATTTTAAAATTCTTTTTCCCCCACAGAACTCATAGAAG**CTGGAGTGGTTCAGTCTCCCAGATATAAGATTATAGAGAAAAAACAGCC**TGTGGCTTTTTGGTGCAATCCTATTTCTGGCCACA**ATACCCTTTACTGGTACCTGCAGAACTTGGGACAGGGCCCGGAGCTTCTGATTCGATATGAGAATGAGGAAGCAGTAGACGATTCACAGTTGCCTAAGGATCGATTTTCTGCAGAGAGGCTCAAAGGAGTAGACTCCACTCTCAAGATCCAGCCTGCAGAGCTTGGGGACTCGGCCGTGTA**TCTCTGTGCCAGCAGCTTAGAcacagtgtagcagaga**cacttccctcctgtgcagaaaaccgcaggactctctcctctctactcagctcacagcagc**ctttccttattcctcatcctcccaggaaag**aagtgagttttcagatatagctaggattcatatagtgggaggaaatgaactatttcttacaacatgagcgctataattgttgcttgaaaatgtgtctcagggatttgaaacac**ttcttggggacaactcaagaa**acctaaagtgacttatcagagattgagtcctcaggggtagtgatggtttatctcccataaaagtatgtgatcatgaaacatgtacaataaacttaataatacagatagtgattataccaagcagcaaacagcaggctccccttatgtagttgtcagcatgtgaatgacttgttacccttagtgacaccccggctgtctagcaaagtctc**aacacacagtatgtgtt**tggaaatctttttccttctgttttatttttggtaaagtatgctaataatgatgttgttaatgatagtgattataatttcaggtatgttattcagccctttttccagttcaaagaaatgcacacaaatggataccccggcaaccaggtgcaaatcttagctccagatgaattctacttctcctgaactctctaactcttctatgtaatcatcccagcccaagaatatatctagacctggaccagtcagttcagaggacatttctacgtgtctgccttgtatattaa**aagttccttagtggattgaaatattccttgtaagaactt**ctgaagcaatcaattgccctctctgataatatacactctgtgtgtctagctccatattcattactctacttctgcctaaactctct**aatactagaaatctttctgaagtatt**ggttagaattacttaaaaagcagcatactggaaattgtctgaaagcactgtcactgtccatttgcccctcctgacttccctcccctccttatccaatc

Uppercase: TRBV11-3-2

Lowercase: Flanking sequence[1000bp]

Red & Bold & Underline: Stem-loop [15]

Blue: Heptamer[37]

Green: Nonamer [1]

id-IGHV3-64[V_gene_segment]

tcagtatttctat**tgagaaaaataatccaaatacataagagcatatactatattttttca**ttcttataaattctagaatataaaagctagtcaaaagacatgtgaaaacatcagtagatttataaagaaatggtagaagaagggaaggagaaaaaccaaaaaaatataaaagagcaagaggaatgctgaggggagttgacttgtcaccttctt**aaaaatagagattttattttt**caaagtttactattgtacagaataaatatgtgaattttcttatctgtcaattaaacctca**taaaatttattacaaaaaaactgaaatttta**gacaaaaagagggtgataggaaggaacaaataaatatgttaaatgtcaaata**tacctaaaaatttatttgtctgacccctagttttctccgtatttttaggta**aatgcagcaaaatcacacaggttgtcgtggcaggaagtggattctgcaaaccacactaggcccgtttatctctgtcctagagttggttaaaagagcaactgaggccagctgtgaggagcataggcccgggtactaggactcactcatgccagatataagcccttag**acacatacatagcccctccatgtgt**gggttcacttttacatctgtacatgaagaaaccactgattcctaaataacataatttatacacataggtaaaaataattaaaaa**tgtgatagttattaagtgtttatcaca**caacaatttcacaataaaacagcattttcccaaatgtaatcattgtcatcgaaatccccaaggacactctcatctgccctgggccctgccctctcctcaggcatctcaccccagagcttgctatatagt**aggagacatgcaaataggtccctccctctcct**gatgaaaaccagcccagccctgactccgcagctctgggagaggagcccccgc**cctgggattcccagg**tgttttcatttggtgatcagcactgaacacagaagagtcatgATGGAGTTTGGGCTGAGCTGGGT**TTTCCTTGTTGCTATTTTTAAAGGTGATTCATGAGGAAA**TAGAGATATTGAGTGTGAGTGGACATGAGTGAG**AGAAACAGTGGATTTGTGTGGCAGTTTCT**GACCTTGGTGTCTCTGTGTTTGCAGGTGTCCAGTGTGAGGTGCAGCTGGTGGAGTCTGGGGAAGGCTTGGTCCAGCCTGGGGGGTCCCTGAGACTCTCCTGTGCAGCCTCTGGATTCACCTTCAGTAGCTATGCTATGCACTGGGTCCGCCAGGCTCCAGGGAAGGGACTGG**AATATGTTTCAGCTATTAGTAGTAATGGGGGTAGCACATATT**ATGCAGACTCTGTGAAGGGCAGATTCACCATCTCCAGAGACAATTCCAAGAACACGCTGTATCTTCAAATGGGCAGCCTGAGAGCTGAGGACATGGCTGTGTATT**ACTGTGCGAGAGAcacagt**gaggagaagttaatgtgggaccatgcagaaacct**ccctgcgggaacgctggggaaagtcatctgcaggg**ggcgctcaggagccactgatcagagtcagccccagcggcaggtgc**agatgaaggctgatttcctgtcacgatgtgggacttcatct**tcttaaagtttctctactgaacctaagttcggaattctgtgattactaggg**tcatttctacatatttttaaaatga**ttgttttaatatgaaaacctat**tctcctatgcagaaaatgcagattgatcctcacagaggaga**tgaaaagttctcaaccatggtcaccactgtcagagtcctg**aggaagctcagggcttcct**ggtgagtcttctcc**agtcagactgaggacaggaactccagaaagaatcctgact**agaactgaggtattgctcctcacaagagaactcaagctgcgg**ggggtctgttcctgcagacccc**tgacccggtggcggatgaataaaacgtacactgacacacagatattctgttttgccagttcagctgagggtccgaggcaac**ttacagactccaagaagaatcctgtaa**agaacggcagcagtggccctgagcagctcgcactccaggcgtgtattgagtatagaattaacaacagaagctctgacaacacacttgtggataattaacatggttaagagagtagttctacgaatgattaaagctcagggactgcggtctaaagtaaataccattagggggcaatacc**ctggttagttaatggaggtagggtaaccag**acttaactggggaggcctctattgtccctagtatttaccctatgcctaacgctctaaggtaagaactggccgccttcagcctgttcaattattacaagctatgta**acctttcggccttctaaaaaggt**atatgactattccctataactttcctt**aatattcccctttaatatt**tccgccaccatcctgagtgaatcccagcaagaa

Uppercase: IGHV3-64

Lowercase: Flanking sequence[1000bp]

Red & Bold & Underline: Stem-loop [23]

Blue: Heptamer[29]

Green: Nonamer [4]

id-IGKV3-11-3[V_gene_segment]

atgcgttagacagagatgggcgggaatgattgaaggtctgagcattttagcacagattaagaccaaatcattaggattttaagagttgtgtacagttagtgaagaaaaagccctagaatttaatttgactgttgataaaacattcttggattagattgaagactcttttctgtgccaagtaagtatatttatgataatgatgatgactgtagtgctaaatatttaatcaataaaaacaaaaataattgccgcatacataatgtcctgagtactactgtaaatgttttatcttattttctttaaactgtctacagcactgtaaggtaggcaccagtat**tgtcacagttacacagatatggaaactgagaca**cagggaagttaagttagttggtcaatttcaagcaatcggcaagccatggagcatctatgtcagggctgccaggacatgtgactgtaaacagaagttttaactttttaactcaaa**gagggtatgtgtctgggttaatggaaagtttcaggaccctc**agaaaacattactaacaagcaaatgaaaggtgtatctggaagattaagttctaacagactcctcatttccatcgatccaataatgcacttagggagatgactgggcatattgaggataggaagagagaaatgaaaacacagccttttatattgttcttaacaggcttgtgccaaacatcatctgggtgaatttaggtgattgaggagaagaaagacataggaatgaaattc**tctgagcacaagggagaagttctacactcaga**ctgagccaacagacttttctggcctgacaaccagggtggcgcaggatgctcagtgcagagaggaagaagcaggtggtctct**gcagctggaagctcagctcccacccagctgc**tttgcatgtccctcccagctgccctaccttccagagcccatatcaatgcctgtgtcagagccctggggaggaactgctcagttaggacccagagggaaccATGGAAGCCCCAGCTCAGCTTCTCTTCCTCCTGCTACTCTGGCTCCCAGGTGAGGGGAACAT**GAGGTGGTTTTGCACATTAGTGAAAACTCTTGCCACCTC**TGCTCAGCAAGAAATATAATT**AAAATTCAAAGTATATCAACAATTTT**GGCTCTACTCAAAGACAGTTGGTTTGATCTTGATTACATGAGTGCATTTCTGTTTTATTTC**CAATTTCAGATACCACCGGAGAAATTG**TGTTGACACAGTCTCCAGCCACCCTGTCTTTGTCTCCAGGGGAAAGAGCCACCCTCTCCTGCAGGGCCAGTCAGAGTGTTAGCAGCTACTT**AGCCTGGTACCAACAGAAACCTGGCCAGGCT**CCCAGGCTCCTCATCTATGATGCATCCAACAGG**GCCACTGGCATCCCAGCCAGGTTCAGTGGC**AGTGGGTCTGGGACAGACTTCACTCTCACCATCAGCAGCCTAGAGCCTGAAGATTTTGCAGTTTATTACTGTCAGCAGCGTAGCAACTGGCCTcccacagtgattccacatgaaacaaaaaccccaacaagaccatcagtgtttactagattattataccagctgcttcctttacagacagctagtgggg**tggccactcagtgttagcatctcagctctatttggccattttggagttcaagttgtcaagtccaaaat**tacttatgttagtccattgcatcgtaccatttcaccgtggctattatgttcaactaaatgcattttagaaggcatctctgtttatggcatcacaaagagtttaataaatcttcggtgcaaaaataaacaacaaacacacatttaaatataaagctgaagtatcaaaacta**tttcagcactctgaaa**attggtgaagtataaaataattaaggatgcatattctttatagaaaaaaaaagtaccagtgctttgagtaaggacagaaaatgtctgtagccttttgcctgtgacagcagccttctacccccagctcaatcagcatgaattacagaactga**agttttaccaatatgagcgtagcaaaaaaaact**agcagcttgctgccaaagtgggtggacttgagtaaagccaaggag**tggaaaaaaaattattcagtgtttcca**gctaaacatgcagaactccataggaaatgaacagaaaaatctcacagctttacaagtccaagatgatgacctgtttgaggcaagtagtac**atctgctgatagtaaatagcagat**tcctggatgagatagacccatattgctgaaataatctctgcacacattcctagtgacctagaagatatagatatgagtgatgagaacaaggagtttccggtgcaaagtaaaaccagaaggattcaagaactag**ctgcattttgcatgcag**tttgctttttaaactacacacagatgggttgacagaagatagatttataacctctagatatttaagcaaaacgtctcaaatcattggtgaccacttcactatgcagatacatgtgcagttcctataaat

Uppercase: IGKV3-11-3

Lowercase: Flanking sequence[1000bp]

Red & Bold & Underline: Stem-loop [16]

Blue: Heptamer[35]

Green: Nonamer [6]

id-TRBV12-2-2[V_gene_segment]

ggggcgcaggggactcagctgtatgcctctgtgccaacagcgaaaatgtggccttaaactgctccttcttctttgttagcaaataacctagttgaccaattc**atagagcagaggttctctcagttctctat**actctacaccccgagaggaagtcagttagtgagatgtaagat**tcttttataagaaaaaga**ccaggtgtaaagattaagaaatgctgagctttgctagaggtaagaataagatgtaaatcaactctggaacatttaatttccaaatctttgtgagaaagacaatcttgtttctgatttattgtgtggatagagaatgtgagagattctgtgctctgggctttgcaaactcaatagagttcaaggatcaataaactcggagagacccttcactgcaaagaagagctttgaaatctaatgagctttcgggacactgattcagctcctgtatctttaacaacgcggttgacatgcttgctcttcctgggtaactcagatgctaacttctaactcatttaaaggcatttgaagaactaggagcacagagtcaccagtgagaggaagaggagagtttgcagagaagctggcttgcaataaggcaatgagttcatctttaaatacttggagtttgaggtgcagatggatatagttggcaggctcctaggtaaggcatgttatggagaaggtgctatgaattgataatatcaaagcaaatctacagggatcctctgcaagtgtgcatctgtatctcagatcaattatagttgacttcagtcctgcctgattcatctcccaaaaatgtagtctgcctgattcatctcccaaaaatgtagcctccgcttaaaggagctttcaagttgggggtggtgggccattcagtgttgtcactaacagatgcatcttgtgggggtaaaatgtcccaaagtatcttttcttgcttatgttcataagggcgctggtctggaatgtgccacatctgttctcactctgccATGGACTCCTGGACCCTCTGTGTGTCCCTTTGTATCCTGGTAGCGAGTGAGTCCTCATGTATTTATCATCCTCATGCTGGGCCTCTGTATAGATGACTTCCTGTATTTTCCTTATTCTGTTTCCTAATTTTGCGTTCTTTTATAGCATGCACAGATGCTGGCATTATCCA**GTCACCCAAGCATGAGGTGAC**AGAAATGGGACAAACAGTGACTCTGAGATGTGAGCCAATTTTTGGCCACAATTTCCTTTTCTGGTACAGAGATACCTTCGTGCAGGGACTGGAATTGCTGAGTTACTTCCGGAGCCGATCTATTATAGATAATGCAGGTATGCCCACAGAGCGATTCTCAGCTGAGAGGCCTGA**TGGATCATTCTCTACTCTGAAGATCCA**GCCTGCAGAGCAGGGGGACTCGGCCGTGTATGTCTGTGCAAGTCGCTTAGCcacagcgctgcagaatctccccctccctgtgcagaaactctggtgcttcctcttttcctcacagctcccagcagtcctgagcaaagtctttcccgcctcaccctccccacaagaataattaagtgggtttggggcatggcaaagacataagatgatacaatatcaacatacaaaatcttgtgtaggagataatgtcgaaaatgcattttgcaaatttctcacaaattgtgtatggggtcataactcacacacaggtaggat**aaaccttaattacccacaccgtaggttt**ccctgccttcctgtatttgaca**ctcccatccaaacagtgggag**ctcttgagaatggctatgtcttgtgaactactctgcccacttcattcccttctcactctaagccatgaacagaaatttcccctacaagtactctttattgtaaatgaaagttcttttctatgattgtggacattcctttataatgctaatttcaatttaccatttatatacggatttttacattaaacttaaagagatcttcagtgtggataatacaggcagtttaatacattagaagtaagctacccctcttggcctacagggatccatttgtttctatcttgcataaagcaagccaggtgctagctaaaggtctgtgtgcactcacaggacactgatgagatctcctccactgcctgttctggaactatcaatatgaatatacacagggtgtctaaaagcacacagggcctggtgtggtggctcatgcctgtaat**ctcagcactttgggaggctgag**gcgggaggaatgcttgaggtggggagttcaagactaggctgggaaacagtgagattttgtctc**taaaatatatatatatttta**aaaattagcctggtatggcagtcttagct**actcaggaggctgagt**tgagagggttgcttgagccctgaagttcgaggctggagtgagccatgatcatcgcaacagagtaagaccccaaatcacaca

Uppercase: TRBV12-2-2

Lowercase: Flanking sequence[1000bp]

Red & Bold & Underline: Stem-loop [9]

Blue: Heptamer[35]

Green: Nonamer [2]

id-TRBV5-1-2[V_gene_segment]

cctgggggttgtattttggagggacatcaatattcagtccactaacaatggcaaaatcttggttactgcg**cagtgggccacatgggggtgctgtggaaccactg**agctccaaatgtctctggggcatgcacagagccaggagcctgggacagggagggttaaggagcagccttgcttgatgtttgtgctcactaaaaaatg**tgatgacggatacataaaactgcaggtgtgcctctcatca**gggagccatgcacttcagctatgcctgggttaggaagcctctgtgagccgaagagggagaatgaagatg**ggatgtcccatgctgggatgggacaacc**tgcatg**tccagggcagggacagatagaggaacaactgcctgga**aggaaaagcaaaggaac**tgtgggggggacatgcagcctaacctcagtccagccccaca**tggtatgtgagagtattaacagcacaaagcagaggtcctggatcgggtaagccaggggcctaacaaggaaggcaggatgcatgggaacccgcagagatgacatcagagtagtgatatcacagcctaacatccgatcagagatgcaatgcccaaaaccccagctctcagaggaccagtatccctcacagggtgacacctgaccagctctgtccc**acctggccatgggctccaggt**acctctgatgggaagacctttgtctcttgggaacaagtgaatccttggcacaggtgggatgtctctgaagcacgcctcatccaaggcaccctcaggctccct**cctgtgttcttgccctagcctctgtctccctccctcacagg**cccagtgga**ttctgctgtgcagaa**cagagagcagtggacctcaggaggcctgcaag**gggaggacataggacagtgacatcacagtatgcccctccc**accaggaaaagcaaggctgagaatttagctctttcccaggaggaccaagccctgagcacagacacagtgctgcctgcccctttgtgccATGGGCTCCAGG**CTGCTCTGTTGGGTGCTGCTTTGTCTCCTGGGAGCAG**GTGAGTCCCTGCA**GACAGGATAGCGCCCCATTCTGAGCCTGTC**CTCACCCCTGTGTCTTCCACTTTACCATGGGGAGGCACCACCAGGCTGTCTCCGGTGCTCATTCTCCATCTGCTTTTCCCACAGGCCCAGTAAAGGC**TGGAGTCACTCAAACTCCA**AGATATCTGATCAAAACGAGAGGACAGCAAGTGACACTGAGCTGCTCCCCTATCTCTGGGCATAGGAGTGTA**TCCTGGTACCAACAGACCCCAGGA**CAGGGCCTTCAGTTCCTCTTTGAATACTTCAGTGAGACACAGAGAAACAAAGGAAACTTCCCTGGTCGAT**TCTCAGGGCGCCAGTTCTCTAACTCTCGCTCTGAGA**TGAATGTGAGCACCTTGGAGCTGGGGGACTCGGCCCTTTATCTTTGCGCCAGCAGCTTG**Gcacagccctacaaagccaaccacattctgtgc**acaaacctccctggcccaatgtggagcaacctcagccctgacatatctgtgagaacctggggactgcagggagaaagaaaggcaatttagaatgctaggatccacagcctggggctttctgcaactcagggctggggaaaacaaggccaagagagcctgctcagaggatgg**tgctgcttccgtaggttgactgcagca**ttaccttctttgccgacactgtaaactacaggagtagaggaaatgtgtgtaatttaacagctggcgcactctttctcaggacaagagatttaattttgtggtcaagattgttgataacagtgtagtgtagaagggagaatcacagggtttgctttatacaacttggggagaagaaattgaggttttgggaaagtcaaggagagagaaaacgaaagatgattgaaaacagagatagagcagaaggtgatgaaaagtggtgttctgtgcaccatgaactctctttcccccacccgaaagttttcaacacctagaaaaaaataaggaaaggtcttatagccaccggaatcacctggtagccggtcccagtcggaggcatcaccaaattgccgcttggccattgtgagagccctggagttgcttctgctgtcttccttcttccagccctgatcttgactcctcaggccacagtcaggtacttcccaaaacacag**aggaaggtaaagtcacccttcttcct**aaaggggcaaatactttccagtttccccccaaataaattctgcaaccctgggattcctttctgacactcacctcccaaatccaattaccttctatgaccaaatgctctttgaacattttctaaaactcagcagtctctacctcctctgcgtttgtccttacagtttccatggcctgggaggacctttgttctcctaagacggggccacatactgatcatctgtcaaggacaggttcacacctgccttcctcttc

Uppercase: TRBV5-1-2

Lowercase: Flanking sequence[1000bp]

Red & Bold & Underline: Stem-loop [17]

Blue: Heptamer[49]

Green: Nonamer [2]

id-IGLV3-9[V_gene_segment]

tagttgctcattaaccttctgttgactt**aattaagtttttaatt**ccttgatttgttttccccagcaactgaagacaacttagtcataataaacattcatttcc**aggggtcttgcagtttgctgacccct**ctcattctctcatggagcaaacattttcttgacctccttatctgtcattggtcctcacagcaccctggggacactgaggcacagcatgtacgacctgccctggagctcacagccaacgaggaaacttaccgcatcc**catggatcttccaacactccatg**tgcacttggcaaccaagaaattttattcttctctagcactgaggagc**tgtgccctggggtggggtctgtacctatccaaggcaca**tgctgtcacctgtagagtgtggatagagggtccttgcattaggttctggaaagtgtcttcaaaaacggatttaggcctacacccaaaatctcctctctggcacaaattattcatgatcaggatacatctagaggcctagaccaaaattcaccacct**ccctcctgagccagaatggaatggaggaggg**gac**tggggaccccagataatttcccca**tcaccactgtgactctatggccacctcctgcaacagttagggataatcacataggaagggcttcatgtatcctaggaaaaacacccagag**tcacttatgagctctaccatgaccaagtga**gcccaggagtcgggttgagcacagcgtggtgtggagatatctgcagagaccgtggaaccagtttgtaaaggagatcctgtaggctgagggtgagagtgcaaggccactcctccccctctctgcatcctttggggctaagcccttctctggaaaccacaaagctcctccagt**agcagcccttgactctgctgatttgcatcacaggctgctctcttcagcaaggggataagagagggctggaaggaacctgcccagcc**tgggcctcaggaagcagcatcgggggtgccgcagccATGGCCTGGACCGCTCTCCTTCTGAGCCTCCTTGCTCACTTTACAGGTGCTGCCCCCAGTGTCCCAGCCACCT**ACCCAGCTCCAAGGCTCTGGGT**CCAGCCTGGCCTGACAGTGATCTCAGCAGGGCCCTGCCTGTGGTGT**GCAGGATGCTCATGATCCTGC**TGCA**GGGGGAGGGGCTGCTGGAGGTGAAATCCCCC**CACACTGTTCTTCTGTGCTCATGGTCC**CCTGAGGACACTTCTATTCCTGAAACTCAGG**CCAGGCAGGTGGGAAGGCATTGTTGGGTTGAGCCTCTCAGTTTCAAGTCTATTCTATTCTCTCCCCTTTTCTTGCAGGTTCTGTGGCCTCCTATGAGCTGACTCA**GCCACTCTCAGTGTCAGTGGC**CCTGGGACAGACGGCCAGGATTACCTGTGGGGGAAACAACATTGGAAGTAAAAATGTGCACTGGT**ACCAGCAGAAGCCAGGCCAGGCCCCTGTGCTGGT**CATCTAT**AGGGATAGCAACCGGCCCTCTGGGATCCCT**GAGCGATTCTCTGGCTCCAACTCGGGGAACACGGCCACCCTGACCATCAGCAGAGCCCAAGCCGGGGATGAGGCTGACTA**TTACTGTCAGGTGTGGGACAGCAGCACTGCAcacagtaa**cacaggcagatgaagaagtgagacacaaaccctttcgccatctatgtcaccctctttctctaattcaaggaggactatgaacaaagccttacgcagatctggttaaattctctttaatctctgcccccagcctgtccctccctccagtccttcaggcggggtctgcagaaagtgcatcagtagttcatttagggtgatgtgatcaggatggactggtgttttggtgactcagctgtgacttgggaaaatagggcctgcatggaagatcagactgaggtagacatccacagttgtctctggatttccataggtggtgacaggccagtccttgccccaaatgtcctagatcagtccccagaactgcccagct**caggctcctgtggtagcctg**tcattctttagagaaacaccaatggagtagacactgt**gggatctgcccagatccc**ctgttctgagcttacccactaagcactactgggagcttctggcaggtcagagctgtgtcctcactgtaaagtgctgttcggtgcagaaaacagactcctccaggtctatgccctttctcagtgaggttagatttcatgactgcc**tgaggccgagttctcgactctgcctca**atataggaagacctgaaaatcctcccagctccacagctctctgtaggaaagactt**gagacctcaatgagggtcagggtctc**ctgcccagggttgcctccctcacctcctttcacatcctctgtgcatgcaattcctcatctcagagtctgcttccagggaacccaattaagatcaccagccatccccaacatgggccatgtgggacttcaggaggccactgtatgctggaaagcacctgtttattcctaaattagcactgtgagtatccacacgcccaatgagatctgagtgtttcttgcatttgcatggaatgagaagggagcctgactcgccaggtgctctggggtctaagggagagttggagtcagatctccc

Uppercase: IGLV3-9

Lowercase: Flanking sequence[1000bp]

Red & Bold & Underline: Stem-loop [22]

Blue: Heptamer[48]

Green: Nonamer [5]

id-IGHV4-30-2-2[V_gene_segment]

ttctggtctccaacaatactcctgcatctgccatctaagtagctggaactaccgttgtgtgccaggaggtttggcttggcttgagtacttattaaaccacacactttttgcaatatttaatgtatagtaataatgtctcaataagactact**acaaatcaatgaatgaataatttgt**tcagtacag**attcatggaaaaatagacactaacatgatgaat**gtctg**acatttatgaaaatacaactgcatgaaatgt**gcttctctttacattcattaggtaaacacaatagtgcatacacatcacaccgtgctttcattacaggaagaaagatctgaaaatgtcactgggg**tgaaccacattgtgctgggcttggttca**gggagcagtcaggcccagtgttgtgaccttcaaccacagaatccttaaaaaataataaaagaggctcccccaaagtccccatcagttcccggactcgctatgtttctggatcgtatcagtgcatccggagctccctggtggctttagtgattccttgcttgccatgctgaggtctccctgtagattatgttgggttttctgaggccgttttgctatttaagacccactccctggcacagagcgattc**cctctaaacctgatagagg**ttctgaactaaaaataacattaagtgaatcctggtgtgtctgaactcaagtgattgttacattaagctgctgttgcaatctgtttcctcacctgggaaaagaggagccaggacatagtgagttgaggccccaggaagataactgaattctcagagggcacagccagcatcctcctccc**agggagagtctaaaagactggggcctccct**catcccttttcacttctccatacagaggcaccacccccatgcaaatctcacttaggcacccac**aggaaaccaccacacatttcct**taaattcag**ggtccagctcacatgggaaatactttctgagagtcctggacc**tcctgcac**aagaacATGAAACACCTGTGGTTCTT**CCTCCTGCTGGTGGCAGCTCCCAGATGTGAGTGTCTCAAGGCTGCAGACATGGAGATATGGGAGGTGCCTC**TGAGCCCAGGGCTCA**CTGTGGGTCTCTCTGTTCACAGGG**GTCCTGTCCCAGCTGCAGCTGCAGGAGTCCGGCTCAGGAC**TGGTGAAGCCTTCACAGACCCTGTCCCTCACCTGCGCTGTCTCTGGTG**GCTCCATCAGCAGTGGTGGTTACTCCTGGAGC**TGGATCCGGCAGCCACCAGGGAAGGGCCTGGAGTGGATTGGGTACATCTATCATAGTGGGAGCACCTACTACAACCCGTCCCTCAAGAGTCGAGTCACCATATCAGTAGACAGGTCCAAGAACCAGTTCTCCCTGAAGCTGAGCTCTGTGACCGCCGCGGACACGGCCGTGTATTACTGTGCCAGAGAcacaatgaggggaggtgagtgtgagcccagacacaaa**cctccctgcagggagg**cggaggggaccggcgcaggtgctgctcaggaccagcagggggcgcgcggggcccacagagcatgaggccgggtcaggagcaggtgcag**ggagggcggggcttcctcatcagctcagtgctctccctcc**tcgccagcacctcagctgtccccaggactcctctttctttattatctgtggttctg**cttcctcacatccttgtggtaggaaaggaaggaggaag**gcaaattttcctcttagagtcaaagtgtcactaattact**aggaactttcctacaagttcct**gaatgtccc**atttttccttcttaattaaaaaaaat**atatattctaatacttctcaccatctcttgatttgtgtcat**cagttgaattgtgctgtctttgaaattcaaatg**ctga**aaccttaaatccaattgatctatattggaattttaaggatggaattaaggtt**aaatgtgatcataagtctgagattctaatgcaatagatctgttgtctttataagaagtggaagagtcaccagagacctctcacttttcccgtgcacgcagagaagaggccatgtggagacatagtggactagaaggtgcaagccaagaagaagccgcaccaagaaccaaccctgccagcacctggaccttggacattcagacttcagaatggtgagaaaatcaatgtttgttgtttaagccacccactcctgtt**gtcttcttatgaagac**ccagacagactaataccacataactctgttagctcctgg**agggagaagcagctccct**gaggctgggcacatctctcagatatccatatgaagtaggcagaaatagtagttctcatataaaaatgtgtc**atggccctgttggccat**ttttttttgacggagtctcgctctgtggccaggctggagtgcagtggtgcaatctcagctgactacaaaccccgtctcctgggt

Uppercase: IGHV4-30-2-2

Lowercase: Flanking sequence[1000bp]

Red & Bold & Underline: Stem-loop [22]

Blue: Heptamer[40]

Green: Nonamer [4]

id-IGKV1-35[V_gene_segment]

ggagctgcacctgagacactaggaacatgctggtagggagcacagcttggagcattacccaggggatatgatctttgtcttcctactttggacacatttgattataagagtctggattgatacagggtttcaagtttatcctaatcaaggctccta**tagttcaatgaagaaacagagaatcttgttgaacta**at**cagatctggcatcactagccagccctttatgcgatgatctg**taagaaatattacacagtgacagaaaaggtggtagaatattctc**cccaaggcaagaataaggtgctatccatttaatccttggg**tattaaagtgaccgatagctctggaaaatggagggtctgtgcgtggtacatgcatgagaaatcatggtattatctttgtgaaactgcagtcaattccttgtccttataccagtaacaacaatggcttaaagttatgacaaact**atttttgaatattgttaaaaat**gcttctgataaagtcttattttcattttaaaaaagacaattcctcactgacag**aaatattgttaacacaacacaaaatattt**ttacactcaccaaatacatttgccagtaacttgaagacccaagagctctactttctaagattttattatgcattccctgcaaacaaggatagtctcctacatactgccaatacaactataaaatgcattactcaatgtagtccttaggttcgtttaaaa**ttttggaaattgtcccaaaa**atacgttttgtaatagtatggccaccaaagagaaatgtattatctgcaggtaagttgatgctaccctgctctgatctgggaccctggggacactgcccctgtgctgagttactgagatgagccagccttgcagctgtgcccagccttccccatcctctgctgatttgcatgtcccagagcacagctcccttcctgaagacttattaattggctggtcacaccccgtgcaggagtcagtttttgtctggacacagcATGGACATGAAGACCCTCATTCAGCTCCCGGGACTTCTGCCATCCTGGTTTCCAGTTAAGAAAAGAAATTACTAGCCTCTTACTCAGCCAGTGTGCTAGGTACTGCCTGGCCATTCAGGGAAGTCTTCTTATAACATGATTAATTGTGTGGATATTTGTTTTTCTGTTTGCAATTTCAGGGACCAGAAGTGACATCCAGTCACCCAGTCTCCATCC**ACACTGTCTGCATCTGTAAGGAGACAGTGT**CACCATCACTTGCCGGGCGAGTCAGAGCATTGGTAGTAATTTAGACTGGTATCAGCAGAAACCAGGGA**AGGCCCCTAAGGGCCT**GATCTATGCTGCATCCAGTTTGCAATCTGGGGCTCCTTCGCGGTTCGGTGGCAGTGGATCTGGGACAGATTTTACTCTCACCATCAGAATCCTGCAGCCTAAAGATGTTGCAACTTATTACTGTCAACAGTATAAAAATTACCCTatcacagtgttacaaaccataacaaaccccccccaggaaagcagacatgtgacgctg**ggctgccccacctgctcttctttgtgcagcc**atctggtgacaacacttctcagactcagcctgagttttgagggttattgggagattacggaagagggg**gccatgaaggtgtctctccaccctaagtctctttcctctcatggc**aatgtctcttcgaatttagtaaagaaaggagattagtctacctgaggagtctgtgttatggggtaattgaaattcttacagtaaaatggaagccacacattccaagcaggaattgttttaactgagtgaattagaatctaaactatagcctttgaaagcctggcatatgttattcatgaagcagatactagagatggggttctctgatgcaaaaatataaatatttattgatctg**tttttaaagatactaagataaaacttaaaaa**tgttgtcactgtatcaaatatattgcatcatatatctataaaaacagtaaaatttgtgtcatttatgaaaaatgcacatatttgagtgtatataacatacatattaatccatata**tatttatttgataacttgttagctataaata**ctaag**tacacatattcatgtgta**gataagattatttaaatatgt**ttatatatatatatataa**gcaacacatgcatacgtatgaaattattgtataaattaaatctgttaagataattccagttactcccatgaaaaaaactgatgtccaggactatatatgttacctgctctggctacagggatttcctttgtaatctttatttctaacactcaccaaatgggtatgtgcacctgaaa**atatcttttagtggaggggtaaaggatat**gaagaaaggta**attttgcctaatgttaaaattaggcaaaat**gttaaaatttcacagtgatgcctgctgggcagagttctaaattgggtgctggaccaaagccacacccctgccatcgtca

Uppercase: IGKV1-35

Lowercase: Flanking sequence[1000bp]

Red & Bold & Underline: Stem-loop [16]

Blue: Heptamer[31]

Green: Nonamer [6]

id-TRBV8-1[V_gene_segment]

tgctaatctttatttttatggtaccattgtcatttggggggacatatttgtttatctttggtaatggaaatgaaattacgttacacttcactttctgtgttttcttcctactttagggtaaatattatttccttctattcttcctattctattcctctctctttagatagttttaagggagaaaatgtcagtgagactgtaactgcagctatggggtaagaattatttagaatgagggtgggatattaagcttgataactcaaaggaacagtcagggttaaaactaatgttgggatcggggttcag**agaagttgcttgtaaaacctgcaggatggcacttct**ggaatagtctcccggctccctctgcagacacttcccagcatcccttgggccatttcccttgcaaaagcaatgatgaaatttcacttattggccacaagatggcactgtggtccactg**ggctctaagagactctggggagagcc**tgggagagcagcctaggaaggaaagggttaagaaaaatcagggctggcatccaatatcatgcagatgttgcagcagttttcagttattgctagactacctacagttatacaacatacaggaaatacctctaatctttaatgagatccacagttgaagaagtttggcctggcagccttggtgtcaagggcaggtgcagacagaggagcaactatctcagaggaaaatgggaaagtgagggg**tgggctgtggcttaagccca**gtgcgtctct**gctgcaccccatcttccctgcagc**tctcac**caggcaacagcctccatgagagtaggtggatcctgcctg**agcccagtcagaatgccccagactaggtgaactctgtaggcatggggagtaacacaacaggcccacctttcacacgtttccaaagtctaggctccaagcctttttttcccctttcctctgcagagtcctgccacttctgaagccttgtcctcactttc**tgcctgcctctcccacagCCCCCATGGAGGCAGGGA**TCAGCCAGATACCAAGATATCACAGACACACAGGGAAAAAGATCATCCTGAAATTTGCTCAGATTAGGAACCATTATTCAGTGTTCTGTTATCAATAAGACCAAGAATAGGGGCTGAGGCTGATCCATTATTCAGGTAGTA**TTGGCAGCATGACCAAAGGCGGTGCCAA**GGAAGGGTACAATGTCTCTGGAAACAAGCTCAAGCATTTTCCCTCAACCCTGGAGTCTACTAGCACCAGCCAGACCTCTGTACCTCTG**TGGCAGTGCATCcgcagccctgcacagcca**gctgctctctgcacaaaaaagggcagccacaggctggaggtgggcactccttcccaaggcctc**tatctcaaccaggagata**aagatgccttttttgaagctcttctcagatgtcccagcaatgggctcccagtctgtgcatcctcctgcagcaggaacttcagtgtgagctgaggtccatcactgtgatgtggatgcacgaatgcaaacaacacttaatgt**tattttactcaattcagaatattctggcagttgcgtaaaata**ggaatagtagctccc**aaaatgaaatttgtgtctcattttctttatgagagaaaagaaaa**aaaaacaga**aagcagctgggcagctttaacctgctt**gagtgatctgggattctacaagtctacaagtggtacaaagaacatcaccttggcctggatgccaatggataccaaacactacaaaaagtattgaaaacaatttataaaagacctaaataaaca**atctacttataggaaatgttagtagat**ccatcgcatagtgccattatacctgacgatgatcaactagtgaaattattat**gtagagattatcagctaagcctctac**ttattacctctgagcatagagcagtgtttgagccttctgagggtatcttaaggaaggaacttaattcttaccattactttggcttggattaaaatccatgagcttgtcgtttttattgtaaaatattacagaataataatgatcaaacgttttattttcctctagcgtttattcctctttagaacgtgcaggttgaagctcacacacatgagcagggttccttaagatgagatccattaactccaggactgggaggtacacagctggcctt**cacaggaagtgcaaaccccaaatgcagtgacccctcctgtg**tctacgctgtgcccaatacccctctgtgaggctgccataaaggcgggcaggctccttgcttggcaaaaaaatcagttcaccaat

Uppercase: TRBV8-1

Lowercase: Flanking sequence[1000bp]

Red & Bold & Underline: Stem-loop [16]

Blue: Heptamer[35]

Green: Nonamer [0]

id-IGHV2-26[V_gene_segment]

aacaacattgtgagttacataatgatcctactttatggccgcggtgcaaatgcttaacttccactaaatcacagagctcagaatggctagcatggaggtgtatatgtatatgtgcattgttctagaaatttcc**aattactgtatattattgtttgcagtaatt**aggtcaatctgaactgacaaaagtctccatggccttataagcacagaga**ccacctccacggcaggtgg**aaatcaaggctgggaacagtctaccagctgctcccctgggattcctccaattaccctgtttaaagcctctgctcactcatgcgct**ggaagagggatgtcagagatgcagtctactcttcc**taaagggctgacagggcataaagacaagactgattatccatggaa**cattcctataggaatg**cacctatg**tgcagacacacaaataagctgtgtgtcttgca**gt**gtgtaaatggctgagatgccatttacac**ttctttac**acaatacatttttaaatgttttgt**tgatgtattctctatcattcaa**aaaaatcatcactttccccctaaacaaagaggattttt**atcagaacactta**tgggaagccccttgctctcccagattccca**ccctcatttctcctgaaggaagaaagaaaaccatctctattgatttccactctcatcctctaggactaaaaccagaaagctgcatgctccctgggtgagccttagagaagaccctgtctgtaggagcaggaa**tctcagagccttggctgaga**ggcatggtcctgaaatgagatggagtcccccatggagatcccacgtggacgcccacacctgagggctcactgctcctcaccacagatgcactccccta**ctgagtcctgagacctgagtgcaccccatagagtaggactcagatgagggaatgcaaatctccaccagctccaccctcctctg**ggttcaaaag**ctgagcacggggcctcgctcag**tgactcctgtgccccaccATGGACACACTTTGCTACACACTCCTGCTG**CTGACCACCCCTTCCTGTGAGTGCTGTGGTCAG**GGACTTCCTCAGAAGTGAAACATCAGTTGTCTCCTTTGTGGGCTTCATCTTCTTATGTCTTCTCCACAGGGGTCTTGTCCCAGGTCACCTTGAAGGAGTCTGGTCCTGTGCTGGTGAAACCCA**CAGAGACCCTCACGCTGACCTGCACCGTCTCTG**GGTTCTCACTCAGCAATGCTAGAATGGGTGTGAGCTGGATCCGTCAGCCC**CCAGGGAAGGCCCTGG**AGTGGCTTGCACACAT**TTTTTCGAATGACGAAAAA**TCCTACAGCACATCTCTGAAGAGCAGGCTCACCATCTCC**AAGGACACCTCCAAAAGCCAGGTGGTCCTT**ACCATGACCAACATGGACC**CTGTGGACACAGCCACATATTACTGTGCACGGATACcacag**agacacagcccaggatgcctcctgtacaagaacctagctgcatctcagtggtgctccctccctacctctgcagaa**caggaaagtgtgactgagatgccatttcctg**c**cagggcttgtgtttcctataccaaccagactcagagccctg**tctgttttcctattctgtactattaaatggcatgttccctgttagtgattcacgcaagcagaggctgtatcctgtttgacaaagattgagcatgaaagattcccattacctg**ggccacatgcatcactgatatgtggcc**acttatttcttgagctcatatccttccaaggggctgaatacaaatatctataacataggccatcttaaactgcaggaagtaatgttggagaagaatgtggagataaaagagcaggatgattaattaaaatccagataccatctttttttgcagaga**aaaatttacactaataaagtaatttt**atgaaacaacaagaaagatggaatgtgtcctcatcatggagtgtctcacttgcagtgcacaaat**aaatttctaaaaattt**tatagggaaggtgtgatagatgaggctgatttccacacagggagtgattaaatacccaggtaagtataaaatccaacacttggaaaggaaaatg**cctcagcttgacatcagaaacccagctcagtaagtctgagg**agcaatgcggaaaagaaatgaaaaattcgggccgggcgcggtggctcacgcctgtaatcccagcactttgggaggccgaggcgggcggatcacgaggtcaggagatcgagaccatcctggctaacacggtgaaaccccgtctctactaaaaatacaaaaaattagccgggcgtggtagcgggcgcctgtagtcccagctactcgggaggctgaggcaggagaatggcgtgaacccgggaggcggagct**tgcagtgagccgagatcgcgccactgca**ctccagcctgggcgacagagcgagactccgtctca

Uppercase: IGHV2-26

Lowercase: Flanking sequence[1000bp]

Red & Bold & Underline: Stem-loop [26]

Blue: Heptamer[44]

Green: Nonamer [3]

id-IGHVIII-25-1-2[V_gene_segment]

gcagggatggcatgagatacggggaataaagaataaagtaactttctccatttggatataagatgtgtccattcactagagagttttctttcagaaggttcccatgtgagccag**tttccttcctgacaaggaaa**tcacccagagga**ttctcataatctcttaagtgagaa**aagttttcctcaaactccagctcagtctagctcacactgtctctaaatgggcaattactttcagacatgtacacacatctgtcccaacgtggactcctccctcagacaagcacacacattgcaatgtggactctcccctcagaaaaccacacatgccccacaatgactccttcttcacataagcatatatgcctgatgctgaactgctgtgtagcaaaattagctcagaatacaagcac**ttatggactctagccataa**caatgtgtagatctgcattttattttaaattctaaatgaacattttgccttcttgttaagaacagtggtttacagctctacatgtagagactacaggcagatatatgtttcctctgggaaaagtttattttcatgtatttactgggttattttgttgataaacattcatctataaagacacctgaaaagtgccctcattagagaaaaagaaagtgtagtttgcagattaactctgagcatccaatcccaggaatcctttga**tctgccctccctgaaatccacaggcaga**ggggggatgatcgctgataagtagcaaa**cacatgtccacaggaagggagacatggaaatgtg**gtcactcccacactcaagaggagcagctcatcccctgtccgtagtggtcctgg**tgaggaaccaccccacacctgtgccttcctca**gtatccacaccatggggtctgtgctgatctgggcttcccttctcatcactctcaatattagtgtccttgcggatcaggctcagctgtggctgctccagatgaggctgttctcagtctgttccctctgtgtttgcagAAGTCCTGTGTGAAGTTCACCGGGGGAGACAGAGGAAATAACGGTGCAGCCGGGGGCTATCT**GAGTCTCTCCTGCAAAGACTC**TGGATTCACCTTCACTGATTGCAGCATAAGCT**TGGTCCAGCAAGCTCCAGGACCA**GGGTTGATGTGGGCAGCAACAGGGAGAAATTGAAGAGGAAGCTCTCAGTGGTGCCCTCCATGAATACAAAGAATCTTCACAGTCCCCAGGACACCCTTACGTGCatggtctcactgatatctttacttcctttatcacttttgttatgtaaatcacaatgaatagtgtattcctcatctattatacatttgttaagtc**ttttttggtgtctttaaaaaaa**ctgataactttatagtatgtaatatccttaagtcctgaaagtgttttttgatgtctaccttgtcttaaatttatacagctactatagcttttcttggttcatgttttct**catatttatcttttctatttgtgaatatgtagagaaactttcttgtacata**gctaatagttgtgtattgcttttttattttggcagtccttaaatattctaattttttatgatactttaagttttagggtacatgtgcacaacatgcaggtttaaacaaacaacctcatcaaaaagtgggcaaaggatatgaacagacacttctcaaaagaagacatttatgcagccaaaaaacacatgaagaaatgctcatca**tcattggccatcagagaaatgcaaatcaaaaccacaatgagatatcatctca**caacagttagaatgacgatcattaaaaagtcaggaaacaacaggtgctggagaggatgtggagaaataagaacact**tttacactgttggtgggactgtaaa**ctagttcaaccattgcggaagtcagtgtggcgattcctcaggtatctagaactagaaataccatttg**acccagccatcccattactgggt**atatacccaaaggattataaatca**tgctgctataaagacacatgcacacgtatgtttatagcagca**ctattcacaatagcaaagacttggaaccaacctaaatgtccagcaacgatagactggattaagaaaatttggcacatatacaccatggaatactatgcagctataaaaaatgatgagttcctgtcctttgtagggacatggatgaaactatctattgcttttttaaatcaactataataaattctatttttttttttgagaaggagtctcgc

Uppercase: IGHVIII-25-1-2

Lowercase: Flanking sequence[1000bp]

Red & Bold & Underline: Stem-loop [16]

Blue: Heptamer[44]

Green: Nonamer [1]

id-IGHV1-58[V_gene_segment]

ggaatggtttcactaattgaaaggatacagaattgtgggagtcactatgttcctatgaataaaaaattcagatttcagtgttaagtaatgttgcctacattgtgtgagtgacagggcagtggtggatctgagagtgtggcaggtgcacagacctagtgagtcagaaatcaatatggaaagatgaggatctatggat**atgaactgaaagtaagtaaacagttcat**gaaattctattaaatggagtaggaaataaaacccaaacttatccaaaacacaaattccttggcgattat**tttgggagcagtgagttcatcaggaaccccaaa**cttctcttacgtcttctgattcctgttgtccatgagatgagaaattcagctctaattgtacatcacagggcaaatctgtaaaccaggagtgtttctattgaggatcatggtggatcaggattccaggcaggtgctggagacactgtctcaggagcgcccagatgatctcagggggac**ctgctggacactcacgtgggacatcagcag**tcactttctcagagtaaccagtgagctgtgct**ggtgcctgatgggactaggatggggtcaaggcacc**ttctcagtgtcatggagagtgattgttccagaaatcatccaggtggtctctacgctaatcaaatatgggttcacagggaggaacatgtgctctgggtgcttggtcttcagtgaaaggacgtctggccaccaaaagtttgtaaatggagcagggcatgcatttcctcaaggaggattagggcttggag**catcagcatcccactcttgtaaggctgatg**tgtcatttaccttccctttcttatcccaaatcagggtcttcagctatgaaatgctctgactcatgaatatgcaaataacctgagatgcactgaggtaaatatggatatttgtcagccctgagagcatcatccagaaaccacatccctccgctagagaagcccctgacggcacagttcctcactATGGACTGGATTTGGAGGATCCTCTTCTTGGTGGGAGCAGCGACAGGCAAGGAGATGCCAAGTCCCAGTGATG**AGGAGGGGATTGAGTCCAGTCAAGGTGGCTTTCATCCACTCCTGTGTTCTCTCCACAGG**TGCCCACTCCCAAATGCAGCTGGTGCAGTCTGGGCCTGAGGTGA**AGAAGCCTGGGACCTCAGTGAAGGTCTCCTGCAAGGCTTCT**GGATTCACCTTTACTAGCTCTGCTGTGCAGTGGGTGCGACAGGCTCGTGGACAACGCCTTGAGTGGATAGGATGGATCGTCGTTGGCAGTGGTAACACAAACTACGCACAGAAGTTCCAGGAAAGAGTCACCATTACCAGGGACATGTCCACAAGCACAGCCTACATGGAGCTGAGCAGCCTGAGATCCGAGGACACGGCCGTGTATT**ACTGTGCGGCAGAcacagt**gtgaaaacccacatcctgagagtgtcagaaacgccaggaaggaggcagctgtactggcatggaggggatgacaaaggttattaga**ttgaagattttcttagaaaacgacttcaa**gtcattaaagaagaggaacaac**ataaatgtgtatttgtgaaattttaattgagagatttttcatacaacatttat**tctgtaagctatttcagggattggaatatgaatcaaattaataaagctgatatagacatcctctgaaggcatcttcgtaaacatcaatttctgaatcagtgttgtaaatattttggaacacagacacaagatca**catttttactctacttttatctctatttttaaaaaatg**ccaaaaagaatcttattttgtgcatgccccattttgaattcccaccgtcaatgcatgatagttcttggttttcctcattcaagttgtcatttgtcattaagagtgttgtgtgttttaaccattctaataggtgagtaacggtatctaatttttatttaaatgcacatgtcccta**aaaaattcatatttaacaatttttatataattttt**ggtgagatgcctctcgtgatatttggttcattttttaaatgcattttttatcagttgtaagtatgcttgcatattgattataaaagtcatttaacaaattaaaataattcatttaacaaatatgcat**cttggaatttttttctccaag**tctgcagttgtcttttactcccttatccctgtgtattgcagaaaaatgtttgtgtgtgtgtgtgcctgtgt**atgtacaaatttagattttaaaaatgtacat**ttttattcatttaaagatca**tgtctttggcagtatatctgaaatctcattgtaaaagaca**caatagtcattattttttccatgtctctaatcttaggaca**caatcaactcatgagtgtttaaccttccctacctgattg**gaggactatccacctgagacatttggaatacttct

Uppercase: IGHV1-58

Lowercase: Flanking sequence[1000bp]

Red & Bold & Underline: Stem-loop [17]

Blue: Heptamer[46]

Green: Nonamer [8]

id-IGLV1-51[V_gene_segment]

tggtctaaacaagctaaagttttaattcttccatctt**caaatacggagcatatttg**ct**tgttttgctctttagcaaaaca**ctgcaaagtcattcgccaggatctcctgcagactggcattcagtgatggagaacagactcctaggtgagatccctggtcacagga**tccagatgactatgactgcaagtctgga**aaagaagagacagaacaaagatttaccagcataaccctctgagaatgtgccccagccatcctggagggaagagctccctattgacccagcactgcagaatccacagaggaaggacagctctggccaccagggagcagccacacgacaagagaggacccggtgtccccacaggtgaggaaatgagtccatggggctggtcagtgctctcgatttcattctgccataac**aggactttgtgtagtcaccacctgagtcct**atacagtgaacagccaaggagacatggacagattcacccctgaggaacccagtagaatgtgaagtctg**ccctgaggtcacgaagaaaagactcaggg**ggctgcctggcc**tggccctgatggagggggcca**tggacagaacaacggggaggcaggggtgtctctgagaggctctggtcaccttgtcataca**atggtggtgtacaatgtcgcaccat**ggacactagggggcgcctgcgcaccattcctgagaagactgggtgtgatgag**agcaggaccagcgccacctgtcctgct**tggt**gccctatgcttagggc**tcacagatgtcaactctccaccccctgggaccacacagccccacccctggcactctctgacatcctcaggcagaggagcttgacccagggcccagggtgggatcagaaagct**ggagggtctgatttgcatggatggaccctcc**t**tctctcagagtataaagaggggcagggagaga**cttggggaagctctgcttcagctgtgagcgcagaaggcaggactcgggacaatcttcatcATGACCTGCTCCCCTCTCCTCCTCACCCTTCTCATTCACTGCACAGGTGCCCAGACACAGGGTCAGGGGAGGGGTCCAGGAAGCCCATGAGGCCCTGCTTTCTCCTTCTCTCTCTAGACCAAGAATCACCGTGTCTGTGTCTCTCCTGCTTCCAGGGTCCTGGGCCCAGTCTGTGTTGACGCAGCCGCCCTCAGTGTCTGCGGCCCCAGGACAGAAGGTCACCATCTCCTGCTCTGGAAGCAGCTCCAACATTGGGAATAATTATGTA**TCCTGGTACCAGCAGCTCCCAGGA**ACAGCCCCCAAACTCCTCATTTATGACAATAATAAGCGACCCTCAGGGATTCCTGACCGATTCTCTGGCTCCAAGTCTGGCACGTCAGCCACCCTGGGCATCACCGGACTCCAGACTGGGGACGAGGCCGATTATTACTGCGGAACATGGGATAGCAGCCTGAGTGCTGGcacagtgctccagcccaatggggaactgagacaagaacccccttcttcctcccccaggagggtgagtgccgccagctgctgctcacgcctgacctgtagcttctgctgctgcaggttctcccatgggccacggggcagccagggccctgcctaggagtggaggctccaccacttttgtcctcagagt**caggaacagggactccaggaatgagaatatcctgctcctg**cagcttgggacacagggtctctgcactgacatcctgggctgaggtggcaggtccagctgtgtcttcacagtccttcctgtgcctgcccatggtgtggggagggagggagtgggggagggggtgtgggcctttgagcttcactagagacctga**gggagcagaggtgctccc**caagacctggactccttccctcaccctcctcccatcactct**cctcactgtagctatggccaggtgagg**gctcagggctacttgaggaccttgctcagcccagggcctccatcccaggcattgtttccaaactctgtaagggctgtctctgaatctgtagaaacagttatttctgt**gtgcccaggactgggttagagggcac**caaccccaccccttgcaaa**ctgctcctcacctcttagcgccccagtgaggagaag**aaagagttagaaatcaggaatctactcagggctcccctctctcccctttcccagcctcccctggctttctctttctacagaaacaccaaggaggcagccaccaaggacgaggggaggcccaggcattataccaggcagcggagccagccagtccc**cccagatggtggagggaggtggtggaggtgtctggg**accagaaaatagtgaggaccctcttactgaagtgaggacgggtggcataaactgaggccc**agatggtggaccagatctcctgagccatct**actcatttcttcgtgtgttccagtttgggtaagaataggctaaaaaaaaaaaaaaaaggctttgaa

Uppercase: IGLV1-51

Lowercase: Flanking sequence[1000bp]

Red & Bold & Underline: Stem-loop [19]

Blue: Heptamer[37]

Green: Nonamer [2]

id-IGKV2D-40[V_gene_segment]

tatttcttatttgttctgtatcattaatctagaagattcttcccttaagacagagtcctctgggccaggcacaatggctcacacctgtaatcccactaccttaggggccaaggcaggcagat**ctcctgaggtcaggag**ttcaagacaagcccagacaacttggtgaatgaaaccccatctctactaaaaacaaaaacagaaacaacaaaaaagaaagagccctctggttaaccttgtatgtgtgagacgattatgatgagatagatcccagattgaacaactggtcaccaggaatttt**aaatttgctgctggaggcacaaaattt**tgtctctctttcctttttcttacactgggctcttggctctaaatgtagagg**ctcacatcattctccctgtgag**gcgcttggacagagagctcttatgctgttcactcaccaggtgccaaggcagagtagattctaatatttgagttgaacattcttgaacagttatcctggaaacagtagataccagacagcccttgaactggctccaggccgctttttatttgcaggctctcagttcagcagtgcttgtggggatgggcctgtttcatactctagattga**ctgggagggaatcaagccagatggcattcacctcccag**agatgtatcctagacacacatttccacattgtcagggttctggtgctttcttacagtcatgccctacacagtgtgtccctacaaaaggtccgaactttcaccttcagatccttcttcccttgattgtgggcaaacttggctgaatctagttctgttttattccaaaggacaatttatatcacattgttcacagaagagacattccccctgccccgtcaaccttttccacaccactgcacccaccaggtgatttgcatattgtcccctagggtggaccttcccttgtgagtctgagataaaagctcagctctatccttgccttgactgatcaggactcctcagttcaccttctcaccATGAGGCTCCCTGCTCAGCTCCTGGGGCTGCTAATGCTCTGGGTCCCTGGTAAGGACAGAAAGAGATGAGGGAGGACAACTGGGTGGGAGGTGAGCT**CTGTGGGCTCCACAG**CT**TCACATGTTTATTCCAATAATGTGA**TAGAGGCACATGGTCTATGCTCCAGGGAATGGAATTCAGGTTTGTCTTATGAATAATCAGGATTCACCTCCAGGGAACGATGACCAGTGCTCTGATTAAGAACTTGAAAAAAAAGAGTTCCCTTGTGGCTAATAAATAATGGGTCTATTTTAGAAAGTCTACTTTTCA**TGATATAAATCAAAACTTTAAAAATGTAACTGTAAATTTATATCA**CAAGAGAA**ATTATGAAAGTTGCTCATAAT**GTATCTATATAAACTTGCACTTCTCTGTTATTATTTCAGGATCCAGTGAGGATATTGTGATGACCCAGACTCCA**CTCTCCCTGCCCGTCACCCCTGGAGAG**CCGGCCTCCATCTCCTGCAGGTCTAGTCAGAGCCTCTTGGATAGTGATGATGGAAACACCTATTTGGACTGGTACCTGCAGAAGCCAGGGCAGTCTCCACAGCTCCTGATCTATACGCTTTCCTATCGGGCCTCTGGAGTCCCAGACAGGT**TCAGTGGCAGTGGGTCAGGCACTGATTTCACACTGAAAATCAG**CAGGGTGGAGGCTGAGGATGTTGGAGTTTATTACTGCATGCAACGTATAGAGTTTCCTt**ccacagtggtacagccctgaacagaaacctccctgctgtgg**tg**ccccagctgctcacatgcactgcttgtctgggg**agcaggtcagcagcgtctctgagtctgcaaaagaggaggctgttggagaatacagggcagggtttgcttctgaggactctgcctgggactacaggtgcatgccactaaacatggctaatttttctatttttttgtagagtcggtgcttcaccatgttgcccagcctgttgtcaaaatcatgggctcaagccacccacctgacttggcctcccaacgtgctggcagtacagtgtgagcca**ctgcggcaggtcagcacccctgtttatgttcctgtcacctgccacag**ccttgactctcataaccaacaggaaaatgaggaggttctagggccctgtgagtaaaaaactgggatgataggga**aaggagaatggaatctcatctgaatcctcctt**ccttgcctacatttgtttaaatttattgagcaaaagg**gccagactactgatcatttctggcaaaacatgttgagtacattttagggtttaacagttttg**ggtacctttcaaa**gaaaatatttggttatatgtaaaattggtattttc**ccactttttaaattcctacttctcctgtttgccattcttctccactccatgagacagtagagacaggattattcactaattctcctctggcggagctggctgaggacaatcagtaagatcttggttgtgagtgtcaaatagattttgtaatttcatagcagacgcaagttcctaatactaaaactctttgattacaattacctcttgctgatagaaaaagggagttcttgaaattttgaaagttggttttaaaaataaaatgcatacactggaagatgcagtatactaaagatgta**gtattttccagggatcactgagaaaatac**aggatgaggtaaacagttttattttccaaagttcagaatttgagattgggcagactgca

Uppercase: IGKV2D-40

Lowercase: Flanking sequence[1000bp]

Red & Bold & Underline: Stem-loop [19]

Blue: Heptamer[37]

Green: Nonamer [5]

id-IGKV2D-30[V_gene_segment]

agtgataaagccatcagctaacttttcatcatgaccaaaggaggctagaaaacagtggaatgacatatgtaaatgctgggaaacaaaagaaaacaaaacctaggattcaatatccagtgaaagtatgcattcaaaataaaagtaaagtaaaaac**atttcttgataaacaaaaatgaagagaaat**aattacttaaagacatgctgtataacaaattctctaggaagtt**cttcagattcacaaaaaatgacagcagacagtaacttgaatctgtag**ggaggaaggaaggcctcaaaagtgctaagggaacaaaaaaagagctagaccaaaccaacatgtttacttccgtttatatgaggtttcatgtagggaaagccaatctatttggacagatgtcagaatgccagtttccttgattgggatggggggaacagctgtttcctgaaaggtaggtacatgaggga**agaatctggagattct**gcag**tattctacagtttaatctcggtggagaata**tatgtaaaactttattcggttgcactttttaacatttctgtcttttactttgtgtgtgtttta**tttaaatttttaaaaaattgaaa**gggccaaatctgaactcttttaaacaaaaatgaacaaaaacataagaattagtaaatatttgtggaaacatggccttattaacaagaagtataaaatgtgcctgggaaagtgctatgaaacaagaaatctgttagggaagacagaaggaaatacttaaatttctcccacatagacagcatagattatatgcctactcatttccctccaaacagagaagatatttaagtcattttgctcacaacagaggctcctaccctccccttggctctttccaccccactacacccaccaggtgatttgcatattatcccttggtgaagactttccttgtgagtctgagataaaagctcagctctaaccttgccttgactgatcaggactcctcagttcaccttctcacaATGAGGCTCCCTGCTCAGCT**CCTGGGGCTGCTAATGCTCTGGGTCCCAGG**TAAGGGTAGAAGGGAGATGAGGGAGGAGAATGGCATGGAACGGTGAGTTCTGGGGCCCCACTGCCTCTAACAACAGTGATCTCTGGGGGTCTCACTACACTCCTATGTGTGTTCCTTTCCTGTATTGG**ACATGCACATGTTGTCCTCCAGAATGGGGCATGT**GATGATCAGATCTGTGAGAGTCAGGAAGATTCAAGAAGAAACAAGGA**TCTGTGCTCTGGGGAAGACTGACACAGA**AAGGGGATGGTGT**GGGGTCTTCTGGAGACCCC**TTTGAGCCTTGGATCCCTTGAGTTCCATTTTGAAACTGT**ATATTTTTGAAATAT**GAACAAATACATATATAGCCTGAGATAAACAACAAATC**AAAATTTATGAAAATTACACATAAACTTT**ATACATAACCTTGCTCTTCTTTCTATTTATTTCAGGATCCAGTGGGGATGTTGTGATGACTCAGTCTCCACTCTCCCTGCCCGTCACCCTTGGACAGCCGGCCTCCATCTCCTGCAGGTCTAGTCAAAGCCTCGTATACAGTGATGGAAACACCTACTTGAATTGGTTTCAGCAGAGGCCAGGCCAATCTCCAAGGCGCCTAATTTATAAGGTTTCTAA**CTGGGACTCTGGGGTCCCAG**ACAGATTCAGCGGCAGTGGGTCAGGCA**CTGATTTCACACTGAAAATCAG**CAGGGTGGAGGCTGAGGATGTTGGGGTTTATTACTGCATGCAAGGTACACACTGGCCTcccacagtggtacagccctgaacaaaaacctccc**tgtggggtggcccagctgcccaca**tgtggtgcttgtctggggagcagctcagcagggtctcagaatctgtgtaagaggaagatgctggagaaccagggaacaattcacag**ctgaggactctggactttgagagcccagtcacacctcag**gcaccactcctttatgccctgccagttgccaccaccttgactgtcataagcagcaggagaatgaggggtccaagtgccctgtgagtaaacaagcaagat**ggaagggagaggagaatgaaagctcaccctaactctccctacc**ttgtgtccatttgttaattaaatgtaatcagcagagcagccaggccattgacacagattgtgactatccatgtgggatacatctttgggtttagcagtttttggcatattgttcagaggacatttgataatatttgatgttggtattttgccagttttctaacttcctgcttctccttttctcccactcccaaaatagagtagagacagcattctataccagttatcctaagcggaagctggctgaggacagtcagt**aaaaatcttgattttggagtatcaaatagattttt**gtaaattcatagaagatacaagatc**ctaatgctaaaactgtttagttagccttag**ttacctcttaatgatagaaaaaaggagtacctgaaattccagaagttgttttcaaaaataaaaagcatatact**ggaaaatgtagagtatataattttcc**ccatagaaaac**tggaaaagtacagaatgatgtgaatgtttttatttcca**aatgttaagaatttaagattgggcagacattagggatgaaagcatgaaggaaacccaggagaggtcagcttcaattaaactacccttggcctttggtaggtagggatgtggatggtggtggagttggcagttcatgatgtggacccaggaggcctgatgtgtttcttgtgaagaatcaca

Uppercase: IGKV2D-30

Lowercase: Flanking sequence[1000bp]

Red & Bold & Underline: Stem-loop [20]

Blue: Heptamer[28]

Green: Nonamer [10]

id-TRGV2[V_gene_segment]

atactttaagtgctaggg**tacatatgcacaacgtgcaggtttgttacatatgta**tacatgtgccatgttggtttgttgcacccatcaacttgtcatctacattaggtatatctcttaatgctatccctcccccctccccccaccccactacaggccccagtgtgtgatgttcccccactctgtgtccaag**tgttctccttgttcaattcccacctacgagtgagaaca**tgcactgtttggatttctgtccttggaatagtttgctcggaatgatggtttccagcttcatc**catgtccttacaaaggacatg**aacgcatccctttttatggccgcatagtattccatggtgtatatgtgccacattttcttaatccagtctatcattgatggacatttgggttggttccaagtctctgctatcgtgaatagtgccacaataaacatatgctttgcgtggtgcctccaagtgtcctcgcagatgccaaa**aggaaggtggttcagttccctaatcagagacttcct**gg**caggacaaggtcctg**gcatcagcattagctgctgcaactctgaa**gggcagggtgtgagcctgctgtgggactgccc**cctcctagggctgggcctgctgactccagggtctgcttcccccacctctgcactccctctgctgcaggcctgagctctccttgcccctggctctcttctgcagagggcctgctctcagctgccacaagagggcgccggagaac**ggacctgagcagagaggcaggggctcctctgcaggtcctcccaggcaccaacccttctgggagg**gaggaaggcttgggagtcctcttagacagatcctcagtcacttccctctgcttgtgtctcag**gaagaccacctcctcctactgtcttc**tgtgctagggatcacttccttgttgagtggaacctgagtttt**aagaggatcttctgctcctctt**catctggtcccttt**ccttccaaggccccagagaggaagg**cATGCAGTGGGCCCTAGCGGTGCTTCTAGCTTTCCTGTCTCCTGGTGAGTGCGCTGCCTACAGAGAGGATCACGGGTTTTGTTTTATTTTCTTCTTTTGCAAGGAGTACCATACTAAGGAATTCCTCATTATATTTTGTGTTGTTCCCATTGCAGCCAGTCAGAAAT**CTTCCAACTTGGAAG**GGAGAACGAAGTCAGTCATCAGGCAGACTGGGTCATCTGCTGAAATCACTTGTGATCTTGCTGAAGGAAGTAACGGCTACATCCACTGGTACCTACACCAGGAGGGGAAGGCCCCACAGCGTCTTCAGTACTA**TGACTCCTACAACTCCAAGGTTGTGTTGGAATCAGGAGTCA**GTCCAGGGAAGTATTATACTTACGCAAGCACAAGGAACAACTTGAGATTGATACTGCGAAATCTAATTGAAAATGACTTTGGGGTCTATT**ACTGTGCCACCTGGGACGGGcacagt**gattcagatccgccctacaccacactgaaaatctgc**cttgtggctgcttctggtacacaag**atagagccgccccctctcatttcctgccaccaaatttaccgtgtgctgaacaagagaaacattgcttaactcctgatctcccctgcaatattacactcccctggcagcagctgcaccctgttccccaccctcccccaggactttcctgaagaccaagccgcca**tctcagctaagcctcagagaagcagcctggctgagagcaaggttctcttagctctcctaggacatgggggaggcccactcactctgcttcctagga**caaatgggtacccctagggtccagctgtaaagcaagatattttggacaacaa**atgggaagggaatttatactaaattgtgtatcccat**catgatccagaaaccgcagctgagactgg**tgcttattccttatgtctataacaatataagca**atactataatgaccactgaaacaccaggcctaggtatgcagttgaacattctctccctctgccctcccttttattctctctctcttttttcttaattctttgcttgcttttattatttttctctctcccttcctccctcgctccatcgctccctccgtccttccctccctctctccctctctccttccctcccttcctctcttctttccttccatccattctccttcgttttccttccttccttctttgcttttctcttgctttcctttttgggtacataatcagaatactttttccacaaaatcatttgaaataatttttctctatgtgttcatgtcaacattgatatagctactttaattgttcagttttcttttttacattttggaaatttgttgtattactttgttttgtttcatatc**tgttttgtatatataaaaca**tttacatggaagtgaagtcaaaattataaaagatgatgcatttgaagattttagcttccattgtagacttctcattcttctt

Uppercase: TRGV2

Lowercase: Flanking sequence[1000bp]

Red & Bold & Underline: Stem-loop [21]

Blue: Heptamer[17]

Green: Nonamer [5]

id-IGHV7-40[V_gene_segment]

taagaaagacagcatgctactagtagaaaaaaatagcccagaaagaaagccaaatatctaaaaccaact**gttgtttgacaaaactgacaaaaatatacactggagaaacaac**cctctattcaataagtggtgccgggaaaattaggtggcttatg**tggaagaatagaacgagacttcca**tatcaccgtagacacaaatta**actgaatatggattcagt**gtttaaa**tttataaactaaaactataaa**aatacttgaataaaatctaaaaagagtcctctggacattggtctaagcaaacaatatacgactaagacttcaaaagcaaatgcaataaaaacagaagtagacaaacagga**tttagttgaactaaa**ggtcttctgcacagcaaaagaaataaccaacatggtgaccagacaatctgcaaatggaaaaaatatctggaatctattcatcttgcaaagggctaatatatagaatctacaagtaactcaaataagtcaactaaaaattacaaataacttcattaaagaatagacataaacagacatttatcaaaagaatacacagaagtggccaacacaaataagaaaatatactcagcatcaccaatcatctgataaatgtaaattagaaacaacatgatacggcatcttccaccagtcagaatg**gctgttattacaaataaaaacagc**aggttttgcagacaaacataggaaaaataatgatttatatatgcttggtgagaatgtaaattagtacaacctccatgaaaaacaacatagaaatttctcaaagaactaaaattagaattaccatttcttccagtaagctgtcccagtaggcatgttcctcccaaacttttatatcagagaatgttgcctgcactcatatgtttatttcaacaccattttcaatagaaaagtcaaataatctaagtgtcaatcagtgg**atgattagataaaatatgatatatgtaaatcat**ggaatactatgcagccagTATGGTATGAATTCAGTGTGACCAGCCCCTGGACAAGGGCTTGAGTGGATGGGATGGATCATCACCTACACTGGGAACCCAACATATACCAACGGCTTCACAGGACGGTTTCTATTCTCCATGGACACCTCTGTCAGCATGGCGTATCTGCAGATCAGCAGCCTAAAGGCTGAGGACACGGCCGTGTATGACTGTATGAGAGAcacagtgtggaaacccacatcccgagggagtcagaaaccccgagggaggaggcagctgtgctgagctgaggcagtggtgcagcagtttctttaacttccatatgatctcattttgcatcatcttctacttttatattagctaagaacttggggtagacaggtgctcctaagagatccttaacttgcccattttgatgggttttccagaagacgtgagaagccactttgttagcaaagcatcccaaatccatgccctgttctagatacatgtgagcccatttcctggtctttgcttaactgacaagctctcatcagtgcacctgggctaatttcacatcaggtagaggaacgcgttataaaggaaagctaatgttgtaatagcaattcctgctt**aaaaaccttcagcttcattgttttt**gtgtaatccatcagcaaattatgttagttcaaggttctcaatgggag**tttctaataaatagaaa**ggatgtatagagcttcccctaattaaaatgaaacaattgtgaacacaacctcggtattcagccatgtctccacccttcacacccttcgccacaaaggaattttcacctctcctggaagctgggttcattttcaaattagttatttttttcaatgtaatatctcaagattatcgtatatgactattttagcagaaagtgaattatgggaacttgaactaaacaactgaaaacaaattcacaactaattaaacaagatgccagaatgtgattggctccaggctttgtaattcagcagtt**catgtacccagactggaaatttacatg**tcttcttgttaccttcacagca**cagtcaactcccattatgtaagaaatggtgactg**cattcccaagggttatgcatagatatgaaaatagactgggtaaggtgag**gagttgattgtttaaattcccctctgaagaagcagcatcaactc**aacaaaccacctctcttcactctgtgactagagctatgtcacagg

Uppercase: IGHV7-40

Lowercase: Flanking sequence[1000bp]

Red & Bold & Underline: Stem-loop [12]

Blue: Heptamer[24]

Green: Nonamer [4]

id-IGHVII-65-1[V_gene_segment]

caccaccatatccggctaatttttgtatttttagtagagacagtgtttcaccacatt**ggtcaggctggtctcaaactactgacc**tcaggtgatccacccgcctcagcctcccaaagtgctgggattacaggtgcaggccaccaagtccggctaatttttgtatttttagtagagatggggtttcacaatattggtctggctggtctcaaactcatgatctcagcctcccaaagtgctgggattacagatgtgagctactgtgcccgg**ccaacatcacctatattaatgttgg**attgaataactacaaagtctaccctaaccaagtgtaccatcaaacttaccattacccagagggaagaatctttaacatgagatcgatttcttaagtgttttaaggt**gcaaaactgaccactgtacaatcacccaattatgattttgc**taatgagttgtagaagttgcatgtacattttggatattaacactttttcagatgcacagcttataattatattctctctttctgtgggt**tggaaattttttcca**tgctttgcagcatctttttgctgtgatgtagttctgcttgttcaattctgcttttattgtctgtgcctttaatttgaaatataggaaatcattcttaatattattttctggg**tgggggaaattttacaattgcaagccatacgtttaagtatttcgcccatttagagtaaattttggtatttaatctaaa**ctaaaattcttaattctttgcatgtgtagattcagtttttctaacaccctcttagggagagactttgattcatccattgtatattgttagttctcatgctgaaaatcagtttgctgtagatatattagtttatttctgagctctctctatatattaatgccaataccatcctgttattatttatggatcaaaatgtgcatatttgtgaatttcaaaat**gttgttcagaacttgtgcatgccaacaac**tgtgtttctcctgcactcttGGGCTAGTGAAGCTCTCACAGACCCTCTCTCTCACCTGTGCTGTCTCTGCATTCTCCATCACAACCAGTG**CTTCCTGCTGGAGCTGCATCCATCACCCCCCGCAAGGAAG**GGACTGGAGCGAATCAG**GTGCACAGGTCATGAGGGAGTGCAC**ATTCCAACCCACTCCTCAAGAGTCCAGTCACTATCTCCAGATCCACATCCAAAAAACAGTGTTTCCTGTAGCTGAGCTACCTGAGCAACAAGTACACAACCATGAATTTTAATACAAAAGAcacaacgaggggaagtcattgtgagcccagatacaaacctccctgcaggggagctcagaaagagcaggaggcactcaggacaccagggaacactctggacacatcaaggcaggtgcaatggggggacaaggggctggagatggggtttggcatcaccatcgtatttcaccactggacacccaccaccgtgtttattctcatgtacctgattctgtgtattattagaaaatggcatttatataaatatatagccatatgtaggtgcatcaagttgtcctctccatcgtatgtggacctttccactaagactctgagtccctgtattta**tatgagcacctcata**aattatggtcaattttgtgggatctctatcttttctttctcccttcttccctccttctctctcactcacaaactcacacacacgcaaaattctacaacttcagttacgtgatgtgttgaagacagtttataatgtgcagcttttccagtttagctg**atgttcatattgctgtgagaataggaacat**tgtgtttctcagctgtgtatttctctaaggtagcagcaacttttt**tttttaatttgacattttttaaaaa**ttttattattattatactttaagttttagggt**acatgtgcacaaagtgcaggtttgttacatatatatacatgtgccatgt**tggtgtgctgcacccattaactcgtcattta**gcattaggtatatctcctaatgc**atttttttaaatacccagatactgaaagcaatccaaatatcaatcagcagcttaatcataaacaaattgtggtaaattcattccctggaaaatcacccactgttacattctggtaccgctggatacgttcaacagcaaaggtaaagtcacaagtacatatgatgagtcaaataagacaaataaatacaagtacacacatacaattcctc**ttttataaattctataaaa**ttaataccaatctaaagttacataaagaaaaccagtagt

Uppercase: IGHVII-65-1

Lowercase: Flanking sequence[1000bp]

Red & Bold & Underline: Stem-loop [15]

Blue: Heptamer[28]

Green: Nonamer [4]

id-TRAV15[V_gene_segment]

ttaagtatagattaaggtttagctttccactctcctgctattgattatattattttactatggttaaggctgagtcaccctttatttcatattgaccttattccatccaattatttgaactgactgtaatggctttcttgtttggtttctttagttcacaggttttatgcac**agaatgaaacattct**tgcccctggtgtaactta**atgtaaattctaagatgaattcacat**gagccttatggagcaggtg**tcagatataatactattatattga**gctatggtgccaataattagatccgtatctgggggctcatttaacaggtttctgcatt**gaaaggagagaaaataacctttc**taggctttccttggggaggaatgctttttggatattcaatttatgacttgtacatacatggtagatgctcaggaaggattatagaagaaaaaaaggaaggacaacaatctggcagacaaacaagcatcactctccaacaaatcagcacctggtcctaccaaggtccttttccatgttacctcttcagacatcaggaggagctgtgttatctgtgcatgggggaatggatcttgtttgtggttggctgggaggactctcagcttcctgtaataacagaactatgaatattcactcagagaaacaagctattggcttggtagtgtttgagggctcccagaccccaaccagagtccgcagtaagtctcatggataaagtggagaagcccttgggagtttgtggagaagcccttgggagttttgttcttgattttctcttggcagctgtgctgtgagttttggggctctggaaatatcattgaaaagagcagtttgtagtatcttctttaaaagtctgcacttggacacaggaaggggagcataacacaccggggcctgttgtggggtggggggacggggtagggatag**cattaggagatatacctaatg**taaatgactagttaatgggtgcagacaccaacatggcgcATGTATACGTATGTAACAAACCTGCGCGTTGTGCACATGTACCCTAGAACGTAAAGTATAATAAAAATAAATAAATAAATAAATAAATAAATAAATAAAATAAAAGTCTGGAAGTTATAGAAGGGTGGTTAGGGCCTCTGTGCTCACCTGTGGGGACTTGAAAGGAGAGTATATCTTCCAAAAGTAGTCAGCCATGAATGAATGTCCTCTGTCTGTGGTTCTCTGTTTAAACAGGGGTGAACAGCCTCCATATTCTGGAGTAGAGTCCTTCATT**CATTCCTGAGTATCCGGGAGGGAATG**CACAACATTCTTAATTGCACTTATGAGGAGAGAACGTTCTCTTAACTTCTACTGGTTCTGGCAGGGTCTGGAAAAGGACTTGTGTCTTTGACCTTAATTCAATCAAGCCAGATGGAGGAGGGAGACAAACATTTTAAAGAAGCGCTTGGAAAAGAGAAGTTTTATAGTGTTTTGAATATGCTGGTCTCTCATCCTGGAGATTCAGGCACCTACTT**CTGTGCTTTGAGGcacag**ggtccccaagcacctgcagcctgtaccaca**acctgcatccgggacccttgacacagccttgccttgcaggt**gggagtgaaggtgttgtctttatatgtagagagaacttctttataaactctgtctccaaacagaagcgaagggtagtgcagaatctttcttatgtgatgaagctg**aatttatattaaatt**aaattgcattttgttatt**attgaaatttcattcttcaat**gtagattgtgctgtgctggttctgggcctgtttgcccaccgatcccttccacagctctgtcctttactctaaagtcgcagaccactgtggcacaggctctgaggccaagaggcaagttttctccgtgctatactctcaccaacagtattttacccatttgcttgtaaattcatgatgtttcttgtatttgtatcatctatactattatttac**ttaattaaatgttaaattaa**cttttatgttgagtagacttcaaaactgtg**taaaatattgttcttatttta**atggatactatcagtgaaa**ttatattttatataa**actatacatctgatgtgcaggctat**gtttgttcctgatacaaac**taaaataacattacatgtataacaattaaaaagaccatctgtgtactgcc**taacatgacctcctatgtta**accacaatttagaaaatactgattcagctcatgtccctcatttaatagatgaaaaaagtagaatcaacagggacagcatcacttactcaaagcatgaattacagcaaaatggaagctaggaatggaacacagacattgtggccccttgtcacttgctttttccatgaaggatgtaaagtaggattcagtgtttgaaaaaaggagtggtatgttg**taattttaaactaaatta**tacttttattacttaattatatgttcattagaataaat**taggagatataaacacccagaaaacatagaacactccta**taatttcataacctagaaatgatag

Uppercase: TRAV15

Lowercase: Flanking sequence[1000bp]

Red & Bold & Underline: Stem-loop [17]

Blue: Heptamer[37]

Green: Nonamer [5]

id-TRBVAOR9-2[V_gene_segment]

ttgaaga**aatatattcactcttatatt**ccttgaagcattattcacacagccaaaatacagaattaacctaggtttccaccaatggatgactagatga**agaatatatgacttatattct**gta**tatgtatatatatattctctatatgtatatgtatatacata**cacacacacacacacaaaaaaaatggaatactactcagccatacagaaaaaggaaatcatataatttgcaacaacatggagacattatgctaagtgaaataatccaggcacagaaagacaaataatgcatggtctcacttattcatggaatccaacaaagtcaaacccacagaagcagagagtagaagggtaattgccaagcactgggatgtggtttgaggaagggaggcatgggaaatagggaaatgttggtcaaaggtacatgttttaatcagaaataagttcaagagatctattatgcagcgtgattactatagttaataatatatttactcttgaaaattgctaagaatgtagattttaaatgttcctaccaaaagaagtcaaaaccaaatggtaagtatgtgaggtaatgcatatgt**taattagcttgatttaatta**ttcc**acaatgtatacatatatcaaaatgtcacaatatcacattgt**ataccataaatatatataatttgtatttgtcaattttaaataagtaaataaagtaatgagatagctttcaactaagc**ttatatagtggtataatataa**aaatttttgtagtaa**caaaattatagtaattttg**gcttatac**tgaaatgtgcattaaacaagtgtttgcataaaatttca**attcaaattactgttcacagcccacagaaagccacgaaaacgtgatttactcttggtatggaaacaaatagcttgtactcctaaggaaaaaagaaaattcctgccttttgatgagaattgtttgagttttatacttactttatggggagctttgctgacctcttcattccacgATAGCTGGATATTCCTCCCCTGCATGTACTTT**TGTTTCCTGGGGGCAGTGAGTCTTGGAAACA**TTTGTGAAATTCTTATTCTGAGTTTGCCTGGCCACAGCCCAGGTTCCTTCACGGACACTTCAGCATCAGGCTCTCTCCTCAGCTTTCTTTCTATCTTCTGTGCCCTCTGCTCATAAACCCCAGCTCATAGAAGCTGAAGTCACCTAGACTCCAAGACACCTGATTGTAGAGACAGGAAAGGAGTTCTCAGGATATGTGCCATAATTTCATACTGGTTTCTACAA**GAATCAGGGTTGGAATTAGAGCTGATTC**ATTATTCAGTTAACATTGACATTGTTGATTAGAGACATATCCCTGTTGAAAATATTTCCCGGCAAAAAACAGAAGTTCCCTTTGGC**TCTGAAATCTGCAAAGCCCTTTCAGA**TGTC**CCTGTGTCCTTGTGCCGTCACTCcacagcactgcacagg**catgtgctcacctcacaaaatggcagtctcaaagggaggagtgtccaaccacaagaggcgccaccctatgctgagaaagaacctttcacaggaggagagaaaagacctttccag**aatctttaacataagatt**ttttgtgtgtg**tgaagtctgacctgagcagcaaccttgcatcagacttca**aaaggcaatatttcctgtacctccttgtg**ctgtacaagtgagtacag**aagtcg**agaaaatgttcctaagtctatggccagttcaaattccatcattttct**atttctgagattaaattatatgtattaagatccttactctgccctttcatgtttattgctcctcggacaagatatcccctcttcaggtcccaaacacattcattgcaatgcctttcctcatccttgtcccagtcctggctctctagcttcttgagacaataacccatcctaaggtatgtggtgacatgcatcttctgcctcactgggatccctaagtggtataagcattcactcaattgctttagctagagtccccagaatatcttcaccctctgctgtcacactcttcttccacattggtccataaccacatctttaacatttctgtttattaaaacctctcaaattcaaccactattttctatcccaaccaatactgccttagctcaggcaaccactgtctttcacctgaattccttcaacactttccaaatatccacctacctgatgtcttactctctgttctaactcagttcattctctgcagctgaaaacacttcctttccattgactttagggttatgaccagtcttctttatatctaaatgttagcctgtaagttctttttgtaaaataaataaataaataaaaatcttccaaactacaaattcttaaaaccctccatgatttggcccttgcatacctatccagtcttgtgaaattcacctactctttacccactttttgcatacttatgctcta

Uppercase: TRBVAOR9-2

Lowercase: Flanking sequence[1000bp]

Red & Bold & Underline: Stem-loop [16]

Blue: Heptamer[27]

Green: Nonamer [7]

id-IGKV1-33-3[V_gene_segment]

cttctgctttcctgaaatttactcctgccttaaaaaacccttacctgcaagccatcagtgaggccaggatttgaatcttagctgcctgattctctttacctgatgccctacaaaaaaacaaacaaacaaacaaacaaacctttactttctcctgctgcaaa**ctcagtgtgaatatctgatctgactgag**ctgagtgagtggactccagttcggttccacagcacaagctctttttatagtggcatttatccac**tcatgagggcagagctctcatga**cttcaacacttcccattaggccccacctcccaat**tctgttgctttgagataaatttaaaaacaga**tgggttttgaaggacacagtgaaaccataacactgcccaaagagagggccattcgagcccctgtggtctgctatgcatgcaggcagagctgccttctccaggttcagaaagagcaatgaggggcatagatggccattggaagtcagcagcagtacagcgaaagggaaaggcatagcctgtagacagggattgcaatcttgatatattttctatttagtttcctttctaaaaacagaataatttcatgctaacagaagtgttgtaagtacagcacaaagtgctttcttccaattaagtgcatattg**tattttttaaaaaata**atctccatctccatacacgaataaaatacattactccatctagtcctcaggaatatttcgaa**ttttgacaattaactcaaaa**aagttctttgtaaccaaacagtctccaggaagaaacacttatttacggacaaatctgtgatgccctggtccgacctgggacactggggacattgctcctatgctgagttactgagaagagccagccctgcagctgtgcccagcctgccctatcccctgctgatttgcatgttcgcagagcacagccccctgccctgaagacttattaataggctggtcgcaccctgtgcaggagtcagtcccaaccaggacacagcATGGACATGAGGGTCCCTGCTCAGCTCCTGGGGCTCCTGCTGCTCTGGCTCTCAGGTAAGGAAGGATAACACTAGGAATTTTCTC**AGCCAGTGTGCTCAGTACAGCCTGGCT**CTTGATGGAAGCCTTCCTATAATATGACTAATAGTATGAATATTTGTGTTTATGTTTCTAATCGCAGGTGCCAGATGTGACATCCAGATGACCCAGTCTCCATCCTCC**CTGTCTGCATCTGTAGGAGACAG**AGTCACCATCACTTGCCAGGCGAGTCAGGACATTAGCAACTATTTAAATTGGTATCAGCAGAAACCAGGGAAAGCCCCTAAGCTCCTGATCTACGATGCATCCAATTTGGAAACAGGG**GTCCCATCAAGGTTCAGTGGAAGTGGATCTGGGAC**AGATTTTACTTTCACCATCAGCAGCCTGCAGCCTGAAGATATTGCAACATATTACTGTCAACAGTATGATAATCTCCCTcccacagtgtaacaagtcataacataaatcacccaggggagcagatgcgtgaggctcagctgtcccagatgccccttctggtgccttcgcctgc**tgagaatgtttctca**aattgcagtcacactttgaagttcactggagagtttttgtaaaagggcc**atgaaggcccacttcat**cgtag**ctgtctttccttgtcctaatccccagtatcatagacag**ggcaatgcctctcctgatttcattgagaagaaatggttacacctgaggggtctgagttgtagcatcag**ttggaattcatgtagcaatagtgagccactctaggtattccaa**gtaggattttttttaaatataagatgtgagaatctaaactacagccttttaaaggtttgcaagtatagtagtcaaagacgcaaatactaga**gaagaggaattctcttc**cggaatccagaatgcatctgatagagaaggtacaactgccaatcatgtggtc**ctcagacctttctgag**aagcccatgggtgggggtgcagatgctctcagctgccta**gaggacttcatcaggtgcttctgcagtcctc**acctcggtccatatgtcttgctgcaggtgttgatggatagtattgaatcctcctcttcttacttctcaatctcagggcaggccccacactgggcaactccaccaaaaaccagagaaggcatagggt**ttctggcaaatgtgcttccagaa**tgatagtgatgatggggaagtgacagctgacatcgtagtgtggtcatgtatctcgactctcaggattttttcagtgaagtgatggcctcagaatacacttggatgtacttccatata**ctatgagtaagtttgaaatcatag**catgaaaatga**tatttagtcatatgataaata**gaactacatggctacattaatcaaaatagcatagtgctggtacaaaaagatacacagacaaatggaacagaatagggaactcagaaataatgctggagaccta

Uppercase: IGKV1-33-3

Lowercase: Flanking sequence[1000bp]

Red & Bold & Underline: Stem-loop [18]

Blue: Heptamer[32]

Green: Nonamer [8]

id-TRAV36DV7[V_gene_segment]

aaaccaagcgatccatcacttcctctcaactgggcaattcagcttttctggaatgcaaatatgtttgtcttcaccttgagatttgaactttctaattagtaattctt**tttcttagtgatcataaagatcaagcaagaaa**gtactagtcttcctagacttgaacactacc**tacatagattatatgtattttactaaaaata**gacattcctagacgatgacaagaaaactatgtaataacatttatggaatggaatgcaaccaaagc**tatactccaagtata**ctagcgaaggccaaactattctaacaaatatatgccaaacgttcagaggcttaatgaacaaagtttgtctttcttgtgaaaatctggagcagatattccaggtcagcagatgactctgccttacatgatctttgaggaactgaggttattctcatttggggactctgctatttctcagagacttaccatcatatgcagggtcatgg**cctggctatgccagg**tttcacttagcaaggagatgaacgagtgggtatggagcaggcatgaccattcccttaagaatcatggacaagaaataatagaattactgactcctaagattat**ggacagctttgggtagcctgtcc**ttca**atggtaagacaattttagaagtatctaccat**ctttcataaaacatgtggacactagttttcagctattctgcaccagctcacaagatggcggtgttctattttgaatgacttccatgggtgattgagtgggagggatttgaatttaaatagaaccaaatgacactttttttaatggttggctgaattgccaaattgctatttccctttttctgagtgtgtgtctgtgtccgtgtgtg**tgcatgtgtgtgtgcacgcacgcgcacacaggca**gacgctaggggtagagtgtgatggttcaaaaagaaaaggaaacgcttaaaggtagtgaatcacgttttgcccaggaaaac**acacttgataactgaaggATGATGAAGTGT**CCACAGGCTTTACTAGCTATCTTTTGGCTTCTACTGAGCTGTAAGTAGGGTTACTACAAAGGGTCAGGGGCAACCAGGAAAC**CTGAGGATTGATCCTCAGAACACTAAAAGGGTAAAGTAGACTTCGTTCTGA**CTTATTGTCTGGGAGCCCTGTGGTTCCAACGGTGAATAATTATCACATTTTTCTACACAGGGGTGAGCAGTGAAGACAAGGTGGTACAAAGCCCTCTATCTCTGGTTGTCCACGAGGGAGACACCGTAACTCTCAATTGCAGTTATGAAGTGACTAACTTTCGAAGCCTACTATGGTACAAGCAGGAAAAGAAAGCTCCCACATTTCTATTTATGCTAACTTCAAGTGGAATTGAAAAGAAGTCAGGAAGACTAAGTAGCATATTAGATAAGAAAGAACTTTTCAGCATCCTGAACATCACAGCCACCCAGACCGGAGACTCGGCCATCTACCT**CTGTGCTGTGGAGGcacag**tgctccctagtcacctgcagcctgtactcaaattctacagctgaggctctgcaactgtaagatggggaa**cttgctacattgagcaag**ccctcaaaaataaactatacggaaaagcagttattggtcaaaatccttgtgcctatgttgacaattacttttgcagtttactgaatcctaactcagagtcatatcgctgcattcactactttctctgtgagttttctatgggattcagcatagctaatatttctagagggtcactcaggatactaactccattgccacaccattatgctctgtggtagaaagcaagcccttt**tatcaggattgtacaggcaactgata**tgtgcactacacattagggaatctacctaacagctatggatgttttagcttctttcccaggacaggcacat**ttataggtaaagttctgcctataa**catttttttctggacctactagcaaagaacaaagggtaggaacaag**caggtatcattacctg**attaaaacaaaaggttttcattctcagttgcattattcttacttactatcttcattttaggattgttgcctatgctaggtacttcattaaagtggtgatagaatttttcaagatat**tacacattaatttgtgta**cttagaaccttaagcagcctacgaaaagaaaggagttagttgaag**taagttatgtgaagctaacataaactta**gcctacttatgttgtgatacactctttcaatgaattatgtaatcatatgt**atggtatttacaaagagcttgcaatggaagctaccat**ttattgaaagcaagtattatgtaccttgt**attttgcaaatatcatctcattgaattctcaaaat**cactttgataaatcatcaatatta**ctttctgttaacatattaggaaatgcagactcagaaag**attaagcccccttccccacctcctaccccagacaaataactataacagaatctgaaatcaggtctatgactcc

Uppercase: TRAV36DV7

Lowercase: Flanking sequence[1000bp]

Red & Bold & Underline: Stem-loop [21]

Blue: Heptamer[25]

Green: Nonamer [3]

id-TRBV6-4[V_gene_segment]

caggacttactggcagatcctttgttcaaacactattaagaatttaaaaatcatgacagcagagtgataaagcaagcgtgggatccttctgaatgcagggctctgtacaccgaagatacagtttttgagctccttttcatttattactaaattaatggcaggaatcctgaacgaagggatattccctatgagacatgacacctcaaataaaa**cgagagcatttttccttatcttgaagtctgctccctctcg**ccaagtccctcactcatcaggacacctttgtgctcacaagtcttcagcagcgctgggtggccaccattgtccacacagagagggcagt**cagcagcatgaggtggttctgcctgctg**tggtctcacc**ccaggaacagaaagcaagactcctgg**atgg**agctgaaggtgctcagct**gga**cttgtcaggagtcccatctgtcagtgaattgacaag**aaacagagcaaaacgtctcctgcaatgttgatgagcctgcccctgggatttggaaacctgataacagagaaaaccaatatagacacaggactttaacaggattatggtcaattaagcaaattggaaaaggatacttgaaggagtatttgggacacaggaatcaaaaacaccagggagacaagaggatttttcctaagattctagactacagcaatacatagataactaacaccagaatattaatgaggaattatatcattgcaggaataaaatttatattgaattacaaactgtttgcacaaaagtcaagaaaaacttgaaagccttatatcaagaagcattcttccc**atccaaacttcagtggtgcatttattttggat**ttgaccctctggggaagggccatggcc**tctctcgacaagaaggttctggggaccaggcagagaga**atgaggtctcaggatgacttccttgacagccctgttcccctttcatcaacacacagacccagaagacctctctgtcttgtagcatctgccATGAGCATCAGGCTCCTGTGCTGTGTGGCCTTTTCTCTCCTGTGGGCAGGTAAGTCCTGGGCAGGGCCCATGTGTAGATTTCAAGACCCAGAGCCTTCCCATTGAGGCTGCAGCATTGGCTTTGTTCTCCTTCTCTGTAGGTCCAGTGACTGCTGGGATCACCCAGGCACCA**ACATCTCAGATCCTGGCAGCAGGACGGAGCATGACACTGAGATGT**ACCCAGGATATGAGACATAATGCCATGTACTGGTATAGACAAGATCTAGGACTGGGGCTAAGGCTCATCCATTATTCAAATACTGCAGGTACCACTGGCAAAGGAGAAGTCCCTGATGGTTATAGTGTCTCCAGAGCAAACACAGATGATTTCCCCCTCACGTTGGCGTCTGCTGTACCCTCTCAGACATCTGTGTACTT**CTGTGCCAGCAGTGACTCcacagtgctgcacag**ccatctcctctctgtacataaatgca**ggggaggctctgccctcctcccc**gaccccagactcaaccatgtccttggcagagttctcagcactgggaatcttggaagcccaatgggtgccgggcagtgtgagcctcagtctgtgccaggtgcctctgcagg**cagtcccagccaggcctggactg**gtc**ccagagcctcagaggtctcttttgtttctctctggtctttcttccaagctatctttttgggatgggaccag**ggctttcccagctcttacttttctactcat**catcctgagtccgaggtccacaggatg**gaacag**gatttgtatttcaaatc**tgtctagactcctgtttctctctgggggccacgttgcttcctctctccagggtttcctccagtacccactctcatgtggtctcacctgtggcccgcctttcccatctgagcagtcacccccgaaggccttgctgggcctctgcacaccacctcccctcgcctttctact**gcagccatgaggctgc**ctct**tctgtgcctccttcctttccaacacaga**gacctcaaaggccatttcctccaccctgggctagtgccttcctttctgcagtggtcagcttctacctgcactt**ctgatctcagcatgatcag**cctctcctgctggaagcactccctcaactctcagctgaggtcaaatttcagtttccctctcataagctcacagaatcatgtgcctcttagtaaagttcagcaccattgcaatttttccaacacttgttttattccttttgtgctccacttaattttaaactcatgaagctgaaggctgagcttgttacatttaccaccagcagtgcctggcaggcagggagtcctcattcatttcacagagaatggatgggtgagtgacaggtgagtgagtgatgagtgggtg**aatgaaccagtaacaggaacacactagattcatt**gtaccctggcagaggtctagagt

Uppercase: TRBV6-4

Lowercase: Flanking sequence[1000bp]

Red & Bold & Underline: Stem-loop [19]

Blue: Heptamer[35]

Green: Nonamer [2]

id-IGKV2-14[V_gene_segment]

acttactctatactaggaagcaataataaaggtattggctaaattttc**atcattactaatagaggtgggatgagagtggaatgat**atatttaaa**tgctgggggagaaaagtcagccacaaatgctatatccagca**aaacaatcatttaaacataaaagcaaggtagaaatgttccttcatacacaaagactaattcattatagcatatgtgccttgtggaaaattcaagagaaaatccttcagaataacagggaatgagagcagacagcaattcaaattgataagaaggaatcaagccctcaaaaaatgaaagaaataagaaagaagctacaaaaaatgtttacctgatatgaagattaagtacaggaaaaaaaaacaattataatagatgtcacagtatcagttacattgattgtgttttgagcagcagttgctgactg**ggaaggtgcataatgaaagcagcaagagaattggaaaccttcc**atattttgatgtg**gatcatgacatgaaatgatc**tggattttgacattcatttgcacccttaagatttgaaccttttgcactgtg**tgtttttccttaaataaaataaaataaaaca**aacctaggctcttttaaataagcctgaaccaagaaataagaatcaccaaatatttatgaaagctcaaccttattaaagagaagtgtaaaacatgcctgagaaagtcctaacgaactagaaattgggtagaggaaatagcaggaaacatctaaatatgtcccttctgccactcaggaaacagattttacctctcttgatatccttaaaa**acataaaatatatttatgt**cattttgctcacaagagaggtccccaccttctccttggctctttccaccccactgcacccacc**aggggatttgcatactgtcccctagggaagaccttccct**tgtgagtctgagataaaagctcagctctaaccttgccttgactgatcaggactcctcagttcaccttcccacaGTGAGGCTCCCTG**CTCAGCTCCCAGGGCTGCTAATGCACTGGGTCCCTGCTGAG**AGCAGAAGGGAAATGAGGGAGGATGATGG**GGTGGGAGGGTGAACTCTGTGGGCCCTGCCACC**TCCCATGTGTGTTCTGTCCTCGTGTTAGATGTGTCTTGTCCTCCAGGATGGGGCATGTGATGTCTAGATCTGTGAGAGTGAGGAAGATTCCAGAAGGAGCAAGGACATGTACTTTAGTGAAAGCTGTGACACAGAAAGAGGGGGATGGGATAGGTGACTTCTAGAGGCTTATTTGTGCTTTGCAAATCTTGGTTCTTTTTAAACC**TGTATGTTTTGGGAGTATTAACCAAAATCATACA**CAAAAAATAATTGAGCAAAACATAAATAACAGACAGAAAATTATTAAAATGGCTCACAATGTTTGTACATAACCTTGCACTTCTCTCTTATTATTTCAGGATCCAATGGAGATGCTGTGTGACCCAGCCTCTACTCTCCCTGCCCGTCACCCCTGGAAAGCCAGCCTCTATCTCCTGTAGATCCAGTCATAGCCCCTTTCACAGTAATGGATACACCTATTTGAATTGGGACCTACAGAAGCCAGGCCAGCCTCCACGGCTCCTGATTGATTTGGTTTCCAACAGGGACCCTGGAGTCCCAGACAGGTTCAGTGGCAGTGGGTCGGGGACAGATTTCATGTTTAAAATAAGGAGGATGGATGCTGAGGATGTTGGGGTTTATTGCTGCCAGCAAAGTACACATTATTCTCCcacaatggtacagtcctgaacacaaacctccctgcttgctgtggcctagctgcccagatgtgttgtttctgtggagagcaggcactgtggattctcttagatgcctaaag**accaagatgttggagaactcagaggacttggt**ac**agctgagggctcatgaccataaatttctcagct**acacctcaggcaccacattttaagttcccatcagctgcagcggcc**tttgcatgacagagtctgcaaaaaggaggaggtacacctcccctttgagcaatgagacacgagagaagagaaagctcaa**tgaaagctcattctaagcctctcttccttccctacattcattcatcaagtaaattcgttctgcgtaacaggcacctaattgagactgattactggcaacgcaaaactaatacattcttttgatttggtttagcagttactagagtatatgtatgttgataaggtttggtaatattaaaacagtttccctcccctcctccacctccccccaccacttttccgaaggcagacccctgaccaggacaccagcaggcactgtattttttcctacttttttcagggagtaccaaaatataactcttcacggtctagatttttaagttactagatatgttgctgcaaatgtatgcaaaaat**aaagacctagtgccaaaagtcttt**ccacagtctcaaccacagcatgttgaagtaaaaagtagttttctgaactccaaagctggg**tttcaacaataaggaccaagtttgaaa**gaataaaggaacataagtatgccatcttaaacagaattacaagat**aatatgtaagccatatttttaccccatgaaaaata**accatctgaagacttggagcagaggtcagtacgtgagagactgg**tgggttgcgtttgagatatgaaccca**gatgcaagttcttatgtatttcctgggaagaatcacagagtcgagagacccctgcctggcattcaaggctgggccctgatgcc

Uppercase: IGKV2-14

Lowercase: Flanking sequence[1000bp]

Red & Bold & Underline: Stem-loop [21]

Blue: Heptamer[29]

Green: Nonamer [5]

id-IGKV1D-8[V_gene_segment]

gagtgcaa**atccatgtccaggttcatatggat**ccagctgatttctccatgtaagttggcaattgcttgataagggattgtctctttcctaaagatg**ttaacagggaggctggtgtctgggtcaggatgatgtccctgataa**aacgtaaaagaagaaagtggcattgttgg**tgcatgacagggacatgctccatgca**gtggtcaccctcaataagagagaagaactttggaaagtaatactcaatgacagaaaagaaggtagacaatgaaggtgcccaaaacaagaataaggtgaagtgaattta**gtctctgggtattaaagagac**ctgtagctcttgataatggtggatgtgtgagtgctgcatgcattgaggaaactcggtatcatctctctgtatctgtagtaaattgc**ttgatcttatagtgataagaacaa**tggcataacaccattacctaatacttacaaatatgtatagcatcatgtcaataaattttatttttaatatttttagaaaggaacaatgttaaagctcacagaaatgttgcaagtataggacaaagtacctccttccctaacccgaatcatatgagagtcttttgaagtcctgagaatcatactgtttaacattttactatgtatttcctacaaacaagatattctaaataatccccatacaccaa**tgaaatacattactccgtcgactcctgaggaatatttca**aattttcaaaaaaatacataaaaaa**tgtttctcataacaaactactctccagtagaaaca**cattcactgcagacaaatttgtgctaccctggtctttcctgggacacctggggacact**gagctggtgctgagttactgagatgagccagctc**tgcagctgtgcccagtcagccccatcccctgctcatttgcatgttcccagagcacaac**ctcctgcactgaagccttattaataggctggccacacttcatgcaggag**tcagacccagtcaggacacagcATGGACATGAGGGTCCCCGCTCAGCT**CCTGGGGCTCCTGCTGCTCTGGCTCCCAGG**TAAGGAAGGAGAACACTAGGAATTTACTCAGCCAATGTGCTCAGTACAGCCTGGCCCTTCAGGGAAATTCTCTTACTACATGATTAATTGTATGGATATTTGTTTTTATGTTTCCAATCTCAGGTG**CCAGATGTGCCATCTGG**ATGACCCA**GTCTCCATCCTTACTCTCTGCATCTACAGGAGAC**AGAGTCACCATCAGTTGTCGGATGAGTCAGGGCATTAGCAGTTATTTAG**CCTGGTATCAGCAAAAACCAGG**GAAAGCCCCTGAGCTCCTGATCTATGCTGCAT**CCACTTTGCAAAGTGG**G**GTCCCATCAAGGTTCAGTGGCAGTGGATCTGGGAC**AGATTTCACTCTCACCATCAGCTGCCTGCAGTCTGAAGATTTTGCAACTTATTACTGTCAACAGTATTATAGTTTCCCTcccacagtg**ttacacacccgaacaaaaacccccagggaagcagatgtgtaa**g**gctgggctgccccagc**tgctcctcctgattccttcattgcctg**agagtgttcctcagatgcagccacactct**gatggtattggtagagggggacctgaaatcacctctgcaacccaattccttttgctttgtaagccccagctgcacagacatagcaatgcctctcctgatgtaataaaggcagagatcttgacacctaaggagtctagtttagggctttggttggaattcaaataacagagaagaaaccactatagatattctaagcaggaattgtcttaatacagaaaattagagtctaaactactgaagtctaaata**aaatgtagagcagaatctctaaattt**aatgttttatttgcaaagaaatgtttgccaaattggacacacaggaaaactcagtggtcttcaatatattggaagtacgaagagaaggttagtgtttt**atgaaaaagggaaaatattaccttttgcat**tttgagaaagttcactggcactagtaagggttgggagctggcaagctcagactgggaagcagtggtggacaaagtgaatcctagaattatatcaagttatctcagaagttgcagttaaatgtgaattcaggttacaagaagccaaagcagtgaaggttgcagagaattttcttactaaaatgccagggattcagtgtacaccctgctgctcaccacacagaaagtcaatctctaaaacaagtattgccaagaaacaggctttaatcaggtactgcagccgt**ggagatgggacaccatttgcgaatgtatctccctgatcaactacaattaagagtttatataacaggga**cgaaatgtgggaaaacaggaatttggggggtgggtaaggaagataatttggtcaacaggaagcaagaggtcagttatgcaatcataatgggtgaaggttctgatgtctcagtgtccggattcagt

Uppercase: IGKV1D-8

Lowercase: Flanking sequence[1000bp]

Red & Bold & Underline: Stem-loop [22]

Blue: Heptamer[23]

Green: Nonamer [2]

id-IGKV1D-17[V_gene_segment]

tgtccaatttcatctggatc**caactgatttgtccatgaacatgggcagttg**cttgataaaagattgagtgtctctttcctaaagatgttagcagggaggctggtgtctgggtcaggatgatgttcccactcactgataaaatgtaaaagcggacagtggcattgatggt**gcatggcagagacatgctccgtgcagtggccatgc**tcactaagagagataaactttggaaaatagtactcaatggcagaaaagaaggtagactataagttgcccaaaacaagaataaggtacagcccattta**gtctctgggtattgaagagac**ctgtcactcttgataatggtgaatgtgtgagtgctgcatgcattgaggaaacatggtatcatctttgtgtatttgtagtaaattgcttgaac**ttatactggtaagaacaatagtataa**caccattacctcatacttac**aaatatatgcatcatcatgtcaatatattt**tatttttaattttttttagaaaggaacaatgataaacttacagaaaagttgcaagtatacaataaatacccccttccctaaccggaattatacaatagtcttttaaagaccttagaattctatcgtctaaaactttactatgtgtttcctacaaaaaaagaatatcctcctatatactccccatacgccaa**tgaaatatgatactccacggactcctaaggaatatttca**aattgtcaaaaaaaatctaaaaaatgtctctcgtaacaaaatagtctccagtagaaacccattctctgcagacaaatctgtgctaccctgatcttacctgggacacctggggacactg**agctgctgctgagttactgagatgagccagccctgcagct**gcgcccagcctgccccatcccctgctcatttgcatgttcccagagcatagcctcctgccctgaagccttattaataggctggacacacttcatggaggaatcagtcccactcaggacacagcATGGACATGAGGGTCCCTGCTCAGCT**CCTGGGGCTCCTGCTGCTCTGGTTCCCAGG**TAAGGATGGAGAACACTAACAGTTTACTCAGCCCAGAGTGCTCAGTACTGCTTTACTGTTCAGGGAAATTCTCTTACAACATGATTAATTGTGTGGACATTTGTTTTTATGTTTCCAATCTCAGGTGCCAGATGTAACATC**CAGATGACCCAGTCTCCATCTG**CCATGTCTGCATCTGTAGGAGACAGAGTCACCATCACTTGTCGGGCGAGGCAGGGCATTAGCAATTATTTAG**CCTGGTTTCAGCAGAAACCAGG**GAAAGTCCCTAAGCACCTGATCTATGCTGCATCCAGTTTGCAAAGTGGG**GTCCCATCAAGGTTCAGCGGCAGTGGATCTGGGAC**AGAATTCACTCTCACAATCAGCAGCCTGCAGCCTGAAGATTTTGCAACTTATTACTGTCTACAGCATAATAGTTACCCTcccacagtgttacacacccgaacataaacccccagggaagaagatgtgtgag**gctgggctgccccagc**tgctcctcctgatgcctccatcggctgagattgttcctcagatgcagccacactctgatggtgttggtagaaggggacatgaagtcacctctgcaccctaattcttttctctttctcagccccaactacacagacatagcaatgcctctccgaatttaatacagacagagatcatgacacctgaggagtctagttta**tggcttcagttggactttatataacagagaagaagcca**ctatagatattctaggcaggaattgtcttaatacagagaattaaagtaaactactgacgtctaaataaaatg**tagagatgaatctcta**aatgtaatgttttatttacacagaaatatttcccaaatggggcatacaggaaaactcagtggtcttcaatatgtggcaagaacaaagaaaaagttagtgtttcatgaaaaagggaaaatattacctatggctctttgagaaagttcattggcactaggaagggttgggagctggcaagctcagactgggaagcagtggtggacaaagtgaatcctagaattatatcaagttatctcagaagttgtggacaaatttgatttcaggttacaagaagccaaagcagtgaaggttgcagagtgttttgttactgaaataccagggattcagcgtagatcctgcagctcaccactcagaaagccaatcactaaggcaagtattgccaaggaaaaggctttaatcaggtgctgtagctgaggagatgggacaccattctcaaatgtatccctgaccaacaaaaattaggggtttatatagcatggaagaaatgtgggaaaacaggaattatggagggttaagggagataatttggtcaacaagaagcaggaggtcagttatgcagtcataatgagtgaagggtctgatgtcccactgtcctgattcagtgatatg

Uppercase: IGKV1D-17

Lowercase: Flanking sequence[1000bp]

Red & Bold & Underline: Stem-loop [14]

Blue: Heptamer[25]

Green: Nonamer [4]

id-IGHV1-17-2[V_gene_segment]

gtttcaagagatggtgtcagcagcatatggtgtcactgaaggagtattctacactagcacacagc**catttcatgctgggctagagaagcctgggggaaatg**atttgaagtctca**gcaaggaacctccttgc**aaggcaggggctgggctgtcgggggcaccgagcaggtgagtcagaaatcaatatgtaaagatgaggacctatggatatgaattgaaaatatataaatacttcaaaaaattccagtaaattgagtccaaaattaaccccaaattattcaaaacacaaattccttgacaattattttgggagcagtgagttcata**aagaattcaaaactactgtttcagcttctgattctt**attgttcct**gagatgagaaaatcatctc**taatcacacatcacacagcaaatctgtaaacaagagtgtttctatgga**agatcctgggggatct**gaacaccaggcaggtgctggagaccctgtttcaggagcgc**ccagcagatctcagagggacctgctgg**tcactcacgtgggacatcagcagtaactttc**tcagtcaccagtcagctgtgctggtgactga**tggacccaggaca**gaaccaaggcacctcctcagtgtcatggagagtgatggttc**cagaaatcatccaggtgctctctgtgcttataaaatgtaggttcactgtgaggagcgtgttctgagggggattgttctgtgaaaggacctctgttcacgagtgttcataaatggagcagggcatgcatttcctcaaacaggaatagcgcttggac**catcagcatctcactcttgtaaatctgatg**tgtcatttatcttccctttcttattattgaccaggctttgcgctatgaaatgctctgtctcatgaatatgcaaataacctgagatccactgaggtgaattt**ggatgtctgtgccctgagagcatcacccaacaaccacatcc**ctcctctatagaagcccctgagagcacagctcctcaccATGGACTG**TACCTGGGGGATCCTCTTCTTGGTGGCAGCTGCCACAGGTA**AGGGGCTCCCAAGTCCTAGTGATGAGGAGGGGATTGAGTCCAGTCAAGGGGGCTTTTATCATCTCCTCCCTTCTCCTCACAGATGTCCATT**CCCAGGTTCAGCTGTTGCAGCCTGGG**GCTGAGGTGAA**GAAGCCTGCGTCCTCAGTGAAGGTCTCCTGCCAGGCTTC**CAGATACACCTTCACCAAATACTTTACACAGTGGGTGCGACAGGGCCCTGGACAAGGGCATAGTGGTTGGGATGCATCAACCCTTACAATGATAACACACACTACGCACAGAAGTTCCGGGGCAGAGTCACCATTACCAGTGACAGGTCCGTGAGCACAGCCTACATGGAGCTGAGCAGCCTGAGATCTGAAGACATGGTCGTGTATTC**CTGTGTGAGAGAcacag**tgcgaaaacccacatcctgaga**gtgtcagaaaccccaggaaggaggcacctgtgctgacac**agaggagatgacaaagattattagattaaag**attttcttagaaaatgacactaagtcattaaagaaaaggaacaatattaatg**tgtatttgagaaattttaattatttgagagatttttcatacagcattt**attctgtaaacaaatttcaatgattacagaat**gaatcaaattaatgaaactaatacagaacttcctctgaaggtatctttgtaaacattaatttcttaatcagtgctgtaagt**attttggaacacagacacaaaat**cacattttaagtctgcatttatgtct**attaaaaatgccaaaaaaatctccttttgtgcatgtagcattttgaat**tcccaccatcaatgcatgatagttcttggttttccacattcatattgtcatgtatccttacgagaattgtgtgttttaaccaatctagtaggtgagtaatgggatctaatttttatttaaatgcacatgtccctccaaaagttcatatttaacaattttcgtataacttctgttgagacgcctctcctgacattcggttcattttaactgcattgcttcatttcgattcat**tgcaagttgacttgca**tattgtttataaaagtcatttcacaaattaaaagaattcatttcacaaatatgt**gacttggaagtattttctccaagtc**tgtggctgtctttattcccttatcagtgtgtattg**cagaaacatatgtgtgtgtatgtgtctg**tgtgtgtatgtgtctgtttgtacaaatttagactcaaaaacatgtaaaattgcattcattcatagatga**tgtatttggcaatatatatgaagtctcattataaaataca**ctaatagtgattattttttccatgtctctcaactcaggcca**caatcaactcatgagtgtttaaacttcatctacttgattg**gaggactatccatt

Uppercase: IGHV1-17-2

Lowercase: Flanking sequence[1000bp]

Red & Bold & Underline: Stem-loop [26]

Blue: Heptamer[47]

Green: Nonamer [6]

id-IGHV5-78[V_gene_segment]

gtgataaagagaagtccttttgcttctgcaccccaggtacaactcctgtgagcaatctggcaaatattttagaaactgtgtagtttgtattgccttgtcttt**tcatttgaatcttttagatgataaggaaatga**gagtggttcacaagataatggggcagacacaggggatagttaagggcctcctccaagcaatgtggatatcataaaatgaatgttaaaaaatgaaagaactaaacaagaaggtgctggggttgacacaaaggacttaaaacagaactaccaggagttgggtgcagcaatggcaatcccttcagttccatcagcaattcagagacctaaacaaatattccctatttatgttgaattggcagactgtagaaatgtagaagacaaaatgaccagaaaaaatgtgtgatttggcatgggtgaaggtctggcaagttaatcaagttgaggactgttgaagccctagggtggtcctgcctttgctgggtgcccagaccataatggcactcatgggaaattgccagcaagtagaaactaggtgttttttttttttttgtagttcttattatcaggcatatgtgcatttgaattttgtcttcatagcatttcctggttctatttctcattttttaaaacacacgtgatttcatctagatttatcaccttcacagggtcacagagaagggtaggaagaagggaggccctggtttggg**tcttgtaggaccatggacaaga**gtgcggagggacaacagggggttgtaagacattattaagaccttattct**gcctctcccaagaggc**tcagttcagccgtttttctgcagttgcggttctgggttataaaccttgtagactcttcccttcaggccagggcggcaactatgcaaattc**aagtgggggcctccccactt**aaacccagggctctcctccacagtgagtctccttcaccacccagctgggatctcagtgctttcttttctgtcctcctcc**aggATGGGGTCAACCGTCATCCT**TTCCCTCGTCCTGGCTGTTCTCCAAGGTCAGTC**CTGCCCAGGGTTTGAGGTCACAGAGAAGAATGGGCAG**AAGGGAGCCCCTGATGCAAATTTTGTGTCTCCCACACAGGTGTCTTTGCCGAGGTGCAGCTGTTGCAGTCTGCAGCAGAGGTGAAAAGACCCGGGGAGTCTCTGAGGATCTCCTGTAAGACTTCTGGATACAGCTTTACCAGCTACTGGATCCACTGGGTGCGCC**AGATGCCCGGGAAAGAACTGGAGTGGATGGGGAGCATCT**ATCCTGGGAACTCTGATACCAGATACAGCCCATCCTTCCAAGGCCACGTCACCATCTCAGCCGACA**GCTCCAGCAGCACCGCCTACCTGCAGTGGAGC**AGCCTGAAGGCCTCGGACGCCGCCATGTATTATTGTGTGAGAGggaccatctaaaaccttccgcggtgcaggtgcagagtgagctgccagacacac**cctccccaggggcctctctattcatccggggagg**aaacactggctgtttgtgtcctcaggagcaaaaaccagagaacaacatgggagcgttcctaacccctaaggc**aactggatgggagacctgacccatccagtt**ctctgagggggctcttgtgtgttctacaaggttgttcatggtgtatattacatggttaacatcaaaaggctgcctaataggcacctcttcaatataatagtcttttaattagtgaaaattttacacagttcatcattgcttgcttgccttcctcccttctgtccgctcttactccctccttcttttattttctacttacttttccaaaatcatttaacccctttttgtaccattaataagttatcttgtttttgttgttgttttccttttaacaatatgcactgaataattca**tctttgtgccaattcgtaaatattttgatataacaaaga**cttctgtcataaatagatttcctgtgagtaatcttgcaaatattttagaacctgtttggttaagaataaa**ttaaaataaatagacaaatttttaa**agacatacaacctatcaaaactg**aaccaccaagaaataaaaaatctgggctgggcgtggtt**gctcatatgtgtattctcagcactttgggagaccgatgcgggtggatcacctcaggtcaggagttcaagatcagcctggccaacatggcaaaaccccgtctctactaaaaatacacagattagctgggtgtggtggtgttcacccgcagtcccatagtcccggctactcgggaggctgaggtaggagaatgacttgaacctgggaggcagaggttgcagaaagccaaaatcgagccactgcactccagactgagcaacagagtgagactccaattaaaaaaaatctgaacagaccaatgagtaaagagatcgagtcagtgatt

Uppercase: IGHV5-78

Lowercase: Flanking sequence[1000bp]

Red & Bold & Underline: Stem-loop [13]

Blue: Heptamer[32]

Green: Nonamer [7]

id-IGLV1-36[V_gene_segment]

ctctggttaaaccaagttaaaattttaattcttcccatctt**caaagactgagcatctttg**ctctttttgttcataagtgaaacactccaaagtcatttgccaggatcccctgcagatgagcatttagtgtacagactcctaggcaagatccctggtcacagtat**ccagatgactatgactgcaaatctgg**ggaagt**agagggagaacaaagatttaccaggaaaaccctct**gggaa**ttcctccccagggaatcctagaggaaagagctccctttctcctcagcgccacaggatccacagaggaaggagagctct**agc**caccaggaaggagccacatgagaggagaggacctggtg**tccaacacaggtggggaggtgagtccatggggctggtccctgctatgggcttgatcctgacat**aggactttatgtagataccacctgagtcct**ctacaatgtacagccaaagagccatggaaagattcactc**ctgaggagtctgaagactcctcag**tagagcgtgaagatagccctgaagtcacagggaggagacacagacaccacttggcctgccttggacctgagagagggggcagtggacagaatggggggaggaagccatgtatctgagaggttctggtcaccttatcatagaatggtttaccttgatctggacaccagagggcgcatggggatcattcctgagaagacaggggtgatcagagcagaaccagccgcct**gccctgcctggtgtcctctgctcagggc**tcacagctgtgtcctccccaccccctgggaccacaaagctccacccctgccaccccctgacatcctcaagccaaggagcc**tgacccagggctcagggtggggtca**caaacctggggggggtctgatttgcatggatgga**ctctccccctctcagagtatgaagaaagggagag**atttgggggaagctcagcttcagctgcgggtagagaagacaggactcaggacaatctccagcATGGCCTGGTCCCCTCTCTTCCTCACCCTCATCACTCACTGTGCAGGTGACAGGATGGGGACCAAGAGAGAGGCCCTGGGAAGCCCATGCGACCCTGCTTTCTCCTCTTGTCTCCTTTTGTCTCTTGTCAATCACCATGTCTGTGTCTCTCTCACTTCCAGGGTCCTGGGCCCAGTCTGTGCTGACTCAGCC**ACCCTCGGTGTCTGAAGCCCCCAGGCAGAGGGT**CACCATCTCCTGTTCTGGAAGCAGCTCCAACATCGGAAATAATGCTGTAAACTGGTACCAGCAGCTCCCAGGAAAGGCTCCCAAACTCCTCATCTATTATGATGATCTGCTGCCCTCAGGGGTCTCTGACCGATTCTCTGGCTCCAAGTCTGGCA**CCTCAGCCTCCCTGGCCATCAGTGGGCTCCAGTCTGAGGATGAGGCTGATTATTACTGTGCAGCATGGGATGACAGCCT**GAATGGTCCcacagtgctccaggccc**agggggaacagagacaagaaccccct**tccttttct**gccaggagggtgagccccggctgctgctgctcaggcctggc**ctgtggcttctgctgctgcagcttcccccatgggtccagggatatccagggccctgcctgagagtggaggctcctcctcccctgagtcctcagagtcaggaacagggtgtccagaaaacaaa**agcagggtgtcctgctccctgtagcttgggacacaggg**tctctgcactgaagtcctgggctgaggtggcaggtccagctgtgtcatcacagacccacttctgttggggaacctgtgtctccatcatcctactttttccatttccaggagtttccagagtggtgtcttcctcccccttctcctcagtgtgaatccctgtgcttcctttctctccagcctattctttttttaataaaactcctttcatgtaggaataatgcacatataataagccacccatatttacaatagggaatgagacaaattttggcatttctatggatccttgaaaacatcaccaccaccg**ggagctatacaacatacaacctgtgaccaaccccaatagctcc**tcatgacccttgtcacctcattccctcccccaacaccctccttggaagttgcagatctgcctctgtcactactgtttggtttctcttttctggaatgtatcagtgttatcaaacattatcttcccatttgttcaggcttctttcactctgcacacttcttttgagattctcgctgttgttgctgcattgtacccatctcgtagagggctccttactatctttgaaggtccttaggccatttttttctgattttatttatgtatttattt**taaagtttgaatttattgtgcatccaaatcaaacttta**ggggcaatgttcatctaaggtgacaggacagtctgtgatggcttgtcgggattcttgcttcttcatctcttgcttctgattctctctata

Uppercase: IGLV1-36

Lowercase: Flanking sequence[1000bp]

Red & Bold & Underline: Stem-loop [20]

Blue: Heptamer[35]

Green: Nonamer [4]

id-TRBV6-3[V_gene_segment]

cagagacggtagtcagtagcgtgaggtggttc**tgcctgctgggatctcaccccaggca**cagaaagaaggagccctgggtggagctgaaagtgctcatctgggtttgtcaggagtcccatctgtcagtgaattcacaagaaacagaacaaaacaactccttcaatgttgatgagactgcccctgggatttggaaagctaataacagagaaaaccaatatagacaaaggattttaaacaggactatcggattatgatcaattaagcaaattagaaaaggatacttgaaggagtatttgggacacaggaatc**aaaaacaccaggaagacatgaggagttttt**ccaagattgtagactatagcaatacttagataaccaataccagtgtattaatgagtaattattagtattattattttgagatggagtcgcactctgtcatccaggctggaa**tgcagtggcatgatctcggcttactgca**acctccgcctccctggttcaagaattctggtgcctccgcctcccgagtagctgggactataggtgcacgtcaggatgcctggataatttctatatttttagtacagatgggatttcaccatgttgg**tcaggctggtctccaactcctggcctgaagtgatctgctctcttcag**cctcccaaagggctaggattacaggcatgggccactatgcacagccaagtaatttgtatcatcatgggaataaaaaatatacattgagttacaaactaattaccaaaagatatttaaaaaatggtagacttgcatcgattcagacaaaggcattcttcccatccaaactgttcactggtgcattgatcttgaatttgaccatctggggaaggggcgtggcctctcctgacagcaaggctctggggcccaggcagggagaatgaggtctcagaatgacgcccttgaaagacgtgttcccttttcaccaatgcacagacccagaggacccctccatcctgcagttcctgccATGAGCCTCGGGCTCCTGTGCTGTGGGGCCTTTTCTCTCCTGTGGGCAGGTGGGTCCTGGGCGGGGCCCCTTGTGTGGATTTCAAGGCCCATCCCCTTTC**CACTGGAGCTGTAGCATCAGCTTTGTCCTTCTCTGCAGGTCCAGTG**AATGCTGGTGTCACTCAGACCCCAAAATTCCGG**GTCCTGAAGACAGGAC**AGAGCATGACACTGCTGTGTGCCCAGGATATGAACCATGAATACATGTACTGGTATCGACAAGACCCAGGCATGGGGCTGAGGCTGATTCATTACT**CAGTTGGTGAGGGTACAACTG**CCAAAGGAGAGGTCCCTGATGGCTACAATGTCTCCAGATTAAAAAAACAGAATTTCCTGCTGGGGTTGGAGTCGGCTGCTCCCTCCCAAACATCTGTGTACTTCTGTGCCAGCAGTTACTCcacagtgctgcacggctgtctcctctctgcacagaaaggcaagggaag**gtgctgccctcctccgcagcac**agattcagcgatgcccttggtcctagcaccgaaaactttggagccccaatgggcccgggcagtgcgagccttcatctgtgccaggtgcctctgcagtcggtctcggccaggcctggatcggtcccagg**acctcagatgtcttccttgttgccctccggtctttcctctgaggt**gtccttttgggctagtgccaggggtttcccagctctgactttcctagtca**ttcacctgagtctgaggcctcctaggtgaaataggatttgtatttca**gatccatctagactcccgtctctacctgggtaccacgtgctttcctctctctatgatttttcctcatgttgaaccctacggtggtctcagctgcagcctgcatttcctatattcacagtcgccctccaaggccttgctgggtttctcctccccgacctccctgcccttctctactccagccatgttgagagactccaaagtccatttcctgtgccctgggctggagccttcctttctgtagtggtcaacacctacctgtgcttcagatctcagcatcatcagccctagtgataatcaccattatgcttcccttggctcccagctgaggtcagtttccctctcataatctctcatggtatcatgtgtctgttaatcagtagaatttagcacagttggaatcttccaagtactttaaaccccatgaaggccaaggctatgcctgggttataaatcagcagctgagcctggcacaaggggagatgcatttcagagaatggatgagtcagtgatggagtgagtgatgagtgg**atggatgaatccataacaggagcaccctagatccat**tttacccttgcggagatctggctgtaagtacaagaactcttacccttttgattgctgggctacgggaggtccttgaaattaatggggagtcgctat

Uppercase: TRBV6-3

Lowercase: Flanking sequence[1000bp]

Red & Bold & Underline: Stem-loop [13]

Blue: Heptamer[30]

Green: Nonamer [4]

id-TRBV4-2-2[V_gene_segment]

ttacacatcagaaaaatgctaacactttcctcctggcccagcacatctccaagttaaaaaaatctctaaattatcaacccatagtttaatcatggtaactcccttctacccccaacatagatttttatgtaactgatttgtaactacaaccttaaccgtacaatcacaattgtcattataaaatactaacaaacatgtgctattaacaatggatggttaaaatcacttagggattatgaatcatttcatttttgtgtttatttattatgtcagtttcaagctgatttccctttcccttgttccacacaccttgggttgttctttttcagcatcctgtgtatgttagctctaagcccagccaaggctcatgcccctccacttcagggacagagggtggctcatccctttcgtgctccctcagctccctccccggtgcctggttgctggttggcatcacagccatgcccttgaaagatgggcgctggtccacactgcatccgtggtcatgtagccatcctccctccacagaaccgagcaaaaagggctccaccgcgcttaggcattgagggaatggacattttgaatggtgctgtcttgtgtgatccctcagaggagaaggagatttcttgaatgggagtggtctgttgctgagag**ggatcctgaaagacacggggacagagacaggggtaggatcc**acataatgaggaagcagctgagaacaaaggagcccctgccagaggaatatccttggggggtaaaaagctggctctgccctttctccccagcccattcatcccagcccagacaattcaaatctgccttctatcaggacctagaaaggatgtaaaacggccgggtataaatatcccctgggtctggggaaactgtcaggagcagtgacatcacaggaaaaaccaccaaccaaggccaaggagaccagagcccagcacctcacccagaggaccccagtcagaggccccatctcagacccgaggctagcATGGGCTGCAGGCTGCTCTGCTGTGCGGTTCTCTGTCTCCTGGGAGCGGGTGAGTTGGGTTCACATCAGCTGTCCTTGAATTCGAAACTTTTTCCTTGTGATTTCAGCAACAAGCCTCCTCCTGGGCTCTGCCTGAATTTTGTCCCTTTCCCCCCGCAGTCCCCATGGAAACGGGAGTTACGCAGACACCAAGACACCTGGTCATGGGAATGACAAATAAGAAGTCTTTGAAATGTGAACAACATCTGGGGCATAACGCTATGTATTGGTACAAGCAAAGTGCTAAGAAGCCACTGGAGCTCATGTTTGTCTACAACTTTAAAGAACAGACTGAAAACAACAGTGTGCCAAGTCGCTTCTCACCTGAATGCCCCAACAGCTCTCACTTATTCCTTCACCTACACACCCTGCAGCCAGAAGACTCGGCCCTGTATCTCTGTGCCAGCAGCCAAG**Acacagccttgcagagtcaccgctttcctgtgt**agaaaccttaggggccctccaggaagctgtgggggccaccaaagccttcgatgaacatttcctacaagagccccgacagaagattcagaacatcatagcacctttaatggcat**aaaatatctagcatatagtagaagcctagctatttt**gatttcctttgctgatctcagacaacctcttccaccacacc**aatgtgcaaagacagatatggacacatt**ctgccaagaaatcttaggttgaggactcagaagtcacagaaatggaaatatatatgcctgataattcatctgataaaattaactagaggatgtagcatacatatctcctcaggca**tttaaaaatcaacattattgatctttaaa**gtatatgaagttaacaggacaccttttaaatgaacatatctgtgatttttgagcaa**catatatctatatatg**caaccccacctctatgaagatgtatctgcatcacccagaaagtccctcactcccctctgcaatgcctctgtctgtcccagcaaccaatgatgtgacttctatcactggagagtcgttttgttggcaaatatttttaatgcaaatgtcaaacctcttctgaataatcctcttgaatacatagctaaatcatcaaggcagggaggagttcctggaggttttatggttttagaagtggttgaagccaggtgagctacctgacccatggtagggacaattctatctggagatggatattgt**gccgaaataacctcaaaaggtcttttcggc**tg**gtagaagctgattctac**aaaatttattcattttcattcgtttattttctcgtcaattcattcaatcaaacaattttatgttacgttttgacaaaagttaatcatctttaaggaaattattttgagttcccatcagac**acacaagcagctgggctttgtgt**tcacacagtgagcatgctggggtcaagcaaaccatgatcacagtg

Uppercase: TRBV4-2-2

Lowercase: Flanking sequence[1000bp]

Red & Bold & Underline: Stem-loop [9]

Blue: Heptamer[30]

Green: Nonamer [3]

id-TRBV17[V_gene_segment]

ctgaccatgccctaagtc**ctgagcattaatttttgctcag**tttcatttacatcattttcaactagtttctcattttctctctattctttttgttttcaatgcagctcccagaaactaaatctatccatttaatattctttttcccaggaaaattcttctcattctatttaaagtgtaatattctagtttaatgggaaagttttttgttgttgttgtttagccttttaaaaatctctactgaaaa**aattttggatatgaaaatt**agccaggcatggtggcgggagcctgtaatcccagctactcaggaggctgaggcaggagaatcgcttgaaacttggaggcggaggt**tgcagtgagctgagactgca**ccactgtgctccagcctgggca**atggagagagactccat**ctcaaaaaaaaaaaaaaaagaaaagaaaagaaaaaattgtgaaatgtggt**ctgagacaacttttctcag**tgtcacctgaggcatgggttaaacaggtatattctaggacaggttcag**aatttcagagtcgaaatt**tcttgttggtggactttaggaatacgtatcattgctcctgagtatttctgttagtgctgaagttctgccccaaggacttcgt**aagattaataacatcgcaaatctt**ctgtagtgctttaaaattttgttcttcatttttttattctcactttcttctctttccttttccacttcctcttatgttttccttaatcttcaactttcctaagcacctgcaagtgggattggagccttgtttaacatcgtccatgtagcaaaataaggatgaggccaaatatttgaaccaaggatccccatctcctatggaaggtgccctgaggttgtgggtgttgctgggga**catgatgtcatggccagatcctacatcatg**cggccaagggaacccagaactttcactgctctttgctactgcacatcagaacccatcgctgggagtgtcttgcactgcctgacctcaccATGGATATCTGGCTCCTCTGCTGGGTGACCCTGTGTCTCTTGGCGGCAGGTGGGTCCAGGTATACTT**AAACATTTGCATAAAGATGTTT**TTGGCTGGGCGTGGTGGCTCACAGCCGTAATCCCACCTTTTTGGGAGGTTGAGGTGAGTAGATCACCAGAGGTCAAGAGTTCGAGACCAGCCTGGTCAACGTGGTGAAACACCTTCT**CTACCAAAAAATACAAAAATTAGCCAGGCGTGGTAG**TGTGCTCCTGTAGTCCCAGCTACTTGGGAGGCTGAGGTGGGAGGATCACTTGAATCTGGGAGGTAG**AGGCTGCAGTGAGCAGAGATCACGACATTTCACTCCAGCCT**GGGCAACACAGAGAGACCCTATCTCAAAAAAAAAAGATGTTTTCTTTGGGCTTCCCTTCACCTTCTATGGCTTCCGTCTTCTTCCACAGGACACTCGGAGCCTGGAGTCAGCCAGACCCCCAGACACAAGGTCACCAACATGGGACAGGAGGTGATTCTGAGGTGCGATCCATCTTCTGGTCACATGTTTGTTCACTGGTACCGACAGAATCTGAGGCAAGAAATGAAGTTGCTGATTTCCTTCCAGTACCAAAACATTGCAGTTGATTCAGGGATGCCCAAGGAACGATTCACAGCTGAAAGACCTAACGGAACGTCTTCCACACTGAAGATCCATCCCGCAGAGCCGAGG**GACTCAGCCGTGTATCTCTACAGTAGCGGTGGcactgcatggctgagtc**agttccctccagggtgcaaaccctctggctgctcttctcccagttgaactccaagaaaacatttgaaaaagcctcttccttatcttcctaccccagaagaaagaagcgagttgattgttgtggctgcagctgctaccgggagagtac**aagaccatgaattaaggtctt**aaatggtcatgatgggcacactggacaatgggctcctgaaactacccaaatacaaaatgagacattc**tgtggatcaggaggaatccaca**tgtttagaaggaagggccccagaccaatttccaagttcagagaccagctatctgaggttgatattacttaccaaaacacaaaaacattcctactgattttatctcaaacacagttctcctgcaaactcttcccccaccaacgtaccccagcagaggcaatgacatgtacatttatggaacagcctgttgactactgcagatccatctcttcaataagaccaactttccctggaggtcagtgactaaaccaaaccgcaaggcagacggcaacacgaccctctta**ggggatggatgccagagaaacactgactaaaatcccc**actcggggactgtcctagcacattcagttcctcacacagctgcccaggggagcgcgacgttttattctgttacttcaaacactcatctgttcctctctctctcttgcaggccctagactcacagacac**tggaattagagtcagaaaaattgctttctaatccca**aaactggttttgtttactccccttcagctgtgaaatcttggcaaatgctcttacttctctgggactcagtgtcctcctcttcaaaa**ttagagacttgctccacttagtctctaa**gatctgata**caaatatgattatttg**acttgctatttatttatttatacaaatattaga**atatttatgtatacatttaacataaatat**atttatatttaataaaaacataaaatttataacattttata

Uppercase: TRBV17

Lowercase: Flanking sequence[1000bp]

Red & Bold & Underline: Stem-loop [19]

Blue: Heptamer[36]

Green: Nonamer [5]

id-IGHV3-47-2[V_gene_segment]

atgtctcatgtctccctaaattgtgtaaaaccaagctgtgtcccaccacactgggcacgtatcat**caggactccctgaggctgtgtcacaggcatgtccttagtcctg**gaaaaatgaacttcctaaatctattgagattagtctcagatactctttggtttataagtatgttttctatctcataacttcaattatctgaaatactaaaggaaatctgcca**aataacacattatcttgtttatctgttatt**gttgttgcaaaa**ataaatatatttatataatttat**gtataattgatcatctttgtacattaccaaactgagtagcagatttgttagtaagacccaatgtaataaaaaattcaaatatcaattagcaacttaatagaa**aaacaaattacgctacatttgttt**gccgaaatgctacca**atttttataaaaaaat**aaacacatataacataaatggtttaa**ttctcagtatttctattgagaa**aaataagccaaatacataagcgaatatactatattatttcattcttataaattctaccaaataaatactagtctaaagaaatatgaaaacatcagtagtttataaagaaatggtagaaaaagggaaggggaaaaaacaaataatataaaagatcaagaggaatgctgaggggagttgacttgttacctca**ttgaaaatagagattattttttcaa**agtttactattgcacatgttaaatatgtg**aattttatcatctgtcagttaaaactcataaaatt**tattacaagtaaacagccgacattttatacaaaaaagggatgataggaagaaacaaataaatacattaaatgt**cagatacaccaaaaacttatctg**cctgacccctagttgtctccgtaatttttggatgaaaaccagcccacccctgaccctgctgctctgggagaggagccccagc**cttgggattcccaag**tgtttgca**ttcagtgatcaggactgaa**cacacaggactCACCAGGGAGTTTGTGCTAAGCTGGGTTTTCCTTGTTGCTATATTAAAATGTGATTCATGGAGAACTAGAGAGATTGAGTGTGAGTTACATGAGTGAGAGAAACAGTGGATATGTTTGGCAATTTCTGACCTTGGTGTCTCTGTGTTTGCAGGTGTCCAGTGTGAGGATCAGCTGGTGGAGTCTGGGGGAGGCTTGGTACAGCCTGGGGGGTCCCTGCGACCCTCCT**GTGCAGCCTCTGGATTCGCCTTCAGTAGCTATGCTCTGCAC**TGGGTTCGCCGGG**CTCCAGGGAAGGGTCTGGAG**TGG**GTATCAGCTATTGGTACTGGTGGTGATAC**ATACTATGCAGACTCCGTGATGGGCCGATTCACCATCTCCAGAGACAACGCCAAGAAGTCCTTGTATCTTCATATGA**ACAGCCTGATAGCTGAGGACATGGCTGTGTATTATTGTGCAAGAGAcacag**tgag**gggaagtcaatgtgagcccagatacaaacttccctgcaggaacgctggaggaaaccagctgcaggg**ggcgctcaggagccactgatcagag**tcagcccaagaggcaggtgcacacagaggctga**tttcctgtcagggtgtgggacttcgtctttttaccatttctctagggaacctctctaagttcagaattctgtgcttaccagtgtcatctctacatgttttttaatgattattttaatatgagaacctattcttacatgcacaaaatgcagatggatgcttacagagatgaaaagtcctcaaccatgg**tcaccaggatcagccttgaggaaactcaggggtgcctggtga**atcttctcc**agtcagactcaggacagaaacctctgtgagattccctgact**agatcagtcttcaggaattttgata**ccagccaatagagaggctgg**gac**agggtcagtgtcatgtagaacctcacaggtttcacgtctgaccct**tctccttctcctgacactaaagtatgcaa**atcagtatcagcactgat**ctgggtccccttttgctcttaaaccattctatttctttttatttgttgttgttcttgcttttcctcgtacttctcttgctccctgtaaaatggggaggtgtttctt**gctgcaaaagccccaagcctcaagcccattccctgcagc**tcaggcggggctcaggctgtggctcctg**cagccacatgggagtggctg**ttggggctttctcttctcccattgctcagcaccctccagtg**tgttgtgtggagactacctgggaacgaatgtggacaaca**ggagtgaaggggatgagcttgcgtggacaaaatgggatgtggatgtgaaatttatcctgtgctgtacaaacaaccacagattcaacttcctcaccagtagtattagaaagagggtgtgaaagttgtcagaataaaaatggaaacacttgtgttaacaccctgacaaatggaacta

Uppercase: IGHV3-47-2

Lowercase: Flanking sequence[1000bp]

Red & Bold & Underline: Stem-loop [27]

Blue: Heptamer[35]

Green: Nonamer [6]

id-IGLV3-24[V_gene_segment]

gctgtgccctgcggtgggacctg**cacctttccaaggtg**tgtgtta**tcatctgcacagtggagatga**agagggaacctgcattgggttctggaaagtaaca**tctaaaaatatatttaga**cctgcaacagaaatctcctctctaa**cataaaatatttatg**atcaggacacactgagaggcctagaccaaaattaagcactg**ccctcctgagccaggatagaatggaggaggg**gactggggaccccagataattccacagttaccagcatgactct**atgatcacctcctggaccagatcaaagatcat**ccc**atatgaaggacttcatat**tccttaggaaatacaaagtcacctctgagctctgctgtgaccaagtgagcccaggaatgaggctgagcacagtgtgatgtggagatgtctgcagagaatatggaatcaattgataaaggagatcctgtgggcaaa**gggtgggagggcaaggccaccc**ctaacctctttgggtcccttggggctgagtccttctggaaaccacaaagctcctccagcag**cagcccctgactctgctgattggcatcacgggctg**t**tctctccaacaaggggttaagagaga**cctgggaggagcctgcccagcctcagacttgagaagcagcattggaggcacttcagccatggcctggcccctcttttccttggtctcctgactcactgcc**caggtgctgcccccagggtctcacccacctg**cccagcc**ccaggcctccaggttcagcctgg**ccatcagtctgagct**cagcagggccctgtgtgtggtgggcaggatgctcgtgaccctgctg**cagggggaggggctggttgggctgaaatgcccccactctgtgctcctgtgctcatgggtgccctgagagctcttttattcctggggctcactccagccctggcaagtagcaagatatcctggggttgagcctctcagtttcaaattttttatctcccctttccttcttgcagGCTCTGTCACC**TCCTGTGAGCTGACACAGCCACCCTCAGTGTCAGTAGCTCCAGGA**CAGATGACTAGGATCACCTGTAGGGAAAACAGTATTGGAAGTAAAAGTGCTTAATGGTACCAGCAGAAGCCAGGACAGGCCCCTGTGTGGGTCATCTATGGGGATAGCAGGCGGCCTTCAGGGATCTCTGAGATTCTCTGGCTCAAACTCAGGGAACAGGACCACCCTGGCCATCAACAGGGCCCAGGCTGGGGACGAGGCTATTACTGTAAGA**TGTGGGACATTAGGACTCCTCATCCcaca**gtgacagaggtag**atggggaagtgagacacaaaccccttccccat**acgtgtcactc**ttgtcctccagcccaggaagactgtggacaa**agccatgagcatgggtttcccaattcacctggatctgagaccccccaggctgccctttccacaggccctccaggcagg**ctctgcagaggggacatcaagactgaatttagggcagag**gtgaccaggatatattgaggtttgggggactcagttgtgatttggggaaatggggccagaatggaaggtcaggccaaggtcaacatccttagttggtctctggatttccatagggggtgaaggccagtccttgggctgaatat**cctgagtcagtttccagaactgcccagctcagg**ctccc**atggcatcctgccat**tctctgaaaagggccctggcatgccaggtgctctgggttctgagggagagttggaatcagatcctcctgcaggaatccagg**gtcaggagaagcagcctcacctgac**acagggaccacagatgcacccccacagggtgagctgc**aggatggatgctgctcatctccaatcctcacatcct**atgcaaaaaatcattcttttcacccctccttcagtgtatcaaa**gttgacacttaaaaagccttaagagcccactcttgtcaac**agaaaacccattcacatctccatcagccacacaatctccacagaatgattcttcaaaaaacgataacagtagacatgaaaaggtatctaaaatgttacataaataattgaacttagattacatatgaaaatgatactatttttggtatgctaggt**aaataaaatatattattt**ggactcctttactttcttctctttgtatttctaatgtggtgactaga**cattgggattccaatg**aggctcacatcatattgtgattgaacagtgctggtctgcacctcttcaggtaacaggggctgtggagtgacctggtgcaggaagccccatctccaatc

Uppercase: IGLV3-24

Lowercase: Flanking sequence[1000bp]

Red & Bold & Underline: Stem-loop [25]

Blue: Heptamer[44]

Green: Nonamer [3]

id-IGLVI-42[V_gene_segment]

cacaatggtgattaaacctggtgctcattcctctggtttgatccacctttatgctcttgtcatttaaagttaatgtaagtagtgttccatttat**ttaaaaggataaaatattttaa**tagttgatcattgcattttgaactgagtaccagtgaagtattctagcttttaggtaatttcaattgtttgctagtcaatgatgttatacgttgttgtcattaaaccattcctggtcatttctgtggattttgtgaggtcagagaaaatgtctgtgcttttcaatggaaagccttgggtacattattcctgacttcccaatgcacaaaatagagaaacacaaatttcctgctcatgttagttagagcataagagattcaggtacgaggaacaatttataccccaaactacatacagtgtcttcaggcatgtattagaagtgtacttctcaaactctctgatacccttagttctgaagaagctcaaagttggctaatacttttatatttatctttttattttttttacctcagccagcactttctcc**tgattcctggggttttctgcatttccagtgaatccagtgaatca**cacaaataggctaactgattcttcccaaactc**cagcactctcccttatgtgctg**atctagtgagg**aatcacatgggaagagagacccatggatt**tctggcggcctgggaagtgagttcatggctggcctcagcttgcaaaccagacacttgtctccttgtttccctttactgttcattgggttcagcagctgtgtcct**tccaggtttcctgga**cagtgatggaccacactctcccaactcccaaccacccgctcctcaaaaagaataaccagactcagcaacttatgtgaggggcacagccaggggtcaggatagattggcattaagtcccctcaattaagagaggccaggga**aggggggctgttccccct**aatccgtgtggctcaggaagcagagctctagagacatctctaccATGGCCTGCACCCCCTCCTCCTCACCCTCCTCAGTCGCGGCACAGGTGGCTGGAGACAGGGAGTCAGGGACTGGCACTGGCAGGA**CCAGGGCTCTGCTTTGCTCCCCTGG**CTCACTGAGCTGCCTCCCTC**ACCCTGTGTCTCTCTCCCGACTCTCAGGGT**CCCGGGCAATCATGGCTCATCAGGTAACCTGCACCGTCTGTCT**GTGGCTAAACCAGCCAC**CATCTCCTGAACTGGAAGCAGCAGCAATAAGGTTCTTGGAATTGTGA**CCTGGTGCCAACAATGACCAGG**AAGTGCCTCCAAGCTTCTGACTCATAGAAGTAAATATCTGCCCGCAGGGATTCAGGACAGACGCTCAGGCTACCAGTCTTGCATG**AAGCCCTTCCTAAGCATCTCTGGGCTT**TAGGCTGAGGACAAGGCTGATCACTCCTGTTGGCTTCAGACAGCCCCCTGGAGGTCCAaacagtgctgcagtctgggaaagtgagatgagaacacgccaggtctcctaggagcatgaccttccaatggcaccacccacaaccaggacagctggtctgtttaccatttgtgtggatgtttcttgatgctgccgctaaccagggcccaagacttagttcggggacagtgatct**agagaatacagctttgttctct**cacagccagccctcagaggaagcccttagcaacagcattgtcagactgcctcaaaaaaatcagagtctttgattccaggaacaggctgccctggggaca**gactcaagatgagtc**agggtaagcagggacagaatccagctgggtctgcctcaggcatgtgcactaacgcaaaggtctgttcatccttattgtctactaaacacctattatgtgccagtctcaggagagtgctcaggatacagagtttacaagac**agacagggttcctgtct**ttacagggataatgcctcacagaagagtgagcaccaacaaatcaatgagaataaggcatattgtgaggtataaaatggttgacatgggccagaagacagtgagcaacaagaagctattaggaggttgtaagcaaggagatacagactctggtttgagattcaggaggcaaagtccagcttctctaaggaagatgaaaaaggaggggacagagtgaagaaagatggtccagttaggacatggtggctttgtacacatgagagatgaggtttctggcaagtaatgtggcagctgagctgaggcctgtgctgtggatcccagacgggagccacctgctaggcctaggctggagccctctgcctctgctcacg**aggaccagcagctgtgtccttacaggtcct**tagggagccacagaacagccccaccctggtccaccccatcccaggctcagcttt**ctaggcaagggtcagagtcagggcctag**ggtgggacttggccagcccaggggaggaggctggtgtgcatgaagagaccctc

Uppercase: IGLVI-42

Lowercase: Flanking sequence[1000bp]

Red & Bold & Underline: Stem-loop [16]

Blue: Heptamer[36]

Green: Nonamer [4]

id-IGHVIII-2-1-2[V_gene_segment]

caggattattttctttccttggttgagaaatatttccccaaacttcagctcag**ttcaggcatacactgtccctgaa**tgggcatttaccct**cagatgggtacacacatctg**tcaacatgtggactcttctgtcagacaaacgcacctttact**cacgtggattcttctctcagacaaacacacatgtccccacgtg**gactctttcctcagactaccacatatgttcttacatttactctttcctcagaaaacagacatttcctcatgtggact**cttgtctcagacaag**caaacatgtctcc**atgtgaactcttcactcacat**aagtacacatatgtccacatttactgtttccttacacaagtacatatatccaatgtcgaattgttctgtggcaaaatgatctcaagataatgataatcataaaccccctccctgacaaggcgtagatctgtatttttttcattgtaacctaactttgccttattgtcaagaacagtagtttgcag**ctctaaatataccaattagag**acaggtgtccattttctctggaaacgtatttttatgttcttactggacatatttgttgataatgtttgctattatgaagataccccaacagtgtccacattagagaataaaaaagagtaatgggcagattaactctgtgcatccagacccagaaatcctttgaccttgacttccctgaaatgtagacacagaggatggatgagcaatgctgagcagtgcacccatgaccacaaaaagaaagacgtggaaatgtgtcccctccacttctcatgaaaggcagctcatcccctgttccctcaggccctggcgaggagccaccccatgtctgtgcccttcctcagtgtccacaccgtggggtctgcattgatctggattcccttctcatccccgtcaatattagtgtccttcgtaaatcaggtccagctgtggcttctcctcacggggctgttctcagtctgttttctgtgttcacggAAGTCCTGTGTGAAGTTTACTGATGGAGTCAGAGGGGGAAAAATTTTACAGCCCAGTGGTGAGACTCTCCTGCAAAGCCTCTGGTTTCACCTTTACTGGTTAC**AGCATGAGCTTGGTCCAGCATGCT**TCACAACAGGGATAGGTGTGGGTGCCAACAGTGAGTGATCAAGTATGAATTCTCAGGGTTACTCTCCATGAGTACAAATAAATTAACAATCTCAAGCAACACCCTTTTAAGTGCAGTCTGCCTTACA**ATGACCAATCTGAAAGCCAAGGACAAGGTCAT**GTATT**ACTGTGAGTGAcacagt**gagggaaaccctgt**gtgagcccagacacaaagctcac**cgcagggagacaggaggggactatgtggtagatgctgctcagaaccaccagggggcactcagaaccatcagggagggtgcacagaaccaccaggaggggctcaggacaccagggggcgctcagaaccaccagggggcactcaggaccatcagggagggtgcacagaaccaccaggaggggctcaggacaccagggggcgctcagaaccaccagggggcactcaggaccatcagggagggtgcacagaaccaccaggaggggctcaggacaccagggggcgctcagaaccaccagggggcactcaggaccatcagggagggtgcacagaaccaccaggaggggctcaggacaccagggggcgctcagaaccaccagggggcactcaggaccatcagggagggtgcacagaaccaccaggaggggctcaggacaccagggggcgctcagaaccaccagggggcactcaggaccatcagggagggtgcacagaaccaccaggaggggctcaggacaccagggggtgctcagaacactaggaggtgctatgaatcactagggggcgctcaggactcaagggagcactcagaaccaccagggatagctcaggacaccagggggcactcagaaccgccagggggcactcaggaccatcagggagggtgcacagaaccaccaggaggggctcaggacaccagggggcgctcagaaccaccagggggcactcaggaccatcagggagggtgcacagaaccaccaggaggggctcaggacaccagggggtgctcagaacactaggaggtgctatgaatcactagggggcgctcaggactcaagggagcactcagaaccaccagggatagctcaggacaccagggggcactcaggaccatcagggagggtgcacagaaccaccaggaggggctcaggacaccagggggcgctcagaaccaccagggg

Uppercase: IGHVIII-2-1-2

Lowercase: Flanking sequence[1000bp]

Red & Bold & Underline: Stem-loop [10]

Blue: Heptamer[31]

Green: Nonamer [3]

id-IGKV2D-36[V_gene_segment]

gaccccaagtctctagcc**acatccaattcttcacagaagctggatgt**atgtgccagggcaaggagaactgggtgggaggtgagt**ctgtgggctccacag**cttcacgtgtttatgaggtaggagttattaagatattattttaggcagatagagaggaaaaaagggatccttgggagacttttcacagctgttgtctagcatgaaagccctgggtcttagccccgggcaggcaaccttttatatgcaaatgcaagccgttagaaactgggtccatccaacatgcacccaacatggtgatttccaccgttgtcctcttgcccttgccccaacatgcgcctggcagcatggttgcccctacatatccccacgtgtgtggaacatcatggcgccctgtatttgcatattaaaaggctagggtgggagggccagttctttcgagggctaccgtgaatgacatgtctggtcaaaccaataccctgagccctgtgcaaatcagacaccgcctcctc**cagccatcaaatatagctggctg**atattacctgaaagagtggttcgtccctcggctttggagcccccctccctctgtctttgtacaggggagcttcatccttctttcttcccccttctttcttgcctattaaactccctgctccttaaaactattctaagtgtgtcagtgtcattttttctcatttgactcgagacaaagaacctggtgtcctctactcatcagagcag**tatcatttattccaacatgata**gagtcacatggtctatgcccagggaaaaga**attcatatttttgtcttatgaat**aatcaaggttcacctccagggaacaatgaacagtactctgattaagatcttgaaaaaaaaaaagagttcccttctggctggt**aaataatgggtttattt**tagaaagtctactttcatgacatgaatcaaaacttgaaaaatgtaactataaatcaatatcataaaagaaatcatgaaagttgttcatAATGGAAAGCCAGCCTTTATCTCCTGCAGGTCTAGTTAGAGCCTCCTGTATAATAATGGAAACACCTATTTGTATTGGTACCTGCAGAAGCCAGGCCAGTCTCCACAGCTCCTGATTTATGAGGTTTCCAACCAGGCCTCCGAATTCTCAGACAGGTTCAGGGGTAATGGGTCAGG**TACTGAGTTTACACTGAAAGTCAGTA**GGACGGAGACTAAGGATGTTGGAGTTTATTAGaaaacttgaacttccatcaatgataaatattccttttg**cctcaagcacatatttgagg**aattttccattgagtagatctaccgataaggtcacatttttctgtctgttttaatctgaatatgtatttatatgat**tctcagaaatgttttttgactatgaaattatatttgtgaga**ccttttttcaagttgttgtttaccatttgataattactgctgaaacgtcatctattaaaaaaaatcggtcatccgttctctaactctcattgaaaggaatttgcctttttaaacccctcagactctttttaagcttttcaattggccttatcttttttttacagattcaatgtattaagtc**aatttattatttatgataaatt**tatttatttattt**atttttactatcacaaaatttaaaaat**cctgtaagttgctatgtcaaacctgcctctagattgcaaacaaaccccacaaaaaagagagtaccattatcttagaaatactagaaaagcaagagtgatagtgtcccattgttaacagggaccccataaggctagaataatgtcccacatgttacctggggcacctgggaggagctgccactgtgctgaatggtgggaaaaagccatctgtgctctgagactggaagcctggccctattcccctatgtggatgtaggagataattggggttataaaggggtaattacggttaaatgaagtcataagggtggggctctgatccaataggattag**tgtccttataaggaca**gaaacc**agagaactcttctct**tttcctttatttctctaactcactgcttccctatggaaaggccatgagaagacatggtgagaaggtgactacctgtaaactaggaag**aaggttctaaccagaaagtgaacctt**g**ctggaacttgatcttggactttccag**gcttgtgaactatgagaataaatttctctggtttaatctacccagcccagggtattttggtgtagcagtgca

Uppercase: IGKV2D-36

Lowercase: Flanking sequence[1000bp]

Red & Bold & Underline: Stem-loop [15]

Blue: Heptamer[19]

Green: Nonamer [5]

id-IGHVII-62-1[V_gene_segment]

cagcaaggccattggtgccttccagagcacagtagtccatcaggaatctcccatgtgtcccagcagcagccttgcctcagtatctccactgtgcacagccattgt**ctgggaggagctcccag**gatgagtgtctttggcacacaccaggtggcgggtgttagagtgcagtacagcagctggctgcctggtctattgggctccctgatgttggagggatttgaggtgcattctcaggcccagcaccctctttgtgaatttttatataaaaaccctgattttacttcattttctcagatgacatagataactaaaaacagaatctgcaaagaaattgtaattttcaactttaccccaaattcattgtttcttaattctgtgcaagatccaaacatatttttgccttccac**atgagaaattgttctgtttaaactgaaatcattttttctcat**attctttctttttgcccaagtacagagatcttgttgaaaataagttgggttctttccacacactaaccctca**cctcccctagagaaagagcagagattgtcctcactctgagtctaggggagg**agctgttcctgtacaattcagagcctgcagagacccccaggtgcagcttcagtgagtcagacatttctccatgtgggcgacctccagtgcctgtgactgcttctcagg**cctaattgtgggttaagaattagg**acactctttaagttatcacatctgaatcctattctgaaaatcaccataaagagagatagttcaatggctattctcctgacataa**gtttctctttattacttggttccaagtatggagaaac**atgtgaccctatatttgtctgaatccaacatcagaatccactgcattactctgggatactcacggatcgaacacaggtgagctctttattctcacagaaatgtatgtagttgggaatttcaacatattgtccagaacgtgtgcctgccaacaactgtgtttctcggtgcactcttGGCCTGGTGAAGCCCTCACAGACCCTCTCCCTCATGTGTGTCATCTCTGCATTCTCCATCACAACCAGTGCTTCCTCCTGGAGCTGCATCCATCAGCCCCTCTCTGGGAGGGAATGGAGTGGATTGGGTGCATAGGTCATGAAGGGAGCACACATTACTCCCCTTTCCTCAAGAGTCCAGTCACCATCCCCAGATCCATGTCCAAAAACAGTTCTTCCTACAGCTGAGCTACATGAGCAACAATCACATAGCCATATATTTTTAAGCAAAAGA**cacagtgagggaatcacagtgtgaactcacaccaaaacctccctgtg**gggatgcataggacagaaggggttactccg**gagcccagggggctc**tcaggacaccaaggggcactcaagaccattgtagaggcacgcaggtagccgggggctctcaggaaccatggaggaaaatcaggacaccaaagggtgctcggtacagcgaggggctcaggacaattgtggggattcagaaagaacaggttcaaggctcagcttcagggcaggtgcagctggtgtgaaaaggggctggatgaggcgttttgtgtcaccatcatgtttcaccaccagacaccctccactacatctg**ttctaatgcatgtgtttgtatgattagaa**aatgatattgatataaatatataaccatagctaggtgtgtctttttctgccttccgtcctcccttcttctctctctctcacacagaaatttacatacacccaccccacaacacacataaatctataact**tttattacctgatgtattcaataaa**cctgattaatgtgttttgcagcttcattgtttatgctgttgtagcaataaaaaccatctgtttcccagctgtgtacttctctaagctgagtagcatctttgttcataatacccagaatgaaaaacaacccaaatgtcaatcatcagcttaactggtaaacacattgtggaaaagtcattcattgacatactacccactactaccatcagctaatgttggctatgctcaaaagcatggttaaattcacaagtacttcagatgagtaaaatgagcccaagtgataaaagtgcatacataggata**caactttcataagttctatagaacaaaaagtaatctaaagttg**caaaaaaaaaatcagtagttcactgtgaatattgtaggagaagggatggtctaggaaggaggaattatgg**aacaagataaaatgttgagggaattgacttgtt**atctatgttgatagtgataatgtc

Uppercase: IGHVII-62-1

Lowercase: Flanking sequence[1000bp]

Red & Bold & Underline: Stem-loop [11]

Blue: Heptamer[42]

Green: Nonamer [7]

id-IGHV3-13[V_gene_segment]

attctcagtatt**tctattgagaaaaataagccaaataga**tgagtacatactatattatttcattcttataaattctagagaataaaaactagtctaaagaaatatgaaaacatcagtacttttataaagaaatggtagaagaaaaggagaaaaacaaaaatatatctataaaagagcaagaggaattctg**aggtgagttgacttgtcacct**tcttgaaaatagagactttctttatcaaagtttactattgtacaggttaaatatgtgaattttatcatctgtcaattaaaactca**taaaatttattacaagtaaacaagtgaaatttta**gacaaaaaagggatgataagaaggaacaaataaatacattaaatgt**cagatacaccaaaaatgtatctg**cctgacgcctagttgtctctgtatttttaggtaaatgcagcaaaatcacacaggttgtcgtggcaggaagtggattctgcaaaccacactaggccgtttatctctgtcctgtagttggttcagagcaactgaggccagctgtgaggcgcataggcccaggtactaggactcactcatgccagatataagccctgagacacgtacatagcccctccatgtgtgggttcacttttacatctgtacatgaagaaaccactgactcctaaataacatcatctatacacataggtaaaaaaattaaaaatatgatagttgttaaatgtttatcgcagaacaatttcaaaataaggcagcattttcccaaatacaatcattgtcatcaaaatcccccaggacgctctcatctactctgcgccctgccttcacctcagatgtcccaccccagagcttgctatatagtaacagacatgcaaatagttgactccctctcctgatgaaaaccagcccagccctgaccctgcagctctgggagtggagccccagc**cttgggattcccaag**tgtt**tgtattcagtgatcaggactgaacaca**caggactcaccATGGAGTTGGGGCTGAGCTGGGTTTTCCTTGTTGCTATATTAGAAGGTG**ATTCATGGAGAACTAGAGATATTGAGTGTGAATGGGCATGAAT**GAGAGAAACAGTGGGTATGTGTGGCAATTTCTGACTTTTGTGTCTCTGTGTTTGCAGGTGTCCAGTGTGAGGTGCAGCTGGTGGAGTCTGGGGGAGGCTTGGTACAGCCTGGGGGGTCCCTGAGACTCTCCTGTGCAGCCTCTGGATTCACCTTCA**GTAGCTACGACATGCACTGGGTCCGCCAAGCTAC**AGGAAAAGGTCTGGAG**TGGGTCTCAGCTATTGGTACTGCTGGTGACCCA**TACTATCCAGGCTCCGTGAAGGGCCGATTCACCATCTCCAGAGAAAATGCCAAGAACTCCTTGTATCTTCAAATGA**ACAGCCTGAGAGCCGGGGACACGGCTGT**GTATT**ACTGTGCAAGAGAcacagt**gaggggaagtcagtatgagcccagacacaaacct**ccctgcagaatgcctgggggaaatcagctgcaggg**ggcgcacaggacccactgatcagagtcatccccagaggcaggtgcagatggaggctggtttcctgtcaggatgtgggacttcatcttt**ttagagtttctctagggaacctctctaa**gttcagaattctgtgcttaccaatgccatccctacatattt**ttaaaatgattattttaa**tatgaaaacctattctcttatgcacgaaacacagactgatgcttacagagatgaaaagccctcaaccattgt**caccaggatcagagtattgaggaaactcaggggtacctggtg**ggtcttctccactcagactcaggacagaaacctcagtgagtttccctgactaggacggtctttaggaattgtgatcgcagccaatagagagtctgggcaaggatcagtgtcatgtagaacttcacaggtttcacttctgacccttctc**ctgacactaaagtatacaacttagtgtcag**cactgatctggggccccttttgctcttagcccattctatttctttttatttgttgttgttgttcttgcttttccttgtagtgttcctgctccctgtaaagtggggatgtggctcttgctgccaaagctccaggtctcaagcccattccctgcagctcaggtggggctcaggct**gtggctcctgcagccac**gtggaagaggctg**atgggactttcctctctcccat**tgctcagcaccctccagtgtgtcacgtggagactcacctgcgaatggaagtggccaacagtagtgaaggggatgagcttgtgtggacaaaatgggatgtggatgtgaaatttatc**ctgtgctgtgcaaagtagcacag**agtgagtcaccttcctcaccagtagtgttagaaagagggtgtgaaagttgtcagaatcaaaatagatccacttgtgttaaaaccctgacaaatgg

Uppercase: IGHV3-13

Lowercase: Flanking sequence[1000bp]

Red & Bold & Underline: Stem-loop [19]

Blue: Heptamer[38]

Green: Nonamer [7]

id-IGHV3-64-2[V_gene_segment]

tcagtatttctat**tgagaaaaataatccaaatacataagagcatatactatattttttca**ttcttataaattctagaatataaaagctagtcaaaagacatgtgaaaacatcagtagatttataaagaaatggtagaagaagggaaggagaaaaaccaaaaaaatataaaagagcaagaggaatgctgaggggagttgacttgtcaccttctt**aaaaatagagattttattttt**caaagtttactattgtacagaataaatatgtgaattttcttatctgtcaattaaacctca**taaaatttattacaaaaaaactgaaatttta**gacaaaaagagggtgataggaaggaacaaataaatatgttaaatgtcaaata**tacctaaaaatttatttgtctgacccctagttttctccgtatttttaggta**aatgcagcaaaatcacacaggttgtcgtggcaggaagtggattctgcaaaccacactaggcccgtttatctctgtcctagagttggttaaaagagcaactgaggccagctgtgaggagcataggcccgggtactaggactcactcatgccagatataagcccttag**acacatacatagcccctccatgtgt**gggttcacttttacatctgtacatgaagaaaccactgattcctaaataacataatttatacacataggtaaaaataattaaaaa**tgtgatagttattaagtgtttatcaca**caacaatttcacaataaaacagcattttcccaaatgtaatcattgtcatcgaaatccccaaggacactctcatctgccctgggccctgccctctcctcaggcatctcaccccagagcttgctatatagt**aggagacatgcaaataggtccctccctctcct**gatgaaaaccagcccagccctgactccgcagctctgggagaggagcccccgc**cctgggattcccagg**tgttttcatttggtgatcagcactgaacacagaagagtcatgATGGAGTTTGGGCTGAGCTGGGT**TTTCCTTGTTGCTATTTTTAAAGGTGATTCATGAGGAAA**TAGAGATATTGAGTGTGAGTGGACATGAGTGAG**AGAAACAGTGGATTTGTGTGGCAGTTTCT**GACCTTGGTGTCTCTGTGTTTGCAGGTGTCCAGTGTGAGGTGCAGCTGGTGGAGTCTGGGGAAGGCTTGGTCCAGCCTGGGGGGTCCCTGAGACTCTCCTGTGCAGCCTCTGGATTCACCTTCAGTAGCTATGCTATGCACTGGGTCCGCCAGGCTCCAGGGAAGGGACTGG**AATATGTTTCAGCTATTAGTAGTAATGGGGGTAGCACATATT**ATGCAGACTCTGTGAAGGGCAGATTCACCATCTCCAGAGACAATTCCAAGAACACGCTGTATCTTCAAATGGGCAGCCTGAGAGCTGAGGACATGGCTGTGTATT**ACTGTGCGAGAGAcacagt**gaggagaagttaatgtgggaccatgcagaaacct**ccctgcgggaacgctggggaaagtcatctgcaggg**ggcgctcaggagccactgatcagagtcagccccagcggcaggtgc**agatgaaggctgatttcctgtcacgatgtgggacttcatct**tcttaaagtttctctactgaacctaagttcggaattctgtgattactaggg**tcatttctacatatttttaaaatga**ttgttttaatatgaaaacctat**tctcctatgcagaaaatgcagattgatcctcacagaggaga**tgaaaagttctcaaccatggtcaccactgtcagagtcctg**aggaagctcagggcttcct**ggtgagtcttctcc**agtcagactgaggacaggaactccagaaagaatcctgact**agaactgaggtattgctcctcacaagagaactcaagctgcgg**ggggtctgttcctgcagacccc**tgacccggtggcggatgaataaaacgtacactgacacacagatattctgttttgccagttcagctgagggtccgaggcaac**ttacagactccaagaagaatcctgtaa**agaacggcagcagtggccctgagcagctcgcactccaggcgtgtattgagtatagaattaacaacagaagctctgacaacacacttgtggataattaacatggttaagagagtagttctacgaatgattaaagctcagggactgcggtctaaagtaaataccattagggggcaatacc**ctggttagttaatggaggtagggtaaccag**acttaactggggaggcctctattgtccctagtatttaccctatgcctaacgctctaaggtaagaactggccgccttcagcctgttcaattattacaagctatgta**acctttcggccttctaaaaaggt**atatgactattccctataactttcctt**aatattcccctttaatatt**tccgccaccatcctgagtgaatcccagcaagaa

Uppercase: IGHV3-64-2

Lowercase: Flanking sequence[1000bp]

Red & Bold & Underline: Stem-loop [23]

Blue: Heptamer[29]

Green: Nonamer [4]

id-IGKV2-23[V_gene_segment]

atgcagtcaaaattttaatcattacatactcacaatcacttctgtggggcctggagatactgcacatacgactgttagcaagacactcactgggacgctgcgttgtgtgatggccccacatacaaacctcaaggaggctcagcctctcaatgcagcaggagcagctggggtacccaggccacacgtccataccaggtgggctcagttagagatggctggagagccttccag**gaagaggccatgaggtttcagtcacaaacactggctcctcttc**tgtgtaaacaggggctagagccctccaggacaattcctagagcctctcgctttctctccaattagtgcactgacaccccacagattctccaggaagtggttgtcatgtcctccctgcaacagccactaaagttccctactgctgtcatgaatgcagggacacttagtcacatcactgggaggcgaccctagtgtatcctgacctcacctgctgccactgatgactttcagggcacctctttctccctttgct**gagtgactctcactc**tcaccaaccatcaggagaatggaaagctgcctgcaatgcatgatgttggctgttgagcaaatcaaagctcacaggagtctcaaacatgtacaccacataataatattttctgataatactatttggacttttcttcctt**tcaattctggaagtaattga**gaatattttttgaactcttagaaacacttagtatatatgtgtagtaggtagtaactagttttgtctactggtttattttgtttgcttgtttcaggccatgatgcggcatgttaaaatactgaagacaaagatacattttagaattaagcatactgtacattggctctttccacaccactgcaaccaccaggggatgtgcatattgtcccttaggaatgaccttcccttgtgagtctgggagtaaagctcagctgtaaccttgcctt**aactgatcaggactcctcagtt**caccttctcacaGTGAGGTTCCCTGCTCAGCTCCTGGGGCTGCTAATGCTTTGGGTTCCTGGTAAGGACAGAGGAGATGAGGGAGGAGAATGGGGTGGGA**GGGTGAGCTCTGGGGGCCCCACTGTCACCC**ATGTGTGTTCTGTC**CACGTGTTGGATGCACGTG**TCTTGTGCTCCAGG**ATAAAATGTATGGTGGCACTTTTAT**ATGTGAAAGAGTGAGGAAGATTCCAGAAAAAGCAAAGACCTGTGCTCTGGTGCAGATTCTGACATAGAAAGAGGAGGGTAGCATAAGTGACTTCCAAAGGGCAACTTGGGCCTTCAAAATGTCTGTTTTTTTTTTAATTA**AATTTTTTTGGTGCATGAATCAAAATT**ACACACACACTCACACACACACACACACACACACACGCGCCGCAA**TACAATTATTTAGCATTAAATAATTGTA**GAGAAATTATGATAATGTCTCATGATTTACATAACATTGTACTTCTTTTTTATATTACTTTAGGATCCGGTGGGAATATTGTGATGACCCAGACTCCACTCTCTCTGCCCGTCACCA**ATGGAGAGCCGGCCTCCAT**CTCCTGCAGGTCTAGTCAGAACCTTTTACATGGTAATGGATACACCTATTTGTATTAGTTCCTGCAGAAGCCAGGCCACTCTCCACAGCTCCTGATCTGTAGGACTTCCAATCAGTTTTCTGCCT**TCCCACACAGGTTCTCCCCAGTGGGa**ggagagagtagaccagtcatccccagatatatcacaggactagtttcaacctttggaagctggtctatatcctatggttaaataggcatttgtgatacgacctgaaatacatttggacaagaacttcactaacaattgagtcactgaagacttacggccctgtgtgacgcaccacataactgtgagtttgcagtggttgcaggtcagggacagattttatgcttaagatcagtagggtggaggct**gaggatcttggctattacaactgccaccacactctacaatatcctc**ccacaatggttcagcaccaaacaaaagcctcctgcttggattgtcccagctgcccaaattagttccttcactgaggagtagacagggtatattctctaaatctatgtaacaggaagatgttggtgaactcaggggattagtatgaagctacacctcaggcatcacacataagatcacttcagcagtcgcagccttagcatgggcagaacctacagaagatgcaagtgccctctgagccaggagacaggaggaaggaggaagggaaaggtgacttagctcatctcaatcctctctcctttgcatacatttgtcaaccag**atgtattcagcctaccagtcacacaactgaggctgatacat**gacaacatagcactggtatattcttggtattgtttggcttagcagttactagtatatatttaat**gggagaatatttggtggtgttaacacattgcttatctccc**tttccccagttgtactttacacttgttcttggcacacattctcctccaggactggagcattcacagggttttatgttactgttcttatgggagtaaaaagaaaaacgattcacattcttgctactgagctaggctgggatgccctgggccaagctgaaaatgtgaaaaataagagtatgaatatttattaagttttatctggatctaagatacttatccatgaaccagtcctgcagctgtgccca

Uppercase: IGKV2-23

Lowercase: Flanking sequence[1000bp]

Red & Bold & Underline: Stem-loop [14]

Blue: Heptamer[40]

Green: Nonamer [5]

id-IGKV1-17[V_gene_segment]

tgccaccctgtgtcttttaagtggggcgtttagcctgttgacattcaaggttaatattgatatgttagatttgataccaccatcatgctgttagcctgttattttgccaacttgtttgtgtggttgctttataatgtcactggtctgtgtacttcagtgtgtttttatagtggctgataacggtgtttcatttccatatttagt**gcttcttccagaagc**ttgtgttaggcaggtctggtggtaacaaatttcctgagcatttgcttgtctga**aaaggatcttatttctccttt**gcttccgaagcttagtttgtccaaatatg**aaattcttagttggaattt**attttctttaagaatgttaaatataggcccccaatctcttctgctgagaggtcttctgttagtctgatggcttccctttgtaggtgatctgttttttctctctagc**tgcatatacttacaaatatatgca**tcatcatgtcaatatattttatttttaattttttttagaaagcaacaatgataaacttacagaaaagttgcaagtatacaataaatacccccttccctaaccggaattatacaatagtcttttaaagaccttagaattctatcgtctaaaactttactatgtgtttcctacaaaaaaagaatatcctcctatatactccccatacgccaa**tgaaatatgatactccaccgactcctgaggaatatttca**aattgtcaaaaaaaatctaaaaaatgtctctcgtaacaaaatagtctccagtagaaacccattctctgcagacaaatctgtgctaccctgatcttacctgggacacctggggacactg**agctgctgctgagttactgagatgagccagccctgcagct**gcgcccagcctgccccatcccctgctcatttgcatgttcccagagca**cagcctcctgccctgaagccttattaataggctg**gacacacttcatgcaggaatcagtcccactcaggacacagcATGGACATGAGGGTCCCCGCTCAGCT**CCTGGGGCTCCTGCTGCTCTGGTTCCCAGG**TAAGGATGGAGAACACT**AGCAGTTTACTCAGCCCAGAGTGCTCAGTACTGCT**TTACTGTTCAGGGAAATTCTCTTACAACATGATTAATTGTGTGGACATTTGTTTTTATGTTTCCAATCTCAGGTGCCAGGTGTGACATCCAGATGACCCAGTCTCCATCCTCC**CTGTCTGCATCTGTAGGAGACAG**AGTCACCATC**ACTTGCCGGGCAAGT**CAGGGCATTAGAAATGATTTAGGCTGGTATCAGCAGAAACCAGGGAAAGCCCCTAAGCGCCTGATCTATGCTGCATCCAGTTTGCAAAGTGGG**GTCCCATCAAGGTTCAGCGGCAGTGGATCTGGGAC**AGAATTCACTCTCACAATCAGCAGCCTGCAGCCTGAAGATTTTGCAACTTATTACTGTCTACAGCATAATAGTTACCCTcccacagtgttacacacccgaacataaacccccagggaagcagatgtgtgag**gctgggctgccccagc**tgctcctc**ctgatgcctccatcag**ctgagattgttcctcagatgcagccacactctgatggtgttggtagaaggggacatgaagtcacctctgcaccctaattcttttctctttctcagccccaactacacagacatagcaatgcctctcctaatttaatacagacagagatcatgacacctgaggagtctagttta**tggcttcagttggactttatataacagagaagaagcca**ctatagatattctaggcaggaattgtcttaatacagagaattaaagtaaactactgacgtctaaataaaatg**tagagatgaatctcta**aatgtaatgttttatttacacagaaatatttcccaaatggggcatacaggaaaactcagtggtcttcaatatgtggcaagaacaaagaaaaagttagtgtttcatgaaaaagggaaaatattacctatggctctttgagaaagttcattggcactaggaagggttgggagctggcaagctcagactgggaagcagtggtggacaaagtgaatcctagaattatatcaagttatctcagaagttgtggacaaatttgatttcaggttacaagaagccaaagcagtgaaggttgcagagtgttttgttactgaaataccagggattcagcgtagatcctgcagctcaccacacagaaagccaatcactaaggcaagtattgccaaggaaaaggctttaatcaggtgctgtagctga**ggagatgggacaccattctcaaatgtatctcc**ctgaccaacaaaaattaggggtttatatagcacggaagaaatgtgggaaaacaggaattatggagggttaagggagataatttggtcaacaagaagcaggaggtcagttatgcagtcataatgagtgaagggtctgatgtcccactgtcctgattcagtgata

Uppercase: IGKV1-17

Lowercase: Flanking sequence[1000bp]

Red & Bold & Underline: Stem-loop [17]

Blue: Heptamer[28]

Green: Nonamer [3]

id-IGHVIII-67-3[V_gene_segment]

cctgtaccccataaatatatacgtgtactatttacccacaaagattcaaaattaaagtgtttatgtgtctaatattctgtaactatattacattaaacctaagaacacactggcttttacacactactgtcatccacatagacattcatagccttc**tttctctgcttgtccataatcccacactccaaaatgagaaa**acctctcctaagatatgacattaatttacttagtgcacaatttcaaaatacacaaatcgtctttaggattgttaaactgacccttgttgaaaatatgtttatcaactagaatacagtgcttatgtggagcttatttatgttttacaatcgcaacatcaaattctttccaagtctcttgtgtccaatatttatgccccaccttcttcagtgtaattagttcaaaccttctgattgttgttagatatttttatacagctgtctttttttgtgtttcatgacctgctaaatactttcgaagttgtatgcatttaggttaacactttgtctcaaaaaatatgagttttcagaaattctttatgccatgtgtccatcattgaatatcataaaaatagtttcattggccg**ggcgtgttggctcacgcc**tgtaatcccagcactttgggaggccgaggtgggcggatcacgaggtcaggagatcgagaccatcctggctaacacaatgaaaccccatctctactaaaaatacaaaaattagccgggagtggtggtgggcacctgtagtcccaactactcgggaggctgaggcaggagaa**tggcgcgaacccgggaggcagagcttgctatgagcagagatcgcgcca**ctgcactccagcctgggtgacagagcaagactccctctcaaaaaaaaaaaatagtttcattgggattaaaaa**gtgtgtgtttcacctaatccacacac**cttctccccaccttcctgggaaacctcaaatgtttactggatctatatttttatctttggcaa**aaCAGCCTGGCTGTT**CTCTTAGCCTCTTTTGCAAAGCATCAAGATTCACCTTCACTGACTACAGCATAAATTGAGCCCAGATGGCTGGGGGACAGAGGCTGGAGTGGGTGGTAACAGTGATTGATTCAAGTGGAAGTTCTCAGTGATATTCTGCATCAGCATAATGAAGATTCACAATTCCCAGGGACACCAATTACCAGCACAGTCTCCCTTAAAATAATCTACTTGGAAGCTGAGGG**GGCTCTCACAGGGGTAGGCAGTGTATTACTGTGAGAGAc**acagcgagggaca**tttctgtgagtccagacagaaa**cctccctgcagggagacaagagaggactttgtgataaatggtgcttaggacaccagggggcactcaggacagcagagggtgctcagcagagcagggtgcgctcaggacatggtgtgtgttgtagggggcattgctcaggatcaccagggggctctcagaaccaccaagggtcactcaggaaaccagggggcgctctggacaccagggggagctcagaaccaccaggtggtactcaggacaccagggggcgctcatgaccagcagg**gcatgcttgggactctatgaggcatgc**aggagaagcaggccgtgctcagggcaacagggggctcacagaaccacctgggggtgctcagaaccactaggggtcactcaggacagcagcaggcattcgggacaccatggggccctcaggacctgcagggggcgctcagaacaccaggggaaggtcagaaagacctgggggcacttggaaccaccagggggcactcaggacagcagggggcgctcaggaccactagggggcgctcaggaccagcagggggtgctcaggacaccatggggcatgcaggagcag**caggctgttctcagggtaacagggggctctcagaacaacctg**g**gggtgctcagaaccaccagggggcaccc**aggacatcatgcggccctcagaacctgcagggggcgctcaggacaccaggggaagctcagaatg**acctgggggcactcagaaccaccaggt**ggcactcaggacagcagggggcgctcagtaccacggggtgcactcaggacagcagggggcgctcaagacacagtgggatgctcaggaccagcaagaggcactcaggaccaccagggggcactcgggggagcaggaggcactcaggacaccagaagaggatcagaaccactcgggggcactcagaaccaccagggggcgctcaggacactgggggcattcagaaccacc

Uppercase: IGHVIII-67-3

Lowercase: Flanking sequence[1000bp]

Red & Bold & Underline: Stem-loop [11]

Blue: Heptamer[32]

Green: Nonamer [4]

id-IGKV1D-39[V_gene_segment]

cctgaacactcacaaacattatggtgggaaaagcagatcagagcattaggcaagttgcattaccttggccttcttcctttggagacaattgatgtggggttctagattgacccagagtttcaagttcatcctgattcagg**cttcaacagctggaggaagaaacagagatgttttttgaag**taaacagatctagcattactaatcaacccttcatactgatgacctatgggaaataatacccaagggcagaaa**aatgggcagaataaggggagccccaaaccaagacgaagctgctgcccatt**gagaccctgggtattacagagacctatagctctgataatggaagatctatgagtggcacaggcgctgaggaatcacagcatcattatcgtgcatctgcagggaattgc**ttgtaaatatactggtaattacaa**atgtttaaggtcactacaaatactttggagtgtattaaatatgcttctgataaagactgtttttc**tcacatgaaacaatgggaaccatgtga**caatcacagaggtgttgttactatagca**aaagggattgttactctccacatcccttt**aagtaacttgaaggcctgatagacccaccctctaagacttcattagacattccctacgaatggttatactctcctgtatactcccaatacaactctaaaatatattattccatatagtccttaggtttgtattaaagtttgacttttttccttcaaaatatctcttgtcacaacagcggctctagagagaaatacattccctccaggcaaatctatgctgcgctggtctgacctgggaccctggggacattgcccctgtgctgagttactaagatgagccagccctgcagctgtgctcagcctgccccatgccctgctgattgatttgcatgttccagagcacagcctcctgccctgaagacttttttatgggctggtcgcaccctgtgcaggagtcagtctcagtcaggacacagcATGGACATGAGGGTCCCCGCTCAGCTCCTGGGGCTCCTGCTACTCTGGCTCCGAGGTAAGGATGGAGAACACTAGGAATTTACTCAGCCAGTGTGCTCAGTACTGACTG**GAACTTCAGGGAAGTTC**TCTGATAACATGATTAATAGTAAGAATATTTGTTTTTATGTTTCCAATCTCAGGTGCCAGATGTGACATCCAGATGACCCAGTCTCCATCCTCC**CTGTCTGCATCTGTAGGAGACAG**AGTCACCATC**ACTTGCCGGGCAAGT**CAGAGCATTAGCAGCTATTTAAATTGGTATCAGCAGAAACCAGGGAAAGCCCCTAAGCTCCTGATCTATGCTGCATCCAGTTTGCAAAGTGGG**GTCCCATCAAGGTTCAGTGGCAGTGGATCTGGGAC**AGATTTCACTCTCACCATCAGCAGTCTGCAACCTGAAGATTTTGCAACTTAC**TACTGTCAACAGAGTTACAGTA**CCCCTcccacagtgttacaagtcataacataaacctccaaggaagcagatgtgtgaggacgagccaccccag**atgctcctcctggtgcctccatctgctgagagcat**ttctcaaactcagtcaggttttg**aaagtcattgggagacttt**tgtagaggggaccagg**gaggctcctctgaactctaagcctcttttgcccctatccccaggagaaaaga**tgtgacaatgcctgtcctgattgaataaggaagagatacaagtccacctgaggagt**ctgtgttatgggataattggaatttacacag**caaaagagaagctattctcggtatttcaaggagaaattgttcaagttgaataaattagagtctaaactagtctttttgaagcctacggtatgttattcgtgaagcagccactagagacaggggattct**caggtgctcctgcagaagtcagagtgcacctg**cccctggtggtatgtgctgagtaccgtgtgatgatcctcagacctgtctgggaagccgagggctggg**gtgctgatgctctcagctgcctgcagcac**gtctccaggtgattctccagtccacaaacaattccacatgttttacttca**gatgtcagagtacatgaatccaccactctgacttc**ccaatctcatgggagtgcctctcattaagcaactctaaagaaaccatagagagaaaaggagttttggaaaatgtgcgtccagaagtgatagtagcgatggggaattgacagctgacaggtcagtaaggttgctctttccacaaggctcaaaactttgccagttacagattgtcccagaatatactcgaatg**tgctatcacgtcttacgagcaactctgggttcatagca**agaaaacttcattaagtcatagatgaaacagaaaaatcaggaaactgtatgaaatacattataaggctgtgtgtggtagctcatgcctgtaattcccagcactttgagaggctaaggcaggagaattgctt

Uppercase: IGKV1D-39

Lowercase: Flanking sequence[1000bp]

Red & Bold & Underline: Stem-loop [19]

Blue: Heptamer[26]

Green: Nonamer [7]

id-IGHV3-69-1[V_gene_segment]

gagacatggtcttgctttgtcatcc**aggctggagctcagtggtgagatcacagctctctgcagcct**cagaccctgggtctccagcaatcctcctccctctgccctatgtatgcaccaccatgctcagcttcagtgcttactacactgcacactttaattatgcaatatttattatacagcaataatgcctcaataagggtatcagaaataagtgaatagataatttgttagataaagattgatggaaagacagacactgacatgagaaatgtatgacactc**aagaaaataaaactgtaggaaacgtgcttttctt**cacatttgttaggtaatcacaacagtgcgtacacatcacaccatattcttgttacagagaaaaggttctgcaaacctcagtaggtgcgacgccctgtgtgctgtgcttagttcagggagcagtcaggctcggtggtgagaagcacaggcccagatacccaggtcaccctgaccaaatgtgaactctagggagattgaacaaccgatctgtgattttgctggtaatttttcatctgttacatggaaataacattgatactacataccatggtttcactgcatatgaaaaaataaaagatgatttgttctaactttaaacatatgcactttctgttgatctactgtacctcaatagaactgttttaaaataaaaattacaaaattataagatttataggttttaaggttttatcacagagcagatttaccataagaaaccacaatttcccaaatgctatcaatatcacaaatctccccaggacactgtcacgtgctctgagcccca**ctctctccaaaggcctctaaccagagag**cttactatatagtaggagacatggaaatagagccctccctctgcttatgaaaaccagcccagccctgaccctgcagctctgggacaggagccccagccctgggattttcaggtgttttcat**ttggtgatcaggactgaacacagaggaccaccAA**GGAGTCATGGCTGAGCTGGCTTTTTCTTGTGGCTATTTTAAAAGGTAATTCATGGAGAAATAGAAAAATTGAGTGTGAGTGGATAAGAGTGAGATAAACAGTGGATTTGTGTGGAAGTTTCTGACCAGGTTGTCTCTTTGTTTGCAGGTGTCCAGTGTGAGGTGCAGCTGGTGGAGTCTGGGGG**AGGCTTGGTAAAGCCT**GGGGGGTCCCTGAGACTCTCCTGTGCAGCCTCTGGATTCACCTTCAGTGACTACTACATGAACTGGGTCCGCCAGG**CTCCAGGGAAGGGGCTGGAG**TGGGTCTCATCCATTAGTAGTA**GTAGTACCATATACTAC**GCAGACTCTGTGAAGGGCCGATTCACCATCTCCAGAGACAACGCCAAGAACTCACTGTATCTGCAAATGA**ACAGCCTGAGAGCCGAGGACACGGCTGT**GTATT**ACTGTGCGAGAGAcacagt**gaggggaagtcagtgtgagcccagacacaaacctccctgcaggggtccccaggaccaccagggggcgcccgggacactgtgcacggggctgtctccagggcaggtgcaggtgctgct**gaggcctggcttccctgtcatggcctgggcggcctc**gttgtcaaatttctccagggaacttctccagatttacaattctgtactgacatttcatgtctct**aaatgcaaaacttttttgttctttttgtattt**ttgtttttgtaacaggaggacacaccctcacct**ccacagaagccacagtgtcactttgggggcagatgatccttctgtgg**tcagcaggatgaaagttccgaggaatctcaggggaacccgaagagtgtttgccagttagactcagggcagcgacctccacaggaatctctgattagaacaggctttgagttctgataggagccaagagagacgctcacccagggtcagagtcctt**aaaaccttgtggtttt**cacagcaatcccccctggtcttgtaaaactgtgtacatatgactcagactgattcacttgaccccctttctgctaatccattttccttctctgcatacttgattctcacagttccctttcttcttctctttcct**gaaaacagaggatgtgttttc**tgtagtcaaaatcccagagat**caggtctgcaggacctg**ggtaggctgaggggactttctcactcaccataggctgatgacactcctgctgtattttgtgtgtggaggtgtttggaaaacgaaatgaacattagtcatgaagggaataatactagttttctccaaagggatgtggatgtagagctgatcttgtgcttctcacactgtcacagaatttat**gctctcccctgtgactttaggagagc**tgaggatggacactccattgtgctgtgagctctggtaatagtaattatagggtctggctaggcagcctaaggtcaatactgctggccttcgggaa

Uppercase: IGHV3-69-1

Lowercase: Flanking sequence[1000bp]

Red & Bold & Underline: Stem-loop [16]

Blue: Heptamer[50]

Green: Nonamer [8]

id-TRAV14DV4[V_gene_segment]

ctactatattaataaataaattatggtaggcaaggcaatgacctgagttgtttgcagtgttgttgagtagtatcactagggcatggtctttaatgtttacacagaaagaaaatttgtcacaatcggtgatcctgtaatcagttgtcctgtatatgggatcgtccagaacacattctcttcagagcggggcatacagaacggcgtggcttgctggggcttatttggagtctgaattgaagtcatggaaaaaatataaattcgaaggtttgataacctaataaccaaa**ctctgtaatctaatatgatctgtatgaacagag**ctggctgttgaaactgaatagaaaagagaatgggacctttaaaacacaacttgatatctgaggggttaattctaaactcttcgtcaaagagagattcgtttttctgctgtttggccccaagattggcaactggataggcattggaattgaggaggaatactgttatatcaatagataagtcaaaagatctgcagagatttgtacaaaccttcttatcataacatg**atgtatttgtactttaggagaaacaacaacatacat**agatgtgaggctagaggattattgtatcttacattgtctgtataattcattacaggtatcaaatatgtgtcaatctcccagttccatcacaggtgcaatgataaaatacagtagt**tggtcaaatgatcatttttcctctcaatgtgacca**tgtttattttcagatttgttaacaaattaagagttggttgatccaaaagaaagtctgcattgggtgag**cttctcagccagtacatagagaagtgtgagatgacacacact**agtttgctatac**caggaaacctctgtttcctg**ttctagggcagcagtagcagcagcacatggcccagtaattcttctctcaccatgccaggttcacttcacagtacagagtcctgaaaataaagaagaaaatttttttttat**ctagaaaaagaaccaaacATGTCACTTTCTAGCCTGCTGAAGGTGGTCACAGCTTCACTGTGGCTAG**GTATGGGTGGCCCAGTGGGAATGGACTTTCTGTATGAACTAAGGACTTTGGAAGTATCTTTATGGAAACTTTGGGAAGGAGGGGTAATGGGGTACAAAAATTGAATTTCTTTGTCCCAATCCACTTTATCCTTAAACCTTCAAATTCTATCTTTTTTCCTTCAGGACCTGGCATTGCCCAGAAGATAACTCAAACCCAACCAGGAATGTTC**GTGCAGGAAAAGGAGGCTGTGACTCTGGACTGCAC**ATATGACACCAGTGATCCAAGTTATGGTCTATTCTGGTACAAGCAGCCCAGCAGTGGGGAAATGATTTTTCTTATTTATCAGGGGTCTTATGACCAGCAAAATGCAACAGAAGGTCGCTACTCATTGAATTTCCAGAAGGCAAGAAAATCCGCCAACCTTGTCATCTCCGCTTCACAACTGGGGGACTCAGCAATGTACTT**CTGTGCAATGAGAGAGGGcacag**tgacagaactgtcggag**ggaggtgtacaaaagccctggggacctgcttgagacctcc**acctgctggagaaccaaggcgggaaatcaacatcacagacaggaagtggctacagctgtggcagcccagatgcagtgttgtaggaaatac**acttaactctaagaaaatacctctttaagt**tcatagaccatgatc**aaagttatcattaactaaaaacttt**cttatccctgaatc**tttggtttaaagtttagacataccaaa**tttgtgactgaagattgaagagaaaaaactgccacaataaa**caatttgccatttggctactaaattg**aactggattcaaaaacatgtgac**tgaaaaatcatttaaaatatctgtgaattttca**tatcaatttgttcagtaaatactggagatct**aaaaaaagttcatgaaaaatacgtattataaaaactatttttggatttcaatttttttttgcaccaaaa**taaacttacacttaaattgctataacatatgtagacaggatctactttgaggcactaagaaa**aataggacagcaatttgaaaagaacccctatt**agagcaacagattctgctaaaa**tgaagcaaaggaaagcttca**catttatggtgaaggttgggtagtagaatggtgaaatcatcaattctctatgaaactttacgaagtcagtgctccaaataaatcagcagttagcaagtggataactcattttaagaagggatgagacaatgttgaaggtaaatcctgcagtggtagaccatccacaccaatttgtttggaaaaagttaaccttatttgtgccctaattgaaatggactgatgattaacagcagaaacaatagccaacaccacagacacctcaatcggcccagcttacac**aattttgactgaaaattaaagttgagcaaacttt**ccattccatgggtgccaaaactattggacctagatcagttgcagacaagatcagagctttcaata

Uppercase: TRAV14DV4

Lowercase: Flanking sequence[1000bp]

Red & Bold & Underline: Stem-loop [22]

Blue: Heptamer[21]

Green: Nonamer [3]

id-IGKV2D-29[V_gene_segment]

aaatttaaatagtgaaatatgatacatttattctaatcctacttccacactacaatttgatattctcaatttaggttactgccatgacgttcaacgttttaaggtcaatatcatttttactcaatttttttcaccaactttgttaaaacatcatatgccagtgcatttcccggaagataatcaggacaagctttgtgt**gcaaaacacctagagctctgttttgc**ttttctgggtgtgtgttctttaccaagtaaatcagatgttgtattaaatcactgaagattacatagcactagaaactattcttttaattgctcccaggatttttgttgtttttttcaggacagtggcagtctctgtgaagtgatttgatgtacgcaccgttattcaggagcagctaaattcaatattaagcacatcactgacaaatgaagaatactcaagttgttttcctattgtaacgatgagttgtgtggtagattcatatgcctgacctgtgatcatctttcaataaaaacctaggaaaatgtcgtccatagtactgctgaccctgagacttaaaaccagggatccagttcttgctcttcctacatagagaatctcattttgaactgtggtgtcatggc**tgtggctgtgccaca**tacaggccagggggagacacgggttcaccctcagagttgacaagaatttggaagccctgacatcctataaaacgttacttgcccaagattgaaactttca**aattcaggtccttcttcctcctctatgatgaatt**aggttttattagtttcctccaagggacactttatatcacattgctcacagagaagacatatctacccccttcaccccccacccaatggctctttccacaccactgcacccaccaggtgatttgcatattgtcccctagggaggaccttcccttgtgagcctgagataaaagctcagctctaaccttgccttgactgatcaggacttctcagttcatcttctcaccATGAGGCTCCCTGCTCAGCTCCTGGGGCTGCTAATGCTCTGGATACCTGGTAAGGATGGAAGGAGATGAGGGAGGAGGAGGGGGTGGGAAGCTGAGCTCTGGCGGCCCCACTGATTCCCGTGTTTATTCTAACCATGTGTTAAAGGAATATGGCCTATGCTCCAGGGAGAGGAATTCATATTTTGCCCTGATGATGATTTGAAAACTCCTAAAAGCAGTGCTCTGAATAATATCTTGAGAAATGAAAG**AACTCTTGTGCCTATTTAATAAAGGGTTCATTTAAAGAGTT**TGTTTTTATGATATGAATACAAATTTGTAAAAATAAAAGATTAGCCATAAATCAATACCATAAGGCAAATCTCAAAAGTTGTTCATTATGCTTTCACATAACCT**TGCACTTCTCTCTCATAATTTCAGGATCCAGTGCA**GATATTGTGATGACCCAGACTCCACTCTCT**CTGTCCGTCACCCCTGGACAG**CCGGCCTCCATCTCCTGCAAGTCTAGTCAGAGCCTCCTGCATAGTGATGGAAAGACCTATTTGTATTGGTACCTGCAGAAGCCAGGCCAGCCTCCACAGCTCCTGATCTATGAAGTTTCCAACCGGTTCTCTGGAGTGCCAGATAGG**TTCAGTGGCAGCGGGTCAGGGACAGATTTCACACTGAA**AA**TCAGCCGGGTGGAGGCTGA**GGATGTTGGGGTTTATTACTGCATGCAAAGTATACAGCTTCCTCCcacagtggtacagaccaatacagaaacctccct**gctggggtgtcccagc**tgctcacttgcactgcttgtctggggagtagctcagcagggact**ctgagtctgcagaagaggaggctgttggagaactcag**tggcagggct**tgctgttgaggactctggcccatgagagtctcagcagca**cctcagtcccacatggtcaaagctttatcagttgccaggctcatcctcaccctgcctgtcttgcccacgtttgccgagtacactcaatcagcataacagccagaacatagatgcagtttagtgacaacacaggtggaatacatgtggcaaatgaccagcttggggtttattttacacaaattaatacttggtgataatatttggaattattgcgaatttggtattttcccactttccttactttctatttcactttactactcacactaaactgcactttccagtgtcatgtgggagaacgtgttctatacaagctgtcctcaaggggagtatggaa**aagaataattagaaaaaaatgtttgatttctgactattctt**agactttgtaaatccatggcagatataagatcttaatgccaaaagattttgatttgcctcaattacgtttttcccagtgaagag**ggagttcttgcaactcc**aaaagtgagcttcagaaataagcagcactggccgggtacagtggctcacgcctgtcatcccagcactttgggaggccgaggtggatggatcacctga**ggtcaggagttcgagaccagcctgacc**aacatggtgaaacccggtctctactgaaaatacaaaat**tagccagggtgtggtggcacacacctgtaatcctggcta**gttgggaggccgatgcaggaaaatcacttgaacctgggaggtggaggttgcagtga**gctggagatagtgccactgcactccagc**ctgggctacaaggcgaaactctgtcaaaaaaaaaaaaagcagcataaactgaaagat

Uppercase: IGKV2D-29

Lowercase: Flanking sequence[1000bp]

Red & Bold & Underline: Stem-loop [16]

Blue: Heptamer[27]

Green: Nonamer [8]

id-IGHVII-30-21-2[V_gene_segment]

aggccattggtgcctcgctgagcacagttgtccatcagggatctcccatgtgtcccaggagcagccatgcctcagtatctccactgtgcacagccactgt**ctgggaggagctcccag**gatgggtgtctttggcacacacaggtgatgggtgttagagtgcagtgcagcagctggctgcctggtctattgggctcctggatattggagggattggaggtgcattctcagggccagcacgctgtttgttaattttttataaaaaccatgtgatttaattcattttctcagatgacatagataattaataacacaatctgc**aaacaatcgtaattttcagctttagcccaagttcattgttt**ctgaattctgtgcaggatccagacgtggtattgcccttctcat**gagaaactgttcgatctaaactgaaaccagttgtttctc**gtatactttggttctccccacgtgcagagatcttgattagagcaagtttggtactttccacacactcaccctca**cctccccagataaagagcagaagttctccttagactgagtctgagggagg**agctgttcctgtaccactcagggcctgcggagacccccaggtgcagcttcactgagtcaggtgtttcactccctgtgattgctgctcaggt**ctaattgtgggctcagaattag**gacagtcttcaggttatcacaggtcaatcatattctaaaaatcatcattatcacacaccgtggtaataattcaatgttcatttttctaaacagcagtttctcttttttatttggttgcaagtctgaggaaaggaacatgtgataatactttttaatctaacctctaaatctactgaattgttctaggagactcacaaatcggacaaagtgagctctttatgctcataaaaaaatgcatgtatttgggaatttcaccgt**gttgtccaggacccgttaacatggacaac**tgtgtttcgcagtgcacttctggctt**gagacatccTCACAGACCCTCTCCCTCACCTGCACTGTCTC**TGGATTCCCCATCATAACCAGTGTTTC**CTTCTAGAATTGTATCTGCTTGCCCCTAGAAG**ATGGACAGGAGTGGATCAGGTGCA**TGGGTTGTGAAGGGAGCACAAATTACAACCCA**CTTCTCAAGAGTCCATATCCGGATCCAAGAAACAGTTCTTACAGCTGAGCTCTGTGCCCAGTGAACACACAACTACGCATTTTGAAGCAAAAGATG**CAATGAAGGGCCTTcattg**tgagcctagacacaaacct**ccctgcaggggtgaataggagcagcaggg**ggcattcggggcagtatgggggcttaggatgattgttaggggtcaggatgagcaggaccaaggcttcgcatcagggcaagtgcaacagggcagaaaaggggctgtagatgtgggttgttctcaccatcatatttcac**caccagacaccctccactacatctcttctaatgtgtctgagtg**ttgatatgattagaaaatggcatttatgtaaatactactatgtacccat**atggaggtgcatccagttgtcctctccat**cttatgtggaccttgtccatcaagcactaagtccctgtaattacttgagtacctcacacattatg**gtcaaaagatgtagggtctccttttgac**atggtctctcctcctgccttctct**ctctgtcacacaaaaacacgtgaactgacacacacacagag**cttcccaactttaattatgtgatgtattgaagcaaattgattagtgtgcagcttttctgctttgcctgctattcatgttatgtaaaaataagaaccatgtttttctcagcttgtcacttctctaagctaagtagcatctttgtttattatacccagagaccaaaagcgatccaactgttattcagcagcttaactggtaaacaaattgtggaaaattcatttactggaataatacccactgttacaatcaagtactgctggatacactcaacaccatgcttaaaataacaagcacctgaataagtaaaataagtcaaacaaatgaaagtgcatacatacgattccacttctataatttctataaagtaaaaatgaacttaaagttacatgaaaagatctgtagttgactggggacatggtacaagaagagaaggtatagaaaggagaaactacaggagagcaaaaggaaatttctaggataattgattttttctctgttagtaaaagtgaggattatgtcactatttgtaa

Uppercase: IGHVII-30-21-2

Lowercase: Flanking sequence[1000bp]

Red & Bold & Underline: Stem-loop [15]

Blue: Heptamer[32]

Green: Nonamer [5]

id-IGHV3-74[V_gene_segment]

atcaagtatttaaattgagtaaaataagccaaacaaataagagtatgtatgattctatttttaaaaattctggaaaatgaaaactgatctaaagtaatataaagaagattagtagtttcctgggaatatgttggcagaagggaaggagaaaggataaggaaatagaaataggaagtagaaggacagaaaagaagttgagggaatttcacttgtccaccttccttataatggtaatagttatgccatgattatcagt**tttacactttaaatatgtaaa**g**tttataatctgtcaatcaaatcttataaa**atgtattatgaggaaacaagttgaaaattagacaatgtaggagtgacagaaagatagatatgagtatgttgaatgtcagagatacctgaaagtttatctacctgaaccctagtt**ctctccatagtttaaggtaaacaggagag**tgcaggaaaatcatccatattctgattaggcagtggcttctgcaaaccacactaggcctggccggctgtgtcctggagttggctaagggaggagtcagggccagtggtgagaagtgcaggcccagataccagaactcactcatcccagacatgagctcttagatacacagagagcccatccatgtgtggatttatcttaca**tctgtaagtagagaacattgactcttacaga**acata**atttacacacataggtaaatctgaaataaggtgatcagtgtgaagattt**tatcacagcacagtttcataataagcacaatttctcaaatcccattgttgtcacccatcttcct**caggacactttcatctgccctgggtcctg**ctctttcttcagg**tgtctcaccccagagcttgatatatagtaggagaca**tgcaaatagggccctcactctgctgaagaaaaccagccctgcagct**ctgggagaggagccccagccctgggattcccag**ctgtttctgcttgctgatcaggactgcacacagagaactcaccATGGAGTTTGGGCTGAGCTGGGTTTTCCTTGTTGCTATTTTAAAAGGTGATTCATGGAGAACTGGAGATATGGAGTGTGAATGGACATGAGTGAGATAAGCAGTGGATGTGTGTGGCAGTTTCTGACCAGGGTGTCTCTGTGTTTGCAGGTGTCCAGTGTGAGGTGCAGCTGGTGGAGTCCGGGGGAGGCTTAGTTCAGCCTGGGGGGTCCCTGAGACTCTCCTGTGCAGCCTCTGGATTCACCTTCAGTAGCTACTGGATGCACTGGGTCCGCCAAGCTCCAGGGAAGGGGCTGGTGTGGGTCTCACGTATTAATAGTGATGGGAGTAGCACAAGCTACGCGGACTCCGTGAAGGGCCGATTCACCATCTCCAGAGACAACGCCAAGAACACGCTGTATCTGCAAATGAACAGTCTGAGAGCCGAGGACACGGCTGTGTATT**ACTGTGCAAGAGAcacagt**gaggggaagtcaatgtgagcccagacacaaacctcgctgcaggggcatctgagaccacgagggggtgtcctgggccctgtgaactgggctgctctccgtggcagcggctggtggtgctaaaggctgattttctctcagcatctggggctgattcatcaagtttcctcagagaacct**ttcagatttacaattctgtacttacgtttaatgtctctgaa**tgtgacactttccttccctgg**tgtgtctttgtttttgtgacaagaggacaca**ttctcacctccacagaagcccgagtgtcactttggggacagaaatgaccctgccctggtcaccagaatcagagtcccgaggaag**cccaggaggacctggg**aagtgttttt**caatcagactcagggcaggcgtctccgtgggaatctctgattg**gaacaggctttgggattcagattgggaccaagagggaggctcacccagggccagggtccttagaatcctga**cagttttcacagtaaccccatcgtcctttaaaactg**aacatctaactcagaactgacccatttggtcctttctctgtaatccattttccttttctctaggcttcattcttacact**tcccttttcaccttcattctgaaaatgga**gggtgtgctt**cctgtggtctaaaccacagg**cctcagatgcattacctggaactcaggtgtggctctggctatggctcctgtggacctg**gcaggctgagggatctttctcattccctggtgcctgc**atgcccctgctgtcttctatgcgtgga**tgcatttgggaaatgca**agtggacactcatagtcgtttcctcaaatgggatactgttgtagagctgatcttgtgcttctcaccctgtcacagagcccccactctcacttgtggatttttggg**agagctgaggatggacactttattgggctgtgagctct**gcatgatggcaatcgtgaggtctgggtgggcacagccagagccaa

Uppercase: IGHV3-74

Lowercase: Flanking sequence[1000bp]

Red & Bold & Underline: Stem-loop [20]

Blue: Heptamer[46]

Green: Nonamer [6]

id-TRAV8-3[V_gene_segment]

gtctatgtatttacaaaatgtcagccatactctttctacacaaaaaacaatctgtacaggcagtaacaaatgaatcaagaacccaaggagagaactggccacatgctcctggaatcaaagaaaataagtattctga**ctctgtaaaagattctcattggccacagagaacctgtcgaatttctcctttctct**tgttttcagagattgcttgtttggtgttggtgtatttccttcag**ccctagaggtttctgtagttgcgtttaccttagctctaggg**gcaaagcctttccatcctgtgttacaccttgtcattagtcatatagaggcactactcccattttaagggtcccgtcttttactctatggccgcacaagcatcttccaagggaaatatctctatcatcctccaccaccaacaaggcaactctgaaccatcaaaaactcatcccgaagcagctagtctgcgtccaacgattc**tcagaaacacagcttaattctga**tatatttttaaaacctgatttctcaatgaggtaacgtattactctatttttacatgacatcttattgctaatggactataggcctaaaacaccgtcatagacttgggcaggaagaaacacatggtgtggatgttttgatacatctcagaaattaggagaaacacactaatgagctctctctgctaaactggttctgcacttggc**ctccagagggcgatgctgcacacactggag**cttttgtttctgttgaagatcaatccactgctcagtttcttcttcctgcagctggttgagttctttccagacaaagacaagtgacaagaattagaggttta**aaaagcaaccagattcatctcagcagctttt**gtagttttaaataagcaagg**agtttctccagcgaaact**tcctcacacctcttggtcttggtctcttcagacactttccttcctgttctctggagatcttgcagaaaagagcctgcagtgtttcccttgttcagccATGCTCCTGGAGCT**TATCCCACTGCTGGGGATA**CATTTTGTCCTGAGTGAGTAAAATTTCTTTATGGTC**TCTAGTTCCACAGGTTCTGACTAGA**AATGCTTGCTTTTTATACTGAGTCTGCACTGCTTTCACTGATAGTACATTGTTTTTCCAGGAACTGCCAGAGCCCAGT**CAGTGACCCAGCCTGACATCCACATCACTG**TCTCTGAAGGAGCCTCACTGGAGTTGAGATGTAACTATTCCTATGGGGCAACACCTTATCTCTTCTGGTATGTCCAGTCCCCCGGCCAAGGCCTCCAGC**TGCTCCTGAAGTACTTTTCAGGAGACA**CTCTGGTTCAAGGCATTAAAGGCTTTGAGGCTGAATTTAAGAGGAGTCAATCTTCCTTCAATCTGAGGAAACCCTCTGTGCATTGGAGTGATGCTGCTGAGTACTTCTGTG**CTGTGGGTGCcacag**tgcctgggtctgcagggggagctgaacacaaactgcctatgctagagaagttttcagagtttcagtatatctccctgtggtgttttcaaagatctccttttttatgatggcgaatgaggcaaagagggctaagccgacccaggcatttctatgtaattctgcagttcctgttatgctccatgtcctgataaattttgtttcagaagctttttgctcattgtagaaactatataataggaaatagtattcttgaatgatggaagagatagtattcttataaca**atcaagtcctccatgacttaccttgat**gttgtctgcattagggggttga**ccatgtgagctgtgacatgg**cctcagagaaaattggttttcatttataatttcatctg**tcatctaatctggaaaatagaagatga**ttttctcatgtttttaccttgcaaatttgttttcaagttggtcattcataggttatttttt**ttagagtttctttttttgcctctaa**ctgtgg**tatttatgtgtctaaata**ttttgtctgagtt**taaataactctattta**ggaagaactgtgagaagttattagtcaacaataaaagagtcattagaatcatgaaggacttagaaaaatgtcaggtgatgttagttccgaca**actatttaagaatagt**gcagaagaaaaatgaagattaaaaattcatcagtgtttctccaaa**tactttttcaccgtgagaaaagta**gtttcctattcttttagtctcagaaatgctgcaaatctgtatcagaaatttgtggctataatccctttcatttttcttctagctactggcatttctattcattgctttcgccctgttgttatatgatttcagctattggaatattaaagatggatt**cttataacattttgataatcaatcatgtgataag**tttagcaatgccaaactgtgctggaaacactgataagaattacaatgcatggtagagagtcactggacccagt

Uppercase: TRAV8-3

Lowercase: Flanking sequence[1000bp]

Red & Bold & Underline: Stem-loop [21]

Blue: Heptamer[29]

Green: Nonamer [5]

id-TRBV10-2[V_gene_segment]

cttctgattctgtgtcctctctcactggagcatgattgtgggaattacccatccagg**ttgcatcaagatatgcaatagataatctattg**tatctgaaataa**caatagcagtgctattg**actggatgtaagtttggtgccaaagcacgtttacacctcctccaatcttctcccatttagtgaatacaagatcttctgaat**ggaaatagaacatttcc**ctgtgaccctagagactccagatagttgtagttctgggtgaacatttcatccacattcttaatgagcctcctcttttcttcataggaagtacagtcacttatgggacttaggtggtgagtgtttgtggcttcacctgaagcaaagggaaaaatcatcctcatgagaacacaccgca**gcctgagcttggatggtgaggctcccatcaggc**gcaatgaggtgctacagtgtgaaacaagac**ttcaaatgttaaagaccttgaatccacaactttgaa**acctttctggatgtcagttgactcttttaaccaagggtatctctgtaggctgaaatacaaatatccagaggttaaacaaagtggaaaat**aattcaacttggagttgaatt**ttgtacagagaa**gtcataaaagtcaccaggactgctatgacaatgctctttagactacagaaatattcataaatgttaaaacagcatt**caaatgaagaatttagcataatgggagtaaaatacgcatagaagaacaa**attcccctattaaaatcagggaat**gagattacacattttaagatttctttcaaaacagatctacaaatttttcccaatctaagcttctcaagtatgtctataataaatgatttgttcctgtggggaagagaacagatgtggcaggttagctctgaggttcaagcatttagcaggga**cagtgacatcatcaagtcactg**agagctcaacttcaatttgcccacag**cagggctgggagacacaagatcctgccctg**gagctgaaATGGGCACCAGGCTCTTCTTCTATGTGGCCCTTTGTCTGCTGTGGGCAGGTGAGGGCTGGTCACAGGAGGGCCTCCTTCCCTGGAAT**TCCCAAGGACTCAATGCAAGCTTTTCTGTTGGGA**TGACAGCATCAGTGTCTGTTGTTCTCTATTACAGGACACAGGGATGCTGGAATCACCCAGAGCCCAAGATACAAGATCACAGAGACAGGAAGGCA**GGTGACCTTGATGTGTCACCAGACTTGGAGCCACAGCTATATGTTCTGGT**ATCGACAAGACCTGGGACATGG**GCTGAGGCTGATCTATTACTCAGC**AGCTGCTGATATTACAGATAAAGGAGAAGTCCCCGATGGCTACGT**TGTCTCCAGATCCAAGACAGAGAATTTCCCCCTCACTCTGGAGTCA**GCTACCCGCTCCCAGACATCTGTGTATTTCTGCGCCAGCAGTGAGTCcacagtgctgcacagctgcctcctctctgcacggaaacggcagttagaaaaactgaggttgcctgtgcacccaagtctgggccccaccctgggacgtctcag**cccccataggagtcacagagtcctgcccaggttggggg**ggtgtct**tagagcaggggccatatgaccctcagtgctgtggctcta**ttcctacccctgtctcagaccatctggagtttgccccaacacactaaccttgctgggtctttc**ttctgcagctctctttttctgcttgcagaa**tggaagagatattttgttgatattcctaattcccttcataatttcaaagccaacatattcttctctcccttatgggatcagtgacttcttcgtgtaaaccttaccgtatcctctactacccttcaactcacagacacaaatcttcccccatgtggtctctgctcctggctgcctttcttct**aagggcaatagccctacaaggtcatctatcatttgccctt**cacctgcttctccatcatcattttgttttttttttttttttttttgagacggagtcttgctctgttgcccaggctggagtgcag**tggcgcgatctcggctcactgcaagcccgcctcccgggttcgcgcca**ttctcctgcctcagcctcccaagtagctgggactacaggtgcctgccacggcgcccggctaattttttgtatttttagtagagatggggtttcacggtgttagccaggatggtctcgatttcctgacctcgttattcacccgccttggcctcccaaagt**gctgggattacaggcgtgagccactgcgcccagc**ctccatcatcattttgcattaagtttcccctggctttctgctttagccacactcccctacattcagttacagcttcgtaggtgtagtgttgttttccaccctggggattgtttatgaataatagattatttctacttcactgagcatatgtgcctccaacttattcccaggcctcc

Uppercase: TRBV10-2

Lowercase: Flanking sequence[1000bp]

Red & Bold & Underline: Stem-loop [23]

Blue: Heptamer[43]

Green: Nonamer [2]

id-IGHV4-4-2[V_gene_segment]

cagaggtgagggctcacagaattcctctctggtttccaggaaaggtaactgcagtaatcttggtaatgagaatatcctccagtgctgacctattatagagtttacatatgaaattttcactgcaattcacaatctactctttcacacagaagtgtacagaggtcaggacacatcctcagggtcacacattgagaatgatgaagatatgtcccacgagtttttcctaaggtctcagaaagaattccaggactcaaaaggtctcagagggcagctcccagtgccttagtt**aaaatggtggctcaggcctgtaatcccagcatttt**gggaggctaaggcaagtggatcacctgaggtcgggacattgagactagcttggccaacatggtgaaaccttatctctactaaaaatataaaaattagacgggggtggttgtgcgtgcctgtactcccagctactcgggaggctgaggcaggagaatcacttgaacccaggaggcggaggttgcggtgagccgagatcgggacactgcactctagcctgggcaaaggagcaaaagttcatctaa**aaaatttattttaatttaaaaatttt**aaaaaaatggcccactccctagaacagagagattccctctaaacatgatggaggtcccgaactatacattaagtgaatcctggtgtgtctgaactcacatgattattacgttaagctgctgttccaatctacttcctcacctgggaaaagaggagccagggcatggctagttgaggccccaggaa**gagaactgagttctc**aaagggcaaagcaagcatcctcatcccagggcgagcctaaaagactggggcctccctcatcccttttcacctctttatacaaaggcaccacctacatgcaaatcctcacttaggcacccac**aggaaaccaccacacatttcct**taaattcagggtccagctcacatgggaaatactttctgagagtcatggacctcctgcac**aagaacATGAAACACCTGTGGTTCTT**CCTCCTCCTGGTGGCAGCTCCCAGATGTGAGTGTCTCAAGGCTGCAGACATGGGGGTATGGGAGGTGCCTCTGATCCCAGGGCTCACTGTGGGTCTCTCTGTTCACAGGG**GTCCTGTCTCAGGTGCAGCTGCAGGAGTCGGGCCCAGGAC**TGGTGAAGCCTTCGGGGACCCTGTCCCTCACCTGCGCTGTCTCTGGTGGCTCCATCAGCAGTAGTAACTGGTGGAGTTGGGTC**CGCCAGCCCCCAGGGAAGGGGCTGGAG**TGGATTGGGGAAATCTATCATAGTGGGAGCACCAACTACAACCCGTCCCTCAAGAGTCGAGTCACCATATCAGTAGACAAGTCCAAGAACCAGTTCTCCCTGAAGCTGAGCTCTGTGACCGCCGCGGACACGGCCGTGTATT**ACTGTGCGAGAGAcacagt**gaggggaggtgagtgtgagcccagacacaaa**cctccctgcagggagg**cggaggggaccggcgcaggtgctgctcagagccagcagggggcgcgcggggcccacagagcaagaggccgggtctggagcaggtgcag**ggagggcggggcttcctcatcagctcagtgctctccctcc**tcgccaggacctcagctgtccccaggcctcctctttctttattatctgtggttctgcttcctcacatcctcgctgcaggcaagaaagaagggagaaatttctctgtttaccattaaggttttaacagtcactggcacctttttttcaacaaattcccacaaggcccattttacctt**cttgaaaaaatatttcaag**gcttcccaccatctcttgattggcgtcatcggctgcatcgtgtcctgctcttaaattcatgtgatgaaacccc**aatcccagtggctctgtatttggagagagggatt**ttaagaaggcaaataagttaactgtggtcataaatgtgggaccctaatccagtaggactcttgtccttatgggaagtggaagacatcaga**ggcctctctctccacatgcacacagaggagaggcc**atgtggggacgcgctgcatgagaaggtggccctttgcaagcctggaagagggctctcaaaaaactaaccctttctgcacccagatcttggacgtcaagactccagaacaataagaaaattaatatttgttatttgaaactacctag**tctgtggtatcttctgatggaaagccaagcagattcacaga**tacaattgtgttagctccgtctcctggagggagaagcagcccaccgaggctgacacatctctca**gattatttttataaagaaaaatagatc**caagatgaaaaccccaccacatttctgctgagtcattcacttagcaggcttctctgagat**cagccctggggggctg**tgttctaggtcatgtctctcactttccactatgagaat

Uppercase: IGHV4-4-2

Lowercase: Flanking sequence[1000bp]

Red & Bold & Underline: Stem-loop [16]

Blue: Heptamer[38]

Green: Nonamer [3]

id-IGLV3-32[V_gene_segment]

ctcccaatattgattccctaataaaatctgggccctgagcatagggctcctcatcttcctactccatatgatgcaagttttcttgaagttatgcttgtaaaacactcaccaatgcatctggcatatattaagggctcaaaagtaggc**caggcatggtggcttatgcctg**taa**tcccaatactttggga**ggccgaggtgggggatcactggagtcca**ggagttccagaccagccaggccaacatggtgaaactcc**atctctactaaaaatacaaaaaaaaaaaaaaaatagctaggcatggtggtgcccttctgtaat**ctcagctattcaggaggctgaggcaggggaatagcttgag**cctgagaggcggaggttgcggtgagcccagatggtgccattgcactccagcctagatgatagagtgagactccatcacaaaaaggaaaaaaaaaaggttgtcattctgcagcccagatcgtgccattgcactccagcctggacgatagagtgagactccaccacacacacacacaaacgcacacacacacaaaaactaacccccgccccgcaaaaatcgaaggtagccattctg**ctgtatgatctattgatacatacaggatgacatttgtcctgt**gggc**accagggggcactgtgacccctggt**ttgggacactaatggctccaagtccctggtagtacttgagtcatttcctttctggccccttctattgagtggaatcctgaaacccacctgactcttctactcttgaaaaattacgcagagaatgtcctcagacatagggtcagacaagaacttcgacatacattttaaagaatggagagcagatttacatccgctgcttaacagggaaccaactaacaaagaattaagaataaaggaagtagagttgcatgaagagactcc**cctttctatgataagaaagg**cctgaaggtcccttcccagctgtggactcagaggcagagctctggggcatttccattATGGCCTGGACCCCTCCCCTGCTCGTCCTCACTCTCTGCACAGGTGCTGCCTCCCAGGGCTCAGCCCCCAGTGGGATCAAGATCAGCCTGGCCCTGACCTTCAACTCAACATAGGGAGTGATGCAGGGTGTGGGGTTCTGGGAATGAGGCCCTCATCCTCAGACTCACCTCTCCTGTCCTCTCTTGTGGGCTCCGTTATTTCCTCTGGGCCAACTCAGGTGCCTGCAGTGTCTGTGGCCTTGGGACAAATGGCCAGGATCACCTGCCAGGGAGACAGCATGGAAGGCTCTTATGAACACTGGT**ACCAGCAGAAGCCAGGCCAGGCCCCCGTGCTGGT**CATCTATGATAGCAGTGACCGGCCCTCAAGGATCCCTGAGCGATTCTCTGGCTCCAAATCAGGCAACACAACCACCCTGACCATCACTGGGGCCCAGGCTGAGGATGAGGCTGATTATTACTATC**AGTTGATAGACAACCATGCTACTCAACT**cac**agtgacacaggcagatggaaaagtgagacacaaaccctttttctgtctgtgtcact**cacttcctccggccccagcaggactgtggacacagccatgggcaggtctggctcagttttcctggatctgagaccccagggtgccctgacctccaagccctccagggagactctgaagagagtgggttaggtcaggagaggacttggggctgg**caggaccagactgtcctg**gtattgtccagagtggctgtgtcttggcttaagatgacctgagtgtaaggatagtcagagggagacagccattgagtggtaatggtccctggatttccttctgtatggtgactggacctaaggcagtgctgcttttctgtggcccaaatgccccagatcattctgaact**tctcaggtttaactgaga**ccctcaagccccagctccatgacatcctcatattctgaacagggag**ggccaccacaatgggcagtgtggtgtcc**atcaagattcccctttcagggctcacccatccgccatacacccaaagctgccgggaggccctggcttctcacagctgctccccaccggaaaat**gccattgattgcaaagaatggc**ctcccccaaaagtttgcccatgtcagagcgtggctcaggtgcaatgactgcttgatgcctcaatcccaaagtcaggccttaatttaggatggccttgaag**tgtgttgcagctgagcactcaattgtaacaca**ttcccactcatgcctcccatacttctttcctgagaatatatccctaaacctcactgcatgg**aattccccctctcagagtaggagtctggggaatt**aaaagatttaaaaacaaaccaaaacagtccaaagggaagagtaacaaagaggagagagttgcacagagagaggactctggagtctgcaaatgttcgtgagcaccgatgagcacatatgtgggaggaaacaacctctggaagagaaaagaaccatccaaagacacttg

Uppercase: IGLV3-32

Lowercase: Flanking sequence[1000bp]

Red & Bold & Underline: Stem-loop [17]

Blue: Heptamer[32]

Green: Nonamer [7]

id-TRBV9-2[V_gene_segment]

tgcactgtctaaa**atttagtaataatgaacctaaat**acaaaatatgaaaacatatgggtgtaattaactgggattagctgtggaatatagggtttccaggctgggtttccttgatagcatcacactgtttaattgtatggcctttataattctggctggtctctaccatcttaacatattatttgcaattatgtaacaattgttt**attttgtaatcaaaat**gttggttcatgtgtcctttgactccattggcta**agggtagatttgaaaatcaaccct**aaattgactcttctgggaaagcatggacggggtgtgtaagagggtgggagtggcaggagtgggggcatctctctttctgatttggacaatttgagcatatcacgtcagagggtatgaatccatgagactgtagactgttggccacttggaggcgctgtggctccacagtgctgcagagaactcagcggaggggtgggagagtggccaagagaggaggactgaaggg**gatgccctggctgcctgctggcatc**agtagagtggatggaattccctgagtggggcctggccatgtagagttaagcacagactatcattgccttgaaagagttggggtgccagaggcctcagtttgcatcaaggagggagtgtctctgctgctgcacgggcagtggagggtgaaggggctggggtggcaagggctggggaatcagtgtcagggcagggacagggcgggcagaggaaagtaaggtgtgatgtgggtatgttggtactgtgttcagtcctcctcacctttagcttctggcctgtgcaagacaggaatagtacatggagtagcagaacctgaggcctgccaaagagtacagtgggctggagtggcctgcaggggaaagt**atgtcagtgtggtgacat**cacaggtcaatgctcctatcaggaagaagggagcttaggaacttcagaatgcttactacagagacaccagccccaagctaggagatcctgccATGGGCTTCAGGCTCCTCTGCTGTGTGGCCTTTTGTCTCCTGGGAGCAGGTGAGTCCTGGGCACAACTTGAAAGTCTCCGATCTT**CATTTCTTGTCCCTGAAATG**CATGTGGGCCAACGATGGCTTCAGCAGGAGGCTTTCTTCTGTGCCTTATGGTTAACTTTTGTCTTCTGACACACAGGCCCAGTGGATTCTGGAGTCACACAAACCCCAAAGCACCTGATCACAGCAACTGGACAGCGAGTGACGCTGAGATGCTCCCCTAGGTCTGGAGACCT**CTCTGTGTACTGGTACCAACAGAG**CCTGGACCAGGGCCTCCAGTTCCTCATTCAGTATTATAATGGAGAAGAGAGAGCAAAAGGAAACATTCTTGAACGATTCTCCGCACAACAGTTCCCTGACTTGCACTCTGAACTAAACCTGAGCTCTCTGGAGCTGGGGGACTCAGCTTTGTATTT**CTGTGCCAGCAGCGTAGcacag**ccctgcatgagcatcagccttctgtgcaataaca**ttcctgccccactcaggaa**gtgacggtgaggggagggctgccagccagaggggctcaggccctg**gagagtggacaggcctttcccaggcactctc**tggtgcttctgtatgtgatctggtttactgctcactgaagctgtgtaagcgaaatattattttacctgtttatatatctgagggacagcagatgtcctggggctcatatttagtagctagctgaagtctattgtttcaacacctctgcactgtccctcctcactccggagatctgtccttttacctggtgtcccctttttgtgtccccaagagcaggatgctaaggtcatatatgaaactgtgatcatgata**acttgttccctgacaagt**ccaatggacacataatttccaggcaaaacatatcca**tctattctttattgtgatttaaaaaaaaaagaaagaaaaga**aaggtgcagttttctgctctgtagccctcctatgggcaaggccaggtcatctcttttatggagactgtatgaggagggagc**tttcctaggttcacagcaaatagcaggaaa**cgttttttatggggcgtgtagtcttcctgtggcggatgcaatctgccatcagtgaaaggacatgaaaaattgttattcattagttaatattttttattcttttttttttttgagatagagtttcactcttgttgcccaggctggag**tgcagtggtgctatcttggctcactgca**acctccac**ctcttgggttcaagag**attctcctgccttagcctc**ccaggtggcctccaccacacctgg**ccagttttgtgtttttagtagaggtggggtttctctttggtggccaggctggtctcaaactcccaatctcaggtgatcacctgccttggcttcccaacgtgctgggattatagagccattgcactaggccaattttttattctttatcaataaatcttttggggcat

Uppercase: TRBV9-2

Lowercase: Flanking sequence[1000bp]

Red & Bold & Underline: Stem-loop [16]

Blue: Heptamer[37]

Green: Nonamer [5]

id-IGKV1-13[V_gene_segment]

ttaaatgaattttgtttgccttttctcttattaatct**tccttttgtcagtttattttcagcaaacatttagaggacaaatgga**gttttcttcctttctcccaatataagcaagttcccttaaaattcaggcggcttacaaagcagcaagaaggtttgtgcacgggctatggcactgtgattt**ggctcccctactcaggcatcagtaaaattttgtggagcc**ctaggctgcagcccactgatgctaatatagttggatccacttcccctgctactgagctaggctgggacagttttgg**gcacattagatatgtgc**gatat**aatgattgcaaatcatt**tc**cagttttgtctggatcaaactg**ctttctccatgtacataggcatcatctctgtggatctgtagtaaattgcttgatcttatagtggtaagaacaatggcataacaccattaccgaatac**tgacatgtatatatagcatcatgtca**ataaattttatttttgattttttttagaaa**ggaacaatgttaaaatcacagaaatgttcc**aagtatagggcaaagtaaccccttccctaaccgggatcatatgagagtcttttggagacctgataatcataccgtctaacattttattatatatttcctacaaacaagaatattctcctaaataatccccatacaccaa**tgaaatacattactccatcaactcctgaggaatatttca**aattgtcaaaaaaaacctaaaaaatgtctctcataataaaatagttcccagtagaaacacattctctggagacaaatttgtgctaccctggtcttacctgggacacctggggacactgaactggtgctgagttactgagatgagccagccctgcagctgtgcccagcctgcccatcctctgctcatttgcatattcccagaacacaacctcctgccctgaagacttcttaataggctggtcacacttcttgcaggagtcagacccactcaggacacagcATGGACATGAGGGTCCCCGCTCAGCT**CCTGGGGCTTCTGCTGCTCTGGCTCCCAGG**TAAGGAAGGAGAACACTAGCAGTT**TACTCAGCCCAGGGGGCTCAGTA**CAGCCTGGCTATTCAGGGAAATTCTCTTACTACATGATTAATTGTGTGGACCATTTGTGTTTATGCTTCCAATCTCAGGTGCCAGATGTGCCATCCAGTTGACCCAGTCTCCATCCTCC**CTGTCTGCATCTGTAGGAGACAG**AGTCACCATC**ACTTGCCGGGCAAGT**CAGGGCATTAGCAGTGCTTTAGCCTGATATCAGCAGAAACCAGGGAAAGCTCCTAAGCTCCTGATCTATGATGCCTCCAGTTTGGAAAGTGGG**GTCCCATCAAGGTTCAGCGGCAGTGGATCTGGGAC**AGATTTCACTCTCACCATCAGCAGCCTGCAGCCTGAAGATTTTGCAACTTATTACTGTCAACAGTTTAATAATTACCCTcacatagtgttacaaacccgaacataaacccc**cagggaagcagatgtgtgagactgggccgccccagctgcttctcctg**atgcctccat**tggctgagagtgttcctcagatgcagcca**cactctgatggtgttggtagaggaggatatgagatcacctctgcatcccaatttctttttcttttctcagccccagctgcacagacattacaatgcctctgctgatttaataaagatagagatcatgacacctgaagagtctagttt**atggctttggttagaattcatataacagagaagaagccat**tatagat**attctaagcaggaatagtcttaatagatagaat**tagagtctaaagtattgaagtct**aaataaaatgtacagataaatttaatgttttattt**gctaagaaatttttgccaaatggggcatacagg**aaaactcaatggtcttcaatatgttggaagagcaaagagtttt**attaaaagggaaattattacctattgttctttgagaaattttgttggctgtagtaagggttgggagctggcaagctcagactggtaagcagtggtggtcaaactgaatcctagaattatattaagttatctcagaagttgtgggtaaatt**tgctttcaggttacaataagccaaagca**gtgaagcttgcagagaattttgt**tactgaaatgccagggattcagta**tagatcctgcggctcaccacacagaaagccaatcactaa**gacaacaagtgttgtc**aaagaacaggctttaatcaggtgctgcagccgaggagacgggacaccattctcaaatgtatc**tccctgacaaaataaattaggcgtttatatagcaggga**agaaatgtggaaaacaggaattagagaggggtaaggaagataatttagtcaacaggaagctggaggtcagttaggcaatcat**aatgggtgaagggtctgatgtctcactgtccccatt**cagtgatatataactttcagctc

Uppercase: IGKV1-13

Lowercase: Flanking sequence[1000bp]

Red & Bold & Underline: Stem-loop [24]

Blue: Heptamer[17]

Green: Nonamer [4]

id-IGKV1-5[V_gene_segment]

atgtccagtttcatctggatccaactgatttctccatgtacatagacaattgcttgataagagattgagtatgtttttcctaaaggtgttaacagggaggctggtgtctgggtcaggatgatgtccccatgcactgataaaaagtataagaagaaagtgtcattgatggtgcatggcagggacatgctccgtgcagtggccaccctcactaagacagatgaactttgggaaataatacccaatggcagaaaagaaggtagactatgaaggtacccaaaacaagaataaggtgcacctcattta**gtctctgggtattaaagagac**ctgcagttcttgatagtggtggatctgtgagtgctgcatgcatggagacaacacggtatcatctttgtatatctgtaataaattgcttgatctaatactagtaagaacaaaggcataacaccattacctaatacttacaaatatatagcatcatgccgatacattttatttttaattttttttagaaaggaacaatgttaaactcacagaaatgttgcaggtatagcacaattacccccttccctacccggaatcttatgag**agtcttttgaagact**tgagaatcctaccatctaacattttactatgtgtttcctacaaacaagaatattctcctaaataatcctgatacaccaatgaaatacattactctatcggctcctgaggaatatttaaaattctcaaaaaaatacctaaa**aattgtttctcataataaaatagtccccagtagaaacacatt**ctctgcagacaaatttgtgctaccctggtcttacctgggacacctggggacact**gagctggtgctgagttactgagatgagccagctc**tgcagctgtgcccagcctgccccatcccctgctcatttgcatgttcccagagcacaacctcctgccctgaagccttattaataggctggtcacactttgtgcaggagtcagacccagtcaggacacagcATGGACATGAGGGTCCCCGCTCAGCT**CCTGGGGCTCCTGCTGCTCTGGCTCCCAGG**TAAGGAAGGAGAACACTAGGAATTTACTCAGCCCAGTGTGCTCAGTACTGCCTGGTTATTCAGGGAAGTCTTCCTATAATATGATCAATAGTATGAATATTTGTGTTTCTATTTCCAATCTCAGGTGCCAAATGTGACATCCAGATGACCCA**GTCTCCTTCCACCCTGTCTGCATCTGTAGGAGAC**AGAGTCACCATCACTTGCCGGGCCAGTCAGAGTATTAGTAGCTGGTTGG**CCTGGTATCAGCAGAAACCAGG**GAAAGCCCCTAAGCTCCTGATCTATAAGGCGTCTAGTTTAGAAAGTGGG**GTCCCATCAAGGTTCAGCGGCAGTGGATCTGGGAC**AGAATTCACTCTCAC**CATCAGCAGCCTGCAGCCTGATG**ATTTTGCAACTTATTACTGCCAACAGTATAATAGTTATTCTcccacagtgttacacacccgaacataaacccccagggaagcagatgtgtgaggcagggctgccc**cagctgctcctcctgatgcctctatcagctgagagtggtcctcagatgcagccacactct**gatggtgttggtagagggggacatggagtcacctctgcaccctaattcttttctctttctcagccccaactgcacagatctagcaatgcctctcctgatttaataaagacagagatcatgacac**ctgaagagtctagtttatggcttcag**ctggactttatataacagagaagaggccactatagatattcaaagcagg**aattgtcttaatacagacaatt**agagtctaaactactgaagtctaaata**aaatgtagagatgaatctctaaattt**aatgttttacgtgcaaagaaatatttgccaaatggggcatacaggaaaactcagtggtcttcaatatgttggaagaacaaagagaaggttagagttttacgaaaaagggaacatgttacctatggctctttgagaaagttcatgggcactaggaagggttgggagctggcaagctcagactgggaagcagtggtggacaaagtgaatcctacagttatatcaagttatctcagaagttgtggataaatttgatttcaggttacaataagccaaaggcagtgaagcttgcagagaattttgtt**actgaaatgccagggattcagt**gtagatcctgctgctcaccacacagaaaaacaatcactaagacaagtattgccaaggaacaggctttaatcaggtgctgcagccga**ggagatgggacaccattctcaaatgtatctccctgaactactaaaatgaggggtatatatatcaggga**agaaatgtgggaaaacaggaattagggaggggtaaggaagatgatttggtcaacaggaagcaggaggtcagttaggcaatcacaatgggtgaagagtctgatgtctcactgtcccgattcagtg

Uppercase: IGKV1-5

Lowercase: Flanking sequence[1000bp]

Red & Bold & Underline: Stem-loop [17]

Blue: Heptamer[24]

Green: Nonamer [4]

id-IGHV3-21[V_gene_segment]

aaatatctaagtatttataccgagtaaaatatgccaaacaaataagaatatatactatgtcattt**catttttataaatgctgatgaataaaaaatg**catctgaagtaaaataatgaggatcagtagttgctgggggaaatggtagaagggaaggggagaggaggaggaatacagcagaataagaggaagtgttgagaagaattcacttgtccactttcttgataatgacgacagttacatcacgtttattaattgtacattttaaa**tatgtgaagtttattatctttcaattaagccacata**aaatgtattacaagcaaatggaaatttagacaagcacggagtaatagaa**agatacaaacatatgttaaatgtcagaagtatctgaaaattagtgtgtctggaccctagttctctccatattttcag**gtgagtgctggtggcaacatcacacatgctcttactacagaaagtgggttctgaaaaccacaccaggcacgtccagctttgtcctggagttggtttagggggatgtcacagccagtgacgaggagcacagggccagataccagcgttcacttatcccagacatgaactcctggatgcatacagagccccctccacttgtgggtttacttccac**ttctgtaaaaggagaaaatattgactcctacagaa**cataatttacacaaatatttacagatgaaatagggtgatcagggcaaagtgtttatcacagcacaatttcataagacagcatattttccaaataccatcgtcagcaaacatctgcagggcaccgtcttattatctgggtacagcctattcctccagcgtcccaccctagagcttgttatatagtaggagatatgcaaatagggacctccctctactgatgaaaaccaaccgaaccctgaccctgcagctctgagagaggagccttagccctggattccaaggcctatccact**tggtgatcagcactgagcaccgaggattcaccA**TGGAACTGGGGCTCCGCTGGG**TTTTCCTTGTTGCTATTTTAGAAGGTGAATCATGGAAAA**GTAGAGAGATTTAGTGTGTGTGGATATGAGTGAG**AGAAACGGTGGATGTGTGTGACAGTTTCT**GACCAATGTCTCTCTGTTTGCAGGTGTCCAGTGTGAGGTGCAGCTGGTGGAGTCTGGGGGAGGCCTGGTCAAGCCTGGGGGGTCCCTGAGACTCTCCTGTGCAGCCTCTGGATTCACCTTCAGTAGCTATAGCATGAACTGGGTCCGCCAGG**CTCCAGGGAAGGGGCTGGAG**TGGGTCTCATCCATTAGTAGTA**GTAGTAGTTACATATACTAC**GCAGACTCAGTGAAGGGCCGATTCACCATCTCCAGAGACAACGCCAAGAACTCACTGTATCTGCAAATGA**ACAGCCTGAGAGCCGAGGACACGGCTGT**GTATT**ACTGTGCGAGAGAcacagt**gaggggaagtcagtgtgagcccagacacaaacctccctgcaggggtccccaggaccaccagggggcgcccgggacactgtgcacggggctgtctccagggcaggtgcaggtgctgctgagggctggcttcctgtcatggcctgcggcggcctcattgtcaaatttccccagggaacttctccagatttacaattctgtactg**acatttcatgtctctaaatgt**aaaacttttt**tgtcctttttgtttctttgtttttgtaacaggaggaca**caccctcacctccacagaa**accacagtgtcactttgggggcagatgatccttctgtggt**cagcaggatgaaagttccgaggaatctcaggggaacccgaagagtgtttgccagttagactcagggcagcgacctccacgggaatctctgattagaacaggctttgagttctgataggagccaagagagacgctcacacagggtcagggtccttaaagcctgatggttttcacagctatcccccctggtcttgtaaaactgtgtacatatgactcagactgattcacttgaccccctttctgctaatccattttccttctctgcatacttgattctcacaattccctttcttctcttcccc**gaaaactgaggatgtgttttc**tgtagtctaaattccagggct**caggtctgcaggacctg**ggtaggctacggggactttctcactcaccattgtctggactctcctgttgtcgtctgtgaacagagatgtttggaagatgaagtggacattagtcatgaagggaataatactagttttctccaatgggatactgatgtagagctgatcttgtgcttctcacactgtcacagagtttggactctcacccgtgactttgagg**agagctgaggatggacactccattgtgctgtgagctct**gggtaacaatagtagggcctggctaggcagcctaaggtcaatactgctggccttcaggaaagacag

Uppercase: IGHV3-21

Lowercase: Flanking sequence[1000bp]

Red & Bold & Underline: Stem-loop [18]

Blue: Heptamer[44]

Green: Nonamer [7]

id-TRBV6-3-2[V_gene_segment]

cagagagggtagtcagtagcgtgaggtggttc**tgcctgctgggatctcaccccaggca**cagaaagaaggagccctgggtggagctgaaagtgctcatctgggtttgtcaggagtcccatctgtcagtgaattcacaagaaacagaacaaaacaactccttcaatgttgatgagactgcccctgggatttggaaagctaataacagagaaaaccaatatagacaaaggattttaaacaggactatcggattatgatcaattaagcaaattagaaaaggatacttgaaggagtatttgggacacaggaatc**aaaaacaccaggaagacatgaggagttttt**ccaagattgtagactatagcaatacttagataaccaataccagtgtattaatgagtaattattagtattattattttgagatggagtcgcactctgtcatcaaggctggaa**tgcagtggcatgatctcggcttactgca**acctccgcctccctggttcaagaattctggtgcctccgtctcccgagtagctgggactataggtgcacgtcaggatgcctggataatttctatatttttagtacagatgggatttcaccatgttgg**tcaggctggtctccaactcctggcctgaagtgatctgctctcttcag**cctcccaaagggctaggattacaggcatgggccactatgcacagccaagtaatttgtatcatcatgggaataaaaaatatacattgagttacaaactaattaccaaaagatatttaaaaaatggtagacttgcatcgattcagacaaaggcattcttcccatccaaactgttcactggtgcattgatcttgaatttgaccatctggggaaggggcgtggcctctcctgacagcaaggctctggggcccaggcagggagaatgaggtctcagaatgacgcccttgaaagacgtgttcccttttcaccaatgcacagacccagaggacccctccatcctgcagttcctgccATGAGCCTCGGGCTCCTGTGCTGTGGGGCCTTTTCTCTCCTGTGGGCAGGTGGGTCCTGGGCGGGGCCCCTTGTGTGGATTTCAAGGCCCATCCCCTTTC**CACTGGAGCTGTAGCATCAGCTTTGTCCTTCTCTGCAGGTCCAGTG**AATGCTGGTGTCACTCAGACCCCAAAATTCCGG**GTCCTGAAGACAGGAC**AGAGCATGACACTGCTGTGTGCCCAGGATATGAACCATGAATACATGTACTGGTATCGACAAGACCCAGGCATGGGGCTGAGGCTGATTCATTACT**CAGTTGGTGAGGGTACAACTG**CCAAAGGAGAGGTCCCTGATGGCTACAATGTCTCCAGATTAAAAAAACAGAATTTCCTGCTGGGGTTGGAGTCGGCTGCTCCCTCCCAAACATCTGTGTACTTCTGTGCCAGCAGTTACTCcacagtgctgcacggctgtctcctctctgcacagaaaggcaagggaag**gtgctgccctcctccgcagcac**agattcagcgatgcccttggtcctagcaccgaaaactttggagccccaatgggcccgggcagtgcgagccttcatctgtgccaggtgcctctgcagtcggtctcggccaggcctggatcggtcccagg**acctcagatgtcttccttgttgccctccggtctttcctctgaggt**gtccttttgggctagtgccaggggtttcccagctctgactttcctagtca**ttcacctgagtctgaggcctcctaggtgaaataggatttgtatttca**gatccatctagactcccgtctctacctgggtaccacgtgctttcctctctctatgatttttcctcatgttgaaccctacggtggtctcagctgcagcctgcatttcctatattcacagtcgccctccaaggccttgctgggtttctcctccccgacctccctgcccttctctactccagccatgttgagagactccaaagtccatttcctgtgccctgggctggagccttcctttctgtagtggtcaacacctacctgtgcttcagatctcagcatcatcagccctagtgataatcaccattatgcttcccttggctcccagctgaggtcagtttccctctcataatctctcatggtatcatgtgtctgttaatcagtagaatttagcacagttggaatcttccaagtactttaaaccccatgaaggccaaggctatgcctgggttataaatcagcagctgagcctggcacaaggggagatgcatttcagagaatggatgagtcagtgatggagtgagtgatgagtgg**atggatgaatccataacaggagcaccctagatccat**tttacccttgcggagatctggctgtaagtacaagaactcttacccttttgattgctgggctacgggaggtccttgaaattaatggggagtcgctat

Uppercase: TRBV6-3-2

Lowercase: Flanking sequence[1000bp]

Red & Bold & Underline: Stem-loop [13]

Blue: Heptamer[30]

Green: Nonamer [4]

id-TRAV8-2[V_gene_segment]

gatagaaaaactgcattaattttttttagtggcagattctcataatttggcacaatatatcagt**ctattctgaacactgacagttggaatag**tactctcactttctctttttaaatataagttatctaaattattttctcctaggttcacaaatattaaacttgttgacctgaaattaggaagtcttttccacactttccacttgacaaagaaaattcttgcct**ttaatcgtttgctttgattaa**ttaatgatatgtgatccaggctgactttaagtatgagtctctgatctaatttatttttt**aaagagtcaactaaaggaataatctcttt**cacccactgtaatgctcacatccagctgcacagggccatatgaagctcagcccagactgagacaactgatcaccacccctcttcttgatgaacgactttactaatatagagtgtctcaatggttttccaaagtctacaacgttgtcttctgtgac**taatttctcagggaagtaaatta**ctcttctatttctacataattctcatttctcatgtatttaaaattaaaaccatatactctacggaagatgacgtcatcacagattcggga**agaaagaaacactttct**gttggttttgatatcacatctctcaaatgag**aaggaaacagttcacttcctt**ggatctgtggttccacctgtgccctccagggggcgacgttgcactaaggaggcatctgtgttcattgccgaccatcctcatccactgagcctcctccctgcagctggctgatgtagctcactgatgtctgtgtagatagggagctgtgacgagagcaagaggtcagaacacatccagactccttaagagaaagcctttctgttttggaaacttttcaaagccagggacttgtccagcccaacctccccattgctcctagctcccgag**gctcaggacccctggcttctgtcctccctgctcagggtcctgcagc**gttgcctctgctcagccATGCTCCTGCTGCTCGTCCCAGTGCTCGAGGTGATTTTTACCCTGGGTGAGTAACATTCCATTCTTTTTCCTCCTTTTGTAAATGTGACTGTGTTGTAATTGCCTCTAAGTA**GACTTCTTAGCGAAGTC**AGCGCTGCTTTTGCTTACACAGCATTGGTGTTGCAGGAGGAACCAGAGCCCAGTCGGTGACCCAGCTTGACAGCCACGTCTCTGTCTCTGAAGGAACCCCGGTGCTGCTGAGGTGCAACTACTCATCTTCTTATTCACCGTCTCTCTTCTGGTATGTGCAACACCCCAACAAAGGACTCCAGCTTCTCCTGAAGTACACATCAGCGGCCACCCTGGTTAAAGGCATCAACGGTTTTGAGGCTGAATTTAAGAAGAGTGAAACCTCCTTCCACCTGACGAAACC**CTCAGCCCATATGAGCGACGCGGCTGAG**TACTT**CTGTGTTGTGAGTGAcacag**tgcttgagactg**caggagagctgaacacaagcctcctg**agatgctg**agactttctgtgactcaagaactcgaccttgaagtct**gttttataatattaaatagcaattccaagtttctgatgcttattgatgtgtgctgtatctttttcatctatttatttttaaactatctatg**actttaaatttaaagt**gtgtttcctttaggtagctttatatataaggtctttctatccactatgtcaatctctggttttgcactggctagtttagataattataatttaatgtatattgatgaaattggattttagtatactatgttattattgtgtttctgttcctcccttattttttgttgttctttttttttgctgacactttttggattatttgaacatttttaacactattttaatc**tatggacttctttggttatatttccata**ttttttgagtggttgct**ctggagattataatacatatgactaactctccag**ggtcttctgtgatgttgtacctctttatgtaaaatgtagaa**atttttgtataatatgtctctttattatacaataat**aattgtct**cttcaacaaacactgaagtttgttgtag**tcgttcatgtatatcaaagctacctatattggaaagctcaccatataatgg**tacaatttttgttttaaataattgta**tggaccgtataaaactcactaggaataataatatttcgtatttaacagctatttactaactctggtatccttcctttgtttctaaagtaagttttcctctgacaccaggagatgt**gtttgaactgatgtcattcccttcaaac**tt**aaaaaactccctttaacatttttt**ctagagtaggtttgctggtaatggtattttttagtcatttgttatctgaagatgtctttattcttaatttgccctttatttgtggaagatatttttactgggtctgtaattctgtgccaaccattttcccccctcacttta

Uppercase: TRAV8-2

Lowercase: Flanking sequence[1000bp]

Red & Bold & Underline: Stem-loop [20]

Blue: Heptamer[24]

Green: Nonamer [1]

id-TRBV7-2-2[V_gene_segment]

aggaaaaaatgaag**tttctagaatattagaaa**gcatataatgttttattgaaacttttctta**ctttcttgtatataactgacgtttcagaaag**agaaaataaggcaaatgatctgaggtgattgtgtatgtgatgctagctgagaatgttcctgagatgtttgaggaccgaaatcctcgttcaagaattcagatgaattgcaagacaagtaggcaggtccctgtgaaactgcagaccacaaatgacaaagaaaagattttaacagcagtcaaaagggaaagataaatatcctccaggggaatagaaagaaagaaatagtcaaagtcaagaaagaggattaaaaataactgttaaatcagaattgagattgataaaaaatacctacaggacaggacaagaaaatatggaaaaccaaaataattctgtgagtttgccaccaaagtatcttttcagagttatagttaacaagggtgggaaatgatcccagaagaaatgtatgagggcagtttgtgagggaacc**catgtgatgggacaatcccgttgggcacatg**tgagtgggggaatggaggaggctggggcatgaatggggatggcagaggggaccctgacttggaggaaagacaatgagctcatcccatggtgcctggtgttgggggcactgttggca**catcctacagaacgtgttcagcagacagaggagcggctgtgggatg**agaaggtgaactcggagatgcagcgtgaggcctccaagtccagaca**gcatgggagcccaaagcgatgtcccatgc**aataacgttgtttacaaggagcttataaatatttaaagtagtcacccaagtgtggtctaatataaatcctgtgttcctgaggtcatgcagattgagaga**ggaagtgatgtcactgtgggaacttcc**gtgtaaggacggggcgtccctcctcctctgctcctgctcacagtgatcctgatctggtaaagctcccatcctgccctgaccctgccATGGGCACCAGGCTCCTCTTCTGGGTGGCCTTCTGTCTCCTGGGGGCAGGTGAGTCCTCAGACACCAACCATTCTCATTGTGTGTGTGTGTGTGTGTGTGTGTGTGTGTGTGTGTGTGTGTGTGTGTGTAGGGGTGGGCTGTGTGCATTCGTGTGATGACTAAAATTATTTTCCTCATTCTGTTCCCAACTTTGTCTCCACAGATCACACAGGAGCTGGAGTCTCCCAGTCCCCCAGTAACAAGGTCACAGAGAAGGGAAAGGATGTAGAGCTCAGGTGTGATCCAATTTCAGGTCATA**CTGCCCTTTACTGGTACCGACAGAGCCTGGGGCAG**GGCCTGGAGTTTTTAATTTACTTCCAAGGCAACAGTGCACCAGACAAATCAGGGCTG**CCCAGTGATCGCTTCTCTGCAGAGAGGACTGGG**GGATCCGTCTCCACTCTGACGATCCAGCGCACACAGCAGGAGGACTCGGCCGTGTATCT**CTGTGCCAGCAGCTTAGCcacagcatggcacag**tcgcctccttcctgctcataaacctcatccttctctctccttgcagctcctagacacccttaacagaggcttctctttgcttctccctccccatgggaaacaagtagatttggacctcagcttgtcctttgggtagaaagagaccacag**atttacccttaagacacagtaactgtaaat**gtgggtggtggaagcaatcgtggcccaccgggctattggagtcctcagagccagctcattgctctaaggcaagc**actgggtgtcttagccttggccttcagggctgggacacacagt**cttctttggggcctgggaggttgctgcccacatagagaagcttcgggcatggctgaggaacaggtgtcttcccctacatgtgttggggcatctgcaaggtctgaa**gcttcatccacaagaacattctttcttctgaagc**ctcttctgttacattggaagttttctgcaatacaagagcaaaccattcttcctgtttgatttaaaagtctttgtggcttttgtggccattacttggtgaatacagccaagataacaataagacttcgtttcttctgcctagtgtaagcaatgcgagttctttattttgttttcatttaccactgctcctgtcctgagagactgaagcatgcgtccactactgctccagtgtcaacttggttccctaggaaatcgggtttctagaacctgaatgctgacaaataagagttgtgtacatgtat**accatgcaacctgcgtttaaaaatgtatgtacatggt**gaagtgactaaatctagctaattaacatatgcagtacctcaatccttatcctttcctgtggtgagaatacttgaaatgtactctgttagcaatattccagaatacaatccactgttactaactttagtcactgtgttgtacaatagatcttttgaacttcttcctcctatttgagtgaaattttgtatcctttaaccaacatctccccagca

Uppercase: TRBV7-2-2

Lowercase: Flanking sequence[1000bp]

Red & Bold & Underline: Stem-loop [13]

Blue: Heptamer[51]

Green: Nonamer [5]

id-TRAV25[V_gene_segment]

ggtgatatttctaaatagctacaaccaaaccttgattgtttttatagaagggcacattgtcaac**caataccatttgaatatggtattg**atatggttct**aatattttaatactttttggaatatt**ttaagtttaaatcgtgacacagtgtatatacaaacacacactttttttttttatcatttaatgaacacctgagtatccacaatttacccgctattagagtaggacatgttggtgaataatagacaaatactttccaatagaaattga**ttatctgttagataa**gactttttttctaattatattaatgtcttccctcatcacgctggcatgtgtaaaagtaacatagtgggatggcagacatgatctc**taaattggataattta**ccagggaaatcagggtgacgaaag**agaaatttgtatttct**tatcgtctgttttgaggtaatgcgttgtctaac**aataaatcacaagtgtctcccacatttatt**aaaaagtctgtgacctaatatgtcccattaaggcactta**taagcatgtgctacatggcacaacatgatgctta**gtgacaatggaagcgacaaaatggaactttctcttttcttcggatgccctagttttaaatcacagactccatttttaagaggcaaattattcctccttgaacaatgtcaccatcagtaaactcttttcttccaatccttgacacacatacataccacctaaattagaaagcagctttattaataaaataatctgttgtcaaatcagca**atgtttgagtccaccacaggggggcagcatcggcaaacat**agactctcctctgagagggtgggcaaaatagtgtgattggttttctcaacttgcagtcacctctttccctttctttccaaaatgcacacatctactttcca**aaatagaccgcaactattt**ctattgatctggaagaagcttgtaccaggcaacccatttaggagaagttggatgaagagggagagggagATGCTACTCATCACATCAATGTTGGTCTTATGGATGCAATTGTCACGTGAGTTTGAGACGTCCCTGACTGCTTCCAGAGAATAAATCCAACAGGATAGAGAAGTATCTGCAGGCAGGGAATGGACTT**TCTGTTTGAGAAACAAGAGTTAAGACAAAGGGAGAGAAACAGA**GAGAAAGAGACAGAATGGATAAAGGTTGAGGGAGATGGGAAGGGAGACGTGGACAGAAAGGGGAAAAAAGAAAAGAGGAGTGAGAGAGACCTGCGTAGCTCAGGAGCTAAATATCTAATCTGATTTTATGTGATGCTTCCCATTGTCTTTCTAAACAGAGGTGAATGGACAACAGGTAATGCAAATTCCTCAGTACCAGCATGTACAAGAAGGAGAGGACTTCACCACGTACTGCAATTCCTCAACTACTTTAAGCAATATACAGTGGTATAAGCAAAGGCCTGGTGGACATCCCGTTTTTTTGATACAGTTAGTGAAGAGTGGAGAAGTGAAGAAGCAGAAAAGACTGACATTTCAGTTTGGAGAAGCAAAAAAGAACAGCTCCCTGCACATCACAGCCACCCAGACT**ACAGATGTAGGAACCTACTTCTGTGCAGGGcacag**tgctccccaagcacctgctgcctgtctccaaatcttgccctgggtcttcaggagcagatcatcctactctccccaaagagcaggcgccagagaaagccaaagtcacaatgtctgtgaggatctcgagcaactcctccccatggcagtaggaatgcatgtagaaattcactctgttaagacctggaagagtttagc**agttttagttcctaaaact**taactaccggttttcttctacacaggtgtctcaaagaattctaattgtcttaggaagaatcggtggattttaacgaattgtgatacagatatttatataaaaagtcatctttcagtgcagt**ggtccccaacatttttggcatcagggacc**ggttttgttgaagacaatttttccacggagagggtggggtgatggtttcgggatgaaactgtttacctcagattatcaggaattagttagattatcttaaggagcgagcagcctagatccctcgcatgcgcggttcgcaatagggttcggg**ctcctatgagaatctaatgccactgctgatctcacaggag**gcggagctctggcagtaatgctcacctgccgctcacttcctgctctgcggccaggttcctaacaggccacagaatggtacagattccctgccgc**caggggtttgggacccctg**tttgggtggtagcagaggtttccaagcagtttctttaaggttttagctcaattgtattgcggaagatt**tgaaaattaaacatttttca**ttatgtacaca**aaacttaaaagaagttt**acgaccaactccctgaagagtcacctctgtagaactcatctgcttcaatgcaacgcagtccaacata**gcataaagattatgc**aggcggtaaccacctttccagagaactgagtgaaaagagctctttttaaaatgaggagtttagacgaaactggaaggtgacccatgaagagatttaggcatggtctctttttcactcttccaactgcaa

Uppercase: TRAV25

Lowercase: Flanking sequence[1000bp]

Red & Bold & Underline: Stem-loop [19]

Blue: Heptamer[29]

Green: Nonamer [1]

id-IGHV3-76[V_gene_segment]

aagaatgttactttggactaaaaccttaaactataaatacaaataaatccatataaattaaatataataacacagaaacaaatgtatgtcactttgatgtttggaaatgcatttatttttgcctgtgcacatatatgtttcattgctgggtgtgtgtgtctgtgtctgtatatatgggtgtgactgtgaagtttcaaaatacattattaacttgtatagttatataaatctaa**gtttttttatgttaaaagatagtaaaaaaac**acacacaaaaatattctagagagcatttgcaataactaaggagtattcaggtgttgtattaataatg**aaaaactagccaaaatgttttt**aaacttttattttaggttcagaagtacttgtgcaagtgttgttttataggcaaattgtgagttctgttggtttgatgtacaga**ttattttgtcaccaaggaaataa**gcatagtacctggtaggtagtttttcaattctcaccctcctcccaccctccacactcaagtaggacctagtgtctgttgttttctttgtgtcaatatgattagctcccacttatacatgag**aacagatggtttgtaattttctgtt**actgtgttactttccttaggataatgacctccagctccatctatgttgctgcaaataacatgatcttattcttcttatgactgtgttgtattccattctgaaatatacataaatcagtaaatgtgattcatcacacacacacaatcatct**cattgatcaatattaactttcaatg**taaatctcccaggaatctttcatcttctcttgaccctcccctctcctcaggcatcccaaaccatagctttctatatattaggatacatgaaaatagggccctctctcagctgatgaaaaccagcccagccctgactctgcagctctgggagacaagccccagccgtgcagtaccaaggtgcttcca**ttcagtgattagcactgaa**cactgaagacatcccATGGAGTTAGGAATGCGCTGGGTTTTCTTTGTTGTTGTTCTAAAAGGCAATCAAGAAGAACTAGAAATATTGTCTGTGAGTGATATGAGTGAG**AGAAACAGTGGATCTGTGTGGCAGTTTCT**GACAAGAATGTCTCTGTGTTTGCAGGTGTCCAATGTGAAGTGTAGCTGGTAGAGTCTGGGGGAGGCTTGGTACAGCTTGGGGGGTCCATGTGACTCTCCTGTGCAGCCTCTGGATACACTTTCAGTAACTATGGCATGCACTAGGTGGACCAAG**CTCCAGGGAAGGGACTGGAG**TGGGTCTTACATTAATGCTAGTGGTGGAGGCATATACTACTCAGACTCTGTGAAGGGCCGGTTGACCATCTCCAGAGAAAACACCAAGAACTCACTGTATCTGCAAATAAACAGTTTCATTGCTGACACCATGGCCGTCTATTACTGTAAGAGAcacagtgaggagaagtcagtgtgagcccagtcacaaacctcctacaggaacgctgggaggaaaatcagctacagggctcactcaaggcccactgatcagagtccactccagagggaggttcacatggaggttaggggctggttttctgtcaggacattggacttcatcatcttcttccattactttctttagggagcctcttcattttagaattctgtgtgtactattg**ttatatatacatatatatatatatatatatataa**aataatcatagtaatgcgaggacatattttctcatgcacaaaatgtaga**ttgacacttacagagaagaaaaattatcaaccaatgtcaa**aaacatcagagtcctggaaaatctta**agtgttctggtgggtcatctctaatcagacccagaacagt**aacctcagtgaaattccatgattcagatactatttttggattctgatcataaccaagacagagtctggaccagggtcagt**gtcatatagaacctcacagcttccatatatgac**ccttctcttgacagtgaaatatgaaattcagtatcagcagtgatttgatttttcttttgctttccgtattctttctttttagttgttgttgtttagttatttcttttgctatatccattttctttttttttagttgtgcctttcacttttggctttcacttttccatttgcttttcctgcttcctgataaagggagatgtaatccctgtggtctaaattccagggctcaagccttttcctgggagctcaggtgggtctcaggat**gtggctcctgcagccac**gt**gaaagaggctgatgggactttctttc**cttctcatgactcaggcattccacggtgttgtgtggagactcatctgagattgcacgtggccattagtagtgaaggggatgagatgtgggtgtggactgatcctgtgctgtgcaaattgtcattagtagtgaaggggatgagatgtggatgtggactgat

Uppercase: IGHV3-76

Lowercase: Flanking sequence[1000bp]

Red & Bold & Underline: Stem-loop [14]

Blue: Heptamer[41]

Green: Nonamer [7]

id-IGKV3-31[V_gene_segment]

ttttagcctgatacaaag**agaagccagggaaagcttct**gagctcctgatctacgatgcatccaatttgcaaacctgggtcccattgcagttatgtggcattggatccaggacagatttgattctcaccattagcatcctccagtct**gaagttgctgcaacttc**ttattattggtcaacagtataaaagtgaccctctcacagtgttacaaacccaataagctccccaag**gaagcagatatgtgagggtgggctgccccagctgcttc**tcctgtttcctccatct**gctgagagtgtttctcagactcagc**cacactctgaaggtcactgagcagttttcatagaagggg**taagggaagtgtctctacaccctta**gtttcttttatcctcctcagccccagcagcagacatggcaatgcctctcctaatttcataga**agagtcattacctatgaggactct**gggttacagcactggtccaggttcatacaacaaaagagaagctattgtaggtaacccaaatagaaagtttttcctaatatgggaaatccatgtctaaattacaacttttcaaagaccagaggatataatggttaggaaaccagaaaggcaaaaggagtgctgggaaatctacttcaagaagcaatggtggccgcgcacagtggttcaggcctgtaatcccagcactttgggaggccgaggcaggcggatcacctgagatcaggagttcaagaccagcctggccaacatggcgaaaccccgtctctactaaaaatacaaaaattagccgggcatagtggtgggtgcctgtaatcctagctactcagaaggctgaggcaggagaatcatttgaaccaggcaggtggtctctgcagtttcgaagcccagcgccatct**gtggctgttatgcatgtctctcccagccac**cctgctgtccagagcccatatcaatccgtgggggtaggtctgtgaaaga**gcagctcagttaggacccagagggaaccATGGAAGCTGC**AGCTCAGCTGCTTTGATTCTTGTTACTCTGGCTCCCAGGTGCAGGAAACACCGGATGGTTCTGCATGTCAGTGAAACTTT**CTCAACCTTGTTGAG**TCCTGTTACCTGGCACACCTGCTGGGAAGGCACAATGATTAAAGCTCAATGTAGATCAATGGTCCTGGATGCACTGGGAAGACAATAGGTATGATGTAGTGTACGTGTGTGACATTTCTGTTTTTATTCCAATTTCAGATACCACCAGAGAACTTGTGATGACAGTCTCCAGCCTCCCTGTCTTTATCTCTAGGGGAAAAAGCCACCCTCACCTACAGAGCCAAGTAGACTATTAGCAGCTCCTTAGCAGGGTATCAGTGGAAACCTGGACAGGCTCTCAGGCTCCTCATCCATGGTGCA**TCCACCAGGACCACCAATGTCCCAGCCTGGTGGA**GTGGCAGTGGGTTCGGGGAAAACTTCAGTCTCATTATCAGCAGGCTGGAGCATGAAGATTTTGCACTTTAA**CACTGTTATCAGCATAGTGGTGGGTATtccacagtg**attccac**aggaaaccaaacctccacaagacagctggtgttttttcct**caagccttctgtttacttatgggaagctactatggtggctgct**tagttattgagagaaaacaatggagacttcaataaata**g**aagtttataccacattcatgcattataaaactt**aatattgtgagtaacataattctcccttaa**ttaattcaatacagtgacaattaa**aatcccagaggacttttgtgaaattgaaatgatgtttcatatatatatagtatatatatatatatatatatatatgtactcacataccacgcactatatata**tatatatatacacacacactatatatggtgtatgtatatatata**gtgatatgtatgtgtataatatatatatagagagtg**tgtgtgtatatatatagatacatacatatcacaca**tgcatttaaaaaggacaaaaatagccattggaatcttgcataagaataacaggtcgg**gctgggcgtggtggatcactcctgtaatcccagc**aatttgggaggctgaggtgggcaggtcacctga**ggtcaggaggttgagactagcctgacc**aatatggagaaaccccatctctactaaaaatacaaaattagtcaggtgtggtggcacatgcctgtaatcccagctactcaggaggctgaggcaggagaatcatttgaacccgggtggtggaggttgcagtgagctgagatcgcaccattgcactccagcctggggaacagagccagaatctatctcaaaaaaacaaaaacaaaaaacaacaacaaaacaaaacaaaaaaagaataacagggcaggcatggtgtctcacgcctataatcccagtactttgagaggctgaggcaagtgtatcgcttgaggtcaggagttagaccaccctggccaacatggtgaaaccttgactctactaaaaaaatacaaaaaattagccaggcatggttgcacattcctgtagtccca

Uppercase: IGKV3-31

Lowercase: Flanking sequence[1000bp]

Red & Bold & Underline: Stem-loop [19]

Blue: Heptamer[36]

Green: Nonamer [1]

id-IGHV1-46[V_gene_segment]

tgtgaagaataggtatgattttactatttcaatggatacagaattgtgggagtcactatattcctatgaacaaaaaattcagatttcagtgttaagtaatgttgcctacattgtgtgagtgacggggcagtggtggatctgagagtgtggtgggtgcacggacataatgattcagaaagcaatatggaaagatgagtatctatggatacgaactga**aagtatgtaaatactt**cacaaaatactaataaacggagttgaatataaaacccataattat**ccaaaacacaaatttcttggaagttattttgg**gaacatgatttcataaagaactccaaactcttgtttcaacttctgactcctcgtttctgtgatataagaaaaccatttccaattatgcatctcagggcaattctgtaaacccagagcatttctgctgaagatcctggggaatcaagacaccgggcaggtgatggagacactgtctcaggtgcgcccaacgaatctcagaggaacctgctggagagtcacgtggaacatctacagtcagtttctcagagtcaacagtgagctgtgttg**gtgcctgaggggaccatgatggggccaaggcac**gtgctcagtgtcgtggacagtgatggtccagaaatgatctagatggtcttgacgctaatgaaatatgggttcagagtgaggagcataatctgtggggacttgttcttcagtgaaaggatcctgtccgcaaac**agaaatggagcaggacatgcatttct**tcaagcaggattagggcttggaccatcagcatcccactcctgtgtggcagatgggacatctatcttctttctcaacctcgatcaggctttgaggtatgaaataatctgtctcatgaatatgcaaataaccttagatctactgaggtaaatatggatacatctgggccctgaaagcatcatccaacaaccacatcccttctctacagaagcctctgagaggaaagttcttcaccATGGACTGG**ACCTGGAGGGTCTTCTGCTTGCTGGCTGTAGCTCCAGGT**AAAGGGCC**AACTGGTTCCAGGGCTGAGGAAGGGATTTTTTCCAGTT**T**AGAGGACTGTCATTCTCTACTGTGTCCTCT**CCGCAGGTGCTCACTCCCAGGTGCAGCTGGTGCAGTCTGGGG**CTGAGGTGAAGAAGCCTGGGGCCTCAGTGAAGGTTTCCTGCAAGGCATCTGGATACACCTTCAC**CAGCTACTATATGCACTGGGTGCGACAGGCCCCTGGACAAGGGCTTGAGTGGATGGGAATAATCAACCCTAGTGGTGGTAGCACAAGCTACGCACAGAAGTTCCAGGGCAGAGTCACCATGACCAGGGACACGTCCACGAGCACAGTCTACATGGAGCTGAGCAGCCTGAGATCTGAGGACACGGCCGTGTATT**ACTGTGCGAGAGAcacagt**gtgagaaaccacat**cctcagagtgtcagaaaccctgagg**gaggag**tcagctgtgctgagctga**gaaaatgacaggggttattcagtttaagactgtttagaaaacgggttatatatttgagaacaaagaacaatagaaacacaatcgaattgtaagagaaatattccattcaagagccaccacataagccaaactgacagagtgggaaaggccacactcagtaaagttgatacaaacataccataaaggtgctactatgaacaagtttttgaattagatgaataaatcatttggagcaaggttatttggtcatatgttaagagtaagcatgattcttacaaagtgggaaaattgtctttcaaatgtttctgtcacttcttaccataaagttcattttagaggttttaggattacagtgaaattgcacagaaggtgtgagaattcccatgaatccctgccccgcacggacaccgcctcctccactacagccatcctgccccacagtcacaaataagtcacaatggatgaatctac**aagaactcttggttctt**tctttttctggtgatcccctaatataacaagcctaaattatcttggaacacccaggtattttcaatggctttctagaagtgatattagtcagagg**gaaagtgagtgaggctattactatttgagcactttc**t**tccaaaatccacaaaatatatgttaatttgga**gtttt**tcttacttctggtttacaatgtcccttcccagagagtaaga**tttttaagcttttagtgaggctggaaaaaaaaatttttaaaaaagagaaataagctttcctgtattaggctgacttatcccagcggcagcaacaagcacagcccagacccaggaaaagtcttaataatattatctaatgtgctctggagactctttcagca**ctccctcaacatagggag**aagaaaaacaaattttcctttgtcttatgatatgagtttatagattcttgttct

Uppercase: IGHV1-46

Lowercase: Flanking sequence[1000bp]

Red & Bold & Underline: Stem-loop [17]

Blue: Heptamer[52]

Green: Nonamer [5]

id-IGHV3-54-2[V_gene_segment]

tgtctacattctcaaataactctgctgagtgaaagtagctgaagaattaagagtgcaattcataaacttctaattgtataaactgcaaaaggtccaactattctaaagtaacagagcagatttgaaatttgtgagaaacgggtgttgaaa**gtaattggctggtgagatgaaattac**agagaagtgacagaaagatttaggggttaacttaattgtacacaacctgattaaagtt**tgcacacatacgttaccattttccaaattgtgca**gtgtagatttg**aattaattattaatt**gtacttaaaaaaagcagtaacaaa**taaacacatgaatatgttta**ctgaggaggaacaaaaaatagatgggtatgaacactggaaacatctcagactcttgaaagtacacaggcttgaacactggttctctccgtatacttccggtaaacggctgaatacactaaaagaaaacagagatgtcctggcaggggtggaatcctgcagacctcactaggtgtgtcccacactgccctggagttgtctcaggggagcagtctcctctagtggtcagaggcacaggctgagataatggggttaactctgtccagctgtgtgactttgaatgcattgtataaacactctgttctgtatgtaatttatcttccttaaaatgcaaca**ttgacacttacattaaatgtattctacaaatatgtcaa**aaagaagatgatgactgctaaatgattatcaaggcacaatcacataatataatgatattttcctgagtgataagatgactaccaatctcg**ggggcactttgtctgctctgagccctgcccc**tcctcaggattcccatcccagagcttgctatacagtaggagacatgcaaataggtttctccctctgctgatgaccagtcctgaccccatagctctgggagagaagcgccag**ccctgggattcccaggg**gtttccatttggtgatcaggactaaagacagaggacccaccATGGAGCTTGGGCTGAGCTGGGTTTTCACTGTTGCTGTTTTAAAAGGTGAACTAGAGAGATTGAGTGTGAATGGATACACTTGAGAGAAACAGTGGATATGTCTGGAACTTTCTGACCA**GGACACCTACAAAGTTTGCAGGTGTCC**AGTGTGAGGTACAGCTGGTGGAGTCTGAAGAAAACCAAAGACAACTTGACAACTTGGGGGATCCCTGAGACTCTCCTGTGCAGACTCTGGATTAACC**TTCAGTAGCTACTGAA**TGAGCTCAGATTCCCAGG**CTCCAGGGAAGGGGCTGGAG**TGAGTAGTAGATATATAGTACGATAGAAGTCAGATATGTTATGCAC**AATCTGTGAAGAGCAGATT**CACCATCTCCAAAGAAAATGCCAAGAACTCACTCTGTTTGCAAATGAACAGTCTGAGAGCAGAGGGCACAGCTGTGTGTTACTGTATGTGAGGCAccaggtaagaagacatcagtgtgaacacagacacagaatttcctgaaataagggaggagtctgggctaaaagggcactcaggacccacagaaaacaggggaagctctagggcaggtgcagatggtcatcatgggctgctttccttgagggtct**gaggcttcctctgcatctaacagtttccctgggagcctc**tctacatttatgcttctgtggccacccctgaggtctctggacattctcatttgttgcaaaggcagatgtaagtattggaggcataaaaatgcacaggaggccagggagtctgtagacattgttaccccagaaggtcaatctcaccactagtgctggaggagggtgggagtttgatgaagctgccctaagtatcctgtggtctaagctaagtccaacgaggccatttgtgcctccctgagcacagttgtccatcagagatgtcccatgtgtcccagcagcagccatgtctcagtgtcttcactgtgcacagccattgt**ctgggaggagctcccag**gatgggtgtctttggcacacaccaggtggcgggtgttagagtgcggtgcagcagctggctgcctgttctattgggctccctga**tgctggagagatgggaggtgcattctcaggtccagca**ccctgtttgtgaatttttatataaaaccatgattttacttcattttctcagatgacatagataattaggaacagaacctgcaaagaaattgtaattttcaactttaccccaaatttattgtttcttaattctgtgtaagatccagacatattattgccttcctca**tgagaaattgttctatttaaaatgaaattagttttttctca**cattctttgtttctgttcaagtacagagatctttattaaagtaagttgggt**tctttccacacactaaccctcacctcccccagagaaaga**gcagagattttcctcactctgagtctaagggaggagctgttcctgcacgactcagagc

Uppercase: IGHV3-54-2

Lowercase: Flanking sequence[1000bp]

Red & Bold & Underline: Stem-loop [16]

Blue: Heptamer[41]

Green: Nonamer [3]

id-IGHV3-15-2[V_gene_segment]

tgatgttcaggaaaataaagc**aaatgaataagaatatgtactatgttattccattttaataaactctgataaattaaa**ttgaatctacagcaatataaagaagaacagaatt**taccatttggggaaatggta**gaagaagggaagaggaaaggaggaggaatatagaagaatgagagggaaattttgtgaattttctggttcaccttggtaactaggatggttacatcaggtttatcaattgtacactttaaatatgtga**agtttattatcagtaaact**g**aaatttataaaattt**attaccagcaaacaaatgaaaacttgcacaagaaataagtgacataaagatagaaaaaatattaaatttcagaaa**cacctaataatttatcttcgtgaaccctagttctcaccatatttttaggtg**aatgctagaatgcagcaaaattacacatgctctcaatacagaaagtgggtttcacaaaccacactaggcatgctcagctctgtcctggagttgggttagggagtaatatagggccagtggatgaggagcacaggcctagatactggggctcactaacctcaggtatgagctcttagatacatacaaagcccctccacgcatgggtttacttccccatctgtaaattgagaaaccattgacccctaaaaatatga**tttacacaaatatgtaaa**aatgtaagagagtgattagtgcaaagtgtttatcacagcacaatttcataacaagacagcaagttttccaaacagcatcattgtcattagattcct**gcagggcatcattaccttatctgggccctgc**cctctgctcaggcatcccaccccagagcttgctatatagtaggtgacatgcaaatagggccctccctctcctgatgaaaaccagcccagtcctgaccctgcagctctgggagaggagccccagc**cttgggattcccaag**tgttttca**ttcagtgatcaggactgaa**cacagaggactcaccATGGAGTTTGGGCTGAGCTGGATTTTCCTTGCTGCTATTTTAAAAGGTGATTTATGGAGAACTAGAGAGATTAAGTGTGAGTGGACGTGAGTGAG**AGAAACAGTGGATATGTGTGGCAGTTTCT**GATCTTAGTGTCTCTGTGTTTGCAGGTGTCCAGTGTGAGGTGCAGCTGGTGGAGTCTGGGGG**AGGCTTGGTAAAGCCT**GGGGGGTCCCTTAGACTCTCCTGTGCAGCCTCTGGATTCACTTTCAGTAAC**GCCTGGATGAGCTGGGTCCGCCAGGCTCCAGGGAAGGGGCTGGAG**TGGGTTGGCCGTATTAAAAGCAAAACTGATGGTGGGACAACAGACTACGCTGCACCCGTGAAAGGCAGATTCACCATCTCAAGAGATGATTCAAAAAACACGCTGTATCTGCAAATGAACAGCCTGAAAACCGAGGACACAGCCGTGTATTACTGTACCACAGAcacagtgag**gggaggtcagtgtgagcccggacacaaacctccc**tgcaggggcgcgcggggccaccagggggcgctcgtgacccactgagggcgggacaggtcccaggagcaggtgccgggagaggtttcctttctcctcagctggaaaagtcaggtttatcttcgcaggactctggagtcttctaggctgtgatattttgttacttatatttattatgaattttatca**ttaatacttaaattttagtaattattaa**cattctacatattattatatttttaagtatatactttcaagaaataaacattcctaattgcat**aataattattgtaacataattatt**gta**acatataattatatcctaaatgcataataattattgt**aacaatcacattctcattttgtatttgaattttcatatttattttactaagatttta**attttaatataaaat**atttttctacaattgtctgttttc**catttttgtgagataaaattagcatatacaaaaatg**cacagattctccaggtacagttagagggtttcaggcaaatgtgcacatacttgtctacagcagctaagtgagggtgaggaacaggtgaggtccatctccccacaaagtgtcctcttagtgcttccagttagttctcacataaggatttttttttatttcaactttataatttagatacagagggttcatgtgtggatttgttacatgggattactgagtgatgctgaggtttggaatacagatttcttcaccccctccctccatggtctagcagtccacagtgtctgttgctcccatatttatgtctatgtgtgctcaatgctgaggtcccacttacaagaatatgtggtgttcagttttattttcctgaattaatttgtttaggattaagaactccagctccatttgttttgctgcaaaggacatgatttcattcttttttatggctgtgtagtattatatattttacatgtaacacattttat

Uppercase: IGHV3-15-2

Lowercase: Flanking sequence[1000bp]

Red & Bold & Underline: Stem-loop [20]

Blue: Heptamer[33]

Green: Nonamer [7]

id-IGLV5-37-2[V_gene_segment]

agcattgtcaaattgcctcaaaaaaaatcagaaagtccttg**gttcctggaacaggctgccctaggaac**agagcccaatcggtcagagcaagcagggacacaactgtgcagaaaacaacaagttttacttagggagataacggttgactgtataaaatgagaaatatgtcacactactcacttgtaaaagtaagtttaccaatgctattcatttccctatatcactacgttgagtaaaaactctaaaatgtatgaaaccactgtagagataacttttgctgaattgctactttaacatgtcatggatcatttatttttcttgcaaaattctcattattcaccgtccttttgagtaacccc**cagctgcccttcaacctagctcctcagctg**acctcttctccatgatgcctaccctgatggcacatgtgggcagttaggtttttctagattttaaaagttatatatgtacatgtctacatcccatggaccaaga**ctcactctgggagtgag**aaatcacaataatcatgcatagccttatggaattagtgtctagcaatcagaccatagaggtgtgaatagagacagtgactaatgagctgcagagatcagctgggccatcaggaggtaagtgtcctctaa**ccagcatgggatgctgg**gggactgccct**atggtccctacacagctgctcagtaaggaccat**tcactcaaagaaatccaagactaggg**gctaagaccccgtgctcactgaggggtacttagc**atctatgtcctcccatggcatggcagagtcccctgagcaggaaggtgtgtatggtttcagtagg**tgctttcttccggagccctctcaaggggaaaagca**aaacttcttcctcctcagtctgcagtgtgagctgagctt**cagccccagagcttagagaggggctg**ggcagtccctggatgaaaacacatttgcatgagcagcc**cctcctctgcagagtatgggaagaaaaggagg**cctggggCAGCCCAGTCCCACTGTGCATGTCAGGCTGTGTCCACCATGGCCTGGACTCCTCTTCTTCTCTTGCTCCTCTCTCACTGCACAGGTAGGGACAGGCCT**CAGAGATCAGGGCCAGCCACCCAACCTGATTCTG**GCTCTTCTGGTAAAGATCCCTGAAAAACCTCACCCTGAACCCTGCCCATCAACCATGAGTGTCTGTGTTTGCAGGTTCCCTCTCCCAGCCTGTGCTGACTCAGCCACCTTCCTCCTCCGCATCTCCTGGAGAATCCGCCAGACTCACCTGCACCTTGCCCAGTGACATCA**ATGTTGGTAGCTACAACAT**ATACTGGTACCAGCAGAAGCCAGGGAGCCCTCCCAGGTATCTCCTGTACTACTACTCAGACTCAGATAAGGGCCAGGGCTCTGGAGTCCCCAGCCGCTTCTCTGGATCCAAAGATGCTTCAGCCAATACAGGGATTTTA**CTCATCTCCGGGCTCCAGTCTGAGGATGAG**GCTGACTATTACTGTATGATTTGGCCAAGCAATGCTTCTCACAGTGACACACACAGATGGGAAAGTGGGACAAAAACCTCACCCTGCTTTAGGTCTTGTTCTTAAAAA**ATTTTAAATTTTAAAAT**AACTGGCCTAGGCACAAACTACATTTGGAAGTACTTTCTAGTTGTAAAAGGCTCTTCTCTCAATTTCCCATCCATTCGTCTGAAGTCTCAGTAAAACAAAACAAAACCAAAGCAACTGATGACCCTGAAGTGTTGGCTGGTTG**TCCACATGTGCTAACACCAGTGCCTGTGGA**GCTGGACTTTTGTTCTTGAAAGAAGCAAAATGAGGCTTTCCTAGCTCACATCATGGTCAGAGCCTGGGGACATCCTAGCAGGTGAACAGTGCCCTGCAGCATGGA**CTCTGCTCTTCCAGCGTACGGGCAGAG**ACCTCCCCTTCCATCCCCAGTAGTCTGTTCTCAGGGCTGTCTGTGGCTCA**CTGTGTCCCAGGAACAAGGCACTCAAATATGAAATGCTGCCTGAGGACACAGGTCAGGGAATTCTAGATATAGTGCTGACCTG**GCTCAGTCCTGGGTACCTGCATTTCAGTGGCAATGTCAGGAGATGGGGTTAGGTATTAGGGTAACTGGGGACCTGAGACCAGTAAACCTCTATGTAGGACATGACAGGAGAGTCAAGGGGAGGCTCCACTGTGGCTGAGACAAGCCAGAGGCTGCTCCTTGGCCACGGGGGAAGTAGGTCCCATGGATGTGACTTTCCTGCAGCTCATCCCACTCAGGACCATGGGGACT**GCTGAGCGTGGGTTTCTTGATCCCTCAGC**GCCTGTGTTCTCTTCTCCTCTCTTACACCATTGGAGCTTCTTGCTGAATCAGGGTCTAGTGG**TGAGTATGTGGAAAGACCACCACATCTCA**AGGTCACCTT**TTTCTGCATGGGGAGCAGCAGAAA**CATTGGGCATTCCCGTGGAAATGGCTACGGAGGCCTCCCAGAAAGAGCCCCCAAGTTTATGACCTAGgacagcagtgatggtccctcaggggtcccagcctgattctcctgctctgggtttatccacacagcctc**ctggagcacctcttggctccagcctgaggactggctgg**ttatcagtgtcacacagctgcatcctgactccatgttaccaagggttcctgttccatgtggaaatgagagaaagatctgccccactcatcctgtctccccatccctcctttatccctgcctgtgaccaacaccagttcatgccaagacttcggctgtc**acttgtgttactgaagcagagcacaagt**t**cagtaaaaataatcccttttcttactg**gtctcccaatgcaaacagaatcaatttgtttctagaagcaaaaggagctgcc**cacacatggtgggtggggatgaggtagggacaagaaagaatccaccctgtgtg**acccagaggcccagaagcctttagtctggttccaaaggtgtctcctcctcttggtgcagttatgctgaccatttcctgagctcagaaactgtgcgagagatgccaagaatgaagaatccagcctgtgtgacccagagacccaaacgtgagacaggaagtctgatgcttaagaaaacgtgggttcagcattctgaaaggtctagatatggcttggcctagagctggtgtagaagagaagcaagagagatgctcagccttaggtggcccaggcacagcggtcctgtcaaggggttgagc**tctctgcaacatggaaatcctgcagaga**catacttgggaggctgtgatcccagaatgacacaaaaatgctaatggaaatgagcaggggattgtttatgggggaatggggagacaggtgtgggataagatgtggtaaggagaatgcagcctgaagaggagactgtccattgtgactcataggaggagtagggccggggcctacagtagtgataaaaggaggacctggaaggacctgggccctgaagacacacagagactggagttttggtctgtctgtg

Uppercase: IGLV5-37-2

Lowercase: Flanking sequence[1000bp]

Red & Bold & Underline: Stem-loop [26]

Blue: Heptamer[52]

Green: Nonamer [6]

id-TRDV2[V_gene_segment]

tttttttcagggcagtctgtt**cccagtcacactggg**gaacttaagaaagcttagatgagtctttcctccactaacctgggaatgtaagtcctgcatccaccctccaatatggaagccacctttgctcttgcctttttgcttaccaggtgactctcttcccagcttctgctgcactctctagttctcattattaatttacctgtttgatttcaaaatttcaaagattgacctttaaac**tcaaaagtggaggtcccaaaaatcctaacctcccctcttttga**agtccatgtaaatttagcccgtcataagttgagatgtagtggtggtttctaagttttttacaatgtttggctttgtgtaatttaacacaggtaacctaagaacccttcagaaaagaagctagatgagctcaataaagatcaggaaactctggctcacatttggaaga**ccatgtttcattggattgctctttaacacatgg**gtaaatcacccttatccaactttgcctagagggagtttgcagtgaacacaacaatcattacaacatcttcagtctttgtgcacatgctgttgatctcctcatttggtggttcttttgattacagctgtaaatgccaataattcttcttggtatga**actatgaacatcaccacatagt**gtctaacatgtacatctaacctgaaactggaacaacaatatcaaatttaacgtctagggaaggaagcagaaaccctgtcaccaattaaataagctgccttaatctcctctcagttggcctaacttacctgaaccgtactattggtcttgaaggtggcactgtcatcccattagcaggcagcaggccctgcgagcctcctgtg**atcagctagatttctctggggaatgaaacagctgctgat**agg**gactgacttcctcagtc**acaagtgtttcacaagtgtgttggttcagaggagggaccaggcagaaggtggttgagaggcagagctgcccctgagtgagccATGCAGAGGATCTCCTCCCTCATCCATCTCTCTCTC**TTCTGGGCAGGTAAGGCAGACCCCAGAA**CTTGGGCCAAGCAAGACTCAGCAGCAAAGAAACATCAAGGCTCACAGGACCTTGTACTCATGGACTGTGCTCCTTACTCAATGACCTCAGCCCCCAGCCCTCACTGAGCCCCTCTCTTGTGTCTTTTCTCTCAGGAGTCATGTCAGCCATTGAG**TTGGTGCCTGAACACCAA**ACAGTGCCTGTGTCAATAGGGGTCCCTGCCACCCTCAGGTGCTCCATGAAAGGAGAAGCGATCGGTAACTACTATATCAACTGGTACAGGAAGACCCAAGGTAACACAATGACTTTCATATACCGAGAAAAGGACATCTATGGCCCTGGTTTCAAAGACAATTTCCAAGGTGACATTGATATTGCAAAGAACCTGGCTGTACTTAAGATACTTGCACCATCAGAGAGAGATGAAGGGTCTTACTACTGTGCCTGTGACACCcaccctgctgcagctctacttctgagcagctcaaaaaccactgaccaggcgcggtggctcacacctgtaatcccagcactttgggaggccgaggtgggtggatcacgaggtcaggagatcgagaccatcctggctaatacggtgaaaccccgtccctactaaaaatacaaaaaaaaaaaattagctggtcgtggtggcgggcacctgcagtccgagctactcaggaggctgaggcaggagaatagcgtgaacatgggaggcggagct**tgcagtgagccgaggtcatgccactgca**ctccagcctgggagacagagcaagagtccatctcaaaaaaaaaaaaaaaagaacattgccatccatgtagagcagcaattccaacaagatcgtgattttctcttgaataaataatttaggaaatagggaccag**agaaagctagtgtttctacctttct**ttcaggctcagccactcagcttgcccgtgccctcccatggtgctcagataggcagcccccagtgggacagaagagaggagagatggagccaaaggaatg**aactgtagcatctactgtgacagtt**ttcttggaggctcataaaaggtatgaccacagagccacagtgcttagagtatagcatagattaaggaggcaccttacaatttacaactatgggtgacatccagtttttccccagccacatgcctctgaatttacatgttct**agaggaaaaggactggcagtggggggaggctcctct**ccctccttcagatcaaatatgtacactacctgtcattttgcccaacatggaaggaaaggctacatctctagcatctctagcatgctctgttgttttgatgattactgactggatattttgttttgctttgttttgttttcttcatgagacggggtcttgctgtgtcccccaggctagag**tgcagtggcgtgatcatagctcactgca**gcctcacactcctgggcccaagctggatatggtgttttg

Uppercase: TRDV2

Lowercase: Flanking sequence[1000bp]

Red & Bold & Underline: Stem-loop [13]

Blue: Heptamer[33]

Green: Nonamer [7]

id-IGKV2D-10[V_gene_segment]

agcaatattaaaatcattggctaaattttc**atcattacaaatggaggtgagaagacagtggaatgat**atttttaaatactgagggagaaaagtcagccacaaattctatatccagaaaaacaatcatttaaatataaaagcaaggtagaaatattccttcatacacaaagattaattcattaatagcatatatgccttgtaagagattcaagagaaaattcttcagaataacagaaaatgacagcagacagcaactcaaattgacaagaaggaatcaaggcctcaaaaaatgaaaaaaattggaaagaagctacaaaaaatgttt**actttacctgatataaagt**ttaagtccatgaataaaacaattataatatatgtcagaatatcatttacattgattgtgttttgagcagcagtttttgactgg**gaaggtgcataatgaaagcagcgagagaattggaaaccttc**tgtattttgatgtg**gatcatgacatgaaatgatc**tggatcttgacattcatttgcacccttaagagttgcaccctttgcactgtgtgttttaccttagataaaataaaataaaataaaataaaataaacaaacctagactcttttaaataaacctaaacaaagaaataagaatcatcaaatatttatgaaagatcaagcttattaaaaagaaacataaaacatgcccgagaaagtcctaaagaactagaagtctagtagaggaaacagcaggaaaatctaaatacgtcccttctgccactcaggcaacacagattttacctctattgatttccttaaaa**acataaaatatatttatgt**cgttttgctcacaagagacgcccccaccttctccttggctctttccaccccactgcacccacc**aggggatttgcatactgtcccct**agggaggactttccttgtgagtctgagataaaagctcagctctacccttgccttgactgatcaggactcctcagttcaccttctcacaGTGAGGCTCCCTGCTCAGCTCCTGGGGCTGCTAATGCTCTGGGTCCCTGGTAAGGGCAGAAGGGAAATGAGGGAGGATGATGGGG**TGGGAGGGTGAACTCTGTGGATCCCGCCACCTCCCA**TGTGTGTCCTGTCCTCGTGTTAGATGTGTCTTGTCCTCCAGGATGGGGCATATGATGTCTAGATCTGTGAGAGTGAGGAAGATTCCAGAAGGAGCAAGGACATGTACTTTAGTGAAAGCTGTGACACAGAAAGAGGGGGATGGGATAGGTGACTTCTAGAGGCCGGCTTTGCCTTGCAAATATTGGTTCTTTTTAAACCTGTATGTTTTGGGAGGATTAATCAAAATCACACACACAAAAATAATTGAGCAAAACATAAATAACAGACAGAAAATGATTAAAATGACTCACAATGTTTGCACATAACCTTGCACTTCTCTCTCATTATTTCATGATCCAATGGAGATGCTGCGAAGACCCAACCTCCA**CTCTCCCTGCCCATCACCCCTGGAGAG**CCAGCCTCCATCTCCTGCAGGTCTAGTCACAGCCCCCTTCACAGTAATGGATACATCTATTTCAATTGGTACCTACAGAAGCCAGGCCAGCCTCCTTGGCTCCCAATCTATTTAGTTTCCAATCA**CGACCCTGGAGTCCCAGACAAGTTCAGTGGCAGTGGGTCG**GGGACAGATTTCATGCTAAAATCAGGAGGATGGATGCTGAGGATGTTGGGGTTTATTGCTGCCAGCAAAGTACACATTATCCTCCcacaatggtagagtcttgaacacaaacctccccacttgctgtggcctagctgcccagatgtgctgtttctgtggagagcaggcactgtggattctcttagatgcctaaagaagaagatgttggagaactcagagggcttggtgcag**ctgagggctcatgaccataaatttctcggctacacctcag**gcatcacattttaaggtcccgtcag**ctgcagcagcctttgcatgacagagtctgcag**tatggaggaggtccacgtgccctctgagcaatgagacac**aagagaagagaaggctcaatgagagctcattctaatcctctcttcctt**ccctacgttcattcatcaactaaattcattctgcataacaggcacctaattgagactgattactggcaacacaaagctaacacattcttttgatttggtttagcagttaccagagtacatgtacattgataagatttggtgatattaaatcagtttccctcccctcctccacctccccccctctactttttgtaaggcagacccttgaccagaacaccagcaagcactgtattttatcctacttttttcagggagaacccaaataaaactcttcacagtctagattttttaattactagatatgttgtggcaaatgta**tgcaaaaataaagacctagttgctaaagtgtttgca**cagtctcactcacagcatattgaagtacaaaagtag**ctttctgaactccaaagctgggtttcaaaaataaggacaaagttcgaaag**aataaaggaacatatgtatgttacttaaacagaatttgaatgaatacaagataatatataagctaactttttaccccatgaaaaattacaatgtgaagactttgagcagagagcagcaagtgagagactg**ctgggttgtgtttgagatatgaacccag**atggaag**ttctcacgtatttcctgggaagaatcacagagttgagaa**gcacctgcctggctttcaaggcagta

Uppercase: IGKV2D-10

Lowercase: Flanking sequence[1000bp]

Red & Bold & Underline: Stem-loop [16]

Blue: Heptamer[38]

Green: Nonamer [10]

id-IGHV3-57[V_gene_segment]

gtggggatatcatgagacat**atgcaacaaagaatgaaggcattttcctcatttgcat**gtaagatgttaccattcactaaagacctttaataa**aaaaaaatttctacgtgatccagtattttttt**ccttctgtcaagcaaataactcataggattcttt**tctttccttggttgagaaaga**ttttctacaagcttcagcacacgtcaggcatacactgtccctgaatgggcatttaccctcagatgggtacacacacctgtcaacatgggggctcttctgtcagacaaacacacctttact**catgtggattcttccctcagacaaacacacatg**tccccacgtggactctttcctcagattacaacatttgtccttacatttactatttcctcagaaaacagacattgcatcatgtggtct**cttgtctcagacaag**caaacatgtctt**cagtcagataagtctgtggatttccacattgactg**tttccttacacaagcacctatatccaatatttaactgttttgt**gagaaatttatcccttttctc**tggaaatgtatttttatgttcttactggacatattttttaataatgtttgctactatgaagatacccgaacactgtccatacgagagaataagaaagagtaataagcagattaaccctgtgcatccagacccaggagtcctttgatcctgc**ccttccgaaatggagacacagaggaagg**atgagcaatgctgagcagtgcacccatgaccacaaaaggaaagacatggcaatgtgtcccctcccctcctcatgaaaggcagctcatcccctgttccttcaggccctggtgaggagcc**accccatgtatattcccttgatcagtgtccacaccatggggt**ctgcactgatctgggcttcccttctcatcaccctcaatattagtgtcccttgtgaatcaggtccagctgcggctgttccacatggggccgttcttccatttcctcagtgtttgcagAAGTCCTGTGTGAAGTTTATTGATGGAGTCAGAGGCAGAAAATTGTACAGCCCAGTGGTTCACTGAGACTCTCCTGCAAAGCCTCTGATTTCACCTTTACTGGCTACAGCATGAGCTTGGTCCAGCAGGCTTCATGACAGGGATTGGTGTGGGTGGAAACAGTGAGTGATCAAGTGGGAGTTCTCAGAGTTACTCTCCATGAGTACAAATAAATTAACAGTCCCAAGCGACACCTTTTCATGTGCAGTCTACCTTACAATGACCAACCTGAAAGCCAAGGACAAGGCTGTGTATTACTGTGAGGG**Acacaggagagggaatatctgtgt**gagcccagacacaaaaatctctgcagagagacaggagggaactgcatggtagatgctcctcataaccacaaaggggcagtcaggaccatcaggaggagctcaggacacctgggggtgctcagaaccatgaggggtgctcaggacatcgggggctctcagaaccatgaggggtgctcaggacatcagggggctctcagaaccaccagggggcgctcaggacacctgggggcgctcagaagcatcaggggtgctcaggatatcagagtgccctcagtaccacgagggggcgctcatgacaccaggggcactc**agaaccaccagggggcgctcagtacacgggggttct**cttaggaagcagctccacatcaggagcctgagaggctgtgatttcgttttaaaccttggtgattcccgacctggtcaagcaaaagtcttccccaggatctctcaccatttcttc**tttgtaaatccatgatttttttttacctacaaa**acattaagttagaacagggatttaattcaacttttaattctgcagattttccgagtaatagtagcaatgttccctcagga**caatttttaaaattg**gttatttatat**attttatttatgtaaaat**ga**tactatcttgaaaatagta**aaatttttaaaatcatacctctaatcccagcactttaggaggttcgggcaggcagatcagttgcgctcaggagttttagaacagcctggcaacacatttaga**tcttttccatacaaaaaga**aaaagaaaaaga**aaatagctaggtgtggtgctacctctggtaccagctattt**gaaagtctgaggtgggaggatcccttaagcctgggaggttgatgctgcagtgagctgtgattgtgccactacactccggcctggttgacagagtgagaccctttcttaaaaatacaaactgtttcaataagtagagctttgtgtctttgtgctgtaataatcagtt

Uppercase: IGHV3-57

Lowercase: Flanking sequence[1000bp]

Red & Bold & Underline: Stem-loop [17]

Blue: Heptamer[46]

Green: Nonamer [5]

id-IGLVI-56[V_gene_segment]

cgagatcaggagatcgagaccatcctggctaacacagtgaaaccctgtctctactaaaaatacaaaaaaattatccaggcgtggtggcaggcacctgtagtcccagctactcgggaggctgaggcaggagaatggcgtgaaccagggaggcggagct**tgcagtgagctgagtttgcgccactgca**ctccagagcctggctgacagagcgagactccatctcaaaaaacaaaacaaaacaaaaaaacactgtgcttgctcgcttacctcctcctcaggtcatttcctgcagtctctaagaacagtccgcctggcttccccatctcctccctgagcgccagcatctggggcgtgaacagcacaatcaaggccacacctgtgagaccctcagtgagtcagcagggct**gtgtcctggtgatgtcaagggacac**ggggtcagcaggggacctgtgaggctgtctcagggggtgagcatgtctggccctgccagctgggaagg**ccctcaggaggttgaggg**gagctgagcacacagtgagctcactgagctgagatgggaagtgactgcagctgtcaggaccttcagggacactcagtctaggatcaggctctctcaccctgtggggcctggttcaggtcaactcctcatcctcctcttcttcctccagacaagagagacattgcctatggctgaggggtcctggaggtgatccgggtgcagggacatatttgaatgtaggaaattctctcccagtgagacctgtagagactaaaagagcgttaggaacattcactgttactgctggagtcttag**ggtgcagatctcttgagcacctgcacc**atggcttggcccctctgctttttgacctcttccctccctgcccagctgaggcacatcctgctctggggagaagggtctatgcggtcctcagcccacctctgctccactccctttctcacagggctcattgttttctctttctctccaccctcctgggtctgtggTCCACATGGCACTGACAGCCTCTTTCACTGACCAAGCCTCCGGGACTGACAATTACAATCACCCACCCTGGAAACAGCCATAATACTGTCAATCAAGGCACAGCTTGGATTCAGGAGCACCTGGGACATGTCCCCAAACCTCAGACCCAGAGGGGGTAACAACCATCCCTCAGGGACCTCAGGGAATTTCCCAGCCCCATGTTAGGCAGTTTGGCCTCCCTGGCCATCTCTGGGCTCCAGGCTGGCGACGGTGCTGATTTTCACTATTTAGCACAGAATGGCAGCCTCGCTGATTAttctctgattatctggatgctctgtgactccttctgtgcatctctgggatcatcattcagactcacctgcaccctgag**cagtaacatcaatgttgtttgctatgacatttactg**gaaacatga**cagggcagggagccttccctccctg**gtatctactgagattctactgagattcaagcagaaccagggctcagggtccccagcctcttctccgtgtgagtgggtgccgtcgccagtgctgaggttttccatatctttggtctcaagcttgaggatgaggctact**catggcacagtagtgcctctcacagtgccatg**gatgggatgttctct**ctaaaatagaagtcttttttttttttag**tttggcccctttgataagtagattctataataattaaaga**aaaaaagtaatgtctttttt**ggagggtagaaacactggtgaaagagaaaccgggaaggacttggca**tggggggtgggtgggcttcaggccacggctctgaccccca**cgaaaggagagaaggaagggggattagtaggaactgccccagcccacaggcaggtcagacgtcagacaaagcctcagccgactgagtgggttctttcccaggggtgcccattagaagtgccctgcttcctgcaggaagacctggctggaggga**ccctggggtcctcattgcctggagccagcccaggg**gcaggggacatgttagtcacttgtggtcctcctgcaggttctcccaaagaggcatctgagtgggacacttccatagccgccaccagcactgctgctgcattccccctcaggacaggggtagtgccttccttccct**ggctcatctacctgagcc**ccatgtgaagtcaaagcttaggt**gtctcagggctttcccccaacaggccctgcagccactgagac**ccttatataccctgagctgcatgctgtggagagagggtctgtggaagaccttggctcttagccatcgactcctctggggaacaccatg

Uppercase: IGLVI-56

Lowercase: Flanking sequence[1000bp]

Red & Bold & Underline: Stem-loop [13]

Blue: Heptamer[41]

Green: Nonamer [3]

id-IGLV5-39[V_gene_segment]

aaatagtataacatctaggacaaattcttgaccaaaaaatatgacagaactgtgcagacaatgacaagttttacttagggagataatggctaacttcataaaatagagaaatacataattgttcactgataaaggtaagtctaccaaatctattagttttcctgtattactaaattgagtcaaatccctataaaacatatgaaaccattgcatatataacttttgctaaattgctgcctttaacatgtcaagaatttttttatttttcctggaaaagtctcattattcaccgtccttttggacagcccccagctgcccttcaacccagctccttagctgatcacttttccataa**tgcccaccctaatggcacatgtgggca**attagatttttctagaatgccctgcggtgatgctgacacctgtaaaagttatatatgtacatgtctgcacatcccacagaccaggactactactgggagtgagaaatcacactagtaattcatagcataatagaagtgtctaggaatcagatcaaggatatgtgaatccagagactgactggtgaatggt**agagaccagctgggttttgaagtctct**gagccatgggaaagtgtgtgtccggggaccagcgtcggatgttgtggggctgccctgtgggtgctacacagctgttcagtaaggaacattcactcaaagaaatccaggccttgggtctgggccctgtgctcactgat**ggacactcagcagctgtgtcc**cctctggcctggcagagttctctgagcaggg**acctgtgtatcgtctcaacaggt**gcttcctcccaggaccttgcaagaaaaaaaacaaaccttcctcctgcttagtctgcagagtgaggtgagctgca**tgcccaggtctcaaggaggggctgggca**gtccctgggtggaaacacatttgcatgagcagcc**cctcctctgcagagggtgggaagaaaaggagg**cctggggcagcccagtcccactGTGGGGTAAGAGGCTGTGTCCACCATGGCCTGGACTCCTCTCCTCCTCCTGCTCCTCTCTCACTGCACAGGTAGGAAAAGGCCTAACAGACCAGGGCCAGCACCTCAGCCTGATTCTGACTCTTCTGGCAAAGATCCCTAAAAAACTTTACCCTCCATTCTGCCTTATCACTCATAAATGCCTGTGTTTCTAGGTTCCCTCTCGCAGCCTG**TGCTGACTCAGCCAACCTCCCTCTCAGCA**TCTCCTGGAGCATCAGCCAGATTCACCTGCACCTTGCGCAGTGGCATCAATGT**TGGTACCTACAGGATATACTGGTACCA**GCAGAAGCCAGG**GAGTCTTCCCCGGTATCTCCTGAGGTACAAATCAGACTC**AGATAAGCAGCAGGGCTCTGGAGTCCCCAGCCGCTTCTCTGGATCCAAAGATGCTTCAACCAATGCAGGCCTTTTA**CTCATCTCTGGGCTCCAGTCTGAAGATGAG**GCTGACTATT**ACTGTGCCATTTGGTACAGCAGCACTTCTCACAGT**GACACACACACATGGGGAAGTGGGACAAAAACCTCACCCTGCTCTCACTCTTGTTCTGTG**ACAATTTTCAAATTGTAAacttaacttgcctatttacaaagtatatttggaagtacttt**cttgtggtaacaggcttttctctccatttccctttcactgcttttgaagtctgagtaaaacaaaaacaaaacaaaaccacctgatgacgctaaactgttggatggttgtccgtatgtgctaaaaaccagtgcctgtggagctgggcctttttctcttgagagaagcaa**aatgtgccttttctagctcacatt**gtagtcagagcctgagaatatcctagcaggtaaatagtccagagacccgctcttccatttccagtagtctgtgctcagggctgtccacggctcactgtgtcccaggatcaaagcactaaaatgtaaaatgcttcatgcggatacagttcagggaggaactccagacataa**acctggcctggcccagtcccaggt**acctgcatttcagtttcaatgtcaggagagctgatgaggtgtgggggtaactggatctgaggccaggaaatcactctgtaggacatgggagaataaacgagtagggggtgggggtctccatcatggcttagacaaagctgaggctgctcctgggccactggtagaagtaggttccat**ggatgagactttcctccagctcatcc**cactcagaatcacgggaacttccgagggtggttttcctgaactcttggggtctgactgtattattttctcttctcttacaccattagctttt**cactgagtcagagtctagtcagtg**caatatctgggtatagaaactgcccaatatttgcttcaagtga**ctcctccccatgaaactcttgaggag**gcctacaacttgatgtcaccttttcctg**catggggagcagcagaagcattgggcagtcccatg**gaaatggctactgagggctcccagggagaacccctctatttgtgacccaggacaacagtgatggaccttcagggatcccagcctgattctcctgctctggctttgtccacgttgcctccatg

Uppercase: IGLV5-39

Lowercase: Flanking sequence[1000bp]

Red & Bold & Underline: Stem-loop [20]

Blue: Heptamer[43]

Green: Nonamer [7]

id-IGHV3-69-1-2[V_gene_segment]

gagacatggtcttgctttgtcatcc**aggctggagctcagtggtgagatcacagctctctgcagcct**cagaccctgggtctccagcaatcctcctccctctgccctatgtatgcaccaccatgctcagcttcagtgcttactacactgcacactttaattatgcaatatttattatacagcaataatgcctcaataagggtatcagaaataagtgaatagataatttgttagataaagattgatggaaagacagacactgacatgagaaatgtatgacactc**aagaaaataaaactgtaggaaacgtgcttttctt**cacatttgttaggtaatcacaacagtgcgtacacatcacaccatattcttgttacagagaaaaggttctgcaaacctcagtaggtgcgacgccctgtgtgctgtgcttagttcagggagcagtcaggctcggtggtgagaagcacaggcccagatacccaggtcaccctgaccaaatgtgaactctagggagattgaacaaccgatctgtgattttgctggtaatttttcatctgttacatggaaataacattgatactacataccatggtttcactgcatatgaaaaaataaaagatgatttgttctaactttaaacatatgcactttctgttgatctactgtacctcaatagaactgttttaaaataaaaattacaaaattataagatttataggttttaaggttttatcacagagcagatttaccataagaaaccacaatttcccaaatgctatcaatatcacaaatctccccaggacactgtcacgtgctctgagcccca**ctctctccaaaggcctctaaccagagag**cttactatatagtaggagacatggaaatagagccctccctctgcttatgaaaaccagcccagccctgaccctgcagctctgggacaggagccccagccctgggattttcaggtgttttcat**ttggtgatcaggactgaacacagaggaccaccAA**GGAGTCATGGCTGAGCTGGCTTTTTCTTGTGGCTATTTTAAAAGGTAATTCATGGAGAAATAGAAAAATTGAGTGTGAGTGGATAAGAGTGAGATAAACAGTGGATTTGTGTGGAAGTTTCTGACCAGGTTGTCTCTTTGTTTGCAGGTGTCCAGTGTGAGGTGCAGCTGGTGGAGTCTGGGGG**AGGCTTGGTAAAGCCT**GGGGGGTCCCTGAGACTCTCCTGTGCAGCCTCTGGATTCACCTTCAGTGACTACTACATGAACTGGGTCCGCCAGG**CTCCAGGGAAGGGGCTGGAG**TGGGTCTCATCCATTAGTAGTA**GTAGTACCATATACTAC**GCAGACTCTGTGAAGGGCCGATTCACCATCTCCAGAGACAACGCCAAGAACTCACTGTATCTGCAAATGA**ACAGCCTGAGAGCCGAGGACACGGCTGT**GTATT**ACTGTGCGAGAGAcacagt**gaggggaagtcagtgtgagcccagacacaaacctccctgcaggggtccccaggaccaccagggggcgcccgggacactgtgcacggggctgtctccagggcaggtgcaggtgctgct**gaggcctggcttccctgtcatggcctgggcggcctc**gttgtcaaatttctccagggaacttctccagatttacaattctgtactgacatttcatgtctct**aaatgcaaaacttttttgttctttttgtattt**ttgtttttgtaacaggaggacacaccctcacct**ccacagaagccacagtgtcactttgggggcagatgatccttctgtgg**tcagcaggatgaaagttccgaggaatctcaggggaacccgaagagtgtttgccagttagactcagggcagcgacctccacaggaatctctgattagaacaggctttgagttctgataggagccaagagagacgctcacccagggtcagagtcctt**aaaaccttgtggtttt**cacagcaatcccccctggtcttgtaaaactgtgtacatatgactcagactgattcacttgaccccctttctgctaatccattttccttctctgcatacttgattctcacagttccctttcttcttctctttcct**gaaaacagaggatgtgttttc**tgtagtcaaaatcccagagat**caggtctgcaggacctg**ggtaggctgaggggactttctcactcaccataggctgatgacactcctgctgtattttgtgtgtggaggtgtttggaaaacgaaatgaacattagtcatgaagggaataatactagttttctccaaagggatgtggatgtagagctgatcttgtgcttctcacactgtcacagaatttat**gctctcccctgtgactttaggagagc**tgaggatggacactccattgtgctgtgagctctggtaatagtaattatagggtctggctaggcagcctaaggtcaatactgctggccttcgggaa

Uppercase: IGHV3-69-1-2

Lowercase: Flanking sequence[1000bp]

Red & Bold & Underline: Stem-loop [16]

Blue: Heptamer[50]

Green: Nonamer [8]

id-IGKV3-7-3[V_gene_segment]

aactgtctacagcactgtaaggcaggtaccactattgtcacagttacacagatatggaaaccgagacacagggaagttaagttacttgattaatttcaagcaatcggcaagccatggagcatctatgtcagggctgccaggacatgtgactgtaaacagaagtttttaactcaaa**gagggtatgtgtctgggttaatggaaagcttcaggaccctc**agaaaacattactaataagcaaatgaaaggtgtatctgggccgggcgcggtggctcatgcctgtaatcccagcactttgggaggccaaggcgggtggatcaccaggtcaggagatcgagaccatcctggctaacatggtgaaaccccgtctctactaaaaatacaaaaaattagctgggcgtggtggcaggcgcttgtaatcccaggcacttgggaggctgaggcaggagaatggcatgaacccgggaggcagagctt**gcagtgagtggagatcgcaccactgc**cctctagcctggaagacagagcgagactccgtctcagaaaaaaaaaaaaaaaatgtctatctggaagattaagttctaacagactcttcatttccatagatccaataatgcacttagggagatgactgggcatactgaggacaggaagagagaaatgaaaacacagccttttatattgttcttaacagacttgtgccaaatatcatacgggtgtatttaggtgattgaagagaagaaaggcacaggagtgaaattatgtgagcacaagggaggagttctacactcagactgagccaacagacttttctggcctgacaaccagggaggcacaggacgctcagtgcagagaggaagaagcaggtggtctct**gcagctggaagctcagctcccacccagctgc**tttgcatgtccctcccagctgccctaccttccagagcccatatcaatgc**ctgggtcagagccctgggaaggaactgctcagttaggacccag**acggaaccATGGAAGCCCCAGCTCAGCTTCTCTTCCTCCTGCTACTCTGGCTCCCAGGTGAGGGGAACAT**GAGGTGGTTTTGCACATCAGTGAAAACTCCTGCCACCTC**TGCTCAGCAAGAAATATAATT**AAAATTCAATGTAGATCAACAATTTT**GGCTCTACTCAAAGACAGCTGGTTTGATCTAGATTACATGAGTGCATTTCTGTTTTATTTCCAATCTTGGATACCACCAGAGAAATTGTAATGACACAGTCTCCACCCACCCTGTCTTTGTCTCCAGGGGAAAGAGTCACCCTCTCCTGCAGGGCCAGTCAGAGTGTTAGCAGCAGCTACTTAACCTGGTATCAGCAGAAACCTGGCCAGGCGCCCAGGCTCCTCATCTATGGTGCATCCACCAGG**GCCACTAGCATCCCAGCCAGGTTCAGTGGC**AGTGGGTCTGGGACAGACTTCACTCTCACCATCAGCAGCCTGCAGCCTGAAGATTTTGCAGTTTATTACTGTCAGCAGGATTATAACTTACCTcccacagtgattcaacatgaaacaaaaacctcaacaagaccatcagtgtttactagattttaccagctgcttcctttacagacagctaatgtggtggccactcagttttagcgtctctgctctatttggacattttgcagttctaaaaaaaaa**tcattgaacaatttggactttgattcttggactctgttcaactga**ggcaccagaatcccaggtttccagaaatagtgactcactgtatgaatccttatatagcctcagtggttcttaactttcccagtagaggtagctcagtgcatgctacactgctccattt**gaattttgcaacattctaagtagtagaaaattctatttatttatccaaatag**ttgactcggtaaaagctgttcatgtgaagatactaccatggctgaataaatcccattctttttctttcttcaggctatcaacatttcag**tggcaaatggttattatggaaacatttgcca**tttaaaagttaactaaattatttcttcaattttctctgtgatgcagtagactgtaaaaagattaaagtttgttaaaataaagtacatattcgataaggaagaaatagattattcctaatgacgtctgcaatgacctagtagaaagagtgatagaagcagttgttttcattatttttgtccaaaacttccttccaaatgggatttcattgatcatattcgtttattaccacctataagacatgttgacattatgtaacatctgatgtgaagcactgaggatacatcctgtctgtattattcttgccaaaaatta**atggtgtgaattgaatcagtagtaaacaccat**atacaaacccaactaggaggacattcttcaacatacctggacagtaaa**cttcaaatgtttgaag**gccatgaaagagaaacaaaagtgaaaaactatcacagatttaaagatattgaggacaggataaccaaataaaatacagaaacctgaattttatcttgtaacat

Uppercase: IGKV3-7-3

Lowercase: Flanking sequence[1000bp]

Red & Bold & Underline: Stem-loop [13]

Blue: Heptamer[39]

Green: Nonamer [7]

id-TRBV12-2[V_gene_segment]

ggggcgcaggggactcagctgtatgcctctgtgccaacagcgaaaatgtggccttaaactgctccttcttctttgttagcaaataacctagttgaccaattc**atagagcagaggttctctcagttctctat**actctacaccccgagaggaagtcagttagtgagatgtaagat**tcttttataagaaaaaga**ccaggtgtaaagattaagaaatgctgagctttgctagaggtaagaataagatgtaaatcaactctggaacatttaatttccaaatctttgtgagaaagacaatcttgtttctgatttattgtgtggatagagaatgtgagagattctgtgctctgggctttgcaaactcaatagagttcaaggatcaataaactcggagagacccttcactgcaaagaagagctttgaaatctaatgagcttccgggacactgattcagctcctgtatctttaacaacgcggttgacatgcttgctcttcctgggtaactcagatgctaacttctaactcatttaaaggcatttgaagaactaggagcacagagtcaccagtgagaggaagaggagagtttgcagagaagctggcttgcaataaggcaatgagttcatctttaaatacttggagtttgaggtgcagatggatatagttggcaggctcctaggtaaggcatgttatggagaaggtgctatgaattgataatatcaaagcaaatctacagggatcctctgcaagtgtgcatctgtatctcagatcaattatagttgacttcagtcctgcctgattcatctcccaaaaatgtagtctgcctgattcatctcccaaaaatgtagcctccgcttaaaggagctttcaagttgggggtggtgggccattcagtgttgtcactaacagatgcatcttgtgggggtaaaatgtcccaaagtatcttttcttgcttatgttcataagggcgctggtctggaatgtgccacatctgttctcactctgccATGGACTCCTGGACCCTCTGTGTGTCCCTTTGTATCCTGGTAGCGAGTGAGTCCTCATGTATTTATCATCCTCATGCTGGGCCTCTGTATAGATGACTTCCTGTATTTTCCTTATTCTGTTTCCTAATTTTGCGTTCTTTTATAGCATGCACAGATGCTGGCATTATCCA**GTCACCCAAGCATGAGGTGAC**AGAAATGGGACAAACAGTGACTCTGAGATGTGAGCCAATTTTTGGCCACAATTTCCTTTTCTGGTACAGAGATACCTTCGTGCAGGGACTGGAATTGCTGAGTTACTTCCGGAGCTGATCTATTATAGATAATGCAGGTATGCCCACAGAGCGATTCTCAGCTGAGAGGCCTGA**TGGATCATTCTCTACTCTGAAGATCCA**GCCTGCAGAGCAGGGGGACTCGGCCGTGTATGTCTGTGCAAGTCGCTTAGCcacagcgctgcagaatctccccctccctgtgcagaaactctggtgcttcctcttttcctcacagctcccagcagtcctgagcaaagtctttcctgcctcaccctccccacaagaataattaagtgggtttggggcatggcaaagacataagatgatacaatatcaacatacaaaatcttgtgtaggagataatgtcgaaaatgcattttgcaaatttctcacaaattgtgtatggggtcataactcacacacaggtaggat**aaaccttaattacccacaccgtaggttt**ccctgccttcctgtatttgaca**ctcccatccaaacagtgggag**ctcttgagaatggctatgtcttgtgaactactctgcccacttcattcccttctcactctaagccatgaacagaaatttcccctacaagtactctttattgtaaatgaaagttcttttctatgattgtggacattcctttataatgctaatttcaatttaccatttatatacggatttttacattaaacttaaagagatcttcagtgtggataatacaggcagtttaatacattagaagtaagctacccctcttggcctacagggatccatttgtttctatcttgcataaagcaagccaggtgctagctaaaggtctgtgtgcactcacaggacactgatgagatctcctccactgcctgttctggaactatcaatatgaatatacacagggtgtctaaaagcacacagggcctggtgtggtggctcatgcctgtaat**ctcagcactttgggaggctgag**gcgggaggaatgcttgaggtggggagttcaagactaggctgggaaacagtgagattttgtctc**taaaatatatatatatttta**aaaattagcctggtatggcagtcttagct**actcaggaggctgagt**tgagagggttgcttgagccctgaagttcgaggctggagtgagccatgatcatcgcaacagagtaagaccccaaatcacaca

Uppercase: TRBV12-2

Lowercase: Flanking sequence[1000bp]

Red & Bold & Underline: Stem-loop [9]

Blue: Heptamer[35]

Green: Nonamer [2]

id-TRBVA[V_gene_segment]

gtactcttgtgttccttgaagcattattcacatagccaaaatacagaattaacctaggtttccaccaagggatgactagatgaagaatatatgatataaattctgtatatgt**atatataaatatatatat**tatttatac**gtatatgtgtatacatacacacatac**acacacacacacaatggaataatactcaaccctacagaaaaaggcaatcatataatttgcaacaacatgtttggacctggagacattatgctaagtgaaataattcaggcacagaaaaacaaataatgcatggtctcacttattcatggaatccaacaaagtcaaacccacagaagcagagagtagaagggtaattgccaagcactgggatggggtttcaggaagggaggcataggaaatagggaaatgttggtcaaaaggtacatgttttaatcagacagaaagaagttctagagacctaatatgcagcatgattactatagttaataatatatgtactcttgaaaattgctaagaatgtagattttaaatgttcctaccaaaaaaaagtcaaaacaaaataagtatgtgagattatgcatatgttagcttgatttaattattccacaa**tgtatacatatattaaaatgtcacaatatcacattgtataca**ataaatatatataatttctatttgtcaattttaaataaataataaagaaatgagatagctttcaactaagc**ttatatagcagtataatataa**aaatttttgtagtaa**caaaattatagtaattttg**gcttatactgaaatgtgcattaaacaagtgtttgcataaaatataattcaaattactattcatagcccacagccatgaaaacatgtgtgccttactcttggtacagaaacaaatagcttttactcctaaggaaaaaagaaaattcctgccttttgatgagaattgtttgagttttatacttattttatggggaactttgctgacctcttcattccacCATAGCTGGTTATTCCTCCCTTGCATGTACTTT**TGTTTCCTGGGGGCAGTGAGTCTTGGAAACA**TTTGGGAAATTCTTATTCTGAGCTTGCCTGCCCACAGCCCAGGTTCCTTCCTGGAGTCTGCAGCACCAGGCTCTCTCCTTGGCTCTCTTTCTGTCTTCTGTCCCC**TCAGCTCATAAACCCCAGCTCGTAGAAGCTGA**AGCCACCTAGACTCTAAGACACCTGATTGCAGAGACAGGAAAGGAGTTCTCAAGATAAGTGCCAAGATTTCATACTGGTTTTCACAAGAATCAGGGTTGGAATTAGAGCCGACTCATTATTCAGTTAACATTGACATTGTTGATTAGAGACATATCCCTATTGAAAATATTTCCTGGCAAAAAATAGAAGTTCTCTTTGGC**TCTGAAATCTGCAACTCCCTTTCAGA**TGTC**CCTGTGTCCTTGTACCGTCACTCcacagcactgcacagg**catgtgctcacctcacaaaatggcagtctcaaagggaggagtgcccacccacaagaggctccacccta**ttctgagaaagaacttctttcagaggaggagagaatagaactttccagaa**ccttttacatgagatttttttttttgggtag**gaagtctgacctgagcagcaaccttgcatcagacttc**aaaaggcaatatttcctatacctccttgtgctctactagtgagtacagaagtcgagaaaatgttcctaagtctatggccagttc**aaatagtctcattttctattt**ctgagattaaattatatgtattaagatccttactctgccctttcaggtttattgctcctcagacaagatatcccctcttcaggtcccaaacacattcattgcaatgcctttcctcatccttgtcccagtcctggctctctagcttcttgagacaataacccaccctaaggtatgtt**gtgacatgcatcttctgtgtcac**tgggatccctaagtggtacaagcattcactcaattgctttagctagagtccccagaatatcttcacctcttctgtcacattcttcttccacattggtcc**ataaacacatctttaacatttctgtttat**taaaacctctcaaattcaaccactattttctatcccaacgaatactgccttagctcaggcaaccactgtcttcgatctgaattacttctttgacctgaattacttcaactcctaccaaatatccacctacctgatgtcttactctcttttctaactcagttcattctctgcagctgaaaacacttcctttccattgactttagggttatgaccagtcttctttatatctaaatgttagcctat**aagttttttcgtaaaaaaaaaaaaaaaaactt**ccaaacta**caaattcttaaaaccctccataatttg**gcccttgcatacctatccagtcttgtgaaattcacctactctttacccactttttg

Uppercase: TRBVA

Lowercase: Flanking sequence[1000bp]

Red & Bold & Underline: Stem-loop [16]

Blue: Heptamer[29]

Green: Nonamer [7]

id-IGHV3-33-3[V_gene_segment]

ggcta**aattctcaaagaatt**gtgctgagtgaaagaaactaaggaattaagagt**aaattttacatgatacatttgtagaaattt**tagaagatgccactattataaattaacatggagaagatttaaatgtttctgagaatatgctattgggagtaatggggatgtgagttaaatttcagaggaataagagaaagatttagggattaattttttcaaaccttgattgaagtgctgagtaaatggttgcaaacataggtctacattt**ttcaaatcattcaccataaatttgaa**tta**tttattaattacactcgaataaa**gcaataaagaaactgatgagataatatttgactgaattgcatcaataaatagatcgatattaacacaaggaatataactgatttccaaaaacatacacatgaaccgtgggtcactctgcatatttaggtaaataacagaaagttgtcataacagatggggaatcctgcagacttcactaggcatgggccatgctgcc**ctggagttgtctcaggggagctgcctcctccag**aggttagagcacaggcccaggtaataggactaaatttttagatgtgttatcttagacacactgcacaactgctgtgttctctatgtaaattatctcctgtaaaatataacattgaagcctgcattaaatatattgtgtaaatatgtaagaataaaagaaagttatgagagctaagtgttaatcaaggcacaagcatataagatataactatattttcctgaatgatggaattactaccagtctcccccaggacactt**catctgccctgagcccagcctctcctcagatgtcccacccagagcttgctatatagtgggggacat**gcaaatagggccctccctctactgatgaaaaccagcccagccctgaccctgcagctctgggagaggagcccagcactagaagtcggcggtgtttccattcggtgatcatcactgaacacagaggactcaccATGGAGTTTGGGCTGAGCTGGGTTTTCCTCGTTGCTCTTTTAAGAGGTGATTCACGGAGAAATAGAGAGACTGAGTGTGAGTGAACATGAGTGAGAA**AAACTGGATTTGTGTGGCAGTTT**CTGATAACGGTGTCCTTCTGTTTGCAGGTGTCCAGTGTCAGGTGCAGCTGGTGGAGTCTGGGGGAGGCGTGGTCCAGCCTGGGGGGTCCCTGAGACTCTCCTGTGCAGCGTCTGGATTCACCTTCAGTAGCTATGGCATGCACTGGGTCCGCCAGG**CTCCAGGCAAGGGGCTGGAG**TGGGTGGCATTTATACGGTATGATGGAAGTAATAAATACTATGCAGACTCCGTGAAGGGCCGATTCACCATCTCCAGAGACAATTCCAAGAACACGCTGTATCTGCAAATGA**ACAGCCTGAGAGCTGAGGACACGGCTGT**GTATT**ACTGTGCGAAAGAcacagt**gaggggaagtcattgtgcgcccagacacaaacct**ccctgcaggaacgctggcgggaaatcagcggcaggg**ggcgctcaggagccactgatcagagtcagccctggaggcaggtgcagatggaggctgtttc**ctgtcagggtgtgggactttgtcttcttctgacagttccccagggaacctcttaaatttagaaaactgt**gcctaacaatgtcttctctatgtatatgaggaccttttctccctggcacaaaatgcagattgacgctgacacggatgaaaattcctcaaccatggtcacaaggatcagagt**cctgagtaacctcagg**gct**tcctggtgattcttctccaatcagacccaggacagggacctccgtgagattccctg**actggaacagtctttatggat**cctggtcacagacaatagagaggctgaaccagg**gtcagcgtcatgtagaacctcacagatttcacgtctgatccttctcctgacacgaaagtatgcaaatcagtatc**agcaccgatctggtgct**tcttttgttcctaatccatttactttattttttcgtcgtttttc**tcctttttccatttgtttttcctgctttttgcaaaagga**agatgttttccctgtgagatgcaggggatgacaattttggga**gatggctggaacatccaatatcctcagggccggccatc**agtaagtgcaggctggaagtctcagaaagagctgaagctgcttaatcaccgtggagttttaccttctccagttctgctctgatggaatcagggccaagcaggttatcaatgataatctacctaacatagagtcaaccgattccagtttcaataacgtctgttaaaaattcacaccaccacctggattactgttttgtcaaat**caatacacggtattg**tccagctaagtagacccaaagacggaccgttgcccatggagaaaaacattaacctgagttctaggttcttacagtgttaaagg

Uppercase: IGHV3-33-3

Lowercase: Flanking sequence[1000bp]

Red & Bold & Underline: Stem-loop [22]

Blue: Heptamer[31]

Green: Nonamer [6]

id-LOC124907873[V_gene_segment]

actgtcctgatctttcccaagttctctcctttatgtgttctccagaaatccagactccagagcacccagggacaggctggtgggcagtgcaactcagaacattactcagtggatatggtattggtcttcttactttagagacaattgatccttaggtaatttcaagtttatcttaatccatgctcctataattggaggaagaaacaagtaacttttttttgaagtaatcagatccggcatcactagtcagcgcttcctgctgataaaccctggggaacatacccaatgacagaaaaggaggtataatatagagatccccaaagcaaaaataagatgcagctcatataatctctgggcattattgaggtctataggtttgcata**aatgtgcaactgtaagtggtacacacatt**aaggacttatagcaccatctttgtgcacctgcagtgaattgcttg**accttacactgataataagaatggtttaaggt**tattatagaatacttacaaatatatttagcattatcttgat**atttttcctttttttttaagaaggaacaat**tacacactcacagaaatgtaactataccacaaagtacccccctccctttcagagtatttttaagacctgctgattccacactctaaaaactttactttgagtttcctacagagaacaatttctcccaaataatccccatacgccaatggaatacattactccctcaagtactcagcagtatttcaaatttcgacaattatcaaaaaatattctt**tgtttcaaattgacccccagcaagaaacagattaaagacaaatctgt**gttgccctggtccgacctgagatcctgaggacactgtccttgtgctgagttactgagatgagccagccctgcagctgtgcccagtgggccccatcccctgctgatttgcatgttcccagagcccagccccctccctgccctgaagacttttttata**ggctggtcacaccctgtgcaggaGTCAGTCCCAGCC**AGGACACAGCATGGAAATGAGGGTCCCCGCTCAGCTCCTGGGGCTCCTGCTGCTCTGGCTCCGAGGTAAGGAAGGAGAACACTAGGAATTTACT**CAGCCAGTGTATTCGGTACTGGCTG**GATTTTCAGGGAGGGCTTCTCATAACGTGATTGATTGTGTGGATGTTTGTTTTTATGCTTCCAACCTCAGGTGCCAGATGTGACATCCAGGTGACCCAGTCTCCATCTTCC**CTGTCTGCGTCTGTAGGAGACAG**AGTCACCATCACCTGCCGGGCAAGTCAGGGCATTAGCAATGGGTTAT**CCTGGTATCAGCAGAAACCAGG**GCAAGCCCCTACGCTCCTGATCTATGCTGCATCCAGTTTGCAGTCGGGG**GTCCCATCTCGGTTCAGTGGCAGTGGATCTGGGAC**AGATTTCACTCTCACCATCAGCAGCCTGCAGCCTGAAGATGTTGCAACTTATTACTGTCTACAGGATTATACTACCCCATTCACAGTGTTACAAGCCATAACAAAAACCCCCCAGAGAAGCAGATGTGTGAGTCTGCACTGTCCTAGCTACTCCTCCTGGTGCCTCCATCTGCTGAGAACATTTCCCAAATTGCATCCA**CACTTTGAAGTTTCTGTAGAAAGTG**ACATGAAGCCTCCTCTATACCTTAACTATCTTTCCTTCACGGCATCCCCAGCGGCACAGACAGGGCAAACCCTCTCCTGATTGCATTAAGAACAGAGATGATTACACCTACGAAGTCTGAGTTATGGTGTG**AGTTGGAATTAATATCATAAAGGAGACACCACTCTTGGAATTCCAAGT**AGAATTTTTCTTCTAAAAACAGGAATGAGAGTCTAAACTACAGTCTTTAAAAGTCTTGATGGCATGAGAATGAAGATGCAAATGCAAAAAAGAGGGATTC**TCTGGATCTCGCCATGAATCCAGA**ATGTATCTGCCACTTCAGAAGGTATAATTGCCAATCATGTGGTCCTCAGACATGTCTGGGAAGCC**CAGGTTTGGGGCTGCTGATGCTCTCTGCCTGGAGAACCTG**CCTGATTTTCTAAGGCACCTGCTTAACTGTTGACCAGCAAGGGTCCTGTTCCAAAGAGACCTAGCCAAAGAGAATGATCAGAGTAAAAGGCCAATTCCTTCCAATGCCTTCCTGGCCCCATTTTCCCTGAATCCTCCCTGTTAATTATAAAAACTCTTTCATGGCTCTTACCTCAGACTCACCGGGCTCCCAACTCTCGGGTTTCCTAGTTCTGCCCTAAACAAGCCTCACACTGTCCTGCCCTTGGAGCTTTACTCTCACTGCTCCCTCAGGG**GTGAGGCTTTTCCCCAGACCCTTGGGTTCCTTCCCTCAC**TCCCAAAGCCAACCCTGTGCTTCCCATCTTAATGATCCCTCTGTATGTGTTCATAGGACACCCTACTCCATGCTTTT**GTCAGAAGCACTTTTCGCCTTCTGAC**TTATGATATAATTTATTTACTTATGATTTTACACTTGATAGGCTGACTCCCTCTAGTGGAATTTGAGCTTCATGGGATCAGGGGCCATTGTCTATTATTTGCATTGCTGCATCCCAGTGTAATGGGAGTACCTGGCAAGCAGTAGGTGCTC**AGAGTGGGGCACTCTTCCCAGGAGTCTGCATGTCTAGAAGTCTGGGA**GGATTGTAAAGAGTGGATTAAACTGCTTGTTATATGGACAAtgttccc**ttttccttttttttattcattttacttacccagtctttaggaaataaggaaaa**ttacaattctcagaacttatatactttgtctcaagtggtattttag**gttgtatttcctgaaaacatacaataataatacaac**attatcatacttaattgatt**atattatactaatat**gtaatgacatgtaatatg**tactatgtaagttataacaatagta**gatttattgactttccatttaaaagacaaaacttaccagatt**atatcaggagtcagataattgatat**ttcaggtttgttggcaaagaaagaaaactgaatacattacttagatacatacataagaaagaaaacacattttcactattgttattaataat**tttaaatggaatgtgatttaaa**gagcttctcagcctgttcaggggaacctcactcaccactacatgttcacaactctgagga**aaacaagaaaaccccagataagattcttgttt**ccatcagtggtgtgaacagaggttttcaaaactgttaaaaggaaaagagtctgagtctgtgtgaaaggctgtgttgtgcagagtggagtgctctatagaagagaagataccacccccatttttcctggccaggtggatgcac**catcatctgtctgcagggagtggaatcacacaatgatg**agtctgagcctggagtctgcttatctttgcag**ctgtggtcactcatgcacccacag**tgcttggcctaacaaagtgatgtcctttactagttcttccaaagaatggaacgttttcagggctcatagaaaagatgtctaagctgcgaaggaaaagtgtgtatagaagatgaatttctgatcaatgacttcagccagg**tgcaatcatgcagcaaggaagatgaatgggcatgttgca**ggaa**gaaagagaaaaagaaaccataggtttttattttttctttc**aagagatttgtactaaacaccatctgtctgagcctaggttattggtgag

Uppercase: LOC124907873

Lowercase: Flanking sequence[1000bp]

Red & Bold & Underline: Stem-loop [29]

Blue: Heptamer[39]

Green: Nonamer [11]

id-TRBV6-1-2[V_gene_segment]

ttcaccatgtttgtcagactggtcttgaactcctgacctcatgatccgcccacctcagccttccaatgtgttgggatta**caggcgtgatccactgcgcctg**gcctttttttttttttttttaattgttactgaattagtggtgagaatcctgaccaagggaatattccctatgaaatatgacacttcaaataaaatgagagcgctttttcttatcctggagcctgcctcttctctcctctccccacatccctcacatgtcaagacacctttg**tgcccacagatcatgggcagcgttgggctgcc**actatcatccacacagagagggcagtcagcagcgtgaggtggttc**tgcctgctgcggtctcaccccaggca**cagaaagcaagagccctgggtgg**agctgaaggtgctcagct**gggcttgtcaggagtcccatctgtcggtgaattgaaaagaaacagagcaaaatgactcctccaatggtgatgagcctgcccctgggatttggaaacttggtaacagagaaaaccaatatagacaaaggattttaaacaggattatggtcaattaagcaaattagaaaaggatacttgaagggggatttgggacacaggagtcaaaaacacctggaagacatgagaagtttccctaagag**tctaaattagagaaatatttaga**taactgacaccagtgtatgaatgaggaattatatcatcacagggataagagttccattgagttacaaactgcttccaaaaaggttaagaaaaactcgtaaggctgtgtcaattcagacaaaggcattcttccc**attcaaacggttcaccggtgcatgaatcttgaat**ttgaccatctggggaaggggcgtggcctctcctgacaggaaggctctggggcccaggcagggagaatgaagtctcagaatgacccacttgagagtcctgttcccctatcaccgatgcacagacccagaagacccctccatcctgtagcacctgccATGAGCATCGGGCTCCTGTGCTGTGTGGCCTTTTCTCTCCTGTGGGCAAGTAAGTC**CTGGGCAGGGCCCCATGTGTGGATTTCAAGGCCCAG**CCTGTTTCCATTGTGG**CTGCAGCATCAGCTTTGTTCTTCTCTGCAG**GTCCAGTGAATGCTGGTGTCACTCAGACCCCAAAATTCCAG**GTCCTGAAGACAGGAC**AGAGCATGACACTGCAGTGTGCCCAGGATATGAACCATAACTCCATGTACTGGTATCGACAAGACCCAGGCATGGGACTGAGGCTGATTTATTACTCAGCTTCTGAGGGTACCACTGACAAAGGAGAAGTCCCCAATGGCTACAATGT**CTCCAGATTAAACAAACGGGAGTTCTCGCTCAGGCTGGAG**TCGGCTGCTCCCTCCCAGACATCTGTGTACTTCTGTGCCAGCAGTGAAGCcacagcgctgcatggccgtctcctctctgcacataaaggc**agggaggctctgccctcctccct**caccccagactcagtgatgc**cctgggcagagttctctgcaccagg**aaacttgaaacccca**tcatcatgggtctgaggcccccaggatga**aacaggatttgtatttcagatccatctagactctcgtctctccctggggaccatgttgcttcttctctctagggtttc**ccccagcccccaccctcatgttgtctctccctgggg**accatgttgcttcttctctctagggtttcccccagcccccaccctcatgttgtctctcctgtggcccacctttcccatctgggcagtcaccctccaaggccttgctgggtctctc**ctcccctcacttccccacacctctccacagcagccatgaggggag**cccctct**tctgtgcctccttccttcccatcacagagacttcaaagtccattttctctg**ccctgggctggaggcttcctttctgcagtggccaactcctacctg**tgcttcacatctcagcatgatcagccctcctgtgggaagca**tgcccttgcctcccagctgagatcaatttttctgttat**aagcactgatggtaacatgtgctt**gttagtcaacagagttgagcacagttggagttttccagttacttgtctggatatttttctgctcatgtcgattttaaactccataaaggggaaggctgtgcttgttacatttaccaacagctgttcctggaatgaagggagacctgtttcacagataatggatgagttaatgaatgacaggctgattgagtgatgagcgggtggatgaaccaataacaggaacactccaggtccgctgtaccctggcagaaatctagagttaactgtgcctgagatgctctgctatgagctcttcaaagggcact**cgcttgttgaacaagcg**ttggaccattcattgtgcaagccagtttgtcagttcatctaggagt

Uppercase: TRBV6-1-2

Lowercase: Flanking sequence[1000bp]

Red & Bold & Underline: Stem-loop [21]

Blue: Heptamer[42]

Green: Nonamer [2]

id-IGHV1-12-2[V_gene_segment]

tatcttaatagtgagaatatgaaggataggcatgttttt**actaattctatgggtacagattagt**tgaagaaactacattcctatgaataagaaattcagatttcagtgttaagtaatgttgcttacattgtgtgagtgacagggcagtggtggatctgagagtgtggcaggtgcacagaccaagtgagtcagaaatcaatatggaaaggtgagggtctgtggatatgaactga**aagtatgtaaatactt**gacaaaatactaataaatggagttcaaaataacccaa**aattgttctaaacacaaattccttgacaatt**actttgggagtagagagttcataatggactccaaactcctgctttatcttctgattcccattgt**ctgtgagatgagaaaatcagctctaattatgcatcacag**ggcaaatctgtaaatcaagagtg**ttcaatagaattgaa**gaccctgggggatcaggacatgaggcaggtgctggagac**agtgtctcaggagcacccagtagatctcagaggtccctcctggacact**catgtgggacataagcatcactttctcagagtcaccaatgagctgtgctggtggctgatgggtccaggaaaagaccaaggcacctgctcagtgtgatggagagtgatggttccaaaaatgatccaggtggtctctatgctaatcaaatgt**aggctcacagtgaggagcct**gttctatatgggcttattcttcagtgaaaggacgtctgtccacaaatgtttgtaaatggagcagggcatgcatttcctcaagcaggattagggcttcgaccgtctgcatctcactcttgttaggctgatgtgtcatttatcttc**cctttcttatcatggattgggctttgagctaagaaagg**ctttgtcttatgaatatgcaaatatactgatatccactgaggtaaatatgttctgtgccctgagagaatcacctgagagaatcccctgagagcacatctcctcATGGGCTGGACCTGCAAGATCCTCTTCTTGGTGGCAGCAGCCACAGGTAAGCAGTTCCCAGGTCCAAGTAATG**AGGAGGGGATTGAGTCCAGTCAAGGGGGCTTTCATCCACTCCTGTGTCCTCCCCACAGG**TGCCCACTCCCAGGTGCAGCTGGTGCAATCTGGGGCTGAGGTGA**AGAAGCCTGGGGCCTCAGTGAAGGTCTCCTGCAAGGCTTCT**GGATACACCTTCACCTACTGCTACTTGCACTGGGTATGACAGGCCCCTGGACAAGGGCTTGAATGGACAGGATtttagttatttgagagatttttcatacaacatttattctgtaagcaaatttcagggattgtagaatgaatcatattaacaaatctgacacagaacttcctctgaatcaatctttgtaaacatcaat**ttctgaatcaatgttgtaaatatttcagaa**cacaagcacaaattcacattttaactctacttttatctctatttaaaaaata**tcaaaaaatctcattttgtgcatgtaacgttttga**atgcccaccatcaatgcatgacatttcttgtttttccacattcatgttaccatt**tatcatcatgagaattgtgagttttagccatgctgata**ggtgagtaatggcatctaa**tttttatttaaatgcacatgtcccaaataaaaaattcgtattaaacaattttt**atataatttttgctgggatgcctttcctgatatgtggttcatttttatctgcattgttttcttttcattagttgtaagtttacttgcatattgattataaaagtcacttaacaaatccaaagaattgatttaacaaatagatgacttggaagtattttctcccagtctgtggttgtctttttctctcttatcagtgggtatttcaaaaaatatatgtgtgtctgtgtg**tctgtgtgtgtgtgtgtgcacaga**tttagacaaaaaacataaaaaattattcattcatagatca**tgtatttggcattatatctgaagtctcattataaaataca**ctaatagtgattatttattccatgttgctaatctcaggcca**caatcaactcatgagtgcttagccttcacctattgattg**gaggactatccacctgagatatttggaatacttccatgaggagatgtgttcttcttcccatcatttctttatttaatcatctattaacatccgtattggtttatggacatctatttcatactctgaagaagatccatgctacattattcattttcttgttcaaatctccacagctttatta

Uppercase: IGHV1-12-2

Lowercase: Flanking sequence[1000bp]

Red & Bold & Underline: Stem-loop [19]

Blue: Heptamer[27]

Green: Nonamer [1]

id-IGHV3-33-2[V_gene_segment]

acattctcaaataactctgctgagtgaaagtagctgaagaattaagagtgcacttcataaacttctaattgtataaactgcagaaggtacaactattctaaagtaacagagcagatttgaaatttgggagaaatgggtgttgaaa**gtaattgggtggtgagatgaaattac**agagaagtgacagaaagatttaggggttaacttaattgtacacaacctgattaaagtttgcacacatacgtt**accattttccaaatggt**gcagtgtagatttg**aattaattattaatt**gtacttaaagcagtaacaaagaaacaaatgaatatgttggttgaggaggagcaaaaaatagatgggtatgaacactcaaaacatctcagactcttg**aaagtacacatgcttgaacactggttttctctgtatacttt**aggtaaagagcagaatacattaaaagaaaacagagatgtcctggcgggggtggaatcctgcaaacctcactaggcgtgtcccacactgccctggagt**tgtctcaggggagcagtctcctctagtggtcagagacacaggcatggataatggggctaatactgtccagctgtgt**gaccttgaacacattgtataaacactgttccatatgtaatttatcttccttaaaatgtaaca**ttgacacttgcattaaatgtattctacaaatatgtcaa**aaagaagatgacgactgctaa**atgattatcaaggcacaatcat**ataatataatgatattttcctgagtgataagatgactaccaatctcccc**cagggcactttgtctgctctgagccctg**cgcctcctcaggattcccatcccagagcttgctatacagtaggagacatgcaaataggtttctccctctgctgatgaaaaccagcccagtcctgaccccacagctctgggagagaagcgccag**ccctgggattcccaggg**gtttctatttggtgatcaggactaaagacagaggacccaccATGGAGCTTGGGCTGAGCTGGGTTTTCACTGTTGCTGTTTTAAAAGGTGAACTAGAGAGATTGAGCGTGAGTGGATACCCTTGAGAGAAATGGTGGATTATGTCTGGGAGTTTCTGACCAGGATGTCTACGAGTTTGCAGGTGTCTAGTGAGAGGTACAGCTCGTGGAGTCCGGAGAGGACCCAAGACAACCTGGGGGATCCTTGAGACTCTCCTGTGCAGACTCTGGATTAACC**TTCAGTAGCTACTGAA**TGAGCTCGGTTTCCCAGG**CTCCAGGGAAGGGGCTGGAG**TGAGTAGTAGATATACAGTGTGATGGAAGTCAGATATGTTATGCCCAATCTGTGAAGAGCAAATTCACCATCTCCAAAGAAAATGCCAAGAACTCACTGTATTTGCAAATGAACAGTCTGAGAGCAGAGGGCACAGCTGTGTGTTACTGTATGTGAGGCAccaggtaagaggacatcagtgtgaacacagacacagtttcctgcaatgacaagggaggaggctgggctaaaaggggcactcaggacccactgaaaacgggcagctctagggcaggtacagatggttatcatgggctgctttccttcagggtctgtggcctcctctgcatctaccagttcccctcggagcctctggacatttatgtttctgtggccacccctgacatctctggattagaaaagtttattataggaagaggaaacattctcatttgtcccaaagcagatgtaagtaatggaagcaaaaaatgcacaggaggccaggcggggctgtagatactgtcaccccagaatgtcaatctcaccactagtactggag**agggaagggagtttgatggaccttccct**aacaaccctgtggtccaagataagtccaggaagaccattggtgcctcccagagcacagttgtccatcagggatgtccaatgcgtcccagcagcagccatgcctcagtgtctccactgtgcacagccactgt**ctgggaggagctcccag**gatgggtgtctttggcacacaccaggtggcgggtgttagagtgcagtgcagcagctggctgcctgttctattgggctccctga**tgctggagagatgggaggtgcattcgcagggccagca**ccctgtttgtgaatttttatataaaaaccctgattttacttcatt**tttgcagatgacatagataactaaaaacagaatctgcaaa**gaaattgtaattttcaactttaccccaaattcattgtttcttaattctgtgcaagatccagacatattattgccttcctcatgataatttattctatgtaaaaccgaaatcattttttctcctactctttgtttctgttcaagtacagagatcttgagcaaagtaagttgggttctttccacacactaaccctcaccttccccagagaatgagcagagattgtcctcagtctgagtctaag

Uppercase: IGHV3-33-2

Lowercase: Flanking sequence[1000bp]

Red & Bold & Underline: Stem-loop [16]

Blue: Heptamer[38]

Green: Nonamer [4]

id-IGHVII-40-1-2[V_gene_segment]

gctactaaatagtaggtgttattcattctttctaactctgtattttgtacccattaaccatatccacctgttctctggcccccactagcctcccagactccagttaccatccttctaccatccatgttcctgagtttaattctgttgatttttagagcacacacataagtaagaacctgcaatgctttttttttcagtgccttgtctttttcacttaacatagtgacctccagttcca**tccatgttgttgcaaatgactggatctcatgga**atacctaattttattcggttctttcatatgacgtagagaagcaagcatgcagtctacaaacaactgtaattttcacatttaccccaaatgaattattacttatttctgtgcagggtccaggcacatagtagcctttctcttgagggactgttaaatttagagtgagatcacttgtttctcatattatttgattcctgtcca**tgtgcagagatcttaagtaatctgcttctttctgcaca**ctaactctgaccttctcgacaaaaagagtagagattttcatgtttcctgagtctggg**gtaggagttagtcctac**acaactcagagcctgcagagacctcccagtgcagttttatagacacaggtgctttctcatggggggtgagacctccacttcctgtgattgctgctcagatctaactgtgatttcagatttaggacaccattaggctctcaaacca**tttctattctaaaaatcagcctcatcatagaaaatggaaataattcaatatccattt**ccctgaccgtatttcatccttgtttttaatttggtgcaagtttattacatatttgcctgaatctaacttcagaatcaactgaattggtctagaaatctcaaaaattgaacagaagtggattttattatttttaccaaaatgtgtgtaggtagggtttccaatgtatcgtccagaacttgtgcatgctaacaactgtttttctcagtcaattatcAGCCTGGTGAAGCCCTTGCAAACCCCCTCACTCACCTGTGCTGCCTCTGGATTCTCTGTCACAAT**CAGTGCTTCCTGtaggcactg**gatggaaagcacaggagtgggtcaggtgcatacg**tgatgagtggaggatgaattccagcccacttatca**tgaattcagacaagcccacatgttcccacatgcactatatctt**cagacaagcagatgtgtctg**cacatgaattcttctttcagacaaacacgtctgtcccctcatggactcttgccaca**cacaagcacacgtggcttcttgtg**gactgtactttcagacaagcacacattgtttcatgtggacatcttcctcccacaaatgcattttcccctgagtagactcttctcccacaagcccat**atgtccgcacatggacat**ttctcacagacaaacacccatttccccatgtagtcccttccctcagacaagcacacgtattcccaggtagactcttcccttagacaagcacagatgttcc**catatgaacccttccatatg**acaagca**cacatgttcccatgtg**aactcccgtataacaagcacacatatttcctcacgggctcatcactcagataggtctgcatgtccacatgtggactctactctcagacaagcccacaggtcctcacatggactcttctctcaaacacccgttctccattgtagactcttcccttagacaagcacgcatt**ttcccatgtgaactcttctgcatgggaa**gcacacatattcccacagggactcttctctgaaacacccattctccattgtagactcttcccttagacaagcacgcattttcccatgtgaactcttccgcatggaaagcacacatattcccacagggattcttcacta**agacaagcatgattgtct**ccactgtgacccttccctcagagaagtaca**cttgtctacatatggactcttctctcagacaag**caaacatgtccc**cacgtggactctttcttctgacaagcacacgtg**tccccgtgttgttataatcattttctcacctctgtctcacaaagttccctgtcctccaaga

Uppercase: IGHVII-40-1-2

Lowercase: Flanking sequence[1000bp]

Red & Bold & Underline: Stem-loop [16]

Blue: Heptamer[23]

Green: Nonamer [2]

id-IGHV3-42[V_gene_segment]

tataagatggtatctcactgtgtctttaaattgcatttttctgatgattaatgtcattgattattttttcatatgctggttgatg**atttttttgtcttttgaaaaat**aaacattgtagtaatttgctcatttttaatgtggttatttgtgggtttttgtgttgcttttgtttcttgtagtttgtaaatattagctctttgtcaga**tacagagtttaaaaatagtttatcccattctgta**cgctttcggttaagttttttgattatttgttttgctgttcagatgctttgcttcttatttgtattaagtccaaatggtctatttcggtttttattgtttacttttaaggttgtagtcatgaattctttgc**ctaggacaatctccagaacaatatttcctag**aatagcatctgcaactttcagactctcaggtctcccagttaagtcttt**tatccatattggcttaattttggata**tagtgagagatacgggtccagttttattctgctgcaaatggctgttcagtttttcctgcacaatttatataacaaggtgacctgttcccagtgtatgcttcttgtctagtttttcacagtcagtttggctgcaggcatttgactttatttatgaaatgtctattctgttccactcatctatggctttctgctagattgttacttgattgttacttgtgtataataatgctactggtttttgtatgcttttttcatttattctaaaactttactgaattaactcatcaattctaggagt**attttgaaagaatattagtttttttaagtacaaaat**catattatcagcaaacac**aaagagtttgacttcctcttt**tccaatttgagtgcctttatttctttctcttgcctaatttctctggctaggatttcaagttccatattgattaagagtggtgaaagtgggcatccttgacgtgttctgagtcttagg**aggaatactttcaacatattcct**attcagtataatgTTGTTTCGGAGTTTGAAGGTGATTTATGGAGAATGAGAGATGTTGAGTGCGAGTGGACATGAGTGAG**AGAAACAGTAGATATGTGTGGCCGTTTCT**GACCAGGGTGTCTCTGTGTTTGCAGGCGTCCAGCGTGAGGCGCAGCTGGTG**GAGTCTGGGGGAGGCTTGGTACAGCCTGGGTGGGTCCCCGAGACTC**TCATTTGCAGCTTCTAGATTCACCTTCAGTGACTTCTGAATGCACTGGATCCGCCAGGCTTCTGGGAAAGGGCTGGAGTGGGTTGGCCGTATTAGAACCAAACGTAACAGTTACACGACAGAATGCGCTGCATCTGTGAAAGGCAGGTTCACCATCTCAAGAGATGATTCAAAGAACACACTGTATCTGCAAGTGAATACCCTGAAAACCG**AGTACACGGCCATCTATTACTGTACT**AGAGAcagtgagggggaggttaacgtaggcccatacacaaatct**ccctgcaggggcgcgcaggg**ccaactgggggcgctc**gggacccactgaggatgggacaggtccc**aggggcgggtgcagggggaggtttcctttctcagctgcaggaggcgggtttgtttttgcaggaatatggagtcttatgaggttttgatattttactatggttatttatcatgattttttaaaattgggatttgtgttttag**taatttttaaatttatatgtaggggtatttttaaaaatta**agttttagggt**acatgtgcacaacgtgcaggtttgttacatatgtatacatgtgccatgt**tggtgtgctgcacccattaactcgtcacttaa**cattaggtatatctcctaatg**ctatccctcccccctccccccacctcacaacaggccccggtgtgtgttccccttcctgtgtccatgtgttctcattgttcaattcccacctatgagtgagagcatgcggtgtttagttttttgtgattgcaatagtttgctgagaatgatggtttccagcttcatc**catgtccctacaaaggacatg**aacttatcattttttatggctgcatagtattccatggtgtatatgtgccacattttcttagtccagtctatcattgttggacatttgggttggttccaagtttttgc**tattgtgaatagtgccacaata**aacatatgtgtgcatgtgtctttatagcagcatgatttataatcctttgggtatat**acccagtaatgggattgctgggt**caaatggttatttctagttcaagatccc**tgtggaatcgccacactgacttccaca**atggttgaactagtttatagtcccaccaacagtgtaaaagtgttccctatttctccacatcctctccagcacctgttgtttcctgaccttttaatgattgccattctaactggtgtgagatggtatctcattgtggttttgatttgcatttct

Uppercase: IGHV3-42

Lowercase: Flanking sequence[1000bp]

Red & Bold & Underline: Stem-loop [19]

Blue: Heptamer[33]

Green: Nonamer [9]

id-TRBV23-1[V_gene_segment]

tgtttttttaagtgatcattctgttggtattttatcaagttattatctatatattcaaagtccattttgtaatgtggagaaaacaacctcactgaacaattaaattagtaaggagaaatctatggaaagatatacattaagctgttaacattacttacctgaaatgaggatgagatggaggaagagaaaagtaagcgaaaaaaaaaatcatcaaaaaatgcacaatttaaaataaatagttacagagagaaaaagcaaaaaagggaggttatgtaaaagaagcactagtaaatgaacaagct**aaaatacaccagaaagagctagaaggtgaagtatttt**tcaaagcagtcactag**tttcagaatcataaagatagcaattacaacagagatctgaaa**tatataaaagataattagataactaatagccagaaggaatgtccagtcttgagttagaaactgatgctcaaattaatg**gcaaaggacgtttttgaaacaactgtgaaactttgc**ataaaaactaaatagtagataagatgaaaatgcatcgatgagaaaggatgatcattaactgaaaaaaagcttatcaaacagaatttcaagaatattgtttactaaaaaaagtgtctatatatctacacatttatatatggggcagagaaggatct**agaaagactcactaaggtgttaacagtgataattgctttct**ggcagtggta**tcatatttttattttgttattttattttctaactttctaaaatatga**gtgtattgcttatattagc**aaaaatagacataacatattttt**aaatgaatacaaaataactgagttatatttcctgagttaggataaaagctttttttccctgatactatgaag**tgtttttgctcaggaacatgaagcaataaca**tgatgattataagtcagtggagggatcacgtcacaagaaggtgcagaaag**gggatgaaaaagcctcatccc**tttgcaacgtcaatgcgatcATGGGCACCAGGCTCCTCGGCTGTGCAGCCCTGTGTCTCCTGGCAGCAGATGCGCCCTGAGCACAAAAGAAAAATCCTTGTCCTGGGCTTGCCAACCAGCAGATCCAAGCTTTGTTTTGTTTTGTTTTGTTTTGTTTTGTTTTGTTTTGTTTTGTTTGTTAAGCTCTTTCCTGGACTCTTTCTACAGCATCTGTTTCTTTCTCTTACAGACTCTTTTCATGCCAAAGTCACACAGACTCCAGGACATTTGGTCAAAGGAAAAGGACAGAAAACAAAGATGGATTGTACCCCCGAAAAAGGACATACTTTTGTTTATTGGTATCAACAGAATCAGAATAAAGAGTTTATGCTTTTGATTTCCTTTCAGAATGAACAAGTTCTTCAAGAAACG**GAGATGCACAAGAAGCGATTCTCATCTC**AATGCCCCAAGAACGCACCCTGCAGCCTGGCAATCCTGTCCTCAGAACCGGGAGACACGGCACTGTATCTCTGCGCCAGCAGTCAATCcacagcactgaaatgtcagttcctcttagcacacaaacttgtcacagacccagctcaggaagcaggtgatgtattaggctggaagggagtaacagaaaataactggagccagcttaagccacagtgtaatttaacacaaggataaagggtgtttagcagaagtcaagaggtgatagaagttctgaaaaggcaccactagggctgggctctagaatggctatctctccagcgctgatttttattcaattttggctgctgaatgcttttctctccctgtac**accatctctttttctccaggcctcgtggtgagaatgagatggt**cactaaagattacaaattcagttgccatcatttccctacactgataaataccaggctggctctttctaagcctcaattcaaaatcccaagagaaaaaaacagatcagtcatgatccagccattaaccttctcacagagcactcagcagtaggcatgtat**gtatatatttaaaatcacataaatattac**ttttccactcagctcaagaaagggaagtgctccgtttttctgtaagtacatttggtcttgttgcttctacatgatagagagtactcatacccatttctt**cagaaatacatagccagattttctccagttttctg**cc**agaattgaaaattct**tgtacaaataccttcatagacttgtatgattgctttataggattatatacccagagcaagtcatgattgctgagacatggaattctatgcatattttatctatctaaataccaccacgggttcttcaggatcactgtatacatctgcaatcccacataaaagtgaatagggtgttttcctacatcctaatatctcctttaatccttatccagtgatctagattgttctcagtctagcaatgt**aaagagattccaatctctgtctctcttt**ctctccaccccccacccctgcaaccctctcacctactgccagaatatcatcagtgtttaccataaagat

Uppercase: TRBV23-1

Lowercase: Flanking sequence[1000bp]

Red & Bold & Underline: Stem-loop [14]

Blue: Heptamer[24]

Green: Nonamer [4]

id-IGKV2-19[V_gene_segment]

atcctcttgtaagaagtccccactccacccaggaagtctagctggcctcacctctcactatcaatgactgaaatgatcattagtagcagtggctttggttaaatgtataccagtagttacatacagggtagaaaaatggtctgaagaggtatattcagcatcaaatgaaaaaattacaccttgtgacccctttctctccctgtatgatcccttaagattgtgctgaaacctcagagcatcaggtcagcctcatattgctccttcctaaatcttggtccctgctcaacctgagcctatgtttggccatccacgggcatccatgtatacaactttgaaggcaaaactccctgcttggagtggctcattagatgaccagggtccagatagcattgccaaggaggctcacagatggcttcatactgagtctctcccctcagcctcccagcatcaccagccctgaaggaatgtccctgttatccgtggcacccagagcac**agtcagccctgccctgact**tcactgcccaattcacagcctcatcgcagattccactgcaaaatagacagtgtgagatctcaaagacaggctgatttgtagtcctgaaaagcaacaacccatcgttccctcgatctctttggtggaatttcaaagttcgtcataagctgaatttctgtacaaacatactctcttaacaatggaaactatttgcaatatgggaaacataaaacatgggaaaagtttaaatctctcaagagcatcttctagtcagattcatattcagcttccctgattctgaactttctgtcttgcggtcagcttcttttggtccatttgtttcaccctttccagctcagatcttctctccctggctacccttagtctcaggttgtccttacgtttgataggcttct**accttatggatgctgtaaggt**tatcctctcatttgctggattcttgtgggaaaaaaaatccctccaagttcacatcctgagacttcagactCCACTCTTCCTTGGAGAGCCATCC**TCCATTTCCTGCAGGTCTGGTCAGAGCCCCTTTCATAGAAATGGA**AACACTTACTTGAATTGGTTTCTGCAGAAGCCAGACCAACCTCTAAGGTGCCTGATTTATGAAGTTTCTAACAGGAAGTCTGGGGTCTCGGGCAGGTTCAGCAGCAGTGGTTCAGGGACAGATTTCATATTGAAAATCAGCAGGGTAGAGGCTGAGGACGTTGGGGTTTATTACTGCCTGCAAGGTACACAAGTGCCTcccacagtggtacaaccctgaatacaaacctccctgcttggagtggcccagctgctcaaata**tgttgtttatctggggagtggcacaaca**gaatctctgctgtataagatgaagatgttgggccgaggcgggcagatcatgaggtcaaaagtttgagaccagcccgaccaacatggtgaaaccccatctctactaaaaatacaaaaattagctggatgtggtggtgcacgcctgtaatcctagttactcaggaggctgaggcaggagaatagcttgaacccgggagatggagcc**tgcagtgagccaagatcacaccactgca**ctccagcctggatgacagaaggagactccatctcaaaaaaaaaaaaaaaaaaaaaaaaaatgaagatgttggtgagctcagggtaaaaggttgcagctgaatgacctgtcccatgggggactcagcagtacatcaggtaaaacccattcatggtcctgtcagctcccacagccttggcatggcataagccaaaggaaaacagagataattcaagtgccttcagagtaagcagacaggactgagggagagcgg**agggaaatctcacactaatcttccct**gccttgcctacattcgacaatgagacttcaaatagcttaatagccagacaagtaacacagattcgtgtcaacacgtgttgaatatctcttgaagtttaggtcttttgtgtatatttttaagagggtagtatttggtagtatttagaaactggtatttttcaacttttccaatttccttcttctcctttttactagttctcttcccattgcaccacataacaaatagtgaaaagagcattctacacaagc**tgtcctcagggagagctggctgaggaca**atcatgaaaaagcttgaatttgcacctccaaatagacttttgtgacgtcatggaagacagaagatcctgatgttaaaactctttcattcatttcaattacttcttgctaataaaaaaggataa

Uppercase: IGKV2-19

Lowercase: Flanking sequence[1000bp]

Red & Bold & Underline: Stem-loop [7]

Blue: Heptamer[23]

Green: Nonamer [6]

id-TRAV27[V_gene_segment]

ttacagttactcagatactacatggctcagcacgcttgttgaggatatcagaaatgataagatggtactttcattattttctattattcttttattctgg**cttttaaatgatcaatgtttaaaag**gatatattttccattttttttaataaggtcacaatactctttctgtaaccggaccccagtccagct**gctctccacttgaaagccaaacagagagc**cgagggtagatgggaaaagcaggtttaatatgagagccagcaaactgagaagatggcaaactagcattctaaagtaccatttta**catttttaaattcaccatgagattttaaaaatg**aaacataggatgggagacatgcaggaattctgcagggtgcagggtttgtgtgtcttgttctcgaggctgtattgggtaattgcgcctctgggggtctggaagttgttatcttgacttcggcctggtggtggtggacaaattgttcaca**agtccccctaagcttgatgatgcctcaatggggact**gcgtggtttgtttctaggtttgactctgggatttcttaagcaagagaataattagacaagcatgcattactggcaaggagtgtccagaggggaagaaatgaagggatgagaggggagggaagggaaggaagaaaaagaaagtgggtgaggtccccagttacatttctactttaatcttgatatttatgcctaccaaatgcaattgggagaaacagagcattattggaaataaatcacttttctatctctccacaggagggtagtatttcaaaaggcaacatacattttgggttggaggaaggggcgggtggagagtgaatgattccattggctgccacactgaggctcctggctttctatggaactcaaagtgggtggcagcaagcactcttctagcccagagaagtctgttccaggacgggctctttcaggagcagctaaagtcaggggccatgtccaccatgtgatagaaagacaagATGGTCCTGAAATTCTCCGTGTCCATTCTTTGGATTCAGTTGGCATGTGAGTTGAAAAGGTTCCCAATTAAATTAACAGAAAATCCCTGATGGCAAGTTGTTTTATGGCAAAGTATGAAAGGAGGAATTATAAAAGAAGGAAGAAAGGGGGAAAGCAAGGGAGGGAGGAGAAAGGAGTAAGGAGGGAGGAAAAATCATTTCAGGTTGGGAAACAATCCCTAATTCCCTTCCCTGGTTCCTTCTGCTCTGTTTCTAAACAGGGGTGAGCACCCAGCTGCTGGAGCAGAGCCCTCAGTTTCTAAGCATCCAAGAGGGAGAAAATCTCACTGTGTACTGCAACTCCTCAAGTGTTTTTTCCAGCTTACAATGGTACA**GACAGGAGCCTGGGGAAGGTCCTGTC**CTCCTGGTGACAGTAGTTACGGGTGGAGAAGTGAAGAAGCTGAAGAGACTAA**CCTTTCAGTTTGGTGATGCAAGAAAGG**ACAGTTCTCTCCACATCACTGCAGCCCAGACTGGTGATACAGGCCTCTACCTCT**GTGCAGGAGcacagtgctcttgaggcacctgctgcctgcac**ccaaaccctgctgccagccccagtca**cgaggctgccacatgcctccagctccgcctcg**cacagcttatggcatgaatagagagaacaacctcagcccctcatggaacca**tttctgctgcctttccagaaa**acattttccacacacactggaggacaagacccctctgctataacacaaatatctggaaatgcagcctgctattaggatcattaacgatatgggcctgcaactccccggggtaggcagtagcaaagagagtcgatatagagggacaaatggaagctatcagacagac**tgtttctgagtgttccacccctgtgtagaaaca**aagcagaaggcaccagggaagggagctagggattctttcccaacctgaaatgagtttttctccctccttccacctttctcaccccctcattctctctttcttccttctttcattaattcttcctttcatagttggcatcagggattttccatctctttaggattttctgttaatttaaatgggaacgccttcagctctcatgctaattgaatccaaaggatcaacatggagaatttcagaatacttcatggttcagattggt**cattccgttttccagaagaccacacccaggaatg**ttcagtacaatggggcagaaatggatggtttcc**cagctttcttgtagtactccagagatgatgaagctg**ctatctttccagactcttacactgcagtaaaatgcagaagtattgtaaaaattagaagaaagttactacctttttggaagaaagataccgatagtacacctcagtagcat**ccaactgtggtttcaaagcaacatcctagttgg**tggttctgaaagtgaccaaacattaactacaatggcctgtatgttcacaggactggcttggggaagagtctttgaatggcaaatgtcaataacttcaagttttcaagcctgaagaagaaagaagttggcag

Uppercase: TRAV27

Lowercase: Flanking sequence[1000bp]

Red & Bold & Underline: Stem-loop [13]

Blue: Heptamer[28]

Green: Nonamer [6]

id-IGHV3-53-2[V_gene_segment]

tctcaaagaattgtgctgagtgaaagaagctaag**gaattaagagtaaattttatataattc**attgtagaaatattagaagatgccactaccataaattaaaatgaagaagacttaaatttttctgagaaaatggtgttgggaatgatgcggatgtgatttaagtttcagaggaataagaaaaagatttagggattaatttaattattcaaaagttgattgaagtgccgagtgaatggctgcaaacatagctctacattt**ttcaaatcattccctataaatttgaa**ttaatt**atttatttttatacttgaataaagcaataacaaagaaataaat**gaatatttttgctaaaatggagcaataaaaagactgatattga**cagaagaaatatgactgacttctg**aaaatacacacacatgagccgtggttctctctacatatttagataaattacagaaagttgtcataactgatggggaatcctgcagacttcactaggcatagtccacactgcc**ctggagttgtctcaggggagctgcctcctccag**tggttagagcacaggcccaggtaataggactcatttttttagatgtgtaattttagacacac**tgcacaactgctgtgttctctgtgca**aattatctcctgtaaaatgtaacattgaaacctgccttaaatatattgtgtaaatatgtaaaaataaaatcagattgtgagagctaaatgctaatcaaggcgcaatcacgt**aatatacaattatatt**ttcctgaatgatggaattaataccaatctcccccaggacactt**catctgcacggagcccggcctctcctcagatgtcccaccccagagcttgctatatagtcggggacat**gcaaatagggccctccctctgctgatgaaaaccagcccagctgaccctgcagctctgggagaggagcccagcactgggattccgaggtgtttccattcggtgatcagcactgaacacagaggactcaccATGGAGTTTTGGCTGAGCTGGGTTTTCCTTGTTGCTATTTCAAAAGGTGATTCATGGAGAACTAGAGATATCGAGTGTGAGTGAACACGAGTGAG**AGAAACAGTGGATATGTGTGGCAGTTTCT**AACCAATGTCTCTGTGTTTGCAGGTG**TCCAGTGTGAGGTGCAGCTGGTGGAGACTGGA**GGAGGCTTGATCCAGCCTGGGGGGTCCCTGAGACTCTCCTGTGCAGCCTCTGGGTTCACCGTCAGTAGCAACTACATGAGCTGGGTCCGCCAGG**CTCCAGGGAAGGGGCTGGAG**TGGGTCTCAGTTATTTATAGCGGTGGTAGCACATACTACGCAGACTCCGTGAAGGGCCGATTCACCATCTCCAGAGACAATTCCAAGAACACGCTGTATCTTCAAATGAACAGCCTGAGAGCCGAGGACACGGCCGTGTATT**ACTGTGCGAGAGAcacagt**gag**gggaggccattgtgcgcccagacacaaacctccctgcaggaacgctggggaaatcagcggcaggg**ggcgctcaggagccactgatcagag**tcagccccggaggcaggtgcagatggaggctga**tttcctgtcaggatgtgggactctgtcttcttctgac**ggttccccagggaacc**tctctaagtttagcattctgtgcctatgaacgtcttctctaagtatttgaaagagattattttaatatgaagagcagttctcactcgcacaaaatgtggattgatgcttactgggatgaaaagtcctcaaacatggtcaccacgatcagagtctgagtgagctcagggcttcctgctgagtctcctcctatcagaccaaggacagggacctcagtgaggttccccgtctagaacagtctttatggatactgattgtgggcggcaagccacccaggtgccgacgcaagagaccgaggacacgagctgttccagtacaataaaatataaaacaagaatagttataccagatatagatcttagat**atgattatatatgaatatcattaatcat**tagttggtagcaattactctttattccaatattataataatcctcactctacaatcataacctaggaaaagccaggccatacagagataggagctgaggggacatagtgagaagtgaccagaagacaagagtgc**gagccttctgttatgcctggacagggcgaccagagggctc**cttggtctagcagtaatgccagcatctgggaagacgcctgttgccaagcggaccatggtctagtggtagactcagtgtcaaggaaaaacacctgctacttagcagacca**ggaaagggagtctccctttcc**ccggggagtttagagaagactctgct**cctccacctcctgtggagg**gcctgatatcagtcagacccgcccgcacttatccggaggcctaacagtctccctgtgatgctgtgcttcagtggcc

Uppercase: IGHV3-53-2

Lowercase: Flanking sequence[1000bp]

Red & Bold & Underline: Stem-loop [21]

Blue: Heptamer[40]

Green: Nonamer [6]

id-IGHVII-28-1-2[V_gene_segment]

gtccaagctaagtccagcaaggccattggtgcctcgctgagcacagttgtccatcagggatctcccatgtgtcccagcagcagccatgcctcaatctctccactgtgcacagccattgt**ctgggaggagctcccag**gatgggtgtctttggcacacacagttgatgggtgttagagtgcagtgcagcagctggctgcctggtctattgggctcctggatattggagggatttgaggtgcattctcagggccagcacgctttgtcaatttttatatgaaaaccatgtgatttaattcattttctcagatgacatagataattaataacacaatctgc**aaacaattataattttcaactttaccccaagttcattgttt**ctgaattctgtgcaggatccagacatggtactgcccttctcat**gagaaactgttcgatctaaactgaaaccagttgtttctc**gtatactttggttctccccacatgcagagatcttgattagagcaagtttggtactttccacgcactcaccctca**cctccccagataaagagcagaagttctccttagactgagtctgagggagg**agctgttcctgtaccactcagggcctgcggagacccccaggtgcagcttcactgagtcaggtgtttcactccctgtgattgctgctcaggt**ctaattgtggctcggaattag**gacagtcttcaggttatcacaggtcaatcatattctaaaaatcatcgttatcacacaccatggtaacaattcaaggttcatttttctaacggcagtttctcttttttatttggttgcaagtttgaggaaaggaacatgtgataatactttttaacctaacctcgaaatctactgaattgttctaggagactcacaaattggacaaagtgagctctttattctcataaaaatgtgtggttttgggaatttcactgtgttgcccagaacctgttaacatcaacaactatgtttctcagcacacttctGGCTTGAGACGTCCTTGCAGACCCTCTCCCTCACCTGCACTGTCTCTGGATTCCCCATCATAACCAGTGTGTCCTGCTAGAATTGTATCTGCTTGCCCCTAGAAGATGGACAGGAGTGGATCAGGT**GCATGGGTTGTGAAGGGAGCACAAATTACAACCCACTGC**TCAAGAGTCCATATCCAGATCCAAGAAACAGTTCTTACAGCTGAGCTCTGTGCCCAGTGAACACACAACTACGCATTTTTAAGCAAAAGAcg**caatgaagggccttcattg**tgagcctagacacaaccct**ccctgcaggggtgaataggagcagcaggg**ggcattcggggcagtatgggggcttaggatgattgttaggggtcaggatgagcaggatcaaggcttcgcatcagggcaagtgcaacagggcagaaaaggggctgtagatgtgggttgttctcaccatcatatttcac**caccagacaccctccactacatctcttctaatgtgtctgagtg**ttgatatgattagaaaatggcatttatgtaaatactaatatgtacccat**atggaggtgcatccagttgtcttctccat**cttatgtggaccttgtccatcaagcactaagtccctgtatttacttgagtacctcacacattatggtcaaaacaggtagggtctccttttggcatggtctctcctcctgccttctctcttctgtcacacaaaaacacgtgaactgacacacacacagagcttcccaactttaattatgtgatgtattgaagcaaattgattagtgtgcagcttttctgctttgcctgctattcatgttatgtaaaaataagaaccatgtttttctcagcttgtcacttctctaagctaagtagcatctttgtttattatacccagagaccaaaagcgatccaactgttattcagcagcttaactggtaaacaaattgtggaaaattcacttactggaatcatacccactgctacaatcaagtaatgctggatacacccaacaccatgcttaaaatagcaagtacttgaataagtaaaataagtcagacaaataaaagtgcatacatacgattccacttctataatttctataaagtaaaaatgaacttaaagttacataaaaagatctgtagttgactggggacatggtacaagaagagaaggtatagaaaggagaaactacag**gagaacaaaaggaaatttctaggataattgatttttttctc**tgttagtaaaagtgagga

Uppercase: IGHVII-28-1-2

Lowercase: Flanking sequence[1000bp]

Red & Bold & Underline: Stem-loop [11]

Blue: Heptamer[32]

Green: Nonamer [3]

id-TRGV8[V_gene_segment]

ttttaccgaa**gtgaaaatctgtgtatgcactactgaattttcac**aattattttactatacttgttttatcacctgtatctccatctatgagtccaggtccactgcagacatcagtcagctttctgaggatacggttgctaagtaaaatatgcaatactcaggaaatttggacg**ttcagataaataacaagtttttagctgaa**gtatgtcccaaatattgcatgacatggcaccctgtattactttaaaaaaaaaactcacacttctacttgaatgaacggaaatcctctaataaagagatcacacacacaggcatacactcccaacacacacacacacacaaactcacagaggatcacacaaacacccgccccacccccacacacccatttaccttagaggagagtccaagacccaccaacatacctgaccctcgatcctgttaataaagcaggttcaggccctctcagagtgaccctgggtgttgcctctagtgtccttgcagatgtgaaaaggatggtggtttagggccctaatcagaagcttcccggcagaatgagggtcttgcagttgccttacttgctgaaactctgaagtgctgggtgtgaacctgctgtgagactaccccctttaaggtctgagcctgttaactccagggtcagcttcccccacatctgcactccctctgctgcaggcct**gagctctctatgcccggagctc**tcttccttagagggcctgctctcagccgccacagagggcgctggagaactggcctga**gcagagaggcagaaactcctctgc**aggtcctcctaggcaccacccagagtcccctaagacagatcctcaatcacttatctctgcctgtgaatcag**gaagaccagctcctcctactgtcttc**tgtgttagggatcacttccttgttgagtgggacctgagttttgagagggtcttctgctcctcttcgtctggtcccttacttccaagaccccagagaggaaggcATGCTGTTGGCTCTAGCTCTGCTTCTAGCTTTCCTGCCTCCTGGTAAGAGTGCTGCCTACAGAGAGGCTCACAGGTTTTATTTTGTTTCGTTTTGTTTAT**TTTCTTCTTTTGCAAGGGATACCATACTAAGAAA**TGCCTCATTACATTTTGTGTTGTTCCCATTGCAGCCAGTCAGAAAT**CTTCCAACTTGGAAG**GGAGAACAAAGTCAGTCACCAGGCCAACTGGGTCATCAGCTGTAATCACTTGTGATCTTCC**TGTAGAAAATGCCGTCTACA**CCCACTGGTACCTACACCAGGAGGGGAAGGCCCCACAGCGTCTTCTGTACTATGACTCCTACAACTCCAGGGTTGTGTTGGAATCAGGAATCAGTCGAGAAAAGTATCATACTTATGCAAGCACAGGGAAGAGCCTTAAATTTATACTGGAAAATCTAATTGAACGTGACTCTGGGGTCTATT**ACTGTGCCACCTGGGATAGGcacagt**gattcagacctgtgctacaccacactgaaaatctgc**cttgtgactactcctggtacacaag**atagaccagcctcatctcatttcctgcccatgaattcactgtattctgtacaaagagaaacacagcttaactcctcatctctcccctaatatcacactcccctggcggcagctgcaccctgttctccaccccaccctcatgagttacctaaagaccaaactacaatctcccagctg**tgtcagccaagcagccttgctgaca**gcaaagttctttcagctcttctatgggct**ggtggagacccattccacc**tgcttcctaag**acagacaaattcctcaaggactggtctgt**aaagggattcatttaggagactatatgggaagactagcttcactcaagcatttatccaattattgtctataggctaaaatagtgc**ttattcattgctcaggataaatgttggaataa**caactttgcacttcagtaagacatctaacgtgtatcagatctgctatcttttggacacactgacatttttttctacataacaaatcatggtactattggtctgcaggagaatgggtaaaaaagtgatagtcataggtacgccaacttgtgggtcatatcataggttttcaaccatttcaagcatccctgccctaatgcaagtatctgaaggatctgacccttgccctgcagcagaattcattccatgtcccctctggatattagactgataatttggatcagg**tctcagcatgggccattgaggaacactacagccctgctgaga**gataaaaaggattatccactcaaggagttcacacctccgggccgacatggagaacgtgcctagaagcagacttggagggtgctcagtggtgggttgtggaatgtaagcctgaagaagcagagtttaattctacgggggtgctttcataatttagtaccctgacagggctcctggtgttggtccttagacactcctgggatgg

Uppercase: TRGV8

Lowercase: Flanking sequence[1000bp]

Red & Bold & Underline: Stem-loop [15]

Blue: Heptamer[20]

Green: Nonamer [9]

id-IGLV5-48[V_gene_segment]

ggacaaatcgtttaccaaaaaatatgacacgactgtgaagaaagtgacgttttacttagggagataatggctaacctcttaaaacagagatata**tgtcacactgtgaca**ggaaagtctacaaaatctcttaatgttcctacatcacaaaattgagtcaaatctctataaaacatatgaaaccattagatgtataacttttgctaaattgctgtctttaacatgtattggatcatttatttttcctggaaaattctcattcttcactgtccttttgaacaaccc**cagctgcccttcaacccagctcctcagctg**acccatttttcatgatgccca**ccctgatggcacatgtgggcaactcgatttttctagaatgccatgcaggg**gtgctgacaactttaaaagttatatatgtacatggctgcacatcccacagaccaagactacttctgggagtgagaaatcacagtaataatgcacagcatgatagaagcag**tgtctgggaatcagaccatggatatgtgaatacagaca**gtgactggag**agctgcagagaccagctgggttctgaagtctctgagct**ttgggaaagtgtgtttcctgggaccagcgtcggatgttgtggggctgccctgtggttctgacacagctgctcagtaaggagcattcactccaagaaatccaggccttgaggctgggccctgtgctcactgat**ggacactcggcagctgtgtcc**cctctggccttgcagagttccctgagcaggg**acctgtgtatcgtctcaacaggt**gcttcctcccaggaccttgcaagaagaaaaataaaccttcctcctgcttagtctgcagagtgaggtgagctgcatc**cccaggtctcaaggaggggcccggcagtccctggg**tggaaacacatttgcatgagcagcc**cctcctctgcaagggtgggaagaaaaggagg**cctggggcagcccagtcccactgtggggtaagaggctgtgtccaccATGGCCTGGACTCCTCTCCTTCTCCTGTTCCTCTCTCACTGCACAGGTAGGAAAAGGCCTCAGAGACCAGGGTCAGCCACACAGCCTGATTCTGACTCTCGTGTCAAAGATCACTAAAAAAAATATTACCTTGGTTTCTGCCTTAAAGCCTATAAATGCCTGTGTTTCCAGGTTCCCTCTCGCAGGCTG**TGCTGACTCAGCCAACTTCCCTCTCAGCA**TCTCCTGGAGCATCAGCCAGACTCACCTGCACCTTGCGCAGTGGCATCAGTGTTGGTAGCTACAGGATATACTGGTACCAGCAGAAGCCAGGGAGTCCTCCCAGGTATCTCCTGAACTACTACTCAGACTCAGATAAGCACCAGGGCTCTGGAGTCCCCAGCCGCTTCTCTGGATCCAAAGATGCTTCGACCAATGCAGGGATTTTATTCATCTCTGGGCTCTAGTCTGAGGATGAGGCTGACTATTACTGTATGATTTGGCACAGCAGTGCTTCTcacagtgacacacacagatggggaagtgggacgaaaacctcaccctgatctgggttcttgttctgtaacagt**ttttaaattttaaaa**taaccagccaagggacaaactaaatttggaaataactttctagttgtaaaaggctcttctctcaatttcccatccattcttctgaagtctcagtaaaacaaaaccaaaccaaagcacctgacgaccctgaagtgttggctggttgtccatatgtgctaacaccactgcctgcggagctggacttttgttcttgaaagaagcaaaatgaggctttcctagctcacatcatggtcagagcct**ggggacatcctagcaggtgaacagtcccc**tacagcat**ggactctgctcttccagtgtacaagcagagacc**tccccttccatccccactagtctgttctcagggctgtctgaggctcactgtgtcccaggaacaaggcactcaaatatgaaatgctgcatgaggaca**caggtcagggaattgcagatatagtgctgacctg**gctcagtactgggta**cctgcatttcagtggaaatgtcagg**agatgggattacgtctgagggtaactgtggacctgaggccggtaaacctctatgtaggacttgagagagtcaaggggaggc**tccaccatggctgaggcaaggctgaggctgctccttggccatggggga**agtaggtcccagggatgtgactttcctgcagctcattccactcaggaccacggggactg**ctgagggtgggtttgctgatccctcag**ggcctgtgttctcttctcctctattgcaccattggaggtt**ctcactgagtcaggttctagtagtgag**tctgtggaacatctgggtacagatactgtttgcttcaagtgactcctccccaggaaactc**ttgagaaggcccccatctcaa**tgtcacctt**tttctgcatggggagcagcagaaa**cattgggcattccctcccagaaagagcccctaatttt

Uppercase: IGLV5-48

Lowercase: Flanking sequence[1000bp]

Red & Bold & Underline: Stem-loop [20]

Blue: Heptamer[51]

Green: Nonamer [6]

id-IGHV3-79[V_gene_segment]

gtagctaaagtattaacagtgcacttcataaa**cttctatttgtataaactgtagaag**gtaaaactattctaaagtaacagagaagagaatttggtgagaattgggtgttgaaagtaatggggtgttgagatgaaattacagagaagtgacagaaagatttagaggttaacttaattgcacacgacttaattga**aaccctgattaaagggttgcacacatacattaacatattccaaattgtgca**gtatatatttgaattaa**ttatttgttaattttgtacttaaaaaagcagtaacaaagaa**acaaatgaatgtgttggttgaggaggagcaaaaaatagatggctattaacacttgaaacatcgctgacttttgaaagtacacatgtatgaacactggttctctctatatatatttcagataaacaacagaatacactaaaagaaaacagagatatcctggcaggggtggaatcctgcaaacctcactaggcatgtcccacactgccctggagttgtctcaggggagcagtcttctctagtggtcagaggcacaggcatggataatgggactaattctgtccagctgtgtgatcttgaccatattgtacagccactctgttctgtgtgtaatgtaaattccttaaaatgtaacattgacacttgcattaaatgtattctgcaaatatgtagaaatgaaataagatgatgactgctaaatagttatcaaggcatgatcacataatataaaatgatattttcctgagtgataagatgactaccaatctcccccaaggcactttgtct**gctctgacccctgcccctcctcaggattcccatcccagagc**ttgctatacagtaggagacatgcaaacaggtttctccctttgttgatgaaaaccagcccagtcctgaccccgtaggtttgggagagaagccccag**ccctgggattcccaggg**gtttccgttttgtgatcaggactaaagacagaggacccatcATGGAGTCTGGGCTGAGCTGGGTTTCACAGTTGTTATTTTAAAAGGTAAACCAGAGAAATAGAGTGTGAGTGAATATATTTGA**GAGAAACAGTGGATATGTCAGGCAGTTTCTC**ACCAGGATCTCTACGAGTTTACGGGTGTCCAGTGTGAGGTACAGCTGGTGGAGTCT**GGAGAGGACCCAAGACAACCTGGGGTTCCCTGAGACTCTCC**TGTGCAGCCTCTGGATTAACC**TTCAGTAGCTACTGAA**TGAGCTCGGTTTCCCAGG**CTCCAGGGAAGGGGCTGGAG**TGAGTAGTAGATATACAGTGTGATGGAAGTCAGATATGTTATGCAC**AATCTGTGACGAGCAGATT**CACCGTCTCCAAAGAAAATGCCAAGAACTCACTGTATTTGCAAATGAACAGTCTGAGAGCAGAGGGCACAGCTGTGTGTTACTGTATGTGAGGCACCAGGTAagaagacatcagtgtgaacacagacacagaggttcctgtaatgataagggaggaggctgggataaagggagcactcaagacccacagaaaacaggggaagctctagggcaggtgcagacggtcgtcatgggctactttccttcagggtctgtggctttctctgcatccaacagttcccctgggagcctcttgacatttatgtttctgtgcccatccctgaggtctctggattagaaaaatttattataagaagtggaaacattctcatttgtcccaaaggcagatgtaagtaatggaggcataaaaatgcacaggaggccagggaggctgtagacactgccaccccagaatgtcaatctcaccactagtgctggagaagagtgggagtttgatggagcttacctaaataccctgggtccaagctaagttcagcgaggccactggtgcctcccagagctcagttgtccatcaggaatctcccatgtgtcccagcagcagccatgcctcagtacttccgctgtgcacagccattgt**ctgggaggagctcccag**gttgggtgtctttggcacacaccaggtgacaggtgttagagtgcagtgcagcagctggctgcctggtctattgggct**ccctgatgttggagagatgggaggtggattctcaggg**ccagcacccagtttgtgaatttttatataaaaaccatgattttacttcattttctcagatgacatagataattaagaacagagtctggtccggacgtggtggctgatgcctgtaatcccagcactttgggaggccaaggcgggcagatcacctgaggtcgagagttcgagaccagcctgaccaatatggaaaaatccatgtctactaacaatacaaaattagc**caggcatggtagcatatgcctg**taatcccagctactagggaggctgaggcaggagaatcgcttgaacccgggaggtgaaggttatggtgagccaagatcacaccattacact

Uppercase: IGHV3-79

Lowercase: Flanking sequence[1000bp]

Red & Bold & Underline: Stem-loop [14]

Blue: Heptamer[30]

Green: Nonamer [4]

id-IGKV1-27-3[V_gene_segment]

atcaagtgccctaaacccagtcactaagaactgttt**tctcttaggagctggaagaga**ttggtgatttggaaatatgcaggtataagaaacagagtagtcacagggatagagggtgacaacttggtttagaggacacctcagcttctgaaggggaatggcttggataagagaaataaaaggcataaataaaattcagggagcacagggaaata**tctagcatgagactgtaggatggcatacagagctaga**atatagctgagaactttcagaagtaaagggagaaaatttatcatgttggctggcccagctgaaagaggtaggaaagaacattcagatatggaggataacaattatgtgtctggagatgggagattagctatgccagatagccagtggcaggacccttccttgctgtggcactattttcaagtattaggttttttttaagttttttatttcttttttaatgagcaaatctatctatctatctatctatctatctatctatctatctatctatctatcattcatgattataccttaatgcatccattgttggtagcactgacaatttacagcactggtggttccagggaattggaccaaaaaggaagtctctctgaccttaatagtactcatctgtatcaaatgcag**gaaacttctaaaatgtcttgagtttc**tagagatgtttttccctagcagaccttgtcataaatagaaagctagcaagagaagcatgtcatgaaacatgaagagagcaaaagaacactccacatat**aggacagtaggctgattctgtcct**gtagc**ctgcagggagaaacacatgctctgcag**actttggacacctgggaggcactgggcctgtgcagtgttattgagataagacatctttgcagctgtgcagatttgcatgtcccacagagcaacgcctactgccctgaacatttatcaataggctggtgacatcctgtgcagaagtctctctcagtcaggacacagcATGGACATGAGGGTCCCTGCTCAGCT**CCTGGGACTCCTGCTGCTCTGGCTCCCAGG**TAAGGAGGGAAACAACAAAAATTTTATTCAGCCAGTGTAGCCACTAATGCCTGGCACTTCAGGAAATTCTTCTTA**GAACATTACTAATCATGTGGATATGTGTTTTTATGTTC**CTAATATCAGATACCAGATGTGACATCCAGATGACCCAGTCTCCATCCTCC**CTGTCTGCATCTGTAGGAGACAG**AGTCACCATCACTTGCCGGGCGAGTCAGGGCATTAGCAATTATTTAG**CCTGGTATCAGCAGAAACCAGG**GAAAGTTCCTAAGCT**CCTGATCTATGCTGCATCCACTTTGCAATCAGG**G**GTCCCATCTCGGTTCAGTGGCAGTGGATCTGGGAC**AGATTTCACTCTCACCATCAGCAGCCTGCAGCCTGAAGATGTTGCAACTTATTACTGTCAAAA**GTATAACAGTGCCCCTcccactgtgatac**aagcccgaacataaaccatggagggaagtagatgtgtgagtctgggctgccccagctgctcctcctggtgccgccctctgctgacagcagttctcagatgcagccaaggtttgaagctcactgggaagttttggtagaaggggtcagggaggcacatttacattctatctctctttatcc**tcagctccatcagctga**tatgcaagtatctctcctgattattattaataaaggacaaagaaaattaaacctaggaggtctagattgcagcaaaagtcagacttatacagaaaaggagaaggtactctacatatttttaaagaattttttttgatacaggggattagagtctaaattatgacctttcaaagatcagacataagt**acatacacatataagtatgt**tcagccaacaacatatcagataa**acatattgcacctatatgt**atactgtaaacatttatgctccgatcccaccttcttccaacactacccagatcaagatccacactcttcccagcatctcaaatatctccccagtttacttccctgtcaattccgtgttccatctatcatcatcctaggcttccactgtattttatatcaatatttattaatttttcctatacttaagctttatataatttcaatcatacagtatttactctttgacgtcttgcttattttgg**tcaaaatatttttga**gcttcatctgcgttgtttctctggcagtaatttgtgcctttttattgaggaacaattttcc**atgaatatgccaaaatatattcat**ttttcaactaattattt**gtttttggtttaagaatattactggtaaaaac**cttaagaacattcttgtgcttgtcttttgtgaacttatgcactcatttctcttgggaatattcctaagaatggaagtgttgagttataaagttagcatagaatttgtcttagtaaacattgacaatagtttactgaaatgctgaacatgtt

Uppercase: IGKV1-27-3

Lowercase: Flanking sequence[1000bp]

Red & Bold & Underline: Stem-loop [18]

Blue: Heptamer[21]

Green: Nonamer [5]

id-TRBV2-2[V_gene_segment]

tctctctctttctttctttctttgtttctttgtttctttctttccttctttcttttttg**agacagagtctcactgtct**ggagtgcagtggcgcgatcttggctcacttccacctccatcttccagattcaagcgattctcctgcctcagcctcccgggtagctggaactacaggcgcccagcaccacgactggctaatttttgtatttttagtagagacagggtttcaccatgttggccaggatggtctcaatctcttaacttcttgatttgcccccctcgggctcccaaagtgctgggattacaggcgtgagccaccgcgcccggcaggtccttttc**tttttctttctttcttttttttttgaaaaa**ctaaagaatacccaatcccc**tcctcaggtggctctatcactgcctcaccctgagga**cagtatttccccatggggtttggatcactgttctctagttccctctccctggggcatgtgcagagtttcaacttcttgttggctgctctcttccactcaggatttaaaacattcacacttgattatttttccttctatctctttaatcctaataccaagttttcttggcattcattatcagccgtggagtctgttg**tacctgaattgttactcacagcaggta**ataagagaacaagacctcctggaatttctgagccagactttgtactcagggttaccttgctgacccccatattttccagcattgctcttattaattaatggccctcagtcctgcacacaccatcagatcccag**gggctgagcacagccc**tgcattgtcaggttctgcaacaacacaaaggctccctcctctctcctatagagggcgctccagggatggtgggtgttgccagagacaccagtaattctgccagaccttgcctgtggggccatgggagctcaaaatgcccct**cctttcctccacaggaccagatgcctgagctaggaaagg**cctcattcctgctgtgatcctgccATGGATACCTGGCTCGTATGCTGGGCAATTTTTAGTCTCTTG**AAAGCAGGTCGATGCTTAGACTCTAGGAAATTCTTGCTTT**GAACTTACCTAAGACAATTCTAAACCATTCTCTTAATCTTCTTCTTTTCTCACAGGACTCACAGA**ACCTGAAGTCACCCAGACTCCCAGCCATCAGGT**CACAC**AGATGGGACAGGAAGTGATCTTGCACTGTGTCCCCATCT**CTAATCACTTATACTTCTATTGGTACAGACAAATCTTGGGG**CAGAAAGTCGAGTTTCTG**GTTTCCTTTTATAATAATGAAATCTCAGAGAAGTCTGAAATATTCGATGATCAATTCTCAGTTGAAAGGCCTGATGGATCAAATTTCACTCTGAAGATCCGGTCCACAAAGCTGGAGGACTCAGCCATGTACTTCTGTGCCAGCAGTGAAGCcacagccttgcaaagacaactccagcctgtgcaaaatccctcaca**gagctgcctccctcccagccgccagctc**cc**acttcctgcctaagaaaaggaagt**ctctggttgggtttgttcttgcag**ctgtctccaaatagacag**atatctgaaattaaattaccaaaggtatcttgggcagaaatgaccagaca**ctgtaggatttgaagaatgtttacagatactacag**agtgtgagtgtgatggtgggctctgaatatgagagaattgtacacaaccaagtaacttatatcttctcctcattcccatttggccatctttattagaattgttt**tgttttagttgcttaggaaaaca**tattttaacttctagggtggcgagatgattcctgagtgggagttcaggtatttccatgagaatatagagggtgctcacagaaatcttcagaagaaggggcttggtacaaagaactcaattatcaatga**ctcttgctcagcatgtggtcaagag**cagaactatctgatgaccctgtgaacctggtgtgatgcatttgggctaatgatgctagctcacaaaataacaagttaagagagtagaaaacatggagtctaatatttcacaccatttcaaggtcaacagagttctagaccagcatgagaatcgccagaataaagcaaggcagtgcaggaacagaattcagcagcaaaagtcaagagacacctagaggcagaggcaggaatgaatttgtcactttcaacttaacctgaagcagccagattaacttcttgagtttcatttaagaaattattcctcttt**taaacatacctattgaagtctaaattttaggtatttta**tatatttcagtt**ctggaatttccatttaaatcttgctacaatttccag**ttgtttgctaaaattgcacattttgtcatttactttgttgactatattaacatctgtcattttaaagccagtttttgataactttaacatctgaatctgttt

Uppercase: TRBV2-2

Lowercase: Flanking sequence[1000bp]

Red & Bold & Underline: Stem-loop [18]

Blue: Heptamer[30]

Green: Nonamer [1]

id-TRGVA[V_gene_segment]

ctctgtctgctggcctgacttagccttcttcctttggagacacacacacactcacacacacgcacgcacacacctggttcttgaaaactgcattagtccctataatttctactttgtgtgaaccccagagatcattt**actggaacccatccagt**ttccaggtgaaacaacaacaccccacatttcttgagcatggtctatgcagctgggattgtgctgcgtgctctgctcacattctcctgtttactgtccatgtggccctatga**cacctctctttacagatgaggaaaaagaggtg**aaacaacttgcccaagactacacagctatcatgttattttagggcactgaaaatctccttctactaggacctctgggtgacacgctccgatgctggctcagacggttggcatgggtttttttagggattgctgttttgtgtttgtcccttcctctggagcaagcagcacactcattgttctctgagtataagatggttacatgaagcagaaatttgtcaccagcctggcctggcttctctcccaactctgattctctttaggtattacagccttggaaaagctacttaaaataacagccttactccttcatttctaaactgaagataacatttgtcctatagagttat**tttaggatttaataaagcaatggccctaaa**ccaattagtgcagtgcaagagtt**ttgctggtgtccaaccaatgattgctattattgctaattataccagcaa**cactctttaagcctttttcagcactctagccattggtcatccctcccttcaccttgtggtcagatttctgtctctcctctggaacttctttcttcctcccccatttctctcatcagttacctccttgcaccaccagggaaatagcagttcaggccctaagctggtcgctaccataactcaggaaagtgtggctcaggtccatagtttgaaacaggtcactctgggctcctctcccccagtcaggatcagagatATGGTGGGCTGCCCTGCCTTCCTTTGGGTCATCCTGTTTC**CAGGTCAGTCTCCTACCTAAGGACCTG**TGTAGAGGTGGCGAGGCTTTGGGACATCCCTACCTCACTGCTCTCTCCCTGA**CAGCTGGCTGGTGGCTCATCAGGCCGGAGCAGCTG**GCCCATGTCCTGGGGCACTAGGGAAGCTTGGTC**ATCCTGCAGTGCGTGGTCCGCACCAGGAT**CAGCTACACCCACTGGTACCAGCAGAAG**GGCCAGGTCCCTGAGGCACTCCACCAGCTGGCC**ATGTCCAAGTTGGATGTGCAGTGGGATT**CCATCCTGAAAGCAGATAAAATCATAGCCAAGGATGG**CAGCAGCTCTATCTTGGCAGTACTGAAGTTGGAGACAGGCATCGAGGGCATGAACTACTGCACAACCTGGGCCCTGcgcagccttgcatgctgccccagccctacacaaaaggactcttcctcccgatccaacaaggccttgggcattttcacttactcttggtcccttgggtttccctgtggcatagaagaaaaaagtcttatcccctgtccctc**acctccccatctatcctggaggt**cccttccccggtcccactacagatgaggctttcagcttgtgcagactcagtgatctatatctcatcgaaaggggtctgtttaaagatgctccagttcagtgtctctgaagcctttgcaggatgacctgtgctcctgaatgcaggtgccac**tagccacctgtggctattgagcatgggaaacataccaagtgcaatag**tccaagctgagatgggctgtgtgtctaaaatcatattttgaagctttagtgttaaaagtgagataactcttataatgatcaaaagttgaaatgattaggtatcttggactaaagccaatagcattaaaactaattttacctgtttct**ctttactttttttaatgcagcaactaaaaaaattaaag**tgtgtgactcacattggtgactcatttatttctgttggacagtgctgttagagactgatcatttaagaatctgaccatctcccctgaagaatgcacaaataaaatattttggaagagatttcaagggtggaggacttgctgagatttaccacaaaatggttgaa**aagctgagactagaagcctactcctgtttccagctt**ccaagctcttggccctgggactgttatcattttgcaatagagtcagcaattagaagcatgggaaacatgact**gcccacttctcagtactttcacccgtgggc**gggctgcataggaactgatgtgagtagacagtcgccatggcagggtgatgcagtcctccctggatgagacatttacc**ttccacttctccttcaacagcaccccagtgtggaa**cacgtgtgtatcccaagcttctctagacttttgttctccta

Uppercase: TRGVA

Lowercase: Flanking sequence[1000bp]

Red & Bold & Underline: Stem-loop [16]

Blue: Heptamer[21]

Green: Nonamer [4]

id-IGKV3-25-3[V_gene_segment]

agaatgataacagttttagacttcgagggtacttcaggaatgggaacacacaagga**acacagtagatatgagctactgtgtctaccgtattagaca**atcctcctgcattttaacttacatttggtgaacttcagatgccagtgggtgttgcaagttgatgacataaacctaaaatcaagtcccgaggtatttcttgtgaagagtcacagagttgaagagaccactgcctggcttcctgagttgagcctgtgccactgcaagtttcatgggaaaatggcgagaaaagctgtttaggtttgtaatcatcaaaggtttagtgaagttcccgtgcaaagagactcaatcccattcaactcagcatgaagagatcttctagaactcattctgcccctatagcagagatgatgtggtcaggtttgtggagatgaacatgcaatgacgtgttaaccaaaaggcccagcagaaagggatagtgatcaatgtccatggttcagacatatacaa**aggatgatttgcacagcactggaggaaaaaacatcct**gagtgttggagatttctgccttcctacccaagctcccgcagcctgtctcctgactcagaactcagcaaatgccacatgaaaaatctggctat**ggttgcacagaaattgccctgtgcaacc**ttcaaagctcagttatgttctctcttcttgctgggtttctctgtctgtg**ccaaatcaaaatttctatccgaacgcacatttgg**taaaaaaaaaaaaaaaaagggcagaaaaacaaaatgtcagatgaacaccaact**caactagaattagttg**tttttttactctagaattttagatcacctagtcctcatttcttccattactttctaatgtcactaataggtaactttgcattttatttgtttttattggcacattgtaat**tgtacatatttatggagtacactttgatgtcttgatatgtaca**tatgttgaaaaatgatcaaatcaggatatttaggATGTGCATCAGCTCATGTATTTACCATTTCTTTATGTTACTCTGGGTCCCAGGTGAGGGGAATATGCGGGGGGGCGTGGGGGGTTCTGTGTGTCAGTGAAAACTCTCTCAACTTTGTTGGGTCCTTCAACCTGGCATTTCCGCTCAGCAAGCACAATAATTAAAGCTCAATGTAAATCAATGGTCTTGACTCTATTGGAAAGACAATGGGT**ATAATCTAGATTACATGTGTGATTAT**TTGTTTTTCTGTTTAAACTGTTTTTTCAA**AAAATAAATTTTATTCTGTATATTTT**AGGTTTGCAATATGATGTTGTGGCATACATATAG**ATAGTAAAATGCTTACTAT**AGTGAAGAGAATGAACATTTATCATCTCACATAGTTACTTTTTCGTCTGTGACAAGAGCAGCTGGAATCTACTTATTTAACAAAAATCTCTAATACAGTACAGTTTTATTAACTATAGCCCTCATTTTCTACAGTAGATCTCTATACTTGCTATTCGTAAATATTTACTACCTTGCATCTTTGGACCTACATCTTCCCATTTCTTCCCATTCCACCCCCGGCCCTTAATAACGACTGTTTCATTCTCTATTTCTGCATAATTTATCTTTTTAGTTATTTAATTTATTTTTAGGTTTACATGTAAGTAAAATCAGGCAATATTTTTATTTCTGTATCTTGCTTTTTTCACTTGGAATACTGTCCTCTAGGTCCATCCACATTACGGCAAATAGAAGAATCTTCTTTTTC**TGCTGGGTGTGGTGGCTCATGCCTGTAATCCCAGCA**CTAGGCGGGTGCATCACCTGACATCAGGAGTTGAATACCAGCCTGGCCAACATGGCAAAACCCTGTCTCTACTAAAAATACAAAAAATACAAAAGAAAATTAGCTGGGCGTGTTGGCATGGGCCTGTAATCCCAACTACTCAGGAAGCTGAGGCAGGAGAATCGCTTGAACCCGGGAGGAGGAGGTTGCAGTGAGCTGAGATAGCATCATTGCACTCCAGCCTGGGCAACAAGAGTGAGACTTCTCTAAAATAAAAATAATAATAATAATAAAAAAGAATCTTCTTGTTTAAGGCTGAATTATATTCCATGGTGTGTGTGTGTGTGTGTGTGTGTGTGTGTGTGTGTGTACATTTATGTATACCACATTTTCTTTATTCATTTATTTGTCCACGGATGCTCAGGTTTTTCTCATTTCTTGTGAATAATGCTGCAATAAACACGGGAGTGTAGGTATCTTTAAAAGAAGGTGGTGATTTCATCTCTTCTGGGTATGTATCCAAAATAGGGTCACTGTTGGGTTATAAGGTGGTTAGGTTTTGAATTTCTTTAGGAACCTCCATACTGTTTTCCATAATGGGTGCACCAATCATCATTCCCACCAACAATGTACAAGTGTTTTATTTTCTTCACACCCTCATCAATATTTATCTCTT**GTCTTTTTTATAATAGCCATCCTAAAGAC**TGTAAGGCGTTTTATTTCTAATCTCAGATT**TCACTGTAGAAACAGTGA**TGACACAG**TCTCCAGCTTCCCTGTCTTTGTCTCTGGAGA**AAAAAGCCACC**CTGACTTGCAGGGCCAGTCAG**TGTTAGCAGCTACTA**AGCCTGGTACCAGAAGAAACCTGAGCGGGTTCCCAGGCT**CCTCATCTATG**GTACAGCCCTGATTTGTGATAGTGGGTCGGGGACAGGGCTTAC**TCTCACCATCGGCAGCCTGGAGCCTGGAGCCTGGAGATTTGCACTTCATCACTGTTATCAGCATAGTAGTTGGTGTCCcatactgattcgacatgcaacaaaaacctccaggagacctaaggtgtttatttgattatactacctgcttcctttttagtcatctgatgtggtgctgctcagttttagcatctctgctttgattggaaattctgaggttctcaaaagtaattccttataatatttatagtttcactcatggatttttttctcagacccaaatgtacagccaggttcaggcacaatttcatggtcaaggccattggatcagactcacatgagtggacgcctctaaaggtcctggccagtgcgataaagtagcagcgacaatgataaagaagaagaattagaaaggcagaattaaaggtataacaattcactgatgaaaggactgtgtgggggaga**aatttctaattgtctacacagaaatt**attagaattaatgagatacatagcaaattgttagattaaagatca**atataaaatatatttatttttatat**aaatgtaacaactgctaaaattaaaattatgaaacacatcatttatagtagcaccaaaccaagaggcattagaaaaaactaacaaaatatactgacagctccagaatataattcaaagaaatcaaagaaaatctaagcaaatggaaggccacagt**actttcaaaggttgaaagt**cccagtattgtaaagatgttcattcttcacagattgatatattaaagataatcgaaatcaaaatttcaaaccgtttttaaatgga**aatgtaaaagctgattctacatatcatattacatt**gttaaggagtttgaataaccacaataaacccgaagaagcccaagaatgaggacttacactacttggatactcgaaccatgctgctgatgtgagttcaaacaaattgcctaatgcaacagaagagagtattctgagaatgactaacacatatagtgacactttttagggacaaaggtgacactgcacaacagagtgaacacaataaaaagctgtccaacatcaataa

Uppercase: IGKV3-25-3

Lowercase: Flanking sequence[1000bp]

Red & Bold & Underline: Stem-loop [21]

Blue: Heptamer[41]

Green: Nonamer [6]

id-IGHV3-29[V_gene_segment]

taagtttgtttttagacgtatgcaatacattttgaggtgaaacctggtggaa**ttttttctaataaattagaaaaaaataaatcatttaattgactattttattcaggttgatttgttt**aatatttgctaaaggccagttctttaagctatgacacataataaatcccaaatggcagtacctcattgtttacttagcttttgtacttatatttctcagaggaagaaccactactgtaaattgtaaatagc**caatacataattgtattg**tatgcaaatctgtgactgtttacagtgtcatctctgagaaacataaagtttatttactatatatatataaagagtttggaaggtggactcctcaccaatttttgaaagagttacagaagggctggcattacttctttaaatgttaaggttcatttt**atgatgtgacatataacccatcat**ggagaatgttcaatgtgttcttgagaaggatgtgtattacgtgactcttggttggaaggttctgtaaatatcattcagataaatttgttcaatagtgttgttcaagttcagaggcacattaggaattttctttccggatttgctatacattatggtcatgaggtattaaggtcttgtgttatttttgtattgttttctatttctttatatctcttacagtttgcttaatgcagttatatttgtatttgtacacatgtgtaaaaacaaaaatcataattgctaaatgagttttatggcacagtcacattataagtaatatttttccaaatgctaccattgccactaaactcctcctggagtctgacgtctgctttgggcactgccttcttctcaggcatcccacactggagcttgctatagaggaggagggagggcccctctcctggtgaaaacaag**cccagacctgaccctgcagctctggg**agaagagccacag**ccctgggattcccaggg**gtttccatttggtaatcaggactgaacacagagaactcactATGGGGTGTGAATTAAGCTGAATTTTTCTTGTTGGTATTTTAAAAGGTA**ACTCATAGGGAACTAGAGTGAGTGAGAGTGAGTGGATATGAGT**GAG**AGAAACAGTGGATATGTTTGCCAGTTTCT**GACCAGGATTTGTTTGTATTTTCAGGTGTTCAGTGTGAGGTGGAGCTGATAGAGCCCACAGAGGACCTGAGACAACCTGGGAAGTTCCTGAGACTCTCCTGTGTAGCCTCTAGATTCGCCTTCAGTAGCTTCTGAATGAGCCCAGTTCACCAGTCTGCAGGCAAGGGGCTGGAGTGAGTAATAGATATAAAAGATGATGGAAGTCAGATACACCATGCAGACTCTGTGAAGGGCAGATTCTCCATCTCCAAAGACAATGCTAAGAAC**TCTCTGTATCTGCAAATGAACAGTCAGAGA**ACTGAGGACATGGCTGTGTATGGCTGTACATAAGGTTccaagtgaggaaacatcggtgtgagt**ccagacacaaaatttcctgcaaaaagaagaaaggagtctgg**gccaaaggggacactcagcactcacaaaacaggt**gcagccccaaggcaggtgcagatggagggagggtaagggctgc**tttccttcaggatctgtggtttcctctgcttctaatatttcccctctgagcctctgtacatttatattttgtgcccaccatgaggtcgctggattagaaaactaatttgaaagaggaaatattctcatatgtcctaaaaacagatgtaagtattggaggcataaaaatgcataggagccgggtgtgtctgtagacactgccacccccaatgccagacccacaactagtgctggagaagggtgggagtttgatggagcttccctgatgaccccgtggtccaagctaagtccagcaaggccattggtgcctcgctgagcacagttgtccatcagggatctcccatgtgtcccagcagcagccatgcctcaatctctccactgtgcacagccattgt**ctgggaggagctcccag**gatgggtgtctttggcacacacagttgatgggtgttagagtgcagtgcagcagctggctgcctggtctattgggctcctggatattggagggatttgaggtgcattctcagggccagcacgctttgtcaatttttatatgaaaaccatgtgatttaattcattttctcagatgacatagataattaataacacaatctgc**aaacaattataattttcaactttaccccaagttcattgttt**ctgaattctgtgcaggatccagacatggtactgcccttctcat**gagaaactgttcgatctaaactgaaaccagttgtttctc**gtatactttggttctccccacatgcagagatcttgattagagcaagtttggtactttccacgcactcaccctcacctccccagataaagagcagaagttctccttagactgagtctgaggga

Uppercase: IGHV3-29

Lowercase: Flanking sequence[1000bp]

Red & Bold & Underline: Stem-loop [14]

Blue: Heptamer[36]

Green: Nonamer [3]

id-IGLV7-35[V_gene_segment]

gctcacccaagcttggtgtccagagggcttttaaggggtggtctgggtgagagtggggttattatgtagaccttgtggatcatcctcaaggctcacttcagtttcc**attgaattaacaataaaacttaaaccttcaat**ctttggttattttct**ttttaaaaattctctgaagtaaattattattaaaa**ttctcacttaagacag**ctgtttatccaatatgaggctcaatataaacag**tttcaaaaagtcagtaggtcattaaaacaatttcttcagtcattcaaaccattcagataaaaattaacaccatgtaatacaaaaattgaattatagtcacagggacaggatttcagcaatggcaagaaaaagtagtgcaaataggaatgagcaaagaatagacaagtaatgggtgtgaattggcctcggaaataccatggtaatgtcaaaaagataaatcaattg**ttattttgttaaataa**gtgaaaggccaggcattctcagtccaggaaagccatcacagggcagtctgagtcttctgcatcttaagaagctcaagaggttcaaacttaataaaccctgcacagttgctcactggttcatgtaaaatgctagagcagttgcttcctcccactcagcaaatccacagcacacagtcaccctggtctggcccttcacagtgtagatgccctcagggcta**agaacctgagtggttct**aaaagtagtaaagatttgcataaagtaccacacaacacaccctctccacatagggagctcagtaggacataaagagccatcagaatccagccccgactctggagcc**aggagtcccttctaatatcagcatcatggcctggactcct**ctccttctccttctccccatgtacctcctcacttgctgcccaggttaatagagatttcaaataccagcctttggagggattcctgtgtctccctttctaattcccaacatgtgtctgttttttgtttcagggtcaaattctCAGGCTGTGATGACTCAGGAACCCTCACTGACTG**TGTCCCCAGGAGGGACA**GTCACTCTCACCTATG**GCTCCAGCACTGGAGC**AGTCAATGGGGGTCATTATCCTTACTGGTTCCAGCAGAAGCTTGGCCAAGCCC**CCAGGACACTGATTTATCATGCAAGCAACAAACACTCCTGG**ACCCCTGCCCAGTTCTCAGGCTCAGTCCTTGGGAGCAAAGCTGCCCAGACACTCTTGGGTGTGCAGCCCGAGAGGTGAAGCTGAGTACTACTGCTTACTGCACCATAGTCGTGCTTGGcacagtgacagactcagaagaggaaccgagacataaacctctctaggcccttgtg**atatgaagatcatat**gatcacgcacaccagctctcaaggcagcctacatgtggaccagccatagaaaggggaaggaaaggatctgaattgattcctatccctgcttaagccctgaagtgaaggaaatgtgagaatgatctgggaagaactggatcaaaaaaaaaatcagaagtttattgctttgtattctaaaaggagcactaacagctggatcagatctaaaggctgaggctaaatgcatttcctccagaaagaagcatcttcaaagtatggg**gtttctgagctaagagtaaagagaagaaac**tctacttttgtatatttctaaagtttgttttattgagttactttcaaagcaatacatgcctattatgtggaatcagaaagtattaaaaatcaccaagttctctgcaaagctaccttatcccagataatcaatagaagtacatttcttcatttggatgcattctctcatttttgttgtcaatattttcacaatggttatcaaaccttgtgctcattcctctggttttatacagctttttgctctgtcatttaatgttaatgtaagtagttttccatttatttgaaagggtaaaatatttgagtagttgctaattttattttt**atttgaataccaatgaaaaattataatttggggtaattcaaat**tgcttgtcagacaaagatgtgacattgggctattattaaaatgctcctggtcttctctgtggatttagtgaagtcaaacaaaatgcctgcaattttctatggaaagccttgagtacattatgcctgacttcacaatgcagaaaatagagaaacaagaatttcctgctcatgtttgctggagcataatagattcaggtacgaggaacaacttatattgaaaactacgtcaatgtctccatgcatgtgttagaacagtacttcccaaactctattgtgc

Uppercase: IGLV7-35

Lowercase: Flanking sequence[1000bp]

Red & Bold & Underline: Stem-loop [12]

Blue: Heptamer[33]

Green: Nonamer [8]

id-IGHVII-67-1[V_gene_segment]

tatatactacag**agatattataactctatatatct**aattcctctttcctatcccatgttttttattttctgtttgtatctatacatgctat**aaaatgtaatattggcatttt**tataattgctttccacacttacataatgaaca**aaatacaaatgcaaaaactgtattt**caacctcatttttccattgttgaccatcttctttgtgcatattttcaggtgttggatctatgtcacatggctattcatctagagactctgctaaaatgtctacggaaagattcatgtgctggaaataggttttctcaggttttattggtccgacagaatatttatttgtctttgattt**attgtctccagagacaat**gtcaagaatatgctatatctgcaaatgggcgatctgtaaaccaagaacacatcagtatatcactgtgcaagaggagcacatgagggaaagccagtgtgtgcccagacataaacctccagggacacatgggcgaattagctgcagggggagcttgggacccactgatgagagtcgaccccagagggaggtgcacagaaaggttaagaactgctttcctgccagtatatttttaaaaaaacaaaacttccatctgacagtttctccagagaacttccctaattttagcattctgtgcctaccaatgccatcactaaataggtttttaaaaaatattgta**atatgaggacatattctcatat**ggacaaaacgcaaattgacatttacagaaatgaaaagtcctcaaccgtggtcgctagaattggagacctgtagcagctcagtgggacctgttgagtcttctccaatcacactcagaacagagaccttagtgcatctccctgactagaatacc**ctttaggtatcgagataacagcctaaag**atggtaaatcatgagagtctcagatgtccagatgacagacatgggtctatgggaatag**tactgtaactgacagta**actaattttccataccgccagATGTCCTATTCAGGAGCAGCTACAGCAGTCATGCCTAG**GTGTGAAGATCACAC**ACTGACCTCACCCATGCTGTCTCTGGCCACTTCATCACAACCAATGCTTAATATTGGACGTGGATCTGCCAGTCCCCGGGGAATGGGTTGAATGGATaaaatgcatggctag**tgctgctggaaacccattcctactgtggcaaatggcagca**tctcttttaaaaggctaaataatattctattctgtatacataccacattgccattatcctttttgtttgtttgtttgtttgtttgtttgagatggagattcactcactctgtcacccaggctggagtgcagtggcgtgatttcaggtcactgtaatccctgccttctgggctcaaacaattctcctgcttcagcctccggagtatctgggattacaggcatgcaccaccactcccggttaatttttgtattttttgtagagatggtgtttcaacatgtttgtcaggctggtctcaaactgtgacctcaaaaaatcaacccacctcagtctttgaaagtgctcagattgcaggcgtgagccatagcatccagccacttttctgcatttttttttcattgacacttagattatttcaatatcttggctcttgtgaatagcgctgcagtaaacataggagtgcagatatctccacaaggtggtgattttttctatcttgggtatattcccagaagggaggttgctggtcatacagtagttctagttctaattgatttaagatcatcttactgctttccatagtggtggagattcagattagtgtagctattatgaaaaacagtttcaatgttggggagtgactgaaaaacaaataatgcttgattatggttctccatgagagtctctaataaacctaatggaagttcagaaagtttcctcaccttgttaagaaaacatttgttttgctcataattctctttcagcccagagttggcagtatgtagaatggaggctgttggctcgtttaaggattctgaagcaaagaacagtaatgagagatgacaatagagaaaaagagagagaaggagatgctgaatcatcatggactcattccacctgcaagcctttcactgtaaagctctattgctaaggatcctgaggattcta

Uppercase: IGHVII-67-1

Lowercase: Flanking sequence[1000bp]

Red & Bold & Underline: Stem-loop [9]

Blue: Heptamer[19]

Green: Nonamer [3]

id-IGHV3-41[V_gene_segment]

gctttgtcatcc**aggctggagctccgtggtgacatcatagctctctgcagcct**cagaccctgggtctccagcaatcctcctccctctgccctatctagctgggactacagctgtgcaccaccatgctcggcttcagtgcttattcaaccagacactttaattatgcagtatttattacgtg**gcaataatgcctcaataagggtattgc**aaataagtggatggataatttgttcaattaagattgatggaaatatagacacttacatgacaaatgtatgacaatcaagaaaataaaactgtaggaaacatactttactttttgttaggtaatcgcaacagtgcatacacatcacaccatgttctcattacagagaaaaggttctgcaaacctcacttggtgtgacctcctgtatgctgggcttggttcagggagaagtcaggaccag**tggttagaagcacgggcccagataacca**gactccctctgaccaaatgtgagtgctggggacattgtacaatccatctgtgttttgctgatattttttcatctgtaacatggaaataacattgatactacataccatggtttctgtgcatatggaaaaataaaagatgataggtgcaaactttaaatacac**acagtttatgtagatcaattgtacctcaataaaactgt**tttaaaataaaaattac**aaaactataaattttataggttt**taagggtttatcatagaacaaacttacaataagaaaccagaatttccaaatgctatcaatatcacaaatctcccccaggacactctcacatgctctgagccccactctctcctcaggcgtcccatcccagagcttg**gcatgtattaggagacatgc**aaatagggccctccctctgcttatgaaaaccagcccagccctgaccctgcagctgtgggagaggagccccagccctgggattttcaggtgctttcattttgtgatcaggactgaacacagaggattcaccATGGAGTCATGGCTGAGCTGGGTTTTTCTTGCCGCTATTTTAAAAGGTAATTCATTGAGAACTATTGAAATTGAGTGTGAGTGGATAAGAGTGAGATAAACAGTGGATTTGTGTGGAAGTTTCTGACCAGGGTTTCTTTTTGTTTGCAGGTGTCCAGTGTGAGGTGCAGCTGGTGGAGTCTGGGGGAGGCTTGGTCCAGCCTGGGGGGTC**CCTGAGACTCTCCTGTGCAGCCTCAGG**ATTCTCCTTTAGTAGCTATGGCATGAGCTGGGTCCGCCAGGCTCCAGGGAAGGGGCTGGACTGAGTGGCACATATCTGGAATGATGGAAGTCAGAAATACTATGCAGACTC**TGTGAAGGGCCGATTCACA**ATCTCCAGAGACAATTCTAAGAGCATGCTCTATCTGCAAATGGACAGTCTGAAAGCTAAGGACACGGCCATGTATTACTGTACCAGACAcagtgagaggaagtccgtgtgagcccagacacaaacctccctgcag**gggcacgcggggccaccagagggtgccc**aggatcccctgaagacagggacagcccaaaggcaggtgcagatggatgtcaagagggtctgtggcttcgtctacatctaactgtttcctggtgagcctctgtatatttatttttctgtgcccactaatgaggttgctaagtttaaacacttttttttttgagatgaagtctcgtgcttgtgccccaggctggagtgcaatggcatgatctcagctcactgcaacatccgcctcatgaattcaagcgattttcctgcctcagcctcccaagtagctgggattccagacaactgccaccacgcccggctaatttttgtatttttagtagagacagggtttcaccatgttggccaggctagtctcgaa**ctcctgacctcaggag**accctcccaccttggcctcccaaagtgctgggattacaggcatgagccactgtgcctagccaaaaaagacttcattattataggaggaaaaattctcatctgcaagtgtcactgatggaagcagaaaaattgcccaggaagtcaggtgaggctgtagacactgtcaacccaggatgcaaatctcaccacaagtgcaggagaaagggcagggggttgatggatcatcctaactaacgtatggtttaagctgagcccagcaagaccctcagtgcctccctaaccacagttgcccttcagggataacccatgtgctcaacagcagccacaccttagtatctccactgtgcagtcattgtctaggacaagctcccaggatgtggggctttggcacaaaccaggtcatggatgtcagagaggggcagctgggtgtctggtccataggctccctg**atgttggagggttgagaggtgcattctctgggccaacat**atactctagggatatttacttcagaacacatacttttatttgttggtttatggttaa

Uppercase: IGHV3-41

Lowercase: Flanking sequence[1000bp]

Red & Bold & Underline: Stem-loop [11]

Blue: Heptamer[39]

Green: Nonamer [11]

id-TRBV27-2[V_gene_segment]

cactgcctggctacaacacgatatccagggacagataccttccatgtacagcaagctgtggagtggcaagaaagcaaagcacgtaccaaaagcaggcaggaag**cagagctggctggtaaggctctg**aggttactgcag**gaagcaggtgttgcctctgcctttcaccagcttcttctggagaagaa**cagagaaggggaaaagaag**aaagatgctccagataccatcttt**ggtggaaacagctgctagctctagcaatgatgactaagtcaat**tttgctaaagagcaaa**gaagctgtgtgtcttggattttaaacagcaatagactagcaaaataatctatgatgtgaatctgagattttacaccctagactcttttaaaatgtagtgactaaaaatatataccagtggataaatgaagtcagggatatacttgaaattttttctgccagagtagaaatgacttcaaaatatatgataatgcagctttcatgtaatcccagctactggggaggctgaggcaggagtattgcttaaacccgggaagcggaggt**tgcagtgagccaagatcatgccactgca**ctccagcctgggt**gacagagtgatactctgtc**tcaaaaaaaaaaaaaaagataatgcagctttcaatattataagctttatgttctcacacttctaagtgataatcaaatgtgttacaccactttataccctgagatccaaaatacataaatatttgacagtttgtacaagactgtattaataagaaaatgtggcacatgtataaagaaaggaatctcacagattaaggaaaat**gacattgattacaaatgtc**ctcagcctgagttggctgtgttgcgtttgta**cacttcagccctctgagctgaagtg**ggagtggtttctctcctgaaaatgtttctgaggcccaaatagctgaag**aggtggagacgttacagaaaccacct**ggagcccccagaac**tggcagacacctgcctgatgctgccA**TGGGCCCCCAGCTCCTTGG**CTATGTGGTCCTTTGCCTTCTAGGAGCAGGTGAGTCCCAGAACACATAG**GCAACTTTTGGCCTTATTTTGTAGGCCTCAGCCCAAGGC**ATCTCCTAAAGGCTTAAGCATTGGAGAT**CCCCTTCAGCGCTGCCTCTAACTCTGTCCCCTTCCTTTACAGGCCCCCTGGAAGCCCAAGTGACCCAGAACCCAAGATACCTCATCACAGTGACTGGAAAGAAGTTAACAGTGACTTGTTCTCAGAATATGAACCATGAGTATATGTCCTGGTATCGACAAGACCCAGGGCTGGGCTTAAGGCAGATCTACTATTCAATGAATGTTGAGGTGACTGATAAGGGAGATGTTCCTGAAGGGTACAAAGTCTCTCGAAAAGAGAAGAGGAATTTCCCCCTGATCCTGGAGTCGCCCAGCCCCAACCAGACCTCTCTGTACTT**CTGTGCCAGCAGTTTATCcacagtgttgcacag**ccagctgctctctgcacaaaaacagagggtagctgcaagaac**aaggagactcctcctt**caggagacccctcaccgaccaacaggataaacttcctccatcatcccacagaagcccttccccacacgccagcctcagggactctgcaatgcagccactctgtggaaccaggggtccatcctctacctgctgtaccaatgaggtttcaaagttgggcaggacaagctcactgagttaggtctagtgccagggtacactctcccatcctttcttcttgtggagcttaaatctttaaattcaactaaactgttgcccaaactgaagactccatttacaatctaatgggtctcaacaatatttggttctgtattccaggtcctatttattctggaggttttgtcacttggaggaacaccatgtttggagttatttatgaaacccagcacctcctattttcttcctctcagcagatggtccaaatgccc**aagccagagatgcaagactgatcttcactctctctctgactt**tctccttccacaccaaatccactaccctatcctgcatttttctaagtactaaattcctcacaaattttactaacttttctaatttcactgatctcacctggatcactttagccaccttccaagtagccaccctacctgcaagcctgcacctgctcccaccca**aaaccactccctcaagcttaaaaccctttggttt**atttctattaactagctaaagtctcaagtttatgttagtttgcaagtctctcagcccctccatgatgtgacctttgcatatcccccacctttctaacactgacttcccttatccttcaactaccacaactgcctgcatctactccagaaacatcccatttctcgcacatccccaaatatgtccactgctgactcactc**ctgggtctttatctacaatggtcactctacccag**aataccatttctctctctggtattcatctgtt

Uppercase: TRBV27-2

Lowercase: Flanking sequence[1000bp]

Red & Bold & Underline: Stem-loop [18]

Blue: Heptamer[31]

Green: Nonamer [3]

id-TRAV18[V_gene_segment]

ccaggggcctaagacactagacacatcttcccacagactgtccttcatcaaagataattttgcccaccagatacagccattcattcattttttccaccaatttttaggggttttttttgagca**tttcctatgagctaaggttctaggaaa**agggctgctgatagtaataggattcaacttctgatgcaggagcactaagtgcagcttgagaaacaggacagaaatgattctaatatgatgtgataaggacagaaaatatggtgggaaaaaggagagcactaactcagctcgagagatggcagaatgttttcacatggaggtgatgcactggttttcaaagaatgaacaggagtcatccaagtgaaagggagtggggagaaggaagggcattctaagcagaagtgctcaccccccggttttgggggaccagtgggaaatatgtcg**agaaaaaaatctcacattttct**tttttttatat**tctcaggatatcctgaga**tgcctaaaactaaattaacacttgttgtgaacacattagatgcagggaatggggagattcaagggaatcatattgctctg**cgccttgaagaaggcg**acaggaatcacccacacgatgaagtcagcagagtgtttgatgctccttccttgagtcccacccttcacctcggacaccgttttctttctgagtctgcaggttaagccgcaccccagggtgaaaatgaacatcattctgagactcagagcaggcgtcagggacagagaggagcctcaagagggcgctgctgccttcagcttgtggtttgacacaaatttcatctttccttccttcgcacgtctctttgttccagcctgtggcctgatcttgtagggcagtttactctgcttaatacg**tgagtcaagattttgtagaagattccgccaaagactca**acaacataaagaaatatatatacctttcggtttggatatctctcaacaaaaccttctactgcttctcagccagccATGCTGTCTGCTTCCTGCTCAGGACTTGTGATCTTGTTGATATTCAG**TGAGTAATGTTTACTCA**AAACTTCTGAGATGTTCTATTCCTTAGCTTTAGGATGACCTTTAACCCCTTCCAAACTGGAGACCATGCCTCTTCTATTGCCACTATAATAAAGTCTTTCCCTTTTTTAGGAAGGACCAGTGGAGACTCGGTTACCCAGACAGAAGGCCCAGTTACCCTCCCTGAGAGGGCAGCTCTGACATTAAACTGCACTTATCAGTCCAGCTATTCAACTTTTCTATTCTGGTATGTCCAGTATCTAAACAAAGAGCCTGAGCTCCTCCTGAAAAGTTCA**GAAAACCAGGAGACGGACAGCAGAGGTTTTC**AGGCCAGTCCTATCAAGAGTGACAGTTCCTTCCACCTGGAGAAGCCCTCGGTGCAGCTGTCGG**ACTCTGCCGTGTACTACTGCGCTCTGAGAGAcagagt**gggagggactgcagcgagagcccagcacaaaccctggggaacgcaggtggggcctgggtgtgagccgctttgggagatgaatgaatatggactcttgttcgctgggaccccaaatg**gagaaaataatctcatatttctc**tgcatcaaagagcacaaaacagaaatgtgggcccctcacacatgggatgtagctcaccacgagaaacttttcctaatattctggagttctgtgagccttgtttgtcctttgtctct**cctgcatgattctgtgagttttacacataaaaatgcagg**ttttgcagggcctgatgc**ttctttgatttctggggctctctttaagaaaaagaaaattaaattataaatttaagtttct**aggttatgcggtagaaaggggccttaaaatcaaagccgaatatacttcatagtaaatctgcatttgaggttacatagagatgatcttctt**taagactgaaatattataatagtctta**gtttgtcctgatactgggaggttccctaactccaagtcctctttcccacggcagcactagaatcactttagctatcattaatatgagtattttcacttctatctttaaagaaataacagctttattgagatataatttgcatac**cttaaacttcattcttttaag**tctacaattcagtggttttagtatattcacagttatccaatcattattgtgatctaattccagaacattttcaccacctcataaggaaactctgtgcctactagcactcacccccattcttttcttaccctagc**cataaatcaaataaaaatattgatttatg**aggaataataacacaaaattttgatgtacccagtcttattgtaa**atacaggacctcttatgaacctgtat**ctagagggatgaaacataagtcccaacatttaagactgcaaagagtttaatgactgtgatgcaggggacaaaagccagtggagattgaactctgctggtattttaa

Uppercase: TRAV18

Lowercase: Flanking sequence[1000bp]

Red & Bold & Underline: Stem-loop [16]

Blue: Heptamer[20]

Green: Nonamer [5]

id-TRBV12-3[V_gene_segment]

ccagtaacacaagctagtctgcaaactaatgcccttttactttgggaagtgcttttgactgcaggggactcagaagcatgcctctg**tgccaacagcaaaaatgtgcccttctcttttgttggca**agtaacttaaccaaccaacccaaaaaaaagatctttc**tctcagctttccataatctctgaga**cgaagtaggtttggagaagtggggttacaggggaaaaagccaggtgttaatgatgaaaaaacattgaacttttctaggggtagtaat**aagatttaattcaagactagaacattttagctgcaaatctt**caagaataag**acaatattatcccctttctgttttattgt**gggactaga**gaatgtgagagaggttacattc**catgggctttgggaatttaatatggttcaaggataaacacacccaggtttttcactgcagagaagagcttcaaatataatcagttttcaggtcatcagctcagctcttgtatccctaacaatgcagttgacatgcgtcttctcagatgtctaactcctaactcactgagggatactttaagtacatataaaggactagaagcaccaagctaccagtgagaggaagaggagagtttgcagagaagctggcttgaaataagacaatgagttcatctttaaatacttgccatttgaggtgcagatggatatagttggcaggctcctatgtaaggcatgttatggagaagctaccgtgaattgataatatcaaaacaaatatccagggagcctctgcaagtgtgcatctctatttcacaccaattatagttgagttaattcctgcctgatt**catctcccagagatg**cagcctcctcttaaagaagttgggggtggtggcccat**tcagtgatgtcactga**cagatgcattctgtggggataaaatgtcacaaaattcatttctttgctcatgctcacagagggcctggtctagaatattccacatctgctctcactctgccATGGACTCCTGGACCTTCTGCTGTGTGTCCCTTTGCATCCTGGTAGCGAGTGAGTCTTCAGAATATTTGCC**ATCATCAGGCTGGGCTTCTGCATGGATGAT**CTCATATATTTTCCTTATTCTGACGCCCAATTCTGTCTTCTTTCATAGAGCATACAGATGCTGGAGTTATCCA**GTCACCCCGCCATGAGGTGAC**AGAGATGGGACAAGAAGTGACTCTGAGATGTAAACCAATTTCAGGCCACAACTCCCTTTTCTGGTACAGACAGACCATGATGCGGGGACTGGAGTTGCTCATTTACTTTAACAACAACGTTCCGATAGATGATTCAGGGATGCCCGAGGATCGATTCTCAGCTAAGATGCCTAATGCATCATTCTCCACTCTGAAGATCCAGCCCTCAGAACCCAGGGACTCA**GCTGTGTACTTCTGTGCCAGCAGTTTAGCcacagc**gctgcagaatcacccctttcctgtgcagaaaacccggtgtttccccttctccttctacctcc**cagcagtcctgggcaaagtctctgctg**ttcctccctccctatgagaaaaaagtggtttgggggtatgaaaaagacagaaaatgagaagggat**caacataggaaaccttatgttg**gtttgaggattacaaa**atgggttttgaggattccttaaaaattgtctctgctcaaaacacat**aggagtaagataaaccttggctactgacactggagatttccctgccctcctgcatttgccatcccatgagaatggtgggggctcttgagaagggctgcattttctgaactgtgaggccctcttcattctctcctaactctaagctgcaaacagaaatttccctcacacgttttctagattgta**aaagaaagttcttcttt**actatgattgtggacgttcctttataatgccaatttcaactttacattacttcaggatttttcactactcctaaagagtgtctcaaatgtggctagagcaagcaggttagtacactagatgtaagctacctggcctggaatctaaggatccatttgtctctgttctgcgtaagatgagccgggtgctggccaaaggctgtgcacactcacagagcactgatgacgcctcctggtaaggacccacactggggtatctaaaagcagacaggcatgtccagtcttctgttgccctgtttcctttctgattatatgtccttaacacacaaatttacattttccttcttatttatatgagaagtttctatacaatacctgcaatccattctgagtggttataatttctgtgtgatattcatatttacatgctgattccttctaaatacctatcatggtatcattgacaactgaggcaaaagacccctatattttgagtgcccaaggccattgaggttttttggagctctgccataagcccaattccactgtgtcattttcc

Uppercase: TRBV12-3

Lowercase: Flanking sequence[1000bp]

Red & Bold & Underline: Stem-loop [14]

Blue: Heptamer[34]

Green: Nonamer [4]

id-TRBV10-3[V_gene_segment]

aatccaggggattgctagtgcaagtcccggaatccaaaggtaggagaatctagaattctgatatccaagggcagtagaagacacatgtctcatctctgagagagaatgcaaattcaccctttctgtgcctttttgtttc**atccaggcccccagctgactggat**attgccatacacattgagagtagatctttcccacacagttcatcaatttacaagccagtctcctccagaaataccctcacaga**cataccaagaagtgttgctgtaccagctatccaggtatg**cc**ttaatctagtcagattaa**cacctaaaattaaccatcaaactaccagcgggaacacacagaggtctgagctctgatgaggaggctcctgtgaggtgaactaatgtgacagagcataaaacaatatacaaattgtattgat**cctgaccttaagactttttgaaaccttcccagatgtcagg**tggctcttttaacagaggccagcactgtaggctgacatccagttatccagagtttgagaaaagataaaatctagtaaaaggtacttagaggcaaatttgaggagatgagttataaaggacaccagactgccacgagaatgtcccttagactacataaatattagaaacaattaaaacagc**atacaaatgaggaattgtat**cctaatgggaataaaatatgcagagaggaatgcattttctttctaataacctgcaaagagcatatgcacttatgtctttaaaaatagtctacaaattattcccattctgagctgttaaattgtatatatgtcc**tataaaatattgttttatataccataaattacatatatgtatata**tttgatttatttttttggagaatgtaaggggtgtggcaggttaactttccagctcaactatttgtcaggggcactgatgtcatcgagtcactgagaacctaagttctatttcc**ccaggcagggctgggagagatgagatcctggcctgg**acctgaaATGGGCACAAGGTTGTTCTTCTATGTGGCCCTTTGTCT**CCTGTGGACAGGTGAGGGCTGGTCACAGG**TGGGCTTCCTTCCCTAGAAT**TCCCAAGGCCTCAATACAAGTCTTTTTCTTGGGA**TT**ACAACATCAGGGTCTGTTGT**TTTCTATTACAGGACACATGGATGCTGGAATCACCCAGAGCCCAAGACACAAGGTCACAGAGACAGGAACACCAGTGACTCTGAGATGTCACCAGACTGAGAACCACCGCTATATGTACTGGTATCGACAAGACCCGGGGCATGGGCTGAGGCTGATCCATTACTCATATGGTGTTAAAGATACTGACAAAGGAGAAGTCTCAGATGGCTATAGTGTCTCTAGATCAAAGACAGAGGATTTCCTCCTCACTCTGGAGTCCGCTACCAGCTCCCAGACATCTGTGTACTTCTGTGCCATCAGTGAGTCcacagtgctgcatggctgcctcctctctgcacgtaaacagcagttagaaagactgaggttgctctgtgtctatccccacccttggaag**tccaggcctccatagaagtcagagggccctggccagcctgga**agccatagagcaggggccttatgaccctcagtgctgacgtccattcctaccccagtctcagaccaactggaggtcaccccaacacacttagcttgccaagtctctc**ttctgcagctctcttttgctgcttgcagaa**aggaaaaggcatcattagttgaggttagcgatgattctcttaaccccaaagcctggactcccttctctcccctgtgggcttcagtgacttcttcatctgcccctttccccacgctccactatccttccactcatagtcatgaaccttcactcctgacctctgttcctggctgcctttcttctatgggccaatagcctcatggggccctctatcatttgctcttcacctacatctccatcatcattttgcacaagtctcccctggctctctctgctccagccatgctcctttatgttcagttatttcctcatatgtgttatgctgtttcccaccctggggacgtgtgtttttacaaattatttcttctacactgaacatatgttcctccctcttattcctaagtctctacttcatttcttacatgaaagtatatttcaggtacatttactcttaactataaacagaaaaacttaaat**ttcactaaaagaaattaaggaaaatatttttatcctttgagtgaa**gacttttttagtggaaacacaaaagaaca**catgccataaaaaataatactggatgggcatg**gtggctcacacctttaattccagcactttgggaggcctaggcgggtggatcacttga**gcccaggagttcaagaccagcctgggc**aacatggcaaaacctcgtctctactaaaaacacataaattagccgggggtagtggcgtttgcctgtaa

Uppercase: TRBV10-3

Lowercase: Flanking sequence[1000bp]

Red & Bold & Underline: Stem-loop [16]

Blue: Heptamer[33]

Green: Nonamer [3]

id-IGKV2-38[V_gene_segment]

cttctctgtcctttctgattatcgttagttttgttttgttttgttttgttttgtttttt**gagacagagtctctctctgtctc**cc**aggttgcagtgcagtggcgcgatctcggctcactgcaagct**ccgccccctgggttcacgctattctcctgcctcagcctcccaagtagctgggactacaggcgcctgccaccacgctcggctattttttttgtatttttaatagagacaaggtttcatcgtgttagccagaatggtctcgatttcctgacctcgtgatctacccg**ctttggcctcccaaag**tcctgggattacaggtgtgagccaccgcgcccagccatgatcattgttagttttaagtctgttgtatctgaaataagaatagcaacccctgctcttttttgctttccatttgtttgatagatctttctttctccctttactttgagcctatgtgtaatcattagaaaccacctgaaagaacaatgaccagtgccttgaataatatcttttaataaagggtcgcttgtacctag**taaatactaagcttattta**tttatttatttatttatttatttatttatttattttgttgcccagtttcgctcttgttgcccaggatggagcgcaatggtgcgatctcggctcaccaaaacctccgcctcccggattcaagtgattctcctgcctcagcctcccaagtagctgggattacaggcataagcccccaacccccaggtgattttatgtatttttagtagagatggggtttttccatgttggtcaggctgaactcccgacctcgggtgatc**cgcctgccgcggaccaagaagtgctgggattacaggcg**tgagccaccacaaccggcctaagctcatttttaaaaacgt**attttttatgatgtgaataaaaaat**cttttaaaagataa**atgataaatccatatcat**aagggaaatcgtgatacttgctcataatatttgtacataactttgtacttcATCTCC**TGCAGGTCTAGTCAGACCCTCCTGCA**TAGCAATGGAAACACCTATTTGTATTGGTAACTATAGAAGCCAGGCGAGTCTCCACAGCTCCTGATCTATGGGGTTTCCAATGGGGCCTCTGGAGTCCCAGACAGGTTCAATAGCAGTGGGTCAGGCACATATTTTAAACTCAAAATTAGCAGGGTGGAGGCTGAGGATATTCGACTTTATTAATACA**TGCAAGCTACATAATATCCTtgca**cagtggtaacgccttgaacacaaacctccttgctggggtgtcccaactgcccacat**gagctgctgtctgggagcagctc**tgcagggtct**ctgagtctgcggaagaggaggctgttggggaactcag**ggaaagggtttgctgctgaagactctggctcatgagagcct**cagctgtgcctcagtcacatatgttaaggtcccatcagctg**ccatggcctggggaagacatcatcaggaggagaattcttgggggccatgagccctgggaacaaccagctatgatga**agggagagtgaatgaaagctcatccctccct**atcttgcctacatttttcaaggacatttac**ttgtcaaaacaaccagaccgtggatgcagattagtgacaacacaagtggaatacgtgttg**gaaatgaccagcttggggtttattttatacaagatgatacttggtcctattgagaaatcatttttttcatttcattctcccaaccttgtatcttttttagcactcgtacactaaactg**ccctctccagtgtcatgctggagaagg**tattctacacaagctgtcctcaggggagtgtggcaaaaaataattattaaaaa**tgtttaggtttgactattcatagacttggtaaaca**catggcaaataaaagatctcaatgccaaaatacttcaattctcctcaattaccttcatcccagtgaaagttcttccagttacaaaagtgagctttgaaaataaacagaatgaactcacagatggacaacatagactcactaaatatgtcacatgaataatgcagaattacataatattttttcttcttaccccag**ttttaagaatttaaaa**agcatttaacctaagtactgttttagaggtgacaaatttagtggtgtgtccccatgagaaaatatttc**aaacaataccaaaattccttgttt**acatattacatcatcgttcctagaaaatatatctaaatcctctattttaggtata

Uppercase: IGKV2-38

Lowercase: Flanking sequence[1000bp]

Red & Bold & Underline: Stem-loop [19]

Blue: Heptamer[14]

Green: Nonamer [4]

id-TRAV2[V_gene_segment]

agccactcac**ctgaatgaagaacatcttttagtattcagaaggcttttccaaaaggcagttatttcctctggatagccttc**atattcctatggaaattgaataaggatgaaaaatttgaacaaaagcaaaggacatgaaatctcttgaagatatgagaagctgtgaacacttgtcttagagggtgcatggagaaattatttttcatgagttcccatgtagtccgctaggtggcggaggttacataaaaaccctactgtat**aaacactgaattgctacacacttgtgtttcctcttctttttcaagtaaaatcccaaggaaac**ctgtagagtataataaaagctggagatagaactttgtagtaagccctatattaataacaataacagtgtcaggttgcaaaatgaggataggtaccatac**caaaagagatgatctccaaatgcagtttttgtttcttttg**tttacag**aaaaattacaactgattttt**ttttccatatttgttccaaagatcaccttgaatgcaaggaatcaaattttgatcagtatatttacacagtaacttttgtaccttaaaaaattcatttcttatctactgcagctactaatatctcac**gctaattattttaaacacattagc**aggtcatcatgtcatatgtgacgggacttagcatactttttgtgtgactatatcactttaatttgtgtgatgaagcagtgccccattaattatgtgactccttctaatacccttagagaagaaaggaagctccttgaatactaaatttgcacaaccacagtgaccttataaaggatgcattcccttttaagattaatttctcagaacaaagcacagtgacgtcattactaagcagcctcacaactgtagcaacccttctaaaggttgtagattctggctgatgatgtcactgacacaaaggaaaaaatgcaaaacaggtagtcttaaataagcattctggtgagaccaactgcattttggccATGGCTTTGCAGAGCACTCTGGGGGCGGTGTGGCTAGGGCTTCTCCTCAACTCTCTCTGGAAGGGTGAGTAGCCTCCTTCACTATTTGGTTTTGTAAAAGTGTATGAAA**TTGTTTTAAAATTTGGACTCATTTTTAATACAA**AAACAAGAGAAGTCAGATCTAACATTCACTGTACCTAAGGGGCAATCAACAAGCCAAATAGTACTATTAAAGTGACAGCCCTTT**TTGTTTTCTGTAGTTGCAGAAAGCAA**GGACCAAGTGTTTCAGCCTT**CCACAGTGGCATCTTCAGAGGGAGCTGTGG**TGGAAATCTTCTGTAATCACTCTGTGTCCAATGCTTACAACTTCTTCTGGTACCTTCACTTCCCGGGATGTGCACCAAGACTCCTTGTTA**AAGGCTCAAAGCCTT**CTCAGCAGGGACGATACAACATGACCTATGAACGGTTCTCTTCATCGCTGCTCATCCTCCAGGTGCGGGAGGCAGATGCTGCTGTTTACTACTGTGCTGTGGAGGAcacagaggcagggaacccatgaagagctgaacagaaacagagatcacagcctttgc**aggaggcaaaacagagatgagcaataactttttcctcct**taattcagtattacccaagc**tttttcatttggtactgctattaggaaaaa**aaattgagtcc**agacatactatatattaacaaatgtct**at**aaaaatatgcttttaattttt**ttttgagacacagtcttgctctgtcatacaagttggagtgcggtggc**atgagcatagctcat**tgtaacctcaaactccagggctcaagggtagctgggactcagccttctgactaggtgagacta**taggcatgtaccaccatgccta**gtgaattttcctcctttttgtagacacgagttctctctgtgtcatccagactggtctcaaactcctggcttcaagtgattctcctgctttggcctccaaaagtgttgggatta**caggtgtgagccaatgcacctg**gaccagaaaatatgtttttaaagggtgattatttcaaatagtcttcctgatatctgaaacacttttgactcattttcttgagatctgattagattatttttgtttttccctcataaaatggctgatgatgaacttgtgatagtagtaagaggcatgtccactcttccattttcttctcttttttaaaaattcagacaggctggagt**gcagtggcatgatcatagctcactgc**ggccttgaacttctggcct**ccagtgatcctcctgccttgacctcccaaagcactgg**gattataggtgtgagccaccaaggccagtcaactctttcattttcttattgctgtctgtgcttaccttttaaatactatctgctaacttcagtctttttgcagcaatattttcaaaacatgtcaatttgtaaaggatggtttagttaacattgggtaatgtaataagaagtgagaaaggaaggaataaatgagtggctgatacctacagccctgcatttgaaat

Uppercase: TRAV2

Lowercase: Flanking sequence[1000bp]

Red & Bold & Underline: Stem-loop [20]

Blue: Heptamer[41]

Green: Nonamer [8]

id-TRBV11-1-2[V_gene_segment]

tcgcccaggctggag**tgcagtggcgcgatctcggctcactgca**agctccacctcccgggttcacgccattctcctgcctcagtctccagagtagctgggactata**ggcatgtgccgccatgcc**cggctaattttttgtatttttagtagagacggggtttcaccgtgttacccaggatggtctcgatctcctgacctcgtgatccacccaccttggcctcccaaagtgctgggactacaggcgtgagccaccatacccggccgatttt**ggatctcttactgtattcacaatgatattttcagggataagagagtcc**ccacccatcttataagatatcagaactttcattacattgatgaccaagaaccataggcagaac**catgatgaagacttggtacacaatgcatcatg**aagcttgataacagaaaagagaaagctggccattgaccttcaccaaaaatgaacccttccaactctctctcagggtattattttacccctttagct**tagagcatgtcagctggttcaacatgcttcta**ggataaacctgtgtcaactaccccagctcccactctgtctaagctcatagcaatgagaaaatggagaacctggtaagacagaaatgcccagagagaaggat**atacagataagacatagaggttactaattgttgtctgtat**tttgagttgcctttgacacattcaagtagaaacatgcagcagaagatggccctggagatgcagtgggtgatccaatcccccaagccctagagatgtgagtaaagacacttggatgaacttacacacagtccatgaatacgctaaagcaacttctagctctgttttagttcataacaactggtttctaagaacttaggcacttgtggagacaatgatgttactgtaggaactaccgtataagg**acaggatgtcccacctcctctgctcctgt**tcacaaggaccctgaactggcaaagctcccatcctgccctgaccctgccATGAGCACCAGGCTTCTCTGCTGGATGGCCCTCTGTCTCCTGGGGGCAGGTGTGTCCTAAGAACACCATGATCATCCCATTGGAAATTCCAGTGATTTCTTCAATCATTTCTGTCTTTCTGTTTTCAAA**TTCTGTCTTTTTCCCCAACAGAA**CTCTCAGAAGCTGAAGTTGCCCAGTCCCCCAGATATAAGATTACAGAG**AAAAGCCAGGCTGTGGCTTTT**TGGTGTGATCCTATTTCTGGCCATGCTACCCTTTACTGGTACCGGCAGATCCTGGGACAGGGCCCGGAGCTTCTGGTTCAATTTCAGGATGAGAGTGTAGTAGATGATTCACAGTTGCCTAAGGATCGATTTTCTGCAGAGAGGCTCAAAGGAGTAGACTCCACTCTCAAGATCCAGCCTGCAGAGCTTGGGGACTCGGCCATGTA**TCTCTGTGCCAGCAGCTTAGCcacagcgttgcagaga**ctttctgtcctgtgcacaaaactccagggctctctccgctctactcagctcacagcagcctttccttattcctcatcctctcagggaagaagtgagttttcagatatagctagg**gttcatatagtgggaggaaatgaac**tattttcttaaaacatgagcgctatagttgttggttgaaaacatgtcttagggatttgaaacacttctgggtgggaattcaagaaaccaaaactgaaagtgatttatcccagacctagtcctcaggggcagtgatggttgttcttctataaaaaatgcacacaatgttgtacgttaattaagaaaataaataaataaacttgaaaaataaaaacatttaatcaccaaaacatgtgtgttaaactttaataatatagatattgattacatgcaagtaacaaacagcaggccctccttctgccgttgtcagcatctgaatgatgtgaatcttatctggctctggagcctcttcttgacctctcactggatgtttcatgcagtcaaatctc**aacagacactgtgtgtttggaaatcttttccctctgtt**tttgtggaaatatg**acaataataataatattgtaatgatgatgatgttgatcatt**tcagctatttcattcagcacactttaaacttcaaagaaatgcacacaaatggttatcctggaaacaattcccaaatcttaactcaaaatgaattttgcttctctggagtgctttaacttttcagtgtaacagacacagcccaacactataattaggcctgggccagctagttcagaggctggtgaagacacttccacctgcctgcctacttccttgtctgcc**cactaaaatagctccttagtg**cattagatattcctgccagaacttcagaagcaatgagttgcccctggtccagttgtctcctctgataatgcactgtgtgtgtccaactctagtctctaccctaacttctgcccagattctc

Uppercase: TRBV11-1-2

Lowercase: Flanking sequence[1000bp]

Red & Bold & Underline: Stem-loop [15]

Blue: Heptamer[41]

Green: Nonamer [3]

id-TRGV10[V_gene_segment]

caacatggtgaaaccccgtctctacaaaaaatacaaaattagccaggcatggtggtgtgcacctgtattcccagttactcaagaggctggggatgaaaggattgcttg**agcctcagaggttgaggctgcagtgagccatgagtgtgccactgca**ctccagggtgggtgacagaacaagatcttgtctcaaaaaatatatgtaaagaaaaaattctccaatgacatgggaagttgctcagaatataagctgaactgaaagaatacagaac**actatacatgtaaatatatctatagt**ccataaaatcatattaatattcatgtatccaaaatgaaaatatctacaagtgagcatatgataatgtgaatagtagtgggcctttgtgatatgaccgtagatgtttttattgttatctttccccattgctctgagtttcttcaatgagcactgttaccagcatgttctacctatcagctgtgtgacttctgtggagttgatgactctgtgcctcattccctcatccaaaaaatgaaaattccaagtacccatattaagtgatcattgtgaagattagaaggtgaaaaacgtttagaaaaaaaaaaggc**tagcaaatagaaaatgctcaataagtatttgcta**ttataaaagttaatgttta**ttaaatatgcaacaatttaa**aatattatgctgaggaaaagtgttcacgaacaactcctaggcaattagaaatccttctcatttccacagaaattaagcaacccccatggccccctcacattcgtgcagagaacaccatcgagattgctagcctgatttgagataagcagggtgagcaggaggggggc**tcagtgctgagcctgcaggaggcactga**agggcccaaaaaaccattttctttctctccccagttcaggccacaccctccagaacctgaccagagtcgtctagccaccaggtccgcactt**tcctgccttctccttgggtacaccgccccgaagacaggA**TGTCACTGCTGGAAGCATTTGCCTTCTCCTCCTGGGCTCCTGAGTTGCTGTCCGAACATTTCAGTCTAGCTGCGTGACCAAAATCATTCTTCAATGTTTCCCTCTTCTACTGTCAGATGTTCATTGGAAATT**CTCCTCTTTTACTTACAGTTGGACTTGGATTATCAAAAGTGGAG**CAGTTCCAGCTATCCATTTCCACGGAAGTCAAGAAAAGTATTGACAT**ACCTTGCAAGATATCGAGCACAAGGT**TTGAAACAGATGTCATTCACTGGTACCGGCAGAAACCAA**ATCAGGCTTTGGAGCACCTGAT**CTATATTGTCTCAACAAAATCCGCAGCTCGACGCAGCATGGGTAAGACAAGCAACAAAGTGGAGGCAAGAAAGAATTCTCAAACTCTCACTTCAATCCTTACCATCAAGTCCGTAGAGAAAGAAGACATGGCCGTTTACTACTGTGCTGCGTGGGATTACaccatactagaactgttgaaacaacatgcacaaaatcccctcccagggtctgtgcccaccacatccttcccaacaggggcaaccacagccagtccccagctgggctcccagactcaggctcgccttctctgaggctcttccggggctttccaggaagctgttatgggttataccttcgttcccacctgatcccaggacttgtgctccagaatctttatttgacactgtttcttaatgcattaaagacttgtctggggactgagagaacataggtcgtgccactgtccatcatatcaccacctagagacttccagtaagtagcaaggtgcagttgagacagggatccttattacgtacagtcaagacaacccccctgcaatataactatcactggcccagtttacaatgaaagaaatgaaagcttagagaactgaggttgcatgatctgccagacaagtcatgtctttgagc**tggtctatgagacca**caccgaacagtttaaggattgcaattctctaagtaaatgctaggatatctgggataaattcgggaaggagtaagtgcccttttttctgggaggggtagggaggagaatttgcctgcatgagatcgtcagaatagagctccaag**tcatcccaggggccattggatga**gacccttcatgttggaataacctttggtcttacctgtaaaggaagagagaattgtatcgtgtgtgtgtgtgtgtgtgtgtgtgtgtgtgtgtaagttctctacagtgttctcaattctttgcatatattagctcctttaaccc**acactacctatagtgt**ctgctctgttgttctcatcaaacatatctaactcctgaaataagtggagtattagttcatgcacttgtccaaatcacagagctgataggtggcaggacagcctagggagggacccaggtaatctggttcctcttgaatatgacactaaactgcttctgaatcacaggagtataaaagaaaagacacatt

Uppercase: TRGV10

Lowercase: Flanking sequence[1000bp]

Red & Bold & Underline: Stem-loop [13]

Blue: Heptamer[28]

Green: Nonamer [4]

id-IGHV1-14[V_gene_segment]

tccagctctcattcttcttggttctctcctgcaggccagcagccctgaagtttagcctggatctcccatgcatccacccatctccttccaagtgcattttaccacagcctccactgtttttgaagcactcttgggctttgtaattctccacactctgttgtaaaggaagtcaggtccttcaagaccagattcgggactctattttatgaccaaatttcagcctcacccctgctcctgagacagag**ctcctagataagattctgcaggtggagattaggag**tgtttttcttcttcaatgtagttgctgagcgctcagtgcagggttggggagaaactttccactttgtcagcatgcagct**cctgctggggtagaccttcttccatagaagcagg**gttgggaaccagggggccaatgtcctcagtgctgctgcacccagggcagagccttcatccatcagtggggctgtggaagaagtgagtctctggtt**ctcagtagctcttgtccagaactgag**cctctgcagcatgttctgtcggccccagtgtcctggcccctaaggagcagcatcctaaaatgggagctagcgtatttgagaataacaacacctacacattcagaagctctttggctttctttccagctaatataatttccttttttt**tgtgtagcaacctgtacaca**cgcatattgatgcatacagacctatgacac**ttttttctcgataagtaaaaaa**ttattgatcactgtgatcttttctccaagttcaccatttccctgaaggtgagcacaggtccttctgcatgtgttcaaacaaaaggcccagagactacctggtaagtgaggtgctcacctggttctggatgtttggtctgtctcctcccctctgttgccccacacaaggtcagcccactctttccaggtccgaagaagagagcacaggtttgtcctgattatatgactcacccagcttctgatgactctcctgttgccagcgtccatgGCCTCAGT**GAAGGTGTCCTGCAAAGCTCTGGATACACCTTC**GCCAGCTACGACATTCACTGTGTGTGACAGGCCCCTGGATAAGGGTTTGAATGGATGGTAGGGAGCTACTCTGGCAATGGTAACACAGGCTATGCACAGAAGTTTCAGGGCAGAGTCACCATGACCAGGGACACGTCCACGAGCACAGCCTACATGGAGCTGAGCAGTCAGAGATCTGAGGACATAGATGTGTACT**ACTGTGCGAGACAcacagt**gtgaaaacccacatcctgagagagtcagaaatcctgagggaggtggcagcagtgctaggcttgagagatgacagggattttatttgctttaaaga**ctttttttagaaagcgaggttaattcattgcagaaaaaag**gaaaatagaaatg**tgtatggactctaattatgtgggaaattttccataca**act**tttgttctctaagcaaaattcagggagtggaaaacaaa**tcaaattaataaacctgataaaagaattcctctgaaaatttagtgtgagcataagtttttgaatgggtgttgtaaatattttggaacacagctgctagatcacattttaactctacacttatctccattatataaaatat**caaaatgttttaatgttttccattttg**tgcaattataatttt**gtgttataatcacattttaacac**tttaactctacacttatctccattatataaaatatcaaaatgttttaatgttttccatttcgtgcaattataattttgtgt**tcatgccagcaatgcatga**tagatcttgttcttccgcatcctcattgccatttggcactaagagtattgtgtattttaatattctaatagattagtagtgatagctcattgctgtttaaatgcacatatttc**taattaaaattttgtatttaatta**ttttatataattgtgatgaagtgtctcgtatggtatttggattattttttattgcattgtttctttttgatcagttgtaagtttc**cttatatacccattatataag**tcactcacaaagttaacaaaaaattgattaacaaatatgtgttttacaagtgtattctccaaattgttgttgtcg**ttttactcccatatcagtgtctgtgggagaaaaa**tatttatatacatatatatgtgtgtgtgc**atatatatatatatatatatatatatatatatatat**agtgtgtgtgtgtgtgtgtgtaaacttagataaaataattatttca

Uppercase: IGHV1-14

Lowercase: Flanking sequence[1000bp]

Red & Bold & Underline: Stem-loop [17]

Blue: Heptamer[36]

Green: Nonamer [7]

id-TRBV11-1[V_gene_segment]

tcgcccaggctggag**tgcagtggcgcgatctcggctcactgca**agctccacctcccgggttcacgccattctcctgcctcagtctccagagtagctgggactata**ggcatgtgccgccatgcc**cggctaattttttgtatttttagtagagacggggtttcaccgtgttacccaggatggtctcgatctcctgacctcgtgatccacccaccttggcctcccaaagtgctgggactacaggcgtgagccaccatacccggccgatttt**ggatctcttactgtattcacaatgatattttcagggataagagagtcc**ccacccatcttataagatatcagaactttcattacattgatgaccaagaaccataggcagaac**catgatgaagacttggtacacaatgcatcatg**aagcttgataacagaaaagagaaagctggccattgaccttcaccaaaaatgaacccttccaactctctctcagggtattattttacccctttagct**tagagcatgtcagctggttcaacatgcttcta**ggataaacctgtgtcaactaccccagctcccactctgtctaagctcatagcaatgagaaaatggagaacctggtaagacagaaatgcccagagagaaggat**atacagataagacatagaggttactaattgttgtctgtat**tttgagttgcctttgacacattcaagtagaaacatgcagcagaagatggccctggagatgcagtgggtgatccaatcccccaagccctagagatgtgagtaaagacacttggatgaacttacacacagtccatgaatacgctaaagcaacttctagctctgttttagttcataacaactggtttctaagaacttaggcacttgtggagacaatgatgttactgtaggaactaccgtataagg**acaggatgtcccacctcctctgctcctgt**tcacaaggaccctgaactggcaaagctcccatcctgccctgaccctgccATGAGCACCAGGCTTCTCTGCTGGATGGCCCTCTGTCTCCTGGGGGCAGGTGTGTCCTAAGAACACCATGATCATCCCATTGGAAATTCCAGTGATTTCTTCAATCATTTCTGTCTTTCTGTTTTCAAA**TTCTGTCTTTTTCCCCAACAGAA**CTCTCAGAAGCTGAAGTTGCCCAGTCCCCCAGATATAAGATTACAGAG**AAAAGCCAGGCTGTGGCTTTT**TGGTGTGATCCTATTTCTGGCCATGCTACCCTTTACTGGTACCGGCAGATCCTGGGACAGGGCCCGGAGCTTCTGGTTCAATTTCAGGATGAGAGTGTAGTAGATGATTCACAGTTGCCTAAGGATCGATTTTCTGCAGAGAGGCTCAAAGGAGTAGACTCCACTCTCAAGATCCAGCCTGCAGAGCTTGGGGACTCGGCCATGTA**TCTCTGTGCCAGCAGCTTAGCcacagcgttgcagaga**ctttctgtcctgtgcacaaaactccagggctctctccgctctactcagctcacagcagcctttccttattcctcatcctctcagggaagaagtgagttttcagatatagctagg**gttcatatagtgggaggaaatgaac**tattttcttaaaacatgagcgctatagttgttggttgaaaacatgtcttagggatttgaaacacttctgggtgggaattcaagaaaccaaaactgaaagtgatttatcccagacctagtcctcaggggcagtgatggttgttcttctataaaaaatgcacacaatgttgtacgttaattaagaaaataaataaataaacttgaaaaataaagacatttaatcaccaaaacatgtgtgttaaactttaataatatagatattgattacatgcaagtaacaaacagcaggccctccttctgccgttgtcagcatctgaatgatgtgaatcttatctggctctggagcctcttcttgacctctcactggatgtttcatgcagtcaaatctc**aacagacactgtgtgtttggaaatcttttccctctgtt**tttgtggaaatatg**acaataataataatattgtaatgatgatgatgttgatcatt**tcagctatttcattcagcacactttaaacttcaaagaaatgcacacaaatggttatcctggaaacaattcccaaatcttaactcaaaatgaattttgcttctctggagtgctttaacttttcagtgtaacagacacagcccaacactataattaggcctgggccagctagttcagaggctggtgaagacacttccacctgcctgcctacttccttgtctgcc**cactaaaatagctccttagtg**cattagatattcctgccagaacttcagaagcaatgagttgcccctggtccagttgtctcctctgataatgcactgtgtgtgtccaactctagtctctaccctaacttctgcccagattctc

Uppercase: TRBV11-1

Lowercase: Flanking sequence[1000bp]

Red & Bold & Underline: Stem-loop [15]

Blue: Heptamer[41]

Green: Nonamer [3]

id-TRBV16[V_gene_segment]

tgatggattggaaaaatggatgtttgagtgcataaggttcacgagtataagaaacaattggtacctcctaactatagattgtcagtttca**gaggatgagaatgaatgtgccctcatcctc**attttaattttgaaagacaatgcagatatccctactatagaatcattttttacatcaccacctatagagattaacaacagtagatgtttccagctccctaagaggccaagaagtattaacccaaggtcagattcaggtaattgatcgtgactcacgaattttcagagatccacaatggttgtttttaaattttcagcagttatgtctttttctctggggagagggtattcaaagctcctcatgaggccattctggactacttttcccctcctgccccctgcccataaaacagtttacagaaataggtgtccagaccgtgggctgtagttcaccaactcc**tgctctagcctaattttgtctcattactagagca**tggccattctgtggatgccagtgaagcctttctggggtttctctccaatgccttttaaagctttttcctacacagacacagtttactgtttagccaaaaaattaagataattc**ctccagacatctggag**ttcttgtgcagcttcctcctctctggtggtctttccttaaaaattctagctgcctcagcccccgctg**attttgtaacctgtttccgcaactcaaaat**ttgccatgctctgttgctctcttccccactgtgtatgtgcagaaattgcccccatgttttggtgttttggcagaatcccagtcctgtgctgtctgctttccagtggctgaatacaattgtttcacatattctcttaatttgtgttacttgtatagagcaggatgctaaaggcaattggattctacaaagtgatcacgtcacagagaagccgccgacagaggtggagagagccacacagatagccagctgcctgtgctgcctgctcttcccctaattctgccATGAGCCCAATATTC**ACCTGCATCACAATCCTTTGTCTGCTGGCTGCAGGT**AAGTCCCTGTTCTGCAGTTGTCAGCTCCCTGCTCTAAGCCTTTCATCCATGTCATCGAACTCCCTCATGGGCTCAGTCTCCAACTCCTGTCTGCTTTCTTTACAGGTTCTCCTGGTGAAGAAGTCGCCCAGACTCCAAAACATCTTGTCAGAGGGGAAGGACAGAAAGCAAAATTATATTGTGCCCCAATAAAAGGACACAGTTAT**GTTTTTTGGTACCAACAGGTCCTGAAAAAC**GAGTTCAAGTTCTTGATTTCCTTCCAGAATGAAAATGTCTTTGATGAAACAGGTATGCCCAAGGAAAGATTTTCAGCTAAGTGCCTCCCAAATTCACCCT**GTAGCCTTGAGATCCAGGCTAC**GAAGCTTGAGGATTCAGCAGTGTATTTTTGTGCCAGCAGCCAATCcacagtgttaaatattagctaatcttaggacacagactcatcacggactcagctcaggaagcaggtggtatactaggttggaaggaaataacagaaactagagctagcttaagccaaaggggaatgtattataaggctaaatgtatgtcccatagaaccacaaagcaagaacacaggaacttcccagaaataaactgcaatgaaaccttagaaccggtggctcgcgcctgtaatcccagagctttgggaggccgaggcgggaggatcacaa**agtcaggagatcaagaccatcctgact**aacacggtgaaaccctgtctctactaaaaaaaaaaaaaaaaaaaaatagccgggcctagtggtgggcgcctgtagtcccagctactcgctactcgggaggctgaggcaggagaatggcgtgaaccgggaggcggagct**tgcagtgagccgagatcgtgccactgca**ctccagcctggacgacagaacgagactccatctcaaaaaaacaaaaaacaaaaaaaaaccttagaaccatggcatggctgattgacaactcattttttatataccttgtcggcctcattcttttattataagttgtc**ttttctgtttcttaggtcttacagtgattagaaaa**ttgcagatttaggtgttagccctccagtcaattgagaccaacgtgactttctctcatgctagattacaaatccagggtgcagggggaaaaaatgtgatcctgctttactcaagaatgggctcgtggaaatccaagggaattgaattattttgaacaagatagtatctgggagtctgtccctctgaccttgtggaccaaggatttcaaggggtggaatccagtcagaacatccaataggcacccatcacaagtgagaagttggggataatacattttctggaatatagagaccttgggaaaagagaagagtttggagaggggatcaaaaccttattatttagaacaatatcagaccacgtaaaga

Uppercase: TRBV16

Lowercase: Flanking sequence[1000bp]

Red & Bold & Underline: Stem-loop [10]

Blue: Heptamer[25]

Green: Nonamer [4]

id-IGKV2D-14[V_gene_segment]

tgacttactctatactaggaagcaataataaaggtattggctaaattttc**atcattactaatagaggtgggatgagagtggaatgat**atatttaaa**tgctggggagaaaagtcagccacaaatgctatatccagca**aaacaatcatttaaacgtaaaagcaaggtagaaatgttccttcatacacaaagactaattcattatagcatatgtgccttgtggaaaattcaagagaaaatccttcagaataacagggaatgagagcagacagcaattcaaattgataagaaggaatcaagccctcaaaaaatgaaagaaataagaaagaagctacaaaaaatgtttacctgatatgaagattaagtacaggaaaaaaaacaattataatagatgtcacagtatcagttacattgattgtgttttgagcagcagttgctgactg**ggaaggtgcataatgaaagcagcaagagaattggaaaccttcc**atattttgatgtg**gatcatgacatgaaatgatc**tggatttcgacattcatttgcacccttaagatttgaaccttttgcactgtg**tgtttttccttaaataaaataaaataaaaca**aacctaggctcttttaaataagcctgaaccaagaaataagaatcaccaaatatttatgaaagctcaaccttattaaagagaagtgtaaaacatgcctgagaaagtcctaacgaactagaaattgggtagaggaaatagcaggaaacatctaaatatgtcccttctgccactcaggaaacagattttacctctcttgatatccttaaaa**acataaaatatatttatgt**cattttgctcacaagagaggtccccaccttctccttggctctttccaccccactgcacccacc**aggggatttgcatgctgtcccctagggaagaccttccct**tgtgagtctgagataaaagctcagctctaaccttgccttgactgatcaggactcctcagttcaccttcccacaGTGAGGCTCCCTG**CTCAGCTCCCTGGGCTGCTAATGCACTGGGTCCCTGCTGAG**AGCAGAAGGGAAATGAGGGAGGATGATGG**GGTGGGAGGGTGAACTCTGTGGGCCCTGCCACC**TCCCATGTGTGTTCTGTCCTCGTGTTAGATGTGTCTTGTCCTCCAGGATGGGGCATGTGATGTCTAGATCTGTGAGAGTGAGGAAGATTCCAGAAGGAGCAAGGACATGTACTTTAGTGAAAGCTGTGACACAGAAAGAGGGGGATGGGATAGGTGATTTCTAGAGGCTTATTTGTGCTTTGCAAATCTTGGTTCTTTTTAAACC**TGTATGTTTTGGGAGTATTAACCAAAATCATACA**CAAAAAATAATTGAGCAAAACATAAATAACAGACAGAAAATTATTAAAATGGCTCACAGTGTTTGTACATAACCTTGCACTTCTCTCTTATTATTTCAGGATCCAATGGAGATGCTGTGTGAACCAGCCTCCACTCTCCCTGCCCATCACCCCTGGAAAGCCAGCCTCTATCTCCTGTAGATCCAGTCATAGCCCCTTTCACAGTAATGGATACACCTATTTGAATTGGGACCTACAGAAGCCAGGCCAGCCTCCACGGCTCCTGATTGATTTGGTTTCCAACAGGGACCCTGGAGTCCCAGACAGGTTCAGTGGCAGTGGGTCGGGGACAGATTTCATGTTTAAAATAAGGAGGATGGATGCTGAGGATGTTGGGGTTTATTGCTGCCAGCAAAGTACACATTATTCTCCcacaatggtacagtcctgaacacaaacctccctgcttgctgtggcctagctgcccagatgtgttgtttctgtggagagcaggcactgtggattctcttagatgcctaaag**accaagatgttggagaactcagaggacttggt**ac**agctgagggctcatgaccataaatttctcagct**acacctcaggcaccacattttaagttcccatcagctgcagcagcc**tttgcatgacagagtctgcaaaaaggaggaggaacacctccccttt**gagcaatgagacac**aagagaagagaaagctcaatgaaagctcattctaagcctctcttcctt**ccctacattcattcatcaagtaaattcgttctgcgtaacaggcacctaattgagactgactactggcaacgcaaaactaatacattcttttgatttggtttagcagttactagagtatatgtatgttgataaggtttggtaatattaaaacagtttccctcccctcctccacctccccccaccacttttccgaaggcagacccctgaccaggacaccagcaggcactgtattttttcctacttttttcagggagtaccaaaatataactcttcacggtctagatttttaagttactagatatgttgctgcaaatgtatgcaaaact**aaagacctagtgccaaaagtcttt**ccacagtctcaaccacagcatgttgaagtaaaaagtagttttctgaactccaaagctggg**tttcaacaataaggaccaagtttgaaa**gaataaaggaacataaatatgccatcttaaacagaattacaagat**aatatgtaagccatatttttaccgcatgaaaaata**accatctgaagacttagagcagaggtcagtacgtgagagactgg**tgggttgcgtttgagatatgaaccca**gatgcaagttcttatgtatttcctgggaagaatcacagagttgagagacccctgcctggcattcaaggctgggccctgacgcc

Uppercase: IGKV2D-14

Lowercase: Flanking sequence[1000bp]

Red & Bold & Underline: Stem-loop [21]

Blue: Heptamer[29]

Green: Nonamer [5]

id-IGKV3-20-3[V_gene_segment]

tagtagtctcagaacagggagagacaga**ctgagtccctccactaactcag**cccaacacctgtggatgagtaggggatgcctgagatcctggggtggatgggaggtggggcactgatctgtcaa**tctgctttttcttcaaggatcaggcagcaga**gacccagaagcttcatgtctttgtaaggctcttccaagccaaca**cagataatttgacaaaacactgtatctg**catcccagacctcaccctgtcacagactgaagtggttgtttcatcctactaaggg**taaactataccagctatacagaataaaaagactggatagttta**gaggatcacccaagaaatagttctttgcctgcatggacaaaaccatcttctgtctttagggaaatggtatcactaccctgaggatttggagcccaggtctcacttatctgtgcagttgtgaaagtcctcacacccacagtgctgaggttaattgaatgttactcttttaatttctgcaaagaatgagacagcttctggaccctcaggaaagatcactaacaagtaaatacaagtatatcc**ggaagataaagttgtaatagactcttcc**tttcaacctgatccatcatgcatttagggagctgactgggcacaagttggagcagaaagagaaaaatgaaaccacagccttctattttgtttctaacagacttgtaccaaacattctgtggctcaatctaggtgatggtgagacaagaggacacaggggttaaattctgtggccgcaggggagaagttctaccctcagactgagccaacggccttttctggcctgatcacctgggcatgg**gctgctgagagcagaaaggggaggcagattgtctctgcagc**tgcaagcccagcacccgccccagctgctttgcatgtccctcccagccgccctgcag**tccagagcccatatcaatgcctgggtcagagctctgga**gaagagctgctcagttaggacccagagggaaccATGGAAACCCCAGCGCAGCTTCTCTTCCTCCTGCTACTCTGGCTCCCAGGTGAGGGGAACATGGGAT**GGTTTTGCATGTCAGTGAAAACC**CTCTCAAGTCCTGTTACCTGGCAACTCTGCTCAGTCAATACAATAATTAAAGCTCAATATAAAGCAATAATTCTGGCTCTTCTGGGAAGACAATGGGTTTGATTTAGATTACATGGGTGACTTTTCTGTTTTATTTCCAATCTCAGATACCACCGGAGAAATTGTGTTGACGCAGTCTCCAGGCACCCTGTCTTTGTCTCCAGGGGAAAGAGCCACCCTCTCCTGCAGGGCCAGTCAGAGTGTTAGCAGCAGCTACTT**AGCCTGGTACCAGCAGAAACCTGGCCAGGCT**CCCAGGCTCCTCATCTATGGTGCATCCAGCAGGGCCACTGGCA**TCCCAGACAGGTTCAGTGGCAGTGGGTCTGGGA**CAGACTTCACTCTCACCATCAGCAGACTGGAGCCTGAAGATTTTGCAGTGTATTACTGTCAGCAGTATGGTAGCTCACCTcccacagtgattcagcttg**aaacaaaaacctctgcaagaccttcattgttt**actagattataccagctgcttcctttacagatagctgctgcaatgacaactcaattttagcatctcttctctgcttgggcattttggggatcttaaaaaagtaatcccttgatatatttttgactctgattcctgcatttttcctcagaccaagatggacagccaggtttaagcacagtttcacagtaatgg**ccactggatcagatttacatcagtgg**atgtcagtaaaggtcccaaccagagccataaggcaacaacaatagcaacaaataatcaaaattggaaaagaa**gaattaaagctgtcataattc**actgatgaaggattgtgtgcagataaaattcaaatttgtctacagagaaactactaaaattgacatgagaaatagaaaatcattagattcaagatcaatttattaattcatagattcaaaaatcaatttcatttctgcataataaaaaatgctaaaaattaacattataaaacaaacacaccatttacaaaaacatcaaagtatcaattatttaaaaaaaatagactaaatacactgacgtctccagaata**ttattttgaaaaataa**aagaaaacctaagtaaat**agaaattcagttcaaagactgaatgtct**cagtactataaaaatgtcaattcttcaaagattaaaata**ttgattacatataagaaaaatcaa**aatcctaaagtatactccaatttaaattaagaagctaatctaaatattatatgggaatgtcaaggatgtagaatagccacagtgaacctgaagaaacaccaaaatgagaac**ttccagtgcctgaatacctggaa**tatagtgtgggtggcagtatggtgatggtgagatcagaagtttaaaaatttgcaaac**gtgcttatttttggaaataatcac**tacgcagatgtagccaaaccctcttcaactgtgccaccagaatctca

Uppercase: IGKV3-20-3

Lowercase: Flanking sequence[1000bp]

Red & Bold & Underline: Stem-loop [18]

Blue: Heptamer[19]

Green: Nonamer [7]

id-TRAV4[V_gene_segment]

actacaaaaaattagatatataatcctacacgactgccttctactagccatggtactctgcgctctgaatcagatgagaagaaagggaaaggaactataatttactgttgactacgtgccacagattgaaaccctgaaatacatattattgttcccattttacagatgaagaaaccaagactcagaggagtgagctgcctgaggtcacaaggtggcagagttgagatttagatttagatctgtctgacacagcctgtgccttttcttcttgttcaccttacctcctccttggttaagtgagtgattccagaccaaatgtcctatcctagctggtcagcagggactaatgagaggtttgttcctctgtttaatatctgttacacagggtctgagaagaaggtattagttgtggacttag**tgggattataaatgatccca**gggaatcaggagatggaatgtccagtccagactgcctcatttgctctttcttggaataggattaggaatccctgtttcttcttggctaaataggggtcagtggagcactgcagggcataaagtaacaagtcaggcaggttacacttagaccatgactgtcaatgagtccatgagtgtcagaccgactacagtacatgaccaccctctcgggctgtggaaagtaatgtaatggggaataggagcccaatgtaatctaccttctcatagctttgcctcatttggaccaa**atctttaagggattagacctctgcaaagat**ctaaaatgaaattcagg**ctgctgtgcttcagtaccagcag**ttttcactccctactcttcttggggctttagctgaactttgaggggagccctgctctgaggagct**gctgccctgctattggctgctgtactggggcagc**ctgagtgacagctgctggtgtgggccctggcag**ttgctgctgggctcattgcagctcagacacagcaa**aagagcctagaacctgggtcctagtttgcacctagaatATGAGGCAAGTGGCGAGAGTGATCGTGTTCCTGACCCTGAGTGAGTTATTTTGGGATGAAGAGGAATGGGATCTGGGCCTGATGATGCTGGAAAGGAATCTGGAACTTTGCCTGCTAGCAGTTGCTCTTTATCCAAGATGTAGAGGGATAGCTTCAGGGTTCCATTTTTCTCCAGGCAGCT**CCTGAGCATTTATGTGAGATGTGTCTCAGG**GCAGCAAAGATTTTCGGGGTTTCTAATCCCTAGGGTCTATCCATGGGAGGGAGATAGTAAACCTGACACTTCTGTGTATGCCTGGGATAAATATTTTTGTACTGAAACATAAATGAGAAGTAAATTGTATGTACATGTATATGGCAGGGAAGTAGAGCGTGGGGATGGATTTGACTGTGAGAGGGAACCGTGGGGCTACTGAGTGGGACCGTCTAACTTACCTTGCTTTCAGGATAGGGGCTAGGATTGTGTTTTACTCCCATAGGTACTTTGAGCCTTGCTAAGACCACCCAGCCCATCTCCATGGACTCATATGAAGGACAAGAAGTGAACATAACC**TGTAGCCACAACAACATTGCTACA**AATGATTATATCA**CGTGGTACCAACAGTTTCCCAGCCAAGGACCACG**ATTTATTATTCAAGGATACAAGACAAAAGTTACAAACGAAGTGGCCTCCCTGTTTATCCCTGCCGACAGAAAGTCCAGCACTCTGAGCCTGCCCCGGGTTTCCCTGAGCGA**CACTGCTGTGTACTACTGCCTCGTGGGTGACAcacagtg**agacagatgggcctgcacctgtgccgttttcctctgtggggtgggagtcacagcc**tagaaagaagtccaaaagtgctttcta**aaatttttattttcaaaaggtatta**gcaaatttatgtattcttctactatttgc**aaaatcaatct**tatttattttttaaata**ggtatttcacttatgtgatctaaaattaaaaaagtataaaagggcatacagcatgatgtcttccttccaccatggtccctgagtactcaattatcaaagtgaggtatctgtggtatttatactattgttgctaagttcactggtactcctcagtt**tgtgtgtatagatagacacaca**cta**catatactttatatg**cataccacgcacagtcaacatgggcaatatgcttcttagcttttgaaacatttagcctaagtatcaggttattatttatagctgcacacattttgaacatttctgtggccatttccaagtatctacatgctggtttccatgcgtcccctgagattagatctctcactctctctgttgcctgtctttctcttttcttttgtcccacttccctattgtggatggtatgtaatggaaaatggaggtgatggtagaggaggagatgggagagaatttg**tttctgatttgcattgtagactatttttgaaacatttaaatcagaaa**tggattctggatcattattagtttttaaagttgggt**tctttgaggctccaaaga**gagaagaataacagaaaatttgctgaaaatcctgcttgggctcctgaggcttcattaggatcatggggtatgtttttgtttgtttgttttttgtgggatgttgtctctgattttgatcacaaggaaagcaatatttgataatttttttattgagaatgagaggatcaggctatgtggcaggaggaggtttccatctagttcatctgagggtttggtgggggagtagtattgtggaaagaaaacaaacatttgtagagta

Uppercase: TRAV4

Lowercase: Flanking sequence[1000bp]

Red & Bold & Underline: Stem-loop [16]

Blue: Heptamer[39]

Green: Nonamer [7]

id-IGHVII-60-1[V_gene_segment]

ggcggggcttcctcatctgctcagtggtctccgtcctcgccagcacctcgctgtcaccagggctcctctttctttattatctgtggttctg**cttcctcacattcttgtgccaggaaagaaacgaggaag**acaaattttcgtctatagttgaagcttcaccaattact**aggaacttgcctacaagttcct**gcatgacccattataacttatcgattaaaaaatatatattctaatgcttctcaccatctcttgatttgtatcatcaac**tgaattgtaccctctttgaaattca**tatgatgaaacctt**aaattcaatggatctatattggaattt**taatgaaataattaaggttaaatgtggtcataattgtaagaccctaatgcaatagacgtgttgtctttataagaagaggaagagacaccagagacctctcacttttcacgtgcaggcagagaagaggccatgtggagacatag**tgcactagaaggtggcccagtgca**agccaggaagaagccgcgccaagaaccagccctgccagcacactaatcttcaacattcagactgcagaattttaagaaaatcaatatttgttgtttaagccacccactcctgttgtcttcttatgaagatcca**gacagactaataccacataactctgttagtgctgtc**ccctggatggagaattagcctcctgaggctgggcacatctctcagatttccacataaaacaggtaaaaaatagtagttctgatataaaaacttgtcatgtccctgttggccaatttctgggcaaggtcttttaaataagccaagtttgcggggaaatggagaccatatgtttgtgggactctaaccgtggaatctactgcattgctctagcagaatcaaaaattgaacaccagtgagctctttattctcattaaaatgtgtgtctttggaaatgttaacattctgtccagaatctgtgctcacaaacaactgtgtttcttagtgcactcttGGCCTGGTGGAGCCCTCGCAGACCCTCTCCCTCACCTGTGCTGTCTCTGGATTTTCCATCACAACCAGTGTTTCCTGCTGGAGCTGGATCCACGA**GTCCACATGGGAAGGACTGGAGTGGAC**CAGGCGCACACGTCATGAAGGGAGCAAAAATTCCCACCCACTCCTTATGAATCCAGTCACCATCTCCAAATTCGGGTCCA**AAAAACACTTGTTTTT**ACAGTGGAGCTA**TGTGAGCAACAAGCTCACA**GCCATGTTTTAAAGAAGAGAcagagtgaggggaccacggtgcgagctcacacccaaaccttcctggaggggtgcacaggacagcaggagtgccgatgatggaaggggg**tggtctggattccagggcacacgcaagacca**ttgtagaggcactcaggtcacccgggggctctcaggaaccatcgaggaaaatcaggacaccagaagagactcggtacagcagggggctcaggaccattgtggggactcagaaacagcaggttctaggctcagcctcagggcaggtgcagatggggtgaaaaggggctggatgaggggttttgtgtcaccatcatatttcaccactagacaccctccactacatgtgttctactgcatatgagt**gtatgattagaaaatcatac**ttctataaatatataaccatacgaaggtgtgtcaagttttcctcttcaacttctatcggccttgttcgtaaggactaattccctgtaattacttgaggacctcataaattgtggtcagttttgtaggtttcctctctttccttgtctccctttccccttattctctctcacacaaatggacataatccccactccacacgcaaagaaatctataactttcattacctgatgtattgaataaaattgattaatgtgcagctttcccatctttgttgttgttcatgctgctgtaagtataagaaccatgagatt**ctcagctgtgtacttctccaagctgag**tggcagctttgcttatcatactgagatggccaacaatccaaatgtcaatcatcagcttaactggtaaacaaattgtggaaaagtcattcattggcatactactgactgtgaccacaatatactaatgttggatacgctcaacagtatggttaaattc**tcatatggtgagtaaaatgagccaaacaaataaaagtacacacatatga**taccacttttatacattctcccatccaagtactaaccaggcccgaccctgcttagcttccgagatcagacgag

Uppercase: IGHVII-60-1

Lowercase: Flanking sequence[1000bp]

Red & Bold & Underline: Stem-loop [13]

Blue: Heptamer[28]

Green: Nonamer [3]

id-IGLVI-63[V_gene_segment]

aactgagggacagtag**gctgggcgcggtggctcacacctgtaatcccagc**actttggcaggccgaggcgggcagaacacgaggtcaggagatcgagaccatcctggctaacacggtgagaccctgtctctactaaaaatacaaaaaattagccggacatggttggggggcacctgtagtcccagctacttgggaggctgaggcaggagaatagcatgaacctgggaggaagaggt**tgcagtgagccgagattgtgccactgca**cttcagcctgggc**gacagagcaagactctgtc**tcaaaaacacaaaaacagacaaacaaacaaagaactgagggacagtatatccccagtggacaccgatccataatcagc**cccagtcactgcacagactggg**tgtatggtgtcctcagtg**cctgctgctcccgtgtaaatgaacagcagg**agatcctagggtattagtgaattatatattcatttccaacgtcacactcaagatgaagtctctacagtatttgaaattgaacttggacagcttcaac**tatttttgtttcaaaaaatattatatatacaaaaata**ccctatttctgaaggtaattggagtttattttacttccaca**ataatactaaacataaatggttagcgtattat**aaagtggtgtgtctaggcatggacagcagaaggcacttggagcctgttcctgagggtctgggggtgttcggagctgagaccagccacttcccctgtctggtgctctct**gttcagggctcacagctgtgatctcctcacccctgaac**ggctcacagcactactcatacttatccctatggcgccctcaggcagaagag**tctgatccagagctcagggtgggatcaga**aaagttcccatttgcatgaggagatcctccctgcctcagagtataaagagagaaaaggagagagttgggaaaagctttacttcaactgtggggcaaaaaggcaggactcaggacaatctcaagaATGGCCTGGTCCCATTTCCTGCTCACCCCCTTTATTCACTGCACAGGTGACTGGA**CATGGGGCCACGGAAGGGGCCCTGGGGAGCCCATG**GAGAGCTGCTTTCTCCTCTTATTTATAGACCCAGAATCACCATGTCTGTGTCTTTCCCACTTTCAGGGCCTTGGGTCCCATCTGCGCTGATTCAGTC**ACCCTCAGTCCCTGGGGCTCTGGGCCAGAGGGT**CACCATCTCCTGCA**CTGGAAGCAGCTCCACTTTGGTTATTGGTGTGATGTACACTGGTTCCAG**CAGCTCCAAGGAATGACCCCCAAACTCCTCATCTATAAAAACAATTGGTGACACCTGTGGGTCCCTGACAATTCTCTGGCTTCAAGTCTGGCAACTCCATTTTTGTGACCATCACTGTGCTA**CAGCCTGAAGATGAGGCTG**ATTATCACTGCCAATTCTACAAAAACAGCCTGAGTGCTTTcacagtgctccgggttgaagtaagtcagaccaaaacacacagtgtgcccagccatgaagctctcccatgcaccccctactctgcagctaagtcaatgt**gttctctcacttgtttgtcctaaaactggggcctgagaac**ttcttcagggtaatttttctagggaatgtgtcctcaccctctcaatatctgccccaaattcactctttggcaacaataccatgtgctcttttctcagaggta**gcagctggtactgaatgccagaggccagctgc**cctgtggtggagtacatatgggtgaatcagtcggctggggcttccacaacaacacaccacagactggagtcttcaccaacagatatgtactgtctcactgttctggagtctgaaaggaccagatgaaggtgccaccccgtttggttttctctttgccctctttcctttgcttccagatggctcttttctcactgtgtcct**cacatggccacccccaggtctgcatgtg**gtttttgtacacatttctt**atgaagtctttagtccttttggattagagactcattcctatgacttcat**tgaatcttaataatcttagtgtccagatctcccaatacagacaaattggtaacttagttataacataaaaaatcagggggccaggctcggtggctcacttctgttaatcccagcactttgggaggctgaggtgggtggatcacgaggtcaggagattgagaccatcctggctaacacggtgaaactccatctttactaaaaatacaaaaaat**tagctgagcatggtggtacatgcctgtagtcccagcta**ctcgggaggctgaggcaggagaatcactggaaccaggaggtag**aggctgcagtgagccaagatcacccactgcactccagcct**gtgtgacagagcaagactccatctcaaaaaaaaaaaaaaattcggggtggatgcaatataatttgtaacaaaggatcaggggtccagggacccagagtc

Uppercase: IGLVI-63

Lowercase: Flanking sequence[1000bp]

Red & Bold & Underline: Stem-loop [19]

Blue: Heptamer[41]

Green: Nonamer [6]

id-IGKV2-18[V_gene_segment]

gggcaacagagcaagactccatctcaaaaaaaaaaagaaaaacaaaaatctatattccatagttccaccttgacagttaatata**cacaaacttcattcagttacactcttaatatttgtg**ccctttactctggggattttgtcaactaaaatgaaactaatttaaaaacctagatgctaaaataaatatgaaaataataaataagaataactggccaggtgtggtggcacatgcctgtaatcccagcactttgggaggtggaggcaggaggatcacttgaagtcaagagttcgagaccggcctggccaacatatgaaaccccatctctactaaaaatacaaaaattaaccaggcatggtggtgcacgccagtaatcccagctagctcgggaggctgaggcaggaaaatcgctttaaccttggaggcggaggttgcagtgagccgagatcctgacactgtactccagtctgggt**gacagagtaagtgaaactctgtc**tcaaaatatatatatatatatttatacacaaaat**aatatatatatttatatatatttatatatatcaaaaatatatatatt**t**atacatatatacatatgtgtatatatatgtat**gtgtgtgtatatatataataaaat**taaataagaataaccaaatattta**tggaaactcagcctattaaagacaagtgtaaaatgtgtctgggaaagtgaaaacaatgtagaaatctcttagaggaaatagaagaacatgattaaatatgtccctttttccactcaggcagcacaaactttattaggttccctctaaacacagaggacacattatgtcactgtgctcacactagatgtccccatactctccttggctctttccaccccactgcacccacc**aggggatttgcatactgtcccct**agggaggaccttccattgtgagtctgagataaaagctcagctgtaaccttgccttgactgatcagaactcctcagctcaccttctcacaATAAGGGTCCCTGCTCAGCTCCTGGGGCTGCTAATGCTTTGGGTTCCTGGTAAGGACAGAAGGGAGATAAGGGAGGAGAATGGAGTGTGAGGGTGAGCTCTGGGGGTCCCACTGCCTGTCCACATGCACATCTTGACCTGCAAGAAAAGGTGTATA**AAGTTTAGAACTGCAAGAGTCAGAAAGAGAAGATTCCTCACTCTTATAGTTCTAAACTT**CATATCCTAGAAGGACAAAGGACTTGTGCTCTCATGAAGAGTGTCACACAGGAAGAGGATAATAGTGTAGGCGACTTCTTGAGTCTCTTTTGTGCCTTGTGAATG**TTGGTTCTTTTTATGCCTGGATGTTTAGGGGTATGAACCAA**AGTCACACAAAAAAATCACTTAGCATAAAGTAGATAACAGAAAGAATATCATTTCAATGGTTCCCAATATTTGTACATAACTTTGCACTTCTCTCACTATTTCAGGATCCA**GTGGAGATATTGTGATGACCCAGACTCCAC**CCTCCCTGCCCGTCAACCCTGGAGAGCCGGCCTCCATCTCT**TGCAGGTCTAGTCAGAGCCTCCTGCA**TAGTAATGGATATACCTATTTGCATTGGTACCTGCAGAAGCCAGGGCAGTCTCCACAGCTCCTGATTTATAGGGTTTCCAATCATCTTTCTGGGGTCCCAGACAGGTTTAGTGGCAGTGGGTCAGGTAG**TGATTTCACACTGAAAATCA**GCTGGGTGGAGGCTGAGGATGTTGGGGTTTATTACTGCATGCAAGCTACACAGTTTCCTaacacagtggtacaacc**ctgaacagaaacctcccttcttgctgtggttcag**ctgcccaaatgtgttgtttatctggaaagcagacactgtctattatcttgggagagtaaagaggaagatgatggagaattcaggagaatacattacagctgagggctcttgaccatcaatatctccgttacatctcaggtaccacaatttagtccccatcagt**tgcaagagtcttggcctggcagaacttgca**ggagaatggagggaggtcaagtgccctctgaacaaccagtcatgttgcaagggaaagctgaatcaaagctcatcgtaatcctctctgccttgcctacgttcattcactaactaaa**ttcattcagcctgacagccacctaaatgaa**acagattagtggcaata**caaaactaatacgttcttggttttg**atttggtttagcagttactagtgtacatgtaccttgacaagatttggtgatactaaaacagtttctctcccatcttctaccttccactttacacttttcttaacatatatctctctctgggacagcagg**aagcacagcattttatcctatttttttttcagggagtgctt**gcgtgtatttacattctagactttgtaactgctagatatattttggtaagtttatgcaaaaa**caaagacctagtgacaaaaatctttg**aacaatcttattcacagtatgttgaaatgaaaagaaaattctatgaaatccaaagctgtgttttaaaaataaatatccaagtttcaaaagacaaaagaaaaaacgactaaaaatgtatgccatagtcaaaaatgtatgccatattcaatggaatctggatt**aatataaaataatatataagcttatattttgccccttgaaaaata**ataatctatagactttgggcagagggcagtaaatgagaatgtgagggcagctgttaactgtgttagtggtggtgtgttggagttgatatatgaacccaagtggaa

Uppercase: IGKV2-18

Lowercase: Flanking sequence[1000bp]

Red & Bold & Underline: Stem-loop [19]

Blue: Heptamer[30]

Green: Nonamer [12]

id-TRBV21OR9-2[V_gene_segment]

gtgtgtcaaaagctactatctgtaaagaccgagttgtatgattccttacctgagatggaagggaggaggggaagaagcaaattgacaagttaaaaaagtaaacagagtgtaaattagttcaaccatgtggaagacagtgtggcgattcctcaaggatctagaaccagaaataccatttg**acccagcaatcccattactgggt**atatacccaaaggattataaatcattctactataaagacacatgcacatgtatgtttactgcagcactgttcacaatagcaaagacttggaaccaacccaaatgcccatcagtgatacactggataaagaaaatgtggcacatatacaccatggaatactatgcagccattaaaaagaatgagtt**catgtcctttgcagggacatg**gatgaagctggaaaccatcattctcagcaaactaacacaagaacagaaaaccaaacactgca**tgttctcactcataagtgggagttgaacaatgagaaca**catggatacagggagggcaacatcatacacggggcctgttgggggatggagggctaggggagggatag**cattaggagaaatacctaatg**tagatgatgggttgatgggtgcagcaaaccaccatggcatgtgtatacctaatgtaacaaaactgcacattctgcacatgtaccccagaacttaaagtataataat**tttttttaataaatgtatgagtgattttacttaaaaaaa**tagcaaaacagtgagtcacagaaaaaaaaaactaaaagcaggctgaaacttacactatttttaacagctaaaacaaagcctgccctcctacattctctgaaattgaaatgattctg**tcccatgtacagagcatctacaaaggctatggga**agtgatcacatcacaagcaggttgctgacgggggcggggggagcgggcggagcgggtgaggaagcctt**acagaaaaagctaccaccacgatttctgtctgagccaaccAGGTGCCTCAGAC**TTCTCTGCTGTGTGGCCCTTTCTTTCTGGGGAGCAGGTAAGTCCCTGTTCT**GAACTGGTGAATTCCAGTTC**CAAGACTCTCTCCTGTGGCTGCAGTATCAGGCTCCCTCCTGGGCTTTTTTTCCTTCCTCCACAGCCTCCATGGACACCAAGGTCACCCAGAGACCTAGA**TTTCTGGTCAAAGCAAATGAACAGAAA**GCAAAGATGGACTGTGTTCCTATAAAAAGACATAGTTATGTTTACTGGTATCATAAGACGCTGGAAGAAGAGCTCAAGTTTTTTATTTA**CTTTCAGAATGAAGAAATTATTCAGAAAG**CAGAAATAATCAATGAGCGATTTTCAGCCCAATGCCCCCAAAACTCACCCTGTACCTTGGAGATCCAGTCCACGGAGTCAGGAGACACAGCACGGTATTTCTGTGCCAACAGCAAAGCcacagtgccgaatgttagcccttcttagaacacaaactcattatggacccagctcaggaaataagtgtatgtcaggttggtacacactataataacagaaagccaacttgaaagacaataaatcagaaggaaaaacttgtaaataacagctctaagtgagcaaaaaacaaagggggaggggtacttaaaacagctgatggggacaaaaccccagggaaggaagcagaaggcataaaaagctcggattcctccgtgttcaggatattagttaagatatactaagttaccttctgtctctcttttcatatgagctttagggatttctaacaatctttatttaaagaatgggattctgtgcttttcacttggtaaagctgtgagttcagaaataagaataagcaaatcaactgtgtctttcaggtagaccctgatcatctgacccagcacatttccttgaactagagacagactctaaagggctttttgactgcttctcacagctcttcctgccatagaagtttagtaggaaccattagcaattcccaccctctggaatgttacaatcacaaaaatttttaaagcacc**tttattgagacataagatacatgtagaatacgtaggcaataaa**catatgcaaccgatgcaacaaaacccaccctcattgtaagtgtatacttcaacaaattttaacacatttttatacccatgtaacc**atgatgacaatcaagatacaaaacattttcatcat**ctcaaaaagtcccccacatctctttattgaggtctcgttaatttctctcagcattgctttgtagtttttattatacagatctcatacatattttcctaaa**ttttattttaataaattataaatgtactcaataaaa**atattctctgccaagatttcacatcagcctgactcagcctaat**tcaatttcatctttacatttttcaattga**catctctacctcagcatattacataaaaataccttctccagaaa

Uppercase: TRBV21OR9-2

Lowercase: Flanking sequence[1000bp]

Red & Bold & Underline: Stem-loop [15]

Blue: Heptamer[36]

Green: Nonamer [7]

id-TRAV23DV6[V_gene_segment]

gtgcttctcatgttattttaacttatatgacttataatacagatcatccctatttcaggggtgctttagaatttatcttcaatttctcccttgtatcttgcctgtgttaatatcaaaagttaaatttattctttatattttcatagttgatcgattttga**agtgtgaataaaatatcgaatgtttattcaaact**caggataagagccaccaaaaatgactgcattacggatggttatagatctatatcttctgcagaatttttctatcctctctttccatttaaagatcacagctcctccagaacagataattccagtatttcctattttctttttc**taggtacatttaaaaaaatatttaatttaccta**atgtgtaatttagaaaaaaatatggagaaaaaataaaattgacttaattgtattatatcatttctgaggatcatctaaaatatggaatgaatacatctgaaaaattgcaacaaaatttaagttctctgaggacaaagggaccatatattctctatgttctgattcatgaccttagcttgt**aatggggtattatgcccattgtgtgccctcaataaactttattaatactcacaat**aaaaaagaaatgccactgtacatgttaatcgtgcggtttcacaagagttcctctcttctaccattgtaaaataaggccctttctattgacatatccagccatcttaaaggtctgaggaaaagttgagggggcttgacagacagaaattctaaactgatgcttatctgtgtgtaaagaaaggattactgattcccaatgaatatatcttcagcaattctaaatttggacaaagtgggg**aagtgcttcctttgacagagacagctttaagtgaaagcactt**gtgaaagggcgggg**cctgctgaaagaattcagttgagggtgaatttacagagtttcagctgg**ttgggaagactggaagaccacctgggctgtcattgagctctggtgccaggaggaATGGACAAGATCTTAGGAGCATCATTTTTAGTTCTGTGGCTTCAACTATGCTGTGAGTTATGGAAGAGAAACTCCAGGATATGTTACTAACTTGGGTGAATGAGAAAATTGGGGTACAGACAAACTCATGCAGGTAGACACTTTAAGAAGAAAATATTATTTTGGGGCCTAATGACATTTTGTTATCTCTCTAAGCAGGGGTGAGTGGCCAACAGAAGGAGAAAAGTGACCAGCAGCAGGTGAAACAAAGTCCTCAATCTTTGATAGTCCAGAAAGGAGGGATTTCAATTATAAACTGTGCTTATGAGAACACTGCGTTTGACTA**CTTTCCATGGTACCAACAATTCCCTGGGAAAG**GCCCTGCATTATTGATAGCCATACGTCCAGATGTGAGTGAAAAGAAA**GAAGGAAGATTCACAATCTCCTTC**AATAAAAGTGCCAAGCAGTTCTCATCGCATATCATGGATTCCCAGCCTGGAGACTCAGCCACCTACTTCTGTGCAGCAAGCAcacagtgctccccaggcacctgaagcctgtacccaaacctgcagttgaggttccagccaaaccccacagtgggagcttacgtaggcagagatgtagcctagttttcatctgcatatgcaagtgataaataaatgtataaacactagtgtggcagtatgagaatggtaattaaagaaaagaaacttagcttaataaatgaccatctctaacaaatgttttattatagttacagtatttgaaattgttacaagcatccag**ccagttttggggcaactgg**gaattttatgccttgctgatgggat**gcaaatcagtattatgacactggaaaatcatttgc**cagtttcttataaattgctcttaacataggaccagaaattccactcctaagt**atttatctataataaat**gaaaatatatcttcacacaagactt**ttgtttacagcagttttaaccataatagccccaagctggaaacaa**cccaaatgtccattagctgg**ggaatagaaaaataaattgtggactattcc**tataatggaatactacagagcaataaaatgcaataagctagtgatacaggtaacacatgacggatctcagaactttatgcttcattgagtgaatggttaaatatactgtggcacatacccacataatggaatatgatt**catcaataaaaaacatgaattattgatg**catgcaacaactttgatgaatctcaagggcattatgttgagtgaacaaaagtcaatctcaaaggc**tatataatgtgtgatttcccttatata**acactatcaatatgtaaaaattacaggacagtggttgccagaagttatgggtggggagtgtgtatatagggagtttatttagggtgatggatggaatagttctgtatcctgattgtagttgttagggaaatctatacatatgataaaatgtcacagaagtatatatcaaaaaaagagtgcatataaaaatgggtgaatccaatagcaaagacttgg

Uppercase: TRAV23DV6

Lowercase: Flanking sequence[1000bp]

Red & Bold & Underline: Stem-loop [15]

Blue: Heptamer[25]

Green: Nonamer [4]

id-TRBV3-2[V_gene_segment]

tgttgacttttgtattttgtcaaagttaaaggagatattttgatcagagtcagagccacaagtaactagcaggaaactgagtgggatggaggttatgccatggtccttgcttccttgcagagctaagaaaccagttgtgagcaccagcgaaaagagacaaagtcagatgaaactgaaatcttcatggga**ctctgtgcccaagaaaaccagcttgacaggattgggcacaagag**ctattccttccatcctacgctgccagagcacacagcacagtttcccccatgagctcttggggagctggaggagggagatagctgtgctgggcggagtcgtattcccatgggaagtccaaaaagtagaggcagggatggaggttttccttctagaaagagtggaagattttgtttacaaacgtacccatacacatgcctttctcccgaagacccaaactttggggaattgcctaaatataacaaggaggttaaatattggacagccaaaaataggaccgaagccaatctcttctgaagtaggtgtagtcactgtgcattagtcagatattcaaatgagcaaagattcaagaggaaaaaaaaaagcattattggaaaaggaagcaaaagatcaagagaagaaaaccagcctgactctgagaggagagcccaaaagggagaggaagggaggaggtcgtcagatcccgggtagggagcagggagggagggggtacaggcacaggaatgactgtcagagacacatcctggaggccacaagtccactctctcaactattcccagaaattccaaaaactcctttaaagaagccagtgcagggcatgaaagggaggagccatgctagaggagatcctgggaatgggggaatgattacag**gctgtgacctcaatggcacagc**acctctcagag**gcagtggaaaccacagcctagtcttctcaccactgcagaccagaatcctgccctgggccttgcctggtctgc**ctgactcggccATGGGC**TGCAGGCTCCTCTGCTATGTGGCCCTCTGCCTCCTGCA**AGCAGGTGAGTCCTGGGCCCAAGTGACAAGATCCTGTTGGAGTCCCCGAGCCTTTTCACCATGACAACAA**CAGGAGGCTACCTCCTG**GGATTTGCCTGAATTCTGCTTCTTTCCTTTACAGATCCACTGGACACAGCCG**TTTCCCAGACTCCAAAATACCTGGTCACACAGATGGGAAA**AAAGGAGTCTCTTAAATGAGAACAAAATCTGGGCCATAATGCTATGTATTGGT**ATAAACAGGACTCTAAGAAATTTCTGAAGACAATGTTTAT**CTACAGTAACAAGGAGCCAATTTTAAATGAAACAGTTCCAAATCGCTTCTCACCTGACTCTCCAGACAAAGTTCATTTAAATCTTCACATCAATTCCCTGGAGCTTGGTGACTCT**GCTGTGTATTTCTGTGCCAGCAGCCAAGAcacagc**cttacagagccactgcatccct**gtgcacaaacctcccggctcagccaggaagctgtgggccgtgtgtgcac**ctgcacccaaggctccagt**ctccattccctgatggcctctgatggag**tttcagtctgtagtacagcctctgccagtgcacctgatgtggcaattctccattttatgtac**acagaactctcagaattctgt**ttatcagctaaagcaggggttcacaacaaagtcagcaaagtatcagagctcaactaacaaaacaggagcctg**tcccatgcactgaaggtctttactgatggga**gaagccatttcttgtctttgtctgctgcacatatgggtccatattttactaatgttatgattataaacacactgtaggcaaaaatggttgatgagcctaagcataaagaagaggggaaggaaagtatttctaagatcaagagacaatttggattatcagtgagtattttgtttgacagagaaatattttgagaggcgtgaaattccatggcaataggtaaactggaaagtgttgaggagaagtcagtggaggcagccttgtgcattgtcggct**ccttctcttcaagaagg**gtgaccagcaagagtttctcccaattctgatgagtgtca**tcttgaactctcctcaaga**gcaggtgacagctggatgacaggggatattgggagacatggaaaggaattttgacacaaatccctagg**cagtggtgagccactg**gtcctactccaagtgaaacaacatgatgaaactgatattttggtacaggaacactggtagtagtatggatcagaaaagagatagatttttgggtggcatgaatgattaggagacttctaaataccctccatgaaagactagaaatcgtttatgtaaagcattgtggacgacataggcttgaggctctggattatgttatctcttggaaagagtgttgggttttattctgggacggttgaatgtaggacttgctaagtt

Uppercase: TRBV3-2

Lowercase: Flanking sequence[1000bp]

Red & Bold & Underline: Stem-loop [16]

Blue: Heptamer[40]

Green: Nonamer [4]

id-TRAV21[V_gene_segment]

tataaaatgctgtggttcaattgaataagagacagaagcttttgcaactcagaaagtttaacccaccttatacttttttgcaagtgtagtaagtaactca**caatacagagctaccataatgtattg**actatctatattgaaagaatatttaaatttttttccacttttttaaccattgcatgcaatgtacagtgctgtaagtgtatttataagagtatttcttaaga**aaaattcctggccaaatggaaatttt**gcattcagta**atatgtaaataacatat**gtagacatttattaattatatttcagctatcagaatatcagagtatccatttcctgacattatcaacaaaactgaatattaatagtatgattctgaactccttttgcaatatgacaagtaagagatgatagaaagggccttttcacaaaaatgcaagttacgggaaaccagtccttaagcactgaagtggcaagaagag**acagcagtcagaggaatctggagaaagttgctgt**aacagtatggaaaacatttatatattttttcttttctaacataagcatttaagaatatatatttctctctatgcact**gatttggctgcattctacaaatc**tcatattcaaacatttttgtttctttaggagaaggaggaggaggtgaaagtggag**gagaaagaagcatttctc**tagcagtactgtgcttggttctgaagtggcagtgttttcatttctttcattccatttccacctcccacattttgcagatctgtgtgtgtgtgtgagagagagagagagagagaggagagagagaggagagagagagagagagagagagagagagagagagagagacttgtctcagattggctgggccaactctgtgtttcctgtagaaatgtttgaacatatgagagagcaaagcagagtgtttatcttgtgagccattctccatatttcagatataagatttcagttctcagtgagtctaagtgacagaaggaATGGAGACCCTCTTGGGCCTGCTTATCCTTTGGCTGCAGCTGCAATGTGAGTTAGAGGGAAATGGAAATGAGCAAAGAGCAGGCTGACTCCAGGACG**TTTCTCCCAGGAAAGAGGGAGAAA**GAAAGGGAGTTGTTATGCCTGTATGGTTTACTGGG**AATTTCTGTGGAAATT**GAAAAAATAAATTTAAAGGTAATTTCTAAGGAATCATTAGCCACTAACCAGTCTATTGCTCTTTTTCTGAACAGGGGTGAGCAGCAAACAGGAGGTGACGCAGATTCCTGCAGCTCTGAGTGTCCCAGAAGGAGAAAACTTGGTTCTCAACTGCAGTTTCACTGATAGCGCTATTTACAACCTCCAGTGGTTTAGGCAGGACCCTGGGAAAGGTCTCACA**TCTCTGTTGCTTATTCAGTCAAGTCAGAGA**GAGCAAACAAGTGGAAGACTTAATGCCTCGCTGGATAAATCATCAGGACGTAGTACTTTATACATTGCAGCTTCTCAGCCTGGTGACTCAGCCACCTACCT**CTGTGCTGTGAGGcacag**tgcacaacaggcacctgcaaccaatacccaaactctatagctggggctctaactgcatgttttatcttgagactgagcaatgtttttgcattaagaggacttctaaattgacactgtctccacacagaagcaatcaaatacacaaaacttgatgtaataatctgaaattaaactatgaattctctgatctttagatttttatcaattttaatat**aaataatagttttattt**caattaatttacatgccatgaagttcagtcttttaaagtgaacaattaagtgttatttagtgtattcacaatgttgtgcagcttttatctctgtgtagtttcaaaaactttttaacatctcaaaaggaaaccacatgtccattaagcagttaccttgcccaccctcctcccagcctctggcagccaccaacctgttttcagtctctatggattttcctattctgtacatttcatataaatggaataatatagtaggtagtggcctttgtgtatggcttctctcacatggcataattttctgaggtttattcaatgtaggaaggaccagagaaagctaaatatgcacatctaaccaatgacatgcttatatcttccccatgcaagagcctccaaccagggcatacct**gaagctttgttagttttcttccactctaaagcttc**cctactcctctgcctatctttaagtctctgccaaacacaagggacggtagctgattcctttggtataccaagctctgaataaattgcctttgcttttttcatttaggtggtcttcacttatatccacagaactattgtaaagacactgagtttttcttccctgccaagttctcacttcaccataagagtcactttgcacatgaaaaattaatgaatcaccataagaaatgtta**aagctggagggcagctt**gtcctccattcagataggtcattgaagcctacatcaatcagatagtacttcttagataagtcagagagatgca

Uppercase: TRAV21

Lowercase: Flanking sequence[1000bp]

Red & Bold & Underline: Stem-loop [13]

Blue: Heptamer[34]

Green: Nonamer [3]

id-IGHVII-1-1-2[V_gene_segment]

ttacac**ttaagaaatttcttaa**tagttttttcctccttaaggtgaactacagtcagataacatacttatcaattgtctctagctcttgtcagaaaaacatatagatgtgtgtgtgcgtgtgtcttggcctttccaatgatgaattaagatgtgcattgagaaggcattcactttatttgacgttaaggaagtaccaagaagacgctctccacagaccctgggaaagccagcagctgcaccccgaggctgtgccaggcagggaacaaggaggcagcaccacctgctg**ggcagggaaaatgtcctcccagtccctgcc**gcttctctgcagaggcacaaagagctgccccttctcctgggccttctcctgggctgatgagattgctccccgatatgccaaatcagggttgtgcatctgaggctctgtctagactctcagctccttcctactcctgcaaagtgaagaaaacaatgcca**aggggtcctggaggcgtctctacccct**g**gagagttttgactctc**ttcaatagtctccactaccctgccctcactccatgtcctccgtttctccctaaagcggtgcccagtctgattgcactgtggcagggataacgaggggccaggacatcaggggagagaagtttctacctgagtcacagcagcggctgccctgcagactcctgaagacacaagaca**catttccatcccagagacccagcgaaatg**caacctcaaactagagacagccagttattttttcttgttctgtcctggagaggccactgagaaagtcgagccccttgttgaggaaaacatgagatctctgtgtgtcgtcctctgcctgatggctgtacctccatgtgagtgtctcagagatttcagaacgggggctgtgggctgtggtgtccgcttgtgactcatctctttgcttcttgtccctgagtgtc**ctgcatcagatgcag**ctactggagtcatgccc**agggctggtgaggtcctcacagacctCTGGGCCTGGACCCAGCAGCCCT**CTGGGAAGGCGCTGGGGCACCTCAGCTCCAGGGGCAGCACACACTTCA**GCCCAGCCTTTCTGGGC**CAACTCTCCATCTGTAGAGACACATCCAAGGCCCAGTTAT**CCCTGCAGCTGAGCTCCGTGATGGCCAAGGGCAGGG**CCGCACATTCCCGTGGGAGACAGAATGGGGACCTCAGCGTGAGCCCAGACACAAACCTCCCTGCAGGGAAGCACAAGACCACCAGGCGGCGCTCCAGACCACACAGCGGCCCCAGAAGCAGGTTTTAGGGGGCGGGGCAGACGTGTCCGCGTTGAGTCAGGTCATTGGTTTTACTTTCCCTGAGCAAACGGCCTCTGCCAAGGACTCACTGCACCTCTCACCTTCACAGTTGTTTTTTTTTTTTTTTTTTAATCACCCTGTAGAGTTTTGCTAGCTAATTTAGATATTGAggagtgcttcatacttccttgggcctctgcttgcagaaacatagcaattgtaaggaggcacgtgggaaag**ccccggctcggtgacccgggg**gatgctgctgtggccctggcaagagggcgtcgggccgcagtaacaaaggtgcagacggctctcagcctgcgcccgcggagtacaacacataagggctgtaacctaacgaaaaaagaatcgcagtgcaactgtcctgcatttgagtttgtgatcagttttgccctttgtctttaacaggttctaacataaaattttgaatgttggttcaagccc**tgtgggtaaaatgcacttacccaca**ttccttaaacaaatagaacactgaggtggaaatgttt**tgaaaaagtagttttca**gacatttggaaacaagcatcacaggatcataacccctgagaaaagaaaaacaaatgaacgaatcctgctattgcctgaaagcagctgccaggacacacggaaaggcttagtgagctgagcggacagagagcagagttcaag**gcagcagcagcccgaggggaggagcaccggggagcaggctgctgtgc**agccaggatgggccggggtggggcggggggagaacagctggagacttgccgcagggagggggatccctcaggtttggggctgagaactgacttatgcctgacttatgcctgcatgaaaagaaactactcgatatcagggggaaatcaccagaaacctgtggacccaaaactacacagagcct**acacaaggaaagcattgtttgtgt**tctcccagccagggtggaaagaccttgagatatgtaaagcttcaagcaatcttccgaagtaatctcgtgagtagtggtgccacattaattcaggactaaaggctgctctgaactgaacctaagaaatgcttcaagtgtagcctggagcccgggtgcagtggctcacacctgtaatcccagcactgtgggaggccgaggcaggcggatcacttgaggtcaggac

Uppercase: IGHVII-1-1-2

Lowercase: Flanking sequence[1000bp]

Red & Bold & Underline: Stem-loop [14]

Blue: Heptamer[39]

Green: Nonamer [5]

id-IGHV3-62[V_gene_segment]

tcaagtaattctgtttagtgaaagaaactaaggagttaagagtaaattttatatgatacatttgtaggaattttagaagatgccattattgtaaattaacacggagaagatttgagtttgtctgacaatacgctgttgggagtaatgtggatgtgagttgaatttcagagaaataaaagaaagatttagggattaatttaattattcaaaacttgattgaagtgctgagcaaatggctgcaaacataggtcaacattt**ttcaaatccttcactataaatttgaattaattacttattaatt**acacttgaataaagcaataacgaagaaaccaataaaataacatttgactaaaatggagcaataaacagatcgatgttaacacaaggaatatgactgacttatgaaaacatgcacatgaaccatggttcactctacgtattttggtaaattacagaaagttgtcataacagatggggaatcctgcagacttcactaggcatggtccacgctgcc**ctggagttgtctcaggggagctgcctcctccag**tggttagagcacaggcccaggtaataggactaatttttttagatgtgtaattttagacacactgcacaactgctgtgttctctgtgtaaattatctcctgtaaaatgtaacattgaaacctgcattaa**acatattgtataaatatgt**aagaataaaataagattatgagagctaaatattaatcaaggcacaagcacat**aatataaaattatatt**ttcctgaatgatacaattattccaatctgtcccaggacactt**catctgccctgagcccagcctctcctcagatgtcccaccacagagcttgctatatagtgggggacat**gcgaatagggccctccctctgctgatgaaaaccagcccagctgaccctgcagctctgggagaggagcccagcactggaagtcggcggtgtttccattccgtgatcagcactgaacacagaggactcaccATGGAGTTTGGGCTGAGCTGGGTTTTCCTTGTTGCTATTTTAAGAGGTGATTCATAGATAAATAGAGATGTTGAGTGGGAGTGGACATGAGTGAG**AGAAACAGTGGATGTGTGTGGCAGTTTCT**GACCTTGGTGTCTTTGTGTTTGCAGGTGTCCAGTGTGAGGTGCAGCTGGTGGAGTCTGGGGAAGGCTTGGTCCAGCCTGGGGGGTCCCTGAGACTCTCCTGTGCAGCCTCTGGATTCACCTTCAGTAGCTCTGCTATGCACTGGGTCCGCCAGGCTCCAAGAAAGGGTTTGTAGTGGGTCTCAGTTATTAGTACAAGTGGTGATACCGTACTCTACACAGACTCTGTGAAGGGCCGATTCACCATCTCCAGAGACAATGCCCAGAATTCACTGTATCTGCAAATGA**ACAGCCTGAGAGCCGACGACATGGCTGT**GTATTACTGTGTGAAAGAcgcagtgagaagtcagtgtgagcccagacacaaacc**tccctgcagggtacctgggacaaccaggga**aag**cctgggacactgtatactgggctgtccccagg**ggcaagtccaggtggtataagcctaggtttcctgtcatggtctagggttgcctttttagcaacttaccccagagaacttctctagatttccaattctgtaata**acatttgatgtcgtctctggctgcaaaatgt**cccctcaactttgtatctttttttttttttttgtaacaggaggacacatcctcaccctgcagaagcctgagtgtcacattggg**ggcagaaatgatctgcc**ttgatctcactgatcactgtcccgaggaacataccccacaggggaccctgatgactccagtaatgtctctgcctcaaaaccattgaagagtccttccttccattagagttgaccacagcacctgggcttcagcacaagccataccacagacgtcacaaagcagcagattga**cacctgatccaggtg**cattttctcacctttagaggctgagagaggggtgtattctcaaggtcaacacactctttgtggatttttacagagaacacctacttttactttatt**tcatttgaagataaatgaacaaatgtgcagctgcaaatga**ttgtaattttcacatttattccaattccaatatttctgaattctgcatgatgtcctggcacagggttgtgttttctacaggtatttttcatc**aaaccaagaaaatgcattttccactttccgtggttt**ttctccatgtgcagaggccttgagtagagcacgtctgactgtgtttgttttcacgctgctgataaagagaccttgagtagatcacgtctgactgtgtttgtctgttttcacgtggctgataaagacataccagagactgggaaacttacaacagaaagagttttattggacttacagttccacgtggctggaaaggcctcacaatcatggtggaa

Uppercase: IGHV3-62

Lowercase: Flanking sequence[1000bp]

Red & Bold & Underline: Stem-loop [16]

Blue: Heptamer[49]

Green: Nonamer [8]

id-IGKV3-15[V_gene_segment]

gagttagacagggatgggtgggaatgattgaaggtgtgagtactttagcacagattaagaccaaatcattaggatttaaagagttgtgtagagttagtgaaggaaaagccttagaattaaatttggctgtggataaaacattcttggattagactgaagactcttttctgtgctaagtaagtatatttatgataatgatgatgactgtagtgctgaatatttaataaataaaaacaaaattaattgccgcatacataatgtcctgaatactattgtaaatgttttatcttattttctttaaactgtctacagcactataaggtaggtaccagtattgtcacagttacacagatatggaaaccgagacacagggaagttaagttacttgatcaatttcaagcaatcggcaagccatggagcatctatgtcagggctgccaggacatgtgactgtaaacagaagtttttcactttttaactcaaa**gagggtatgtggctgggttaatggaaagcttcaggaccctc**agaaaacattactaacaagcaaatgaaaggtgtatctggaagattaagttttaacagactcttcatttccatcgatccaataatgcacttagggagatgactgggcatattgaggataggaagagagaagtgaaaacacagctttttatattgttcttaacaggcttgtgccaaaca**tcttctgggtggatttaggtgattgaggagaaga**aagacacaggagcgaaattc**tctgagcacaagggaggagttctacactcaga**ctgagccaacagacttttctggcctgacaaccagggcggcgcaggatgctcagtgcagagaggaag**aagcaggtggtctttgcagctgaaagctcagctcccaccccagctgctt**tgcatgtccctcccagctgccctacctt**ccagagcccatatcaatgcctgggtcagagctctgg**ggaggaactgctcagttaggacccagacggaaccATGGAAGCCCCAGCGCAGCTTCTCTTCCTCCTGCTACTCTGGCTCCCAGGTGAGGGGAATAT**GAGGTGGTTTTGCACATCAGTGAAAACTCCTGCCACCTC**TGCTCAGCAAGAAATATAATT**AAAATTCAATGTAGATCAACAATTTT**GGCTCTACTTAAAGACAGTGGGTTTGATTTTGATTACATGAGTGCATTTCTGTTTTATTTCCAATTTCAGATACCA**CTGGAGAAATAGTGATGACGCAGTCTCCAG**CCACCCTGTCTGTGTCTCCAGGGGAAAGAGCCACCCTCTCCTGCAGGGCCAGTCAGAGTGTTAGCAGCAACTT**AGCCTGGTACCAGCAGAAACCTGGCCAGGCT**CCCAGGCTCCTCATCTATGGTGCATCCACCAGG**GCCACTGGTATCCCAGCCAGGTTCAGTGGC**AGTGGGTCTGGGACAGAGTTCACTCTCACCATCAGCAGCCTGCAGTCTGAAGATTTTGCAGT**TTATTACTGTCAGCAGTATAATAA**CTGGCCTcccacagtgattcaacatgaaacaaaaacctcaagaagaccatcagtgtttactagattataccagctgcttcctttacagacagctagtgtgg**tggccactcagttttagcatctctgctctatttggccattttggagctcaagttctcaagtccaaaat**tacttatgttagtccattacatcataccatttcagtgtggctattacattcatttaaacgcatttcagaaggcatctctgtttatggcatcacaaagagtttaataaatctgtgcaagaataaacaacaaacacacctataaatataaagttgaaatatcaaaacta**tttcagcactctgaaa**attggcaaagcataaaataattaaggatgcatattctttatagaaaaaaaaagtactagtgctttgagtaaggacagaaaatgtctgtagccttttgcctgtgacagcacccttctattcccagctcagtcaatatgaattgcagaactgg**agttttacccatgtaaggatagcaaataaaactggcagcttgctgcca**aagtgggtggacttgagtgaagccaaggagtgggaaataattcttcagtgtttccagctaaacagggagaacaccataggaaatgaacagaaaaagcccacagccttgctagtccaagatgatgccttgtttgaggcaagtagtac**atctgctgatagtaaatagcagat**tcctggataagatagacccacattgctgaaataatctctgcacatattcctagtgacctagaagttatagatatgagtgatgagaactaggagtttccggtgcaaagtaaaaccagagggaggtaagaactag**ctgcattttgcatgcag**tttgctttttaaactacacacagatgggttgacagaagatagatttataacctccagatatttgagcaaaatgtctcagatcattggtgaccacttcgctatgcagatacatgtgtagttcctataaaatcaaact

Uppercase: IGKV3-15

Lowercase: Flanking sequence[1000bp]

Red & Bold & Underline: Stem-loop [18]

Blue: Heptamer[31]

Green: Nonamer [5]

id-IGKV2-36[V_gene_segment]

gaccccaagtctctagcc**acatccagttcttcacagaagctggatgt**atgtgccagggcaaggagaactgggtgggaggtgagt**ctgtgggctccacag**cttcacgtgtttatgaggtaggagttattaagaaattattttaggcagatagagaggaaaaaagggatccttgggagacttttcacagctgttgtctagc**ataaaagccctgggtcttagccccgggcaggcaaccttttat**atgcaaatgcaagccgttagaaactgggtccatccaacatgcacccaacatggtgatttccaccgttgtcctcttgcccttgccccaacatgcgcctggcagcatggttgcccctacatatccccacgtgtgtggaacatcatggcgccctgtatttgcatattaaaaggctagggtgggagggccagttctttcgagggctaccgtgaatgacatgtctggtcaaaccaataccctgagccctgtgcaaatcagacaccgcctcctccagacatcaaatatagctggctgatattacctgaaagagtggttcgtccctcggctttggagcccccctccctctgtctttgtacaggggagcttcatccttctttcttcccccttctttcttgcctattaaactccctgctccttaaaactattctaagtgtgtcagtgtcattttttctaatttgactcgagacaaagaacctggtgtcctctactcatcagagcag**tatcatttattccaacatgata**gagtcacatggtctatgcccagggaaaaga**attcatatttttgtcttatgaat**aatcaaggttcacctccagggaacaataaacagtactctgattaagatcttgaaaaaaaaaaagagttcccttctggctggt**aaataatgggtttattt**tagaaagtctactttcatgacatgaatcaaaacttgaaaaatgtaactataaatcaatatcataaaagaaatcatgaaagttgttcatAATGGAAAGCCAGCCTTTGTCTCCTGCAGGTCTAGTTAGAGCCTCCTGTATAATAATGGAAACACCTATTTGTATTGGTACCTGCAGAAGCCAGGCCAGTCTCCACAGCTCCTGATTTATGAGGTTTCCAACCAAGCCTCCGAATTCTCAGACAGGTTCAGGGGTAATGGGTCAGG**TACTGAGTTTACACTGAAAGTCAGTA**GGACGGAGACTAAGGATGTTGGAGTTTATTAGaaaacttgaacttccatcaatgataaatattccttttg**cctcaagcacatatttgagg**aattttccattgagtagatctaccgataaggtcacatttttctgtctgttttaatctgaatatgtatttatatgat**tctcagaaatgttttttgactatgaaattatatttgtgaga**ccttttttcaagttgttgtttaccatttgataattactgctgaaacgtcatctattaaaaaaaatcggtcatccgttctctaactctcattgaaaggaatttgcctttttaaacccctcagactctttttaagcttttcaattggccttatcttttttttacagattcaatgtattaagtc**aatttattatttatgataaatt**tatttatttattt**atttttactatcacaaaatttaaaaat**cctgtaagttgctatgtcaaacctgcctctagattgcaaacaaaccccacaaaaaagagagtaccattatcttagaaatactagaaaagcaagagtgatagtgtcccattgttaacagggaccccataaggctagaataatgtcccacatgttacctggggcacctgggaggagctgccactgtgctgaatggtgggaaaaagccatctgtgctctgagactggaagcctggccctattcccctatgtggatgtaggagataattggggttataaaggggtaattacggttaaatgaagtcataagggtggggctctgatccaataggattag**tgtccttataaggaca**gaaacc**agagaactcttctct**tttcctttatttctctaactcactgcttccctatggaaaggccatgagaagacatggtgagaaggtgactacctgtaaactaggaag**aaggttctaaccagaaagtgaacctt**g**ctggaacttgatcttggactttccag**gcttgtgaactatgagaataaatttctctggtttaatctacccagcccagggtattttggtgtagcagtgca

Uppercase: IGKV2-36

Lowercase: Flanking sequence[1000bp]

Red & Bold & Underline: Stem-loop [15]

Blue: Heptamer[19]

Green: Nonamer [5]

id-IGHV3-75[V_gene_segment]

atacacacaacaaactaaaatatctaggtatttatgctgagtaaaataagccaaaaaaataagtatatatgctatattattccatttctataaattctggaaaatgaaaactaacctacaataatttaatgaagaacagttgttgcctgaggaaatggtagaagaagggaaagggataggaggaaatgtggaacaggagaaaaaattgaggcgaattcacttgtccactttc**ttgataatggtgatagttatgtcatgtttatcaa**tg**tatactttaagtata**tgaagtttattatctctcaattagaccttataaag**tttattacaagcaaacaaataaa**aaattagacaaaaagaactgatagaaaatg**ggtaacatatgttaaatgccagttacctgaaaatttatctgcgtgaaccttagttctctccataattttaggt**gaacgctggagtgtaacagagtcacaatattttcattaaggcagtaagttctgcaagccacactagacatgtcccgctctgtcctggagttagttcagggaacacagt**gatgaggaacacgggctcagataccggaactcactcatc**tcagacatgagcacttagacacatacataggcactctgtgtgtgggtttactttcacatccgtacatggagaaacattgagtctcacagacataatttacacacatatttaaaatataataaggtgtttga**tgctgtttatcacagca**caatttcctcgtaaaagtgcaaatcttctaaataccataagttaccaaactcctgcaggacaccatcatcctctctcggcaatgccttctcctcacgtgtcccaccctagagttttttatgtagttggaaatatgcaaaggaccctccttctttggtgaaaaccagccctgccctcaccctacagttctgggatgggagacccagccctggaatttcaaggttttt**ccacatggtgattagcactgaacagagaactcaccATGGGG**TTTGTGCTGACTTGGGATTTACTTGTTACTATTTTAAAAGTAGATTCAGGGTGAACTAGAG**ATTTTGAGTGTGAGTGGATAACAGTTAAGGAAACACTGAAAAT**ATGTGCCAGTTTCTGACCAGGATGCCCCTGTGTTTTTGGGTGTCCAGTGTGAGGCGCAGTTGTTGGAGCCTGGGGGAGACTTGGTACAGCCTGGCGCGGTGGGGGTGGGGGTCCCTGAGACTGTCCTGTGCAGCCTCTGGATTTGCCTTCAGTAGCCACTGGATGCACTGGGCCTGCCAGG**CTCCAGGGAACAGACTGGAG**TGGGTCTCATGTATTAGTACTGATGGGAGTAGCATACGCTATGCACA**TTCTGGGAAGTTCTGATTCCCCATCTCCAGAA**ACAATGCCAAGAACACACTCTTATCTGGAAATGAACAGTTTGAGAGCTGAGTACACAGCCACGTATTACTGTGCTGGAGAcgcagtgaagggaagtcaggttgagcccagacacaaacctctctgcagaaacacatgaagtaaatcagctacgggggcgctcaggacccactgatcagattcaacctcagaggcagatgcagatacagggtaggagctggttttctcatagtgtttgtaacttccctctccaactgacagtttcccagagaacctcactaatttagagttctgtgcctaccaaaaaaatctccaaatatacaccttttaaaaggtagtaatataaaggcctagtctcacatgcacaaaatgcagatcaccacttccaggcatgagaagttctcaaccacggtcaccaagatcagagt**cctgaggaagctcagg**ggtgccttgtgggtcttctccaatcacattcgggacaggaaactaag**tttaattccctgattaaa**aaaggctataattccctgatttaaaacgtttcattccctgattaaaaaa**ggctggaatttttctgattccagcc**aagaaagaggctgggccagagtcagtgtcacgtacaagctcacaggtttcacgtctgccccttctc**ctgatatgagactatgcaaagcagtatcag**cactcatctgctgcactttttgctgtcaatccattttctttcttttcagctgttgtcactttttcgt**tttcttttctgctccctgaaaaagaaa**gatgtggtccatgcggtctaaattccaaggctcaagctcttccgctggagctcaggtggggctcaggctgtggctcctgcagccgcatgggagaggctgatggggctttcttctt**tccccagtgcttgggaccctccaccgtgtttcatgggga**ctcatgtggaaatttgagtggcccaggggaactgaagggcataagcttgtttggtgagagtgaaacgtgactgtggaattgattctttgttgtgcaaaccagaagaaa**ttttgaattttgtcagaatcaaaa**cggagtcacttttgttaaaact

Uppercase: IGHV3-75

Lowercase: Flanking sequence[1000bp]

Red & Bold & Underline: Stem-loop [18]

Blue: Heptamer[35]

Green: Nonamer [4]

id-TRBVB[V_gene_segment]

aggttgctgcagac**cagcagggattcactgctg**ataactcaagataccttccaccagtaacttggagatgacaccaagagacaacacaggcaaaggaaagtgtattaattctgttttaggtgtagaaactggagtgattggaaaataattcctaaaagaaattgaaaagtcaaattgtcaagcagtaagttcaccatcacatttaccataaagcatgtagagaaatggaaatatttacacatcatggttgagagtgcaggatttgcagtcagacagtttctggcacttagtagctttagaaagtgacttaagccctctaagccctcagttttcacatctataaaatgaagtaaataatataagcctctcagaagtggggtgagggttgaatgaaacaatatgtgagtagtttaacacagagactggcactgacttaagaaagtattgtgataaaagcagcagtggagaaaatattttgaatagctgtatgcatggcaatcctttaacctctttattca**tggcaactcatttaattgcca**caacaaatatgtaaggtaagtacttttattatttccattgttatggactgagttgtgctaccttcttcatttcatatgttgatgtcctaagtcccagtg**ccccataatgtgactgactatatttggagatgggg**tctttacatcgataattaagttaaaattaggtc**attagggtgggccctaat**tcaatatgactg**gtgtccttatcaggaggttaggacac**agacatgcaaagagggaagaccagaaagaaggtatagaaaaaaaaaaattgttatctacaagccaaggagagaggcctcagaagaaaccaactcaacaaattttaaaagatgtattttggagaacaaaactgttagaatatatttagaacagaaaaaaaccatttcttctagaactaagttggtctaaaagtgcatatcttggtaaggctctgtgaggaaatgacctaatctggagagAAGCTGCTGGGACACTCATCTGTGTGTGG**CCTGTGGCTCACAGG**GGCAGGTATTCTCAGAACGCACATGGGATAATCACCATCCTGGTGTGTCCAGGCTCTGGGAACAGGCTCTTCCTCGTGGCTTTCACATGAGTTTCCTGGTGGGCTATGTAGTCACTTTTCTGCATCCTTCTTTTAAAGGCTCATGAATGTCAAAGTAACACAGACCCTGAGATGAGGCAGGAAAGTTGTATCGGAATGTTTTCAGACTATCAACCAGACCAAACGTTCTGGAATCCATAAGATCCAGGGCTGGGGATAAATTTAATCTATTAATTATTCTTCTGAAGTTGGCAGCTTTGAAAAAGATGCCCTGAGTCAGCATAGTCATTCCAACCAAAAAGAAAAAA**GGGAAAAAATAATCATCTTTCCC**AACCCTTAAATCGTCCAGCACTAACCAGACTCTGAGACCCTCTGCAGCAGCAGCCTATCAGTGCAGCCACATCCTCTCTGAGCGGATATGACAAACCCCAGGGGTGAAGCGACCTAACCTATGAGCCgccacacacactcaagatgccccagacaccctgcactccgatcttactcgttcctttactgttttcatcctaattgccctcttacacatttgaccacacatttttggtcttggtgg**ttgtattagttttctgttgctactatacaa**attgccacccatttagtggcttaaaacaatgcaaatgtatcaccttacagttccgtagttcagaaaccaacataggtatc**acccagccaaaatcaaggtgtcggcagatctgggt**tcctttctgaaggctccaggggagaatcgattttcttaccttatgggttgttttcctctatcttcaaaggcagcacgttctgttgctctgaccctacttgcgtactcacagctccctctaaacactgccgggaaaggctctccacgtttaaggacctgcgt**gattacattggacccacctgggtaatc**cacgaactgcttcctatctcaaggtcggctgattagcagccctaattgcacctgcagctttaattcccatttgccatctaactgaacatccacagctgttacgggttgcagggattattagcacctggacatctttaaggaaccattcagcccaccactccgatgttccactagagggcactaacatattcctgtaatataaacattaaaattcagataatggtgggagggaaagtttga**taaaattgaatactgtattatcaatttta**aaaacttgatatataaatatgtctcattggggtctttcaaaagctaatcatatagtacaagaaaatacataaactggagtgactctcagaactaaaatggcctgagcgtaagtgaaat**agaatttgaagtgaggtgagaattct**gcagtctattcctgggctcaaaatcacatagaaaaaaacaaaaaaaggttgctttatatttaaagggacaaggcgtaggttctcttacaggggtttttaggaaatggttcatcagtttagtaggtgttagcttaattg

Uppercase: TRBVB

Lowercase: Flanking sequence[1000bp]

Red & Bold & Underline: Stem-loop [12]

Blue: Heptamer[18]

Green: Nonamer [4]

id-IGHVII-46-1[V_gene_segment]

agaagctatttagg**aagatgtaatgggaaggcatctt**t**aacattctgttgaatgtt**tccacatggcacttcaatcccctaaaatactttgctgtatcagcatgtgatgaatcggagtataatattgaaggtaaaatgggaaatacatgggcttttgatgaatcaagtcataggatgagaatgtctgtgtccttgaggaaggagaccatgggttgtaagttctagtggaggtacctttggcaaagggagttcatgagtttctgaaccttatgctactttta**atttaaggaagcaatttatgatgtatgtttacttaaat**ctttccagagacatttgtcagtaaggacagggtagcatttgtgtaatagtgatgacttagagagttattttgtaattgctcctgtaaggtatgcacattgctcacttgatacagaagttcaaatgtcacaggtggaaaaacaggaataaaggtgggcagatcacaaggtaaggatcacaaggttaaccatcctggctaacacggtgaaaccccatcgctactaaaaaatacaaaaaattagccaggcatggtggcgggcacctgtagtcccagctactcaggaggctgaggcgggagaatagcgtgagcccgggaggcagagcttgcagtgagccgagatcacaccactgaacttcagcctgggcaacagagggagactccatctcaaaaaaaaaaaaaaagaaaacaggaataaagcaaattttatgaactgtcatgactgtagtttttggcaaggacattatcatgtcaacctgtaaataaacagacaacaataaaacatatcaaatccatggggagtctgacctgtgtccctctgtcttataagcacaaggctttg**ccacatccaaaatattattcaggctccagggtataaaatgcttttggactgtgg**aaggtaacagctctcccctcaggcaggggtaaggtatctggggaatgcagagttgtgttcataaggaagATGTCATTATCGTGTCTTCTCCTGTGCCTGGTAGCAGGTCCTGAAGGTGAGCGTCTCAGATGTCAGACATGGGTCTATCGGGTTAGGATAGGTACATGTGACT**GACAGGGACTGATTCTCCATGTAATCACATGCCCTGTC**CCAGGAGCAGCTGCAGGAGTCAGCCCTGGACCTGAATAGCACACACTTACCCTCTGCCTCACCTACACTGTTACTGGCCACTCCGTCACAACCAGTCCTTACTAGTGGACCTGGATCTGCCGGCTCTCAGGGAGGGGCT**GCAATGGATaaaatccattgc**tagtggtggtgggagt**ccatttgtcttgtggaaaatgg**cagcattt**ccttattttataagg**cataataatgctatgttgtgtacacataccacattgtctttatccatttgcccattgacagacacttagtttccatatcttggctgttgtgaatacagctgcaataatcacaggagcgcaggtatcttcacaaggcggtaatttcatctctttgggtatatttccataagctgcatcactggtcatatggtatttctgttttaattcatttaggagccaccacactgttttgcctaatggtaatgatgggaatacagaatgtcatagccgctacgaagaacagttttagatttgaggtataatccaaaagcaaataatgtttgatcagggttctcatatgaggctctaataaataagtccagaaagttttcccatctt**gggcaaggatgagtttgccc**ctaattgtctttgaggaagaagattggcagaacgtgagatggagtctgttggcaggattcaggattcagtaataactagtaatggcacagaaaaatagagagatagagacatacagagaaagagagaatatgaatctcataagaggaaaatttgctaaatatcagcattgggttttcagtcatagacacatttgtatgagccaggaaccatggagtcaagtgaggagtg**gagaatatgttcagtccaaaaatcagcatattctc**agaggcacccattgccccatcacacaggtggagaattttggaaaccagtgaagtgcgagttcacattaggtgattaagctac**catgtttcgataaactttcattaacatg**caaagtacttccatagataactgatgcacaaatatacgagatgcctttttgcgtttatggatgtctaga**gaaaaataagtgagaaaatttttc**caggttgcagagatctgtttaagttgcagattccataggagagtgtctttgaatgaatactg

Uppercase: IGHVII-46-1

Lowercase: Flanking sequence[1000bp]

Red & Bold & Underline: Stem-loop [12]

Blue: Heptamer[30]

Green: Nonamer [3]

id-IGHV1-69D[V_gene_segment]

caatgggcacataactgtgggagacactatattcctatgaagaaggtattcagacttcagagataagtaaagtttcctacgttgtgtttgtgacttggaagcagtggattgaagagtgtgataggtgcccagaccaagcggaacagaaatcaacatgtaaagatgatgatctatggatatgatctaaaaccatgaaaatacttcaaataattctatttaatgc**agtttgaaataaaacacaaact**tattcaaaataca**aattacttggtaatt**attttgggagcta**tgagttcatcaagaaactca**aattcctatttctacttcaacccctgattcctactgtcaatgggagggaagtctcagaaccaatcacacatcagacggcaaatctgtcaaccaagagt**ctttcccttgaaggcctgggaggtcaggaccctcaggaaag**tgctggggactctgtcttgg**gagtgtccagcagatctcagaactctccatgggtcctgctggacactc**atgtag**ggtaacgagtggccaccttttcagtgttacc**agtgagctctgagtgttcctaatgggaccaggatgggtctaggtgcctgctcaatgtcagagacagcaatggtcccacaaaa**aacccaggtaatctttaggccaataaaatgtgggtt**cacagtgaggagtgcatcctggggttggggtttgttctgcagcgggaagagcgctgtgcacagaaagcttagaaatggggcaagagatgcttttcctcaggcaggatttagggcttggtctctcagcatcccacacttgtacagctgatgtggcatctgtgttttctttctcatcgtagatcaggctttgagctgtgaaataccctgcctcatgcatatgcaaataacctgaggtcttctgagataaatatagatatattggtgccctgagagcatcacataacaaccacattcctcctctaaagaagcccctgggagcacagctcatcaccATGGACTGGACCTGGAGGTTCCTCTTTGTGGTGGCAGCAGCTACAGGTAAGGGG**CTTCCTAGTCCTAAGGCTGAGGAAG**GGATCCTGGTTTAGTTAAAGAGGATTTTATTCACC**CCTGTGTCCTCTCCACAGG**TGTCCAGTCCCAGGTGCAGCTGGTGCAGTCTGGGGCTGAGGTGA**AGAAGCCTGGGTCCTCGGTGAAGGTCTCCTGCAAGGCTTCT**GGAGGCACCTTCAGCAGCTATGCTATCAGCTGGGTGCGACAGGCCCCTGGACAAGGGCTTGAGTGGATGGGAGGGATCATCCCTATCTTTGGTACAGCAAACTACGCACAGAAGTTCCAGGGCAGAGTCACGATTACCGCGGACGAATCCACGAGCACAGCCTACATGGAGCTGAGCAGCCTGAGATCTGAGGACACGGCCGTGTATT**ACTGTGCGAGAGAcacagt**gtgaaaacccacatcctgagagtgtcagaaaccctgagggagaaggcagctgtgccgggctgaggagatgacagggtttattaggtttaaggctgtttacaaaatgggttatatatttgagaaaaaaagaacagtagaaacaagtacatactcctctaattttaagataattattccattcaagagtcgtaatataagccagattcac**agagtggaaaaggccacactct**ataaagttgatacaaacattccatgaaggtgctactgtgaacaagttttcaaattggatgaat**acatgatttggagcaaggttatttgatcatgt**ggtgagac**taagaatgattctta**aaaagtgccaaaagtttccttcaaatgtttctgtcactccttatcataaagtttattttacagcagttttaggattacaaagaaattgcacaggaggcgtgagaattcccatgactccctgccctacacaggcacagcctcctccactacgaccatcctgcaccgcagtcacaaatcagttacaatggaggaatctccaaggacgcttggttctttctttttctggtgatctcctaatataacaagcctaagtatctcaagattccacggttttttcagtgttttctagaactgatattagtcagagggaaagtgggtaaggctattactatttgaactctttcttccaaaatccacaaaatatatattaatttagagctt**atcttacttctggtttacaatgctccttcccagacagtaagat**ttttttaag**cttttagaggccagttcatgtctctaaaag**accagaaagttctggggaatcccataatgaacatcctttcatgagaattggagaccctggcaatgagagactccatgtataatgccctagagttggattagatgctctgtaagctcttgggtggtgagtctgagtaagggggtttgtgcagcaaacgcaaacacatgcatgggacccaggtag

Uppercase: IGHV1-69D

Lowercase: Flanking sequence[1000bp]

Red & Bold & Underline: Stem-loop [16]

Blue: Heptamer[47]

Green: Nonamer [5]

id-IGHVIII-38-1-2[V_gene_segment]

aaaatttatagagcttcttcattgtggatg**aatatttgcttcccttctccaaatttggaaatatt**ttttgct**attattttgaataat**ttatctcttctctctctctctctctctctctctctctctctctctctctctccctctctctctctctctctctctctctctctctctctctctctccccctctctcccttcactaccccttttgtcttgtctttgtataactcctcaaatgcgtagttgagtgtttagtgtctttcctggcccataagtcttgaactcttttttatttctttcacttaaatttttcttctctgtctctactatttcaaatgatctttctttgagtttactgtttcttctattacttgatctagtttggtctaaaatcttgttaaaatttttggctgtattcttcaggatcaatatttttgtttagtcatc**ttttaaatgttttatctttattaaaa**ttctcacattgttcttgcatgctgccactagctcattgaacatgtttctaacatttactttaaagtttctgtcaggtaatccatgttcttccttttcattgatgtcagtttatggaccttttggttgttcttctgtttggaccatgttcccctgcttcttcattttccttgactgtctttg**ttgttatctgttctttataacaa**aagccatcatatacttcttcaaagactaggctcatagagatcaccctcaccaatcagcccagacagtaattgtgggtctctcaaaacttgataggattcaaacttctgtctgttcttagtggcccctaggttttagaggatgatagattgtgccaggactcacaggtaagggaggtagaaactatgtactcatgaccatggggtctgtgctgatctgagcttctcttgtcatcactctcaatattcaacttccc**cgtggatcaggctcagctgtggctgctccacg**tggggctgttctcagtctgttgcctctgtgtgtgcagAAGTCCTCTGTGAAGTTAACTAGTGGAGTCAGACAGGAAAATACTACAGACCAAGAATTCTCAGAC**TGTTCTGCAAAGCCCCTGGATTCACTGAAAAAAGAACA**AGTTTGGTCCAGCAGGATTCATGGCAAGTGTTGGTGAGGGAGATAACAGTAATTCAAGTGGAAGTTCTCAATGGGACTCGCCTTCAGTACAAAGAAGATT**AACAGTCCTCAGAGACACTGTT**CAGAAGATTCTCTTTTAAGATAATAAAACTGAGAGCCCAAGACAAGTCTGTGTATT**ACTGTGAGGGAcacagt**gtggggacatctatgtgagcacagacacaaa**agtccctgcagggagacaggaggggact**gcatgttagttgctgctcagaaccaccagggggcattcaggacaccagtaggcactcagaaccaccaaggagagctcagaatcaccagggggcactcaggaaactggcaggtgctcagaaccaccaaggggcgctcaagatcccagcgggagctcataaccaccagggggcgctcaggacactagggggctcagaaccactagggg**gcgcccaggaccaccaggtctctcaagacaggagggcgc**actcaggacactagggggcgctcagaaccatcagggggcaatcaggacaccagagggacttcagaaccaccagggggcgctgagaaccaccagggggtgctcaggacatgaccggccctcaggaccaccaggggatgttcagaacaccaggaagcactcaggacaccagagagcgatcagaacaccagggggcactcagatcctccagggggcactcaggacaacagg**gagctctcaggacaccagggtgagctc**agtaaaccagtgttcgctcagaaccacc**agggagcactgaggacaccaccgctccct**taggaggcagctccacatcagg**tccctggatccgggcaggga**gggcggttccttttggatcttgccactaaccttttgggagttttcctgcttcctttgtggtttc**aagaatcatttgcagattctt**ctcatgtacaaagctctgctttcttggattatgtaatgtttttggctttggatgctaccagaattacgttgtactgtgagagga**ttcattcgtggtgtgtgcaatagtgaatgaa**agctccaatgttaggggtggctttgaaagctacgttaggggtggctgagggcagttagcaga**aaatgatcatcactatagaaggctactcattt**ctttgcacatttgcataaacaattgtagtttatgccctaaaa

Uppercase: IGHVIII-38-1-2

Lowercase: Flanking sequence[1000bp]

Red & Bold & Underline: Stem-loop [16]

Blue: Heptamer[22]

Green: Nonamer [6]

id-TRBV14[V_gene_segment]

attttaggaccaatgaggaggcccatatgactgaaaaattaaggaagggagtggtaggctttatttccaagcggtaatggggcaggatggatcatttaagaccttgatggccttaagtgctttgttgttttgttttcttat**tccaaataaaactgggagctagtagatgatttgga**gatgagtgacatgatctagcatgttctagagggctgagtttgagagctagttcagaagtaggtgaaagcagggagatcagttaggaatctattgcagtaatccaggaaaaagatgatggtggccttgtcctgagggttagcaaagaaggtg**atgaaaaaatggatggattctgagtgtattttcat**gatagagccagttgaatttcctaatggattggctatagggtgggatagaaaaagaggatttgggaaccactctaaggttttcagctggaaggctgttggtggagtaaagttgaggataatctttcatgtttagagcatgctgtttgagagtggtggggggaaaatgttgattggagtgcattagagagataatgggaaaagagaatttggaggtaagtattgatcattagcaatagcaacaatagctaccgtagattaaaatgccccacagtctatggggatgtgatacatgagggaagacagggccagatgggatgaggtaggatccagacatcagactcaggagctacgagtggtatatataaggttaacacctagtcaaatgcataaacaggttgattttaattgatgtcagtttttctcc**attgccccctctagaggcaat**cttcttcagagaaccctggctaggtcttctatgtttc**atgtccgtagagggagctcctgagactgtggacat**tggctaatatgctgatgtcactggaggccacatcttacagggccaagagacagatttgctttcctttttctcatgcttgtaagctccttcat**ctggaaatgtgatttacctgggtcctgccATGGTTTCCAG**GCTTCTCAGTTTAGTGTCCCTTTGTCTCCTGGGAGCAAGTGAGTCTTCAGGTACTTAAATATCTGTGCTGTACCCTATCCCAGTCTATTCATGTCATGTATTCTGTTTTTGTCTCTTCCCACAGAGCACATAGAAGCTGGAGTTACTCAGTTCCCCAGCCACAGCGTAATAGAGAAGGGCCAGACTGTGACTCTGAGATGTGACCCAATTTCTGGACATGATAATCTTTATTGGTATCGACGTGTTATGGGAAA**AGAAATAAAATTTCT**GTTACATTTTGTGAAAGAGTCTAAACAGGATGAGTCCGGTATGCCCAACAATCGATTCTTAGCTGAAAGGACTGGAGGGACGTATTCTACTCTGAAGGTGCAGCCTGCAGAACTGGAGGATTCTGGAGTTTATTTCTGTGCCAGCAGCCAAGAcacagtgcttcacagtcgtgcccttgctgtgcaaaaccatagccttctcctctcaactcacagctgcccaaaag**gaaggctttccctgtgccttctcccccaagggaggggaga**taaagaaccagaattaactcatgaaatacaag**agtattccaagaagatttgggtgaaaatact**tgtggtttgggggatctctgaagttttttaaacagaactcaaactatgtgtttctcatatttatttttattttttaaagtgtatgttgccctggttataaagcagacttcttgcctatcttgctgctctgaaaggtttgtgactctgactgtgaggccacatctttatatcccttcttatagtaagagaaacaagtctttctgatgaaaataaaaatagtgaataaaaatgcatttactactggcttttaggatgctcaatgatgatgctagcaatggtggtagtggtgacgttggtaatggtggtgatgagtttaggttgcttttgtctagcatttataaatttccaa**ccactttaaaacaatctcattttagagtgg**aatagctaagtccattatcagaggttaatgctaattttctgttcacatagggtctgag**tttttaatgtaaaaa**a**tatagttagcactata**aatagatatctgtcagccgatacagaacattttttaagatcacttccagttgctgc**aataggaatatgtattcctatt**ttttttcttttttctttttctttttttgagacgaagtcttgctctgtcgcccaggctggag**tgcagtggcaaaatctcagctcactgca**acttctgcctcctgggtttaagcgattctcctgcctcagcctaccaagtagctgggattacaggtgcctgccaccatgcctggctatttgttgtatttttagtagagacgggatttcaccatgttggccaggctggtcttgaactcctgacctcaagtgatctgcctgcctcagcctcccaaagtgc

Uppercase: TRBV14

Lowercase: Flanking sequence[1000bp]

Red & Bold & Underline: Stem-loop [14]

Blue: Heptamer[33]

Green: Nonamer [6]

id-IGHV1-18[V_gene_segment]

agtccc**cccaaacccctgaagagtttttggg**ttcaccgtgtgtccaatgattcagtgcctcttgagctccaggaaagggctccccagtgatgcatgagatctttcttggggtctccctgcagagttcactgggattcctaaggccaattcagtatttcaaaagatggtgtgagaagcacaggctgtcactaaaggagaattctgagccagggca**cagccactttatacttggctg**gggacactggtaggaatatactctgtgagatcagacaggaacctccttgcaggggcagggcagggctgcag**ggggcgctcaggacacacagagcacaggcttccgccccagagcaggtgaaggaggctgggg**aggggttcctctcagggcctgggacttcctttaaaaaatctaaaataagtatttcacaaggactgccgatgtttatataaatatcctattcaattgtgagcatttatgaaactcgatg**ttgtaatgagaaccacttttacaa**tgggaatttcaaacttccctagacatcttaatagtaagcagctggaggtcaggaggagatcctttcttataaataa**gtgcaatttttggagaaacacactcattcccaaaatagcac**attcacatattaaggtctagaaatgattcgagttgcccctgagacagtcaaatgtgggttctaagtgaggtgcgtgtcctgggg**gagcttgttctccagtgggggaagctctgtcaacacagag**ttcagggatgggtaggggatgcgtggcctctaacaggattacggcttgaaccctcagcttctacaattgtgtcgtccatgtgtcatgtatttgctctttctcatcctgggtcaggaattgggctattaaatagcatccttcatgaatatgcaaataactgaggtgaatatagatatctgtgtgccctgagagcatcacccaaaaaccacacccctccttgggagaatcccctagatcacagctcctcaccATGGACTGGACCTGGAGCATCCTTTTCTTGGTGGCAGCAGCAACAGGTAACGGACTCCCCAGTCCCAGGGCTGAGAGAGAAACCAGGCCAGTCATGTGAGACTTCACCCACT**CCTGTGTCCTCTCCACAGG**TGCCCACTCCCAGGT**TCAGCTGGTGCAGTCTGGAGCTGA**GGTGA**AGAAGCCTGGGGCCTCAGTGAAGGTCTCCTGCAAGGCTTCT**GGTTACACCTTTA**CCAGCTACGGTATCAGCTGG**GTGCGACAGGCCCCTGGACAAGGGCTTGAGTGGATGGGATGGATCAGCGCTTACAATGGTAACACAAACTATGCACAGAAGCTCCAGGGCAGAGTCACCATGACCACAGACACATCCACGAGCACAGCCTACATGGAGCTGAGGAGCCTGAGATCTGACGACACGGCCGTGTATT**ACTGTGCGAGAGAcacagt**gtgaaaacccacat**cctgagggtttcagaaaccccagg**gaggaggcagctgcactgaatttgaggagattacagggcttacaatgtttaaagttgtttagaaaatgagctgagcaattgaggaatgtgagtaatg**gaaacatggatgcactctatataggaaatgtttc**tttcaacagtcaccctatatgcaaaattcagaatggtaaaggcagcaatcagtgaggctgacgcaaatattcccatggaggccttgtgcagacatacgttttaaaatcagatagataaataatttggaacaagattgctggtaacgtggctaagactaaatatgattcctaaaaactggccaaaatctattccaaattgtctctgccactccttttacataaatgta**ttaaaaagtagttttaa**gaccacagcaaaattggacagaaggtgcagagagttctcatgtgcccctgcttcaccatgcacagcctttcccactgtcaccatcctgccccagagtcatcaa**taagttacaatggatgaactta**catgggcggattggttctttcctcttctggtggtctcttggcataccaagcccaaattatcttgaagcaccataggttctactgtaatgcctaaccttgttttttactctagtttgctactttaaattttccctttttttgtctccttaa**ttgccagccatgtttcccatatgaatagactctccctggctgggaa**agctgggcaaactccatttgaccttttgatttataagacattaagggctccttacccaacccccttcttcaaggaattaacctgtgtaagcagatcctcagcatttcaaaggagcccaattaactgataaggtactggaacaaacaatgtatgaagttcccaggatttttctcaaagagataacaacataaagccttgagttcatgcccagcatagcatcatatctaactataatgaaggatttagagccctgcacctgg

Uppercase: IGHV1-18

Lowercase: Flanking sequence[1000bp]

Red & Bold & Underline: Stem-loop [18]

Blue: Heptamer[42]

Green: Nonamer [6]

id-IGHV4-59[V_gene_segment]

agggtatgttcatgcaattcaatggaggagaaagtc**tttcagagacagatggatctgaaa**tgataaatatgtgggtaaggact**ctggacttgagtgtcattgtccag**ccatgttt**cacaagtgtgtcctgtcagggaaggatcagagttccttgtg**ctctcagagggaaggggtcacagagttcctctctggttcccaggaaaggtaatcgcactaatct**tcatgatcttcatga**gactatcctccagtgctgacctgttatagagtttttgtctgaagttctcactgcaatccccaatctacatattttcaatcagaagtgtttagaggccaggacacatcttcaaggtcacacatt**gagaaggatgtagatatgtcccactaccttctc**ctgagatctcagacagaatcccagatttcaaaaggacacagaaggacagctctcaggtgcttttaaaaaatgacccacttccagggacagggagcttccctataaccatggtggatgttctgaactacaataaacattggatggatccaggattgtttgaagtcactgtcattattacattcagctgctgtttcaatgtgtctgaagtagtaaatgacaatttagatgacaatttatatgaatcttcaagggtagaacaatattgaccatattccaaaatctgtccttgatccatgatcacactcatctcccagaccaggtccttcagcacgtctctttacctgaaagaagaggactctgggcttggagaggggagaccccaagaagacaactgagttctcaaagggcacagccagcatcctactcccagggcgagcccaaaagactggggcctccctcctcctttttcacctctccatacaaaggcaccacccacatgcaaatcctcacttaagcacccac**aggaaaccaccacacatttcct**taaattcaggttccagctcacatgggaaatactttctgagagtcctggacctcctgtgc**aagaacATGAAACATCTGTGGTTCTT**CCTTCTCCTGGTGGCAGCTCCCAGATGTGAGTATCTCA**GGGATCCAGACATGGGGATATGGGAGGTGCCTCTGATCCC**AGGGCTCACTGTGGGTCTCTCTGTTCACAGGG**GTCCTGTCCCAGGTGCAGCTGCAGGAGTCGGGCCCAGGAC**TGGTGAAGCCTTCGGAGACCCTGTCCCTCACCTGCACTGTCTCTGGTG**GCTCCATCAGTAGTTACTACTGGAGC**TGGATCCGGCAGCCCCCAGGGAAGGGACTGGAGTGGATTGGGTATATCTATTACAGTGGGAGCACCAACTACAACCCCTCCCTCAAGAGTCGAGTCACCATATCAGTAGACACGTCCAAGAACCAGTTCTCCCTGAAGCTGAGCTCTGTGACCGCTGCGGACACGGCCGTGTATT**ACTGTGCGAGAGAcacagt**gaggggaggtgagtgtgagcccagacaaaaa**cctccgtgcagggaggcggagg**ggaccggcgcaggtgctgctcagcgccagcagggggcgcgcggggcccacagagcaggaggcccggtcaggagcaggtgcag**ggagggcggggcttcctcatctgctcagtggtctccctcc**tcgccagcacctcagctgtccccaggggtcctctttctttattatctgtggttc**tgcttcctcacattcttgtgccaagaaagaaatgaggaagaca**aattttcgtctgtagttgaagtttcaccaattact**aggaactttcctagaagttcct**gcatggcccattatagcttactgattaaaaaatatatattctaacgcttctcagcatctcttgatttgtgtcatcaac**tgaattgtgccctctttgaaattca**tatgcagaaacctt**aaattcaattgatgtatattggaattt**taatgaaataattaaggttaaatgtggtcataagtgtaagactctaattcaacagacgtgtcgtctttataagaagaggaagagacaccagagacctctcacttttcacgtgcaggcagagaagaggccatgtggagacgtaa**tgcactagaaggtggcccagtgca**agccaggaagaagcctcaccaagaaccaaccctgccagaac**attgatcttcaacattcagactgcagaattttaagaaaatcaat**atttgttgtttaagccacccactcctgttgtcttcttatgaagatccagacagactaataccacataactctgttagcgctgtcccctggatgcagaat**cagcccgctggggctg**ggcacatctctcagatttccacataaagtaggcaaaaaatagtagttctgatataaaaatttgtcatgtccctgttggccaatttctgggcaaggtcttttaaag**aagccctgggggctt**tgtcacaaaagttgccttttatcatttattagga

Uppercase: IGHV4-59

Lowercase: Flanking sequence[1000bp]

Red & Bold & Underline: Stem-loop [21]

Blue: Heptamer[41]

Green: Nonamer [6]

id-IGHV2-70-2[V_gene_segment]

gaattcagccaagacagagccccatttaggagcaggtgcatcaaggaaccagccagagaacctctgactacaatc**tagggttgtcttcttttgcttcacagcatccaacccta**aggagaccacccactatatcatc**tgtaactcagaactttcttgttaca**aaggaaaacacaaatgaatataactctgcagttaacttcttgaacctggattgatgagtttccatggcctcatgagcacagagaccacatccagtgcaggtgcagagca**aggctgcaacagtcagcct**gctgctccctcaggagtcttccaa**ttcccttgtccaaagcatccgctcattcatgggctgagaaaagggaa**gtcattcatgcagtctactctctccacagaactgacggggcacaaggacaacatcgatttttcatggaacatgcctctaggaatgcagc**tgtgtgcacacacactgctaagcacaca**cttctttacataattacttgtaactgtattttcttatttattctctccaatttttttacacaaattcatcacttttccccata**atcaaagaggattttgat**cagaatgcttgtggggagccccttgccccccagatgcccaccatcattattcttgaagggaggaggaacggcagctctcttgatttctactctaatcctctaggactaaaaccagaaggttgcatgtccggtgcgggagcatcgaagaagatcctgtctgtagaagcaggagcgtcaagacttgactgagagccatggtgctgaaatgagatagattccctgatggagagcacacgtggacccccacacctgagggctcactgctcctcaccacaga**tgcactcccctactgagtcctgagacctgagtgca**ccccatagagtagggctcagatg**aggggatgcaaatctccaccagctccaccctcccct**gggttcaaaagacgaggacagggcctcgctcagtgaatcctgctccccac**cATGGACATGCTTTGTTCCATG**CTCCTGCTACTGACTGTCCCGTCTTGTGAGCGCTGTGGTCAGGTCGTACTTCAGAAGCAAAAAATCTATTCTCTCCTTTGTGGGCTTCATCTTCTTATGTCTTCTCCACAGGGGTCTTATCCCAGGTCACCTTGAAGGAGTCTGGTCCTGCGCTGGTGAAACCCACACAGACCCTCACGCTGACCTGCACCTTCTCTGGGTTCTCACTCAGCACTACTGGAATGCGTGTGAGCTGGATCCGTCGGCCC**CCAGGGAAGGCCCTGG**AGTGGCTTGCACGCATTGATTGGGATGATGATAAATACTGCAGCACATCTCTGAAGACCAGGCTCACCATCTCC**AAGGACACCTCCAAAAACCAGGTGGTCCTT**ACAATGACCAACATGGACC**CTGTGGACACAGCCACGTATTACTGTGCACGGATACcacag**agacacagcccagggcgcctcctgtacaagaacccaggctgcttctcagtggtgctccctccccacctctgcagaacaggatagtgtggctgagatgccatttcctgccagggcctgcgtttcccatccccatctgactcagagccttgttttcctccctcttctttactaataaatggcatgtcccctgttagtggttcgtgcaagcagaagctgtatcctgtttgacaaagattcagcatgaaaggtcctgctacctaaa**aaaaaaatagacagatgagacttaattaacctaaataattttttt**cacaacaacagagtgaatacgcaatttacagaatgacagaaaacttttgcacactttgcctgtgacagggaactaatatgaagaatttgcaaggaactcaaacaactctacaacaacaacagcaacaagaaccaaataaccccgtt**aaaatgagcaaagaacatgagtagacatttt**caaaagaacacatagaaatggataataaatatataaacaatgctcaacatcactaaccatcagggaaatgcaaattgaaaccacaataagatatcatcttccac**cagtcacaatgactg**ttactaaaaactcaaataatatcagatgttgctgaggatgggaaataaaggcaactcttagacattgttgatgaggatgtagacgagtacaacctctgtggaaaatggtatggagatttcccagaaaactagaaatagaactgccatttggtccagcaatcccactactgtgtaactacccaaagga**aaataaaccattattt**caaaaagatacccaccttctatgcttaccataaaactactctcaatagcacatatgtcaaactgagtgtctgccaacagatgattttataaaagaatatatcatgtatgcacaattcaatactagtcagccacaataaggaatgaaactgtgtcttttgcagcaagatgcatagaagtgggggacaata

Uppercase: IGHV2-70-2

Lowercase: Flanking sequence[1000bp]

Red & Bold & Underline: Stem-loop [16]

Blue: Heptamer[39]

Green: Nonamer [5]

id-IGLVI-38[V_gene_segment]

ccaggctgtggtctcaactcactgcaaccttcacctcctgactcaagcaatcttctcagctcagcatcccaagtagctgtgactatgggcacgcaccaccacaccctgctaatatcttgtattttt**ggtagacatggggtttccccatgttacc**caggctggtcttgaatttctgagctcaagtgatccaccctcctcagcctcccaaagtcctggaattacaggcatgagccgctgtgcccagcctattttttctcttttacgagcctctctgattccctagttctgaagtagcccagagttggctgattctcctattttgcctcagctagcacttcctccttctcctttttttttttctccttctgatccctggagtcctctgcatttccactgaatccagtgaaccacacaagcatgctcatttattctccccaaactccaccactctgccttatgtgctgatccagtgagaatcaagaaggaagagagacccatggatttctgtgcagcctggaaagtcagttcatggctggacttggctgggacaccagacacctgtctccctgtctctatttagtattcatagggcccagcagctgcatcctcccagggtgcctgaagagtcactgaccatgctctcctacccagcctcaagaagccacccagtcctcgagaagtataatcagactcagcatcttatgtgaagggaacaaccat**ggggacaggttagacttgcattgagtcccc**tcctctaagaagggccagggaaaaggggctgttccccctaatctgtgtggctcaggaa**gtagagctccagagacttctctac**catggcctgaatccctctgctgctcac**cctcctcagtctctgcacaggagg**ctggagat**agaaaggcagggactggcactgggaggaccagggctctgccttgct**cccccggctcaccgagccacttcccacaccctgtgtctctgt**cctgactcttagggtcctgggCAGTCAGG**GCTCACT**TGGTGGCCTGCACAGTTTGTCTGTGACTAAACCAGCCACCA**TCTCCTGCACTGGAAACAGCAGCAATAACGTCCTTGGAATTGTGTCCTGGTACCAACAATGAACAGGAAGTGTCCTTAAACCTCTGATCT**GTGGATGTAAATATCCAC**CTGCAGAT**CCAGAACTGATTCTTGAGTGATCAGTCTGG**CAAGGAGGCCTTCCTGAGCATATCTGGGCTCCAGGCTGAGGACAAGGCTGATCATTAACGTTGGATTTGGACAGTTCTCTGGAGGC**CCCcacggtgctgcagtctgtgggg**aagtgagatgggaacatttcaggtctcctaggagcacgacccgacagcaccacctacaactgggaaaactgggctgtttaccatttgtgttcatgtttgcttgggctggtgctagccagggcccaggtctcaggtcagggacagtgatttagagaatgcagctttgtttcctcagagtcagccctcagagaaagcccatagcaacagcattgtcaaattgcctcaaaaaaaatcagaaagtccttg**gttcctggaacaggctgccctaggaac**agagcccaaatcggtcagagcaagcagggacacaactgtgcagaaaacaacaagttttacttagggagataacggttgactgtataaaatgagaaatatgtcacactactcacttgtaaaagtaagtttaccaatgctattcatttccctatatcactacgttgagtaaaaactctaaaatgtatgaaaccactgtagagataacttttgctgaattgctactttaacatgtcatggatcatttatttttcttgcaaaattctcattattcaccgtccttttgagtaacccc**cagctgcccttcaacctagctcctcagctg**acctcttctccatgatgcctaccctgatggcacatgtgggcagttaggtttttctagattttaaaagttatatatgtacatgtctacatcccatggaccaaga**ctcactctgggagtgag**aaatcacaataatcatgcatagccttatggaattagtgtctagcaatcagaccatagaggtgtgaatagagacagtgactaatgagctgcagagatcagctgggccatcaggaggtaagtgtcctctaa**ccagcatgggatgctgg**gggactgccct**atggtccctacacagctgctcagtaaggaccat**tcactcaaagaaatccaagactaggg**gctaagaccccgtgctcactgaggggtacttagc**atctatgtcctcc

Uppercase: IGLVI-38

Lowercase: Flanking sequence[1000bp]

Red & Bold & Underline: Stem-loop [16]

Blue: Heptamer[33]

Green: Nonamer [2]

id-IGHVIII-16-1[V_gene_segment]

aacccagataattcttt**tctttccttggttgagaaaga**ttttccccaaacttcagctcag**ttcaggcacacactgtccctgaa**tgggcat**ttaccctcagatgggtaa**cacacctgtcaacatggggactcttctgtcagacaaacacacctttactcacgtggattcttccctcagaccaacacacctgtccccacatggactctttcctcagaccaccacat**atgtccttacatttactctttcctctagtaatgacat**ttcctcaagtggactcttgtct**cagacaaacaaacatgtctg**catgtgaactcttcactctgataagtacaaatatttccacagtgactgtttcctgacacaagcacatatatccaatgttgaactgttttatggcaaaatgatctcaagataataattataaaatccctcctgacaaggtgtagatctgcatttttttattgtaccttaactttgcct**tattgtcaagaacaata**gtttgcagctctatatgcaccgattagagattggtgtccattttctctggaaatgtatttttatattcttactggatgtatttttttgataatgtttgctactgtgaagatacctgaacagggtccacactagaaaataaaaaagactaatcggcagattaaccctgtgcatccagacccacgagtcctttgatcctgcccccctgaaatggagacacagaggacagatgagcaatgctgagcggtgcacccaagaccacaaaaagaaagacaaggaaatgtgtcccctcccctcctcatgaaaggcagctcatcccctgttccttcaggccctggtgaggagccaa**cccatgtctgttcccttcctcggtgtccacaccatggg**gtctgccctgatttgggcttcccttctcatcaccctcaatattagtgtcccttgtgaatcaggtccagctgcggctgctccacatggggccgttcttccatttcctctgtgtttgcagAAGTCCTGTGTGAAATTTACTGATAGAGTCAGGGGGGAAAAATTGTACAGCCCAGTGGTTCACTGAGACTCTCATGCAAAGCCTCTGATTTCACCTTTACTGGCTACAGCATGAGCTTGGTCCAGCAGGCTTCATGACAGGGATTGGTGTGGGTGGAAACAGTGAGTGATCAAGTGGGAGTTCTCAGAGTTACTCTCCATGAGTACAAATAAATTAACAGTCCCAAGTGACACCTTTTCATGTGCAGTCTACCTTAAAGGGACCAAACTGAAAGTCAAGGACAAGGCCTTGTAATACTGTGAGAGAcacaggagagggaatatctgcgtgagcccagacagaaaaatctctgcaggaagacaggagggagctgcatggtagatgctcctcagaaccaccagggcaccttggggacaacctgggggcactcagaaccaccagggtgtgcttaggaccacgggacactcaggatattagggggtgctcaggatcacgagggggcgctcatgacaccaggggacactcagaatcacaagggggcgctcaggacacgagggtt**tgcctgggaccaccagcaggca**ctcaggtcaccagggggtgctcaggaccaccagggggcgctcaggacaccagggggcgctcagaaccaccagggggcgctcaggacacctgggggcgctcagaagcatcaggggtgctcaggatatcagagtgccctcagtaccacgagggggcgctcatgacaccaggggcactc**agaaccaccagggggcgctcagtacaggggggttct**cttaggaggcagctccaaatcagaagcctgagaggctgtggttttcttttaaaccttggtgattcccgacctggtcaagcaaaagtcttccccaggatctctcaccatttcttccttgtaagtccatgattacttttacctacaaaacattaacttagaacaggaatttaattcaacttttaatgctgcatattttccaagtaatactagcaatgatctctcaggacaatttttacaataggttatttata**tattttcttgattaaaaata**a**tactattattaaaatagta**aagttataaaaatcacacctgtaatcccagcactttgggaggctaaggcaggcagatcacttgagctcaggacttttagaacagcctggcagcacagtgaggtcttttctatatatatatataaaataataataataataataaatacctgggtatggttccacctgtggtagcagctacttgaaaggctgaggcaggaggatcccttaagcctgggaggtt

Uppercase: IGHVIII-16-1

Lowercase: Flanking sequence[1000bp]

Red & Bold & Underline: Stem-loop [11]

Blue: Heptamer[35]

Green: Nonamer [5]

id-IGHV3-57-2[V_gene_segment]

gtggggatatcatgagacat**atgcaacaaagaatgaaggcattttcctcatttgcat**gtaagatgttaccattcactaaagacctttaataa**aaaaaaatttctacgtgatccagtattttttt**ccttctgtcaagcaaataactcataggattcttt**tctttccttggttgagaaaga**ttttctacaagcttcagcacacgtcaggcatacactgtccctgaatgggcatttaccctcagatgggtacacacacctgtcaacatgggggctcttctgtcagacaaacacacctttact**catgtggattcttccctcagacaaacacacatg**tccccacgtggactctttcctcagattacaacatttgtccttacatttactatttcctcagaaaacagacattgcatcatgtggtct**cttgtctcagacaag**caaacatgtctt**cagtcagataagtctgtggatttccacattgactg**tttccttacacaagcacctatatccaatatttaactgttttgt**gagaaatttatcccttttctc**tggaaatgtatttttatgttcttactggacatattttttaataatgtttgctactatgaagatacccgaacactgtccatacgagagaataagaaagagtaataagcagattaaccctgtgcatccagacccaggagtcctttgatcctgc**ccttccgaaatggagacacagaggaagg**atgagcaatgctgagcagtgcacccatgaccacaaaaggaaagacatggcaatgtgtcccctcccctcctcatgaaaggcagctcatcccctgttccttcaggccctggtgaggagcc**accccatgtatattcccttgatcagtgtccacaccatggggt**ctgcactgatctgggcttcccttctcatcaccctcaatattagtgtcccttgtgaatcaggtccagctgcggctgttccacatggggccgttcttccatttcctcagtgtttgcagAAGTCCTGTGTGAAGTTTATTGATGGAGTCAGAGGCAGAAAATTGTACAGCCCAGTGGTTCACTGAGACTCTCCTGCAAAGCCTCTGATTTCACCTTTACTGGCTACAGCATGAGCTTGGTCCAGCAGGCTTCATGACAGGGATTGGTGTGGGTGGAAACAGTGAGTGATCAAGTGGGAGTTCTCAGAGTTACTCTCCATGAGTACAAATAAATTAACAGTCCCAAGCGACACCTTTTCATGTGCAGTCTACCTTACAATGACCAACCTGAAAGCCAAGGACAAGGCTGTGTATTACTGTGAGGG**Acacaggagagggaatatctgtgt**gagcccagacacaaaaatctctgcagagagacaggagggaactgcatggtagatgctcctcataaccacaaaggggcagtcaggaccatcaggaggagctcaggacacctgggggtgctcagaaccatgaggggtgctcaggacatcgggggctctcagaaccatgaggggtgctcaggacatcagggggctctcagaaccaccagggggcgctcaggacacctgggggcgctcagaagcatcaggggtgctcaggatatcagagtgccctcagtaccacgagggggcgctcatgacaccaggggcactc**agaaccaccagggggcgctcagtacacgggggttct**cttaggaagcagctccacatcaggagcctgagaggctgtgatttcgttttaaaccttggtgattcccgacctggtcaagcaaaagtcttccccaggatctctcaccatttcttc**tttgtaaatccatgatttttttttacctacaaa**acattaagttagaacagggatttaattcaacttttaattctgcagattttccgagtaatagtagcaatgttccctcagga**caatttttaaaattg**gttatttatat**attttatttatgtaaaat**ga**tactatcttgaaaatagta**aaatttttaaaatcatacctctaatcccagcactttaggaggttcgggcaggcagatcagttgcgctcaggagttttagaacagcctggcaacacatttaga**tcttttccatacaaaaaga**aaaagaaaaaga**aaatagctaggtgtggtgctacctctggtaccagctattt**gaaagtctgaggtgggaggatcccttaagcctgggaggttgatgctgcagtgagctgtgattgtgccactacactccggcctggttgacagagtgagaccctttcttaaaaatacaaactgtttcaataagtagagctttgtgtctttgtgctgtaataatcagtt

Uppercase: IGHV3-57-2

Lowercase: Flanking sequence[1000bp]

Red & Bold & Underline: Stem-loop [17]

Blue: Heptamer[46]

Green: Nonamer [5]

id-TRAV8-6[V_gene_segment]

tgcttatttggaacttctctcttattttcttcattaacctacttagtggtgtatcaatcttgtttattcttcctgaaagacaacttttggttttattgatcctctgtatggattttcaagtctcaattttgttcagttcttttctgattttgtttatttattttcttctgctagcttttggctttgtttgttcttcttt**ttctagtccttctaattgtgatgctagaa**tgctcacttgagatctttctaatttcttgatataagtatttagcagtataaactttcctattaacacctttagctgcaaacc**aaagactctggaattttgtcttt**ctgttttcattaatttttgctctactttgttgtttacccaaaagtcattgaggagcaagttgtttaatttccgtgtaactgtgtggttttgagagatctccttggtaatgatttccatttttattacactgtggtccaagattatgcttggtatgattttggtatttttgaatttattgcaatttgctttatggcttcaatt**tctcaataattttgaga**ttaatatcatgtactatacagaaggtaacatcatcacagattcaggaggaaagaaacgctttctgttgaagttttgatatcatgtctgaaatgagaaggaaacagcttgctcccttggatccgtggttcctcctgttgcctctagagg**gcgatgttgcatcaagtaggcatctgttttcatcgc**agaccatcctcatccattgagcctcctccctgtagctggctgatgtagctcgttgatgtctgtgtggatagggagctgtgacgagagcaagaggtcagaacacatccaggctccttaagggaaagcctctttctgtttctgaaacttttcaaagccagggacttgtccaatccaacctcctcacagttcctagctcctgaggctcagcgcccttggcttctgtccgcccagcttaaggtcct**gcagcattgccactgctcagccATGCTCCTGCTGC**TCGTCCCAGCGTTCCAGGTGATTTTTACCCTGGGTGAGTAACATTCCATTCTTTTTCCTTCCACAGAAGTGACTGTGTTGTAACTGAATCTAAATTGAGACACATTTTTTACTCAAACAGCATTGATGTTGCAGGAGGAACCAGAGCCCAGTCTGTGACCCAGCTTGACAGCCAAGTCCCTGTCTTTGAAGAAGCCCCTGTGGAGCTGAGGTGCAACTACTCATCGTCTGTTTCAGTGTATCTCTTCTGGTATGTGCAATACCCCAACCAAGGACTCCAGCTTC**TCCTGAAGTATTTATCAGGA**TCCACCCTGGTTGAAAGCATCAACGGTTTTGAGGCTGAATTTAACAAGAGTCAAACTTCCTTCCACTTGAGGAAACCCTCAGTCCATATAAGCGACACGGCTGAGTACTTCTGTGCTGTGAGTGAcacagtgcctgagactg**caggagagctgaacacaaacctcctg**agatgctgagactttctgtgactcaagaactcaacctgtggagctttcaagagggtcccttttttc**tgtgcccgttttgaaggcaca**tagggcagggtgggctgagcccttggtagggtttgtctccggaataattgctccaatgtcaatgtcatgccacattcagatatgctttgtcccacatggaggtttcctgatgaa**tttcttcagtggggacctcctccacactctgaagaaa**gcacatcataaggtgtcagaactcc**gaaataaaaggcaacaacatttttatacaaaattatttc**tagcaaccattatgattagtattgcctcaggtgttaacttgtgaataaccgatgtaattgattagaatgttagctaaaagctgtgcaggatggattctcaattagggtaaccca**tacagcacagattctcaatcagtgtgctgta**gattctcaacctagatttgaagcttaacttctagcacatacataatatcccactgtgtggtattgacatatttaaagtaaattagacataatat**caagatttttaatcttg**tgagcattatgcaaaacaggaaattaaacataaatcttaatacaatattgaaaaacta**agctgtcaagttgtgatcttttttaaaacagct**attccagtgatttc**caagttacaaacttg**agggataaaaatgtaaccaatataaagatatcaaataccctagattcaactac**tttaaaaactactttaaa**actccgaggcattttattttaagagattacacagatttattatataaaattacacaacaaaaaattccagttgcattatcagagagtatatgaattctaatgggcatatccaagaccagggtaccctattacttactctgaaataattatatcgaaaaaccaacatgttaagatattttatgtacatgattattactccaaattttta

Uppercase: TRAV8-6

Lowercase: Flanking sequence[1000bp]

Red & Bold & Underline: Stem-loop [15]

Blue: Heptamer[30]

Green: Nonamer [8]

id-TRBV6-9[V_gene_segment]

acctggaggaagggtgttggctgggtctaggagaagatgatgagctcagaaaatagacaaggagaaaccccaggggaaaa**cttgtgaaaccccggaggaatcacaag**atatgttatttgagctcctt**ttaaattttatttaa**tactaaattaatggcaggaatcctgcacaaagggatattccctatgaaacatgacatctcagataaaaacagagcattcttccttaccctaaagtctgccctctctccccaagtccctcatgcatcaggtcacctttg**tgcccataagtcatgggca**gcgct**gtgtggtcaacatcatccacac**ctagaggacagtcagcaaagtgaggtggttc**tgcctgctgtggtctcaccccaggca**tggaaagcaagagccctgggagcagcggaaggtacacagctgg**gtttctcaggagtctcatctgtcagtgaattgacaagaaac**agagcaaaacgactcctccaatgttgatgagcctgcccctgggatttggaaacttgataacagagaaaaccaatatagacaaaggattttaaacaggattatggtcaattaagcaaattagaaaaggatacttgaaggagta**tttgggacacagaagtcaaaaacactaggaagacatgaggtgtgtccctaa**gactctagactacagcactgtgtagataactaacacttaactattaattatatcattgaagaaataaaatttacattgaattacaaactgtttccaaaaggtcatgaaaatcttgtagacttgtttcaaatcacacaaatgtgttcttctcattctcagctcttcactggtgcatttattttggatttgatcatctggggaagaggtgtggcctctcctgaaaggagggctctgggcccaggc**agggagactgaggtctcagaatgactccct**tgagagtcctgttcccctttcatcaatgcacagacccagaagacccctacgtcctgcagcccctgccATGAGCATCGGGCTCCTGTGCTGTGTGGCCTTTTCTCTCCTGTGGGCAGGTGGGTCCTGGGCAGAGCCCCTTGCGTGGATGTCAAGGCCCATCCCCTTTCCACTGGGG**CTGCAGCATCAGCTTTGTTCTTCTCTGCAG**GTCCAGTGAATGCTGGTGTCACTCAGACCCCAAAATTCCACATCCTGAAGACAGGACAGAGCATGACACTGCAGTGTGC**CCAGGATATGAACCATGGATACTTGTCCTGG**TATCGACAAGACCCAGGCATGGGGCTGAGGCGCATTCATTACTCAGTTGCTGCTGGTATCACTGACAAAGGAGAAGTCCCCGATGGCTACAATGTATCCAGATCAAACACAGAGGATTTCCCGCTCAGGCTGGAGTCAGCTGCTCCCTCCCAGACATCTGTATACTTCTGTGCCAGCAGTTATTCcacagcgctgcaagcctgtctcctctctgcacataaaggcacagaggctctgccctcctcccacccaagactcaaggatgc**cctgggcagagttctctgcaccagg**aaccttggaaccc**agagtggccccaagtggccaggacagtatgagtctcactct**atgccaggtg**ccacttcaggcagtctcagccaggcctggaagtgg**tcctaggttctcagatgtctcctttgttgctctctctggtatatcctccaagctgtcctttttgtgtagggccagggcttccccagctcctactttcctactcatt**atcctgaatccaaggtgcccaggatgaaactggatttgtatgtcagattcatc**tatactcccatctctccctggtgaccctgttgcttcctctctctatggtttcccccagcccccatcctcacgtgatctctcctgtggcccacctttcccatctgggtagtcaccctccaagtccttgctgggtctctcctcccctcac**ctccccgcccctttctactgcagccattaggggag**cccctgt**tctgtgcctccttacttcccatcacaga**gacttcaaagtccatttcctctgccctgggctggagccttccttcctctaatggccagctcctacctgtgcttcagatctcagcatgatccgcccctcctgcgggaagcacaccctcacctccctgctgagatcaactttcctctcataagctctcacagaatcatatgtctgttggtaacctttagcacagttggactttctcaggtacttatctgattatttttgtgctccacccaattttttttttttttttta**gacagagtcttgctctgtc**accaagctggaa**tgcagtggcaccatctcggttcactgca**acctctgactctttggttcaagcaattctcctacctcagactcccgagc**agctgggattacaggcacacaccaccacacccagct**aattcttgtttgtattt

Uppercase: TRBV6-9

Lowercase: Flanking sequence[1000bp]

Red & Bold & Underline: Stem-loop [20]

Blue: Heptamer[34]

Green: Nonamer [4]

id-IGKV1-37-3[V_gene_segment]

aaaccaaaaataagatgcagcttatataatctctgggtattgaggtggcttacaggtttaaataatggtgaatctgtgagtgataccggccttgaggacttgtggtttcatttttatgcatctgtagtgaattgcgtgaccatacagtcatgtaagaatggcttaaggtcattacaaaatactttcaaatatatttagcattatcttgtaaaattgtctcttttttttaagaaggaac**catttcactctcacagaaatg**ttataagtatagaacaaagtaccctctcctttgcagagtatattcaagacttgatgttcccaccctctaaaactttactgagtttcctacaaataaaaatattcccctggctggacgtggtggctcatgcctgtaatcccagcactttgggaggctgaggcgagggggtcacctgaggttaggagtttgagaccagcttggccaacatggtgaaaccccatttctactactaaaaatacaaaattagcctggcatggtggtgcgtacctgtagtcccacctaccctggaggctgaggcaggagaaatcgatggtacctggcaggtggaggt**tgcagtgaggtgagatcgtgccactgca**ctccagcctaggt**gacagaacgagactctgtc**tcaaaaaaaaaaaaccttaattaatccccatatacaaatagaaaacattacttcatcgagtactcaagagtatttcaaatat**taacaattattaaaaaaaaaagttctttgtta**caaaacggccccaagaggaaacatgttcttacagacaaatctgtgctaccctcctctgacttgggacactggggacacga**gggactggctcagagatgagccagcccc**gcagctgtgcccagcctgccccaacccctgctgatttgcatgttcccagagcacagccccctgccctgaagtcttcttaacaggctggtcacaccccgtgcagcagtcagtcccagtcaggacacagcATGGACATGAGGGTCCCCGCTCAGCT**CCTGGGGCTCCTACTGCTCTGGGTCCCAGG**TAAGGAAGGAGAACACTAGGAATTTACTCAGCTGGTGCGCTCAGTATAGCCTGGCTCTTCAGGGAGGTCTTCTTATAACATGATGGATTGAATGGATGTTTGTTTTTATATTTCCAATCTCAGGTGCCAGATGTGACATCCAGTTGACCCAGTCTCCATCCTCC**CTGTCTGCATCTGTAGGAGACAG**AGTCACCATCACTTGCCGGGTGAGTCAGGGCATTAGCAGTTATTTAAATTGGTATCGGCAGAAACCAGGGAAAGTTCCTAAGCTCCTGATCTATAGTGCATCCAATTTGCAATCTGGA**GTCCCATCTCGGTTCAGTGGCAGTGGATCTGGGAC**AGATTTCACTCTCACTATCAGCAGCCTGCAGCCTGAAGATGTTGCAACTTATTACGGTCAACGGACTTACAATGCCCCTcccacagtgttacaaggcataacataaaccccccaaggaagcagatgta**tggggctggcctgcccca**gatactcctcctactgcctccagctgctcagagggtttctcatattccagtcaagctttgaaagtcactgaaaagttttgatggaaggggccatgaagactctgaaactgcctttgcaaaaa**tcataactgagaaaattatga**cggtggaagatatt**acacagccagctctgtgt**gaattactctttcactactgctattgccgtgtcttgataaatcgactctgtctaagcagcaggcaaggtgaaccccttgggcgattacaactcctctataccttaactgtttttccttctcatcccctgcagcacaaacatggcaatactgttcctgatttcattaagaaaagagatgattacacctgagtcttggctatggtgtgagttggaatcaataccacaaaggaaaacctactcttggaattccaagtctatttttttttttctaaatacacgcaatgagagtctaaactac**agcctctgggaggct**tgggagcatggtactcaaagatgcaaataattagaaaacgat**tctctggatctccccaagaatccagaga**gcatctggtgcagaagctataattgccaatcatgtggtcctcagacatgtctgggaagcccatgttcggggctgttgatgctctgcctggaggaccttcctgattttctatggcatctgcttacaattgaccagccccctattcaaagcttgccgtgttgcatctcaactcttgtccgtgggaccccaagtctctagcc**acatccagttcttcacagaagctggatgt**atgtgccagggcaaggagaactgggtgggaggtgagt**ctgtgggctccacag**cttcacgtgtttatgaggtaggagttattaagaaattattttaggcagatagagaggaaaaaagggatccttgggagacttttcacagctgttgt

Uppercase: IGKV1-37-3

Lowercase: Flanking sequence[1000bp]

Red & Bold & Underline: Stem-loop [15]

Blue: Heptamer[29]

Green: Nonamer [4]

id-IGHV3-19-2[V_gene_segment]

gagaaagctgttgcttttacgccaatcaatc**tggagtcataaaagatacactcca**aaaagttcaaaaaaatctagataggagccaacaagaacgagaaaataacacagcctggtatcaaagcatgttcaattggaacccctggttaactactttaatcactggtttagctggacccatcatcatcatattattgagtttaatttttggcccttgtatattaaattggttccttgattttgtaaaacaacgtatagcgtctgtcaaacttatgtatctaagaactcaatataacccccttgttgtaactgaa**gaatcaacgacttgattcccctaaaacacaagtggggaa**atgaaatgcctaacggtgtttttactttaactcgttactttgaattttgtcctgcttgtctctttaatcacctaaccttgcttctcatgtaaataagactctctctagctaggaaagccagacaaactccaattgac**cccttagtttacaagacactaagggctcctcacccaaccccctttcgtgaggag**ttggcctgggtaaacagatcctcagcatttcaaaggagcccaattaactgataaggtacccacaccaacaatgtatgaagttcccaggaattttctccaagagataacaacatgaaaccttgagtttgtgtccggcatagaccctatatctaattataataaaagatttagaaccttgcacctggtaccgttgctcttcttgtaaccatttgtcttttaagttgtttatcactctgtaactattttgattattttgattcttgcatgt**ttttacttctgtaaaa**ttattacatttgagtccctctccccttcctaaacctaggtataaaatttactcgagccccttcctcgtggccgagagaattttgagcatgagctgtctctttggcagccggcttaataaaggactcttaattcgtctcaaagtgtggcgttttcttaactcacctgggtacaacaGTGCAGCTGGTGGAGTCTGGGGGAGGCTTGGTAGAGCCTGGGGGGTCCCTGAGACTCTCCTGTGCAGCCTCTGGATTCACCTTCAGTAACAGTGACATGAACTGGGTCCGCCAGG**CTCCAGGAAAGGGGCTGGAG**TGGGTATCGGGTGTTAGTTGGAATGGCAGTAGGACGCACTATGCAGACTCTGTGAAGGGCCGATTCATCATCTCCAGAGACAATTCCAGGAACTTCCTGTATCAGCAAATGA**ACAGCCTGAGGCCCGAGGACATGGCTGT**GTATTACTGTGTGAGAAAcactgtgagaggacggaagtgtgagcccagacacaaacct**cctgcaggaacgttgggggaaatcagctgcagg**gggcgctcaagacccactcatcagagtc**aaccccagagcaggtgcacatggaggctggggtt**tgtttc**ctgtcaggatttgggacttcctctgcttctgacagtttctctagggaaact**ctttaattttagatttctgtgcccaccaatgtcatctctacatttttttaatcattgtaatatgataggaggacttattctcacatgcacaatatgtatattgccacctacgggaatgaaaac**tcctcaaccatggtcaccagcatcagagtcgtgagga**agctcaggggtgcctgg**tgagtcttctccagtcagactca**ggacagtaacct**caaggggattcccttg**tgagaactcacaca**ctttcatgagaacagcaccaggagttggttctaaaccattcatgaaag**acccactccatgacccagtcacctcccaccaggtcaccccttcacaattgaggattataatacaacatga**gatttggggcaggacacaaatc**caaaccatatcagatacacattgtgaaatgctcatgatggtcagggtaatttgtatttctatcacctcacagcgctaccattttattatttttttaaattaactgcatgagtgagtgtctgatgagaacacctcagatataccctttcagcaacaatcatttttataatacagtattaactataggaccattgctgtacgttagatctccagaactcatccaacctgcacaactgaaactgtacaatttaacaaacatcacccgatttccctctcctcccaggtcctgagaccctctattctgctctctactttcaagagcttgaatattttagatcccacatgtaaatgagatc**atgcagcatttgtctttctgcat**ctggcttattccactcagcatcatgtcctccaggcccatccgtgttgt

Uppercase: IGHV3-19-2

Lowercase: Flanking sequence[1000bp]

Red & Bold & Underline: Stem-loop [18]

Blue: Heptamer[25]

Green: Nonamer [5]

id-IGHVII-33-1-2[V_gene_segment]

aggaagaccattggtgcctcccagagcacagttgtccatcagggatgtccaatgcgtcccagcagcagccatgcctcagtgtctccactgtgcacagccactgt**ctgggaggagctcccag**gatgggtgtctttggcacacaccaggtggcgggtgttagagtgcagtgcagcagctggctgcctgttctattgggctccctga**tgctggagagatgggaggtgcattcgcagggccagca**ccctgtttgtgaatttttatataaaaaccctgattttacttcatt**tttgcagatgacatagataactaaaaacagaatctgcaaa**gaaattgtaattttcaactttaccccaaattcattgtttcttaattctgtgcaagatccagacatattattgccttcctcatgataatttattctatgtaaaaccgaaatcattttttctcctactctttgtttctgttcaagtacagagatcttgagcaaagtaagttgggttctttccacacactaaccctcaccttccccagagaatgagcagagattgtcctcagtctgagtctaagggaggagctgtt**tctgcacaactcagagcctgcaga**gacccccaggtgcagcttcagtgagtcagacatttctccatgtgggcgacctccagtgcctgtgactgctgctcaggcctaattgttggttaagtattacgacacccttta**ggtgatcacatctcaagcctattctgaaaatcacc**atgaacagagatagttcaatggccattctcctgacataagtttctctttat**tacttggttccaagta**tggagaaaaatgtgaccctatatttgtctgaatccaacatcagcatccacggcattacgctaggaga**ctcactaattgagcacaagtgag**ctcgttattctcatagaaatgtaagtatttggaaatttcagcgtgttctccagaacctgtggctgccaacaactgtatttctcggtgcactcttGGCCTGGTGAAGCCTTCACAGAACCTCTCCCTCACCTGTGCCATCTCTGCATTCTCCATCACAACCAGTGTTTCCTTCTGGAGCTGCATCCATCAGCGCCCCCATGGGAGGTACTGGAGTGGATCGGGTGCATAGGTCATGGAGGGAGCACAGATTACTCCCCTCTTCTCAAGAGTCCAGTCACCATCTCCAGATCCATGTCCAAAAAGTACTTCTTCTTACAGGTGAA**CTATGTGAGCAACAAACACATAG**CCATGTATTTTAGAGCAAAAGA**cacagtgaggaaaccacagtgtgaactcatacccaagcctccctgtg**ggggtgcataggacagccagggttactcaggacaccaggctccctcaggacaccaaggggcactcaagaccattgtagaggcatgcaggtagctgggggctctcaggaaccatgggggaaaatcaggacaccaaagggtgcttggtacagcagggggctcaggattattgtggggattcagaaagagcaggttcaaggctcaacctcagggcatgtgcagctggtgtgaaaaggggctggatgaggcgttttgtgtcaccatcatgtttcaccaccagacaccctccactacctcta**ttctaatgcatgtgtttgtatgattagaa**aatgatattgatataaatatataaccatagctaggtgtgtcaagttgtcctctccatcttatatcagccttgtctgtaaggactaattccc**cataattacttgagaacctcataaattgtggtcaattatg**taggattcctctctttttctgccttccttcctccctccttctctctctcacacagaaacttatatacagccaccccacaacacacataaatcta**taacttttattacctgatgtattgataagtaatctaaagtta**cataaaaatcagtagttcactgtcaatattgtaggagaagggaaggtctaggaaggaggaattacag**aacaagaggatattttgagggtaattgacttgtt**atctatgtggataatgataatgtctatgaccatatttgtaaaattgaacacttcgtgtggagactattatttttttatttaaccccatgccattattttgccgtattctag**catttctctaaaaatacagaaatg**ttcagcactcattcatgtgtatattcaggagtttctgacttttcatgtattttatttatctctgtctaattgctttgataccaaattttacctagtataattagtactacctttagcactgatt

Uppercase: IGHVII-33-1-2

Lowercase: Flanking sequence[1000bp]

Red & Bold & Underline: Stem-loop [14]

Blue: Heptamer[29]

Green: Nonamer [5]

id-IGKV2-26[V_gene_segment]

cagtcaggttggaaagtaagtccagggatatgtagggttaaataaaacccatctgatgagaatttatagtttgtagggcatgactccccagactccttagataggaactgagacaaggaaaaaaaaaatcagagcttagtccttaccttgataaacacaaatgaaaagaaattattacttgtagacattcaataaatta**tctgaaatcaatttaagaaaacattgttcaga**atcacatg**aaattatagaggacagtaattt**gaatccacaagcagaaataaaggcctcgaaatagtaaagaactgggaaaaaaatagacacaatgataaaaaaaagtttacttcaatttctatgatattca**ctaccaagtacaaacaaactatcgtggtag**atgtcagaaagacagttactttggttgggatggggaggcagttattgaatgggaaggtgtacgagagaatcatccagaaattt**tgaaacattctatagtttca**gctggatag**tgaatatacataaaactttattca**tt**ccacatttaggatttctgccctttatatgtgg**tttatttcaatgaaaaaaaaatagaaaagccaagccta**gattcttttaaagaaaattggaggcaagaaaaaagaatc**agccaatatttatggaaactcaactttattaaagacaagtgcaaaatgtgcctggaaaagtgctaaagaactgttgaaaaacaggtggaaagaaaaggaaatattataatatgtctcttctccaactgaagcaacacagattttatcttggttagttttccttgaaacacgcaggatattttatgtccccttgcccacaagagaagccctcatcctcctgttgttgcgcttagcatcactgcacccaccgggggatttgcatatt**gtcctctagggaggac**cttcccttgtgggtctgagataaaagctcagctctaacccttaccttgactgatcaggactcctcagttcaccttctcactATGAGGCTCCCTGCTCAGCTCTTGGGGCTGCTAATGCTCTGGGTCCCTGGTAAGGACAGAAGGAGATGAGGGAGGAGAATGGG**GTGGGAAGGTAAGCCTGGGGGTCCCAC**TGCCTTCCATGTGTGTTCTGCCCTGCCCATGTGTTAGATGTACAGGTCTTGTTCTCCAGGATGGGGAATGTGAGGTTTAAATCTGTGAGAGTGAGGACAATTCAAAAAGAAGCAAGGAC**CTGTGTGATCTGGTGAAGATTGTCACACAG**AG**AAAGGGAGGTGGTGTAGGTGATTTCTAGAATCCCCTTT**GCGGCTTGCAAATTTGGAATATGTTTAGTGTATAAATACAAACAACAAAAAATTATATAGCCTGAAATAAAAAATGAAAATTTATGATAAATGACACATGATATTTGTACATATCCTTCCACTTCTTTCTATTTATTTTAGGATCCAGTGCAGAGATTGTGATGACCCAGACTCCA**CTCTCCTTGTCTATCACCCCTGGAGAG**CAGGCCTCCATGTCC**TGCAGGTCTAGTCAGAGCCTCCTGCA**TAGTGATGGATACACCTATTTGTATTGGTTTCTGCAGAAAGCCAGGCCAGTCTCCACAGCTCCTGATCTATGAAGTTTCCAACCGGTTCTCTGGAGTGCCAGATAGG**TTCAGTGGCAGCGGGTCAGGGACAGATTTCACACTGAA**AA**TCAGCCGGGTGGAGGCTGAGGATGTTGGAGTTTATTACTGCATGCAAGATGCACAAGATCCTC**Ccacagtgctacagccctgagcacaaacctctctgcttgaagtggccca**gctgctcacatgtgatgtttgtctggggagcagc**acagcaagtt**ctctgagtctgtgcatgaggaaaatgttggacaactcagag**gagtagattggtctcatgaattccttggttacatctcaggaactactcatttgtgcagctgcaaaagccttgacatggcaaaagcagtaggagaatggggaaggtcccatggagctgatgctgattcatgatgcaactgaataaaagctcattcgaatcctcctttcttacatttc**tcaattgtagttaatcagcataagtgccaggcaattgatacaaatgttggcaacacatgctgagtgcctcctcatttgtat**gtatttcatccatttttgtatattttaataggataattatttggtcatatttagaaattggtattttccaccttccaagctttctactttccttacttattgcttctcttcctatgtaataaagatactgtagaca**acattttacactagttgtaaaatgt**ggctccccatttatacgggtagctggttgggggcaattag**caaaaatcttggtttttg**agtgcctaaatagacttttgtaaattaatgtcagatacaagatccaaatgctaaaactctttgatttgccttgattacttcttactgatagatcataaaa**ggaattcttgaaattcc**aaaagttgggttttaaaaataaaagcatacactgg**aatatacagggcatagaattcaccctatatatt**gctagaagaatagagaatgataacagttttagacttcgagggtacttcaggaatgggaacacacaagga**acacagtagatatgagctactgtgtctaccgtattagaca**atcctcctgcattttaacttacatttggtgaacttcagatgccagtgggtgttgcaagttgatgacataaacctaaaatcaagtcccgaggtatttc

Uppercase: IGKV2-26

Lowercase: Flanking sequence[1000bp]

Red & Bold & Underline: Stem-loop [26]

Blue: Heptamer[23]

Green: Nonamer [9]

id-IGHV3-47[V_gene_segment]

atgtctcatgtctccctaaattgtgtaaaaccaagctgtgtcccaccacactgggcacgtatcat**caggactccctgaggctgtgtcacaggcatgtccttagtcctg**gaaaaatgaacttcctaaatctattgagattagtctcagatactctttggtttataagtatgttttctatctcataacttcaattatctgaaatactaaaggaaatctgcca**aataacacattatcttgtttatctgttatt**gttgttgcaaaa**ataaatatatttatataatttat**gtataattgatcatctttgtacattaccaaactgagtagcagatttgttagtaagacccaatgtaataaaaaattcaaatatcaattagcaacttaatagaa**aaacaaattacgctacatttgttt**gccgaaatgctacca**atttttataaaaaaat**aaacacatataacataaatggtttaa**ttctcagtatttctattgagaa**aaataagccaaatacataagcgaatatactatattatttcattcttataaattctaccaaataaatactagtctaaagaaatatgaaaacatcagtagtttataaagaaatggtagaaaaagggaaggggaaaaaacaaataatataaaagatcaagaggaatgctgaggggagttgacttgttacctca**ttgaaaatagagattattttttcaa**agtttactattgcacatgttaaatatgtg**aattttatcatctgtcagttaaaactcataaaatt**tattacaagtaaacagccgacattttatacaaaaaagggatgataggaagaaacaaataaatacattaaatgt**cagatacaccaaaaacttatctg**cctgacccctagttgtctccgtaatttttggatgaaaaccagcccacccctgaccctgctgctctgggagaggagccccagc**cttgggattcccaag**tgtttgca**ttcagtgatcaggactgaa**cacacaggactCACCAGGGAGTTTGTGCTAAGCTGGGTTTTCCTTGTTGCTATATTAAAATGTGATTCATGGAGAACTAGAGAGATTGAGTGTGAGTTACATGAGTGAGAGAAACAGTGGATATGTTTGGCAATTTCTGACCTTGGTGTCTCTGTGTTTGCAGGTGTCCAGTGTGAGGATCAGCTGGTGGAGTCTGGGGGAGGCTTGGTACAGCCTGGGGGGTCCCTGCGACCCTCCT**GTGCAGCCTCTGGATTCGCCTTCAGTAGCTATGCTCTGCAC**TGGGTTCGCCGGG**CTCCAGGGAAGGGTCTGGAG**TGG**GTATCAGCTATTGGTACTGGTGGTGATAC**ATACTATGCAGACTCCGTGATGGGCCGATTCACCATCTCCAGAGACAACGCCAAGAAGTCCTTGTATCTTCATATGA**ACAGCCTGATAGCTGAGGACATGGCTGTGTATTATTGTGCAAGAGAcacag**tgag**gggaagtcaatgtgagcccagatacaaacttccctgcaggaacgctggaggaaaccagctgcaggg**ggcgctcaggagccactgatcagag**tcagcccaagaggcaggtgcacacagaggctga**tttcctgtcagggtgtgggacttcgtctttttaccatttctctagggaacctctctaagttcagaattctgtgcttaccagtgtcatctctacatgttttttaatgattattttaatatgagaacctattcttacatgcacaaaatgcagatggatgcttacagagatgaaaagtcctcaaccatgg**tcaccaggatcagccttgaggaaactcaggggtgcctggtga**atcttctcc**agtcagactcaggacagaaacctctgtgagattccctgact**agatcagtcttcaggaattttgata**ccagccaatagagaggctgg**gac**agggtcagtgtcatgtagaacctcacaggtttcacgtctgaccct**tctccttctcctgacactaaagtatgcaa**atcagtatcagcactgat**ctgggtccccttttgctcttaaaccattctatttctttttatttgttgttgttcttgcttttcctcgtacttctcttgctccctgtaaaatggggaggtgtttctt**gctgcaaaagccccaagcctcaagcccattccctgcagc**tcaggcggggctcaggctgtggctcctg**cagccacatgggagtggctg**ttggggctttctcttctcccattgctcagcaccctccagtg**tgttgtgtggagactacctgggaacgaatgtggacaaca**ggagtgaaggggatgagcttgcgtggacaaaatgggatgtggatgtgaaatttatcctgtgctgtacaaacaaccacagattcaacttcctcaccagtagtattagaaagagggtgtgaaagttgtcagaataaaaatggaaacacttgtgttaacaccctgacaaatggaacta

Uppercase: IGHV3-47

Lowercase: Flanking sequence[1000bp]

Red & Bold & Underline: Stem-loop [27]

Blue: Heptamer[35]

Green: Nonamer [6]

id-IGLV2-18[V_gene_segment]

tgtcgcagggttgagagcagccagctttaggttgccctgaggatgcgctcagatgggaaaaagcgagaaagcaggaaacgcgtct**ctgtgacctgacagataaatcacag**taatggaaataagtctttcaataattcaaacattcacaataaatatcactctgatgcacacagaaaggcctggtaacaaactttccctttgacctgctggctcatatccccatggatcacctgtgtgcagagtggactctctggtccccat**ttctcaggctgagaa**ctagaggctgggaaggacaagtggc**ctgctcatgagtgggaagggcgggattgggagcag**cagactcgggctcaggactctggccctggtcagtgctccttgttccctccacagggtccct**cccacatgtgcccgtgtggatgtggg**tgcactgcctggggtggctgatgtgctccagggattcaatgcggcagatgcttcagcaccaaaggtagggagtagggatgggacgaggctgcagctgtggagaccagggacccatgatgcccagcagaggtttccacggggtgggtagccccatttcctgacagtagagacagcctagaggtgggtgccaagcaga**gggcactgtgccatgtgccc**aggcctggagggccacagagacacaggcgccacacagcagagacactgagagccaggagctca**ctcaggtaagggatgtcagcagatctttctctcctgag**caaatcaggtacaaaaaaattaagatcctgccatcagaatagacaaacagggactttaatctcctcggctgagccccgctgtccagggaagaagaagtctctgagcccaggcccaggtgagggtggagtgaggagaggagctcagggtgcagatttgcataaaggccccgccct**cctctgaggcagagg**ggataagacagggctggggtcaggcccagtgctgggatctcaggaggcagctctctcggaatatctccaccATGGCCTGGGCTCTGCTCCT**CCTCACCCTCCTCACTCAGGGCACAGGTGAGG**CCTCCAGGGAAGGGGCTTCGGGGACCTCTGGGCTGATCCTTAACTCCTGCTCCTCAGGCTCACCTGGGCCCAGCACTGACTTACTAAAATGTGTTTCTTCCTTTTTCCAGGATCCTGGGCTCAGTCTGCCCTGACTCAGCCTCCCTCCGTGTCCGGGTCTCCTGGACAGTCAGTCACCATCTCCTGCACTGGAACCAGCAGTGACGTTGGTAGTTATAACCGTGTCTCCTGGTACCAGCAGCCCCCAGGCACAGCCCCCAAACTCATGATTTATGAGGTCAGTAATCGGCCCTCAGGGGTCCCTGATCGCTTCTCTGGGTCCAAGTCTGGCAACACGGCCTCCCTGACCATCTCTGGGCTCCAGGCTGAGGACGAGGCTGATTATTACTGCAGCTTATATACAAGCAGCAGCACTTTCcacagaggtcca**agttcctggggaactgagaccaaaacctgccctgggctctcag**gccccctccttgctctgaagatgcttcctcgcccagtgcaagcggcttcctgcagtgcggccttgagaattcttctctctcagctccttccctttccaccatgaattccaaaag**gaaacccgctctgtggtttc**tcatccaggac**agggacagcttcctgatgcttgtgcgctgtggtccct**caatgtgcagctcttcccagctcttcaaatgcaggga**cagtgacaaggagctgcctgagtggtgcagtcactg**cttttttca**gggatgtcttcaccctacatgcatcaccatccc**ctacactgtggg**tagaattttagcaactacattcta**atggttatcgccacaactttgatcttaaataacagtgcagcgaacatccctacgcagactcctttgagt**tcctgtgtgaatatgaccacagga**ttcatttctaaaagtgaaattgcaggtcagaaagattgtgtgtttgtaattttcacccattgttgtcagattcccctccagagggattgtaccaattttcaatgccagctggaagtacttgttcagagtacattgttgaaagtggaa**atttttgccagcttatcagacagaaaaat**gttaccttgttacagttttatttttgagtctcctgttctgcttacattttggccctttcagaaatagttcagtactttttct**ttttttaggatttagtaaactttataaaaaa**tatccagataaggagaattttagtgtttgtgggcaggcacagtggctcacacctgtaatcc**caagcacttcgaaagagtaagacaggtgaattgcttg**ttcccaggagtttgagattagtctgggccacacagcaaaccctcatctctattaaattaatgcaaata**aaaataaattttaaaaagaattatttt**agtctttgtggcccaagaatcaactttagggatatta

Uppercase: IGLV2-18

Lowercase: Flanking sequence[1000bp]

Red & Bold & Underline: Stem-loop [20]

Blue: Heptamer[51]

Green: Nonamer [2]

id-TRBV29-1-2[V_gene_segment]

ctctgttacttactcttggattatatgaacttggataggttatttaaggtctctaggtcttatctgtaaaatttcctaatccgtaaaaatagtgtgatgataataaatagcctagccaaaaaaatagacttaaataaatgtatgaaaagtagagtaagtgctcaaaaacttagct**ctgcttttatgcaaaaatgaactattttaagcag**ttacatgatacgagaaatcagcttgataatg**gaaaaagatcataccattgtggtcaagttttacacaaacatgatctgtttc**gtggcctagactgagttgaattagaatagaagaaaccatattacaagtattcaggaattaagcaaagacaaaaaaaaaat**agttatgactgtgataact**gaaacaagggaagagatataagagacaatgcattta**atttgtgaattgacaaat**aaggtaggtgaggaaaaaagaggcttatggaa**ttcttgaacctaacatgcaagaa**gagcagggatgccatttgttgggagaggggaattgaggaataagagcccgtttattggtgtgtgtacgtgttgggggaggaaggagttcagtttgggacaaccatgaagacccagcaggcagctgaaatactgtaatgaagttcccttatattctaacatattacttttagttctttgaagtcaattctgcaccccactttttgctctacacccagacagtagcacagattggtgttcagtaatgacagatggaaaagtgatgggagaatggtggtgagtggctggtgcatgctgtgagacttgtgcatgaagactcatcctgagtgaagagccacacggttccaccagggggaagactgatttttgcatatcatgcagaaactcaaccacctgttagaggagggaaacagagtgggaaggaactacgctgtaggaaggaaagatgaacttgagtttcacttcttagtgccttttctcaggggagaggccatcacttgaagATGCTGAGTCTTCTGCTCCTTCTCCTGGGACTAGGTATGAGCCTTGTCTTTTGAGGAGTCTGGGGTTAATGTTAACACTGGGTGGAAAGGTGGAAAGACTTTCTCTCCAGAGAGTCTGAAGCCACCAGACAGAAGATGAAGGGGATATGTTAGATTAAAGAACCATAACAGAGTAATTGTTGGCACAATAA**AAAGGAGGCAGTTCCCTGGGGTCCTTT**GGGAGCTGAGGTGGTATTGGAGAGATAAAGCAGAGACCTTCCTGCCTGGGGAGATGTCTGACCCCAAGCTTGTCGGTCCTCATGAATATCTCTCTGGAACTAGGCTCTGTGTTCAGTGCTGTCATCTCTCAAAAGCCAAG**CAGGGATATCTGTCAACGTGGAACCTCCCTG**ACGATCCAGTGTCAAGTCGATAGCCAAGTCACCATGATGTTCTGGTACCGTCAGCAACCTGGA**CAGAGCCTGACACTGATCGCAACTGCAAATCAGGGCTCTG**AGGCCACATATGAGAGTGGATTTGTCATTGACAAGTTTCCCATCAGCCGCCCAAACCTAACATTCTCAACTCTGACTGTGAGCAACATGAGCCCTGAAGACAGCAGCATATATCTCTGCAGCGTTGAAGAcacagtgcggggca**cagatcaaagatctg**agcaagaacctcagctctctcctacccagctcctctcacacgagcctgaaggccctgccaaggtgggacagaaggaggaaaccacagcttttgggcag**acacagctgtttctgtgt**ttgtgggtgggcatgggtatgaatggatagactggggatgaggagagcatctgtaggactagctggagcgttctatccacaggcaggaagagtggctcctggaaagctgagggtaaa**atttcttctagaagggtgctcagaaat**acccaataaaaacaatcagatacagtttccttcaagtggcggggtcagtaat**agcctttaactgttttatctcaaggct**ctttttagctggttttctatcataatatattttggaagaactttagtcatcttgaaacaaagctggtacattatattaatgacatctggttgttctcagtgaataggaatgataaagccctctagatgcttcaatatgacaatatgacacatgcatcccaggccaatataaaatctgcaaagttcaggaaccttctatactggtgagttttggagtggtccagatgtaagaagtaagctgc**tgcaccttagatcatgtaccacggagaaagaggtgca**atgcttagtctttggactttgaattaaactttggatatactgcatttgattgtgctactctatcccatttactatgtaatctgtaaagctggggccagtgcaagagagggttctgtggcagaagaagcccatgatgaaagcttcctg**ctgcttgggccatggactaagcag**acccagttgttcttgaagagcctatggtagatactgatgccctgtggagctcctgacattccccaatacgagaatcacaaggcagtgtggaacaggtttgctgtgccctgatttgcacttcagaccattgcatcttgggctatacttttgttgctttccagatcccagagagaaagaatgctc

Uppercase: TRBV29-1-2

Lowercase: Flanking sequence[1000bp]

Red & Bold & Underline: Stem-loop [14]

Blue: Heptamer[27]

Green: Nonamer [4]

id-IGHVII-44-2[V_gene_segment]

tgcacctctctctccctccccactccagcagttcacagtgcctattgtttccatggtcatgtc**cacatgtgctcaacctttagcactcactttcaagtgagaacatgtg**atattttgtttcctttttctgtgttaacttgtttacgactatgacctccagctgcatccatgttgctgcaaaggacatgattttgtttctttt**gtgacagagtctcactctctcac**caggcaggag**tgcagtgacgcgatctcagcttactgca**acctctaagtccctggttcaagtgattctcctgcctcagcctcccggg**tagctgggattacaggcatgtgccactatgcccagctaaattttgtatttttagc**agagagggggtttcactatattggccaggatgatcttgatctcctgacctcatgatctgtctgcctcagcctcccaaagtgctgggattataggtgtgagcc**accacacttttttttatgaatgtgtggtattttatgattttatattgcatacatacca**ttttattcatctaatacatcattgattggcacctagattgattccatatctttgctgttgtgaatagtgtggcaagaggcatgagtgcatgtgcattttttggtagaatgatttattttactttggctatatacctaatgatgggattgctgggtc**aaacagtagctctgttt**caagttctttgagaattctctaaactgctttgcacagtgctgaactaatttacatttccaccacccatgtatgtgttcccttttcttcacttcccacccagcagctgttttttgtttaacttttttcatattagctattatgactggtctgagatggtatttcattgttgtttttatttgcatttctctgataattaatgatgttggaaatttttttcatgtctgttggccactgagattgacaggcttacaagtgtcttagtccagcagttttcggtaacaggctatccactttcattttCTCTTCCTCACATGCCCCATTTCTGGGTTGTCCATCCCAACCAACAGTTACTGCTGGAGCTGAATCCACTAGCCCGCTTCCATGGGCTGTCAGGAACACATTGCAAGGGTGACACACACGATGATCC**ATCTCTGCAGAGCCAACTCTCCTTCTCCAGAGAT**TCATCCAAGAAACAATTTTGACTATACCTGAGCTCTGTGACATCTGAGGACATG**GTTTGTATTACTGTGCAAGACAaac**agtgag**aggaagtcaatgtgagtccagacataaaccttcct**gctgagaacaatggaaag**cttttcttctaagataaggaataagaaaag**aatgcccagtcttaataattctaatcagcatattgatgatagttttaaccataga**aatttgagaaagaaaaaaatagtaaaaggcatacaaatt**gaggggaat**aagttaaattgtttctgtttacagacaatataactt**tataaattccaaaaaaataaaaaattctgaaaaaacagccactgaaaactaacaaac**aaatttagtaacattgcagtatacaaaattt**atatggaaaactcagtagtgttactaaaaactaaaaattaactatttaaaaatcataagacaatgtcagttgcatttacaatattaaataacttacttagtggccatgcacagtagtttacacctgtaattcttgtactttgggaggtaggggagaatcacttgaggctaggagtttgagaccagcctgggcaacacagcaagaccaatatctatttctaaaaa**tcacttagaaaaaatgttggtcaaataagtga**aaatctgtgcattgaagactatatgacattgatgaaataaataaatctatatgcaaat**atatgggaaactattccatat**tcacagattgaagataataatactgttaacatgtctatactacccaaagcaatctacagatccaaagcaatttctatacaaatgacattttcaacaaactactttttaaatgataaaattgaatcacaaaagaccccagatgtccaaagtaaccttgaatacaaacatcaaagcggggggaggtttcctagtacctgaactgcaaatatgctatgaaactacaataataaaactgcatgctgctgactttctaaataaactttaatattctgaattgaagttatttgacaatttcttctatcattttatctttactaaagtgttgcatgagtgtttatctagacag

Uppercase: IGHVII-44-2

Lowercase: Flanking sequence[1000bp]

Red & Bold & Underline: Stem-loop [17]

Blue: Heptamer[25]

Green: Nonamer [5]

id-IGHV3-72-2[V_gene_segment]

tggaccatccctctccttagctgggtggttgcagactgcaagggaccctggtgaaacgagggagatgtccgctttattagtgtgttctcgtattgctagaaagaaatacctgaaac**tgagtaattcacatagaaaataggtttaatttactca**tgattctgctggctgtatgggaagcatgatgaaaaatctgttcagcttctggagaggcctcaggaaatttacaatgatagtgaaaggcaaaaggggagcagccactttatatggtgggagcaggagcaaagaaagagtgagggtggaggtcccacacactgttaaacaaccaaatctcatgagaactcatttactatcatagcaaggggaccgtgctaaccattattcataaatccggacccacgatacaaccaccgggccccgcctcaaacattagggaagcaaaatcacatacagttgttatgacag**gtgggacatcctgaaaacctctctaggcatgtcccac**acggcc**ctggagctgtctcaggggagcagtctcctccag**tgtttagacgcacaggcacggataatagggctaagtctggccag**atgtgtgatattcaacacat**tgcacaactgctctgttctgtatgtaatttatcttctctacaaatgtaacattgatggttgcattaaatatattctgcaaatatgtaaaaataaaataagatga**tgattgctaaatgattatcaaggcacaatca**ca**taatccgaagttacaatttcctgagagataggatta**ctgcccatgcttaccaggacactcacatct**gctctgggcactgccttctcctcaggcgtcccaccccagagc**ttgctatatagtaggagacatgcaaatagggtcctccctctgctgatgaaaaccagcccagccctgaccctgcagctctgagagcggagccccagccccagaattcccaggtgttttcatt**tggtgatcagcactgaacacagaggactcaccA**TGGAGTTTGGGCTGAGCTGGGTTTT**CCTTGTTGTTATTTTACAAGG**TGATTTATGGAGAACTAGAGATGTTAAGTGTGAGTGGACGTGAGTGAG**AGAAACAGTGGATTTGTGTGACAGTTTCT**GACCAGGGTGTCTCTGTGTTTGCAGGTGTCCAGTGTGAGGTGCAGCTGGTGGAGTCTGGGGGAGGCTTGGTCCAGCCTGGAGGGTCCCTGAGACTCTCCTGTGCAGCCTCTGGATTCACCTTCAGTGACCACTACATGGACTGGGTCCGCCAGG**CTCCAGGGAAGGGGCTGGAG**TGGGTTGGCCGTACTAGAAACAAAGCTAACAGTTACAC**CACAGAATACGCCGCGTCTGTG**AAAGGCAGATTCACCATCTCAAGAGATGATTCAAAGAACTCACTGTATCTGCAAATGAACAGCCTGAAAACCGAGGACACGGCCGTGTATTACTGTGCTAGAGAcacagcgag**gggaggtcagtgtgagcccggacacaaacctccc**tgcaggggcgcgtggggccaccagggggcggtcgggaaccactgaggacgg**gacaggtcccaggagcaggtgcagggggcggtttcctgtc**agctgcaggaggcgggtttgtttttgcaggactctggagccttatgaggttgcactattttattatggttatttatcatgtgttttttattgatatttgtgttttagtaatttttacatttatatgtaggg**gtattttttaaaaatac**ataacttcaagaaataattcttcctagtaatttacacctattcctctaaagttttattaacacctgttgatatcagcaactacatagctatgcaacattaatttacatctataaacatatgtgtgaatacatgaaactatgcatacatgtgtagtc**atttatatttaacattataataaaataataaattaaataaaat**aattgaattaaattaaaagaattaaacttaatgatgaactaaatataaattaga**gtaatccataattgatttcaatccttctctatggtttac**ataaatcgatgtctacttgtaagcttaagtatagtgaattggtcattttaaaatagc**caagaacaaatttcaaatgttcttg**tcaccaaaaaatgataaggatttgaagatttgaggtgatatatatgtcaattagctcgattcaattattccacattgcattcataaatcataacatagct**ttgtaccccatcagtatatggggtacaa**cc**aaaatttctcaatttccaatgaaatttt**aattatatattttttaatttgatgcctctctttggatcaggccacctcctcaggattacagtgctgttcagttttctcaacatgctgttgtaccagataagcacaaaacatttaatttcattatgctttgctttagtttttcaaaacaacataaaggtaataattttcccaacaaacatattacag

Uppercase: IGHV3-72-2

Lowercase: Flanking sequence[1000bp]

Red & Bold & Underline: Stem-loop [20]

Blue: Heptamer[30]

Green: Nonamer [8]

id-TRBV26-2[V_gene_segment]

aaaggaaaaccaaaaccaagccgaattgaatactcaaaaagttacagcaccttactattgtacttatttttttggatgctaacccaaagccaggactgaaataatctaaccaggacctccctagggtaggattcaaaccgatgaagctgcacaaaaa**taggactctaatctacagtccta**gaccaaagacaatttaaggtgtaagcacatattaacaaggtattgactccaaaagacatcttatccagaagaatttctttcctcaaaagagaaagtagcactcaaagggcctgaaatgtcatgacagagaataaggacctcacagctaagcctctggataaactggtctaagtagcaccactcaggg**tcagtgaagaaactttaattcccactga**ggggctacagtgccttggcaggtgttggcctcacattgggcacctat**atttgttatcaaacaaat**gggtttgcacacagaagccaataactatggcaccagcttttgagaaaagaaaaggctttattgaaacaccagtcagcaaaaagaaaggagtgggtt**caaatctgtttccatgatttg**gggtctagggcaggttttaagggatcagagagcaagggaaaggatttaggaatgttgatatggtagggtctgattagaaggcttcaaatatgactatctatggtaagatatgttaagaaacattttagcccgggatcttctgggccagtggaccgctcacta**ctgaaagaggtctggctttcag**gttccagtcatgtcttggtcttcctgtttctaaggggaggaatcttcggttctgggtattgttagaggtcaaagcttttactactgcacatgcctgg**gctacatgatttgcagttatgtagc**tgttggcagtgtgggagatgagtgttgtttctctggccaggaaataaatgaaaagtgctggttatcagggactaaattcatca**cagcaccaagcaacatcctgcttttggtgctg**ccATGAGCAACAGGCTTCTCTGCTGTGTGATCATTTGTCTCCTAAGAGCAGGTGAGTCCTGGGCACAGGTAAGGAATCCCTATTACTGAATTCACAGGATCTAACTATAAAATGTTTCTTCATAAAACTCACATTAGTCTCCTCTTTCCCAGGCCCTGTCTCTATTTGATCATCCATTTTTCCCCCCAACAGGCCTCAAGGATGCTGTAGTTACACAAT**TCCCAAGACACAGAATCATTGGGA**CAGGAAAGGA**ATTCATTCTACAGTGTTCCCAGAATATGAAT**CATGTTACAATGTACTGGTATCGACAGGACCCAGGACTTGGACTGAAGCTGGTCTATTATTCACCTGGCACTGGGAGCACTGAAAAAGGAGATATCTCTGAGGGGTATCATGTTTCTTGAAATACTATAGCATCTTTTCCCCTGACCCTGAAGTCTGCCAGCACCAACCAGACATCTGTGTATCT**CTATGCCAGCAGTTCATCcatagcactgcatag**ccatatcctctccacacaaaaaaaggtgcataccaaa**gaggaaaagcctgccctcaaaattcctc**accccgaataagagaagtttcctcacaagtattgacaaaaagagccaaactctgtaaaatatttgaagagatttattc**tgagccaaacatgagtgagtgtggccagaggcagaggctca**agagggcgtaagaac**atgtgtccaaggtgctcaggctacagcttggttttttggagacat**cagacatcagtcaatacatgtaagctgtacattgtttcaatttggaaaggtgggacaactcaaagagggggcttccaggtcataggtggattcagagatttttcaggctggcagttggttgaaagagttaagttatcatctaaagatttgaaatcaatataaaggagtgtctgagttgtggggacaaaggttcttattatgcaga**tgaagcctccagggagcaggcttca**gagagaagagattgtgtttcttaatcagacttttaaaaggtgccagactc**ataattctctcctggatcaggggaaaaaacctgaaaaggaaagaattat**ctacagaatgcagattttcctcatgatagacagctttgtagggccatttaaaaatatgtcaa**aaaatatactttggggtaaaatatttt**gatttttttcagggcctgctgtgagtcatgttggtgtcttattgct**acaaaaggtctgttttgt**cacttgtaaggtctccattctaatgttaatgctggttggttgtgcctgaattctaaagtgaagagggtataatgaggcatgtctgaccccacttcccactaag**acctgaactggtttttcaggt**taactttggaatgctcttggccaagagaaggggtccacacagttggttgtggggcttgggattttatt**tttggtttacacaagattataccaaa**gcccttccctaaatttgatatgccctgcttccctctttcca

Uppercase: TRBV26-2

Lowercase: Flanking sequence[1000bp]

Red & Bold & Underline: Stem-loop [19]

Blue: Heptamer[19]

Green: Nonamer [4]

id-TRBV22-1-2[V_gene_segment]

gcagataacctcagaaatagaacaacatttgtctaccaaaacaaaagcattcattaagaaatcagctaggatcctatttttatttactaaaactacttggaagaacccatgttatacaatgaatttacactcatacaaatttctactcagttgga**aaatgagactttagtcagctggtacaaccctatgaccataaatgtctcattt**cacttgatacagtattgacttcctaacttcctgtgaattagaacttcaggtccagaaagtaataagagcaaacgcaaatctagtgt**ttcttatgtactaagaa**ctgttctaaatgctttatttatgtaatgttcatcacaggttttgggtagttacttattatc**tcaattttacagataaagtgaaattga**agagat**gctctgtaacttttccaaggttacatagc**tgcgaaatggcagagacaggactaaagcccatgccgtgtcccaagtccctgctctaatccattgtcacaagggaaggggcgcttcagacactttgctattcctaaaattgcaggagtagcccccaccccatctcta**ctgtattaattatttatagtgaatacag**agggtgtagagaatcggttctttaaaaaacaaaaacaaaaacaaaaccactaaacttccaaaccccttcttgtgttgagttgagcacaggtgcatggcactctgtgtatgactctgaccaggaagtgagggattggctcttctgaccatgcaggcaggcacttgtccatt**tagaagggatataaaaagcttcta**taagcagaagaggaggtgcatgtgacttgatcgctctaattcttagtctcaccatctgcaaaatcagctgtgaaaatcagtcatcacgtagcaatgacaaattctttatattaagcccc**ggggaggggctagatctcagccccttccc**ttggcctctctc**tctctcttagagcctgtgtctgtaacttcagagaga**ttttgtcgctgccATGGGGAGCTGGGTCCTCTGCTATGTGACCCTGTGTCTCCTGGGAGCAGGTGAGCTCAGCACACATCCAGGTGGTCAATTCCTATTCTGG**GCTTGCCAGGCCCTTCCCTGGCCAGC**ACCTCCCTTCCAGGCATGATCTCCAGCCTTTGTCTCTTGGACCCTTGGATGCTGACATCTATCAGATGCCAT**TCCAGCTCACTGGGGCTGGA**TGGGATGTGACTCTGGAGTGGAAACGGAATTTGAGACACAATGACATGTACTGCTACTGGTACTGGCAGGACCCAAAGCAAAATCTGAGACTGATCTATTACTCAAGGGTTGAAAAGGATATTCAGAGAGGAGATCTAACTGAAGGCTACGTGTCTGCCAAGAG**GAGAAGGGGCTATTTCTTCTC**AGGGTGAAGTTGGCCCACACCAGCCAAACAGCTTTGTACTTCTGTCCTGGGAGCGCACc**acaatggaagcacaaccattgt**ctctctgtgcggaaatgtgtcctcaccctacagcccccaccacatcctctagcttaattttttcatttttaatattttcttgagattttactatgtcctgtta**cttttgcaaacacaaaag**atacaaaactgaaaagaaaatagatcctttatccaaggagctaagagtctagcaataaagtagttataacagaatggaaaatataatgtataatactaaaatcagaagtcatttgggagctttagggaggacatgcccaattctg**tctgattgaatcaga**gaaaacaaagaggaagtcacattagaacttcaccagctagacctggttctgagtagggcattctgggcaaagagaacagcttttgcaaaggcatattaaagagtttgtcactttggggaacagtttacaaagagaaagaagatgatcaatttatatgaatcaaaagaaatggaatacatttgaggaagtagacagaaaggagtttttgaaaaagttgagtttttggttccatgaatacaataagtatgaaagtcctttgtaggctgttgctcaaaagagagagctgaagcttcattgtcatcagcagacaagcaaacaatacaatttttc**aatatatatatatatt**ttcctcatgtaactttgaaatgatactatttttccattttcctcatataactttgatccttaacaaagggcacagtttcaaacaatatgaaatcatgttagcagtagcagttcccatttgggataatagggttcaactgtctatggaacaatgagataaaaactagaagcaggcaactgcatagacaagatgtgaaattgagatgccataaggctggactagaggttgagattctgcagatttcagcatgtaaataattgaactatggggacagaaaattccaaccaaatatagatttagatcacagagataaaaggcatggaagttaagactgtggtattctggttcct

Uppercase: TRBV22-1-2

Lowercase: Flanking sequence[1000bp]

Red & Bold & Underline: Stem-loop [15]

Blue: Heptamer[26]

Green: Nonamer [7]

id-IGLVIV-65[V_gene_segment]

tcacatccatctttcagggtcctgctcccagactcttctgacttatctgttcactgtgctgggttggcagggaaaagtcatctgcagcagaaaaagtgacaaaatggagatacatagtgtgtcttggccaaaaatcagccaacagatgagagagatctaggat**tgtgagaagttctgccgttccccagcacctcaca**ctatttgatgtctcaatgtttgtacatatcactatat**tttctagaagatgcactacatctagaaa**tagtgtttgctttgtttgtccccaagcaggaccccactctccactcagagccagttcacagttgccttcttccctggcccctgtggaaggagctgctcaggtgtttgctcccagagccctccgcattttccccttactgcggcctcctcac**aatgaacaattgtcatgtttttcatttctcctgttcctgttattctagaaat**ccctcaatcaaaaaggatggttatgaatataagttacctgagacagagcaggggtgtggtccatggatgctgtgacctgacagagacccagcactcaggccaactccactcacatgattctgact**gaaagactgaagtgtgcagcagggggaaatctgggtctttc**caagctctgaacagtgg**tgcccacatatctgttatccctgggca**atggagcctgcactcactgaggaagataccacagctgtgtcttctctgggctgggtagagtctcctactcacaggtctgtggaaggtgcaggagctgcttcctcacttactcctcccgtgggacaaacgctgaatctcacccaccaacccaccatatctaaagaatcatgctcagacttcagctcttcctagggtggaacaaagattacca**ggaggatacacaaatttgtgtggacagatcctcc**cttt**tcaggccactggagaagataaaagaggcctga**gttaagcagcttcctctgtggggccacaggaggcggtgtccaccATGGCCTGGACCCCTCTGCTCCTCATGTTCCTCTCTCATTGCTCAGGTGAGAACTGGTTTTAGGTGTCTAGGGTTAGCTCCTTAGCTTGCGTCAGCCTCTCTGGCTCAGATCCCCAAGACATGTATCTCTAGTTCCTGCCCCAGCCCCCATGAGTGTCTGTGCTTGCAGGTTCCCTCTCCCAGCCTGTGCTGACTCAGATGACCTCACTCTCTGCATCTCCAGGAGCATGAGCCAGACTTACCTGCACCCTGAGCAG**TGGCTTCAGAGTTGGTAACTTTGCAATATAGTGGTCAGCTGAAGCCA**GGCAGCCGTCCCAGGTGTCTCCTGAGATACTACTCAGACTCAAATATCACTCAGCACCCTGGGGTTCCCATCCACTTCTCTGGATCCAATGATACATTAGCCAATGCAGGGATTCTGTACATTCCTGGGCTGAAGCCTGAGGGTGAGGCTATTACTGTTGTACGTGTCACAGCAGCTCCAAGTcagcactccagacccactgg**gaggttacaaaaacctc**ttctctgatctcctggcctggtgtagtcactcctgctggtggctctaataaagtctatctcactgggtgacttatattttagatcttcaaccagcacctgattcttt**ttttcagacaagtagatctgaaaa**tgtgactttattttcttcaattccactttcatattgattcatgctcataccttctcatttc**tttctggttttcacaggcagaaa**tcccttgctcagagacaaaaattctctaaattgagactttggaggagaccaaaagagagtggttcgggccatgcccggctcagagtctaggagtgactcagagatgcttctggtcccagagcttcaatctaggcattgg**atacaataccagcattcacaatggaattgtat**tctctatattagcaacaaaccatcacagattttgaaataaaaagaatataatttataataacatcaatataa**aatatttaagggtaaatatt**aaaaaatggaaagtcttaaactgaaaactataaaacgttgagagtcattacaaaaaaaccttaaaataactgatattccttgtttgtagttggaaaactcaa**tgtgattataatttttataatgaca**attctttccaaagttatctaaggaatcattgtaaggtcttcagtaaaaataggtttgcagggaatgaggagataaaaaaatctcaacaaagaaatgaaaactgtaacagaaaacatcaatgaaacaacagacctgggataatccttctcagctgccatgggatctaactgtatttaagaaaggggtgagtaggaaggcggctgccaaatccaggaacacagatgtcattttgagtgaccaaggtgagccaagggaggacagctttagtcagaagagcatagaccaggagcagaactcacatatctgaagctgatgggcagagtgtgggtgcaataggcctggcattgggtgctgctga

Uppercase: IGLVIV-65

Lowercase: Flanking sequence[1000bp]

Red & Bold & Underline: Stem-loop [15]

Blue: Heptamer[32]

Green: Nonamer [2]

id-TRBV16-3[V_gene_segment]

tgatggattggaaaaatggatgtttgagtgcataaggttcacgagtataagaaacaattggtacctcctaactatagattgtcagtttca**gaggatgagaatgaatgtgccctcatcctc**attttaattttgaaagacaatgcagatatccctactatagaatcattttttacatcaccacctatagagattaacaacagtagatgtttccagctccctaagaggccaagaagtattaacccaaggtcagattcaggtaattgatcgtgactcacgaattttcagagatccacaatggttgtttttaaattttcagcagttatgtctttttctctggggagagggtattcaaagctcctcatgaggccattctggactacttttcccctcctgccccctgcccataaaacagtttacagaaataggtgtccagaccgtgggctgtagttcaccaactcc**tgctctagcctaattttgtctcattactagagca**tggccattctgtggatgccagtgaagcctttctggggtttctctccaatgccttttaaagctttttcctacacagacacagtttactgtttagccaaaaaattaagataattc**ctccagacatctggag**ttcttgtgcagcttcctcctctctggtggtctttccttaaaaattctagctgcctcagcccccgctg**attttgtaacctgtttccgcaactcaaaat**ttgccatgctctgttgctctcttccccattgtgtatgtgcagaaattgcccccatgttttggtgttttggcagaatcccagtcctgtgctgtctgctttccagtggctgaatacaattgtttcacatattctcttaatttgtgttacttgtatagagcaggatgctaaaggcaattggattctacaaagtgatcacgtcacagagaagccgccgacagaggtggagagagccacacagatagccagctgcctgtgctgcctgctcttcccctaattctgccATGAGCCCAATATTC**ACCTGCATCACAATCCTTTGTCTGCTGGCTGCAGGT**AAGTCCCTGTTCTGCAGTTGTCAGCTCCCTGCTCTAAGCCTTTCATCCATGTCATCGAACTCCCTCATGGGCTCAGTCTCCAACTCCTGTCTGCTTTCTTTACAGGTTCTCCTGGTGAAGAAGTCGCCCAGACTCCAAAACATCTTGTCAGAGGGGAAGGACAGAAAGCAAAATTATATTGTGCCCCAATAAAAGGACACAGTTAT**GTTTTTTGGTACCAACAGGTCCTGAAAAAC**GAGTTCAAGTTCTTGATTTCCTTCCAGAATGAAAATGTCTTTGATGAAACAGGTATGCCCAAGGAAAGATTTTCAGCTAAGTGCCTCCCAAATTCACCCT**GTAGCCTTGAGATCCAGGCTAC**GAAGCTTGAGGATTCAGCAGTGTATTTTTGTGCCAGCAGCCAATCcacagtgttaaatattagctaatcttaggacacagactcatcacggactcagctcaggaagcaggtggtatactaggttggaaggaaataacagaaactagagctagcttaagccaaaggggaatgtattataaggctaaatgtatgtcccatagaaccacaaagcaagaacacaggaacttcccagaaataaactgcaatgaaaccttagaaccggtggctggcgcctgtaatcccagagctttgggaggccgaggcgggaggatcacaa**agtcaggagatcaagaccatcctgact**aacacggtgaaaccctgtctctactaaaaaaaaaaaaaaaaaaatagccgggcctagtggtgggcgcctgtagtcccagctactcgctactcgggaggctgaggcaggagaatggcgtgaaccgggaggcggagct**tgcagtgagccgagatcgtgccactgca**ctccagcctggacgacagaacgagactccatctcaaaaaaacaaaaaacaaaaaaaaaccttagaaccatggcatggctgattgacaactcattttttatataccttgtcggcctcattcttttattataagttgtc**ttttctgtttcttaggtcttacagtgattagaaaa**ttgcagatttaggtgttagccctccagtcaattgagaccaacgtgactttctctcatgctagattacaaatccagggtgcagggggaaaaaatgtgatcctgctttactcaagaatgggctcgtggaaatccaagggaattgaattattttgaacaagatagtatctgggagtctgtccctctgaccttgtggaccaaggatttcaaggggtggaatccagtcagaacatccaataggcacccatcacaagtgagaagttggggataatacattttctggaatatagagaccttgggaaaagagaagagtttggagaggggatcaaaaccttattatttagaacaatatcagaccacgtaaagagt

Uppercase: TRBV16-3

Lowercase: Flanking sequence[1000bp]

Red & Bold & Underline: Stem-loop [10]

Blue: Heptamer[24]

Green: Nonamer [4]

id-IGKV3D-11[V_gene_segment]

gcgttagacagagatgggcgggaatgattgaaggtctgagcattttagtacagattaagaccaaatcattaggattttaagagatgtgtacagttggtgaagaaaaagccctagaatttaatttgactgttgataaaacattcttggattagactgaagactcttttctgtgctaagtaagtatatttatgataatgatgatgactgtagtgctaaatatttaatcaataaaaacaaaattaactgccgcatacataatgtcctgaatactattgtaaatgttttatcttattttctttaaactgtctacagcactgtaaggtaggcaccagtat**tgtcacagttacacagatatggaaactgagaca**tagggaagttaagttacttgatcaattgcaagcaatcggcaagccatggagcatctatgtcagggctgccaggacatgtgactgtaaacagaagtttttcactttttaactcaaa**gagggtatgtgtctgggttaatggaaagcttcaggaccctc**agaaaacattactaacaagcaaatgaaaggtgtatctggaagattaagttctaacagactcttcatttccatcgatccaataatgcacttagggagatgactgggcatattgaggataggaagacagaaatgaaaacacagccttttatattgttcttaacaggcttgtgccaaacatcatctgggtgaatttaggtgattgaggagaagaaagacataggaatgaaattc**tctgagcacaagggaggagttctacactcaga**ctgagccaacagacttttctggcctgacaaccagggcggcgcaggatgctcagtgcagagaggaagaaacaggtggtctct**gcagctggaagctcagctcccaccccagctgc**tttgcatgtccctcccagctgccctacctt**ccagagcccatatcaatgcctgggtcagagctctgg**ggaggaactgctcagttaggacccagacggaaccATGGAAGCCCCAGCGCAGCTTCTCTTCCTCCTGCTACTCTGGCTCCCAGGTGAGGGGAATAT**GAGGTGGTTTTGCACATCAGTGAAAACTCCTGCCACCTC**TGCTCAGCAAGAAATATAATTAAAATT**CAAAATAGATCAACAATTTTG**GCTCTACTCAAAGACAGTGGGTTTGATTTTGATTACATGAGTGCATTTCTGTTTTATTTC**CAATTTCAGATACCACCGGAGAAATTG**TGTTGACACAGTCTCCAGCCACCCTGTCTTTGTCTCCAGGGGAAAGAGCCACCCTCTCCTGCAGGGCCAGTCAGAGTGTTAGCAGCTACTT**AGCCTGGTACCAGCAGAAACCTGGCCAGGCT**CCCAGGCTCCTCATCTATGATGCATCCAA**CAGGGCCACTGGCATCCCAGCCAGGTTCAGTGGCAGTGGGCCTG**GGACAGACTTCACTCTCACCATCAGCAGCCTAGAGCCTGAAGATTTTGCAGTTTATTACTGTCAGCAGCGTAGCAACTGGCATcccacagtgattccacatgaaacaaaaaccccaacaagaccatcagtgtttactagattattataccagctgcttcctttacagacagctagtgggg**tggccactcagtgttagcatctcagctctatttggccattttggagttcaagttgtcaagtccaaaat**tacttatgttagtccattgcatcataccatttcagtgtggctattatgttcaactaaatgcattttagaaggcatctctgtttatggcatcacaaagagttcaataaatcttctgtgcaaaaataaacaacaaacacacatttaaatataaagctgaagtatcaaaacta**tttcagcactctgaaa**attggtgaagtataaaataattaaggatgcatattctttgtagaaaaaaaaagtaccagtgctttgagtaaggacagaaaatgtctgtagccttttgcctgtgacagcagccttctacccccagctcaatcagcatgaattacagaactgg**agttttaccaatatgagcgtagcaaataaaact**agcagcttgctgccaaagggggtggacttgagtaaagccaaggag**tggaaaaaaattcttcagtgtttcca**gctaaacatgcagaactccataggaaatgaacagaaaaatctcacagctttacaagtccaagacaatgacctgtttgaggcaagtagtac**atctgctgatagtaaatagcagat**tcctggataagataggcccatattgctgaaataatctctgcacacattcctagtgacctagaagatatagatatgagtgatgagaacaaggagtctccggtgcaaagtaaaaccagaaggaggtaagaactag**ctgcattttgtatgcag**tttgctttttaaactacacacagatgggttgacagaagatagatttataacctccagat**atttgagcaaaacgtctcaaat**cattggtgaccacatcactatgcagatacatgtgcagttcctataaatt

Uppercase: IGKV3D-11

Lowercase: Flanking sequence[1000bp]

Red & Bold & Underline: Stem-loop [18]

Blue: Heptamer[32]

Green: Nonamer [6]

id-IGHVII-15-1[V_gene_segment]

taacctcaatgggattccctgactagaatggcttttagggatttcgattacagccaagagagaggctggaccaggttcag**tgtcatgtaggacctcacagattttatgtatgaca**tttctcctgacaataaagtatgcaaatgagtatcagcactgatctggtgcttcctgactttcatttttagtggttctcgcttttccatttgattttcctgctttctggataaaaggatgttgtccctgtggtctaaatcctggggctcaagccctttccctggagcttaggtggggctcaggctgtgactcct**gcagccattgggagaggctgc**tgagactttcttctctctcattgttcaggatactccactctgttttctggag**acgcgtctgagaacgcgt**gtggccattagtaatgagggcacaagcttctttggtcaaaatggggtgaggatgtggaattgatcctgtgctgtgtaaactgtcacagagtcaccttcttcaccagtagtgttagaagagcttgtaaaatttgtcggaatccaactggagtcccctgtgttcaaaccctgacaaatggagctggggaaggccatggatggaagc**ttctcatgcacatacccctgagaa**caagacctagctttataacactccaaaaaaccacaaattgaacaaaggcaaccacaacaataaaagaattacttatatccctgagaacaagagctagctttataacactcagaataaccacaacaacaaaataattacttatatccctaacaacaagagctagctttttaacactcggaaaggccacaatcttgcacaaggcaaccacaacaatgaaagaattacttctgtgaaaatatctgcacagcaactgcctatccaaccttacactgatgtcaaccttgttattgatgcttctagcc**caggatcgtcctcatcaaaatgtcattcagatcctg**cgta**tgaaaaaaattgtgtttttca**gtgcactcatGTTCTAGTAAAGCCCT**CAGAGACCCTCTCCTTCACCTGTGCTGTCTCTG**GATTTCCAATCATAACCAGTACTTCCTCCTGGAGCTGTATCTGCCAGCCCCCAGGGAAGAAGCTGAAGTGGGTCAG**GTGTGTAGGTCACGAGGAAACACAC**AGTGCAACCCGCTTCTCAAGAGTCCAGTCACCACCTCCAGATCCACATTCAGAAAACAGTTTTTCCTACAGCTGAGCTACCCGCACAATGAGTAAACCACCATGGATTTTTATACAAAAGAcacagaag**gggaggtcattgtgaggccagacacaaacctccc**tgcagggaagctcaggacaccagggggtgctcagacaccaagggctctcaggacacatcaaggcaggtgcaagaggg**gaaaaaaggtgctggaggtggggtttttc**atcactgtcatattttactctccactatatttat**tctaatgtatattattttattattaga**caatg**atatttatataaatat**atagccatacgtaggtgcaccaagttgtcctttccatcttatgtggaccttgtccattaggacgaagtccctgtatgtatttgagcacctcataagttatggttaattatgtaggatctctcactttttctgtgtcccttcctccctcctctctcacacacaaactgacacccacaaagagttctacaactttaattacctgatgcgttgaagaaaattt**actgaagtgcagctttttcagt**ttaactgctgttcatgttgatgtaggaataagaacattgtctttctcaactgtgtagttctctaagctgagtagcacctttatttataatacccagatactgaaagcaatccaaatattgatcagcaggttaagcagtaaacactttgttctaaattcgttcacttgaataatacccactgttcaaacaagtacggctggaaatgctcaacaagggtaaattcacaagta**cttatgacgagtaacataag**ccaaagaaatacaagtacatacatacagttccac**ttttataaattttataaaa**tgaaaactaatctaaagttacataatgaaaaccagtagttggccgtggacatggtatgagaagggaaggtgtagcaagtggaaattacaggacaacgagaggaaattttgaggataattttttctgtattgagaaaggttatggttatgtcattatttgtcaaattgtacacattaagtgaagattattactttctaatttcgacttcttaaaatactacaaatt

Uppercase: IGHVII-15-1

Lowercase: Flanking sequence[1000bp]

Red & Bold & Underline: Stem-loop [15]

Blue: Heptamer[38]

Green: Nonamer [6]

id-IGHV3-72[V_gene_segment]

tggaccatccctctccttagctgggtggttgcagactgcaagggaccctggtgaaacgagggagatgtccgctttattagtgtgttctcgtattgctagaaagaaatacctgaaac**tgagtaattcacatagaaaataggtttaatttactca**tgattctgctggctgtatgggaagcatgatgaaaaatctgttcagcttctggagaggcctcaggaaatttacaatgatagtgaaaggcaaaaggggagcagccactttatatggtgggagcaggagcaaagaaagagtgagggtggaggtcccacacactgttaaacaaccaaatctcatgagaactcatttactatcatagcaaggggaccgtgctaaccattattcataaatccggacccacgatacaaccaccgggccccgcctcaaacattagggaagcaaaatcacatacagttgttatgacag**gtgggacatcctgaaaacctctctaggcatgtcccac**acggcc**ctggagctgtctcaggggagcagtctcctccag**tgtttagacgcacaggcacggataatagggctaagtctggccag**atgtgtgatattcaacacat**tgcacaactgctctgttctgtatgtaatttatcttctctacaaatgtaacattgatggttgcattaaatatattctgcaaatatgtaaaaataaaataagatga**tgattgctaaatgattatcaaggcacaatca**ca**taatccgaagttacaatttcctgagagataggatta**ctgcccatgcttaccaggacactcacatct**gctctgggcactgccttctcctcaggcgtcccaccccagagc**ttgctatatagtaggagacatgcaaatagggtcctccctctgctgatgaaaaccagcccagccctgaccctgcagctctgagagcggagccccagccccagaattcccaggtgttttcatt**tggtgatcagcactgaacacagaggactcaccA**TGGAGTTTGGGCTGAGCTGGGTTTT**CCTTGTTGTTATTTTACAAGG**TGATTTATGGAGAACTAGAGATGTTAAGTGTGAGTGGACGTGAGTGAG**AGAAACAGTGGATTTGTGTGACAGTTTCT**GACCAGGGTGTCTCTGTGTTTGCAGGTGTCCAGTGTGAGGTGCAGCTGGTGGAGTCTGGGGGAGGCTTGGTCCAGCCTGGAGGGTCCCTGAGACTCTCCTGTGCAGCCTCTGGATTCACCTTCAGTGACCACTACATGGACTGGGTCCGCCAGG**CTCCAGGGAAGGGGCTGGAG**TGGGTTGGCCGTACTAGAAACAAAGCTAACAGTTACAC**CACAGAATACGCCGCGTCTGTG**AAAGGCAGATTCACCATCTCAAGAGATGATTCAAAGAACTCACTGTATCTGCAAATGAACAGCCTGAAAACCGAGGACACGGCCGTGTATTACTGTGCTAGAGAcacagcgag**gggaggtcagtgtgagcccggacacaaacctccc**tgcaggggcgcgtggggccaccagggggcggtcgggaaccactgaggacgg**gacaggtcccaggagcaggtgcagggggcggtttcctgtc**agctgcaggaggcgggtttgtttttgcaggactctggagccttatgaggttgcactattttattatggttatttatcatgtgttttttattgatatttgtgttttagtaatttttacatttatatgtaggg**gtattttttaaaaatac**ataacttcaagaaataattcttcctagtaatttacacctattcctctaaagttttattaacacctgttgatatcagcaactacatagctatgcaacattaatttacatctataaacatatgtgtgaatacatgaaactatgcatacatgtgtagtc**atttatatttaacattataataaaataataaattaaataaaat**aattgaattaaattaaaagaattaaacttaatgatgaactaaatataaattaga**gtaatccataattgatttcaatccttctctatggtttac**ataaatcgatgtctacttgtaagcttaagtatagtgaattggtcattttaaaatagc**caagaacaaatttcaaatgttcttg**tcaccaaaaaatgataaggatttgaagatttgaggtgatatatatgtcaattagctcgattcaattattccacattgcattcataaatcataacatagct**ttgtaccccatcagtatatggggtacaa**cc**aaaatttctcaatttccaatgaaatttt**aattatatattttttaatttgatgcctctctttggatcaggccacctcctcaggattacagtgctgttcagttttctcaacatgctgttgtaccagataagcacaaaacatttaatttcattatgctttgctttagtttttcaaaacaacataaaggtaataattttcccaacaaacatattacag

Uppercase: IGHV3-72

Lowercase: Flanking sequence[1000bp]

Red & Bold & Underline: Stem-loop [20]

Blue: Heptamer[30]

Green: Nonamer [8]

id-IGLV2-5[V_gene_segment]

catgtatgaggtttcctcagcctgag**ccacagctgcctcaaggacaccaaccctgtgg**gaca**ctcctctgggaaaagcaaacaagtgccaagaggag**catctgaagctctggtgtgggagatgt**cctccaccctgggtcaggggagtctggagg**ggatgagaagggaccaggggcctgggattgagctgtgaagggaaccaaaaggcaggagggacagggcaggggctgtcagctatgactcag**gggaggttcctgggcctcaggatcctccc**tctgag**gccaccagggggcgggggtggc**acatgcct**ggacctgggaggtcc**ctgctgggcttcaccctgggtgggtcc**taggagctccttcctccta**agtccccctaaaga**gacagaggcattctggggtcctaaatctgtc**atgcccccataaatgcatttctacgagggccaataaatgaactccaggtttatccaagcagcagcttcaggcgtctgcagacacagagcggggaggaattagccaacctgaggcaccctagaagggctgaagggggctgaaggggactgaagggtccctgtggggcctgtggtcctggggaggggaga**gctggggtgtctcccagc**cactctgggccctgtcctgacacttctcccacaaagaagggaagggaaa**tcctgggaccccacagccagga**ccaaccgtgaaccacaggacaggaaggacagggacccccaaggctggctcc**atttcccaggcactgtcatgggctgagtctcaggaaat**ccaagtcaaggagtttcaatccccctggctgagccccactgtccaaggaaacagaagtctacgggcccaggcccaggtgagggtggggtaagaagaggagcttaggatgcagatttgcatggaggccccgccctcctctgaggcatcagggtaagacaaggctgggggcaggcccagtgctggggtctcaggaggcagcgctctggggacgtctccaccATGGCCTGGGCTCTGCTCCTCC**TCACCTCCTCACTCAGGGCACAGGTGA**CGCCTCCAGGGAAGGGGCCTCGGGGACCCTTGGGCTGATCCTTGGTCTCCTGCTCCTCAGGCTCACCTGGGCCCAGCACTGACTCACTAGAGTGTGTTTCTCCCTCTTTCCAGGCTCTTGGGCCCAGTCTGCCCTGATTCAGCCTCCCTCCGTGTCCGGGTCTCCTGGACAGTCAGTCACCATCTCCTGCACTGGAACCAGCA**GTGATGTTGGGAGTTATGACTATGTCTCCTGGTACCAACAGCAC**CCAGGCACAGTCCCCAAACCCATGATCTACAATGTCAATACTCAGCCCTCAGGGGTCCCTGATCGTTTCTCTGGCTCCAAGTCTGGCAATACGGCCTCCATGACCATCTCTGGACTCCAGGCTGAGGACGAGGCTGATTATTAGTGCTGCTCATATACAAGCAGTGCCACTTAACcacagtggtccaagttcttggggaa**ctgagacgaaaacctgccctgggctctcag**gctccctttttgctctgaagatgtttcctcacccagtgcaacgggcttcctgaagcacagccttgagaattcttctccctcagcaactctcttttcccaccatgaaatccaaag**gaaacctgctctgtggtttc**tcatccagga**cagggacagcttccttttgcttgtgtgttgtggtccctg**agtgggtgcaactcttcctagctttttaaattatgggagggtgacaatgagctccctgactggtgcagtccctgctgttttcaggaacatcctcatcctaaatgcatctgaatctcccactgtgtgcagaccaa**tctggacagatgttattagggggagtttccaga**agccacatcttactcaactctgtatccaccacactctggtgaagatgcttttttggtgcccatcctcttgggtaaagtccactctctactcatgacagccatgtgtcccatcttctgaacttgggaaattgccacaccaaaccctcaggctcagggcccacataggagctcagaaagacgcactgggaaatggagaagagatctgcatccacccctcagtggagagacacctaactagataataagataaggaagtaggtttgcatgatgagtctccccttcttatgataagagagacctagagacccccttctcaggtgtgcgc**tcagaggcagaactctga**agcatctccaccatggcctggacccctttcctgctcaccctgctcactctctgcaca**ggtgctgcttcccagggctcaacccccacagcacc**agggaacagcctggccctgactttcagct**catcacggcagtgatg**cagggtatggcactctgagaatgagaccctcatcttcagactcacctctcctgtcctcgcttgcaggttttgtggccttctctaagctgactcagctgcctgaagtgtctgtggc

Uppercase: IGLV2-5

Lowercase: Flanking sequence[1000bp]

Red & Bold & Underline: Stem-loop [20]

Blue: Heptamer[44]

Green: Nonamer [3]

id-IGHV3-33-2-2[V_gene_segment]

acattctcaaataactctgctgagtgaaagtagctgaagaattaagagtgcacttcataaacttctaattgtataaactgcagaaggtacaactattctaaagtaacagagcagatttgaaatttgggagaaatgggtgttgaaa**gtaattgggtggtgagatgaaattac**agagaagtgacagaaagatttaggggttaacttaattgtacacaacctgattaaagtttgcacacatacgtt**accattttccaaatggt**gcagtgtagatttg**aattaattattaatt**gtacttaaagcagtaacaaagaaacaaatgaatatgttggttgaggaggagcaaaaaatagatgggtatgaacactcaaaacatctcagactcttg**aaagtacacatgcttgaacactggttttctctgtatacttt**aggtaaagagcagaatacattaaaagaaaacagagatgtcctggcgggggtggaatcctgcaaacctcactaggcgtgtcccacactgccctggagt**tgtctcaggggagcagtctcctctagtggtcagagacacaggcatggataatggggctaatactgtccagctgtgt**gaccttgaacacattgtataaacactgttccatatgtaatttatcttccttaaaatgtaaca**ttgacacttgcattaaatgtattctacaaatatgtcaa**aaagaagatgacgactgctaa**atgattatcaaggcacaatcat**ataatataatgatattttcctgagtgataagatgactaccaatctcccc**cagggcactttgtctgctctgagccctg**cgcctcctcaggattcccatcccagagcttgctatacagtaggagacatgcaaataggtttctccctctgctgatgaaaaccagcccagtcctgaccccacagctctgggagagaagcgccag**ccctgggattcccaggg**gtttctatttggtgatcaggactaaagacagaggacccaccATGGAGCTTGGGCTGAGCTGGGTTTTCACTGTTGCTGTTTTAAAAGGTGAACTAGAGAGATTGAGCGTGAGTGGATACCCTTGAGAGAAATGGTGGATTATGTCTGGGAGTTTCTGACCAGGATGTCTACGAGTTTGCAGGTGTCTAGTGAGAGGTACAGCTCGTGGAGTCCGGAGAGGACCCAAGACAACCTGGGGGATCCTTGAGACTCTCCTGTGCAGACTCTGGATTAACC**TTCAGTAGCTACTGAA**TGAGCTCGGTTTCCCAGG**CTCCAGGGAAGGGGCTGGAG**TGAGTAGTAGATATACAGTGTGATGGAAGTCAGATATGTTATGCCCAATCTGTGAAGAGCAAATTCACCATCTCCAAAGAAAATGCCAAGAACTCACTGTATTTGCAAATGAACAGTCTGAGAGCAGAGGGCACAGCTGTGTGTTACTGTATGTGAGGCAccaggtaagaggacatcagtgtgaacacagacacagtttcctgcaatgacaagggaggaggctgggctaaaaggggcactcaggacccactgaaaacgggcagctctagggcaggtacagatggttatcatgggctgctttccttcagggtctgtggcctcctctgcatctaccagttcccctcggagcctctggacatttatgtttctgtggccacccctgacatctctggattagaaaagtttattataggaagaggaaacattctcatttgtcccaaagcagatgtaagtaatggaagcaaaaaatgcacaggaggccaggcggggctgtagatactgtcaccccagaatgtcaatctcaccactagtactggag**agggaagggagtttgatggaccttccct**aacaaccctgtggtccaagataagtccaggaagaccattggtgcctcccagagcacagttgtccatcagggatgtccaatgcgtcccagcagcagccatgcctcagtgtctccactgtgcacagccactgt**ctgggaggagctcccag**gatgggtgtctttggcacacaccaggtggcgggtgttagagtgcagtgcagcagctggctgcctgttctattgggctccctga**tgctggagagatgggaggtgcattcgcagggccagca**ccctgtttgtgaatttttatataaaaaccctgattttacttcatt**tttgcagatgacatagataactaaaaacagaatctgcaaa**gaaattgtaattttcaactttaccccaaattcattgtttcttaattctgtgcaagatccagacatattattgccttcctcatgataatttattctatgtaaaaccgaaatcattttttctcctactctttgtttctgttcaagtacagagatcttgagcaaagtaagttgggttctttccacacactaaccctcaccttccccagagaatgagcagagattgtcctcagtctgagtctaag

Uppercase: IGHV3-33-2-2

Lowercase: Flanking sequence[1000bp]

Red & Bold & Underline: Stem-loop [16]

Blue: Heptamer[38]

Green: Nonamer [4]

id-IGLV3-2[V_gene_segment]

acacgattgatggccaagtcggttgagccacagtgacacagatg**aaggggaagtgaggccaaaacctcccacctcccctt**tctc**ctcagggactcagagtcaacagccctgggttcctctccagtaccctgag**acctctgactctcctccctccaagacacccctaggcagaagcagatgcaggccatgactgaggccattttgaaaactcatctttctgtttcctccacaacagcctgtggggaccatgctgggagagaccagagcagcccaggcaggaaaaggaccttgagagagtggctagg**atgagatgaccgtgtcccagtgtctcat**tgtgccctgtgctctattaccaggctctgccccaggaaaggacatggatcttgacctgtcagccatcatttatgagagttcaa**tttttacattttttaaaatataaaaa**ctacaatctccaagatgctaggagcaatttcactgagtctcatcatggaattctgacctcccccagcatgtcccctgttcctacaatctcctgttcaagaccaactcaactccattacccagaccagctgctggatgatcaagacagggctgtgagagcctaagaatctttaaggtccacgatgagaacacgtcaa**gaaggatttatgttggaacttccttc**cccatggatattcacatcaacagagaagatatctgagttcttgcctacacctatgaatacctaagacatgaagtgtgtgtgtgcata**tggtgtgggcagcacttcctcactcaacacca**gag**agtgtctgaggacact**gggtacagacatggagacagggcatatttttatacactcaattaagccctcgtccttctctgctcaccaaggagccacccaacaaggccacagcaggaagagaagttga**tttccatgaagagccttccctttctaagagtcctggaaa**cccactgtctagttgtggaatcagaggcagagctctggggcatctccaccATGGCCTGGACCCTCCCCTGCTCACCCTCCCGACTCTCTGCATAGGTGCTTCCTCCCAGGGAATAAGCCCCATGGAACCAGG**GGTCAGCCTGGCCCTGACC**TTGAGCTCAGTGCAGGGAGTGATGCAGGGTGTGGGGCTATGGGAA**TGAGACCCTAATTCTCAGTCTCA**CGTCTCCTGTCTCCTCT**TGCAGGCTGTGTGGCCTCCTCTGAGCTGACTCAGCTGCCTGCA**GTGTCTGTGGCTTTGGGACGTACGGCCAGGATCATCTGCCAGGGAGATAGCATAGGTGACTCTGATACAAACTCG**TACCAGCAGAAGCCAGGTCAGGTCCCTGTGCTGGTA**ATTTATGGTGACAGCAACCAGCCCTCAGTGATTCCTGAGCAATTTTCTGACTGCATATCAGAGGACATGGCCACCTTGATTATTAATGGGGCACAGGATGGAAACAAGGCTATTACTGTCGCTCGGAACAGCACTGCTTCT**CATCTcacagtgacacagcatacagtgaagtgagatg**caaaccccttcccatctatgtcattctctcccttcaaccccaagaggactgtggtcacagccatcagcaggtctggcccagttcattcagatttgagacccccaggctgcccttccctccagtcctccaggcaggttcattagatggtaaatcaggggtggatacacaactattaaatatcat**catgttccaatttttcatgaagtagaggaatgattgaacatg**ttaagtagaagcacagaataaccaaaaaaggactaattgagcttgtgtacagctaataattgcaatgtctgtcactttaaaaacatactggatgtgattaacaacagattagattatgtaaaacaaaagattagcaaacttgatcatgtagcactagaaaatattcaaaatgcagcacataaagaaaaaacaatgacaaaaatgaaaaagcattatcggatattcataaagaaaatgaaggtgagttataggacaacttcaaacagcctgaggttgactaggagtaatggttcacgcctataatcccagcactttgggaggtctaggagggaggatcttttaagactaggaattcaagaccagactaggcaacatagctctataatttaaaaaaaaaaaaaaaacttaaaaattaactgggcttcgtggcttgcacctgtagtcgtacctacttataaacctgaggcagtaagattggtagagcccagaaattca**agactgcagtgagtgagctatgatcatgccactggactgcagcct**gggcaacagagcaagttcctgtctctaatatagggtctcaatctgtcattcaggctggagtg**cagtggtgcaatcatggatcactgcagcactg**acttcccaggcccaagtgatcctcccagctcagcctcccaag**tagctgcgaatactggcgtgcaccaccatgcccagcta**atttttgttttttctggagagactgggtc

Uppercase: IGLV3-2

Lowercase: Flanking sequence[1000bp]

Red & Bold & Underline: Stem-loop [17]

Blue: Heptamer[47]

Green: Nonamer [5]

id-IGHV3-22[V_gene_segment]

ttgctttgtcatcc**aggctggagctcagtggtgagatcacagctctctgcagcct**cagaccctgggtctccaatcctcctccctctgccctatgtagctgggactacagttgtgcaccaccatgctcggcttc**agtgtgtattcaactgcacact**ttaattatgcaatatttattatata**gcaataatgcctcaataagggtattgc**aaataagtgaatatataatttgttagataaagactgatggaaagatagacacgaacatgagagatgtatgacacttaagaaaataaaactgtgggaaacatgctgttctttacaattgttaggtaatcacaaaagagcatacacatcacaccatgttccca**ttacagagaaatggttctgtaa**acctcactaggtgcaacccctgtgtgctgggcttggttcagggagaagtcaggtccagtggtgagaagcacagg**cccagatgcccaggtcactctgaccaaatgtgagctctggg**gacactgtacaacccatctgtgcttctgctggtaatttctcatttgtaacatggaaataacattgatactacataccatggtttctctgcatgtgtaaaaataaaagattattggtgtgaacttcaaaaatatgcagtttatgtagatctattgtacctcaataaaacttttaaaataaaaattacaaaattataagatttataggttttaagggtttaccacagaacaaacttacaataagtaaccacaattcccaaatgctatcaatatcacaaatctcccccaggacgct**ctcacatgctgtgag**ccccactctctcctcaggcctctcaccccagagcttactatatagtaggagacatgcaaatagagccctccctctgctgatgaaaaccagcccagccctgaccctgcagctctgggagaggagccccagccctgggattttcagatgttttcatt**tggtgatcaggactgaacacagaggactcaccA**TGGAGTCATGGCTGAGCTGGGTTTTTCTTGCCGCTATTTTAAAAGGTAATTCATTGAGAACTATTGAAATTGAGTGTGAGCGGATAAGAGTGAG**AGAAACAGTGGATACGTGTGGCAGTTTCT**GACCAGGGTTTCTTTTTGTTTGCAGGTGTCCAGTGTGAGGTGCAGCTGGTGGAGTCTGGGGGAGGCTTGGTACAGCCTGGGGGGTCCCTGAGACTCTCCTGTGCAGCCTCTGGATTCACCTTCAGTTACTACTACATGAGCGGGGTCCGCCAGGCTCCCGGGAAGGGGCTGGAATGGGTAGGTTT**CATTAGAAACAAAGCTAATG**GTGGGACAACAGAATAGACCACGTC**TGTGAAAGGCAGATTCACA**ATCTCAAGAGATGATTCCAAAAGCATCACCTATCTGCAAATGAAGAGCCTGAAAACCGAGGACACGGCCGTGTATTACTGTTCCAGAGAcacagtgag**gggaggtcagtgtgagcccggacacaaacctccc**tgcagggccgcgcggggccaccagggggcgctagggacccactgagggcgggacaggagcaggtgccgggagaggtttcctttctcctcagctggaaaagtcaggtttgtgtttgccggactctggagctttctaggctgtgacgttttattacttgtgtttattatgaatttattatcgttagta**tttaaattttagtaatttaaa**aattatatatatatgtttacactttaatgaaataagcattcctatttgcaccgattcttccagagttttattaacatttgttgacgtcagcagctacgaagctatagggacataaatttataaccatagaaagatgtataggccaggcgcggtggctcacgcatgtaatcccagcacgttggaagccataggtgggctgatcacttgaggcca**ggagttcgagaccagcctgggcaaaatagtgaaactcc**atctctactaaaaatgcaaaaatcagctgtgcgtattggcatgtgcctttaatcccagctactcggtaggctgaagcaggagaatcacttgaacacggaaggcagagat**tgcagtgagctaaatggcgccactgca**ctccagcctgggtgacagagcaagattcgatctcaaaaaaacaacaaaaaacaaaaacaaaacaaagaaagaaaaaatatataaatacaaag**acatatgcatatatgt**gtaggaattcataataaacattacaataaagtaattctaaaaatatgtcctaaagaatcaaacttaatgatgagctaaatgtaaattatttgagtatttca**caattgattttggttatttttaattgtttacatcaattg**atttatatttgttc**atttaactactgttaaataatggttatttcaaatgtggttgtcacaataaataataagaatttgaa**ttgataaatgataattatcttaatcatttcttaat

Uppercase: IGHV3-22

Lowercase: Flanking sequence[1000bp]

Red & Bold & Underline: Stem-loop [19]

Blue: Heptamer[37]

Green: Nonamer [8]

id-IGLV3-17[V_gene_segment]

ccagcaggtctttggtatctcattaaggtagcactagcaaatgaaaacaaaagggaccccaaagaccttgggtgccagagaaggaggagatggaggatggtgcaggaggcggggcagggaggggctggaagttcaggctctgaggtgcatctcttgggtgcaatcctgactccactcactgctgtctggggacttggggaagaatcattaacct**cccaatattaattccataataaagattggg**tct**tgagcatagggctgctca**tcctcctattccatatgatgcaaatcttcttgaggtcatgcttgtaaaacactcagcaatgtatctggcatacactatggcc**tcagaagtggttattctga**attttctagtgatacatagagaatggcattgttcctgtgggcagaggaggcactgtggaccttggttt**gaggactcagaaagacaccaagtcctc**tggtagtgctcaagtgcgaagagcgtcacagatgcttcctcccaggccccttctattgggtaaaatttactctccactcctgacacccatctggtccttcttctgaccttagcacattatcagtttgaagacttaaggctcagggctcagatgggagctaggacaggcacactggg**gaatgcggagtagatttgcattc**actgctcatcagggagcccacccaacaaggatacaagaataaaagaagtagatttgcatgaagag**cctctccttcctatgataagagagg**gctgaaggtccctccccgctgtgggctcagagacagagctctggagcatctccaccgtggcctggacccctcccctgctcaccctcttcactctctgaactgcctcccagggctcagcccccaaaggaccagagatcagcctggac**ctgagcttcagctcag**cacagggagtgatggagagtgtggggtctgggaatgaccctcatcctcataattacctcttctgtcctctcttataggctttgtggctTCCCGTAA**GCTGACTCCAGAGCCAGCAGTGTCTGTGGACAGTCAGC**AAGGGTAACCTGCCAGGGAGAAAACCTAAGAAGCCA**TGGTACAAACTGGTACCA**GCAGAACCCTGCCAGATGCCTGTGCAGGTCATACATGGTAACAACAACCAGCCCTCAGGGATCCCAGACCGATTCTCTGGCTCCAA**GTCAGGAACACAGCCACCCTGAC**CATCACTGGGGCTCAGGTTGAACATGAAGCTGACTATTACCGTCACTCATGGGACAACAGTGGTACTCATCTcaca**gtgacacagacagattggaaagtgagatctaaagaccttcactgtctgtatcac**cctctttct**ccagccatagcaggactgagcagggctgg**cccgggtcacctggatcgaagcccttcaggctgcccttccctccagccctcccggcagactgcagagagtaggtcaggataggatttggggctggcagaaccaggctgtcctgctgttatttggaatggatgtaccttggctcaagatgacctgagtggaaggacagtaagagggagacagctactgagtgctcatggtcactgaatttcctcctgtgtgttgactgaacccaagacagtgcctctactctgtggcccaaataccctggattatttctctgaactgcccatgtttaactgagaccctcaagccccagctttgtgcatcctgatgttctgaacatggaagcctttcataatgtgcactgtgacatccttcagattcccatttcaggacttatgcacccaccatccacccagggctgcagggagttcctggcttctcacagctcttccccaccagaaactgccatgcagtacagagaatagctttgctcaaagaat**tgcccatttcaaaatgtgtttcatgggca**gtgactccctgattgccttaatctgaaagtcatgctttaatttaagaccctgaatatca**aattgaagtgccttctagctgaacgctcaatt**gtaaa**acattcccagtcacacactccacacttctttcctgagaatgt**atcctgaaaccctactgcatgg**aatcctccctctcagagtaggatt**gcagggaactcaaaaattaaaaacaaacaaacaaaaaacagtctagagggaaaagtattgaagaggagagag**agagttgcacaaggagacaactct**gaagtctgcaaatattcatgcactgaagagcgcttgtgtgtacagaaactccctctgaaagagaaagacccttgagggaacactccaaaga

Uppercase: IGLV3-17

Lowercase: Flanking sequence[1000bp]

Red & Bold & Underline: Stem-loop [17]

Blue: Heptamer[39]

Green: Nonamer [1]

id-IGHV3-19[V_gene_segment]

gagaaagctgttgcttttacgccaatcaatc**tggagtcataaaagatacactcca**aaaagttcaaaaaaatctagataggagccaacaagaacgagaaaataacacagcctggtatcaaagcatgttcaattggaacccctggttaactactttaatcactggtttagctggacccatcatcatcatattattgagtttaatttttggcccttgtatattaaattggttccttgattttgtaaaacaacgtatagcgtctgtcaaacttatgtatctaagaactcaatataacccccttgttgtaactgaa**gaatcaacgacttgattcccctaaaacacaagtggggaa**atgaaatgcctaacggtgtttttactttaactcgttactttgaattttgtcctgcttgtctctttaatcacctaaccttgcttctcatgtaaataagactctctctagctaggaaagccagacaaactccaattgac**cccttagtttacaagacactaagggctcctcacccaaccccctttcgtgaggag**ttggcctgggtaaacagatcctcagcatttcaaaggagcccaattaactgataaggtacccacaccaacaatgtatgaagttcccaggaattttctccaagagataacaacatgaaaccttgagtttgtgtccggcatagaccctatatctaattataataaaagatttagaaccttgcacctggtaccgttgctcttcttgtaaccatttgtcttttaagttgtttatcactctgtaactattttgattattttgattcttgcatgt**ttttacttctgtaaaa**ttattacatttgagtccctctccccttcctaaacctaggtataaaatttactcgagccccttcctcgtggccgagagaattttgagcatgagctgtctctttggcagccggcttaataaaggactcttaattcgtctcaaagtgtggcgttttcttaactcacctgggtacaacaGTGCAGCTGGTGGAGTCTGGGGGAGGCTTGGTAGAGCCTGGGGGGTCCCTGAGACTCTCCTGTGCAGCCTCTGGATTCACCTTCAGTAACAGTGACATGAACTGGGTCCGCCAGG**CTCCAGGAAAGGGGCTGGAG**TGGGTATCGGGTGTTAGTTGGAATGGCAGTAGGACGCACTATGCAGACTCTGTGAAGGGCCGATTCATCATCTCCAGAGACAATTCCAGGAACTTCCTGTATCAGCAAATGA**ACAGCCTGAGGCCCGAGGACATGGCTGT**GTATTACTGTGTGAGAAAcactgtgagaggacggaagtgtgagcccagacacaaacct**cctgcaggaacgttgggggaaatcagctgcagg**gggcgctcaagacccactcatcagagtc**aaccccagagcaggtgcacatggaggctggggtt**tgtttc**ctgtcaggatttgggacttcctctgcttctgacagtttctctagggaaact**ctttaattttagatttctgtgcccaccaatgtcatctctacatttttttaatcattgtaatatgataggaggacttattctcacatgcacaatatgtatattgccacctacgggaatgaaaac**tcctcaaccatggtcaccagcatcagagtcgtgagga**agctcaggggtgcctgg**tgagtcttctccagtcagactca**ggacagtaacct**caaggggattcccttg**tgagaactcacaca**ctttcatgagaacagcaccaggagttggttctaaaccattcatgaaag**acccactccatgacccagtcacctcccaccaggtcaccccttcacaattgaggattataatacaacatga**gatttggggcaggacacaaatc**caaaccatatcagatacacattgtgaaatgctcatgatggtcagggtaatttgtatttctatcacctcacagcgctaccattttattatttttttaaattaactgcatgagtgagtgtctgatgagaacacctcagatataccctttcagcaacaatcatttttataatacagtattaactataggaccattgctgtacgttagatctccagaactcatccaacctgcacaactgaaactgtacaatttaacaaacatcacccgatttccctctcctcccaggtcctgagaccctctattctgctctctactttcaagagcttgaatattttagatcccacatgtaaatgagatc**atgcagcatttgtctttctgcat**ctggcttattccactcagcatcatgtcctccaggcccatccgtgttgt

Uppercase: IGHV3-19

Lowercase: Flanking sequence[1000bp]

Red & Bold & Underline: Stem-loop [18]

Blue: Heptamer[25]

Green: Nonamer [5]

id-IGHVIII-44-2[V_gene_segment]

ataaacgcactaaattcacgagtacatataagtaaaataagccaaaaaaaaaaaactacatcctgtatgatgacacttttatacattctacagaatagaaactgaagtgacatgaaggtcgggagttgaatggagaaaaggtgagagaaaagagggaggaaggcgttgcaagagaataag**agaaaatgttgagggtaattgatttgttttct**atcttaataatggtgatggttacatcaataatttccaaattgtacattttgtgtgttaatt**agaattttatattattcaagctattacaaataaatacctgtataaatttct**tagggacggtgtaagggagatggataaatcacataaaaagtcagatatttctgaatgtacaatgaataacccttagctctatctctattttttggtaattctaaaatgcagcagttttttttaatgtttttattacaagaagggtttcttcaaaccacaccaggtatgcgtaactctgggataaagttggctaa**gggagctgtgggatccggtggagaggagcacaggtccc**ga**tcctggagcccattcttgtcacatgtgagctccagga**cacatctcatagctccctcatgcttctggg**tttacattcacatctgtaaa**tggaggaccaatgggcatctgcctctaacaatgtattttcgtgtgttaagataatgtatgctaaatgtttaccacagcacaatcttttactaagaaattacattttttccaaatattatcattgtcttcaagccccaccacccactagaaaacttcatctgctctgtgccttgccctgtcctcagacatcctgcccaagagacttctatatagtaggagacatgcaaatagttccctccctctgctgatgaaaaccagcccagccttgaccctgcagct**ctgggagaggagctcaagtgccaggattcccag**gtgttttcact**tggtgatcagaacttaacacagaggactcaccA**TGTTGTTTGGGCTGAGCTGGGCTTTCCTTGTTACTATTTTAAGAGGTGATTCATGAAGAACTACAGATATTGTTTGTGAGTGGATATTAGAGAAACAGTGGATATGTGTGGCAGTTGCTGACCAGGATTTCTCTGTGTTTGCAGGTGTGCAGTATGAGGTGCAGCTGGTAGAGTCTtttttttttttttttttcactttttagcgaacatccatgggttacaaaataatgggttggcttttcttccaacactttacagacaccatcaattttcctcttgcttataaggttttaaccagaagaatgctgtcatcatctttcctgttcttttagaaggaatgccccctcaactcatctccacttgtctgcatgtatttctatttgtctttggttttcagcagttttaataagattcacctaaatgtgtgtgtgtgtgtcgaggggtgttatgctattgttctgtgttctctgagatgcatggattcaccgtttactctgtctccatttttgtgaaaacaattagaaaaaaagtcagtgtgagcctagaaacaagcctccctgaagtgggcactggaccacctgggggc**gctcaggacccactgagc**acaagagccagccccagggcaggtgcagaggggtgttaaggtctggtttcctgtcagccctgtggcttcctctccataaaacagtttcctttgtggcacatctctggattccttatcctttttttcctgtgaagtctgaagaagaaacatttgtcgtaacaa**gagaaaaactttctc**atatgcaccaaaggcagagtcacctacagtcatttactcctgtttctgaatgtcaataaggtgtcaatgcttctgaagttaatcagctaaatctataaaaggtgcggtgtttaactcagcatggcagcccagctcaacagaactccaagggccagtgagcaagcaggcaggataaagtgcatgctgggcattggggcagagggagttagcatccagtgcaagagaagaaagcccccgtggtggtcattgtcaggactccaagcccacagttccaattgtaggtgatactgggcaaaggaagagagaccccaccaatggtt**agtgtggatttcgagtctgatggttccacact**cacactccaggtgaatatgaaaacatttattaactctattttt

Uppercase: IGHVIII-44-2

Lowercase: Flanking sequence[1000bp]

Red & Bold & Underline: Stem-loop [10]

Blue: Heptamer[22]

Green: Nonamer [3]

id-IGHVIII-5-1[V_gene_segment]

attattttgtctgaaattaacatagctattcaagttttctttggtttatattttcatggtctatatttttccatcttatttcttctctttgtgaatatgttaagaagttttcttggagaaagataattgctgggtcttttttttttttagtcaactatgaggaattattttttaatgatattacccttttgtccaataatggttgttttgtcttttt**acaataatatgtattgt**ttctattaatttattatttaattattattgatgaata**tcttattggttaccataaggataacaataaga**catttctataa**ttgaattctaattcaaaaatgtagtacttctttttcttttttatttttttgagacggagtctca**ctctgtcgcccaggctggagtgcagtggcacaatctcagatcactgtaagctctgcctcctgggttctcgccattctcctgcctcagcctcccaagtagctgggactacaggcacctgccaccatgcccggataattttttgtatctttag**tagagacaggctttcaccgtgttagccaggatggtctcta**tcttctgaccttgtgatc**tgcctgacctggcctcccaaagtactgggattacaggca**tgagccaccgtgcccagcctacccctttatat**atagaacagagatattataactatatatgtctaattcctcttttctat**cccttttttattttatcagtttgtaattacataagc**tataaaaatgtaatattgacatttttata**attgctttacacatttatgtaataaagaaaacaca**aatgcaaaaactaaatttaaacctcatttttttcatt**gttgacaatctttatcttttgg**atacattcaggtattggatgtat**atcatatggctgctcatctagaaactctgctaaaacgtctatggcaagattaatatgctgggaatagattttctcaggatttgtttgtccgacagagtatttat**ttgtctttGATTTATCATCTCAAGAGACAA**TGTCAAGAAGATGCTGTTTCTGCAAATGGGCAATCTGCAAACCAAGGACACGTCACTACATTACTGTGCAAGAGAAGcacatgaggaaaggccggtgtgagacacaaacctccaggaacacctgggctaatgagctgcagggggtgctcaggacccactgatcagtcaaccacagaggggagtgcaaaggttaggactgctttcctgcaagcattgaaaaaagaagaaaaaaactccgtctgacagtttctccaaggaacttctgtaattttagaattctgtgcctaccaatgacatctccaaatatttttttaa**aatattgtgttttgaggacatactctcacatggacaaaacacaaatt**gacacagagatgaaaagccctcaaccatagtcaccaaaattggagacatgtagaagctcactgggacctgttgagtcttctccaatcacactcagaaca**gagaccttagtgggtctc**ccttagagtagtcattaaggatggtgataacagcctaaagatggtagaccatggtcagtgtcatatagaatatggggaccctcacaagttttttgtctgacccttctcctgacactaaattatgcaaattaataacactgatctggtgcttcttttgattctaatttattttatttttagttgtcgttctca**cttttcctttggattttcctgctccctggaaaag**gtaaatgtggtctccgtgacctcaattc**aagggctgaagccctt**tccctgtagctcagctggggctcaggctgtggctactgcagccatgtggaagaggctgaagggactttcttcactctccttgctcaggaccatccactgtattgtgtataggcttctctggaaatgcaagtgg**ccatttgtagtgaaagaaatatgtttgtctggttaaaatgg**gaggtgg**atgtagagttaattggctgctacat**aaactgtccttctccaccagtgcttttaggatgagattgtgaaatttgtaagaatcaaaatggagtcacatatgttaaaaccctgacaaatggattcaggaagtgtagggagaattcttaca

Uppercase: IGHVIII-5-1

Lowercase: Flanking sequence[1000bp]

Red & Bold & Underline: Stem-loop [18]

Blue: Heptamer[17]

Green: Nonamer [3]

id-TRBV6-6-2[V_gene_segment]

tctagaaccatctggggctgtcttcagatcggacaccagagttaagagagcctgtgtaccctacag**cactgcagatagaagcatcatggtgttggcagtg**gggaggtactagagtgtgtaagtggttagtggcctacaaccagaagacctgagttttgagaaaattattcccatcacaagatgtagcagagacagtgatatggatatactagac**cagagagctctgtgaggagctctgtg**aggagct**gtggcctaatcatgcatcaggccac**cgttgtgtccataggtcatgggcagtgctgggtggccaccatcatccatacagagagggcagtcagcagcgtgaggtggttctc**tgcctgctgtggtctcaccccaggca**tggaaagcaagagccctgggtgg**agctgaaggtgctcagct**ggg**cttgtcaggagtctcatctgtcagtggattgacaag**aaacagagcaaaatgattcctccaatgttgatgagcctgcccgtgggatctggaaagctaataacagagaaaaccaatatagacaaaggattttaaacaggattatggtcaattaagcaaattagaaaatgatacttgaaggagtatttgggacgcagtagtcaaaaacaccaggaagacatgaggaattgccctaagactctagactacagcactatgtagataactaacaccagactattaattatatcattgaaaaaataaaatttacagtgaattacaaactgtttaccaaaggttatgaaaatcttgtagacttgttttaattcagacaaatgttttcttctcattgtcagctgttctctggcgcatttattttggattgaaccatctgggga**agaggcgtggcctct**cctgacaggaaggctctggggcccaggcagggagaatgaggtctcagaatgacttccttgagagtcctgttcccctttcatcaatgcacagatacagaagacccctccgtcctggagcacctgccATGAGCATCAGCCTCCTGTGCTGTGCAGCCTTTCCTCTCCTGTGGGCAGGTGGGTCCTGGACAGGGCCCCTTGCATAGATTTCAAGGCC**CAGCCCCTTTCCATTGGGGCTGCAGCATCAGCTGTTTCCTTCTCTGCAG**GTCCAGTGAATGCTGGTGTCACTCAGACCCCAAAATTCCGCATCCTGAAGATAGGACAGAGCATGACACTGCAGTGTGCCCAGGATATGAACCATAACTACATGTACTGGTATCGACAAGACCCAGGCATGGGGCTGAAGCTGATTTATTATTCAGTTGGTGCTGGTATCACTGATAAAGGAGAAGTCCCGAATGGCTACAACGT**CTCCAGATCAACCACAGAGTATTTCCCGCTCAGGCTGGAG**TTGGCTGCTCCCTCCCAGACATCTGTGTACTTCTGTGCCAGCAGTTACTCcacagcgctacaagaccatctcctctctgcacataaaggca**gggtggctctgccctcctcccccaccc**aagactcagggatgacctgggcagagttttctgcaatggggaccttggaaccccgagtggccctaagtggcccggacagtatgagcctcagtctgtgccaggtgctcctgcagccatctcagccaggcctggactggtcccaggtcctcagatgtctcctttgttgctctctctggtctatcctctgagctctccttttggggttgggccagggcttccccagctcctacttttctactca**tcatcctgactccgaggcccccaggatga**aacaggatttgtatttcagatccatctaggctgtcctctccctggtgagcatgctgcttcctctatctagagttttcccctcccccaccccacccatggtctgtcctgtggcctatctttcccatctgggcagtcaccttccatcccccgc**ctccccgcccctctctactgcagccatggggggag**cacctcttctgagactccttccttcccatcacagagacttcaaaggccatttcctctgccctggatggagacttcctttctctagtggccagctcctacctgtgctt**cagatctcagaatgatctg**ccctcccgcaggaagcaatccctcaccgcccagctgagatcaattttcctctcataagctctcatagaatcatatgtctcttagtaacccttaggacaattaggctttctcagatacttgtctgatgatttttgtgctctacccaattttttttttttttctgagacagtctcactctgtcaccaggctggagtgcggtggcatgatttccgcttactgcaacctctgactctgattcaagtgattctcctgcctcagtctcccgag**tagctgggattacaggcatgtgccaccatgctcagcta**attcttttttgtatttttagtagagatgcggtttcaccatgttgg

Uppercase: TRBV6-6-2

Lowercase: Flanking sequence[1000bp]

Red & Bold & Underline: Stem-loop [15]

Blue: Heptamer[35]

Green: Nonamer [4]

id-TRAV7[V_gene_segment]

cactggacacagaggtttcgagctggaaaagtgacaccctaagtgtcctgtgacaatcctacagaaaagttgtattgctttctagagtacatttccaaatacgtattcctaacattcactttgatatggctacgggaggccttgcattgtgtgttattttctttttcttttttttttttttttaagatggagtttcgctctgtcatccaggatggag**tgcagtggcacgatctcagcttactgca**accttcgcctcccgggttcaagcaattctcctgcctcagcctcctgag**tagctgggactacaggtgcatgctgccatacccagcta**agtgttgtatttttagtagagacggggtttcgctatgttggcctggcttgtcttgaacttctgacctcatgatccacctgcctcggcctcccaaagtgctgggattacaggcgtgagccaccacgtctggccactgtgtgttatttggaacctccgtgtacctatgttagctttccttggctttatcagttgagattgttgcattttgaagagaacatatatccagttcttctgttttgcccatctgctccataatcactggtgatagttttgctctgagatagatcaggcctcaatgaacagattagtaaaagaaaaacacaggagaaagtcatgatgatcacattatagaggcttataaaataagaac**aattcctccaatttagaagttggaggaatt**tttgcacgtggtgttagcagatgtgcacagcgttaacatttctttaataattttgcttatagccacaaggtgacaggggttctctttcattt**gtaaactatgtctttcttatacctaggaagtatgtttac**gttagaagagcatattgctgattggttgacagacagacaggc**tgtttcatttattagattaaaagaaacaaacaataaaagctttgtttg**gctacataattttatgcagcatttttggtatcaagacaaagtatcaggATGGAGAAGATGCGGAGACCTGTCCTAATTATATTTTGTCTATGTCTTGGCTGTAAGTTGAGGGTTCTAAGAACTGGGGACCCCAGGAGACATTTATTC**AAGTCCTTTTGGGGAGATGGGGATGTAGTCTGGACTT**ACTTGTCATTGCTTGTTTGAGATTAAGAAATAAAATTATGAAAGGTCTAAATTAAAATGTACATATTGTACCTGATGTCTTTCTGAATAGGGGCAAATGG**AGAAAACCAGGTGGAGCACAGCCCTCATTTTCT**GGGACCCCAGCAGGGAGACGTTGCCTCCATGAGCTGCACGTACTCTGTCAGTCGTTTTAACAATTTGCAGTGGTACAGGCAA**AATACAGGGATGGGTCCCAAACACCTATTATCCATGTATT**CAGCTGGATATGAGAAGCAGAAAGGAAGACTAAATGCTACATTACTGAAGAATGGAAGCAGCTTGTACATTACAGCCGTGCAGCCTGAAGATTCAGCCACCTATTT**CTGTGCTGTAGATGcacag**tactccctaggcacctgcaacctgtatccaaacatgcagctgggtagaagtaccataacagaagcatcagcaataggggccctgagcctgagtagacgtgaagaactaaggcatgagtatgaccagagaggggaggagataaagggtgctgttcatttgactcatgtatcttttatcaggccaccctt**cacaaagaccgtctttgtg**ccttatttgaattgatcaatcctgacccttcctatcttcagcagaatcctttagtgcccaggtcagtcttctagcatagagagcacttgttctcttctcaggcctggtggaaacacatgtcaaagaaatttgaagactgttgtttttctactgtaaaatctgacaaaggaactacaatggttggaatcatatgtgaaaaatgctgtag**cagtggttttcatttactcacccactg**aacaagtatctctcagctcctgatatgtgccaggtactctttagtcattgggatgcattggtattgaacatagcagataagccatcatagagctcat**gtttcctaagtttctgggatacacaaaggggaaac**caagtctgcctaggatttagaggttcccgagaaattgcagcaagagtttagtttcatcttcatcaatcatttttctcatggaaacatggttaaagattggggacagctgaatgctgcttgaatctgtgggtcaagaattggcttcacatgttgaggtattgtctgccagagttgagga**gaaatgccaggtggttttcggtatcatctggccatttccaaacaagtctggaaa**gtgaagtgcta**caggctgtcataaataaaacaaatctataaagcctg**accc**tttatttaatggaattgtggagttcaaataaa**atatcaaatacga**tgctattgatgaatagca**ctaaacattggac**attattttaaataat**ttgtagagtagctttctgttactgtagattctagatgctcttttggg

Uppercase: TRAV7

Lowercase: Flanking sequence[1000bp]

Red & Bold & Underline: Stem-loop [19]

Blue: Heptamer[27]

Green: Nonamer [1]

id-IGHV3-38[V_gene_segment]

cagaagttaaactgtgc**tttcttatagctagacatcttatactctaaagaaa**ata**aaatcaatgacatgattt**ggctctgtgtccccaccccaatctcatcttggattgt**aataattcccacatgtcaagtggaaaaaggtgggaataatt**gaat**catggaggctgttttctccatg**ctgttctcctgttattctcacaagatctgatggttgtataaggggcctccctcttcccttggcacttctctcttgcagccatgtgaagaagcacgtgtttgctaccccttccatcatgattgtaaatttctt**gtggctttctcagccac**gcagaactgtaagacaattaaaccttttttctttataatttacccagcctcaggtatgtccttatagcagcatgaaaacggacaaatgtaagcaacttaattatatacaaaaactcttttatacctgcctactgtgaaaggaaaataagtcttgagaccgcaaaatcaccaagctaaatggaagagtcaaactggtgttacaggagatagaa**agaaattattttggtagatagttaggatgacagaatttct**ggcaaaaacatttcttctaacaaaaatcagcttggcaattacttattttctattcacacagagttcaaagaaatcacttctaacaaaaag**cagcctgaaagatcaggctg**taaaacatataagcaactctgccacagagagggtgtttctgggtgtaatcaccaaacttcacatacatacactgggccctagtaaaaacagtgggccttaatgagcacattcctttcct**tttctgggggcactataacatcagaaa**gctgg**aaacttgcacaagttt**aggatgcctgcacttgcaaggaggtacctgggacctagcatggaaaatcctcttcccctttttagcact**ctgggagtggagccccagccttgggattcccag**gtgtttccc**ttcagtgatcaggactgaa**cacacacaactcatcATGCAGTTTGTGCTGAGCTGGGTTTTCCTTGTTGGTATTTTAAAAGGTGATTCATGGAGAACTACAGATGTTGAGTGTGAGTGGACATGAGTGAGCAAAACAGTGGGTTTGTGTGGCAGTTTCTGACCTTGGTGTCTCTGTGTTTGCAGGTGTCCAGTGTGAGGTGCAGCTGGTGGAGTCTGGGGGAGGCTTGGTACAGCCTAGGGGGTCCCTGAGACTCTCCTGTGCAGCCTCTGGATTCACCGTCAGTAGCAATGAGATGAGCTGGATCCGCCAGG**CTCCAGGGAAGGGGCTGGAG**TGGGTCTCATCCATTAGTGGTGGTAGCACATACTACGCAGACTCCAGGAAGGGCAGATTCACCATCTCCAGAGACAATTCCAAGAACACGCTGTATCTTCAAATGAACAACCTGAGAGCTGAGGGCACGGCC**GTGTATTACTGTGCCAGATAtacac**agaggggaagtcattgtgcgcccagacacaaacct**ccctgcaggaacgctggggggaaatcagcggcaggg**ggcgctcaggagccactgatcagagtcagtcccggaggcaggtgcagatggaggctgatttcctgtcaggatgtgggacttcatcttcttccaacagtttctctaatgaacctccctaattttagaattctgtggttcctaatgtcatctcta**catattttcaaaagatcattttaatatg**aggacataacctctcatgtaccaaatgcacattgatgcttacaaagatgaaaagttctcaaccattgtcaccaggatcgcagtcctgaggaagctcacgggtgtctgat**gagtctcctccattcaggcccaggacagaaacctcagggagactc**ctggactagaacggcagggattctgatcacagccaatagagagcctgggcc**agggtcagtgtcctgtagaagctcacagggttcacatctgaccct**tctcctgaccctaaagccaatccgcatcagcactgatctggtgctccttttgctcccaatacatgttctttctttggagtgtttgttctcccttttttatttgct**tttctttcttcctgaaaaagaaa**cacatggtctctatgctctacactccagggctcaaggcattttcttagaactcaggcaaggctcaggcttggctactccagcccacgt**gggagaggctgatgggatttccttctctccc**catattctcagggccctcctctgtgttgtgtgtagactcatctgggaatgcaattggccgttaatagtgaaggggataaactcatttgatcaaaatgggatgtggatgtggaattaaccctgttctatgcacactgtcagagtcatcttcttcagaagtagtgttagaaagagcttgtgaactttatcagaatcaaaatggatccacttgtgttaaaaccctaatgaatgaagctggggaaggcca

Uppercase: IGHV3-38

Lowercase: Flanking sequence[1000bp]

Red & Bold & Underline: Stem-loop [19]

Blue: Heptamer[36]

Green: Nonamer [14]

id-TRBV11-2[V_gene_segment]

gttttgcgggtttggtggaatatctctgagtgcacctcactgaacacatatatcctcattcccacctccctggacgtgcatctccttgtgttattgtcttatgccaactctgaggagcgccatcccctaactc**cttttatttcttattactcaggcaaggaataaaag**tgcgtttgtga**ctaatgttgccccattag**aaacaaaaaagagataatgaccactactgttcaactgtaatttacactagaggtatagctataaggttatttttagatttcagcaagtctctttgattttgcatttcttacaatatatactgatattttcagggataagagaattccccacctatcttcgataaaaatgagaacttactttcattgcactggtgacccaaaactacaggcagaac**catgatgaaggcttggtacacaatgtatcatg**aatgat**aacagtaaagaggaaactgttcatcggccttcactaaaaatgaac**ccttccaactctgtgcccaggttattgt**tttatccctttagcttacagcgcgtcagcttctaggataaa**cttgtgtcaattaccccagcttctaagtgcataggacaaggaagatagagaacctagtaaaagagaaacacccaaagagaaggataattgctgataagacaaagaggtcactaactgatatctc**tgtttcgagttgccttcaactcgaaaca**tccagcagaggatgggcctagagatggagtaggagatccagtccccaagccctagagatgtgagtgaggacacttggctgaacttacacagtctgtgaatacggtaaagcaacttcgtgctctcttttagcttataataattggtttctaacaatgtaggcatttgtggaggcaatgatgtcactgtgggaactgccatgagaggacagggatgtccctcctc**ctttgctgttgctcacagtgaccctgattgggcaaag**ctcccatccttccctgaccctgccATGGGCACCAGGCTCCTCTGCTGGGCGGCCCTCTGTCTCCTGGGAGCAGGTGAGTCCTAGGAACACCATGATCACATTGGATCTCTCTGATTATTTCAATCATTTCCTTCTGTTTTCAAA**TTCTGTCTTTTTCCTTCTCAGAACTCACAGAA**GCTGGAGTTGCCCAGTCTCCCAGATATAAGATTATAGAGAAAAGGCAGAGTGTGGCTTTTTGGTGCAATCCTATATCTGGCCATGCTACCCTTTACTGGTACCAGCAGATCCTGGGACAGGGCCCAAAGC**TTCTGATTCAGTTTCAGAA**TAACGGTGTAGTGGATGATTCACAGTTGCCTAAGGATCGATTTTCTGCAGAGAGGCTCAAAGGAGTAGACTCCACT**CTCAAGATCCAACCTGCAAAGCTTGAG**GACTCGGCCGTGTA**TCTCTGTGCCAGCAGCTTAGAcacagtgtagcagaga**cacttccctcctgtgcagaaaaccagaaaaccgcaggactctctcctctctactcagctcacagcagcct**ttccttattcctcatcctcccaaggaa**gaagtgagttttcagatatagctaggactcatatagtgggaggaaataaactttttcttttcttttctttttttctttttgtttttgag**atggagtcttgctccat**tgcccaggctg**gagtgcagtggtgcgatctgggctcactgcagtc**tctgcctcctgagttcaagtggttgtcctgcctcagcctccccag**tagctgggatgacaggcatgtgccaccatgcccagcta**atttctgtatttttagtagagacgggattttgatatgttggccaggctgatttcgaacttctgacctcaagtgattcgcccttctcggcatctcaaagcgctggaattacaagcttgagccactgtgcctggctggagataaactatttcttaaaacatgaaggctgcagttgttatttgaaaatatgtctaaggaatctgaaacacctatcggtgagaattcaagaaaccaaaactgaaagtgacttatcacagacttagtcttctggggtactaatagttgttcttctatttaaaaaatgcatacaa**tattttatatcaattaaaaaata**cgtaaatttgaaaaataaaaacatgcagtcaataaacatgtgtgataaactttaataatacagataa**tgattgtatgtgagtaacaatca**gcaggttttctctctgcagttgtcagcatatgaataatttgaatcctatccttggtgacaccctggctctggagcctcctctagacctctcattggatgtattatgcaaccaaatctc**aacaggtactatgtgtttggaaatctttttcccctgtt**tttgcggaaatatgctaataagagtaatgttaaagatgatgatgttgataatttcaactatttcattcagcaccc

Uppercase: TRBV11-2

Lowercase: Flanking sequence[1000bp]

Red & Bold & Underline: Stem-loop [19]

Blue: Heptamer[37]

Green: Nonamer [4]

id-TRAV12-3[V_gene_segment]

tttaacataatgacttccagttccatccatgttgctgcaaaggacacgatttcatttatttttatacctgaatagtattccattgtgtaactatacaacattttctttatccattcaccagtagatagacactttgttgatttaatatcttt**actattgagaatagt**gctacaataaacatgaggaggcaggtatgccttcaaaacacttctttcctttggataaatactcagtagttggattgctggattgtatggtagttctattactacatttttgagaaatttccctactgttttccagagtggctgtactaatttacattcctaccaacaatgtttgagagttccattttctatgaatcctcatcacat**ctattattttctatatttttaataatag**cgattctaactgggataggatgatatctcattgtggttttgattcgcatttccttgatgattagtgatgttgagcatgttttaaatacctattggctatttgtcttcttttgagaaatgtcaatttagatcttctg**cccatttttaaatggg**attgtttgacttttttttttgctgttgagttgtttgagttccttgtatattctgaatttaact**aaagtgtcatattttcacttt**tagcagttcacataacacaggtctgtagttggaatacatctttgatcactttcatgagt**cctcaagatggcagtgtcttctcagggaccttgagg**caagctctcatgtgctcctgcggtggggaggggaggggatgca**tgtgcacttgcaagtgcaca**cacacacatgaataagagcaagacagagggagagatgagggaggagcttaatgatggagcagaggtgtta**aaaagaacatcctttttctaattggtaggacagatttctttt**atgattcctacagcag**aaaaatgagaaacgtttgttattattttt**ttttcgtgtttaaggtttgaatcctcagtgaaccagggcagaaaagaATGATGAAATCCTTGAGAGTTTTACTGGTGATCCTGTGGCTTCAGTTAAGCTGTGAGTTTTGGGCATCTCTGTAGAAACATAATGTATATTGTTTGCAATTATTGTGTATTCTAGGCCTGTGGGTAGAAACATGAGTGGTTTTTTCTCTAACAAAGGGAGACTGGGGAGAGTGGGGGGCCCATGAAAAGATGATTTAAATTC**TGGGTACGAAGTGTCCCCTACCCA**ATCTGTCTTCTCTGTCTGACTGCTGTCTTTTTTCACAGGGGTTTGGAGCCAACAGAAGGAGGTGGAGCAGGATCCTGGACCACTCAGTGTTCCAGAGGGAGCCATTGTTTCTCTCAACTGCACTTACAGCAACAGTGCTTTTC**AATACTTCATGTGGTACAGACAGTATT**CCAGAAAAGGCCCTGAGTTGCTGATGTACACATACTCCAGTGGTAACAAAGAAGATGGAAGGTTTACAGCACAGGTCGATAAATCCAGCAAGTATATCTCCTTGTTCATCAGAGACTCACAGCCCAGTGATTCAGCCACCTACCT**CTGTGCAATGAGCGcacag**tgctccccagacacttgcagtctgtacccaaacctgctgggccccaggaatgcctgatgtagagcttagactgcaaggcaatgaagttctatttgttctctagatgcaaatgaaaactaaaagcagtagtattgaaggtttgattaatttgggacat**attttcataattctgaaaat**tattgagaccccaaag**gaaaataaaagacaccatttggtctacttgaaatattttc**tagtcttctctttccactttagttttgaaattttttttttgctgactttgtggctcatgtctgtaatctcaacacttttggaggcatagggaagaggatcacttaaacccaagaa**tttgagactagcctgggcaacacagggagaccttgtctcaaa**aaaagaaaaaaaatcgccaagagtggtgacatgcacctgtagtcccagcaacttgggagggaagaggtgggaggatttcttgagtcggagaggtggaggctgcaatgagctgtgatcacaggactg**cactccagcctggacaatggagtg**ataccctgtctcaaaaaacaagcaaaaacaaaacacaaaacaacaacaaaacaca**aataaaatatttttatt**gaggtgttattgacacacaaaaaggtgcacatattcactttatacagacaatataatcgtgccataaacatatgtaccatttctaaaaatttgcttcctccctggttt**ctttttattaagttattattgctatacaaaaag**gtacatttatttaatccatgcccagtctacttttatgccacatctttactttatgagattggttttatgattctttttatcaaagacaagaaaatatttccatacaataacagatatgctctcatggttcctaagacattttgctctttccccaaggcatttgatagacat**tgctttaagaatgaaacagcaaagca**gaaatgcaaagagaagaaaaatcataagcagactaatttgaaaat

Uppercase: TRAV12-3

Lowercase: Flanking sequence[1000bp]

Red & Bold & Underline: Stem-loop [18]

Blue: Heptamer[40]

Green: Nonamer [3]

id-TRBV5-1[V_gene_segment]

cctgggggttgtattttggagggacatcaatattcagtccactaacaatggcaaaatcttggttactgcg**cagtgggccacatgggggtgctgtggaaccactg**agctccaaatgtctctggggcatgcacagagccaggagcctgggacagggagggttaaggagcagccttgcttgatgtttgtgctcactaaaaaatg**tgatgacggatacataaaactgcaggtgtgcctctcatca**gggagccatgcacttcagctatgcctgggttaggaagcctctgtgagccgaagagggagaatgaagatg**ggatgtcccatgctgggatgggacaacc**tgcatg**tccagggcagggacagatagaggaacaactgcctgga**aggaaaagcaaaggaac**tgtgggggggacatgcagcctaacctcagtccagccccaca**tggtatgtgagagtattaacagcacaaagcagaggtcctggatcgggtaagccaggggcctaacaaggaaggcaggatgcatgggaacccgcagagatgac**atcagagaagtgatatcacagcctaacatctgat**cagagatgcaatgcccaaaaccccagctctcagaggaccagtatccctcacagggtgacacctgaccagctctgtccc**acctggccatgggctccaggt**acctctgatgggaagacctttgtctcttgggaacaagtgaatccttggcacaggtgggatgtctctgaagcacgcctcatccaaggcaccctcaggctccct**cctgtgttcttgccctagcctctgtctccctccctcacagg**cccagtgga**ttctgctgtgcagaa**cagagagcagtggacctcaggaggcctgcaag**gggaggacataggacagtgacatcacagtatgcccctccc**accaggaaaagcaaggctgagaatttagctctttcccaggaggaccaagccctgagcacagacacagtgctgcctgcccctttgtgccATGGGCTCCAGG**CTGCTCTGTTGGGTGCTGCTTTGTCTCCTGGGAGCAG**GTGAGTCCCTGCA**GACAGGATAGCGCCCCATTCTGAGCCTGTC**CTCACCCCTGTGTCTTCCACTTTACCATGGGGAGGCACCACCAGGCTGTCTCCGGTGCTCATTCTCCATCTGCTTTTCCCACAGGCCCAGTAAAGGC**TGGAGTCACTCAAACTCCA**AGATATCTGATCAAAACGAGAGGACAGCAAGTGACACTGAGCTGCTCCCCTATCTCTGGGCATAGGAGTGTA**TCCTGGTACCAACAGACCCCAGGA**CAGGGCCTTCAGTTCCTCTTTGAATACTTCAGTGAGACACAGAGAAACAAAGGAAACTTCCCTGGTCGAT**TCTCAGGGCGCCAGTTCTCTAACTCTCGCTCTGAGA**TGAATGTGAGCACCTTGGAGCTGGGGGACTCGGCCCTTTATCTTTGCGCCAGCAGCTTG**Gcacagccctacaaagccaaccacattctgtgc**acaaacctccctggcccaatgtggagcaacctcagccctgacatatctgtgagaacctggggactgcagggagaaagaaaggcaatttagaatgctaggatccacagcctggggctttctgcaa**cttggctggggaaaacaaggccaag**agagcctgctcagaggatgg**tgctgcttccgtaggttgactgcagca**ttaccttctttgccgacactgtgaactacaggagtagaggaaatgtgtgtaatttaacagctggcgcactctttctcaggacaagagatttaattttgtggtcaagattgttgataacagtgtagtgtagaagggagaatcacagggtttgctttatacaacttggggagaagaaattgaggttttgggaaagtcaaggagagagaaaacgaaagatgattgaaaacagagatagagcagaaggtgatgaaaagtggtgttctgtgcaccatgaactctctttcccccacccgaaagttttcaacacctagaaaaaaataaggaaaggtcttatagccaccggaatcacctggtagccggtcccagtcggaggcatcaccaaattgccgcttggccattgtgagagccctggagttgcttctgctgtcttccttcttccagccctgatcttgactcctcaggccacagtcaggtacttcccaaaacacag**aggaaggtaaagtcacccttcttcct**aaaggggcaaatactttccagtttccccccaaataaattctgcaaccctgggattcctttctgacactcacctcccaaatccaattaccttctatgaccaaatgctctttgaacattttctaaaactcagcagtctctacatcctctgcgtttgtccttacagtttccatggcctgggaggacctttgttctcctaagacggggccacatactgatcatctgtcaaggacaggctcacacctgccttcctcttcac

Uppercase: TRBV5-1

Lowercase: Flanking sequence[1000bp]

Red & Bold & Underline: Stem-loop [19]

Blue: Heptamer[51]

Green: Nonamer [2]

id-IGKV2-4-3[V_gene_segment]

aaattttttgaaaagattatgcaatctgctgtgctggtggagtgttctcgaagtgtcaattatgtc**cagatggttgattgtgctgtccaattcatctg**catctgcactggcctgagaggcaggactgcttgtgctacaggctgaacctgctgcagagccctttttttgccccgggtcccactcaagg**ctggctgactcgtgtatcagtcggtcagtgagccag**tacagtattccaaggtacagaatttctt**gcctctagaatccaaagaggcctaccaaggcct**tcctcccttcaaagacaacgacatgcattagatacaat**ggaatggaatatgcattcc**tcttcccgtgaactctaattttctccttgtttacgaagcacacattacagctagcttattcactggggcatcatttag**tgccatttaatggca**g**ctgtagagccaagtgcccagaaataccagctacag**tgaaatacaagctgagggaaatcagccttgcaaccacagccaaataaccctggaactgtttcaagttcatcctgatacttagagacattctatcaatgagcaccagagttagtgagagctagttcctcctcagaattagctgggtggagccaagaagaaagcctattcaccacctaaagattgatacacatcttcatcatcacaa**tctccattgcatcagcaatgaaga**gtgacaggctgcagtcacagtgtcttccacattgattcatggtatttcagaatgtcacccttctaagccaggcaacaccaatattttattttggtttcatcaacacaaaagatattctgtgtcactttgcttacataagaagcccactcttcgaccccaactggatatttcttcactgcttcactcacc**aggggatttgcatgttgtcccctagggaagaccttccct**tgtgagtctgagataaaagctcagctctaaccttgccttgactgatcaggactcctcagttcaccttctcacaGTGAGGCTCCCTGCTCAGCTCCTGGGGCTGCTAATGCTCTGGGTCCCTGGTAAGGACAGAAGAGAGGAGAATGGGACTGGAGGGTGAGCTCTGTGG**AATCTGCAGCCTCCCATTTGTTGTTCTGTCTACATGTCAGATGCAGATT**TCTTGTCCTCCAGCAAAGGGAATTTAATATTCAGACCTGTAAGAACTAAGAACGAGCCACAAGGAAAAAGAACTTCACTGTCGAGAACTTCAAAA**ACAAAGAGGCCACTTTGT**GCCTTTAATTATTGGATTATTTTTGGATATTGGATAAATGTAGTGTATGAATCTAATCAAACACAAAAAAGTAGCCTAACGTATACAATAAAACTAAATTATAAAAATGTCTCACATTGTTTGTATGTAACCTTGTACTTCTCACTGTTATTATTTTAGGATCCAGTGGGGATATTGTGATGACCCAGCATCT**GCTCTCCCTGCCCATCCCTCTGGGAGAGC**CGGCCTCCATCTCC**TGCAGGTCTAGTCAGAGCCTCCTGCA**TAGTGATGGAAACACCTATTTGGATTGGTACCTGC**AGAAGCCAGGCCAGTCTCCACAGCTTCT**TATCTACACAATTTCTAACAAATTCTATGGAGTCCCAAACAAGTTCAGTGGCAGCAGGTCAGGGACAGGTTTCACACTTAAATT**CAGCAAAGTGGAGGCTGAGGATGTTGGGGTTTATTGCTG**TGAA**CAGGGTCTGCAAGGTCCTcacacagtggtaaaaccctg**aacacaaacctccct**acttgggatggcccagccatccacaagt**gtttgcacgtggactgtctgcatggcagattctgagttggcttcacaggtagatgttagatgactgcagggaataatttgcaggaagatccagactgttctggccccatcactgtaacagccttatttcacatggctcaagcagaagaatgaaggtgatcaaagt**gctctctgagcaactagacatgacagaagagagagc**tacctggaaagtgcattctgatcctccctgtgtttcctgcatttgtcaatgagatttagtcagcataacagcaagaaaattgaggcctactcatggctacaaaggttgggtatatgttaggattagcaggttttgttatatttttaacacgataacgtttggtaatattcagaaattggtattttcccactttctaacttcctacctctccattagtggttataccaaacctccacttacatacctgtgattgcattctacacaagctatactcagggggagtgtgactcacagcaattagtgaaa**atgtttagtttgcaaacat**ctaaatagactttgtcaattcatagtggagataacattttatcccaaaagtctttgatttgcctcaactg**ccttttgtgagagaaaagg**agagctcttagaattgcaagtggattttgaaaataaaagtataagccatttggtgacaactgcagaggagagtgcccacaaattt**cagctgtacaatactgatacttccagctg**gggtaaatcagccatccaaccacagccaggatgttactaggcagtccagaaacagttccagggcattttcacataacttttttttcatctttatacagtatctgagctagtgctttagttaggaggctct**aaaatttgacattataatttt**cattacttctgctgtatgtacctccatctagtccttatgtcaatacaaccccaggac

Uppercase: IGKV2-4-3

Lowercase: Flanking sequence[1000bp]

Red & Bold & Underline: Stem-loop [23]

Blue: Heptamer[47]

Green: Nonamer [6]

id-IGHV3-30[V_gene_segment]

ggcta**aattctcaaagaatt**gtgctgagtgaaagaaactaaggaatgaagagt**aaattttatatgatacatttgtagaaattt**tagaagatgccactattataaattaacatggagaagatttaaatgtttctgagaatatgctattgggagtaatggggatgtgagttaaatttcagaggaataagagaaagatttagggattaatttattcaaaccttgattgaagtgctgagtaaatggttgcaaacataggtctacattt**ttcaaatcattcaccataaatttgaa**tta**tttattaattacactcgaataaa**gcaataaagaaactgatgagataatatttgactgaattgcagcaataaatagatcgatattaacacaaggaatataactgacttccaaaaacatacacatgaaccgtggttcactctgcgtatttaggtaaataacagaaagttgtcataacagatggggaatcctgcagacttcactaggcatgggccatgctgcc**ctggagttgtctcaggggagctgcctcctccag**aggttagagcacaggcccaggtaataggactaaatttttagatgtgttatcttagacacactgcacaactgctgtgttctctatgtaaattatctcctgtaaaatataacattgaagcctgcattaaatatattgtgtaaatatgtaagaataaaagaaagttatgagagctaagtgttaatcaaggcacaagcatataagatataactatattttcctgaatgatggaattactaccagtctcccccaggacactt**catctgccctgagcccagcctctcctcagatgtcccacccagagcttgctatatagtgggggacat**gcaaatagggccctccctctactgatgaaaaccagcccagccctgaccctgcagctctgggagaggagcccagcactagaagtcggcggtgtttccattcggtgatcagcactgaacacagaggactcaccATGGAGTTTGGGCTGAGCTGGGTTTTCCTCGTTGCTCTTTTAAGAGGTGATTCATGGAGAAATAGAGAGACTGAGTGTGAGTGAACATGAGTG**AGAAAAACTGGATTTGTGTGGCATTTTCT**GATAACGGTGTCCTTCTGTTTGCAGGTGTCCAGTGTCAGGTGCAGCTGGTGGAGTCTGGGGGAGGCGTGGTCCAGCCTGGGAGGTCCCTGAGACTCTCCTGTGCAGCCTCTGGATTCACCTTCAGTAGCTATGGCATGCACTGGGTCCGCCAGG**CTCCAGGCAAGGGGCTGGAG**TGGGTGGCAGTTATATCATATGATGGAAGTAATAAATACTATGCAGACTCCGTGAAGGGCCGATTCACCATCTCCAGAGACAATTCCAAGAACACGCTGTATCTGCAAATGA**ACAGCCTGAGAGCTGAGGACACGGCTGT**GTATT**ACTGTGCGAAAGAcacagt**gaggggaagtcattgtgcgcccagacacaaacct**ccctgcaggaacgctggcgggaaatcagcggcaggg**ggcgctcaggagccactgatcagagtcagccctggaggcaggtgcagatggaggctgtttc**ctgtcaggatgtgggactttgtcttcttctgacagttccccagggaacctcttaaatttagaaaactgt**gcctaacaatgtcttctctatgcatatgaggaccttttctccctggcacaaaatgcagattgacgctgacacggatgaaaattcctcaaccatggtcacaaggatcagagt**cctgagtaacctcagg**gct**tcctggtgagtcttctccaatcagacccaggacagggacctccgtgagattccctg**actggaacagtctttatggat**cctggtcacagacaatagagaggctgaaccagg**gtcagcgtcatgtagaacctcacagatttcacgtctgatccttctcctgacacgaaagtatgcaaatcagtatc**agcaccgatctggtgct**tcttttgttcctaatccatttactttattttttcgtcgtttttc**tcctttttccatttgtttttcctgctttttgcaaaagga**agatgttttccctgtgagatgcaggggatgacaattttggga**gatggctggaacatccaatatcctcagggccggccatc**agtaagtgcaggctggaagtctcagaaagagctgaagctgcttaatcaccgtggagttttaccttctccagttctgctctgatggaatcagggccaagcaggttatcaatgataatctacctaacatagagtcaaccgattccagtttcaataacgtctgttaaaaattcacaccaccacctggattagtgttttgtcaaat**caatacacagtattg**tccagctaagtagacccaaagacggaccgttgcccatggagaaaaacattaacctgagttctaggttcttacagtgttaaagg

Uppercase: IGHV3-30

Lowercase: Flanking sequence[1000bp]

Red & Bold & Underline: Stem-loop [22]

Blue: Heptamer[30]

Green: Nonamer [6]

id-LOC105379603[V_gene_segment]

ctactgaatttgaaccata**atttttccctaaaaat**cctagttggggtgcttgatgttgagagctaccacgtatcaggctcacctgaggacagactctggttaaaccaagttaaaattttaattcttcccatctt**caaagactgagcatctttg**ctctttttgttcataagtgaaacactccaaagtcatttgccaggatcccctgcagatgagcatttagtgtacagactcctaggcaagatccctggtcacagtat**ccagatgactatgactgcaaatctgg**ggaagt**agagggagaacaaagatttaccaggaaaaccctct**gggaa**ttcctccccagggaatcctagaggaaagagctccctttctcctcagcgccacaggatccacagaggaaggagagctct**agc**caccaggaaggagccacatgagaggagaggacctggtg**tccaacacaggtggggaggtgagtccatggggctggtccctgctatgggcttgatcctgacat**aggactttatgtagataccacctgagtcct**ctacaatgtacagccaaagagccatggaaagattcactc**ctgaggagtctgaagactcctcag**tagagcgtgaagatagccctgaagtcacagggaggagacacagacaccacttggcctgccttggacctgagagagggggcagtggacagaatggggggaggaagccatgtatctgagaggttctggtcaccttatcatagaatggtttaccttgatctggacaccagagggcgcatggggatcattcctgagaagacaggggtgatcagagcagaaccagccgcct**gccctgcctggtgtcctctgctcagggc**tcacagctgtgtcctccccaccccctgggaccacaaagctccacccctgccaccccctgacatcctcaagccaaggagcc**tgacccagggctcagggtggggtca**caaacctggggggggtctgatttgcatggatgga**ctcTCCCCCTCTCAGAGTATGAAGAAAGGGAGAG**ATTTGGGGGAAGCTCAGCTTCAGCTGCGGGTAGAGAAGACAGGACTCAGGACAATCTCCAGCATGGCCTGGTCCCCTCTCTTCCTCACCCTCATCACTCACTGTGCAGGTGACAGGATGGGGACCAAGAGAGAGGCCCTGGGAAGCCCATGCGACCCTGCTTTCTCCTCTTGTCTCCTTTTGTCTCTTGTCAATCACCATGTCTGTGTCTCTCTCACTTCCAGGGTCCTGGGCCCAGTCTGTGCTGACTCAGCC**ACCCTCGGTGTCTGAAGCCCCCAGGCAGAGGGT**CACCATCTCCTGTTCTGGAAGCAGCTCCAACATCGGAAATAATGCTGTAAACTGGTACCAGCAGCTCCCAGGAAAGGCTCCCAAACTCCTCATCTATTATGATGATCTGCTGCCCTCAGGGGTCTCTGACCGATTCTCTGGCTCCAAGTCTGGCA**CCTCAGCCTCCCTGGCCATCAGTGGGCTCCAGTCTGAGGATGAGGCTGATTATTACTGTGCAGCATGGGATGACAGCCT**GAATGGTCCCACAGTGCTCCAGGCCC**AGGGGGAACAGAGACAAGAACCCCCT**TCCTTTTCT**GCCAGGAGGGTGAGCCCCGGCTGCTGCTGCTCAGGCCTGGC**CTGTGGCTTCTGCTGCTGCAGCTTCCCCCATGGGTCCAGGGATATCCAGGGCCCTGCCTGAGAGTGGAGGCTCCTCctcccctgagtcctcagagtcaggaacagggtgtccagaaaacaaa**agcagggtgtcctgctccctgtagcttgggacacaggg**tctctgcactgaagtcctgggctgaggtggcaggtccagctgtgtcatcacagacccacttctgttggggaacctgtgtctccatcatcctactttttccatttccaggagtttccagagtggtgtcttcctcccccttctcctcagtgtgaatccctgtgcttcctttctctccagcctattctttttttaataaaactcctttcatgtaggaataatgcacatataataagccacccatatttacaatagggaatgagacaaattttggcatttctatggatccttgaaaacatcaccaccaccg**ggagctatacaacatacaacctgtgaccaaccccaatagctcc**tcatgacccttgtcacctcattccctcccccaacaccctccttggaagttgcagatctgcctctgtcactactgtttggtttctcttttctggaatgtatcagtgttatcaaacattatcttcccatttgttcaggcttctttcactctgcacacttcttttgagattctcgctgttgttgctgcattgtacccatctcgtagagggctccttactatctttgaaggtccttaggccatttttttctgattttatttatgtatttattt**taaagtttgaatttattgtgcatccaaatcaaacttta**ggggcaatgttcatctaaggtgacaggacagtctgtgatggcttgtcgggattcttgcttcttcatctcttgcttctgattctctctatagtggaagaggttctcgtctttcatttatttatatatttatttatttactgatttatttgggtttttaagggttttagtgatttttctctttttggttatttttcaatttttttattttaataggtttttgggggacaagtgttgctttgtcacataaataagttcgtta

Uppercase: LOC105379603

Lowercase: Flanking sequence[1000bp]

Red & Bold & Underline: Stem-loop [21]

Blue: Heptamer[37]

Green: Nonamer [6]

id-IGLVI-20[V_gene_segment]

ttatggaatgaaaagggagcctgacatgccaggtgctctgggttgagggattgttggagtcagatct**ccctgcaggaaagcccggggcaggg**gga**gcagcctcacccctcacaggaaccacagatacacccacaaggtgagctgcaggatggatgctgcccacctccaccctccacatcct**ctgtaaatgttgctcctt**tctacaactccaaccagatatgtagatgtggcgaactacgtaaaatacggatcattcatcacatc**aaaacccactgcaggacaccctggtcaacaaagaacccaatcacatccccatcaactacatagtttccaaattttccatctccagaaaaataacaataacaatat**acatgaaaatcgatgtaatttatctcatacataatttcatgt**tgataacgtgaaaatgatagtattttgctctactg**aaataaataaaatatatatatatatctgaatttattt**gtcttatcttttcatatttaatgtggtgactagacactagggggtcacaggaggatcgtgtcataccactatgggacagagctcctcacaactctttcaggtgacaggtactgttgagtaacctgctgcaagcatccccatctccaccagaccatataagtgtgaacccaggaagaggcactggaacaatagagagaaaaacctgcttgtgcagaagacggtgcccttgagccctgctcctgctccatcctacgggtgccacattcatctcatggtgtaatatttcgtgc**cctgcctgagcttatgaccgaggggatatggcagg**tctgactgtgtggttactggtgtctcatgaggttctggatgtaacaaagccctcgaatatagaagaggttgttttcaaaaggaaataattatctactgcacatgacatagacttgttgctaaatcccatgcgtctacactaggattctcct**ctgaagccttgccttaagcacaaggtttcag**ttcctatgtcCAGTTCCTCTATTATGGTAGAGTCTGCTAGTTTCTCTGGCCCAAT**AGCAGGACACTCACTCCCCCACCTGCACCTGCT**GCAGAGCCTTTCTACTCTTGGCCCCAAAACACTGGG**TGACACAGTTCTCAGACCCATGATTTATAGTGTCA**GTATTCAGGCCTCAGGGGTCCCTGATGGCTTCTCTGGCTCCAAGTCTGGAAACA**CAGCCTCCATGACCATCTCTGGGTTCCAGGCTG**AGGATGAGGCTGATTATTACTGCAACTCACATAGGAGAGGTGGCACTTTCcaccgtggtccaagttcatggggaattgagacccaaacctgccctgggctctcagcctctctcttgttctgaagatgcttcctcaccctg**tgcaaggggcttcttgca**gcactgccttgagaatttcccctctcccagctcctc**tcctttctcaccaggaagtccaaaagga**aacctgctctgtgatttctcatccaggacagtgacagcttcctgatgcttgtgtgctgtggtccctgaatgtgcaactcttcctagctcttcaaatg**caggcacatagtgagaaaagctgcctg**actggtgcattcactgctgttttta**aggatgtcctcacccaaaatgcatcct**cctcccaaattgtgaagaacaatctggacagaggtcattacagggagtttcaagaaactg**catcttattcaattgtgtccaccatggtctggtaaagatg**gccctcctggatggactattcctctgcatgtctgtcctgaagcagtgaccactgtgagaagatctgaacatgtttgtgaggtattaaggacgagaggaaactgttgtttttattattcttttgtt**tttgtttttgaaacaaa**cttttgctttgtcgccaggctggagtgcagtggcagaatcttggcttactgaaatctcagcc**tcccaggctctaccaatgctccctgcctcagcctcccgaagagctggga**taacaggtgaccaccaccatgcctggctgatttttgtatatttagtagagacgggatttcaccatgttggccaggctggtcttgaactcctgatctcaggtgatgcacccacctcggcctctcaaagggatgggaatacacacaggagc**cactgcatctggcagtg**ttttttttatttttgctcctcctctttgcctcaatacctcaggttgctgagctg**gggagattttgcgtgacaggctcagtgctccctcaaaatcctccc**gtctcaattcgctggggccctgtcctggaaactc

Uppercase: IGLVI-20

Lowercase: Flanking sequence[1000bp]

Red & Bold & Underline: Stem-loop [21]

Blue: Heptamer[42]

Green: Nonamer [6]

id-LOC102724999[V_gene_segment]

cccacaacactcactactgaatttgaaccaaaatcttctcctagaaatcctcgttggcggctcgatagtaagagctgccatgtatcaggctcacctgaggacagacgctggttaaaccaag**ttgaagttttaattcttcccatcttcaa**agac**tgagcatctttgctctttttgctca**taagtgaaacactccaaagtcatttgccaggatcccctgtagatgggcattcagtgtacagactcctaggcaagatccctggtcacagtatccaaatgactttgactgcaaacctggggaagtagagagagaacaaagatttaccagaaaaaccctctgggaatg**tcttcccaaggcaccctagagggaagagctccctttctcttcagctccacaggatccacagaggaaggagagctct**agccagcaggaaggagccacatgagaggagaggaccaggtgtcccacacaggtggggaagtgagtccatggggctggtccccgctctgggcttgatcctgacata**agaggactctgtgtagacaccacctgagtcctct**gcaatgtacagccacagagccatggacagattcactcctgaggagtccagtagagcgcgaagactgctttgaggtcacagcgggaagacacaggtgtcacttgagctggcctggacctgagagagggggcagtggacagaacagaggggaggcagtggtgtgaatgagaggctctggtcaccttatcatagaatggtttaccttggctggacaccaggggacgctgggggaccatttctgagaagacgcgcggggggcggg**ggtgggcggtgatgggagcacaaccggccacc**tgttgtgcctgttgtcctctgctcagggctcacaactgtgttctccccaccctctgggaccacagagctccacccctgccacaccctgacatcctcaaggcaaggagcc**tgacccagggctcagggtggggtca**caaagctgggggggtCTGATTTGCATGGATGGAC**TCTCCCCCTCTCAGAGTATGAAGAGAGGGAGAGA**TCTGGGGGAAGCTCAGCTTCAGCTGTGGGTAGAGAAGACAGGACTCAGGACAATCTCCAGCATGGCCAGCTTCCCTCTCCTCCTCACCCTCCTCACTCACTGTGCAGGTGACAGGATGGGGACCAAGAAAG**GGGCCCTGGGAAGCCCATGGGGCCC**TGCTTTCTCCTCTTGTCTCCTTTTGTCTCTTGTCAATCACCATGTCTGTGTCTCTCTCACTTCCAGGGTCCTGGGCCCAGTCTGTGCTGACTCAGCC**ACCCTCAGCGTCTGGGACCCCCGGGCAGAGGGT**CACCATCTCTTGTTCTGGAAGCAGCTCCAACATCGGAAGTAATACTGTAAACTGGTACCAGCAGCTCCCAGGAACGGCCCCCAAACTCCTCATCTATAGTAATAATCAGCGGCCCTCAGGGGTCCCTGACCGATTCTCTGGCTCCAAGTCTGGCA**CCTCAGCCTCCCTGGCCATCAGTGGGCTCCAGTCTGAGGATGAGGCTGATTATTACTGTGCAGCATGGGATGACAGCCT**GAATGGTCCCACAGTGCTCCAGGCCAATGGGGAACTGAGACAAGAACCCCCTT**CCTCCTCTGTCAGGAGG**GTGAGCCC**CAGCAGCTGCTGCTCAAGCCTGGCCTGTGGCTTCTGCTGCTG**CAGCTTCCTTCATGGGTCCAGGGGCATCCAGGGCCCTGCCTGAgagtggaggctcctcctccccttcagtcctcagagtcaggaacaggacatccaggaaacagaacatcctcctccctgcagcttaggacacagggtctctgcactgaagtcctgggct**gaggtggcaggtccagttgtgtcatctcagacccacctc**tgatggggaacccgtgtctccgtcatcctccttttcccatttccaggagtttccagagtggtgtcttcctccccctgctcctcagtgtgaatccctgtgcttcctttctctccagcctattcttttttaataaaactcctttcatgtaggaactatgcacatctaataaactacccatatttataacaggaaattagacaaattttggcatttttctggttccttgaaaacatcaccatcc**ccgggagctattgtgaccatcccgg**cagctcctcatgatgcttgtctcctccctccctcccctaatggcctctcaggaagccacagatctgcctctctcactactgtttggtttttttttctggaatgtatcagtgttatcaaacgttatcttcccatttgttcaggcttctttcactctgcacacttgctttgagattatccctgctgtcgctgaattgcacccat**cccttagagggctcattactatctttaaggg**tccttaggccatttttttctgattttattcatgtatttcctttaaa**gtttgaatttattgttcatcaaac**aaactctatgggtgatgttcatctaaggtgacaggacagtccgtgatggtttgtcaggattcttgcttcttcagttcttttcttggctctctctatagtggaagaggttctcatcttttatttatttatttatttattttggggtttttttaagggttttagcgattttccctttttaattattt**ttaaaatttgttattttattttaa**taggtttttggagaacaagtgttgcttggttacatatataagttcgttagtggtgattcctgagat

Uppercase: LOC102724999

Lowercase: Flanking sequence[1000bp]

Red & Bold & Underline: Stem-loop [19]

Blue: Heptamer[38]

Green: Nonamer [3]

id-IGHVIII-47-1[V_gene_segment]

ctgagaaaaagaaaataacctcctgcatgcacttctcaagtctctgcttgtatcctgtttgctt**taggtctctaggggaaaaagtcagatacctggaccta**gtgtcagtgtaggaggcacatcctaaaagctacaaactaggaagaagggaagatgggtgttattggaatagtggatatggagtgggctttcttctgaaagtatatgcacctgctggtattctcagatgcaacattcaactgcaagagcccattgaagaaacaaggcactcccaaatctcctgtaagattctgtattcatttcagataagcccacatgtccccgcattaactttttc**ttcagacaagcacatacatctgaa**actgaacagctatgtggcaaaataagctcaggatagaagt**aattgtggactccagctctgacaatt**tgtagatctgcatttttaaaattctaactgaagactttgctttattgtagaggacagtggtttacagctcgaattgcacagcctacaggcagatgtccatttcctctgggcaaggtttatttttatttgtttactgtacttatttgttgacaaacattgatactataaagataccctaacagcgtccacatgaaagaaaaagaaagagcaatggacacatcaaccctgaacatccagtcccaggaatcctttgaccctgccctccttgcaacccagagatatagattggatgagccctgctgagcagaacaca**catgtccccaggagaaagacatg**gaaatgaggcccctccctgctaatgaaaagcagctcttcccctcttctcctgcaggtcctgg**tgaggagccacccaatatctgtgcccttcctca**gtgtccacaccatcgggtcta**tgatgatctgggcttcacttgtcatca**ctctcaatattgaggttccccgttaaacagactgagtgaactgtggctgctccacgtgggggctgttctcagtctgttgcttctgtgcttgcagAGGTCCCCTGTGAAGTTAATTACTGGAGTCTCTCAGAGAAATACTACAGACCAAGAATTCTCAGACTTTTCTGGAAACCCTGTGGATTCACTTTCACTGAAAACAGCATAAGCTTGGTCCAGCAGGCTTCATGACAGGGGTGGGTGTAGGTGATAACATCAGTAATTCAAGTGGAAGTTCTCAGTGGGACTCTCCTTGAGTACAAAGAAGATTAACAGTCCTCAGAGACACGCTTTTCAGATGATTCTCTCTTAAGATGATTAACCTGAGAGCTCAGGAAAATTCCGTTTATTACTGTGAGGGAcacggtgaggggacatctgtgtgagctcagacacaaacctgcc**tgcagggagacacaaacctccctgca**tggtagatgcttctcagaaccaccagggggtgcacaggaaaccagaaggtgctcaggacaccagggggtgctcagaaccaccatgactcactcaagacgccaggggcgctcagaaccaccgtggggcactcaggacaccagggggactcaggaaactagggggtactcagaaccaccaggcggtgctcaggacaccagggagctctcagaaccatctgggggcactgaggaaaccaggggactcaaacactagggtgtgctctgaagcaccagcggaccctcaggactgcagggggcgctcaggacactagggaacactcagaaccaccaggaagcgctcagaaccaccagggggc**cctcaggacatcagagcgtgctgagg**acctccgggggcgctcaggaccttcagagagtgatcagaacaccagagggcgctcagaacaccaggaggtgctcaggacagcaaggggctctcaggacactagggtgtgctcagtaaaccaggggtccctcacaaccaccagggggcactcaggacaccaggggatgctcaggacaccaggggccactgaggacaccaccgctcccttagcaggcagctccacatcaggcccctgggttggggcaggaagggtgttttccttttggatcttgccact**aaactcttgggagttt**ttctccttcctttgtggtttcaagaaacattggtagattcttctcaggtataaagctctgctttcttggattatgtaatgtttttggtttcggatgttaccagaattacactgcactgtgagagga**ttcattcctcgtgtgtgcaatagtgaatgaa**agctccaatgttaggggtggctttgaaagctacgttaggggtggctgagggcagttagcaggaaatgatcatcactatagaaggctactcatttctttg

Uppercase: IGHVIII-47-1

Lowercase: Flanking sequence[1000bp]

Red & Bold & Underline: Stem-loop [10]

Blue: Heptamer[32]

Green: Nonamer [5]

id-IGKV1-9[V_gene_segment]

aaagaaaatgtggtacatatgcaacatggaatacttaatatacagctatgaaaaatgagatcatgtcttttgcaacactgtggatggtgctggaggccattatcctaagcaaactaacacaaagattcaagtttatgctaatcaaggctcctgtagtagaagaagaaaaaaaattgttttaaagtaatcagatttagcattgctagtcagacttttatgctgatggaccatgggaaataattcacaatgacagaaaagggggtagaatatgggggtccccaagcagagaataaaatgaatcccattaaaactcgagcattaaagagacttatagctctggacaatgcaggaactgtggatgtcatatgctttaaggaatcccaatatcatcttctttgtcaatctgcag**taaacctcttcagcttagactggtaataacattggttta**gggccattac**aaaatgcttttgagaacatttt**acttgctcatgactaag**tgttcttttttacttaaaaaaagatca**atttcatgcttacaaaaatgtagtatgtatgtcacaaagtatcgctcccaactggaacaatttcacagtgtgttgaagacctgatagccccactctctaagactttattaagaattctctacaaacaaggatattctcctacttatgcccaatacggcattgaaatatattattccatctagttctcacaaccacttcgagatttgccaataaatgctcaaaaatgtacttggtaacaaaatatcctttaggaagaaacattctctgcaggcaaatctaggtgccctggtctgacctgggacactggggacactgcccctgtgctgagttactgagataagccagccacgcagctgtatccagcctgccccaccccctgccgatttgcttgttcccagagcaccaccccctgccctaaagacttcttaataggctggtcacacctgtgcaggagtcagtcccagtcaggacacagcATGGACATGAGGGTCCCCGCTCAGCT**CCTGGGGCTCCTGCTGCTCTGGCTCCCAGG**TAAGGAAGGA**GAACACTAGGAATTTACTCAGCCCAGTGTGTTC**CGTACAGCCTGGCTCTTGAGGGAAGTTCTCTTACAACATGATTAATTCTATGGACATTTGTGTTTATATTTCCAATCTCAGGTGCCAGATGTGACATCCAGTTGACCCAGTCTCCATCCTTC**CTGTCTGCATCTGTAGGAGACAG**AGTCACCATCACTTGCCGGGCCAGTCAGGGCATTAGCAGTTATTTAG**CCTGGTATCAGCAAAAACCAGG**GAAAGCCCCTAAGCTCCTGATCTATGCTGCAT**CCACTTTGCAAAGTGG**G**GTCCCATCAAGGTTCAGCGGCAGTGGATCTGGGAC**AGAATTCACTCTCACAATCAGCAGCCTGCAGCCTGAAGATTTTGCAACTTATTACTGTCAACAGCTTAATAGTTACCCTcccacagtgttacaaacctgaacataaacccccagggaagcagacatgtgaggccgggctgccc**cagctgctcctcctgattcctccatcagctgagagtgttcctcagatgcaggcacactct**gatggtgttggtagagg**gggatgtgaagtcacctctgcatccc**aatttctttttctttctcagcaccaggtgcacagacataacagttcctctcctgatttaaaaaaggcagggatcatgacacctgaggagtctagttta**tggcttcagttggaattcaagtaacagagaagaagcca**ctatagatattctaagcaggaattgtcttgatacagagaaatagagtataaactatggaagtct**aaataaaaatatagagatgaatctcaaatttcatgttttattt**gctaagaaatatttgctaaatggggcatacagaaaaactcaatggtcttcaatatgttgaaagaacaaagagaagcttagagtttcatgaaaaaaggaaaatgttacctattgctctttgagaaagtttattggctccagaaaattttgggagctggcaagctcagaatggtaagcactgatggacaaa**ctgaatcctagaattatgttaaattattcag**aagttggggttgaatttgatttccggtt**acaagaggccaaagcagtgaagcttgcagagaatcttgt**tactgaaatgccagggatttggtgtagatcctgctgctcaccacatataaaaccaatcactaagacaacaagtattgccaagaaacaggctttaatcaggtgctgcagctgtggaaatgggaccccattctcaaatgtatc**tccctgacaaactaaaattagggggttatatagcaggga**agaaatgtgggcaaacaggaattagggaggggtaatgaagttaattttgtcaacaggaagcagaaggt**cagtgaggcaatcacaatgtgtgaagggtctgacgtctcactg**tcctgattcagt

Uppercase: IGKV1-9

Lowercase: Flanking sequence[1000bp]

Red & Bold & Underline: Stem-loop [18]

Blue: Heptamer[27]

Green: Nonamer [4]

id-TRBV6-5-2[V_gene_segment]

acaacttattagcatcccgttgg**ttcaaaatatcataagaatttgaa**gaacatgacaatgttatactgactgtggcgtcctaataagtgtgggactctgtgcaccaagatatgttatttga**attcctttttattgattactaaattaatggcaggaat**cccacacaaagggatactccctatgaaacatgacaactcaaattaaaaaagagcactcttccttactctaaagtctgccctctctccccgagtccctcacgcataaggccacatttgtgcccctaggtcatgggcagcgctggttggccaccatcgtccacatgtaga**aggcagtcagcaacgtgaagtggttctgcct**actgcagtctcaccccagacatggaaagcaagagccctgggtgg**agctgaaggtgctcagct**gggt**ttgtcagaagtctcatctgtcagtgagttgacaa**gaaacagagcaaaacgactcctccaatgttgacgagcctgcccctgggatttggaaacttcataacagaaaaaaccaatatagacaaaggattttaaacaggattaaggtcaattaagcaaattagaaaaggacacttgaaggagtatttgggacaggggaatcaaaagcaccaggaagacatgaggagtttccctaagactctagactacagcactagcactatgtagataactaacactaaaatattaattatatcattgaaaaaataaaatttacagtgaattacaaactgtttccaaaaggtcatgaaaatcttgtagacttgtttcaattcagacagatgtgttcttctcattctcagctgttcactggtgcatttattttggatttgaccatctggggaatgggtgtggcctctcctggcc**tctccctccctggggcccaggcagggaga**atgtctcagaatgacttccttgagagtcctgctcccctttcatcaatgcacagatacagaagacccctccgtcatgcagcatctgccATGAGCATCGGCCTCCTGTGCTGTGCAGCCTTGTCTCTCCTGTGGGCAGGTGGGTC**CTGGGCAGAGCCCCTTGTGTGAATTTCAAGGCCCAG**CGCCTTCCCATTGGGG**CTGCAGCATCAGCTTTGTTCTCCTTCTCTGCAG**GTCCAGTGAATGCTGGTGTCACTCAGACCCCAAAATTCCAG**GTCCTGAAGACAGGAC**AGAGCATGACACTGCAGTGTGC**CCAGGATATGAACCATGAATACATGTCCTGG**TATCGACAAGACCCAGGCATGGGGCTGAGGCTGATTCATTACTCAGTTGGTGCTGGTATCACTGACCAAGGAGAAGTCCCCAATGGCTACAATGTCTCCAGATCAACCACAGAGGATTTCCCGCTCAGGCTGCTGTCGGCTGCTCCCTCCCAGACATCTGTGTACTTCTGTGCCAGCAGTTACTCcacagcgctacaaggccgtctcctctctgcacataaaggca**gggaggttctgccctcctccc**ccacccaagactcagggatgc**cctgggcagagatctctgcgccagg**aaccttggaacccacagtggccccaagtggcctggacagtatgagccttgctctgtgccaggtgcctctgcaggcatctcagccaggcctggactggtcccaggtcctcagatgtctcctttgttgctttctctggtctatccttggagctttccttttggggtggggccagggctttcccagctcctacttttctactca**ttatcctgagtccgaggcccccaggataa**aacaggatttgcatttcagatccatctagactcccgtctctccctggtgaccctgttgcttcctctctctaatgtttcccccagcccccaccctcatgttttctctcctgtggcccacctttcccatctgggcagtcaccctccaaggcgttgctgggtctctcctccccccacctcctcg**cccctctctactgcagccatgagggg**agcccctgt**tctgtgcctccttccttcccatcacaga**gacttcaaaggccatttcctctgcccttgcatggagccttcctttctgcagtggccagctcctacctgtccttcatatctcagcatgatcaacccctcttgtgggaagcgctccctcccctcccatctgaaatcaactttcctctcataagctgtcacagattcatgtgtctgttagtaacctttagcacagcttggctttctcagatacttgtctggttatttttgtactccacccaatttttaagcccatgagagctaaggctgtgcctgctgcatttaccaacattggtgtctggcatgtagggagacccatttcacaaagaatggatgagtgaatgacaggctgagtgagtgatgagtgggtggataaaccaataataaaaacacactacatctactgtaacctggcagagatc

Uppercase: TRBV6-5-2

Lowercase: Flanking sequence[1000bp]

Red & Bold & Underline: Stem-loop [15]

Blue: Heptamer[36]

Green: Nonamer [3]

id-IGLVVI-25-1[V_gene_segment]

ggggatctgatgggataggtggcatttaggcaggtattgggtgaagaatagtgatatacacaagaaactttcaaagagaaaaacacacggaagttggaacacagaagagaacataatatgtctgagaaataaggaaagtccaga**ctcactggagctgagtgag**aacaagaaaaatggcataagccatgtatttctctctggatatctcatcaaaatattgtctcctccccaacacaccacccatgtgaattat**aaaaatgcccgtttttaattttt**gtttgcatgtcgtgacagagtcaagcacgtgaatcaagctgagtttgatgagtccagggctgttaagctccagacccaggtggcctcagaaggacttggtggaacaagggcttgttgtcaaaagcagaggaagtgagattttccattgtggcttagaccagagattcttaaattccagtggacacacaaggccccttaaggatctggttaagatgcagatgcagcctcagcagttctggatgtggccccaaattctctgtgtccggcaactcccaggtgaggctgctgctgc**tggtcctcccaggacca**cactttgagtggctgaactcttat**tcacccacctcactcaagccacgggtga**gactcacacaaggagggcaccttctagaagcacaggtcacttctcactcctccccttgtgtcttagatcctttcctgttactataaagaag**tacccaaggctgggta**atttatacaga**aaataaaagttttattt**ggcagacagttccaatggctgcagagttcaaaatgtggcatctgctcttgttgaagacctcaggctgcttcctgtcatggtgaaaggtgaaggcgagccagtgtgtgctgaggtcaatgaggggaaggaagcaagagagaaagggaagataccagtctcatttaaataagcagctcttctggaaacagaggaaattcactcaccccaagtgagggcattaatctattcATGAGAGATCCACTCCTATCACCCTAAAAGCTCCCATTAATTCCCACCCCCAACACTGGGGATTCAATTTCAGCATAAGATTTGCAAGGGATGAAGATCAAAGCCATAGCACCCTGAGGCTTGGCCACACACTTGCTCATCTCCACATTTGTCCCTTCCT**TCAGGACCGGGATCCTGA**CTGCTCTGACTCAGTCTCCGTCTACCTCATGGACACCTGGCAGTCGGTCATCATCGGTT**TTGCTGAATCAGCAGCAA**CATTGGGGGTAACAATTG**TGTCTCCTGGTTCCACCAGAGACA**GAGTAGAGA**CCCACACAGCTGAAGTGTGGG**GTCAAATATTAAAACTAAGAGACATAAGATTGATTCTCAGGCTCCAAGTCTGACAAGTGAGCTTCTTTGAGACTCCCTGGGATCCCAG**CAGTGACACTGATCACTG**TTGCTGTTCCACACATCCCAAGTGATGAGGAagtaagaccaaaaccctcctgagattcctgacttgtgtcctgacaccaggtctgttcttccctcccctagaataaaacatctcttaagcacaaggctgaagaaatgtggcctcctcctttctcccacagctggtcaaagtggatacctgagagaattggctcactgaggcctcttgggaacaaaggctgggttcctgcctgtgatctgtgtactgagacacattccggacctgcccaggaaaggcaagtatagatgatgtttagttgtcacagagtccaagtagggacaattcagcggtgtcttcttgggacccctgcctctggctattttcatctacttt**ataatgggccagggccattat**ttccaacttgtcatatccttttt**ctcctttcctcaccaggtaaggag**gtggaggacagttgatgagggctggcttatgagttaccccaggatgcaaatgactttgaagtcaagcccatgtggggtca**agctcagtctataaccagggtcttttgagct**ctggaggagccaggcatcccagccatca**aagaggtcttgcatttgctaataaatcctctt**cttccaacccacagcttatcttggctgcctaccctcatctctgtcaggaagcaatccttcgttgtttaacaccaaattattcttacttggtcagtttaaaagtcccttcctcagaaaggtcatctgtggcctgaagacaaagtgagaaggctttgctgcattttcatttggaacctt**acattgccccttctgcaatgt**tctaaccacacatggcacatatttacaggacagcgtgataattcacttttgaaaatacagaatgtgctcatg**gaacaggacaatgtcatacttttggggcctgatgagcatgtccttttc**acaggaccccccagagctgcccatgcatccagaaggtaccattctactctatgcaaaacgatggggcatttctatattgcagtttcttagggttccttagaa

Uppercase: IGLVVI-25-1

Lowercase: Flanking sequence[1000bp]

Red & Bold & Underline: Stem-loop [17]

Blue: Heptamer[26]

Green: Nonamer [1]

id-TRAV22[V_gene_segment]

attatttctaaactgt**tgaaagccatcttttctttagtattcttctgtccctttca**tgtaactagaaccccctaccctcaaccaaaaataaataaataaatagataaataaataagtaaataaataaaagagagagagagaaaaggaagaacatggaaacagaaggtaatgataggagaaagtggaaggaaaaaaggaagacagaaaaaaaaagaaaaaagatttactggactgtaagataagctataaaggaagtttgtataaaataagaattagaaaaataaaacatattgcccatccagggtaatacgt**agaaaaaaatttgaattgatcttattttttct**ctcacaatttacagtcttactagggtcatttcttataacaatatttaccacaattcttgattatttcagaggagtcctaatccacacttttcatatattc**taataataaaaattatta**catatata**atacataacatgtat**gttcttattactttacatgactggaatgactttcacctaaattggctaataaagaccttcttgtcagaa**cacattcttgttttgggggtttaaaatgtg**gacactcattctgtgatcctctccaggctcgaattagtattacagttgaggcacgttgtcctcccgaggaggcccctccctttccttctcacaggatctttcacattctttactatatccagaattattagaaaacatccctcaggttacggaataatgacctttctttttcccttccactccaccccttt**ccaccccggctctcagccaggtttaagtcggcgggtgg**ggcttcctgtcacagaacgctgtttaaaaactcagtagcatctgggcgtgagcaggctttgtcgggtggagctgattggttgcaggagcagcaacagttccagagccaagtcatgacaccgacctccccaaggtttagttaaatatatcttatggtgaaaatgcccggagcaagaaggcaaagcatcATGAAGAGGATATTGGGAGCTCTGCTGGGGCTCTTGAGTGCCCAGGTTTGCTGTGAGTTGGGGCTGTCCAAGGTGGGAATCTGAAAGGATTTAGCAGCTGCAGCCTTGTTGAGGGGAGGAAGGGCAAGGAGTTAAAAAGTCATGCAGTCTTTCTATTCACACAACCTGTGGGGGCATTTTCAGGATTCTTCAGATGTTAAACTGCCCATCACCTATGACTTTTTCTCCATTTCTGACCAGGTGTGAGAGGAATACAAGTGGAGCAGAGTCCTCCAGACCTGATTCTCCAGGAGGGAGCCAATTCCACGCTGCGGTGCAATTTTTCTGACTCTGTGAACAATTTGCAGTGGTTTCATCAAAACCCTTGGGGACAGCTCATCAAC**CTGTTTTACATTCCCTCAGGGACAAAACAG**AATGGAAGATTAAGCGCCACGACTGTCGCTACGGAACGCTACAGCTTATTGTACATTTCCTCTTCCCAGA**CCACAGACTCAGGCGTTTATTTCTGTGCTGTGG**AGCcacagtgctccccaggcacctgcggcctgtacacaaaccctcatccgggctcggttcctctaccagtaacaaccacatcacgaggccaccgcagcagcattttgcacagcttaatattccttttgtacatag**gtagagttttaaccgtagattctttttggctctac**aggtttgagactttggtctttattttcccttttcaatacacacctcctcctaccctataaatatctaacttggaaggagtagagtagctgataagaggacagaatcaaaataaatatttaagtgacagtatgcaggactccaaaagggtaaaaagtaaaaagaataaaaagaaaagaaacaccattccaataagcatgttgttagacattagttagacagtgatccagcttagttgtgtgtctcaagcacattcaggtctacctcaccacattatgtaccaacagacaagggcaaatatttgcatactatcccactagtgcagcaaagtaaaatttctttaaatattcacatattgataaatagctgtggaga**aaaaactttaaaatgttatgttcccatatgtgttttt**caggcttgttgtataacctagaatatgatctatcctggaaaatataaccatgcatgattgagaagaatgtgtaatctgatcttg**ggtagatagatgtctgttaggtctacc**tggcttttgtgttgttcaagtcttctatttccttgttggtattctatttgtcctatctattattgaaagtgaggtattgagtcttcacctgttattcttgaattttctgtttttcccttcaattctgtcagtttttgtgcatcatgtattttgggatatgtcattaggtgcatatatg**tttataactgatatattcttaatggatgcccttttatcattataaa**atttccctttttatctctagtaacattttttggtttaaagtatactttgtctaataataatgtagtgactccagtcatatcttttcct

Uppercase: TRAV22

Lowercase: Flanking sequence[1000bp]

Red & Bold & Underline: Stem-loop [12]

Blue: Heptamer[21]

Green: Nonamer [7]

id-TRAV1-1[V_gene_segment]

tgatctgcctgccttggcctcccaaagtggtgggattacaggtgtgagccactgctcccagctcttttttcctgttatacctcttttctttcctttagttttttaaaaaattacataatcaaacatg**tctattttaacattaaccaataga**gggatgtaccaaaaaaaattaactcaactcactgcaacccactgcaacccctgacataaccaatgttagtagtttattgagtatatcctcacacttttaaaaatgtatgcatatgtacataagttt**atgataaaaatatcattcaatactcatcactctgcaacttacttttgaat**atattaaagattatttctatattagctgttgtaagcacacttaaatggtaggtaaatttccttgtctttctagcttccaaaatatatatgacacacaaacaaacaatatttagtatatgcacacacacactgcatctctaattgatcctggatttcattttgttgtatcatgagaaagaaaactgttagtatg**gtcaaattgattagttttgac**tttgccttatgttcccatttgttttctctgttctttacatgttcgatgttcaccataatcacttggattaaaatgtgtggattagtttt**tggagatagggacctcaccatgttgcttaggctggtctcca**gttcctggc**ctcaagggattcttctacctcagcgtcttgag**t**agctgggattacaggcataagccactgtgcccagct**taaaacctgtggatttatcagtagaaaatgttcatgtaaagatactcctgtaagagaaaccatagctgctccagtggaaggaagcttaaactcatcccttcaagaaagaagctcctccctttgtatttctactgggttttgcatccggactgatcttccttccctcacccacatgaagtgtctaccttctgcagactacagtggctcaggaaccggggatgcagtgccaggctcatggtatcctgcagcagATGTGGGGAGCTTTCCTTCTCTATGTTTCCATGAAGATGGGAGGTGAGTCTCAATCTAATAGTAAATGCTGCTAGGAATTTTCAAAACAATTTCCTTTCAGCTAAATTATTGCAAATTTTGACATTTGTAATGAGAGTATTTCCTGAATATGCATTTTCCTAACGTGGTGCTAATTGTCCTCCTGTTACTATTGCTGCTGCTGTTACTGCAACCATTTATTTCAGTCTAAGAAATTCTCCCATCAATGGCAGTTCTTTTGTGACCACATGGAAGCATCATTTAAAAAATTATTCCAATAGTTTTTGGAGGAAA**CATCATTTTTAATAATGATG**GGGCTTCTGGGGGTGCTGCCCTAGTAACAATCATGTATCTTGTCATAGGCACTGCAGGACAAAGCCTTGAGCAGCCCTCTGAAGTGACAGCTGTGGAAGGAGCCATTGTCCAGATAAACTGCAC**GTACCAGACATCTGGGTTTTATGGGCTGTCCTGGTAC**CAGCAACATGATGGCGGAGCACCCACATTTCTTTCTTACAATGCTCTGGATGGTTTGGAGGAGACAGGTCGTTTTTCTTCATTCCTTAGTCGCTCTGATAGTTATGGTTACCTCCTTCTACAGGAGCTCCAGATGAAAGACTCTGCCTCTTACTTCTGCGCTGTGAGAGA**cacagtgactatgaggcctccttaactgtg**ccaaaattcaaaagacaatcagtggagtacaggtgggcttga**gaagttctagaacttc**ctgagtgtatctttgcttaccgtctaattttaaacttgtttagaatatgtagcatagggctctgagacggaaccccattattttctgaaccattgattgctaa**agagaaggataaactattttctct**ctctctttcttctttagtttcttcttttcctcttcctcctgcttatttatgcctccagtatcctgtggttccacgtatggaagatggaatttacttttggaggaagctttttata**caagggaaggatcccttg**gtgagtgctttagtttatgcagtagtaagttacattaatattcatgtatccttgggaggctctga**aatttgtgaagtaaatgcaaatt**cctcagcttgagatagtatgatttcatttaattatgtcaaatatatcctcatttttgtatttacctgagtgataatgttgatattatttagcattttttggtggatttatccaacatattggagagtgtgggggctgtggacttagagggtga**agagaagatcatggagagctcttctct**gtgaaacagaagagtgaggtcatgcgatgaggtgggggtcaagtgtgga**aggaattggtcagtagggaattcct**gtaaaagcactggataatggacagaacagtaatataaaat**attcagtttaaagattttcttttactgaat**tgtaatttttgtgtgctatccta**aatgtttttgaagtttgttaaacatt**acagcctacatagtttgtccttagagggcaatgcagaatttcttccccataggtgaattattttttggcttagtacatttatgtatgtatgtatgtatgtatgtatgtatgtatgtatgtatgtatgatggagtctcgctctgtcacccaggctggagtgcagtggcccaatctcggctcactgcaagctc

Uppercase: TRAV1-1

Lowercase: Flanking sequence[1000bp]

Red & Bold & Underline: Stem-loop [18]

Blue: Heptamer[23]

Green: Nonamer [5]

id-TRBV8-2[V_gene_segment]

attatcatcctaatcttttgctaatctttatgtttatggtaacattgattgttattttgggggacatg**tatttcatctttggtaatgaaatgaaata**actatgcacttcaattcctgtatgttctgcctacttaagagtaaatccaattttcttccattcctcccattctatccca**ctccctttatatgatagttttaagggag**aaaatattaattagactctaactgcagctatgtgataagagttatttagaatgagggtgggatattaagctttgagtaaagcaatagtcagggttagaactagtgttggaatcagggtttagggaagttgctcataaaacctgcaggatgacacttctagaatagtctctcactcactctgcagacacttcccagcatcccttgggc**cattgcaaaagcaatg**atgaaacttcacgtattggccacaagatggcactgtggtccactgggatctaagggactctggggagagcctgggagagcagcctaggaagggaagggttaagaaaaattagggctggaatccaatattatgcagatgttgcagcagttttcagtcattgctaggctacctacagctatgcaagatgcaggaaatacctctaatctttaatgagatccacagttgaagaattttggcctggcagccttggtgtcaagcgcaggtgcaagcagaggagcaactgcctcaaaggaacgtgggaaagctaggggcgggctgtgccctaagctcagtatgtctctgctgcaccccatcttccctgcagacctcgccgggcaacagccttcatgagaggaggtggatcctgcctgagaccagtcagaaggccctggaccagctggactccgtaggcatggggggtgacaccacaggcccaccagactattccctagggagcagcctccgagccccagctcacagtcc**tgaggatcactgcacttagttgggccagcccctcctca**ctctgaccctaccATGAACCCCAAACTCTTCTGTGTGACCCTTTGTCTCCTGGGAGCAGATAAGTTCTGGAAACAGCTGCA**AAACCCTGTCATGGGTTT**GCAATGTCTGGGCTACAAGCCTTTTTTGTCCTTTCTTCTGCAGAGTCCCCCTATTTCTTAAGTCTTGTCTTCACTTTCTGTCTGCCTCT**CCAGCAGGCTCTATTGATGCTGG**GATCACCCAGATGCCAAGATAT**CACATTGTACAGAAGAAAGAGATGATCCTGGAATGTG**CTCAGGTTAGGAACAGTGTTCTGATATCGACAAGACCCAAGACGGGGGCTGAAGCTTATCCACTATTCAGGCAGTGGTCACAGCAGGACCAAAGTTGATGTCACAGAGGGGTACTGTGTCTCTTGAAACAAGCTTGAGCATTTCCCCAATCCTGGCATCCACCAGCACCAGCCAGACCTATCTGTACCACTGTGGCAGCACATCcgcagccctgcacagccag**ctgccctctgcacaaaaagggcag**tcacaggctggaggtgggcactccttatggaagcccgtgtctcaaccagaagaaaaagctgccctttctgaagctcttcccagacttcccagcaatgggatctcattctatgtgtgctcctgcagcataaaacttagagtgagctgggtctgtcactgtagtgcggatgtgtaaatgcaaacaacgtaatgttat**tttgctcaatttggaatattctggtaattgtgcaaa**ggaatggtagcttctaacaagaaattcatatttcac**tttcttttcaggagagaagaaa**aaagcaga**aagcagctgggaagctttagcctgctt**gagtgatctgggattctcaggtagcatacataacatcaccttggctttggttcctttagataccaagcatgtgttctgtggaggacaagaaaatccattttttttgttataaagttataaacacacaagagaacaatccagttagggagaatgttaataaatacaacaaactctgccattagacctgaa**gataatcagctagtagaattatc**aaagagattgtaagactcttactgagtacctgtttgaggtcctctgaggatagatgaaggaaggaacttaattcttaccatcactgtggcttggattagaatccatgagttccttaacttgtttttcttgtaaa**atattataaaataatat**agaacacacattttattttcctgtagcctgtagtcct**ctttagcacctaaagggagaagcgcatatcccggtgcaaaag**ggagctggttgcttgcttggcaaaaaaatgaattcaccaattggccaatctgctgaaagtcaaaatgaaaggctaatatatgagactgatgacagttcatggttcttggtataacttcaacagcgagtgagcatttttttctttgtctatagctttctttcagccatcgatcctcagttgtgtcttagcct

Uppercase: TRBV8-2

Lowercase: Flanking sequence[1000bp]

Red & Bold & Underline: Stem-loop [14]

Blue: Heptamer[23]

Green: Nonamer [0]

id-TRBV29-1[V_gene_segment]

actctgttacttactcttggattatatgaacttggataggttatttaaggtctctaggtcttatctgtaaaatttcctaatccgtaaaaatagtgtgatgataataaatagcctagccaaaaaaatagacttaaataaatgtatgaaaagtagagtaagtgctcaaaaacttagct**ctgcttttatgcaaaaataaactattttaagcag**ttacatgatacgagaaatcagcttgataatg**gaaaaagatcataccattgtggtcaagttttacacaaacatgatctgtttc**gtggcctagactgagttgaattagaatagaagaaaccatattacaagtattcaggaattaagcaaagacaaaaaaaaat**agttatgactgtgataact**gaaacaagggaagagatataagagacaatgcattta**atttgtgaattgacaaat**aaggtaggtgaggaaaaaagaggcttatggaa**ttcttgaacctaacatgcaagaa**gagcagggatgcaatttattgggagaggggaattgaggaataagagcccgtttattggtgtgtgtacgtgttgggggaggaaggagttcagtttgggacaaccatgaagacccagcagacagctgaaatactgtaatgaagttcccttatattctaacatattacttttagttctttgaagtcaattctgcaccccactttttgctctacacccagacagtagcacagattggtgttcagtaatgacagatggaaaagtgatgggagaatggtggtgagtggctggtgcatgctgtgagacttgtgcatgaagactcatcctgagtgaagagccacacggttccaccagggggaagactgatttttgcatatcatgcagaaactcaaccacctgttagaggagggaaacagagtgggaaggaactacgctgtaggaaggaaagatgaacttgagtttcacttcttagtgccttttctcaggggagaggccatcacttgaagATGCTGAGTCTTCTGCTCCTTCTCCTGGGACTAGGTATGAGCCTTGTCTTTTGAGGAGTCTGGGGTTAATGTTAACACTGGGTGGAAAGGTGGAAAGACTTTCTCTCCAGAGAGTCTGAAGCCACCAGACAGAAGATGAAGGGGATATGTTAGATTAAAGAACCATAACAGAGTAATTGTTGGCACAATAA**AAAGGAGGCAGTTCCCTGGGGTCCTTT**GGGAGCTGAGGTGGTATTGGAGAGATAAAGCAGAGACCTTCCTGCCTGGGGAGATGTCTGACCCCAAGCTTGTCGGTCCTCATGAATATCTCTCTGGAACTAGGCTCTGTGTTCAGTGCTGTCATCTCTCAAAAGCCAAG**CAGGGATATCTGTCAACGTGGAACCTCCCTG**ACGATCCAGTGTCAAGTCGATAGCCAAGTCACCATGATGTTCTGGTACCGTCAGCAACCTGGA**CAGAGCCTGACACTGATCGCAACTGCAAATCAGGGCTCTG**AGGCCACATATGAGAGTGGATTTGTCATTGACAAGTTTCCCATCAGCCGCCCAAACCTAACATTCTCAACTCTGACTGTGAGCAACATGAGCCCTGAAGACAGCAGCATATATCTCTGCAGCGTTGAAGAcacagtgcggggca**cagatcaaagatctg**agcaagaacctcagctctctcctacccagctcctctcacacgagcctgaaggccctgccaaggtgggacagaaggaggaaaccacagcttttgggcag**acacagctgtttctgtgt**ttgtgggtgggcatgggtatgaatggatagactggggatgaggagagcatctgtaggactagctggagcgttctatccacaggcaggaagagtggctcctggaaagctgagggtaaa**atttcttctagaagggtgctcagaaat**acccaataaaaacaatcagatacagtttccttcaagtggcggggtcagtaat**agcctttaactgttttatctcaaggct**ctttttagctggttttctatcataatatattttggaagaactttagtcatcttgaaacaaagctggtacattatattaatgacatctggttgttctcagtgaataggaatgataaagccctctagatgcttcaatatgacaatatgacacatgcatcccaggccaatataaaatctgcaaagttcaggaaccttctatactggtgagttttggagtggtccagatgtaagaagtaagctgc**tgcaccttagatcatgtaccacggagaaagaggtgca**atgcttagtctttggactttgaattaaactttggatatactgcatttgattgtgctactctatcccatttactatgtaatctgtaaagctggggccagtgcaagagagggttctgtggcagaagaagcccatgatgaaagcttcctg**ctgcttgggccatggactaagcag**acccagttgttcttgaagagcctatggtagatactgatgccctgtggagctcctgacattccccaatacgagaatcacaaggcagtgtggaacaggtttgctgtgccctgatttgcacttcagaccattgcatcttgggctatacttttgttgctttccagatcccagagagaaagaatgctc

Uppercase: TRBV29-1

Lowercase: Flanking sequence[1000bp]

Red & Bold & Underline: Stem-loop [14]

Blue: Heptamer[27]

Green: Nonamer [4]

id-IGHV3-32[V_gene_segment]

tttttagacatatgcaatacattttgaggtgaaacctggtggaattttttcc**aataaattagtaaaaataaatcatttaattgactattttatt**caggttgatttgtttaatatttgctaaaggccagttctttaagctatgacacataataaatcccaaatggcagtacctcattgtttacttagcttttgtacttatatttttcagaggaagaaccactactgtaaattgtaaatagc**caatacataattgtattg**tatgcaaatctgtgactgtttacagtgtcatctctgagaaacataaagtttatttactatatatatataaagagtttggaaggtggactccaccaatttttgaaagagttacagaagggctgctattacttctttaaatgttaaggttcattttatgatgtaacagataacctatcatggagaatgttcaatgtgttcttgagaaggatgtgtattacgtgactcttggttggaaggttctgtaaatatcattcagataaatttgttcaatagtgttgttcaagttcagaggcacattgagaattttctttctggatttgctaaacattatggtcatgaggtattaaggtcttgtgttatttttgtattgttttctatttctttatatctcttacagtttgcttaatgcagttatatttgtatttgtacacatgtgtaaaaacaaaaatcataattgctaaatgagttttatggcacagtcacattataagtaatatttttccaaatgctaccattgccactaaagtcctcctggagtctgacgtctgctctgggcactgccttcttctcaggcgtcccacactggagcttgctatagaggaggaggcatgaaaacagggccctccctctcctggtgaaatcaag**cccagacctgaccctgcagctctggg**agaacagccacag**ccctgggattcccaggg**gtttccatttggtaatcaggactgaacacagagaactcactATGGGGTGTGAATTAAGCTGAATTTTTCTTGTTGGTATTTTAAAAGGTGACTCCTAGGGAACTAGAGTGAGTGAGAGTGAGTGGATATGAGTGAG**AGAAACAGTGGATATGTTTGCCAGTTTCT**GACCAGGATGTGTGTGTATTTTCAGGTGTTCAGTGTGAGGTGGAGCTGATAGAGTCCATAGAGGACCTGAGACAACCTGGGAAGTTCCTGAGACTCTCCTGTGTAGCCTCTAGATTCGCCTTCAGTAGCTTCTGAATGAGCCGAGTTCACCAGT**CTCCAGGCAAGGGGCTGGAG**TGAGTAATAGATATAAAAGATGATGGAAGTCAGATACACCATGCAGACTCTGTGAAGGGCAGATTCTCCATCTCCAAAGACAATGCTAAGAA**CTCTCTGTATCTGCAAATGAACACTCAGAGAG**CTGAGGACGTGGCCGTGTATGGCTATACATAAGGTCccaagtg**aggaaatatcggtgtgagtccagacacaacatttcct**gcaaaaagaagaaaggagtctgggccgaaggggacactcagcactcacaaaacaggt**gcagccccaaggcaggtgcagatggagggagggtaagggctgc**tttccttcagggtctgtggtttcctctgcttctaatatttcccctctgaacctctgtatatttatgttttgtgcccaccatgaggtccctggattagaaaactaatttaaaagaggaaatattctcatatgtc**ccaaaaacagatgtaagttttgg**aggcataaaaatgcataggagccgcgtgagtctgtagacactgccaccccacaatgccagacccaaaactagtgctggagaagggtgggagtttgatggagcttccctgatgaccccgtggtccaagctaagtccagcaaggccattggtgcctcgctgagcacagttgtccatcagggatctcccatgtgtcccagcagcagccatgcctcagtatctccactgtgcacagccatcgt**ctgggaggagctcccag**gatgggtgtctttggcacacacaggtgatgggtgttagagtgcagtgcagcagctggctgcctggtctattgggctcctggatattggagggattggaggtgcattctcaaggccagcacgctgtttgttaatttttgtataaaaaccatgtgatttaattcattttctcagatgacatagataattaataacacaatctgc**aaacaattgtaattttcaactttaccccaagttcattgttt**cttaattctgtgcaggatccagacatggtattgcccttctcat**gagaaattgttcgacctaaactgaaaccagttgtttctc**gtatactttggttctccccacatgcagagatcttgattagagcaagtttggtactttccacgcactcaccctcacctccccagataaagagcagaagttctccttagactgagtctgag

Uppercase: IGHV3-32

Lowercase: Flanking sequence[1000bp]

Red & Bold & Underline: Stem-loop [13]

Blue: Heptamer[39]

Green: Nonamer [5]

id-IGHV3-37[V_gene_segment]

aactaaattcacaagtacatataagtaaaataagccaaaaaaaaaaaactacatcccgtatgatga**cacttttatacattctacagaatagaaactaaagtg**atatgaaggttgggagttgaatggagaaaaggtgagcgaaaagagggaggacggaggagttgcaagagaacaag**agaaaatgttgagggtaattgatctgttttct**atcttaataatggttatggttacatcaataatttctaaattgtacattttgtgtgttaattataattttatattattcaagctattacaaataaatacctgtataaatttcttagggacggtttaagggagatggataaatcacataaaaagtcagatatttctgaatgtacaatgaataacccttagctctatctttattttttggtaattctaaaatgcagcagatttttttaatgtttttattacaagaagggtttcttcaaaccacaccaggtgtgtgtaactctgggctaaagttggttaagggagcagtgggatccggtggagaggagcacaggcctcgatcctggagcccattcttgtcacatgtgagctcctggacacatctcatagctccctcatgcttctgcg**tttacattctcatctgtaaa**tggaggaccaatgggcatctgcctctaacaatgtattttcatgtgttaagatgatgtatgttaagtgtttactatagcacaatcttttaataagaaattacatttttttccaaatactatcattgtcttcaagccccaccacccaccagaaaacttaatatgctctgtgccctgccctgtcctcagacatccagcccaagagacttttatatagtaggagatatgcaaatagttccctccctctgctgatgatgaaaaccagcccagccgtgaccgtgcagctctgggagacgagctcaagtgccaggattc**ccaagtgttttcacttggtgatcagaacttaacacagaggaatcaccA**TGTTGTTTGGGCTGAGCTGGGCTTTCCTTTTTACTATTTTAAGAGGTGATACATGAAGAACTACAGATATTGTTCGTGAGTGGATATTAG**AGAAACAGTGGATATGTGTGGCAGTTTCT**GACCAGGATGTCTCTGTGTTTGCAGGTGTGCAGTATGAGGTG**CAGCTGGTAGAGTCTGGGGGAGACTTGGTACAGCTG**TGGTGGGGTTGGGGGTGTCCTGAGACTCTCATGTGCAGCCTGCGGATTCATCTTGAGAAGCAATGACATGAACTGGGCCCACCGGGCTTCAAGAAAGGGGCTGGAATGGGTCTCATACATTAGTGCTAGTGGTGGTAGCCTATACTATGCAGACACTGAAGGGTAGATTCACCATCTCTAGAGACAATGGCAAGAACATGCTGTACTTGCAAATGAACAGTCTGAGAGATGAGGACTCGGCTGTGTGAGAGAcatggtgaggggaaatcagtatgagcccagccagaaacct**ccctgcaggaaccctggggtggggggaaatcagctgcaggg**ggcactcaggacccactgatcagaatcaacccca**gaaggaggtgcacatggagattaggggctggattccttc**caggatgtgggacttcaacttcttctgacagtttccccagggaaccgctctaagtttagaattctgtgtcatctctatgtat**ttaaaaaagatcattttaatgtgaggaactaactattctcacat**gcacaaaatgcagattcacactcacagagatgaaaagtcctcaactgtggtcactaggatcagagt**cctgaggaatctcagg**cgt**tcctggtgagtcttctccaatcagacccagga**cagacacctcagtgagattccatgactagaacaggctttatggattttaaaaaaaatagactgagcc**agggtcagtgtcatgtagaacctcacaggtttcacatctgaccct**tctcctgacactaaagtgtgcaaatcagcatcaatactgatctggtgctgcttttgctcccaatctgtgttctttcttgagttgtccc**tttttcatttgcttttcctgctccctgaaaaa**agaaagatgttgtctctgtggtctaaattccagggctcaagcccttttcttggagctcaggtgggtctcaggtt**gtggctcctgcagccac**gctggggaggctg**atgggactttcttctctcccat**tgctcagggcc**ctccactatattgtgtggag**actcatctcagaatagatgtggccagtgggagctgaaggggacaagcttgtttggtcaaaatgagatacaaatatggaattcaccctgtgctgtacaaactgtcacagagtc**actttctgaaacaatagtgttagaaagaacttgagaaagt**tgtcagaatcaaaattgggtcacttgtgttaaaactctgctgaatgaagctgggaaaggccg

Uppercase: IGHV3-37

Lowercase: Flanking sequence[1000bp]

Red & Bold & Underline: Stem-loop [19]

Blue: Heptamer[30]

Green: Nonamer [6]

id-IGKV3-15-3[V_gene_segment]

gagttagacagggatgggtgggaatgattgaaggtgtgagtactttagcacagattaagaccaaatcattaggatttaaagagttgtgtagagttagtgaaggaaaagccttagaattaaatttggctgtggataaaacattcttggattagactgaagactcttttctgtgctaagtaagtatatttatgataatgatgatgactgtagtgctgaatatttaataaataaaaacaaaattaattgccgcatacataatgtcctgaatactattgtaaatgttttatcttattttctttaaactgtctacagcactataaggtaggtaccagtattgtcacagttacacagatatggaaaccgagacacagggaagttaagttacttgatcaatttcaagcaatcggcaagccatggagcatctatgtcagggctgccaggacatgtgactgtaaacagaagtttttcactttttaactcaaa**gagggtatgtggctgggttaatggaaagcttcaggaccctc**agaaaacattactaacaagcaaatgaaaggtgtatctggaagattaagttttaacagactcttcatttccatcgatccaataatgcacttagggagatgactgggcatattgaggataggaagagagaagtgaaaacacagctttttatattgttcttaacaggcttgtgccaaaca**tcttctgggtggatttaggtgattgaggagaaga**aagacacaggagcgaaattc**tctgagcacaagggaggagttctacactcaga**ctgagccaacagacttttctggcctgacaaccagggcggcgcaggatgctcagtgcagagaggaag**aagcaggtggtctttgcagctgaaagctcagctcccaccccagctgctt**tgcatgtccctcccagctgccctacctt**ccagagcccatatcaatgcctgggtcagagctctgg**ggaggaactgctcagttaggacccagacggaaccATGGAAGCCCCAGCGCAGCTTCTCTTCCTCCTGCTACTCTGGCTCCCAGGTGAGGGGAATAT**GAGGTGGTTTTGCACATCAGTGAAAACTCCTGCCACCTC**TGCTCAGCAAGAAATATAATT**AAAATTCAATGTAGATCAACAATTTT**GGCTCTACTTAAAGACAGTGGGTTTGATTTTGATTACATGAGTGCATTTCTGTTTTATTTCCAATTTCAGATACCA**CTGGAGAAATAGTGATGACGCAGTCTCCAG**CCACCCTGTCTGTGTCTCCAGGGGAAAGAGCCACCCTCTCCTGCAGGGCCAGTCAGAGTGTTAGCAGCAACTT**AGCCTGGTACCAGCAGAAACCTGGCCAGGCT**CCCAGGCTCCTCATCTATGGTGCATCCACCAGG**GCCACTGGTATCCCAGCCAGGTTCAGTGGC**AGTGGGTCTGGGACAGAGTTCACTCTCACCATCAGCAGCCTGCAGTCTGAAGATTTTGCAGT**TTATTACTGTCAGCAGTATAATAA**CTGGCCTcccacagtgattcaacatgaaacaaaaacctcaagaagaccatcagtgtttactagattataccagctgcttcctttacagacagctagtgtgg**tggccactcagttttagcatctctgctctatttggccattttggagctcaagttctcaagtccaaaat**tacttatgttagtccattacatcataccatttcagtgtggctattacattcatttaaacgcatttcagaaggcatctctgtttatggcatcacaaagagtttaataaatctgtgcaagaataaacaacaaacacacctataaatataaagttgaaatatcaaaacta**tttcagcactctgaaa**attggcaaagcataaaataattaaggatgcatattctttatagaaaaaaaaagtactagtgctttgagtaaggacagaaaatgtctgtagccttttgcctgtgacagcacccttctattcccagctcagtcaatatgaattgcagaactgg**agttttacccatgtaaggatagcaaataaaactggcagcttgctgcca**aagtgggtggacttgagtgaagccaaggagtgggaaataattcttcagtgtttccagctaaacagggagaacaccataggaaatgaacagaaaaagcccacagccttgctagtccaagatgatgccttgtttgaggcaagtagtac**atctgctgatagtaaatagcagat**tcctggataagatagacccacattgctgaaataatctctgcacatattcctagtgacctagaagttatagatatgagtgatgagaactaggagtttccggtgcaaagtaaaaccagagggaggtaagaactag**ctgcattttgcatgcag**tttgctttttaaactacacacagatgggttgacagaagatagatttataacctccagatatttgagcaaaatgtctcagatcattggtgaccacttcgctatgcagatacatgtgtagttcctataaaatcaaact

Uppercase: IGKV3-15-3

Lowercase: Flanking sequence[1000bp]

Red & Bold & Underline: Stem-loop [18]

Blue: Heptamer[31]

Green: Nonamer [5]

id-TRBV7-1[V_gene_segment]

atggaggaggctggggcatcaatggggatggcacaggggactctgacttgcaggaaagacaatgagctcaccttttggtgctttgtgttggcggagctgttgacacatcctagagaacatgcccagcagacagaggagcggctgtgggatgaggagataaactcagagatgcagcgtgaggcctccgggtccagacagcatgagagcccaaagcgatgatacatgcattgatgttgttaaaaaggatttttttttttttttga**gacagattctcgctctgtc**gcccaggctggagtacagtggcgcaatctcggctcactgcaagctccgcctcccgggttcaagccattctcctgcctcagcctcccaagtagcttggactagaggcgcccgccacaacgcctggctcattttttgtatttttagaagagacagggtttcaccatattagccaggatggtctcgatctcctgacctcatgatccacctgcctccgc**ctcccaaagtgctgggctcacgcctgtaatcccagcactttgggag**accaaggtgggcggatcacttgaggtcatgagtttgagaccagccggggcaatgtggtgaaaccccgtctctactaaaaatacaaaaattagcctggtatggtggtgtgagtctgtaatcccagctactcaggaggctgaggcaggaggatcacttgaacttgggaggcagagttt**gcagtgagccgagatcacactgc**tgca**ctccattctgggcaatggag**caaggctctgtctcaaagaacaaacaagcaaaaagcaaggaactcataaatattt**aaaggaggcatttaagtaggtcctaaaataaatccttt**gttcctgtcattgcgtggattgagagaggatgt**gatgtcactatgggaccttctgtgtgggaacaaggacatc**cctcctcctctgctcctactcacagtgactctgatctggtaaagctcccatcctgccctgaccctgccATGGGCACAAGGCTCCTCTGCTGGGCAGCCATATGTCTCCTGGGGGCAGGTGATTCCTCAGATGCCAAGCAGTCTCCTGTGTGTGTGTGTGTGTGTGTGTATGTGTGTGAGATGTGTGTGTGAGAGAGAGAGAGAGAGAGCTGACTATAGTTGTTTTTCTCATTCAGTTCCCAATTTCTGTCTCCACAGATCACACAGGTGCTGGAGTCTCCCAGTCCCTGAGACACAAGGTAGCAAAGAAGGGAAAGGATGTAGCTCTCAGATATGATCCAATTTCAGGTCATAATGCCCTTTATTGGTACCGACAGAGCCTGGGGCAGGGCCTGGAGTTTCCAATTTACTTCCAAGGCAAGGATGCAGCAGACAAATCGGGGCTTCCCCGTGATCGGTTCTCTGCACAGAGGTCTGAGGGATCCATCTCCACTCTGAAGTTCCAGCGC**ACACAGCAGGGGGACTTGGCTGTGT**ATCTCTGTGCCAGCAGCTCAGCcacagcactactgctccagtgtcagcttggttccctaggaaatggggtttctagaacctgaatgctgacaaataagagttgtatatgtgtataccatgcaacctgcgtttaaaaatgtatgtacatagtgcaatgactaaatctagctagttaacatatgcagtacctcaatccttatcctttccagtggtgagaatacttaaaatgtactct**gttagcattttcccagaatacaatccactgttgctaac**tctagtcatttgttgtataatagatctttgtaacatcttcgtcttatctgagtgagattttgtat**ctgttgaccaacatcttcccaacag**tccatctctactccagcctttgct**aattactgctctagtctttgaatttgaagtaatt**tccctgaggtctttagctcaacaagatgggggaattttctct**ataatgatgcttacattat**ttttgctttgtttttgtttcgttattttatgtcttgcaatatagtgtttattgta**aatataaataagtatatt**gtaataaaaatccttcacctctctttgggtaaagctacagttcacatattctaaatttcatgctgtgaccaacgctgacataattatggatcatgggtttctgggagtcctcagagacagccccatacaccaagtttaagatagaatattccagacccagc**caggacagaagtgcatggtcctg**catagctccttggaaaattatagttccccaaattcagttattggatattgtggtgccgcagacaccaaacgtctttctctagcaaatgatggtccttggcagggattgttctgaacccattacaattttgccatcatcaaa**tcactccttgctgttaccttgtgtctcctgggagtga**ggacgccccgggcacagatggaaattccctgaccttcggatgctatttcaaggacttcctaagaccttgtgtccatctttttccacctttatccacatgactcct

Uppercase: TRBV7-1

Lowercase: Flanking sequence[1000bp]

Red & Bold & Underline: Stem-loop [14]

Blue: Heptamer[38]

Green: Nonamer [4]

id-IGHV3OR16-8[V_gene_segment]

aaggcatgcctcag**tcacataagaatgtga**ggacattatcaaataattgtgctgagtgtaggaagctaaa**gaattcacagtaaatctcctgtgatttc**atttgtataaattgtagaaaatgcaactattctaaattaacatggagaagatctgaatttttctgagaaaagtgtggtgagagtaacaagatggtgaaataaaattacagggaagtgagagaaaa**aaattagaggttaattt**aattgctaatagcatgattgaagtgctgattcaaaggctgcacacatataccaacattttc**caaattgtacactataaatttgaattcactatggattgaatttttgaataaagcagtaacaaaaa**aatgagtatattggctgaggaagaggaagagatgaatattgacacttgaataatcacggact**cctgaaaatacacacatgtgaacactgattgcatatttcagg**taaacactagaaaaagcaaagtcacataaagttgttatgacag**gtgggacatcctgaaaacctcactaggcatgtcccac**atcgcc**ctggagctgtctcaggggagcagtctcctccag**tgtttagaggcacagacacaggtaatagggctaactctggccag**atgtgtgatattggacacat**tgcacaactgctctgttatgtatgtaattcatcttctctacaaatgtaacattgacacttgcactgaatatattctgcaaatgtgtaaacattaaataaga**tgatgactgctaattgatcatca**aggcacaatcacataatctgaagttatattttcctgagagataggattacctccaatgttttctgggatgctctcatcttctctgggcactgccctctcctcagctgtcccaccacagagcttgctatatagtaggagacatgcaaatagggccctctgtctgctgataaaaaccagcccagccctgaccctgcagc**tctgggagaagagccccagccccaga**ATTCCCAGGAATTTCCATC**TGGTGATCAGCACTGAACACAGAGGACTCACCA**TGGAGTTTGGGCTGAACTGGGTTTTCCTTGTTGCTATTATAAAAGGTGATTTATGGCGAACTAGAGACACTGAGAGGACGTGAGTGAGATAAGCAGTGAATATATGTGGCAGTTTCTGACCAGGTTGTCTCTGTGTTTGCAGGTGCCCAGTGTGAGGTGCAGCTGGTGGAGTCTGGGGGAGGCTTGGTACAGCCTGGGGGGTCCCTGAGACTGTCCTGTCCAGCCTCTGGATTCACCTTCAGTAACCACTACATGAGCTGGGTCCGCCAGG**CTCCAGGGAAGGGACTGGAG**TGGGTTTCATACATTAGTGGTGATAGTGGTTACACAAACTACGCAGACTCTGTGAAGGGCCGATTCACCATCTCCAGGGACAACGCCAATAACTCACCGTATCTGCAAATGAACAGCCTGAGAGCTGAGGACACGGCTGTGTATTA**CTGTGTGAAACACACAG**CGAGGGGAAGTCAGTGAgagcccaggcacaaacctccctgaaggggtcccagagaccaccagggggcgccaggacactgtgcacggggctgtctccagggcaggtgcaggtgctgctgagggctggctttctgtcatgg**cctggggcagcctcatcgtcaaatttccccagg**gaacttctccagatttacaattctatatta**acatttgatgtctctaaatgt**aaaacgtttttttttg**ttcctttgtttttgtaacaagaggaaacaccctcacctccacagaagccagggtgt**cactttgggggcagaaataatcctttcatggtcaggatgagagt**cctgaggaatctcagg**gaaacctggagagtgttttccaattagactcagag**cagagacctccatgggaatctctg**attagaacaggccttgagctctgatgggagccaagagagaggctcacccagggtcagggtcctt**aaaacctgatggtttt**cacagctatcccctctcaacttgtaaaactgtgcccatctgactcagactgattcagctgaccctctttctgctgatccattttccatctctgtagacttgattctcacagttccctttcttcttctcttccct**gaaaacagaagatgtgttttc**tgtagtcaaaattccagggcttgggtctgcaggacctgggtaggctgaggggactttctcactcaccattgtctggacactcctgttgtcttctgtgcatggaggcatttggaaaatgtagtggacattagccatgaagggaataatactagttttctccaatgggatattgatgtagagctgatcttgtgcttctcatactatctgagtttggactctcacctgtgactttgaga**agagctgaggatgggcactccattgtgctgtgagctct**gggtaacaataattgtagaatctggctaggcagtttaaggtcaatactactggccttcgggaaagacaggctggaattcctgggaagatctgca

Uppercase: IGHV3OR16-8

Lowercase: Flanking sequence[1000bp]

Red & Bold & Underline: Stem-loop [24]

Blue: Heptamer[41]

Green: Nonamer [9]

id-IGHV3-6-2[V_gene_segment]

acaaggaaaagtatctaagtatttatgct**gagtaaaataagccagacaaataagaatatgtactatattactc**catttatgtaatttatttcaaatgaaaaaaatctaaagtaatatacaaaagatgagtagttacctgggaaaagggtagaataaaggaaggggaaatgtggaaacatacagaagaacaagagaaaatgttgaggggaactcacttgtccacctcagta**atgatgacgcttacatcat**gttttcaattc**tacactttaaatacgtgaagtgta**ttatctgccaattaacactcacaaaatttattgcaagcaaacaaatgaaaaattagaccaaaagcggtggtataaagatacaaaatatccattaaaagtcagaaatatctgagaatttacctcctgaaccctagttctctccttatttttaggtgaatgctagagtgccacaaaaatcactcaagttctcaatacaggaagtgggttccacaaagcacactaggcatgtccaattcttacctggagttggttcagggagtaactggggccactgatgtggagcacaggccaagctaccagggctcattcatctcagacatgagaca**cacacagcccctcctccatgtgtg**ggtttatttcaca**tctgtaaatggagaaaccattgacttctacaga**acataa**tttgcacaaatatgtaaaaatataatagtgcaaaatgtttatcacaacacatttt**cataataagacagt**gtatttcccaaatac**cataattgtcacccaactcctgcggggccctgtcatttta**tctggggtctgccgtctcctcagggttcccaccccaga**gctctctgtgtagcaggagacacgcaaatagggccctccccatgctgatgaaaaccagctaaaccctgaccctgcagctctggcagaggagcctcagccccgggattcccaggtgtttctgcttggtcaacactgaacatacaggattcaccATGGAGTTGGGGCTGAGGTGGGTTTTCCTTGCTGCTATTTTAAAAGGTGATTTATGGAGAACTAGAGCTATTGAGTGTGAATGGACATAAGTGAGCGAAACAGTGGATATGTGTGGCAGTTTCTTACCAGGATGTCTCTGTGTTTGCAGGTGTCCAGTGTGAGATGCAGCTGGTAGAGTCTGGAGCAAACTTGACAAAGCCTGGGTGTC**CCAGAGACTCTCCTGTGCAGCCTCTGG**ATTCACCTTCAGTAGCCATAGCACGC**ACTGGGTCCCCCAGGCTCCAGGGAAGGGTCTGCAGTGGGTCCCAGT**TATTAGTGGTAGTG**GTAGTACCATGTACTAC**GCAGACTCTGTGAAGGGCCGATTCAC**CATTTCCAGAGACAATACGAAAAACTCACTGTATCTGCAAATG**AACAGACTGAGGGCAGAGGATGCAGCTGCATATGACTCTGTGAGAGAtacggtaaggagaagtcagtgtgagcccagacacaaacctcccttcagggtacctgggacaaccagggaaag**cctgggacactgtgcactgtgctgaccccagg**ggcaagtgcaggtgctacaagggctgggtttcctgtcttgatctggggctgcgttgttaccaaattacccgagggaacttctctagatttgcaattctgtactaacatttgatgtctgtgactgctaaacttccctcacatatttgttttttgtttgtttgttttgttttgttttttgtaacaggaagacccatcctcacccctacagaagcccgagtgtcaccttgggggcagaaatgaccctgccttggtcacattgatcagtgtcctgaggaagcccagtggaacctggggagtgttttccagtcaggctcaggg**cagagacctccgtggaaatctctg**attagaataggtttgggattcagacttgggccaagagggagtctcacccatgggagggtccttaaatcctg**acagtttttacagctgttctccctcctctggcaaaactgt**gcacatctgactcagaactgatccagctggcccttacttgctaatcaatttttcttctctctacacttcattctcatagttcccttttcttcccttcc**cagaaaacagagagtgtgttttctg**tggtctaaatctcagggctcaggcctgttacctggaactcaggtgtgg**cccaggctgtggctcctgtgagacctggg**gaggctgaggggactttcccactcaccattgcctggacccgcctgctgtcttctgtgcatggaggcatttgggaaatgaagtggacattagtcactgaggggataatggtagctttctcaaatgagatactgatgtagagctgatcttctgtttctcacact**gtcacagaatttcctccttcatctgtgac**tttggggaaagctaaggatggacactctattgtgctgtgagctctgggtgac

Uppercase: IGHV3-6-2

Lowercase: Flanking sequence[1000bp]

Red & Bold & Underline: Stem-loop [19]

Blue: Heptamer[42]

Green: Nonamer [4]

id-IGHVII-51-2-2[V_gene_segment]

accatgtggtctaaactaagtccaactaag**tccctgagctctgggtgcccatcaggga**tccgccatgtgcccggcagccgcgtgcctttttgtctcctctgcgcccaatcactgtctgtgatgagcttccaggatgtgtgtgtttggcacaaaccaggtgatggacgtcagacagcagcagctggtgcccggatcatgggctccctaatgctggaggaatgagaggtgcattctcagggacaagacattgttgatggatttttatgtagaaaccacgattttac**ttgcttctctcaggagacatagaagaagcaa**ccatgcagtcagcaaataattataatttccacaattaccccaaatggttaatccttaattctgcgcagggtcccaccgtagagtcacctttctcatgaggaatggttgaatctagaatgagtccacatgattttcatatttttggcttctgtccatgttcagagagttagagtagagtaagtttggacctttccacacactaagcctcacctcccccacagaaagagcagagacttcaactattcctgagtgtggggtaggggctggtc**ctgcacacctcagagcctgcagagactgccacgtgcag**tgttatagacttgggtgttttctcattcaggaggggtgacctccacggcctccgat**ttctgctaagatataactgtgagtgcagaa**ttaggacactattaggctatcatgcctcacttctattctgaaaatcaccctctttatagtaaaagaaaacaattcaatgtccatgcccctgaaagtcatttctttttaatttggttgcgagtttaccacatagtgcctgaatttattctcagaatctactgaattgtttataaatctcacaaattgaacaaaagtgaattcttcattcttactaaaatgtgtgtatgtaggaatttcagtgtgttttccagaagccatgcacaccaaccactgtgtttcatggtcatttattGGCCT**GGTGAGGCCCTCACAGACCCCCTCCCTCACC**TGTGCTGTCTCTGGATTCTCCATCACAACTAGTGCTTCCTGCTGGAGCTGGATCCGCCAGTCCTCAGTCAAGGGAGTGGATCAGGGCATAGGTCACGAGGGAGCACAAATTCTAACCCACTCCTCATGAGCTCAGTCACCATCTCCAGATCCACGTCCAAGAACCAAATTTTCTTTTAGCTGAGTTCTGTGACCAACAATGCCACAACCTTGTATTACTGTGAGAGGaacagaagagatgtcagtgtgatcccagacacaaacttccctggagaggggcccaggaccaccaaagagcactcaggctccatgaaaacagggcccaagctggagaacgggtttcctgtcaccctcacgtttcaccattagatactctacactatgcctatgctgaggtgtgtgtttaatataattagaaaatggtatttatatacgtgtataatcatatctagtgtaagaatttgtagtttccaccttatgtgaccattaaagacaacatctctcacgttattcaaggaccttat**aaagtataatacttt**tacataagatttatctccttttttgtttctccttctgtctatattctctctgtctctctcttcctccctctttctctctctctctccatctctctctctctgtctctctctcacacacaaatacacacacatgcaatcgttgacagtatataggctaattaagaatatattcccatatgatttaaaaataacagttttacaaatatttgcc**tttaatttttcaaagaaaattaaa**tccaaatgccagtgattttttatgtgactttt**aaaatttaaattgttaatttt**tgttttcatcaatgcttgtatgtgtgcttttgtattctttcagattatttggtgcct**tcacagaattaaacatgtgaaactctccacttgaatttctgtgactcaggactggtgagcaaaaagttagaagtcac**tctcctgatccttccctacagctgcagattcctgaaggtaatgacctcacagtagaccaaggccctcaaaggtgaccttgagttccagcacatcttgggaggccaagcggatgtttgtgaaagaaatagtggctatgtgacagtttctaaccagaatatctctgtgttttcaggtatccagggtgaggcccagcttacagagtctgggggagacttggtacacttagaggggcccctgaggctctcctgtgcagcctcttggttcac

Uppercase: IGHVII-51-2-2

Lowercase: Flanking sequence[1000bp]

Red & Bold & Underline: Stem-loop [10]

Blue: Heptamer[25]

Green: Nonamer [6]

id-IGKV2-14-3[V_gene_segment]

acttactctatactaggaagcaataataaaggtattggctaaattttc**atcattactaatagaggtgggatgagagtggaatgat**atatttaaa**tgctgggggagaaaagtcagccacaaatgctatatccagca**aaacaatcatttaaacataaaagcaaggtagaaatgttccttcatacacaaagactaattcattatagcatatgtgccttgtggaaaattcaagagaaaatccttcagaataacagggaatgagagcagacagcaattcaaattgataagaaggaatcaagccctcaaaaaatgaaagaaataagaaagaagctacaaaaaatgtttacctgatatgaagattaagtacaggaaaaaaaaacaattataatagatgtcacagtatcagttacattgattgtgttttgagcagcagttgctgactg**ggaaggtgcataatgaaagcagcaagagaattggaaaccttcc**atattttgatgtg**gatcatgacatgaaatgatc**tggattttgacattcatttgcacccttaagatttgaaccttttgcactgtg**tgtttttccttaaataaaataaaataaaaca**aacctaggctcttttaaataagcctgaaccaagaaataagaatcaccaaatatttatgaaagctcaaccttattaaagagaagtgtaaaacatgcctgagaaagtcctaacgaactagaaattgggtagaggaaatagcaggaaacatctaaatatgtcccttctgccactcaggaaacagattttacctctcttgatatccttaaaa**acataaaatatatttatgt**cattttgctcacaagagaggtccccaccttctccttggctctttccaccccactgcacccacc**aggggatttgcatactgtcccctagggaagaccttccct**tgtgagtctgagataaaagctcagctctaaccttgccttgactgatcaggactcctcagttcaccttcccacaGTGAGGCTCCCTG**CTCAGCTCCCAGGGCTGCTAATGCACTGGGTCCCTGCTGAG**AGCAGAAGGGAAATGAGGGAGGATGATGG**GGTGGGAGGGTGAACTCTGTGGGCCCTGCCACC**TCCCATGTGTGTTCTGTCCTCGTGTTAGATGTGTCTTGTCCTCCAGGATGGGGCATGTGATGTCTAGATCTGTGAGAGTGAGGAAGATTCCAGAAGGAGCAAGGACATGTACTTTAGTGAAAGCTGTGACACAGAAAGAGGGGGATGGGATAGGTGACTTCTAGAGGCTTATTTGTGCTTTGCAAATCTTGGTTCTTTTTAAACC**TGTATGTTTTGGGAGTATTAACCAAAATCATACA**CAAAAAATAATTGAGCAAAACATAAATAACAGACAGAAAATTATTAAAATGGCTCACAATGTTTGTACATAACCTTGCACTTCTCTCTTATTATTTCAGGATCCAATGGAGATGCTGTGTGACCCAGCCTCTACTCTCCCTGCCCGTCACCCCTGGAAAGCCAGCCTCTATCTCCTGTAGATCCAGTCATAGCCCCTTTCACAGTAATGGATACACCTATTTGAATTGGGACCTACAGAAGCCAGGCCAGCCTCCACGGCTCCTGATTGATTTGGTTTCCAACAGGGACCCTGGAGTCCCAGACAGGTTCAGTGGCAGTGGGTCGGGGACAGATTTCATGTTTAAAATAAGGAGGATGGATGCTGAGGATGTTGGGGTTTATTGCTGCCAGCAAAGTACACATTATTCTCCcacaatggtacagtcctgaacacaaacctccctgcttgctgtggcctagctgcccagatgtgttgtttctgtggagagcaggcactgtggattctcttagatgcctaaag**accaagatgttggagaactcagaggacttggt**ac**agctgagggctcatgaccataaatttctcagct**acacctcaggcaccacattttaagttcccatcagctgcagcggcc**tttgcatgacagagtctgcaaaaaggaggaggtacacctcccctttgagcaatgagacacgagagaagagaaagctcaa**tgaaagctcattctaagcctctcttccttccctacattcattcatcaagtaaattcgttctgcgtaacaggcacctaattgagactgattactggcaacgcaaaactaatacattcttttgatttggtttagcagttactagagtatatgtatgttgataaggtttggtaatattaaaacagtttccctcccctcctccacctccccccaccacttttccgaaggcagacccctgaccaggacaccagcaggcactgtattttttcctacttttttcagggagtaccaaaatataactcttcacggtctagatttttaagttactagatatgttgctgcaaatgtatgcaaaaat**aaagacctagtgccaaaagtcttt**ccacagtctcaaccacagcatgttgaagtaaaaagtagttttctgaactccaaagctggg**tttcaacaataaggaccaagtttgaaa**gaataaaggaacataagtatgccatcttaaacagaattacaagat**aatatgtaagccatatttttaccccatgaaaaata**accatctgaagacttggagcagaggtcagtacgtgagagactgg**tgggttgcgtttgagatatgaaccca**gatgcaagttcttatgtatttcctgggaagaatcacagagtcgagagacccctgcctggcattcaaggctgggccctgatgcc

Uppercase: IGKV2-14-3

Lowercase: Flanking sequence[1000bp]

Red & Bold & Underline: Stem-loop [21]

Blue: Heptamer[29]

Green: Nonamer [5]

id-IGHV1-18-2[V_gene_segment]

agtccc**cccaaacccctgaagagtttttggg**ttcaccgtgtgtccaatgattcagtgcctcttgagctccaggaaagggctccccagtgatgcatgagatctttcttggggtctccctgcagagttcactgggattcctaaggccaattcagtatttcaaaagatggtgtgagaagcacaggctgtcactaaaggagaattctgagccagggca**cagccactttatacttggctg**gggacactggtaggaatatactctgtgagatcagacaggaacctccttgcaggggcagggcagggctgcag**ggggcgctcaggacacacagagcacaggcttccgccccagagcaggtgaaggaggctgggg**aggggttcctctcagggcctgggacttcctttaaaaaatctaaaataagtatttcacaaggactgccgatgtttatataaatatcctattcaattgtgagcatttatgaaactcgatg**ttgtaatgagaaccacttttacaa**tgggaatttcaaacttccctagacatcttaatagtaagcagctggaggtcaggaggagatcctttcttataaataa**gtgcaatttttggagaaacacactcattcccaaaatagcac**attcacatattaaggtctagaaatgattcgagttgcccctgagacagtcaaatgtgggttctaagtgaggtgcgtgtcctgggg**gagcttgttctccagtgggggaagctctgtcaacacagag**ttcagggatgggtaggggatgcgtggcctctaacaggattacggcttgaaccctcagcttctacaattgtgtcgtccatgtgtcatgtatttgctctttctcatcctgggtcaggaattgggctattaaatagcatccttcatgaatatgcaaataactgaggtgaatatagatatctgtgtgccctgagagcatcacccaaaaaccacacccctccttgggagaatcccctagatcacagctcctcaccATGGACTGGACCTGGAGCATCCTTTTCTTGGTGGCAGCAGCAACAGGTAACGGACTCCCCAGTCCCAGGGCTGAGAGAGAAACCAGGCCAGTCATGTGAGACTTCACCCACT**CCTGTGTCCTCTCCACAGG**TGCCCACTCCCAGGT**TCAGCTGGTGCAGTCTGGAGCTGA**GGTGA**AGAAGCCTGGGGCCTCAGTGAAGGTCTCCTGCAAGGCTTCT**GGTTACACCTTTA**CCAGCTACGGTATCAGCTGG**GTGCGACAGGCCCCTGGACAAGGGCTTGAGTGGATGGGATGGATCAGCGCTTACAATGGTAACACAAACTATGCACAGAAGCTCCAGGGCAGAGTCACCATGACCACAGACACATCCACGAGCACAGCCTACATGGAGCTGAGGAGCCTGAGATCTGACGACACGGCCGTGTATT**ACTGTGCGAGAGAcacagt**gtgaaaacccacat**cctgagggtttcagaaaccccagg**gaggaggcagctgcactgaatttgaggagattacagggcttacaatgtttaaagttgtttagaaaatgagctgagcaattgaggaatgtgagtaatg**gaaacatggatgcactctatataggaaatgtttc**tttcaacagtcaccctatatgcaaaattcagaatggtaaaggcagcaatcagtgaggctgacgcaaatattcccatggaggccttgtgcagacatacgttttaaaatcagatagataaataatttggaacaagattgctggtaacgtggctaagactaaatatgattcctaaaaactggccaaaatctattccaaattgtctctgccactccttttacataaatgta**ttaaaaagtagttttaa**gaccacagcaaaattggacagaaggtgcagagagttctcatgtgcccctgcttcaccatgcacagcctttcccactgtcaccatcctgccccagagtcatcaa**taagttacaatggatgaactta**catgggcggattggttctttcctcttctggtggtctcttggcataccaagcccaaattatcttgaagcaccataggttctactgtaatgcctaaccttgttttttactctagtttgctactttaaattttccctttttttgtctccttaa**ttgccagccatgtttcccatatgaatagactctccctggctgggaa**agctgggcaaactccatttgaccttttgatttataagacattaagggctccttacccaacccccttcttcaaggaattaacctgtgtaagcagatcctcagcatttcaaaggagcccaattaactgataaggtactggaacaaacaatgtatgaagttcccaggatttttctcaaagagataacaacataaagccttgagttcatgcccagcatagcatcatatctaactataatgaaggatttagagccctgcacctgg

Uppercase: IGHV1-18-2

Lowercase: Flanking sequence[1000bp]

Red & Bold & Underline: Stem-loop [18]

Blue: Heptamer[42]

Green: Nonamer [6]

id-IGKV3-7[V_gene_segment]

aactgtctacagcactgtaaggcaggtaccactattgtcacagttacacagatatggaaaccgagacacagggaagttaagttacttgattaatttcaagcaatcggcaagccatggagcatctatgtcagggctgccaggacatgtgactgtaaacagaagtttttaactcaaa**gagggtatgtgtctgggttaatggaaagcttcaggaccctc**agaaaacattactaataagcaaatgaaaggtgtatctgggccgggcgcggtggctcatgcctgtaatcccagcactttgggaggccaaggcgggtggatcaccaggtcaggagatcgagaccatcctggctaacatggtgaaaccccgtctctactaaaaatacaaaaaattagctgggcgtggtggcaggcgcttgtaatcccaggcacttgggaggctgaggcaggagaatggcatgaacccgggaggcagagctt**gcagtgagtggagatcgcaccactgc**cctctagcctggaagacagagcgagactccgtctcagaaaaaaaaaaaaaaaatgtctatctggaagattaagttctaacagactcttcatttccatagatccaataatgcacttagggagatgactgggcatactgaggacaggaagagagaaatgaaaacacagccttttatattgttcttaacagacttgtgccaaatatcatacgggtgtatttaggtgattgaagagaagaaaggcacaggagtgaaattatgtgagcacaagggaggagttctacactcagactgagccaacagacttttctggcctgacaaccagggaggcacaggacgctcagtgcagagaggaagaagcaggtggtctct**gcagctggaagctcagctcccacccagctgc**tttgcatgtccctcccagctgccctaccttccagagcccatatcaatgc**ctgggtcagagccctgggaaggaactgctcagttaggacccag**acggaaccATGGAAGCCCCAGCTCAGCTTCTCTTCCTCCTGCTACTCTGGCTCCCAGGTGAGGGGAACAT**GAGGTGGTTTTGCACATCAGTGAAAACTCCTGCCACCTC**TGCTCAGCAAGAAATATAATT**AAAATTCAATGTAGATCAACAATTTT**GGCTCTACTCAAAGACAGCTGGTTTGATCTAGATTACATGAGTGCATTTCTGTTTTATTTCCAATCTTGGATACCACCAGAGAAATTGTAATGACACAGTCTCCACCCACCCTGTCTTTGTCTCCAGGGGAAAGAGTCACCCTCTCCTGCAGGGCCAGTCAGAGTGTTAGCAGCAGCTACTTAACCTGGTATCAGCAGAAACCTGGCCAGGCGCCCAGGCTCCTCATCTATGGTGCATCCACCAGG**GCCACTAGCATCCCAGCCAGGTTCAGTGGC**AGTGGGTCTGGGACAGACTTCACTCTCACCATCAGCAGCCTGCAGCCTGAAGATTTTGCAGTTTATTACTGTCAGCAGGATTATAACTTACCTcccacagtgattcaacatgaaacaaaaacctcaacaagaccatcagtgtttactagattttaccagctgcttcctttacagacagctaatgtggtggccactcagttttagcgtctctgctctatttggacattttgcagttctaaaaaaaaa**tcattgaacaatttggactttgattcttggactctgttcaactga**ggcaccagaatcccaggtttccagaaatagtgactcactgtatgaatccttatatagcctcagtggttcttaactttcccagtagaggtagctcagtgcatgctacactgctccattt**gaattttgcaacattctaagtagtagaaaattctatttatttatccaaatag**ttgactcggtaaaagctgttcatgtgaagatactaccatggctgaataaatcccattctttttctttcttcaggctatcaacatttcag**tggcaaatggttattatggaaacatttgcca**tttaaaagttaactaaattatttcttcaattttctctgtgatgcagtagactgtaaaaagattaaagtttgttaaaataaagtacatattcgataaggaagaaatagattattcctaatgacgtctgcaatgacctagtagaaagagtgatagaagcagttgttttcattatttttgtccaaaacttccttccaaatgggatttcattgatcatattcgtttattaccacctataagacatgttgacattatgtaacatctgatgtgaagcactgaggatacatcctgtctgtattattcttgccaaaaatta**atggtgtgaattgaatcagtagtaaacaccat**atacaaacccaactaggaggacattcttcaacatacctggacagtaaa**cttcaaatgtttgaag**gccatgaaagagaaacaaaagtgaaaaactatcacagatttaaagatattgaggacaggataaccaaataaaatacagaaacctgaattttatcttgtaacat

Uppercase: IGKV3-7

Lowercase: Flanking sequence[1000bp]

Red & Bold & Underline: Stem-loop [13]

Blue: Heptamer[39]

Green: Nonamer [7]

id-IGKV1-33[V_gene_segment]

cttctgctttcctgaaatttactcctgccttaaaaaacccttacctgcaagccatcagtgaggccaggatttgaatcttagctgcctgattctctttacctgatgccctacaaaaaaacaaacaaacaaacaaacaaacctttactttctcctgctgcaaa**ctcagtgtgaatatctgatctgactgag**ctgagtgagtggactccagttcggttccacagcacaagctctttttatagtggcatttatccac**tcatgagggcagagctctcatga**cttcaacacttcccattaggccccacctcccaat**tctgttgctttgagataaatttaaaaacaga**tgggttttgaaggacacagtgaaaccataacactgcccaaagagagggccattcgagcccctgtggtctgctatgcatgcaggcagagctgccttctccaggttcagaaagagcaatgaggggcatagatggccattggaagtcagcagcagtacagcgaaagggaaaggcatagcctgtagacagggattgcaatcttgatatattttctatttagtttcctttctaaaaacagaataatttcatgctaacagaagtgttgtaagtacagcacaaagtgctttcttccaattaagtgcatattg**tattttttaaaaaata**atctccatctccatacacgaataaaatacattactccatctagtcctcaggaatatttcgaa**ttttgacaattaactcaaaa**aagttctttgtaaccaaacagtctccaggaagaaacacttatttacggacaaatctgtgatgccctggtccgacctgggacactggggacattgctcctatgctgagttactgagaagagccagccctgcagctgtgcccagcctgccctatcccctgctgatttgcatgttcgcagagcacagccccctgccctgaagacttattaataggctggtcgcaccctgtgcaggagtcagtcccaaccaggacacagcATGGACATGAGGGTCCCTGCTCAGCTCCTGGGGCTCCTGCTGCTCTGGCTCTCAGGTAAGGAAGGATAACACTAGGAATTTTCTC**AGCCAGTGTGCTCAGTACAGCCTGGCT**CTTGATGGAAGCCTTCCTATAATATGACTAATAGTATGAATATTTGTGTTTATGTTTCTAATCGCAGGTGCCAGATGTGACATCCAGATGACCCAGTCTCCATCCTCC**CTGTCTGCATCTGTAGGAGACAG**AGTCACCATCACTTGCCAGGCGAGTCAGGACATTAGCAACTATTTAAATTGGTATCAGCAGAAACCAGGGAAAGCCCCTAAGCTCCTGATCTACGATGCATCCAATTTGGAAACAGGG**GTCCCATCAAGGTTCAGTGGAAGTGGATCTGGGAC**AGATTTTACTTTCACCATCAGCAGCCTGCAGCCTGAAGATATTGCAACATATTACTGTCAACAGTATGATAATCTCCCTcccacagtgtaacaagtcataacataaatcacccaggggagcagatgcgtgaggctcagctgtcccagatgccccttctggtgccttcgcctgc**tgagaatgtttctca**aattgcagtcacactttgaagttcactggagagtttttgtaaaagggcc**atgaaggcccacttcat**cgtag**ctgtctttccttgtcctaatccccagtatcatagacag**ggcaatgcctctcctgatttcattgagaagaaatggttacacctgaggggtctgagttgtagcatcag**ttggaattcatgtagcaatagtgagccactctaggtattccaa**gtaggattttttttaaatataagatgtgagaatctaaactacagccttttaaaggtttgcaagtatagtagtcaaagacgcaaatactaga**gaagaggaattctcttc**cggaatccagaatgcatctgatagagaaggtacaactgccaatcatgtggtc**ctcagacctttctgag**aagcccatgggtgggggtgcagatgctctcagctgccta**gaggacttcatcaggtgcttctgcagtcctc**acctcggtccatatgtcttgctgcaggtgttgatggatagtattgaatcctcctcttcttacttctcaatctcagggcaggccccacactgggcaactccaccaaaaaccagagaaggcatagggt**ttctggcaaatgtgcttccagaa**tgatagtgatgatggggaagtgacagctgacatcgtagtgtggtcatgtatctcgactctcaggattttttcagtgaagtgatggcctcagaatacacttggatgtacttccatata**ctatgagtaagtttgaaatcatag**catgaaaatga**tatttagtcatatgataaata**gaactacatggctacattaatcaaaatagcatagtgctggtacaaaaagatacacagacaaatggaacagaatagggaactcagaaataatgctggagaccta

Uppercase: IGKV1-33

Lowercase: Flanking sequence[1000bp]

Red & Bold & Underline: Stem-loop [18]

Blue: Heptamer[32]

Green: Nonamer [8]

id-IGHV1-58-2[V_gene_segment]

ggaatggtttcactaattgaaaggatacagaattgtgggagtcactatgttcctatgaataaaaaattcagatttcagtgttaagtaatgttgcctacattgtgtgagtgacagggcagtggtggatctgagagtgtggcaggtgcacagacctagtgagtcagaaatcaatatggaaagatgaggatctatggat**atgaactgaaagtaagtaaacagttcat**gaaattctattaaatggagtaggaaataaaacccaaacttatccaaaacacaaattccttggcgattat**tttgggagcagtgagttcatcaggaaccccaaa**cttctcttacgtcttctgattcctgttgtccatgagatgagaaattcagctctaattgtacatcacagggcaaatctgtaaaccaggagtgtttctattgaggatcatggtggatcaggattccaggcaggtgctggagacactgtctcaggagcgcccagatgatctcagggggac**ctgctggacactcacgtgggacatcagcag**tcactttctcagagtaaccagtgagctgtgct**ggtgcctgatgggactaggatggggtcaaggcacc**ttctcagtgtcatggagagtgattgttccagaaatcatccaggtggtctctacgctaatcaaatatgggttcacagggaggaacatgtgctctgggtgcttggtcttcagtgaaaggacgtctggccaccaaaagtttgtaaatggagcagggcatgcatttcctcaaggaggattagggcttggag**catcagcatcccactcttgtaaggctgatg**tgtcatttaccttccctttcttatcccaaatcagggtcttcagctatgaaatgctctgactcatgaatatgcaaataacctgagatgcactgaggtaaatatggatatttgtcagccctgagagcatcatccagaaaccacatccctccgctagagaagcccctgacggcacagttcctcactATGGACTGGATTTGGAGGATCCTCTTCTTGGTGGGAGCAGCGACAGGCAAGGAGATGCCAAGTCCCAGTGATG**AGGAGGGGATTGAGTCCAGTCAAGGTGGCTTTCATCCACTCCTGTGTTCTCTCCACAGG**TGCCCACTCCCAAATGCAGCTGGTGCAGTCTGGGCCTGAGGTGA**AGAAGCCTGGGACCTCAGTGAAGGTCTCCTGCAAGGCTTCT**GGATTCACCTTTACTAGCTCTGCTGTGCAGTGGGTGCGACAGGCTCGTGGACAACGCCTTGAGTGGATAGGATGGATCGTCGTTGGCAGTGGTAACACAAACTACGCACAGAAGTTCCAGGAAAGAGTCACCATTACCAGGGACATGTCCACAAGCACAGCCTACATGGAGCTGAGCAGCCTGAGATCCGAGGACACGGCCGTGTATT**ACTGTGCGGCAGAcacagt**gtgaaaacccacatcctgagagtgtcagaaacgccaggaaggaggcagctgtactggcatggaggggatgacaaaggttattaga**ttgaagattttcttagaaaacgacttcaa**gtcattaaagaagaggaacaac**ataaatgtgtatttgtgaaattttaattgagagatttttcatacaacatttat**tctgtaagctatttcagggattggaatatgaatcaaattaataaagctgatatagacatcctctgaaggcatcttcgtaaacatcaatttctgaatcagtgttgtaaatattttggaacacagacacaagatca**catttttactctacttttatctctatttttaaaaaatg**ccaaaaagaatcttattttgtgcatgccccattttgaattcccaccgtcaatgcatgatagttcttggttttcctcattcaagttgtcatttgtcattaagagtgttgtgtgttttaaccattctaataggtgagtaacggtatctaatttttatttaaatgcacatgtcccta**aaaaattcatatttaacaatttttatataattttt**ggtgagatgcctctcgtgatatttggttcattttttaaatgcattttttatcagttgtaagtatgcttgcatattgattataaaagtcatttaacaaattaaaataattcatttaacaaatatgcat**cttggaatttttttctccaag**tctgcagttgtcttttactcccttatccctgtgtattgcagaaaaatgtttgtgtgtgtgtgtgcctgtgt**atgtacaaatttagattttaaaaatgtacat**ttttattcatttaaagatca**tgtctttggcagtatatctgaaatctcattgtaaaagaca**caatagtcattattttttccatgtctctaatcttaggaca**caatcaactcatgagtgtttaaccttccctacctgattg**gaggactatccacctgagacatttggaatacttct

Uppercase: IGHV1-58-2

Lowercase: Flanking sequence[1000bp]

Red & Bold & Underline: Stem-loop [17]

Blue: Heptamer[46]

Green: Nonamer [8]

id-IGKV2-26-3[V_gene_segment]

cagtcaggttggaaagtaagtccagggatatgtagggttaaataaaacccatctgatgagaatttatagtttgtagggcatgactccccagactccttagataggaactgagacaaggaaaaaaaaaatcagagcttagtccttaccttgataaacacaaatgaaaagaaattattacttgtagacattcaataaatta**tctgaaatcaatttaagaaaacattgttcaga**atcacatg**aaattatagaggacagtaattt**gaatccacaagcagaaataaaggcctcgaaatagtaaagaactgggaaaaaaatagacacaatgataaaaaaaagtttacttcaatttctatgatattca**ctaccaagtacaaacaaactatcgtggtag**atgtcagaaagacagttactttggttgggatggggaggcagttattgaatgggaaggtgtacgagagaatcatccagaaattt**tgaaacattctatagtttca**gctggatag**tgaatatacataaaactttattca**tt**ccacatttaggatttctgccctttatatgtgg**tttatttcaatgaaaaaaaaatagaaaagccaagccta**gattcttttaaagaaaattggaggcaagaaaaaagaatc**agccaatatttatggaaactcaactttattaaagacaagtgcaaaatgtgcctggaaaagtgctaaagaactgttgaaaaacaggtggaaagaaaaggaaatattataatatgtctcttctccaactgaagcaacacagattttatcttggttagttttccttgaaacacgcaggatattttatgtccccttgcccacaagagaagccctcatcctcctgttgttgcgcttagcatcactgcacccaccgggggatttgcatatt**gtcctctagggaggac**cttcccttgtgggtctgagataaaagctcagctctaacccttaccttgactgatcaggactcctcagttcaccttctcactATGAGGCTCCCTGCTCAGCTCTTGGGGCTGCTAATGCTCTGGGTCCCTGGTAAGGACAGAAGGAGATGAGGGAGGAGAATGGG**GTGGGAAGGTAAGCCTGGGGGTCCCAC**TGCCTTCCATGTGTGTTCTGCCCTGCCCATGTGTTAGATGTACAGGTCTTGTTCTCCAGGATGGGGAATGTGAGGTTTAAATCTGTGAGAGTGAGGACAATTCAAAAAGAAGCAAGGAC**CTGTGTGATCTGGTGAAGATTGTCACACAG**AG**AAAGGGAGGTGGTGTAGGTGATTTCTAGAATCCCCTTT**GCGGCTTGCAAATTTGGAATATGTTTAGTGTATAAATACAAACAACAAAAAATTATATAGCCTGAAATAAAAAATGAAAATTTATGATAAATGACACATGATATTTGTACATATCCTTCCACTTCTTTCTATTTATTTTAGGATCCAGTGCAGAGATTGTGATGACCCAGACTCCA**CTCTCCTTGTCTATCACCCCTGGAGAG**CAGGCCTCCATGTCC**TGCAGGTCTAGTCAGAGCCTCCTGCA**TAGTGATGGATACACCTATTTGTATTGGTTTCTGCAGAAAGCCAGGCCAGTCTCCACAGCTCCTGATCTATGAAGTTTCCAACCGGTTCTCTGGAGTGCCAGATAGG**TTCAGTGGCAGCGGGTCAGGGACAGATTTCACACTGAA**AA**TCAGCCGGGTGGAGGCTGAGGATGTTGGAGTTTATTACTGCATGCAAGATGCACAAGATCCTC**Ccacagtgctacagccctgagcacaaacctctctgcttgaagtggccca**gctgctcacatgtgatgtttgtctggggagcagc**acagcaagtt**ctctgagtctgtgcatgaggaaaatgttggacaactcagag**gagtagattggtctcatgaattccttggttacatctcaggaactactcatttgtgcagctgcaaaagccttgacatggcaaaagcagtaggagaatggggaaggtcccatggagctgatgctgattcatgatgcaactgaataaaagctcattcgaatcctcctttcttacatttc**tcaattgtagttaatcagcataagtgccaggcaattgatacaaatgttggcaacacatgctgagtgcctcctcatttgtat**gtatttcatccatttttgtatattttaataggataattatttggtcatatttagaaattggtattttccaccttccaagctttctactttccttacttattgcttctcttcctatgtaataaagatactgtagaca**acattttacactagttgtaaaatgt**ggctccccatttatacgggtagctggttgggggcaattag**caaaaatcttggtttttg**agtgcctaaatagacttttgtaaattaatgtcagatacaagatccaaatgctaaaactctttgatttgccttgattacttcttactgatagatcataaaa**ggaattcttgaaattcc**aaaagttgggttttaaaaataaaagcatacactgg**aatatacagggcatagaattcaccctatatatt**gctagaagaatagagaatgataacagttttagacttcgagggtacttcaggaatgggaacacacaagga**acacagtagatatgagctactgtgtctaccgtattagaca**atcctcctgcattttaacttacatttggtgaacttcagatgccagtgggtgttgcaagttgatgacataaacctaaaatcaagtcccgaggtatttc

Uppercase: IGKV2-26-3

Lowercase: Flanking sequence[1000bp]

Red & Bold & Underline: Stem-loop [26]

Blue: Heptamer[23]

Green: Nonamer [9]

id-IGHV3-7-2[V_gene_segment]

acatttaccacacttgacttcaccaattcagtatctatgtatttttttctcttataatagcctacatttttatatgccagactctgcctcagttcaactcgtacccatgcctttttactctctgcaagaacttgtctctccctaaatttcggatttgatgattatcttaaaacctccagta**tttaaagtatattttaaa**attggcaaaattccatctcctgtgtttatctgttatgttgatgctgtaaaacagtaagtaaaatatactcctcatctatgtacattttgaagctgagttgcaggttttttggtaagacccagagtcacagagaattcaaatattgttaagctgcttaatagaaaaacaaattatggtaaatgtgttcactggaatactacccatg**atttataataaataaat**gcctgacacacagaacagcagcaaaaccacacatgctcttattacagaaagtggcttctgaaaaccacaccgggcatgtacagctttgtcctggagttggtttagggggatgtcagagccagtgacgagaagcacagggccagatggcagcgttcactcat**cccagacatgagctcctgggtgcatacagagcccccccatgtgtggg**tttacttccact**tctgtaaaaggagaaaatactgactcctacaga**gcataatttac**acattttttaaaaaatgt**aatagggtgatcagggcaaagtgtttatcacagcacaatttcataagacagcatattttccaaataccatcattgtcagcaaacttctgcagagcaccgtcttcttatatgggtacagcctattcctccagcatcccactagagcttcttatatagtaggagacatgcaaatagggccctccctctactgatgaaaaccaacccaaccctgaccctgcaggtctcagagaggagccttagccctggactccaaggcctttccact**tggtgatcagcactgagcacagaggactcaccA**TGGAGTTGGGGCTGAGCTGG**GTTTTCCTTGTTGCTATTTTAGAAGGTGATTCATGGAAAAC**TAGGAAGATTGAGTGTGTGTGGATATGAGTGTG**AGAAACAGTGGATTTGTGTGGCAGTTTCT**GACCTTGGTGTCTCTTTGTTTGCAGGTGTCCAGTGTGAGGTGCAGCTGGTGGAGTCTGGGGGAGGCTTGGTCCAGCCTGGGGGGTCCCTGAGACTCTCCTGTGCAGCCTCTGGATTCACCTTTAGTAGCTATTGGATGAGCTGGGTCCGCCAGG**CTCCAGGGAAGGGGCTGGAG**TGGGTGGCCAACATAAAGCAAGATGGAAGTGAGAAATACTATGTGGACTCTGTGAAGGGCCGATTCACCATCTCCAGAGACAACGCCAAGAACTCACTGTATCTGCAAATGAACAGCCTGAGAGCCGAGGACACGGCCGTGTATT**ACTGTGCGAGAGAcacagt**gaggggaagtcagtgtgagcccagacacaaacctccctgcaggggtcccttgggaccaccagggggcgacagggcattgagcacggggctgtctccagggcaggtgcaggtgctgctgagggctggcttcctgtcgcggt**ctggggctgcctcgttgtcaaatttccccag**gaacttctccagatttacaattctgtactgacatttcatgtctctaaatgcaatactttttttgtcctttttgtttctttgtttttttgcaacaggagtacatatcctcagctccacagaagccagggtgtcactttgggggcagaaataatcctttcgtggtcaacaggatgagagtcctgaggaatcccagggaaacctggagagtgttttccaattagactcagggcagagacctccatgggaatttctgattagaacgggctttgagctctgatgggagccaagaga**caggctcacccagggtcggggtccttaaagcctg**atggttttcacagctatcccccctcgtcttgtaaactcagactgattcagttgaccctttttctgctaatccattttgcttctctgtaggtttgattctcacagttcgctttcttcttctc**ttccctgagaacaaacgatgtgttttctgttttcagggaa**aatcccagggct**caggtctgcaggacctg**ggtaggctgaggagactttctcactcaccattgtctggacactcctgttgtcttctgtgcatggaggcatttgggaaatgaagtggacattagtcattaagggaataatactagttttctccaattggatattgatgtagagctgaacttgtgcttctcacactgtcacagagtttggaccctcacctataactttgaggagagttgatgatggatagtccattgtgggtgagctctgggtaacagtaattgtagggtctggctaggcagcctaaggtcaatacttctggacctcaggaaagacaggc

Uppercase: IGHV3-7-2

Lowercase: Flanking sequence[1000bp]

Red & Bold & Underline: Stem-loop [14]

Blue: Heptamer[37]

Green: Nonamer [5]

id-IGHV3-7[V_gene_segment]

acatttaccacacttgacttcaccaattcagtatctatgtatttttttctcttataatagcctacatttttatatgccagactctgcctcagttcaactcgtacccatgcctttttactctctgcaagaacttgtctctccctaaatttcggatttgatgattatcttaaaacctccagta**tttaaagtatattttaaa**attggcaaaattccatctcctgtgtttatctgttatgttgatgctgtaaaacagtaagtaaaatatactcctcatctatgtacattttgaagctgagttgcaggttttttggtaagacccagagtcacagagaattcaaatattgttaagctgcttaatagaaaaacaaattatggtaaatgtgttcactggaatactacccatg**atttataataaataaat**gcctgacacacagaacagcagcaaaaccacacatgctcttattacagaaagtggcttctgaaaaccacaccgggcatgtacagctttgtcctggagttggtttagggggatgtcagagccagtgacgagaagcacagggccagatggcagcgttcactcat**cccagacatgagctcctgggtgcatacagagcccccccatgtgtggg**tttacttccact**tctgtaaaaggagaaaatactgactcctacaga**gcataatttac**acattttttaaaaaatgt**aatagggtgatcagggcaaagtgtttatcacagcacaatttcataagacagcatattttccaaataccatcattgtcagcaaacttctgcagagcaccgtcttcttatatgggtacagcctattcctccagcatcccactagagcttcttatatagtaggagacatgcaaatagggccctccctctactgatgaaaaccaacccaaccctgaccctgcaggtctcagagaggagccttagccctggactccaaggcctttccact**tggtgatcagcactgagcacagaggactcaccA**TGGAGTTGGGGCTGAGCTGG**GTTTTCCTTGTTGCTATTTTAGAAGGTGATTCATGGAAAAC**TAGGAAGATTGAGTGTGTGTGGATATGAGTGTG**AGAAACAGTGGATTTGTGTGGCAGTTTCT**GACCTTGGTGTCTCTTTGTTTGCAGGTGTCCAGTGTGAGGTGCAGCTGGTGGAGTCTGGGGGAGGCTTGGTCCAGCCTGGGGGGTCCCTGAGACTCTCCTGTGCAGCCTCTGGATTCACCTTTAGTAGCTATTGGATGAGCTGGGTCCGCCAGG**CTCCAGGGAAGGGGCTGGAG**TGGGTGGCCAACATAAAGCAAGATGGAAGTGAGAAATACTATGTGGACTCTGTGAAGGGCCGATTCACCATCTCCAGAGACAACGCCAAGAACTCACTGTATCTGCAAATGAACAGCCTGAGAGCCGAGGACACGGCCGTGTATT**ACTGTGCGAGAGAcacagt**gaggggaagtcagtgtgagcccagacacaaacctccctgcaggggtcccttgggaccaccagggggcgacagggcattgagcacggggctgtctccagggcaggtgcaggtgctgctgagggctggcttcctgtcgcggt**ctggggctgcctcgttgtcaaatttccccag**gaacttctccagatttacaattctgtactgacatttcatgtctctaaatgcaatactttttttgtcctttttgtttctttgtttttttgcaacaggagtacatatcctcagctccacagaagccagggtgtcactttgggggcagaaataatcctttcgtggtcaacaggatgagagtcctgaggaatcccagggaaacctggagagtgttttccaattagactcagggcagagacctccatgggaatttctgattagaacgggctttgagctctgatgggagccaagaga**caggctcacccagggtcggggtccttaaagcctg**atggttttcacagctatcccccctcgtcttgtaaactcagactgattcagttgaccctttttctgctaatccattttgcttctctgtaggtttgattctcacagttcgctttcttcttctc**ttccctgagaacaaacgatgtgttttctgttttcagggaa**aatcccagggct**caggtctgcaggacctg**ggtaggctgaggagactttctcactcaccattgtctggacactcctgttgtcttctgtgcatggaggcatttgggaaatgaagtggacattagtcattaagggaataatactagttttctccaattggatattgatgtagagctgaacttgtgcttctcacactgtcacagagtttggaccctcacctataactttgaggagagttgatgatggatagtccattgtgggtgagctctgggtaacagtaattgtagggtctggctaggcagcctaaggtcaatacttctggacctcaggaaagacaggc

Uppercase: IGHV3-7

Lowercase: Flanking sequence[1000bp]

Red & Bold & Underline: Stem-loop [14]

Blue: Heptamer[37]

Green: Nonamer [5]

id-IGLVVI-22-1[V_gene_segment]

aaaatggcataagccatgtatttctctctggatatctcattaaaatattgtctcctccccaacacaccacccatgtgaattat**aaaaatgcccgttttaaattttt**gtttgcatgtcgtgacagagtcaagcacgtgaatcaagctgagtttgatgagtccagggctgttaagccccagacccaggtggcctcagaaggacttggtggaacgagggcttgttgtcaaaagcagaggaagtgagattttccattgtggctgagaccagtttcttaaattccagtggacacacaaggccccttaaggatcggttaagatgcagatgcagcctcagcagttctggatgtggccccaaattctctgtgtctggcaactcccaggtgaggctgctgctgc**tggtcctcccaggacca**cactttgagtggctgaactcttat**tcacccacctcactcaagccacgggtga**gactcacacaaggagagcatattctagaagcacaggtcacttctcactcctccccttgtgtcttagatccttttctgttactataaagaag**tacccaaggctgggta**atttatacaga**aaataaaagttttattt**ggcagacagttccaatggctgcagagttcaaaatgtggcatctgctcttgttgaagacctcaggctgcttcctgtcatggtgaaaggtgaaggggagccagtgtgtgcagatgtcaatgggggagaggaagcaagagagaagacaaagatgccaggctctttttaataaccagctctttcggaaacaggggaattcactcaccccaagtgagggcattaatctattcatgagagatccactcccatcaccctaaaagctcccattagttcc**cacctccaatactggggatccaatttcaacataagatttggaggtg**acaaagatcaaagccatagcaccctgaggtttggccacacacttgctcatctccacatttgtccctcccc**tcaggactagggtcCTGA**CTGCTCTGACTTAGTCTCCCTCCACCTTATGGGCACCTGGCAGTCGGTCATCATCAGCT**TTGCTGAATCAGCAGCAA**CATTGAGGGTAAAAATTGTGTCTCCTGGTTCCACCAGAAATAAAGTAGAG**CCCCACACAGCTGAGGTGTGGGG**TCAAAAATTAAAACTAAGAGACATAAGACTCATTCTCAGGCTCCAAGTCTGGCCAGTGAGCTTCTTTGAGACTCCCTGGGATCCCAGCAGTGACACTG**ATCACTATTGCTGTCCCACACATCCCAAGTGAT**GAGGAagtaagaccaaaaccctcctgagattcctggcttgtgtcctgacactggggctgttgggattcctgtctttccttcaa**gattgttcaaataagcaccgacaatc**acttc**catgtgagatatgaggagagaataattccacatg**aaccaaatccacgaagcctacactgaagcctgccaagttctctgcagtccagcacgatgaagggagcattttcatgggaacttttgggaatatgaggggagccaggctcagctatgggttccagagacaaagctcctggatgtttccccctggccttggtccatcacctgctacctctcctcactctttccacagatgctgccccaggttaagcccccacaggactctaagcctattcctgaccctgagctcagcccaagtgatgct**tcaggagatgggatcctga**gaatgagcgccaacgtctctttaacctg**aggtgtgaggtcatgagacacct**gttggccataagacacctgtccccca**cctgctcaggagacagtggagcagg**caacagtgcagagagaaccagcaggagccaggccaggccctgcatccaaagcaggccccaatcccacagattctccagctc**cagcctggggtacagccccaactttctgtaggctg**aaatgggcttctcaggatgggcaggccatgatgggggatggcag**tgtgtccacaggacaca**gccctgctgcctgcctggccctcatgcccactccttccccaccctggttaatctccacccaaaagggcccaggctcagaactcaggacacctggatcagctcctggacttgcccccactgctttctgttcccctgagccacttagtctcttttggcctcagtgttacctgctgtgaagtgaggaggcttatactgaactgaggaagc**tcttttcttcattcaaaattaaaaga**cagaaacaataaagtcatgtgaattaaagccttctaaggggatgtcatttaatgggctc

Uppercase: IGLVVI-22-1

Lowercase: Flanking sequence[1000bp]

Red & Bold & Underline: Stem-loop [18]

Blue: Heptamer[31]

Green: Nonamer [1]

id-TRBV7-4[V_gene_segment]

tggatgtttcccatatattaga**atatgcaaatttttcataaattaccatttttttttgcatat**gattgcagtttcaaatagataaaataggtagaatgttatgaggtgatatttatgagatgttagctgacaatt**atttagaggtgtttaaggacctaaat**tcagagtttccagaatcaaaatgaatctcaagtaagacaaatagtcaagtcatagggaaattgtataacatgagggacaaagagaagattttgaaagcagtcagaaagaaaagacagatacctcccataggaatagatggagaacaacagtaagtcaagatagtggttaaaaataaccgttagc**tcaatcgtcattctccttcattagaggtgagattga**ttaaaaaaaaaacttttatacgtggaatatgaaaaaccaccactgtgagtttgctaccaagttatcttttctgaattacagttaacaaaggtgggaaatgatcccaaagaaaggcatgagggcagttcatgagggaacgcatgtgaggggacagtcctgttgggcacgtgtgactgggggaatggaggaggctggggcatgaatggggatggcagaggggaccctgacttgcaggaaagacaatgagctcacccttttgtgttctatgttaggggcactgttggtacatcctacaggacatgcctagcagacagaagagcagctgtgggctggcaagatgaactcagagatgcagcgtgaggcctccggg**tccagacagatctgga**gcccaaagcaatgagccatgcattgatgttgttaaaaaggagcttataaatatttaaagcagcacccaaatgtgttctaatagaaatgctggggccctgaggtcctggggattgacagaggaagtgatgtcactgtgggaaccgccctgtggagacaaggac**gtcactcatcctctgctcctgctcatagtgac**actgacctggtaaaacccccgtcctggcctgaccctgccATGGGCACCAGGCTCCTCTGCTGGGTGGTCCTGGGTTTCCTAGGGACAGGTGAGTCCTTAGAACACAAAGTAGTTTCAGTTTTTTTCTGTGTGTAGGCTTGTGTGTGTGTGTGCGATGACTACAAATATTTTCCTTATTCTGTTGCCAAATTCTATTTCCACAGATCACACAGGTGCTGGAGTC**TCCCAGTCCCCAAGGTACAAAGTCGCAAAGAGGGGACGGGA**TGTAGCTCTCAGGTGTGATTCAATTTCGGGTCATGTAACCCTTTATTGGTACCGACAGACCCTGGGGCAGGGCTCAGAGGTTCTGACTTACTCCCAGAGTGATGCTCAACGAGACAAATCAGGGCGGCCCAGTGGTCGGTTCTCTGCAGAGAGGCCTGAGAGATCCGTCTCCACTCTGAAGATCCAGCGCACAGAGCAGGGGGACTCAGCTGTGTATCT**CTGTGCCAGCAGCTTAGCcacagcgtggcacag**tcgcctcctttctgctcacaaacctcatccttctctctccttgcagctcctacaaacccttaacagaggcctctctttgctccttacttttcatgggaaaaaggtagatttggacctcagctgtcctttgggtagaaagagaccgcagattcattcctgaaacacagtgagtgcaaatgtaggtggtgaaaacaat**cagagcccactgcgctctg**ggagtcctccgagccagctcactgttccaagcaaggagtcagtgtcttagccttggccttcagggcagacatgcatcttctataggtcttggaggctgctgtgctgcccacatccatgaggttgtcataggcaggaaacacgcctcttcttcatatgttggggcatttggaaggtctgaggctacatccccaggaacatctttcttcagaagcctgttctacccctgtcaccttggaagtttcttcaacaaaatatcaaacatctcttcctataaagtaaacatctttgcaaatttgtggtccttattaataaatacaatggtgataacaataagacttcacttcttctgcctactttaagccacgtgtatcctttattttgtttgcatttgccat**tgctactgtcctgatagacagaagca**tgcatcaccaccgctgcccgttcaccttgatt**ccctcaggaaatctaatttctagactctgaggg**ttttcattgctgtcaaactcatttgatttgaaatcattttcctgaggcctttaactcaagaagtgttttatt**tataatattgaatatatttcttttttttatttttcatattata**ttttattataca**gtacttgttataaatagaagtacaatgattatattgta**atagaatcttcaactcgtctgtgggtgctgctgcagtttgtatctatgaaagcgaatgcactggtcagagctgatgtgattatggatcatgggtttcagggagtcttcggcatcagac

Uppercase: TRBV7-4

Lowercase: Flanking sequence[1000bp]

Red & Bold & Underline: Stem-loop [13]

Blue: Heptamer[39]

Green: Nonamer [5]

id-TRBV26[V_gene_segment]

aaggaaaaccaaaaccaagccgaattgaatactcaaaaagttacagcaccttactattgtacttatttttttggatgctaacccaaagccaggactgaaataatctaaccaggacctccctagggtaggattcaaacccatgaagctgcacaaaaa**taggactctaatctacagtccta**gaccaaagacaatttaaggtgtaagcacatattaacaaggtattgactccaaaagacatcttatccagaagcaatttcttccctcaaaagagaaagtagcactcaaagggcctgaaatgtcatgacagagaataaggacctcacagctaagcctctggataaactggtctaggtagcaccactcaggg**tcagtgaagaaactttaattcccactga**ggggctacagtgccttggcaggtgttggcctcacattgggcacctat**atttgttatcaaacaaat**gggtttgcacacagaagccaataactatggcaccagcttttgagaaaagaaaaggctttattgaaacaccagtcagcaaaaagaaaggagtgagtt**caaatctgtttccatgatttg**gggtctagggcaggttttaagggatcagagagcaagggaaaggatttaggaatgttgatatggtagggtatgattagaaggcttcaaatatgactatctatggtaagatatgttaagaaacattttagcctgggatcttctgggccagtggaccgctcacta**ctgaaagaggtctggctttcag**gttccagtcatgtcttggtcttcctgtttctaaggggaggaatcttcggttctgggtattgttagaggtcaaagcttttactactgcacatgcctgg**gctacatgatttgcagttatgtagc**tgttggcagtgtgggagatgagtgttgtttctctggccaggaaataaatgaaaagtgctggttatcagggactaaattcatcacaa**caccaagcaacatcctgcttttggtg**ctgccATGAGCAACAGGCTTCTCTGCTGTGTGATCATTTGTCTCCTAAGAGCAGGTGAGTCCTGGGCACAGGTAAGGAATCCCTATTACTGAATTCACAGGATCTAACTATAAAATGTTTCTTCATAAAACTCACATTAGTCTCCTCTTTCCCAGGCCCTGTCTCTATTTGATCATCCATTTTTCCCCCCAACAGGCCTCAAAGATGCTGTAGTTACACAAT**TCCCAAGACACAGAATCATTGGGA**CAGGAAAGGA**ATTCATTCTACAGTGTTCCCAGAATATGAAT**CATGTTACAATGTACTGGTATCGACAGGACCCAGGACTTGGACTGAAGCTGGTCTATTATTCACCTGGCACTGGGAGCACTGAAAAAGGAGATATCTCTGAGGGGTATCATGTTTCTTGAAATACTATAGCATCTTTTCCCCTGACCCTGAAGTCTGCCAGCACCAACCAGACATCTGTGTATCT**CTATGCCAGCAGTTCATCcatagcactgcatag**ccatatcctctccacacaaaaaaaggtgcataccaaa**gaggaaaagcctgccctcaaaattcctc**accccgaataagagaagtttcctcacaagtattgacaaaaagagccaaactctgtaaaatatttgaagagatttattc**tgagccaaacatgagtgagtgtggccagaggcagaggctca**agagggcgtaagaac**atgtgtccaaggtgctcaggctacagcttggttttttggagacat**cagacatcagtcaatacatgtaagctgtacattgtttcaatttggaaaggtgggacaactcaaagagggggcttccaggtcataggtggattcagagatttttcaggctggcagttggttgaaagagttaagttatcatctaaagatttgaaatcaatataaaggagtgtctgagttgtggggacaaaggttcttgttatgcaga**tgaagcctccagggagcaggcttca**gagagaagagattgtgtttcttaatcagacttttaaaaggtgccagactc**ataattctctcctggatcaggggaaaaaacctgaaaaggaaagaattat**ctacagaatgcagattttcctcatgatagacagctttgtagggccatttaaaaatatgtcaa**aaaatatactttggggtaaaatatttt**gatttttttcagggcctgctgtgagtcatgttggtgtcttattgct**acaaaaggtctgttttgt**cacttgtaaggtctccattctaatgttaatgctggttggttgtgcctgaattctaaagtgaagagggtataatgaggcatgtctgaccccacttcccactaag**acctgaactggtttttcaggt**taactttggaatgctcttggccaagagaaggggtccacacagttggttgtggggcttgggattttatt**tttggtttacacaagattataccaaa**gcccttccctaaatttgatatgccctgcttccctctttcca

Uppercase: TRBV26

Lowercase: Flanking sequence[1000bp]

Red & Bold & Underline: Stem-loop [19]

Blue: Heptamer[20]

Green: Nonamer [5]

id-TRBV4-1-2[V_gene_segment]

tcttcattctcccctggcctttca**aacccttctctgctgctcagggtt**gtgagatgtggaacccatctggatgctgacactgggtcggtga**gtggccaaatactgttttggccac**aggcaggagctgtgctctggtgaaacctctaaggcagggacctggggaagggatagagaataagaggagag**atgggccaggagggatcttgcccat**ggggagccaaccgtgcctcaactgagagcatggctgtactcagcccaagcacactaac**atcagcttcaagctgat**ttccctttcctttgctccacacaccttgtgttgttctttttcagcatcctgtgtatgttagctctaagcccagccaagccccatgccctccacttcagggacagagggtggctcatcccttttgtgctccctcagctccctccccagtgcctggtggctggttggcaacacagtcatgcccttgaatgatgggagctggtgcac**actgcatccgtgttcatgcagt**caccctccctccacagagccgggcagaaagggctccaccacacttagacattgagggaagggacgttttgaatggcactgttttatgtgatcccgcaggacagaaggagatttcttgaatgggagtggtctgttgttgagag**ggatcctgaaagacagggagacaaagatagagggaggatcc**acataattaggaagcagctgagagcaaaggagcccctgccagaggaatatccttgaggggtaaaaagccggctctgccctttctccccagccgattcatcctagcccagcaaattcaaatctaccttctatcagaacttagaaa**ggatgtaaagcagtcaggaagaaacatcc**cctgggtctggggaaactatcaggagcagtgacatcacaagaaaaaccaccaaccagggccaaggagaccagagcccagcacctcgcccaaaggaccccagtcagaggccccatctcagacccgaggctagcATGGGCTGCAGG**CTGCTCTGCTGTGCGGTTCTCTGTCTCCTGGGAGCAG**GTGAGTTAGGTTGAAATTGTCTGTCCTTGGACTCCAACCCTTTTCCTTGTGGCTG**CAGGAACAACCCTCTTCCTG**GGCTCTGCCTGAATTTTGTCCCTTTCCTCCTACAGTTCCCATAGACACTGAAGTTACCCAGACACCAAAACACCTGGTCATGGGAATGACAAATAAGAAGTCTTTGAAATGTGAACAACATATGGGGCACAGGGCTATGTATTGGTACAAGCAGAAAGCTAAGAAGCCACCGGAGCTCATGTTTGTCTACAGCTATGAGAAACTCTCTATAAATGAAAGTGTGCGAAGTCGCTTCTCACCTGAATGCCCCAACAGCTCTCTCTTAAACCTTCACCTACACGCCCTGCAGCCAGAAGACTCAGCCCTGTAT**CTCTGCGCCAGCAGCCAAGAcacagccttgcagag**tcaccgct**ttcctgtgcagaaaccttcggggcctgccaggaa**g**ccgtgggggccacgg**a**gggctcgggtgaacatttcctccaagagccc**cgaagaagcttcagaacatcatagcacctgctaattcatccatgtggcaactttacatcctatgacatatttagagtgtgggtttccttttgcctgagtgtgactctgccccgtcaactgatttgaaaaaagagaaacatattcagatctagt**ttaaaaaaaaaaaattttttaa**tcatgaaataggccgggcatggcggctcatgcttgtaatcccagcactttgggaggccaagacgagcggatcatgaggtcaggagtttgagaccagcctggccaacatggtgaaaccccgtctctactaaaaatacaaaaatc**agctgggtgtggtggcatatgcctgtaatcccagct**actcaggaggcaggagaatcacttgaactcgggaggtggaggttgcggtgagccgagaccgtgacattgcactccagcctgggc**gacagagtgagattctgtc**tcaaaaaaaaaaaaaaattcctgaaataaagcaagcatgctgtaag**gattgtgtagcttaaaattttataagaaccctctaaacaatc**acctgtgtcaccacagaggcacaccctgcactgggtcagacactgtcctgtgaggagctgtgttctcctgacacttcatgtccagtggtttccatgactcttctttggggccaagggttccctttcatcataggttgttgtcatgcagagtcaagcaacactga**gcactgggaaatcagagaaggaacttccaggcaagggtcccagtgc**agaagccagctagagaggacagtagtagactcaccctttgggaaatatgtttgagaaatatgtatctattttcaccagttctattactctatattccagcttgtctattctctaatttctcatttgtacactgaaagcta

Uppercase: TRBV4-1-2

Lowercase: Flanking sequence[1000bp]

Red & Bold & Underline: Stem-loop [18]

Blue: Heptamer[38]

Green: Nonamer [3]

id-IGLVIV-64[V_gene_segment]

ttggctgcct**tcaggctctttgttgagcctgacaccaaaggatgtctttctaaagtgtcagtttcatctattctgcccccagttcacaacctgaaac**aacctccgatagtgaatcaagccaagcccctcactctaaagcacagagacatgccatgaattcctaaggggatgacagaatacatcattgatgagaagattgaacaagtgaagtgtgagtggcaagtcaggtgag**gacatttgaatccaatctggcatttaaaatgtc**tcttcatttccaaagtcctatctctccctagtaatgttgtttgactgctagtcccagagcaggaccccacatcccagtgacagccaagtcacagtcgcctccttccctggtccctgtggaaagagctgctc**agggctttgctcccagagccct**ctgcattcccaacttcctgcagcctcctttcattggatgggggtgatagtttgcacatgcctttctagtaatccaagaaaaac**ttactgaaaaagtctgtttgttcatgtcagtaa**tcaggggcatgaccagggcatggcccataaatgcctctggactgatactcaccaggctcccaggccaacaccacatacatgatttccactgagagaatgcagtctggacagagagcagcagggtacactgtaggtctgcccagtgggcatccacagctgtgccaacccctcccaccccacaaattcctcagggct**gggcagtggaccctgccc**tcactgagcctgactcagcagctctgtcctccc**tgggtgtggtcccagcaccca**cttcctccca**ggtctggcctgccagactcccccctccaccagaccctaggacttgaataaatacatagggt**aaggtagaggcaggc**aatgtgggggacacatt**tgggtccataactcctcctctgtgaggtcagtggaggggataagagg**cctgaggcagcccactacggctgtggacctcaggaggactgtgttgaccATGGCCTGGGCCCCTCTCCTG**CTGTTTCTCTCTCAATTCTCAGGTAGGCACAGGCCTCCCATGTCCAGGGCATGTCCCTCAGGCTTCACCAGTCCCCTCTGGCTAAAATCTTACAGCAACTTTATGCTGGTAGGTGGCCCAACTCCCATGTGCATCTGTGTCTGCAGGTTCCCTCTCCCAGCCTGTGCTGACTCAGCCGCCCTCCCTCTCTGCACCTCCGGATCAGCCAGACTCTCCTGCACCCTGAGCAGTGGCTTCAGTGCTGATTTCTACT**GGATATACTGGTACCAGCATAAACCAGGGAGCCTCCCCAGTATCTCC**TGAGCCACTACCAAAATTCACTTCATGACCAGGGCTCCGGGGTCCCCAGGCACTTCTCTGGATTAATGGAAGGCCGGTCCAATAAAGGGCTCTTGCTCATATCTGATCTCCAGTCTGAGGATGAGGCTTACTATTACTGTATGATCGAGCACAGCAGAGCTTCTCATGCTGACACAcacaggtggggaagtgggacaaaatctcagcctgc**tcagagtcttgttctctga**tgaaatttagatcttaaaataacttatatcacttgtgtgggatgagtgagatatcccgagctcacaggaactgtcgatga**acagagtcgagctctgt**aaaatattgaagacatttattccaagccaaatatcagtgaccaatggcccacaacgcagcc**ctcaggagatcctgag**aacatgtaccccaggtggtcaggctgcaacttggttttatacattttaaggagacatgagacatcaatcaacacatgcgagatgtacgttgg**ttgagtccaggaaggcagcacaactcaa**agtagtggggctccaaagatcg**tctttttgataattggttaaaaga**gttaaattattgtctaaagacttagaaccaaagg**aatgcctgggttaaattaaggcatt**gtagtgaccaaggttttgttgctcagaggaagcctccaggtagcaggctgcagagagaatagattgtaaatctttcttatcagatgtaaaaagtctgttctctcagaaattccaaaaatgaggaggcatatctggctccccgttcccaagatgacctgaactctctttcagattaactttggaatgcccttggccaaaaggaggggtctgttcagatggatgggagtcttagaattttatttttggtttacaatcctgtaacacatcctgtaatttgctgcccagaggtcattaatggcctggaagaaataatgtgccagctgcagtgttgtccacagaaagact**tactcattctgctgccctgagta**gtgatatcagaaagatccatctgccaacagtcagaatgcc**cttccaggagggttgcacagtaccaggggtgctggaag**acc**tgggatctgaggcaagcagggggcactgttggatggatccca**agatgacaggattaccaggccacacaaggattaaacaagagacagtaacagtccac

Uppercase: IGLVIV-64

Lowercase: Flanking sequence[1000bp]

Red & Bold & Underline: Stem-loop [23]

Blue: Heptamer[36]

Green: Nonamer [1]

id-IGHVII-51-2[V_gene_segment]

accatgtggtctaaactaagtccaactaag**tccctgagctctgggtgcccatcaggga**tccgccatgtgcccggcagccgcgtgcctttttgtctcctctgcgcccaatcactgtctgtgatgagcttccaggatgtgtgtgtttggcacaaaccaggtgatggacgtcagacagcagcagctggtgcccggatcatgggctccctaatgctggaggaatgagaggtgcattctcagggacaagacattgttgatggatttttatgtagaaaccacgattttac**ttgcttctctcaggagacatagaagaagcaa**ccatgcagtcagcaaataattataatttccacaattaccccaaatggttaatccttaattctgcgcagggtcccaccgtagagtcacctttctcatgaggaatggttgaatctagaatgagtccacatgattttcatatttttggcttctgtccatgttcagagagttagagtagagtaagtttggacctttccacacactaagcctcacctcccccacagaaagagcagagacttcaactattcctgagtgtggggtaggggctggtc**ctgcacacctcagagcctgcagagactgccacgtgcag**tgttatagacttgggtgttttctcattcaggaggggtgacctccacggcctccgat**ttctgctaagatataactgtgagtgcagaa**ttaggacactattaggctatcatgcctcacttctattctgaaaatcaccctctttatagtaaaagaaaacaattcaatgtccatgcccctgaaagtcatttctttttaatttggttgcgagtttaccacatagtgcctgaatttattctcagaatctactgaattgtttataaatctcacaaattgaacaaaagtgaattcttcattcttactaaaatgtgtgtatgtaggaatttcagtgtgttttccagaagccatgcacaccaaccactgtgtttcatggtcatttattGGCCT**GGTGAGGCCCTCACAGACCCCCTCCCTCACC**TGTGCTGTCTCTGGATTCTCCATCACAACTAGTGCTTCCTGCTGGAGCTGGATCCGCCAGTCCTCAGTCAAGGGAGTGGATCAGGGCATAGGTCACGAGGGAGCACAAATTCTAACCCACTCCTCATGAGCTCAGTCACCATCTCCAGATCCACGTCCAAGAACCAAATTTTCTTTTAGCTGAGTTCTGTGACCAACAATGCCACAACCTTGTATTACTGTGAGAGGaacagaagagatgtcagtgtgatcccagacacaaacttccctggagaggggcccaggaccaccaaagagcactcaggctccatgaaaacagggcccaagctggagaacgggtttcctgtcaccctcacgtttcaccattagatactctacactatgcctatgctgaggtgtgtgtttaatataattagaaaatggtatttatatacgtgtataatcatatctagtgtaagaatttgtagtttccaccttatgtgaccattaaagacaacatctctcacgttattcaaggaccttat**aaagtataatacttt**tacataagatttatctccttttttgtttctccttctgtctatattctctctgtctctctcttcctccctctttctctctctctctccatctctctctctctgtctctctctcacacacaaatacacacacatgcaatcgttgacagtatataggctaattaagaatatattcccatatgatttaaaaataacagttttacaaatatttgcc**tttaatttttcaaagaaaattaaa**tccaaatgccagtgattttttatgtgactttt**aaaatttaaattgttaatttt**tgttttcatcaatgcttgtatgtgtgcttttgtattctttcagattatttggtgcct**tcacagaattaaacatgtgaaactctccacttgaatttctgtgactcaggactggtgagcaaaaagttagaagtcac**tctcctgatccttccctacagctgcagattcctgaaggtaatgacctcacagtagaccaaggccctcaaaggtgaccttgagttccagcacatcttgggaggccaagcggatgtttgtgaaagaaatagtggctatgtgacagtttctaaccagaatatctctgtgttttcaggtatccagggtgaggcccagcttacagagtctgggggagacttggtacacttagaggggcccctgaggctctcctgtgcagcctcttggttcac

Uppercase: IGHVII-51-2

Lowercase: Flanking sequence[1000bp]

Red & Bold & Underline: Stem-loop [10]

Blue: Heptamer[25]

Green: Nonamer [6]

id-IGLV6-57[V_gene_segment]

gaaattacctgacagatatattgaaatcagccaccataaaatcctgttgatgagcaatttcaggcattcttgaaacaaatgaaaaaggataaattctcagaaaaaatagaagtcataaaatcaaactaaatataaatacaaataaaataaatgaaaaatttaaaaatacctctctggagagactcaagaacaagtaactatgaaactaaaatccacgcattagagaagacacttaacactgt**taactagagaagcacaaaccatagtta**tcacagcccagcagggcgagcaatcagtctgctcagaagtgagaggcctgaaaaaaaactcctggtgtccctgatggtgaaatctggcccctgatcccagggttcct**ccctcttcgtgctcagaggg**ctagacctagcagtgaagggttcccaacctcaccctatgtggcccctccagaaggcagagcccattcttgctgatattcccgatctccaataacacaaagtcaaagagaatgtcttaaccagagtactgagtcttggcctatctgaacatatttcaaagtttaaatgtacagtgagcataggatgctgagcagaggccccctggacagtggctggagttgggagcagtgcaggatggaagg**acaggtcaggggccctgagcacctgt**acctgcacctgcacctgcccag**ggtggacatgactcccacatccacc**agagggcgctgtggtctaagcaactaatcagagctccccagatcctctgctgttttccctccacctgctggtgtttctgtgccctccagatggtgttgctcctcccccttctatgaccacttttcagccataaccttgtctgaggtcaaaggagggggttgtgcaacctccagaaagggagaaatttgcatggagccctaccactctgaggatacgcgtgacagataagaagggctggtgggatcagtcctggtggtagctcaggaagcagagcc**tggagcatctccactATGGCCTGGGCTCCA**CTACTTCTCACCCTCCTCGCTCACTGCACAGGTGGCTGCCTGCAAGGAATTCAGGGAGCGTTCCTGGATGTCACCTGGGCTGATGATCTGTTCCTCCTGCCTGGGAACCAGTCTTCATCTCTCCCCACTGATCTCTGTGTTGCTCTCTTCTTGCAGGTTCTTGGGCCAATTTTATGCTGACTCAGCCCCACTCTGTGTCGGAGTCTCCGGGGAAGACGGTAACCATCTCCTGCACCGGCAGCAGTGGCAGCATTGCCAGCAACTATGTG**CAGTGGTACCAGCAGCGCCCGGGCAGTGCCCCCACCACTG**TGATCTATGAGGATAACCAAAGACCCTCTGGGGTCCCTGATCGGTTCTCTGGCTCCATCGACAGCTCCTCCAACTCTGCCTCCCTCACCATCTCTGGACTG**AAGACTGAGGACGAGGCTGACTACTACTGTCAGTCTT**ATGATAGCAGCAATCAcacagtgctccagaccca**tggggaagtgagacagaaactcccca**gagcatctctacctgggccagtctcagcctgtctccacc**ggagagggtagctctcc**catctctcctgtctaagtgcttgaggtgaactttgcttatgacttcctcttttccttttcaaaataccaatctgtttttccacaccttgatggctgattttctgatatgtgcacgatcaattgtgggataaatgtagtttataatcgggtctctgagccacaccatgattgatgttctggttggggctgccttgcacagtataagttattgacttgcatctctggcctctacccttagatactattaacaccctcccctcatttgtgacaacaagactgtcttgagacactgccaca**tgtcccatgcttggggaca**aaatcagcccaggttgagatactcagttaaagagcatgtcttcagctactaagaagcttggctttgaaatgtgtttatgccattctgctgagcatgaggctggttttctaatttttgtttctggttcccatctatgaga**caaaacaataacaccaaacttcaatgttttg**atggttgaatggtttttccaagcaacaagaagtcagagattgtgctacttagaggtgtagaagccctaaccagatcctgtaacacagagtttagctctgactaggacagcacacaatgccttccatcaatctccaacccacagcccgcaggctgcatgcagcccagaaaggctttgaatgtggcccaacacaaatttgtaaactt**tcataaaacattatga**tttttctgcaattttct**atagctcaccagctattgatactaacagttagtatca**gttttatgtgtgttccgagacaattcttcttccattgtggcccagggaaat**caaaagatgggacacccctgctttgcacttttg**tacaatgcacatggaaatataatagagccttcttccttctaaggtcatgtgactc

Uppercase: IGLV6-57

Lowercase: Flanking sequence[1000bp]

Red & Bold & Underline: Stem-loop [15]

Blue: Heptamer[37]

Green: Nonamer [3]

id-TRAV13-2[V_gene_segment]

gtttcatttttggagcctgcagcctgatgttttctgagaatttctccttgtttt**ttctgcactttgcagaa**tacaagccacatagcaataggaaaaggtttcttaaaacatgccagatttctccttgcaaaccagcccaatctatgggtcaaactttcctattgttggagtatttaataactcatggaacttgttcccttcctcctttatcattgta**ctttaattctttaccttttattaaag**ccatcct**taccaaagaaagattccacatttcgactgttggta**ctgtcaagtctctgggaatagaaagtcatggggaagaccagtcttggagaagcagagcctgagataatatgcacgtgaaatattgagtagtcctctctctctt**actcttatctttggcttacatattcctttaagaaagagt**ctttctccataacaaa**caaatcatgatgaccgatttg**actgcaagggaatgtattacttttgtttgtccaggacagtgaatcatggtgttttcctgatttcaaacatgtaaacttggtgca**tcattgtataaacacctgcaacatctacctctttctaactttttaacaatga**ctcattagtttgcataa**atatcaatcttgatat**ctacattcttactgtgaacttcatctcagagtgaagactttatttgtttgacctattgctccttttgtgcatctgagtccctcctagggttatagaaaa**acattcaaactgcaagtttagtagctacgaatgt**gaccacaagatggcagtgcttctctgctgatgcagaacacacagaggtttcattttagtggattctgattaaataggtttatggcagaagtagagcctttttctattggctgagtagacaatgtgagaaaccactatgattgttagaggaaagaaaggaatacccgatgatggaagtagctcttatggctggagattgcaggtttatgactgatcctatttgggaagaacaatgATGGCAGGCATTCGAGCTTTATTTATGTACTTGTGGCTGCAGCTGGACTGTAAGTTTAGAGTTTTGGGTAACACAGTATATTAGTTGATGATTTTGAGACATCCAGGGGGAAGTCTTGCTTTATTTAGGCTTCCTCTAGTTACTAT**AGAAAAAGAAAATTCTGGAGAATAATAACTTGATAAACCTTTCTCTTTTTCT**TTCTGCACAGGGGTGAGCAGAGGAGAGAGTGTGGGGCTGCATCTTCCTACCCTGAGTGTCCAGGAGGGTGACAACTCTATTATCAACTGTGCTTATTCAAACAGCGCCTCAGACTACTTCATTTGGTACAAGCAAGAATCTGGAAAAGGTCCTCAATTCATTATAGACATTCGTTCAAATATGGACAAAAGGCAAGGCCAAAGAGTCACCGTTTTATTGAATAAGACAGTGAAACATCTCTCTCTGCAAATTGCAGCTACTCAACCTGGAGACTCAGCTGTCTACTTTTGTGCAGAGAATAcacattgctttccaggcatctgtaaccatcacccaaacctgagatgggaggtgaagcagcatccctttcctttgcaataaattttagttatagcacttgtcattttgtttgttcataagtggaaactaacatatagatattaaaaccatgtaatgaggttagattcaaagtgacccagaaaggtttagagcattttgttggccagttgatgcttaatggattccgtttttcaaaaaaatatatggtacgcatttaagttaacaacatgaagttttgacacacatgtacatagtaaagtaaagagtcagctaaagagacgcatagggcaaggtatgggaaaaatggccctgagcttccatgccttctctggctctgccacccaccacatgttcagcaatcaggaggttctgctcaagggatccttacagagatttagtgatgttggattttgtaatctagtcttcaaaggtgactttacatttgc**tattcagacttctctcctatgaata**accccttagtgtcaatattgcaatccctattatggata**aattttaacttttagttttcaaaaatt**gcttttgttatctttttcttattgtaaaacattacatgccaaaaccagatagtgtaaaatataaaaaatacaaaattgataattctgtacccagataggttttgct**aatattaggttggtgcaaaagtaagtgctgtttttgccattaaaagtcatggcaaaaataacaattacttttgcaccaacctaatatt**tagatttgtttcttgatagagtcctattataattaggatcatgattttttcacctattcttatatctcgaattgagcttttctcacttcgtcctctcctcaaacacatgctttttaatagctccgtgagaaattttcaccagtcacagttatttttattttcttactctctgtatctgtggtaaaatttggcagtcccttctacctgctgtttcttgttgggaaaacgtgat

Uppercase: TRAV13-2

Lowercase: Flanking sequence[1000bp]

Red & Bold & Underline: Stem-loop [12]

Blue: Heptamer[23]

Green: Nonamer [5]

id-IGKV7-3-3[V_gene_segment]

atcacaagtaacagaatagagtcctggagtggagctcatagagaagtgatggttggcatcatgttccctgacttttcatgttccttgtagccttgtgttgg**tgtctgcacatttgaagatgcagtcccttcttccagaca**ttttaagtt**ggttttgttgggggaaaacc**ttcacctatggatgggtgagagtgccagctgactggtgtggacaccatatcttgctgtaggagaagatggacttccttagccccagcttccagtggcaaggcagatggaaccagtgggagttgggggatggttattctccactctgaataggatccactacaatgactcttctgcctctgagggcctctggtggatacagcattggtttcaggactaagcagatggagagaccctgccttatccacaatgaggcaagtt**ccttgattagtttaggttaccagtcttcatcaagg**tttcatgctcacccccttcatcctttccagatcatgataacatttatagacttgttttaaataccacttactcagtggaacaaacttctccttccca**tatgaaccagtttcata**aaacccacccaacttgctttctctttctgacaatgataacttaaaatg**cattattattgataatg**atataaaatatgaa**ttttgaaaaatcaaaa**gtatttctgaaataacttatatccaaatggcagggcagccccagtacactcctccacatggcatctcaagtagactggcaggcaaagatccaatagaaaaggtcaccaaatcccttgggacatgcttgagctaccctaggaaatgttttgttagccccaactcccactgggtgaaataagagcaggtgtcccc**tttgagcatcaactcaaa**cttttctgagctgatttgcatgttgggccattagacaacggtc**caagcttctcagagcatccttaaagacagcttg**tcagggcccctgaggagcagcctctagttcagctctcagagATAGGGTCCGGGGCTCCTTTGCTGTGGATCCTCCTGCTCTGGGCTCCAAGTGAGGGGGTATAGGGAAACATCGGGCAAACTCATGTGGCCATCATTGTTTTTTGTGTCTCTGGATTCTTGGTTGTTGGATTTAGGGCACTCATTGTAGGTTGTAAGTTCCCCAGTTATCTTTTGTTTTCTGGTTACTCAAGATGTTGCATATGACTGTGGTTACAAGTGTCACCAATGCATGGCCTACATGGGGACAGTCTCAGTATATCTGATATAATTTATCTGTCTTTATCATTTC**AGGCTGCAATGGGGACATTGTGCTGACCCAGTCTCCAGCCT**CCTTGGCCGTGTCTCCAGGACAGAGGGCCACCATCACCTGCAGAGCCAGTGAGAGTGTCAGTTTCTTGGGAATAAACTTAATTCACTGGTATCAGCAGAAACCAGGACAACCTCCTAAACTCCTGATTTACCAAGCATCCAATAAAGACACTGG**GGTCCCAGCCAGGTTCAGCGGCAGTGGGTCTGGGACC**GATTTCACCCTCACAATTAATCCTGTGGAAGCTAATGATACTGC**AAATTATTACTGTCTGCAGAGTAAGAATTT**TCCTcccacagtgctttaggtctaaacaaaaacctccccaggcagctgctcc**ctgaggctcaaatccctcag**atgtggctttttatgcaggtccatcagcctgctgtcataggcttgtttgaacaaaggggtcaaattatcaaagagaaaatccttaaattgatgtgtgtgtgtgtgtgtgcatgcatgtgtgtcacagccagttttataatttaataatttctttactgtatcaggacctctc**agaagttttctacttct**caatgtatattttttatgtcaaagacccttaccaatagaagaa**ttttaaacattaagaaaactttgtagtcattttaaaa**aatgctctgaatagtcaaaattattagct**ctatatctttcagatataggtgaccttctcacct**gaattttga**cttttctccacttttatttgctgaacccacatgaaaag**gct**tttctagttatttagaaa**cctaagacttgtgaggtagttatctctgcttatttaatttagatgtctctatctcggtgactttcccaaaagttagattctcagctctagttttaataaattgtcctttactcttcatgcagtgttcatgtcctttttcttttctttttttttttttttaa**acagagtctcactctgt**cacccaggctggagggcaatggcacaatcttggctcacggcagtctccacctcccaggttgaagtgattctcctgcctcaatgtctc**aagtagctgggattacaggtgcctgccaccacgcccagctaatt**tctgtatttttagtagagatggggtttcaccatgttggccagcctggtcacgaactcctgacctcatgatccacctgccttggcctcccaaagtgctgggattacaggtcatgtcctttttctatgatattttccatgtccattgttaaggttttttttctcatcttctctttttgcttttctaattgctctaacactacaggcagagtggactccagaaatg

Uppercase: IGKV7-3-3

Lowercase: Flanking sequence[1000bp]

Red & Bold & Underline: Stem-loop [20]

Blue: Heptamer[31]

Green: Nonamer [7]

id-IGHV3-48[V_gene_segment]

tatgttgagtaaaataagctaaacaaataagaatatatactatgtaattt**catttttataaattctgataaataaaaatg**catctgaagtaaaataatgaagataagtagttgcctggggtaatggtagaagaagggagggggagaggaggaggaatacagcagaacaaggggaaatgttgagaagaattcacttgtccactttcttggt**aatgatagcagttacatcatt**tttattagttgtacattttaaatatgtgaagtttattatctttcaattaagcctcataaaatgtcttacaagcaaacaaatggaaacttagacaaggaaagagtaatagaaagacagaaaaaataagttcaatgtcagaagtac**ctgaaaattaatgtgcctggatcctagttctctccatattttcag**aagagtgctggagggcagcaaaaccacacatgctcttattacggaatgtgggttctgataaaaacactagac**acatccagctttgtcctggagttggtttagggggatgt**cagagacagtgatgaagagcacagggccagataccggggttcactcat**cccagacatgagctccgagacgcatacagagcccccccatgtgtggg**tttacttccac**ttctgtaaatggagaaaatattgtctcctacagaa**catagtttacatgaatatttaaaatgaaatagggtgattagtgcaaagtgtttatcacagcacaatttcataataagacagcatattttccaaatgcaatcattgccagcaaacttctacagggcaccgtcgtcttatctgggtacagcctactcctcaagggtcccaccctagagcttgctatatagtaggagatatgcaaatagggccctccctctactgatgaaaaccaacccaaccctgaccctgcagctctcagagagg**tgccttagccctggattccaaggca**tttccact**tggtgatcagcactgaacacagaggactcaccA**TGGAGTTGGGGCTGTGCTGG**GTTTTCCTTGTTGCTATTTTAGAAGGTGATTCATGGAAAAC**TAGAGAGATTTAGTGTGTGTGGATATGAGTGAG**AGAAACAGTGGATATGTGTGGCAGTTTCT**GACCTTGGTGTCTCTTTGTTTGCAGGTGTCCAGTGTGAGGTGCAGCTGGTGGAGTCTGGGGGAGGCTTGGTACAGCCTGGAGGGTCCCTGAGACTCTCCTGTGCAGCCTCTGGATTCACCTTCAGTAGTTATGAAATGAACTGGGTCCGCCAGG**CTCCAGGGAAGGGGCTGGAG**TGGGTTTCATACATTAGTAGTAGTG**GTAGTACCATATACTAC**GCAGACTCTGTGAAGGGCCGATTCACCATCTCCAGAGACAACGCCAAGAACTCACTGTATCTGCAAATG**AACAGCCTGAGAGCCGAGGACACGGCTGTT**TATT**ACTGTGCGAGAGAcacagt**gaggggaagtcagtgtgagcccagacacaaacctccctgcaggggtcccttgggaccaccagggggc**gacagcacactgagcacggggctgtc**tccagggcaggtgcaggtgctgctgagggctggcttcctgtgatggcctggggcggcctcattgtcaaatttccccggggaacttctccagatttacaatcctgtactaatatttgatgtctctaaatgcaaccttttttttcctttttgtgt**ctgttttttttttttaaaacag**gaggacacatcctcacctccacagaagccacagtgtcactttgggggcagaaataatcctttcgtggtcaacagggtgagagttttgaggaat**cccagggaaacctggg**gaatgttttccaattagactcaggg**cagagacctccatgggaatctctg**attagaacaggctttgagttctgatgggagccaagagagaggctcacccagggtcagggttctt**aaaacctgatggttttcacagcaatcccccttcatcttgtgaaa**ctgggcacatctgactcagactgattcagttgaccctctttctgctaatccattttccttcccagtagacttgattctcacagatccctttcttcttctctttcct**gaaaacagaggatgtgttttc**tgtagtctaaattccaaggctcaggtccgcaggagctgggtaggctgaggggtcttcctcactcactattgcctggaaaatcctgctgtcttctgtgcatggaggcatttggaaaatgaagcaggcattagtcatgaagggaataatactagttttctccaatgggatgttgatgtagagctgatcttatgcttctcacact**gtcacaaagtttggactctcacctgtgac**tttgaggagagcttgtgataccttatcttgttttaatatgaatagactctcccttagctcaggaagctggaacagactccatttggctccttcatttgtaag

Uppercase: IGHV3-48

Lowercase: Flanking sequence[1000bp]

Red & Bold & Underline: Stem-loop [22]

Blue: Heptamer[34]

Green: Nonamer [7]

id-TRBV5-6-2[V_gene_segment]

gaattgggaggagcagactagcaaaagtctagattcctgaaaacagagcataaagggcaattctggtgaaggctcagggggaaatgagg**aacaaggtattgaaaatcgaagtaaatgccatccttgtt**gtaagtagcaaaaaccttggcaaaattgtgtcctgttctagaactttatggaatataaaaattatgagccatttgctagtatatctgctgaaagaaatatctaagaggcaaagcattcaggctactgtgtgactacttttagccaccaggaaatttaacccagcaagaagggagccatgggaatagattttgcaaaccagcacagatggtgactctacctccctctgctgtcctgtctcccataagccaaactctgtgctgtgagctgttaaagtccaagaattatttctggggaa**tctgtactcttaataatttacaga**caacacaggtctgctttggatctgatcagacagactaaatcttggggactgtaccagtggccactgagaaaaggggcccgg**aatgttgtcttggatagaaaaatataataagaaacatt**ggtgtgaatctagcatcagaaagatgacgtgaggacagcaaggaagagctggaaactgaggtttactcacaggttcctcaccctccgctgacgggcaggtgtgtga**gctccagcatggagc**accacagcactaggtgggagaacagtgatagggtgatggggcagcctgtgagctggggca**gtgtaggcagaggaggaactgtatcaccacagaaacctctgccttcac**acatccctccagctcgg**caggacaggtagagagtccagtgtcctg**gagcactagacctaaggaaggctgcatg**gggaggacaaaggacagtgacatcacaggatacccctccc**atcaggaaaatcaaggcccagaactcactcggctcttccccaggagaaccaagccctgaatcagatgcagtgcttcctgtccctctgtgccATGGGCCCCGGGCTCCTCTGCTGGGCACTGCTTTGTCTCCTGGGAGCAGGTGA**GTCCTGGGCACAGGAC**AGCAGCCCCATTCTCAGCTTTCCCACCCCTGT**GTCCTCCACTTTACCTTGGGGAGGAC**CTCCAGGCTGTC**CCTGTGCTCATCCTCCATCTGCTTTTCCCACAGG**CTTAGTGGACGCTGGAGTCACCCAAAGTCCCACACACCTGATCAAAACGAGAGGACAGCAAGTGACTCTGAGATGCTCTCCTAAGTCTGGGCATGACACTGTGTCCTGGTACCAACAGGCCCTGGGTCAGGGGCCCCAGTTTATCTTTCAGTATTATGAGGAGGAAGAGAGACAGAGAGGCAACTTCCCTGATCGATTCTCAGGTCACCAGTTCCCTAACTATAGCTCTGAGCTGAATGTGAACGCCTTGTTGCTGGGGGACTCGGCCCTCTATCT**CTGTGCCAGCAGCTTGGcacag**cccatcagagtcactgacgttctgtatataaacttcctgccttagctttgccttgagagctgcaggccccacccagatttcactccttcaagggaagcttttagttgtttggaaggcatgtcttgtgtcctactgagggcagagctctcccaaccaataaagcccaagtttcctgtgtcctgagtgtgcccacttctgt**gctgcatcttcttgcagc**ttgtcacctcagtacaagagtgactgctgagccccagatgtgtgcttgattatttgcatttcttacatagcttaatgcctttcaacaatcttgcacatcaaacattcttattcttcctttacagatagaaggctcagg**gacattgagtcattttccccaatgtc**tcttggcttgtaaggatcagaagtaggaaacaaactagtccatccattttccacctacccccctgctccatcatcatcttctgcatc**ctggtcagtaaatcagagccctcacactgccgtctagtgaccag**cagggccgcagtagaggctgaaatgtcttcaggggtcattactatggccttttcattagctcctttgag**gaatgttaaatttcacacattttaacaatc**atttatatgcattccctttgtctttccccagtctgcagcttatattttcatttttatgg**agtttttggtgcacaagagcttaatttaatgcaacaaaaact**aaaaacaa**ttttgtttgtgtgtagacaaaa**accactttttcatataaatgtctaagcatatttttctc**aacttattctactaaaatttaagtt**tggattttcatcactaagaataaatgtacctgaaatacatctcatgtaagaaataagatacagacctaggaatgggagggaaggacatatattcagtgaagaaacaatagcttgtcaaaaagagccctaagtggacacctggagctgacggtggagaggtaggtggagggcaaatgagcatgaactttgt

Uppercase: TRBV5-6-2

Lowercase: Flanking sequence[1000bp]

Red & Bold & Underline: Stem-loop [18]

Blue: Heptamer[28]

Green: Nonamer [8]

id-IGHV3-36[V_gene_segment]

tgtatttctattttgggtagagtttttatggtgaaaccttgaatttttttaatgttattttctgtctgagaagttaatgggaccttcttcgtattta**atttggcataccaaat**gtattgatttgaggattttgaatcatctgtgcat**ctcaaaaataaatttgag**ttggtcatggtgtatgatccttctaaagtcctgtggaacttagttgactattatttggtgtgtggagggggctcatttatgtctactaggaatattggtctgtagtttacttttttgtgaccttgtctattactgaaaatacggtaatttgggccttatagaaagactggaaggttggcatctcaccaatttttgaaaaagttagagaatggttggcattacttctttaaatgttgagagtcctttttttacttaacatgtgacctatcatggagaatatttagtctttgcttcggaagcatacatattacactgctcttggttggaaggttctgtagatgtctattaggtcaatttgttcaatagtgttgttcaagtccactgtcttc**ttaagaattctcttaatgatgttctatccatcatt**gaaagtgaggtgctaaagtcttctcctaatttttattattttctatttatctctccatatctcttaaggtttgcttaatgcagttat**atttatttttgtacaaatatgtaaaaaataaat**gaaataataattgctaagtaattttaatggcaaaatcacataataagaaattatattttcccaaatgctgtcattgacactaagctcctccaggaatctcacatct**gctctggacacggctctcctggtgtcctaccccagagc**ttgctatagaggaggagacatgcaaatagggccctcactgtcctgatgaaaaccagccttgcctgcgtctacgggagaagagccccagtccagaagtaccaggggtttccatttggtggtcaggtctctgaacacagaggactcactATGGAGTTTGGGCTGAGCTGGGTTTTCCTTGTTGCTAATGTAAAAGGTGACTCATGGAGAACTAGAGATATTGAGTGTGAGTGGACACAAGTGAGAGAAACAGTGGATATGTGTGGCAGGTTCTGACCAGGGTGTCTGTGTGTGTTTGCAGGTGTCCAGTGTGAGGTGCAGCTGGTGGAGTCTTTGGGAGGCTTGTTATAGCCTGGGGGTCCCTGAGACTTTCTTTTGCAGCCTCTGGATTCACCTTT**AGTACCTTTATTAGGTACT**GGATGAGCTGGGTCCATCAGGCTCCTGGGAAAGGGCTGGAGTAGGTCTCATTTATGAGTTGTTGTGTAGGTAGCACAAGCTATGCAGACTCTGTGAAGGGTCGATTCACCCTCTCCAGAGATGATGCCAAGAAATCACTGTATCTGCAAATGAACAGCGTCAGAGCCGAGGATAGGTCTGTGTATTACTGTGGTGGcattgtgtgcatcccttgtttaggtacatgcagagatgctgct**ttggtgtgttcaggggctcctgttttggggacaccaa**ttttggagtttgcagtatccttgagtccagtacgttcatggtggcattgggtctggggtgtgcagtggcaatgagtccaggatacatgcagctgatgccgta**tcttgtgaggacaaga**acaactacaaaacgat**cctattaataaatagg**caaatgacttgaatacacatttttcaaaaaatacatacaaatgacatatagatatataaaatgttaaatatgagtaataatttttaaaattcaaatgtaatctacaataaattatcaccccatacctgttagaatagctatttttaaagaagatgaaagataaccagtattggggcagaagtggagtaaagggagcctatatatattgttactgaa**aatgtcaattagtacagacatt**atgga**gaatactatggaggttcctcaaaaaataaaaatagaactaccatagtttc**caacaagcccactactggctatgtatccacaggaagtgcaatcagtatgtcaaagagacatctgtgcccccacgttcattgcaggaatatttatgatagtaatgatgtgggaacatcctaaattaccataaatggatgaattactaaagaaaatgtagtatgtacccaatggagtaatgttcatcctaaaataagaagggaatcctgcatttgcaacagcaggcatgactctagaggaccatatgtgaagttccacaggcaggcgcagaagacaagtaattcgcgatttcactgtagtttgtttaaatggtagactcacagaagtagagtggagtgttggttaccaggggctggaggggtggactggaaaaggcgagattttggtcaaagggtgaaatgttccagttagacagaaggagaaagttacagtgttctaatgcacaagatggaaaatatggctaataatgcattatatat

Uppercase: IGHV3-36

Lowercase: Flanking sequence[1000bp]

Red & Bold & Underline: Stem-loop [12]

Blue: Heptamer[32]

Green: Nonamer [2]

id-TRBV27[V_gene_segment]

cactgcctggctacaacacgatatccagggacagataccttccatgtacagcaagctgtggagtggcaagaaagcaaagcacgtaccaaaagcaggcaggaag**cagagctggctggtaaggctctg**aggttactgcag**gaagcaggtgttgcctctgcctttcaccagcttcttctggagaagaa**cagagaaggggaaaagaag**aaagatgctccagataccatcttt**ggtggaaacagctgctagctctagcaatgatgactaagtcaat**tttgctaaagagcaaa**gaagctgtgtgtcttggattttaaacagcaatagactagcaaaataatctatgatgtgaatctgagattttacaccctagactcttttaaaatgtagtgactaaaaatatataccagtggataaatgaagtcagggatatacttgaaattttttctgccagagtagaaatgacttcaaaatatatgataatgcagctttcatgtaatcccagctactggggaggctgaggcaggagtattgcttaaacccgggaagcggaggt**tgcagtgagccaagatcatgccactgca**ctccagcctgggt**gacagagtgatactctgtc**tcaaaaaaaaaaaaaaagataatgcagctttcaatattataagctttatgttctcacacttctaagtgataatcaaatgtgttacaccactttataccctgagatccaaaatacataaatatttgacagtttgtacaagactgtattaataagaaaatgtggcacatgtataaagaaaggaatctcacagattaaggaaaat**gacattgattacaaatgtc**ctcagcctgagttggctgtgttgcgtttgta**cacttcagccctctgagctgaagtg**ggagtggtttctctcctgaaaatgtttctgaggcccaaatagctgaag**aggtggagacgttacagaaaccacct**ggagcccccagaac**tggcagacacctgcctgatgctgccA**TGGGCCCCCAGCTCCTTGG**CTATGTGGTCCTTTGCCTTCTAGGAGCAGGTGAGTCCCAGAACACATAG**GCAACTTTTGGCCTTATTTTGTAGGCCTCAGCCCAAGGC**ATCTCCTAAAGGCTTAAGCATTGGAGAT**CCCCTTCAGCGCTGCCTCTAACTCTGTCCCCTTCCTTTACAGGCCCCCTGGAAGCCCAAGTGACCCAGAACCCAAGATACCTCATCACAGTGACTGGAAAGAAGTTAACAGTGACTTGTTCTCAGAATATGAACCATGAGTATATGTCCTGGTATCGACAAGACCCAGGGCTGGGCTTAAGGCAGATCTACTATTCAATGAATGTTGAGGTGACTGATAAGGGAGATGTTCCTGAAGGGTACAAAGTCTCTCGAAAAGAGAAGAGGAATTTCCCCCTGATCCTGGAGTCGCCCAGCCCCAACCAGACCTCTCTGTACTT**CTGTGCCAGCAGTTTATCcacagtgttgcacag**ccagctgctctctgcacaaaaacagagggtagctgcaagaac**aaggagactcctcctt**caggagacccctcaccgaccaacaggataaacttcctccatcatcccacagaagcccttccccacacgccagcctcagggactctgcaatgcagccactctgtggaaccaggggtccatcctctacctgctgtaccaatgaggtttcaaagttgggcaggacaagctcactgagttaggtctagtgccagggtacactctcccatcctttcttcttgtggagcttaaatctttaaattcaactaaactgttgcccaaactgaagactccatttacaatctaatgggtctcaacaatatttggttctgtattccaggtcctatttattctggaggttttgtcacttggaggaacaccatgtttggagttatttatgaaacccagcacctcctattttcttcctctcagcagatggtccaaatgccc**aagccagagatgcaagactgatcttcactctctctctgactt**tctccttccacaccaaatccactaccctatcctgcatttttctaagtactaaattcctcacaaattttactaacttttctaatttcactgatctcacctggatcactttagccaccttccaagtagccaccctacctgcaagcctgcacctgctcccaccca**aaaccactccctcaagcttaaaaccctttggttt**atttctattaactagctaaagtctcaagtttatgttagtttgcaagtctctcagcccctccatgatgtgacctttgcatatcccccacctttctaacactgacttcccttatccttcaactaccacaactgcctgcatctactccagaaacatcccatttctcgcacatccccaaatatgtccactgctgactcactc**ctgggtctttatctacaatggtcactctacccag**aataccatttctctctctggtattcatctgtt

Uppercase: TRBV27

Lowercase: Flanking sequence[1000bp]

Red & Bold & Underline: Stem-loop [18]

Blue: Heptamer[31]

Green: Nonamer [3]

id-IGKV2-4[V_gene_segment]

aaattttttgaaaagattatgcaatctgctgtgctggtggagtgttctcgaagtgtcaattatgtc**cagatggttgattgtgctgtccaattcatctg**catctgcactggcctgagaggcaggactgcttgtgctacaggctgaacctgctgcagagccctttttttgccccgggtcccactcaagg**ctggctgactcgtgtatcagtcggtcagtgagccag**tacagtattccaaggtacagaatttctt**gcctctagaatccaaagaggcctaccaaggcct**tcctcccttcaaagacaacgacatgcattagatacaat**ggaatggaatatgcattcc**tcttcccgtgaactctaattttctccttgtttacgaagcacacattacagctagcttattcactggggcatcatttag**tgccatttaatggca**g**ctgtagagccaagtgcccagaaataccagctacag**tgaaatacaagctgagggaaatcagccttgcaaccacagccaaataaccctggaactgtttcaagttcatcctgatacttagagacattctatcaatgagcaccagagttagtgagagctagttcctcctcagaattagctgggtggagccaagaagaaagcctattcaccacctaaagattgatacacatcttcatcatcacaa**tctccattgcatcagcaatgaaga**gtgacaggctgcagtcacagtgtcttccacattgattcatggtatttcagaatgtcacccttctaagccaggcaacaccaatattttattttggtttcatcaacacaaaagatattctgtgtcactttgcttacataagaagcccactcttcgaccccaactggatatttcttcactgcttcactcacc**aggggatttgcatgttgtcccctagggaagaccttccct**tgtgagtctgagataaaagctcagctctaaccttgccttgactgatcaggactcctcagttcaccttctcacaGTGAGGCTCCCTGCTCAGCTCCTGGGGCTGCTAATGCTCTGGGTCCCTGGTAAGGACAGAAGAGAGGAGAATGGGACTGGAGGGTGAGCTCTGTGG**AATCTGCAGCCTCCCATTTGTTGTTCTGTCTACATGTCAGATGCAGATT**TCTTGTCCTCCAGCAAAGGGAATTTAATATTCAGACCTGTAAGAACTAAGAACGAGCCACAAGGAAAAAGAACTTCACTGTCGAGAACTTCAAAA**ACAAAGAGGCCACTTTGT**GCCTTTAATTATTGGATTATTTTTGGATATTGGATAAATGTAGTGTATGAATCTAATCAAACACAAAAAAGTAGCCTAACGTATACAATAAAACTAAATTATAAAAATGTCTCACATTGTTTGTATGTAACCTTGTACTTCTCACTGTTATTATTTTAGGATCCAGTGGGGATATTGTGATGACCCAGCATCT**GCTCTCCCTGCCCATCCCTCTGGGAGAGC**CGGCCTCCATCTCC**TGCAGGTCTAGTCAGAGCCTCCTGCA**TAGTGATGGAAACACCTATTTGGATTGGTACCTGC**AGAAGCCAGGCCAGTCTCCACAGCTTCT**TATCTACACAATTTCTAACAAATTCTATGGAGTCCCAAACAAGTTCAGTGGCAGCAGGTCAGGGACAGGTTTCACACTTAAATT**CAGCAAAGTGGAGGCTGAGGATGTTGGGGTTTATTGCTG**TGAA**CAGGGTCTGCAAGGTCCTcacacagtggtaaaaccctg**aacacaaacctccct**acttgggatggcccagccatccacaagt**gtttgcacgtggactgtctgcatggcagattctgagttggcttcacaggtagatgttagatgactgcagggaataatttgcaggaagatccagactgttctggccccatcactgtaacagccttatttcacatggctcaagcagaagaatgaaggtgatcaaagt**gctctctgagcaactagacatgacagaagagagagc**tacctggaaagtgcattctgatcctccctgtgtttcctgcatttgtcaatgagatttagtcagcataacagcaagaaaattgaggcctactcatggctacaaaggttgggtatatgttaggattagcaggttttgttatatttttaacacgataacgtttggtaatattcagaaattggtattttcccactttctaacttcctacctctccattagtggttataccaaacctccacttacatacctgtgattgcattctacacaagctatactcagggggagtgtgactcacagcaattagtgaaa**atgtttagtttgcaaacat**ctaaatagactttgtcaattcatagtggagataacattttatcccaaaagtctttgatttgcctcaactg**ccttttgtgagagaaaagg**agagctcttagaattgcaagtggattttgaaaataaaagtataagccatttggtgacaactgcagaggagagtgcccacaaattt**cagctgtacaatactgatacttccagctg**gggtaaatcagccatccaaccacagccaggatgttactaggcagtccagaaacagttccagggcattttcacataacttttttttcatctttatacagtatctgagctagtgctttagttaggaggctct**aaaatttgacattataatttt**cattacttctgctgtatgtacctccatctagtccttatgtcaatacaaccccaggac

Uppercase: IGKV2-4

Lowercase: Flanking sequence[1000bp]

Red & Bold & Underline: Stem-loop [23]

Blue: Heptamer[47]

Green: Nonamer [6]

id-IGKV2-24[V_gene_segment]

aaactgaatgcaatatgggatccagaaattttatcctaaaacattaaaagtaaaaatggtaaatacatgtatcagtggaaagctcagtgaaattcaaatgtagattgtaactt**cattaataatagtggattaaccattaatg**ttaaagctatttgaagtactaga**aaaatcagtttaaaatgatttt**atattcagcaaaactatccttaaagaaaagaaaagaagccgtgactagcatatatctcctataagaaactcaagaagaaatccttcagaattcagaatcacagtaaacgacaatgaacagta**atttaaatccatgaaattaaat**gaaagcttcataaatatacttacctcaattcatatgttgttgatgttcacgaaaactgaatctttgtgatagatatcagagttgcagttcccttggtaggttagaggcagaagctattgactagaaagg**tgaatgaaggcagcatgtggagaatttcaaatcattca**tatttgtatctgggtagtgaatgtgagtactttatttggctgagcagcgaacatgtttgcactttactcagggcacaatttattttgatttataaaattaacagcaaaccaagaccctttcaacacacatgaagaaaaaaataagaagcaccaaatatttacagaaactcagccgtattaaagagaagtgtaataagcactgggaaaatactaggaagtaaaaaaattgacagtaaacacagtaaacatagaaatatatcctgtcccaatcaggctgcatagattgttatttctgccagttttttctcaagcatacaaaatatgttgttcataggaaaggcccccatacccctgcacatatcatgttatttctacaccactgcacccaccaggggatttgcatattgtcccccagggaggaccttcccttgcaagtctgagataaaagctcagcaccaaccttgacttaactaattaggactcctcaggtcaccttctcacaATGAGGCTCCTTGCTCAGCTTCTGGGGCTGCTAATGCTCTGGGTCCCTGGTGAGGACAGAAGAGAGATGAGGGAGGAGAATGGGG**TGGGAGGGTGAACTCTGGGGGCCCCATTGCCTCCCA**TGTGTGTTCTGTCCTCATGTTAGATGTGTACGTCTTGTACTCCAGGATGGGGCTTGTAACTTTTATATCTGCGTGAGTAAGGCATGTGAGGTTTAGATCTGTAAGAATGAGGAAGATTCCAGAAGGAACAAAGACCAGTGCTCCGGTGAAGACTCTAACAGAGAAAGAGGGAATGGTAGAGGAAACTTCTAGCAC**TCAAAGCACTCTGCTGTGCTTTGAAAATATGTTTTTATTTT**GAAATTATATATTACTAGGGTCTG**AATCAAATTATAAAAATTGATT**TAGCCTGAAAT**AAATAACAGAAGAAAAATTATTT**TAAAATTGTGCTTAAAGTTTCTACATAACCTTGCACTTCTCTCTCATTATTTCAGGATCCAGTGGGGATATTGTGATGACCCAGACTCCACTCTCCTCACCTGTCACCCTTGGACAGCCGGCCTCCATCTCCTGCAGGTCTAGTCAAAGCCTCGTACACAGTGATGGAAACACCTACTTGAGTTGGCTTCAGCAGAGGCCAGGCCAGCCTCCAAGACTCCTAATTTATAAGATTTCTAACCGGTTCTCTGGGGTCCCAGACAGA**TTCAGTGGCAGTGGGGCAGGGACAGATTTCACACTGAA**AATCAGCAGGGTGGAAGCTGAGGATGTCGGGGTTTATTACTGCATGCAAGCTACACAATTTCCTcacacagtggtacagctctgaacaaaaacctcccgctggagtggcccag**ctgctcaagtgtgttgtttctctggggagcag**ttgaacagaatctctatctgtatgagataaacatgttggagaactcagggcaacaggttgcat**ctgagggttctgtcccatgggtgcctcag**ttgtacgtcaggcaaaacccgttcacagccctgtcagctgcaacagccttggcatggcataagccataggaaaccagaggtgatcccagtgcctgcacaggtaatagactgccctg**agggagagcttaagaaaatcctattccaatcctccct**gccttgcctgcattgggaaataagacttaaagaggtaaataaccagacaagtaacccagatttgttgcaac**acttcttgaatatatcttgaggtttagaagt**ttaaagtctacatttaagaggataatatgtggtaatatccaaaaattgaacttttcaactttcctaacttcttatttttctctttcaccaccta**tcttcccaccacatattgatggtggaaaga**gccttccgcacaagctgtcatcatgaggagctggatgagggcaattagtgaaaatcttggatttcagcctcagaatggacttttgtaaattggtgagagatag**aaaatatgaatgctaaaattatttt**attcacttcaattgtgtcttgctgacagaaaaggatagtttttgaaatttcagaagttg**agtttcataaacagaaact**taaa**ctataagacataggttatag**aatttacctcatagaacactgaaataacacagaatgatatgcgatttctttccccaaaatgtaagagt**ttgaagacagtgggcagacttcaa**gaatgggagaattaatggaagatagtggaggtcaactatggcccaataacctgctctttgacttacattaggtacagttgtggatgacagtgactgttgggggttggtgatataaactcagaaagg

Uppercase: IGKV2-24

Lowercase: Flanking sequence[1000bp]

Red & Bold & Underline: Stem-loop [19]

Blue: Heptamer[31]

Green: Nonamer [8]

id-IGHVII-78-1[V_gene_segment]

agagtctggtccggacgtggtggctgatgcctgtaatcccagcactttgggaggccaaggcgggcagatcacctgaggtcgagagttcgagaccagcctgaccaatatggaaaaatccatgtctactaacaatacaaaattagc**caggcatggtagcatatgcctg**taatcccagctactagggaggctgaggcaggagaatcgcttgaacccgggaggtgaaggtt**atggtgagccaagatcacaccat**tacactccagcctgggcaacaagagtgagactccatcaaaaaaaaaaaacaacaaaaaaaaaacagactttgcaaaataatgttcaaccttatcccaaattcattgtttcttaattctgtgcagcatccagacatattattaccttcctcatgatgaattgttctgttt**aaactgagatcagttt**tttctcatattctttggtttctgtct**aagaacagagatcttgattacagtaaggttggttctt**tcctcatactaaccctca**cctcccccagagaaagagcagagattgtcctcactctgagtctgagggagg**agctgttcctgtacaactcagagcctgcagagacccccaggtgcagcttcagtgagtcagacatttctccatgtgggt**gacctccactgccagtgattgctgctgaggtctaattgtgggttaataattag**gacacccttcaggttgtcacatcccaatcctattctgaacatcaccattatcatagacagagatagttcaatgcccattctactgacataag**tttctctttataatttggttccaactatggagaaa**aacgtgaccctatatttgtc**tgaatctaacctcagaatctgctgcattgctctaggagattca**caaattgaacacaagtgagctgttt**attctcataaaaatgtacatatttgagaat**ttcaatgtattgcccagaacctgtgcatgccaacaactgtgtttctcagcgcactctTGGCCTGGTGAAGCCCT**CACAGACCCTCTCCCTTACCTGCGCTGTGTCTGTG**TTCCCCATGATAACGAGGTCTTCCTGCTGAAGCTGGATCCATCAGCCCCCCAGGGAAGGGAATGGAGT**GGATTGGGTGCATAGGTCATGAAGGGAGCACACATTACCACCCACTCC**TCAAGTGTCCAGTCACCATCCCCAGATCCGTGTCCAAAAAAGCAGTTCTTCCTACAGCTGAGC**TACATGAGCAACAAGCACATAGCCATGTA**TTTTTAAGCCAAAGA**cacagtgagggaatcacagtgtgagctcacacccaaacctccctgtg**ggggtgcataggacagaaggggttgctcaggtccccagggggctctcaggacaccaaggggcactcaagaccattgtagaggcacgcaggtcagccgggggctctcaggaaccatcgaggaaaatcaggacaccagagggtgctcggtacagcagggggctcaggaccattgtgaggactcagaaagagcaggttcaaggctcagccttagggcaggtgaagctgggatgaaaaggggctggatgaggggttttgtgtcaccatcatatttcaccactagacaccctccactacatctgttctaatgcatgttctaacacatgtgagtgtttgtatgactagaaaatgatattttggggccaggtgcagtgactcatgcctgtaatcccagcactttgggaggctgaggcgggcagatcacctgaggtca**ggagttcaagactagcctggccaacatgatgaaactcc**gtttctactaaaaatacaaatattagccaggcccagtggcacacgcctgtaatcccagctactcgggaggctgaggcaggagaatcgcttgaacctgggaggcagaggttgcgttgagccaagatgtgccactgcactccagcctgggcaacagagcgagactccattaaaaaaaagagagaaaaaaagaaaatgatat**ttatatcaatatataa**ccatagctaggtgtgtcaagttgtcctctccatctatgtcagccttgtccataaggactaattccctgtaatttcttgaggacctcataaactgtggtcaattatgtaatattcctctcttt**tttctgccttccttcctgccttctctctctcacacagaaa**ctgacgtacacccaccccacaacacacataaatctatacct**tttattacctgatgtatttaataaa**cttgattaatgtgcagcttttcaagcttagttgcttgtgagg

Uppercase: IGHVII-78-1

Lowercase: Flanking sequence[1000bp]

Red & Bold & Underline: Stem-loop [18]

Blue: Heptamer[26]

Green: Nonamer [5]

id-TRAV35[V_gene_segment]

atcctgggatgctcattcatacttagtaggatacaaatctgagattcaagcaaagaaaaatggctccagttcctgtttttaaatacttcttcacttcactctcaactaagcctgatagcccagaatttcatgttatttattcaatatcttca**gaaaaatagtcttttttccacaaaagtggttggaaa**ggagtggacacctgactattccaggtcaaatgggaaatctgagagtataactgcaatttattcaattttactttgcattttctctataaagtctcctacccctaccctctctctgattttctgaggtaccaggtatcctactcactaagacttctggtgcacctgaatttttgttggcatttgcttctgtgggaccttagaatttgcctttcttcagttcatttaagtcagttaccacttgttcattcacctgcattttccaaaacatctcacatgctgttatcctttgcctgttctatgtgtcctttggggttcataacttttaaaaacatttttactctcatttcaccaggtactcagaaaggaatagag**aagtatgtttccaattcaccatactt**aat**tatcagtctagccttcactgata**tttcagagcaattataaggtgcccaggggcagagtttccatctgtcttgttcatgtctccatatttctctaatatatagtgaatactcattaagtgtgttcatcaagtgaaacggagaaatagaagaaactgaaaaagtggcatgaaaggacaaaaatttagcatttcaccatatgggggcagtgtctctattcat**agctaaccttcaaggttagct**caaaaccctgggaaaacatctggtaagagatccgattggctgatatctggaacagagatttaaatttcgtggtagtgtttttagtgtcactctaagcccaagagagtttcttgaagcaaaaaaaaaaaaaaaaaaaaaacccattcaggaaataattctttgctgataaggATGCTCCTTGAACATTTATTAATAATCTTGTGGATGCAGCTGACATGTGAGTCTTGAGAGTTTCTGACTCTTCCATAGCGAATTCTAAAGGGAAGGAAATTCTGTGAGGGCAGGAGGTGAGGCTTTATCAGAGACTATGAAAAGAGGAAAACAGAAGGAAAAAAGCTAGAGAAGGAGGGAGGAAAGGAAAGACACAGATGCAAGGCTGGGAAGTTTGGTGATATAGTGTCCAACACAGCTCTGAAA**AAGGTGATCTAAAGATAACCTTAACCACCTT**CTTTTCTCTTTCTAAACAGGGGTCAGTGGTCAACAGCTGAATCAGAGTCCTCAATCTATGTTTACCCAGGAAGGAGAAGATGTCTCCATGAACTGCACTTCTTCAAGCATATTTAACACCTGACTATGGTACAAG**CAGGACCCTGGGGAAGGTCCTG**TCCTCTTGAT**AGCCTTATATAAGGCT**GGTGAATTGACCTCAAATGGAAGACTGACTGCTCAGTTTGGTATAACCAGAAAGGACAGCTTCCTGAATATCTCAGCATCCATACCTAGTGATGTAGGCATCTACTT**CTGTGCTGGGCAGcacag**tgctccccaaacacctgcagcctgtactcaaacttgcagctggaactctagtctctatgctgccttcagctcttagtcctct**tggcatgaaatgtgattatgcatgcca**cctttgccactgccattctgtcaccacaaactctcctttatccaaggatttgtcctc**caaagattgtgctctttg**cttgatgaacttgcccctctcaattcatttttagtgtatgtaatagattttcttagttatttctgccacgatatgaggagaattgatcaacacacagttacagtagaag**attttaataagtatctcttttaaaat**ggatgatatcaataaagtaaaaatgagagtccaga**aataaatccataaatctatagccaatttatt**ccaacaagatcattcaatgg**gaaaagaatagtcttttc**gaca**agtggtgctgggaccact**ggatagccacagttggatctctacttcacatatgtacaaaagttaactcaaatggattaaagacctaaatgtaagatctagaagtacaaacacttagatgaaaa**catagttgtaaatcttggccaccttggactatg**caacaatctcttatatacaacatcaaaaaaagcaactgaaggaaaaatagataaattgggcattatcagaatttaaat**ctttgtgcatcaaaattcgcaaaatattccacaaag**gaaaatgaaaagataacccacataatagaagaatatatttgcaagtcatgtttatctgataaggatctaatatcaagaatatatattttaaactcttacaatttaacactaagacaaatgccaattttaaaacggtca**aatgatttaaaccatcatt**gctccaaagaagacacacaaatggccaaaaagcatgaaaaaaaagatgctcaacattattat**tcattgggaaaatgcaaattaaaccacaatga**gctgccacatcacacccactaggatgttcataatttttcaaaaaggaa

Uppercase: TRAV35

Lowercase: Flanking sequence[1000bp]

Red & Bold & Underline: Stem-loop [19]

Blue: Heptamer[16]

Green: Nonamer [0]

id-TRBV1-2[V_gene_segment]

tttacttcggcgtttcaaatgccttcttgttttgtgcattcccctgcggaatccctttgtattctattctctcaatgctgtatcaggaggactgagggcagatggctgacatgagggcaatgtgcctc**tttcacctccaggaggagctacgactttagtgaaa**ttcctgagtcatgtggagggagaagagaggcaggagggaggaaaggactaacgaaggaatcaaatgctctggtatcagggtttgctccttgttgtagttgagagaccatgtgtggacaacttgaagacactaaagtccgttccacctggacctgcatctctgcagtccctctgctcatgcccttagggcctcctgttattctagtctgtgttcaggtcctgcatcctaagtaggtgagatgctccctgggggtaggggtcgcagattatttctctcatgtcccctc**gagcacccactgaaaggtgctc**cctccctagttggcattaccttactgggtcttaaaatgatgcacagctggctccagggaagggctccactgagctaggtgaggtgtcctcctggaattcactga**gatgagggaggggagctggagtgtgctcatc**ctgggtccaagacaggcatcgggaaggcatctgcccaaagggaaggggtctgtgtgttagggaggaggggagccataagtagaaagaggaaggggagacccattcattcgttgtgggaagggcaggcagctgctaagaaaaaagcaactgtctaaagaacccgccctgcacacctggccctgagaagctagtctaaaccca**cctcttgaggtgccagtgccaagcttggaaaggaaagagg**aagtgtgagctgtagacactaatagtgacaccaacaggagcagaga**cttcccaagcagcccctgtctcagggccagggaag**cacacccagacgacaagga**cacagagcagggagacacagggtccccctgcctgtg**ccccgggtgaccctgccATGGGCTGAAGTCTCCACTGTGGTGTGGTCCATTGTCTCAGGTGAGTCCTGGGCACAGGTGGGACATTTCTGTCCTTAAATTTTTTGCTTTTTTCATGGAACTGC**TTCAGAAGACTCTGTCCTAGGCTTAGTCTGAA**TTTGGCTTCTTATTTTCATAGGCTCCATGGATACTGGAATTACCCAGACACCAAAATACCTGGTCACAGCAATGGGGAGTAAAAGGACAATGAAACGTGAGCATCTGGGACATGATTCTATGTATTGGTACAGACAGAAAGCTAAGAAATCCCTGGAGTTCA**TGTTTTACTACAACTGTAAGGAATTCATTGAAAACA**AGACTGTGCCAAATCACTTCACACCTGAATGCCCTGACAGCTCTCGCTTATACCTTCATGTGGTCGCACTGCAGCAAGAAGACTCAGCTGCGTAT**CTCTGCACCAGCAGCCAAGAcacagccctgcagag**tcaccgcctccctgtgcacaaacctc**ctggatctaatcagaaaaccgtgggggcaatgcatccag**ctgagcctcagcactcagttcagcattctgtaagacctcagcagacatctcagatcataatactgtcatttattggatgt**ggcaatgctaaattgcc**cagactcctgcagcctgtggcctgagtttgtcttgaatgagactgtgcctg**aacttgaatgtaggacagatgccgtcaagtt**taagtttttccaagtatctttcagtcggtttgtgcctctgggaaatgttgt**gtgaggtatttaacactctggcctcac**caatccaccttccaatatggactgttttttgacttttcttttgactcaagttagaccctggatctcagccctcatcccagctccaaatatgtatggtcagaccttttcttttcttttcctttctttttttttttttttcttttctttccctcccttcctccctccctccctcctttctctctctctctctctttctttctttctttgtttctttgtttctttctttccttctttcttttttg**agacagagtctcactgtct**ggagtgcagtggcgcgatcttggctcacttccacctccatcttccagattcaagcgattctcctgcctcagcctcccgggtagctggaactacaggcgcccagcaccacgactggctaatttttgtatttttagtagagacagggtttcaccatgttggccaggatggtctcaatctcttaacttcttgatttgcccccctcgggctcccaaagtgctgggattacaggcgtgagccaccgcgcccggcaggtccttttc**tttttctttctttcttttttttttgaaaaa**ctaaagaatacccaatcccc**tcctcaggtggctctatcactgcctcaccctgagga**cagtatttccccatggggtttggatcactgttctctagttccctctccctggggca

Uppercase: TRBV1-2

Lowercase: Flanking sequence[1000bp]

Red & Bold & Underline: Stem-loop [16]

Blue: Heptamer[39]

Green: Nonamer [3]

id-IGLV10-54[V_gene_segment]

atgctagccagttggaagtaatacaacttcttatgcgacacgaataaaaatataattatcagttgaaaaacaataacaacaaactcccaccaaattaaacaaaatcaaaagctctcaagaaaaaaacaatgtgccacttattttaaagcagaaacatgcataacgaattcttcaagcttatataacgggctttttggaaagtagactggaggcacattatttcacataaatgg**gtgtcttaggaagagatggagacac**gtataatgg**ctaaaaaatgaaataaatccatagagaggtttttag**tgtctagtgataaaagtactaataatattacgtaattaaattccaaaaaattaatacagaaaaatattcttgatttgggggtggtaacagatgattaagtaaagcgagtctac**atgagctgctgatgcagtggatgacaagctctctgcagctcat**tgttagtgagaagggcactgctttgccagaaaccccttttatttagattt**ctcatggtccctgggtccctctccaacaccatgag**cagcttcagaagaggatcaggataaccagtgactctctggctaccatggcccagttgtgtccgagaggtaagaaaggtttgggatcttggacagagtgggacagtgggagaaattcctgagatggccccaaccagcctggaaaccactcagaacctgcattgaagtttagccatcagggtgctcttgagactcggtttgggctgcaacacagacctgtgggga**caggggtcaccataagccaggccacctccacccctg**ggtcagctcatctcctccctctctgcttcagatctttgagtaaccagcacatggaggcccgggaaaagtcagagagcaagatcttatttaaatgctgtgactccacccccatgaaggctttgggagctataaaaaa**gagacttaggggagcccagtctc**caaacagagcttcagcaagcatagtgggaatctgcaccATGCCCTGGGCTCTGCTCCTCCTGACCCTCCTCACT**CACTCTGCAGGTGAGAGTG**GACCTTACCCAGGGATCTGCACCCACCTCTGCTCCAGCTTC**TCCACTCCCTGGCTCAGTGGA**CTCTGATCCTGCTCTCACATTCCTTTCTGTCCCCTCTACAGTGTCAGTGGTCCAGGCAGGGCTGACTCAGCCACCCTCGGTGTCCAAGGGCTTGAGACAGACCGCCACACTCACCTGCACTGGGAACAGCAACATTGTTGGCAACCAAGGAGCAGCTTGGCTGCAGCAGCACCAGGGCCACCCTCCCAAACTCCTATCCTACAGGAATAACAACCGGCCCTCAGGGATCT**CAGAGAGATTCTCTG**CATCCA**GGTCAGGAAACACAGCCTCCCTGACC**ATTACTGGACTC**CAGCCTGAGGACGAGGCTG**ACTATTACTGCTCAGCATTGGACAGCAGCCTCAGTGCTCAcacagtgcctcaggccagtgggga**agtgagataaaaactcaagagctccctcggcctcact**gaacaggcctcacagagcactgtttaaactggaccacccaaaagacaagggatgcattcac**aatagtgtacactatt**taggactgttaaaaataggaacatgaagtgaaattagcaagcctgccttattaacaaatgtgtgtagggaacatcattgt**ttatatattagtagttacatatatataa**ta**ctagtgcatgaacatacatatgcactag**caattagtaaaaaagaaacactaggtttaactaaggaaatatttattattccagagttttcttatccattccatttgaagatgctgggatcaagtcaccgagggtgctgctctctggccttgagaatccactc**agcagggccaccagcagcccctcccttctgtggcctgct**catgccctgtcttctcattagaatcccct**tccctgctgtcacaggga**tccctggactcctccagat**ccaaatatgtccaatacatttggttgaggcctcctgggccccacattactgtgtaatctcaac**acccacacactcaaa**acagtgttcccatgcgtattcttctttcgactcatcactgt**cctctaacagccatgggcagtgtgtggagagagcaggtgaagttggagctttggggtacagttgacagagatgttgacattttgagaaaaa**gaaaacacctatatagctgttttc**aggataattttccatcaagataaaaatccacgcatgagag**catcactcttccagtgatg**aatgcatggaggatccttctcactctattgttgggtagtgacacctgggaaacttactgtgttatttagtgtgcaga**cagctggtaaaccactcagctg**actctagggaagttgaccaaacagtgtccccactgatgtgtgacttttctccacccatgaggccccactaccctgtcacctcctgagtccacct

Uppercase: IGLV10-54

Lowercase: Flanking sequence[1000bp]

Red & Bold & Underline: Stem-loop [23]

Blue: Heptamer[39]

Green: Nonamer [4]

id-TRBV5-7-2[V_gene_segment]

tgtgtcctgttctaggactttatggaatataaaattatgagccattagctaggatatctgctgaaagaaatatctaagctgcaaagcattcaggccactgtgtgactacttttaggcaccaggcaatttaacccagcaagaaggcagccaagggaatagattttgcaaaccagcacagatggtgacactacctccctctgctgtcctgtctcccataagccaaactcactctgaagttagagaaaagaggagacagttaacatgatacactggagtcagcctgtgaaggcagggtgga**tctggagggattaacataaactttccaga**gcactacatataggatggccttgagggtgttgagtccaaagctggctaggtctgtggtgtgagctgataaggccctagatttatttctggggaatctgcactcttaataatttacagacaacacaggtctgctttggatctgatcagatggactaaatcttggggcctctgtaccgctggccactcaggaaatgggttgggaatgttgcctgggacaggaaaatatagtaagaaaccttggtgtgaatctagcttcagaaagatgatgtgaggagagcaaggaagagctggaaact**gatggtttaatcacaggctcctcaccatc**tgctgataggcaggtgtgcga**gctccagcatggagc**accgcagcactaggtcagaggagggtgatagggtgatggggcagcctgtgagctggggcagtt**aggcagaggagcaactgtatcaccacagaagcttctgcct**tcacccatccctccagctcggcaggacaggtagagagtccagtgtccgtggagcactagacctaaggaaggctgcatagggaagacacaagacagtgacatcacagcatacccctcccatcaggaaaatcaaggcccagaactcactcagctctttcccaggaggaccaagccctgaatcaggtgcagtgctgcctgccccactgtgccATGGGCCCCGGGCTCCTCTGCTGGGTGCTGCTTTGTCCCCTAGGAGAAGGTGAGTCCCGGG**CACAGGACAGCTGCTCCATTCTCAGCTTTCCCACCCCTGTGTCCTCCACTTTACCTTGGGGAGGAC**CTCCAGGCTGTCTCCAGTGCTCATTATACATCTGCTTTTCCCACAGGCCCAGTGGACGCTGGAGTCACCCAAAGTCCCACACACCTGATCAAAACGAGAGGACAGCACGTGACTCTGAGATGCTCTCCTATCTCTGGGCACACCAGTGTGTCCTCGTACCAACAGGCCCTGGGTCAGGGGCCCCAGTTTA**TCTTTCAGTATTATGAGAAAGA**AGAGAGAGGAAGAGGAAACTTCCCTGATCAATTCTCAGGTCACCAGTTCCCTAACTATAGCTCTGAGCTGAATGTGAACGCCTTGTTGCTAGGGGACTCGGCCCTCTATCT**CTGTGCCAGCAGCTTGGcacag**cccagcagagtcactgacattctgtatataaacttccgccttagctttgacttgagaactgcaggccccacccaggtttcactccttcaagggaagcttttagttgtttggaaggcatgtcttgtgtcctactgagggcagacctttcccaaccaatagagcccaggtttcctgtgccctgagtgtgcctgcttctgtgctgcaacttcttgtagttt**gtcacttcctggataacttcagtagaagagtgac**tgttgagccccagatacgtgctagattcttcgtatttgttacatagcttaagggctttccacactcttgcacatcaaacat**ttctattcttcctttatagatagaa**ggctcagggacattgagtcattttccccagtgtctcttggcttgtaaggatcagaagtaggaaacaaactagtccatccattttccacctacccccctgctccatcatcaccttctgcatc**ctggtcagtaagtcagagccctcgcactgccctctagtgaccag**cagggctgcagtagacgctcagatgccttcaggggtcattactatggcctcttcattagcaactttgaggaatgttaaatttaacactttttaaaaatcatttatatgcattccctttgtctttccccagtctgcagcttatgttttcacttttatggtgtt**tttttgatgcacaagagtttaatttaatgcaacaaaaa**ctaaaatcaattttgtttgtgtg**tagacaaaaaccgctttcttatataaatgtcta**agcgtagttttatca**atttattctagtaaaatttaagcttggattttcaccactaagaataaat**gtacctgaagta**tatcttatgtaagaaataagata**gagacctaggaatgggagggaaggacatatgttcagtgaagaagtaatagcctgtcaaagagagccctaaagtagacccccagagctgatggtggagaggtaagtggagggcaaatg

Uppercase: TRBV5-7-2

Lowercase: Flanking sequence[1000bp]

Red & Bold & Underline: Stem-loop [15]

Blue: Heptamer[20]

Green: Nonamer [4]

id-TRGV4[V_gene_segment]

tactttaagtgctagggtacatttgcacaacgtgcaggtttgttacatatgtatacatgtgccatgttggtttgttgcacccatcaacttgtcatcta**cattaggtatatctcctaatg**ctatccctcccccctgcccccaccccactacaggccccagtgcgtgatgttcccccactctgtgtccaag**tgttctcattgttcaattcccacctacgagcgagaaca**tgcactgtttggatttctgtccttggaatagtttgctcggaatgatggtttccagcttcatc**catgtccttacaaaggacatg**aacgcatccctttttatggccgcatagtattccatggtgtatatgtgtcacattttcttaatccagtctatcactgatggacatttgggttggttccaagtctctgctatcgtgaatagtgccgcaataaacacacgctttgcgtggtgcctccaagtgtccttgcagatgccaaa**aggaaggtggttcagttccctaatcagagacttcct**gg**caggacaaggtcctg**gcatcagcattagctgctgcaactctgaa**gggcagggtgtgagcctgctgtgggactgccc**cctcctagggctgggcct**gctgactccagggtcagc**ttccgctacatctgcattccctctgctgcaggcctgagctctcctggcccctggctctcttctgcagagggcctgctctcagctgccacaagagggcgccggagaac**ggacctgagcaagagaggcaggagctcctctgcaggtcctcccaggcaccaatccttctgggagg**gaggaaggcttgggagtcctcttagacagatcctcagtcacttctctctgcttgtgtctcag**gaagaccagctcctcctactgtcttc**tgtgctagggatcacttccttgttgagtggggcctgagtttt**aagaggatcttctgctcctctt**catctggtcccttt**ccttccaaggcctcagagaggaagg**cATGCAGTGGGCCCTAGCGGTGCTTCTAGCTTTCCTGTCTCCTGGTGAGTGCGCTGCCTACAGAGAGGATCATGGGTTTTGTTTTCTTTATTTTCTTCTTTTGCAAGGATTGCCATACTAAGGAATTCCTCATTATATTTTGTGTTGTTCCCATTGCAGCCAGTCAGAAAT**CTTCCAACTTGGAAG**GGAGAACGAAGTCAGTCATCAGGCAGACTGGGTCATCTGCTGAAATCACTTGTGATCTTGCTGAAGGAAGTACCGGCTACATCCACTGGTACCTACACCAGGAGGGGAAGGCCCCACAGCGTCTTCTGTACTATGACTCCTACACCTCCAGCGTTGTGTTGGAATCAGGAATCAGCCCAGGGAAGTATGATACTTACGGAAGCACAAGGAAGAACTTGAGAATGATACTGCGAAATCTTATTGAAAATGACTCTGGAGTCTATT**ACTGTGCCACCTGGGATGGGcacagt**gattcagatccgccctacaccacactgaaaacctgc**cttgtggctgcttctggtacacaag**atagagccgccccctctcatttcctgccaccaaatttaccgtgtgctgaacaagagaaacattgcttaactcctgatctcccctgcaatatcacactcccctggcagcagctgcaccctgttccccaccctcccccaggacttccctgaagaccaagccacca**tctcagctaaacctcagagaagcagcctggctgagagcaaggttctcttagctctc**ctaggacatggc**ggaagcccactcactctgcttcc**taggacaaatgggtacctctagggtccagctgtgaagcaagatattttggacaacaaatgagaagggtatttatactaaattgtgtatcccatcatgatccagaaatcgcagctgagactgg**tgcttattccttatatctataacaatgtaagca**atactataatgaccactgaaacaccaggcctaggtatgcagttgaacattctctccctctgccctcccttttattctctctctctttttttaattctttgcttgcttttattatttttctctctctcccttcctccttccctccgtccttccctccctccttccctccctctctccctccctccttccctcccttcctctcttctttccttccatccattctccttcgttttccttccttccttctttgcttttctcttgctttcctttttgggtacataatcagaatactttttccacaaaatcatttgaaataatttttctctatgtgttcatgtcaacattgatatagctactttaattgttcagttttcttttttacattttggaaatttgttgtattactttgttttgtttcatatc**tgttttgcatatataaaaca**tttacatggaagtgaagtc**aaaatcataaaagatgatgcatttgaagatttt**agcttccattgtagacttctcattcttctt

Uppercase: TRGV4

Lowercase: Flanking sequence[1000bp]

Red & Bold & Underline: Stem-loop [21]

Blue: Heptamer[20]

Green: Nonamer [2]

id-IGLV3-25[V_gene_segment]

agcctcattgaggagacactctccagagggatggggattctcacctgcctcaggccacagctctgaccccgga**cctgtccatgctcagtgacagg**acaataaccaggaacctggctggagtaaccagggaccacccatcctctggagctcacatgctgtgcagagaccatgaccctgggaccctgggaccctgggaccctgggaccctgggaccctggaaccgtgac**agaatccatggggcctaaagttagattct**cagtccactatggttttctgctgctttttgcaagtttacataaaacattgatgagaaaaacataaataaataacagacgagctgagggtgccaattgcttatctcttgtggcaatgagggaatggtatagaaggagcagcaccatggctgaaatgagaccagagg**ccctgttgactccacaggg**gtgtgtggttgcttaaatgacccaactattcccactgaataatcagcaagtgtcccaatgacacagcacagcatttagatgaaatgaacaagaggggactgagtctg**agggaccagggggaggaagatcatgggtccct**gggccacctgc**tgctgagtcaggtgtcagca**tgatctctcggcagggcccagctcagccccatggccaaac**tcctggaaggtccacagctctgctgagcctggcccagga**cagaccccaggtcaacgaggt**ttgggacagtggtcatgaagtcaattcccaa**gaaggtgactgcgatcacagggcttcaaggctgcagctgattttctaaccagagatcctgcttcttcttcttcctcttgtcgcctgtgctgcccccacagctggtttgggtgacatctctccaggaggagtcccagaggaagtaaatttgcataaacaccaaacactgactaccctaaaaagcctgagagagaataagagaggcctggggagcctagctgtgctgtggg**tccaggaggcagaactctgggtgtctcaccATGGCCTGGA**TCCCTCTACTTCTCCCCCTCCTCACTCTCTGCACAGGTGCTGTCCCCAGGCCCTGCTCCAGGCCCTGCTCCAGTCTTATTCCCCACAGATCCCAAGTTGAGCCTGCCCTGAATCCCGAGCAAAG**CCCAGACACAGCCTCTGGG**TGCGACTCCTGGGAATGGGTCCTTTGTCTTCAAGCCCCCTCTCTTGTTCTTCCTTGCAGGCTCTGAGGCCTCCTATGAG**CTGACACAGCCACCCTCGGTGTCAG**TGTCCCCAGGACAGACGGCCAGGATCACCTGCTCTGGAGATGCATTGCCAAAGCAATATGCTTATTGGT**ACCAGCAGAAGCCAGGCCAGGCCCCTGTGCTGGT**GATATATAAAGACAGTGAGAGGCCCTCAGGGA**TCCCTGAGCGATTCTCTGGCTCCAGCTCAGGGA**CAACAGTCACGTTGACCATCAGTGGAGTCCAGGCAGAAGACGAGGCTGACTATTACTGTCAATCAGCAGACAGCAGTGGTACTTATCCcacagtgacacaagcagacagggaagtgagacataaacccctccccctctatttcatcctct**tcctcctgccccaggagga**ctgtggacaaagccatgagtaggcttggccaaattcacccaaatctgaacccccaggctgcccttccctccagccctccaggcaggcactgcagagtagtcagtctggagtggatttgt**gcctggcaggccaggc**tgtcctgtgtttctttgcctggggtttgagttgtcgatggagggtctgactggaaagacagacacagggagacatccatgcagg**catgtccaaagggacatg**gggaactctcaggtctcagtt**gaatgcagtcattgttgtatcctctgggaaagcattc**ctctagtgcatctaagacctccagacccacagagcctgagatcaggaggagagaatgttaaatttttgctaatggaccactgaagacatttttgagtggagccactacaatttactccttgattctggactcatggatcctgactgtgggagaaacagcaccatctattgggcactgatgcacagaacatgccctatcctggaagacattgtccagccctacacatacttgccacttgcctggtgcagt**aatggtgtctccaagcagtcattaaaccatt**catataagccatatgtctcagaatgtttgggacaagagtaagaatcttgattgccagaagaaaaggcccatagccctttatttgcagtgaaatgaattccttggtcagaacctcctgtggaataccatcatgatgggtgcgtcattctgtaatccagagacaacagttttggcagaagcattgctgtcaatgaaagcaaatcctctg**tagaataaatgtcaattcta**gtaagaaaaaagagctattcctgccatgatggaagagtgcaaacgaaattaatattaagcaggcccctggctgttttccacagagaaagtcatcctatagggatctctgtgttagtc

Uppercase: IGLV3-25

Lowercase: Flanking sequence[1000bp]

Red & Bold & Underline: Stem-loop [18]

Blue: Heptamer[53]

Green: Nonamer [4]

id-TRBV7-5-2[V_gene_segment]

ggaagaaaaaaatggatgtttcccatatattaga**atatgcatatttttcataatttcccattttttgttgcatat**gattgaattttcagacagataaaataggtagaatgttatgaagtgatatttatgagatgttagctgagaagttttttgaggtgtttaaggacctaaattcagagtttccagaatcacaatgaatctcaaataaggcagtcaagtcataagcaaattgtagaaca**tgacggacaaagagaagattttgaaagccgtca**gaaagaaaagacaaatacgtcccataggaatagatggagaacaacagtaagttaagacagtggtaaaaataaccattagctcaa**taatcattctccttcatttaaggtaagattgatta**aaaaaaaataacttttagaaggggaacatgaaaaaatgactgctgtgagtttgctatcaagg**tttcttttctgaattacagttaagaaa**ggtgggaaatgatcccagaagaaaggcatga**gggcagttcgtgaggtaacccaagtgatggggctgccc**cgttgggcacgtgtgactgagggaatggaggaggctggggcataaatggggatggcagaggggaccctgaattgcaggatagacaatgagctaatgccttggtgccttgttttggggatgctgttggca**catcctacaagacatgcccagcagacagaggagtggctgtaggatg**agaagatgaactcagagatgcagtgtgaggcctct**ggctccagacagcacgggagcc**caaagcaatgagccaggcattgatgttgttaaaaaggagcttataaatatttaaagtgtgttctaatataaatactgtgaccctgacatcctggggattgagagaggaagtgatgttactgtgggaactgccctgtggagacaaggacattcctcgtcctctgctcctgctcacagtgacactgatctggtaaagcccccatcctggcctgaccctgccATGGGCACCAGGCTCCTCTGCTGGGTGGTCCTGGGTTTCCTAGGGACAGGTGAGTCCTCAGAACATTAAGTAGTTTTATTTTTTCTCTGTGTGTAGGTGTGTGTGTGTTTGTGTGTGTGTGTGTGCGATGACTACAGATGTTTTCCTTATTCTGTTGCCAAATTCTATTTCCACAGATCACACAGGTGCTGGAGTCTCCCA**GTCCCCAAGGTACGAAGTCACACAGAGGGGAC**AGGATGTAGCTCCCAGGTGTGATCCAATTTCGGGTCAGGTAACCCTTTATTGGTACCGACAGACCCTGGGGCAGGGCCAA**GAGTTTCTGACTTCCTTCCAGGATGAAACTC**AACAAGATAAATCAGGGCTGCTCAGTGATCAATTCTCCACAGAGAGGTCTGA**GGATCTTTCTCCACCTGAAGATCC**AGCGCACAGAGCAAGGGCGACTCGGCTGTGTATCT**CTGTGCCAGAAGCTTAGccacagtgtggcacag**tcgtgtccttcctgctcacaaacctcatccttctctctccttgcatctcctagagacccttaatagaggcctatctttgatcctcacttttcgtgggaaataagtagatttggacatcggctgtcctttgggtagaaagagaccacagattcattcctgaaacacagtgactgcaaatgtaggtggtgaaaacaatcacgtcccactgccctctaggagtgc**ttggagccagctcactgctccaa**acggagtgggtgtcttagccttggccttcagggcagacatgcatcttctataggtcttggaggctgctgtgttacccacatatatgaggttgtcaagg**gcaggaaacatgctcttctcctgc**atattttggggcatctggaaggtctgaggctacatccc**caggaatatctttcttctaaagcctcttctattcctg**tcaccttggaaatttttgcaacaaaatatcgaacctctcttcctgttta**aagtaaaggtctttgcaactttcgtggtctttactt**gataaatacaatcatggtaacaataagacttcatttcttctgcctactttaagccacttgtatcctttattttatttccatttgccat**tgctactgtcctgatagacagaagca**tgcattcaccactgctgcctgttcacatcgattccctcaggaaatctgatttttagactctgagtgttttcattgttgtccaactcatttgatttgaaataattttaccgaggcctttaactcaagaagtgttttatttataatattga**atctattcctttttattttattttttataatagatagtactatatagtact**tgttataaatagaagtacaatgattatattgcaatagaatcttccacctatctgtgggtgctgctgcagtgtgtatctgtgaaagcgaatgcactggtcagagctgatgtgattatggatcgtgggtttctgggagtccttgg

Uppercase: TRBV7-5-2

Lowercase: Flanking sequence[1000bp]

Red & Bold & Underline: Stem-loop [18]

Blue: Heptamer[37]

Green: Nonamer [5]

id-TRAV12-2[V_gene_segment]

aggttaaacactgcgctttacctcagtgccgggcttggctctcttccaagtgtaccttcctttctttcctattctaaagccttttaaaataaatttccacccttgctctgaaaaaaaaaaaaaaagaaaggctatgtactgaaggcaggtttatctctattataccaacccatgtaagtactgttttgacatgcatgtgtcattggtgcaagatgcaatggctattcccaagcccctctattccctggtcttgttaaaaaatgttgctaaggctgtgagctagaaccatgcttgttaagctagattttaggtcattctg**ttttgaaagaattattttgaattacaagaaacttctttaaaaa**ttggagataacagaatggtatagggatgtgtggcctgacatgaagagcta**tgcaatgtgaaatgctccttgaattgca**agaatctcacctaaatttttataatagaatacttcatgactgttatttcagtgtaggaaagtggctattagaaacaactgctcttactcatacaacaattgccatttgatatcatacacatacacacataaacatccccacagacattaacttggagcatttgcttctcaaatttttaactgatgtgtcagatttttaaagtgacttgtttacataataaggctctacattttgagtatatgtttgatcactatcatgattccacaagatggcagtgtgttcttagggac**cttgaggacagctctcaag**tgctcgtgtgtgtgtgtgtgtgtgtgtgtgtgtgtgtgtgtgtgtgagagagagagagagaggaggaggaggaggaggaggaaggatgaggggaggggcttagtggtgaattaggggtgttaaaaagagcatcatttttttgaactggtaaagcagattctttttatgatttttaaagtagaaatatccattctaggtgcatt**ttttaagggtttaaaa**tttgaatcctcagtgaaccagggcagagaagaATGATGAAATCCTTGAGAGTTTTACTAGTGATCCTGTGGCTTCAGTTGAGCTGTGAGTTTGGGGCATCTTTATATAGGAAAATAATGT**ACAGTATCTGGAGTTACTGT**CTATTCTAGATCTATAGGTAGAAGCATGATTTTTTTTCTAACTAAGGGAAAGTAGGGGCACCAGTGAAAAGAGGATTTAAATTCTGGGGAGTAAGCATCTATTACCCAGCCAGTCTTCTCTGTCTGACCACTGTCTTTTTCACAGGGGTTTGGAGCCAACAGAAGGAGGTGGAGCAGAAT**TCTGGACCCCTCAGTGTTCCAGA**GGGAGCCATTGCCTCTCTCAACTGCACTTACAGTGACCGAGG**TTCCCAGTCCTTCTTCTGGTACAGACAATATTCTGGGAA**AAGCCCTGAGTTGATAATGTTCATATACTCCAATGGTGACAAAGAAGATGGAAGGTTTACAGCACAGCTCAATAAAGCCAGCCAGTATGTT**TCTCTGCTCATCAGAGA**CTCCCAGCCCAGTGATTCAGCCACCTACCTCTGTGCCGTGAACAcacagtgctccccagacacctgcagtctgtacccaaacctgccatgccccaggaatgcctgatgtagagcttagactgcagggtag**tgaaactccccttgctctctagtttca**agtggaaattataagaaccagtatggaggattaattaattaggg**aagtattttcataattctgaaaataatt**gaaaccctga**aaaaaataaaaacaaaaatcttggtctggttgacatatttttt**agtcctgtttttccactatagttttaaatatttttttaattgaggtctgactgataaacaaaaaa**gtatgtgtatttactttatacatac**agcacaaactatgccataacatatccataatttctaaaagtttacttcctccct**ccttttttaaaaattgaggtatgattgctatataaaaagg**cacacatatttaatgtatacatcccgatgctcagtctagttttatgccacctctttatgagattggttttatggtttttaaattacataacaagaaaatattttcacacaatgacagatattctctcatagttcccaggacaccttcgtctttccttctttagttactcaaagtgtttgaaagtcattaatttgggaatgcaacagcaaaacaggaatgcaaagcagagaaaaactcacaaacaaa**tgacttcaaaaattctacaagtcaaaaattctttttttttttttttttga**gatggagtctggctctgtc**gcccaggctggagtgcagtggcgcgatctgggc**tcactgtaagctccgccttccaggttcacgccattctcctg**tcccagcctcccgagtagctggga**atacaggcgcctgccaccacaccggccaccatgcccggctaatgttttgtatttttttttttagtagaaacgtggtttcaccgtgttagccaggatggtctcgatctcttgacctcgtgatctgcccgcctcagcctcccaaagtgctgggattacaggtgtgag

Uppercase: TRAV12-2

Lowercase: Flanking sequence[1000bp]

Red & Bold & Underline: Stem-loop [17]

Blue: Heptamer[44]

Green: Nonamer [4]

id-TRGV6[V_gene_segment]

agaggtggtggtaatctgtgcatgccctactgaattttcacaattatattactgtacttgttttgtccccttatttccatctgtgaatccaggtgcactgcagatattagtctgctttctgagggtagggatgacaaataaaatatacaatacccagggaatttggaagttcagatatataacaagtttttagttcaagtatgttccaaatatttcatgacatggcacccagtactactttaaaaacaaacaagcaaacaaacaaacttacactcctactttaatgaacaaaaattgtctaataaagagatcacacccacaggattacaggcacccccccccccgccccccgccacacacacacacatttaccttagagcagagtccaaggcccaccaacacacatgatcctcagtccttttaataaagaacattcaaccctcctcagagtggccttgggcagttgcctccaagtgtccttgcaga**tgtgaaaaggaagttggttcaca**gccctaatcagaggcttcctgg**caggacaaggtcctg**gcatcagcattagctgctgcaactctgaa**gggcagggtgtgagcctgctgtgggactgccc**cctcctagggctgggcct**gctgactccaggggtcagc**tttccccatatctgcactccctctgctggaggcctgaactctccatgcccctggctctcttctgcagagggcctgctctcagctgccacagagggtgctgcagaactggcctga**gcagagaggcaggagctcctctgc**aggtc**ctcccaggcaccacccttctggaaggaaggcgtgggag**tcccctaagacaca**tcctcagtcacttctccctgcttgtgactcagga**agaccagccttccccactgccttctgtgctagggatcacttccttgttgagtgggacctgagttttgagaggatcttctgctcctgttca**tctgggccctttctttcaagaccccaga**gacgaaggcATGTGGTGGGCCCTAGCCCTGCTTCTAGCTTTCCTGTCTCTTGGTAAGTGTGCTGCCTACAGAGAGGCTCACAGGTTGGGTTTTGTTTTGGATTTTTCTTCTTTAGCAAGGGACGCCATACTAAGGAATACCTCATTATATTTTATGTTGTTCCCATTGCAGCCAGTCAGATATCTACTAACTTGGAAGCGAAAATAAAGTCAGGCACCAG**GCAGATGGGGTCATCTGC**TGTAATCACCTGTGATCTTCC**TGTAGAAAATGCCTTCTACA**TCCACTGGTACCTACGCCAGGAGGGGAAGGCCCCACAGCATCTTCTGTACTATGACATCTGCAACTCCAGGGATATGTTGGAATCAGGAGTCAGTCAGGAAAGCATGATACTTATGGAAGTAGAAGGATAAGCTGGAAATTTATACCTCCAAAACTAAATGAAAATGCCTCTGGGGTCTATTACTGTGCCACCTAGGACAGGcagagtg**attcagacctgctctacaccacactgaat**a**tctgtcttgtggctgcttctggtacacaagagaga**gccgtcccctctcatttcctgccctgaatttactgtattctgtacaaagagaaacagcttaactcctgatctcccccctaatattacactctcctgtcagcagctgtaccctgttccccaccctcccccaggactttcctgaagactaagctgccatctcagctaagcctcagctaagcagcctg**gctgagagcaaggttttctcagc**tctcctaggacatgggggaggcccactcactctgtttcttagggtccagctgtgaagcgagatattttggacagcaaatgggaagggtatttatactgaatcatgtatccaatcatggtccagagatcgcagctgagagtgctgcttactccttatgtctataacaatataggcaatattataaataccactaaaacactaggcctagatatgcagttaaacactctctccctctgccctcccttttattctctctcctttcttacttccttgcttgcttttattattttttcttccctccctccctccctccctctctccctcccttccttccttccttttttcatttcttccttccttgcttttctctctctttctttcctttttgggtacataatccgaat**attttttcaacaaaatcatttgaaataat**ttttctttacatgttcatgccaacattgatatagctagtttatttgttcagttttcattttaaaattttagacatttgttgtattattttgttttgtttcatatttgttttgcat**atgtaaactatttacat**ggaactaaagtcataattttaacgaagatgcatttggagattttaacttacattctagatttctttttctctctcttttttttttttttttttttttgagacggattctcagtctgtcactcaggctggagtgcagtggagcaatctcggctcac

Uppercase: TRGV6

Lowercase: Flanking sequence[1000bp]

Red & Bold & Underline: Stem-loop [15]

Blue: Heptamer[17]

Green: Nonamer [1]

id-TRAV6[V_gene_segment]

cttctatctgtccatccatctgttcagctgtcattgatctctccatggaatggcataaatgcatctaaggaagtgactacaatacatccatcctggcattaaaaagtagaaaagaaaggatt**tctgatatatataatcaga**gggctgctgtat**tttaaaaataattttaaa**c**accctctcagagggt**attctaggggtaaaaaacccttcactc**gtttgaattctcgctgaatttaaac**tgaatgtctacttattatagcccaacaagataaag**aaaaaaggccttttgaagtcaaagcaatactttttt**ttcacctgtgggaaggcctattccctagatctgggaaagtttcctagaattcagccatacattttctgtgttttacattatctagagcacatgacatctgcaactgtggcaagaaggacagagtagttcaactgggcaatgaaatggagaacttggaaaacttgatcatttgacctgtatccagaattaatattgtcaattccatgttatgtactgagactcaattagcattcttctgg**acttgttacttacaagt**gtggtctgagaaccaactgtatcagcatcaccagtaaacttgttaaaatgtggaatatccagccccacctcagacctaatgaaacagaattcacagcttaacaaaatccccaggtgatttgcatacataattaagttttagaagccctgacctgggagcacgttccattattataacaactttctgaacacaagagggcagtatttcctaccccctcagagcctgtctcagat**gtgaaccgaagttcac**tgcatagctggattaggccagtatgtgtaaggggctgaacaggcttgccattgattggctggat**aggaaggccagaacttccttctaggggtagaag**aaccccagtaacacctatcaaactaaacagaatggctttttggctgagaagcctgggtctacatttcaggccacatttggggagacgaATGGAGTCATTCCTGGGAGGTGTTTTGCTGATTTTGTGGCTTCAAGTGGACTGTGAGTTGGAGATTTCCTGTTGATGGAGACAGAGAGATCCAGGGCTGCAGTGCACAGGATGAAGGGGAGGGGCTAGGATCCATTCTTCCCCATCTTTGGTTATATCTGGGGTTGTGGGAGGAAAATAGACTCAACAGTGTCAGTGTAGTGACTAACCCTCCTACTTTTGGTGTTGTGTTTCTGCCTAGGGGTGA**AGAGCCAAAAGATAGAACAGAATTCCGAGGCTCT**GAACATTCAGGAGGGTAAAACGGCCACCCTGACCTGCAACTATACAAACTATTCTCCAGCATACTTACAGTGGTACCGACAAGATCCAGGAAGAGGC**CCTGTTTTCTTGCTACTCATACGTGAAAATGAGAAAGAAAAAAGG**AAAGAAAGACTGAAGGTCACCTTTGATACCACCCTTAAACAGAGTTTGTTTCATATCACAGCCTCCCAGCCTGCAGACTCAGCTACCTACCTCTGTGCTCTAGACAcacagtagtgccctggcagctgcttcctgcacccaaactctgctaactctcacaatcagagctcatg**gctgtgctgtctcccaaaggctaatcacagc**tcctgacagaatgggggggtgttag**ttgaaagggcaattttcaa**aacaaacaaacaaatg**ctctgaagaattcctcagag**aaaggagggaggacttgcttttaaaaaatgcataaattctgcttagtttttgtgactagcctagagcagcgtgataaatttatggagttaaacacattacacttaagcattaaaaca**tggattcttctagatgaaaaaataatcca**aacatttttgagtcccttcccaaactgaacactgaataatgatgatgatgatgataatga**taaaaaattgaccttttta**aaaattgttcatcttatgaatggataaccatgctatattggaaaac**atgccataagtcatggcat**tgcagagataaatttagagccgttgtttactatgtatgtggtatcagtgagtaggtcaatatacaagtcttctttttcatttgtttcttaagagacagggtcacactctgtcacccaggctggaggctggagtgcaatagcacaatcacagctcactgcagaccacaactcctgggcttaagtgatcctcctgcctcagcctcctgagtagctggaattacaagcacacaccaccatgcctgggttgtacaa**ctcttttaagaaagag**aacttacaccttctcatgttaggcatgggttctgagttgaggacttctccgttattattgcctgcattgccatttctatcaatttctgtcatctatgtattaccttatttcttgatctatttttagattattgccaaaacagacagatcccataggtcatg**catgttcagttactaacatg**gcagaatctgtgggttttgactaactctgttttaggctagggctgatatttctgtctgctttgggccacatacagtcactgcc

Uppercase: TRAV6

Lowercase: Flanking sequence[1000bp]

Red & Bold & Underline: Stem-loop [19]

Blue: Heptamer[40]

Green: Nonamer [4]

id-TRBV10-1-2[V_gene_segment]

tgggcctcctccctgtctccacttctgattctgtgtcctctctcactggagcatgattgtgggaattacccatccaggttgcatcaagatg**tgcaatagatagcctattgtatctgaaataactattgcagtgctatggactacatgtgagttttgcaccatagcac**gtttacaaatccatcaaccttc**tcccatttggtgaatataagatctgctggatggga**atagaggattttcctgtgaccctagagactccaggtaggtgtagttcttggtgaacatttcatccatattcctaatgagcctcctcttttcttcacaggaagtacagtcactcaggggacttagatggggca**gtgtttgtggcttcacctgaagcaaagggagaaaacac**ccccatgagaatacacagcagcctgagcttggatggtgacattcccatcaggtgtaatgaggtgctacagcatgtaacaagtc**ttcaagcggtcatgaccttgaa**cctggaacttggaacttggagacatttctggatgtcatttaaccgagggtatccc**tgtatgctggcataca**agtgtccagaggttaaacgaagaggaaaat**aattcaacttggagttgaatt**ttgtgcaggaa**gtcataaaagtcaccaggactgctatgac**aatgttctttagcctacagaaatattcataaaagtgaaaatagtattcaaataaagaatttctcatgatggaaataaaatatacatagaagaacaaattcccctattaaaatcagg**gaaagagcttaggagtttattagatttctttc**aaaacagacctaa**aaattattcaactctgttataaataattt**gttc**ctgcggggaagagaaggggcatggcaggttagctccgcag**ttcaagcattcagcaggaa**cagtgacatcatcaagtcactg**agagcccaacttcagtctgcccacag**cagggctgggagacacaagatcctgccctg**gagctgaaATGGGCACGAGGCTCTTCTTCTATGTGGCCCTTTGTCTGCTGTGGGCAGGTGAGGGCTGGTCACAGGAGGGCCTCCTTCCCTGGAAT**TCCCAAGGCCTCAGTGCAGGCTTTTCTGTTGGGA**TGACAGCA**TCAGCATCTGTTGTTCTCTATTACAGGACACAGGGATGCTGA**AATCACCCAGAGCCCAAGACACAAGATCACAGAGACAGGAAGGCA**GGTGACCTTGGCGTGTCACCAGACTTGGAACCACAACAATATGTTCTGGT**ATCGACAAGACCTGGGACATGGGCTGAGGCTGATCCATTACTCATATGGTGTTCAC**GACACTAACAAAGGAGAAGTCTCAGATGGCTACAGTGTC**TCTAGATCAAACACAGAGGACCTCCCCCTCACTCTGGAGTCTGCTGCCTCCTCCCAGACATCTGTATATTTCTGCGCCAGCAGTGAGTCcacagtgctgcacagctgcctcctctctgcacataaagggcagttagaatgactgaggttg**cctgtgctcccaagtcccagccttcacagg**agtcggagagccctggctagcctgggggccatagagcaggggccatataaaccttgatgttgaggtccattcctaccccagtctcagaccaactggaggtcaccccaacacccttagcttctgcagttccctcttt**ccttttgtaaaatggaaaagg**catctttagtcgagattcctaatttcgttgaaaatttgaaacccaccatattcttctctaccctaagggctgggtgacttttccacataccctttcccctaccctctactgcctttcaactcccaatcataaatctttgcccatgtggtctctgctcctgggg**gccctttattctaagggc**aatagacccacgaggtcctatatcattagccatttacgtacctcttcatcatcattttgcaccatgttttccctggctctctgctccagccacactcccttacatgcagttatttcttcatatgtgtctgttgttttccaccctggggattgtttatataaattctgtcttctgcactgagcatacataccacccacttagtcctacatctctacttcatttcttaaatta**aagtgtatttcaggcacactt**gctcttaattgtaaaaagcaaaaaagaaagtttactagaaaaatttaagaaaagcattttatgcattcagttgagggatgacatttttagaggagacaaaaagccacaagccaaaagaacaacattcataacgctagttgcattaaattgtttcaagccaaaacacaccataaaaaataaagttaagctacagcctgggaaaggcatgaggaatgcatacaaatgatgaaaaatcagtgtgacagagaac**attcagtgtaacagagaacattcagaacatcactgaat**acctgtagctcctccatggtttctggtctcc

Uppercase: TRBV10-1-2

Lowercase: Flanking sequence[1000bp]

Red & Bold & Underline: Stem-loop [23]

Blue: Heptamer[44]

Green: Nonamer [5]

id-IGKV3D-25[V_gene_segment]

aagaatagagaatgataacagttttagacttcgagggtacttcaggaatgggaacacacaagga**acacagtagatatgagctactgtgtctaccgtattagaca**atcctcctgcattttaacttacatttggtgaacttcagatgccagtgggtgttgcaagttgatgacataaacctaaaatcaagtcccgatgtatttcttgtgaagagtcacagagttgaagagaccactgcctggcttcctgagttgagcctgtgccactgcaagtttcatgggaaaatggtgagaaaaggtgtttaggtttgtaatcatcaaaggtttagtgaagttcccgtgcaaaaagactcaatcccattcaactcagcatgaagagatcttctagaactcattctgcccctatagcagagatgatgtggtcaggtttgtggagatgaacatgcagtgacgtgttaaccaaaaggcccagcagaaagggatagtgatcaatgtccatggttgagacatatacaa**aggatgatttgcacagcactggaggaaaaaacatcct**gagtgttggagatttctgccttcctacccaaactcccacagcctgtctcctgactcagaactcagcaaatgccacatgaaaaatctggctat**ggttgcacagaaattgccctgtgcaacc**ttcaaagctcagttatgttctctcttcttgctgggtttctctgtctgtg**ccaaatcaaaatttctatccgaacgcacatttgg**taaaaaaaaaaggcagaaaaacaaaatgtcagatgaacaccaact**caactagaattagttg**tttttttactctagaattttagatcgcctagtcctcatttcttccattactttctaatgtcactaataggtaactttgcattttatttgtttttattggcacattgtaat**tgtacatatttatggagtacactttgatgtcttgatatgtaca**tatgttgaaaaatgatcaaatcaggatatttaggATGTGCATCAGCTCATGTATTTATCATTTCTTTATGTTACTCTGGGTCCCAGGTGAGGGGAATATGCGGGGGGTTGTGGGGGGTTCTGTGTGTCAGTGAAAACTCTCTCAACTTTGTTGGGTCCTTCAACCTGGCATCTCTGCTCAGCAAGCACAATAATTAAAGCTCAATGTAGATCAATGGTCTTGACTCTATTGGAAAGACAATGGGT**ATAATCTAGATTACATGTGTGATTAT**TTGTTTTTCTGTTTAAACTGTTTTTTCAA**AAAATAAATTTTATTCTGTATATTTT**AGGTTTGCAATATGATGTTGTGGCATACATATAG**ATAGTAAAATGCTTACTAT**AGTGAAGAGAATGAACATTTATCATCTCACATAGTTACTTTTTCGTCTGCGACAAGAGCAGCTGGCAGCTGGAATCTACTTATTTAACAAAAATCTCTAATACAGTACAGTTTTATTAACTATAGCCCTCATTTTCTAC**AGTAGATCTCTATACTCGTTATTCATAAATATCTACT**ACCTTGCATCTTTGGACCTACATCTTCCCATTTCTTCCCGTTCCACCCCCGGCCCTTAATAACGACTGTTTCATTCTCTATTTCTGCATAATTTATCT**TTTTACTTATTTAATTTATTTTTAGGTTTACATGTAAGTAAAA**TCAGGCAATATTTTTATTTCTGTGTCTCGCTTTTTTCACTTAGAATACTGTCCTCTAGGTCCATCCATATTACGGCAAATAGAAGAATCTTATTTTTCG**GCTGGGTGTGGTGGCTCATGCCCGTAATCCCAGC**ACTAGGCGGGTGCATCACCTGACATCAGGAGTTGAATACCAGCCTGGCCAACATGGCAAAACCCTGTCTCTACTAAAAATACAAAAAATACAAAAGAAAATTAGCTGGGCGTGTTGGCATGGGCCTGTAATCCCAACTACTCAGGAAGCTGAGGCAGGAGAATCGCTT**CAACCTGGGAGGAGGAGGTTG**CAGTGAGCTGAGATAGCATCATTGCACTCCAGCCTGGGCGACAAGAGTGAGACTTCTCTAAAGTAAAAATAATAATAATAATAATAAAGAATCTTCTTGTTTAAGGCTGAATTATATTCCATTGTGTGTGTGTGTGTGTGTGTGCATTTATGTATACCACATTTTCTTTATTCATTTATTTGTCCACGGATGCTCAGGTTTTTCTCATTTCTTGTGAATAATGCTGCAATAAACACGGGAGTGTAGGTATCTTTAAAAGAAGGTGGTGATTTCATCTCTTCTGTGTATGTATCCAAAATAGGGTCACTGTTGGGTTATAAGGTGGTTAGGTTTTGAATTTCTTTAGGAACCTCCATACTGTTTTCCATAATGGGTGCACCAATCATCATTCCCACCAACAATGTACAAGTGTTTTATTTTCTTCACACCCTCATCAATATTTATCTCTT**GTCTTTTTTATAATAGCCATCCTAAAGAC**TGTAAGGCATTTTATTTCTAATCTCAGATT**TCACTGTAGAAACAGTGA**TGACACAG**TCTCCAGCTTCCCTGTCTTTGTCTCTGGAGA**AAAAAGCCACC**CTGACTTGCAGGGCCAGTCAG**TGTTAGCAGCTACTA**AGCCTGGTACCAGAAGAAACCTGAGCGGGCTCCCAGGCT**CCTCATCTATG**GTACAGCCCTGATTTGTGATAGTGGGTCAGGACAGGGCTTAC**TCTCACCATCGGCAGGCTGGAGCCTGAAGATTTGCACTTCATCACTGTTATCAGCATAGTAGTTGGTGTCCcatactgattcaacatgcaacaaaaacctccaggagacctaaggtgtttatttgattataccacctgcttcctttttagtcatctgatgtggtgctgctcagttttagcatctctgctttgattggaaattctgaggttctcaaaagtaattccttataatatttatagtttcactgatggatttttttctcagacccaaatgcacagccaggttcaggcacaatttcatggtcaaggccattggatcagactcacatgagtggacgcctctaaaggtcctggccagtgcgataaagtagcaacgacaatgataaagaagaagaattagaaaggcagaattaaaggtataacaattcactgatgaaagga**ctgtgtgggggagaaatttcaaattgtctacacag**aaattattagaatcaatgagatacatagtaaatagattaaagatca**atataaaaatatatttatttttatat**aaatgtaacaactgctaaaaattaaaattatgaaacacatcatttatagtagcatcaaaccaagaggcattagaaaaaactaacaaaatatactgacagctccagaatataattcaaagaaatcaaagaaaatctaagccaatggaaggccatagt**actttcaaaggttgaaagt**cccagtattgtaaagatgttcattcttcacagattgatatattaaagataatcgaaatcaaaattcaaactgtttgtaaatgga**aatgtaaaagctgattctaaatatcatattacatt**gttaaggagtttgaataaccacaataaacccgaagaagcccaagaatgaggacttacactacttggatactcgaaccgtgctgctgatgtgagttcaaacaaattgcctaatgcaacagaagagggtattctgagaatgactaacacatatagtgacactttttagggacaaaggtgacactgcacaacagagtgaaaacaataaaaagctgtccaacatcaataatca

Uppercase: IGKV3D-25

Lowercase: Flanking sequence[1000bp]

Red & Bold & Underline: Stem-loop [24]

Blue: Heptamer[45]

Green: Nonamer [6]

id-TRBV12-1[V_gene_segment]

ccttaggatcggaaagtactctggggtgcaggggactcagcagtatgcctctgtgccagcagcaaaaatgtggccctaaactgctccttcttggttagcaattaacccagttgaccaaccc**acagagcagaggcttctctcagctctgt**acactctacaccctgagaggaagtcggcttctgagatgcaagactcttttataaggaaaagtccagttgtaaagattaagaaactttgagctttgctagaggtaaaaataggatgtaagtcaactctggagcatttaatttccaaatgtttatgaaaaagacaatcttcttcctgttttattgtatggatagagaatgagagagattctgtactctgggctttgcaaactcactagagttcaagaataaatacactcagagaggtctttcactgcaaagaagagctttgaaatcgaatgaactttcaggatatcagttcggctcctgtatctctaacaacgcagttgacatgcttgcccttcctagggatctcaggggccaactcctaactcacttgtggataattgaagtgcattcaaaggagtaggagcacagagacatcagggagaggaataggagagtttgcagagaatctggcttgctgtaagacaatgag**ttcatcttgaaataattggagtgtgaggtgcagatgaa**tatagttggcaagatcctaggtaagatatactatagagaaggctttatgaattggtaatatcaag**acaaatccacagagagcctccgtaggtgtgcatttgt**gtctcagatcaattatagttcagtcctgcctgatt**catctcccagagatg**cag**cctccacttaaataaggagctttcaaattggagg**tggtggcccat**tcagtgacgtcactga**cagatgcatcttgtgtggataaaatgtcacaaaattaatttctttgttcatgctcacagaggccctggtctggaatgttccacttctgctctcactctgccATGGGCTCTTGGACCCTCTGTGTGTCCCTTTATATCCTGGTAGCGAGTGAGTTCTCCGATATTTATTATGGTCATGCTGGACCTCTGTCTAGATGACTCCTTATATTTTCCTTATTCTTTCCCCCAATTCTGTCTTCTTTTATAGCACACACAGATGCTGGTGTTATCCAGTCACCCAGGCACAAAGTGACAGAGATGGGACAATCAGTAACTCTGAGATGCGAACCAATTTCAGGCCACAATGATCTTCTCTGGTACAGACAGACCTTTGTGCAGGGACTGGAATTGCTGAATTACTTCTGCAGCTGGACCCTCGTAGATGACTCAGGAGTGTCCAAGGATTGATTCTCAGCACAGATGCCTGATGTATCATTCTCCACTCTGAGGATCCAGCCCATGGAACCCAGGGACTTGGGCCTATATTTCTGTGCCAGCAGCTTTGCcacagcactgcagaatctccccatctctgtgcagaaaccctggtgcttcctcttctccccacagctctcagcagtcgtcagcaaagtctttcctgctctctgctcaccatggctcacgcctataatcccagcactttgggaggccgaggtgggtggatcatgaagtcaggagatcgagaccatcctggctaacacagtgaaaccccgtctccactaaaaatacaaaaaattagccgggcacggtggcaagcgcctgtagtcccagctactcgagaggctgaggcaggagaatggcgtgaacccgggaggcggagct**tgcagtaagccgagatcgtgtcactgca**cttcagcctgggcaacagagcaagactccatctcaaaaataaaataaaataaaaattaagtgggtttggagaatgacaaagacagaagatggtaggatattaacataaaaaaaatcatgtgtaggggataatgtagaaaatgtgt**tttgtgaatttctcacaaa**ttgtctaaggagtcagaattcacaggcatgtaagat**aaaccttaattacccccactgtaggttt**ccttgtctccaggtatttgccatccgcattagcaagctgagagctcctgagaatggctatgtcttttgaactagtctgccctctgcattctattttcaccctaagctgtgaacagaaatttcccccacaagttttcttgattgt**aaaagaaagttctttt**ctatggcattccttataatgttaatttcaacttaccatttatatagggattttgcattgcacttaaagagtaaacctcctgtgggtaatacaggcagtgaaataggttagatgtaagctgctcctcctggactctagggatctatttgtctttatcctgcttaaggagagccaagttctagctgaagg**cctgtgccaactcacagg**acactgacgtggtctcttccactgcctgcttggagctatccacatgtatccacacaggctgt

Uppercase: TRBV12-1

Lowercase: Flanking sequence[1000bp]

Red & Bold & Underline: Stem-loop [11]

Blue: Heptamer[44]

Green: Nonamer [4]

id-IGHV3-32-2[V_gene_segment]

gacttatgcaatacattttgaggtgaagcttggtggaa**ttttttctaataaattagaaaaaaataaatcatttaattgactattttattcaggttgatttgttt**aatatttgctaaaggccagttctttaagctatgacacataataaatcccaaatggcagtacctcattgtttacttagcttttgtacttatatttctcagaggaagaaccactactgtaaattgtaaatagc**caatacatatttgtattg**tatgcaaaactgtga**ctgtttacagtgtcatctctgagaaacag**ataaagtttatttactatatacatataaagagtttggaaggtggactcctcaccaatttttgaaagagttacagaagggctggtattacttctttaaatgttaaggttcatttt**atgatgtaacatataacgcatcat**ggagaatgttcaatgtgtgcttgagaaggatgggtattacgtgactcttggttggaaggttctgtaaatatcattcagatacatttgttcaatagtgttgttcaagttcagaggcacattaagaatttactttctggatttgctaaacattatggtcatgaggtattaaggtcttgtgttatttttgtattgttttctatttctttatatctcttacagtttgcttaatgcagttatatttgtatttgtacacatgtgtaaaaacaaaaatcataattgctaaatgagttttatggcacagtcacattataagtaatattttcccgaatgctaccattgccactaaactcctcctggagtctgacatctgctctgggcactgccttcttctcaggcgtcccacactggagcttgctatagaggaggaggcatgaaaacagggccctccctctcctggtgaaatcaag**cccagacctgaccctgcagctctggg**agaacagccacag**ccctgggattcccaggg**ggtttccatttggtaatcaggactgaacacagagaactcactATGGGGTGTGAATTAAGCTGAATTTTTCTTGTTGGTATTTTAAAAGGTG**ACTCATAGGGAACTAGAGTGAGTGAGAGTGAGCGGATATGAGT**GAG**AGAAACAGTGGATATGTTTGCCAGTTTCT**GACCAGGATGTGTGTGTATTTTCAGGTGTTCAGTGTGAGGTGGAGCTGATAGAGTCCATAGAGGACCTGAGACAACCTGGGAAGTTCCTGAGACTCTCCTGTGTAGCCTCTAGATTCGCCTTCAGTAGCTTCTGAATGAGCCGAGTTCACCAGT**CTCCAGGCAAGGGGCTGGAG**TGAGTAATAGATATAAAAGATGATGGAAGTCAGATACACCATGCAGACTCTGTGAAGGGCAGATTCTCCATCTCCAAAGACAATGCTAAGAA**CTCTCTGTATCTGCAAATGAACAGTCAGAGAG**CTGAGGACATGGACGTGTATGGCTGTACATAAGGTCCCAAGTGAGAGCTGAGGACATGGCTGTGCATGGCTGTACATAAGGTCccaagtgagcaaacatcggtgtgagt**ccagacacaacacttcctgcaaaaacaagaaaggagtctgg**gccgaaggggacactcagcactcacaaaacaggt**gcagccccaaggcaggtgcagatggagggagggtaagggctgc**tttccttcagggtctgtggtttcctctgcttctaatatttcacctctgagcctctgtacatttatgttttgtgcccaccatgaggtccctggattagaaaactaatttaaaagaggaaatattctcatatgtcccaaaagcagatgtaagtattggaggcataaaaatgcataggagctgagtgtgtctgtagacactgccaccccacaatgccagacccaaaactagtgctggagaagggtgggagtttgatggagcttccctgatgaccccgtggtccaagctaagtccagcaaggccattggtgcctcgctgagcacagttgtccatcagggatctcccatgtgtcccaggagcagccatgcctcagtatctccactgtgcacagccactgt**ctgggaggagctcccag**gatgggtgtctttggcacacacaggtgatgggtgttagagtgcagtgcagcagctggctgcctggtctattgggctcctggatattggagggattggaggtgcattctcagggccagcacgctgtttgttaattttttataaaaaccatgtgatttaattcattttctcagatgacatagataattaataacacaatctgc**aaacaatcgtaattttcagctttagcccaagttcattgttt**ctgaattctgtgcaggatccagacgtggtattgcccttctcat**gagaaactgttcgatctaaactgaaaccagttgtttctc**gtatactttggttctccccacgtgcagagatcttgattagagcaagtttggtactttccacacactcaccctcacctccccagataaagagcagaagttctccttagactgagtctgagg

Uppercase: IGHV3-32-2

Lowercase: Flanking sequence[1000bp]

Red & Bold & Underline: Stem-loop [16]

Blue: Heptamer[39]

Green: Nonamer [2]

id-TRBV6-6[V_gene_segment]

agtctagaaccatctggggctgtcttcagatcggacaccagagttaagagagcctgtgtaccctacag**cactgcagatagaagcatcatggtgttggcagtg**gggaggtactagagtgtgtaagtggttagtggcctacaaccagaagacctgagttttgagaaaattattcccatcacaagatgtagcagagacagtgatatggatatactagac**cagagagctctgtgaggagctctgtg**aggagct**gtggcctaatcatgcatcaggccac**gtttgtgtccataggtcatgggctgtgctgggtggccaccatcatccatacagagagggcagtcagcagcgtgaggtggttc**tgcctgctgtggtctcaccccaggca**tggaaagcaagagccctgggtgg**agctgaaggtgctcagct**ggg**cttgtcaggagtctcatctgtcagtggattgacaag**aaacagagcaaaatgattcctccaatgttgatgagcctgcccgtgggatctggaaagctaataacagagaaaaccaatatagacaaaggattttaaacaggattatggtcaattaagcaaattagaaaatgatacttgaaggagtatttgggacgcagtagtcaaaaacaccaggaagacatgaggaattgccctaagactctagactacagcactatgtagataactaacaccagactattaattatatcattgaaaaaataaaatttacagtgaattacaaactgtttaccaaaggttatgaaaatcttgtagacttgttttaattcagacaaatgttttcttctcattgtcagctgttctctggcgcatttattttggattgaaccatctgggga**agaggcgtggcctct**cctgacaggaaggctctggggcccaggcagggagaatgaggtctcagaatgacttccttgagagtcctgttcccctttcatcaatgcacagatacagaagacccctccgtcctggagcacctgccATGAGCATCAGCCTCCTGTGCTGTGCAGCCTTTCCTCTCCTGTGGGCAGGTGGGTCCTGGACAGGGCCCCTTGCATAGATTTCAAGGCC**CAGCCCCTTTCCATTGGGGCTGCAGCATCAGCTGTTTCCTTCTCTGCAG**GTCCAGTGAATGCTGGTGTCACTCAGACCCCAAAATTCCGCATCCTGAAGATAGGACAGAGCATGACACTGCAGTGTGCCCAGGATATGAACCATAACTACATGTACTGGTATCGACAAGACCCAGGCATGGGGCTGAAGCTGATTTATTATTCAGTTGGTGCTGGTATCACTGACAAAGGAGAAGTCCCGAATGGCTACAACGT**CTCCAGATCAACCACAGAGGATTTCCCGCTCAGGCTGGAG**TTGGCTGCTCCCTCCCAGACATCTGTGTACTTCTGTGCCAGCAGTTACTCcacagcgctacaagaccatctcctctctgcacataaaggca**gggtggctctgccctcctcccccaccc**aagactcagggatgacctgggcagagttttctgcaatggggaccttggaaccccgagtggccctaagtggcccggacagtatgagcctcagtctgtgccaggtgctcctgcagccatctcagccaggcctggactggtcccaggtcctcagatgtctcctttgttgctctctctggtctatcctctgagctctccttttggggttgggccagggcttccccagctcctacttttctactca**tcatcctgactccgaggcccccaggatga**aacaggatttgtatttcagatccatctaggctgtcctctccctggtgagcatgctgcttcctctatctagagttttcccctcccccaccccacccatggtctgtcctgtggcctatctttcccatctgggcagtcaccttccatcccccgc**ctccccgcccctctctactgcagccatggggggag**cacctcttctgagactccttccttcccatcacagagacttcaaaggccatttcctctgccctggatggagacttcctttctctagtggccagctcctacctgtgctt**cagatctcagaatgatctg**ccctcccgcaggaagcaatccctcaccgcccagctgagatcaattttcctctcataagctctcatagaatcatatgtctcttagtaacccttaggacaattaggctttctcagatacttgtctgatgatttttgtgctctacccaatttttttttttttctgagacagtctcactctgtcaccaggctggag**tgcagtggcatgatttccgcttactgca**acctctgactctgattcaagtgattctcctgcctcagtctcccgag**tagctgggattacaggcatgtgccaccatgctcagcta**attcttttttgtatttttagtagagatgcggtttcaccatgttggc

Uppercase: TRBV6-6

Lowercase: Flanking sequence[1000bp]

Red & Bold & Underline: Stem-loop [16]

Blue: Heptamer[36]

Green: Nonamer [5]

id-IGKV1-39-3[V_gene_segment]

ctgaacactcacaaacattatggtgggaaaagcagatcagagcattaggcaagttgcattaccttggccttcttcctttggagacaattgatgtggggttctagattgacccagagtttcaagtttatcctgattcagg**cttcaacagctggaggaagaaacagagatgttttttgaag**taaacagatctagcattactaatcaacccttcatactgatgacctatgggaaataatacccaagggcagaaa**aatgggcagaataaggggagccccaaaccaagacgaagctgctgcccatt**gagaccctgggtattacagagacctatagctctggataatggaagatctatgagtggcacaggcgctgaggaatcacagcatcattatcgtgcatctgcagggaattgc**ttgtaaatatactggtaattacaa**atgtttaaggtcactacaaatactttggagtgtattaaatatgcttctgataaagactgtttttc**tcacatgaaacaatgggaaccatgtga**caatcacagaggtgttgttactatagca**aaagggattgttactctccacatcccttt**aagtaacttgaaggcctgatagacccaccctctaagacttcattagacattccctacgaatggttatactctcctgtatactcccaatacaactctaaaatatattattccatatagtccttaggtttgtattaaagtttgacttttttccttcaaaatatctcttgtcacaacagcggctctagagagaaatacattccctccaggcaaatctatgctgcgctggtctgacctgggaccctggggacattgcccctgtgctgagttactaagatgagccagccctgcagctgtgctcagcctgccccatgccctgctgattgatttgcatgttccagagca**cagccccctgccctgaagacttttttatgggctg**gtcgcaccctgtgcaggagtcagtctcagtcaggacacagcATGGACATGAGGGTCCCCGCTCAGCTCCTGGGGCTCCTGCTACTCTGGCTCCGAGGTAAGGATGGAGAACACTAGGAATTTACTCAGCCAGTGTGCTCAGTACTGACTG**GAACTTCAGGGAAGTTC**TCTGATAACATGATTAATAGTAAGAATATTTGTTTTTATGTTTCCAATCTCAGGTGCCAGATGTGACATCCAGATGACCCAGTCTCCATCCTCC**CTGTCTGCATCTGTAGGAGACAG**AGTCACCATC**ACTTGCCGGGCAAGT**CAGAGCATTAGCAGCTATTTAAATTGGTATCAGCAGAAACCAGGGAAAGCCCCTAAGCTCCTGATCTATGCTGCATCCAGTTTGCAAAGTGGG**GTCCCATCAAGGTTCAGTGGCAGTGGATCTGGGAC**AGATTTCACTCTCACCATCAGCAGTCTGCAACCTGAAGATTTTGCAACTTAC**TACTGTCAACAGAGTTACAGTA**CCCCTcccacagtgttacaagtcataacataaacctccaaggaagcagatgtgtgaggacgagccaccccag**atgctcctcctggtgcctccatctgctgagagcat**ttctcaaactcagtcaggttttg**aaagtcattgggagacttt**tgtagaggggaccagg**gaggctcctctgaactctaagcctcttttgcccctatccccaggagaaaaga**tgtgacaatgcctgtcctgattgaataaggaagagatacaagtccacctgaggagt**ctgtgttatgggataattggaatttacacag**caaaagagaagctattctcggtatttcaaggagaaattgttcaagttgaataaattagagtctaaactagtctttttgaagcctacggtatgttattcgtgaagcagccactagagacaggggattct**caggtgctcctgcagaagtcagagtgcacctg**cccctggtggtatgtgctgagtaccgtgtgatgatcctcagacctgtctgggaagccgagggctggg**gtgctgatgctctcagctgcctgcagcac**gtctccaggtgattctccagtccacaaacaattccacatgttttacttca**gatgtcagagtacatgaatccaccactctgacttc**ccaatctcatgggagtgcctctcattaagcaactctaaagaaaccatagagagaaaaggagttttggaaaatgtgcgtccagaagtgatagtagcgatggggaattgacagctgacaggtcagtaaggttgctctttccacaaggctcaaaactttgccagttacagattgtcccagaatatactcgaatg**tgctatcacgtcttacgagcaactctgggttcatagca**agaaaacttcattaagtcatagatgaaacagaaaaatcaggaaactgtatgaaatacattataaggctgtgtgtggtagctcatgcctgtaattcccagcactttgagaggctaaggcaggagaattgctt

Uppercase: IGKV1-39-3

Lowercase: Flanking sequence[1000bp]

Red & Bold & Underline: Stem-loop [20]

Blue: Heptamer[26]

Green: Nonamer [7]

id-TRBV8-1-2[V_gene_segment]

tgctaatctttatttttatggtaccattgtcatttggggggacatatttgtttatctttggtaatg**gaaatgaaattacgttacatttcactttc**tgtgttttcttcctactttagggtaaatattatttccttctattcttcctattctattcctctctctttagatagttttaagggagaaaatgtcagtgagactgtaactgcagctatggggtaagaattatttagaatgagggtgggatattaagcttgataactcaaaggaacagtcagggttaaaactaatgttgggatcggggttcag**agaagttgcttgtaaaacctgcaggatggcacttct**ggaatagtctcccggctccctctgcagacacttcccagcatcccttgggccatttcccttgcaaaagcaatgatgaaatttcacttattggccacaagatggcactgtggtccactgggctctaagagactctggggagaacctgggagagcagcctaggaaggaaagggttaagaaaaatcagggctggcatccaatatcatgcagatgttgcagcagttttcagttattgctagactacctacagttatacaacatacaggaaatacctctaatctttaatgagatccacagttgaagaagtttggcctggcagccttggtgtcaagggcaggtgcagacagaggagcaactatctcagaggaaaatgggaaagtgagggg**tgggctgtggcttaagccca**gtgcgtctct**gctgcaccccatcttccctgcagc**tctcac**caggcaacagcctccatgagagtaggtggatcctgcctg**agcccagtcagaatgccccggactaggtgaactctgtaggcatggggagtaacacaacaggcccacctttcacacgtttccaaagtctgggctccaagcctttttttcccctttcctctgcagagtcctgccacttctgaagccttgtcctcactttc**tgcctgcctctcccacagCCCCCATGGAGGCAGGGA**TCAGCCAGATACCAAGATATCACAGACACACAGGGAAAAAGATCATCCTGAAATTTGCTCAGATTAGGAACCATTATTCAGTGTTCTGTTATCAATAAGACCAAGAATAGGGGCTGAGGCTGATCCATTATTCAGGTAGTA**TTGGCAGCATGACCAAAGGCGGTGCCAA**GGAAGGGTACAATGTCTCTGGAAACAAGCTCAAGCATTTTCCCTCAACCCTGGAGTCTACTAGCACCAGCCAGACCTCTGTACCTCTG**TGGCAGTGCATCcgcagccctgcacagcca**actgctctctgcacaaaaaagggcagccacaggctggaggtgggcactccttcccaaggcctc**tatctcaaccaggagata**aagatgccttttttgaagctcttctcagatgtcccagcaatgggctcccagtctgtgcatcctcctgcagcaggaacttcagtgtgagctgaggtccatcactgtgatgtggatgcacgaatgcaaacaacacttaatgt**tattttactcaattcagaatattctggcagttgcgtaaaata**ggaatagtagctccc**aaaatgaaatttgtgtctcattttctttatgagagaaaagaaaa**aaaaacaga**aagcagctgggcagctttaacctgctt**gagtgatctgggattctacaagtctacaagtggtacaaagaacatcaccttggcctggattccaatggataccaaacactacaaaaagtattgaaaacaatttataaaagacctaagtaaaca**atctacttataggaaatgttagtagat**ccatcgcatagtgccattatacctgacgatgatcaactagtgaaattattat**gtagagattatcagctaagcctctac**ttattacctctgagcatagagcagtgtttgagccttctgagggtatcttaaggaaggaacttaattcttaccattactttggcttggattaaaatccatgagcttgtcgtttttattgtaaaatattacagaataataatgatcaaacgttttattttcctctagcgtttattcctctttagaacgtgcaggttgaagctcacacacatgagcagggttccttaagatgagatccattaactccaggactgggaggtacacagctggcctt**cacaggaagtgcaaaccccaaatgcagtgacccctcctgtg**tctacgctgtgcccaatacccctctgtgaggctgccataaaggcgggcaggctccttgcttggcaaaaaaatcagttcaccaat

Uppercase: TRBV8-1-2

Lowercase: Flanking sequence[1000bp]

Red & Bold & Underline: Stem-loop [16]

Blue: Heptamer[35]

Green: Nonamer [0]

id-IGLV5-37[V_gene_segment]

tggttcctggaacaggctgccctaggaacagagcccaaatcggtcagagcaagcagggacacaactgtgcagaaaacaacaagttttacttagggagataacggttgactgtataaaatgagaaatatgtcacactactcacttgtaaaagtaagtttaccaatgctattcatttccctatatcactacgttgagtaaaaactctaaaatgtatgaaaccactgtagagataacttttgctgaattgctactttaacatgtcatggatcatttatttttcttgcaaaattctcattattcaccgtccttttgagtaacccc**cagctgcccttcaacctagctcctcagctg**acctcttctccatgatgcctaccctgatggcacatgtgggcagttaggtttttctagattttaaaagttatatatgtacatgtctacatcccatggaccaaga**ctcactctgggagtgag**aaatcacaataatcatgcatagccttatggaattagtgtctagcaatcagaccatagaggtgtgaatagagacagtgactaatgagctgcagagatcagctgggccatcaggaggtaagtgtcctctaa**ccagcatgggatgctgg**gggactgccct**atggtccctacacagctgctcagtaaggaccat**tcactcaaagaaatccaagactaggg**gctaagaccccgtgctcactgaggggtacttagc**atctatgtcctcccatggcatggcagagtcccctgagcaggaaggtgtgtatggtttcagtagg**tgctttcttccggagccctctcaaggggaaaagca**aaacttcttcctcctcagtctgcagtgtgagctgagctt**cagccccagagcttagagaggggctg**ggcagtccctggatgaaaacacatttgcatgagcagcc**cctcctctgcagagtatgggaagaaaaggagg**cctggggcagcccagtcccactgtgcatgtcaggctgtgtccaccATGGCCTGGACTCCTCTTCTTCTCTTGCTCCTCTCTCACTGCACAGGTAGGGACAGGCCT**CAGAGATCAGGGCCAGCCACCCAACCTGATTCTG**GCTCTTCTGGTAAAGATCCCTGAAAAACCTCACCCTGAACCCTGCCCATCAACCATGAGTGTCTGTGTTTGCAGGTTCCCTCTCCCAGCCTGTGCTGACTCAGCCACCTTCCTCCTCCGCATCTCCTGGAGAATCCGCCAGACTCACCTGCACCTTGCCCAGTGACATCA**ATGTTGGTAGCTACAACAT**ATACTGGTACCAGCAGAAGCCAGGGAGCCCTCCCAGGTATCTCCTGTACTACTACTCAGACTCAGATAAGGGCCAGGGCTCTGGAGTCCCCAGCCGCTTCTCTGGATCCAAAGATGCTTCAGCCAATACAGGGATTTTA**CTCATCTCCGGGCTCCAGTCTGAGGATGAG**GCTGACTATTACTGTATGATTTGGCCAAGCAATGCTTCTcacagtgacacacacagatggggaagtgggacaaaaacctcaccctgctttaggtcttgttcttaaaaa**attttaaattttaaaat**aactggcctaggcacaaactacatttggaagtactttctagttgtaaaaggctcttctctcaatttcccatccattcgtctgaagtctcagtaaaacaaaacaaaaccaaagcaactg**atgacactgaagtcttggctggttgtcat**atgtgctaacaccactgcctgtggagctggacttttgttcttgaaagaagcaaaatgaggctttcctagctcacatcatggtcagagcctggggacatcctagcaggtgaacagtgccctgcagcatgga**ctctgctcttccagcgtacgggcagag**acctccccttccatccccagtagtctgttctcagggctgtctgtggctca**ctgtgtcccaggaacaaggcactcaaatatgaaatgctgcctgaggacacaggtcagggaattctagatatagtgctgacctg**gctcagtcctgggtacctgcatttcagtggcaatgtcaggagatggggttaggtattagggtaactggggacctgagaccagtaaacctctatgtaggacatgacaggagagtcaaggggaggctccactgtggctgagacaagccagaggctgctccttggccacgggggaagtaggtcccatggatgtgactttcctgcagctcatcccactcaggaccatggggact**gctgagcgtgggtttcttgatccctcagc**gcctgtgttctcttctcctctcttacaccattggagcttcttgctgaatcagggtctagtgg**tgagtatgtggaaagaccaccacatctca**aggtcacctt**tttctgcatggggagcagcagaaa**cattgggcattcccgtggaaatggctacggaggcctcccagaaagagcccccaagtttatgacctaggacagcagtgatggtccctcaggggt

Uppercase: IGLV5-37

Lowercase: Flanking sequence[1000bp]

Red & Bold & Underline: Stem-loop [19]

Blue: Heptamer[30]

Green: Nonamer [4]

id-IGKV7-3[V_gene_segment]

atcacaagtaacagaatagagtcctggagtggagctcatagagaagtgatggttggcatcatgttccctgacttttcatgttccttgtagccttgtgttgg**tgtctgcacatttgaagatgcagtcccttcttccagaca**ttttaagtt**ggttttgttgggggaaaacc**ttcacctatggatgggtgagagtgccagctgactggtgtggacaccatatcttgctgtaggagaagatggacttccttagccccagcttccagtggcaaggcagatggaaccagtgggagttgggggatggttattctccactctgaataggatccactacaatgactcttctgcctctgagggcctctggtggatacagcattggtttcaggactaagcagatggagagaccctgccttatccacaatgaggcaagtt**ccttgattagtttaggttaccagtcttcatcaagg**tttcatgctcacccccttcatcctttccagatcatgataacatttatagacttgttttaaataccacttactcagtggaacaaacttctccttccca**tatgaaccagtttcata**aaacccacccaacttgctttctctttctgacaatgataacttaaaatg**cattattattgataatg**atataaaatatgaa**ttttgaaaaatcaaaa**gtatttctgaaataacttatatccaaatggcagggcagccccagtacactcctccacatggcatctcaagtagactggcaggcaaagatccaatagaaaaggtcaccaaatcccttgggacatgcttgagctaccctaggaaatgttttgttagccccaactcccactgggtgaaataagagcaggtgtcccc**tttgagcatcaactcaaa**cttttctgagctgatttgcatgttgggccattagacaacggtc**caagcttctcagagcatccttaaagacagcttg**tcagggcccctgaggagcagcctctagttcagctctcagagATAGGGTCCGGGGCTCCTTTGCTGTGGATCCTCCTGCTCTGGGCTCCAAGTGAGGGGGTATAGGGAAACATCGGGCAAACTCATGTGGCCATCATTGTTTTTTGTGTCTCTGGATTCTTGGTTGTTGGATTTAGGGCACTCATTGTAGGTTGTAAGTTCCCCAGTTATCTTTTGTTTTCTGGTTACTCAAGATGTTGCATATGACTGTGGTTACAAGTGTCACCAATGCATGGCCTACATGGGGACAGTCTCAGTATATCTGATATAATTTATCTGTCTTTATCATTTC**AGGCTGCAATGGGGACATTGTGCTGACCCAGTCTCCAGCCT**CCTTGGCCGTGTCTCCAGGACAGAGGGCCACCATCACCTGCAGAGCCAGTGAGAGTGTCAGTTTCTTGGGAATAAACTTAATTCACTGGTATCAGCAGAAACCAGGACAACCTCCTAAACTCCTGATTTACCAAGCATCCAATAAAGACACTGG**GGTCCCAGCCAGGTTCAGCGGCAGTGGGTCTGGGACC**GATTTCACCCTCACAATTAATCCTGTGGAAGCTAATGATACTGC**AAATTATTACTGTCTGCAGAGTAAGAATTT**TCCTcccacagtgctttaggtctaaacaaaaacctccccaggcagctgctcc**ctgaggctcaaatccctcag**atgtggctttttatgcaggtccatcagcctgctgtcataggcttgtttgaacaaaggggtcaaattatcaaagagaaaatccttaaattgatgtgtgtgtgtgtgtgtgcatgcatgtgtgtcacagccagttttataatttaataatttctttactgtatcaggacctctc**agaagttttctacttct**caatgtatattttttatgtcaaagacccttaccaatagaagaa**ttttaaacattaagaaaactttgtagtcattttaaaa**aatgctctgaatagtcaaaattattagct**ctatatctttcagatataggtgaccttctcacct**gaattttga**cttttctccacttttatttgctgaacccacatgaaaag**gct**tttctagttatttagaaa**cctaagacttgtgaggtagttatctctgcttatttaatttagatgtctctatctcggtgactttcccaaaagttagattctcagctctagttttaataaattgtcctttactcttcatgcagtgttcatgtcctttttcttttctttttttttttttttaa**acagagtctcactctgt**cacccaggctggagggcaatggcacaatcttggctcacggcagtctccacctcccaggttgaagtgattctcctgcctcaatgtctc**aagtagctgggattacaggtgcctgccaccacgcccagctaatt**tctgtatttttagtagagatggggtttcaccatgttggccagcctggtcacgaactcctgacctcatgatccacctgccttggcctcccaaagtgctgggattacaggtcatgtcctttttctatgatattttccatgtccattgttaaggttttttttctcatcttctctttttgcttttctaattgctctaacactacaggcagagtggactccagaaatg

Uppercase: IGKV7-3

Lowercase: Flanking sequence[1000bp]

Red & Bold & Underline: Stem-loop [20]

Blue: Heptamer[31]

Green: Nonamer [7]

id-IGLV3-19[V_gene_segment]

tggactgatgtcctcctaggaagaagaaatttgcacacacagaaatgaggcaccagaggtgagcgtgcagagaaaagaccaggtgaggattcagcaaggaggtagcaacctgcaagccaaggagagagtcctcaggggaaacc**aaacccactaccacctttatcttgggttt**tccagcttcagaactgtgagaaaatatgtttctgccatttcggtcactaattctttcctatct**tcttgtgggagctctagcaaaaacaaga**gggaccccaaagaccttggatgagggagaaggaggagatggagcagggtgcaggaggcggtgcaggaaggggctggaaggtcgggctctgaggtgcatctcctgggtggaatcttgactccactccctattgtctggaggacttgggaaaaacatttaacctcctaatattcactcactaataaa**gatgggcttgaagcacaaggctccccatc**atcctattctata**ttacaaaagtcttcttgaggtaacacttgtaa**aactctcgctaatgcatctggcatgtattatggactcataagtagcccttctgagtgatctagtgatgtgcagaaaatggcattcatgctgtgtgcaccagggggcactgtgaggtttagtctgaggcccctaatgagtccaagcccctagtaatgctcaagggcgaagagcctgactgttgcttcctatgaggccccttctagtgggtaaatctgaaaatgcacttggc**ccttcttctgatcttgagaaattactcagagaagg**ccatcaggctcagggctcagacaagaaccaggacaaatgttttagggaatggagaacagatttgcatccactgctcaccagagccacctaacgacgacacaagaataaa**ggaagtagatttgcatgaagagacttcc**cttcctatgataagagaggcctggaggttcctccttagctgtgggctcagaagcagagttctggggtgtctccaccATGGCCTGGACCCCTCTCTGGCTCACTCTCCTCACTCTTTGCATAGGTGCTGCCTCCCAGGGCTCAACCCCATATTATCATGCTAGCTGTGCCAACCTGGCCCTGAGCTTCGGCTCAACACAGGGAGTAGTGTAGGGTGTGGGACTCTAGGCGTGAAACCCTTATCCTCACCTCTTCTGTCCTCTTTTGCAGGTTCTGTGGTTTCTTCTGAGCTGACTCAGGACCCTGCTG**TGTCTGTGGCCTTGGGACAGACA**GTCAGGATCACATGCCAAGGAGACAGCCTCAGAAGCTATTATGCAAGCTGGTACCAGCAGAAGCCAGGACAGGCCCCTGTACTTGTCATCTATGGTAAAAACAACCGGCCCTCAGGGATCCCAGACCGATTCTCTGGCTCCAGCTCAGGAAACACAGCTTCCTTGACCATCACTGGGGCTCAGGCGGAAGATGAGGCTGACTATTACTGTAACTCCCGGGACAGCAGTGGTAACCATCTcacat**tgacacagacagatggggaagtgagacagaaaccccttcactatctgtgtca**tcctctctct**ccagccccagcaggactgtgaacaaagccataaacagggctgg**cccagttcacctgcatctgagacctccaggctgcctttccctccattcctccaggtaggctctgcagaaggtgggtcaggatgatgtgaggcttgaaggaccaggctgttctgctgttgtttagactgagtgtgccttgcctcaagatgacctgaatggaag**gaccacaagaaggagatgggcagctgttgagtggtc**atgatccctggagttcctc**ttgtgtggtgactggatctaacacaa**tgcctgtactttgtggcccaaatgccctggattattggtctgaactccccatgtttaactgagaccctcaagccccagctccatgcatcctgatgttctgaacaggaaagtcttttctagtgagcactgtggcatcccttggcatcccagcttcacacagcaattccccaccaaaaaatgcctttgagtgcaaagaacggcttcatgcaaagatgtgcctccttcagaat**gtagttcaggagcaataactac**ctgatgccttaatcccaaagccatattttaatttagaatgaccttgaaatgccttctagctgagcagtcagttgtaacacattcccagtcacgcctcccacacttc**tttcctgacagtatgtcatgaaa**ccctactgca**tggaatcctgcctcttagagtaggattcca**gggatctcaaaaattgtttagaaatgaacaaaaacagtctagttggaagagtatcaaagaggaggcagttgcaca**tgcagagaactctgaggtctgca**aatatacatgaactgatgagcacttgtgtgtaaagaaacacctccgaaagagaaaagaaccacgcaaagactcttgagggaacacctcacagagctcactcccacacacacccacccacgcactcaca

Uppercase: IGLV3-19

Lowercase: Flanking sequence[1000bp]

Red & Bold & Underline: Stem-loop [15]

Blue: Heptamer[35]

Green: Nonamer [5]

id-TRGV5[V_gene_segment]

acagaggtggtgataatctgcctatgccttactgaattttcacaataatattactatactttttttgtcccctatatttccaactgtgaatccaggtgcactgcagatattagtctgctttctgagggtagggatgacaaataaaatatacaataccgggggaattggaagtacagaaaaataacaagtttttagtttaagtatgttccaaatattgcatgacatggcacccagtactactttaaaaacaaacaagcaaacaaacaaattcacactcctactttaatgaacaaa**aatcctctaataaggaggtcacacacacaggatt**acaggcacaacccctgcccccccaacacacacacacacatatttaccttagagcagactccgggacacaccaacacacatgatcctcagtccttttaataaagcacgttcaacccttctcagagtgaccttgagtggttgcctccatgagtctgcagatgcgaaaaggaacttggttcagggccctaatcagaggcttcctgtcaggacaaggttctggcatcagcattacctgctgcaactctgaa**gggcagggtgggagcctgctatgggactgccc**cctccaaggtgtgagcctgttaattccagggtcagcttacccaacatctgcagtccctcagct**ggaggcctgagctcttaatgcccctgcctcc**cttctgccctgggcctgctctcaactgccacaagatggcgcagaagaaagggcct**gaggagatgcgggagctcctc**tgcagg**tcctcccaggcaccaccccttctgggagga**aggcttgggagtcccctaagacacatcttc**agtcacttctctctgcctgtgact**cag**gaagaccagctcctcctactgtcttc**tgtgctagggatcacttccttgttgagtggggccggagttttaagaggatcttctgctcctcctcatctggtcccttt**ccttccaaggcccccgagaggaagg**cATGCGGTGGGCCCTACTGGTGCTTCTAGCTTTCCTGTCTCCTGGTGAGTACGCTGCCTACAGAGAGGCTCACAGGTTGGGTTTTGTTTTGTTTACTTCTTTTGAAAGGGGTGCCATACAAAGGAATACCTCATTAAATTTTGTGTTGTTCCCATTGCAGCCAGTCAGAAAT**CTTCCAACTTGGAAG**GGGGAACGAAGTCAGTCACGAGGCCGACT**AGGTCATCTGCTGAAATCACTTGTGACCT**TACTGTAATAAAT**GCCTTCTACATCCACTGGTACCTACACCAGGAGGGGAAGGC**CCCACAGCGTCTTCTGTACTATGACGTC**TCCAACTCAAAGGATGTGTTGGA**ATCAGGACTCAGTCCAGGAAAGTATTATACTCATACACCCAGGAGGTGGAGCTGGATATTGATACTACGAAATCTAATTGAAAATGATTCTGGGGTCTATT**ACTGTGCCACCTGGGACAGGcacagt**gattcagacctgtcctacaccacactgaaaatctgccttgtggctgcctctggttc**acaggatagagccgccccctctcatttcctgt**caccaaatttactgtattctgaacaagagaaagacagcttaactcctgatctccctcctaatatcacactgtcctggcagcagccgcaccctgttccccacccctcccccacaactttcctgaagatcaagctgccatct**ccaggcctcagctaagcagcctggctgagagcaaggttctctcagc**tctcctaggacatgggggaggcccactcactctgcttcctatgacacacaggtacaactagggtccagctgtgaagcgagatattttggacagcaaatgggaagggtatttatactgaatcatgtatccagtcatggtccagagatcacagctgagagtggtgct**tattccttatgtctataacaacataagcaata**ctataatgaccactaaaacactaggcccaggtatgcagttgaacatggtctccctctgccctcccttttattctctctctcttttttcttacttctttgcttgtttttatttttctctccctccctaactccctccctccatccatcctttcctcccttcctcccttgtttccttccatccattctccttccttgttttccttccttccttccttgcttttctctctctttctttcctttttgggtacataatcagaatactttttccacaaaatcatttgatataatttttctctatttgttcatgccaacatttatacagctagtttaattgttcagttttcattttt**acattttggaaatgt**gttgtattactttgttttgcttcatatctgttttgtatatataaaacgtttacatgaaact**gaagccaaaattataaaagatgatgcatttgaagattttagcttc**cattgtagatttttcgttcctcttttctccctgacactataggtaacaatat

Uppercase: TRGV5

Lowercase: Flanking sequence[1000bp]

Red & Bold & Underline: Stem-loop [19]

Blue: Heptamer[21]

Green: Nonamer [6]

id-IGHV3-25-2[V_gene_segment]

gctgagtggaggaagctaaagaattcaccatcaat**ttcatgtgatttcatttgtataaatcacagaa**gatgcaatattctaaattaacat**agagaagatctgaatttctct**gaaaatagggtgggcagagtaacaacgtggtgaaataaaattatagac**aagttagataaaaaaaattagaggttaactt**aattgctaaaacatg**attgaagtgctgattcaat**gactgcacacatatatcaacattttccaaatgctgcactatcaatttgat**ttcattattgattgtttttttaataaagcagtaacaaaaatgaa**tatattggctgagaaagagcaagaaatggataaataccgacacttgaataatcaaggact**cctgaaaatacacacatgtgaacactgattgtagatttcagg**taaacactagaaaaggcaaaattacacaaagttgttatgacagga**gggacatcctgaaaacctctctaggcatgtccc**acaccacc**ctggagctgtctcaggggagcagtctcctccag**tgtttagaggcacaggcacgcataatagggctaactctggccag**atgtgtgatattggacacat**tgcacaactgctctgttctgtatgtaatttatcttctctacaaatgtaacattgacagttgcattaaatatattctgcaaatatgtaaaagttaaataagctgatgtctgctaaatgattaccaaggcacaatcacataatctgaagttatattttcc**agagtgataggattacctccagagttttccaggacactct**catctgctccgggcactgccttctcctcagccgtcccacccctgagcttgctatatagtaggagacatggaaatagggcccactctctgctgataaaaaccagcccagccccgaccctgcagct**ctgggagaggagcccagcccccgaattcccag**gtgttttcatctggtga**tcagcaccgaacacagaggactcaccATGGAGTTTGTGCTGA**GCTGGGTTTTCCTTGTTGCTATTTTAAAACGTGATCTATAGAGAACTAGAGATATTGAGTATGAATGGATATGAGTG**AGAAACAGTGGATACGTGTGGCAGTTTCT**GACCGGGGTGTCTCTGTGTTTGCAGGTATCCAGTGTGAGATGCAGCTGGTG**GAGTCTGGGGGAGGCTTGGCAAAGCCTGCGTGGTCCCCGAGACTC**TCCTGTGCAGCCTCTCAATTCACCTTCAGTAGCTACTACATGAACTGTGTCCGCCAGG**CTCCAGGGAATGGGCTGGAG**TTGGTTGGACAAGTTAATCCTAATGGGGGTAGCACATACCTCATAGACTCCGGTAAGGACCGATTCAATACCTCCAGAGATAACGCCAAGAACACACTTCATCTGCAAATGAACAGCCTGAAAACCGAGGACACGGCCCTGTATTAGTGTACCAGAGAcacagtgaggggaggtcagtgtgagcccagacacaaa**cctccctgcagggccatgcgggtggtttcctttctcagctgcaggagg**cgggcttattgttgcaggactctggagacttattaggttgtgatattttactatggttatttatcatgaacttttttattgggaattgtgttttattaatttttattttatatgtaaggt**tatttttttaaaata**cataccttcaagaaataattcttcctaataatttgcacttgttctttttagagttttgttaacatctgttgatatcagctactacatagctatagggacattaatttacatctatag**acatatgtgtaaatacacaaacttatacatatatgt**ggtcttttatatt**taatgttataataaaattattataacaatta**taaaatacatcctaaagaatgaaacatatatagtaatccatattcctttcaatcattgcctatagtttacataaattgatgtctatttgtaaccttaaacatagtgtattggtcattttcaaacatccaagaacaaatt**tcaaatgtccttgtcacacacacaaaaataaaaatttga**ggatttgaggtgatatatatgtcaatcagctcgattcaattattccatattgcattcataaatcataacatagctttattcactataaatacatacaacc**aaaatttctcaattttcaatgaaatttt**aattatacatattttatctgtcctaggtcatgaatgtt**tttctccctgtccggatgtgattaagattgtcctgagaaa**caaattcagcctc**ctgcctcctgaaggcttcaggaagcag**ctcctagatggggtgaagcaggcaacccgtgtgtgtccacaggacccggagcctctctctccttggattaggccccctcctcaggattacagggctcttcagttttctcaacatgctgttataccagagaagcacaaaacacttaatttcattatattt

Uppercase: IGHV3-25-2

Lowercase: Flanking sequence[1000bp]

Red & Bold & Underline: Stem-loop [23]

Blue: Heptamer[25]

Green: Nonamer [8]

id-IGHV4-55[V_gene_segment]

atgttcatgcaattgaatcgaggagaaagtct**ttcagagacagatggatctgaa**ctggtaaatatgtgtgtaaggactctgggcttaagtgtcattgtccagccatgtttc**acaggtgtgacctgt**cagggaagaaccagagttccttgtgttctcagaggggaggggtcccagaagtcctctctggttcccaggaaaggtaattgcattaatcttggtgatgagact**atcatccagtgatgat**gtactatagagtttatgtttgaagttgacactgctatcgcaatctacatcttttcacacagaagtgtttagaggtcaggccacatcttcaggatcccacatt**gagaaggacagagatatattccactaccttctcctgagatctcaggcagaaacccaaatttcaaaaggtctcag**aagggcagctctcaggggctatttaaaaataacccacttcgtgggacagggagcatccttctaaccatgatggatgttctgaactacaataaacattgcatggatccagggtctgaattcactgtgattattacactccactgctgtttcaatgtgtctgaaggggtaaatgacaatttagatgacctgggtgtgtggtttgttttatataaatcttcaaggatagaacagcattgaacctattccaaaatctgtccctgatcca**agatcacactgatct**cccagaccagcatcttcagcacatttccctacctggaagaagaggactatgggcttggtaaggggaggccacaggaa**gagaactgagttctc**agagggcacagccagcttcctactcccagggcgagcccaaaagactggggcctccctcctcccttttcacctgtccatacaaagtcaccgcccacatgcaaatcctcacttaggcacctac**aggaaaccagcacacatttcct**taaatttgggatccagctcacatgggaaatactttctgagactcatgggcctcctgcac**aagaacATGAAACACCTGTGGTTCTT**CCTCCTGCTGGTGGCAGCTCCCAGATGTGAGTGCCTCAGGGATCCAGACCTGAAGAT**ATGAGATGCTGCCTCTCAT**CCCAGGGCTCACCGTGGTTCTCTCTGTTCACAGGG**GTCCTGTCCCAGGTGCAGCTGCAGGAGTCGGGCCCAGGAC**TGGTGAAGCCTTCGGAGACCCTGTCCCTCATCTGCGCTGTCTCTGGTGACTCCATCAGCAGTGGTAACTGGTGAATCTGGGTC**CGCCAGCCCCCAGGGAAGGGGCTGGAGTGGATTGGGGAAATCCA**TCATAGTGGGAGCACCTACTACAACCCGTCCCTCAAGAGTCGAATCACCATGTCCGTAGACACGTCCAAGAACCAGTTCTACCTGAAGCTGAGCTCTGTGACCGCCGCGGACACGGCCGTGTATT**ACTGTGCGAGATAcacagt**gag**gggaggtgagtgtgagcccagacacaaacctccc**tacagataggcagagggggcgggcacaggtgctgctcaggaccaacagggggcgcgcgaggccacagagcccgaggccgggtcaggagcaggtgcag**ggagggcggggcttcctcatcagctcagtgatctccctcc**tcgccagcactcagatgtccccagggctcctgttt**ctttattgtctgtggttctgcttcctcacatccttgtggcagacaagaaag**gaggaagacaatttttctgtttactgttgaggtttcaccaattact**aggaactttcctacaagttcct**gcgtgactcattttaccgtata**tgtgtgtgtgtataaatatatacacatacacacacaca**cc**atatatatacaccatatatat**tatatagatacacaccatatatatt**atatatatacacaccatatatat**tatatatacacacc**atatatattatatatatacacacgatatatataatatatat**atacac**accatatgtatattatatataaacacaccgtatatatatatggt**tctcaccatctcttgatttgtgtcatcaatggaattgtgcctatttgaaattcatttacccaaaccttaaatccaatggatctataccggaattttaatgatgtaattaaggttaaatgtggtcaaagtgtgagaccctaattcaataaaccagttgtctttataagaagaggaagagacacccgagacctctcacttttcgc**gtgcacacagagaagaagccgtgaggagacgtagtgcac**tagaaggtggccctgtgcaagccaggaagaagccgcgctaagaaccaatttttacagctccttgatcttccacattcagactgcagaattgtaagaaaatcaatatttgttgtttaacccacccactcctgttgtcttcttatgaagatccaaacagactgataccacgtaattctgttagctctggt

Uppercase: IGHV4-55

Lowercase: Flanking sequence[1000bp]

Red & Bold & Underline: Stem-loop [24]

Blue: Heptamer[42]

Green: Nonamer [8]

id-IGLV7-43-2[V_gene_segment]

gtata**tatgtaacatacata**ctgacaaataaaatagaactgagtgcctagaaaaaaaactcttagttatacagtcaaataattttaaacaaggttgtc**aagacttttcagtggaaacagtacagtctt**tcacaaatatagacgttctgggaaataatactgaatggcttttgctattgttgtttttgaatggtctcaa**atgttctctcaaagatgctacagaacat**agcttattgattttgtaagttaaaaggctgagtaagactgtgagagactccctgctcatcatcattcttcttccacttccagtgactactcttatattctgtgccataagccaacagctactgaccccaggatcttttaatccttctgtaaaactgtcattaaa**aaaaaattgtcgaaagtaaatttttt**aaaaaattctcacttaaaccagctgtttatccaatataaggctcaacacaaatagttacagaaagccagtaggtcattaaaacaattaatttagtcattcaaataaaaatgaacaccatgtaacactaatatcaaattatagtcacaggggcagaatttcagcaacagcaagagaaagtagcacaaataggaatgagcaaagaatagacaattaatgagtgtgaattggcctggaaaatattgtggcaatgtccaaagataaatccacagttatttgtgaaacaagtgaaagaccacgcattctcagtccaggaaagccaccagggggtggtctgggtctc**ttggactgtaagaagctcaagaggtccaa**acttcatgaaccctgcacaggtgctcattgactcatgtaaatgttagagcagctgcttcctcccacaggacaaatccacagcacacagcctccctgggctggcccctcacagtgtagctgctctcaggcctgtggacccaggtggtgataaaggcagtaaagatttgcataaagcagcacacagcacaccccctccatggagagagctcaatagGAGATAAAGAGCCATCAGAATCCAGCCCCAGCTCTGGCGCCAGGGGTCCCTTCCAATATCAGCACCATGGCCTGGACTC**CTCTCTTTCTGTTCCTCCTCACTTGCTGCCCAGGTTAAGAGAG**ATTTCAAATACCAGCCTTTGGAGGGATCCTTCTGTCTGCCCTTCTAATTTCTAACATGTGTCTGTTTTTTGTTTCAGGGTCCAATTCTCAGACTGTGGTGACTCAGGAGCCCTCACTGACTG**TGTCCCCAGGAGGGACA**GTCACTCTCACCTGTGCTTCCAG**CACTGGAGCAGTCACCAGTG**GTTACTATCCAAACTGGTTCCAGCAGAAACCTGGACAAGCACCCAGGGCACTGATTTATAGTACAAGCAACAAACACTCCTGGACCCCTGCCCGGTTCTCAGGCTCCCTCCTTGGGGGCAAAGCTGC**CCTGACACTGTCAGG**TGTG**CAGCCTGAGGACGAGGCTGAGTATTACTGCCTGCTCTACTATGGTGGTGCTCAGC**ACAGTGACAGACTCATAAGAGGAACCAAGACATAAACCTCCCTCGGCCCTtgtgatgtggagattgtgtgatcatacacaccagctctcaagacagcctacatgtggaccagcc**atagaaaggggaaggaaagggtctgaattgatttctat**ccctccttgtgccctgaagtggaggaaatgtgagagtgatttgcagtaattgaatgagacaaagcaaaagttatt**tgttttatatgaaaaaaaaaca**gaaacagcaggatcagatctaaaggctgagtctaaatgcatttcctccagacagaagcttcttcaaacgatgggctttctgagctaagagca**aagaaaataaactctccacgggtatattattaaagtttattttatt**gagttactttcaaagcaatccatgactattatataaagtcagaaagtattaaaaatcaccaagttctctgctaagctaccttatcccatgcaatcaaaataagtacttttcttcatttggatgcattttttatttctgtttttaatatttccacaatggtgattaaacctggtgctcattcctctggtttgatccacctttatgctcttgtcatttaaagttaatgtaagtagtgttccatttat**ttaaaaggataaaatattttaa**tagttgatcattgcattttgaactgagtaccagtgaagtattctagcttttaggtaatttcaattgtttgctagtcaatgatgttatacgttgttgtcattaaaccattcctggtcatttctgtggattttgtgaggtcagagaaaatgtctgtgcttttcaatggaaagccttgggtacattattcctgacttcccaatgcacaaaatagagaaacacaaatttcctgctcatgttagttagagcataagagattcaggtacgaggaacaatttataccccaaactacatacagtgtcttcaggcatgtattagaagtgtacttctcaaactctctgatacccttagttctgaagaagctcaaagttggctaatacttttatatttatcttttta

Uppercase: IGLV7-43-2

Lowercase: Flanking sequence[1000bp]

Red & Bold & Underline: Stem-loop [15]

Blue: Heptamer[37]

Green: Nonamer [9]

id-IGLV3-30[V_gene_segment]

cccccatgtaggaaac**tcctggggtcagagacagtctttctcgtgctgaattcccagga**tggaatattgccggatcagagtgggtctcaataaatactggctgggtggatat**gctcctgcctttgtttttggggcaggagc**caccactgcctcatgtttttctacacctggagccatgaaacttggtgtctgaggtctgcggcactatatgcagagcc**ttggtaagatgcactggaccacttaccaa**cccagtgacagtagagggatgtcttaggtgggccctacccattttcctccaataggacgtgctcataatttcatgggtaaccgagcccct**tatgcctggtattgaggcata**taaggccaaataaagaagaggaattcctctcttggaaccagcagtccatggacaagggaatcagggacctgagcagcaggttctgaatccacacctga**tctggggcaggacccaga**acctatcaaagtcttctgctgttaactatagaatgggctcacaaaatgcagacagaatgatctgtgaactccacc**tggcctggctagctgagaaccaaggcca**agacatagtttaaggtaaagatgttggcccagactttgggaccagcaagtgaaggagaattctgtgggctgaggtgggtctgtagggtcctcccctcccctctctgtgtcccttggggctgagcccttctctggaaactacaaagctactccagtagcagcccctgattctgctgacttgcatcacgggccgc**tttctccatcaaggggataagagaggcctgggagaaa**cctacccgccctgggccacaggaagcagcatcagggatgtctcagtcatggactggacctttctcagcctcctcgctcactgcacaggtgctgccgccagagtcccagccacccgc**ccagccccgggctgtgggaccaggctgg**ccttgattgtgagctcagcagggctctgcctgtggtggaaaggatgctcATGACCTTGCTGCAGGGTGAGGGGCTGGTAAGGTTGAAATCTTCCTGTACTC**CTGTGCTCATGGGCACTCTGAGGGGCACAG**TCAGGCTCCCTTTCATCCCAGAAGCTCTGTCCCAGCCAGGCAGGTGCAAAGGAATC**CTGAGGATCAGCTCCTCAG**CTTCAAGCCTTTTCTCCCTCATCTCCTGCAGGTTTCATGGCCACATATGAGCTGTCTCAGCCACCCTCAGTGTCAGTGGCCCCATTACAGATGGCCAGGATCACCTGCTGTGGAAATACTGTTGGAGGTGGATATGTTTATTGGTACCAGCAGAAGTCAGGCCAGACCCCTGTGCCGGTCATCTATAAAGACAGCAAGTGGCCCTCAAGGACCCCTGAGTGATTCTCTGGCTCCAACTTGTGGAACACAGCCACTCTGACCATTAGTGGGGCCCAGGCCAGGGACGAGGCTATTACTGTAGCACCTATGATGGCTGAGGGAGCAGCAGGCAGTGGCTcacagtgaaacagacagatggggatgtgggacaggaacctctaatataactaggctcaatatctc**acatcctctctaagccaggatgt**ccaacattttggcttctctgggccacattggaagaagaaaaattgtcttgggccacacataaatacattaacactaacaatagctgatgag**cttaaaaaattgcaaaaaaatctcataaagttttaag**aaagtttacaaatttatgtag**ggccacgttcaaagccatcctaggctgcatgtggcctgcaggctgcaggc**tggacaagcttgctctaagcagtgaccatgactttggtctcagttcactcaggtg**gtttctcatggtagccaaagcagcaggagaaac**tgtcggatttctatttaaccatgaat**tttacaattacattcattgtaaa**tccaaacatcttgtccctttattacctgggccctacagagactagtgcacaggttcatgggataagaaa**tgtctccagagagcttggtttctgcactagagaca**gtcgtttccctc**cctttcaacacagagtgtggcaggaaagg**aacc**agttctctgagctaaccctgagaact**tgggcttgagaaaaagaaagcattttgaaccagtaggattgtgcccacagaatgagatgtgctact**ctggcccagcccaactcctttcatttcaggagctctgtctgggccag**gcaggtgcaatggaatcctagggatcagctcctcagtttcaagccttttcttctctgggtgtctgagaacaggagttgtcttgcac**gtgcacatgagaaagaagtgcac**acacgactctcaggtgtgagcagcaaagacagaacaaacctttccagaaggggcaaatttcattaagatgacagggccttgttacacaaccccacatgtctggactgggatctcctattggatcttgccata**aatgcaaggtttcttctcccatcataaattcctgcatt**gcagtagac

Uppercase: IGLV3-30

Lowercase: Flanking sequence[1000bp]

Red & Bold & Underline: Stem-loop [22]

Blue: Heptamer[35]

Green: Nonamer [4]

id-TRAV16[V_gene_segment]

catgacccatagttttctgctcttaatcccggagcttctgtctacttgaccccagatttatatctgaccccaaaatgggacacctcacttgagctctctcaacacagcttgactgggagctggagaaaatctttggttaaggaccagggattggggactctttcaatctaccatttttctagaatttccgtctttctgagccaaatgattgttcctttttttatggccttcttttcttttcttttttctttttttttttttaaagtgtatagaactaaaatcccaaagggccatttcctgtgtggggaaggtcagaatcaatgatggtggacggaaaataaaataacttgatctggcaatgtttaaacatttcactaaaatatctgtg**tagaaatttaaatttcta**tgaagtgtaatcgatcaatattttttcttatgttttttgactgttatgc**ttagaaaatccatcaccattctaaatttattttaaaaattt**tt**tattttacttaaaata**ttaaatgttacgtcctttgaactttatttgataacaggtattaaggtag**gcatcctaacttatctgttaacttccaaatgattgttaggattgcacattttcccccattttaactaagtttccaaatgt**atttgtatatttgaactttgtattctacttcattaagccctatatgtggattttaaaaataacattctattgcaattagtatagcatcacacttttactgaggagg**aaattctcattctctgaattt**ga**aggcttcctgaagcct**gagttagctgcagtattgtatttggcctctagagagagatgttacacaaattgaatttg**atagccaaaaacaacctgatctttttgaagaatagcttttggctat**acatgccgggtaaactcatccacctctctgtcagagtgtctatgtggctgaatcgtttccagaaaagacctccagaaaatagcttcctgtttctccacaggtcagacATGAAGCCCACCCTCATCTCAGTGCTTGTGATAATATTTATACTCAGTAAGTAAAGCCTCTTTTTGTTCTATTTTTCTATGG**TATTCTGATTTAGAATA**ACTTTAATGAATATTATCCCCAGTCTGTAAGACTTTTCCTGATCCAGAATTGGTGTTATAGGAGGAACAAGAGCC**CAGAGAGTGACTCAGCCCGAGAAGCTCCTCTCTG**TCTTTAAAGGGGCCCCAGTGGAGCTGAAGTGCAACTATTCCTATTCTGGGAGTCCTGAACTCTTCTGGTATGTCCAGTACTCCAGACAACGCCTCCAGTTACTCTTGAGACACATCTCTAGAGAGAGCATCAAAGGCTTCACTGCTGACCTTAACAAAGGCGAGACATCTTTCCACCTGAAGAAACCATTTGCTCAAGAGGAAGACTCAGCCATGTAT**TACTGTGCTCTAAGTGGcacagta**gctggttttgcaaggaagcagaacacaaaccctttaa**atacaggaaatatttctttgcaaactctctgtat**ggccacagcagggcattctttctccagaaattaatattgagtttatctcgtagatattatatcatcagtcatcagttagctatagattaacctcat**aattttgtccaaaaaatt**ctcattatttaccctgc**ttaaattaaatttaaaaatagtaaatgtatttatttt**gtctagtagtactggaaac**atactttcgaaataaaaactatattgcactgcataaaaagtat**tcttttctttgttc**agaggtactgtccaggcttccatggtgcagtcctacctct**tcaactcctctgctgccccacgtgatccccac**atgtggttgtcaccacat**acctgagcatagttcctcaaagtgctctgctgtttcatggcttatgattgttcatctttctatatctattacccttctttccatgagtttcatagactcctcctatctcaattggtgtccagccaggaaaaaagaagcccagtgaag**tttaatataaagatttattaaa**taataataagtaattaactactaatgggtaaaaaaacctctaaggaatataggaatgcacatgtaagcagaag**ttcctgtctctaggactaaggtagaatcccaggaa**aggaacaga**ttggcataagcatgacccctcctgccaa**ggctatgattcagatctcattgagagtgtgtagctggagcttgctgcatagtggagtagtttgctgaggtgccagagaaaaatactgacaagcagaaaaatacctaccaggattacaacacaattcatgggaagcccgcaaacgagtgctgctgaacttgcttgtggtggctgtgaaacttgccaagaaaccacctgtcgggcttgctggaggggagtgccactggtgtcctacaggccaacgaagcatagtggacaagaagtgtggaaaacat

Uppercase: TRAV16

Lowercase: Flanking sequence[1000bp]

Red & Bold & Underline: Stem-loop [22]

Blue: Heptamer[27]

Green: Nonamer [6]

id-TRBV5-2-2[V_gene_segment]

tcttattcacattggcctaaaagaagtaggagagtgtgtggatctcggggcttttgacattttacatttggggtaaggatgtttgtagggtgctgtgacctttggtagagaaaagcaatcaatatgaaatagtccagcaaggaggcagccaggagacacacaaaccgggacagttggttaccctctctctctctctctctcctgcca**ttgtctctcataagacaa**actcactctggagtacacaaaaagagatgcagttactataatccattggcatcaacctctgaa**ggcaggatggatctggagggataaacataacctgcc**cagaggatgactcatagggtggccttgtgggtgctgaatcccaagtggttaggtctgtgctgtgagctgataaagccccag**aattatttatggggaatctatactcctaataatt**tacagacaacacagatccactttggacttgatcagatggacaaaatcttggggacttgacatcactgcgcacacaggggaggagctgggattgttgtccaggaaaggaaaatataaggagaaaccttggtttgaatttggtgtcagacatatgatgagaggacagccaggaagatctggaacatgaagatttaattacaggccactaactctcagctgacgggcaggtctgtgagttccagaatggagcactgcagcagtaggtggggaggaggctgagagggc**gatgaggcagtctgagtgctgagggcagtgcagtcagagaagcaactgcctcatc**acagaagcttctgcccttactcagcccccttgctctgcaggatgaggagggagttccatgggtagggaccactggacctaaggaagcctgaagg**ggaaggaccacaggacagtgacatcacaggatatccttcc**tatcaggaaaagtgag**gctcagaactcagctcttcctgggaggaccaagccctgagcacaggtgcagtgctgcctgctccgctgtgc**cATGGGCTCCGGACTCCTCTGCTGGACGCTGCTTTGTTTCCTGGGAGCAGGTGAATCCTGTGGACAGGACAGCACCCCTATTCTCAGCTTGCCCACCCCTGTGTCCCCCACTTCACCATGGGGAGGCACAAGTTCATT**CTCCACCTATTTTTTCCTCAGGCCCAGTGGAG**GCTGGAATCACCCAAGCTCCAAGACACCTGATCAAAACAAGAGACCAGCAAGTGACACTGAGATGCTCCCCTGCCTCTGGGCATAACTGTGTGTCCTGGTACCTACGAACTCCAAGTCAGCCCCTCTAGTTATTGTTACAATA**TTGTAATAGGTTACAA**AGAGCAAAAGGAAACTTGCCTAATTGATTCTCAGCTCACCACGTCCATAACTATTA**CTGAGTCAAACACGGAGCTAGGGGACTCAG**CCCTGTATCTCTGTGTCAGCAATTTGAtgcaggcctgcagagccaagaacattctgtgtacaaacatccctgccccagtgtggagaacttcagccctaacatatctgtgagaacttgaggactgtagtgggaaagaaaagcagtttcaggaagctggatcct**aaaacctggggtttt**ctgccagttaggagtggggataacaaggccaagagggtctgctcaggaggtggtgaaaactaccttcttcgccaaacgctgta**cactatagggatagtg**gaagtgtttgtaatttaatag**ctggtgccctctttacagggcagcag**atttaatttcagggtcaagattcttgataatagtgtggaagggaaaatcatagggtttgctttatataattcagggagaagaaagtgagatttggggaaattctatgaaagagaaaatgatcaaagatagttgaaaacagagatagagcagaaggtaatgaaa**catggtgatctgtgcaccatg**agctcccttctcctcccacagtttttgatatccagaaaatacaaagaaaggtcttata**ccaactgacatcacctggggactggtcctagttgg**ggatatcacaaatagcagcctgcctgatgtgagggccttggggttggttctgacatcttctttctcgacttctggatctagtcttgcctgctcgaggcacagtcaaatacttcccaaaacacaaaggaggtcaggtcagtcttggtcctaaaggggcaaatgctgtccgttttctcccaagataaaattctgcatccctgggattcctctctgagactcagtgcccagattc**aattaccttctgtaatt**aaacagttctttgaacattttctaaaactcatcagcgttggtatcatctgcatttctgctatttccttggcctggaaacttttgttctctccagacacatctagcccctgctcatctgtcaaggatggtctcacatctgccttactctttgcttcttaccctgacattccctggaact

Uppercase: TRBV5-2-2

Lowercase: Flanking sequence[1000bp]

Red & Bold & Underline: Stem-loop [16]

Blue: Heptamer[41]

Green: Nonamer [6]

id-IGLV3-15[V_gene_segment]

gtggtgggacccgcacctgtccaaggtgtgcattatcacctgtacagtggagacgaagagggaacttgcattgggttctggaaagtact**gtctaaaaatagatttagac**ctacaccagaaatgtcctct**ctaacataaaatatttatgatag**gacacattagaggcctagaccaaaatgaaatacct**ccctccagagccaagatagaatggaggaggg**gactggggaccccagataattccacagttaccagcatgact**ctatggtcacctcccagaccaggtcaaaacccatcccatag**gagggacttcatattc**ctttgtaaatacagacaaag**tcacctctgagctctgctgtgaccaagggagcccaggagtgaggctgagcacagtgtgatgtggggatgtgtgcagagaatatggaatcaattcataaaggagatcctgtgggcaaagggtgggagggcaaggtcacccctaacctctttgggtccctt**ggggctgtgcccttctcgaaatcacaaggctcctccagcagcagcccc**tgactctgctgattggcatcatggaccct**tctctccaacaaggggttgagaga**cctgggaggagcctgcccagcctcagacttgagaagcagcattggaggcacttcagccatggcctggcccctcttttccttggtctcctgactcactgca**caggtgctgcccccagggtctcacccacctg**cccagcc**ccaggcctccaggttcagcctgg**ccgtcagtctgagct**cagcagggccctgtgtgtggtgggcaggatgctcgtgaccctgctg**cagggggaggggctggttggggtgaaatgcccccactctgtgctcctgtgctcataggtgccctgagagctcttttattcctggggctcactccagccctggcaagtagccagatatccttgggttgagaccctcagtttcaagttttttctgtccccttttctttccttgtaggctctgtcACCTATGAGCTGACACAGTCACCCTCAGCGTCAGTGGCCCCAGGACAGATGACCAGGATCACCTGTGGGGAAAGCAGTATTGGAAGTAAAAGTGCTCAATGGTACCAGCAGAAGCCAGCCAGGCCCGTGTGTGGGTCATCTATGGGGATAGCGGGCGGCCTTCAGGGATCTCTGAGAGATTCTCTGGCTCCAACTTGGGGAACGTGG**CCACCCTGACCATCAACAGGACCCAGGGTGG**GGACAAGGCTATTACTGTAAGA**TGTGGGACATTAGCACTCCTCATCCcaca**gtgacacagggagatgaaggagcaaaacacaaaaacctcttccattggtgtcactc**ttgtcctccagcccaggaagacagtggacaa**agccatgagtgtgtgttgcccagttaacctggatgtgagacccccaggctgccctttccacaggccctccaggcaggctctgcagagggggcatcaagagtgaatttagggctgagatgaccaggatatattgaggtgtggaggaatcagttgtgatttggggaagtggggcctggttggaagatcagaccaaggtcgacatgcttagttgggctctggatttccatagggggtgaaggctagtccttgggctgaatatcctgagtcagtttccagaactgcccagctcacgttcccgtggcatcctgccattctcttcaggaaaaccagtggagtggacactgtgggatccacccatattccctcttctgagctcacccacccagcactgctggt**agctctggcagctcagagct**gtgccctcactgagaagggctgtcagcagagaaaactgtctcctccaggtttatgccccttcttagcaaggtgtggtttcataactgcctgaggtgaagctctcaagactccctttcaata**tgggagaaacagaaaatcctccca**gcttcacagctccttattgggatggtagaagactcaatgtgggtcagagtgtccttgtgcccggtgttgcctccctcacttag**tttcacacctactgtgaaa**gcaaagctccatctcagagtctgcttccaggcatcccaaaataacataaccagtcatcccgaacgtgagccatcaggaacctcaggaggccacagtataccagaaagcaccagtttattcctaatgatcattgtgagtttg**caaatgcccaatgagatctgagtgtttcttgcatttg**tatggaatgaaaagggccctggcatgccaggcactgtggggtctgagggagagttggagtcagatctc

Uppercase: IGLV3-15

Lowercase: Flanking sequence[1000bp]

Red & Bold & Underline: Stem-loop [17]

Blue: Heptamer[45]

Green: Nonamer [5]

id-TRBV28[V_gene_segment]

acaaatatgtagtaagaatactatgatcacaaggaacaaag**acatggtggactttggagaaatccatgt**gagacttccttatgttctcttcca**tgagagagcacactctca**ccccaacaatgaaaatgcagcaacgtgagaacaacctttctgcccagggaagctcattagagac**tcagcacctccaatactttcagtggctggtgttaggtgctga**atcgtgtcattaaacattcatatgttgaaatcccagcctccagtactaagaatatgactatgtttggagataaggattttacagaggtggttaaggttaaatgaggtcctcactaatatgactgggatccttataagaagaaattaggacacaaacatgcatagagggaagacaatgtgcagacacagggggaagccatctacaagccaaggagcgaggctccacgagaaaccaaccatgctgacaccccgatcttggatgtctagtctccag**atgtatgagaaaatacat**ttctgttgcttaaatgacctagtctgtggtgacttatggcagccctcgcaaaataattcagctgatcacattggcaccctctgcctatcatgcagcataattccagacttgacgtaaagcaagtgttcagcctaacccacgtggtttgtgcaagcagtctagacacactaagccatgcttaccagttactatagtgggaacactcccaaaattcaagttcgcagatgcaaatcaagagccaaccatgcaagcagtcccttctaaggccagcagtctcaggcccg**ctgtgtcaactcttttctatacgtgcacatacacacag**tgacacttccatgagcacaagggatgtccttcactctctagcacc**tctagaccctttctagtctaga**caatggctgagggattatgag**agtgggccgggacaatgacatcacagaccatcaacccact**gcctggtcctgggagaagacctattctttcttcaaagcagccATGGGAATCAGGCTCCTCTGTCGTGTGGCCTTTTGTTTCCTGGCTGTAGGTGAGTCCTGGGAGTTGCTGGGTCTGGGGAGTTGGTGGCAACCCCTGTCCTACTATTTTGAGGTCTTTTATACCAAGCTCTTTTCTCCTCTCCTCCTTGAT**GGCCTGTTTTTCACCCTTGCTTTCTGTCTCTCACAGGCC**TCGTAGATGTGAAAGTAACCCAGAGCTCGAGATATCTAGTCAAAAGGACGGGAGAGAAAGTTTTTCTGGAATGTGTCCAGGATATGGACCATGAAAATATGTTCTGGTATCGACAAGACCCAGGTCTGGGGCTACGGCTGATCTATTTCTCATATGATGTTAAAATGAAAGAAAAAGGAGATATTCCTGAGGGGTACAGTGTCTCTAGA**GAGAAGAAGGAGCGCTTCTC**CCTGATTCTGGAGTCCGCCAGCACCAACCAGACATCTATGTACCT**CTGTGCCAGCAGTTTATGcacag**cgcagcacagccgcatc**ctctctgcacaaaaagagcggacgtaagagag**aaggggccctaactcagggctggtgctggctccgatggcacattcgtgctaaatagaaaaaaagcggccactaaggagaaactagctccataagcacacttctttgcaaggagagaatgaggtattatttttcacccctttacaagtcag**ggactgttctggatttaattcccccagtcc**gatgtgttgtaaattcacgagtcccaacaaggaggaaaagtcctctcgcatattcacagtgtaaccctgttctaccatttcgttaatcaattttaccttcccttttctatcctcgtctctcttaactgtgcattttctactttgcttagtgttttgctcttggttgacatttcactagaggacattaacatattctaaattctttcgatgatttttactgtgcctgtgctgattcaagagaaagtttgtggaat**ttgaattttgttcattcaa**tttagaaagagaaaaactctgattaaaacaaattc**ttccaggttctccgatcaatcatattttgctggaa**aaaattctgatcacagaaggtttaaactctgctgcagaaggtggtaggaatttggttataactgaactcacaatcttctggaaaaagtgtgtatttgtctgtctttaagaataaaacatatgctcttctgtcagtaaaaattaacagtgctttggaaatacaaataaatgtaaagggtattagaaatcccaaactgcagagcacttgcacaattattattggcaagacaaaacaggcatctgaaaacgtattttgattcagtgcta**aatttttgtgtggaagaagtcagagaaaaaaattgaatgaatgtgtatcacaatattcaa**gattaagtttgtgtgagataggacttgagctga**ttttgaatacaagtaaatgagggaagagaaaccatcaaaa**ggaagaaataccctgcaaatgttt

Uppercase: TRBV28

Lowercase: Flanking sequence[1000bp]

Red & Bold & Underline: Stem-loop [17]

Blue: Heptamer[27]

Green: Nonamer [3]

id-TRBV12-1-2[V_gene_segment]

ccttaggatcggaaagtactctggggtgcaggggactcagcagtatgcctctgtgccagcagcaaaaatgtggccctaaactgctccttcttggttagcaattaacccagttgaccaaccc**acagagcagaggcttctctcagctctgt**acactctacaccctgagaggaagtcggcttctgagatgcaagactcttttataaggaaaagtccagttgtaaagattaagaaactttgagctttgctagaggtaaaaataggatgtaagtcaactctggagcatttaatttccaaatgtttatgaaaaagacaatcttcttcctgttttattgtatggatagagaatgagagagattctgtactctgggctttgcaaactcactagagttcaagaataaatacactcagagaggtctttcactgcaaagaagagctttgaaatcgaatgaactttcaggatatcagttcggctcctgtatctctaacaacgcagttgacatgcttgcccttcctagggatctcaggggccaactcctaactcacttgtggataattgaagtgcattcaaaggagtaggagcacagagacatcagggagaggaataggagagtttgcagagaatctggcttgctgtaagacaatgag**ttcatcttgaaataattggagtgtgaggtgcagatgaa**tatagttggcaagatcctaggtaagatatactatagagaaggctttatgaattggtaatatcaag**acaaatccacagagagcctccgtaggtgtgcatttgt**gtctcagatcaattatagttcagtcctgcctgatt**catctcccagagatg**cag**cctccacttaaataaggagctttcaaattggagg**tggtggcccat**tcagtgacgtcactga**cagatgcatcttgtgtggataaaatgtcacaaaattaatttctttgttcatgctcacagaggccctggtctggaatgttccacttctgctctcactctgccATGGGCTCTTGGACCCTCTGTGTGTCCCTTTATATCCTGGTAGCGAGTGAGTTCTCCGATATTTATTATGGTCATGCTGGACCTCTGTCTAGATGACTCCTTATATTTTCCTTATTCTTTCCCCCAATTCTGTCTTCTTTTATAGCACACACAGATGCTGGTGTTATCCAGTCACCCAGGCACAAAGTGACAGAGATGGGACAATCAGTAACTCTGAGATGCGAACCAATTTCAGGCCACAATGATCTTCTCTGGTACAGACAGACCTTTGTGCAGGGACTGGAATTGCTGAATTACTTCTGCAGCTGGACCCTCGTAGATGACTCAGGAGTGTCCAAGGATTGATTCTCAGCACAGATGCCTGATGTATCATTCTCCACTCTGAGGATCCAGCCCATGGAACCCAGGGACTTGGGCCTATATTTCTGTGCCAGCAGCTTTGCcacagcactgcagaatctccccatctctgtgcagaaaccctggtgcttcctcttctccccacagctctcagcagtcgtcagcaaagtctttcctgctctctgctcaccatggctcacgcctataatcccagcactttgggaggccgaggtgggtggatcatgaagtcaggagatcgagaccatcctggctaacacagtgaaaccccgtctccactaaaaatacaaaaaattagccgggcacggtggcaagcgcctgtagtcccagctactcgagaggctgaggcaggagaatggcgtgaacccgggaggcggagct**tgcagtaagccgagatcgtgccactgca**cttcagcctgggcaacagagcaagactccatctcaaaaataaaataaaataaaaattaagtgggtttggagaatgacaaagacagaagatggtaggatattaacataaaaaaaatcatgtgtaggggataatgtagaaaatgtgt**tttgtgaatttctcacaaa**ttgtctaaggagtcagaattcacaggcatgtaagat**aaaccttaattacccccactgtaggttt**ccttgtctccaggtatttgccatccgcattagcaagctgagagctcctgagaatggctatgtcttttgaactagtctgccctctgcattctattttcaccctaagctgtgaacagaaatttcccccacaagttttcttgattgt**aaaagaaagttctttt**ctatggcattccttataatgttaatttcaacttaccatttatatagggattttgcattgcacttaaagagtaaacctcctgtgggtaatacaggcagtgaaataggttagatgtaagctgctcctcctggactctagggatctatttgtctttatcctgcttaaggagagccaagttctagctgaagg**cctgtgccaactcacagg**acactgacgtggtctcttccactgcctgcttggagctatccacatgtatccacacaggctgt

Uppercase: TRBV12-1-2

Lowercase: Flanking sequence[1000bp]

Red & Bold & Underline: Stem-loop [11]

Blue: Heptamer[43]

Green: Nonamer [4]

id-TRBV30[V_gene_segment]

tgtgttttttgtggctcgtgatcccttcagcccctaccttccaccccatgccattgtaaatctcccaatctttgctgcagacatagggagtcaaaaaaaaaaaaaaaaaaaaaaaaaaaaaaaagccaacatatgtaagagtgcc**tgcctgcagtgcaccaggca**aaatgcatctttcaacaatttttagatattgtcatcatt**attttcactttatagataagaaaat**tgagaatcagggtttgcttgagagttcccaagccagcaagaagcagagctgtgaaatgaacccaaatccatatgattccaagtctctgtgatgtttcctattactcatattgcatttagaggataaattagatct**agttggcccattagagacacatggagcagattccaact**ttgatcagttcaggagttaattatgcctgtccaggtctcaaaatctgtctgcc**agggatgaagattatgcaggcgatccct**ttggctgtcatagaagcttcctgccctcagaactgagcatttctgctttgattaccccaaaa**tacctgaatctcttgagccctatcctttcatctcccaggta**acatgtctagctc**tagtttccaatttcagataaacta**aaatcttttttcatccttctatcagtatcacagtttctctgaagctttcccaggcttctgatgtctcccgggggagctggagag**ttacctttagtttacaggtaaataatgagtcattaaatatatttac**ataccaggatgtcagcaggaatgcaaagaagataacaagtgcagagtatagctggttcccagggaggctgggggtgattcaccacactcttaaaagaagactagcctcatttcctccccagctcatggggtaggattcc**tctgctctggcagcaga**tctcccagagggagcagcctgaccacatcactggcccagaagaggaggcgtctgtcccccagactagctgaaggaaaggctggcttggatgATGCTCTGCTCTCTCCTTGCCCTTCTCCT**GGGCACTTTCTTTGGTGGGTGCCC**TCCTCTCTGGCAAGCCTCTTTTCTCATCCTCACCCTCACCCCAATCCCGTGAATGCAGAGTCTGAGCTGTTGGTTTGACACTCTTGAGTCTATTCCTGATTGTCTGGAACAGATATGGGAATTTAGCCTTAGGGGCTGATCTGATGATTCTTAGTTCTCTCTATCTCCTGAGAAGGAGTTCGTTCTGTTGGTCTTCCCTGGC**CCAGGGATGCTTTGCATCCTTGG**CTCCCCCAGTCTACTCCCCATGATGGATTCCAGGTGCCTGTGTCATGCAAGAGCTTGAGCTGCCCCTCTGTCTGGGGAAGGCAAGCCAGGCTGGGGTGCTCCTCTGTAAATGGCACTTTATCTTTCCCACAGGGGTCAGATCTCAGACTATTCATCAATGGCCAGCGACCCTGGTGCAGCCTGTGGGCAGCCCGCTCTCTCTGGAGTGCACTGTGGAGGGAACATCAAACCCCAACCTATACTGGTACCGACAGGCTGCAGGCAGGGGCCTCCAGCTGCTCTTCTACTCCGTTGGTATTGGCCAGATCAGCTCTGAGGTGCCCCAGAATCTCTCAGCCTCCAGACCCCAGGACCGGCAGTTCATCCTGAGTTCTA**AGAAGCTCCTTCTCAGTGACTCTGGCTTCT**ATCTCTGTGCCTGGAGTGTcacactgagct**gggtggggcagacatctgtgcaaaaaccccaccc**tctcctgag**ccctaaccatactccccaggggccttcacttaggg**actgggtggaggatatttgtaagtaggtttctgactgctggccaatgtgaaagcaatggaataaagcgagaattctacaaaagactctactaa**gttctcagggagaac**aaaaaattgtatg**tgtgcagaacctgtgatttgcctgcaca**tagtcaagttctcaat**gtatggatgtcccggcccaggctaccatac**tccagccct**caaggtgtgctataccttg**cctgattttctagtcttccaaattcttctatgtgtgccgtggggagggggtttgtgcatatatggctgagagaaaggatatacctgtggcaggaggtgagctggtgaacaaacgggaggatgccaac**ccaggttacttaattggttaatttcatcacctgg**ag**agtctcctgtcattgatctttctgctgagact**agaaattatggggaggcttagacctgaggctttgagtcaggcaaagaggcaagatgttcaggagactgttctcattcgaggcagaagtgtggttttgttagtataggtaacatgcacttaggccagacgtggtagctcatgcctgtaatcccagcactttgggaggctgaggcaggaggatcacctgaggtcaggagtttgagaccagcctggccaatatggtgaaacctgtctctactaaaagtacaaaaattaaccgggcgtggtggcaggcgtctgtaatactagctacttgggaggctgaggcaagagaatcacttgaacctgggaggtggaggt**tgcagtgagccaagatcatgccactgca**ctctagcctgggt**gacagagcgagactctgtc**tcaaaaaaaaaaaaaataaaaataagcttgcacttagaaattttggtgatgtttgaggttttaaaaaatgtcctgttgtactatcttgtagccc

Uppercase: TRBV30

Lowercase: Flanking sequence[1000bp]

Red & Bold & Underline: Stem-loop [22]

Blue: Heptamer[19]

Green: Nonamer [5]

id-IGLV5-45-2[V_gene_segment]

tttttttttttttttagacagaatctcgctttgtcgcccaggctgggg**tgcagtggcccaatatcagctcactgca**agctccacctcccgggttcccgggttcacgccattctcctgcatcagcctcccgag**tagctgggactacaggcgcctgccaccacgcccagcta**atatgttgtatttttagtagagacggggtttcaccgcgttagccaggatggtcttgatctactgacctcgtgatccgcccgccttggcttccc**aaaggaattgctgcctttaacatgtcattgatccttt**atttttcctggaaatttttcattattcaccatctttttgaacaaccc**cagctgcccttcaacccagctcctcagctg**accccttttccatga**tgcccactctgatgacatatgtgggca**aatagatttttctaaaatgccctgttggggtgctgacacctataaaagttatgtatgtaca**ggtctgcacatcccacagacc**aagactacttctgggagtgagaaatcgtagtaataatgcatagcatgatagaagcag**tgtctgggaatcagaccatggatatgtgaatacagaca**gtgactaaag**agctgcagagaccagctgggttctgaagtctctgagct**ttgggaaagtgtgtttcctgggaccagcgtcagatgttgtggggctgccctgtggttccgacacagc**tgctcagtaatgagca**ttcactcagagaaatccaggccttgaggctgggccctgtgctcactgat**ggacactcggcagctgtgtcc**cctctggcctggcagagttccctgagcaggg**acctgtgtatggtctcaacaggt**gcttcctcccagggccttgcaaaaagaaaagcaaaccttcctcct**gctcagtctgcagagagagctgagc**tgcatc**cccaggtctcaaggaggggcccggcagtccctggg**tggaaacacatttgcatgagcaacccctcctctgcaa**cAGTGGGAAGAAAAGGAGGCCTGGGGCAGCCCAGTCCCACTG**CGGGGGTAAGAGGTTGTGTCCACCATGGCCTGGACTCCTCTCCTCCTCCTGTTCCTCTCTCACTGCACAGGTAGGAATAGACTTCAGAGACCAGGGTCAGCCACCCAGCCTGATTCTGACTCTTCTGGCAAAGATCCCTGAAAAACTTTACCCTGGTTTCTGCCTTAGCACCCATTAATGTCTGTGTTTCCAGGTTCCCTCTCGCAGGCTGTGCTGACTCAGCCGTCTTCCCTCTCTGCATCTCCTGGAGCATCAGCCAGTCTCACCTGCACCTTGCGCAGTGGCATCAATGTTGGTACCTA**CAGGATATACTGGTACCAGCAGAAGCCAGGGAGTCCTCCCCAGTATCTCCTG**AGGTACAAATCAGACTCAGATAAGCAGCAGGGCTCTGGAGTCCCCAGCCGCTTCTCTGGATCCAAAGATGCTTCGGCCAATGCAGGGATTTTA**CTCATCTCTGGGCTCCAGTCTGAGGATGAG**GCTGACTATTACTGTATGATTTGGCACAGCAGCGCTTCTCACAGTGACACACACAGATGGGGAAGTGGGACAAAAACCTCACCCTGCTCTGGGTCTTGCTCTGTACCAAT**TTTTAAATTTTAAAA**TAACTGGCCTAGGcacaaactatatttggaagtactttctagttgtaaaaggctcttctctcaattttctatccatttttctgaagtctcagtaaaacaaaacaaaacaaaacaaaaccacctgatgaccctgaagtgttggctggttg**tccacatgtgctaacaccagtgcctgtgga**gctggacgtttttttcttgaaagaagcaaaatgaggctttcctagctcacatcatggtcagagcctggggacaacatagcaggtgaacagtgccctgcagcatgga**ctctgctcttccagcgtacgggcagag**acctccccttccatccccagtagtctgttctcagggctgtctgaggctcactgtgtcccaggaacaaggcactcaaatatgaaatgctgcctgaggact**caggtcagggaattctggatatagtgctgacctg**gctcagtcctgggtacctgcatttcagtggcaatgtcaggagatggcgttagatattagggtaactgcgggcccgatgccagtaaacctctatgtaggacatgacagaatcaaggggaggc**tccaccatggctgagacaagccagaggctgctccttggccatggggga**agtagatcccatggatgtgactcctgtagctcatcccactcaggaccacggggactg**ctgagggtgggtttcctgatccctcag**ggcctgactgtgttctcttctcctctcttacaccattggagcttct**cactgactcaggtctagtggtgagtctgtg**gaatatctggatacagaaactgcccaatatttgcttcaagtgactcctccctgggaaact**cttgaggaggcccccacctcaag**gtcatctttttc**tccatggggagcagcagaaacattggccattcccatgga**aatggctatgaggcctcccagagagaacccctaagtttatgacctaggacaatagtgatggtccctcaggggtcccagcctgcttctcctgatctgggtttatcc

Uppercase: IGLV5-45-2

Lowercase: Flanking sequence[1000bp]

Red & Bold & Underline: Stem-loop [25]

Blue: Heptamer[40]

Green: Nonamer [7]

id-IGLV11-55[V_gene_segment]

cccaatctcctgtctcaggttctagaacttgtcctggaacttccccagagctgagtctctaataggagagctgatgacaggcacagccgcggtcaa**gccctctagccagcagatggaggagaatcagcctgagggccctgtgcatcctccctgctgcacacctgggccc**atagccccctccaccctccatatgctcctgcagcccccacctcagtgtctgcctcagtgtctcctcccagcaccttc**ctcaggaagggcctgag**atcttgcatggtgtgtttccccttggtgaactcctagtcacagcagcatcctgagtaagagcctcctggtgaagagatcgtcttggtgctcca**ccctggcattgtgtcctcagctccaggg**caggtcacatgctttcctggaccaagattacatgcagtccacccctgctgg**cccctgccccacagagggaggcccggggtcagggg**tggcaccaagaccacagctccactgatccctgtaggaattcttcctgcatgtttgaaagacaagagttacagaaacaaagcagagggaagattctggatcattgcaaaaggagaaatttgcctgcagaccagcagggggcgcggtgagaacttctctgtgtatttctgcaggacaggtgctcacagctggggagccctcaactga**cagaggcccagggctgctccttcacttcctcctgggactctg**cagccttgtctcatctgtgcctggcagagtcctctacacatgaccctgtgtgtggtcccagcagctgctgctttctctggtcccacccgtggggcaaactcttgcttcatgccagcctccctgtccaagggtctggtccctgcgagtgcacctgcgggagaggcaatggggaaacacatttgcatggtcagcccctcctctgtgagcccagtgacgctgggataaga**gaggcctggggcagcccacccctaggtctgtggcctc**agaggcagctgtgtccactATGGCCCTGACTCCTCTCCTCCTCCTGCTCCTCTCTCACTGCACAGGTAGGGACAGGGCTCAGAGCCCAGGGTGGTCCCCAGCCT**GATCTGTCCCTCATGGCTCAGATC**CCTCAGCAGCTGCGCCCTGACCCTGCTCCTCACTGTGCTGTGTCTGTGTCTGCAGGTTCCCTCTCCCGGCCCGTGCTGACTCAGCCGCCCTCCCTGTCTGCATCCCCGGGAGCAACAGCCAGACTCCCCTGCACCCTGAGCAGTGACCTCAGTGTTGGTGGTAAAAACATGTTCTGGTACCAGCAGAAGCTAGGGAGCTCTCCCAGGTTATTCCTGTATCACTACTCAGACTCAGACAAGCAGCT**GGGACCTGGGGTCCC**CAGTCGAGTCTCTGGCTCCAAGGAGACCTCAAGTAACACAGCGTTTTTGCTCATCTCTGGGCTCCAGCCTGAGGACGAGGCCGATTATTACTGCCAGGTGTACGAAAGTAGTGCTAATcacagtgagacagatgaggaagtcggacaaaaaccaaggttttaagcttgtcattttt**actgaactggttaagaacttcagt**ggttaataaaatcacattaaatactggattgttgtt**aaaaatggtggcacattataactctctcccattttt**ttcacataatgcccatagtgactctaagcagctaaaatcaaaacacaaaaacttataccaatactaagaaatactaaca**ttactaaccaatttccactagtaa**aagagttgtgaaacagctgaaatacatccagatggcataatacataataccacta**tatacaagagttgtata**aaattatgtatcatgtatatataaaattatgtatcatgtatatgtataatatatgaaataa**aatgtcagcagacatt**accgtcttcccaccagtagccagcaattctgccctgcctctagaataaatggttaccatgggttgaataggtgaacagccctggcagctctgcaggatgactctcactgtttgttattgttttatttttaagattgcttttctttcctttttttttttgaggcagagtcttgctgtcagcgaggctggagtgcag**tgcagtgagctgggctcactgca**acctaagcctccctggttcaaacgattctcctgcctcagcctcctgagtagctgggatacaggcgtgcgccaccatgcctggctaaattttttgtattttttagtagagacggggtttcaccatgttg**gtcaggctggtctggaactcctgac**ttcgggatccgccgtcctcggcctcccaaagtgctgggattacaggcgtgaaccaccgtgcccgacctcttttttttttttttttttttttggttttgttagtttagttttgttttatctctttaacttga**cagagatgatccatgtggatctctg**catgggtctgggagctg**caaggctcctggtgcatggcccagtcccgcctgccttg**gcagggctcctc

Uppercase: IGLV11-55

Lowercase: Flanking sequence[1000bp]

Red & Bold & Underline: Stem-loop [18]

Blue: Heptamer[59]

Green: Nonamer [4]

id-TRBV5-6[V_gene_segment]

gaattgggaggagcagactagcaaaagtctagattcctgaaaacagagcataaagggcaattctggtgaaggctcagggggaaatgagg**aacaaggtattgaaaatcgaagtaaatgccatccttgtt**gtaagtagcaaaaaccttggcaaaattgtgtcctgttctagaactttatggaatataaaaattatgagccatttgctagtatatctgctgaaagaaatatctaagaggcaaagcattcaggctactgtgtgactacttttagccaccaggaaatttaacccagcaagaagggagccatgggaatagattttgcaaaccagcacagatggtgactctacctccctctgctgtcctgtctcccataagccaaactctgtgctgtgagctgttaaagtccaagaattatttctggggaa**tctgtactcttaataatttacaga**caacacaggtctgctttggatctgatcagacagactaaatcttggggactgtaccagtggccactgagaaaaggggctcgg**aatgttgtcttggatagaaaaatataataagaaacatt**ggtgtgaatctagcatcagaaagatgacgtgaggacagcaaggaagagctggaaactgaggtttactcacaggctcctcaccctccgctgatgggcaggtgtgtga**gctccagcatggagc**accacagcactaggtgggagaacggtgatagggtgatggggcagcctgtgagctggggca**gtgtaggcagaggaggaactgtatcaccacagaaacttctgccttcac**acatccctccagctagg**caggacaggtagagagtccagtgtcctg**gagcactagacctaaggaaggctgcatg**gggaggacaaaggacagtgacatcacaggatacccctccc**atcaggaaaatcaaggcccagaactcactcggctcttccccaggagaaccaagccctgaatcagatgcagtgcttcctgtccctctgtgccATGGGCCCCGGGCTCCTCTGCTGGGCACTGCTTTGTCTCCTGGGAGCAGGTGA**GTCCTGGGCACAGGAC**AGCAGCCCCATTCTCAGCTTTCCCACCCCTGT**GTCCTCCACTTTACCTTGGGGAGGAC**CTCCAGGCTGTC**CCTGTGCTCATCCTCCATCTGCTTTTCCCACAGG**CTTAGTGGACGCTGGAGTCACCCAAAGTCCCACACACCTGATCAAAACGAGAGGACAGCAAGTGACTCTGAGATGCTCTCCTAAGTCTGGGCATGACACTGTGTCCTGGTACCAACAGGCCCTGGGTCAGGGGCCCCAGTTTATCTTTCAGTATTATGAGGAGGAAGAGAGACAGAGAGGCAACTTCCCTGATCGATTCTCAGGTCACCAGTTCCCTAACTATAGCTCTGAGCTGAATGTGAACGCCTTGTTGCTGGGGGACTCGGCCCTCTATCT**CTGTGCCAGCAGCTTGGcacag**cccatcagagtcactgacgttctgtatataaacttcctgccttagctttgccttgagagctgcaggccccacccagatttcactccttcaagggaagcttttagttgtttggaaggcatgtcttgtgtcctactgagggcagagctctcccaaccaataaagcccaagtttcctgtgtcctgagtgtgcccacttctgt**gctgcatcttcttgcagc**ttgtcacctcagtacaagagtgactgctgagccccagatgtgtgcttgattatttgcatttcttacatagcttaatgcctttcaacaatcttgcacatcaaacattcttattcttcctttacagatagaaggctcagg**gacattgagtcattttccccaatgtc**tcttggcttgtaaggatcagaagtaggaaacaaactagtccatccattttccacctacccccctgctccatcatcatcttctgcatc**ctggtcagtaaatcagagccctcacactgccgtctagtgaccag**cagggccgcagtagaggctgaaatgtcttcaggggtcattactatgtccttttcattagctcctttgag**gaatgttaaatttcacacattttaacaatc**atttatatgcattccctttgtctttccccagtctgcagcttatattttcatttttatgg**agtttttggtgcacaagagcttaatttaatgcaacaaaaact**aaaaacaa**ttttgtttgtgtgtagacaaaa**accactttttcatataaatgtctaagcatatttttctc**aacttattctactaaaatttaagtt**tggattttcaccactaagaataaatgtacctgaaatacatctcatgtaagaaataagatacagacctaggaatgggagggaaggacatatattcagtgaagaaacaatagcttgtcaaaaagagccctaagtggacacctggagctgacggtggagaggtaggtggagggcaaatgagcatgaactttgt

Uppercase: TRBV5-6

Lowercase: Flanking sequence[1000bp]

Red & Bold & Underline: Stem-loop [18]

Blue: Heptamer[27]

Green: Nonamer [8]

id-IGHV3-74-2[V_gene_segment]

atcaagtatttaaattgagtaaaataagccaaacaaataagagtatgtatgattctatttttaaaaattctggaaaatgaaaactgatctaaagtaatataaagaagattagtagtttcctgggaatatgttggcagaagggaaggagaaaggataaggaaatagaaataggaagtagaaggacagaaaagaagttgagggaatttcacttgtccaccttccttataatggtaatagttatgccatgattatcagt**tttacactttaaatatgtaaa**g**tttataatctgtcaatcaaatcttataaa**atgtattatgaggaaacaagttgaaaattagacaatgtaggagtgacagaaagatagatatgagtatgttgaatgtcagagatacctgaaagtttatctacctgaaccctagtt**ctctccatagtttaaggtaaacaggagag**tgcaggaaaatcatccatattctgattaggcagtggcttctgcaaaccacactaggcctggccggctgtgtcctggagttggctaagggaggagtcagggccagtggtgagaagtgcaggcccagataccagaactcactcatcccagacatgagctcttagatacacagagagcccatccatgtgtggatttatcttaca**tctgtaagtagagaacattgactcttacaga**acata**atttacacacataggtaaatctgaaataaggtgatcagtgtgaagattt**tatcacagcacagtttcataataagcacaatttctcaaatcccattgttgtcacccatcttcct**caggacactttcatctgccctgggtcctg**ctctttcttcagg**tgtctcaccccagagcttgatatatagtaggagaca**tgcaaatagggccctcactctgctgaagaaaaccagccctgcagct**ctgggagaggagccccagccctgggattcccag**ctgtttctgcttgctgatcaggactgcacacagagaactcaccATGGAGTTTGGGCTGAGCTGGGTTTTCCTTGTTGCTATTTTAAAAGGTGATTCATGGAGAACTGGAGATATGGAGTGTGAATGGACATGAGTGAGATAAGCAGTGGATGTGTGTGGCAGTTTCTGACCAGGGTGTCTCTGTGTTTGCAGGTGTCCAGTGTGAGGTGCAGCTGGTGGAGTCCGGGGGAGGCTTAGTTCAGCCTGGGGGGTCCCTGAGACTCTCCTGTGCAGCCTCTGGATTCACCTTCAGTAGCTACTGGATGCACTGGGTCCGCCAAGCTCCAGGGAAGGGGCTGGTGTGGGTCTCACGTATTAATAGTGATGGGAGTAGCACAAGCTACGCGGACTCCGTGAAGGGCCGATTCACCATCTCCAGAGACAACGCCAAGAACACGCTGTATCTGCAAATGAACAGTCTGAGAGCCGAGGACACGGCTGTGTATT**ACTGTGCAAGAGAcacagt**gaggggaagtcaatgtgagcccagacacaaacctcgctgcaggggcatctgagaccacgagggggtgtcctgggccctgtgaactgggctgctctccgtggcagcggctggtggtgctaaaggctgattttctctcagcatctggggctgattcatcaagtttcctcagagaacct**ttcagatttacaattctgtacttacgtttaatgtctctgaa**tgtgacactttccttccctgg**tgtgtctttgtttttgtgacaagaggacaca**ttctcacctccacagaagcccgagtgtcactttggggacagaaatgaccctgccctggtcaccagaatcagagtcccgaggaag**cccaggaggacctggg**aagtgttttt**caatcagactcagggcaggcgtctccgtgggaatctctgattg**gaacaggctttgggattcagattgggaccaagagggaggctcacccagggccagggtccttagaatcctga**cagttttcacagtaaccccatcgtcctttaaaactg**aacatctaactcagaactgacccatttggtcctttctctgtaatccattttccttttctctaggcttcattcttacact**tcccttttcaccttcattctgaaaatgga**gggtgtgctt**cctgtggtctaaaccacagg**cctcagatgcattacctggaactcaggtgtggctctggctatggctcctgtggacctg**gcaggctgagggatctttctcattccctggtgcctgc**atgcccctgctgtcttctatgcgtgga**tgcatttgggaaatgca**agtggacactcatagtcgtttcctcaaatgggatactgttgtagagctgatcttgtgcttctcaccctgtcacagagcccccactctcacttgtggatttttggg**agagctgaggatggacactttattgggctgtgagctct**gcatgatggcaatcgtgaggtctgggtgggcacagccagagccaa

Uppercase: IGHV3-74-2

Lowercase: Flanking sequence[1000bp]

Red & Bold & Underline: Stem-loop [20]

Blue: Heptamer[46]

Green: Nonamer [6]

id-IGKV6-21[V_gene_segment]

tgtttattttgggggcatttcaggatcttgctcaagaactgtagagatttttttctgtgactcttttttggtgcttgcatggaggtttacagagtttcctcatctaatatagattatctagcaccaggcaatgtgctggatctcatggctgaagtgacagaggcatttgcatta**aaactcaaacttactacagaatattttctttctcagagttt**attcataaaagacagccttccaagttagctgataaa**tgggttggtatagtaaaccca**agtgcaaaatgcattgtcaacactctaggatggcttaaccagtaatgtgcttcattgctagtggttggaagtacaaggtgcaattatttttccttactttggaggggataagccagcatgactcatactccttttataaacacttgacatcttctctaatgtgacaagcccttgatgttttggggcgtgcatcccaccctctagagcacatgtgttttcacaagaaattcagagttcttacaatgtccagctcatcacgtctaattaccatgatgt**catcaatatagtgttgatgctttgtggaacgttcacaaagc**tttttcagcctacattgtgacagagagcaggagagttaacatagccctgggacgagactgaggatgtgagctgttattcaccccagataactgcagactctcccagagatggcgatggactctgccttcactctgcagctgtgccctggggtcttggtcaagccctgccagagcctcagcggagctcgtctgcaggtgccagcagagggcgcttcacacccctcatggaaggggccgggagggcgctctcctggcaacagtgatttctgtttatttaaaccagcaggacatccccataatttgcatgtatcgttcctcctatatgtga**agaggccctgcctct**cggtatcttaaaagaggttctttctctgggatgtggcatgagcaaaactgacaagtcaaggcaggaagATGTTGCCATCACAACTCATTGGGTTTCTGCTGCTCTGGGTTCCAGGTG**AGAATATTTCCACAAACCTAGGCGGAGATATTCT**TTCAATCTGTAATTTCTTTCATTGGGGACTCTGCAATAGGTGATTTTTGGCTTGA**TTTTAAAATCCTAATTTTAAAA**ATGTAATGCATATTCTTTCTTCATGTCTAGCAAGATTAAAGGTGATTTTCATACACAGATATTTATGTTGTACTGATGTTTGCTGTATATTTTCAGCCTCCAGGGGTGAAATTGTGCTGACTCAGTCTC**CAGACTTTCAGTCTGTGACTCCAAAGGAGAAAGTCAC**CATCACCTGCCGGGCCAGTCAGAGCATTGGTAGTAGCTTACACTGGTACCAGCAGAAACCAGATCAGTCTCCAAAGCTCCTCATCAAGTATGCTTCCCAGTCCTTCTCAGGGGTCCCCTCGAGGTTCAGTGGCAGTGGATCTGGGACAGATTTCACCCTCACCATCAATAGCCTGGAAGCTGAAGATGCTGCAACGTATTACTGTCATCAGAGTAGTAGTTTACCTcacactgtgttacaacccagaacaaaaacta**gttcagcctggctgaac**agagaaactgggtgataccctagaatacttctgattgttacaggtgctttggaggcaatgagttaaccaatacaatgaagtctggctcacc**cagcagagaggaaactagagtcactgctg**catactttcatctttttaaaaatgaattattttatttatttcaatagtttttgggggtataggtggtttttagttacatggataagttctttagtggtgatgtctgagattttggtggacctgttacctgagcagtgcatactgtgcccaatatgttgtcttctagccttcacctccccttctatccttcctccccagtccccaaagtccattatatcattcttacgcctttgcatcctcatagcttagctcccacttatagat**gaaaacatataggttttc**cattcctgagttacttcatttagaataatagcctccagcttcatccatgttgctgcaaagttcattattttgttctgttctgttttatggctgagaagtattt**cgtggtgtatatacaccacg**ttttctttatccactcgttgcttgattggcacttatggtggttccatatttttgaaatggagaaatgtgctggaat**aaacatgcatgtgcatgttt**ctttttcctataataactttt**tttttctttgggtagataagaaaaa**taagtactgggattgctggactgaatggtatttctacttttagttctttaaggaatctccatactgtttttcatagtggttgtactag**tttacattcccaccagctgtgtaaa**agtgttccctcttcaccacatccacgccaatatctattattttttgacattttaattatggccattcttgcatgagtaaggtggtatttcaaggctatggttaccaaaacagcatggttctagtataaaaataggcacatagatcaatggaacacaatagagaacacagaaa

Uppercase: IGKV6-21

Lowercase: Flanking sequence[1000bp]

Red & Bold & Underline: Stem-loop [16]

Blue: Heptamer[26]

Green: Nonamer [8]

id-IGHV1-68-2[V_gene_segment]

ggaagcagagtt**tgcagtgagctgagattgtgccactgca**ctccagcttgggcaacaaagcgagattccatctccaaaaaaaaaattcagatttcagtgttaagtaaagttgcctacattgtgtgagtgacagggcagtggtggatccgagagtgtggatctgagagtgagtcagaaatcagcatgtaaagatgaggatctatgca**catgaactgaaagtatgtaaacagttcatg**aaattctaataaatccagtaggaaataaaacccaaacttatccaaaacacaaattcccttgaaattattatgggagcat**gagttcataaagaactc**ctaactcctgtttcaacttctgaatcctagtgtccatgacataagaaaatcatctccaattatgcatcacagggcaaatctgtaaactaagagtttttctgttgacga**tcctggggaatcaggacaccagga**aggtgctggagaaactgtctcaggagcgccccagggatctcagaggaacgtgctggccactcacgtgggacatcagcgtcacttgctcagagtcatcagtgagctg**tgctggtgtctgacgggtccagca**tagggccaaggcacctgctctgtgtcatggaccgagatggtccccagaatgatccaagtggtctctgtgctaatctaatgtagtttcacagtgagaggccgttctgagggggcttcttcttcagtgaaaggacct**ctgtccacaaatattcctaaatggacag**gggcatgcatttcctcaagcaggattagggcttg**gaccatcagcatctcactcttgcaaggctgatgtgtc**atttgtcttccctttcttatcatggatcaggctttgagctatgaaatgccctgtctcatgaatatgtaaatacctgagatccactgaggtaaatatggtctgtgccctgagagcttcacccaacaatcacatcccgcctctagagaatcccccgagagcatggctcctcaccATGGACAGGACCTAGAGCTAATCTTCCTGGTGGCAGCAGCTACAGGTAAGGTGCTCCCAAGTCCCAGTGATGAGAAGGGGATTGAGTACAGTCAAGGAGGCTTTCATCCACTCCTGTGTCCCACCCTCCAATGGGTGTCTGCTCCCAGGTGCAGCTGGGGCAGT**CTGAGGCTGAGGTAAAGAAGCCTGGGGCCTCAGTGAAGGTCTCCTGCAAGGCTTCCGGATACACCTTCACT**TGCTGCTCCTTGCACTGGTTGCAACAGGCCCCTGGACAAGGGCTTGAAAGGATGAGATGGATCACACTTTACAATGGTAACACCAACTATGCAAAGAAGTTCCAGGGCAGAGTCACCATTAC**CAGGGACATGTCCCTG**AGGACAGCCTACATAGAGCTGAGCAGCCTGAGATCTGAGGACTCGGCT**GTGTATTACTGGGCAAGATACAC**GGTGCGAGAACCCACA**TCCTGAGAGAGTCAGGA**ACCccagggaggaggcagctgtgctggcatggaggagatgacaaagattattagattgaagactttcttagaaaatagcattaagt**catttacgaaaaggaacaatataaatg**tgtatttg**agaaattgtaattatttgagagatttct**tatgcaacatttattctgtaagca**aattctagggattggagaattaatcaaattaat**aaagctgacat**agaaattcctctgaaggtatctttgtaaacatcaatttct**gaatcagtgttgt**aaatgttttggaacacagacacaagatcacattt**taactctacttttgtctctataa**aaaatgccaaaaagagtctcatttt**gaccatgtacctctttgaattcccaccatcaatgaatgattgttcttgggtttccacattatattaccatttatcattatgagaattgtgtgtttta**accattataataggtgagtaatggt**atctaatttttgcttaaatgcacatgcccctaat**aaaattcatatttaacaattttcatataatttt**tggtgagatgcctctcctgatatttggttcattttcaatagcactgttttcttttgattagttgtaagtttacttgcatattgattataaaag**ttaattaacaaattaa**aagaattcatttaacaaatatgtgacttggaagt**tgcaggggtttgtcctgca**caccctgacacaatgacgaatgaataaagtgcactgacacaaagatattctgctttgccagctcgactgagcatctgggccacttagtcacagccaccgcctcgatcagtcagggagacttacatttattcagtaaagattaattgacaaaggccgtgagtaaacaccactagaaggtaattgacattgtggacctcttgagtggaaagcaattaagcacccgcggtagatcaaaggttagtcttaggaacacatgagtaaacaagctagttagataagctcctcacattcctttgtttc

Uppercase: IGHV1-68-2

Lowercase: Flanking sequence[1000bp]

Red & Bold & Underline: Stem-loop [23]

Blue: Heptamer[45]

Green: Nonamer [4]

id-IGLV1-47[V_gene_segment]

caagttaaaattttaattcttccc**atgttcaaagactgaacatctttgctctttttgctcataagtgaaacactccaaaga**catt**tgccaggatcccccacagactggca**ttcagtgcttggaggacagactcctaggcaagatccctggtcacagtatccggatgactatgactgcaaatctggggaagtagagggagaacaaagatttaccagaaaaa**ccctctgggaattcctccccagggcatcccagaggg**aagagctccctttctcctcagcgcagcaggatccacagaggaagtggag**ccctggccaccaggg**aggagtcacatgagaggagaggacccggtgtccaacactggtggggaagtgagtccatggggccggtccccgctctgggcttgatcctgaaataaa**aggactttgtgtagacatcacctgagtcct**ctacaatgtacagccaaagagccatagacagattcactcctgaggagtccagtagagcgtgaagactgccctgaggtcacaggagggagacacaggtgccaccttactgggccgggacttgggagcgggggcagtggccagaaaacagggcggggcggaggcagcggtgaatctcagaggctctggtcacctgatcatagaacggtttgccttagtctggacactagggggcgcttggggaccatttctgagaagacttggggtaatcagagcagaac**caggcacctgccctgcctg**atgtcctctgctcagggctggcagctgtgtcctgtgtcctccccaccccctgggaccacaaagctccacccctgccacaccctgacatactcaagcccaggagcc**tgacccagggctcagggtggggtca**aaaaccggggggatctgatttgcatggatggac**tctccccctctcagagtatgaagagagggagaga**tctgggggaagctcagcttcagctgtggtagagaagacaggattcaggacaatctccagcATGGCCGGCTTCCCTCTCCTCCTCACCCTCCTCACTCACTGTGCAGGTGACAGGATGGGGACCAAGAGAG**GGGCCCTGGGAAGCCCATGGGGCCC**TGCTTTCTCCTCTTGTCTCCTTTCGTCTCTTGTCAATCACCATGTCTGTGTCTCTCTCACTTCCAGGGTCCTGGGCCCAGTCTGTGCTGACTCAGCC**ACCCTCAGCGTCTGGGACCCCCGGGCAGAGGGT**CACCATCTCTTGTTCTGGAAGCAGCTCCAACATCGGAAGTAATTATGTATACTGGTACCAGCAGCTCCCAGGAACGGCCCCCAAACTCCTCATCTATAGTAATAATCAGCGGCCCTCAGGGGTCCCTGACCGATTCTCTGGCTCCAAGTCTGGCA**CCTCAGCCTCCCTGGCCATCAGTGGGCTCCGGTCCGAGGATGAGGCTGATTATTACTGTGCAGCATGGGATGACAGCCT**GAGTGGTCCcacagtgctccaggccc**agggggaactgagacaagaaccccct**t**cctcctctgtcaggagg**gtgagccccggcagctgctgctcaggcccggcctgtggcttctgctgctgtagcttcccccatgggtccaagggcatccagggccctgcctgagagtggaggctcctcctccccttcatcctcagagtcatgaacaggctgtc**caggaaacagaacatcctgcttcctg**caacttgggacacagggtctctgcactgaagtcctggact**gaggtggcaggtacagctgtgtcgtcacagacccacctc**tg**atgggaaacctgtgtctctgtcatcctccttttcccat**ttccaggagtttccagagtggtgtcttcctccccctgctcctcagtgtgaatccctgtgcttcctttctctccagccaattctttttttaataaaactcctttcatgtaggaataatgcatatttaataaaccacccagatttataataggaaattagacaaatgttgacattttttgatccttgaatacatcaccaccaacacgcagctatgcaacctgtgaccaaccccagcaacccctcatgaccccttgtcatctccctccctcccttccccagtgctatcctgtgcagccacagatctgtctctgtcactattgtttgctttccattttttggaatgtatcagtgttatcagacattatcttcccatttgttctggcttttttcacactgcaaactccttt**tgggattcttcctgttctcgctgaattgcacccatccca**tagagggctcattactacctttgagggcctttaggttgattattcctgattttgtttatttattttaa**agtttgaatttatcgtccatcaaaatcaaact**cttggagttatgttcatctaaggtgacaggagagtctgtgatgtcttgtcgggattcttgcttcttcatctcttgcctctcgttctgtctatattggaagaggttc

Uppercase: IGLV1-47

Lowercase: Flanking sequence[1000bp]

Red & Bold & Underline: Stem-loop [20]

Blue: Heptamer[35]

Green: Nonamer [2]

id-TRBV7-9-2[V_gene_segment]

atatgaaagagggtgcccatgaagttttaagagttaagttaggaatgaaaatgagaaattaattgctcataatgcagactgttaggaaccccaaggattaattcaactccctattttttaattgtactactcccatcacaatgtcaagggtaaaacagttctgactttttctaaaggctatggaacatttattgatttcagaggtgacttagaaacacaggaggtgtcgtgataagtagtagagacaaaaaagaaaaagtattgagattaatgaattcatatggggaattgtttaactgttctttttttaacctttagaatgagttgaagatcccaacttagcttggtgcctgctggaggcaacgtttcttcccttgcccctctgagctgt**gaattgaaactctttcgggaaaaagacgcggcaattc**tctccacttcccaaacacctcccactcctacccagaccgtggatgggcaggaaatgcaggaacagagccagaaacaggagatctccaaggaaggttga**cagtcagcactgggatcgtctgtgtaaagtgctgctg**aagcagccaggtggcatgtccagccgacaatgcgaaaggaaaaagtgagaagacttcccgaaggcggagggtggaatgcgggcag**cagccccctggagggctg**agtggggaaaacaaaatggacctca**cagaagctgtgtgtgtggaaacccacttctg**acttatcacttgt**catgaattctatgcttcatg**gtgttacaccgtttattgtttctgatgagtgacagtaattattttctttcttgctggtacataataaagtggtgcacatcagagttgctgccatcttagacttaactcatcagtatcaggtgatcctgaggctcagtgatgtcactgtgggaactgctctgtggcgacaaggacgtccctcatcctctgctcctgctcacagtgaccctgatctggtaaagctcccatcctgccctgaccctgccATGGGCACCAGCCTCCTCTGCTGGATGGCCCTGTGTCTCCTGGGGGCAGGTGAGTCCTCAGAACACCAAGCAATCTCATTGTGTCTGTGTATGTCTGTGTGTGTGTGCGTGTGTGTGTGTGTGTGTGTGATGACTACAATTGTTTTCCTCCTGTTCCCAACTT**GTATCTCCACAGATCACGCAGATAC**TGGAGTCTCCCAGGACCCCAGACACAA**GATCACAAAGAGGGGACAGAATGTAACTTTCAGGTGTGATC**CAATTTCTGAACACAACCGCCTTTATTGGTACCGACAGACCCTGGGGCAGGGCCCAGAGTTTCTGACTTACTTCCAGAATGAAGCTCAACTAGAAAAATCAAGGCTGCTCAGTGATCGGTTCTCTGCAGAGAGGCCTAAGGGATCTT**TCTCCACCTTGGAGA**TCCAGCGCACAGAGCAGGGGGACTCGGCCATGTATCT**CTGTGCCAGCAGCTTAGCcacagcatggcacag**tcgcctccttcctgctcacaaaccctcaggcacttacttctccttc**cagctctcagaagccctgaacaaaggagctg**ccctgctctttcctcagcaaggagaatgaatgcatttggaactgcaggtgttc**ttctgatactaggaggtcagaa**aataacctctgaaatacaggaacagggaatactgggtagtaataattttgacttatggatttctgggattccttatatatagttcaaatttccataattaggatataacagag**cttagtctcatggatagtgactaag**taaatattctcttatagaactatgaagtttcagcacatttatattaaactactgttaccacatgtcaccaactcagacctataatctaccaaggagtggggaagccaaacgcaaacactgctaaggccatttacagctaccacccttggaag**aaaattggaatctttagtaagaatttccatcaaaatcaatttt**ttagaaataactattacccacaataatgcacattaactcatgttcatacattgctttttttccttctgaaaatcttgtagtca**ttctatgtttaatttccttcttctccagtctaacttaaaaaaatagaa**atacaaattgagattgtttcataagataaaccccacctgggaaggtgttaaaaggacacaattcaaagaaaacctgaataaaattatatcaaccacgtagga**ttctttgaggtttccaaagaa**gatgtaccagtttacattcccacgagaggtatatgagttttggatggaatttttttttttctctctctctctctgtag**attagtttttccctctactaat**agtttctcttgacaaacagaagttcttaagtgacaatcattccatgtttctcccatttcagttagtgttttgtattttggtgtccactttaggaaatccttgcttacttcaaaatgatcatgttttcttacatttccttctaaaat

Uppercase: TRBV7-9-2

Lowercase: Flanking sequence[1000bp]

Red & Bold & Underline: Stem-loop [16]

Blue: Heptamer[39]

Green: Nonamer [4]

id-IGLV1-44[V_gene_segment]

aaccaag**ttgaagttttaattcttcccatcttcaa**agac**tgagcatctttgctctttttgctca**taagtgaaacactccaaagtcatttgccaggatcccctgtagatgggcattcagtgtacagactcctaggcaagatccctggtcacagtatccaaatgactttgactgcaaacctggggaagtagagagagaacaaagatttaccagaaaaaccctctgggaatg**tcttcccaaggcaccctagagggaagagctccctttctcttcagctccacaggatccacagaggaaggagagctct**agccagcaggaaggagccacatgagaggagaggaccaggtgtcccacacaggtggggaagtgagtccatggggctggtccccgctctgggcttgatcctgacata**agaggactctgtgtagacaccacctgagtcctct**gcaatgtacagccacagagccatggacagattcactcctgaggagtccagtagagcgcgaagactgccttgaggtcacagcgggaagacacaggtgtcacttgagctggcctggacctgagagagggggcagtggacagaacagaggggaggcagtggtgtgaatgagaggctctggtcaccttatcatagaatggtttaccttggctggacaccaggggacgctgggggaccatttctgagaagacgcgcggggggcggg**ggtgggcggtgatgggagcacaaccggccacc**tgttgtgcctgttgtcctctgctcagggctcacaactgtgttctccccaccctctgggaccacagagctccacccctgccacaccctgacatcctcaaggcaaggagcc**tgacccagggctcagggtggggtca**caaagctgggggggtctgatttgcatggatggac**tctccccctctcagagtatgaagagagggagaga**tctgggggaagctcagcttcagctgtgggtagagaagacaggactcaggacaatctccagcATGGCCAGCTTCCCTCTCCTCCTCACCCTCCTCACTCACTGTGCAGGTGACAGGATGGGGACCAAGAAAG**GGGCCCTGGGAAGCCCATGGGGCCC**TGCTTTCTCCTCTTGTCTCCTTTTGTCTCTTGTCAATCACCATGTCTGTGTCTCTCTCACTTCCAGGGTCCTGGGCCCAGTCTGTGCTGACTCAGCC**ACCCTCAGCGTCTGGGACCCCCGGGCAGAGGGT**CACCATCTCTTGTTCTGGAAGCAGCTCCAACATCGGAAGTAATACTGTAAACTGGTACCAGCAGCTCCCAGGAACGGCCCCCAAACTCCTCATCTATAGTAATAATCAGCGGCCCTCAGGGGTCCCTGACCGATTCTCTGGCTCCAAGTCTGGCA**CCTCAGCCTCCCTGGCCATCAGTGGGCTCCAGTCTGAGGATGAGGCTGATTATTACTGTGCAGCATGGGATGACAGCCT**GAATGGTCCcacagtgctccaggccaatggggaactgagacaagaaccccctt**cctcctctgtcaggagg**gtgagccc**cagcagctgctgctcaggcctggcctgtggcttctgctgctg**cagcttccttcatgggtccaggggcatccagggccctgcctgagagtggaggctcctcctccccttcagtcctcagagtcaggaacaggacatccaggaaacagaacatcctcctccctgcagcttaggacacagggtctctgcactgaagtcctgggct**gaggtggcaggtccagttgtgtcatctcagacccacctc**tgatggggaacccgtgtctccgtcatcctccttttcccatttccaggagtttccagagtggtgtcttcctccccctgctcctcagtgtgaatccctgtgcttcctttctctccagcctattcttttttaataaaactcctttcatgtaggaactatgcacatctaataaactacccatatttataacaggaaattagacaaattttggcatttttctggttccttgaaaacatcaccatcc**ccgggagctattgtgaccatcccgg**cagctcctcatgatgcttgtctcctccctccctcccctaatggcctctcaggaagccacagatctgcctctctcactactgtttggtttttttttctggaatgtatcagtgttatcaaacgttatcttcccatttgttcaggcttctttcactctgcacacttgctttgagattatccctgctgtcgctgaattgcacccat**cccttagagggctcattactatctttaaggg**tccttaggccatttttttctgattttattcatgtatttcctttaaa**gtttgaatttattgttcatcaaac**aaactctatgggtgatgttcatctaaggtgacaggacagtccgtgatggtttgtcaggattcttgcttcttcagttcttttcttggctctctctatagtggaagaggttctcatcttttat

Uppercase: IGLV1-44

Lowercase: Flanking sequence[1000bp]

Red & Bold & Underline: Stem-loop [18]

Blue: Heptamer[36]

Green: Nonamer [1]

id-TRAV29DV5[V_gene_segment]

ttatgtgttatcagtccttgctaaaaat**aatttcaaaaaacattttggaatt**taatctca**gctgaaatcactagttgccctactaattcagc**attaaaagtacaattgattcttcttccttcctccatcatagtaaaccatggggaaatacattacctgtggaccaagagt**gaagataagcagctatgttgcctttttggcatcttc**tggtaaatttcatgaatatattgtctatagactttacaggattatgtcaccgtttttctcttttgaaaatacagaacacact**aaatgatctcattcaagttccaatagagataattt**cagttcagaaggtgatctaatacagaagataaattttgttgtgctgatttgtgtttttgagagaggtacccagaat**ttaaatttaatttaa**ttatgttgtgctatttatcatatagaa**aatatgggcatatttgaagaccatatt**agaattcaattgcactaaagagtaagctacgaaacttctttgtgttctaaattcatatcattgtattaaataaaacatcatacccatatgcgtcccc**aataagtatcaatccttatt**aatgtttttaaacaagatgtctgccttagtatgccagttgacaaggtgtcagtcatttgaccttatgtaattctctaattttcagctatcattttctactattgctgtgtgtgtgtgtgttctct**aaaaacttttccttctctttcaatctcttgttttt**taact**ttttatggagaaggcattataaaa**tcatctttaggaaatacagaaacacatttacagaaactttctacatttgaactaaagggggcagtgcttcctttcataaagataaataagcttagagc**tccacacaggttgttcatatgtaaaatgaagggtctgtgga**aggacatgaataaagcacaggaggttgaagt**cagatttgcagctttctaggcaggagataagacaatctg**catcttca**caggagggATGGCCATGCTCCTG**GGGGCATCAGTGCTGATTCTGTGGCTTCAGCCAGACTGTGAGTTGTGCATGGGAGGTTTGAATATAGTAAGAAAAGCTACAAGGAACACTACAAGGCTGGGAGATAATTGGAGAAGTTTTGTTTTGTTTTCCGGATCTTGATGGTTTCCTGTGGGGACATTAAAAATAAAACTGAAAA**TCTGTTATTCTCTTTCCAAACAGGGGTAAACAGTCAACAGA**AGAATGATGACCAGCAAGTTAAGCA**AAATTCACCATCCCTGAGCGTCCAGGAAGGAAGAATTT**CTATTCTGAACTGTGACTATACTAACAGCATGTTTGATTATTTCCTATGGTACAAAAAATACCCTGCTGAAGGTCCTACATTCCTGATATCTATAAG**TTCCATTAAGGATAAAAATGAAGATGGAA**GATTCACTGTTTTCTTAAACAAAAGTGCCAAGCACCTCTCTCTGCACATTGTGCCCTCCCAGCCTGGAGACTCTGCAGTGTACTT**CTGTGCAGCAAGCGcacag**tgctctccaggcacctgcagcccgtactcaaacctgctttggggactcagactgggagacacatagactcgcttccatttacacatgccaatatgagagattatgctttgaagtaggtgaggcaagttgagcctaaagaaagactgatacatgacaaaaccttacaaataagttaagaaagttccctgagtttt**agagtgtttaagagtctaaccactct**ccttttctactttcagaaagaggttgctgatttagct**aaagcataaatctctagtgatgcttt**gtttctgctagcttctggaatgcgggtgccctgtctcccttgtatactagggggcaaggatgtgtgtttggaatgggaaaaccaaggtcctactgcaggttcttccagtgatgaacatttc**ctgagatgtcttttcttttagtttttctcag**tgtacaggcatttctgagcaaagaattgcatgtgtgaatactaacttatgtatatctgcatatctataaatatttctatatgtagcataaccatctgtatctatattaagttaaacattcgttc**ttagcaatgtctccaatgctaa**atcattaccacgtggatcacactagcctcctccccttacttatccataaattcccactacaggggtgcacagcctggctctcactg**ttaacaaaaagttattcagtaatacttgttaa**aacacggtgaggcagaatttatttaggaccatcatgataggtacagagac**cactgcaacaggattttgcagtg**agggagaagaattggattcaactccgaatacagcatagacaagtggagatttctagccaaggggcagattaggggtcagtggataggaaattactaagaggaaacattagaaccaagggatagtctggctagaccagccaaacaggatccttgcagaagacaggtcagggtgatcagacatcaatcccctgggggatgatagaggatgaggagcccagttgga

Uppercase: TRAV29DV5

Lowercase: Flanking sequence[1000bp]

Red & Bold & Underline: Stem-loop [22]

Blue: Heptamer[38]

Green: Nonamer [4]

id-LOC102725003[V_gene_segment]

ggaatggttgatcctaaaagctaccacgtgtcaggctcacttaagtacagactctggtcaaaacaagctgaagttttaactcttccatctt**caaatactgagcatatttg**ct**tgttttgctctttagcaaaaca**ctgcaaagtcatttgccaggatcccccgcagattggcattcagtgatggagaacagactcctaggtgagatccctg**gtcacaggatcccgatgactgtgac**tgcaagtctggaaaagaagagacagaacaaagatttaccagcataaccctctgagaatgtgccccaggtatcctggagggaagagctccctatcgacccagcactgcagaatccacagaggaaggacagctctggccaccagggagcagccacacgagaggagaggacccggtgtccccacaggtgaggaaatgagtccatggggctggtcactgctctcgatttcattctgacataag**aggactttgtgtagtcaccacctgagtcct**atacagtgaacagccaaggagacatggacagattcacccctgaggaacccagtagaatgtgaagtctg**ccctgagctcacgaaaaaaaagactcaggg**ggctgcctggcc**tggccctgatggagggggcca**tggacagaacaacggggagtgaggggtgtctctgagaggctctggtcaccttgttatacaatga**tggtgtacatgaccaaaatggacacca**ggggatgcctgcgtaccattactgagaagactgggtgtgatgag**agcaggaccagcgccacctgtcctgct**tggt**gccctatgcttagggc**ttacagatgtcaactctgcaccccctgggaccacatagccccacccctggcactctctgacatcc**tcaggcagaggagcctga**cccagggcccaaggtgggatcagaaagct**ggagggtctgatttgcatggttggaccctcc**t**tctctcagagtataaagaggggcagggagaga**cttggggAAGCTCTGCTTCAGCTGTGAGCACAGAAGGCAGGACTCGGGACAATCTTCATCATGACCTGCTCCCCTCTCCTCCTCACCCTTCTCATTCACTGCACAGGTGCCCAGACACAGGGTCAGGGGAGGGGTCCAGGAAGCCCATGAGGCCCTGCTTTCTCCTTCTCTCTCTAGACCAAGAATCACTGTGTCTGTGTCTCTCCTGCTTCCACGGTCCTGGGCCCAGTCTGTGTTGACGCAGCCGCCTTCAGTGTCTGCGGCCCCAGGACAGAAGGTCACCATCTCCTGCTCTGGAAGCAGCTCCGACATGGGGAATTATGCGGTA**TCCTGGTACCAGCAGCTCCCAGGA**ACAGCCCCCAAACTCCTCATCTATGAAAATAATAAGCGACCCTCAGGGATTCCTGACCGATTCTCTGGCTCCAAGTCTGGCACCTCAGCCACCCTGGGCATCACT**GGCCTCTGGCCTGAGGACGAGGCC**GATTATTACTGCTTAGCATGGGATACCAGCCCGAGAGCTTGCACAGTGCTCCAGGCCAATGGGGAACTGAGACAAGAACCCCCTT**CCTCCTCCGCCAGGAGG**GTGAGTGCCTGCAGCTGCTGCTCACACCTGACCTGTAGCTTCTGCTGCTGTAGCTTCCCCCATGGGCCTCGGGGCAGCCAGGGCCTTGCCTAGGAGTGGAGGCTCCACCACTTTTGTCCTCAGAGTCAGGAACAGGGACCCCAGGAGACAGAATATCCTGCTCCTCAGCTTGGGACACAGGGTCTCTGCACTGAAATCGTGGGCTGAGGTGGCAGGTCCA**ACTGTGTCTTCACAGT**C**CTTCCTGTGCCTGCCCATGGTGTGGGGACGGAGTGAGGAAG**TGTGGGCCATTGAGCTTCACTAGAGGCCTGA**GGGAGCAGAGGTGCTCCC**GAAGACCTGGACTCCTTCCGTCACCCTCCTCCCATCACTCT**CCTCACTGTAGCTGTGGCCAGGTGAGG**GCTCAGGGCTACTTGAGGACCTTGCTCAGCCCAGGGCCTCCATCCCAGGCATTGTTTCCAAACTCTGTAAGGGCTGTCTCTGAATCTG**TAGAAACATTTATTTCTA**T**GTGCCCAGGACTGGGTTAGAGGGCAC**CAACCCCACCCCTTGCAAACTGCTCATCACCTCTCAGTGAAACAATGAGGAGAAGAAAGAGTTAGAAATCAGAAGCCTCC**CCAAGGCTCCCCTCTCTCCTCTCAACCAGCCTCCCCTTGGCTTCCTCCTTCCACAGAAATAAAGAGGAGGCAGCCA**CAGACGGCAAGGGAAGGTCCAGGCATTATACCAGGCAGTGGAGCCAGCCAGTCCC**CCCAGATGGTGGAGGGAGGTGGTGGAGGTGTCTGGG**ACCAGAAAATAGTGAGGACCCTCTTACTAAAGTGAGGAGGGGTGGCATAAACTGA**GGCTCAGATGGTGGACCAGATCTCCTGAGCC**ATCTACTCATTTCTTCTTGAGTTCCAGTTTCAGTAAGAATAGGATAAAAAAAAAAGTCTGAAGAGGTTCCTTTCCTCAGATAAGAATTCGTAGAGGATTATTGCCCACTCAGGCATTAAG**GGGAGGCTGAGCCTCCC**TCTCAGGCCAATAGAA**CTGGGGGCTTTCATGGGAATTTGATTGTACTGAAGCTCCCAG**ACTCTGACATCTAAGACTTATGCTCTCGTAGCAGAGCCCAGCACTGATACCAAAATGGATAAAGGCTGATTGATAAGCCTGGAATATCCTAGCTGACACCGACTTGTGGAAAAGAGAATCAAGTTAGAATATTGATGTGATAAGGTGGCATCGCACAGTGAGGAGAGGAGAGCTTGTGTGGGGGAGGGTAGAGGAGCCATTAGAGGGTTGGACAAAGAGGACACAGAATCTGAGGTAAGTTTGTGAGGAAAACTCCAGATGCTTCAGAGAAGATGGACAAGGAGGTAGGCAGTGTGAAGGCCATGGGACCAGCTCTAGTCACCGGGAGTGAGTGGTGAGATTCAGCACCAGTGTCTGGGGAGGAGGTACTGAAGCCTGGGTGAGGGGAGATGCTCAAGGCTGGCCTTAGGACATGAGTGGGGCACAGTGATCACACTGCTGGGGGGTCCTTCAAGTGCAAGGGGGCTACAGGAAGGCTCGAGC**TCATTGTCACAATGA**AAACTTAGACAACAGCTCCCCAGACTTGTTCAAAGGTGTCACAGGACT**TTTTCCTCTCCACATGGCAACGGGGAGAGGAAAA**CTTATCTCTGAGATATATAAGATGTTTGTTCTCCATAACCTCTCCTGTCCCCTGGCTGAATTCCTATTTGGAAACAAAATGACTCTCCAGTTTTACATATGACATCCTCACTAAAAAAAAAGTTCCAGAATGAGAAAGATCTGCCCTCCCTGACCCACTGGTGAGGAATCTCAGCCTTCAGGACAACCAGACGAAATCGTTCATGCCATCT**CCCATGACAGCATCATGGG**GACCAGGGAGCATATCTTATGG**GTTTGGGCCACACACTGCCTGTCCCAAAC**ATTGCAGCTGGGGACACTTATATGCTGAACTCATGGCTGGGAGATCTGGGGTCTAGACATGACAGGAAGAGGAAAAAGACCAGGAGCTCTGCCTAAGTAGCCCCAGTGTGTGAATCACTCTACCCTCCACCCTCAACTTCAGTTCGTGATGGGTGTTGTCTTGCGAAAACTGACAGCTGGGCTGGGACAC**AGGCAAGAGAGACTCTGACCCACAAATCTCCATCGTGGGCTCTCTGGCCT**TTCTGATATTCTTTTTGCACATCAATTCTGGGCCAAGACACATCCATGCTCATCAAGAAGGCATTGATAGGAGAATCAGGTAGCCTCCAGAGAGACAGGAAGCAGATAAACACCCTCATGTTGGCCTCCTAAGCTCAGAGGAGAAAAAAAGGGAAAGAAGTTCAAGAAGCCACACTAGAAGCCACTGATCACCATCACCTCGGCACCTGTCCTCCATCTGCCCCTCTTTCAAGTAAGGACCAGACTCAGG**GGACAGCACCTGCACCTCCCTTTCTGTCC**ATGCACTGTGTAGGGAGCCTCCTTCTGCTCTCATCTCAAGGCGACTGCTCCGCCAGGTCAGCCGTGTCACTGCTGAGAGTCTTACCTCAGTCCTGGTCACAGGGTCTTGGATCAGCAGCAACCTGGCTTTGATTCAGCTGTGACCCTGCCCCTTCAGGACACAAAGCCGATGATTAAAGCCTAACTGAACTCTCCTAGATTTCTCAAACCTTAGATTCTCCTTCTCTGTGTCCCTCTAGAAAGTGCCACCTCTACC**AAATAATTTCATGTTATTT**CCTCAGGTCTAGAGCTATTTGATTACCAAAATTCTATACTCACTCTTCTGCTCTCATTCCTTGTCATGGTCCATTTTCCACTCAGGAGCTAATGATGCCAACCCTGTATCTACAGGCCATTTTCACAAAAAAAAAAAAGAAATGAGAGGGTTCCTTAGATAAATACAGGGTTAGCAAAGAAAAGTTTCATTACCCCTTA**GGAATTACTCCAAGATCTAAGAAAATTAATTCC**AGAGCTTCAAGAGTTTACTCCTCAGAAACAACAGCATGAATACATTAGGTAGTGAATGTTCTTGGTCTTCTCAATGAATAAC**CTCTGAAAGGTTCAGAG**TATACGTTTCAAAAAGATAAGTGCATAACTTTCACAAAAATGTAATCTTAACTTATGTCATCAATTTTAATTGGTATATATATTTCTGGAAAAAAAAGCAGTATGATGGTAAAAGAAATGTAAAAAAGCCTTTGTAAAACCCAGTGCTTACAGTCCACCAGGATAGCACCTGTGA**AGCAAATTTTCTTTGCT**GAAGATACACCGCCTTTTAGAATAATAGCAAGATAGTTTGAATTTGTCTTCTTCTCTCACAACAATCTAAAAATTAACATGATCCCATTTATATGAAATATCTAGAAATATATCTGTACATGTGGACATTCCCATCGCACACCTCCCATCCAGGCTGTCCTAAAGTGGCATCTCCTCTTCTTCTGCTCATGTTCTCCCCCTCTCCTCACCTCTCCTGCTGACACGTGGCTCCTCCCTGCCTTCCTCTCCAGTAGCCCCAGCAGATCAGGCCCAACCAGTCAACAAGCTCCTGGACTTGGGGAACAAAAGTACATCTTCTTGGATTTCTGAGGACTTGCCCTGCCAGGGTGTCTTTCTCAATCCAATCTCTCCTTCCCGGCAGTCCACTTCTCCTAGATCCTGTAATTTATCTAATCTTATTCATCTGTATCTACTTCGTGGACTTCCTTACATCTTC**ACCTGAACACCAGGAATCTTCAGGT**CATGCCCATATGCCCTTGGTTCTCCTGTAGTCCCTTCTTCCTTCTTCCCCTGTGATGCCACCAAGTC**GAAGTCTCTTCTCAGACCCTGGAGTGCAGCAGGGACTTC**TCTGGTCTCTGCCCCAGGGATTAAGGGGGTGGAAGAGTGGACTCCTGTGCCCTTCCTATGCCTGTGTTTTCATCAGGGTTAGCTGCAAAAGGTAGGGATTAGGCTGTCTAGGTAGGGATTATGGCATTAGGTAGGGATTATGGCATTTCCAGA**GCTGAGCTCTATGCCCAGCAAAAGATACGTGCTCAGC**AACTACTTGACAGAGACCTACCTCTTATCACATGGGACACATTATACTGGACCCAGGATGGGGCTGAACTGGGAGAGGTAGCTGTGGACCTGCTCCTCGCTCTGCCCTCACCCTAAGAGCTGATGGGGCATCCA**CCAGGAAGAGCATTCCTGG**GCAGGGAGAGGGCCATGAGCAGCCTTGTCCTGTATGTGAGACACAGCTCAGCCCGGTCACAAGATCCAGATGACTATGACTGCAATTCAGGGGAAGTAGAGAATGAAGATTGACCAGAATAA**CCCTCTGGGAATGTGCCCTGGGCCTCCTAGAGGGAAGAGCTTCCCT**TCTATTCAGCGCTGCAGGATCCACAGAGGAAGGAGAG**CCCTGGCCACCAGGG**AGGAGCCACATGAGAGGACCCGGTGTCCCCACAAATGGGAACGTGAGTCCATGAAAGTCGGTCCCTGCTCTGGGTTTGATTGTGACCTAT**GAGGACATTGTGTAGTCACCACCTGAGTCCTC**TACAATAAACAGTGAGAGCGCCACGGACAGATTCACTCCCGAGGAGCCCAGTAAAGTGGGAGGACTGAGGTCACAGGGAGAAGACACAGGCACCACCTGGCCTGGCCCTGACAGGGGGCAGTGGACAGAACAGTGCGGAGGCAGTGGGGTCTCTAGAGGCTCTGGTCACCTTGTCATACCATGGTTTGTCTCGGTCTGGACACTAGAGGGCACTTGGGGACCATTCCTGAGAAGACTGAGGTTGATCA**GAGCAGAACCAGCCACCTGTCCTGCCTAGTGTCCTCTGCTC**AGGGCTCACAGCTGCGTCCTTTACACCCGTGGGACCACACAGCCCCACCCCTGCCACCCCCTGACATCCTCAGGCATAGGGACCTGACCCAGGGCCCAGGGTGGGATTAGAAAGCTGGGGGTCTGATTTGCATGGATGGACCCTCCCACTCTCAGAGTATGAAGAGGGGCAGGGAGAGATTTGGGGAGGCTCTGCTTCAGCTGTGGGCACAAGAGGCAGCACTCAGGACAATCTCCAGCATGGCCTGGTCTCCTCTCCTCCTCACTCTCCTCGCTCACTGCACAGGTGACTGGATACAGGT**CCAGGGGAGGGGCCCTGG**GAAGCCTATGGATTCTTGCTTTCTCCTGTTGTCTCTAGAAGCCGAATAATGATGCCTGTGTCTCTCCCACTTCCAGGGTCCTGGGCCCAGTCTGTGCTGACGCAGCCGCCCTCAGTGTCTGGGGC**CCCAGGGCAGAGGGTCACCATCTCCTGCACTGGG**AGCAGCTCCAACATCGGGGCAGGTTATGATGTACACTGGTACCAGCAGCTTCCAGGAACAGCCCCCAAACTCCTCATCTATGGTAACAGC**AATCGGCCCTCAGGGGTCCCTGACCGATT**CTCTGGCTCCAAGTCTGG**CACCTCAGCCTCCCTGGCCATCACTGGGCTCCAGGCTGAGGATGAGGCTGATTATTACTGCCAGTCCTATGACAGCAGCCT**GAGTGGTTCCACAGTGCTCCAGGCCCGGGGGGAACTGAGACAAGAACCCACTT**CCTCCTCTGCCAGGAGG**GTGAGCCCCGGCAGCTGCTGCTCAGGCCTGGCCTGTGGCTTCTGCGGCTGCTGCTTCCCTCATGG**TCCCAGGGGCATCCAGGGCCCTGCCTGGGA**GTGGAGGCTCCTCCTCCCCTTCTGTCCTCAGAGTCAGGAACAGGGAATCCAGGAAGCAGAATATCCCTCTCCCTTAACTTGGGACACAAAGTCTCTACACTGAagttctggactc**aggtggcagatccagttgtgtcacctcagacccacct**ctgatagggaacccatgtattcattattctcctttccccaattccaggagtttccggagtggtgtcttcctctccctgctcctcaatgtgagtccctgtgtttcttttctcccaggcctattcttttttttttttttaacaaaactcctttcatgaaggaaaaatgcacacctaataaacca**gctgaatttatagcattaaattcggc**ccattttggcatttttctggatctttgaaaatatcaccaccactgagctatgcaacctctgactaaccccagcagcccttcatgatgctttgtcatctccttcactccccgaatgccctcctgggcagtcaggaatctgcctttgccactactgtatggtttccattttctgaaacttac**tagtgttatcaaacacta**tcttcccatttgttcgagcttctttcactctgcacacttcttttcagattctccctgctgttgctgaattgcatcccattctacagagggctcattacaacctttgagggtctaggacattctcttctgattctatttatttacttatttattttaa**agtttgaattcattgtccatcaaatcaaact**ctacgggtgatgttcatctaaggtgacaggagtcagtgagggcttctcaggattcttgcttcttcatctcttccctct**aattctttctataatgggagaatt**tcttatctttttaaagtttttctatttattattttatgttggc**ttttaaggtttttagtgttttttaaaa**tttttttatttcaataggtttttgggaaacagatgttgttttgttacataaataagtttgttagt**ggtgatttctgggattttggtgcactcatcacc**caagcagggaacactgtacccaatgtgtagtcttttatccctcaccccatcccaccctttctcctgagtccccaaagtccgttgtat

Uppercase: LOC102725003

Lowercase: Flanking sequence[1000bp]

Red & Bold & Underline: Stem-loop [60]

Blue: Heptamer[108]

Green: Nonamer [10]

id-IGHV3-52[V_gene_segment]

tcctactctgtgacccaggcaggactgcagtggtgggatcatagttcaatgtagcttccaacttctggtcaccaacgatcctcctgcatctgccatctaagt**agctgaaactacagttgtgtgccaccaggctcagct**tgagtactt**attaaatcaaacactttatgcaatatttaat**gtatggcaataatgtctcattgagagtatt**acaaataaatgaatggataatttgt**tcagtacagattgatggaaaatagacactaacatgaggaatgtctgaca**tttatgaacatacaactgcataaa**a**tgtgttctcttacattcattaggtaaacaca**atagtgcatacacatcaaaccatgctttcattacaggaaggaagttctgaaaatgtcactgggggtgacccacgctgtgctgggcttggttcgggggcagtcaggcccggtggtgagaagcacagg**cccagatacccaggcttactctgcaaatgtgagctctggg**gacattgtaccacccatctgtgcttctgctggtaat**tttccatctgtaacgtggaaa**taacattgatac**tacataccgtgatttctccacatatgta**aaaataaaataagatgattgctgctaagtttaaataagggcagttttcataggtccattgtacctcaataaaatta**ctttaaaataaaaattacaaatacagttgtaggtttaaag**agtttatcacagaacaaacttataataagaaactatattttcaaaaattgtatcaatatctctaaactcccccaggacacactcacct**gctctgggctctccactctcctcaggattcccaccccagagc**ttgctatatagtaggagacatgcaaacagagccaaacctctgctgatgaaaagcagcccagccctgaccctgcagct**ctgggagaggagccccagctccaggattcccag**gtctttccatttagtcttcagggctgagcacagaggactcaccATGGAGTCTGGGCTGAGCTGGGTTTTCCTTGTTGCTATTTTGAAAGGTGATTCATGGGGAATGAGTTGAATGTAAGTGAATATGAGTGAG**AGAAACAGTGGATGTGTGCGGCAGTTTCT**GACCAGGGTGTCTCTGTGTTTGCAGGTGTCCAGTGTGAGGTGCAGCTGGTGGAGTCTGGGTGAGGCTTGGTACAGCCTGGAGGGTCCCTGAGACTCTCCTGTGCAGCCTCTGGATTCACCTTCAGTAGCTCCTGGATGCACTGGGTCTGCCAGGCTCCGGAGAAGGGGCTGGAGTGGGTGGCCGACATAAAGTGTGACGGAAGTGAGAAATACTATGTAGACTCTGTGAAGGGCCGATTGACCATCTCCAGAGACAATGCCAAGAACTCCCTCTATCTGCAAGTGAACAGCCTGAGAGCTGAGGACATGACCGTGTATT**ACTGTGTGAGAGGcacagt**gagg**ggaggtcagtgtgagcccagacacaaacctcc**tgcaggggcatctggagccacaagggggcgctcaggatacacagaggacaggg**gcagccccagggcaggtgcaggtggaggtcaagggctgc**tctccttcagggtctgtggcttcctctcatctaacagttccgcagggagcctcttgtatttacagtgatgtgctactgaggtttctaagtttgtaaagtttattactacaggaggaaccactatcaaacgcccttaaggcaggtgtcactaatggagaaaggaaagtgcacaggaggctgggtgaggctgtggacactgtctgcctatgattcaagtttcacaagcagtgacggagaaatgggagtttgatggagctccctaactaccatgtggtctaaactaagtccaactaag**tccctgagctctgggtgcccatcaggga**tccgccatgtgcccggcagccgcgtgcctttttgtctcctctgcgcccaatcactgtctgtgatgagcttccaggatgtgtgtgtttggcacaaaccaggtgatggacgtcagacagcagcagctggtgcccggatcatgggctccctaatgctggaggaatgagaggtgcattctcagggacaagacattgttgatggatttttatgtagaaaccacgattttac**ttgcttctctcaggagacatagaagaagcaa**ccatgcagtcagcaaataattataatttccacaattaccccaaatggttaatccttaattctgcgcagggtcccaccgtagagtcacctttctcatgaggaatggttgaatctagaatgagtccacatgattttcatatttttggcttctgtccatgttcagagagttagagtagagtaagtttggacctttccacacactaagcctcacctcccccacagaaagagcagagacttcaactattcctgagtgtggggtaggggctggtcctgcacacctcagagcc

Uppercase: IGHV3-52

Lowercase: Flanking sequence[1000bp]

Red & Bold & Underline: Stem-loop [17]

Blue: Heptamer[29]

Green: Nonamer [9]

id-TRAV8-1[V_gene_segment]

cttctcctggacaatgagtctcaggagttccccaccagaacaccttgtgacccccgcccctgcccgcaagagaaaaccccctttaactgtaattttccactacctacccaaatcctataaaactgccccaccccatctccctttgctgactc**ctttttaggacttagtctgcctgcacccaggtgattaaaaag**ctttattgctcacacaaaacctgtttggtggtctcttca**cagatgcgcataacactagacttttattttgtacatctg**atcttatttttaattttaccaaagtaggtatctccacatcctaccgcaacctcctctcaacctgatgctcactgcaacaagaccatttaaagctcagtcagaagagttgatctaactatttttgcttaaaggatgtctccacagcttttatacaaagatactgagggactccaaaatctgctagattcattgtgagatctttttct**ctgtaaattcctcctctcttttacag**tcctcatttttcattgaggctaggattaatgtaatatactacagagggcaacacagtacagattagggaagaaagaaatgttcactggaagtttttgacatgtctccaaacaaggaaagcacaggttcacttcttgtggggagtctgtatcctcagtcctgaatctggcctccagagggtgacgttgcacaagggacagatggctgtttccaatgatgatcaacaaacactttctgctcagccccttcttcctgcagctggcttactgaagcaccgtcactggagt**tctttgaagatgggacaaatggagagaatcaaaga**ttgaaaagcacttaagattatctcagcagattttgtattcttaccaaaacaagagacttgcctagcccaaccttcctcacgctcgctattctcaagacctgggttccagccactttcctactggccccgaggagaatttccaaagagacgcctgcagtgtttccacagctcagccATGCTCCTGTTGCTCATACCAGTGCTGGGGATGATTTTTGCCCTGAGTGAGTAACATTCTATTATGGTCTCTAGTTCCACAGAAGTAACTGTTTTCTGATTCAAATCTTAGTAGAAATACTTTTCATAGACAAGTCTGCACTATTTTCACTGATACAACATTGATTTTTTCAGGAGATGCCAGAGCCCAGTCTGTGAGCCAGCATAACCACCACGTAATTCTCTCTGAAGCAGCCTCACTGGAGTTGGGATGCAACTATTCCTATGGTGGAACTGTTAATCTCTTCTGGTATGTCCAGTACCCTGGTCAACACCTTCAGCTTCTCCTCAAGTACTTTTCAGGGGATCCACTGGTTAAAGGCATCAAGGGCTTTGAGGCT**GAATTTATAAAGAGTAAATTC**TCCTTTAATCTGAGGAAACCCTCTGTGCAGTGGAGTGACACAGCTGAGTACTTCTGTGCCGTGAATGCcacagtgtctgggactgcaaagggagctgaacacaaacttcctaaggtgctagggagaataactgcctctgaaagattttggattctgtcacagtagaaaccatgatgttagtattt**ttaaaaaaatttatttttaa**tttttatggatacatagtaggtgaatatgtttatggagtacatgagatattttgatacaggcattcaatgcataacaatgacatcacgtaaaatggggtatccatcccctgaaacatttatcctttgtgttacaaacaacccaatta**tacacttttagttatttaaaaatgta**caattaaattattattgactgtagctaccctgttgtgctatcaaatactaggtcttatttattctttctatttttttgtacctgttaaacatccctcctctccccgaccctcccactatcgttcccagcctctggaaaccatccttatcttctactctctatcactatgagttcgattgttttgatttttaaatcccacagataagtgagaacatgcaatgtttgtctttcttgcctggcttatttcacttagtgtaatgatctccagttccatc**catgctgttgctctaacccaaaagcatg**tgagacgggtctcaatcaatttagaaagtttatttcccaaggttaacaacatatctgtgacacagcctcaggaggtcctgacaacatgtgcccaaggtagctggggaacaggttgcttatatacagtttagggaaacataatacatcaatcaatacatgtaagatttacagtggtt**tgatctggaatggcagtacaacttgaaggggagcttccagatca**taggtagattaaaaaattttctgattggcaattgtttatcaatagaaagcaatatctggggtataataaggggt**tgtggaggccaaagttttatcatacagatgaagcctccaca**tagcaggcttcagagagaatagattgtaaatgtttttatcagactgaaaata

Uppercase: TRAV8-1

Lowercase: Flanking sequence[1000bp]

Red & Bold & Underline: Stem-loop [10]

Blue: Heptamer[27]

Green: Nonamer [2]

id-TRAV5[V_gene_segment]

tctgtagtctgatgtttcctgagatgtttcctcttttatgtcaatactttgcaataagtaagccacatatccgtaatttcagatgatctgttcttccagaggaaaatatttcagagtgcatgtcagatttttcatcacaatctataggctaaactctttcctttcattagaccaatattattaactcatggaagcctttctccctccttttatcttttattgtataatttagtcattttgaag**aaaacaacactcatgtttt**gacctctggcactgtcaat**ttcctgggaatggcaagaacaggaa**caaggc**caggcttggaggagcagagcctg**agatagcatgcatctgatatatttagtgaacctgtcttactccaatttttgtgcttaaatgctagcttcagaaagagttttcctctgtgagtgctgaaacaggatggctaatttggctgattgattttctggccaattcaaaaaatagtagagggataaggaggaaacgttttatacctcagtagctttgtagactgaatcatggcattttcttaatttcaagaagactgtcattaataaacacgaaaaacctctaacttcttcaaatatatttaacgtgactcagtagtgtgccgagaagatgcaatggaattctaaaacctacgtttgcactttttctgtctgctaatccacataatttctgactggaggtacacttggaatttactttatttggccttctgctgtttgatgtgccagaatctcccacagatacaggataattggtttagcgtttggggatctggccacaaggtg**gcagagcttctctgc**tcatgtagaatgtacgaaaggatccttttggttga**cttctgtgagtaggctgacggcagaag**cagatgcct**ctgtggacagtttaagaaaccacag**tgctttggaggaaagg**aagagatacttgataatatagctctctt**ggctggagattgcaggtcccagtggggagaacaATGAAGACATTTGCTGGATTTTCGTTCCTGTTTTTGTGGCTGCAGCTGGACTGTGAGTCGAGAGCTTTTGGGGAACAGAGGGTTTAGTAAATACTCATTAGAAGTCTGAGGAGGAGACTCATCTGGTCTTTTCCAGAATGTCTGAAGTTACTACAGTGAAAAAAATAAAAAAGCAAACTCCAGAGAATGATGGCCTGATGATCTTGTCTGACTTTTCCTTTTGCACAGGTATGAGTAGAGGAGAGGATGTGGAGCAGAGTCTTTTCCTGAGTGTCCGAGAGGGAGACAGCTCCGTTATAAACTGCACTTACACAGACAGCTCCTCCACCTA**CTTATACTGGTATAAG**CAAGAAC**CTGGAGCAGGTCTCCAG**TTGCTGACGT**ATATTTTTTCAAATAT**GGACATGAAACAAGACCAAAGACTCACTGTTCTATTGAATAAAAAGGATAAACATCTGTCTCTGCGCATTGCAGACACCCAGACTGGGGACTCAGCTATCTACTTCTGTGCAGAGAGTAcacattgcttctcaggcacctgtatcctgtacccaaacctgcacctgggactaaagccacactctatttcctt**tacctttaagtcagggattttgctgtaaggtatttttaatgtacggacattacaaa**acatcctaaaaggtcagtt**tcaaaatgacttaatgattttcagaacattttga**ttaaactctttttgggagttaatgatttcttagttgatatcctggtcttagg**gttgactttatgggtgccataaagtcatc**tgaatgatttctcccct**accaaggacatcttggt**gtcagagaatcctctccttgatggagtcttttatcttttaacattag**tatttattaactttttctaactataaata**tattaagtgcacggtataggaaaaaacaggaaattataaaatagaaaatcaaaatgtcttataatcctatcatacacataattcatagctaacatttagtattccttcttcattaaaaaaaaaattaggatcgcttttcctgtggcagcagctggggtgagaagagcagtggctctctcctctctctgccatggcatgtgcttgcccactgatgtcagcgtactctgagaagggagagtcatctggcaaaaatgtcactttgtctgccatattcaaggctcctattcgaccagatatt**gtgaactttgttcac**gccaacttgtgcaaaaacaaca**gacagccctatgctgtc**agtgaatcagcaggtcatcaaaccagtgttgagtcttggagtactggcagagctgcagcttgaattcccagagttcgaagtggtgggactcaccattctggccagggagcttttgg**aaacatgtgtcgtggaggctgaatgttt**gcaccaaccaaaacctggctccattggcatagtagagtgaatacaaccccaaaacgatatgtcatctcttctgccctggctgcctcagccttaccagcactggttatgtctaagggtcatcgtattgagga

Uppercase: TRAV5

Lowercase: Flanking sequence[1000bp]

Red & Bold & Underline: Stem-loop [19]

Blue: Heptamer[24]

Green: Nonamer [6]

id-IGHV4-28-2[V_gene_segment]

tccaacttctggtctccaacaatactcctgcatctgccatctaagtagctggaactaccgttgtttgccagcaggcttggcttgagtacttattaaaccgcacactttttgcaatatttaatgtatagtaataatgtctcaataagactatt**acaaatcaatgaatgaataatttgt**tctgtacagatttatggaaaaatagacactaacatgatgaatgactg**acatttatgaaaatacagctgcatgaaatgt**gcttctctttacattcattaggtaaacacaatagtgcatacacatcacaccgtgctttcattacaggaagaaagatctgaaaatgtcactgggg**tgaaccatattgtgctgggcttggttca**ggaagcagtcaggcccagtgttgtgaccttcaaccacagaatcctaaaaaataataaaagaggctcccccaaagtccccatcagttcctggactcgccatgtgtctgggccgtatcagtgcatccggagctccctggtggctttagggattccttgcttgccatgctgaggtctcgctatagattatgttgggttttctgaggccgttttgctatttaagacccactccctggcacagagcgattc**cctctaaacctgatagagg**ttctgaactacaaataacgttaagtgaatcctggtgtgtctgaactcacatgattgttacattaagctgctgttgcaatctgtttcctcacctgggaaaagaggagccaggacatagtgagttgaggccccaggaagataactgaattctcagagggcacaaccagcatcctccttcc**agggagagcctaaaagactggggcctccct**catcccttttcacctctccatacagaggcaccacccacatgcaaatctcacttaggcacccaagggaaaccatcacac**atttccttaaattcagggtcctgctcacatgggaaat**actttctgagagtcctggacctcctgtgc**aagaacATGAAACACCTGTGGTTCTT**CCTCCTGCTGGTGGCAGCTCCCAGATGTGAGTGTCTCAAGGCTGCAGACATGGAGATATGGGAGGTGCCTC**TGAGCCCAGGGCTCA**CTGTGGGTCTCTCTGTTCACAGGG**GTCCTGTCCCAGGTACAGCTGCAGGAGTCGGGCCCAGGAC**TGGTGAAGCCTTCGGACACCCTGTCCCTCACCTGCGCTGTCTCTG**GTTACTCCATCAGCAGTAGTAAC**TGGT**GGGGCTGGATCCGGCAGCCCC**CAGGGAAGGGACTGGAGTGGATTGGGTACATCTATTATAGTGGGAGCACCTACTACAACCCGTCCCTCAAGAGTCGAGTCACCATGTCAGTAGACACGTCCAAGAACCAGTTCTCCCTGAAGCTGAGCTCTGTGACCGCCGTGGACACGGCCGTGTATT**ACTGTGCGAGAAAcacagt**gaggggaggtgagtgtgagcccagacacaaa**cctccctgcagggagg**cggaggagccggcgcaggtg**ctgctcaggacgagcag**ggggcgcgcggggcccacagagcaggaggccgggtcaggagcaggtgcag**ggagggcggggcttcctcatcagctcagtgctctccctcc**tccccagcacctcagctgtccccaggactcctctttctttattatctgtggttctacttcctcacatccttgctgcaggttttaacagtcactggcacccttttttcaacaaattcccacaaggcccattttaccttcttgacaagcaaaatattctaaggcttctcactatcccttgattggcgtcatctactgcatcgtgtctttctcttaaattcatgtgatgaaaccccaatcccagtgg**ctctgtattggacagag**ggattttaagagggcaaataagttaactgtggccataaatgtgggaccctaatccagtaggactcttgtccttatgagaagtggaagacatcagaggcctctctct**ccacatgcacacagaggagagtccatgtgg**ggatgcgctgcatgagaaggtggccctttgcaagcctggaagaggcctctcaaaaaacttaccctttctgcacccagatcttggacgtcaagactccagaacaataagaaaattaatatttgttatttgaaatagcctagtctgtggtatcttctcatggaaagccaagtagattcagagattcaattgtgttagctccgtctcctgcatggagtagcagcccaccgaggctggacacatctctcaaatta**ttttcataaagaaaaatagatccaagatgaaaa**ccccaccacattt**ctgctgagccattcacttagcagcag**tctc**tgagatcagccctgggggtatgttctaggtcgtgtatctca**ctttccactatgagaatccatgtagataatgagaaccagccagatggagtcctga

Uppercase: IGHV4-28-2

Lowercase: Flanking sequence[1000bp]

Red & Bold & Underline: Stem-loop [20]

Blue: Heptamer[39]

Green: Nonamer [3]

id-TRBV28-2[V_gene_segment]

acaaatatgtagtaagaatactatgatcacaaggaacaaag**acatggtggactttggagaaatccatgt**gagacttccttatgttctcttcca**tgagagagcacactctca**ccccaacaatgaaaatgcagcaacgtgagaacaacctttctgcccagggaagctcattagagac**tcagcacctccaatactttcagtggctggtgttaggtgctga**atcgtgtcattaaacattcatatgttgaaatcccagcctccagtactaagaatatgactatgtttggagataaggattttacagaggtggttaaggttaaatgaggtcctcactaatatgactgggatccttataagaagaaattaggacacaaacatgcatagagggaagacaatgtgcagacacagggggaagccatctacaagccaaggagcgaggctccacgagaaaccaaccatgctgacaccccgatcttggatgtctagtctccag**atgtatgagaaaatacat**ttctgttgcttaaatgacctagtctgtggtgacttatggcagccctcgcaaaataattcagctgatcacattggcaccctctgcctatcatgcagcataattccagacttgacgtaaagcaagtgttcagcctaacccacgtggtttgtgcaagcagtctagacacactaagccatgcttaccagttactatagtgggaacactcccaaaattcaagttcgcagatgcaaatcaagagccaaccatgcaagcagtcccttctaaggccagcagtctcaggcccg**ctgtgtcaactcttttctatacgtgcacatacacacag**tgacacttccatgagcacaagggatgtccttcactctctagcacc**tctagaccctttctagtctaga**caatggctgagggattatgag**agtgggccgggacaatgacatcacagaccatcaacccact**gcctggtcctgggagaagacctattctttcttcaaagcagccATGGGAATCAGGCTCCTCTGTCGTGTGGCCTTTTGTTTCCTGGCTGTAGGTGAGTCCTGGGAGTTGCTGGGTCTGGGGAGTTGGTGGCAACCCCTGTCCTACTATTTTGAGGTCTTTTATACCAAGCTCTTTTCTCCTCTCCTCCTTGAT**GGCCTGTTTTTCACCCTTGCTTTCTGTCTCTCACAGGCC**TCGTAGATGTGAAAGTAACCCAGAGCTCGAGATATCTAGTCAAAAGGACGGGAGAGAAAGTTTTTCTGGAATGTGTCCAGGATATGGACCATGAAAATATGTTCTGGTATCGACAAGACCCAGGTCTGGGGCTACGGCTGATCTATTTCTCATATGATGTTAAAATGAAAGAAAAAGGAGATATTCCTGAGGGGTACAGTGTCTCTAGA**GAGAAGAAGGAGCGCTTCTC**CCTGATTCTGGAGTCCGCCAGCACCAACCAGACATCTATGTACCT**CTGTGCCAGCAGTTTATGcacag**cgcagcacagctgcatc**ctctctgcacaaaaagagcggacgtaagagag**aaggggccctaactcagggctggtgctggctccgatggcacattcgtgctaaatagaaaaaaagcggccactaaggagaaactagctccataagcacacttctttgcaaggagagaatgaggtattatttttcacccctttacaagtcag**ggactgttctggatttaattcccccagtcc**gatgtgttgtaaattcacaagtcccaacaaggaggaaaagtcctctcgcatattcacagtgtaaccctgttctaccatttcgttaatcaattttaccttcccttttctatcctcgtctctcttaactgtgcattttctactttgcttagtgttttgctcttggttgacatttcactagaggacattaacatattctaaattctttcgatgatttttactgtgcctgtgctgattcaagagaaagtttgtggaat**ttgaattttgttcattcaa**tttagaaagagaaaaactctgattaaaacaaattc**ttccaggttctccgatcaatcatattttgctggaa**aaaattctgatcacagaaggtttaaactctgctgcagaaggtggtaggaatttggttataactgaactcacaatcttctggaaaaagtgtgtatttgtctgtctttaagaataaaacatatgctcttctgtcagtaaaaattaacagtgctttggaaatacaaataaatgtaaagggtattagaaatcccaaactgcagagcacttgcacaattattattggcaagacaaaacaggcatctgaaaacgtattttgattcagtgcta**aatttttgtgtggaagaagtcagagaaaaaaattgaatgaatgtgtatcacaatattcaa**gattaagtttgtgtgagataggacttgagctga**ttttgaatacaagtaaatgagggaagagaaaccatcaaaa**ggaagaaataccctgcaaatgttt

Uppercase: TRBV28-2

Lowercase: Flanking sequence[1000bp]

Red & Bold & Underline: Stem-loop [17]

Blue: Heptamer[27]

Green: Nonamer [3]

id-IGHV5-51[V_gene_segment]

gtagctgatcccaacctcaagtaggatgttcttgatgagacagcttccctggtgcactggatatcagatcctgtatagtttcagaacttgtgtagtttgtgttgctttggctttttctctggatcttttaggtgctaatgaaatgagagtgcttcacaaaaaaatggggcagacaaaggggatagttaagggcctcctccaagcaatgtggatatcataaaataaatgttaaaaaatgaaagaactaaacaagaag**gtgctggggttgacacaaaggacttaaaacagcac**taccaggagttgggtgcagcaatgggcatcccttcaggtccatcagcaagtcagagacctaaacaaatattccctattcatgttgaattggcagagtgtaaaaatctagaaga**caaaatggccaggaaaaatgtgcgattttg**catgggtgaaggtctggcaagttaatcatcttgaggactgttgaggtcctgaggtggtcctgcccttgcttgatgcccaggccgtagtccaactcacacgaaattgccagggaatagacagtagtttttttagtagttcttgttatcaggcataagtgcatttgaattttctcttcatggcctttcctggcactatttctcattttttttaacacacatagtttcaactagatttatcaccttc**acagggtcacagagaagggtggaagaagggaggccctgt**atgggtctcgaagaaacatggaaaagagtggagagggacaatagcagggt**gtaaggaattattgagaccttac**tctgcccctcccaggaggctcaggccagcctttttctgcatttgaggttctgggttataaacgctgtagactcctcccttcagggcagggtgacaactatgcaaatgc**aagtgggggcctccccactt**aaacccagggctcccctcca**cagtgagtctccctcactg**cccagctgggatctcagggcttcattttctgtcctccaccatcATGGGGTCAACCGCCATCCTCGCCCTCCTCCTGGCTGTTCTCCAAGGTCAGTCCTGCCGAGGGCTTGAGGTCACAGAGGAGAACGGGTGGAAAGGAGCCCCTGATTCAAATTTTGTGTCTCCCCCACAGGAGTCTGTGCCGAGGTGCAGCTGGTGCAGTCTGGAGCAGAGGTGAAAAAGCCCGGGGAGTCTCTGAAGATCTCCTGTAAGGGTTCTGGATACAGCTTTACCAGCTACTGGATCGGCTGGGTGCGCCAGATGCCCGGGAAAGGCCTGGAGTGGATGGGGATCATCTATCC**TGGTGACTCTGATACCAGATACAGCCCGTCCTTCCAAGGCCAGGTCACCA**TCTCAGCCGACAAGTCCATCAGCACCGCCTACCTGCAGTGGAGCAGCCTGAAGGCCTCGGACACCGCCATGTATT**ACTGTGCGAGACAcacagt**gagagaaaccagccccgagcccgtctaaaaccctccacaccgcaggtgcagaatgagctgctagagactca**ctccccaggggcctctctattcatccggggag**gaaacactggctgtttgtgtcctcaggagcaagaaccagagaacaatgtgggagggttcccag**cccctaaggcaactgtatagggg**acctgaccatgggaggtggattctctgacggggctcttgtgtgttctacaaggttgttcatggtgtatattagatggttaacatcaaaaggctgcctaacaggcacctctccaatatgacagtattttaattagtgaaaattttacacagttcatcattgcttgcttgccttcctccctcctgtccactctcactcactccttcttttattttctacttaattttacaaaatcatttaacccctttttgaactattaataggttatctttgtttggtgattgttttcctttcaataatatgtactgaataattcatctttgtgccaattcataagtattctggtgtaataaagacttctttca**taaaaattggataaattaaaataaagataaattttta**aaaacatacgatctatcaaaactgaaccataaagaaataaaaactctgggttgggtgtgtttgctcattcctgtaatcccagcactttgggaggccatggccggtggatcacctgaggtcaggagttcgatatcggtctggcaaac**gtggagaaacgctgtctccac**taaaaatacaaaaattagctggacatggtggtgctcgcctgtagtcccagatacttgggagcctaaggccagagaagagattgaacccgggaggcagaggttgaaataagccgaaatctagccactgcattccagactgggcaacagagtgagactccatcccgaaaaaaaaaactgaacagacctatgagtaaagagattgagtcagtgatttttcaaacatctcaaatcaaagaaa

Uppercase: IGHV5-51

Lowercase: Flanking sequence[1000bp]

Red & Bold & Underline: Stem-loop [12]

Blue: Heptamer[29]

Green: Nonamer [6]

id-TRBV15-2[V_gene_segment]

aaaagaaataatatccttaggaaggccaccatggaattcattctatctgaacaatccttacataatatcatttcttataatatgtcctcaacatcgtgctctgcatatatttctccaccatctctaaaaatcctcttagcccaggcctcctcaaactttaatgtgcatgtaaattatgaccttgttaaaatgcagattctaatccaatcagtcagggttggagctaaggttccctgtattttgaacaagagtccgtccaagtgatgccaaagctgccagtccatggaccacaatgagcagccaagctctagatg**acacctacaataattacaggtgt**aaggcccctcttctcaagtatcattttgcacagttcctggactaggtc**agatacagaccctacttaccacatgtatct**ctgtattccaagtattacgtaatcatcagttttcctaaatctaggcaccaatattttaacatatattttcatggtataa**atagaaattaattttctat**agtaaatacactaatac**ctaatcacattcatgcatctttaatgtgagtag**caaaaagccaaggcactatcttaatggttattgagtgagactttagcaaatactatctttatggagaaacctctgggctgctttttaggatgtgagaaaagtgaggtaaagtgaggagaactggatgaacacgggaca**gagacagggacaggggcaaatatggggacacctgtctc**aaggaagcagcaaatgatatagaaaataaatatctgttccatccctgttccagacaagcccatgtacctgccaagtaggaagctgtgtatcacattgcaacaaggaatgaccccggccctggtaaagtcaacagcaacagtcatcacaggccaatctgcctatcagggactggagactctctaaactcccacctctcaacc**caggaatcagagcctgagacagacagatgcttcattcctg**tatggggtggtattcctgccATGGGTCCTGGGCTTCTCCACTGGATGGCCCTTTGTCTCCTTGGAACAGGTGAGTACCGGGCAGAAAGGAAATCTTTGAGCAAAGCTATCTTGTCCTCAGTCTGCACCTTTCATTCACAGCAGTAACACTGTTCTCCTTAACTCTGACTCCAAATTTGTCTTCTTTCTCTACAGGTCATGGGGATGCCATGGTCATCCAGAACCCAAGATACCAGGTTACCCAGTTTGGAAAGCCAGTGACCCTGAGTTGTTCTCAGACTTTGAACCATAACGTCATGTACTGGTAC**CAGCAGAAGTCAAGTCAGGCCCCAAAGCTGCTG**TTCCACTACTATGACAAAGATTTTAACAATGAAGCAGACACCCCTGATAACTTCCAATCCAGGAGGCCGAACACTTCTTTCTGCTTTCTTGACATCCGCTCACCAGGCCTGGGGGACGCAGCCATGTAC**CTGTGTGCCACCAGCAGAGAcacagagctgcagtgcttcctgctctctgt**tcataaacctcattgtttcccagatccaggtgctttctctaggacttctccctcaccacctcttacaacaataggaagtgggttggtggctgtcaatatctgtagacagaagttgagcacaaaccaataaaaaccattgacagtt**atgccaagagtggaaaagtggttacacatgcctggcat**tcagttggtggtattttcccaggatcacatttagaacccatgctgtccttttcagaagacaaataagaactttttctttctttttttctttgttttaa**tctaagtcagaggcttaga**aatataagaggtgatatgtgaagaattatggacacccaatgtctataaagtccagtctctggcattcgatgaactcttataggatttaatcctcctggagaatccacatgtttttagcctagacctggagcaactacctcattaatgggagaatgaaagcacaagtatcccaagataaaaa**tttctcagaggtaaaaccacattggagagaaa**aaaagagaagaccatacattaatc**tgctcatatctggagctggaggtactattgagatgagca**aatggaaaccataaatacataaagaaatgatattgaaaatagagtacatttggagatgaagatcatggagccatctgtatagaagagagtaataatgtcattaaggtgcataagagtgccagcaaatgtcaataagaaaaactacaggagggaagaagatgggagatcctgctccccttaacaaagataaataaaatgctgagtgagaaatattgcaacctcataaattgtcacatggaagtatgtat**ttataaattaccttataa**ttaacatccctcaatgagaatgatattctgagaaaaaaaagcagaaaatatagttatatggctatctaaaaatgtaaaattttgcacattaaattaagataatctaggaaaaatatgacaagacctat

Uppercase: TRBV15-2

Lowercase: Flanking sequence[1000bp]

Red & Bold & Underline: Stem-loop [14]

Blue: Heptamer[24]

Green: Nonamer [2]

id-IGLV4-60[V_gene_segment]

gagctagaactagaaataccatttg**acccagccattccattactgggt**atatacccaaaggactataaatcatgctgctataaagacacatgcacacgtatgtt**tattgtggcactattcacaata**gcaaagacttggaaccaacccaaatgtccaacaatgatagactggattaagaaaacgtggcacatatacaccatggaatactatgcagccataaaaaaggatgagtt**catgtcctttgtagggacatg**gatgaaattggaaattatcattctcagtaaactattgcaagaacaaaaaaccaaacaccgcata**ttctcactcataggtgggaattgaacaatgagaa**aacatggacacaggaaggggaacatcacactctggggactgttgtgtggtggggggaggggggaggga**tagcattgggagatatacctaatgcta**gatgatgagttagcgggtgcagcacactagcatggcacatgtatacatatgtaactaacctgcacattgtgcacatgtaccctaaaacttaaagtataataataattaaaaaaaaaaaaagagagttaaaggctga**gatgaggtaggaggctgccctgaggaagggtctcatc**ctctggcatctggtcacagacacgatgggcctgggcctgggctgcagagagcacaatggggctgctggtgggggctctgtgcggctctcagctgggaaataaagtccatgaactttgttttgcctcacggctcactggggccagcagctgggtcctctctgggt**gcttggggagcttcaggccaagc**tcagcccaggctgactcctcctagtagtgtcacccagccacagaggctgtacacaggcccagggagagttaggttaagcaggggaagaggagcacatttgcatgaagggcccct**ccctctcctttgaggctagaggg**tggataagagagacctgcagcgtggctgcctcagcagagctctggggagtctgcacCATGGCCTGGACCCCACTCCTCCTCCTCTTCCCTC**TCCTCCTCCACTGCACAGGTCAGGAGGA**CCCTCAGCATCCTCATGCCCCAGCTCACTGACACCATCTCCCAAACTCATACCAGAAATGTTGTTTGCTCTTGTCCTTCCTTCAGGCCATAATGAGCGTCTCTGTTTTCAGGGTCTCTCTCCCAGCCTGTGCTGACTCAATCATCCTCTGCCTCTGCTTCCCTGGGATCCTCGGTCAAGCTCACCTGCACTCTGAGCAGTGGGCACAGTAGCTACATCATCGCATGGCATCAGCAGCAGCCAGGGAAGGCCCCTCGGTACTTGATGAAGCTTGAAGGTAGTGGAAGCTACAACAAGGGGAGCGGAGTTCCTGATCGCTTCTCAGGCTCCAGCTCTGGGGCTGACCGCTACCTCACCATCTCCAACCTCCAGTTTGAGGATGAGGCTGATTA**TTACTGTGAGACCTGGGACAGTAA**CACTCAcacagtgatacaggcaga**tgaggaagtgggacaaaatcctca**acctgctgaggctattgttcagtgacaatttttaattttaaaacattttctgtatgtaaaaaatctatctggatgcatcctctgatttagaacaattttactcacaatttcccattta**atgtttcagaagtctcagaagaaattaaaacat**aaacaagaac**ataagcctcctgatgacactgagatgtctgcttat**ccttatgggctgataccaaagtctgtgaagctcacc**ttttctcttgaaagaaaa**aaagaatgagccttccct**tgagcaaatcttgctca**gagcccaaacttca**agcccacaaatggacagtcccctgcagcatgggct**cttgttcttcaaggggtcaatgaggagacctccctttcccaccccaggtgtctgtagtcagggctgcctgatgttccctgcatcctgggactaggcactcaggtgtgtaataccccagagga**caggcctgggagtaattccaaacacagagctggcctg**attcactttcaggttccttt**attttaagttaaaat**gtcaggatagaaagaagttgggtatggggttattcatgaaactgaggccagaagatgtctttgtgggctgtgagagggtgac**atggtggctccaccat**tgctaggggaatgttgagtgagctcctaagaagtttggggaaatcggtcttg**tgagtgtggcttcccttcagctcattccactcagaaacatgaggacacccatgggtggatttttctga**actcacaaacactgacttgtgtcctcttttttcttatcccattggagatattcagtaatctactctgtgagatggggtctacaggagatcatggagaagaaactcttcaacctgtgcaagtgaaacttccttgtgtgaagctcttgatgtgtccagattccaaggtcaccttctcctgcacaggggtcagcagaaacattgaatattaccat

Uppercase: IGLV4-60

Lowercase: Flanking sequence[1000bp]

Red & Bold & Underline: Stem-loop [21]

Blue: Heptamer[41]

Green: Nonamer [1]

id-IGLV8-61[V_gene_segment]

ctcatcctcctcctccactgcatgggtt**aggatggtccccagcatcct**gatcccccaggtcactgacatcagctcccaaactcatgccaggaatgtccttccctcttcttccttcaggccataatgggtatctatttttccagcatctttctcccaatctgtgctgatttagccaccctctgtatctgcctccctgggacaaagggtcaaactcaccagcactctgagcaagagctacagcagctgccacatagactatccccagcagagctcagtgatggacaccaggttcacaatgccagtgggcagtggtgctgtttactgcatccagtggggatgga**gtggttgtccaggcacagacagatggacaatcac**acagcaggaacaaatgtac**attctgaaatttaacaaagacagaat**aagggactcttggggaatagaggtctcaattgcagacatagtgagtgcctcaaaaattagagaactg**gggccatctctgtcatggccc**ggattcctctctccttcatcctcctcagtcatcaaagagattccattagggggcaatcaaaggatgctggggcctgctactagctcccaaaaaacaccaca**agcatttcagcagggagaaaagaaaatgct**cacaaggatggggttagatttagttataaaccagattgtgtggcaggagggaggacatatcccccatcctttggccactccaaaattgggagagctaatctggggaaaggcaccaacacctattcattagtgtctcggtcctgtgtgatcatgggtctgtgtgagttctcagcagctgcctctctcaaggtctccccacaggacagcagggtgaaacctggggcaggaagagaggggaagctgatttgcataaagaactttttttatcctattgggt**ctggggaggcatgaaaaggccctgaggaaaacaaaccccag**ctgggaagcctgagaacacttagccttcatgagtgtccccaccATGGCCTGGATGATGCTTCTCCTCGGACTCCTTGCTTATGGATCAGGTCAGGGGAAGGGACTCTATCCCTGGGGGACCACAGAAAACAGGGTCCAGGTTACTCTCATCCTCATGATCATAACTGTGTCTCTCCTGTTCGTTTTAGGAGTGGATTCTCAGACTGTGGTGACCCAGGAGCCATCGTTCTCAG**TGTCCCCTGGAGGGACA**GTCACACTCACTTGTGGCTTGAGCTCTGGCTCAGTCTCTACTAGTTACTACCCCAGCTGGTACCAGCAGACCCCAGGCCAGGCTCCACGCACGCTCATCTACAGCACAAACACTCGCTCTTCTGGGGTCCCTGATCGCTTCTCTGGCTCCATCCTTGGGAACAAAGCTGCCCTCACCATCACGGGGGCCCAGGCAGATGATGAATCTGATTATT**ACTGTGTGCTGTATATGGGTAGTGGCATTTCcacagt**gatttaaacctatgaggaagtgcaactaaaacctctttatatactgagaacagttcagcccttacagacaggagggaaagtgagagggtggaaatggtcaacacggtgagtgaggagtctcctcgccccagtctctgacgcaggggttcactgccaggtgcctcacccagtctctccccgggtgtgtccaccttcccagtgtggcttttccatggctgtgcttcgt**ctccactgagaataatgtggag**ggagaattatctacaaaatatataagctcacagtgagtcacagacaccccagtgctccaggaagacgcttcttattcaaagagaattaacaggcatatgctcactcattaatttgaccatggaggctgactgaatattttatttatcactgaagtgggag**cattttcaaggggcttgttaaaaatg**taattctctaacctaaacctactaagtcagaaacttgtgttggtgacagatagcctgtatttaacaagctgtacatgagatttttattagcgttaatgtttgggaagaacctac**ttttattatcataaaa**ttattcataaggatctccatttataatgatgtcaggttaatattccataatcctactctcctaaattcccattcaaagcccccaaatactcagaaacagattttaaatctcaagccatagcaaaaattataacttgatttatgcagacagagtagggaggaatttctaaaaccgtaagttcctaggtgccaggtgtgctgactcatgcttataatcacaatactttgggaggttgagacaagaggactgcttcaagccaggagattgagaccagtgtcttccacatagcgagagtctatcaccacgaaagaaaaaaaaaagatataagtcccttcaggaatttccaaaaacttgtttccttagggtatatttagtggtccatatgaacatctgttactgaaaaccatcccgtagaggactggcttccaga

Uppercase: IGLV8-61

Lowercase: Flanking sequence[1000bp]

Red & Bold & Underline: Stem-loop [11]

Blue: Heptamer[41]

Green: Nonamer [1]

id-IGKV2-28-3[V_gene_segment]

gtggatccagttcgtgctcttcttacatagagaatttccactcgaattgtgaactcatggctgtggctgcaccacttacaggcccaggggacactcagatctcacttggtagttgacgaaaaactggaagtcctgatagga**cctcattccacatgatgagg**aagactgtgggaagagcttttgttaacaccattcag**aaaaacatagtgcaaagttagttttt**gttc**tttctatataattatcctagaaa**agtcttcccttaataaatccttctggtt**aatctggcatatgtgagatgattatgatgggatatataccagatt**gaacaattggtcaccaggaattttatattcactgcct**gaggaataaattgtttcccactttcctc**ttacctgcactgggctcttgaatctaaatatagagacccacattattttccctatgaggcccttggacagagcgctcttatggggctcactcaccaggtgccaggggagggtagattccaacacttgctatgaacattcttgaacagttatcctggaaaccgcagataccagaccac**tcttgaactggctcaaga**ctatgttttatttgtaggctgtctgctcatcagtgcttgt**aggaaagggtaacgtttcct**tttttagattagctgggagggagccaaga**agaatggcattcatccatattcattct**agacatatctcta**cattgttagggttgttatgctttcctagagttgcatatcctatacaatg**ggacctacccaagatccaaactgtcacagtcagatccttcctcccattttatatcacattgctcacaggagagacatatcccctgcccgcctgccccattgactctttccacaccactgcatgcaccaggggatttgcatattgtcccacagggaggaccttcccttgtgagtctgagataaaagctcagctgtaactgtgccttgactgatcaggactcctcagttcaccttctcacaATGAGGCTCCCTGCTCAGCTCCTGGGGCTGCTAATGCTCTGGGTCTCTGGTAAGAAAAGAAGGAGATGAGGAAGGAGAATAGGGTGGGAGGGTGAGCTCTAGGGCTCCACAGCATCCCATGATCCCATGTTTAGTCTTACCCTGTGTTAGAGGAGTATAATCTGTGCTGTAGAAAAGGGAACTTGATATTTTGCTCTGTGAATAATTAGAAGCCTCATAAGAAATATGACGTCTGGTGCTCTGATTAAGATCTTCAAAATATAAAGGTCTCTTATACTTTACAAAAATTGAATTCATTTTAGAATGTGTATTTTTATGGCATAAATCACT**ATTTTTTAAAATTAAGTTTAAATAAAT**GACATAAGATAAATTATGAA**AATTGCTCATTAGGTTTGTACATAACTTTGCAATT**CATTATTTCAGGATCCAGTGGGGATATTGTGATGACTCAGTCTCCA**CTCTCCCTGCCCGTCACCCCTGGAGAG**CCGGCCTCCATCTCC**TGCAGGTCTAGTCAGAGCCTCCTGCA**TAGTAATGGATACAACTATTTGGATTGGTACCTGCAGAAGCCAGGGCAGTCTCCACAGCTCCTGATCTATTTGGGTTCTAATCGGGCCTCCGGGGTCCCTGACAGG**TTCAGTGGCAGTGGATCAGGCACAGATTTTACACTGAA**AATCAGCAGAGTGGAGGCTGAGGATGTTGGGGTTTATTACTGCATGCAAGCTCTACAAACTCCTcccacagtggta**caacccctaacagaaacctcctcctggggttg**cctagttgctcacatgt**gctgcttgtctggagagcagc**tcagcagggtctctgagtctgcagaagaggaggctgttg**gagacctcagggcagaggttgctgctgaggactc**tggctcatgatagcctcagctgtacttcagtcccacatgttaaggccccattaggtgaaaaataaa**tgattccaaaaactgagatgaaataccaaggagaatca**gagtacaattaaggctgttacaaagaagcctcaaaatatggtggactaaatgtgacatggtttctgtgtctgttgcctgacagtgcagaggcaggtgggtggctttggtggtgtcggtggctcttctccaagaggtcgctcagatgggcaggagacacgaccaccctcagacacagccttcctccttcctcagagtcacttgcccccatgcccatcctcgacagcatgaagtggaacgactagagagaaagctgt**tttcttctaaagaccaaagaaa**tctagagtttcccatcagggataaatgttcttttactttaagcacgcattggagaaattttccattcagtggatctgctgataaacccactttgttttgtttgtttgttttgttgtgttttcagatggagtctctctctgtcacccaggctagagc**gcagtgacatgatctcggctcactgc**aaactgcctcctgggttcaagagattctcatgcctcagcctccggagcagctgggattacaggcacccaccaccattccaggctaatttttgtatttttagtagagacagggtttcaccatgttggccagggtggtgtcgaactcctgatctgaagcgatccacctgccttagcctcccaaagtgctgggattataggcgtgagccactgcacctggcagttatttgctcttatatgataatgtaaaattattttagaaaataattttttgagtatatagtt

Uppercase: IGKV2-28-3

Lowercase: Flanking sequence[1000bp]

Red & Bold & Underline: Stem-loop [20]

Blue: Heptamer[41]

Green: Nonamer [7]

id-IGHVIII-47-1-2[V_gene_segment]

ctgagaaaaagaaaataacctcctgcatgcacttctcaagtctctgcttgtatcctgtttgctt**taggtctctaggggaaaaagtcagatacctggaccta**gtgtcagtgtaggaggcacatcctaaaagctacaaactaggaagaagggaagatgggtgttattggaatagtggatatggagtgggctttcttctgaaagtatatgcacctgctggtattctcagatgcaacattcaactgcaagagcccattgaagaaacaaggcactcccaaatctcctgtaagattctgtattcatttcagataagcccacatgtccccgcattaactttttc**ttcagacaagcacatacatctgaa**actgaacagctatgtggcaaaataagctcaggatagaagt**aattgtggactccagctctgacaatt**tgtagatctgcatttttaaaattctaactgaagactttgctttattgtagaggacagtggtttacagctcgaattgcacagcctacaggcagatgtccatttcctctgggcaaggtttatttttatttgtttactgtacttatttgttgacaaacattgatactataaagataccctaacagcgtccacatgaaagaaaaagaaagagcaatggacacatcaaccctgaacatccagtcccaggaatcctttgaccctgccctccttgcaacccagagatatagattggatgagccctgctgagcagaacaca**catgtccccaggagaaagacatg**gaaatgaggcccctccctgctaatgaaaagcagctcttcccctcttctcctgcaggtcctgg**tgaggagccacccaatatctgtgcccttcctca**gtgtccacaccatcgggtcta**tgatgatctgggcttcacttgtcatca**ctctcaatattgaggttccccgttaaacagactgagtgaactgtggctgctccacgtgggggctgttctcagtctgttgcttctgtgcttgcagAGGTCCCCTGTGAAGTTAATTACTGGAGTCTCTCAGAGAAATACTACAGACCAAGAATTCTCAGACTTTTCTGGAAACCCTGTGGATTCACTTTCACTGAAAACAGCATAAGCTTGGTCCAGCAGGCTTCATGACAGGGGTGGGTGTAGGTGATAACATCAGTAATTCAAGTGGAAGTTCTCAGTGGGACTCTCCTTGAGTACAAAGAAGATTAACAGTCCTCAGAGACACGCTTTTCAGATGATTCTCTCTTAAGATGATTAACCTGAGAGCTCAGGAAAATTCCGTTTATTACTGTGAGGGAcacggtgaggggacatctgtgtgagctcagacacaaacctgcc**tgcagggagacacaaacctccctgca**tggtagatgcttctcagaaccaccagggggtgcacaggaaaccagaaggtgctcaggacaccagggggtgctcagaaccaccatgactcactcaagacgccaggggcgctcagaaccaccgtggggcactcaggacaccagggggactcaggaaactagggggtactcagaaccaccaggcggtgctcaggacaccagggagctctcagaaccatctgggggcactgaggaaaccaggggactcaaacactagggtgtgctctgaagcaccagcggaccctcaggactgcagggggcgctcaggacactagggaacactcagaaccaccaggaagcgctcagaaccaccagggggc**cctcaggacatcagagcgtgctgagg**acctccgggggcgctcaggaccttcagagagtgatcagaacaccagagggcgctcagaacaccaggaggtgctcaggacagcaaggggctctcaggacactagggtgtgctcagtaaaccaggggtccctcacaaccaccagggggcactcaggacaccaggggatgctcaggacaccaggggccactgaggacaccaccgctcccttagcaggcagctccacatcaggcccctgggttggggcaggaagggtgttttccttttggatcttgccact**aaactcttgggagttt**ttctccttcctttgtggtttcaagaaacattggtagattcttctcaggtataaagctctgctttcttggattatgtaatgtttttggtttcggatgttaccagaattacactgcactgtgagagga**ttcattcctcgtgtgtgcaatagtgaatgaa**agctccaatgttaggggtggctttgaaagctacgttaggggtggctgagggcagttagcaggaaatgatcatcactatagaaggctactcatttctttg

Uppercase: IGHVIII-47-1-2

Lowercase: Flanking sequence[1000bp]

Red & Bold & Underline: Stem-loop [10]

Blue: Heptamer[32]

Green: Nonamer [5]

id-IGLV3-13[V_gene_segment]

ataatgcactttgtataaccattagagagtctctatttgcacatcaaaatgtggatgcttaacgttaacattggaaatagtattttaatatgcatatgagatgaattttctttagactatttttgtgaattatctttcttaataaggacatactcttctcttagaacaatttaacttgtgatattttttctattttctgtatcagttctgttttttctaaacactggcaaagatatcctatggctactg**attaataaattagattaat**gtgtatt**taacttaggtaagtta**gttcaagagtggacagatgtgatttccccacatatcctatggctactg**attaataaattcaattaat**gggtatt**taacttaggtaagtta**gttcaagagtggacagatgtgatttccccacatatcctatggctactg**attaataaattcaattaat**gggtatt**taacttaggtaagtta**gttcaagagtggacagatgtgatttctccacatatacctcacagatgttgctggtgcttactcactcttctcttctaatcttagcaaataaagtgggaaatttgacattcctaaacacccagatgacaacaggagagaggaggtgaagccccaaagctgttattctttaaatcctcaggatgaatcaccctggagaatcacccacacttcagttggatgatacgcaaatgagagtttggtcttagagtggtgctgagttagttcaggatttgggtcactgggatggggtgaatgtgttttacaagtgaaaaatacatgagttttggggtctgcagggtagagagttcttggctgaattgtgtcccccagatctgggttgagtgacatcccaccaagaggagtccaggagaagtaaatttgtgtaaacaccaaacatgcactgccccact**aggcctgagaggaaataagcgaggcct**gggaagcccacctgtgctttgggttcaggaggcagagctctgggtgtcttaccATGGCCTGGACCCCCTCTCTGCCTCTCCTTCCTCACCCTCTACAAAGGTGATGCCCCCAACCCTGCCTTAGGCTCAGCCCTTACAGGATCCTGAGCTGGTCCTGCCCTGA**ACCCTGAGCTCAGCCTAGGCATAGCCTCAGGGT**GACACCACTGGAAATGTGTTTGTTATCTTCAAGCCCCCTCTCCTTTCCTCTCCTGCAGGCTCTGTGGCCTCCTATGAG**CTGACACAGCCACCCGCGGTGTCAG**TGTCCCCAGGACAGACAGCCAGGATCAGCTGCTCTGGAGATGTACTGAGGGATAATTATGCTGACTGGTACCCGCAAAAGCCAGGCCAGGCCCCTGTGCTGGTGATATATAAAGATGGTGAGCGGCCCTCTGGAA**TCCCTGAGCGATTCTCTGGGTCCACCTCAGGGA**ACACAACCGCCCTGACCATTAGCAGGGTCCTGACCAAAGGCGGGGCTGACTATTACTGTTTTTCTGGTGATTAGAACAATCTcacagtgacacaggcagatgggg**aagtgagacacaaatccatttttcatctgtctcatctt**ttcctccagcaccaggatgactgtgggccaagccatgagc**aggtctggccaagttctactggatatgagacct**ccaggccatccttttgttcagccctacaggttggtgctgcagagggtagagcaggagtggatttagggctgctataatcagaattttcttgtgttttcaggcccctgaagtgacttggggaaataagaacgggatggaagatgagagagatgtagagaa**caatcacttggccttagatttccatacgatggtgattg**ggtcactccttccaaatatcttacatcagtccccaaaaatgccc**agctcagctcaacacctgagct**t**ccaggcttccatggcagcctgg**gattttgaacaggaaaagcagaggaacag**gaaaagcactgtggcatctgcctagatatgcttttc**tgagctcacccaccctcaactgctgggagcttctagaagcttagagctgtgccctcacaggaaagtgctattgtttccagaaaacttcctttctcatattta**ggctcagttttgagagttttggtttgatgactgcctgagcc**aagaaccc**aaggccccgccttaatacaggatgacactgaaaggcctt**ccagctccagacctcccattgtggatgcctgaggcctcagttgcaatcttatgatgggtcagggtctccttctggtgggtgatgcttctctcactcctttcagattctctgtgcatgaaaatgtttatctcaaagtctactttcagggaacccaaactaagacggccagctgtcaccaaaatga**gccaccatagacctgaggtggc**cactataccctggaatgcatcagcgaatttcctaaatttccactgtgagcatccaaatgcccaacgatttctgagagatccttgtattcttataaaatgaaaagggatcctgaaatgcc

Uppercase: IGLV3-13

Lowercase: Flanking sequence[1000bp]

Red & Bold & Underline: Stem-loop [19]

Blue: Heptamer[30]

Green: Nonamer [2]

id-IGHVIII-2-1[V_gene_segment]

caggattattttctttccttggttgagaaatatttccccaaacttcagctcag**ttcaggcatacactgtccctgaa**tgggcatttaccct**cagatgggtacacacatctg**tcaacatgtggactcttctgtcagacaaacgcacctttact**cacgtggattcttctctcagacaaacacacatgtccccacgtg**gactctttcctcagactaccacatatgttcttacatttactctttcctcagaaaacagacatttcctcatgtggact**cttgtctcagacaag**caaacatgtctcc**atgtgaactcttcactcacat**aagtacacatatgtccacatttactgtttccttacacaagtacatatatccaatgtcgaattgttctgtggcaaaatgatctcaagataatgataatcataaaccccctccctgacaaggcgtagatctgtatttttttcattgtaacctaactttgccttattgtcaagaacagtagtttgcag**ctctaaatataccaattagag**acaggtgtccattttctctggaaacgtatttttatgttcttactggacatatttgttgataatgtttgctattatgaagataccccaacagtgtccacattagagaataaaaaagagtaatgggcagattaactctgtgcatccagacccagaaatcctttgaccttgacttccctgaaatgtagacacagaggatggatgagcaatgctgagcagtgcacccatgaccacaaaaagaaagacgtggaaatgtgtcccctccacttctcatgaaaggcagctcatcccctgttccctcaggccctggcgaggagccaccccatgtctgtgcccttcctcagtgtccacaccgtggggtctgcattgatctggattcccttctcatccccgtcaatattagtgtccttcgtaaatcaggtccagctgtggcttctcctcacggggctgttctcagtctgttttctgtgttcacggAAGTCCTGTGTGAAGTTTACTGATGGAGTCAGAGGGGGAAAAATTTTACAGCCCAGTGGTGAGACTCTCCTGCAAAGCCTCTGGTTTCACCTTTACTGGTTAC**AGCATGAGCTTGGTCCAGCATGCT**TCACAACAGGGATAGGTGTGGGTGCCAACAGTGAGTGATCAAGTATGAATTCTCAGGGTTACTCTCCATGAGTACAAATAAATTAACAATCTCAAGCAACACCCTTTTAAGTGCAGTCTGCCTTACA**ATGACCAATCTGAAAGCCAAGGACAAGGTCAT**GTATT**ACTGTGAGTGAcacagt**gagggaaaccctgt**gtgagcccagacacaaagctcac**cgcagggagacaggaggggactatgtggtagatgctgctcagaaccaccagggggcactcagaaccatcagggagggtgcacagaaccaccaggaggggctcaggacaccagggggcgctcagaaccaccagggggcactcaggaccatcagggagggtgcacagaaccaccaggaggggctcaggacaccagggggcgctcagaaccaccagggggcactcaggaccatcagggagggtgcacagaaccaccaggaggggctcaggacaccagggggcgctcagaaccaccagggggcactcaggaccatcagggagggtgcacagaaccaccaggaggggctcaggacaccagggggcgctcagaaccaccagggggcactcaggaccatcagggagggtgcacagaaccaccaggaggggctcaggacaccagggggcgctcagaaccaccagggggcactcaggaccatcagggagggtgcacagaaccaccaggaggggctcaggacaccagggggtgctcagaacactaggaggtgctatgaatcactagggggcgctcaggactcaagggagcactcagaaccaccagggatagctcaggacaccagggggcactcagaaccgccagggggcactcaggaccatcagggagggtgcacagaaccaccaggaggggctcaggacaccagggggcgctcagaaccaccagggggcactcaggaccatcagggagggtgcacagaaccaccaggaggggctcaggacaccagggggtgctcagaacactaggaggtgctatgaatcactagggggcgctcaggactcaagggagcactcagaaccaccagggatagctcaggacaccagggggcactcaggaccatcagggagggtgcacagaaccaccaggaggggctcaggacaccagggggcgctcagaaccaccagggg

Uppercase: IGHVIII-2-1

Lowercase: Flanking sequence[1000bp]

Red & Bold & Underline: Stem-loop [10]

Blue: Heptamer[31]

Green: Nonamer [3]

id-IGHV3-64D-2[V_gene_segment]

ttaactctcagcatttctattgagaaaaataagccaaatagttaagatatatactatattacttcattcttataaattctagaaaataaaagctagttgaaagaaatatgaaaacatcagtagttttataaagaaatggtagaagaaaggaaggagaaaaaacaaaaaaaataagagagcaagaggaatcctgagggaagttgacttgtcaccttc**ttgaaaatagagattttttttcaa**tgtttactattgtacaggttaaatatgtgaattttgttatc**tgtcaattaaaacgtatgaaatttattacaagtaaacaattgaca**ttttagacaaaataggaaatacagcaaggaacaa**ataaatatgttaaatgtcagatatacctaaatatttat**ctacctgaatcttgtctccacatttgtaagtaaacacagcaaaatcagacaggttctcgtgacaggaagtggattctgcaaaccacacttggcccgtttatctctgtcctgtagttggttcagag**caactgaggccagttg**tgaggagcataggcccaggtactaggactcactcatgccagatataagcccttag**acacatacatagcccctccatgtgt**gggttcacttttacgtctgtacatgaagaaaccactgattcctaaataacataatttatacacatagg**taaacataattaaaaatgtgatagttattaaatgttta**tcatggaacaatttcacaataagtcagcattttcccaaatacaatcattgtcatcgaaatccccaaggacactctcatctgccctgggccctgccctctcctcaggcgtcccaccccagagcttgctatatagt**aggagacatgcaaataggtccctccttctcct**gatgaaaaccagcccagccctgactccgcagct**ctgggagaggagccccaggcccgggattcccag**gtgtttcca**ttcagtgatcagcactgaa**gacagaagactcatcATGGAGTTCTGGCTGAGCTGG**GTTCTCCTTGTTGCCATTTTAAAAGGTAATTCATGGAGAAC**TAGAGATAGTGTGAGTGGACGTGAGTGAG**AGAAACAGTGGATATGTGTGGCAGTTTCT**GACCTTGGTGTCTCTTTGTTTGCAGATGTCCAGTGTGAGGTGCAGCTGGTGGAGTCTGGGGGAGGCTTGGTCCAGCCTGGGGGGTCCCTGAGACTCTCCTGTTCAGCCTCTGGATTCACCTTCAGTAGCTATGCTATGCACTGGGTCCGCCAGGCTCCAGGGAAGGGACTGGAATATGTTTCAGCTATTA**GTAGTAATGGGGGTAGCACATACTAC**GCAGACTCCGTGAAGGGCAGATTCACCATCTCCAGAGACAATTCCAAGAACACGCTGTATCTTCAAATGAGCAGTCTGAGAGCTGAGGACACGGCTGTGTATT**ACTGTGTGAAAGAcacagt**gaggggaagtcagtgtgagcccagacacaaacctcc**ctgcaggaacgctggagaaaaacagctgcagtgggcgctcaggacccactgatcagcgtcaacctcagaggcaggtgcacaaggaggctgat**ttcctgtcacgatgtgggacttcatctctttacagtttctctagtgaacctctctaagttcagaactctgtgcttactaatgtcatctgtacatatttt**ttaaaatgattattttaa**tatgaaaacctat**tctcctgtgtacaaaatgcggattgatccttacagaggaga**tgaaaactcctcaaccatggtcaccatgatcagagttctgaggaaactaggggcttcctggtgagtcttctccaatcaaattcaggacagga**acctcagaaagattcctgactagaactgaggt**attgctcctcacaagagagctctagctccgg**ggggtctgttcctgcagacccc**tgattcggtggtgtatgaataaaatgtacactgacacacagatactctgtttttccagtgcagctgagggtccgaggccac**ttacagactccaagaagaatcctgtaa**agaatggcagcagtggccctgagcagctcgcactccaggcgtgtatttagtatagaattaacaacagaagctctgagtcaacacacttgtggataattaacatggttaagagagtagttctacgaatgagtaaagctcagggactgtggtctgaagtaaataccattagggggcaatattcttggtcgacctcccgctgagagggtcttctggcttaaaggtt**agttaatggggtagggtaaccagacttaact**ggggaagcctctgttgtccctagtatttaccctacaacctaatgctctaaggtaagaaccggccatcttcagcctgttcagttattgcaagctatgt**aaacttccggccttccaaaaaagttt**gtgactatttcctataactttcgttaatatttc

Uppercase: IGHV3-64D-2

Lowercase: Flanking sequence[1000bp]

Red & Bold & Underline: Stem-loop [23]

Blue: Heptamer[30]

Green: Nonamer [2]

id-TRBV6-2[V_gene_segment]

cagagagggtagtcagtagcgttaggtggttctgcttg**ctggggtctcaccccag**gcacagaaagcaggagccctgggtggagctgaaagtgctcatctgggtttgtcaggagtcccatctgtcagtgaattcacaagaaacagaacaaaacaactccttcaatgttgatgagactgcccctgggatttggaaagctaataacagagaaaaccaatatagacaaaggattttaaacaggactatcggattatgatcaattaagcaaattagaaaaggatacttgaaggagtatttgggacacaggaatc**aaaaacaccaggaagacatgaggagttttt**ccaagattgtagactatagcaatacttagataaccaataccagtgtattaatgagtaattattagtattattattttgagatggagtcgcactctgtcatcaaggctggaa**tgcagtggcatgatctcggcttactgca**acctccgcctccctggttcaagaattctggtgcctccgcctcccgagtagctgggactataggtgcacgtcaggatgcctggataatttctatatttttagtacagatgggatttcaccatgttgg**tcaggctggtctccaactcctggcctgaagtgatctgctctcttcag**cctcccaaagggctaggattacaggcatgggccactatgcacagccaagtaatttgtatcatcatgggaataaaaaatatacattgagttacaaactaattaccaaaagatatttaaaaaatggtagacttgcatcgattcagacaaaggcattcttcccatccaaactgttcactggtgcattgatcttgaatttgaccatctggggaaggggcgtggcctctcctgacagcaaggctctggggcccaggcagggagaatgaggtctcagaatgacgcccttgaaagacgtgttcccttttcaccaatgcacagacccagaggacccctccatcctgcagttcctgccATGAGCCTCGGGCTCCTGTGCTGTGGGGCCTTTTCTCTCCTGTGGGCAGGTGGGTCCTGGGCGGGGCCCCTTGTGTGGATTTCAAGGCCCATCCCCTTTC**CACTGGAGCTGTAGCATCAGCTTTGTCCTTCTCTGCAGGTCCAGTG**AATGCTGGTGTCACTCAGACCCCAAAATTCCGG**GTCCTGAAGACAGGAC**AGAGCATGACACTGCTGTGTGCCCAGGATATGAACCATGAATACATGTACTGGTATCGACAAGACCCAGGCATGGGGCTGAGGCTGATTCATTACT**CAGTTGGTGAGGGTACAACTG**CCAAAGGAGAGGTCCCTGATGGCTACAATGTCTCCAGATTAAAAAAACAGAATTTCCTGCTGGGGTTGGAGTCGGCTGCTCCCTCCCAAACATCTGTGTACTTCTGTGCCAGCAGTTACTCcacagtgctgcacggctgtctcctctctgcacagaaaggcaagggaag**gtgctgccctcctccgcagcac**agattcagcgatgcccttggtcctagcaccgaaaactttggagccccaatgggcccgggcagtgcgagccttcatctgtgccaggtgcctctgcagtcggtctcggccaggcctggatcggtcccagg**acctcagatgtcttccttgttgccctccggtctttcctctgaggt**gtccttttgggctagtgccaggggtttcccagctctgactttcctagtca**ttcacctgagtctgaggcctcctaggtgaaataggatttgtatttca**gatccatctagactcccgtctctacctgggtaccacgtgctttcctctctctatgatttttcctcatgttgaaccctacggtggtctcagctgcagcctgcatttcctatattcacagtcgccctccaaggccttgctgggtttctcctccccgacctccctgcccttctctactccagccatgttgagagactccaaagtccatttcctgtgccctgggctggagccttcctttctgtagtggtcaacacctacctgtgcttcagatctcagcatcatcagccctagtgataatcaccattatgcttcccttggctcccagctgaggtcagtttccctctcataatctctcatggtatcatgtgtctgttaatcagtagaatttagcacagttggaatcttccaagtactttaaaccccatgaaggccaaggctatgcctgggttataaatcagcagctgagcctggcacaaggggagatgcatttcagagaatggatgagtcagtgatggagtgagtgatgagtgg**atggatgaatccataacaggagcaccctagatccat**tttacccttgcggagatctggctgtaagtacaagaactcttacccttttgattgctgggctacgggaggtccttgaaattaatggggagtcgctat

Uppercase: TRBV6-2

Lowercase: Flanking sequence[1000bp]

Red & Bold & Underline: Stem-loop [13]

Blue: Heptamer[30]

Green: Nonamer [3]

id-IGHVIII-67-4[V_gene_segment]

aaaggattcttt**tctttcattggttgagaaaga**ttttcccctaacttcagcttag**ttcaggcatacgctgacctgaa**tgggcatttaccctcagatgggtacacacctgtcaatatgtggactcttctgtcagacagacacagcttcact**catgtggattcttccctcaaacacaaatgtccccacatg**gactatttcctcagactaccacatatgtccttacatttacactttcctcagaaaacagacatttcctcatgtggactcttgtctcagataagcaaacatgtctcagtgtgaatgaagtcttcactcagataagtacacatatttcaacactgactgtttcctgacacaagcacatatgtccaatgttaaactgtcttgcgacaaaatgatctcaagataatgataattataaactccaaccctgaaaatctgtagatctgcattttgtctattgtaacataacttcatctcattgtcagaaacagtcgtttgcagctataaatgcactgattacagtcag**atttccattttctctggaaat**gtatttcttatgttcttactggactaatttgttgataatgtttgctcacatgaagatacctgaacagtgtccacattagagaataaaaaagagcaatgggcagattaaccctgtgcatccagacgcaggaa**ccatttgactctgccttccctgaaatgg**agacacagaggatggatgagcaacgctgagtggtgcacctacgaccacaaagagaaagacctggaaatatgtcccatcccctcctcatgaaaggcagctcatcccctgttccttcaggccctggtgaggagccaccccatatctgtgtctttct**tcagtgttcacactatggagtctgcactga**tctgggtttcccttctcatcaccctccatattagtgtcccttgtaaatcaggtccagctgtggctgctcatagggttgttctcagtctatttcctctgtg**ttcatagAAGTCCTGTATGAA**GTTCAGTGGTGGAGTCAGAGGGGTAAACGTAGTACAGCCCAGTGGTTCACTGAGACTTTCTTGCAAAGCGTCTGGATTCACCTTTTCTGGCAAC**AGCCTGAGGTTGGTCCAGCAGGCT**TCACAACAGGGATTGTGGTGGCTGGCAACAGTGAGTCAACAAGTGGGAGTGCTCAGGTTTACTCTTCATGAGTACAAATAAATTAACTGGTCCAGCGACACCCTTTCACGTGCACTCTACCTTACAATGACTAACCTGAAAGCCAAGGACAAGGTTGTGTAATA**CTGTGAGCTTcacag**gagagagattatctgcacaagcccagacacaaaaatctgcagggagacaggagggaactgcatggtagatgctgctcagaagcaccagggggcactcaacacaagggggcgctcaggacaccaggcggcactcaggatacagctgggggcgctcaggacaccagggggccactcaggatacagcggggggtgctcggatccaccagggggctcttagggcaccagggggcattcaggaccactagggagccctcagggccaccaggaggacctcaggacacgtcagcgtgctggagccaccgggggggggggggggggggggcgctgaggaaaccaagagacgctctgaaccgcgaggggaaggaagctcaggaccacgaggggacgctcaggacaccagggggcgctcagaacaccagtgggcactcagaacaggaagaatctcttagaaagcag**ctccacatcaggatcctgggaggctgtggag**aaatgggaacactcataaactgttgagcatttaaattagtacaaccactatagagaacagcttagagtttcctcaaaacctgacgatagagttatcatataatctggcaatcctgctgctggtatatacccctaaaaaggaaatgtttttgcactaccatgtctgtaatagcactgttaaagc**actgttcacaacagt**caaaatttgaaagcaacctatgtgccgattaacacatgaatggataaataaaatatggtacatataggatatgggttaatatcatgccataaaa**aaatgagattttgtcattt**gcagcaacatggatgaaactggagttcactatattaattgaaatgagcccagaacagaaagacaaaa**ttcacatgttcccatttatttgtgaa**agctaaaaattaagacaattgaactcgtggagatagagagggaaaagatggtcaccagaagctgggaaggtctatggtagatggggattattggggatggtcaa

Uppercase: IGHVIII-67-4

Lowercase: Flanking sequence[1000bp]

Red & Bold & Underline: Stem-loop [13]

Blue: Heptamer[37]

Green: Nonamer [6]

id-IGHVIII-22-2-2[V_gene_segment]

ctaacttttagaggttttattatgagatcttgaattttttaaatgttattttctatgtctggtgagaagttaa**ctgatgcttttcctttattctcttaatttaccacatcag**atttattgatttgaagatattgattcatcagcgcatctcaaaaataatttggagcttgtcatggtgtacaatccttttaaagtcttatagaacttagttgactatgaattgggagtagcttgtttacatatactaagaacatttgtcagaagtttggttttcttcttttgccttgtctattactggtaatacagtgattggagcctcttagaaagagattagaaggtggggacctcacggatt**ttttaaaaagttagagaacggttgtcatcacttttttaaa**cgctgagagtcatttgttatctaatatataacccatcatgaggaatatctctagtgtgcttgggaagcatgcatattacgctgctgttggttggaatgttccgtagatgtctgttagggcagtttgttcgagagtgcttaagtccacgggcttcctaagaatttttctgtggatgctctgtacattattgaaggcgaggttttgaagtctcctaatttttttatagttttctatttctccttttatacatcttaaagtttcataaatgtagttatatttctagttttt**caaatatgtaaaagagaaataaaaatatttg**cttagtaattttaatga**gaaaatcacataacaagaaattatattttc**caaatgctgccatcaacactaaactcctccaggagtctcacatctgccctgggctctgctct**ctcctcaggcatcccacctcagcatttgctgcagaggaggggacgtgaaaatagggccctcc**ctttcctgatggaaaccatcgcagccctgagcctacagctctgggagaagagccccagtccagcattaccaggagttttgatttgacaatcagctctgaacacacatggctcaccATAGAGTTAGGGCTGAGCTGGGTTTTCTTGTTGCTATTTTAAAAGGCAAATAATGGAGAACTAGAGATATTAAGCATGAGTGGATGTGAGTGAAAATACAGTGATTATGTGTGGCAGGTTCGGACCAAAATGTCCCTATGTTTGTAGGTGTCTAGTGTGAGGTGCAGGTGGTGGAGTGTAGGGGAGAGacaagaggcagaaaagtgtgcaggaggccgggtgaggctgtagacactgtcagcccactatgccaa**tcccaccacgagtgctggagaaggtggga**gtctgatgaagcttactaacaaccctgtggtccaagctaattccatcaaggccattggttcttcccagagcatagctgtccatcgggaatctcccatgtgcccagcagcagccacgcgttagcaacttcactgtgcacagtcattgtttgggaggagctccaggatgtgtctctttgtcaaaaactgggtgatgg**tgtcagagagctgtgccgggtgcctggtccacgtgctccctgaca**ttggaggtataggaggtgcgttctcagggtggaaacaccttttatggaaatgtacataataaatcacgattttc**cttggttctcttagacgacatggagaagcaag**catgcaccctgcaaacaattgtaatttcctgctttaccccaagttcatgatttcttgtaaactttgatttctgtccaagtacagagatactgattagagtgagtttggttctttccacacactaaccctca**cctcccccagatagagcacattgtcctcactctgagtctgagggagg**agctgttcctgtacaactcagggactgcagagacccccgtgtgcagctttgcagagtcagatctttcctcatgtgaggcgacctccactgcccgtgattgctgctcaggtctaattgtggattcgggattatgacaccctttgggctatcacaggtcaatcttattctgaaaatcaccgtaattata**gacagaggtaataatccagtacccattcttctgtc**cgcagtttgtc**tttcttatttggttgaaagtttgaagaaa**aaatgtgaccatatattcatctaattctaacctctgaatctgctatttgctctcggggacctacaaattgcaaaacagtgagttcttcattttcatcaaaatgtgcaaatttgtgaatt

Uppercase: IGHVIII-22-2-2

Lowercase: Flanking sequence[1000bp]

Red & Bold & Underline: Stem-loop [12]

Blue: Heptamer[27]

Green: Nonamer [4]

id-IGHV4-34-2[V_gene_segment]

ttcttgcaattcagcggaggagaaattct**ttcagagacagatggatctgaa**ttggtaaatatgtgggtacgaattctgggcttgagtgtcattgtccagccatgtttc**acaggtgtgacctgt**cagggaagaaccagagttccttgttctctcagagggtagagctcacagaggtcctctctggttcccaggaaaggtaatttcactaatcttggtgatgagactatcctccagtgctgatgtactatagagttttcatctgaagctgtcactgctatccccaatgtacatcttttcacacagaaatgtttagaggtcaggccatattctcagggttacacatt**gagaaggatggagatatattctactaccttctc**c**tgagatctcacacacaatctca**aatttcaaaaggtctcagaagggcagctctcaggtactatttaaaaataacccacttcctgggacaggtagcatccttctaaccatgatggatgttctgaactacagtacacattgcatggatccaggtttgtctcaattcactgtgattattacactcagcagctgtttcaatatgtctgaaggggtaaatgacaatttaggtgacctgggtgtatggttggtgttatatgaatctttaaatgtagaacagtattaactgtattccaaaatctgtctttgatccatgatcacacttgtctcccagaccagctccttcagcacatttcctacctggaagaagaggactctgggtttggtgaggggaggccacaggaa**gagaactgagttctc**agagggcacagccagcatacacctcccagggtgagcccaaaagactggggcctccctcatccctttttacctatccatacaaaggcaccacccacatgcaaatcctcacttaggcacccac**aggaaatgactacacatttcct**taaattcagggtccagctcacatgggaagtgctttctgagagtcatggacctcctgcac**aagaacATGAAACACCTGTGGTTCTT**CCTCCTCCTGGTGGCAGCTCCCAGATGTGAGTGTCTCAGGAATGCGGATATGAAGATATGAGATGCTGCCTCTGATCCCAGGGCTCACTGTGGGTTTCTCTGTTCACAGGG**GTCCTGTCCCAGGTGCAGCTACAGCAGTGGGGCGCAGGAC**TGTTGAAGCCTTCGGAGACCCTGTCCCTCACCTGCGCTGTCTATGGTGGGTCCTTCAGTGGTTACTACTGGAGCTGGATC**CGCCAGCCCCCAGGGAAGGGGCTGGAG**TGGATTGGGGAAATCAATCATAGTGGAAGCACCAACTACAACCCGTCCCTCAAGAGTCGAGTCACCATATCAGTAGACACGTCCAAGAACCAGTTCTCCCTGAAGCTGAGCTCTGTGACCGCCGCGGACACGGCTGTGTATT**ACTGTGCGAGAGGcacagt**gag**gggaggtgagtgtgagcccagacaaaaacctccc**tgcaggtaggcagagggggcgggcgcaggtactgctcaagaccagcaggtggcgcgcggcgcccacagatcccgaggccgggtccggagcaggtgcaaggagggcggggcttcctcaacagctcagtggtctgtctcctcgccagcacctcagatgtccccaggactctctttctttattatctgtggttctg**cttcctcacatccttgtggcagggaagaaaggaggaag**acaatttttctgtttactgttgaggtttcaccaattactagaaactttcctacaagttcctgcatgactcattttgccatatatggattctcaccatctcttgatttgtttcatcaatcgaattgtgccctatttgaaattaacttactgaaaccttaaatccaatggatctatactggaattttaatgatgtaattgaggttaaatgtggtcaaagtgtgagaccctaatgcaataaaccgttgtctttataagaagaggaagagacaccagagacctctcacttttcac**gtgcacacagagaagaggccatgtggagacgtagtgcac**tagaaggtggccctgtgcaagccaggaagaagccatactaagaaccaattttgccagctccttgatcttcgacattcagactgtagaattgtaagaaaatcaatatttgttgtttaagccacccaatcctgtt**gtcttcttacgaagac**ccaaactgactaataccacctaactctgttagctctgtctcctggagggagaagcagccccctgaggctgggcactgcatctctcagatttccacgtgaagtaggcaaaaatagtagttctcatat**aaaaatgtgtcatggctctgttggccattttt**gagcatggtctctgaaaccagccctggtgtgtgtgtgtaacaa**atgtcccttatcttttatttggacat**aacaaatagacgataggtac

Uppercase: IGHV4-34-2

Lowercase: Flanking sequence[1000bp]

Red & Bold & Underline: Stem-loop [16]

Blue: Heptamer[40]

Green: Nonamer [5]

id-LOC107987384[V_gene_segment]

gcggcgggggcaaaga**gccgcggcgacggggtcaaaaagccgcggc**agagaaagccacggcggcgggggcaaaaagccgcggcggcggaggtagaaagccgcaaaaa**gccgcggcggcggggacaaaaagccgcagcggc**aaaaagctgcgtcggcggggggcataaagccggggcgggagaaacctgcggcggcggcaaaaacc**tgcggcggcggccaaaagccgca**aaaagccgcggcggcgggatcaaaaagccagctgctcccctgggattcctccaattaccctgtttaaagcctctgctcactcatgcgct**ggaagagggatgtcagagatgcagtctactcttcc**taaagggctgacagggcataaagacaagattgattatccatggaa**cattcctacaggaatg**cacctatg**tgcagacacactaataagcagtgtgtcttgca**gt**gtgtaaatggctgaaatgctatttacac**ttctttacacaataca**tttttaaatgttttgttgatgtattctctatcatttaaaaaaatcatcactttccccctaaacaaagaggattttt**atcagaaactta**tgggaagccccttgctctaccagattccca**ccctcatttctcctgaaggaagaaagaaaaccatctctattgatttctcctctcatcctctaggactaaaaccagaaagctgcatgctgcctgggtgagacttagagaagaccctgtctgtaggagcaggaatctcagagccttggctgaaaggcatggtcctgaaatgagatggagtcccccatgg**agagcacacgtggaagtccacacctgagggctcactgctct**tcaccacagatgcactccccta**ctgagtcctgagacctgagtgcaccccatagagtaggactcagatgaggggatgcaaatctccaccagctccaccctcctctg**ggttcaaaagccgagaatggggcctcgctcagtgactcctgtcccccactATGGACACGTTTTGCTCCACACTCCTGCTG**CTGACCACCCCTTCCTGTGAGTGTTGTGGTCAG**GGACTTCCTCAGAAGTGAAACATCAGTTCTCTCCTTTGTGGGCTTCATCTTCTTATGTCTTCTCCACAGGGGTCTTGTCCCAGGTCACCTTGAAGGAGTCTGGTCCTGCGCTGGTGA**AACCCACAGAGACCCTCACGCTGACCTGCACTCTCTCTGGGTT**CTCACTCAGCACTTCTGGAATGGGTATGAGCTGGATCCGTCAGCCC**CCAGGGAAGGCCCTGG**AGTGGCTTGCTCACAT**TTTTTTGAATGACAAAAAA**TCCTACAGCACGTCTCTGAAGAACAGGCTCATCATCTCC**AAGGACACCTCCAAAAGCCAGGTGGTCCTT**ACCATGACCAACATGGACCCTGTGGACACAGCCACGTATTA**CTGTGCATGGAGAGCACAGAGACACAGCCCAGGATGCCTCCTGTACAAGAACCCAGGCTGCGTCTC**AGTGGTGCTCCCTCCCTACCTCTGCAGAA**CAGGAAAGTGTGACTGAGATGCCATTTCCTG**C**CAGGGCTTGTGTTTCCTATACCAACCAGACTCAGAGCCCTG**TCTGTTTTCCTATTCTGTATTATTAAATGTCATGCTCCCTGTTAGTGATCCATGTAAGCAGAGGCTGTATCCTGTTTGACAAAGATTGAGCATCTAAGAGTCCCATTACCTC**GGCCACATGCATCACTGATATGTGGCC**ACTTATTTCTTGAGCTCATATCCTTCCAAGGGTCTGAATACAAATATCTATAACATATG**ACATTCTTAAACTGCAGGAAATAGTGTTGGAGAAGAATGT**GGAGATAAAAGAGCAGGATGATTAATTAAAATCCAGATACCATCTTTTCTTGCAGAGACAAATTTACACTAATAAAGTAATTTTATGAAACAACAAGAAAGACAGAATGTGTCCTCATCATGGAGTGTCTCACTTGCAGTGCACAAAT**AAATTTCTAGAAATTT**TATAGGGAAGGTGTGATAGATGAGGCTGATTTCCACACAGGGGGTGATTAAATACCCAGGTAAGTATAAAATCCAACACTTGGAAAGTAAAATG**CCTCAGCTTGACATCAGAAACCCATCTCAGTAAGTCTGAGG**AGCAATGTG**GAAAAGAAATGAAAAATTCGACAACACTTTTC**TTATGAaattcctagctcaagtttgaacaggactataaatcatgctgctataaagacacatgcacacgtatgtttattgcggcactattcacaataccaaagacttggaaccaacccaaatgtccaacaatgatagactggattaagaaaatgtggcacatatacaccatggaatactatgcagccataaaaaatgatgaatt**catgtcctttgtagagacatg**gatgaaactggaaaccatcattctcagtaaactatcacaaggacaaaaaaccgaacaccacatgttctcactcataggtgggaattgaacaatgagaccacatggacacaggaaggggatcacactctggggactgttgtggggtggggggaggggggagggac**agcattaggagatatacctaatgct**aaatgatgagttaatgggtgcagcacaccaacatggctcatgga**tacatatgtaactaacctgcacattgtacacatgta**ccctaaaacttaaagtataataattaaaaaaaaaaaaagaaaacagaattgtccacaattttaggaaagccatcgcatgacaacaaaccactacaatgataggagca**tcactgagtcaacggagttcaagcagtga**gcagatcagtattgaggccccaggaaaatcaaggctcttgggtacattttgcagaacccagaattgagccacataatctatggccaagtgatctttgacaaagttaacaaaaattaaccaacgtatacaccagggaaaggacacctcattcaataaagggtgctggggaaattggatagctatatgcacatgaattaaactggacacccaccctcaacatgtaaataattaacagaaaatagattaaaggtttaaatgtaagacctcaaactgtaaacgtattcaaataaaacaccaggacaactcttctggacactggcctaggtaaataatttacgactaagaactcaaaagcacaagccaaaaaccaaaagaca

Uppercase: LOC107987384

Lowercase: Flanking sequence[1000bp]

Red & Bold & Underline: Stem-loop [31]

Blue: Heptamer[52]

Green: Nonamer [7]

id-IGHV1-67[V_gene_segment]

aaataaaggattcg**ccaaaagtcttgaaaattttgg**tgctgactatgtgtccactgattcaatgcatcctgagttccagggaagggaatccccaattctttcatagagttgactgggtctcctaaggtcaatgttttcaaacgatggtgtcagtaacatatggtgtcactgaaagagcattttaaa**ctagggcatggccacttcctatagccctag**agacactgagaggtaaa**tgccctgtgagcccagatggaaacctccatgcagggca**gaggcagtgctgcagggggcgcccaggacccacccagaacaggctccagccccagagctggtgcacaggaggctgcagagggagtctctcacaaggactgagtgttactttactgaaaaccaa**aaaattataacatgctaaataagaatttt**atgaggactatttatgtatatt**taaataccctaatgtattta**gaaataaataaatcaataattcagtgagaaccacatttaaaaggcaaaattattactgctttgagatattttactagtaaaaagataaagaacaagaatgg**ttttattaaataaaa**atgtacacatttcagagacacactggtcccgagagtaagactgcggagagcaaatcctggaaagggcggaggctgtccatgtgccatgggaacttgggctcacgctgagaaccatgtccggtgtgagtcagcttcttagcggagaaattc**tcccttcacaaatttccgggcatataaaggga**aacatgtcattaaataagaatgaagatttgtacctcagcatcccacagttgtgttgtccatgtgagatctattttctctttctcgtgctggatcaggtgtaatgctatgaagtagtagtcctcatgaatacgcaaatcacctgaggtgaacactacagataactctgtgccctgagagcatcacccaataaccacatccctcctctagagaagcccctgagagcacagctcctcaccATGGAGTGGACTTGGAGG**ATCCTAATTTTGGTGGTCGTAGCTGCAGGTAGGAT**AATTCTCAGTCCCCAGGACTAAGG**TGACTGGGTCCAGTCA**AAGGGGGTTTTATCCACTCCCGTGTCCTCTCCACAGGTGCCCAGTCCCAGGTGCAGCTGGTGCAGTCTGGGGATGAGATGAAGAAGGCTGGGGCATCAGTGA**AAGTCTCCTGCAAGACTT**GTGGATA**CACCTACCTTCACCAGTTACTCTATGCACTAGGTG**CGCCAGGCCCATGCACAAGGGCTTGAGTGGATGGGAAGGATGTGCCCTAGTGATGGCAGCATAAGCTACGCAGAGAAGTTCCAGGGCAGAGTCACCATGACCAGGGACACATCCACGAGCACAGCCTACATGGAGCTGAGCAGCCTGAGATCTGAAGACACGGCCATGTATT**ACTGTGGGAGAGAcacagt**gtgaaaactcatatcctgagagtgtcagtaaccctgagggaggaagcagctgtcccagttttcaggatatgacaggatttatggggtttaatgttgtttagaaaataggttatataattgaggaaaagaagaaatagaaagatatatgcattctaattatataagaaatattcttttcaaatctcaccctataagcaaaattaacagagtgggaaaagcagcaatcaatcaagctgaaacaaacattcccatggaggatttgtgggggcagacgttttaaaattgaatgggtaaat**catttagggcaagattgcttgatccaatactaagactaaatg**taatttctgag**aaactgcctatctttttatattaaatttattataaagcagttt**tagggttacagcagatgtaaaaagtagaaacagagagctcccatgtacccctgcccaacacatacacagccctccccatgatcagctccctgaaacagagtcctaactgcgttatgtgggatgaacctacatggacacactagttctttcctttcctggtggtcccccggtataacaaacataaattatt**ttgaagcgccacgggttcttcaa**gtgggttactgggaatgatgccagttagagggaaagtgggtggggcc**attcctttttgcactctttcttcaggaat**ccataaaatgcacattgacttagagctcatctgactcctggttttctatgcc**ccttcccagagggtaagttctccaagcgtgggaggaagg**tccatttcactaaaaggccagaagcttctggtgaattccataatgcacaaccttttttgagaagtggagactttgg**acatgagggatttcatgt**atgatgccctggagttggatt**aaatacagcgtaagctcttgggggtattt**ctggaggaggccctttctgcaggaaattagaatacatgcaggggatccaggcaggaagaatcacttcctttgacatcg

Uppercase: IGHV1-67

Lowercase: Flanking sequence[1000bp]

Red & Bold & Underline: Stem-loop [19]

Blue: Heptamer[33]

Green: Nonamer [2]

id-IGKV1-9-3[V_gene_segment]

aaagaaaatgtggtacatatgcaacatggaatacttaatatacagctatgaaaaatgagatcatgtcttttgcaacactgtggatggtgctggaggccattatcctaagcaaactaacacaaagattcaagtttatgctaatcaaggctcctgtagtagaagaagaaaaaaaattgttttaaagtaatcagatttagcattgctagtcagacttttatgctgatggaccatgggaaataattcacaatgacagaaaagggggtagaatatgggggtccccaagcagagaataaaatgaatcccattaaaactcgagcattaaagagacttatagctctggacaatgcaggaactgtggatgtcatatgctttaaggaatcccaatatcatcttctttgtcaatctgcag**taaacctcttcagcttagactggtaataacattggttta**gggccattac**aaaatgcttttgagaacatttt**acttgctcatgactaag**tgttcttttttacttaaaaaaagatca**atttcatgcttacaaaaatgtagtatgtatgtcacaaagtatcgctcccaactggaacaatttcacagtgtgttgaagacctgatagccccactctctaagactttattaagaattctctacaaacaaggatattctcctacttatgcccaatacggcattgaaatatattattccatctagttctcacaaccacttcgagatttgccaataaatgctcaaaaatgtacttggtaacaaaatatcctttaggaagaaacattctctgcaggcaaatctaggtgccctggtctgacctgggacactggggacactgcccctgtgctgagttactgagataagccagccacgcagctgtatccagcctgccccaccccctgccgatttgcttgttcccagagcaccaccccctgccctaaagacttcttaataggctggtcacacctgtgcaggagtcagtcccagtcaggacacagcATGGACATGAGGGTCCCCGCTCAGCT**CCTGGGGCTCCTGCTGCTCTGGCTCCCAGG**TAAGGAAGGA**GAACACTAGGAATTTACTCAGCCCAGTGTGTTC**CGTACAGCCTGGCTCTTGAGGGAAGTTCTCTTACAACATGATTAATTCTATGGACATTTGTGTTTATATTTCCAATCTCAGGTGCCAGATGTGACATCCAGTTGACCCAGTCTCCATCCTTC**CTGTCTGCATCTGTAGGAGACAG**AGTCACCATCACTTGCCGGGCCAGTCAGGGCATTAGCAGTTATTTAG**CCTGGTATCAGCAAAAACCAGG**GAAAGCCCCTAAGCTCCTGATCTATGCTGCAT**CCACTTTGCAAAGTGG**G**GTCCCATCAAGGTTCAGCGGCAGTGGATCTGGGAC**AGAATTCACTCTCACAATCAGCAGCCTGCAGCCTGAAGATTTTGCAACTTATTACTGTCAACAGCTTAATAGTTACCCTcccacagtgttacaaacctgaacataaacccccagggaagcagacatgtgaggccgggctgccc**cagctgctcctcctgattcctccatcagctgagagtgttcctcagatgcaggcacactct**gatggtgttggtagagg**gggatgtgaagtcacctctgcatccc**aatttctttttctttctcagcaccaggtgcacagacataacagttcctctcctgatttaaaaaaggcagggatcatgacacctgaggagtctagttta**tggcttcagttggaattcaagtaacagagaagaagcca**ctatagatattctaagcaggaattgtcttgatacagagaaatagagtataaactatggaagtct**aaataaaaatatagagatgaatctcaaatttcatgttttattt**gctaagaaatatttgctaaatggggcatacagaaaaactcaatggtcttcaatatgttgaaagaacaaagagaagcttagagtttcatgaaaaaaggaaaatgttacctattgctctttgagaaagtttattggctccagaaaattttgggagctggcaagctcagaatggtaagcactgatggacaaa**ctgaatcctagaattatgttaaattattcag**aagttggggttgaatttgatttccggtt**acaagaggccaaagcagtgaagcttgcagagaatcttgt**tactgaaatgccagggatttggtgtagatcctgctgctcaccacatataaaaccaatcactaagacaacaagtattgccaagaaacaggctttaatcaggtgctgcagctgtggaaatgggaccccattctcaaatgtatc**tccctgacaaactaaaattagggggttatatagcaggga**agaaatgtgggcaaacaggaattagggaggggtaatgaagttaattttgtcaacaggaagcagaaggt**cagtgaggcaatcacaatgtgtgaagggtctgacgtctcactg**tcctgattcagt

Uppercase: IGKV1-9-3

Lowercase: Flanking sequence[1000bp]

Red & Bold & Underline: Stem-loop [18]

Blue: Heptamer[27]

Green: Nonamer [4]

id-IGHVIII-5-1-2[V_gene_segment]

attattttgtctgaaattaacatagctattcaagttttctttggtttatattttcatggtctatatttttccatcttatttcttctctttgtgaatatgttaagaagttttcttggagaaagataattgctgggtcttttttttttttagtcaactatgaggaattattttttaatgatattacccttttgtccaataatggttgttttgtcttttt**acaataatatgtattgt**ttctattaatttattatttaattattattgatgaata**tcttattggttaccataaggataacaataaga**catttctataa**ttgaattctaattcaaaaatgtagtacttctttttcttttttatttttttgagacggagtctca**ctctgtcgcccaggctggagtgcagtggcacaatctcagatcactgtaagctctgcctcctgggttctcgccattctcctgcctcagcctcccaagtagctgggactacaggcacctgccaccatgcccggataattttttgtatctttag**tagagacaggctttcaccgtgttagccaggatggtctcta**tcttctgaccttgtgatc**tgcctgacctggcctcccaaagtactgggattacaggca**tgagccaccgtgcccagcctacccctttatat**atagaacagagatattataactatatatgtctaattcctcttttctat**cccttttttattttatcagtttgtaattacataagc**tataaaaatgtaatattgacatttttata**attgctttacacatttatgtaataaagaaaacaca**aatgcaaaaactaaatttaaacctcatttttttcatt**gttgacaatctttatcttttgg**atacattcaggtattggatgtat**atcatatggctgctcatctagaaactctgctaaaacgtctatggcaagattaatatgctgggaatagattttctcaggatttgtttgtccgacagagtatttat**ttgtctttGATTTATCATCTCAAGAGACAA**TGTCAAGAAGATGCTGTTTCTGCAAATGGGCAATCTGCAAACCAAGGACACGTCACTACATTACTGTGCAAGAGAAGcacatgaggaaaggccggtgtgagacacaaacctccaggaacacctgggctaatgagctgcagggggtgctcaggacccactgatcagtcaaccacagaggggagtgcaaaggttaggactgctttcctgcaagcattgaaaaaagaagaaaaaaactccgtctgacagtttctccaaggaacttctgtaattttagaattctgtgcctaccaatgacatctccaaatatttttttaa**aatattgtgttttgaggacatactctcacatggacaaaacacaaatt**gacacagagatgaaaagccctcaaccatagtcaccaaaattggagacatgtagaagctcactgggacctgttgagtcttctccaatcacactcagaaca**gagaccttagtgggtctc**ccttagagtagtcattaaggatggtgataacagcctaaagatggtagaccatggtcagtgtcatatagaatatggggaccctcacaagttttttgtctgacccttctcctgacactaaattatgcaaattaataacactgatctggtgcttcttttgattctaatttattttatttttagttgtcgttctca**cttttcctttggattttcctgctccctggaaaag**gtaaatgtggtctccgtgacctcaattc**aagggctgaagccctt**tccctgtagctcagctggggctcaggctgtggctactgcagccatgtggaagaggctgaagggactttcttcactctccttgctcaggaccatccactgtattgtgtataggcttctctggaaatgcaagtgg**ccatttgtagtgaaagaaatatgtttgtctggttaaaatgg**gaggtgg**atgtagagttaattggctgctacat**aaactgtccttctccaccagtgcttttaggatgagattgtgaaatttgtaagaatcaaaatggagtcacatatgttaaaaccctgacaaatggattcaggaagtgtagggagaattcttaca

Uppercase: IGHVIII-5-1-2

Lowercase: Flanking sequence[1000bp]

Red & Bold & Underline: Stem-loop [18]

Blue: Heptamer[17]

Green: Nonamer [3]

id-IGHV3-20[V_gene_segment]

aattatgtagaatcactttctttttcagtctcatttcctccctctttctctctctcacacacactgacaaacacgctggttgctataactttaattacctgatgtgttaaaaaaaattgattcatttggaacttaaccagtttagctgttgttcacgctgttatgtaaaaaagatccctaaatgacttctacctggtaagaggagggaaggcgttggaaggtgaaattgcag**gagaaaaagaaaatcgtgaaacggattgatttgttttctc**tatgaatggtgattaggtcaatatttttcaaacggtgaacatattgtgtgaaga**ttatttgtcaatattatctcattaaaactattacaaataa**aag**tgtataatttggtaaaaaaaatgaagagagaaataaaaataataca**agaaaagtcatgaactcctgactccacacgtgagttaaccctca**gtttttctatgttacttagaaaaac**act**aagatacagccaaagaatatcatgatatctt**tataagaa**gagggttttgtaaccctc**actcgaaatttctagctctgtcctagagttaatttagggagcagtcagatccagctgtgagaaacacgggccagacagtgagactggcttttaccag**atgtgagctcttagacatgtcacat**ggcccttccgtgcttgggg**gtttaccttcacatctgtaaatgaaggaaac**tttgactcccacagaacataattcatgtgcatctataagtagaaatgctaatgagaattaattatttatgaagtataatcactcgcatccactctgggacacagcctactctgaggcatcccttccagaagtcactatatagtaggagacatgcaaatggggtcctccctctgccgatgaaaaccagcccagccctgaccctgcagctctgggagaggagccccagccctgagattcccacgtgtttcca**ttcagtgatcagcactgaa**cacagaggactcgccATGGAGTTTGGGCTGAGCTGGGTTTTCCTTGTTGCTATTTTAAAAGGTGATTCATGGATCAATAGAGATGTTGAGTGTGAGTGAACACGAGTGAG**AGAAACAGTGGATTTGTGTGGCAGTTTCT**GACCAGGGTGTCTCTGTGTTTGCAGGTGTCCAGTGTGAGGTGCAGCTGGTGGAGTCTGGGGGAGGTGTGGTACGGCCTGGGGGGTCCCTGAGACTCTCCTTTGCAGCCTCTGGATTCACCTTTGATGATTATGGCATGAGCTGGGTCCGCCAAG**CTCCAGGGAAGGGGCTGGAG**TGGGTCTCTGGTATTAATTGGAATGGTGGTAGCACAGGTTATGCAGACTCTGTGAAGGGCCGATTCACCATCTCCAGAGACAACGCCAAGAACTCCCTGTATCTGCAAATGAACAGTCTGAGAGCCGAGGACACGGCCTTGTA**TCACTGTGCGAGAGAcacagtga**ggggaagccagtgagagcccagacacaaacgtcc**ctgcaggaagacaggaggggcctgggctgcag**agggcgctcaggacacattaagaacacagtaaactcagggaccaggtgcccagggaggttaagggctggtttcctttcagagacttagctgtttctccatctaacagtttccccagaaccctgcctagatttgtgatctgtatctgctcaaactatctgtgtcac**cttcctcacctgtgactttgaggaag**cttagtgtggacattccagtgggtggagagatcttggtgacagcaattgtgtgttctgtctaggcatgtctatggctggccatcaggaacgacaggctggaattcttggaaagaggcacacctgccatccatcaggaaattttgttgtcttctgttctgctagaattaaatcaggcacgcccaggttaactactactatcttcctagcttgagaaaattaatggtgggcttgaataacacctatatgaaatcatcagagcaacacctagattcg**agtctcatttaataattgagact**atgttctagccaaggagacacataaaactggtgattgccgtggtcagttcatatcatatatttgtcagtagaatctggctggtattatgaacagctttg**ccaacacatgtgttagtgttgg**tcttcggagaagcacattctatgaatggattaggtttatgtaca**gtttcctaaggaaccggtccacgcttttgagaggaaac**gcagagcgggtggaaggaggctggggaagctgaaggcatatgagggaagttggtcc**ctgagtgaaggagagagggaggataactcagtggaacttccttaaactgtgctgttctaggaaggtcca**gcaagtcactgtatct**gagtcatgtcaaagtttcccatcggggacccccgtgactc**ccagcagtggctctactcagatcagcgcggagctcattgttctcctgagagtgg

Uppercase: IGHV3-20

Lowercase: Flanking sequence[1000bp]

Red & Bold & Underline: Stem-loop [20]

Blue: Heptamer[31]

Green: Nonamer [5]

id-IGHVIII-67-2[V_gene_segment]

gttgtcatgttcctttattctcttctggcaaatcacagtttgtgtttctgaagttttctagtttaatattttatagataaaaatt**tatcaagtagattgata**gtcctatttaggacacttatatctttacttatttg**aacactgcttgtcaatcacaatgagtgtt**gatttctccgtcttctatacatgttattatattttctg**ttccaaatccattcattaagtatttttaaatgtattttggaa**agttgatcc**tattactttatgtaata**tttctcttgagtcccaaaggtcttctttttatgattattttgt**cttaaattaacatagctatttaag**ttttcttttggtttatattttcatgatatatatttttccatctttattttttct**atttctgaatatgtaaagaaat**tttcttggagataaata**atagttgggtgtttttttttagtcaactat**g**aagaattattttttaatgatgttactcttttgtccaattattctt**gttttaatttgtattttt**acaataaagtttattgt**ttctattaacttattatttaatcattctgtgactaacatt**ttatgggttaccataa**ggataacaataaaaagtttctataa**ttgaattctaattcaa**aaatttaattcccattcatatatactacag**agatattataactctatatatct**aattcctctttcctatcccatgttttttattttctgtttgtatctatacatgctat**aaaatgtaatattggcatttt**tataattgctttccacacttacataatgaaca**aaatacaaatgcaaaaactgtattt**caacctcatttttccattgttgaccatcttctttgtgcatattttcaggtgttggatctatgtcacatggctattcatctagagactctgctaaaatgtctacggaaagattcatgtgctggaaataggttttctcaggttttattggtccgacagaatatttatttgtctttGATTT**ATTGTCTCCAGAGACAAT**GTCAAGAATATGCTATATCTGCAAATGGGCGATCTGTAAACCAAGAACACATCAGTATATCACTGTGCAAGAGGAGcacatgagggaaagccagtgtgtgcccagacataaacctccagggacacatgggcgaattagctgcagggggagcttgggacccactgatgagagtcgaccccagagggaggtgcacagaaaggttaagaactgctttcctgccagtatatttttaaaaaaacaaaacttccatctgacagtttctccagagaacttccctaattttagcattctgtgcctaccaatgccatcactaaataggtttttaaaaaatattgta**atatgaggacatattctcatat**ggacaaaacgcaaattgacatttacagaaatgaaaagtcctcaaccgtggtcgctagaattggagacctgtagcagctcagtgggacctgttgagtcttctccaatcacactcagaacagagaccttagtgcatctccctgactagaatacc**ctttaggtatcgagataacagcctaaag**atggtaaatcatgagagtctcagatgtccagatgacagacatgggtctatgggaatag**tactgtaactgacagta**actaattttccataccgccagatgtcctattcaggagcagctacagcagtcatgcctag**gtgtgaagatcacac**actgacctcacccatgctgtctctggccacttcatcacaaccaatgcttaatattggacgtggatctgccagtccccggggaatgggttgaatggataaaatgcatggctag**tgctgctggaaacccattcctactgtggcaaatggcagca**tctcttttaaaaggctaaataatattctattctgtatacataccacattgccattatcctttttgtttgtttgtttgtttgtttgtttgagatggagattcactcactctgtcacccaggctggagtgcagtggcgtgatttcaggtcactgtaatccctgccttctgggctcaaacaattctcctgcttcagcctccggagtatctgggattacaggcatgcaccaccactcccg

Uppercase: IGHVIII-67-2

Lowercase: Flanking sequence[1000bp]

Red & Bold & Underline: Stem-loop [20]

Blue: Heptamer[18]

Green: Nonamer [3]

id-IGLV3-26[V_gene_segment]

gctctttaggccagcaggtctttggtatctcattaaggtagcactagcaaatgaaaacaaaagggaccccaaagaccttggatgcgggaggaggaggagatggagcagggtgcaggaggtggggcagggaggggctggaagttcaggctctgaggtcatctcctgggtggaatcctgactccactcactgctgtctggggacttggggaaaaac**aattaaccgcccaatattaatt**ccatagtaaagatcgggccttgagcacaggattcctcatcttccttttttatatgatgaaagtcttcttgaggtcatgcttgtaaaacactcagcaatgtatctggcatacactatggtctcagaagtggttattatgaattttctagtgatatgtagaggatgacattggtcctgtgggcagagggggcactgtggac**cttggtttgaggactctgaaagacaccaag**tccgctggtagtgctcaagagcgaag**agcatcacagatgct**tcctcccaggccccttctattgggtaaaatttactctccactcctgacacccatctggtccttcttctgaccttagcacattatcagtttgaagacttaaggctcagggctcagatgggagctaggacaggcacactggggaatgaggagtagatttgcattcactgctcatcagagagcccacccaacaaggatacaagaataaaagaagtagatttgcatgaaga**gcctctccttcccatgataagagaggc**ctgaaggtccctccccagccatgggctcagagacagagctatgtggcatctccaccgtggcctggacccctcccttgctcaccctcctccgtctctgtgctgcctcccagggctcagcccccaaaggaccagagatcagactggcc**ctgagcttcagctcag**cacagggagtgatgcagagtgtggggctctgggaatgaccctcatcctcataattaactctcctgtcctctcttgtagGCTCTGTGGCCTCCTCCATGCTGACTCAGGAGCCAGTAGGGTCTGTGG**CCCTGGGACAGTCAGCCAGGG**TAACC**TGCCAGGGAGAAAACCTAAGAAGCCATGGTACAAACTGGCA**CCAGCAGAAGCCAGGCTAGGCTCCTGTGCAGGTCATACATCGACAAAATAACCAGCCCTCAGGGCTCCCAGACCAATTCTCTGGCTCCAA**GTCAGGAACACAGCCACCCTGAC**CATCACTGGGGCTCAGGTTGAACATGAAGCTGACTATTACCATCACTCATGGGACAGCAGTGCTACTCACCTcacaa**tgacacagacagattgggaagtgagatctaaagaccttcactgtctgtgtca**ccctcttcctccagccatagcaggactgtggagacagccatgagcagggctggcctgggtcacctgcatcgaagtcctccaagctgcccttccccccagctctccaggtagtctctgcagagaatgagtcaggataggatttggggctggcagaaccaggctgtcctgctgttatttggaatggatgtacct**tggctcaagatgacctgagccatcttactggaaggacttactggaaggacggtaaga**gggagacagctcctgagtgctcatggtcactgaatttcctcctgtgtgttgactgaacccaatgcagtgcctctactctgtggcccaaataccctggattatttttctgaactgcccgtgtttaactgagaccctcaagccccagctttgtgcatcctgatgttctgaacatggaagcctt**tcacagtgtgcactgtga**catccctcagattcccgtttcaggacttatgcacccaccatcctc**ccagggctgcagggaggccctgg**cttctcacagctcttccccaccagaaactgccatgcagtacagagaatagctttgctcaaaga**attgcccatttcagagtatgtttcgggggcaatgactccctgatgccttaatctgaaagtcat**gctttaatttaagaccctgaatatca**aattgaagtgccttctagctgaacactcaatt**gtaaaacattcccagtcacacactccacacttctttcctgagaatatatcctgaaaccctattgca**tggaatcctccctctcagagtaggattcca**gggaacttaaaaattaaaaaaaaaaacaacacaccagtctagagggaaaaatattgaagaggag**agagttagggagttgcacaaggaaagaactct**gaagtctgcaaatatgcatgcactgaagagcacttgtgtgtaaagaaactc

Uppercase: IGLV3-26

Lowercase: Flanking sequence[1000bp]

Red & Bold & Underline: Stem-loop [18]

Blue: Heptamer[41]

Green: Nonamer [0]

id-IGHV1-67-2[V_gene_segment]

aaataaaggattcg**ccaaaagtcttgaaaattttgg**tgctgactatgtgtccactgattcaatgcatcctgagttccagggaagggaatccccaattctttcatagagttgactgggtctcctaaggtcaatgttttcaaacgatggtgtcagtaacatatggtgtcactgaaagagcattttaaa**ctagggcatggccacttcctatagccctag**agacactgagaggtaaa**tgccctgtgagcccagatggaaacctccatgcagggca**gaggcagtgctgcagggggcgcccaggacccacccagaacaggctccagccccagagctggtgcacaggaggctgcagagggagtctctcacaaggactgagtgttactttactgaaaaccaa**aaaattataacatgctaaataagaatttt**atgaggactatttatgtatatt**taaataccctaatgtattta**gaaataaataaatcaataattcagtgagaaccacatttaaaaggcaaaattattactgctttgagatattttactagtaaaaagataaagaacaagaatgg**ttttattaaataaaa**atgtacacatttcagagacacactggtcccgagagtaagactgcggagagcaaatcctggaaagggcggaggctgtccatgtgccatgggaacttgggctcacgctgagaaccatgtccggtgtgagtcagcttcttagcggagaaattc**tcccttcacaaatttccgggcatataaaggga**aacatgtcattaaataagaatgaagatttgtacctcagcatcccacagttgtgttgtccatgtgagatctattttctctttctcgtgctggatcaggtgtaatgctatgaagtagtagtcctcatgaatacgcaaatcacctgaggtgaacactacagataactctgtgccctgagagcatcacccaataaccacatccctcctctagagaagcccctgagagcacagctcctcaccATGGAGTGGACTTGGAGG**ATCCTAATTTTGGTGGTCGTAGCTGCAGGTAGGAT**AATTCTCAGTCCCCAGGACTAAGG**TGACTGGGTCCAGTCA**AAGGGGGTTTTATCCACTCCCGTGTCCTCTCCACAGGTGCCCAGTCCCAGGTGCAGCTGGTGCAGTCTGGGGATGAGATGAAGAAGGCTGGGGCATCAGTGA**AAGTCTCCTGCAAGACTT**GTGGATA**CACCTACCTTCACCAGTTACTCTATGCACTAGGTG**CGCCAGGCCCATGCACAAGGGCTTGAGTGGATGGGAAGGATGTGCCCTAGTGATGGCAGCATAAGCTACGCAGAGAAGTTCCAGGGCAGAGTCACCATGACCAGGGACACATCCACGAGCACAGCCTACATGGAGCTGAGCAGCCTGAGATCTGAAGACACGGCCATGTATT**ACTGTGGGAGAGAcacagt**gtgaaaactcatatcctgagagtgtcagtaaccctgagggaggaagcagctgtcccagttttcaggatatgacaggatttatggggtttaatgttgtttagaaaataggttatataattgaggaaaagaagaaatagaaagatatatgcattctaattatataagaaatattcttttcaaatctcaccctataagcaaaattaacagagtgggaaaagcagcaatcaatcaagctgaaacaaacattcccatggaggatttgtgggggcagacgttttaaaattgaatgggtaaat**catttagggcaagattgcttgatccaatactaagactaaatg**taatttctgag**aaactgcctatctttttatattaaatttattataaagcagttt**tagggttacagcagatgtaaaaagtagaaacagagagctcccatgtacccctgcccaacacatacacagccctccccatgatcagctccctgaaacagagtcctaactgcgttatgtgggatgaacctacatggacacactagttctttcctttcctggtggtcccccggtataacaaacataaattatt**ttgaagcgccacgggttcttcaa**gtgggttactgggaatgatgccagttagagggaaagtgggtggggcc**attcctttttgcactctttcttcaggaat**ccataaaatgcacattgacttagagctcatctgactcctggttttctatgcc**ccttcccagagggtaagttctccaagcgtgggaggaagg**tccatttcactaaaaggccagaagcttctggtgaattccataatgcacaaccttttttgagaagtggagactttgg**acatgagggatttcatgt**atgatgccctggagttggatt**aaatacagcgtaagctcttgggggtattt**ctggaggaggccctttctgcaggaaattagaatacatgcaggggatccaggcaggaagaatcacttcctttgacatcg

Uppercase: IGHV1-67-2

Lowercase: Flanking sequence[1000bp]

Red & Bold & Underline: Stem-loop [19]

Blue: Heptamer[33]

Green: Nonamer [2]

id-TRBV5-3-2[V_gene_segment]

cttattcacattggcctgaaataagtaggagaggatgtggatcccaaagcatcaggtactgtagctctaaggcaaggctgcctataaggtgctgtggactttggtagggaaaagtaacctatataaaataatgaagcaagaggagagccagggcatagatgaatcagaacaagtggtgcccctacctccctctactctcctgccattgtctcccattagccaaactcactctggagttagagaagaagagagccagttaacataatacactgaggtcagcctctgaaggcagggtg**gatctggagaaattaacgtaaattatccagatc**actacgcataagatggccttgagggtgctgaatcccaaactgtctggctaggtctgtgctgtgagctgataaagccctagtgttatttctggggaa**tctgtactcttaataatttacaga**caaccaaggtctgctttggatctgatcagatagactaaatcttggggactttgcaccactggccactcaagggaggggctggaaat**attttctgaggacaaaaaaatagaattagaaaat**ttggttgaatctagcgtcagagagatgacattaggacagccagggagagctggaacaagaaggtttaatcacaggctcctcaccctctgctgatgggcaggtgtgtga**gctccagcatggagc**acgacagcactaggtgggaggagggtgagagggtgatggggcagcctgggagctggagcagtgta**ggcagaggagcaactgtctcatcacagaagcttctgcc**ctcacccaacccttctgctggacagggagggagttcaggttgtggggagtactggacctcaggaagcatttatagggaggacacaggacagtgacatcacaggatatcccacccatcaggaaaatcaaggcccagaactcacttggcccttccccaggaggaccaagccctgaatcagatgcagtgctgcctgccccactgtgccATGGGCCCCGGG**CTCCTCTGCTGGGAACTGCTTTATCTCCTGGGAGCAGATGAG**TCCTGTGCACAGAACAGCAGCCCCATTCTCAGATTTCCCACCCCTGT**GTCCTCCACTTTACACTGGGGAGGAC**CTCCAGGCTGTCTGCTGTGCTCAT**CCTCCATCTGCTTTTCCCACAGGCCCAGTGGAGG**CTGGAGTCACCCAAAGTCCCACACACCTGATCAAAACGAGAGGACAGCAAGTGACTCTGAGATGCTCTCCTATCTCTGGGCACAGCAGTGTGTCCTGGTACCAACA**GGCCCCGGGTCAGGGGCC**CCAGTTTATCTTTGAATATGCTAATGAGTTAAGGAGATCAGAAGGAAACTTCCCTAATCGAT**TCTCAGGGCGCCAGTTCCATGACTGTTGCTCTGAGA**TGAATGTGAGTGCCTTGGAGCTGGGGGACTCGGCCCTGTATCT**CTGTGCCAGAAGCTTGGcacag**ccctg**cagagtcactggaactctg**tgcactaatctctctgcttccgtgt**acagcagtctcagaccagacagctgt**gagaacctggggccttcagggggaaagataaacaatttcaggactctgaacaaggctggcggaaacaaggccaggaggatatgctcagaggatggtgctgcttcaggaggttcacggcaaaattactttctttgtcggatgctgtacactatagtgatagagtagctggcactctcctttttagaaaaagaaattcattttggtgtcaggattgttcgacagatgtgtagtatagaaggtaaaatctcaagggtttgctttatataattaagggagaagaaattgagatttggtttggagaaacttgaggagagagaacatgattaaagattattaaataaatagaagtagagctaccagatgataaaaaattgtgttctctgcaacataacctgtcaaagttttcaa**ttgctagaaaatacgcagcaaggccttgtagcaa**ctggcatcaccccgaggccactctatgatggtcatcaccacgtagccagttgccctgtgggagggccctggagctggttccaatgtcttccttctggacttatatctcaattctcaaaacagaggaattcatttcactgctgatggtaaacagataaaggctctcccatttccccaa**agatacattctgcttcccttgtatct**ctttctgacacttaccgcccagatccattaccttctgtgatcaaacacctcttcgaatatttcctaaaactctgcagcctttacattctctgcatttgtgctcagtatttccatggcctggaaggaccttctgtctcccaagaaacagccagaccctgcttatcggtcaaggatgggctcacacgtgcttcctctttgcttcttaccatgacattccccagcattgtactataccaccatctttattattttcataggaaaagccaagtattaat

Uppercase: TRBV5-3-2

Lowercase: Flanking sequence[1000bp]

Red & Bold & Underline: Stem-loop [15]

Blue: Heptamer[36]

Green: Nonamer [1]

id-IGHVII-53-1-2[V_gene_segment]

gccatttgtgcctccctgagcacagttgtccatcagagatgtcccatgtgtcccagcagcagccatgtctcagtgtcttcactgtgcacagccattgt**ctgggaggagctcccag**gatgggtgtctttggcacacaccaggtggcgggtgttagagtgcggtgcagcagctggctgcctgttctattgggctccctga**tgctggagagatgggaggtgcattctcaggtccagca**ccctgtttgtgaatttttatataaaaccatgattttacttcattttctcagatgacatagataattaggaacagaacctgcaaagaaattgtaattttcaactttaccccaaatttattgtttcttaattctgtgtaagatccagacatattattgccttcctca**tgagaaattgttctatttaaaatgaaattagttttttctca**cattctttgtttctgttcaagtacagagatctttattaaagtaagttgggt**tctttccacacactaaccctcacctcccccagagaaaga**gcagagattttcctca**ctctgagtctaagggaggagctgttcctgcacgactcagag**cctgcagagacccccccccgccaggtgcagcttcagtgagtcaggtatttctccttgtgggtgacctccaccgccagtgattgctgctcaggtctaattgtgggttaagcattaggacaccc**ttcaggtgatcacatctcagtcttattctgaaaatcaccatgaa**cagagatagttc**aatgtctattctcctgacatt**ag**tttctctttattatttggttccaagtatggagaaa**aatg**tgacaataaatttgtca**gaatctaacctcagtatccactgcattactctaggagactcacaaattgaacacaaatgagccctttattctcataaaagtgtatgtatttgggaatttcaatgtgttctccagaacctgtgcatgccaacaactgtgtttctcagtgcccacttGGCCTGGTGAAGCCCTCACAGACCCTCTCCCTCACCTGTGCTGTCTCTGGATTCCCCATCACAACCAGTG**CTTCCTGCTGTAGCTGCATTCATAAACCCCCCAGGAAG**GGACTGGAGTGAATCCG**GTGCACAGGTCATGAGGGAGTGCAC**ATTCCAACCCACTCCTCAAGAGTCCAGTCACCATCTCCAGATCCATGTCCAAAAAGCAGTTCTTCCTACAGCCGAGCTAAGTGAGCCGCAAGCACACAGCCATGTATTTTTAACAAAAGAcacagtaaggtaaccacagtgggaactcacacccaaacctc**cctgtgggggtgcacagg**acagccacagttactcaggaccccaggattcctcaggacaccaaggggcactcaaggccattgtagatgccctcaggtagccaagggttctcaggaaacatggaggaaaaccaggaccccaaaaggtgctccgtacagcaggggactcaggacaattgtggggactcagagcaggctcaaagctcagcttcagggcaggtgcagctggggttgaaaggggctggatgaggggttttgtgacaccatcatatttcaccac**tagacacactccactttgtctattctaacgcatgagagtgtatgattagaa**aatgatatttatataaatacataaccatagttagctgtgtcaagttgtcctcttgctaggtgtccatagctaggtgcatcagccttgtccataaggactaattccccgcaattactgga**gaatctcataaattgtggtcaattatgtcagattc**ctctctttttctgccttcctttctcccttcttctctctctctcacacagaaacttacatacacccaccccacaacacaccaaaatctataact**tttattacctgatatattcaataaa**cctgattaatgtgcagcttttccagcttcgttatttatgctgttgtaacaataagaacaatgtgtttcctagctgtgtacttctctaagctgagtagcatct**ttgtttataatacctagaattaaaaacaa**cccaaaagtcaatcaccagcttaactggtaaacaaattgaggaaaagtcattcattgacatactatccactactaccatcaactaatgttgggtacactcaacagcatggttaaat**tcacaagtacttgtga**tgagtaaaatgagccaaagtaacaaaagtgcatacataagatacaactttcataaattctatagaacaaaaagtaatctaaagttacataaaaatc

Uppercase: IGHVII-53-1-2

Lowercase: Flanking sequence[1000bp]

Red & Bold & Underline: Stem-loop [18]

Blue: Heptamer[42]

Green: Nonamer [5]

id-TRBV7-3[V_gene_segment]

aatatattctatagaatatattatatac**aatatataaatatatatatt**tcccatt**tttcttatagataattgaagtttcaggaaaagaaa**gtagagggaataagtgattgtgatgagacattagc**tgagaattttccagaggtggttcaggacctaaattctcaatttcaagaattcaaatgaatatgaaat**aagataaatgtttatattataatgagattgtggaacataagtgacaaatagatgattctaaaagctgtcaaaaagaaaagataaatctctcctagtggaatagactgaaaaaatcaatgaaagaaagaggatcagaaataaccatttagtaggaattataatccccattcattaaacactgagatggaggaaaatatatttacaag**tggaaaaattaaaaaaaaagactatgagttttcca**ataaggtatcttttctgaattatagtcaacaaaggtgggaagtggtcccagaagaaaggcatgaggtgcagcttgtgagggaacccatgtgttgggacagccccgttgggcacgtgtgactgggggatggaggaggctggggcatcaatggggatggcataggggaccctgacttgcaggaaagacaatgagctcatcccttggtgccttgtgttgggggcgctgttggcacatcctagagaacatgcccagcagacagaggagcgcctgtgtggtgaggagataaactcagaaatgcagcatgaggcctgtagctccagacagctctagagcacaaagtgaagaccgatgcattgatgttgtttaaagggagctaataaatatctaaagcagtcatccaagtgtgttctaatataaatcctgtgttcctgaggttgtggggattgagagaggaagtgatgtcactgtgggtactgttctgtgtcaggacaaggacgtccctcctcctctgctcctgctcacagtgaccctgatctggtaaagctcccatcctgccctgactctgtcATGGGCACCAGGCTCCTCTGCTGGGCAGCCCTGTGCCTCCTGGGGGCAGGTGAGTCCT**CAGACACCAACCAGTCTCATTCTGTGTGTCTG**TATGTGTGTGTGTGTGTGTGTGTGT**GATGAGTACAACTGTTTTCCTCATC**CTGTTCTCAACTTGTGTCTCCGCAGATCACACAGGTGCTGGAGTCTCCCAGACCCCCAGTAACAAGGTCACAGAGAAGGGAAAAGATGTAGAGCTCAGGTGTGATCCAATTTCAGGTCATA**CTGCCCTTTACTGGTACCGACAAAGCCTGGGGCAG**GGCCCAGAGTTTCTAATTTACTTCCAAGGCACGGGTGCGGCAGATGA**CTCAGGGCTGCCCAAAGATCGGTTCTTTGCAGTCAGGCCTGAG**GGATCCGTCTCTACTCTGAAGATCCAGCGCACAGAGCAGGGGGACTCAGCCGCGTATCTCCGTGCCAGCAGCTTAACcacagcatgacacaatcgcctccttcctgctcataaacctcctcctctctctccttgcttccttatgatactattttgcaccaggggatcctcatctcacaccactccactgcctcttccagaatcattgcttgcatgtccgctgctcagatcagcagatgtgcgttgtatatgttaatcattttctgtggaaggctcgttgctggtcttcat**tatccatccttattttccacattggata**ctgctcccttcaagattacttgttgcctacaaaagatcctactttttacat**tcaaattttactttttattaatacgtatttga**tgccatcttgaaggcaatctcattatcatcctaatcttttgctaatctttatgtttatggtaacattgattgttattttgggggacatg**tatttcatctttggtaatgaaatgaaata**actatgcacttcaattcctgtatgttctgcctacttaagagtaaatccaattttcttccattcctcccattctatccca**ctccctttatatgatagttttaagggag**aaaatattaattagactctaactgcagctatgtgataagagttatttagaatgagggtgggatattaagctttgagtaaagcaatagtcagggttagaactagtgttggaatcagggtttagggaagttgctcataaaacctgcaggatgacacttctagaatagtctctcactcactctgcagacacttcccagcatcccttgggc**cattgcaaaagcaatg**atgaaacttcacgtattggccacaagatggcactgtggtccactgggatctaagggactctggggagagcctgggagagcagcctaggaagggaagggttaagaaaaattagggctggaatccaatattatgcagatgttgcagcagttttcagtcattgctaggctacctacagctatgcaagatgcaggaaatacctctaatctttaatgagatccacagttgaag

Uppercase: TRBV7-3

Lowercase: Flanking sequence[1000bp]

Red & Bold & Underline: Stem-loop [14]

Blue: Heptamer[28]

Green: Nonamer [4]

id-IGLV1-40[V_gene_segment]

cttatcacatgggacacattatactggacccaggatggggctgaactgggagaggtagctgtggacctgctcctcgctctgccctcaccctaagagctgatggggcatcca**ccaggaagagcattcctgg**gcagggagagggccatgagcagccttgtcctgtatgtgagacacagctcagcccggtcacaagatccagatgactatgactgcaattcaggggaagtagagaatgaagattgaccagaataa**ccctctgggaatgtgccctgggcctcctagagggaagagcttccct**tctattcagcgctgcaggatccacagaggaaggagag**ccctggccaccaggg**aggagccacatgagaggacccggtgtccccacaaatgggaacgtgagtccatgaaagtcggtccctgctctgggtttgattgtgacctat**gaggacattgtgtagtcaccacctgagtcctc**tacaataaacagtgagagcgccacggacagattcactcccgaggagcccagtaaagtgggaggactgaggtcacagggagaagacacaggcaccacctggcctggccctgacagggggcagtggacagaacagtgcggaggcagtggggtctctagaggctctggtcaccttgtcataccatggtttgtctcggtctggacactagagggcacttggggatcattcctgagaagactgaggttgatca**gagcagaaccagccacctgtcctgcctagtgtcctctgctc**agggctcacagctgcgtcctttacacccgtgggaccacacagccccacccctgccaccccctgacatcctcaggcatagggacctgacccagggcccagggtgggattagaaagctgggggtctgatttgcatggatggaccctcccactctcagagtatgaagaggggcagggagagatttggggaggctctgcttcagctgtgggcacaagaggcagcactcaggacaatctccagcATGGCCTGGTCTCCTCTCCTCCTCACTCTCCTCGCTCACTGCACAGGTGACTGGATACAGGT**CCAGGGGAGGGGCCCTGG**GAAGCCTATGGATTCTTGCTTTCTCCTGTTGTCTCTAGAAGCCGAATAATGATGCCTGTGTCTCTCCCACTTCCAGGGTCCTGGGCCCAGTCTGTGCTGACGCAGCCGCCCTCAGTGTCTGGGGC**CCCAGGGCAGAGGGTCACCATCTCCTGCACTGGG**AGCAGCTCCAACATCGGGGCAGGTTATGATGTACACTGGTACCAGCAGCTTCCAGGAACAGCCCCCAAACTCCTCATCTATGGTAACAGC**AATCGGCCCTCAGGGGTCCCTGACCGATT**CTCTGGCTCCAAGTCTGG**CACCTCAGCCTCCCTGGCCATCACTGGGCTCCAGGCTGAGGATGAGGCTGATTATTACTGCCAGTCCTATGACAGCAGCCT**GAGTGGTTCcacagtgctccaggcccggggggaactgagacaagaacccactt**cctcctctgccaggagg**gtgagccccggcagctgctgctcaggcctggcctgtggcttctgcggctgctgcttccctcatgg**tcccaggggcatccagggccctgcctggga**gtggaggctcctcctccccttctgtcctcagagtcaggaacagggaatccaggaagcagaatatccctctcccttaacttgggacacaaagtctctacactgaagttctggactc**aggtggcagatccagttgtgtcacctcagacccacct**ctgatagggaacccatgtattcattattctcctttccccaattccaggagtttccggagtggtgtcttcctctccctgctcctcaatgtgagtccctgtgtttcttttctcccaggcctattctttttttttttttttaacaaaactcctttcatgaaggaaaaatgcacacctaataaacca**gctgaatttatagcattaaattcggc**ccattttggcatttttctggatctttgaaaatatcaccaccactgagctatgcaacctctgactaaccccagcagcccttcatgatgctttgtcatctccttcactccccgaatgccc**tcctgggcagccagga**atctgcctttgccactactgtatggtttccattttctgaaacttac**tagtgttatcaaacacta**tcttcccatttgttcgagcttctttcactctgcacacttcttttcagattctccctgctgttgctgaattgcatcccattctacagagggctcattacaacctttgagggtctaggacattctcttctgattctatttatttacttatttattttaa**agtttgaattcattgtccatcaaatcaaact**ctacgggtgatgttcatctaaggtgacaggagtcagtgagggcttctcaggattcttgcttcttcatctcttccctctaattctttctataatggga

Uppercase: IGLV1-40

Lowercase: Flanking sequence[1000bp]

Red & Bold & Underline: Stem-loop [18]

Blue: Heptamer[40]

Green: Nonamer [1]

id-TRBV7-6-2[V_gene_segment]

cagctgagtacccaggatatcatacatcccagcatcttctaacaagggaaaattcttcacatgccagccttcttcctgcaccacattgactaacc**ctggaaacttcgcctttccag**aaaaaagctgtcttctcttggcccagtctagcccagtaggtttaattcatgatttatgtgaccttttaa**tttactcagaggagaacatatgatttcagagtaaa**gaatat**tcattttactttgtataagttggacctaaacagtaaactga**gtacttctcttccatcagtgtcttaaattagacaacggtaaaacaaatctttgatgcaagaactttaggtcctggtgtttttcatatggaagatcccagctatccagtccagggcatggaatagatgagcgttaataagtcactctatgcctgagatggttttctattggcatcttttctgagttacagttaacaaaggtgggaaatgatcccagaagaaaggcacgagggcagttcttgagggaatccatgtgatagga**cagtcattgggcacgtgtgactg**ggggaatggaggaggctggagcatgaatggggatggcactggggaccctgacttgcaggaaatgcaatgagctcaccactttgtgccctatgttaggggcggtgttggtgcatcctagagtaaatgtccagcagacaaaggagcaagaagcggctgtgggatgacaagataaactcagagatacagcatgagacctccggg**tccagacagctctgga**gcccaaggcgatgagccatgcattgatgttgttaaaaaggagctgataaatatttaaagcagcacccaactgtgttctaatagaaatgctgtgatcctgaggtcctggggattgagagaggaagt**gatgtcactgtgggaactgccctgtggagacaaggacatc**cctcatcctctgctcctgctcacagtgacactgatctggtaaagccctcatcctggcctgaccctgccATGGGCACCAGTCTCCTATGCTGGGTGGTCCTGGGTTTCCTAGGGACAGGTGAGTCCTCAGAACACAAAGTAGTTTCATTTTTTTTTCTGTGTGTAGGCGTGTGGGCGTGTGTGTGTGTGTGTGTGTGTGTGTGTGTGTGTGTGTGCTGACGACAAATGTTTTCCTTATTCTGTTGCCAAATTCTATTTCCACAGATCACACAGGTGCTGGAGTCTCCCAGTCTCCCAGGTACAAAGTCACAAAGAGGGGACAGGATGTAGCTCTCAGGTGTGATCCAATTTCGGGTCATGTATCCCTTTATTGGTACCGACA**GGCCCTGGGGCAGGGCC**CAGAGTTTCTGACTTACTTCAATTATGAAGCCCAACAAGACAAATCAGGGCTGCCCAATGATCGGTTCTCTGCAGAGAGGCCTGAGGGATCCGTCTCCACTCTGACGATCCAGCGCACAGAGCAGCGGGACTCGGCCATGTATC**GCTGTGCCAGCAGCTTAGCcacagc**atggcatagtcgcctccttcctgttcacaaacctcatccttctctctccttgcacctcctagagacccttaacagaggcctctctttgctcctcacttttgatgggaaagaagtagatttggacatcggctgtcctttgggtagaaagagaccacagattcattcctgaaacacagtgactgcaaatgtaggtggtgaaaacaatcacgtcccact**gccctctaggagggc**tcggagccagctcactgctcgaaacagggagtgggtgtcttagccttggccttcagggcagacatgcatcttctataggtcttgggggctgctgtg**ctgcccacatacatgaggttgtcatgggcag**gaaacatgcctctttttcatatgttggggcatctggaaggtct**gaggctacatccccagtaacatctttcttctgaagcctc**ttctatccctgtcaccttggaagtttctgcaacaaaata**tcaaacctctcttcctgtttga**agtaaaggtctgtgcaacttttgtggtccttacgtgataaaagcaatcatgataacaataagacttcatttcttctgtctactttaagccaagagtatccttcattttgtttccatttgccattgctgctgttctgatacacacaagcatgcattcaccactgctgccggttcaccttgattcgctcagcaaatctgattattagactctcgagtgttttcattgctgtc**aaactcatttgatttgaaataagttt**cctgaggccttgaactcaagaggtgttttacttataatattgaatctattcctttttattttaattttcacat**tatatattgttatatagtacttgttataaatagaagtac**aatgattatattgcagtagaatcttccacctatctgtgggtgatgctgcagtttgtatccatgaaagtgaatgcactggtcagagatcatgtgatcatggatcatgggtttctgagagtcctcag

Uppercase: TRBV7-6-2

Lowercase: Flanking sequence[1000bp]

Red & Bold & Underline: Stem-loop [15]

Blue: Heptamer[38]

Green: Nonamer [6]

id-IGHVII-22-1-2[V_gene_segment]

ctatgccaa**tcccaccacgagtgctggagaaggtggga**gtctgatgaagcttactaacaaccctgtggtccaagctaattccatcaaggccattggttcttcccagagcatagctgtccatcgggaatctcccatgtgcccagcagcagccacgcgttagcaacttcactgtgcacagtcattgtttgggaggagctccaggatgtgtctctttgtcaaaaactgggtgatgg**tgtcagagagctgtgccgggtgcctggtccacgtgctccctgaca**ttggaggtataggaggtgcgttctcagggtggaaacaccttttatggaaatgtacataataaatcacgattttc**cttggttctcttagacgacatggagaagcaag**catgcaccctgcaaacaattgtaatttcctgctttaccccaagttcatgatttcttgtaaactttgatttctgtccaagtacagagatactgattagagtgagtttggttctttccacacactaaccctca**cctcccccagatagagcacattgtcctcactctgagtctgagggagg**agctgttcctgtacaactcagggactgcagagacccccgtgtgcagctttgcagagtcagatctttcctcatgtgaggcgacctccactgcccgtgattgctgctcaggtctaattgtggattcgggattatgacaccctttgggctatcacaggtcaatcttattctgaaaatcaccgtaattata**gacagaggtaataatccagtacccattcttctgtc**cgcagtttgtc**tttcttatttggttgaaagtttgaagaaa**aaatgtgaccatatattcatctaattctaacctctgaatctgctatttgctctcggggacctacaaattgcaaaacagtgagttctt**cattttcatcaaaatg**tgcaaatttgtgaatttcaacatgttgttcagaacctgtgcatgccgacaactgtgattctcggtgcactcctGGCCTGGTGAAGCCCTCACAGATCCTCCCCCTGGCCTGTGGCATCTCTGCTTCTCCATTACAACCAGTGCTTCCTGCTGGAGCTGGATCCGTCACCCCCCGGGGAAGGGACTGGAGCGAATCAG**GTGCACAGGTCATGAGGGAGTGCAC**AATCCAACCCACTCCTCAAGAGTCCAGTCACCATCTCCAGATCCACATCCAAAAAACAGTTTCTCCTACAGCTGAGCTACCTTAACAAGGAGTACACAACCATGATTTTTATACAAAAGAcacagcgaggggaagccattgtgcgctcag**aacactctacaaattttcctccctagtgtt**ttaccaaaactg**gtatatatttcagatactgaaatatttac**aacctacgttattatgctaaga**attctaagttagaagttagaat**gataattgtctcttttttttccttttttttttttttttttttgatactgagtctcactgttgttgcctgggctggagttcaatggtgcaatctcggctca**ctgcaacctctgcatcccaggttgcag**caattctcctgc**ctcaaccccccaagtagttgag**attacaggcgcctgccacgatgcccggctaatttctgtatttttaatagagacggggtttcaccatgttggcctggctggtctcaaactcctgacctcaggtgatccaaccgcctcagcctcccaaagtgctggaattacaggcaggagccactgcacctggccaa**ttatcctcttttcatagatggataa**actaacctaggcctttgaaaataaacc**cttatctgagagtgaaaagataag**ccata**gatttggagagtttgcttgcaaatc**aa**atatttggaaaaggacttttattacaaatat**acaaattaccttacaattgaacaacaaaaccacacaatttaattttaaaatgggcaaagacctgaatagaaagtgcatctaacaataagtataaaaagtgacca**actggattttcattaggtaaatgtatgtttaatattgaaaaaatcctagt**aacctaatagaatgggtaaaatacacaatatagacaataccaaatgttgacgaggaagt**tgaaaaacaggaattttca**cccattgctcctgagaatgctgaattggcacatagacaactacagttaa**taatttattgtgtatttcaaatta**gctagaagat**aacatttagaatgtt**cttaccacaaagaagtagtcaatgattgagatgaaggttgaaggttgtcccagttatcc

Uppercase: IGHVII-22-1-2

Lowercase: Flanking sequence[1000bp]

Red & Bold & Underline: Stem-loop [21]

Blue: Heptamer[30]

Green: Nonamer [5]

id-IGHV1-45[V_gene_segment]

atcctgtgtgagagcagcacttgagtgatttcaataacaacttcccaggagaatcagctgaaaactacttgtcccattttccatacagatataacccc**tctattttcctgaagaaataga**aagagctgaatgctgaatacactgaatgtctgctggttttgcaagtttgtgactatatcactttctaatttctgacctgtgcagaccactgtacagacttttctcactggtgggaccagccttccagatgtcaaatataact**gagcttcttcatataaaagtcaacacaagctc**ctcatggt**ttcagtgctcactgaa**tggagttggaaataaaacccaca**attatccataacacaaattccttggaagttattttgggataat**gagttcataacctgtagaccaagagtccaagagtgtttctattgaaga**gcctgggggatcaagacaccaggc**aggtgatgcagacactgtctaaagagtgcccagcggctctcagagggacctactggatactcacgtgggacatcagcaatcactttctcagagtcaccagtgagctgtgctggttcctgaagggtccaggatagggccaaggcacctgctctgtgtcgtggagagtgattgttccagaaat**catagaggtggtctctatg**cttataaaatctatgttcacagtgagaagtctgttctgagagggcttattcttcagtgaaaggacctctgcccacaaatgttcataaatggagcagggcattcatttc**ctcaagcaggatcagggcttgag**t**catcagcatctcactcttgcaaggctgatg**tgtcgtttgtcttccctttcttatcatcgaccaggctttgagctatgaaatgccctgtctcatcaatatgcaaataacctgagatcgactgaggtaaatatggatatgtctgtgccctgagagcatcacccaacaaccacatccctcctctagagaatcccctgaaagcacagctcctcaccATGGACTGGACCTGGAGAATCCTCTTCTTGGTGGCAGCAGTCACAGGTAAGGGGCTCCCAAGTCCCAGTGATG**AGGAGGGGATTGAGTCCAGTCAAGGTGGCTTTTATCCACTCCT**GTGTCCCCTCCACAGATGCCTACT**CCCAGATGCAGCTGGTGCAGTCTGGG**G**CTGAGGTGAAGAAGACTGGGTCCTCAGTGAAGGTTTCCTGCAAGGCTTCCGGATACACCTTCAC**CTACCGCTACCTGCACTGGGTGCGACAGGCCCCCGGACAAGCGCTTGAGTGGATGGGATGGATCACACCTTTCAATGGTAACACCAACTACGCACAGAAATTCCAGGACAGAGTCACCATTACCAGGGACAGGTCTATGAGCACAGCCTACATGGAGCTGAGCAGCCTGAGATCTGAGGACACAGCCATGTATT**ACTGTGCAAGATAcacagt**gtgaaaacccacatcctgagaccgtcagaaaccccaaggaggaggcagcttcactgaatgaggaggttacagggcttacgatgtttaaagttgttcagaaaataggctaagcaattgaggaatatgagtaa**tagaaatatgtatgcactctatacaggaaatatttcta**ataattgtcaccctatatgcaaaattcgcagagaggtaaaagcagaaatcagtcaagctgatgcaaagttccccacgtaggctttgtgcagatgtaagttctaaaatcagatagataaataatttggagcaagattgcttgataacatggctaatgctgaatatgattcctaaaaactggccaaaatatattccaatttgtctctgccacctctcttacataaaatgta**ttaaaaagtagttttaa**gaccacagcaaaattgaacagaaggtgcagagatttcctatgtgcccctgcttcacacatgcacagccttccccactgtcaccatcctgccccagagtcatcaa**taagttacaatggatgaactta**catggatggattggttctttcctcttccggcggtcccttggcataccaagtctaaactatcttgaagcacaacaggttcttcgagtgggttcctgggaatgaagccagttagaggaaaagtgggtggggctattccta**ttaggagtcttttttagaagactcataa**aatgtatatgttcctatagattctgtgactcctgacttagtatcccttcccagacggtaagcttcctaaatgtttagaggcagatccatatctatggaaagaaagcaagttctagtgaatcccataaggaatgtcctttaatgagaagtggagaccttggtcatgaggcacatcatgtatgattttctataattcctttagattcactgtaagcttttgagggtg**tttctggatgaggccctttgtacagaaa**atgaaaactcaggcatgagttccaggcacaacca

Uppercase: IGHV1-45

Lowercase: Flanking sequence[1000bp]

Red & Bold & Underline: Stem-loop [18]

Blue: Heptamer[42]

Green: Nonamer [5]

id-IGHV3-6[V_gene_segment]

acaaggaaaagtatctaagtatttatgct**gagtaaaataagccagacaaataagaatatgtactatattactc**catttatgtaatttatttcaaatgaaaaaaatctaaagtaatatacaaaagatgagtagttacctgggaaaagggtagaataaaggaaggggaaatgtggaaacatacagaagaacaagagaaaatgttgaggggaactcacttgtccacctcagta**atgatgacgcttacatcat**gttttcaattc**tacactttaaatacgtgaagtgta**ttatctgccaattaacactcacaaaatttattgcaagcaaacaaatgaaaaattagaccaaaagcggtggtataaagatacaaaatatccattaaaagtcagaaatatctgagaatttacctcctgaaccctagttctctccttatttttaggtgaatgctagagtgccacaaaaatcactcaagttctcaatacaggaagtgggttccacaaagcacactaggcatgtccaattcttacctggagttggttcagggagtaactggggccactgatgtggagcacaggccaagctaccagggctcattcatctcagacatgagaca**cacacagcccctcctccatgtgtg**ggtttatttcaca**tctgtaaatggagaaaccattgacttctacaga**acataa**tttgcacaaatatgtaaaaatataatagtgcaaaatgtttatcacaacacatttt**cataataagacagt**gtatttcccaaatac**cataattgtcacccaactcctgcggggccctgtcatttta**tctggggtctgccgtctcctcagggttcccaccccaga**gctctctgtgtagcaggagacacgcaaatagggccctccccatgctgatgaaaaccagctaaaccctgaccctgcagctctggcagaggagcctcagccccgggattcccaggtgtttctgcttggtcaacactgaacatacaggattcaccATGGAGTTGGGGCTGAGGTGGGTTTTCCTTGCTGCTATTTTAAAAGGTGATTTATGGAGAACTAGAGCTATTGAGTGTGAATGGACATAAGTGAGCGAAACAGTGGATATGTGTGGCAGTTTCTTACCAGGATGTCTCTGTGTTTGCAGGTGTCCAGTGTGAGATGCAGCTGGTAGAGTCTGGAGCAAACTTGACAAAGCCTGGGTGTC**CCAGAGACTCTCCTGTGCAGCCTCTGG**ATTCACCTTCAGTAGCCATAGCACGC**ACTGGGTCCCCCAGGCTCCAGGGAAGGGTCTGCAGTGGGTCCCAGT**TATTAGTGGTAGTG**GTAGTACCATGTACTAC**GCAGACTCTGTGAAGGGCCGATTCAC**CATTTCCAGAGACAATACGAAAAACTCACTGTATCTGCAAATG**AACAGACTGAGGGCAGAGGATGCAGCTGCATATGACTCTGTGAGAGAtacggtaaggagaagtcagtgtgagcccagacacaaacctcccttcagggtacctgggacaaccagggaaag**cctgggacactgtgcactgtgctgaccccagg**ggcaagtgcaggtgctacaagggctgggtttcctgtcttgatctggggctgcgttgttaccaaattacccgagggaacttctctagatttgcaattctgtactaacatttgatgtctgtgactgctaaacttccctcacatatttgttttttgtttgtttgttttgttttgttttttgtaacaggaagacccatcctcacccctacagaagcccgagtgtcaccttgggggcagaaatgaccctgccttggtcacattgatcagtgtcctgaggaagcccagtggaacctggggagtgttttccagtcaggctcaggg**cagagacctccgtggaaatctctg**attagaataggtttgggattcagacttgggccaagagggagtctcacccatgggagggtccttaaatcctg**acagtttttacagctgttctccctcctctggcaaaactgt**gcacatctgactcagaactgatccagctggcccttacttgctaatcaatttttcttctctctacacttcattctcatagttcccttttcttcccttcc**cagaaaacagagagtgtgttttctg**tggtctaaatctcagggctcaggcctgttacctggaactcaggtgtgg**cccaggctgtggctcctgtgagacctggg**gaggctgaggggactttcccactcaccattgcctggacccgcctgctgtcttctgtgcatggaggcatttgggaaatgaagtggacattagtcactgaggggataatggtagctttctcaaatgagatactgatgtagagctgatcttctgtttctcacact**gtcacagaatttcctccttcatctgtgac**tttggggaaagctaaggatggacactctattgtgctgtgagctctgggtgac

Uppercase: IGHV3-6

Lowercase: Flanking sequence[1000bp]

Red & Bold & Underline: Stem-loop [19]

Blue: Heptamer[42]

Green: Nonamer [4]

id-IGHV3-16[V_gene_segment]

gtgaggtgacatggggtatatgtgaggtgacccccctgacgtgtg**aggtgacctgagcacatgtgaggtcacct**aagg**cacatctgagttgacctgggggatgtg**tgaggtgacctagggaaatgtatgagctgacaaagggcatgtgtgcagtgacctgggtgcatgtgaggtgacctggtgcatgtgttagtttacctgggggatgtgtgaggtgaacaggggcaggtgtgaagtgcctgaggaatgcatgaggtgacctggtgcatgtgtgtggtgacctagcagatttacgaggtgacctgcgggatgtgtgaggtgaatagggacttgtgtgaggtggccacaggaatgtgtgaggtgacctgggggatgtgtaaggtgacattggggatgtgtgaggtgactcagggcacatgtgaggtgacctgggggatgtg**caaggtcaccttgggtgagtgtgaggtgaccttg**gggatgtgcaagg**tcacactgtgtgagtgtga**ggtcatctgggatatttcaaacttcgtaaggaaagcctgcaacagtatatggcactaaaagcttcacactccaacaccgttaatgcgcacatcctcaaggaactctaagtgtttccaggttagcttaaatgccatggagcgacacctgc**cccaggcacactcatatatgcatcctggg**tctccctatcaacccctatcctcaaattttcagtttatgtttgctccatg**atatcaactctgatat**gttgaaggtttttttttttcttttatttgtagttgttcaggtttgctatttcactctcattactctgggctcagtcctctcctcaggtgtcccacttcagagcttgctatataataggagacatgcaaataggaccctcc**cttttctgatgaaaag**cagcgcagccctgaccctgcagccctgggagagaagccccagccctgggattctcaggtgtttctattgggtcaacagcaataaa**caaattaccATGGAATTTG**GGCTGAGCTGGGTTTTTCTTGCTGGTATTTTAAAAGGTGATTCATGGAGAACTAAGGATATTGAGTGAGTGGACATGAGTGAG**AGAAACAGTGGATATGTGTGGCAGTTTCT**GACCAGGGTGTCTCTGTGTTTGCAGGTGTCCAGTGTGAGGTGCAGCTGGTGGAGTCTGGGGGAGGCTTGGTACAGCCTGGGGGGTCCCTGAGACTCTCCTGTGCAGCCTCTGGATTCACCTTCAGTAACAGTGACATGAACTGGGCCCGCAAGG**CTCCAGGAAAGGGGCTGGAG**TGGGTATCGGGTGTTAGTTGGAATGGCAGTAGGACGCACTATGTGGACTCCGTGAAGCGCCGATTCATCATCTCCAGAGACAATTCCAGGAACTCCCTGTATCTGCAAAAGAACAGACGGAGAGCCGAGGACATGGCTGTGTATTACTGTGTGAGAAAtcctgtgaggggacacaagtgcgagcccagacacaa**acctcctgcaggaacactgggcgaaatcagctgcagggggt**gctcagggcccactcatcagagtc**aaccccagagcaggtgcacatggaggctggggtt**tgtttc**ctgtcaggatttgggacttcctctgcttctgacagtttccctagggaaact**ctttaattttagatttctgtgcccaccaatgtcatctctacatatatttttttaaatcattataatacgaggactcattctcacatgcacaatatgtatattgccacctacgggaatgaaaac**tcctcaaccatggtcaccagcatcagagttgtgagga**agctcaggggtgcctgg**tgagtcttctccagtcagactca**ggacagtaacctcaatgggattccctgactagaatggcttttagggatttcgattacagccaagagagaggctggaccaggttcag**tgtcatgtaggacctcacagattttatgtatgaca**tttctcctgacaataaagtatgcaaatgagtatcagcactgatctggtgcttcctgactttcatttttagtggttctcgcttttccatttgattttcctgctttctggataaaaggatgttgtccctgtggtctaaatcctggggctcaagccctttccctggagcttaggtggggctcaggctgtgactcct**gcagccattgggagaggctgc**tgagactttcttctctctcattgttcaggatactccactctgttttctggag**acgcgtctgagaacgcgt**gtggccattagtaatgagggcacaagcttctttggtcaaaatggggtgaggatgtggaattgatcctgtgctgtgtaaactgtcacagagtcaccttcttcaccagtagtgttagaagagcttgtaaaatttgtcggaatccaactggagtcccctgtgttcaaaccctgacaaatggagctggggaaggccatggatggaag

Uppercase: IGHV3-16

Lowercase: Flanking sequence[1000bp]

Red & Bold & Underline: Stem-loop [19]

Blue: Heptamer[36]

Green: Nonamer [4]

id-IGHVIII-25-1[V_gene_segment]

gcagggatggcatgagatacggggaataaagaataaagtaactttctccatttggatataagatgtgtccattcactagagagttttctttcagaaggttcccatgtgagccag**tttccttcctgacaaggaaa**tcacccagagga**ttctcataatctcttaagtgagaa**aagttttcctcaaactccagctcagtctagctcacactgtctctaaatgggcaattactttcagacatgtacacacatctgtcccaacgtggactcctccctcagacaagcacacacattgcaatgtggactctcccctcagaaaaccacacatgccccacaatgactccttcttcacataagcatatatgcctgatgctgaactgctgtgtagcaaaattagctcagaatacaagcac**ttatggactctagccataa**caatgtgtagatctgcattttattttaaattctaaatgaacattttgccttcttgttaagaacagtggtttacagctctacatgtagagactacaggcagatatatgtttcctctgggaaaagtttattttcatgtatttactgggttattttgttgataaacattcatctataaagacacctgaaaagtgccctcattagagaaaaagaaagtgtagtttgcagattaactctgagcatccaatcccaggaatcctttga**tctgccctccctgaaatccacaggcaga**ggggggatgatcgctgataagtagcaaa**cacatgtccacaggaagggagacatggaaatgtg**gtcactcccacactcaagaggagcagctcatcccctgtccgtagtggtcctgg**tgaggaaccaccccacacctgtgccttcctca**gtatccacaccatggggtctgtgctgatctgggcttcccttctcatcactctcaatattagtgtccttgcggatcaggctcagctgtggctgctccagatgaggctgttctcagtctgttccctctgtgtttgcagAAGTCCTGTGTGAAGTTCACCGGGGGAGACAGAGGAAATAACGGTGCAGCCGGGGGCTATCT**GAGTCTCTCCTGCAAAGACTC**TGGATTCACCTTCACTGATTGCAGCATAAGCT**TGGTCCAGCAAGCTCCAGGACCA**GGGTTGATGTGGGCAGCAACAGGGAGAAATTGAAGAGGAAGCTCTCAGTGGTGCCCTCCATGAATACAAAGAATCTTCACAGTCCCCAGGACACCCTTACGTGCatggtctcactgatatctttacttcctttatcacttttgttatgtaaatcacaatgaatagtgtattcctcatctattatacatttgttaagtc**ttttttggtgtctttaaaaaaa**ctgataactttatagtatgtaatatccttaagtcctgaaagtgttttttgatgtctaccttgtcttaaatttatacagctactatagcttttcttggttcatgttttct**catatttatcttttctatttgtgaatatgtagagaaactttcttgtacata**gctaatagttgtgtattgcttttttattttggcagtccttaaatattctaattttttatgatactttaagttttagggtacatgtgcacaacatgcaggtttaaacaaacaacctcatcaaaaagtgggcaaaggatatgaacagacacttctcaaaagaagacatttatgcagccaaaaaacacatgaagaaatgctcatca**tcattggccatcagagaaatgcaaatcaaaaccacaatgagatatcatctca**caacagttagaatgacgatcattaaaaagtcaggaaacaacaggtgctggagaggatgtggagaaataagaacact**tttacactgttggtgggactgtaaa**ctagttcaaccattgcggaagtcagtgtggcgattcctcaggtatctagaactagaaataccatttg**acccagccatcccattactgggt**atatacccaaaggattataaatca**tgctgctataaagacacatgcacacgtatgtttatagcagca**ctattcacaatagcaaagacttggaaccaacctaaatgtccagcaacgatagactggattaagaaaatttggcacatatacaccatggaatactatgcagctataaaaaatgatgagttcctgtcctttgtagggacatggatgaaactatctattgcttttttaaatcaactataataaattctatttttttttttgagaaggagtctcgc

Uppercase: IGHVIII-25-1

Lowercase: Flanking sequence[1000bp]

Red & Bold & Underline: Stem-loop [16]

Blue: Heptamer[44]

Green: Nonamer [1]

id-IGHV1-14-2[V_gene_segment]

tccagctctcattcttcttggttctctcctgcaggccagcagccctgaagtttagcctggatctcccatgcatccacccatctccttccaagtgcattttaccacagcctccactgtttttgaagcactcttgggctttgtaattctccacactctgttgtaaaggaagtcaggtccttcaagaccagattcgggactctattttatgaccaaatttcagcctcacccctgctcctgagacagag**ctcctagataagattctgcaggtggagattaggag**tgtttttcttcttcaatgtagttgctgagcgctcagtgcagggttggggagaaactttccactttgtcagcatgcagct**cctgctggggtagaccttcttccatagaagcagg**gttgggaaccagggggccaatgtcctcagtgctgctgcacccagggcagagccttcatccatcagtggggctgtggaagaagtgagtctctggtt**ctcagtagctcttgtccagaactgag**cctctgcagcatgttctgtcggccccagtgtcctggcccctaaggagcagcatcctaaaatgggagctagcgtatttgagaataacaacacctacacattcagaagctctttggctttctttccagctaatataatttccttttttt**tgtgtagcaacctgtacaca**cgcatattgatgcatacagacctatgacac**ttttttctcgataagtaaaaaa**ttattgatcactgtgatcttttctccaagttcaccatttccctgaaggtgagcacaggtccttctgcatgtgttcaaacaaaaggcccagagactacctggtaagtgaggtgctcacctggttctggatgtttggtctgtctcctcccctctgttgccccacacaaggtcagcccactctttccaggtccgaagaagagagcacaggtttgtcctgattatatgactcacccagcttctgatgactctcctgttgccagcgtccatgGCCTCAGT**GAAGGTGTCCTGCAAAGCTCTGGATACACCTTC**GCCAGCTACGACATTCACTGTGTGTGACAGGCCCCTGGATAAGGGTTTGAATGGATGGTAGGGAGCTACTCTGGCAATGGTAACACAGGCTATGCACAGAAGTTTCAGGGCAGAGTCACCATGACCAGGGACACGTCCACGAGCACAGCCTACATGGAGCTGAGCAGTCAGAGATCTGAGGACATAGATGTGTACT**ACTGTGCGAGACAcacagt**gtgaaaacccacatcctgagagagtcagaaatcctgagggaggtggcagcagtgctaggcttgagagatgacagggattttatttgctttaaaga**ctttttttagaaagcgaggttaattcattgcagaaaaaag**gaaaatagaaatg**tgtatggactctaattatgtgggaaattttccataca**act**tttgttctctaagcaaaattcagggagtggaaaacaaa**tcaaattaataaacctgataaaagaattcctctgaaaatttagtgtgagcataagtttttgaatgggtgttgtaaatattttggaacacagctgctagatcacattttaactctacacttatctccattatataaaatat**caaaatgttttaatgttttccattttg**tgcaattataatttt**gtgttataatcacattttaacac**tttaactctacacttatctccattatataaaatatcaaaatgttttaatgttttccatttcgtgcaattataattttgtgt**tcatgccagcaatgcatga**tagatcttgttcttccgcatcctcattgccatttggcactaagagtattgtgtattttaatattctaatagattagtagtgatagctcattgctgtttaaatgcacatatttc**taattaaaattttgtatttaatta**ttttatataattgtgatgaagtgtctcgtatggtatttggattattttttattgcattgtttctttttgatcagttgtaagtttc**cttatatacccattatataag**tcactcacaaagttaacaaaaaattgattaacaaatatgtgttttacaagtgtattctccaaattgttgttgtcg**ttttactcccatatcagtgtctgtgggagaaaaa**tatttatatacatatatatgtgtgtgtgc**atatatatatatatatatatatatatatatatatat**agtgtgtgtgtgtgtgtgtgtaaacttagataaaataattatttca

Uppercase: IGHV1-14-2

Lowercase: Flanking sequence[1000bp]

Red & Bold & Underline: Stem-loop [17]

Blue: Heptamer[36]

Green: Nonamer [7]

id-IGKV2-10-3[V_gene_segment]

aaatcattggctaaattttcaccattacaaatggaggtgagaagacagtggaatgatatttttaaatactgagccacaaattctcagccacaaattctatatccagaaaaacaatcgtttaaatataaaagcaaggtagaaatattccttcatacacaaagattaattcattaatagcatatatgccttgtaagagattcaagagaaaattcttcagaataacagagaatgacagcagacagcaactcaaattgacaagaaggaaccaaggcctcaaaaaatgaaaaaagttagaaagaagctacaaaaaatgtttactttacctgatataaagattaagtccaggactaaaacaattatatgtcagaatatcatttacattgattgtgttttgagcagcagtttttgactgg**gaaggtgcataatgaaagcagcgagagaattggaaaccttc**tgcgtttctatgtg**gatcatgatatgaaatgatc**tggatcttgacattcatttgcacccttaagagttgcaccctttgcactgtgtgttttaccttagataaaataaaataaaataaaataaaataaaataaaataaaataaacaaacctagactcttttaaataaacctaaacaaagaaataagaatcatcaaatatttatgaaagatcaagcttattaaaaagaaacataaaacatgcccgagaaagtcctaaagaactagaagtctagtagaggaaacagcaggaaaatctaaatacgtcccttctgccactcaggcaacacagattttacctctattgatttccttaaaa**acataaaatatatttatgt**cattttgctcacaagagacgcccccaccttctccttggctctttccaccccactgcacccacc**aggggatttgcatactgtcccct**agggaggactttccttgtgagtctgagacgaaagctcactctaaccttgccttgactgatcaggactcctcagttcaccttctcacaGTGAGGCTCCCTGCTCAGCTCCTGGGGCTGCTAATGCTCTGGGTCCCTGGTAAGGGCAGAAGGGAAATGAGGGAGGATGATGGGG**TGGGAGGGTGAACTCTGTGGACCCCGCCACCTCCCA**TGTGTGTCCTGTCCTCGTGTTAGATGTGTCTTGTCCTCCAGGATGGGGCATGTGATGTCTAGATCTGTGAGAGTGAGGAAGATTCCAGAAGGAGCAAGGATATGTAAATATTGGTTCTTTTTAAATCTGTATGTTTTGGGAGGATTAATCAAAATCACACACACAAAAATAATTGAGCAAAACATAAATAACAGACAGAAAATGATTAAAATGACTCACAATGTTTGCACATAACCTTGCACTTCTCTCTCATTATTTCATGATCCAATGGAGATGCTGTGATGACCCAACCTCCA**CTCTCCCTGCCCATCACCCCTGGAGAG**CCAGCCTCCATCTCCTGCAGGTCTAGTCACAGCCCCCTTCACAGTAATGGATACATCTATTTCAACTGGTACCTACAGAAGACAGGCCAGCCTCCTTGGCTCCCAATCTATTTGGTTTCCAATCACGACCCTGGAGTCCCAGACAGGTTCAGTGGCAGTGGGTTGGGGACA**GATTTCATGCTGAAATC**AGGAGGATGGATGCTGAGGATGTTGGGGTTTATTGCTGCCAGCAAAGTACACATTATCCTCCcacaatggtatagtcttgaacacaaacccccccacttgctgtggcctagctgcccagatgtgctgtttctgtggagagcaggcactgtggattctcttagatgcctaaag**accaagatgttggagaactcagaggacttggt**gcag**ctgagggctcatgaccataaatttcttggctacacctcag**gcaccacattttaaggtcccgtcag**ctgcagcagcctttgcatgacagagtctgcag**tatggaggaggtccacgtgccctctgagcaatgagacac**aagagaagagaaggctcaatgagagctcattctaatcctctcttcctt**ccctatgttcattcatcaactaaattcattctgcataacaggcacctaattgagactgattactggcaacacaaaactaacacattcttttgatttggtttagcagtt**accagagtacatgtacattgataagatctggt**gatattaaatcagtttccctcccctcctccacctcccccctccactttttgtaaggccgacccttgaccagaacagcagcaagcactgtattttatcctacttttttcagggagaagccaaataaaactcttcacagtctagattttttaattactagatatgttgtggcaaatgta**tgcaaaaataaagacctagttgctaaagtgtttgca**cagtctcactcacagcatattgaagtacaaaagtag**ctttctgaactccaaagctgggtttcaaaaataaggacaaagttcgaaag**aataaaggaacatatgtatgttatttaaacagaatttgaatgaatacaagataatatataagctaactttttaccccatgaaaaattacaatgtgaagattttgagcagagggcagcaagtgagagactgg**tgggttgtgtttgagatatgaaccca**gatggaag**ttctcacgtatttcctgggaagaatcacagagttgagaa**gcgcctgcctggctttcaaggcagtaa

Uppercase: IGKV2-10-3

Lowercase: Flanking sequence[1000bp]

Red & Bold & Underline: Stem-loop [16]

Blue: Heptamer[35]

Green: Nonamer [8]

id-IGHV3-60[V_gene_segment]

aaaaatatctaagtgtttatgctgagtaaaatacgccagacaaataag**aatatgtaccatatt**actccatttatgcaat**ttcttttaagtgaaaaagaa**tctaaagcaatatccagaagatcagtagttacctggaaaaagggtagaccaaaggaaggggaaaggaggaaacttacagaagaacaagagaaaatgttgaggggagttcacttgtccagcttggaa**atgatgggttacatcat**gttgatcaattgcacactttaaatatgtgaagtctattatctgccaattaacactcgcaaaatttattgcaagcagacaaatgaaaaattagacaaagaaaggatggtataaagatagaaaatatatattaaatgtcagaaatgtctgagaatttaactcctgaccctagttccgtccttatttttaggtgaatggtagcgtgcaccaaaatcacacacattctcagtacaggaagtgggttccacaaagcacacgaggtatgtccaattcttaccaagatttggttcagggagtaacagt**gatgaggaatcacaggcccagataccggggctcactcatc**tcagacatgacctcgtggacacacacttagcccctc**ctccatgtgtaggttgacttccacatatgtaaatggag**aaaccattgactcctacagaacataatttacagaaatatacaaaagataaaatagtgcaaatacttatcacaacaaaatttcctaataagacagt**gtattttccaaatac**cgtaattgtcacccaactcctgtggggccgtgtcatttta**tctggggtctgccgtctcctcaggattcccaccccaga**gctctctatgtagtaggagacaagcaaatagggccctccctctgctgctgaaaatcagccaaatcctgaccctgcagct**ctgggagaggagcccccgccccgggattcccag**ctgtctccacttggtcatgaacactgaacacagaagacacaccATGGAGTCTGGGCTGAGCTGGATTTTCCTTGTTGCAGTTTTAAAAGGTGATTTATGGAGAATGAGACACACTGAGTGTGACTGGACATAAGTGAG**AGAAACAGTGGATTTGTGTGGCAGTTTCT**GACCAGGGTGTCTCCGTGTTTGCAGGTGTCCAGTGTGAGGTGCAGCTGGTGGAGTCTGGGGGAGGCTTAGTAAAGACTGGAGGGGTCTCTGAGACTCTCCTGTGCAGCCTCTGGATTCACCTTCAGTAGCTCTGCTATGCACTGGGTCCACCAGGCTCCAGGAAAGGGTTTGGAGTGGGTCTCAGTTATTAGTACAAGTGGTGATACCGTACTCTACACAGACTCTGTGAAGGGCTGATTCACCATCTCT**AGAGACAATGCCCAGAATTCACTGTCTCT**GCAAATGAACAGCCTGAGAGCCGAGGG**CACAGTTGTGTACTACTGTG**TGAAAGAcgcagtgagaagtcagtgtgagcccagacacaaacctcctgcagggtacctgggacaatcagggaaagcctgggacactgtatactgggctgtc**cccaggggcaagtccaggtgatataagcctggg**tttcctgtcatgatctagggttgcattgttagcaaattaccccagggaccatctctagatttccaattctgtaataacatttgatgtcgtctctgactgcacaatgtcccctcaactttgtatcttttttttttttgtaacaggaggacacatcctcaccctgcagaagcctgagtgtcactttgggggcagaaatgacctgcct**tgatcacattgatca**ctgtcctgaggaaaataccccacaggggaccccgatgactccagcaaaggctctgcctcaaaaccattgaagagtccttcctttcattagaattgaccacagcacctgggcttcagcacaagccataccacagacgtcacaaagcagcagcttga**cacctgatccaggtg**cattttctc**acctttagaagctgagagaggggtgtattctcaaggt**caacacactctttgtg**ggtttttacacagaaaacc**tgcttttactttat**ttcatttgaagataaatgaacaaatgtgcatctataaatgattgtaattttcacatttattc**caaatccaatatttctgaattctgcataatgtcctggcacaaggttgtg**ttttctataggtatttttcaacagaccaagaaaa**tgcattttccaatttccctgtttttcctccatgtgcagagaccttgagtagagcacgtctgactgtgtttgtctgttttcacgctgctgatgaagacataccagagacgaggaaacttacaatagaaagagttttattggacttacag**ttccacgtggctggaaaggtctcacaatcatggtggaa**ggtgaaaggcacatctgacatggcggcagacaagagaagagagcttgtgctgggtct

Uppercase: IGHV3-60

Lowercase: Flanking sequence[1000bp]

Red & Bold & Underline: Stem-loop [20]

Blue: Heptamer[46]

Green: Nonamer [5]

id-TRAV33[V_gene_segment]

attttttctttagttgctggtgctttatatgtcatatttttaaaaacattgcctaatccaaggtcacagatagttatatctatgttttctaagagtttcatagtttcagtttgtacattgaggttattgctccacttcaagttaatttttgaatatggtgagaagtagggatgaaagttcattcttttgcatgtgaataataagacatttggagtccccaaccaaaagaaagctgggtagctatactaatatcagataaaattgacattaaaccaggaattattactagagacacagagcaacacatcaaaatgattgagtacgatctacccagaacaaattctatttttatatatacctaatggcgtagcttcaaaatatatcaaacataaatcagtagagttgaccagtagaattaaattggtaga**atttattgattaaaagatcaataaat**aattatacaaccatcatgtcaggcattaacacacatctctcagtagaaaagtattgaaattatgaccaacacaactaatatcctttacctcacatacatagattgaatactttacctcaaaagaacagattacacaaggaatatttagca**aaattgactatattctgaaacataaagtaagccttaatcaattt**caaagaacttaaatcattcagaatagtcatagtctctgaccatatggtattaagcaaaaattcaataacaaaatgattattaaactttattcctgtaccttgctagatataaggttaatttttcaaatgtattgccacagaaaacaccaagacaaaa**cagagatttggggtagggtagatttttcagtctctg**aatagagcaatgtaacatactacttgcacactttatttcctgtttgggggcagaaacactgaacttctccagggactttgcactcagg**cactcagagaagcattttgtttccaggcttgagtg**aggaagca**aatttcaaccataagaagaaatt**ctaccATGCTCTGCCCTGGCCTGCTGTGGGCATTCGTGGTCCCCTTTGGCTTCAGTACCTATGGGCGGCCCCAAACATGGGAC**TGCAGCTTCCTCGAGAGCTGGGCTGCA**CAAATTGGAACTGGTGGAGTATTAAGTAAGAGGTAAGGGAAGTGGGGAAATGAAACACTATTTAATTATTTTCTTTGTCCTTTTGCCATAAATTAATCTAGTCCTCATGACCCATCTGAGACATTTAAGGGAACCAACTCTGTTTTCTCCTTTTTCCCATAGGATCCAGCATGGCTCAGAAAGTAACCCAAGTTCAGACCACAGTAACTAGGCAGAAAGGAGTAGCTGTGACCTTGGACTGCATGTTTGAAACCAGATAGAATTCGTACACTTTATACTGGTACAAGCAACAAGCAACCTCCCAGTGAAGAGATGGTTTTCCTTATTCATCAGGGTTATTCTAAGTCAAATGCAAAGCC**TGTGAACTTTGAAAAAAAGAAAAAGTTCATCA**ACCTCACCATCAATTCCTTAAAACTGACTCAGCCAAG**TACTTCTGTGCTCTCAGGAATCCCacagaagta**gaaatgacagtggaagataaacaaaaaccttagcactccataaaggaagccacctgctcaggagcttagggaaaatacatgaagcacagacaggaagaaggc**acattagtggcacagttgagaccttataatgt**aaacattcctctgtaaaaaaaaagtttccttattgagtttatatataaagcagcattctttcttgcgatttccaactattacgtttctatcttgggttaaagtaaatcacaaagtg**tattagtcagggttctctagagagacagaactaata**ggagagttgtatgtatgaaggggagtttattaaggagtattgactcacacaatcgaaaggtgaagtctcacattaggccgtctgcaagctgaggagcaacgaagatagtcggagtcccaaaacctcaaaagtagggaagccgacagtgcagccttccatttgcggtcaaaggtccgagagctcctggcaaaccactgctgtaagcccaagagtccaaaagctgaagaacttggagcctgatgtttgagggcaagaagcatctggcatgggagaaagatgaagga**tggaagacttaacccgtcaagtccttcca**cattcctctgcctgcttttatcctagttgagttggcaactgattagatggtg**cccacccagattaagggtggg**tcagcctctcccagtgcagactcaactattaatttcctttggcaacaccctcacagacacacccaggaacaatactttgcatccttcaatccaatcaagttgacactcactattaaccatcatacaaagtgctttttaaaatcgttgctgaacaataagattaaatgaacctatgtatttcatgaaattcagaag**cagtttgctgattctcatcttaaactg**ttgaatttcc**agatttgaagtagcactataaaccattttatcttccaatct**catgttgattttagcagtttttataaatattatcattttatttatt

Uppercase: TRAV33

Lowercase: Flanking sequence[1000bp]

Red & Bold & Underline: Stem-loop [14]

Blue: Heptamer[16]

Green: Nonamer [2]

id-IGHVII-26-2[V_gene_segment]

gaattgatgtgtgtgtgtgtgtgtgtgtgtcct**caggcctgttgggcctgttgagaacacagcagggctctatgtacaacagg**aaagcaatgttttctctataccatctattt**tcttataaagtgttcataggtaatcaaaaggaaatgaatcacagtttataaga**tgttgtggactggactgatgttgtcactcaaggctactatgggggctccaggaaaagaaacctagaaaaggataagactgtgagcaaagaagcaaaacctgggaagcagcagggagagggtgagcaggggtgactcaactggggagggataacaccacttaccacagagggtgaaaactccgtgctcagcaaagaggagaacttactcagcatttcccctaatcctccctccagaagcgcgactgccaggaaagttcacaatgtatc**atctccagctgggaacccaaaagaaacaggagat**tttagggcgtaggtcctgggcacaaggtgtttccctggttcctaccccttccctctctgtccagcaaaactgcttgtttctatcacttgccttcctgtccttggggtaagaatggatgctttgacactcagcagagggatgctctttcagaggaggaatgagtttgtgttatctgatgtctgagcaggacactcacctctgggaagaggctccaagacagaaggtcgccgtggttcct**gccccttatatgtgaggggc**tcttgacacatgcacacttttcctcaaactctgtttatac**cccagcaacaatgctggg**tgttgtcagcctctctcctccccagatactgctctatcatcagcttgagcacagtttccccatgaaaggcattattctctctctcaagaagatcaactttttacatgtcttcgatggatgacagtttttctgtgtgagaatctagaaataagaaatggggtaatggggaca**agtgagctctctgtgtgactcact**aggaagtccactgttattccagGT**GTCCTGTCCCAGGTGCAGCTGCAGGAGTCAGGTCCAGGAC**CCATGAGGCCCT**CAGACACTGCCCTCTGCCTGTGCTGTGTCTGGATTCTTTTTTTTACCATGGATATTATTTGAGATGAATCCA**CCATCCCGTGGGCAACAGGCTGGAATGGCTGGGGCACAGTTACTAGACTGGATCATCAAGGAATACACATCATGCACCCTCCACCCAGGTCCATGTCCCCATCAACAGTGACTCAACCAAGAGCCAGTTCTCTGTGAAG**CTCAGCTCCATGACCACCTAGGACACGGCTGAG**TATTACTGTGAAAGActcagtgaggaggtgtccttatga**gccctgacacaaacctgtcagggc**acttaggacctccaggaagactcaagaccaccaaggggactcacgaccactggggaagggcaggttgcagtaagcacaggg**ccagccccagagcaggtgcaggtggagatgagaggctgg**tttactgccagggtctggggctgcttctccatttagatgtttccttcagagaccctccct**tatttcatgatgctctgcttagctgtaagtctctgaaata**tcacagttttgttgtaccaggaggaaaccttctctcaggtgctaaatgtagaacaatctctctgccagtggttgccagggtcacagccctgtggaagctgaagggtacccagtgagtcttctccagtcacattcaggaca**ggaaccttagtggggttccctgaggagaacagtctttaggaattctaacctcag**ccaagagaggctgggccagggtc**agtgtcaggtagaacctcacaggatttacgtctgacccttctcctgacact**aaagcatgcaa**atcagtatcagcactgat**ctggggccccttttgctcctagcccattctatttcttgttagttgtatttgttgttgattttccttttgctgttcctgatccctgtaaagtggagatgtggttcctgctgtaaaaattccagggctcaagccctttccctgcagctcaggtggggctcagactgtggctcctgcaaccacgt**gggagaggctgatgggactatcttctctccc**gttgctcagaaccctccagtgtgttgtgtggagactcacctgggaatgcaagtggccaatagttgtgaaaggtatgagcttgtttggtcaaagtggaatgtggatgtggaatttat**cctgtgttgtgcaaactaacacagg**ttcaccttcctcacctgtagtgttagaaagagtgtgaaagttgtaagaatcaaaacagagccacttgtgttaacaccctga

Uppercase: IGHVII-26-2

Lowercase: Flanking sequence[1000bp]

Red & Bold & Underline: Stem-loop [20]

Blue: Heptamer[41]

Green: Nonamer [4]

id-IGLVI-68[V_gene_segment]

ggaatcagaagcctacctagggctcccctctctcctctcaaaattgcacttggtgtcatattgccctagaatttatgcctggtgctggcaaatatttttaaatttattttttaatcttttccttttctgcaatcaa**atttcttgaggggctgagaagaagaaat**agg**cccagagagaagcaaggcagtctggg**tgcaggctctagacagcccagggagaagaaagagggagca**gctccctgtgtcctggagggagc**cctgtccccattgggcttcctgctctgatgtcccctagt**ggaggctcctcagggagtccagcctcc**cagaactgccctataaacctcacagtgagctaagcaagtatatccaggaaggagggtctgtggagatgtgagactaaggtgtgttatggcctaaacactgcccatctccatccttcttggggatgacaaagtgaacagtggaggagacaggatacatgggtgcatctggacctcaagcagtaggctgttctgtggtctgtccctaaatcttgtgctggagaccccatacgagagccactccctgaccctgggctggagccctctaccccctgctcacaagaccagca**gctgtttccttataggcccccagggagccacagaacagc**atcaaccaagaccatcctcttcatgtgtcagcttttcagggagggatcgaagtcatggcccaggaaagggcttggacagcc**caggggaggaggctggttggtatgaaggatgccctcctcaccctg**agctgctggagggagtacaagggacttggagagcccaggctcagttgatgatcctcaggatgtagagctatgtatgggtctccactgtggtctctcccccgccttctccttcactgcacaggtggctagggctcagtgtcagggctggtgatcaaaagaccaaggtttatgcatgtgtctctgtccactgacccacctctcaccctgtgtctgtcttaatgtccagCGTCCTGAGCCCAGCCTGTGCTGACCC**AGCCACCTGCAGTGCATGCATCTGTGGGGCAGGTGGGT**ACCAGCTCCTGCACTAGAAGCAGCAGCAATGTTGGGCACG**GGCATGTGCTCTGGTACAAACATGCC**TCCGAGAGACCCAGAGCTCCACATATATAAAAGCAGTCATTGACCCTTAGGGGCATAGGCCCAATGCTGAGGCTCCAGGTTGGAGAACATGGCCTCTCTGAGCATCTCTGGACTCCAGGCAGAGGAAAAGGCTGATTTTTATTCTCAGCTTGGGACACAAGCACCAAGGCTCAggcagtacttcaggccagtggggaagtgggagaaaaagctgctgcccatccagcaatggagcttctctgtgcagcccccacttcttgggcaagtcagctgattaacgttgcttttcatttgtttagtaaaaatttagattttaaaactcactccagtgaacgcagtgtggagaatgtgtttctgtcctctcaattctaccctcacacccaacccttggcagtaacaaattgtc**gtattgtctcatattaagagaaatccctcaatac**caagatcaggct**gtcctggaaaccaaatccataatgggtcaaaacaggac**cagggtcattgtgtctgactcaggatagtcgactcccaaaggtctgtttatccacctcattcactgagaagctactatgcaccagcctcagatccagggattcaaggtacatgatgaacaagagaggaaggggccctgtcctcatggagctgacactgagtggtggaacagataaggcaatcaagtaaaaatgtcaatgcgacaatatgatgtcatcacacagtgaggagagctcagttgggggtagaggagccatcagaaggttgggcaaggaagacacaggatctgactcaagtttatgaagaaaagtccagatgcctcaaagaagatgaacaaggagacagcaagcatgaaggccacggggccagttggggtcattggagagaaaggtcaggtctagcctgtgtttgtaagaggagggactgaagcctgggtgaggagagagggctcaggaatagccttggaaagtggagggagtggatgcgaaacccctgctgagg**gggcctccatgagcaatgaggctgggggaggccc**tgagctaatcaccatgatggaaactca**gacaacagcctcccagcctctttcaaagttgtc**acaggacgcttttctctccaaatggcaaggggggaatcccactatgagttttatttgagatgtctcttctccatgtcctgtcctgtccc

Uppercase: IGLVI-68

Lowercase: Flanking sequence[1000bp]

Red & Bold & Underline: Stem-loop [12]

Blue: Heptamer[39]

Green: Nonamer [0]

id-TRBV14-2[V_gene_segment]

attttaggaccaatgaggaggcccatatgactgaaaaattaaggaagggagtggtagactttatttccaagcggtaatggggcaggatggatcatttaagaccttgatggccttaagtgctttgttgttttgttttcttat**tccaaataaaactgggagctagtagatgatttgga**gatgagtgacatgatctagcatgttctagagggctgagtttgagagctagttcagaagtaggtgaaagcagggagatcagttaggaatctattgcagtaatccaggaaaaagatgatggtggccttgtcctgagggttagcaaagaaggtg**atgaaaaaatggatggattctgagtgtattttcat**gatagagccagttgaatttcctaatggattggctatagggtgggagagaaaaagaggatttgggaaccactctaaggttttcagctggaaggctgttggtggagtaaagttgaggataatctttcatgtttagagcatgctgtttgagagtggtggggggaaaatgttgattggagtgcattagagagataatgggaaaagagaatttggaggtaagtattgatcattagcaatagcaacaatagctaccgtagattaaaatgccccacagtctatggggatgtgatacatgagggaagacagggccagatgggatgaggtaggatccagacatcagactcaggagctacgagtggtatatataaggttaacacctagtcaaatgcataaacaggttgattttaattgatgtcagtttttctcc**attgccccctctagaggcaat**cttcttcagagaaccctggctaggtcttctatgtttc**atgtccgtagagggagctcctgagactgtggacat**tggctaatatgctgatgtcactggaggccacatcttacagggccaagagacagatttgctttcctttttctcatgcttgtaagctccttcat**ctggaaatgtgatttacctgggtcctgccATGGTTTCCAG**GCTTCTCAGTTTAGTGTCCCTTTGTCTCCTGGGAGCAAGTGAGTCTTCAGGTACTTAAATATCTGTGCTGTACCCTATCCCAGTCTATTCATGTCATGTATTCTGTTTTTGTCTCTTCCCACAGAGCACATAGAAGCTGGAGTTACTCAGTTCCCCAGCCACAGCGTAATAGAGAAGGGCCAGACTGTGACTCTGAGATGTGACCCAATTTCTGGACATGATAATCTTTATTGGTATCGACGTGTTATGGGAAA**AGAAATAAAATTTCT**GTTACATTTTGTGAAAGAGTCTAAACAGGATGAGTCCGGTATGCCCAACAATCGATTCTTAGCTGAAAGGACTGGAGGGACGTATTCTACTCTGAAGGTGCAGCCTGCAGAACTGGAGGATTCTGGAGTTTATTTCTGTGCCAGCAGCCAAGAcacagtgcttcacagtcgtgcccttgctgtgcaaaaccatagccttctcctctcaactcacagctgcccaaaag**gaaggctttccctgtgccttctcccccaagggaggggaga**taaagaaccagaattaactcatgaaatacaag**agtattccaagaagatttgggtgaaaatact**tgtggtttgggggatctctgaagttttttaaacagaactcaaactatgtgtttctcatatttatttttattttttaaagtgtatgttgccctggttataaagcagacttcttgcctatcttgctgctctgaaaggtttgtgactctgactgtgaggccacatctttatatcccttcttatagtaagagaaacaagtctttctgatgaaaataaaaatagtgaataaaaatgcatttactactggcttttaggatgctcaatgatgatgctagcaatggtggtagtggtgacgttggtaatggtggtgatgagtttaggttgcttttgtctagcatttataaatttccaa**ccactttaaaacaatctcattttagagtgg**aatagctaagtccattatcagaggttaatgctaattttctgttcacatagggtctgag**tttttaatgtaaaaa**a**tatagttagcactata**aatagatatctgtcagccgatacagaacattttttaagatcacttccagttgctgc**aataggaatatgtattcctatt**ttttttcttttttctttttctttttttgagacgaagtcttgctctgtcgcccaggctggag**tgcagtggcaaaatctcagctcactgca**acttctgcctcctgggtttaagcgattctcctgcctcagcctaccaagtagctgggattacaggtgcctgccaccatgcctggctatttgttgtatttttagtagagacgggatttcaccatgttggccaggctggtcttgaactcctgacctcaagtgatctgcctgcctcagcctcccaaagtgc

Uppercase: TRBV14-2

Lowercase: Flanking sequence[1000bp]

Red & Bold & Underline: Stem-loop [14]

Blue: Heptamer[33]

Green: Nonamer [6]

id-IGKV1D-12[V_gene_segment]

gtgtccagtttcatctggatccaagtgatttctccatgtacataggc**aactgcttgataagagattcagtgtctctttcctaaagcagtt**aacagggaggctggtgtctgggtcaggatgatttccccaatcactgataaaaagtaaaagaggaaagtgtcattg**atggtgcatggcagggacatgctccat**gcagtggtcaccctcagtaagagagatgaactttgggaaataatattgaatggcagaaaagaaggtagactatgaaggtgcccaaaacaagaataaggtgcagcccatttagtgtctgagtactatagagacctgtcgc**tcttgataattgtggatctgtgactgctgcatgcatcaaga**aaacacggtatcatctttgtgtatctatagtacatagtttgatcgcatactggtaagaacaatggcataacaccattacctgatacttacaaatgtatgtaccatcatgtcaataaattttatttttaattttttttacataggaacaatgttaaactcacagaaatgttgcatg**tatatgacaaatacccccttccctaaccggaatcatata**ag**agtcttttgaagact**tgagaattgtaccgtctaacattttactatgtgtttcctacaaacaacaatattctcctaaataatccccatacaccaa**tgaaatacattagtccatcaactcctgaggaatatttca**aatcgtcaaaaagaaaccaaaaaatgttactcacaacaaaatagttcccagtagaaacacattctctgcagacaaatttgtgctaccctggtcttacctgggacacctggggacactgaactggtgctgagttactgagatgagccagccctgcagctgtgcccagcctgccccatcctctgctcatttgcatattcccagaacacaacctcctgcctgaagacttcttaataggctggtcacacttcttgcaggagtcagacccactcaggacacagcATGGACATGATGGTCCCCGCTCAGCT**CCTGGGGCTCCTGCTGCTCTGGTTCCCAGG**TAAGAAAGGAGAACACTAGGAATT**TACTCAGCCCAGTGTGCTGAGTA**CTGCTTTACTATTCAGGGAACTTCTCTTACAGCATGATTAATTGTGTGGACATTTGTTTTTATGTTTCCAATCTCAGGTTC**CAGATGCGACATCCAGATGACCCAGTCTCCATCTTCCGTGTCTGCATCTG**TAGGAGACAGAGTCACCATCACTTGTCGGGCGAGTCAGGGTATTAGCAGCTGGTTAG**CCTGGTATCAGCAGAAACCAGG**GAAAGCCCCTAAGCTCCTGATCTATGCTGCATCCAGTTTGCAAAGTGGG**GTCCCATCAAGGTTCAGCGGCAGTGGATCTGGGAC**AGATTTCACTCTCACCATCAGCAGCCTGC**AGCCTGAAGATTTTGCAACTTACTATTGTCAACAGGCT**AACAGTTTCCCTtccacagtgttaccaacccgaacataaacccccagggaagcagatgtgtga**agctgggctgccccagct**gctcctcctgatgcctccattggctg**agagtgttgctcagatgcagccacactct**gatggtgttggtagaggggtacgtgaaatcgcctctgcaccctaattcttttctctttctcagccccaactgcacagacatagcaatgcatctcctgatttgataaatacagagatcatgacacttgaggagtctagttta**tggcttcagcttgaattcatataacacagaagaagcca**ctatagatattctaagcaggaatcgtcttaatacagagaattaaagtaaactactaaagtcaaaata**aaatgtagagatgaatctctaaattt**aagttttatttgcaaagaaatatttgccaggtggggcatacaggaaaactcagtggtcttcaagatattggaagaacgaagagaaaattag**cattttatgaaaaagggaaaatg**ttacctgtggctctttgagaaagttcattggctctaggaagggttgggagctggcaagctcagactggtaagcagtggtgggcaaaatgaatcctagaattatatcaagttatctcagaagttgtggacaaatttgatttcaggttacaataagccaaagcagtgaaggttgcagagaattttgtt**actgaaatgccagggattcagt**gaagaccctgcagcttaccccacagaaagccaatcactaagacaacaagcattgccaaggaacaggctttaatcaggtgctgcagccga**ggagatgcgataccattctcaaatgtatctccctgaccaactgaaattaggagtttatatagcaggga**agaaatgtgggaaaacaggaattagggaggggtaaggaagataatttggtcaacaggaagcagga**ggtcagtgaggcaatcataatgggtgaaggttctgatgtctcactgtcc**caattcagtga

Uppercase: IGKV1D-12

Lowercase: Flanking sequence[1000bp]

Red & Bold & Underline: Stem-loop [21]

Blue: Heptamer[21]

Green: Nonamer [5]

id-IGHV1-3-2[V_gene_segment]

ttcatccacaggaaccgacaagagattaggtgttctcccaaagcccctggaagctcctggactcactgtgagtgtggactgatccagtgcttccggagcgccacagaaggggctccctggtggtttcatagaatccttgcttggggtgtttctgcagagttcactggctttttcctacgaccaatttactattgcaagcgatggcgtcagcagcacatggtgtcacgtgtcactaaaggagcattctgagccaggacacagccacttcatactggggggaaaatgctctggg**agcccagacaggagcctctctgcagtgcaagggctgggct**gcagggggcgctcaaggcccacccagcacaggctccagccccagagcaggtgcacaggaggctgcggatggagtccccttaagatctgtgtcattcttctaaaaaaatctaaaataagtatttgacaaaaactgctgaagagtccataaatatcctattcaattgcaagaatttatcaatttacactggagatttctaaccctgctacatatcttaatagtaagcatctggaggtcaattaagctt**ttattttatataaataa**gtgcaacttttggagaaacacactcatcccccaaataacacattcatgta**ttaaagcctagaaatactttaa**attacctctgagctattcaagtgtgggttcccagtgaagtcctgttctagggaaaattg**ttctccagtgggagaa**gctctgtcaacatacagcttagggatatggcagggcacacatggcct**ctaaggggattatagcttgaacccttag**catcctcctgttgggtaatccatgtgtcatctctccactctttctcatgctgtgttaggtatgaaatagcatcgctcatgaatatgcaaataactgagatgactatagatatctttgtgccctgagagcatcacccaacaaccacatccctcctcagaagcccccagagcacaacgcctcaccATGGACTGGACCTGGAGGATCCTCTTTTTGG**TGGCAGCAGCCACAGGTAAGGGGCTGCCA**AATCCCAGTGAGGAGGAAGGGATCGAGGCCAGTCAAGGGGAATTCCATCCACTCCTGTGTCTTCTCTACAGGTGCCCACTCCCAGGTCCAGCTTGTGCAGTCTGGGGCTGAGGTGAAGAAGCCTGGGGCCTC**AGTGAAGGTTTCCTGCAAGGCTTCTGGATACACCTTCACT**AGCTATGCTATGCATTGGGTGCGCCAGGCCCCCGGACAAAGGCTTGAGTGGATGGGATGGATCAACGCTGGCAATGGTAACACAAAATATTCACAGAAGTTCCAGGGCAGAGTCACCATTACCAGGGACACATCCGCGAGCACAGCCTACATGGAGCTGAGCAGCCTGAGATCTGAAGACACGGCTGTGTATT**ACTGTGCGAGAGAcacagt**gtgaaaacccacatcctgagagtgtcagaaaccccaggggggaagcagctgtgctggcatggaggaaatgacaaagattattagattgaagactttctcagaaaatgatattaagtcattaaggaaaaggaacaatataaacatgtatttgagaaattttaattatttgagagatttttcatacaatatttattctgcaagcaaatttcagggattgaattaataaaactga**tacagaacttcctctgta**ggtatctgtgtaaacatcaatttctgaatcagtgttgtaaatattttggaacacacacacaaatcacattttatctctacttttatctctatt**attaaaaatgccaaaaaaactcattttgtgcatgtagcattttgaat**tcccaccatcaatgcatgatagttcttggttttccacattcatattgccatttatcattatgagaattgtgtgtttta**accattctaataggtgagtaatggt**atctaattttta**gttaaatgcacatttccgtaataaaaattcacatttaac**aattttcatataatttttgccgagatgcctcttctcatatttggttcatttttaactgcattgttttcttttgattagttgtaagtttacttgcatattgattataaaatcatttaacaaattaaaagaattcatttaacaaatatgt**gacttggaagtattttctccaagtc**tgcggctgtcttttactcccttatcagtatgtattgcagaaaagtgtgtgtgtgtgtgtgtgtgtgtttatacaaatttag**attttaaaaatgtaaaat**gttattcatccacagatcatgtctttggtattatatctgaaatctcattataaaatac**agtaatatcaattact**ttttccatgtctctaatctcaggcta**caatcatctcatgagtgtttaagcttcacctacttgattg**gaggactatcaacctaacatatttggaatacttctgta

Uppercase: IGHV1-3-2

Lowercase: Flanking sequence[1000bp]

Red & Bold & Underline: Stem-loop [16]

Blue: Heptamer[35]

Green: Nonamer [6]

id-IGHV7-34-1-2[V_gene_segment]

taatcaactgtaggaccctaaaggaaataagagtctcccctcaacccctgaagagtgtttgggttcaccgtgtgtccaatgattctgtgcctcttgag**ctccaggaaagggctccctggtgatgcatgagatcttttcttggag**tctctctgcagagttcactgggtttcctaaaggcaattcactatttcaaaagatggtgtgaggagcatgtggtgtccctaaaggagaattctgagccagggcacaaccactttata**ctgagctggatacactggtaggaatatactctgtcagctcag**atagaaacct**ccctgcatggttggggctgggctgcagggggcgctccggatacacccagcacaggctcccgccccagagcaggtgcacaggaggctgggg**agaggttcc**tcccagggcctggga**cttcctttaaaaatatctaaaataagtatttcacaaagactgctga**agtttgtataaatatctattcaattgtgagcatttatcaaact**ggatg**ttgtaatgagaaccacttttacaa**tggggatttcaaactctgctggaggtcaggaagagatcctttcttataaataaatgcaatttttggataaacacagtcattccctaaataacgcattcacatattatggtctagaaatgatgcaagttgaccctgagacagtcaaatgtggtttcaaagtgaggtgctgtccttgag**gagcttgttctccagtgggggaagctctgtcaacacagag**ttcaggg**atgtgtaggggacacat**ggcctctaacaggattacggcttgaaccctcagcttctacagttgtgtcacccatgtgtctgtttctcatactgggtcaggaattgggctattaaatagcatccttcatgaatatgcaattaactgaggtgactatagtatctccgttccctgagagcctcacccaacaaccacacccctcctctggagaagcccctagatcacagctcctcaccATGGACTGGACCTGAAGGATCCTCTTCTTGATGGCAGCAGCAACAGGTAAGGGGCTCCCC**AGTCTCAGGGCTGAGGAAGAAACCAGGCCAGTCATGTGAGACT**TCACCCACTCTTGTGTCCACTCCACAGGTGCCCACTCCCTGCAGCTGGTGCAGTCTGGGC**CTGAGGTGAAGAAGCCTGGGGCCTCAGTGAAGGTCTCCTATAAGTCTTCTGGTTACACCTTCAC**CATCTATGGTATGAATTGGGTATGATAGACCCCTGGACAGGGCTTTGAGTGGAT**GTGATGGATCATCAC**CTACACTGGGAACCCAACGTATACCCACGGCTTCACAGGATGGTTTGTCTTCTCCATGGACACGTCTGTCAGCACGG**CGTGTCTTCAGATCAGCAGCCTAAAGGCTGAGGACACG**GCCGAGTATT**ACTGTGCGAAGTAcacagt**gtggaaacccacatcccgagagtttcagaaagcctgaggaaggaggcagctgtgctgagctgaggcagtggtacagcagttttctgaacttccatagtatctcattttgcattgagttccgctttaatattagccaagaatatgggatagacgggtgctcctaagagatccttaacttgcccattttgatgggttttcccaaagacgtgagaagccacttttttcgcaaagcatcccaaagccatgccctgctccagaaacacgtgtatccatttcctggtctttgattaactgacaaactctcatcagcgcacctgggctaatttcacatcaggtagaaatatgtgctttaaagcaaggctaacgttgtaatagcaattcctgcttaataaccttcagcattgttgttgtgtgctccatc**aactaattacgttagtt**caaggttctcaatgggagtttctaataaatataagggatgtatagaagttcccctaattaaaacaattgtgaagacaacctcagtgttcaaccatatttcaacccttcaccacaaaggaactttcatctctcctggaagttgggttcattttcaaattagtttttttattttaatatctcaagattattgtatgtgactattttagcagaaagtgaattatgggaacttgaactaaccaactgaaaat**acattcagaactaattaaacaagatgccagaatgt**gattggctccaggcattttaaattcaacaggttatgtaaccaggctttaaatttgcacatcttcgtgttaccttcatgacacagtcaactcccattatgtaagaaatggtgagtgcattcccaagggtcttgcacagttataaaaatagacttgatgaggtgaggagttgtttaaattcccctctgaagaagcagcatcaacccaacaaaccactctcttccct**ctgtgactagagctctgtcacag**gccacatggacctaaatccttgatgg

Uppercase: IGHV7-34-1-2

Lowercase: Flanking sequence[1000bp]

Red & Bold & Underline: Stem-loop [20]

Blue: Heptamer[37]

Green: Nonamer [3]

id-IGHVIII-76-1-2[V_gene_segment]

ggggcgtgaagcaggggcttgctgagggttcttgggcccttggtcagagaatgaagtgtgcactgttggcttttctggttttgagacttctggacttagactgagccactaccagtttctctctcttcccaagcttttagaaagcctatcatggaactttcattttgt**aattttgtgagaaaatt**ctaataaactatatttcctatatacatatgttatatt**agttctatttctctagagaact**ctgggtaatacagctttgatacaaaaaaca**acattgttgtattcacaaaaatgt**atcctaagataagccaacacataaaagtgcataaagtatgattctatttctaggaattataaaaaatgaaaatgaatctaaagtaacaaaaaaagatcagtagttgcctaaggacaga**ttggaaaattatatttttccaaatgctacaaatgatactctcacctactatgagcatt**gctctctcctcaggaatttcacccaagaa**ccgactctatagtcgg**agacatgcaaacaggttcctgttcatgtacgaactcttcctaacactttcttttcatgtacttccttgacttatttgttgatgaaggttgataccataaaaataactaaaccaggctcacattagagaaaaagaagagagtgatggaaagattaagccggagcatccagtttcagtgattctttgaccctgccctccctgaagtctagaggcagagggatgagccctgctgagcagttcacacttgtccacagggagaaagaggtgtaaatgtgccccccatccatgctcataaaatggcc**catggacagcttatcccctgtccatgaaggggatgccacctgatgtcggtgcccttc**ctcagtgtccacaccttagggtctctgctaatctggggttctcttcttattctcagaggtagggtccctgtggatcaagcctagctgtggctgcttctcagcctgttccctctgtgtttgtaaGAGTCCTACAAGAAGTTCACTGGTGGAATCTGGAAGAGAAATGATACAGCCTAACTGTTC**TCAGAGCCTTTTTTACAAAGCCTCTGA**ATTTACCTTCATTGACTCTAGCATGAGCTCAGCCCAGCAGGCTTCAGGACAGGGGTTGGTGTGGGAGGCAAAAGCGAGGGATTCAAGTGGAATTTGTTAGTGGTACCCTCCATCAATACAAAGAAAAATCATAATCCTCAGGGACACCCTTGTCAGCACAGTCTCCCTCAAAATGACCAACCTGAGAGCCGAGGAGAAGGCCATGTATTACTGTGAGAGAcacaggagatatccgtgtggcaacctaacacaggggacacctgta**tttgtgtctgagcccagacacaaa**cctccctgcagggagacaggaggggaccgtgtgacagacactgctcagaaccaccaaggggagctcagaaccagcaggaggcagtggggacacaaaggaagactcaggagccaccagggggcgctcaga**accacctgggggtggt**gaggacaccagggggcgctgaggacacaaggggaggatcaggacaccagggtcggctcaggacgcgaggaggtgctcaggacaccagggtctgctcagaaccaccagggggcactcaggaaccaccaggagaggctcagccaccaccagggggcactcaggacatcaggagctgctcagaatcaccagacggcg**gctgggcgcggtggctcacgtctgtaatcccagc**actttgggaggccgaggcgggcggatcacgaggtcaggagatcgagaccatcccggctaaaacggtgaaaccccgtctctactaaaaatacaaaaaattagc**caggcgtagtggcgggcgcctg**tagtcccagctacttgggaggctgaggcaggagaatggcgtgaacccgg**gtggcggagcttgcagtgagccgagatcccgccac**tgcactccagcctggggtgacagagcgagactccgtctcaaaaaaaaaaaaaaaaaaaaaaagaatcaccagacggcgcacaggacatgaggtggcgcacaggacatgaggtggcactcaggacacaagggggagctcaga**accacctgggggtggt**gaggacaccaggggatgctgaggacacaaggggaggatcaggacaccagggtctgctcagcaaccaccagggggcgctcaggacatcaggagctgctcagaatcaccagacgtcgctcaggacatgaggtggcactcaggacacaagggggagctcagaaccaccagggggcgctcaggacatgaggtggcgc

Uppercase: IGHVIII-76-1-2

Lowercase: Flanking sequence[1000bp]

Red & Bold & Underline: Stem-loop [15]

Blue: Heptamer[28]

Green: Nonamer [6]

id-TRAV8-4[V_gene_segment]

tagatgggcctctgttaatgttctttttagtgtgagatataatttggcatgatgtatcagactgtaggagctttgacaattggaacatgattctttttaaacatgaagtacctaagttgttttttcctagagttacaaatatcgaagttattgacctgaaaccaaggggcctctttaaca**tttttgcttaacaaaaa**aaattttgctttatttgtttgctttgattaattaattagtgacatgtgatccatgctgattttaagtatgagactcaggtctagtttgt**ttatttttttaagaatcaaataa**ggaataatctctttcacccaccgtaatgctgacattcagaaacacagggccatatgaaactcagcccagactgaggcaactgatcacctcccttctccttgatgaatggttttacaaaataatatagagtgtctcaatggtattccaaagtctagaatgttgtcttctgtgactagttgctc**agtaaagtaaattagtcttctatttttacattact**ctcatttctcatgtatttaggattaaaaccatgtactctacagaaggtgacgtcatcacagatttggga**agaaagaaacactttct**gttggttttgataccacatttctcaaatgagaagcaaacagttcacttccttggatccgtggtttcgcctgtggccttcagggggcgacgttgcactaaggaggcatctgtgttcattgccgaccatcctcatccactgagcctcctccctgcagctggctgatgtagctcactggtgtctgtgtagatagggagctgtgatgagaacaagaggtcagaacacatccaggctccttaagagaaagcctttctttaaccatt**tttgaaacccttcaaaggcagagacttgtccagcctaacctgcct**gctgctcctagctcctgaggctc**agggcccttggcttctgtccgctctgctcagggccct**ccagcgtggccactgctcagccATGCTCCTGCTGCTCGTCCCAGTGCTCGAGGTGATTTTTACCCTGGGTTAGTAACATTCCATTACCTTTCTTCCTTTTGTAAATGTGACTGTGTTGTAATTTTTTCTGAATAGAGACATCTTTTCTTCATAGCAAAGTCTGTGCTGCCTTTACTTACACAGCATTGGTGTTGCAGGAGGAACCAGAGCCCAGTCGGTGACCCAGCTTGGCAGCCACGTCTCTGTCTCTGAAGGAGCCCTGGTTCTGCTGAGGTGCAACTACTCATCGTCTGTTCCACCATATCTCTTCTGGTATGTGCAATACCCCAACCAAGGACTCCAGCTTCTCCTGAAGTACACATCAGCGGCCACCCTGGTTAAAGGCATCAACGGTTTTGAGGCTGAATTTAAGAAGAGTGAAACCTCCTTCCACCTGACGAAACC**CTCAGCCCATATGAGCGACGCGGCTGAG**TACTTCTGTGCTGTGAGTGAcacagtgcttgagactg**caggagaggtgaacataaacctcctg**agatgctg**agactttctgtgactcaagaactcaaccttgaagtct**gttttataatattaaatagcatttccaagtttctgatgcttattgatgtatgctgtatctttttcatctatttatt**tttaaactatctgtgactttaaa**gtgtgtttcctgtaggtagcat**aaaagaccatctttt**ctttttatctactttgttaatctctgtttttgcattgtgaagtttagataattaaaatggaatataaatatcaatcaggttggatttaagtatacttcattattgtgtttcacccatgttcctttttttgttctttttttctgactccttttggattatttaaatacttttcaatttacattttaatttatttatgaactttttggctataactctgtactttttgagtagttgctctgaagatggcaatgtacatgcctaaatatccagggtcttatgtgagcaaatactgtacctccttatgtgaaatgtcaaaatcttacacaacgtctctttatctaccaacactaaagtttgttattgttgttcatatatattagtcacatacattgaaaacttcaccaggcctggcacagtggctcatgtttgtaa**tcccaacactttggga**ggcagagatgggaggattgcttgaactcaggg**gtttcagaccatcctgaaac**aaagtgagacccccatctctatgaaatataaaaaaaatagctgggtatgatggcactcatctgtagcctcaactactcggcaggttgaagcattatgatagcttgagcccaggagttcgaggctgcagttagctaccatcatgccactgtattccagcttaagcaatagagcgagatccttgtctca**atttttttttttaaaaat**cacaatatatagaaaacttcac**catataatgttataatttttgctttcaacaattatatg**cactttataaa

Uppercase: TRAV8-4

Lowercase: Flanking sequence[1000bp]

Red & Bold & Underline: Stem-loop [16]

Blue: Heptamer[29]

Green: Nonamer [1]

id-TRBV7-7[V_gene_segment]

cagatcagctgagtacccaggatatcacacatcccagcatcttctaacgaggaaaaatacttcacgtaccagcctccttcctgcaccacattgactaacc**ctggaaacttagcctttccag**aaaaaagctctgtcttctcttggcccagtctagcctggtaggtttaattcatgatttatgtgaccttttaatttactcaaaggagaacgtatgatttcagggtaaagaatattcattttatttttataagttggacctaaacagtaaactgagtacttttcttccatcagcgtcttaaattagacaacggtaaaacaaatctttgatgcta**gaacttcagatcctagtgtttttcacatggaagttc**ccagttacccagtctagggcatggaatagatgagcattaataagtcactgtatgcctgagatggttttctatgggcatcttttctgaattacggttaacaagggtgggaaatgatcccagaagaaaggcatgagggcagttcttgagggaacccatgtgatgggacagtctcattgggcacgtgtgactgggggaatggaggaagctggggcatgattgggatggcacaggggaccctgacttgcgggaaagacaatgagctcaaccctttgtgccctatgttaggggcgctgttgttg**catcctacaggacatgcccagcaggcaggggagcggctgtaggatg**agaagatgaactcagagatgcagtgtgaggt**ctccaggcacagacggctctggag**cacaaaccaatgagccatgaattgatgttgttaaaacggaggttataaatatttaaagtgtcacccaagtgtgttctaatagaaatgctgtgaccctgaggtcctgaggattgagagaggaagt**gatgtcactgtgggaactgccctgtggagacaaggacatc**cctcatcctccgctcctgctcacagtgacactgatctggtaaagcccccatcctggtctgacactgtcATGGGTACCAGTCTCCTATGCTGGGTGGTCCTGGGTTTCCTAGGGACAGGTGAGTCCTCAAAACACAAAGTAGTTTCATATTTTTTCTGTATGTAGGTGTGTGTGTGTATGCATGTGTGTCTGTGTGTGTGTGTGTGTGTGTGTGTGTGTGTGAGATGACTACAAATGTTTTCCTTATTCTGTTGCCAGATTCTGTTTCCACAGATCACACAGGTGCTGGAGTCTCCCAGTCTCCCAGGTACAAAGTCACAAAGAGGGGACAGGATGTAACTCTCAGGTGTGATCCAATTTCGAGTCATGCAACCCTTTATTGGTATCAACA**GGCCCTGGGGCAGGGCC**CAGAGTTTCTGACTTACTTCAATTATGAAGCTCAACCAGACAAATCAGGGCTGCCCAGTGATCGGTTCTCTGCAGAGAGGCCTGAGGGATCCATCTCCACTCTGACGATTCAGCGCACAGAGCAGCGGGACTCAGCCATGTATCG**CTGTGCCAGCAGCTTAGCcacagcatggcacag**tcgcctccttcctgttcacaaacctcatccttctctctctttgcagctcccagagatcattaacagaggcct**ctctttgctcctcacttttcctgggaaagag**gtagatttggacctcggttgtcctttgggtggaaagagaccacagattcattcctgaaacacagtgactgcaaatgtaagtggtgaaaacaatcacgtcccactgcactctaggagggctcggagccagctcactgctccaaacaaggagtgggtgtcttagccttggccttcagagcagacatgcatcttctataggtcttggaggctgctgtgttgcccacatacatgaggttgtcataggcaggaaacatgcctcttcttcatatgttggggcatctggaaggtctgaggctacatccccg**agaagatctttcttct**gaagcctcttctatccctgtcaccttggaagttttctgccacaaaata**tcaaacctctcttcctgtttga**agtaaaggtctttgcaaatttgtggtccttacttgataaatacaatcatggtaacaataacacttcatttcttctgcctactttaagccaagagtatcctttattttatttccatttgccat**tgctgctgtcctgatagacagaagca**tgcatcaccaccgctgccacttcaccttgatt**cactcaggaaatctgattttcagactctgagtg**ttttgattgttgtccaactcatttgatttgaaataattctcccaaggcctttaactcaaaaagtgttttatttataatattgaatctattcctttt**tattatattttttcataatatatagtgttatata**gcacttgttagaaatagaagtacaatgattatattgcaatagaatcttccatctgtctgtgggtgatgctgcagtttgtatctatgaaatgg**aatgcactggtcagagctgatgtgattatgcatt**atgggtttcggggagtcctcagc

Uppercase: TRBV7-7

Lowercase: Flanking sequence[1000bp]

Red & Bold & Underline: Stem-loop [15]

Blue: Heptamer[41]

Green: Nonamer [7]

id-IGHVIII-76-1[V_gene_segment]

ggggcgtgaagcaggggcttgctgagggttcttgggcccttggtcagagaatgaagtgtgcactgttggcttttctggttttgagacttctggacttagactgagccactaccagtttctctctcttcccaagcttttagaaagcctatcatggaactttcattttgt**aattttgtgagaaaatt**ctaataaactatatttcctatatacatatgttatatt**agttctatttctctagagaact**ctgggtaatacagctttgatacaaaaaaca**acattgttgtattcacaaaaatgt**atcctaagataagccaacacataaaagtgcataaagtatgattctatttctaggaattataaaaaatgaaaatgaatctaaagtaacaaaaaaagatcagtagttgcctaaggacaga**ttggaaaattatatttttccaaatgctacaaatgatactctcacctactatgagcatt**gctctctcctcaggaatttcacccaagaa**ccgactctatagtcgg**agacatgcaaacaggttcctgttcatgtacgaactcttcctaacactttcttttcatgtacttccttgacttatttgttgatgaaggttgataccataaaaataactaaaccaggctcacattagagaaaaagaagagagtgatggaaagattaagccggagcatccagtttcagtgattctttgaccctgccctccctgaagtctagaggcagagggatgagccctgctgagcagttcacacttgtccacagggagaaagaggtgtaaatgtgccccccatccatgctcataaaatggcc**catggacagcttatcccctgtccatgaaggggatgccacctgatgtcggtgcccttc**ctcagtgtccacaccttagggtctctgctaatctggggttctcttcttattctcagaggtagggtccctgtggatcaagcctagctgtggctgcttctcagcctgttccctctgtgtttgtaaGAGTCCTACAAGAAGTTCACTGGTGGAATCTGGAAGAGAAATGATACAGCCTAACTGTTC**TCAGAGCCTTTTTTACAAAGCCTCTGA**ATTTACCTTCATTGACTCTAGCATGAGCTCAGCCCAGCAGGCTTCAGGACAGGGGTTGGTGTGGGAGGCAAAAGCGAGGGATTCAAGTGGAATTTGTTAGTGGTACCCTCCATCAATACAAAGAAAAATCATAATCCTCAGGGACACCCTTGTCAGCACAGTCTCCCTCAAAATGACCAACCTGAGAGCCGAGGAGAAGGCCATGTATTACTGTGAGAGAcacaggagatatccgtgtggcaacctaacacaggggacacctgta**tttgtgtctgagcccagacacaaa**cctccctgcagggagacaggaggggaccgtgtgacagacactgctcagaaccaccaaggggagctcagaaccagcaggaggcagtggggacacaaaggaagactcaggagccaccagggggcgctcaga**accacctgggggtggt**gaggacaccagggggcgctgaggacacaaggggaggatcaggacaccagggtcggctcaggacgcgaggaggtgctcaggacaccagggtctgctcagaaccaccagggggcactcaggaaccaccaggagaggctcagccaccaccagggggcactcaggacatcaggagctgctcagaatcaccagacggcg**gctgggcgcggtggctcacgtctgtaatcccagc**actttgggaggccgaggcgggcggatcacgaggtcaggagatcgagaccatcccggctaaaacggtgaaaccccgtctctactaaaaatacaaaaaattagc**caggcgtagtggcgggcgcctg**tagtcccagctacttgggaggctgaggcaggagaatggcgtgaacccgg**gtggcggagcttgcagtgagccgagatcccgccac**tgcactccagcctggggtgacagagcgagactccgtctcaaaaaaaaaaaaaaaaaaaaaaagaatcaccagacggcgcacaggacatgaggtggcgcacaggacatgaggtggcactcaggacacaagggggagctcaga**accacctgggggtggt**gaggacaccaggggatgctgaggacacaaggggaggatcaggacaccagggtctgctcagcaaccaccagggggcgctcaggacatcaggagctgctcagaatcaccagacgtcgctcaggacatgaggtggcactcaggacacaagggggagctcagaaccaccagggggcgctcaggacatgaggtggcgc

Uppercase: IGHVIII-76-1

Lowercase: Flanking sequence[1000bp]

Red & Bold & Underline: Stem-loop [15]

Blue: Heptamer[28]

Green: Nonamer [6]

id-TRBV24-1[V_gene_segment]

cacttgggctcctgcatctgtccctctgtgtgttccctttcccctcaggggtttgag**cagttgcagcaactg**aaaaggtgagcaacactcctgtcgcacgtccttccaggggggtcagggaactcttcctttttcaccaggacgagcacagatctgaagcagcaaagagccagttgtttattacattgaacaatgtttcacactatcaccataactgttctgtacttgcaatctgctgcattaaatattagctctttcctcctaaagtcaggaataatttgtgttagaataaaatgaattctttaatgaacaactg**tattttcagctaattcaaaata**ggaatgaaggcgttaagtacttctttgaagcacatacccacattaaactaacatctttctacaataagccaatatttttaataaagctctgtgcccagaaaacattgaccagaaaagcaggagccactaaaaaataggagcaatttgaaacaaagttaggaatcctgtccttcttacaacaaaaaagtctcaaactttacagagaatgtattcaaattgcatagtttaccaagaactgagttagaataaatgattaaatgcagaaaaaaaggagaagagttataatgtcctatcattcaagtgagactagacttcctttattcg**aaattcaatttggttttgaattt**tctccagcctgttttatcaggaggatgatgtcaccttgaacaactgaaatggatatgccctgagaaaggggagagatacaactctctggtggttgccagtaaagcgccagaaaaacagaagagaagcaagtagaagaatttcagcatctgctaagaacatataatacataagccaacatggcatctttctttaagacccagtacct**tgggagggacaatgacatcacttcctgaatcctccca**gttttcta**tttccatgccctgcttccctcaacatccagagctggaaa**cacctccatcctgcctcttcatgccATGGCCTCCCTGCTCTTCTTCTGTGGGGCCTTTTATCTCCTGGGAACAGGTGAGTTTGGAACACAGATGGGGAAATCACTGCCTTAAATTTTCCAGGTTCTGAATTAAGCC**TATCCTCAGAGATTGCAGCACGAGGATA**TTTACCAGTTTCTGTCTTCAATTTCTGTCTTATTTCCCACAGGGTCCATGGATGCTGATGTTACCCAGACCCCAAGGAATAGGATCACAAAGACAGGAAAGAGGATTATGCTGGAATGTTCTCAGACTAAGGGTCATGATAGAATGTACTGGTATCGACAAGACCCAGGACTGGGCCTACAGTTGATCTATTA**CTCCTTTGATGTCAAAGATATAAACAAAGGAGAGATCTCTGATGGATACAGTGTCTCTCGACAGGCACAGGCTAAATTCTCCCTGTC**CCTAGAGTCTGCCATCCCCAACCAGACAGCTCTTTACTT**CTGTGCCACCAGTGATTTGcacag**tgcttcttggccacctgctctctacacagaaagacagacacatgggtgagttgtttgctctgaagggtacctggatgtgggttgtgggatgtggggtgtttagagctttcagtggtcttaggtagtgtgagctaagggccactttggatcaatgtccccaagccatgtgatgactctgaaagcacaggctacactgaatcaatctcccctgtctatttatttttcctagggagctagacgaatggtgacctttcaggagaaggaacttgagcatttgacaga**agctcatgattttcaacaagagcatttcttatgagct**gaatacaatgcagtttttatagtccttttgccattatttccacccactttgcaacatcccacactttgttctgttcccactgacagctcctttatcctgtctaacctttctctgccccagttcttccaagctcagctccaccatttagctctataattgattgcctttctacta**aaaacaaaataactagcttgtttt**ctaaaacttcactgtaagtaa**atttaattgtcattaaat**ctgactcagaagttaaagttcaagtaacagagactgtgatttacaaaagtaatttttcatctactaccccacactttgatctctgctgagattcattttaacctgcaggtg**cagatagacttgcattattatctg**gatgtact**gaatttcaaaaaataaaattc**cctgtccccagtatgaggtcaaagctaggagcttgga**agtgccccttgtttccaggcact**agaataattccctggttcccctgctgtggggattcttttaggagccagtgaggcccaggcacaaaaataattgatcaaggactgccagagcttgacgtaaaagtcttcctgagaatcaagttgatgtcttcctcatggatttagtcttctt**gtcccataagcctcaattatttcagcaaacgaatccatgggac**ttggtgactgttcagaaatga

Uppercase: TRBV24-1

Lowercase: Flanking sequence[1000bp]

Red & Bold & Underline: Stem-loop [17]

Blue: Heptamer[23]

Green: Nonamer [5]

id-IGHV5-51-2[V_gene_segment]

gtagctgatcccaacctcaagtaggatgttcttgatgagacagcttccctggtgcactggatatcagatcctgtatagtttcagaacttgtgtagtttgtgttgctttggctttttctctggatcttttaggtgctaatgaaatgagagtgcttcacaaaaaaatggggcagacaaaggggatagttaagggcctcctccaagcaatgtggatatcataaaataaatgttaaaaaatgaaagaactaaacaagaag**gtgctggggttgacacaaaggacttaaaacagcac**taccaggagttgggtgcagcaatgggcatcccttcaggtccatcagcaagtcagagacctaaacaaatattccctattcatgttgaattggcagagtgtaaaaatctagaaga**caaaatggccaggaaaaatgtgcgattttg**catgggtgaaggtctggcaagttaatcatcttgaggactgttgaggtcctgaggtggtcctgcccttgcttgatgcccaggccgtagtccaactcacacgaaattgccagggaatagacagtagtttttttagtagttcttgttatcaggcataagtgcatttgaattttctcttcatggcctttcctggcactatttctcattttttttaacacacatagtttcaactagatttatcaccttc**acagggtcacagagaagggtggaagaagggaggccctgt**atgggtctcgaagaaacatggaaaagagtggagagggacaatagcagggt**gtaaggaattattgagaccttac**tctgcccctcccaggaggctcaggccagcctttttctgcatttgaggttctgggttataaacgctgtagactcctcccttcagggcagggtgacaactatgcaaatgc**aagtgggggcctccccactt**aaacccagggctcccctcca**cagtgagtctccctcactg**cccagctgggatctcagggcttcattttctgtcctccaccatcATGGGGTCAACCGCCATCCTCGCCCTCCTCCTGGCTGTTCTCCAAGGTCAGTCCTGCCGAGGGCTTGAGGTCACAGAGGAGAACGGGTGGAAAGGAGCCCCTGATTCAAATTTTGTGTCTCCCCCACAGGAGTCTGTGCCGAGGTGCAGCTGGTGCAGTCTGGAGCAGAGGTGAAAAAGCCCGGGGAGTCTCTGAAGATCTCCTGTAAGGGTTCTGGATACAGCTTTACCAGCTACTGGATCGGCTGGGTGCGCCAGATGCCCGGGAAAGGCCTGGAGTGGATGGGGATCATCTATCC**TGGTGACTCTGATACCAGATACAGCCCGTCCTTCCAAGGCCAGGTCACCA**TCTCAGCCGACAAGTCCATCAGCACCGCCTACCTGCAGTGGAGCAGCCTGAAGGCCTCGGACACCGCCATGTATT**ACTGTGCGAGACAcacagt**gagagaaaccagccccgagcccgtctaaaaccctccacaccgcaggtgcagaatgagctgctagagactca**ctccccaggggcctctctattcatccggggag**gaaacactggctgtttgtgtcctcaggagcaagaaccagagaacaatgtgggagggttcccag**cccctaaggcaactgtatagggg**acctgaccatgggaggtggattctctgacggggctcttgtgtgttctacaaggttgttcatggtgtatattagatggttaacatcaaaaggctgcctaacaggcacctctccaatatgacagtattttaattagtgaaaattttacacagttcatcattgcttgcttgccttcctccctcctgtccactctcactcactccttcttttattttctacttaattttacaaaatcatttaacccctttttgaactattaataggttatctttgtttggtgattgttttcctttcaataatatgtactgaataattcatctttgtgccaattcataagtattctggtgtaataaagacttctttca**taaaaattggataaattaaaataaagataaattttta**aaaacatacgatctatcaaaactgaaccataaagaaataaaaactctgggttgggtgtgtttgctcattcctgtaatcccagcactttgggaggccatggccggtggatcacctgaggtcaggagttcgatatcggtctggcaaac**gtggagaaacgctgtctccac**taaaaatacaaaaattagctggacatggtggtgctcgcctgtagtcccagatacttgggagcctaaggccagagaagagattgaacccgggaggcagaggttgaaataagccgaaatctagccactgcattccagactgggcaacagagtgagactccatcccgaaaaaaaaaactgaacagacctatgagtaaagagattgagtcagtgatttttcaaacatctcaaatcaaagaaa

Uppercase: IGHV5-51-2

Lowercase: Flanking sequence[1000bp]

Red & Bold & Underline: Stem-loop [12]

Blue: Heptamer[29]

Green: Nonamer [6]

id-LOC124905154[V_gene_segment]

cacagaactatttgggagaacagaggacagtgatacacta**gggaagtaaaacacaccccttccc**cttgcattggtttcctgttgctgctgtaacaaattaccacaaccttactgctacaca**taatacaactgtatta**ccttacatttctggaggtcagaagtctcaatgaagtaaaatcaaggagtaatagggctctattcagtctaggcttcaagagagagaatccaatatcgagcattccatctttctgatgttcccacattcctagcagcatggcccctacctccatcactccagtttccccgtccgttgtcccaggtcctctctggctgttaccttcctccctccctattgtaaggacccttgtgattatgatggtctcacccagataat**tcaggatactctcctga**ccccaaaattctcaaccatgtctgccaagttatttttgacatattcataagtaatgatcatagattccagatattaggacaatgatgtctttagtgggtgtattatttattccacaaacaactctcatcatccacacaatggtcttcccctaaggtagaataaaaatatcacaaggcagatttatgaggcgatcga**cctagaaaaaacctgagactctagg**actgtctgatgtgtggatgtcaaatcctgggagattctgagtctctgctctatgtggactctatgttgtgtagccatttgtggaaggcttctg**tgattttgtgacctagagaaaatgaatctctgctaaaatca**aatctaagaaagattggcaa**agggaatttaaagatttcctaaatttttggaatttccct**aggcattaaaacaggagaagtggcaataattcaaaccaatgatgccctccaagaatgaggatttttccaatgcattaggttgggtcccctcagtgagaaggatgccaaagattcgcatgcaggcagtgtatttacaaagtgcgggaaacaagcaagtgagcaagggaggggaggaGGGAAAGGGAAAGTGAAAGGTG**CCTCAGAAGGAGCCACCTCTGAGG**ATGACGAGAGCTCAAGCCCACATAGAAACACAGGAAAAATGCCTCTGTTATTCCACCTGAGAGGTG**AGGGAGCTGGGGGATGTGTACACCTCCCT**TGTCATCACTGATTGACAGCCGTCCTAGGGGATGTTAATTCCAGGCC**ATGAGGTCTGCCTCAT**TTGCAGCCTGAGCTGCTTCCCCAGGTTCAGATAGAGCAGTGAAGGGGAGAAAGGGCCATAGAGAGTCGGCTGAAGTATAATGACTAGAATCCCCAAGGCGTAGTAACAATGACTGCAAAAATTATGCACAAAGAAAAAGCGCATTTGA**ATCCAGAGATGTATCTCTCTGAATCTGGAT**ATATGGATCCCGGCAGCCTGTTCAGTAGCCATTTCCCAGAAATCCAGTCCTCTGGAAAAGCAGCAGGAGGTTTGTGCACAGGCTGC**ACTACCTTGGTCTGGCCACTGGTAGT**CGTGCA**TGAGAACTACTCCCTGGAGTATTTCTCA**GTCCACTGACACTGACGTAAT**TGGCTCCACTTCCCCTGCTGTTGAGCCA**GGCCGACACACCCTGGACAAAGGCATCTGTGTGAAGTATTGAGGTTCAAATCAGTGCTTAAGATATGTTTGGATGCAAAATACTTTTTCATCTACATGGGCAGTGTCTTGGCAGAAGATGGAGATTCTCTCTAAATGGATGTGAGACAGGGTGGCTGGCATCTGGGTCAGGATGTTGCCCTGG**TGCATGGCAAGAACATGCA**TT**GGGCAGCAGCTGCCC**TCGCTAAGGAGAGAGGTTCACTGAC**CTGGCTTTTCCCCCCTCACTTGCTCTCCAGAAAGCCAG**ACTCTAGGGCAAATGCTCCTGAGACCCCAAGAACAGGCTGGTGGGGAGCGCAGCTCAGAGCATTACTCAGGGGATGTGGCCTTTGTCATCCTACTTTGAAACAATTGACTATTT**GAGCCTAGATTGATAGAGGGCTTCAAGTTGATTTTAATCCAGGCTC**CTATAGTCAGCGAGTGAAACAGAGTTTTTAGTTGAAATAATGAGACCTGGTATTACTAGTCAGC**TCTCCATGCTGGAGA**ACCATAAGAAATTATACCAAAGGCAGGAAAGGGGATAGAATATGGGGATCATCACACCAAGAATAAGGTGCAGCCCATTTAGCCCC**TGGGTCTTAAAGAGACCCATAGCTCTGGATAATTAATGGCAGATCTATG**CGTGACACAGTTATCATCTTTGTGCATCTTCAGAGA**ATTGTTTTTCCTTTTACTCCTAGGAACAAT**GTCTTAAGTTTGTTAG**TAAATTCTATTGAATTTA**TTAAAGATGCTTCTGATAAA**TTCTTTTTATATTCATTTCAAAAAAGAA**GAAATTTCACACTGACAGAGACATTGTTATTATAGCACTAAATACTTTTACACTCTTCAAATTCCTTTGAGACTAACTGAAATTTCTGACAGCCCCACATTCTACAACTTTATTGTAAATTTTCTGCCAAAAATGATGCTTCCCTATACACTCCTAATACAAGTA**TAAATATATTATTTA**ATCTAGTCTTAGGTTGATTTAAAATTTTGAAAATTCACTCCAAAAATATGTTCTGTAACCGTATGGCCACCAATGAGAAATGTATTCTTTCAAGGTAAATCTGTGCTGCCCTGGTCTGACCTGGGACTCTGGGGATACTGCGCCCCTGTGCTGAGTTACTGAAATGAGCCAGCCCTGCAGCTGTGCTCAGCCTGCCCCATCCCCTGCTGATTTCCCTGTTCCTAGAGCACAGCCCCCTGCCCTGAAGACTTTTTATAGGCTGGTCACACCCGGTGCAGGAGTCAGTCCCAGTCAGGACACAGCACAGACGTGA**GAGCCCCCACTCAGCTCCTGGGGCTC**CTGGTGCTCTGGCTGCCAGGTAAGGAAGGAGAACACTAGGATTATACTTGGTCAGTGTGCTCAGTACTGTCTGGAACTTCAGGGAAGTCCTCTGATAACATGATTAATTGCAAGAATATTTGTTTTTATGTTTCCAACTTCAGGTGCCAGATGTGACATCCAGATGACCCAGTCTCCATCCTCC**CTGTCTGCATCTGTAGGAGACAG**AGTCACCATCACTTGCCGGGCGAGTCAGGGCATTAGCAATAATTTAAATTGGTATCAGCAGAAACCAGGGAAAACTCCTAAGCTCCTGATCTATGCTGCATCCAGTCTGCAAAGTGGGATTCCCTCTCGGTTCAGTGACAGTGGATCTGGGACAGATTACACTCTCACCATCAGCAGCCTGCAGCCTGAAGATTTTGCAACTTAT**TACTGTCAACAGAGTGACAGTA**TCCCTCCCACAGTGTTACAAGTCATAACATAAACCCCAAGGAAGCAGatgcgtgaggctgggctgccccaatgctccttctggtgcctctatctgctgagggaagttctcaaactcagtcaggtttggaaagtcatagggagattttcct**agaggaggccagggaggttcctct**gaaccctaagcctctttcaccctcatccccagcagaaaagacatgacaatgactgtcctgactgaataaagaagagagataagtccagctgaggtgtctgtgttatgggataatcggaatttgtacagcaaaagagaagctattctcagtatttcaaggagcaa**ttattcaagttgaataa**attagagtctaaaccacagtctttctgaagcctatggagtgttattcatgaagcaggtactagacacaggggattct**caggtgctacttcagaagccagggttcacctg**cccctgatggtatgtgctgaacaccgtgtgatgatcctcagtcctg**tctgggaagcccaga**gctgggg**gtgctgatgctctcagctgcctgcagcac**atctccaggtgattctccagtcc**acacctaacagcatgtgttttacttcaggtgt**cagtgtacatgaatccaccactctgacttcccaatctcatg**acagtaattagttgtaacttactgt**gacctcatggagcaactctaaagaaaccagagagagaaaaggagttttggaaaatgtgctcccagaagtgatagtagtgatggggaattgacagctgacggggaagtaaggtgactctttccacaaggctcaac**attttgccagttacgagttgttgcaaaat**acatttgaatgtgctttcaattattaacagtttggggtcatagctgaaaaactttattaagtcacagataaaatgggaaaatcaggaa**attgtatgaaatatacaat**aacactgtgtgtgatggctcagg**tctgtaatcctgtgatagttaatacaga**ttgtcaacttgattacattgaaggatgtaagcattgctcctggg

Uppercase: LOC124905154

Lowercase: Flanking sequence[1000bp]

Red & Bold & Underline: Stem-loop [37]

Blue: Heptamer[46]

Green: Nonamer [5]

id-IGKV1-13-3[V_gene_segment]

ttaaatgaattttgtttgccttttctcttattaatct**tccttttgtcagtttattttcagcaaacatttagaggacaaatgga**gttttcttcctttctcccaatataagcaagttcccttaaaattcaggcggcttacaaagcagcaagaaggtttgtgcacgggctatggcactgtgattt**ggctcccctactcaggcatcagtaaaattttgtggagcc**ctaggctgcagcccactgatgctaatatagttggatccacttcccctgctactgagctaggctgggacagttttgg**gcacattagatatgtgc**gatat**aatgattgcaaatcatt**tc**cagttttgtctggatcaaactg**ctttctccatgtacataggcatcatctctgtggatctgtagtaaattgcttgatcttatagtggtaagaacaatggcataacaccattaccgaatac**tgacatgtatatatagcatcatgtca**ataaattttatttttgattttttttagaaa**ggaacaatgttaaaatcacagaaatgttcc**aagtatagggcaaagtaaccccttccctaaccgggatcatatgagagtcttttggagacctgataatcataccgtctaacattttattatatatttcctacaaacaagaatattctcctaaataatccccatacaccaa**tgaaatacattactccatcaactcctgaggaatatttca**aattgtcaaaaaaaacctaaaaaatgtctctcataataaaatagttcccagtagaaacacattctctggagacaaatttgtgctaccctggtcttacctgggacacctggggacactgaactggtgctgagttactgagatgagccagccctgcagctgtgcccagcctgcccatcctctgctcatttgcatattcccagaacacaacctcctgccctgaagacttcttaataggctggtcacacttcttgcaggagtcagacccactcaggacacagcATGGACATGAGGGTCCCCGCTCAGCT**CCTGGGGCTTCTGCTGCTCTGGCTCCCAGG**TAAGGAAGGAGAACACTAGCAGTT**TACTCAGCCCAGGGGGCTCAGTA**CAGCCTGGCTATTCAGGGAAATTCTCTTACTACATGATTAATTGTGTGGACCATTTGTGTTTATGCTTCCAATCTCAGGTGCCAGATGTGCCATCCAGTTGACCCAGTCTCCATCCTCC**CTGTCTGCATCTGTAGGAGACAG**AGTCACCATC**ACTTGCCGGGCAAGT**CAGGGCATTAGCAGTGCTTTAGCCTGATATCAGCAGAAACCAGGGAAAGCTCCTAAGCTCCTGATCTATGATGCCTCCAGTTTGGAAAGTGGG**GTCCCATCAAGGTTCAGCGGCAGTGGATCTGGGAC**AGATTTCACTCTCACCATCAGCAGCCTGCAGCCTGAAGATTTTGCAACTTATTACTGTCAACAGTTTAATAATTACCCTcacatagtgttacaaacccgaacataaacccc**cagggaagcagatgtgtgagactgggccgccccagctgcttctcctg**atgcctccat**tggctgagagtgttcctcagatgcagcca**cactctgatggtgttggtagaggaggatatgagatcacctctgcatcccaatttctttttcttttctcagccccagctgcacagacattacaatgcctctgctgatttaataaagatagagatcatgacacctgaagagtctagttt**atggctttggttagaattcatataacagagaagaagccat**tatagat**attctaagcaggaatagtcttaatagatagaat**tagagtctaaagtattgaagtct**aaataaaatgtacagataaatttaatgttttattt**gctaagaaatttttgccaaatggggcatacagg**aaaactcaatggtcttcaatatgttggaagagcaaagagtttt**attaaaagggaaattattacctattgttctttgagaaattttgttggctgtagtaagggttgggagctggcaagctcagactggtaagcagtggtggtcaaactgaatcctagaattatattaagttatctcagaagttgtgggtaaatt**tgctttcaggttacaataagccaaagca**gtgaagcttgcagagaattttgt**tactgaaatgccagggattcagta**tagatcctgcggctcaccacacagaaagccaatcactaa**gacaacaagtgttgtc**aaagaacaggctttaatcaggtgctgcagccgaggagacgggacaccattctcaaatgtatc**tccctgacaaaataaattaggcgtttatatagcaggga**agaaatgtggaaaacaggaattagagaggggtaaggaagataatttagtcaacaggaagctggaggtcagttaggcaatcat**aatgggtgaagggtctgatgtctcactgtccccatt**cagtgatatataactttcagctc

Uppercase: IGKV1-13-3

Lowercase: Flanking sequence[1000bp]

Red & Bold & Underline: Stem-loop [24]

Blue: Heptamer[17]

Green: Nonamer [4]

id-IGKV2-30-3[V_gene_segment]

cagtgataaagccatcagctaacttttcatcgtgcccaaaggaggctagaaaacagcggaatgacatatgtaaatgcagggaaacaaaacaaaacaaaacctaggattcaatatccagtgaaagtatgcattcaaaataaaagtaaagtaaaaac**atttcttgataaacaaaaatgaagagaaat**aattacttaaagacatgctgtataacaatttatctaggaaatt**cttcagattcacaaaaaatgacagcagacagtaacttgaatctgtaag**gaggaaggaaggcctcaaaagtactaagggaacaaaaaaagagctagaccaaaccaacatgtttacttccgtttatatgaggtttcgtatagggaaagccaatctatttggacagatgtcagaatgccagtttccttgattgggatgg**ggaaacagctgtttcc**tgaaaggtaggtacatgaggga**agaatctggagattct**gcag**tattctacagtttaatctcggtggagaata**tatgtaaaactttattcggttgcactttttaacatttctgtcttttactttgtgtgtgtttta**tttaaatttttaaaaaattgaaa**gggccaaatctgaactcttttaaacaaaaatgaacaaaaacataagaattagtaaatatttgtggaaacatggccttattaacaagaagtataaaatgtgcctgggagagtactatgaaacaagaaatctgttagggaagacagaaggaaatacttaaatttctccaacatagacagcatagattttatgcctattcgtttccctccaaacagagaagatatttaagtcattttgctcacaagagaggctcctaccctccccttggctctttccaccccactgcacccaccaggtgatttgcatattatcccttagtgaagactttccttgtgagtctgagataaaagctcagctctacccttgccttgactgatcaggactcctcagttcaccttctcacaATGAGGCTCCCTGCTCAGCT**CCTGGGGCTGCTAATGCTCTGGGTCCCAGG**TAAGGGTAGAAGGGAGATGAGGGAGGAGAATGGCATGGAACGGTGAGTTCTGGGGCCCCACTGCCTCTAACAACAGTGATCTCTGGGGGTCTCACTACACTCCTATGTGTGTTCCTTTCCTGTATTGG**ACATGCACATGTTGTCCTCCAGAGTGGGGCATGT**GATGATCAGATCTGTGAGAGTGAGGAAGATTCAAGCAGAAACAAGGA**TCTGTGCTCTGGGGAAGACTGACACAGA**AAGGGGATGGTGT**GGGGTCTTCTGGAGACCCC**TTTGAGCCTTGGATCCCTTGAGTTCCATTTTGAAACTG**TGTATTTTTGAAATATGAACAAATACA**TATATAGCCTGAAATAAACAACAAATC**AAAATTTATGAAAATTACACATAAACTTT**ATACATAACCTTGCTCTTCTTTCTATTTATTTCAGGATCCAGTGGGGATGTTGTGATGACTCAGTCTCCACTCTCCCTGCCCGTCACCCTTGGACAGCCGGCCTCCATCTCCTGCAGGTCTAGTCAAAGCCTCGTATACAGTGATGGAAACACCTACTTGAATTGGTTTCAGCAGAGGCCAGGCCAATCTCCAAGGCGCCTAATTTATAAGGTTTCTAACCGGGACTCTGGGGTCCCAGACAGATTCAGCGGCAGTGGGTCAGGCA**CTGATTTCACACTGAAAATCAG**CAGGGTGGAGGCTGAGGATGTTGGGGTTTATTACTGCATGCAAGGTACACACTGGCCTc**ccacagtggtacagccctgaacaaaaacctccctgtgg**agtggccc**agctgcccacatgtggtgcttgtctggggagcagct**cagcagggtctcagaatctgtgtaagaggaagatgctggagaaccagggaacaattcacat**ctgaggactctggactttgagagcccagccacacctcag**gcaccactcctttatgccctgccagttgccaccaccttgactgtcataagcagcaggagaatgaggggtccaagtgccctgtgagtaaacaagcaagatggaagggggaggagaatgaaagctcaccctaactctccctaccttgtgtccatttgt**taattaaatgtaatta**gcagagcagccaggccattgacacagattgtgac**tatccatgttggata**catctttgggtttagcagtttttggcatattgttcagaggacatttgataatatttaatgttggtattttgccagttttctaacttcctgcttccccttttctcccactcccaaaatagagtagagacagcattctataccagttatcctaagcggaagctggctgaggacagtcagt**aaaaatcttgattttggagtatcaaatagattttt**gtaaattcatagaagatacaagatc**ctaatgctaaaactgtttagttagccttag**ttacctcttaatgatagaaaaaaggagtacctgaaattccagaagttgttttcaaaaataaaaagc**atatactggaaaatgtagagtatat**aat**ttttcccatagaaaactggaaaa**gtacagaatgatgtgaacgtttttatttccaaatgttaagaatttaagattgggcagacattagggatgaaagcatgaaggaaacccaggagaggtcagcttcaattaaactacccttggcctttggtaggtagggatgtggatggtggtggagttggcagttcatgatgtggacccaggaggcctgatgtgtttcttgtgaagaatcaca

Uppercase: IGKV2-30-3

Lowercase: Flanking sequence[1000bp]

Red & Bold & Underline: Stem-loop [22]

Blue: Heptamer[27]

Green: Nonamer [11]

id-IGKV2-30[V_gene_segment]

cagtgataaagccatcagctaacttttcatcgtgcccaaaggaggctagaaaacagcggaatgacatatgtaaatgcagggaaacaaaacaaaacaaaacctaggattcaatatccagtgaaagtatgcattcaaaataaaagtaaagtaaaaac**atttcttgataaacaaaaatgaagagaaat**aattacttaaagacatgctgtataacaatttatctaggaaatt**cttcagattcacaaaaaatgacagcagacagtaacttgaatctgtaag**gaggaaggaaggcctcaaaagtactaagggaacaaaaaaagagctagaccaaaccaacatgtttacttccgtttatatgaggtttcgtatagggaaagccaatctatttggacagatgtcagaatgccagtttccttgattgggatgg**ggaaacagctgtttcc**tgaaaggtaggtacatgaggga**agaatctggagattct**gcag**tattctacagtttaatctcggtggagaata**tatgtaaaactttattcggttgcactttttaacatttctgtcttttactttgtgtgtgtttta**tttaaatttttaaaaaattgaaa**gggccaaatctgaactcttttaaacaaaaatgaacaaaaacataagaattagtaaatatttgtggaaacatggccttattaacaagaagtataaaatgtgcctgggagagtactatgaaacaagaaatctgttagggaagacagaaggaaatacttaaatttctccaacatagacagcatagattttatgcctattcgtttccctccaaacagagaagatatttaagtcattttgctcacaagagaggctcctaccctccccttggctctttccaccccactgcacccaccaggtgatttgcatattatcccttagtgaagactttccttgtgagtctgagataaaagctcagctctacccttgccttgactgatcaggactcctcagttcaccttctcacaATGAGGCTCCCTGCTCAGCT**CCTGGGGCTGCTAATGCTCTGGGTCCCAGG**TAAGGGTAGAAGGGAGATGAGGGAGGAGAATGGCATGGAACGGTGAGTTCTGGGGCCCCACTGCCTCTAACAACAGTGATCTCTGGGGGTCTCACTACACTCCTATGTGTGTTCCTTTCCTGTATTGG**ACATGCACATGTTGTCCTCCAGAGTGGGGCATGT**GATGATCAGATCTGTGAGAGTGAGGAAGATTCAAGCAGAAACAAGGA**TCTGTGCTCTGGGGAAGACTGACACAGA**AAGGGGATGGTGT**GGGGTCTTCTGGAGACCCC**TTTGAGCCTTGGATCCCTTGAGTTCCATTTTGAAACTG**TGTATTTTTGAAATATGAACAAATACA**TATATAGCCTGAAATAAACAACAAATC**AAAATTTATGAAAATTACACATAAACTTT**ATACATAACCTTGCTCTTCTTTCTATTTATTTCAGGATCCAGTGGGGATGTTGTGATGACTCAGTCTCCACTCTCCCTGCCCGTCACCCTTGGACAGCCGGCCTCCATCTCCTGCAGGTCTAGTCAAAGCCTCGTATACAGTGATGGAAACACCTACTTGAATTGGTTTCAGCAGAGGCCAGGCCAATCTCCAAGGCGCCTAATTTATAAGGTTTCTAACCGGGACTCTGGGGTCCCAGACAGATTCAGCGGCAGTGGGTCAGGCA**CTGATTTCACACTGAAAATCAG**CAGGGTGGAGGCTGAGGATGTTGGGGTTTATTACTGCATGCAAGGTACACACTGGCCTc**ccacagtggtacagccctgaacaaaaacctccctgtgg**agtggccc**agctgcccacatgtggtgcttgtctggggagcagct**cagcagggtctcagaatctgtgtaagaggaagatgctggagaaccagggaacaattcacat**ctgaggactctggactttgagagcccagccacacctcag**gcaccactcctttatgccctgccagttgccaccaccttgactgtcataagcagcaggagaatgaggggtccaagtgccctgtgagtaaacaagcaagatggaagggggaggagaatgaaagctcaccctaactctccctaccttgtgtccatttgt**taattaaatgtaatta**gcagagcagccaggccattgacacagattgtgac**tatccatgttggata**catctttgggtttagcagtttttggcatattgttcagaggacatttgataatatttaatgttggtattttgccagttttctaacttcctgcttccccttttctcccactcccaaaatagagtagagacagcattctataccagttatcctaagcggaagctggctgaggacagtcagt**aaaaatcttgattttggagtatcaaatagattttt**gtaaattcatagaagatacaagatc**ctaatgctaaaactgtttagttagccttag**ttacctcttaatgatagaaaaaaggagtacctgaaattccagaagttgttttcaaaaataaaaagc**atatactggaaaatgtagagtatat**aat**ttttcccatagaaaactggaaaa**gtacagaatgatgtgaacgtttttatttccaaatgttaagaatttaagattgggcagacattagggatgaaagcatgaaggaaacccaggagaggtcagcttcaattaaactacccttggcctttggtaggtagggatgtggatggtggtggagttggcagttcatgatgtggacccaggaggcctgatgtgtttcttgtgaagaatcaca

Uppercase: IGKV2-30

Lowercase: Flanking sequence[1000bp]

Red & Bold & Underline: Stem-loop [22]

Blue: Heptamer[27]

Green: Nonamer [11]

id-IGHVII-15-1-2[V_gene_segment]

taacctcaatgggattccctgactagaatggcttttagggatttcgattacagccaagagagaggctggaccaggttcag**tgtcatgtaggacctcacagattttatgtatgaca**tttctcctgacaataaagtatgcaaatgagtatcagcactgatctggtgcttcctgactttcatttttagtggttctcgcttttccatttgattttcctgctttctggataaaaggatgttgtccctgtggtctaaatcctggggctcaagccctttccctggagcttaggtggggctcaggctgtgactcct**gcagccattgggagaggctgc**tgagactttcttctctctcattgttcaggatactccactctgttttctggag**acgcgtctgagaacgcgt**gtggccattagtaatgagggcacaagcttctttggtcaaaatggggtgaggatgtggaattgatcctgtgctgtgtaaactgtcacagagtcaccttcttcaccagtagtgttagaagagcttgtaaaatttgtcggaatccaactggagtcccctgtgttcaaaccctgacaaatggagctggggaaggccatggatggaagc**ttctcatgcacatacccctgagaa**caagacctagctttataacactccaaaaaaccacaaattgaacaaaggcaaccacaacaataaaagaattacttatatccctgagaacaagagctagctttataacactcagaataaccacaacaacaaaataattacttatatccctaacaacaagagctagctttttaacactcggaaaggccacaatcttgcacaaggcaaccacaacaatgaaagaattacttctgtgaaaatatctgcacagcaactgcctatccaaccttacactgatgtcaaccttgttattgatgcttctagcc**caggatcgtcctcatcaaaatgtcattcagatcctg**cgta**tgaaaaaaattgtgtttttca**gtgcactcatGTTCTAGTAAAGCCCT**CAGAGACCCTCTCCTTCACCTGTGCTGTCTCTG**GATTTCCAATCATAACCAGTACTTCCTCCTGGAGCTGTATCTGCCAGCCCCCAGGGAAGAAGCTGAAGTGGGTCAG**GTGTGTAGGTCACGAGGAAACACAC**AGTGCAACCCGCTTCTCAAGAGTCCAGTCACCACCTCCAGATCCACATTCAGAAAACAGTTTTTCCTACAGCTGAGCTACCCGCACAATGAGTAAACCACCATGGATTTTTATACAAAAGAcacagaag**gggaggtcattgtgaggccagacacaaacctccc**tgcagggaagctcaggacaccagggggtgctcagacaccaagggctctcaggacacatcaaggcaggtgcaagaggg**gaaaaaaggtgctggaggtggggtttttc**atcactgtcatattttactctccactatatttat**tctaatgtatattattttattattaga**caatg**atatttatataaatat**atagccatacgtaggtgcaccaagttgtcctttccatcttatgtggaccttgtccattaggacgaagtccctgtatgtatttgagcacctcataagttatggttaattatgtaggatctctcactttttctgtgtcccttcctccctcctctctcacacacaaactgacacccacaaagagttctacaactttaattacctgatgcgttgaagaaaattt**actgaagtgcagctttttcagt**ttaactgctgttcatgttgatgtaggaataagaacattgtctttctcaactgtgtagttctctaagctgagtagcacctttatttataatacccagatactgaaagcaatccaaatattgatcagcaggttaagcagtaaacactttgttctaaattcgttcacttgaataatacccactgttcaaacaagtacggctggaaatgctcaacaagggtaaattcacaagta**cttatgacgagtaacataag**ccaaagaaatacaagtacatacatacagttccac**ttttataaattttataaaa**tgaaaactaatctaaagttacataatgaaaaccagtagttggccgtggacatggtatgagaagggaaggtgtagcaagtggaaattacaggacaacgagaggaaattttgaggataattttttctgtattgagaaaggttatggttatgtcattatttgtcaaattgtacacattaagtgaagattattactttctaatttcgacttcttaaaatactacaaatt

Uppercase: IGHVII-15-1-2

Lowercase: Flanking sequence[1000bp]

Red & Bold & Underline: Stem-loop [15]

Blue: Heptamer[38]

Green: Nonamer [6]

id-IGHV1-69D-2[V_gene_segment]

aatgggcacataactgtgggagatactatattcctatgaagaaggtattcagacttcagagataagtaatgtttcctacattgtgtttgtgacttggaagcagtggattgaagagtgtgataggtgcccagaccaagcagaacagaaatcagcatgtaaagatgatgatctatggatatgatctaaaaccatgtaaatacttcaaataattctatttaatgc**agtttgaaataaaacacaaact**tattcaaaataca**aattacttggtaatt**attttgggagcta**tgagttcaccaagaaactca**aattcctatttctatttcaacccctgattcctactgtcaatgggagggaagtctcagaaccaatcacacatcagacggcaaatctgtcaaccaagagtctttccactgaaggacctgggaggtcaggaccctcaggaaagtgctggggaccctgtcttgg**gagtgcccagcagatctcagaactctccatgggccctgctggacactc**atgtgg**ggtaaccagtgccaaccttttcagtgttacc**agtgagctctgagtgttcctaatgggaccaggatgggtctaggtgcctgctcaatgtcagagacagcaatggtcccacaaac**aacccaggtaatctttaggccaataaaatgtgggtt**cacagtgaggagtgcatcctggggttggggtttgttcttcagcgggaagagtgctgtgcacagaaagcttagaaatggggcaggggatgcgtttcctcaggcaggatttagggcttggtctctcagcatcccacacttgtacagctgatgtggcatctgtgttttctttctcatcctagatcaggctttgagctgtgaaataccctgcctcatgaatatgcaaataatctgaggtcttctgagataaatatagatatattggtgccctgagagcatcacataacaaccagattcctcctctaaagaagcccctgggagcacagctcatcaccATGGACTGGACCTGGAGGTTCCTCTTTGTGGTGGCAGCAGCTACAGGTAAGGGG**CTTCCTAGTCCTAAGGCTGAGGAAG**GGATCCTGGTTTAGTTAAAGAGGATTTTATTCACC**CCTGTGTCCTGTCCACAGG**TGTCCAGTCCCAGGTCCAGCTGGTGCAGTCTGGGGCTGAGGTGA**AGAAGCCTGGGTCCTCGGTGAAGGTCTCCTGCAAGGCTTCT**GGAGGCACCTTCAGCAGCTATGCTATCAGCTGGGTGCGACAGGCCCCTGGACAAGGGCTTGAGTGGATGGGAGGGATCATCCCTATCTTTGGTACAGCAAACTACGCACAGAAGTTCCAGGGCAGAGTCACGATTACCGCGGACGAATCCACGAGCACAGCCTACATGGAGCTGAGCAGCCTGAGATCTGAGGACACGGCCGTGTATT**ACTGTGCGAGAGAcacagt**gtgaaaacccacatcctgagagtgtcagaaaccctgagggagaaggcagctgtgccgggctgaggagatgacaggggttattaggtttaaggctgtttacaaaatgggttatatatttgagaaaaaaagaacagtagaaacaagtacatactctaattttaagataaatattccattcaagagtcgtaatataagccaaattcac**agagtggaaaaggccacactct**ataacgttgatacaaacattccatgaaggtgctactgtgaacaagttttcaaattggatgaat**acatgatttggagcaaggttatttgatcatgt**ggtgagac**taagaatgattctta**aaaagtgccaaaagtttccttcaaatgtttctgtcactccttatcataaagtttattttacagcagttttaggattacaaagaaattgcacaggaggcgtgagaattcccacgactccctgccctacacaggcacagcctcctccactacgaccatcctgcaccgcagtcacaaatcagttacaatggaggaatctccaaggacgcttggttctttctttttctggtgatctcctaatataacaagcctaagtatctcaagattccacggtttttcagtgttttctagaactgatattagtcagagggaaagtgggtaaggctattactatttgaactctttcttccaaaatccacaaaatatatattaatttagagctt**atcttacttctggtttacaatgctccttcccagagagtaagat**ttttttaag**cttttagaggccagttcatgtctctaaaag**accagaaagttctggggaatcccataatgaacatcctt**tcattggaattggagaccctggcaatga**gagactccatgtataatgccctagagttggattagatgctctgtaagctcttgggtggtgagtctgagtaagggggtttgtgcagcaaacgcaaacacatgcatgggacccaggtaggaac

Uppercase: IGHV1-69D-2

Lowercase: Flanking sequence[1000bp]

Red & Bold & Underline: Stem-loop [16]

Blue: Heptamer[48]

Green: Nonamer [5]

id-TRAV10[V_gene_segment]

tactttgcttatacaactataatataccaactttctttttattgtttcttgtataactatttccattctttcaacttctctgagctttaaatttcaaacttaaaataaatcctcatatatctacactctcaactgttcataagcttcttatttcactga**gaaaatagaaacaatcagaagagaattttc**aaaaacttcaacacttaccagtcacttgcatctatgctcaaatactctgtctttcctctcattagaatgtcatcccctgtgtgcagggactttgcttattctcagcacttatcatcatacctgaaaatgtaggcatttgaaaaataattgttgaattaaaaatgaataataaaaataactctgtcattaccagaacaaaactatcatttattcattccc**tcaataaaaatttattatatgcctatttattga**caggcct**tattctagcaactgacaatgtagcagagaata**gaacagataaaagtccctatcctaagggagcttatatttcagttgaa**ggctaagttcatgttagcc**actacaataagcctttacatgaatcatctcttcaatctttcaaaaattgt**taggtaagtgttatcataccta**ttttcagatgagaaagctaaggctcaactctcctatttaatatcacatagagccagaatgcagagcttcaaatatatgtctaaattactaatctactgt**gctgatatagacatgaagggctttatcctctatatcatgc**tgcaaataggaggaaaaaagcttgatcaaaagttctaagcaattgtagctctgaccccaagagggcagtatttgcattcgccccgcctgcaggtctgaatgtgtagtatgtgtaacatagcgaagcctggtctctgattggctagggataccctctgagaaaccctttggttagtgttaaaaaaaaagagaagatgttgaatacacaagtcaacttctgggagcagatctctgcagaataaaaATGAAAAAGCATCTGACGACCTTCTTGGTGATTTTGTGGCTTTATTTTTATAGTAAGTTAGTGGAGAGTTTTAGCAAAGTTAACATAGTAAGTCAAGGGAATATTTTCTTAGGATTTGGGGTGAAGATGTAGTCCACTGGGATAGGAGGAAGGG**AGAAAATGGGACCATTTTCT**GTTTCACTTGGGATTCATTGTGGGGAGTAAGGAAGGGATGCAAATTAGACAGTATTAATACTGGATGATGGATCTCTTGAATCTCACTTTTCTTTCTGAATAGGGGGGAATGGCAAAAACCAAGTGGAGCAGAGTCCTCAGTCCCTGATCATCCTGGAGGGAAAGAACTGCACTCTTCAATGCAATTATACAGTGAGCCCCTTCAGCAACTTAAGGTGGTATAAGCAAGATACGGGGAGAGGTCCTGTTTCCCTGACAATCAT**GACTTTCAGTGAGAACACAAAGTC**GAACGGAAGATATACAGCAACTCTGGA**TGCAGACACAAAGCAAAGCTCTCTGCA**CATCACAGCCTCCCAGCTCAGCGATTCAGCCTCCTACATCTGTGTGGTGAGCGcactgtgctccacaggcacttgaagccagtatgcaaacctgcacctggaggttatcaaggaggcataggagttagagtagaccgttattttttatgcagaatatgatttcactagtgaatatttatctctccgacatgatttgatttgag**cttgagcctaaagttctacttaggctcaag**tagaatacatgtcctcaagcgaggataaagacattctcactcaacacacaaagttaattgctacatctgggtgttcccaagggcaagacagt**ctgtcatgctgtgttactattgctgaagtagatgacag**tgttgtttggaagggagagactagcttggtttttgatctggacacctaccagcagaatgtcaactaagtgaaagagcaacaagtttatgtggctgatggaaatttttttttttttttttttttttttttgagatggagtctcactcagtcgcccaggctggag**tgcagtggcgtgatttcggctcactgca**agctccacctcccgggttcatgccattctcctgcttcagcctcctgagtagctgggactacaggcgcccaccaccacgcccggctaatttttttgtatttttagtagagatggggtttcaccgtgtta**gccaggatggtctcgatctcctgacctcgtgatccacccatcttggc**ctcccaaagtgctgggattacaggcgtgagccactgctcctggccggaaatatcttgagaagtttccttcccctgaaagacattcagttgtcatttcatggctattcaag**ttctcagaaagcatcctgtgagaa**taaatctgctcttcccatttagctatgctccagatcaaggcttttgtccaggtgctgcatcaactctttattat**cttccatcactgctcctttagaggacactggaaggccaggggatcctggc**aga**cacagtcactctcaggcagcagaacaactgtg**ccaggcttttcaaccatcagtgttctagaaa

Uppercase: TRAV10

Lowercase: Flanking sequence[1000bp]

Red & Bold & Underline: Stem-loop [17]

Blue: Heptamer[36]

Green: Nonamer [8]

id-IGKV2-36-2[V_gene_segment]

gaccccaagtctctagcc**acatccagttcttcacagaagctggatgt**atgtgccagggcaaggagaactgggtgggaggtgagt**ctgtgggctccacag**cttcacgtgtttatgaggtaggagttattaagaaattattttaggcagatagagaggaaaaaagggatccttgggagacttttcacagctgttgtctagc**ataaaagccctgggtcttagccccgggcaggcaaccttttat**atgcaaatgcaagccgttagaaactgggtccatccaacatgcacccaacatggtgatttccaccgttgtcctcttgcccttgccccaacatgcgcctggcagcatggttgcccctacatatccccacgtgtgtggaacatcatggcgccctgtatttgcatattaaaaggctagggtgggagggccagttctttcgagggctaccgtgaatgacatgtctggtcaaaccaataccctgagccctgtgcaaatcagacaccgcctcctccagacatcaaatatagctggctgatattacctgaaagagtggttcgtccctcggctttggagcccccctccctctgtctttgtacaggggagcttcatccttctttcttcccccttctttcttgcctattaaactccctgctccttaaaactattctaagtgtgtcagtgtcattttttctaatttgactcgagacaaagaacctggtgtcctctactcatcagagcag**tatcatttattccaacatgata**gagtcacatggtctatgcccagggaaaaga**attcatatttttgtcttatgaat**aatcaaggttcacctccagggaacaataaacagtactctgattaagatcttgaaaaaaaaaaagagttcccttctggctggt**aaataatgggtttattt**tagaaagtctactttcatgacatgaatcaaaacttgaaaaatgtaactataaatcaatatcataaaagaaatcatgaaagttgttcatAATGGAAAGCCAGCCTTTGTCTCCTGCAGGTCTAGTTAGAGCCTCCTGTATAATAATGGAAACACCTATTTGTATTGGTACCTGCAGAAGCCAGGCCAGTCTCCACAGCTCCTGATTTATGAGGTTTCCAACCAAGCCTCCGAATTCTCAGACAGGTTCAGGGGTAATGGGTCAGG**TACTGAGTTTACACTGAAAGTCAGTA**GGACGGAGACTAAGGATGTTGGAGTTTATTAGaaaacttgaacttccatcaatgataaatattccttttg**cctcaagcacatatttgagg**aattttccattgagtagatctaccgataaggtcacatttttctgtctgttttaatctgaatatgtatttatatgat**tctcagaaatgttttttgactatgaaattatatttgtgaga**ccttttttcaagttgttgtttaccatttgataattactgctgaaacgtcatctattaaaaaaaatcggtcatccgttctctaactctcattgaaaggaatttgcctttttaaacccctcagactctttttaagcttttcaattggccttatcttttttttacagattcaatgtattaagtc**aatttattatttatgataaatt**tatttatttattt**atttttactatcacaaaatttaaaaat**cctgtaagttgctatgtcaaacctgcctctagattgcaaacaaaccccacaaaaaagagagtaccattatcttagaaatactagaaaagcaagagtgatagtgtcccattgttaacagggaccccataaggctagaataatgtcccacatgttacctggggcacctgggaggagctgccactgtgctgaatggtgggaaaaagccatctgtgctctgagactggaagcctggccctattcccctatgtggatgtaggagataattggggttataaaggggtaattacggttaaatgaagtcataagggtggggctctgatccaataggattag**tgtccttataaggaca**gaaacc**agagaactcttctct**tttcctttatttctctaactcactgcttccctatggaaaggccatgagaagacatggtgagaaggtgactacctgtaaactaggaag**aaggttctaaccagaaagtgaacctt**g**ctggaacttgatcttggactttccag**gcttgtgaactatgagaataaatttctctggtttaatctacccagcccagggtattttggtgtagcagtgca

Uppercase: IGKV2-36-2

Lowercase: Flanking sequence[1000bp]

Red & Bold & Underline: Stem-loop [15]

Blue: Heptamer[19]

Green: Nonamer [5]

id-IGHV7-34-1[V_gene_segment]

taatcaactgtaggaccctaaaggaaataagagtctcccctcaacccctgaagagtgtttgggttcaccgtgtgtccaatgattctgtgcctcttgag**ctccaggaaagggctccctggtgatgcatgagatcttttcttggag**tctctctgcagagttcactgggtttcctaaaggcaattcactatttcaaaagatggtgtgaggagcatgtggtgtccctaaaggagaattctgagccagggcacaaccactttata**ctgagctggatacactggtaggaatatactctgtcagctcag**atagaaacct**ccctgcatggttggggctgggctgcagggggcgctccggatacacccagcacaggctcccgccccagagcaggtgcacaggaggctgggg**agaggttcc**tcccagggcctggga**cttcctttaaaaatatctaaaataagtatttcacaaagactgctga**agtttgtataaatatctattcaattgtgagcatttatcaaact**ggatg**ttgtaatgagaaccacttttacaa**tggggatttcaaactctgctggaggtcaggaagagatcctttcttataaataaatgcaatttttggataaacacagtcattccctaaataacgcattcacatattatggtctagaaatgatgcaagttgaccctgagacagtcaaatgtggtttcaaagtgaggtgctgtccttgag**gagcttgttctccagtgggggaagctctgtcaacacagag**ttcaggg**atgtgtaggggacacat**ggcctctaacaggattacggcttgaaccctcagcttctacagttgtgtcacccatgtgtctgtttctcatactgggtcaggaattgggctattaaatagcatccttcatgaatatgcaattaactgaggtgactatagtatctccgttccctgagagcctcacccaacaaccacacccctcctctggagaagcccctagatcacagctcctcaccATGGACTGGACCTGAAGGATCCTCTTCTTGATGGCAGCAGCAACAGGTAAGGGGCTCCCC**AGTCTCAGGGCTGAGGAAGAAACCAGGCCAGTCATGTGAGACT**TCACCCACTCTTGTGTCCACTCCACAGGTGCCCACTCCCTGCAGCTGGTGCAGTCTGGGC**CTGAGGTGAAGAAGCCTGGGGCCTCAGTGAAGGTCTCCTATAAGTCTTCTGGTTACACCTTCAC**CATCTATGGTATGAATTGGGTATGATAGACCCCTGGACAGGGCTTTGAGTGGAT**GTGATGGATCATCAC**CTACACTGGGAACCCAACGTATACCCACGGCTTCACAGGATGGTTTGTCTTCTCCATGGACACGTCTGTCAGCACGG**CGTGTCTTCAGATCAGCAGCCTAAAGGCTGAGGACACG**GCCGAGTATT**ACTGTGCGAAGTAcacagt**gtggaaacccacatcccgagagtttcagaaagcctgaggaaggaggcagctgtgctgagctgaggcagtggtacagcagttttctgaacttccatagtatctcattttgcattgagttccgctttaatattagccaagaatatgggatagacgggtgctcctaagagatccttaacttgcccattttgatgggttttcccaaagacgtgagaagccacttttttcgcaaagcatcccaaagccatgccctgctccagaaacacgtgtatccatttcctggtctttgattaactgacaaactctcatcagcgcacctgggctaatttcacatcaggtagaaatatgtgctttaaagcaaggctaacgttgtaatagcaattcctgcttaataaccttcagcattgttgttgtgtgctccatc**aactaattacgttagtt**caaggttctcaatgggagtttctaataaatataagggatgtatagaagttcccctaattaaaacaattgtgaagacaacctcagtgttcaaccatatttcaacccttcaccacaaaggaactttcatctctcctggaagttgggttcattttcaaattagtttttttattttaatatctcaagattattgtatgtgactattttagcagaaagtgaattatgggaacttgaactaaccaactgaaaat**acattcagaactaattaaacaagatgccagaatgt**gattggctccaggcattttaaattcaacaggttatgtaaccaggctttaaatttgcacatcttcgtgttaccttcatgacacagtcaactcccattatgtaagaaatggtgagtgcattcccaagggtcttgcacagttataaaaatagacttgatgaggtgaggagttgtttaaattcccctctgaagaagcagcatcaacccaacaaaccactctcttccct**ctgtgactagagctctgtcacag**gccacatggacctaaatccttgatgg

Uppercase: IGHV7-34-1

Lowercase: Flanking sequence[1000bp]

Red & Bold & Underline: Stem-loop [20]

Blue: Heptamer[37]

Green: Nonamer [3]

id-IGKV5-2-3[V_gene_segment]

gctcacaaattatgttgttggttctgttttccagtaatacatatcagataccagcaatgctgaagtttggaaactttctagaatgctttttgtttcattaggaagagtcctccacctccatgattccctttgccttctag**aagaataaaatgggaaaaacagaattttattatt**aaaaaaaaaaacccagaagagtaacatcactttccactcaaactgactacactaatctaatttctcattctttatctcttttacactaaggtatacctgtaattttcagagttc**cattttcaaagactgcttatagtttttaaaaatg**tgcatttatggtatagaaagaaaatcattttacgactaccaggttccactgaaaagtaattaatatactgtagtcaaaa**taaaataaaaatttta**atacatgaaaacaactaccgaaatgttatgaaattatagtttagtagaactaacaagtgcattaatgcaaaagaaaagtagggctcagtaatcaggg**aaccaagtgtgcattgtaaaagtgcagcctctctaacactgggtt**tcatcacaagtaacagaacaggatgcctgatgcagggaaaaaagaaaggcaattgt**tggcaatgtcaggaatattgcca**tggcgtctaccactggtgggttactgagcatcagaaccacagaagaggtaacactgaagcctgcagaataatgagtcttgttgcagaaatgcaaagaaaaagaaagatgataaatgcacacccagg**aggttacgttaattttaacct**tc**tgagtgtgcacagctgtgaagataaaactatgcacagctca**tgggttaggtttcacacagtctgccatttaaacactccctatgtcaccatcatgtaccatataatttgcatatttgtcctttatataaccctataatatgtttgtggccataaaatctgtgctgtcaaactgattaggaactgactaccacctgcaggtcagggccaaggttATGGGGTCCCAGGTTCACCTCCTCAGCTTCCTCCTCCTTTGGATCTCTGGTAAGAGAAACACTTCCTCTCCTCTGTGCCACCAAGTCCCCTGCATATCCACA**AAAATAATATATTTT**CATAAGGAATTGATTTTCCTCATTCTCTGCAAATATGATGCATTTGATTTATGTTTTTTACTTTGCTCCATAATCAGATACCAGGGCAGAAACGACACTCACGCAGTCTCCAGCATTCATGTCAGCGACTC**CAGGAGACAAAGTCAACATCTCCTG**CAAAGCCAGCCAAGACATTGATGATGATATGAACTGGTACCAACAGAAACCAGGAGAAGCTGCTATTTTCATTATTCAAGAAGCTACTACTCTCGTTCCTGGAATCCCACCTCGATTCAGTGGCAGCGGGTATGGAA**CAGATTTTACCCTCACAATTAATAACATAGAATCTG**AGGATGCTGCATATTACTTCTGTCTACAACATGATAATTTCCCTctcacagtgatacaccctgttacaaaaacctccaagttctctcagtgggattgcctcagctgctgctgaaaatgttcagaatgtggaactgggtggccctgtggtgtttataaggcttctccatgcccatgcaactcacttcctttttctttttgtttctgtagtttctgctgcccccatgctgtgcctgacacacttgatatggcaaaaaccaggacagcctttctcacgttgtccagctgtgccaacaatttcacttccaagcagaggtaaattctcatggccaactccctgtcttttgccacactcatcatttccacctccaagaaaaacaaaaccaaaaaccaaaatgcatctagcttagattccaaatagtagcattaaaggaaattgtccctccctataagcattctcaccctcatccacacagtcatcgtgactcttccctaacccactgagtgctcaccagtcctcttcagtcctccctccagtgatggggaaaagcgagtgatagttatgagagcaacatgt**aacctgggtgcagactcttactagtcaggtt**tgcctcagatttcaagtatttatctttcttttattttccattttttttcttaattatatggatcagtaagatgtctcatttcttttttttaagttacttgttcttttaacacaattcaattctataattccaaattt**aaagagataccctgtatgtcttt**ggcatgtgttatatctggacaagttttgggttatccccttcatgccactgctcaaatcttacctgaaaccccacatatttttccaaaaacctgaaaactgtgtagcggttgaggaaattaaatttggcaaatcacaatacttgaggaaacatgtatctttttatcttcactctctaatgttca**ggtgatgctgtttaattgagcatttgctatttagaatcacc**cctctgaaacttgtgggaaaacagcaccagcaaggcagtcttcatgttttgacc

Uppercase: IGKV5-2-3

Lowercase: Flanking sequence[1000bp]

Red & Bold & Underline: Stem-loop [13]

Blue: Heptamer[33]

Green: Nonamer [10]

id-IGHV1-24[V_gene_segment]

gtctggaccgtatcagtggatccggagctc**cagggaaggggctccctg**atggcttttatgattccttgcttgccgtgctgaggtctccctgtagattatgttgggttttctaaggcctattttctattgtaagaggtggtgggggaaacaatttctgtcactgaaagagcattctgagccagggcacagccccttcctactgagcttgagattctgggaggaaaaagttctctgacaggaggaaa**aagttcccagacaggaactt**ccctgcagggcaggag**ctgagcggcagggggcgctcag**ggagcacccagcacagga**cccagccctggggcaggtgcacaggaggctggg**gcagggttttctctcaggaattgaatgttc**cttatttcaaagcaataatgacctaacatttaaataag**aatttagcaagtactgatgtgcctgtaagtattttattgtatacgcagcctatacctaacccaataatttaatgcaaaccggatttaaaagg**agaaatgtctaggtctttcaaatgtatttct**agttaggaattgaggggtggttttattaattcaatgggtgttactgtccggagacacactcatcccagaagttagatgtgcagagggcaa**ggctcaggaaaagttcaggtagacagtgagcc**atatgaacagggaatcacattgaggaccatgtcctgtgagattctgggtttctgtgagaggagttctgtcttcatggacttctgggcatgtcagaggacaaatattaaacaaagttcagggcagggagctcagcatcccactgtggcgtggtccacatgtcacctatcttcttc**ctcagggtggcgtggcctgag**ctatgaaaaacctgcctcatgaatatgcaaatgcactgctgtctaccgaggtaaatacagatctttccttgcccagagaccatcacacaacagccacatccctcccctacagaagcccccagagcgcagcacctcaccATGGACTGCACCTGG**AGGATCCTCTTCTTGGTGGCAGCAGCTACAGGCAAGAGAATCCT**TAGTTCCAGGGCT**GATGAGGGGACTGGGTCCAGTTAAGTGGTGTCTCATC**CACTCCTCTGTCCTCTCCACAGGCACCCACGCCCAGGTCCAGCTGGTACAGTCTGGGG**CTGAGGTGAAGAAGCCTGGGGCCTCAGTGAAGGTCTCCTGCAAGGTTTCCGGATACACCCTCACTGA**ATTATCCATGCACTGGGTGCGACAGGCTCCTGGAAAAGGGCTTGAGTGGATGGGAGGTTTTGATCCTGAAGATGGTGAAACAATCTACGCACAGAAGTTCCAGGGCAGAGTCACCATGACCGAGGACACATCTACAGACACAGCCTACATGGAGCTGAGCAGCCTGAGATCTGAGGACACGGCCGTGTATT**ACTGTGCAACAGAcacagt**gtgaaaacccacatcctgagagcgtcagaaaccctgaggaatgaggcagctgtgctgaggctgaggagatgacag**gatttatgaagtttaaatctgtttagaaaatgggttgagtaattgagtaaaacag**caatggaaagatgtgtacactgtcattatgtaggaaatagtattttcaactgtcaccctataagtaacactcacagagtgggaaaggcagccatcaatcaagctgatgcaaacattcccatgggggctttgtggggacatacat**tttataattggatctataaa**tcattagaaacaggattgcttgatctcatggtaagactaaatat**aatttccaagaagtggccgaaatt**tcttccaaaatgtctttgccct**tccttttacactgaatttattttaaagga**gttttaggatcacaacaagtttgagtagaagaaacagagttcccatgcactcctgccccccgacgctcagccttctctgctatcaacaccctgcaccagtgtcataaatcggttacagaggatgaatctccacagacacattggttgtttccttttctggtggtccctggtataacaagcctcaactat**cttgaagcaccccaggttcaagtggattcttcaag**tgggttactgggaatgatgccaggtaga**aggaaagtgggtgggaccattcctctttgcactctttcct**caaggatccataaaatgtacatttacatggagctcctctttctcctggttttatatgcctcttccccaatggtaaggtttcccagtgtttacagcgggggtccccagtccccgtgctgtggatccgttaaggtgtgcagtgcgttaggaaccgggccacacagctggagatgagtgggggcgagccagcatcgctgcctgagctctgtctcctgtcggatcagctgtgcattagattctcctacaagccgaattcactttttttatatatatactttaattctagggta

Uppercase: IGHV1-24

Lowercase: Flanking sequence[1000bp]

Red & Bold & Underline: Stem-loop [20]

Blue: Heptamer[53]

Green: Nonamer [5]

id-IGHV1-3[V_gene_segment]

ttcatccacaggaaccgacaagagattaggtgttctcccaaagcccctggaagctcctggactcactgtgagtgtggactgatccagtgcttccggagcgccacagaaggggctccctggtggtttcatagaatccttgcttggggtgtttctgcagagttcactggctttttcctacgaccaatttactattgcaagcgatggcgtcagcagcacatggtgtcacgtgtcactaaaggagcattctgagccaggacacagccacttcatactggggggaaaatgctctggg**agcccagacaggagcctctctgcagtgcaagggctgggct**gcagggggcgctcaaggcccacccagcacaggctccagccccagagcaggtgcacaggaggctgcggatggagtccccttaagatctgtgtcattcttctaaaaaaatctaaaataagtatttgacaaaaactgctgaagagtccataaatatcctattcaattgcaagaatttatcaatttacactggagatttctaaccctgctacatatcttaatagtaagcatctggaggtcaattaagctt**ttattttatataaataa**gtgcaacttttggagaaacacactcatcccccaaataacacattcatgta**ttaaagcctagaaatactttaa**attacctctgagctattcaagtgtgggttcccagtgaagtcctgttctagggaaaattg**ttctccagtgggagaa**gctctgtcaacatacagcttagggatatggcagggcacacatggcct**ctaaggggattatagcttgaacccttag**catcctcctgttgggtaatccatgtgtcatctctccactctttctcatgctgtgttaggtatgaaatagcatcgctcatgaatatgcaaataactgagatgactatagatatctttgtgccctgagagcatcacccaacaaccacatccctcctcagaagcccccagagcacaacgcctcaccATGGACTGGACCTGGAGGATCCTCTTTTTGG**TGGCAGCAGCCACAGGTAAGGGGCTGCCA**AATCCCAGTGAGGAGGAAGGGATCGAGGCCAGTCAAGGGGAATTCCATCCACTCCTGTGTCTTCTCTACAGGTGCCCACTCCCAGGTCCAGCTTGTGCAGTCTGGGGCTGAGGTGAAGAAGCCTGGGGCCTC**AGTGAAGGTTTCCTGCAAGGCTTCTGGATACACCTTCACT**AGCTATGCTATGCATTGGGTGCGCCAGGCCCCCGGACAAAGGCTTGAGTGGATGGGATGGATCAACGCTGGCAATGGTAACACAAAATATTCACAGAAGTTCCAGGGCAGAGTCACCATTACCAGGGACACATCCGCGAGCACAGCCTACATGGAGCTGAGCAGCCTGAGATCTGAAGACACGGCTGTGTATT**ACTGTGCGAGAGAcacagt**gtgaaaacccacatcctgagagtgtcagaaaccccaggggggaagcagctgtgctggcatggaggaaatgacaaagattattagattgaagactttctcagaaaatgatattaagtcattaaggaaaaggaacaatataaacatgtatttgagaaattttaattatttgagagatttttcatacaatatttattctgcaagcaaatttcagggattgaattaataaaactga**tacagaacttcctctgta**ggtatctgtgtaaacatcaatttctgaatcagtgttgtaaatattttggaacacacacacaaatcacattttatctctacttttatctctatt**attaaaaatgccaaaaaaactcattttgtgcatgtagcattttgaat**tcccaccatcaatgcatgatagttcttggttttccacattcatattgccatttatcattatgagaattgtgtgtttta**accattctaataggtgagtaatggt**atctaattttta**gttaaatgcacatttccgtaataaaaattcacatttaac**aattttcatataatttttgccgagatgcctcttctcatatttggttcatttttaactgcattgttttcttttgattagttgtaagtttacttgcatattgattataaaatcatttaacaaattaaaagaattcatttaacaaatatgt**gacttggaagtattttctccaagtc**tgcggctgtcttttactcccttatcagtatgtattgcagaaaagtgtgtgtgtgtgtgtgtgtgtgtttatacaaatttag**attttaaaaatgtaaaat**gttattcatccacagatcatgtctttggtattatatctgaaatctcattataaaatac**agtaatatcaattact**ttttccatgtctctaatctcaggcta**caatcatctcatgagtgtttaagcttcacctacttgattg**gaggactatcaacctaacatatttggaatacttctgta

Uppercase: IGHV1-3

Lowercase: Flanking sequence[1000bp]

Red & Bold & Underline: Stem-loop [16]

Blue: Heptamer[35]

Green: Nonamer [6]

id-IGKV1D-35[V_gene_segment]

ggagctgcacctgagacactaggaacatgctggtagggagcacagcttggagcattacccaggggatatgatctttgtcttcctactttggacacatttgattataagagtctggattgatacagggtttcaagtttatcctaatcaaggctccta**tagttcaatgaagaaacagagaatcttgttgaacta**at**cagatctggcatcactagccagccctttatgcgatgatctg**taagaaatattacacagtgacagaaaaggtggtagaatattctc**cccaaggcaagaataaggtgctatccatttaatccttggg**tattaaagtgaccgatagctctggaaaatggagggtctgtgcgtggtacatgcatgagaaatcatggtattatctttgtgaaactgcagtcaattccttgtccttataccagtaacaacaatggcttaaagttatgacaaact**atttttgaatattgttaaaaat**gcttctgataaagtcttattttcattttaaaaaagacaattcctcactgacag**aaatattgttaacacaacacaaaatattt**ttacactcaccaaatacatttgccagtaacttgaagacccaagagctctactttctaagattttattatgcattccctgcaaacaaggatagtctcctacatactgccaatacaactataaaatgcattactcaatgtagtccttaggttcgtttaaaa**ttttggaaattgtcccaaaa**atacgttttgtaatagtatggccaccaaagagaaatgtattatctgcaggtaagttgatgctaccctgctctgatctgggaccctggggacactgcccctgtgctgagttactgagatgagccagccttgcagctgtgcccagccttccccatcctctgctgatttgcatgtcccagagcacagctcccttcctgaagacttattaattggctggtcacaccccgtgcaggagtcagtttttgtctggacacagcATGGACATGAAGACCCTCATTCAGCTCCCGGGACTTCTGCCATCCTGGTTTCCAGTTAAGAAAAGAAATTACTAGCCTCTTACTCAGCCAGTGTGCTAGGTACTGCCTGGCCATTCAGGGAAGTCTTCTTATAACATGATTAATTGTGTGGATATTTGTTTTTCTGTTTGCAATTTCAGGGACCAGAAGTGACATCCAGTCACCCAGTCTCCATCCACA**CTGTCTGCATCTGTAAGGAGACAG**CGTCACCATCACTTGCCGGGCGAGTCAGAGCATTGGTAGTAATTTAGACTGGTATCAGCAGAAACCAGGGA**AGGCCCCTAAGGGCCT**GATCTATGCTGCATCCAGTTTGCAATCTGGGGCTCCTTCGCGGTTCGGTGGCAGTGGATCTGGGACAGATTTTACTCTCACCATCAGAATCCTGCAGCCTAAAGATGTTGCAACTTATTACTGTCAACAGTATAAAAATTACCCTatcacagtgttacaaaccataacaaaccccccccaggaaagcagacatgtgacgctg**ggctgccccacctgctcttctttgtgcagcc**atctggtgacaacacttctcagactcagcctgagttttgagggttattgggagattacggaagagggg**gccatgaaggtgtctctccaccctaagtctctttcctctcatggc**aatgtctcttcgaatttagtaaagaaaggagattagtctacctgaggagtctgtgttatggggtaattgaaattcttacagtaaaatggaagccacacattccaagcaggaattgttttaactgagtgaattagaatctaaactatagccttcgaaaacctggcatatgttattcatgaagcagatactagagatggggttctctgatgcaaaaatataaatatttattgatctg**tttttaaagatactaagataaaacttaaaaa**tgttgtcactgtatcaaatatattgcatcatatatctataaaaacagtaaaatttgtgtcatttatggaaaatgcacatatttgagtgtatataacatacatattaatccatata**tatttatttgataacttgttagctataaata**c**tatgtacacatattcatgtgtagata**agattatttaaatatgt**ttatatatatatatatatataa**gcaacacatgcatacgtatgaaattattgtataaattaaatctgttaagataattccagttactcccatgaaaaaaactgatgtccaggactatatatgttacctgctctggctacagggatttcctttgtaatctttatttctaacactcaccaaatgggtatgtgcacctgaaa**atatcttttagtggaggggtaaaggatat**gaagaaaggta**attttgcctaatgttaaaattaggcaaaat**gttaaaatttcacaatgatgcctgctgggcagagttctaaattgggtgctggaccaaagccacacccctgccatc

Uppercase: IGKV1D-35

Lowercase: Flanking sequence[1000bp]

Red & Bold & Underline: Stem-loop [16]

Blue: Heptamer[30]

Green: Nonamer [6]

id-IGHVII-20-1[V_gene_segment]

gtctctaacccacagggatgaagatgccgcggtgtgtctgtgtgtctgtaa**tgcagctttaaagaggagtgaagcactgacataggctgca**acttggatgagccttgaaaacattgggtgactgaactgagggagacacggaagtctacatccctaattgtcccattgacacgaagcacgcagagcaggaaaatcccagatcacaggaggtcaggggaag**ggaaacacagagtgaccgcttaatggaggtgaggtttcc**ttttctgtgatgaaaatgttcccaaaccaaatagtggaaatggttgcacagcattgtgaatgtgtaaagtatcactgacccatac**acttaccaagggctgaaatggtaagt**tttatgttatgtgtatctcaacacaata**aaaatgagtgccacatttt**gtttagcatatcaatgacaataacagttcaggtaaatgctggattcctcatcacaaccagtgcttcctgatggagctggattcaggggagggact**ggagtgggttgggtgcacaggtcatgaaggcagcaaaaattccaacccactcc**tcaagagtccagtcaccacctccagatctatgtccaaaaaacagctcttcctatggctgagctacatgag**caacaagcaaataaccatgttttttgttgttg**ttgtttagttttgttttgctttgtttttagacagagtctcgctgtgtcacccaggctg**gagtgcagtggtgcgatctcagctcactgcaatc**tctgcctcctgagttcaagtgattctcctgc**ctcagcctccctagtagctgag**actacaggcactcactaccacgcccagctaatatttgtatttttagtagagacagggtttcactatgttggccaggctgttcttgaactcttgacttcgtgatccacccgcctcagcctcccaaagtgccgggattacaggcgtgagccaccgcgctcagccacaaccatgtatttttaagcaaaagaCACAGTGAGGGGACATCAGTGTGAGCCCAGACACAAACCtccctatgcgggttcacaggacagcatggggtgctgaggacagaggtgggcactcaggaaccagcagggaaacccagggggcgcttggcacttcatgggggctcaggaccattgtgggctcagtggtcaggcaggctcaaggctcagcct**cagggcaggtgcagcaggcggggaaaggccctg**gagactgggtttagtgtcaccttctcattgcgccactggacaccctccactacatctattctaatgtgtatgagtacttatgagtttagaaaataacattgatattataaatctatagccatatgtgggtgcatcaagttaccctcttcaacctatgtggaccctgttcatcaggaataagtccctgcatttgaggaccttataaatcaataattatgtagaatcactttctttttcagtctcatttcctccctctttctctctctcacacacactgacaaacacgctggttgctataactttaattacctgatgtgttaaaaaaaattgattcatttggaacttaaccagtttagctgttgttcacgctgttatgtaaaaaagatccctaaatgacttctacctggtaagaggagggaaggcgttggaaggtgaaattgcag**gagaaaaagaaaatcgtgaaacggattgatttgttttctc**tatgaatggtgattaggtcaatatttttcaaacggtgaacatattgtgtgaaga**ttatttgtcaatattatctcattaaaactattacaaataa**aag**tgtataatttggtaaaaaaaatgaagagagaaataaaaataataca**agaaaagtcatgaactcctgactccacacgtgagttaaccctca**gtttttctatgttacttagaaaaac**act**aagatacagccaaagaatatcatgatatctt**tataagaa**gagggttttgtaaccctc**actcgaaatttctagctctgtcctagagttaatttagggagcagtcagatccagctgtgaga

Uppercase: IGHVII-20-1

Lowercase: Flanking sequence[1000bp]

Red & Bold & Underline: Stem-loop [15]

Blue: Heptamer[33]

Green: Nonamer [4]

id-TRGV11[V_gene_segment]

ccctgtgtccatgtgttctcatcattaagctcccacttataaatgcggac**atgcagtgtttggctttctgttcctgcat**tagtttgctgaggataatggctttcagctcatccgttttcctgt**gaaggacatgatctccttc**cattttatggctgcatagtactcc**atggtatatatgtaccat**aatttctttatccaatctatcattgatgggcatttgggttaataccatgtctttgctaccgt**gaatagtgctgcaatgaacaataccaagctattc**tatttccaaccataaatatgttgagaaaaattcaaaggaaaagtaaatgatgggtgccttttgttgtcaatttttatatctcatttt**actcagattttccttaacggatgactgctactcacctgagt**tcttatgtttaccagctctgtgtttttg**aatgagtcactcaaactctgtttgccgttgtcctcatt**tgacacatggagataatcataaaacttactgcacaggactgttgagaggatcaagtaaaataatataagaaaacatcta**gacaagtagcaagttgtaaatggatattagatgcttgtc**aagtttcttactgcag**ttctgaattaacagctaattacagaa**agatagtcattgggacagaacactgcaggtttatcttcacattttcaaag**ccatatttctgcaatatgg**tactgctggggaatgggagagtgggcaagagaacatccctgccccacccctagaggtacacattcatactaggtatttctttccagatagttaacccacttacccgatctgcccagcactcatgggatgcaggaaaccctggaactgctcataactgtagcagctcatgaagaaccctaaggaaccacctcatctagtacttctgagagacctcaacagccccgccttctgccactgacttgagacctccagtgaacagggttaccttctccctggggaaccagactggcagaaagcATGCCACTGGTAGTAGCTGTTATCTTCTTCTCCCTCTGGGTTTGTGAGTTGCCCCCTAATCATTTCAGTC**TACAGAAGAGATTGAAGTCATTCTGTA**TTATTACCTCTTCACGTATCGTTGGCTTATGGGAATTTCTCTTCACTTGCAATTGCACTTGGGCAGTTGGAACAACCTGAAATATCTATTTCCAGACCAGCAAATAAGAGTGCCCACATAT**CTTGGAAGGCATCCATCCAAG**GCTTTAGCAGTAAAATCATACACTGGTACTGGCAGAAACCAAACAAAGGCTTAGAATATTTATTACATGTCTTCTTGA**CAATCTCTGCTCAAGATTG**CTCAGGTGGGAAGACTAAGAAACTTGAGGTAAGTAAAAATGCTCACACTTCCACTTCCACTTTGAAAATAAAGTTCTTAGAGAAAGAAGATGAGGTGGTGTACCACT**GTGCCTGCTGGATTAGGCAC**cacagtgttagagttgtcaagataaccta**cacagaaactatctccgagtctgtg**cctgt**ccacatccttctccatgtgggcaaccacagcggtttgctcagctgggtgccca**gccggagcctgccctc**cctgcagctcttctggaatgttgcagg**aagacatctta**gattctgctcccttggtgtgcagaatc**acattggtgattcatggtgtccaaacgggaaaagttttcttgttccctctcagggcgtgcgataaggatgtggctcccttcttggtgcccctctcctcaaacctctagggggagcatgcacacgtgcaggt**tgtggggctccgaccccaca**gcagtgtctaggggtgaatgcttataactcctg**aggccccagtgggcct**gtgttacaaggtgtccttttagtttggctgtccataggtgacttgctttagccagctcaattagaccccctgccttattgcaaggacggagggctttctgtatccctgggtttcttgccttggtgtactgaaagaattggataacatgtgggcttggagaatgagtgcaagttttttatttttgtttttgttttgagacggagtctcgctttgtcgcctgggctggagtgcagtggcgcaatctcggctcacgcaagctccgcctcccgggttcacgccattctcctgggcagtctcccgagtagctgggactacaggcgccccccaccatgcctggctaattttttgtatttttagtagagacggggtttcaccgtgttagccaggattgtctcgaactcctgacctcgtgatctgcccacctcggcctcccaaagtgctgggattacaggtgtgagccaccgcgcctggccgagtgcaaggttttattgagtggaagtagctcttagcagatgggggagccagacgggagatggagtgggaaggtggttttcccctggagttgactgctcagcggccaggctttcctccaaccaccccgccaaac

Uppercase: TRGV11

Lowercase: Flanking sequence[1000bp]

Red & Bold & Underline: Stem-loop [20]

Blue: Heptamer[28]

Green: Nonamer [9]

id-IGKV5-2[V_gene_segment]

gctcacaaattatgttgttggttctgttttccagtaatacatatcagataccagcaatgctgaagtttggaaactttctagaatgctttttgtttcattaggaagagtcctccacctccatgattccctttgccttctag**aagaataaaatgggaaaaacagaattttattatt**aaaaaaaaaaacccagaagagtaacatcactttccactcaaactgactacactaatctaatttctcattctttatctcttttacactaaggtatacctgtaattttcagagttc**cattttcaaagactgcttatagtttttaaaaatg**tgcatttatggtatagaaagaaaatcattttacgactaccaggttccactgaaaagtaattaatatactgtagtcaaaa**taaaataaaaatttta**atacatgaaaacaactaccgaaatgttatgaaattatagtttagtagaactaacaagtgcattaatgcaaaagaaaagtagggctcagtaatcaggg**aaccaagtgtgcattgtaaaagtgcagcctctctaacactgggtt**tcatcacaagtaacagaacaggatgcctgatgcagggaaaaaagaaaggcaattgt**tggcaatgtcaggaatattgcca**tggcgtctaccactggtgggttactgagcatcagaaccacagaagaggtaacactgaagcctgcagaataatgagtcttgttgcagaaatgcaaagaaaaagaaagatgataaatgcacacccagg**aggttacgttaattttaacct**tc**tgagtgtgcacagctgtgaagataaaactatgcacagctca**tgggttaggtttcacacagtctgccatttaaacactccctatgtcaccatcatgtaccatataatttgcatatttgtcctttatataaccctataatatgtttgtggccataaaatctgtgctgtcaaactgattaggaactgactaccacctgcaggtcagggccaaggttATGGGGTCCCAGGTTCACCTCCTCAGCTTCCTCCTCCTTTGGATCTCTGGTAAGAGAAACACTTCCTCTCCTCTGTGCCACCAAGTCCCCTGCATATCCACA**AAAATAATATATTTT**CATAAGGAATTGATTTTCCTCATTCTCTGCAAATATGATGCATTTGATTTATGTTTTTTACTTTGCTCCATAATCAGATACCAGGGCAGAAACGACACTCACGCAGTCTCCAGCATTCATGTCAGCGACTC**CAGGAGACAAAGTCAACATCTCCTG**CAAAGCCAGCCAAGACATTGATGATGATATGAACTGGTACCAACAGAAACCAGGAGAAGCTGCTATTTTCATTATTCAAGAAGCTACTACTCTCGTTCCTGGAATCCCACCTCGATTCAGTGGCAGCGGGTATGGAA**CAGATTTTACCCTCACAATTAATAACATAGAATCTG**AGGATGCTGCATATTACTTCTGTCTACAACATGATAATTTCCCTctcacagtgatacaccctgttacaaaaacctccaagttctctcagtgggattgcctcagctgctgctgaaaatgttcagaatgtggaactgggtggccctgtggtgtttataaggcttctccatgcccatgcaactcacttcctttttctttttgtttctgtagtttctgctgcccccatgctgtgcctgacacacttgatatggcaaaaaccaggacagcctttctcacgttgtccagctgtgccaacaatttcacttccaagcagaggtaaattctcatggccaactccctgtcttttgccacactcatcatttccacctccaagaaaaacaaaaccaaaaaccaaaatgcatctagcttagattccaaatagtagcattaaaggaaattgtccctccctataagcattctcaccctcatccacacagtcatcgtgactcttccctaacccactgagtgctcaccagtcctcttcagtcctccctccagtgatggggaaaagcgagtgatagttatgagagcaacatgt**aacctgggtgcagactcttactagtcaggtt**tgcctcagatttcaagtatttatctttcttttattttccattttttttcttaattatatggatcagtaagatgtctcatttcttttttttaagttacttgttcttttaacacaattcaattctataattccaaattt**aaagagataccctgtatgtcttt**ggcatgtgttatatctggacaagttttgggttatccccttcatgccactgctcaaatcttacctgaaaccccacatatttttccaaaaacctgaaaactgtgtagcggttgaggaaattaaatttggcaaatcacaatacttgaggaaacatgtatctttttatcttcactctctaatgttca**ggtgatgctgtttaattgagcatttgctatttagaatcacc**cctctgaaacttgtgggaaaacagcaccagcaaggcagtcttcatgttttgacc

Uppercase: IGKV5-2

Lowercase: Flanking sequence[1000bp]

Red & Bold & Underline: Stem-loop [13]

Blue: Heptamer[33]

Green: Nonamer [10]

id-IGHV1-69-2-2[V_gene_segment]

gactcgccatgtgtctggaccatatcagtgcatctggagct**ccagggaaggggctccctgg**tggatttagtgattccttgcttcctgtgccaaggtctccctgtgggatatgttaggtttcctaaggtctattttctagtgtaagaagtgatgtgagaagtagttgctgtcactgaaggagcattcttagccagggcacagccaggtcata**ctgggcttgagatgctgggagaaaaatgctctgtgagcccag**aaaagaactt**ccctgcagggcaggagctgagctgcaggg**ggcgctcagga**tgcacccagcacaggatccagctctggagcaggtgca**caggaggctgtggaggggttttctctcagggattgaatattc**cttatttcaaagccataatgacataaaatttaaataag**aatttagcaagtactgatgtgtctttagtattctcttgcatacacagaggctaccaaacccaataacttaacgcaaaac**acatttaaaaggagaaatgtctaggcctttcaaatgt**atttatagttaggaattgaggggtggttttattaattcaatgggtgttactctccggagacacactcatcccagaagttagatgtgcagagggcaaggcccaggaaaagttcagattgtcagtgagccataggaacaagaaa**tcacagtgaggacaatgccctgtga**gattctgggtttctgtaagaggagttctgtcttcatggacttctaagtgtgtcagaggacaaatatcattaaacaaagttcagggcagggagctcagtgtcccactgtggcatggtccgtgtgtcacctatcttcttccccagggtggggtggcctaagctatgaaatacctgcctcatgaatatgcaaatgcagtggtgtctaccgaggtaaatacagatctgtccttgcccagagagcatcacacaacagccacatccctcccctacagaagcccccagagagcagcacctcaccATGGACTGCACCTGGAGGATCCTCCTCTTGGTGGCAGCAGCTACAGGTAAGAGAATCCTTAGTTCCAGGGCT**GATGAGGGGACTGGGTCCAGTTAAGTGGTGTCTCATC**CACTCCTTTCTCTTCTCCACAGGCACCCACGCCGAGGTCCAGCTGGTACAGTCTGGGGCTGAGGTGAAGAAGCCTGGGGCTACAGTGAAAATCTCCTGCAAGGTTTCTGGATACACCTTCACCGACTACTACATGCACTGGGTGCAACAGGCCCCTGGAAAAGGGCTTGAGTGGATGGGACTTGTTGATCCTGAAGATGGTGAAACAATATACGCAGAGAAGTTCCAGGGCAGAGTCACCATAACCGCGGACACGTCTACAGACACAGCCTACATGGAGCTGAGCAGCCTGAGATCTGAGGACACGGCCGTGTATT**ACTGTGCAACAGAcacagt**gtgaaaacccacatgctgagagtgtcagaaaccctgaggaaggaggcagctgtgctggggctgaaaagatgacagcatttatgaggtttaaagtgtttagaagatgggttaagtcacagagtaaaagtagaaatagaaagatgtatacattccaattatataggaaataatcttttcaactttcaccctgtaagtaacattcacagagtgggaaaggcagcaatcaatggagtggatgcaaacattcctttgtgaggacataaattttaaaatcgaatggataaatcatttggagcaggattgcttgatcacatggtaagactaaatataatttctaaaaagtgacca**acatttctaacaaaatgt**ctttgccactccttttacattaattttattttaaaacatttttaggattgcagcaaatttgagtagaataaacagcattcccatgtactcctgcccaagacatgcacagacttctccatgatcaacaccctgcgctgaagtaataaataacttgcaacagatgaacccacatggacaccttgattgtttcctttcctggtggcccctggtataacaagcctaaattaccttgaagcaccccaggttcttctagtgggttacaggaaatgatgccaggtagagggaaagtgagtggggatgatcctctttgcactgtttcctcaagaatccataaaatgtacattgatttagagctcaactaactcctggttttctatgcccctttccagaaggtaaggtctccaagaatttacagcaggggatctcctcaagagtccataaaatgtacagcggtttagagctcgtctgactcctggttttatatgcccctaccaagaggataaggtctccaagcacttacggc**aggggtctccaacccct**aggcagtgaacccgtactggtca**gcagcctgatagaaagcaggctgc**acagcaggagatgaacagggggtgagcatcactgcctgagc

Uppercase: IGHV1-69-2-2

Lowercase: Flanking sequence[1000bp]

Red & Bold & Underline: Stem-loop [12]

Blue: Heptamer[54]

Green: Nonamer [9]

id-IGKV1-17-3[V_gene_segment]

tgccaccctgtgtcttttaagtggggcgtttagcctgttgacattcaaggttaatattgatatgttagatttgataccaccatcatgctgttagcctgttattttgccaacttgtttgtgtggttgctttataatgtcactggtctgtgtacttcagtgtgtttttatagtggctgataacggtgtttcatttccatatttagt**gcttcttccagaagc**ttgtgttaggcaggtctggtggtaacaaatttcctgagcatttgcttgtctga**aaaggatcttatttctccttt**gcttccgaagcttagtttgtccaaatatg**aaattcttagttggaattt**attttctttaagaatgttaaatataggcccccaatctcttctgctgagaggtcttctgttagtctgatggcttccctttgtaggtgatctgttttttctctctagc**tgcatatacttacaaatatatgca**tcatcatgtcaatatattttatttttaattttttttagaaagcaacaatgataaacttacagaaaagttgcaagtatacaataaatacccccttccctaaccggaattatacaatagtcttttaaagaccttagaattctatcgtctaaaactttactatgtgtttcctacaaaaaaagaatatcctcctatatactccccatacgccaa**tgaaatatgatactccaccgactcctgaggaatatttca**aattgtcaaaaaaaatctaaaaaatgtctctcgtaacaaaatagtctccagtagaaacccattctctgcagacaaatctgtgctaccctgatcttacctgggacacctggggacactg**agctgctgctgagttactgagatgagccagccctgcagct**gcgcccagcctgccccatcccctgctcatttgcatgttcccagagca**cagcctcctgccctgaagccttattaataggctg**gacacacttcatgcaggaatcagtcccactcaggacacagcATGGACATGAGGGTCCCCGCTCAGCT**CCTGGGGCTCCTGCTGCTCTGGTTCCCAGG**TAAGGATGGAGAACACT**AGCAGTTTACTCAGCCCAGAGTGCTCAGTACTGCT**TTACTGTTCAGGGAAATTCTCTTACAACATGATTAATTGTGTGGACATTTGTTTTTATGTTTCCAATCTCAGGTGCCAGGTGTGACATCCAGATGACCCAGTCTCCATCCTCC**CTGTCTGCATCTGTAGGAGACAG**AGTCACCATC**ACTTGCCGGGCAAGT**CAGGGCATTAGAAATGATTTAGGCTGGTATCAGCAGAAACCAGGGAAAGCCCCTAAGCGCCTGATCTATGCTGCATCCAGTTTGCAAAGTGGG**GTCCCATCAAGGTTCAGCGGCAGTGGATCTGGGAC**AGAATTCACTCTCACAATCAGCAGCCTGCAGCCTGAAGATTTTGCAACTTATTACTGTCTACAGCATAATAGTTACCCTcccacagtgttacacacccgaacataaacccccagggaagcagatgtgtgag**gctgggctgccccagc**tgctcctc**ctgatgcctccatcag**ctgagattgttcctcagatgcagccacactctgatggtgttggtagaaggggacatgaagtcacctctgcaccctaattcttttctctttctcagccccaactacacagacatagcaatgcctctcctaatttaatacagacagagatcatgacacctgaggagtctagttta**tggcttcagttggactttatataacagagaagaagcca**ctatagatattctaggcaggaattgtcttaatacagagaattaaagtaaactactgacgtctaaataaaatg**tagagatgaatctcta**aatgtaatgttttatttacacagaaatatttcccaaatggggcatacaggaaaactcagtggtcttcaatatgtggcaagaacaaagaaaaagttagtgtttcatgaaaaagggaaaatattacctatggctctttgagaaagttcattggcactaggaagggttgggagctggcaagctcagactgggaagcagtggtggacaaagtgaatcctagaattatatcaagttatctcagaagttgtggacaaatttgatttcaggttacaagaagccaaagcagtgaaggttgcagagtgttttgttactgaaataccagggattcagcgtagatcctgcagctcaccacacagaaagccaatcactaaggcaagtattgccaaggaaaaggctttaatcaggtgctgtagctga**ggagatgggacaccattctcaaatgtatctcc**ctgaccaacaaaaattaggggtttatatagcacggaagaaatgtgggaaaacaggaattatggagggttaagggagataatttggtcaacaagaagcaggaggtcagttatgcagtcataatgagtgaagggtctgatgtcccactgtcctgattcagtgata

Uppercase: IGKV1-17-3

Lowercase: Flanking sequence[1000bp]

Red & Bold & Underline: Stem-loop [17]

Blue: Heptamer[28]

Green: Nonamer [3]

id-IGHVIII-5-2[V_gene_segment]

ctgaactcgtggagatagagagtgaaaggatggtcaccagaggctgggaaggtctatggtagatggggattattggggatgctcaatggatgaaaaaagtggttagaaataa**caaataatacatagtatttg**ataacacaacagactaagtataatccttaataacataattatagattgtaaaataactaaaagtgtataaatggaatgttggtaactcaaagaattaatacttgaaaggatgtaggaaccattcat**cataatgtgagtattatg**cagttcatgctatatcaaagtcaggtaccccataaatatacacatataccatttacccacaaaaattcaaaattaaagtatttatgtatctc**atcttctgtatctatattacattaaacctaagaagat**actggtttcttcacactaccaagatccacatagacatccgtagccttctttctttgcctgtccataatctgacattgaaaaatgaggaaacctttcctatcatatgacattaatttacttagtg**cacaatttcaaaatacacaaattgtg**ttagaattattaaactgacccttgttgaaaatatgtttatcgactagaataccgtgcttatatgcagcttatttatattttacaatcacaacatcaaattatttccaaagatacttgtttctgatattcatgcccccctttctgcagtgaaattagtacaaacctgtttatcattgttagatatttttatacagctttcattttattctgtgtttcatgaccagctaaatacttttgaagttgtatgcatttaggttaacactttgtctcaaaaaattatgagttttcacaaattatttatgccatgtgtttatcattgaa**caccatgaaaatagtttcattgtg**atta**aaaagtgtgtgtttcacctaatccacacaccttttccccagattcctggaaaa**ccacaaaggtttactggatcta**tatttttgtctttggcaaaatAGCCTGGCTATT**CTCTTAGCTTCTTCTGCAAAGCATCAAGATTCACCTTCACTGACTGCAGCATTTATTGGGCCCTG**ATGGCTGGAGGAGAGGGGCTGGAGTGGGTGGTAGCCAT**GAGTGACCCAAGTGGAAGTTCTCAGTGGTGCTCTGCTCCAGCACAAAGAAGATTCACAGTTCCTGGGGACAACACTTAACATCACAATCTCCCTTAAAATTATCTACTGGAAAGCTGAGGAGTAGGCAGTGTATT**ACTGTGAGAGAcacagt**gagggacatatgggtgagtctagacgcaaacttccctgcaggaagacaagaggggactttgtggtaaatggtgctcagaaccactaggggacactcagaaccactaggagacactcaggatagcagaaggtgctcaggatgccgcagggtgctcaaaacactaggggtgctcaagaccagcagggggcactcaggacaccaggagacacttaggacatcagggggcgctcaggacaccagggggcgctcagcacaagcaggggccacagaatgacaagggtgcattccagaccaccaggaaacactcaggaccaacaagagattctcagaaaccaccaggtgtaacaggaccacctaaaggcccacaagacaaccaggagacactca**tggccactgaagaacattgaggacaccaggggttcagggcca**acagggagtgctcggggccaccatggggctttcagtaaccacgagggaatcaatcactcaggacaccacggttttcttaggaggcagctccacatcaggtgcctggggaaggt**gacgtttgcttttagacctagctgattcctgagctggtcaagcaaaagtc**ttcccacgatctctcactatttcttccttgtaacccatggtttctttcacatacaaatcattaacttaggataggaattcaattcaacttttaaccttgcatatttttagaataataatagcaatagtaattcaatgtgttttgataataagaaattgtgtatgcctaattcaaactctggttccacgcatgggtttttgtttcttgtgctgtgccagtcacactacagtcaatgcttttctaataataactcaacaatgtcacggtgttaagtttcctttcattttgtctgccattt**gtggagatgaaacaccactttcacgggtcagttcctccac**tttggttgggttctggtcttctgctcttcacactgtttctccaccttcccttctgtcaaatcct

Uppercase: IGHVIII-5-2

Lowercase: Flanking sequence[1000bp]

Red & Bold & Underline: Stem-loop [14]

Blue: Heptamer[31]

Green: Nonamer [6]

id-IGHV4-34[V_gene_segment]

ttcttgcaattcagcggaggagaaattct**ttcagagacagatggatctgaa**ttggtaaatatgtgggtacgaattctgggcttgagtgtcattgtccagccatgtttc**acaggtgtgacctgt**cagggaagaaccagagttccttgttctctcagagggtagagctcacagaggtcctctctggttcccaggaaaggtaatttcactaatcttggtgatgagactatcctccagtgctgatgtactatagagttttcatctgaagctgtcactgctatccccaatgtacatcttttcacacagaaatgtttagaggtcaggccatattctcagggttacacatt**gagaaggatggagatatattctactaccttctc**c**tgagatctcacacacaatctca**aatttcaaaaggtctcagaagggcagctctcaggtactatttaaaaataacccacttcctgggacaggtagcatccttctaaccatgatggatgttctgaactacagtacacattgcatggatccaggtttgtctcaattcactgtgattattacactcagcagctgtttcaatatgtctgaaggggtaaatgacaatttaggtgacctgggtgtatggttggtgttatatgaatctttaaatgtagaacagtattaactgtattccaaaatctgtctttgatccatgatcacacttgtctcccagaccagctccttcagcacatttcctacctggaagaagaggactctgggtttggtgaggggaggccacaggaa**gagaactgagttctc**agagggcacagccagcatacacctcccagggtgagcccaaaagactggggcctccctcatccctttttacctatccatacaaaggcaccacccacatgcaaatcctcacttaggcacccac**aggaaatgactacacatttcct**taaattcagggtccagctcacatgggaagtgctttctgagagtcatggacctcctgcac**aagaacATGAAACACCTGTGGTTCTT**CCTCCTCCTGGTGGCAGCTCCCAGATGTGAGTGTCTCAGGAATGCGGATATGAAGATATGAGATGCTGCCTCTGATCCCAGGGCTCACTGTGGGTTTCTCTGTTCACAGGG**GTCCTGTCCCAGGTGCAGCTACAGCAGTGGGGCGCAGGAC**TGTTGAAGCCTTCGGAGACCCTGTCCCTCACCTGCGCTGTCTATGGTGGGTCCTTCAGTGGTTACTACTGGAGCTGGATC**CGCCAGCCCCCAGGGAAGGGGCTGGAG**TGGATTGGGGAAATCAATCATAGTGGAAGCACCAACTACAACCCGTCCCTCAAGAGTCGAGTCACCATATCAGTAGACACGTCCAAGAACCAGTTCTCCCTGAAGCTGAGCTCTGTGACCGCCGCGGACACGGCTGTGTATT**ACTGTGCGAGAGGcacagt**gag**gggaggtgagtgtgagcccagacaaaaacctccc**tgcaggtaggcagagggggcgggcgcaggtactgctcaagaccagcaggtggcgcgcggcgcccacagatcccgaggccgggtccggagcaggtgcaaggagggcggggcttcctcaacagctcagtggtctgtctcctcgccagcacctcagatgtccccaggactctctttctttattatctgtggttctg**cttcctcacatccttgtggcagggaagaaaggaggaag**acaatttttctgtttactgttgaggtttcaccaattactagaaactttcctacaagttcctgcatgactcattttgccatatatggattctcaccatctcttgatttgtttcatcaatcgaattgtgccctatttgaaattaacttactgaaaccttaaatccaatggatctatactggaattttaatgatgtaattgaggttaaatgtggtcaaagtgtgagaccctaatgcaataaaccgttgtctttataagaagaggaagagacaccagagacctctcacttttcac**gtgcacacagagaagaggccatgtggagacgtagtgcac**tagaaggtggccctgtgcaagccaggaagaagccatactaagaaccaattttgccagctccttgatcttcgacattcagactgtagaattgtaagaaaatcaatatttgttgtttaagccacccaatcctgtt**gtcttcttacgaagac**ccaaactgactaataccacctaactctgttagctctgtctcctggagggagaagcagccccctgaggctgggcactgcatctctcagatttccacgtgaagtaggcaaaaatagtagttctcatat**aaaaatgtgtcatggctctgttggccattttt**gagcatggtctctgaaaccagccctggtgtgtgtgtgtaacaa**atgtcccttatcttttatttggacat**aacaaatagacgataggtac

Uppercase: IGHV4-34

Lowercase: Flanking sequence[1000bp]

Red & Bold & Underline: Stem-loop [16]

Blue: Heptamer[40]

Green: Nonamer [5]

id-IGKV2-28[V_gene_segment]

gtggatccagttcgtgctcttcttacatagagaatttccactcgaattgtgaactcatggctgtggctgcaccacttacaggcccaggggacactcagatctcacttggtagttgacgaaaaactggaagtcctgatagga**cctcattccacatgatgagg**aagactgtgggaagagcttttgttaacaccattcag**aaaaacatagtgcaaagttagttttt**gttc**tttctatataattatcctagaaa**agtcttcccttaataaatccttctggtt**aatctggcatatgtgagatgattatgatgggatatataccagatt**gaacaattggtcaccaggaattttatattcactgcct**gaggaataaattgtttcccactttcctc**ttacctgcactgggctcttgaatctaaatatagagacccacattattttccctatgaggcccttggacagagcgctcttatggggctcactcaccaggtgccaggggagggtagattccaacacttgctatgaacattcttgaacagttatcctggaaaccgcagataccagaccac**tcttgaactggctcaaga**ctatgttttatttgtaggctgtctgctcatcagtgcttgt**aggaaagggtaacgtttcct**tttttagattagctgggagggagccaaga**agaatggcattcatccatattcattct**agacatatctcta**cattgttagggttgttatgctttcctagagttgcatatcctatacaatg**ggacctacccaagatccaaactgtcacagtcagatccttcctcccattttatatcacattgctcacaggagagacatatcccctgcccgcctgccccattgactctttccacaccactgcatgcaccaggggatttgcatattgtcccacagggaggaccttcccttgtgagtctgagataaaagctcagctgtaactgtgccttgactgatcaggactcctcagttcaccttctcacaATGAGGCTCCCTGCTCAGCTCCTGGGGCTGCTAATGCTCTGGGTCTCTGGTAAGAAAAGAAGGAGATGAGGAAGGAGAATAGGGTGGGAGGGTGAGCTCTAGGGCTCCACAGCATCCCATGATCCCATGTTTAGTCTTACCCTGTGTTAGAGGAGTATAATCTGTGCTGTAGAAAAGGGAACTTGATATTTTGCTCTGTGAATAATTAGAAGCCTCATAAGAAATATGACGTCTGGTGCTCTGATTAAGATCTTCAAAATATAAAGGTCTCTTATACTTTACAAAAATTGAATTCATTTTAGAATGTGTATTTTTATGGCATAAATCACT**ATTTTTTAAAATTAAGTTTAAATAAAT**GACATAAGATAAATTATGAA**AATTGCTCATTAGGTTTGTACATAACTTTGCAATT**CATTATTTCAGGATCCAGTGGGGATATTGTGATGACTCAGTCTCCA**CTCTCCCTGCCCGTCACCCCTGGAGAG**CCGGCCTCCATCTCC**TGCAGGTCTAGTCAGAGCCTCCTGCA**TAGTAATGGATACAACTATTTGGATTGGTACCTGCAGAAGCCAGGGCAGTCTCCACAGCTCCTGATCTATTTGGGTTCTAATCGGGCCTCCGGGGTCCCTGACAGG**TTCAGTGGCAGTGGATCAGGCACAGATTTTACACTGAA**AATCAGCAGAGTGGAGGCTGAGGATGTTGGGGTTTATTACTGCATGCAAGCTCTACAAACTCCTcccacagtggta**caacccctaacagaaacctcctcctggggttg**cctagttgctcacatgt**gctgcttgtctggagagcagc**tcagcagggtctctgagtctgcagaagaggaggctgttg**gagacctcagggcagaggttgctgctgaggactc**tggctcatgatagcctcagctgtacttcagtcccacatgttaaggccccattaggtgaaaaataaa**tgattccaaaaactgagatgaaataccaaggagaatca**gagtacaattaaggctgttacaaagaagcctcaaaatatggtggactaaatgtgacatggtttctgtgtctgttgcctgacagtgcagaggcaggtgggtggctttggtggtgtcggtggctcttctccaagaggtcgctcagatgggcaggagacacgaccaccctcagacacagccttcctccttcctcagagtcacttgcccccatgcccatcctcgacagcatgaagtggaacgactagagagaaagctgt**tttcttctaaagaccaaagaaa**tctagagtttcccatcagggataaatgttcttttactttaagcacgcattggagaaattttccattcagtggatctgctgataaacccactttgttttgtttgtttgttttgttgtgttttcagatggagtctctctctgtcacccaggctagagc**gcagtgacatgatctcggctcactgc**aaactgcctcctgggttcaagagattctcatgcctcagcctccggagcagctgggattacaggcacccaccaccattccaggctaatttttgtatttttagtagagacagggtttcaccatgttggccagggtggtgtcgaactcctgatctgaagcgatccacctgccttagcctcccaaagtgctgggattataggcgtgagccactgcacctggcagttatttgctcttatatgataatgtaaaattattttagaaaataattttttgagtatatagtt

Uppercase: IGKV2-28

Lowercase: Flanking sequence[1000bp]

Red & Bold & Underline: Stem-loop [20]

Blue: Heptamer[41]

Green: Nonamer [7]

id-IGLV2-33[V_gene_segment]

cccagcactttgggagactgaggagggtggatcataaggtcagaagatcgagaccatcctggccaacgtggtgaaaccccgtctctactaaaaatacaaaaat**tagctgggcgtggtggcttatgcctgtaatctcagcta**gtcgcgaacctgaggcaggagaactgcttgaaccagggagttggaggttgcagtgagccgagatggcgccaccgtgttccagcctggtgacagcgcgagactccatctcaaaaaaaaa**aaagtccctgaatctcaaaaaaaagaatgtgacttt**atgattaactatgaacagctgctcattaataaaaagccattaaaaaaaaaaaagaaatcccaagagctacttacccttgaatatctctaaaggt**actgggaagcagtgctccaaatctgttgtggcccagt**gaattcctttttagaagcaccattaatggtcctagttcaacatgtcaggccccagctttaaac**cactgcaggggcaggggatgtagcagtg**cttagaacccaccttagctgagaggaccaagatggttgggcaggggatgagggaaatgagaagttccagcactgtctgaagtctctgggaggaggagtgattgaactttcccacccggcctgggagcagcctgtcctcacattcatccctcacagcacaaaggagaagtcctggtttcctcagcctggagctcctgcaatagtgagggttaggatatctctggactggcctgatgtcctgggcactctcatgggttcagtctcagga**aatccatgcctaggctggatt**aataccctcagctgagcctcacagggaagcagaagtctctgagcccaggcccaggtgagggtggggtgaggagaggagctcaggacacagatttgcatggag**gccctgccctcctctgaggcggggggataagacagggc**tagaggcaggcccggtgctggggtctcaaggcagcgctctcgggacatctccaccATGGCCTGGGCTCTGCTCCTCCTCACCCTCCTCACTCAGGGCACAGGTGACACCTCCAGGGAAATGGCCTTGGGGACCTCTGAGCTGATGCTTGGTCTTCTGCTCCTGCTCCTCA**GGGTCACTGGACCCAGTACTGACCC**AGTAGAGTGTGTTTCTCCCTCTTTCCAGGGTCCTGGGCCCAATCTGCCCTGACTCAGCCTCCTTTTGTGTCCGGGGCTCCTGGACAGTCGGTCACCATCTCCTGCACTGGAACCAGCAGTGACGTTGGGGATTATGATCATGTCTTCTGGTACCAAAAGCGTCTCAGCACTACCTCCAGACTCCTGATTTACAATGTCAATACTCGGCCTTCAGGGATCTCTGACCTCTTCTCAGGCTCCAAGTCTGGCAACATGGCTTCCCTGACCATCTCTGGGCTCAAGTCCGAGGTTGAGGCTAATTATCACTGCAGCTTATATTCAAGTAGTTACACTTTCcatagtggtcca**agttcctagggaactgagaccaaaacctgccctgggctctcag**gctctgtctttgctctgaagatgcttcctca**ccctgtgccaggggcttcctgcaacaggg**ccttgagaattcacttctctgtcagctcctc**tccttttccatcaattcccaaagga**aacctcctctcttgtttactctccgggatatgacagcttcttcttcactcgtatgacggatgtcctctctgaattggaaac**tacttctagctctttaacaggaagta**catacc**ggcagggaaatgcctgcc**tagtgcagtccccatacttctccggatgatcctcactttattatttatttttatttttgtatgcactaaagtcgtcaatgtaaatccttacgtcgaatgcctccacttcccacattgtgtagcacagcctggattcagttcttcagaaatgacgtcctcttcaatgataaaggaccactgctctgtgaacaccccactctagtgaagatagcccaccttgctgggttattcatgt**acatcttttgtggaagcagtaaccactgtgtaaagctccaaagatgttttaaagcaatgaaaaca**aagaggaagaagtgtgtgtgtgtgtgtgtgtgtgtgtgtgtgtgtgtgtgtgtgtagggtccatcctgtttgccctaatatctcaggttgccgagctagggagtatttgagtgacagacttggtgccctctcaggatcctcctcctctcacatcactgagtccctgtcctggaaacaccctacaggtggacaatctccccatagtatggaagtttccaaaatggctccacaggg**aagagttgagctgaacacacccactctt**tctcatgggcatcaaatatatctaatttttctggggaaatcatcaatttctagctggaatcctgattaatcgacagactgtgagaattaaaaattttaaaaatattcagacatatccagagtccagtatttaaatt

Uppercase: IGLV2-33

Lowercase: Flanking sequence[1000bp]

Red & Bold & Underline: Stem-loop [16]

Blue: Heptamer[35]

Green: Nonamer [4]

id-IGHV3-49[V_gene_segment]

ggtaaaatatccatttatgctaactaaaataagccaaacaaatgagaatgtatactatgttat**ttcatttttataaattctgaaaaattaaaatgaa**tttgcagcaatgtaaagatcagtagttgccagggaaatggtagaagaaagaaaggaaaaggagaaagaatacagaagaacaaaaggaaatgttaagaattctcttgtccaac**ttgataaggatgacggttacatcatttttatcaa**ctgtaatctttaaatatgtg**aaagtttattatctgtaaactaaacttataaacttt**attacaagcaaaaattgaaagttagacaagaaggagtgatagaaagagaaaatgtata**ttaaatttcagaaatatttaa**gaatgtatctgcctgaaccctagttctcaccatatctttaggtgaatgctaaaatgcagcaaaatcacgcatgttctcactacagaaagtgggttctacaaaccacactcggcacatttagctttgtcctggagttggttcagggagttattggggcc**agtgatgaggagcacaggccaagataccagcgatcact**tatcccaaacatgagctctaacat**acacacttagtcccttttccgtgtgt**ggtttacttccacatctgtacatggagagaccactgactgacaaaata**taatttatacaaatatgtaaaatta**aatagggtgatcagttcaaggtgtttatcacagcataattttacaataagacagcatatttcccaaataccatcattgtcaccaaactccttcaaggcacagtcatctta**tctgggccccgtcctctcctcaggtgtcccaccccaga**gcttggtatatagtaggagacatgcaaataaggccctccctctgctgatgaaaatgagcccagccctgaccctgcagct**ctgggagaggagccccagccgtgagattcccag**gagtttccac**ttggtgatcagcactgaacacagaccaccaa**ccATGGAGTTTGGGCTTAGCTGGGTTTTCCTTGTTGCTATTTTAAAAGGTAATTCATGGTGTACTAGAGATACTGAGTGTGAGGGGACATGAGTGGT**AGAAACAGTGGATATGTGTGGCAGTTTCT**GACCTTGGTGTTTCTGTGTTTGCAGGTGTCCAATGTGAGGTGCAGCTGGTGGAGTCTGGGGGAGGCTTG**GTACAGCCAGGGCGGTCCCTGAGACTCTCCTGTAC**AGCTTCTGGATTCACCTTTGGTGATTATGCTATGAGCTGGGTCCGCCAGG**CTCCAGGGAAGGGGCTGGAG**TGGGTAGGTTTCATTAGAAGCAAAGCTTATGGTGGGACAACAGAATACGCCGCGTCTGTGAAAGGCAGATTCACCATCTCAAGAGATGATTCCAAAAGCATCGCCTATCTGCAAATGAACAGCCTGAAAACCGAGGACACAGCCGTGTATTACTGTACTAGAGAcacagtgag**gggaggtcagtgtgagcccagacacaaacctccc**tgcaggggcgcacagagccaccagggggcgcta**gggaccgactgagtacgggacaggtccc**aggagcaggtgcagggggaggtttcct**ttttccttggctggaaaa**gtcacctttatcttcccaggactcgagccttctaggctgtgatattttattacttgtatttactgttcatttattatcattagtttttaaattttggtaatttttacaactctatggatatatttttaagtgtatactttcaag**aaataaacattcctaattattt**gcactgattctcccagagttttattaacatttgttgacatcagcaactacatagctatagggacaaacacttttaacgatagacagttgtttaggcctgaaaccccgtttatactaaaaatttacaaaaattagcctggcgtggtgaggggcgcctgtaatcccagctactcgggaggctgagcaaggagaattgcttgaatccgggaagcggaagtcacagt**gagtggagtggactgccactacactc**cagcctggcgacagagcgagactttgtatgaaagaaagaaaagaaaaggaagaaaggaagaaaggaaggaaggaaggaaggaaggaaggaaggaaggaaggaaggaaggaaggaaaagagagaaagaaagagagaggaaggaaggaaggaaggaaggaaggaaggaaggaaaagagagaaagaaagagagaggaaggaaggaaggaagaaaggaaggaaggaagaaaagaaggaaagaaagaaaaaagatatataaacacgcaga**cctatgcatataaccatagg**gatttatattaaacattacaataaaataattctaaaaatgtgtcctaaggaatcaaacataatgatgaagtaaa**tataaatgttagagtaatttata**attgatttgttatttttaatcgtttacttgaatttatttctatttgttcattta

Uppercase: IGHV3-49

Lowercase: Flanking sequence[1000bp]

Red & Bold & Underline: Stem-loop [20]

Blue: Heptamer[21]

Green: Nonamer [3]

id-IGHV4-59-2[V_gene_segment]

agggtatgttcatgcaattcaatggaggagaaagtc**tttcagagacagatggatctgaaa**tgataaatatgtgggtaaggact**ctggacttgagtgtcattgtccag**ccatgttt**cacaagtgtgtcctgtcagggaaggatcagagttccttgtg**ctctcagagggaaggggtcacagagttcctctctggttcccaggaaaggtaatcgcactaatct**tcatgatcttcatga**gactatcctccagtgctgacctgttatagagtttttgtctgaagttctcactgcaatccccaatctacatattttcaatcagaagtgtttagaggccaggacacatcttcaaggtcacacatt**gagaaggatgtagatatgtcccactaccttctc**ctgagatctcagacagaatcccagatttcaaaaggacacagaaggacagctctcaggtgcttttaaaaaatgacccacttccagggacagggagcttccctataaccatggtggatgttctgaactacaataaacattggatggatccaggattgtttgaagtcactgtcattattacattcagctgctgtttcaatgtgtctgaagtagtaaatgacaatttagatgacaatttatatgaatcttcaagggtagaacaatattgaccatattccaaaatctgtccttgatccatgatcacactcatctcccagaccaggtccttcagcacgtctctttacctgaaagaagaggactctgggcttggagaggggagaccccaagaagacaactgagttctcaaagggcacagccagcatcctactcccagggcgagcccaaaagactggggcctccctcctcctttttcacctctccatacaaaggcaccacccacatgcaaatcctcacttaagcacccac**aggaaaccaccacacatttcct**taaattcaggttccagctcacatgggaaatactttctgagagtcctggacctcctgtgc**aagaacATGAAACATCTGTGGTTCTT**CCTTCTCCTGGTGGCAGCTCCCAGATGTGAGTATCTCA**GGGATCCAGACATGGGGATATGGGAGGTGCCTCTGATCCC**AGGGCTCACTGTGGGTCTCTCTGTTCACAGGG**GTCCTGTCCCAGGTGCAGCTGCAGGAGTCGGGCCCAGGAC**TGGTGAAGCCTTCGGAGACCCTGTCCCTCACCTGCACTGTCTCTGGTG**GCTCCATCAGTAGTTACTACTGGAGC**TGGATCCGGCAGCCCCCAGGGAAGGGACTGGAGTGGATTGGGTATATCTATTACAGTGGGAGCACCAACTACAACCCCTCCCTCAAGAGTCGAGTCACCATATCAGTAGACACGTCCAAGAACCAGTTCTCCCTGAAGCTGAGCTCTGTGACCGCTGCGGACACGGCCGTGTATT**ACTGTGCGAGAGAcacagt**gaggggaggtgagtgtgagcccagacaaaaa**cctccgtgcagggaggcggagg**ggaccggcgcaggtgctgctcagcgccagcagggggcgcgcggggcccacagagcaggaggcccggtcaggagcaggtgcag**ggagggcggggcttcctcatctgctcagtggtctccctcc**tcgccagcacctcagctgtccccaggggtcctctttctttattatctgtggttc**tgcttcctcacattcttgtgccaagaaagaaatgaggaagaca**aattttcgtctgtagttgaagtttcaccaattact**aggaactttcctagaagttcct**gcatggcccattatagcttactgattaaaaaatatatattctaacgcttctcagcatctcttgatttgtgtcatcaac**tgaattgtgccctctttgaaattca**tatgcagaaacctt**aaattcaattgatgtatattggaattt**taatgaaataattaaggttaaatgtggtcataagtgtaagactctaattcaacagacgtgtcgtctttataagaagaggaagagacaccagagacctctcacttttcacgtgcaggcagagaagaggccatgtggagacgtaa**tgcactagaaggtggcccagtgca**agccaggaagaagcctcaccaagaaccaaccctgccagaac**attgatcttcaacattcagactgcagaattttaagaaaatcaat**atttgttgtttaagccacccactcctgttgtcttcttatgaagatccagacagactaataccacataactctgttagcgctgtcccctggatgcagaat**cagcccgctggggctg**ggcacatctctcagatttccacataaagtaggcaaaaaatagtagttctgatataaaaatttgtcatgtccctgttggccaatttctgggcaaggtcttttaaag**aagccctgggggctt**tgtcacaaaagttgccttttatcatttattagga

Uppercase: IGHV4-59-2

Lowercase: Flanking sequence[1000bp]

Red & Bold & Underline: Stem-loop [21]

Blue: Heptamer[41]

Green: Nonamer [6]

id-IGKV1-8[V_gene_segment]

caaatccatgtccagtttcatctggatccagctgatttctccatgtaagttggcaattgcttgataagggattgtctctttcctaaagatg**ttaacagggaggatggtgtctgggtcaggatgatgtccctgataa**aacgtaaaagaagaaagtggcattgttgg**tgcatgacagggacatgctccatgca**gtggtcaccctcaataagagagaagaactttggaaagtaatactcaatgacagaaaagaaggtagacaatgaaggtgcccaaaacaagaataaggtgaagtgaattta**gtctctgggtattaaagagac**ctgtagctcttgataatggtggatgtgtgagtgctgcatgcattgaggaaactcggtatcatctctctgtatctgtagtaaattgcttgatcttatagtggtaagaacaatggcataacaccattacctaatacttacaaatatgtatagcatcatgtcaataaattttat**ttttaatatttttagaaaggaacaatgttaaaa**ctcacagaaatgttgaagtataggacaaagtacctccttccctaacccgaatcatatgagagtcttttgaagtcctgagaatcatactgtttaacattttactatgtatttcctacaaacaagatattccaaataatccccatgtgccaa**tgaaatacattactccgtcgactcctgaggaatatttca**aattttcaaaaaaatacataaaaaa**tgtttctcataacaaactactctccagtagaaaca**cattcaccacagacaaatttgtgctaccctggtctttcctgggacacctggggatactgagctggtgctgagttactgagatgagccagccctgcagctgtgcccagtcagccccatcccctgctcatttgcatgttcc**cagggcacaacctcctgccctg**aagacttattaataggctggacacacttcatgcaggagtca**gaccctgtcaggacacagcatagacATGAGGGTC**CCCGCTCAGCT**CCTGGGGCTCCTGCTGCTCTGGCTCCCAGG**TAAGGAAGGAGAACACTAGGAATTTACTCAGCCCAGTGTGCTTGGTACAGCCTGGCCCTTCAGGGAAGTTCTCTTACAACATGATTAATTGTATGGACATTTGTTTTTATGTTTCCAATCTCAGGTGCCAGATGTGCCATCCGGATGACCCA**GTCTCCATCCTCATTCTCTGCATCTACAGGAGAC**AGAGTCACCATCACTTGTCGGGCGAGTCAGGGTATTAGCAGTTATTTAG**CCTGGTATCAGCAAAAACCAGG**GAAAGCCCCTAAGCTCCTGATCTATGCTGCAT**CCACTTTGCAAAGTGG**G**GTCCCATCAAGGTTCAGCGGCAGTGGATCTGGGAC**AGATTTCACTCTCACCATCAGCTGCCTGCAGTCTGAAGATTTTGCAACTTATTACTGTCAACAGTATTATAGTTACCCTcccacagtgttacacacccaaacaaaaacccccagggaagcaaatgtgtgag**gctgggctgccccagc**tgctcctcctgatgcctccatcgcctg**agagtgttcctcagatgcaaccacactct**gatggtgttggtagagggggacctgaaatcacctctgcaacccaattcctttttctttgtaagccccagctgcacagacataacaatgcctctcctgatttaataaaggcagagatcttgacacctaaggagtctagtttagggctttggttggaattcaaataacagagaagaaaccactatagatattctaagcaggaattgtcttaatacagaaaattagagtctaaactactgaagtctaaata**aaatgtagagcggaatctctaaattt**catgttttatttgcaaagaaatgtttgccaaattggacacacaggaaaactcagtggtcttcaatatattggaagtacgaagagaaggttagtgtttt**atgaaaaagggaaaatattaccttttgcat**tttcagaaagttcactggcactagtaagggttgggagctggcaagctcagactggtaagcagtggtggacaa**agtgaattaggattcact**aattatatcaattatatcaagttgtctcagaagttgcagttaaatgtgaattcaggttacaagaagccaaagcagtgaaggttgcagagaattttcttactaaaatgccaaggattcattgtacaccctgctgctcaccacgcagaaagccaatctctaaaacaagtattgccaagaaacaggctttaatcaggtactgcagccga**ggagatgggacaccatttgcgaatgtatctccctgaccaactaaaattaagactttatataacaggga**caaaatgtgggaaaacaggaatttgggaggggtaaggaagataatttggtcaacaggaagcaagaggtcagttatgcaatcataatgggtgaaggttctgatgtctcagt

Uppercase: IGKV1-8

Lowercase: Flanking sequence[1000bp]

Red & Bold & Underline: Stem-loop [20]

Blue: Heptamer[15]

Green: Nonamer [2]

id-IGHVIII-26-1[V_gene_segment]

aagc**aaataacccacaggattattttctttccttgattgagaaaga**ttttccccaaacttcagctcagttcaggcatatactgtccctaaatgggcatttaccctcagatgagtccacacacctgtcaacacgtggagtcttctgtcagacaaacacacctttactg**atgtggattcttccctcagacaaaaacacatgtccccacatggacactttcctcagactaccatgt**atgtccttacatttacgctttcctcagaaaacagacttttcctcatgtggact**cttgtctcagacaag**caaatatgtctccatgtgaactcttcactcagataagtacacatatgtccacatttactgtttccttacacaggcacatatattcaatgttgaattgttttgtggcaaaatgatctcaagataatggtaattataaatcccctccctaataaagtgtagatctgtattt**tttttgttgtaacctaactttgccttattgtcaaaaa**cagtaatttgcag**ctctaaatgcaccagttagag**acaggtgtccattttctctggaaatgtatttttattttcttactggacatatttgttgataatgttgctactatgaagatacctgaacagtatccacattaaagaataaaaaagagtaatgggcagaataaccctgtgcatccagacccagaaatcctttgaccctgccctcccgaaatgtagacacagaggatggatgagcagtgctgagcagtgtacccatgaccacaaacatgaagacatggaaatgtgtccccactcctcctcatgaaaggcagctcatgccctcttccttcaggccctggtatggagcc**accccatgtctctgcccttccttagtgtccacaccatggggt**ctgcactgatctggattcccttctcatccccctcaatattagtgtgccttgtgaatcaggtccagctgtggctgttctcagtctgtttcctctgtattcacagAAGTCCTGTGTGATGTTTACTGATGGAGTCAGAAGGAGAAAAATTTAACAGCCCAGCAGTTCACTGAGACTGTCCTGCAAAGCCTCTGGTTTCGGGTTTACTGGGTGCAGCATGAGCTTGGTCCAGCACACTTCACAACAGGAATTGGTGTGGGTAGCAACAGTGAGAGATCATATGGGAGTTCTAAGGGTTACTCTCCATGAGTACAGATAAATCAACATTCCCAAGTGACACC**CTTTCAAGTGCAGTCTACCTTACAAGGACCAACCTGAAAG**CCAAGGGCAAGGCCGTATATTACAGTGAGGGAcacaggggagggaagatc**tgtctgagcccagaca**caaaaatctctgcagagagacaggagggaactgcgtggtagatgctgctcagaaccaccatggggtggccaggaaacaaggggtgcttaggacctctagggagcgctcaggacgccaggggggtgctcagaaccactaggggacgctcaggacactaggtggcgctcaagacaacactactgcactcatgtcaccacggggtactcagaaccaccagggggcgctcagaaccaccagggggcgctcagaaccaccaggtggcgctcagcaccaccagggggcgctcaggacaccagggggcgctcagaaacactcggggacgctcaggacaagagggggtactcagaaccaacagggggcgctcaggaccaccaatgggtgctcagaaccacaagggggcgctcaggacataggtggtgctaaagacaacaattctgggctcaggtcaccagggggcgctc**agaaccaccaggggacgctcaggacatgcaggttct**cttaggaggcagctgcacatcaggagcttcagaggctgtg**atttcctttcagaccctggaaat**tcctgacctggtcaagcaaaagtcttccccaggatgtctgatttttccttttaattccatgatttcttttacctacaaaacactaacttagaacagggatgtaatgcaactttt**attgctgcgtattttctgagtaatactagcaat**gatctctcgggac**aattattaaaataggttatttatatattttaattatt**aaaaataa**tactattattaaaatagta**aaattttttaaatcacacttgtaatcccagcacttggggagactgaggcagacagattacttgagctcaggagttttagaacagcctggcaacatggtgagatgctttctatacataaaagaagaaa**tagctgagattggtgcaaccagcta**cttaaaaggctgaggagggaggattcctta

Uppercase: IGHVIII-26-1

Lowercase: Flanking sequence[1000bp]

Red & Bold & Underline: Stem-loop [16]

Blue: Heptamer[34]

Green: Nonamer [5]

id-IGHVII-65-1-2[V_gene_segment]

caccaccatatccggctaatttttgtatttttagtagagacagtgtttcaccacatt**ggtcaggctggtctcaaactactgacc**tcaggtgatccacccgcctcagcctcccaaagtgctgggattacaggtgcaggccaccaagtccggctaatttttgtatttttagtagagatggggtttcacaatattggtctggctggtctcaaactcatgatctcagcctcccaaagtgctgggattacagatgtgagctactgtgcccgg**ccaacatcacctatattaatgttgg**attgaataactacaaagtctaccctaaccaagtgtaccatcaaacttaccattacccagagggaagaatctttaacatgagatcgatttcttaagtgttttaaggt**gcaaaactgaccactgtacaatcacccaattatgattttgc**taatgagttgtagaagttgcatgtacattttggatattaacactttttcagatgcacagcttataattatattctctctttctgtgggt**tggaaattttttcca**tgctttgcagcatctttttgctgtgatgtagttctgcttgttcaattctgcttttattgtctgtgcctttaatttgaaatataggaaatcattcttaatattattttctggg**tgggggaaattttacaattgcaagccatacgtttaagtatttcgcccatttagagtaaattttggtatttaatctaaa**ctaaaattcttaattctttgcatgtgtagattcagtttttctaacaccctcttagggagagactttgattcatccattgtatattgttagttctcatgctgaaaatcagtttgctgtagatatattagtttatttctgagctctctctatatattaatgccaataccatcctgttattatttatggatcaaaatgtgcatatttgtgaatttcaaaat**gttgttcagaacttgtgcatgccaacaac**tgtgtttctcctgcactcttGGGCTAGTGAAGCTCTCACAGACCCTCTCTCTCACCTGTGCTGTCTCTGCATTCTCCATCACAACCAGTG**CTTCCTGCTGGAGCTGCATCCATCACCCCCCGCAAGGAAG**GGACTGGAGCGAATCAG**GTGCACAGGTCATGAGGGAGTGCAC**ATTCCAACCCACTCCTCAAGAGTCCAGTCACTATCTCCAGATCCACATCCAAAAAACAGTGTTTCCTGTAGCTGAGCTACCTGAGCAACAAGTACACAACCATGAATTTTAATACAAAAGAcacaacgaggggaagtcattgtgagcccagatacaaacctccctgcaggggagctcagaaagagcaggaggcactcaggacaccagggaacactctggacacatcaaggcaggtgcaatggggggacaaggggctggagatggggtttggcatcaccatcgtatttcaccactggacacccaccaccgtgtttattctcatgtacctgattctgtgtattattagaaaatggcatttatataaatatatagccatatgtaggtgcatcaagttgtcctctccatcgtatgtggacctttccactaagactctgagtccctgtattta**tatgagcacctcata**aattatggtcaattttgtgggatctctatcttttctttctcccttcttccctccttctctctcactcacaaactcacacacacgcaaaattctacaacttcagttacgtgatgtgttgaagacagtttataatgtgcagcttttccagtttagctg**atgttcatattgctgtgagaataggaacat**tgtgtttctcagctgtgtatttctctaaggtagcagcaacttttt**tttttaatttgacattttttaaaaa**ttttattattattatactttaagttttagggt**acatgtgcacaaagtgcaggtttgttacatatatatacatgtgccatgt**tggtgtgctgcacccattaactcgtcattta**gcattaggtatatctcctaatgc**atttttttaaatacccagatactgaaagcaatccaaatatcaatcagcagcttaatcataaacaaattgtggtaaattcattccctggaaaatcacccactgttacattctggtaccgctggatacgttcaacagcaaaggtaaagtcacaagtacatatgatgagtcaaataagacaaataaatacaagtacacacatacaattcctc**ttttataaattctataaaa**ttaataccaatctaaagttacataaagaaaaccagtagt

Uppercase: IGHVII-65-1-2

Lowercase: Flanking sequence[1000bp]

Red & Bold & Underline: Stem-loop [15]

Blue: Heptamer[28]

Green: Nonamer [4]

id-IGHVII-30-21[V_gene_segment]

ggccattggtgcctcgctgagcacagttgtccatcagggatctcccatgtgtcccagcagcagccatgcctcagtatctccactgtgcacagccatcgt**ctgggaggagctcccag**gatgggtgtctttggcacacacaggtgatgggtgttagagtgcagtgcagcagctggctgcctggtctattgggctcctggatattggagggattggaggtgcattctcaaggccagcacgctgtttgttaatttttgtataaaaaccatgtgatttaattcattttctcagatgacatagataattaataacacaatctgc**aaacaattgtaattttcaactttaccccaagttcattgttt**cttaattctgtgcaggatccagacatggtattgcccttctcat**gagaaattgttcgacctaaactgaaaccagttgtttctc**gtatactttggttctccccacatgcagagatcttgattagagcaagtttggtactttccacgcactcaccctca**cctccccagataaagagcagaagttctccttagactgagtctgagggagg**agctgttcctgtaccactcagggcctgcggagacccccaggtgcagcttcactgagtcaggtgtttcactccctgtgattgctgctcaggt**ctaattgtgggctcagaattag**gacagtcttcaggttatcacaggtcaatcatattctaaaaatcatcattatcacacaccgtggtaataattcaatgttcatttttctaaacagcagtttctcttttttatttggttgcaagtctgaggaaaggaacatgtgataatactttttaatctaacctctaaatctactgaattgttctaggagactcacaaatcggacaaagtgagctctttatgctcataaaaaaatgcatgtatttgggaatttcaccgt**gttgtccaggacccgttaacatggacaac**tgtgtttcgcagtgcacttctggctt**gagacatccTCACAGACCCTCTCCCTCACCTGCACTGTCTC**TGGATTCCCCATCATAACCAGTGTTTC**CTTCTAGAATTGTATCTGCTTGCCCCTAGAAG**ATGGACAGGAGTGGATCAGGTGCA**TGGGTTGTGAAGGGAGCACAAATTACAACCCA**CTTCTCAAGAGTCCATATCCGGATCCAAGAAACAGTTCTTACAGCTGAGCTCTGTGCCCAGTGAACACACAACTACGCATTTTGAAGCAAAAGATG**CAATGAAGGGCCTTcattg**tgagcctagacacaaacct**ccctgcaggggtgaataggagcagcaggg**ggcattcggggcagtatgggggcttaggatgattgttaggggtcaggatgagcaggaccaaggcttcgcatcagggcaagtgcaacagggcagaaaaggggctgtagatgtgggttgttctcaccatcatatttcac**caccagacaccctccactacatctcttctaatgtgtctgagtg**ttgatatgattagaaaatggcatttatgtaaatactactatgtacccat**atggaggtgcatccagttgtcctctccat**cttatgtggaccttgtccatcaagcactaagtccctgtaattacttgagtacctcacacattatg**gtcaaaagatgtagggtctccttttgac**atggtctctcctcctgccttctct**ctctgtcacacaaaaacacgtgaactgacacacacacagag**cttcccaactttaattatgtgatgtattgaagcaaattgattagtgtgcagcttttctgctttgcctgctattcatgttatgtaaaaataagaaccatgtttttctcagcttgtcacttctctaagctaagtagcatctttgtttattatacccagagaccaaaagcgatccaactgttattcagcagcttaactggtaaacaaattgtggaaaattcatttactggaataatacccactgttacaatcaagtactgctggatacactcaacaccatgcttaaaataacaagcacctgaataagtaaaataagtcaaacaaatgaaagtgcatacatacgattccacttctataatttctataaagtaaaaatgaacttaaagttacatgaaaagatctgtagttgactggggacatggtacaagaagagaaggtatagaaaggagaaactacaggagagcaaaaggaaatttctaggataattgattttttctctgttagtaaaagtgaggattatgtcactatttgtaa

Uppercase: IGHVII-30-21

Lowercase: Flanking sequence[1000bp]

Red & Bold & Underline: Stem-loop [15]

Blue: Heptamer[31]

Green: Nonamer [5]

id-TRGV7[V_gene_segment]

cttatctatactcccaattccaattccttgtaaacatactttgtaaagtcctgtaagatcctgtctcctttgccatgacgctgcaaggtcataaagtagat**aaaacctaagttgcaattccggtttt**cctcaagatctaagac**atgttacaaatggttaattgcctttgtttctcgctttggtaacat**cttcccgcctcaggtatttcccgccttga**agagtttaaaaggcaatcctataatctaactct**ggctacccattctggaccccctccatgctttggaagctttgtactttcactctgctcaataaagcctacagctttttctctctctcggtccgtgtttctatcactcgctgcggtcagccgccacaccaattctatggcgtggctaggcaa**gaaccttaggtgttacactgttaataaagcaggttc**aggccttctcagagtggccctgggtattttgcctccaagtgtccttgcgaatgtgaaaaggaaagcagttgaaagaccaaatcaa**gcttcctgacaggacaaggccctggaagc**agcatcaactgctgaaactctgaa**gggcagggtgtgagcctgctgtggactgccc**cgtcctatgg**ccctggccagctggcgccaggg**tcagctttccccacatctgcactccctctgctggaggcctgagctctccatgcccctggctctcttctgcagagggcctg**ctctctgctgccacaagatggcgctggagaaggggcctgagcagagag**gcaggagctcctctgcagg**tcctcccagacaccacccctctgggagga**aggctggggagtcccctaagacagatcctccatcacttctccctgcctgtgactcaggaagaccagctcctcctactgccttctgtgctagggaacacttcctgtttgagtgggacctgagttttgaaagcgtcttctgctcctcttcatctggtcccttt**ccttccaagaccccagagaggaagg**cATGCGGTGGGCCCCAGCCCTGCTTCTAGCTTTCCT**GCCTCTTGGTAAGTGTGCTGCCTACAGAGAGGC**TGATGGGTTTTATTTTGTTCTGTTTTGTTTTCTCATTTTGCAAGGGGTGCCGTTCTAAGGAGTTCCTCATTATATTTTGTGCTGTTCCCATTGCAGCCAGTCAGAAATCTTCCAACTTGCAAGGGAGAAGGAAGTCAGTCACCAGGCCAGCTGGGTCATCTGCTGTAATCACTTGTGATCTTACTGTAATAAATACCTTCTACATCCACTGGTACCTGCACCAGGCGGGGAAGGCCCCACAGCATCTTCCATACTATGACCCCTACTACTCCAGGGTTGTGTTGGAATCAAGAATCAGTAGAGGAAAGTATTTTACTTATGCAAGCATGAGGAGGAGCTGGAAATTGATACTGCAAAATCTAATTGAAAATGATTCTGGATCTATT**ACTGTGCCACCTGGGACAGGcacagt**gattcacacctgccctacaccacactgaaaatctgc**cttgtgactgcttctggtacacaag**atagaccagccccctctcatttgttgccaccgaatttactgtattctgtacaaagagaaacacagcataattcctgatctctcccctaatatcacactctcctggcagcagctgcaccttgttccccaccctcccccaggacttt**cctgaagacaagctgtcatcttcaggcctcagccaagcagcctggctgagagc**aagactctctcagctc**tccgaggacatgggggaggcccactcactctgcttcctcgga**ccaacaggtacacctagg**gtccaactgcgaagcgagatattttggac**agcaaatgggaaggatatttatactgaatcatatatctagagatcacagctgagagtg**ttgcttattctttatgtctataacaatataagcaa**tagtataattaccactataacactaagcctaggtatgcagttgaacattgtctccctctgctttcctttttgtctctctctctactttcttggttacttgctatttttctacacatccctttttctggcctcccctccctccctccctagctccctccctagcttgccttccttccttccttccttccttcctttcttcctccctctctccttcccttccttacttccttcctttctttcttcctccctctctccttcccttccttacttccttcctttcttcctccctccttctcttacttccttctttttttcctttcttccttttctctctctttctttcttttttgggcacacaatcagaatacagaatcattctctatatgttcatgccaactttgatacagctagtttattcgttcagttttcatttttatattttggaaatctgttgtattattttgatttgtat**atgtaaaacctttacat**ggaactgaagtcaaaattataaaacaagatcaatttagagattttagc

Uppercase: TRGV7

Lowercase: Flanking sequence[1000bp]

Red & Bold & Underline: Stem-loop [19]

Blue: Heptamer[16]

Green: Nonamer [6]

id-IGHV1-69-2[V_gene_segment]

gactcgccatgtgtctggaccatatcagtgcatctggagct**ccagggaaggggctccctgg**tggatttagtgattccttgcttcctgtgccaaggtctccctgtgggatatgttaggtttcctaaggtctattttctagtgtaagaagtgatgtgagaagtagttgctgtcactgaaggagcattcttagccagggcacagccaggtcata**ctgggcttgagatgctgggagaaaaatgctctgtgagcccag**aaaagaactt**ccctgcagggcaggagctgagctgcaggg**ggcgctcagga**tgcacccagcacaggatccagctctggagcaggtgca**caggaggctgtggaggggttttctctcagggattgaatattc**cttatttcaaagccataatgacataaaatttaaataag**aatttagcaagtactgatgtgtctttagtattctcttgcatacacagaggctaccaaacccaataacttaacgcaaaac**acatttaaaaggagaaatgtctaggcctttcaaatgt**atttatagttaggaattgaggggtggttttattaattcaatgggtgttactctccggagacacactcatcccagaagttagatgtgcagagggcaaggcccaggaaaagttcagattgtcagtgagccataggaacaagaaa**tcacagtgaggacaatgccctgtga**gattctgggtttctgtaagaggagttctgtcttcatggacttctaagtgtgtcagaggacaaatatcattaaacaaagttcagggcagggagctcagtgtcccactgtggcatggtccgtgtgtcacctatcttcttccccagggtggggtggcctaagctatgaaatacctgcctcatgaatatgcaaatgcagtggtgtctaccgaggtaaatacagatctgtccttgcccagagagcatcacacaacagccacatccctcccctacagaagcccccagagagcagcacctcaccATGGACTGCACCTGGAGGATCCTCCTCTTGGTGGCAGCAGCTACAGGTAAGAGAATCCTTAGTTCCAGGGCT**GATGAGGGGACTGGGTCCAGTTAAGTGGTGTCTCATC**CACTCCTTTCTCTTCTCCACAGGCACCCACGCCGAGGTCCAGCTGGTACAGTCTGGGGCTGAGGTGAAGAAGCCTGGGGCTACAGTGAAAATCTCCTGCAAGGTTTCTGGATACACCTTCACCGACTACTACATGCACTGGGTGCAACAGGCCCCTGGAAAAGGGCTTGAGTGGATGGGACTTGTTGATCCTGAAGATGGTGAAACAATATACGCAGAGAAGTTCCAGGGCAGAGTCACCATAACCGCGGACACGTCTACAGACACAGCCTACATGGAGCTGAGCAGCCTGAGATCTGAGGACACGGCCGTGTATT**ACTGTGCAACAGAcacagt**gtgaaaacccacatgctgagagtgtcagaaaccctgaggaaggaggcagctgtgctggggctgaaaagatgacagcatttatgaggtttaaagtgtttagaagatgggttaagtcacagagtaaaagtagaaatagaaagatgtatacattccaattatataggaaataatcttttcaactttcaccctgtaagtaacattcacagagtgggaaaggcagcaatcaatggagtggatgcaaacattcctttgtgaggacataaattttaaaatcgaatggataaatcatttggagcaggattgcttgatcacatggtaagactaaatataatttctaaaaagtgacca**acatttctaacaaaatgt**ctttgccactccttttacattaattttattttaaaacatttttaggattgcagcaaatttgagtagaataaacagcattcccatgtactcctgcccaagacatgcacagacttctccatgatcaacaccctgcgctgaagtaataaataacttgcaacagatgaacccacatggacaccttgattgtttcctttcctggtggcccctggtataacaagcctaaattaccttgaagcaccccaggttcttctagtgggttacaggaaatgatgccaggtagagggaaagtgagtggggatgatcctctttgcactgtttcctcaagaatccataaaatgtacattgatttagagctcaactaactcctggttttctatgcccctttccagaaggtaaggtctccaagaatttacagcaggggatctcctcaagagtccataaaatgtacagcggtttagagctcgtctgactcctggttttatatgcccctaccaagaggataaggtctccaagcacttacggc**aggggtctccaacccct**aggcagtgaacccgtactggtca**gcagcctgatagaaagcaggctgc**acagcaggagatgaacagggggtgagcatcactgcctgagc

Uppercase: IGHV1-69-2

Lowercase: Flanking sequence[1000bp]

Red & Bold & Underline: Stem-loop [12]

Blue: Heptamer[54]

Green: Nonamer [9]

id-IGHVII-49-1-2[V_gene_segment]

cacaacagtgctggagaaggtgggagtc**tgatggagcttcccaacaaccctgtggtctaagctaagtccatca**aggccgctggttcctcctggaacacagctgtccatcaggaatcccccatgtgcccagcagcagccacgcgttagcatcttcactgtgcacagtcattgtttgggaggagctccaggatgggtctctttgtcaaaaaccgggtgatggt**gtcagagagctgtgccgggtgcctggtccaggggctccctgac**gttggaggtataggaggtgcgttctcagggtgggaacaccttttatggaaatgtacagaagaaaacatgattttccttggtcctctcagatgacatggagaa**gcaagcgcgcagcttgc**aaacaattgtaatgtccttgtaatttcctactttatcccaagttcattgattcttgtaaaccttggtttctgtccaagtacagagatattgattagagtgagtttggttctttccacacactaaccctca**cctcccccagataaagagcacattgtccttactctgagtctgagggagg**agctgttc**ctgtacaactaagggcctgcagagacccccgtgtacag**ctttgcagagtcagatctttccacatgtgaggcgacctccacggcccatgattgctgctcaggtctaattgtggattcaggattaggacatcctttaggctatcacaggtcaatcagattctgaaaaatcactgttatcatagacagaggtaataattcaatacccactcccctgacagcagattctctttcttatttggttgaaagtttg**aagaaaataagtgaccatttattattctt**attctaactttagtatccact**atttgttctaggggattcacaaat**ttcaaaacactgagctcttcattctcatgaaaatgtgcatatttgtgaatttcaacgtgttgttcagaacctgtgcatgccgacatctgtatttctcgtgcgttcttGGCCTGGCGAAGCCCT**CACAGACCCTCTCCCTCATCTGTG**CTGTCTCTGCTTCTCCATCACAACCAGTGCTTCCTGCT**GGAGCTGGATCCCTCAGCTCC**CCAGGGAAGGGACTGGAGTGAATCAGGTGCACAGGTCATGAGGGAGAACACAACGCAACCCACGCCTCAAGAGTCCAGTCACCATCTCCAGATCCACATCCAAAACACAGTTTCTTCTACAGCTGAGCTACCTGAGCAACGAGTACACAACCATGAATTTTTACACAAAAGAtacagcaaggggaagtcattgtgagcccagaaacaaa**cctccctgcagggaagctcaggaactgcgggagg**ccctcgggacaccagggggcgctcaggacacacatcaaggcaggtgcaagaggaaaaggtgctggagatggggtttggcatcatcatcatatttcactgacacccgccactgtgtttattctcatgtacgtgattctttgtattattagaaaatggcgtttatgtaaataaataaccatatgtaggtgcatcaagtcgtcctctcccttttttttttttttttttttttttgagtcctgctctgtcaccc**aggctggagtgcagtggcgcgatctcggctcactccaacct**ccgcctcccgagttcaagcgattctcctgactcagcctcccgagtagctggaattacaggtgtgcaccaccgcgcctggctaagttttgtatttttagtagagaccggatttcgccatattggccaggctggcctcaaactcatgatctcagttgatccgcccgccttggcctcccacagtgctgggattacaggtgtgagccaccgcgcccggcccgtcctctccatcttatgtggaacttttccattaagactctgagtccctg**gatttatttgagaacctcataaatc**atggtcaattatgtctctttctcctctttctcccttcttcactcctctctagctcacaaactgaaacacacacagaattttctaa**cttcaattacctgatgtgttgaag**acaattgattaaagcgcagcttttccagttcagccactgttcatattgttgtaagaataagaacgttgtgtttctc**agctgcgtgcttctctaagatgagtagcagct**tttttttaataatacccagaggctgaaagcaatctgaataccaatcatcagcttaatcagtaaacacaaggtggtaaattcgctcactggaaaaacaccgactgttacaatcaggtaaggctggatactc

Uppercase: IGHVII-49-1-2

Lowercase: Flanking sequence[1000bp]

Red & Bold & Underline: Stem-loop [14]

Blue: Heptamer[37]

Green: Nonamer [7]

id-IGHV1-69[V_gene_segment]

aatgggcacataactgtgggagatactatattcctatgaagaaggtattcagacttcagagataagtaatgtttcctacattgtgcttgtgacttggaagcagtggattgaagagtgtgataagtgcccagaccaagcagaacagaaatcagcatgtaaagatgatgatctatggatatgatctaaaaccatgtaaatacttcaaataattctatttaatgc**agtttgaaataaaacacaaact**tattcaaaataca**aattacttggtaatt**attttgggagcta**tgagttcaccaagaaactca**aattcctatttctatttcaacccctgattcctactgtcaatgggagggaagtctcagaaccaatcacacatcagacggcaaatctgtcaaccaagagtctttccactgaaggacctgggaggtcaggaccctcaggaaagtgctggggaccctgtcttgg**gagtgcccagcagatctcagaactctccatgggtcctgctggacactc**atgtag**ggtaacgagtggccaccttttcagtgttacc**agtgagctctgagtgttcctaatgggaccaggatgggtctaggtgcctgctcaatgtcagagacagcaatggtcccacaaaa**aacccaggtaatctttaggccaataaaatgtgggtt**cacagtgaggagtgcatcctggggttggggtttgttctgcagcgggaagagcgctgtgcacagaaagcttagaaatggggcaagagatgcttttcctcaggcaggatttagggcttggtctctcagcatcccacacttgtacagctgatgtggcatctgtgttttctttctcatcctagatcaggctttgagctgtgaaataccctgcctcatgcatatgcaaataacctgaggtcttctgagataaatatagatatattggtgccctgagagcatcacataacaaccacattcctcctctgaagaagcccctgggagcacagctcatcaccATGGACTGGACCTGGAGGTTCCTCTTTGTGGTGGCAGCAGCTACAGGTAAGGGG**CTTCCTAGTCCTAAGGCTGAGGAAG**GGATCCTGGTTTAGTTAAAGAGGATTTTATTCACC**CCTGTGTCCTCTCCACAGG**TGTCCAGTCCCAGGTGCAGCTGGTGCAGTCTGGGGCTGAGGTGA**AGAAGCCTGGGTCCTCGGTGAAGGTCTCCTGCAAGGCTTCT**GGAGGCACCTTCAGCAGCTATGCTATCAGCTGGGTGCGACAGGCCCCTGGACAAGGGCTTGAGTGGATGGGAGGGATCATCCCTATCTTTGGTACAGCAAACTACGCACAGAAGTTCCAGGGCAGAGTCACGATTACCGCGGACAAATCCACGAGCACAGCCTACATGGAGCTGAGCAGCCTGAGATCTGAGGACACGGCCGTGTATT**ACTGTGCGAGAGAcacagt**gtgaaaacccacatcctgagagtgtcagaaaccctgagggagaaggcagctgtgccgggctgaggagatgacaggggttattaggtttaaggctgtttacaaaatgggttatatatttgagaaaaaaagaacagtagaaacaagtacatactctaattttaagataaatattccattcaagagtcgtaatataagccaaattcac**agagtggaaaaggccacactct**ataacgttgatacaaacattccatgaaggtgctactgtgaacaagttttcaaattggatgaat**acatgatttggagcaaggttatttgatcatgt**ggtgagac**taagaatgattctta**aaaagtgccaaaagtttccttcaaatgtttctgtcactccttatcataaagtttattttacagcagttttaggattacaaagaaattgcacaggaggcgtgagaattcccatgactccctgccctacacaggcacagcctcctccactacgaccatcctgcaccgcagtcacaaatcagttacaatggaggaatctccaaggacgcttggttctttctttttctggtgatctcctaatataacaagcctaagtatctcaagattccacggttttttcagtgttttctagaactgatattagtcagagggaaagtgggtaaggctattactatttgaactctttcttccaaaatccacaaaatatatattaatttagagctt**atcttacttctggtttacaatgctccttcccagagagtaagat**ttttaag**cttttagaggccagttcatgtctctaaaag**accagaaagttctggggaatcccataatgaacatcctt**tcattggaattggagaccctggcaatga**gagattccatgtataatgccctagagttggattaggtgcactgtgagctcttgggtggtgagtctgagtaaggtggtttgtgcagcaaatgcaaacacattcatgggacccaggtaggaaca

Uppercase: IGHV1-69

Lowercase: Flanking sequence[1000bp]

Red & Bold & Underline: Stem-loop [16]

Blue: Heptamer[50]

Green: Nonamer [5]

id-IGHV7-4-1-2[V_gene_segment]

aaaggaactgacaagagattaggtgttttcccaaagtccctgggagttcctggactcactatgagtgtggactgatccagtgcttccggagct**ccagggaaggggctccctgg**tggtttcatagaatccttgcttggggtgtttctgcagagttcactggctttcctacggccaatttactattgtaagagatggtgtcagcagtacatggtgtcacgtgtcactaaaggagcattccgagccaggacacagccacttcatacttgggggaaaatgctctggg**agcccagacaggagcctctctgcagtgcaagggctgggct**gcaggggacgctcaaagcccacccagtacaagtttcag**ccccagagcaggtgcacaggaggctgggg**agggattcctctcgagatctgtgtcattctttaaaaaatctaaaatacatatttgacaactgaacattctataaatatcctattcaattgcgagcatttatcaaacttgatgttgtaatgagaaccacttttacactggagatttctaacccttctagatatcttaatagtatgcagctggaggttaaggaaactacttttcttttctataaataagtgaaacttttggagaaacacactcatcccccaaa**taatacattcatgtatta**aagcctagaaatgcttatattacctctgagccatttgagtgtgggttcccagtgaagtcctgttctagggaaaattgttagccaatgggag**aagctctgtcaacatagagctt**agggatatggcagggctcacatggcctctaaggggattacagcttgaaccttgagcatcctccgtgtgtcatctctctgctctttctcatgcaatatcaggtatgaaataggatcactcatgaatatgcaaataactgaggtaaatatagctatctttgggccctgagagcatcacccaacaaccacacccctcctaagaagaagcccctagaccacagctccacaccATGGACTGGACCTGGAGGATCCTCTTCTTGGTGGCAGCAGCAACAGGTAAGGGGCTCCCCAGTCACTGGGCTGAGGGAGAAACCAGCACAGTCAAGTGAGACTTCATGCACTCCCATCTCCTCTCCACAGGTGCCCACTCCCAGGTGCAGCTGGTGCAATCTGGGTCTGAGTTGAAGAAGCCTGGGGCCTC**AGTGAAGGTTTCCTGCAAGGCTTCTGGATACACCTTCACT**AGCTATGCTATGAATTGGGTGCGACAGGCCCCTGGACAAGGGCTTGAGTGGATGGGATGGATCAACACCAACACTGGGAACCCAACGTATGCCCAGGGCTTCACAGGACGGTTTGTCTTCTCCTTGGACACCTCTGTCAGCACGGCATATCTGCAGATCTGCAGCCTAAAGGCTGAGGACACTGCCGTGTATT**ACTGTGCGAGAGAcacagt**gtggaaa**cccacatcctgagagtgtcaaaaaccctgtggg**aggaggcagctgtgctgagctgaggcagtgacagggacaacgtggctgcaccctttgtagggttataaacatacatatatatgcacccaacagtggagtacccagatatatcaagcaaacattattagagctaaacagagagatagacccaatacagtaaaagctggagactttaacaccccaccctcagcattggacagatcttccagacagaaaattaacaatgaaatcttagactaaatgtacatgacagaccaaatggacataatggatattgacagaacacttcatccaatggctgtagagtgcacagtcttttcctaaac**atatggatcagtctcaaggatagaccatat**agtagaccagaaaattagtcattttttaatt**gagaagatatcaagtaccttctc**tgatcataatggaagaaaactaaattaataagaggaatcttggaaaactatgcaaacacatggaaattaaacaatatgctccctaattaaacaatgtgctcccaaattgaaccacatgctccaaaatgggttaataaagacataaaaaattttaaaatttattgaaagaaatgataaagaagacaaagcatatgaaaacctatgaaatacagtgaaagcagtattaagaggaaagtttatagctataagcact**tactttaaaaaagta**gacaaccataaaataaacatcctaggaatgtcttttaaaaaatagaaaagcaagagcaaatcaaacccaaacacagtaaaaaaaaaaaaaggaaagatcaaaacgttaatacataaaattgaaataaagaaaacaatacaaaagatggatcaaatgaaaagctgttttttaaaaagataaaagacatctacaaacctttaaccagactaagaaaaaaagaaagaagaccaaagtaaatgaaatcagagagaaaaatggaaatattacaaccaatgg

Uppercase: IGHV7-4-1-2

Lowercase: Flanking sequence[1000bp]

Red & Bold & Underline: Stem-loop [11]

Blue: Heptamer[42]

Green: Nonamer [1]

id-IGHVII-30-1[V_gene_segment]

ccagcaagaccattggtgcctcccagagcacagttgtccatcagggatgtccaatgcttcccagcagcagccatgcctcagtgtctccactgtgcacagccactgt**ctgggaggagctcccag**gatgggtgtctttggcacacaccaggtggcgggtgttaagagtgcagtgcagcagctggctgcctggtctgttgggcctg**atgctggagagatgggaggtgcattcttggggccagcat**gctgtttgtgaatttttatataaaaactctgattttacttcattttc**gcagatgacatagataactaaaaacagaatctgc**aaagaaattgtaattttcaactttaccccaaattcattgtttcttaattctgtgcaagatccagacatattattgccttcctcatgataatttattctatgtaaaaccgaaatcattttttctcctactctttgtttctgttcaagtacagagatcttgagcaaagtaagttgggttctttccacacactaaccctcaccttccccagagaatgagcagagattgtcctcagtctgagtctaagggaggagctgtt**tctgcacaactcagagcctgcaga**gacccccaggtgcagcttcagtgagtcagacatttctccatgtgggcgacctccagtgcctgtgactgctgctcagg**cctaattgttggttaagcattagg**acactcttta**ggtgatcacatctcaagcctattctgaaaatcacc**atgatcagagatagttcaatggccattctcctaacataagtttctctttat**tacttggttccaagta**tggagaaaaatgtgaccctatatttgtctgaatccaacatcagcatcaactgcattatgctaggaga**ctcactaattgagcacaagtgag**ctcgttattctcatagaaatgtaagtatttggaaatttcagtgtgttctccagaacctgtggctgccaacaactgtatttctcggtgcactcttGGCCTGGTGAAGCCTTCACAGAACCTCTCCCTCACCTGTGCCATCTCTGCATTCTCCATCACAACCAGTGTTTCCTTCTGGAGCTGCATCCATCAGCGCCCCCATGGGAGGTACTGGAGTGGATCGGGTGCATAGGTCATGGAGGGAGCACAAATTACTCCCCTCTTCTCAAGAGTCCAGTCACCATCTCCAGATCCATGTCCAAAAAG**TAGTTCTTCTTACAGCTGAACTATGTGAGGAACAAACACATAG**CCATGTATTTTAGAGCAAAAGA**cacagtgaggaaaccacagtgtgaactcatacccaagcctccctgtg**ggggtgcataggacagccagggttactcaggacaccaggctccctcaggacaccaaggggcactcaagaccattgtagaggcatgcaggtagctgggggctctcaggaaccatgggggaaaatcaggacaccaaagggtgcttggtacagcggggggctcaggattattgtggcgattcagaaagagcaggttcaaggctcaacctcagggcatgtgcagctggtgtgaaaaggagctggatgaggcgttttgtgtcaccatcatgtttcaccaccagacaccctccactacctcta**ttctaatgcatgtgtttgtatgattagaa**aatgatattgatataaatatataaccatagctaggtgtgtcaagttgtcctctccatcttatatcagccttgtctgtaaggactaattccc**cataattacttgagaacctcataaattgtggtcaattatg**taggattcctctctttttctgccttccttcctccctcattctctctctctcacacagaaacttatatacagccaccccacaacacacataaatcta**taacttttattacctgatgtattgataagtaatctaaagtta**cataaaaatcagtagttcactgtcaatattgtaggagaagggaaggtctaggaaggaggaattacag**aacaagaggatattttgagggtaattgacttgtt**atctatgtggataatgataatgtctatgaccatatttgtaaaattgaacacttcgtgtggagactattatttttttatttaaccccatgccattattttgccgtattctag**catttctctaaaaatacagaaatg**ttcagcactcattcatgtgtatattcaggagtttctgacttttcatgtattttatttatctctgtctaattgctttgataccaaattttacctagtataattagtactacctttagcactga

Uppercase: IGHVII-30-1

Lowercase: Flanking sequence[1000bp]

Red & Bold & Underline: Stem-loop [16]

Blue: Heptamer[28]

Green: Nonamer [4]

id-TRBV24OR9-2[V_gene_segment]

tctgcacacttgggctcctgcatctgtccgtctgcgtgttccctttcccct**caggggtttgagcccctg**cagcaaccgaagaggtga**gcaacactcctgttgc**acgtccttccaggggggtcagggaactctcctttttcaccaggacaagcacagatctgaagcagcaaagagccagttgtttattacattggacaatatctcacactatcactataactgttctgtgcttgcaatctgctgcattaaatattaaactcttgcctcctaaagtcagg**aataatttatttagaataaaatgaattatt**taatgaaccactgtattttcagctaattcaaaataggaatgaaggtgttaa**gtgcttctttgaagcac**atacccacattaaactaacacctttctacaataaaccaatatttttaataaagctctgtgcccagaaaacactgaccagaaaagcaggagccactaaa**aaattggagcaattt**gaaacaaagttaggaatcctgt**cattcttaaaacaaaaaagtctcaaactttacagagaatg**tattcaaattgcatagtttaccaagaaccgagttagaataaatgattaagcacagaaaaaaaaggagaacaattatattgtcctatcattcaagtgagactagactccctttattc**aaaattcaacttggttttgaatttt**ctccagcctgctttatcaggaggatgatgtctccttgaacaactgaaatagatatgccttgagaaaggggagagatacaa**ctctctggtggttgtcagtaaagcaccagagag**agaagagaagcaagtacaggaatttcagcatctgctaagaacatctaatacataagccaacatggcatctttaagacccagtaccttggaagggacaatgacatcacttcctgaatcctcctagttttcta**tttccatgccctgcttccctcaacatccagagctggaaa**cacctccatcattcctcttcatgccATGGCCTCCCTGCTCCTCTTCTGTGTGGCCTTTTGTCTCCTGGGAACAGGTGAGTT**TGGAAAACAGATGGGGAAATCACTGCCTTAAATTTTCCA**GGTTCTGGATTAAGCCTATCCATAGAGATTGCAGCT**TGAAGACATTTACCAGTCTCTGTCTTCA**GTTTCTGTCTTATTTCTCCACAGGGTCCATGGATGCTGATGTTATCCAGACCCCAAGGAATAGGATCACAAAGACAGGAAAGAGGATTATGCTGGCATGTTCTCAGACTAAGGGTCATGATGGAATGTACTGGTATCGACAAGACCCAGGACTGGGGCTACGGTTGATCTATTGCTCCTTTGATGTCAAAATATATAAACAA**AAGAGAGATCTCTGATGGATACAGTGTCTCTT**GACAGGAA**CAGGCTAAATTCTCCCTGTCCCTAGAGCCTG**CCACCCCCAACCAGACAGCTTCTAGGTTACTTCAGTGCCACCAGTGATTTCcacagtgcttcttggccacctgctctctacacagaaagacagacacatgggtgagttgtttgctctgaagggtacctggatgtggtttgtgggatgtggggtgtttagagctttcagtggttttaggtagtgtgagctaagggccactttggatcaatgtcctcaagccatgtgatgaatctgaaaccacagaatcaatctcccctattttattcttcctagggagctagcccaatggtgacctttcaggagaatgaacttgagcatttgacagaagctcatgattttcaacaagagcatttcttataagctcaatacaacgtagtttttatagtctctttgccattatctccacctgctttgcaatatcccacactttgttctgttcccacagacaactcttttatcctgtctaacctttctctgccccagttcttccaagctcagctccaccatttagctctatatttgattgcctttctacta**aaaacaaaataactagcttgtttt**ctaaaacttcactgtaagtaa**atttaattatcattaaat**ctgactcagaagctaaagttcaagtatcagagactgatttacaaaagtaatttttcatctactaccccacactttcatctctgctgagattcattttaacgtgcagatgcagacagacttgcaatattatctggatgtact**gaatttcaaaaaataaaattc**cctgtccccagtatgaggtcaaagctaggagcttggag**gtgcctgttttctttccaggcac**tggaataat**tccctggttcccctgccgtgggagttcttttaggagccaggga**ggcccaggcacattg**atcaaggactgccagagcttgat**gtaaaagtcttcctgagagtcaagttgatgtcgtcctcatgggttttgtcttcttgtcccacaagcctcaattatttcagcaaaagaatccatgggacttggtgactgttcagaaagaaagaatccaacttgttccct

Uppercase: TRBV24OR9-2

Lowercase: Flanking sequence[1000bp]

Red & Bold & Underline: Stem-loop [19]

Blue: Heptamer[18]

Green: Nonamer [8]

id-TRBV4-1[V_gene_segment]

tcttcattctcccctggcctttca**aacccttctctgctgctcagggtt**gtgagatgtggaacccatctggatgctgacactgggtcggtga**gtggccaaatactgttttggccac**aggcaggagctgtgctctggtgaaacctctaaggcagggacctggggaagggatagagaataagaggagag**atgggccaggagggatcttgcccat**ggggagccaaccgtacctcaactgagagcatggctgtactcagctcaagcacactaac**atcagcttcaagctgat**ttccctttcctttgctccacacaccttgtgttgttctttttcagcatcctgtgtatgttagctctaagcccagccaagccccatgccctccacttcagggacagagggtggctcatcccttttgtgctccctcagctccctccccagtgcctggtggctggttggcaacacagtcatgcccttgaatgatgggagctggtgcac**actgcatccgtgttcatgcagt**caccctccctccacagagccgggcagaaagggctccaccacacttagacattgagggaagggacgttttgaatggcactgttttatgtgatcccgcaggacagaaggagatttcttgaatgggagtggtctgttgttgagag**ggatcctgaaagacagggagacaaagatagagggaggatcc**acataattaggaagcagctgagagcaaaggagcccctgccagaggaatatccttggggggtaaaaagccggctctgccctttctccccagccgattcatcctagcccagcaaattcaaatctaccttctatcagaacttagaaa**ggatgtaaagcagtcaggaagaaacatcc**cctgggtctggggaaactatcaggagcagtgacatcacaagaaaaaccaccaagcagggccaaggagaccagagcccagcacctcgcccaaaggaccccagtcagaggccccatctcagacccgaggctagcATGGGCTGCAGG**CTGCTCTGCTGTGCGGTTCTCTGTCTCCTGGGAGCAG**GTGAGTTAGGTTGAAATTGTCTGTCCTTGGACTCCAACCCTTTTCCTTGTGGCTG**CAGGAACAACCCTCTTCCTG**GGCTCTGCCTGAATTTTGTCCCTTTCCTCCTACAGTTCCCATAGACACTGAAGTTACCCAGACACCAAAACACCTGGTCATGGGAATGACAAATAAGAAGTCTTTGAAATGTGAACAACATATGGGGCACAGGGCTATGTATTGGTACAAGCAGAAAGCTAAGAAGCCACCGGAGCTCATGTTTGTCTACAGCTATGAGAAACTCTCTATAAATGAAAGTGTGCCAAGTCGCTTCTCACCTGAATGCCCCAACAGCTCTCTCTTAAACCTTCACCTACACGCCCTGCAGCCAGAAGACTCAGCCCTGTAT**CTCTGCGCCAGCAGCCAAGAcacagccttgcagag**tcaccgct**ttcctgtgcagaaaccttcggggcctgccaggaa**g**ccgtgggggccacgg**a**gggctcgggtgaacatttcctccaagagccc**cgaagaagcttcagaacatcatagcacctgctaattcatccatgtggcaactttacatcctatgacatatttagagtgtgggtttcctttcgcctgagtgtgactgccccgtcaactgatttgaaaaaagagaaacatattcagatctagt**ttaaaaaaaaaaatttttttaa**tcatgtaataggccgggcatggcggctcatgcttgtaatcccagcactttgggaggccaagacgagcggatcatgaggtcaggagtttgagaccagcctggccaacatggtgaaaccccgtctctactaaaaatacaaaaatc**agctgggtgtggtggcatatgcctgtaatcccagct**actcaggaggcaggagaatcacttgaactcgggaggtgg**aggttgcggtgagccgagaccgtgacattgcactccaacct**gggc**gacagagtgagattctgtc**tcaaaaaaaaaaaaaaattcctgaaataaagcaagcatgctgtaag**gattgtgtagcttaaaattttataagaaccctctaaacaatc**acctgtgtcaccacagaggcacaccctgcactgggtcagacactgtcctgtgaggagctgtgttctcctgacacttcatgtccagtggtttccatgactcttctttggggccaagggttccctttcatcataggttgttgtcatgcagagtcaagcaacactga**gcactgggaaatcagagaaggaacttccaggcaagggtcccagtgc**agaagccagctagagaggacagtagtagactcaccctttgggaaatatgtttgagaaatatgtatctattttcaccagttctattactctatattccagcttgtctattctctaatttctcatttgtacactgaaagctaga

Uppercase: TRBV4-1

Lowercase: Flanking sequence[1000bp]

Red & Bold & Underline: Stem-loop [19]

Blue: Heptamer[38]

Green: Nonamer [3]

id-IGKV1-6-3[V_gene_segment]

cgttcatcatggtacagttaacccttgaataacttgggtttgaaatttgcagatacacttatattttgttctgtgtgtgtcacctgtaagcaaatgcttaataagtagtaaatatattttctctattttatggtttccttaataacatttttttctttagc**atactttattggaataatagagtatataatatatataacattaaaaaacatgtta**atcaactatttatttt**attgctaaggcttccaatcaaaagcaat**ctattagtacttaagttagggatgcaaaaattctacacaggtgcagcccatttagtctctgggtattaaaaagacctgtagctcttgataatggtggggctgttagtgccacatgcattgaagaagcacggcatcatctttgtgtatctgtagtaa**attgtttgatcttatactggtaagaacaat**ggcataacaccattacctaatacttacaaatatatatagcatcatgtcaataaattttatttttaattttttagaaaggaacaatgttaaactcacagaaatgtttcaagtatagcacaattacccccttccctaaatggaatcttataacagtcttttgaagacctgagaatcgtaccatctaatatgtactgtgtttcctacaaacaagaatattctccaaaataatccccatacaccaatgaaatatattaccccattgactctcgaggaatattgtcaaaaaatatctttaaaa**tgtttctcataacaaaatagtccccagtagaaaca**cattctctgcagacaaatttgtgctaccctggtcttacctgggacacctggggacactgagccggtgctgagttactgagatgagccagccctgcagctgtgcccagcctgccccttcccctgctcatttgcatgttcccagagcacagtctcctgacctgaagacttattaacaggctgatcacaccctgtgcaggagtcagacccactcaggacacagcATGGACATGAGGGTCCCCGCTCAGCT**CCTGGGGCTCCTGCTGCTCTGGCTCCCAGG**TAAGGATGGAGAACACTGGCAGTTTACTCAGCCCAGGGTGCTCAGCACAGCCTGGCTATTCAGGGAAATTCTCTTACTACATGATTAATTGTGTGGACCATTTGTTTTTGTGTTTCCAATCTCAGGTGCCAGATGTGCCATCCAGATGACCCA**GTCTCCATCCTCCCTGTCTGCATCTGTTGGAGAC**AGAGTCACCATC**ACTTGCCGGGCAAGT**CAGGGCATTAGAAATGATTTAGGCTGGTATCAGCAGAAACCAGGGAAAGCCCCTAAGCTCCTGATCTATGCTGCATCCAGTTTACAAAGTGGGGTCCCATCAAGGTTCAGCGGCAGTGGATCTGGCACAGATTTCACTCTCACCATCAGCAGCCTGCAGCCTGAAGATTTTGCAACTTATTACTGTCTACAAGATTACAATTACCCTcccacagtgttacacacccgaacagaaacccccagggaagcagatgtgtgagactgggctgccccagctgctcctcctgatgcctccat**tggctgagattgttcctcagatgcagcca**cactctgatggtgttggtagagggggacacgaagtcacctctgcaccctaattcttttctctttctcagccccaactgcacagttctagcaatgcctctcttgatttaataaagacagagatcatgacacctgaagagtctagttta**tggcttcagttggaattcgtataacagagaagaagccactatagatatttgaagcagatatagt**cttaatacagagaattagagtaaactactgaagtctaaata**aaatgtagagatgaatctctaaattt**aatgttttatttgcaaagaaatatttgccaaatggggcatacaggaaaactcagtggtcttcaatatgttggaagaacaaagagaaatttag**cattttatgaaaaagggaaaatg**ttacctatggctcttttgagaaagctcattggcactaggaagggttgggagctggcaagctcagactggtaagcagtggtggacaaagtgaatcctagaattatatcaagttatctcagaa**gttttggataaatttgatttcaggttacaataaaccaaaac**ggtgaagcttgcagagaattttgttactgaaatgccagggattcggtgtagatcctgcagctcgtcacacagaaagccaatcactaagacaagtattttcaaggaacaggctttaatcaggtgctgcagctgaggagatgggatgccattctcaaatgtatctctctgaccaactgaaattaggggtttatataacaaggaagaaatgtggaaaaacaggagttagggagggttaagaaagataattgggtcaacaggaagcaggaggtcagttaggcaatcataatgggtgaagggtctgatgtctcactgtcccgattcagtgat

Uppercase: IGKV1-6-3

Lowercase: Flanking sequence[1000bp]

Red & Bold & Underline: Stem-loop [15]

Blue: Heptamer[27]

Green: Nonamer [6]

id-IGHVII-62-1-2[V_gene_segment]

cagcaaggccattggtgccttccagagcacagtagtccatcaggaatctcccatgtgtcccagcagcagccttgcctcagtatctccactgtgcacagccattgt**ctgggaggagctcccag**gatgagtgtctttggcacacaccaggtggcgggtgttagagtgcagtacagcagctggctgcctggtctattgggctccctgatgttggagggatttgaggtgcattctcaggcccagcaccctctttgtgaatttttatataaaaaccctgattttacttcattttctcagatgacatagataactaaaaacagaatctgcaaagaaattgtaattttcaactttaccccaaattcattgtttcttaattctgtgcaagatccaaacatatttttgccttccac**atgagaaattgttctgtttaaactgaaatcattttttctcat**attctttctttttgcccaagtacagagatcttgttgaaaataagttgggttctttccacacactaaccctca**cctcccctagagaaagagcagagattgtcctcactctgagtctaggggagg**agctgttcctgtacaattcagagcctgcagagacccccaggtgcagcttcagtgagtcagacatttctccatgtgggcgacctccagtgcctgtgactgcttctcagg**cctaattgtgggttaagaattagg**acactctttaagttatcacatctgaatcctattctgaaaatcaccataaagagagatagttcaatggctattctcctgacataa**gtttctctttattacttggttccaagtatggagaaac**atgtgaccctatatttgtctgaatccaacatcagaatccactgcattactctgggatactcacggatcgaacacaggtgagctctttattctcacagaaatgtatgtagttgggaatttcaacatattgtccagaacgtgtgcctgccaacaactgtgtttctcggtgcactcttGGCCTGGTGAAGCCCTCACAGACCCTCTCCCTCATGTGTGTCATCTCTGCATTCTCCATCACAACCAGTGCTTCCTCCTGGAGCTGCATCCATCAGCCCCTCTCTGGGAGGGAATGGAGTGGATTGGGTGCATAGGTCATGAAGGGAGCACACATTACTCCCCTTTCCTCAAGAGTCCAGTCACCATCCCCAGATCCATGTCCAAAAACAGTTCTTCCTACAGCTGAGCTACATGAGCAACAATCACATAGCCATATATTTTTAAGCAAAAGA**cacagtgagggaatcacagtgtgaactcacaccaaaacctccctgtg**gggatgcataggacagaaggggttactccg**gagcccagggggctc**tcaggacaccaaggggcactcaagaccattgtagaggcacgcaggtagccgggggctctcaggaaccatggaggaaaatcaggacaccaaagggtgctcggtacagcgaggggctcaggacaattgtggggattcagaaagaacaggttcaaggctcagcttcagggcaggtgcagctggtgtgaaaaggggctggatgaggcgttttgtgtcaccatcatgtttcaccaccagacaccctccactacatctg**ttctaatgcatgtgtttgtatgattagaa**aatgatattgatataaatatataaccatagctaggtgtgtctttttctgccttccgtcctcccttcttctctctctctcacacagaaatttacatacacccaccccacaacacacataaatctataact**tttattacctgatgtattcaataaa**cctgattaatgtgttttgcagcttcattgtttatgctgttgtagcaataaaaaccatctgtttcccagctgtgtacttctctaagctgagtagcatctttgttcataatacccagaatgaaaaacaacccaaatgtcaatcatcagcttaactggtaaacacattgtggaaaagtcattcattgacatactacccactactaccatcagctaatgttggctatgctcaaaagcatggttaaattcacaagtacttcagatgagtaaaatgagcccaagtgataaaagtgcatacataggata**caactttcataagttctatagaacaaaaagtaatctaaagttg**caaaaaaaaaatcagtagttcactgtgaatattgtaggagaagggatggtctaggaaggaggaattatgg**aacaagataaaatgttgagggaattgacttgtt**atctatgttgatagtgataatgtc

Uppercase: IGHVII-62-1-2

Lowercase: Flanking sequence[1000bp]

Red & Bold & Underline: Stem-loop [11]

Blue: Heptamer[42]

Green: Nonamer [7]

id-IGKV3D-20[V_gene_segment]

tagtctcagaac**agggacagacagactgagtccct**ccactaactcaaaccaacccctgtggatgagtaggggatccctgagatcctggggtggatgggaggtggggca**ctgatctgtcaatctgttttttcttcaaggatcag**gcagcagagacccagaagcttcatgtctttgtaaggctcttccaagccacca**cagataatttgacaaaacactgtatctg**catcccagacctcaccctgtcacggactgaagtggttgtttcatcctactaagggtaaaccataccagctatacaggataaaaagactggatagattagaggatcacccaaggaatagttctttgcctgcatggacaaaaccatcttctgtctttagggaaatggtatcactaccctgaggatttggagcccaggtatcacttatctgtgcagttgtgaaagtcctcacactcacagtgctgaggttaattgaatgttactcttttaatttctgcaaagaatgagacagcctctggaccctcaggaaagattgctaacaagtaaatacaagtatatcc**ggaagataaagtagtaatagactcttcc**tttcaacctgatccatcatgcatttagggaactgactgggcacaagttggagcagaaagagaaaaatgaaaccacagccttctattttgtttctaacagacttgtaccaaacattctgtggctgaatctaagtgatggtgagacaagaggacacaggggttaaattctgtggccgcaggggagaagttctaccc**tcaggctgagccaacggccttttctggcctga**tcacctgggcatgggctgctgagtg**cagagaggggaggcagatggtctctg**a**agctggaagcccagcacccgcccccccagct**gctttgcatgtccctcccagccgccctgcagt**ccagagcccatatcaatgcctgggtcagagctctgg**ggaggaactgctcagttaggacccagagggaaccATGGAAACCCCAGCGCAGCTTCTCTTCCTCCTGCTACTCTGGCTCCCAGGTGAGGGGAACATGGGAT**GGTTTTGCATGTCAGTGAAAACC**CTCTCAAGTCCTGTTACCTGGCAACTCTGCTGAATCAATACAATAATTAAAGCTCAATATAAAGCAATAATTCTGGCTCTTCTGGGAAGACAGTGGGTTTGATTTAGATTACATGGGTGACTTTTCTATTTTATTTCCAATCTCAGATACCACCGGAGAAATTGTGTTGACGCAGTCTCCAGCCACCCTGTCTTTGTCTCCAGGGGAAAGAGCCACCCTCTCCTGCGGGGCCAGTCAGAGTGTTAGCAGCAGCTACTTAGCCTGGTACCAGCAGAAACCTG**GCCTGGCGCCCAGGC**TCCTCATCTATGATGCATCCAGCAGGGCCACTGGCA**TCCCAGACAGGTTCAGTGGCAGTGGGTCTGGGA**CAGACTTCACTCTCACCATCAGCAGACTGGAGCCTGAAGATTTTGCAGTGTATTACTGTCAGCAGTATGGTAGCTCACCTcccacagtgattcagcttg**aaacaaaaacctctgcaagaccttcattgttt**actagattataccagctgcttcctttacagatagctgctgcaatgacaactcaattttagcatctcttctctgcttgggcattttggggatcttaaaaaagtaatcccttgatatatttagactctgattcctgcattattcctcagaccaagatggacagccaggtttaaagcacagtttcacagtaatgg**ccactggatcagatttacgtcagtgg**atgtcaataaaggtcccaaccagagccataaggcaacaacaatagcaacaaataatcaaaattggaaaagaa**gaattaaagctgtcataattc**actgatgaaggattgtgtgcagataaaattcaaatttgtctacagagaaactattaaaattgacatgagaaatagaaaatcgttagattcaagatcaatttattaattcgtagattcaaaaatcaa**tttcatttctgcataatgaaa**aaaatgctaaaaattaacattataaaacaaacacaccttttacaaaaacatcaaagtatcaattatttaaaaaaaatagactaaatacactgacgtctccagaata**ttattttgaaaaataa**aagaaaacctaagtaaat**agaaattcagttcaaagactgaatgtct**cagtactataaaaatgtcaattcttcgaagattaaaata**ttgattacatataagaaaaatcaa**aatcctaaagtatactccaatttaaattaagaagctaatctaaatattatatgggaatgtcaaggatgtagaatggccacagtgaacctgaagaaacaccaaaatgagaacttacagtgcctgaatacctggaatatagtgtgggtgacagtatggtgatggtgagatcagaagattaaaaatttgcgaacgtgc**ttatttttggaaataa**ttactatgcagatgtagccaaaccctcttcaactgtgccgccagaatct

Uppercase: IGKV3D-20

Lowercase: Flanking sequence[1000bp]

Red & Bold & Underline: Stem-loop [19]

Blue: Heptamer[22]

Green: Nonamer [7]

id-TRBV6-8[V_gene_segment]

ggagggtgttggctgggtcaaggagaagatgatgagctcagaaaatagacaaggagaaaccccagtggaaagcttgtaaaactctggatgaatcacaagatatgttattagagctcctttttattactaaattttatttattactaaattaatggcaggaaccctgcacaaagggatattccctatgaaagatggcatctcaaataaaaacagagcattcttccttaccctaaagtctgccctttctccccaagtccctcatgcatcaggccacctttg**tgcccacagatcatgggca**gcgctgggtggccatcatcatccacacagagagggcagtcagcagcatgagctggttctgcctgatat**ggtctcaccctaggcatggaaagcaagagacc**tgcgttg**agctgaaggtgctcagct**gggt**ttgtcaggagtcccatctgtcagtgaattgacaa**gaaacagagcaaaacgactcctgcaatgttgagaagcctgcacctaggatctggaaacttgataacagagaaaaccaatatagacaaaggattttaaacaggattatggtcaattaagcaaattagaaaaggatacttgaaggagtatttgggatacaggaatcaaaaacaccaggaagacatgagaagtttccctaagagtctagactacagcactgtgtagataactaacatcag**aatattaattatattattgaagaattaaaatt**tacattgaattacaaactctttacaaaaggtcatgaaaatcttgtagacttgtttcaattcagacaaatgtgttcttctcattctcagctgttcactggtgcatttatcttgaatttgaccatctggggaatgggcgtggcctctcctgactggaaggctctggggcccaggc**agggagaatgatgtctcagaatgactccct**tgagagtcctgttccccttttatcaatgcacagacccagaagacccctccgtcctgcagcccctgccATGAGCCTCGGGCTCCTGTGCTGTGCGGCCTTTTCTCTCCTGTGGGCAGGTGGGTCCTGGGCAGGGCCCTTGCGTGGATTTCAAGGCCCATCCCCTTTCCACTGGGG**CTGCAGCATCAGTTTTGTTCTTCTCTGCAG**GTCCCGTGAATGCTGGTGTCACTCAGACCCCAAAATTCCACATCCTGAAGACAGGACAGAGCATGACACTGCAGTGTGC**CCAGGATATGAACCATGGATACATGTCCTGG**TATCGACAAGACCCAGGCATGGG**GCTGAGACTGATTTACTACTCAGC**TGCTGCTGGTACTACTGACAAAGAAGTCCCCAATGGCTACAATGTCTCTAGATTAAACACAGAGGATTTCCCACTCAGGCTGGTGTCGGCTGCTCCCTCCCAGACATCTGTGTACTTGTGTGCCAGCAGTTACTCcacagcgctacaaggttgtctcctctctgcacataaaggcagggaggctctgccctccaccccgacctgagactcagggatgacctgggcagagtattctgcaacgggaaccttggaacccgaagtggccccaagtggcccagacagtatgagcctcgctctgtgccaggtgcctctgtaggcatctcagccaggcctggactggtcccaggtcctcatatgtctcctttgttgctctctcctacaagctctacttttggggttggggccagggcttccccagctcctacttttctactca**tcatcctgagtccaaggcccccaggatga**a**acaggatttgtatttcagatccatctagactcctgt**ctctccctggtgaccatgttgcttcctctctagggtttcccccagcccccaccctcctgtggcccacctttcccacctccgcagtcatcctccaccccccacttccctg**cccctctctactgcagccatgagggg**agcacc**tctctgtgactccttccttcccgtcacagaga**cttcaaaggccattttctttgccctgggccagagccttcctttctctagtggccagctccttcctgtgcttcagatctcagtatgatcagcccctcctgcgggaagcactccctcccctcccaggtaagatcaactttcctctcataagctctcaatgaatcatatgtctgttagcaacctttagcacagttgggctttctcagatgcttgtctgattatttttgtgctccacccaatttttttttttttaagacacagtct**cactccgtcaccaggctggagtgcagtggcatgatctctgctcactgca**acctctgatactctgattcaagtgattctcctgcctcagcctcccgag**tagctgggattacagacatgtgccaccacgcccagcta**attcttgtttgtatttttagtagagatggggttttaccatgttggccaggatggtctcgatct

Uppercase: TRBV6-8

Lowercase: Flanking sequence[1000bp]

Red & Bold & Underline: Stem-loop [16]

Blue: Heptamer[33]

Green: Nonamer [3]

id-IGHV3-50-2[V_gene_segment]

aggggtgctaagttgacaagaggtgggcttttgatgcttgatactaggtgtcaacttgattggattaaaggatgcctacatggctgggaaagtattgtttcttggtgtgt**ctctgggggttttgccagag**gaggctgacatttgagtcaatggactggaaatttgagtttgtcatgttgcatgatccttttaaagtcttatagaacttagttgactattattttggtgtagttcatttgtataaatatttctttgaagtttaattttcttcctttgtgtttgtctgtcactggcaacacggtaattgtagccttttagaaatagcttagaaggttggcgactcatcattt**tttaaaaaagttagagaacagttggcatgacttttttaaa**ttctgagagccattttttatctaacatataggctatcatgaagaatatttatagtgtgcttgagaagcatgcgtattacactgcgcttggttggaatgttctgtagatgtctattaggtcagtttgttcaatagtgttgtttaagtccatgggcttcctaagaatttttttatggatgttctatacattattgagggtgaggtgttgaagtctcctcctttcttattgttttctatttctcacttcatatctcttaaactcttgttaatgcggtta**tatttctatttttataactatgtaaaatagaaataaaataataattgcttagtaatttta**a**tgggaaaatcacataataagaaattatattttccca**aatgctgccatcaccactaaactcctccaggagtctcacatctgctctgggctctgctctctcctaaggggtcccacatcagagtttgctacagagg**aggagacaagaaaatagggccctccctctcct**gatgaaaaccagccctgccctgaccctgcaactctgggagaagatctctagtccagaattaccaggagtttggatttgatgatcagctctgtacaaacatggctcaccATAGAGTTAGGGCTGAGCTGGGTTTCCCTTGT**CATTATTTTAAAAGGCGAATAATG**GAGAACTTGAGATATGGAGTGTGAGTGGATATGAGTGAAAAAACAGTGATTCTGTGTGGCAGGTTCTGACTCAGATGTCTCTGTGCTTGTAGGTGTCTAGTGTGGGGTGCAGATGGTGGAGTCTTGGGGAGAGTT**GGCACAAGCTGAATGTGCC**TGAGACTCTGCCGTGCATCCTCTGAATCCACCTTCTGTAGCTACTAGATCAGCTGAATCTGCCAGGCTCCAGGAAAGGGGCTGCAGTGAGTAGTAGATATAATGTACGATGGAAGTCAGACATAATATGCAGACTCTGTGAAGGTCAGATTCACCATCTCCAAAGACAATGCCAAGCACAGGTTGTATCTGCAAATGAACAGTCTGAGAGCTGAGAATATGGCTCTGTATTATTGAGTCAAAGGTAccaaatgaagggacatcagtgtgaacccagacacaaaatttc**ctgcagggaggagggaggaggctgggctgcag**tgggcactcagcacacacaaaaggcagggacagttccaggggcaggtacaggtgcaggtgaaggcaaaggtctactttccttccagatctgtggattcctctgcatccaacagttcgcctgggcctctgtctttatggatctgcgcctaccactgatgtctctgggttagtaaagtttgctactataggaggaaacattgtcatttgtcagaaaggcgaataatggagaacaaaagatattgagtgtgagtggatatgagtgaaaaaacagtgattatgtgtggcaggttctgaccaaaatgtctctgtgtttgtaagtgtctagtgtgaggtgcaggtggtggagtgtaggggagaggcaagaggcagaaaagcgcgcaggaggccgggtgaggctgtagacattgtcagctcactatgccaatctcacaacagtgctggagaaggtgggagtc**tgatggagcttcccaacaaccctgtggtctaagctaagtccatca**aggccgctggttcctcctggaacacagctgtccatcaggaatcccccatgtgcccagcagcagccacgcgttagcatcttcactgtgcacagtcattgtttgggaggagctccaggatgggtctctttgtcaaaaaccgggtgatggt**gtcagagagctgtgccgggtgcctggtccaggggctccctgac**gttggaggtataggaggtgcgttctcagggtgggaacaccttttatggaaatgtacagaagaaaacatgattttccttggtcctctcagatgacatggagaa**gcaagcgcgcagcttgc**aaacaattgtaatgtccttgtaatttcctactttatcccaagttcattgattcttgtaaaccttggtttctgtccaagtacagagatattgattagagtgagtttggttctttcca

Uppercase: IGHV3-50-2

Lowercase: Flanking sequence[1000bp]

Red & Bold & Underline: Stem-loop [12]

Blue: Heptamer[33]

Green: Nonamer [7]

id-TRBV7-4-2[V_gene_segment]

atggatgtttcccatatattaga**atatgcaaatttttcataaattaccattttttttgcatat**gattgcagtttcaaatagataaaataggtagaatgttatgaggtgatatttatgagatgttagctgacaatt**atttagaggtgtttaaggacctaaat**tcagagtttccagaatcaaaatgaatctcaagtaagacaaatagtcaagtcatagggaaattgtataacatgagggacaaagagaagattttgaaagcagtcagaaagaaaagacagatacctcccataggaatagatggagaacaacagtaagtcaagatag**tggttaaaaataacca**ttagc**tcaatcgtcattctccttcattagaggtgagattga**ttaaaaaaaaaacttttatacgtggaatatgaaaaaccaccactgtgagtttgctaccaagttatcttttctgaattacagttaacaaaggtgggaaatgatcccaaagaaaggcatgagggcagttcatgagggaacgcatgtgaggggacagtcctgttgggcacgtgtgactgggggaatggaggaggctggggcatgaatggggatggcagaggggaccctgacttgcaggaaagacaatgagctcacccttttgtgttctatgttaggggcactgttggtgcatcctacaggacatgcctagcagacagaagagcagctgtgggctggcaagatgaactcagagatgcagcgtgaggcctccggg**tccagacagatctgga**gcccaaagcaatgagccatgcattgatgttgttaaaaaggagcttataaatatttaaagcagcacccaaatgtgttctaatagaaatgctggggccctgaggtcctggggattgacagaggaagtgatgtcactgtgggaaccgccctgtggagacaaggac**gtcactcatcctctgctcctgctcatagtgac**actgacctggtaaaacccccgtcctggcctgaccctgccATGGGCACCAGGCTCCTCTGCTGGGTGGTCCTGGGTTTCCTAGGGACAGGTGAGTCCTTAGAACACAAAGTAGTTTCAGTTTTTTTCTGTGTGTAGGCTTGTGTGTGTGTGTGCGATGACTACAAATATTTTCCTTATTCTGTTGCCAAATTCTATTTCCACAGATCACACAGGTGCTGGAGTC**TCCCAGTCCCCAAGGTACAAAGTCGCAAAGAGGGGACGGGA**TGTAGCTCTCAGGTGTGATTCAATTTCGGGTCATGTAACCCTTTATTGGTACCGACAGACCCTGGGGCAGGGCTCAGAGGTTCTGACTTACTCCCAGAGTGATGCTCAACGAGACAAATCAGGGCGGCCCAGTGGTCGGTTCTCTGCAGAGAGGCCTGAGAGATCCGTCTCCACTCTGAAGATCCAGTGCACAGAGCAGGGGGACTCAGCTGTGTATCT**CTGTGCCAGCAGCTTAGCcacagcgtggcacag**tcgcctcctttctgctcacaaacctcatccttctctctccttgcagctcctacaaacccttaacagaggcctctctttgctccttacttttcatgggaaaaaggtagatttggacctcagctgtcctttgggtagaaagagaccgcagattcattcctgaaacacagtgagtgcaaatgtaggtggtgaaaacaat**cagagcccactgcgctctg**ggagtcctccgagccagctcactgttccaagcaaggagtcagtgtcttagccttggccttcagggcagacatgcatcttctataggtcttggaggctgctgtgctgcccacatccatgaggttgtcataggcaggaaacacgcctcttcttcatatgttggggcatttggaaggtctgaggctacatccccaggaacatctttcttcagaagcctgttctacccctgtcaccttggaagtttcttcaacaaaatatcaaacatctcttcctataaagtaaacgtctttgcaaatttgtggtccttattaataaatacaatggtgataacaataagacttcacttcttctgcctactttaagccacgtgtatcctttattttgtttgcatttgccattgctactgtcctgatagacagaagcgtgcattcaccaccgctgcccattcaccttgatt**ccctcaggaaatctaatttctagactctgaggg**ttttcattgctgtcaaactcatttgatttgaaatcattttcctgaggcctttaactcaagaagtgttttatt**tataatattgaatatatttctttttcttttatttttcatattata**ttttattatata**gtacttgttataaatagaagtacaatgattatattgta**atagaatcttcaactcgtctgtgggtgctgctgcagtttgtatctatgaaagcgaatgcactggtcagagctgatgtgattatggatcatgggtttcagggagtcttcggcatca

Uppercase: TRBV7-4-2

Lowercase: Flanking sequence[1000bp]

Red & Bold & Underline: Stem-loop [13]

Blue: Heptamer[39]

Green: Nonamer [5]

id-IGKV2-24-3[V_gene_segment]

aaactgaatgcaatatgggatccagaaattttatcctaaaacattaaaagtaaaaatggtaaatacatgtatcagtggaaagctcagtgaaattcaaatgtagattgtaactt**cattaataatagtggattaaccattaatg**ttaaagctatttgaagtactaga**aaaatcagtttaaaatgatttt**atattcagcaaaactatccttaaagaaaagaaaagaagccgtgactagcatatatctcctataagaaactcaagaagaaatccttcagaattcagaatcacagtaaacgacaatgaacagta**atttaaatccatgaaattaaat**gaaagcttcataaatatacttacctcaattcatatgttgttgatgttcacgaaaactgaatctttgtgatagatatcagagttgcagttcccttggtaggttagaggcagaagctattgactagaaagg**tgaatgaaggcagcatgtggagaatttcaaatcattca**tatttgtatctgggtagtgaatgtgagtactttatttggctgagcagcgaacatgtttgcactttactcagggcacaatttattttgatttataaaattaacagcaaaccaagaccctttcaacacacatgaagaaaaaaataagaagcaccaaatatttacagaaactcagccgtattaaagagaagtgtaataagcactgggaaaatactaggaagtaaaaaaattgacagtaaacacagtaaacatagaaatatatcctgtcccaatcaggctgcatagattgttatttctgccagttttttctcaagcatacaaaatatgttgttcataggaaaggcccccatacccctgcacatatcatgttatttctacaccactgcacccaccaggggatttgcatattgtcccccagggaggaccttcccttgcaagtctgagataaaagctcagcaccaaccttgacttaactaattaggactcctcaggtcaccttctcacaATGAGGCTCCTTGCTCAGCTTCTGGGGCTGCTAATGCTCTGGGTCCCTGGTGAGGACAGAAGAGAGATGAGGGAGGAGAATGGGG**TGGGAGGGTGAACTCTGGGGGCCCCATTGCCTCCCA**TGTGTGTTCTGTCCTCATGTTAGATGTGTACGTCTTGTACTCCAGGATGGGGCTTGTAACTTTTATATCTGCGTGAGTAAGGCATGTGAGGTTTAGATCTGTAAGAATGAGGAAGATTCCAGAAGGAACAAAGACCAGTGCTCCGGTGAAGACTCTAACAGAGAAAGAGGGAATGGTAGAGGAAACTTCTAGCAC**TCAAAGCACTCTGCTGTGCTTTGAAAATATGTTTTTATTTT**GAAATTATATATTACTAGGGTCTG**AATCAAATTATAAAAATTGATT**TAGCCTGAAAT**AAATAACAGAAGAAAAATTATTT**TAAAATTGTGCTTAAAGTTTCTACATAACCTTGCACTTCTCTCTCATTATTTCAGGATCCAGTGGGGATATTGTGATGACCCAGACTCCACTCTCCTCACCTGTCACCCTTGGACAGCCGGCCTCCATCTCCTGCAGGTCTAGTCAAAGCCTCGTACACAGTGATGGAAACACCTACTTGAGTTGGCTTCAGCAGAGGCCAGGCCAGCCTCCAAGACTCCTAATTTATAAGATTTCTAACCGGTTCTCTGGGGTCCCAGACAGA**TTCAGTGGCAGTGGGGCAGGGACAGATTTCACACTGAA**AATCAGCAGGGTGGAAGCTGAGGATGTCGGGGTTTATTACTGCATGCAAGCTACACAATTTCCTcacacagtggtacagctctgaacaaaaacctcccgctggagtggcccag**ctgctcaagtgtgttgtttctctggggagcag**ttgaacagaatctctatctgtatgagataaacatgttggagaactcagggcaacaggttgcat**ctgagggttctgtcccatgggtgcctcag**ttgtacgtcaggcaaaacccgttcacagccctgtcagctgcaacagccttggcatggcataagccataggaaaccagaggtgatcccagtgcctgcacaggtaatagactgccctg**agggagagcttaagaaaatcctattccaatcctccct**gccttgcctgcattgggaaataagacttaaagaggtaaataaccagacaagtaacccagatttgttgcaac**acttcttgaatatatcttgaggtttagaagt**ttaaagtctacatttaagaggataatatgtggtaatatccaaaaattgaacttttcaactttcctaacttcttatttttctctttcaccaccta**tcttcccaccacatattgatggtggaaaga**gccttccgcacaagctgtcatcatgaggagctggatgagggcaattagtgaaaatcttggatttcagcctcagaatggacttttgtaaattggtgagagatag**aaaatatgaatgctaaaattatttt**attcacttcaattgtgtcttgctgacagaaaaggatagtttttgaaatttcagaagttg**agtttcataaacagaaact**taaa**ctataagacataggttatag**aatttacctcatagaacactgaaataacacagaatgatatgcgatttctttccccaaaatgtaagagt**ttgaagacagtgggcagacttcaa**gaatgggagaattaatggaagatagtggaggtcaactatggcccaataacctgctctttgacttacattaggtacagttgtggatgacagtgactgttgggggttggtgatataaactcagaaagg

Uppercase: IGKV2-24-3

Lowercase: Flanking sequence[1000bp]

Red & Bold & Underline: Stem-loop [19]

Blue: Heptamer[31]

Green: Nonamer [8]

id-TRBV4-2[V_gene_segment]

ttacacatcagaaaaatgctaacactttcctcctggcccagcacatctccaagttaaaaaaatctctaaattatcaacccatagtttaatcatggtaactcccttctacccccaacatagatttttatgtaactgatttgtaactacaaccttaaccgtacaatcacaattgtcattataaaatactaacaaacatgtgctattaacaatggatggttaaaatcacttagggattatgaatcatttcatttttgtgtttatttattatgtcagtttcaagctgatttccctttcccttgttccacacaccttgggttgttctttttcagcatcctgtgtatgttagctctaagcccagccaaggctcatgcccctccacttcagggacagagggtggctcatccctttcgtgctccctcagctccctccccggtgcctggttgctggttggcatcacagccatgcccttgaaagatgggcgctggtccacactgcatccgtggtcatgtagccaccctccctccacagaaccgagcaaaaagggctccaccgcgcttaggcattgagggaatggacattttgaatggtgctgtcttgtgtgatccctcagaggagaaggagatttcttgaatgggagtggtctgttgctgagag**ggatcctgaaagacacggggacagagacaggggtaggatcc**acataatgaggaagcagctgagaacaaaggagcccctgccagaggaatatccttggggggtaaaaagctggctctgccctttctccccagcccattcatcccagcccagacaattcaaatctgccttctatcaggacctagaaaggatgtaaaacggccgggtataaatatcccctgggtctggggaaactgtcaggagcagtgacatcacaggaaaaaccaccaaccaaggccaaggagaccagagcccagcacctcacccagaggaccccagtcagaggccccatctcagacccgaggctagcATGGGCTGCAGGCTGCTCTGCTGTGCGGTTCTCTGTCTCCTGGGAGCGGGTGAGTTGGGTTCACATCAGCTGTCCTTGAATTCGAAACTTTTTCCTTGTGATTTCAGCAACAAGCCTCCTCCTGGGCTCTGCCTGAATTTTGTCCCTTTCCCCCCGCAGTCCCCATGGAAACGGGAGTTACGCAGACACCAAGACACCTGGTCATGGGAATGACAAATAAGAAGTCTTTGAAATGTGAACAACATCTGGGGCATAACGCTATGTATTGGTACAAGCAAAGTGCTAAGAAGCCACTGGAGCTCATGTTTGTCTACAACTTTAAAGAACAGACTGAAAACAACAGTGTGCCAAGTCGCTTCTCACCTGAATGCCCCAACAGCTCTCACTTATTCCTTCACCTACACACCCTGCAGCCAGAAGACTCGGCCCTGTATCTCTGTGCCAGCAGCCAAGAcacagccttgcagagtcaccgct**ttcctgtgcagaaaccttaggggccctccaggaa**gctgtgggggccaccaaagccttcgatgaacatttcctacaagagccccggcagaagattcagaacatcatagcacctttaatggcat**aaaatatctagcatatagtagaagcctagctatttt**gatttcctttgctgatctcagacaacctcttccaccacacc**aatgtgcaaagacagatatggacacatt**ctgccaagaaatcttaggttgaggactcagaagtcacagaaatggaaatatatatgcctgataattcatctgataaaattaactagaggatgtagcatacatatctcctcaggca**tttaaaaatcaacattattgatctttaaa**gtatatgaagttaacaggacaccttttaaatgaacatatctgtgatttttgagcaa**catatatctatatatg**caaccccacctctatgaagatgtatctgcatcacccagaaagtccctcactcccctctgcaatgcctctgtctgtcccagcaaccaatgatgtgacttctatcactggagagtcgttttgttggcaaatatttttaatgcaaatgtcaaacctcttctgaataatcctcttgaatacatagctaaatcatcaaggcagggaggagttcctggaggttttatggttttagaagtggttgaagctaggtgagctacctgacccatggtagggacaattctatctggagatggatgttgtg**ccgaaataacctcaaaaggtcttttcgg**atg**gtagaagctgattctac**aaaatttattcattttcattcgtttattttctcgtcaattcattcaatcaaataattttatgttacgttttgacaaaagttaatcatctttaaggaaattattttgagttcccatcagacacacatgcagctgggctttgtgttcacacagtgagcatgctggggtcaagcaaaccatgatcacagtg

Uppercase: TRBV4-2

Lowercase: Flanking sequence[1000bp]

Red & Bold & Underline: Stem-loop [8]

Blue: Heptamer[30]

Green: Nonamer [5]

id-IGHVII-44-2-2[V_gene_segment]

tgcacctctctctccctccccactccagcagttcacagtgcctattgtttccatggtcatgtc**cacatgtgctcaacctttagcactcactttcaagtgagaacatgtg**atattttgtttcctttttctgtgttaacttgtttacgactatgacctccagctgcatccatgttgctgcaaaggacatgattttgtttctttt**gtgacagagtctcactctctcac**caggcaggag**tgcagtgacgcgatctcagcttactgca**acctctaagtccctggttcaagtgattctcctgcctcagcctcccggg**tagctgggattacaggcatgtgccactatgcccagctaaattttgtatttttagc**agagagggggtttcactatattggccaggatgatcttgatctcctgacctcatgatctgtctgcctcagcctcccaaagtgctgggattataggtgtgagcc**accacacttttttttatgaatgtgtggtattttatgattttatattgcatacatacca**ttttattcatctaatacatcattgattggcacctagattgattccatatctttgctgttgtgaatagtgtggcaagaggcatgagtgcatgtgcattttttggtagaatgatttattttactttggctatatacctaatgatgggattgctgggtc**aaacagtagctctgttt**caagttctttgagaattctctaaactgctttgcacagtgctgaactaatttacatttccaccacccatgtatgtgttcccttttcttcacttcccacccagcagctgttttttgtttaacttttttcatattagctattatgactggtctgagatggtatttcattgttgtttttatttgcatttctctgataattaatgatgttggaaatttttttcatgtctgttggccactgagattgacaggcttacaagtgtcttagtccagcagttttcggtaacaggctatccactttcattttCTCTTCCTCACATGCCCCATTTCTGGGTTGTCCATCCCAACCAACAGTTACTGCTGGAGCTGAATCCACTAGCCCGCTTCCATGGGCTGTCAGGAACACATTGCAAGGGTGACACACACGATGATCC**ATCTCTGCAGAGCCAACTCTCCTTCTCCAGAGAT**TCATCCAAGAAACAATTTTGACTATACCTGAGCTCTGTGACATCTGAGGACATG**GTTTGTATTACTGTGCAAGACAaac**agtgag**aggaagtcaatgtgagtccagacataaaccttcct**gctgagaacaatggaaag**cttttcttctaagataaggaataagaaaag**aatgcccagtcttaataattctaatcagcatattgatgatagttttaaccataga**aatttgagaaagaaaaaaatagtaaaaggcatacaaatt**gaggggaat**aagttaaattgtttctgtttacagacaatataactt**tataaattccaaaaaaataaaaaattctgaaaaaacagccactgaaaactaacaaac**aaatttagtaacattgcagtatacaaaattt**atatggaaaactcagtagtgttactaaaaactaaaaattaactatttaaaaatcataagacaatgtcagttgcatttacaatattaaataacttacttagtggccatgcacagtagtttacacctgtaattcttgtactttgggaggtaggggagaatcacttgaggctaggagtttgagaccagcctgggcaacacagcaagaccaatatctatttctaaaaa**tcacttagaaaaaatgttggtcaaataagtga**aaatctgtgcattgaagactatatgacattgatgaaataaataaatctatatgcaaat**atatgggaaactattccatat**tcacagattgaagataataatactgttaacatgtctatactacccaaagcaatctacagatccaaagcaatttctatacaaatgacattttcaacaaactactttttaaatgataaaattgaatcacaaaagaccccagatgtccaaagtaaccttgaatacaaacatcaaagcggggggaggtttcctagtacctgaactgcaaatatgctatgaaactacaataataaaactgcatgctgctgactttctaaataaactttaatattctgaattgaagttatttgacaatttcttctatcattttatctttactaaagtgttgcatgagtgtttatctagacag

Uppercase: IGHVII-44-2-2

Lowercase: Flanking sequence[1000bp]

Red & Bold & Underline: Stem-loop [17]

Blue: Heptamer[25]

Green: Nonamer [5]

id-TRBV12-5-2[V_gene_segment]

atccactctatgaacagaattcttactactatttataccctaaaaagcaggccattttctagagtttcaccccctcaccagaaagaatttatttgtgagtattggttgacttactataacaagagactcaagtgttaatcattaaaggaacctgttagtgctttcttagaagtaagaggaagatcacatttaactggagagtgccttattgtggttttcaaggaaggtcaaagctcttacttatcactgctcaatcttcacccatgcccaaagaattaacataattttgaaccttacc**ctggagaagcgaaatgggaccttgtcctccag**aaatgcagaagacactgtgctgtgggctgggggagtcacggagcacgagagtatccatctccagccctca**ctggaaataaaatctgcacagttctatctctttccag**ggcagccaccttgttctctgcatgaggagcatggaggt**ccagatgcttccaggagggatctgg**ggcaaagctctaactaagcaacacattattttgattttgtgtgtctgggatataaggatggccggtaaggggaa**caataaagttcttgagatttcacaagtaatttattg**atgtccagaatttgaggtgctgatggtacacctaagtggcaatatgcaccaggcagatcaatatgtgacgcgtcagagaaagcagaatgggtgatgtgatgtgcaatgccacagaagcactgcagccaggagaggtgacagctaatgggga**tgtttggagtctttgagtgaaccaaaca**catcccagagtaattgtaatttatttcagtcaatcttctgtacagacttagcattcacctttggaggaaggtcctttgagcagggacagagatggtgatgtcactgacagtcccccttttactctgggtgagaggtctagaatcctcagctcctgtattcgtgcccacaagggcctcatctaggtgaaggctccacctgccccaccctgccATGGCCACCAGGCTCCTCTGCTGTGTGGTTCTTTGTCTCCTGGGAGAAGGTGAGTCCCCACAAATAAAGCACCTGCATTTTTGGATATTGCCAGTTATGATTCCAATTATGTTTCTTATTCTGTCCCCAAATTCTATCTCTTTTCACAGAGCTTATAGATGCTAGA**GTCACCCAGACACCAAGGGACAAGGTGAC**AGAGATGGGACAAGAAGTAACAATGAGATGTCAGCCAATTTTAGGCCACAATACTGTTTTCTGGTACAGACAGACCATGATGCAAGGACTGGAGTTGCTGGCTTACTTCCGCAACCGGGCTCCTCTAGATGATTCGGGGATGCCGAAGGATCGATTCTCAGCAGAGATGCCTGATGCAACTTTAGCCACTCTGAAGATCCAGCCCTCAGAACCCAGGGACTCA**GCTGTGTATTTTTGTGCTAGTGGTTTGGTcacagc**gctgcagaatcacctgctccctgtgcagaaaccctggtgcttcctcttctcctccagtacccagcagctctcagcagcctttcttgctcctcccctagcacaggaagtacataggtttcgtgttccacatgtccctaggcaaggcaagaacaggtcataaggacacatcacgttaggaaacttttggtaggaagtcagtgggtgtgatggttcctgggattccacacatacttctcacaagggtgcctgagtccaagtttgagggcgagtgtcacagactggagctttcaagtgattgagtgagcttaaaac**tgtgcagccattgcaca**tggatgcttatctcctttgctgctttccgttatggcttctttgccatttcttttctcttctaccttaaaactttcaacatct**ttgatgacttccaacatcaa**tattggcttcccagttttttcacttgctcacttcaaatgatcatttccttcaccttgtgtcagatattcatttccactttctttttctagattccaatgac**aaattatttgaattacccccaaatagttt**cctcaagcatcccattgtctcatgcgggcttctcttttcatctctggtactacttctgcagcaattcttcaaccccatcaggaccactaattgacttctcctgcaaatattgcaccatatatcatctcctcatgtactatattccttatctgacttagactatgtgttccaaggtgattgtgaaattttgcatatgtcttcagctcttttctttttctgtttcttacctggaaaataaaatggggccta**ctctgtgcccctttcaatgacacagag**aacataaaatgtcccttacccttgtcaagcagtattataacgatacagtgtgtcttgaggtctgtaagtccctctatgcatataaaattatatacttttcttct**attaaaataatttgttaccttttaat**gagttcatcagtttcatctttcaatta

Uppercase: TRBV12-5-2

Lowercase: Flanking sequence[1000bp]

Red & Bold & Underline: Stem-loop [12]

Blue: Heptamer[28]

Green: Nonamer [5]

id-IGHV8-51-1[V_gene_segment]

agtcaccatctccagatccacgtccaagaaccaaattttcttttagctgagttctgtgaccaacaatgccacaaccttgtattactgtgagaggaacagaagagatgtcagtgtgatcccagacacaaacttccctggagaggggcccaggaccaccaaagagcactcaggctccatgaaaacagggcccaagctggagaacgggtttcctgtcaccctcacgtttcaccattagatactctacactatgcctatgctgaggtgtgtgtttaatataattagaaaatggtatttatatacgtgtataatcatatctagtgtaagaatttgtagtttccaccttatgtgaccattaaagacaacatctctcacgttattcaaggaccttat**aaagtataatacttt**tacataagatttatctccttttttgtttctccttctgtctatattctctctgtctctctcttcctccctctttctctctctctctccatctctctctctctgtctctctctcacacacaaatacacacacatgcaatcgttgacagtatataggctaattaagaatatattcccatatgatttaaaaataacagttttacaaatatttgcc**tttaatttttcaaagaaaattaaa**tccaaatgccagtgattttttatgtgactttt**aaaatttaaattgttaatttt**tgttttcatcaatgcttgtatgtgtgcttttgtattctttcagattatttggtgcct**tcacagaattaaacatgtgaaactctccacttgaatttctgtgactcaggactggtgagcaaaaagttagaagtcac**tctcctgatccttccctacagctgcagattcctgaaggtaatgacctcacagtagaccaaggccctcaaaggtgaccttgagttccagcacatcttgggaggccaagcggatgtttgtgaaagaaatagtggctatgtgacagtttctaaccagaatatctctgtgttttcagGTATCCAGGGTGAGGCCCAGCTTACAGAGTCTGGGGGAGACTTGGTACACTTAGAGGGGCCCCT**GAGGCTCTCCTGTGCAGCCTC**TTGGTTCACCTTCAGTATCTATGAGATTCACTGGGTTTGCCAGGCCTCAGGGAAGGGGCTGGAATGGGTTGCAGTTATATGGCGTGGTGAAAGTCATCAATACAATGCAGACTATGTTAGGGGCAGACTCACCACTTCCAGAGACAACACCAAGTACATGCTGTACATGCAAATGATCAGCCTGAGAACCCAGAACATGGCAGCATTTAACTGTGCAGGAAAcatcgtgatgggaagtccacgtgggctcagagacagactgccatgcaggacacagggggtggcttggctgaagggggcactcagcacccacagaagacagg**agcagcccagggcaggggcaggtggagttcaagggctgct**ttcctgtcaggttctgtggcttcttctgcatcaaatgcgttctcctgggagcctctctatatttatggttctctgcctaccactgaggcctctggattggaaaagattactactagaaaaaaaaattctcatatacccctgaagaatacatcattaattgaggcagaagatgtcacaggaggccagggaggctgtggaaactgtgcgacgtggatgcacatctgacaacaaagacataaaaaatcagggaacactgataaaacctcctaattatgccatggccctctctaagcctagtaaagccattgatgccttcctgggccaaacccacccatcagggatcactcctgtgtcccagcagcagacatgccttagaatctccactgtatgtaatcactgt**ctgggaggagctcccag**ggcatgtgtctttgcattaaccaggtgatgggtgtcagagaacagcagctggatgcctggtctatgggttccctgatgttgaagaaatgggaggtgcagtctcagagccaatacactgtttatgcatttttatatagaaaccatgat**ttttcttgcttctctcagatgacacagaggatcaagaaaa**cagtctgtaaacaattgtaatgtccccatttcttctgaataaatgcattatttcttaattctgagtgggggtttgtcagaccatgcccttctcctaagcaattgttcaatccaggctg**aaaccacatggggatcacatactttggttt**ggttcaggtgcagaggtcttcagtagagcaaatttggtccttcccacacagtaagcctcaccttcctcagagaaagagctgaaattgatccaactttgtctgaggtg

Uppercase: IGHV8-51-1

Lowercase: Flanking sequence[1000bp]

Red & Bold & Underline: Stem-loop [10]

Blue: Heptamer[35]

Green: Nonamer [4]

id-IGHVII-20-1-2[V_gene_segment]

gtctctaacccacagggatgaagatgccgcggtgtgtctgtgtgtctgtaa**tgcagctttaaagaggagtgaagcactgacataggctgca**acttggatgagccttgaaaacattgggtgactgaactgagggagacacggaagtctacatccctaattgtcccattgacacgaagcacgcagagcaggaaaatcccagatcacaggaggtcaggggaag**ggaaacacagagtgaccgcttaatggaggtgaggtttcc**ttttctgtgatgaaaatgttcccaaaccaaatagtggaaatggttgcacagcattgtgaatgtgtaaagtatcactgacccatac**acttaccaagggctgaaatggtaagt**tttatgttatgtgtatctcaacacaata**aaaatgagtgccacatttt**gtttagcatatcaatgacaataacagttcaggtaaatgctggattcctcatcacaaccagtgcttcctgatggagctggattcaggggagggact**ggagtgggttgggtgcacaggtcatgaaggcagcaaaaattccaacccactcc**tcaagagtccagtcaccacctccagatctatgtccaaaaaacagctcttcctatggctgagctacatgag**caacaagcaaataaccatgttttttgttgttg**ttgtttagttttgttttgctttgtttttagacagagtctcgctgtgtcacccaggctg**gagtgcagtggtgcgatctcagctcactgcaatc**tctgcctcctgagttcaagtgattctcctgc**ctcagcctccctagtagctgag**actacaggcactcactaccacgcccagctaatatttgtatttttagtagagacagggtttcactatgttggccaggctgttcttgaactcttgacttcgtgatccacccgcctcagcctcccaaagtgccgggattacaggcgtgagccaccgcgctcagccacaaccatgtatttttaagcaaaagaCACAGTGAGGGGACATCAGTGTGAGCCCAGACACAAACCtccctatgcgggttcacaggacagcatggggtgctgaggacagaggtgggcactcaggaaccagcagggaaacccagggggcgcttggcacttcatgggggctcaggaccattgtgggctcagtggtcaggcaggctcaaggctcagcct**cagggcaggtgcagcaggcggggaaaggccctg**gagactgggtttagtgtcaccttctcattgcgccactggacaccctccactacatctattctaatgtgtatgagtacttatgagtttagaaaataacattgatattataaatctatagccatatgtgggtgcatcaagttaccctcttcaacctatgtggaccctgttcatcaggaataagtccctgcatttgaggaccttataaatcaataattatgtagaatcactttctttttcagtctcatttcctccctctttctctctctcacacacactgacaaacacgctggttgctataactttaattacctgatgtgttaaaaaaaattgattcatttggaacttaaccagtttagctgttgttcacgctgttatgtaaaaaagatccctaaatgacttctacctggtaagaggagggaaggcgttggaaggtgaaattgcag**gagaaaaagaaaatcgtgaaacggattgatttgttttctc**tatgaatggtgattaggtcaatatttttcaaacggtgaacatattgtgtgaaga**ttatttgtcaatattatctcattaaaactattacaaataa**aag**tgtataatttggtaaaaaaaatgaagagagaaataaaaataataca**agaaaagtcatgaactcctgactccacacgtgagttaaccctca**gtttttctatgttacttagaaaaac**act**aagatacagccaaagaatatcatgatatctt**tataagaa**gagggttttgtaaccctc**actcgaaatttctagctctgtcctagagttaatttagggagcagtcagatccagctgtgaga

Uppercase: IGHVII-20-1-2

Lowercase: Flanking sequence[1000bp]

Red & Bold & Underline: Stem-loop [15]

Blue: Heptamer[33]

Green: Nonamer [4]

id-IGHV2-5-2[V_gene_segment]

aaaattggagacatgtagaagctcactgggacctgttgagtcttctccaatcacactcagaaca**gagaccttagtgggtctc**ccttagagtagtcattaaggatggtgataacagcctaaagatggtagaccatggtcagtgtcatatagaatatggggaccctcacaagttttttgtctgacccttctcctgacactaaattatgcaaattaataacactgatctggtgcttcttttgattctaatttattttatttttagttgtcgttctca**cttttcctttggattttcctgctccctggaaaag**gtaaatgtggtctccgtgacctcaattc**aagggctgaagccctt**tccctgtagctcagctggggctcaggctgtggctactgcagccatgtggaagaggctgaagggactttcttcactctccttgctcaggaccatccactgtattgtgtataggcttctctggaaatgcaagtgg**ccatttgtagtgaaagaaatatgtttgtctggttaaaatgg**gaggtgg**atgtagagttaattggctgctacat**aaactgtccttctccaccagtgcttttaggatgagattgtgaaatttgtaagaatcaaaatggagtcacatatgttaaaaccctgacaaatggattcaggaagtgtagggagaattcttacacacatatccctgacaa**caagaactatcataaaatagttcttg**caaaaagaccaacatgacctcataatcatgacttctgcaaagacttctactcag**aatctacttgcccagccttagatt**aatgccatctgaattacactgatcatgttactatcactgctcctcaccacagatgcaacaccctcctgagtcctgaaacctgactccatcccatagagtagggca**cagatgaggggaatgcaaatctccaccagctccaccctcctctg**ggttgaaaaagccga**gcacaggtcccagctcagtgactcctgtgc**cccaccATGGACACACTTTGCTCCACGCTCCTGCTG**CTGACCATCCCTTCATGTGAGTGCTGTGGTCAG**GGACTCCTTCACGGGTGAAACATCAGTTTTCTTGTTTGTGGGCTTCATCTTCTTATGCTTTCTCCACAGGGGTCTTGTCCCAGATCACCTTGAAGGAGTCTGGTCCTACGCTGGTGAAACCCACACAGACCCTCACGCTGACCTGCACCTTCTCTGGGTTCTCACTCAGCACTAGTGGAGTGGGTGT**GGGCTGGATCCGTCAGCCC**CCAGGAAAGGCCCTGGAGTGGCTTGCACTCATTTATTGGGATGATGATAAGCGCTACAGCCCATCTCTGAAGAGCAGGCTCACCATCACC**AAGGACACCTCCAAAAACCAGGTGGTCCTT**ACAATGACCAACATGGACCCTGTGGACACAGCCACATATTACTGTGCACACAGACcacaaagacacagcccagggcacctcctgtacaaaaacccaggctgcttctcattggtgctccctccccacctctgcagaaca**ggaaagtgcagctgagatacgttttcctgccagggcctgcatttcc**catccccattagactcagagccctgtcttcctccttcttctttaataataaatggcatgactcctgttaatagttcatagaagcagaagctgagtcctgtttgtcaaacattcag**catgaaatgttcatg**ttacctgggccagatgcatcactggtatgt**ggccgccagtttaatgggctcatactactccaagcggcc**gaatacaaaacagcagaacatatgcctttattaaactgcaggaaatgacactggagaaaaatgtggggataagggagcaggaggaattgtgaaaattcagataatagct**tttctcgtagagagaaa**ggtgcgtta**ataaaataattttat**aaaacaataaaacagatggaatgtgtccccatcacggagtgtctcacagtaacagtgcagaaataaatttctaaaaatgtcgtagggtaggtgtgatagacaaggcttatttccacacagggagcccataaatacccatgtaattaaaaatccagctcttagaaagggagtgcctcaacttgacatcagaaacccacgtcagtgagtctgaggagcaatgtggaa**aagaaatggaaaattcaacaatacctttctt**atgaaaatcctagctcac**gtttcaacatgaaac**ataactgtctacaattttaggaaaaccatcccatggcaacaagtcactacactcaaaaggtactgagctaacagagttccagcagtgagcaggtcagtgctgaaatccccggagaatcaaggctctcaggtacgttgtacaaaacccagaactgagccacacaatctatggtcaagtgatctttgacaaagtcaacaaaaatatgcacggggaagaggccacgctcttcatgc

Uppercase: IGHV2-5-2

Lowercase: Flanking sequence[1000bp]

Red & Bold & Underline: Stem-loop [19]

Blue: Heptamer[39]

Green: Nonamer [6]

id-IGHV2-26-2[V_gene_segment]

aacaacattgtgagttacataatgatcctactttatggccgcggtgcaaatgcttaacttccactaaatcacagagctcagaatggctagcatggaggtgtatatgtatatgtgcattgttctagaaatttcc**aattactgtatattattgtttgcagtaatt**aggtcaatctgaactgacaaaagtctccatggccttataagcacagaga**ccacctccacggcaggtgg**aaatcaaggctgggaacagtctaccagctgctcccctgggattcctccaattaccctgtttaaagcctctgctcactcatgcgct**ggaagagggatgtcagagatgcagtctactcttcc**taaagggctgacagggcataaagacaagactgattatccatggaa**cattcctataggaatg**cacctatg**tgcagacacacaaataagctgtgtgtcttgca**gt**gtgtaaatggctgagatgccatttacac**ttctttac**acaatacatttttaaatgttttgt**tgatgtattctctatcattcaa**aaaaatcatcactttccccctaaacaaagaggattttt**atcagaacactta**tgggaagccccttgctctcccagattccca**ccctcatttctcctgaaggaagaaagaaaaccatctctattgatttccactctcatcctctaggactaaaaccagaaagctgcatgctccctgggtgagccttagagaagaccctgtctgtaggagcaggaa**tctcagagccttggctgaga**ggcatggtcctgaaatgagatggagtcccccatggagatcccacgtggacgcccacacctgagggctcactgctcctcaccacagatgcactccccta**ctgagtcctgagacctgagtgcaccccatagagtaggactcagatgagggaatgcaaatctccaccagctccaccctcctctg**ggttcaaaag**ctgagcacggggcctcgctcag**tgactcctgtgccccaccATGGACACACTTTGCTACACACTCCTGCTG**CTGACCACCCCTTCCTGTGAGTGCTGTGGTCAG**GGACTTCCTCAGAAGTGAAACATCAGTTGTCTCCTTTGTGGGCTTCATCTTCTTATGTCTTCTCCACAGGGGTCTTGTCCCAGGTCACCTTGAAGGAGTCTGGTCCTGTGCTGGTGAAACCCA**CAGAGACCCTCACGCTGACCTGCACCGTCTCTG**GGTTCTCACTCAGCAATGCTAGAATGGGTGTGAGCTGGATCCGTCAGCCC**CCAGGGAAGGCCCTGG**AGTGGCTTGCACACAT**TTTTTCGAATGACGAAAAA**TCCTACAGCACATCTCTGAAGAGCAGGCTCACCATCTCC**AAGGACACCTCCAAAAGCCAGGTGGTCCTT**ACCATGACCAACATGGACC**CTGTGGACACAGCCACATATTACTGTGCACGGATACcacag**agacacagcccaggatgcctcctgtacaagaacctagctgcatctcagtggtgctccctccctacctctgcagaa**caggaaagtgtgactgagatgccatttcctg**c**cagggcttgtgtttcctataccaaccagactcagagccctg**tctgttttcctattctgtactattaaatggcatgttccctgttagtgattcacgcaagcagaggctgtatcctgtttgacaaagattgagcatgaaagattcccattacctg**ggccacatgcatcactgatatgtggcc**acttatttcttgagctcatatccttccaaggggctgaatacaaatatctataacataggccatcttaaactgcaggaagtaatgttggagaagaatgtggagataaaagagcaggatgattaattaaaatccagataccatctttttttgcagaga**aaaatttacactaataaagtaatttt**atgaaacaacaagaaagatggaatgtgtcctcatcatggagtgtctcacttgcagtgcacaaat**aaatttctaaaaattt**tatagggaaggtgtgatagatgaggctgatttccacacagggagtgattaaatacccaggtaagtataaaatccaacacttggaaaggaaaatg**cctcagcttgacatcagaaacccagctcagtaagtctgagg**agcaatgcggaaaagaaatgaaaaattcgggccgggcgcggtggctcacgcctgtaatcccagcactttgggaggccgaggcgggcggatcacgaggtcaggagatcgagaccatcctggctaacacggtgaaaccccgtctctactaaaaatacaaaaaattagccgggcgtggtagcgggcgcctgtagtcccagctactcgggaggctgaggcaggagaatggcgtgaacccgggaggcggagct**tgcagtgagccgagatcgcgccactgca**ctccagcctgggcgacagagcgagactccgtctca

Uppercase: IGHV2-26-2

Lowercase: Flanking sequence[1000bp]

Red & Bold & Underline: Stem-loop [26]

Blue: Heptamer[44]

Green: Nonamer [3]

id-IGHV3-25[V_gene_segment]

gctgagtggaggaagctaaagaattcaccatcaat**ttcatgtgatttcatttgtataaatcacagaa**gatgcaatattctaaattaacat**agagaagatctgaatttctct**gaaaatagggtgggcagagtaacaacgtggtgaaataaaattatagac**aagttagataaaaaaaattagaggttaactt**aattgctaaaacatg**attgaagtgctgattcaat**gactgcacacatatatcaacattttccaaatgctgcactatcaatttgat**ttcattattgattgtttttttaataaagcagtaacaaaaatgaa**tatattggctgagaaagagcaagaaatggataaataccgacacttgaataatcaaggact**cctgaaaatacacacatgtgaacactgattgtagatttcagg**taaacactagaaaaggcaaaattacacaaagttgttatgacagga**gggacatcctgaaaacctctctaggcatgtccc**acaccacc**ctggagctgtctcaggggagcagtctcctccag**tgtttagaggcacaggcacgcataatagggctaactctggccag**atgtgtgatattggacacat**tgcacaactgctctgttctgtatgtaatttatcttctctacaaatgtaacattgacagttgcattaaatatattctgcaaatatgtaaaagttaaataagctgatgtctgctaaatgattaccaaggcacaatcacataatctgaagttatattttcc**agagtgataggattacctccagagttttccaggacactct**catctgctccgggcactgccttctcctcagccgtcccacccctgagcttgctatatagtaggagacatggaaatagggcccactctctgctgataaaaaccagcccagccccgaccctgcagct**ctgggagaggagcccagcccccgaattcccag**gtgttttcatctggtga**tcagcaccgaacacagaggactcaccATGGAGTTTGTGCTGA**GCTGGGTTTTCCTTGTTGCTATTTTAAAACGTGATCTATAGAGAACTAGAGATATTGAGTATGAATGGATATGAGTG**AGAAACAGTGGATACGTGTGGCAGTTTCT**GACCGGGGTGTCTCTGTGTTTGCAGGTATCCAGTGTGAGATGCAGCTGGTG**GAGTCTGGGGGAGGCTTGGCAAAGCCTGCGTGGTCCCCGAGACTC**TCCTGTGCAGCCTCTCAATTCACCTTCAGTAGCTACTACATGAACTGTGTCCGCCAGG**CTCCAGGGAATGGGCTGGAG**TTGGTTGGACAAGTTAATCCTAATGGGGGTAGCACATACCTCATAGACTCCGGTAAGGACCGATTCAATACCTCCAGAGATAACGCCAAGAACACACTTCATCTGCAAATGAACAGCCTGAAAACCGAGGACACGGCCCTGTATTAGTGTACCAGAGAcacagtgaggggaggtcagtgtgagcccagacacaaa**cctccctgcagggccatgcgggtggtttcctttctcagctgcaggagg**cgggcttattgttgcaggactctggagacttattaggttgtgatattttactatggttatttatcatgaacttttttattgggaattgtgttttattaatttttattttatatgtaaggt**tatttttttaaaata**cataccttcaagaaataattcttcctaataatttgcacttgttctttttagagttttgttaacatctgttgatatcagctactacatagctatagggacattaatttacatctatag**acatatgtgtaaatacacaaacttatacatatatgt**ggtcttttatatt**taatgttataataaaattattataacaatta**taaaatacatcctaaagaatgaaacatatatagtaatccatattcctttcaatcattgcctatagtttacataaattgatgtctatttgtaaccttaaacatagtgtattggtcattttcaaacatccaagaacaaatt**tcaaatgtccttgtcacacacacaaaaataaaaatttga**ggatttgaggtgatatatatgtcaatcagctcgattcaattattccatattgcattcataaatcataacatagctttattcactataaatacatacaacc**aaaatttctcaattttcaatgaaatttt**aattatacatattttatctgtcctaggtcatgaatgtt**tttctccctgtccggatgtgattaagattgtcctgagaaa**caaattcagcctc**ctgcctcctgaaggcttcaggaagcag**ctcctagatggggtgaagcaggcaacccgtgtgtgtccacaggacccggagcctctctctccttggattaggccccctcctcaggattacagggctcttcagttttctcaacatgctgttataccagagaagcacaaaacacttaatttcattatattt

Uppercase: IGHV3-25

Lowercase: Flanking sequence[1000bp]

Red & Bold & Underline: Stem-loop [23]

Blue: Heptamer[25]

Green: Nonamer [8]

id-IGHV3-48-2[V_gene_segment]

tatgttgagtaaaataagctaaacaaataagaatatatactatgtaattt**catttttataaattctgataaataaaaatg**catctgaagtaaaataatgaagataagtagttgcctggggtaatggtagaagaagggagggggagaggaggaggaatacagcagaacaaggggaaatgttgagaagaattcacttgtccactttcttggt**aatgatagcagttacatcatt**tttattagttgtacattttaaatatgtgaagtttattatctttcaattaagcctcataaaatgtcttacaagcaaacaaatggaaacttagacaaggaaagagtaatagaaagacagaaaaaataagttcaatgtcagaagtac**ctgaaaattaatgtgcctggatcctagttctctccatattttcag**aagagtgctggagggcagcaaaaccacacatgctcttattacggaatgtgggttctgataaaaacactagac**acatccagctttgtcctggagttggtttagggggatgt**cagagacagtgatgaagagcacagggccagataccggggttcactcat**cccagacatgagctccgagacgcatacagagcccccccatgtgtggg**tttacttccac**ttctgtaaatggagaaaatattgtctcctacagaa**catagtttacatgaatatttaaaatgaaatagggtgattagtgcaaagtgtttatcacagcacaatttcataataagacagcatattttccaaatgcaatcattgccagcaaacttctacagggcaccgtcgtcttatctgggtacagcctactcctcaagggtcccaccctagagcttgctatatagtaggagatatgcaaatagggccctccctctactgatgaaaaccaacccaaccctgaccctgcagctctcagagagg**tgccttagccctggattccaaggca**tttccact**tggtgatcagcactgaacacagaggactcaccA**TGGAGTTGGGGCTGTGCTGG**GTTTTCCTTGTTGCTATTTTAGAAGGTGATTCATGGAAAAC**TAGAGAGATTTAGTGTGTGTGGATATGAGTGAG**AGAAACAGTGGATATGTGTGGCAGTTTCT**GACCTTGGTGTCTCTTTGTTTGCAGGTGTCCAGTGTGAGGTGCAGCTGGTGGAGTCTGGGGGAGGCTTGGTACAGCCTGGAGGGTCCCTGAGACTCTCCTGTGCAGCCTCTGGATTCACCTTCAGTAGTTATGAAATGAACTGGGTCCGCCAGG**CTCCAGGGAAGGGGCTGGAG**TGGGTTTCATACATTAGTAGTAGTG**GTAGTACCATATACTAC**GCAGACTCTGTGAAGGGCCGATTCACCATCTCCAGAGACAACGCCAAGAACTCACTGTATCTGCAAATG**AACAGCCTGAGAGCCGAGGACACGGCTGTT**TATT**ACTGTGCGAGAGAcacagt**gaggggaagtcagtgtgagcccagacacaaacctccctgcaggggtcccttgggaccaccagggggc**gacagcacactgagcacggggctgtc**tccagggcaggtgcaggtgctgctgagggctggcttcctgtgatggcctggggcggcctcattgtcaaatttccccggggaacttctccagatttacaatcctgtactaatatttgatgtctctaaatgcaaccttttttttcctttttgtgt**ctgttttttttttttaaaacag**gaggacacatcctcacctccacagaagccacagtgtcactttgggggcagaaataatcctttcgtggtcaacagggtgagagttttgaggaat**cccagggaaacctggg**gaatgttttccaattagactcaggg**cagagacctccatgggaatctctg**attagaacaggctttgagttctgatgggagccaagagagaggctcacccagggtcagggttctt**aaaacctgatggttttcacagcaatcccccttcatcttgtgaaa**ctgggcacatctgactcagactgattcagttgaccctctttctgctaatccattttccttcccagtagacttgattctcacagatccctttcttcttctctttcct**gaaaacagaggatgtgttttc**tgtagtctaaattccaaggctcaggtccgcaggagctgggtaggctgaggggtcttcctcactcactattgcctggaaaatcctgctgtcttctgtgcatggaggcatttggaaaatgaagcaggcattagtcatgaagggaataatactagttttctccaatgggatgttgatgtagagctgatcttatgcttctcacact**gtcacaaagtttggactctcacctgtgac**tttgaggagagcttgtgataccttatcttgttttaatatgaatagactctcccttagctcaggaagctggaacagactccatttggctccttcatttgtaag

Uppercase: IGHV3-48-2

Lowercase: Flanking sequence[1000bp]

Red & Bold & Underline: Stem-loop [22]

Blue: Heptamer[34]

Green: Nonamer [7]

id-IGKV3D-15[V_gene_segment]

gcgttagacagggatgggtgggaatgattgaaggtgtgagtactttagcacagattaagaccaaatcattaggatttaaagagttgtgtagagttagtgaaggaaaagccttagaattaaatttggctgtggataaaacattcttggattagactgaagactcttttctgtgctaagtaagtatatttatgataatgatgatgactgtagtgctaaatatttaataaataaaaacaaaattaattgccacatacataatgtcctgaatactattgtaaatgttttatcttattttctttaaactgtctacagcactataaggtaggtaccagtattgtcacagttacacagatatggaaaccgagacacagggaagttaagttacttgatcaatttcaagcaatcggcaagccatggagcatctatgtcagggctgccaggacatgtgactgtaaacagaagtttttcactttttaactcaaa**gagggtatgtggctgggttaatggaaagcttcaggaccctc**agaaaacattactaacaagcaaatgaaaggtgtatctggaagattaagttctaacagattcttcatttccatcgatccaataatgcacttagggagatgactgggcatattgaggataggaagagagaagtgaaaacacagctttttatattgttcttaacaggcttgtgccaaaca**tcttctgggtggatttaggtgattgaggagaaga**aagacacaggagcgaaattc**tctgagcacaagggaggagttctacactcaga**ctgagccaacagacttttctgacctgacaaccagggcggcgcaggatgctcagtgcagagaggaag**aagcaggtggtctttgcatctggaagctcagctcccaccccagctgctt**tgcatgtccctcccagctgccctaccttccagggcccatatcaatgc**ctgggtcagagctctggggaggaactgctcagttaggacccag**acggaaccATGGAAGCCCCAGCGCAGCTTCTCTTCCTCCTGCTACTCTGGCTCCCAGGTGAGGGGAATAT**GAGGTGGTTTTGCACATCAGTGAAAACTCCTGCCACCTC**TGCTCAGCAAGAAATATAATT**AAAATTCAATGTAGATCAACAATTTT**GGCTCTACTTAAAGACAGTGGGTTTGATTTTGATTACATGAGTGCATTTCTGTTTTATTTCCAATTTCAGATACCA**CTGGAGAAATAGTGATGACGCAGTCTCCAG**CCACCCTGTCTGTGTCTCCAGGGGAAAGAGCCACCCTCTCCTGCAGGGCCAGTCAGAGTGTTAGCAGCAACTT**AGCCTGGTACCAGCAGAAACCTGGCCAGGCT**CCCAGGCTCCTCATCTATGGTGCATCCATCAGG**GCCACTGGCATCCCAGCCAGGTTCAGTGGC**AGTGGGTCTGGGACAGAGTTCACTCTCACCATCAGCATCCTGCAGTCTGAAGATTTTGCAGT**TTATTACTGTCAGCAGTATAATAA**CTGGCCTcccacagtgattcaacatgaaacaaaaacctcaagaagaccatcagtgtttactagattataccagctgcttcctttacagacagctagtgtgg**tggccactcagttttagcatctctgctctatttggccattttggagctcaagttctcaagtccaaaat**tacttatgttattccattacatcataccatttcagtgtggctattacattcatttaaacgcatttcagaaggcatctctgtttatggcatcacaaagagtttaataaatctgtgcaagaataaacaacaaacacacctataaatataaa**gctgaaatatcaaaactatttcagc**actctgaaaattggcaaagcat**aaaataattaaggatgcatattttttatagaaaaaaa**aagtactagtgctttgagtaaggacagaaaatgtctgtagccttttgcctgtgacagcacccttctattcccagctcagtcaatatgaattgcagaactgg**agttttacccatgtaaggatagcaaataaaactggcagcttgctgcca**aagtgggtggacttgagtaaagccaaggagtgggaaataattcttcagtgtttccagctaaacagggagaacaccataggaaatgaacagaaaaagcccacagccttgctagtccaagatgatgccttgtttgaggcaagtagtac**atctgctgatagtaaatagcagat**tcctggataagatagacccacattgctgaaataatctctgcacatattcctagtgacctagaagttatagatatgagtgatgagaactaggagtttccggtgcaaagtaaaaccagagggaggtaagaactag**ctgcattttgcatgcag**tttgctttttaaactacacacagatgggttgacagaagatagatttataacctccagatatttgagcaaaatgtctcagatcattggtgaccacttcgctatgcagatacatgtgtagttcctataaaatcaaact

Uppercase: IGKV3D-15

Lowercase: Flanking sequence[1000bp]

Red & Bold & Underline: Stem-loop [20]

Blue: Heptamer[32]

Green: Nonamer [5]

id-TRBV3-1[V_gene_segment]

aaaggggactgtggtcctcaacattagataattcacaaaacgcgaaggcattttctgtcgtcacattgccatcaggtgggtagggccca**aggatgccgccaagcatcct**ccctacacaggacagctccccacaacaaacaatcacctgatccaacatggcagtagtgctgaaattgagaaaccccgcaataagcttcggttgtaaactttgtaccaagaaggagaattctctgcagacaactcaaggggcaatacc**gtcaactacttttgttttctctgactgccgcgttgttgac**cttcatcgccattctttcccagcacatggtgtctgcatattttccagcagcctcctgtttcctcataaatctgccccttggacagataagaacatcacagggagcagggctgcagttcactcccaataataacgcctcagcagggtttgctctgcagtagcatcacagtcatgtggagtaggtggctatggctggt**attcaggtgaaatctcaggccaatctctcctgaat**caaggggaccactgatcattaatgagaccttcaagtgagcaaagattcaagagagcaaattaatgtcctgcgctattgtgaagggacgcaaaggatccagagaggaaaccagctgactctgagagaagggcctgaaaaggagaggaagggaggaggtggtcagatccccgggaaggagcagggagggagggggctgctggc**ccaggaatgactgtcagagacacatcctgg**aggtcacaattccaccctctcaattattcccagaaattccaaaaactcctttaaaggagcaagtgcagggcatgaaagggaggagccatgctagaggagaccctgggaatgggggaatgattacaggctgtgacctcactggcgcagcacctctcagcg**gcagtggaaaccacagcctagtcctctcaccactgcagaccagaatcctgccctgggccttgcctggtctgc**ctcactctgccATGGGCTGCAGGCTCCTCTGCTGTGTGGTCTTCTGCCTCCTCCAAGCAGGTGAGTCCCGGGCCCAGGTGACATGATCCTATTGGAGTCCCTAAGCCTTTTCACCATGACAACAACAGCAGGCCGTCTCCTAGGATTTGCCTGAATTCTGCTTCTTTCCTTTGCAGGTCCCTTGGACACAGCT**GTTTCCCAGACTCCAAAATACCTGGTCACACAGATGGGAAAC**GACAAGTCCATTAAATGTGAACAAAATCTGGGCCATGATACTATGTATTGGTATAAACAGGACTCTAAGAAATTTCTGAAGATAATGTTTAGCTACAATAATAAGGAGCTCATTATAAATGAAACAGTTCCAAATCGCTTCTCACCTAAATCTCCAGACA**AAGCTCACTTAAATCTTCACATCAATTCCCTGGAGCTT**GGTGACTCT**GCTGTGTATTTCTGTGCCAGCAGCCAAGAcacagc**cctgcaaagtcactgcatccctgtgcacaaacctcccggctcagccaggaagc**tgcagggcagcgtgtgcacctgca**cccagggctccagt**ctccattccctgatggcctctgatggag**tttcagtctgtagtacagccagctagtgcacccagtggagaagtcctccatcttatgcacacaaaagtctcacagaattgtttatcagctagagcagggcttcacaacaaagccactaaagtattagggctccacaaacaaaaaagagcctgtcctatgcactgatgatcttaactgatgggggaagctctttcttttctttctgtgtggcacatattggtacgtattttatataacattatgaaaaaaacacgattgctgaaaatggttgatgagcacaagcataaaggagggggaaggaaatatttccaagatcaagagacaattctgatgatcagtgagtactttattcaacagaaaaatatttggagaggcatgaaattccatggcaataggtaaactggaaagtgttgaggagatgccggtggaggtagccttgtgcgttgtgggct**ccttctcttcaagaagg**gtgaccagcaagtgtctctcccaattctgatgaatatca**tcttgaactgtcctcaaga**gcaggcaatagccagatgactggggatattgggagacctgaaaggactttggacacaaatctctaggcagtggtgagccattggtcctactccaagtgaaacaacatgatgaaactgatattttggtagaggaacactggtagtagtatggatcagaaaggagatagatttttgggtggcatgaatgattaggagac**ttctaaatatcctacatgaaagattagaa**agggttgatgtgaagcattgtggacaatataggcttgaggctctgcattatgttatttcttggaaatgtattgggttttattctgggagggttgaatgtaggacttgctagggtctatttca

Uppercase: TRBV3-1

Lowercase: Flanking sequence[1000bp]

Red & Bold & Underline: Stem-loop [14]

Blue: Heptamer[48]

Green: Nonamer [6]

id-TRBV7-7-2[V_gene_segment]

cagatcagctgagtacccaggatatcacacatcccagcatcttctaacgaggaaaaatacttcacgtaccagcctccttcctgcaccacattgactaacc**ctggaaacttagcctttccag**aaaaaagctctgtcttctctcggcccagtctagcctggtaggtttaattcatgatttatgtgaccttttaatttactcaaaggagaacgtatgatttcagggtaaagaatattcattttatttttataagttggacctaaacagtaaactgagtacttttcttccatcagcgtcttaaattagacaacggtaaaacaaatctttgatgcta**gaacttcagatcctagtgtttttcacatggaagttc**ccagttacccagtctagggcatggaatagatgagcattaataagtcactgtatgcctgagatggttttctatgggcatcttttctgaattacggttaacaagggtgggaaatgatcccagaagaaaggcatgagggcagttcttgagggaacccatgtgatgggacagtctcattgggcacgtgtgactgggggaatggaggaagctggggcatgattgggatggcacaggggaccctgacttgcgggaaagacaatgagctcaaccctttgtgccctatgttaggggcgctgttgttg**catcctacaggacatgcccagcaggcaggggagcagctgtaggatg**agaagatgaactcagagatgcagtgtgaggt**ctccaggcacagacggctctggag**cacaaaccaatgagccatgaattgatgttgttaaaacggaggttataaatatttaaagtgtcacccaagtgtgttctaatagaaatgctgtgaccctgaggtcctgaggattgagagaggaagt**gatgtcactgtgggaactgccctgtggagacaaggacatc**cctcatcctccgctcctgctcacagtgacactgatctggtaaagcccccatcctggtctgacactgtcATGGGTACCAGTCTCCTATGCTGGGTGGTCCTGGGTTTCCTAGGGACAGGTGAGTCCTCAAAACACAAAGTAGTTTCATATTTTTTCTGTATGTAGGTGTGTGTGTGTATGCATGTGTGTCTGTGTGTGTGTGTGTGTGTGTGTGTGTGTGTGAGATGACTACAAATGTTTTCCTTATTCTGTTGCCAGATTCTGTTTCCACAGATCACACAGGTGCTGGAGTCTCCCAGTCTCCCAGGTACAAAGTCACAAAGAGGGGACAGGATGTAACTCTCAGGTGTGATCCAATTTCGAGTCATGCAACCCTTTATTGGTATCAACA**GGCCCTGGGGCAGGGCC**CAGAGTTTCTGACTTACTTCAATTATGAAGCTCAACCAGACAAATCAGGGCTGCCCAGTGATCGGTTCTCTGCAGAGAGGCCTGAGGGATCCATCTCCACTCTGACGATTCAGCGCACAGAGCAGCGGGACTCAGCCATGTATCG**CTGTGCCAGCAGCTTAGCcacagcatggcacag**tcgcctccttcctgttcacaaacctcatccttctctctctttgcagctcccagagatcattaacagaggcct**ctctttgctcctcacttttcctgggaaagag**gtagatttggacctcggttgtcctttgggtggaaagagaccacagattcattcctgaaacacagtgactgcaaatgtaagtggtgaaaacaatcacgtcccactgcactctaggagggctcggagccagctcactgctccaaacaaggagtgggtgtcttagccttggccttcagagcagacatgcatcttctataggtcttggaggctgctgtgttgcccacatacatgaggttgtcataggcaggaaacatgcctcttcttcatatgttggggcatctggaaggtctgaggctacatccccg**agaagatctttcttct**gaagcctcttctatccctgtcaccttggaagttttctgccacaaaata**tcaaacctctcttcctgtttga**agtaaaggtctttgcaaatttgtggtccttacttgataaatacaatcatggtaacaataacacttcatttcttctgcctactttaagccaagagtatcctttattttatttccatttgccat**tgctgctgtcctgatagacagaagca**tgcatcaccaccgctgccacttcaccttgatt**cactcaggaaatctgattttcagactctgagtg**ttttgattgttgtccaactcatttgatttgaaataattctcccaaggcctttaactcaaaaagagttttatttataatattgaatctattcctttt**tattatattttttcataatatatagtgttatata**gcacttgttagaaatagaagtacaatgattatattgcaatagaatcttccatctgtctgtgggtgatgctgcagtttgtatctatgaaatgg**aatgcactggtcagagctgatgtgattatgcatt**atgggtttcggggagtcctcagc

Uppercase: TRBV7-7-2

Lowercase: Flanking sequence[1000bp]

Red & Bold & Underline: Stem-loop [15]

Blue: Heptamer[41]

Green: Nonamer [7]

id-IGHV3-20-2[V_gene_segment]

aattatgtagaatcactttctttttcagtctcatttcctccctctttctctctctcacacacactgacaaacacgctggttgctataactttaattacctgatgtgttaaaaaaaattgattcatttggaacttaaccagtttagctgttgttcacgctgttatgtaaaaaagatccctaaatgacttctacctggtaagaggagggaaggcgttggaaggtgaaattgcag**gagaaaaagaaaatcgtgaaacggattgatttgttttctc**tatgaatggtgattaggtcaatatttttcaaacggtgaacatattgtgtgaaga**ttatttgtcaatattatctcattaaaactattacaaataa**aag**tgtataatttggtaaaaaaaatgaagagagaaataaaaataataca**agaaaagtcatgaactcctgactccacacgtgagttaaccctca**gtttttctatgttacttagaaaaac**act**aagatacagccaaagaatatcatgatatctt**tataagaa**gagggttttgtaaccctc**actcgaaatttctagctctgtcctagagttaatttagggagcagtcagatccagctgtgagaaacacgggccagacagtgagactggcttttaccag**atgtgagctcttagacatgtcacat**ggcccttccgtgcttgggg**gtttaccttcacatctgtaaatgaaggaaac**tttgactcccacagaacataattcatgtgcatctataagtagaaatgctaatgagaattaattatttatgaagtataatcactcgcatccactctgggacacagcctactctgaggcatcccttccagaagtcactatatagtaggagacatgcaaatggggtcctccctctgccgatgaaaaccagcccagccctgaccctgcagctctgggagaggagccccagccctgagattcccacgtgtttcca**ttcagtgatcagcactgaa**cacagaggactcgccATGGAGTTTGGGCTGAGCTGGGTTTTCCTTGTTGCTATTTTAAAAGGTGATTCATGGATCAATAGAGATGTTGAGTGTGAGTGAACACGAGTGAG**AGAAACAGTGGATTTGTGTGGCAGTTTCT**GACCAGGGTGTCTCTGTGTTTGCAGGTGTCCAGTGTGAGGTGCAGCTGGTGGAGTCTGGGGGAGGTGTGGTACGGCCTGGGGGGTCCCTGAGACTCTCCTTTGCAGCCTCTGGATTCACCTTTGATGATTATGGCATGAGCTGGGTCCGCCAAG**CTCCAGGGAAGGGGCTGGAG**TGGGTCTCTGGTATTAATTGGAATGGTGGTAGCACAGGTTATGCAGACTCTGTGAAGGGCCGATTCACCATCTCCAGAGACAACGCCAAGAACTCCCTGTATCTGCAAATGAACAGTCTGAGAGCCGAGGACACGGCCTTGTA**TCACTGTGCGAGAGAcacagtga**ggggaagccagtgagagcccagacacaaacgtcc**ctgcaggaagacaggaggggcctgggctgcag**agggcgctcaggacacattaagaacacagtaaactcagggaccaggtgcccagggaggttaagggctggtttcctttcagagacttagctgtttctccatctaacagtttccccagaaccctgcctagatttgtgatctgtatctgctcaaactatctgtgtcac**cttcctcacctgtgactttgaggaag**cttagtgtggacattccagtgggtggagagatcttggtgacagcaattgtgtgttctgtctaggcatgtctatggctggccatcaggaacgacaggctggaattcttggaaagaggcacacctgccatccatcaggaaattttgttgtcttctgttctgctagaattaaatcaggcacgcccaggttaactactactatcttcctagcttgagaaaattaatggtgggcttgaataacacctatatgaaatcatcagagcaacacctagattcg**agtctcatttaataattgagact**atgttctagccaaggagacacataaaactggtgattgccgtggtcagttcatatcatatatttgtcagtagaatctggctggtattatgaacagctttg**ccaacacatgtgttagtgttgg**tcttcggagaagcacattctatgaatggattaggtttatgtaca**gtttcctaaggaaccggtccacgcttttgagaggaaac**gcagagcgggtggaaggaggctggggaagctgaaggcatatgagggaagttggtcc**ctgagtgaaggagagagggaggataactcagtggaacttccttaaactgtgctgttctaggaaggtcca**gcaagtcactgtatct**gagtcatgtcaaagtttcccatcggggacccccgtgactc**ccagcagtggctctactcagatcagcgcggagctcattgttctcctgagagtgg

Uppercase: IGHV3-20-2

Lowercase: Flanking sequence[1000bp]

Red & Bold & Underline: Stem-loop [20]

Blue: Heptamer[31]

Green: Nonamer [5]

id-IGHV3-30-2[V_gene_segment]

aataactgtgctgagtgaaagtagcta**acaaattaacagtgcacttcataaacttctatttgt**ataaactgcagaaggtacagctattctaaagcaacagagaagattacatattggtgggaactgggttttgaaagtaatggggtggtgagatgaaattacagagaagtgacacaaagatttaggtgtgaattgtacacaaattaattgaagctctgattaaagggttgcacacatgttaccattttccaaattatgcagtatatatttg**aattaattattaatt**gtatttaaataaagcagtaacaaagaaacaaatgaatacatttattgaggtggagccaaaactaaatgggaatgaacactcgaaacatctctgactcttga**aaatacacaagcgtgaacactggttctctctatgtattt**caggtcaatagcagaatacactaaaagaaaacagagatgtcctggcaggggtggaatcctgcagacctcactaggcatgtctcacactgccctggagttgtctcaggggagcagtctcctctagtggtcagaggcacaggccgagataatggggctaactctgtccagctgtgtgaccttgaatgcattgtataaacactctgttctctgtgtgatttatcttccttaaaatgtgaca**ttgacacttgcattaaatgtattctacaaatatgtcaa**aaagatgatgactgctaa**atgattatcaaggcacaatcat**ataatataatgatattttcctgagtgataagatgactaccaatctcccc**cagggcactttgtctgctctgagccctg**cacctcctcaggattcccatcccagagcttgctatacagtaggagacatgcaaataggtttctccctctgctgatgaaaaccagcccagtcctgaccgcacagctctgggagagaagcgccag**ccctgggattcccaggg**gtttctatttggtgatcaggactaaagacagaggacccaccATGGAGCTTGGGCTGAGCTGGGTTTTCACTGTTGCTGTTTTAAAAGGTGAACTAGAGAGATTGAGCGTGAGTGGATACCCTTGAGAGAAATGGTGGATTATGTCTGGGAGTTTCTGACCAGGATGTCTACGAGTTTGCAGGTGTCTAGTGAGAGGTACAGCTCGTGGAGTCCGGAGAGGACCCAAGACAACCTGGGGGATCCCTGAGACTCTCCTGTGCAGACTCTGGATTAA**CCTTCAGTAGCTACTGAAGG**AACTCGGTTTCCCAGG**CTCCAGGGAAGGGGCTGGAG**TGAGTAGTAGATATACAGTGTGATGGAAGTCAGATATGTTATGCATAA**TCTTTGAAGAGCAAATTCACCATCTCCAAAGA**AAATGCCAAGAACTCACTGTATTTGCTAATGAACAGTCTGAGAGCAGCGGGCACAGCTGTGTGTTACTGTATGTGAGGCAccaggtaagaagacatcagtgtgaacacagacacagtttcctgcaatgataagggaggaggctgggctaaaaggggcactcaggacccactgaaaacgggcagctctagggcaggtacagatggttatcatgggctgctttccttcggggtctgtggcttcctctgcatctaccagttcccctcggagcctctggacatttatgtttctgtggccacccctgacatctctggattagaaaagtttattataggaagaggaaacattttcatttgtcccaaagcagatgtaagtaatggaagcaaaaaatgcacaggaggccaggtggggctgtagatactgtcaccccagaatgtcaatctcaccactagtactggagaggcatgggagtttgatggagcttccctaacaactctgtcttccaagataagtccagcaagaccattggtgcctcccagagcacagttgtccatcagggatgtccaatgcttcccagcagcagccatgcctcagtgtctccactgtgcacagccactgt**ctgggaggagctcccag**gatgggtgtctttggcacacaccaggtggcgggtgttaagagtgcagtgcagcagctggctgcctggtctgttgggcctg**atgctggagagatgggaggtgcattcttggggccagcat**gctgtttgtgaatttttatataaaaactctgattttacttcattttc**gcagatgacatagataactaaaaacagaatctgc**aaagaaattgtaattttcaactttaccccaaattcattgtttcttaattctgtgcaagatccagacatattattgccttcctcatgataatttattctatgtaaaaccgaaatcattttttctcctactctttgtttctgttcaagtacagagatcttgagcaaagtaagttgggttctttccacacactaaccctcaccttccccagagaatgagcagagattgtcctcagtctgagtctaaggg

Uppercase: IGHV3-30-2

Lowercase: Flanking sequence[1000bp]

Red & Bold & Underline: Stem-loop [13]

Blue: Heptamer[35]

Green: Nonamer [3]

id-IGHV3-71-2[V_gene_segment]

ttgtcatcc**aggctggagctcagtggtgagatcatagctctctgcagcct**cagaccctgcgtctccaccaatcttcctccctctgccatacgtagctgggactactggtatgtaagacaatacttggcttcag**tgcttattaaaatgcacactttaattaagca**gtatttgttatacagcaatcatgcctcaataagagtattgcaaataagtggatagataatttgttcaataaagattgatggaaagatagatgctaacatg**agaaaatgcatgacacccaagaaaataacactgtaggaaacatgcttttct**ttacaattgttaggtaatcacaacagagcatacacatcacaccatgttctcgttacagagaaaaggttctgcaaacctcactagctgcaacccctgtgtgctgggcttggttcagggagaagtcaggtccagtggtgagaagcacaggcccagataaccaa**gctcactctgaccgaatgtgagc**actgggtacattgtacaacccatctgtgcttctgctggtaatttttcatctgt**aacatggaaatgacattgatactatataccatgtt**tcctctgcatatgtaaaaataaaaggtgattggtgctaactttaaatatacacagtttatgtacattgattgcacctcaatacaacagttttaaatagatattacaacattataagttttataggtttaatattttatcacagaacaaactttcaataagaaaccacaatttccaaatgctatcaatatcacaaatctcccccaggacactctcacatgctctgagccccactctctcctcaggcgtcccatcccagagcttg**gcatgtagtaggagacatgc**aaatagagccctccctcggcttatgaaaaccagcccagccctgaccctgcagcactggaagaggagccccagccctgggattttcaggtgttttcatt**tggtgatcaggactgaacacagaggactcaccA**TGGAGTCATGGCTGAGCTGGGTTTTTCTTGCCGCTATTTTAAAAGGTAATTCATTGAGAACTATTGAAATTGAGTGTGAGTGGATAAGAGTGAG**AGAAACAGTGGATTTGTGTGGCAGTTTCT**GACCAGGGTTTCTTTTTGTTTGCAGGTGTCCAGTGTGAGGTGCAGCTGGTGGAGTCTGGGGGAGGCTTGGTCCAGCCTGGGGGGTCCCTGAGACTCTCCTGTGCAGCCTCTGGATTCACCTTCAGTGACTACTACATGAGCTGGGTCCGCCAGGCTCCCGGGAAGGGGCTGGAGTGGGTAGGTTT**CATTAGAAACAAAGCTAATG**GTGGGACAACAGAATAGACCACGTC**TGTGAAAGGCAGATTCACA**ATCTCAAGAGATGATTCCAAAAGCATCACCTATCTGCAAATGA**ACAGCCTGAGAGCCGAGGACATGGCTGT**GTATT**ACTGTGCGAGAGAcacagt**gag**gggaggtcagtgtgagcccagacacaaacctccc**tgcaggggtccccatgaccaccagggggcgcccgggacactgtgcacggggctgtctccagggcaggtgcaggtgctgctgagggctggcttcctgtcgcggt**ctggggcggcctcattgtcaaatttccccag**ggaacttctccagatttacaattctgtactgacatttcatgtctctaaatgcaaaacttttttgttctttctgtatttttgtttttgtaacaggaggacacattctcgcctcc**acagaagccacagtgtcactttgggggcagatgatccttctgt**agtcagcaggatgaaagtgcagaggaatctcagtggaacctgggaagtcttttccaattagactcaggg**cagagacctccatgggaatctctg**attagaacaggctttgagttctgataggagccaagagagaggctcacccagggtcagagtcctt**aaaacctgacggttttcacagctatctcacctcgtcttgaaaa**actgtgcacatctgactcacactgattcagttgaccctatttctgctaatctattttccttctctgcagacttgattctcacagttccctttcttcttctctttcct**gaaaacagaggatgtgttttc**tgtagtcaaaatcccagagat**caggtctgcaggacctg**ggtaggctgaggggactttctcactcaccataggctgatgacactcctgctgtattttgtgtgtggaggtgtttggaaaacgaagtgaacattagtcatgaagggaataatactagttttctccaaagggatgtggatgtagagctgatcttgtgcttctcacactgtcacagaatttat**gctctcccctgtgactttaggagagc**tgaggatggacactccattgtgctgtgagctctggtaatagtaattatagggtctggctaggcagcctaaggtcaatactgctggccttcaggaa

Uppercase: IGHV3-71-2

Lowercase: Flanking sequence[1000bp]

Red & Bold & Underline: Stem-loop [21]

Blue: Heptamer[44]

Green: Nonamer [9]

id-TRBV5-4-2[V_gene_segment]

gggaggagcagactagcaaaagcctagactcctgaaaacagagcattaagggcaattctggtgaagtgtcagggggaaatgagggacaagctatcggaaattgaagcaaataccatccttgttataaatagcaaaaaccttggcaaaattgtgtcctgttctaggactttatggaatataaaattatgagccattcactaggatatcaactgaaagaaatatctaagcggcaaagcattcaggctactgtgtgactactttcgggcaccaggaaatttcacccagcaagaagggagccatgggaatagattttgcaaaccagcacagatggtaaccctacctccctctgctgtcctgtctcccataagccaaaccctgtgctgtgagctcttaaaatcctagaattatttctggggaa**tctatactcttaataatttataga**caacccaggtctgctttggatctgatcagacggactaaatcttggggactctgcaccactggccactgaagaaaggggctggg**aatgttgatgggacaggaaaatataataagaaacatt**ggtgtgaatctagtatcagaaagatgatgtgaggacaacaaggaagagctggaat**gtgagggtttaatcacaggctcctcac**cctccgctgatgggtaggtgtgtta**gctccagcatggagc**accaccgcactaggtgggggaagggtgatagggtgatggggcagcctgtgagctggggcaacgt**aggcagagaaggaactgtgtcaccacagaaacttctgcct**tcacccatcccttcagctctgcaggacaggtagagactccaggatcatccactgagcactggacataaggaaggctgcatggggaggactcaggacagtgacatcacaggatacccctcctattaggaaaatcaaggcccagaattcactcggctcttccccaggaggaccaagccctgaatcaggtgcagtgctgcct**ggcccactgtgccATGGGCC**CTGGGCTCCTCTGCTGGGCGCTGCTTTGTCTCCTGGGAGCAGGTGA**GTCCTGGGCACAGGAC**AGCAGCCCCATTCTCAGCTTTCCCACCCAT**GTCCTCCACTTTACCTTGGGGAGGAC**CTCCAGGCTTTCTGCTGTGCTCATCCTCCATCTGCTTTTCCCACAGGCTCAGTGGACGCTGGAGTCACCCAAAGTCCCACACACCTGATCAAAACGAGAGGACAGCAAGTGACT**CTGAGATGCTCTTCTCAG**TCTGGGCACAACACTGTGTCCTGGTACCAACAGGCCCTGGGTCAGGGGCCCCAGTTTATCTTTCAGTATTATAGGGAGGAA**GAGAATGGCAGAGGAAACTTCCCTCCTAGATTCTC**AGGTCTCCAGTTCCCTAATGATAGCTCTGAGCTGAATGTGAATGCCTTGGAGCTGGACGACTCGGCCCTGTATCT**CTGTGCCAGCAGCTTGGcacag**cccggcagagtctctgacattctatacataaacttcctgccttagctttgacttgagagc**tgcaggccccacccaggtttcactcctgca**agggaagcttttagttgtatggaaggcatgtgttgtgtcctactgagcacagagctctcccaaacagagcccaggtttcctatgccctgagtgtgcccgcttctgtgcagcatcttcttgcagctt**gtcacttcctgggtaacttcagtagaagagtgac**tgcggagccccagatgtgtgctagattctttgaatttgttatatagcttaaagtctttcaacaaccttgcacatcaatcattcttattcttccttgacagatggaagtctcagggacattcagtcattttccccagtgtctcctggcttgtaaggaccagaagtaggaaacaaagtagtgcatccattttccacctagccccctgctccatcatcaccttctgtgtcctggtcagtaagtcagagccctcaaact**gccatctagtgaccagcagggcctcactagaggc**ttacatgtcttcaggggtcattactctggcctcttcattagaaactttgaggaatgttaaatttaacactttttaaaaatcatttatatgcattccctttgtctatccccagtctgcagcctacactttcatttttatggtgttttacacaagtttaatttaatgcaacaataactaaaaacaa**ttttgtttctgtgtagacaaaa**actgctttctcatataaatgtctaatgatatttttctcaatttattctactaaaatttaagtttggattttcaccaataagaataaatgaacctgaaatatatctcatgtgagaaataagatagagaagatagagacctaggaattggagggaagcacatatgttcagtgaagaagcaatcgtctttcaaaaagagctctaaagtggacacctcgtgctgacagtggagagataattggaaggcaaatta

Uppercase: TRBV5-4-2

Lowercase: Flanking sequence[1000bp]

Red & Bold & Underline: Stem-loop [15]

Blue: Heptamer[33]

Green: Nonamer [6]

id-TRBV5-5[V_gene_segment]

aattgtgtcctgttctaggactttctggaataaaaatattatgagcta**ttagctaggatatctactgaaggaaatatctaagcagctaa**gcattcaggctattgtggactactttcaggcaccaggaaatttaacccagcaagaaaggagccaagggaatagattttgcaaaccagcacagatggtgacactacctacctctgctgtcctgtctccaataagacaaactcgctctggagttagagaaaagaggagccagttaacataatacactggggtcagcttctgaaggcagggtggatctgaagggaataacataaactgtccagagcactacgtataggatgaccttgagggtgctgagtcccaagctggctaggtctgtgctatgagctgagaaggccctagaattatttctggggaatctgaactcttgataatttacagacaacacaggtctgctttgattctgatcagatggactaaatcttggggactctgcaccgctggcca**ctcaggaaatgggttgggaatgttgcctgag**acaggaaaatatgataagaaacatggtgtgaatctagcatcagaaagatgatgtgaggacagcaaggaagaactgcaacttgaggtttaatcacaggctcct**catcctcccctgatgggcaggtgtgtgagctccaggatg**gagtaccacagcactaggtgggagaagggtgatagggtggtggggctgcctgtgagctggggcagtgtaggtagaggagcaactgtatcaccacagaagcttctgccttcacacattcctccagctctgcaggacaggttgagtccagggtccgtagtgcgctagacttaaggaaggctgcatg**gggaggacacaggacagtgacatcacaggatacccctccc**atcaggaaaatcaaggctcagaactcactgggttcttccccaggaggaccaagccctgaatcaggtgcagtgctgcctgccccactgtgccATGGGCCCTGGGCTCCTCTGCTGGGTGCTGCTTTGTCTCCTGGGAGCAGGTGA**GTCCTGGGCACAGGAC**AGCAGCCCCATTCTCAGCTTTTCCACCCGTGTCCTCCACTTTACCATGGAGAAGACCTCCAGGCTGTCTCCTGAGCTCATCCTCCATCTGCTTTTCCCACAGGCCCAGTGGACGCTGGAGTCACCCAAAGTCCCACACACCTGATCAAAACGAGAGGACAGCAAGTGACTCTGAGATGCTCTCCTATCTCTGGGCACAAGAGTGTGTCCTGGTACCAACAGGTCCTGGGTCAGGGGCCCCAGTTTA**TCTTTCAGTATTATGAGAAAGA**AGAGAGAGGAAGAGGAAACTTCCCTGATCGATTC**TCAGCTCGCCAGTTCCCTAACTATAGCTCTGAGCTGA**ATGTGAACGCCTTGTTGCTGGGGGACTCGGCCCTGTATCT**CTGTGCCAGCAGCTTGGcacag**cc**cggcagaatcactgacattctgtatataaacttcctgccg**taactttgacttgagagctgcaggccccacccaggtttcactccttcaagggaagcttttagttgcttggaaggcatgtcttgtgtcctactgagggcagacctttcccaaccaatagagcccaggtttcctgtgccctgagtgtgcctgcttctgt**gctgcatcttcttgcagc**ttgtcccttcctgggtaacttcagtataagagtgactgctgagcgcttggtgtgtgctagatttttgtatatgttacatatcttaaagcttttccacaaccttgcaaatcaaacattctcatccttcctttacagatgggaggctcagggacattgagtcattttccccggtgtctcctggcttgtaaggatcagaagtgggaaacaaactagtccatccattttccacctacccccctgtttcatcaccaccttctgtatcctgttcagtaaaccagagccctcacactgccctctagtgaccagcagggccgcagcagaggctcagatgtcttcagggggttattactatggcctcctaattagcaattttgaggaatgttaaatttcacactttttaaaaaatcatttatatgcattccctttatctttcccatgtctgtagcttacattttcatttttatggtgttttttgatgtacaagagtttaatttaatgcc**acaaaaatgaaaaacaattttgt**ttg**tgtgtagacaaaaacttctttctcatataaatgtctaaaca**tatttttcttaatttattctagtaaaatttaagtttggcttttctccattaagaagaaatgtacctgaaatatatctcacgtcagaaataacatagagacctaggaatgggagggaagcacatatgttcagtgaagaagcaacagcctgtcaaaaagatcccaaaatggacaccctgcgctgacggtggagaggtagctggagggtaaatg

Uppercase: TRBV5-5

Lowercase: Flanking sequence[1000bp]

Red & Bold & Underline: Stem-loop [12]

Blue: Heptamer[20]

Green: Nonamer [3]

id-IGLV3-16[V_gene_segment]

gggtgggtgtggtggggcatttgggaccctggccctggatgggcatccccagcctccttcaggagacactcttcagagggacagggcttctcacctgcctcaggccacagctctgaccatggtgctgtccac**gctcagtgacaggacactgagc**aggggcctggctggagtaaccagggaccacccaa**tctctggagctcacatgctgtgcagaga**ccatgaccctgggacccttacagagaccatggacctaacgttagattctcagtccagtgtggttttctgctgctttttgcaagtttacataaaatattgatgagaaaaacatgaataaataacagacaagttgacggtgctgattgctaatctcttgtggcaatgaggggtgatgtagcaggagcagcaccacgactgaaatgagaccagatgccatgttgactccacaggggtgtgtggttgcttaaatgacccaaatattcccactgaataatcagcaagtgtcctaatgacacagcacagcatttagatgaaatgaacaagaggggactgagtctg**agggaccaggaggaggaagatcatgggtccct**gggc**cacctgctgctgagtcaggtg**tcatcatgatctctccacagagcccagctcagccccatggccaacctcctggaaggtccacagctct**gctgagcctggcccaggacagaccccaggtcagc**gaggtttgggacagtggtcatgaggttaattcccacaaaggtctatgatcacagggcttgaaggctgcagctgattttctaaccagagaccctgcttgttcctcctgttgcctgtgctgccctcacagctggtttgagagacatctctccaggaagagccacagaggatgtaaatttgcataaacgcttaacactgactgcccccgaaagcctgagagggaataagagaggcctgcggagcccaactgttctgtggg**tccaggaggcacagctctgggaatctcaccATGGCCTGGA**TCCCTCTCCTGCTCCCCCTCCTCACTCTCTGCACAGGTGCTGACCCCAGGCCCTTCCC**CAGGCTCAGTCCCCACAGATTCCAAGTTGAGCCTG**ACCTGAATCCTGAGCAAAG**CCCAGACACAGCCTCTGGG**TGGGACTCCTGGAAATGGGTCCTTTGTCTTCAAGCCCCCTCTCTTGTTCTTCCTTGCAGGCTCTGAGGCCTCCTATGAG**CTGACACAGCCACCCTCGGTGTCAG**TGTCCCTAGGACAGATGGCCAGGATCACCTGCTCTGGAG**AAGCATTGCCAAAAAAATATGCTT**ATTGGT**ACCAGCAGAAGCCAGGCCAGTTCCCTGTGCTGGT**GATATATAAAGACAGCGAGAGGCCCTCAGGGA**TCCCTGAGCGATTCTCTGGCTCCAGCTCAGGGA**CAATAGTCACATTGACCATCAGTGGAGTCCAGGCAGAAGACGAGGCTGACTATTA**CTGTCTATCAGCAGACAG**CAGTGGTACTTATCCcacagtgacacaaggagacagggaagtgaaacataaacccctccccctctatttcatccccttccttctgccccaggaggactgtggacaaagccatgagcaggcttggc**caaattcacccaaatttg**aacccccaggctgcccttccctccagccctccaggcaggcgctgcaaaggagtcagtctgcagtggatttgtgcctggcaggcctggcagtcctgtgtttctttgcctggggtttgagttgttgatggagggtctgactggaaagacagacacagggag**acatccatgcaggcatgtccagagggatgt**ggggaactctcaggtctcagttcagtgcagtcattgttgtatcctctgggaaagcattcctccagtgcacctaa**gacctctagacccacagagcctgaggtc**gggatgagagaatgtcaaatttttgctgatggatcactaaaggcatttttgagtggagccactacaatttactccttgattctggactcgtggatcctgactgtgggagaaacagcaccatcta**ttgggcactgatgcacagcacatgccccatcctggaagacattgcccaa**ccctacacatacttgccacttgcctggtgcagtaatggagtctccatgcagtcattaaaccattcatataagccacatgtctcagaatgttggggacaagagtaagaatcttgaatgccaggagaaaaggcccatagccctttatttgcggtgaaacgagttccttggtcagaacctcctgtggaataccatcgtgatgggtgcatcattctgtaatccagagataacagttttggcagaagcattgctgtcaatgaaagcaaatcctctgtacatgccaatgctagtaaggaaaa**agagctgtctctcccatgatggaagagctct**aatgaaattaatatcaagcaggcccctggctgttttccacagagaaggtcatcctatagggatgcctgtgttggtctctgc

Uppercase: IGLV3-16

Lowercase: Flanking sequence[1000bp]

Red & Bold & Underline: Stem-loop [18]

Blue: Heptamer[56]

Green: Nonamer [2]

id-IGHV3-76-2[V_gene_segment]

aagaatgttactttggactaaaaccttaaactataaatacaaataaatccatataaattaaatataataacacagaaacaaatgtatgtcactttgatgtttggaaatgcatttatttttgcctgtgcacatatatgtttcattgctgggtgtgtgtgtctgtgtctgtatatatgggtgtgactgtgaagtttcaaaatacattattaacttgtatagttatataaatctaa**gtttttttatgttaaaagatagtaaaaaaac**acacacaaaaatattctagagagcatttgcaataactaaggagtattcaggtgttgtattaataatg**aaaaactagccaaaatgttttt**aaacttttattttaggttcagaagtacttgtgcaagtgttgttttataggcaaattgtgagttctgttggtttgatgtacaga**ttattttgtcaccaaggaaataa**gcatagtacctggtaggtagtttttcaattctcaccctcctcccaccctccacactcaagtaggacctagtgtctgttgttttctttgtgtcaatatgattagctcccacttatacatgag**aacagatggtttgtaattttctgtt**actgtgttactttccttaggataatgacctccagctccatctatgttgctgcaaataacatgatcttattcttcttatgactgtgttgtattccattctgaaatatacataaatcagtaaatgtgattcatcacacacacacaatcatct**cattgatcaatattaactttcaatg**taaatctcccaggaatctttcatcttctcttgaccctcccctctcctcaggcatcccaaaccatagctttctatatattaggatacatgaaaatagggccctctctcagctgatgaaaaccagcccagccctgactctgcagctctgggagacaagccccagccgtgcagtaccaaggtgcttcca**ttcagtgattagcactgaa**cactgaagacatcccATGGAGTTAGGAATGCGCTGGGTTTTCTTTGTTGTTGTTCTAAAAGGCAATCAAGAAGAACTAGAAATATTGTCTGTGAGTGATATGAGTGAG**AGAAACAGTGGATCTGTGTGGCAGTTTCT**GACAAGAATGTCTCTGTGTTTGCAGGTGTCCAATGTGAAGTGTAGCTGGTAGAGTCTGGGGGAGGCTTGGTACAGCTTGGGGGGTCCATGTGACTCTCCTGTGCAGCCTCTGGATACACTTTCAGTAACTATGGCATGCACTAGGTGGACCAAG**CTCCAGGGAAGGGACTGGAG**TGGGTCTTACATTAATGCTAGTGGTGGAGGCATATACTACTCAGACTCTGTGAAGGGCCGGTTGACCATCTCCAGAGAAAACACCAAGAACTCACTGTATCTGCAAATAAACAGTTTCATTGCTGACACCATGGCCGTCTATTACTGTAAGAGAcacagtgaggagaagtcagtgtgagcccagtcacaaacctcctacaggaacgctgggaggaaaatcagctacagggctcactcaaggcccactgatcagagtccactccagagggaggttcacatggaggttaggggctggttttctgtcaggacattggacttcatcatcttcttccattactttctttagggagcctcttcattttagaattctgtgtgtactattg**ttatatatacatatatatatatatatatatataa**aataatcatagtaatgcgaggacatattttctcatgcacaaaatgtaga**ttgacacttacagagaagaaaaattatcaaccaatgtcaa**aaacatcagagtcctggaaaatctta**agtgttctggtgggtcatctctaatcagacccagaacagt**aacctcagtgaaattccatgattcagatactatttttggattctgatcataaccaagacagagtctggaccagggtcagt**gtcatatagaacctcacagcttccatatatgac**ccttctcttgacagtgaaatatgaaattcagtatcagcagtgatttgatttttcttttgctttccgtattctttctttttagttgttgttgtttagttatttcttttgctatatccattttctttttttttagttgtgcctttcacttttggctttcacttttccatttgcttttcctgcttcctgataaagggagatgtaatccctgtggtctaaattccagggctcaagccttttcctgggagctcaggtgggtctcaggat**gtggctcctgcagccac**gt**gaaagaggctgatgggactttctttc**cttctcatgactcaggcattccacggtgttgtgtggagactcatctgagattgcacgtggccattagtagtgaaggggatgagatgtgggtgtggactgatcctgtgctgtgcaaattgtcattagtagtgaaggggatgagatgtggatgtggactgat

Uppercase: IGHV3-76-2

Lowercase: Flanking sequence[1000bp]

Red & Bold & Underline: Stem-loop [14]

Blue: Heptamer[41]

Green: Nonamer [7]

id-TRGV1[V_gene_segment]

aatcccagctacttgggaggctgaggcaggagaattgcttgaaccccggaggtggagat**tgcagtgagctgagattgtgccactgca**ctccagtctgagacacagagcgagactctatctatctaaagaaaaaaaaaaagaactttcattcttaagctctgtggtattctaaaatgtttgt**acagaaatggcaaggtaaatgcttgattattttcttttctgt**tttcagaatgagatggtgtct**tagcaacctccagaggagactttattttgcta**tattatgtttttc**aagaatttgtgtaggtgttattctt**ttggtgcttaaaatgtcctaccgatggttagtgagagccactgccgttgtttttatttgagtagagtccaaggcaaaccaacatgcctgaccctcagtcctgttaataaagcaggctcaagccttctcagaatggccctgggtggac**ttgcaaatgtggacttgcaa**atgtgaaa**aggaaggtggttcagggccctaatcagaagcttcct**ggcaggaccaggttctggcagctgcgttggctgctgaaactctgaagtgcaggatgtgagcctgctgtgggactgccctctcccaagggccatgaccagctagcgccagggtcagcttcc**cccacatctgcactccctctgctgtggg**cctgagcaatccatgcccctggctctctcctgaagagggcctgctctcagctgccacaagagggcgatggagaacgggcctga**gcagagaggcaggagctcctctgc**agg**tcctcccaggaaccaccgcttctgggagga**aggcttgggagtcccctaagacaca**tcctcagtcacttctccctgcctgtgactcaggaagaccagctcctcttactgtcttc**tgtgctagggatcacttccttgttgagtggggcctgagtttt**aagaggatcttctgctcctctt**catctggtcccttt**ccttccaaggccccagagaggaagg**cATGCGGTGGGCCCTAGCGGTGCTTCTAGCTTTCCTGTCTCCTGGTGAGTGCGCTGCCTACAGAGAGGATCACGGGTTTTGTTTTGTTTTGTTAT**TTTCTTCTTTTGCAAGGAGCGACATACTAAGAAA**TGCCTCATTATATTTTGTGTTGTTCCCATTGCAGCCAGTCAGATAT**CTTCCAACTTGGAAG**GGAGAACGA**AGTCAGTCACCAGGCTGACT**GGGTCATCTGCTGAAATCACCTGTGATCT**TCCTGGAGCAAGTACCTTATACATCCACTGGTACCTGCACCAGGA**GGGGAAGGCCCCACAGTGTCTTCTGTACTATG**AACCCTACTACTCCAGGGTT**GTGCTGGAATCAGGAATCACTCCAGGAAAGTATGACACTGGAAGCACAAGGAGCAATTGGAATTTGAGACTGCAAAATCTAATTAAAAATGATTCTGGGTTCTATT**ACTGTGCCACCTGGGACAGGcacagt**gattcagacactgaaaatctgcctgtggttgcttctggtacacaagatagaccagccaactctcatttcctgccctgaatttactgtattctgtacaaagagaaacacagcttaactcctgatctcccctgcaatatcacactcccctggcagcagctgcaccctgttccccaccctcccccaggactttcctgaagaccaagctgccacctccaagcctcagctaagcagcctg**gctgagagcaagtttctctcagc**tctcctagaacatggggaaggcccactcactctgcttcctaggacagacaggt**acagctagggtccagctgt**gaagcaaga**cattttggacaaaaatg**ggaagggtattcatactgaatcatacatccaataatggtccagagatcgcagctgagagtgg**tgcttattccttatgtctataacaatataagca**atactataatgaccactaaaacactaggccaatatatgcagttgaacattctctctctctgccctcccttttattctcgctctcctttcttacttccatgcttgcttttatcattttctctccctccctcccttcctcctttccttccatcaccccaccctccttcctttctttccttccttccttccttgcttttctctctttctctcct**ttttgtgtacataatcacaatactttttccacaaaa**tcatttg**aaatagtttttctctattt**gttcatgccaacattgatatagctgctttaattgttcagtttcatttttatattttggaaatttgtgtattattttgttttgtttcatatt**tgttttgtatacgtaaaaca**tttagttggaactgaagtcaaaattataaaagatgacacatttgaagattttagcttccattgtagacttctcattcctcttttctccctgacactataggtaacaatatttttatccgttttcttgaactttttcactctgtcatagcgcagtatgta

Uppercase: TRGV1

Lowercase: Flanking sequence[1000bp]

Red & Bold & Underline: Stem-loop [26]

Blue: Heptamer[17]

Green: Nonamer [4]

id-IGHVIII-67-4-2[V_gene_segment]

aaaggattcttt**tctttcattggttgagaaaga**ttttcccctaacttcagcttag**ttcaggcatacgctgacctgaa**tgggcatttaccctcagatgggtacacacctgtcaatatgtggactcttctgtcagacagacacagcttcact**catgtggattcttccctcaaacacaaatgtccccacatg**gactatttcctcagactaccacatatgtccttacatttacactttcctcagaaaacagacatttcctcatgtggactcttgtctcagataagcaaacatgtctcagtgtgaatgaagtcttcactcagataagtacacatatttcaacactgactgtttcctgacacaagcacatatgtccaatgttaaactgtcttgcgacaaaatgatctcaagataatgataattataaactccaaccctgaaaatctgtagatctgcattttgtctattgtaacataacttcatctcattgtcagaaacagtcgtttgcagctataaatgcactgattacagtcag**atttccattttctctggaaat**gtatttcttatgttcttactggactaatttgttgataatgtttgctcacatgaagatacctgaacagtgtccacattagagaataaaaaagagcaatgggcagattaaccctgtgcatccagacgcaggaa**ccatttgactctgccttccctgaaatgg**agacacagaggatggatgagcaacgctgagtggtgcacctacgaccacaaagagaaagacctggaaatatgtcccatcccctcctcatgaaaggcagctcatcccctgttccttcaggccctggtgaggagccaccccatatctgtgtctttct**tcagtgttcacactatggagtctgcactga**tctgggtttcccttctcatcaccctccatattagtgtcccttgtaaatcaggtccagctgtggctgctcatagggttgttctcagtctatttcctctgtg**ttcatagAAGTCCTGTATGAA**GTTCAGTGGTGGAGTCAGAGGGGTAAACGTAGTACAGCCCAGTGGTTCACTGAGACTTTCTTGCAAAGCGTCTGGATTCACCTTTTCTGGCAAC**AGCCTGAGGTTGGTCCAGCAGGCT**TCACAACAGGGATTGTGGTGGCTGGCAACAGTGAGTCAACAAGTGGGAGTGCTCAGGTTTACTCTTCATGAGTACAAATAAATTAACTGGTCCAGCGACACCCTTTCACGTGCACTCTACCTTACAATGACTAACCTGAAAGCCAAGGACAAGGTTGTGTAATA**CTGTGAGCTTcacag**gagagagattatctgcacaagcccagacacaaaaatctgcagggagacaggagggaactgcatggtagatgctgctcagaagcaccagggggcactcaacacaagggggcgctcaggacaccaggcggcactcaggatacagctgggggcgctcaggacaccagggggccactcaggatacagcggggggtgctcggatccaccagggggctcttagggcaccagggggcattcaggaccactagggagccctcagggccaccaggaggacctcaggacacgtcagcgtgctggagccaccgggggggggggggggggggggcgctgaggaaaccaagagacgctctgaaccgcgaggggaaggaagctcaggaccacgaggggacgctcaggacaccagggggcgctcagaacaccagtgggcactcagaacaggaagaatctcttagaaagcag**ctccacatcaggatcctgggaggctgtggag**aaatgggaacactcataaactgttgagcatttaaattagtacaaccactatagagaacagcttagagtttcctcaaaacctgacgatagagttatcatataatctggcaatcctgctgctggtatatacccctaaaaaggaaatgtttttgcactaccatgtctgtaatagcactgttaaagc**actgttcacaacagt**caaaatttgaaagcaacctatgtgccgattaacacatgaatggataaataaaatatggtacatataggatatgggttaatatcatgccataaaa**aaatgagattttgtcattt**gcagcaacatggatgaaactggagttcactatattaattgaaatgagcccagaacagaaagacaaaa**ttcacatgttcccatttatttgtgaa**agctaaaaattaagacaattgaactcgtggagatagagagggaaaagatggtcaccagaagctgggaaggtctatggtagatggggattattggggatggtcaa

Uppercase: IGHVIII-67-4-2

Lowercase: Flanking sequence[1000bp]

Red & Bold & Underline: Stem-loop [13]

Blue: Heptamer[37]

Green: Nonamer [6]

id-IGHV1-46-2[V_gene_segment]

tgtgaagaataggtatgattttactatttcaatggatacagaattgtgggagtcactatattcctatgaacaaaaaattcagatttcagtgttaagtaatgttgcctacattgtgtgagtgacggggcagtggtggatctgagagtgtggtgggtgcacggacataatgattcagaaagcaatatggaaagatgagtatctatggatacgaactga**aagtatgtaaatactt**cacaaaatactaataaacggagttgaatataaaacccataattat**ccaaaacacaaatttcttggaagttattttgg**gaacatgatttcataaagaactccaaactcttgtttcaacttctgactcctcgtttctgtgatataagaaaaccatttccaattatgcatctcagggcaattctgtaaacccagagcatttctgctgaagatcctggggaatcaagacaccgggcaggtgatggagacactgtctcaggtgcgcccaacgaatctcagaggaacctgctggagagtcacgtggaacatctacagtcagtttctcagagtcaacagtgagctgtgttg**gtgcctgaggggaccatgatggggccaaggcac**gtgctcagtgtcgtggacagtgatggtccagaaatgatctagatggtcttgacgctaatgaaatatgggttcagagtgaggagcataatctgtggggacttgttcttcagtgaaaggatcctgtccgcaaac**agaaatggagcaggacatgcatttct**tcaagcaggattagggcttggaccatcagcatcccactcctgtgtggcagatgggacatctatcttctttctcaacctcgatcaggctttgaggtatgaaataatctgtctcatgaatatgcaaataaccttagatctactgaggtaaatatggatacatctgggccctgaaagcatcatccaacaaccacatcccttctctacagaagcctctgagaggaaagttcttcaccATGGACTGG**ACCTGGAGGGTCTTCTGCTTGCTGGCTGTAGCTCCAGGT**AAAGGGCC**AACTGGTTCCAGGGCTGAGGAAGGGATTTTTTCCAGTT**T**AGAGGACTGTCATTCTCTACTGTGTCCTCT**CCGCAGGTGCTCACTCCCAGGTGCAGCTGGTGCAGTCTGGGG**CTGAGGTGAAGAAGCCTGGGGCCTCAGTGAAGGTTTCCTGCAAGGCATCTGGATACACCTTCAC**CAGCTACTATATGCACTGGGTGCGACAGGCCCCTGGACAAGGGCTTGAGTGGATGGGAATAATCAACCCTAGTGGTGGTAGCACAAGCTACGCACAGAAGTTCCAGGGCAGAGTCACCATGACCAGGGACACGTCCACGAGCACAGTCTACATGGAGCTGAGCAGCCTGAGATCTGAGGACACGGCCGTGTATT**ACTGTGCGAGAGAcacagt**gtgagaaaccacat**cctcagagtgtcagaaaccctgagg**gaggag**tcagctgtgctgagctga**gaaaatgacaggggttattcagtttaagactgtttagaaaacgggttatatatttgagaacaaagaacaatagaaacacaatcgaattgtaagagaaatattccattcaagagccaccacataagccaaactgacagagtgggaaaggccacactcagtaaagttgatacaaacataccataaaggtgctactatgaacaagtttttgaattagatgaataaatcatttggagcaaggttatttggtcatatgttaagagtaagcatgattcttacaaagtgggaaaattgtctttcaaatgtttctgtcacttcttaccataaagttcattttagaggttttaggattacagtgaaattgcacagaaggtgtgagaattcccatgaatccctgccccgcacggacaccgcctcctccactacagccatcctgccccacagtcacaaataagtcacaatggatgaatctac**aagaactcttggttctt**tctttttctggtgatcccctaatataacaagcctaaattatcttggaacacccaggtattttcaatggctttctagaagtgatattagtcagagg**gaaagtgagtgaggctattactatttgagcactttc**t**tccaaaatccacaaaatatatgttaatttgga**gtttt**tcttacttctggtttacaatgtcccttcccagagagtaaga**tttttaagcttttagtgaggctggaaaaaaaaatttttaaaaaagagaaataagctttcctgtattaggctgacttatcccagcggcagcaacaagcacagcccagacccaggaaaagtcttaataatattatctaatgtgctctggagactctttcagca**ctccctcaacatagggag**aagaaaaacaaattttcctttgtcttatgatatgagtttatagattcttgttct

Uppercase: IGHV1-46-2

Lowercase: Flanking sequence[1000bp]

Red & Bold & Underline: Stem-loop [17]

Blue: Heptamer[52]

Green: Nonamer [5]

id-IGHV3-43[V_gene_segment]

gtggaaaagtcgtttattgggatagtacccactactacaatcagtaaatgttggatacgatcaacagcatggttcaattcacaagta**catatgatgagtatagtgagccaaagcacatacatatg**attccgttttataaactgtacaaagtgaatactcattggaaattacataacaaagatcactaactga**cttctccatagtaagagaag**cgaaggtataggagggagaaattgtgagaga**caaagagaaaattgagaggcgaattgatttgttttctctgtg**aatggtcattaggtcaatgtttgtcaaatggtga**acatattgtgtgaagattatatgt**ctgtattacttcattaaagctattataaataaaagtctaatgtggtagaaaaagatgaagagagaaataaaaataatacaagaaaagtcatgaactcctgaatgaattaaccctta**gtttttctctattacttataaaaac**accaagatacagccaaataatatcacgatatcattataagaagagtgttttgtaaacctcactgggaatttatagctctttcctagagttaattttgggaacagttggatccaattgtgagaaatgcaggctggacactgagactggctcttatgag**atgtgagctcttgtctatgtcacat**ggtccttccatacttggggg**tttacattcacatctgtaaa**tgaaggaaacattgactctcaaagaacatatttcatgtgcatgtaaaagtatgaatgctagtgagaattaattacttatgaagtataatcacccacatccactcttggacacagcccactctgaggcatctgttacagaactcattatatagtaggagacatgcaaatagggtcctccctctgctgatgaaaaccagcccagccctgaccctgcagct**ctgggagaggagccccagccctgagattcccag**gtgtttccattcggtgatcagcactgaacacagagaacgcaccATGGAGTTTGGACTGAGCTGGGTTTTCCTTGTTGCTATTTTAAAAGGTGATTCATGGATAAATAGAGATGTTGAGTGTGAGTGAACATGAGTGAGAGAAACAGTGGATATGTGTGGCAGTGTCTGACCAGGGTGTCTCTGTGTTTGCAGGTGTCCAGTGTGAAGTGCAGCTGGTGGAGTCTGGGGGAGTCGTGGTACAGCCTGGGGGGTCCCTGAGACTCTCCTGTGCAGCCTCTGGATTCACCTTTGATGATTATACCATGCACTGGGTCCGTCAAGCTCCGGGGAAGGGTCTGGAGTGGGTCTCTCTTATTAGTTGGGATGGTGGTAGCACATACTATGCAGACTCTGTGAAGGGCCGATTCACCATCTCCAGAGACAACAGCAAAAACTCCCTGTATCTGCAAATGAACAGTCTGAGAACTGAGGACACCGCCTTGTATT**ACTGTGCAAAAGATAcacagt**gaggggaagtcagcgagagcccagacaaaaaccttg**ctgcaggaagacaggaggggcctgggctgcag**aggccactcaagacaca**ctgagcatagggttaactctgggacaagttgctcag**gaaggttaagagctggtttcctttcagagtcttcacaatttctccatctaacagtttccccaggaaccctgtctagatctgtgatctggatctgctgaaactgcctgtgtcaccttcctcacctgtgactttgggggagctgattgtggacactccagtgtgtgggatttcttggtgacagcaattgtgtcttctgtctaggcatgtctagggctggccatcaggaag**ggcaggctggaatttttggaaagaggcgcacctgcc**atccaccaggaaattttgttgtcttttgttc**tgctagaattaaatcagacacaccaaggttaactagca**ctatcttcctagcttgagaaact**tgatggcaggcttgaataacacctgtatgaagccatca**gagcaacaactagattaatgtc**tgctagaattaaatcaggcacaccaaggttaactagca**ctatcttcctagctcgagaaacttgatagcaggcttgaataacacttgtatgaaacc**atcagagcaacacctagaatagtgtctgat**ttaataattgagactatggtctagccaaggagacac**ataaaacatgatttccatggttggttcatattttat**atttattaatagaatctggctggtattataaacagctttgccaaaatatatgtgttagtgttggtcttcggagaagcaaactctgtgactggattagcttcg**tgtacagtttccttcgtgtaca**gtttccctgtccactcttctgagaggaaatgcagaggtggtggaaggggtctggggtagctgaatgcatgtgagggaagtcggtccctgagtgaaggagaaagggaggaggactgggtggaactttcctaaacttc

Uppercase: IGHV3-43

Lowercase: Flanking sequence[1000bp]

Red & Bold & Underline: Stem-loop [18]

Blue: Heptamer[30]

Green: Nonamer [3]

id-IGKV3-34-3[V_gene_segment]

tcagataataaatatttaggtttggtgaggcaagatgcaaaattgaatatattatgtgagtacttagagaagagagaaaacaaatttctgcaatgtctattaatgaactcagatggaatgtaacccaaagagcttctcagcctgtccagtgaactgtcacttaccaccaaatgctca**caattttaaggaacacaaggaaaccccaaattg**aattattgtttagatcagaaacatgagcagaggtttccaaaattattaaagtgagaaaggcctgagtctgattgaatggatgtgttgtgcaggatggggtggcatagaaagtagggcatcacccaaggaattgttcctggcctgctggacagcattgtcgtctgtttgcagggaatggaattacacacactgaagggtctgagcccggggcctgctcatctgtgcagccgtgatgatcatcacacccacagtgctgggcctaattgcgtgttgccctttaatagtttttacaaagaatgaggaggcttcagggctcataggaaagatggctaagctgcaaaggaaaagtgtgtgtggaagatgaacttctgatgaactagttttcagccaggtacattcatgcaccaatggagatgactc**ggttctttggagaaataaggagaggaaaagaacc**acaggcttctattttattcccaacaggtttgtactaaataccatctggctgggtctaggtgactagtgagacgggaagaaacaggggctatagtctgtgaccacagggaagaagttctacca**tcaggctgagccaatggacttttctggcctga**ccacctgggcaggggctgctgag**tgcagagaggaggaggcagggggtctctgca**gctggaagcccagcacccactccaactactttgcatatccttccagccactctgctgtccagagcccatatcaatgcctgggtcag**agctctcgggaagagct**gctcggttaggacccagagggaaccATGGAAGTTCCAGCTCAGCTCCTCTCCTTTCTGGTACTCTGGCTCCCAGGTGAGGGGAACACGGGATGATTCTGCATGCCAATGAAACTCTGTCAATGTTATTGGGTCCTGTGACCTGCTCAGCAAGAACAATAATTAATATTCAATGTACATCAATGA**TCCCAGCTGTACTGGGA**AGACAGTGGATTTG**ATCTAGATTGCACAATTGACTTTCCTGTTTTATTCCAATCTCAGAT**ACCACTGGAGAAGTTGTGCTGACATGGTCCCCAATCACCCTGTGTTTGTCCCCAGGGGAAGGAGCCACCCTCACCTGAAAGGCCAGTCAGATCATTGACAACTATTC**AGCCTGGGACCAGTAGAAACCTGTTCTGGTTCCCAGGCT**CCTTATTTATAGGGCATCTACCAAGGCTACCAGCATCCCAGCCCGGTTCAGTGGTGGTGGGCCTGAGGCAGACTTTACCCCAACCATCAACAGCCTAGACCCTGAAGATGTCACAATTTTATTACCCTCATCAGTACAGCAGTGGGTGTCCcacagtaattcaacat**gaaacaaaaactttcacaaaaccattgattttttttttc**taaaacc**agcagctttatgggctgcagctatgatggctgct**cagttttagcaactgtgcctctatttggaaattttgaaattctgaaaagtaactcgttgtcaagatggtgactccaatg**tctgtatttttctcagacttatttacaga**gttaggttttgaatcaattttactgcaaggccttcagacagact**cacctgcatggatgcctttaaagacacaggtg**ccgctgggcacagtggctcactctttttttttccaatagagtagaacaaacatagcttttcatttaataaattttcacttaatggcagcattttcacgaacaagtaaaggccaccgagtgaaggttaggtgcacgtgagtaaataagacaattttgggtcccaattcagtaggggaaagagccatgaaggaagaataaccacacaataaaagtgcgtagaataatttctgtagttgacaaattgtgtaaaaagcaggaatggtctgtggtatctggaggctgaggccatatatttgtgctgatcagggaaggccccagggtcatactgaagattcatctgagtggtgaggaggcagccaggtgaccacaaaaacagcaaagaccacgggatcattatgggaagggcagggaccagtgtgtactgaaaaagtcagcaagtgctaaact**aatcagatgccttctctgatt**tttattcccgta**aagataaaacttactgtaatatctt**acatcatacaccttggtctgggtcactcccatggcatagaatgttccctggttcagaagttggaagcccaggtctttcctttgttgacagtttgcaggctccccagagcccttccctgcc**actcaagcccttatcactgggtcagtctctgtctttgagt**tgggatcaccataggcaggggcagcactagacactggacatggggctctggccct

Uppercase: IGKV3-34-3

Lowercase: Flanking sequence[1000bp]

Red & Bold & Underline: Stem-loop [15]

Blue: Heptamer[22]

Green: Nonamer [7]

id-TRBV24-1-2[V_gene_segment]

cacttgggctcctgcatctgtccctctgtgtgttccctttcccctcaggggtttgag**cagttgcagcaactg**aaaaggtgagcaacactcctgtcgcacgtccttccaggggggtcagggaactcttcctttttcaccaggacgagcacagatctgaagcagcaaagagccagttgtttattacattgaacaatgtttcacactatcaccataactgttctgtacttgcaatctgctgcattaaatattagctctttcctcctaaagtcaggaataatttgtgttagaataaaatgaattctttaatgaacaactg**tattttcagctaattcaaaata**ggaatgaaggcgttaagtacttctttgaagcacatacccacattaaactaacatctttctacaataagccaatatttttaataaagctctgtgcccagaaaacattgaccagaaaagcaggagccactaaaaaataggagcaatttgaaacaaagttaggaatcctgtccttcttacaacaaaaaagtctcaaactttacagagaatgtattcaaattgcatagtttaccaagaactgagttagaataaatgattaaatgcagaaaaaaaggagaagagttataatgtcctatcattcaagtgagactagacttcctttattcg**aaattcaatttggttttgaattt**tctccagcctgttttatcaggaggatgatgtcaccttgaacaactgaaatggatatgccctgagaaaggggagagatacaactctctggtggttgccagtaaagcgccagaaaaacagaagagaagcaagtagaagaatttcagcatctgctaagaacatataatacataagccaacatggcatctttctttaagacccagtacct**tgggagggacaatgacatcacttcctgaatcctccca**gttttcta**tttccatgccctgcttccctcaacatccagagctggaaa**cacctccatcctgcctcttcatgccATGGCCTCCCTGCTCTTCTTCTGTGGGGCCTTTTATCTCCTGGGAACAGGTGAGTTTGGAACACAGATGGGGAAATCACTGCCTTAAATTTTCCAGGTTCTGAATTAAGCC**TATCCTCAGAGATTGCAGCACGAGGATA**TTTACCAGTTTCTGTCTTCAATTTCTGTCTTATTTCCCACAGGGTCCATGGATGCTGATGTTACCCAGACCCCAAGGAATAGGATCACAAAGACAGGAAAGAGGATTATGCTGGAATGTTCTCAGACTAAGGGTCATGATAGAATGTACTGGTATCGACAAGACCCAGGACTGGGCCTACGGTTGATCTATTA**CTCCTTTGATGTCAAAGATATAAACAAAGGAGAGATCTCTGATGGATACAGTGTCTCTCGACAGGCACAGGCTAAATTCTCCCTGTC**CCTAGAGTCTGCCATCCCCAACCAGACAGCTCTTTACTT**CTGTGCCACCAGTGATTTGcacag**tgcttcttggccacctgctctctacacagaaagacagacacatgggtgagttgtttgctctgaagggtacctggatgtgggttgtgggatgtggggtgtttagagctttcagtggtcttaggtagtgtgagctaagggccactttggatcaatgtccccaagccatgtgatgactctgaaagcacaggctacactgaatcaatctcccctgtctatttatttttcctagggagctagacgaatggtgacctttcaggagaaggaacttgagcatttgacaga**agctcatgattttcaacaagagcatttcttatgagct**gaatacaatgcagtttttatagtccttttgccattatttccacccactttgcaacatcccacactttgttctgttcccactgacagctcctttatcctgtctaacctttctctgccccagttcttccaagctcagctccaccatttagctctataattgattgcctttctacta**aaaacaaaataactagcttgtttt**ctaaaacttcactgtaagtaa**atttaattgtcattaaat**ctgactcagaagttaaagttcaagtaacagagactgtgatttacaaaagtaatttttcatctactaccccacactttgatctctgctgagattcattttaacctgcaggtg**cagatagacttgcattattatctg**gatgtact**gaatttcaaaaaataaaattc**cctgtccccagtatgaggtcaaagctaggagcttgga**agtgccccttgtttccaggcact**agaataattccctggttcccctgctgtggggattcttttaggagccagtgaggcccaggcacaaaaataattgatcaaggactgccagagcttgacgtaaaagtcttcctgagaatcaagttgatgtcttcctcatggatttagtcttctt**gtcccataagcctcaattatttcagcaaacgaatccatgggac**ttggtgactgttcagaaatga

Uppercase: TRBV24-1-2

Lowercase: Flanking sequence[1000bp]

Red & Bold & Underline: Stem-loop [17]

Blue: Heptamer[23]

Green: Nonamer [5]

id-TRBV12-4[V_gene_segment]

aacacaagctagtctgcaaactaatgccctttaacattggcaagtgctttcaactgcatgggactcagaagcatgcctctgtgccaagagcaaaaatatgcccttttatggcaagtaagttaattgatcaactcaaaaaacaaacattt**ctctcagctttccagaacctctgagag**gaagtaggtttggagaagtgggattaaggggttggcagggggccggaggaaaggtcaggtgttaatgatgaaaaaacattgaacttttctt**ttctaggggtagtaataagatataattcaagactagaa**cattgtagctgcaaatcttcaaa**ataagacaatcttat**cctctgttttattgtgggaataga**gaatggagagaagttccattc**cacgggctttgggaattcaatatggttcaaggataaataaacccaggttttttctactgcagagaagagcttcaaatataatcagttctcagggcatcagctcagttcctctatccctaacaatgcagttgacat**gcatcttctcagatgc**ctaactcctaactcgctgatggataatttaggtacacatagaggactagaaacaccaagccaccagtgagaggaagaggagagtttgcagagaagctggcttgcaataagacaatgagttcatctttaattacctgcaatttgaggagatatatatagttggcaagctcctaggtaaggcatgttatggagaagctaccgtgaattgataatatcaaaataaatcttaagggagcctctgcatgtgtgcatttgtatctcagatctgctatagttgagttaattcctgcctgatt**catctcccagagatg**cagcctcctcttaaagttgggggtggtggcccat**tcagtgatgtcactga**cagatgcattctgtggggataaaatgtcacaaaattcatttctttgctcatgttcacagagggcctggtctggaatattccacatctgctctcactctgccATGGGCTCCTGGACCCTCTGCTGTGTGTCCCTTTGCATCCTGGTAGCAAGTGAGTCTTCAGAACATTTACC**ATCATCAGGCTGGGCTTCTGCATGGATGAT**CTCATATATTTTCCTTATTCTGACGCCCAATTCTGTCTTCCTTCATAGAGCACACAGATGCTGGAGTTATCCA**GTCACCCCGGCACGAGGTGAC**AGAGATGGGACAAGAAGTGACTCTGAGATGTAAACCAATTTCAGGACACGACTACCTTTTCTGGTACAGACAGACCATGATGCGGGGACTGGAGTTGCTCATTTACTTTAACAACAACGTTCCGATAGATGATTCAGGGATGCCCGAGGATCGATTCTCAGCTAAGATGCCTAATGCATCATTCTCCACTCTGAAGATCCAGCCCTCAGAACCCAGGGACTCA**GCTGTGTACTTCTGTGCCAGCAGTTTAGCcacagc**gctgcagaatcacccctttcctgtgc**agaaaccctggtgtttct**ccttctccttctacctcccagcagtcctgggcaaagtctttcctgttcctccctccccatgagaaaaagtggttttgggttgtgacaaagacagaaaatgaggtt**tcaacataggaaaccttatgttga**tttgaggattatgaa**atgggttttgaggattccttaaaaaattgtctctgctcaaaacacat**aggagtaagataaaccttggctactgacactggagatttccctgccctcctgcatttgccatcccatgagaatggtgggagctcccgagaaaggctgcattttctgaactgtgaggccctcttcattctctcctaactctaagctgcaaacagaaatttccctcacacgttttctagattgta**aaagaaagttcttcttt**actatgattgtggacattcctttataatgccaatttcaactttacatttacttcaggatttttcactactcctaaagagtgtctcaaatgtggctagagcaagcaggttagtacactagatgtaagctacctggcctggaatctaaggatccatttgtctctgttctgcgtaagatgagccgggtgctggccaaaggctttgcacactcacagagcactgatgacgcctcctggtaaggacccacactggggtatctaaaagcagacaggcatgtccagtcttctgttgccctgtttcctttctgattacatgtccttaacacacaaatttacatttttcttcttatttatatgagaagtttctatacaatacctgcaatccattctgagtggttataatttctgtgtgatattcatatttacatactgattccttctaaatccctatcatggt**atcattgacaagtgaggcgaatgat**ccctgtattttg**agtgcccaaggcact**tgaggttttttggagttctgccatatgccaaattccactttgtcattttccca

Uppercase: TRBV12-4

Lowercase: Flanking sequence[1000bp]

Red & Bold & Underline: Stem-loop [16]

Blue: Heptamer[25]

Green: Nonamer [3]

id-IGHV1-2[V_gene_segment]

ctccctttgtttccagtgatccagtgcatctagacctccaggaagtggactccctggtgattttagcgattcttctcttggagccaccctgaagaggacattgggtttccaaaggcccattcactatttcaagaggtggtgtcatcagctcatgttgtcactgaaggagcattctgagccaggtcacagtcacttcctagtgagct**acagaggctgagagaaaaatgctctgt**gagacccaatgggaa**gctccctgcagtgcaaggtctgggtggcagggagc**gctagggcctcgcccagcacaggctgcagccctggagcaggtgcaagggaggctg**gggaggggttcctccc**agggtctgatgtcttccttttctcggacaaacatgctttaataagttaaacaagactttagt**aaagactattgatgtgtcttt**gtgtctttcagtatacagttctatttgtaggatttatctaacctaacaagtcaatgagaatcacatgtaaaaggagaaatttctaggattttcagatatcttaataggtaggagatggagaaaagggatggttttattaattcagtgcttgccaatcttaacagagacagtagtaagacatgcagaaagcaaagcccagaaaagtatgaaggtgtcaaagtgccatttaagtatgggttcacttggaggaccatgttctgcgggaacttgttttcagcagacaatctattttagcagagttctgggcatacaaggggacacacatcattaaacaaggat**tgggacagggacttcagcgtccca**ctgttgcatggcccataaattatgtgtgttctctttctcatcttggatcaagtctagagctatgaaatagtatccctcatgaatatgcaaataacctgag**atttactgaagtaaat**acagatctgtcctgtgccctgagagcatcacccagcaaccacatctgtcctctagagaatcccctgagagctccgttcctcaccATGGACTGGACCTGGA**GGATCCTCTTCTTGGTGGCAGCAGCCACAGGTAAGAGGCTCC**CTAGTCCC**AGTGATGAGAAAGAGATTGAGTCCAGTCCAGGGAGATCTCATCCACT**TCTGTGTTCTCTCCACAGGAGCCCACTCCCAGGTGCAGCTGGTGCAGTCTGGGGCTGAGGTGA**AGAAGCCTGGGGCCTCAGTGAAGGTCTCCTGCAAGGCTTCT**GGATACACCTTCACCGGCTACTATATGCACTGGGTGCGACAGGCCCCTGGACAAGGGCTTGAGTGGATGGGATGGATCAACCCTAACAGTGGTGGCACAAACTATGCACAGAAGTTTCAGGGCTGGGTCACCATGACCAGGGACACGTCCATCAGCA**CAGCCTACATGGAGCTGAGCAGGCTG**AGATCTGACGACACGGCCGTGTATT**ACTGTGCGAGAGAcacagt**gtgaaaacccacatcctgagggtgtcagaaa**ccccagggaggaggcagctgtgctgggg**ctgagaaatgaaagggattact**atttttaatgttgtttacagtatgtcattaataaat**tgaaaaaaagtaacaatagaagtatatactctaattatatgggaactttgttttttcagttttttcatttttttttttttttttggtttgttt**gtgacagagtctcactctgccac**ccaggctggagtgtaacggcacaatctcagctcactacaacctccacctcccaggttcaagcaattctcctgcctcggcctccagagtagttgggattacaggcacccgccaccatgccc**ggtgaatttttgtatttttagtagagacggggtttcacc**atgttagctaggctggtctcaaactgctgatctcaggtgatctaccctcctcagcctcccaaagtcctgggatta**caggcgtgagccactgcgcctg**gccc**aattatatgggaattgtttatataatt**atcaccctataagcaaaattcatggaggaggaaaagctctactgaagaaagctgataccggcattcccatgaaagtatctgtgtagaagtaagtattaaaatcagttga**ataggcaaggcatggtggctcacgcctat**aatcccagcactttgggagaccgaggcaggtggatcacaaggtaaggagttcaagatcagcctgcccaagatggtgaaaccccttctctactaaaaatacaaagaat**tagctgggcgtggtggtgggtgcctgtaatcgcagcta**ttcgggaggctgaggcagagaattgcttgaacctgggaggtgaaggt**tgcagtgagccgagatcacgccactgca**ctccagcctgggcgacagagtgaaactccatctcaaaacaaaacaaaacaaaacaaaacaaaaaaacagttgaataaagtaccttagagtcatctgttcaattaacatgtttaact

Uppercase: IGHV1-2

Lowercase: Flanking sequence[1000bp]

Red & Bold & Underline: Stem-loop [20]

Blue: Heptamer[43]

Green: Nonamer [4]

id-IGLVIV-59[V_gene_segment]

ccttctatgtcattgccttgaacaaccccttcatgccactcaagtcagcacaactgtttcagcatctgtggtggtgccatagtgacgtatgcaggtcatttagctttttctagaatgtactcttggggcactggcacc**ttaaaaggttataaaatttttaa**acagttataaacgaacatatttgctcctctcatagatcaagactacttacaggagtgagatactggctaggcatctagtagaccacgtcatcatcacaggatataggatt**caggacacagatgtgtatacagaggaatagtgtcctg**gtgagctg**cagagatcagctgggctctctggtctctg**agccatggggaggtgagtgt**cctgctatcagcagg**ctgcactgtgggacttgcctgtggtgtctgcacagctgctctttaagaaccattcattccaaggagcctaggtctaggggctgggccttgtgctcattgatggggactcggcagctgtcttctctggtctggcagagt**ctgctgaacagggacctgtgtatggtctcagcag**gtgatatctcctagggcccttccaaggagggaaagcaaatcatcttcctcagtctgcagtgtgagctgggctccagccccagggctcagga**agggactgggcagtccct**gcatagaaacacatttgcatgagcagcc**cctcctctgctgggggttgggaagaaaaggagg**cctggggcagctcaggctcactg**tggtggtcaggggctgtgtccacca**tggccagaacttctctcctcctcctgatcctctttcactgctggcctgaacttctcttc**tccttatgctcctctctcactgcacaggtaagga**caggcctcggagatcaggactcagttttc**agcctgcatcagtccctcaggct**caggaacccatgacaaatttcatcctcatcttaaccctgtctctt**atttgtgtctgtgtttacaaat**tccctctcccagcttgtgCTGACTCAGCTGTGTCCTCCCCCCCACTTCCTGTATCTCTGGGAGCATCTTCCAGACTCACCTGCACCCTGAGCAGTGGCTGCAGTGTTGACATTTATCTCATATCCCGCTGTCAGCAAAAGCCAGTGAGCCCTTCCCAGTATCTCCTGATCTACCTCTCAAAAGTCAAGTAAAACTAGGGCTCTGGAGTCTCCAGCTGCTTCTCTAAATCCAAAGATGCCTTGGCCAGTGCAGGCAATTTGCTCAT**CTCTGGGGTCCAGCCAGAG**GACAAGACTATCTGTTCTATCTATTACTGTCAGACCTGGGATATTGATACTTCAGTTAcacaggcagatgagaaagtgagacgaaactcagcctactaagaatggaactatggctctttttccaattgtcaaataattttcacatacacaaactattttggaagtagctactgattcacaatggttctctcaatgtcccattcaatgtttctgagtcctcaggcaaactaacaacaaaacaatgataaaa**catatgacaaaacatatg**ataacactgactctttccctgctgaaccacatgtaata**aaaccagaggagctggttt**aataaaacctgaggagctga**cttttcttttaaaagaaaag**atgacagtcttttggagtctttttcttggtcaaaatttcctagaccatcccagcaggtggacagtcccctacagaagacacgat**cctgggaggaatcccaggtacagagctggcctgcttcacttgtgggtacctg**ccacttcaattgtgaaatcagggaggggagagtaagcatagggggtaggtgaggatcaggaaaggggcaaggccactacatttcattaaccacctttggactccagtgcaagatagaggctgatgatcccatcacacaggtgactacaacctccttgtcatcaacagctcctgccc**atggggaagcgaaatgaaaatctatccccat**ccctctcttcttacatatccctgtcctgcccctgccggtgaccaacaccagctcaggttgaattttgttttgtcacttgtgctattaaagaaaataactccttctcttactggtctcctagttcagacagaatcagtttgtttccagaaatagaaaacgctgctcacacacatgagtgtaaagtaaggatgacaaaaagaaaaaatccagg**ctatgtggcccagagacccaggtgccctctgattggcctcatag**atgtcccctcctcctggcccaggtttgcttgtcattccttaagctctggaaccttgaactacatgggttctgtcttcatagttgcagaagaaaac

Uppercase: IGLVIV-59

Lowercase: Flanking sequence[1000bp]

Red & Bold & Underline: Stem-loop [19]

Blue: Heptamer[30]

Green: Nonamer [3]

id-IGHVII-28-1[V_gene_segment]

gtccaagctaagtccagcaaggccattggtgcctcgctgagcacagttgtccatcagggatctcccatgtgtcccagcagcagccatgcctcaatctctccactgtgcacagccattgt**ctgggaggagctcccag**gatgggtgtctttggcacacacagttgatgggtgttagagtgcagtgcagcagctggctgcctggtctattgggctcctggatattggagggatttgaggtgcattctcagggccagcacgctttgtcaatttttatatgaaaaccatgtgatttaattcattttctcagatgacatagataattaataacacaatctgc**aaacaattataattttcaactttaccccaagttcattgttt**ctgaattctgtgcaggatccagacatggtactgcccttctcat**gagaaactgttcgatctaaactgaaaccagttgtttctc**gtatactttggttctccccacatgcagagatcttgattagagcaagtttggtactttccacgcactcaccctca**cctccccagataaagagcagaagttctccttagactgagtctgagggagg**agctgttcctgtaccactcagggcctgcggagacccccaggtgcagcttcactgagtcaggtgtttcactccctgtgattgctgctcaggt**ctaattgtggctcggaattag**gacagtcttcaggttatcacaggtcaatcatattctaaaaatcatcgttatcacacaccatggtaacaattcaaggttcatttttctaacggcagtttctcttttttatttggttgcaagtttgaggaaaggaacatgtgataatactttttaacctaacctcgaaatctactgaattgttctaggagactcacaaattggacaaagtgagctctttattctcataaaaatgtgtggttttgggaatttcactgtgttgcccagaacctgttaacatcaacaactatgtttctcagcacacttctGGCTTGAGACGTCCTTGCAGACCCTCTCCCTCACCTGCACTGTCTCTGGATTCCCCATCATAACCAGTGTGTCCTGCTAGAATTGTATCTGCTTGCCCCTAGAAGATGGACAGGAGTGGATCAGGT**GCATGGGTTGTGAAGGGAGCACAAATTACAACCCACTGC**TCAAGAGTCCATATCCAGATCCAAGAAACAGTTCTTACAGCTGAGCTCTGTGCCCAGTGAACACACAACTACGCATTTTTAAGCAAAAGAcg**caatgaagggccttcattg**tgagcctagacacaaccct**ccctgcaggggtgaataggagcagcaggg**ggcattcggggcagtatgggggcttaggatgattgttaggggtcaggatgagcaggatcaaggcttcgcatcagggcaagtgcaacagggcagaaaaggggctgtagatgtgggttgttctcaccatcatatttcac**caccagacaccctccactacatctcttctaatgtgtctgagtg**ttgatatgattagaaaatggcatttatgtaaatactaatatgtacccat**atggaggtgcatccagttgtcttctccat**cttatgtggaccttgtccatcaagcactaagtccctgtatttacttgagtacctcacacattatggtcaaaacaggtagggtctccttttggcatggtctctcctcctgccttctctcttctgtcacacaaaaacacgtgaactgacacacacacagagcttcccaactttaattatgtgatgtattgaagcaaattgattagtgtgcagcttttctgctttgcctgctattcatgttatgtaaaaataagaaccatgtttttctcagcttgtcacttctctaagctaagtagcatctttgtttattatacccagagaccaaaagcgatccaactgttattcagcagcttaactggtaaacaaattgtggaaaattcacttactggaatcatacccactgctacaatcaagtaatgctggatacacccaacaccatgcttaaaatagcaagtacttgaataagtaaaataagtcagacaaataaaagtgcatacatacgattccacttctataatttctataaagtaaaaatgaacttaaagttacataaaaagatctgtagttgactggggacatggtacaagaagagaaggtatagaaaggagaaactacag**gagaacaaaaggaaatttctaggataattgatttttttctc**tgttagtaaaagtgagga

Uppercase: IGHVII-28-1

Lowercase: Flanking sequence[1000bp]

Red & Bold & Underline: Stem-loop [11]

Blue: Heptamer[32]

Green: Nonamer [3]

id-TRBV12-4-2[V_gene_segment]

agaaacacaagctagtctgcaaactaatgccctttaacattggcaagtgctttcaactgcatgggactcagaagcatgcctctgtgccaagagcaaaaatatgcccttttatggcaagtaagttaattgatcaactcaaaaaacaaacattt**ctctcagctttccagaacctctgagag**gaagtaggtttggagaaggattaaggggttggcagggggccggaggaaaggtcaggtgttaatgatgaaaaaacattgaacttttctt**ttctaggggtagtaataagatataattcaagactagaa**cattgtagctgcaaatcttcaaa**ataagacaatcttat**cctctgttttattgtgggaataga**gaatggagagaagttccattc**cacgggctttgggaattcaatatggttcaaggataaataaacccaggttttttctactgcagagaagagcttcaaatataatcagttctcagggcatcagctcagttcctctatccctaacaatgcagttgacat**gcatcttctcagatgc**ctaactcctaactcgctgatggataatttaggtacacatagaggactagaaacaccaagccaccagtgagaggaagaggagagtttgcagagaagctggcttgcaataagacaatgagttcatctttaattacctgcaatttgaggagatatatatagttggcaagctcctaggtaaggcatgttatggagaagctaccgtgaattgataatatcaaaataaatcttaagggagcctctgcatgtgtgcatttgtatctcagatctgctatagttgagttaattcctgcctgatt**catctcccagagatg**cagcctcctcttaaagttgggggtggtggcccat**tcagtgatgtcactga**cagatgcattctgtggggataaaatgtcacaaaattcatttctttgctcatgttcacagagggcctggtctggaatattccacatctgctctcactctgccATGGACTCCTGGACCCTCTGCTGTGTGTCCCTTTGCATCCTGGTAGCAAGTGAGTCTTCAGAACATTTACC**ATCATCAGGCTGGGCTTCTGCATGGATGAT**CTCATATATTTTCCTTATTCTGACGCCCAATTCTGTCTTCTTTCATAGAGCACACAGATGCTGGAGTTATCCA**GTCACCCCGGCACGAGGTGAC**AGAGATGGGACAAGAAGTGACTCTGAGATGTAAACCAATTTCAGGACACGACTACCTTTTCTGGTACAGACAGACCATGATGCGGGGACTGGAGTTGCTCATTTACTTTAACAACAACGTTCCGATAGATGATTCAGGGATGCCCGAGGATCGATTCTCAGCTAAGATGCCTAATGCATCATTCTCCACTCTGAAGATCCAGCCCTCAGAACCCAGGGACTCA**GCTGTGTACTTCTGTGCCAGCAGTTTAGCcacagc**gctgcagaatcacccctttcctgtgcagaaaacccggtgtttccccttctccttctacctcc**cagcagtcctgggcaaagtctctgctg**ttcctccctccctatgagaaaaaagtggtttgggggtgtgaaaaagacagaaaatgag**aaggtttcaacataggaaacctt**atgttgatttgaggattatgaa**atgggttttgaggattccttaaaaaattgtctctgctcaaaacacat**aggagtaagataaaccttggctactgacactggagatttccctgccctcctgcatttgccatcccatgagaatggtgggagctcccgagaaaggctgcattttctgaactgtgaggccctcttcattctctcctaactctaagctgcaaacagaaatttccctcacacgttttctagattgta**aaagaaagttcttcttt**actatgattgtggacattcctttataatgccaatttcaactttacatttacttcaggatttttcactactcctaaagagtgtctcaaatgtggctagagcaagcaggttagtacactagatgtaagctacctggcctggaatctaaggatccatttgtctctgttctgcgtaagatgagccgggtgctggccaaaggctgtgcacactcacagagcactgatgacgcctcctggtaaggacccacactggggtatctaaaagcagacaggcatgtccagtcttctgttgccctgtttcctttctgattatatgtccttaacacacaaatttacattttccttcttatttatatgagaagtttctatacaatacctgcaatccattctgagtggttataatttctgtgtgatattcatatttacatgctgattccttctaaatacctatcatggt**atcattgacaagtgaggcgaatgat**ccctgtattttg**agtgcccaaggcact**tgaggttttttggagttctgccatatgccaaattccactttgtcatttt

Uppercase: TRBV12-4-2

Lowercase: Flanking sequence[1000bp]

Red & Bold & Underline: Stem-loop [16]

Blue: Heptamer[27]

Green: Nonamer [3]

id-IGLV1-41[V_gene_segment]

ctggtcaaaacaagctgaagttttaactcttccatctt**caaatactgagcatatttg**ct**tgttttgctctttagcaaaaca**ctgcaaagtcatttgccaggatcccccgcagattggcattcagtgatggagaacagactcctaggtgagatccctggtcacaggatcccgatgactatgactgcaagtctggaaaagaagagacagaacaaagatttaccagcataaccctctgagaatgtgccccaggcatcccggagggaagagctccctatcgacccaggactgcagaatccacagaggaaggagagctctggccaccagggagcagccacacgagaggagaggacccagtgcccccacaggtgaggaaatgagtccatggggctggtcactgctctcgatttcattctgacataag**aggactttgtgtagtcaccacctgagtcct**atacagtgaacagccaaggagacatggacagattcacccctgaggaacccagtagaatgtgaagtctg**ccctgagctcacgaaaaaaatgactcaggg**ggctgcctggcc**tggccctgatggagggggcca**tggacagaacaacggggagtgaggggtgtctctgagaggctctggtcaccttgttatacaatgg**tggtgtacatgaccaaaatggacacca**ggggatgcctgcgtaccattactgagaagactgggtgtgatgag**agcaggaccagcgccacctgtcctgct**tggt**gccctatgcttagggc**ttacagatgtcaactctgcaccccctgggaccacatagccccacccctggcactctctgacatcc**tcaggcagaggagcctga**cccagggcccaaggtgggatcagaaagct**ggagggtctgatttgcatggatggaccctcc**t**tctctcagagtataaagaggggcagggagaga**cttggggaagctctgcttcagctgtgagcacagaaggcaggactcgggacaatcttcatcATGACCTGCTCCCCTCTCCTCCTCACCCTTCTCATTCACTGCACAGGTGCCCAGACACAGGGTCAGGGGAGGGGTCCAGGAAGCCCATGAGGCCCTGCTTTCTCCTTCTCTCTCTAGACCAAGAATC**ACCGTGTCTGTGTCTCTCCTGCTTCCACGGT**CCTGGGCCCAGTCTGTGTTGACGCAGCCGCCTTCAGTGTCTGCGGCCCCAGGACAGAAGGTCACCATCTCCTGCTCTGGAAGCAGCTCCGACATGGGGAATTATGCGGTA**TCCTGGTACCAGCAGCTCCCAGGA**ACAGCCCCCAAACTCCTCATCTATGAAAATAATAAGCGACCCTCAGGGATTCCTGACCGATTCTCTGGCTCCAAGTCTGGCA**CCTCAGCCACCCTGGGCATCACTGGCCTCTGGCCTGAGG**ACTAGGCCGATTATTACTGCTTAGCATGGGATACCAGCCTGAGAGCTTGcacagtgctccaggccaatggggaactgagacaagaaccctctt**cctcctccgccaggagg**gtgagtgcctgcagctgctgctcacacctgacctgtagcttctgctgctgtagcttcccccatgggcctcggggcatccagggccttgcctaggagtggaggctccaccacttttgtcctcagagtcaggaacagggaccccaggagacagaatatcctgctcctcagcttgggacacagggtctctgcactgaaatcgtgggctgaggtggcaggtcca**actgtgtcttcacagt**c**cttcctgtgcctgcccatggtgtggggacggagtgaggaag**tgtgggccattgagcttcactagaggcctga**gggagcagaggtgctccc**gaagacctggactccttccgtcaccctcctcccatcactct**cctcactgtagctgtggccaggtgagg**gctcagggctacttgaggaccttgctcaacccagggcctccatcccaggcattgtttccaaactctgtaagggctgtctctgaatctg**tagaaacatttatttcta**t**gtgcccaggactgggttagagggcac**caaccccaccccttgcaaactgctcatcacctctcagtgaaacaatgaggagaagaaagagttagaaatcagaagcctcc**ccaaggctcccctctctcctctcaaccagcctccccttggcttcctccttccacagaaataaagaggaggcagcca**cagaaggcaagggaaggtccaggcattataccaggcagtggagccagccagtccc**cccagatggtggagggaggtggtggaggtgtctggg**accagaaaatagtgaggaccctcttactaaagtgaggaggggtggcataaactga**ggctcagatggtggaccagatctcctgagcc**atctactcatttcttcttgagtttcagtttcagtaagaataggataaaaaaaaagtctgaagaggttcctttcctcag

Uppercase: IGLV1-41

Lowercase: Flanking sequence[1000bp]

Red & Bold & Underline: Stem-loop [25]

Blue: Heptamer[34]

Green: Nonamer [2]

id-TRBV13-2[V_gene_segment]

atgtgtttatttctccagcttagatgattgtttcactgatagcatgtcactgtcagctgcctccaggaattatctttggttaaag**ggaatgctttgcccaggtctcaccccattcc**tatggcagctggcatttaatgactggctgatatctcccttcctacaccattatttgatga**aatttaaaatttaaatt**aaaatgaaaaatttttctttcttgtttaaaa**agacatttgctttatgtct**ccctcaccaacctgtgcaaaatttgagtttgaggcaggtaaacctgaaatcagcaagagctgtgctgtgtcctgagagatcctgagggtctaattt**aggaagcacccagtctccctagtcagggtccttagagcttcct**catagtgactacacactggccacttgggggcactgtggatccactgagtgggtcactgatagcg**ctgtctgagagagagagggttaagggagacag**tcttgatttcgtctctgtatcagggatattaccagatgaaccagaggtactgctggcacaggaagagttaatcatagtct**tatcagatcaactgata**tgcaacatgtatttgtcatgacaggaagccatgaactgaagccttgtgtggcctctctagcaacctaggtttgatggggagtgtgatgagggtttcagcagggcagcctggataacccaagccaggtggagataaaggag**catctgcctcagatg**aaaatctcagaagctgtgcggatggctgtctgcccggaagcctgccccctctcgctcagcacagct**agcttcccctgctctgcaggaagct**ggacaggatgggggaaagcctgagttagctgagcta**gtcctggagtacagggtgcctatgggagcctacaggac**gatgacatcagaacagtgacatcacagtaaaaacctccaaaaacggtgaggaggagcaaaag**ccctgctttctcaccccaggagaccagcaacctgagcaggg**agATGCTTAGTCCTGACCTGCCTGACTCT**GCCTGGAACACCAGGC**TCCTCTGCCGTGTCATGCTTTGTCTCCTGGGAGCAGGTGAGTCCTGAG**TACAGGTGGGACATCCCTGTA**TCCACAGTGTTCAATTGTTGCTGAAGTGTCAAACTCTCCCG**AGCTGAGTCTTCAGCT**TCTGTCTCCTTCCTCCACAGGTTCAGTGGCTGCTGGAGTCATCCAGTCCCCAAGACATCTGATCAAAGAAAAGAGGGAAACAGCCACTCTGAAATGCTATCCTATCCC**TAGACACGACACTGTCTA**CTGGTACCAGCA**GGGTCCAGGTCAGGACCC**CCAGTTCCTCATTTCGTTTTATGAAAAGATGCAGAGCGATAAAGGAAGCATCCCTGATCGATTCTCAGCTCAA**CAGTTCAGTGACTATCATTCTGAACTG**AACATGAGCTCCTTGGAGCTGGGGGACTCAGCCCTGTACTT**CTGTGCCAGCAGCTTAGGcacag**ccctggagaattactggctttctgtacccaaac**cctcctatctcacttgaggatgtaatagggagaaggagg**tg**ggggctgccacacaactttagccaagcccc**agagatgcttctattcttttctaacattttcccctgccctgct**gagctcagtgagagctc**ctgcacttgtgggctccagacccactggaagttctcacatcttagccagtactttttaattcctagcaagtggcgggagcttctactctgtgcc**aacataggggtatatgtt**ttagtgtgtttcctgttgccatacag**aatacctgagactgggtatt**ttata**aagaaatgaaatttgtttctt**acagctctggaggctgggaagtcc**aaggtcaaggacccacatctagtgaggacctt**cttgctggtga**ggactctgcagagtcc**ccaggtggcatagggcattacatatgagggggctcatgaaagatggtcaaactggcttttataacagacccaacctcataactaactaattccttctataaccca**ttaatccatgaatggattaa**cctttatgagggcagagtccttaagatccagtcaccttccaaaggtcctacctctcaacgctgctgcattgggaaccaagttttcaatatataaattatttaggaacacagtcaaatcatagcagtacacattgcgggtttttttttttttttcttttt**tgagacagggtctca**ctctgtcacccaggctgaagtgcagtggtgctgtgatcatggctcattgcaaccttgagcgcgtgggttcaagcaatcctcctgcctcagcggcctgagtagctatgactaca**gggacacatcactacaaggtgtgtccc**cagttcctcagttcctacaaggtgatgtgtccccttgcctggctaattttttttaatttttatttttgtagagatgaggccttgtcatgttgtccagattagtctcaaactcctcaaatgatcct

Uppercase: TRBV13-2

Lowercase: Flanking sequence[1000bp]

Red & Bold & Underline: Stem-loop [28]

Blue: Heptamer[43]

Green: Nonamer [3]

id-IGHVII-22-1[V_gene_segment]

ctatgccaa**tcccaccacgagtgctggagaaggtggga**gtctgatgaagcttactaacaaccctgtggtccaagctaattccatcaaggccattggttcttcccagagcatagctgtccatcgggaatctcccatgtgcccagcagcagccacgcgttagcaacttcactgtgcacagtcattgtttgggaggagctccaggatgtgtctctttgtcaaaaactgggtgatgg**tgtcagagagctgtgccgggtgcctggtccacgtgctccctgaca**ttggaggtataggaggtgcgttctcagggtggaaacaccttttatggaaatgtacataataaatcacgattttc**cttggttctcttagacgacatggagaagcaag**catgcaccctgcaaacaattgtaatttcctgctttaccccaagttcatgatttcttgtaaactttgatttctgtccaagtacagagatactgattagagtgagtttggttctttccacacactaaccctca**cctcccccagatagagcacattgtcctcactctgagtctgagggagg**agctgttcctgtacaactcagggactgcagagacccccgtgtgcagctttgcagagtcagatctttcctcatgtgaggcgacctccactgcccgtgattgctgctcaggtctaattgtggattcgggattatgacaccctttgggctatcacaggtcaatcttattctgaaaatcaccgtaattata**gacagaggtaataatccagtacccattcttctgtc**cgcagtttgtc**tttcttatttggttgaaagtttgaagaaa**aaatgtgaccatatattcatctaattctaacctctgaatctgctatttgctctcggggacctacaaattgcaaaacagtgagttctt**cattttcatcaaaatg**tgcaaatttgtgaatttcaacatgttgttcagaacctgtgcatgccgacaactgtgattctcggtgcactcctGGCCTGGTGAAGCCCTCACAGATCCTCCCCCTGGCCTGTGGCATCTCTGCTTCTCCATTACAACCAGTGCTTCCTGCTGGAGCTGGATCCGTCACCCCCCGGGGAAGGGACTGGAGCGAATCAG**GTGCACAGGTCATGAGGGAGTGCAC**AATCCAACCCACTCCTCAAGAGTCCAGTCACCATCTCCAGATCCACATCCAAAAAACAGTTTCTCCTACAGCTGAGCTACCTTAACAAGGAGTACACAACCATGATTTTTATACAAAAGAcacagcgaggggaagccattgtgcgctcag**aacactctacaaattttcctccctagtgtt**ttaccaaaactg**gtatatatttcagatactgaaatatttac**aacctacgttattatgctaaga**attctaagttagaagttagaat**gataattgtctcttttttttccttttttttttttttttttttgatactgagtctcactgttgttgcctgggctggagttcaatggtgcaatctcggctca**ctgcaacctctgcatcccaggttgcag**caattctcctgc**ctcaaccccccaagtagttgag**attacaggcgcctgccacgatgcccggctaatttctgtatttttaatagagacggggtttcaccatgttggcctggctggtctcaaactcctgacctcaggtgatccaaccgcctcagcctcccaaagtgctggaattacaggcaggagccactgcacctggccaa**ttatcctcttttcatagatggataa**actaacctaggcctttgaaaataaacc**cttatctgagagtgaaaagataag**ccata**gatttggagagtttgcttgcaaatc**aa**atatttggaaaaggacttttattacaaatat**acaaattaccttacaattgaacaacaaaaccacacaatttaattttaaaatgggcaaagacctgaatagaaagtgcatctaacaataagtataaaaagtgacca**actggattttcattaggtaaatgtatgtttaatattgaaaaaatcctagt**aacctaatagaatgggtaaaatacacaatatagacaataccaaatgttgacgaggaagt**tgaaaaacaggaattttca**cccattgctcctgagaatgctgaattggcacatagacaactacagttaa**taatttattgtgtatttcaaatta**gctagaagat**aacatttagaatgtt**cttaccacaaagaagtagtcaatgattgagatgaaggttgaaggttgtcccagttatcc

Uppercase: IGHVII-22-1

Lowercase: Flanking sequence[1000bp]

Red & Bold & Underline: Stem-loop [21]

Blue: Heptamer[30]

Green: Nonamer [5]

id-IGHVII-46-1-2[V_gene_segment]

agaagctatttagg**aagatgtaatgggaaggcatctt**t**aacattctgttgaatgtt**tccacatggcacttcaatcccctaaaatactttgctgtatcagcatgtgatgaatcggagtataatattgaaggtaaaatgggaaatacatgggcttttgatgaatcaagtcataggatgagaatgtctgtgtccttgaggaaggagaccatgggttgtaagttctagtggaggtacctttggcaaagggagttcatgagtttctgaaccttatgctactttta**atttaaggaagcaatttatgatgtatgtttacttaaat**ctttccagagacatttgtcagtaaggacagggtagcatttgtgtaatagtgatgacttagagagttattttgtaattgctcctgtaaggtatgcacattgctcacttgatacagaagttcaaatgtcacaggtggaaaaacaggaataaaggtgggcagatcacaaggtaaggatcacaaggttaaccatcctggctaacacggtgaaaccccatcgctactaaaaaatacaaaaaattagccaggcatggtggcgggcacctgtagtcccagctactcaggaggctgaggcgggagaatagcgtgagcccgggaggcagagcttgcagtgagccgagatcacaccactgaacttcagcctgggcaacagagggagactccatctcaaaaaaaaaaaaaaagaaaacaggaataaagcaaattttatgaactgtcatgactgtagtttttggcaaggacattatcatgtcaacctgtaaataaacagacaacaataaaacatatcaaatccatggggagtctgacctgtgtccctctgtcttataagcacaaggctttg**ccacatccaaaatattattcaggctccagggtataaaatgcttttggactgtgg**aaggtaacagctctcccctcaggcaggggtaaggtatctggggaatgcagagttgtgttcataaggaagATGTCATTATCGTGTCTTCTCCTGTGCCTGGTAGCAGGTCCTGAAGGTGAGCGTCTCAGATGTCAGACATGGGTCTATCGGGTTAGGATAGGTACATGTGACT**GACAGGGACTGATTCTCCATGTAATCACATGCCCTGTC**CCAGGAGCAGCTGCAGGAGTCAGCCCTGGACCTGAATAGCACACACTTACCCTCTGCCTCACCTACACTGTTACTGGCCACTCCGTCACAACCAGTCCTTACTAGTGGACCTGGATCTGCCGGCTCTCAGGGAGGGGCT**GCAATGGATaaaatccattgc**tagtggtggtgggagt**ccatttgtcttgtggaaaatgg**cagcattt**ccttattttataagg**cataataatgctatgttgtgtacacataccacattgtctttatccatttgcccattgacagacacttagtttccatatcttggctgttgtgaatacagctgcaataatcacaggagcgcaggtatcttcacaaggcggtaatttcatctctttgggtatatttccataagctgcatcactggtcatatggtatttctgttttaattcatttaggagccaccacactgttttgcctaatggtaatgatgggaatacagaatgtcatagccgctacgaagaacagttttagatttgaggtataatccaaaagcaaataatgtttgatcagggttctcatatgaggctctaataaataagtccagaaagttttcccatctt**gggcaaggatgagtttgccc**ctaattgtctttgaggaagaagattggcagaacgtgagatggagtctgttggcaggattcaggattcagtaataactagtaatggcacagaaaaatagagagatagagacatacagagaaagagagaatatgaatctcataagaggaaaatttgctaaatatcagcattgggttttcagtcatagacacatttgtatgagccaggaaccatggagtcaagtgaggagtg**gagaatatgttcagtccaaaaatcagcatattctc**agaggcacccattgccccatcacacaggtggagaattttggaaaccagtgaagtgcgagttcacattaggtgattaagctac**catgtttcgataaactttcattaacatg**caaagtacttccatagataactgatgcacaaatatacgagatgcctttttgcgtttatggatgtctaga**gaaaaataagtgagaaaatttttc**caggttgcagagatctgtttaagttgcagattccataggagagtgtctttgaatgaatactg

Uppercase: IGHVII-46-1-2

Lowercase: Flanking sequence[1000bp]

Red & Bold & Underline: Stem-loop [12]

Blue: Heptamer[30]

Green: Nonamer [3]

id-TRAV1-2[V_gene_segment]

tgggattacaggcgtaagccactgtgcccggcagtaactaatgttaatagcttttttctcatttcttcctgtatattcgtacatgtacatacacctacagacatacaaattatgtttcttttgaactttaaattaatatctgatcatctgttgggagacaaatttagatggttgaaaactctctgtttcttccttattgtcacaaacaccaccaccaccactaccaccaacacacacacacacacacacacacacacacacacacacacacaccccacctctaattgcaacct**tgtttccatttcttcctgtcttgagaagagaaaca**cttatggggaaattgaatactttgtctttttttttttttttttttttttttt**tgagacggagtctca**ctttgtcgcccaggctggag**tgcagtggcgcgatctctgctcactgca**aactccgcctcccgggttcccgccattctcctgcctcagcctctcgag**tagctgggactacaggcgcccgccacagcgcccagcta**atttttttgtattttttggtagagacggggtttcaccgtgttagccaggatggtctcgatctcctgacctcgtgatccgcccacctcggcctcccaaagcgctgggattaaaggcgtgagccaccgcgcccggcctactttgtcttatgttattcccatttgccgtctctgttccttatacattatgctttttcaactttaccagaatcacttggattaaaacccgtggatttctcagtaggaaatgttcatgtgaagacac**ttctgtagtaacagaa**c**ctacagctgctcctgtag**aaggaagttgaaagtcatcccttcaaga**aaggggctcctcccctt**gtaattctactgggttttgcatccagactgagtttccttccctcacccacatgaagtgtctaccttctgcagactccaatggctcaggaactgggaatgcagtgccaggctcgtggtatcctgcagcagATGTGGGGAGTTTTCCTTCTTTATGTTTCCATGAAGATGGGAGGTAAGTCTCAATCTAATAGCAAATGCTGCTGGGAGTTTTCAAAACGATTTTCTTTCAGCTAAATGATTGCCATTTCTGACACTCGTAACAAAAGTGTTTCTTAATAGATATTTTCTTTATGCGGTGCTAATT**GTTATTGGATATCAAATAATAAC**GACTTTGC**TGCCATTTATTTCTAAGAATCTCTCCTTAATGGCA**ATTCTTTTGTGACCACATGGAGACATCGTTTTTAATGATAATGGGTTTCCAGGAGCGCTGCCCTAGTAACAGTCATGTATCTTGTCTTAGGCACTACAGGACAAAACATTGACCAGCCCACTGAGATGACAGCTACGGAAGGTGCCATTGTCCAGATCAACTGCAC**GTACCAGACATCTGGGTTCAACGGGCTGTTCTGGTACCAGCAACATGCTGG**CGAAGCACCTACATTTCTGTCTTACAATGTTCTGGATGGTTTGGAGGAGAAAGGTCGTTTTTCTTCATTCCTTAGTCGGTCTAAAGGGTACAGTTAC**CTCCTTTTGAAGGAG**CTCCAGATGAAAGACTCTGCCTCTTACCTCTGTGCTGTGAGAGAcacggtgactatgaggcctctttagctgcaccaaaattcaaaaggc**aaccacagcagcgagaagctgtatttcctgagtgtatgcctgctgtgagtt**aagactggggactttggaaccagactgcctggatttgagtttgctactctctagttgggtggtcttaagtcagtcactatttttttttcttttggtttatttccttatccagaaagtgagaataatgagtatctact**tcacaagcctgttgtga**gttatgactctctgtatcatctatatctctatctatagctatatctagtgttactcagagaagtagtggcacacag**taaactttatttaagttta**gttatttcaattctagtcttgaattttcactaagtattacgtggtaggacctctgagactggaac**tcagtatttctctacccttggtttgtggaaggtgaaatcctga**ccactatttgtatcttgtggatctaaagc**tccacctaatgatggagtttgtcgttcaggtgga**ttgttgctttggaaggatcactgtggtgatactcattttctttggggggactccgaactttggaaatccaaatgcaaatgcttcagagtgagatgttaag**tagcctgattaattacgaaaggcta**tcttcattatgtgcatcctctataaataaatgtaatcttaacatattttggaggtttataactcagactagaagtgaagtttaggggcttcagacttggggagcatgaacataatcaggggagtctatagaagatgggaaaaagaatgaacataaacaatttggtggcaaatctgggtgaggcaggagagccagtaggagattcctatgacaagatgcagtcagtaggcattaagcacaataaaaatttacagtaaatgttaagattttcatttcctgctttataattcccttatgttctcttcaaatcttttaatgaagtctgttaaatgctacaacttgctgtcagctaggtttag

Uppercase: TRAV1-2

Lowercase: Flanking sequence[1000bp]

Red & Bold & Underline: Stem-loop [18]

Blue: Heptamer[31]

Green: Nonamer [3]

id-IGLV3-31[V_gene_segment]

atctgc**cagagggagcattaggatgtcctcacctctg**tccccagaactcaggaagggcagtacacctgactaaagagtagagggttgtgaaaagagggacctagggcactggataagcaggaggagaaggagcactgggtgggaggtagagaaaggagggggctgaatggttaggctctgaggtgtgtctcctggatggagtcttggattccactcactaccatctgagaactttgagaaaaccgtgtaacctcctgatgttgggtctcttgtcag**tagacatcagtgctttcctgatgtata**gggttactgtagaattataatgaatgcctggttgaggtgacatgtctaatacaccctccaaagcatcaggcatatattaagtactcataagcaggcattctgtgtgaactagtcatacatggaagatgacattccccgtgtgggcacagtaggatgctgtggactttggttt**ggggaccctaataacttcaagtcccc**tggt**agcacccaagtgtaaaaggcaaagggcatgactggtgct**tcctgtgaggcctcttctatttggtaaaattcactctccactcatcgcacccacctggcctttattctaaacttggaaaattgctaacatgaaggtctcagcctcagggctcagataagaactcagacacatcaatagagaggatggctgaggtgcttggccgag**ctcagtgaacaccccagaacaggaactgag**gagtgtatgcatgtgttggggg**cagcactttctcactcatccagaaagtgtg**aggaatctgtacagaactggggacagggcccatttttatacccttctataagcccagcccttgtctgctcactagggagccattcaccaaggccacaggag**ggaaggaagtagatttgcatgaagagcatccccttcc**tatgttaagagaggcctggaggccccttcccagctgtggactcagaggcagagctctggggcatctccaccATGGCCTGGACCCCTCCCCTGCTCACCCTCCTCACTCTCTGCACAGGTGCTGCCTCCTAGGGCTCAGCCCCCACAGGACCAAGAATCAGCCTGGCCCTGACATTCAGCTCAGCACAGGGAGTAATGCAGGGTGTGGGGCTCTGGAAATGAGACCCTTATCTTCAGACTCACCTCTTCTCTCTTCTCT**TGCAGGCTCCGTGGTTTCCTCTGAGCTGAGTCAGGAGCCTGCA**GTGTCTGTGGCCTTGGGATAGACAGCCAGGATCACCTGCCAGGGAGACAGCATAGAAGACTCCGTTGTAAACTGGTACAAGCAGAAGCCAAGCCAGGCCCCTGGGCTGGTCATCTAACTTAACAGTGTCCAGTCTTCAGGGATTCCTAAGAAATTCTCTGGCTCCAGC**TCAGGGAACATGGCCACCCTGA**CCATCACTGGGATTCAGGTTGAAGACAAGGCTGACTATTACTGTCAGTCATGGGACAGCAGTCGTACTCATTCcgcggtgacacaggcagaaggggaagtgagacaaaaaccttttctccatctgtcacactctttcttcagctccaggaaaaaactgtggacaaattcatgagcaggtcttgccctgttcacccagatat**gagaccttgaggctgcccttccctccaggtctc**caggtaggctatgaaaagggtgaatcaggagtcaatacacacacaattattataatt**ttattcttgtttgaagaagaagtagaggaataa**ttgaacatgtaaagtgaaaacatggaaaaccgaaaaagaggaattattgaacttgtaaagatgaaaactacaatgtctgaaatttagaaaatacactgggtgagtttgacaacagaaaaaagattagctgcaaaaaaaaagattagctgcaaaaaaaaagattagcaaattgtataatacagcagtagaaattgtccaagatgaaacacatatgggaaaaataattatggaaattaatgatagaaaggaacagagtatcagtg**agttgtaggacaacttcaaacagcctaaagtatgtgtaattagtgtttgaa**atagaaatgggtggaggagaaatagaa**aaaagtatttgcataaataatggccaaatatttttcacctttggtgaaaa**ccataaactgaaatatccaagaagctcaacaaaattcaagtataggagacatgaaggaaacaactccaacataaatcataatcaaattaataaaatccagtgatgaagagtcacctaaataataatgctaagagcagttctcattaaaagcaatgcaagaaagaagacggcggagcaatgtcttgaaaacact**aaaagaagaatcctgtcaaccttgaattctttt**tcttttcttttcattttctttctttctttttcttttcttttcttttctttttttctttattttttttttttgagacagggtcttactctgtcacccaggctggagtgcagtggcacaatctcagttcacaac

Uppercase: IGLV3-31

Lowercase: Flanking sequence[1000bp]

Red & Bold & Underline: Stem-loop [16]

Blue: Heptamer[28]

Green: Nonamer [4]

id-IGHV1-17[V_gene_segment]

gtttcaagagatggtgtcagcagcatatggtgtcactgaaggagtattctacactagcacacagc**catttcatgctgggctagagaagcctgggggaaatg**atttgaagtctca**gcaaggaacctccttgc**aaggcaggggctgggctgtcgggggcaccgagcaggtgagtcagaaatcaatatgtaaagatgaggacctatggatatgaattgaaaatatataaatacttcaaaaaattccagtaaattgagtccaaaattaaccccaaattattcaaaacacaaattccttgacaattattttgggagcagtgagttcata**aagaattcaaaactactgtttcagcttctgattctt**attgttcct**gagatgagaaaatcatctc**taatcacacatcacacagcaaatctgtaaacaagagtgtttctatgga**agatcctgggggatct**gaacaccaggcaggtgctggagaccctgtttcaggagcgc**ccagcagatctcagagggacctgctgg**tcactcacgtgggacatcagcagtaactttc**tcagtcaccagtcagctgtgctggtgactga**tggacccaggaca**gaaccaaggcacctcctcagtgtcatggagagtgatggttc**cagaaatcatccaggtgctctctgtgcttataaaatgtaggttcactgtgaggagcgtgttctgagggggattgttctgtgaaaggacctctgttcacgagtgttcataaatggagcagggcatgcatttcctcaaacaggaatagcgcttggac**catcagcatctcactcttgtaaatctgatg**tgtcatttatcttccctttcttattattgaccaggctttgcgctatgaaatgctctgtctcatgaatatgcaaataacctgagatccactgaggtgaattt**ggatgtctgtgccctgagagcatcacccaacaaccacatcc**ctcctctatagaagcccctgagagcacagctcctcaccATGGACTG**TACCTGGGGGATCCTCTTCTTGGTGGCAGCTGCCACAGGTA**AGGGGCTCCCAAGTCCTAGTGATGAGGAGGGGATTGAGTCCAGTCAAGGGGGCTTTTATCATCTCCTCCCTTCTCCTCACAGATGTCCATT**CCCAGGTTCAGCTGTTGCAGCCTGGG**GCTGAGGTGAA**GAAGCCTGCGTCCTCAGTGAAGGTCTCCTGCCAGGCTTC**CAGATACACCTTCACCAAATACTTTACACAGTGGGTGCGACAGGGCCCTGGACAAGGGCATAGTGGTTGGGATGCATCAACCCTTACAATGATAACACACACTACGCACAGAAGTTCCGGGGCAGAGTCACCATTACCAGTGACAGGTCCGTGAGCACAGCCTACATGGAGCTGAGCAGCCTGAGATCTGAAGACATGGTCGTGTATTC**CTGTGTGAGAGAcacag**tgcgaaaacccacatcctgaga**gtgtcagaaaccccaggaaggaggcacctgtgctgacac**agaggagatgacaaagattattagattaaag**attttcttagaaaatgacactaagtcattaaagaaaaggaacaatattaatg**tgtatttgagaaattttaattatttgagagatttttcatacagcattt**attctgtaaacaaatttcaatgattacagaat**gaatcaaattaatgaaactaatacagaacttcctctgaaggtatctttgtaaacattaatttcttaatcagtgctgtaagt**attttggaacacagacacaaaat**cacattttaagtctgcatttatgtct**attaaaaatgccaaaaaaatctccttttgtgcatgtagcattttgaat**tcccaccatcaatgcatgatagttcttggttttccacattcatattgtcatgtatccttacgagaattgtgtgttttaaccaatctagtaggtgagtaatgggatctaatttttatttaaatgcacatgtccctccaaaagttcatatttaacaattttcgtataacttctgttgagacgcctctcctgacattcggttcattttaactgcattgcttcatttcgattcat**tgcaagttgacttgca**tattgtttataaaagtcatttcacaaattaaaagaattcatttcacaaatatgt**gacttggaagtattttctccaagtc**tgtggctgtctttattcccttatcagtgtgtattg**cagaaacatatgtgtgtgtatgtgtctg**tgtgtgtatgtgtctgtttgtacaaatttagactcaaaaacatgtaaaattgcattcattcatagatga**tgtatttggcaatatatatgaagtctcattataaaataca**ctaatagtgattattttttccatgtctctcaactcaggcca**caatcaactcatgagtgtttaaacttcatctacttgattg**gaggactatccatt

Uppercase: IGHV1-17

Lowercase: Flanking sequence[1000bp]

Red & Bold & Underline: Stem-loop [26]

Blue: Heptamer[47]

Green: Nonamer [6]

id-TRBV7-2[V_gene_segment]

aggaaaaaatgaag**tttctagaatattagaaa**gcatataatgttttattgaaacttttctta**ctttcttgtatataactgacgtttcagaaag**agaaaataaggcaaatgatctgaggtgattgtgtatgtgatgctagctgagaatgttcctgagatgtttgaggaccgaaatcctcgttcaagaattcagatgaattgcaagacaagtaggcaggtccctgtgaaactgcagaccacaaatgacaaagaaaagattttaacagcagtcaaaagggaaagataaatatcctccaggggaatagaaagaaagaaatagtcaaagtcaagaaagaggattaaaaataactgttaaatcagaattgagattgataaaaaatacctacaggacaggacaagaaaatatggaaaaccaaaataattctgtgagtttgccaccaaagtatcttttcagagttatagttaacaagggtgggaaatgatcccagaagaaatgtatgagggcagtttgtgagggaacc**catgtgatgggacaatcccgttgggcacatg**tgagtgggggaatggaggaggctggggcatgaatggggatggcagaggggaccctgacttggaggaaagacaatgagctcatcccatggtgcctggtgttgggggcactgttggca**catcctacagaacgtgctcagcagacagaggagcggctgtgggatg**agaaggtgaactcggagatgcagcgtgaggcctccaagtccagaca**gcatgggagcccaaagcgatgtcccatgc**aataacgttgtttacaaggagcttataaatatttaaagtagtcacccaagtgtggtctaatataaatcctgtgttcctgaggtcatgcagattgagaga**ggaagtgatgtcactgtgggaacttcc**gtgtaaggacggggcgtccctcctcctctgctcctgctcacagtgatcctgatctggtaaagctcccatcctgccctgaccctgccATGGGCACCAGGCTCCTCTTCTGGGTGGCCTTCTGTCTCCTGGGGGCAGGTGAGTCCTCAGACACCAACCATTCTCATTGTGTGTGTGTGTGTGTGTGTGTGTGTGTGTGTGTGTGTGTGTAGGGGTGGGCTGTGTGCATTCGTGTGATGACTAAAATTATTTTCCTCATTCTGTTCCCAACTTTGTCTCCACAGATCACACAGGAGCTGGAGTCTCCCAGTCCCCCAGTAACAAGGTCACAGAGAAGGGAAAGGATGTAGAGCTCAGGTGTGATCCAATTTCAGGTCATA**CTGCCCTTTACTGGTACCGACAGAGCCTGGGGCAG**GGCCTGGAGTTTTTAATTTACTTCCAAGGCAACAGTGCACCAGACAAATCAGGGCTG**CCCAGTGATCGCTTCTCTGCAGAGAGGACTGGG**GGATCCGTCTCCACTCTGACGATCCAGCGCACACAGCAGGAGGACTCGGCCGTGTATCT**CTGTGCCAGCAGCTTAGCcacagcatggcacag**tcgcctccttcctgctcataaacctcatccttctctctccttgcagctcctagacacccttaacagaggcttctctttgcttctccctccccatgggaaacaagtagatttggacctcagcttgtcctttgggtagaaagagaccacag**atttacccttaagacacagtaactgtaaat**gtgggtggtggaagcaatcgtggcccaccgggctattggagtcctcagagccagctcattgctctaaggcaagc**actgggtgtcttagccttggccttcagggctgggacacacagt**cttctttggggcctgggaggttgctgcccacatagagaagcttcgggcatggctgaggaacaggtgtcttcccctacatgtgttggggcatctgcaaggtctgaa**gcttcatccacaagaacattctttcttctgaagc**ctcttctgttacattggaagttttctgcaatacaagagcaaaccattcttcctgtttgatttaaaagtctttgtggcttttgtggccattacttggtgaatacagccaagataacaataagacttcgtttcttctgcctagtgtaagcaatgcgagttctttattttgttttcgtttaccactgctcctgtcctgagagactgaagcatgcgtccactactgctccagtgtcaacttggttccctaggaaatcgggtttctagaacctgaatgctgacaaataagagttgtgtacatgtat**accatgcaacctgcgtttaaaaatgtatgtacatggt**gaagtgactaaatctagctaattaacatatgcagtacctcaatccttatcctttcctgtggtgagaatacttgaaatgtactctgttagcaatattccagaatacaatccactgttactaactttagtcactgtgttgtacaatagatcttttgaacttcttcctcctatttgagtgaaattttgtatcctttaaccaacatctccccagca

Uppercase: TRBV7-2

Lowercase: Flanking sequence[1000bp]

Red & Bold & Underline: Stem-loop [13]

Blue: Heptamer[51]

Green: Nonamer [5]

id-IGHV7-81[V_gene_segment]

agtccc**cccaaacccctgaagagtgtttggg**ttcaccatgtgtccaatgattcagtgcctctcgagctccaggaaacggctccctggtgatgcgtgagatcttttcttggggtgtccctgcagagttcgctgggtttcctaaggctgattcactatttcaaaagatggtgtgagaagcatatggtgtaaataaagcagaattctgagccagggcacagccactttatactgggctagagacactggtaggaatatactctgtcagctcagatagaaacct**ccctgcagggtgggggcagggctgcagggggcgctcaggacacatcgagcacagtcttccgccccagagcaggtgcacatgaggctgggg**agaggttcctctcagggcctgggacttcctttaaaaatatctaaaataagtatttcacaaggactgctgat**gtttgtataaatatcctattcaattgtgagcatttatcaaac**tggatgttgtaatgagaaccacttttataatggcgatttcaaact**ctgctagttatcttaataatagcag**ctggaggtcaggaagagattattacttataaataagtgcaat**ttttggagagacacactcattcccaaaa**taacgcattcacatattaaggtctagaaatggttcacgttgcccctgagacattcaaatgtgggttcaaagtgaggtgctgtcctcggggagttgttccccagtggaggaag**ctctgtcaacacagag**ttcagggatgggtaggggatgcatggcctctaacaggattacgactcgaaccctcagcttctacaattgtgtcgtccgtgtgtcatggattttctctttctcatactgggtcaggaattggtctattaaatagcatccttcatgaatatgcaaataactgaggggaatatagtatctctgtaccctgaaagcatcacccaacaacaacatccctccttgggagaatcccctagagcacagctcctcaccATGGACTGGACCTGGAGCATCCTCTTCTTGGTGGCAGCAGCAACAGGTAAGGGGCTCCCCAGTCTCAGGGCTGAGGAAGAAACCAGGCCACTCAAGTGAGGCTTTACCCACC**CCTGTGTCCTCTCCACAGG**TACCTACTCCCAGGTGCAGCTGGTGCAGTCTGGCCATGAGGTGAAGCAGCCTGGGGCCTCAGTGAAGGTCTCCTGCAAGGCTTCTGGTTACAGTTTCACCACCTATGGTATGAATTGGGTGCCACAGGCCCCTGGACAAGGGCTTGAGTGGATGGGATGGTTCAACACCTACACTGGGAACCCAACATATGCCCAGGGCTTCACAGGACGGTTTGTCTTCTCCATGGACACCTCTGCCAGCACAGCATACCTGCAGATCAGCAGCCTAAAGGCTGAGG**ACATGGCCATGTATTACTGTGCGAGATAcaccatgt**ggaaa**cccacatcctgagagtgtcagaaatcctgatgtggg**aggcagctgtgctgagctgaggcagtgatgcagcagtttccttaacttccatcttatctcattttgcatcgacatccacctttatattagccaagaatgtggaatagacgggtgctcataagagatccttaacttgcccattttaatgggttttccagaagatgtgagaagccactttgttagcaaaacatgccaaagccatgccctgctccagacacatgtgagcccatttcctgctctttgcttaactgacaagctctcatcagtgcacctgggctaatttcacatcaggtacaggaatatgttctaaaggaaagctaattttataatagcaattcctgcttaataaccttcagcttcattgtttttgtgtaatctatcaacaaattatgttagttcaaggttctcaatgggagtttctaataaatagaagggatgtatagaagttcccctaattaaaacaattgtgaacacaatcttggtattcagctgtgtctccacccttcttaccattcaccacaaagtaattctcacttctggaagctgggttcattttcaaattagttttttttttcattttaatatctcaaggttattgtatgtgactattttagcagaaat**tgaattttgggaaattgaactaaccaactgaaaataaattca**caactaaataacaagatgccagaatataattggctccaggctttgttaattcagcagtttatctacctaggcattatatttacacatcttcatgctatttttacagtcaactc**ccattatgtaagaaatggtcatacaccatttcttacatatgg**taagatatctacgatggatattcagaactacgataaacattggatgcatccaggtttgtctgaactcactgagattattacactcagctgctgtttcaatgtgtctgaaggtataaatgacaattcgaatgac

Uppercase: IGHV7-81

Lowercase: Flanking sequence[1000bp]

Red & Bold & Underline: Stem-loop [13]

Blue: Heptamer[34]

Green: Nonamer [5]

id-IGHV3-73[V_gene_segment]

tctctctaaagggcgtgaccagatttgcagtagaatgagaatagtttcaaccctaaagttcatctaaaatgttagtggttt**tcatggtaagtaattaatagaaacctgacagctccatga**cagcaatatagaaatcaatagtccactcattttaacagagagaattagaaaacctataatttaagaataaccagctgggcactgtggctctgcctgcagttgaatttacttgggagtttgaggtgagaggatcacttgagcccaggatttcaagcgcagcctggaaacatagctagacttaatttcttgaaaaaaatcagtgctattttggggtcagaacaatcctcaaaaatggtccccggaaagaagcccaaatttaattgcaccagattgttgaacgatttatgctacaaaacagtgtcacatgtcagaaatgagcacagtgtaacatctacatggtttgttaagagacacaaatggtcaagtagaacaatcaggtaattag**gctgtcgaaggcaacactgccaaatgacagc**atttccgtggaaactgcgtgtacatccaggactgcac**ctgtgaacgatgacatcatacccttcacag**tgtcgaggaaagagacatcactcaaacagacaagccaagggacttcagaaaatataaggggaaatacagtgtgcaaatatgtaaaaaatgcaataagatgattactcctaaatgaatatcaagacacaatcacataatatgaaattaaattttcctgaatgataggattactaccaatcaccccccaggacaccctcatctactctgtgcacagccttctcgtcaggcgtcccagcccagaccttgctatgtagcagaagacatgcaaataagacccccctttttgctgatgaaaagcagcccagccctgaccctgcagct**ctgggagaggagctccagccttgggattcccag**ct**gtctccactcggtgatcggcactgaatacaggagac**tcaccATGGAGTTTGGGCTGAGCTGGGTTTTCCTTGTTGCTATTTTAAAAGGTGATTCATGGGGAACTAGAGATACTGAGTGTGAGTGGACATGAGTGAGAGAAACAGTGGACGTGTGTGGCACTTTCTGACCAGGGTGTCTCTGTGTTTGCAGGTGTCCAGTGTGAGGTGCAGCTGGTGGAGTCCGGGGGAGGCTTGGTCCAGCCTGGGGGGTCCCTGAAACTCTCCTGTGCAGCCTCTGGGTTCACCTTCAGTGGCTCTGCTATGCACTGGGTCCGCCAGGCTTCCGGGAAAGGGCTGGAGTGGGTTGGCCGTATTAGAAGCAAAGCTAACAGTTACGCGACAGCATATGCTGCGTC**GGTGAAAGGCAGGTTCACC**ATCTCCAGAGATGATTCAAAGAACACGGCGTATCTGCAAATGAACAGCCTGAAAACCGAGGACACGGCCGTGTATTACTGTACTAGACAcacagtgag**gggaggtcagtgtgagcccggacacaaacctccc**tgcaggggcgcgcggggctaccagggggcgctcgggactcactgagggcgggacaggtcccaggaacaggtgcagcggtaggttttct**ttctcctcagctggagaa**gtcaggtttgtgttttcagaactctggagtcttacaggttgctacattttcatacagttattagtatgtatttattatcattggtattta**agttttaataattttaaaccttttatgtagtgttattttttaaaact**gtttactttcatttgcagttattcttccagagtttcattaacatctatt**gctatgagcaactacatagc**tatgagagcataa**atttacacctgtagacgtaggtctaaat**gccacaaacctgtgcataaatgtatagtgacttatatttaacattataataagataatttttaaaatatatcctaaacgatcaaacttactgatgaacta**aatataaattatcagagtaatgcataattgatttcaataattttatatt**gtttatattaattaatatctatttctttactgaaacataatatattggtcatttcaaaatagctacgatacatttcaaatggccttgatgctaataatgaaaagattttgaggtgattaatatgc**taattagttatatttaatta**ttccatattgtattaatatatcaaaacattgctttgtacctcataaatagatacagtcgtaatttgtcaattttcaagacagttatttaatccatcctgggtcataatctttttttcccctgtccatggctgattggattgtcctgataaacccatccatac**gcctgcctcccgcaggcttctgagacgggctccatgaaggccagaagc**aggcaccctgtgagagtccacacgacctggagcctccctctccttggattaggccatctcctcgggatcgcagggctcttcattatcctcacccccttgt

Uppercase: IGHV3-73

Lowercase: Flanking sequence[1000bp]

Red & Bold & Underline: Stem-loop [15]

Blue: Heptamer[32]

Green: Nonamer [6]

id-IGLV4-3[V_gene_segment]

tctctaagatattaggtaggaaatattctcccgccactcgtatttattaaagttaaaatgcttcagtatctaatgtgacagttgtccaaattcaatacccttcaccttgttagaaatggaatcctgatcctaacttgtctggagaggaggctggcatgaggagcatgaccacacatctggatgtccccagtgtcactgctaacacacgctcttttgttacttgccgtgaagggagcagagggagcatccacacctttgtccaaaggacttggaggacagtgcacctgattaaaaaatgcagggctgtgataagtgggttccgggtcctggatgagggaagagaagatgaggaggcatggggtaggaggcagggcagggaggagctgtaag**gtcaggctctgagtatgggtgaaatcctgac**tccactcactactctctgggaggatttgataaatcttgtaacctcctgatgttggttctcccatcaataaactaaaggagaggtcgtgtcttaagcgtatgcttgctgtggggctactgtgaatgtgtggttgaggtggtgcctgcaaaacactctccaaagggcctggcacgtattaatgtctcaaaggagtcattctgcttaagctcgt**ggaggatgactttcctcc**tgtgtgcactagagggccttgttttgaggaccccagtgacttccctggtagggctcaagcacaaagggcatgactgatgcttcctctgaggccccttctattgggtaaaattcatgctccactcatgacacccacctggcccttcttctgaccttgagaaattgccaatatgaagccctcagactcagagctcagacagtacctcagacagacatgctgtaaagtggagagcagatttgcattcaaagctcc**tcccttcttgagcttcaagaagagga**taagacaggagctcatagtgtccaggttccactgggcagtctcgaatagagctcttggaagtccctccaaccATGGCCTGGGTCTCCTTCTACCTACTGCCCTTCATTTTCTCCACAGGTCAGAACATCCCAGGGAATTCA**GGGAAATGTTTTCACTGCTATTTTCCC**ATGAGCACCAGTCCTCAGGGGCATTCTTTCCAGTTCTTCTGTGCATTCAGCATCATTCATGACATTCTGTTTACAGGTCTCTGTGCTCTGCCTGTGCTGACTCAGCCCCCGTCTGCATCTGCCTTGCTGGGAGCCTCGATCAA**GCTCACCTGCACCCTAAGCAGTGAGC**ACAGCACCTACACCATCGAATGGTATCAACAGAGACCAGGGAGGTCCCCCCAGTATATAATGAAGGTTAAGAGTGATGGCAGCCACAGCAAGGGGGACGGGATCCCCGATCGCTTCATGGGCTCCAGTTCTGGGGCTGACCGCTACCTCACCTTCTCCAACCTCCAGTCTGACGATGAGGCTGAGTATCACTGTGGAGAGAGCCACACGATTGATGGCCAAGTCGGTTGAGCcacagtgacacagatg**aaggggaagtgaggccaaaacctcccacctcccctt**tctc**ctcagggactcagagtcaacagccctgggttcctctccagtaccctgag**acctctgactctcctccctccaagacacccctaggcagaagcagatgcaggccatgactgaggccattttgaaaactcatctttctgtttcctccacaacagcctgtggggaccatgctgggagagaccagagcagcccaggcaggaaaaggaccttgagagagtggctagg**atgagatgaccgtgtcccagtgtctcat**tgtgccctgtgctctattaccaggctctgccccaggaaaggacatggatcttgacctgtcagccatcatttatgagagttcaa**tttttacattttttaaaatataaaaa**ctacaatctccaagatgctaggagcaatttcactgagtctcatcatggaattctgacctcccccagcatgtcccctgttcctacaatctcctgttcaagaccaactcaactccattacccagaccagctgctggatgatcaagacagggctgtgagagcctaagaatctttaaggtccacgatgagaacacgtcaa**gaaggatttatgttggaacttccttc**cccatggatattcacatcaacagagaagatatctgagttcttgcctacacctatgaatacctaagacatgaagtgtgtgtgtgcata**tggtgtgggcagcacttcctcactcaacacca**gag**agtgtctgaggacact**gggtacagacatggagacagggcatatttttatacactcaattaagccctcgtccttctctgctcaccaaggagccacccaacaaggccacagcaggaagagaagttga**tttccatgaagagccttccctttctaagagtcctggaaa**cccactgtctagttgtggaatcagaggcagagctctggggcatctccaccatggcctggaccctcccctgctcaccct

Uppercase: IGLV4-3

Lowercase: Flanking sequence[1000bp]

Red & Bold & Underline: Stem-loop [13]

Blue: Heptamer[44]

Green: Nonamer [1]

id-IGKV2-29-3[V_gene_segment]

aaatttaaatagtgaaatatgatacatttattctaatcctacttccacactacaatttgatattctcaatttaggttactgccatgacgttcaacgttttaaggtcaatatcatttttactcaatttttttcaccaactttgttaaaacatcatatgccagtgcatttcccggaagataatcaggacaagctttgtgt**gcaaaacacctagagctctgttttgc**ttttctgggtgtgtgttctttaccaagtaaatcagatgttgtattaaatcactgaagattacatagcactagaaactattcttttaattgctcccaggatttttgttgtttttttcaggacagtggcagtctctgtgaagtgatttgatgtacgcaccgttattcaggagcagctaaattcaatattaagcacatcactgacaaatgaagaatactcaagttgttttcctattgtaacgatgagttgtgtggtagattcatatgcctgacctgtgatcatctttcaataaaaacctaggaaaatgtcgtccatagtactgctgaccctgagacttaaaaccagggatccagttcttgctcttcctacatagagaatctcattttgaactgtggtgtcatggc**tgtggctgtgccaca**tacaggccagggggagacaagggttcaccctcagagttgacaagaatttggaagccctgacatcctataaaacgttacttgcccaagattgaaactttca**aattcaggtccttcttcctcctctatgatgaatt**aggttttattagtttcctccaagggacactttatatcacattgctcacagagaagacatatctacccccttcaccccccacccaatggctctttccacaccactgcacccaccaggtgatttgcatattgtcccctagggaggaccttcccttgtgagcctgagataaaagctcagctctaaccttgccttgactgatcaggacttctcagttcatcttctcaccATGAGGCTCCCTGCTCAGCTCCTGGGGCTGCTAATGCTCTGGATCCCTGGTAAGGATGGAAGGAGATGAGGGAGGAGGAGGGGGTGGGAGGCTGAGCTCTGGCGGCCCCACTGATTCCCGTGTTTATTCTAACCATGTGTTAAAGGAATATGCCCTATGCTCCAGGGAGAGGAATTCATATTTTGCCCTGATGATGATTTGAAAACTCCTAAAAGCAGTGCTCTGAATAATATCTTGAG**AAATGAAAGAACTCTTGTGCCTATTTAATAAAGGGTTCATTT**AAAA**TTTTGTTTTTATGATATGAATACAAATTTGTAAAAATAAAA**GATTAGCCATAAATCAATATCATAAGGCAAATCTCAAAAGTTGTTCATTATGCTTTCACATAACCTTGCACTTCTCTCTCATAATTTCAGGATCCAGTGCGGATATTGTGATGACCCAGACTCCACTCTCT**CTGTCCGTCACCCCTGGACAG**CCGGCCTCCATCTCCTGCAAGTCTAGTCAGAGCCTCCTGCATAGTGATGGAAAGACCTATTTGTATTGGTACCTGCAGAAGCCAGGCCAGTCTCCACAGCTCCTGATCTATGAAGTTTCCAGCCGGTTCTCTGGAGTGCCAGATAGG**TTCAGTGGCAGCGGGTCAGGGACAGATTTCACACTGAA**AA**TCAGCCGGGTGGAGGCTGA**GGATGTTGGGGTTTATTACTGAATGCAAGGTATACACCTTCCTcccacagtggtacagaccaatacagaaacctccct**gctggggtgtcccagc**tgctcacttgcactgcttgtctggggagtagctcagcagggact**ctgagtctgcagaagaggagactgttggagaactcag**gggcagggct**tgctgttgaggactctggcccatgagagtctcagcagca**cctcagtcccacatggtcaaagctttatcagttgccaggctcatcctcaccctgcctgtcttgcccacgtttgccgagtacactcaatcagcataacagccagaacatagatgcagtttagtgacaacacaggtggaatacatgtggcaaatgaccagcttggggtttattttacacaaattaatacttggtgataatatttggaattattgcgaatttggtattttcccactttccttactttctatttcactttactactcacactaaactgcactttccagtgtcatgtgggagaacgtgttctatacaagctgtcctcagggggagtatgga**taagaataattagaaaaaaaatgtttgatttctgactattctta**gactttgtaaatccatggcagatat**aaatcttaatgccaaaagattt**tgatttgcctcaattacgtttttcccagtgaagag**ggagttcttgcaactcc**aaaagtgagcttcagaaataagcagcactggccgggtacagtggctcacgcctgtcattccagcactttgggaggccgaggtggatggatcacctga**ggtcaggagttcgagaccagcctgacc**aacatggtgaaacccggtctctactgaaaatacaaaattagccagggcgtggtggcacacacctgtaatcccggctagttgggaggccgatgcaggaaaatcacttgaacctgggaggtggaggttgcagtga**gctggagatagtgccactgcactccagc**ctgggctacaaggcgaaattctgtcaaaaaaaaaaaaaaaaaaagcagcataaac

Uppercase: IGKV2-29-3

Lowercase: Flanking sequence[1000bp]

Red & Bold & Underline: Stem-loop [16]

Blue: Heptamer[25]

Green: Nonamer [8]

id-TRBVB-2[V_gene_segment]

aactgagg**ttgctgcagaccagcaa**ggattcactgctgataactcaagataccttccaccagtaacttggagatgacaccaagagacaacacaggcaaaggaaagtgtattaattctgttttaggtgtagaaactggagtgattggaaaataattcctaaaagaaattgaaaagtcaaattgtcaagcagtaagttcaccatcacatttaccataaagcatgtagagaaatggaaatatttacacatcatagttgagagtgcaggatttgcagtcagacagtttctggcacttagtagctttagaaagtgacttaagccctctaagccctcagttttcacatctataaaatgaagtaaataatataagcctctcagaagtggggtgagggttgaatgaaacaatatgtgagtagtttaacacagagactggcactgacttaagaaagtattgtgataaaagcagcagtggagaaaatattttgaatagctgtatgcatggcaatcctttaacctctttattca**tggcaactcatttaattgcca**caacaaatatgtaagtacttttattatttccattgttatggactgagttgtgctaccttcttcatttcatatgttgatgtcctaagtcccagtg**ccccataatgtgactgactatatttggagatgggg**tctttacatcgataattaagttaaaattaggtc**attagggtgggccctaat**tcaatatgactg**gtgtccttatcaggaggttaggacac**agacatgcaaagagggaagaccagaaagaaggtatagaaaaaaaaaaattgttatctacaagccaaggagagaggcctcagaagaaaccaactcaacaaattttaaaagatgtattttggagaacaaaactgttagaatatatttagaacagaaaaaaaccatttcttctagaactaagttggtctaaaagtgcatatcttggtaaggctctgtgaggaaatgacctaatctggagagAAGCTGCTGGGACACTCGTCTGTGTGTGG**CCTGTGGCTCACAGG**GGCAGGTATTCTCAGAACGCACATGGGATAATCACCATCCTGGTGTGTCCAGGCTCTGGGAACAGGCTCTTCCTCGTGGCTTTCACATGAGTTTCCTGGTGGGCTATGTAGTCACTTTTCTGCATCCTTCTTTTAAAGGCTCATGAATGTCAAAGTAACACAGACCCTGAGATGAGGCAGGAAAGTTGTATCGGAATGTTTTCAGACTATCAACCAGACCAAACGTTCTGGAATCCATAAGATCCAGGGCTGGGGATAAATTTAATCTATTAATTATTCTTCTGAAGTTGGCAGCTTTGAAAAAGATGCCCTGAGTCAGCATAGTCATTCTAACCAAAAAGAAAAAA**GGGAAAAAATAATCATCTTTCCC**AACCCTTAAATCGTCCAGCACTAACCAGACTCTGAGACCCTCTGCAGCAGCAGCCTATCAGTGCAGCCACATCCTCTCTGAGCGGATATGACAAACCCCAGGGTTGAAGCGACCTAACCTATGAGCCgccacacacactcaagatgccccagacaccctgcactccgatcttactcgttcctttactgttttcatcctaattgccctcttacacatttgaccacacatttttggtcttggtgg**ttgtattagttttctgttgctactatacaa**attgccacccatttagtggcttaaaacaatgcaaatgtatcaccttacagttccgtagttcagaaaccaacataggtatc**acccagccaaaatcaaggtgtcggcagatctgggt**tcctttctgaaggctccaggggagaatcgattttcttaccttatgggttgttttcctctatcttcaaaggcagcacgttctgttgctctgaccctacttgcatactcacagctccctctaaacactgccgggaaaggctctccacgtttaaggacctgcgt**gattacattggacccacctgggtaatc**cacgaactgcttcctatctcaaggtcggctgattagcagccctaattgcacctgcagctttaattcccatttgccatctaactgaacatccacagctgttacgggttgcagggattattagcacctggacatctttaaggaaccattca**gccctccactccgatgttccactagagggc**actaacatattcctgtaatataaacattaaaattcagataatggtgggagggaaagtttga**taaaattgaatactgtattatcaatttta**aaaacttgatatataaatatgtctcattggggtctttcaaaagctaatcatatagtacaagaaaatacataaactggagtgactctcagaactaaaatggcctgagcgtaagtgaaat**agaatttgaagtgaggtgagaattct**gcagtctattcctgggctcaaaatcacatagaaaaaaacaaaaaaaggttgctttatatttaaagggacaaggcataggttctcttacaggggtttttaggaaatggttcatcagtttagtaggtgttagcttaattg

Uppercase: TRBVB-2

Lowercase: Flanking sequence[1000bp]

Red & Bold & Underline: Stem-loop [13]

Blue: Heptamer[18]

Green: Nonamer [4]

id-IGKV1-22[V_gene_segment]

ggacaagaacttcactaacaattgagtcactgaagacttacggccctgtgtgacgcaccacataactgtgagtttgcagtggttgcaggtcagggacagattttatgcttaagatcagtagggtggaggct**gaggatcttggctattacaactgccaccacactctacaatatcctc**ccacaatggttcagcaccaaacaaaagcctcctgcttggattgtcccagctgcccaaattagttccttcactgaggagtagacagggtatattctctaaatctatgtaacaggaagatgttggtgaactcaggggattagtatgaagctacacctcaggcatcacacataagatcacttcagcagtcgcagccttagcatgggcagaacctacagaagatgcaagtgccctctgagccaggagacaggaggaaggaggaagggaaaggtgacttagctcatctcaatcctctctcctttgcatacatttgtcaaccag**atgtattcagcctaccagtcacacaactgaggctgatacat**gacaacatagcactggtatattcttggtattgtttggcttagcagttactagtatatatttaat**gggagaatatttggtggtgttaacacattgcttatctccc**tttccccagttgtactttacacttgttcttggcacacattctcctccaggactggagcattcacagggttttatgttactgttcttatgggagtaaaaagaaaaacgattcacattcttgctactgagctaggctgggatgccctgggccaagctgaaaatgtgaaaaataagagtatgaatatttattaagttttatctggatctaagatacttatccatgaaccagtcctgcagctgtgcccagcctgctccattccctgctgatttgcatgttcccagagcacaaccccctgttctgaagacttcttaataggctggtcacaccctgtgcaggagtcagtctcagtcaggacacagcATGGACATGAGGGTCCCCACTCAGCTCCAGGGGCTCCTGCTGCTCTGGCTCCCAGGTACGGATGGAGAACACTAGGAATTTACTCAGCCAATGTGCTCAGTACAGCCTGGCCTTTCAGGGAAATCATCTTATAAATAGTTGTGTGGATTATTTGTTTTTATGTCCCAG**GAGTCAGATGTAATTTCCAGATGACTC**AGTCTCCATCCTCCCTGACTGCATCTGTAGGAGAGAGAGTCACCATCACTTGCTGGGCGAGTCAGGGCATTTGCAATTATTTAAGCTAGTATCAGTAGAAACTAGAGAATCCTCCTAAGCTCCTGATCTATGCTGCATCCAGTTTGCAATCTGGGGTCCCGTCACGGTTCAGTGGCAGTAGGTCTGGGACACATTTCACACATTCTCACCATCAGGAGCCTGCAACCTGAAGATGTTATAACTTAT**TGCTGTCTATAGACTTACAGCA**GCCATcctagagtgttacaggtcataaaataaaccc**ccagggaagcagaagtatgactcatggctgccccaggtgcttccactgg**tgcctccatctg**ctgagagtgtttctcag**gtgcagccaagatttaaaggtttttgtaggaatggccagaagtctcatctgcattctaactctttttcttcctgcttagccccagcagcacagacatgacactatctctcctgatttaataagggatagcatttacgatacctgaagaatctgtgttattgcatccatctgggtcatagattaaaagagaaaccactctacagattgccagaaggcattgttttaatacagggaattagagttgaatatacaaaactgggagtgtggtagttagggaagctgacactagaaacacgggagt**ctctggaggtctgccagaagccagag**ttcatcagccgctaaaggcatgggctatctaaccatatagtcttctttgtctaggaagtccgtatgcgaagatgctgatgctatcagttgttgcagcacctcaccaggtgattctccagtccttatctcagtgaacatgtttgcctaccggtgtcaaagaatattgaatcgccttcttcttaccttcaaatatgatgagaggtcttctctttgagtaactctacaagaaaccatagagggtttaatgtgtttcaggaaaggtgcttttagaaatcatggtgaatatgaggaattacagccaag**tgagataagtatttcccaaaatctca**gaattttccaggtatggggtggcttcag**aatacatttggatgttcttacatgtatt**attagaaagtttggtattattgcaagaaaattttattaagtcgtaaagtaaaagaaaaaaatgacaac**attgcttgaaatacatagcaat**cctttgacaaatgaaaaaaaaattgacaaaacaaacaagaacacctataggtgcacgtagcatactttttccttaatataagagcactt

Uppercase: IGKV1-22

Lowercase: Flanking sequence[1000bp]

Red & Bold & Underline: Stem-loop [11]

Blue: Heptamer[35]

Green: Nonamer [5]

id-IGHVII-26-2-2[V_gene_segment]

gaattgatgtgtgtgtgtgtgtgtgtgtgtcct**caggcctgttgggcctgttgagaacacagcagggctctatgtacaacagg**aaagcaatgttttctctataccatctattt**tcttataaagtgttcataggtaatcaaaaggaaatgaatcacagtttataaga**tgttgtggactggactgatgttgtcactcaaggctactatgggggctccaggaaaagaaacctagaaaaggataagactgtgagcaaagaagcaaaacctgggaagcagcagggagagggtgagcaggggtgactcaactggggagggataacaccacttaccacagagggtgaaaactccgtgctcagcaaagaggagaacttactcagcatttcccctaatcctccctccagaagcgcgactgccaggaaagttcacaatgtatc**atctccagctgggaacccaaaagaaacaggagat**tttagggcgtaggtcctgggcacaaggtgtttccctggttcctaccccttccctctctgtccagcaaaactgcttgtttctatcacttgccttcctgtccttggggtaagaatggatgctttgacactcagcagagggatgctctttcagaggaggaatgagtttgtgttatctgatgtctgagcaggacactcacctctgggaagaggctccaagacagaaggtcgccgtggttcct**gccccttatatgtgaggggc**tcttgacacatgcacacttttcctcaaactctgtttatac**cccagcaacaatgctggg**tgttgtcagcctctctcctccccagatactgctctatcatcagcttgagcacagtttccccatgaaaggcattattctctctctcaagaagatcaactttttacatgtcttcgatggatgacagtttttctgtgtgagaatctagaaataagaaatggggtaatggggaca**agtgagctctctgtgtgactcact**aggaagtccactgttattccagGT**GTCCTGTCCCAGGTGCAGCTGCAGGAGTCAGGTCCAGGAC**CCATGAGGCCCT**CAGACACTGCCCTCTGCCTGTGCTGTGTCTGGATTCTTTTTTTTACCATGGATATTATTTGAGATGAATCCA**CCATCCCGTGGGCAACAGGCTGGAATGGCTGGGGCACAGTTACTAGACTGGATCATCAAGGAATACACATCATGCACCCTCCACCCAGGTCCATGTCCCCATCAACAGTGACTCAACCAAGAGCCAGTTCTCTGTGAAG**CTCAGCTCCATGACCACCTAGGACACGGCTGAG**TATTACTGTGAAAGActcagtgaggaggtgtccttatga**gccctgacacaaacctgtcagggc**acttaggacctccaggaagactcaagaccaccaaggggactcacgaccactggggaagggcaggttgcagtaagcacaggg**ccagccccagagcaggtgcaggtggagatgagaggctgg**tttactgccagggtctggggctgcttctccatttagatgtttccttcagagaccctccct**tatttcatgatgctctgcttagctgtaagtctctgaaata**tcacagttttgttgtaccaggaggaaaccttctctcaggtgctaaatgtagaacaatctctctgccagtggttgccagggtcacagccctgtggaagctgaagggtacccagtgagtcttctccagtcacattcaggaca**ggaaccttagtggggttccctgaggagaacagtctttaggaattctaacctcag**ccaagagaggctgggccagggtc**agtgtcaggtagaacctcacaggatttacgtctgacccttctcctgacact**aaagcatgcaa**atcagtatcagcactgat**ctggggccccttttgctcctagcccattctatttcttgttagttgtatttgttgttgattttccttttgctgttcctgatccctgtaaagtggagatgtggttcctgctgtaaaaattccagggctcaagccctttccctgcagctcaggtggggctcagactgtggctcctgcaaccacgt**gggagaggctgatgggactatcttctctccc**gttgctcagaaccctccagtgtgttgtgtggagactcacctgggaatgcaagtggccaatagttgtgaaaggtatgagcttgtttggtcaaagtggaatgtggatgtggaatttat**cctgtgttgtgcaaactaacacagg**ttcaccttcctcacctgtagtgttagaaagagtgtgaaagttgtaagaatcaaaacagagccacttgtgttaacaccctga

Uppercase: IGHVII-26-2-2

Lowercase: Flanking sequence[1000bp]

Red & Bold & Underline: Stem-loop [20]

Blue: Heptamer[41]

Green: Nonamer [4]

id-TRBV5-2[V_gene_segment]

gctcttattcacattggcctaaaataagtaggagagtgtgtggatctcggggcttttgacattttacatttggggtaaggatgtttgtagggtgctgtgacctttggtagagaaaagcaatcaatatgaaatagtccagcaaggaggcagccaggagacacacaaaccgggacagttggttaccctctctctctctactctcctgcca**ttgtctctcataagacaa**actcactctggagtacacaaaaagagatgcagttactataatccattggcatcaacctctgaa**ggcaggatggatctggagggataaacataacctgcc**cagaggatgactcatagggtggccttgtgggtgctgaatcccaagtggttaggtctgtgctgtgagctgataaagccccag**aattatttatggggaatctatactcctaataatt**tacagacaacacagatccactttggacttgatcagatggacaaaatcttggggacttgacatcactgcgcacacaggggaggagctgggattgttgtccaggaaaggaaaatataaggagaaaccttggtttgaatttggtgtcagacatatgatgagaggacagccaggaagatctggaacatgaagatttaattacaggccactaactctcagctgacgggcaggtctgtgagttccagaatggagcactgcagcagtaggtggggaggaggctgagagggc**gatgaggcagtctgagtgctgagggcagtgcagtcagagaagcaactgcctcatc**acagaagcttctgccctta**ctcatccccttgctctgcaggatgag**gagggagttccatgggtagggaccactggacctaaggaagcctgaagg**ggaaggaccacaggacagtgacatcacaggatatccttcc**tatcaggaaaagtgag**gctcagaactcagctcttcctgggaggaccaagccctgagcacaggtgcagtgctgcctgctccgctgtgc**cATGGGCTCCGGACTCCTCTGCTGGACGCTGCTTTGTTTCCTGGGAGCAGGTGAATCCTGTGGACAGGACAGCACCCCTATTCTCAGCTTGCCCACCCCTGTGTCCCCCACTTCACCATGGGGAGGCACAAGTTCATT**CTCCACCTATTTTTTCCTCAGGCCCAGTGGAG**GCTGGAATCACCCAAGCTCCAAGACACCTGATCAAAACAAGAGACCAGCAAGTGACACTGAGATGCTCCCCTGCCTCTGGGCATAACTGTGTGTCCTGGTACCTACGAACTCCAAGTCAGCCCCTCTAGTTATTGTTACAATA**TTGTAATAGGTTACAA**AGAGCAAAAGGAAACTTGCCTAATTGATTCTCAGCTCACCACGTCCATAACTATTA**CTGAGTCAAACACGGAGCTAGGGGACTCAG**CCCTGTATCTCTGTGCCAGCAACTTGAtgcaggcctgcagagccaagaacattctgtgtacaaacatccctgccccagtgtggagaacttcagccctaacatatctgtgagaacttgaggactgtagtgggaaagaaaagcagtttcaggaagctggatcct**aaaacctggggtttt**ctgccagttaggagtggggataacaaggccaagagggtctgctcaggaggtggtgaaaactaccttcttcgccaaacgctgta**cactatagggatagtg**gaagtgtttgtaatttaatag**ctggtgccctctttacagggcagcag**atttaatttcagggtcaagattcttgataatagtgtggaagggaaaatcatagggtttgctttatataattcagggagaagaaagtgagatttggggaaattctatgaaagagaaaatgatcaaagatagttgaaaatagagatagagcagaaggtaatgaaa**catggtgatctgtgcaccatg**agctcccttctcctcccacagtttttgatatccagaaaatacaaagaaaggtcttatagcaactgacatcacctggggactggtcctagttggggatatcacaaatagcagcctgcctgatgtgagggccttggggttggttctgacatcttctttctcgacttctggatctagtcttgcctgctcgaggcacagtcaaatacttcccaaaacacaaaggaggtcaggtcagtcttggtcctaaaggggcaaatgctgtccgttttctcccaagataaaattctgcatccctgggattcctctctgagactcagtgcccagattc**aattaccttctgtaatt**aaacagttctttgaacattttctaaaactcatcagcgttggtatcatctgcatttctgctatttccttggcctggaaacttttgttctctccagacacatctagcccctgctcatctgtcaaggatggtctcacatctgccttactctttgcttcttaccctgacattccctggaact

Uppercase: TRBV5-2

Lowercase: Flanking sequence[1000bp]

Red & Bold & Underline: Stem-loop [16]

Blue: Heptamer[41]

Green: Nonamer [6]

id-IGLVVII-41-1[V_gene_segment]

ttcttggaattgtga**cctggtgccaacaatgaccagg**aagtgcctccaagcttctgactcatagaagtaaatatctgcccgcagggattcaggacagacgctcaggctaccagtcttgcatg**aagcccttcctaagcatctctgggctt**taggctgaggacaaggctgatcactcctgttggcttcagacagccccctggaggtccaaacagtgctgcagtctgggaaagtgagatgagaacacgccaggtctcctaggagcatgaccttccaatggcaccacccacaaccaggacagctggtctgtttaccatttgtgtggatgtttcttgatgctgccgctaaccagggcccaagacttagttcggggacagtgatct**agagaatacagctttgttctct**cacagccagccctcagaggaagcccttagcaacagcattgtcagactgcctcaaaaaaatcagagtctttgattccaggaacaggctgccctggggaca**gactcaagatgagtc**agggtaagcagggacagaatccagctgggtctgcctcaggcatgtgcactaacgcaaaggtctgttcatccttattgtctactaaacacctattatgtgccagtctcaggagagtgctcaggatacagagtttacaagac**agacagggttcctgtct**ttacagggataatgcctcacagaagagtgagcaccaacaaatcaatgagaataaggcatattgtgaggtataaaatggttgacatgggccagaagacagtgagcaacaagaagctattaggaggttgtaagcaaggagatacagactctggtttgagattcaggaggcaaagtccagcttctctaaggaagatgaaaaaggaggggacagagtgaagaaagatggtccagttaggacatggtggctttgtacacatgagagatgaggtttctggcaagtaatgtggcagctgagctgaggcctgtgctgtggatcccagacgggagcCACCTGCTAGGCCTAGGCTGGAGCCCTCTGCCTCTGCTCACG**AGGACCAGCAGCTGTGTCCTTACAGGTCCT**TAGGGAGCCACAGAACAGCCCCACCCTGGTCCACCCCATCCCAGGCTCAGCTTT**CTAGGCAAGGGTCAGAGTCAGGGCCTAG**GGTGGGACTTGG**CCAGCCCAGGGGAGGAGGCTGG**TGTGC**ATGAAGAGACCCTCCTTCAT**CCTGGGCTGCTGGCGAAAGTAATAGGGACTGGGGGAGCTCAAGCCCAGCTGAGTGTCATCAGGATGCAGAGCTGTGTGTGGGTCCCcaccaaggcctgagctcctccctttctcccccttctcattcactgcacaggtaactagggctagggatcagggctgtgggtgaatggaacgaggaccctgcatgctgccctctcaactgacccacctctctcaccctctgttagtccaggtgtcaaggttctgggcccagtctgtgctgactcagtggtgtgaagagttcctgtgggtggggtaaactgccgccatctatcacacttgaagcagaagcaaggctgggtgtgggcctgtgtcctggtatcagcagtgtccccaaactcctcatctatgacagcagtcattggtctgcacaggtcccagcgaagtt**ctcaggtcccaggttggggatcatggcatccctgag**ca**tctctgggctccaggcagaga**atgaggctgattctttcgctcatcttgagacccaaggatccatgattaggcagtgctgcaggccagtgggtagctgggaggaaaaccttctgtctccccagtaatgtgtcttccctcttcag**cctccaccccttcagcacaagtttctggggaggagg**gtctgaagcccgggtgagaaaaagggtcaaggctggccttaggacctggatgggcacagtgaccccctgctgagagg**gccttcatgagcaagaaggc**tgggggaggtcctgagttgatttttacaatggaaacagacaccagctccccatccttgttcaaaagcatctcaggaccttttcctctgcagatggcaagggggagaatctccctctgagatacatgagaattcccttcccatgacctgtcctgtcacctggatgaattcgt**atttggaaaccaaat**gactccccagtgccaggcattacatcctcacagaaaacctgccagaatgagcaagatcttccctccctgacccactggccaggaatctcggacatcag**gatggctggactagatcactcatgccatc**tcccctgacatcatcatggttaccggggg

Uppercase: IGLVVII-41-1

Lowercase: Flanking sequence[1000bp]

Red & Bold & Underline: Stem-loop [15]

Blue: Heptamer[33]

Green: Nonamer [4]

id-IGHVII-67-1-2[V_gene_segment]

tatatactacag**agatattataactctatatatct**aattcctctttcctatcccatgttttttattttctgtttgtatctatacatgctat**aaaatgtaatattggcatttt**tataattgctttccacacttacataatgaaca**aaatacaaatgcaaaaactgtattt**caacctcatttttccattgttgaccatcttctttgtgcatattttcaggtgttggatctatgtcacatggctattcatctagagactctgctaaaatgtctacggaaagattcatgtgctggaaataggttttctcaggttttattggtccgacagaatatttatttgtctttgattt**attgtctccagagacaat**gtcaagaatatgctatatctgcaaatgggcgatctgtaaaccaagaacacatcagtatatcactgtgcaagaggagcacatgagggaaagccagtgtgtgcccagacataaacctccagggacacatgggcgaattagctgcagggggagcttgggacccactgatgagagtcgaccccagagggaggtgcacagaaaggttaagaactgctttcctgccagtatatttttaaaaaaacaaaacttccatctgacagtttctccagagaacttccctaattttagcattctgtgcctaccaatgccatcactaaataggtttttaaaaaatattgta**atatgaggacatattctcatat**ggacaaaacgcaaattgacatttacagaaatgaaaagtcctcaaccgtggtcgctagaattggagacctgtagcagctcagtgggacctgttgagtcttctccaatcacactcagaacagagaccttagtgcatctccctgactagaatacc**ctttaggtatcgagataacagcctaaag**atggtaaatcatgagagtctcagatgtccagatgacagacatgggtctatgggaatag**tactgtaactgacagta**actaattttccataccgccagATGTCCTATTCAGGAGCAGCTACAGCAGTCATGCCTAG**GTGTGAAGATCACAC**ACTGACCTCACCCATGCTGTCTCTGGCCACTTCATCACAACCAATGCTTAATATTGGACGTGGATCTGCCAGTCCCCGGGGAATGGGTTGAATGGATaaaatgcatggctag**tgctgctggaaacccattcctactgtggcaaatggcagca**tctcttttaaaaggctaaataatattctattctgtatacataccacattgccattatcctttttgtttgtttgtttgtttgtttgtttgagatggagattcactcactctgtcacccaggctggagtgcagtggcgtgatttcaggtcactgtaatccctgccttctgggctcaaacaattctcctgcttcagcctccggagtatctgggattacaggcatgcaccaccactcccggttaatttttgtattttttgtagagatggtgtttcaacatgtttgtcaggctggtctcaaactgtgacctcaaaaaatcaacccacctcagtctttgaaagtgctcagattgcaggcgtgagccatagcatccagccacttttctgcatttttttttcattgacacttagattatttcaatatcttggctcttgtgaatagcgctgcagtaaacataggagtgcagatatctccacaaggtggtgattttttctatcttgggtatattcccagaagggaggttgctggtcatacagtagttctagttctaattgatttaagatcatcttactgctttccatagtggtggagattcagattagtgtagctattatgaaaaacagtttcaatgttggggagtgactgaaaaacaaataatgcttgattatggttctccatgagagtctctaataaacctaatggaagttcagaaagtttcctcaccttgttaagaaaacatttgttttgctcataattctctttcagcccagagttggcagtatgtagaatggaggctgttggctcgtttaaggattctgaagcaaagaacagtaatgagagatgacaatagagaaaaagagagagaaggagatgctgaatcatcatggactcattccacctgcaagcctttcactgtaaagctctattgctaaggatcctgaggattcta

Uppercase: IGHVII-67-1-2

Lowercase: Flanking sequence[1000bp]

Red & Bold & Underline: Stem-loop [9]

Blue: Heptamer[19]

Green: Nonamer [3]

id-IGLV2-34[V_gene_segment]

gcctggctaatttttgtatttttagtaaaagt**ggagttttgccatgttggccagtctggtcttgaactcc**tgccctcaagcgatcctcctgcctcagtctcccaaactgctgggataacag**gcgtgagctatcacgc**tcggtctgaaatctttagataagtatgaagtctatataaaaaaaaaaggctcttactaggctagaaatggcccattttagaagctttcattttccacctcacactcatctctctatagatcctctgcatgggcagtgggtgggccctg**tggtcctcagttctcaggctgagaacca**gaggctgaaaaggacaagtcacctgctcatgagtgggaagggcaggattg**ggagcagcggactgtggctccattgctcc**ttgtttcctccacaggcccctcccacgtgtgcccctgaatgacctagaatgactaatgtgctcaagggatttgatgctgcagatgcttcggcatgaagggtgggggtggggatgggacaaggctgcagctgaagagatgggagaccaacaacacacagcaggggtttcc**atggggtgggcacccccat**ttcctcacagtttagagagcctggaggtg**ggtgccaagcagagggcacc**atggcgtggcccaggcctggaggttcacagagacacaggcacaacacagcagagacactgagggccaggagctcactcagatgagggacttcagcagatctttctctcttgaggaaattaggtacaaaggaatcaagttcctaccatcagaatagacaacaagttttaatctcctcagctgagccccgctgtccagggaagcagaagtctctgagcccggcccaggtgagagtggggtgaggagaggagctc**agggtgcagatttgcatgaaggccccaccctcctctgaggcagagg**ggataagacaagtctgggggtaggccc**agcgctggggtctcaggaggcagcgct**ctctcgggacgtctccaccATGGCCTGGGCTCTGCTCCTTCTCACCCTCCTCACTCAGGGCACAGGTGACGCCTCCAGGGAAGGGGCCTTGGGGACCTCTGGGCTGATCTTTGGTCTCCTGCTCCTCAGACTCACCTGGGCCCAGCACTGACTCACCAGAGTGTATTTCTCCCTCTTTCCAGGGTCCTGGGCCCAGTCTGTTCTGACTCAGCCTCGCTC**AGTGTCCAGGTCTCCTGGACAGT**AGGTTACTATCTTCTGCACTGGAACCAGCAGTGACATTGGGGGTTATGACCTTGTCTCCTGGTGCCAGTAGCACCCAGGCAAAGCCCCCAAACTCATGATTTATGATGTCGGTAATTGGCCCTCAGGGGCCCCTGGTTGCTTCTCTGGCTCCAAGTCTGGCAACACGGCCTCCCTGACCATCTCTGGGCTCCAGGCTGAGGACGAGGCTGATTATTACTGCAGCTCATATGCAGGCAGCTACAATTTCcacagtggtccaagttcatgcgtaat**gagaccaaaacctgccctggtctc**tcaggctctctctcgct**atgaaggtgcttccttaccctgtgcagaagagggcttcat**gcaacatggccttgaga**atttcatccactctcagcccctctcccctccaacaatgaaat**gcaaaagaaacatgctctctggtta**attgtctagggacaat**ggcagcttcttcttttcctgtgtgatatggtcctgcatggtgactctttccaacttttcaatggcagggacagagcaataggatccatctgctcagtaccatttctgcagttttctatgaaaccctcacattaaatgcctccatctcccacactgtgctgactggcctgggtctgtgctccagaaataacgt**cttactcaactccatatccaccacgctgtagtaag**gatggctgtctttgctgggttatttgttcctctgcaggtctcttcccaaagcaacaccaactgagagaagatctgaagatgttgagagagtgagc**ataaaagtaaagggttttat**ttatttgctttcttttttttttaactttgtttttatttatttttttgatacagagttttgctcttgttgcccaggctggagtgcaat**ggtgcaatctcagctcactgcacc**ctctgcctcccgggttcaagcgattctcctgcctcagcctcctgag**tagctgggactacaggcacgcgccaccatgcccagcta**atgtagagatggggtttcaccatgttggcctggctggtcttgaactcctggcctcaggtgatctgcccacctcagcctcccgaagtgctggcattacagacatgagctaccatgcctggccttgtttttattttttagagttcctttgtctttgacccgatgtctatctatgggggcttagccagggcacatttgagtgat**tgttctttgcgagaaca**ttgttctttgcagatcattaaacccctgtcctggaaac

Uppercase: IGLV2-34

Lowercase: Flanking sequence[1000bp]

Red & Bold & Underline: Stem-loop [19]

Blue: Heptamer[40]

Green: Nonamer [3]

id-IGKV4-1[V_gene_segment]

tttagatatggcaactctcattcattttgacaagaaaaatctaaagctcataaaccctgaatcctatatgcttactctcacaaaaatctctaatgtcctgctgggatttatccacagtttagattagacctggaatacatatggtcatgcaacaatgatcttagaacaggactttaacttggctttaggaactgaggctgagagtaatagaattgatttttttgtgtgtgtgtgaagctcctattataataatgagaatactttgattcactcagttaaagttttcccctgatttattgtgtacatacaatgaaggatcaagaaagagaaatttttaaatggaagcattagccagacaagtttgacctcacagtttt**actaggggatatatcacctagt**tttggatctatttctaacatcttaacattgtgaaaagagtc**ttgggaaactggttaaatcccaa**agaatgctgcaataggaggttggcccttatgagttatttaatatcttgagctgccttcggaaaatgttgctgagcaggcattgaagagtatcgataaaatttattgagaatttgtttattatgattaacagaggtaaaagccag**tatattactgattaatata**ggtaaaaggcagtt**aagaaattgggaatgctttctcttctgctttctt**ctacgatgcacaaggcgtttcacatttatgcccctatgaa**aattactaggctgtcctagtcatt**agatctttcagcagtttgtagttttagagcttctaagttgacttctgtcttttctattcatacaattacacattctgtgatgatatttttggctcttgatttacattgggtactttcacaacccactgctcatgaaatttgcttttgtactcactggttgtttttgcataggcccctccaggccacgaccagctgtttggattttataaacgggccgtttgcattgtgaactgagctacaacaggcaggcaggggcagcaagATGGTGTTGCAGACCCAGGTCTTCATTTCTCTGTTGCTCTGGATCTCTGGTGAGGAATTAAAAAGTGCCACAGTCTTTTCAGAGTAATATCTGTGTAGAAATAAAAAAAATTAAGATATAG**TTGGAAATAATGACTATTTCCAA**TATGGATCCAATTATCTGCTGACTTATAATACTACTAGAAAGCAAATTTAAATGACATATTTCAATTATATCTGAGACAGCGTGTATAAGTTTATGTATAATCATTGTCCATTACTGACTACAGGTGCCTACGGGGACATCGTGATGACCCAGTCTCCAGACTCCCTGGCTGTGTCTCTGGGCGAGAGGGCCACCATCAACTGCAAGTCCAGCCAGAGTGTTTTATACAGCTCCAACAATAAGAACTACTTAGCTTGGTACCAGCAGAAACCAGGACAGCCTCCTAAGCTGCTCATTTACTGGGCATCTACCCGGGAATCCGGG**GTCCCTGACCGATTCAGTGGCAGCGGGTCTGGGAC**AGATTTCACTCTCACCATCAGCAGCCTGCAGGCTGAAGATGTGGCAGTTTATTACTGTCAGCAATATTATAGTACTCCTcccacagtgcttcagcctcgaacacaaacctcctccccatacgctgggccagtaggtctttgctgcagcagctgcttcctctgcacacagc**ccccaacatgcatgcttcctctgtgtgttgggg**aggtcac**tctcttgatttattcgttggagggtttgcagggcccaggattaaattaagaga**cttgac**ttttgctggatctctttttgtagaagattattaaagcaaaa**tgttgtaaagatcccttagagacattgtcaggagtttttgtgttgcaggaacctgcatgtttcacatggacacatcacatgaccgagccaaatagatttatctttactctctagaacagggtccactcagttttacgcacagatgggtcaatt**ctttcctctatgtgcctccttgactctggaaag**tttctacttggattcctaattctcttttttctttcttcatctgcaatccttcccaatattaaaccttgatccttcctattcatagcctcccacatagaaggtagtgtccctctgtagctttttccttccagagaccctgtttctttccatcatattaatactttcataggcaattttctaattctttattactct**agaaatatgtgacagcatttct**cattgccttttacaatagtttgccaactttcttcttatttatcatttttccatttttaattttcatggtatacagtagaaagattatatatacacacacacacacacacat**atatatatacacatatgtatatatat**ttatggggtacctgagatgtttagatacaggcatgcaatgtgtaataatcacatca**tggagaatggggtctcca**gccccttaagcatttatcctttgtgttacaaacaatccaattacactcttttagttatttaaaaatgtatagttaagttattattgactaaagtcaccctgttgtactagcaaatagtaggacttactcattctttctaagtattttttgtatgg

Uppercase: IGKV4-1

Lowercase: Flanking sequence[1000bp]

Red & Bold & Underline: Stem-loop [14]

Blue: Heptamer[25]

Green: Nonamer [10]

id-IGLV2-14[V_gene_segment]

taatgtatgttattat**atttaaaaggaacttaaat**tcattaat**atttattttaatatgaatcattaaaaaaat**ttgtcataggcaatggggaaggaggaaaagatattaaaatt**attttgaggaaaaaaccaaaat**tgtatttctcgaatgtctgagtgtcttcagtgt**ctcagggttgacagcggtcagctttaggttgtcctgag**gatgtactccattgggggaaagcaagaaagcatgaaacacacatct**ctgtgacctgacacataaatcacag**taaaatggaagtaaatcttttaataaatcaaacactcacaataaatatcactctggtgaacacagaaatgcctgttaacaaactttccctttcccctgctggctcatatccccatggatcacctgtgtgcagagtggactctgtggtccccat**ttctcaggctgagaa**ccagaggctgggaaggacaagtcacctgcttgtgagtggcaggggaaggattgggagcagtggactcggtctcaggactgcggccctcaggaccaatgccacctagcagaggtttccacggggcgggctgccccatttcctgacagtagggacaacctggaggtgggtgccaagcaga**gggcactgcagggtgtgccc**aggcctggagggtcacagagacacaggcaccacacagcagagacactgagggccagaagatca**ctcaggtaagggacttcagcagatctttctcctgag**caaatcaggtacaaaaaaattaagatcctgccatcagaatagataaacaaggactttaatctcagctgagctccactatccaaggaagcagaagcctctgagcccaggcccaggtgagggtggggtgaagggaggagctcaaggtggagatttgcatgcaggccccacc**ctcctctttggcagaggag**ataagacagggctgggggcaggcccagtgctggggtctcaggaggcagcgctctcaggacatctccaccATGGCCTGGGCTCTGCTGCTCCTCACCCTCCTCACTCAGGGCACAGGTGACGCCTCCAGGGAAGGGGCTTCAGGGACCTCTGGGCTGATCCTTGGTCTCCTGCTCCTCAGGCTCACCGGGGCCCAGCACTGACTCACTGGCATGTGTTTCTCCCTCTTTCCAGGGTCCTGGGCCCAGTCTGCCCTGACTCAGCCTGCCTCCGTGTCTGGGTCTCCTGGACAGTCGATCACCATCTCCTGCACTGGAACCAGCAGTGAC**GTTGGTGGTTATAACTATGTCTCCTGGTACCAAC**AGCACCCAGGCAAAGCCCCCAAACTCATGATTTATGAGGTCAGTAATCGGCCCTCAGGGGTTTCTAATCGCTTCTCTGGCTCCAAGTCTGGCAACACGGCCTCCCTGACCATCTCTGGGCTCCAGGCTGAGGACGAGGCTGATTATTACTGCAGCTCATATACAAGCAGCAGCACTCTCcacagtggtcca**agttcctggggaactgagaccaaaacctgccctgggctctcag**gccccctccttgctctgaagatgcttcctcacccagtgcaagcggcttcctgcagtgcggccttgagaattcttctctctcagctccttccctttccaccatgaattccaaaag**gaaacccgctctgtggtttc**tcatccagga**cagggacagcttcctgatgcttgtgtgccgcggtccctg**aatgtgcaactcttgttagctcttcaaatgcaggga**cagtgacaaggagcttcctgattgatgcagtcactg**cttttttcagggatgtcttcaccctacatgcatcaccatctcccacactgtgggtagaattttagcaactacattctgatggttatcgccacaactttgatcttaaataacaatgcagcgaacatccctgtgcagcctcctttgagt**tcctgtgtgaatatgaccacagga**ttcatttctaaaagtgaaattgagggtcagaaagatgtgtgtgtgtaattttcacccgttgttgtcagattcccctccagagggattctaccaattttcaatgccagccggaagtacttgttcagagtacattgttgaaagtggaagtttttgccaacctgtc**agactcaaaaatgatacctgttatggttttatttttgagtct**cctgttctgcttaagtttgggccttttc**agaaatagttcagtactttttcttttttaggatttgggaaactttttcttaaatatccagataagaa**gaattttaatcttcgtggccggacaccgtggctcacgcctgtaatcccaagtgtgttggaagggtaggatgggtcaattgcttgctcccaggagttgagattagtccgggccacacagtgaactctcatctctattaaattaaagaacataaaaataaattttaaaaggaagtattttagtctttgtggctcaagaatcaactttggggatattccata

Uppercase: IGLV2-14

Lowercase: Flanking sequence[1000bp]

Red & Bold & Underline: Stem-loop [19]

Blue: Heptamer[46]

Green: Nonamer [3]

id-IGHVII-49-1[V_gene_segment]

cacaacagtgctggagaaggtgggagtc**tgatggagcttcccaacaaccctgtggtctaagctaagtccatca**aggccgctggttcctcctggaacacagctgtccatcaggaatcccccatgtgcccagcagcagccacgcgttagcatcttcactgtgcacagtcattgtttgggaggagctccaggatgggtctctttgtcaaaaaccgggtgatggt**gtcagagagctgtgccgggtgcctggtccaggggctccctgac**gttggaggtataggaggtgcgttctcagggtgggaacaccttttatggaaatgtacagaagaaaacatgattttccttggtcctctcagatgacatggagaa**gcaagcgcgcagcttgc**aaacaattgtaatgtccttgtaatttcctactttatcccaagttcattgattcttgtaaaccttggtttctgtccaagtacagagatattgattagagtgagtttggttctttccacacactaaccctca**cctcccccagataaagagcacattgtccttactctgagtctgagggagg**agctgttc**ctgtacaactaagggcctgcagagacccccgtgtacag**ctttgcagagtcagatctttccacatgtgaggcgacctccacggcccatgattgctgctcaggtctaattgtggattcaggattaggacatcctttaggctatcacaggtcaatcagattctgaaaaatcactgttatcatagacagaggtaataattcaatacccactcccctgacagcagattctctttcttatttggttgaaagtttg**aagaaaataagtgaccatttattattctt**attctaactttagtatccact**atttgttctaggggattcacaaat**ttcaaaacactgagctcttcattctcatgaaaatgtgcatatttgtgaatttcaacgtgttgttcagaacctgtgcatgccgacatctgtatttctcgtgcgttcttGGCCTGGCGAAGCCCT**CACAGACCCTCTCCCTCATCTGTG**CTGTCTCTGCTTCTCCATCACAACCAGTGCTTCCTGCT**GGAGCTGGATCCCTCAGCTCC**CCAGGGAAGGGACTGGAGTGAATCAGGTGCACAGGTCATGAGGGAGAACACAACGCAACCCACGCCTCAAGAGTCCAGTCACCATCTCCAGATCCACATCCAAAACACAGTTTCTTCTACAGCTGAGCTACCTGAGCAACGAGTACACAACCATGAATTTTTACACAAAAGAtacagcaaggggaagtcattgtgagcccagaaacaaa**cctccctgcagggaagctcaggaactgcgggagg**ccctcgggacaccagggggcgctcaggacacacatcaaggcaggtgcaagaggaaaaggtgctggagatggggtttggcatcatcatcatatttcactgacacccgccactgtgtttattctcatgtacgtgattctttgtattattagaaaatggcgtttatgtaaataaataaccatatgtaggtgcatcaagtcgtcctctcccttttttttttttttttttttttttgagtcctgctctgtcaccc**aggctggagtgcagtggcgcgatctcggctcactccaacct**ccgcctcccgagttcaagcgattctcctgactcagcctcccgagtagctggaattacaggtgtgcaccaccgcgcctggctaagttttgtatttttagtagagaccggatttcgccatattggccaggctggcctcaaactcatgatctcagttgatccgcccgccttggcctcccacagtgctgggattacaggtgtgagccaccgcgcccggcccgtcctctccatcttatgtggaacttttccattaagactctgagtccctg**gatttatttgagaacctcataaatc**atggtcaattatgtctctttctcctctttctcccttcttcactcctctctagctcacaaactgaaacacacacagaattttctaa**cttcaattacctgatgtgttgaag**acaattgattaaagcgcagcttttccagttcagccactgttcatattgttgtaagaataagaacgttgtgtttctc**agctgcgtgcttctctaagatgagtagcagct**tttttttaataatacccagaggctgaaagcaatctgaataccaatcatcagcttaatcagtaaacacaaggtggtaaattcgctcactggaaaaacaccgactgttacaatcaggtaaggctggatactc

Uppercase: IGHVII-49-1

Lowercase: Flanking sequence[1000bp]

Red & Bold & Underline: Stem-loop [14]

Blue: Heptamer[37]

Green: Nonamer [7]

id-IGLV3-4[V_gene_segment]

tctcctgcactggaaccagca**gtgatgttgggagttatgactatgtctcctggtaccaacagcac**ccaggcacagtccccaaacccatgatctacaatgtcaatactcagccctcaggggtccctgatcgtttctctggctccaagtctggcaatacggcctccatgaccatctctggactccaggctgaggacgaggctgattattagtgctgctcatatacaagcagtgccacttaaccacagtggtccaagttcttggggaa**ctgagacgaaaacctgccctgggctctcag**gctccctttttgctctgaagatgtttcctcacccagtgcaacgggcttcctgaagcacagccttgagaattcttctccctcagcaactctcttttcccaccatgaaatccaaag**gaaacctgctctgtggtttc**tcatccagga**cagggacagcttccttttgcttgtgtgttgtggtccctg**agtgggtgcaactcttcctagctttttaaattatgggagggtgacaatgagctccctgactggtgcagtccctgctgttttcaggaacatcctcatcctaaatgcatctgaatctcccactgtgtgcagaccaa**tctggacagatgttattagggggagtttccaga**agccacatcttactcaactctgtatccaccacactctggtgaagatgcttttttggtgcccatcctcttgggtaaagtccactctctactcatgacagccatgtgtcccatcttctgaacttgggaaattgccacaccaaaccctcaggctcagggcccacataggagctcagaaagacgcactgggaaatggagaagagatctgcatccacccctcagtggagagacacctaactagataataagataaggaagtaggtttgcatgatgagtctccccttcttatgataagagagacctagagacccccttctcaggtgtgcgc**tcagaggcagaactctga**agcatctccaccATGGCCTGGACCCCTTTCCTGCTCACCCTGCTCACTCTCTGCACA**GGTGCTGCTTCCCAGGGCTCAACCCCCACAGCACC**AGGGAACAGCCTGGCCCTGACTTTCAGCT**CATCACGGCAGTGATG**CAGGGTATGGCACTCTGAGAATGAGACCCTCATCTTCAGACTCACCTCTCCTGTCCTCGCTTGCAGGTTTTGTGGCCTTCTCTAAGCTGACTCAGCTGCCTGAAGTGTCTGTGGCTTTGGGACAGATGGCCAGGATCACCTGCCATGGAGAGGACTTAGAAAGCTATTATGCACACTGGTACCAACACAAGTCAGTCCAAACCCCTAATC**TCATTGTATACAATGA**TAGAAAGCTGT**GCTCAGAGATCACTGAGC**GATTCTCTGGTTCCTGCTCAGGGGGAACAGCCACACTGACCATTACTGGGGCTCACGTTGAAGACGAGGCTATTTT**TGTTTTTCTGGAGATAAAAACA**CATTcatagtgacacag**acagatagggaagtgaggcacaaaccccatcaccatctgt**atcactctctttcttaagccccaggagtattgtggacaaagtcgtgagcagggttagcccagctcag**ctggatgtgtgaccccgaggcacctttcactctagtcatccag**gaaggctctacagagggtgggtcaggagaaaatttggggctgaaaactaagctgtcctggtgttgtttagctaggtatgaattggcttaagatgacc**tgactggtaggacagtca**cagcaagataaccattgagtaatcatgttgcctggatttccttttgtttaatgaccaaacctaaggtagctctctggtccaaaatgccctggattagtcccctggactcctcattctttaactgagacctttgatcccagctgtcatggcatcatgatgtcgtgaacagggaagtccattgtaatggcaactgtgacatcaggggtcatttacccactatctac**ccaaggccgcagggaatccttgg**cttctcacagctttttctcactggaaaatgccattgagtgcagagtactgtgtcttccaaggatgtgccccctccataatgtggtttgggagcaatgattgcctgatgcctagattccaaagccatgctttaattcaggatgaccttgaagtgcctcctgctgagtgctcaattgtaacacat**tcccagaagagcctcccacacttattttctggga**ctatatacccgaaccctactgcatggaattcaacctctcagagtaactttccaagaaacccaaaaattaaaaactaaacaacaacaagaaaacaatatagagagaagagtatcaaagaaaagcaagttgtacaaagagagaactctggagtcttcaaatattaataagcatcaagcagcacatgtgtggacggaaactccattttcagacaaacagcagtccaaagacactcgcgggaacactccacagagcattctt

Uppercase: IGLV3-4

Lowercase: Flanking sequence[1000bp]

Red & Bold & Underline: Stem-loop [16]

Blue: Heptamer[35]

Green: Nonamer [4]

id-IGLV9-49[V_gene_segment]

aatgatagctcactgcatccttgaactcttgggtctaggcaagcctcctgcctcagactcctgagtagctgggactacaggtgcaagccaccatgccaggcaattctatcacattgtgaatatacttaacaccactgaattttacagttaaaagtgatttaaatggcaaattttatgttgt**gtgtattctatgacaattaaaaatacac**ttgttcaaatgtcaa**aatatatatgatgttatatt**cataaaattttccttttcacacttatatatttatcctattctaactcattctcatcaattaaaaaaagtactgtaattactggtc**attatgtactaaattttccataat**tcacattgaagccttaacccccaatattactgtatttggagacagagcattacagagggtaattaagatgaaatgagatc**ataaggaccggcatccttat**aagaggagattagaaaatggacaacataaaatg**agatgaggacagagtgagaaggtggtcatct**acaagtgtggacacatcttcagtttgtgtggagtaacagttgaagggct**cgcccatcctataatgaaagttggaacaggtgccttgggcg**ggctccaggattaactgatctagtcactgcatcccactgtgggaccccagagggaaagaagcactgtggagattcctgagacattagaaa**tggagcctgagaaatttccaggacacccacatgggctcca**ggtgaagcaggggcccagaaggccttgtctgatccctgagtgtgacgcagcacctggccactagggggaggcaaactctgtgaatcctaagggacctgagctcttgctctgagccctgcccttgttaatcaccagccctgggagggagccgatttgcatggaatgtgatgtccctgcccacttcttca**ctgagggaataagaggctttagggcctcag**gctcagctgagagactgaagaacccagcattgcagcagctccaccATGGCCTGGGCTCCTCTGCTCCT**CACCCTCCTCAGTCTCCTCACAGGTCAGGGTG**GGCAGTG**GGCTGGGCCCCCAAAGGGACCCCCACCTCCCAGCC**TCCATCTCCCCATCCCTGCTCTTCCTCCTCCAACAGCTCAT**CAGCCACCCACCAACAGGAGCCCTCATGGGTGTCTG**TGTTTCCAGGGTCCCTCTCCCAGCCTGTGCTGACTCAGCCACCTTCTGCATCAGCCTCCCTGGGAGCCTCGGTCACACTCACCTGCACCCTGAGCAGCGGCTACAGTAATTATAAAGTGGACTGGTACCAGCAGAGACCAGGGAAGGGCCCCCGGTTTGTGATGCGAGTGGGCACTGGTGGGATTGTGGGATCCAAGGGGGATGGCATCCCTGATCGCTTCTCAGTCTTGGGCTCAGGCCTGAATCGGTACCTGACCATCAAGAACATCCAGGAAGAGGATGAGAGTGACTA**CCACTGTGGGGCAGACCATGGCAGTGG**GAGCAACTTCGTGTAACCcacagtgacacaggcagaggggaagtgagacaaaaacctcccggctgatctgctctgtccacacttccttcccactggcctgactcacatttcagagggctgtctcttaactctgtatttaaagttttcaccttcccagacattttggagttctaattcacagcatatgttccctgtgtgtaaatattttctcaagagcttttgaaacattgaagtttgcacaccaacatttgtggtttttgccactcacctgtcccctgatgggattgaagtttccaggttgtggaactgctcacaggtgcgtttatcaggaaatcaccgtcaccagctcccacgacctttgtccatattcaacatttgtgagctgcctattcaatgatgggaactgacaatttccctggagacacaaggccttcaccctgactcacagtgtcacataatcgacgtggacttcagccatctccagtcacctccatgtgcacagagctctcatggtacatgtgcttggctgccctagaaagagagaatggacacacaccaaagatagtagaatgtgagcctgctctatatctgag**attctgaaagaggcccccactgccttcacagaat**gaattccttcacatgcagggagagagctctggtggaatcactctgcaatgctttctgatgacaaatacctggattgaggtcaaatttcacaggttaataacccttttcaagactacctccactacagactccagttcccaggcattaagcctgaggacccttgtagacaaccaacatggggctgtgaccagcaggagcaatgggtaactgtcccaggccctgcct**tggggctgctcacagccgggtgagcccca**ctggccacaggctggcagaggctga**gctgctccctgtgctcgctgatgaaactcagcagc**ctagtgctctcagggtccagcaaagcctcttgggcagggtcctgtgcctgttcccaacagccgctccttcgaggacc

Uppercase: IGLV9-49

Lowercase: Flanking sequence[1000bp]

Red & Bold & Underline: Stem-loop [15]

Blue: Heptamer[47]

Green: Nonamer [8]

id-IGHV3-63-2[V_gene_segment]

tgttagagaccagttctttacactatggcatgtaataaattccaaatggcagtacgtcattgtttacttagcttttgtgcttata**tttttcagaggaaaaa**a**aatactgtaaattgtaaatacccaatacctaacagtatt**gtatgcaa**atctgtgactgttggcagtgtcatctctgagaaacagat**aaagtttatttactatatataaaaaaagagtttggaaggtggactcctcaccaatttttgaaagagttagagaagcattagcattaattct**ttaaatgttaagactaattttatgatgtaacatataa**cctatcatggagaatgttcaatgggtgcttgagaaggatgtgtattacatggctcttggttggaaggttctgtaaatgtctttcaggtaaatttgttcaatactgttgttcaagttcagaggcttattaagaattttctttctggatttgctatacattattctaagtgaggtattaaggttttcccttatttt**tatatagttttctatttctctctatata**tatatcttaaagtttgcttaatgcattta**tatttgtatttgtactgtgtaaaaagtaaaataaaata**ataattgcttagtgagtttcatggcacaatcacgttatgaataatcatattttcccaaatgctgtcattgccactaactcctccagatacatggtatattatcca**attctgtgagttggaattatttcgttgctttgcagaat**ctttttttgttaatctagttccacttgttcaattatgctcctccgggagtctcacatctgctctgggcactgccttcttctcaggcatcccacactggagcttgcta**tagaggaggaggcatgaaaacagggccctccctcta**ctggtgaaaacaag**cccagacctggccctgcagctctggg**agaagagccacagcactgggattcccaggggtttccatgtggtaatcagggctgaacacag**agctcactATGGGGTTTGAGCT**AACCAGAATTTTTCTTGTTGCTATTTTAAAAGGTGACTCATAGAGAAATAGAGTGAGTGAGAGTGAGTGGATATAAGTGAGAAAAACAGTAGATGTGTTTGGCAGTTTCTGACCAGGACGTTTGTGTATTTTCAGGTGTTCAGTGTGAGGTGGAGCTGATAGAGTCCATAGAGGGCCTGAGACAACTTGGGAAGTTCCTGAGACTCTCCTGTGTAGCCTCTGGATTCACC**TTCAGTAGCTACTGAA**TGAGCTGGGTCAATGAGACTCTAGGGAAGGGGCTGGAGGGAGTAATAGATGTAAAATATGATGGAAGTCAGATATACCATGCAGACTCTGTGAAGGGCAGATTCACCATCTCCAAAGACAATGCTAAGAACTCACCGTATCTGCAAACGAACAGTCTGAGAGCTGAGGACATGACCATGCATGGCTGTACATAAGGTTCCAAGTG**AGGaaacatcggtgtgagtccagacacaaaatttcct**gcagaaagaagaaaggattctgggccgaaggggacactcagcactcacaaaacaggtggagccccagggcaggtacaga**aaagcagtcaagggctgctgtccttcaggatctgtgcttt**cctctgcatgtagcagttcccctcgaatcctctgcacttttatgtttctgtgcccaccatgaggtccctggattacaaaactttaatttaaaagaggaaacattcttatatgtcccaaaaacaaaagtaagtattagaggcacaaaagtgcacaggcggccaggtgaggctgtagacactgccaccccgc**aatgccagtctcacaactagcactggagagtagtgggcatt**cagtggagcttcctacctatcctgtggtccgagctaagtccagcaaggccattggtgccttccagagcacagtagtccatcaggaatctcccatgtgtcccagcagcagccttgcctcagtatctccactgtgcacagccattgt**ctgggaggagctcccag**gatgagtgtctttggcacacaccaggtggcgggtgttagagtgcagtacagcagctggctgcctggtctattgggctccctgatgttggagggatttgaggtgcattctcaggcccagcaccctctttgtgaatttttatataaaaaccctgattttacttcattttctcagatgacatagataactaaaaacagaatctgcaaagaaattgtaattttcaactttaccccaaattcattgtttcttaattctgtgcaagatccaaacatatttttgccttccac**atgagaaattgttctgtttaaactgaaatcattttttctcat**attctttctttttgcccaagtacagagatcttgttgaaaataagttgggt**tctttccacacactaaccctcacctcccctagagaaaga**gcagagattgtcctcactctgagtctaggggaggagct

Uppercase: IGHV3-63-2

Lowercase: Flanking sequence[1000bp]

Red & Bold & Underline: Stem-loop [17]

Blue: Heptamer[31]

Green: Nonamer [4]

id-IGHV3-42-2[V_gene_segment]

tataagatggtatctcactgtgtctttaaattgcatttttctgatgattaatgtcattgattattttttcatatgctggttgatg**atttttttgtcttttgaaaaat**aaacattgtagtaatttgctcatttttaatgtggttatttgtgggtttttgtgttgcttttgtttcttgtagtttgtaaatattagctctttgtcaga**tacagagtttaaaaatagtttatcccattctgta**cgctttcggttaagttttttgattatttgttttgctgttcagatgctttgcttcttatttgtattaagtccaaatggtctatttcggtttttattgtttacttttaaggttgtagtcatgaattctttgc**ctaggacaatctccagaacaatatttcctag**aatagcatctgcaactttcagactctcaggtctcccagttaagtcttt**tatccatattggcttaattttggata**tagtgagagatacgggtccagttttattctgctgcaaatggctgttcagtttttcctgcacaatttatataacaaggtgacctgttcccagtgtatgcttcttgtctagtttttcacagtcagtttggctgcaggcatttgactttatttatgaaatgtctattctgttccactcatctatggctttctgctagattgttacttgattgttacttgtgtataataatgctactggtttttgtatgcttttttcatttattctaaaactttactgaattaactcatcaattctaggagt**attttgaaagaatattagtttttttaagtacaaaat**catattatcagcaaacac**aaagagtttgacttcctcttt**tccaatttgagtgcctttatttctttctcttgcctaatttctctggctaggatttcaagttccatattgattaagagtggtgaaagtgggcatccttgacgtgttctgagtcttagg**aggaatactttcaacatattcct**attcagtataatgTTGTTTCGGAGTTTGAAGGTGATTTATGGAGAATGAGAGATGTTGAGTGCGAGTGGACATGAGTGAG**AGAAACAGTAGATATGTGTGGCCGTTTCT**GACCAGGGTGTCTCTGTGTTTGCAGGCGTCCAGCGTGAGGCGCAGCTGGTG**GAGTCTGGGGGAGGCTTGGTACAGCCTGGGTGGGTCCCCGAGACTC**TCATTTGCAGCTTCTAGATTCACCTTCAGTGACTTCTGAATGCACTGGATCCGCCAGGCTTCTGGGAAAGGGCTGGAGTGGGTTGGCCGTATTAGAACCAAACGTAACAGTTACACGACAGAATGCGCTGCATCTGTGAAAGGCAGGTTCACCATCTCAAGAGATGATTCAAAGAACACACTGTATCTGCAAGTGAATACCCTGAAAACCG**AGTACACGGCCATCTATTACTGTACT**AGAGAcagtgagggggaggttaacgtaggcccatacacaaatct**ccctgcaggggcgcgcaggg**ccaactgggggcgctc**gggacccactgaggatgggacaggtccc**aggggcgggtgcagggggaggtttcctttctcagctgcaggaggcgggtttgtttttgcaggaatatggagtcttatgaggttttgatattttactatggttatttatcatgattttttaaaattgggatttgtgttttag**taatttttaaatttatatgtaggggtatttttaaaaatta**agttttagggt**acatgtgcacaacgtgcaggtttgttacatatgtatacatgtgccatgt**tggtgtgctgcacccattaactcgtcacttaa**cattaggtatatctcctaatg**ctatccctcccccctccccccacctcacaacaggccccggtgtgtgttccccttcctgtgtccatgtgttctcattgttcaattcccacctatgagtgagagcatgcggtgtttagttttttgtgattgcaatagtttgctgagaatgatggtttccagcttcatc**catgtccctacaaaggacatg**aacttatcattttttatggctgcatagtattccatggtgtatatgtgccacattttcttagtccagtctatcattgttggacatttgggttggttccaagtttttgc**tattgtgaatagtgccacaata**aacatatgtgtgcatgtgtctttatagcagcatgatttataatcctttgggtatat**acccagtaatgggattgctgggt**caaatggttatttctagttcaagatccc**tgtggaatcgccacactgacttccaca**atggttgaactagtttatagtcccaccaacagtgtaaaagtgttccctatttctccacatcctctccagcacctgttgtttcctgaccttttaatgattgccattctaactggtgtgagatggtatctcattgtggttttgatttgcatttct

Uppercase: IGHV3-42-2

Lowercase: Flanking sequence[1000bp]

Red & Bold & Underline: Stem-loop [19]

Blue: Heptamer[33]

Green: Nonamer [9]

id-IGKV2-29[V_gene_segment]

aaatttaaatagtgaaatatgatacatttattctaatcctacttccacactacaatttgatattctcaatttaggttactgccatgacgttcaacgttttaaggtcaatatcatttttactcaatttttttcaccaactttgttaaaacatcatatgccagtgcatttcccggaagataatcaggacaagctttgtgt**gcaaaacacctagagctctgttttgc**ttttctgggtgtgtgttctttaccaagtaaatcagatgttgtattaaatcactgaagattacatagcactagaaactattcttttaattgctcccaggatttttgttgtttttttcaggacagtggcagtctctgtgaagtgatttgatgtacgcaccgttattcaggagcagctaaattcaatattaagcacatcactgacaaatgaagaatactcaagttgttttcctattgtaacgatgagttgtgtggtagattcatatgcctgacctgtgatcatctttcaataaaaacctaggaaaatgtcgtccatagtactgctgaccctgagacttaaaaccagggatccagttcttgctcttcctacatagagaatctcattttgaactgtggtgtcatggc**tgtggctgtgccaca**tacaggccagggggagacaagggttcaccctcagagttgacaagaatttggaagccctgacatcctataaaacgttacttgcccaagattgaaactttca**aattcaggtccttcttcctcctctatgatgaatt**aggttttattagtttcctccaagggacactttatatcacattgctcacagagaagacatatctacccccttcaccccccacccaatggctctttccacaccactgcacccaccaggtgatttgcatattgtcccctagggaggaccttcccttgtgagcctgagataaaagctcagctctaaccttgccttgactgatcaggacttctcagttcatcttctcaccATGAGGCTCCCTGCTCAGCTCCTGGGGCTGCTAATGCTCTGGATCCCTGGTAAGGATGGAAGGAGATGAGGGAGGAGGAGGGGGTGGGAGGCTGAGCTCTGGCGGCCCCACTGATTCCCGTGTTTATTCTAACCATGTGTTAAAGGAATATGCCCTATGCTCCAGGGAGAGGAATTCATATTTTGCCCTGATGATGATTTGAAAACTCCTAAAAGCAGTGCTCTGAATAATATCTTGAG**AAATGAAAGAACTCTTGTGCCTATTTAATAAAGGGTTCATTT**AAAA**TTTTGTTTTTATGATATGAATACAAATTTGTAAAAATAAAA**GATTAGCCATAAATCAATATCATAAGGCAAATCTCAAAAGTTGTTCATTATGCTTTCACATAACCTTGCACTTCTCTCTCATAATTTCAGGATCCAGTGCGGATATTGTGATGACCCAGACTCCACTCTCT**CTGTCCGTCACCCCTGGACAG**CCGGCCTCCATCTCCTGCAAGTCTAGTCAGAGCCTCCTGCATAGTGATGGAAAGACCTATTTGTATTGGTACCTGCAGAAGCCAGGCCAGTCTCCACAGCTCCTGATCTATGAAGTTTCCAGCCGGTTCTCTGGAGTGCCAGATAGG**TTCAGTGGCAGCGGGTCAGGGACAGATTTCACACTGAA**AA**TCAGCCGGGTGGAGGCTGA**GGATGTTGGGGTTTATTACTGAATGCAAGGTATACACCTTCCTcccacagtggtacagaccaatacagaaacctccct**gctggggtgtcccagc**tgctcacttgcactgcttgtctggggagtagctcagcagggact**ctgagtctgcagaagaggagactgttggagaactcag**gggcagggct**tgctgttgaggactctggcccatgagagtctcagcagca**cctcagtcccacatggtcaaagctttatcagttgccaggctcatcctcaccctgcctgtcttgcccacgtttgccgagtacactcaatcagcataacagccagaacatagatgcagtttagtgacaacacaggtggaatacatgtggcaaatgaccagcttggggtttattttacacaaattaatacttggtgataatatttggaattattgcgaatttggtattttcccactttccttactttctatttcactttactactcacactaaactgcactttccagtgtcatgtgggagaacgtgttctatacaagctgtcctcagggggagtatgga**taagaataattagaaaaaaaatgtttgatttctgactattctta**gactttgtaaatccatggcagatat**aaatcttaatgccaaaagattt**tgatttgcctcaattacgtttttcccagtgaagag**ggagttcttgcaactcc**aaaagtgagcttcagaaataagcagcactggccgggtacagtggctcacgcctgtcattccagcactttgggaggccgaggtggatggatcacctga**ggtcaggagttcgagaccagcctgacc**aacatggtgaaacccggtctctactgaaaatacaaaattagccagggcgtggtggcacacacctgtaatcccggctagttgggaggccgatgcaggaaaatcacttgaacctgggaggtggaggttgcagtga**gctggagatagtgccactgcactccagc**ctgggctacaaggcgaaattctgtcaaaaaaaaaaaaaaaaaaagcagcataaac

Uppercase: IGKV2-29

Lowercase: Flanking sequence[1000bp]

Red & Bold & Underline: Stem-loop [16]

Blue: Heptamer[25]

Green: Nonamer [8]

id-TRAV38-2DV8[V_gene_segment]

aaatgaaactagtagttctctgtattgtgagtttt**catttttgctctgttgcctcttctataaaatg**cagactaattcctgacc**tttcctccctcaaagaagaatgtgtgcatgaggaaa**aaatcagatgccaatgaaaagctgtcagatgtgcaacacagaatgttcccaaatgaattttatgttgattgttctgagtcttctttcacaaagtcattaacagaagctcacaaaagcccttgccaccagtattgtagagctccttgacacagccaaaatcaggta**ttctttttttaaaaaagaa**agtcttcagcaaattacagtataattagatatcaaaagtctggtttcaattttctctagactatctgaatgttattgttgttgtcattatttttcatcatttctcaaagtattaaaatacaatttggtcaaacaataatataaccaatacctgt**attatcataacccagagataat**tttctgatattggcttcaagacagaggagggaaagaaaaaaaggaaagaaggaaggaagaaaaaaaaaaggaaaaaagaggagtaaggggaagggaaagagaaggaaagacgggaaggaaagagagagaaagagagagagaaaggaaagacaataaaagcttatagagaaaactgaagtcacagcattagcttggcctgtcttttccccacccatctcattctcctatccaaattatgtctttaaaaaacaatgtatcagctccctttcctcttcatg**tttattcccaaataaa**ttacagttccaacaacatcttttttcgttaaatattatggtccaagttttattctctgaagcaaaagacaaaaagtatcgaagagctttccctaagggagacatcataaacagcccacctgacatcccatttcctgttctgaagcagctacggcaccagtgcagctgatactcaaggttcagatcagaagaggaggcttctcaccctgcagcagggacctgtgagcATGGCATGCCCTGGCTTCCTGTGGGCACTTGTGATCTCCACCTGTCTTGGTAAGGAAGGCGCATGTATTCACCTTAACTAGTATTCAATAGAACAGAGAAGTGGGAAGAA**AGATTAATAGTGCAAGAATCTAATCT**CTTCACTGGGCCCAGCTATTAGGGGCAAGAAAGCTGATGGATTTTGTCAGAGGTTTGGATCAAGAGGAAGAAAGGCGGGTGGCAATGGGAACAGGCT**TGTGAGTCTGTTTCTGTCTCTCACA**TGGTATCCCAGTGAAGTCCTTTCTCTCCTCTTTCCCACAGAATTTAGCATGGCT**CAGACAGTCACTCAGTCTCAACCAGAGATGTCTGTGCAGGAGGCAGAGACCGTGACCCTGAGCTGCACA**TATGACACCAGTGAGAGTGATTATTATTTATTCTGGTACAAGCAGCCTCCCAGCAGGCAGATGATTCTCGTTATTCGCCAAGAAGCTTATAAGCAACAGAATGCA**ACAGAGAATCGTTTCTCTGT**GAACTTCCAGAAAGCAGCCAAATCCTTCAGTCTCAAGATCTCAGACTCACAGCTGGGGGATGCCGCGATGTATTT**CTGTGCTTATAGGAGCGcacag**tgagacaagcaacagggagaggcttacagaaacctcagacctcagcatctgtgcaaaggtcagggggaagtgttcagggttataggaacataaagaggaaataattgtcttccctaaagaaaaaaaaatgtatctgactataggaaacacagaggggatgtctataagctgtaatttttgaaatagtaataaaataatgaatccacacatgccttatagattttcaaagtaccttcacatgttaatccattaaacctgcataagcctgtgtgttacataggtcagatcagatat**tctaaggatcactgctatccagagaatcttaga**gcttgacc**taaagctatattcttttagaattatatatttttgcttta**ttattttaattactatctatctgtacatcccatgatcatatcaaagtcaatatatccaaaactgaatttcttaccatctccacaaacctttattttttcttcatttttttctcctcagccttgctcaagattcaggaacacaataaaagtttgctattgctgggtttccagtttggggaatttcttcaccctcgactttccagcctgggctatcaga**tttttatgcataccttattacattctaattataaaatggtgcaaaaaaa**aacactacatcatgtcagtctcactgtttcccctctatccgtcattctttctgactttaacctggcttaatcctgtatttcttaacgttctttaaagcatcatttgt**cttgacataagaaagtaaggaaagaaagcaaacgactgtcaag**taggatttcatgtactgctagtactcaccagcaaacgctaaaaaaaaaaaaaaaagaggaagttaacttctggtaatgctataattgcttctggtaaggcaacctagcaggtggtgacaattttgataaacctggtccaacag**ggagaccaagcagctgccatggtccaagtctcc**acccctatgtcttctgagaattgggagat

Uppercase: TRAV38-2DV8

Lowercase: Flanking sequence[1000bp]

Red & Bold & Underline: Stem-loop [16]

Blue: Heptamer[23]

Green: Nonamer [9]

id-TRAV9-2[V_gene_segment]

accatcagggaagtttttaaaaaatgaaaccctttgaatcttggaaattgtggtttgaaatgttggaaatcgtgattcctaaagttagtagtcttgtcccaacatctgtaattttaaatgactcacaagttaattcagttctcacacaaccaggttttagaacctcgattatcagccgatgtatctttatcaatttgtaaagcaagagaagaggaaagttctaactcatgcttcag**tctgttgatgcaaatctacatgtggattccctacatcatcaga**ggaaattgattaaactttggcaataaaatgttggcatatgaccacagtagaaagcgtccttgaggccagatctccagattatagggttaacacttgttgcctgattgtcataagcaagcaacttgtcttggagtcattgaaatcgtgagggaataagctattaatctgaggtcatatcttatcagtaaatgtcagtcactgctctacatgttaggatacagtgctgaaaaacatagactcagtaacagcaagacttcttaacattccctctgatttcaactcaaaacccttgtctgacatc**ccattgaattataccactggcctgcaatggatacgtaaaaatattggcctgtattaaaatatcca**agaaaatggaactctcacctacagaaatactctaggatgctttgctaaagatc**ttttgtttcaggtagtgtccaaagaacaaaa**ttgacaaaatagaccatagagatgacaacatctgcccttcat**atttacaaggaccacctgagactggcatgggccttataaat**gaggtcagggttgaggacctatttctgacttttggaaggagtccaggaaaacattcatcccagttcagcccctgtccaccagagggagtggcttcctaacacaaactcatttcctgctatcactgtaggtttcactgtgatttcttcatgttaaggatcaagaccattatttgggtaacacactaaagATGAACTATTCTCCAGGCTT**AGTATCTCTGATACT**CTTACTGCTTGGTTAGTGAGATGCCTTTTAAAATTTGGTCTCTGTCTTCTATCAAGCCAAAAAAATCTTTAAC**CATAATTTGACCTAAGAGCATTATG**CTCAAGGCTAAACAGATCTCTCTCTCTCTCTTTTTTTTTTTTTCCCTAGGAAGAACCCGTGGAGATTCAGTGACCCAGAT**GGAAGGGCCAGTGACTCTCTCAGAAGAGGCCTTCC**TGACTATAAACTGCACGTACACAGCCA**CAGGATACCCTTCCCTTTTCTGGTATGTCCAATATCCTG**GAGAAGGTCTACAGCTCCTCCTGAAAGCCACGAAGGCTGATGACAAGGGAAGCAACAAAGGTTTTGAAGCCACATACCGT**AAAGAAACCACTTCTTT**C**CACTTGGAGAAAGGCTCAGTTCAAGTG**TCAG**ACTCAGCGGTGTACTTCTGTGCTCTGAGT**GAcacagtgacagggactgcagggggagctgagcacaaactctg**agcagcacggggggcccggctgct**gagtgtcagcc**actgtgatccactctggttctagcacagt**ctgtggttgtttgtgcctcagtttctctgattgatgcagtaatcatacacatggccagaggagaaaaacaagaaatgaacatccccaaacccagctcttggttagtgaaattaaaaacagacattgacaccaaatagttcctcatccagtgccaaagtaatttcaaggttaaac**cacagaaacaatgaagctgtctctgtg**atcatccttgcagacaaagttcaaccc**ccacctcttgttccaggtcactctggtagatattggaggtgg**atgtgttctattacatttt**tacagagccaaatatgccagagaaggtggcgtctgta**caatgtctctgtaacagagtacagcaggagagatgacgtgagagccacagctcctcaattcagttcaaatgtctatgcaggattcttaatagagaaagaagggaagagaagaaaggaggagggacagaatgtcacatgtaaaaaaataatacggagaagaaagcaacaattgtcagagagagggaagttttcttttttggtaagccagatggtagacaaaggctgatgttccaaaaagaaagggaaggatcaaaatcattattgaagaaatgactcatttgtaaatcataagtattc**cttgcttgttacattgatcaattggctttctaacaacaag**tccttaagtgagttagttttgcagtgccaatggaagattaagcatacagatttatatttgtatccctgtgaaatcctattaaagacacagtaaagaaattgaaaagggcataaaaccacaaaaagaatagaaaaggaggtgacagcagatggtgataccaatacaattttagaaggcaggagtgtatggagaaatgaaaacttgcatagcagaccagataaagaagaaacc

Uppercase: TRAV9-2

Lowercase: Flanking sequence[1000bp]

Red & Bold & Underline: Stem-loop [18]

Blue: Heptamer[32]

Green: Nonamer [3]

id-TRAV30[V_gene_segment]

tcatttgatacctaatatgtgtcttagacttgaaaacaatagatgcaaatgagaagaat**aaggcattgtttctgcctt**cagggagca**tgtaatattgcagagtgtattaca**atatagtataataaaagctatcataggagaagcgtagaatataaccacctaacagtgg**catttaatttggaaacttaaatg**taagggaaagctgcctacgggaagtagtatctaggttaaga**tctgtaaaacagttagccaaaaatggagtcagagacagggtattacaga**aaaagataaatagaatgtacaaagacatgaagtcaatatttaggttaagagaatgacattaaactaattcaggatggccagag**catacaactcaaaatgtatg**tcagtaaatataaaactaaagcaataagcagaaaccactttgtgaaaataattaaaagccattataagatgtttgtacattacattatcttgaagacaatgatgagcagtgaaagaaatctgcacaagtaaataatataatgaaatgtcactgtttgaaagattactatgagttcacatacttatata**ttatatccatgtgtgggcatgtatataa**aatata**tatatgtagtaaaatacatata**atacattttcccttgatacttcttttttctcttaattctggagtaattctttctgtaacacagtatatt**gataggcccttcttatagcaagaaaaaaacctatc**ctgtatgaaaacttactcaagaaaagccaaaatccacagtttaatcaacattcctcagtctggccacaggggagagctgtttaaaaagtattctaggtagatgcatttagttctctgcagttaaactgttgtgatgtgtctctgattggttgcccccaaa**gttgcaatatcctggaagcaattgcaacaggcctcattctgagttcaaagcaactcctgtt**aaggaagcccattcagaagctgactggatattctggcaggccaaggATGGAGACTCT**CCTGAAAGTGCTTTCAGG**CACCTTGTTGTGGCAGTTGACCTGTGAGTACACGATTCAGTCTGGGAGAGGTTGTGATATGACAGCATCTTAATGACAATAGCTTTCTGTCTTGGGAAGGGCCCTTTTAAGAACCAGACCTAAGAAGTGAGGAAAATGTACCGTAATTATGTGGTGATGCTTAATGTTTTAAATAA**GGAAGGAAGAGAGAAAATCACCCTTCC**CTTCACGAGATTGAGCATCTCATTCGATTGTCTTTCTGATCAGGGGTGAGAAGCCAACAACCAGTGCAGAGTCCTCAAGCCGTGATCCTCCGAGAAGGGGAAGATGCTGTCATCAACTGCAGTTCCTCCAAGGCTTTATATT**CTGTACACTGGTACAG**GCAGAAGCATGGTGAAGCACCCGTCTTCCTGATGATATTACTGAAGGGTGGAGAACAGAAGGGTCATGACAAAATAT**CTGCTTCATTTAATGAAAAAAAGCAG**CAAAGCTCCCTGTACCTTACGGCCTCCCAGCTCAGTTACTCAGGAACCTACTTCTGCGGCACAGAGAcacagtgatacccaggcctccaagacctgtactcaaacctaaagctgagccgcagatgctcccctagcatagatgcccaccacaggagtatggggaacttaccagaaggttcatccatgataaaagagaaacctgaaggaaattcat**ttgcttcaacatataacacaaaaagcaa**gccatggccttaagaccatatctaatttggcattgttttcttgataattatttaattgctttgttggtattatatgatagataacttagattcagagaaaattacatagattt**cagtttcaaaaactg**gagtgaattcctatttgactgaagctatacatataaactccttttggatctctgcattcctatctgaaagtgaaaagtaacctctctttcttctcctaggtacccacagctt**gacaataggacagcaaaaaaaaaaaaaaaaaaaattgtc**tggttctttcttggacgtgaggattgtacacgg**taggtatttagattgttaatcttgttaccta**tc**tggaaaggtcactagcattattttttcca**aggttttaaatatctgttttggggaaatggcatttagaagtacttcaattttatatttgccttgccttctcttctcattgaggtttattctgattgatttccagctttgcccaagggaccctgagttatcttaccagtttgcccaatggtcaagatccttcctcaattgttcctttcacatcaacccagcacaacactgcatgtaggaaac**actcaacaagttttaattgagt**catatgttgataagaattgagaatggccaggtgtggtggctcacgtctataatcccagcactttggaaggctgaggcgggtggatcacttgaggccaggagttcaagaccagcttggccaacatggcgaaagcctaactctactaaaaatataaaaattagc**caggtgtggtgacacacacctg**taatcccagctacttgagagattgaggcatgaga

Uppercase: TRAV30

Lowercase: Flanking sequence[1000bp]

Red & Bold & Underline: Stem-loop [21]

Blue: Heptamer[31]

Green: Nonamer [8]

id-TRBV6-8-2[V_gene_segment]

ggagggtgttggctgggtcaaggagaagatgatgagctcagaaaatagacagggagaaaccccagtggaaagcttgtaaaactctggatgaatcacaagatatgttattagagctcctttttattactaaattttatttattactaaattaatggcaggaaccctgcacaaagggatattccctatgaaagatggcatctcaaataaaaacagagcattcttccttaccctaaagtctgccctttctccccaagtccctcatgcatcaggccacctttg**tgcccacagatcatgggca**gcgctgggtggccatcatcatccacacagagagggcagtcagcagcatgagctggttctgcctgatat**ggtctcaccctaggcatggaaagcaagagacc**tgcgttg**agctgaaggtgctcagct**gggt**ttgtcaggagtcccatctgtcagtgaattgacaa**gaaacagagcaaaacgactcctgcaatgttgagaagcctgcacctaggatctggaaacttgataacagag**aaaaccaatatagacaaaggtttt**taaacaggattatggtcaattaagcaaattagaaaaggatacttgaaggagtatttgggatacaggaatcaaaaacaccaggaagacatgagaagtttccctaagagtctagactacagcactgtgtagataactaacatcag**aatattaattatattattgaagaattaaaatt**tacattgaattacaaactctttacaaaaggtcatgaaaatcttgtagacttgtttcaattcagacaaatgtgttcttctcattctcagctgttcactggtgcatttatcttgaatttgaccatctggggaatgggcgtggcctctcctgactggaaggctctggggcccaggc**agggagaatgatgtctcagaatgactccct**tgagagtcctgttccccttttatcaatgcacagacccagaagacccctccgtcctgcagcccctgccATGAGCCTCGGGCTCCTGTGCTGTGCGGCCTTTTCTCTCCTGTGGGCAGGTGGGTCCTGGGCAGGGCCCTTGCGTGGATTTCAAGGCCCATCCCCTTTCCACTGGGG**CTGCAGCATCAGTTTTGTTCTTCTCTGCAG**GTCCAGTGAATGCTGGTGTCACTCAGACCCCAAAATTCCACATCCTGAAGACAGGACAGAGCATGACACTGCAGTGTGC**CCAGGATATGAACCATGGATACATGTCCTGG**TATCGACAAGACCCAGGCATGGG**GCTGAGACTGATTTACTACTCAGC**TGCTGCTGGTACTACTGACAAAGAAGTCCCCAATGGCTACAATGTCTCTAGATTAAACACAGAGGATTTCCCACTCAGGCTGGTGTCGGCTGCTCCCTCCCGGACATCTGTGTACTTGTGTGCCAGCAGTTACTCcacagcgctacaaggttgtctcctctctgcacataaaggcagggaggctctgccctccaccccgacctgagactcagggatgacctgggcagagtattctgcaacgggaaccttggaacccgaagtggccccaagtggcccagacagtatgagcctcgctctgtgccaggtgcctctgtaggcatctcagccaggcctggactggtcccaggtcctcatatgtctcctttgttgctctctcctacaagctctacttttggggttggggccagggcttccccagctcctacttttctactca**tcatcctgagtccaaggcccccaggatga**a**acaggatttgtatttcagatccatctagactcctgt**ctctccctggtgaccatgttgcttcctctctagggtttcccccagcccccaccctcctgtggcccacctttcccacctccgcagtcatcctccaccccccacttccctg**cccctctctactgcagccatgagggg**agcacc**tctctgtgactccttccttcccgtcacagaga**cttcaaaggccattttctttgccctgggccagagccttcctttctctagtggccagctccttcctgtgcttcagatctcagtatgatcagcccctcctgcgggaagcactccctcccctcccaggtaagatcaactttcctctcataagctctcaatgaatcatatgtctgttagcaacctttagcacagttgggctttctcagatgcttgtctgattatttttgtgctccacccaatttttttttttttaagacacagtct**cactccgtcaccaggctggagtgcagtggcatgatctctgctcactgca**acctctgatactctgattcaagtgattctcctgcctcagcctccc**aagtagctgggattacaggcacacaccaccacgcccagctaatt**cttgtttgtatttttagtagagatggggttttaccatgttggccaggatggtctcgatct

Uppercase: TRBV6-8-2

Lowercase: Flanking sequence[1000bp]

Red & Bold & Underline: Stem-loop [17]

Blue: Heptamer[34]

Green: Nonamer [3]

id-IGKV2-23-3[V_gene_segment]

atgcagtcaaaattttaatcattacatactcacaatcacttctgtggggcctggagatactgcacatacgactgttagcaagacactcactgggacgctgcgttgtgtgatggccccacatacaaacctcaaggaggctcagcctctcaatgcagcaggagcagctggggtacccaggccacacgtccataccaggtgggctcagttagagatggctggagagccttccag**gaagaggccatgaggtttcagtcacaaacactggctcctcttc**tgtgtaaacaggggctagagccctccaggacaattcctagagcctctcgctttctctccaattagtgcactgacaccccacagattctccaggaagtggttgtcatgtcctccctgcaacagccactaaagttccctactgctgtcatgaatgcagggacacttagtcacatcactgggaggcgaccctagtgtatcctgacctcacctgctgccactgatgactttcagggcacctctttctccctttgct**gagtgactctcactc**tcaccaaccatcaggagaatggaaagctgcctgcaatgcatgatgttggctgttgagcaaatcaaagctcacaggagtctcaaacatgtacaccacataataatattttctgataatactatttggacttttcttcctt**tcaattctggaagtaattga**gaatattttttgaactcttagaaacacttagtatatatgtgtagtaggtagtaactagttttgtctactggtttattttgtttgcttgtttcaggccatgatgcggcatgttaaaatactgaagacaaagatacattttagaattaagcatactgtacattggctctttccacaccactgcaaccaccaggggatgtgcatattgtcccttaggaatgaccttcccttgtgagtctgggagtaaagctcagctgtaaccttgcctt**aactgatcaggactcctcagtt**caccttctcacaGTGAGGTTCCCTGCTCAGCTCCTGGGGCTGCTAATGCTTTGGGTTCCTGGTAAGGACAGAGGAGATGAGGGAGGAGAATGGGGTGGGA**GGGTGAGCTCTGGGGGCCCCACTGTCACCC**ATGTGTGTTCTGTC**CACGTGTTGGATGCACGTG**TCTTGTGCTCCAGG**ATAAAATGTATGGTGGCACTTTTAT**ATGTGAAAGAGTGAGGAAGATTCCAGAAAAAGCAAAGACCTGTGCTCTGGTGCAGATTCTGACATAGAAAGAGGAGGGTAGCATAAGTGACTTCCAAAGGGCAACTTGGGCCTTCAAAATGTCTGTTTTTTTTTTAATTA**AATTTTTTTGGTGCATGAATCAAAATT**ACACACACACTCACACACACACACACACACACACACGCGCCGCAA**TACAATTATTTAGCATTAAATAATTGTA**GAGAAATTATGATAATGTCTCATGATTTACATAACATTGTACTTCTTTTTTATATTACTTTAGGATCCGGTGGGAATATTGTGATGACCCAGACTCCACTCTCTCTGCCCGTCACCA**ATGGAGAGCCGGCCTCCAT**CTCCTGCAGGTCTAGTCAGAACCTTTTACATGGTAATGGATACACCTATTTGTATTAGTTCCTGCAGAAGCCAGGCCACTCTCCACAGCTCCTGATCTGTAGGACTTCCAATCAGTTTTCTGCCT**TCCCACACAGGTTCTCCCCAGTGGGa**ggagagagtagaccagtcatccccagatatatcacaggactagtttcaacctttggaagctggtctatatcctatggttaaataggcatttgtgatacgacctgaaatacatttggacaagaacttcactaacaattgagtcactgaagacttacggccctgtgtgacgcaccacataactgtgagtttgcagtggttgcaggtcagggacagattttatgcttaagatcagtagggtggaggct**gaggatcttggctattacaactgccaccacactctacaatatcctc**ccacaatggttcagcaccaaacaaaagcctcctgcttggattgtcccagctgcccaaattagttccttcactgaggagtagacagggtatattctctaaatctatgtaacaggaagatgttggtgaactcaggggattagtatgaagctacacctcaggcatcacacataagatcacttcagcagtcgcagccttagcatgggcagaacctacagaagatgcaagtgccctctgagccaggagacaggaggaaggaggaagggaaaggtgacttagctcatctcaatcctctctcctttgcatacatttgtcaaccag**atgtattcagcctaccagtcacacaactgaggctgatacat**gacaacatagcactggtatattcttggtattgtttggcttagcagttactagtatatatttaat**gggagaatatttggtggtgttaacacattgcttatctccc**tttccccagttgtactttacacttgttcttggcacacattctcctccaggactggagcattcacagggttttatgttactgttcttatgggagtaaaaagaaaaacgattcacattcttgctactgagctaggctgggatgccctgggccaagctgaaaatgtgaaaaataagagtatgaatatttattaagttttatctggatctaagatacttatccatgaaccagtcctgcagctgtgccca

Uppercase: IGKV2-23-3

Lowercase: Flanking sequence[1000bp]

Red & Bold & Underline: Stem-loop [14]

Blue: Heptamer[40]

Green: Nonamer [5]

id-TRBV7-5[V_gene_segment]

agggaagaaaaaaatggatgtttcccatatattaga**atatgcatatttttcataatttcccattttttgttgcatat**gattgaattttcagacagataaaataggtagaatgttatgaagtgatatttatgagatgttagctgagaagttttttgaggtgtttaaggacctaaattcagagtttccagaatcacaatgaatctcaaataaggcagtcaagtcataagcaaattgtagaacatgacggacaaagagaagattttgaaagccatcagaaagaaaagacaaatacgtcccataggaatagatggagaacaacagtaagttaagacagtggtaaaaataaccattagctcaattatcattctccttcatttaaggtaagattgattaaaaaaaataacttttagaaggggaacatgaaaaatgaccgctgtgagtttgctatcaagg**tttcttttctgaattacagttaagaaa**ggtgggaaatgatcccagaagaaaggcatga**gggcagttcgtgaggtaacccaagtgatggggctgccc**cgttgggcacgtgtgactgagggaatggaggaggctggggcataaatggggatggcagaggggaccctgaattgcaggatagacaatgagctaatgccttggtgccttgttttggggatgctgttggca**catcctacaagacatgcccagcagacagaggagtggctgtaggatg**agaagatgaactcagagatgcagtgtgaggcctct**ggctccagacagcacgggagcc**caaagcaatgagccaggcattgatgttgttaaaaaggagcttataaatatttaaagtgtgttctaatataaatactgtgaccctgacatcctggggattgagagaggaagtgatgttactgtgggaactgccctgtggagacaaggacattcctcgtcctctgctcctgctcagagtgacactgatctggtaaagcccccatcctggcctgaccctgccATGGGCACCAGGCTCCTCTGCTGGGTGGTCCTGGGTTTCCTAGGGACAGGTGAGTCCTCAGAACATTAAGTAGTTTTATTTTTTCTCTGTGTGTAGGTGTGTGTGTGTTTGTGTGTGTGTGTGTGCGATGACTACAGATGTTTTCCTTATTCTGTTGCCAAATTCTATTTCCACAGATCACACAGGTGCTGGAGTCTCCCA**GTCCCCAAGGTACGAAGTCACACAGAGGGGAC**AGGATGTAGCTCCCAGGTGTGATCCAATTTCGGGTCAGGTAACCCTTTATTGGTACCGACAGACCCTGGGGCAGGGCCAA**GAGTTTCTGACTTCCTTCCAGGATGAAACTC**AACAAGATAAATCAGGGCTGCTCAGTGATCAATTCTCCACAGAGAGGTCTGA**GGATCTTTCTCCACCTGAAGATCC**AGCG**CACAGAGCAAGGGCGACTCGGCTGTGTATCTCTGTG**TCAGAAGCTTAGccacagtgtggcacagtcgtgtccttcctgctcacaaacctcatccttctctctccttgcacctcctagagacccttaatagaggcctatctttgatcctcacttttcgtgggaaataagtagaattggacatcggctgtcctttgggtagaaagagaccacagattcattcctgaaacacagtgactgcaaatgtaggtggtgaaaacaatcacgtcccactgccctctaggagtgc**ttggagccagctcactgctccaa**acggagtgggtgtcttagccttggccttcagggcagacatgcatcttctataggtcttggaggctgctgtgttacccacatatatgaggttgtcaagg**gcaggaaacatgctcttctcctgc**atattttggggcatctggaaggtctgaggctacatccc**caggaatatctttcttctaaagcctcttctattcctg**tcaccttggaaatttttgcaacaaaatatcgaacctctcttcctgttta**aagtaaaggtctttgcaactttcgtggtctttactt**gataaatacaatcatggtaacaataagacttcatttcttctgcctactttaagccacttgtatcctttattttatttccatttgccat**tgctactgtcctgatagacagaagca**tgcattcaccactgctgcctgttcacatcgattccctcaggaaatctgatttttagactctgagtgttttcattgttgtccaactcatttgatttgaaataattttaccgaggcctttaactcaagaagtgttttatttataatattga**atctattcctttttattttatttttcataatagatagtactatatagtact**tgttataaatagaagtacaatgattatattgcaatagaatcttccacctatctgtgggtgctgctgcagtgtgtatctgtgaaagcgaatgcactggtcagagctgatgtgattatggatcgtgggtttctgggagtccttgg

Uppercase: TRBV7-5

Lowercase: Flanking sequence[1000bp]

Red & Bold & Underline: Stem-loop [16]

Blue: Heptamer[36]

Green: Nonamer [5]

id-IGHV3-33[V_gene_segment]

ggcta**aattctcaaagaatt**gtgctgagtgaaagaaactaaggaattaagagt**aaattttacatgatacatttgtagaaattt**tagaagatgccactattataaattaacatggagaagatttaaatgtttctgagaatatgctattgggagtaatggggatgtgagttaaatttcagaggaataagagaaagatttagggattaattttttcaaaccttgattgaagtgctgagtaaatggttgcaaacataggtctacattt**ttcaaatcattcaccataaatttgaa**tta**tttattaattacactcgaataaa**gcaataaagaaactgatgagataatatttgactgaattgcatcaataaatagatcgatattaacacaaggaatataactgatttccaaaaacatacacatgaaccatggttcactctgcgtatttagataaattacagaaagttgtcataacagatggggaatcctgcagacttcactaggcatgggccatgctgcc**ctggagttgtctcaggggagctgcctcctccag**aggttagagcacaggcccaggtaataggactaaatttttagatgtgttatcttagacacactgcacaactgctgtgttctctatgtaaattatctcctgtaaaatataacattgaagcctgcattaaatatattgtgtaaatatgtaagaataaaagaaagttatgagagctaagtgttaatcaaggcacaagcatataagatataactatattttcctgaatgatggaattactaccagtctcccccaggacactt**catctgccctgagcccagcctctcctcagatgtcccacccagagcttgctatatagtgggggacat**gcaaatagggccctccctctactgatgaaaaccagcccagccctgaccctgcagctctgggagaggagcccagcactagaagtcggcggtgtttccattcggtgatcagcactgaacacagaggactcaccATGGAGTTTGGGCTGAGCTGGGTTTTCCTCGTTGCTCTTTTAAGAGGTGATTCATGGAGAAATAGAGAGACTGAGTGTGAGTGAACATGAGTG**AGAAAAACTGGATTTGTGTGGCATTTTCT**GATAACGGTGTCCTTCTGTTTGCAGGTGTCCAGTGTCAGGTGCAGCTGGTGGAGTCTGGGGGAGGCGTGGTCCAGCCTGGGAGGTCCCTGAGACTCTCCTGTGCAGCGTCTGGATTCACCTTCAGTAGCTATGGCATGCACTGGGTCCGCCAGG**CTCCAGGCAAGGGGCTGGAG**TGGGTGGCAGTTATATGGTATGATGGAAGTAATAAATACTATGCAGACTCCGTGAAGGGCCGATTCACCATCTCCAGAGACAATTCCAAGAACACGCTGTATCTGCAAATGA**ACAGCCTGAGAGCCGAGGACACGGCTGT**GTATT**ACTGTGCGAGAGAcacagt**gaggggaggtcattgtgcgcccagacacaaacct**ccctgcaggaacgctggggggaaatcagctgcaggg**ggcgctcaggagccactgatcagagtcagccctggaggcaggtgcagatggaggctgtttc**ctgtcaggatgtgggactttgtcttcttctgacagttccccagggaacctcttaaatttagaaaactgt**gcctaacaatgtcttctctatgcatatgaggaccttttctccctggcacaaaatgcagattgacgctgacacggatgaaaattcctcaaccatggtcacaaggatcagagt**cctgagtaacctcagg**gct**tcctggtgattcttctccaatcagacccaggacagggacctccgtgagattccctg**actggaacagtctttatggat**cctggtcacagacaatagagaggctgaaccagg**gtcagcgtcatgtagaacgtcacagatttcacgtctgatccttctcctgacacgaaagtatgcaaatcagtatc**agcaccgatctggtgct**tcttttgttcctaatccatttactttcttttttcgtcgtttttc**tcctttttccatttgtttttcctgctttttgcaaaagga**agatgttttccctgtgagatgcaggggatgacaattttgggagatggctggaacatccaatatcctcagggccgaccatcagtaagtgcaggctagaagtctcagaaagagctgaagctgcttaatcaccgtggagttttaccttctccagttctgctctgatggaatcagggccaagcaggttatcaatgataatctacctaacatagagtcaactgattccagtttcaataacgtctgttaaaaattcacaccaccacctggattactgttttgtcaaat**caatacacagtattg**tccagctaagtagacccaaagacggaccgttgcccatggagaaaaacattaacctgagttctaggttcttacagtgttaaagg

Uppercase: IGHV3-33

Lowercase: Flanking sequence[1000bp]

Red & Bold & Underline: Stem-loop [21]

Blue: Heptamer[29]

Green: Nonamer [6]

id-TRBV12-5[V_gene_segment]

atccactctatgaacagaattcttactactatttataccctaaaaagcaggccattttctagagtttcaccccctcaccagaaagaatttatttgtgagtattggttgacttactataacaagagactcaagtgttaatcattaaaggaacctgttagtgctttcttagaagtaagaggaagatcacatttaactggagagtgccttattgtggttttcaaggaaggtcaaagctcttacttatcactgctcaatcttcacccatgcccaaagaattaacataattttgaaccttacc**ctggagaagcgaaatgggaccttgtcctccag**aaatgcagaagacactgtgctgtgggctgggggagtcacggagcacgagagtatccatctccagccctca**ctggaaataaaatctgcacagttctatctctttccag**ggcagccaccttgttctctgcatgaggagcatggaggtccagatgcttccaggagggatatggggcaaagctctaactaagcaacacattattttgattttgtgtgtctgggatataaggatggccggtaaggggaa**caataaagttcttgagatttcacaagtaatttattg**atgtccagaatttgaggtgctgatggtacacctaagtggcaatatgcaccaggcagatcaatatgtgacgcgtcagagaaagcagaatgggtgatgtgatgtgcaatgccacagaagcactgcagccaggagaggtgacagctaatgggga**tgtttggagtctttgagtgaaccaaaca**catcccagagtaattgtaatttatttcagtcaatcttctgtacagacttagcattcacctttggaggaaggtcctttgagcagggacagagatggtgatgtcactgacagtcccccttttactctgggtgagaggtctagaatcctcagctcctgtattcgtgcccacaagggcctcatctaggtgaaggctccacctgccccaccctgccATGGCCACCAGGCTCCTCTGCTGTGTGGTTCTTTGTCTCCTGGGAGAAGGTGAGTCCCCACAAATAAAGCACCTGCATTTTTGGATATTGCCAGTTATGATTCCAATTATGTTTCTTATTCTGTCCCCAAATTCTATCTCTTTTCACAGAGCTTATAGATGCTAGA**GTCACCCAGACACCAAGGCACAAGGTGAC**AGAGATGGGACAAGAAGTAACAATGAGATGTCAGCCAATTTTAGGCCACAATACTGTTTTCTGGTACAGACAGACCATGATGCAAGGACTGGAGTTGCTGGCTTACTTCCGCAACCGGGCTCCTCTAGATGATTCGGGGATGCCGAAGGATCGATTCTCAGCAGAGATGCCTGATGCAACTTTAGCCACTCTGAAGATCCAGCCCTCAGAACCCAGGGACTCA**GCTGTGTATTTTTGTGCTAGTGGTTTGGTcacagc**gctgcagaatcacctgctccctgtgcagaaaccctggtgcttcctcttctcctccagtacccagcagctctcagcagcctttcttgctcctcccctagcacaggaagtacataggtttcgtgttccacatgtccctaggcaaggcaagaacaggtcataaggacacatcacgttaggaaacttttggtaggaagtcagtgggtgtgatggttcctgggattccacacatacttctcacaagggtgcctgagtccaagtttgagggcgagtgtcacagactggagctttcaagtgattgagtgagcttaaaac**tgtgcagccattgcaca**tggatgcttatctcctttgctgctttccgttatggcttctttgccatttcttttctcttctaccttaaaactttcaacatct**ttgatgacttccaacatcaa**tattggcttcccagttttttcacttgctcacttcaaatgatcatttccttcaccttgtgtcagatattcatttccactttctttttctagattccaatgac**aaattatttgaattacccccaaatagttt**cctcaagcatcccattgtctcatgcgggcttctcttttcatctctggtactacttctgcagcaattcttcaaccccatcaggaccactaattgacttctcctgcaaatattgcaccatatatcatctcctcatgtactatattccttatctgacttagactatgtgttccaaggtgattgtgaaattttgcatatgtcttcagctcttttctttttctgtttcttacctggaaaataaaatggggccta**ctctgtgcccctttcaatgacacagag**aacataaaatgtcccttacccttgtcaagcagtattataacgatacagtgtgtcttgaggtctgtaagtccctctatgcatataaaattatatacttttcttct**attaaaataatttgttaccttttaat**gagttcatcagtttcatctttcaatta

Uppercase: TRBV12-5

Lowercase: Flanking sequence[1000bp]

Red & Bold & Underline: Stem-loop [11]

Blue: Heptamer[28]

Green: Nonamer [5]

id-IGHV3-71[V_gene_segment]

ttgtcatcc**aggctggagctcagtggtgagatcatagctctctgcagcct**cagaccctgcgtctccaccaatcttcctccctctgccatacgtagctgggactactggtatgtaagacaatacttggcttcag**tgcttattaaaatgcacactttaattaagca**gtatttgttatacagcaatcatgcctcaataagagtattgcaaataagtggatagataatttgttcaataaagattgatggaaagatagatgctaacatg**agaaaatgcatgacacccaagaaaataacactgtaggaaacatgcttttct**ttacaattgttaggtaatcacaacagagcatacacatcacaccatgttctcgttacagagaaaaggttctgcaaacctcactagctgcaacccctgtgtgctgggcttggttcagggagaagtcaggtccagtggtgagaagcacaggcccagataaccaa**gctcactctgaccgaatgtgagc**cctgggtacattgtacaacccatctgtgcttctgctggtaatttttcatctgt**aacatggaaatgacattgatactatataccatgtt**tcctctgcatatgtaaaaataaaaggtgattggtgctaactttaaatatacacagtttatgtacattgattgcacctcaatacaacagttttaaatagatattacaacattataagttttataggtttaatattttatcacagaacaaactttcaataagaaaccacaatttccaaatgctatcaatatcacaaatctcccccaggacactctcacatgctctgagccccactctctcctcaggcgtcccatcccagagcttg**gcatgtagtaggagacatgc**aaatagagccctccctcggcttatgaaaaccagcccagccctgaccctgcagcactggaagaggagccccagccctgggattttcaggtgttttcatt**tggtgatcaggactgaacacagaggactcaccA**TGGAGTCATGGCTGAGCTGGGTTTTTCTTGCCGCTATTTTAAAAGGTAATTCATTGAGAACTATTGAAATTGAGTGTGAGTGGATAAGAGTGAG**AGAAACAGTGGATTTGTGTGGCAGTTTCT**GACCAGGGTTTCTTTTTGTTTGCAGGTGTCCAGTGTGAGGTGCAGCTGGTGGAGTCCGGGGGAGGCTTGGTCCAGCCTGGGGGGTCCCTGAGACTCTCCTGTGCAGCCTCTGGATTCACCTTCAGTGACTACTACATGAGCTGGGTCCGCCAGGCTCCCGGGAAGGGGCTGGAGTGGGTAGGTTT**CATTAGAAACAAAGCTAATG**GTGGGACAACAGAATAGACCACGTC**TGTGAAAGGCAGATTCACA**ATCTCAAGAGATGATTCCAAAAGCATCACCTATCTGCAAATGAACAGCCTGAGAGCCGAGGACACGGCCGTGTATT**ACTGTGCGAGAGAcacagt**gaggggaagtcagtgtgagcccggacacaaacc**tccctgcaggggtccccaggaccaccagggggcgccaggga**cactgtgcacggggctgtctccagggcaggtgcaggtgctgctgagggctggcttcctgtgatggt**ctggggctgcctcattgtcaaatttccccag**ggaacttctccagatttacaatcctgtactaacatttcatgtctctaaatgcaaaacttttttgttctttctgtatttttgtttttgtaacaggaggacacattctcgcctcc**acagaagccacagtgtcactttgggggcagatgatccttctgt**agtcagcaggatgaaagtgcagaggaatctcagtggaacctgggaagtcttttccagttagactcaggg**cagagacctccatgggaatctctg**attagaacaggctttgagttctgataggagccaagagagaggctcacccagggtcagagtcctt**aaaacctgacggttttcacagctatctcacctcgtcttgaaaa**actgtgcacatctgactcacactaattcagttgaccttatttctgctaatctattttccttctctgcagacttgattctcacagttccctttcttcttctcttgcct**gaaaacagaggatgtgttttc**tgtagtcaaaatcccagagat**caggtctgcaggacctg**ggtaggctgaggggactttctcac**tcaccatagcctggtga**cactcctgctgtattttgtatgtggaggtgtttggaaaacgaagtgaacattagtcatgaagggaataatactagttttctccaaagggatgtggatgtagagctgatcttgtgcttctcacactgtcacagaatttat**gctctcccctgtgactttaggagagc**tgaggatggacactccattgtgctgtgagctctggtaatagtaattatagggtctcgctaggcagcctaaggtcaatactgctggccttcgggaa

Uppercase: IGHV3-71

Lowercase: Flanking sequence[1000bp]

Red & Bold & Underline: Stem-loop [21]

Blue: Heptamer[48]

Green: Nonamer [8]

id-IGHV1-69-3[V_gene_segment]

aatgggcacataactgtgggagatactatattcctatgaagaaggtattcagacttcagagataagtaatgtttcctacattgtgcttgtgacttggaagcagtggattgaagagtgtgataagtgcccagaccaagcagaacagaaatcagcatgtaaagatgatgatctatggatatgatctaaaaccatgtaaatacttcaaataattctatttaatgc**agtttgaaataaaacacaaact**tattcaaaataca**aattacttggtaatt**attttgggagcta**tgagttcaccaagaaactca**aattcctatttctatttcaacccctgattcctactgtcaatgggagggaagtctcagaaccaatcacacatcagacggcaaatctgtcaaccaagagtctttccactgaaggacctgggaggtcaggaccctcaggaaagtgctggggaccctgtcttgg**gagtgcccagcagatctcagaactctccatgggtcctgctggacactc**atgtag**ggtaacgagtggccaccttttcagtgttacc**agtgagctctgagtgttcctaatgggaccaggatgggtctaggtgcctgctcaatgtcagagacagcaatggtcccacaaaa**aacccaggtaatctttaggccaataaaatgtgggtt**cacagtgaggagtgcatcctggggttggggtttgttctgcagcgggaagagcgctgtgcacagaaagcttagaaatggggcaagagatgcttttcctcaggcaggatttagggcttggtctctcagcatcccacacttgtacagctgatgtggcatctgtgttttctttctcatcctagatcaggctttgagctgtgaaataccctgcctcatgcatatgcaaataacctgaggtcttctgagataaatatagatatattggtgccctgagagcatcacataacaaccacattcctcctctgaagaagcccctgggagcacagctcatcaccATGGACTGGACCTGGAGGTTCCTCTTTGTGGTGGCAGCAGCTACAGGTAAGGGG**CTTCCTAGTCCTAAGGCTGAGGAAG**GGATCCTGGTTTAGTTAAAGAGGATTTTATTCACC**CCTGTGTCCTCTCCACAGG**TGTCCAGTCCCAGGTGCAGCTGGTGCAGTCTGGGGCTGAGGTGA**AGAAGCCTGGGTCCTCGGTGAAGGTCTCCTGCAAGGCTTCT**GGAGGCACCTTCAGCAGCTATGCTATCAGCTGGGTGCGACAGGCCCCTGGACAAGGGCTTGAGTGGATGGGAGGGATCATCCCTATCTTTGGTACAGCAAACTACGCACAGAAGTTCCAGGGCAGAGTCACGATTACCGCGGACAAATCCACGAGCACAGCCTACATGGAGCTGAGCAGCCTGAGATCTGAGGACACGGCCGTGTATT**ACTGTGCGAGAGAcacagt**gtgaaaacccacatcctgagagtgtcagaaaccctgagggagaaggcagctgtgccgggctgaggagatgacaggggttattaggtttaaggctgtttacaaaatgggttatatatttgagaaaaaaagaacagtagaaacaagtacatactctaattttaagataaatattccattcaagagtcgtaatataagccaaattcac**agagtggaaaaggccacactct**ataacgttgatacaaacattccatgaaggtgctactgtgaacaagttttcaaattggatgaat**acatgatttggagcaaggttatttgatcatgt**ggtgagac**taagaatgattctta**aaaagtgccaaaagtttccttcaaatgtttctgtcactccttatcataaagtttattttacagcagttttaggattacaaagaaattgcacaggaggcgtgagaattcccatgactccctgccctacacaggcacagcctcctccactacgaccatcctgcaccgcagtcacaaatcagttacaatggaggaatctccaaggacgcttggttctttctttttctggtgatctcctaatataacaagcctaagtatctcaagattccacggttttttcagtgttttctagaactgatattagtcagagggaaagtgggtaaggctattactatttgaactctttcttccaaaatccacaaaatatatattaatttagagctt**atcttacttctggtttacaatgctccttcccagagagtaagat**ttttaag**cttttagaggccagttcatgtctctaaaag**accagaaagttctggggaatcccataatgaacatcctt**tcattggaattggagaccctggcaatga**gagattccatgtataatgccctagagttggattaggtgcactgtgagctcttgggtggtgagtctgagtaaggtggtttgtgcagcaaatgcaaacacattcatgggacccaggtaggaaca

Uppercase: IGHV1-69-3

Lowercase: Flanking sequence[1000bp]

Red & Bold & Underline: Stem-loop [16]

Blue: Heptamer[50]

Green: Nonamer [5]

id-TRGV5P[V_gene_segment]

ttcttacagaggtggtgataatc**tgtgaatgccttactgaattttcaca**ataatattactgtacttgatttgtccccttattttcatctgtgaatccaggtgcactgcagatattagtctgctttctgagggtagggatgacaaataaaatatacaatacgcagagaattggaagttcagataaataacaagtttttagtttaagtatgttccaaatgttgcatgacatggcacccagtactactttaaaaacaaacaagcaaacaaacaaactcacattcctacttaatgaacaaa**aatcctctaataaggaggtcacacacacaggatt**acaggcacaactcaccccccccaacacacacacacatttaccttagagcagagtccaagacacaccaacacacatgatcctcagtccttttaatttaaagcacgttcaacccttctcagagtggccttgagtggttgcctccatgtgtctgcagatgcgaaaaggaacttggttcagggccctaatcagaggcttcctgt**caggacaaggtcctg**gcatcagcattagctgctgcaactctgaa**gggcagggtaggagcctgttgtgggactgccc**cctccaaggtctgagcctgttaattccagggtcagcttcccctaaatctgcagtccctctgct**ggaggcctgaactctcaatgcccctgcctcc**cttctgccctgggcctgctctcagctgccacaagagggcgccgaagaaagggcct**gaggagacacaggagctcctc**tgcagg**tcctcccaggaaccaccccttctgggagga**aggcttgggagtcc**cctaagacacatcctcagtcacttctctctgcctgtgtctcagg**aataccagctcctcctactgtcttctgtgctagggatcacttccttgttgagtgggacctgagttttgagaggatcttctactccttttcatctggtcccttt**ccttccaaggccccagaaaggaagg**cATGCGGTGAGCCCTAGCGGTGCTTCTAGCTTTCCTGTATCCTGGTGAGTGTGCTGCCTACAGAGAGGCTCACAGGTTGGGTTTTGTTTTGTTTTCTTCTTTTGAAAGGGGTGCCATACAAAGGAATACCTCATTATATTTTATGTTTTTCCCATTGCAGCCAGTCAGAAAT**CTTCCAACTTGGAAG**GGAGAATGAAGTCAGTCACCAGGCCGACTGGGTCATCTGCTGAAATCACTTGTGACCTTACTGTAATAAATGCCGTCTACATCCACTGGTACCTACAGCAGGAGGGGAAGACCCCACAGCATCTTCTGCACTATGATGTC**TCCAACTCAAGGGATGTGTTGGA**ATCAGGTCTCAGTCTTGGAAAGTATTATACTCATACACCGAGGAGGTGGAGCTGGAATTTGAGACTGCAAAATCTAATTGAAAATGATTCTGGGGTCTATT**ACTGTGCCACCTGGGGCAGGcacagt**gattcagatctgccctacaccacactgaacatctgc**cttgtggctgcctctggtacacaag**atagagccgccccctctcatttcctgccaccaaatttactgtattctgaacaagagaaagacagcttaactcctgatctccctcctaatatcacactgtcctggcagcagctgcaccctgttccctacccctcccccacaactttcctgaagatcaagctgccatctccgggcctcagttaagcagcctg**gctgagagtaaggttctctcagc**tctcctaggacatgggggaggcccactcactctgcttcctatgacagacaggtacaactagggtccagctgcgaatctagatattttggacagcaaataggaaggttatttatactgaatcatttatccagtcatggtccagagatcacagctgagagtgg**tgcttattccttatgtctataacaatataagca**atagtataatgaccactaaaacactaggcctaggtatgcagttgaacattctctccctctgccctcccttttattctctctctcttttttcttacttctttgcttgtttttatttttctctccctccctaactccctccctccatccatcctttcctcccttcctcccttgtttccttccatccattctccttccttgttttcctccttccttccttgtttttctctctctttctttcctttttgggtacataatgagaatactttttccacaaaatcatttgaaataatttttctctatttgttcatgccaacatttatacagctagtttaactgttcagttttcatttttacattttggaaatttgttgtattactttgttttgtttcaaatctgttttgtatatataaaatatttacatggaactgaagtc**aaaatcataaaaggtgatgcatttgaagatttt**agcttccattgtagatttcttgttcctcttttctccctgacactacaggtaacaatatt

Uppercase: TRGV5P

Lowercase: Flanking sequence[1000bp]

Red & Bold & Underline: Stem-loop [16]

Blue: Heptamer[22]

Green: Nonamer [4]

id-IGHV3-65[V_gene_segment]

gaattacatatgttaagtaaaataagccaa**acaaataagaatatatactgcattgatttgt**ataaattttaataaactaaatgactcagccgcaatgtatagctcaggagttgccagagaaatggtagaagaatgaaagggaaaaggaggaagaatgtagaagaacaaaaggaaatgttgagaattctcttgtccacc**ttgataacaatgatggttacattatatttatcaa**ttgtacaatttaaatatgtgaa**agtttattatctgtaaact**aaaactctcaaaatttattacaagcaaacaaatggaaacttagacaaagaaggagtgatagaaagatagaaaaaatgtatattaaatttcagaaataccta**agaacttatctgcctgaaccctagttct**caccatatttttaggtgaatgctagaatgcagcaaaatcacacgtgttctcactacagaaagtgggttccacaaccacactaggcacgcccagctctgtcctggagttggctcagggagtaattggggccagt**gatgaggagcacaggctcaggtactggggcttactcatc**cagacgtgagctcttagacacatacttagccc**tttctccatgcgtggttgacttccacatctgtacatggagaaa**caattgactccgacaaaacataatttgcacaaatatgtaaaaataaaataggatgatgaattcaaaaagtttatcacagcataattttataataagacagcatattttctgagtaccatcgttgtcaccaaattcttgcagggcacagtcatttta**tctgggtactgccttctcgtcaggcttcccaccccaga**gcttgctatatagtaggagacatgcaaatagggccatctctctgctgaagaaaaccagcccagccctgactctgcagctctgggagaggggccccagc**cctgggatgcccaggtgtttccacttggtgatcagcactgaacacc**gactaccaaccatATGAAGTCTGGGCTGAGCCGGTTTTTCCTTGTTGCTATTTTAAAAGATAATTAATGGTGAACTAGAGATACTTAGTGTGAGGGGACATGAGTGAG**AGAAACAGTGGATAGTGTGGCAGTTTCT**GATCACGATGTCTCTGGAGGTGTCCAGTGTGAGGTGCAGCTGGTGGAGTCTGGGGGAGGCTTGGTACAGCCTGGGGGGTCCCTGAGACTCTCCTGTGCAGCCTCTGGATTCACCTTCTGTTAACTACTGCATGCAGTGGATCTGCCGGG**CTCCAGGAAAGGGGCTGGAG**TGGGTAGGTTTCACTAAAAACAAAACTAATCGTGGAACAACAGAATACGCCGCGTCTGTGAAAGGCAGATTCACCATCTCAA**GCGATGATTCCAAAAGCATCGC**CTATCTGCAAATGAACAGCCTGAAAACCGAGGACACGGCCGTGTATTACTATACCAGAGACACAGTGAG**GGGAGGTCAGTGTGAGCCCAGACACAAACCtccc**tgcaggggcgcgcggggccaccgggggcggtcaaaacccactgaggacaggacaggtcccaggagcagatgctggggtagttttcctttctcatcagctggagaagtcaggtttgtatttgcaggactctgcagcattctaggctgtgacattttatcacttgtatttattatgactttattatcgttagtattta**aattttagtaattattaaaatt**atatat**aagtacattttaaatatgtactt**tgaagatataaatattcctaattatttgcactgattcttccagagttttattaacatttgttgacatcagcaactacatagctatagggacataaatttataaacatagaaatatgtatcggcaagacatggtggtctgtgcctgtaaccc**cagcttgggaagctg**aggcagaagtactgcttgaacccagaaggcggaggttgcaatgagccgtgatcgcgccactgcactccagccgggcaactgagactccatctcaaaaaactaaataaataaaaagaaagaaagaaaaaaagatgtacaaa**tacacagacctatgcatatatgtgta**ggcattcatattaaacattaagataaaataattctaaaaacatgtcctgaagaatcaaaattaatgatgaactaaatata**aattattagagtaattcataatt**gattttggttatttttaattgtttacatgttgtgattttacacaaaatattagtcatttatcc**tgataaaacaggtcaaaatgttatatca**agagtttatttactttgtatta**ctaataatataaaatgttccacatatttttag**ccatgttttacttatctgtgaaatgtgtt**tattaatttttcattttaata**tgtcacctttatctttctaatttcctggatttaattccactacagctgaatgcattttaataattattgtaatcacatttacattttttatttgaattttcatatatattttattagga

Uppercase: IGHV3-65

Lowercase: Flanking sequence[1000bp]

Red & Bold & Underline: Stem-loop [21]

Blue: Heptamer[29]

Green: Nonamer [4]

id-IGHV3-64D[V_gene_segment]

ttaactctcagcatttctattgagaaaaataagccaaatagttaagatatatactatattacttcattcttataaattctagaaaataaaagctagttgaaagaaatatgaaaacatcagtagttttataaagaaatggtagaagaaaggaaggagaaaaaacaaaaaaaataagagagcaagaggaatcctgagggaagttgacttgtcaccttc**ttgaaaatagagattttttttcaa**tgtttactattgtacaggttaaatatgtgaattttgttatc**tgtcaattaaaacgtatgaaatttattacaagtaaacaattgaca**ttttagacaaaataggaaatacagcaaggaacaa**ataaatatgttaaatgtcagatatacctaaatatttat**ctacctgaatcttgtctccacatttgtaagtaaacacagcaaaatcagacaggttctcgtgacaggaagtggattctgcaaaccacacttggcccgtttatctctgtcctgtagttggttcagag**caactgaggccagttg**tgaggagcataggcccaggtactaggactcactcatgccagatataagcccttag**acacatacatagcccctccatgtgt**gggttcacttttacgtctgtacatgaagaaaccactgattcctaaataacataatttatacacatagg**taaacataattaaaaatgtgatagttattaaatgttta**tcatggaacaatttcacaataagtcagcattttcccaaatacaatcattgtcatcgaaatccccaaggacactctcatctgccctgggccctgccctctcctcaggcgtcccaccccagagcttgctatatagt**aggagacatgcaaataggtccctccttctcct**gatgaaaaccagcccagccctgactccgcagct**ctgggagaggagccccaggcccgggattcccag**gtgtttcca**ttcagtgatcagcactgaa**gacagaagactcatcATGGAGTTCTGGCTGAGCTGG**GTTCTCCTTGTTGCCATTTTAAAAGGTAATTCATGGAGAAC**TAGAGATAGTGTGAGTGGACGTGAGTGAG**AGAAACAGTGGATATGTGTGGCAGTTTCT**GACCTTGGTGTCTCTTTGTTTGCAGATGTCCAGTGTGAGGTGCAGCTGGTGGAGTCTGGGGGAGGCTTGGTCCAGCCTGGGGGGTCCCTGAGACTCTCCTGTTCAGCCTCTGGATTCACCTTCAGTAGCTATGCTATGCACTGGGTCCGCCAGGCTCCAGGGAAGGGACTGGAATATGTTTCAGCTATTA**GTAGTAATGGGGGTAGCACATACTAC**GCAGACTCCGTGAAGGGCAGATTCACCATCTCCAGAGACAATTCCAAGAACACGCTGTATCTTCAAATGAGCAGTCTGAGAGCTGAGGACACGGCTGTGTATT**ACTGTGTGAAAGAcacagt**gaggggaagtcagtgtgagcccagacacaaacctcc**ctgcaggaacgctggagaaaaacagctgcagtgggcgctcaggacccactgatcagcgtcaacctcagaggcaggtgcacaaggaggctgat**ttcctgtcacgatgtgggacttcatctctttacagtttctctagtgaacctctctaagttcagaactctgtgcttactaatgtcatctgtacatatttt**ttaaaatgattattttaa**tatgaaaacctat**tctcctgtgtacaaaatgcggattgatccttacagaggaga**tgaaaactcctcaaccatggtcaccatgatcagagttctgaggaaactaggggcttcctggtgagtcttctccaatcaaattcaggacagga**acctcagaaagattcctgactagaactgaggt**attgctcctcacaagagagctctagctccgg**ggggtctgttcctgcagacccc**tgattcggtggtgtatgaataaaatgtacactgacacacagatactctgtttttccagtgcagctgagggtccgaggccac**ttacagactccaagaagaatcctgtaa**agaatggcagcagtggccctgagcagctcgcactccaggcgtgtatttagtatagaattaacaacagaagctctgagtcaacacacttgtggataattaacatggttaagagagtagttctacgaatgagtaaagctcagggactgtggtctgaagtaaataccattagggggcaatattcttggtcgacctcccgctgagagggtcttctggcttaaaggtt**agttaatggggtagggtaaccagacttaact**ggggaagcctctgttgtccctagtatttaccctacaacctaatgctctaaggtaagaaccggccatcttcagcctgttcagttattgcaagctatgt**aaacttccggccttccaaaaaagttt**gtgactatttcctataactttcgttaatatttc

Uppercase: IGHV3-64D

Lowercase: Flanking sequence[1000bp]

Red & Bold & Underline: Stem-loop [23]

Blue: Heptamer[30]

Green: Nonamer [2]

id-IGKV3-11[V_gene_segment]

atgcgttagacagagatgggcgggaatgattgaaggtctgagcattttagcacagattaagaccaaatcattaggattttaagagttgtgtacagttagtgaagaaaaagccctagaatttaatttgactgttgataaaacattcttggattagattgaagactcttttctgtgccaagtaagtatatttatgataatgatgatgactgtagtgctaaatatttaatcaataaaaacaaaaataattgccgcatacataatgtcctgagtactactgtaaatgttttatcttattttctttaaactgtctacagcactgtaaggtaggcaccagtat**tgtcacagttacacagatatggaaactgagaca**cagggaagttaagttagttggtcaatttcaagcaatcggcaagccatggagcatctatgtcagggctgccaggacatgtgactgtaaacagaagttttaactttttaactcaaa**gagggtatgtgtctgggttaatggaaagtttcaggaccctc**agaaaacattactaacaagcaaatgaaaggtgtatctggaagattaagttctaacagactcctcatttccatcgatccaataatgcacttagggagatgactgggcatattgaggataggaagagagaaatgaaaacacagccttttatattgttcttaacaggcttgtgccaaacatcatctgggtgaatttaggtgattgaggagaagaaagacataggaatgaaattc**tctgagcacaagggagaagttctacactcaga**ctgagccaacagacttttctggcctgacaaccagggtggcgcaggatgctcagtgcagagaggaagaagcaggtggtctct**gcagctggaagctcagctcccacccagctgc**tttgcatgtccctcccagctgccctaccttccagagcccatatcaatgcctgtgtcagagccctggggaggaactgctcagttaggacccagagggaaccATGGAAGCCCCAGCTCAGCTTCTCTTCCTCCTGCTACTCTGGCTCCCAGGTGAGGGGAACAT**GAGGTGGTTTTGCACATTAGTGAAAACTCTTGCCACCTC**TGCTCAGCAAGAAATATAATT**AAAATTCAAAGTATATCAACAATTTT**GGCTCTACTCAAAGACAGTTGGTTTGATCTTGATTACATGAGTGCATTTCTGTTTTATTTC**CAATTTCAGATACCACCGGAGAAATTG**TGTTGACACAGTCTCCAGCCACCCTGTCTTTGTCTCCAGGGGAAAGAGCCACCCTCTCCTGCAGGGCCAGTCAGAGTGTTAGCAGCTACTT**AGCCTGGTACCAACAGAAACCTGGCCAGGCT**CCCAGGCTCCTCATCTATGATGCATCCAACAGG**GCCACTGGCATCCCAGCCAGGTTCAGTGGC**AGTGGGTCTGGGACAGACTTCACTCTCACCATCAGCAGCCTAGAGCCTGAAGATTTTGCAGTTTATTACTGTCAGCAGCGTAGCAACTGGCCTcccacagtgattccacatgaaacaaaaaccccaacaagaccatcagtgtttactagattattataccagctgcttcctttacagacagctagtgggg**tggccactcagtgttagcatctcagctctatttggccattttggagttcaagttgtcaagtccaaaat**tacttatgttagtccattgcatcgtaccatttcaccgtggctattatgttcaactaaatgcattttagaaggcatctctgtttatggcatcacaaagagtttaataaatcttcggtgcaaaaataaacaacaaacacacatttaaatataaagctgaagtatcaaaacta**tttcagcactctgaaa**attggtgaagtataaaataattaaggatgcatattctttatagaaaaaaaaagtaccagtgctttgagtaaggacagaaaatgtctgtagccttttgcctgtgacagcagccttctacccccagctcaatcagcatgaattacagaactga**agttttaccaatatgagcgtagcaaaaaaaact**agcagcttgctgccaaagtgggtggacttgagtaaagccaaggag**tggaaaaaaaattattcagtgtttcca**gctaaacatgcagaactccataggaaatgaacagaaaaatctcacagctttacaagtccaagatgatgacctgtttgaggcaagtagtac**atctgctgatagtaaatagcagat**tcctggatgagatagacccatattgctgaaataatctctgcacacattcctagtgacctagaagatatagatatgagtgatgagaacaaggagtttccggtgcaaagtaaaaccagaaggattcaagaactag**ctgcattttgcatgcag**tttgctttttaaactacacacagatgggttgacagaagatagatttataacctctagatatttaagcaaaacgtctcaaatcattggtgaccacttcactatgcagatacatgtgcagttcctataaat

Uppercase: IGKV3-11

Lowercase: Flanking sequence[1000bp]

Red & Bold & Underline: Stem-loop [16]

Blue: Heptamer[35]

Green: Nonamer [6]

id-IGKV3D-31[V_gene_segment]

ttttagcctgatacaaag**agaagccagggaaagcttct**gagctcctgatctacgatgcatccaatttgcaaacctgggtcccattgcagttatgtggcattggatccaggacagatttgattctcaccattagcatcctccagtct**gaagttgctgcaacttc**ttattattggtcaacagtataaaagtgaccctctcacagtgttacaaacccaataagctccccaag**gaagcagatatgtgagggtgggctgccccagctgcttc**tcctgtttcctccatct**gctgagagtgtttctcagactcagc**cacactctgaaggtcactgagcagttttcatagaagcgg**taagggaagtgtctctacaccctta**gtttcttttatcctcctcagccccagcagcagacatggcaatgcctctcctaatttcataga**agagtcattacctatgaggactct**gggttacagcactggtccaggttcatacaacaaaagagaagctattgtaggtaacccaaatagaaagtttttcctaatatgggaaatccatgtctaaattacaacttttcaaagaccagaggatataatggttaggaaaccagaaaggcaaaaggagtgctgggaaatctacttcaagaagcaatggtggccgcgcacggtggttcaggcctgtaatcccagcactttgggaggccgaggcaggcggatcacctgagatcaggagttcaagaccagcctggccaacatggtgaaaccccgtctctactaaaaatacaaaaattagccgggcatagtggtgggtgcctgtaatcctagctactcagaaggctgaggcaggagaatcatttgaaccaggcaggtggtctctgcagtttcgaagcccagcgccatct**gtggctgttatgcatgtctctcccagccac**cctgctgtccagagcccatatcaatccatgggggtaggtctgtgaaaga**gcagctcagttaggacccagagggaaccATGGAAGCTGC**AGCTCAGCTGCTCTGATTCTTGTTACTCTGGCTCCCAGGTGCAGGAAACATCGGATGGTTCTGCATGTCAGTGAAACTTT**CTCAACCTTGTTGAG**TCCTGTTACCTGGCACACCTGCTGGGAAGGCACAATGATTAAAGCTCAATGTAGATCAATGGTCCTGGATGCACTGGGAAGACAATAGGTATGATGTAGTGTACATGTGTGACATTTCTGTTTTTATTCCAATTTCAGATACCACCAGAGAACTTGTGATGACAGTCTCCAGCCTCCCTGTCTTTATCTCTAGGGGAAAAAGCCACCCTCACCTACAGAGCCAAGTAGACTATTAGCAGCTCCTTAGCAGGGTATCAGTGGAAACCTGGACAGGCTCTCAGGCTCCTCATCCATGGTGCA**TCCACCAGGACCACCAATGTCCCAGCCTGGTGGA**GTGGCAGTGGGTTCGGGGAAAACTTCAGTCTCATTATCAGCAGGCTGGAGCATGAAGATTTTGCACTTTAA**CACTGTTATCAGCATAGTGGTGGGTATtccacagtg**attccac**aggaaaccaaacctccacaagacagctggtgttttttcct**caagccttctgtttacttatgggaagctactatggtggctgct**tagttattgagagaaaacaatggagacttcaataaata**g**aagtttataccacattcatgcattataaaactt**aatattgtgagtaacataattctcccttaa**ttaattcaatacagtgacaattaa**aatcccagaggacttttgtgaaattgaaatgatgttt**catatatatatagtatatatatatatatatatatatatatatatatatatatg**tactcacataccacacactatatata**tatatatatacacacacactatatatggtgtatgtatatatata**gtgatatgtatgtgtataatatatatatagagagtg**tgtgtgtatatatatagatacatacatatcacaca**tgcatttaaaaaggacaaaaatagccattggaatcttgcataagaataacaggtcaggctgggcgtggtggatcactcctgtaatcccagtaatttgggaggctgaggtgggcaggtcacctga**ggtcaggaggttgagactagcctgacc**aacatggagaaaccccatctctactaaaaatacaaaattagtcaggtgtggtggcacatgcctgtaatcccagctactcaggaggctgaggcaggagaatcatttgaacctgggtggtggaggttgcagtgagctgagatcgcaccattgcactccagcctggggaacagagccagaatctatctcaaaaaaacaaaaacaaaaaacaacaacaaaacaaaacaaaaaaagaataacagggcaggcatggtgtctcacgcctataatcccagtactttgagaggctgaggcaagtgtatcgcttgaggtcaggagttagaccaccctggccaacatggtgaaaccttgactctactaaaaaaatacaaaaaattagccaggcatggttgcaca

Uppercase: IGKV3D-31

Lowercase: Flanking sequence[1000bp]

Red & Bold & Underline: Stem-loop [19]

Blue: Heptamer[38]

Green: Nonamer [1]

id-TRBV30-2[V_gene_segment]

ttttttgtggctcgtgatcccttcagcccctaccttccaccccatgccattgtaaatctcccaatctttgctgcagacatagggagtcaaaaaaaaaaaaaaaaaaaaaaaaaaaaaaaaaaaagccaacatatgtaagagtgcc**tgcctgcagtgcaccaggca**aaatgcatctttcaacaatttttagatattgtcatcatt**attttcactttatagataagaaaat**tgagaatcagggtttgcttgagagttcccaagccagcaagaagcagagctgtgaaatgaacccaaatccatatgattccaagtctctgtgatgtttcctattactcatattgcatttagaggataaattagatct**agttggcccattagagacacatggagcagattccaact**ttgatcagttcaggagttaattatgcctgtccaggtctcaaaatctgtctgcc**agggatgaagattatgcaggcgatccct**ttggctgtcatagaagcttcctgccctcagaactgagcatttctgctttgattaccccaaaa**tacctgaatctcttgagccctatcctttcatctcccaggta**acatgtctagctc**tagtttccaatttcagataaacta**aaatcttttttcatccttctatcagtatcacagtttctctgaagctttcccaggcttctgatgtctcccgggggagctggagag**ttacctttagtttacaggtaaataatgagtcattaaatatatttac**ataccaggatgtcagcaggaatgcaaagaagataacaagtgcagagtatagctggttcccagggaggctgggggtgattcaccacactcttaaaagaagactagcctcatttcctccccagctcatggggtaggattcc**tctgctctggcagcaga**tctcccagagggagcagcctgaccacatcactggcccagaagaggaggcgtctgtcccccagactagctgaaggaaaggctggcttggatgATGCTCTGCTCTCTCCTTGCCCTTCTCCT**GGGCACTTTCTTTGGTGGGTGCCC**TCCTCTCTGGCAAGCCTCTTTTCTCATCCTCACCCTCACCCCAATCCCGTGAATGCAGAGTCTGAGCTGTTGGTTTGACACTCTTGAGTCTATTCCTGATTGTCTGGAACAGATATGGGAATTTAGCCTTAGGGGCTGATCTGATGATTCTTAGTTCTCTCTATCTCCTGAGAAGGAGTTCGTTCTGTTGGTCTTCCCTGGC**CCAGGGATGCTTTGCATCCTTGG**CTCCCCCAGTCTACTCCCCATGATGGATTCCAGGTGCCTGTGTCATGCAAGAGCTTGAGCTGCCCCTCTGTCTGGGGAAGGCAAGCCAGGCTGGGGTGCTCCTCTGTAAATGGCACTTTATCTTTCCCACAGGGGTCAGATCTCAGACTATTCATCAATGGCCAGCGACCCTGGTGCAGCCTGTGGGCAGCCCGCTCTCTCTGGAGTGCACTGTGGAGGGAACATCAAACCCCAACCTATACTGGTACCGACAGGCTGCAGGCAGGGGCCTCCAGCTGCTCTTCTACTCCGTTGGTATTGGCCAGATCAGCTCTGAGGTGCCCCAGAATCTCTCAGCCTCCAGACCCCAGGACCGGCAGTTCATCCTGAGTTCTA**AGAAGCTCCTCCTCAGTGACTCTGGCTTCT**ATCTCTGTGCCTGGAGTGTcacactgagct**gggtggggcagacatctgtgcaaaaaccccaccc**tctcctgag**ccctaaccatactccccaggggccttcacttaggg**actgggtggaggatatttgtaagtaggtttctgactgctggccaatgtgaaagcaatggaataaagcgagaattctacaaaagactctactaa**gttctcagggagaac**aaaaaattgtatg**tgtgcagaacctgtgatttgcctgcaca**tagtcaagttctcaat**gtatggatgtcccggcccaggctaccatac**tccagccct**caaggtgtgctataccttg**cctgattttctagtcttccaaattcttctatgtgtgccgtggggagggggtttgtgcatatatggctgagagaaaggatatacctgtggcaggaggtgagctggtgaacaaacgggaggatgccaac**ccaggttacttaattgattaatttcatcacctgg**ag**agtctcctgtcattgatctttctgctgagact**agaaattatggggaggcttagacctgaggctttgagtcaggcaaagaggcaagatgttcaggagactgttctcattcgaggcagaagtgtggttttgttagtataggtaacatgcacttaggccagacgtggtagctcatgcctgtaatcccagcactttgggaggctgaggcaggaggatcacctgaggtcaggagtttgagaccagcctggccaatatggtgaaacctgtctctactaaaagtacaaaaattaaccgggcgtggtggcaggcgtctgtaatactagctacttgggaggctgaggcaagagaatcacttgaacctgggaggtggaggt**tgcagtgagccaagatcatgccactgca**ctctagcctgggt**gacagagcgagactctgtc**tcaaaagaaaaaaaaataaaaataaacttgcacttagaaattttggtgatgtttgaggttttaaaaaatgtcctgttgtactatcttgtagccc

Uppercase: TRBV30-2

Lowercase: Flanking sequence[1000bp]

Red & Bold & Underline: Stem-loop [22]

Blue: Heptamer[19]

Green: Nonamer [5]

id-IGLV5-52[V_gene_segment]

ttctgatgagcctctccaaaaggaggcaatcagatatgtgtttctctcagtgagcagaggggagactttcaataga**ctgggaggcaggtttgctctaagcagttcccag**gttgagtggggtcccaagatttgttttcctttcacactgtcttctcctgagccccccaaacccttccaacctctggccatgactgagttccacagctgcttttgtactatcaggtatcttttagccatactttatttaccaattatctgtgttaggacattgttgcattgctacaaagaaa**tttgttgggaaatttacaaacaaa**agaggttgaattggcttatggttctgcaggctgtacaagagtcatggtgcctccatttgcttggcttctgaggacacctcaaggggattttacttggtggaacatgaagtgggagcagacatctctcatggcagggtgggagcaagaaagagagtaaggcaggaggttccacacatctaaacaatcagatcttgcaagaactcactcactgtcatgaggacagcaccaagcctcgcggaatccgcttcaatgacccaaccacctcccaccaggtgccacatgcaacattggggattacgtttcaacaggagattgagagagg**acacacatccacactatatcaggaggtgtgt**gccttgcaaccatcaggggatgctgcaggactgccctatggtccctacacagctgcttagtaagaaccattcactccaaggaacctgtgccctgtgctcaatgat**ggggactcaggagctgtgtcctctctggcctggcagagtcccc**taagca**gagacctgtgtatggtctc**ggcaggtgcttcctcccagggccctcccgagggggaaaccaaaccttcctcctcctcagt**ctggagtgtgagctaagcttcagctccag**ggctcagggaggggctgggcagtccttggatggaaacatatttgcatgagcagccc**ctcctctggcagaggATGGAGAAGAAGAGGAG**ACCTGGGGCAGCTCAGGCCCACTGTTAGGGCTCAGGGGCTGTGTCCACCATGGCCTGGACTCTTCTCCTTCTCGTGCTCCTCTCTCACTGCACAGGTAGGGAAAGTCCTTATAAACTGAGTCTCAGTGTCCAACCTACACCATCCCCTGTGGCTCAGACCTACAAGAAGCTTTACCCTGGGAACTGCCTTATCACCCATGATGTCTGTGTTTTCAGGTTCCCTCTCCCAGCCTGTGCTGACTCAGCCATCTTCCCATTCTGCATCTTCTGGAGCATCAGTCAGACTCACCTGCATGCTGAGCAGTGGCTTCAG**TGTTGGGGACTTCTGGATAAGGTGGTACCAACA**AAAGCCAGGGAACCCTCCCCGGTATCTCCTGTACTACCACTCAG**ACTCCAATAAGGGCCAAGGCTCTGGAGT**TCCCAGCCGCTTCTCTGGATCCAACGATGCATCAGCCAATGCAGGGATTCTGCGTATCTCTGGGCTC**CAGCCTGAGGATGAGGCTG**ACTATT**ACTGTGGTACATGGCACAGCAACTCTAAGACTCAcacagt**gctccagacccatgaggaagtaagacaaaaccctcccctctactctcctggtctagtgaaatca**cccctgctggtggctctgaccaaatctagctcagggg**gtgacatctgttgtctcatcgacagcactggagctcccacggagaagtaggtctaacaaagtcctaccttgctcgcatttcacaggcagacatcccttgctcaggggtaaaatctctcagtgttgacattgaagaaggcaagagggctgggttggggttcatctcagctcagagtctagaagtga**ctcagagatgctgctggtctgag**agttttaatatgagccttggg**atacaatacagggatgcagatgggaattgtat**tctctttgtcagtaacgaaccatcagtaattgaaaagtcaatacattgagaatagcataaagcatgaa**atactttaggataagtat**gacaaaaattgt**tgttttacacactagaagtcctaaaaca**ttgtttag**acacaggtcttaaagaagacctgagt**aaatgcacagataatcctttttttttttttttttttttgaggcagagtttcactcttgttgcccaggctggagtgcaatggcatgatcttggatctcggctcaccacaacctccacctcctgggttcaagcgattctcctgcctcagcctcccgagtagctgggattacaggcatgtgccaccacgcctggctaatttttgtatttttagtagagatggggtttctccatgttggtcaggctggtctcgaacacctgatctccggtgatctgcccgcctcggcctcctgaagt**gctgggattactggcgtgagccaatgcgcccagc**ccagataacccttttttttttttttttttttgagcaggagtctcgctctgtcgccc**aggatgcagtgcagtggcgcgatcttggctcactgcaacct**ctgcctcccggttgaagtgattctcctgcctcagcctccctagtagct

Uppercase: IGLV5-52

Lowercase: Flanking sequence[1000bp]

Red & Bold & Underline: Stem-loop [19]

Blue: Heptamer[38]

Green: Nonamer [2]

id-IGKV1-12[V_gene_segment]

ctgtccagtttcatctggatccaagtgatttctccatgtacataggc**aactgcttgataagagattcagtgtctctttcctaaagcagtt**aacagggaggctgttgcctgggtcaggatgatttccccaatcactgataaaaagtaaaagaggaaagtgtcattg**atggtgcatggcagggacatgctccat**gcagtggtcaccc**tcagtaagagagatgaactttgggaaataatactga**atggcagaaaagaaggtagactatgaaggtgcccaaaacaagaataaggtgcaacccatttagtgtctgagtactatagagacctgtcgctcttgataattgtggatctgtgactgctgcatgcatcgagaaaacacggtatcatctttgtgtatctatagtacatagtttgatctcatactggtaagaacaatggcataacaccattacctgatacttacaaatgtatgtaccatcatgtcaataaattttatttttaatttttttacataggaacaatgttaaactcacagaaatgttgcatgtatacgacaaatacccccttccctagccggaatcatataag**agtcttttgaagact**tgagaattgtaccgtctaacattttactatgtgtttcctacaaacaacaatattctcctaaataatccccatacaccaa**tgaaatacattagtccatcaactcctgaggaatatttca**aatcgtcaaaaagaaaccaaaaaatgttactcacaacaaaatagttcccagtagaaacacattctctgcagacaaatttgtgctaccctggtcttacctgggacacctggggacactgagctggtgctgagttactgagatgagccagccctgcagctgcgcccagcctgccccatcccctgctcatttgcatgttcccagagcacagtctcctgacctgaagacttattaacaggctgatcacaccctgtgcaggagtcagacccagtcaggacacagcATGGACATGAGGGTCCCCGCTCAGCT**CCTGGGGCTCCTGCTGCTCTGGTTCCCAGG**TAAGAAAGGAGAACACTAGGATTATACTCGGTCAGTGTGCTGAGTACTGCTTTACTATTCAGGGAACTTCTCTTACAGCATGATTAATTGTGTGGACATTTGTTTTTATGTTTCCAATCTCAGGTTC**CAGATGCGACATCCAGATGACCCAGTCTCCATCTTCCGTGTCTGCATCTG**TAGGAGACAGAGTCACCATCACTTGTCGGGCGAGTCAGGGTATTAGCAGCTGGTTAG**CCTGGTATCAGCAGAAACCAGG**GAAAGCCCCTAAGCTCCTGATCTATGCTGCATCCAGTTTGCAAAGTGGG**GTCCCATCAAGGTTCAGCGGCAGTGGATCTGGGAC**AGATTTCACTCTCACCATCAGCAGCCTGC**AGCCTGAAGATTTTGCAACTTACTATTGTCAACAGGCT**AACAGTTTCCCTcccacagtgttaccaacccgaacataaacccccagggaagcagatgtgtga**agctgggctgccccagct**gctcctcctgatgcctccattggctg**agagtgttgctcagatgcagccacactct**gatggtgttggtagaggggtacgtgaaatcgcctctgcaccctaattcttttctctttctcagccccaactgcacagacatagcaatgcatctcctgatttgataaatacagagatcatgacacttgaggagtctagttta**tggcttcagcttgaattcatataacacagaagaagcca**ctatagatattctaagcaggaattgtcttaatacagagaattaaagtaaactactaaagtcaaaata**aaatgtagagatgaatctctaaattt**aagttttatttgcaaagaaatatttgccaggtggggcatacaggaaaactcagtggtcttcaagatattggaagaacgaagagaaaattag**cattttatgaaaaagggaaaatg**ttacctgtggctctttgagaaagttcattggcactaggaagggttgggagctggcaagctcagactggtaagcagtggtggacaaaatgaatcctagaattatatcaagttatctcagaagttgtggacaaatttgatttcaggttacaataagccaaagcagtgaaggttgcagagaattttgtt**actgaaatgccagggattcagt**gaagactctgcagcttaccccacagaaagccaatcactaagacaacaagcattgccaaggaacaggctttaatcaggtgctgcagccta**ggagatgtgataccattctcaaatgtatctccctgaccaactgaaattaggagtttatatagcaggga**agaaatgtgggaaaacaggaattagggaggggtaaggaagataatttggtcaacaggaagcagga**ggtcagtgaggcaatcataatgggtgaaggttctgatgtctcactgtcc**cgattcagtga

Uppercase: IGKV1-12

Lowercase: Flanking sequence[1000bp]

Red & Bold & Underline: Stem-loop [19]

Blue: Heptamer[25]

Green: Nonamer [5]

id-IGHVII-53-1[V_gene_segment]

gccatttgtgcctccctgagcacagttgtccatcagagatgtcccatgtgtcccagcagcagccatgtctcagtgtcttcactgtgcacagccattgt**ctgggaggagctcccag**gatgggtgtctttggcacacaccaggtggcgggtgttagagtgcggtgcagcagctggctgcctgttctattgggctccctga**tgctggagagatgggaggtgcattctcaggtccagca**ccctgtttgtgaatttttatataaaaccatgattttacttcattttctcagatgacatagataattaggaacagaacctgcaaagaaattgtaattttcaactttaccccaaatttattgtttcttaattctgtgtaagatccagacatattattgccttcctca**tgagaaattgttctatttaaaatgaaattagttttttctca**cattctttgtttctgttcaagtacagagatctttattaaagtaagttgggt**tctttccacacactaaccctcacctcccccagagaaaga**gcagagattttcctca**ctctgagtctaagggaggagctgttcctgcacgactcagag**cctgcagagacccccccccgccaggtgcagcttcagtgagtcaggtatttctccttgtgggtgacctccaccgccagtgattgctgctcaggtctaattgtgggttaagcattaggacaccc**ttcaggtgatcacatctcagtcttattctgaaaatcaccatgaa**cagagatagttc**aatgtctattctcctgacatt**ag**tttctctttattatttggttccaagtatggagaaa**aatg**tgacaataaatttgtca**gaatctaacctcagtatccactgcattactctaggagactcacaaattgaacacaaatgagccctttattctcataaaagtgtatgtatttgggaatttcaatgtgttctccagaacctgtgcatgccaacaactgtgtttctcagtgcccacttGGCCTGGTGAAGCCCTCACAGACCCTCTCCCTCACCTGTGCTGTCTCTGGATTCCCCATCACAACCAGTG**CTTCCTGCTGTAGCTGCATTCATAAACCCCCCAGGAAG**GGACTGGAGTGAATCCG**GTGCACAGGTCATGAGGGAGTGCAC**ATTCCAACCCACTCCTCAAGAGTCCAGTCACCATCTCCAGATCCATGTCCAAAAAGCAGTTCTTCCTACAGCCGAGCTAAGTGAGCCGCAAGCACACAGCCATGTATTTTTAACAAAAGAcacagtaaggtaaccacagtgggaactcacacccaaacctc**cctgtgggggtgcacagg**acagccacagttactcaggaccccaggattcctcaggacaccaaggggcactcaaggccattgtagatgccctcaggtagccaagggttctcaggaaacatggaggaaaaccaggaccccaaaaggtgctccgtacagcaggggactcaggacaattgtggggactcagagcaggctcaaagctcagcttcagggcaggtgcagctggggttgaaaggggctggatgaggggttttgtgacaccatcatatttcaccac**tagacacactccactttgtctattctaacgcatgagagtgtatgattagaa**aatgatatttatataaatacataaccatagttagctgtgtcaagttgtcctcttgctaggtgtccatagctaggtgcatcagccttgtccataaggactaattccccgcaattactgga**gaatctcataaattgtggtcaattatgtcagattc**ctctctttttctgccttcctttctcccttcttctctctctctcacacagaaacttacatacacccaccccacaacacaccaaaatctataact**tttattacctgatatattcaataaa**cctgattaatgtgcagcttttccagcttcgttatttatgctgttgtaacaataagaacaatgtgtttcctagctgtgtacttctctaagctgagtagcatct**ttgtttataatacctagaattaaaaacaa**cccaaaagtcaatcaccagcttaactggtaaacaaattgaggaaaagtcattcattgacatactatccactactaccatcaactaatgttgggtacactcaacagcatggttaaat**tcacaagtacttgtga**tgagtaaaatgagccaaagtaacaaaagtgcatacataagatacaactttcataaattctatagaacaaaaagtaatctaaagttacataaaaatc

Uppercase: IGHVII-53-1

Lowercase: Flanking sequence[1000bp]

Red & Bold & Underline: Stem-loop [18]

Blue: Heptamer[42]

Green: Nonamer [5]

id-IGHV3-49-2[V_gene_segment]

ggtaaaatatccatttatgctaactaaaataagccaaacaaatgagaatgtatactatgttat**ttcatttttataaattctgaaaaattaaaatgaa**tttgcagcaatgtaaagatcagtagttgccagggaaatggtagaagaaagaaaggaaaaggagaaagaatacagaagaacaaaaggaaatgttaagaattctcttgtccaac**ttgataaggatgacggttacatcatttttatcaa**ctgtaatctttaaatatgtg**aaagtttattatctgtaaactaaacttataaacttt**attacaagcaaaaattgaaagttagacaagaaggagtgatagaaagagaaaatgtata**ttaaatttcagaaatatttaa**gaatgtatctgcctgaaccctagttctcaccatatctttaggtgaatgctaaaatgcagcaaaatcacgcatgttctcactacagaaagtgggttctacaaaccacactcggcacatttagctttgtcctggagttggttcagggagttattggggcc**agtgatgaggagcacaggccaagataccagcgatcact**tatcccaaacatgagctctaacat**acacacttagtcccttttccgtgtgt**ggtttacttccacatctgtacatggagagaccactgactgacaaaata**taatttatacaaatatgtaaaatta**aatagggtgatcagttcaaggtgtttatcacagcataattttacaataagacagcatatttcccaaataccatcattgtcaccaaactccttcaaggcacagtcatctta**tctgggccccgtcctctcctcaggtgtcccaccccaga**gcttggtatatagtaggagacatgcaaataaggccctccctctgctgatgaaaatgagcccagccctgaccctgcagct**ctgggagaggagccccagccgtgagattcccag**gagtttccac**ttggtgatcagcactgaacacagaccaccaa**ccATGGAGTTTGGGCTTAGCTGGGTTTTCCTTGTTGCTATTTTAAAAGGTAATTCATGGTGTACTAGAGATACTGAGTGTGAGGGGACATGAGTGGT**AGAAACAGTGGATATGTGTGGCAGTTTCT**GACCTTGGTGTTTCTGTGTTTGCAGGTGTCCAATGTGAGGTGCAGCTGGTGGAGTCTGGGGGAGGCTTG**GTACAGCCAGGGCGGTCCCTGAGACTCTCCTGTAC**AGCTTCTGGATTCACCTTTGGTGATTATGCTATGAGCTGGGTCCGCCAGG**CTCCAGGGAAGGGGCTGGAG**TGGGTAGGTTTCATTAGAAGCAAAGCTTATGGTGGGACAACAGAATACGCCGCGTCTGTGAAAGGCAGATTCACCATCTCAAGAGATGATTCCAAAAGCATCGCCTATCTGCAAATGAACAGCCTGAAAACCGAGGACACAGCCGTGTATTACTGTACTAGAGAcacagtgag**gggaggtcagtgtgagcccagacacaaacctccc**tgcaggggcgcacagagccaccagggggcgcta**gggaccgactgagtacgggacaggtccc**aggagcaggtgcagggggaggtttcct**ttttccttggctggaaaa**gtcacctttatcttcccaggactcgagccttctaggctgtgatattttattacttgtatttactgttcatttattatcattagtttttaaattttggtaatttttacaactctatggatatatttttaagtgtatactttcaag**aaataaacattcctaattattt**gcactgattctcccagagttttattaacatttgttgacatcagcaactacatagctatagggacaaacacttttaacgatagacagttgtttaggcctgaaaccccgtttatactaaaaatttacaaaaattagcctggcgtggtgaggggcgcctgtaatcccagctactcgggaggctgagcaaggagaattgcttgaatccgggaagcggaagtcacagt**gagtggagtggactgccactacactc**cagcctggcgacagagcgagactttgtatgaaagaaagaaaagaaaaggaagaaaggaagaaaggaaggaaggaaggaaggaaggaaggaaggaaggaaggaaggaaggaaggaaaagagagaaagaaagagagaggaaggaaggaaggaaggaaggaaggaaggaaggaaaagagagaaagaaagagagaggaaggaaggaaggaagaaaggaaggaaggaagaaaagaaggaaagaaagaaaaaagatatataaacacgcaga**cctatgcatataaccatagg**gatttatattaaacattacaataaaataattctaaaaatgtgtcctaaggaatcaaacataatgatgaagtaaa**tataaatgttagagtaatttata**attgatttgttatttttaatcgtttacttgaatttatttctatttgttcattta

Uppercase: IGHV3-49-2

Lowercase: Flanking sequence[1000bp]

Red & Bold & Underline: Stem-loop [20]

Blue: Heptamer[21]

Green: Nonamer [3]

id-IGLV3-6[V_gene_segment]

ccaaataacctaaagttagtttctcagtcaggtctgggtttctggtgctttttacacatttatatgaaacactgatgagaaaaaataaagaatcaaagagcagaggatggcaatttctagtctattagagagatggggggtgtcatacaaggag**caactcaggtgagaaacgagactagactgtgtaatgaatccacatgagttg**tggctgctcaaaggaccaaaccatggtaatggaaatcaag**tttaggatataagtatcctaaa**gacacagaaccacattctaattaaactaccaggacaggaccaagtgag**tctgggaaacccaga**gctgacttggaggaggaggaagatgatgggttcctggcccacctgc**tgctgagtcaggtgtcattgatgatctctcagca**cggcccagctcagccccat**ggccaggctcctggaaggtccacagccctgctgagcctggcc**taggacacagcacaggccaggatggtataaaacagtggtcatgtggccaaactccaggaggtgactgtgatcacaggccttcaaggatggagtctgatcttctgaccagagaagcagataaagtgagctgggacccacttcttcttcctcagtctgctgtgcttcccacagagctggtttgagtga**catccttcagtgaggagtcccagaggatg**ccgatttgcatcaaacactcac**tgcctcaaacggcctgagaggca**ataagagagagctgg**tgagcccagctgtgctgtgggctca**ggaggcagagctctgggtgtctcaccatggcctggatccctctcctgctccccctcctcactctctgcacaggtgctgcccccaagcccttccttcag**gctcagaacccataggatcctgagc**tgggcctgcccaaacatgagttca**tcccaggcacaacctcagggtgggaccccctggga**acagattcatcatttacaagcctcctctcctgtcctctcttgc**aagcTCCTATGAGCTT**ACACAGCCACCCTCAGTGTCAGTGTCACCAGGACAGGCAGCCATGATCACCTGCTCTTGAGATAACCTCAAAGATGAGTATGTTTA**CTGGTTCTGGCAGAAGCCAGACCAG**GCCCATACTGGTGATATATGAAGGCAGCAAGCGGCCCTCAGGAATTTCTGATTTTCTGAGTCCAGC**TCAGGGAACATGGCCACCCTGA**CCATCAGCAGGGCTCAGACTGAGGACGAGGCTGACTATTACTGTCACAGGTACAATAGAAACAGTGATGAGCCcacagtgacacagg**cagattaggaagtgagacacaaaccccttcccaatctg**tgtcaccctctttct**ccagccccaggatggctgtggacaaagccatgagcagggctgg**tccagttcacccagttctgtgacccccaggctg**ctcttccctccagccctccaggcaggttgtgaagag**ggtgagttaagaagagatttgaggctgtgtgacaggc**tatcctggggtttttctggccaaggata**aagctggtgaaagagagcctgaatggacgggcagacagagtggcccatgcacgtagtggcaaaagacctgggtttccacatgccagcccttcaggcaggctctgcagagagtgggttaggaggagatttgaggctgtgtggacaggctgttttggggttttctaacccaggataaagctggtgaaagagggcctg**gatgaaaggacacacaaagcgtatctttcatc**tagtggtaaaggatgtgagttccatatggtgaagcctgggtcaccccttggcccagacgtcctaggttagtcctcagcatttccctagcacttaagttcctgaatctcctggcagccttatattcaaaacggtgtaagctggagtggacgctgtg**gggtccacctagactccacgttaaggtcgaccc**acacactctcagagcctacgagtggatacccatgctgtggggcacatgtcacctgacagagtggctgaccacccctgatctcttctttctgaccacttgggtgtttggcacta**tgtccatcatggaca**ctctatgggctattaacctatgattcaaagtttcttctgtgtggtgcccgtgcccaaagaccacccacaggcttctgcccatactccacttcccccatttccaagtttttctcttgcagtccctggaacacctgcccaggtcatttcccactgccctaagcccctatctgccctggcatccggccactcacctttcctcacaaaatgaatgac

Uppercase: IGLV3-6

Lowercase: Flanking sequence[1000bp]

Red & Bold & Underline: Stem-loop [20]

Blue: Heptamer[52]

Green: Nonamer [6]

id-IGLVI-70[V_gene_segment]

aacttttgggtcatcttcttgatctttccataaaactctaacccttctattcttaacctcccccattcccaatttcaccctgatgtgcacatcctacctggtgcctagtcatcccggtggcccctctctgggctgtatctagggagagtggtgaccttggtcatcttcctcaggtccaagtatccata**agagaagcagacttttctct**ctttcataatgatggtctcttcttgaataatttccctgtgcattttcattaagggtctcaccccatcaagggtgtgatgaggacagtgatgagggtcacggagcccggcctcccacggggggatgtggggtatgatgagagagagaagcaggtgccctcaggcccatatggaggctgg**ggagggttgttaaccctcctgccctcctggccactgggtgcagga**catgttctctctgaggctcactcacctatggccccaaccacacaaaaatctaaatggggaagaagcagaggatgacttcatgctgaggaggcccatcttgcagcgagaaa**tgtgctgtggtgagcacctgcagcaca**aagagcctggggtgggggtggaggggcgtg**agcgctgcctctgtgtggggatggtgtgcgcaaggcagtgct**gaggaccttcaagccagtcaggaatgaacctgggctctgagacagtggagggaagtgtggcctgtccctgagggtctgtggg**ggtttcaactgggaaacc**aggccctggacccctacc**ctgggccctctgctctgtgctttctggggcccag**cagctgggacctccc**aggagcagaagaggctcct**ctagccccaccccacactcaactttccccaagttcaggcccagggatagcagggggaggggtgtgatttgcatgaagggtgctctgtgcc**ctcttaggaggagaggataagag**agactctgtgcagt**tctgctcagctgtgggcttaggaagcaga**gcctggggcatctccaccATGGCCTGGACCCCTCTCCTCCTCCAGCTTCTCACCCTCTGCTCAGGTGACTGCCTGTGGAATGCCAAAGTGATTATTGGGGACACATGGGATGACTTTTCTCTTATATTTTAACATTGTGGGGTGGGTAGTGAACCCAGACTCACCTCTCTGTGCCTGCCTCCTCTGTTCCAGGG**TCCTGGGCACAGTCTGCGCTGACCCAGGA**AGCCTCGGTGTCAGGGACCGTGGGACAGAAGGTCACCCTCTCCTGTACTGGAA**ACAGCAACAACGTTGGAAGTTATGCTGT**GGGCTGGTACCAACAGATTTCTCACGGTGCTCCCAAAACTGTGATGTTTGGAAATTCTCTGCCCTCAGGGATCCCTGACCGCTTCTCTGGCTCAAAGTCTGGGACCACAGCCTCCCTGACTATCTCGGGCCTCTAGCCTGAGGAC**GAGGCTGATTATTACTGTTCAACATGGGACTACAGCCTC**AGTGCTCAcacagtgctgcaggcacatggggaaccgagacaaaaacctgcccttggcctgtcccgaggctgatcactccatacttgcctatgacaaacaaagagggt**gcctgtggctgatcgtacagtttaagcaagggaggaagtgagactcagccacaggcccctgcaaagggtcagcgggcct**tgttcccacattgtctctatttgaaa**aagtgagataaatttcactt**cgtataaaatgtacaattaaattgtcttattgtgtttacaga**attatgcatctatcaccataat**ctaattttaaaacgttttcatcaccccacaaaaaaacccccacaattagcagccacccttcaccctttctccccagccactggtcattgctaaagactttctgtaaagaacatgcatttaaagacctaagaaactcagtgatgccaaacagctaggtttttcaaatgcccaagcatgtaataatagattactgaaaccaaagataaagacaca**attatgaaagcagacagaaagctttataat**atatacagaaaatcaacaattcaaacaatgagaggtattttatgagtatacattgaggccaaatggcagtgttat**gttcttacaaagtcaaaagaaagaaagaaagaac**tgttgagtcataatgctatatccaggtaaagatt**ttttttttttaaaaaa**ggtaaaaaataagactttgtc**agatgaagaaaacctgaaagaattcatct**aaaaaaaatcaaaacaaatcaactctaagagaaaattttcaggtggaagataaatgataatggaaacattgaacccaaagaatgcaggaagaataaccaataaatatagtagtaaaagaccaaatttttagctctacagttctggaaaatatgc**atatcaaaagctaagttatagcatttttgatat**aatttctacttatgaaaaatat**tatatatacatatatacacatatata**catatatatacacacacacacacac

Uppercase: IGLVI-70

Lowercase: Flanking sequence[1000bp]

Red & Bold & Underline: Stem-loop [23]

Blue: Heptamer[41]

Green: Nonamer [4]

id-IGKV1-16-3[V_gene_segment]

tctggaac**caactgatttctccatgtacctgggcagttg**cttgataaaagattgagtgcctctttccaaaagatg**ttaacagggatcttggtgtctgggtcaggatgatgtccctgataa**aaagtaaaagaagaaagtgtcacct**atggtgcatggcagggacatgctccat**gcagtgatcaccctcactaagagagatgaatgttgggaaataatacttaatggcagaaaagaaggtagactatgaaggtgcccaaaacaggaataaggtgcaaaacaggaataaggtgcagcccattta**gtctctggatattaaagagac**ctagagctcttgataatggtggatctgtaattgctgcatgcattgaggaaacacggtatcaacttagtgtatctgaagtaaattgc**ttgatcttacagtggtaagaacaa**tggcataacaccattacctaacatttacaaatatatgtagcatcatgtgaataaattttatttttaatttttttttagaaaggagcaatgttaaactcacagaaatgttgcaagtatatgataagtaccaccttccctaacagaaatcacatgagagtcttttgaagacctgaaaa**ttgtagggtctaaaattttactatgtgtttcctacaa**aaaagaatattctcctaaataatccccatacaccaa**tgaaatacattgctccatcgactcccaaggaatatttca**aattgtcaaaaagaaaccataaaa**tgtttcccataacaaaatagtccccagtagaaaca**cattctctgcagacaaatttgagctaccttgatcttacctgggacacgtggggacact**gagctggtgctgagttactgagatgcgccagctc**tgcagctgtgcccagcctgccccatcccctgctcatttgcatgttcccagagcacaacctcctgccctgaagccttattaataggctggtcagactttgtgcaggaatcagacccagtcaggacacagcATGGACATGAGAGTCCTCGCTCAGCT**CCTGGGGCTCCTGCTGCTCTGTTTCCCAGG**TAAGGATGGAGAACACT**AGCAGTTTACTCAGCCCAGGGTGCTCAGTACTGCT**TTACTATTCAGGGAAATTCTCTTACAACATGATTAATTGTGTGGACATTTGTTTTTATGTTTCCAATCTCAGGTGCCAGATGTGACATCCAGATGACCCAGTCTCCATCCTCA**CTGTCTGCATCTGTAGGAGACAG**AGTCACCATCACTTGTCGGGCGAGTCAGGGCATTAGCAATTATTTAG**CCTGGTTTCAGCAGAAACCAGGGAAAGCCCCTAAGTCCCTG**ATCTATGCTGCATCCAGTTTGCAAAGTGGG**GTCCCATCAAAGTTCAGCGGCAGTGGATCTGGGAC**AGATTTCACTCTCACCATCAGCAGCCTGCAGCCTGAAGATTTTGCAACTTATTACTGCCAACAGTATAATAGTTACCCTcccacagtgttacacacccaaacataaacccccagggaagcagatgtgtgag**gctgggctgccccagc**agcttctc**ctgatgcctccatcag**ctgagagtgttcctcagatgcagccacgctctgatggtgttggtagatggggacatgaagtcacctctgcaccctaattcttttctctttctcagcaccaactgcacagacatagcaatgcttctccttatttaataaaaacagagatcatgacacctgaggagtctagttta**tggcttcagttggaaatcataatgcagaagaagcca**ctataga**tattctaagcaggaatagtcttaatacagagaattagaata**aactgctgaagtctaaata**aaatgtagagatgaatctctaaattt**aatgttttatttgaagagaaatatttgccaaatgaggcatacaggaaaactcagtggtcttcagtacgttggaagaacaaagaaaaagtcagtgttccatgaaaaagggaaatattacctattgcactttgagaaagttcattggcactaggaagggttgggagctggcaagctcagactggtaaacagtggtggacaaagtgaatcctagagttatatcaagttatctcagaaattgtggataaatttgatttcaggttacaataagccaaagcactgaaggttgcagagaattttcttactgaaataccagggacttggtgtagatcctgcagctcaccacacagaaagccaatctcgaagacaacaagtattgccaaggaacaggctttaatcaggtgctgcagccgaggagatgggacgccattctcaattgtatctctctgacaaaattaattaggggtttatataggagggaagaaatgttgaaaacaggaattagggaggggtaaggaagataatttggtcaagaagaagcaggaggtcagttatgcaatc**acagtgggtgaaggttctgatgtctcactgt**cccaattcagtgatatg

Uppercase: IGKV1-16-3

Lowercase: Flanking sequence[1000bp]

Red & Bold & Underline: Stem-loop [21]

Blue: Heptamer[25]

Green: Nonamer [3]

id-IGHV3-52-2[V_gene_segment]

tcctactctgtgacccaggcaggactgcagtggtgggatcatagttcaatgtagcttccaacttctggtcaccaacgatcctcctgcatctgccatctaagt**agctgaaactacagttgtgtgccaccaggctcagct**tgagtactt**attaaatcaaacactttatgcaatatttaat**gtatggcaataatgtctcattgagagtatt**acaaataaatgaatggataatttgt**tcagtacagattgatggaaaatagacactaacatgaggaatgtctgaca**tttatgaacatacaactgcataaa**a**tgtgttctcttacattcattaggtaaacaca**atagtgcatacacatcaaaccatgctttcattacaggaaggaagttctgaaaatgtcactgggggtgacccacgctgtgctgggcttggttcgggggcagtcaggcccggtggtgagaagcacagg**cccagatacccaggcttactctgcaaatgtgagctctggg**gacattgtaccacccatctgtgcttctgctggtaat**tttccatctgtaacgtggaaa**taacattgatac**tacataccgtgatttctccacatatgta**aaaataaaataagatgattgctgctaagtttaaataagggcagttttcataggtccattgtacctcaataaaatta**ctttaaaataaaaattacaaatacagttgtaggtttaaag**agtttatcacagaacaaacttataataagaaactatattttcaaaaattgtatcaatatctctaaactcccccaggacacactcacct**gctctgggctctccactctcctcaggattcccaccccagagc**ttgctatatagtaggagacatgcaaacagagccaaacctctgctgatgaaaagcagcccagccctgaccctgcagct**ctgggagaggagccccagctccaggattcccag**gtctttccatttagtcttcagggctgagcacagaggactcaccATGGAGTCTGGGCTGAGCTGGGTTTTCCTTGTTGCTATTTTGAAAGGTGATTCATGGGGAATGAGTTGAATGTAAGTGAATATGAGTGAG**AGAAACAGTGGATGTGTGCGGCAGTTTCT**GACCAGGGTGTCTCTGTGTTTGCAGGTGTCCAGTGTGAGGTGCAGCTGGTGGAGTCTGGGTGAGGCTTGGTACAGCCTGGAGGGTCCCTGAGACTCTCCTGTGCAGCCTCTGGATTCACCTTCAGTAGCTCCTGGATGCACTGGGTCTGCCAGGCTCCGGAGAAGGGGCTGGAGTGGGTGGCCGACATAAAGTGTGACGGAAGTGAGAAATACTATGTAGACTCTGTGAAGGGCCGATTGACCATCTCCAGAGACAATGCCAAGAACTCCCTCTATCTGCAAGTGAACAGCCTGAGAGCTGAGGACATGACCGTGTATT**ACTGTGTGAGAGGcacagt**gagg**ggaggtcagtgtgagcccagacacaaacctcc**tgcaggggcatctggagccacaagggggcgctcaggatacacagaggacaggg**gcagccccagggcaggtgcaggtggaggtcaagggctgc**tctccttcagggtctgtggcttcctctcatctaacagttccgcagggagcctcttgtatttacagtgatgtgctactgaggtttctaagtttgtaaagtttattactacaggaggaaccactatcaaacgcccttaaggcaggtgtcactaatggagaaaggaaagtgcacaggaggctgggtgaggctgtggacactgtctgcctatgattcaagtttcacaagcagtgacggagaaatgggagtttgatggagctccctaactaccatgtggtctaaactaagtccaactaag**tccctgagctctgggtgcccatcaggga**tccgccatgtgcccggcagccgcgtgcctttttgtctcctctgcgcccaatcactgtctgtgatgagcttccaggatgtgtgtgtttggcacaaaccaggtgatggacgtcagacagcagcagctggtgcccggatcatgggctccctaatgctggaggaatgagaggtgcattctcagggacaagacattgttgatggatttttatgtagaaaccacgattttac**ttgcttctctcaggagacatagaagaagcaa**ccatgcagtcagcaaataattataatttccacaattaccccaaatggttaatccttaattctgcgcagggtcccaccgtagagtcacctttctcatgaggaatggttgaatctagaatgagtccacatgattttcatatttttggcttctgtccatgttcagagagttagagtagagtaagtttggacctttccacacactaagcctcacctcccccacagaaagagcagagacttcaactattcctgagtgtggggtaggggctggtcctgcacacctcagagcc

Uppercase: IGHV3-52-2

Lowercase: Flanking sequence[1000bp]

Red & Bold & Underline: Stem-loop [17]

Blue: Heptamer[29]

Green: Nonamer [9]

id-TRBV6-4-2[V_gene_segment]

catgacttactggcagatcctttgttcaaacactattaagaatttaaaaatcatgacagcagagtgataaagcaagcgtgggatccttctgaatgcagggctctgtacaccgaagatacagtttttgagctccttttcatttattactaaattaatggcaggaatcctgaacgaagggatattccctatgagacatgacacctcaaataaaa**cgagagcatttttccttatcttgaagtctgctccctctcg**ccaagtccctcactcatcaggacacctttgtgctcacaagtcttcagcagcgctgggtggccaccattgtccacacagagagggcagt**cagcagcatgaggtggttctgcctgctg**tggtctcacc**ccaggaacagaaagcaagactcctgg**atgg**agctgaaggtgctcagct**gga**cttgtcaggagtcccatctgtcagtgaattgacaag**aaacagagcaaaactactcctgcaatgttgatgagcctgcccctgggatttggaaacctgataacagagaaaaccaatatagacacaggactttaacaggattatggtcaattaagcagattggaaaaggatacttgaagga**gtgtttgggacacaggaatcaaaaacac**cagggagacaagaggatttttcctaagattctagactacagcaatacatagataactaacaccagaatattaatgaggaattatatcattgcaggaataaaatttatgttgaattacaaactgtttgcacaaaagtcaagaaaaacttgaaagccttatatcaagaagcattcttccc**atccaaacttcagtggtgcatttattttggat**ttgaccctctggggaaggggcatggcc**tctctcgacaagaaggttctggggaccaggcagagaga**atgaggtctcaggatgacttccttgacagccctgttcccctttcatcaacacacagacccagaagacctctctgtcttgtagcatctgccATGAGAATCAGGCTCCTGTGCTGTGTGGCCTTTTCTCTCCTGTGGGCAGGTAAGTCCTGGGCAGGGCCCATGTGTAGATTTCAAGACCCAGAGCCTTCCCATTGAGGCTGCAGCATTGGCTTTGTTCTCCTTCTCTGTAGGTCCAGTGATTGCTGGGATCACCCAGGCACCA**ACATCTCAGATCCTGGCAGCAGGACGGCGCATGACACTGAGATGT**ACCCAGGATATGAGACATAATGCCATGTACTGGTATAGACAAGATCTAGGACTGGGGCTAAGGCTCATCCATTATTCAAATACTGCAGGTACCACTGGCAAAGGAGAAGTCCCTGATGGTTATAGTGTCTCCAGAGCAAACACAGATGATTTCCCCCTCACGTTGGCGTCTGCTGTACCCTCTCAGACATCTGTGTACTT**CTGTGCCAGCAGTGACTCcacagtgctgcacag**ccatctcctctctgtacataaatgca**ggggaggctctgccctcctcccc**gaccccagactcaaccatgtccttggcagagttctcagcactgggaatcttggaagcccaatgggtgccgggcagtgtgagcctcagtctgtgccaggtgcctctgcagg**cagtcccagccaggcctggactg**gtc**ccagagcctcagaggtctcttttgttgctctctggtctttcttccaagctatctttttgggatgggaccag**ggctttcccagctcttacttttctactcat**catcctgagtctgaggtccacaggatg**gaacag**gatttgtatttcaaatc**tgtctagactcctgtttctctctgggggccacgttgcttcctctctccagggtttcctccagtacccactctcatgtggtctcacctgtggcccgcctttcccatctgagcagtcacccccgaaggccttgctgggcctctgcacaccacctcccctcgcctttctact**gcagccatgaggctgc**ctct**tctgtgcctccttcctttccaacacaga**gacctcaaaggccatttcctccgccctgggctagtgccttcctttctgcagtggtcagcttctacctgcactt**ctgatctcagcatgatcag**cctctcctgctggaagcactccctcaactctcagctgaggtcaaatttcagtttccctctcataagctcacagaatcatgtgcctcttagtaaagttcagcaccgttgcaatttttccaacacttgttttattccttttgtgctccacttaattttaaactcatgaagctgaaggctgagcttgttacattcaccaccagcagtgcctggcaggcagggagtc**ctcattcatttcacagagaatggatgag**tgagtgacaggtgagtgagtgatgagtgggtg**aatgaaccagtaacaggaacacactagattcatt**gtaccctggcagaggtctagagt

Uppercase: TRBV6-4-2

Lowercase: Flanking sequence[1000bp]

Red & Bold & Underline: Stem-loop [21]

Blue: Heptamer[34]

Green: Nonamer [2]

id-IGKV1D-33[V_gene_segment]

cttctgctttcctgaaatttactcctgccttaaaaaacccttacctgcaagccatcagtgaggccaggatttgaatcttagctgcctgattctctttacctgatgccctacaaaaaaacaaacaaacaaacaaacaaacctttactttctcctgctgcaaactcaatgtgaatatctgatctgactgagctgagtgagtggactccagttcggttccacagcacaagctctgtttatagtggcatttatccac**tcatgagggcagagctctcatga**cttcaacacttcccattaggccccacctcccaat**tctgttgctttgagataaatttaaaaacaga**tgggttttgaaggacacagtgaaaccataacactgcccaaagagagggccattcgagcccctgtggtctgctatgcatgcaggcagagctgccttctccaggttcagaaagagcaatgaggggcatagatggccattggaagtcagcagcagtacagcgaaagggaaaggcatagcctgtagacagggattgcaatcttgatatattttctatttagtttcctttctaaaaacagaataatttcatgctaacagaagtgttgtaagtacagcacaaagtgctttcttccaattaagtgcatattg**tattttttaaaaaata**atctccatctccatacacgaataaaatacattactccatctagtcctcaggaatatttcaaa**ttttgacaattaactcaaaa**aagttctttgtaaccaaacagcctccaggaagaaacacttatttacggacaaatctgtgatgccctggtccgacctgggacactggggacattgctcctatgctgagttactgagaagagccagccctgcagctgtgcccagcctgccctatcccctgctgatttgcatgttcgcagagcacagccccctgccctgaagacttattaataggctggtcgcaccctgtgcaggagtcagtcccaaccaggacacagcATGGACATGAGGGTCCCTGCTCAGCTCCTGGGGCTCCTGCTGCTCTGGCTCTCAGGTAAGGAAGGATAACACTATGAATTTTCTC**AGCCAGTGGGCTCAGTACAGCCTGGCT**CTTGATGGAAGCCTTCCTATAATATGACTAATAGTATGAATATTTGTGTTTATGTTTCTAATCGCAGGTGCCAGATGTGACATCCAGATGACCCAGTCTCCATCCTCC**CTGTCTGCATCTGTAGGAGACAG**AGTCACCATCACTTGCCAGGCGAGTCAGGACATTAGCAACTATTTAAATTGGTATCAGCAGAAACCAGGGAAAGCCCCTAAGCTCCTGATCTACGATGCATCCAATTTGGAAACAGGG**GTCCCATCAAGGTTCAGTGGAAGTGGATCTGGGAC**AGATTTTACTTTCACCATCAGCAGCCTGCAGCCTGAAGATATTGCAACATATTACTGTCAACAGTATGATAATCTCCCTcccacagtgtaacaagtcataacataaatcacccaggggagcagatgcgtgaggctcagctgtcccagatgccccttctggtgccttcgcctgc**tgagaatgtttctca**aattgcagtcacactttgaagttcactggagagtttttgtaaaagggcc**atgaaggcccacttcat**cgtag**ctgtctttccttgtcctaatccccagtatcatagacag**ggcaatgcctctcctgatttcattgagaagaaatggttacacctgaggggtctgagttgtagcatcag**ttggaattcatgtagcaatagtgagccactctaggtattccaa**gtaggattttttttaaatacaagatgtgagaatctaaactacagccttttaaaggtttgcaagtatagtagtcaaagacgcaaatactag**agaagaggaattctcttctggaatccagaa**tgcatctgatagagaaggtacaactgccaatcatgtggtc**ctcagacctttctgag**aagcccatgggtgggggtgcagatgctctcagctgccta**gaggacttcatcaggtgcttctgcagtcctc**acctcggtccatatgtcttgctgcaggtgttgatggatagtattgaatcctcctcttcttacttctcaatctcagggcaggccccacactgggcaactccaccaaaaaccagagaaggcatagggt**ttctggcaaatgtgcttccagaa**taatagtgatgatggggaagtgacagctgacatcgtagtgtggtcatgtatctcgactctcaggattttttcagtgaagtgatggcctcagaatacacttggatgtacttccatata**ctatgagtaagtttgaaatcatag**catgaaaatga**tatttagtcatatgataaata**gaactacatggctacattaatcaaaatagcatagtgctggtacaaaaagatacacagacaaatggaacagaatagggaactcagaaataatgctggagaccta

Uppercase: IGKV1D-33

Lowercase: Flanking sequence[1000bp]

Red & Bold & Underline: Stem-loop [18]

Blue: Heptamer[34]

Green: Nonamer [8]

id-TRAV39[V_gene_segment]

tcctaaagaaataaaagccttccttccctcccactctccctccatcaatggt**ccacagagttgagagaacacaactgtggtaggcatctacca**ttttgttcac**ttagaatggcaactcattcctggtaaaatataaattctaa**ccatgtgttatgaatggagactgctatcttacttatgaatgggctgatga**atccacccagaggtggat**gcctgacccagctgaaccaatcatgttatgtatacaccactctagtcacaaattgcccccctggatgggaacctggctcagactgaagcaatcaaagccttacctcaatatttttaaatctgggatcagaaaggatcctcagt**atttctttggtggctacactgtgagatgaatcccaaccaaaaaat**gtttacaatcatgcctcctgctttgtggtggaatctggtctgaggaatgaaac**taagacacgaatgaaaggaaaagtctta**aaggattaagttccctggcaccctggatgcctgaagtccagatacacttctgctcttgccaaggttattggagtcagtgcatttctcttcttgcctaagcacat**ttgagttacatttctgtcacttgtaacatcaa**tgagacgttaaagagttaattcactaggaaatttttaggaacaccttgctggcttacaagaagcatcaattctacgaattactttc**tcaggaatttctctaaatatcctga**agagggcactgtttccattcataacttctgtccaaacccacccccattcttttgtttttttttaatttttctttttatttcttttttttctttttcttttttttttatgaaagggagggactgagtaaattgcaggatgtgattctaattggttggaacatcttttgaaatcgtgtttctgtagagaaagaaaaactaccatatttggatagccctggccaactttcaaggctcctaaatctgagttttcagtgaactgg**acagaaaaaaaaaATGAAGAAGCTACTAGCAATGATTCTGT**GGCTTCAACTAGACCGTGAGCTGGGGGTTCATTGAAAAGGGAGCCATGGGAGGAAAGGAATTGTCCATACAATGTTTGGGGGTAGAGACAAGGTTCAATGCGACTCATTTGGGTTCCCTCGGGAGGAACAGGATTATTGGGGTAACCAGTGAATGCTTCCTTCTGAAATGTTCTCTTTGGACAGGGTTAAGTGGAGAGCTGAAAGT**GGAACAAAACCCTCTGTTCC**TGAGCATGCAGGAGGGAAAAAACTATACCATCTACTGCAATTATTCAACCACTTCAGA**CAGACTGTATTGGTACAGGCAGGATCCTGGGAAAAGTCTG**GAATCTCTGTTTGTGTTGCTATCAAATGGAGCAGTGAAGCAGGAGGGACGATTAATGGCCTCACTTGATACCAAAGCCCGTCTCAGCACCCTCCACATCACAGCTGCCGTGCATGACCTCTCTGCCACCTACTTCTGTGCCGTGGACAcacagtgctcccctgacg**ccaccagtctgtacccaaacctgcagctggtgg**gcccactcctcctgcaggaactatgactgtgaggcttcgttcactgtctgtgcatttctttctgcaaggaaaattctgagatgggtgaaaaaagatgctgttttatcttcttaactgtggaatattttttaaaaggaaaatgtattcaggctgagggaagattcctaaccatgtatttcctgagaatatgaggtctcattctttactgcctcacagtcagaatttgttagtcttccattatggaataattttacatgtagtgcaactgtc**acaaataattcctatgaaatatggaatcattatttgt**tttatatatgaatgtaacatatg**gcatttaccaatgaaagaaagaataaatgc**tttataaaagaatggtctatgtgaggttccttacctagtgtttcctgctaaaata**ctgactacagtagtcag**ataatatccgtgtgatctaagtacctgtttatctcaatgtgtggagagtctctgggaaatgtatcctttggctggatttactcgaaaactatctgagggtccttgccaaccagaagaatattatattttttcttgtattatttgccagtttcctcttctcctccctccatttttttttgtaattgttttgctacatggtagaatttatattaaactttatttgcccattgcagactagataaaatgaatttcactttcagga**aaattgtcttataagaaatccccaattt**gaatgttatacaaaaataattgaaaataagttttcctgtcccactaatgagataatcagtaccttcagatatcaattgcatca**tattttcatcccagacttctaggaaaata**ataaacttgattcagtcaggcttttaagtgacaatatttcagaaattaccattacaatgtaaaagctgaccttaacatttcaacatttgtaccttcctaagcaacagcaccttta

Uppercase: TRAV39

Lowercase: Flanking sequence[1000bp]

Red & Bold & Underline: Stem-loop [17]

Blue: Heptamer[26]

Green: Nonamer [4]

id-IGLV3-22[V_gene_segment]

tgcatgaggcatcaccagcctccttcaggagatgctgtccagaggaacagggcttctcacctgcctcaggccacagctctcaccacggggctgtcctttgatcagtggcaggatctgcatggctggagggactactgaccacatgctctctggagctcatgtcttgtgtggagaccatgacctggggaccctcagaaagaccaagcagcctaaagttagattttcagtccagtgtgattctctggttcttttcgcaagtttacgtaaaacgttgatgagaaaaagtgaataaataaaggaagagttgaggatggcaattgctggtctac**tgtggggctgggggataccatggaaagagcagcaccaca**tgtcaaatgagactagagattgggatgcatccgaaatatgtgtggttgctcaaatgacaaaactcttccaactgcaatgattcataagtgtcctacagacacagtacatcatgtacataacattaccaggatggggccaagtgactcgggggtatccggagcagggatgaaagaggag**aaagatgatgcatcttt**ggtgtacttgctgctgagtcaggcgtcatcagtgatttctcaacagggcccagctcagccccaggccaggctcctggaaggtccata**gtcctgctcaatgcagcccaggac**aggccacatgacacagaggtttgagacagggaccatgttgtcaatccccaggacggtgactgtgatcacag**gccttggaggccaaggc**tgattttctgactggagaagcagacgaaggaagctggggattcaccccttcattttaagtttctcatgctgcccccagagctggcttgagggatatcccgccatgaggagtcccagagaagtaaatttgcataaacaccaaggatgcactgccccagtgggcctgagaggaaataagagaggcctggga**agcccagctgtgctgtgggct**caggaggcagagctgtgggtgtctcaccATGGCATGGGCCACACTCCTGCTCCCACTCCTCAACCTCTACACAGGTGCTGCCCCCAGACCCTGCCCCAGGCTCAGCCCTCCTAAGCCCCTGGTCTTACCCTGAA**CCCTGAGCTCAGCCCAGGCATAGCCTCAGGG**CGATACTACTGGAATGGGTTTGTTATCTTCAAGCCCCCTCTCTTGTCCTCTCTTGCAGGCTCTGTTGCCTCCTATGAG**CTGACACAGCTACCCTCGGTGTCAG**TGT**CCCCAGGACAGACAGCCAGGATCACCTGCTCTGGAGATGTACTGGGG**GAAAATTATGCTGACTGGTACCAGCAGAAGCCAGGCCAGGCCCCTGAGTTGGTGATATACGAAGATAGTGAGCGGTACCCTGGAATCCCTGAACGATTCTCTGGGTCCACCTCAGGGAACACGACC**ACCCTGACCATCAGCAGGGT**CCTGACCGAAGACGAGGCTGACTATTACTGTTTGTCTGGGGATGAGGACAATCCctcagtgacacaggcagatggggaagtgag**acacaaactcctttttcatctgtgttgtgttcttcctccagcaccaggaaga**ctgtggaccaagcaatgagc**aggtctggccaagttctactggatctgagacct**ccaggccatccttttttccagctgtccaggttggctctgcagagggtggagcaggagtggatttaaggctgctataatcagaatttatggtgttttggggcctcatgagtgacttggggaaataagaactggatggaagatgaaagagatgtagaga**accatcacttgaccttagatttccatacgatggt**gattgggtcactccttcttcaaaatatcctacatcagtcccgaaaaatacccacctc**agctcaacacctgagct**t**ccaggcttccatggcagcctgg**gattttgaacaggaaaagccaaggaggggacactgtggcatctgcctcaatacccttt**tctgagctcacccaacctcaactgctgggagcttctaggagctcaga**gctgtgccctcactg**ggaagtgctattagctccaaagaaaacttcc**ttcctcatatttaggcccagttttgagagttttggtttgatgactgcctgagccaagaaccc**aaggccctgtcctaatagaggatgacactgaaaggcctt**ccagatccagacatcccagtatggatgcctgaggcctcagttgcaaccctattgtgggtcagggtctccttctggtgggcgatgcctccctcactccttccagatcctctgtgcatgcacatctttttctcagagtctacttccaggaaacccaaactaagatggctggctatcaccaaaataaactaccatagacctgagg**tggaaacaccagcgaatttcccaaatttcca**ttgtgaacatccaaatgcccatttatatctgagtgttccttgtattcttataaaatgaaaatggagcctgacataccaggtttttggagtcc

Uppercase: IGLV3-22

Lowercase: Flanking sequence[1000bp]

Red & Bold & Underline: Stem-loop [19]

Blue: Heptamer[36]

Green: Nonamer [4]

id-TRBV5-3[V_gene_segment]

cttattcacattggcctgaaataagtaggagaggatgtggatcccaaagcatcaggtactgtagctctaaggcaaggctgcctataaggtgctgtggactttggtagggaaaagtaacctatataaaataatgaagcaagaggatagccagggcatagatgaatcagaacaagtggtgcccctacctccctctactctcctgccattgtctcccattagccaaactcactctggagttagagaagaagagagccagttaacataatacactgaggtcagcctctgaaggcagggtg**gatctggagaaattaacgtaaattatccagatc**actacgcataagatggccttgagggtgctgaatcccaaactgtctggctaggtctgtgctgtgagctgataaagccctagtattatttctggggaa**tctgtactcttaataatttacaga**caaccaaggtctgctttggatctgatcagatagactaaatcttggggactttgcaccactggccactcaagggaggggctggaaat**attttctgaggacaaaaaaatagaattagaaaat**ttggttgaatctagcgtcagagagatgacattaggacagccagggagagctggaacaagaaggtttaatcacaggctcctcaccctctgctgatgggcaggtgtgtga**gctccagcatggagc**acgacagcactaggtgggaggagggtgagagggtgatggggcagcctgggagctggagcagtgta**ggcagaggagcaactgtctcatcacagaagcttctgcc**ctcacccaacccttctgctggacagggagggagttcaggttgtggggagtactggacctcaggaagcatttatagggaggacacaggacagtgacatcacaggataccccacccatcaggaaaatcaaggcccagaactcacttggcccttccccaggaggaccaagccctgaatcagatgcagtgctgcctgccccactgtgccATGGGCCCTGGG**CTCCTCTGCTGGGAACTGCTTTATCTCCTGGGAGCAGATGAG**TCCTGTGCACAGAACAGCAGCCCCATTCTCAGATTTCCCACCCCTGT**GTCCTCCACTTTACACTGGGGAGGAC**CTCCAGGCTGTCTGCTGTGCTCAT**CCTCCATCTGCTTTTCCCACAGGCCCAGTGGAGG**CTGGAGTCACCCAAAGTCCCACACACCTGATCAAAACGAGAGGACAGCAAGTGACTCTGAGATGCTCTCCTATCTCTGGGCACAGCAGTGTGTCCTGGTACCAACA**GGCCCCGGGTCAGGGGCC**CCAGTTTATCTTTGAATATGCTAATGAGTTAAGGAGATCAGAAGGAAACTTCCCTAATCGAT**TCTCAGGGCGCCAGTTCCATGACTATTGCTCTGAGA**TGAATGTGAGTGCCTTGGAGCTGGGGGACTCGGCCCTGTATCT**CTGTGCCAGAAGCTTGGcacag**ccctg**cagagtcactggaactctg**tgcactaatctctctgcttccgtgt**acagcagtctcagaccagacagctgt**gagtacctggggccttcagggggaaagataaacaatttcaggactctgaacaaggctggcggaaacaaggccaggaggatatgctcagaggatggtgctgcttcaggaggttcacggcaaaattactttctttgtcggatgctgtacactatagtgatagagtagctggcactctcctttttagaaaaagaaattcattttggtgtcaggattgttcgacagatgtgtagtatagaaggtaaaatctcaagggtttgctttatataattaagggagaagaaattgagatttggtttggagaaacttgaggagagagaacatgattaaagattattaaataaatagaagtagagctaccagatgataaaaaattgtgttctctgcaacataacctgtcaaagttttcaa**ttgctagaaaatacgcagcaaggccttgtagcaa**ctggcatcaccccgaggccactctatgatggtcatcaccacgtagccagttgccctgtgggagggccctggagctggttccaatgtcttccttctggacttatatctcaattctcaaaacagaggaattcatttcactgctgatggtaaacagataaaggctctcccatttccccaa**agatacattctgcttcccttgtatct**ctttctgacacttaccgcccagatccattaccttctgtgatcaaacacctctttgaatatttcctaaaactctgcagcctttacattctctgcatttgtgctcagtatttccatggcctggaaggaccttctgtctcccaagaaacagccagaccctgcttatcggtcaaggatgggctcacacctgcttcctctttgcttcttaccatgacattccccagcattgtactataccaccatctttattattttcataggaaaagccaagtattaat

Uppercase: TRBV5-3

Lowercase: Flanking sequence[1000bp]

Red & Bold & Underline: Stem-loop [15]

Blue: Heptamer[37]

Green: Nonamer [1]

id-TRBV20-1-2[V_gene_segment]

cctagaaactattttcactgtccagtacttagtatgacctccattacaat**agaataaatttaattattct**cacaacctcaacagtgattaacacgg**aaacatacaagctaaattgagaaatattattagaatgttt**agaattaagtccctttatatatgagggaaattgtc**atgagatacacatgaggtaactatctcatttagaaaatga**tgaagtggcttcaacatgattatgaagaaacattggacttccagccga**ggcactggcctgtgctttatcaatatttatggtagtgcc**aacctgtgggtctagtgattaacaaccatgagttgtcttggtgttattactttgaggatatgaaacagaagcagcaaacagactccaatgccattcattacctaaacctacactcc**tttccttataggaaa**ctcacttaagtcattacaatgctttatgtctcagtgtatctttaataaatttggaagttgaattgggaagtctttaggccactttaggcaatgtatgggttttttgttttcttttgatatggggtctcactctgtcacccaggctggagtgcagtggcatgatttcagctcattgcaacctctgcctcctgggttcaagcaattctctccctcagcctcctgagtagctgggattacaggtgtccaccaccacgcctggctaatttttgtgtttttagtagagatgggttttcaccatgttggccagattggtcttgaactcctaaccttgtgatccacccgcctcggactcccaaagttctggaat**tacaggcgtgagccacctcacccgacctgta**tggttttttgacaggcagagatggacgtaacatcagtcatgggcaaagattaccaccagggggcagactagggcatccttgggattctgtgatcagtcatccctcctcgctggtgaatggaggcagtggtcacaactctccccagagaaggtggtgtgaggccatcacggaagATGCTGCTGCTTCTGCTGCTTCTGGGG**CCAGGTATAAGCCTCCTTCTACCTGG**GAGCTTGGGTGGGCATGTGCGTGTGTTGGCATGGTCAAGTGGTGGCCAGCAGGGTTGCAGTGTGGATTGTTTATGCTCATCGAAGGGAGAGGGAGAGGCCCTGCTCTCTAGAGGTGTAAATGGTAAGGTGAAAGCCGCCGGTCAGAGGAGATGGGGGATTATGGCCCTAGGGAGATGACGGGAAGATTGCACAAAACAAACAGGACTCTCCAG**GAGCTGGGAGCACAGGGAGGGAGTGAGGCTCAGCTC**TGCCTGGCGTCCTGTCTGACTCGGCTCCCACTGGGCTCTCCTCTCTCTCTGGCTTCTGTCTCAGCAGGCTCCGGGCTTGGTGCTGTCGTCTCTCAACATCCGAGCTGGGTTATCTGTAAGAGTGGAACCTCTGTGAAGATCGAGTGCCGTTCCCTGGACTTTCAGGCCACAACTATGTTTTGGTATCGTCAGTTCCCGAAACAGAGTCTCATGCTGATGGCAACTTCCAATGAGGGCTCCAAGGCCACATACGAGCAAGGCGTCGAGAA**GGACAAGTTTCTCATCAACCATGCAAGCCTGACCTTGTCC**ACTCTGACAGTGACCAGTGCCCATCCTGAAGACAGCAGCTTCTACATCTGCAGTGCTAGAGAcacagcgccaggaggggatcagacaccgcggcaagaacccctgcagctgccctccgccccagcgggccccctgagtgctgagaggggaagcgtggagaatggaaaaccacagctttcctgactgagacatctgggagtgtgtgtgcagggggtggggcgggggggagggggagaaaaagaaaagaagcactgtgagtgtggagtatggaggaggtgtagtgtttgagacccacgaaatggcaatagagttgggcctaaagtggtcggcggacatggaaagttccctg**aaaattataaaacctggaatttt**gccctgactctgccacattagtcatgtattcttgaatag**atctacacataatcctagaggtagat**atgaagacagttgtataatcactgtgtatttcctataataaacttcagcgattggctttcttgtcatcatgctgggctctcaagacaaagataaccaagaggcagccaagtagagagttatgcatgaatacgttaccttaggggaggatagcaacagcattgaatagcattcactccaaatttgtgttctgtcctccttagaccaggccaccaaaataaaatcctatc**acatttcctgtaaatgt**tgaacgccatcatgttctccatggtgtttcggttgatgaatgaacagagtaagactcggtacaaagggaggggagcaccctctatgcctaggtacacaccatacggaatgtatcgctagggaaggtgctggtgagctggctcctgggaaaggtgtcaaggggaagagggggcagaggtacctagaaggaagacctc**agacaacatggcttgtct**acagagttggaaagttctgccagtagttagaagacagtaagtggaacagtgaaaatcctttcctaggagaaaccaaacatttacattgaccacaaatctgtatgaaccaacagaaaatgagcaaaacatatgaccatacatttaatagaataaacatatatgacta

Uppercase: TRBV20-1-2

Lowercase: Flanking sequence[1000bp]

Red & Bold & Underline: Stem-loop [14]

Blue: Heptamer[38]

Green: Nonamer [4]

id-TRBV5-7[V_gene_segment]

tgtgtcctgttctaggactttatggaatataaaattatgagccattagctaggatatctgctgaaagaaatatctaagctgcaaagcattcaggccactgtgtgactacttttaggcaccaggcaatttaacccagcaagaaggcagccaagggaatagattttgcaaaccagcacagatggtgacactacctccctctgctgtcctgtctcccataagccaaactcactctgaagttagagaaaagaggagacagttaacatgatacactggagtcagcctgtgaaggcagggtgga**tctggagggattaacataaactttccaga**gcactacatataggatggccttgagggtgttgagtccaaagctggctaggtctgtggtgtgagctgataaggccctagatttatttctggggaatctgcactcttaataatttacagacaacacaggtctgctttggatctgatcagatggactaaatcttggggcctctgtaccgctggccactcaggaaatgggttgggaatgttgcctgggacaggaaaatatagtaagaaaccttggtgtgaatctagcttcagaaagatgatgtgaggagagcaaggaagagctggaaact**gatggtttaatcacaggctcctcaccatc**tgctgataggcaggtgtgcga**gctccagcatggagc**accgcagcactaggtcagaggagggtgatagggtgatggggcagcctgtgagctggggcagtt**aggcagaggagcaactgtatcaccacagaagcttctgcct**tcacccatccctccagctcggcaggacaggtagagagtccagtgtccgtggagcactagacctaaggaaggctgcatagggaagacacaagacagtgacatcacagcatacccctcccatcaggaaaatcaaggcccagaactcactcagctctttcccaggaggaccaagccctgaatcaggtgcagtgctgcctgccccactgtgccATGGGCCCCGGGCTCCTCTGCTGGGTGCTGCTTTGTCCCCTAGGAGAAGGTGAGTCCCGGG**CACAGGACAGCTGCTCCATTCTCAGCTTTCCCACCCCTGTGTCCTCCACTTTACCTTGGGGAGGAC**CTCCAGGCTGTCTCCAGTGCTCATTATACATCTGCTTTTCCCACAGGCCCAGTGGACGCTGGAGTCACCCAAAGTCCCACACACCTGATCAAAACGAGAGGACAGCACGTGACTCTGAGATGCTCTCCTATCTCTGGGCACACCAGTGTGTCCTCGTACCAACAGGCCCTGGGTCAGGGGCCCCAGTTTA**TCTTTCAGTATTATGAGAAAGA**AGAGAGAGGAAGAGGAAACTTCCCTGATCAATTCTCAGGTCACCAGTTCCCTAACTATAGCTCTGAGCTGAATGTGAACGCCTTGTTGCTAGGGGACTCGGCCCTCTATCT**CTGTGCCAGCAGCTTGGcacag**cccagcagagtcactgacattctgtatataaacttccgccttagctttgacttgagaactgcaggccccacccaggtttcactccttcaagggaagcttttagttgtttggaaggcatgtcttgtgtcctactgagggcagacctttcccaaccaatagagcccaggtttcctgtgccctgagtgtgcctgcttctgtgctgcaacttcttgtagttt**gtcacttcctggataacttcagtagaagagtgac**tgttgagccccagatacgtgctagattcttcgtatttgttacatagcttaagggctttccacactcttgcacatcaaacat**ttctattcttcctttatagatagaa**ggctcagggacattgagtcattttccccagtgtctcttggcttgtaaggatcagaagtaggaaacaaactagtccatccattttccacctacccccctgctccatcatcaccttctgcatc**ctggtcagtaagtcagagccctcgcactgccctctagtgaccag**cagggctgcagtagacgctcagatgccttcaggggtcattactatggcctcttcattagcaactttgaggaatgttaaatttaacactttttaaaaatcatttatatgcattccctttgtctttccccagtctgcagcttatgttttcacttttatggtgtt**tttttgatgcacaagagtttaatttaatgcaacaaaaa**ctaaaatcaattttgtttgtgtg**tagacaaaaaccgctttcttatataaatgtcta**agcgtagttttatca**atttattctagtaaaatttaagcttggattttcaccactaagaataaat**gtacctgaagta**tatcttatgtaagaaataagata**gagacctaggaatgggagggaaggacatatgttcagtgaagaagtaatagcctgtcaaagagagccctaaagtagacccccagagctgatggtggagaggtaagtggagggcaaatg

Uppercase: TRBV5-7

Lowercase: Flanking sequence[1000bp]

Red & Bold & Underline: Stem-loop [15]

Blue: Heptamer[20]

Green: Nonamer [4]

id-IGHV4-55-2[V_gene_segment]

atgttcatgcaattgaatcgaggagaaagtct**ttcagagacagatggatctgaa**ctggtaaatatgtgtgtaaggactctgggcttaagtgtcattgtccagccatgtttc**acaggtgtgacctgt**cagggaagaaccagagttccttgtgttctcagaggggaggggtcccagaagtcctctctggttcccaggaaaggtaattgcattaatcttggtgatgagact**atcatccagtgatgat**gtactatagagtttatgtttgaagttgacactgctatcgcaatctacatcttttcacacagaagtgtttagaggtcaggccacatcttcaggatcccacatt**gagaaggacagagatatattccactaccttctcctgagatctcaggcagaaacccaaatttcaaaaggtctcag**aagggcagctctcaggggctatttaaaaataacccacttcgtgggacagggagcatccttctaaccatgatggatgttctgaactacaataaacattgcatggatccagggtctgaattcactgtgattattacactccactgctgtttcaatgtgtctgaaggggtaaatgacaatttagatgacctgggtgtgtggtttgttttatataaatcttcaaggatagaacagcattgaacctattccaaaatctgtccctgatcca**agatcacactgatct**cccagaccagcatcttcagcacatttccctacctggaagaagaggactatgggcttggtaaggggaggccacaggaa**gagaactgagttctc**agagggcacagccagcttcctactcccagggcgagcccaaaagactggggcctccctcctcccttttcacctgtccatacaaagtcaccgcccacatgcaaatcctcacttaggcacctac**aggaaaccagcacacatttcct**taaatttgggatccagctcacatgggaaatactttctgagactcatgggcctcctgcac**aagaacATGAAACACCTGTGGTTCTT**CCTCCTGCTGGTGGCAGCTCCCAGATGTGAGTGCCTCAGGGATCCAGACCTGAAGAT**ATGAGATGCTGCCTCTCAT**CCCAGGGCTCACCGTGGTTCTCTCTGTTCACAGGG**GTCCTGTCCCAGGTGCAGCTGCAGGAGTCGGGCCCAGGAC**TGGTGAAGCCTTCGGAGACCCTGTCCCTCATCTGCGCTGTCTCTGGTGACTCCATCAGCAGTGGTAACTGGTGAATCTGGGTC**CGCCAGCCCCCAGGGAAGGGGCTGGAGTGGATTGGGGAAATCCA**TCATAGTGGGAGCACCTACTACAACCCGTCCCTCAAGAGTCGAATCACCATGTCCGTAGACACGTCCAAGAACCAGTTCTACCTGAAGCTGAGCTCTGTGACCGCCGCGGACACGGCCGTGTATT**ACTGTGCGAGATAcacagt**gag**gggaggtgagtgtgagcccagacacaaacctccc**tacagataggcagagggggcgggcacaggtgctgctcaggaccaacagggggcgcgcgaggccacagagcccgaggccgggtcaggagcaggtgcag**ggagggcggggcttcctcatcagctcagtgatctccctcc**tcgccagcactcagatgtccccagggctcctgttt**ctttattgtctgtggttctgcttcctcacatccttgtggcagacaagaaag**gaggaagacaatttttctgtttactgttgaggtttcaccaattact**aggaactttcctacaagttcct**gcgtgactcattttaccgtata**tgtgtgtgtgtataaatatatacacatacacacacaca**cc**atatatatacaccatatatat**tatatagatacacaccatatatatt**atatatatacacaccatatatat**tatatatacacacc**atatatattatatatatacacacgatatatataatatatat**atacac**accatatgtatattatatataaacacaccgtatatatatatggt**tctcaccatctcttgatttgtgtcatcaatggaattgtgcctatttgaaattcatttacccaaaccttaaatccaatggatctataccggaattttaatgatgtaattaaggttaaatgtggtcaaagtgtgagaccctaattcaataaaccagttgtctttataagaagaggaagagacacccgagacctctcacttttcgc**gtgcacacagagaagaagccgtgaggagacgtagtgcac**tagaaggtggccctgtgcaagccaggaagaagccgcgctaagaaccaatttttacagctccttgatcttccacattcagactgcagaattgtaagaaaatcaatatttgttgtttaacccacccactcctgttgtcttcttatgaagatccaaacagactgataccacgtaattctgttagctctggt

Uppercase: IGHV4-55-2

Lowercase: Flanking sequence[1000bp]

Red & Bold & Underline: Stem-loop [24]

Blue: Heptamer[42]

Green: Nonamer [8]

id-IGHVIII-11-1-2[V_gene_segment]

aagaagatccatgctacattattcattttcttgttcaaatctccacagctttattaggtgctgggacctcatttagtttggatcctgcatccttacagcaaagctcatccttttgtttttgaacacttccctgtttcctgatattacgatagattctaagcttgtttctttatcacctttttcatacatagaattagccatttatataaagattgcttgtttctg**attttaaagagtagtgttaaaat**aaaatattgtgataatgggtatgtgtgttgttaatgtggtataagtacttctgggacctctcaaccttctgttctagcaaatgtgcatgtt**tatatgaatcatgtcatgtccatata**tccaatgttgaactatgttgcgaaaagtgatctcaa**gataatggtaattatc**aaccccctccctaacaaggtgtagatctgcatttttttttcattgtaactcaactttgcctcatggtcaggaacagtggtttccagctctaaatgtactgattacagcgagatgtccgttttccctggaaatgtatttttaagttcttactggatgtatttgttgataatgttttctactatgaagatacctgaacagtgtccgcactagagagtaagaaagagtaatgggcagattaaccctgtgcatccagacccagga**atcctctgatcctgccctccctgaaacggagacacagaggat**ggatgagcaatgctgagcggtgcacccaagaccacaaaaagaaagacatggaaatatgtcccctcccctcctcaagaaaggcagctcatcccctgttccctcaggcccgggcgaggagccaccccatgtctcttcccttcc**tcagtgtccacactgtggggtctgcactga**tctgggcttcccttctcatcaccctcgtattagtggcccttgtgaatcaggtccagctgtggctgctgctcatggggctgctctcagtctgctctctctgtgtttgcagAAGTCCTGTGTGAAGTTTACTGATGGAGTCAGAGGGAGAAAGATTGTACAGCCCAGCGGTTCACTCAGCCACTC**CTGCAAAGACTCTGGATCACCTTCACTGATTGCAG**CATAAGCT**TGGTCCAGCAGGCTCCAGGACCA**GAGTTGGCGTGGGCAGCAACAGGGAGAAATTCAAGAGGAAGTTCTTAcatgcacccttacgtgcacggtctcactgagatctttacttcctttatcacgtttgttctgtaaatcacaacgaatggtgcattcttcatctattatacacttgttaagtct**ttttttggcatcttttaaaaaa**ctggtaactttatcctgtgtaatatccctgttaagtcctaaaagtcttttttgatgtctattttttcttaactttacacagctactatagatttattttggttcatatttttgtaatccatgttttctcatctttagtttttacatttgtgaatacgtacagtaattttctaatacatagcttc**tagttggggcttgctttttaaaatcaacta**taataagctctatt**tttaaactaatattatttttctgttatattgtttaaa**ttagcatttcacaatgatgtttatttctctactaacatataatctaattcacttttataaatattatattggttaccataagttt**tacaataagaattgta**tataattagattctataagaattgtatataattggattctaatgcagatactgtgatggccttgatatgaagaacagagacagtgtaactgtgtgctttgaattcctccttctcactgctctttcttgtttattcccagtttgcacttacatatactataaaacataatatataaattttttattcctttcatagttatataatatagcaattacaaagataaaaactgcacttcatcttgattttcccattttgtacagtcttaatttcttcgtatagatttatgttttgaatgtatatcacacggctactagcagagaaagtttgttaaaatgtgcactgaagcatgaatgtgctgacaataaattgtctcaagatctttttgtaatgacagaattttatttgcctttcacttttaatgaaaatgt**aatgcacgtagaattccagtttgcatt**tcacttgtaatttattttcttgtgtttattattttgttcatggag

Uppercase: IGHVIII-11-1-2

Lowercase: Flanking sequence[1000bp]

Red & Bold & Underline: Stem-loop [12]

Blue: Heptamer[40]

Green: Nonamer [4]

id-IGKV2D-18[V_gene_segment]

ggcgacacagcaagactccatctcaaaaaaaaaaaaaaaatc**tatattccatagttccaccttgacagtgaatatacacaaatttcattcagttacacgcttaatatttgtg**ccctttactctggg**gattttatcaactaaaatc**aaactaatttaaaaacctagatgctaaaataaatatgaaaataataaataagaataactggccaggtgtggtggcacatgcctgtaatcccagcactttgggaggtcgaggcaggaggatcacttgaagtcaagagtttgagaccggcctggccaatatatgaaaccccatctctactaaaaatacaaaaattaaccaggtatggtggtgcatgccagtaatcccagctagctcgggaggctgaggcaagagaatcgctttaacctgggaggcagaggttgcagtgagccgagatcctgacactgtactccagtctgggt**gacagagtaagtgaaactctgtc**tca**aaatatatacatatatattt**atacaca**aaataatatatatttttatatatagatatatatatcaaaaatatatatatatttatacatatatacatatgtgtatatatatgtat**gtgtgtgtgtatatatataataaaattaaacaaataagaataaccaaatatttatggaaactcagcctattaaagacaagtgtaaaatgtgtctgggaaagtgaaaacaatgtagaaatctcttagaggaaatagaagaacatgattaaatatgtccctttttccactcaggcagcacaaactttattaggttccctccaaacacagaggacacattatgtcactgtgctcacactagatgtccccatactctccttggctctttccaccccactgcacccacc**aggggatttgcatactgtcccct**agggaggaccttccattgtgagtctgagataaaagctcagctgtaaccttgccttgactgatcagaactcctcagctcaccttctcacaATAAGGTTCCCTGCTCAGCTCCTGGGGCTGCTAATGCTTTGGGTTCCTGGTAAGGACAGAAGGGAGATAAGGGAGGAGAATGGAGTGTGAGGGTGAGCTCTGGGGGCCCCACTGCCTGTCCACATGCACATCTTGACCTGCAAGAAAAGGTGTATA**AAGTTTAGAACTGCAAGAGTCAGAAAGAGAAGATTCCTCACTCTTATAGTTCTAAACTT**CATATCCTAGAAGGACAAAGGACTTGTGCTCTCATGAAGAGTGTCACACAGGAAGAGGATAATAGTGTAGGTGACTTCTTGAGTCTCTTTTGTGCCTTGTGAATG**TTGGTTCTTTTTATGCCTGGATGTTTAGGGGTATGAACCAA**AGTCACACAAAAAAATCACTTAGCATAAAGTAGATAACAGAAAGAATATCATTTCAATGGTTCCCAATATTTGTACATAACCTTGCACTTCTCTCACTACTTCAGGATCCA**GTGGAGATATTGTGATGACCCAGACTCCAC**CCTCCCTGCCCGTCAACCCTGGAGAGCCGGCCTCCATCTCC**TGCAGGTCTAGTCAAAGCCTCCTGCA**TAGTAATGGATATACCTATTTGCATTGGTACCCGCAGAAGCCAGGGCAATCTCCACAGCTCCTGATTTATAGGGTTTCCAGTCGTTTTTCTGGGGTCCCAGACAGGTTTAGTGGCAGTGGGTCAGGCAG**TGATTTCACACTGAAAATCA**GCTGGGTGGAGGCTGAGGATGTTGGGGTTTATTACTGCATGCAAGCTACACAGTTTCCTaacacagtggtacaacc**ctgaacagaaacctcccttcttgctgtggttcag**ctgcccaaatgtgttgtttatctggaaagcagacactgtcgattatcttgggagagtaaagaggaagatgatggagaattcaggagaatacattacagctgagggctcttgaccatcaatatctcggttacatctcaggtaccacaatttagtccccatcagt**tgcaagagtcttggcctggcagaacttgca**ggagaatggagggaggtcaagtgcctctgaacaaccagtcatgttgcaagggaaagctgaatcaaagctcatcgtaatcctctctgccttgcctaagttcattcactaactaaa**ttcattcagcctgacagccacctaaatgaa**acagattagtggcaata**caaaactaatatatttttggttttg**atttggtttagcagttactagtgtacatgtaccttgacaagatttggtgatactaaaacagtttctctcccatcttctaccttccactttacacttttcttaacatacatctctctctgggacagcagg**aagcacagcattttatcctatttttttttcagggagtgctt**gcgtgtatttacattctagactttgtaactgctagata**tgttttggtaagtttatgcaaaaacaaagacctagtgacaaaaatctttg**aacaatcttattcacagtatgttgaaatgaaaagaaaattctatgaaatccaaagctgtgttttaaaaataaatatccaagtttcaaaagacaaaagaaaaaaggactaaaaatgtagccatagtcaaaaatgtatgccatattcaatggaatctggattaatataaaataatatataagcttatgttttgccccttgaaaaataataatctatagactttgggcagagggcagtaaatgagaatgtgagggcagctgttaactatgttagtggtggtgtgttggagttgatatatgaacccaagtgaaagc

Uppercase: IGKV2D-18

Lowercase: Flanking sequence[1000bp]

Red & Bold & Underline: Stem-loop [20]

Blue: Heptamer[32]

Green: Nonamer [9]

id-IGKV3D-7[V_gene_segment]

actgtaaggcaggtaccactattgtcacagttacacagatatggaaaccgagacacagggaagttaagttacttgattaatttcaagcaatcgggaagccatggagcatctatgtcagggctgccaggacatgtgactgtaaacagaagtttttaactcaaa**gagggtatgtatctgggttaatggaaagcttcaggaccctc**agaaaacattactaataagcaaatgaaaggtgtatctgggccgggcgcggtggctcatgcctgtaatcccagcactttgggaggccaaggcgggtggatcaccaggtcaggagatcgagaccatcctggctaacacggtgaaaccccgtctctactaaaaatacaaaaaattagttgggcatggtggcaggcgcctgtagtcccagccactcgggaggctgaggcaggagaatggcgtgaacctgggaggcagagctt**gcagtgagtggagatcgcaccactgc**cctctagcctggaagacagag**tgagactccgtctca**aaaaaaaaaaaaaaaaaaaaaaaaaaaaaaaaaaaaatgtgtatctggaagattaagttctaacagactcttcatttccatcgatccaataatgcacttagggagatgactgggcatattgaggacaggaagagagaaatgaaaacacagccttttatattgttcttaacagacttgtgccaaacattatacgggtgtatttaggtgattgaagagaagaaaggcacaggagtgaaattctgtgagcacaagggaggagttctacactcagactgagccaacagacttttctgacctgacaaccaaggcggcgcaggatgctcagtgcagagaggaagaagcaggtggtatct**gcagctggaagcccagctcccaccccagctgc**tttgcatgtccctcccagctgccctacctt**ccagggcccatatcaatgcctgggtcagagccctgg**ggaggaactgctcagttaggacccagATGGAACCATGGAAGCCCCAGCACAGCTTCTTCTTCCTCCTGCTACTCTGGCTCCCAGGTGAGGGGAATATGAGGTGGTTTTGCACAT**CAGTGAAAACTCCATCAGGAGTTTTCTCTG**CTCAGCAAGAAATATAATTAAAATTCAA**AGTAGATGAACAATTTTGGCTCTACT**CAAAGACAGCTGGTTTGATCTAGATTACATGAGTGCATTTCTGTTTTATTTCCAATCTCAGATACCACCGGAGAAATTGTAATGACACAGTCTCCAGCCACCCTGTCTTTGTCTCCAGGGGAAAGAGCCACCCTCTCCTGCAGGGCCAGTCAGAGTGTTAGCAGCAGCTACTTATCCTGGTACCAGCAGAAA**CCTGGGCAGGCTCCCAGG**CTCCTCATCTATGGTGCATCCACCAGG**GCCACTGGCATCCCAGCCAGGTTCAGTGGC**AGTGGGTCTGGGACAGACTTCACTCTCACCATCAGCAGCCTGCAGCCTGAAGATTTTGCAGTTTATTACTGTCAGCAGGATTATAACTTACCTcccacagtgattcaacatgaaacaaaaacctcaacaagaccatcagtgtttactagattttaccagctgcttcctttacagacagctaatgtggtggccactcagttttagcgtctctgctctatttggacattttgcagttctaaaaaaaaatcattgaataatttggacttt**gattcttggactcttttcaactgaggcaccagaatc**ccaggtttccagaaatagtgactcactgtatgaatccttatatagcctcagtggttcttaactttcccagtagaggtagctcagtgcatgctacactgctccattt**gaattttgcaacattctaagtagtagaaaattctatttatttatccaaatag**ttgactcagtaaaagctgttcatgtgaagatactaccatggctgaataaatcccattctttttctttcttcaggctatcaacatttcag**tggcaaatggttattatggaaacatttgcca**tttaaaagtgaactaaattatttcttcaattttcgctgtgatgcagtagactgtaaaaagattaaagtttgttaaaataaagtacatattcgataaggaagaaacagattattcctaatgacgtctgcaatgacctagtagaaagagtgatagaagcagttgttttcattatttttgtccaaaacttcctttcaaatgggatttcattgatcatattcatttattaccacctataagacatgttgacattatgtaacatctgatgtgaagcgctgaggatacatctgtctgtattattcttgccaaaaattaatggtgtgaattaaatcagtagtaaacatcatatacaaacccaactaggaggacattcttcaacatacctggacagtaaa**cttcaaatgtttgaag**gccatgaaagagaaacaaaagtgaaaaactatcacagatttaaagatattaaggacaggataaccaaataaaatacagaaacctgaattttatcttgtaacata

Uppercase: IGKV3D-7

Lowercase: Flanking sequence[1000bp]

Red & Bold & Underline: Stem-loop [14]

Blue: Heptamer[35]

Green: Nonamer [7]

id-TRBV1[V_gene_segment]

tttacttcggcgtttcaaatgccttcttgttttgtgcattcccctgcggaatccctttgtattctattctctcaatgctgtatcaggaggactgagggcagatggctgacatgagggcaatgtgcctc**tttcacctccaggaggagctacgactttagtgaaa**ttcctgagtcatgtggagggagaagagaggcaggagggaggaaaggactaacgaaggaatcaaatgctctggtatcagggtttgctccttgttgtagttgagagaccatgtgtggacaacttgaagacactaaagtccattccacctggacctgcatctctgcagtccctctgctcatgcccttagggcctcctgttattctagtctgtgttcaggtcctgcatcctaagtaggtgagatgctccctgggggtaggggtcgcagattatttctctcatgtcccctcg**agcacccactgaaaggtgct**tcctccctagttggcattaccttactgggtcttaaaatgatgcacagctggctccagggaagggctccactgagctaggtgaggtgtcctcctggaattcactga**gatgagggaggggagctggagtgtgctcatc**ctgggtccaagacaggcatcgggaaggcatctgcccaaagggaaggggtctgtgtgttagggaggaggggagccataagtagaaagaggaaggggagacccattcattcgttgtgggaagggcaggcagctgctaagaaaaaagcaactgtctaaagaacccgccctgcacacctggccctgagaagctagtctaaaccca**cctcttgaggtgccagtgccaagcttggaaaggaaagagg**aagtgtgagctgtagacactaatagtgacaccaacaggagcagaga**cttcccaagcagcccctgtctcagggccagggaag**cacacccagacgacaagga**cacagagcagggagacacagggtccccctgcctgtg**ccccgggtgaccctgccATGGGCTGAAGTCTCCACTGTGGTGTGGTCCATTGTCTCAGGTGAGTCCTGGGCACAGGTGGGACATTTCTGTCCTTAAATTTTTTGCTTTTTTCATGGAACTGC**TTCAGAAGATTCTGTCCTAGGCTTAGTCTGAA**TTTGGCTTCTTATTTTCATAGGCTCCATGGATACTGGAATTACCCAGACACCAAAATACCTGGTCACAGCAATGGGGAGTAAAAGGACAATGAAACGTGAGCATCTGGGACATGATTCTATGTATTGGTACAGACAGAAAGCTAAGAAATCCCTGGAGTTCA**TGTTTTACTACAACTGTAAGGAATTCATTGAAAACA**AGACTGTGCCAAATCACTTCACACCTGAATGCCCTGACAGCTCTCGCTTATACCTTCATGTGGTCGCACTGCAGCAAGAAGACTCAGCTGCGTAT**CTCTGCACCAGCAGCCAAGAcacagccctgcagag**tcaccgcctccctgtgcacaaacctc**ctggatctaatcagaaaaccgtgggggcaacgcatccag**ctgagcctcagcactcggttcagcattctgtaagacctcagcagacatctcagatcataatactgtcatttattggatgt**ggcaatgctaaattgcc**cagactcctgcagcctgtggcctgagtttgtcttgaatgagactgtgcctg**aacttgaatgtaggacagatgccgtcaagtt**taagtttttccaagtatctttcagtcggtttgtgcctctgggaaatgttgt**gtgaggtatttaacactctggcctcac**caatccaccttccaatatggactgttttttgacttttcttttgactcaagttagaccctggatctcagccctcatcccagctccaaatatgtatggtcagaccttttcttttcttttcctttcttttttttttttttcttttctttccctcccttcctccctccctccctcctttctttctctctctctctctctctttctttctttctttctttctttctttgtttctttgtttctttctttccttctttcttttttg**agacagagtctcactgtct**ggagtgcagtggcgcgatcttggctcacttccacctccatcttccagattcaagcgattctcctgcctcagcctcccgggtagctggaactacaggcgcccagcaccacgactggctaatttttgtatttttagtagagacagggtttcaccatgttggccaggatggtctcaatctcttaacttcttgatttgcccccctcgggctcccaaagtgctgggattacaggcgtgagccaccgcgcccggcaggtccttttc**tttttctttctttcttttttttttgaaaaa**ctaaagaatacccaatcccc**tcctcaggtggctctatcactgcctcaccctgagga**cagtatttccccatggggtttggatcactgttctctagt

Uppercase: TRBV1

Lowercase: Flanking sequence[1000bp]

Red & Bold & Underline: Stem-loop [16]

Blue: Heptamer[40]

Green: Nonamer [3]

id-IGKV6D-41[V_gene_segment]

tggtcattgcttggtcttctcta**aaatcccagggattt**atgccgtgagttttattggagtttgcaagagagttcatgaagtattcccatctaatgcagatgcatctagcaccgaacaatgttctggatcacatggctcaagtgacagagctacttgtgtcaaagcccaaaaccttcctcaaaatcttct**ctttttctgagcccactccaaaaag**cagccttccaagttagcttataagtgcggtggtggagtagacccaagtgtgggatttgttatgtcaatactctaagcaggccttgccaacactgtgctgcattctcagtgattggagctgcaaagtacaattatttgtccttacct**tgaaggggatgagccaacgctactcatgttccttca**ctaaatccctgataccttctccaatgtgacaggtccttaacttttgtagggttgaatcccac**actctggtcttcatgtgctttcactgaaaatccagagt**ccttattattttctgcttatcctatctagttac**cacgatgacatcaatatcgtg**ttgtatggagtgatgctttgtgaagcgtccacaggatcaaagtttcttcagcctcattgtgaatgtgttgtcaggaaagttaagatagccctgtgccgagactgaggatgtgagctgctgttcaccccagatgactgcagagcctcccagagactgcgatggactcttccttcactctgaggctgggccctgggtctggtcaagccctgccggggcctcagcggcgctcatctgcaggtgccagcagagggcgcctctcacat**tccccgaaggggcgggga**gggcgctctcctgacaacagtgatgtctgtgtatttaaaacaacaggacattcccgtaatttgcatgtaaccttcctcccacatctgaagtggtcct**gcctctcagtatcttataagaggc**tttttctcctagatgtgt**cttcagcaaaactgaag**tcaaaacactgagATGGTGTCCCCGTTGCAATTCCTGCGGCTTCTGCTCCTCTGGGTTCCAGGTGAGAAT**ATTTAGAAAAAGCTAAAACTAATTCTTTGAACCATTAATTTTCTTAAT**TAGGAACCTGGCACCATATGGAACTTGGCTTGTTTTTAAATGTGATTTTTTTTTAAGTAATGCGTATTCTTTCATCTTGTGCTACTAGATTAGTGGTGATTTCATTAAGCAGATGCTTATATTGTGCTAATGTTTGCTGTATGGTTTCAGCCTCCAGGGGTGATGTTGTGATGACACAGTCTCCAGCTTTCCTCTCT**GTGACTCCAGGGGAGAAAGTCAC**CATCACCTGCCAGGCCAGTGAAGGCATTGGCAACTACTTATACTGGTACCAGCAGAAACCAGATCAAGCCCCAAAGCTCCTCATCAAGTATGCTTCCCAGTCCATCTCAGGGGTCCCCTCGAGGTTCAGTGGCAGTGGATCTGGGACAGATTTCACCTTTACCATCAGTAGCCTGGAAGCTGAAGATGCTGCAACATATTACTGTCAGCAGGGCAATAAGCACCCTCAcactgtgctacaacccaaaacaaaaattagctcagcctggcggaacagagaaactgaacaataccccgtttttatgatccttgcaggtgcagttggggaaataatttaccaaataccatcacatatggcccattcagcagagctgacacccaagagtcagtgcctcacactttcatctttacccacagtaactttc**aaactaggagaagagttgttgaagattacttctcctgtttctttccataggtgtacagaaac**tgtcctttacaagttggtaaaaactgtggattattcttcagggctacagtaattcatttatgacaaaatcactttaaagggtaaagattaggaaacaatcaggtgatggcctgga**gctcttttgacagaagagc**catggactcaggtttcagtcattgtgggttcaaaggaatgagacattgttcctatttccaaagcggctctgtgaagtctgacatt**tgttaaatcagattaaca**tttaaggttcactttcactgaatatttgggttttgaaacatgaaactggtccatcaagaacagggttataaatatctagctatagctttggaaatggggactatcccagaaacaatatttacctggaataaaaaaggaatcagcgtcttttgttgccagtcttagaagcaggacagggctacaagccaaagaaggttgaaatcagcaactcttaattcctccataagaactgcttcaccaaaatctgagtgtaccaaagcaaaaactacaaagagaaagtagcttcaggaactcaagtaagatg**taaaaacaaaatggacatttgatttta**tatcttgtttgacatatgaaacacagggagatcacccatgatccttggaattttcttttctgcacatcaaaagcattgtctgtctcctgtttttccacctctattatccaaattgtgtttg**aaaaaatgtatttttt**aaattttcattattcttgattaaaagaccagtatctt

Uppercase: IGKV6D-41

Lowercase: Flanking sequence[1000bp]

Red & Bold & Underline: Stem-loop [16]

Blue: Heptamer[42]

Green: Nonamer [8]

id-TRBV22-1[V_gene_segment]

gcagataacctcagaaatagaacaacatttgtctaccaaaacaaaagcattcattaagaaatcagctaggatcctatttttatttactaaaactacttggaagaacccatgttatacaatgaatttacactcatacaaatttctactcagttgga**aaatgagactttagtcagctggtacaaccctatgaccataaatgtctcattt**cacttgatacagtattgacttcctaacttcctgtgaattagaacttcaggtccagaaagtaataagagcaaacgcaaatctagtgt**ttcttatgtactaagaa**ctgttctaaatgctttatttatgtaatgttcatcacaggttttgggtagttacttattatc**tcaattttacagataaagtgaaattga**agagat**gctctgtaacttttccaaggttacatagc**tgcgaaatggcagagacaggactaaagcccatgccgtgtcccaagtccctgctctaatccattgtcacaagggaaggggcgcttcagacactttgctattcctaaaattgcaggagtagcccccaccccatctcta**ctgtattaattatttatagtgaatacag**agggtgtagagaatcggttctttaaaaaacaaaaacaaaaacaaaaccactaaacttccaaaccccttcttgtgttgagttgagcacaggtgcatggcactctgtgtatgactctgaccaggaagtgagggattggctcttctgaccatgcaggcaggcacttgtccatt**tagaagggatataaaaagcttcta**taagcagaagaggaggtgcatgtgacttgatcgctctaattcttagtctcaccatctgcaaaatcagctgtgaaaatcagtcatcacgtagcaatgacaaattctttatattaagtccc**ggggaggggctagatctcagccccttccc**ttggcctctctc**tctctcttagagcctgtgtctgtaacttcagagaga**ttttgtcgctgccATGGGGAGCTGGGTCCTCTGCTATGTGACCCTGTGTCTCCTGGGAGCAGGTGAGCTCAGCACACATCCAGGTGGTCAATTCCTATTCTGG**GCTTGCCAGGCCCTTCCCTGGCCAGC**ACCTCCCTTCCAGGCATGATCTCCAGCCTTTGTCTCTTGGACCCTTGGATGCTGACATCTATCAGATGCCAT**TCCAGCTCACTGGGGCTGGA**TGGGATGTGACTCTGGAGTGGAAACGGAATTTGAGACACAATGACATGTACTGCTACTGGTACTGGCAGGACCCAAAGCAAAATCTGAGACTGATCTATTACTCAAGGGTTGAAAAGGATATTCAGAGAGGAGATCTAACTGAAGGCTACGTGTCTGCCAAGAG**GAGAAGGGGCTATTTCTTCTC**AGGGTGAAGTTGGCCCACACCAGCCAAACAGCTTTGTACTTCTGTCCTGGGAGCGCACc**acaatggaagcacaaccattgt**ctctctgtgcggaaatgtgtcctcaccctacagcccccaccacatcctctagcttaattttttcatttttaatattttcttgagattttactatgtcctgtta**cttttgcaaacacaaaag**atacaaaactgaaaagaaaatagatcctttatccaaggagctaaaagtctagcaataaagtagttagaatggaaaatataatgtataatactaaaatcagaagtcatttgggagctttagggaggacatgcccaattctg**tctgattgaatcaga**gaaaacaaagaggaagtcacattagaacttcaccagctagacctggttctgagtagggcattctgggcaaagagaacagcttttgcaaaggcatattaaagagtttgtcactttggggaacagtttacaaagagaaagaagatgatcaatttatatgaatcaaaagaaatggaatacatttgaggaagtagacagaaaggagtttttgaaaaagttgagtttttggttccgtgaatacaataagtatgaaagtcctttgtaggctgttgctcaaaagagagagctgaagcttcattgtcatcagcagacaagcaaacaatacaatttttc**aatatatatatatatt**ttcctcatgtaact**ttgaaatgatactatttttcaa**ttttcctcatataactttgatccttaacaaagggcacagtttcaaacaatatgaaatcatgttagcagtagcagttcccatttgggataatagggttcaactgtctatggaacaatgagataaaaactagaagcaggcaactgcatagacaagatgtgaaattgagatgccataaggctggactagaggttgagactctgcagatttcagcatgtaaataattgaactatggggacagaaaattccaaccaaatatagatttagatcacagagataaaaggcatggaagttaagactgtggtattctggttccttatgg

Uppercase: TRBV22-1

Lowercase: Flanking sequence[1000bp]

Red & Bold & Underline: Stem-loop [16]

Blue: Heptamer[25]

Green: Nonamer [7]

id-IGKV2D-26[V_gene_segment]

cagtcaggttggaaagtaagtccaaggatatgtagggttaaataaaacccatctgatgagaatttatagtttgtagggcatgactccccagactccttagataggaactgagacaaggaaaaaaaaaatcagagcttagtccttaccttgataaacacaaatgaaaagaaaatattacttgtagatattcaa**taaattatctgaaatcaattta**agaaaacattgctcagaatcacatg**aaattatagaggacagtaattt**gaatccacaaacagaaataaaggcctcaaaatagtaaagaactgggaaaaaaatagacacaatgataaaaaaatgtttacttcaatttctatgatattca**ctaccaagtacaaacaaactattgtggtag**atgtcagaaagacagttactttggttgggatggggaggcagttattgaatgggaaggtgtacgagagaatc**atccagaaattttgaaatattctatagtttcagctggat**ag**tgaatatacataaaactttattca**tt**ccacatttaggatttctgccctttatatgtgg**tttatttcaatgaaaaaaaaatagaaaagccaaaccta**gattcttttaaagaaaattggaggcaagaaaaaagaatc**agccaatatttatggaaactcaactttattaaagacaagtgcaaaatgtgcctggaaaagtgctaaagaactgttgaaaaacaggtggaaagaaaaggaaatattataatatgtctcttctccaactgaagcaacacagattttatcttggttagttttccttgaaacacgcaggatattttatgtccccttgcccacaagagaagccctcatcctcctgttgttgcgcttagcatcactgcacccaccgggggatttgcatatt**gtcctctagggaggac**cttcccttgtgggtctgagataaaagctcagctctaacccttaccttgactgatcaggactcctcagttcaccttctcactATGAGGCTCCCTGCTCAGCTCTTGGGGCTGCTAATGCTCTGGGTCCCTGGTAAGGACAGAAGGAGATGAGGGAGGAGAA**TGGGGTGGGAAGGTAAGCCTGGGGACCCCA**CTGCCTTCCATGTGTGTTCTGCCCTGCCCATGTGTTAGATGTACAGGTCTTGTTCTCCAGGATGGGGAATGTGAGGTTTAAATCTGTGAGAGTGAGGACGATTCAAAAAGAAGCAAGGAC**CTGTGTGCTCTGGTGAATATCGTCACACAG**AG**AAAGGGAGGTGGTGTAGGTGACTTCTAGAATCCCCTTTGCAGCTTGCAAA**TTTGGAATATGTTTAGTGTATAAATACAAACAACAAAAAATTATATAGCCTGAAATAAAAAATGAAAATTTATGATAAATGACACATGATATTTGTACATATCCTTCCACTTCTTTCTATCTATTTTAGGATCCAGTGCAGAGATTGTGATGACCCAGACTCCA**CTCTCCTTGTCTATCACCCCTGGAGAG**CAGGCCTCCATGTCC**TGCAGGTCTAGTCAGAGCCTCCTGCA**TAGTGATGGATACACCTATTTGTATTGGTTTCTGCAGAAAGCCAGGCCAGTCTCCACGCTCCTGATCTATGAAGTTTCCAACCGGTTCTCTGGAGTGCCAGATAGG**TTCAGTGGCAGCGGGTCAGGGACAGATTTCACACTGAA**AA**TCAGCCGGGTGGAGGCTGAGGATTTTGGAGTTTATTACTGCATGCAAGATGCACAAGATCCTC**Ccacagtgctacagccctgagcacaaacctctctgcttgaagtggccca**gctgctcacatgtgatgtttgtctggggagcagc**acagcaagtt**ctctgagtctgtgcatgaggaaaatgttggacaactcagag**gagtagattggtctcatgaattccttggttacatctcaggaactactcatttgtgtagctgcaaaagccttgacatggcaaaagcagtaggagaatggggaaggtcccatggagctgatgctgattcatgatgcaactgaataaaagctcattcgaatcctcctttcttacatttc**tcaattgtagttaatcagcataagtgccaggcaattgatacaaatgttggcaacacatgctgagtgcctcctcatttgtat**gtatttcatccatttttatatattttaataggataattatttggtcatatttagaaattggtattttccaccttccaagctttctattttccttacttattcct**tatcttcctatgtaataaagata**ctgtagaca**acattttacactagttgtaaaatgt**ggctccccatttatacgggtagctggttgggggcaattag**caaaaatcttggtttttg**agtgcctagacttttgtaaattaatgtcagatacaagatccaaatgctaaaactctttgatttgccttgattacttcttactgatagatcataaaa**ggaattcttgaaattcc**aaaagttgggttttaaaaataaaagcatacactgg**aatatacagggcatagaattcaccctatatatt**gctagaagaatagagaatgataacagttttagacttcgagggtacttcaggaatgggaacacacaagga**acacagtagatatgagctactgtgtctaccgtattagaca**atcctcctgcattttaacttacatttggtgaacttcagatgccagtgggtgttgcaagttgatgacataaacctaaaatcaagtcccgatgtatttcttgt

Uppercase: IGKV2D-26

Lowercase: Flanking sequence[1000bp]

Red & Bold & Underline: Stem-loop [28]

Blue: Heptamer[22]

Green: Nonamer [8]

id-IGHV3-62-2[V_gene_segment]

tcaagtaattctgtttagtgaaagaaactaaggagttaagagtaaattttatatgatacatttgtaggaattttagaagatgccattattgtaaattaacacggagaagatttgagtttgtctgacaatacgctgttgggagtaatgtggatgtgagttgaatttcagagaaataaaagaaagatttagggattaatttaattattcaaaacttgattgaagtgctgagcaaatggctgcaaacataggtcaacattt**ttcaaatccttcactataaatttgaattaattacttattaatt**acacttgaataaagcaataacgaagaaaccaataaaataacatttgactaaaatggagcaataaacagatcgatgttaacacaaggaatatgactgacttatgaaaacatgcacatgaaccatggttcactctacgtattttggtaaattacagaaagttgtcataacagatggggaatcctgcagacttcactaggcatggtccacgctgcc**ctggagttgtctcaggggagctgcctcctccag**tggttagagcacaggcccaggtaataggactaatttttttagatgtgtaattttagacacactgcacaactgctgtgttctctgtgtaaattatctcctgtaaaatgtaacattgaaacctgcattaa**acatattgtataaatatgt**aagaataaaataagattatgagagctaaatattaatcaaggcacaagcacat**aatataaaattatatt**ttcctgaatgatacaattattccaatctgtcccaggacactt**catctgccctgagcccagcctctcctcagatgtcccaccacagagcttgctatatagtgggggacat**gcgaatagggccctccctctgctgatgaaaaccagcccagctgaccctgcagctctgggagaggagcccagcactggaagtcggcggtgtttccattccgtgatcagcactgaacacagaggactcaccATGGAGTTTGGGCTGAGCTGGGTTTTCCTTGTTGCTATTTTAAGAGGTGATTCATAGATAAATAGAGATGTTGAGTGGGAGTGGACATGAGTGAG**AGAAACAGTGGATGTGTGTGGCAGTTTCT**GACCTTGGTGTCTTTGTGTTTGCAGGTGTCCAGTGTGAGGTGCAGCTGGTGGAGTCTGGGGAAGGCTTGGTCCAGCCTGGGGGGTCCCTGAGACTCTCCTGTGCAGCCTCTGGATTCACCTTCAGTAGCTCTGCTATGCACTGGGTCCGCCAGGCTCCAAGAAAGGGTTTGTAGTGGGTCTCAGTTATTAGTACAAGTGGTGATACCGTACTCTACACAGACTCTGTGAAGGGCCGATTCACCATCTCCAGAGACAATGCCCAGAATTCACTGTATCTGCAAATGA**ACAGCCTGAGAGCCGACGACATGGCTGT**GTATTACTGTGTGAAAGAcgcagtgagaagtcagtgtgagcccagacacaaacc**tccctgcagggtacctgggacaaccaggga**aag**cctgggacactgtatactgggctgtccccagg**ggcaagtccaggtggtataagcctaggtttcctgtcatggtctagggttgcctttttagcaacttaccccagagaacttctctagatttccaattctgtaata**acatttgatgtcgtctctggctgcaaaatgt**cccctcaactttgtatctttttttttttttttgtaacaggaggacacatcctcaccctgcagaagcctgagtgtcacattggg**ggcagaaatgatctgcc**ttgatctcactgatcactgtcccgaggaacataccccacaggggaccctgatgactccagtaatgtctctgcctcaaaaccattgaagagtccttccttccattagagttgaccacagcacctgggcttcagcacaagccataccacagacgtcacaaagcagcagattga**cacctgatccaggtg**cattttctcacctttagaggctgagagaggggtgtattctcaaggtcaacacactctttgtggatttttacagagaacacctacttttactttatt**tcatttgaagataaatgaacaaatgtgcagctgcaaatga**ttgtaattttcacatttattccaattccaatatttctgaattctgcatgatgtcctggcacagggttgtgttttctacaggtatttttcatc**aaaccaagaaaatgcattttccactttccgtggttt**ttctccatgtgcagaggccttgagtagagcacgtctgactgtgtttgttttcacgctgctgataaagagaccttgagtagatcacgtctgactgtgtttgtctgttttcacgtggctgataaagacataccagagactgggaaacttacaacagaaagagttttattggacttacagttccacgtggctggaaaggcctcacaatcatggtggaa

Uppercase: IGHV3-62-2

Lowercase: Flanking sequence[1000bp]

Red & Bold & Underline: Stem-loop [16]

Blue: Heptamer[49]

Green: Nonamer [8]

id-TRDV1[V_gene_segment]

catggaagattaacatctaaattccatacaatcacgtaacagcatacaaaaaattcagccaccaacttctgtggtattgacatgcacaaagttcttgcaaccttcagataacacgttctgtgggagtatattatagcattgtatcatttgcatcacagagagatcttgtatgcttatcatgacaaacaatgtcttatgcttatcattaaaaattttaaaaatgcccagccacagaatgtaacaagtgagactgaatactttggatcaatgggatagaaaatgtgatatcaggacaattctaggacacatgatctaagaagtataattctacttgtgaaggcaacag**gactgaatatactgatcctcctcagtc**tagtaaataaaacttagtgtcagattt**tgaaatgattacattttattgctgttataatttca**taaaga**gataaaaaactgagcagcttctgaaagtcgagaaggactgatttttatc**atttgattttgaagtagctttacttattatagaacttaactagggccctagacctcagtaaggaggaagaaagagatgcagtaattgtttagaataggactgcctttcacagagaaagtcaaatatctgagtatttttcttttaaatttaattagtttctgatggcaaaaagtaaatatatgctaaatgaataatatcaatgtatgtcaactggttcaactgcagttcccctcaatacaaattgcttttcttatatcccttttctaggttc**tgtctatcctaatgactagggaaagacaactagaca**aagtaaagattcaggggtgagccagacatttttcctcatctgtgcatagccatgtagtctatgaacatgagactcaat**ttcctgtttggggagatagcatcagaacccaggaa**ctacatatacacaccgcagggccctggtaggcactgaactttgagcttcaggca**gcacaactcacatttgtgc**aaagagctacat**gccacATGCTGTTCTCCAGCCTGCTGTGTGTATTTGTGGC**CTTCAGCTACTCTGGTATGGATGGTTCCAAGTGTGAGTAAAAATGTTAAGGATGGAAGGATTCCAAACAGGCATATGGT**ATTTCTCATGGACTTGAGGAGGAAAGGAGAAAT**TGTAAAAAAGCGAGCATATTATGATTTTAGTTTGGTTTGTCTCTGGGTCCTAAATTAGTCTAGCTCTTACGATATTTTATTCACTATATCAAAGGATCAGCTCTGTTTTCTGATTTTTCCCACAGGATCAAGTGTGGCCCAGAAGGTTACTCAAGCCCAGTCATCAGTATCCATGCCAGTGAGGAAAGCAGTCACCCTGAACTGCCTGTATGAAAC**AAGTTGGTGGTCATATTATATTTTTTGGTACAAGCAACTT**CCCAGCAAAGAGATGATTTTCCTTATTCGCCAGGGTTCTGATGA**ACAGAATGCAAAAAGTGGTCGCTATTCTGT**CAACTTCAAGAAAGCAGCGAAATCCGTCGCCTTAACCATTTCAGCCTTACAGCTAGAAGATTCAGCAAAGTACTTTTGTGCTCTTGGGGAACTcacagtgtttgaagtgatagtaaaagcaaaacaaaaaccctagggctcaataagagaacccctctactccccatcctttgctacaggagccaatctgaaatgcacacctgcagatctcaggcaagagagggtagtgactgtggtctgtggatttaatctgtggtctggatcagatgatttaa**ttccaagtaaaacctccttctatctcacttggaa**acaggtgttcagagcaactttcttctaatacttcaatttttacttaaaatagatcatacagaaaatgtgtaaagttgtctaaagttggaa**agacttccacataataacttaaagtggcagtct**cacttaacgatgctcacagcagaaatctgggtctagttatcttgaatcaccataaaaatcatttccttagccctttcctcttaagctttatgtcagttaggatacaatcagggaagcagaaccatcgtgaaagatgtagaataagattaattataaggattagacctcatgcaattgctggaacgggtggctctcggaagttacttagttgtttatgt**ttttatgaaattgtaacaggaataaaa**tgtaatcttttcaaaatctgacactgagttttatgtactaggctgaagaggatcaggacattcagtcctattggcttcacaagtagaatagtacttcttagatcatgtgtcctagaattctcctgataccacatttctatcccattaatccaaactattcagtctcacttgttaaattctgtggctgggcatgtttttcatgtgtaaaataaagtcgtgataagcatgc**cagatttctgtgatacgatgccataatctg**ctcttactcctatgcaaagctgtttgctgaatttactctaggtcagctagactcacatttgggaaggagagttgaacatggacaagagcatgcaaaagctgtgatttgtaaagacattttggaacccatggtgacaggtggaacccacgaggacaaa

Uppercase: TRDV1

Lowercase: Flanking sequence[1000bp]

Red & Bold & Underline: Stem-loop [14]

Blue: Heptamer[33]

Green: Nonamer [1]

id-IGHV3-23[V_gene_segment]

gctttgtcgtcc**aggctggagctccgtggtgagatcatagctctctgcagcct**cagaccctgggtctccagcaatcctcctccctctgccctatgtagctgggactacaggggtgcaccaccatgctcggcttcagtgcgtattaaaccgcacacttcaattatgcagtatttattatatagcaataatgcctcaataagggtatt**acaaataagtggataatttgt**taaagattgatggaaagatagataccaacatgagaaatgtatgacactc**aagaaaataaaactgtaggaaacttgcttttctt**tatatttgttaggtaatcaccacagtgtgtacacatcacaccatgttcccattacagagaaa**aggttctgcgaacct**cacgagctgtgacccctgtgtgctgggcttggttcagggagaagtcaggtccagtggtgagaagcacaggcccagatg**cccaggctcactctgaccaaatgtgagcactggg**gacattgtaaaacccacctgtgcttttgctgataatttttcatctttaacatggaaataatattgatactatataccatggtttctctgcgtatgtaaaaataaaagatgattggtgctaactttaaaaatatg**cagtttatgtagatctatggtacctcaataaaactg**ttttaaaa**taaaaattacaaaattataagattttta**ggttttaaggtttaagtttatcacaaaacaaactgacaatag**gaaagcacaatttcccaatgctttc**aatatcacagatctccccgaggacattctgacatgctctgagccccactatctccaaaggcctctcaccccagagcttactatatagtaggagatatgcaaatagagccctccgtctgctgatgaaaaccagcccagccctgaccctgcagctctgagagaggagcccagccctgggattttcaggtgttttcatt**tggtgatcaggactgaacagagagaactcaccA**TGGAGTTTGGGCTGAGCTGGCTTTTTCTTGTGGCTATTTTAAAAGGTAATTCATGGAGAAATAGAAAAATTGAGTGTGAATGGATAAGAGTGAG**AGAAACAGTGGATACGTGTGGCAGTTTCT**GACCAGGGTTTCTTTTTGTTTGCAGGTGTCCAGTGTGAGGTGCAGCTGGTGGAGTCTGGGGGAGGCTTGGTACAGCCTGGGGGGTCCCTGAGACTCTCCTGTGCAGCCTCTGGATTCACCTTTAGCAGCTATGCCATGAGCTGGGTCCGCCAGG**CTCCAGGGAAGGGGCTGGAG**TGGGTCTCAGCTATTAGTGGTAGTGGTGGTAGCACATACTACGCAGACTCCGTGAAGGGCCGGTTCACCATCTCCAGAGACAATTCCAAGAACACGCTGTATCTGCAAATGAACAGCCTGAGAGCCGAGGACACGGCCGTATATT**ACTGTGCGAAAGAcacagt**gaggggaagtcattgtgagcccagacacaaacctccctgcaggaacgatgggggggaa**atcagcggcagggggcgctcaggacccgctgat**cagagtcatccccagaggcaggtgcagatggaggctgtttc**ctgtcagggtgtgggacttcatcttcttctgacag**tttctctagtgaacctctctaacctcagaattctgtgcttactaatgtcatctctacgtatttt**ttaaaagatcattttaa**tatgagcacctattctcacacgcaccaaatgcagattgacgcttacagagatgaaaagttctcaaccatg**gtcaccaggatgagagtcctgaggaaactcaggggtgcctggtgac**tcttccgcaatcagtcctgggacagaaacctcagggacagcctttatgagttttgatcacatctaacagagaggatgggccagggccagagtcatgta**gaacctcacaggttc**cacgtctgacccttctcctgacactaaagccaatcagcatcagcactgacctggtgctccttttgctcccaatccatgttctttctttggagagtttcttctcccttttttatttgcttttc**ctgtttcctgaaaaagaaacag**atggtccccttggtccacattccagggctcaaggcattttcttggagctcaggtggggctcaggct**gtggctcctgcagccac**gt**gggagaggctgatgggacttccttctctccc**cattttctcaggaccctccactgtgttgtgtgtaggctcatctgggaatgcagttgaccattaatactgaaggggacaaactcatttgatcaaaatgggatgtggatgtggagttaaccctgttccgtgcacagtgtcagagtcgtctgcttcagaagtagtgttagaaagagctggtgaaatttatcagaatcaagatggagccacttgtgttaaagccctaatgaatgaagctggggaaggccatgaaggagggt

Uppercase: IGHV3-23

Lowercase: Flanking sequence[1000bp]

Red & Bold & Underline: Stem-loop [20]

Blue: Heptamer[42]

Green: Nonamer [6]

id-LOC107985528[V_gene_segment]

ctctagggcagatgctcctgagaccccaggaacaggctggtggggagcgcagctcagagcattactcaggggatgtggcctttgtcatcctactttgaaacaattgactatttgggcctagattgatacagggcttcaagttgattttaatccaggctcctatagtcagcgcgtgaaacagagattttagttgaaataatgagacctggtattactagtcagc**tctccatgctggaga**accataagaaattataccaaaggcaggaaaggggatagaatatggggatcatcacaccaagaataaggtgcagcccatttagcccc**tgggtcttaaagagacccatagctctggataatggcagatctatg**cgtgacacagttatcatctttgtgcatcttcagagaattgtttttccttttactcctagggacaatgtcttaagtttgttag**taaattctattgaattta**ttaaagatgcttctgataaattctttttatattcatttcaaacaagaagcaatttcacactgacattgttattatagcactaaatacttttacactcatcaaattcctttgagactaactgaaatttctgacagccccacattctacaactttattgtaaattttctgccaaaaatgatgctttcctatacactcctaatacaagta**taaatatattattta**atctagtcttaggttgatttaaaattttgaaaattcactccaaa**catatgttctgtaaccatatg**gccaccaatgagaaatgcattctttcaaggtaaatctgtgctgccctggtttgacctgggactctggggatactgcgcccctgtgctgagttactgagatgagccagccctgcagctgtgctcagcctgccccatcccctgctgatttgcctgttcctagagcacagccccctgccctgaagactttttataggctggtcacacccggtgcaggagtcagccccagtcaggacacagcacagacATGAGGGCCCCCACTCAGCTCCTGGGGCTCCTGGTGCTCTGGCTGCCAGGTAAGGAAGGAGAACACTAGGATTATACTCGGTCAGTGTGCTCAGTACTGTCTGGAACTTCAGGGAAGTCCTCTGATAACATGATTAATTGCGACAATATTTGTTTTTATGTTTCCAACTTCAGGTGCCAGATGTGACATCCAGATGACCCAGTCTCCATCCTCC**CTGTCTGCATCTGTAGGAGACAG**AGTCACCATCACTTGCCGGGCGAGTCAGGGCATTAGCAATAATTTAAATTGGTATCAGCAGAAACCAGGGAAAACTCCTAAGCTCCTGATCTATGCTGCATCCAGTCTGCAAAGTGGGATTCCCTCTCGGTTCAGTGACAGTGGATCTGGGGCAGACTACACTCTCACCATCCGCAGCCTGCAGCCTGAAGATTTTGCAACTTAT**TACTGTCAACAGAGTGACAGTA**CCCCTCCCACAGCGTTACAAGTCATAACATAATCCCCAAGGAAGCAGATGTGTGAGGCTGGGCTGCCCCAATGCTCCTTCCGGTGCCTCTATCTGCTGAGGGAAGTTCTCAAACTCAGTCAGGTTTGGAAAGTCATTGGGAGATTTTCCT**AGAGGAGGCCAGGGAGGTTCCTCT**GAACCCTAAGCCTCTTTCACCCTCATCCCCAGCAGAAAAGATGTGACAATGCCTGTCCTGACTGAATAAAGAAGAGAGATAAGTCCAActgaggagtctgtgttatgggataatcggaatttgtacagcaaaagagaagctattctcagtatttcaaggagaaa**ttattcaagttgaataa**attagagcctaaaccacagtctttccgaagcctatggaatgttattcatgaagcaggtactagacacaggggattct**caggtgctacttcagaagccagggtgcacctg**cccctggtggtatgtgctgaacaccgtgtgatgatcct**cagtcctgtctgggaagcccaggtcttcccagacaggaatg**ctttgataaatccactgctaggtaggtaattcttctgtaacattaataattcggttttacttttggatcctaaagtggaactacttagaaaatctgcaaaaatcatgcaaagaaa**atcatgtaaaatcatgat**ctttacttaggcacttacaggtgaaatgacgaggtatctggaattgcattaaaatagatcaaggtgaaaaagactgacaaaaacattaactgttaaaacttggggctatatatcaaagacattagtttctctatgagtatatttcaaattttcctcaataaatt**ttttaaaaattcaatttaaaa**ttcaagccacaaataaatgctttttaaaaatttcttagattatacaaatagtgccataacatcaagaataatcccaaatcattccatatacttcaaaagatacaaaaccaggctgggtgcggtggctcacgtctgtaa**tcccaacactttggga**ggccgaggtgggcggatcacctgaggttgtgagtttgaaaccagtctgaccaacatggagaaaccccgtctctaccaaaaatacaaaat**tagctgggtgtggtggtgcatgcctgtaatcccagcta**ctggggaggctgaggcaggagaatcacttgaactcgggaagtggaggttgcagtgagccgagattgcgccattgcactccagcctgggcgacaagagcaaaactccatctcaaaaaaaaa

Uppercase: LOC107985528

Lowercase: Flanking sequence[1000bp]

Red & Bold & Underline: Stem-loop [16]

Blue: Heptamer[23]

Green: Nonamer [4]

id-IGHV4-28[V_gene_segment]

tccaacttctggtctccaacaatactcctgcatctgccatctaagtagctggaactaccgttgtttgccagcaggcttggcttgagtacttattaaaccgcacactttttgcaatatttaatgtatagtaataatgtctcaataagactatt**acaaatcaatgaatgaataatttgt**tctgtacagatttatggaaaaatagacactaacatgatgaatgactg**acatttatgaaaatacagctgcatgaaatgt**gcttctctttacattcattaggtaaacacaatagtgcatacacatcacaccgtgctttcattacaggaagaaagatctgaaaatgtcactgggg**tgaaccatattgtgctgggcttggttca**ggaagcagtcaggcccagtgttgtgaccttcaaccacagaatcctaaaaaataataaaagaggctcccccaaagtccccatcagttcctggactcgccatgtgtctgggccgtatcagtgcatccggagctccctggtggctttagggattccttgcttgccatgctgaggtctcgctatagattatgttgggttttctgaggccgttttgctatttaagacccactccctggcacagagcgattc**cctctaaacctgatagagg**ttctgaactacaaataacgttaagtgaatcctggtgtgtctgaactcacatgattgttacattaagctgctgttgcaatctgtttcctcacctgggaaaagaggagccaggacatagtgagttgaggccccaggaagataactgaattctcagagggcacaaccagcatcctccttcc**agggagagcctaaaagactggggcctccct**catcccttttcacctctccatacagaggcaccacccacatgcaaatctcacttaggcacccaagggaaaccatcacac**atttccttaaattcagggtcctgctcacatgggaaat**actttctgagagtcctggacctcctgtgc**aagaacATGAAACACCTGTGGTTCTT**CCTCCTGCTGGTGGCAGCTCCCAGATGTGAGTGTCTCAAGGCTGCAGACATGGAGATATGGGAGGTGCCTC**TGAGCCCAGGGCTCA**CTGTGGGTCTCTCTGTTCACAGGG**GTCCTGTCCCAGGTACAGCTGCAGGAGTCGGGCCCAGGAC**TGGTGAAGCCTTCGGACACCCTGTCCCTCACCTGCGCTGTCTCTG**GTTACTCCATCAGCAGTAGTAAC**TGGT**GGGGCTGGATCCGGCAGCCCC**CAGGGAAGGGACTGGAGTGGATTGGGTACATCTATTATAGTGGGAGCACCTACTACAACCCGTCCCTCAAGAGTCGAGTCACCATGTCAGTAGACACGTCCAAGAACCAGTTCTCCCTGAAGCTGAGCTCTGTGACCGCCGTGGACACGGCCGTGTATT**ACTGTGCGAGAAAcacagt**gaggggaggtgagtgtgagcccagacacaaa**cctccctgcagggagg**cggaggagccggcgcaggtg**ctgctcaggacgagcag**ggggcgcgcggggcccacagagcaggaggccgggtcaggagcaggtgcag**ggagggcggggcttcctcatcagctcagtgctctccctcc**tccccagcacctcagctgtccccaggactcctctttctttattatctgtggttctacttcctcacatccttgctgcaggttttaacagtcactggcacccttttttcaacaaattcccacaaggcccattttaccttcttgacaagcaaaatattctaaggcttctcactatcccttgattggcgtcatctactgcatcgtgtctttctcttaaattcatgtgatgaaaccccaatcccagtgg**ctctgtattggacagag**ggattttaagagggcaaataagttaactgtggccataaatgtgggaccctaatccagtaggactcttgtccttatgagaagtggaagacatcagaggcctctctct**ccacatgcacacagaggagagtccatgtgg**ggatgcgctgcatgagaaggtggccctttgcaagcctggaagaggcctctcaaaaaacttaccctttctgcacccagatcttggacgtcaagactccagaacaataagaaaattaatatttgttatttgaaatagcctagtctgtggtatcttctcatggaaagccaagtagattcagagattcaattgtgttagctccgtctcctgcatggagtagcagcccaccgaggctggacacatctctcaaatta**ttttcataaagaaaaatagatccaagatgaaaa**ccccaccacattt**ctgctgagccattcacttagcagcag**tctc**tgagatcagccctgggggtatgttctaggtcgtgtatctca**ctttccactatgagaatccatgtagataatgagaaccagccagatggagtcctga

Uppercase: IGHV4-28

Lowercase: Flanking sequence[1000bp]

Red & Bold & Underline: Stem-loop [20]

Blue: Heptamer[39]

Green: Nonamer [3]

id-IGHV1-12[V_gene_segment]

tatcttaatagtgagaatatgaaggataggcatgttttt**actaattctatgggtacagattagt**tgaagaaactacattcctatgaataagaaattcagatttcagtgttaagtaatgttgcttacattgtgtgagtgacagggcagtggtggatctgagagtgtggcaggtgcacagaccaagtgagtcagaaatcaatatggaaaggtgagggtctgtggatatgaactga**aagtatgtaaatactt**gacaaaatactaataaatggagttcaaaataacccaa**aattgttctaaacacaaattccttgacaatt**actttgggagtagagagttcataatggactccaaactcctgctttatcttctgattcccattgt**ctgtgagatgagaaaatcagctctaattatgcatcacag**ggcaaatctgtaaatcaagagtg**ttcaatagaattgaa**gaccctgggggatcaggacatgaggcaggtgctggagac**agtgtctcaggagcacccagtagatctcagaggtccctcctggacact**catgtgggacataagcatcactttctcagagtcaccaatgagctgtgctggtggctgatgggtccaggaaaagaccaaggcacctgctcagtgtgatggagagtgatggttccaaaaatgatccaggtggtctctatgctaatcaaatgt**aggctcacagtgaggagcct**gttctatatgggcttattcttcagtgaaaggacgtctgtccacaaatgtttgtaaatggagcagggcatgcatttcctcaagcaggattagggcttcgaccgtctgcatctcactcttgttaggctgatgtgtcatttatcttc**cctttcttatcatggattgggctttgagctaagaaagg**ctttgtcttatgaatatgcaaatatactgatatccactgaggtaaatatgttctgtgccctgagagaatcacctgagagaatcccctgagagcacatctcctcATGGGCTGGACCTGCAAGATCCTCTTCTTGGTGGCAGCAGCCACAGGTAAGCAGTTCCCAGGTCCAAGTAATG**AGGAGGGGATTGAGTCCAGTCAAGGGGGCTTTCATCCACTCCTGTGTCCTCCCCACAGG**TGCCCACTCCCAGGTGCAGCTGGTGCAATCTGGGGCTGAGGTGA**AGAAGCCTGGGGCCTCAGTGAAGGTCTCCTGCAAGGCTTCT**GGATACACCTTCACCTACTGCTACTTGCACTGGGTATGACAGGCCCCTGGACAAGGGCTTGAATGGACAGGATtttagttatttgagagatttttcatacaacatttattctgtaagcaaatttcagggattgtagaatgaatcatattaacaaatctgacacagaacttcctctgaatcaatctttgtaaacatcaat**ttctgaatcaatgttgtaaatatttcagaa**cacaagcacaaattcacattttaactctacttttatctctatttaaaaaata**tcaaaaaatctcattttgtgcatgtaacgttttga**atgcccaccatcaatgcatgacatttcttgtttttccacattcatgttaccatt**tatcatcatgagaattgtgagttttagccatgctgata**ggtgagtaatggcatctaa**tttttatttaaatgcacatgtcccaaataaaaaattcgtattaaacaattttt**atataatttttgctgggatgcctttcctgatatgtggttcatttttatctgcattgttttcttttcattagttgtaagtttacttgcatattgattataaaagtcacttaacaaatccaaagaattgatttaacaaatagatgacttggaagtattttctcccagtctgtggttgtctttttctctcttatcagtgggtatttcaaaaaatatatgtgtgtctgtgtg**tctgtgtgtgtgtgtgtgcacaga**tttagacaaaaaacataaaaaattattcattcatagatca**tgtatttggcattatatctgaagtctcattataaaataca**ctaatagtgattatttattccatgttgctaatctcaggcca**caatcaactcatgagtgcttagccttcacctattgattg**gaggactatccacctgagatatttggaatacttccatgaggagatgtgttcttcttcccatcatttctttatttaatcatctattaacatccgtattggtttatggacatctatttcatactctgaagaagatccatgctacattattcattttcttgttcaaatctccacagctttatta

Uppercase: IGHV1-12

Lowercase: Flanking sequence[1000bp]

Red & Bold & Underline: Stem-loop [19]

Blue: Heptamer[27]

Green: Nonamer [1]

id-IGHV3-30-3[V_gene_segment]

ggcta**aattctcaaagaatt**gtgctgagtgaaagaaactaaggaatgaagagt**aaattttatatgatacatttgtagaaattt**tagaagatgccactattataaattaacatggagaagatttaaatgtttctgagaatatgctattgggagtaatggggatgtgagttaaatttcagaggaataagagaaagatttagggattaatttattcaaaccttgattgaagtgctgagtaaatggttgcaaacataggtctacattt**ttcaaatcattcaccataaatttgaa**tta**tttattaattacactcgaataaa**gcaataaagaaactgatgagataatatttgactgaattgcagcaataaatagatcgatattaacacaaggaatataactgacttccaaaaacatacacatgaaccgtggttcactctgcgtatttaggtaaataacagaaagttgtcataacagatggggaatcctgcagacttcactaggcatgggccatgctgcc**ctggagttgtctcaggggagctgcctcctccag**aggttagagcacaggcccaggtaataggactaaatttttagatgtgttatcttagacacactgcacaactgctgtgttctctatgtaaattatctcctgtaaaatataacattgaagcctgcattaaatatattgtgtaaatatgtaagaataaaagaaagttatgagagctaagtgttaatcaaggcacaagcatataagatataactatattttcctgaatgatggaattactaccagtctcccccaggacactt**catctgccctgagcccagcctctcctcagatgtcccacccagagcttgctatatagtgggggacat**gcaaatagggccctccctctactgatgaaaaccagcccagccctgaccctgcagctctgggagaggagcccagcactagaagtcggcggtgtttccattcggtgatcagcactgaacacagaggactcaccATGGAGTTTGGGCTGAGCTGGGTTTTCCTCGTTGCTCTTTTAAGAGGTGATTCATGGAGAAATAGAGAGACTGAGTGTGAGTGAACATGAGTG**AGAAAAACTGGATTTGTGTGGCATTTTCT**GATAACGGTGTCCTTCTGTTTGCAGGTGTCCAGTGTCAGGTGCAGCTGGTGGAGTCTGGGGGAGGCGTGGTCCAGCCTGGGAGGTCCCTGAGACTCTCCTGTGCAGCCTCTGGATTCACCTTCAGTAGCTATGGCATGCACTGGGTCCGCCAGG**CTCCAGGCAAGGGGCTGGAG**TGGGTGGCAGTTATATCATATGATGGAAGTAATAAATACTATGCAGACTCCGTGAAGGGCCGATTCACCATCTCCAGAGACAATTCCAAGAACACGCTGTATCTGCAAATGA**ACAGCCTGAGAGCTGAGGACACGGCTGT**GTATT**ACTGTGCGAAAGAcacagt**gaggggaagtcattgtgcgcccagacacaaacct**ccctgcaggaacgctggcgggaaatcagcggcaggg**ggcgctcaggagccactgatcagagtcagccctggaggcaggtgcagatggaggctgtttc**ctgtcaggatgtgggactttgtcttcttctgacagttccccagggaacctcttaaatttagaaaactgt**gcctaacaatgtcttctctatgcatatgaggaccttttctccctggcacaaaatgcagattgacgctgacacggatgaaaattcctcaaccatggtcacaaggatcagagt**cctgagtaacctcagg**gct**tcctggtgagtcttctccaatcagacccaggacagggacctccgtgagattccctg**actggaacagtctttatggat**cctggtcacagacaatagagaggctgaaccagg**gtcagcgtcatgtagaacctcacagatttcacgtctgatccttctcctgacacgaaagtatgcaaatcagtatc**agcaccgatctggtgct**tcttttgttcctaatccatttactttattttttcgtcgtttttc**tcctttttccatttgtttttcctgctttttgcaaaagga**agatgttttccctgtgagatgcaggggatgacaattttggga**gatggctggaacatccaatatcctcagggccggccatc**agtaagtgcaggctggaagtctcagaaagagctgaagctgcttaatcaccgtggagttttaccttctccagttctgctctgatggaatcagggccaagcaggttatcaatgataatctacctaacatagagtcaaccgattccagtttcaataacgtctgttaaaaattcacaccaccacctggattagtgttttgtcaaat**caatacacagtattg**tccagctaagtagacccaaagacggaccgttgcccatggagaaaaacattaacctgagttctaggttcttacagtgttaaagg

Uppercase: IGHV3-30-3

Lowercase: Flanking sequence[1000bp]

Red & Bold & Underline: Stem-loop [22]

Blue: Heptamer[30]

Green: Nonamer [6]

id-IGKV3-25[V_gene_segment]

agaatgataacagttttagacttcgagggtacttcaggaatgggaacacacaagga**acacagtagatatgagctactgtgtctaccgtattagaca**atcctcctgcattttaacttacatttggtgaacttcagatgccagtgggtgttgcaagttgatgacataaacctaaaatcaagtcccgaggtatttcttgtgaagagtcacagagttgaagagaccactgcctggcttcctgagttgagcctgtgccactgcaagtttcatgggaaaatggcgagaaaagctgtttaggtttgtaatcatcaaaggtttagtgaagttcccgtgcaaagagactcaatcccattcaactcagcatgaagagatcttctagaactcattctgcccctatagcagagatgatgtggtcaggtttgtggagatgaacatgcaatgacgtgttaaccaaaaggcccagcagaaagggatagtgatcaatgtccatggttcagacatatacaa**aggatgatttgcacagcactggaggaaaaaacatcct**gagtgttggagatttctgccttcctacccaagctcccgcagcctgtctcctgactcagaactcagcaaatgccacatgaaaaatctggctat**ggttgcacagaaattgccctgtgcaacc**ttcaaagctcagttatgttctctcttcttgctgggtttctctgtctgtg**ccaaatcaaaatttctatccgaacgcacatttgg**taaaaaaaaaaaaaaaaagggcagaaaaacaaaatgtcagatgaacaccaact**caactagaattagttg**tttttttactctagaattttagatcacctagtcctcatttcttccattactttctaatgtcactaataggtaactttgcattttatttgtttttattggcacattgtaat**tgtacatatttatggagtacactttgatgtcttgatatgtaca**tatgttgaaaaatgatcaaatcaggatatttaggATGTGCATCAGCTCATGTATTTACCATTTCTTTATGTTACTCTGGGTCCCAGGTGAGGGGAATATGCGGGGGGGCGTGGGGGGTTCTGTGTGTCAGTGAAAACTCTCTCAACTTTGTTGGGTCCTTCAACCTGGCATTTCCGCTCAGCAAGCACAATAATTAAAGCTCAATGTAAATCAATGGTCTTGACTCTATTGGAAAGACAATGGGT**ATAATCTAGATTACATGTGTGATTAT**TTGTTTTTCTGTTTAAACTGTTTTTTCAA**AAAATAAATTTTATTCTGTATATTTT**AGGTTTGCAATATGATGTTGTGGCATACATATAG**ATAGTAAAATGCTTACTAT**AGTGAAGAGAATGAACATTTATCATCTCACATAGTTACTTTTTCGTCTGTGACAAGAGCAGCTGGAATCTACTTATTTAACAAAAATCTCTAATACAGTACAGTTTTATTAACTATAGCCCTCATTTTCTACAGTAGATCTCTATACTTGCTATTCGTAAATATTTACTACCTTGCATCTTTGGACCTACATCTTCCCATTTCTTCCCATTCCACCCCCGGCCCTTAATAACGACTGTTTCATTCTCTATTTCTGCATAATTTATCTTTTTAGTTATTTAATTTATTTTTAGGTTTACATGTAAGTAAAATCAGGCAATATTTTTATTTCTGTATCTTGCTTTTTTCACTTGGAATACTGTCCTCTAGGTCCATCCACATTACGGCAAATAGAAGAATCTTCTTTTTC**TGCTGGGTGTGGTGGCTCATGCCTGTAATCCCAGCA**CTAGGCGGGTGCATCACCTGACATCAGGAGTTGAATACCAGCCTGGCCAACATGGCAAAACCCTGTCTCTACTAAAAATACAAAAAATACAAAAGAAAATTAGCTGGGCGTGTTGGCATGGGCCTGTAATCCCAACTACTCAGGAAGCTGAGGCAGGAGAATCGCTTGAACCCGGGAGGAGGAGGTTGCAGTGAGCTGAGATAGCATCATTGCACTCCAGCCTGGGCAACAAGAGTGAGACTTCTCTAAAATAAAAATAATAATAATAATAAAAAAGAATCTTCTTGTTTAAGGCTGAATTATATTCCATGGTGTGTGTGTGTGTGTGTGTGTGTGTGTGTGTGTGTGTACATTTATGTATACCACATTTTCTTTATTCATTTATTTGTCCACGGATGCTCAGGTTTTTCTCATTTCTTGTGAATAATGCTGCAATAAACACGGGAGTGTAGGTATCTTTAAAAGAAGGTGGTGATTTCATCTCTTCTGGGTATGTATCCAAAATAGGGTCACTGTTGGGTTATAAGGTGGTTAGGTTTTGAATTTCTTTAGGAACCTCCATACTGTTTTCCATAATGGGTGCACCAATCATCATTCCCACCAACAATGTACAAGTGTTTTATTTTCTTCACACCCTCATCAATATTTATCTCTT**GTCTTTTTTATAATAGCCATCCTAAAGAC**TGTAAGGCGTTTTATTTCTAATCTCAGATT**TCACTGTAGAAACAGTGA**TGACACAG**TCTCCAGCTTCCCTGTCTTTGTCTCTGGAGA**AAAAAGCCACC**CTGACTTGCAGGGCCAGTCAG**TGTTAGCAGCTACTA**AGCCTGGTACCAGAAGAAACCTGAGCGGGTTCCCAGGCT**CCTCATCTATG**GTACAGCCCTGATTTGTGATAGTGGGTCGGGGACAGGGCTTAC**TCTCACCATCGGCAGCCTGGAGCCTGGAGCCTGGAGATTTGCACTTCATCACTGTTATCAGCATAGTAGTTGGTGTCCcatactgattcgacatgcaacaaaaacctccaggagacctaaggtgtttatttgattatactacctgcttcctttttagtcatctgatgtggtgctgctcagttttagcatctctgctttgattggaaattctgaggttctcaaaagtaattccttataatatttatagtttcactcatggatttttttctcagacccaaatgtacagccaggttcaggcacaatttcatggtcaaggccattggatcagactcacatgagtggacgcctctaaaggtcctggccagtgcgataaagtagcagcgacaatgataaagaagaagaattagaaaggcagaattaaaggtataacaattcactgatgaaaggactgtgtgggggaga**aatttctaattgtctacacagaaatt**attagaattaatgagatacatagcaaattgttagattaaagatca**atataaaatatatttatttttatat**aaatgtaacaactgctaaaattaaaattatgaaacacatcatttatagtagcaccaaaccaagaggcattagaaaaaactaacaaaatatactgacagctccagaatataattcaaagaaatcaaagaaaatctaagcaaatggaaggccacagt**actttcaaaggttgaaagt**cccagtattgtaaagatgttcattcttcacagattgatatattaaagataatcgaaatcaaaatttcaaaccgtttttaaatgga**aatgtaaaagctgattctacatatcatattacatt**gttaaggagtttgaataaccacaataaacccgaagaagcccaagaatgaggacttacactacttggatactcgaaccatgctgctgatgtgagttcaaacaaattgcctaatgcaacagaagagagtattctgagaatgactaacacatatagtgacactttttagggacaaaggtgacactgcacaacagagtgaacacaataaaaagctgtccaacatcaataa

Uppercase: IGKV3-25

Lowercase: Flanking sequence[1000bp]

Red & Bold & Underline: Stem-loop [21]

Blue: Heptamer[41]

Green: Nonamer [6]
