## Supplementary Data 2 for "Adaptive immunity: from CRISPR to CRIHSP?"

id-IGHD3-3-2[D_gene_segment]

ctgctggcagctcctggggcctgatgtggagcaggcacagagccgtatccccccgaggacatatacccccaaggacggcacagttggtacattccggagacaagcaactcagccacactcccaggccagagcccgagagggacgcccatgcacagggaggcagagcccagctcctccacagccagcagcac**ctgtgcaggggccgccatctggcaggcacag**agcatgggctgggaggaggggcagggacaccaggcagggttggcaccaactgaaaattacagaagtctcatacatctacctcagccttgcctgacctgggcctcacctgacctggacctcacctggcctggacctcacctggcctagacctcacctctgggcttcacctgagctcggcctcacctgacttggaccttgcctgtcctgagctcacatgatctgggcctcacctgacctgggtttcacctgacctgggcttcacctgacctgggcctcatctgacctgggcctcactggcctggacctcacct**ggcctgggcttcacctggcctcaggcc**tcatctgcacctgctccaggtcttgctggaa**cctcagtagcactgagg**ctgcaggggctcatccagggttgcagaatgactctagaacctcccacatctcagctttctgggtggaggcacctggtggcccagggaatataaaaagcctgaatgatgcctgcgtga**tttgggggcaatttataaacccaaa**aggacatggccatgcagcgggtagggacaatacagacagatatcagcctgaaatggagcctcagggcac**aggtgggcacggacactgtccacct**aagccaggggcagacccgagtgtccccgcagtagacctgagagcgctgggcccacagcctcccctcggtgccctgctacctcctcaggtcagccctggacatcccgggtttccccaggcctggcggtaggtttggggtgagg**tctgtgtcactgtgGTATTACGATTTTTGGAGTGGTTATTATACCcacagtgtcacaga**gtccatcaaaaacccatccctgggaaccttctg**ccacagccctccctgtgg**ggcaccgctgcgtgccatgttaggattttgactgaggacacag**caccatgggtatggtg**gctaccgcagcagtgcag**cctgtgacccaaacacacagg**gcagcaggcacaacagacaagcccacaagtgaccaccctgagctcctgcctgccagccctggagaccatgaaacagatggccaggattatcccataggtcagc**cagacctcagtccaacaggtctg**catcgctgctgccctccaataccagtccggatggggacag**ggccggcccacattaccatttgctgccatccggcc**aacagtcccagaagcccctccctcaaggctgggccacatgtgtggaccctgag**agccccccatgtctgagtaggggcaccaggaaggtggggct**ggccctgtgcactgtcactgcccctgtggtccctggcctgcctggccctgacacctgggcctc**tcctgggtcatttccaagacagaagacattcccagga**cagctggagctgggagtccatcatcctgcctggccatcctgagtcctgcgcctttccaaacctcacccgggaagccaacagaggaatcacctcccacaggc**agagacaaagaccttccagaaatctctgtctct**ctccccagtgggcaccctcttccagggcagtcctcagtgatatcacagtgggaacccacatctggatcgggactgcccccagaacacaagatggcccacagggacagccccacagcccagcccttcccagacccctaaaaggcgtcccaccccctgcatctgccccagggctcaaactc**caggaggactgactcctgcacaccctcctg**ccagacatcacctcagcccctcctgga**agggacaggagcgcgcaagggtgagtcagaccctcctgccct**cgatggcaggcggagaagattcagaaaggt

Uppercase: IGHD3-3-2

Lowercase: Flanking sequence[1000bp]

Red & Bold & Underline: Stem-loop [16]

Blue: Heptamer[47]

Green: Nonamer [2]

id-IGHD4-4[D_gene_segment]

gtggctaccgcagcagtgcag**cctgtgacccaaacacacagg**gcagcaggcacaacagacaagcccacaagtgaccaccctgagctcctgcctgccagccctggagaccatgaaacagatggccaggattatcccataggtcagc**cagacctcagtccaacaggtctg**catcgctgctgccctccaataccagtccggatggggacag**ggccggcccacattaccatttgctgccatccggcc**aacagtcccagaagcccctccctcaaggctgggccacatgtgtggaccctgag**agccccccatgtctgagtaggggcaccaggaaggtggggct**ggccctgtgcactgtcactgcccctgtggtccctggcctgcctggccctgacacctgggcctc**tcctgggtcatttccaagacagaagacattcccagga**cagctggagctgggagtccatcatcctgcctggccatcctgagtcctgcgcctttccaaacctcacccgggaagccaacagaggaatcacctcccacaggc**agagacaaagaccttccagaaatctctgtctct**ctccccagtgggcaccctcttccagggcagtcctcagtgatatcacagtgggaacccacatctggatcgggactgcccccagaacacaagatggcccacagggacagccccacagcccagcccttcccagacccctaaaaggcgtcccaccccctgcatctgccccagggctcaaactc**caggaggactgactcctgcacaccctcctg**ccagacatcacctcagcccctcctgga**agggacaggagcgcgcaagggtgagtcagaccctcctgccct**cgatggcaggcggagaagattcagaaaggtctgagatccccaggacgcagcaccactgtcaatgggggc**cccagacgcctggaccagggcctgcgtgggaaaggcctctggg**cacactcaggggctttttgtgaagggtcctcct**actgtgTGACTACAGTAACTACcacagt**gatgaacccagcagcaaaaactgaccggactcccaaggtttatgcacacttctccgctcagagctctccaggatcagaagagccgggcccaagggtttctgcccagaccctcggcctctagggacatcttggccatgacagcccatgggctggtgccccacacatcgtctgccttcaaacaagggcttcagagggctctgaggtgacctcactgatgaccacaggtgccctggccccttccccgccagctgcaccagaccccgtcctgacagatgccccgattccaacagcca**attcctggggccaggaat**cgctgtagacaccagcctccttccaacacctcttgccaattgcctg**gattcccatcccggttggaatc**aagaggacagcatcccccaggctcccaacaggcaggactcccacaccctcctctgagaggccgctgtgttccgtagggccaggctgcagacagtccccctcacctgccactagacaaatgcctgctgtagatgtcccc**acctggaaaagaccactcatggagcccccagccccaggt**acagccatagagagagtctctgaggcccctaagaagtagccatgcccagttctgccgggaccctcggccaggctgacaggagtggacgctggagctgg**gcccacactgggccacataggagctcaccagtgagggc**aggagagcacatgccggggagcacc**cagcctcctgctgaccagaggcccgtcccagagcccaggaggctg**cagaggcctctccagggggacactgtgcatgtctggtccctgagca**gccccccatgtccccagtcctgggggc**ccctggcacagctgtctggaccctctctattccctgggaagctcctc**ctgacagccccgcctccagttccaggtgtggttattgtcagggggtgtcagactgtggtggatacagctatggttaccacagt**ggtgctgcccatagcagcaaccaggccaa

Uppercase: IGHD4-4

Lowercase: Flanking sequence[1000bp]

Red & Bold & Underline: Stem-loop [18]

Blue: Heptamer[41]

Green: Nonamer [6]

id-IGHD2-15[D_gene_segment]

tctcccccctgtgcccacaccctacctcctcctgcccacaactctaactcttcttctcctggagcccctgagccatggcattgaccctgccctcccaccacccacagcccatgccctcaccttcctcctggccactccgaccccgccccctctcaggccaagccctggtatttccaggacaaaggctcacccaagtcttt**cccaggcaggcctggg**ctcttgccctcacttcccggttacacgggagcct**cctgtgcacagaagcagggagctcagcccttccacagg**cagaaggcactgaaagaaatcggcctccagcaccttgacacacgtccccccgtgtctctcactgcccgcacctgc**agggaggctccgcactccct**ctaaagacaagggatccaggcagcagcatcacgggagaatgcagggctcccagaca**tcccagtcctctcacaggcctctcctggga**agagacct**gcagccaccaccaaacagccacagaggctgc**tggatagtaactgagtcaatgaccgacctggagggcaggggagcagtgagccggagcccataccatagggacagagaccagccgctgacatcccgagctcctcaatggtggccccataacacacctaggaaacataacacacccacagccccacctggaacagggcagagactgctgagcccccagcacc**agccccaagaaacaccaggcaacagtatcagagggggct**cccgagaaagagaggaggggagatctccttcaccatcaaatgcttcccttgaccaaaaacagggtccacgcaactcccccaggacaaaggag**gagccccctatacagcactgggctcagagtcctctctgag**acaccctgagtttcagacaacaacccgctggaatgcacagtctcagcaggagaacagaccaaagccagcaaaagggacctcggtgacaccagtagggacaggaggattttgtgggggctc**gtgtcactgtgAGGATATTGTAGTGGTGGTAGCTGCTACTCCcacagtgacac**agacccattcccaaagccctactgcaaacacacccactcctggggctgaggggctgggggagcgtctgggaagtagggtccaggggtgtctatcaatgtccaaaatgcaccagactgcccgccaaacaccaccccaccagccagcgagcagggtaaacagaaaatgagaggctctgggaag**cttgcacaggccccaaggaaagagctttggcgggtgtgcaag**aggggatgcaggcagagcctgagcagggccttttgctgtttctgctttcctgtgcagagagttccataaactggtgttcaagatcagtggctgggaatgagcccaggagggcagtctgtgggaagagcacagggaaggaggagcagccgctatcctacactgtcatctttcaaaagtttgccttgtgaccacactattgcatcatgggatgcttaagagctgatgtagacacagctaaagagagaatcagtgagatgaatttgcagcatagatctgaataaactctccagaatgtggagcagtacagaagcaaacacacagaaag**tgcctgatgcaaggacaaagttcagtgggcaccttcaggca**ttgctgctgggcacagacactctgaaaagccctggcaggatctccctgcgacaaagcagaacc**ctcaggcaatgccagccccagagccctccctgag**agcgtcatggggaaagatgtgcagaacagctgattatcatagactcaaactgagaacagagcaaacgtccatctgaagaacagtcaaataagcaatggtaggttcatgcaatgcaaacccagacagccaggggacaacagtagagggctacaggcggctttgcggttgagttcatgacaatgctgagtaattggagtaacagaggaaagcccaaaaaatacttttaatgtgatttcttctaaataaaatttacaccaggcaaaatgaactgtcttcttaagggataaactttcccctggaaaa

Uppercase: IGHD2-15

Lowercase: Flanking sequence[1000bp]

Red & Bold & Underline: Stem-loop [12]

Blue: Heptamer[32]

Green: Nonamer [4]

id-IGHD5-12[D_gene_segment]

ggccacactcgggctttttgtgaagggccctcctgctgtgtgactacagtaactaccatagtgatgaacccagtggcaaaaactggctggaaacccaggggctgtgtgcacgcctcagcttggagctctccagga**gcacaagagccgggcccaaggatttgtgcccagaccctcagcctctagggacacctgggc**catctcagcctgggctggtgccctgcacaccatcttcctccaaataggggcttcagagggctctgaggtgacctcactcatgaccacaggtgacctggcccttccctgccagctataccagaccctgtcttgacagatgccccgattccaacagccaattcctgggaccctgaatagctgtagacaccagcctcattccagtacctcctgccaattgcctggattcccat**cctggctggaatcaagaaggcagcatccgccagg**ctcccaacaggcaggactcccgcacaccctcctctgagaggccgctgtgttccg**cagggccaggccctg**gacagttcccctcacctgccactagagaaacacctgccattgtcgtcccc**acctggaaaagaccactcgtggagcccccagccccaggt**acagctgtagagagagtcctcgaggcccctaagaaggagccatgcccagttctgccgggaccctcggccaggccgacaggagtggacgctggagctgg**gcccacactgggccacataggagctcaccagtgagggc**aggagagcacatgccggggagcacc**cagcctcctgctgaccagaggcctgccccagagcccaggaggctg**cagaggcctctccagggagacactgtgcatgtctggtacctaagca**gccccccacgtccccagtcctgggggc**ccctggctca**gctgtctggaccctccctgttccctgggaagctcctcctgacagc**cccgcctccagttccaggtgtggttattgtcaggcgatgtcag**actgtgGTGGATATAGTGGCTACGATTACcacagt**ggtgccgcccata**gcagcaaccaggccaagtagacaggcccctgctgc**gc**agccccaggcatccacttcacctgcttctcctggggct**ctcaaggctgctgtctgtcctctggccctctgtggggagggttccctcagtgggaggtctgtgctccagggcagggatgattgagatagaaatcaaaggctggcagggaaaggcagcttcccgccctgagaggtgcaggcagcaccacggagccacggagtcacagagccacggagcccccattgtgggcatttgagagtgctgtgcccccggcaggcccagccct**gatggggaagcctgtcccatc**ccacagcccgggtcccacg**ggcagcgggcacagaagctgcc**aggttgtcctctatgatcctcatccctccagcagcatcccctccacagtggggaaactgaggcttggagcaccacccggccccctggaaatgaggctgtgagcccag**acagtgggcccagagcactgt**gagtaccccggcagtac**ctggctgcagggatcagccag**agatgccaaaccctgagtgaccagcctacaggaggatccggccccacccaggccactcgattaatgctcaaccccctgccctggagacctcttccagtaccaccagcagctcagcttctcagggcctcatccctgcaaggaaggtcaagggctgggcctgccagaaacacag**caccctccctagccctggctaagacagggtg**ggcagacggctgtggacgggacatattgctggggcatttctcactgtcacttctgggtggtagctctgacaaaaacgcagaccctgccaaaatccccactgcctcccgctaggggctgg**cctggaatcctgctgtcctaggaggctgctgacctccagg**atggctccgtccccagttccagggcgagagcaga**tcccaggcaggctgtaggctggga**ggccacccctgcccttgccggggttgaatgca

Uppercase: IGHD5-12

Lowercase: Flanking sequence[1000bp]

Red & Bold & Underline: Stem-loop [19]

Blue: Heptamer[46]

Green: Nonamer [4]

id-IGHD6-19-2[D_gene_segment]

ttacccaggacccagccctgcccctcctcccctctgctctcctctcatcaccccatgggaatccggtatccccaggaagccatcaggaagggctgaaggaggaagcggggccgtgcaccaccgggc**aggaggctccgtcttcgtgaacccagggaagtgccagcctcct**agagggtatggtccac**cctgcctggggctcccaccgtggcagg**ctgcggggaaggaccagggacggtgt**gggggagggctcagggccctgcgggtgctcctccatcttcggtgagcctccccc**ttcacccaccgtcccgcccacctcctctccaccctggctgcacgtcttccacaccatcctgagtcctacctacaccagagccagcaaagccagtgcagacaaaggctggggtgcaggggggctgccagggcagcttcggggagggaaggatggaggg**aggggaggtcagtgaagaggcccccttcccctgggtccaggatcctcctctgggaccc**ccggatccca**tcccctcctggctctgggaggagaagcaggatggga**gaatctgtgcgggaccctctcacagtggaatatccccacagc**ggctcaggccagacccaaaagcccctcagtgagcc**ctccactgcagtcctgggc**ctgggtagcagcccctcccacagaggacagacccag**caccccgaagaag**tcctgccagggggagctcagagccatgaaagagcagga**tatggggtccccgatacaggcacagacctcagctccatccaggcccaccgggacccaccatgggaggaacacctgtctccgggttgtgaggtagctgg**cctctgtctcggaccccactccagacaccagacagagg**ggcaggccccccaaaaccagggttgagggatgatccgtcaaggcagacaagaccaaggggcactgaccccagcaagggaaggctcccaa**acagacgaggaggtttctgaagctgtctgt**atcacagtgGGGTATAGCAGTGGCTGGTACcacagtgacactcgccaggccagaaaccccgtcccaagtcagcggaagcagagagagcagggaggacacgtttaggatctgaggccgcacctgacacccagggcagcagacgtctcccctccagggcaccctccaccgtcctgcgtttcttcaagaataggggcggcct**gagggggtccagggccaggcgataggtcccctc**taccccaaggaggagcca**ggcaggacccgagcaccgtccccattgaggctgacctgcccagacgggcctgggc**ccaccc**cacacaccggggcggaatgtgtg**ca**ggccccagtctctgtgggtgttccgctagctggggcc**cccagtgctcaccccacacctaaagcgagccccagcctccagagccccctaagcattccccgcccagcagcccagcccctgcccccacccaggaggccccagagctcagggcgcctggtcggattctgaacagccccgagtcacagtgggtataactggaacgaccaccgtgagaaaaactgtgtccaaaactgactcctggcagcagtcggaggccccgccagagaggggagcagccggcctgaacccatgtcctgccggttcccatgacccccagcacccagagccccacggtgtccccgttggataatgaggacaagggctgggggctccggtggtttgcggcagggacttgatcacatccttctgctgtggccccattgcctctggctggagttgaccc**ttctgacaagtgtcctcagaa**agacagggatcaccggcacctcccaatatcaaccccaggcagcacagacacaaaccccacat**ccagagccaactccaggagcagagacaccccaacactctgg**gggaccccaaccgtgataactccccactggaatccgccccagagtctaccaggaccaa**aggccctgccctgtctctgtccctcactcagggcct**cctgcagggcgagcgcttgggagcagactcggtctt

Uppercase: IGHD6-19-2

Lowercase: Flanking sequence[1000bp]

Red & Bold & Underline: Stem-loop [19]

Blue: Heptamer[29]

Green: Nonamer [8]

id-IGHD1-1-2[D_gene_segment]

atgcaggaatgactgggccacacccctcccgtgcacgccccctcctgccctgcaccccacagcccagccccccgtgctggatgccccccc**acagcagaggtgctgt**tctgtgatcccctgggaaagacgccctcaacctccaccctgtcccacggcccaaggaagacaagacacaggccctctcctcacagtctccccacctggctcctgctgggaccctcaaggtgtgaacagggaggatggttg**tctgggtggcccctaggagcccaga**tcttcactccacagaccccaacccaagcacccccttctgcagggcccagctcatccccctcctcctccctctgctctcctctcgtcgcctctacgggaaatccgggactcagcagtaaccctc**aggaagcagggcccaggcgccgtttaataggaggcttcct**cacaatgaaacttttagaaagccttgactacaatgatgaccttggtgtg**gctgtgaacactgtcagctcccacagc**tgctgcagcaaaaaatgtccatagacagggtgggggcccggggtcgtc**tgctgtcctgctcagcccacagca**cgcatggaggatctgaggtgccacacctgacgcccaggccagaacatgcc**tccctccagggtgacctgccatgtcctgcattgctggaggga**caggggcagcctatgagga**tctggggccaggagatgaatcctattaacccaga**ggaaaactaacaggacccaagcaccctccccgttgaagctgacct**gcccagaggggcctgggc**ccaccccacac**accggggcggaatgtgtacaggccccggt**ctctgtgggtgttccgctaa**ctggggctcccagtgctcaccccacaactaaagcgagccccag**cctccagagcccccgaaggagatgccgcccacaagcccagcccccatccaggaggccccagagctcagggcgc**cggggcagattctgaacagccccg**ag**tcacggtgGGTACAACTGGAACGACcaccgtga**gaaaaactgtgtccaaaa**ctctctcctggcccctgctggaggccgcgccagagag**gg**gagcagccgccccgaacctaggtcctgctc**agctcacacgacccccagcagccagagca**cagtggagtccccactg**aaccccactgaatggtgaggacggggaccagggctccag**ggggtcatggaaggggctggacccc**atcctactgctatggtcccagtgctcctggccagaactgaccctaccaccgacaagagtccctcagggaaacgggggtcactggcacctcccagcatcaaccccaggcagcacaggcataaaccccacatccagagccgactccaggagcagagaca**ccccagtaccctgggg**gacaccgaccctgatgactccccactggaatccaccccagagtccaccaggaccaaagac**cccgccccggtctctgtccctcactcaggacctgctgcggggcggg**ccatgagaccagactcgggcttagggaacacca**ctgtggccccaacctcgaccaggccacag**gcccttccttcctgccctgcggca**gcacagactttggggtctgtgc**agagaggaatcacagaggccccaggctgaggtgg**tgggggtggaagaccccca**ggaggtggcccacttcccttcctcccagctggaacccaccatgaccttcttaagataggggtgtcatccgaggcaggtcctccatggagctcccttcaggctcctccctggtcctcactaggcctcagtcccgg**ctgtgggaatgcagccaccacag**gcacaccaggcagcccagacccagc**cagcctgcagtgcccaagcccacattctggagcagagcaggctg**tgtctgggagagtctgggctccccaccgccccccgcacaccccacccacccctgtccaggccctatgcaggagggtcagagccccccatggggtatggacttagggtctcactcacgcggctcccctcc

Uppercase: IGHD1-1-2

Lowercase: Flanking sequence[1000bp]

Red & Bold & Underline: Stem-loop [23]

Blue: Heptamer[32]

Green: Nonamer [1]

id-IGHD3-22[D_gene_segment]

gctgtatatccccaaggaaggtacagtcagtgcattccagagagaagcaactcagccaca**ctccctggccagaacccaagatgcacacccatgcacagggag**gcagagcccagcacctccgcagccaccaccacctgcgcacgggccaccaccttgcaggcacagag**tgggtgctgagaggaggggcagggacaccaggcagggtgagcaccca**gagaaaactgcagaagcctcacacatccacctcagcctcccctgacctggacctcacctggcctgggcctcacctgacctggacctcacctggcctgggcttcacctggcctgggcttcacctgacctggacctcacctggcctcgggcctcacctggcctgggcttcacctggcctgggcttcacctgacctggacctcacctggcctgggcctcacctgacctggacctcacctggcctgggcttcacctggcctgggcttcacctggcctgggcttcacctgacctggacctcacctggcctgggcttcacctgacctggacctcacctggcctcgggcctcacctgcacct**gctccaggtcttgctggagc**ctgagtagcactgaggctgtagggactcatccagggttggggaatgactctgcaactctcccacatctgacctttctgggtggaggcacctggtggcccagggaatataaaaagccccagaatgatgcctgtgtgatttgggggcaatttatgaacccgaaaggacatggccatggggtgggtagggacagtagggacagatgtcag**cctgaggtgaagcctcagg**acac**aggtgggcatggacagtgtccacct**aagcg**agggacagacccgagtgtccct**gcagtagacctgagagcgct**gggcccacagcctcccctcggggccc**tgctgcctcctcaggtcagccctggacatcccgggtttccccaggcctggcggtaggtttgaagtgagg**tctgtgtcactgtgGTATTACTATGATAGTAGTGGTTATTACTACcacagtgtcacaga**gtccatcaaaaactcatgcctgggagcctcccaccacagccctccctgcgggggaccgctgcat**gccgtgttaggattttgatcgaggacacggc**gc**catgggtatggtggctaccacagcagtgcagcccatg**acccaaacacacggggcagcagaaacaatggacaggcccacaagtgaccatgatgggctccagcccaccagccccagagaccatgaaacagatggccaaggtcaccctacaggtcatccagatctggctccaaggggtctgcatcgctgctgccctcccaacgccaaac**cagatggagacagggccggccccatagcaccatctgctgccgtccacccagcag**tcccggaagcccctccctgaacgctgggccacgtgtgtgaaccctgcg**agccccccatgtcagagtaggggcagcaggagggcggggct**ggccctgtgcactgtcactgcccctgtggtccctggcctgcctggccctgacacctgagcctctcctgggtcatttccaagacattcccag**ggacagccggagctgggagtcgctcatcctgcctggctgtcctgagtcctgctcatttccagacctcaccagggaagccaacagaggactca**cctcacacagtcagagacaatgaaccttccagaaatccctgtttctctccccagtg**agagaaaccctcttccagggtttctct**tctctcccaccctcttccaggacagtcctcagcagcatcacagcgggaacgcacatctggatcaggacggcccccagaacacgcgatggcccatggggacagcccagcccttcccagacccctaaaaggtatccccaccttgcacctgccccagggctcaaactc**caggaggcctgactcctgcacaccctcctg**ccagatatcacctcagc**cccctcctggagggg**acaggagcccgggagggtgagtcagacccacctgccctcaatggcag

Uppercase: IGHD3-22

Lowercase: Flanking sequence[1000bp]

Red & Bold & Underline: Stem-loop [18]

Blue: Heptamer[31]

Green: Nonamer [1]

id-IGHD5-24-2[D_gene_segment]

tcaggggctttttgtgaagggccctcctgctgtgtgactacggtggtaactcccacagtgatgaaaccagcagcaaaaactgaccggactcgcagggtttatgcacacttctcggctcggagctctccaggagcacaagagcc**aggcccgagggtttgtgcccagaccctcggcct**ctagggacacccgggccatcttagccgatgggctgatgccctgcacaccgtgtgctgccaaacaggggcttcagagggctctgaggtgacttcactcatgaccacaggtgccctggtcccttcactgccagctgcaccagaccctgttccgagagat**gccccagttccaaaagccaattcctggggc**cgggaattactgtagacaccagcctcattccagtacctcctgccaattgcctggattcccat**cctggctggaatcaagagggcagcatccgccagg**ctcccaacaggcaggactcccacacaccctcttctgagaggccgctgtgttccgcagggccaggccgcagacagttcccctcacctgcccatgtagaaacacctgccattgtcgtcccc**acctggcaaagaccacttgtggagcccccagccccaggt**acagctgtagagagagtcctcgaggcccctaagaaggagccatgcccagttctgctgggaccctcggccaggccgacaggagtggacgctggagctgg**gcccacactgggccacataggagctcaccagtgagggc**aggagagcacatgccggggagcacc**cagcctcctgctgaccagagacccgtcccagagcccaggaggctg**cagaggcctctccagggggacacagggcatgtctggtccctgagca**gcccccaggctctctagcactgggggc**ccctggcaca**gctgtctggaccctccctgttccctgggaagctcctcctgacagc**cccgcctccagttccaggtgtggttattgtcagggggtgccaggcc**gtgGTAGAGATGGCTACAATTACcac**agtggtgccgcccatagcagcaaccaggccaagtagacagacccctgccacgc**agccccaggcctccagctcacctgcttctcctggggct**ctcaaggctgctgtctgccctctggccctctgtggggagggttcc**ctcagtgggaggtctgtgctccagggcagggatgactgag**atagaaatcaaaggctggcagggaaaggcagcttcccgccctgagaggtgcaggcagcaccacagagccatggagtcacagagccacggagcccccagtgtgggcgtgtgagggt**gctgggctcccggcaggcccagc**cctgatggggaagc**ctgccccgtcccacagcccaggtccccaggggcag**cag**gcacagaagctgccaagctgtgc**tctacgatcctcatccctccagcagcatccactccacagtggggaaactga**gccttggagaaccacccagccccctggaaacaaggc**ggggagcccag**acagtgggcccagagcactgt**gtgtatcctggcactaggtgcagggaccacccggagatccccatcactgagtggccagcctgcagaaggacccaaccccaaccaggccgcttgattaagctccatccccctgt**cctgggaacctcttcccagcgccaccaacagctcggcttcccaggccctcatccctccaaggaaggccaaaggctgggcctgccaggggcacagtaccctcccttgccctggc**taagacagggtgggcagacggctgcagataggacatattgctggggcatcttgctctgtgactactgggtactggctctcaacgcagaccctaccaaaatccccactgcctcccctgctaggggctggcctggtct**cctcctgctgtcctaggagg**ctgctgacctccaggatggcttctgtccccagttctagggccagagcaga**tcccaggcaggctgtaggctggga**ggccacccctgtccttgccgaggttcagtgcag

Uppercase: IGHD5-24-2

Lowercase: Flanking sequence[1000bp]

Red & Bold & Underline: Stem-loop [21]

Blue: Heptamer[42]

Green: Nonamer [5]

id-IGHD2-15-2[D_gene_segment]

tctcccccctgtgcccacaccctacctcctcctgcccacaactctaactcttcttctcctggagcccctgagccatggcattgaccctgccctcccaccacccacagcccatgccctcaccttcctcctggccactccgaccccgccccctctcaggccaagccctggtatttccaggacaaaggctcacccaagtcttt**cccaggcaggcctggg**ctcttgccctcacttcccggttacacgggagcct**cctgtgcacagaagcagggagctcagcccttccacagg**cagaaggcactgaaagaaatcggcctccagcaccttgacacacgtccccccgtgtctctcactgcccgcacctgc**agggaggctccgcactccct**ctaaagacaagggatccaggcagcagcatcacgggagaatgcagggctcccagaca**tcccagtcctctcacaggcctctcctggga**agagacct**gcagccaccaccaaacagccacagaggctgc**tggatagtaactgagtcaatgaccgacctggagggcaggggagcagtgagccggagcccataccatagggacagagaccagccgctgacatcccgagctcctcaatggtggccccataacacacctaggaaacataacacacccacagccccacctggaacagggcagagactgctgagcccccagcacc**agccccaagaaacaccaggcaacagtatcagagggggct**cccgagaaagagaggaggggagatctccttcaccatcaaatgcttcccttgaccaaaaacagggtccacgcaactcccccaggacaaaggag**gagccccctatacagcactgggctcagagtcctctctgag**acaccctgagtttcagacaacaacccgctggaatgcacagtctcagcaggagaacagaccaaagccagcaaaagggacctcggtgacaccagtagggacaggaggattttgtgggggctc**gtgtcactgtgAGGATATTGTAGTGGTGGTAGCTGCTACTCCcacagtgacac**agacccattcccaaagccctactgcaaacacacccactcctggggctgaggggctgggggagcgtctgggaagtagggtccaggggtgtctatcaatgtccaaaatgcaccagactgcccgccaaacaccaccccaccagccagcgagcagggtaaacagaaaatgagaggctctgggaag**cttgcacaggccccaaggaaagagctttggcgggtgtgcaag**aggggatgcaggcagagcctgagcagggccttttgctgtttctgctttcctgtgcagagagttccataaactggtgttcaagatcagtggctgggaatgagcccaggagggcagtctgtgggaagagcacagggaaggaggagcagccgctatcctacactgtcatctttcaaaagtttgccttgtgaccacactattgcatcatgggatgcttaagagctgatgtagacacagctaaagagagaatcagtgagatgaatttgcagcatagatctgaataaactctccagaatgtggagcagtacagaagcaaacacacagaaag**tgcctgatgcaaggacaaagttcagtgggcaccttcaggca**ttgctgctgggcacagacactctgaaaagccctggcaggatctccctgcgacaaagcagaacc**ctcaggcaatgccagccccagagccctccctgag**agcgtcatggggaaagatgtgcagaacagctgattatcatagactcaaactgagaacagagcaaacgtccatctgaagaacagtcaaataagcaatggtaggttcatgcaatgcaaacccagacagccaggggacaacagtagagggctacaggcggctttgcggttgagttcatgacaatgctgagtaattggagtaacagaggaaagcccaaaaaatacttttaatgtgatttcttctaaataaaatttacaccaggcaaaatgaactgtcttcttaagggataaactttcccctggaaaa

Uppercase: IGHD2-15-2

Lowercase: Flanking sequence[1000bp]

Red & Bold & Underline: Stem-loop [12]

Blue: Heptamer[32]

Green: Nonamer [4]

id-IGHD5-24[D_gene_segment]

tcaggggctttttgtgaagggccctcctgctgtgtgactacggtggtaactcccacagtgatgaaaccagcagcaaaaactgaccggactcgcagggtttatgcacacttctcggctcggagctctccaggagcacaagagcc**aggcccgagggtttgtgcccagaccctcggcct**ctagggacacccgggccatcttagccgatgggctgatgccctgcacaccgtgtgctgccaaacaggggcttcagagggctctgaggtgacttcactcatgaccacaggtgccctggtcccttcactgccagctgcaccagaccctgttccgagagat**gccccagttccaaaagccaattcctggggc**cgggaattactgtagacaccagcctcattccagtacctcctgccaattgcctggattcccat**cctggctggaatcaagagggcagcatccgccagg**ctcccaacaggcaggactcccacacaccctcttctgagaggccgctgtgttccgcagggccaggccgcagacagttcccctcacctgcccatgtagaaacacctgccattgtcgtcccc**acctggcaaagaccacttgtggagcccccagccccaggt**acagctgtagagagagtcctcgaggcccctaagaaggagccatgcccagttctgctgggaccctcggccaggccgacaggagtggacgctggagctgg**gcccacactgggccacataggagctcaccagtgagggc**aggagagcacatgccggggagcacc**cagcctcctgctgaccagagacccgtcccagagcccaggaggctg**cagaggcctctccagggggacacagggcatgtctggtccctgagca**gcccccaggctctctagcactgggggc**ccctggcaca**gctgtctggaccctccctgttccctgggaagctcctcctgacagc**cccgcctccagttccaggtgtggttattgtcagggggtgccaggcc**gtgGTAGAGATGGCTACAATTACcac**agtggtgccgcccatagcagcaaccaggccaagtagacagacccctgccacgc**agccccaggcctccagctcacctgcttctcctggggct**ctcaaggctgctgtctgccctctggccctctgtggggagggttcc**ctcagtgggaggtctgtgctccagggcagggatgactgag**atagaaatcaaaggctggcagggaaaggcagcttcccgccctgagaggtgcaggcagcaccacagagccatggagtcacagagccacggagcccccagtgtgggcgtgtgagggt**gctgggctcccggcaggcccagc**cctgatggggaagc**ctgccccgtcccacagcccaggtccccaggggcag**cag**gcacagaagctgccaagctgtgc**tctacgatcctcatccctccagcagcatccactccacagtggggaaactga**gccttggagaaccacccagccccctggaaacaaggc**ggggagcccag**acagtgggcccagagcactgt**gtgtatcctggcactaggtgcagggaccacccggagatccccatcactgagtggccagcctgcagaaggacccaaccccaaccaggccgcttgattaagctccatccccctgt**cctgggaacctcttcccagcgccaccaacagctcggcttcccaggccctcatccctccaaggaaggccaaaggctgggcctgccaggggcacagtaccctcccttgccctggc**taagacagggtgggcagacggctgcagataggacatattgctggggcatcttgctctgtgactactgggtactggctctcaacgcagaccctaccaaaatccccactgcctcccctgctaggggctggcctggtct**cctcctgctgtcctaggagg**ctgctgacctccaggatggcttctgtccccagttctagggccagagcaga**tcccaggcaggctgtaggctggga**ggccacccctgtccttgccgaggttcagtgcag

Uppercase: IGHD5-24

Lowercase: Flanking sequence[1000bp]

Red & Bold & Underline: Stem-loop [21]

Blue: Heptamer[42]

Green: Nonamer [5]

id-IGHD3-16-2[D_gene_segment]

atgatacagac**atacatttagtacatgagacatcgatgatgtat**ccccaaagaaatgactttaaagagaaaaggcctgatgtgtggtggcactcacctccctgggatccccggacaggttgcaggcacactgtgtggcagggcaggctggtacatgctggcagctcctggggcctgatgtggagcaagcgcagggctgtatacccccaaggatggcacagtcagtgaattccagagagaagcagctcagccacactgcccaggcagagcccgagagggacgcccacgcacagggaggcagagcccagctcctccacagccacca**ccacctgtgcacgggccaccaccttgcaggcacagagtgggtgctgagaggaggggcagggacaccaggcagggtgagcaccca**gagaaaactgcagaagcctcacacatccacctcagcctcccctgacctggacctcacctggtctggacctcacctggcctgggcctcacctgacctggacctcacctggcctgggcttcacct**gacctggacctcacctggcctccggcctcacctgcacctgctccaggtc**ttgctggaacctgagtagcactgaggctgcagaagctcatccagggttggggaatgactctggaactctcccacatctgacctttctgggtggaggcatctggtggccctgggaatataaaaagccccagaatggtgcctgcgtgatttgggggcaatttatgaacccgaaaggacatggccatggggtgggtagggacatagggacagatgccag**cctgaggtggagcctcagg**acacagttggacgcggacactatccacataagcgagggacagacccgagtgttcctgcagtagacctgagagcgctgggcccacagcctcccctcggtgccctgctgcctcctcaggtcagccctggacatcccgggtttccccaggccagatggtaggtttgaagtgaggtctgtgtcactgtgGTATTATGATTACGTTTGGGGGAGTTATCGTTATACCcacagcatcacacggtccatcagaaacccatgccacagccctccccgcaggggaccgccgcgtgccatgttacgattttgatcgaggacacagcgc**catgggtatggtggctaccacagcagtgcagcccatg**acccaaaca**cacagggcagcaggcacaatggacaggcctgtg**agtgaccatgctgggctccagcccgccagccccggagaccatgaaacagatggccaaggtc**accccacagttcagccagacatggctccgtggggt**ctgcatcgctgctgccctctaacaccagccc**agatggggacaaggccaaccccacattaccatct**cctgctgtccacccagtggtcccagaagcccctccctcatggctgagccacatgtgtgaaccctgagagcaccccatgtcagagtaggggcagcagaagggcggggctggccctgtgcactgtccctgcacccatggtccctcgcctgcctggccctgacacctgagcc**tcttctgagtcatttctaagatagaaga**cattcccgg**ggacagccggagctgggcgtcgctcatcccgcccggccgtcctgagtcctgcttgtttccagacctcaccagggaagccaacagaggactca**cctcacacagtca**gagacaaagaaccttccagaaatccctgtctc**actccccagtgggcaccttcttccaggacattcctcggtcgcatcacagcaggcacccacatctggatcaggacggcccccagaacacaagatggcccatggggacagccccacaacccaggccttcccagacccctaaaaggcgtcccaccccctgcacctgccccagggctaaaaatccaggaggcttgactcccgca**taccctccagccagacatcacctcagccccctcctggagggga**caggagcccgggagggtgagtcagaccca**cctgccctcgatggcagg**cggggaagattcagaaaggcctgagatcccc

Uppercase: IGHD3-16-2

Lowercase: Flanking sequence[1000bp]

Red & Bold & Underline: Stem-loop [15]

Blue: Heptamer[42]

Green: Nonamer [3]

id-IGHD6-6[D_gene_segment]

acccagccctgcccctcc**tcccatctgctctcctctcatcaccccatggga**atccagaatccccaggaagccatcaggaagggctgagggaggaagtggggccactgcaccaccaggc**aggaggctccgtctttgtgaacccagggaggtgccagcctcct**agagggtatggtccac**cctgcctatggctcccacagtggcagg**ctgcagggaaggaccagggacggtgtggggga**gggctcagggccccgcgggtgctccatcttggatgagccc**atctctctcacccacggactcacccacctcctctccaccctggccacacgtcgtccacaccatcctaagtcccacctacaccagagccggcacagccagtgcagacagaggctggggtgcaggggggccgccagggcagctttggggagggaaggatggagga**aggggagttcagtgaagaggcccccctcccctgggtccaggatcctcctctgggaccc**ccgga**tcccatcccctccaggctctgggaggagaagcaggatggga**gaatctgtgcgggaccctctcacagtggaatacctccacagc**ggctcaggcaagacccaaaagcccctcagtgagcc**ctccactgcagtcctgggc**ctgggtagcagcccctcccacagaggatgaacccag**caccccgaggatg**tcctgccagggggagctcagagccatgaaggagcagga**tatgggacccccgatacaggcacagacctcagctccattcaggactgccacgtcctgccctgggaggaacccctttctctagtccctgcaggc**caggaggcagctgactcctg**acttggacgcctattccagacaccagacagaggggcaggccccccagaaccagggatgaggacgccccgtcaaggccagaaaagaccaagttgtgctgagcccagcaagggaaggtccccaaacaaaccaggaagtttctgaaggtgtct**gtgtcacagtgGAGTATAGCAGCTCGTCCcacagtgacac**tcgccaggccagaaaccccatcccaagtcagcggaatgcagagagagcagggaggacatgtttaggatctgaggccgcacctgacacccaggccagcagacgtctcctgtc**catggcaccctgccatg**tcctgcatttctggaagaacaagggcaggctgaagggggtccaggaccaggagatgggtcccctctacccagagaaggagcca**ggcaggacacaagccccctccccattgaggctgacctgcccagagggtcctgggc**ccaccc**cacacaccggggcggaatgtgtg**caggcctcggtctctgtgggtgttccgcta**gctggggctcacagtgctcaccccacacctaaaacgagccacagc**ctcagagcccctgaaggagaccccgcccacaagcccagcccccacccaggaggccccagagcacagggcgccccgtcggattctgaacagccccgag**tcacagtgggtataactggaactaccactgtga**gaaaagcttcgtccaaaacggtctcctggccacagtcggaggccccgccagagaggggagcagccaccccaaacccatgttctgcc**ggctcccatgaccccgtgcacctggagcc**ccacagtgtccccactggatgggaggacaagggccgggggctccggcgggtcggggcaggggcttgatggcttccttctgccgtgg**ctccagtgcccctggctggag**ttgacccttctgacaagtgtcctcagagagtcagggatcagtggcacctcccaacatcaaccccacgcagcccaggcacaaaccccacat**ccagggccaactccaggaacagagacaccccaataccctgg**gggaccccaaccctgatgactcccgtcccatctctgtccctcacttggggcctgctgcggggcgagcacttgggagcaaac**tcaggcttaggggacaccactgtgggcctga**cctcgagcaggccacagacccttc

Uppercase: IGHD6-6

Lowercase: Flanking sequence[1000bp]

Red & Bold & Underline: Stem-loop [22]

Blue: Heptamer[38]

Green: Nonamer [7]

id-IGHD4-11[D_gene_segment]

acagccctccccatggggccctgctgcctcctcaggtcagccccggacatcccgggtttccccaggctgggcggtaggtttggggtgagg**tctgtgtcactgtggtattactatggttcggggagttattataaccacagtgtcacaga**gtccatcaaaaacccatccctgggagcctcccgccacagccctccctgcaggggaccggtacgtgccatgttaggattttgatcgaggagacag**caccatgggtatggtg**gctaccacagcagtgcagcctgtgacccaaacccgcagggcagcaggcacgatggacaggcccgtgactgaccacgctggg**ctccagcctgccagccctggag**atcatgaaacagatggccaaggtcaccctacaggtcatccagatctggctccgaggggtctgcatcgctgctgccctcccaacgccagtccaaatgggacagggacggcctcacagcaccatctgctgccatcaggccagcgatcccagaagcccctccctcaaggctgggccacatgtgtggacactgagagccctcatgtctgagtaggggcaccaggaggg**aggggctggccctgtgcactgtccctgcccct**gtggtccctggcctgcctggccctgacacctgagcctctcctgggtcatttccaagacagaagacattcctgg**ggacagccggagctgggcgtcgctcatcctgcccggccgtcc**tgagtcctgctcatttccagacctcaccggggaagccaacagaggactcgcctcccacattcagagacaaagaaccttccagaaatccctgcctctctccccagtggacaccctcttccaggacagtcctcagtggcatc**acagcggcctgagatccccaggacgcagcaccgctgt**caataggggccccaaatgcctggaccagggcctgcgtgggaaaggtctctggccacactcgggctttttgtgaagggccctcctgctgtgTGACTACAGTAACTACcatagtgatgaacccagtggcaaaaactggctggaaacccaggggctgtgtgcacgcctcagcttggagctctccagga**gcacaagagccgggcccaaggatttgtgcccagaccctcagcctctagggacacctgggc**catctcagcctgggctggtgccctgcacaccatcttcctccaaataggggcttcagagggctctgaggtgacctcactcatgaccacaggtgacctggcccttccctgccagctataccagaccctgtcttgacagatgccccgattccaacagccaattcctgggaccctgaatagctgtagacaccagcctcattccagtacctcctgccaattgcctggattcccat**cctggctggaatcaagaaggcagcatccgccagg**ctcccaacaggcaggactcccgcacaccctcctctgagaggccgctgtgttccg**cagggccaggccctg**gacagttcccctcacctgccactagagaaacacctgccattgtcgtcccc**acctggaaaagaccactcgtggagcccccagccccaggt**acagctgtagagagagtcctcgaggcccctaagaaggagccatgcccagttctgccgggaccctcggccaggccgacaggagtggacgctggagctgg**gcccacactgggccacataggagctcaccagtgagggc**aggagagcacatgccggggagcacc**cagcctcctgctgaccagaggcctgccccagagcccaggaggctg**cagaggcctctccagggagacactgtgcatgtctggtacctaagca**gccccccacgtccccagtcctgggggc**ccctggctca**gctgtctggaccctccctgttccctgggaagctcctcctgacagc**cccgcctccagttccaggtgtggttattgtcaggcgatgtcag**actgtggtggatatagtggctacgattaccacagt**ggtgccgcccatagcagcaaccaggcc

Uppercase: IGHD4-11

Lowercase: Flanking sequence[1000bp]

Red & Bold & Underline: Stem-loop [16]

Blue: Heptamer[46]

Green: Nonamer [8]

id-IGHD2-2-2[D_gene_segment]

agtttctaccctctgtgcctaccccctgcctcctcctgcccacaactcgagctcttcctctcctggggcccctgagccatggcactgaccgtgcactcccacccccacactgcccatgccctcaccttcctcctggacactctgaccctgctcccctcttggacccagccctggtatttccaggacaaaggctcacccaagtcttccccatgcaggcccttgccctcactgcccggttacacggcagcct**cctgtgcacagaagcagggagctcagcccttccacagg**cagaaggcactgaaagaaatcggcctccagcaccctgatgcacgtccgcctgtgtctctcactgcccgcacctg**cagggaggctcggcactccctg**taaagacgagggatccaggcagcaacatca**tgggagaatgcagggctccca**gacagcccagccctctc**gcaggcctctcctgggaagagacctgc**agccaccactgaacagccacggagcccgctggatagtaactgagtcagtgaccgacctggagggcaggggagcagtgaaccggagcccagaccatagggacagagaccagccgctgacatcccgagcccctcactggcggccccagaacaccgcgtggaaacagaacagacccacattcccacctggaa**cagggcagacactgctgagcccccagcaccagccctg**agaaacaccaggcaacggcatcagagggggctcctgagaaaga**aaggaggggaggtctcctt**caccagcaagtacttcccttgaccaaaaacagggtccacgcaactcccccaggacaaaggag**gagccccctgtacagcactgggctc**agagtcctctccaacacaccctgagtttcagacaaaaaccccctggaaatcatagtatcagcaggagaactagccagagacagcaagaggggactcagtgactcccgcggggacaggaggattttgtgggggctc**gtgtcactgtgAGGATATTGTAGTAGTACCAGCTGCTATGCCcacagtgacac**agccccattcccaaagccctgctgtaaacgcttccacttctggagctgaggggctggggggagcgtctgggaagtagggcctaggggtggccatcaatgcccaaaacgcaccagactcccccccagacatcaccccactggccagtgagcagagtaaacagaaaatgagaagcagctgggaag**cttgcacaggccccaaggaaagagctttggcgggtgtgcaag**aggggatgcgggcagagcctgagcagggccttttgctgtttctgctttcctgtgcagatagttccataaactggtgttcaagatcgatggctgggagtgagcccaggag**gacagtgtgggaagggcacagggaaggagaagcagccgctatcctacactgtc**atctttcaagagtttgccctgtgcccacaatgctgcatcatgggatgcttaacagctgatgtagacacagctaaagagagaatcagtgaaatggatttgcagcacagatctgaataaattctccagaatgtggagccacacagaagcaagcacaaggaaagt**gcctgatgcaagggcaaagtacagtgtgtaccttcaggc**tgggcacagacactctgaaaagccttggcaggaactc**cctgcaacaaagcagagccctgcagg**caatgccagctccagagccctccctgagagcctcatgggcaaagatgtgcagaacata**tgtttgtcatagccccaaactgagaatgaagcaaacagccatctgaaggaaaacaggcaaataaacgatggc**aggttcatgaaatgcaaacccagacagccagaaggacaacagtgagggttacaggtgactctgtggttgagttcatgacaatgctgagtaattggagtaacaaag**gaaagtccaaaaaatactttc**aatgtgatttcttctaaataaaatttacagccggcaaaatgaactatcttcttaagggataaactttccactaggaaaacta

Uppercase: IGHD2-2-2

Lowercase: Flanking sequence[1000bp]

Red & Bold & Underline: Stem-loop [15]

Blue: Heptamer[47]

Green: Nonamer [5]

id-IGHD3-9-2[D_gene_segment]

gaggacggcacagtcagtgaattccagagagaagcaactcagccacactccccaggcagagcccgagagggacgcccacgcacagggaggcagagcccagcacctccgcagccagca**ccacctgtgcacgggccaccaccttgcaggcacagagtgggtgctgagaggaggggcagggacaccaggcagggtgagcaccca**gagaaaactgcagacgcctcacacatccacctcagcctcccctgacctggacctcactggcctgggcctcacttaacctgggcttcacctgaccttggcctcacctgacttggacctcgcctgtcccaagctttacctgacctgggcctcaactcacctgaacgtctcctgacctgggtttaacctgtcctggaactcacctggccttggcttcccctgacctggacctcatctggcctgggcttcacctggcctgggcctcacctgacctggacctcatctggcctggacctcacctggcctggacttcacctggcctgggcttcacctgacctggacctcacctggcctcgggcctcacctgcacct**gctccaggtcttgctggagc**ctgagtagcactgagggtgcagaagctcatccagggttggggaatgactctagaagtctcccacatctgacctttctgggtggaggcagctggtggccctgggaatataaaaatctccagaatgatgactctgtgatttgtgggcaacttatgaacccgaaaggacatggccatggggtgggtagggacatagggacagatgccag**cctgaggtggagcctcagg**acacaggtgggcac**ggacactatccacataagcgagggatagacccgagtgtcc**ccacagcagacctgagagcgctgggcccacagcctcccctcagagccctgctgcctcctccggtcagccctggacatcccaggtttccccaggcctgccggtaggtttagaatgagg**tctgtgtcactgtgGTATTACGATATTTTGACTGGTTATTATAACcacagtgtcacaga**gtccatcaaaaa**cccatgcctggaagcttcccgccacagccctccccatggg**gccctgctgcctcctcaggtcagccccggacatcccgggtttccccaggctgggcggtaggtttggggtgagg**tctgtgtcactgtggtattactatggttcggggagttattataaccacagtgtcacaga**gtccatcaaaaacccatccctgggagcctcccgccacagccctccctgcaggggaccggtacgtgccatgttaggattttgatcgaggagacag**caccatgggtatggtg**gctaccacagcagtgcagcctgtgacccaaacccgcagggcagcaggcacgatggacaggcccgtgactgaccacgctggg**ctccagcctgccagccctggag**atcatgaaacagatggccaaggtcaccctacaggtcatccagatctggctccgaggggtctgcatcgctgctgccctcccaacgccagtccaaatgggacagggacggcctcacagcaccatctgctgccatcaggccagcgatcccagaagcccctccctcaaggctgggccacatgtgtggacactgagagccctcatgtctgagtaggggcaccaggaggg**aggggctggccctgtgcactgtccctgcccct**gtggtccctggcctgcctggccctgacacctgagcctctcctgggtcatttccaagacagaagacattcctgg**ggacagccggagctgggcgtcgctcatcctgcccggccgtcc**tgagtcctgctcatttccagacctcaccggggaagccaacagaggactcgcctcccacattcagagacaaagaaccttccagaaatccctgcctctctccccagtggacaccctcttccaggacagtcctcagtggcatc**acagcggcctgagatccccaggacgcagcaccgctgt**caataggggccccaaatgcctggaccagggcctgcgtggga

Uppercase: IGHD3-9-2

Lowercase: Flanking sequence[1000bp]

Red & Bold & Underline: Stem-loop [13]

Blue: Heptamer[52]

Green: Nonamer [5]

id-TRBD1[D_gene_segment]

tcagtttctgttgtatctttacagtgtttc**tgtatacaaaatgtataca**aaatcaagtaaatacaaaagtataattcttcttttcccccctctgttagacaaaggtggcatctagttatgctattctgagact**ttctcttttctgggttaataatgtatcttagagaa**ttttccatagcaatataaaatattttgattctttttttagcattgaatattactccattgtgtgaacatatcacagtttatccagtcccttactgatgggcatataattcaattttgtcagtcaaattagtgaaaagtgtattttgatgaagtttttttaattttatttttcttattgtgattttttttcccttatagttaggtaccatttggatttatttttctctgaacaatttactcttcacctatttttattgggctgctggattattttcctattgatttttaggagt**tcaccacatgttgaggagattagccctgatggtga**gaactaggaatatttttcccagattgttatgtatattttgattctgtttatattttaccatgcgagtgtgtcatcattatgtcgttgtttgtcttttttttttttttttct**ggattttaagtcatagcttaaaaccc**tccgagtgacg**cacagcctcggggcagggccagggtagtgatgggggctgtg**gcttctctataaggacatgccccaacgtgacaacagcttggagaggggtgggtactggagaagaccagccccttcgccaaacagccttacaaagacatccagctctaaggagctcaaaacatcctgaggacagtgcctggaggtgagaaggaagcccccggcctggtccataccccaccaccaacttgcataatggggggtgatgtcacccaccctccactcccctcaaaggagcagctgctctggtggtct**ctcccaggctctgggggcggacccatgggag**gggctgtttttgtacaaagctgtaa**cattgtgGGGACAGGGGGCcacaatg**attcaactctacgggaaacctttacaaaaacct**ctctggcggtcccaactcccagag**tcctcttctttcctcctgggtcacaggtcttaatgcaatttggttcagaatgcctctgcctcactcctgatcacatgtcagaccaagactgtggacaaggacaggcccagatgagaactaaagcttccc**aggcagagagaggtcagacataagaagactgcct**caggaacctcacaagtggaggactcagggagggtcccaatccccaaaaattgagacaaagtcaggtggaaggttcatcggaggtgaccagctctccagaggactcgggaagaagt**caggggtatctatagatggagtcacaggttctgggcccctg**ccatcctctgcaggccatgcactttccctttcgatggaccctcacagagggagcatctgaatggggca**tcctttgaaaaagga**acc**taggaccctgtggatggactctgtcattctccatggtccta**aaaagcaaaagtcaaagtgttcttctgtgtaatacccataaagcaca**ggaggagatttcttagctcactgtcctcc**atcctagccagggccctctcccctctctatgccttcaatgtgattttcaccttgacccctgtcactgtgtgaacactgaagctttctttggacaaggcaccagactcacagttgtaggtaagacatttttcaggttcttttgcagatccgtcacagggaaaagtgggtccacagtgtcccttttagagtggctatattcttatgtgctaactatggctacaccttcggttcggggaccaggttaaccgttgtaggtaaggctgggggtctctaggaggggtgcgatgagggaggactctgtc**ctgggaaatgtcaaagagaacagagatcccag**ctcccggagccagactgagggagacgtcatgtcatgtcccgggattgagttcaggggaggctccctgtgagggcgaatcc

Uppercase: TRBD1

Lowercase: Flanking sequence[1000bp]

Red & Bold & Underline: Stem-loop [14]

Blue: Heptamer[38]

Green: Nonamer [7]

id-IGHD6-13-2[D_gene_segment]

cagacagtgggcccagagcactgtgagtaccccggcagtac**ctggctgcagggatcagccag**agatgccaaaccctgagtgaccagcctacaggaggatccggccccacccaggccactcgattaatgctcaaccccctgccctggagacctcttccagtaccaccagcagctcagcttctcagggcctcatccctgcaaggaaggtcaagggctgggcctgccagaaacacag**caccctccctagccctggctaagacagggtg**ggcagacggctgtggacgggacatattgctggggcatttctcactgtcacttctgggtggtagctctgacaaaaacgcagaccctgccaaaatccccactgcctcccgctaggggctgg**cctggaatcctgctgtcctaggaggctgctgacctccagg**atggctccgtccccagttccagggcgagagcaga**tcccaggcaggctgtaggctggga**ggccacc**cctgcccttgccggggttgaatgcaggtgcccaaggcagg**aaat**ggcatgagcacagggatgaccgggacatgcc**ccaccagagtgcgccccttcctgctctgcaccctgcaccccccaggccagcccacgacgtccaacaactgggc**ctgggtggcagccccacccag**a**caggacagacccagcaccctgaggaggtcctg**ccagggggagctaagagccatgaaggagcaagatatggggcccccgatacaggcacagatgtcagctccatccaggaccacccagcccacaccctgagaggaacgtctgtctccagcctctgcaggtcgggaggcagctgacccctgacttggacccctattccagacaccagacagaggcgcaggccccccagaaccagggttgagggacgccccgtcaaagccagacaaaaccaaggggtgttgagcccagcaagggaaggcccccaa**acagaccaggaggtttctgaaggtgtctgtgtcacagtgGGGTATAGCAGCAGCTGGTACcacagtgacac**tcacccagccagaaaccccattccaagtcagcggaagcagagagagcagggaggacacgtttaggatctgagactgcacctgacacccaggccagcagacgtctcccctccagggcaccccaccctgtcctgcatttctgcaagatcaggggcggcctgagggggggtctagggtgaggagatgggtcccctgtacaccaaggaggagtta**ggcaggtcccgagcactctccccattgaggctgacctgcccagagagtcctgggc**ccaccc**cacacaccggggcggaatgtgtg**caggcctcggtctctgtgggtgttccgcta**gctggggctcacagtgctcaccccacacctaaaatgagccacagc**ctccggagcccccgcaggagaccccgcccacaagcccagcccccacccaggaggccccagagctcagggcgccccgtcggattccgaacagccccgagtcacagcgggtataaccggaacca**ccactgtcagaatagctacgtcaaaaactgtccagtgg**ccactgccggaggccccgccagagagggcagcagccactctgatcccatgtcctgcc**ggctcccatgacccccagcacgcggagcc**ccacagtg**tccccactggatgggaggacaagagctgggga**ttccggcgggtcggggcaggggcttgatcgcatccttctgccgtgg**ctccagtgcccctggctggag**ttgacccttctgacaagtgtcctcagagagacaggcatcaccggcgcctcccaacatcaaccccaggcagcacaggcacaaaccccacatccagagccaactccaggagcagagacaccccaataccctgggggaccccgaccctgatgacttcccactggaattcgccgtagagtccaccaggaccaaagaccctgcctctgcctctgtccctcactcaggacctgctgccgggcgaggccttgggagcagacttgggcttaggg

Uppercase: IGHD6-13-2

Lowercase: Flanking sequence[1000bp]

Red & Bold & Underline: Stem-loop [18]

Blue: Heptamer[28]

Green: Nonamer [8]

id-IGHD3-10-2[D_gene_segment]

ggcagggtgagcacccagagaaaactgcagacgcctcacacatccacctcagcctcccctgacctggacctcactggcctgggcctcacttaacctgggcttcacctgaccttggcctcacctgacttggacctcgcctgtcccaagctttacctgacctgggcctcaactcacctgaacgtctcctgacctgggtttaacctgtcctggaactcacctggccttggcttcccctgacctggacctcatctggcctgggcttcacctggcctgggcctcacctgacctggacctcatctggcctggacctcacctggcctggacttcacctggcctgggcttcacctgacctggacctcacctggcctcgggcctcacctgcacct**gctccaggtcttgctggagc**ctgagtagcactgagggtgcagaagctcatccagggttggggaatgactctagaagtctcccacatctgacctttctgggtggaggcagctggtggccctgggaatataaaaatctccagaatgatgactctgtgatttgtgggcaacttatgaacccgaaaggacatggccatggggtgggtagggacatagggacagatgccag**cctgaggtggagcctcagg**acacaggtgggcac**ggacactatccacataagcgagggatagacccgagtgtcc**ccacagcagacctgagagcgctgggcccacagcctcccctcagagccctgctgcctcctccggtcagccctggacatcccaggtttccccaggcctgccggtaggtttagaatgagg**tctgtgtcactgtggtattacgatattttgactggttattataaccacagtgtcacaga**gtccatcaaaaa**cccatgcctggaagcttcccgccacagccctccccatggg**gccctgctgcctcctcaggtcagccccggacatcccgggtttccccaggctgggcggtaggtttggggtgagg**tctgtgtcactgtgGTATTACTATGGTTCGGGGAGTTATTATAACcacagtgtcacaga**gtccatcaaaaacccatccctgggagcctcccgccacagccctccctgcaggggaccggtacgtgccatgttaggattttgatcgaggagacag**caccatgggtatggtg**gctaccacagcagtgcagcctgtgacccaaacccgcagggcagcaggcacgatggacaggcccgtgactgaccacgctggg**ctccagcctgccagccctggag**atcatgaaacagatggccaaggtcaccctacaggtcatccagatctggctccgaggggtctgcatcgctgctgccctcccaacgccagtccaaatgggacagggacggcctcacagcaccatctgctgccatcaggccagcgatcccagaagcccctccctcaaggctgggccacatgtgtggacactgagagccctcatgtctgagtaggggcaccaggaggg**aggggctggccctgtgcactgtccctgcccct**gtggtccctggcctgcctggccctgacacctgagcctctcctgggtcatttccaagacagaagacattcctgg**ggacagccggagctgggcgtcgctcatcctgcccggccgtcc**tgagtcctgctcatttccagacctcaccggggaagccaacagaggactcgcctcccacattcagagacaaagaaccttccagaaatccctgcctctctccccagtggacaccctcttccaggacagtcctcagtggcatc**acagcggcctgagatccccaggacgcagcaccgctgt**caataggggccccaaatgcctggaccagggcctgcgtgggaaaggtctctggccacactcgggctttttgtgaagggccctcctgctgtgtgactacagtaactaccatagtgatgaacccagtggcaaaaactggctggaaacccaggggctgtgtgcacgcctcagcttggagctctccaggagcacaagagccgggcccaaggatttgtgcccagaccctca

Uppercase: IGHD3-10-2

Lowercase: Flanking sequence[1000bp]

Red & Bold & Underline: Stem-loop [11]

Blue: Heptamer[52]

Green: Nonamer [7]

id-IGHD2-8-2[D_gene_segment]

gtgctcctcctgcccacagctcgagctcttcctctcctagggcccctgagggatggcattgaccgtgccctcgcacccacacactgcccatgccctcacattcctcctggccactccagccccactcccctctcaggcctggctctggtatttctgggacaaagccttacccaagtctttcccatgcaggcctgggcccttaccctca**ctgcccggttacagggcag**cct**cctgtgcacagaagcagggagctcagcccttccacagg**cagaaggcactgaaagaaatcggcctccagcgcctt**gacacacgtctgcctgtgtc**tctcactgcccgcacctgc**agggaggctcggcactccct**ctaaagacgagggatccaggcagcagcatcacaggagaatgcagggctaccagaca**tcccagtcctctcacaggcctctcctggga**agagacctgaagacgcccagtcaacggagtctaacaccaaacctccctggaggccgatgggtagtaacggagtcattgccagacctggaggcaggggagcagtgagcccgagcccacaccatagggccagaggacagccactgacatcccaagccactcactggtggtcccacaacaccccatggaaagaggacagacccacagtcccacctggaccagggcagagactgctgagacccagcaccagaaccaaccaagaaacaccaggcaacagcatcagagggggctctggcagaacagaggaggggaggtctccttcaccagcaggcgcttcccttgaccgaagacaggatccatgcaactcccccaggacaaaggag**gagccccttgttcagcactgggctc**agagtcctctccaagacacccagagtttcagacaaaaaccccctggaatgcacagtctcagcaggagagccagccagagccagcaagatggggctcagtgacacccgcagggacaggaggattttgtgggggctc**gtgtcactgtgAGGATATTGTACTAATGGTGTATGCTATACCcacagtgacac**agccccattcccaaagccctactgcaaacgcattccacttctggggctgaggggctgggggagcgtctgggaaatagggctcaggggtgtccatcaatgcccaaaacgcaccagactcccctccatacatcacacccaccagccagcgagcagagtaaacagaaaatgagaa**gcaagctggggaagcttgc**acaggccccaaggaaagagctttggcgggtgtgtaagaggggatgcgggcagagcctgagcagggccttttgctgtttctgctttcctgtgcagagagttccataaactggtgttcgagatcaatggctgggagtgagcccaggaggacagcgtgggaagagcacagggaaggaggagcagccgctatcctacactgtcatctttcgaaagtttgccttgtgcccacactgctgcatcatgggatgcttaacagctgatgtagacacagctaaagagagaatcagtgagatggatttgcagcacagatctgaataaattctccagaatgtggagcagcacagaagcaagcacacagaaag**tgcctgatgcaaggacaaagttcagtgggcaccttcaggca**ttgctgctgggcacagacactctgaaaagccctggcaggaactccctgtgacaaagcagaacc**ctcaggcaatgccagccccagagccctccctgag**agcctcatgggcaaagatgtgcacaacaggtgtttctcatagccccaaactgagagcaaagcaaacgtccatctgaaggagaacaggcaaataaacgatggcaggttcatgaaatgcaaacccagacagccacaagcacaaaagtacagggttataagcgactctggttgagttcatgacaatgctgagtaa**ttggagtaacaaagtaaactccaa**aaaatactttcaatgtgatttcttctaaataaaatttacaccctgcaaaatgaactgtcttcttaagggatacatttcccagtt

Uppercase: IGHD2-8-2

Lowercase: Flanking sequence[1000bp]

Red & Bold & Underline: Stem-loop [11]

Blue: Heptamer[44]

Green: Nonamer [4]

id-IGHD3-22-2[D_gene_segment]

gctgtatatccccaaggaaggtacagtcagtgcattccagagagaagcaactcagccaca**ctccctggccagaacccaagatgcacacccatgcacagggag**gcagagcccagcacctccgcagccaccaccacctgcgcacgggccaccaccttgcaggcacagag**tgggtgctgagaggaggggcagggacaccaggcagggtgagcaccca**gagaaaactgcagaagcctcacacatccacctcagcctcccctgacctggacctcacctggcctgggcctcacctgacctggacctcacctggcctgggcttcacctggcctgggcttcacctgacctggacctcacctggcctcgggcctcacctggcctgggcttcacctggcctgggcttcacctgacctggacctcacctggcctgggcctcacctgacctggacctcacctggcctgggcttcacctggcctgggcttcacctggcctgggcttcacctgacctggacctcacctggcctgggcttcacctgacctggacctcacctggcctcgggcctcacctgcacct**gctccaggtcttgctggagc**ctgagtagcactgaggctgtagggactcatccagggttggggaatgactctgcaactctcccacatctgacctttctgggtggaggcacctggtggcccagggaatataaaaagccccagaatgatgcctgtgtgatttgggggcaatttatgaacccgaaaggacatggccatggggtgggtagggacagtagggacagatgtcag**cctgaggtgaagcctcagg**acac**aggtgggcatggacagtgtccacct**aagcg**agggacagacccgagtgtccct**gcagtagacctgagagcgct**gggcccacagcctcccctcggggccc**tgctgcctcctcaggtcagccctggacatcccgggtttccccaggcctggcggtaggtttgaagtgagg**tctgtgtcactgtgGTATTACTATGATAGTAGTGGTTATTACTACcacagtgtcacaga**gtccatcaaaaactcatgcctgggagcctcccaccacagccctccctgcgggggaccgctgcat**gccgtgttaggattttgatcgaggacacggc**gc**catgggtatggtggctaccacagcagtgcagcccatg**acccaaacacacggggcagcagaaacaatggacaggcccacaagtgaccatgatgggctccagcccaccagccccagagaccatgaaacagatggccaaggtcaccctacaggtcatccagatctggctccaaggggtctgcatcgctgctgccctcccaacgccaaac**cagatggagacagggccggccccatagcaccatctgctgccgtccacccagcag**tcccggaagcccctccctgaacgctgggccacgtgtgtgaaccctgcg**agccccccatgtcagagtaggggcagcaggagggcggggct**ggccctgtgcactgtcactgcccctgtggtccctggcctgcctggccctgacacctgagcctctcctgggtcatttccaagacattcccag**ggacagccggagctgggagtcgctcatcctgcctggctgtcctgagtcctgctcatttccagacctcaccagggaagccaacagaggactca**cctcacacagtcagagacaatgaaccttccagaaatccctgtttctctccccagtg**agagaaaccctcttccagggtttctct**tctctcccaccctcttccaggacagtcctcagcagcatcacagcgggaacgcacatctggatcaggacggcccccagaacacgcgatggcccatggggacagcccagcccttcccagacccctaaaaggtatccccaccttgcacctgccccagggctcaaactc**caggaggcctgactcctgcacaccctcctg**ccagatatcacctcagc**cccctcctggagggg**acaggagcccgggagggtgagtcagacccacctgccctcaatggcag

Uppercase: IGHD3-22-2

Lowercase: Flanking sequence[1000bp]

Red & Bold & Underline: Stem-loop [18]

Blue: Heptamer[31]

Green: Nonamer [1]

id-TRBD2-2[D_gene_segment]

acgcgtgtatatgtttatatttacacacatgtacacacatccactcatta**aacacatatccctgtgatgtgtt**gttatcaacttccaagaaaattaagatacccaaattacagtccccaagaagcgta**tctgttcacagagcaactcaatcatgaacaga**aagagccaaacgcaagtccatgctttgaaagagcaacatgaactgcttactaaaatgcagtcagagagtggccctcaat**ttctcactgggataatcagtaaagagtctgagaa**ggtggcaatctcttcaaaggaagaattggtctcagttgttcactctagagaatggccatgctggtcagaggcatgaacaaaagccctaatgacatggctaggtacctctgggtaggcccgtttagctggttgcaaggttcagaaatgggagtgctcaaaattagcgctggaaaggcaggttggtgcatgccactctgggaataaacttac**ctccatgcaaccagcatggag**cctgcacatggtggatgttcactaaacacctgtggagtaaatgaagaatatggagttcagacatc**gttcaggaagcaactgaac**aaagggaaattatagaggtttctggattgtttgtcctcctgtcataaggtgccatcaactgctctgtggattttcctatgagctgcctgccacccctcgctcctcccacccacttcactataaatgccagtctgagca**ggtgggcacagtgagccccacc**agggagacccagtgacatagatggtctgctcagggtgatgcatgttccaaggagggacctctctgccccccaccattaccatcactgtgactttccccaagcccttcccattttaattcactgcctttgtcttttccaagccccacacagtcagactaacctctgccacctgcgcttcctgccgctgcccagtggttgggggagggggactagcagggaggaaacatttttgtatcatggtgtaacattgtgGGGACTAGCGGGGGGGcacgatgattcaggtagaggaggtgcttttacaaaaaaccctgatgcagtaagcatc**cccacccagctcagggaatgcagctaccaggtggg**aagagttctctggggctggtcccagctgtggtcttgcagggtcccccaacccagcgagcacctgtccatctccctgtccagactcggcttccaaggaataagaaggccaagacagc**aaagtgggattatcactcagcacttt**taataaaacttgttcttgacaaagtacttgcacatgcattatttattaagaactgatgaaaaccctgag**ggaaagatattgtcccatctttcc**aatgaggaaactgagatcagaggttacaggtcatataactaggaaacggcaaggtctagcctgcaatatcgcccagctccagccgttccagtaccaccaatgccccttcagatttca**aatccactgtgttgtcccccagccaagtggatt**ctcctctgcaaattggtggtggcctcatgcaagatccaggttaccgtgtccagctaactcgagacaggaaaagataggctcaggaaagagaggaagggtgtgccctctgtctgtgctaagggaggtg**gggaaggagaaggaattctgggcagccccttccc**actgtgctcctacaatgagcagttcttcgggccagggacacggctcaccgtgctaggtaagaagggggctccaggtgggagagagggtgagcagcccagcctgcacgaccccagaaccctgttcttaggggagtggacactgggcaatccagggccctcctcgagggaagcggggtttgcgccagggtccccagggctgtgcgaacaccggggagctgttttttggagaaggctctaggctgaccgtactgggtaaggaggcggctggggctccggagagctccgagagggcgggat**gggcagaggtaagcagctgccc**cactctgagaggggctgtgctgagaggcgctgctgggcgtctg

Uppercase: TRBD2-2

Lowercase: Flanking sequence[1000bp]

Red & Bold & Underline: Stem-loop [12]

Blue: Heptamer[40]

Green: Nonamer [8]

id-IGHD2-8[D_gene_segment]

gtgctcctcctgcccacagctcgagctcttcctctcctagggcccctgagggatggcattgaccgtgccctcgcacccacacactgcccatgccctcacattcctcctggccactccagccccactcccctctcaggcctggctctggtatttctgggacaaagccttacccaagtctttcccatgcaggcctgggcccttaccctca**ctgcccggttacagggcag**cct**cctgtgcacagaagcagggagctcagcccttccacagg**cagaaggcactgaaagaaatcggcctccagcgcctt**gacacacgtctgcctgtgtc**tctcactgcccgcacctgc**agggaggctcggcactccct**ctaaagacgagggatccaggcagcagcatcacaggagaatgcagggctaccagaca**tcccagtcctctcacaggcctctcctggga**agagacctgaagacgcccagtcaacggagtctaacaccaaacctccctggaggccgatgggtagtaacggagtcattgccagacctggaggcaggggagcagtgagcccgagcccacaccatagggccagaggacagccactgacatcccaagccactcactggtggtcccacaacaccccatggaaagaggacagacccacagtcccacctggaccagggcagagactgctgagacccagcaccagaaccaaccaagaaacaccaggcaacagcatcagagggggctctggcagaacagaggaggggaggtctccttcaccagcaggcgcttcccttgaccgaagacaggatccatgcaactcccccaggacaaaggag**gagccccttgttcagcactgggctc**agagtcctctccaagacacccagagtttcagacaaaaaccccctggaatgcacagtctcagcaggagagccagccagagccagcaagatggggctcagtgacacccgcagggacaggaggattttgtgggggctc**gtgtcactgtgAGGATATTGTACTAATGGTGTATGCTATACCcacagtgacac**agccccattcccaaagccctactgcaaacgcattccacttctggggctgaggggctgggggagcgtctgggaaatagggctcaggggtgtccatcaatgcccaaaacgcaccagactcccctccatacatcacacccaccagccagcgagcagagtaaacagaaaatgagaa**gcaagctggggaagcttgc**acaggccccaaggaaagagctttggcgggtgtgtaagaggggatgcgggcagagcctgagcagggccttttgctgtttctgctttcctgtgcagagagttccataaactggtgttcgagatcaatggctgggagtgagcccaggaggacagcgtgggaagagcacagggaaggaggagcagccgctatcctacactgtcatctttcgaaagtttgccttgtgcccacactgctgcatcatgggatgcttaacagctgatgtagacacagctaaagagagaatcagtgagatggatttgcagcacagatctgaataaattctccagaatgtggagcagcacagaagcaagcacacagaaag**tgcctgatgcaaggacaaagttcagtgggcaccttcaggca**ttgctgctgggcacagacactctgaaaagccctggcaggaactccctgtgacaaagcagaacc**ctcaggcaatgccagccccagagccctccctgag**agcctcatgggcaaagatgtgcacaacaggtgtttctcatagccccaaactgagagcaaagcaaacgtccatctgaaggagaacaggcaaataaacgatggcaggttcatgaaatgcaaacccagacagccacaagcacaaaagtacagggttataagcgactctggttgagttcatgacaatgctgagtaa**ttggagtaacaaagtaaactccaa**aaaatactttcaatgtgatttcttctaaataaaatttacaccctgcaaaatgaactgtcttcttaagggatacatttcccagtt

Uppercase: IGHD2-8

Lowercase: Flanking sequence[1000bp]

Red & Bold & Underline: Stem-loop [11]

Blue: Heptamer[44]

Green: Nonamer [4]

id-IGHD1-14[D_gene_segment]

aatgcaggtgcccaaggcaggaaat**ggcatgagcacagggatgaccgggacatgcc**ccaccagagtgcgccccttcctgctctgcaccctgcaccccccaggccagcccacgacgtccaacaactgggc**ctgggtggcagccccacccag**a**caggacagacccagcaccctgaggaggtcctg**ccagggggagctaagagccatgaaggagcaagatatggggcccccgatacaggcacagatgtcagctccatccaggaccacccagcccacaccctgagaggaacgtctgtctccagcctctgcaggtcgggaggcagctgacccctgacttggacccctattccagacaccagacagaggcgcaggccccccagaaccagggttgagggacgccccgtcaaagccagacaaaaccaaggggtgttgagcccagcaagggaaggcccccaa**acagaccaggaggtttctgaaggtgtctgtgtcacagtggggtatagcagcagctggtaccacagtgacac**tcacccagccagaaaccccattccaagtcagcggaagcagagagagcagggaggacacgtttaggatctgagactgcacctgacacccaggccagcagacgtctcccctccagggcaccccaccctgtcctgcatttctgcaagatcaggggcggcctgagggggggtctagggtgaggagatgggtcccctgtacaccaaggaggagtta**ggcaggtcccgagcactctccccattgaggctgacctgcccagagagtcctgggc**ccaccc**cacacaccggggcggaatgtgtg**caggcctcggtctctgtgggtgttccgcta**gctggggctcacagtgctcaccccacacctaaaatgagccacagc**ctccggagcccccgcaggagaccccgcccacaagcccagcccccacccaggaggccccagagctcagggcgccccgtcggattccgaacagccccgagtcacagcgGGTATAACCGGAACCA**Ccactgtcagaatagctacgtcaaaaactgtccagtgg**ccactgccggaggccccgccagagagggcagcagccactctgatcccatgtcctgcc**ggctcccatgacccccagcacgcggagcc**ccacagtg**tccccactggatgggaggacaagagctgggga**ttccggcgggtcggggcaggggcttgatcgcatccttctgccgtgg**ctccagtgcccctggctggag**ttgacccttctgacaagtgtcctcagagagacaggcatcaccggcgcctcccaacatcaaccccaggcagcacaggcacaaaccccacatccagagccaactccaggagcagagacaccccaataccctgggggaccccgaccctgatgacttcccactggaattcgccgtagagtccaccaggaccaaagaccctgcctctgcctctgtccctcactcaggacctgctgccgggcgaggccttgggagcagacttgggcttaggggacaccagtgtgaccccgaccttgaccaggacgcagacctttccttcctt**tcctggggcagcacagactttggggtctgggccagga**ggaacttctggcaggtcgccaagcacaga**ggccacaggctgaggtggccctggaaagacctccagg**aggtggccactccccttcctcccagctggaccccatgtcctccccaagataagggtgcca**tccaaggcaggtgctccttgga**gccccattcagactcctccctggaccccactgggcctcagtcccagctctggggatgaagccaccacaagcacaccaggcagcccaggcccagc**caccctgcagtgcccaagcacacactctggagcagagcagggtg**cctctgggaggggctgagctccccaccccacccccacctgcacaccccacccacccctgcccagc**ggctctgcaggagggtcagagcc**ccacatggggtatggacttagggtctcactcacgtggctccca

Uppercase: IGHD1-14

Lowercase: Flanking sequence[1000bp]

Red & Bold & Underline: Stem-loop [19]

Blue: Heptamer[21]

Green: Nonamer [8]

id-IGHD6-19[D_gene_segment]

ttacccaggacccagccctgcccctcctcccctctgctctcctctcatcaccccatgggaatccggtatccccaggaagccatcaggaagggctgaaggaggaagcggggccgtgcaccaccgggc**aggaggctccgtcttcgtgaacccagggaagtgccagcctcct**agagggtatggtccac**cctgcctggggctcccaccgtggcagg**ctgcggggaaggaccagggacggtgt**gggggagggctcagggccctgcgggtgctcctccatcttcggtgagcctccccc**ttcacccaccgtcccgcccacctcctctccaccctggctgcacgtcttccacaccatcctgagtcctacctacaccagagccagcaaagccagtgcagacaaaggctggggtgcaggggggctgccagggcagcttcggggagggaaggatggaggg**aggggaggtcagtgaagaggcccccttcccctgggtccaggatcctcctctgggaccc**ccggatccca**tcccctcctggctctgggaggagaagcaggatggga**gaatctgtgcgggaccctctcacagtggaatatccccacagc**ggctcaggccagacccaaaagcccctcagtgagcc**ctccactgcagtcctgggc**ctgggtagcagcccctcccacagaggacagacccag**caccccgaagaag**tcctgccagggggagctcagagccatgaaagagcagga**tatggggtccccgatacaggcacagacctcagctccatccaggcccaccgggacccaccatgggaggaacacctgtctccgggttgtgaggtagctgg**cctctgtctcggaccccactccagacaccagacagagg**ggcaggccccccaaaaccagggttgagggatgatccgtcaaggcagacaagaccaaggggcactgaccccagcaagggaaggctcccaa**acagacgaggaggtttctgaagctgtctgt**atcacagtgGGGTATAGCAGTGGCTGGTACcacagtgacactcgccaggccagaaaccccgtcccaagtcagcggaagcagagagagcagggaggacacgtttaggatctgaggccgcacctgacacccagggcagcagacgtctcccctccagggcaccctccaccgtcctgcgtttcttcaagaataggggcggcct**gagggggtccagggccaggcgataggtcccctc**taccccaaggaggagcca**ggcaggacccgagcaccgtccccattgaggctgacctgcccagacgggcctgggc**ccaccc**cacacaccggggcggaatgtgtg**ca**ggccccagtctctgtgggtgttccgctagctggggcc**cccagtgctcaccccacacctaaagcgagccccagcctccagagccccctaagcattccccgcccagcagcccagcccctgcccccacccaggaggccccagagctcagggcgcctggtcggattctgaacagccccgagtcacagtgggtataactggaacgaccaccgtgagaaaaactgtgtccaaaactgactcctggcagcagtcggaggccccgccagagaggggagcagccggcctgaacccatgtcctgccggttcccatgacccccagcacccagagccccacggtgtccccgttggataatgaggacaagggctgggggctccggtggtttgcggcagggacttgatcacatccttctgctgtggccccattgcctctggctggagttgaccc**ttctgacaagtgtcctcagaa**agacagggatcaccggcacctcccaatatcaaccccaggcagcacagacacaaaccccacat**ccagagccaactccaggagcagagacaccccaacactctgg**gggaccccaaccgtgataactccccactggaatccgccccagagtctaccaggaccaa**aggccctgccctgtctctgtccctcactcagggcct**cctgcagggcgagcgcttgggagcagactcggtctt

Uppercase: IGHD6-19

Lowercase: Flanking sequence[1000bp]

Red & Bold & Underline: Stem-loop [19]

Blue: Heptamer[29]

Green: Nonamer [8]

id-IGHD3-16[D_gene_segment]

atgatacagac**atacatttagtacatgagacatcgatgatgtat**ccccaaagaaatgactttaaagagaaaaggcctgatgtgtggtggcactcacctccctgggatccccggacaggttgcaggcacactgtgtggcagggcaggctggtacatgctggcagctcctggggcctgatgtggagcaagcgcagggctgtatacccccaaggatggcacagtcagtgaattccagagagaagcagctcagccacactgcccaggcagagcccgagagggacgcccacgcacagggaggcagagcccagctcctccacagccacca**ccacctgtgcacgggccaccaccttgcaggcacagagtgggtgctgagaggaggggcagggacaccaggcagggtgagcaccca**gagaaaactgcagaagcctcacacatccacctcagcctcccctgacctggacctcacctggtctggacctcacctggcctgggcctcacctgacctggacctcacctggcctgggcttcacct**gacctggacctcacctggcctccggcctcacctgcacctgctccaggtc**ttgctggaacctgagtagcactgaggctgcagaagctcatccagggttggggaatgactctggaactctcccacatctgacctttctgggtggaggcatctggtggccctgggaatataaaaagccccagaatggtgcctgcgtgatttgggggcaatttatgaacccgaaaggacatggccatggggtgggtagggacatagggacagatgccag**cctgaggtggagcctcagg**acacagttggacgcggacactatccacataagcgagggacagacccgagtgttcctgcagtagacctgagagcgctgggcccacagcctcccctcggtgccctgctgcctcctcaggtcagccctggacatcccgggtttccccaggccagatggtaggtttgaagtgaggtctgtgtcactgtgGTATTATGATTACGTTTGGGGGAGTTATCGTTATACCcacagcatcacacggtccatcagaaacccatgccacagccctccccgcaggggaccgccgcgtgccatgttacgattttgatcgaggacacagcgc**catgggtatggtggctaccacagcagtgcagcccatg**acccaaaca**cacagggcagcaggcacaatggacaggcctgtg**agtgaccatgctgggctccagcccgccagccccggagaccatgaaacagatggccaaggtc**accccacagttcagccagacatggctccgtggggt**ctgcatcgctgctgccctctaacaccagccc**agatggggacaaggccaaccccacattaccatct**cctgctgtccacccagtggtcccagaagcccctccctcatggctgagccacatgtgtgaaccctgagagcaccccatgtcagagtaggggcagcagaagggcggggctggccctgtgcactgtccctgcacccatggtccctcgcctgcctggccctgacacctgagcc**tcttctgagtcatttctaagatagaaga**cattcccgg**ggacagccggagctgggcgtcgctcatcccgcccggccgtcctgagtcctgcttgtttccagacctcaccagggaagccaacagaggactca**cctcacacagtca**gagacaaagaaccttccagaaatccctgtctc**actccccagtgggcaccttcttccaggacattcctcggtcgcatcacagcaggcacccacatctggatcaggacggcccccagaacacaagatggcccatggggacagccccacaacccaggccttcccagacccctaaaaggcgtcccaccccctgcacctgccccagggctaaaaatccaggaggcttgactcccgca**taccctccagccagacatcacctcagccccctcctggagggga**caggagcccgggagggtgagtcagaccca**cctgccctcgatggcagg**cggggaagattcagaaaggcctgagatcccc

Uppercase: IGHD3-16

Lowercase: Flanking sequence[1000bp]

Red & Bold & Underline: Stem-loop [15]

Blue: Heptamer[42]

Green: Nonamer [3]

id-IGHD5-12-2[D_gene_segment]

ggccacactcgggctttttgtgaagggccctcctgctgtgtgactacagtaactaccatagtgatgaacccagtggcaaaaactggctggaaacccaggggctgtgtgcacgcctcagcttggagctctccagga**gcacaagagccgggcccaaggatttgtgcccagaccctcagcctctagggacacctgggc**catctcagcctgggctggtgccctgcacaccatcttcctccaaataggggcttcagagggctctgaggtgacctcactcatgaccacaggtgacctggcccttccctgccagctataccagaccctgtcttgacagatgccccgattccaacagccaattcctgggaccctgaatagctgtagacaccagcctcattccagtacctcctgccaattgcctggattcccat**cctggctggaatcaagaaggcagcatccgccagg**ctcccaacaggcaggactcccgcacaccctcctctgagaggccgctgtgttccg**cagggccaggccctg**gacagttcccctcacctgccactagagaaacacctgccattgtcgtcccc**acctggaaaagaccactcgtggagcccccagccccaggt**acagctgtagagagagtcctcgaggcccctaagaaggagccatgcccagttctgccgggaccctcggccaggccgacaggagtggacgctggagctgg**gcccacactgggccacataggagctcaccagtgagggc**aggagagcacatgccggggagcacc**cagcctcctgctgaccagaggcctgccccagagcccaggaggctg**cagaggcctctccagggagacactgtgcatgtctggtacctaagca**gccccccacgtccccagtcctgggggc**ccctggctca**gctgtctggaccctccctgttccctgggaagctcctcctgacagc**cccgcctccagttccaggtgtggttattgtcaggcgatgtcag**actgtgGTGGATATAGTGGCTACGATTACcacagt**ggtgccgcccata**gcagcaaccaggccaagtagacaggcccctgctgc**gc**agccccaggcatccacttcacctgcttctcctggggct**ctcaaggctgctgtctgtcctctggccctctgtggggagggttccctcagtgggaggtctgtgctccagggcagggatgattgagatagaaatcaaaggctggcagggaaaggcagcttcccgccctgagaggtgcaggcagcaccacggagccacggagtcacagagccacggagcccccattgtgggcatttgagagtgctgtgcccccggcaggcccagccct**gatggggaagcctgtcccatc**ccacagcccgggtcccacg**ggcagcgggcacagaagctgcc**aggttgtcctctatgatcctcatccctccagcagcatcccctccacagtggggaaactgaggcttggagcaccacccggccccctggaaatgaggctgtgagcccag**acagtgggcccagagcactgt**gagtaccccggcagtac**ctggctgcagggatcagccag**agatgccaaaccctgagtgaccagcctacaggaggatccggccccacccaggccactcgattaatgctcaaccccctgccctggagacctcttccagtaccaccagcagctcagcttctcagggcctcatccctgcaaggaaggtcaagggctgggcctgccagaaacacag**caccctccctagccctggctaagacagggtg**ggcagacggctgtggacgggacatattgctggggcatttctcactgtcacttctgggtggtagctctgacaaaaacgcagaccctgccaaaatccccactgcctcccgctaggggctgg**cctggaatcctgctgtcctaggaggctgctgacctccagg**atggctccgtccccagttccagggcgagagcaga**tcccaggcaggctgtaggctggga**ggccacccctgcccttgccggggttgaatgca

Uppercase: IGHD5-12-2

Lowercase: Flanking sequence[1000bp]

Red & Bold & Underline: Stem-loop [19]

Blue: Heptamer[46]

Green: Nonamer [4]

id-TRDD1[D_gene_segment]

gttggcagggagggtcacaggtgagcattgtattttaatagcaataatagccataagctaaacactgcagcaaccacagcagatatacaaatgg**tatatatacaaatatacaattatatttgtataaata**tatataaacatataataaagtaagctcctatt**ttcttaattagaagattcaaagttaagaa**ttttgtccaacttgaatatatgaaactttagggttatagagaaattagccaattctactttagatgttttaatgtattaatacagttaaacacacatattttaaaattac**atattatataaaataatat**atcaattttgaatactgctatttccttttaataattcataaaataactttaaattcttcacaggcatttcaaatcaatgtatttgttgccactagagccacttttttatcctcaaatatttttccagagtatatgaaactcactttgctaaattgaaaattaagaggcacattagattta**gaaacataattttaaattgtttc**aatgatggcacattattaatacctactttgagacacaatacatactcacataatattagaatcattgttttgccatttatcctgtaagtgaatattcatgatctccacaacaaaacatttac**tttgatagatttgtttaaacaatatcaaa**cacttgcagtcttgaaggcactggaagacccaaaaaatagttactacgtttct**gttaaactacttagtcatctttttaac**taattctatcagctgactgccataagcatccatgagctagcattgattgagatcattccaatgtgtcctcactaaactgt**acttaactgttcaaagccaaataattaagt**gaaagtatggtaatactgtttttcataattattgtaaggaatcaaatataagggatttcttcttctctgaggatcatgaaccttactccatgttcaaatagatatagtattttttcacagtaaattccccaaagtgGAAATAGTcactcaaacgaatacttttactcttttgacattaaccaaatggcttccaa**taaaaatcaaacaatttatattttta**ccaacatcttgcccacattgggagtgtcaacattttgagcaaaagctttaattaaatatccatgcaaaaaatggtttgttaatactttacagttttattactagagggttaaaatcctttttcaagtctgataatcaatgattaactttcttcatttgtccttcacccatttgttttttaggttgatggtgttttacttattgatttgtgtaattataataa**ttttgtgtctgagttttacagcatttaaccacaaaa**acagcattggtgaaaggagtttcaggggtattgtggatggcagcgggtggtgatggcaaagtgccaaggaaagggaaaaaggaagaagagggtttttatactgatgtgtttcattgtgccttcctaccacacaggttggagtgcattaagcctttgtccaaaaacacccagccgtgacccgctatgtatgtctcagcattgggaagagtcctctgagtgtcatgg**gaaaataatatatgagttttattgcccctgtgtcccaattattttc**tgtttactccatgttggtctacagcaatagcttcagaacataaacaaatatcaccaaaaacatcagtgatgaacaattccaaccacacatttcaatgtttactaaatatacccgtggcactgtaaaaaagaatataatggaagagtaaaggaattgggactgatcgtgactgagaacctgtatagtaggctacacactagctatatgatgaaccaagaagttgaacccaaatctaactaacttacataatcccttcctttaaagaacttaatgtccagtgcaaagcaagtgttcaaaaaagtgtttgttgtgaattgaattcatttcattggaatacagcatgcttgcaaagtcccaactttttaaaataatgtgtttctaaagaccactggtct

Uppercase: TRDD1

Lowercase: Flanking sequence[1000bp]

Red & Bold & Underline: Stem-loop [10]

Blue: Heptamer[26]

Green: Nonamer [7]

id-IGHD1-14-2[D_gene_segment]

aatgcaggtgcccaaggcaggaaat**ggcatgagcacagggatgaccgggacatgcc**ccaccagagtgcgccccttcctgctctgcaccctgcaccccccaggccagcccacgacgtccaacaactgggc**ctgggtggcagccccacccag**a**caggacagacccagcaccctgaggaggtcctg**ccagggggagctaagagccatgaaggagcaagatatggggcccccgatacaggcacagatgtcagctccatccaggaccacccagcccacaccctgagaggaacgtctgtctccagcctctgcaggtcgggaggcagctgacccctgacttggacccctattccagacaccagacagaggcgcaggccccccagaaccagggttgagggacgccccgtcaaagccagacaaaaccaaggggtgttgagcccagcaagggaaggcccccaa**acagaccaggaggtttctgaaggtgtctgtgtcacagtggggtatagcagcagctggtaccacagtgacac**tcacccagccagaaaccccattccaagtcagcggaagcagagagagcagggaggacacgtttaggatctgagactgcacctgacacccaggccagcagacgtctcccctccagggcaccccaccctgtcctgcatttctgcaagatcaggggcggcctgagggggggtctagggtgaggagatgggtcccctgtacaccaaggaggagtta**ggcaggtcccgagcactctccccattgaggctgacctgcccagagagtcctgggc**ccaccc**cacacaccggggcggaatgtgtg**caggcctcggtctctgtgggtgttccgcta**gctggggctcacagtgctcaccccacacctaaaatgagccacagc**ctccggagcccccgcaggagaccccgcccacaagcccagcccccacccaggaggccccagagctcagggcgccccgtcggattccgaacagccccgagtcacagcgGGTATAACCGGAACCA**Ccactgtcagaatagctacgtcaaaaactgtccagtgg**ccactgccggaggccccgccagagagggcagcagccactctgatcccatgtcctgcc**ggctcccatgacccccagcacgcggagcc**ccacagtg**tccccactggatgggaggacaagagctgggga**ttccggcgggtcggggcaggggcttgatcgcatccttctgccgtgg**ctccagtgcccctggctggag**ttgacccttctgacaagtgtcctcagagagacaggcatcaccggcgcctcccaacatcaaccccaggcagcacaggcacaaaccccacatccagagccaactccaggagcagagacaccccaataccctgggggaccccgaccctgatgacttcccactggaattcgccgtagagtccaccaggaccaaagaccctgcctctgcctctgtccctcactcaggacctgctgccgggcgaggccttgggagcagacttgggcttaggggacaccagtgtgaccccgaccttgaccaggacgcagacctttccttcctt**tcctggggcagcacagactttggggtctgggccagga**ggaacttctggcaggtcgccaagcacaga**ggccacaggctgaggtggccctggaaagacctccagg**aggtggccactccccttcctcccagctggaccccatgtcctccccaagataagggtgcca**tccaaggcaggtgctccttgga**gccccattcagactcctccctggaccccactgggcctcagtcccagctctggggatgaagccaccacaagcacaccaggcagcccaggcccagc**caccctgcagtgcccaagcacacactctggagcagagcagggtg**cctctgggaggggctgagctccccaccccacccccacctgcacaccccacccacccctgcccagc**ggctctgcaggagggtcagagcc**ccacatggggtatggacttagggtctcactcacgtggctccca

Uppercase: IGHD1-14-2

Lowercase: Flanking sequence[1000bp]

Red & Bold & Underline: Stem-loop [19]

Blue: Heptamer[21]

Green: Nonamer [8]

id-IGHD4-17[D_gene_segment]

ctaccacagcagtgcagcccatgacccaaaca**cacagggcagcaggcacaatggacaggcctgtg**agtgaccatgctgggctccagcccgccagccccggagaccatgaaacagatggccaaggtc**accccacagttcagccagacatggctccgtggggt**ctgcatcgctgctgccctctaacaccagccc**agatggggacaaggccaaccccacattaccatct**cctgctgtccacccagtggtcccagaagcccctccctcatggctgagccacatgtgtgaaccctgagagcaccccatgtcagagtaggggcagcagaagggcggggctggccctgtgcactgtccctgcacccatggtccctcgcctgcctggccctgacacctgagcc**tcttctgagtcatttctaagatagaaga**cattcccgg**ggacagccggagctgggcgtcgctcatcccgcccggccgtcctgagtcctgcttgtttccagacctcaccagggaagccaacagaggactca**cctcacacagtca**gagacaaagaaccttccagaaatccctgtctc**actccccagtgggcaccttcttccaggacattcctcggtcgcatcacagcaggcacccacatctggatcaggacggcccccagaacacaagatggcccatggggacagccccacaacccaggccttcccagacccctaaaaggcgtcccaccccctgcacctgccccagggctaaaaatccaggaggcttgactcccgca**taccctccagccagacatcacctcagccccctcctggagggga**caggagcccgggagggtgagtcaga**cccacctgccctcgatggcaggcggg**gaagattcagaaaggcctgagatccccaggacgcagcaccactgtcaatgggggccccagacgcctggaccagggcctgcgtgggaaaggccgctgggcacactc**aggggctttttgtgaaggcccct**cct**actgtgTGACTACGGTGACTACcacagt**gatgaaactagcagcaaaaactggccggacacccagggaccatgcacacttctcagcttggagctctccaggaccagaagagtcaggtct**gagggtttgtagccagaccctc**ggcctctagggacaccctggccatcacagcggatgggctggtgccccacatgccatctgctccaaacaggggcttcagagggctctgaggtgacttcactcatgaccacaggtgccctggccccttccccgccagctacaccgaaccctgtcccaacagct**gccccagttccaacagccaattcctggggc**ccagaattgctgtagacaccagcctcgttccagcacctcctgccaattgcctggattcacat**cctggctggaatcaagagggcagcatccgccagg**ctcccaacaggcaggactcccgcacaccctcctctgagaggccgctgtgttccg**cagggccaggccctg**gacagttcccctcacctgccactagagaaacacctgccattgtcgtcccc**acctggaaaagaccactcgtggagcccccagccccaggt**acagctgtagagagactccccgagggatctaagaaggagccatgcgcagttctgccgggaccctcggccaggccgacaggagtggacactggagctgg**gcccacactgggccacataggagctcaccagtgagggc**aggagagcacatgccggggagcacc**cagcctcctgctgaccagaggcccgtcccagagcccaggaggctg**cagaggcctctccagggggacactgtgcatgtctggtccctgagca**gccccccacgtccccagtcctgggggc**ccctggcacagctgtctggaccctccctcttccctgggaagctcctc**ctgacagccccgcctccagttccaggtgtggttattgtcagggggtgtcagactgtggtggatacagctatggttaccacagt**ggtgctgcccatagcagcaaccaggcca

Uppercase: IGHD4-17

Lowercase: Flanking sequence[1000bp]

Red & Bold & Underline: Stem-loop [21]

Blue: Heptamer[37]

Green: Nonamer [5]

id-IGHD1-26-2[D_gene_segment]

atcccctccaggctctgggaggagaagcaggatgggagaatctgtgcgggaccctctcacagtggaatacctccacagc**ggctcaggccagatacaaaagcccctcagtgagcc**ctccactgca**gtgctgggcctgggggcagcccctcccacagaggacagacccagcac**cccgaagaagtcctgccagggggagctcagagccatgaaggagcaagatatggggaccccaatactggcacagacctcagctccatccaggcccaccaggacccaccatgggtggaacacctgtctccggcccctgctggctgtgaggcagctgg**cctctgtctcggacccccattccagacaccagacagagggacaggccccccagaaccagtgttgagggacacccctgtcc**agggcagccaagtccaagaggcgcgctgagcccagcaagggaaggcccccaaacaaaccaggaggtttctgaagctgtct**gtgtcacagtcgggtatagcagcggctaccacaatgacac**tgggcaggacagaaaccccatcccaagtcagccgaaggcagagagagcaggcaggacacatttaggatctgaggccacacctgacactcaagccaacagatgtctcccctccagggcgcc**ctgccctgttcagtgttcctgagaaaacaggggcag**cct**gaggggatccagggccaggagatgggtcccctc**taccccgaggaggagccaggcgggaatcccagc**cccctccccattgaggccatcctgcccagagggg**cccggacccaccc**cacacacccaggcagaatgtgtg**caggcctcaggctctgtgggtgccgctagctggggctgccagtc**ctcaccccacacctaaggtgag**ccacagccgccagagcctccacaggagaccccacgcagcagcccagcccct**acccaggaggccccagagctcagggcgcctgggt**ggattctgaacagccccgagtcacggtgGGTATAGTGGGAGCTACTACcactgtgagaaaagctatgtccaaaactgtctcccggccactgctggaggcccagccagagaagggaccagccgcccgaacatacgaccttcccagccctcatgacccccagcacttggagctccacagtgtccccattggatggtgaggacgggggccggggccatctgcacctcccaacatcacccccaggcagcacaggcacaaaccccaaatccagagccgacaccaggaacacagacaccccaataccctgggggaccctggccctggtgacttcccactgg**gatccacccccgtgtccacctggatc**aaagaccccaccgctgtctctgt**ccctcactcagggcctgctgaggg**gcgggtgctttggagcagactcaggtttaggggccaccattgtggggcccaacctcgaccaggacacagatttttctttcctgccctggggcaacacagacttt**ggggtctgtgcagggaggaccttctggaaagtcaccaagcacagagccc**tgactgaggtggtctcaggaag**acccccaggagggggt**ttgtgccccttcctctcatgtggacccc**atgccccccaagataggggcat**catgcagggcaggtcctccatgcagccaccactaggcaactccctggcgccggtccccactgcgcctc**catcccggctctggggatg**cagccaccatggccacaccaggcagcccgggtccagcaaccctgcagtgcccaagcccttggcaggattcccagaggctggagcccacccctcctcatccccccacacctgcacacacacacctaccccctgcccagtccccctccaggagggt**tggagccgcccatagggtgggcgctcca**ggtctcactcactcgcttcccttcctgggcaaaggagcctc**gtgccccggtcccccctgacggcgctgggcac**aggtgtgggtactgggccccagggctcctccagccccagctgccctgctctccctgg

Uppercase: IGHD1-26-2

Lowercase: Flanking sequence[1000bp]

Red & Bold & Underline: Stem-loop [19]

Blue: Heptamer[36]

Green: Nonamer [10]

id-IGHD2-21-2[D_gene_segment]

ctgtgcccaccccctaacccctcctgcccacaacttgagttcttcctctcctggagcccttgagccatggcactgaccctacactcccacccacacactgcccatgccatcaccttcctcctggacactctgaccccgctcccctccctctcagacccggccctggtatttccaggacaaaggctcacccaagtcttccccatgcaggcccttgccctcactgcctggttacacgggagcct**cctgtgcgcagaagcagggagctcagctcttccacagg**cag**aaggcactgaaagaaatcggcctccagtgccttgacacacgtccgcctgtgtc**tctcactgcctgcacctgc**agggaggctccgcactccct**ctaaagatgagggatccaggcagcaacatcacgggagaatgc**agggctcccagacagcccagccct**ctc**gcaggcctctcctgggaagagacctgc**agccaccactgaacagccacgg**aggtcgctggatagtaaccgagtcagtgaccgacct**ggagggcaggggagcagtgaaccggagcccataccatagggacagagaccagccgctaacatcccgagcccctcactggcggccccagaacaccccgtggaaagagaacagacccacagtcccacctggaacagggcagacactgctgagcccccagcacc**agccccaagaaacactaggcaacagcatcagagggggct**cctgagaaagagaggaggggaggtctccttcaccatcaaatgcttcccttgaccaaaaacagggtccacgcaactcccccaggacaaaggag**gagccccctgtacagcactgggctcagagtcctctctgag**acaggctcagtttcagacaacaacccgctggaatgcacagtctcagcaggagagccaggccagagccagcaag**aggagactcggtgacaccagtctcct**gtagggacaggaggattttgtg**ggggttcgtgtcactgtgAGCATATTGTGGTGGTGACTGCTATTCCcacagtgacacaacccc**attcctaaagccctactgcaaacgcacccactcctgggactgaggggctgggggagcatctgggaagtatggcctaggggtgtccatcaatgcccaaaatgcaccagactctccccaagacatcaccccaccagccagtgagcagagtaaacagaaaatgagaagcagctgggaag**cttgcacaggccccaaggaaagagctttggcaggtgtgcaag**aggggatgtgggcagagcctcagcagggccttttgctgtttctgctttcctgtgcagagagttccataaactggtattcaagatcaatggctgggagtgagcccaggag**gacagtgtgggaagagcacagggaaggaggagcagccgctatcctacactgtc**atcttttgaaagtttgccctgtgcccacaatgctgcatcatgggatgcttaacagctgatgtagacacagctaaagagagaatcagtgaaatgcatttgcagcacagatctgaataaatcctccagaatgtggagcagcacagaagcaagcacacagaaag**tgcctgatgccaaggcaaagttcagtgggcaccttcaggca**ttgctgctgggcacagacactctgaaaagcactggcaggaac**tgcctgtgacaaagcagaaccctcaggca**atgccagccctagagcccttcctgagaacctcatgggcaaagatgtgcagaacagctgtttgtcat**agccccaaactatggggct**ggacaaagcaaacgtccatctgaaggagaacagacaaataaacgatggcaggttcatgaaatacaaactaggacagccagaggacaacagtagagagctacaggcggctttgcggttgagttcatgacaatgctgagtaattggagtaacagaggaaagcccaaaaaatacttttaatgtgatttcttctaaataaaatttacacccggcaaaatgaactatcttcttaagggataaactttccc

Uppercase: IGHD2-21-2

Lowercase: Flanking sequence[1000bp]

Red & Bold & Underline: Stem-loop [17]

Blue: Heptamer[39]

Green: Nonamer [5]

id-IGHD1-1[D_gene_segment]

atgcaggaatgactgggccacacccctcccgtgcacgccccctcctgccctgcaccccacagcccagccccccgtgctggatgccccccc**acagcagaggtgctgt**tctgtgatcccctgggaaagacgccctcaacctccaccctgtcccacggcccaaggaagacaagacacaggccctctcctcacagtctccccacctggctcctgctgggaccctcaaggtgtgaacagggaggatggttg**tctgggtggcccctaggagcccaga**tcttcactccacagaccccaacccaagcacccccttctgcagggcccagctcatccccctcctcctccctctgctctcctctcgtcgcctctacgggaaatccgggactcagcagtaaccctc**aggaagcagggcccaggcgccgtttaataggaggcttcct**cacaatgaaacttttagaaagccttgactacaatgatgaccttggtgtg**gctgtgaacactgtcagctcccacagc**tgctgcagcaaaaaatgtccatagacagggtgggggcccggggtcgtc**tgctgtcctgctcagcccacagca**cgcatggaggatctgaggtgccacacctgacgcccaggccagaacatgcc**tccctccagggtgacctgccatgtcctgcattgctggaggga**caggggcagcctatgagga**tctggggccaggagatgaatcctattaacccaga**ggaaaactaacaggacccaagcaccctccccgttgaagctgacct**gcccagaggggcctgggc**ccaccccacac**accggggcggaatgtgtacaggccccggt**ctctgtgggtgttccgctaa**ctggggctcccagtgctcaccccacaactaaagcgagccccag**cctccagagcccccgaaggagatgccgcccacaagcccagcccccatccaggaggccccagagctcagggcgc**cggggcagattctgaacagccccg**ag**tcacggtgGGTACAACTGGAACGACcaccgtga**gaaaaactgtgtccaaaa**ctctctcctggcccctgctggaggccgcgccagagag**gg**gagcagccgccccgaacctaggtcctgctc**agctcacacgacccccagcagccagagca**cagtggagtccccactg**aaccccactgaatggtgaggacggggaccagggctccag**ggggtcatggaaggggctggacccc**atcctactgctatggtcccagtgctcctggccagaactgaccctaccaccgacaagagtccctcagggaaacgggggtcactggcacctcccagcatcaaccccaggcagcacaggcataaaccccacatccagagccgactccaggagcagagaca**ccccagtaccctgggg**gacaccgaccctgatgactccccactggaatccaccccagagtccaccaggaccaaagac**cccgccccggtctctgtccctcactcaggacctgctgcggggcggg**ccatgagaccagactcgggcttagggaacacca**ctgtggccccaacctcgaccaggccacag**gcccttccttcctgccctgcggca**gcacagactttggggtctgtgc**agagaggaatcacagaggccccaggctgaggtgg**tgggggtggaagaccccca**ggaggtggcccacttcccttcctcccagctggaacccaccatgaccttcttaagataggggtgtcatccgaggcaggtcctccatggagctcccttcaggctcctccctggtcctcactaggcctcagtcccgg**ctgtgggaatgcagccaccacag**gcacaccaggcagcccagacccagc**cagcctgcagtgcccaagcccacattctggagcagagcaggctg**tgtctgggagagtctgggctccccaccgccccccgcacaccccacccacccctgtccaggccctatgcaggagggtcagagccccccatggggtatggacttagggtctcactcacgcggctcccctcc

Uppercase: IGHD1-1

Lowercase: Flanking sequence[1000bp]

Red & Bold & Underline: Stem-loop [23]

Blue: Heptamer[32]

Green: Nonamer [1]

id-IGHD2-2[D_gene_segment]

agtttctaccctctgtgcctaccccctgcctcctcctgcccacaactcgagctcttcctctcctggggcccctgagccatggcactgaccgtgcactcccacccccacactgcccatgccctcaccttcctcctggacactctgaccctgctcccctcttggacccagccctggtatttccaggacaaaggctcacccaagtcttccccatgcaggcccttgccctcactgcccggttacacggcagcct**cctgtgcacagaagcagggagctcagcccttccacagg**cagaaggcactgaaagaaatcggcctccagcaccctgatgcacgtccgcctgtgtctctcactgcccgcacctg**cagggaggctcggcactccctg**taaagacgagggatccaggcagcaacatca**tgggagaatgcagggctccca**gacagcccagccctctc**gcaggcctctcctgggaagagacctgc**agccaccactgaacagccacggagcccgctggatagtaactgagtcagtgaccgacctggagggcaggggagcagtgaaccggagcccagaccatagggacagagaccagccgctgacatcccgagcccctcactggcggccccagaacaccgcgtggaaacagaacagacccacattcccacctggaa**cagggcagacactgctgagcccccagcaccagccctg**agaaacaccaggcaacggcatcagagggggctcctgagaaaga**aaggaggggaggtctcctt**caccagcaagtacttcccttgaccaaaaacagggtccacgcaactcccccaggacaaaggag**gagccccctgtacagcactgggctc**agagtcctctccaacacaccctgagtttcagacaaaaaccccctggaaatcatagtatcagcaggagaactagccagagacagcaagaggggactcagtgactcccgcggggacaggaggattttgtgggggctc**gtgtcactgtgAGGATATTGTAGTAGTACCAGCTGCTATGCCcacagtgacac**agccccattcccaaagccctgctgtaaacgcttccacttctggagctgaggggctggggggagcgtctgggaagtagggcctaggggtggccatcaatgcccaaaacgcaccagactcccccccagacatcaccccactggccagtgagcagagtaaacagaaaatgagaagcagctgggaag**cttgcacaggccccaaggaaagagctttggcgggtgtgcaag**aggggatgcgggcagagcctgagcagggccttttgctgtttctgctttcctgtgcagatagttccataaactggtgttcaagatcgatggctgggagtgagcccaggag**gacagtgtgggaagggcacagggaaggagaagcagccgctatcctacactgtc**atctttcaagagtttgccctgtgcccacaatgctgcatcatgggatgcttaacagctgatgtagacacagctaaagagagaatcagtgaaatggatttgcagcacagatctgaataaattctccagaatgtggagccacacagaagcaagcacaaggaaagt**gcctgatgcaagggcaaagtacagtgtgtaccttcaggc**tgggcacagacactctgaaaagccttggcaggaactc**cctgcaacaaagcagagccctgcagg**caatgccagctccagagccctccctgagagcctcatgggcaaagatgtgcagaacata**tgtttgtcatagccccaaactgagaatgaagcaaacagccatctgaaggaaaacaggcaaataaacgatggc**aggttcatgaaatgcaaacccagacagccagaaggacaacagtgagggttacaggtgactctgtggttgagttcatgacaatgctgagtaattggagtaacaaag**gaaagtccaaaaaatactttc**aatgtgatttcttctaaataaaatttacagccggcaaaatgaactatcttcttaagggataaactttccactaggaaaacta

Uppercase: IGHD2-2

Lowercase: Flanking sequence[1000bp]

Red & Bold & Underline: Stem-loop [15]

Blue: Heptamer[47]

Green: Nonamer [5]

id-TRDD3[D_gene_segment]

caacaacaacaaaattcctactcttctttcatggttaaagtttaataaggtcactttcgatgagcctatcacctatcatattgctttgttttgacagcactagaacagagatcaaaggcaa**acaagaccagtgtcttgt**gagccctgagg**agaggttttgagaaggcaatgactgggtcaaaactct**gaggtcaaacaagattaaaactgaaaatcatccgttgaatttggccagagctgctgcctctgattgctcaggttgtgccttcacaagggcatctggcagagaggagaagtggggctaaaattcaatgcgtgctctgccgccagccaaggccctgatgtggcgttgcatcttccggaaagggaaggccttttataaaccacacaaaagtgcaatctgttcgccaagcttgaacccaaggggctatctctacattcacccagcagggctgccttttctaaatcgcacaaagcaccatatgggctactggcctgaatttagcaactgggaggaggtcaatggaggggacacaattttaaa**tttacatgaaaaccctttgtaaa**ctgtataacactattcaagtgtaaagcaggaagaggtgattcagaagcttagcaggtgacaggagaaattacactgaccacaagccttttctgcctaacccacagcatgtagaatctccatgagacgtttaagtacccgacaaactcctcagtgggaaaccatttgagggcagactggccagaaagcacagatgg**gaatggatgtgattatttctaaaagtccattc**tagtcacaaaccccaaggcagatctagccag**tggggcagaagagcccca**gacagaagcacctgagccagcttgg**cctgacctaactgtcagg**accctttgatcttgctggagcttgacttggagaaaacatctggttctggggattctcaggggccatatagtgtgaaaccgaggggaagtttttgtaaagctc**tgtagcactgtgACTGGGGGATACGcacagtgctaca**aaacctacagagacctgtacaaaaactgcaggggcaaaagtgccatttccctgggatatcctcaccctgggtcccatgcctcaggagacaaacacagc**aagcagcttccctccctgctt**tggggcctggaagggatagcaggaagttgactggaccagggagatgaccacagctgctgacctctcactcactgctgttcttccttgggtgaaactggcatttctacattttcttacagcacatttggggaatacaaaaaggcctttcttaaaaactattcttgtcttgttttcatgttgattctattgcaaaagagagtta**tatgagccacctcata**cggaatttctaaattcaaacctctagagagatttacccaagtgctttgctttgcagtttgggaggatggatttgaagagagattgatttttttgtaggcaatcaccggccacagttgctcattctaaagctgactgctctgtaaatcacccagtgcttcatgccaccctttctcctcttgctgtgccacacgttatct**gcctttaaagcagcagcactggtgtctgtaaaggc**cttaaccctggagtagtcatggagccaagacccacccctttgacagtgccagctttccaacacagagagctgagtatgggtctaggaagtgagagcaatgtaaaacaatagaaagcaacagttcagagcactgcatcaagtgtactgtgctggaaaggtccgccataggaaatatggtcctccatactcctcagacaacagccttccgaaagcaaacctgtccctacctgcagatgattaaccatctatgaaccggctgggtaagcaacaagtgcc**atctttcatggagctgagccttaaagat**cctccagtcctaaagctgacgggaagaaggtaggtgggagcagcgctgaggtttttggaacgtcctcaagtgctgtgacaccgataaactcatctttggaaaaggaacccgtgt

Uppercase: TRDD3

Lowercase: Flanking sequence[1000bp]

Red & Bold & Underline: Stem-loop [11]

Blue: Heptamer[22]

Green: Nonamer [8]

id-IGHD3-9[D_gene_segment]

gaggacggcacagtcagtgaattccagagagaagcaactcagccacactccccaggcagagcccgagagggacgcccacgcacagggaggcagagcccagcacctccgcagccagca**ccacctgtgcacgggccaccaccttgcaggcacagagtgggtgctgagaggaggggcagggacaccaggcagggtgagcaccca**gagaaaactgcagacgcctcacacatccacctcagcctcccctgacctggacctcactggcctgggcctcacttaacctgggcttcacctgaccttggcctcacctgacttggacctcgcctgtcccaagctttacctgacctgggcctcaactcacctgaacgtctcctgacctgggtttaacctgtcctggaactcacctggccttggcttcccctgacctggacctcatctggcctgggcttcacctggcctgggcctcacctgacctggacctcatctggcctggacctcacctggcctggacttcacctggcctgggcttcacctgacctggacctcacctggcctcgggcctcacctgcacct**gctccaggtcttgctggagc**ctgagtagcactgagggtgcagaagctcatccagggttggggaatgactctagaagtctcccacatctgacctttctgggtggaggcagctggtggccctgggaatataaaaatctccagaatgatgactctgtgatttgtgggcaacttatgaacccgaaaggacatggccatggggtgggtagggacatagggacagatgccag**cctgaggtggagcctcagg**acacaggtgggcac**ggacactatccacataagcgagggatagacccgagtgtcc**ccacagcagacctgagagcgctgggcccacagcctcccctcagagccctgctgcctcctccggtcagccctggacatcccaggtttccccaggcctgccggtaggtttagaatgagg**tctgtgtcactgtgGTATTACGATATTTTGACTGGTTATTATAACcacagtgtcacaga**gtccatcaaaaa**cccatgcctggaagcttcccgccacagccctccccatggg**gccctgctgcctcctcaggtcagccccggacatcccgggtttccccaggctgggcggtaggtttggggtgagg**tctgtgtcactgtggtattactatggttcggggagttattataaccacagtgtcacaga**gtccatcaaaaacccatccctgggagcctcccgccacagccctccctgcaggggaccggtacgtgccatgttaggattttgatcgaggagacag**caccatgggtatggtg**gctaccacagcagtgcagcctgtgacccaaacccgcagggcagcaggcacgatggacaggcccgtgactgaccacgctggg**ctccagcctgccagccctggag**atcatgaaacagatggccaaggtcaccctacaggtcatccagatctggctccgaggggtctgcatcgctgctgccctcccaacgccagtccaaatgggacagggacggcctcacagcaccatctgctgccatcaggccagcgatcccagaagcccctccctcaaggctgggccacatgtgtggacactgagagccctcatgtctgagtaggggcaccaggaggg**aggggctggccctgtgcactgtccctgcccct**gtggtccctggcctgcctggccctgacacctgagcctctcctgggtcatttccaagacagaagacattcctgg**ggacagccggagctgggcgtcgctcatcctgcccggccgtcc**tgagtcctgctcatttccagacctcaccggggaagccaacagaggactcgcctcccacattcagagacaaagaaccttccagaaatccctgcctctctccccagtggacaccctcttccaggacagtcctcagtggcatc**acagcggcctgagatccccaggacgcagcaccgctgt**caataggggccccaaatgcctggaccagggcctgcgtggga

Uppercase: IGHD3-9

Lowercase: Flanking sequence[1000bp]

Red & Bold & Underline: Stem-loop [13]

Blue: Heptamer[52]

Green: Nonamer [5]

id-IGHD4-23-2[D_gene_segment]

cccatgacccaaacacacggggcagcagaaacaatggacaggcccacaagtgaccatgatgggctccagcccaccagccccagagaccatgaaacagatggccaaggtcaccctacaggtcatccagatctggctccaaggggtctgcatcgctgctgccctcccaacgccaaac**cagatggagacagggccggccccatagcaccatctgctgccgtccacccagcag**tcccggaagcccctccctgaacgctgggccacgtgtgtgaaccctgcg**agccccccatgtcagagtaggggcagcaggagggcggggct**ggccctgtgcactgtcactgcccctgtggtccctggcctgcctggccctgacacctgagcctctcctgggtcatttccaagacattcccag**ggacagccggagctgggagtcgctcatcctgcctggctgtcctgagtcctgctcatttccagacctcaccagggaagccaacagaggactca**cctcacacagtcagagacaatgaaccttccagaaatccctgtttctctccccagtg**agagaaaccctcttccagggtttctct**tctctcccaccctcttccaggacagtcctcagcagcatcacagcgggaacgcacatctggatcaggacggcccccagaacacgcgatggcccatggggacagcccagcccttcccagacccctaaaaggtatccccaccttgcacctgccccagggctcaaactc**caggaggcctgactcctgcacaccctcctg**ccagatatcacctcagc**cccctcctggagggg**acaggagcccgggagggtgagtcaga**cccacctgccctcaatggcaggcggg**gaagattcagaaaggcctgagatccccaggacgcagcaccactgtcaatgggggcc**ccagacgcctggaccagggcctgtgtgggaaaggcctctgg**ccacactcaggggctttttgtgaagggccctcctgctgtgTGACTACGGTGGTAACTCCcacagtgatgaaaccagcagcaaaaactgaccggactcgcagggtttatgcacacttctcggctcggagctctccaggagcacaagagcc**aggcccgagggtttgtgcccagaccctcggcct**ctagggacacccgggccatcttagccgatgggctgatgccctgcacaccgtgtgctgccaaacaggggcttcagagggctctgaggtgacttcactcatgaccacaggtgccctggtcccttcactgccagctgcaccagaccctgttccgagagat**gccccagttccaaaagccaattcctggggc**cgggaattactgtagacaccagcctcattccagtacctcctgccaattgcctggattcccat**cctggctggaatcaagagggcagcatccgccagg**ctcccaacaggcaggactcccacacaccctcttctgagaggccgctgtgttccgcagggccaggccgcagacagttcccctcacctgcccatgtagaaacacctgccattgtcgtcccc**acctggcaaagaccacttgtggagcccccagccccaggt**acagctgtagagagagtcctcgaggcccctaagaaggagccatgcccagttctgctgggaccctcggccaggccgacaggagtggacgctggagctgg**gcccacactgggccacataggagctcaccagtgagggc**aggagagcacatgccggggagcacc**cagcctcctgctgaccagagacccgtcccagagcccaggaggctg**cagaggcctctccagggggacacagggcatgtctggtccctgagca**gcccccaggctctctagcactgggggc**ccctggcaca**gctgtctggaccctccctgttccctgggaagctcctcctgacagc**cccgcctccagttccaggtgtggttattgtcagggggtgccaggcc**gtggtagagatggctacaattaccac**agtggtgccgcccatagcagcaaccaggcc

Uppercase: IGHD4-23-2

Lowercase: Flanking sequence[1000bp]

Red & Bold & Underline: Stem-loop [19]

Blue: Heptamer[34]

Green: Nonamer [5]

id-IGHD5-18[D_gene_segment]

acactc**aggggctttttgtgaaggcccct**cct**actgtgtgactacggtgactaccacagt**gatgaaactagcagcaaaaactggccggacacccagggaccatgcacacttctcagcttggagctctccaggaccagaagagtcaggtct**gagggtttgtagccagaccctc**ggcctctagggacaccctggccatcacagcggatgggctggtgccccacatgccatctgctccaaacaggggcttcagagggctctgaggtgacttcactcatgaccacaggtgccctggccccttccccgccagctacaccgaaccctgtcccaacagct**gccccagttccaacagccaattcctggggc**ccagaattgctgtagacaccagcctcgttccagcacctcctgccaattgcctggattcacat**cctggctggaatcaagagggcagcatccgccagg**ctcccaacaggcaggactcccgcacaccctcctctgagaggccgctgtgttccg**cagggccaggccctg**gacagttcccctcacctgccactagagaaacacctgccattgtcgtcccc**acctggaaaagaccactcgtggagcccccagccccaggt**acagctgtagagagactccccgagggatctaagaaggagccatgcgcagttctgccgggaccctcggccaggccgacaggagtggacactggagctgg**gcccacactgggccacataggagctcaccagtgagggc**aggagagcacatgccggggagcacc**cagcctcctgctgaccagaggcccgtcccagagcccaggaggctg**cagaggcctctccagggggacactgtgcatgtctggtccctgagca**gccccccacgtccccagtcctgggggc**ccctggcacagctgtctggaccctccctcttccctgggaagctcctc**ctgacagccccgcctccagttccaggtgtggttattgtcagggggtgtcagactgtgGTGGATACAGCTATGGTTACcacagt**gg**tgctgcccatagcagca**accaggccaagtagacaggcccctgctgtgc**agccccaggcctccagctcacctgcttctcctggggct**ctcaaggtcactgttgtctgtactctgccctctgtggggagggttccctcagtgggaggtctgttctcaacatcccagggcctcatgtctgcacggaaggccaatggatgggcaacctcacatgccgcggctaagatagggtgggcagcctggcgggggacagtacatactgctggggtgtctgtcactgtgcctagtggggcactggctcccaaacaacgcagtcctcgccaaaatccccacagcctcccctgctaggggctggcctgatctcctgcagtcctaggaggctgctgacctccagaatgtctccgtccccagttccagggcgagagcaga**tcccaggccggctgcagactggga**ggccaccccctccttcccagggttcactggaggtgaccaaggtaggaaatggccttaacaca**gggatgactgcgccatccc**cca**acagagtcagccccctcctgctctgt**accccgcaccccccaggccagtccacgaaaa**ccagggccccacatcagagtcactgcctggcccggccctgg**ggcggacccctcag**cccccaccctgtctagaggacttggggggacaggacacaggccctctccttatggttccccc**acctgcctccggccgggacccttggggtgtggacagaaaggacacctgcctaattggcccccaggaacccagaacttctctccagggaccccagcccgagcacccccttacccaggacccagccctgcccctcctcccctctgctctcctctcatcaccccatgggaatccggtatccccaggaagccatcaggaagggctgaaggaggaagcggggccgtgcaccaccgggc**aggaggctccgtcttcgtgaacccagggaagtgccagcctcct**agagggtat

Uppercase: IGHD5-18

Lowercase: Flanking sequence[1000bp]

Red & Bold & Underline: Stem-loop [21]

Blue: Heptamer[38]

Green: Nonamer [4]

id-IGHD5-5-2[D_gene_segment]

cacactcaggggctttttgtgaagggtcctcct**actgtgtgactacagtaactaccacagt**gatgaacccagcagcaaaaactgaccggactcccaaggtttatgcacacttctccgctcagagctctccaggatcagaagagccgggcccaagggtttctgcccagaccctcggcctctagggacatcttggccatgacagcccatgggctggtgccccacacatcgtctgccttcaaacaagggcttcagagggctctgaggtgacctcactgatgaccacaggtgccctggccccttccccgccagctgcaccagaccccgtcctgacagatgccccgattccaacagcca**attcctggggccaggaat**cgctgtagacaccagcctccttccaacacctcttgccaattgcctg**gattcccatcccggttggaatc**aagaggacagcatcccccaggctcccaacaggcaggactcccacaccctcctctgagaggccgctgtgttccgtagggccaggctgcagacagtccccctcacctgccactagacaaatgcctgctgtagatgtcccc**acctggaaaagaccactcatggagcccccagccccaggt**acagccatagagagagtctctgaggcccctaagaagtagccatgcccagttctgccgggaccctcggccaggctgacaggagtggacgctggagctgg**gcccacactgggccacataggagctcaccagtgagggc**aggagagcacatgccggggagcacc**cagcctcctgctgaccagaggcccgtcccagagcccaggaggctg**cagaggcctctccagggggacactgtgcatgtctggtccctgagca**gccccccatgtccccagtcctgggggc**ccctggcacagctgtctggaccctctctattccctgggaagctcctc**ctgacagccccgcctccagttccaggtgtggttattgtcagggggtgtcagactgtgGTGGATACAGCTATGGTTACcacagt**gg**tgctgcccatagcagca**accaggccaagtagacaggcccctgctgtgc**agccccaggcctccagctcacctgcttctcctggggct**ctcaaggctgctgttttctgcact**ctcccctctgtggggag**ggttccctcagtgggagatctgttctcaacatcccagggcctcattcctgcaaggaaggccaatggatgggcaacctcacatgccgcggctaagatagggtgggcagcctggcggggacaggacatcctgctggggtatctgtcactgtgcctagtggggcactggctcccaaacaacgcagtcctcgccaaaatccccacggcctcccccgctaggggctggcctgatctcctgcagtcctaggaggctgctgacctccagaatggctccgtccccagttccagggcgagagcaga**tcccaggccggctgcagactggga**ggccaccccctccttcccagggttcactgcaggtgaccagggcaggaa**atggcctgaacacagggataaccgggccat**ccccca**acagagtccaccccctcctgctctgt**accccgcacccccaaggccagcccatgacatccgacaaccccacaccagagtcactgcccggtgctgccctagggaggacccctcagcccccac**cctgtctagaggactggggaggacagg**acacgccctctccttatggttcccccacctggctctggctgggacccttggggtgtggacagaaaggacgcttgcctgattggcccccaggagcccagaacttctctccagggaccccagcccgagcacccccttacccaggacccagccctgcccctcc**tcccatctgctctcctctcatcaccccatggga**atccagaatccccaggaagccatcaggaagggctgagggaggaagtggggccactgcaccaccaggc**aggaggctccgtctttgtgaacccagggaggtgccagcctcct**agagggtatg

Uppercase: IGHD5-5-2

Lowercase: Flanking sequence[1000bp]

Red & Bold & Underline: Stem-loop [18]

Blue: Heptamer[34]

Green: Nonamer [7]

id-IGHD4-17-2[D_gene_segment]

ctaccacagcagtgcagcccatgacccaaaca**cacagggcagcaggcacaatggacaggcctgtg**agtgaccatgctgggctccagcccgccagccccggagaccatgaaacagatggccaaggtc**accccacagttcagccagacatggctccgtggggt**ctgcatcgctgctgccctctaacaccagccc**agatggggacaaggccaaccccacattaccatct**cctgctgtccacccagtggtcccagaagcccctccctcatggctgagccacatgtgtgaaccctgagagcaccccatgtcagagtaggggcagcagaagggcggggctggccctgtgcactgtccctgcacccatggtccctcgcctgcctggccctgacacctgagcc**tcttctgagtcatttctaagatagaaga**cattcccgg**ggacagccggagctgggcgtcgctcatcccgcccggccgtcctgagtcctgcttgtttccagacctcaccagggaagccaacagaggactca**cctcacacagtca**gagacaaagaaccttccagaaatccctgtctc**actccccagtgggcaccttcttccaggacattcctcggtcgcatcacagcaggcacccacatctggatcaggacggcccccagaacacaagatggcccatggggacagccccacaacccaggccttcccagacccctaaaaggcgtcccaccccctgcacctgccccagggctaaaaatccaggaggcttgactcccgca**taccctccagccagacatcacctcagccccctcctggagggga**caggagcccgggagggtgagtcaga**cccacctgccctcgatggcaggcggg**gaagattcagaaaggcctgagatccccaggacgcagcaccactgtcaatgggggccccagacgcctggaccagggcctgcgtgggaaaggccgctgggcacactc**aggggctttttgtgaaggcccct**cct**actgtgTGACTACGGTGACTACcacagt**gatgaaactagcagcaaaaactggccggacacccagggaccatgcacacttctcagcttggagctctccaggaccagaagagtcaggtct**gagggtttgtagccagaccctc**ggcctctagggacaccctggccatcacagcggatgggctggtgccccacatgccatctgctccaaacaggggcttcagagggctctgaggtgacttcactcatgaccacaggtgccctggccccttccccgccagctacaccgaaccctgtcccaacagct**gccccagttccaacagccaattcctggggc**ccagaattgctgtagacaccagcctcgttccagcacctcctgccaattgcctggattcacat**cctggctggaatcaagagggcagcatccgccagg**ctcccaacaggcaggactcccgcacaccctcctctgagaggccgctgtgttccg**cagggccaggccctg**gacagttcccctcacctgccactagagaaacacctgccattgtcgtcccc**acctggaaaagaccactcgtggagcccccagccccaggt**acagctgtagagagactccccgagggatctaagaaggagccatgcgcagttctgccgggaccctcggccaggccgacaggagtggacactggagctgg**gcccacactgggccacataggagctcaccagtgagggc**aggagagcacatgccggggagcacc**cagcctcctgctgaccagaggcccgtcccagagcccaggaggctg**cagaggcctctccagggggacactgtgcatgtctggtccctgagca**gccccccacgtccccagtcctgggggc**ccctggcacagctgtctggaccctccctcttccctgggaagctcctc**ctgacagccccgcctccagttccaggtgtggttattgtcagggggtgtcagactgtggtggatacagctatggttaccacagt**ggtgctgcccatagcagcaaccaggcca

Uppercase: IGHD4-17-2

Lowercase: Flanking sequence[1000bp]

Red & Bold & Underline: Stem-loop [21]

Blue: Heptamer[37]

Green: Nonamer [5]

id-IGHD2-21[D_gene_segment]

ctgtgcccaccccctaacccctcctgcccacaacttgagttcttcctctcctggagcccttgagccatggcactgaccctacactcccacccacacactgcccatgccatcaccttcctcctggacactctgaccccgctcccctccctctcagacccggccctggtatttccaggacaaaggctcacccaagtcttccccatgcaggcccttgccctcactgcctggttacacgggagcct**cctgtgcgcagaagcagggagctcagctcttccacagg**cag**aaggcactgaaagaaatcggcctccagtgccttgacacacgtccgcctgtgtc**tctcactgcctgcacctgc**agggaggctccgcactccct**ctaaagatgagggatccaggcagcaacatcacgggagaatgc**agggctcccagacagcccagccct**ctc**gcaggcctctcctgggaagagacctgc**agccaccactgaacagccacgg**aggtcgctggatagtaaccgagtcagtgaccgacct**ggagggcaggggagcagtgaaccggagcccataccatagggacagagaccagccgctaacatcccgagcccctcactggcggccccagaacaccccgtggaaagagaacagacccacagtcccacctggaacagggcagacactgctgagcccccagcacc**agccccaagaaacactaggcaacagcatcagagggggct**cctgagaaagagaggaggggaggtctccttcaccatcaaatgcttcccttgaccaaaaacagggtccacgcaactcccccaggacaaaggag**gagccccctgtacagcactgggctcagagtcctctctgag**acaggctcagtttcagacaacaacccgctggaatgcacagtctcagcaggagagccaggccagagccagcaag**aggagactcggtgacaccagtctcct**gtagggacaggaggattttgtg**ggggttcgtgtcactgtgAGCATATTGTGGTGGTGACTGCTATTCCcacagtgacacaacccc**attcctaaagccctactgcaaacgcacccactcctgggactgaggggctgggggagcatctgggaagtatggcctaggggtgtccatcaatgcccaaaatgcaccagactctccccaagacatcaccccaccagccagtgagcagagtaaacagaaaatgagaagcagctgggaag**cttgcacaggccccaaggaaagagctttggcaggtgtgcaag**aggggatgtgggcagagcctcagcagggccttttgctgtttctgctttcctgtgcagagagttccataaactggtattcaagatcaatggctgggagtgagcccaggag**gacagtgtgggaagagcacagggaaggaggagcagccgctatcctacactgtc**atcttttgaaagtttgccctgtgcccacaatgctgcatcatgggatgcttaacagctgatgtagacacagctaaagagagaatcagtgaaatgcatttgcagcacagatctgaataaatcctccagaatgtggagcagcacagaagcaagcacacagaaag**tgcctgatgccaaggcaaagttcagtgggcaccttcaggca**ttgctgctgggcacagacactctgaaaagcactggcaggaac**tgcctgtgacaaagcagaaccctcaggca**atgccagccctagagcccttcctgagaacctcatgggcaaagatgtgcagaacagctgtttgtcat**agccccaaactatggggct**ggacaaagcaaacgtccatctgaaggagaacagacaaataaacgatggcaggttcatgaaatacaaactaggacagccagaggacaacagtagagagctacaggcggctttgcggttgagttcatgacaatgctgagtaattggagtaacagaggaaagcccaaaaaatacttttaatgtgatttcttctaaataaaatttacacccggcaaaatgaactatcttcttaagggataaactttccc

Uppercase: IGHD2-21

Lowercase: Flanking sequence[1000bp]

Red & Bold & Underline: Stem-loop [17]

Blue: Heptamer[39]

Green: Nonamer [5]

id-IGHD7-27-2[D_gene_segment]

gtgtcctccaacgacaggtcccagcctcccagcctttgccttgcctgttcctctccctggaactctgccccgacacagaccctccccagcaagccc**gcaggggcacctcccctgc**ccccagacaccctgtgcccgtcagttcatccccagcagaggccctcaccaggcacacccccatgctcacacctggccgcagg**cctcagcctccctgagg**gccccacccagcccgcgtctggccagtggtgcgtgcaa**agcccctcacccagactcggcggaaggcagccagtgcaggcctggggaggggct**ctccttagaccaccttgcac**cttccctggcacccaccatgggaag**ag**ctgagactcactgaggaccagctgaggctcag**agaagggacccagcactggtggacacgcagggagcccacgccagggcgccgtggtgagt**gaggcccagtgccacccactgaggcctc**ccgttcagtgggacgacggtgaacaggtggaaccaaccaggcaacccccgccgggccccacagacgggatcaga**gcaggaaaggcttcctgc**ccctgcaggccagcgaggagccc**tggcgggggccatggccctccaggcgaggaggctcccctggccaccgcca**cccgggcctctctgctgctgggaaaacaagtcagaaagcaagtggatgagaggtggcgtgacagacccagcttcagatctgctctaatttacaaaagaaaaggaaaaacacacttggcagccttcagcactctaatgattcttaacagcagcaaattattggcacaagactccagagtgactggcagggttgagggctgggg**tctcccgcgtgttttggggctaacagcggaagggaga**gcactggcaaaggtgctgggggcccctggacccgacccgccctggagaccgcagccacatcagcccccagccccacaggccccctaccagccgcagggttttggctgagctgagaac**cactgtgCTAACTGGGGAcacagtg**attggcagct**ctacaaaaaccatgctcccccgggaccccgggctgtgggtttctgtag**cccctggctcagggctgactcaccgtggctgaatacttccagcactggggccagggcaccctggtcaccgtctcctcaggtgagtctgctgtctggggatagcggggagccaggtgtactgggccaggcaagggctttggcttcagacttggggacaggtgctcagcaaaggaggtcggcaggagggcggagggtgtgtttttgtatgggagaagcaggagggcagaggctgtgctactggtacttcgatctctggggccgtggcaccctggtcactgtctcctcaggtgagtcccactgcagccccctcccagtcttctctgtccaggcac**caggccaggtatctggggtctgcagccggcctgggtctggcctg**aggccacaccagctgccatccctggggtctccgccatgggctgcatgccagagccctgctgtcacttagccctggggcca**gctggagcccccaaggacaggcagggaccccgctgggcttcagc**cccgtcagggaccctccacaggtagcaagcaggccgagggcagggacgggaaggagaagttgtgggcagagcctgggctggggctgggcgctggctgttcatgtgccggggaccaggcctgcgctttagtgtggctaca**agtgcttggagcact**gggg**ccagggcagcccggccaccgtctccctgg**gaacgtcacccctccctgcctgggtctcagcccgggggtctgtgtggctggggacagggacgccggctgcctctgctctgtgcttgggccatgtgacccattcgagtg**tcctgcacgggcacaggtttatgtctgggcaggaacagggactgtgtccctgt**gtgatgcttttgatatctggggccaagggacaatggtcaccgtctcttcaggtaagatggctttccttctgcctcctttctctgggc

Uppercase: IGHD7-27-2

Lowercase: Flanking sequence[1000bp]

Red & Bold & Underline: Stem-loop [17]

Blue: Heptamer[34]

Green: Nonamer [6]

id-IGHD1-7-2[D_gene_segment]

ccatcccctccaggctctgggaggagaagcaggatgggagaatctgtgcgggaccctctcacagtggaatacctccacagc**ggctcaggcaagacccaaaagcccctcagtgagcc**ctccactgcagtcctgggc**ctgggtagcagcccctcccacagaggatgaacccag**caccccgaggatg**tcctgccagggggagctcagagccatgaaggagcagga**tatgggacccccgatacaggcacagacctcagctccattcaggactgccacgtcctgccctgggaggaacccctttctctagtccctgcaggc**caggaggcagctgactcctg**acttggacgcctattccagacaccagacagaggggcaggccccccagaaccagggatgaggacgccccgtcaaggccagaaaagaccaagttgtgctgagcccagcaagggaaggtccccaaacaaaccaggaagtttctgaaggtgtct**gtgtcacagtggagtatagcagctcgtcccacagtgacac**tcgccaggccagaaaccccatcccaagtcagcggaatgcagagagagcagggaggacatgtttaggatctgaggccgcacctgacacccaggccagcagacgtctcctgtc**catggcaccctgccatg**tcctgcatttctggaagaacaagggcaggctgaagggggtccaggaccaggagatgggtcccctctacccagagaaggagcca**ggcaggacacaagccccctccccattgaggctgacctgcccagagggtcctgggc**ccaccc**cacacaccggggcggaatgtgtg**caggcctcggtctctgtgggtgttccgcta**gctggggctcacagtgctcaccccacacctaaaacgagccacagc**ctcagagcccctgaaggagaccccgcccacaagcccagcccccacccaggaggccccagagcacagggcgccccgtcggattctgaacagccccgag**tcacagtgGGTATAACTGGAACTACcactgtga**gaaaagcttcgtccaaaacggtctcctggccacagtcggaggccccgccagagaggggagcagccaccccaaacccatgttctgcc**ggctcccatgaccccgtgcacctggagcc**ccacagtgtccccactggatgggaggacaagggccgggggctccggcgggtcggggcaggggcttgatggcttccttctgccgtgg**ctccagtgcccctggctggag**ttgacccttctgacaagtgtcctcagagagtcagggatcagtggcacctcccaacatcaaccccacgcagcccaggcacaaaccccacat**ccagggccaactccaggaacagagacaccccaataccctgg**gggaccccaaccctgatgactcccgtcccatctctgtccctcacttggggcctgctgcggggcgagcacttgggagcaaactcaggcttaggggacacca**ctgtgggcctgacctcgagcaggccacag**acccttc**cctcctgccctggtgcagcacagactttggggtctgggcagggagg**aacttctggcaggtcaccaagcacagagcccccaggctgaggtggccccagggggaaccccagcaggtggcccactacccttcctcccagctggaccccatgtcttccccaagataggggtgccatccaaggcaggtcctccat**ggagcccccttcaggctcc**tctccagaccccactgggcctcagtccccactctaggaatgcagccaccacgggcacaccaggcagcccaggcccagc**caccctgcagtgcccaagcccacaccctggaggagagcagggtg**cgtctgggaggggctgggctccccacccccacc**cccacctgcacaccccacccacccttgcccgggccccctgcaggaggg**tcagagcccccatgggatatggacttagggtctcactcacgcacctcccctcctgggagaaggggtctcatgcccagatccccccagcagc

Uppercase: IGHD1-7-2

Lowercase: Flanking sequence[1000bp]

Red & Bold & Underline: Stem-loop [19]

Blue: Heptamer[33]

Green: Nonamer [7]

id-IGHD6-6-2[D_gene_segment]

acccagccctgcccctcc**tcccatctgctctcctctcatcaccccatggga**atccagaatccccaggaagccatcaggaagggctgagggaggaagtggggccactgcaccaccaggc**aggaggctccgtctttgtgaacccagggaggtgccagcctcct**agagggtatggtccac**cctgcctatggctcccacagtggcagg**ctgcagggaaggaccagggacggtgtggggga**gggctcagggccccgcgggtgctccatcttggatgagccc**atctctctcacccacggactcacccacctcctctccaccctggccacacgtcgtccacaccatcctaagtcccacctacaccagagccggcacagccagtgcagacagaggctggggtgcaggggggccgccagggcagctttggggagggaaggatggagga**aggggagttcagtgaagaggcccccctcccctgggtccaggatcctcctctgggaccc**ccgga**tcccatcccctccaggctctgggaggagaagcaggatggga**gaatctgtgcgggaccctctcacagtggaatacctccacagc**ggctcaggcaagacccaaaagcccctcagtgagcc**ctccactgcagtcctgggc**ctgggtagcagcccctcccacagaggatgaacccag**caccccgaggatg**tcctgccagggggagctcagagccatgaaggagcagga**tatgggacccccgatacaggcacagacctcagctccattcaggactgccacgtcctgccctgggaggaacccctttctctagtccctgcaggc**caggaggcagctgactcctg**acttggacgcctattccagacaccagacagaggggcaggccccccagaaccagggatgaggacgccccgtcaaggccagaaaagaccaagttgtgctgagcccagcaagggaaggtccccaaacaaaccaggaagtttctgaaggtgtct**gtgtcacagtgGAGTATAGCAGCTCGTCCcacagtgacac**tcgccaggccagaaaccccatcccaagtcagcggaatgcagagagagcagggaggacatgtttaggatctgaggccgcacctgacacccaggccagcagacgtctcctgtc**catggcaccctgccatg**tcctgcatttctggaagaacaagggcaggctgaagggggtccaggaccaggagatgggtcccctctacccagagaaggagcca**ggcaggacacaagccccctccccattgaggctgacctgcccagagggtcctgggc**ccaccc**cacacaccggggcggaatgtgtg**caggcctcggtctctgtgggtgttccgcta**gctggggctcacagtgctcaccccacacctaaaacgagccacagc**ctcagagcccctgaaggagaccccgcccacaagcccagcccccacccaggaggccccagagcacagggcgccccgtcggattctgaacagccccgag**tcacagtgggtataactggaactaccactgtga**gaaaagcttcgtccaaaacggtctcctggccacagtcggaggccccgccagagaggggagcagccaccccaaacccatgttctgcc**ggctcccatgaccccgtgcacctggagcc**ccacagtgtccccactggatgggaggacaagggccgggggctccggcgggtcggggcaggggcttgatggcttccttctgccgtgg**ctccagtgcccctggctggag**ttgacccttctgacaagtgtcctcagagagtcagggatcagtggcacctcccaacatcaaccccacgcagcccaggcacaaaccccacat**ccagggccaactccaggaacagagacaccccaataccctgg**gggaccccaaccctgatgactcccgtcccatctctgtccctcacttggggcctgctgcggggcgagcacttgggagcaaac**tcaggcttaggggacaccactgtgggcctga**cctcgagcaggccacagacccttc

Uppercase: IGHD6-6-2

Lowercase: Flanking sequence[1000bp]

Red & Bold & Underline: Stem-loop [22]

Blue: Heptamer[38]

Green: Nonamer [7]

id-IGHD6-13[D_gene_segment]

cagacagtgggcccagagcactgtgagtaccccggcagtac**ctggctgcagggatcagccag**agatgccaaaccctgagtgaccagcctacaggaggatccggccccacccaggccactcgattaatgctcaaccccctgccctggagacctcttccagtaccaccagcagctcagcttctcagggcctcatccctgcaaggaaggtcaagggctgggcctgccagaaacacag**caccctccctagccctggctaagacagggtg**ggcagacggctgtggacgggacatattgctggggcatttctcactgtcacttctgggtggtagctctgacaaaaacgcagaccctgccaaaatccccactgcctcccgctaggggctgg**cctggaatcctgctgtcctaggaggctgctgacctccagg**atggctccgtccccagttccagggcgagagcaga**tcccaggcaggctgtaggctggga**ggccacc**cctgcccttgccggggttgaatgcaggtgcccaaggcagg**aaat**ggcatgagcacagggatgaccgggacatgcc**ccaccagagtgcgccccttcctgctctgcaccctgcaccccccaggccagcccacgacgtccaacaactgggc**ctgggtggcagccccacccag**a**caggacagacccagcaccctgaggaggtcctg**ccagggggagctaagagccatgaaggagcaagatatggggcccccgatacaggcacagatgtcagctccatccaggaccacccagcccacaccctgagaggaacgtctgtctccagcctctgcaggtcgggaggcagctgacccctgacttggacccctattccagacaccagacagaggcgcaggccccccagaaccagggttgagggacgccccgtcaaagccagacaaaaccaaggggtgttgagcccagcaagggaaggcccccaa**acagaccaggaggtttctgaaggtgtctgtgtcacagtgGGGTATAGCAGCAGCTGGTACcacagtgacac**tcacccagccagaaaccccattccaagtcagcggaagcagagagagcagggaggacacgtttaggatctgagactgcacctgacacccaggccagcagacgtctcccctccagggcaccccaccctgtcctgcatttctgcaagatcaggggcggcctgagggggggtctagggtgaggagatgggtcccctgtacaccaaggaggagtta**ggcaggtcccgagcactctccccattgaggctgacctgcccagagagtcctgggc**ccaccc**cacacaccggggcggaatgtgtg**caggcctcggtctctgtgggtgttccgcta**gctggggctcacagtgctcaccccacacctaaaatgagccacagc**ctccggagcccccgcaggagaccccgcccacaagcccagcccccacccaggaggccccagagctcagggcgccccgtcggattccgaacagccccgagtcacagcgggtataaccggaacca**ccactgtcagaatagctacgtcaaaaactgtccagtgg**ccactgccggaggccccgccagagagggcagcagccactctgatcccatgtcctgcc**ggctcccatgacccccagcacgcggagcc**ccacagtg**tccccactggatgggaggacaagagctgggga**ttccggcgggtcggggcaggggcttgatcgcatccttctgccgtgg**ctccagtgcccctggctggag**ttgacccttctgacaagtgtcctcagagagacaggcatcaccggcgcctcccaacatcaaccccaggcagcacaggcacaaaccccacatccagagccaactccaggagcagagacaccccaataccctgggggaccccgaccctgatgacttcccactggaattcgccgtagagtccaccaggaccaaagaccctgcctctgcctctgtccctcactcaggacctgctgccgggcgaggccttgggagcagacttgggcttaggg

Uppercase: IGHD6-13

Lowercase: Flanking sequence[1000bp]

Red & Bold & Underline: Stem-loop [18]

Blue: Heptamer[28]

Green: Nonamer [8]

id-IGHD1-20-2[D_gene_segment]

tcccatcccctcctggctctgggaggagaagcaggatgggagaatctgtgcgggaccctctcacagtggaatatccccacagc**ggctcaggccagacccaaaagcccctcagtgagcc**ctccactgcagtcctgggc**ctgggtagcagcccctcccacagaggacagacccag**caccccgaagaag**tcctgccagggggagctcagagccatgaaagagcagga**tatggggtccccgatacaggcacagacctcagctccatccaggcccaccgggacccaccatgggaggaacacctgtctccgggttgtgaggtagctgg**cctctgtctcggaccccactccagacaccagacagagg**ggcaggccccccaaaaccagggttgagggatgatccgtcaaggcagacaagaccaaggggcactgaccccagcaagggaaggctcccaa**acagacgaggaggtttctgaagctgtctgt**atcacagtggggtatagcagtggctggtaccacagtgacactcgccaggccagaaaccccgtcccaagtcagcggaagcagagagagcagggaggacacgtttaggatctgaggccgcacctgacacccagggcagcagacgtctcccctccagggcaccctccaccgtcctgcgtttcttcaagaataggggcggcct**gagggggtccagggccaggcgataggtcccctc**taccccaaggaggagcca**ggcaggacccgagcaccgtccccattgaggctgacctgcccagacgggcctgggc**ccaccc**cacacaccggggcggaatgtgtg**ca**ggccccagtctctgtgggtgttccgctagctggggcc**cccagtgctcaccccacacctaaagcgagccccagcctccagagccccctaagcattccccgcccagcagcccagcccctgcccccacccaggaggccccagagctcagggcgcctggtcggattctgaacagccccgagtcacagtgGGTATAACTGGAACGACcaccgtgagaaaaactgtgtccaaaactgactcctggcagcagtcggaggccccgccagagaggggagcagccggcctgaacccatgtcctgccggttcccatgacccccagcacccagagccccacggtgtccccgttggataatgaggacaagggctgggggctccggtggtttgcggcagggacttgatcacatccttctgctgtggccccattgcctctggctggagttgaccc**ttctgacaagtgtcctcagaa**agacagggatcaccggcacctcccaatatcaaccccaggcagcacagacacaaaccccacat**ccagagccaactccaggagcagagacaccccaacactctgg**gggaccccaaccgtgataactccccactggaatccgccccagagtctaccaggaccaa**aggccctgccctgtctctgtccctcactcagggcct**cctgcagggcgagcgcttgggagcagactcggtcttaggggacaccactgtgggccccaactttgatgaggccactgacccttccttcctttcctggggcagcacagactttggggtctgggcagggaagaactactggctggtggccaatcacagagcccccaggccgag**gtggccccaagaaggccctcaggaggtggccac**tccacttcctcccagctggaccccaggtcctccccaagataggggtgccatccaaggcaggtcctccatggagcccccttcagactcctcccgggaccccactggacctcagtccctgctctgggaatgcagccaccacaagcacaccaggaagcccaggcccagc**caccctgcagtgggcaagcccacactctggagcagagcagggtg**cgtct**gggaggggctaacctccc**caccccccaccccccatctgcacacagccacctaccactgcccagaccctctgcaggagggccaagccaccatggggtatggacttagggtctcactcacgtgcctc

Uppercase: IGHD1-20-2

Lowercase: Flanking sequence[1000bp]

Red & Bold & Underline: Stem-loop [16]

Blue: Heptamer[33]

Green: Nonamer [8]

id-IGHD5-18-2[D_gene_segment]

acactc**aggggctttttgtgaaggcccct**cct**actgtgtgactacggtgactaccacagt**gatgaaactagcagcaaaaactggccggacacccagggaccatgcacacttctcagcttggagctctccaggaccagaagagtcaggtct**gagggtttgtagccagaccctc**ggcctctagggacaccctggccatcacagcggatgggctggtgccccacatgccatctgctccaaacaggggcttcagagggctctgaggtgacttcactcatgaccacaggtgccctggccccttccccgccagctacaccgaaccctgtcccaacagct**gccccagttccaacagccaattcctggggc**ccagaattgctgtagacaccagcctcgttccagcacctcctgccaattgcctggattcacat**cctggctggaatcaagagggcagcatccgccagg**ctcccaacaggcaggactcccgcacaccctcctctgagaggccgctgtgttccg**cagggccaggccctg**gacagttcccctcacctgccactagagaaacacctgccattgtcgtcccc**acctggaaaagaccactcgtggagcccccagccccaggt**acagctgtagagagactccccgagggatctaagaaggagccatgcgcagttctgccgggaccctcggccaggccgacaggagtggacactggagctgg**gcccacactgggccacataggagctcaccagtgagggc**aggagagcacatgccggggagcacc**cagcctcctgctgaccagaggcccgtcccagagcccaggaggctg**cagaggcctctccagggggacactgtgcatgtctggtccctgagca**gccccccacgtccccagtcctgggggc**ccctggcacagctgtctggaccctccctcttccctgggaagctcctc**ctgacagccccgcctccagttccaggtgtggttattgtcagggggtgtcagactgtgGTGGATACAGCTATGGTTACcacagt**gg**tgctgcccatagcagca**accaggccaagtagacaggcccctgctgtgc**agccccaggcctccagctcacctgcttctcctggggct**ctcaaggtcactgttgtctgtactctgccctctgtggggagggttccctcagtgggaggtctgttctcaacatcccagggcctcatgtctgcacggaaggccaatggatgggcaacctcacatgccgcggctaagatagggtgggcagcctggcgggggacagtacatactgctggggtgtctgtcactgtgcctagtggggcactggctcccaaacaacgcagtcctcgccaaaatccccacagcctcccctgctaggggctggcctgatctcctgcagtcctaggaggctgctgacctccagaatgtctccgtccccagttccagggcgagagcaga**tcccaggccggctgcagactggga**ggccaccccctccttcccagggttcactggaggtgaccaaggtaggaaatggccttaacaca**gggatgactgcgccatccc**cca**acagagtcagccccctcctgctctgt**accccgcaccccccaggccagtccacgaaaa**ccagggccccacatcagagtcactgcctggcccggccctgg**ggcggacccctcag**cccccaccctgtctagaggacttggggggacaggacacaggccctctccttatggttccccc**acctgcctccggccgggacccttggggtgtggacagaaaggacacctgcctaattggcccccaggaacccagaacttctctccagggaccccagcccgagcacccccttacccaggacccagccctgcccctcctcccctctgctctcctctcatcaccccatgggaatccggtatccccaggaagccatcaggaagggctgaaggaggaagcggggccgtgcaccaccgggc**aggaggctccgtcttcgtgaacccagggaagtgccagcctcct**agagggtat

Uppercase: IGHD5-18-2

Lowercase: Flanking sequence[1000bp]

Red & Bold & Underline: Stem-loop [21]

Blue: Heptamer[38]

Green: Nonamer [4]

id-TRBD2[D_gene_segment]

acgcgtgtatatgtttatatttacacacatgtacacacatccactcatta**aacacatatccctgtgatgtgtt**gttatcaacttccaagaaaattaagatacccaaattacagtccccaagaagcgta**tctgttcacagagcaactcaatcatgaacaga**aagagccaaacgcaagtccatgctttgaaagagcaacatgaactgcttactaaaatgcagtcagagagtggccctcaat**ttctcactgggataatcagtaaagagtctgagaa**ggtggcaatctcttcaaaggaagaattggtctcagttgttcactctagagaatggccatgctggtcagaggcatgaacaaaagccctaatgacatggctaggtacctctgggtaggcccgtttagctggttgcaaggttcagaaatgggagtgctcaaaattagcgctggaaaggcaggttggtgcatgccactctgggaataaacttac**ctccatgcaaccagcatggag**cctgcacatggtggatgttcactaaacacctgtggagtaaatgaagaatatggagttcagacatc**gttcaggaagcaactgaac**aaagggaaattatagaggtttctggattgtttgtcctcctgtcataaggtgccatcaactgctctgtggattttcctatgagctgcctgccacccctcgctcctcccacccacttcactataaatgccagtctgagca**ggtgggcacagtgagccccacc**agggagacccagtgacatagatggtctgctcagggtgatgcatgttccaaggagggacctctctgccccccaccattaccatcactgtgactttccccaagcccttcccattttaattcactgcctttgtcttttccaagccccacacagtcagactaacctctgccacctgcgcttcctgccgctgcccagtggttgggggagggggactagcagggaggaaacatttttgtatcatggtgtaacattgtgGGGACTAGCGGGAGGGcacgatgattcaggtagaggaggtgcttttacaaaaaaccctgatgcagtaagcatc**cccacccagctcagggaatgcagctaccaggtggg**aagagttctctggggctggtcccagctgtggtcttgcagggtcccccaacccagcgagcacctgtccatctccctgtccagactcggcttccaaggaataagaaggccaagacagc**aaagtgggattatcactcagcacttt**taataaaacttgttcttgacaaagtacttgcacatgcattatttattaagaactgatgaaaaccctgag**ggaaagatattgtcccatctttcc**aatgaggaaactgagatcagaggttacaggtcatataactaggaaacggcaaggtctagcctgcaatatcgcccagctccagccgttccagtaccaccaatgccccttcagatttca**aatccactgtgttgtcccccagccaagtggatt**ctcctctgcaaattggtggtggcctcatgcaagatccaggttaccgtgtccagctaactcgagacaggaaaagataggctcaggaaagagaggaagggtgtgccctctgtctgtgctaagggaggtg**gggaaggagaaggaattctgggcagccccttccc**actgtgctcctacaatgagcagttcttcgggccagggacacggctcaccgtgctaggtaagaagggggctccaggtgggagagagggtgagcagcccagcctgcacgaccccagaaccctgttcttaggggagtggacactgggcaatccagggccctcctcgagggaagcggggtttgcgccagggtccccagggctgtgcgaacaccggggagctgttttttggagaaggctctaggctgaccgtactgggtaaggaggcggttggggctccggagagctccgagagggcgggat**gggcagaggtaagcagctgccc**cactctgagaggggctgtgctgagaggcgctgctgggcgtctg

Uppercase: TRBD2

Lowercase: Flanking sequence[1000bp]

Red & Bold & Underline: Stem-loop [12]

Blue: Heptamer[40]

Green: Nonamer [8]

id-IGHD3-10[D_gene_segment]

ggcagggtgagcacccagagaaaactgcagacgcctcacacatccacctcagcctcccctgacctggacctcactggcctgggcctcacttaacctgggcttcacctgaccttggcctcacctgacttggacctcgcctgtcccaagctttacctgacctgggcctcaactcacctgaacgtctcctgacctgggtttaacctgtcctggaactcacctggccttggcttcccctgacctggacctcatctggcctgggcttcacctggcctgggcctcacctgacctggacctcatctggcctggacctcacctggcctggacttcacctggcctgggcttcacctgacctggacctcacctggcctcgggcctcacctgcacct**gctccaggtcttgctggagc**ctgagtagcactgagggtgcagaagctcatccagggttggggaatgactctagaagtctcccacatctgacctttctgggtggaggcagctggtggccctgggaatataaaaatctccagaatgatgactctgtgatttgtgggcaacttatgaacccgaaaggacatggccatggggtgggtagggacatagggacagatgccag**cctgaggtggagcctcagg**acacaggtgggcac**ggacactatccacataagcgagggatagacccgagtgtcc**ccacagcagacctgagagcgctgggcccacagcctcccctcagagccctgctgcctcctccggtcagccctggacatcccaggtttccccaggcctgccggtaggtttagaatgagg**tctgtgtcactgtggtattacgatattttgactggttattataaccacagtgtcacaga**gtccatcaaaaa**cccatgcctggaagcttcccgccacagccctccccatggg**gccctgctgcctcctcaggtcagccccggacatcccgggtttccccaggctgggcggtaggtttggggtgagg**tctgtgtcactgtgGTATTACTATGGTTCGGGGAGTTATTATAACcacagtgtcacaga**gtccatcaaaaacccatccctgggagcctcccgccacagccctccctgcaggggaccggtacgtgccatgttaggattttgatcgaggagacag**caccatgggtatggtg**gctaccacagcagtgcagcctgtgacccaaacccgcagggcagcaggcacgatggacaggcccgtgactgaccacgctggg**ctccagcctgccagccctggag**atcatgaaacagatggccaaggtcaccctacaggtcatccagatctggctccgaggggtctgcatcgctgctgccctcccaacgccagtccaaatgggacagggacggcctcacagcaccatctgctgccatcaggccagcgatcccagaagcccctccctcaaggctgggccacatgtgtggacactgagagccctcatgtctgagtaggggcaccaggaggg**aggggctggccctgtgcactgtccctgcccct**gtggtccctggcctgcctggccctgacacctgagcctctcctgggtcatttccaagacagaagacattcctgg**ggacagccggagctgggcgtcgctcatcctgcccggccgtcc**tgagtcctgctcatttccagacctcaccggggaagccaacagaggactcgcctcccacattcagagacaaagaaccttccagaaatccctgcctctctccccagtggacaccctcttccaggacagtcctcagtggcatc**acagcggcctgagatccccaggacgcagcaccgctgt**caataggggccccaaatgcctggaccagggcctgcgtgggaaaggtctctggccacactcgggctttttgtgaagggccctcctgctgtgtgactacagtaactaccatagtgatgaacccagtggcaaaaactggctggaaacccaggggctgtgtgcacgcctcagcttggagctctccaggagcacaagagccgggcccaaggatttgtgcccagaccctca

Uppercase: IGHD3-10

Lowercase: Flanking sequence[1000bp]

Red & Bold & Underline: Stem-loop [11]

Blue: Heptamer[52]

Green: Nonamer [7]

id-IGHD6-25-2[D_gene_segment]

cagccctgcccctcctcccctctgctctcctctcatcactccatgggaatccagaatccccaggaagccatcaggaagggctgaaggaggaagcggggccgctgcaccaccgggc**aggaggctccgtcttcgtgaacccagggaagtgccagcctcct**agagggtatggtccac**cctgcctggggctcccaccgtggcagg**ctgcggggaaggaccagggacggtgtgg**gggagggctcaggtccctgcaggtgctccatcttggatgagcccatccc**tctcacccaccgacccgcccacctcctctccaccctggccacacgtcgtccacaccatcctgagtcccacctacaccagagccggcagagccagtgcagacagaggctggggtgcaggggggccgccagggcagctttggggagggaggaatggagga**aggggaggtcagtgaagaggcccccctcccct**gggtcta**ggatccacctttgggacccccggatcccatcccctccaggctctgggaggagaagcaggatggga**gaatctgtgcgggaccctctcacagtggaatacctccacagc**ggctcaggccagatacaaaagcccctcagtgagcc**ctccactgca**gtgctgggcctgggggcagcccctcccacagaggacagacccagcac**cccgaagaagtcctgccagggggagctcagagccatgaaggagcaagatatggggaccccaatactggcacagacctcagctccatccaggcccaccaggacccaccatgggtggaacacctgtctccggcccctgctggctgtgaggcagctgg**cctctgtctcggacccccattccagacaccagacagagggacaggccccccagaaccagtgttgagggacacccctgtcc**agggcagccaagtccaagaggcgcgctgagcccagcaagggaaggcccccaaacaaaccaggaggtttctgaagctgtct**gtgtcacagtcGGGTATAGCAGCGGCTACcacaatgacac**tgggcaggacagaaaccccatcccaagtcagccgaaggcagagagagcaggcaggacacatttaggatctgaggccacacctgacactcaagccaacagatgtctcccctccagggcgcc**ctgccctgttcagtgttcctgagaaaacaggggcag**cct**gaggggatccagggccaggagatgggtcccctc**taccccgaggaggagccaggcgggaatcccagc**cccctccccattgaggccatcctgcccagagggg**cccggacccaccc**cacacacccaggcagaatgtgtg**caggcctcaggctctgtgggtgccgctagctggggctgccagtc**ctcaccccacacctaaggtgag**ccacagccgccagagcctccacaggagaccccacgcagcagcccagcccct**acccaggaggccccagagctcagggcgcctgggt**ggattctgaacagccccgagtcacggtgggtatagtgggagctactaccactgtgagaaaagctatgtccaaaactgtctcccggccactgctggaggcccagccagagaagggaccagccgcccgaacatacgaccttcccagccctcatgacccccagcacttggagctccacagtgtccccattggatggtgaggacgggggccggggccatctgcacctcccaacatcacccccaggcagcacaggcacaaaccccaaatccagagccgacaccaggaacacagacaccccaataccctgggggaccctggccctggtgacttcccactgg**gatccacccccgtgtccacctggatc**aaagaccccaccgctgtctctgt**ccctcactcagggcctgctgaggg**gcgggtgctttggagcagactcaggtttaggggccaccattgtggggcccaacctcgaccaggacacagatttttctttcctgccctggggcaacacagactttggggtctgtgcagggaggaccttctggaaag

Uppercase: IGHD6-25-2

Lowercase: Flanking sequence[1000bp]

Red & Bold & Underline: Stem-loop [19]

Blue: Heptamer[34]

Green: Nonamer [9]

id-TRDD2[D_gene_segment]

cactttgctaaattgaaaattaagaggcacattagattta**gaaacataattttaaattgtttc**aatgatggcacattattaatacctactttgagacacaatacatactcacataatattagaatcattgttttgccatttatcctgtaagtgaatattcatgatctccacaacaaaacatttac**tttgatagatttgtttaaacaatatcaaa**cacttgcagtcttgaaggcactggaagacccaaaaaatagttactacgtttct**gttaaactacttagtcatctttttaac**taattctatcagctgactgccataagcatccatgagctagcattgattgagatcattccaatgtgtcctcactaaactgt**acttaactgttcaaagccaaataattaagt**gaaagtatggtaatactgtttttcataattattgtaaggaatcaaatataagggatttcttcttctctgaggatcatgaaccttactccatgttcaaatagatatagtattttttcacagtaaattccccaaagtggaaatagtcactcaaacgaatacttttactcttttgacattaaccaaatggcttccaa**taaaaatcaaacaatttatattttta**ccaacatcttgcccacattgggagtgtcaacattttgagcaaaagctttaattaaatatccatgcaaaaaatggtttgttaatactttacagttttattactagagggttaaaatcctttttcaagtctgataatcaatgattaactttcttcatttgtccttcacccatttgttttttaggttgatggtgttttacttattgatttgtgtaattataataa**ttttgtgtctgagttttacagcatttaaccacaaaa**acagcattggtgaaaggagtttcaggggtattgtggatggcagcgggtggtgatggcaaagtgccaaggaaagggaaaaaggaagaagagggtttttatactgatgtgtttcattgtgCCTTCCTACcacacaggttggagtgcattaagcctttgtccaaaaacacccagccgtgacccgctatgtatgtctcagcattgggaagagtcctctgagtgtcatgg**gaaaataatatatgagttttattgcccctgtgtcccaattattttc**tgtttactccatgttggtctacagcaatagcttcagaacataaacaaatatcaccaaaaacatcagtgatgaacaattccaaccacacatttcaatgtttactaaatatacccgtggcactgtaaaaaagaatataatggaagagtaaaggaattgggactgatcgtgactgagaacctgtatagtaggctacacactagctatatgatgaaccaagaagttgaacccaaatctaactaacttacataatcccttcctttaaagaacttaatgtccagtgcaaagcaagtgttcaaaaaagtgtttgttgtgaattgaattcatttcattggaatacagcatgcttgcaaagtcccaactttttaaaataat**gtgtttctaaagaccactggtctttagaaagac**caatgaccaaaga**ccttagaccattggtctaagg**accatctggaagcagatccccaaggtgaaattctgacgttgaattcccaaatctaacaggtcatgtgccttgttctgatgagaaaactgtcgaataagggagtagcagtggcatgaaggttgttagtaggtagcataggccaaatggcccagtgatgtgatgaaaagtgcactcctccaaataagaccctagttactagaaagtcacaagtaggtaaacagatgccgggagggacagtttttgttggataaagtcagcagagaaaggcttttggaggagatgaatcctgagctggtgtctacagatagtgacagacttagtgcgactaggaatattatggagaataaaagaatctccctctctaactaatgccaccaagcagtcccttcaccccagagcaaaggcataacctatctt

Uppercase: TRDD2

Lowercase: Flanking sequence[1000bp]

Red & Bold & Underline: Stem-loop [9]

Blue: Heptamer[27]

Green: Nonamer [7]

id-IGHD7-27[D_gene_segment]

gtgtcctccaacgacaggtcccagcctcccagcctttgccttgcctgttcctctccctggaactctgccccgacacagaccctccccagcaagccc**gcaggggcacctcccctgc**ccccagacaccctgtgcccgtcagttcatccccagcagaggccctcaccaggcacacccccatgctcacacctggccgcagg**cctcagcctccctgagg**gccccacccagcccgcgtctggccagtggtgcgtgcaa**agcccctcacccagactcggcggaaggcagccagtgcaggcctggggaggggct**ctccttagaccaccttgcac**cttccctggcacccaccatgggaag**ag**ctgagactcactgaggaccagctgaggctcag**agaagggacccagcactggtggacacgcagggagcccacgccagggcgccgtggtgagt**gaggcccagtgccacccactgaggcctc**ccgttcagtgggacgacggtgaacaggtggaaccaaccaggcaacccccgccgggccccacagacgggatcaga**gcaggaaaggcttcctgc**ccctgcaggccagcgaggagccc**tggcgggggccatggccctccaggcgaggaggctcccctggccaccgcca**cccgggcctctctgctgctgggaaaacaagtcagaaagcaagtggatgagaggtggcgtgacagacccagcttcagatctgctctaatttacaaaagaaaaggaaaaacacacttggcagccttcagcactctaatgattcttaacagcagcaaattattggcacaagactccagagtgactggcagggttgagggctgggg**tctcccgcgtgttttggggctaacagcggaagggaga**gcactggcaaaggtgctgggggcccctggacccgacccgccctggagaccgcagccacatcagcccccagccccacaggccccctaccagccgcagggttttggctgagctgagaac**cactgtgCTAACTGGGGAcacagtg**attggcagct**ctacaaaaaccatgctcccccgggaccccgggctgtgggtttctgtag**cccctggctcagggctgactcaccgtggctgaatacttccagcactggggccagggcaccctggtcaccgtctcctcaggtgagtctgctgtctggggatagcggggagccaggtgtactgggccaggcaagggctttggcttcagacttggggacaggtgctcagcaaaggaggtcggcaggagggcggagggtgtgtttttgtatgggagaagcaggagggcagaggctgtgctactggtacttcgatctctggggccgtggcaccctggtcactgtctcctcaggtgagtcccactgcagccccctcccagtcttctctgtccaggcac**caggccaggtatctggggtctgcagccggcctgggtctggcctg**aggccacaccagctgccatccctggggtctccgccatgggctgcatgccagagccctgctgtcacttagccctggggcca**gctggagcccccaaggacaggcagggaccccgctgggcttcagc**cccgtcagggaccctccacaggtagcaagcaggccgagggcagggacgggaaggagaagttgtgggcagagcctgggctggggctgggcgctggctgttcatgtgccggggaccaggcctgcgctttagtgtggctaca**agtgcttggagcact**gggg**ccagggcagcccggccaccgtctccctgg**gaacgtcacccctccctgcctgggtctcagcccgggggtctgtgtggctggggacagggacgccggctgcctctgctctgtgcttgggccatgtgacccattcgagtg**tcctgcacgggcacaggtttatgtctgggcaggaacagggactgtgtccctgt**gtgatgcttttgatatctggggccaagggacaatggtcaccgtctcttcaggtaagatggctttccttctgcctcctttctctgggc

Uppercase: IGHD7-27

Lowercase: Flanking sequence[1000bp]

Red & Bold & Underline: Stem-loop [17]

Blue: Heptamer[34]

Green: Nonamer [6]

id-IGHD4-23[D_gene_segment]

cccatgacccaaacacacggggcagcagaaacaatggacaggcccacaagtgaccatgatgggctccagcccaccagccccagagaccatgaaacagatggccaaggtcaccctacaggtcatccagatctggctccaaggggtctgcatcgctgctgccctcccaacgccaaac**cagatggagacagggccggccccatagcaccatctgctgccgtccacccagcag**tcccggaagcccctccctgaacgctgggccacgtgtgtgaaccctgcg**agccccccatgtcagagtaggggcagcaggagggcggggct**ggccctgtgcactgtcactgcccctgtggtccctggcctgcctggccctgacacctgagcctctcctgggtcatttccaagacattcccag**ggacagccggagctgggagtcgctcatcctgcctggctgtcctgagtcctgctcatttccagacctcaccagggaagccaacagaggactca**cctcacacagtcagagacaatgaaccttccagaaatccctgtttctctccccagtg**agagaaaccctcttccagggtttctct**tctctcccaccctcttccaggacagtcctcagcagcatcacagcgggaacgcacatctggatcaggacggcccccagaacacgcgatggcccatggggacagcccagcccttcccagacccctaaaaggtatccccaccttgcacctgccccagggctcaaactc**caggaggcctgactcctgcacaccctcctg**ccagatatcacctcagc**cccctcctggagggg**acaggagcccgggagggtgagtcaga**cccacctgccctcaatggcaggcggg**gaagattcagaaaggcctgagatccccaggacgcagcaccactgtcaatgggggcc**ccagacgcctggaccagggcctgtgtgggaaaggcctctgg**ccacactcaggggctttttgtgaagggccctcctgctgtgTGACTACGGTGGTAACTCCcacagtgatgaaaccagcagcaaaaactgaccggactcgcagggtttatgcacacttctcggctcggagctctccaggagcacaagagcc**aggcccgagggtttgtgcccagaccctcggcct**ctagggacacccgggccatcttagccgatgggctgatgccctgcacaccgtgtgctgccaaacaggggcttcagagggctctgaggtgacttcactcatgaccacaggtgccctggtcccttcactgccagctgcaccagaccctgttccgagagat**gccccagttccaaaagccaattcctggggc**cgggaattactgtagacaccagcctcattccagtacctcctgccaattgcctggattcccat**cctggctggaatcaagagggcagcatccgccagg**ctcccaacaggcaggactcccacacaccctcttctgagaggccgctgtgttccgcagggccaggccgcagacagttcccctcacctgcccatgtagaaacacctgccattgtcgtcccc**acctggcaaagaccacttgtggagcccccagccccaggt**acagctgtagagagagtcctcgaggcccctaagaaggagccatgcccagttctgctgggaccctcggccaggccgacaggagtggacgctggagctgg**gcccacactgggccacataggagctcaccagtgagggc**aggagagcacatgccggggagcacc**cagcctcctgctgaccagagacccgtcccagagcccaggaggctg**cagaggcctctccagggggacacagggcatgtctggtccctgagca**gcccccaggctctctagcactgggggc**ccctggcaca**gctgtctggaccctccctgttccctgggaagctcctcctgacagc**cccgcctccagttccaggtgtggttattgtcagggggtgccaggcc**gtggtagagatggctacaattaccac**agtggtgccgcccatagcagcaaccaggcc

Uppercase: IGHD4-23

Lowercase: Flanking sequence[1000bp]

Red & Bold & Underline: Stem-loop [19]

Blue: Heptamer[34]

Green: Nonamer [5]

id-IGHD4-4-2[D_gene_segment]

gtggctaccgcagcagtgcag**cctgtgacccaaacacacagg**gcagcaggcacaacagacaagcccacaagtgaccaccctgagctcctgcctgccagccctggagaccatgaaacagatggccaggattatcccataggtcagc**cagacctcagtccaacaggtctg**catcgctgctgccctccaataccagtccggatggggacag**ggccggcccacattaccatttgctgccatccggcc**aacagtcccagaagcccctccctcaaggctgggccacatgtgtggaccctgag**agccccccatgtctgagtaggggcaccaggaaggtggggct**ggccctgtgcactgtcactgcccctgtggtccctggcctgcctggccctgacacctgggcctc**tcctgggtcatttccaagacagaagacattcccagga**cagctggagctgggagtccatcatcctgcctggccatcctgagtcctgcgcctttccaaacctcacccgggaagccaacagaggaatcacctcccacaggc**agagacaaagaccttccagaaatctctgtctct**ctccccagtgggcaccctcttccagggcagtcctcagtgatatcacagtgggaacccacatctggatcgggactgcccccagaacacaagatggcccacagggacagccccacagcccagcccttcccagacccctaaaaggcgtcccaccccctgcatctgccccagggctcaaactc**caggaggactgactcctgcacaccctcctg**ccagacatcacctcagcccctcctgga**agggacaggagcgcgcaagggtgagtcagaccctcctgccct**cgatggcaggcggagaagattcagaaaggtctgagatccccaggacgcagcaccactgtcaatgggggc**cccagacgcctggaccagggcctgcgtgggaaaggcctctggg**cacactcaggggctttttgtgaagggtcctcct**actgtgTGACTACAGTAACTACcacagt**gatgaacccagcagcaaaaactgaccggactcccaaggtttatgcacacttctccgctcagagctctccaggatcagaagagccgggcccaagggtttctgcccagaccctcggcctctagggacatcttggccatgacagcccatgggctggtgccccacacatcgtctgccttcaaacaagggcttcagagggctctgaggtgacctcactgatgaccacaggtgccctggccccttccccgccagctgcaccagaccccgtcctgacagatgccccgattccaacagcca**attcctggggccaggaat**cgctgtagacaccagcctccttccaacacctcttgccaattgcctg**gattcccatcccggttggaatc**aagaggacagcatcccccaggctcccaacaggcaggactcccacaccctcctctgagaggccgctgtgttccgtagggccaggctgcagacagtccccctcacctgccactagacaaatgcctgctgtagatgtcccc**acctggaaaagaccactcatggagcccccagccccaggt**acagccatagagagagtctctgaggcccctaagaagtagccatgcccagttctgccgggaccctcggccaggctgacaggagtggacgctggagctgg**gcccacactgggccacataggagctcaccagtgagggc**aggagagcacatgccggggagcacc**cagcctcctgctgaccagaggcccgtcccagagcccaggaggctg**cagaggcctctccagggggacactgtgcatgtctggtccctgagca**gccccccatgtccccagtcctgggggc**ccctggcacagctgtctggaccctctctattccctgggaagctcctc**ctgacagccccgcctccagttccaggtgtggttattgtcagggggtgtcagactgtggtggatacagctatggttaccacagt**ggtgctgcccatagcagcaaccaggccaa

Uppercase: IGHD4-4-2

Lowercase: Flanking sequence[1000bp]

Red & Bold & Underline: Stem-loop [18]

Blue: Heptamer[41]

Green: Nonamer [6]

id-IGHD3-3[D_gene_segment]

ctgctggcagctcctggggcctgatgtggagcaggcacagagccgtatccccccgaggacatatacccccaaggacggcacagttggtacattccggagacaagcaactcagccacactcccaggccagagcccgagagggacgcccatgcacagggaggcagagcccagctcctccacagccagcagcac**ctgtgcaggggccgccatctggcaggcacag**agcatgggctgggaggaggggcagggacaccaggcagggttggcaccaactgaaaattacagaagtctcatacatctacctcagccttgcctgacctgggcctcacctgacctggacctcacctggcctggacctcacctggcctagacctcacctctgggcttcacctgagctcggcctcacctgacttggaccttgcctgtcctgagctcacatgatctgggcctcacctgacctgggtttcacctgacctgggcttcacctgacctgggcctcatctgacctgggcctcactggcctggacctcacct**ggcctgggcttcacctggcctcaggcc**tcatctgcacctgctccaggtcttgctggaa**cctcagtagcactgagg**ctgcaggggctcatccagggttgcagaatgactctagaacctcccacatctcagctttctgggtggaggcacctggtggcccagggaatataaaaagcctgaatgatgcctgcgtga**tttgggggcaatttataaacccaaa**aggacatggccatgcagcgggtagggacaatacagacagatatcagcctgaaatggagcctcagggcac**aggtgggcacggacactgtccacct**aagccaggggcagacccgagtgtccccgcagtagacctgagagcgctgggcccacagcctcccctcggtgccctgctacctcctcaggtcagccctggacatcccgggtttccccaggcctggcggtaggtttggggtgagg**tctgtgtcactgtgGTATTACGATTTTTGGAGTGGTTATTATACCcacagtgtcacaga**gtccatcaaaaacccatccctgggaaccttctg**ccacagccctccctgtgg**ggcaccgctgcgtgccatgttaggattttgactgaggacacag**caccatgggtatggtg**gctaccgcagcagtgcag**cctgtgacccaaacacacagg**gcagcaggcacaacagacaagcccacaagtgaccaccctgagctcctgcctgccagccctggagaccatgaaacagatggccaggattatcccataggtcagc**cagacctcagtccaacaggtctg**catcgctgctgccctccaataccagtccggatggggacag**ggccggcccacattaccatttgctgccatccggcc**aacagtcccagaagcccctccctcaaggctgggccacatgtgtggaccctgag**agccccccatgtctgagtaggggcaccaggaaggtggggct**ggccctgtgcactgtcactgcccctgtggtccctggcctgcctggccctgacacctgggcctc**tcctgggtcatttccaagacagaagacattcccagga**cagctggagctgggagtccatcatcctgcctggccatcctgagtcctgcgcctttccaaacctcacccgggaagccaacagaggaatcacctcccacaggc**agagacaaagaccttccagaaatctctgtctct**ctccccagtgggcaccctcttccagggcagtcctcagtgatatcacagtgggaacccacatctggatcgggactgcccccagaacacaagatggcccacagggacagccccacagcccagcccttcccagacccctaaaaggcgtcccaccccctgcatctgccccagggctcaaactc**caggaggactgactcctgcacaccctcctg**ccagacatcacctcagcccctcctgga**agggacaggagcgcgcaagggtgagtcagaccctcctgccct**cgatggcaggcggagaagattcagaaaggt

Uppercase: IGHD3-3

Lowercase: Flanking sequence[1000bp]

Red & Bold & Underline: Stem-loop [16]

Blue: Heptamer[47]

Green: Nonamer [2]

id-IGHD6-25[D_gene_segment]

cagccctgcccctcctcccctctgctctcctctcatcactccatgggaatccagaatccccaggaagccatcaggaagggctgaaggaggaagcggggccgctgcaccaccgggc**aggaggctccgtcttcgtgaacccagggaagtgccagcctcct**agagggtatggtccac**cctgcctggggctcccaccgtggcagg**ctgcggggaaggaccagggacggtgtgg**gggagggctcaggtccctgcaggtgctccatcttggatgagcccatccc**tctcacccaccgacccgcccacctcctctccaccctggccacacgtcgtccacaccatcctgagtcccacctacaccagagccggcagagccagtgcagacagaggctggggtgcaggggggccgccagggcagctttggggagggaggaatggagga**aggggaggtcagtgaagaggcccccctcccct**gggtcta**ggatccacctttgggacccccggatcccatcccctccaggctctgggaggagaagcaggatggga**gaatctgtgcgggaccctctcacagtggaatacctccacagc**ggctcaggccagatacaaaagcccctcagtgagcc**ctccactgca**gtgctgggcctgggggcagcccctcccacagaggacagacccagcac**cccgaagaagtcctgccagggggagctcagagccatgaaggagcaagatatggggaccccaatactggcacagacctcagctccatccaggcccaccaggacccaccatgggtggaacacctgtctccggcccctgctggctgtgaggcagctgg**cctctgtctcggacccccattccagacaccagacagagggacaggccccccagaaccagtgttgagggacacccctgtcc**agggcagccaagtccaagaggcgcgctgagcccagcaagggaaggcccccaaacaaaccaggaggtttctgaagctgtct**gtgtcacagtcGGGTATAGCAGCGGCTACcacaatgacac**tgggcaggacagaaaccccatcccaagtcagccgaaggcagagagagcaggcaggacacatttaggatctgaggccacacctgacactcaagccaacagatgtctcccctccagggcgcc**ctgccctgttcagtgttcctgagaaaacaggggcag**cct**gaggggatccagggccaggagatgggtcccctc**taccccgaggaggagccaggcgggaatcccagc**cccctccccattgaggccatcctgcccagagggg**cccggacccaccc**cacacacccaggcagaatgtgtg**caggcctcaggctctgtgggtgccgctagctggggctgccagtc**ctcaccccacacctaaggtgag**ccacagccgccagagcctccacaggagaccccacgcagcagcccagcccct**acccaggaggccccagagctcagggcgcctgggt**ggattctgaacagccccgagtcacggtgggtatagtgggagctactaccactgtgagaaaagctatgtccaaaactgtctcccggccactgctggaggcccagccagagaagggaccagccgcccgaacatacgaccttcccagccctcatgacccccagcacttggagctccacagtgtccccattggatggtgaggacgggggccggggccatctgcacctcccaacatcacccccaggcagcacaggcacaaaccccaaatccagagccgacaccaggaacacagacaccccaataccctgggggaccctggccctggtgacttcccactgg**gatccacccccgtgtccacctggatc**aaagaccccaccgctgtctctgt**ccctcactcagggcctgctgaggg**gcgggtgctttggagcagactcaggtttaggggccaccattgtggggcccaacctcgaccaggacacagatttttctttcctgccctggggcaacacagactttggggtctgtgcagggaggaccttctggaaag

Uppercase: IGHD6-25

Lowercase: Flanking sequence[1000bp]

Red & Bold & Underline: Stem-loop [19]

Blue: Heptamer[34]

Green: Nonamer [9]

id-IGHD5-5[D_gene_segment]

cacactcaggggctttttgtgaagggtcctcct**actgtgtgactacagtaactaccacagt**gatgaacccagcagcaaaaactgaccggactcccaaggtttatgcacacttctccgctcagagctctccaggatcagaagagccgggcccaagggtttctgcccagaccctcggcctctagggacatcttggccatgacagcccatgggctggtgccccacacatcgtctgccttcaaacaagggcttcagagggctctgaggtgacctcactgatgaccacaggtgccctggccccttccccgccagctgcaccagaccccgtcctgacagatgccccgattccaacagcca**attcctggggccaggaat**cgctgtagacaccagcctccttccaacacctcttgccaattgcctg**gattcccatcccggttggaatc**aagaggacagcatcccccaggctcccaacaggcaggactcccacaccctcctctgagaggccgctgtgttccgtagggccaggctgcagacagtccccctcacctgccactagacaaatgcctgctgtagatgtcccc**acctggaaaagaccactcatggagcccccagccccaggt**acagccatagagagagtctctgaggcccctaagaagtagccatgcccagttctgccgggaccctcggccaggctgacaggagtggacgctggagctgg**gcccacactgggccacataggagctcaccagtgagggc**aggagagcacatgccggggagcacc**cagcctcctgctgaccagaggcccgtcccagagcccaggaggctg**cagaggcctctccagggggacactgtgcatgtctggtccctgagca**gccccccatgtccccagtcctgggggc**ccctggcacagctgtctggaccctctctattccctgggaagctcctc**ctgacagccccgcctccagttccaggtgtggttattgtcagggggtgtcagactgtgGTGGATACAGCTATGGTTACcacagt**gg**tgctgcccatagcagca**accaggccaagtagacaggcccctgctgtgc**agccccaggcctccagctcacctgcttctcctggggct**ctcaaggctgctgttttctgcact**ctcccctctgtggggag**ggttccctcagtgggagatctgttctcaacatcccagggcctcattcctgcaaggaaggccaatggatgggcaacctcacatgccgcggctaagatagggtgggcagcctggcggggacaggacatcctgctggggtatctgtcactgtgcctagtggggcactggctcccaaacaacgcagtcctcgccaaaatccccacggcctcccccgctaggggctggcctgatctcctgcagtcctaggaggctgctgacctccagaatggctccgtccccagttccagggcgagagcaga**tcccaggccggctgcagactggga**ggccaccccctccttcccagggttcactgcaggtgaccagggcaggaa**atggcctgaacacagggataaccgggccat**ccccca**acagagtccaccccctcctgctctgt**accccgcacccccaaggccagcccatgacatccgacaaccccacaccagagtcactgcccggtgctgccctagggaggacccctcagcccccac**cctgtctagaggactggggaggacagg**acacgccctctccttatggttcccccacctggctctggctgggacccttggggtgtggacagaaaggacgcttgcctgattggcccccaggagcccagaacttctctccagggaccccagcccgagcacccccttacccaggacccagccctgcccctcc**tcccatctgctctcctctcatcaccccatggga**atccagaatccccaggaagccatcaggaagggctgagggaggaagtggggccactgcaccaccaggc**aggaggctccgtctttgtgaacccagggaggtgccagcctcct**agagggtatg

Uppercase: IGHD5-5

Lowercase: Flanking sequence[1000bp]

Red & Bold & Underline: Stem-loop [18]

Blue: Heptamer[34]

Green: Nonamer [7]

id-IGHD1-20[D_gene_segment]

tcccatcccctcctggctctgggaggagaagcaggatgggagaatctgtgcgggaccctctcacagtggaatatccccacagc**ggctcaggccagacccaaaagcccctcagtgagcc**ctccactgcagtcctgggc**ctgggtagcagcccctcccacagaggacagacccag**caccccgaagaag**tcctgccagggggagctcagagccatgaaagagcagga**tatggggtccccgatacaggcacagacctcagctccatccaggcccaccgggacccaccatgggaggaacacctgtctccgggttgtgaggtagctgg**cctctgtctcggaccccactccagacaccagacagagg**ggcaggccccccaaaaccagggttgagggatgatccgtcaaggcagacaagaccaaggggcactgaccccagcaagggaaggctcccaa**acagacgaggaggtttctgaagctgtctgt**atcacagtggggtatagcagtggctggtaccacagtgacactcgccaggccagaaaccccgtcccaagtcagcggaagcagagagagcagggaggacacgtttaggatctgaggccgcacctgacacccagggcagcagacgtctcccctccagggcaccctccaccgtcctgcgtttcttcaagaataggggcggcct**gagggggtccagggccaggcgataggtcccctc**taccccaaggaggagcca**ggcaggacccgagcaccgtccccattgaggctgacctgcccagacgggcctgggc**ccaccc**cacacaccggggcggaatgtgtg**ca**ggccccagtctctgtgggtgttccgctagctggggcc**cccagtgctcaccccacacctaaagcgagccccagcctccagagccccctaagcattccccgcccagcagcccagcccctgcccccacccaggaggccccagagctcagggcgcctggtcggattctgaacagccccgagtcacagtgGGTATAACTGGAACGACcaccgtgagaaaaactgtgtccaaaactgactcctggcagcagtcggaggccccgccagagaggggagcagccggcctgaacccatgtcctgccggttcccatgacccccagcacccagagccccacggtgtccccgttggataatgaggacaagggctgggggctccggtggtttgcggcagggacttgatcacatccttctgctgtggccccattgcctctggctggagttgaccc**ttctgacaagtgtcctcagaa**agacagggatcaccggcacctcccaatatcaaccccaggcagcacagacacaaaccccacat**ccagagccaactccaggagcagagacaccccaacactctgg**gggaccccaaccgtgataactccccactggaatccgccccagagtctaccaggaccaa**aggccctgccctgtctctgtccctcactcagggcct**cctgcagggcgagcgcttgggagcagactcggtcttaggggacaccactgtgggccccaactttgatgaggccactgacccttccttcctttcctggggcagcacagactttggggtctgggcagggaagaactactggctggtggccaatcacagagcccccaggccgag**gtggccccaagaaggccctcaggaggtggccac**tccacttcctcccagctggaccccaggtcctccccaagataggggtgccatccaaggcaggtcctccatggagcccccttcagactcctcccgggaccccactggacctcagtccctgctctgggaatgcagccaccacaagcacaccaggaagcccaggcccagc**caccctgcagtgggcaagcccacactctggagcagagcagggtg**cgtct**gggaggggctaacctccc**caccccccaccccccatctgcacacagccacctaccactgcccagaccctctgcaggagggccaagccaccatggggtatggacttagggtctcactcacgtgcctc

Uppercase: IGHD1-20

Lowercase: Flanking sequence[1000bp]

Red & Bold & Underline: Stem-loop [16]

Blue: Heptamer[33]

Green: Nonamer [8]

id-TRBD1-2[D_gene_segment]

gtcagtttctgttgtatctttacagtgtttc**tgtatacaaaatgtataca**aaatcaagtaaatacaaaagtataattcttcttttcccccctctgttagacaaaggtgtcatctagttatgctattctgagact**ttctcttttctgggttaataatgtatcttagagaa**ttttccatagcaatataaaatattttgattctttttttagcattgaatattactccattgtgtgaacatatcacagtttatccagtcccttactgatgggcatataattcaattttgtcagtcaaattagtgaaaagtgtattttgatgaagtttttttaattttatttttcttattgtgattttttttcccttatagttaggtaccatttggatttatttttctctgaacaatttactcttcacctatttttattgggctgctggattattttcctattgatttttaggagt**tcaccacatgttgaggagattagccctgatggtga**gaactaggaatatttttcccagattgttatgtatattttgattctgtttatattttaccatgcgagtgtgtcatcattatgtcgttgtttgtctttttttttttttttct**ggattttaagtcatagcttaaaaccc**tccgagtgacg**cacagcctcggggcagggccagggtagtgatgggggctgtg**gcttctctataaggacatgccccaacgtgacaacagcttggagaggggtgggtactggagaagaccagccccttcgccaaacagccttacaaagacatccagctctaaggagctcaaaacatcctgaggacagtgcctggaggtgagaaggaagcccccggcctggtccataccccaccaccaacttgcataatggggggtgatgtcacccaccctccactcccctcaaaggagcagctgctctggtggtct**ctcccaggctctgggggcggacccatgggag**gggctgtttttgtacaaagctgtaa**cattgtgGGGACAGGGGGCcacaatg**attcaactctacgggaaacctttacaaaaacct**ctctggcggtcccaactcccagag**tcctcttctttcctcctgggtcacaggtcttaatgcaatttggttcagaatgcctctgcctcactcctgatcacatgtcagaccaagactgtggacaaggacaggcccagatgagaactaaagcttccc**aggcagagagaggtcagacataagaagactgcct**caggaacctcacaagtggaggactcagggagggtcccaatccccaaaaattgagacaaagtcaggtggaaggttcatcggaggtgaccagctctccagaggactcgggaagaagt**caggggtatctatagatggagtcacaggttctgggcccctg**ccatcctctgcaggccatgcactttccctttcgatggaccctcacagagggagcatctgaatggggca**tcctttgaaaaagga**acc**taggaccctgtggatggactctgtcattctccatggtccta**aaaagcaaaagtcaaagtgttcttctgtgtaatacccataaagcaca**ggaggagatttcttagctcactgtcctcc**atcctagccagggccctctcccctctctatgccttcaatgtgattttcaccttgacccctgtcactgtgtgaacactgaagctttctttggacaaggcaccagactcacagttgtaggtaagacatttttcaggttcttttgcagatccgtcacagggaaaagtgggtccacagtgtcccttttagagtggctatattcttatgtgctaactatggctacaccttcggttcggggaccaggttaaccgttgtaggtaaggctgggggtctctaggaggggtgcgatgagggaggactctgtc**ctgggaaatgtcaaagagaacagagatcccag**ctcccggagccagactgagggagacgtcatgtcatgtcccgggattgagttcaggggaggctccctgtgagggcgaatcc

Uppercase: TRBD1-2

Lowercase: Flanking sequence[1000bp]

Red & Bold & Underline: Stem-loop [14]

Blue: Heptamer[38]

Green: Nonamer [7]

id-IGHD4-11-2[D_gene_segment]

acagccctccccatggggccctgctgcctcctcaggtcagccccggacatcccgggtttccccaggctgggcggtaggtttggggtgagg**tctgtgtcactgtggtattactatggttcggggagttattataaccacagtgtcacaga**gtccatcaaaaacccatccctgggagcctcccgccacagccctccctgcaggggaccggtacgtgccatgttaggattttgatcgaggagacag**caccatgggtatggtg**gctaccacagcagtgcagcctgtgacccaaacccgcagggcagcaggcacgatggacaggcccgtgactgaccacgctggg**ctccagcctgccagccctggag**atcatgaaacagatggccaaggtcaccctacaggtcatccagatctggctccgaggggtctgcatcgctgctgccctcccaacgccagtccaaatgggacagggacggcctcacagcaccatctgctgccatcaggccagcgatcccagaagcccctccctcaaggctgggccacatgtgtggacactgagagccctcatgtctgagtaggggcaccaggaggg**aggggctggccctgtgcactgtccctgcccct**gtggtccctggcctgcctggccctgacacctgagcctctcctgggtcatttccaagacagaagacattcctgg**ggacagccggagctgggcgtcgctcatcctgcccggccgtcc**tgagtcctgctcatttccagacctcaccggggaagccaacagaggactcgcctcccacattcagagacaaagaaccttccagaaatccctgcctctctccccagtggacaccctcttccaggacagtcctcagtggcatc**acagcggcctgagatccccaggacgcagcaccgctgt**caataggggccccaaatgcctggaccagggcctgcgtgggaaaggtctctggccacactcgggctttttgtgaagggccctcctgctgtgTGACTACAGTAACTACcatagtgatgaacccagtggcaaaaactggctggaaacccaggggctgtgtgcacgcctcagcttggagctctccagga**gcacaagagccgggcccaaggatttgtgcccagaccctcagcctctagggacacctgggc**catctcagcctgggctggtgccctgcacaccatcttcctccaaataggggcttcagagggctctgaggtgacctcactcatgaccacaggtgacctggcccttccctgccagctataccagaccctgtcttgacagatgccccgattccaacagccaattcctgggaccctgaatagctgtagacaccagcctcattccagtacctcctgccaattgcctggattcccat**cctggctggaatcaagaaggcagcatccgccagg**ctcccaacaggcaggactcccgcacaccctcctctgagaggccgctgtgttccg**cagggccaggccctg**gacagttcccctcacctgccactagagaaacacctgccattgtcgtcccc**acctggaaaagaccactcgtggagcccccagccccaggt**acagctgtagagagagtcctcgaggcccctaagaaggagccatgcccagttctgccgggaccctcggccaggccgacaggagtggacgctggagctgg**gcccacactgggccacataggagctcaccagtgagggc**aggagagcacatgccggggagcacc**cagcctcctgctgaccagaggcctgccccagagcccaggaggctg**cagaggcctctccagggagacactgtgcatgtctggtacctaagca**gccccccacgtccccagtcctgggggc**ccctggctca**gctgtctggaccctccctgttccctgggaagctcctcctgacagc**cccgcctccagttccaggtgtggttattgtcaggcgatgtcag**actgtggtggatatagtggctacgattaccacagt**ggtgccgcccatagcagcaaccaggcc

Uppercase: IGHD4-11-2

Lowercase: Flanking sequence[1000bp]

Red & Bold & Underline: Stem-loop [16]

Blue: Heptamer[46]

Green: Nonamer [8]

id-IGHD1-26[D_gene_segment]

atcccctccaggctctgggaggagaagcaggatgggagaatctgtgcgggaccctctcacagtggaatacctccacagc**ggctcaggccagatacaaaagcccctcagtgagcc**ctccactgca**gtgctgggcctgggggcagcccctcccacagaggacagacccagcac**cccgaagaagtcctgccagggggagctcagagccatgaaggagcaagatatggggaccccaatactggcacagacctcagctccatccaggcccaccaggacccaccatgggtggaacacctgtctccggcccctgctggctgtgaggcagctgg**cctctgtctcggacccccattccagacaccagacagagggacaggccccccagaaccagtgttgagggacacccctgtcc**agggcagccaagtccaagaggcgcgctgagcccagcaagggaaggcccccaaacaaaccaggaggtttctgaagctgtct**gtgtcacagtcgggtatagcagcggctaccacaatgacac**tgggcaggacagaaaccccatcccaagtcagccgaaggcagagagagcaggcaggacacatttaggatctgaggccacacctgacactcaagccaacagatgtctcccctccagggcgcc**ctgccctgttcagtgttcctgagaaaacaggggcag**cct**gaggggatccagggccaggagatgggtcccctc**taccccgaggaggagccaggcgggaatcccagc**cccctccccattgaggccatcctgcccagagggg**cccggacccaccc**cacacacccaggcagaatgtgtg**caggcctcaggctctgtgggtgccgctagctggggctgccagtc**ctcaccccacacctaaggtgag**ccacagccgccagagcctccacaggagaccccacgcagcagcccagcccct**acccaggaggccccagagctcagggcgcctgggt**ggattctgaacagccccgagtcacggtgGGTATAGTGGGAGCTACTACcactgtgagaaaagctatgtccaaaactgtctcccggccactgctggaggcccagccagagaagggaccagccgcccgaacatacgaccttcccagccctcatgacccccagcacttggagctccacagtgtccccattggatggtgaggacgggggccggggccatctgcacctcccaacatcacccccaggcagcacaggcacaaaccccaaatccagagccgacaccaggaacacagacaccccaataccctgggggaccctggccctggtgacttcccactgg**gatccacccccgtgtccacctggatc**aaagaccccaccgctgtctctgt**ccctcactcagggcctgctgaggg**gcgggtgctttggagcagactcaggtttaggggccaccattgtggggcccaacctcgaccaggacacagatttttctttcctgccctggggcaacacagacttt**ggggtctgtgcagggaggaccttctggaaagtcaccaagcacagagccc**tgactgaggtggtctcaggaag**acccccaggagggggt**ttgtgccccttcctctcatgtggacccc**atgccccccaagataggggcat**catgcagggcaggtcctccatgcagccaccactaggcaactccctggcgccggtccccactgcgcctc**catcccggctctggggatg**cagccaccatggccacaccaggcagcccgggtccagcaaccctgcagtgcccaagcccttggcaggattcccagaggctggagcccacccctcctcatccccccacacctgcacacacacacctaccccctgcccagtccccctccaggagggt**tggagccgcccatagggtgggcgctcca**ggtctcactcactcgcttcccttcctgggcaaaggagcctc**gtgccccggtcccccctgacggcgctgggcac**aggtgtgggtactgggccccagggctcctccagccccagctgccctgctctccctgg

Uppercase: IGHD1-26

Lowercase: Flanking sequence[1000bp]

Red & Bold & Underline: Stem-loop [19]

Blue: Heptamer[36]

Green: Nonamer [10]

id-IGHD1-7[D_gene_segment]

ccatcccctccaggctctgggaggagaagcaggatgggagaatctgtgcgggaccctctcacagtggaatacctccacagc**ggctcaggcaagacccaaaagcccctcagtgagcc**ctccactgcagtcctgggc**ctgggtagcagcccctcccacagaggatgaacccag**caccccgaggatg**tcctgccagggggagctcagagccatgaaggagcagga**tatgggacccccgatacaggcacagacctcagctccattcaggactgccacgtcctgccctgggaggaacccctttctctagtccctgcaggc**caggaggcagctgactcctg**acttggacgcctattccagacaccagacagaggggcaggccccccagaaccagggatgaggacgccccgtcaaggccagaaaagaccaagttgtgctgagcccagcaagggaaggtccccaaacaaaccaggaagtttctgaaggtgtct**gtgtcacagtggagtatagcagctcgtcccacagtgacac**tcgccaggccagaaaccccatcccaagtcagcggaatgcagagagagcagggaggacatgtttaggatctgaggccgcacctgacacccaggccagcagacgtctcctgtc**catggcaccctgccatg**tcctgcatttctggaagaacaagggcaggctgaagggggtccaggaccaggagatgggtcccctctacccagagaaggagcca**ggcaggacacaagccccctccccattgaggctgacctgcccagagggtcctgggc**ccaccc**cacacaccggggcggaatgtgtg**caggcctcggtctctgtgggtgttccgcta**gctggggctcacagtgctcaccccacacctaaaacgagccacagc**ctcagagcccctgaaggagaccccgcccacaagcccagcccccacccaggaggccccagagcacagggcgccccgtcggattctgaacagccccgag**tcacagtgGGTATAACTGGAACTACcactgtga**gaaaagcttcgtccaaaacggtctcctggccacagtcggaggccccgccagagaggggagcagccaccccaaacccatgttctgcc**ggctcccatgaccccgtgcacctggagcc**ccacagtgtccccactggatgggaggacaagggccgggggctccggcgggtcggggcaggggcttgatggcttccttctgccgtgg**ctccagtgcccctggctggag**ttgacccttctgacaagtgtcctcagagagtcagggatcagtggcacctcccaacatcaaccccacgcagcccaggcacaaaccccacat**ccagggccaactccaggaacagagacaccccaataccctgg**gggaccccaaccctgatgactcccgtcccatctctgtccctcacttggggcctgctgcggggcgagcacttgggagcaaactcaggcttaggggacacca**ctgtgggcctgacctcgagcaggccacag**acccttc**cctcctgccctggtgcagcacagactttggggtctgggcagggagg**aacttctggcaggtcaccaagcacagagcccccaggctgaggtggccccagggggaaccccagcaggtggcccactacccttcctcccagctggaccccatgtcttccccaagataggggtgccatccaaggcaggtcctccat**ggagcccccttcaggctcc**tctccagaccccactgggcctcagtccccactctaggaatgcagccaccacgggcacaccaggcagcccaggcccagc**caccctgcagtgcccaagcccacaccctggaggagagcagggtg**cgtctgggaggggctgggctccccacccccacc**cccacctgcacaccccacccacccttgcccgggccccctgcaggaggg**tcagagcccccatgggatatggacttagggtctcactcacgcacctcccctcctgggagaaggggtctcatgcccagatccccccagcagc

Uppercase: IGHD1-7

Lowercase: Flanking sequence[1000bp]

Red & Bold & Underline: Stem-loop [19]

Blue: Heptamer[33]

Green: Nonamer [7]
