## Supplementary Data 3 for "Adaptive immunity: from CRISPR to CRIHSP?"

id-TRAJ13[J_gene_segment]

aaataagtttcagagttgtctagagaatatcacttggactgttctcaggctttccactacatctattgtaccactatggtgccctgaaaagaggctttctcaggcttaattccatggcttagtctttattccagtatcaaaaagggggaatcccagccagagctcccatgagggaggatagctgcatgctaaccacattaatctattatcaaggtaactcggtcatttttgtcaggcagcacagtgctgtgatttatagcacattcatctttgggagtgggacaagattatcagtaaaacctggtaagtaggcaatatgtcactaaagtaggaggcttaatgtggctactgagacccactaaacttactgcagtatttggaaggcccaagtgtcaagaaattaatggtttatgcagacttaagtggattccaatgaaggaagaaatgttaaagtaatggcacagaggatagaagagctagctgtgaaaaaaaatagccatgtggatgaaccaaacgcaacagactgaaagagcctagagagttgatcctaaagaaaaag**gcagaaagtctggctgagcacttccagttctgc**tgacctggcttgattagtgaatctgggccacagtatctttatctat**aaaatgaaaacatttaatatcacctcagattttttgttttaatttt**gtgggtacataataggtgtatatatttattttaaagataagctaatatatgtgaaagtattttgtaaattcttaaatgctatccacatgtaaaatatta**tttgtttatagaacaaa**aaaaaactatgtcagatttag**aattcttttaacatgagaatt**tgtaggttcttagtgaaatagccatttacatgtgatctgtagccctaggcattggtaaaaagcactttctcttctactgaaatatagtggctcaaatgctaccc**aaatgaaaagggcaggggaagggaagtcattt**tgtaaaggcaggcattacagtgTGAA**TTCTGGGGGTTACCAGAA**AGTTACCTTTGGAACTGGAACAAAGCTCCAAGTCATCCCAAgtgagtccaatttcctatgctttcctcttccttgtgttgtcttctctcagaccgtaacatttggagcacat**aaagagatattatgggcctgctcttt**tctggctg**tttcaggagactgaaa**aggattgta**taagaacatcctttagcatgttctta**gtattgttttgtccagtgtgtgtttcttcttaatatcaaaaaaaactaagacattgcctgtgaagccaatggaaaacagtagatttgcaaagaaaagaactaaagagaagacaagtggtactttttaccaagc**ttaaagcagagtgggtgtcctttaa**tcagagggatgcttggttctgggggctggatag**gactcaggtaagtaacaaggctaccactgagtc**actg**agcttctaaaatgagttgactcgggggagcaggaagct**gccactcatgatg**atctgaatgctattccgtgaagtttcagat**tgattgaagttaaaattaggggtgacactatgtcaggaatctaaaaattgataactgtaatggaaatgaaagaagggctagctacatctgttggataaatggagttgaaaatcatgaatttgatggcgacagacccagagagttgggcaggtagaaacttcatttcaagtgatatgcagtaaataccccaaatttgagtcaaatggcctgtgtaaaaaggaagtgaaaatgtaacccgtatctccttgtaggtgaaaaggccatatctctaggtcttaggtatgaaaaggatgtgtgggcttctgg**gtgtttttgactgactaagaaacac**tgtgggatggatagcagctataaattgatcttcgggagtgg**gaccagactgctggtc**aggcctggtaagtaaggtgtcagagaggcaacagaaagattgagggtaaaatgtcttcatgtctcaggaaatgattgtataatgcaaaatgagctggagttttttaggagcactgttactagagttctgatgtctggttctagcaaa

Uppercase: TRAJ13

Lowercase: Flanking sequence[1000bp]

Red & Bold & Underline: Stem-loop [15]

Blue: Heptamer[32]

Green: Nonamer [5]

id-TRBJ1-4-2[J_gene_segment]

acccaggcttcccagaggctctgagcagtcacagctgagcccagggtgatggggcagaag**agggaaggggagggggcctctcctcatagttccct**gagatagcccagagaaagcccggtgggtaatgaatgagccacaacacctctccatctatctgcttcactg**acagaggttctctgt**agattcttcgtatattcctgtgctggattttataggaggccactctgtgtctctttttgtcacctgcctgagtcttgggca**agctctggaagggaacacagagtactggaagcagagct**gctgtccctgtgagggaa**gagttcccatgaactc**ccaac**ctctgcctgaatcccagctgtgctcagcagag**actggggggttttgaagtggccctgggaggctgtgctctggaaacaccatatattttggagagggaagttggctcactgttgtaggtgagtaagtcaaggctggacagctgggaacttgcaaaaaggggctggaatccagacggagcctttgtctctagtgcttaggtgaaagtgtatttttgtcaggaaggcctatgaggcagat**gaggaggggatagcctccctctcctc**tcgactattttgtagactgcctgtgccaagttaggttcccctactgagagatgggtagactcagcttggaaggggtcaccttgaacatctcctgtctccttgaagggtgccggtcacggccatgacagataaaagagcctctgaccttaccaccacggtcctaccgtttctc**tccctcacacagaaaggagaaggtcacagaagaggga**acttgggggatcacacggggcctaattggtctgctgaccaccgcattttgggttgtaccattgtctacccctctacccaccagggctaaaattctactaaggaacaggagaggacctggcaggtggacttggggaggcag**gagtggaaggcagcaggtcgcggttttccttccagtc**tttaatgttgtgCAACTAATGAAAAACTGTTTTTTGGCAGTGGAACCCAGCTCTCTGTCTTGGgtatgtaaaagacttctttcgggatagtgtatcataaggtcggagttccaggaggaccccttgcgggagggcagaaactgagaacacagccaagaaaagctcataaaatgtgggtcagtggagtgtgtggtggggccccaagagttctgtgtgtaagcagcttctggaaggaagggcccacaccagctcctctggggtttgccacactcatgatgcactgtgtagcaatcagccccagcattttggtgatgggactcgactctccatcctaggtaagttgcagaatcagggtggtatggccattgtcccttgaaggcagagttctctgcttctcctcccggtgctggtgaggcagattgagtaaaat**ctcttaccccatggggtaagag**ctgtgcctgtgcctgcgttccctttggtgtgtcttggttgactcctctatttctcttctctaagtcttcagtccataatctgcctcctcactcccttcttggctcatcctccctcttatgtgcatggctctgcctctcctaagcctcttcctcttgcgccttatgctgcacagtatgcttaggcctttttcctaacagaatccctttggtccagagccatgaatccaggcagagaaaggcagccatcctgctgtcagggagctaagacttgccctctgactggagatcgccgggtgggttttatctaagcctctgcagctgtgctcctataattcacccctccactttgggaacgggaccaggctcactgtgacaggtatgggggctccactcttgactcgggggtgcctgggtttgactgcaatgatcagttgctgggaagggaattgagt**gtaagaacggaggtcagggtcaccccttcttac**ctggagcactgtgccctctcctcccctccctggagctcttccagcttgttgctctgctgtgttgcctgcagttcctcagctgtagagctccttgcttagtcttcagggctgtgtgtttctttgctc

Uppercase: TRBJ1-4-2

Lowercase: Flanking sequence[1000bp]

Red & Bold & Underline: Stem-loop [10]

Blue: Heptamer[47]

Green: Nonamer [4]

id-TRAJ52[J_gene_segment]

tttgatcatcaccacatcatgaaactgc**ctggattttgctcttcacattttttgtgacatccagacacagccagggtgactcccaggattcctgtggttgtgtctgttggcaacaga**agcattatttggtcaagggatgaa**atttacaggaaccccaagtaaat**gtgcaaaagcctgcccctaatagagacctcacttaaccctaccactgtggtcaccgggctgctctg**tgaggtcagaacacctca**ggtgagactgtggagtaaccaga**tgtcctggttttcctgggtttcaggggttttccaggaca**aagg**tcattctttgctaagcccagaatga**tccccaggatagttaatcaccgtaagaagg**cacagataaaaacttttttctgtg**agctactacttacatttcaccaaattcaagactgacaaaaaaactacgttgtgtgaaagctcagttacatgtgcccatcatttttatttcttatgctaggg**catgaagggaagctcaggagatcctgtttcatg**aggctacagtctaaatctcatctaattctagtcattctccaaaaagctgaacactctcctcttgtctccttgtaatcaattcattgtcatcagaaatgtgtgacacctcgaggggaggggaggacgtgtcttgaaaactgatcagagaaattgtcctgaaaatgatgctgtaaaatggaaatgagacctttagaaagagagatgctgaacatgagaaatcagtaagcctcaagacaaaaaccagaggaaatggagagagatgggaggcaggaagccggacctcgggggacgcagcaggtcagtgtcccaggaccagaggtggaaatggtcttccagcctttggtgggatgaaactgagcctcacagacccaggcccctttgggtgagtttattctccccagaattaaaagaggaggaagtctgcgagcttcagaagtccccctagggttcttgtaaaggcctccagtgcagtgCTAATGCTGGTGGTACTAGCTATGGAAAGCTGACATT**TGGACAAGGGACCATCTTGACTGTCCA**TCCAAgtaagtgtaacaagacacagcagtatactggaaattctggaatgtcaccccagtgtcaggttttaaagaaccagatattgcttaaagttaagtggtgtcctc**tgaggtcctttctcttctttaacctca**atattcacagattttcttttggaccctgagttgttaggtctgtctccctgctgcgtgtacagctgaccttgaccct**ggaaagtcagaatctttcc**tgctttcagtgtcttctctggaggtgatgccctaacttttttgag**ggggaggtcatcctttccaaatgactcccc**tgagtaacctgttctaatgtttagtagcattccctgtcagaagtccttccttacctcaatcttaaattattcttggtatagccgaggccgtttccaccagcatcatccacaataaagaggaaaattgagagcggagtttgaataggtatgtggtatttggttcagaatgggttttaaatgaaagtggagagggaaaaaatgaaacctgaatgggaatctgaggtcttatttcaaatccctctctgtttcctgtctgtcatctcc**ttttgcttctatagatgcaaaa**ctacagagagagaggcggggagagaaaggatagacagaaataaacatccactgtgtccattccctgtcactgaaaccaccaggcaagggattacaattaattccaaagagtttcagatacacggcttaatacctttctccttgtggtgacttgtatcctggctgataattagagcagatagaaagatttggctttggtgtgtttgccgctgcaccccgctgggaagagacagaaacttgactgttttttaagattcataaagttccttctgtcagtcgttgtaaaa**ctccctgaagcagggag**atgcgtgacagctatgagaagctgatatttggaaaggagacatgacta**actgtgaagccaagcaagctggaaagacctaaactcacagt**gttccttatgtacttttgctccc

Uppercase: TRAJ52

Lowercase: Flanking sequence[1000bp]

Red & Bold & Underline: Stem-loop [16]

Blue: Heptamer[24]

Green: Nonamer [5]

id-TRAJ34[J_gene_segment]

cattgcgggtccggcactcaagtgattgttttaccacgtaagtatatcttttctcatttctgtgggctgttatgtccgtaaatcatatagaacagttcttcttatgcacacacacacacacacacccattttccagtcactgaggagaaacagtctctgggtctacctacacagccag**tttgaccattaagatggtcaaa**ttcatttttcaagataattttaactggttccatt**taatgctgccaacagccaagggcatta**gcatgtcgtgggaacctaatctatgttcactcactgctgtgggaggacagtcatttcttcttctcccctggcagtcagacttcagaatccagctgaccttttccatgcctgtctcttctggcagattaagtg**gttatttccactaactttcttgtggaaaaac**acacaagacagatatattctccttcagtttttactaaga**aagttttgtggttatcattttgacaataaactt**cagctagcatagtccctactattcatgttaataatatcttgacttgaaccatcatattttccccacatcattcacaatgcatttcaatgctgtaacccactgcaccatccccagcccttttatcttcacaacgttctctaataaggtaaacaggatcatacatcatcaaacatatgtcacagacagaaaagctgacacacagacaagttaaataatttaagtcaaaccaaaaaaaaatcataactgggattagagcttgagattccctattttcttacagtgaactgtcttgagctgtcacttaatgcattgaaaacataataagcacaggccataggcat**tttcagaacatgaaacttgactgaaa**caaaaactgactgtttgattgctttggtgttttaaatgtatcc**ccaaatctcctttgctattgatagtcatttgg**tgccagtgacagggaagagcagaggggcttaggaggtttttgtagatctcagtatcactgtgTCTTATAACACCGACAAGCTCATCTTTGGGACTGGGACCAGATTACAAGTCTTTCCAAgtaagtactagaaaccaaggagccattttgaaaaggtgttttctttttctttttcttttttttttaaatgaagttttgctctttgttgcgcaggctggagtgcagtggcatgatctcggctcactacaacctccacctcccgggttcaagcgattctcctgtctcagcctcccgagtagctgggattacaggcacctaccaccacgcctggctaatttttgtatttttagtagaaacagtgtttcaccatgttggccaggctggtcttgaactcctgacctcaagtgatctgcctgcctc**ggcctgccaaagattacaggcc**actgcgcccagccttaaaaggtgttttctataaacaggaacgttctcacagcactgcaggccacccttaccaactccgagctctgtaggggcctttagcacctatagctgcagaaccttctataaaccaattaatctatatgtgaacaatttaacctatatgtgaacaccatggtactgagtaaaatcttaccagcagtgaaaagaaggcactagagcgtcagatgcattatgactctaaacaaatgatctgt**ttccctcttttgcatttttatttaaaatatcagggaa**ggtgaagcaagttcaacttctccaatttgaaaactgtttctgaattatgcttcccttgtatgcagagagacctagattgactttggatgtttagtcattcg**atttaatcaaatgtaattaaat**gagcaaataatctccaatagccaatattcctatgttgtttcattgttttatgtgctttgctctagactttttgtctgggctttgtctctaataggatccccggaaggacagtgaaggtttttgttaaggtttttgtgtctgtgtggatagcaactatcagttaatctggggcgctgggaccaagctaattataaagccaggtaagtctcagagatgtgactgcacgggagaggagacactagttgaataatgcacaaagtgtagc

Uppercase: TRAJ34

Lowercase: Flanking sequence[1000bp]

Red & Bold & Underline: Stem-loop [9]

Blue: Heptamer[36]

Green: Nonamer [6]

id-TRBJ1-3-2[J_gene_segment]

tttccctttcgatggaccctcacagagggagcatctgaatggggca**tcctttgaaaaagga**acc**taggaccctgtggatggactctgtcattctccatggtccta**aaaagcaaaagtcaaagtgttcttctgtgtaatacccataaagcaca**ggaggagatttcttagctcactgtcctcc**atcctagccagggccctctcccctctctatgccttcaatgtgattttcaccttgacccctgtcactgtgtgaacactgaagctttctttggacaaggcaccagactcacagttgtaggtaagacatttttcaggttcttttgcagatccgtcacagggaaaagtgggtccacagtgtcccttttagagtggctatattcttatgtgctaactatggctacaccttcggttcggggaccaggttaaccgttgtaggtaaggctgggggtctctaggaggggtgcgatgagggaggactctgtc**ctgggaaatgtcaaagagaacagagatcccag**ctcccggagccagactgagggagacgtcatgtcatgtcccgggattgagttcaggggaggctccctgtgagggcgaatccacccaggcttcccagaggctctgagcagtcacagctgagcccagggtgatggggcagaag**agggaaggggagggggcctctcctcatagttccct**gagatagcccagagaaagcccggtgggtaatgaatgagccacaacacctctccatctatctgcttcactg**acagaggttctctgt**agattcttcgtatattcctgtgctggattttataggaggccactctgtgtctctttttgtcacctgcctgagtcttgggca**agctctggaagggaacacagagtactggaagcagagct**gctgtccctgtgagggaa**gagttcccatgaactc**ccaac**ctctgcctgaatcccagctgtgctcagcagag**actggggggttttgaagtggccctgggaggctgtgCTCTGGAAACACCATATATTTTGGAGAGGGAAGTTGGCTCACTGTTGTAGgtgagtaagtcaaggctggacagctgggaacttgcaaaaaggggctggaatccagacggagcctttgtctctagtgcttaggtgaaagtgtatttttgtcaggaaggcctatgaggcagat**gaggaggggatagcctccctctcctc**tcgactattttgtagactgcctgtgccaagttaggttcccctactgagagatgggtagactcagcttggaaggggtcaccttgaacatctcctgtctccttgaagggtgccggtcacggccatgacagataaaagagcctctgaccttaccaccacggtcctaccgtttctc**tccctcacacagaaaggagaaggtcacagaagaggga**acttgggggatcacacggggcctaattggtctgctgaccaccgcattttgggttgtaccattgtctacccctctacccaccagggctaaaattctactaaggaacaggagaggacctggcaggtggacttggggaggcag**gagtggaaggcagcaggtcgcggttttccttccagtc**tttaatgttgtgcaactaatgaaaaactgttttttggcagtggaacccagctctctgtcttgggtatgtaaaagacttctttcgggatagtgtatcataaggtcggagttccaggaggaccccttgcgggagggcagaaactgagaacacagccaagaaaagctcataaaatgtgggtcagtggagtgtgtggtggggccccaagagttctgtgtgtaagcagcttctggaaggaagggcccacaccagctcctctggggtttgccacactcatgatgcactgtgtagcaatcagccccagcattttggtgatgggactcgactctccatcctaggtaagttgcagaatcagggtggtatggccattgtcccttgaaggcagagttctctgcttctcctcccggtgctggtgaggcagattgagtaaaatctcttaccccatggggtaagagctgtgcctgtgcctg

Uppercase: TRBJ1-3-2

Lowercase: Flanking sequence[1000bp]

Red & Bold & Underline: Stem-loop [12]

Blue: Heptamer[53]

Green: Nonamer [6]

id-TRAJ2[J_gene_segment]

tgta**agcaacaaagacccaaggagccttttagaatgttgct**ctgggaccccagtggcttccagcagattagcatcaagaaggttatctcaaagaccttacccacagtgggggtacagcagtgcttccaagataatctttggatcagggaccagactcagcatccggccaagtaagtagaatgaagcaggagagcaagggaggac**ggacaactatttcttctttgtcc**aaaatgccaacttgaacccaggtattttcctctgtgggctccaaaatggtttcttccacccatccccagaaaatcctcccatccagctctcggtgctctgcagacatcatccttaggctgggaccttaggaaccatgaggcaggcaatgagcgagtactcctgaaccttccacaaacatatccaagatactgctgcctggaattgaggtttttgttctagggaattattggtttattgaaatcatatgaagtcacaacctcagcccagggaatcacagcagaatggaacaggaggagaa**cttttgtatcagaagcgaaatttcagtgcaaaag**agcagaaggtttaggcct**ggcgtggtggctcacgcc**tgtaatcctagcactttgggaggccgaggcagggagatcacaaggtcaggagatcaagaccatcctggctaacacagtgaaaccctatctctactaaaaatacaaaaattagccgggcgtgatggcacgcgcctgt**agtctcagctactcgggagact**gaggcaggagaattgttgaacccgggaggcagagattgcagtgagctgagatcgcatcattgcactccagcctgggcaacagagcaagact**ccttctcaaaaaaacaaaaaaaaaaaaaaaaaagaagg**tctaacccttaggagtgtgattatcctgttctcctgccttgtgggggagatcagtgtttctcttatttaagtaatag**gtaggtcctgtgagtttgtgcaatggtgtcacctac**ggtaTGAATACTGGAGGAACAATTGATAAACTCACATTTGGGAAAGGGACCCATGTATTCATTATATCTGgtgagtcatcccaggtggcaccacgtgcaaccccatgggccagtgtcactaatcctttctctggagatatcacttattactatggtgaggcttgctgtagatgttgtaactaattttcttacagaggtctgggaagggaaaagcattactatctatcttgaatattcatgtttctctag**gtcaaacacattaaaatttgac**tttaatcattcaatgggtattgtaaaatgccttctatgtgactatcactctataaaatgttagactgagtatgaagtgtgagatagattcctgtcctgtcctcaagtggtataaaaactagacaaaggtactgaactattgtaaattaagcagccagaaaactattttagt**attgaccagttgatgatgtcaat**ggacacaatagagattcagaggggatagggtgctggctgagaagttggcctagactgagaagtttcctgtggtacaaaggattcattgagccctgaaggatggataagatctgtatgggcagagaaaggagagaagggaagttctgggcgtagggaacgacaagaaagaaggcatgatcttgggaataatcaaggcacatgcaaagtagcctaagtatacatctgataataaaattggttgaaaagtagtcagagaagatgt**ctttttaggcatggaaaaag**gaaatactagagcattcaacagaatacagaaattagggccagggccagccattgggaaactgagaatccgatttagagatgcagactagaagtgaaggtgagagcagccagctatggtgccgcagacct**cccctctccttcctcagtgggctctgagagggg**tcatcccaca**ccttagaggaggagaaacctaagg**gattctgtaatagagacacggggcatggtatgaaagtattacctcccagttgcaatttggcaaaggaaccagagtttccacttctccccgtacgtctgcccatgcccacagtttcctgatgctcact

Uppercase: TRAJ2

Lowercase: Flanking sequence[1000bp]

Red & Bold & Underline: Stem-loop [12]

Blue: Heptamer[21]

Green: Nonamer [2]

id-IGHJ2P[J_gene_segment]

tcagatctgctctaatttacaaaagaaaaggaaaaacacacttggcagccttcagcactctaatgattcttaacagcagcaaattattggcacaagactccagagtgactggcagggttgagggctgggg**tctcccgcgtgttttggggctaacagcggaagggaga**gcactggcaaaggtgctgggggcccctggacccgacccgccctggagaccgcagccacatcagcccccagccccacaggccccctaccagccgcagggttttggctgagctgagaac**cactgtgctaactggggacacagtg**attggcagct**ctacaaaaaccatgctcccccgggaccccgggctgtgggtttctgtag**cccctggctcagggctgactcaccgtggctgaatacttccagcactggggccagggcaccctggtcaccgtctcctcaggtgagtctgctgtctggggatagcggggagccaggtgtactgggccaggcaagggctttggcttcagacttggggacaggtgctcagcaaaggaggtcggcaggagggcggagggtgtgtttttgtatgggagaagcaggagggcagaggctgtgctactggtacttcgatctctggggccgtggcaccctggtcactgtctcctcaggtgagtcccactgcagccccctcccagtcttctctgtccaggcac**caggccaggtatctggggtctgcagccggcctgggtctggcctg**aggccacaccagctgccatccctggggtctccgccatgggctgcatgccagagccctgctgtcacttagccctggggcca**gctggagcccccaaggacaggcagggaccccgctgggcttcagc**cccgtcagggaccctccacaggtagcaagcaggccgagggcagggacgggaaggagaagttgtgggcagagcctgggctggggctgggcgctggctgttcatgtgccggggaccaggcctgcgctttagtgtgGCTACA**AGTGCTTGGAGCACT**GGGG**CCAGGGCAGCCCGGCCACCGTCTCCCTGG**GAACGTcacccctccctgcctgggtctcagcccgggggtctgtgtggctggggacagggacgccggctgcctctgctctgtgcttgggccatgtgacccattcgagtg**tcctgcacgggcacaggtttatgtctgggcaggaacagggactgtgtccctgt**gtgatgcttttgatatctggggccaagggacaatggtcaccgtctcttcaggtaagatggctttccttctgcctcctttctctggg**cccagcgtcctctgtcctggagctggg**agataatgtccgggggctccttggtctgcgctgggccatgtggggccctccggggctccttctccggctgtttgggaccacgttcagcagaaggcctttctttgggaactgg**gactctgctgctggggcaaagggtgggcagagtc**atgcttgtgctggggacaaaatgaccttgggacacggggctggctgccacggccggcccgggacagtcggagagtcaggtttttgtgcaccccttaatggggcctcccacaatgtgactactttgactactggggccagggaaccctggtcaccgtctcctcaggtgagtcctcacaacctctctcctgctttaactctgaagggttttgctgcatttttggggggaaataagcgtgctgggtctcctgccaaga**gagccccggagcagcctggggggctcaggaggatgccctgag**gcaacagcggccacacagacgaggggcaa**gggctccagatgctccttcctcctgagccc**agcagcacgggtctctctgtgg**ccagggccaccctgg**gcctctggggtccaatgtccaacaacc**cccgggccctccccggg**ctcagtctgagagggtcccagggacttagcgggg**tgccagttcttgcctggggtcctggca**tt**gttgtcacaatgtgacaac**tggttcgacccctggggccagggaaccctggtcaccgtctcctcaggtgagtcctcaccaccccctc

Uppercase: IGHJ2P

Lowercase: Flanking sequence[1000bp]

Red & Bold & Underline: Stem-loop [18]

Blue: Heptamer[37]

Green: Nonamer [6]

id-TRAJ3[J_gene_segment]

taaagcaccatctgattgtgtgttttctggtggctacaataagctgatttttggagcagggaccaggctggctgtacacccatgtgagtatgaccctg**caagtgaccagtgcaaaaagaactgaccacttg**tctatgagaaaacaggtgatgatctatagcaaacttggggatatattgagaagcactcattttccattcctacaagctctcttggtgacttaaaatgttcccatttctcatatgaaacacagaagtgccaagagccaattgcgtaaatgaaaatctgagcagaataatttatagaaacataaaaagagttccaaataatgagtgctttcaagtgaagctgataatatcggcaccagataatctgaacattcaggaaattagctaagtgcccttggggg**aaagagatgagttaatggcaaacacaaattctcttt**tattgcagcatgtagctaaattccctggattggctaataaacatggtatggggatccctactcatagaatcttcctctctggag**ttaggagagttttcagcttcctcctaa**gggtaaaagtggcccagacccctaactgaacggcaag**tataccacatgcaattacaaggtata**gaggcagcaatgtagagggaggcccacagacctggggaaaatctgggctctactctcactggctgttttcaataaagcctccactttctcagcagcaaaatgactataactacatcataataatatgagataatggccagaatgtaaaaagtgatcaataagtagtagcttttctaaaatattattataattattattattgaggctttgcttatactaaggtttacactatcataaagggaaaggagaatc**gcttccttgggaaataaaagatgtaagcaacaaagacccaaggagc**cttttagaatgttgctctgggaccccagtggcttccagcagattagcatcaagaaggttatctcaaagaccttacccacagtgGGGGTACAGCAGTGCTTCCAAGATAATCTTTGGATCAGGGACCAGACTCAGCATCCGGCCAAgtaagtagaatgaagcaggagagcaagggaggac**ggacaactatttcttctttgtcc**aaaatgccaacttgaacccaggtattttcctctgtgggctccaaaatggtttcttccacccatccccagaaaatcctcccatccagctctcggtgctctgcagacatcatccttaggctgggaccttaggaaccatgaggcaggcaatgagcgagtactcctgaaccttccacaaacatatccaagatactgctgcctggaattgaggtttttgttctagggaattattggtttattgaaatcatatgaagtcacaacctcagcccagggaatcacagcagaatggaacaggaggagaa**cttttgtatcagaagcgaaatttcagtgcaaaag**agcagaaggtttaggcct**ggcgtggtggctcacgcc**tgtaatcctagcactttgggaggccgaggcagggagatcacaaggtcaggagatcaagaccatcctggctaacacagtgaaaccctatctctactaaaaatacaaaaattagccgggcgtgatggcacgcgcctgt**agtctcagctactcgggagact**gaggcaggagaattgttgaacccgggaggcagagattgcagtgagctgagatcgcatcattgcactccagcctgggcaacagagcaagact**ccttctcaaaaaaacaaaaaaaaaaaaaaaaaagaagg**tctaacccttaggagtgtgattatcctgttctcctgccttgtgggggagatcagtgtttctcttatttaagtaatag**gtaggtcctgtgagtttgtgcaatggtgtcacctac**ggtatgaatactggaggaacaattgataaactcacatttgggaaagggacccatgtattcattatatctggtgagtcatcccaggtggcaccacgtgcaaccccatgggccagtgtcactaatcctttctctggagatatcacttattactatggtgaggcttgctgtagatgt

Uppercase: TRAJ3

Lowercase: Flanking sequence[1000bp]

Red & Bold & Underline: Stem-loop [11]

Blue: Heptamer[16]

Green: Nonamer [3]

id-TRGJP1[J_gene_segment]

gaattttctattaagaagaaaccctttccaaacataattttcacc**cagccccattgtcctctccattgaattgggggctg**tgagctgctgtgtatggacaggtaccctctgtga**cctcattcatatcttcctgaatgagg**actcaggggacttaggctggagcagggtggatctggagtggacatggaggttgtcaaggcttgtcctcctcagcccttgcaagtgaacacctcatctgtactcaggagatcacacagctgaccccggcttaagatggcagatttttcaacatattagataactggatttgttttttttggtcatgcctcagtataattaaagaatgtattatttcatgggagatagaattctacaa**atttcttctattattccaggagaaat**aataagtcattgcagaacaca**ttttttcatttggcattgccttggtaaagcatcatgttgccaaatgataaaa**gtggctt**tttgaaagtggagaccgttgttcaaa**tttctcaagtatatctgcattgacacatttgcattttcagttactaaagagcttcctcctttcagaagtatc**ttcccaaattgctagcagagaggctctggtgtttgtgggaa**cagaagggcttttc**ctttgatgtaatctcaaag**cagtttcaacacaattgaacccctggaaaaaaaaaagacaattctctgaata**cttttcctggtaatttagaaaag**cagtgggttgacccattaaacctgctgttttctcttcctgcttttttttttccatttcccttttctgttggtgcttttcaagaattactgccttagaagaaaacaggcaatttatgaggaaaaattacaaactatcacatgtcacaaaacctatattcaaggacttccaaaaaagccagaagatgaaattgctagttcaaagttgttggattgctagtcgtgtcccgaggatcagaaggctgagatttttgtagaagcttagaccggtgtgATACCACTGGTTGGTTCAAGATATTTGCTGAAGGGACTAAGCTCATAGTAACTTCACCTGgtaagtaacttttctttct**gtttttattccagtaatgaaaaac**tgata**gatgatttttagaaaaaaatgatc**aaccttatctgaatatgtcacattcctggcctcagtatacgaacagcaattttgcagattagctgaatatggaa**acaaaacgattttgt**acatgatatcttcccttccctgtctgtaagcggattaaaattctactttaagaatacacacacacacacacacacacacacacacacaaaagatagactctggcaagaaattaaaatgttttgaggcccatgtctatacaccattagtgtaagtccttggtccaagttacttttttgttggtcttttaagtgagagac**ataaaaatggagttttat**taattacttcttctctgtattttgagatataacatagtgctcccttcttctgctagcctgccaaacccatgttattgattttgatctcatcctgagaaatagtatagaaaaaaatta**cttcttatagtaaattactatagaag**gtcaatacatcccacttattagtgttctagtttctttgagggaggctgaagtcagtcaaaatttatgcaattgggactcttgttatatcacctggaaatggcccggtttaatagaatttgtaagttcaaccagccatgtcagctccaaggtc**aatgaaaagagtcttcatcattttacatcttcatt**tgttcattcatcctctcagcaaatatttat**caaaaacagtttttg**ttctggatgctagagataaaagggcaaagaagacacagtctcaacactcaaaag**ctgccagtctggcag**ataagtattctagattctggcttggttgggccctgtccacgttctttggcctgtggctgccggagaagcagaaggtcttgggttcatgagctcattttctttcttaatttgtgtagcagttattgcttataagatctatgtattgcgtttggtgggaggatttgataaaatgacccaat

Uppercase: TRGJP1

Lowercase: Flanking sequence[1000bp]

Red & Bold & Underline: Stem-loop [16]

Blue: Heptamer[17]

Green: Nonamer [5]

id-TRAJ11[J_gene_segment]

aaggctaccactgagtcactg**agcttctaaaatgagttgactcgggggagcaggaagct**gccactcatgatg**atctgaatgctattccgtgaagtttcagat**tgattgaagttaaaattaggggtgacactatgtcaggaatctaaaaattgataactgtaatggaaatgaaagaagggctagctacatctgttggataaatggagttgaaaatcatgaatttgatggcgacagacccagagagttgggcaggtagaaacttcatttcaagtgatatgcagtaaataccccaaatttgagtcaaatggcctgtgtaaaaaggaagtgaaaatgtaacccgtatctccttgtaggtgaaaaggccatatctctaggtcttaggtatgaaaaggatgtgtgggcttctgg**gtgtttttgactgactaagaaacac**tgtgggatggatagcagctataaattgatcttcgggagtgg**gaccagactgctggtc**aggcctggtaagtaaggtgtcagagaggcaacagaaagattgagggtaaaatgtcttcatgtctcaggaaatgattgtataatgcaaaatgagctggagttttttaggagcactgttactagagttctgatgtct**ggttctagcaaaagaacc**ctaatatagctgtaatccctctctagaaggaacaaggaagaggattcttctctgaaagctcctcctaagatttatccatcctctgccaaagtgtcttgtggatccaggatacttacatctttgttagcagtgctggcccaacctgctcagcctcagatcacctatc**cctcatttgtaaaatgagg**ataatatttaagctcatggcacagttgaagcccaaatgaggtagtggaatatatgcagattattaatccatttggctcataaatgaaggttttctttccttttccctaagtagatcatgggttgaaatacctaacact**gcaaatcatttttgtatggggatttgc**tatagtgTGAATTCAGGATACAGCACCCTCACCTTTGGGAAGGGGACTATGCTTCTAGTCTCTCCAGgtacatgttgaccccatcccacccatgttttccccctatctggtttaaggcttccatatgtattgcgtgttatcctcatggatttcatcatccttgttttattatcaatgttctgtgaatttaagattgagcctccatgg**actcttcatttaaaaatgaaaatagctaatagaagagt**tggaaataacagtagaactaattcactaggtcaggatggagaagggagtaataccctagacaattaggacaaacgtggtttttccagaaatagactcacttcctgtttaaagcctagactgtggtctctccccgggcacccttcacattcctctaaccctccatatcccaaatt**tagctgtggaatcttagacaatctgtgacctatgcagcta**cagaa**atcttttcttactgagaatatctgcctatttgaaccaaaagat**ccttgagaatagacagtctctaatattccatgaagtgtctgatatggtccttttacaaaagtaaatacttgataaattcttgcttaattgaagttaaacacccaaagaaatcaggtttccaggccaaagggaagaagaaaattaag**caaatgacacagtacatttg**agcgtgtagtggggaggaagaaatgctagattgggattcaaatagagccatgttt**tggtctctgctgccaactagcaatgtaacttagacca**gatctcttgcctctaagcctcagtttctttatctgtaaatggggaagtgggtccaatggcctctctggcctcttagtac**aaaaagtctatgaatgtatcccatttggggcacttttt**tctcctaggagta**actcaatagctattttaaagcttgagt**gttttctaactcattagagcacatcaagaagaggggtgaagtgacaaaagggagtttattgtgaggcatcaaacactgtgatactcacgggaggaggaaacaaactcacctttgggacaggcactcagctaaaagtggaactcag

Uppercase: TRAJ11

Lowercase: Flanking sequence[1000bp]

Red & Bold & Underline: Stem-loop [14]

Blue: Heptamer[24]

Green: Nonamer [4]

id-TRBJ2-6[J_gene_segment]

taccgtgtccagctaactcgagacaggaaaagataggctcaggaaagagaggaagggtgtgccctctgtctgtgctaagggaggtg**gggaaggagaaggaattctgggcagccccttccc**actgtgctcctacaatgagcagttcttcgggccagggacacggctcaccgtgctaggtaagaagggggctccaggtgggagagagggtgagcagcccagcctgcacgaccccagaaccctgttcttaggggagtggacactgggcaatccagggccctcctcgagggaagcggggtttgcgccagggtccccagggctgtgcgaacaccggggagctgttttttggagaaggctctaggctgaccgtactgggtaaggaggcggttggggctccggagagctccgagagggcgggat**gggcagaggtaagcagctgccc**cactctgagaggggctgtgctgagaggcgctgctgggcgtctgggcggaggactcctggttctgggtgctgg**gagagcgatggggctctcagcggtgggaaggacccgagctgag**tctgggacagcagagcgggcagcaccggtttttgtcctgggcctccaggctgtgagcacagatacgcagtattttggcccaggcacccggctgacagtgctcggta**agcgggggctcccgct**gaagccccggaactggggagggggcg**ccccgggacgccgggg**gcgtcgcagggccagtttctgtgccgcgtctcggggctgtgagccaaaaacattcagtacttcggcgccggga**cccggctctcagtgctgggtaagctggggccgccggg**ggaccggggacgagactgcgctcgggtttttgtgcggggctcgggggccgtgaccaagagacccagtacttcgggccaggcacgcggctcctggtgctcggtgagcgcgggctgctggggcgcgggcgcgggcggcttgggtctggtttttgcggggagtccccgggctgtgCTCTGGGGCCAACGTCCTGACTTTCGGGGCCGGCAGCAGGCTGACCGTGCTGGgtgagttttcgcgggaccacccgggcggcgggattcaggtggaaggcggcggctgcttcgcggcacccggtccggccctgtgctgggaga**cctgggctgggtccccagg**gtgggcaggagctcggggagccttagaggtttgcatgcgggggtgcacctccgtgctcctacgagcagtacttcgggccgggcaccaggctcacggtcacaggtgagattcgggcgtctccccaccttccagcccct**cggtccccggagtcggagggtggaccg**gagctgg**aggagctgggtgtccggggtcagctctgcaaggtcacctccccgctcct**ggggaaagactggggaagagggagggggtggggagg**tgctcagagtccggaaagctgagca**gagggcgaggccacttttaatcttttttctggggtgtttagagagaaggtgaacgatggaggagaggatttgttaggactctgggagaggcgagactggagaggacgaagggaaatcctggtttggggaatgggtaggagtgggggtaactgctattcgtaggcaaaaagagctgagcaggctgggaacagcgcgggtgggcaagggtcagcactgcgggcaggcgggtgggtgttagggggcagaaatcctgcagccgagggtgcagtagaacacagaagaaaaagcctgccaaacaaaagtggaacagagaagccaaaaagggagatgaacatgagtcagtgaagaaaagaatgaaagtttactgtttagcagtgtggatctctaatccgacttaaaactccttgttcccgattcctattcctcctaagccagagatccctgggtccagggtgagggcacggcattcatgcttacccacgggctggtcaacaaagaggtgctgacctgagagtagggcacataacctcagccactggggtacacttaccacccccgcccccgtgtagctccctcccctatcctgaaatctcccttagcacactaagta

Uppercase: TRBJ2-6

Lowercase: Flanking sequence[1000bp]

Red & Bold & Underline: Stem-loop [11]

Blue: Heptamer[28]

Green: Nonamer [11]

id-TRAJ28[J_gene_segment]

ccacccgcctcagcctcccaaagt**gctgggattacaggcgtgagccaccgtgcccagc**cccaaaagccatttttaaatacattgcaggtttcta**tttaatgttgttattcattttgatttgcccaagtaatacattaaa**atttctcattgtaaaccattcaa**attacagataaagtccaggcacggtggctcacacctgtaat**cccagcactttagaaggctgaggtgggcaaatcacctgaggtcaggagtttgagaccagtctggccaacatgatgaaatcccatctctactaaaaatacgaaaagtagtcaatcgtggtggcaggcacctataatcccagctactcaggaggctgaggcaggataatcacttgaacccgggaggttgaggttgccatgagccgagatcgtgccactgcactccagcctgggtgacagagtgagactccatctcaaaaaaaaaaaaaaaattacagataaaaccagcattcaccttggccacttcc**ttcagtatcagtttcctcctccccaaaagtgaccactgaa**accagcttggtgtgttgtgatgagtttctctgtggaatatgtttatgtgaaatcccatcttcaaccaggaaagcttgagagggaaagggctcagactgcagtttgcaggatttcaattagatataaggaacagttgattagatgtaagagctgttagcaaaggaaaaccttataaaaactaggagctcttaataactagacaggtctttggtcagacaacacactgactgaggacaggaggatggattcgatgacctctgaaagtccagccaggactctggaggactctgaggaatggtctgttccatagcctgcctctgtaatgcccttctctcttgcctattgtctggttgttgttacagtctgagcttttgtcagagctgctcct**atgctgtgagtggtctgattttctcagcat**ctctggggtttttgcaaagcaaggaaactctgtgCATACTCTGGGGCTGGGAGTTACCAACTCACTTTCGGGAAGGGGACCAAACTCTCGGTCATACCAAgtaagttcttctttctggctaattattcttcccgagaagcctgtcttccatcatgcagaagctgtc**aaaacacaggtggtgtttt**ctttgcttggtttgtgttgggtggttagtaatatcagttggaaaacaaggttattaatgcacatattccctggggcattgtattcggacatttaatatccatgtagtctctccctgtgaaaatatgtgaacctccacgaaaagaaaggtccaaggaaagtagcagggaaaaggcaggaattggatgaaaatagccaaagagcctgcaggaatataaacaaggcaggatcccaggagacagagcagtagccactttgagtgaatttcc**caggaggtgctcctg**ccaaggcccataccttcaaggaaaattaaggcaaatagaattgggctggggagttgctacttattagtattcctcccacgttctaacctaattataaggaggttgttttggccatgggcagtcatctcaggttttgttttcctgctttcctccc**taacctccacctgtcttcctagaggcctgagtcaaggtta**ttgcaatagcactaaagactgtgtaacaccaatgcaggcaaatcaacctttggggatgggactacgctcactgtgaagccaagtaagttgtgttcttctttgcctaggccttcaggggcaatcaa**tcaaaccattagtttga**aaaagactttaatcctatgcatctggttgggctctttattaatgttctttccccaggcca**aagagagttggttctctt**ccctgctttaaaatgagatatgag**tgcatgtatgcacacacgcatgcccacatgca**gactcttgctctagctcatggtaagggcttctcaggagcatatacaacattttgaaagaaatagagaaacaaacaataatgagccaatggagct**gtcaggaaggttctccaccacccctgac**caggcttcccagcaggatctctggccatataggatgtgctt

Uppercase: TRAJ28

Lowercase: Flanking sequence[1000bp]

Red & Bold & Underline: Stem-loop [12]

Blue: Heptamer[22]

Green: Nonamer [4]

id-TRAJ56[J_gene_segment]

agcagttgctgt**tttccccactttacaagtgaggaaa**ctgaggtgtag**caagttaggaaacttg**tcc**tgagttttaaactca**agcctacctgccccaaagcctgagctcttttaaat**taactgcaactgctgaccagtta**gattaactaattacttaattaataagataacaacctaacaggctgtaaggtaaaaaagtaatcagatttgttctataggtcccctcccttttcgggaatagctatacaattgtataattttatgcttccc**ctgggagagtgggtggagccctggctcccag**cccatgatggaagggtcttggcagtatttgtaaagcagtctgtgggggtgtaactcagggcggatctgaaaagctggtctttggaaagggaacgaaactgacagtaaacccatgtaagtctgaataatgcttccaaatttctccctggaaccctgatttccaaattttccattctgttttataacccaggtccaaacca**cagcagtcccattaatggattccagtgcaaaacaactgctg**gtgtattcctactacacgcaggtctctgttgtttgccctcatgt**gctattttatctaatagc**tgagaatgacagtaccaatggagcactgagttaaggagtcagactgttcgaaggcttcattcttcgttataattggtgag**attttccatgggactaagagaaaat**tgattaactctctgagcctctattttcctcctct**gtagaatgggggaggcagttcctgttcccattttaccacagccggcgtgctgtg**aggggatgttggtacatttccaataaggggacaatgagtgt**gaagagaaaggaaggccccaggtggtccgtaaactcttc**caccagtcccacactataaacagctggttttatcagggggattcttggatgacaagtaagcacttaagtaaatatcaaggggagtctgggcaactgagtttttgtagatcctcgtgtcattgtgTTATACTGGAGCCAATAGTAAGCTGACATTTGGAAAAGGAATAACTCTGAGTGTTAGACCAGg**tatgttttaatgaatgttatttgtttccaaacata**agccaccatccttagaaattca**gtgaaagataaccgaatctcctgcccagttattagcatctttcac**catgggtctttctggagaaaatgacaatgtgggcagcccctgactgcagcccctttgggactgtttctttaacacctttaagtacttgggaatgttcagtgtgtttttgttaatgttggagatatgtgtctgacaaatggaatctgaattgaagttttagtgtgtaggggcagaaagcatttagaaaggacaaaagaaggacagattagactaaaatacataccaatggctgggagtatgcaatgcaacccaatccaaaaggaaacagtgagatgtagcctgctgattaaacaactgagccagcactccgtgtcagctgacttgtcttcccaaagcttccatgttggtgcaattaggaaaagaattgtctaatccccataactcaaaatcttggagccaagctaaattgggtaaagccgtgtaagattttctggtactgacactgactacaagctgatatttagggaaaagttaagattgaaagcaaatattcaaattag**tcagaaagaccacgaacttctga**aaacaagccccagggtgctgttaactgctgcatttctaattgggtcctcatggaacattttca**tctctgccttgtctgagctttctgcaccagaga**agccccttttgatacccactcacttctgagtccctcattgaaaaggtggcaactaagcttagagaaggattccctaatccaaat**gcaatacttgggggaatattttgagtattgc**tccaccattctcaaaacaaacct**aatttcagttaacttgaatgtagctgaaatt**ttgttccatgagggaatctctgctttctatgataatgcaagctcatttcaaacaattg**tttgggatttaacttttgctcataattcccaaa**agataccaactattcaat

Uppercase: TRAJ56

Lowercase: Flanking sequence[1000bp]

Red & Bold & Underline: Stem-loop [18]

Blue: Heptamer[22]

Green: Nonamer [4]

id-TRAJ15[J_gene_segment]

agacaagtaataagcttttttggaaatctaagtggactgccaaatgttctcgatttatccatattgcatggggagcagagcattcttcatgccttctccatcctcacttctgccaggtagcagtttgcat**tgttaacagggaagcagaatggaagcttaaca**gagaagcagtatggaagacgtctccttagcccagtggtcagttgca**tcagagaagtctgctctga**aatgcgaacactttctc**ttgcagctggcagccccctgcaa**cctcagtgccctccatggccagggccaaagcttccaggtgcccagataacatcccaaagtcactctgagagaagag**agaaaatcggtcttactttaccttttctaaccaggaatttacatggttag**ttttacatctaacacttcagcagggagaagacatgatcta**agtgtccatgtggacact**ctgtgtgactcggatccaggatctgctctttcagcccctccatttatcgcaaaagggatgatgattaattcctccatgcctagacttcactttagacatcacctcagagaggcctgtcttacccagccaatctaacaagccatcaaaca**cagccactgcctggctg**ctctctaccctgttatttagcctgattttaattcacacactttcatcacttaacattatactatctgtgtatttattgtttgccttcttctagaccg**tagacaccaggaggacggggacagtgtcta**agtggctctcaccgtatccctagtgccggcacaatgtctttacacatcggaggtgttcagtgaacagttactgagtgaataaaggacttagcaccaactgccgctctttttc**atgaaattacagagctttgctatttcat**acccaaatactaaaacctactgagatttttgcaaatttccatcatttatagttatatccaaggtggatttgagtgagcaggtac**atgaggtatttgcagggcctcat**ttcactgtgCCAACCAGGCAGGAACTGCTCTGATCTTTGGGAAGGGAACCACC**TTATCAGTGAGTTCCAgtaagtacctgataa**ttattgatcatagtgcttgtacttatcctgtagtcatatattgcacaggtgaccaaatgcctcctatttctttgagttaaggaatattgacatttgatgaggaaagccatttaaacacatgggaaaattggatgttgtctctcacccctaatactccaaccatagcccctgtcctattccatgctc**ttaaaaaaattttaa**ttgatagaagaaagaaagattatctacaaggatatgagcaatttgctaaacagtggtaaaacgagattatgtacaggtgttgattcctaggtaaaaatgggggtcagagctgctatcctttggctagtaaaacaagattacttggtgaaattatgtaaatctcctcgaaagaaataagtttcagagttgtctagagaatatcacttggactgttctcaggctttccactacatctattgtaccactatggtgccctgaaaagaggctttctcaggcttaattccatggcttagtctttattccagtatcaaaaagggggaatcccagccagagctcccatgagggaggatagctgcatgctaaccacattaatctattatcaaggtaactcggtcatttttgtcaggcagcacagtgctgtgatttatagcacattcatctttgggagtgggacaagattatcagtaaaacctggtaagtaggcaatatgtcactaaagtaggaggcttaatgtggctactgagacccactaaacttactgcagtatttggaaggcccaagtgtcaagaaattaatggtttatgcagacttaagtggattccaatgaaggaagaaatgttaaagtaatggcacagaggatagaagagctagctgtgaaaaaaaatagccatgtggatgaaccaaacgcaacagactgaaagagcctagagagttgatcctaaagaaaaag**gcagaaagtctggctgagcacttccagttctgc**tgacctggcttgattagtgaa

Uppercase: TRAJ15

Lowercase: Flanking sequence[1000bp]

Red & Bold & Underline: Stem-loop [13]

Blue: Heptamer[35]

Green: Nonamer [1]

id-TRAJ33[J_gene_segment]

ggtgccagtgacagggaagagcagaggggcttaggaggtttttgtagatctcagtatcactgtgtcttataacaccgacaagctcatctttgggactgggaccagattacaagtctttccaagtaagtactagaaaccaaggagccattttgaaaaggtgttttctttttctttttcttttttttttaaatgaagttttgctctttgttgcgcaggctggagtgcagtggcatgatctcggctcactacaacctccacctcccgggttcaagcgattctcctgtctcagcctcccgagtagctgggattacaggcacctaccaccacgcctggctaatttttgtatttttagtagaaacagtgtttcaccatgttggccaggctggtcttgaactcctgacctcaagtgatctgcctgcctc**ggcctgccaaagattacaggcc**actgcgcccagccttaaaaggtgttttctataaacaggaacgttctcacagcactgcaggccacccttaccaactccgagctctgtaggggcctttagcacctatagctgcagaaccttctataaaccaattaatctatatgtgaacaatttaacctatatgtgaacaccatggtactgagtaaaatcttaccagcagtgaaaagaaggcactagagcgtcagatgcattatgactctaaacaaatgatctgt**ttccctcttttgcatttttatttaaaatatcagggaa**ggtgaagcaagttcaacttctccaatttgaaaactgtttctgaattatgcttcccttgtatgcagagagacctagattgactttggatgtttagtcattcg**atttaatcaaatgtaattaaat**gagcaaataatctccaatagccaatattcctatgttgtttcattgttttatgtgctttgctctagactttttgtctgggctttgtctctaataggatccccggaaggacagtgaaggtttttgttaaggtttttgtgtctgtgTGGATAGCAACTATCAGTTAATCTGGGGCGCTGGGACCAAGCTAATTATAAAGCCAGgtaagtctcagagatgtgactgcacgggagaggagacactagttgaataatgcacaaagtgtagcatgcagattatatttttaagaacaagtcagcctgc**tggagacaatgcactcaacccaaatggggtctcca**ctcccagtgagaagcatgtcagccgctaactcttttgtttggtctagtagcttccggaatgaataatgcttaagtagccctctaagagagcagagccaggttgtt**agggaatatctaattccct**ctgatatggttaaaactctttgccaagggcagaaccagcctcattccattggaagagcaaattcagagaaaaaggaaagggtcacacagccaaagacggtaaaatgttcaaaggtgaaaaatagccaagtctgcagctctcctgacatccttgcctcaggctgctgtcccacgtg**gagagcagatgcctgcaaagctctc**ggtcacttgctggagtcactcagggttctgggccttgggatgtaacttaaa**tgacagtgtcagggcactgtca**gaggaatatgaaggctgcagcgatgcctgcaagtcagcttttgctacactgggactgaggtgttctaagcctggaaagacacaaagctgtcctcatcccaacctgccgtgcccctccttgagggttagtgtaaggctctgaaggactgtgtgaattatggtggtgctacaaacaagctcatctttggaactggcactctgcttgctgtccagccaagtacgtaagtagtggcatgtgtcaggtggattctgtgtccatggcaagtaggaagcga**cagccacccttaggtggaaaggatggctg**aaggttgactttgttcactgctgtcatctcttatctgccgtgatatagcaggctgtcaaaattcccattctcctggatggcacaccacagtcaggggaggggaaacatgctaattttatgataaaccccaggggagaaaaagactactgtgggaaaatttagttg

Uppercase: TRAJ33

Lowercase: Flanking sequence[1000bp]

Red & Bold & Underline: Stem-loop [8]

Blue: Heptamer[35]

Green: Nonamer [6]

id-TRGJP[J_gene_segment]

ttgcagtgctaagtcgcataactcagagctgcctcagattctctgttgagtagcacacgatttaatttgatgaaactgtatagatattttcaagtcaaaagtagcagtctttcagca**atgaatcatcattaagaattcat**attttctaacaactcatcagtattctatcagtgagaactggattc**taaaaagaacttactttattttta**ttttttattttttgagatggagtctcactctgtcgcccagctggaa**tgcagtggcatcatctctgctcactgca**acctccgcttcccagtttcaagagattcttctgcttcggcctcctgagtagctaggattacaggctcctgccactacgcccaactaatgttttgggttttttttgtatttttagtagagatggggtttcaccatgttgaccaggctggttttcaactcctgagctcaagtgatccgcttgcctcagcctcccaaagtgctaggattacagggatgagccactgtgcctggccaaaaagaagttattttagaatcaatctagatacccatccctccatac**atatgaatcccaaaatacctataaatcatat**tgaatatctatttattaaatgtcaaacataggaatcctattattgtctctctttgaaaattaaggaaatattaactctaaat**tacacttaagacaaagtgta**tacaagaaaactttttagggaagatttagaagattcaggcaaaatcatgagaaaaggtaaaatcagactctctaatgtatgggatacaaggagagaatgtacatagcccaaagtgcccagggcaaggtgaggtcagttcttaaattcctgattacattcagatccagtgtgattttgtttgacctttgtcttgacttaacgtcagcagggccaatttttatgtatttatgtaaatattgaaaaaaatgttggcacaattttaagacaaacacaaagggagattcttataaaggcttctcaggtggTGGGCAAGAGTTGGGCAAAAAAATCAAGGTATTTGGTCCCGGAACAAAGCTTATCATTACAGgtaagtt**ttctttaaattttgcaatgtaaagaa**gggatgggaggctggcaggcaggagctggctcagaattctaggattccctccttggtcactaatcatggtgaaatctccagagaggtcaggtgacttggttcatgccttaggagtcagaacttttcctgccctagcatgggagacattactaaggagcttagaccaactgcaaatctctaaaggatggaggccttatg**ttgttccattggtatctctctgttgaaagttttgtgaacaa**gtttagttttgagatt**ttatttcttcagttctaaaaataa**ccaatggaaaaaattaaaaaaaaaaaaagtaaaatcagctaccctcc**ttatttggtcaccaaaagatgaaatatttgataatttgaccaaagaa**tggttaatatcaaatgttaaaatatttctcggtgtgaccatatttagaagtaacataagctgcatttattgttatttaaattggtccaatgagtttgtttattatttgttagcttaagtttaaagatgtaattttgcctgg**atgagagaaaacatctcat**aaagatactcttagtctgttagatcca**ctgaaatgaaagttttcag**agaattttcacaaagttgataaagcatcgga**aaaaatgaaagcagttttacattttt**aattccttagtggttgagatcttatgataaaatgaacttataagttaaaaagtgaccttccg**attctcttttatccaattgacttaatgagaat**tgtagcaat**caaattcataatttg**taattttctgagaagtgatttttcaggaaagtgcggtgcaagaaaaggtatggactttgtttctcgactcctggagctagcactttctatgtaactcccttttaggaagtgtattttgatttaccagttgttcaaattataaaaatgctggttcattatagaaaaattggaaaacacaaaaataaacaaaaaattataatttgatattgaccc

Uppercase: TRGJP

Lowercase: Flanking sequence[1000bp]

Red & Bold & Underline: Stem-loop [14]

Blue: Heptamer[12]

Green: Nonamer [1]

id-TRBJ1-5[J_gene_segment]

gaacacagagtactggaagcagagctgctgtccctgtgagggaa**gagttcccatgaactc**ccaac**ctctgcctgaatcccagctgtgctcagcagag**actggggggttttgaagtggccctgggaggctgtgctctggaaacaccatatattttggagagggaagttggctcactgttgtaggtgagtaagtcaaggctggacagctgggaacttgcaaaaaggggctggaatccagacggagcctttgtctctagtgcttaggtgaaagtgtatttttgtcaggaaggcctatgaggcagat**gaggaggggatagcctccctctcctc**tccactattttgtagactgcctgtgccaagttaggttcccctactgagagatgggtagactcagcttggaaggggtcaccttgaacatctcctgtctccttgaagggtgccggtcacggccatgacagataaaagagcctctgaccttaccaccacggtcctaccgtttctc**tccctcacacagaaaggagaaggtcacagaagaggga**acttgggggatcacacggggcctaattggtctgctgaccaccgcattttgggttgtaccattgtctacccctctacccaccagggttaaaattctactaaggaacaggagaggacctggcaggtggacttggggaggcag**gagtggaaggcagcaggtcgcggttttccttccagtc**tttaatgttgtgcaactaatgaaaaactgttttttggcagtggaacccagctctctgtcttgggtatgtaaaagacttctttcgggatagtgtatcataaggtcggagttccaggaggaccccttgcgggagggcagaaactgagaacacagccaagaaaagctcataaaatgtgggtcagtggagtgtgtggtggggccccaagagttctgtgtgtaagcagcttctggaaggaagggcccacaccagctcctctggggtttgccacactcatgatgcactgtgTAGCAATCAGCCCCAGCATTTTGGTGATGGGACTCGACTCTCCATCCTAGgtaagttgcagaatcagggtggtatggccattgtcccttgaaggcagagttctctgcttctcctcccggtgctggtgaggcagattgagtaaaat**ctcttaccccatggggtaagag**ctgtgcctgtgcctgcgttccctttggtgtgtcttggttgactcctctatttctcttctctaagtcttcagtccataatctgcctcctcactcccttcttggctcatcctccctcttatgtgcatggctctgcctctcctaagcctcttcctcttgcgccttatgctgcacagtatgcttaggcctttttcctaacagaatccctttggtccagagccatgaatccaggcagagaaaggcagccatcctgctgtcagggagctaagacttgccctctgactggagatcgccgggtgggttttatctaagcctctgcagctgtgctcctataattcacccctccactttgggaatgggaccaggctcactgtgacaggtatgggggctccactcttgactcgggggtgcctgggtttgactgcaatgatcagttgctgggaagggaattgagt**gtaagaacggaggtcagggtcaccccttcttac**ctggagcactgtgccctctcctcccctccctggagctcttccagcttgttgctctgctgtgttgcctgcagttcctcagctgtagagctccttgcttagtcttcagggctgtgtgtttctttgctcttcttttcattgttttctgggactcttctcatctctactttcttagtggatgtattgttttactttcccttttttaaattgcatcttctccattttttccttcccattctaactccacttctgcattgttgactccttttggtgactagctctgtcttctatgttaagattctccccactgccagcctccagcacagaactctgctcatgtcttcatctccctccttctttctttctctaccagtcttagaagatgcatctatgtcttcctg

Uppercase: TRBJ1-5

Lowercase: Flanking sequence[1000bp]

Red & Bold & Underline: Stem-loop [7]

Blue: Heptamer[38]

Green: Nonamer [4]

id-TRBJ2-7-2[J_gene_segment]

agcctgcacgaccccagaaccctgttcttaggggagtggacactgggcaatccagggccctcctcgagggaagcggggtttgcgccagggtccccagggctgtgcgaacaccggggagctgttttttggagaaggctctaggctgaccgtactgggtaaggaggcggctggggctccggagagctccgagagggcgggat**gggcagaggtaagcagctgccc**cactctgagaggggctgtgctgagaggcgctgctgggcgtctgggcggaggactcctggttctgggtgctgg**gagagcgatggggctctcagcggtgggaaggacccgagctgag**tctgggacagcagagcgggcagcaccggtttttgtcctgggcctccaggctgtgagcacagatacgcagtattttggcccaggcacccggctgacagtgctcggta**agcgggggctcccgct**gaagcccgggaactggggagggggcg**ccccgggacgccgggg**gcgtcgcagggccagtttctgtgccgcgtctcggggctgtgagccaaaaacattcagtacttcggcgccggga**cccggctctcagtgctgggtaagctggggccgccggg**ggaccggggacgagactgcgctcgggtttttgtgcggggctcgggggccgtgaccaagagacccagtacttcgggccaggcacgcggctcctggtgctcggtgagcgcgggctgctggggcgcgggcgcgggcggcttgggtctggtttttgcggggagtccccgggctgtgctctggggccaacgtcctgactttcggggccggcagcaggctgaccgtgctgggtgagttttcgcgggaccacccgggcggcgggattcaggtggaaggcggcggctgcttcgcggcacccggtccggccctgtgctgggaga**cctgggctgggtccccagg**gtgggcaggagctcggggagccttagaggtttgcatgcggggatgcacctccgtgCTCCTACGAGCAGTACGTCGGGCCGGGCACCAGGCTCACGGTCACAGgtgagattcgggcgtctccccaccttccagcccct**cggtccccggagtcggggggtggaccg**gagctgg**aggagctgggtgtccggggtcagctctgcaaggtcacctccccgctcct**gggaaaagactggggaagagggagggggtggggagg**tgctcagagtccggaaagctgagca**gagggcgaggccacttttaatcttttttctggggtgtttagagagaaggtgaacgatggaggagaggatttgttaggactctgggagaggcgagactggagaggacgaagggaaatcctggtttggggaatgggtaggagtgggggtaactgctattcgtaggcaaaaagagctgagcaggctgggaacagcgcgggtgggcaagggtcagcactgcgggcaggcgggtgggtgttagggggcagaaatcctgcagccgagggtgcagtagaacacagaagaaaaagcctgccaaacaaaagtggaacagagaagccaaaaagggagatgaacatgagtcagtgaagaaaagaatgaaagtttactgtttagcagtgtggatctctaatccgacttaaaactccttgttcccgattcctattcctcctaagccagagatccctgggtccagggtgagggcacggcattcatgcttacccacgggctggtcaacaaagaggtgctgacctgagagtagggcacataacctcagccactggggtacacttaccacccccgcccccgtgtagctccctcccctatcctgaaatctcccttagcacactaagtattctaggttaaacagcccagatgttcagggagttcattcgccacaaacacacactaaaatg**cagacaatttgcctgtgagatgaggaaaattctctg**gaaga**tttaggccctgagagctgaaaagggaccctaaa**cattacctggtgacaactgccctgaggccagagaagagaactcacaatattggtatattaaccggtaccatttgta

Uppercase: TRBJ2-7-2

Lowercase: Flanking sequence[1000bp]

Red & Bold & Underline: Stem-loop [12]

Blue: Heptamer[25]

Green: Nonamer [10]

id-TRAJ22[J_gene_segment]

atggatgagaaaactgaggctcagggaagagacagatcccggcccca**cagccagtctatggctg**agctagaactagattttaggtctctaatttgccaaacctgtcagttggc**tcaaacttgaagtttga**agagtccgcaagattcatctgacaaatattttcatgtacttgccctctgccaggctcgttgctgaaaccagggatccaacaggaagcaaactcagtgtgatattgccttcatgaaacttacattctggaggtggattccatgtttcttcccagatatgacaatgcttaccctgatttctcttgggcatggctctcttatactatcatcacttccttaggaaatactaaaactaatttttgctgcagtttaaagtccttgagcagataactaacacacatac**cactttagtcaggagaagggaaatgcccaaaagtg**aagtagaaaactaggaatttgt**ctgataattgattattataaagttattttatcag**tgtgaaatgagtggccaaattaactggacaatgccagctctgtacctacctcactttgattctatagactcagtctcaggaggattcaagaattcagacagttctctgatgacataatccagtgatgcccacaacagaggaggcgtattctctattctggactttgtgagtctccttggatgaaaagattcaagtgctgttttgacacctaaaatcagaga**gtatttgctctatacatatatcgaaaatac**acagttaatgtttagaactttgtagtggaaagtggaaagcaaaggtcccctaaaagaaaatgggattgtaaagacgaaggagggttagtttaggatttgtagaaagtca**gtttggctactccagaccaaac**ctagattaacagtgatgcaggcctaattcataaagg**aagcactgccagctctttattcagtgctt**tgccaacctggcctgtttgatctggtttttgttgttgagcaaatcatagtgTTTCTTCTGGTTCTGCAAGGCAACTGACCTTTGGATCTGGGACAC**AATTGACTGTTTTACCTGgtaggctgcctcaatt**a**aatactatttgcactgatttactaatctacaaatgtatt**ctgtacatgtaacttaatgggcctgaggttgaagatggggagagacagcaacaatgcatactagaaagggtaatgcgagcagagaattatttatcagagaccaagaagagaaagagaagctttttgctaaggcttttagtaataggagttaatg**aaaatgaatagaaaaaaatttcatttt**aacttagaagaattgctgaataatgggaacagattccataaggggaacctctgcttctgaaaatatttgcactaggaccagcctacgtgctacaaaaactgtctcacaccaagtctagttggtcacttacatttttagcctattctagtagctatttctaaataagtctttaaacatagaggtgcctat**atttaagatatttaaat**gaggctctatttctatagctctctggtgcccttc**cagcaaagcattgctg**taacacattgcttatcatttctacaagtgagaacaagtaagcccctggccctccaaaattgcatgagtattataattacttggcttagattgaagtcacttctgtgttacttctcataactgtgttcatttgttgagatctgtttaactccaaactgtgtgggcgtctaactgttggcactaaggtttcttttctcttgaaattgccagcaaaatattttacccataaataatgtttcatctagacttgcaaatgacaactaacactgcaggctctttttttttcttttgctttgtttc**tttttttctgaagggaatatgcaagcaggaagcaaaaaaaaa**aaaaaagccaaaatgtacagtttgactgtggggtctgggaggagagtgtcaatgtggatgctaaaa**tatacatggttgtata**atgtaggtactgtcacagagaggctatag**tgatcttattagatca**tacatacagagcaccactggtcagctggcgttgctcggg

Uppercase: TRAJ22

Lowercase: Flanking sequence[1000bp]

Red & Bold & Underline: Stem-loop [15]

Blue: Heptamer[22]

Green: Nonamer [3]

id-TRBJ2-2[J_gene_segment]

ccattttaattcactgcctttgtcttttccaagccccacacagtcagactaacctctgccacctgcgcttcctgccgctgcccagtggttgggggagggggactagcagggaggaaacatttttgtatcatggtgtaacattgtggggactagcgggagggcacgatgattcaggtagaggaggtgcttttacaaaaaaccctgatgcagtaagcatc**cccacccagctcagggaatgcagctaccaggtggg**aagagttctctggggctggtcccagctgtggtcttgcagggtcccccaacccagcgagcacctgtccatctccctgtccagactcggcttccaaggaataagaaggccaagacagc**aaagtgggattatcactcagcacttt**taataaaacttgttcttgacaaagtacttgcacatgcattatttattaagaactgatgaaaaccctgag**ggaaagatattgtcccatctttcc**aatgaggaaactgagatcagaggttacaggtcatataactaggaaacggcaaggtctagcctgcaatatcgcccagctccagccgttccagtaccaccaatgccccttcagatttca**aatccactgtgttgtcccccagccaagtggatt**ctcctctgcaaattggtggtggcctcatgcaagatccaggttaccgtgtccagctaactcgagacaggaaaagataggctcaggaaagagaggaagggtgtgccctctgtctgtgctaagggaggtg**gggaaggagaaggaattctgggcagccccttccc**actgtgctcctacaatgagcagttcttcgggccagggacacggctcaccgtgctaggtaagaagggggctccaggtgggagagagggtgagcagcccagcctgcacgaccccagaaccctgttcttaggggagtggacactgggcaatccagggccctcctcgagggaagcggggtttgcgccagggtccccagggctgtgCGAACACCGGGGAGCTGTTTTTTGGAGAAGGCTCTAGGCTGACCGTACTGGgtaaggaggcggttggggctccggagagctccgagagggcgggat**gggcagaggtaagcagctgccc**cactctgagaggggctgtgctgagaggcgctgctgggcgtctgggcggaggactcctggttctgggtgctgg**gagagcgatggggctctcagcggtgggaaggacccgagctgag**tctgggacagcagagcgggcagcaccggtttttgtcctgggcctccaggctgtgagcacagatacgcagtattttggcccaggcacccggctgacagtgctcggta**agcgggggctcccgct**gaagccccggaactggggagggggcg**ccccgggacgccgggg**gcgtcgcagggccagtttctgtgccgcgtctcggggctgtgagccaaaaacattcagtacttcggcgccggga**cccggctctcagtgctgggtaagctggggccgccggg**ggaccggggacgagactgcgctcgggtttttgtgcggggctcgggggccgtgaccaagagacccagtacttcgggccaggcacgcggctcctggtgctcggtgagcgcgggctgctggggcgcgggcgcgggcggcttgggtctggtttttgcggggagtccccgggctgtgctctggggccaacgtcctgactttcggggccggcagcaggctgaccgtgctgggtgagttttcgcgggaccacccgggcggcgggattcaggtggaaggcggcggctgcttcgcggcacccggtccggccctgtgctgggaga**cctgggctgggtccccagg**gtgggcaggagctcggggagccttagaggtttgcatgcgggggtgcacctccgtgctcctacgagcagtacttcgggccgggcaccaggctcacggtcacaggtgagattcgggcgtctccccaccttccagcccct**cggtccccggagtcggagggtggaccggagctggaggagctgggtgtccggggtcagctc**tgcaaggtcacct

Uppercase: TRBJ2-2

Lowercase: Flanking sequence[1000bp]

Red & Bold & Underline: Stem-loop [14]

Blue: Heptamer[39]

Green: Nonamer [12]

id-TRAJ16[J_gene_segment]

gaaaggcaaggtaatatgatcaatacatgaacgtgtgcaaaataaatcattaaggcattcaactagaagccaaaaaacagtatggaccccggctcagccactcactcactaggcagccatgagaagatcaccaagctctcccagcctcagttaacctaaaatgtggagattattattccagccttgactacttcaaaggatgctatgaaattcagatgagataatatatgtgaaatctccttgaaaaagatgggtggtcaaatgattcatgatttttttttttttttttttttttttttgagacggagtctcgctctgtcgcccaggctggag**tgcagtggcgggatctcggctcactgca**agctccgcctcccgggttcacgccattctcctgcctcagcctcccaagtagctgggaccacaggcgcccgccactacgcccggctaattttttgtatttttagtagagacgaggtttcaccgttttagccgggatggtctcgatctcctgacctcgtgatccgcccgcctcggcctcccaaagtgctgggattacaggcgtgagccaccgcgcccggcctcatgatttttaatagttagattttatgcaaaatttcatttgaggg**taaacttaaaggtgaacaagttta**atggactttggccatggcagagaaatggttaggtttggatataaggtggacctctctaattataagaaaaaaaaggaacatgagaattggtgaacaagaccttccctggagatcttcagaaa**cctgaggtcactcagg**agagaaatgtgtgaagttgagagcttacagta**aatggtctctgaagtttcccaccatt**ccaaagagtgcgtgagcttaaaatttttcgtgagtttagctaaatatgtgcttagtgttagattaggtttcaa**ccaagcaaaagaaccctgggagaagtactctgcttgg**aaatgaagcatcctttggtttttgtggtacaatagatcactgtgGGTTTTCAGATGGCCAGAAGCTGCTCTTTGCAAGGGGGACCATGTTAAAGGTGGATCTTAgtaagtattatt**actaatgaattcttaattgattagt**ttttgagacaagtaataagcttttttggaaatctaagtggactgccaaatgttctcgatttatccatattgcatggggagcagagcattcttcatgccttctccatcctcacttctgccaggtagcagtttgcat**tgttaacagggaagcagaatggaagcttaaca**gagaagcagtatggaagacgtctccttagcccagtggtcagttgca**tcagagaagtctgctctga**aatgcgaacactttctc**ttgcagctggcagccccctgcaa**cctcagtgccctccatggccagggccaaagcttccaggtgcccagataacatcccaaagtcactctgagagaagag**agaaaatcggtcttactttaccttttctaaccaggaatttacatggttag**ttttacatctaacacttcagcagggagaagacatgatcta**agtgtccatgtggacact**ctgtgtgactcggatccaggatctgctctttcagcccctccatttatcgcaaaagggatgatgattaattcctccatgcctagacttcactttagacatcacctcagagaggcctgtcttacccagccaatctaacaagccatcaaaca**cagccactgcctggctg**ctctctaccctgttatttagcctgattttaattcacacactttcatcacttaacattatactatctgtgtatttattgtttgccttcttctagaccg**tagacaccaggaggacggggacagtgtcta**agtggctctcaccgtatccctagtgccggcacaatgtctttacacatcggaggtgttcagtgaacagttactgagtgaataaaggacttagcaccaactgccgctctttttc**atgaaattacagagctttgctatttcat**acccaaatactaaaacctactgagatttttgcaaatttccatcatttatagttatatccaaggtggatttgagt

Uppercase: TRAJ16

Lowercase: Flanking sequence[1000bp]

Red & Bold & Underline: Stem-loop [15]

Blue: Heptamer[18]

Green: Nonamer [4]

id-TRAJ59[J_gene_segment]

ttatcgttgccttaagggagctggggggttgtcaacggg**ccagaggtgggatgaaaaatgacaacagatttacctctgg**gaccgggacatggttaaccacagcggccctgggtaagtagcttagcttcagaagaaaatgtgcccaacagcatgggtaacctaaaacaccgggcaatccaaatattcttttatgattggctttagcatgtattttattcttttgtagggcaggtttatctcaccaattatatattttcttaactga**cctgtaaaatctacagg**ggaaaagtattttaagaattatatgtttctgcaattaggctcccagcagtcaacaaagaagtggta**ctttttgtctttccagtgatgaaaaag**ggaacctggcatcc**ctggtggcccaccag**cgtctccttccctggcctaggtcagaacaagccgtaaatcagcaggccgttatcttcttataaatctgtag**agcagggtggacaacaaaaggcagcctgct**aggttttcagaacatgagttccttgtgtagccagagaacctg**ggaccatcctgacgtggctggtcc**tgctgtcctc**acagcctgaatcccaggctgt**atgaataggagaggttcaagtccagatgactgttcacgatgctggtcccccttgcatccctaatcatgctagagacatgaccagggtctgagaggaggaagttacagcacagcaccaacaggggcttttggtaaagggcctgggcactatgtgaagatcacctagatgctcaactttgggaaggggactgagttaattgtgagcctgggtgag**tacctcaactccagaggta**gctttagcggaac**ccctctatacctaacacctggcaatcagaggg**ctgaaacacttggccagataatgaattctcttatccggtgggaaaaggctgtaaagatcaaaccacctttcctatgggtaagcaagagtctgtagtttatgtaaaggcagcagc**tcctgtgGGAAGGAAGGAAACAGGA**AATTTACATTTGGAATGGGGACGCAAGTGAGAGTGAagctatcttt**aaaccaaaggtgtcaggttatttggttt**ggtttttgaattatctggaagttccaaagaaagaacacttctccctgaggatttgattgcaaaattctgacttcaaacttctaaaaagatcaaatgttaaatcagatagtaggcttggaaaactctatttctctatgta**aaaagtagagaactactttt**cttttgtttgatcatt**ttattttgtttaggaaataa**gaggattagataccctggtggtgagtggggagggcagggacatttgcacagcatttctggataaattagatccaaataatcaaatcaaagactctcagctcagaaagt**aattaaagatctcctcacccacctctttaatt**ttgcaaatgaagacagtgaaactcagagaggttatgaacttgctcaaggtcacacaactgatcctgatatcaaggtccagggcaaagccaagacagctcatagttctcaccagcccagac**ccaagggagaaaaaaacatcatagtccttgg**ccacggcagcatcatttgcatcccaaacattctttacccaagacttaat**gaactcaaaaagaaatcctgagttc**caaagggaaataaaatcattctgcgtctgttgaaaaaaaaagcaagctaaagtggaacaataattgaacttgatattaggggaaaggtgcagccatctgcagactgagagaagggtgaaaaaaacaaaatgaaaatgctagtctt**gtgttcagacaatgattatgaccatagaacac**tacctataagttctgtagagctaatttctttcccagtggggttga**tattctatgatagattgcatcagcggtgtgattgttttggaccatagaata**cgtatacagaattggcttgttgagccttctggtgtcttgaagtgaaaatcatatctctggacatatctgtcaggcctgaagccctaggctactggaaagaaagataccatgacttatccttcctctaaa

Uppercase: TRAJ59

Lowercase: Flanking sequence[1000bp]

Red & Bold & Underline: Stem-loop [18]

Blue: Heptamer[18]

Green: Nonamer [3]

id-TRAJ6[J_gene_segment]

gtggaggggccccagccctttactcccaccctgagtttggtggagccacacagctctttgctaagtgaccctcaaacccagtgtgagataagctgagtgttgacagtgcccacctgctagggaaccaagtactgagttcctgcccca**gctcaggactgtggcgcaggccatgacctccctgagc**ctcagttttcatatctctgaagtagacagcatggtctctgtttcctcccatctttcaaaaattctatggaaattattgtgactttttt**ctctgtaattgtagggtttcacagag**gcagggccttgtctgagtcaaggtcattagaaata**aaggaatccaagggctttggatgcctt**ggcctaattctaggtggtaaagagggaaggatgaaagcacaaatcacaggaagtgc**ctaaatcctgcttggatttag**tgttgcctcaaacagggatatgcagaaatccaaccccagggtctcccataggaagaagactaggagtgggaagtgccaaagagctctggaaaggaaaagagaacaaggagcactctgagaaaaggaggtcagaacagcttc**aggaaaaacgtcacataaaatttcct**ttgctctctgtcctcgcagcactttcagaagctccttgcatctgactaaagagtagcagagccagaccctcagaatgtctg**ccccaagcaagcagagtgtggtgtttgtattgggg**tgctggg**agcagacatggggagttgctgccctgcttctctgct**gatgggaaggtggtattttcccctgggtacacgggggccaaagagctctccagtcccc**agcccctggctcccgcccctgggctccagcactcgggggct**ctaaggacagcagtaggagactttctacaactcagagagtattcaggagggtttcccaggatg**ttgcccaataccatggggcaa**ttggtagagggaggtcacatgaaacaggttttatcaaaggctgtcctcactgtgTGCATCAGGAGGAAGCTACATACCTACATTTGGAAGAGGAACCAGCCTTATTGTTCATCCGTgtaagtattatagaaatgatcaagggaaattttgcagacagattatattatggaaggatactgtagtagtgagagtttgtgttgtgatctattccatcattaaaggagtcctttgttgcctgggagtaactatcagtgtcaggtctgatgagatgattgaggctgtgccggacaaggg**ttttgcacaatgatttcagaggacaaatccccaagttgtgaaaaa**agacttcactcttggttaggtttctaaaacagaactttcttcttggcaaccaaggggtctactctgccccctcactcctatgtctcttccacctgagactctgtcaccacctcccctagaatccgtgagataccttcccctgaattagagcataccagccagggtgctgaggcactctgtggtactggaatggatggtacagggatgcgtttctg**ggcttctggctgcaagaagcc**tcatcctttcccccagtgtaaagcattgctgggaatcatcccattatgagctactatttactgaatgcctgacacgtaccagtggccattctcttgtctttta**tgtaatcctcattcagtgattaca**cacatcctaataataggtagccgatattttgcccattttacaggtgaggaaacagagg**cttgggataagtaacttatccaag**attatatagccacgaatgacagag**ctagagtttaactcaaatctgtttgactctag**ggccttgttattaaccgtgccaccaccatgcctcctgcattc**tgtgaaggaagacttcaca**cct**caaaggccatctgtttggcctttg**gtttgcccatcctgagccttcttaccatagcgaccatggcatgg**tgactcagcacttctgaagagatcagcagagtca**ttgggttgggtggcagaatacaggtatggcagggagggaaggagaaacttagggggactgtttattgcgcatacagttagagagaaagaaaaatgtatcaacaaaccata

Uppercase: TRAJ6

Lowercase: Flanking sequence[1000bp]

Red & Bold & Underline: Stem-loop [17]

Blue: Heptamer[34]

Green: Nonamer [2]

id-TRAJ9[J_gene_segment]

gccctgctctttcttttattttattttattttatttatttatttatttatttctga**gacagagtcttgctctgtc**accc**aggctggagtgcagtgcagtctcagctcactgcagcct**ctgcctcccaggttcaagcgattcgattctcctgcctcagcctcccaagtagctggggttacaggcatgcgccaccacacctgaaagccctgctctttctactgcatgatgtcg**tagagcttgtgctcta**gtttacttagaaatgagaggtgatgccaatcacagtgactggtgaaatgcttggaatttatcttttctactttagatattgctgtgagttagagagggtgcttcgggaaatcttgacatggatgccccaagtgtctaaagagttctcctcccctttcaatgacaaaacaaagtacagagttgctgggagcatttaccccaat**agggagaaaagagaggggagctggctgcagccactccct**tgttctgactgtccctttccttcgctgtttcttgtattttccccaggcagaccagccaccaatataactgatctacaagagcctgactcctttgtgagttatagagaaatgcagggtgaaggcactttgcacatagaactttagctctgagagacagcctcaaaatatctagttcacccctctttattttcccaataaggaaattaagacccagagaggttggctgaattgtctaaggtcagtgcgaactagggctgggactagaacctggagatcctggtcctggggccatattctgtgactgctgtttggtgaactaaacttctgataggtaggaagtgtgagttatgctcataaacaggaaaaacaaataggttttacaaggggagtataaaaatgtcctccttattactagcaccttagtataggctccctagacctgctccccagctgtggtgta**aaaatggaaggagaaggaaatggcccatttt**gtcgcagtgcaaatcactgtgGGAAATACTGGAGGCTTCAAAACTATCTTTGGAGCAGGAACAAGACTATTTGTTAAAGCAAgtaagttccatgaaataacctgatttatattacagttgaagaacatctgttttcattgatttttttctttttagggggaaggggacatagttttgtctgccagttaaattagaagatacattggcaataataatgttgcactgtacagtatccatgggcatcactcttgcaagaaacatgtataataacaaacagttaagtgtggatggataaaacttccctaggtcctatctagaactgaaactctgtaattatattaataacttttttaacttgaatattagtgacatttttatataacccaaggtat**acaaagcagaatcttcagttgggttctatctgatttgt**tgtcatcttccaatggtcttaggaaaattaaggttaaatagtaagattagatgcctcctaccactttggtacatg**catgcagacatagaaacacccttagtaactaacttgcatg**tgacttggcaatcctaggctatggcccagtgacagggtcccattttgtacagagttatgtcagagtgtgaacacaggctttcagaaacttgtatttggaactggcacc**tgacttctggtcagtccaagtaagtca**aatctgcagaaatgtgtagccctatcctcaatattgggcatatatggatataattattctggtct**ataatagattattat**tactactattttgttcttattaatcataacaatgctgctaatttcaaatatacattttctgccatattattatttttacctttaaggagaattattgtcaagatatcaggag**ggaaatatttgaatttcc**tttgagtcaagtttcttatttgttcgtcagttcccacctctctgctgagggtcatatgaacctcacgatgttttggtttttataccgccttcttattatctctcttagaaatgcttccttggtttttccttgcctaagcacttagattttgcagctcacatttacccagagtatacttagcttttaacctct

Uppercase: TRAJ9

Lowercase: Flanking sequence[1000bp]

Red & Bold & Underline: Stem-loop [10]

Blue: Heptamer[16]

Green: Nonamer [3]

id-TRBJ2-3-2[J_gene_segment]

tgcagggtcccccaacccagcgagcacctgtccatctccctgtccagactcggcttccaaggaataagaaggccaagacagc**aaagtgggattatcactcagcacttt**taataaaacttgttcttgacaaagtacttgcacatgcattatttattaagaactgatgaaaaccctgag**ggaaagatattgtcccatctttcc**aatgaggaaactgagatcagaggttacaggtcatataactaggaaacggcaaggtctagcctgcaatatcgcccagctccagccgttccagtaccaccaatgccccttcagatttca**aatccactgtgttgtcccccagccaagtggatt**ctcctctgcaaattggtggtggcctcatgcaagatccaggttaccgtgtccagctaactcgagacaggaaaagataggctcaggaaagagaggaagggtgtgccctctgtctgtgctaagggaggtg**gggaaggagaaggaattctgggcagccccttccc**actgtgctcctacaatgagcagttcttcgggccagggacacggctcaccgtgctaggtaagaagggggctccaggtgggagagagggtgagcagcccagcctgcacgaccccagaaccctgttcttaggggagtggacactgggcaatccagggccctcctcgagggaagcggggtttgcgccagggtccccagggctgtgcgaacaccggggagctgttttttggagaaggctctaggctgaccgtactgggtaaggaggcggctggggctccggagagctccgagagggcgggat**gggcagaggtaagcagctgccc**cactctgagaggggctgtgctgagaggcgctgctgggcgtctgggcggaggactcctggttctgggtgctgg**gagagcgatggggctctcagcggtgggaaggacccgagctgag**tctgggacagcagagcgggcagcaccggtttttgtcctgggcctccaggctgtgAGCACAGATACGCAGTATTTTGGCCCAGGCACCCGGCTGACAGTGCTCGgta**agcgggggctcccgct**gaagcccgggaactggggagggggcg**ccccgggacgccgggg**gcgtcgcagggccagtttctgtgccgcgtctcggggctgtgagccaaaaacattcagtacttcggcgccggga**cccggctctcagtgctgggtaagctggggccgccggg**ggaccggggacgagactgcgctcgggtttttgtgcggggctcgggggccgtgaccaagagacccagtacttcgggccaggcacgcggctcctggtgctcggtgagcgcgggctgctggggcgcgggcgcgggcggcttgggtctggtttttgcggggagtccccgggctgtgctctggggccaacgtcctgactttcggggccggcagcaggctgaccgtgctgggtgagttttcgcgggaccacccgggcggcgggattcaggtggaaggcggcggctgcttcgcggcacccggtccggccctgtgctgggaga**cctgggctgggtccccagg**gtgggcaggagctcggggagccttagaggtttgcatgcggggatgcacctccgtgctcctacgagcagtacgtcgggccgggcaccaggctcacggtcacaggtgagattcgggcgtctccccaccttccagcccct**cggtccccggagtcggggggtggaccg**gagctgg**aggagctgggtgtccggggtcagctctgcaaggtcacctccccgctcct**gggaaaagactggggaagagggagggggtggggagg**tgctcagagtccggaaagctgagca**gagggcgaggccacttttaatcttttttctggggtgtttagagagaaggtgaacgatggaggagaggatttgttaggactctgggagaggcgagactggagaggacgaagggaaatcctggtttggggaatgggtaggagtgggggtaactgctattcgtaggcaaaaagagctgagcaggctgggaacagcgcgggtgggcaagggtcagcac

Uppercase: TRBJ2-3-2

Lowercase: Flanking sequence[1000bp]

Red & Bold & Underline: Stem-loop [14]

Blue: Heptamer[27]

Green: Nonamer [12]

id-IGHJ3P-2[J_gene_segment]

ttctccggctgtttgggaccacgttcagcagaaggcctttctttgggaactgg**gactctgctgctggggcaaagggtgggcagagtc**atgcttgtgctggggacaaaatgaccttgggacacggggctggctgccacggccggcccgggacagtcggagagtcaggtttttgtgcaccccttaatggggcctcccacaatgtgactactttgactactggggccagggaaccctggtcaccgtctcctcaggtgagtcctcacaacctctctcctgctttaactctgaagggttttgctgcatttttggggggaaataagcgtgctgggtctcctgccaaga**gagccccggagcagcctggggggctcaggaggatgccctgag**gcaacagcggccacacagacgaggggcaa**gggctccagatgctccttcctcctgagccc**agcagcacgggtctctctgtgg**ccagggccaccctgg**gcctctggggtccaatgtccaacaacc**cccgggccctccccggg**ctcagtctgagagggtcccagggacttagcgggg**tgccagttcttgcctggggtcctggca**tt**gttgtcacaatgtgacaac**tggttcgacccctggggccagggaaccctggtcaccgtctcctcaggtgagtcctcaccaccccctct**ctgagtccacttagggagactcag**cttgccagggtctcagggtcagagtcttggaggcattttggaggtcaggaaagaaa**gctggggagagggacccttcgaatgggaacccagc**ctgtcctccccaagtccggccacagatgtcggcagctggggggctccttcggctggtctggggtgacctctctccgcttcacctggagcattctcaggggctgtcgtgatgattgcgtggtgggactctgtcccgctccaa**ggcacccgctctctgggacgggtgcc**ccccggggtttttggactcctgggggtgacttagcagccgtctgCTTGCAGTTGGACTTCCCAGGCCGACAGTGG**TCTGGCTTCTGAGGGGTCAGgccaga**atgtggggtacgtgggaggccagcagagggttccatgagaagggcaggacagggccacggacagtcagcttccatgtgacgcccggagacagaaggtctctgggtggctgggtttttgtggggtgaggatggacattctgccattgtgattactactactactactacatggacgtctggggcaaagggaccacggtcaccgtctcctcaggtaagaatggccactctagggcctttgttttctgctactgcctgtggggtttcctgagcattgcaggttggt**cctcggggcatgttccgagg**ggacctgggcggactggccaggaggggatgggcactggggtgccttgaggatctgggagcctctgtgg**attttccgatgcctttggaaaat**gggactcaggttgggtgcgtctgatggagtaactgagcctgggggcttggggagccacatttggacgagatgcctgaacaaaccaggggtcttagtgatggctgaggaatgtgtctcaggagcggtgtctgtaggactgcaaga**tcgctgcacagcagcga**atcgtgaaatattttctttagaattatgaggtgcgctgtgtgtcaacctgcatcttaaattctttat**tggctggaaagagaactgtcggagtgggtgaatccagcca**ggagggacgcgtagccccggtcttgatgagagcagggttgggggcaggggtagcccagaaacggtggctgcc**gtcctgacaggggcttagggaggctccaggac**ctcagtgccttgaagctggtttccatgagaaaaggattgtttatcttaggaggcatgcttactgttaaaagacaggatatgtttgaagtggcttctgagaaaaatggttaagaaaattatg**acttaaaaatgtgagagattttcaagt**atattaatttttttaactgtccaagtatttgaaattcttatcatttgattaacacccatgagtgatatgtgtctggaatt

Uppercase: IGHJ3P-2

Lowercase: Flanking sequence[1000bp]

Red & Bold & Underline: Stem-loop [18]

Blue: Heptamer[24]

Green: Nonamer [5]

id-TRAJ14[J_gene_segment]

gccttcttctagaccg**tagacaccaggaggacggggacagtgtcta**agtggctctcaccgtatccctagtgccggcacaatgtctttacacatcggaggtgttcagtgaacagttactgagtgaataaaggacttagcaccaactgccgctctttttc**atgaaattacagagctttgctatttcat**acccaaatactaaaacctactgagatttttgcaaatttccatcatttatagttatatccaaggtggatttgagtgagcaggtac**atgaggtatttgcagggcctcat**ttcactgtgccaaccaggcaggaactgctctgatctttgggaagggaaccacc**ttatcagtgagttccagtaagtacctgataa**ttattgatcatagtgcttgtacttatcctgtagtcatatattgcacaggtgaccaaatgcctcctatttctttgagttaaggaatattgacatttgatgaggaaagccatttaaacacatgggaaaattggatgttgtctctcacccctaatactccaaccatagcccctgtcctattccatgctc**ttaaaaaaattttaa**ttgatagaagaaagaaagattatctacaaggatatgagcaatttgctaaacagtggtaaaacgagattatgtacaggtgttgattcctaggtaaaaatgggggtcagagctgctatcctttggctagtaaaacaagattacttggtgaaattatgtaaatctcctcgaaagaaataagtttcagagttgtctagagaatatcacttggactgttctcaggctttccactacatctattgtaccactatggtgccctgaaaagaggctttctcaggcttaattccatggcttagtctttattccagtatcaaaaagggggaatcccagccagagctcccatgagggaggatagctgcatgctaaccacattaatctattatcaaggtaactcggtcatttttgtcaggcagcacagtgctgtgATTTATAGCACATTCATCTTTGGGAGTGGGACAAGATTATCAGTAAAACCTGgtaagtaggcaatatgtcactaaagtaggaggcttaatgtggctactgagacccactaaacttactgcagtatttggaaggcccaagtgtcaagaaattaatggtttatgcagacttaagtggattccaatgaaggaagaaatgttaaagtaatggcacagaggatagaagagctagctgtgaaaaaaaatagccatgtggatgaaccaaacgcaacagactgaaagagcctagagagttgatcctaaagaaaaag**gcagaaagtctggctgagcacttccagttctgc**tgacctggcttgattagtgaatctgggccacagtatctttatctat**aaaatgaaaacatttaatatcacctcagattttttgttttaatttt**gtgggtacataataggtgtatatatttattttaaagataagctaatatatgtgaaagtattttgtaaattcttaaatgctatccacatgtaaaatatta**tttgtttatagaacaaa**aaaaaactatgtcagatttag**aattcttttaacatgagaatt**tgtaggttcttagtgaaatagccatttacatgtgatctgtagccctaggcattggtaaaaagcactttctcttctactgaaatatagtggctcaaatgctaccc**aaatgaaaagggcaggggaagggaagtcattt**tgtaaaggcaggcattacagtgtgaa**ttctgggggttaccagaa**agttacctttggaactggaacaaagctccaagtcatcccaagtgagtccaatttcctatgctttcctcttccttgtgttgtcttctctcagaccgtaacatttggagcacat**aaagagatattatgggcctgctcttt**tctggctg**tttcaggagactgaaa**aggattgta**taagaacatcctttagcatgttctta**gtattgttttgtccagtgtgtgtttcttcttaatatcaaaaaaaactaagacattgcctgtgaagccaatggaaaacagtagat

Uppercase: TRAJ14

Lowercase: Flanking sequence[1000bp]

Red & Bold & Underline: Stem-loop [14]

Blue: Heptamer[31]

Green: Nonamer [2]

id-TRAJ1[J_gene_segment]

cctgtgagtttgtgcaatggtgtcacctacggtatgaatactggaggaacaattgataaactcacatttgggaaagggacccatgtattcattatatctggtgagtcatcccaggtggcaccacgtgcaaccccatgggccagtgtcactaatcctttctctggagatatcacttattactatggtgaggcttgctgtagatgttgtaactaattttcttacagaggtctgggaagggaaaagcattactatctatcttgaatattcatgtttctctag**gtcaaacacattaaaatttgac**tttaatcattcaatgggtattgtaaaatgccttctatgtgactatcactctataaaatgttagactgagtatgaagtgtgagatagattcctgtcctgtcctcaagtggtataaaaactagacaaaggtactgaactattgtaaattaagcagccagaaaactattttagt**attgaccagttgatgatgtcaat**ggacacaatagagattcagaggggatagggtgctggctgagaagttggcctagactgagaagtttcctgtggtacaaaggattcattgagccctgaaggatggataagatctgtatgggcagagaaaggagagaagggaagttctgggcgtagggaacgacaagaaagaaggcatgatcttgggaataatcaaggcacatgcaaagtagcctaagtatacatctgataataaaattggttgaaaagtagtcagagaagatgt**ctttttaggcatggaaaaag**gaaatactagagcattcaacagaatacagaaattagggccagggccagccattgggaaactgagaatccgatttagagatgcagactagaagtgaaggtgagagcagccagctatggtgccgcagacct**cccctctccttcctcagtgggctctgagagggg**tcatcccaca**ccttagaggaggagaaacctaagg**gattctgtaatagagacacggggcatgGTATGAAAGTATTACCTCCCAGTTGCAATTTGGCAAAGGAACCAGAGTTTCCACTTCTCCCCgtacgtctgcccatgcccacagtttcctgatgctcactaaagcctcggt**gggacccagagtgactgtcactaattctgatttctgggtccc**tagtgcccaaacacgggggacagatttaatggtaaggaagctttcaatcactgctgtgtccctagggatcta**aagcactagagcacatgtgctt**ctgcagttcattttgaatttaaaggacagcttaggatctagaatagctgaatttccacctcaaaacattggttccgtcttgccaagcctaccttctgatatcatcagtgatgggatgtgtttttcttactagggtagaataggatgtctctccccaaaggactctggcagacagacccctaaacacctccaaattaaaagcggcaaagagataaggttgaactagacgtacatggggataaaaagtataaaaggtacatgggaatgaaaggataaaaaggctaaaaaaattaagtacctctaactcagcccctgttgccatttctcagagtcttgtgttctgtggcattgcgctttctagaccaacagtgtccaatagaactttctgtggcaatggaaatgtcctgtcaatctgcactgtcccata**caatagccaccagctacatgtggctattg**agctcttgaaatgaagtttccatttttaa**ttgaaaacattttatttcacattgactaatttttatttcaa**cagccacatgtagctagagactattataccagacagagcagcctagatcttctccagtctgacacccaccagccccaggac**ttgagtgagtgtttaaccaggactcaa**agttgggtttctgccccacaaggccaccccctttcctctttaaagccaacctgcatctggtggcccctgatcccctgccttgaggatcggcacttccagactcctctccccctctgcagtgctgtccagtacccccactgatgactaacaatcagggggatgtgttggtagagctaatggct

Uppercase: TRAJ1

Lowercase: Flanking sequence[1000bp]

Red & Bold & Underline: Stem-loop [10]

Blue: Heptamer[16]

Green: Nonamer [2]

id-IGHJ6[J_gene_segment]

tactttgactactggggccagggaaccctggtcaccgtctcctcaggtgagtcctcacaacctctctcctgctttaactctgaagggttttgctgcatttttggggggaaataagcgtgctgggtctcctgccaaga**gagccccggagcagcctggggggctcaggaggatgccctgag**gcaacagcggccacacagacgaggggcaa**gggctccagatgctccttcctcctgagccc**agcagcacgggtctctctgtgg**ccagggccaccctgg**gcctctggggtccaatgtccaacaacc**cccgggccctccccggg**ctcagtctgagagggtcccagggacttagcgggg**tgccagttcttgcctggggtcctggca**tt**gttgtcacaatgtgacaac**tggttcgacccctggggccagggaaccctggtcaccgtctcctcaggtgagtcctcaccaccccctct**ctgagtccacttagggagactcag**cttgccagggtctcagggtcagagtcttggaggcattttggaggtcaggaaagaaa**gctggggagagggacccttcgaatgggaacccagc**ctgtcctccccaagtccggccacagatgtcggcagctggggggctccttcggctggtctggggtgacctctctccgcttcacctggagcattctcaggggctgtcgtgatgattgcgtggtgggactctgtcccgctccaa**ggcacccgctctctgggacgggtgcc**ccccggggtttttggactcctgggggtgacttagcagccgtctgcttgcagttggacttcccaggccgacagtgg**tctggcttctgaggggtcaggccaga**atgtggggtacgtgggaggccagcagagggttccatgagaagggcaggacagggccacggacagtcagcttccatgtgacgcccggagacagaaggtctctgggtggctgggtttttgtggggtgaggatggacattctgccattgtgATTACTACTACTACTACTACATGGACGTCTGGGGCAAAGGGACCACGGTCACCGTCTCCTCAGGTaagaatggccactctagggcctttgttttctgctactgcctgtggggtttcctgagcattgcaggttggt**cctcggggcatgttccgagg**ggacctgggcggactggccaggaggggatgggcactggggtgccttgaggatctgggagcctctgtgg**attttccgatgcctttggaaaat**gggactcaggttgggtgcgtctgatggagtaactgagcctgggggcttggggagccacatttggacgagatgcctgaacaaaccaggggtcttagtgatggctgaggaatgtgtctcaggagcggtgtctgtaggactgcaaga**tcgctgcacagcagcga**atcgtgaaatattttctttagaattatgaggtgcgctgtgtgtcaacctgcatcttaaattctttat**tggctggaaagagaactgtcggagtgggtgaatccagcca**ggagggacgcgtagccccggtcttgatgagagcagggttgggggcaggggtagcccagaaacggtggctgcc**gtcctgacaggggcttagggaggctccaggac**ctcagtgccttgaagctggtttccatgagaaaaggattgtttatcttaggaggcatgcttactgttaaaagacaggatatgtttgaagtggcttctgagaaaaatggttaagaaaattatg**acttaaaaatgtgagagattttcaagt**atattaatttttttaactgtccaagtatttgaaattcttatcatttgattaacacccatgagtgatatgtgtctggaattgaggccaaagcaagctcagctaagaaatactagcacagtgctgtcggccccgatgcgggactgcgttttgaccatcataaatcaagtttatttttttaattaattgagcgaagctggaagcagatgatgaattagagtcaagatggctgcatgggggtctccggcacccacagcaggtggcaggaagcaggtcaccgcgagagtctattttaggaagca

Uppercase: IGHJ6

Lowercase: Flanking sequence[1000bp]

Red & Bold & Underline: Stem-loop [17]

Blue: Heptamer[21]

Green: Nonamer [4]

id-TRBJ2-5[J_gene_segment]

cccagctccagccgttccagtaccaccaatgccccttcagatttca**aatccactgtgttgtcccccagccaagtggatt**ctcctctgcaaattggtggtggcctcatgcaagatccaggttaccgtgtccagctaactcgagacaggaaaagataggctcaggaaagagaggaagggtgtgccctctgtctgtgctaagggaggtg**gggaaggagaaggaattctgggcagccccttccc**actgtgctcctacaatgagcagttcttcgggccagggacacggctcaccgtgctaggtaagaagggggctccaggtgggagagagggtgagcagcccagcctgcacgaccccagaaccctgttcttaggggagtggacactgggcaatccagggccctcctcgagggaagcggggtttgcgccagggtccccagggctgtgcgaacaccggggagctgttttttggagaaggctctaggctgaccgtactgggtaaggaggcggttggggctccggagagctccgagagggcgggat**gggcagaggtaagcagctgccc**cactctgagaggggctgtgctgagaggcgctgctgggcgtctgggcggaggactcctggttctgggtgctgg**gagagcgatggggctctcagcggtgggaaggacccgagctgag**tctgggacagcagagcgggcagcaccggtttttgtcctgggcctccaggctgtgagcacagatacgcagtattttggcccaggcacccggctgacagtgctcggta**agcgggggctcccgct**gaagccccggaactggggagggggcg**ccccgggacgccgggg**gcgtcgcagggccagtttctgtgccgcgtctcggggctgtgagccaaaaacattcagtacttcggcgccggga**cccggctctcagtgctgggtaagctggggccgccggg**ggaccggggacgagactgcgctcgggtttttgtgcggggctcgggggccgtgACCAAGAGACCCAGTACTTCGGGCCAGGCACGCGGCTCCTGGTGCTCGgtgagcgcgggctgctggggcgcgggcgcgggcggcttgggtctggtttttgcggggagtccccgggctgtgctctggggccaacgtcctgactttcggggccggcagcaggctgaccgtgctgggtgagttttcgcgggaccacccgggcggcgggattcaggtggaaggcggcggctgcttcgcggcacccggtccggccctgtgctgggaga**cctgggctgggtccccagg**gtgggcaggagctcggggagccttagaggtttgcatgcgggggtgcacctccgtgctcctacgagcagtacttcgggccgggcaccaggctcacggtcacaggtgagattcgggcgtctccccaccttccagcccct**cggtccccggagtcggagggtggaccg**gagctgg**aggagctgggtgtccggggtcagctctgcaaggtcacctccccgctcct**ggggaaagactggggaagagggagggggtggggagg**tgctcagagtccggaaagctgagca**gagggcgaggccacttttaatcttttttctggggtgtttagagagaaggtgaacgatggaggagaggatttgttaggactctgggagaggcgagactggagaggacgaagggaaatcctggtttggggaatgggtaggagtgggggtaactgctattcgtaggcaaaaagagctgagcaggctgggaacagcgcgggtgggcaagggtcagcactgcgggcaggcgggtgggtgttagggggcagaaatcctgcagccgagggtgcagtagaacacagaagaaaaagcctgccaaacaaaagtggaacagagaagccaaaaagggagatgaacatgagtcagtgaagaaaagaatgaaagtttactgtttagcagtgtggatctctaatccgacttaaaactccttgttcccgattcctattcctcctaagccagagatccctgggtccagggtgagggcacggcattcatgcttacccacgggc

Uppercase: TRBJ2-5

Lowercase: Flanking sequence[1000bp]

Red & Bold & Underline: Stem-loop [12]

Blue: Heptamer[29]

Green: Nonamer [11]

id-IGKJ5[J_gene_segment]

gggagtttttgtataggagggaagttaagaggaaccattgtgtgtgcagttttggccaggggaccaagctggagatcaaacgtaagtacttttttccactgattcttcactgttgctaattagtttactttgtgttcctttgtgtggattttcattagtcggatgccagggatctaacaaacttcattcccaggttaggtacagaggaggggaaattgttccacaggacgctagcttgtggctaatttttaagatttctaaatcaaaataacttcattgggggaaagaggcttgctgagctttcagggaggtttttgtaaagggaaaagttaagac**gaatcactgtgattcactttcggccctgggaccaaagtg**gatatcaaacgtaagtacatctgtctcaattattcgtgagattttagtgccattgtatcatttgtgcaagttttgtgatattttggttgaataaac**ctggtgacccagaagtaaatagcaggacaccag**aaaatgaacttaaaaagctgagcaaatagacgaatcattgggtttgagaggagaataggattcatgggggaaatggggaagaaatagctagatttttctctgaacaagcagcctatctcatatgattggcttcaagagaggtttttgt**tgaggggaaagggtgagatccctca**ctgtggctcactttcggcggagggaccaaggtggagatcaaacgtaagtgcactttcctaatgctttttcttataaggttttaaatttggagcgtttttgtgtttgagatattagctcaggtcaattccaaagagtaccagattctttcaaaaagtc**agatgagtaagggatagaaaattagttcatct**taaggaacagccaagcgctagccagttaagtgaggcatctcaattgcaagattttctctgcatcggtcaggttagtgata**ttaacagcgaaaagagatttttgttaa**ggggaaagtaattaagttaacactgtgGATCACCTTCGGCCAAGGGACACGACTGGAGATTAAACgtaagtaatttttcactattgtcttctgaaatttgggtctgatggccagtattgacttttagaggcttaaataggagtttggtaaagattgg**taaatgagggcattta**agatttgccatgggttgcaaaagttaaactcagcttcaaaaatggatttggagaaaaaaagattaaattgctctaaactgaatgacacaaagtaaaaaaaaaaagtgtaactaaaaaggaacccttgtatttctaaggagcaaaagtaaatttatttttgttcactcttgccaaatattgtattggttgttgctgattatgcatgatacagaaaagtggaaaaatacattttttagtctttctcccttttgtttgataaattattttgtcagacaacaataaaaatcaatagcacgccctaagaaaaatcagggaaaagtgaagtgtacctatttgctatgtagaagaggcagcttacttgaaaatcagcagcaatgttgtttttagagtctgtaataagtaataaactcaaaaagacacattctataggaataagggcttcacagatagagctcattttttaaaaatccaatttgtacattagactaaacgtgaaattatctcttattgtaatggtggaaaggtggttattcccaaaagctcaatctcaaagaaatgtgtttaaatgaaaaaaagtaaataattgcattttttaatgaccgtgggtctgtgaaa**aaaataggaaatatttt**aaagagtatgttctttcattatcctctgttattacttgtctacatttttattctgccaagaaggccgtggc**accgcgagctgtagacagagccgcggt**ctttctcgattgagtggcttt**ggtggccatgccacc**gcgctcttggggcagccgccttgccgctagtggccgtggccaccctgtgtctgcccgattgatgctgccgtagccagctttcctgatgcacagtgatacaaataatgccactaagggaaagagaac

Uppercase: IGKJ5

Lowercase: Flanking sequence[1000bp]

Red & Bold & Underline: Stem-loop [10]

Blue: Heptamer[24]

Green: Nonamer [7]

id-TRAJ8[J_gene_segment]

gagaaatgcagggtgaaggcactttgcacatagaactttagctctgagagacagcctcaaaatatctagttcacccctctttattttcccaataaggaaattaagacccagagaggttggctgaattgtctaaggtcagtgcgaactagggctgggactagaacctggagatcctggtcctggggccatattctgtgactgctgtttggtgaactaaacttctgataggtaggaagtgtgagttatgctcataaacaggaaaaacaaataggttttacaaggggagtataaaaatgtcctccttattactagcaccttagtataggctccctagacctgctccccagctgtggtgta**aaaatggaaggagaaggaaatggcccatttt**gtcgcagtgcaaatcactgtgggaaatactggaggcttcaaaactatctttggagcaggaacaagactatttgttaaagcaagtaagttccatgaaataacctgatttatattacagttgaagaacatctgttttcattgatttttttctttttagggggaaggggacatagttttgtctgccagttaaattagaagatacattggcaataataatgttgcactgtacagtatccatgggcatcactcttgcaagaaacatgtataataacaaacagttaagtgtggatggataaaacttccctaggtcctatctagaactgaaactctgtaattatattaataacttttttaacttgaatattagtgacatttttatataacccaaggtat**acaaagcagaatcttcagttgggttctatctgatttgt**tgtcatcttccaatggtcttaggaaaattaaggttaaatagtaagattagatgcctcctaccactttggtacatg**catgcagacatagaaacacccttagtaactaacttgcatg**tgacttggcaatcctaggctatggcccagtgacagggtcccattttgtacagagttatgtcagagtgTGAACACAGGCTTTCAGAAACTTGTATTTGGAACTGGCACC**TGACTTCTGGTCAGTCCAAgtaagtca**aatctgcagaaatgtgtagccctatcctcaatattgggcatatatggatataattattctggtct**ataatagattattat**tactactattttgttcttattaatcataacaatgctgctaatttcaaatatacattttctgccatattattatttttacctttaaggagaattattgtcaagatatcaggag**ggaaatatttgaatttcc**tttgagtcaagtttcttatttgttcgtcagttcccacctctctgctgagggtcatatgaacctcacgatgttttggtttttataccgccttcttattatctctcttagaaatgcttccttggtttttccttgcctaagcacttagattttgcagctcacatttacccagagtatacttagcttttaacctctcccaaaactggcctccctgtcccaataattatgcacagtaaaagtcagttcaattcagtaaatgattattgagtatcatcaacaaaccagggaatcatcatt**ggttttaaaagaaagaaacaaagaaaagaatttttaaaaccctaagctatcttagg**ct**tctgatctcaagaacttccagtcttatcaga**gaaactaagcactcataataaaaagtctaacaacaatttatatgtgtatgtgccaagataggggggtgggtttcctgagggctggagacaggacaggtatgaccatgcacagccgcattctttggcagagaacacacatgattcccccaaatatgtttctggaaacggaattgagtctgttctgtgctgagatctttgactttcttgaggacaaaagcccccttggcagttagcatccttgagcgggggtgaggacagaaacaggaatgagtcaggcgt**gagtcacccaagactctcactcagccccaagaggatggagtgactc**tgagtcatgccgaggcatgtctttagcatgtgctccagcacagtgtgcatttatggcaaggtgaaaaatacatt

Uppercase: TRAJ8

Lowercase: Flanking sequence[1000bp]

Red & Bold & Underline: Stem-loop [10]

Blue: Heptamer[17]

Green: Nonamer [4]

id-TRBJ1-1[J_gene_segment]

gtagtgatgggggctgtggcttctctataaggacatgccccaacgtgacaacagcttggagaggggtgggtactggagaagaccagccccttcgccaaacagccttacaaagacatccagctctaaggagctcaaaacatcctgaggacagtgcctggaggtgagaaggaagcccccggcctggtccataccccaccaccaacttgcataatggggggtgatgtcacccaccctccactcccctcaaaggagcagctgctctggtggtct**ctcccaggctctgggggcggacccatgggag**gggctgtttttgtacaaagctgtaa**cattgtggggacagggggccacaatg**attcaactctacgggaaacctttacaaaaacct**ctctggcggtcccaactcccagag**tcctcttctttcctcctgggtcacaggtcttaatgcaatttggttcagaatgcctctgcctcactcctgatcacatgtcagaccaagactgtggacaaggacaggcccagatgagaactaaagcttccc**aggcagagagaggtcagacataagaagactgcct**caggaacctcacaagtggaggactcagggagggtcccaatccccaaaaattgagacaaagtcaggtggaaggttcatcggaggtgaccagctctccagaggactcgggaagaagt**caggggtatctatagatggagtcacaggttctgggcccctg**ccatcctctgcaggccatgcactttccctttcgatggaccctcacagagggagcatctgaatggggca**tcctttgaaaaagga**acc**taggaccctgtggatggactctgtcattctccatggtccta**aaaagcaaaagtcaaagtgttcttctgtgtaatacccataaagcaca**ggaggagatttcttagctcactgtcctcc**atcctagccagggccctctcccctctctatgccttcaatgtgattttcaccttgacccctgtcactgtgTGAACACTGAAGCTTTCTTTGGACAAGGCACCAGACTCACAGTTGTAGgtaagacatttttcaggttcttttgcagatccgtcacagggaaaagtgggtccacagtgtcccttttagagtggctatattcttatgtgctaactatggctacaccttcggttcggggaccaggttaaccgttgtaggtaaggctgggggtctctaggaggggtgcgatgagggaggactctgtc**ctgggaaatgtcaaagagaacagagatcccag**ctcccggagccagactgagggagacgtcatgtcatgtcccgggattgagttcaggggaggctccctgtgagggcgaatccacccaggcttcccagaggctctgagcagtcacagctgagcccagggtgatggggcagaag**agggaaggggagggggcctctcctcatagttccct**gagatagcccagagaaagcccggtgggtaatgaatgagccacaacacctctccatctatctgcttcactg**acagaggttctctgt**agattcttcgtatattcctgtgctggattttataggaggccactctgtgtctctttttgtcacctgcctgagtcttgggca**agctctggaagggaacacagagtactggaagcagagct**gctgtccctgtgagggaa**gagttcccatgaactc**ccaac**ctctgcctgaatcccagctgtgctcagcagag**actggggggttttgaagtggccctgggaggctgtgctctggaaacaccatatattttggagagggaagttggctcactgttgtaggtgagtaagtcaaggctggacagctgggaacttgcaaaaaggggctggaatccagacggagcctttgtctctagtgcttaggtgaaagtgtatttttgtcaggaaggcctatgaggcagat**gaggaggggatagcctccctctcctc**tccactattttgtagactgcctgtgccaagttaggttcccctactgagagatgggtagactcagcttggaaggggtcaccttgaacatctcctgtctcctt

Uppercase: TRBJ1-1

Lowercase: Flanking sequence[1000bp]

Red & Bold & Underline: Stem-loop [15]

Blue: Heptamer[49]

Green: Nonamer [7]

id-TRBJ1-3[J_gene_segment]

tttccctttcgatggaccctcacagagggagcatctgaatggggca**tcctttgaaaaagga**acc**taggaccctgtggatggactctgtcattctccatggtccta**aaaagcaaaagtcaaagtgttcttctgtgtaatacccataaagcaca**ggaggagatttcttagctcactgtcctcc**atcctagccagggccctctcccctctctatgccttcaatgtgattttcaccttgacccctgtcactgtgtgaacactgaagctttctttggacaaggcaccagactcacagttgtaggtaagacatttttcaggttcttttgcagatccgtcacagggaaaagtgggtccacagtgtcccttttagagtggctatattcttatgtgctaactatggctacaccttcggttcggggaccaggttaaccgttgtaggtaaggctgggggtctctaggaggggtgcgatgagggaggactctgtc**ctgggaaatgtcaaagagaacagagatcccag**ctcccggagccagactgagggagacgtcatgtcatgtcccgggattgagttcaggggaggctccctgtgagggcgaatccacccaggcttcccagaggctctgagcagtcacagctgagcccagggtgatggggcagaag**agggaaggggagggggcctctcctcatagttccct**gagatagcccagagaaagcccggtgggtaatgaatgagccacaacacctctccatctatctgcttcactg**acagaggttctctgt**agattcttcgtatattcctgtgctggattttataggaggccactctgtgtctctttttgtcacctgcctgagtcttgggca**agctctggaagggaacacagagtactggaagcagagct**gctgtccctgtgagggaa**gagttcccatgaactc**ccaac**ctctgcctgaatcccagctgtgctcagcagag**actggggggttttgaagtggccctgggaggctgtgCTCTGGAAACACCATATATTTTGGAGAGGGAAGTTGGCTCACTGTTGTAGgtgagtaagtcaaggctggacagctgggaacttgcaaaaaggggctggaatccagacggagcctttgtctctagtgcttaggtgaaagtgtatttttgtcaggaaggcctatgaggcagat**gaggaggggatagcctccctctcctc**tccactattttgtagactgcctgtgccaagttaggttcccctactgagagatgggtagactcagcttggaaggggtcaccttgaacatctcctgtctccttgaagggtgccggtcacggccatgacagataaaagagcctctgaccttaccaccacggtcctaccgtttctc**tccctcacacagaaaggagaaggtcacagaagaggga**acttgggggatcacacggggcctaattggtctgctgaccaccgcattttgggttgtaccattgtctacccctctacccaccagggttaaaattctactaaggaacaggagaggacctggcaggtggacttggggaggcag**gagtggaaggcagcaggtcgcggttttccttccagtc**tttaatgttgtgcaactaatgaaaaactgttttttggcagtggaacccagctctctgtcttgggtatgtaaaagacttctttcgggatagtgtatcataaggtcggagttccaggaggaccccttgcgggagggcagaaactgagaacacagccaagaaaagctcataaaatgtgggtcagtggagtgtgtggtggggccccaagagttctgtgtgtaagcagcttctggaaggaagggcccacaccagctcctctggggtttgccacactcatgatgcactgtgtagcaatcagccccagcattttggtgatgggactcgactctccatcctaggtaagttgcagaatcagggtggtatggccattgtcccttgaaggcagagttctctgcttctcctcccggtgctggtgaggcagattgagtaaaatctcttaccccatggggtaagagctgtgcctgtgcctg

Uppercase: TRBJ1-3

Lowercase: Flanking sequence[1000bp]

Red & Bold & Underline: Stem-loop [12]

Blue: Heptamer[53]

Green: Nonamer [6]

id-TRDJ3[J_gene_segment]

actccattcgcttctctggttttcttctcctgcagtggtgaaatgtaatatgatgatgtgaagctcaggtaggggatgtgaaaattttcaaatcaaaccccaagtccttaaagctt**tgacagtgaataatggccctacacaactgtca**ttctcttttgcctggctctagcccaacgatcttgatttttggtgatgaggcagagtaattcacatggtctttatgttacattgcacatgatgactatata**ctcctgaaacttatatctggatccagagtcaggag**aaattataaaacaaaaataagcaatgcaaagcaagcactggcttctctgacc**tgtgtccatgacaca**cagcaccagtgggctc**cctgagcagtgaccaatttcttgttcaaccctcagg**tccttagaaagcctttcgccaagctcagggaattagaaagcaggttcgttattattgtgagactcagtctgagtgatgggggaaattaggtttctaatcaaaattctgtacttgtgacccaagagaatggaaccagagtataataagaatggaaactctttctataaccctcctctcctcaacctaacatttgttgtaagaaaggtcaagttaagtgccccatccctaccctcaagctattgtgttcttcttggatgcccctctcaataaactggaagttagattttccatcatgaaggcagctatttgtcacctgccaccctcttctcttgctctgtctcagccctgccctagtatctcttggatggggatatcaccctcacatctctatcaag**cagttcaactattattagcttcaaagggaactgagactctgtgtactcagccccttggtctca**tcaagagcagctttgt**agttctctgagctgtggggtctctaggctgagaact**gagg**ctggggaggcagggcacagatgttacagctcaggccccag**ggccagctccaggctagttacctgtgaggcactgtcataatgtgCTCCTGGGACACCCGACAGATGTTTTTCGGAACTGGCATCAAACTCTTCGTGGAGCCCCgtgagttgatctttttcctatatttctgggataatttgagtcctggcactggggctgcaatccagtttgcattataaattataactagtaaatgaaattataacaaggagacagagtattacagatgtgaaataggccagagtgaacacactagactgagtcacttaagtagctctgtaactctattta**ttttaatatatatactctatattaaaa**caatagctcccatttatt**tctgaaatcttcaga**gcagcccaaaaaacacaaagaaatctagaagaactacactaagagttggagcattcaggtgcaagggagacagtgggagggatacttcagccccaattcctcaggaagctgcttcatctctgcacatccctttggctaagctaactagtatgcgtgcctccttgcaaactgaaaacaagtgtcctacagcttggaagattctggagaaaggtagttagcatggtcagagtaattaggtagaacaaacttttatttcagggtttaagttaaatcccatattggcagaaagtcttgaaattctgagtgacatttattacagttgttgcaattcttaggggatcagtaggactagctatgagaatctaaaacataggtgacttctaagtttataaattaagaaagactgttcaggaatggtacacacttttttggtattgtttaccagcacataaacttagcaatgtattggtgttcataaaaagagaaagtctgacaataaagcaattttccaaagaaataacacttgaaacacatgttgcaatgtaattatatttgtggactaagaaaaaaagactacaagttcttc**aaagataaacgattaaatcttt**aatctaaattctggactacaatttcatacagcttcacgtctgggtttaagtcaaatacaaaatagactaggatgcatatggcaactatattgagtgcactccagtttttaaaaaggtttaattcagtcagacacccatttctt

Uppercase: TRDJ3

Lowercase: Flanking sequence[1000bp]

Red & Bold & Underline: Stem-loop [11]

Blue: Heptamer[17]

Green: Nonamer [2]

id-TRAJ48[J_gene_segment]

caccggtaaccagttctattttgggacagggacaag**tttgacggtcattccaagtaagtcaaa**gaaaattttccatcaccattgtgttgagcaaaccctttaaactgcagaaagagctgtcaaagtctgctactctatttctctgcttctaggtataactttatctggaccagagtaaagaggcccccgacccaccccga**gaatgatttatgcttggacaggtctttatttcattc**cttgaactatctatctgtctatctatctatccagctatctatgtatctatctatctatcatctatctcagaaatagttgtttttaacagttgattctgtgcctcacccaaattgtcattaactaaatgcagatgtgctgccacactgataaacttcattggaaaaaaagaaaaaaccaaacc**cagagaatgttccttgaacattcctg**aagtgggggggcaggcga**tgtcacttgcagtgaca**gctggcctctctagctcacccccacatctgtgccaaccagagaagctgggccctttgtagcacaagcctcaggagca**gagaatggagggacattctc**aactgagaggagagggactaggaggctagatgggaaatgaggtgactctacagaaaatcgaatatgggatggtataaaaggatgagaaagatggcagggggtccacgatcgctcttgtatttagacctgggggatgcactgatagataccacctggagagagaaagagga**tttctgtttccctctgaagaaaaatgtggcagaaa**aatcaaaacagatcgcacatgtcaagcctctcctaattacagcctttcattctggtttctctcagctgttccagcaggaaatggtaggaaagggtcattggtgcctttgatgttgtggaagcattgcaggaaggagatt**tgcaagaaacggggcaggggcttgca**gtagagtctggagagtttagaatgatggtttttgcaatgacttagaacactgtgTATCTAACTTTGGAAATGAGAAATTAACCTTTGGGACTGGAACAAGACTCACCATCATACCCAgtaagttcttcatccttggtcaggaaatc**agcctgcataagattctggggatcatagtagagatgcaggct**tagagaattttcatctgtccttgttcataactggtagagacaggtggcagccaaatatctgagttttccgggggaactccagtcctgactatttaattcaaaactagaaaatatgattcacaattctgtttttgctgctgccgtcaagtgctaatgtggcat**gtgaagagaattcaggcagaatgactgcatcttcac**tttcatcttgtgtgataattttagctttgagtactgcaagtgtttgggagaaatggtagaaacatagaatatcaaagagacaagcaaaccaagagacccagagaggcaggcagacggaacgtcaggcagtataattttacaaaactaattcagtgttgtgtctctctctgcaaatggcaaggtatcgggaaaggtgctaagagctagtgctccgggtggggcaggattatctaaatagaaaactaagcactgatctcagggggaaagtttaaac**ttggaaatctctgggtttggttttttgtttttttccaa**tgaagtttatcagtgtggcagcacatctgcatttaattaatgacaatttgggtgaggcacagaattaactgttaaaaacaactatttctatgcagccataaaaaatgatgagtt**catgtcctttgtagggacatg**gatgaaactagaaatcatcattctcagtaaactatcgcaagaacaaaaaaccaaacaccgcatattctcactcataggtgggaattgaacaatgagatcacatggacacaggaaggggaacatcacactctggggactgttgcggggtggggggagtggggaggga**tagcattgggagatatacctaatgcta**aatgacgagttagtgggtgcagcacaccagcatg**gcacatgtatacatatgtaactaacctgcacaatgtgc**ataagtaccctaaaactta

Uppercase: TRAJ48

Lowercase: Flanking sequence[1000bp]

Red & Bold & Underline: Stem-loop [13]

Blue: Heptamer[20]

Green: Nonamer [1]

id-TRBJ2-2-2[J_gene_segment]

ccattttaattcactgcctttgtcttttccaagccccacacagtcagactaacctctgccacctgcgcttcctgccgctgcccagtggttgggggagggggactagcagggaggaaacatttttgtatcatggtgtaacattgtggggactagcgggggggcacgatgattcaggtagaggaggtgcttttacaaaaaaccctgatgcagtaagcatc**cccacccagctcagggaatgcagctaccaggtggg**aagagttctctggggctggtcccagctgtggtcttgcagggtcccccaacccagcgagcacctgtccatctccctgtccagactcggcttccaaggaataagaaggccaagacagc**aaagtgggattatcactcagcacttt**taataaaacttgttcttgacaaagtacttgcacatgcattatttattaagaactgatgaaaaccctgag**ggaaagatattgtcccatctttcc**aatgaggaaactgagatcagaggttacaggtcatataactaggaaacggcaaggtctagcctgcaatatcgcccagctccagccgttccagtaccaccaatgccccttcagatttca**aatccactgtgttgtcccccagccaagtggatt**ctcctctgcaaattggtggtggcctcatgcaagatccaggttaccgtgtccagctaactcgagacaggaaaagataggctcaggaaagagaggaagggtgtgccctctgtctgtgctaagggaggtg**gggaaggagaaggaattctgggcagccccttccc**actgtgctcctacaatgagcagttcttcgggccagggacacggctcaccgtgctaggtaagaagggggctccaggtgggagagagggtgagcagcccagcctgcacgaccccagaaccctgttcttaggggagtggacactgggcaatccagggccctcctcgagggaagcggggtttgcgccagggtccccagggctgtgCGAACACCGGGGAGCTGTTTTTTGGAGAAGGCTCTAGGCTGACCGTACTGGgtaaggaggcggctggggctccggagagctccgagagggcgggat**gggcagaggtaagcagctgccc**cactctgagaggggctgtgctgagaggcgctgctgggcgtctgggcggaggactcctggttctgggtgctgg**gagagcgatggggctctcagcggtgggaaggacccgagctgag**tctgggacagcagagcgggcagcaccggtttttgtcctgggcctccaggctgtgagcacagatacgcagtattttggcccaggcacccggctgacagtgctcggta**agcgggggctcccgct**gaagcccgggaactggggagggggcg**ccccgggacgccgggg**gcgtcgcagggccagtttctgtgccgcgtctcggggctgtgagccaaaaacattcagtacttcggcgccggga**cccggctctcagtgctgggtaagctggggccgccggg**ggaccggggacgagactgcgctcgggtttttgtgcggggctcgggggccgtgaccaagagacccagtacttcgggccaggcacgcggctcctggtgctcggtgagcgcgggctgctggggcgcgggcgcgggcggcttgggtctggtttttgcggggagtccccgggctgtgctctggggccaacgtcctgactttcggggccggcagcaggctgaccgtgctgggtgagttttcgcgggaccacccgggcggcgggattcaggtggaaggcggcggctgcttcgcggcacccggtccggccctgtgctgggaga**cctgggctgggtccccagg**gtgggcaggagctcggggagccttagaggtttgcatgcggggatgcacctccgtgctcctacgagcagtacgtcgggccgggcaccaggctcacggtcacaggtgagattcgggcgtctccccaccttccagcccct**cggtccccggagtcggggggtggaccggagctggaggagctgggtgtccggggtcagctc**tgcaaggtcacct

Uppercase: TRBJ2-2-2

Lowercase: Flanking sequence[1000bp]

Red & Bold & Underline: Stem-loop [14]

Blue: Heptamer[39]

Green: Nonamer [12]

id-TRAJ10[J_gene_segment]

tagtgtgaattcaggatacagcaccctcacctttgggaaggggactatgcttctagtctctccaggtacatgttgaccccatcccacccatgttttccccctatctggtttaaggcttccatatgtattgcgtgttatcctcatggatttcatcatccttgttttattatcaatgttctgtgaatttaagattgagcctccatgg**actcttcatttaaaaatgaaaatagctaatagaagagt**tggaaataacagtagaactaattcactaggtcaggatggagaagggagtaataccctagacaattaggacaaacgtggtttttccagaaatagactcacttcctgtttaaagcctagactgtggtctctccccgggcacccttcacattcctctaaccctccatatcccaaatt**tagctgtggaatcttagacaatctgtgacctatgcagcta**cagaa**atcttttcttactgagaatatctgcctatttgaaccaaaagat**ccttgagaatagacagtctctaatattccatgaagtgtctgatatggtccttttacaaaagtaaatacttgataaattcttgcttaattgaagttaaacacccaaagaaatcaggtttccaggccaaagggaagaagaaaattaag**caaatgacacagtacatttg**agcgtgtagtggggaggaagaaatgctagattgggattcaaatagagccatgttt**tggtctctgctgccaactagcaatgtaacttagacca**gatctcttgcctctaagcctcagtttctttatctgtaaatggggaagtgggtccaatggcctctctggcctcttagtac**aaaaagtctatgaatgtatcccatttggggcacttttt**tctcctaggagta**actcaatagctattttaaagcttgagt**gttttctaactcattagagcacatcaagaagaggggtgaagtgacaaaagggagtttattgtgaggcatcaaacactgtgATACTCACGGGAGGAGGAAACAAACTCACCTTTGGGACAGGCACTCAGCTAAAAGTGGAACTCAgtaagtatgagattctatggtaattacaaaatgtattctgg**tgacaataagtggaagaaaaggtgatgtaatttttgtca**ttgtatcttatttttaatctattactgaagcattaagaattgtgacatccagataaattagtgaaagtattcaaaggctatataagcatgctttcgttagggtggaatttagctaaccactaagactttttccaagatatgagattttcagctctcttcaacgtgagagttt**ttaatactagtattaa**gctaaacgatgctcaatggaagaaaagagctagaagagcagaaaattgcaccgtcatgatcaaatagatcctttctactt**ccactgccatggactcagtgg**tgcatttagaagagggtattcaaggtcattttagccatgccccaaaaaaaccactcccaattcattccagagattgtgcccccttatgctaaacatcatccaagaagagtcagtgaagtacagagaaaggaacccagggcttcttgtttcggtccagagttctaggtttgatttggtcccaactggatgtgtgatattggaacgagtcattgcccctctctgcacctcagactcctccagtgga**tgagggggtcaaagccctca**aaatctgttgttctgggatttcgctattataccacccgagagtttataggactcgagagtatggaatcttgtgagtatttcataaacaaattctatgaccctcaggaaatagccc**tgttaatcaacaacctaacttaaca**tagaaataataaccaccatctcagtccctaccatgtttagatacgatgctagatgcatgacatgtagaattgttcatctgaaataactctgtgaggtaaggggaagtatcttattttacaaaagaagaaactgaggctcagagaaggaagtgcccaagatcccaaagctagtaagtaaagggcctaaatttaaacttgggtctgcctgcctgactaaaaagccctgc

Uppercase: TRAJ10

Lowercase: Flanking sequence[1000bp]

Red & Bold & Underline: Stem-loop [12]

Blue: Heptamer[14]

Green: Nonamer [2]

id-TRAJ43[J_gene_segment]

actgtgtgttcctttaaaacaggggatactgcagaggggatatcctggctggatgtagtccttccttccttccagtgggaggtgatctcactgaggacacatacctgcattcagtcagttcttgtgctatttttatacctgttacggaggctttcctttccacctgtgaaaacagaacctctctgctccaagcttcttcagtctttattaccctatgtatttcctttagaacagtggtttgggaattaatt**tatcatcaggactttgtctctgatgata**tcactctgggcaacaaaatgaaagggctctcaaatagagaacagagagtcctgaatactcggataacttttccagagtgatatgtgttatgcgtaatgggatgaagtgggctagaatgaccttccaatgactattttggagacacac**taaggaaacttgcttaacatcctatatcctta**acttgttagaacattggtgaagtaaagcaaaatcttgaccctctaagaacataaacaaatgccaactcctctcccccaaagcactgcca**tgtttcattatgtgcaattatgaaaca**aatacatga**taatttatatgaaatta**catagatgagtagtttgagtattttgacccatgagtaagatgcaaattgtagaagttctcaggtagt**tcttcataaaaagttgtccctgaagacaaatagtgtacctgggcataatttgt**ttgcagtgttggtttatgcaccatttggggacctgtttgaaagctgattatgtcccagtgtgccaatttggggtctcaattaactgtgctcagtgcactgataagtgcccatatttacagtctagaagcatgacaaaatgaccccatttctgtaggaaaaggagaatggaactgtgtgtggcgtggtacttgtcagtgctgctcccccttgtccctgccatgctgggaggtggctgcaagttttccccaggaggtttttgttaga**gcatgtattactgtgACAATAACAATGACATGC**GCTTTGGAGCAGGGACCAGACTGACAGTAAAACCAAgtaagttgggggaatgggtcaatcttaaaagctgacctgagtgagcaatggtgctgttctg**ggcagtttctgtggggttaagacactgcc**actcccaagaggatctaagtacctggtttctttgtgcctgagaccctgagctggttggaagtctgttagttgatgaattctggcaatcttaaggattaggtcaa**gaagaagtagaaaatttcactggtttcttc**attggcttggctctggatatataaaaatgctaaatataagccgggtgcggtagctcactcctgtaatcccagcactttgggaggccgaggcgggaagatcatgaggtcaagagatgcagaccatcctggccaacatggtgaaaccccatctctactaaaaatacaaaaaat**tagctgggcatggtggcatgcgccggtagtcccagcta**ctcaggaggctgaggcaggagaatcgcttgaacccggaaggcggaggttgccgtgggccgagatcgtgccactgcactccagcctggcgacagagcgagactacatctcaaaaaaagaaagaaaaagaaatatggctgagaaagtggcatggtgctcct**ggtccctaaagtagcccctgggacc**tcaatgttgacctcctgtgctgggtgggatgtttctgcccaaccttgcta**ttgtgcctctggcacaa**gcattctgtcagaacccagtgtcctaggaacaaaacaggggatttacgatcaagttcagatctatctgcacagccagctgttaattttagagaactgtcctg**accttgcaaaggaactggaaacagaggctgcaaggt**ggggagacttccacta**gaggggaaggtgatctcacttgctcagccttcccctc**catccttcccacctgttgattattgtaaagccccataggactgtgtgaattatggaggaagccaaggaaatctcatctttggaaaaggcactaaactctctgttaaaccaagtaagtgttgggga

Uppercase: TRAJ43

Lowercase: Flanking sequence[1000bp]

Red & Bold & Underline: Stem-loop [14]

Blue: Heptamer[35]

Green: Nonamer [3]

id-IGHJ1[J_gene_segment]

cacctcccctgcccccagacaccctgtgcccgtcagttcatccccagcagaggccctcaccaggcacacccccatgctcacacctggccgcagg**cctcagcctccctgagg**gccccacccagcccgcgtctggccagtggtgcgtgcaa**agcccctcacccagactcggcggaaggcagccagtgcaggcctggggaggggct**ctccttagaccaccttgcac**cttccctggcacccaccatgggaag**ag**ctgagactcactgaggaccagctgaggctcag**agaagggacccagcactggtggacacgcagggagcccacgccagggcgccgtggtgagt**gaggcccagtgccacccactgaggcctc**ccgttcagtgggacgacggtgaacaggtggaaccaaccaggcaacccccgccgggccccacagacgggatcaga**gcaggaaaggcttcctgc**ccctgcaggccagcgaggagccc**tggcgggggccatggccctccaggcgaggaggctcccctggccaccgcca**cccgggcctctctgctgctgggaaaacaagtcagaaagcaagtggatgagaggtggcgtgacagacccagcttcagatctgctctaatttacaaaagaaaaggaaaaacacacttggcagccttcagcactctaatgattcttaacagcagcaaattattggcacaagactccagagtgactggcagggttgagggctgggg**tctcccgcgtgttttggggctaacagcggaagggaga**gcactggcaaaggtgctgggggcccctggacccgacccgccctggagaccgcagccacatcagcccccagccccacaggccccctaccagccgcagggttttggctgagctgagaac**cactgtgctaactggggacacagtg**attggcagct**ctacaaaaaccatgctcccccgggaccccgggctgtgggtttctgtag**cccctggctcagggctgactcaccgtgGCTGAATACTTCCAGCACTGGGGCCAGGGCACCCTGGTCACCGTCTCCTCAGGTgagtctgctgtctggggatagcggggagccaggtgtactgggccaggcaagggctttggcttcagacttggggacaggtgctcagcaaaggaggtcggcaggagggcggagggtgtgtttttgtatgggagaagcaggagggcagaggctgtgctactggtacttcgatctctggggccgtggcaccctggtcactgtctcctcaggtgagtcccactgcagccccctcccagtcttctctgtccaggcac**caggccaggtatctggggtctgcagccggcctgggtctggcctg**aggccacaccagctgccatccctggggtctccgccatgggctgcatgccagagccctgctgtcacttagccctggggcca**gctggagcccccaaggacaggcagggaccccgctgggcttcagc**cccgtcagggaccctccacaggtagcaagcaggccgagggcagggacgggaaggagaagttgtgggcagagcctgggctggggctgggcgctggctgttcatgtgccggggaccaggcctgcgctttagtgtggctaca**agtgcttggagcact**gggg**ccagggcagcccggccaccgtctccctgg**gaacgtcacccctccctgcctgggtctcagcccgggggtctgtgtggctggggacagggacgccggctgcctctgctctgtgcttgggccatgtgacccattcgagtg**tcctgcacgggcacaggtttatgtctgggcaggaacagggactgtgtccctgt**gtgatgcttttgatatctggggccaagggacaatggtcaccgtctcttcaggtaagatggctttccttctgcctcctttctctggg**cccagcgtcctctgtcctggagctggg**agataatgtccgggggctccttggtctgcgctgggccatgtggggccctccggggct**ccttctccggctgtttgggaccacgttcagcagaagg**cctttctttgggaactgggactctgc

Uppercase: IGHJ1

Lowercase: Flanking sequence[1000bp]

Red & Bold & Underline: Stem-loop [18]

Blue: Heptamer[34]

Green: Nonamer [6]

id-TRGJ2[J_gene_segment]

gattagaaaaaggc**tcacttgttttgaaaaacacaagtga**agaatttccatacactgacactgactgcatatattagactttagccaatattc**cctttgattttcttaacacggcaggtaaggcaaagg**ataacctattaaagattgggtgtgtggaaggcatgtttcttgtgatgatggggacgatggctttgagaatcccagagcaaagtggaatgcaaacagaggaactgagaaattattcttcctgcttaattgctatggatttaactgccactccaaaatgctgaattttttttagtaagggcaatgcttggtcctatagggttaaaatgtcatgtcaaggcacacaatcatagcaaacagattgccaatcataacaatgacaccatattcatcataatctcttatatttccacagcatttttttttgaggcagggtcttcccctgttgcccagtcgggag**tgcagtggtgtgatcaaggctcactgca**gcctcgaactcctaggctcaagtgaccctcctgcctcagcctctcgag**tagctgggcgtacagtcgtgcacatcatgctcagcta**atgctttttgtatttttagtaaatatggggtctcactagatactgggttgcccaggctggcttcaaaattctgagctcaagcaatcctcccacctcagcctcccaaagtgctggga**ttataggcacgagccactgcactgggacctataa**cattttaatcaaattgcttttttatatcttgtttcattttggtctcacttcagtgttggtagc**gatgttgaactgattttgaaacatc**ac**tgtttttagacaaataaaaca**ccaaaagctttaagttatttgatttgtggagcaacagaacttgttatgagcaaaatga**accaggactggaaccctggtcttttgagaatcccagacca**ccagaatttgaagaactcagggaaactgaattagagtttttgatatggactgaatcactgtgGAATTATTATAAGAAACTCTTTGGCAGTGGAACAACACT**TGTTGTCACAGgtaagtatcggaagaatacaacatttccaaggtaatagagggaaggcaggaaatg**attaaactggaataatgtaataatgtttagaaaaaagaggaattggatgg**ggatttgatgtagaaatcc**taggagagactttaaaacaaatgctcatactaaaagagaacatagataacatggcacagatatcataataggatttggctttgtgcatatacggacttccataaagggctcatatgtaatgtatgaaatgatctcattaaaatgtctggccctgagaccaaatgtattatgacaggttagtttgtgatagactctataaagcggacacatgtt**catctctgatggtcaaagagatg**t**tgactcttgttgtgaaggagcccgggtgttggagtca**ggtccacctggaactctggctccactactcactgcttagtgaaattgagtgattattctctggcctcagttttctaatccataaaatgggataacagtatgaattcagcagggttgtataagaattgcacaacatagtgtgaattaagtacttggcacattgtccaacccaaaataggtgcccaacaaatgttttctggattcacatgtaaagagacaatgggatctacta**tggagagttgctctcagtccatctaatttacagagcagcaattctcca**gaggattcagctgtacttctaactgctcaaatggaaggttatcttaacatcagctcacagacaaaaattgaacttatggattcgtttcttgttagcagactttttaatcacgtggc**aaaaacattgtattaagatgtctgttttt**tatttttgttgttctatgtgctttacttaatcctttatcttaattgatatcatttctaacaccaacatattggtcctaagatttatagccaattagtttagggttctgttcatatgtctcaggaaaaaaaggttaaaatcttaccaaaaatagtcaagatatcaaatcaattaaatcc

Uppercase: TRGJ2

Lowercase: Flanking sequence[1000bp]

Red & Bold & Underline: Stem-loop [16]

Blue: Heptamer[18]

Green: Nonamer [2]

id-TRAJ60[J_gene_segment]

cactcctgtgggtaccgggttaataggaaactgacatttggagccaacactagaggaatcatgaaactcagcaagtaatatttggcag**aatttttttttctatctgaaaatt**atcagtgagagattctaatgtgccttaacaaacaggaacaaaagatgagtgtttaatacaattcaatttaacaaatatttattaagagcctacagttattcccatggatcatctgagtcagtttccgaggaaacattatcgttgccttaagggagctggggggttgtcaacggg**ccagaggtgggatgaaaaatgacaacagatttacctctgg**gaccgggacatggttaaccacagcggccctgggtaagtagcttagcttcagaagaaaatgtgcccaacagcatgggtaacctaaaacaccgggcaatccaaatattcttttatgattggctttagcatgtattttattcttttgtagggcaggtttatctcaccaattatatattttcttaactga**cctgtaaaatctacagg**ggaaaagtattttaagaattatatgtttctgcaattaggctcccagcagtcaacaaagaagtggta**ctttttgtctttccagtgatgaaaaag**ggaacctggcatcc**ctggtggcccaccag**cgtctccttccctggcctaggtcagaacaagccgtaaatcagcaggccgttatcttcttataaatctgtag**agcagggtggacaacaaaaggcagcctgct**aggttttcagaacatgagttccttgtgtagccagagaacctg**ggaccatcctgacgtggctggtcc**tgctgtcctc**acagcctgaatcccaggctgt**atgaataggagaggttcaagtccagatgactgttcacgatgctggtcccccttgcatccctaatcatgctagagacatgaccagggtctgagaggaggaagttacagcacagcaccaacaggggcttttggtaaagggcctgggcactatgTGAAGATCACCTAGATGCTCAACTTTGGGAAGGGGACTGAGTTAATTGTGAGCCTGGgtgag**tacctcaactccagaggta**gctttagcggaac**ccctctatacctaacacctggcaatcagaggg**ctgaaacacttggccagataatgaattctcttatccggtgggaaaaggctgtaaagatcaaaccacctttcctatgggtaagcaagagtctgtagtttatgtaaaggcagcagc**tcctgtgggaaggaaggaaacagga**aatttacatttggaatggggacgcaagtgagagtgaagctatcttt**aaaccaaaggtgtcaggttatttggttt**ggtttttgaattatctggaagttccaaagaaagaacacttctccctgaggatttgattgcaaaattctgacttcaaacttctaaaaagatcaaatgttaaatcagatagtaggcttggaaaactctatttctctatgta**aaaagtagagaactactttt**cttttgtttgatcatt**ttattttgtttaggaaataa**gaggattagataccctggtggtgagtggggagggcagggacatttgcacagcatttctggataaattagatccaaataatcaaatcaaagactctcagctcagaaagt**aattaaagatctcctcacccacctctttaatt**ttgcaaatgaagacagtgaaactcagagaggttatgaacttgctcaaggtcacacaactgatcctgatatcaaggtccagggcaaagccaagacagctcatagttctcaccagcccagac**ccaagggagaaaaaaacatcatagtccttgg**ccacggcagcatcatttgcatcccaaacattctttacccaagacttaat**gaactcaaaaagaaatcctgagttc**caaagggaaataaaatcattctgcgtctgttgaaaaaaaaagcaagctaaagtggaacaataattgaacttgatattaggggaaaggtgcagccatctgcagactgagagaagggtgaaaaaaacaaaatgaaaatgctagtcttgtgttcagacaat

Uppercase: TRAJ60

Lowercase: Flanking sequence[1000bp]

Red & Bold & Underline: Stem-loop [17]

Blue: Heptamer[23]

Green: Nonamer [1]

id-IGHJ3[J_gene_segment]

cgcagccacatcagcccccagccccacaggccccctaccagccgcagggttttggctgagctgagaac**cactgtgctaactggggacacagtg**attggcagct**ctacaaaaaccatgctcccccgggaccccgggctgtgggtttctgtag**cccctggctcagggctgactcaccgtggctgaatacttccagcactggggccagggcaccctggtcaccgtctcctcaggtgagtctgctgtctggggatagcggggagccaggtgtactgggccaggcaagggctttggcttcagacttggggacaggtgctcagcaaaggaggtcggcaggagggcggagggtgtgtttttgtatgggagaagcaggagggcagaggctgtgctactggtacttcgatctctggggccgtggcaccctggtcactgtctcctcaggtgagtcccactgcagccccctcccagtcttctctgtccaggcac**caggccaggtatctggggtctgcagccggcctgggtctggcctg**aggccacaccagctgccatccctggggtctccgccatgggctgcatgccagagccctgctgtcacttagccctggggcca**gctggagcccccaaggacaggcagggaccccgctgggcttcagc**cccgtcagggaccctccacaggtagcaagcaggccgagggcagggacgggaaggagaagttgtgggcagagcctgggctggggctgggcgctggctgttcatgtgccggggaccaggcctgcgctttagtgtggctaca**agtgcttggagcact**gggg**ccagggcagcccggccaccgtctccctgg**gaacgtcacccctccctgcctgggtctcagcccgggggtctgtgtggctggggacagggacgccggctgcctctgctctgtgcttgggccatgtgacccattcgagtg**tcctgcacgggcacaggtttatgtctgggcaggaacagggactgtgtccctgt**gTGATGCTTTTGATATCTGGGGCCAAGGGACAATGGTCACCGTCTCTTCAGGTaagatggctttccttctgcctcctttctctggg**cccagcgtcctctgtcctggagctggg**agataatgtccgggggctccttggtctgcgctgggccatgtggggccctccggggctccttctccggctgtttgggaccacgttcagcagaaggcctttctttgggaactgg**gactctgctgctggggcaaagggtgggcagagtc**atgcttgtgctggggacaaaatgaccttgggacacggggctggctgccacggccggcccgggacagtcggagagtcaggtttttgtgcaccccttaatggggcctcccacaatgtgactactttgactactggggccagggaaccctggtcaccgtctcctcaggtgagtcctcacaacctctctcctgctttaactctgaagggttttgctgcatttttggggggaaataagcgtgctgggtctcctgccaaga**gagccccggagcagcctggggggctcaggaggatgccctgag**gcaacagcggccacacagacgaggggcaa**gggctccagatgctccttcctcctgagccc**agcagcacgggtctctctgtgg**ccagggccaccctgg**gcctctggggtccaatgtccaacaacc**cccgggccctccccggg**ctcagtctgagagggtcccagggacttagcgggg**tgccagttcttgcctggggtcctggca**tt**gttgtcacaatgtgacaac**tggttcgacccctggggccagggaaccctggtcaccgtctcctcaggtgagtcctcaccaccccctct**ctgagtccacttagggagactcag**cttgccagggtctcagggtcagagtcttggaggcattttggaggtcaggaaagaaa**gctggggagagggacccttcgaatgggaacccagc**ctgtcctccccaagtccggccacagatgtcggcagctggggggctccttcggctggtctggggtgacctctctccgcttcacctggagcatt

Uppercase: IGHJ3

Lowercase: Flanking sequence[1000bp]

Red & Bold & Underline: Stem-loop [19]

Blue: Heptamer[35]

Green: Nonamer [6]

id-TRAJ41[J_gene_segment]

tgtcagaacccagtgtcctaggaacaaaacaggggatttacgatcaagttcagatctatctgcacagccagctgttaattttagagaactgtcctg**accttgcaaaggaactggaaacagaggctgcaaggt**ggggagacttccacta**gaggggaaggtgatctcacttgctcagccttcccctc**catccttcccacctgttgattattgtaaagccccataggactgtgtgaattatggaggaagccaaggaaatctcatctttggaaaaggcactaaactctctgttaaaccaagtaagtgttggggatt**caaagtcctgatttatcatcagtactttg**tcactctgggcaacagaatgaaagggctctcaaatagagaacagagagtcctgaatactcagataacttttccagagtgatatgtgttatgtgtaatgggatgaagtgggctagaatgaccttccaatgactgttttggagacacactaaggaaacttgcttaacatcctatacccttaacttgttagaacattggtgaagtgaaaacaaaagcagtaagtgcaactaaagtgcaatgcagagtgagtgagaagagacttaccaactcctttctctgtgatatcttccccccaccttcacccttgaacccc**acagaattgtttctgt**ccatgcccctggctgttcccagtccaga**ccagcacgtcgttctgctgg**acataggaaaactccagacttcataacatgtcacagggccccctccctcagatggcagatctcaatattgatca**cccacatcactgtggg**gagtggtgaccttgtcagcaatggtctcatggaagagggaagctttatttacaccatcaagggcccatggatgacccacagtctgtgtgactgctgtgtgattggttttcagacattagctttaataggaatcataggagaagggacatggtggctactgcaaggggttttttgtttagggagaacgcactgtgGAACTCAAATTCCGGGTATGCACTCAACTTCGGCAAAGGCACCTCGCTGTTGGTCACACCCCgtgagtttttgtggtttactaattgtcctctctggaaagaaatccaatgggacctgttgaaacacagctgaatttaattgctatgcttagcatgcagttgttaactatgtctgatgtgtgagcaagatatgaatacatgtttccctggaggctggatttgg**ttatcaggtctcggggcagtttgataa**attgtactaatgctgcaatcactgtttttcaaaggtcca**caaagcacgttgtggctttg**ggaaaggcagagataagaagcaaagctttgtgatagagacagaaacaaggccatgaaaagggaagctaccaaagcaatggcatagccaaggaagtgtgtc**ctcaacagataagtggcaaggaccctgttgag**ttgatgcttgtgttgtctggtagaattaaaaaataagatgagtgggctgggcacagtggctcatgcctgtaattctatcactttgggaggctgaggcaggtggatagcttgaggtcaagagttcaagaccagcctgatcaacataatgaaaccccatctctactaaaaatacaaaaattagccgggcgtggtggcgggcacctgtaattccagctacttgggaagctaagg**caagagaatctcttg**aacccgggaggcagagat**tgcagtgagcctagatcgtggcactgca**ttccagcctgggtgatagagcgagactccgtctcaaaaacaaaaagaaagaaaaacagaaaaataaggcgagtggactgaggactccagtgaagggagggcagcaggagttccaagcttggcatggctttcttcctctacgagaatagcagaacaatgtaact**ttctctagagagagaacagagcccttgatttatcacatcaatgcttctgtt**taggtagcccccttcctcaccgtagggacaaagcaggcccccacatggacttcacccccttaactaggctggccaagccaaccctatgggagaggcagagaatcccaggactgca

Uppercase: TRAJ41

Lowercase: Flanking sequence[1000bp]

Red & Bold & Underline: Stem-loop [13]

Blue: Heptamer[42]

Green: Nonamer [4]

id-TRAJ24[J_gene_segment]

taaaggtgttgagcagaccaaggcccagtacctcgtcaccttcttcatccttgaaggagctttaagaggtgtggaggggaagaaacccaccaggaccccaccaataagcccagccttgagacccctccactctgtcagacttgaataagaacca**tctgagaacaaggtcttcctcaga**aggggactccagcatagtcatccccatttgatagattttgaaactcagggcggggcagagtggctcatgcctgtaatcccagcactttgggaagctg**aggcaggtggatcacttgaaggtcaggagttcgagacctgcct**ggccaacatggtgaaagcccgtctctactaaaaataaaaaaaat**tagctgggtgtggtggcacgcacctgtaatcccagcta**ctcagaaggctgaggcaggagaatcgcttgaacccgggaggcggaggttgcagtgagccaagatcatgccattgcactccagcctgggcaacaaaagcgaaacttcatctcaaacaaacaaacaagctcaggcaagtagaggccagtgattaacactcttccaataggaacagatcccaagcaacctgcagtagtttttccagtaggctcgtgaactcaaaacaccaagttattt**aaatgacagggcaaccgttgtgatcagctgtcattt**gagtagtgctgaggactgggtctgaccttgagtaaacgcctaggctggtgggtttctatccctgcagcatctaaagcggacgcccaggcttcactgaccccagcagtctggctttctgcccttgccgcccaggaggtgcagctcttggcacacaccatccttagtgtcttgaaagaagagaaattaaaagagaaagggggaaaagcttatatctcatatcatgcagttgcctcattttgtagcaagattgt**ttccccatgagcagtttgtcttcattcagcagctgtcttctctggggaa**gccattttgtagaggtgt**ttgtcacagtgTGACAA**CTGACAGCTGGGGGAAATTCCAGTTTGGAGCAGGGACCCAGGTTGTGGTCACCCCAGgtaagccccattccctggagcctcacctgccctt**agtatttggcatgccctgcatgccaaatatt**tctagccgagactatgagaaacacatctgaaaggaagccatttccctgaacagcagtacccaaagccattaagcaaaaatgagagatcagactggtgttaaatagaaacgtttttggataaatagtaattctcattattctctatcagtctcctgtgtgaatac**ccttatttcccttataagg**aaaaaagttacaataagcaggatagccatgattcatgctgtggtgtgggagcaatctgctgtggatttgggggtggaggattagaaatgtgttcaggcagactgga**tgtgtttttgacaggatatgtaacaca**gtgtgatttataaccagggaggaaagcttatcttcggacagggaacggagttatctgtgaaacccagtaagtataaaattgtatccctggattaagcaatgtctgtggattaagggctgatttagacccaattatacaccatatgaagaatgtttagagagggggatgagcatcaaataggaagccaatcgtcgtcctggctaggatctagcatctcagtgcaaaatgggctatgtaagtgtgcctctgggaattgctcctgaatatatgtgttgggactaaaatgtatgactggctacttgttgtattggctgggatcatatccctggatttgtgcagtaattggtgacaagatttctaattccttaacaaaccttctaaggcactattctatttggctaagattatgtcaatttatagaaggaaagaaccttgtaaataatgatcattaaccaaaagtaatgtgattagtctctctcaaagtctgaaaaacc**acttttttttaaaaagt**ttactcatgactactgagggacacttcctgtttctgagactttcagccaaaaattcgtacagcgtagcctcacggagcagagagaaccttgacaacattcctttttga

Uppercase: TRAJ24

Lowercase: Flanking sequence[1000bp]

Red & Bold & Underline: Stem-loop [10]

Blue: Heptamer[22]

Green: Nonamer [2]

id-TRAJ32[J_gene_segment]

agcaagttcaacttctccaatttgaaaactgtttctgaattatgcttcccttgtatgcagagagacctagattgactttggatgtttagtcattcg**atttaatcaaatgtaattaaat**gagcaaataatctccaatagccaatattcctatgttgtttcattgttttatgtgctttgctctagactttttgtctgggctttgtctctaataggatccccggaaggacagtgaaggtttttgttaaggtttttgtgtctgtgtggatagcaactatcagttaatctggggcgctgggaccaagctaattataaagccaggtaagtctcagagatgtgactgcacgggagaggagacactagttgaataatgcacaaagtgtagcatgcagattatatttttaagaacaagtcagcctgc**tggagacaatgcactcaacccaaatggggtctcca**ctcccagtgagaagcatgtcagccgctaactcttttgtttggtctagtagcttccggaatgaataatgcttaagtagccctctaagagagcagagccaggttgtt**agggaatatctaattccct**ctgatatggttaaaactctttgccaagggcagaaccagcctcattccattggaagagcaaattcagagaaaaaggaaagggtcacacagccaaagacggtaaaatgttcaaaggtgaaaaatagccaagtctgcagctctcctgacatccttgcctcaggctgctgtcccacgtg**gagagcagatgcctgcaaagctctc**ggtcacttgctggagtcactcagggttctgggccttgggatgtaacttaaa**tgacagtgtcagggcactgtca**gaggaatatgaaggctgcagcgatgcctgcaagtcagcttttgctacactgggactgaggtgttctaagcctggaaagacacaaagctgtcctcatcccaacctgccgtgcccctccttgagggttagtgtaaggctctgaaggactgtgTGAATTATGGTGGTGCTACAAACAAGCTCATCTTTGGAACTGGCACTCTGCTTGCTGTCCAGCCAAgtacgtaagtagtggcatgtgtcaggtggattctgtgtccatggcaagtaggaagcga**cagccacccttaggtggaaaggatggctg**aaggttgactttgttcactgctgtcatctcttatctgccgtgatatagcaggctgtcaaaattcccattctcctggatggcacaccacagtcaggggaggggaaacatgctaattttatgataaaccccaggggagaaaaagactactgtgggaaa**atttagttgagaaaatcaaattgttctcaactaaat**gctgcaggctttatgcttctgtgaacccagtttcccagttattcattgaccctgaacatacgtttacctgtgcaaatacaatctctcctaacctatttatacaattatgtgtctgccggtccaaatgccttcttctataggccatgtttttatt**tctcttccaaaagaga**ctccagagccatccttgggaagagtgctgagaagtcaggcaggcctaactcccaggggtcagggaggggaaagagaag**gctaagggtaacttagc**tcagtactagaactgtgatccacgtatctgtcactctatttcctacgtttgcttcccttttgttttctctagaataagttctttttgtttgctctggacacccactgctcccaggatgaaaggagagaaatgagatcagttttgaacacttcctcttgaaatataaagaatcaacaagttacagtcatgttggggacttcttctctctccaagcttaaatttctattacgtaagccttacttttaaccaagaaagttagtgacatggtatcactgtctgggatgctcacccctctctcttttctccagatactccgtgtatact**aaagctcagccaatgctcagcttt**gt**gataaaggagctgttgtgtgtctttatc**tctgcctttaggatacagtcttttttataacagaaagtgttcacttctcccatttatttcatggctggactctctacc

Uppercase: TRAJ32

Lowercase: Flanking sequence[1000bp]

Red & Bold & Underline: Stem-loop [11]

Blue: Heptamer[34]

Green: Nonamer [5]

id-TRAJ55[J_gene_segment]

tcttatgcctaaaacagcataagagagaaaagcatttccacgctaacctgtgtgttcactggttaacaaaggtctaaa**gagcaggtgagtcagctgctc**catttccaaact**ggactggacatttacagtcc**tgcagatactaaaaatgataatcaggaaaagcacacacaattctactttttcttataattcccatctttcccaaagaactgttataaattctctcccaaaggtgttatattcatccttcttttgttgcctctatttgtgatgtgtactgaggtgaaatatttgttgagattcaaaaaccccatgagatttaacttaagctttccctgtcagagagattctaagagctaaaagatgggaagctggtat**ttgggaactagattaaaaagaaaatccaaattaatcccaa**ggagcaaggctcaatagatagttatttagacataacatgtccaccctgcaagcctagttgctcagttgactgggctgcccctatcttagtcttgcctggaagagaaataagttgtttgccccatacctgtcttgagaaatgtcagcctccagacctgaaagcttcttgtgatttggtttatttctgtagagtttattattatacacagtgttcttgaagtaatagagaagtgcattagaagctcctgcaaatggaaatgaaatgtaaatttagattgtaaatacatgactgcggtaaatagatagcaagaaaaaggatgaagagatt**agaaatttgccacatttct**ctaaccctaaagggatgtttctataaatagtttaaaactgttgaagacgaagggggaggagga**aggagaacagaggcaaacagattttctcct**aatcctttccatttggcaaaatt**agaaatggtttttcaattaaatttccctgagcagaggaagaaaccatgtct**gtttccacactaaaattcctgtgggtgggagtctctagagttgatttggaggatggatccctgttagtgACAAGTGCTGGTAATGCTCCTGTTGGGGAAAGGGGATGAGTACAAAAATAAATCCAAgtaagtgtggagggacaagaagatctcacagtgcaggatttcccctggattttctgcattgccttttcacctttcctgtcttagcacgacaaattaggtcccagatgagcaggccctcgcattcaaaccggaaattttaaggaggaagccagattaactttactcgggagacctagtagactctgactttaaagat**tcttttaatccagctgtttttagggagagctctgtaaaaga**tgaaagaaatg**ttttaaaaaaaaattaaaa**ttctgtggggtgaaaacaaagatgttaaatatttgattggcaaggcaactggaaaatctggaccatgtctacaactgctaaaggaggctttgtgaaagagaaaatgagcagcccaaggagatcctgtcctaaacttctctggccagtgaaattcgggccattc**tctgccacagccctggactgctaggagggcaga**tcatatgtcttcctcagtggggagaggtgggccctcgctggcagtttctgtaaagcctcgtgctgtggtgtaattcagggagcccagaagctggtatttggccaaggaaccaggctgactatcaacccaagtaagtatgacagggtgaagctacatgcagctgagtacagtcttttcctttctagaccgtgtcctgcaagctctccttgagggacgtatactcattttgcattgtcctttgtagagaagcagaccaggaaagacaggaaagccctcaaatttccacttttaaacacctccctgtaa**aaactgtctcgcttccctccccttctacaatcagttt**ctagtaaatcagaatccggtgaattgatatgcaatttcaacgaaaaaaaaagcagagaaatagttaccccaacaagtgcaaaaagtagaaactatctgagtaccagccaagggaagataattaggaaagaaaaaaaaagaaagaaatggattactggaaccatgatggcagcttagtcacaaagaaaatgataa

Uppercase: TRAJ55

Lowercase: Flanking sequence[1000bp]

Red & Bold & Underline: Stem-loop [10]

Blue: Heptamer[18]

Green: Nonamer [4]

id-IGHJ1P-2[J_gene_segment]

tgcccgcacggtgcctga**gggggccttcttgggcagcgcctaagcaagccccc**agcacccttcggccccttca**aggcacacaggccccctttccacccagcctcaggaaaccacctgtgtcct**ccaacgacaggtcccagcctcccagcctttgccttgcctgttcctctccctggaactctgccccgacacagaccctccccagcaagccc**gcaggggcacctcccctgc**ccccagacaccctgtgcccgtcagttcatccccagcagaggccctcaccaggcacacccccatgctcacacctggccgcagg**cctcagcctccctgagg**gccccacccagcccgcgtctggccagtggtgcgtgcaa**agcccctcacccagactcggcggaaggcagccagtgcaggcctggggaggggct**ctccttagaccaccttgcac**cttccctggcacccaccatgggaag**ag**ctgagactcactgaggaccagctgaggctcag**agaagggacccagcactggtggacacgcagggagcccacgccagggcgccgtggtgagt**gaggcccagtgccacccactgaggcctc**ccgttcagtgggacgacggtgaacaggtggaaccaaccaggcaacccccgccgggccccacagacgggatcaga**gcaggaaaggcttcctgc**ccctgcaggccagcgaggagccc**tggcgggggccatggccctccaggcgaggaggctcccctggccaccgcca**cccgggcctctctgctgctgggaaaacaagtcagaaagcaagtggatgagaggtggcgtgacagacccagcttcagatctgctctaatttacaaaagaaaaggaaaaacacacttggcagccttcagcactctaatgattcttaacagcagcaaattattggcacaagactccagagtgactggcagggttgagggctgggg**tctcccgcgtgttttggggctaacagcggaagggaga**gcactggcAAAGGTGCTGGGGGCCCCTGGACCCGACCCGCCCTGGAGACCGCAGCCACATCAgcccccagccccacaggccccctaccagccgcagggttttggctgagctgagaac**cactgtgctaactggggacacagtg**attggcagct**ctacaaaaaccatgctcccccgggaccccgggctgtgggtttctgtag**cccctggctcagggctgactcaccgtggctgaatacttccagcactggggccagggcaccctggtcaccgtctcctcaggtgagtctgctgtctggggatagcggggagccaggtgtactgggccaggcaagggctttggcttcagacttggggacaggtgctcagcaaaggaggtcggcaggagggcggagggtgtgtttttgtatgggagaagcaggagggcagaggctgtgctactggtacttcgatctctggggccgtggcaccctggtcactgtctcctcaggtgagtcccactgcagccccctcccagtcttctctgtccaggcac**caggccaggtatctggggtctgcagccggcctgggtctggcctg**aggccacaccagctgccatccctggggtctccgccatgggctgcatgccagagccctgctgtcacttagccctggggcca**gctggagcccccaaggacaggcagggaccccgctgggcttcagc**cccgtcagggaccctccacaggtagcaagcaggccgagggcagggacgggaaggagaagttgtgggcagagcctgggctggggctgggcgctggctgttcatgtgccggggaccaggcctgcgctttagtgtggctaca**agtgcttggagcact**gggg**ccagggcagcccggccaccgtctccctgg**gaacgtcacccctccctgcctgggtctcagcccgggggtctgtgtggctggggacagggacgccggctgcctctgctctgtgcttgggccatgtgacccattcgagtg**tcctgcacgggcacaggtttatgtctgggcagga**acagggactgtgtccctgtgtgatgcttttgat

Uppercase: IGHJ1P-2

Lowercase: Flanking sequence[1000bp]

Red & Bold & Underline: Stem-loop [18]

Blue: Heptamer[37]

Green: Nonamer [6]

id-TRBJ2-1-2[J_gene_segment]

ttcctatgagctgcctgccacccctcgctcctcccacccacttcactataaatgccagtctgagca**ggtgggcacagtgagccccacc**agggagacccagtgacatagatggtctgctcagggtgatgcatgttccaaggagggacctctctgccccccaccattaccatcactgtgactttccccaagcccttcccattttaattcactgcctttgtcttttccaagccccacacagtcagactaacctctgccacctgcgcttcctgccgctgcccagtggttgggggagggggactagcagggaggaaacatttttgtatcatggtgtaacattgtggggactagcgggggggcacgatgattcaggtagaggaggtgcttttacaaaaaaccctgatgcagtaagcatc**cccacccagctcagggaatgcagctaccaggtggg**aagagttctctggggctggtcccagctgtggtcttgcagggtcccccaacccagcgagcacctgtccatctccctgtccagactcggcttccaaggaataagaaggccaagacagc**aaagtgggattatcactcagcacttt**taataaaacttgttcttgacaaagtacttgcacatgcattatttattaagaactgatgaaaaccctgag**ggaaagatattgtcccatctttcc**aatgaggaaactgagatcagaggttacaggtcatataactaggaaacggcaaggtctagcctgcaatatcgcccagctccagccgttccagtaccaccaatgccccttcagatttca**aatccactgtgttgtcccccagccaagtggatt**ctcctctgcaaattggtggtggcctcatgcaagatccaggttaccgtgtccagctaactcgagacaggaaaagataggctcaggaaagagaggaagggtgtgccctctgtctgtgctaagggaggtg**gggaaggagaaggaattctgggcagccccttccc**actgtgCTCCTACAATGAGCAGTTCTTCGGGCCAGGGACACGGCTCACCGTGCTAGgtaagaagggggctccaggtgggagagagggtgagcagcccagcctgcacgaccccagaaccctgttcttaggggagtggacactgggcaatccagggccctcctcgagggaagcggggtttgcgccagggtccccagggctgtgcgaacaccggggagctgttttttggagaaggctctaggctgaccgtactgggtaaggaggcggctggggctccggagagctccgagagggcgggat**gggcagaggtaagcagctgccc**cactctgagaggggctgtgctgagaggcgctgctgggcgtctgggcggaggactcctggttctgggtgctgg**gagagcgatggggctctcagcggtgggaaggacccgagctgag**tctgggacagcagagcgggcagcaccggtttttgtcctgggcctccaggctgtgagcacagatacgcagtattttggcccaggcacccggctgacagtgctcggta**agcgggggctcccgct**gaagcccgggaactggggagggggcg**ccccgggacgccgggg**gcgtcgcagggccagtttctgtgccgcgtctcggggctgtgagccaaaaacattcagtacttcggcgccggga**cccggctctcagtgctgggtaagctggggccgccggg**ggaccggggacgagactgcgctcgggtttttgtgcggggctcgggggccgtgaccaagagacccagtacttcgggccaggcacgcggctcctggtgctcggtgagcgcgggctgctggggcgcgggcgcgggcggcttgggtctggtttttgcggggagtccccgggctgtgctctggggccaacgtcctgactttcggggccggcagcaggctgaccgtgctgggtgagttttcgcgggaccacccgggcggcgggattcaggtggaaggcggcggctgcttcgcggcacccggtccggccctgtgctgggagacctgggctgggtccccagggtgggcaggagctc

Uppercase: TRBJ2-1-2

Lowercase: Flanking sequence[1000bp]

Red & Bold & Underline: Stem-loop [12]

Blue: Heptamer[44]

Green: Nonamer [11]

id-IGKJ2[J_gene_segment]

taagagctgagctcttcctgtgctgtgaaaacagacaaaccaaccaagtaaagtctacttttctactctattagtcttcactttggtttcgtataccatctggagctacatttcaaaatgcatttcaaagttatgagccttaagttgatatatatttagtctacctttttttaaataacattgcagcaaaggagaagataaaatagtaagacaacc**tgtaattattactcattgagaagctgatgatttccataattaca**ctaaatgaagtttatcctttgcaaaagcccccccagcccaccccaaaagaaagtac**aaaaaaactggccattttttt**taattgcttgtttttctttgtaattaacattcagtctactttctaaaaaataaataaataataagcagtccagatgtggcaagttgctaaagaaaggaaccatcaggccatagacgtaaatatattctcttcttggatt**ttaggtctcacctaa**gaaaataaacacatgctatgtcaga**gaagcctcagggcttc**cacacctgctcgaaaagggagttgagcttcagcagctgacccaggactctgttcccctttggtgagaagggtttttgttcagcaagacaatggagagctctcactgtggtggacgttcggccaagggaccaaggtggaaatcaaacgtgagtagaatttaaactttgcttcctcagttgtctgtgtcttctgttccctgtgtctatgaagtgatctataaggtgactctgcaatcagcctctgatatccttcagggaaaagataaagataagtctgtagtcaaactcgagaattgattgcacattttctttgaagagcaagcaagattcagtcattgggtgagaataacttgtctaagtaatagcttcagaaatgtcctggg**gaacataacatgttc**tggacagagccttggtcaattgtcagaaagggagtttttgtataggagggaagttaagaggaaccattgtgTGTGCAGTTTTGGCCAGGGGACCAAGCTGGAGATCAAACgtaagtacttttttccactgattcttcactgttgctaattagtttactttgtgttcctttgtgtggattttcattagtcggatgccagggatctaacaaacttcattcccaggttaggtacagaggaggggaaattgttccacaggacgctagcttgtggctaatttttaagatttctaaatcaaaataacttcattgggggaaagaggcttgctgagctttcagggaggtttttgtaaagggaaaagttaagac**gaatcactgtgattcactttcggccctgggaccaaagtg**gatatcaaacgtaagtacatctgtctcaattattcgtgagattttagtgccattgtatcatttgtgcaagttttgtgatattttggttgaataaac**ctggtgacccagaagtaaatagcaggacaccag**aaaatgaacttaaaaagctgagcaaatagacgaatcattgggtttgagaggagaataggattcatgggggaaatggggaagaaatagctagatttttctctgaacaagcagcctatctcatatgattggcttcaagagaggtttttgt**tgaggggaaagggtgagatccctca**ctgtggctcactttcggcggagggaccaaggtggagatcaaacgtaagtgcactttcctaatgctttttcttataaggttttaaatttggagcgtttttgtgtttgagatattagctcaggtcaattccaaagagtaccagattctttcaaaaagtc**agatgagtaagggatagaaaattagttcatct**taaggaacagccaagcgctagccagttaagtgaggcatctcaattgcaagattttctctgcatcggtcaggttagtgata**ttaacagcgaaaagagatttttgttaa**ggggaaagtaattaagttaacactgtggatcaccttcggccaagggacacgactggagattaaacgtaagtaatttttcactattgtcttctgaaatttgggtctgat

Uppercase: IGKJ2

Lowercase: Flanking sequence[1000bp]

Red & Bold & Underline: Stem-loop [11]

Blue: Heptamer[24]

Green: Nonamer [7]

id-TRAJ29[J_gene_segment]

taactcccactgagttcaaactgctgagagtgttcagtaggaaagtttctaagacatgtgggtggcctgtaggtcctgggctcttctcttgcagaggtccagtc**ctcagcctttccacagctgag**ccaaataccaaattctcccctccagggctggaaaagttgctaccagtcctctgctacccaaatataaattatttttttattgtttattaagaagagtaaaagaatcgtgacactttgtccagaacctattttccagtcttttcccccaggcttgtccctggtaacctctgtttggactcattgttaagcccagcgtgatttttggctccaactaaattgattttggaaatgactccagcatacctatgacccctctccttccaccctccataaaagatttatttttctttatttggcctctgct**gagattcctggctttgatgccccaggatctc**tcatttcccttgttccagggataagtgag**agatgactttttaaagagctactttcatct**tagagacaacaaagagagtatgcca**gcaagagcaacatcttgc**attgagccttctcactctatgccattttccaaatctttttctttatttaggataagtgaccactcctttttatttctagtacagcaaagagtacatcatgatgtcagaaacagggatttcctttggaatgtttttcacaggctaacaataagctagaagtctgcaagcaattcagaaatgcatccc**tgagctacaactccatgtatggaagctca**tcagcagggtagacaggcaaagcagaaacattttaattataccatactttgtagagattgggggagaagtgggcaaatgcgtgctaaggaaaaacaaaaactatggttaagtggggagatttctagatggttctaacagggaagaagaccaacaagaggaaacttccagg**aaaaccaccaaggccaggcattcagggtttt**tgttatggaggaaatcactgtgGGAATTCAGGAAACACACC**TCTTGTCTTTGGAAAGGGCACAAGA**CTTTCTGTGATTGCAAgtaagtgtttct**agccatccttgattttgatcagcaatggct**tcttcccttgaattatttttcagtgtacctagaatgcttttgcccccaagaaaggtttggaaggagctgggtcattagcattgcgcaggaaaattacaggttattctgttataattcgaaagccaactggacagtcatg**aatgcacacagatctgggccctgcatt**gcaccctggcctagttgcctccagcttgttctgaccgtaatcctggcccagtttcctcctctgat**cccaggagacaaaagccctggg**cagagaaccagactacttagcctgggggaagcac**aaaaaggcttgtctgagatccataagccttttt**tttttttctttttcttttttttttttttttttttttga**gacagagtctcgctctgtc**gcccaggctggag**tgcagtggtgcgatctcggctcactgca**agctccgcctcccgggttcacaccattctcctgcctcagcctccctagtagctgggactacaggtgctcaccaccacgtccggctaattttttgtatttttttttagtagagacagggtttcaccgcgttggccaggatgatctcgatctcccgacctcgtgatccacccgcctcagcctcccaaagt**gctgggattacaggcgtgagccaccgtgcccagc**cccaaaagccatttttaaatacattgcaggtttcta**tttaatgttgttattcattttgatttgcccaagtaatacattaaa**atttctcattgtaaaccattcaa**attacagataaagtccaggcacggtggctcacacctgtaat**cccagcactttagaaggctgaggtgggcaaatcacctgaggtcaggagtttgagaccagtctggccaacatgatgaaatcccatctctactaaaaatacgaaaagtagtcaatcgtggtggcaggcacctataatcccagctactcaggaggctgaggcaggataatcacttg

Uppercase: TRAJ29

Lowercase: Flanking sequence[1000bp]

Red & Bold & Underline: Stem-loop [16]

Blue: Heptamer[15]

Green: Nonamer [4]

id-TRBJ2-6-2[J_gene_segment]

taccgtgtccagctaactcgagacaggaaaagataggctcaggaaagagaggaagggtgtgccctctgtctgtgctaagggaggtg**gggaaggagaaggaattctgggcagccccttccc**actgtgctcctacaatgagcagttcttcgggccagggacacggctcaccgtgctaggtaagaagggggctccaggtgggagagagggtgagcagcccagcctgcacgaccccagaaccctgttcttaggggagtggacactgggcaatccagggccctcctcgagggaagcggggtttgcgccagggtccccagggctgtgcgaacaccggggagctgttttttggagaaggctctaggctgaccgtactgggtaaggaggcggctggggctccggagagctccgagagggcgggat**gggcagaggtaagcagctgccc**cactctgagaggggctgtgctgagaggcgctgctgggcgtctgggcggaggactcctggttctgggtgctgg**gagagcgatggggctctcagcggtgggaaggacccgagctgag**tctgggacagcagagcgggcagcaccggtttttgtcctgggcctccaggctgtgagcacagatacgcagtattttggcccaggcacccggctgacagtgctcggta**agcgggggctcccgct**gaagcccgggaactggggagggggcg**ccccgggacgccgggg**gcgtcgcagggccagtttctgtgccgcgtctcggggctgtgagccaaaaacattcagtacttcggcgccggga**cccggctctcagtgctgggtaagctggggccgccggg**ggaccggggacgagactgcgctcgggtttttgtgcggggctcgggggccgtgaccaagagacccagtacttcgggccaggcacgcggctcctggtgctcggtgagcgcgggctgctggggcgcgggcgcgggcggcttgggtctggtttttgcggggagtccccgggctgtgCTCTGGGGCCAACGTCCTGACTTTCGGGGCCGGCAGCAGGCTGACCGTGCTGGgtgagttttcgcgggaccacccgggcggcgggattcaggtggaaggcggcggctgcttcgcggcacccggtccggccctgtgctgggaga**cctgggctgggtccccagg**gtgggcaggagctcggggagccttagaggtttgcatgcggggatgcacctccgtgctcctacgagcagtacgtcgggccgggcaccaggctcacggtcacaggtgagattcgggcgtctccccaccttccagcccct**cggtccccggagtcggggggtggaccg**gagctgg**aggagctgggtgtccggggtcagctctgcaaggtcacctccccgctcct**gggaaaagactggggaagagggagggggtggggagg**tgctcagagtccggaaagctgagca**gagggcgaggccacttttaatcttttttctggggtgtttagagagaaggtgaacgatggaggagaggatttgttaggactctgggagaggcgagactggagaggacgaagggaaatcctggtttggggaatgggtaggagtgggggtaactgctattcgtaggcaaaaagagctgagcaggctgggaacagcgcgggtgggcaagggtcagcactgcgggcaggcgggtgggtgttagggggcagaaatcctgcagccgagggtgcagtagaacacagaagaaaaagcctgccaaacaaaagtggaacagagaagccaaaaagggagatgaacatgagtcagtgaagaaaagaatgaaagtttactgtttagcagtgtggatctctaatccgacttaaaactccttgttcccgattcctattcctcctaagccagagatccctgggtccagggtgagggcacggcattcatgcttacccacgggctggtcaacaaagaggtgctgacctgagagtagggcacataacctcagccactggggtacacttaccacccccgcccccgtgtagctccctcccctatcctgaaatctcccttagcacactaagta

Uppercase: TRBJ2-6-2

Lowercase: Flanking sequence[1000bp]

Red & Bold & Underline: Stem-loop [11]

Blue: Heptamer[28]

Green: Nonamer [11]

id-TRAJ46[J_gene_segment]

ataacaagttcgttttccaagccgtgcttccagaagctgttagtggttggtagttgctaagggttttgaaactggaggac**atcatagcagagtggaattatttggcacaaggggctatgat**t**agaattaatgagatgaaagaaaattct**ggctagaagactagaaaaatccggaaagaaaagatacagtaaaccttgggggagttgtagcatttatcctagtccaagcagcaaagagcagacagaagccaaattccaataaacaattggcctttttacttttacaagtttctgcaattatcacaatgtcttcagcttcttgggcaaatatcaacaggtcctaaatgcaaagtggaacagtcctctgtttccagaaccgagtagcttagccccgccccacacaccacccatgctcaaccactgtttttgtagagg**agtttgacgctgtgtggaatatggaaacaaact**ggtctttggcgcaggaaccattctgagagtcaagtcctgtgagtataaaacacactcaagtcccaacctgggtttcattgcatattaaataaactttttctggctgggaagtatttgattctaaagaaaggaaaag**tgaaattccaagcgcaacgcaagcatttca**ttagtgtgattcattaagaatttccatgggttgccttcgagagcgttaatcacatccattcactggtgagacccacagtttgaaaattgagttaaatttgaaaaaaaaattgtcaccatctctgtagaccacagtgcttgctgtttgggtgtctagacttccaaaatggaacctcaatctgtagtttcgagcttggagatttatcccgagcccgtgacttagagatgtgtaagaaaagctgc**acactgctgggccttgtagggcatttgtgggatcagtgt**catgaaatagcctgggtagacaggttggtgagg**agacactggtgacaggtgtct**gccctgtttctgtaaa**gctgctgacagccgtgAGAAGAAAAGCAGC**GGAGACAAGCTGACTTTTGGGACCGGGACTCGTTTAGCAGTTAGGCCCAgtaagtctgagcagaaagtaagatatttatgccttttcctattatttgttctagcctgacatttgagttgtcctcctttggattccagatcaacaaaccatagtgtcttttcatactccttttaatatttgtggctaagggccctccgttcttcccatctgtcctacatgagggtcctgtggccaggtccacattcataaagaagccaacag**tgatgcagttggttggcagtgtgtcaagaaactgcagtca**tcatctccagggcccttactctggttatgtgtgtttcaagcttttatgccagcaacagaggagagggagaagtgaccagtagcaatg**accagggctattagctgtacctggt**gtctgtgggtttcagagcagctgtggaggttacgggaggttcagggcatgagtttttctggcagaggagtttatgtaaagggttggcccagagtgtgtattcaggaggaggtgctgacggactcacctttggcaaagggactcatctaatcatccagccctgtaagtgcttttgcctgggaggtggggttcaaaatgcagtcctcatggggacatttctaaaccttggctcatgtgcttctg**gtgccactaccatttttggcac**tatcctact**tacatgggaggttttgagccatgta**gacaaggaggtgatgcagtttgtgctttggaggactcctgagatctgccctttggcaactgtgtttgagtaggagtccctcattggaaaatctggatgccagcagacaaaagatatgagcttaggatgggtcacaaacaaatggaggccactttgccagcagcacctttgcaacgaaaattgctttcataggaaatgtg**gaaatggctgtgtagatagagatttggccatttc**tccttttctgtctttataccctcttctctgcctgacttagggaccttaaattagagcagttattaatcatttgatttctcggccccatcactcatcaagtatgac

Uppercase: TRAJ46

Lowercase: Flanking sequence[1000bp]

Red & Bold & Underline: Stem-loop [12]

Blue: Heptamer[30]

Green: Nonamer [2]

id-TRBJ2-3[J_gene_segment]

tgcagggtcccccaacccagcgagcacctgtccatctccctgtccagactcggcttccaaggaataagaaggccaagacagc**aaagtgggattatcactcagcacttt**taataaaacttgttcttgacaaagtacttgcacatgcattatttattaagaactgatgaaaaccctgag**ggaaagatattgtcccatctttcc**aatgaggaaactgagatcagaggttacaggtcatataactaggaaacggcaaggtctagcctgcaatatcgcccagctccagccgttccagtaccaccaatgccccttcagatttca**aatccactgtgttgtcccccagccaagtggatt**ctcctctgcaaattggtggtggcctcatgcaagatccaggttaccgtgtccagctaactcgagacaggaaaagataggctcaggaaagagaggaagggtgtgccctctgtctgtgctaagggaggtg**gggaaggagaaggaattctgggcagccccttccc**actgtgctcctacaatgagcagttcttcgggccagggacacggctcaccgtgctaggtaagaagggggctccaggtgggagagagggtgagcagcccagcctgcacgaccccagaaccctgttcttaggggagtggacactgggcaatccagggccctcctcgagggaagcggggtttgcgccagggtccccagggctgtgcgaacaccggggagctgttttttggagaaggctctaggctgaccgtactgggtaaggaggcggttggggctccggagagctccgagagggcgggat**gggcagaggtaagcagctgccc**cactctgagaggggctgtgctgagaggcgctgctgggcgtctgggcggaggactcctggttctgggtgctgg**gagagcgatggggctctcagcggtgggaaggacccgagctgag**tctgggacagcagagcgggcagcaccggtttttgtcctgggcctccaggctgtgAGCACAGATACGCAGTATTTTGGCCCAGGCACCCGGCTGACAGTGCTCGgta**agcgggggctcccgct**gaagccccggaactggggagggggcg**ccccgggacgccgggg**gcgtcgcagggccagtttctgtgccgcgtctcggggctgtgagccaaaaacattcagtacttcggcgccggga**cccggctctcagtgctgggtaagctggggccgccggg**ggaccggggacgagactgcgctcgggtttttgtgcggggctcgggggccgtgaccaagagacccagtacttcgggccaggcacgcggctcctggtgctcggtgagcgcgggctgctggggcgcgggcgcgggcggcttgggtctggtttttgcggggagtccccgggctgtgctctggggccaacgtcctgactttcggggccggcagcaggctgaccgtgctgggtgagttttcgcgggaccacccgggcggcgggattcaggtggaaggcggcggctgcttcgcggcacccggtccggccctgtgctgggaga**cctgggctgggtccccagg**gtgggcaggagctcggggagccttagaggtttgcatgcgggggtgcacctccgtgctcctacgagcagtacttcgggccgggcaccaggctcacggtcacaggtgagattcgggcgtctccccaccttccagcccct**cggtccccggagtcggagggtggaccg**gagctgg**aggagctgggtgtccggggtcagctctgcaaggtcacctccccgctcct**ggggaaagactggggaagagggagggggtggggagg**tgctcagagtccggaaagctgagca**gagggcgaggccacttttaatcttttttctggggtgtttagagagaaggtgaacgatggaggagaggatttgttaggactctgggagaggcgagactggagaggacgaagggaaatcctggtttggggaatgggtaggagtgggggtaactgctattcgtaggcaaaaagagctgagcaggctgggaacagcgcgggtgggcaagggtcagcac

Uppercase: TRBJ2-3

Lowercase: Flanking sequence[1000bp]

Red & Bold & Underline: Stem-loop [14]

Blue: Heptamer[27]

Green: Nonamer [12]

id-IGLJ2[J_gene_segment]

ccttccagggtgaataattaatgtcttctctcatggtgaactctaggattcaagccatctaatgtttttgaa**gccactgtcattatatttaattgatgatgacaggtggc**caccaatgatgaatattttcccagggggagtctccctaagtggctttagacttcctcacatggccccaggggattaaatggctcctgattactcagag**gataagaggttctgtcttatc**atgttcctttcttatttgtcttatgtgtctttcctgccccaggcctgggatcccccactgatctcccttcccttagtgagaggtggtatttggagaccacattctggaggctcccttatgtcccccatttgaaaaagacaacggcagccaccaccccagctgtcccacccaacatgaggccagattcggggtgcagggatgctcccaaggttaccctaacagatgtgactggcacttcatattgggaccagcc**aggcctcactgaccaggcct**atccaactagaactactccagaa**ggtggggctgaaacccacc**aaggttcccagaacactgcactctagggcaatcagcctctgcatgggagg**agaggggcaccctct**gcaccaccccatggtgttacca**aaagttgaaccatgggttggttcaacttt**gcagagaagagaccacctaacccatctgtggaaattcactccttagcgatactgatgctccctaagaaattcaatcctgggcctgagtgatggttggtgcaaaaaacaaattcaa**gatcccagtgtcctccagaagcctggatttccagggatc**ctgctgtgagtcacaggacgtcaccggtccccttctctttgtgggttgagtgtg**ggggccatgtggactccctcatgagcagatgccaccagggccactggcccc**agcttcctccttcacagctgcagtgggggctggggctggggcatcccagggagggtttttgtatgagcctgtgtcacagtgTGTGGTATTCGGCGGAGGGACCAAGCTGACCGTCCTAGgtgagtctcttctcccctctccttccccgctcttgggacaatttctgctgtttttgtttgtttctgtatcttgtctcaacttgtggtcagcctttctccctgcatcccaggcctg**agcaaggacctctgccctccctgttcagacccttgct**tgcctcagcaggtcactacaaccacttcacctctgaccacaggggcaggggactagatagaatgacctactgagcctcgtctgtctgtctgtctgtctgtctctctgtttgtctctctgtctctctgtttgtctctctgactgtctgacaggcgcaggctgggtctctaagccttgtt**ctgttctggcctcctcagtctgggttcttgtcggaacag**ctttgtccttgggttacctgggttccatctcctggggaattgggaacaaggggtctgagggaggcacctcctgggagactttagaaggacccagtgccctcggggctgatgctcgggaatcacagagctgggacccagagccaggatccagacccagaatgaggtaggaggtggaggggctg**ccctgggcgtctgggggctgccaggg**actgagc**cctgagccagcctgagactcagg**aaaccccgtcaggagggagaagggagaagcagactctggacaccagaaagccaggggaagg**gtcacaaaaggagtggatgtgac**ggaagggcgggctcctgggtctcttcagaacatat**cccctgtgcccagggg**gatcagaggggcagagtccactgcgtgaaag**ccccactgctatgaccaggtagccgggacgtgggg**tggatgccagaaaagactccacggaataaga**gagagcccaggacagcaggcaggctctc**cgatccccccaggcccttgccccatacacgggctccagaacacacatttggctggaacagcctgagggaccaaaaggccccagtatcccacagagctgaggagccaggccagaaaagtaaccccagagttcgc

Uppercase: IGLJ2

Lowercase: Flanking sequence[1000bp]

Red & Bold & Underline: Stem-loop [16]

Blue: Heptamer[28]

Green: Nonamer [4]

id-TRAJ40[J_gene_segment]

atgggagaggcagagaatcccaggactgcagagggcctccccatgtccactagattatctgtaataaaacagaatatttatccaagttgtgaagctgtgagggagagggtaatcgtgatagattataaaaagcaatattgaaaattatacaagctgtctc**cacacttgcagaattcagtgtg**ttttttgttgttattgggtttt**tttctgttagaattacagaaacaggggtgctttcctgcaggcaccctgttt**gcaa**aaatgtctgccatctcacattt**cagtagtatttcatcaacaatcttctgcacacgaaccagctgtgacaataaggtaatggggct**gatttgttaacaaatc**gagtttatacatttcaa**tgagtggcctagtcaaaacactca**gcttgaagcatccaacactgtccattgctttccccctccctgaaaccaccctcaaaggccaaaacactttctttttgtagcatccatgatggctttatcttttaaaggt**ggattttactcttgataatcaaaatcc**tttcagaaccaaggctattaaatagaggagagaatcgac**tagttactgatactgttttggatccaaaaatagatgtaacta**taaaataatgtcatgggttt**gtttatgtggctcataaac**tggctctcggggggaatcctagatggaatatttcagagggttcaacagtcatttagatgcttaagtcctcatatgttcactaaatcagac**tctttaattaataaacatgcataaaga**tagcatgagattgaagaacttttgcaaactcggacccaatagaagcagtgaagactttctgtatgaattaagatataatccgaaataaattacttgtagaaaaatacgaggtcaaggcctttcaatggggaaagtgcatgtaatgccccttttattcatggagagtaaagctgcttccataaatggtaacattatgttggt**ttatgtagagacacataa**cactgtgACTACCTCAGGAACCTACAAATACATCTTTGGAACAGGCACCAGGCTGAAGGTTTTAGCAAgtgagtattacagacaagacaggctgtattttaaaggttgttggtgttcatgtttgagggacaagggaaggtaggacaagggaaatacccatgtaaactgtgcaagctaaaatcactgctatctgaagcggcaattctgggtccctctactctggacaggaggggcgaaggagagctctgaggcttacc**atgatggacttactgccgatttcataagaattgtatccatcat**tagcaaaatgtttatgacttgaaggcctcaaga**aactcctgttaacacaacaggagtt**ctgtgttctttattcttttaaaactatattctcactttacatttttcactcatttttctgcctttttttatgcgtctctcctctgtgcaactcaatgcttcatctgggcattcatatt**ttgagaaatcatctcaa**gaggaaatcatgt**ccagatcctagcatgtcactcaaagctctctgaaatctgg**gccctttctattctccaaatttattcccactgattccctttcatgaatgctctgctccaggaaaactaggtatctccctcccttgaatatttacacatgcccacacctgtaccctgtgtccatacatcagacatcctggacccaccacctaaaacaggcttgcacaggtgacaaaatttggcctcatcatctctcagccttgagacccagcacaggcatcaccagatttag**tttcataggaagagcatgaaaaatcatatcattttt**taattaagcaggccaaggtaagagatccatcagacaa**agaagcaaagtaggaaggagagagatggcttct**gtaatgggtttgcaaaggagagctctgtccctgaaatagcactaatataatccaagtaggagatttacaggctgcatgtttcttggtttccctttggagtgctccattcctaacaccagttgttaagcccctggtcctgaatgtggccgggggtgaggaagtgggaggagaccatgctggccact

Uppercase: TRAJ40

Lowercase: Flanking sequence[1000bp]

Red & Bold & Underline: Stem-loop [18]

Blue: Heptamer[20]

Green: Nonamer [4]

id-TRAJ42[J_gene_segment]

tttttgttaga**gcatgtattactgtgacaataacaatgacatgc**gctttggagcagggaccagactgacagtaaaaccaagtaagttgggggaatgggtcaatcttaaaagctgacctgagtgagcaatggtgctgttctg**ggcagtttctgtggggttaagacactgcc**actcccaagaggatctaagtacctggtttctttgtgcctgagaccctgagctggttggaagtctgttagttgatgaattctggcaatcttaaggattaggtcaa**gaagaagtagaaaatttcactggtttcttc**attggcttggctctggatatataaaaatgctaaatataagccgggtgcggtagctcactcctgtaatcccagcactttgggaggccgaggcgggaagatcatgaggtcaagagatgcagaccatcctggccaacatggtgaaaccccatctctactaaaaatacaaaaaat**tagctgggcatggtggcatgcgccggtagtcccagcta**ctcaggaggctgaggcaggagaatcgcttgaacccggaaggcggaggttgccgtgggccgagatcgtgccactgcactccagcctggcgacagagcgagactacatctcaaaaaaagaaagaaaaagaaatatggctgagaaagtggcatggtgctcct**ggtccctaaagtagcccctgggacc**tcaatgttgacctcctgtgctgggtgggatgtttctgcccaaccttgcta**ttgtgcctctggcacaa**gcattctgtcagaacccagtgtcctaggaacaaaacaggggatttacgatcaagttcagatctatctgcacagccagctgttaattttagagaactgtcctg**accttgcaaaggaactggaaacagaggctgcaaggt**ggggagacttccacta**gaggggaaggtgatctcacttgctcagccttcccctc**catccttcccacctgttgattattgtaaagccccataggactgtgTGAATTATGGAGGAAGCCAAGGAAATCTCATCTTTGGAAAAGGCACTAAACTCTCTGTTAAACCAAgtaagtgttggggatt**caaagtcctgatttatcatcagtactttg**tcactctgggcaacagaatgaaagggctctcaaatagagaacagagagtcctgaatactcagataacttttccagagtgatatgtgttatgtgtaatgggatgaagtgggctagaatgaccttccaatgactgttttggagacacactaaggaaacttgcttaacatcctatacccttaacttgttagaacattggtgaagtgaaaacaaaagcagtaagtgcaactaaagtgcaatgcagagtgagtgagaagagacttaccaactcctttctctgtgatatcttccccccaccttcacccttgaacccc**acagaattgtttctgt**ccatgcccctggctgttcccagtccaga**ccagcacgtcgttctgctgg**acataggaaaactccagacttcataacatgtcacagggccccctccctcagatggcagatctcaatattgatca**cccacatcactgtggg**gagtggtgaccttgtcagcaatggtctcatggaagagggaagctttatttacaccatcaagggcccatggatgacccacagtctgtgtgactgctgtgtgattggttttcagacattagctttaataggaatcataggagaagggacatggtggctactgcaaggggttttttgtttagggagaacgcactgtggaactcaaattccgggtatgcactcaacttcggcaaaggcacctcgctgttggtcacaccccgtgagtttttgtggtttactaattgtcctctctggaaagaaatccaatgggacctgttgaaacacagctgaatttaattgctatgcttagcatgcagttgttaactatgtctgatgtgtgagcaagatatgaatacatgtttccctggaggctggatttgg**ttatcaggtctcggggcagtttgataa**attgtactaatgctgcaatcactgtttttcaaaggtccacaaagca

Uppercase: TRAJ42

Lowercase: Flanking sequence[1000bp]

Red & Bold & Underline: Stem-loop [13]

Blue: Heptamer[46]

Green: Nonamer [3]

id-TRDJ1[J_gene_segment]

tttgtaaagctc**tgtagcactgtgactgggggatacgcacagtgctaca**aaacctacagagacctgtacaaaaactgcaggggcaaaagtgccatttccctgggatatcctcaccctgggtcccatgcctcaggagacaaacacagc**aagcagcttccctccctgctt**tggggcctggaagggatagcaggaagttgactggaccagggagatgaccacagctgctgacctctcactcactgctgttcttccttgggtgaaactggcatttctacattttcttacagcacatttggggaatacaaaaaggcctttcttaaaaactattcttgtcttgttttcatgttgattctattgcaaaagagagtta**tatgagccacctcata**cggaatttctaaattcaaacctctagagagatttacccaagtgctttgctttgcagtttgggaggatggatttgaagagagattgatttttttgtaggcaatcaccggccacagttgctcattctaaagctgactgctctgtaaatcacccagtgcttcatgccaccctttctcctcttgctgtgccacacgttatct**gcctttaaagcagcagcactggtgtctgtaaaggc**cttaaccctggagtagtcatggagccaagacccacccctttgacagtgccagctttccaacacagagagctgagtatgggtctaggaagtgagagcaatgtaaaacaatagaaagcaacagttcagagcactgcatcaagtgtactgtgctggaaaggtccgccataggaaatatggtcctccatactcctcagacaacagccttccgaaagcaaacctgtccctacctgcagatgattaaccatctatgaaccggctgggtaagcaacaagtgcc**atctttcatggagctgagccttaaagat**cctccagtcctaaagctgacgggaagaaggtaggtgggagcagcgctgaggtttttggaacgtcctcaagtgctgtgACACCGATAAACTCATCTTTGGAAAAGGAACCCGTGTGACTGTGGAACCAAgtaagtaactcatttatttatctgaagtttaaggttaaggcatcctccatctaaggaggcagaaataatcctgaaatgggaaatgggtgaaatagctagcatttaggaggactcctgggaagaggtgaaatatggttaatcctttccataggagaggagcagaaggtgtctgtaaaaaaagagtttggggcatatgaagggacccttctggctcagaggaaacagagaactctgcccagtccatcgaatttgggtcaggctgtagaaaccacagagtcattggcccaactccctgagcag**agtcagaatggacacagaatgaagctgact**tgagtccaggtggttttaccaacgtgattccaaac**gttctgacctcccttatgccaaacacatcctcctggaagagctcagaac**tgaatccgtctcctgggcatggagggtggggcgagcgcaagtagtccctggtgttgcgatatttctctttagaagaaaataacctgtttcaga**catttgtttctaattgtttgagctgcaaatg**aaaactaggttactagtgagaaatatgggacat**tctggttcccactgagaccaga**ttccaagcaggacaaagtgaatgtgtagtgcttccaactaagaaatgctttgaagtctccaggacacagccaggagagttccctaccactcaccagtgttcatcgtttttgcaaagagtatatggtgccagtcacttattaacagttatgcaagaatcaatgctaaggt**tgttgaaggattgtgctcaaca**cagaaggcatctttggtgtcctttcaatgacatttgaattcgaccg**tgtcttgctaaaaaataaaataaaataagaca**gtggtctccacctggcagcagattccagttagatagtacatattaatcttaagtaagaaaatgtaataaaccaatacaaaaacccatttcgaaggtcctagggaagtctagcaataggggtcagtggtcccaaatctg

Uppercase: TRDJ1

Lowercase: Flanking sequence[1000bp]

Red & Bold & Underline: Stem-loop [11]

Blue: Heptamer[26]

Green: Nonamer [5]

id-TRAJ51[J_gene_segment]

ttcagaagtccccctagggttcttgtaaaggcctccagtgcagtgctaatgctggtggtactagctatggaaagctgacatt**tggacaagggaccatcttgactgtcca**tccaagtaagtgtaacaagacacagcagtatactggaaattctggaatgtcaccccagtgtcaggttttaaagaaccagatattgcttaaagttaagtggtgtcctc**tgaggtcctttctcttctttaacctca**atattcacagattttcttttggaccctgagttgttaggtctgtctccctgctgcgtgtacagctgaccttgaccct**ggaaagtcagaatctttcc**tgctttcagtgtcttctctggaggtgatgccctaacttttttgag**ggggaggtcatcctttccaaatgactcccc**tgagtaacctgttctaatgtttagtagcattccctgtcagaagtccttccttacctcaatcttaaattattcttggtatagccgaggccgtttccaccagcatcatccacaataaagaggaaaattgagagcggagtttgaataggtatgtggtatttggttcagaatgggttttaaatgaaagtggagagggaaaaaatgaaacctgaatgggaatctgaggtcttatttcaaatccctctctgtttcctgtctgtcatctcc**ttttgcttctatagatgcaaaa**ctacagagagagaggcggggagagaaaggatagacagaaataaacatccactgtgtccattccctgtcactgaaaccaccaggcaagggattacaattaattccaaagagtttcagatacacggcttaatacctttctccttgtggtgacttgtatcctggctgataattagagcagatagaaagatttggctttggtgtgtttgccgctgcaccccgctgggaagagacagaaacttgactgttttttaagattcataaagttccttctgtcagtcgttgtaaaa**ctccctgaagcagggAG**ATGCGTGACAGCTATGAGAAGCTGATATTTGGAAAGGAGACATGACTA**ACTGTGAAGCCAAgcaagctggaaagacctaaactcacagt**gttccttatgtacttttgctcccttccccgtttcaggccttttggctgtttcctgtagggattctaagagagtcaaatggaaatggattgaagagagtctgaagccccgtgggtgctcagaaatgataaatgaaaggaataagagacagaaagggaaacaaagttgaaaagcaggaatatgtctttcactgcagcttgataagactgagatgtgaggtgggtgtcagagaagggcatgatcgctccagctgctattgcagaacactctgttttcctaggatctctatcaagggcagtgtcctcttatgccaaagggatgagtcaagtccctatatttctggagtagctgccc**ttgtgcaggcatagttggtcaggcagacagcacaa**tcccagctcagtgtgtatggaggaggccacctgacattgccagctgtatctgctccattcagtggcagcat**aaatccataccaggtgggtgtggattt**ggaatatgccagtgccactacaaggctgaatgttgagttgttgtgatcatcttgaatgtgtgagatgcctccaagtgaagtctggttctaatgagcagggtgcaggttggaccaatattgcccattttctgttgagtttctccaagcccagtagtgtgggagccaacattgagagactgacatttgagctagggatcctattgactgaaatatgccaaaggga**tctgaaaggtcaaaactgccttcaga**gctcagcctttgttgttatctctctggcagttgtcatttcagcaaaatgcatccacttagaaagctcagctcttcattaggcagacaaatgagaacaacatattgaccagagataaggagttgggagcccaga**tctcagcctctgaga**gcctggcatttccagttctttaatcacgtgagccagtgagcatatgctatctcagagaaagcagatcaatggtgcctgaggacctggtgt

Uppercase: TRAJ51

Lowercase: Flanking sequence[1000bp]

Red & Bold & Underline: Stem-loop [11]

Blue: Heptamer[22]

Green: Nonamer [2]

id-IGHJ5[J_gene_segment]

cgctttagtgtggctaca**agtgcttggagcact**gggg**ccagggcagcccggccaccgtctccctgg**gaacgtcacccctccctgcctgggtctcagcccgggggtctgtgtggctggggacagggacgccggctgcctctgctctgtgcttgggccatgtgacccattcgagtg**tcctgcacgggcacaggtttatgtctgggcaggaacagggactgtgtccctgt**gtgatgcttttgatatctggggccaagggacaatggtcaccgtctcttcaggtaagatggctttccttctgcctcctttctctggg**cccagcgtcctctgtcctggagctggg**agataatgtccgggggctccttggtctgcgctgggccatgtggggccctccggggctccttctccggctgtttgggaccacgttcagcagaaggcctttctttgggaactgg**gactctgctgctggggcaaagggtgggcagagtc**atgcttgtgctggggacaaaatgaccttgggacacggggctggctgccacggccggcccgggacagtcggagagtcaggtttttgtgcaccccttaatggggcctcccacaatgtgactactttgactactggggccagggaaccctggtcaccgtctcctcaggtgagtcctcacaacctctctcctgctttaactctgaagggttttgctgcatttttggggggaaataagcgtgctgggtctcctgccaaga**gagccccggagcagcctggggggctcaggaggatgccctgag**gcaacagcggccacacagacgaggggcaa**gggctccagatgctccttcctcctgagccc**agcagcacgggtctctctgtgg**ccagggccaccctgg**gcctctggggtccaatgtccaacaacc**cccgggccctccccggg**ctcagtctgagagggtcccagggacttagcgggg**tgccagttcttgcctggggtcctggca**tt**gttgtcacaatgtgACAAC**TGGTTCGACCCCTGGGGCCAGGGAACCCTGGTCACCGTCTCCTCAGGTgagtcctcaccaccccctct**ctgagtccacttagggagactcag**cttgccagggtctcagggtcagagtcttggaggcattttggaggtcaggaaagaaa**gctggggagagggacccttcgaatgggaacccagc**ctgtcctccccaagtccggccacagatgtcggcagctggggggctccttcggctggtctggggtgacctctctccgcttcacctggagcattctcaggggctgtcgtgatgattgcgtggtgggactctgtcccgctccaa**ggcacccgctctctgggacgggtgcc**ccccggggtttttggactcctgggggtgacttagcagccgtctgcttgcagttggacttcccaggccgacagtgg**tctggcttctgaggggtcaggccaga**atgtggggtacgtgggaggccagcagagggttccatgagaagggcaggacagggccacggacagtcagcttccatgtgacgcccggagacagaaggtctctgggtggctgggtttttgtggggtgaggatggacattctgccattgtgattactactactactactacatggacgtctggggcaaagggaccacggtcaccgtctcctcaggtaagaatggccactctagggcctttgttttctgctactgcctgtggggtttcctgagcattgcaggttggt**cctcggggcatgttccgagg**ggacctgggcggactggccaggaggggatgggcactggggtgccttgaggatctgggagcctctgtgg**attttccgatgcctttggaaaat**gggactcaggttgggtgcgtctgatggagtaactgagcctgggggcttggggagccacatttggacgagatgcctgaacaaaccaggggtcttagtgatggctgaggaatgtgtctcaggagcggtgtctgtaggactgcaaga**tcgctgcacagcagcga**atcgtgaaatattttctttagaattatgaggtgcgctgtgtg

Uppercase: IGHJ5

Lowercase: Flanking sequence[1000bp]

Red & Bold & Underline: Stem-loop [20]

Blue: Heptamer[29]

Green: Nonamer [5]

id-TRBJ2-2P-2[J_gene_segment]

acattgtggggactagcgggggggcacgatgattcaggtagaggaggtgcttttacaaaaaaccctgatgcagtaagcatc**cccacccagctcagggaatgcagctaccaggtggg**aagagttctctggggctggtcccagctgtggtcttgcagggtcccccaacccagcgagcacctgtccatctccctgtccagactcggcttccaaggaataagaaggccaagacagc**aaagtgggattatcactcagcacttt**taataaaacttgttcttgacaaagtacttgcacatgcattatttattaagaactgatgaaaaccctgag**ggaaagatattgtcccatctttcc**aatgaggaaactgagatcagaggttacaggtcatataactaggaaacggcaaggtctagcctgcaatatcgcccagctccagccgttccagtaccaccaatgccccttcagatttca**aatccactgtgttgtcccccagccaagtggatt**ctcctctgcaaattggtggtggcctcatgcaagatccaggttaccgtgtccagctaactcgagacaggaaaagataggctcaggaaagagaggaagggtgtgccctctgtctgtgctaagggaggtg**gggaaggagaaggaattctgggcagccccttccc**actgtgctcctacaatgagcagttcttcgggccagggacacggctcaccgtgctaggtaagaagggggctccaggtgggagagagggtgagcagcccagcctgcacgaccccagaaccctgttcttaggggagtggacactgggcaatccagggccctcctcgagggaagcggggtttgcgccagggtccccagggctgtgcgaacaccggggagctgttttttggagaaggctctaggctgaccgtactgggtaaggaggcggctggggctccggagagctccgagagggcgggat**gggcagaggtaagcagctgccc**cactctgagaggggctgtgCTGAGAGGCGCTGCTGGGCGTCTGGGCGGAGGACTCCTGGTTCTGGgtgctgg**gagagcgatggggctctcagcggtgggaaggacccgagctgag**tctgggacagcagagcgggcagcaccggtttttgtcctgggcctccaggctgtgagcacagatacgcagtattttggcccaggcacccggctgacagtgctcggta**agcgggggctcccgct**gaagcccgggaactggggagggggcg**ccccgggacgccgggg**gcgtcgcagggccagtttctgtgccgcgtctcggggctgtgagccaaaaacattcagtacttcggcgccggga**cccggctctcagtgctgggtaagctggggccgccggg**ggaccggggacgagactgcgctcgggtttttgtgcggggctcgggggccgtgaccaagagacccagtacttcgggccaggcacgcggctcctggtgctcggtgagcgcgggctgctggggcgcgggcgcgggcggcttgggtctggtttttgcggggagtccccgggctgtgctctggggccaacgtcctgactttcggggccggcagcaggctgaccgtgctgggtgagttttcgcgggaccacccgggcggcgggattcaggtggaaggcggcggctgcttcgcggcacccggtccggccctgtgctgggaga**cctgggctgggtccccagg**gtgggcaggagctcggggagccttagaggtttgcatgcggggatgcacctccgtgctcctacgagcagtacgtcgggccgggcaccaggctcacggtcacaggtgagattcgggcgtctccccaccttccagcccct**cggtccccggagtcggggggtggaccg**gagctgg**aggagctgggtgtccggggtcagctctgcaaggtcacctccccgctcct**gggaaaagactggggaagagggagggggtggggagg**tgctcagagtccggaaagctgagca**gagggcgaggccacttttaatcttttttctggggtgtttagagagaaggtgaacgatggag

Uppercase: TRBJ2-2P-2

Lowercase: Flanking sequence[1000bp]

Red & Bold & Underline: Stem-loop [15]

Blue: Heptamer[33]

Green: Nonamer [11]

id-TRAJ20[J_gene_segment]

acagaggacacaagaatgaacccaagcagaaaactaaagaaaagaaatcaataaaaccactgaagaaaaccagttgcccttatcttgcctgtctctgataatgaatgtgccttccc**tgctcattcctcactttggttctcaaattttccagggaattagca**aagaactggtagggttgggttagtcttggtctaatttggttagtcatctctggataacaagtctcctttttaggaagtgccaggggattttttgtaatgccaataaacatggtgtacaacttcaacaaa**ttttactttggatctgggaccaaactcaatgtaaaa**ccaagtaagttatagttgcct**agaagaaaaagttaccaacacattatgctaaattcttct**tttcagtctgtatttcagttcttttattttgcattttgggtccccacccatgatattttattagtccttattcatatgtcacttgaacagatatgtagtaaacaaatctcatgatcgagtagacatgctgaaactgccaatgttctcaaggtgctattctttataaggaaaatgc**ccaaattttaactgttgctactgaacaaaagagcatttgg**atataatttaaagcacccctgaatgtccaggatttcagagaggagacatgacaaattaaaattatatgtattatttaaagatgaaaaga**attagatcctcttgaaaatactctaat**aactaacatttcttcaacattcgcaatatgcaaagcacctttcataccttgactcatctgaccctcctaacaaccctctgggatagatattacctatctccgcttccacagatgaggaaacagtgtgaaaggtaaaactaaagcaaagaggttgaaatagttacccaagggctcaccgcagatgaggggcagagctaga**ttaagagcctggcctgcctgtcccctagcccactctcttaa**ccatggtcttggtgaggtttgtgtagggcgacctcgc**actgtgGTTCTAACGACTACAAGCTCAGCTTTGGAGCCGGAACCACAGT**AACTGTAAGAGCAAgtaagtaagaaagaaaagtccagaataattttaagcaaaatggtgggtaggtttttcagcaatttcacctaggaagtgcaatgtcaagaactaaa**ttctaagagctttcccagtttgtgttagaa**ccataatttttctacctcacacgtctccctcgccttctctgtcacctcagaacagctcctcctaaggcatgacttcacaatggtacatttgttggtggcgcagtctttgtgg**tcagataaaaactgagctaattatctga**aattatctcaggtctctaagtgagagagttaactcctttcaaatatttgaagaactacgatgtagaccaggtaaggggtgagtgtggaggccaaatcaaaacgagcaggtgcaagatgtgccccaacctcccctcaaacattgttgaacaagtggaactgccctgcagtggaaaggctgctttgtgaagtatggaacttttctt**ccttagatatcttcatgccagacccctaagg**ttaaccatgagatgttgcagaaggcacttctatgccattttagcta**ttagattgaatcacctctgggctccctttcaaatctaa**atgctaagatcccatgactcaagcctggaggattaataggtgagctcagacttgtttcttctatttttactattttgatagccaaaga**aataattcatcaaattatt**cataagctaaagcctacttgggatttttacacctagaagatggtggggtatgatttcccagtgcagtaaatgagaaaacaataggagacatcaaggaggaaaaaaagaaggaagagataaagggaatgtcttaagggaggctcagaggttgaatgaaggaaatgaggtgattttgcagaggacagatgtggctatcaaagattttacaatttcacc**tttggaaagggatccaaa**cataatgtcactccaagtaagtgagcagccttttgtactcgaaaatagggccaggggagcaaagtttcttccaat

Uppercase: TRAJ20

Lowercase: Flanking sequence[1000bp]

Red & Bold & Underline: Stem-loop [13]

Blue: Heptamer[13]

Green: Nonamer [5]

id-TRAJ25[J_gene_segment]

aggcagtttatttcaaagaaaagaggttgagggcaaggtgggagttaagtcacacatctaaatctgggaaaggaagcattaaacagaagacggaggggctcggctctccacctccaccaggattcagaccacaggacatggctaatactgcaagaagggagttaggttagtggt**aaaacctttctaataggtttt**tgaacagtgaaaatgagctgtcaggagtagttgccagggcactttctctagagagtttttgaaaggagatagacaccctgctatcagaggtgattggagtgcagc**cctgcctcgaggcagg**aggagaaatgagagaacctcttaaagcctctttccagccccca**gagcctgtctccatgaaaaaggcccaggctc**attaatgcagagctgcatcctccaggacacagggggttgctgggttgtta**agtttggaacctttgccggcttagtggttaccaaact**ccctttgttctcaagacttgacttgataaatgggc**tggctttatccctttaaagccaaaaacacaggttctgagtgctgtgttgtttg**tggtgttgagatgcccaggctggagggaggaagctctagggggtttttgctgagcccagaaacactgtggggataactatggtcagaattttgtctttggtcccggaaccagattgtccgtgctgccctgtaagtacagttaagtggagatagaaaatgagtccagtgcttgatgtggggagaagctgcagggtcttgaggccaggagtccaccgtgacctcaggagtggtgatggagaagaaggttaaaaaaggaatgagaaacagaaccataaggacaatacagaggaagggttc**attttcgggaaggaaaatcacaaggccgtgtgatt**aacgctggaggaaatgcactcctttggagggtcccttctcaggagggctttgtgtcggggaaagctgagctgttaggtttttgatgctgagataatcactatgCAGAAGGACAAGGCTTCTCCTTTATCTTTGGGAAGGG**GACAAGGCTGCTTGTC**AAGCCAAgtaagtg**acatataatttatatgt**gctgaatatgattatctcc**aagaggaaaacttgccctctt**ttccacgggtg**cctccttgggtcaggagg**acatta**aaggtgttgagcagaccaaggcccagtacctcgtcacctt**cttcatccttgaaggagctttaagaggtgtggaggggaagaaacccaccaggaccccaccaataagcccagccttgagacccctccactctgtcagacttgaataagaacca**tctgagaacaaggtcttcctcaga**aggggactccagcatagtcatccccatttgatagattttgaaactcagggcggggcagagtggctcatgcctgtaatcccagcactttgggaagctg**aggcaggtggatcacttgaaggtcaggagttcgagacctgcct**ggccaacatggtgaaagcccgtctctactaaaaataaaaaaaat**tagctgggtgtggtggcacgcacctgtaatcccagcta**ctcagaaggctgaggcaggagaatcgcttgaacccgggaggcggaggttgcagtgagccaagatcatgccattgcactccagcctgggcaacaaaagcgaaacttcatctcaaacaaacaaacaagctcaggcaagtagaggccagtgattaacactcttccaataggaacagatcccaagcaacctgcagtagtttttccagtaggctcgtgaactcaaaacaccaagttattt**aaatgacagggcaaccgttgtgatcagctgtcattt**gagtagtgctgaggactgggtctgaccttgagtaaacgcctaggctggtgggtttctatccctgcagcatctaaagcggacgcccaggcttcactgaccccagcagtctggctttctgcccttgccgcccaggaggtgcagctcttggcacacaccatccttagtgtcttgaaagaagagaaattaaaagagaaagggggaaaagcttatatctcatatcatgcagttgcct

Uppercase: TRAJ25

Lowercase: Flanking sequence[1000bp]

Red & Bold & Underline: Stem-loop [17]

Blue: Heptamer[25]

Green: Nonamer [7]

id-IGHJ4[J_gene_segment]

cagaggctgtgctactggtacttcgatctctggggccgtggcaccctggtcactgtctcctcaggtgagtcccactgcagccccctcccagtcttctctgtccaggcac**caggccaggtatctggggtctgcagccggcctgggtctggcctg**aggccacaccagctgccatccctggggtctccgccatgggctgcatgccagagccctgctgtcacttagccctggggcca**gctggagcccccaaggacaggcagggaccccgctgggcttcagc**cccgtcagggaccctccacaggtagcaagcaggccgagggcagggacgggaaggagaagttgtgggcagagcctgggctggggctgggcgctggctgttcatgtgccggggaccaggcctgcgctttagtgtggctaca**agtgcttggagcact**gggg**ccagggcagcccggccaccgtctccctgg**gaacgtcacccctccctgcctgggtctcagcccgggggtctgtgtggctggggacagggacgccggctgcctctgctctgtgcttgggccatgtgacccattcgagtg**tcctgcacgggcacaggtttatgtctgggcaggaacagggactgtgtccctgt**gtgatgcttttgatatctggggccaagggacaatggtcaccgtctcttcaggtaagatggctttccttctgcctcctttctctggg**cccagcgtcctctgtcctggagctggg**agataatgtccgggggctccttggtctgcgctgggccatgtggggccctccggggctccttctccggctgtttgggaccacgttcagcagaaggcctttctttgggaactgg**gactctgctgctggggcaaagggtgggcagagtc**atgcttgtgctggggacaaaatgaccttgggacacggggctggctgccacggccggcccgggacagtcggagagtcaggtttttgtgcaccccttaatggggcctcccacaatgtgACTACTTTGACTACTGGGGCCAGGGAACCCTGGTCACCGTCTCCTCAGGTgagtcctcacaacctctctcctgctttaactctgaagggttttgctgcatttttggggggaaataagcgtgctgggtctcctgccaaga**gagccccggagcagcctggggggctcaggaggatgccctgag**gcaacagcggccacacagacgaggggcaa**gggctccagatgctccttcctcctgagccc**agcagcacgggtctctctgtgg**ccagggccaccctgg**gcctctggggtccaatgtccaacaacc**cccgggccctccccggg**ctcagtctgagagggtcccagggacttagcgggg**tgccagttcttgcctggggtcctggca**tt**gttgtcacaatgtgacaac**tggttcgacccctggggccagggaaccctggtcaccgtctcctcaggtgagtcctcaccaccccctct**ctgagtccacttagggagactcag**cttgccagggtctcagggtcagagtcttggaggcattttggaggtcaggaaagaaa**gctggggagagggacccttcgaatgggaacccagc**ctgtcctccccaagtccggccacagatgtcggcagctggggggctccttcggctggtctggggtgacctctctccgcttcacctggagcattctcaggggctgtcgtgatgattgcgtggtgggactctgtcccgctccaa**ggcacccgctctctgggacgggtgcc**ccccggggtttttggactcctgggggtgacttagcagccgtctgcttgcagttggacttcccaggccgacagtgg**tctggcttctgaggggtcaggccaga**atgtggggtacgtgggaggccagcagagggttccatgagaagggcaggacagggccacggacagtcagcttccatgtgacgcccggagacagaaggtctctgggtggctgggtttttgtggggtgaggatggacattctgccattgtgattactactactactactacatggacgtctggggcaaagggaccacgg

Uppercase: IGHJ4

Lowercase: Flanking sequence[1000bp]

Red & Bold & Underline: Stem-loop [19]

Blue: Heptamer[31]

Green: Nonamer [4]

id-TRAJ7[J_gene_segment]

ctggcctccctgtcccaataattatgcacagtaaaagtcagttcaattcagtaaatgattattgagtatcatcaacaaaccagggaatcatcatt**ggttttaaaagaaagaaacaaagaaaagaatttttaaaaccctaagctatcttagg**ct**tctgatctcaagaacttccagtcttatcaga**gaaactaagcactcataataaaaagtctaacaacaatttatatgtgtatgtgccaagataggggggtgggtttcctgagggctggagacaggacaggtatgaccatgcacagccgcattctttggcagagaacacacatgattcccccaaatatgtttctggaaacggaattgagtctgttctgtgctgagatctttgactttcttgaggacaaaagcccccttggcagttagcatccttgagcgggggtgaggacagaaacaggaatgagtcaggcgt**gagtcacccaagactctcactcagccccaagaggatggagtgactc**tgagtcatgccgaggcatgtctttagcatgtgctccagcacag**tgtgcatttatggcaaggtgaaaaatacatttttgcaca**tgtcactaggaaaatggtaccttcttctcaggcatggcagatgaagtaggaaaatgcccgcctgctctctggtatctttagaaatactcacacccatccagcctatgtaggatacaatgtttttcatgaagttttatgaagattctgtttttcccaaaggagaaatagccaaagctgtgtttattgatagctgtgatgaggaaaatgaaggccaacaggtagcagtggggtctggtgctgtttataccctttagggagg**ttccttcaagtcttcataccacccattacaaggaa**aactttagcatccgcctttcagggtttccagatcttagagctactgtgaaggggatgtgtcttcagagaaggggtgggc**aaaaccaggaggtttt**tgtaatacacttacacagtgTGACTATGGGAACAACAGACTCGCTTTTGGGAAGGGGAACCAAGTGGTGGTCATACCAAgtaagtgagctgggatc**ctcctgcacaaatggccacagccacccccatccctaccctgtgctggag**agcctcttatcatatttccatgggagcgggggaaggacatgccattcatcagtgctatatcgaagaaccttaggcggccactgtcccaaatctttgcctgtccctggtttctttcactctccccttctctgcatctccactgaccacactcccattctcatgtaagctgcccctgatcttccctgaggctggatttcagattgaacaaaggaccacaaaagtgagagctggaggggaccgtggggacatcttgctcatgttgcagagggacctgatgccagggtctagcatctgtccaagaccgtgctgatggtcagtgcacagcagtggaggggccccagccctttactcccaccctgagtttggtggagccacacagctctttgctaagtgaccctcaaacccagtgtgagataagctgagtgttgacagtgcccacctgctagggaaccaagtactgagttcctgcccca**gctcaggactgtggcgcaggccatgacctccctgagc**ctcagttttcatatctctgaagtagacagcatggtctctgtttcctcccatctttcaaaaattctatggaaattattgtgactttttt**ctctgtaattgtagggtttcacagag**gcagggccttgtctgagtcaaggtcattagaaata**aaggaatccaagggctttggatgcctt**ggcctaattctaggtggtaaagagggaaggatgaaagcacaaatcacaggaagtgc**ctaaatcctgcttggatttag**tgttgcctcaaacagggatatgcagaaatccaaccccagggtctcccataggaagaagactaggagtgggaagtgccaaagagctctggaaaggaaaagagaacaaggagcactctgagaaaaggaggtcagaacagcttcaggaaaaacgtcacataaaatttcctttgc

Uppercase: TRAJ7

Lowercase: Flanking sequence[1000bp]

Red & Bold & Underline: Stem-loop [12]

Blue: Heptamer[37]

Green: Nonamer [3]

id-TRAJ21[J_gene_segment]

acattgcttatcatttctacaagtgagaacaagtaagcccctggccctccaaaattgcatgagtattataattacttggcttagattgaagtcacttctgtgttacttctcataactgtgttcatttgttgagatctgtttaactccaaactgtgtgggcgtctaactgttggcactaaggtttcttttctcttgaaattgccagcaaaatattttacccataaataatgtttcatctagacttgcaaatgacaactaacactgcaggctctttttttttcttttgctttgtttc**tttttttctgaagggaatatgcaagcaggaagcaaaaaaaaa**aaaaaagccaaaatgtacagtttgactgtggggtctgggaggagagtgtcaatgtggatgctaaaa**tatacatggttgtata**atgtaggtactgtcacagagaggctatag**tgatcttattagatca**tacatacagagcaccactggtcagctggcgttgctcggggtgtccttaagaaaagctcaggtctagttgtcaagtgcaacccagattcagatacagagtctcagagagaagcttccagatcaccctggagctaaatgcagaatatgaatt**ttccccttagttaagtctacctgaggggaa**ttttcatttataattaggaaaagagaaaggtaaagaacatttacttgca**gtgtcctgtttcaaatattgtgtgcattgagcacagaggacac**aagaatgaacccaagcagaaaactaaagaaaagaaatcaataaaaccactgaagaaaaccagttgcccttatcttgcctgtctctgataatgaatgtgccttccc**tgctcattcctcactttggttctcaaattttccagggaattagca**aagaactggtagggttgggttagtcttggtctaatttggttagtcatctctggataacaagtctcctttttaggaagtgccaggggattttttgtaatgccaataaacatggtgTACAACTTCAACAAA**TTTTACTTTGGATCTGGGACCAAACTCAATGTAAAA**CCAAgtaagttatagttgcct**agaagaaaaagttaccaacacattatgctaaattcttct**tttcagtctgtatttcagttcttttattttgcattttgggtccccacccatgatattttattagtccttattcatatgtcacttgaacagatatgtagtaaacaaatctcatgatcgagtagacatgctgaaactgccaatgttctcaaggtgctattctttataaggaaaatgc**ccaaattttaactgttgctactgaacaaaagagcatttgg**atataatttaaagcacccctgaatgtccaggatttcagagaggagacatgacaaattaaaattatatgtattatttaaagatgaaaaga**attagatcctcttgaaaatactctaat**aactaacatttcttcaacattcgcaatatgcaaagcacctttcataccttgactcatctgaccctcctaacaaccctctgggatagatattacctatctccgcttccacagatgaggaaacagtgtgaaaggtaaaactaaagcaaagaggttgaaatagttacccaagggctcaccgcagatgaggggcagagctaga**ttaagagcctggcctgcctgtcccctagcccactctcttaa**ccatggtcttggtgaggtttgtgtagggcgacctcgc**actgtggttctaacgactacaagctcagctttggagccggaaccacagt**aactgtaagagcaagtaagtaagaaagaaaagtccagaataattttaagcaaaatggtgggtaggtttttcagcaatttcacctaggaagtgcaatgtcaagaactaaa**ttctaagagctttcccagtttgtgttagaa**ccataatttttctacctcacacgtctccctcgccttctctgtcacctcagaacagctcctcctaaggcatgacttcacaatggtacatttgttggtggcgcagtctttgtggtcagataaaaactgagctaattatctgaaattatct

Uppercase: TRAJ21

Lowercase: Flanking sequence[1000bp]

Red & Bold & Underline: Stem-loop [13]

Blue: Heptamer[18]

Green: Nonamer [6]

id-TRAJ38[J_gene_segment]

ctctgattgcttttgctttggtgctc**tggacataaactgttcctactttgtcca**agtgaggtatataattttcagaacgctggagaattattatctaaaacaccagtggatacgctacaattttggcattcagtattctgggcttgtgcaggaaaac**tctttctttagaacatgtagaaaga**ctatcctttagttatccagtgtggaaacagtcctgaaatccaaactgatactgcgatcctgtgccaggttattactgtgacaattacttaatattctctgtgctcacgatatcactccttccagaagatctttagggaaaacttagttcagtttctaggaggtttttgctcagccgaagatcactgtgtgaataataatgcaggcaacatgctcacctttggagggggaacaaggttaatggtcaaaccccgtgagtatctctgctgaatccataatgaatgctctaatttcaaaaggaagccgtagcactgagctctgttttctgttttctctacttaattttatcttttatattaattctaatggttacatgttatcg**atacatggcatatgtat**atatgaaaatggacatacatgtaactgtctggcgtgggaacaaatgtgctggagaaaaacagatttcttgtcctttttgtcttaaccatttgg**gtcagaaggtctgac**tgagtagaacatgggctgggtgagcactgcagtgggggaatccttct**ccagcagctttgtcagaattatgctctttgctgg**cctggtcagaaggcagctttgtcaggag**gtggaaggatgaactagttaacctcagaagagcctttccac**tccgaccatgctgggaatagaccgccataataaacagtcttttccgctgcctctttagccaacgtcccagctggaggagcacgccggccagatttggggattgaaagggatttcctgatacacattggcctggtcggttttggtaaagctttctatgactgtgTAATGCTGGCAACAACCGTAAGCTGATTTGGGGATTGGGAACAAGCCTGGCAGTAAATCCGAgtgagtcttcgtgttaactctgtcaaacctgtctgt**gcagtttgaaatatctcagaaactgc**ctccttcttttttagccctgaggtttaagatatctagttatggctagtgttttcttcctgtccaccaaagctgaacccttgcctacccctcaatcaaaataactagatagagggatccatatgagcagggtaggaaggccattggagtgttcagctgttttcttagagagcaaatacattcgtgcattccaaaaaggcctgtgtgggagcactgaatattacagaaggtgagtgtgtagcacatttcatggccgctttagtctgcactatgccaacctcaccgtgggccattggtaaatactgttgctttctggccaaagacaggtcaagtgctgggaactctcgggactcacagtcacgctagtgaccgtgctagagcacatctccccactgtggattatccagaatgactcaaacttgcgatataactcctgcatctctcagttctctttggcttagagccttattgtggatttgctagaggtaacaggaaaggaactgatttcatcctaattctagcctcttctgagaagttattctctgcaagagtcttcaggagcatcttacagaccacctacactattcagaattttgttccgcacaccacattgtgtt**cccagcagtgtttgctgtggctttggctggg**gccaggtcctgagtcagttctggagctccaggcttggcaaagctggataagggcaaggggagttgtacagcagaaccaaagggcttcaggaaagacatctttgatatagacattaatgaatttctaaatgggcttctatatgctaacctgataagactctctggggagaaggtcatgctttcctttgcagctctaaatataagacacatgcagaaaaaaaaaaagaatgggaaatgtctgttacagtgccaaaatgaagataatcagtataaatattagatttgtcttaata

Uppercase: TRAJ38

Lowercase: Flanking sequence[1000bp]

Red & Bold & Underline: Stem-loop [8]

Blue: Heptamer[32]

Green: Nonamer [2]

id-IGLJ5[J_gene_segment]

atctggtgcccagccatcctccagggcgcacagcacaatgtagtaccggagtgagctctagcgtgtgaggacatctgacatgtgggctccactgcagatatactgaattgcaatgacaatgcggctacaaaacataaacatttacccactgggcgcctcctcaggtggcatctgattttctcccattgccc**caggagcttccatggctcctg**atttctcggag**gatgagaggttctgtctcatc**atgtccctttcctgccccaggcctgggatcccgcactgacc**tcacctcccttagcagaaggtga**tatttggagaccacactcgggagctcctttatgtccctcacatttgaataaggcagtggcagccactaccccacctcacccaccaaaatgagaccaggttgaggggtgcaggagatccttccattttaccctggaggatgg**ggctggcatttccagtggggaccagcc**aggcctcactggccaggcccatcccaactaggaca**agcccagggaaggctgggct**gaggctcctggagtcacagataggttcatgg**gaagcttcccaagacaccgcactctagggtaaccagcttc**ttcctggagggagagggcactctctgcatcaccccagggcgtcaccaagcagtcagtgtcgagtcagctccac**cagggagaccatttatccctg**accatgggagttcactcctagtgacacagtgccctccaataaactcatccccatggctgcatgatggttggtgggaaaaccaaatccactgtcctccaggaaccaggatttctaggg**atcctgctggtcacaggat**gtcagctgtccccttctctctgtgggggtgagt**gtggcagccgtgtgaactccctcatgagcagatgccac**caggg**gctgtggcctcagcttcctccatcacagc**tgcagcgggggttgggggtagaggcgtccagagagggtttttgtatgagcctgtgtcacagcaCTGGGTGTTTGGTGAGGGGACCGAGCTGACCGTCCTAGatgagtct**tttccccctccttccctggtctccccaaggtactgggaaa**ttttctgctgcttttgttcttttctgtatcttgtgttgacctgtggtgatgctttctctctggagcctaggccctggtcaaggacctctcccctccctgtttagacccttacctcagtgggtcaccaagaccccttcacctctgacctcagatgtagggcactagactggatgacctactgagactcatctgtctgtctgtctgccagagccaggctgcttccctaaaacttgctcagttctgtcctcccccacctgggcttctgtctaacgaactttgtgcaagggaaactgaggccccatctcatgagggagagggaacaaggggctcgaaggagtgaccacctggtggactttagaaggacctgaaaccctcagagccaagataggggaatgaaaactcagagtctcagggccc**agtcccctggactgtgggact**ctggatccaggctgggaacaaggtaggaggtgcaggggcctctccaggtttctg**tgggctcccagggagagagccctgagctggcctgggaccca**tgaagccctgtcaggagggacgggaaggctctggacatgaaggagccaggtgaagtgtcacgaaaggccatggcattcagggaggtggctgatgggtctctgtgggaggcacccctagaagcaggaacccctgagttcaccgacaggcatatcccaaggcagaaaaactgtagattggccctaaacacagagagactctaacacagactccacagacaaagagcccaggacagacaga**cagtgggacttgggtgagcaaaggccctgactccactg**cagaagatccaggagagacggat**gtgggtacaaacaagagctcttacgtgagagacccac**tctcccccaacccagagcagctgtgtcaggtgagaaaagtttccagagtgaatctaacaagaggctcacagagctcagagaac

Uppercase: IGLJ5

Lowercase: Flanking sequence[1000bp]

Red & Bold & Underline: Stem-loop [15]

Blue: Heptamer[30]

Green: Nonamer [3]

id-TRAJ57[J_gene_segment]

gcttggctaaacaatgctcacggatgtgcacgtgtgcagtgggatgcagctttttctaacctctcacaaaatcacgcaacccctt**caccaaactgcttggtg**gctcttctgtgccctcacagcccttctagtttagcagttagaggaagacaagtgcccaagggctttctcatttttaggtcattgtggaaagaggaacgttggtgatatgtttgcctgcttgggaggaaaataataatgatgatttactcattggtttctatatgctagttgatgtgctgcattctttctcttatcacatttaatccttataccaggtctgtgagttaggtgctattattatcctcattt**tatagaagagatatatttatattttatatttctata**gaagcacagagatgttgaataacttgttcaagccgatgaagtaagaaagagaacctggagtttaattcttggcactcagattccagagctaaaagcttttcagtgaggggattatgt**ctagagggaatgtccctgatgctctag**tctctaaggttgctaagataatcgctaaagaactgcgctctcagcaagtaaagccaggggttggtcaacaggaccacggtgacccagttacatattcttaatccttacaacatcatcaagcagttgctgt**tttccccactttacaagtgaggaaa**ctgaggtgtag**caagttaggaaacttg**tcc**tgagttttaaactca**agcctacctgccccaaagcctgagctcttttaaat**taactgcaactgctgaccagtta**gattaactaattacttaattaataagataacaacctaacaggctgtaaggtaaaaaagtaatcagatttgttctataggtcccctcccttttcgggaatagctatacaattgtataattttatgcttccc**ctgggagagtgggtggagccctggctcccag**cccatgatggaagggtcttggcagtatttgtaaagcagtctgtgggggtgTAACTCAGGGCGGATCTGAAAAGCTGGTCTTTGGAAAGGGAACGAAACTGACAGTAAACCCATgtaagtctgaataatgcttccaaatttctccctggaaccctgatttccaaattttccattctgttttataacccaggtccaaacca**cagcagtcccattaatggattccagtgcaaaacaactgctg**gtgtattcctactacacgcaggtctctgttgtttgccctcatgt**gctattttatctaatagc**tgagaatgacagtaccaatggagcactgagttaaggagtcagactgttcgaaggcttcattcttcgttataattggtgag**attttccatgggactaagagaaaat**tgattaactctctgagcctctattttcctcctct**gtagaatgggggaggcagttcctgttcccattttaccacagccggcgtgctgtg**aggggatgttggtacatttccaataaggggacaatgagtgt**gaagagaaaggaaggccccaggtggtccgtaaactcttc**caccagtcccacactataaacagctggttttatcagggggattcttggatgacaagtaagcacttaagtaaatatcaaggggagtctgggcaactgagtttttgtagatcctcgtgtcattgtgttatactggagccaatagtaagctgacatttggaaaaggaataactctgagtgttagaccagg**tatgttttaatgaatgttatttgtttccaaacata**agccaccatccttagaaattca**gtgaaagataaccgaatctcctgcccagttattagcatctttcac**catgggtctttctggagaaaatgacaatgtgggcagcccctgactgcagcccctttgggactgtttctttaacacctttaagtacttgggaatgttcagtgtgtttttgttaatgttggagatatgtgtctgacaaatggaatctgaattgaagttttagtgtgtaggggcagaaagcatttagaaaggacaaaagaaggacagattagactaaaatacataccaatggctgggagtatgcaatgcaac

Uppercase: TRAJ57

Lowercase: Flanking sequence[1000bp]

Red & Bold & Underline: Stem-loop [16]

Blue: Heptamer[25]

Green: Nonamer [2]

id-IGKJ1[J_gene_segment]

aattcaggtcacctgctcagggctaatct**gagagaaggtctctc**ttcagttgaattttgaaagacaattagcagttcacaagctaacccaggtggacaaagatg**ttcccaagcagagggagtgcttgtgaa**agctggaggccatagaaaaactctaaggagtgtagggaggtgggagtaatgtatggaaggggtggagatggaaggttaagagagatacaaggctgcaaaaatggagctggactcaaaagaaaat**actgaaaaggtcttcagtgttgttgatgagattactatggaaacact**atggaacactgggactccatggcagctccaaagatggcatgcgcctggtcc**agctcagtaagagctgagct**cttcctgtgctgtgaaaacagacaaaccaaccaagtaaagtctacttttctactctattagtcttcactttggtttcgtataccatctggagctacatttcaaaatgcatttcaaagttatgagccttaagttgatatatatttagtctacctttttttaaataacattgcagcaaaggagaagataaaatagtaagacaacc**tgtaattattactcattgagaagctgatgatttccataattaca**ctaaatgaagtttatcctttgcaaaagcccccccagcccaccccaaaagaaagtac**aaaaaaactggccattttttt**taattgcttgtttttctttgtaattaacattcagtctactttctaaaaaataaataaataataagcagtccagatgtggcaagttgctaaagaaaggaaccatcaggccatagacgtaaatatattctcttcttggatt**ttaggtctcacctaa**gaaaataaacacatgctatgtcaga**gaagcctcagggcttc**cacacctgctcgaaaagggagttgagcttcagcagctgacccaggactctgttcccctttggtgagaagggtttttgttcagcaagacaatggagagctctcactgtgGTGGACGTTCGGCCAAGGGACCAAGGTGGAAATCAAACgtgagtagaatttaaactttgcttcctcagttgtctgtgtcttctgttccctgtgtctatgaagtgatctataaggtgactctgcaatcagcctctgatatccttcagggaaaagataaagataagtctgtagtcaaactcgagaattgattgcacattttctttgaagagcaagcaagattcagtcattgggtgagaataacttgtctaagtaatagcttcagaaatgtcctggg**gaacataacatgttc**tggacagagccttggtcaattgtcagaaagggagtttttgtataggagggaagttaagaggaaccattgtgtgtgcagttttggccaggggaccaagctggagatcaaacgtaagtacttttttccactgattcttcactgttgctaattagtttactttgtgttcctttgtgtggattttcattagtcggatgccagggatctaacaaacttcattcccaggttaggtacagaggaggggaaattgttccacaggacgctagcttgtggctaatttttaagatttctaaatcaaaataacttcattgggggaaagaggcttgctgagctttcagggaggtttttgtaaagggaaaagttaagac**gaatcactgtgattcactttcggccctgggaccaaagtg**gatatcaaacgtaagtacatctgtctcaattattcgtgagattttagtgccattgtatcatttgtgcaagttttgtgatattttggttgaataaac**ctggtgacccagaagtaaatagcaggacaccag**aaaatgaacttaaaaagctgagcaaatagacgaatcattgggtttgagaggagaataggattcatgggggaaatggggaagaaatagctagatttttctctgaacaagcagcctatctcatatgattggcttcaagagaggtttttgt**tgaggggaaagggtgagatccctca**ctgtggctcactttcggcggagggaccaaggtggagatcaaac

Uppercase: IGKJ1

Lowercase: Flanking sequence[1000bp]

Red & Bold & Underline: Stem-loop [14]

Blue: Heptamer[24]

Green: Nonamer [5]

id-IGHJ3-2[J_gene_segment]

cgcagccacatcagcccccagccccacaggccccctaccagccgcagggttttggctgagctgagaac**cactgtgctaactggggacacagtg**attggcagct**ctacaaaaaccatgctcccccgggaccccgggctgtgggtttctgtag**cccctggctcagggctgactcaccgtggctgaatacttccagcactggggccagggcaccctggtcaccgtctcctcaggtgagtctgctgtctggggatagcggggagccaggtgtactgggccaggcaagggctttggcttcagacttggggacaggtgctcagcaaaggaggtcggcaggagggcggagggtgtgtttttgtatgggagaagcaggagggcagaggctgtgctactggtacttcgatctctggggccgtggcaccctggtcactgtctcctcaggtgagtcccactgcagccccctcccagtcttctctgtccaggcac**caggccaggtatctggggtctgcagccggcctgggtctggcctg**aggccacaccagctgccatccctggggtctccgccatgggctgcatgccagagccctgctgtcacttagccctggggcca**gctggagcccccaaggacaggcagggaccccgctgggcttcagc**cccgtcagggaccctccacaggtagcaagcaggccgagggcagggacgggaaggagaagttgtgggcagagcctgggctggggctgggcgctggctgttcatgtgccggggaccaggcctgcgctttagtgtggctaca**agtgcttggagcact**gggg**ccagggcagcccggccaccgtctccctgg**gaacgtcacccctccctgcctgggtctcagcccgggggtctgtgtggctggggacagggacgccggctgcctctgctctgtgcttgggccatgtgacccattcgagtg**tcctgcacgggcacaggtttatgtctgggcaggaacagggactgtgtccctgt**gTGATGCTTTTGATATCTGGGGCCAAGGGACAATGGTCACCGTCTCTTCAGGTaagatggctttccttctgcctcctttctctggg**cccagcgtcctctgtcctggagctggg**agataatgtccgggggctccttggtctgcgctgggccatgtggggccctccggggctccttctccggctgtttgggaccacgttcagcagaaggcctttctttgggaactgg**gactctgctgctggggcaaagggtgggcagagtc**atgcttgtgctggggacaaaatgaccttgggacacggggctggctgccacggccggcccgggacagtcggagagtcaggtttttgtgcaccccttaatggggcctcccacaatgtgactactttgactactggggccagggaaccctggtcaccgtctcctcaggtgagtcctcacaacctctctcctgctttaactctgaagggttttgctgcatttttggggggaaataagcgtgctgggtctcctgccaaga**gagccccggagcagcctggggggctcaggaggatgccctgag**gcaacagcggccacacagacgaggggcaa**gggctccagatgctccttcctcctgagccc**agcagcacgggtctctctgtgg**ccagggccaccctgg**gcctctggggtccaatgtccaacaacc**cccgggccctccccggg**ctcagtctgagagggtcccagggacttagcgggg**tgccagttcttgcctggggtcctggca**tt**gttgtcacaatgtgacaac**tggttcgacccctggggccagggaaccctggtcaccgtctcctcaggtgagtcctcaccaccccctct**ctgagtccacttagggagactcag**cttgccagggtctcagggtcagagtcttggaggcattttggaggtcaggaaagaaa**gctggggagagggacccttcgaatgggaacccagc**ctgtcctccccaagtccggccacagatgtcggcagctggggggctccttcggctggtctggggtgacctctctccgcttcacctggagcatt

Uppercase: IGHJ3-2

Lowercase: Flanking sequence[1000bp]

Red & Bold & Underline: Stem-loop [19]

Blue: Heptamer[35]

Green: Nonamer [6]

id-IGLJ4[J_gene_segment]

accatgttgcctgccatctcgtgaagatgaacaattatttcatggtgagctcaaagttatgttactgtatgtgactcacttgagtccaccatggttctatt**tcattgatgatgacaatga**cccaccgtggcccactcagtgcctcttctggtggccccaggatc**ctcctgaaggaacccaggag**acctcgatggctttccgctctctgttcacaatctatcctg**ggcacatctttctcctgccttgtgcc**tggaattgcccattaaccccaagtggactagtccccataactgggaggtgggatttagtgaccacacttggggtgcttctcacacagcccttttgagtcagacactccagacatacccagaaatgaga**caagaccctgaaagggtaacaggggcttg**cttccaacttctccctggaggttgaggctggcatttcatactaaaacctagtgagacccatcccaaactaagacaacacaaggaggacggaagtgagacgccctggagttgtggttgtggtcacgttggagcttcccatgactgctgactct**ggggcaagctgcccc**tcctctaaggcactcactggggacacctgaggacgcctcctgctcttaccctgtagtcacaccaagagatc**agggttacaacaaccct**atagagaatccctgtccccttccatgtca**cttcactccttcgtgaag**caaatgccctcaaggagctcattcccattcctgggtcacagtcacctggaaaacctgatccagacaccaacctcctcaggcctcgccatttccagacgtcccgttactgcatacgcttggtcgactgtcccatctcagcttgagaagggcaggcaggtgtgtggactctgctgagcaaatgccttccaggggcagtggtctggcttcctgcaccatagcttcaggtgggggatggggagggggagttaggggccccagggaagagtttttgtatgaacctgtgtcaccgcaTTTTGTATTTGGTGGAGGAACCCAGCTGATCATTTTAGatgagtctcttcttccctttctttccctgccaagttggtgacaattttattctgatttcgatctttgtctgtgacttgccacagcctgtggtcagggtttcctttgggacctcggtcct**gggaggctgatctctctcctccc**tattcagacccctgtatgcctcagctggtcactgagacaccttcatct**cctctgaccccagagg**cagggagctccaagacaaggccacactggtgtgtctcatgagtgacttctacccgagagccatgacagtggcctggaagatagatggcatcaccatcaccc**agggtgtggagaccaccacaccct**ccaaacagagcaacaagtatgcggccagcagctacctaagactggcacccgacagtggaagtcccacaacctctacagctgccaggtcacgcatgaaaggaa**cactgtggagaagacagtg**gcccctgcagaatgttct**taggtccccgaccctcacctacccacgggggccta**gagctgcaggatcagggcatgtgtctcccctcccactccaagtcatccagcccttctccctgcacccagtaaccctcaataaatatcctcattgtcaacc**agaaatcctgctgtctgtcttcatttct**tatctcatatttagtttgcaacctccttaaattctaagcaaggatgaggaaaatccaggtgcccagtttatcgggtgagaagtccatggtggtgccatcaccaggaacttgtggaaaggtctgggaatggaaactcacaggtgaatttcacagattttcacaatacagggtggctaagtaaagacacttacaagtcctgcaatagggaaacaggaagtccagaatcct**gctcaccatcccagccaacttagtgagc**cctaggatgctctgcaagatactggtgttcacgtcgctagctctggaaagtggggtgaggctggggcacacgggtgatcagttatgatcagatgggcttagggtgaggttc

Uppercase: IGLJ4

Lowercase: Flanking sequence[1000bp]

Red & Bold & Underline: Stem-loop [14]

Blue: Heptamer[29]

Green: Nonamer [4]

id-TRBJ1-5-2[J_gene_segment]

gaacacagagtactggaagcagagctgctgtccctgtgagggaa**gagttcccatgaactc**ccaac**ctctgcctgaatcccagctgtgctcagcagag**actggggggttttgaagtggccctgggaggctgtgctctggaaacaccatatattttggagagggaagttggctcactgttgtaggtgagtaagtcaaggctggacagctgggaacttgcaaaaaggggctggaatccagacggagcctttgtctctagtgcttaggtgaaagtgtatttttgtcaggaaggcctatgaggcagat**gaggaggggatagcctccctctcctc**tcgactattttgtagactgcctgtgccaagttaggttcccctactgagagatgggtagactcagcttggaaggggtcaccttgaacatctcctgtctccttgaagggtgccggtcacggccatgacagataaaagagcctctgaccttaccaccacggtcctaccgtttctc**tccctcacacagaaaggagaaggtcacagaagaggga**acttgggggatcacacggggcctaattggtctgctgaccaccgcattttgggttgtaccattgtctacccctctacccaccagggctaaaattctactaaggaacaggagaggacctggcaggtggacttggggaggcag**gagtggaaggcagcaggtcgcggttttccttccagtc**tttaatgttgtgcaactaatgaaaaactgttttttggcagtggaacccagctctctgtcttgggtatgtaaaagacttctttcgggatagtgtatcataaggtcggagttccaggaggaccccttgcgggagggcagaaactgagaacacagccaagaaaagctcataaaatgtgggtcagtggagtgtgtggtggggccccaagagttctgtgtgtaagcagcttctggaaggaagggcccacaccagctcctctggggtttgccacactcatgatgcactgtgTAGCAATCAGCCCCAGCATTTTGGTGATGGGACTCGACTCTCCATCCTAGgtaagttgcagaatcagggtggtatggccattgtcccttgaaggcagagttctctgcttctcctcccggtgctggtgaggcagattgagtaaaat**ctcttaccccatggggtaagag**ctgtgcctgtgcctgcgttccctttggtgtgtcttggttgactcctctatttctcttctctaagtcttcagtccataatctgcctcctcactcccttcttggctcatcctccctcttatgtgcatggctctgcctctcctaagcctcttcctcttgcgccttatgctgcacagtatgcttaggcctttttcctaacagaatccctttggtccagagccatgaatccaggcagagaaaggcagccatcctgctgtcagggagctaagacttgccctctgactggagatcgccgggtgggttttatctaagcctctgcagctgtgctcctataattcacccctccactttgggaacgggaccaggctcactgtgacaggtatgggggctccactcttgactcgggggtgcctgggtttgactgcaatgatcagttgctgggaagggaattgagt**gtaagaacggaggtcagggtcaccccttcttac**ctggagcactgtgccctctcctcccctccctggagctcttccagcttgttgctctgctgtgttgcctgcagttcctcagctgtagagctccttgcttagtcttcagggctgtgtgtttctttgctcttcttttcattgttttctgggactcttctcatctctactttcttagtggatgtattgttttactttcccttttttaaattgcatcttctccattttttccttcccattctaactccacttctgcattgttgactccttttggtgactagctctgtcttctatgttaagattctccccactgccagcctccagcacagaactctgctcatgtcttcatctccctccttctttctttctctaccagtcttagaagatgcatctatgtcttcctg

Uppercase: TRBJ1-5-2

Lowercase: Flanking sequence[1000bp]

Red & Bold & Underline: Stem-loop [7]

Blue: Heptamer[38]

Green: Nonamer [4]

id-TRAJ18[J_gene_segment]

tatctcaggtctctaagtgagagagttaactcctttcaaatatttgaagaactacgatgtagaccaggtaaggggtgagtgtggaggccaaatcaaaacgagcaggtgcaagatgtgccccaacctcccctcaaacattgttgaacaagtggaactgccctgcagtggaaaggctgctttgtgaagtatggaacttttctt**ccttagatatcttcatgccagacccctaagg**ttaaccatgagatgttgcagaaggcacttctatgccattttagcta**ttagattgaatcacctctgggctccctttcaaatctaa**atgctaagatcccatgactcaagcctggaggattaataggtgagctcagacttgtttcttctatttttactattttgatagccaaaga**aataattcatcaaattatt**cataagctaaagcctacttgggatttttacacctagaagatggtggggtatgatttcccagtgcagtaaatgagaaaacaataggagacatcaaggaggaaaaaaagaaggaagagataaagggaatgtcttaagggaggctcagaggttgaatgaaggaaatgaggtgattttgcagaggacagatgtggctatcaaagattttacaatttcacc**tttggaaagggatccaaa**cataatgtcactccaagtaagtgagcagccttttgtactcgaaaatagggccaggggagcaaagtttcttccaatttaaacacactcaaaaggatgtgtaattg**ctttctgatgggagggaactctgaaggtagaaag**actattgttaccacagatcacttgtccctggagatatagcatctgaggactgactgtctagcttagactgctgtctgcacaagaagaagacttacagatgcttaaggaggatggaggcaaattttcaaccctgtactccaaagctgaggggagaggggatgggaaacattagggctgggttcatgtaaaggggaccagcattgtgCCGACAGAGGCTCAACCCTGGGGAGGCTATACTTTGGAAGAGGAACTCAGTTGACTGTCTGGCCTGgtgagtgagtcgctttctattccaggaaaatattactgtggagaaattaaaaggggagatgaattaactc**ctttagtcttgaaactaaag**agatatgtgtacttcccctcctggaggaccccagtccctaggtagattaggacgaggaagcagagggagacaaaggcgatggaaagtcccttttaggaacccaggaatcagagagaacttgatggtccccaccaaaggcaaagaagggaggtcccactaaaacatgtgagcatcctaaaacacctgctctggtccacccacaatatttggtgaaggacatggccctccacccagaatgggcaagcaaactagaccagctttctaaggtgtgtttttcttgggtcctgtgactcttggccgtctccctgtcagaggctccatctgtcgctgcactctttcttattgagaatggcctctccgtggttctctgtaaactttccccaagaagcctgtttccatgcttcctcagcacttagcctcaccgatggatgggctctagagtagctggagggagtctgggtctgaa**actttcactgagaaagt**aaagttgatccgcagtatccagtggatatggcagctgg**gaggacctaggactgaagtactcgtcctc**tctttggggcctttcctgggcactcgattgaatacagaaaccctgttagcaagtgcatgcatgtatatgagtgtgtt**cattcagcatttccattctgaatg**tatgaatagcctta**ctctgaagactccctggagcttacagggctttctgttttcagag**aaaattgcctttgtgagacaaaaatggccaagtgggcccctgaatgaggttatggttgagagctattaatattatccttattcacagagcaagatccgaattcccaaagcagagtagacaataactatgtgaagggcttagacccaaatatgtgcccacttgcaagcattccagctctatgaagaggatacagtgctagga

Uppercase: TRAJ18

Lowercase: Flanking sequence[1000bp]

Red & Bold & Underline: Stem-loop [10]

Blue: Heptamer[16]

Green: Nonamer [5]

id-TRAJ49[J_gene_segment]

taatgctgaatcttacctga**gatgccttttgaagtggcatc**aagactgggctctaaaatctttaagagtcacctgtcagttattgtaa**aggtttggatggctgtgtgaaaacct**cctacgacaaggtgatatttggg**ccagggacaagcttatcagtcattccaagtaagtgtccctgg**ggtgctgcctgtggagtgcgctggggcta**acagtctcatacattagggcttaaatgactgt**gcagatggcatcgtggttgaggacacaaaactgagagaatcgagatacattgactctgatcagaaaaaaaggtcacggaatagcaaacataggtttagtccttaaaaggtagcatagataacatgggacttgaacgtggcaatttgaaaaacaacctgtggtctatcagcacgttctcactgcacacgggggataaacaggggtgggggaataagcaatggacattgcatgataggggtgctagggaa**atgaggcaggtcaggaagaaggaaggtgatcagagcctcat**acggagttaatctagagaagatgaattgttccataagtttcacaaagaaataaagtccaaagatataataagaag**tcctgattaaaatccacttcagaatcatatgatcatcagga**gagaaattaagtc**ttaaaagaaaaagaagtttttaa**a**tgtaggtccataaaacaaatggacctaca**cctttaattagacagcaagcagcacaagatgcacgttagccttccttttaatcccctctcaggaggaatcagtccctc**cagctgcacatggtcacagctg**ctctaagccctggagctccttcccagtaatacccttgct**cctgtgcctgcacagg**gaaaggagatgcctgcctgcctgcctgcctgcctgcctaccggtctgccttttggtccgccgcttggtggggccccaggtgtaacctctggctgtctcttctgcctggtttttgttgagcttcctatcacagtgGAACACCGGTAACCAGTTCTATTTTGGGACAGGGACAAG**TTTGACGGTCATTCCAAgtaagtcaaa**gaaaattttccatcaccattgtgttgagcaaaccctttaaactgcagaaagagctgtcaaagtctgctactctatttctctgcttctaggtataactttatctggaccagagtaaagaggcccccgacccaccccga**gaatgatttatgcttggacaggtctttatttcattc**cttgaactatctatctgtctatctatctatccagctatctatgtatctatctatctatcatctatctcagaaatagttgtttttaacagttgattctgtgcctcacccaaattgtcattaactaaatgcagatgtgctgccacactgataaacttcattggaaaaaaagaaaaaaccaaacc**cagagaatgttccttgaacattcctg**aagtgggggggcaggcga**tgtcacttgcagtgaca**gctggcctctctagctcacccccacatctgtgccaaccagagaagctgggccctttgtagcacaagcctcaggagca**gagaatggagggacattctc**aactgagaggagagggactaggaggctagatgggaaatgaggtgactctacagaaaatcgaatatgggatggtataaaaggatgagaaagatggcagggggtccacgatcgctcttgtatttagacctgggggatgcactgatagataccacctggagagagaaagagga**tttctgtttccctctgaagaaaaatgtggcagaaa**aatcaaaacagatcgcacatgtcaagcctctcctaattacagcctttcattctggtttctctcagctgttccagcaggaaatggtaggaaagggtcattggtgcctttgatgttgtggaagcattgcaggaaggagatt**tgcaagaaacggggcaggggcttgca**gtagagtctggagagtttagaatgatggtttttgcaatgacttagaacactgtgtatctaactttggaaatgagaaattaacctttgggactggaacaagactcacc

Uppercase: TRAJ49

Lowercase: Flanking sequence[1000bp]

Red & Bold & Underline: Stem-loop [17]

Blue: Heptamer[26]

Green: Nonamer [2]

id-TRAJ30[J_gene_segment]

gctggagtcaccacattgactgt**ctgttattaaaccctataacag**gttataattaagattatggaatgactaacccaaggagc**tttaaaataggatagcctagttttaaa**cggttcagccttgggaaatgctggaagacaaggaaatgagatagatgactccataggtttgttttacgtctggtgaacatatccatcccatgttaattttttttaacgactatgataaagagctt**ctttggggtcaggtagaacccaaag**gaggaagagaaaatgggacttggttctttaagtgtcaaacaaggatccagccagggttgagggcaatggcataaggcttccctctcttggtggtgagagaccacatccatggactgtggaaggagaacattttctaaaagacggggatcccttgggctaaaatgaatgatgctcatgacccccagctagggcagggctataatgagtacagcaaacataaggcaaggtatcctc**agaatttggtgacagggcaaattct**agaacttaaggcttagagccagggtgaagggaagactctgtaatgcagttccttctcaggtggaaatccagatatgtgccctaagtgacctattcctgccccatttttgtttgggctctgaaaagtggattcctagctcagtggctgtctgccaggtatcttttgccttcagtaacagaattatgtttgatattggaaaggagtttgttgtattgttggctttataattcactctcagtgt**caaatgaggtcattcaacttcctctcgcacatttg**caataagcagcatgaaattcaagcagaacaactagaagacacagagaaagatggctgagctgataatgaggcctgtttattacaggagaagagacaatataggaatctgttaagtcccatgaggttgaaatatccaccaaaaaatatgcagctgtcccc**caaaaaaaagtcactaggagtttttg**ttatggtcccaatcacagtgTGAACAG**AGATGACAAGATCATCT**TTGGAAAAGGGACACGACTTCATATTCTCCCCAgtaagtgctgtttatgtgattttctgacattaactcccactgagttcaaactgctgagagtgttcagtaggaaagtttctaagacatgtgggtggcctgtaggtcctgggctcttctcttgcagaggtccagtc**ctcagcctttccacagctgag**ccaaataccaaattctcccctccagggctggaaaagttgctaccagtcctctgctacccaaatataaattatttttttattgtttattaagaagagtaaaagaatcgtgacactttgtccagaacctattttccagtcttttcccccaggcttgtccctggtaacctctgtttggactcattgttaagcccagcgtgatttttggctccaactaaattgattttggaaatgactccagcatacctatgacccctctccttccaccctccataaaagatttatttttctttatttggcctctgct**gagattcctggctttgatgccccaggatctc**tcatttcccttgttccagggataagtgag**agatgactttttaaagagctactttcatct**tagagacaacaaagagagtatgcca**gcaagagcaacatcttgc**attgagccttctcactctatgccattttccaaatctttttctttatttaggataagtgaccactcctttttatttctagtacagcaaagagtacatcatgatgtcagaaacagggatttcctttggaatgtttttcacaggctaacaataagctagaagtctgcaagcaattcagaaatgcatccc**tgagctacaactccatgtatggaagctca**tcagcagggtagacaggcaaagcagaaacattttaattataccatactttgtagagattgggggagaagtgggcaaatgcgtgctaaggaaaaacaaaaactatggttaagtggggagatttctagatggttctaacagggaagaagaccaacaagaggaaacttccaggaaaaccaccaaggccaggcattc

Uppercase: TRAJ30

Lowercase: Flanking sequence[1000bp]

Red & Bold & Underline: Stem-loop [12]

Blue: Heptamer[21]

Green: Nonamer [5]

id-TRAJ23[J_gene_segment]

ttgcagtgagccaagatcatgccattgcactccagcctgggcaacaaaagcgaaacttcatctcaaacaaacaaacaagctcaggcaagtagaggccagtgattaacactcttccaataggaacagatcccaagcaacctgcagtagtttttccagtaggctcgtgaactcaaaacaccaagttattt**aaatgacagggcaaccgttgtgatcagctgtcattt**gagtagtgctgaggactgggtctgaccttgagtaaacgcctaggctggtgggtttctatccctgcagcatctaaagcggacgcccaggcttcactgaccccagcagtctggctttctgcccttgccgcccaggaggtgcagctcttggcacacaccatccttagtgtcttgaaagaagagaaattaaaagagaaagggggaaaagcttatatctcatatcatgcagttgcctcattttgtagcaagattgt**ttccccatgagcagtttgtcttcattcagcagctgtcttctctggggaa**gccattttgtagaggtgt**ttgtcacagtgtgacaa**ctgacagctgggggaaattccagtttggagcagggacccaggttgtggtcaccccaggtaagccccattccctggagcctcacctgccctt**agtatttggcatgccctgcatgccaaatatt**tctagccgagactatgagaaacacatctgaaaggaagccatttccctgaacagcagtacccaaagccattaagcaaaaatgagagatcagactggtgttaaatagaaacgtttttggataaatagtaattctcattattctctatcagtctcctgtgtgaatac**ccttatttcccttataagg**aaaaaagttacaataagcaggatagccatgattcatgctgtggtgtgggagcaatctgctgtggatttgggggtggaggattagaaatgtgttcaggcagactgga**tgtgtttttgacaggatatgtaacaca**gtgTGATTTATAACCAGGGAGGAAAGCTTATCTTCGGACAGGGAACGGAGTTATCTGTGAAACCCAgtaagtataaaattgtatccctggattaagcaatgtctgtggattaagggctgatttagacccaattatacaccatatgaagaatgtttagagagggggatgagcatcaaataggaagccaatcgtcgtcctggctaggatctagcatctcagtgcaaaatgggctatgtaagtgtgcctctgggaattgctcctgaatatatgtgttgggactaaaatgtatgactggctacttgttgtattggctgggatcatatccctggatttgtgcagtaattggtgacaagatttctaattccttaacaaaccttctaaggcactattctatttggctaagattatgtcaatttatagaaggaaagaaccttgtaaataatgatcattaaccaaaagtaatgtgattagtctctctcaaagtctgaaaaacc**acttttttttaaaaagt**ttactcatgactactgagggacacttcctgtttctgagactttcagccaaaaattcgtacagcgtagcctcacggagcagagagaaccttgacaacattcctttttgac**tttctccatggatgagaaa**actgaggctcagggaagagacagatcccggcccca**cagccagtctatggctg**agctagaactagattttaggtctctaatttgccaaacctgtcagttggc**tcaaacttgaagtttga**agagtccgcaagattcatctgacaaatattttcatgtacttgccctctgccaggctcgttgctgaaaccagggatccaacaggaagcaaactcagtgtgatattgccttcatgaaacttacattctggaggtggattccatgtttcttcccagatatgacaatgcttaccctgatttctcttgggcatggctctcttatactatcatcacttccttaggaaatactaaaactaatttttgctgcagtttaaagtccttgagcagataactaacacacataccactttagtcaggagaagggaaatgccc

Uppercase: TRAJ23

Lowercase: Flanking sequence[1000bp]

Red & Bold & Underline: Stem-loop [10]

Blue: Heptamer[23]

Green: Nonamer [2]

id-TRBJ2-7[J_gene_segment]

agcctgcacgaccccagaaccctgttcttaggggagtggacactgggcaatccagggccctcctcgagggaagcggggtttgcgccagggtccccagggctgtgcgaacaccggggagctgttttttggagaaggctctaggctgaccgtactgggtaaggaggcggttggggctccggagagctccgagagggcgggat**gggcagaggtaagcagctgccc**cactctgagaggggctgtgctgagaggcgctgctgggcgtctgggcggaggactcctggttctgggtgctgg**gagagcgatggggctctcagcggtgggaaggacccgagctgag**tctgggacagcagagcgggcagcaccggtttttgtcctgggcctccaggctgtgagcacagatacgcagtattttggcccaggcacccggctgacagtgctcggta**agcgggggctcccgct**gaagccccggaactggggagggggcg**ccccgggacgccgggg**gcgtcgcagggccagtttctgtgccgcgtctcggggctgtgagccaaaaacattcagtacttcggcgccggga**cccggctctcagtgctgggtaagctggggccgccggg**ggaccggggacgagactgcgctcgggtttttgtgcggggctcgggggccgtgaccaagagacccagtacttcgggccaggcacgcggctcctggtgctcggtgagcgcgggctgctggggcgcgggcgcgggcggcttgggtctggtttttgcggggagtccccgggctgtgctctggggccaacgtcctgactttcggggccggcagcaggctgaccgtgctgggtgagttttcgcgggaccacccgggcggcgggattcaggtggaaggcggcggctgcttcgcggcacccggtccggccctgtgctgggaga**cctgggctgggtccccagg**gtgggcaggagctcggggagccttagaggtttgcatgcgggggtgcacctccgtgCTCCTACGAGCAGTACTTCGGGCCGGGCACCAGGCTCACGGTCACAGgtgagattcgggcgtctccccaccttccagcccct**cggtccccggagtcggagggtggaccg**gagctgg**aggagctgggtgtccggggtcagctctgcaaggtcacctccccgctcct**ggggaaagactggggaagagggagggggtggggagg**tgctcagagtccggaaagctgagca**gagggcgaggccacttttaatcttttttctggggtgtttagagagaaggtgaacgatggaggagaggatttgttaggactctgggagaggcgagactggagaggacgaagggaaatcctggtttggggaatgggtaggagtgggggtaactgctattcgtaggcaaaaagagctgagcaggctgggaacagcgcgggtgggcaagggtcagcactgcgggcaggcgggtgggtgttagggggcagaaatcctgcagccgagggtgcagtagaacacagaagaaaaagcctgccaaacaaaagtggaacagagaagccaaaaagggagatgaacatgagtcagtgaagaaaagaatgaaagtttactgtttagcagtgtggatctctaatccgacttaaaactccttgttcccgattcctattcctcctaagccagagatccctgggtccagggtgagggcacggcattcatgcttacccacgggctggtcaacaaagaggtgctgacctgagagtagggcacataacctcagccactggggtacacttaccacccccgcccccgtgtagctccctcccctatcctgaaatctcccttagcacactaagtattctaggttaaacagcccagatgttcagggagttcattcgccacaaacacacattaaaatg**cagacaatttgcctgtgagatgaggaaaattctctg**gaaga**tttaggccctgagagctgaaaagggaccctaaa**cattacctggtgacaactgccctgaggccagagaagagaactcacaatattggtatattaaccggtaccatttgta

Uppercase: TRBJ2-7

Lowercase: Flanking sequence[1000bp]

Red & Bold & Underline: Stem-loop [12]

Blue: Heptamer[26]

Green: Nonamer [10]

id-TRAJ35[J_gene_segment]

aaaacaaatgctgttgtgctttttcccagtctgtctcctctccacac**agaaagtttccctttct**actaaaca**gcatctcttcacacgaagaaaagatgc**c**cagcaagccatttgacttattgctg**ctgctagtttgcatttatatgactcctttccttagaagggatgaagacactttatacgacttattagctaaacagattgacagcctaattaggaagagagaaacagattctgagtatctttttgtggctgctgttcctaccccctcgtgtagaacatttggtcagatggctctttgagacagaaataagacagggacgtggtctctgcttgtagtgtgggacgtctcagctgcacctaagaggttgtcacaggagtactaaataggtcagggcaactgacacagccaggaaaatttctttcccctagaacttgtgtgtgctcaccaagcaaagcgtgactaattttgacatccattgttagtgtgtgtctacccttcttgcctttgttttcagacccttcctctctagatccttttccttccatctccaactcctgtatgtaaatccctgctctctcctttgaattcctgactgctatgcccatgcccagcaccggcacccccacgcacactgctgcctgcctctgtaaccatcagctgtatttc**tgaggactttcctagctccttgagtgatcctca**aatggctcagggccctgagggaatctttgctgagggttggccccagcagagccgctgcagaggtagaggtggccaccaagggacacagactgcctgcatgaaggctggagctgggcccaggatgaggaaaagcctcaggaaggaggggctgacacgaaataaggaataccatggcattcatgagatgtgcgtctgaatcctctctcttgcctgagaagctttagcttccaccttgagacacaaaacatgtggttatgaagagatgacaaggtttttgtaaaagaatgagccattgtgGATAGGCTTTGGGAATGTGCTGCATTGCGGGTCCGGCACTCAAGTGATTGTTTTACCACgtaagtatatcttttctcatttctgtgggctgttatgtccgtaaatcatatagaacagttcttcttatgcacacacacacacacacacccattttccagtcactgaggagaaacagtctctgggtctacctacacagccag**tttgaccattaagatggtcaaa**ttcatttttcaagataattttaactggttccatt**taatgctgccaacagccaagggcatta**gcatgtcgtgggaacctaatctatgttcactcactgctgtgggaggacagtcatttcttcttctcccctggcagtcagacttcagaatccagctgaccttttccatgcctgtctcttctggcagattaagtg**gttatttccactaactttcttgtggaaaaac**acacaagacagatatattctccttcagtttttactaaga**aagttttgtggttatcattttgacaataaactt**cagctagcatagtccctactattcatgttaataatatcttgacttgaaccatcatattttccccacatcattcacaatgcatttcaatgctgtaacccactgcaccatccccagcccttttatcttcacaacgttctctaataaggtaaacaggatcatacatcatcaaacatatgtcacagacagaaaagctgacacacagacaagttaaataatttaagtcaaaccaaaaaaaaatcataactgggattagagcttgagattccctattttcttacagtgaactgtcttgagctgtcacttaatgcattgaaaacataataagcacaggccataggcat**tttcagaacatgaaacttgactgaaa**caaaaactgactgtttgattgctttggtgttttaaatgtatcc**ccaaatctcctttgctattgatagtcatttgg**tgccagtgacagggaagagcagaggggcttaggaggtttttgtagatctcagtatcactgtgtcttataacaccgacaagctcatctttgggactggga

Uppercase: TRAJ35

Lowercase: Flanking sequence[1000bp]

Red & Bold & Underline: Stem-loop [10]

Blue: Heptamer[31]

Green: Nonamer [6]

id-TRAJ61[J_gene_segment]

tttggtaaagggcaaatttttccatttctgtcttttccatggaccatttcccctgctatcaattgctgttgcctgcagcctaagaaataaaatttttaaataaatgagaacgttctataattctctctaaaatcataaacaatgtgtcaaggccaaacacttccct**ggtgtgatttattcctatgtggactcatccttgtataattagcaggaataatcacacc**ctgagccagctggcagagcccctggggacagagaccacccaaagaaggt**ccttgttctcctgagtcacataaagggagtcacacaagg**cgggagtg**gaactcaggtgtccggagttc**tagtcccagctctgcttctacctatctacatgaccttgagcaaatttatcccgcagagcttcag**cattttcatctctaaaatg**agggcattggcccaggcagctgttaaatttccttccaagtct**cacattctaggatgctaatgtg**taaaattatagaactgacccaagagttatt**agttcatatccccctgtgcatgttgcttgaact**cctcattgcccagcaggaggaaaagggaaaacatgtttcttcattggggggtagcataatttcctggttgactatgtgtgactcatcagagcctggcggtccaagcccatatgccatttgaaatccggttattattacatctgggagagagaaaagtgctgaaaacagcctttgggacactatcgatttggccaaaaaaaaaagaagaagaagaagaaggctctgtctagtgt**gataacattttgttatc**ttattcattgtcttcatccctgaaatacactctgctctctcctatctctgctctgaaaggcagaa**agagggcagccctct**ccaaggcaaaatggggctcctgtgggga**acagaggggtgcctctgt**caacaaaggtgatgccacatccctttcaaccatgctgacacctctggtttttgtaaaggtgcccactcctgtgGGTACCGGGTTAATAGGAAACTGACATTTGGAGCCAACACTAGAGGAATCATGAAACTCAgcaagtaatatttggcag**aatttttttttctatctgaaaatt**atcagtgagagattctaatgtgccttaacaaacaggaacaaaagatgagtgtttaatacaattcaatttaacaaatatttattaagagcctacagttattcccatggatcatctgagtcagtttccgaggaaacattatcgttgccttaagggagctggggggttgtcaacggg**ccagaggtgggatgaaaaatgacaacagatttacctctgg**gaccgggacatggttaaccacagcggccctgggtaagtagcttagcttcagaagaaaatgtgcccaacagcatgggtaacctaaaacaccgggcaatccaaatattcttttatgattggctttagcatgtattttattcttttgtagggcaggtttatctcaccaattatatattttcttaactga**cctgtaaaatctacagg**ggaaaagtattttaagaattatatgtttctgcaattaggctcccagcagtcaacaaagaagtggta**ctttttgtctttccagtgatgaaaaag**ggaacctggcatcc**ctggtggcccaccag**cgtctccttccctggcctaggtcagaacaagccgtaaatcagcaggccgttatcttcttataaatctgtag**agcagggtggacaacaaaaggcagcctgct**aggttttcagaacatgagttccttgtgtagccagagaacctg**ggaccatcctgacgtggctggtcc**tgctgtcctc**acagcctgaatcccaggctgt**atgaataggagaggttcaagtccagatgactgttcacgatgctggtcccccttgcatccctaatcatgctagagacatgaccagggtctgagaggaggaagttacagcacagcaccaacaggggcttttggtaaagggcctgggcactatgtgaagatcacctagatgctcaactttgggaaggggactgagttaattgtgagcctgggtgagtacctcaa

Uppercase: TRAJ61

Lowercase: Flanking sequence[1000bp]

Red & Bold & Underline: Stem-loop [17]

Blue: Heptamer[27]

Green: Nonamer [1]

id-TRBJ2-2P[J_gene_segment]

acattgtggggactagcgggagggcacgatgattcaggtagaggaggtgcttttacaaaaaaccctgatgcagtaagcatc**cccacccagctcagggaatgcagctaccaggtggg**aagagttctctggggctggtcccagctgtggtcttgcagggtcccccaacccagcgagcacctgtccatctccctgtccagactcggcttccaaggaataagaaggccaagacagc**aaagtgggattatcactcagcacttt**taataaaacttgttcttgacaaagtacttgcacatgcattatttattaagaactgatgaaaaccctgag**ggaaagatattgtcccatctttcc**aatgaggaaactgagatcagaggttacaggtcatataactaggaaacggcaaggtctagcctgcaatatcgcccagctccagccgttccagtaccaccaatgccccttcagatttca**aatccactgtgttgtcccccagccaagtggatt**ctcctctgcaaattggtggtggcctcatgcaagatccaggttaccgtgtccagctaactcgagacaggaaaagataggctcaggaaagagaggaagggtgtgccctctgtctgtgctaagggaggtg**gggaaggagaaggaattctgggcagccccttccc**actgtgctcctacaatgagcagttcttcgggccagggacacggctcaccgtgctaggtaagaagggggctccaggtgggagagagggtgagcagcccagcctgcacgaccccagaaccctgttcttaggggagtggacactgggcaatccagggccctcctcgagggaagcggggtttgcgccagggtccccagggctgtgcgaacaccggggagctgttttttggagaaggctctaggctgaccgtactgggtaaggaggcggttggggctccggagagctccgagagggcgggat**gggcagaggtaagcagctgccc**cactctgagaggggctgtgCTGAGAGGCGCTGCTGGGCGTCTGGGCGGAGGACTCCTGGTTCTGGgtgctgg**gagagcgatggggctctcagcggtgggaaggacccgagctgag**tctgggacagcagagcgggcagcaccggtttttgtcctgggcctccaggctgtgagcacagatacgcagtattttggcccaggcacccggctgacagtgctcggta**agcgggggctcccgct**gaagccccggaactggggagggggcg**ccccgggacgccgggg**gcgtcgcagggccagtttctgtgccgcgtctcggggctgtgagccaaaaacattcagtacttcggcgccggga**cccggctctcagtgctgggtaagctggggccgccggg**ggaccggggacgagactgcgctcgggtttttgtgcggggctcgggggccgtgaccaagagacccagtacttcgggccaggcacgcggctcctggtgctcggtgagcgcgggctgctggggcgcgggcgcgggcggcttgggtctggtttttgcggggagtccccgggctgtgctctggggccaacgtcctgactttcggggccggcagcaggctgaccgtgctgggtgagttttcgcgggaccacccgggcggcgggattcaggtggaaggcggcggctgcttcgcggcacccggtccggccctgtgctgggaga**cctgggctgggtccccagg**gtgggcaggagctcggggagccttagaggtttgcatgcgggggtgcacctccgtgctcctacgagcagtacttcgggccgggcaccaggctcacggtcacaggtgagattcgggcgtctccccaccttccagcccct**cggtccccggagtcggagggtggaccg**gagctgg**aggagctgggtgtccggggtcagctctgcaaggtcacctccccgctcct**ggggaaagactggggaagagggagggggtggggagg**tgctcagagtccggaaagctgagca**gagggcgaggccacttttaatcttttttctggggtgtttagagagaaggtgaacgatggag

Uppercase: TRBJ2-2P

Lowercase: Flanking sequence[1000bp]

Red & Bold & Underline: Stem-loop [15]

Blue: Heptamer[33]

Green: Nonamer [11]

id-IGHJ2[J_gene_segment]

ttagaccaccttgcac**cttccctggcacccaccatgggaag**ag**ctgagactcactgaggaccagctgaggctcag**agaagggacccagcactggtggacacgcagggagcccacgccagggcgccgtggtgagt**gaggcccagtgccacccactgaggcctc**ccgttcagtgggacgacggtgaacaggtggaaccaaccaggcaacccccgccgggccccacagacgggatcaga**gcaggaaaggcttcctgc**ccctgcaggccagcgaggagccc**tggcgggggccatggccctccaggcgaggaggctcccctggccaccgcca**cccgggcctctctgctgctgggaaaacaagtcagaaagcaagtggatgagaggtggcgtgacagacccagcttcagatctgctctaatttacaaaagaaaaggaaaaacacacttggcagccttcagcactctaatgattcttaacagcagcaaattattggcacaagactccagagtgactggcagggttgagggctgggg**tctcccgcgtgttttggggctaacagcggaagggaga**gcactggcaaaggtgctgggggcccctggacccgacccgccctggagaccgcagccacatcagcccccagccccacaggccccctaccagccgcagggttttggctgagctgagaac**cactgtgctaactggggacacagtg**attggcagct**ctacaaaaaccatgctcccccgggaccccgggctgtgggtttctgtag**cccctggctcagggctgactcaccgtggctgaatacttccagcactggggccagggcaccctggtcaccgtctcctcaggtgagtctgctgtctggggatagcggggagccaggtgtactgggccaggcaagggctttggcttcagacttggggacaggtgctcagcaaaggaggtcggcaggagggcggagggtgtgtttttgtatgggagaagcaggagggcagaggctgtgCTACTGGTACTTCGATCTCTGGGGCCGTGGCACCCTGGTCACTGTCTCCTCAGGTgagtcccactgcagccccctcccagtcttctctgtccaggcac**caggccaggtatctggggtctgcagccggcctgggtctggcctg**aggccacaccagctgccatccctggggtctccgccatgggctgcatgccagagccctgctgtcacttagccctggggcca**gctggagcccccaaggacaggcagggaccccgctgggcttcagc**cccgtcagggaccctccacaggtagcaagcaggccgagggcagggacgggaaggagaagttgtgggcagagcctgggctggggctgggcgctggctgttcatgtgccggggaccaggcctgcgctttagtgtggctaca**agtgcttggagcact**gggg**ccagggcagcccggccaccgtctccctgg**gaacgtcacccctccctgcctgggtctcagcccgggggtctgtgtggctggggacagggacgccggctgcctctgctctgtgcttgggccatgtgacccattcgagtg**tcctgcacgggcacaggtttatgtctgggcaggaacagggactgtgtccctgt**gtgatgcttttgatatctggggccaagggacaatggtcaccgtctcttcaggtaagatggctttccttctgcctcctttctctggg**cccagcgtcctctgtcctggagctggg**agataatgtccgggggctccttggtctgcgctgggccatgtggggccctccggggctccttctccggctgtttgggaccacgttcagcagaaggcctttctttgggaactgg**gactctgctgctggggcaaagggtgggcagagtc**atgcttgtgctggggacaaaatgaccttgggacacggggctggctgccacggccggcccgggacagtcggagagtcaggtttttgtgcaccccttaatggggcctcccacaatgtgactactttgactactggggccagggaaccctggtcaccgtctcctcaggtgagtcctcacaacctc

Uppercase: IGHJ2

Lowercase: Flanking sequence[1000bp]

Red & Bold & Underline: Stem-loop [16]

Blue: Heptamer[38]

Green: Nonamer [7]

id-IGHJ5-2[J_gene_segment]

cgctttagtgtggctaca**agtgcttggagcact**gggg**ccagggcagcccggccaccgtctccctgg**gaacgtcacccctccctgcctgggtctcagcccgggggtctgtgtggctggggacagggacgccggctgcctctgctctgtgcttgggccatgtgacccattcgagtg**tcctgcacgggcacaggtttatgtctgggcaggaacagggactgtgtccctgt**gtgatgcttttgatatctggggccaagggacaatggtcaccgtctcttcaggtaagatggctttccttctgcctcctttctctggg**cccagcgtcctctgtcctggagctggg**agataatgtccgggggctccttggtctgcgctgggccatgtggggccctccggggctccttctccggctgtttgggaccacgttcagcagaaggcctttctttgggaactgg**gactctgctgctggggcaaagggtgggcagagtc**atgcttgtgctggggacaaaatgaccttgggacacggggctggctgccacggccggcccgggacagtcggagagtcaggtttttgtgcaccccttaatggggcctcccacaatgtgactactttgactactggggccagggaaccctggtcaccgtctcctcaggtgagtcctcacaacctctctcctgctttaactctgaagggttttgctgcatttttggggggaaataagcgtgctgggtctcctgccaaga**gagccccggagcagcctggggggctcaggaggatgccctgag**gcaacagcggccacacagacgaggggcaa**gggctccagatgctccttcctcctgagccc**agcagcacgggtctctctgtgg**ccagggccaccctgg**gcctctggggtccaatgtccaacaacc**cccgggccctccccggg**ctcagtctgagagggtcccagggacttagcgggg**tgccagttcttgcctggggtcctggca**tt**gttgtcacaatgtgACAAC**TGGTTCGACCCCTGGGGCCAGGGAACCCTGGTCACCGTCTCCTCAGGTgagtcctcaccaccccctct**ctgagtccacttagggagactcag**cttgccagggtctcagggtcagagtcttggaggcattttggaggtcaggaaagaaa**gctggggagagggacccttcgaatgggaacccagc**ctgtcctccccaagtccggccacagatgtcggcagctggggggctccttcggctggtctggggtgacctctctccgcttcacctggagcattctcaggggctgtcgtgatgattgcgtggtgggactctgtcccgctccaa**ggcacccgctctctgggacgggtgcc**ccccggggtttttggactcctgggggtgacttagcagccgtctgcttgcagttggacttcccaggccgacagtgg**tctggcttctgaggggtcaggccaga**atgtggggtacgtgggaggccagcagagggttccatgagaagggcaggacagggccacggacagtcagcttccatgtgacgcccggagacagaaggtctctgggtggctgggtttttgtggggtgaggatggacattctgccattgtgattactactactactactacatggacgtctggggcaaagggaccacggtcaccgtctcctcaggtaagaatggccactctagggcctttgttttctgctactgcctgtggggtttcctgagcattgcaggttggt**cctcggggcatgttccgagg**ggacctgggcggactggccaggaggggatgggcactggggtgccttgaggatctgggagcctctgtgg**attttccgatgcctttggaaaat**gggactcaggttgggtgcgtctgatggagtaactgagcctgggggcttggggagccacatttggacgagatgcctgaacaaaccaggggtcttagtgatggctgaggaatgtgtctcaggagcggtgtctgtaggactgcaaga**tcgctgcacagcagcga**atcgtgaaatattttctttagaattatgaggtgcgctgtgtg

Uppercase: IGHJ5-2

Lowercase: Flanking sequence[1000bp]

Red & Bold & Underline: Stem-loop [20]

Blue: Heptamer[29]

Green: Nonamer [5]

id-IGHJ4-2[J_gene_segment]

cagaggctgtgctactggtacttcgatctctggggccgtggcaccctggtcactgtctcctcaggtgagtcccactgcagccccctcccagtcttctctgtccaggcac**caggccaggtatctggggtctgcagccggcctgggtctggcctg**aggccacaccagctgccatccctggggtctccgccatgggctgcatgccagagccctgctgtcacttagccctggggcca**gctggagcccccaaggacaggcagggaccccgctgggcttcagc**cccgtcagggaccctccacaggtagcaagcaggccgagggcagggacgggaaggagaagttgtgggcagagcctgggctggggctgggcgctggctgttcatgtgccggggaccaggcctgcgctttagtgtggctaca**agtgcttggagcact**gggg**ccagggcagcccggccaccgtctccctgg**gaacgtcacccctccctgcctgggtctcagcccgggggtctgtgtggctggggacagggacgccggctgcctctgctctgtgcttgggccatgtgacccattcgagtg**tcctgcacgggcacaggtttatgtctgggcaggaacagggactgtgtccctgt**gtgatgcttttgatatctggggccaagggacaatggtcaccgtctcttcaggtaagatggctttccttctgcctcctttctctggg**cccagcgtcctctgtcctggagctggg**agataatgtccgggggctccttggtctgcgctgggccatgtggggccctccggggctccttctccggctgtttgggaccacgttcagcagaaggcctttctttgggaactgg**gactctgctgctggggcaaagggtgggcagagtc**atgcttgtgctggggacaaaatgaccttgggacacggggctggctgccacggccggcccgggacagtcggagagtcaggtttttgtgcaccccttaatggggcctcccacaatgtgACTACTTTGACTACTGGGGCCAGGGAACCCTGGTCACCGTCTCCTCAGGTgagtcctcacaacctctctcctgctttaactctgaagggttttgctgcatttttggggggaaataagcgtgctgggtctcctgccaaga**gagccccggagcagcctggggggctcaggaggatgccctgag**gcaacagcggccacacagacgaggggcaa**gggctccagatgctccttcctcctgagccc**agcagcacgggtctctctgtgg**ccagggccaccctgg**gcctctggggtccaatgtccaacaacc**cccgggccctccccggg**ctcagtctgagagggtcccagggacttagcgggg**tgccagttcttgcctggggtcctggca**tt**gttgtcacaatgtgacaac**tggttcgacccctggggccagggaaccctggtcaccgtctcctcaggtgagtcctcaccaccccctct**ctgagtccacttagggagactcag**cttgccagggtctcagggtcagagtcttggaggcattttggaggtcaggaaagaaa**gctggggagagggacccttcgaatgggaacccagc**ctgtcctccccaagtccggccacagatgtcggcagctggggggctccttcggctggtctggggtgacctctctccgcttcacctggagcattctcaggggctgtcgtgatgattgcgtggtgggactctgtcccgctccaa**ggcacccgctctctgggacgggtgcc**ccccggggtttttggactcctgggggtgacttagcagccgtctgcttgcagttggacttcccaggccgacagtgg**tctggcttctgaggggtcaggccaga**atgtggggtacgtgggaggccagcagagggttccatgagaagggcaggacagggccacggacagtcagcttccatgtgacgcccggagacagaaggtctctgggtggctgggtttttgtggggtgaggatggacattctgccattgtgattactactactactactacatggacgtctggggcaaagggaccacgg

Uppercase: IGHJ4-2

Lowercase: Flanking sequence[1000bp]

Red & Bold & Underline: Stem-loop [19]

Blue: Heptamer[31]

Green: Nonamer [4]

id-IGHJ2P-2[J_gene_segment]

tcagatctgctctaatttacaaaagaaaaggaaaaacacacttggcagccttcagcactctaatgattcttaacagcagcaaattattggcacaagactccagagtgactggcagggttgagggctgggg**tctcccgcgtgttttggggctaacagcggaagggaga**gcactggcaaaggtgctgggggcccctggacccgacccgccctggagaccgcagccacatcagcccccagccccacaggccccctaccagccgcagggttttggctgagctgagaac**cactgtgctaactggggacacagtg**attggcagct**ctacaaaaaccatgctcccccgggaccccgggctgtgggtttctgtag**cccctggctcagggctgactcaccgtggctgaatacttccagcactggggccagggcaccctggtcaccgtctcctcaggtgagtctgctgtctggggatagcggggagccaggtgtactgggccaggcaagggctttggcttcagacttggggacaggtgctcagcaaaggaggtcggcaggagggcggagggtgtgtttttgtatgggagaagcaggagggcagaggctgtgctactggtacttcgatctctggggccgtggcaccctggtcactgtctcctcaggtgagtcccactgcagccccctcccagtcttctctgtccaggcac**caggccaggtatctggggtctgcagccggcctgggtctggcctg**aggccacaccagctgccatccctggggtctccgccatgggctgcatgccagagccctgctgtcacttagccctggggcca**gctggagcccccaaggacaggcagggaccccgctgggcttcagc**cccgtcagggaccctccacaggtagcaagcaggccgagggcagggacgggaaggagaagttgtgggcagagcctgggctggggctgggcgctggctgttcatgtgccggggaccaggcctgcgctttagtgtgGCTACA**AGTGCTTGGAGCACT**GGGG**CCAGGGCAGCCCGGCCACCGTCTCCCTGG**GAACGTcacccctccctgcctgggtctcagcccgggggtctgtgtggctggggacagggacgccggctgcctctgctctgtgcttgggccatgtgacccattcgagtg**tcctgcacgggcacaggtttatgtctgggcaggaacagggactgtgtccctgt**gtgatgcttttgatatctggggccaagggacaatggtcaccgtctcttcaggtaagatggctttccttctgcctcctttctctggg**cccagcgtcctctgtcctggagctggg**agataatgtccgggggctccttggtctgcgctgggccatgtggggccctccggggctccttctccggctgtttgggaccacgttcagcagaaggcctttctttgggaactgg**gactctgctgctggggcaaagggtgggcagagtc**atgcttgtgctggggacaaaatgaccttgggacacggggctggctgccacggccggcccgggacagtcggagagtcaggtttttgtgcaccccttaatggggcctcccacaatgtgactactttgactactggggccagggaaccctggtcaccgtctcctcaggtgagtcctcacaacctctctcctgctttaactctgaagggttttgctgcatttttggggggaaataagcgtgctgggtctcctgccaaga**gagccccggagcagcctggggggctcaggaggatgccctgag**gcaacagcggccacacagacgaggggcaa**gggctccagatgctccttcctcctgagccc**agcagcacgggtctctctgtgg**ccagggccaccctgg**gcctctggggtccaatgtccaacaacc**cccgggccctccccggg**ctcagtctgagagggtcccagggacttagcgggg**tgccagttcttgcctggggtcctggca**tt**gttgtcacaatgtgacaac**tggttcgacccctggggccagggaaccctggtcaccgtctcctcaggtgagtcctcaccaccccctc

Uppercase: IGHJ2P-2

Lowercase: Flanking sequence[1000bp]

Red & Bold & Underline: Stem-loop [18]

Blue: Heptamer[37]

Green: Nonamer [6]

id-TRAJ26[J_gene_segment]

gcagagagctgcatagctgggatcctcaaggctacacacacacgcttggagcagggatgcggttcacaggatggcagggca**aaggttgcagaactgaaacctt**gtacttccccatcaccaaaatgaccccagcatgatttcctttaaggtgtcccga**gccccttcccaaaagacaggctccccatgtggaggggc**aatgacagagggaagatggaaactttatgggaagggcgatctatcccaaggttgtccttctcctc**ttcagggtagtgaagctcatgagaaatatggctcctgaa**aatagccactaaatgaaactctccaacctggaggtagaggatcagtaaagtctgcagaggcaaaatgaggcagtttatttcaaagaaaagaggttgagggcaaggtgggagttaagtcacacatctaaatctgggaaaggaagcattaaacagaagacggaggggctcggctctccacctccaccaggattcagaccacaggacatggctaatactgcaagaagggagttaggttagtggt**aaaacctttctaataggtttt**tgaacagtgaaaatgagctgtcaggagtagttgccagggcactttctctagagagtttttgaaaggagatagacaccctgctatcagaggtgattggagtgcagc**cctgcctcgaggcagg**aggagaaatgagagaacctcttaaagcctctttccagccccca**gagcctgtctccatgaaaaaggcccaggctc**attaatgcagagctgcatcctccaggacacagggggttgctgggttgtta**agtttggaacctttgccggcttagtggttaccaaact**ccctttgttctcaagacttgacttgataaatgggc**tggctttatccctttaaagccaaaaacacaggttctgagtgctgtgttgtttg**tggtgttgagatgcccaggctggagggaggaagctctagggggtttttgctgagcccagaaacactgtgGGGATAACTATGGTCAGAATTTTGTCTTTGGTCCCGGAACCAGATTGTCCGTGCTGCCCTgtaagtacagttaagtggagatagaaaatgagtccagtgcttgatgtggggagaagctgcagggtcttgaggccaggagtccaccgtgacctcaggagtggtgatggagaagaaggttaaaaaaggaatgagaaacagaaccataaggacaatacagaggaagggttc**attttcgggaaggaaaatcacaaggccgtgtgatt**aacgctggaggaaatgcactcctttggagggtcccttctcaggagggctttgtgtcggggaaagctgagctgttaggtttttgatgctgagataatcactatgcagaaggacaaggcttctcctttatctttgggaaggg**gacaaggctgcttgtc**aagccaagtaagtg**acatataatttatatgt**gctgaatatgattatctcc**aagaggaaaacttgccctctt**ttccacgggtg**cctccttgggtcaggagg**acatta**aaggtgttgagcagaccaaggcccagtacctcgtcacctt**cttcatccttgaaggagctttaagaggtgtggaggggaagaaacccaccaggaccccaccaataagcccagccttgagacccctccactctgtcagacttgaataagaacca**tctgagaacaaggtcttcctcaga**aggggactccagcatagtcatccccatttgatagattttgaaactcagggcggggcagagtggctcatgcctgtaatcccagcactttgggaagctg**aggcaggtggatcacttgaaggtcaggagttcgagacctgcct**ggccaacatggtgaaagcccgtctctactaaaaataaaaaaaat**tagctgggtgtggtggcacgcacctgtaatcccagcta**ctcagaaggctgaggcaggagaatcgcttgaacccgggaggcggaggttgcagtgagccaagatcatgccattgcactccagcctgggcaacaaaagcgaaacttcatctcaaacaaacaaacaagctcaggcaagt

Uppercase: TRAJ26

Lowercase: Flanking sequence[1000bp]

Red & Bold & Underline: Stem-loop [19]

Blue: Heptamer[25]

Green: Nonamer [7]

id-IGHJ6-2[J_gene_segment]

tactttgactactggggccagggaaccctggtcaccgtctcctcaggtgagtcctcacaacctctctcctgctttaactctgaagggttttgctgcatttttggggggaaataagcgtgctgggtctcctgccaaga**gagccccggagcagcctggggggctcaggaggatgccctgag**gcaacagcggccacacagacgaggggcaa**gggctccagatgctccttcctcctgagccc**agcagcacgggtctctctgtgg**ccagggccaccctgg**gcctctggggtccaatgtccaacaacc**cccgggccctccccggg**ctcagtctgagagggtcccagggacttagcgggg**tgccagttcttgcctggggtcctggca**tt**gttgtcacaatgtgacaac**tggttcgacccctggggccagggaaccctggtcaccgtctcctcaggtgagtcctcaccaccccctct**ctgagtccacttagggagactcag**cttgccagggtctcagggtcagagtcttggaggcattttggaggtcaggaaagaaa**gctggggagagggacccttcgaatgggaacccagc**ctgtcctccccaagtccggccacagatgtcggcagctggggggctccttcggctggtctggggtgacctctctccgcttcacctggagcattctcaggggctgtcgtgatgattgcgtggtgggactctgtcccgctccaa**ggcacccgctctctgggacgggtgcc**ccccggggtttttggactcctgggggtgacttagcagccgtctgcttgcagttggacttcccaggccgacagtgg**tctggcttctgaggggtcaggccaga**atgtggggtacgtgggaggccagcagagggttccatgagaagggcaggacagggccacggacagtcagcttccatgtgacgcccggagacagaaggtctctgggtggctgggtttttgtggggtgaggatggacattctgccattgtgATTACTACTACTACTACTACATGGACGTCTGGGGCAAAGGGACCACGGTCACCGTCTCCTCAGGTaagaatggccactctagggcctttgttttctgctactgcctgtggggtttcctgagcattgcaggttggt**cctcggggcatgttccgagg**ggacctgggcggactggccaggaggggatgggcactggggtgccttgaggatctgggagcctctgtgg**attttccgatgcctttggaaaat**gggactcaggttgggtgcgtctgatggagtaactgagcctgggggcttggggagccacatttggacgagatgcctgaacaaaccaggggtcttagtgatggctgaggaatgtgtctcaggagcggtgtctgtaggactgcaaga**tcgctgcacagcagcga**atcgtgaaatattttctttagaattatgaggtgcgctgtgtgtcaacctgcatcttaaattctttat**tggctggaaagagaactgtcggagtgggtgaatccagcca**ggagggacgcgtagccccggtcttgatgagagcagggttgggggcaggggtagcccagaaacggtggctgcc**gtcctgacaggggcttagggaggctccaggac**ctcagtgccttgaagctggtttccatgagaaaaggattgtttatcttaggaggcatgcttactgttaaaagacaggatatgtttgaagtggcttctgagaaaaatggttaagaaaattatg**acttaaaaatgtgagagattttcaagt**atattaatttttttaactgtccaagtatttgaaattcttatcatttgattaacacccatgagtgatatgtgtctggaattgaggccaaagcaagctcagctaagaaatactagcacagtgctgtcggccccgatgcgggactgcgttttgaccatcataaatcaagtttatttttttaattaattgagcgaagctggaagcagatgatgaattagagtcaagatggctgcatgggggtctccggcacccacagcaggtggcaggaagcaggtcaccgcgagagtctattttaggaagca

Uppercase: IGHJ6-2

Lowercase: Flanking sequence[1000bp]

Red & Bold & Underline: Stem-loop [17]

Blue: Heptamer[21]

Green: Nonamer [4]

id-IGHJ1P[J_gene_segment]

tgcccgcacggtgcctga**gggggccttcttgggcagcgcctaagcaagccccc**agcacccttcggccccttca**aggcacacaggccccctttccacccagcctcaggaaaccacctgtgtcct**ccaacgacaggtcccagcctcccagcctttgccttgcctgttcctctccctggaactctgccccgacacagaccctccccagcaagccc**gcaggggcacctcccctgc**ccccagacaccctgtgcccgtcagttcatccccagcagaggccctcaccaggcacacccccatgctcacacctggccgcagg**cctcagcctccctgagg**gccccacccagcccgcgtctggccagtggtgcgtgcaa**agcccctcacccagactcggcggaaggcagccagtgcaggcctggggaggggct**ctccttagaccaccttgcac**cttccctggcacccaccatgggaag**ag**ctgagactcactgaggaccagctgaggctcag**agaagggacccagcactggtggacacgcagggagcccacgccagggcgccgtggtgagt**gaggcccagtgccacccactgaggcctc**ccgttcagtgggacgacggtgaacaggtggaaccaaccaggcaacccccgccgggccccacagacgggatcaga**gcaggaaaggcttcctgc**ccctgcaggccagcgaggagccc**tggcgggggccatggccctccaggcgaggaggctcccctggccaccgcca**cccgggcctctctgctgctgggaaaacaagtcagaaagcaagtggatgagaggtggcgtgacagacccagcttcagatctgctctaatttacaaaagaaaaggaaaaacacacttggcagccttcagcactctaatgattcttaacagcagcaaattattggcacaagactccagagtgactggcagggttgagggctgggg**tctcccgcgtgttttggggctaacagcggaagggaga**gcactggcAAAGGTGCTGGGGGCCCCTGGACCCGACCCGCCCTGGAGACCGCAGCCACATCAgcccccagccccacaggccccctaccagccgcagggttttggctgagctgagaac**cactgtgctaactggggacacagtg**attggcagct**ctacaaaaaccatgctcccccgggaccccgggctgtgggtttctgtag**cccctggctcagggctgactcaccgtggctgaatacttccagcactggggccagggcaccctggtcaccgtctcctcaggtgagtctgctgtctggggatagcggggagccaggtgtactgggccaggcaagggctttggcttcagacttggggacaggtgctcagcaaaggaggtcggcaggagggcggagggtgtgtttttgtatgggagaagcaggagggcagaggctgtgctactggtacttcgatctctggggccgtggcaccctggtcactgtctcctcaggtgagtcccactgcagccccctcccagtcttctctgtccaggcac**caggccaggtatctggggtctgcagccggcctgggtctggcctg**aggccacaccagctgccatccctggggtctccgccatgggctgcatgccagagccctgctgtcacttagccctggggcca**gctggagcccccaaggacaggcagggaccccgctgggcttcagc**cccgtcagggaccctccacaggtagcaagcaggccgagggcagggacgggaaggagaagttgtgggcagagcctgggctggggctgggcgctggctgttcatgtgccggggaccaggcctgcgctttagtgtggctaca**agtgcttggagcact**gggg**ccagggcagcccggccaccgtctccctgg**gaacgtcacccctccctgcctgggtctcagcccgggggtctgtgtggctggggacagggacgccggctgcctctgctctgtgcttgggccatgtgacccattcgagtg**tcctgcacgggcacaggtttatgtctgggcagga**acagggactgtgtccctgtgtgatgcttttgat

Uppercase: IGHJ1P

Lowercase: Flanking sequence[1000bp]

Red & Bold & Underline: Stem-loop [18]

Blue: Heptamer[37]

Green: Nonamer [6]

id-TRBJ1-4[J_gene_segment]

acccaggcttcccagaggctctgagcagtcacagctgagcccagggtgatggggcagaag**agggaaggggagggggcctctcctcatagttccct**gagatagcccagagaaagcccggtgggtaatgaatgagccacaacacctctccatctatctgcttcactg**acagaggttctctgt**agattcttcgtatattcctgtgctggattttataggaggccactctgtgtctctttttgtcacctgcctgagtcttgggca**agctctggaagggaacacagagtactggaagcagagct**gctgtccctgtgagggaa**gagttcccatgaactc**ccaac**ctctgcctgaatcccagctgtgctcagcagag**actggggggttttgaagtggccctgggaggctgtgctctggaaacaccatatattttggagagggaagttggctcactgttgtaggtgagtaagtcaaggctggacagctgggaacttgcaaaaaggggctggaatccagacggagcctttgtctctagtgcttaggtgaaagtgtatttttgtcaggaaggcctatgaggcagat**gaggaggggatagcctccctctcctc**tccactattttgtagactgcctgtgccaagttaggttcccctactgagagatgggtagactcagcttggaaggggtcaccttgaacatctcctgtctccttgaagggtgccggtcacggccatgacagataaaagagcctctgaccttaccaccacggtcctaccgtttctc**tccctcacacagaaaggagaaggtcacagaagaggga**acttgggggatcacacggggcctaattggtctgctgaccaccgcattttgggttgtaccattgtctacccctctacccaccagggttaaaattctactaaggaacaggagaggacctggcaggtggacttggggaggcag**gagtggaaggcagcaggtcgcggttttccttccagtc**tttaatgttgtgCAACTAATGAAAAACTGTTTTTTGGCAGTGGAACCCAGCTCTCTGTCTTGGgtatgtaaaagacttctttcgggatagtgtatcataaggtcggagttccaggaggaccccttgcgggagggcagaaactgagaacacagccaagaaaagctcataaaatgtgggtcagtggagtgtgtggtggggccccaagagttctgtgtgtaagcagcttctggaaggaagggcccacaccagctcctctggggtttgccacactcatgatgcactgtgtagcaatcagccccagcattttggtgatgggactcgactctccatcctaggtaagttgcagaatcagggtggtatggccattgtcccttgaaggcagagttctctgcttctcctcccggtgctggtgaggcagattgagtaaaat**ctcttaccccatggggtaagag**ctgtgcctgtgcctgcgttccctttggtgtgtcttggttgactcctctatttctcttctctaagtcttcagtccataatctgcctcctcactcccttcttggctcatcctccctcttatgtgcatggctctgcctctcctaagcctcttcctcttgcgccttatgctgcacagtatgcttaggcctttttcctaacagaatccctttggtccagagccatgaatccaggcagagaaaggcagccatcctgctgtcagggagctaagacttgccctctgactggagatcgccgggtgggttttatctaagcctctgcagctgtgctcctataattcacccctccactttgggaatgggaccaggctcactgtgacaggtatgggggctccactcttgactcgggggtgcctgggtttgactgcaatgatcagttgctgggaagggaattgagt**gtaagaacggaggtcagggtcaccccttcttac**ctggagcactgtgccctctcctcccctccctggagctcttccagcttgttgctctgctgtgttgcctgcagttcctcagctgtagagctccttgcttagtcttcagggctgtgtgtttctttgctc

Uppercase: TRBJ1-4

Lowercase: Flanking sequence[1000bp]

Red & Bold & Underline: Stem-loop [10]

Blue: Heptamer[47]

Green: Nonamer [4]

id-TRAJ39[J_gene_segment]

tagcactaatataatccaagtaggagatttacaggctgcatgtttcttggtttccctttggagtgctccattcctaacaccagttgttaagcccctggtcctgaatgtggccgggggtgaggaagtgggaggagaccatgctggccactcagtggtggtgcccatggagtttcgggctcagactttctgtgagatacctcaaaattgagagaagctcaaaaggatcatgcagcaaaaatgggagaagtcaccttcttagtcataagaggattatttctgattagtccttgagctgtgttatgtcctggggaagggacagtaagagtagaaaagtctctcct**aggacaaggataggaagtggtgaaaagtccttttcct**ggtgactagcaaagcaggggatgaaaagatctggagttggagtccagggtggattccagggcagacaactgcataggactgtctcgagttctgtggattgaaaaggttgggcatcccttcagatgttcctgagctgacctcagtcagaggtgcgtggctccagcacccagaa**ggtgtgaccagcacacc**aacccctcctcccaggtgggccgtgacagcttccctcaatccctcccactctgacaacagtctctctgattgcttttgctttggtgctc**tggacataaactgttcctactttgtcca**agtgaggtatataattttcagaacgctggagaattattatctaaaacaccagtggatacgctacaattttggcattcagtattctgggcttgtgcaggaaaac**tctttctttagaacatgtagaaaga**ctatcctttagttatccagtgtggaaacagtcctgaaatccaaactgatactgcgatcctgtgccaggttattactgtgacaattacttaatattctctgtgctcacgatatcactccttccagaagatctttagggaaaacttagttcagtttctaggaggtttttgctcagccgaagatcactgtgTGAATAATAATGCAGGCAACATGCTCACCTTTGGAGGGGGAACAAGGTTAATGGTCAAACCCCgtgagtatctctgctgaatccataatgaatgctctaatttcaaaaggaagccgtagcactgagctctgttttctgttttctctacttaattttatcttttatattaattctaatggttacatgttatcg**atacatggcatatgtat**atatgaaaatggacatacatgtaactgtctggcgtgggaacaaatgtgctggagaaaaacagatttcttgtcctttttgtcttaaccatttgg**gtcagaaggtctgac**tgagtagaacatgggctgggtgagcactgcagtgggggaatccttct**ccagcagctttgtcagaattatgctctttgctgg**cctggtcagaaggcagctttgtcaggag**gtggaaggatgaactagttaacctcagaagagcctttccac**tccgaccatgctgggaatagaccgccataataaacagtcttttccgctgcctctttagccaacgtcccagctggaggagcacgccggccagatttggggattgaaagggatttcctgatacacattggcctggtcggttttggtaaagctttctatgactgtgtaatgctggcaacaaccgtaagctgatttggggattgggaacaagcctggcagtaaatccgagtgagtcttcgtgttaactctgtcaaacctgtctgt**gcagtttgaaatatctcagaaactgc**ctccttcttttttagccctgaggtttaagatatctagttatggctagtgttttcttcctgtccaccaaagctgaacccttgcctacccctcaatcaaaataactagatagagggatccatatgagcagggtaggaaggccattggagtgttcagctgttttcttagagagcaaatacattcgtgcattccaaaaaggcctgtgtgggagcactgaatattacagaaggtgagtgtgtagcacatttcatggccgctttagtctgcactatgccaacctcaccgtgggccattggtaaatactgttgctt

Uppercase: TRAJ39

Lowercase: Flanking sequence[1000bp]

Red & Bold & Underline: Stem-loop [9]

Blue: Heptamer[30]

Green: Nonamer [3]

id-TRDJ4[J_gene_segment]

tagtgcaat**tacattccccacgcttctagtaaaatgta**tcttctctgagtttagccctaataaacattctaataattacatcgagtctctctagttcacagatgctgtatctaatcttataacctataatacggaagatgttatc**ctcacttcgtagagaaaccagataaagctcagtgag**tttaaggaacttggccaagattacacacttggaaagggacagagt**caaaactaggttttg**gatcttctgacatccttctactttacaacgactaggaaccacccttcatgttagtgattctccaactatcatctacagtcttttaaggattgtaaactgatccgaaaaatccatattccctccttggtttt**tatgagtgacatacaggcaatctcata**gattctgatgttcacgtataacagaaattaatttaactgtcacttctaagtgtgagagtgtcattgaagcttgctaaaatttgatatgcatgaatgtgttattaaaatggatggaaaaacttaattctattttgtactaagcagtttggctgctgcacaatacttggcaatgatgcaaaactctcagatatgaaataaatttcccctaaaaggttggaatgcatggtatatataacctgctaggtccctgcctcacctaagtgtagcccgaa**ggtggagaaccactgctctatgaccaacgttccacc**cttctttcctcttgatcaaaccaaaaaacatctttccagtctatattcattctctcttatttgagcacaaattctgtcagattgtaatatgggcaaaagcttggatttctatttgatgctagagtcacctcttgttccatcctgaaggacaacctctaagacggaggagaagaggctaggccagatgggacagaaaa**tcacatgggatgaatatcatgtga**ctt**ggcaactggggctagggttgcc**gtaaagggaaaagcaaggtagtttttggattaggtatgtcagctgtgCCAGACCCCTGATCTTTGGCAAAGGAACCTATCTGGAGGTACAACAACgtaagtaacctacttttccttcttatgaccactatgtttggcaattacatttgttaaatatttttctaagtcaccaggttaggaagcagtgctccagtaggtaatatagcaagggatggcttgaggaacaggaactgagtttccgataaaatccaggatggcgtctgtgcttcagtgagccgtcttgatccattgtccttcctagagagtttctattccctctacctggctccagaaagctaccctcctcaacatctatgcatccaccatgtacagatctaggccaccactgtgatcagactcacatgacagcacttcggggggtggacacaaggtgaaatgaggagcacgctgaaccagtaaatcagaggccaagttccaatctgtcc**aatgcattaactgacaatgcatt**agctgacaaactctgtgacttaactgcactg**aactatagtttccctatagtt**agatgaaggatttccaagctgtcttccagcccttgcccactatgatcctgattatccccagaccaattcttatgaaca**attgggaggaaaagcaaggaagcccaat**ggtgtgactgtcctattacccagtaggaattctgtgatttaatggaaaaaaaaaaaaaggcactggaa**agggagtgaaaactctccct**atgcatcccttaattagcccttgtgtgatcacagaggtttggaaaaatgatgccttgaggctcctttcagtcctaccatctattactctggttcaggggctggaaactggaaat**tgagtgaagcccaactcagccactca**cttaaccacctttacagtgcctttgtctgtgagatggatctaaaaatatcacctcatagagtttatttcaataactttgttgaagtagta**taaacaccagctgttta**agtaggaaagtatttatacaaatcatagtcatctgtgattattatagtcctttggatagaccccaagtcttttgcatcaatggctcttgctccaa

Uppercase: TRDJ4

Lowercase: Flanking sequence[1000bp]

Red & Bold & Underline: Stem-loop [13]

Blue: Heptamer[24]

Green: Nonamer [1]

id-IGHJ2-2[J_gene_segment]

ttagaccaccttgcac**cttccctggcacccaccatgggaag**ag**ctgagactcactgaggaccagctgaggctcag**agaagggacccagcactggtggacacgcagggagcccacgccagggcgccgtggtgagt**gaggcccagtgccacccactgaggcctc**ccgttcagtgggacgacggtgaacaggtggaaccaaccaggcaacccccgccgggccccacagacgggatcaga**gcaggaaaggcttcctgc**ccctgcaggccagcgaggagccc**tggcgggggccatggccctccaggcgaggaggctcccctggccaccgcca**cccgggcctctctgctgctgggaaaacaagtcagaaagcaagtggatgagaggtggcgtgacagacccagcttcagatctgctctaatttacaaaagaaaaggaaaaacacacttggcagccttcagcactctaatgattcttaacagcagcaaattattggcacaagactccagagtgactggcagggttgagggctgggg**tctcccgcgtgttttggggctaacagcggaagggaga**gcactggcaaaggtgctgggggcccctggacccgacccgccctggagaccgcagccacatcagcccccagccccacaggccccctaccagccgcagggttttggctgagctgagaac**cactgtgctaactggggacacagtg**attggcagct**ctacaaaaaccatgctcccccgggaccccgggctgtgggtttctgtag**cccctggctcagggctgactcaccgtggctgaatacttccagcactggggccagggcaccctggtcaccgtctcctcaggtgagtctgctgtctggggatagcggggagccaggtgtactgggccaggcaagggctttggcttcagacttggggacaggtgctcagcaaaggaggtcggcaggagggcggagggtgtgtttttgtatgggagaagcaggagggcagaggctgtgCTACTGGTACTTCGATCTCTGGGGCCGTGGCACCCTGGTCACTGTCTCCTCAGGTgagtcccactgcagccccctcccagtcttctctgtccaggcac**caggccaggtatctggggtctgcagccggcctgggtctggcctg**aggccacaccagctgccatccctggggtctccgccatgggctgcatgccagagccctgctgtcacttagccctggggcca**gctggagcccccaaggacaggcagggaccccgctgggcttcagc**cccgtcagggaccctccacaggtagcaagcaggccgagggcagggacgggaaggagaagttgtgggcagagcctgggctggggctgggcgctggctgttcatgtgccggggaccaggcctgcgctttagtgtggctaca**agtgcttggagcact**gggg**ccagggcagcccggccaccgtctccctgg**gaacgtcacccctccctgcctgggtctcagcccgggggtctgtgtggctggggacagggacgccggctgcctctgctctgtgcttgggccatgtgacccattcgagtg**tcctgcacgggcacaggtttatgtctgggcaggaacagggactgtgtccctgt**gtgatgcttttgatatctggggccaagggacaatggtcaccgtctcttcaggtaagatggctttccttctgcctcctttctctggg**cccagcgtcctctgtcctggagctggg**agataatgtccgggggctccttggtctgcgctgggccatgtggggccctccggggctccttctccggctgtttgggaccacgttcagcagaaggcctttctttgggaactgg**gactctgctgctggggcaaagggtgggcagagtc**atgcttgtgctggggacaaaatgaccttgggacacggggctggctgccacggccggcccgggacagtcggagagtcaggtttttgtgcaccccttaatggggcctcccacaatgtgactactttgactactggggccagggaaccctggtcaccgtctcctcaggtgagtcctcacaacctc

Uppercase: IGHJ2-2

Lowercase: Flanking sequence[1000bp]

Red & Bold & Underline: Stem-loop [16]

Blue: Heptamer[38]

Green: Nonamer [7]

id-TRAJ5[J_gene_segment]

ttgttgcctgggagtaactatcagtgtcaggtctgatgagatgattgaggctgtgccggacaaggg**ttttgcacaatgatttcagaggacaaatccccaagttgtgaaaaa**agacttcactcttggttaggtttctaaaacagaactttcttcttggcaaccaaggggtctactctgccccctcactcctatgtctcttccacctgagactctgtcaccacctcccctagaatccgtgagataccttcccctgaattagagcataccagccagggtgctgaggcactctgtggtactggaatggatggtacagggatgcgtttctg**ggcttctggctgcaagaagcc**tcatcctttcccccagtgtaaagcattgctgggaatcatcccattatgagctactatttactgaatgcctgacacgtaccagtggccattctcttgtctttta**tgtaatcctcattcagtgattaca**cacatcctaataataggtagccgatattttgcccattttacaggtgaggaaacagagg**cttgggataagtaacttatccaag**attatatagccacgaatgacagag**ctagagtttaactcaaatctgtttgactctag**ggccttgttattaaccgtgccaccaccatgcctcctgcattc**tgtgaaggaagacttcaca**cct**caaaggccatctgtttggcctttg**gtttgcccatcctgagccttcttaccatagcgaccatggcatgg**tgactcagcacttctgaagagatcagcagagtca**ttgggttgggtggcagaatacaggtatggcagggagggaaggagaaacttagggggactgtttattgcgcatacagttagagagaaagaaaaatgtatcaacaaaccatagaataggctgcttaaaagtttttctcccaagaaactgtgagtgtatgagggtcatggg**cagctgcccccagctg**gcagagggcaggattttgtactgtgatgtaccagggtgTGGACACGGGCAGGAGA**GCACTTACTTTTGGGAGTGGAACAAGACTCCAAGTGC**AACCAAgtaagtacccaaacttaggctctggccaaagacac**agaaagcccctactctgctttct**tagatgatgaatctgtacttcctgaactaatttttcatgtttctttatgaggcttgaatatcttgaaattttaattcctagccactagcttaatagtctgttttctaacggtagtattt**gccaatgggaattggc**catttcatcattgcagagacagattcctttgagacaaaagatctttatcagaaaagaaactgtgactgattatgccaaaagatttatttttgttccatttagttttcatgaaaggtgatgaattatgcttcccaaagactctgggcattggacacactt**tctcaaaactgcacatccaattgaga**gatataggaagaacttagagcctaaataatgccataaaaataataaaaagtttgtaaaaaggaaccatatagaaattaaagaagaaatgaaatagtcttaatgaaagaagataacagaactgcattagaaacgaatgg**aaatatttagagggtttatgtattcaacaaatattt**attggacaattattttattccaagcactatgttaagtgtgaggaagggaatgggagggaagggaagggaaggaaaggaaagaaaggagaggaagtagaggaaggaccgggagggagggaagaaggaagaaaaggaggaaaggagggaaggagggaggttctctttcatccagggaaggagggagagtctctgccatccgggagctcagagtgcaaacagacaactatagcctattgtcaccacgtcatagtggaggtacatgcaagttgcaatggaagcacaacagaagacatgactagcctccaattctcattgcctggtacatacacacatcctccactataccttaagctccatgagggcaggaccctttttcatcctcacagctctacagctcagcacaatgcctggcacataggtgtcaccaaacaaatgattgtggggaaaa

Uppercase: TRAJ5

Lowercase: Flanking sequence[1000bp]

Red & Bold & Underline: Stem-loop [14]

Blue: Heptamer[27]

Green: Nonamer [3]

id-IGHJ3P[J_gene_segment]

ttctccggctgtttgggaccacgttcagcagaaggcctttctttgggaactgg**gactctgctgctggggcaaagggtgggcagagtc**atgcttgtgctggggacaaaatgaccttgggacacggggctggctgccacggccggcccgggacagtcggagagtcaggtttttgtgcaccccttaatggggcctcccacaatgtgactactttgactactggggccagggaaccctggtcaccgtctcctcaggtgagtcctcacaacctctctcctgctttaactctgaagggttttgctgcatttttggggggaaataagcgtgctgggtctcctgccaaga**gagccccggagcagcctggggggctcaggaggatgccctgag**gcaacagcggccacacagacgaggggcaa**gggctccagatgctccttcctcctgagccc**agcagcacgggtctctctgtgg**ccagggccaccctgg**gcctctggggtccaatgtccaacaacc**cccgggccctccccggg**ctcagtctgagagggtcccagggacttagcgggg**tgccagttcttgcctggggtcctggca**tt**gttgtcacaatgtgacaac**tggttcgacccctggggccagggaaccctggtcaccgtctcctcaggtgagtcctcaccaccccctct**ctgagtccacttagggagactcag**cttgccagggtctcagggtcagagtcttggaggcattttggaggtcaggaaagaaa**gctggggagagggacccttcgaatgggaacccagc**ctgtcctccccaagtccggccacagatgtcggcagctggggggctccttcggctggtctggggtgacctctctccgcttcacctggagcattctcaggggctgtcgtgatgattgcgtggtgggactctgtcccgctccaa**ggcacccgctctctgggacgggtgcc**ccccggggtttttggactcctgggggtgacttagcagccgtctgCTTGCAGTTGGACTTCCCAGGCCGACAGTGG**TCTGGCTTCTGAGGGGTCAGgccaga**atgtggggtacgtgggaggccagcagagggttccatgagaagggcaggacagggccacggacagtcagcttccatgtgacgcccggagacagaaggtctctgggtggctgggtttttgtggggtgaggatggacattctgccattgtgattactactactactactacatggacgtctggggcaaagggaccacggtcaccgtctcctcaggtaagaatggccactctagggcctttgttttctgctactgcctgtggggtttcctgagcattgcaggttggt**cctcggggcatgttccgagg**ggacctgggcggactggccaggaggggatgggcactggggtgccttgaggatctgggagcctctgtgg**attttccgatgcctttggaaaat**gggactcaggttgggtgcgtctgatggagtaactgagcctgggggcttggggagccacatttggacgagatgcctgaacaaaccaggggtcttagtgatggctgaggaatgtgtctcaggagcggtgtctgtaggactgcaaga**tcgctgcacagcagcga**atcgtgaaatattttctttagaattatgaggtgcgctgtgtgtcaacctgcatcttaaattctttat**tggctggaaagagaactgtcggagtgggtgaatccagcca**ggagggacgcgtagccccggtcttgatgagagcagggttgggggcaggggtagcccagaaacggtggctgcc**gtcctgacaggggcttagggaggctccaggac**ctcagtgccttgaagctggtttccatgagaaaaggattgtttatcttaggaggcatgcttactgttaaaagacaggatatgtttgaagtggcttctgagaaaaatggttaagaaaattatg**acttaaaaatgtgagagattttcaagt**atattaatttttttaactgtccaagtatttgaaattcttatcatttgattaacacccatgagtgatatgtgtctggaatt

Uppercase: IGHJ3P

Lowercase: Flanking sequence[1000bp]

Red & Bold & Underline: Stem-loop [18]

Blue: Heptamer[24]

Green: Nonamer [5]

id-TRAJ53[J_gene_segment]

ttaaaaaaaaattaaaattctgtggggtgaaaacaaagatgttaaatatttgattggcaaggcaactggaaaatctggaccatgtctacaactgctaaaggaggctttgtgaaagagaaaatgagcagcccaaggagatcctgtcctaaacttctctggccagtgaaattcgggccattc**tctgccacagccctggactgctaggagggcaga**tcatatgtcttcctcagtggggagaggtgggccctcgctggcagtttctgtaaagcctcgtgctgtggtgtaattcagggagcccagaagctggtatttggccaaggaaccaggctgactatcaacccaagtaagtatgacagggtgaagctacatgcagctgagtacagtcttttcctttctagaccgtgtcctgcaagctctccttgagggacgtatactcattttgcattgtcctttgtagagaagcagaccaggaaagacaggaaagccctcaaatttccacttttaaacacctccctgtaa**aaactgtctcgcttccctccccttctacaatcagttt**ctagtaaatcagaatccggtgaattgatatgcaatttcaacgaaaaaaaaagcagagaaatagttaccccaacaagtgcaaaaagtagaaactatctgagtaccagccaagggaagataattaggaaagaaaaaaaaagaaagaaatggattactggaaccatgatggcagcttagtcacaaagaaaatgataaaaaaaaattaaaattaaatttcaaaaactttaaag**atatatacatatatatatatctatatatatatgatatat**tgtcctctggaactccagcactcacctacagcagctgcattctggacttattatcaaataacctataaacctgtagcttaatgaagagttgagcaatcgtgactatattgacccccaggacccttctgcaaagagcagcttctgttcctgtttctgtaaagccttctgtggctgtgAGAATAGTGGAGGTAGCAACTATAAACTGACATTTGGAAAAGGAACTCTCTTAACCGTGAATCCAAgtaa**gtttgaagggagtgggggaagggggaattcaaac**acttctgatttaattacttgcccctcca**aaacattccagcttaggttccaaatctaatgttt**gtgctggggggatatggtgcccatcagagggattagaactccactgtacaaagactaaattactttcagcccaaaacattcctatcattcatcttaaacaagattattggtgcaaggagaaaaactgaatacagaa**tcctgagggatccagctaagggtcagga**aatgtttgccgaaaagataaagaggaaattggcatatcgtgactacgcaaatgcagtagaccaaatataaattgtcctgacttagatactaacactcctcttaaatttatcgcctgaaataatggaaccctgcaacttcattaactaatgctgaaaattctataagcaggctccttttc**ttcttttaccttcactttacctctctgaaaagaa**ctgttctcaatccccagtctggtaactgaggtccagtc**agttaactgtctggttaact**caggcagagcatttaaccccactttttctttggttaccaaaat**agtggattcagcctccact**ttacttagttcacctgagaaggggaaaattcatctgtcctctgagtcgt**gagaatagagcagtacattctc**tccttcaaatatacacccacccttccctaatggtaatgtgtgagtacttagaaaagtttttaaatgttagatataaaagtaaggcacagtggttgctaatggcacttcttgaaaatctgcgctgctggtggacttctgaaagtggtgcttataaactaaggctggagaagactgacttgtaaaattcagataaaatgaaaatgggaaaaacttatcaaagggccccggtctcctaagtgccctgatgt**tgagtaaaggccaaatgccccagttttacttactca**gatctgctttctgtgattaagaagagatgcacactagagagaggcact

Uppercase: TRAJ53

Lowercase: Flanking sequence[1000bp]

Red & Bold & Underline: Stem-loop [11]

Blue: Heptamer[17]

Green: Nonamer [4]

id-TRAJ31[J_gene_segment]

atccacgtatctgtcactctatttcctacgtttgcttcccttttgttttctctagaataagttctttttgtttgctctggacacccactgctcccaggatgaaaggagagaaatgagatcagttttgaacacttcctcttgaaatataaagaatcaacaagttacagtcatgttggggacttcttctctctccaagcttaaatttctattacgtaagccttacttttaaccaagaaagttagtgacatggtatcactgtctgggatgctcacccctctctcttttctccagatactccgtgtatact**aaagctcagccaatgctcagcttt**gt**gataaaggagctgttgtgtgtctttatc**tctgcctttaggatacagtcttttttataacagaaagtgttcacttctcccatttatttcatggctggactctctaccccttcctggctgaaacagcatatgggtccatggtggaacttctttgggaaaaaaaaaaaatagaactgtttgacttttgccactcttccctcatgaaatgttttttaggctatcaaataagcattattacacattgagaaggctttgaaaaaacttgatcactgagtctggaggtgagctgggatggctctgtagctggaaaatatccctgtgggtgtaggaatgtaccgtcccttcaaact**tgtgtgtttagggctgagctcacaca**tacgacagcaagagttgttaacaaattagattcagctttggcagttttatgcctag**aaatgcaaaagaattttaaaagctcataagtttggcattt**ttaaaggagacattgtttgctttatgttgcagagccaatttttcttcctatcataaaaggccagaaacacagagaaagcaaaaaaaaaaaaggaaaaaaaaaaaaaacacaaggagagtctaactgccccctcaccacagtgctatgtgtttggtagggt**tttactatgggtttcagtaaa**ggcaggaagtgctgtgGGAATAACAATGCCAGACTCATGTTTGGAGATGGAACTCAGCTGGTGGTGAAGCCCAgtaagtggccatgttttattga**tatttgaccaaacaaata**aatcccgtgaagttagtgg**agatttaatttaatatgtaaacaaatct**acttcttgaaaaatgacttttgtgaatacatgaaacactcgtg**attattgagactagagcaataat**gataaa**taaaggcaaaatgagaccaccccaaatcaccttta**acattgtggcttagccctctttcttctag**agagatatggggccatgagctatttacagccgcagtatctct**taaactatgttctgcaatgacttgtgtgtttactctgattttagccacagaatttgttttccattct**attaaaaatgagaatctatctcattttaaat**gctactaatatttata**aggtaccttctttgtctcttgtacct**gcatatatattcatttcaaaaagatctcataagaaacccaaacattgtcgcat**tttgacaaatgtccacatctagtcccgtaatgtcaaa**tttcactaaaaacacagcctgactatgcaaatgcagtggaagaaaaaaacaattcagaaatggattgacactcccaggtggctttataaacacctaaaaggaaatccagagtgaaataaaatggcaggtatgataacacgatttttttttaatgatcaattcctccatggggaataaaagtttttaaac**ttacctccttgaaaaggagggaa**ataggagtt**caggagcctggcattcccaggtgtgctcctg**gggtgtctgtt**tctccacattgccattttggaga**caataggacccatccaaatgtccaaccaaaaaactaatattaagaggctggagtcaccacattgactgt**ctgttattaaaccctataacag**gttataattaagattatggaatgactaacccaaggagc**tttaaaataggatagcctagttttaaa**cggttcagccttgggaaatgctggaagacaaggaaatgagatagatgactccataggtttgtt

Uppercase: TRAJ31

Lowercase: Flanking sequence[1000bp]

Red & Bold & Underline: Stem-loop [18]

Blue: Heptamer[30]

Green: Nonamer [1]

id-TRAJ4[J_gene_segment]

tataccttaagctccatgagggcaggaccctttttcatcctcacagctctacagctcagcacaatgcctggcacataggtgtcaccaaacaaatgattgtggggaaaaaaatgccttctggctagatgagttatgatttatacattaacaaatattc**tctaacagagagagttgttaga**cctcaaaaccaggagattgcagaatccatccagatgtctgtggtgggagaaggcccatctctcagggaagagcacctctgtatttatctgttcatatgtgtgcttcatcaaagcctcttc**tcagccatggttcaggctga**atttataaagcttgaaacctatcgtatttctcaagaatcagagatcagaggtggaaggtttacaatatctcgataagtaagcagtacaaatgagttctagcagtaaaaaagaaagaggtctaatgataaatatatgaagcctgtctccacttgagagacctcct**agaaaagatttgttctaaggaaacttttct**ccattaggagcagaatcat**cttggaaatatacagctccaag**ttgtgttagaaaggtgtgtacaaagaagaccaaaaattcacatttggaataaggcattaaagttgccattgtttctagtagggaaaccctgcaaagtgttggggcatcgtagatcctgcatgttgacgcctgcaactaatctgccccggagctgttccgacagtggctggagcatttgctgcacagttgcataggtgttcaacaattgccaaaataaaatgaagctatttcgtga**tgattagaaagaaataatcagcatcaatgaggcctctgatgc**a**tcttcaggagtgagtaggtggaatgaaga**cctggtaggtgagact**tttctctcttgtcagagaaa**gtttgacttgggggagcctggaaaacagggacaggagatggagtgccccagccaggcatgccctgcccaggtcagagttcttgtaaagcaccatctgattgtgTGTTTTCTGGTGGCTACAATAAGCTGATTTTTGGAGCAGGGACCAGGCTGGCTGTACACCCATgtgagtatgaccctg**caagtgaccagtgcaaaaagaactgaccacttg**tctatgagaaaacaggtgatgatctatagcaaacttggggatatattgagaagcactcattttccattcctacaagctctcttggtgacttaaaatgttcccatttctcatatgaaacacagaagtgccaagagccaattgcgtaaatgaaaatctgagcagaataatttatagaaacataaaaagagttccaaataatgagtgctttcaagtgaagctgataatatcggcaccagataatctgaacattcaggaaattagctaagtgcccttggggg**aaagagatgagttaatggcaaacacaaattctcttt**tattgcagcatgtagctaaattccctggattggctaataaacatggtatggggatccctactcatagaatcttcctctctggag**ttaggagagttttcagcttcctcctaa**gggtaaaagtggcccagacccctaactgaacggcaag**tataccacatgcaattacaaggtata**gaggcagcaatgtagagggaggcccacagacctggggaaaatctgggctctactctcactggctgttttcaataaagcctccactttctcagcagcaaaatgactataactacatcataataatatgagataatggccagaatgtaaaaagtgatcaataagtagtagcttttctaaaatattattataattattattattgaggctttgcttatactaaggtttacactatcataaagggaaaggagaatc**gcttccttgggaaataaaagatgtaagcaacaaagacccaaggagc**cttttagaatgttgctctgggaccccagtggcttccagcagattagcatcaagaaggttatctcaaagaccttacccacagtgggggtacagcagtgcttccaagataatctttggatcagggaccagactcagcatccggccaagtaagtagaatgaagcaggag

Uppercase: TRAJ4

Lowercase: Flanking sequence[1000bp]

Red & Bold & Underline: Stem-loop [13]

Blue: Heptamer[10]

Green: Nonamer [2]

id-TRGJ1[J_gene_segment]

aagttgtgctgctgattagaaaaaggc**tcacttgttttgaaaaacacaagtga**agaattcccatacactgactgcatatattagactttagccaatgttc**cctttgactttcttaacacggtaggtaaggcaaagg**ataacctattaaagattgggcgtgtg**aaagccatgtttcttgtgatgatggggacgatggcttt**gagaatcccagagcaaagtggaatgcaaaca**gaggaactgagaaattattcctc**ctgcttaattgctatggatttaactgccactccaaaatgctgaattttttttagtaagggcaatgcttggtcctatagggttaaaatgtcatgtcaaggcacacaatca**tagcaaacagattgcta**atcataacaatgacatcatcatcatcataatctcttatatttccatagcatttatttttgaggcagggtcttcccctgttgcccaggcaggag**tgcagtggtgtgatcaattctcactgca**gcctcgaactcctaggctcaagtgaccctcctgcctcagcttctcgag**tagctgggcctacagtcgtgcacatcatgctcagcta**atgctttttgtatttttagtgaatatggggtctcactgtgttgcccaggctggcctcaaaattctgagctcaagcaatcctcccacctcagcctcccaaagtgctggaa**ttataggcacaaaccactgcactgggacctataa**cattttaatcaaattgcttttttatatcttgtttcattttggtctcacttcagtgctggtagc**gatgttgaactgattttgaaacatc**ac**tgtttttagacaaataaaaca**ccaaaagctttaagttatttgatttgtggagcaacagaacttgttatgagcaaaatga**accaggactggaaccctggtcttttgagaatcccagacca**ccagaatttgaagaactcagggaaactgaattagagtttttgatatggactgaatcactgtgGAATTATTATAAGAAACTCTTTGGCAGTGGAACAACACT**TGTTGTCACAGgtaagtatcggaagaatacaacatttccaaggtaatagagggaaagcaggaaattattaaactggaataatgtaataat**gtttagaaaaaagaggaattggatggggatttgatgtagaaaacttaggagagactttaaaacaaacgctcatactaaaagagaacatagataacat**tgcacagatatcataataagatttggctttgtgca**tatacggacttccataaagggctcatatgtaatgtatgaaatgatctcattaaaatgtctggccctgagaccaaatgtattatgacaggtaagtttgtgatagactctacagagtggacacatattcatctctgatggtcaaagacatgtttactcttgttgtgaaggagcccgggtgttggagtcaggtccacctggaactctggctccac**cactcactgcttagtgaaattgagtg**attattctctggcctcagttttctaatccataaaatgggataacagtatgaattcagcagggttgtataagaattgcacaacatagtgtgaattaagtacttggcacattgtccaacccaaaataggtgcccaacaaatgttttctggattcacatgtaaagagacaatgggatctacta**tggagagttgctctcagtccatctaatttacagagcagcaattctcca**gaggattcagctgtacttctaactgctcaaatggaaggttatcaacatcagctcacacacgcaaaaattgaacttatggattcgtttcttgttagcaggctttttaatcacgtggc**aaaaacattgtattaagatgtctgttttt**tatttttgttgttctatgtgctttacttaatcctttatcttaattgatatcatttctaacaccaacatattggtcctaagatttatagccaattagtttagggttctgttcatatgtctcaggaaaaaaaggttaaaatcttaccaaaaatggtcaagatatcaaatcaattaaatcc

Uppercase: TRGJ1

Lowercase: Flanking sequence[1000bp]

Red & Bold & Underline: Stem-loop [19]

Blue: Heptamer[21]

Green: Nonamer [3]

id-TRDJ2[J_gene_segment]

gcattaactgacaatgcattagctgacaaactctgtgacttaactgcactg**aactatagtttccctatagtt**agatgaaggatttccaagctgtcttccagcccttgcccactatgatcctgattatccccagaccaattcttatgaaca**attgggaggaaaagcaaggaagcccaat**ggtgtgactgtcctattacccagtaggaattctgtgatttaatggaaaaaaaaaaaaaggcactggaa**agggagtgaaaactctccct**atgcatcccttaattagcccttgtgtgatcacagaggtttggaaaaatgatgccttgaggctcctttcagtcctaccatctattactctggttcaggggctggaaactggaaat**tgagtgaagcccaactcagccactca**cttaaccacctttacagtgcctttgtctgtgagatggatctaaaaatatcacctcatagagtttatttcaataactttgttgaagtagta**taaacaccagctgttta**agtaggaaagtatttatacaaatcatagtcatctgtgattattatagtcctttggatagaccccaagtcttttgcat**caatggctcttgctccaaccattg**tgtccttccacctctcgctgaggcacagtcctgttgtgattgatttagttgaggcttgttgttacgtttctgcctacaacggccatttggcaattctcagcaaaatatgtgaaaatgttatcctgaagcaagatctgggagtaagagatggcaggttgaggagataatagcagaaaatctaaggacttgcctaagtgtcaaaactcagaagatagcagacatcaaaaaagctgccaaaatataatgctcagatctcagaaacccttgcaaagctctgtctccagctacacccattctgctccacccacccacttccaaacagatgaattagactgacagggagcaaataggtagcaaggtttttcgtaatgatgcctgtggtagtgCTTTGACAGCACAACTCTTCTTTGGAAAGGGAACACAACTCATCGTGGAACCAGgtaagttatgcattttactacagctcagtgtgtatatttctttagggttgttgtagcgtagtcctgaatgaatggggcttcttgggaagtaaagagttccaaattagaaacccat**tgtttgttgcaaaca**gtaggggtaaaatttgggagcaaatgcattagatttctcttaagaaaatattctgtatatgcttggaaactccattctgaggcttggtattcataag**ttttttacaactactgaaaaaaagccctaggtcaggctttt**agttttgggctccctagaaccaggaaaaaggcaagctgtggttgaggagggtctctccagtatttatgagggataaaagtcatggtcacagtgtaggaagagggaaggttagaaagaaaaaagtattacgaaagaaggagttttaaaa**agaggatctttctctcctct**ctctctctctctctctctctctctctctggataagaagccaatcttgcccttgtcacttttt**tctgaacttgcattaaaccaacataaatggttcaga**ttaaatggttctcaaggtcccatcctggcctgggtctgctgagagcagttatgagctcacctctcctgttagc**cctgaaactgggaactttgcacaatggttcagg**agcctgtatccaatgtagaaacacagccaccaa**ctccaggacaatagagaagtcgtggtcccctactggagacattgtctactggtagcctactccaatgt**ttggaaatccaggggctatattctggtagtttgggtagctgccaacagtcaatcactaagaccagtccagagaatagccttcagacaccttccaaaatgccaagtccattgggagagccattacctcttacatgattaggtattgattttg**tttattcagaggaaataaa**taagtagggaatggtacatgaatgtggctaggctatctgaccaaaagtccaaaaagtgactggtgctgtcaacatggtctctgcctctc

Uppercase: TRDJ2

Lowercase: Flanking sequence[1000bp]

Red & Bold & Underline: Stem-loop [15]

Blue: Heptamer[24]

Green: Nonamer [3]

id-IGHJ1-2[J_gene_segment]

cacctcccctgcccccagacaccctgtgcccgtcagttcatccccagcagaggccctcaccaggcacacccccatgctcacacctggccgcagg**cctcagcctccctgagg**gccccacccagcccgcgtctggccagtggtgcgtgcaa**agcccctcacccagactcggcggaaggcagccagtgcaggcctggggaggggct**ctccttagaccaccttgcac**cttccctggcacccaccatgggaag**ag**ctgagactcactgaggaccagctgaggctcag**agaagggacccagcactggtggacacgcagggagcccacgccagggcgccgtggtgagt**gaggcccagtgccacccactgaggcctc**ccgttcagtgggacgacggtgaacaggtggaaccaaccaggcaacccccgccgggccccacagacgggatcaga**gcaggaaaggcttcctgc**ccctgcaggccagcgaggagccc**tggcgggggccatggccctccaggcgaggaggctcccctggccaccgcca**cccgggcctctctgctgctgggaaaacaagtcagaaagcaagtggatgagaggtggcgtgacagacccagcttcagatctgctctaatttacaaaagaaaaggaaaaacacacttggcagccttcagcactctaatgattcttaacagcagcaaattattggcacaagactccagagtgactggcagggttgagggctgggg**tctcccgcgtgttttggggctaacagcggaagggaga**gcactggcaaaggtgctgggggcccctggacccgacccgccctggagaccgcagccacatcagcccccagccccacaggccccctaccagccgcagggttttggctgagctgagaac**cactgtgctaactggggacacagtg**attggcagct**ctacaaaaaccatgctcccccgggaccccgggctgtgggtttctgtag**cccctggctcagggctgactcaccgtgGCTGAATACTTCCAGCACTGGGGCCAGGGCACCCTGGTCACCGTCTCCTCAGGTgagtctgctgtctggggatagcggggagccaggtgtactgggccaggcaagggctttggcttcagacttggggacaggtgctcagcaaaggaggtcggcaggagggcggagggtgtgtttttgtatgggagaagcaggagggcagaggctgtgctactggtacttcgatctctggggccgtggcaccctggtcactgtctcctcaggtgagtcccactgcagccccctcccagtcttctctgtccaggcac**caggccaggtatctggggtctgcagccggcctgggtctggcctg**aggccacaccagctgccatccctggggtctccgccatgggctgcatgccagagccctgctgtcacttagccctggggcca**gctggagcccccaaggacaggcagggaccccgctgggcttcagc**cccgtcagggaccctccacaggtagcaagcaggccgagggcagggacgggaaggagaagttgtgggcagagcctgggctggggctgggcgctggctgttcatgtgccggggaccaggcctgcgctttagtgtggctaca**agtgcttggagcact**gggg**ccagggcagcccggccaccgtctccctgg**gaacgtcacccctccctgcctgggtctcagcccgggggtctgtgtggctggggacagggacgccggctgcctctgctctgtgcttgggccatgtgacccattcgagtg**tcctgcacgggcacaggtttatgtctgggcaggaacagggactgtgtccctgt**gtgatgcttttgatatctggggccaagggacaatggtcaccgtctcttcaggtaagatggctttccttctgcctcctttctctggg**cccagcgtcctctgtcctggagctggg**agataatgtccgggggctccttggtctgcgctgggccatgtggggccctccggggct**ccttctccggctgtttgggaccacgttcagcagaagg**cctttctttgggaactgggactctgc

Uppercase: IGHJ1-2

Lowercase: Flanking sequence[1000bp]

Red & Bold & Underline: Stem-loop [18]

Blue: Heptamer[34]

Green: Nonamer [6]

id-TRAJ19[J_gene_segment]

ccctagcccactctcttaaccatggtcttggtgaggtttgtgtagggcgacctcgc**actgtggttctaacgactacaagctcagctttggagccggaaccacagt**aactgtaagagcaagtaagtaagaaagaaaagtccagaataattttaagcaaaatggtgggtaggtttttcagcaatttcacctaggaagtgcaatgtcaagaactaaa**ttctaagagctttcccagtttgtgttagaa**ccataatttttctacctcacacgtctccctcgccttctctgtcacctcagaacagctcctcctaaggcatgacttcacaatggtacatttgttggtggcgcagtctttgtgg**tcagataaaaactgagctaattatctga**aattatctcaggtctctaagtgagagagttaactcctttcaaatatttgaagaactacgatgtagaccaggtaaggggtgagtgtggaggccaaatcaaaacgagcaggtgcaagatgtgccccaacctcccctcaaacattgttgaacaagtggaactgccctgcagtggaaaggctgctttgtgaagtatggaacttttctt**ccttagatatcttcatgccagacccctaagg**ttaaccatgagatgttgcagaaggcacttctatgccattttagcta**ttagattgaatcacctctgggctccctttcaaatctaa**atgctaagatcccatgactcaagcctggaggattaataggtgagctcagacttgtttcttctatttttactattttgatagccaaaga**aataattcatcaaattatt**cataagctaaagcctacttgggatttttacacctagaagatggtggggtatgatttcccagtgcagtaaatgagaaaacaataggagacatcaaggaggaaaaaaagaaggaagagataaagggaatgtcttaagggaggctcagaggttgaatgaaggaaatgaggtgattttgcagaggacagatgtgGCTATCAAAGATTTTACAATTTCACC**TTTGGAAAGGGATCCAAA**CATAATGTCACTCCAAgtaagtgagcagccttttgtactcgaaaatagggccaggggagcaaagtttcttccaatttaaacacactcaaaaggatgtgtaattg**ctttctgatgggagggaactctgaaggtagaaag**actattgttaccacagatcacttgtccctggagatatagcatctgaggactgactgtctagcttagactgctgtctgcacaagaagaagacttacagatgcttaaggaggatggaggcaaattttcaaccctgtactccaaagctgaggggagaggggatgggaaacattagggctgggttcatgtaaaggggaccagcattgtgccgacagaggctcaaccctggggaggctatactttggaagaggaactcagttgactgtctggcctggtgagtgagtcgctttctattccaggaaaatattactgtggagaaattaaaaggggagatgaattaactc**ctttagtcttgaaactaaag**agatatgtgtacttcccctcctggaggaccccagtccctaggtagattaggacgaggaagcagagggagacaaaggcgatggaaagtcccttttaggaacccaggaatcagagagaacttgatggtccccaccaaaggcaaagaagggaggtcccactaaaacatgtgagcatcctaaaacacctgctctggtccacccacaatatttggtgaaggacatggccctccacccagaatgggcaagcaaactagaccagctttctaaggtgtgtttttcttgggtcctgtgactcttggccgtctccctgtcagaggctccatctgtcgctgcactctttcttattgagaatggcctctccgtggttctctgtaaactttccccaagaagcctgtttccatgcttcctcagcacttagcctcaccgatggatgggctctagagtagctggagggagtctgggtctgaa**actttcactgagaaagt**aaagttgatccgcagtatccagtggatatggcag

Uppercase: TRAJ19

Lowercase: Flanking sequence[1000bp]

Red & Bold & Underline: Stem-loop [10]

Blue: Heptamer[18]

Green: Nonamer [4]

id-TRBJ1-1-2[J_gene_segment]

gtagtgatgggggctgtggcttctctataaggacatgccccaacgtgacaacagcttggagaggggtgggtactggagaagaccagccccttcgccaaacagccttacaaagacatccagctctaaggagctcaaaacatcctgaggacagtgcctggaggtgagaaggaagcccccggcctggtccataccccaccaccaacttgcataatggggggtgatgtcacccaccctccactcccctcaaaggagcagctgctctggtggtct**ctcccaggctctgggggcggacccatgggag**gggctgtttttgtacaaagctgtaa**cattgtggggacagggggccacaatg**attcaactctacgggaaacctttacaaaaacct**ctctggcggtcccaactcccagag**tcctcttctttcctcctgggtcacaggtcttaatgcaatttggttcagaatgcctctgcctcactcctgatcacatgtcagaccaagactgtggacaaggacaggcccagatgagaactaaagcttccc**aggcagagagaggtcagacataagaagactgcct**caggaacctcacaagtggaggactcagggagggtcccaatccccaaaaattgagacaaagtcaggtggaaggttcatcggaggtgaccagctctccagaggactcgggaagaagt**caggggtatctatagatggagtcacaggttctgggcccctg**ccatcctctgcaggccatgcactttccctttcgatggaccctcacagagggagcatctgaatggggca**tcctttgaaaaagga**acc**taggaccctgtggatggactctgtcattctccatggtccta**aaaagcaaaagtcaaagtgttcttctgtgtaatacccataaagcaca**ggaggagatttcttagctcactgtcctcc**atcctagccagggccctctcccctctctatgccttcaatgtgattttcaccttgacccctgtcactgtgTGAACACTGAAGCTTTCTTTGGACAAGGCACCAGACTCACAGTTGTAGgtaagacatttttcaggttcttttgcagatccgtcacagggaaaagtgggtccacagtgtcccttttagagtggctatattcttatgtgctaactatggctacaccttcggttcggggaccaggttaaccgttgtaggtaaggctgggggtctctaggaggggtgcgatgagggaggactctgtc**ctgggaaatgtcaaagagaacagagatcccag**ctcccggagccagactgagggagacgtcatgtcatgtcccgggattgagttcaggggaggctccctgtgagggcgaatccacccaggcttcccagaggctctgagcagtcacagctgagcccagggtgatggggcagaag**agggaaggggagggggcctctcctcatagttccct**gagatagcccagagaaagcccggtgggtaatgaatgagccacaacacctctccatctatctgcttcactg**acagaggttctctgt**agattcttcgtatattcctgtgctggattttataggaggccactctgtgtctctttttgtcacctgcctgagtcttgggca**agctctggaagggaacacagagtactggaagcagagct**gctgtccctgtgagggaa**gagttcccatgaactc**ccaac**ctctgcctgaatcccagctgtgctcagcagag**actggggggttttgaagtggccctgggaggctgtgctctggaaacaccatatattttggagagggaagttggctcactgttgtaggtgagtaagtcaaggctggacagctgggaacttgcaaaaaggggctggaatccagacggagcctttgtctctagtgcttaggtgaaagtgtatttttgtcaggaaggcctatgaggcagat**gaggaggggatagcctccctctcctc**tcgactattttgtagactgcctgtgccaagttaggttcccctactgagagatgggtagactcagcttggaaggggtcaccttgaacatctcctgtctcctt

Uppercase: TRBJ1-1-2

Lowercase: Flanking sequence[1000bp]

Red & Bold & Underline: Stem-loop [15]

Blue: Heptamer[49]

Green: Nonamer [7]

id-TRAJ36[J_gene_segment]

aagaaggtgatttttcataataatatactaaagtagaatccacccttgtgaccaccagaattattcaagtacagtcttccaaggacaaaatgggtagcatggaaatgat**atgtttttttggggggaaaaaacat**ggggagagagagagaaaggggactattagttgtctggattgttaggaaacatggaagacagcacttcttgttgtaaggtttggcaaaggttatcaccaagtagaagtccaaatgtgtgtaacatacccctggcaaacgttatagaagaaggaa**gctggggccgggcacggtggctcacgcctgtaatcccagc**actttgggaggccgaggcgggtggatcacgaggtcaggagttcgagaccatcctggctaacacagtgaaaccccgtctctactaaaaaaaaatacaaaaaattagc**caggcgtggcgggcgcctgtagtcccaactactacag**aggctgaggcaggagaatggcgtgaacctgggaggtggagtttgtgccactgctgaga**tcgcgccactgcactccagcgtgcgcga**cagagtgagactccatctcaaaaaaagaagaaggaagctgggagcagtgtctctcacctgtaat**ctcagcattttggaaggctgag**gcaggaggatcacttgaggccaggagttcaagaccaccctgggcaacacagtgagacccccatctctac**aaaaaataaataaataaaatttttt**taaaaatagccaggcatggtagcatgcacctgtcatcctagctactgaagaggttgaggcaggaggatcacttgtgcccaggaggtcaaggttacagtgagccaagaacacaccactgcactccagcctggacaacaaagtaagatcctgtctctaaaaata**tatatttaaaaatata**aaataaataagttaaagaaggaggaaggagatgaagaaaatttggcttagtaggaaagcgtttttgtactgggcagaaacactgtgTCAAACTGGGGCAAACAACCTCTTCTTTGGGACTGGAACGAGACTCACCGTTATTCCCTgtaagtccttacctcttgacaaaaaagctcttagtctgtaatgacaagttctcacatcccagacacctaatgaatttctaaatgagcttccatgtgttaaactgataaaactctctagggagaaggtcacccttacctttgcggctctaaatgtaagacatgttgtaagaaaaattgggaactgttaattgcagtgccaaaatgaagacagtgattacaaatattgggctagttttaatgttagatttataaaatataagcctttccaaagaaccatctcagggtgcatttattagtttgtgtctatacagcccaattgtccagttactttgattaatgttggtagctgatgcagatatttagggacttccttgagaaaacttctttgtgctcaaaattgcatgaattgaatggcacattacctaaataattgccaaatctgtttcactttgtgctgtatagactcagcctaataaaaacaaatgctgttgtgctttttcccagtctgtctcctctccacac**agaaagtttccctttct**actaaaca**gcatctcttcacacgaagaaaagatgc**c**cagcaagccatttgacttattgctg**ctgctagtttgcatttatatgactcctttccttagaagggatgaagacactttatacgacttattagctaaacagattgacagcctaattaggaagagagaaacagattctgagtatctttttgtggctgctgttcctaccccctcgtgtagaacatttggtcagatggctctttgagacagaaataagacagggacgtggtctctgcttgtagtgtgggacgtctcagctgcacctaagaggttgtcacaggagtactaaataggtcagggcaactgacacagccaggaaaatttctttcccctagaacttgtgtgtgctcaccaagcaaagcgtgactaattttgacatccattgttagtgtgtgtctacccttcttgcctttgttttcagacccttc

Uppercase: TRAJ36

Lowercase: Flanking sequence[1000bp]

Red & Bold & Underline: Stem-loop [11]

Blue: Heptamer[27]

Green: Nonamer [4]

id-IGKJ4[J_gene_segment]

gtggacgttcggccaagggaccaaggtggaaatcaaacgtgagtagaatttaaactttgcttcctcagttgtctgtgtcttctgttccctgtgtctatgaagtgatctataaggtgactctgcaatcagcctctgatatccttcagggaaaagataaagataagtctgtagtcaaactcgagaattgattgcacattttctttgaagagcaagcaagattcagtcattgggtgagaataacttgtctaagtaatagcttcagaaatgtcctggg**gaacataacatgttc**tggacagagccttggtcaattgtcagaaagggagtttttgtataggagggaagttaagaggaaccattgtgtgtgcagttttggccaggggaccaagctggagatcaaacgtaagtacttttttccactgattcttcactgttgctaattagtttactttgtgttcctttgtgtggattttcattagtcggatgccagggatctaacaaacttcattcccaggttaggtacagaggaggggaaattgttccacaggacgctagcttgtggctaatttttaagatttctaaatcaaaataacttcattgggggaaagaggcttgctgagctttcagggaggtttttgtaaagggaaaagttaagac**gaatcactgtgattcactttcggccctgggaccaaagtg**gatatcaaacgtaagtacatctgtctcaattattcgtgagattttagtgccattgtatcatttgtgcaagttttgtgatattttggttgaataaac**ctggtgacccagaagtaaatagcaggacaccag**aaaatgaacttaaaaagctgagcaaatagacgaatcattgggtttgagaggagaataggattcatgggggaaatggggaagaaatagctagatttttctctgaacaagcagcctatctcatatgattggcttcaagagaggtttttgt**tgaggggaaagggtgagatccctca**ctgtgGCTCACTTTCGGCGGAGGGACCAAGGTGGAGATCAAACgtaagtgcactttcctaatgctttttcttataaggttttaaatttggagcgtttttgtgtttgagatattagctcaggtcaattccaaagagtaccagattctttcaaaaagtc**agatgagtaagggatagaaaattagttcatct**taaggaacagccaagcgctagccagttaagtgaggcatctcaattgcaagattttctctgcatcggtcaggttagtgata**ttaacagcgaaaagagatttttgttaa**ggggaaagtaattaagttaacactgtggatcaccttcggccaagggacacgactggagattaaacgtaagtaatttttcactattgtcttctgaaatttgggtctgatggccagtattgacttttagaggcttaaataggagtttggtaaagattgg**taaatgagggcattta**agatttgccatgggttgcaaaagttaaactcagcttcaaaaatggatttggagaaaaaaagattaaattgctctaaactgaatgacacaaagtaaaaaaaaaaagtgtaactaaaaaggaacccttgtatttctaaggagcaaaagtaaatttatttttgttcactcttgccaaatattgtattggttgttgctgattatgcatgatacagaaaagtggaaaaatacattttttagtctttctcccttttgtttgataaattattttgtcagacaacaataaaaatcaatagcacgccctaagaaaaatcagggaaaagtgaagtgtacctatttgctatgtagaagaggcagcttacttgaaaatcagcagcaatgttgtttttagagtctgtaataagtaataaactcaaaaagacacattctataggaataagggcttcacagatagagctcattttttaaaaatccaatttgtacattagactaaacgtgaaattatctcttattgtaatggtggaaaggtggttattcccaaaagctcaatctcaaagaaatgtgtttaaatgaaaaaa

Uppercase: IGKJ4

Lowercase: Flanking sequence[1000bp]

Red & Bold & Underline: Stem-loop [8]

Blue: Heptamer[19]

Green: Nonamer [7]

id-TRAJ45[J_gene_segment]

tcaagtcccaacctgggtttcattgcatattaaataaactttttctggctgggaagtatttgattctaaagaaaggaaaag**tgaaattccaagcgcaacgcaagcatttca**ttagtgtgattcattaagaatttccatgggttgccttcgagagcgttaatcacatccattcactggtgagacccacagtttgaaaattgagttaaatttgaaaaaaaaattgtcaccatctctgtagaccacagtgcttgctgtttgggtgtctagacttccaaaatggaacctcaatctgtagtttcgagcttggagatttatcccgagcccgtgacttagagatgtgtaagaaaagctgc**acactgctgggccttgtagggcatttgtgggatcagtgt**catgaaatagcctgggtagacaggttggtgagg**agacactggtgacaggtgtct**gccctgtttctgtaaa**gctgctgacagccgtgagaagaaaagcagc**ggagacaagctgacttttgggaccgggactcgtttagcagttaggcccagtaagtctgagcagaaagtaagatatttatgccttttcctattatttgttctagcctgacatttgagttgtcctcctttggattccagatcaacaaaccatagtgtcttttcatactccttttaatatttgtggctaagggccctccgttcttcccatctgtcctacatgagggtcctgtggccaggtccacattcataaagaagccaacag**tgatgcagttggttggcagtgtgtcaagaaactgcagtca**tcatctccagggcccttactctggttatgtgtgtttcaagcttttatgccagcaacagaggagagggagaagtgaccagtagcaatg**accagggctattagctgtacctggt**gtctgtgggtttcagagcagctgtggaggttacgggaggttcagggcatgagtttttctggcagaggagtttatgtaaagggttggcccagagtgTGTATTCAGGAGGAGGTGCTGACGGACTCACCTTTGGCAAAGGGACTCATCTAATCATCCAGCCCTgtaagtgcttttgcctgggaggtggggttcaaaatgcagtcctcatggggacatttctaaaccttggctcatgtgcttctg**gtgccactaccatttttggcac**tatcctact**tacatgggaggttttgagccatgta**gacaaggaggtgatgcagtttgtgctttggaggactcctgagatctgccctttggcaactgtgtttgagtaggagtccctcattggaaaatctggatgccagcagacaaaagatatgagcttaggatgggtcacaaacaaatggaggccactttgccagcagcacctttgcaacgaaaattgctttcataggaaatgtg**gaaatggctgtgtagatagagatttggccatttc**tccttttctgtctttataccctcttctctgcctgacttagggaccttaaattagagcagttattaatcatttgatttctcggccccatcactcatcaagtatgacaagagggattcattaagataaaacccagaaacagaaaggacaaagaattcaattctcaatgtcttca**gagttgaacttgacaactc**ctcagccttaaccatctagtcagcaccccatgaccaactgaggtcaaagagggcaaccagcaggtaatgttagtcaagtgagtatctggccagagcccagctgccctcataggaattaatactgtggagtttgggaaagggaaatgcatctgtgagggctcacccaggctgcccatctgggagagatcatggggagagttctgagtgagggagtgaacaccagatggaaagtcaagcaacaggtttctgttatgaagcatctcacagtgtaaataccggcactgccagtaaactcacctttgggactggaacaagacttcaggtcacgctcggtaggtaacagaaaccgtgaggtaacttaactgtgtgttcctttaaaacaggggatactgcagaggggatatcctggctggatgtagtccttccttccttccagtgg

Uppercase: TRAJ45

Lowercase: Flanking sequence[1000bp]

Red & Bold & Underline: Stem-loop [10]

Blue: Heptamer[28]

Green: Nonamer [5]

id-TRGJP2[J_gene_segment]

agataactaaatttttttttttggttatgccatagtttaaataaagaatgtattacttcatgggacatagaattctacaaatttcttctgctattcc**aggaaaaatgataagtcattgtggaaaacatttcctcatttggcattgccttggtaaagcatcgtgttgccaaatga**tacaagtggctt**tttgaaagtggagacctttgttcaaa**tttctcaagtatatctgtgttgacacatttgcattttcagttactaaagagcttcctcctttcagaagtatc**ttcccaaattactaacaggccctggtgtttgtgggaa**cagaagggcttttc**ctttgatgtaatctcaaag**cagtttcaacacaattgaacccctggaaattaaaaaagaaaaaaaacgacaattctctgaa**tacttttcctggtaatttagaaaagta**gtgcattgacccactg**agcctgctcttttttctctctatcgcccaggct**ggagtgcaatggcacgatctcggctcactgcaacctctgcctccctggttcaagcaattctcctgcctcagcctcccgagtagctgtgactacaggcacacgccaccacgcccggataatatttttttgtatgttagtagagacggggtttcactgtgttgccaaagctggtcttgaactc**ctgagttcaggcaatccacctgactcag**cctctgaaagtgctaggattacaggcgtgacccactgcgcccggcctttctttttttgttgttgttggttttttatcttaacatttcttttttctgttga**tgcttttcaagaattactgccttagaagaaagca**ggcaatttatgaggaaaaattacaaactatcacatgtcacaaaacctatattcaaggacttccaaaaaagccagaagatgaaattgctagttcaaagttgttggattgctagtcatgtcatgaggatcagaaggttgagatttttgtagaagcttagaccagtgtgATAGTAGTGATTGGATCAAGACGTTTGCAAAAGGGACTAGGCTCATAGTAACTTCGCCTGgtaagtaattttttttctgtttttattccagtaatg**aaaaactgatagatgttttt**tagaaaaaaatgatcaaccttacctgaatatgtcacattcctggcctcagtatacgaacagcaatttttcagggtagctgaatgcctggcacttggtaggtgatagaaatattatttcctcttccagaattcaagttactacccaagagaggtctttatacactcaggaatacagataaatgtatcaatccccatctattcaacagctcacttcagaagaaatgtcactattcataaaat**gctttttatttctaaaagcctaattgtacctgcttctaaaagcaattag**aatgttttacgtattatctcaaatatgatctgcctgaatacaaggtgatttttc**attttccagagaaaat**ttatacaagtttaa**gaaaatatgtattttc**ccctcccccaacttgaatatgtaaatttatagggatacaagtaaaaaagtgaaatgtctaattataacattcaatttc**ttaactggatcatacctcaaagttaa**agattatatcaatac**caatatcagttgagaaatattg**ccttttttgcccttatatttcattaaaatttttgtttt**atgcttcacatgcatgtaaagcat**ctatctcagagcaactttctaaggcatctcaggtctcagtgtccattttggattcctcaaatgatgatgtccctgaatattttactctaagaaacaatgatccactcac**aaaataaaaggaaattatttt**ttccgatttattctgttattctacacttgtatgaagtaagtaagaactacagggactttagctccaagtaataatgaccaaatccatgataaatttctttgctttttttccttttaccagtgctatccccaaaactagcattcattgagttcatttcatgtcggtatctcctccagttagtccttctggatcaaagcagagtattagaagaacattttttaa

Uppercase: TRGJP2

Lowercase: Flanking sequence[1000bp]

Red & Bold & Underline: Stem-loop [18]

Blue: Heptamer[22]

Green: Nonamer [4]

id-IGKJ3[J_gene_segment]

aaagaaagtac**aaaaaaactggccattttttt**taattgcttgtttttctttgtaattaacattcagtctactttctaaaaaataaataaataataagcagtccagatgtggcaagttgctaaagaaaggaaccatcaggccatagacgtaaatatattctcttcttggatt**ttaggtctcacctaa**gaaaataaacacatgctatgtcaga**gaagcctcagggcttc**cacacctgctcgaaaagggagttgagcttcagcagctgacccaggactctgttcccctttggtgagaagggtttttgttcagcaagacaatggagagctctcactgtggtggacgttcggccaagggaccaaggtggaaatcaaacgtgagtagaatttaaactttgcttcctcagttgtctgtgtcttctgttccctgtgtctatgaagtgatctataaggtgactctgcaatcagcctctgatatccttcagggaaaagataaagataagtctgtagtcaaactcgagaattgattgcacattttctttgaagagcaagcaagattcagtcattgggtgagaataacttgtctaagtaatagcttcagaaatgtcctggg**gaacataacatgttc**tggacagagccttggtcaattgtcagaaagggagtttttgtataggagggaagttaagaggaaccattgtgtgtgcagttttggccaggggaccaagctggagatcaaacgtaagtacttttttccactgattcttcactgttgctaattagtttactttgtgttcctttgtgtggattttcattagtcggatgccagggatctaacaaacttcattcccaggttaggtacagaggaggggaaattgttccacaggacgctagcttgtggctaatttttaagatttctaaatcaaaataacttcattgggggaaagaggcttgctgagctttcagggaggtttttgtaaagggaaaagttaagac**gaatcactgtgATTCACTTTCGGCCCTGGGACCAAAGTG**GATATCAAACgtaagtacatctgtctcaattattcgtgagattttagtgccattgtatcatttgtgcaagttttgtgatattttggttgaataaac**ctggtgacccagaagtaaatagcaggacaccag**aaaatgaacttaaaaagctgagcaaatagacgaatcattgggtttgagaggagaataggattcatgggggaaatggggaagaaatagctagatttttctctgaacaagcagcctatctcatatgattggcttcaagagaggtttttgt**tgaggggaaagggtgagatccctca**ctgtggctcactttcggcggagggaccaaggtggagatcaaacgtaagtgcactttcctaatgctttttcttataaggttttaaatttggagcgtttttgtgtttgagatattagctcaggtcaattccaaagagtaccagattctttcaaaaagtc**agatgagtaagggatagaaaattagttcatct**taaggaacagccaagcgctagccagttaagtgaggcatctcaattgcaagattttctctgcatcggtcaggttagtgata**ttaacagcgaaaagagatttttgttaa**ggggaaagtaattaagttaacactgtggatcaccttcggccaagggacacgactggagattaaacgtaagtaatttttcactattgtcttctgaaatttgggtctgatggccagtattgacttttagaggcttaaataggagtttggtaaagattgg**taaatgagggcattta**agatttgccatgggttgcaaaagttaaactcagcttcaaaaatggatttggagaaaaaaagattaaattgctctaaactgaatgacacaaagtaaaaaaaaaaagtgtaactaaaaaggaacccttgtatttctaaggagcaaaagtaaatttatttttgttcactcttgccaaatattgtattggttgttgctgattatgcatgatacagaaaagtggaaaaatacattttttagtct

Uppercase: IGKJ3

Lowercase: Flanking sequence[1000bp]

Red & Bold & Underline: Stem-loop [11]

Blue: Heptamer[22]

Green: Nonamer [7]

id-TRAJ27[J_gene_segment]

ttgcaggatttcaattagatataaggaacagttgattagatgtaagagctgttagcaaaggaaaaccttataaaaactaggagctcttaataactagacaggtctttggtcagacaacacactgactgaggacaggaggatggattcgatgacctctgaaagtccagccaggactctggaggactctgaggaatggtctgttccatagcctgcctctgtaatgcccttctctcttgcctattgtctggttgttgttacagtctgagcttttgtcagagctgctcct**atgctgtgagtggtctgattttctcagcat**ctctggggtttttgcaaagcaaggaaactctgtgcatactctggggctgggagttaccaactcactttcgggaaggggaccaaactctcggtcataccaagtaagttcttctttctggctaattattcttcccgagaagcctgtcttccatcatgcagaagctgtc**aaaacacaggtggtgtttt**ctttgcttggtttgtgttgggtggttagtaatatcagttggaaaacaaggttattaatgcacatattccctggggcattgtattcggacatttaatatccatgtagtctctccctgtgaaaatatgtgaacctccacgaaaagaaaggtccaaggaaagtagcagggaaaaggcaggaattggatgaaaatagccaaagagcctgcaggaatataaacaaggcaggatcccaggagacagagcagtagccactttgagtgaatttcc**caggaggtgctcctg**ccaaggcccataccttcaaggaaaattaaggcaaatagaattgggctggggagttgctacttattagtattcctcccacgttctaacctaattataaggaggttgttttggccatgggcagtcatctcaggttttgttttcctgctttcctccc**taacctccacctgtcttcctagaggcctgagtcaaggtta**ttgcaatagcactaaagactgtgTAACACCAATGCAGGCAAATCAACCTTTGGGGATGGGACTACGCTCACTGTGAAGCCAAgtaagttgtgttcttctttgcctaggccttcaggggcaatcaa**tcaaaccattagtttga**aaaagactttaatcctatgcatctggttgggctctttattaatgttctttccccaggcca**aagagagttggttctctt**ccctgctttaaaatgagatatgag**tgcatgtatgcacacacgcatgcccacatgca**gactcttgctctagctcatggtaagggcttctcaggagcatatacaacattttgaaagaaatagagaaacaaacaataatgagccaatggagct**gtcaggaaggttctccaccacccctgac**caggcttcccagcaggatctctggcc**atataggatgtgcttgcactatat**cggatggtctaggatgggaaatatctctctgtaattaattccaaaatt**tggcaacccttattgcca**gaaagtacttctcatatctaagtttgatctccctgctgcgatttaaatatctacttgaaaagaaccactagaactctgtacatctggattgagaatcaagtaactgccatagaatatgaggctgagcactaagcccacctctgcagcacctctgacgtgtcatgggtcccctctggacattttgagctcaagtaataatggattaagcaggatagcacaagctttggaaacagaaaaacctgaattcaagtccttgctccactacttacctagctgtgtgaccttgaacaagttacttaatctctctgagcctcagtttccccatccacaaaatgaagataaaaatattcctacctcgttgagtcatcgtaaagatgaaataatacagctaaagtagttagcatagtgctcagaacatagaaaaagc**tcattaaaccctcgctattaataataatga**taaacatcatctctaaccc**acagcagagcctttgtttggcccctggggctcttctgt**caaggttctgttgtgcatcagcagcagagctggaattggggggagggtggagaa

Uppercase: TRAJ27

Lowercase: Flanking sequence[1000bp]

Red & Bold & Underline: Stem-loop [12]

Blue: Heptamer[23]

Green: Nonamer [5]

id-IGLJ6[J_gene_segment]

ggggaggcgcttgctgtggggaatactgttggggactgcatggtgccaggagtgtgtcccaggtgtgagctcttcaggtgatgggcccagaccaggatgcaggtccaggctgtgtgtgtccaggc**cccacactgcacatgcccatccactgtggg**tgccccccgtgcc**ggtgacactccagagcacaggctgtgtgggtcacc**cgtgcatgtaggggactgcagag**cacacacatgtgggtgtgtgtg**taagcc**gggggagctgaagtggacagcagagatccccc**acacgtagagctcctggaggaaaccacactgccagtgaggcctatgacctcccttgagaagctctcctctg**ggccattgcagctctaatccagcacacatggcc**tataggtaggaagctgtccccatgggttcaggaggcggacttgggcagtgatctgtgagctgctccttcaggacagctgggccaaggatgctggagct**cagaccttgcagtggggtctg**tgtccctgtggtcaacgtcattcttcccacaacccccaattcccattccaagcccatcccatcataacctccagtgctgaggat**gtggggggcatcaaggaccctccccac**cattcccaacttctcaaagaggtcccagctgcaccatgtcagtggcgtcacccagccgctcacctccccttcctccctg**gcattcctgacaccagctgatccaggccacccaactcctcagaaatgc**aattacctgggagacaattccacacacagacctgtcaacccttcccatgactgaggtggatgaacccctaagcccccaaggaagtgatattcaggtcagtagaaggtgacccccttcaccccacctatggctcacccacccatgagaaaagggggctggaccctgggccttgtgagcagctgcagggggttgggggggggtggcctgggtcgggtgtatcaggagggtttgtgtgcagggttatatcacagtgTAATGTGTTCGGCAGTGGCACCAAGGTGACCGTCCTCGgtgagtccccttttctattcttttgggtctagggtgagatctggggagacttttctgtcctttctgttctctctagggtagaggtctcaatctctctggggtctgccaccattgccttttttcctggccccaaattcctccagcctgtcccttcttaggcacctggtggggcatgatgggaagatcccagtgacctcttaatgctccctgctcaggcctgaacctgccgctgtcact**cagggggcattctcccctg**ggctctgggatgttctgctccctcgcacagagtctccagtccccaaagaccagcagagcaggggctcaggctggggcaccagggccatgggacag**gggaaggatgctggaaaaagttcagcttccc**agggttctgggtcccaactatggggctgctttaaacaccaagaagggaggcctttgactggggacttggggaaatgaagggggacaaggatggaggaagaatgtcctgtgaggtggcgccagggtgctgggtccctcctcctcccccgggacaggcaggctgccatggacagggtggttctcaggacactcagtccaaggttagagcctccccatcccacccaaaaagagagacccccaaagcagatgctgagggaggcactcctggtgggcgcagg**tgacagggacctgtcaggacagacatttgtcct**aggacagccaga**tctcccaacgacagggaga**ccccatagagcagacacagcgccaggctcagaacagaaaatatacctcacatgcaagccctccatc**tcctggactcccagga**cccggctcccaggactgacatcccct**ccccaccaaggggcctctgtgggaaagtgggg**cagagactgcaatgatggtgctgggggatatgtgaggaagaaattcacttgtcaaaagggaggagaacctaaaaacaggacacagatgccctcgtgaacagacccagaggagggacctggcaggggatgttcagaca

Uppercase: IGLJ6

Lowercase: Flanking sequence[1000bp]

Red & Bold & Underline: Stem-loop [15]

Blue: Heptamer[31]

Green: Nonamer [2]

id-TRBJ2-5-2[J_gene_segment]

cccagctccagccgttccagtaccaccaatgccccttcagatttca**aatccactgtgttgtcccccagccaagtggatt**ctcctctgcaaattggtggtggcctcatgcaagatccaggttaccgtgtccagctaactcgagacaggaaaagataggctcaggaaagagaggaagggtgtgccctctgtctgtgctaagggaggtg**gggaaggagaaggaattctgggcagccccttccc**actgtgctcctacaatgagcagttcttcgggccagggacacggctcaccgtgctaggtaagaagggggctccaggtgggagagagggtgagcagcccagcctgcacgaccccagaaccctgttcttaggggagtggacactgggcaatccagggccctcctcgagggaagcggggtttgcgccagggtccccagggctgtgcgaacaccggggagctgttttttggagaaggctctaggctgaccgtactgggtaaggaggcggctggggctccggagagctccgagagggcgggat**gggcagaggtaagcagctgccc**cactctgagaggggctgtgctgagaggcgctgctgggcgtctgggcggaggactcctggttctgggtgctgg**gagagcgatggggctctcagcggtgggaaggacccgagctgag**tctgggacagcagagcgggcagcaccggtttttgtcctgggcctccaggctgtgagcacagatacgcagtattttggcccaggcacccggctgacagtgctcggta**agcgggggctcccgct**gaagcccgggaactggggagggggcg**ccccgggacgccgggg**gcgtcgcagggccagtttctgtgccgcgtctcggggctgtgagccaaaaacattcagtacttcggcgccggga**cccggctctcagtgctgggtaagctggggccgccggg**ggaccggggacgagactgcgctcgggtttttgtgcggggctcgggggccgtgACCAAGAGACCCAGTACTTCGGGCCAGGCACGCGGCTCCTGGTGCTCGgtgagcgcgggctgctggggcgcgggcgcgggcggcttgggtctggtttttgcggggagtccccgggctgtgctctggggccaacgtcctgactttcggggccggcagcaggctgaccgtgctgggtgagttttcgcgggaccacccgggcggcgggattcaggtggaaggcggcggctgcttcgcggcacccggtccggccctgtgctgggaga**cctgggctgggtccccagg**gtgggcaggagctcggggagccttagaggtttgcatgcggggatgcacctccgtgctcctacgagcagtacgtcgggccgggcaccaggctcacggtcacaggtgagattcgggcgtctccccaccttccagcccct**cggtccccggagtcggggggtggaccg**gagctgg**aggagctgggtgtccggggtcagctctgcaaggtcacctccccgctcct**gggaaaagactggggaagagggagggggtggggagg**tgctcagagtccggaaagctgagca**gagggcgaggccacttttaatcttttttctggggtgtttagagagaaggtgaacgatggaggagaggatttgttaggactctgggagaggcgagactggagaggacgaagggaaatcctggtttggggaatgggtaggagtgggggtaactgctattcgtaggcaaaaagagctgagcaggctgggaacagcgcgggtgggcaagggtcagcactgcgggcaggcgggtgggtgttagggggcagaaatcctgcagccgagggtgcagtagaacacagaagaaaaagcctgccaaacaaaagtggaacagagaagccaaaaagggagatgaacatgagtcagtgaagaaaagaatgaaagtttactgtttagcagtgtggatctctaatccgacttaaaactccttgttcccgattcctattcctcctaagccagagatccctgggtccagggtgagggcacggcattcatgcttacccacgggc

Uppercase: TRBJ2-5-2

Lowercase: Flanking sequence[1000bp]

Red & Bold & Underline: Stem-loop [12]

Blue: Heptamer[29]

Green: Nonamer [11]

id-TRBJ2-4[J_gene_segment]

tattaagaactgatgaaaaccctgag**ggaaagatattgtcccatctttcc**aatgaggaaactgagatcagaggttacaggtcatataactaggaaacggcaaggtctagcctgcaatatcgcccagctccagccgttccagtaccaccaatgccccttcagatttca**aatccactgtgttgtcccccagccaagtggatt**ctcctctgcaaattggtggtggcctcatgcaagatccaggttaccgtgtccagctaactcgagacaggaaaagataggctcaggaaagagaggaagggtgtgccctctgtctgtgctaagggaggtg**gggaaggagaaggaattctgggcagccccttccc**actgtgctcctacaatgagcagttcttcgggccagggacacggctcaccgtgctaggtaagaagggggctccaggtgggagagagggtgagcagcccagcctgcacgaccccagaaccctgttcttaggggagtggacactgggcaatccagggccctcctcgagggaagcggggtttgcgccagggtccccagggctgtgcgaacaccggggagctgttttttggagaaggctctaggctgaccgtactgggtaaggaggcggttggggctccggagagctccgagagggcgggat**gggcagaggtaagcagctgccc**cactctgagaggggctgtgctgagaggcgctgctgggcgtctgggcggaggactcctggttctgggtgctgg**gagagcgatggggctctcagcggtgggaaggacccgagctgag**tctgggacagcagagcgggcagcaccggtttttgtcctgggcctccaggctgtgagcacagatacgcagtattttggcccaggcacccggctgacagtgctcggta**agcgggggctcccgct**gaagccccggaactggggagggggcg**ccccgggacgccgggg**gcgtcgcagggccagtttctgtgccgcgtctcggggctgtgAGCCAAAAACATTCAGTACTTCGGCGCCGGGA**CCCGGCTCTCAGTGCTGGgtaagctggggccgccggg**ggaccggggacgagactgcgctcgggtttttgtgcggggctcgggggccgtgaccaagagacccagtacttcgggccaggcacgcggctcctggtgctcggtgagcgcgggctgctggggcgcgggcgcgggcggcttgggtctggtttttgcggggagtccccgggctgtgctctggggccaacgtcctgactttcggggccggcagcaggctgaccgtgctgggtgagttttcgcgggaccacccgggcggcgggattcaggtggaaggcggcggctgcttcgcggcacccggtccggccctgtgctgggaga**cctgggctgggtccccagg**gtgggcaggagctcggggagccttagaggtttgcatgcgggggtgcacctccgtgctcctacgagcagtacttcgggccgggcaccaggctcacggtcacaggtgagattcgggcgtctccccaccttccagcccct**cggtccccggagtcggagggtggaccg**gagctgg**aggagctgggtgtccggggtcagctctgcaaggtcacctccccgctcct**ggggaaagactggggaagagggagggggtggggagg**tgctcagagtccggaaagctgagca**gagggcgaggccacttttaatcttttttctggggtgtttagagagaaggtgaacgatggaggagaggatttgttaggactctgggagaggcgagactggagaggacgaagggaaatcctggtttggggaatgggtaggagtgggggtaactgctattcgtaggcaaaaagagctgagcaggctgggaacagcgcgggtgggcaagggtcagcactgcgggcaggcgggtgggtgttagggggcagaaatcctgcagccgagggtgcagtagaacacagaagaaaaagcctgccaaacaaaagtggaacagagaagccaaaaagggagatgaacatgagtcagtgaagaaaagaatgaaagtttact

Uppercase: TRBJ2-4

Lowercase: Flanking sequence[1000bp]

Red & Bold & Underline: Stem-loop [13]

Blue: Heptamer[29]

Green: Nonamer [11]

id-TRBJ1-2-2[J_gene_segment]

catcctgaggacagtgcctggaggtgagaaggaagcccccggcctggtccataccccaccaccaacttgcataatggggggtgatgtcacccaccctccactcccctcaaaggagcagctgctctggtggtct**ctcccaggctctgggggcggacccatgggag**gggctgtttttgtacaaagctgtaa**cattgtggggacagggggccacaatg**attcaactctacgggaaacctttacaaaaacct**ctctggcggtcccaactcccagag**tcctcttctttcctcctgggtcacaggtcttaatgcaatttggttcagaatgcctctgcctcactcctgatcacatgtcagaccaagactgtggacaaggacaggcccagatgagaactaaagcttccc**aggcagagagaggtcagacataagaagactgcct**caggaacctcacaagtggaggactcagggagggtcccaatccccaaaaattgagacaaagtcaggtggaaggttcatcggaggtgaccagctctccagaggactcgggaagaagt**caggggtatctatagatggagtcacaggttctgggcccctg**ccatcctctgcaggccatgcactttccctttcgatggaccctcacagagggagcatctgaatggggca**tcctttgaaaaagga**acc**taggaccctgtggatggactctgtcattctccatggtccta**aaaagcaaaagtcaaagtgttcttctgtgtaatacccataaagcaca**ggaggagatttcttagctcactgtcctcc**atcctagccagggccctctcccctctctatgccttcaatgtgattttcaccttgacccctgtcactgtgtgaacactgaagctttctttggacaaggcaccagactcacagttgtaggtaagacatttttcaggttcttttgcagatccgtcacagggaaaagtgggtccacagtgtcccttttagagtggctatattcttatgtgCTAACTATGGCTACACCTTCGGTTCGGGGACCAGGTTAACCGTTGTAGgtaaggctgggggtctctaggaggggtgcgatgagggaggactctgtc**ctgggaaatgtcaaagagaacagagatcccag**ctcccggagccagactgagggagacgtcatgtcatgtcccgggattgagttcaggggaggctccctgtgagggcgaatccacccaggcttcccagaggctctgagcagtcacagctgagcccagggtgatggggcagaag**agggaaggggagggggcctctcctcatagttccct**gagatagcccagagaaagcccggtgggtaatgaatgagccacaacacctctccatctatctgcttcactg**acagaggttctctgt**agattcttcgtatattcctgtgctggattttataggaggccactctgtgtctctttttgtcacctgcctgagtcttgggca**agctctggaagggaacacagagtactggaagcagagct**gctgtccctgtgagggaa**gagttcccatgaactc**ccaac**ctctgcctgaatcccagctgtgctcagcagag**actggggggttttgaagtggccctgggaggctgtgctctggaaacaccatatattttggagagggaagttggctcactgttgtaggtgagtaagtcaaggctggacagctgggaacttgcaaaaaggggctggaatccagacggagcctttgtctctagtgcttaggtgaaagtgtatttttgtcaggaaggcctatgaggcagat**gaggaggggatagcctccctctcctc**tcgactattttgtagactgcctgtgccaagttaggttcccctactgagagatgggtagactcagcttggaaggggtcaccttgaacatctcctgtctccttgaagggtgccggtcacggccatgacagataaaagagcctctgaccttaccaccacggtcctaccgtttctc**tccctcacacagaaaggagaaggtcacagaagaggga**acttgggggatcacacggggcctaattgg

Uppercase: TRBJ1-2-2

Lowercase: Flanking sequence[1000bp]

Red & Bold & Underline: Stem-loop [16]

Blue: Heptamer[50]

Green: Nonamer [7]

id-TRBJ1-6[J_gene_segment]

taccgtttctc**tccctcacacagaaaggagaaggtcacagaagaggga**acttgggggatcacacggggcctaattggtctgctgaccaccgcattttgggttgtaccattgtctacccctctacccaccagggttaaaattctactaaggaacaggagaggacctggcaggtggacttggggaggcag**gagtggaaggcagcaggtcgcggttttccttccagtc**tttaatgttgtgcaactaatgaaaaactgttttttggcagtggaacccagctctctgtcttgggtatgtaaaagacttctttcgggatagtgtatcataaggtcggagttccaggaggaccccttgcgggagggcagaaactgagaacacagccaagaaaagctcataaaatgtgggtcagtggagtgtgtggtggggccccaagagttctgtgtgtaagcagcttctggaaggaagggcccacaccagctcctctggggtttgccacactcatgatgcactgtgtagcaatcagccccagcattttggtgatgggactcgactctccatcctaggtaagttgcagaatcagggtggtatggccattgtcccttgaaggcagagttctctgcttctcctcccggtgctggtgaggcagattgagtaaaat**ctcttaccccatggggtaagag**ctgtgcctgtgcctgcgttccctttggtgtgtcttggttgactcctctatttctcttctctaagtcttcagtccataatctgcctcctcactcccttcttggctcatcctccctcttatgtgcatggctctgcctctcctaagcctcttcctcttgcgccttatgctgcacagtatgcttaggcctttttcctaacagaatccctttggtccagagccatgaatccaggcagagaaaggcagccatcctgctgtcagggagctaagacttgccctctgactggagatcgccgggtgggttttatctaagcctctgcagctgtgCTCCTATAATTCACCCCTCCACTTTGGGAATGGGACCAGGCTCACTGTGACAGgtatgggggctccactcttgactcgggggtgcctgggtttgactgcaatgatcagttgctgggaagggaattgagt**gtaagaacggaggtcagggtcaccccttcttac**ctggagcactgtgccctctcctcccctccctggagctcttccagcttgttgctctgctgtgttgcctgcagttcctcagctgtagagctccttgcttagtcttcagggctgtgtgtttctttgctcttcttttcattgttttctgggactcttctcatctctactttcttagtggatgtattgttttactttcccttttttaaattgcatcttctccattttttccttcccattctaactccacttctgcattgttgactccttttggtgactagctctgtcttctatgttaagattctccccactgccagcctccagcacagaactctgctcatgtcttcatctccctccttctttctttctctaccagtcttagaagatgcatctatgtcttcctgaggtagtttgaaggttcatgagccaggcatgaccaggttggggagacaggtggtttcagggttgctcttgaggcctgagggcagaag**tccctgtcacagcattgggcgagctgcaggga**gtctctgaggtgcctgtgtttggcaggtgtttgggagataggtctgaagagagtgttacaactggaaactgaca**ttatcttagcaacagataa**ggtat**aaagaagacaaaatgggtaggagactctattttcttt**ttttctgagacggtgtctagctctattgcccaggctggag**tgcagtggcacgatctcggctcactgca**acctccgcctcctgggtttgagcgattctcctcccttggcctcccg**agtagctgggattacaggtgcctgccaccaaacccagctact**ttttgcatttttagtagagatagggtttcactgtgttagccgggctgttctcaaactcctgacctcgtgatccgcctgccttg

Uppercase: TRBJ1-6

Lowercase: Flanking sequence[1000bp]

Red & Bold & Underline: Stem-loop [9]

Blue: Heptamer[32]

Green: Nonamer [4]

id-TRBJ1-2[J_gene_segment]

catcctgaggacagtgcctggaggtgagaaggaagcccccggcctggtccataccccaccaccaacttgcataatggggggtgatgtcacccaccctccactcccctcaaaggagcagctgctctggtggtct**ctcccaggctctgggggcggacccatgggag**gggctgtttttgtacaaagctgtaa**cattgtggggacagggggccacaatg**attcaactctacgggaaacctttacaaaaacct**ctctggcggtcccaactcccagag**tcctcttctttcctcctgggtcacaggtcttaatgcaatttggttcagaatgcctctgcctcactcctgatcacatgtcagaccaagactgtggacaaggacaggcccagatgagaactaaagcttccc**aggcagagagaggtcagacataagaagactgcct**caggaacctcacaagtggaggactcagggagggtcccaatccccaaaaattgagacaaagtcaggtggaaggttcatcggaggtgaccagctctccagaggactcgggaagaagt**caggggtatctatagatggagtcacaggttctgggcccctg**ccatcctctgcaggccatgcactttccctttcgatggaccctcacagagggagcatctgaatggggca**tcctttgaaaaagga**acc**taggaccctgtggatggactctgtcattctccatggtccta**aaaagcaaaagtcaaagtgttcttctgtgtaatacccataaagcaca**ggaggagatttcttagctcactgtcctcc**atcctagccagggccctctcccctctctatgccttcaatgtgattttcaccttgacccctgtcactgtgtgaacactgaagctttctttggacaaggcaccagactcacagttgtaggtaagacatttttcaggttcttttgcagatccgtcacagggaaaagtgggtccacagtgtcccttttagagtggctatattcttatgtgCTAACTATGGCTACACCTTCGGTTCGGGGACCAGGTTAACCGTTGTAGgtaaggctgggggtctctaggaggggtgcgatgagggaggactctgtc**ctgggaaatgtcaaagagaacagagatcccag**ctcccggagccagactgagggagacgtcatgtcatgtcccgggattgagttcaggggaggctccctgtgagggcgaatccacccaggcttcccagaggctctgagcagtcacagctgagcccagggtgatggggcagaag**agggaaggggagggggcctctcctcatagttccct**gagatagcccagagaaagcccggtgggtaatgaatgagccacaacacctctccatctatctgcttcactg**acagaggttctctgt**agattcttcgtatattcctgtgctggattttataggaggccactctgtgtctctttttgtcacctgcctgagtcttgggca**agctctggaagggaacacagagtactggaagcagagct**gctgtccctgtgagggaa**gagttcccatgaactc**ccaac**ctctgcctgaatcccagctgtgctcagcagag**actggggggttttgaagtggccctgggaggctgtgctctggaaacaccatatattttggagagggaagttggctcactgttgtaggtgagtaagtcaaggctggacagctgggaacttgcaaaaaggggctggaatccagacggagcctttgtctctagtgcttaggtgaaagtgtatttttgtcaggaaggcctatgaggcagat**gaggaggggatagcctccctctcctc**tccactattttgtagactgcctgtgccaagttaggttcccctactgagagatgggtagactcagcttggaaggggtcaccttgaacatctcctgtctccttgaagggtgccggtcacggccatgacagataaaagagcctctgaccttaccaccacggtcctaccgtttctc**tccctcacacagaaaggagaaggtcacagaagaggga**acttgggggatcacacggggcctaattgg

Uppercase: TRBJ1-2

Lowercase: Flanking sequence[1000bp]

Red & Bold & Underline: Stem-loop [16]

Blue: Heptamer[50]

Green: Nonamer [7]

id-TRAJ50[J_gene_segment]

agtcaagtccctatatttctggagtagctgccc**ttgtgcaggcatagttggtcaggcagacagcacaa**tcccagctcagtgtgtatggaggaggccacctgacattgccagctgtatctgctccattcagtggcagcat**aaatccataccaggtgggtgtggattt**ggaatatgccagtgccactacaaggctgaatgttgagttgttgtgatcatcttgaatgtgtgagatgcctccaagtgaagtctggttctaatgagcagggtgcaggttggaccaatattgcccattttctgttgagtttctccaagcccagtagtgtgggagccaacattgagagactgacatttgagctagggatcctattgactgaaatatgccaaaggga**tctgaaaggtcaaaactgccttcaga**gctcagcctttgttgttatctctctggcagttgtcatttcagcaaaatgcatccacttagaaagctcagctcttcattaggcagacaaatgagaacaacatattgaccagagataaggagttgggagcccaga**tctcagcctctgaga**gcctggcatttccagttctttaatcacgtgagccagtgagcatatgctatctcagagaaagcagatcaatggtgcctgaggacctggtgtcaccaagctagggcttagagacagaaatgagatgcagatggggatgttgtgtgaattggggaataatagagaagtgaggaggagacagggagttcaatgatgctttttaaaaccagcagcatgtgagaaattgtaatgacacaaaggtgaagaagtatgtattgcatataatcctagcaagtttcctaccattcccaacagaatctagcctcactgtggctccaactcacagatcctat**aggtaatgctgaatcttacct**ga**gatgccttttgaagtggcatc**aagactgggctctaaaatctttaagagtcacctgtcagttattgtaa**aggtttggatggctgtgTGAAAACCT**CCTACGACAAGGTGATATTTGGG**CCAGGGACAAGCTTATCAGTCATTCCAAgtaagtgtccctgg**ggtgctgcctgtggagtgcgctggggcta**acagtctcatacattagggcttaaatgactgt**gcagatggcatcgtggttgaggacacaaaactgagagaatcgagatacattgactctgatcagaaaaaaaggtcacggaatagcaaacataggtttagtccttaaaaggtagcatagataacatgggacttgaacgtggcaatttgaaaaacaacctgtggtctatcagcacgttctcactgcacacgggggataaacaggggtgggggaataagcaatggacattgcatgataggggtgctagggaa**atgaggcaggtcaggaagaaggaaggtgatcagagcctcat**acggagttaatctagagaagatgaattgttccataagtttcacaaagaaataaagtccaaagatataataagaag**tcctgattaaaatccacttcagaatcatatgatcatcagga**gagaaattaagtc**ttaaaagaaaaagaagtttttaa**a**tgtaggtccataaaacaaatggacctaca**cctttaattagacagcaagcagcacaagatgcacgttagccttccttttaatcccctctcaggaggaatcagtccctc**cagctgcacatggtcacagctg**ctctaagccctggagctccttcccagtaatacccttgct**cctgtgcctgcacagg**gaaaggagatgcctgcctgcctgcctgcctgcctgcctaccggtctgccttttggtccgccgcttggtggggccccaggtgtaacctctggctgtctcttctgcctggtttttgttgagcttcctatcacagtggaacaccggtaaccagttctattttgggacagggacaag**tttgacggtcattccaagtaagtcaaa**gaaaattttccatcaccattgtgttgagcaaaccctttaaactgcagaaagagctgtcaaagtctgctactctatttctctgcttctaggtataacttt

Uppercase: TRAJ50

Lowercase: Flanking sequence[1000bp]

Red & Bold & Underline: Stem-loop [16]

Blue: Heptamer[34]

Green: Nonamer [2]

id-IGLJ1[J_gene_segment]

gcagagaaaattaggagactacagagagcagaacccagggtggggatctgggagtcagcagttgggcatgggcctggtagaaagggaagccaaggaggaggagagggggcagtctcagacaccaaggaggggagagtgactagaaagaaaaccttcttgcagagacataggggatggggaagaactgcagactgaactggggcaaaggactgttggccttaaccagagagatttgagggagagatgaggctgagagccaggggatcctgccatgtcccagcataaaaacagtacctgacacagatgggtgcttgggagctgttgtcggatgaatgagtggacagatgcatggatggacggatggatggaaggatgatagattgatggacaaacagatgaacagatgaatagctggatggacaactggatggatgggtagacagaa**tgatctcagagatca**gaaaaagcttcatgcactaagtgggactgaaccgcgtctccatgggtagaaagcagaggaatctccacttgagtcaggaatgacccagtgctctcaatccagggaga**aagccagcctggctt**cactggggacacttgtgtgggggactcagaggccctttaaatgaggccagacgaggttggacaggtccaagccaactcagcactcctctgccacactgcacaggaggggatgtgtcactcagggagttg**ctgggacctatgggtcccag**tgttgtcatcagcaccgacagcctcagagaggaaagacacacactggggtaac**tccaaggctgtgtgtggcacttgccttgga**cagcagacaggcacagggacacctctag**ggggctggccacccccctgcctcatgtctaggtcccagcccc**gcccactgcaaccctgtgcccgtcatgcccagcaggctc**ctgctccagcccagcccccagagagcagaccccaggtgctggccccgggggttttggtctg**agcctcagtcactgtgTTATGTCTTCGGAACTGGGACCAAGGTCACCGTCCTAGgtaagtggctctcaacctttcccagcctgtctcaccctctgctgtccctggaaaatctgttttctctctctggggcttcctcccctctgtcctcccagccttaagcactgacccttacctttctccatggggcctggaggaggtgcattagtctccgggtaaccggcaggaagggcctccacagtgggagcagccggatgcagcctggtcccggggcctgagctgggattgggcagggtcagggctcctcctctcttccagggcagatgtctgagtgagggacagaggctggttctgatgaggggccctgcag**tgtccttagggacattgcccagtgactcctggggtcaaggaca**gaggctgctggggtgggcc**tgggagctgctgagtctcatagtctaggggagcagcccca**agaacagctgagggtctaggctgagga**ctggatgccaatccag**cctgggagggccacacggcct**ggtgacacagaggtcacc**ccaaggggagaccaatgg**agggcacagagagggctctgggtctaggctgcagctctgtggcct**gtgctgggtcatgaggacatggggacacagagggacgggtgagactgggtgaggtgccagaatccaaccctcccaggacagtcaccagaaaggagacagtctcttagggcagagatgtgtctgtccctggagccccgtcacctctggggcccagtgtctctctgttcacggatcggcctcctgccttcctcaaagggcatgttagactcaggaaatgaccagaggggagtgaatgaggggtgcagagaactccatggctaccaggtgaagtttggggtcatcacaggctgctggggtggg**cctgggggctgctgagtctcatagtctgtgggagcagccccagg**aacagctgaggtgaagggttctgtggtcgggcttgtggagacaggaaacatctcagagcctcagaggagccctgaggcttgtctaggtggagccca

Uppercase: IGLJ1

Lowercase: Flanking sequence[1000bp]

Red & Bold & Underline: Stem-loop [13]

Blue: Heptamer[23]

Green: Nonamer [3]

id-TRBJ1-6-2[J_gene_segment]

taccgtttctc**tccctcacacagaaaggagaaggtcacagaagaggga**acttgggggatcacacggggcctaattggtctgctgaccaccgcattttgggttgtaccattgtctacccctctacccaccagggctaaaattctactaaggaacaggagaggacctggcaggtggacttggggaggcag**gagtggaaggcagcaggtcgcggttttccttccagtc**tttaatgttgtgcaactaatgaaaaactgttttttggcagtggaacccagctctctgtcttgggtatgtaaaagacttctttcgggatagtgtatcataaggtcggagttccaggaggaccccttgcgggagggcagaaactgagaacacagccaagaaaagctcataaaatgtgggtcagtggagtgtgtggtggggccccaagagttctgtgtgtaagcagcttctggaaggaagggcccacaccagctcctctggggtttgccacactcatgatgcactgtgtagcaatcagccccagcattttggtgatgggactcgactctccatcctaggtaagttgcagaatcagggtggtatggccattgtcccttgaaggcagagttctctgcttctcctcccggtgctggtgaggcagattgagtaaaat**ctcttaccccatggggtaagag**ctgtgcctgtgcctgcgttccctttggtgtgtcttggttgactcctctatttctcttctctaagtcttcagtccataatctgcctcctcactcccttcttggctcatcctccctcttatgtgcatggctctgcctctcctaagcctcttcctcttgcgccttatgctgcacagtatgcttaggcctttttcctaacagaatccctttggtccagagccatgaatccaggcagagaaaggcagccatcctgctgtcagggagctaagacttgccctctgactggagatcgccgggtgggttttatctaagcctctgcagctgtgCTCCTATAATTCACCCCTCCACTTTGGGAACGGGACCAGGCTCACTGTGACAGgtatgggggctccactcttgactcgggggtgcctgggtttgactgcaatgatcagttgctgggaagggaattgagt**gtaagaacggaggtcagggtcaccccttcttac**ctggagcactgtgccctctcctcccctccctggagctcttccagcttgttgctctgctgtgttgcctgcagttcctcagctgtagagctccttgcttagtcttcagggctgtgtgtttctttgctcttcttttcattgttttctgggactcttctcatctctactttcttagtggatgtattgttttactttcccttttttaaattgcatcttctccattttttccttcccattctaactccacttctgcattgttgactccttttggtgactagctctgtcttctatgttaagattctccccactgccagcctccagcacagaactctgctcatgtcttcatctccctccttctttctttctctaccagtcttagaagatgcatctatgtcttcctgaggtagtttgaaggttcatgagccaggcatgaccaggttggggagacaggtggtttcagggttgctcttgaggcctgagggcagaag**tccctgtcacagcattgggcgagctgcaggga**gtctctgaggtgcctgtgtttggcaggtgtttgggagataggtctgaagagagtgttacaactggaaactgaca**ttatcttagcaacagataa**ggtat**aaagaagacaaaatgggtaggagactctattttcttt**ttttctgagacggtgtctagctctattgcccaggctggag**tgcagtggcacgatctcggctcactgca**acctccgcctcctgggtttgagcgattctcctcccttggcctcccg**agtagctgggattacaggtgcctgccaccaaacccagctact**ttttgcatttttagtagagatagggtttcactgtgttagccgggctgttctcaaactcctgacctcgtgatccgcctgccttg

Uppercase: TRBJ1-6-2

Lowercase: Flanking sequence[1000bp]

Red & Bold & Underline: Stem-loop [9]

Blue: Heptamer[32]

Green: Nonamer [4]

id-TRAJ47[J_gene_segment]

aagcatgcttgcaacagcgagaactggaagtatggaacctttctgtcttaaccactcagttgcctagaagaatcttt**tttctattttgctgcattagaaa**ttaaaggcatgagtattgaggatttgggtaggtcagacttttatagtgacaatcacattacaaagcacctcttagctaggttggataaagcacctttaaacatcaaagtc**tttttcttctttctggatgagatagaaaaa**tggtctctcctcactctgtatctaaggacagggggctcagagagttccagacttacaaagtgagactgcagcagagctgagccatgggacct**ggagggacccagtgacggttagtctgctccctcc**cacgggtgctatagtagccacaaccgttcaggaaaccgtccccaggtgagaacgtaacacacgcagggaaaatcccactgctactaagagcaactcagcataagaatgcattacccagcaagggacagtcaga**ggggctctaaaccagagcccc**aaatatttgaaaactttatgtggaagccagctataacaagttcgttttccaagccgtgcttccagaagctgttagtggttggtagttgctaagggttttgaaactggaggac**atcatagcagagtggaattatttggcacaaggggctatgat**t**agaattaatgagatgaaagaaaattct**ggctagaagactagaaaaatccggaaagaaaagatacagtaaaccttgggggagttgtagcatttatcctagtccaagcagcaaagagcagacagaagccaaattccaataaacaattggcctttttacttttacaagtttctgcaattatcacaatgtcttcagcttcttgggcaaatatcaacaggtcctaaatgcaaagtggaacagtcctctgtttccagaaccgagtagcttagccccgccccacacaccacccatgctcaaccactgtttttgtagagg**agtttgacgctgtgTGGAATATGGAAACAAACT**GGTCTTTGGCGCAGGAACCATTCTGAGAGTCAAGTCCTgtgagtataaaacacactcaagtcccaacctgggtttcattgcatattaaataaactttttctggctgggaagtatttgattctaaagaaaggaaaag**tgaaattccaagcgcaacgcaagcatttca**ttagtgtgattcattaagaatttccatgggttgccttcgagagcgttaatcacatccattcactggtgagacccacagtttgaaaattgagttaaatttgaaaaaaaaattgtcaccatctctgtagaccacagtgcttgctgtttgggtgtctagacttccaaaatggaacctcaatctgtagtttcgagcttggagatttatcccgagcccgtgacttagagatgtgtaagaaaagctgc**acactgctgggccttgtagggcatttgtgggatcagtgt**catgaaatagcctgggtagacaggttggtgagg**agacactggtgacaggtgtct**gccctgtttctgtaaa**gctgctgacagccgtgagaagaaaagcagc**ggagacaagctgacttttgggaccgggactcgtttagcagttaggcccagtaagtctgagcagaaagtaagatatttatgccttttcctattatttgttctagcctgacatttgagttgtcctcctttggattccagatcaacaaaccatagtgtcttttcatactccttttaatatttgtggctaagggccctccgttcttcccatctgtcctacatgagggtcctgtggccaggtccacattcataaagaagccaacag**tgatgcagttggttggcagtgtgtcaagaaactgcagtca**tcatctccagggcccttactctggttatgtgtgtttcaagcttttatgccagcaacagaggagagggagaagtgaccagtagcaatg**accagggctattagctgtacctggt**gtctgtgggtttcagagcagctgtggaggttacgggaggttcagggcatgagtttttctggcagaggagtttatgtaa

Uppercase: TRAJ47

Lowercase: Flanking sequence[1000bp]

Red & Bold & Underline: Stem-loop [13]

Blue: Heptamer[29]

Green: Nonamer [2]

id-TRAJ54[J_gene_segment]

ttggtttatttctgtagagtttattattatacacagtgttcttgaagtaatagagaagtgcattagaagctcctgcaaatggaaatgaaatgtaaatttagattgtaaatacatgactgcggtaaatagatagcaagaaaaaggatgaagagatt**agaaatttgccacatttct**ctaaccctaaagggatgtttctataaatagtttaaaactgttgaagacgaagggggaggagga**aggagaacagaggcaaacagattttctcct**aatcctttccatttggcaaaatt**agaaatggtttttcaattaaatttccctgagcagaggaagaaaccatgtct**gtttccacactaaaattcctgtgggtgggagtctctagagttgatttggaggatggatccctgttagtgacaagtgctggtaatgctcctgttggggaaaggggatgagtacaaaaataaatccaagtaagtgtggagggacaagaagatctcacagtgcaggatttcccctggattttctgcattgccttttcacctttcctgtcttagcacgacaaattaggtcccagatgagcaggccctcgcattcaaaccggaaattttaaggaggaagccagattaactttactcgggagacctagtagactctgactttaaagat**tcttttaatccagctgtttttagggagagctctgtaaaaga**tgaaagaaatg**ttttaaaaaaaaattaaaa**ttctgtggggtgaaaacaaagatgttaaatatttgattggcaaggcaactggaaaatctggaccatgtctacaactgctaaaggaggctttgtgaaagagaaaatgagcagcccaaggagatcctgtcctaaacttctctggccagtgaaattcgggccattc**tctgccacagccctggactgctaggagggcaga**tcatatgtcttcctcagtggggagaggtgggccctcgctggcagtttctgtaaagcctcgtgctgtggtgTAATTCAGGGAGCCCAGAAGCTGGTATTTGGCCAAGGAACCAGGCTGACTATCAACCCAAgtaagtatgacagggtgaagctacatgcagctgagtacagtcttttcctttctagaccgtgtcctgcaagctctccttgagggacgtatactcattttgcattgtcctttgtagagaagcagaccaggaaagacaggaaagccctcaaatttccacttttaaacacctccctgtaa**aaactgtctcgcttccctccccttctacaatcagttt**ctagtaaatcagaatccggtgaattgatatgcaatttcaacgaaaaaaaaagcagagaaatagttaccccaacaagtgcaaaaagtagaaactatctgagtaccagccaagggaagataattaggaaagaaaaaaaaagaaagaaatggattactggaaccatgatggcagcttagtcacaaagaaaatgataaaaaaaaattaaaattaaatttcaaaaactttaaag**atatatacatatatatatatctatatatatatgatatat**tgtcctctggaactccagcactcacctacagcagctgcattctggacttattatcaaataacctataaacctgtagcttaatgaagagttgagcaatcgtgactatattgacccccaggacccttctgcaaagagcagcttctgttcctgtttctgtaaagccttctgtggctgtgagaatagtggaggtagcaactataaactgacatttggaaaaggaactctcttaaccgtgaatccaagtaa**gtttgaagggagtgggggaagggggaattcaaac**acttctgatttaattacttgcccctcca**aaacattccagcttaggttccaaatctaatgttt**gtgctggggggatatggtgcccatcagagggattagaactccactgtacaaagactaaattactttcagcccaaaacattcctatcattcatcttaaacaagattattggtgcaaggagaaaaactgaatacagaatcctgagggatccagctaagggtcaggaaatgtttgccgaa

Uppercase: TRAJ54

Lowercase: Flanking sequence[1000bp]

Red & Bold & Underline: Stem-loop [10]

Blue: Heptamer[21]

Green: Nonamer [4]

id-TRBJ2-1[J_gene_segment]

ttcctatgagctgcctgccacccctcgctcctcccacccacttcactataaatgccagtctgagca**ggtgggcacagtgagccccacc**agggagacccagtgacatagatggtctgctcagggtgatgcatgttccaaggagggacctctctgccccccaccattaccatcactgtgactttccccaagcccttcccattttaattcactgcctttgtcttttccaagccccacacagtcagactaacctctgccacctgcgcttcctgccgctgcccagtggttgggggagggggactagcagggaggaaacatttttgtatcatggtgtaacattgtggggactagcgggagggcacgatgattcaggtagaggaggtgcttttacaaaaaaccctgatgcagtaagcatc**cccacccagctcagggaatgcagctaccaggtggg**aagagttctctggggctggtcccagctgtggtcttgcagggtcccccaacccagcgagcacctgtccatctccctgtccagactcggcttccaaggaataagaaggccaagacagc**aaagtgggattatcactcagcacttt**taataaaacttgttcttgacaaagtacttgcacatgcattatttattaagaactgatgaaaaccctgag**ggaaagatattgtcccatctttcc**aatgaggaaactgagatcagaggttacaggtcatataactaggaaacggcaaggtctagcctgcaatatcgcccagctccagccgttccagtaccaccaatgccccttcagatttca**aatccactgtgttgtcccccagccaagtggatt**ctcctctgcaaattggtggtggcctcatgcaagatccaggttaccgtgtccagctaactcgagacaggaaaagataggctcaggaaagagaggaagggtgtgccctctgtctgtgctaagggaggtg**gggaaggagaaggaattctgggcagccccttccc**actgtgCTCCTACAATGAGCAGTTCTTCGGGCCAGGGACACGGCTCACCGTGCTAGgtaagaagggggctccaggtgggagagagggtgagcagcccagcctgcacgaccccagaaccctgttcttaggggagtggacactgggcaatccagggccctcctcgagggaagcggggtttgcgccagggtccccagggctgtgcgaacaccggggagctgttttttggagaaggctctaggctgaccgtactgggtaaggaggcggttggggctccggagagctccgagagggcgggat**gggcagaggtaagcagctgccc**cactctgagaggggctgtgctgagaggcgctgctgggcgtctgggcggaggactcctggttctgggtgctgg**gagagcgatggggctctcagcggtgggaaggacccgagctgag**tctgggacagcagagcgggcagcaccggtttttgtcctgggcctccaggctgtgagcacagatacgcagtattttggcccaggcacccggctgacagtgctcggta**agcgggggctcccgct**gaagccccggaactggggagggggcg**ccccgggacgccgggg**gcgtcgcagggccagtttctgtgccgcgtctcggggctgtgagccaaaaacattcagtacttcggcgccggga**cccggctctcagtgctgggtaagctggggccgccggg**ggaccggggacgagactgcgctcgggtttttgtgcggggctcgggggccgtgaccaagagacccagtacttcgggccaggcacgcggctcctggtgctcggtgagcgcgggctgctggggcgcgggcgcgggcggcttgggtctggtttttgcggggagtccccgggctgtgctctggggccaacgtcctgactttcggggccggcagcaggctgaccgtgctgggtgagttttcgcgggaccacccgggcggcgggattcaggtggaaggcggcggctgcttcgcggcacccggtccggccctgtgctgggagacctgggctgggtccccagggtgggcaggagctc

Uppercase: TRBJ2-1

Lowercase: Flanking sequence[1000bp]

Red & Bold & Underline: Stem-loop [12]

Blue: Heptamer[44]

Green: Nonamer [11]

id-IGLJ3[J_gene_segment]

cttccagggtgaacaattcatgtcttctctcatggtgaactctaggattcaagccatctaatgcttttgaa**gccactgtcattatatttaattgatgatgacaggtggc**caccaatgatgaatattttcccagggggagtctccccaagtggcttcagacttcctcacatggccccaggggattaaatggctcctgattactcagag**gataagaggttctgtcttatc**atgttcctttcttatttgtcttatgtgtctttcctgccccaggcctgggatcccccactgatctcccttcccttagtgagaggtgatatttggagaccacattctggaggctccctcatgtcccccatttgaaaaagacaacgg**cagctaccaccctagctg**tcccaccaaacatgaggccagattcaggggtgcagggatgctcccaaggttaccctaacagatgtgactggcatttcatattgggaccagcc**aggcctcactgaccaggcct**atccaactagaactactccagaa**ggtggggctgaaacccacc**aaggttcccagaacactgcactctagggcaatcagcctctgcatgggaggaaaggagcaccctctgcaccaccccatggtgttacca**aaagttgaaccatgggttggttcaacttt**gcagagaagagaccacctatcccatctgtggaaattcactccttagcgacactaatgccctctaataaattcaatcctgggcctgagtgatggttggtgcaaaaaacaaattcaa**gatcccagtgtcctccagaagcctggatttccagggatcctgctgtgggtcacaggat**gtcaccggtcccctctctctgtgggttgagtgtgggggccatgtggactccctcatgagcagatgccaccaggaccactggt**cccagcttcctccttcacagctgcagtgggggctggg**gctaggggcatcccagggagggtttttgtatgagcctgtgtcacagtgTTGGGTGTTCGGCGGAGGGACCAAGCTGACCGTCCTAGgtgagtctcttctcccctctccttccccgctcttgggacaatttctgctgtttttgtttgtttctgtatcttgtctcaacttgtggtcagcctttctccctgcatcccaggcctg**agcaaggacctctgccctccctgttcagacccttgct**tgcctcagcaggtcattacaaccacttcacctctgaccgcaggggcaggggactagatagaatgacctactgagcctcgtctgtctgtctgtctgtctctctgtctgtctgtctgtctctctgtttgtctctctgtctgtctgacaggcgcaggctgggtctctaagccttgtt**ctgttctggcctcctcagtctgggttcttgtcggaacag**ctttgcccttgggttacctgggttccatctcctggggaattgggaacaaggggtctgagggaggcacctcctgggagactttagaaggacccagtgccctcggggctgatgctcgggaatcacagagctgggacccagagccaggatccagacccagaatgaggtaggaggtggaggggctg**ccctgggcgtctgggggctgccaggg**actgagc**cctgagccagcctgagactcagg**aaaccccgtcaggagggagaagggagaagcagactctggacaccagaaagccaggggaagg**gtcacaaaaggagtggatgtgac**ggaagggcgggctcctgggtctcttcagaacatat**cccctgtgcccagggg**gatcagaggggcagagtccactgcgtgaaag**ccccactgctatgaccaggtagccgggacgtgggg**tggatgccagaaaagactccacggaataaga**gagagcccaggacagcaggcaggctctc**cgatccccccaggcccttgccccatacacgggctccagaacacacatttggctggaacagcctgagggaccaaaaggccccagtatcccacagagctgaggagccaggccagaaaagtaaccccagagttcgctgtg

Uppercase: IGLJ3

Lowercase: Flanking sequence[1000bp]

Red & Bold & Underline: Stem-loop [17]

Blue: Heptamer[29]

Green: Nonamer [3]

id-IGLJ7[J_gene_segment]

ctccttaggggcccaccctgttcctgactcccaggaccctgccctctacccccacatctatctctgaagcaggcacagactggaccctgaccctctgg**acccaggtcactcaccccacaggccagctgggt**acccctggtcccacatgtgtcaccagtgtcccagttatactcaagtcccctgggggataagggtcaactctactctctctcacccctaagtaaccagcccaaaactgaccacacctcaagtacctaatgtccagttatccacggatggtcagtggggctggggaggtcaaagtcaatcatgagattccagagtggtgccatagacgtgcctgtaaaccaattggtctttttaagagctgtatgtgggatctaagaaccgggttgatgattgt**cccaggaggtagcacaggatggggggcctggg**ctatgagacaaacacacatacataca**tgttcacaaacataaacacacatgaaca**cctcataggcacacaggtattcacaaacaaacatgcctgcatgcaaatacatacacacacactctcctgtgctatttagaatcattaccacatgggcatcccactaggggctgtgctgggcatccaggaccctcccagggaactcggatctccataggggaccagctccaccttccctgtggcataaacaggattctcacttcttccatggtgacagtctccaaaggagctgagccgggacacatggcttcctccaggacagagagtagcaggggacccagatgtgatgtgcagacatgcctcctccctctcccctctccctctgggggtgaagggatccctgacttccagtaaagtgtgcatccatgcatg**gaggggccatccccctc**accctcctatcggccagacctgcttcatgacagggccactggacctgggcgcccaagctgctgcagcggggaacgggtcgggtgtgtcaggaggggtttgtgtgtggggctgtgtcactgtgTGCTGTGTTCGGAGGAGGCACCCAGCTGACCGTCCTCGgtaagtctccccgcttctctcctctttgagatcccaagttaaacacggggagtttttccctttcctgtctgtcgaaggctaaggtctaagcctgtctggaggtctggaatctttgcccctccttgcctgggctcctgccctcttctgtgattctgtcctctgtgggtcccagttacggggctgcattaaacacagtgacaggaggcctttgactgaggacttggagagatgggggaggaaatggcaggaggacaaagatagaggaagaatattccgtgagaaggtggccccacagcgctgggtcacacgccatcccccaagacaggcagga**caccacagacagggtggtg**ggtctcagaaaactcaggccctaaacgtggatgcttaccaa**ttcctccactggaggaa**gacctcagagcagatgcccaggacagggacttctggtagggacggtgactgggacgggtgcctgtttgtcagggaaaacccactggagagtcagatcccccagataacttctcacgacatggagactctttcgaacagacaaagctccacgttcagctcagggagtaaaaaaaaaatgcctcaaatggaggcctttgatctactggaatccagcccccaggactgacaccctgtctcaccag**gcagcccagaggggtctctgcagggaggtggggtgggggctgc**aatgatggcaccagggagatgt**gtgggtaagaaacccac**tccctgtgagagagaagagcctgaacccaggaccaacagctgccctgcatgaagagatgagaacaaggggaactggtaggaggtgttcagacagacacccccaagatagacaaatacccagggtgagatgtggtcctggactccatcccatccagtgtggagccagcaccggtgggggtctataggtgatggaaaatatgaaaaagagacagatccaaga**gggggtctgtgaccccc**aagagtgggggcaactcccatctgacagcaa

Uppercase: IGLJ7

Lowercase: Flanking sequence[1000bp]

Red & Bold & Underline: Stem-loop [9]

Blue: Heptamer[31]

Green: Nonamer [2]

id-TRBJ2-4-2[J_gene_segment]

tattaagaactgatgaaaaccctgag**ggaaagatattgtcccatctttcc**aatgaggaaactgagatcagaggttacaggtcatataactaggaaacggcaaggtctagcctgcaatatcgcccagctccagccgttccagtaccaccaatgccccttcagatttca**aatccactgtgttgtcccccagccaagtggatt**ctcctctgcaaattggtggtggcctcatgcaagatccaggttaccgtgtccagctaactcgagacaggaaaagataggctcaggaaagagaggaagggtgtgccctctgtctgtgctaagggaggtg**gggaaggagaaggaattctgggcagccccttccc**actgtgctcctacaatgagcagttcttcgggccagggacacggctcaccgtgctaggtaagaagggggctccaggtgggagagagggtgagcagcccagcctgcacgaccccagaaccctgttcttaggggagtggacactgggcaatccagggccctcctcgagggaagcggggtttgcgccagggtccccagggctgtgcgaacaccggggagctgttttttggagaaggctctaggctgaccgtactgggtaaggaggcggctggggctccggagagctccgagagggcgggat**gggcagaggtaagcagctgccc**cactctgagaggggctgtgctgagaggcgctgctgggcgtctgggcggaggactcctggttctgggtgctgg**gagagcgatggggctctcagcggtgggaaggacccgagctgag**tctgggacagcagagcgggcagcaccggtttttgtcctgggcctccaggctgtgagcacagatacgcagtattttggcccaggcacccggctgacagtgctcggta**agcgggggctcccgct**gaagcccgggaactggggagggggcg**ccccgggacgccgggg**gcgtcgcagggccagtttctgtgccgcgtctcggggctgtgAGCCAAAAACATTCAGTACTTCGGCGCCGGGA**CCCGGCTCTCAGTGCTGGgtaagctggggccgccggg**ggaccggggacgagactgcgctcgggtttttgtgcggggctcgggggccgtgaccaagagacccagtacttcgggccaggcacgcggctcctggtgctcggtgagcgcgggctgctggggcgcgggcgcgggcggcttgggtctggtttttgcggggagtccccgggctgtgctctggggccaacgtcctgactttcggggccggcagcaggctgaccgtgctgggtgagttttcgcgggaccacccgggcggcgggattcaggtggaaggcggcggctgcttcgcggcacccggtccggccctgtgctgggaga**cctgggctgggtccccagg**gtgggcaggagctcggggagccttagaggtttgcatgcggggatgcacctccgtgctcctacgagcagtacgtcgggccgggcaccaggctcacggtcacaggtgagattcgggcgtctccccaccttccagcccct**cggtccccggagtcggggggtggaccg**gagctgg**aggagctgggtgtccggggtcagctctgcaaggtcacctccccgctcct**gggaaaagactggggaagagggagggggtggggagg**tgctcagagtccggaaagctgagca**gagggcgaggccacttttaatcttttttctggggtgtttagagagaaggtgaacgatggaggagaggatttgttaggactctgggagaggcgagactggagaggacgaagggaaatcctggtttggggaatgggtaggagtgggggtaactgctattcgtaggcaaaaagagctgagcaggctgggaacagcgcgggtgggcaagggtcagcactgcgggcaggcgggtgggtgttagggggcagaaatcctgcagccgagggtgcagtagaacacagaagaaaaagcctgccaaacaaaagtggaacagagaagccaaaaagggagatgaacatgagtcagtgaagaaaagaatgaaagtttact

Uppercase: TRBJ2-4-2

Lowercase: Flanking sequence[1000bp]

Red & Bold & Underline: Stem-loop [13]

Blue: Heptamer[29]

Green: Nonamer [11]

id-TRAJ44[J_gene_segment]

gtacctggtgtctgtgggtttcagagcagctgtggaggttacgggaggttcagggcatgagtttttctggcagaggagtttatgtaaagggttggcccagagtgtgtattcaggaggaggtgctgacggactcacctttggcaaagggactcatctaatcatccagccctgtaagtgcttttgcctgggaggtggggttcaaaatgcagtcctcatggggacatttctaaaccttggctcatgtgcttctg**gtgccactaccatttttggcac**tatcctact**tacatgggaggttttgagccatgta**gacaaggaggtgatgcagtttgtgctttggaggactcctgagatctgccctttggcaactgtgtttgagtaggagtccctcattggaaaatctggatgccagcagacaaaagatatgagcttaggatgggtcacaaacaaatggaggccactttgccagcagcacctttgcaacgaaaattgctttcataggaaatgtg**gaaatggctgtgtagatagagatttggccatttc**tccttttctgtctttataccctcttctctgcctgacttagggaccttaaattagagcagttattaatcatttgatttctcggccccatcactcatcaagtatgacaagagggattcattaagataaaacccagaaacagaaaggacaaagaattcaattctcaatgtcttca**gagttgaacttgacaactc**ctcagccttaaccatctagtcagcaccccatgaccaactgaggtcaaagagggcaaccagcaggtaatgttagtcaagtgagtatctggccagagcccagctgccctcataggaattaatactgtggagtttgggaaagggaaatgcatctgtgagggctcacccaggctgcccatctgggagagatcatggggagagttctgagtgagggagtgaacaccagatggaaagtcaagcaacaggtttctgttatgaagcatctcacagtgTAAATACCGGCACTGCCAGTAAACTCACCTTTGGGACTGGAACAAGACTTCAGGTCACGCTCGgtaggtaacagaaaccgtgaggtaacttaactgtgtgttcctttaaaacaggggatactgcagaggggatatcctggctggatgtagtccttccttccttccagtgggaggtgatctcactgaggacacatacctgcattcagtcagttcttgtgctatttttatacctgttacggaggctttcctttccacctgtgaaaacagaacctctctgctccaagcttcttcagtctttattaccctatgtatttcctttagaacagtggtttgggaattaatt**tatcatcaggactttgtctctgatgata**tcactctgggcaacaaaatgaaagggctctcaaatagagaacagagagtcctgaatactcggataacttttccagagtgatatgtgttatgcgtaatgggatgaagtgggctagaatgaccttccaatgactattttggagacacac**taaggaaacttgcttaacatcctatatcctta**acttgttagaacattggtgaagtaaagcaaaatcttgaccctctaagaacataaacaaatgccaactcctctcccccaaagcactgcca**tgtttcattatgtgcaattatgaaaca**aatacatga**taatttatatgaaatta**catagatgagtagtttgagtattttgacccatgagtaagatgcaaattgtagaagttctcaggtagt**tcttcataaaaagttgtccctgaagacaaatagtgtacctgggcataatttgt**ttgcagtgttggtttatgcaccatttggggacctgtttgaaagctgattatgtcccagtgtgccaatttggggtctcaattaactgtgctcagtgcactgataagtgcccatatttacagtctagaagcatgacaaaatgaccccatttctgtaggaaaaggagaatggaactgtgtgtggcgtggtacttgtcagtgctgctcccccttgtccctgccatgctgggaggtggctgcaagttttccccagg

Uppercase: TRAJ44

Lowercase: Flanking sequence[1000bp]

Red & Bold & Underline: Stem-loop [10]

Blue: Heptamer[33]

Green: Nonamer [5]

id-TRAJ17[J_gene_segment]

taggtagattaggacgaggaagcagagggagacaaaggcgatggaaagtcccttttaggaacccaggaatcagagagaacttgatggtccccaccaaaggcaaagaagggaggtcccactaaaacatgtgagcatcctaaaacacctgctctggtccacccacaatatttggtgaaggacatggccctccacccagaatgggcaagcaaactagaccagctttctaaggtgtgtttttcttgggtcctgtgactcttggccgtctccctgtcagaggctccatctgtcgctgcactctttcttattgagaatggcctctccgtggttctctgtaaactttccccaagaagcctgtttccatgcttcctcagcacttagcctcaccgatggatgggctctagagtagctggagggagtctgggtctgaa**actttcactgagaaagt**aaagttgatccgcagtatccagtggatatggcagctgg**gaggacctaggactgaagtactcgtcctc**tctttggggcctttcctgggcactcgattgaatacagaaaccctgttagcaagtgcatgcatgtatatgagtgtgtt**cattcagcatttccattctgaatg**tatgaatagcctta**ctctgaagactccctggagcttacagggctttctgttttcagag**aaaattgcctttgtgagacaaaaatggccaagtgggcccctgaatgaggttatggttgagagctattaatattatccttattcacagagcaagatccgaattcccaaagcagagtagacaataactatgtgaagggcttagacccaaatatgtgcccacttgcaagcattccagctctatgaagaggatacagtgctaggagataggattaccctgcacattgcaaaccctccacttttccaa**gaccatagaaaatgtcctttgttgagcaccaactgagtgttattttcttggtc**cctgtggtttttgctgggccttaaatcattgtgTGATCAAAGCTGCAGGCAACAAGCTAACTTTTGGAGGAGGAACCAGGGTGCTAGTTAAACCAAgtgagtactggggcttgacccac**aattgagccttgtcatcaatttgcaattcaatgtgccgtattggcccaaatt**aatgtctcaaacttgtctttaaaatgttt**catgttgctcttatgcataaacatg**cacttatgtacaaattcagcatggataaaggatattgtcatcttgacattttctagaatctattttcttaagagaaggcacatagtaaaaggtacttttcttatttcttcacatttcttaactgattttcccaagaggcagaggaacttggacgtggcgtttctgtc**tattcagagtagttgatttgaata**tggttacaatagcagagcaaggacttacaatttctaaggttcaaaatattgctgtctgtaccagcctgcagcctctataaccattattctgatttatttcctctggccctcatttttcccaattaatgtaactgaaatattggtaaccattatcttccggttcacagaaagtgtgtgaggattaaaaagtcgctgctctgaaaaggaattgagctggagggttttcttttgggggattattgttttgttgatggtattgtctctttcattgagggagagaataagaaaggaggcatacttgaaaggcaaggtaatatgatcaatacatgaacgtgtgcaaaataaatcattaaggcattcaactagaagccaaaaaacagtatggaccccggctcagccactcactcactaggcagccatgagaagatcaccaagctctcccagcctcagttaacctaaaatgtggagattattattccagccttgactacttcaaaggatgctatgaaattcagatgagataatatatgtgaaatctccttgaaaaagatgggtggtcaaatgattcatgatttttttttttttttttttttttttttgagacggagtctcgctctgtcgcccaggctggag**tgcagtggcgggatctcggctcactgca**agctccgcctcccgggttc

Uppercase: TRAJ17

Lowercase: Flanking sequence[1000bp]

Red & Bold & Underline: Stem-loop [10]

Blue: Heptamer[22]

Green: Nonamer [5]

id-TRAJ58[J_gene_segment]

aattctgacttcaaacttctaaaaagatcaaatgttaaatcagatagtaggcttggaaaactctatttctctatgta**aaaagtagagaactactttt**cttttgtttgatcatt**ttattttgtttaggaaataa**gaggattagataccctggtggtgagtggggagggcagggacatttgcacagcatttctggataaattagatccaaataatcaaatcaaagactctcagctcagaaagt**aattaaagatctcctcacccacctctttaatt**ttgcaaatgaagacagtgaaactcagagaggttatgaacttgctcaaggtcacacaactgatcctgatatcaaggtccagggcaaagccaagacagctcatagttctcaccagcccagac**ccaagggagaaaaaaacatcatagtccttgg**ccacggcagcatcatttgcatcccaaacattctttacccaagacttaat**gaactcaaaaagaaatcctgagttc**caaagggaaataaaatcattctgcgtctgttgaaaaaaaaagcaagctaaagtggaacaataattgaacttgatattaggggaaaggtgcagccatctgcagactgagagaagggtgaaaaaaacaaaatgaaaatgctagtctt**gtgttcagacaatgattatgaccatagaacac**tacctataagttctgtagagctaatttctttcccagtggggttga**tattctatgatagattgcatcagcggtgtgattgttttggaccatagaata**cgtatacagaattggcttgttgagccttctggtgtcttgaagtgaaaatcatatctctggacatatctgtcaggcctgaagccctaggctactggaaagaaagataccatgacttatccttcctctaaagcttagaagttggagaaaaattctagttcttgggaagctctgcaaagcagggttcgttaagctgat**gccacaggtttttgcaaagcccctcagcacagtgTTTAAGAAACCAGTGGC**TCTAGGTTGACCTTTGGGGAAGGAACACAGCTCACAGTGAATCCTGgtaagtggaggggagcattgaatcctctgcctgaatgtatccattttatgccaatgaattgagatatggcttcttgccccatctctgccacattcaagtcctgcttggctaaacaatgctcacggatgtgcacgtgtgcagtgggatgcagctttttctaacctctcacaaaatcacgcaacccctt**caccaaactgcttggtg**gctcttctgtgccctcacagcccttctagtttagcagttagaggaagacaagtgcccaagggctttctcatttttaggtcattgtggaaagaggaacgttggtgatatgtttgcctgcttgggaggaaaataataatgatgatttactcattggtttctatatgctagttgatgtgctgcattctttctcttatcacatttaatccttataccaggtctgtgagttaggtgctattattatcctcattt**tatagaagagatatatttatattttatatttctata**gaagcacagagatgttgaataacttgttcaagccgatgaagtaagaaagagaacctggagtttaattcttggcactcagattccagagctaaaagcttttcagtgaggggattatgt**ctagagggaatgtccctgatgctctag**tctctaaggttgctaagataatcgctaaagaactgcgctctcagcaagtaaagccaggggttggtcaacaggaccacggtgacccagttacatattcttaatccttacaacatcatcaagcagttgctgt**tttccccactttacaagtgaggaaa**ctgaggtgtag**caagttaggaaacttg**tcc**tgagttttaaactca**agcctacctgccccaaagcctgagctcttttaaat**taactgcaactgctgaccagtta**gattaactaattacttaattaataagataacaacctaacaggctgtaaggtaaaaaagtaatcagatttgttctataggtcccctcccttttcgggaatagctatacaa

Uppercase: TRAJ58

Lowercase: Flanking sequence[1000bp]

Red & Bold & Underline: Stem-loop [15]

Blue: Heptamer[18]

Green: Nonamer [3]

id-TRAJ37[J_gene_segment]

gattatccagaatgactcaaacttgcgatataactcctgcatctctcagttctctttggcttagagccttattgtggatttgctagaggtaacaggaaaggaactgatttcatcctaattctagcctcttctgagaagttattctctgcaagagtcttcaggagcatcttacagaccacctacactattcagaattttgttccgcacaccacattgtgtt**cccagcagtgtttgctgtggctttggctggg**gccaggtcctgagtcagttctggagctccaggcttggcaaagctggataagggcaaggggagttgtacagcagaaccaaagggcttcaggaaagacatctttgatatagacattaatgaatttctaaatgggcttctatatgctaacctgataagactctctggggagaaggtcatgctttcctttgcagctctaaatataagacacatgcagaaaaaaaaaaagaatgggaaatgtctgttacagtgccaaaatgaagataatcagtata**aatattagatttgtcttaatatt**aagtttatgaaatgtaagtctttcagaagaactatctcaggat**atatttattaggtcaatgaaatat**tttatccatgtgactggtatacacacttcaaaggagaacccaaaatggcaggcaggggagggtggctaatgaaaggtatggcatagtagttaagaatatgaattttggaataaaactgtttgggtaaatcctaattcagaaggttacttaatctctttatgcctcttttttctcatctgtaaaatggggatggtaatagtacc**tagctcactgagatgttacgagatttaaagagcta**atacctataatggactttgcataatggcagcccgattgaaagttttcagttaatatgtttttattgctactatgtcaggttgctggagggaaggaagaa**aaatggaaatgatccccattt**atagtttttgtaaagtacagcattagagtgTGGCTCTAGCAACACAGGCAAACTAATCTTTGGGCAAGGGACAACTTTACAAGTAAAACCAGgtaggtctggatgtttctaagtgaagtgcagttatccactctgttc**gttttttgtttaatcttctctgtctcaacttctaaccaaaaac**t**cttactcttaggttatattttccttttataaaagtaag**tagaatgtgtgtaaagtctggttgatatgatcctttctaactcacttggtttcagctagtggcaaaaagaagggtgtattggatctggaattcttaggttaaggctgagagaatgtgagtttggctgacttggaagaaataaaaacatcaaatgtgggtggttcacacaaaagaaggtgatttttcataataatatactaaagtagaatccacccttgtgaccaccagaattattcaagtacagtcttccaaggacaaaatgggtagcatggaaatgat**atgtttttttggggggaaaaaacat**ggggagagagagagaaaggggactattagttgtctggattgttaggaaacatggaagacagcacttcttgttgtaaggtttggcaaaggttatcaccaagtagaagtccaaatgtgtgtaacatacccctggcaaacgttatagaagaaggaa**gctggggccgggcacggtggctcacgcctgtaatcccagc**actttgggaggccgaggcgggtggatcacgaggtcaggagttcgagaccatcctggctaacacagtgaaaccccgtctctactaaaaaaaaatacaaaaaattagc**caggcgtggcgggcgcctgtagtcccaactactacag**aggctgaggcaggagaatggcgtgaacctgggaggtggagtttgtgccactgctgaga**tcgcgccactgcactccagcgtgcgcga**cagagtgagactccatctcaaaaaaagaagaaggaagctgggagcagtgtctctcacctgtaat**ctcagcattttggaaggctgag**gcaggaggatcacttgaggccaggagttcaagaccaccctgggcaacacagtgagacccc

Uppercase: TRAJ37

Lowercase: Flanking sequence[1000bp]

Red & Bold & Underline: Stem-loop [13]

Blue: Heptamer[23]

Green: Nonamer [3]

id-TRAJ12[J_gene_segment]

ccatttacatgtgatctgtagccctaggcattggtaaaaagcactttctcttctactgaaatatagtggctcaaatgctaccc**aaatgaaaagggcaggggaagggaagtcattt**tgtaaaggcaggcattacagtgtgaa**ttctgggggttaccagaa**agttacctttggaactggaacaaagctccaagtcatcccaagtgagtccaatttcctatgctttcctcttccttgtgttgtcttctctcagaccgtaacatttggagcacat**aaagagatattatgggcctgctcttt**tctggctg**tttcaggagactgaaa**aggattgta**taagaacatcctttagcatgttctta**gtattgttttgtccagtgtgtgtttcttcttaatatcaaaaaaaactaagacattgcctgtgaagccaatggaaaacagtagatttgcaaagaaaagaactaaagagaagacaagtggtactttttaccaagc**ttaaagcagagtgggtgtcctttaa**tcagagggatgcttggttctgggggctggatag**gactcaggtaagtaacaaggctaccactgagtc**actg**agcttctaaaatgagttgactcgggggagcaggaagct**gccactcatgatg**atctgaatgctattccgtgaagtttcagat**tgattgaagttaaaattaggggtgacactatgtcaggaatctaaaaattgataactgtaatggaaatgaaagaagggctagctacatctgttggataaatggagttgaaaatcatgaatttgatggcgacagacccagagagttgggcaggtagaaacttcatttcaagtgatatgcagtaaataccccaaatttgagtcaaatggcctgtgtaaaaaggaagtgaaaatgtaacccgtatctccttgtaggtgaaaaggccatatctctaggtcttaggtatgaaaaggatgtgtgggcttctgg**gtgtttttgactgactaagaaacac**tgtgGGATGGATAGCAGCTATAAATTGATCTTCGGGAGTGG**GACCAGACTGCTGGTC**AGGCCTGgtaagtaaggtgtcagagaggcaacagaaagattgagggtaaaatgtcttcatgtctcaggaaatgattgtataatgcaaaatgagctggagttttttaggagcactgttactagagttctgatgtct**ggttctagcaaaagaacc**ctaatatagctgtaatccctctctagaaggaacaaggaagaggattcttctctgaaagctcctcctaagatttatccatcctctgccaaagtgtcttgtggatccaggatacttacatctttgttagcagtgctggcccaacctgctcagcctcagatcacctatc**cctcatttgtaaaatgagg**ataatatttaagctcatggcacagttgaagcccaaatgaggtagtggaatatatgcagattattaatccatttggctcataaatgaaggttttctttccttttccctaagtagatcatgggttgaaatacctaacact**gcaaatcatttttgtatggggatttgc**tatagtgtgaattcaggatacagcaccctcacctttgggaaggggactatgcttctagtctctccaggtacatgttgaccccatcccacccatgttttccccctatctggtttaaggcttccatatgtattgcgtgttatcctcatggatttcatcatccttgttttattatcaatgttctgtgaatttaagattgagcctccatgg**actcttcatttaaaaatgaaaatagctaatagaagagt**tggaaataacagtagaactaattcactaggtcaggatggagaagggagtaataccctagacaattaggacaaacgtggtttttccagaaatagactcacttcctgtttaaagcctagactgtggtctctccccgggcacccttcacattcctctaaccctccatatcccaaatt**tagctgtggaatcttagacaatctgtgacctatgcagcta**cagaaatcttttcttactgagaatatctgcctatttgaaccaaaa

Uppercase: TRAJ12

Lowercase: Flanking sequence[1000bp]

Red & Bold & Underline: Stem-loop [16]

Blue: Heptamer[27]

Green: Nonamer [5]
