## Supplementary Data 4 for "Adaptive immunity: from CRISPR to CRIHSP?"

id-IGHG3-4[C_gene_segment]

aatggggcctccctgtggcctgggggtcctggcaccacgcagggtggggagggccaagggcaggtgcaaggctcctacctg**tgctggggggcctgggttgagcccagca**gggaccttgccgggggaagctctggagagagggaggaggtgggctggtggctgagaaggccaggccagggctgggagggtgacggtgtggtgactga**gcctccagaagtaatgcaggacactgggaggc**agggggcatccaggcactcagggccctgacctgggctgctgcacactggggctaaggggaaaggaggggagaggctgaggaggaggctcccggggcgatattccaaggcagggggttccggggccctggggctgaagggcgccgaccctatgcagtgtctggc**ccctctgctgcacagaagaaaagggccttggagggcagaggg**caggctatgaccag**ggccctgggcaagtcaggcccactcactagcggagggcc**acgctggggcggcagggtcaggagcttcaggggactcaggggacccacgagaagccatctgagaacagtgtccactggtcaagccaggcacccataaaaggctggagtggggccaatgggcatgagccgtccctgaggtggcaccgatggccagagctgaggccaagctagaggccctggactgtgctgactcc**cggcagacacagagcgctgacctggctgccg**agccccgcctcctagg**ctgcaggggtgcctgcag**aagggcaccacagggccaccggtcctgcaagctttctggggcgggccgg**gcctgaccttggctttggggcagggagggggctaaggtgaggc**aggtggcgccagccaggcgcacacccaatgcccgtgagcccagacactggaccctgcctgga**ccctcgtggatagacaagaaccgaggg**gcctctgcgccctgggcccagctctgtcccacaccgcagtcacatggcgccatctctcttgcagCTTCCACCAAGGGCCCATCGGTCTTCCCCCTGGCGCCCTGCTCCAGGAGCACCTCTGGGGGCACAGCGGCCCTGGGCTGCCTGGTCAAGGACTACTTCCCAGAACCGGTGACGGTGTCGTGGAACTCAGGCGCCCTGACCAGCGGCGTGCACACCTTCCCGGCTGTCCTACAGTCCTCAGGACTCTACTCCCTCAGCAGCGTGGTGACC**GTGCCCTCCAGCAGCTTGGGCAC**CCAGACCTACACCTGCAACGTGAATCACAAGCCCAGCAA**CACCAAGGTGGACAAGAGAGTTGGTG**AGAGGCCAGCGCAGGGAGGGAGGGTGTCTGCTGGAAG**CCAGGCTCAGCCCTCCTGCCTGG**ACGCATCCCGGCTGTGCAGTCCCAGCCCAGGGCACCAAGGCAGGCCCCGTCTGACTCCTCACCCGGAGGCCTCTGCCCGCCCCACTCATGCTCAGGGAGAGGGTCTTCTGGCTTTTTCCACCAGGCTCCGGGCAGGCACAGGCTGGA**TGCCCCTACCCCAGGCCCTTCACACACAGGGGCA**GGTGCTGCGCTCAGAGCTGCCAAGAGCCATATCCAGGAGGACCCTGCCCCTGACCTAAGCCCACCCCAAAGGCCAAACTCTCTACTCACTCAG**CTCAGATACCTTCTCTCTTCCCAGATCTGAG**TAACTCCCAATCTTCTCTCTGCAGAGCTCAAAACCCCACTTGGTGACACAACTCACACATGCCCACGGTGCCCAGGTAAGCCAGCCCAGGCCTCGCCCTCCAGCTCAAGGCGGGACAAGAGCCCTAGAGT**GGCCTGAGTCCAGGGACAGGCC**CCAGCAGGGTGCTGACGCATCCACCTCCATCCCAGATCCCCGTAACTCCCAATCTTCTCTCTGCAGAGCCCAAATCTTGTGACACACCTCCCCCGTGCCCACGGTGCCCAGGTAAGCCAGCCCAGGCCTCGCCCTCCAGCTCAAGGCAGGACAAGAGCCCTAGAGTGGCCTGAGTCCAGGGACAGGCCCCAGCAGGGTGCTGACGCGTCCACCTCCATCCCAGATCCCCGTAACTCCCAATCTTCTCTCTGCAGAGCCCAAATCTTGTGACACACCTCCCCCATGCCCACGGTGCCCAGGTAAGCCAGCCCAGGCCTCGCCCTCCAGCTCAAGGCGGGACAAGAGCCCTAGAGTGGCCTGAGTCCAGGGACAGGCCCCAGCAGGGTGCTGACGCATCCACCTCCATCCCAGATCCCCGTAACTCCCAATCTTCTCTCTGCAGAGCCCAAATCTTGTGACACACCTCCCCCGTGCCCAAGGTGCCCAGGTAAGCCAGCCCAGGCCTCGCCCTCCAGCTCAAGGCAGGACAGGTGCCCTAGAGTGGCCTGCATCCAGGGACAGGTCCCAGTCG**GGTGCTGACACATCTGCCTCCATCTCTTCCTCAGCACC**TGAACTCCTGGGAGGACCGTCAGTCTTCCTCTTCCCCCCAAAACCCAAGGATACCCTTATGATTTCCCGGACCCCTGAGG**TCACGTGCGTGGTGGTGGACGTGA**GCCACGAAGACCCCGAGGTCCAGTTCAAGTGGTACGTGGACGGCGTGGAGGTGCATAATGCCAAGACAAAGCCGCGGGAGGAGCAGTACAACAGCACGTTCCGTGTGGTCAGCGTCCTCACC**GTCCTGCACCAGGAC**TGGCTGAACGGCAAGGAGTACAAGTGCAAGGTCTCCAACAAAGCCCTCCCAGCCCCCATCGAGAAAACCATCTCCAAAACCAAAGGTGGGACCCGCGGGGTATGAGGGCCACATGGA**CAGAGGCCAGCTTGACCCACCCTCTG**CCCTGGGAGTGACCGCTGTGCCAACCT**CTGTCCCTACAGGACAG**CCCCGAGAACCACAGGTGTACACCCTGCCCCCATCCCGGGAGGAGATGACCAAGAAC**CAGGTCAGCCTGACCTG**CCTGGTCAAAGGCTTCTACCCCAGCGACATCGCCGTGGAGTGGGAGAGCAGCGGGCAGCCGGAGAACAACTACAACACCACGCCTCCCATGCTGGACTCCGACGGCTCCTTCTTCCTCTACAGCAAGCTCACCGTGGACAAGAGCAGGTGGCAGCAGGGGAACATCTT**CTCATGCTCCGTGATGCATGAGGCTCTGCACAACCGCTTCACGCAGAAGAGCCTC**TCCCTGTCTCCGGGTAAATGAgtgcgacggccggcaagcccccgctccccgggctct**cggggtcgcgcgaggatgcttggcacgtaccccg**tgtacatacttcccgggcacccagcatggaaataaagcacccagcgctgccctgggcccctgcgagactgtgatggttctttccacggg**tcaggccgagtctgaggcctga**gtggcatgagggaggcagagcgggtcccactgtccccacact**ggcccaggctgtgcaggtgtgcctgggcc**gcctagggtggggctcagccaggggctgccctcggcagggtgggggatttgccag**cgtggccctccctccagcagcagctgccctgggctgggccacg**ggaagccctaggagcccctgg**ggacagacacacagcccctgcctctgtaggagactgtcc**tgtcctgtgagcgccctgtcctccgaccc**gcatgcccactcgggggcatgc**ctagtccatgtgcgtagggacaggccctccctcacccatctacccccacggcactaacccctggcagccctgcccagcctcgcacccgcatggggacacaaccgactccggggacatgcactctcgggccctgtggagagactggtccagatgcccacacacacactcagcccagacccgttcaacaaaccccgcactgaggt**tggccggccacacggcca**ccacacacacacgtgcacgcctcacacacggagcctcacccgggcgaaccgcacagcacccagaccagagcaaggtcctcgcacacgtgaacactcctcggacacaggcccccacgagccccacgcggcacctcaaggcccacgagccgctcggcagcttctccacatgctgaccagctcagacaaacccagccctcctctcacaaggtgcccctgcagccgccacacacacacaggcccccacacacaggggaacacacgccacgtcgcgtccctggcactggcccacttcccaatacagcccttccctgcagctgg

Uppercase: IGHG3-4

Lowercase: Flanking sequence[1000bp]

Red & Bold & Underline: Stem-loop [29]

Blue: Heptamer[56]

Green: Nonamer [1]

id-IGHG1[C_gene_segment]

gagagaaatggggcctccctgtggcctgggggtcctggcaccatgcagggtggggagggccaagggcaggtgcaaggctcctacctg**tgctggggggcctgggttgagcccagca**gggaccttgccgggggaagctctggagagagggaggaggtgggctggtggccgagaaggccaggccagggctgggagggtgacggtgtggtgactga**gcctccagaagtaatgcaggacactgggaggc**agggggcatccaggcactcagggccctgacctgggctgctgcacactggggctaaggggaaaggaggggagaggctgaggaggaggctccaggaggctattccaaggcagggggttccggggccctggggctgaagggcgccgaccctatgcagtgtctggc**ccctctgctgcacagaagaaaagggccttggagggcagaggg**caggctatgaccag**ggccctgggcaagtcaggccaactcactaggggagggcc**acgctggggcggcagggtcagg**ggcttcagggggctcgggggacccacgagaagcc**atctgagaacagtgtccactggtcaagccaggcacccataaaaggctggagtggggccaatgggcatgagccgtccctgaggtggcaccgatggccagagctgaggccaagctagaggccctggactgtgctgactcc**cggcagacacagagcgctgacctggctgccg**agccccgcctcctagg**ctgcaggggtgcctgcag**aagggcaccacagggccaccggtcctgcaagctttctggggcaggccgggcctgaccttggctttggggcagggggtgggctaaggtgacgcaggtggcgccagccaggcgcacacccaatgcccgtgagcccagacactggacgctgaacctcgcggacagttaagaa**cccaggggcctctgcgccctggg**cccagctctgtcccacaccgcggtcacatggcaccacctctcttgcagCCTCCACCAAGGGCCCATCGGTCTTCCCCCTGGCACCCTCCTCCAAGAGCACCTCTGGGGGCACA**GCAGCCCTGGGCTGC**CTGGTCAAGGACTACTTCCCCGAACCGGTGACGGTGTCGTGGAACTCAGGCGCCCTGACCAGCGGCGTGCACACCTTCCCGGCTGTCCTACAGTCCTCAGGACTCTACTCCCTCAGCAGCGTGGTGACC**GTGCCCTCCAGCAGCTTGGGCAC**CCAGACCTACATCTGCAACGTGAATCACAAGCCCAGCAA**CACCAAGGTGGACAAGAAAGTTGGTG**AGAGG**CCAGCACAGGGAGGGAGGGTGTCTGCTGG**AAGCCAGGCTCAGCGCTC**CTGCCTGGACGCATCCCGGCTATGCAGCCCCAGTCCAGGGCAG**CA**AGGCAGGCCCCGTCTGCCT**CTTCACCCGGAGGCCTCTGCCCGCCCCACTCATGCTCAGGGAGAGGGTCTTCTGGCTTTTTCCCCAGGCTCTGGGCAGGCACAGGCTAGG**TGCCCCTAACCCAGGCCCTGCACACAAAGGGGCAGGTGCTGGGCTCAGACCTGCC**AAGAGCCATATCCGGGAGGACCCTGCCCCTGACCTAAGCCCACCCCAAAGGCCAAACTCTCCACTCCCTCAGCTCGGACACCTTCTCTCCTCCCAGATTCCAGTAACTCCCAATCTTCTCTCTGCAGAGCCCAAATCTTGTGACAAAACTCACACATGCCCACCGTGCCCAGGTAAGCCAGCCCAGGCCTCGCCCTCCAGCTCAAGGCGGGACAGGTGCCCTAGAGTAGCCTGCATCCAGGGACAGGCCCCAGCCG**GGTGCTGACACGTCCACCTCCATCTCTTCCTCAGCACC**TGAACTCC**TGGGGGGACCGTCAGTCTTCCTCTTCCCCCCA**AAACCCAAGGACACCCTCATGATCTCCCGGACCCCTGAGGTCACATGCGTGGTGGTGGACGTGAGCCACGAAGACCCTGAGGTCAAGTTCAACTGGTACGTGGACGGCGTGGAGGTGCATAATGCCAAGACAAAGCCGCGGGAGGAGCAGTACAACAGCACGTACCGTGTGGTCAGCGTCCTCACC**GTCCTGCACCAGGAC**TGGCTGAATGGCAAGGAGTACAAGTGCAAGGTCTCCAACAAAGCCCTCCCAGCCCCCATCGAGAAAACCATCTCCAAAGCCAAAGGTGGGACCCGTGGGGTGCGAGGGCCACATGGA**CAGAGGCCGGCTCGGCCCACCCTCTGCCCTGAGAGTGACCGCTGTACCAACCTCTGTCCCTACAGGGCAG**CCCCGAGAACCACAGGTGTACACCCTGCCCCCATCCCGGGATGA**GCTGACCAAGAACCAGGTCAGC**CTGACCTGCCTGGTCAAAGGCTTCTATCCCAGCGACATCGCCGTGGAGTGGGAGAGCAATGGGCAGCCGGAGAACAACTACAAGACCACGCCTCCCGTGCTGGACTCCGACGGCTCCTTCTTCCTCTACAGCAAGCTCACCGTGGACAAGAGCAGGTGGCAGCAGGGGAACGTCTT**CTCATGCTCCGTGATGCATGAGGCTCTGCACAACCACTACACACAGAAGAGCCTC**TCCCTGTCTCCGGGTAAATGAgtgccacggccggcaagcccccgctccccaggctct**cggggtcgcgcgaggatgcttggcacgtaccccg**tgtacatacttcccaggcacccagcatggaaataaagcacccagcgcttccctgggcccctgcgagactgtgatggttctttccacggg**tcaggccgagtctgaggcctga**gtggcatgagggaggcagagtgggtcccactgtccccacact**ggcccaggctgtgcaggtgtgcctgggcc**gcctagggtggggctcagccaggggctgccctcggcagggtgggggatttgccag**cgtggccctccctccagcagcagctgccctgggctgggccacg**agaagccctaggagcccctgg**ggacagacacacagcccctgcctctgtaggagactgtcc**tgttctgtgagcgccctgtcctccgaccc**gcatgcccactcgggggcatgc**ctagtccatgtgcgtagggacaggccctccctcacccatctacccccacggcactaacccctggcagccctgcccagcctcgcacccgcatggggacacaaccgactccggggacatgcactctcgggccctgtggagagactggtccagatgcccacacacacactcagcccagacccgttcaacaaaccccgcactgaggt**tggccggccacacggcca**ccacacacacacgtgcacgcctcacacacggagcctcacccgggcgaaccgcacagcacccagaccagagcaaggtcctcgcacacgtgaacactcctcggacacaggcccccacgagccccacgcggcacctcaaggcccacgagccgctcggcagcttctccacatgctgacctgctcagacaaacccagccctcctctcacaaggtg**cccctgcagccgccacacacacacagggg**atcacacaccacgtcacgtccctggccctggcccacttcccagtgccgcccttccctgcagctggggtcacatgaggtg

Uppercase: IGHG1

Lowercase: Flanking sequence[1000bp]

Red & Bold & Underline: Stem-loop [32]

Blue: Heptamer[47]

Green: Nonamer [0]

id-IGLC4[C_gene_segment]

cccattaaccccaagtggactagtccccataactgggaggtgggatttagtgaccacacttggggtgcttctcacacagcccttttgagtcagacactccagacatacccagaaatgaga**caagaccctgaaagggtaacaggggcttg**cttccaacttctccctggaggttgaggctggcatttcatactaaaacctagtgagacccatcccaaactaagacaacacaaggaggacggaagtgagacgccctggagttgtggttgtggtcacgttggagcttcccatgactgctgactct**ggggcaagctgcccc**tcctctaaggcactcactggggacacctgaggacgcctcctgctcttaccctgtagtcacaccaagagatc**agggttacaacaaccct**atagagaatccctgtccccttccatgtca**cttcactccttcgtgaag**caaatgccctcaaggagctcattcccattcctgggtcacagtcacctggaaaacctgatccagacaccaacctcctcaggcctcgccatttccagacgtcccgttactgcatacgcttggtcgactgtcccatctcagcttgagaagggcaggcaggtgtgtggactctgctgagcaaatgccttccaggggcagtggtctggcttcctgcaccatagcttcaggtgggggatggggagggggagttaggggccccagggaagagtttttgtatgaacctgtgtcaccgcattttgtatttggtggaggaacccagctgatcattttagatgagtctcttcttccctttctttccctgccaagttggtgacaattttattctgatttcgatctttgtctgtgacttgccacagcctgtggtcagggtttcctttgggacctcggtcct**gggaggctgatctctctcctccc**tattcagacccctgtatgcctcagctggtcactgagacaccttcatct**cctctgaccccagagg**cagggagctccaagACAAGGCCACACTGGTGTGTCTCATGAGTGACTTCTACCCGAGAGCCATGACAGTGGCCTGGAAGATAGATGGCATCACCATCACCC**AGGGTGTGGAGACCACCACACCCT**CCAAACAGAGCAACAAGTATGCGGCCAGCAGCTACCTAAGACTGGCACCCGACAGTGGAAGTCCCACAACCTCTACAGCTGCCAGGTCACGCATGAAAGGAA**CACTGTGGAGAAGACAGTG**GCCCCTGCAGAATGTTCT**TAGgtccccgaccctcacctacccacgggggccta**gagctgcaggatcagggcatgtgtctcccctcccactccaagtcatccagcccttctccctgcacccagtaaccctcaataaatatcctcattgtcaacc**agaaatcctgctgtctgtcttcatttct**tatctcatatttagtttgcaacctccttaaattctaagcaaggatgaggaaaatccaggtgcccagtttatcgggtgagaagtccatggtggtgccatcaccaggaacttgtggaaaggtctgggaatggaaactcacaggtgaatttcacagattttcacaatacagggtggctaagtaaagacacttacaagtcctgcaatagggaaacaggaagtccagaatcct**gctcaccatcccagccaacttagtgagc**cctaggatgctctgcaagatactggtgttcacgtcgctagctctggaaagtggggtgaggctggggcacacggg**tgatcagttatgatca**gatgggcttagggtgaggttcaaagttaaccagcacgtggctgagatctcaaccatgaagttcccaattctaaagtcaggctctggggtggagtgagtatgtgcttggtgtgtggctgagcctgtgatggtcagctcgtgtgaggggaggactcctgtggactgagacaaatgagcaaagacaccatcccagg**cacagaacgggcatcccatggttgtcggggagagtctgtg**tcagagtctcattctggactagagtcaaggctgggtcacgca**aggtcagcacagggtgaacatgacct**aggggctatctataggcaaagtcaggctttcacgggatctcaactgccccaaacacccccatcccaccaggccccactccctctgtcactcacgttgttccgtcccctcaccccctgcaccatggtgcaccggcagcctcactcagagacaccctcatcccggggtccctgacagtgggcaatttggtccctt

Uppercase: IGLC4

Lowercase: Flanking sequence[1000bp]

Red & Bold & Underline: Stem-loop [14]

Blue: Heptamer[30]

Green: Nonamer [4]

id-IGHD[C_gene_segment]

atggggagaagaggagggtcatccagaatttgggaaagcagggcgacagtttct**gccccaagggagaagggaaggaggatggggc**caccgccacaccagatgaccttgcgtaccaggccaaagaacgggaacacctggc**cccacctgagcagcaatagtcagtgtggtggtggg**cagacatgggtggaggcagggggtgagtagaaggttagactaagacggagcacctggggcctcca**gggacccaggcaagaaccctgcacttgctcagctgccctgggtacc**caggtctccaggaaagtgaggctgagagccaagcc**cagcaggcagccacacattctgggacctgccaccctacagcctgctg**t**ccatgagtaacaccccttacaaggggccaggtcagctcatgg**gtttat**cccaggcagcagagcccctggg**gccaggaatcaggga**gaggagcatccaatcccaccagctcctc**cggagccactca**gagggccagacgcatggccctc**cacagggaccccatggcccctgcagggcagctgaggacccgtggctggg**agcctgggcaggagggtcatacagccctaggcccgttgctcccaggct**tgagt**gcccctcctcccctcaggggc**ccaagggaagtgggttccagagaggttgggggcagcagggaaggtggaggtcccaggaatgc**ccagaggggcaccaaagcctctggagggaagacccctcc**cttccaggagctctcggcaacaagagcccagggtccacaaagccacaggtcccactcggttattctgactcacaacacaggagcggcagcaggggcattcgtgttcacgggccacttggtcagccccgctcaccctgggcactcctcctgggccccttttccctgccttccctgtc**accctgctgccagggt**cctctgccctgccctgccccttgtcctcagagcctccagcctcagactcccactgtgtctgtcttccagCACCCACCAAGGCTCCGGATGTGTTCCCCATCATATCAGGGTGCAGACACCCAAAGGATAACAGCCCTGTGGTCCTGGCATGCTTGATAACTGG**GTACCACCCAACGTCCGTGACTGTCACCTGGTAC**ATGGGGACACAGAGCCAGCCCCAGAGAACCTTCCCTGAGATACAAAGACGGGACAGCTACTACATGACAAGCAGCCAGCTCTCCACCCCCCTCCAGCAGTGGCGCCAAGGCGAGTACAAATGCGTGGTCCAGCACACC**GCCAGCAAGAGTAAGAAGGAGATCTTCCGCTGGC**CAGGTAGGTCGCACCGGAGATCACCCAGAAGGGCCCCCCAGGACCCCCAGCACCTTCCACTCAGGGCCTGACCACAAAGACAGAAGCAAGGGCTGGGCTGTGAGGCAACCCCCACCTCCCCCTCAGAGCACGTTCCTCCCCCTTCACCCTGTATCCACCCCTCCGGACCCTCCCCATCTCAGTCCCTCCGCTCCCTCTCTCTGAGGCCCATCTCCCAATACCCAGATCACTTTCCTTCCAGACCCTTCCCTCAGTGTGCACGGAGGCAGCTTGCCCAGCAAAGGTGACTGTCTAGTGGGCTTCCCACAGCCAAGCTCCCACCCCATGCTGCGGCCCCTCCCTTCTTCCTGCTTGGCTGCCTGTGCCCCCCACCTGCCTGTCCACAACCCAGCCTCTGGTACATCCATGCCCTCTGCCCTCAGCCTCACCTGCACTTTTCCTTGGATTTCAGAGTCTCCAAAGGCACAGGCCTCCTCAG**TGCCCACTGCACAACCCCAAGCAGAGGGCA**GCCTCGCCAAGGCAACCACAGCCCCAGCCACCACCCGTAACACAGGTGAGAAGCCCCTTCCCTGCACACTCCACCCCCACCCACCTGCTCATTCCTCAGCCGCCTCCTCCAGGCAGCCCTTCATAA**CTCCTTGTCTGAGTCTCCAAGTCACACTTTGGTAAGGAG**AGGGACACTGAACGGACCTCTAACAAACACCTACTGCCAGCCAGCCCC**AGTCTGGGGGCCAGCAGATGCCAAACAACCAGCAGACT**CCCAGA**GCAGACCTGGGCCGGCTCCCTGGCCCATGGACCCAGCTCTGC**CTC**GCTGAGCTGAGGCATGGGCTCTCAGCGCAGC**CTCACATAGAGCCACCCTGCCGAGGCAGTCCGGCTTGCAGACTCACA**GGTCACTTGGGCCGCAGCAGCCCCTCCCCGTGACC**CTCGCCTCCCGCCCGCCCCAGCCTGGCTCTCTCCAAGTGTTGGATCTTGGTGGCCAGCCTGCTTCTCACCCTCA**CCCTGCCTGCCACCTCAGAATGGCAGGG**GAAAGAGGGCCCTCACCAAGAACTTTATCTGAGAAGTCTGAGGCTTGTGACTCTGACCTGCCTGAGATGTCCATGTGGCCGGGGGGACGGGTTCAGTGTTCGGGAGAACTCGGGTACGTGCCTGACTTTCTCTGAGTAGGGCAGGAAGCTGTTAGGAGAAGCAGCAGTGAGGTGGGCTGGACCAACAGGCAGAATGACTGTCCCTCAGCCACCCTCTGGGATGTGGGTCAAGCTCTGACAAAGG**CATGGCACAGCCATGGTGGCCCCTGCTTGGATGAGTGGCCAC**GGTGCCCTCACCCTGGGCCAGAATCTGCCTCCACTCTGCAGGTGCAGAAACACGACATTCCCGTCTCTAAACACACCTAGCTCCTAGGCTTGGGGTGGGCCTATCAAATGCAGGGAGATGGACACAGCACAAGGGCCAGAGCTTCCCATGAGAAAGGTGAGGGCAGCTGCTCCCTGACCCGGGCATCTGCACTTGTCCC**TCTCCACCCTCCTCATGGGCAGTGGAGA**CTCAGCAACAAAACAAGTTGAGTGCATTAGCAGCCAGCTCTGG**AGCCAAGTCACTCACCCCACGGCCTTGGCTGCTGGTGGAGGGGCCTTCCCCTGGGCAGCC**TCCAAGAAGACAGCCAAGTGCTCTTACTCAGACCACGGCGCTGCTTCCTGGCACCTCGATTTCCCACAACAACATGGGGTGCAGACAGGCTAG**GGCCCCCTGCCCTGGGGCC**TGGACGGCATCCAGTTAAAGATGACCCTTCACGGGCGGTGC**CTGAGGTGTGCTGACCTCAG**CAGCTAAGCCCTCAGGTCTGGTCTGCACTGCCCCACCTGGAGGACCCAACTGACCCAGACACAGCCAGGGTTATGGCATGACCCCGTGGACGGTGACCCAC**AGGCCAGATGCAGCCGGGGGCTGTTTTGTGTGGCCT**AGAAATGTCTTTACAGTTGTAGTGGGATGGAGGAGGAAGAGGAAGAGAGGAGGGGAGAGGAAAGCAGGGAAGGGGAAAAAGAGGAGTTCAATGCAACCCCAAAAGCCAGAACAGTTTTGAGCTGAAAGAACAAGGCAGGAAACATCCCAGTACCTGACTTCAAAACATACTATAAAGCAGTTGTAATCAAAACAGGATCATAAAAACAGACACACAGACCCATGGAACAGAAAAGCGAGCCCAGAAATAAATCTACATGCTTGCAGT**CCATTGATTTTCAACAAAGGCACCAGGAAAACACAATGG**GGAGAGGA**CAGTTTCCTCAATAAATAGTGCTGGGGAAACTG**GATATCCATGTGCAGACTAATGAAACTACACAAAAATCAATTGAAAACAGTCTAGGCCAGGCGCGGTGGCTCATGCCGGTAATCCCAGCACTTTGGGAGGCCGAGACAGGCGGATCACCTGAGGTCAGGAGTTC**GAGACCAGCTTGGCCAACATGGCGAAACCCGGTCTC**CACTAAAAATACAAAAATTAGCACATGGTGGCCTACGTCTGTTATCCCAGCTTTTCAGGAGGCTGAGGCAGGAGAATCGCTTGAATCCGGGAGGTGAAGGTTGCAGGGAGCCAAGATTGCGCCACTGCATTCCAGCCTGGGCAATGGAGCGAGACTGTCTCAAAAAAAAAAAAAAAAAAAAGAAAAGAAAACAGTCTAAAGGTTTAACTGAACAGATAAAGCTACTAGAAGAAAACATAGGGGGAAAACTCCATGACATTAGTCTGAGCAACGAT**TTTTGGATATGATCCCAAAA**GCTCAGGCAGCACTAGTCACAAAAGCCAAGATACAGAACCAACCTAAGCACCCCTCAGCAGATGCACAGGTAAAGAAAATGTGGTACGTATGGGGCACAATGGAATACGATTCAGCCTTTAAAAACAGTGAAATTCTGTCATTGGCAACAATGTAGATGAACCTGAAGGA**CACTTATGCTAAGTG**AAATAAGCCAGGCACAGAAGGAGCAATACTGCATGATTGCACTTACATCTGGCAGGTTAAAAAGGCAAACTCTTAGAGGCAGACAGTAGAGAGGTGGTGCCAGGGAGCGGGCACTGGTGGCTGGGGAGATGTTGGTCAAAGGGCACAAAACTGCAGTTGGGAGGAATTAGTTCAGGACATCCCTTGTACATGGGGACAGTGGTTAGTAACAACGGATTGTATCCTTGAAAACCGCTAAGAAAATAGTTTTTAAGTGTTCTTGACACAAAAAGTGACACGTATGTGAGATACTGCATGGTCATTAGCTGGATTTAGCCATTC**CACAATGTACACATATTTCAAACATTGTG**TTGTATATGATAAACATGTAT**AATTTTTGTCAATTAAAAATT**TTTAGGAAGAGGAGGAGAAGAGAAGAAGAAGGAGAAGGAGAAAGAGGAACAAGAAGAGAGAGAGACAAAGACACCAGGTTTTTTCTGACCCCTGGGCTATCAAAACACCTATTGCCCAATAACTAGTTGGCCGTTGGTGCCCTAAACTATTGAAGCGATTGCTGTTATGTGGATGGGCCCCGGACACTTAGAAACTCGTGACCCCTGA**GGACCCCCACGAGGACAGTCAGGGTCC**CCCCGAACTCAGGGAGCACTGAGGAAGGAGCTCTTAGAGGCGT**GGGGCCCCTCAGGCCCCTCAGAGGGCTCTGCCACATGGGTCAGGGGCAGGCTGAGGGG**GAGTCCCAGGCTCCATGCCCAGCCTCTGTGCCTCTGACCAGGGTGTCCCCCACACCGCCTCCTCC**CCAGTGCCCTCCACTGGCCACACCTGGCCAG**AAGCTGGGGAGAGGAGAGCACAGTGGTTAAGTCAGTCCCTGCAGGGAGACGGCACCAGAAAAACCTGGC**CTGTGGATGAGTCCCGGCCTGGCAGCCACAGAGCAGAGAGCTCTG**GAAGCAACGAAGGCCCGAGTCTGCTCAGGGAAGAGCGGGCAGCAGCCCCAGGGCCGGACAGTGACCAAGAGTGGCACCGCCCATGGCTCAACGGGTCTTTGCC**CACAGATCCCCCAGCCCCTGGAGACAGGGTCTGTGTGCCTGGCCGTGCAGGCAGGCACCACA**CTCAGGGGGAGGCCACTGTGGAGCTCTGTGCAGAGCCCCGGG**CGGGAGCCTACTGCTCCCG**AAGGTCCGGCCACAGCTGCTCTCGTTTGCTCTCCCCTGCAGAGTGTCCGAGCCACACCCAGCCTCTTGGCGTCTACCTGCTAACCCCTGCAGTGCAGGACCT**GTGGCTCCGGGACAAAGCCACCTTCACCTGCTTCGTGGTGGGCAGTGACCTGAAGG**ATGCTCACCTGACCTGGGAGGTGGCTGGGAAGGTCCCCACAGGGGGCGTGGAGGAAGGGCTGCTGGAGCGGCACAGCA**ACGGCTCCCAGAGCCAGCACAGCCGT**CTGACCCTGCCC**AGGTCCTTGTGGAACGCGGGGACCT**CCGTCACCTGCACACTGAACCATCCCAGCCTCCCACCCCAGAGGTTGATGGCGCTGAGAGAACCCGGTGAGCCTGGCTCCCAGGTGGGGAGACGAGGGTGCCCACAGCCTGCTGACCCCTACGCCTGCCCCAGGGCC**ATGACCCCAGCTGGGCCCCAGCAGCACCGGTCAT**CCTCCACAGGAAAGGAGAAGGGAGGCACCAGCACC**CTGGCCGGCCCCACTTCTCTCCCAGTGCCCCCGTGGCCAG**AGCCTGACAGCCTCCCCCACCTCCCCGCAGCTGCGCAGGCACCCGTCAAGCTTTCCCTGAACCTGCTG**GCCTCGTCTGACCCTCCCGAGGC**GGCCTCGTGGCTCCTGTGTGAGGTGTCTGGCTTCTCGCCCCCCAACA**TCCTCCTGATGTGGCTGGAGGA**CCAGCGTGAGGTGAACACTTCTGGGTTTGCCCCCGCACGCCCCCCTCCACA**GCCCAGGAGCACCACGTTCTGGGC**CTGGAGTGTGCTGCGTGTCCCAGCCCCGCCCAGCCCTCAGCCAGCCACCTACACGTGTGTGGTCAGCCACGAGGACTCCCGGACTCTGCTCAACGCCAGCCGGAGCCTAGAAGTCAGCTGTGAGTCACCCCCAGGCCCAGGGTTGGGACGGGGACTCTGAGGGGGGCCATAAGGAGCTGGAATCCATACTAGGCAGGGGTGGGCACTGGGC**AGGGGCGGGGCTAGGCTGTCCTGGGCACACAGGCCCCT**TCTCGGTGTCCGG**CAGGAGCACAGACTTCCCAGTACTCCTG**GGCCATGGATGTCCCAGCGTCCATCCTTGCTGTCCACACCACGTGCTGGCCCAGGCTGGCTGGCACAGTGTAAGAGGTGGATACAACCCCTCGCCGTGCCCTGAGGAGTGGCGGTTTCCTCCCAAGACATTCCCCACGGCTGGGTGCTGGGCACAGGCCTTCCCTGGT**GTGACCGTGAATGTGGTCAC**CCTGAACAGCTGCCCTCTCTGGGGACAT**CTGACTGTCCAAGACCACAGTCAG**CA**CCTCTGGGAGCCAGAGGGGTCTCCAGAGACCCC**CAGATGTCAGGCTTGGGCTCAGTGCCCAGCGAAAGGTCAGCCCCACACATGCCCATAATGGGCGCCCACCCAGAGTGACAGCCCCCAGCCTCCTGCCAGGCCCACCCTTTTCCGCCCCCTTGAGGCATGGCACACAGACCAGTGCG**CCCACTGCCCGAGCATGGCCCCAGTGGG**ATGTGGTGGCCACGAGGGGCTGTACACA**CAGCAGGAGGCTGTCCGCCCTGCTCAGGGCCTGCTG**CCTATGCCCCAGCTGTCCAGCCAAGGGAGGCATGGAAGGGCCCCTGGTGTAAGCTGGAGCCAGGCACCCAGGCCCCCGGCCACCCTGCAGAGCCAAGGAAAGGAAGACACCCAAGTCAACAAGGGGCAGGGCTGAGGGCTGTCCCAGGCTCTTTTGGCCCGAG**GGGCTGCCAGCAGCCC**TGACCCGGCATGGGCCTTCCCCAGAAGCGACC**CTGTGAGGTGGCCTCACAG**AGAACCCCCTCTGAGGACAGTGTCTGACCCTGCCTGCCTCACACAGATGGGCCCCACAGCAGTGGGCAACCTGGGGGGCAGCAGCCCAACCTGACC**CTGCAGGGACTGCCCCCTGCAG**CAGCAGCTGCTTCTCAGTCCCCCAACCTCCCTGTCCCCGCCAGAGGGTCTTCCCCGAAGCT**GCAGCCCCAACCCATGGCTGC**CCACCTGGAACCGGGACTCCCTGTCCACTGCCCCCTCCCCTTCG**GGGCCCCATCTGTGCTGGGGCCC**AGGTTCGGCCTACAGATTCCCATCATTGCCATGGCCTCCTGACCTTGCCTATCCACCCCCAACCACCGGCTCCATGCTGACCCTCCCCCAGGCTCCCACGCCCAGCTGGCCGGCCATCCCCAGGCACAGACAGTCTGGGA**TCTCACAGGTTAGCCTGGACCATCCACCTGGCCAGACCTGGGAGA**GGCTGGAAGCT**GCCCTGCCACCATGCTCCAGGGC**CCCAGGTTGCAGTACTATGGGGTGA**GGGTGTGTGTGCACACCC**GTGTGTACCTAGGATATCCGAGTGTACC**CTTGTGCCCCCAAGCACAAG**TCTCCCTCCCAGGCAGTGAGGCCCAGATGGTGCAGTGGTTAGAGCTGAGGCTTATCCCACAGAGAACCCTGGCGCCTTGGTCAAGGAAGCCCCTATGCCTTTCTTGCCTCGATTTCCCCTCTTGT**CTGCTGAGCCAGCAG**GGGCCACGTCCTGGGCTGCTGTGAGGAGGAAGTGAGTTGGTGCTAGGAGGGGCTCCTGTGTGTGCATGGGCGGGA**GGGGTGCAGGTATCTGAGCACCCC**GGTCTCCACTTGAGAGAGCAGGGCAGGAGCTCCCTGACCCACCCAGACTACACACGCTGTGTCCACGTGTCTCACATTATCTGTGGCAGAGGATCCGGCTTCTTTCTCAATTTCCAGTTCTTC**ACAAAGCAATGCCTTTGT**AAAATGCAATAAGAAATACTAGAAAAATGATATGAACAGAAAGACACGCCGATTTTTTGTTATTAGATGTAACAGACCATGGCCCCATGAAATGAtcccggaccagatccgtccacacccgccactcagcagctctggccgagctcacagtacaaccacaataaactcttgttgaatgaactct**aggaagtctgtgacgtggctggttcttgtcaatgcttcct**gcctgcccacaggctcttcctcgtggatggggctgtgcttgccatggaagcgtttttcccggcctaggcttgccttgggccccactgccgtctccagctggagatgaccttctatacacacatttgctcatgacagacccttgcttagcccccttccatggctccctcctgctgct**gggataaaatcaccttgcctggatatccc**ctcctgggcccctttccaccctccttagtcagcacccccagttcagggcacctgctttccccgctgcggagaagccactctctccttgctgcccggctgtgtctt**gccttccacaccttgtcacagtggccacttcctaaggaaggc**ctccctgtgtgcaggtgtgcagaagtgccccagc**ctcccgtcacctttgtcacgggag**cccaatccatgagagtctatggttctgtctgt**ctgccccactcagggcag**cgacaagtccaggcggggaggacacagtaggcagagatttgtcgaggggacatatgagcaagagggtgaggctgggagctccctggagataaccacgcctcctgggaagactcgccgtcatttcag**ctccacgctgtgcgggggtgggtggag**gggtagcctggccctcatgaccagggagcttctcactcagcccccgttcctccccagacctggccatgacccccctgatccctcagagcaaggatgagaacagcgatgactacacgacctttgatgatgtgggcagcct**gtggaccaccctgtccac**gtttgtggccctcttcatcctcaccctcctctacagcggcattgtcactttcatcaaggtcaggggagcggccaggctctcagtgaccctcggggtgggtg

Uppercase: IGHD

Lowercase: Flanking sequence[1000bp]

Red & Bold & Underline: Stem-loop [83]

Blue: Heptamer[146]

Green: Nonamer [10]

id-IGHGP-2[C_gene_segment]

aaatatata**tttttcaaggtgaaaaa**aaaaaaagaaaaacctgccataatgaagagcagaccaatattccagaaaactgtcact**ttaacagagaagaccaaattctagtttcacatgaactgttaa**t**attaaagctaattttaat**taaaccttataaataattccatccag**gctgggcaaggtggcttacacctgtagtcccagc**attccaggagggtggatcacttaa**gcccaggagtttgagatcagcctgggc**aacatggggaagccctgtctctacaaaaaatacaaaaaaaaaattggccaggcatggtggcacgtgcccgtagtcccagctactt**gggaggctgaggtgggaggattgcttgagccctcccacctcaacctccc**tcagcct**cagtgctgagtgccactgcactg**ctgcctaggcaacacagtgggaccctgtctcaaaaataaataaataaataaatccggccaggtgcagtggctcatgcctgtaa**tcccaacactttggga**ggcctaggcgggtggatcatgaggtcaggagttcgagaccagcctggccaacgtggtgaaacccctgtctctactaaaactataaaaat**tagctgggcgtggtggtgggcacctgtaatcccagcta**ctctggaggctgaggcaggagaatcgcttgaacccaagaggcagaggttgcagtgaaccgagatcacaccattgcactccagcttaggcaacaagagtgaaactctgtctcaaaaaaataaataaataaatccaatcacagccgcctttgaccacataagatcccttttccacaatccttttacaactttttatttgtttgttttttgttttgttttgttttttgtttttgtttttgttttttttgagacaaagcactgacctggctgccgagccccgccccctagg**ctgcaggggtgcctgcag**aagggcaccacagggccaccagtcctgcaagctttctggggcaggccggGCCTGACTTTGGCTTTGGGGCAGGGAGGGGGCTAAGGTGAGGCAGGTGGCACCAGCCAGGTGCACACTCAATGCCCGTGAGCCCAGACACTGGACCCTGCCTGGA**CCCTCGCGGATAGACAAGAACCGAGGG**GCCTCTGCACCCTGGGCCCAGCTCTGTCCCACACCGCGGTCACATGGCACCACCTCTCTTGCAGCCTCCACCAAGGGCCCATCGGTCTTCCCCCTG**GTGCCCTCCTCCAGGAGCGTCTCTGAGGGCAC**AGCGGCCCTGGGCTGCCTGGTCAAGGACTACTTCCCCGAACCGGTGACGGTGTCGTGGAACTCAGGGGCCCTGACCAGAAGCGTGCACACCTTCCCGGCTGTCCTACAGTCCTCAGGACTCTACTCCCTCAGCAGCGTGGTGACC**GTGCCCTCCAGCAGCTTGGGCAC**CCAGACCTACACCTGCAACGTAGATCACAAGCCCAGCAA**CACCAAGGTGGACAAGACAGTTGGTG**AGAGG**CCAGCACAGGGAGGGAGGGTGTCTGCTGG**AAG**CCAGGCTCAGCCCTCTTGCCTGG**ACGTACCCCGGCTGTGCAGCCCCAGTCCAGGGCAGCAAGGCAGGCCCCATCTGTCTCCTCACCCGGAGGCCTCTGCCCGCCCCACTCATGCTCAGGGAGAGGGTCTTCTGGCTTTTTCCACCAGGCTCCAGGCAGCCACAGGCTGGAA**GCCCCTACCCCAGGCCCTGCGCACAAAGGGGC**AGGTGCTGCACTTAGACTGGCCAAGAGCCATATCCGGGAAGACCCTGCCCCTGACCTAAGCCCACCCCAAAGGCCAAGATCTCCACTCCCTCAG**CTCAGACACCTCTCCTCCCAGATCTGAG**TAACTCCCAATCTTCTCTCTGCAGAGCCCAAAACCCCATGTTGTGACACAACTCA**CACATGCCCACCATGTG**CAAGTAAGCCAGCCCAGGCCTCGCCCTCCAGCTCAAGGCGGGACAGGTGCCCTAGAGTAGCCTGCGTCCAGGGACAGGCCCCAACCGGG**TGCTGACACGTCCGCCTCCATCTCTTCCTCAGCA**ACTGAACCCC**TGGGGGGACCGTCAGTCTTCCTCTTCCCCCCA**AAACCCAAGGATACCCTCATGATCTCCCGGACCCCTGAGG**TCACGTGCGTGGTGGTGGACGTGA**GCCACGAAGACCCTGAGGTCAAGTTCAACTGGTACGTGGACGGCGTGGAGGTGCATAATGCCAAGACAAAGCCGTGGGAGGAGCAGTACAACAGCACGTACCATGTGGTCAGCGTCCTCACCGTCGTGCACCAGAACTGGCTGAACGGCAAGGAGTACAAGTGCAAGGTCTCCAACAAAGGCCTCCCAGCCCCCATCGAGAAAACCATCTCCAAAACCAAAGGTGGGACCCACGGAGCGCGAAGGCCACGTGGA**CAGAGGCCGGCTTGGCCCACCCTCTG**CCCTGGGAGTGACCGCTGTACCAACCTCTGTCCCTACA**GGGCAGCCCCGAGAACCACAGGTGTACACCCTGCCC**CCATCCCAGAAGATGACCAAGAAC**CAGGTCACCCTGACCTG**CCTGGTCAAAGGCTTCTACCCCAGCGACATCGCCGTGGAGTGGGAGAGCAATGGGCAGCCGGAGAACAACTACAAGACCACGCCTCCCATGCTGGACTCCAACGGCTCCTTCTTCCTCTATAGCAAGCTCACCGTGGACAAGAGCAGGTGGCAGCAGGGGAACGTCTT**CTCATGCTCCGTGATGCATGAG**GGTCTGCAGAACCACTACACGCAGAAGAGCCTCTCCCTGTCCCCGGGGTAAatga**gtgcgacggccggcaagcccccgctccccgggctctcgcggtcgcac**gaggatgcttggcacgtaccccgtctacatacttcccaggcacccagcatggaaataaagcacccaccactgccctgggcccctgcgagactgtgatggttctttccacggg**tcaggccgagtctgaggcctga**gtggcatgagggaggcagagcgggtcccactgtccccacact**ggcccaggctgtgcaggtgtgcctgggcc**gcctagggtggggctcagccaggggctgccctcggcagggtgggggatttgccagcgtggccctccctccagcagcacctgccctgggctgagccacgagaagccctaggagcccctgg**ggacagacacacagcccctgcctctgtaggagactgtcc**tgttctgtgagcgccctgtcctccgaccccc**catgcccactcgggggcatg**cctagtccatgtgcgtagggacaggccctccctcacccatctacccccacggcactaacccctggcagccctgcccagcctcgcacccgcatggggacacaaccgactccggggacatgcactctcgggccctatggagggactggtccagatgcccacacacacactcagcccagacccgttcaacaaaccccgcactgaggt**tggccggccacacggcca**ccacacacacacgtgcacgcctcacacacggagcctcacccgggcgaactgcacagcacccagaccagagcaaggtcctcgcacacgtgaacactcctcggacacaggcccctacaagccccatgcggcacctcaaggcccacgagcctctcggcagcttctccacttgctgaccagctcagacaaacccagtcctcctctcacaaagtg**cccctgcagccgccacacacacacagggg**atcacacaccacgtcacgtccctggccctggcccacttcccaatacagcccttccctgctcctggggtcacatg

Uppercase: IGHGP-2

Lowercase: Flanking sequence[1000bp]

Red & Bold & Underline: Stem-loop [33]

Blue: Heptamer[50]

Green: Nonamer [2]

id-IGHG3-3[C_gene_segment]

aatggggcctccctgtggcctgggggtcctggcaccacgcagggtggggagggccaagggcaggtgcaaggctcctacctg**tgctggggggcctgggttgagcccagca**gggaccttgccgggggaagctctggagagagggaggaggtgggctggtggctgagaaggccaggccagggctgggagggtgacggtgtggtgactga**gcctccagaagtaatgcaggacactgggaggc**agggggcatccaggcactcagggccctgacctgggctgctgcacactggggctaaggggaaaggaggggagaggctgaggaggaggctcccggggcgatattccaaggcagggggttccggggccctggggctgaagggcgccgaccctatgcagtgtctggc**ccctctgctgcacagaagaaaagggccttggagggcagaggg**caggctatgaccag**ggccctgggcaagtcaggcccactcactagcggagggcc**acgctggggcggcagggtcaggagcttcaggggactcaggggacccacgagaagccatctgagaacagtgtccactggtcaagccaggcacccataaaaggctggagtggggccaatgggcatgagccgtccctgaggtggcaccgatggccagagctgaggccaagctagaggccctggactgtgctgactcc**cggcagacacagagcgctgacctggctgccg**agccccgcctcctagg**ctgcaggggtgcctgcag**aagggcaccacagggccaccggtcctgcaagctttctggggcgggccgg**gcctgaccttggctttggggcagggagggggctaaggtgaggc**aggtggcgccagccaggcgcacacccaatgcccgtgagcccagacactggaccctgcctgga**ccctcgtggatagacaagaaccgaggg**gcctctgcgccctgggcccagctctgtcccacaccgcagtcacatggcgccatctctcttgcagCTTCCACCAAGGGCCCATCGGTCTTCCCCCTGGCGCCCTGCTCCAGGAGCACCTCTGGGGGCACAGCGGCCCTGGGCTGCCTGGTCAAGGACTACTTCCCAGAACCGGTGACGGTGTCGTGGAACTCAGGCGCCCTGACCAGCGGCGTGCACACCTTCCCGGCTGTCCTACAGTCCTCAGGACTCTACTCCCTCAGCAGCGTGGTGACC**GTGCCCTCCAGCAGCTTGGGCAC**CCAGACCTACACCTGCAACGTGAATCACAAGCCCAGCAA**CACCAAGGTGGACAAGAGAGTTGGTG**AGAGGCCAGCGCAGGGAGGGAGGGTGTCTGCTGGAAG**CCAGGCTCAGCCCTCCTGCCTGG**ACGCATCCCGGCTGTGCAGTCCCAGCCCAGGGCACCAAGGCAGGCCCCGTCTGACTCCTCACCCGGAGGCCTCTGCCCGCCCCACTCATGCTCAGGGAGAGGGTCTTCTGGCTTTTTCCACCAGGCTCCGGGCAGGCACAGGCTGGA**TGCCCCTACCCCAGGCCCTTCACACACAGGGGCA**GGTGCTGCGCTCAGAGCTGCCAAGAGCCATATCCAGGAGGACCCTGCCCCTGACCTAAGCCCACCCCAAAGGCCAAACTCTCTACTCACTCAG**CTCAGATACCTTCTCTCTTCCCAGATCTGAG**TAACTCCCAATCTTCTCTCTGCAGAGCTCAAAACCCCACTTGGTGACACAACTCACACATGCCCACGGTGCCCAGGTAAGCCAGCCCAGGCCTCGCCCTCCAGCTCAAGGCGGGACAAGAGCCCTAGAGT**GGCCTGAGTCCAGGGACAGGCC**CCAGCAGGGTGCTGACGCATCCACCTCCATCCCAGATCCCCGTAACTCCCAATCTTCTCTCTGCAGAGCCCAAATCTTGTGACACACCTCCCCCGTGCCCACGGTGCCCAGGTAAGCCAGCCCAGGCCTCGCCCTCCAGCTCAAGGCAGGACAAGAGCCCTAGAGTGGCCTGAGTCCAGGGACAGGCCCCAGCAGGGTGCTGACGCGTCCACCTCCATCCCAGATCCCCGTAACTCCCAATCTTCTCTCTGCAGAGCCCAAATCTTGTGACACACCTCCCCCATGCCCACGGTGCCCAGGTAAGCCAGCCCAGGCCTCGCCCTCCAGCTCAAGGCGGGACAAGAGCCCTAGAGTGGCCTGAGTCCAGGGACAGGCCCCAGCAGGGTGCTGACGCATCCACCTCCATCCCAGATCCCCGTAACTCCCAATCTTCTCTCTGCAGAGCCCAAATCTTGTGACACACCTCCCCCGTGCCCAAGGTGCCCAGGTAAGCCAGCCCAGGCCTCGCCCTCCAGCTCAAGGCAGGACAGGTGCCCTAGAGTGGCCTGCATCCAGGGACAGGTCCCAGTCG**GGTGCTGACACATCTGCCTCCATCTCTTCCTCAGCACC**TGAACTCCTGGGAGGACCGTCAGTCTTCCTCTTCCCCCCAAAACCCAAGGATACCCTTATGATTTCCCGGACCCCTGAGG**TCACGTGCGTGGTGGTGGACGTGA**GCCACGAAGACCCCGAGGTCCAGTTCAAGTGGTACGTGGACGGCGTGGAGGTGCATAATGCCAAGACAAAGCCGCGGGAGGAGCAGTACAACAGCACGTTCCGTGTGGTCAGCGTCCTCACC**GTCCTGCACCAGGAC**TGGCTGAACGGCAAGGAGTACAAGTGCAAGGTCTCCAACAAAGCCCTCCCAGCCCCCATCGAGAAAACCATCTCCAAAACCAAAGGTGGGACCCGCGGGGTATGAGGGCCACATGGA**CAGAGGCCAGCTTGACCCACCCTCTG**CCCTGGGAGTGACCGCTGTGCCAACCT**CTGTCCCTACAGGACAG**CCCCGAGAACCACAGGTGTACACCCTGCCCCCATCCCGGGAGGAGATGACCAAGAAC**CAGGTCAGCCTGACCTG**CCTGGTCAAAGGCTTCTACCCCAGCGACATCGCCGTGGAGTGGGAGAGCAGCGGGCAGCCGGAGAACAACTACAACACCACGCCTCCCATGCTGGACTCCGACGGCTCCTTCTTCCTCTACAGCAAGCTCACCGTGGACAAGAGCAGGTGGCAGCAGGGGAACATCTT**CTCATGCTCCGTGATGCATGAGGCTCTGCACAACCGCTTCACGCAGAAGAGCCTC**TCCCTGTCTCCGGGTAAATGAGTGCGACGGCCGGCAAGCCCCCGCTCCCCGGGCTCT**CGGGGTCGCGCGAGGATGCTTGGCACGTACCCCG**TGTACATACTTCCCGGGCACCCAGCATGGAAATAAAGCACCCAGCGCTGCCCTGGGCCCCTGCGAGACTGTGATGGTTCTTTCCACGGG**TCAGGCCGAGTCTGAGGCCTGA**GTGGCATGAGGGAGGCAGAGCGGGTCCCACTGTCCCCACACT**GGCCCAGGCTGTGCAGGTGTGCCTGGGCC**GCCTAGGGTGGGGCTCAGCCAGGGGCTGCCCTCGGCAGGGTGGGGGATTTGCCAG**CGTGGCCCTCCCTCCAGCAGCAGCTGCCCTGGGCTGGGCCACG**GGAAGCCCTAGGAGCCCCTGG**GGACAGACACACAGCCCCTGCCTCTGTAGGAGACTGTCC**TGTCCTGTGAGCGCCCTGTCCTCCGACCC**GCATGCCCACTCGGGGGCATGC**CTAGTCCATGTGCGTAGGGACAGGCCCTCCCTCACCCATCTACCCCCACGGCACTAACCCCTGGCAGCCCTGCCCAGCCTCGCACCCGCATGGGGACACAACCGACTCCGGGGACATGCACTCTCGGGCCCTGTGGAGAGACTGGTCCAGATGCCCACACACACACTCAGCCCAGACCCGTTCAACAAACCCCGCACTGAGGT**TGGCCGGCCACACGGCCA**CCACACACACACGTGCACGCCTCACACACGGAGCCTCACCCGGGCGAACCGCACAGCACCCAGACCAGAGCAAGGTCCTCGCACACGTGAACACTCCTCGGACACAGGCCCCCACGAGCCCCACGCGGCACCTCAAGGCCCACGAGCCGCTCGGCAGCTTCTCCACATGCTGACCAGCTCAGACAAACCCAGCCCTCCTCTCACAAGGTGCCCCTGCAGCCGCCACACACACACAGGCCCCCACACACAGGGGAACACACGCCACGTCGCGTCCCTGGCACTGGCCCACTTCCCAATACAGCCCTTCCCTGCAGCTGGGGTCACATGAGGTGTGGGCTTCACCATCCTCCTGCCCTCTGGGCCTCAGGGAGGGACACGGGAGACGGGGAGTGGGTCCTGCTGAGGGCCAGGTCGCTATCTAGGGCCGGGTGTGTGGCTGAGTCCCGGGGCCAAAGCTGGTGCCCAGGGCGGGCAGCTGTGGGGAGCTGACCTCAGGACACTGTTGGCCCATCCCGGCCGGGCCCTACATCCTGGGTCCTGCCACAGAGGGAATCACCCCCAGAGGCCCGAGCCCAGCAGGACACAGCACTGACCACCCTCTTCCTGTCCAGAGCTGCAACTGGAGGAGAGCTGTGCGGAGGCGCAGGACGGGGAGCTGGACGGGCTGTGGACGACCATCACCATCTTCATCACACTCTTCCTGTTAAGCGTGTGCTACAGTGCCACCGTCACCTTCT**TCAAGGTCGGCCGCACGTTGTCCCCAGCTGTCCTTGA**CATTGTCCTCCATGCTGTCACACACTGTCCCTGACACTGTCCCCAGGCTGTCCCCACCTGTCCCTGACACTGTCCCCCACGCTCTCACAAACTGTCCCTCACACTGTCCCCCATGCTGTCACAAACTGTCACTGACACTGTCCCCCATGCTATCCCCACCTGTCCCTGACACTGTCCCTGACACTGTCTCTCATGCTGTCCCCACTCATCTGCGACACTGTACCCCACGCTGTCCCCACTTGTCCTCAACAATGTCCCCCATGCTGTCCCCACCTGTCCCTGATGCTGTCCCCCACACTGTCCCAATCTGTCCCCACCACTCTCCCCCACGCTGTCCCCACCTGTCCCTGACACTGTCCCCCATGCCATCCCCATCTGTCCCGACAATGTCCCCAGGGTGTCCCCAGCTGTCCCTGATGCTGTCCCCCACACTGTCCCCACCTCTCCCTGACGCTGTCCCCCACGTGGTCCCCACTTGTCCCTGATGCTGTCCCCCACACTGTCCCCACCTGTCCCTGACACTGTCCCCCATGCCATCCCCATCTGTCCCGACAATGTCCCTATGGTGTCCCCAGCTGTCCCTGATGCTGTCCCCCACACTGTCCCCACCTGTCCCTGACGCTGTCCCCCACACTGTCCCCACCTCCCCCTGACACTGTCCCCCACACTGTCCCCACCTCTCCCTAACACTGTCCCACACACTGTCCCCTCCTGTCCCCAACACTTTCCCCCATGCTGTCCCCACCAGTCCCCAACACTGTACACCATGCTTTTCCCACCTGTCCCCAACACTGTCCCCCATGCTGTCCCCTCCTGTCCCCAACAATGTCCCCCATGCTGTTTCCTCCTGTCCCCAACACTGTCCGCCACTCTGTTTCCTCCTTTCCCTGACACTGTCCCCCACTCTGTCCCCACCTGTAGCCAACACTATCCCCTACGCTGTCTCCACCTGTCCCTGATGCTGTCCCCCACACTGTCCCCACTCCTCCCTGACACTGTCCCCTATGCTGTCCCCACCGGTTCCTAACACTGTCCCCCACACTGTCCCTACCTGTCCCCGACACTTTCTCCCATGCTGTTCCCACGTGTCTCCAACACTGTCCCCCACACAGTCTCCACCTGTCCCTGACACTGTCCCCCATGCTGTCCTCACCCATCTCTGACACTGTACACATACTGTCCCCACCTGTCCCTGATGCTGTCCTCCATGATGTCCCCACCTCTCCCTGACACTGTCACCCATGCTGTCCCCACCTGCCCCTGACACTCTCCTCCACGCTGTTCTCACCTGTCCCCAACACTCTCCCCCACACTGTCTCCACCTGTCCCTGACACTGTCCTCCACGCTGTCCCCACCTATCCCTGACACTGTCCCCCATGCTGTCCTCACCTGTCCCCAACACTCTCCTCCACACTGTCCTCACCTGTCCCCAACACTCTCCCCCCACACTGTCTCAACCTGTCCCTGACACTGTCCCCCATGCTGTCCTCACCTGTCCCTGACACTGTCCCCCATGCTGTCCTCACCTGTCTCTGACACTGTCCCCCGTGCTGTCCCCACCTGACACTATCTTCTGTGCTGTCCACATGCTGTTGCTGCCCTGGCTCTGCTCTCCATGT**CCAGGCCTCAGAGCAGGCAGTGGTGAGGCCCTGG**CACATGGGTGGCATGAGGGGCCGGATAGGCCTCAGGGGCAGGGCTGTGGCCTGGGTGGCCTG**AGGGGTGAGCAGGCCTCGGGGGCAGGGCTGTGGCCTCGCTCACCCCT**GTGCTGTGCCTTGCCTACAGGTGAAGTGGATCTTCTCCTCGGTGGTGGACCTGAAGCAGACCATCATCCCCGACTATAGGAACATGATTGGGCAGGGGGCCTAGggccaccctctgcggggtgtccagggccacccagatcccacacacgagccgtgggccatgctcagccaccacccaggccacaactgcccccgacctcaccgccctcaaccccatggctctctgtctttgcagtcgccctctgagccctgacacgccccccttccagaccctgtgcatagc**aggtctaccccagacct**ccgctgcttggtgcatgcagggagct**ggggaccaggtgtcccc**tcagcaggatgtccctgccctccagaccgcca**gatgctcacacaaaaggaggcagtgaccagcatc**cgaggcccccacccagg**caggagctggccctggagccaaccccgtccacgccagcctcctg**aacacaggcgtggtttccagatggtgagtgggagcatcagccgccaaggtag**ggaagccacagcaccatcaggccctgttggggaggcttcc**gagagctgc**gaaggctcactcagacggccttc**ctcccagcccgcagccagccagcctc**cattccgggcactcccgtgaactcctgacatgaggaatg**aggttgttctgatttcaagcaaagaacgctgctctctggctcctgggaacagtctcggtg**ccagcaccaccccttggctgcctgcctacactgctgg**attctcgggtggaactggacccgcagggacagccag**ccccagagtccgcactgggg**agagaaggggccaggcccaggacactgccacctcccacccactccagtccaccgagatcactcagagaagagcctgggccatgtggccactgcaggagccccacagtgcaagagtgaggatagcccaaggaaggg**ctgggcatctgcccag**acaggcctcccagag**aaggctggtgaccaggtcccaggcgggcaagactcagcctt**ggtg**gggcctgaggacagaggaggccc**aggagcatcggggagagaggtggagggacaccgggagagccaggagcgtggacacag

Uppercase: IGHG3-3

Lowercase: Flanking sequence[1000bp]

Red & Bold & Underline: Stem-loop [44]

Blue: Heptamer[124]

Green: Nonamer [2]

id-IGHE-2[C_gene_segment]

tccagctttgctgagctaaactggaccgggctaaattgatctggactgaccattctcacctggctaagaggagctgagtcagaagcaagctggttgagctggctggactgaaataagagtttgctgcctgcaaggggaggtcctgggctgacctgggccaggctgaaccaggctggcttagagtgaacttcagagggcgactcccccggtaggccagtctcagctgaacttggctgtcccggtgggcagagcggggctggatactgtgattttgggggtacctagagcagacttcaagaccaagctaaactgggctccaggggcaggatgggctggggacttgggactccaggccaggggcgaagggccacgctgtacagaccgcactatctgggccagggttctgtggtgggagggactgactgcctggggcatcagggcaagtcttccc**gccctcccctagaggtcaggggtgggc**agagcaccatgggggtctggcaggtcaggtgagggctgctgtgatggggagatccaggcttggcactcaagagcccgaggagctgagaccacagccttggggggttggggtcagggttggagggcaggcagaccatccaccat**gagcccagagagagtttgaagggggagggctctggggtcccaggccccatggggtccctggg**tttcagcctaggggcatggcccagtgtctctgctcctgagtgcccaccgtgcagcacttgcaggggg**aggctggggtcatcctggaggcaccccccttcctgagcccagcct**gatgatagtggctgagcaacagcttctggtgggggaatgggggccctgggagccgcc**ctgggcctggggattgtggggaaaaaggcccag**aatgagcctggccatctggatccctgccacggggtccccagctcc**cccatccaggccccccaggcctgatggg**cgctggcct**gaggctggcactgactaggttctgtcctcacagCCTC**CACACAGAGCCCATCCGTCTTCCCCTTGACCCGCTGCTGCAAAAACATTCCCTCCAATGCCACCTCCGTGACTCTGGGCTGCCTGGCCACGGGCTACTTCCC**GGAGCCGGTGATGGTGACCTGGGACACAGGCTCC**CTCAACGGGACAACTATGACCTTACCAGCCACCACCCTCACGCTCTCTGGTCACTATGCCAC**CATCAGCTTGCTGACCGTCTCGGGTGCGTGGGCCAAGCAGATG**TTCACCTGCCGTGTGGCACACACTCCATCGTCCACAGACTGGGTCGACAACAAAACCTTCAGCGGTAAGAGAGGGCCAAGCTCAGAGACCACAGTTCCCAGGAG**TGCCAGGCTGAGGGCTGGCA**GAGTGGGCAGGGGTTGAGGGGG**TGGGTGGGCTCAAACGTGGGAACACCCA**GCATGCCTGGGGACCCGGGCCAGGACGTGGGGGCAAGAGGAGGGCACAC**AGAGCTCAGAGAGGCCAACAACCCTCATGACCACCAGCTCT**CCCCCAGTCTGCTCCAGGGA**CTTCACCCCGCCCACCGTGAAG**ATCTTACAGTCGTCCTGCGACGGCGGCGGGCACTTCCCCCCGACCATCCAGCTCCTGTGCCTCGTCTCTGGGTACACCCCAGGGACTATCAACATCACCTGGCTGGAGGACGGGCAGGTCATGGACGTGGACTTGTCCACCGCCTCTACCACGCAG**GAGGGTGAGCTGGCCTCCACACAAAGCGAGCTCACCCTCAGCCAGAAGCACTGGCTG**TCAGACCGCACCTACACCTGCCAGGTCACCTATCAAGGTCACACCTTTGAGGACAGCACCAAGAAGTGTGCAGGTACGTT**CCCACCTGCCCTGGTGGCCGCCACGGAGGCCAGAGAAGAGGGGCGGGTGGG**CCTCACACAGCCCTCCGGTGTACCACAGATTCCAACCCGAGAGGGGTGAGCGCCTACCTAAGCCGGCCCAGCCCGTTCGACCTGTTCATCCGCAAGTCGCCCACGATCACCTGTCTGGTGGTGGACCTGGCACCCAGCAAG**GGGACCGTGAACCTGACCTGGTCCC**GGGCCAGTGGGAAGCCTGTGAACCACTCCACCAGAAAGGAGGAGAAGCAGCGCAATGGCACGTTAACCGTCACGTCCACCCTGCCGGTGGGCACCCGAGACTGGATCGAGGGGGAGACCTACCAGTGCAGGGTGACCCACCCCCACCTGCCCAGGGCCCTCATGCGGTCCACGACCAAGACCAGCGGTGAGCCATGGGCAGGCCGGGGTCGTGGGGGAAGGGAGGGAGCGAGTGAGCGGGGCCCGGGCTGACCCCACGTCTGGCCACAGGCCCGCGTGCTG**CCCCGGAAGTCTATGCGTTTGCGACGCCGGAGTGGCCGGGG**AGCCGGGACAAGCGCACCCTCGCCTGCCTGATCCAGAACTTCATGCCTGAGGACATCTCGGTGCAGTG**GCTGCACAACGAGGTGCAGC**TCCCGGACGCCCGGCACAGCACGACGCAGCCCCGCAAGACCAAGGGCTCCGGCTTCTTCGTCTTCAGCCGCCTGGAGGTGACCAGGGCCGAATGGGAGCAGAAAGATGAGTTCATCTGCCGTGCAGTCCATGAGGCAGCGAGCCCCTC**ACAGACCGTCCAGCGAGCGGTGTCTGT**AAATCCCGGTAAATGAcgtactcctgcctccctc**cctcccagggctccatccagctgtgcagtggggagg**actggccagaccttctgtccactgttgcaatgaccccaggaagctacccccaataaactgtgcctgctcagagccccaggtacacccattcttgggagcgggcagggctgtgggcaggtgcatcttggcacagaggaatgggccccccaggaggggcagtgggaggaggtgggcagggctgagtccccccaggagaggcggtgggaggaggtgggcagggctgaggtgccactcatccatctgccttcgtgtcagggttatttgtcaaacagcatatctgcagggactcatcacagctaccccgggccctctctgcccccact**ctgggtctaccccctccaaggagtccaaagacccag**gggaggtcctcagggaaggggcaaggg**agcccccacagccctctctcttgggggct**tggcttctacccccctggacaggagcccctgcacccccaggtatagatgggcacacaggcccctccaggtggaaaaacagccctaagtgaaaccccca**cacagacacacacgacccgacagccctcgcccaagtctgtg**ccactggcgttcgcctctctgccctgtcccgccttgccgagtcct**ggccccagcaccggggcc**ggtggagccgagcccactcacaccccgcagcctccgccaccctgccctgtgggcacaccaggcccaggtcagcagccaggccccctctcctactgccccccaccgccccttggtccatcctgaatcggcctccaggggatcgccagcctcacacacccagtctcgcccactcacgcctcactcaaggcaca**gctgtgcacacactaggccccatagcaactccacagc**accctgtaccaccaccagggcgccatagacacccca**cacgtggtcacacgtg**gcccacactccgcctctcacgctgcctccagccaggctactgccaagcc

Uppercase: IGHE-2

Lowercase: Flanking sequence[1000bp]

Red & Bold & Underline: Stem-loop [27]

Blue: Heptamer[58]

Green: Nonamer [1]

id-IGHG2-2[C_gene_segment]

aaatggggcctccctgtggcctgggggtcctggcaccacgcagggtggggagggccaagggcaggtgcaaggctcctacctg**tgctggggggcctgggttgagcccagca**gggaccttgccgggggaagctctggagagagggaggaggtgggctggtggccgagaaggccaggccagggctgggagggtgaggttgtggtgactga**gcctccagaagtaatgcaggacactgggaggc**agggggcatccaggcactcagggccctgacctgggctgctgcacactggggctaaggggaaaggaggggagaggctgaggaggaggctccaggaggctattccaaggcagggggttccggggccctggggctgaagggcgccgaccctatgcagtgtctggc**ccctctgctgcacagaagaaaagggccttggagggcagaggg**caggctatgaccag**ggccctgggcaagtcaggcccactcactagcggagggcc**acgctggggcggcagggtcaggagcttcaggggactcgggggacccacgagaagccatctgagaacagtgtccactggtcaagccaggcacccataaaaggctggagtggggccaatgggcatgagccgtccctgaggtggcaccgatggccagagctgaggccaagctagaggccctggactgtgctgactcc**cggcaggcacagagcgctgacctggctgccg**agccccgcctcctagg**ctgcaggggtgcctgcag**aagggcaccacagggccaccggtcctgcaagctttctggggcaggccgggcctgactttggctttggggcagggagggggctaaggtgacgcaggtggcgccagccaggtgcacacccaatgcccgtgagcccagacactggaccctgcctgga**ccctcgcagatagacaagaaccgaggg**gcctctgcgccctgggcccagctctgtcccacaccgcggtcacatggcaccacctctcttgcagCCTCCACCAAGGGCCCATCGGTCTTCCCCCTGGCGCCCTGCTCCAGGAGCACCTCCGAGAGCACAGCGGCCCTGGGCTGCCTGGTCAAGGACTACTTCCCCGAACCGGTGACGGTGTCGTGGAACTCAGGCGCTCTGACCAGCGGCGTGCACACCTTCCCGGCTGTCCTACAGTCCTCAGGACTCTACTCCCTCAGCAGCGTGGTGACCGTGCCCTCCAGCAACTTCGGCACCCAGACCTACACCTGCAACGTAGATCACAAGCCCAGCAA**CACCAAGGTGGACAAGACAGTTGGTG**AGAGGCCAGCTCAGGGAGGGAGGGTGTCTGCTGGAAG**CCAGGCTCAGCCCTCCTGCCTGG**ACGCACCCCGGCTGTGCAGCCCCAGCCCAGGGCAGCAAGGCAGGCCCCATCTGTCTCCTCACCCGGAGGCCTCTGCCCGCCCCACTCATGCTCAGGGAGAGGGTCTTCTGGCTTTTTCCACCAGGCTCCAGGCAGGCACAGGCTGGG**TGCCCCTACCCCAGGCCCTTCACACACAGGGGCAGGTGCTTGGCTCAGACCTGCC**AAAAGCCATATCCGGGAGGACCCTGCCCCTGACCTAAGCCGACCCCAAAGGCCAAACTGTCCACTCCCTCAG**CTCGGACACCTTCTCTCCTCCCAGATCCGAG**TAACTCCCAATCTTCTCTCTGCAGAGCGCAAATGTTGTGTCGAGTGCCCACCGTGCCCAGGTAAGCCAGCCCAGGCCTCGCCCTCCAGCTCAAGGCGGGACAGGTGCCCTAGAGTAGCCTGCATCCAGGGACAGACCCCAGCTG**GGTGCTGACACGTCCACCTCCATCTCTTCCTCAGCACC**ACCTGTGGCAGGACCGTCAGTCTTCCTCTTCCCCCCAAAACCCAAGGACACCCTCATGATCTCCCGGACCCCTGAGG**TCACGTGCGTGGTGGTGGACGTGA**GCCACGAAGACCCCGAGGTCCAGTTCAACTGGTACGTGGACGGCGTGGAGGTGCATAATGCCAAGACAAAGCCACGGGAGGAGCAGTTCAACAGCACGTTCCGTGTGGTCAGCGTCCTCACCGTCGTGCACCAGGACTGGCTGAACGGCAAGGAGTACAAGTGCAAGGTCTCCAACAAAGGCCTCCCAGCCCCCATCGAGAAAACCATCTCCAAAACCAAAGGTGGGACCCGCGGGGTATGAGGGCCACATGGA**CAGAGGCCGGCTCGGCCCACCCTCTG**CCCTGGGAGTGACCGCTGTGCCAACCTCTGTCCCTACA**GGGCAGCCCCGAGAACCACAGGTGTACACCCTGCCC**CCATCCCGGGAGGAGATGACCAAGAAC**CAGGTCAGCCTGACCTG**CCTGGTCAAAGGCTTCTACCCCAGCGACA**TCTCCGTGGAGTGGGAGAGCAATGGGCAGCCGGAGA**ACAACTACAAGACCACACCTCCCATGCTGGACTCCGACGGCTCCTTCTTCCTCTACAGCAAGCTCACCGTGGACAAGAGCAGGTGGCAGCAGGGGAACGTCTT**CTCATGCTCCGTGATGCATGAGGCTCTGCACAACCACTACACACAGAAGAGCCTC**TCCCTGTCTCCGGGTAAATGAgtgccacggccggcaagcccccgctccccaggctct**cggggtcgcgcgaggatgcttggcacgtaccccg**tctacatacttcccgggcacccagcatggaaataaagcacccagcgctgccctgggcccctgcgagactgtgatggttctttccgtggg**tcaggccgagtctgaggcctga**gtggcatgagggaggcagagcgggttccactgtccccacact**ggcccaggctgtgcaggtgtgcctgggcc**gcctagggtggggctcagccaggggctgccctcggcagggtgggggatttgccag**cgtggccctccctccagcagcagctgccctgggctgggccacg**ggaagccctaggagcccctgg**ggacagacacacagcccctgcctctgtaggagactgtcc**tgtcctgtgagcgccctgtcctccgacctc**catgcccactcgggggcatg**cctagtccatgtgcgtagggacaggccctccctcacccatctacccccacggcactaacccctggctgccctgcccagcctcgcacccgcatggggacacaaccgactccggggacatgcactctcgggccctgtggagggactggtccagatgcccacacacaca**ctcagcccagacccgttcaacaaaccccgcgctgag**gt**tggccggccacacggcca**ccacacacacacgtgcacgcctcacacacggagcctcacccgggcgaaccgcacagcacccagaccagagcaaggtcctcgcacacgtgaacactcctcagacacaggcccccacgagccccacgcggcacctcaaggcccacgagccgctcggcagcttctccacatgctgacctgctcagacaaacccagccctcctctcacaaggtgcccctgcagccgccacacacacacaggcccccacacacaggggaacacacgccacgtcgcgtccctggcactggcccacttcccaatgccgcccttccctgcagctga

Uppercase: IGHG2-2

Lowercase: Flanking sequence[1000bp]

Red & Bold & Underline: Stem-loop [28]

Blue: Heptamer[53]

Green: Nonamer [2]

id-TRBC2-2[C_gene_segment]

aatttacctgtcatccctaagaatctacaaaggagatgctcaggacagaaactgtatcaacacaactagtagcaagaagttac**tctgatgatatcaga**tgtttatttgggaaacttgctagtagagaaagctacatataatatttggatgcaa**agggacacagaaggttgaagagtccct**aattttgaaataagggaagatgactaactgtctgagctgagaaaactcaggggtacctggaggcagaggaatggataagatgac**ttcatgcaccacaaaaagaaaaaacctcacattctcatgaa**cgcactgtaaaaccaaaggatgtcctcatatgaatgcaaaaaataggccatctgtaaatccaaagaaagcccccagatccaaaatgtctccctcatcccagattccccttcattcctgagcaccttagatttggtataaataacctgcttgggagggggctttttgaattcgtacataatttaaccttcacacag**tttctgcaaagtcagaatggtgattattacctcacatgcagaaa**aaagtgataggaatttctgtcttaaaagtcttgttggtggacaaaggaagttctaggatttggatctt**gtttttttgggttccaatcccttgctccagttaaaaaac**taccacataaaatggtgagaagtaggtaggcaagtttttattgatagagaggaaatcaaataatggcaatgaggagacatcacctggaat**gttaggcagtgcctaac**tgggggatggacagacaatgggcagtgccaacccatagggtggatacaaaagacaggcaaggaaggggtagaaccatcaaagaggaataggctggtgaccccaaagcaagg**aggacctagtaacataattgtgcttcattatggtcct**ttcccggccttctctctcacacatacacagagcccctaccaggaccagacagctctcagagcaaccctagccccattacctcttccctttccagAGGACCTGAAAAACGTGTTCCCACCCGAGGTCGCTGTGTTTGAGCCATCAGAAGCAGAGATCTCC**CACACCCAAAAGGCCACACTGGTGTG**CCTGGCCACAGGCTTCTACCCCGACCACGTGGAGCTGAGCTGGTGGGTGAATGGGAAGGAGGTGCACAGTGGGGTCAGCACAGACCCGCAGCCCCTCAAGGAGCAGCCCGCCCTCAATGACTCCAGATACTGCCTGAGCAGCCGCCTGAGGGTCTCGGCCACCTTCTGGCAGAACCCCCGCAACCACTTCCGCTGTCAAGTCCAGTTCTACGGGCTCTCGGAGAATGACGAGTGGACCCAGGATAGGGCCAAACCTGTCACCCAGATCGTCAGCGCCGAGGCCTGGGGTAGAGCAGGTGAGTGGGGCCTGGGGAGATGCCTGGAGGAGATTAGGTGAGACCA**GCTACCAGGGAAAATGGAAAGATCCAGGTAGC**GGACAAGACTAGATCCAGAAGAAAGCCAGAGTGGACAAGGTGGGATGATCAAGGTTCACAGGGTCAGCAAAGCACGGTGTGCACTTCCCCCACCAAGAAGCATAGAGGCTGAATGGAGCACCTCAAGCTCATTCTTCCTTCAGATCCTGACACCTTAGAGCTAAGCTTTCAAGTCTCCCTGAGGACCAGCCATACAGCTCAGCATCTGAGTGGTGTGCATCCCATTCTCTTCTGGGGTCCTGGTTTCCTAAGATCATAGTGACCACTTCGCTGGCACTGGAGCAGCATGAGGGAGACAGAACCAGGGCTATCAAAGGAGGCTGACTTTGTACTATCTGATATGCATGTGTTTGTGGCCTGTGAGTCTGTGA**TGTAAGGCTCAATGTCCTTACA**AAGCAGCATTCTCTCATCCATTTTTCTTCCCCTGTTTTCTTTCAGACTGTGGCTTCACCTCCGGTAAGTGAGTCTCTCCTTTTTCTCTCTATCTTTCGCCGTCTCTGCTCTCGAACCAGGGCATGGAGAATCCACGGA**CACAGGGGCGTGAGGGAGGCCAGAGCCACCTGTG**CACAGGTACCTACATG**CTCTGTTCTTGTCAACAGAG**TCTTACCAGCAAGGGGTCCTGTCTGCCACCATCCTCTATGAGATCTTGCTAGGGAAGGCCACCTTGTATGCCGTGCTGGTCAGTGCCCTCGTGCTGATGGCCATGGTAAGGAGGAGGGTGGGATAGGGCAGATGATGGGGGCAGGGGATGGAACATCACACATGGGCATAAAGGAATCTCAGAGCCAGAGCACAGCCTAATATATCCTATCACCTCAATGAAACCATAATGAAGCCAGACTGGGGAGAAAATGCAGGGAATATCACAGAATGCATCATGGGAGGATGGAGACAACCAGCGAGCCCTACTCAAATTAGGCCTCAGAGCCCGCCTCCCCTGCCCTACTCCTGCTGTGCCATAGCCCCTGAAACCCTGAAAATGTTC**TCTCTTCCACAGGTCAAGAGA**AAGGATTCCAGAGGCTAGctccaaaaccatcccaggtcattcttcatcctcacccaggattctcctgtacctgctcccaatctgtgttcctaaaagtgattctcactctgcttctcatctcctacttacatgaatacttctctcttttttctgtttccctgaagattgagctcccaacccccaagtacgaaataggctaaaccaataaaaaattgtgtgttgggcctggttgcatttcaggagtgtctgtggagttctgctcatcactgacctatcttctgattt**agggaaagcagcattccct**tggacatctgaagtgacagccctctttctctccacccaatgctgctttctcctgttcatcctg**atggaagtcctcaaacaccatttccatacccaggcattctgggt**ccccactggagggttagtctgaagggcaatggctgggctttggaaaaccagcaagttgaggacagagaggaaggcacacagcaaaccataagcccttacccagtgcaggacagaggatgcgggcagacctatgggttacaatgtctggtcatttcccaattccagattaaactgtcacctgttttacctttagttttattagtttgtagtcttaacacctccagcttctcttgtttcaggatttgggcttaaaattgagtgct**actctgcatgtctagtttgaaatactagagaaggcagagt**tgagacaactgatatgtaaagcctggggaagagtgatttctcaggagcgagacacactaagtcaggagcaatgggatatagggcccagtgggggctgaagtgctatgttcagagtagcccttccaatgggcttcttcgtttgatggatggaaaccaaaccactccaaacacaaggtgt**taactgctcctacttgggcaaagacagtta**tcctgtcaaggtaaattctgcatacaggctgaatgcattgtggtaaaacactacatggaggaagaggaggaatgggattaaagaaaaggaggccta

Uppercase: TRBC2-2

Lowercase: Flanking sequence[1000bp]

Red & Bold & Underline: Stem-loop [18]

Blue: Heptamer[35]

Green: Nonamer [5]

id-IGKC[C_gene_segment]

ctgcttattttccagtgatcacattattttgctaccatggttattttatacaattatctgaaaaaaattagttatgaagattaaaagagaagaaaatattaaacataagagattcagtctttcatgttgaactgcttggttaacagtgaagt**tagttttaaaaaaaaaaaaaacta**tttctgttatcagctgacttctccctatctgttgacttctcccagcaaaagattcttattttacattttaactactgctctc**ccacccaacgggtggaatcccccagagggggatttcca**agaggccacctggcagttgctgagggtca**gaagtgaagctagccacttc**ctcttaggcaggtggccaagattacagttgacctctcctggtatggctgaaaattgctgcatatggttacaggccttgaggcctttgggagggcttagagagttgctggaacagtcagaaggtggaggggctgacaccacccaggc**gcagaggcagggctcagggcctgctctgc**agggaggttttagcccagcccagccaaagtaacccccgggagcctgttatcccagcacagtcctggaagaggcacaggggaaataaaagcggacggaggctttccttgactcagccgctgcctggtcttc**ttcagacctgttctgaa**ttctaaactctgagggggtcggatgacgtggccattctttgcctaaagcattgagtttactgcaaggtcagaaaagcatgcaaagccctcagaatggctgcaaagagctccaacaaaacaatttagaactttattaaggaatagggggaagctaggaagaaactcaaaacatcaagattttaaatacgcttcttggtctccttgctataattatctgggataagcatgctgttttctgtctgtccctaaca**tgccctgtgattatccgcaaacaacacacccaagggca**gaactttgttacttaaacaccatcctgtttgcttctttcctcagGAACTGTGGCTGCACCATCTGTCTTCATCTTCCCGCCATCTGATGA**GCAGTTGAAATCTGGAACTGC**CTCTGTTGTGTGCCTGCTGAATAACTTCTATCCCAGAGAGGCCAAAGTACAGTGGAAGGTGGATAACGCCCTCCAATCGGGTAACTCCCAGGAGAGTGTCACAGAGCAGGACAGCAAGGACAGCACCTACAGC**CTCAGCAGCACCCTGACGCTGAG**CAAAGCAGACTACGAGAAACACAAAGTCTACGCCTGCGAAGTCACCCATCAGGGCCTGAGCTCGCCCGTCACAAAGAGCTTCAACAGGGGAGAGTGTTAGagggagaagtgcccccacctgctcctcagttccagcctgaccccctcccatcctttggcctctgaccctttttccacaggggacctacccctattgcggtcctccagctcatctttcacctcacccccctc**ctcctccttggctttaattatgctaatgttggaggag**aatgaataaataaagtgaatctttgcacctgtggtttctctctttcctcatttaataattattatctgttgttttaccaactactcaatttctcttataagggactaaatatgtagtcatcctaaggcgcataaccatttataaaaatcatccttcattctattttaccctatcatcctctgcaagacagtcctccctcaaacccacaagccttctgtcctcacagtcccctgggccatggtaggagaga**cttgcttccttgttttcccctcctcagcaag**ccctcatagtcctttttaagggtgacaggtcttacagtcatatatcct**ttgattcaattccctgagaatcaa**ccaaagcaaatttttcaaaagaagaaacctgctataaagagaatcattcattgcaacatgatataaaataacaacacaataaaagcaattaaa**taaacaaacaatagggaaatgtttaagttcatcatggtacttagacttaa**tggaatgtcatgccttatttacatttttaaacaggtactgagggactcctgtctgccaagggccgtattgagtactttccacaacctaa**tttaatccacactatactgtgagattaaaaacattcattaaaatgtt**gcaaaggttctataaa**gctgagagacaaatatattctataactcagc**aatcccacttctagatgactgagtgtccccacccaccaaaaaactatgcaagaatgttcaaagcagctttatttacaaaagccaaaaattggaaatagcccgattgtccaacaatagaatgagttattaaactgtggtatgtttatacattag

Uppercase: IGKC

Lowercase: Flanking sequence[1000bp]

Red & Bold & Underline: Stem-loop [17]

Blue: Heptamer[22]

Green: Nonamer [1]

id-TRAC[C_gene_segment]

catgctaatcctccggcaaacctctgttt**cctcctcaaaaggcaggagg**tcggaaagaataaacaatgagagtcacattaaaaacacaaaatcctacggaaatactgaagaatgagtctcagcactaaggaaaagcctccagcagctcctgctttctgagggtgaaggatagacgctgtggctctgcatgactcactagcactctatcacggccatattctggcagggtcagtggctccaactaacatttgtttggtactttacagtttattaaatagatgtttatatggagaagctctcatttctttctcagaagagcctggctaggaaggtggatgaggcaccatattcattttgcaggtgaaattcctgagatgtaaggagctgctgtgacttgctcaaggccttatatcgagtaaacggtagtgctggggcttagacgcaggtgttctgatttatagttcaaaacc**tctatcaatgagagagcaatctcctggtaatgtgataga**tttcccaacttaatgccaacataccataaacctcccattctgctaatgcccagcctaagttggggagaccactccagattccaagatgtacagtttgctttgctgggcctttttcccatgcctgcctttactctgccagagttatattgctggggttttgaagaagatcctattaaataaaagaataagcagtattattaagtagccctgcatttcaggtttccttgagtggcaggccaggcctggccgtgaacgttcactga**aatcatggcctcttggccaagatt**gatagcttgtgcctgtccctgagtcccagtccatcacgagcagctggtttctaagatgctatttcccgtataaagcatgagaccgtgact**tgccagccccacagagccccgcccttgtccatcactggca**tctggactccagcctgggttggggca**aagagggaaatgagatcatgtcctaaccctgatcctctt**gtcccacagATATCCAGAACCCTGACCCTGCCGTGTACCAGCTGAGAGACTCTAAATCCAGTGACAAGTCTGTCTGCCTATTCACCGATTTTGATTCTCAAACAAATGTGTCACAAAGTAAGGATTCTGATGTGTATATCACAGACAAAACTGTGCTAGACATGAGGTCTATGGACTTCAAGAGCAACAGTGCTGTGGCCTGGAGCAACAAATCTGACTTTGCATGTGCAAACGCCTTCAACAACAGCATTATTCCAGAAGACACCTTCTTCCCCAGCCCAGGTAAGGGCAGCTTTGGTGCCTTCGCAGGCTGTTTCCTTGCTTCAGGAATGGCCAGGTTCTGCCCAGAGCTCTGGTCAATGATGTCTAAAACTCCTCTGA**TTGGTGGTCTCGGCCTTATCCATTGCCACCAA**AACCCTCTTTTTACTAAGAAACAGTGAGCCTTGTTCTGGCAGTCCAGAGAATGACACGGGAAAAAAGCAGATGAAGAGAAGGTGGCAGGAGAGGGCACGTGGCCCAGC**CTCAGTCTCTCCAACTGAG**TTCCTGCCTGCCTGCCTTTGCTCAGACTGTTTGCCCCTTACTGCTCTTCTAGGCCTCATTCTAAGCCCCTTCTCCAAGTTGCCTCTCCTTATTTCTCCCTGTCTGCCAAAAAATCTTTCCCAGCTCACTAAGTCAGTCTCACGCAGTCACTCATTAACCCACCAATCACTGATTGTGCCGGCACATGAATGCACCAGGTGTTGAAGTGGAGGAATTAAAAAGTCAGATGAGGGGTGTGCCCAGAGGAAGCACCATTCTAGTTGGGGGAGCCCATCTGTCAGCTGGGAAAAGTCCAAATAACTTCAGATTGGAATG**TGTTTTAACTCAGGGTTGAGAAAACA**GCTACCTTCAGGACAAAAGTCAGGGA**AGGGCTCTCTGAAGAAATGCTACTTGAAGATACCAGCCCT**ACCAAGGGCAGGGAGAGGACCCTATAGAGGCCTGGGACAGGAGCTCAATGAGAAAGGAGAAGAGCAGCAGGCATGAGTTGAATGAAGGAGGCAGGGCCGGGTCACAGGGCCTTCTAGGCCATGAGAGGGTAGACAGTATTCTAAGGACGCCAGAAAGCTGTTGATCGGCTTCAAGCAGGGGAGGGACACCTAATTTGCTTTTCTTTTTTTTTTTTTTTTTTTTTTTTTTTTTTGAGATGGAGTTTTGCTCTTGTTGCCCAGGCTGGAGTGCAATGGTGCATCTTGGCTCACTGCAACCTCCGCCTCCCAGGTTCAAGTGATTCTCCTGC**CTCAGCCTCCCGAGTAGCTGAG**ATTA**CAGGCACCCGCCACCATGCCTG**GCTAATTTTTTGTATTTTTAGTAGAGACAGGGTTTCACTATGTTGGCCAGGCTGGTCTCGAACTCCTGACCTCAGGTGATCCACCCGCTTCAGCCTCCCAAAGTGCTGGGATTACAGGCGTGAGCCACCACACCCGGCCTGCTTTTCTTAAAGATCAATCTGAGTGCTGTACGGAGAGTGGGTTGTAAGCCAAGAGTAGAAGCAGAAAGGGAGCAGTTGCAGCAGAGAGATGATGGAGGCCTGGGCAGGGTGGTGGCAGGGAGGTAACCAACACCATTCAGGTTTCAAAGGTAGAACCATGCAGGGATGAGAAAGCAAAGAGGGGATCAAGGAAGG**CAGCTGGATTTTGGCCTGAGCAGCTG**AGTCAATGATAGTGCCGTTTACTAAGAAGAAACCAAGGAAAAAATTTGGGGTGCAGGGATCAAAACTTTTTGGAACATATGAAAGTACGTGTTTATACTCTTTATGGCCCTTGTCACTATGTATGCCTCGCTGCCTCCATTGGACTCTAGAATGAAGCCAGGCAAGAGCAGGGTCTATGTGTGATGGC**ACATGTGGCCAGGGTCATGCAACATGT**ACTTTGTACAAACAGTGTATATTGAGTAAATAGAAATGGTGTCCAGGAGCCGAGGTATCGGT**CCTGCCAGGGCCAGGGGCTCTCCCTAGCAGG**TGCTCATATGCTGTAAGTTCCCTCCAGATCTCTCCACAAGGAGGCATGGAAAGGCTGTAGTTGTTCACCTGCCCAAGAACTAGGAGGTCTGGGGTGGGAGAGTCAGCCTGCTCTGGATGCTGAAAGAATGTCTGTTTTTCCTTTTAGAAAGTTCCTGTGATGTCAAGCTGGTCGAGAAAAGCTTTGAAACAGGTAAGACAGGGGTCTAGCCTGGGTTTGCACAGG**ATTGCGGAAGTGATGAACCCGCAAT**AACCCTGCCTGGATGAGGGAGTGGGAAGAAATTAGTAGATGTGGGAATGAATGATGAGGAATGGAAACAGCGGTTCAAGACCTGCCCAGAGCTGGGTGGGGTCTCTCCTGAATCCCTCTCACCATCTCTGACTTTCCATTCTAAGCACTTTGAGGATGAGTTTCTAGCTTCAATAGACCAAGGACTCTCTCCTAGGCCTCTGTATTCCTTTCAACAGCTCCACTGTCAAGAGAGCCAGAGAGAGCTTCTGGGTGGCCCAGCTGTGAAATTTCTGAGTCCC**TTAGGGATAGCCCTAA**ACGAACCAGATCATCCTGAGGACAGCCAAGAGGTTTTGCCTTCTTTCAAGACAAGCAACAGTACTCACATAGGCTGTGGGCAATGGTCCTGTCTCTCAAGAATCCCCTGCCACTCCTCACACCCACCCTGGGCCCATATTCATTTCCATTTGAGTTGTT**CTTATTGAGTCATCCTTCCTGTGGTAGCGGAACTCACTAAG**GGGCCCATCTGGACCCGAGGTATTGTGATGATAAA**TTCTGAGCACCTACCCCATCCCCAGAAGGGCTCAGAA**ATAAAATAAGAGCCAAGTCTAGTCG**GTGTTTCCTGTCTTGAAACAC**AATACTGTTGGCCCTGGAAGAATGCACAGAATCTGTTTGTAAGGGGATATGCACAGAAGCTGCAAGGGACAGGAGGTGCAGGAGCT**GCAGGCCTCCCCCACCCAGCCTGC**TCTGCCTTGGGGAAAACCGTGGGTGTGTCCTG**CAGGCCATGCAGGCCTG**GGACATGCAAGCCCATAACCGCTGTGGCCTCTT**GGTTTTACAGATACGAACCTAAACTTTCAAAACC**TGTCAGTGATTGGGTTCCGAATCCTCCTCCTGAAAGTGGCCGGGTTTAATCTGCTCATGACGCTGCGGCTGTGGT**CCAGCTGAGGTGAGGGGCCTTGAAGCTGG**GAGTGGGGTTTAGGGACGCGGGTCTCTGGGTGCATCCTAAGCTCTGAGAGCAAACCTCCCTGCAGGGTCTTGCTTTTAAGTCCAAAGCCTGAGCCCACCAAACTCTCCTACTTCTTCCTGTTACAAATTCCTCTTGTGCAATAATAATGGC**CTGAAACGCTGTAAAATATCCTCATTTCAG**CCGCCTCAGTTGCACTTCTCCCCTATGAGGTAGGAAGAACAGTTGTTTAGAAACGAAGAAACTGAGGCCCCACAGCTAATGAGTGGAGGAAGAGAGACACTTGT**GTACACCACATGCCTTGTGTTGTAC**TTCTCTCACCGTGTAACCTCCTCATGTCCTCTCTCCCCAGTACGGCTCTCTTAGCTCAGTAGAAAGAAGACATTACACTCATATTACACCCCAATCCTGGCTAGAGTCTCCGCACCCTCCTCCCCCAGGGTCCCCAGTCGTCTTGCTGACAACTGCATCCTGTTCCATCACCATCAAAAAAAAACTCCAG**GCTGGGTGCGGGGGCTCACACCTGTAATCCCAGC**ACTTTGGGAGGCAGAGGCAGGAGGAGCACAGGAGCTGGAGACCAGCCTGGGCAACACAGGGAGACCCCGCCTCTACAAAAAGTGAAAAAATTAAC**CAGGTGTGGTGCTGCACACCTG**TAGTCCCAGCTACTTAAGAGGCTGAGATGGGAGGATCGCTTGAGCCCTGGAATGTTGAGGCTACAATGAGCTGTGATTGCGTCACTGCACTCCAGCCTGGAAGACAAAGCAAGATCCTGTCTCAAATAATAAAAAAAATAAGAACTCCAGGGTACATTTGCTCCTAGAACTCTACCACATAGCCCCAAACAGAGCCATCACCATCACATCCCTAACAGTCCTGGGTCTTCCTCAGTGTCCAGCCTGACTTCTGTTCTTCCTCATTCCAGATCTGCAAGATTGTAAGACAGCCTGTGCTCCCTCGCTCCTTCCTCTGCATTGCCCCTCTTC**TCCCTCTCCAAACAGAGGGA**ACTCTCCTACCCCCAAG**GAGGTGAAAGCTGCTACCACCTC**TGTGCCCCCCCGGCAATGCCACCAACTGGATCCTACCCGAATTTATGATTAAGATTGCTGAAGAGCTGCCAAACACTGCTGCCACCCCCTCTGTTCCCTTATTGCTGCTTGTCACTGCCTGACATTCACGGCAGAGGCA**AGGCTGCTGCAGCCT**CCCCTGGCTGTGCACATTCCCTCCTGCTCCCCAGAGACTGCCTCCGCCATCCCACAGATGATGGATCTTCAGTGGGTTCTCTTGGGCTCTAGGTCCTGCAGAATGTTGTGAGGGGTTTATTTTTTTTTAATAGTGTTCATAAAGAAATACATAGTATTCTTCTTCTCAAGACGTGGGGGGAAATTATCTCATTATCGAGGCCCTGCTATGCTGTGTATCTGGGCGTGTTGTATGTCCTGCTGCCGATGCCTTCattaaaatgatttggaagagcagagactgtgcctctgtttgactgggtttggtaggagtcattttctgcttgctggtgatcactagctgggcagagaaaaaccaaggcatttgtctatgatgctgtccaggaagcctcattcaacaagctgcctaagtcaacctcttcttggaataacctctaaaagcttccgcttagcaggctatgctgagggccaggaaaacccacctaccagcttggacccctcctctcccactctcatgccacgccacggga**ccacccataacaggagcccacacacatgggtgg**cagtgacctgcggcagacagggaccacacagcaagtgtccccaaaatgccacccactgt**ctcctgccctccaggag**catttcctttgcctctcctctcagactgggtttccactgaaactgtgcattgtctcacaaattcgt**ggctggggaccacccaccactctgctgcctgatccagcc**ccacgccagccctttgaggtgcccaagctgacaccaggagcaaggttgagaggaagctgtgaccccagcaggactttatgttcccaccatcccggatgtgagaatgaggaaaaaaggagatgagctgtctccccacaagcccagagatttgaccgaggagagtagaggcctcgagctctcacctaagagaaaagacatggggcttcctggggtccacagctcactgcgctctccctcctgagactcctgctgccagagcacc**tttccccagggtcatggatgctgagggaaa**cacaacttagagaccactccaccatccacccagcaagcacagctaccaagacccaaagctgaggcttaccaatgcccagggtgggagggggttccatccctgaataactccatggttcccctatgcgtctgaccatcccagccagaaatacataaatcatctcagctacaattcaggcctgcttcttttcatagggatgaagctacaggttgagtatc

Uppercase: TRAC

Lowercase: Flanking sequence[1000bp]

Red & Bold & Underline: Stem-loop [34]

Blue: Heptamer[83]

Green: Nonamer [12]

id-IGHEP1[C_gene_segment]

tgtcacccccaggacagggacagccaacccagagccgggagggagggtggggaggcggca**gcagggagctgtcctgagctccactgc**gcaactggctgatcttggcaagtccgagctgggtggactgaggggggcttggctgagtggactagactgagacgggcctaacagactgagctgaggcgagctgggtgggctgagagggctac**cctgtcccttagaggacagg**tggccaagctgggctgtcctgagccagggcgatcggggctggcccgggccaggcgggtttagctgagttgagtgagtggactgggtagagggaaatgagctaggctcagctgagctaggcttgagctgggttatcctaagccctaaggtggactgagctgggctgagctggacttatctggggag**cagggcaaagtcaggctgagctgaggtggcctgccctg**ggtg**gtccaggattgagttaagctgaattaggctgacctggac**ttgactggacttggttgaaataagctgggccgacacaggagtagggacaagctacagttctctacttaggataaaatgggtgctcgtggactatccgggctgaaggagaccaagctggggtatta**cctgctgagcttacctgacctggcctgagttcagcagg**gctgcgctgagctggacagacctg**agccaagcttagctggttgggctgagtaagctgggct**gagctaaatgggattgagctgaggagggctaggctgggggagagacctgacgacggacagggttaaaagctggagtgagcaggccttaaattattgaa**ctaaattgggctggggtgatctgaatttag**ctgggatgagctgggctgggctgaactgtgcccacgtgaactgggctaaactaggctcgcctgagtgga**ctcagctgggttggtctcaactgggttcagctgag**ctgggctcggctagactacactgggttcagctgacactacactgggttcAACCCGAGAGGGGTGAGCGCCTACCTAAGCCGGCCCAGCCCGTTCGACCTGTTCATCCGCAAGTCGCCCACGATCACCTGTCTGGTGGTGGACCTGGCACCCAGCAAGTGGACCGTGAACCTGACCTGGTCCCGGGCCAGTGGGAAGCCTGTGAACCACTCCACCAGAAAGGAGGAGAAGCAGCGCAATGGCACGTTAACCGTCACGTCCACCCTGCCGGTGGGCACCCGAGACTGGATCGAAGGGGAGACCTACCAGTGCAGGGTGACCCACCCCCACCTGCCCAGGGCCCTCGTGCGGTCCACGACCAAGACCAGCGGTGAGCCACGGGCAGGCCGGGGTCGTGGGGGGAGGGAGGGAGCGAGTGAGCG**GGGCCTGGGCTGACCCCACGTCTGGCCACAGGCCC**GCGTGCTGCCCCGGAAGTCTATGCGTTTGCGACGCCGGAGTGGCTGGGGAGCCGGGACAAGCGCACCCTCACCTGCCTGATCCAGAACTTCATGCCTGAGGACATCTCGGTGCAGTG**GCTGCACAACGAGGTGCAGC**TCCCGGACGCCCGGCACAGCACGACGCAGCCCCGCAAGACCAAGGGCTCCGGCTTCTTCATCTTCAGCCGCCTGGAGGTGACCAGGGCCGAGTGGGAGCAGAAAGATGAGTTCATCTGCCGTGCAGTCCATGAGGCAGCGAGCCCCTC**ACAGACCGTCCAGCGAGCGGTGTCTGT**AAATCCCGGTAAATGAcgtactcctgcctccct**ccctcccagggctccgtccagctgtgcagtggggaggg**ctggccagaccttctgtc**cactgttgcaacgaccccaggaagctacccccaataaacagtg**cctgctcagagcccagggtacacccgttcttgggagcgggcagggctgtgggcaggtgcatcttggcacagaggaatgggccccccaggaggggcagtgggaggaggtgggcagggctgagtccccccaggagaggtggtgggaggaggtgggcagggctgaggtgccactcatccatctgccttcgtgtcagggttatttgtcaaacagcgtatctgcagggactcatcacagctaccccgggccctctctgcccccactctcggtctaccccctccaaggagtccaaagacccaggggaggtcctcagggaaggggcaaggg**agccccgacagccctctctcttgggggct**tggcttctacccccctggacaggagcccctgcacccccaggtatagatgggcacacaggcccctccaggtagaaaaacagccctaagtgaaaccccca**cacagacacacacgacccgacagccctcgcccaagtctgtg**ccactggcgttcgcctctctgccctgtcccaccttgccgagtc**ctggccccagcaccggggccag**tggagccgagcccactcacaccccgcagcctccgccaccccgccctgtgggcacaccaggcccaggtcagcagccaggccccctctcctactgccccccaccgccccttggtccatcctgaatcggcctccaggggatcgccagcctcacacacccagtctcgcccactcacgcctcactcaaggcaca**gctgtgcacacactaggccccatagcaactccacagc**accctgtaccaccaccagggcgccatagacacccca**cacgtggtcacacgtg**gcccacactccgcctcccaccctgcctccagcgaggctactgccaagcc

Uppercase: IGHEP1

Lowercase: Flanking sequence[1000bp]

Red & Bold & Underline: Stem-loop [18]

Blue: Heptamer[27]

Green: Nonamer [0]

id-IGHM[C_gene_segment]

tgggctatactgggcttagctgggctgggctatactgggcttagctgggctgggctatactgggcttagctgggctgg**gctgagctgagatggtcttaggtggtctgagctcagc**taggctgggctgagctggtctg**agctcatctgagttgggctgagct**gagcttggctttgctgagctggggtggggtgggctgggctggattgagctggcctgggctgggatgaactggattgagctggcctgggctgggatgaactggaggacatggcactgggccaatcttcatgatcttgttggacatagatggatagcctcagctgagtctacactgcgttccccatcacactcaccctccctatactcact**cccaggcctgggttgtctgcctggg**gagacttcagggtagctggagtgtgactgagctggg**ggcagcagaagctgggctggagggactctattggctgcc**tgcggggtgtgtggctccaggcttcacattcaggtatgcaacctgggccctccagctgcatgtgctgggagctgagtgtgtg**cagcacctacgtgctg**atgcctcgggggaaagcaggcctggtccacccaaacctgagccctcagccat**tctgagcagggagccaggggcagtcaggcctcaga**gtgcagcagggcagccagctgaatggtggcagggatggctcagcctgctccaggagaccccaggtctgtccaggtgttcagtgctgggccctgcagcaggatgggc**tgaggcctgcagccccagcagccttggacaaagacctgaggcctca**ccacggccccgccacccctgatagccatgacagtctgggctttggaggcctgcaggtgggctcggccttggtggggcagccacagcgggacgcaagtagtgagggcactcagaacgccactcagccccgacaggcagggcac**gaggaggcagctcctc**acc**ctccctttctcttttgtcctgcgggtcctcagGGAG**TGCATCCGCCCCAACCCTTTTCCCCCTCGTCTCCTGTGAGAATTCCCCGTCGGATACGAGCAGCGTGGCCGTTGGCTGCCTCGCACAGGACTTCCTTCCCGACTCCATCACTTTCTCCTGGAAATACAAGAACAACTCTGACATCAGCAGCACCCGGGGCTTCCCATCAGTCCTGAGAGGGGGCAAGTACGCAGCCACCTCACAGGTGCTGCTGCCTTCCAAGGACGTCATGCAGGGCACAGACGAACACGTGGTGTGCAAAGTCCAGCACCCCAACGGCAACAAAGAAAAGAACGTGCCTCTTCCAGGTGAGGGCCGGGCCCAGCCACCGGGACAGAGAGGGAGCCGAAGGGGGCGGGAGTGGCGGGCACCGGGCTGACACGTGTCC**CTCACTGCAGTGATTGCTGAGCTGCCTCCCAAAGTGAG**CGTCTTCGTCCCACCCCGCGACGGCTTCTTCGGCAACCCCCGCAAGTCCAAGCTC**ATCTGCCAGGCCACGGGTTTCAGTCCCCGGCAGAT**TCAGGTGTCCTGGCTGCGCGAGGGGAAGCAGGTGGGGTCTGGCGTCACCACGGACCAGGTGCAGGCTGAGGCCAAAGAGTCTGGGCCCACGACCTACAAGGTGACCAGCACACTGACCATCAAAGAGAGCGACTGGCTCGGCCAGAGCATGTTCACCTGCCGCGTGGATCACAGGGGCCTGACCTTCCAGCAGAATGCGTCCTCCATGTGTGTCCCCGGTGAGTGACCTGTCCCCAGGGGCAGCACCCACCGACACACAGGGGTCCACTCGGGTCTGGCATTCGCCACCCCGGATGCAGCCATCTACTCCCTGAG**CCTTGGCTTCCCAGAGCGGCCAAGG**G**CAGGGGCTCGGGCGGCAGGACCCCTG**GGCTC**GGCAGAGGCAGTTGCTACTCTTTGGGTGGGAACCATGCCTCCGCC**CACATCCACACCTGCCCCACCTCTGACTCCCTTC**TCTTGACTCCAGATCAAGA**CACAGCCATCCGGGTCTTCGCCATCCCCCCATCCTTTGCCAGCATCTTCCTCACCAAGTCCACCAAGTTGACCTGCC**TGGTCACAGACCTGACCACCTATGACAGCGTGACCA**TCTCCTGGACCCGCCAGAATGGCGAAGCTGTGAAAACCCACACCAACATCTCCGAGAGCCACCCCAATGCCACTTTCAGCGCCGTGGGTGAGGCCAGCATCTGCGAGGATGACTGGAATTCCGGGGAGAGGTTCACGTGCACCGTGACCCACACAGACCTGCCCTCGCCACTGAAGCAGACCATCTCCCGGCCCAAGGGTAGGCCCCACTCTTGCCCCTCTTCCTGCACTCCCTGGGACCTCCCTTGGCCTCTGGGGCATGGTGGAAAGCACCCCTCAC**TCCCCCGTTGTCTGGGCAACTGGGGA**AAAGGGGACTCAACC**CCAGCCCACAGGCTGG**TCCCCCCACTGCCCCGCCCTCACCACCATCTCTGTTCACAGG**GGTGGCCCTGCACAGGCCCGATGTCTACTTGCTGCCACC**AGCCCGGGAGCAGCTGAACCTGCGGGAGTCGGCCACCATCACGTGCCTGGTGACGGGCTTCTCTCCCGCGGACGTCTTCGTGCAGTGGATGCAGAGGGGGCAGCCCTTGTCCCCGGAGAAGTATGTGACCAGCGCCCCAATGCCTGAGCCCCAGGCCCCAGGCCGGTACTTCGCCCACAGCATCCTGA**CCGTGTCCGAAGAGGAATGGAACACGG**GGGAGACCTACACCTGCGTGGTGGCCCATGAGGCCCTGCCCAACAGGGTCACCGAGAGGACCGTGGACAAGTCCACCGGTAAACCCACCCTGTACAACGTGTCCCTGGTCATGTCCGACACAGCTGGCACCTGCTACTGACCCTGCTGGCCTGCCCACAGGCTCGGGGCGGCTGGCCGCTCTGTGTGTGC**ATGCAAACTAACCGTGTCAACGGGGTGAGATGTTGCAT**CTTATAAAATTAGAAATAAAAAGATCCATTCAAAAGATACTGGTCCTGAGTGCACGATGCTCTGGCCTACTGGGGCGGCGGCTGTGCTGCACCCACCCTGCGCCTCCCCTGCAGAACACCTTCCTCCACAGCCCCCACCCCTGCCTCACCCACCTGCGTGCCTCAGTGGCTTCTAGAAACCCCTGAATTCCCTGCAGCTGCTCACAGCAGGCTGACCTCAGACTTGCCATTCCTCCTACTGCTTCCAGAAAGAAAGCTGAAAGCAAGGCCACACGTATACAGGCAGCACACAGG**CATGTGTGGATACACATG**GACAGACACGGACACACACAAACACATGGACACACAGAGACGTGCTAACCCATGGGCACA**CACATACACAGACATGGACCCACACACAAACATATGTG**GACACACATGTACAAACATGCACAGGCACACAAAGAGAACACTGACTACAGGCACACACACACACGGGCACACACATGGATATGTGCACACATGGACACATACA**TGTGCAGGACATGCACA**CACACAGACACACTAGCACAGAGGCATACACACACAGACACACACATTCACAAACAC**ACATGTGCATGCAAACACACACACATGT**ACAGACACAAGTACATGGACACATGCACACCCAGAGACACACTGACACAGACACACAGGAGCATGTGATACACTAACACGTGGACACACACGTCTACCCACAGGCACACAACAGATGGACACGCGTACACAGACATGCACACACCCACAGGCACAACACGTGCGCATGCCGGCCGGCCCCCGCCAACATTCTCCCAGGGCCCTGCCGGATACTCTGTCCCTGCAGCAGTTTGCTCCCTGCGCTGTGCTGGCACCGGGGCTTTGGGCCCAGGCTCTGCTTGTCCTTCTGTCTCTGCTTGGAGGTGCTGCCATGGCACCCAGCTTGTGCTCTG**CCTGGGGAGCGGAGGCCCCAGG**GATAGCATGTGACCC**CTGCTGAGGCCAGGCTCCTGATGAAGGCAGCAG**ATAGCCCCCACACCCACCGGTGAGCAGAACCAGAGCCTGTGCCATGTGCTGAGAGCAGGCAGTGACTAAGCATATGGGCCCAG**AGGGCAGAGTGGCTGCCCT**GGGCAGCTGCTCCTCTTAGCAGGAGGCCTCAGGAGATGAGCTAGAGCAAGTCTGCCCCTGCAAATACCA**CCTGCTCCCCAACCCACAGCAGGGAGCAGG**CGAGGTCAGACAGCAGCAGCCCGGGAAGGACCGAGCCCCAGCAGGGAAGGCAGGGCCCGAGTGAGGTCTCCACACCCAACGCACAGTGCTGTCTCTAACTGGGGCCACCTCCGAGTCCCCGCCACACTCTT**GGCCCTTTGGAGTCCTGGGCTCCAGGTGTCTCCCAAGGGCC**CATCTGTGCAG**GGGATGCAACCCCCCGAATGTCCTCATCCC**ACTGTGGAGCTCAGGTCTCTGTCTGCTCCCTGGG**TCCTGGCAGGGTAGGACAAGTCCGCCAGGA**TGTCCCCATGCA**GACTCTGCTCCAAGAGGGAGCTGGAGAGTC**AGGGCCTTGGTGAGGGAGTCAGGATCGGGTTCCCCCCAGCTCAGTCCTCCCACCTGCCAGCCCCCACAGCACAGG**GCAGGGCCACACCCCCTGC**TTCCCCCTCCAGGAGAGTCAGGACATGCTGGCCGCTGCTCCGCTGGGGCCCCGCCCTCCAGCCCCCACCTTGGTCTGTGTGCTGCATCCCCCACGCTCTCTCTGCCACCCCA**GGACTCTGAGGAAAAGACCTCAGAGTCC**CAGCCCTGCCCAGTCTCGGCCTGTGCCCCCGCTGCATCAGGCTTTCAGGG**GCCCAGCCCATGCCCTGGGC**AGTGCCCGAGCCCCCCTGCACTTGCTCTC**CCCACCCCTGGGTGCAGCACAGCCTAGGGGCCAAGGGTGGG**CCTAGAGGATGGGCCCCGGGGGGGCTTT**GCTGGGTGCCACCCCAGC**CTGACCCTAT**TCCCCCGTGCTGTGTCTCCTGCAGAGGGGGA**GGTGAGCGCCGACGAGGAGGGCTTTGAGAACCTGTGGGCCACCGCCTCCACCTTCATCGTCCTCTTCCTCCTGAGCCTCTTCTACAGTACCACCGTCACCTTGTTCAAGGTAGCACGGCTGTGGCACAGGGAGGAGGGTGCAGGGCGAGTGTGGG**GCCCAGGGAGCAGCCTGGGC**TGGACGTCTAGCCCGGAGGCCCCCACACCACCCCACTGGGTCATCTCTGCCCCGGCTCCCTTCCCGACCACGGGGAAAG**CATTTCACACTGTCTCTGTTGCCTGTAGGTGAAATG**Atcccaacagaagaacatcggagaccagagagaggaactcaaaggggcgctgcctccgggtctggggtcctggcctgcgtggcctgttggcacgtgtttctcttccccgcccggcctccagt**tgtgtgctctcacaca**ggcttccttctcgaccggcaggggctggctggcttgcaggccacgaggtgggctctaccccacactgctttgctgtgtatacgcttgttgccctgaaataaatatgcacattttatccatgaaactgctttctggtgagggtttttgtttctttttcaaaactttcctgctacatgggcatctcaagggggaccccagttccaaagggagctgtggaaaaga**gcgctgggtcagccagcgc**agggggtttcgaggaaagccacgtgcccaggaaaggggccgcagaagcaggtgggccagactcagactcggggcatgcccagcctgatggaaggaaggggactgagcaggagagggttccaggcctggtcctccaagcaca**gcctgaattgagagactggggctcaggc**ctcgggggcctctgtgtgtgctccacatgcctacaactgcccgggtcaccttgccacccttcccagcaagcccagacagttcttggccttgccccaaaccttcatgatgtgtggtgcacgccaccgggatccaggaggtgcaggctgagccctcgagagcatgtgggcctcaccgggatccaggag**gtgcaggctgagccctcgagagcatgtgggcctgtctgcac**agtgtgggggccttgcactccacaggagcacaggggtggggtagcagtcgcgcccgtggcaggggagtggaagttggagcaaaagtttcaggtgaacgagtgtcttagtctaattgggcagctatcacaaaacactatcggctgcaaagcttgagcaacagacattt**gcctcccacagcgctggaggc**tgaaagtctaagatcaaggcgccggcaacttcaacgtc

Uppercase: IGHM

Lowercase: Flanking sequence[1000bp]

Red & Bold & Underline: Stem-loop [46]

Blue: Heptamer[100]

Green: Nonamer [4]

id-IGHG1-2[C_gene_segment]

gagagaaatggggcctccctgtggcctgggggtcctggcaccatgcagggtggggagggccaagggcaggtgcaaggctcctacctg**tgctggggggcctgggttgagcccagca**gggaccttgccgggggaagctctggagagagggaggaggtgggctggtggccgagaaggccaggccagggctgggagggtgacggtgtggtgactga**gcctccagaagtaatgcaggacactgggaggc**agggggcatccaggcactcagggccctgacctgggctgctgcacactggggctaaggggaaaggaggggagaggctgaggaggaggctccaggaggctattccaaggcagggggttccggggccctggggctgaagggcgccgaccctatgcagtgtctggc**ccctctgctgcacagaagaaaagggccttggagggcagaggg**caggctatgaccag**ggccctgggcaagtcaggccaactcactaggggagggcc**acgctggggcggcagggtcagg**ggcttcagggggctcgggggacccacgagaagcc**atctgagaacagtgtccactggtcaagccaggcacccataaaaggctggagtggggccaatgggcatgagccgtccctgaggtggcaccgatggccagagctgaggccaagctagaggccctggactgtgctgactcc**cggcagacacagagcgctgacctggctgccg**agccccgcctcctagg**ctgcaggggtgcctgcag**aagggcaccacagggccaccggtcctgcaagctttctggggcaggccgggcctgaccttggctttggggcagggggtgggctaaggtgacgcaggtggcgccagccaggcgcacacccaatgcccgtgagcccagacactggacgctgaacctcgcggacagttaagaa**cccaggggcctctgcgccctggg**cccagctctgtcccacaccgcggtcacatggcaccacctctcttgcagCCTCCACCAAGGGCCCATCGGTCTTCCCCCTGGCACCCTCCTCCAAGAGCACCTCTGGGGGCACA**GCAGCCCTGGGCTGC**CTGGTCAAGGACTACTTCCCCGAACCGGTGACGGTGTCGTGGAACTCAGGCGCCCTGACCAGCGGCGTGCACACCTTCCCGGCTGTCCTACAGTCCTCAGGACTCTACTCCCTCAGCAGCGTGGTGACC**GTGCCCTCCAGCAGCTTGGGCAC**CCAGACCTACATCTGCAACGTGAATCACAAGCCCAGCAA**CACCAAGGTGGACAAGAAAGTTGGTG**AGAGG**CCAGCACAGGGAGGGAGGGTGTCTGCTGG**AAGCCAGGCTCAGCGCTC**CTGCCTGGACGCATCCCGGCTATGCAGCCCCAGTCCAGGGCAG**CA**AGGCAGGCCCCGTCTGCCT**CTTCACCCGGAGGCCTCTGCCCGCCCCACTCATGCTCAGGGAGAGGGTCTTCTGGCTTTTTCCCCAGGCTCTGGGCAGGCACAGGCTAGG**TGCCCCTAACCCAGGCCCTGCACACAAAGGGGCAGGTGCTGGGCTCAGACCTGCC**AAGAGCCATATCCGGGAGGACCCTGCCCCTGACCTAAGCCCACCCCAAAGGCCAAACTCTCCACTCCCTCAGCTCGGACACCTTCTCTCCTCCCAGATTCCAGTAACTCCCAATCTTCTCTCTGCAGAGCCCAAATCTTGTGACAAAACTCACACATGCCCACCGTGCCCAGGTAAGCCAGCCCAGGCCTCGCCCTCCAGCTCAAGGCGGGACAGGTGCCCTAGAGTAGCCTGCATCCAGGGACAGGCCCCAGCCG**GGTGCTGACACGTCCACCTCCATCTCTTCCTCAGCACC**TGAACTCC**TGGGGGGACCGTCAGTCTTCCTCTTCCCCCCA**AAACCCAAGGACACCCTCATGATCTCCCGGACCCCTGAGGTCACATGCGTGGTGGTGGACGTGAGCCACGAAGACCCTGAGGTCAAGTTCAACTGGTACGTGGACGGCGTGGAGGTGCATAATGCCAAGACAAAGCCGCGGGAGGAGCAGTACAACAGCACGTACCGTGTGGTCAGCGTCCTCACC**GTCCTGCACCAGGAC**TGGCTGAATGGCAAGGAGTACAAGTGCAAGGTCTCCAACAAAGCCCTCCCAGCCCCCATCGAGAAAACCATCTCCAAAGCCAAAGGTGGGACCCGTGGGGTGCGAGGGCCACATGGA**CAGAGGCCGGCTCGGCCCACCCTCTGCCCTGAGAGTGACCGCTGTACCAACCTCTGTCCCTACAGGGCAG**CCCCGAGAACCACAGGTGTACACCCTGCCCCCATCCCGGGATGA**GCTGACCAAGAACCAGGTCAGC**CTGACCTGCCTGGTCAAAGGCTTCTATCCCAGCGACATCGCCGTGGAGTGGGAGAGCAATGGGCAGCCGGAGAACAACTACAAGACCACGCCTCCCGTGCTGGACTCCGACGGCTCCTTCTTCCTCTACAGCAAGCTCACCGTGGACAAGAGCAGGTGGCAGCAGGGGAACGTCTT**CTCATGCTCCGTGATGCATGAGGCTCTGCACAACCACTACACACAGAAGAGCCTC**TCCCTGTCTCCGGGTAAATGAgtgccacggccggcaagcccccgctccccaggctct**cggggtcgcgcgaggatgcttggcacgtaccccg**tgtacatacttcccaggcacccagcatggaaataaagcacccagcgcttccctgggcccctgcgagactgtgatggttctttccacggg**tcaggccgagtctgaggcctga**gtggcatgagggaggcagagtgggtcccactgtccccacact**ggcccaggctgtgcaggtgtgcctgggcc**gcctagggtggggctcagccaggggctgccctcggcagggtgggggatttgccag**cgtggccctccctccagcagcagctgccctgggctgggccacg**agaagccctaggagcccctgg**ggacagacacacagcccctgcctctgtaggagactgtcc**tgttctgtgagcgccctgtcctccgaccc**gcatgcccactcgggggcatgc**ctagtccatgtgcgtagggacaggccctccctcacccatctacccccacggcactaacccctggcagccctgcccagcctcgcacccgcatggggacacaaccgactccggggacatgcactctcgggccctgtggagagactggtccagatgcccacacacacactcagcccagacccgttcaacaaaccccgcactgaggt**tggccggccacacggcca**ccacacacacacgtgcacgcctcacacacggagcctcacccgggcgaaccgcacagcacccagaccagagcaaggtcctcgcacacgtgaacactcctcggacacaggcccccacgagccccacgcggcacctcaaggcccacgagccgctcggcagcttctccacatgctgacctgctcagacaaacccagccctcctctcacaaggtg**cccctgcagccgccacacacacacagggg**atcacacaccacgtcacgtccctggccctggcccacttcccagtgccgcccttccctgcagctggggtcacatgaggtg

Uppercase: IGHG1-2

Lowercase: Flanking sequence[1000bp]

Red & Bold & Underline: Stem-loop [32]

Blue: Heptamer[47]

Green: Nonamer [0]

id-IGHM-3[C_gene_segment]

tgggctatactgggcttagctgggctgggctatactgggcttagctgggctgggctatactgggcttagctgggctgg**gctgagctgagatggtcttaggtggtctgagctcagc**taggctgggctgagctggtctg**agctcatctgagttgggctgagct**gagcttggctttgctgagctggggtggggtgggctgggctggattgagctggcctgggctgggatgaactggattgagctggcctgggctgggatgaactggaggacatggcactgggccaatcttcatgatcttgttggacatagatggatagcctcagctgagtctacactgcgttccccatcacactcaccctccctatactcact**cccaggcctgggttgtctgcctggg**gagacttcagggtagctggagtgtgactgagctggg**ggcagcagaagctgggctggagggactctattggctgcc**tgcggggtgtgtggctccaggcttcacattcaggtatgcaacctgggccctccagctgcatgtgctgggagctgagtgtgtg**cagcacctacgtgctg**atgcctcgggggaaagcaggcctggtccacccaaacctgagccctcagccat**tctgagcagggagccaggggcagtcaggcctcaga**gtgcagcagggcagccagctgaatggtggcagggatggctcagcctgctccaggagaccccaggtctgtccaggtgttcagtgctgggccctgcagcaggatgggc**tgaggcctgcagccccagcagccttggacaaagacctgaggcctca**ccacggccccgccacccctgatagccatgacagtctgggctttggaggcctgcaggtgggctcggccttggtggggcagccacagcgggacgcaagtagtgagggcactcagaacgccactcagccccgacaggcagggcac**gaggaggcagctcctc**acc**ctccctttctcttttgtcctgcgggtcctcagGGAG**TGCATCCGCCCCAACCCTTTTCCCCCTCGTCTCCTGTGAGAATTCCCCGTCGGATACGAGCAGCGTGGCCGTTGGCTGCCTCGCACAGGACTTCCTTCCCGACTCCATCACTTTCTCCTGGAAATACAAGAACAACTCTGACATCAGCAGCACCCGGGGCTTCCCATCAGTCCTGAGAGGGGGCAAGTACGCAGCCACCTCACAGGTGCTGCTGCCTTCCAAGGACGTCATGCAGGGCACAGACGAACACGTGGTGTGCAAAGTCCAGCACCCCAACGGCAACAAAGAAAAGAACGTGCCTCTTCCAGGTGAGGGCCGGGCCCAGCCACCGGGACAGAGAGGGAGCCGAAGGGGGCGGGAGTGGCGGGCACCGGGCTGACACGTGTCC**CTCACTGCAGTGATTGCTGAGCTGCCTCCCAAAGTGAG**CGTCTTCGTCCCACCCCGCGACGGCTTCTTCGGCAACCCCCGCAAGTCCAAGCTC**ATCTGCCAGGCCACGGGTTTCAGTCCCCGGCAGAT**TCAGGTGTCCTGGCTGCGCGAGGGGAAGCAGGTGGGGTCTGGCGTCACCACGGACCAGGTGCAGGCTGAGGCCAAAGAGTCTGGGCCCACGACCTACAAGGTGACCAGCACACTGACCATCAAAGAGAGCGACTGGCTCGGCCAGAGCATGTTCACCTGCCGCGTGGATCACAGGGGCCTGACCTTCCAGCAGAATGCGTCCTCCATGTGTGTCCCCGGTGAGTGACCTGTCCCCAGGGGCAGCACCCACCGACACACAGGGGTCCACTCGGGTCTGGCATTCGCCACCCCGGATGCAGCCATCTACTCCCTGAG**CCTTGGCTTCCCAGAGCGGCCAAGG**G**CAGGGGCTCGGGCGGCAGGACCCCTG**GGCTC**GGCAGAGGCAGTTGCTACTCTTTGGGTGGGAACCATGCCTCCGCC**CACATCCACACCTGCCCCACCTCTGACTCCCTTC**TCTTGACTCCAGATCAAGA**CACAGCCATCCGGGTCTTCGCCATCCCCCCATCCTTTGCCAGCATCTTCCTCACCAAGTCCACCAAGTTGACCTGCC**TGGTCACAGACCTGACCACCTATGACAGCGTGACCA**TCTCCTGGACCCGCCAGAATGGCGAAGCTGTGAAAACCCACACCAACATCTCCGAGAGCCACCCCAATGCCACTTTCAGCGCCGTGGGTGAGGCCAGCATCTGCGAGGATGACTGGAATTCCGGGGAGAGGTTCACGTGCACCGTGACCCACACAGACCTGCCCTCGCCACTGAAGCAGACCATCTCCCGGCCCAAGGGTAGGCCCCACTCTTGCCCCTCTTCCTGCACTCCCTGGGACCTCCCTTGGCCTCTGGGGCATGGTGGAAAGCACCCCTCAC**TCCCCCGTTGTCTGGGCAACTGGGGA**AAAGGGGACTCAACC**CCAGCCCACAGGCTGG**TCCCCCCACTGCCCCGCCCTCACCACCATCTCTGTTCACAGG**GGTGGCCCTGCACAGGCCCGATGTCTACTTGCTGCCACC**AGCCCGGGAGCAGCTGAACCTGCGGGAGTCGGCCACCATCACGTGCCTGGTGACGGGCTTCTCTCCCGCGGACGTCTTCGTGCAGTGGATGCAGAGGGGGCAGCCCTTGTCCCCGGAGAAGTATGTGACCAGCGCCCCAATGCCTGAGCCCCAGGCCCCAGGCCGGTACTTCGCCCACAGCATCCTGA**CCGTGTCCGAAGAGGAATGGAACACGG**GGGAGACCTACACCTGCGTGGTGGCCCATGAGGCCCTGCCCAACAGGGTCACCGAGAGGACCGTGGACAAGTCCACCGGTAAACCCACCCTGTACAACGTGTCCCTGGTCATGTCCGACACAGCTGGCACCTGCTACTGACCCTGCTGGCCTGCCCACAGGCTCGGGGCGGCTGGCCGCTCTGTGTGTGC**ATGCAAACTAACCGTGTCAACGGGGTGAGATGTTGCAT**CTTATAAAATTAGAAATAAAAAGATCCATTCAAAAGATACTGGTCCTGAGTGCACGATGCTCTGGCCTACTGGGGCGGCGGCTGTGCTGCACCCACCCTGCGCCTCCCCTGCAGAACACCTTCCTCCACAGCCCCCACCCCTGCCTCACCCACCTGCGTGCCTCAGTGGCTTCTAGAAACCCCTGAATTCCCTGCAGCTGCTCACAGCAGGCTGACCTCAGACTTGCCATTCCTCCTACTGCTTCCAGAAAGAAAGCTGAAAGCAAGGCCACACGTATACAGGCAGCACACAGG**CATGTGTGGATACACATG**GACAGACACGGACACACACAAACACATGGACACACAGAGACGTGCTAACCCATGGGCACA**CACATACACAGACATGGACCCACACACAAACATATGTG**GACACACATGTACAAACATGCACAGGCACACAAAGAGAACACTGACTACAGGCACACACACACACGGGCACACACATGGATATGTGCACACATGGACACATACA**TGTGCAGGACATGCACA**CACACAGACACACTAGCACAGAGGCATACACACACAGACACACACATTCACAAACAC**ACATGTGCATGCAAACACACACACATGT**ACAGACACAAGTACATGGACACATGCACACCCAGAGACACACTGACACAGACACACAGGAGCATGTGATACACTAACACGTGGACACACACGTCTACCCACAGGCACACAACAGATGGACACGCGTACACAGACATGCACACACCCACAGGCACAACACGTGCGCATGCCGGCCGGCCCCCGCCAACATTCTCCCAGGGCCCTGCCGGATACTCTGTCCCTGCAGCAGTTTGCTCCCTGCGCTGTGCTGGCACCGGGGCTTTGGGCCCAGGCTCTGCTTGTCCTTCTGTCTCTGCTTGGAGGTGCTGCCATGGCACCCAGCTTGTGCTCTG**CCTGGGGAGCGGAGGCCCCAGG**GATAGCATGTGACCC**CTGCTGAGGCCAGGCTCCTGATGAAGGCAGCAG**ATAGCCCCCACACCCACCGGTGAGCAGAACCAGAGCCTGTGCCATGTGCTGAGAGCAGGCAGTGACTAAGCATATGGGCCCAG**AGGGCAGAGTGGCTGCCCT**GGGCAGCTGCTCCTCTTAGCAGGAGGCCTCAGGAGATGAGCTAGAGCAAGTCTGCCCCTGCAAATACCA**CCTGCTCCCCAACCCACAGCAGGGAGCAGG**CGAGGTCAGACAGCAGCAGCCCGGGAAGGACCGAGCCCCAGCAGGGAAGGCAGGGCCCGAGTGAGGTCTCCACACCCAACGCACAGTGCTGTCTCTAACTGGGGCCACCTCCGAGTCCCCGCCACACTCTT**GGCCCTTTGGAGTCCTGGGCTCCAGGTGTCTCCCAAGGGCC**CATCTGTGCAG**GGGATGCAACCCCCCGAATGTCCTCATCCC**ACTGTGGAGCTCAGGTCTCTGTCTGCTCCCTGGG**TCCTGGCAGGGTAGGACAAGTCCGCCAGGA**TGTCCCCATGCA**GACTCTGCTCCAAGAGGGAGCTGGAGAGTC**AGGGCCTTGGTGAGGGAGTCAGGATCGGGTTCCCCCCAGCTCAGTCCTCCCACCTGCCAGCCCCCACAGCACAGG**GCAGGGCCACACCCCCTGC**TTCCCCCTCCAGGAGAGTCAGGACATGCTGGCCGCTGCTCCGCTGGGGCCCCGCCCTCCAGCCCCCACCTTGGTCTGTGTGCTGCATCCCCCACGCTCTCTCTGCCACCCCA**GGACTCTGAGGAAAAGACCTCAGAGTCC**CAGCCCTGCCCAGTCTCGGCCTGTGCCCCCGCTGCATCAGGCTTTCAGGG**GCCCAGCCCATGCCCTGGGC**AGTGCCCGAGCCCCCCTGCACTTGCTCTC**CCCACCCCTGGGTGCAGCACAGCCTAGGGGCCAAGGGTGGG**CCTAGAGGATGGGCCCCGGGGGGGCTTT**GCTGGGTGCCACCCCAGC**CTGACCCTAT**TCCCCCGTGCTGTGTCTCCTGCAGAGGGGGA**GGTGAGCGCCGACGAGGAGGGCTTTGAGAACCTGTGGGCCACCGCCTCCACCTTCATCGTCCTCTTCCTCCTGAGCCTCTTCTACAGTACCACCGTCACCTTGTTCAAGGTAGCACGGCTGTGGCACAGGGAGGAGGGTGCAGGGCGAGTGTGGG**GCCCAGGGAGCAGCCTGGGC**TGGACGTCTAGCCCGGAGGCCCCCACACCACCCCACTGGGTCATCTCTGCCCCGGCTCCCTTCCCGACCACGGGGAAAG**CATTTCACACTGTCTCTGTTGCCTGTAGGTGAAATG**Atcccaacagaagaacatcggagaccagagagaggaactcaaaggggcgctgcctccgggtctggggtcctggcctgcgtggcctgttggcacgtgtttctcttccccgcccggcctccagt**tgtgtgctctcacaca**ggcttccttctcgaccggcaggggctggctggcttgcaggccacgaggtgggctctaccccacactgctttgctgtgtatacgcttgttgccctgaaataaatatgcacattttatccatgaaactgctttctggtgagggtttttgtttctttttcaaaactttcctgctacatgggcatctcaagggggaccccagttccaaagggagctgtggaaaaga**gcgctgggtcagccagcgc**agggggtttcgaggaaagccacgtgcccaggaaaggggccgcagaagcaggtgggccagactcagactcggggcatgcccagcctgatggaaggaaggggactgagcaggagagggttccaggcctggtcctccaagcaca**gcctgaattgagagactggggctcaggc**ctcgggggcctctgtgtgtgctccacatgcctacaactgcccgggtcaccttgccacccttcccagcaagcccagacagttcttggccttgccccaaaccttcatgatgtgtggtgcacgccaccgggatccaggaggtgcaggctgagccctcgagagcatgtgggcctcaccgggatccaggag**gtgcaggctgagccctcgagagcatgtgggcctgtctgcac**agtgtgggggccttgcactccacaggagcacaggggtggggtagcagtcgcgcccgtggcaggggagtggaagttggagcaaaagtttcaggtgaacgagtgtcttagtctaattgggcagctatcacaaaacactatcggctgcaaagcttgagcaacagacattt**gcctcccacagcgctggaggc**tgaaagtctaagatcaaggcgccggcaacttcaacgtc

Uppercase: IGHM-3

Lowercase: Flanking sequence[1000bp]

Red & Bold & Underline: Stem-loop [46]

Blue: Heptamer[100]

Green: Nonamer [4]

id-IGLC6[C_gene_segment]

accagggccatgggacag**gggaaggatgctggaaaaagttcagcttccc**agggttctgggtcccaactatggggctgctttaaacaccaagaagggaggcctttgactggggacttggggaaatgaagggggacaaggatggaggaagaatgtcctgtgaggtggcgccagggtgctgggtccctcctcctcccccgggacaggcaggctgccatggacagggtggttctcaggacactcagtccaaggttagagcctccccatcccacccaaaaagagagacccccaaagcagatgctgagggaggcactcctggtgggcgcagg**tgacagggacctgtcaggacagacatttgtcct**aggacagccaga**tctcccaacgacagggaga**ccccatagagcagacacagcgccaggctcagaacagaaaatatacctcacatgcaagccctccatc**tcctggactcccagga**cccggctcccaggactgacatcccct**ccccaccaaggggcctctgtgggaaagtgggg**cagagactgcaatgatggtgctgggggatatgtgaggaagaaattcacttgtcaaaagggaggagaacctaaaaacaggacacagatgccctcg**tgaacagacccagaggagggacctggcaggggatgttca**gacagagacgtcccccaaggaaagacgagggcccagggctgaacc**cagccagcgtcggggagcataggtgaggtgagagctggctg**ataaggagg**ggagaatctggagaggaggctggagcagaagaggcctgattctcctcagtcctcagatgctgaggag**tttcccatcaggaggtaagcatccccgccaccgccaagttgacctcagtacagcaag**ggcccagcctgaggtccctatcctgggcc**ttagtccttcac**ccacctgaaaactgaggccaggggctccccaggtgg**acaccaggact**ctgaccccctgcccctcatccaccccgcagGTCAG**CCCAAGGCTGCCCCATCGGTCACTCTGTTCCCGCCCTCCTCTGAGGAGCTTCAAGCCAACAAGGCCACACTGGTGTGCCTGATCAGTGACTTCTACCCGGGAGCTGTGAAAGTGGCCTGGAAGGCAGATGGCAGCCCCGTCAACACGGGAGTGGAGACCACCACACCCTCCAAACAGAGCAACAACAAGTACGCGGCCAGCAGCTAGCTACCTGAGCCTGACGCCTGAGCAGTGGAAGTCCCACAGAAGCTACAGTTGCCAGGTCACGCATGAAGGGAGCACCGTGGAGAAGACAGTGGCCCCTGCAGAATGCTCTT**AGgcccccgaccctcaccccacccacaggggcct**ggagctgcaggttccca**ggggagggggtctctctcccc**atcccaagtcatccagcccttcttcctacactcaataaaccctcaataaatatcctcagtcaaccagaaaccctgggttttgtgttttgcttctgtttatacattttctaccctaaattggtccaacctgcaagatgaacaaaattcttttagccttgtgggaatgaagtctggattgtttttgaggggtagttccaccatcacctagaaaggcccagagaatattccagaacagagcctt**gggcagcaggccccaggactgccc**ctttcccatacgtgctgtgtttgtgtggacatgtccacctccagcggcctggggcgaccgctccttggttccccatggcccagaagaacgagtaccctccacggatcatcaggcctc**acccctggacagcctccctctctcaaggggt**cc**ctccttctgaggcggaaggag**cctctccagtcca**tgggcccttggtctcccctactccttaggggccca**ccctgttcctgactcccaggaccctgccctctacccccacatctatctctgaagcaggcacagactggaccctgaccctctgg**acccaggtcactcaccccacaggccagctgggt**acccctggtcccacatgtgtcaccagtgtcccagttatactcaagtcccctgggggataagggtcaactctactctctctcacccctaagtaaccagcccaaaactgaccacacctcaagtacctaatgtccagttatccacggatggtcagtggggctggggaggtcaaagtcaatcatgagattccagagtggtgccatagacgtgcctgtaaaccaattggtctttttaagagctgtatgtgggatctaagaaccgggttgatgattgt**cccaggaggtagcacaggatggggggcctggg**ctatgagacaaacacacatacataca

Uppercase: IGLC6

Lowercase: Flanking sequence[1000bp]

Red & Bold & Underline: Stem-loop [21]

Blue: Heptamer[23]

Green: Nonamer [3]

id-IGHA1[C_gene_segment]

gggctgagctgtactaagctggcctgggctgggctgagctgtactgagctggcctgggctgggttgacctgggctgagctggactaagctgggctgacctgggctgggatgggatgggctaggatgacttgggctggactgggcgggactgagctggactggcctgggctgagctgggctgggctggactgagctggactggcctgggctgggctgggcgggatgggctgaggtggctgctaatgtgggaaagaggccgtgggttgagtgtgattccacctgcagagccctgagcccagctgtgttcttaggggttctgagggccacgcagctctgttgcaccatgattctgtcttctctctt**gcccactgcctgaaggaaatttggagtgggc**tgggcccagagctcccctgtatag**caggccctgtcctggagggcctg**gcagggacatggcttag**cctgttggcctctagtcccgagacctcataggccacagg**ggtccactgtggcttgtttgggcctggggtggggctcatggagtggtgggtgttggactgagactctgaccagggacaggggg**atggggtcacagccaagccactccacccctaccccat**gcacacagcactcagagcccagaccctctcctaagagcccccaccaaaatcctctctaggggcaggggatagagcaagacatgtcccccacccagagcaggggctgcggtcagggagctcaggggactcagccactccatggcagagccctgtttaatacaacttgtgtctgggatggcctgaatcagagaccctatctaaggagcatgttcagaaaccatgttgctgggatcagacagcagggtccaactgcaggcctgtggtgcaggagctgtgtgaccatggggctgtcaccag**gcctctctgtgctgggttcctccagtatagaggagaggc**agtatagaggagagggccgcgtcctcacagtgcattctgtgttccagCATCCCCGACCAGCCCCAAGGTCTTCCCGCTGAGCCTCTGCAGCACCCAGCCAGATGGGAACGTGGTCATCGCCTGCCTGGTCCAGGGCTTCTTCCCCCAGGAGCCACTCAGTGTGACCTGGAGCGAAAGCGGACAGGGCGTGACCGCCAGAAACTTCCCACCCAGCCAGGATGCCTCCGGGGACCTGTACACCACGAGCAGCCAGCTGACCC**TGCCGGCCACACAGTGCCTAGCCGGCA**AGTCCGTGACATGCCACGTGAAGCACTACACGAATCCCAGCCAGGATGTGACTGTGC**CCTGCCCAGGTCAGAGGGCAGG**CTGGGGAGTGGGGCGGGGCCACCCCGTCGTGCCCTGACACTGCGCCTGCACCCGTGTTCCCCACAGGGAGCCGCCCCTTCACTCACACCAGAGTGGACCGCGGGCCGAGCCCCAGGAGGTGGTGGTGGACAGGCCAGGAGGGGCGAGGCGGGGGCATGGGGAAGTATGTGCTGACCAGCTCAGGCCATCTCTCCACTCCAGTTCCCTCAACTCCACCTACCCCATCTCCCTCAACTCCACCTACCCCATCTCCCTCATGCTGCCACCCCCGACTGTCACTGCACCGACCGGCCCTCGAGGACCTGCTCTT**AGGTTCAGAAGCGAACCT**CACGTGCACACTGACCGG**CCTGAGAGATGCCTCAGGTGTCACCTTCACCTG**GACGCCCTCAAGTGGGAAGAGCGCTGTTCAAGGACCACCTGAGCGTGACCTCTGTGGCTGCTACAGCGTGTCCAGTGTCCTGCCGGGCTGTGCCGAGCCATGGAACCATGGGAAGACCTTCACTTGCACTGCTGCCTACCCCGAGTCCAAGACCCCGCTAACCGCCACCCTCTCAAAATCCGGTGGGTCCAGACCCTGCTCGGGGCCCTGCTCAGTGCTCTGGTTTGCAAAGCATATTCCTGGCCTGCCTCCTCCCTCCCAATCCTGGGCTCCAGTGCTCATGCCAAGTACAGAGGGAAACTGAGGCAGGCTGAGGGGCCAGGACACAGCCCAGGGTGCCCACCAGAGCAGA**GGGGCTCTCTCATCCCCTGCCCAGCCCC**CTGACCTGGCTCTCTACCCTCCAGGAAACACATTCCGGCCCGAGGTCCACCTGCTGCCGCCGCCGTCGGAGGAGCTGGCCCTGAACGAGCTGGTGACGCTGACGTGCCTGGCACGCGGCTTCAGCCCCAAGGATGTGCTGGTTCGCTGGCTGCAGGGGTCACAGGAGCTGCCCCGCGAGAAGTACCTGACTTGGGCATCCCGGCAGGAGCCCAGCCAGGGCACCACCACCTTCGCTGTGACCAGCATACTGCGCGTGGCAGCCGAGGACTGGA**AGAAGGGGGACACCTTCT**CCTGCATGGTGGGCCACGAGGCCCTGCCGCTGGCCTTCACACAGAAGACCATCGACCGCTTGGCGGGTAAACCCACCCATGTCAATGTGTCTGTTGTCATGGCGGAGGTGGACGGCACCTGCTACTGAgccgcccgcctgtccccacccctgaataaactccatgctcccccaagcagccccacgcttccatccggcgcctgtctgtccatcctcagggtctcagcacttgggaaagggcc**agggcatggacagggaagaataccccctgccct**cagcctc**ggggggcccctggcacccccc**tgagcctttccaccctggtgtgagtgtgagttgtgagtgtgagagtgtgtggtgcaggaggcctcgctggtgtgagatcttaggtctgccaaggcaggcacagcccaggatgggttctgagagatgca**catgccccggacagttctgagtgagcagtggcatg**gccgtttgtccctgagagagccgcctctggctgtagctgggagggaatagggagggtaaaaggagcaggctagccaagaaaggcgcaggtagtggcaggagcggcgagggagtgaggggctggactcca**gggccccactgggaggacaagctccaggagggccc**ca**ccaccctagtgggtgg**gcctcaggacgtcccactgacgcatgcagg**aaggggcacctcccctt**aaccacactgctctgtacggggcacgtgggcacaggtgcacactcacactcacatatacgcctgagccctgcaggagcggaacgttcacagcccagacccagttccagaaaa**gccaggggagtcccctcccaagcccccaagctcagcctgctcccctaggc**ccctctggcttccctgtgtttccactgtgcacagatcaggcaccaactccacagacccctc**ccaggcagcccctgctccctgcctgg**ccaagtctcccatcccttcctaagcccaactaggacccaaagcatagacagggaggggccacgtggggtggcatcagaa**gcaggccagtgagacagggcctgc**ccagggccctctgcatgcctctggcttctg**cctggggctcccagg**agtgtaagaacagtcccacaaccactgtggggacacc

Uppercase: IGHA1

Lowercase: Flanking sequence[1000bp]

Red & Bold & Underline: Stem-loop [22]

Blue: Heptamer[63]

Green: Nonamer [3]

id-TRBC2[C_gene_segment]

aatttacctgtcatccctaagaatctacaaaggagatgctcaggacagaaactgtatcaacacaactagtagcaagaagttac**tctgatgatatcaga**tgtttatttgggaaacttgctagtagagaaagctacatataatatttggatgcaa**agggacacagaaggttgaagagtccct**aattttgaaataagggaagatgactaactgtctgagctgagaaaactcaggggtacctggaggcagaggaatggataagatgacttcatgcaccacaaaaagaaaaaacctca**cattctcatgaacgcactgtaaaaccaaaggatgtcctcatgtgaatg**caaaaaataggccatctgtaaatccaaagaaagcccccagatctaaaatgtctccctcatcccagattccccttcattcctgagcaccttagatttggtataaataacctgcttgggagggggctttttgaattcgtacataatttaaccttcacacag**tttctgcaaagtcagaatggtgattattacctcacatgcagaaa**aaagtgataggaatttctgtcttaaaagtcttgttggtggacaaaggaagttctaggatttggatctt**gtttttttgggttccaatcccttgctccagttaaaaaac**taccacataaaatggtgagaagtaggtaggcaagtttttattgatagagaggaaatcaaataatggcaatgaggagacatcacctggaat**gttaggcagtgcctaac**tgggggatggacagacaatgggcagtgccaacccatagggtggatacaaaagacaggcaaggaaggggtagaaccatcaaagaggaataggctggtgaccccaaagcaagg**aggacctagtaacataattgtgcttcattatggtcct**ttcccggccttctctctcacacatacacagagcccctaccaggaccagacagctcttagagcaaccctagccccattacctcttccctttccagAGGACCTGAAAAACGTGTTCCCACCCAAGGTCGCTGTGTTTGAGCCATCAGAAGCAGAGATCTCC**CACACCCAAAAGGCCACACTGGTGTG**CCTGGCCACAGGCTTCTACCCCGACCACGTGGAGCTGAGCTGGTGGGTGAATGGGAAGGAGGTGCACAGTGGGGTCAGCACAGACCCGCAGCCCCTCAAGGAGCAGCCCGCCCTCAATGACTCCAGATACTGCCTGAGCAGCCGCCTGAGGGTCTCGGCCACCTTCTGGCAGAACCCCCGCAACCACTTCCGCTGTCAAGTCCAGTTCTACGGGCTCTCGGAGAATGACGAGTGGACCCAGGATAGGGCCAAACCTGTCACCCAGATCGTCAGCGCCGAGGCCTGGGGTAGAGCAGGTGAGTGGGGCCTGGGGAGATGCCTGGAGGAGATTAGGTGAGACCA**GCTACCAGGGAAAATGGAAAGATCCAGGTAGC**GGACAAGACTAGATCCAGAAGAAAGCCAGAGTGGACAAGGTGGGATGATCAAGGTTCACAGGGTCAGCAAAGCACGGTGTGCACTTCCCCCACCAAGAAGCATAGAGGCTGAATGGAGCACCTCAAGCTCATTCTTCCTTCAGATCCTGACACCTTAGAGCTAAGCTTTCAAGTCTCCCTGAGGACCAGCCATACAGCTCAGCATCTGAGTGGTGTGCATCCCATTCTCTTCTGGGGTCCTGGTTTCCTAAGATCATAGTGACCACTTCGCTGGCACTGGAGCAGCATGAGGGAGACAGAACCAGGGCTATCAAAGGAGGCTGACTTTGTACTATCTGATATGCATGTGTTTGTGGCCTGTGAGTCTGTGA**TGTAAGGCTCAATGTCCTTACA**AAGCAGCATTCTCTCATCCATTTTTCTTCCCCTGTTTTCTTTCAGACTGTGGCTTCACCTCCGGTAAGTGAGTCTCTCCTTTTTCTCTCTATCTTTCGCCGTCTCTGCTCTCGAACCAGGGCATGGAGAATCCACGGA**CACAGGGGTGTGAGGGAGGCCAGAGCCACCTGTG**CACAGGTACCTACATG**CTCTGTTCTTGTCAACAGAG**TCTTACCAGCAAGGGGTCCTGTCTGCCACCATCCTCTATGAGATCTTGCTAGGGAAGGCCACCTTGTATGCCGTGCTGGTCAGTGCCCTCGTGCTGATGGCCATGGTAAGGAGGAGGGTGGGATAGGGCAGATGATGGGGGCAGGGGATGGAACATCACACATGGGCATAAAGGAATCTCAGAGCCAGAGCACAGCCTAATATATCCTATCACCTCAATGAAACCATAATGAAGCCAGACTGGGGAGAAAATGCAGGGAATATCACAGAATGCATCATGGGAGGATGGAGACAACCAGCGAGCCCTACTCAAATTAGGCCTCAGAGCCCGCCTCCCCTGCCCTACTCCTGCTGTGCCATAGCCCCTGAAACCCTGAAAATGTTC**TCTCTTCCACAGGTCAAGAGA**AAGGATTCCAGAGGCTAGctccaaaaccatcccaggtcattcttcatcctcacccaggattctcctgtacctgctcccaatctgtgttcctaaaagtgattctcactctgcttctcatctcctacttacatgaatacttctctcttttttctgtttccctgaagattgagctcccaacccccaagtacgaaataggctaaaccaataaaaaattgtgtgttgggcctggttgcatttcaggagtgtctgtggagttctgctcatcactgacctatcttctgattt**agggaaagcagcattccct**tggacatctgaagtgacagccctctttctctccacccaatgctgctttctcctgttcatcctg**atggaagtcctcaaacaccatttccatacccaggcattctgggt**ccccactggagggttagtctgaagggcaatggctgggctttggaaaaccagcaagatgaggacagagaggaaggcacacagcaaaccataagcccttacccagtgcaggacagaggatgcgggcagacctatgggttacaatgtctggtcatttcccaattccagattaaactgtcacctgttttacctttagttttattagtttgtagtcttaacacctccagcttctcttgtttcaggatttgggcttaaaattgagtgct**actctgcatgtctagtttgaaatactagagaaggcagagt**tgagacaactgatatgtaaagcctggggaagagtgatttctcaggagcgagacacactaagtcaggagcaatgggatatagggcccagtgggggctgaagtgctatgttcagagtagcccttccaatgggcttcttcgtttgatggatggaaaccaaaccactccaaacacaaggtgt**taactgctcctacttgggcaaagacagtta**tcctgtcaaggtaaattctgcatacaggctgaatgcattgtggtaaaacactacatggaggaagaggaggaatgggattaaagaaaaggaggccta

Uppercase: TRBC2

Lowercase: Flanking sequence[1000bp]

Red & Bold & Underline: Stem-loop [18]

Blue: Heptamer[36]

Green: Nonamer [5]

id-TRGC1[C_gene_segment]

tagacagtatgtacatgcggaagtagattctctttaaaacaagtgactgtgtatgttaaaaataaaagtgaacaaaatggctaggcatggtggctcacacctataatcccaggatttgggaggctgaggcaggcagatcacttgagcgcaggagttttaaaccagcctggtcaatatggtgaaacatgttcctagaaaaaaaatattagccaggcatggtgatgcatacctgtagtcccagctacttgggaggctgaggtgggaggatcacttgagcctgggaagtcgagactgcagtgagctatgatcttgccgctgcattccaacctggacgacagagcaagccccagtctcaacaacaacaaaaattttgacattgattcatatgggaaataagataaataataagataaatatggtgcatcgcagaatcagttaaatgaagagtgggagtacacgaaattgcagaacatgagaatgtgtcatatttggccaaagaacataagttacaaaggatggaaaagggaagtgggaaaaggaccaaagagtctcatctgtagagagatcattaaggttttctgaccttccactttcccctgccatccaccttgaaaacctgcttcactatgatgaaacaaaagagattaaaaaataaaataaatatgctcatgaactttggaagccctggtaggaggcagttaaaaatcacactcatcacagcatgtgcagaataaacaaaggccaggttttcgtccagcatctgacatttgagagctgtgacttttggcaagttatttaatgtcttgattctcttttcaacatctgtaaaataagcacaataataagtactgtgcagcctatcctggatgaaaggccgcagtggacaccagctcaatggcatcttctctttttatggttatgtactaggccactccaaactgtgcaatgtgtgtgtttctctaatgattcttttaaactcatatttcatttctccccatagATAAACAACTTGATGCAGATGTTTCCCCCAAGCCCACTATTTTTCTTCCTTCAATTGCTGAA**ACAAAGCTCCAGAAGGCTGGAACATACCTTTGT**CTTCTTGAGAAATTTTTCCCTGATGTTATTAAGATACATTGGCAAGAAAAGAAGAGCAACACGATTCTGGGATCCCAGGAGGGGAACACCATGAAGACTAACGACACATACATGAAATTTAGCTGGTTAACGGTGCCAGAAAAGTCACTGGACAAAGAACACAGATGTATCGTCAGACATGAGAATAATAAAAACGGAGTTGATCAAGAAATTATCTTTCCTCCAATAAAGACAGGTATGTGTTTACGCATATCATCTGTCAGAACACTTCTT**TGAAAGTGAATGCTGCATTTTTTCCTTTCA**GTATTAATGAAAAACAAACATAAATCTTTCTTAAATATTGTTACATTTAATG**GTAGCATAAATGCCCTGCTAC**TTTTCTATAGAATTAAAATGGTATAGGTTTTGGAGAAAACAAAATTGAAAAAGTTACTGAAGGTTTGTCAGCCTCAGCTCCATTATCCAAAATAAGAAAGTCACGTGCTGGTTTTTAGG**GTTGTTAGATGGATTAAAGAAACAACATACACAGAAGCATCTAGCAAC**GTGACACGTGGTAAACGC**TCAAAAAGTGTTCTCCCTTCTTTTGA**TGACTTTACTTGATCAGGAAATAACATATATATGTCTTTCAGGAATGTTCTGCCCAAGCAGGAGAGTCACTCACCTCAATCTTGCTACCCACAAAGTTTAACCTAAAAACAACGGGTTCATTGTTGACAAAATGATGTTTATCTGTTGTTGACAGAATGATGTTTATCTAAAAACAGTTCCAATTTTCTATTT**CCTTTGCTGAGACACAAAGG**GGAGGCAAATGTGCAAAGCTTGAGGGTAGTCTTACCACTGTGCTTAAGTGTTCTGATTTTTCTAGTGATCAGGGCAAAATAAAAAGTATAGTAAGTTCCAAGGCAGTGAATATTATACAGGAGAGAAGTTACAGTTTTATAATGTGTTTTCCTTTACACTAAATTCTAAAAGTAAAAAGTCTTTTTTTTTTTTTGACAGAGTTTCACTCTTGTTGCCCAAGCAGGTGTGCTATGGTATGATCTCAGCTCACTGCAACCTCCACCTCCCGGGTTCAAGTGATTCTCTTACTT**CAGCCTCCCGACAGGCTG**GGATTGCAGGCGCCTGCCACCACACCTGGCTAATTTTTGTGTTTTTAGTAGAGATGGGGTTTCACCATGTTGGCCAGGCTGGTCTCAAATTCCTGACCTCAAGTGATCCATCCACCTCGGCCTCCAAGT**GCTGGGATTATGGGCGTCAGCCACTGTGCCCAGC**CTAAAAGTAAAATGTCTTTCATGAGCTTCCCAAGGCAGCTACGTTAAGGAGGACACTTCTCTTAATGTCATTCTACAGTAGATTTCTAATGCTCTTTCTTGGAAGTTTGTTTTTCTGAGAAAAGCTAAAA**ATATAACATGGAAGTGATCATATTATAT**AATCAATGAAGTGCTTTTCAAGGAGATAAAACTAATCTGGTCCACACTTGCAACCAACCTTGATTGAGAGAGAGAGAGAACTCAGGATACACTTGAAGATTTTATTATGGGGAACAGTTACTTTATTCTTTTTACCTCAATCAATGCATGGAAATAAGTGATAGTCATT**TTCATTTATCTTTTAATAAATGAA**GTCACCATGAGGAAAATAAAAAGACATTGAAAACCCATTAAAGTCAGCCCTTAAAGATATTTGGACATGCAGACTTGATAACTAACGTTTGCATT**CTTGAGACTTACCCAAAACCCATACCTCAAG**TCCAAGTTTTTAGAATTCATGAAATAAAGATCTCAGTGAGTGCATAAAATTGCGCACCAGAATCATATCCGTATAGACAAGAACACATCTACTAGAAAAATAATAAACC**AACACACCAATGCAACTGTGTT**TTCTTCTGTTTTAAAGTATGTTGTCTTTGTATGCATGTTTGCTTCTTCCTTTTTTTTTTTAACATCACAGATAAATTCAA**CTCTCACCTCAGGTTTTATTGAGAG**AACTGTCAATGTGACTTGGCCTCTG**TCTTTCTAGTCCCAGAAAGA**ATTGCACTGAAATCTGAGCTCCTGTAATAAAAACAACCATTTGCTGAGAGTAATTAACATACTGAAAGAGATTTTCTTAGAGTACACAATGGTGACATTATATTGCCTCTTTATAAATAACTTTCTATCTATTTCTGTGGATTATTCCTACAAAGTACTTTTCATATGTCCAATTTCTTTTCTTCCCCTACAACTACTGTCTGAATACTGGCTCTGCTATTTGCTGATATGATTCTCGGCAAGTTGCCTGCACTTTTTAAACTTTATTTCCTCATTCAGAACATGGGGCCATACATAATACAACTCACTTCAGTGTTATTGGGGAATTAAACAAA**AAATGCATGGGAAGCATTT**AACATAGTGCCTGACACAATAATGAGTACTCAGTAGATGTTAGCTTTTATTAATATTGTTGTTGTTATGTCCAGAAACACTATACCTCCAGA**AAATCATGGGTACTTGCTGGGGACATTGGGGATATGCATGATTT**GGAAAAGAATGACTGCTTTTTTT**GCTTAGATGAGAAATTTTTCTAAGC**CAGACTCCTTCAAATATGTAAGATTCTGTTGTGGATTCAAGGACTGAAAGAATTCTTGGCCGAGTGTGGTGGCTTATCCCTGTAATCCCAGCATTTTGTGAGGACAAGGCAGGAAGATTGCTTGAGTCCAGGAGTTTGAAACCAGCCTGCGCAACATGGCGAAACCCTGTCTCTACAAAAAATACAAACAT**TAGCTCGGAGTGAGTGCTGACATGTGCCTGTACTCCCAGCTA**CTCAGAAGGCTGAGATGGGAGGATCTCATGAGCCTGGGGAGTTTGAGGCTTCAGTGAGCCGTGATGACACCGTACTATACTCCACTCCAGCCTGGGTGACAGTGAGACCCTGCCTCAAAAAACAAACAAACAAACAAACAAAACAAAATTAATCTTTT**TGCTGATGTCATGTCAGCA**GTGTGTGTTGAAGGCTGTAAAGCAGCCAT**TTGTTCAGTTTATTTTTCCATTGAACAAGTATTTATCAAAAACATACTT**TGTGGCAGTCACTATGCTAGGAGCTATGAATACAGAAGGAAAAGTAAATGCTCTTGGATACTACACTCCAGTTGTGATAAAAAAGAAAAAATGTATTCTTCACCAACTTCAACATCTTGATGTGCAAAAACATAATACATG**AATTAGATCTACCTAATT**ACACAGAATTAGACCAATTGTTTCTGGAATTGTGGGCTCATATTTTTAATAACTGTCCTCCTGCCTCTCTGTCGACAGGTTTTATAAATATTCATTTAATTACACACACACACACGAACAATTGACTAGTACTTGCTCTCATTCTTCTAGATGTCATCACAATGGATCCCAAAGACAATTGTT**CAAAAGATGCAAATGGTAAGCTTTTG**TGTTTTTCCCTTCCTCCTGATCATTTTGTTTTGAACTTCTCTGGCTTGA**AAAATCAGGGAATGGATTTT**G**CTAGGTTGGATGCTGCAGAATGGACCTAG**TGATATTTTAAATTAGTCCCTCATTTTCTAGGAGTTGTATTAACAAACCTAACTACTGCTTTGGGGTATGAGATG**ACTGTAAATTAGAGAGGGTACAGT**GGTATAGTGATATGCTTTTAATTATTTCAA**AAAAAAGATTTTATTCATTCATGTGTCTTTTTT**CTTTTTCTTTTCTTTTTTTTTTTTTTTTG**GACAGAGTCTTGCTCTGTC**ACCCAGGCTGGAGTGCGGTGGCAGTATCTCAGCTCACCACAACCTCCGCCTCCCGGCTTCAAGTGATTCTCCTGCCTCAGCTTCTCGAGTAGCTGGGACTACAGGCGCGTGCCACCATGCCCGGCTAATTTTTGTATTTTTAGTAGAGTTGGGGTTTCACCATGTTGGCCAGGATGGCCTCGAATTTGTGACCTCGTGATCTGCCCCCTCGCCCTCCCGAACTGTTGGGATTACAGGCGTGAGTCACTGTGCCCGGCCTCCTGTCCTGTCTTTTGTTTAATGACTGGGAAA**AACATGATACCATGTT**GCTTCTCGAGTTGTTTTGTTTTAGTCTTTGGTCTTTGCTAGTAGCTAATAACACGAACTAGTGTTTATCAAGTGCTTTTT**ACACAGAAGGGCTTGGGCTGTGT**TCTGCATTTTCTTGTTTAACCCTCTTAAAACTCCTATAAAATGGTACATATTTTTCTCCCAATTTACAGTCCCTTTAAAGCAAATAATTATAA**AAATCCCTATACATGTCACACAGCTAGATCTGGGATTT**CAAATCAGGCCATCAAACAAAGAGTTTATGTACTTAGTAAGTTTTCTGTTCTTTTTCTACAATAGAGTCAGATAGCAAGAAATTACCAAGCCAGGAACCTGAAACAAAACGGACATCATGTGGGGCTGGGTGGGTGCATGGGCTTTGCAGACTGGACTTTCACTCCAGCTCTTTTAATGATTAGGTGTAAGTGACCTACATTTTGTGAGCAACAGTTTTCTCATCAGCCAACAAAGAATAATTACACCAGATTCACAGTTATTGAAGAGATAAAGGCATGAATG**TGAGATGTCTGGCATAGGGCATCTCATTTAGCAGACACAGAATGAG**TACTTGTTTCTGGCTTTTTCTCTCTACATATGCACAAAGAATGCGACTAGAAGCATGGGCTCTAGCCCTGCTCAACTTTCCTCTATTTCCAATACCAAGGGGCTCTGACTTAGGCTGCCACACCAGGCAAGGAGGGCAGTACCACCTCACTTGACCAAGGGCAGGGAGTCACGGACACATCACTTCTTGAGATCCTTTTCCACACCAAGGACTG**ATGTTTCTGGAATTCTCACTTTATGAAGACAAAACAT**ATAAATGGAAATTTTCTCAGGTAGAGACTCACTCTTGTAGCTCATTGAGTAGGCACTAGTGGTCCACCCCCACTGTCTTTACTTATTCCTTGACATCACATATCTCTTGCAAAACCTCAAATAATATTAAATGCAATCACCCAATAATAGCATAGCCATAATTAGAGGCATTTAGGAAAGACAGGTGAGTGTGCCACAACTACCTAACACATCAGCAAATCTGGATTAACCACTTTCTTTGATTTTCCACAATGCAACCTTACTTTTTAATAGTTGGGAATGTTCTAAGTGAATTTAGCAGAGGTTGTTAATCAACTTGAAAGCTGAATTCTGACTTGTCTGACTCTTGGTGGTGCTGGTAGCAGTAGATGTTTACTTTTAGGTTTTGGTGGTGGTGGAATATCACTTCAACGTAAATCATCAGAAATAAGTATTTGTGAACCCCTCTCGCATTAATGTATCTTATTCTGTAAAAAGAACATGTGCAATTTCTCTTAGATACACTACTGCTGCAGCTCACAAACACCTCTGCATATTACATGTACCTCCTCCTGCTCCTCAAGAGTGTGGTCTATTTTGCCATCATCACCTGCTGTCTGCTTAGAAGAACGGCTTTCTGCTGCAATGGAGAGAAATCATAAcagacggtggcacaaggaggccatcttt**tcctcatcggttattgtccctagaagcgtcttctgagga**tctagttgggctttctttctgggtttgggccatttcagttctcatgtgtgtactattctatcattattgtataacggttttcaaaccagtgggcac**acagagaacctcactctgt**aataacaatgaggaatagccacggcgatctccagcaccaatctctccatgttttccacagctcctccagccaacccaaatagcgcctgctatagtgtagacatcctgcggcttctagccttgtccctctcttagtgttctttaatcagataactgcctggaagcctttcattttacacgccctgaagcagtcttctttgctagttgaattatgtggtgtgtttttccgtaataagcaaaataaatttaaaaaaatgaaaagttgacttttgtccatggtattttaa**ttggatgacatcaaattgaacatccaa**ggtaagaaacagcatggcaattgggctgtggaattctgtattggttgt**aagaatggtccaacaccccatttctaattctt**tccctgagatcgtggttatcacaccttctaagaagaactacaaccaaatgaaggagctcatgtgacttctgtttgaaaggtcaccagagtcagattcattcggtttaggacattccagtggctataggacactatctactgtgacgcgtaccgtgtgagctcagctctagagtgtttcacagacactgtgtttcctgatcctcacgatacccccatgaga**gctgccctaaaagcagagaggcagc**gtgatggagaggttcagcacatgctctctgat**cccaggaatcctggg**tatggtgtttcgtatctgtgtgacctcaggt**gagttccaggaactc**tatgtgccataatctcctcatgtaaaatgaagttataatgccccgtttcctggagttatgtggattagatgagttaatgacacctgg

Uppercase: TRGC1

Lowercase: Flanking sequence[1000bp]

Red & Bold & Underline: Stem-loop [41]

Blue: Heptamer[99]

Green: Nonamer [9]

id-PCDHACT-2[C_gene_segment]

gggccacctcaatctccgcccatgaaaacgcatctagaggag**tgtcacaagtttttcacagtgaca**tttttgcttactgatacaagacagtgatggtgactgatgatgtcccagtgatttctgagtagcttctaaccagcacataactcccccaacagtctttaagtctttaggtgcccatattttcctctttgttctcccccttcagactgagagtttgtagagagagggcaacagatcttttcaatacacaactaatgcaaaatgtatcaggtttttcttggatttcagctactccctgttaaacaatcagagcttagtgaacagtgattcagtgaggagggaaagcactcaggaaggagcaggaacaagtacaagtttatgaaaaggattagctagcaaaacaaggt**caaactctgcaatagtttg**ttttcctctccctagtatccctcttcaatcagaaaagagactgttatcagttgctggtgttatgactgggcacatccgccctgggtcaaatatg**ctgcagtctgcaaagccagcagcagattgcag**tcctctgcagtccagccaggccagcagaacttgtgtagccatgtgccctgttatagctgtaatactgaattgggaatgttcccttaatggggcacttgagggc**aaaagcagtgaaagctttt**cctttctcaaagcagac**tgttcttcccgtagtgttttaagaaca**cagacatgtattgggcaaggcaaagccaaaggtggcctttacaagattattaaatctggtcttc**cagggtatctaatctgtgtgaggaccctg**atgaactaattttcttctaaagtgctatatatgtagatatcatcatagagttacacatgaaatggctcattcaatacttttttagatgcctggaaatatttaagggagtaactaatcaattagcagcattcctgggagaacattgtcttgtcattttaacagaaaactctctttgtgattttgcagCCACGACAGCCCAACCCTGACTGGCGTTACTCTGCCTCCCTGAGAGCAGGCATGCACAGGTATGTATTTCCCTCCTCATTCACTCAGAAGTAACCTTAACTTGGTATGGCTCAGATAAACTGCATCTCCATAGGCCAGAAGCAGCTGTCAAAACTAAAAAGCTTTAGGTACTTTGCCAGGAAAATGCAATTATTTTGTCCCCATGTTTATTCCTTGAAAGATCGCAAATGGTCAG**TGCCAGATGCTTATCAAGTGCTGGCA**TATAAGAGTCCTCTGTAAAATCACAGAAACAGGCTGCTATGTATTTTCTCCCATCAAAATTTCTACAGGGAAGTAATTTCAACCTCCTTCATCCCTCTCTACCTATGCTTTCTTTTCCTCCTTTAAAAACTGTAATTAATACTCATGCTTTGAGACTTGGG**TACATTGTGCAATGTA**TACATACATGTTGTCTACCTTGTTTTTTTTTTAATCTCACATTGGCTATTACATCCTATTACCCTCAATAATTGATTGCTATTGTTGTTTGTGTTCACACCTATTAGAGCCTCCTCATCTTTCCCATCTGTTGCTATCTTATTGTCATCAATG**ACATGGTTCTTCAGAAGATGAGCCATGT**AAAGGGCTCCAAATCTAGCTTACTTTAAATTAACCTAGAGTAACGGTATTAGTCTAAGACTCAGATTAAATATAATTTTGCTTTCTCTACTTTGTCTCTCTGACAGTAATTGATTAACTACCATTATTTCTGGAGGTGATCCAGTATCCATGCCATGGGGC**CAAATAAAAGATTCATTATTTG**CGAATGTCTTTGGAAACCAAATGGGAAGGACCAAGAAACAAATGATCACAACTATCAAAAGGATTTAATTTTAAAGAAGAAATAATCTTCAAACTTAAGCCCCTCAAAATATCTGGGCAACTATCAAGTGAATATTCACCAAACTTAGATCAGTTCGTAAAGAGAAAGCCTACAAAGTATGTGTAGAG**TTAATGTGAAATTAGTTTTAGCCCATTAA**AATGCATTAGATTGAAATAAATTAACATACTCTCAAGCATTACAAATGAG**CAGAAATCATTACATTGGGTGCTATTTCTGATTCAGAAGCAATCAG**TGAAGGGCTGAAGATTAGTAGTTGGCTTGGTAAGATGTCACATTGGAACCTGGGTCATAATTTTAGGCCAGAAACATTCATGCATATACCAGAATATTAGGTATCAGAAGAAATTCTTTATATTTATTAGAG**ACCAACTTGTGCTTTTGCCTGCATCTGAGCTGTTGGT**GGAGACATGCAATGGGTAAAAGCATGGTTTACAGTAC**CAACTCTTGAAAAGTACCAAAGCTATGAGTTG**TGCCTTAAAAACTACATTTGAAATAAAACATTAAAACATAACTTCCTGGACTGGGCGCGGTGGCTCACACCTGTAATCCCAGCACTTTGGGAGGCCGAGGTGGGCAGATCATGAGGTCAAGAGATCGAGACCATCCTGGACAACACGGTGAAACCCTGTCTCTACTAAAAATACAAAAAT**TAGCTGGGCGTGATGGCATGTGCCTGTAGTTCCAGCTA**CTAGGGAGGCTGAGGCAGGAGAATCGCTTGAACCCGGGAGGCGGAAGT**TGCAGTGAGCCAAGATCGAGCCACTGCA**CTCCAGCCTGGCGACAGTGCGAGACTCTGTCTCAGAAGAATAAATAAATGAATAAAATAACATAACTTCCTTATCCCA**TTTTCAAATTGAAAA**AAAAAAGCCAAATGTGCTCCTATTCGGGTTTCAATTAAGATATTATGAGATTTGAGTAGGGTAAGAAATAAAATAAAAATTGAA**ATTAAAATGCCATTTCTTTTTTGCATTGTAAT**ACATTGAACATATTAAATGAGTTGTGAACCTAAATAATACTAATCTTTTTCGTATGTGTGCTTGGGTGTTCTCGGTCTTTCCAGTCTTGGACATCATGTAACTATTCTTTAAAAAATTCTGCTTTGAGCTGAGCTGGCTCCAGGATAGTTACACCTTCATGAATCTGACTGAGCCCACACAATTTGCTAGTAGGATCCAGGAACACTTGAAGGCTGTTAATATTTGGGGAAAAAAAACAGATAATTCTAGAGTGTAGACAAGGGGAAGAATAGTAAAAGGTCAGAGTTTAATGAGTGAATTTCTACTGGATATGTTGTTTGAAGTCAAAGAGTGAGA**AAACATTGAACTTATATGTTGCCTTCCCTCTAATAGTTCAAGTTT**GCCTGCTCTGTTGCCTCATATAACCCCTTTAGTCAGT**AGTCTAAATTTTATTTTAGAAATTTAACT**TTCAAGATAAGCAAATGTCTAGTTTAAAAGGGTCCTCTAGTCTAGGTGTAGTGGCTCATGCCTGTAATCCCAGCACTTTGGGAGGGTGAGGCAGGTGGATCACTTGAGGTCAGGAGTTCAAGACCAGCCTGGTCAATATGGTGAAAACCTGTCTCTACTAAAAATACAAAAATGAGC**CAGGCATGGTGGCGGGTGCCTG**TAGTCCCAGCTACTTGGGAGGCTGAGGCAGGAGAATTGCTTGAACCTGGGAGGCAGAGGTTGCAGTGAGCTGAGATCGTGCCACTGTACTCCAGCCTGAGTGACAGAGTGAGGCTTTGTCTCAAAAATAAATAAAATAAAA**CAAAATGTTCCTCTAATTTTG**ATGAGGGTTTTCTTGGACATTTTCTCTTAGGATCCCACTTATTTCTTCCTTCCTTTCTTCCTTCCTCCCTTCCATCATTCATTCATTCATTCATTCATTCATCCAACAAATATTTGAGAGATTAATATGAGTT**AGTATTAGACATACATAAATGAATACT**GCACCATGTTCTCTTTTCCCTTGAACAGTTTATGTTCTATCTCTGCTTGCCTCTAAAGGTCTCCCAGTTTGTATCTCACTCCCAGCAATGTTTTATGCTGAATTAATCTCTTCTGAGCGGGGATCTGTGAGTGGTGGTGATCAAGTTTCTCTAGTCTCAGGAAATATAGGGTGGGTCATCTATGCATAAAAGATATAGAAAGAGTAAAATAGAAAATAAGGTTAAGAATTAATTAGATAGCCA**AATTGGAACAATACTCCAATT**ATC**AGAAAATATTTTAGTGTGTTTTCT**TCTTTAGAGTAGAGAACCTAGGAACAAGAGAACCTGCAAGAGA**GGCTTGGAACTTTTGAGAACAAGCC**CTCCTCATCTGCTTCAGTATCGAGATGTTAAAATGGCTTAGTCCTCTGATGGGCTTCCTGTTAGATTTAGTGAGCGCCACATGGCGTTAATAAAAAACAGAATTGCCATAAAGATAGAACATGTGTGTTCCTGGAATAGTATAGCAGGCAATAAGTAAGTCAGCAATGCTTCTGCAGTTTATGCAGGGTGA**CTGCTCAGCAGTAATTGCTTCAGTTCAAGCATGAGCAG**AATGTGTTAGCTGCAGCCCTGGCTTCATAGTTGTAAGCAATTTCTGAGGGTGGAAGAAGAGATGGGAAAGAATTTATGATCTAACCGTTATCTGGGTCTGTGTGTTTATTCAGCTCTGTGCACCTAGAGGAGGCTGGCATTCTACGGGCTGGTCCAGGAGGGCCTGATCAGCAGTGGCCAACAGTATCCAGTGCAACACCAGGTAAAGAGCTGGGGTCTCTCCATTCTTTCTTGGTTTCTGGAAAGTGATCAGATGACCTACTTTTGTAAGATCAGGAATGTTGATGGCTCTTTTTCTTTTATATTTTTGTTATTCCCTTTTTTCCATACATACATGATTTCCTTACATATATGATTATTTTGATTTTATACCTAATGCTCTTCAGGAGTTGAAAAAGGATAACAAGGAAAGTGTGTGTGCACGCATGTGTGCATGTGTGTGTGTGTGTATGAAGTTTTTGGGGTTTGTTTGTTTGTTTAAATCAGGTACCTTTCAAATGCTTAGGTCATCCAAGCCATGCAGAGAAGATCTGGTGGCCTTATGCACAGAGATGACACTGTTAACAAAGATCTTTTGGTTGAAGACTATGGAAACCCACCCAAAGTAGTAAGGAAAGAAAGAGAAAAAGAAGGAAAGAAAGAAAAAGGAAGGAAGGAAGGAAGATGGTTTCTCATGGAAGTGGAAAATTATCGGAACCAAGGCATTGTTTTGAGTTCAAGTC**AAAGTCAATCTGCTTCTCTCTGCACATCAACAACATTCTGCAGACTGACTTT**TAGTGCC**TTGGCATGCATGTTGCCAA**ACATGACCGCCTCACAATTTCTTAGTTTAGAGGGATAATAGGGACTA**TTTCCTAATCCAAACTTTCAGGAAA**GAGAACCTGCTAAGTTGTGTAAAAAACCTAATGGCTGG**GTGAGTATAGGAAAATTGCTTAAACTCAC**ATTTGCTTGGTATCCTTATTCTTGCCCTATCACTAAAGCAGGGTCATGTAATAGAAATATGGCTTTGGAGGCCCATCGCTGTGGCAGTTTTCAGAAAGGGAAGATTAAGTGTTGGTAGAGACCACAAATTGTGTCTACTCTAATCCTCTATTAATAGAACATCATGATGATAATAGTAGTTACTAATTATTAAGCCATAATATGCCTAGACACTGTGCCAAGTACATTGTATGTGTGGTCTCATTTATTCCTTAAGTCAAGCTTGCAAGGGATTTATTACATCTATTCTACATATGAGAAACTGGAGAGGCAGAGAGATTAAGAAATGTACCCAAGTTCACGTAGCTTGTAAGTGGCAGAGGGTAGGATTCAAACCCAGATGTGTTTAGTTTCAAATCCAAGTATATCTAGCACTTATATTCATAACATGGCTGGCTTGCAATAATCCATTCAAATTCAAATACATATCTACATACATAACAGATGACAGAATGTGTGTGTAAAAGGTTTTTTCCCAAA**AATAATCAGATGCCTTTCAAATTATT**AAGTAACATGCAGCTAAGGGCCCATTTTTACTGGCAACTTTAAGGGCATTCGTTGATTCTAATCAGCCAGGATTTGCTATTTATGGATGTTGCACAA**TTCAACTAAAAGTCACATTTGTCCAAAAAATATACGAGTTGAA**GCAATTCATTAGAGAGCTAATATTGCCAGATTGCAAGGGGAAAAACATAAAATAGTTCATTGACAAATCTGTACCCTCAGTGCCAACGATGGAGTGAAGAAATGATGGAGGAGGAAGTGGTTTTAGACTGCCAAGTGTTGCAGGATGTGGAGGCATC**TGGGAAGGTGAGAACTTCCCA**AAGAAGCCACG**TGAAATCATGACTTTCTACCTTGCCTTTATTTCA**GAGTTTTTCTTGCTCTTATGGAGGCATTGTAGGTCGACCTGGTAAGCCACAAACTAACTTTGAATACATTCTCCCTCCCATTGGTGATGCTGGTTGGTGTG**TATTCCTAGGCAAATGTGGAATA**GGAACC**ATGTATTGATATTTCATACAT**CTGGCCAAGTCCCTCTTTCAGATTCAAAAAATGTTGAGAAC**CTATCTTTTTTACAGAGATAG**AGAAGGGGATCTCCCTTGTTCCCTTTCTTACTGTCCCAGCCCCTCTTGTATAACCCATTTTATCCAGAACTGTGCCTGGCTGCTGATGCATGAGT**CACAGTCTTCATGGACTGTG**CTGGATAGAGCTTACATCTTCCAACTACTCCATGGCAACCTAATCATACTTTTCAATACATACCTCTGCATCAGTGGTGTAAAGTTAAAGGGATTCTCTGCCTTCTCCCTGTCCTTCTGGTACTTTTAGGTTTTTAGGACTCAATATATGTTCTGCACTGCTTGGAGGGAATATGGCATAAAGATTAAGATTATGATTTAGAGTCAGATTTGAGTTGAATTCTAATCCCAAGCTTACTTGCTGGGTGAGCATAGACAAACTGCCTGAATTCATATTTTCTTAATTACCCTTTCTGTAAATTGGGTGTAGTAATAATAATAACACCTATTTTATTGAGTTACCATGAGAACTAAAGGAGAAAAAAAGAACTGAGCATAGTGCTTGACATATAGTTAATAAATGTCTAATCTTTTTTTTTTGA**GACAGAGTCTCGCTCTGTC**CCCCAGGCTGGAGTCCAGTGGCACGATATCGGCTCACAGCAACCTCTGCCTCCTGGGTTCAAGTGATTCTCCTGC**CTCAGCCTCCTGAGTAGCTGAG**ACTACAGGCGTGTGCCACCAAGCCTGGCTAATTTTTTGTGTTTTTAGTAAAGACGGGGTTTCACCGTGTTAGCCAGGATTGTCTCAATCTCCTGACCTCGTAATCCGCCTGCCTCGGTC**TCCCAAAGTGTTGGGA**TTA**CAGGCGTGAGCCACCGCGCCTG**GCCTAATCTTCTTACTCTTTTTTCTTTCTGGACTACTTTTCTGCAATCTATGATATAGTGTTGGCTGATAGCCTGGTGGCCAGAATTCAGTAGTTCTCATTTGCAGGCCCAG**ATATAGACCCTCTGAGGTTATCTGGGTCTATAT**AATCCAGTCACCCCAACTGTTCCCCTGGAAATGGAGTGAGGAGGATTTATTAGTTGCTGCCTGAAGAAAAGGGAAATGCTCCAAAAAATTTGGTTGTTTCCAGACTCAAATAGAGCCTGCCTTTCATTGATTCTGTTGCCCTTAAAGCTTCACGGTGAAGATGCAGTTGCTTCCAAA**AGGCTTCTTTCTGGTGCCTAAGCCT**CCTTATACTTGCTTCAGAGCCCTTTCCGTGAACCAGCTGTGTATTGCTCTTCTCATCCCAACACTTGCAATGGCTGAATAAAGGAAGTGGGGCCTGCCTTACGCTAATCCTCGTTCATATGTGTTTCTTAAAGTTATTTTTCCTTCACTGATGAATTCCTTTTTTTTTTTTTTTTTTTTGAGACAGTCTCGCTCTGTCGCCCAGGCTGGAGTGCAGTGGCACGATCTCAGCTCACAACAAGCTCTGCCTCCCGGGTTCATGCCACTCTCCTGCCTCAGCCTCCTGAGTAGCTGGGACTACAGGCGCCCGCCACCACTCCCGTCTAATTTTTTGTATTTTTAGTAGAGCCGGGGTTTCACTGTGTTAGCCAGGATGGTCTCAATCTCCTGACCTCGTGATCCGCCCACCTCAGCCTCCCAAAGTGCTGGGATTA**CAGGCGTGAGCCACTGCGCCTG**GCCTCACTGATGAA**TTCTTTTGCTTTTTAAAGAA**ACTGTTCATTTATTTTCACAGTCTGCAAAAGCCTAAGATAAA**GATGCCACACTCTGAAAGGATCAACAAGGGCATC**ACCAAGTAATGTTTTCTGCAGGATAAACAAG**TCAGGCATTAAATTGGTTAATCCTGA**TTACTGGCCCCTTTCTCTAGCCTCCCCTCTGTGTGAGCAGACCCGGACCACAGGCTTTCTTATTTCCTTTCAGCTTCCCTTGAGACTGAGCAGAGAGAGAAAATTAGCTAAATCAGGAATGCAGGGAATACAGTTGCAGCCTCTTCTTCAGATGGAGGAATGCGTTTTGGGGGGAGGGACATTAAAGGGCCAGTCG**CTCATGTTACAGCTCTTTTTAACTTCATGAG**TACTAATGCCCTGAAGAGGTTTTAATGAA**TGCCCTCTTGTGATCAGTTCCTAGGGCA**ATCCCAGGTTATAAAAGGACTGCCCCTGCCTGTGAGGGAACTG**GCCTGGCTTCAGTGGGCCAGGC**TGCTTTGTTATCTGTTATTGGTTTTTCCAGCTCCTCTTTCTACATTTCAAGGGATCTCAGGCCTTACCTAAGGCAACAGTACATTAGTTTTAGAGTGGGAGATGCTCACAGTTTTCAGAAGAGTTCAGAAAGTTTCAAAACACACAGCACTGCAGAAGATAACATTATAGCTTCTCAAGACCCCAGGGGATCTGGGACTAAACAGTGAAAGATTAATTAGGTAGCGGAAGCC**ACTAAGGCAGTGAGTCTTAGT**TA**GAGAACTTTGGTTTAGACAATGGTTCTC**AAAGGGGCAGCAACACCAACAATACCCGGAAACTTGTTAGAAATGCAAATTCTTAGGCCCTATCCTAGACTAATGAATCAGAAATTCTCAGGATGGAGGCTGGGTGTGGTCGCTCA**TGCCTGTAATTCCAGCACTTTGGGAGGCCAAGGCAGGCA**GATCACTTGAGGTCATGAGTTCGAGACCAGCCTGGTCAACATGGCGAAACCCCATCTCTACTAAAGTTACAAAAATG**AGCTGGGCGTGGTGGCAGGTGCCTGTAATCCCAGCT**ACTCGGGAGGCTGAGGCAGGAGAATTGCTTGAACTCGGGAGGTGGAGGT**TGCAGTGAGCTGAGATTGCACCACTGCA**CTCCAGCCTGGGTGACAGAGTGAGACTCCATCTCAAAAAAAAAAAAAATAATAAATAAAGAAAGAAAGACATTCTCAGGAATGGGACCCGGCAGTCTATGTTTTAACAAGCCTTCTATGTGATACCAATGTACTGTGAAGTTTTAAGAACTGGTCTAAGGTAAATATTCCTGAGGTTGTCTTATATCATTACAGGGTCAGAATGCATGCAAGGAAGCCATCTGTTTATGGTTCTTGTGAGAAGCAGGGGGCCTTTCCCCATGCCCGAGAGATAATTGTTAAGAGCTCAAGCTTGGGAGTCAGTGACCCTTTCTGAATTCTACCTCTGCCACTC**AGTAATTGTATGTTCCTGGGACCATTACT**TAACTTTCCTG**ATTCTCAGTTTCTTTCTCTATAAAATGGGGAGAAT**AGTGGTGTCTACCTTATAGGGTTCTTGGAAGAATTAGATGAGATAATGCACACATATTGCAGAATCTGAAGAACAATCAGGGTTTAGT**AAATAGTAGCTATTT**TTAAATGATTTTTCCAGGTATGAGTCTATCCTACAGCTTCAAAATTTAGACCCAGGTTGTTCTGAGTA**TTCTCTGTGGGAAGAATCTGCTATAGAGAA**GATTTTTTTAAAGTGCCTGTCTCTTTGTTTCCTTAGGGGATTGCTTTTGCCCTG**ATTTGCCACATCTCTTTACTCTGTGGAAAAT**GGACAGTTTATGTGCCCTAGTTTTATATGGGGATTTATATTCTTAATTGTCTCAAGGATTCTTACCTGTCTGACAAAACCAACTCC**CCATGGAAAGACTCCATGG**AGACTCCATCTCTGATCCTTCCCCAGAAAGAAAGCAT**GATTCTTAAGTTTTTTAGAATC**TGTTTAGGAGCACTGTCAA**CATGAATTTTTCTATTTCATG**AGTGAGTGCAGCCTCGGGCCTTGTTGGAGAATTTAGAAAGCATGCTGTTCCACTAACCTGTTCAACCTCAACTTCTGCCGT**TGTCATAGCAATGACA**GTCCTGGGAGGTGTGTGCATATCCTTATTAGGAAAAAAAAATGAGATCAGGGATCTATGTGAGTGGGGCAGCTCCCGCCTGTGAGTATCCTTC**GCTGTCACCTGCCATCTGACAGC**CCAGGAGTGCCAGCTTGGCTTGGCTTTCTCTACCCGAG**GAAAGTAAGTCCTTTTAAGATGCACTTTTACTTTC**TGGGGTTGTGAAACTCATTGTGTTTGCCAGAGTTCTCTTCGCAGCTTATGTAAAGAATTTGTTTGTTTTGGATTGACCTGAAGGGAGGAAGCAAGGGTGTGGGA**AGGGGAATTAGCATCCCCT**ACCTAGGAGAAGCCATGAAGCTTACTTAAGTCTCTGCTGGCTCCATCCATTCATGACTTTCTTCATCTTCTTCTTGGGAAAACACTCTGTACCTTCCA**CTTTTAATGGTCATGTAAATAAAAG**ACTAGAATGGAGATGTCCTGGTTTTCTGAAAT**TAAACATTTTTGTTTA**TGAATAGACTCTAAGATAATTCTTTCCCTAGACGCTCTGTATTTTCTTGGACCTCTCATTTGCCCCATAGTAAT**TATTCTAGGATTGGTGGCCTGGGCAGAATA**CAG**TCATGGTTAAGACCATGA**TACATAATAGAAGAATTTCTTCCGTAACAGTAGCCCCAAACAGAGATCACGTGTCTCCTGAAACC**TATCCAGTCTCATGTGCACTCAGATGTACGATCCTGGATA**TAACATTTTAGAGGGTGTAGTGAGGAGAAT**AGAACAACACTGTTCT**CTTAGGCTGCAGTTTCCTTCAAGCCCCACGATGGAGAGAAGCAGGCTAATGCAGGGTAAGGAAATAGAGACTTATTATAGTTTCTTCAGTTATCATTGATTCCTTTAATATGCCAGCTACATTGAGGCAATACAGTCTGGCCTCTGAAAACCTTGTCAGGAGAAACAACTTTTGAATATATCTTAGAAAAATAAGACACTTTATCCCTTCTTTTGTTTAAGAGTGTTGCATAAAACAGGAGAGTTTCTGAATTACCCTCCCTTCTAGCTTTCTTTATACACCATTCTTTGTAGCTGAAGTTTTAAGCCCCTTGTCAAAAGGGAGATTCAAGTTTCTGCTGGGGACATCTCAGTGTCACAAAGGGCCAAGGAAGTAGGGTCTCACCAACTTTGCCACCTGACTCAGCTCAA**AGGTGATAGGTCACCT**GTGTGACTGAGAGCTTTGTGGGAAAGATGATATGGAGGGTGGAGAGTCTGCACTTCACTCTGTGGGAAAAATTGCTGAGGTTGTTTAGAATTGTTAGGCTTGGAACTCCTCCTGGGAGGTGCTTGAACAAGAAAGACCTGCTTATAACCTGAGTTGAGGGCAGAGGAGGAAGATAGTTTGTAATTCCTTTACGTTTTGTGGCTCCGGCAAGCCCTGTCCTCAGCCTCACTGACACAAGTAAACTAAAA**ATGAAAGTCTGTCCTAGTGACAGCAAGGGTCTTTCAT**GGCAATATTTTAAAAGAAACTCTGCCCAGATTTCAAAGGAACGTGAAAATTTTATCTTCAGAGGCAGTCAGCTTTGCAGTTGAAAAGGCCATTGCCTAATCTGGAGAAACATATTCAGTTCAAGCACTGGCTAGAGACTAGAGGCCCCCAGGAAAGGGCCATAAGATTGGTCACTGCCAGGAAGCCTGTCAATGAGTGTGTGGACTGGAGAGGAATCTTTCTCTGCCTCAGCACTTGGAGTCTCCGTTATTCACAATCAGAAAAAGAGGGAGAGCAGAGATAGATGACATTCCACGTTTTTCTCGGTGAGACCAAGAGCTTCCTCAGCAACAGCCCTGCCTGAAATTGTCTATCTCCAAGGGCGTGGGCTAGGCAAAGGATGGGAGAAGTCAGTCCTGAAGATGGTAATACTTAAAGGAAATAAGGGGGACAAAGGATGCTTGTGCTACTATTGAAACAGGAAATTGAGAGCTCTGTAGAAGTCAGACTCAAGAGGCATAATAATAGAAAGTTAGGGTATGAAAAGGTGACTTTTAAGAACCAAATGTGGACCCGAAAGAGAACAAAGAGAAGTTTATTGTAACTAGCGTTGTATTCCTTGTCTGCTGGATACCAA**ACAATGTACCCCGGGTCTTGAGATTATCGATGCCATTGT**CTGTAAAAAACCAACCAAACAAACTTTAAAAATAAGTAAAGCCTGCCCTGTACAGAAAATCTAATAAAGCAAATTTGT**TGGTGATGTTCCAAGAAGGTTCACCA**GCTTGGGCTTTCAACCAGCATTGACCCAATCTTGTGTCCAGAGCTGTTGCTGTAGGTGGTATCATGTATCAAGCTGAATAGTGAGGTGACATTCCCCAAGTCTTCTCTCTTGTCTTTTTTGACTGTGCATCATAGATACTGTATTACCCAAGAACACACACCTGTTCGATCTCTTTTCCTATAATCACCTGGAGTCAGATGAATACCGTAAAGGTCTGTGGTAAGACTAGAAGCCTCAGCCAGTATGAATGATTTACATTAGATGCACACCAAGCTGCTTTCGGAGAGTCCAGATTTATGGGGATAAGAGCATCACT**AGGTATATCAACAGCCCTAGGGTGGATACCT**TTGAGCCTGTAAATTTGGCTCTGTGTTGGACGATGAACCATGGAATACAGAAGAAGCTCCCTTCTCTCAGCTAAAGCCTATTAGCAAAAATAGAGCCCTGAGGACATTGTATTTTAAGTGTATGTTCCAGTGAGTGGGTCGTTGCTGGGTCAAGTTTTATAAGTCTTTAAATCCCATACTGTTGCCTCCTGGACTGTAGAAATAG**TCTCTTAGAACAAGAGA**AAGGAAGACAATAGCTACCATTTATTGAGAACTGTGATAAGCACTTTACATATGTTGCATACTTAACTTTGAGAGAAGTACATTATTATTCCCATGTTTCAGATAAAAAAATTAAAGCTCA**TAAGAATTAAGTGGCTTTCTTA**CATTCACACAGCTAGTAAAAATGTTTTTTGCTCACCATTGTATTCC**CATTGTCAAGCAGGATGCCTAGCACACAATG**ATGCTCAATAAATTTTGTTAAATTCATTAATAAATGCAGTGGTAAAGGCAGTATTTGAACTCATGTCTGCTTGCTGCTGAACTCTATACACTTAACACATTACTATTATCTTGTCCCGCATGATACATGAAGGGAATAGCTTGGTAACTTGGAAAAGACTATTTACTGTCTGAGTTCTAGGCCAACTGTAGGGTGTCTGTATTTATTTCCCATTATGCTATAACAAATTAACCATAAACTTCATGGCTTAAAACAACACAATGTATTATCTTACAGTTCTGGAAGTCAGAAGTCCA**CAGTGAGTTTCACTG**GCTAAAATCAAGGTGTGTTCTTGCCACTATATAACAAGGATCCACTTTCCTTTAGTTTCCAATAACAGGCTCCTTATTTCTACCTGAGCCCTCACTGGCAGCACCTTTAATGCCATATTTGTAATAACAATCTGTTCATGACAATTTAGGTTTGCTCCTCACTTTTTTCTGAGTCCTCT**CTAGTAGAGCCATTAATATGCATATTGCTACTAG**CAGCCTGTTCAAGGTAATGTAGGCTCTTTCTATCATGCTCTTCAAAATTCTTCTAGACTCTGCCCATTTCCCAATCTCAAGGCCACTTCCGCATTTTTAGGCATTTATAACGGCAGAACCGCACTTCCAGATACCAAAATCTGTATTAGTTTCCTAGGGCTGCCATAACAAATTAACACAAAATCCCTTTTGCCATGTAACATAACATATTCACAAGTTCTAAAATTAGAATGCAGTCATTTTGGTGGGGCCATTAT**TCTGCCTACCACCCGGTCTTTCATGTTCAGGCAGA**GGTGGCTCTGTGTGTGTGTGTGTGTGTGTGTGTGTGTGTGTGTGTAAGTTAATTTCAGTAGAGAATGAGCTAGAGTAGAGAAAGAGAATTAAGGTGAGATGTGTCATGGAAGAACAGTGACTGATGATGCTAACTTTGCTCAATCAAGAAGTGTGATCCATTTAAATCATGCTTTCAGTGATCTATCCAATCAGATAAACTACTCTC**CCTTCCTGGGAAGAGTAAGGAAGG**AAGTAGAGCTAAAGATGAAAGTTCTTTTCGTAGCACATCCCTGCAAAGGATGGGAGTATTGTTTTGGTGGGCATTCCCTTTTCTAAGAGCAAAGATGGAAATGTGGAGAGAGGAGAAAAATGGTTCCCATTACATTATTGTGTTCTGGA**CTTAGAGATATTGTTCTGCCCCAGGTCTAAG**AATATTGTTCC**AAGAGTTGGGAGCAGGCAGCAGAGGACAGTACTCTT**TAGATCACCCAGAGGCCAATCTGTAAGGATTCAATCCTGGGGCATGGTAGCTAATGGAAAAGTCACTGTCACAAGTGATGCCAGGAGATGGGGCAACACATATTGCCTGCCTTGTATGCATCTAGCGTCTGTGTCTAAGTGAAAGCAGATTATATCCAGAACTTCCCTGGAAAATAGAAGAAAGAGCCTGATTGGTGTGTATGTGTGTGTTAGGGGTTGAGG**GGTGGGTAATGTTCCTGCCAGTATTCGTAAATCCCACC**TGCTTTTCCTGTGATGCTTCCTGTGTTGGGGATAGGAGGGTGGGATGGTTCAGGGTCAATTTTATGGATCCATATGTATCTAAGGATGTGTTTGTTTTTTGGGAACATTTTGTATAACAACCAGTTTCACTCTAGCTCCCTCCTTGATATTCTAAACAAGCTGGATATCTCCAAAGCTTAGGCCC**TCTCCATACTTACAGCAGCTCTGACTGGAGA**TTAAATCACCCCTTATACTGCTGACAATGATTAGGCCATGGGACCCATGAAAGAGCCTCCCCAGAGTCCAGCATCCCCTGTGGGCTCCGTGTGTCATCAAGATGTCAATCCAA**CTGCTGTTCCTGCAGTCTGCAATCAGCAG**GGCTATGTTTATTCAGCGGTCAGTGTCACATCAATTTCCTTCTGTTGCAACAAGTATAAATGGATTCTAAATATTTGCCTTTGGGAAATTT**CTTTAGAGGGAAACTCACTAAAGCTAATTTTTTTAGCTTT**TTGGTATGTTCTCTCTGAATCTGGGGATTTAGATATATAAATTAGCTTCTTTGGTCTTTTCTTGCCCAGGGTCACAACCTTGCCTCCAGGATAATACCTTCTA**AGTGTTTGTGATTTGAAGGGCACTACGAAGATCCTCACAAAAACT**ACCTCCCAGCCGGGTCCCTGAAACTCCATCATTAACCACCTTCATCAGCATTTCTCTTTTAAAATCCTTATTCATTCCTGTTCTCCTTCTTTCTTTCTCACACACACACACACACACACACACACACACACACACACACACACACGGGGAGAGAGAGACAGAGAGAGAGAGAGATAGAGAAGTGAAGTATATAGTATCCTTTCTAGGGATGCTTTTCTTGGCTTGGCTCCA**ATAGAGAATTTTGTTGGGACCCTCTAT**ATATAGTCGTGT**ACCAAATAATGACACTTTGGT**TAACAATGGACCACATATGTGACAGCTGTCCCATAAGATTGTAATACCATATTTTTACTGTACCTTTTCTATGTTTAGA**TACACAAATACTTGCCATTATATTACAATTGCGCAGTATTTTGTA**CAGTAACATGCTGTGCAGGTTTGTAGCCTAGGAGCAA**TAGGCTATACCATATAGCCTA**GGTATGTAGTAGGCTATGCCATCTAGGTTTGTGTAAGTACGCTCTATGATGTCCATACAACAA**AAATGCCTAGTGATGCATTT**CTCAGAACATATCCCCATTGTTAAGTGATACATGACTGTAGTCATCATCAAACATTTATTAAGCACTTAGGTCAGGCCAGGCTCTGTTCTAGGTGATGCAGATATAATATTAAATATAACACAGTCCTGCCCTTATAAATCTAATGGTGAAGGGAGA**AATGTAAAGAAATATGATGGAAATATTACATTGTCATGTGATAAGGGCGACAAT**AATG**CCCTACGTAGGTAGGG**TCAACGAAGGGGAAGGAAGCTGGGACAAACCACCAGGGCCTGTTGGTCCAGAATGGGACCCAGGGTCTGTCTATGTTATAATCAATTCAAACCCTAGGTAAATAAGGTAAGCTAGGCTGCCTTTCTTGAGACAGTCCCCAGATTGTTTTCACAGGGCCCAAACACTCTCAGCAACCATGAACAGTGGATGTTGTGGGACCTCAGAAGATGGAATTGGAAATTCAGGGGACTAAGTCAGAGATTATTTGACATTTAATACTGGATATTGAGGCCGGGCGCAGTGGCTCACACCTGTGATCCCAGCACTTTGGGAGGCTGAGGCGGGTGGATTGCCTGAA**CCCAGGAGTTTGAGAACAGCCTGGG**CAACATGGCAAAACCCCATCTCTACAAAAAAAATATAAAAATTAGCCGGGTGTGGTGGCATGCACTTGTAGT**CTCAGCTACTTGGGAGGCTGAG**GCACGAGAGTCACTTGAACCTGGAAGGTGGAGGT**TGCAGTGAGCTGAGATCACACTACTGCA**CTCCAGCCTGGGTGACAGAA**TGAGACCCTGTCTCA**AAAAAAAATTTAAAATACTGGGTATTGCAGGATGGCAAGTAGGATGTGGAGAGGAAGAGAATTCCAGGCAAAGGAAACAATGCGTGCATGCAAAAACAAAATACAAAAAACTGAATTGTGTAGTCTGGGTTAATAAGATGAGTGCATTTGATGGATGTGGTAGAATAAAGAGATGTTAGGAAATGAGATCAGAAATGTAGACTGGAATCAGATTTTGTAGGACCTTAAACATCCTGCTAAATAATTTGTAGTTTCATTTGTTGGCCATGGAGACCCACTAGAAGGTTAGTTGGTTTGTTTATATTT**AGGAAAGAAGATTCTTTTAAGTTTCCT**TAACTCTACTGAATTTTCAAAAATCAACCTACAGCAATCCAAACTTATCTGTGGAGATACATTCCAAGACCCTCAGTGGATGCCTGAAACCTCATTTAGTACTGACCTTTATATACATTATGTTCTT**TCATAGGTGCACCTATGA**TAAAGTTTAATTTATAAATTAG**GCACAATACTCTTGTGC**TTTGGGGCCA**GTATTAAGTAAAATAAGGGTACTTGAATAC**AAGCCCTTTGATACCAAAACAGCCAGTCGGATACCAAGACACTTTCTAAGTGAACTAACAGGTGAGTAGTGTAGACGGCATGGATAGGTCGGACAGAGGGATGATTCACGTACTGGGCAGGATAAAGCGGGATGGCTCGAGATTTCATCACATTACTCAGAACAGCTTGCAATTTAAAACTTATGAATTGTTTATTTTTGAAATTTTTCATTTAATATTTTCAGACTAAGGTTGACTGTGGGTAACTGAAGCCTCAGAAATCAAACCTCAAATAAGGGGCGATCACTGTACTGAATAATATGTCAAGATGAGCTATGAGCTTTTAACTTAGTGTTTTTCTTCTCCCAAACCAAATATTGGAAACTTTGGAGTGTTTAGAAAAGGAGAATCGAAAAGGGAAGTAAATGCAGCATTTTTTTTAATTGTTAAATAAAGGGCTGGGCTTGGAAATGTTCTTAGGTAATCCTGGTGGATTTGGTATATCATGTCTGAGAAAATGTGGAGAGTGAAGGGCAGGTTTAAAATTTGTCTCATGTTAGTTTCTGAAGGGAGATAGCCTGGCATTGCACAGTGCCTCACATTTCTATCGTATCACTTAATAATTGTGCAACCTGTGCTTCAGATTTATCAATGTATAAAAGAGGAGAGACCATAGGCTGTTGTAAGTATTAAATGACATAATGCACGCAAAATGCTTAGA**ACCATGCTTAGTTCATGGT**AGCCACTCAAAAATGTCGTCAGAACTATTACTACTCAGTTTAATGCTCCT**AGCACTTAACACAGTGCT**TGGCACACAGTAAAGAGGATTCAAGATGTTTTTGCTAGATTACATGGAATGGTATGGAAGTGTGCAGGGTGGTGTAGAGGGTTCTCCCATTATCTTGCCTTCCTTATACTGCAATGTGCTGACAGCACCTCCATTTTATACCCTCAATGCAAGGATATCACTACTTGTCTAGAAATGGTTGCTCAAGGCTGAGAAGTGA**CAGCAACACAGAGCCTGGGATTGCTG**CCAAATTTCAAGAAGCACAGATTGTAACAAAGTAAGGGGGGAGGGTAGAGAAGAAAAGTTTGAAAACCCAACCAAAGTCAGAAGCCATTTTGTTGTCGG**CTGAAATAATGCCTCATAGAGTTTCAG**GTGGGGCAGGCAGTGTGAAAACTTTGGGAATAGTCAGTGTCAAGTTGTGGTGCTAAGGGAGGAACCCAGTAGTGCTCAGGC**CTGGGCAAGTCTGGCTTCTTGGGGCCCAGCATTTAGAAAGGCCCTGTGTGTTTTGATATTATGCTG**TCACTATCTTGAAGTTCTTGATAATTTGTTAACAAGGGCCGCCTCCCCCCATTTTCATTTGGCAAATTTCACTGGTT**CCTGCAAATTATGTAGTTAATCCTGGGTGCAGG**CTAGTTTTTGAACCATGTTGGGCTAGTA**TTCTTCTGAACCTCTATCTCTTTGATTTAATTACAGAAGAA**ACAAAAGTCATAACAAATTATATCTGTGCAGGACTAAAATAAAATGAGTAGTGCCTCACTCCCTACATCCAATCATGCTTTCCAAAAGTAACCACTACCAATAATTTAGCGTGCATCTTCAGAATTCTTTATATGCCTACATTTTAATG**TATTTTTAAATAAAATA**GAATTACATTGTTGAAGTAAATATTA**AAAAATATAGCATTTTT**GCATAGAAATATTTATTTCCCTCAATCCCACTCCCCTTTGGTGAAGCAACCACTGTTAACAATTTTGTGTGTATCCTCCAATATTTTTTTGAAATTAATGAACTTTACTTTTTAGAGCAGTTTTAGGTTCACAGGAAAGTTGAGTAGAA**AGTGCAGAGTTCTCATGCACT**CCTGCTCTCCCACATACACAATGCCCCCACACCCCCGCCACAGTGACATCCTGCCCAGAGTGGTACATTCATGACAATTGATGAAACTATATTGACATCATTATTGAAACTATCATTACCACCCAACTTCCATGTTTACTTTAGGGTTCACTCTTCCAAATATTTCTTGCATCACTTAACAATGGGGATACACTGAGAAATGTGTCTTTAGGCAGTTTTTTCGTTGTACAA**ATATCATAGAATGTACTTACATAAACCTAGATGATAT**AGCATACTACACACTTAGGCTCTATCGTATGGCCTATTTCTCCTAGGCTACAAACCTGTATATCATGTTACTATACTGAATACTGTAGGCAATTTT**TACACAATGATAAGTATTTGTGTA**TCTCAACATACCTAAACGCAGAAAAAGTACAATAAAAATACATTATTATAATCTTACAGGACAACTGTCATATGTGTGGTCCGTTGTTGACTGAAACATGATTATGGGGCGCATGACTATATAGAGATTTTCAAAAAGCAAAATGGGATAATGCAATATGTATTATTATACAGCTTGTGTGTGTGTGTGTGTGTGTGTGTGTGTATGTTAACAAACACCTTTCTACGTCAGTACATATAGATTTGCCACAATCTTCTTGAGTAAGGATATAGCACTATGTTGTATACCTATATTATAATTTATCCAATTTGCTGTTGGTATCTATGTTTTCTAAACAATACAATATACATTCTTATACATATTTCTTATGCATGCTTGCTAGT**ATTTATACAGGATAAAT**TTCAAGAAGTAGAATTACAGAATCATAGGATATAAAAATATCAAATTGGCAAATTGGCATTCACGTACCTGTGGTTGGACTGCACTTGTTACATACTTCCCTCTGAGCCTTCCATCCCCACCTCGAGCTAGTGCTAT**ATGTCACTTAACTCAGTGACAT**CATCATCAACAGACTTAG**CCTCTGCAGTTGAAAATGCTAGAGCAAACAGAGG**AGAAAATTTACTTGTGAATCATAATAGCTAACCTGTACTGAACAGTTAAGCCATGTGCCAAGTATTATTCTAAGCACTTTACAAGTATTAACTCCTTTAATCTGTATAACCACACCCAGAAATTTGTGCATTATTATACTCATTTTACTGCTAAGAAAACTGACACCCATAGGATTAAA**TCAGATCACACATTTAGTAAGGGCACCAGGATCTGA**AGCAGGATTGTTTGACTTCTGAGTCTGTGCTCTTAACCACTGCACACACCGTCTCTAGAAAGTTTAATGCCATCTTTATGCCAAAGTTTATCTACCTGGTTTATCCTTTAACACTATTATTGTATTTATTCATTTACTTGTCTTTTCCTCCCACTGTGCTGTAACTCTTTGAGAACAGGGTACTTGTCCTAATATTATTCATATTTATATCCCTAATTCCTAATTTAATGTCTAATTTATTGTAAATAAGTTTTTTTTTTTTGAGACAGAGTTTTGCTCTTGTTGCCCAGGCTGGAGTGCAATGGCACAATCTCAGCTCACTGCAACCTCTGCCTCCTGGGTTCAAGTGATTCTCCTGCCTCAGCCTCCCAAGTAGCTGGGATTACAAGCTTGCACCACCACGCTAGGCTAATTTTGTATTTTTAGAAGAGACACAGTTTCACTATGTT**GGTCAGGCTGGTCTGGAACCCCTGACC**TCAGGTGATCTGCCTGCCTTGGCCTCCCAAAGTGCTGGATTACTGGTGTGAGCCACATGCCTGGCCTTGTAAATAATAAGTTTAGTTGAATAAATAACAATGCCTCCGGGAGGTAGCTATTATATCCATT**TTACAGATGAAGAAACTGTAA**GTCAATGTTAATCAAATAGTATCCTAGAAAATAGTAGAAAGCAGAATGCTGGAGCTGAGATTTGAACCCAAGACTTTTGATACTTCGTCCAGTGTGCTTTCCACCATGCCTAAGTAGTCTCCTTCACTTCCTCCTTCAGAGGGCTATGGAGAGTAACCTAGCAACCATTTCTAAGCTGGAAAATGTCACAGCCGGAAGTCTTCAGTCCCCTAGAAGGAAAAGAGCCTGATGGGGAGAGGGTCCTTGGAAAAGAGAATTTCTGGAAGTACTACATTTGAGAATAGGTGGTTAAAGTGGGAGTTGGATTAGTAAAGGAATACTGCATAGTAGTAATACCCATTCTTTATTTCAGAATTTCTTCATGACAGACATTGATCTGTGTGATTTATAAGCCTTGTCTCTTTTTAATGTTCACAATCCCATTAGATGGTTATTGTTATCTTTGTTTTGCATATGAAAGAATGGGAGCTAAGAGGCCAAGTACCTTG**CCTCTTATTCGTTCAACGAATAAGTGG**TGAAGCAGATTCCAACTCAAGTCTATGTTACCCAAGAACCTACATTTTAAGCATTTTGT**TACCCCCTGGATATGACAGCCAATGAAGAGGGGGTA**TTTTGAGAAGAGACTATAA**AGGGAAATTGCCTTCCCT**ACA**TCCTGGGGGACCTTATCAACCAGGA**AACAAGGTAGAGAAAACTGTGCAGCC**TGAGCCCTGCTGGGTTGCGGGGGGCTCA**CAGAAAGAAGAAATGTGATTTTTTTTTAGCTAACTACGGAGACCAGCCATTCCAATGTTTGAATCTGGGTTCGCAGCACATGATGTCTTTATACTCTTAACCTAGAAATGGCAGAGTTATTTTGGGCACAAAGCAAGAGCTGTGGCTTTAAAAATATGCCAAGTGTATTTATCTCTTCCGCTCCAAGATTACTGAAAATTAGCCCAGCTGTAGCTTGGGACACCAAACAGCCAAAAAATCTTCTTCCAGCTCAACTCATTCCACCCAAAGAGGGTGAAA**TACCCAGGAGTAGCAGCTCTAGCGGCCTCTGGGTA**GTGGTATTA**GATTGGCCTCCCCATTGCTAAGCCTGACATCCAATC**ACACACACACCACTCTCCCAGCTGCTCTGTAGATCACAATGCTAGGCCTGTGAATGGAGCTCAACTCCGTCTCTTCCCTCATCCCCAAGGCTTCACAAGAAGTAAACAAAACAAAC**AAAAACTATTTGATTATTGAGCCAAGGAGTCAATGTGAGAATAGTTTTT**CACCTTCATTATCAAATGCCTGTGTGGAGCTGAATGTGGTGGCTGACACCTGTGATCCTAGCACTTTGGGGAGGCAGAGGTGGGAATATTGCTTGAGGCCAGGAGTTCAAGACCAGCTTGGGCAACACAGCAAGGCCCTATCTCTCTCTCTGTCTCTCTCTGTCTCTCTCTCTCTCTCTCTCTCTCTCTCTCTCTC**TATATATATATATATATATATATATA**TTTTTTTTTTTTTTTTTTTTTTTGAGACAGAGTTTTGCTCTTGTTGCCCAAGCTGGAGTGCAATGGTAAGATCTCGGCTCACTGCAACCTCCTTCTCCAGGGTTCAAGTGATTCTCATGCCTCAAACTCCCGAG**TAGCTGGGATTACAGGTGCCTGCCACCATGCCCAGCTA**ATTTTTGTATTTTTAGTAGAGACAGGGTTTCACCATGTTGGGCAGGCTGGTCTCGAACTCCTGACCT**CAGGTGATCCACCTG**CCTCTGCCTCCCAAAGTGCTGGGATTA**CAGGCATGAGCCACTGTGCCTG**GCCCCTATCTCTGTATATTAAAAAAAAAAAATCTTAGCCAGGCATAGTGGCGCACACCTGTATTCCTAGCTACTCAGAAGGCTGAGGTGGGAGGATCACTTGAGTCCAGGAGATCCAGTCTGGCAGTCAGCCATGACCTCACCATTGTACTCCAACCTGGGCAACAGATATAGACGCTGTCTCTAAAAAAAAAAATCCTGTGTGATTTAGGAC**AAATTATCTTGCTGTAATTT**AAGCCTTCATATTCCTTTTTTATA**AAATAAATATGCTTTAAATATGTATTT**AGATATACACTTATAATAATAGC**TTTAATTAATTGTAATTAAA**TCATTTAAAATTAAATACATGCAAAATACCCCATAAACTGTCTAATCCATAATAAGCAATCAATAGTCCTTAAACAAATGAATCTTCTGTTGCTCTGATCTTAAT**AAGTAAAATTTAACGAGTTTTTACTT**TGCACCATGCACTTATGCTATGTGCTGTATAAGGATAGTTACATTTAATCTATAAATCAACCCTATGGGGCGGGCACTCTTATGAACTTTTTTTACGGATGAAAACTGAGGCCCAGAGATATGAAATAATTTGCCAA**ACATCACCATAATTTACATGATGT**AGCCAGGTTTAAGCCTACTATTCTGATTACAGAGCCTAAGATCCTACAGAGAACAGGGAGTATATTTTTACATTCCCTTGTTTCCA**TGGAAAAGTCTTTCCA**CTGTCAATTGAAGGACTAAGCAGCAGCGGGGAAGTGCTGGAAATGCTAGCCCAGGTGGGCTTAGCTTTCTCCCTCTCTCTCTGATCCTGG**ACAGGATTTGGGTTTCAGACCACGATTCTCCTGT**GTTTTGTGGGCTGTGATTACTCAGATTAGGTTTGCCCAATTCATAAAGGACTGGCGGGGGTTGGGGGTGAAGGTGTGGGAGAAGGCGGAGCTTGTCCAGTCGGTCCAACAAACCCCACAGATGGCGAAATAGGGGGCGGGGAAGGAGCTTGAGATATTTTACAACCTGGGCTGTTTCAGTGGTTGATGGCGGATGGTTTTTGCCTTTTGGTCAGCCTGAGTATATACTATCATTCTACAATCGGCCAAATTCTGACAGAGAGGGAGACAGAGAGAGAGGTTGATTAAATTGATGCCCAAAACCAAGAAGGAGCAAAGAAATCGGGGCTGTTTGAAAAAGACTGCAGTGGCTGACTTGGGTGGTGAGCGGAAATAAGGAGGAGGGAGAGGCGGGGTGTCTCTGCGCGGAAAGCCTGGAAGTTCACTTGCAAACACAGAATCTGCACAGCCTTCGGTGCCCTGACTCTTTCCTGGGCATCCAGAGGCAGCAGCAGCCGCCAGCGCCAAAGAACGAGCAGTCCAGGGGCTGGGCCGGAAAC**GGCTATAATCATTTAATAGCC**TTTGCCGGCTGCACTGACTTAGCAGAGTGGGCGGTAGGCAGGCTCCAGAGTGCTGTCTGGCAAGATAGTCCCCGGCTTTAATCAAAATGATGGGTTTTCTGGA**AGCTCTTTATTAACCTCAGCACTGAAATCCCAGAGCT**GGTAACAAAGGGATGAATGGGGAGCAAAGGGGCGGGGCCGAAACC**TGGAGGCCGGGCTGCATCCGCACCCCTTCCCCCACCTCCA**CTCCTTTCAACTCATTCTGGCTTAGGGCTCCTGCTGGCATCTCTGTGCCTCCCAAATAGTAGTAACAAAACAGGCAATAACCATAATAATTGGCACATT**TGTATAACGCTTTAGCATTTTCAAAGGATGACCTTGTTATACA**GCTCAGGATCT**GAGTCTCCTAGCTGGAGTCAGACTC**CCTAGATTTGAGTCCCAGGATCCACCAGTTAGTGACCATGTGACCATTAGTCCTTAGTCCCCTCATCTATAAAACAGACACATAACACAGACAGAGATACTAGAAATGTCTACTTCATAGGGCTCTTGTCAGGACCAAAACCTATACGACGTGCAGATTTGTTTAACTCTGTTGCACACACATAGGAAGGGCACAGTAAATGTTAA**CTACATCGACAACCCTGTGATGTAG**ACAGGAAGGGGATCATCTCCATTTTATGGATGAGGAAACTGAGGCTCAGAGATATTTATGTAAATGGCCC**AGGACTATGCACTAGTGAGAGAACCAGGGCAAGTCCT**CAAGAACAGAAAGAGAACAAAGCAAGATGAAGAGAAGATCAGAAAAGTGAACACCCTCCAACACCCTCCAGCACATGCCCCTCTGAGAGCTTTCCTGAAAATAGTATCCTTGGTCAAAATGAGTTTATGTTCTATTAGGGAAAGTCAGTGTAAATCAGAAAAGCCCTCCTGGCCTTAACAGAACAGTTC**CAAATGCAGATGAGGGTGAGTTTAACAGGCCTGGCATTTG**CTTCACAATCTTCTGGCCCTTTCCTGGCATTTGCCTTGGCAAAGACTCTGACCTGATCAATCCTAGTC**CCTGAGGCTCAACTCCATCAACTCAGG**CAGCCAGGGTTAGTTTAGCATGAAAGAGGAAAGCTGGAAATTTTGCCAAAAAGATTCC**TGGGCAGTGCCTAAGGGAAGTGCCCA**AATTGGATTATAGGATTACATGAAGTGGCCAGCTACTTCCAGAGGGCAGGGTTTTTTGTTTGTTTGCTCTGTCACCCAGGCTGGAGTGCAGTGGTGCAATCTCAGCTCACTGAAACCTCCGCCTCCCGGGTTCGAGTGATTCTTATGCCTCAGCCTCCCAAGTAGATGAAATTACAGGCGTGCACCACCACAGTCTCGCTAATTTTTATAGTTTTAGTAGAGATGGGGTTTCACCATGTTCCCTAGGCTGGT**CTTGAACTCCTGGCTTCAAG**TGATCCACCTGCCTCAGACTCCCAAAGTGCTGGGATTTTAGATGTGAGCCACCATTCCCGGCCCCAGAGGGCAGTTTTTAAAGTAGAAGTGAAGATCTGTATGCATTTA**TTAAAATATATGTTGTTCTGATTTTAA**AATATATCCCTACCAATTGTGAAGATATCGGCTAATCCAATAAAAAAGCAAA**ACCATATTTTGTATAATTATGGT**CGTATTCTGTTAAATAAACTTTTAAATCCTATTGGGTTGTAATCTGTAGTCTGGGAAGGGCTCTGCCTAACGA**TTCAGACCCCTACCAGATTTATATGTGTCTGAA**ATCCTCCTTCTAGCCTCTTAACAGGCCCATTCATTTGGGTCTTGACTGCAT**CCTCAGTCTTGTCTTTGCCTGAGG**GTGAGCTAGGACCCTGGGAGCATAAGGGAGGGGACTTGCCGGAGATCCTAGAAGGGAAAGGAGGCAGCACTGAAAGAAGAAACATTTCCTTGATTGATCATTTGCTGATACTGGCCCCTGGTTGGGGTTAGGGGTAGAAATGTGCTTCTTCCGTTTTCATCTTCTTCAAAAGGGAGTATCTCCATGATTCTGCCCCAAAGGCATGACATTTTATAGGCAAAGCCAGCCAAGTGTCTTGAGTGCTCTTAACCAAAAGGAATTCAGATTAGGAGGTCAGTCAGTGTTTGTCAGATCCTTAAGCTTTTGTTTTACTGGAATGAGTCACAGTCTACATGCTGCTGAGCTCTCAGAGATGTTTGTTTTATACAACAACCAGAACAGGTGAGGCACA**GTGGTCTGTGAGGGACTGGAGAGACCAC**ACCTTGTTCTGCTCTGCGGAAGCTGGACATGTGG**AGGAGGCACCTGACTAGATCTCTGCCTCCTGGAGTTTAGGCTGAGTCATATAGAACCAGGA**AGCTCAGTGACATGTTAAAGTGGATTAACTC**TTGGCAGTCCTGCTGTGAGGGGCTCCCTCTGCCAA**TACACACAGCCATTAATGTCCTTTAACTGTGCAAGATGAACATATCTCTGTGT**TCTGCATCTCCAGATGACAGTGCTTAGGCCTCATGCAGA**GTCCTGGTTATGGTTGAAGAAAAATTTCCATTTTGGGCTCTGAG**GGCAACTAAAGGACTGTGGAGTGGTGCTGAAACCCAGTTTTAGGTGCC**GAATCAGAGGTTTTTAAATACATTTCTCTCTTT**TGTCTCAGTCTCTTAGAGACA**GGACCTGTATTTTAAAGGTAAACAGGCAGATTCTGGCTGAGCTCAATTGCAGA**TTTGATTAACTTAGATAGATCAAA**GTCATTAGTCTCA**GAGAAAAAATTTGTTTCTC**ATCCCTAAAGTGCTATTGTGTCAGCTCTGCCAGGGTGTAAGGAAAAGGGTTTGGA**AGAGAGAAGAAAATTTCTTACCCTCTCT**CGTCACTGCCTGCTGTAAATTTGCAGTATCATTATCTTGGTCTCCTAAGCCCTGAGAGCTGCATTATATATAATTGCCCATATGTGATACCATTTACTGAATTCTTGCATTGTGCTAAGGATCAAGCTTAGTACTTTACATGCATTACATCATTGTCTTTAGGGACCCCAAAAGTGCCTCATTGGAGAGTTGTTTGGTCTGCTAAAGAGTGGGTACCACTTTCACTTCTCACATCAGTCCTTTCATTCATTCATCCAGCAAGTATTTACTAAACACCTGTTATGGGCCAGGAATTGTGCTAGGCACTGGTGATACATTGGTGAATAAACAGATACATTTTTTGCCTTTGTGAATCTTACAGTGGTAGAGAATAATAG**GCCAAGGGGGTGCTGCTTGGC**TAGGAATGTCATAGAATGCCTTTCCAAGGAGGTTACATTTAATACAGACTTGAAGAGTGAGGAGTCATGCTAAGAATGGATGAGAGGCTCACGCCTGTAATCCCAGCACTTTGGGAGGCCGAGACGGGCGGATCATGAGGTCAGGAGATCGAGACCATCCTGGCTAACACGGTGAAACCCCGTCTCTACTAAAAATACAAAAATTAGCCGGGCATGGTGGCGCGTGCCTGTAGTCCCAGCTACACAGGAGGCTGAGGCAGGAGAATGGCGTGAACCCGGGAGGCGGAGCT**TGCAGTGAGTCGAGATCGCGCCACTGCA**CTCCAGCCTGGGCGACAGAGCGAAACTCCGTCTCAAAAAAAAAAAAAAAAAAAAAAAAAAAAAAGAATGGATGAGAAATCATTCAAGACAAAAGCAAACCAGATGTGTAAAGATCCTGAGGCAAAAACAACTCCAAGGAGCCAGGTATGGTGGCCTGTAGTCCCAGCCACTTGGGAGGCTGAGACAGGAGGGTCGATTGAGTCCAGGAGTTTGAGGTTACATCAAGCAATGATTGCACCACTGCACTTCAGCCTGGTTGACAGAGTGAGAACCTATCTCTAA**CAAACAACAATAAAAAAACAATTCCAAAGAGTGTTTG**AGGATCTGAAAGAAGGCCTGTATGGTTGGAGTATAATAGTAAGAGGGAGATTTGTAGATGAGAGTGGAGAAGAAATAAAAATCAGATTATTGAAGGGACTTATGTATCATGGTAAGGAGTTTTTTTTTTTTTTCTCAAGGCAGTAGAAAGCTTAAGCAGAGGAGTG**ATATAATCTGATTTATAT**TTTTAAAAGAGTTTGCTATATGTATGTTATGCCTCATTTTTTTTTAAATTTTTTATTTTTAGATGGAGTCTTGCTCTGTTGCCCAGGCTGGAC**TGCAGTGGCACGATCTCAGCTCACTGCA**AGCTCCACTTCCCGGGTTCATGCCATTCTCCTGCCTCAGCCTCCTGAGTAGCTGGGACTATAGGCGCCCACCACCACGCCCGGCTAAGTTTTTTCTATTTTTTAGTAGAGACGCGGTTTCACTGTGTTAGCCAGGATGGTCTCAATCTCCTGACCTCGAGATCTGCCTGTCTCGGCCTCCCAAAGTGCTGGGATTACATGTGTGAGCCACCGCGCCTGGCTGTTATACATCAGTTTTTAAAGAGAGACATTAAGAAATAAAGATGACTCTGGCTACTGTGTGGAGGGTTGGAGAGAAGCAAAAGTGGAAGCAGACTGAATAAGGAGACTATTGCTGCAATTCATATAAGAGATGATGGTGTCCTGAAAGAGTGGTGGCAGTGAAAATTGGGAGAAGAGGACAGAG**TCAAGATATATTTAGGAGGCAGAAATGACAGGTCTTGA**TGATGTATTATAAATGGAGAATGAAGAATAGAATAGAGAAAAATGAATAATTAGCTTTGAACTTTCTGGCTTGAGAAATGGG**GTAAATGGGGTGTAATTTAC**TGAAACTGGAAAGATTTAGAAAGGAATAGATTCGAGG**AATCAAGAGTTTGATT**TCAGATTTCTTCAGTTTGGGATGCCCATGAGATTTCCAACTGGGGATACCAGATAGGCAGTTATACATGAGATTGGAGCACAGA**GAGATGTGGGCTTAAAATATAAGTCTGCATCTC**ATCAGCTTATTCATATGGTATTTATATCTATGGATATGGCTGAGATCCCAAAAGGAGAGAGTGAAGAGAAAATAGAGAAGAGAGTCTAGGGCCAAACCCAAGGAAGCTTCAACACACAGATGAGGAGACTGACAAAGGAAACTGTCAAAGAACAGTCAGAGAGAAAGGTCAGGAGAATGTTTTGATGGTGGGGGCCAGAATATGCTGTCCCAAAATAAGAAGGATTGTTGAGCTGAAGGCAAGTTAAAAGAAGC**AGATACAGGCTGGGTGCAGTGGGCTCATGCCTGTAATCT**TAGCATTTTGGGAGGCTAAAGTGGACAGATTGCCTGAGCTCAGGAGTTCGAGACCAGCCTGGGCAACATGATGGAACACCATCTCTACTAAAATACAAAAAAAAAAAAAAAAAAAAAAATTAGC**CAGGCATGGTGGCATGTGCCTG**TAGTCCCAGCTACTCAGGAGGCTGAGGCACGAGAATTACTTGGACCTAGGAGGCAGAGAC**TGCAGTGAGCTGATATCTCGCCACTGCA**CTCCAGCTTGGGCAACAGAGCAAGGCTCTATCTCAAATTTAAAAAATAATAATCATGATAATAATAAAAGAAGCAGATATAGGAGAGGTCTTCTGCCCTCCCTGTATTTGCCTAAAAAACCATAAATTTACAAAGACAAAAGTTATCCTACTTCCAC**CTCCCTCCTCTGCCTCCCACCACCAGGGAG**AACAAAGGTTAACCACTGAAGATAACTTTGGACTCTTATTGGCCTGGAAATGGTACTGCTTTACAAATTAGCCTTTATCTGCCATTCATTTGCCTTCCCCCAAGTAGCTACCCATTAGAGACTCAAAGTCCTTAAAGGTCTTTTCCTTTGTCTTACACTTCTTTAAAAATTTATTGTTCTTTGTTGAAGATGCTATATAAGCTGGAATTCTAAGCCACCTTTTTGAGAACTGCTCATTCTCTGGGTGTCTGT**CATGTATATATGAAATGTACATG**TTAATAAACTTCTGTTTCCTTTTTTTTCTTGTTAATCTGCCTTTTGTAACAGGGGTCCAGTCCATCTAAGAACTTATTGGGG**TTATTCTACATGACAAAGAACTCATGAGAATAA**GTGTTTCAAGAAGGGGATGACAAATGCTGCTGAGAGGTTTGATAAGATGAGGACTAAAATGTGACCATTGATTGTGGAAGTAATATAATGAACTTGACATGAGTTATGTTAGAAA**ATTACTGCTCAGGGACACCAAATTAGAGGAGACTGAGAAGTAAT**AGGAAATTGAAATAGCAGTTGTACCCAACCC**TAAACTGTAATTGAACATAAACAGTTTA**TTTGATGGAGCTTTTCACGTGTCAACCAAAGGAGCAGTGTTAGATACATAA**ACATTGCTATGATGTCAGAGTTCCAATGT**CATGGCCACTGGGATCACTTTGCCCAGACAGACCATTACCCTTCCTCTCCAGCTC**TGACTTCTACTCTTCTAGAAGTCA**CTGGTATACTTTATGGTGTGTCTTCCAATCAGACTCTTGGGAATGTCTTTTATTGAAAACTCCTGTGGATGCATAATCATTTTCCCAAGACTCAGGGCAGATTTCACCTCCTCTGGAACCCCATTAACTTTCTCAAGTATAATTAACAATTTCTTCTTCTGGAGTCCCACAATACTTTACATATACTTTAGTTATTGCATGTATTAAGTTGGTT**TCTAGTTGCTTGAGTGTATGTCCAACTAGA**CTGTGAGCACACTGAGGGAAGGAATTTGGCTTGGAAAGTAAAATTGGCTCCCAGTGTTTTATCTAAACAAATAGGCTCAATTTACAACAGGATTCCATCCTCTTACGCTGTGTATTCCCATCTCTTCCATGCTGCATCCCACCACCCTTCAATGTTATTACAATA**AAATTAAGATCAAATAATTT**ATTCAGCTAATTTTTCTTGTTTTGGACAAAATAGACATCCTAATACATTTATAGCCAAAGTTTAATCTAGACACTAAAAGGAGCATATTTTGCCTTTAGGCTCTGTATTTCCCATCCCGTTCCCAGTCAAGATAATCACAACTGTA**TTCTTTAGGAAATTCTTCACAAAGAA**CCAAACATATGTTACTATGAAACTAGATTCTTGGTATCCTGGTGAAAAAACCTGCCCTCTTGCTTGTCTGTAAATACTGGCCTTGGCTGGGTGTGGTAGCTCATGTCTATGATTCCAACACTTTGGGAGGCCAAGGTGGGAGGATCTCTTGAGCTTCAGTGTTTGAGACCAGCCTGGGCAACATAGTGAGATCCCATTTCTATAAAATTTTTTTTAAAAAGCTAGCCTGGCATGGGAGCTTGTGCCTGTAGTTCCAGCTACTTGGGAGGCTAAGATGGGAGGATTGATTGAGCACAGGAGGTCG**AGGCTGCAGTGACTGCATTTCAGCCT**GGGTGACAGAGGGAAATCCTGTCTCAAAAAAATTAAACAAATAAATAAATAAGTAAACACTTGCCTTGCCCTCAGACTTGAACAAACAACTCGTGATTTTTCTGGGGAGGTTCAGCCTGCCTATGCAGTACTCCTGTACTCTACCAGCAGTGTGGCATCAAGAGCATGTGTTGACCCTGTTAATGATTTGTAATGTTTTGTCTTTCAGAACCAGAGGCAGGAGAAGTGTCCCCTCCAGTCGGTGCGGGTGTCAACAGCAACAGCTGGACCTTTAAATACGGACCA**GGCAACCCCAAACAATCCGGTCCCGGTGAGTTGCC**CGACAAATTCATTATCCCAGGAT**CTCCTGCAATCATCTCCATCCGGCAGGAG**CCTACTAACAGCCAAATTGACAAAAGTGACTTCATAACCTTCGGCAAAAAGGAGGAGACCAAGAAAAAGAAGAAAAAGAAGAAGGGTAACAAGACCCAGGAGAAAAAAGAGAAAGGGAACAGCACGACTGACAACAGTGACCAGTGAGGTCCTCAAATGGAAACAAGCCACTTAGCCAGTTTTTGTAATAATGGCAAATCTCTCC**CATGTAGCAATTCCCTGCTCCTTTTTCCTATCTACATG**AGCCCTCTTAGAGACCTCAGAAATCTGCAGAAAGTTCCCTGTGTCTGTCTAGAACGCATTTAACAGGTTTTGTCGTAAAAGCTTTACTAAGTCTGGTGTTAACTCTTTCTCTCCACTCTGGCTTGTTTTCAGAACCTAAAAAGCAGACCCAAGTTTCCTT**TCTCCTCCGCCGCAAAGGAGA**GGCTTCCCAGCCCCGCCAGTGAGAGGTTGGACTCTCTGCCCTGTGCTCCGGGGATCCTGTCTTGATGACACTTGCAGGGCAGGCTGAAAAGTTTTGAGATTGAGCAGCTTGGGAGTTTGTGGCCACTGGGTATGTGTGGCTACCGCGGGTATGCGAGTGCCAGATATTGGCTGAGACGAGCCAGCTTAGACTAATTGGTACAAGGAAGGCAAGAAAACAAAGACAAATAAACAGCGGAAGTTATCAGTATGGAGGGGAAGTGTAAACTTAAAGGGACCAGACTTTCTAAATCTTACAACTCAAGAGGTGGCAGCCACCCTCTAGGAGACAAAACTACCCCCACTGACAAGG**CTTTAGGAGACCCTAAAG**TCTGTTGGCTGTGACGTCATTATACCTAAAATCTGCATCATACCTGCAAGCCAACAGTTCAGTGTTTTAACAGAGAACCACCCTG**GGAAACAGAAGCAGATCTGATGTGTTTCC**TATACATGTCCTGTGCTCACTTTATTAAAAATTCTTTTGCACACAATGTTTAT**GAAAAGGCCAGATCCTTTTC**CAATACTTATGCAAAAGCAAAAGAAAACCCCGACACCTCACCTTTCGCTGTTTGTTGTTTCAT**AGATTTATTTAAAAAAAGAGAAAGTCTATAGCTATAAATCT**TTAAAGAGAAATATGAATACAATTCCCCTAAACTCTCCTCAAAAGAGAATTCAGTCTACAGCCATTTAAATGATCATTGCTGC**TACAGAAGTGCTTTAAGAGAATTGCCTGAAACATCTGTA**TTATATCGGCCACCTGCCAATCACAGCTTTACTCTTTCAGGTCACTCTGGGGCTGCCTCTTGCATGTATTACTAAATAAAATGATCTCTCTTTCTCTCTCTCTCTCTCTTTTCTAAGAAACAATTATGTGCACTTTGATACACAACCTTCTCTAACCAACTATATATCAAGACCCAAAAATTGA**AGAAAAATATTGTTTTCT**CATACAGTGAGCAGATTTTTCAATCTACTAATTCTGTGACTTGTCTTGGTGTGCTAGCCTACACCTTCTCTTTGGTTTAGTTTTCCTTTTCTATAACACTCTGAATTGCTAATCTTACTAACACCTATGATGTTACCTGAAATCAATCTCCCATATGTATGCTGTATGCTATGCTAAGACTCCTGAAATATACTTACTCTGTGCTTGTGTATGTGAATGTTAATGCAACTATTACCTAGAGTGAACTTTAAGCTTTATTGTTGAATGTAATTCCATTATATTTCCTTTTGTACACCTGTGAAAAAGTGGAGTAGTGTTTTTTTAACCATTGTTAATCAGCTTTTGTGTAT**GAAAGACACAGTAAAATTTCTTTC**TTAAATCAAGATACTGGTGATTCAAGGAATTTTATTTATGGTCCAGCCAAGAGCCATCTCGTGCCAAGACTTCTGCTGGCAAGGGAATGGATAAAGCTGTTTTGTTCTAGTAACAATTT**TGGAATGAATACTGACAATATTCCA**TGAGGGTGTGCAAGCACAAATTTTACCAATCTGACCTCTTTGAAGTTGCAGAATGCTTTGAAATTCTAATGGTATCTGAAATATCAGCTCATAGAAAG**TAACAAAATTTGCTGTCACCTTAAATAAGACATTTTAATTTTGTTA**TAATGTACAATTTAGAAGTTTGATTAA**TTATATTATCTATTTAGGCATTAATATAA**AAGAGGTAGGAGTCTGTT**ATTTAAAAAAAGCATTAAAT**TTAAAAAAAAACTGTCTTGTCTACTTTTAGCTTCATTCTCCCATATTTTGAAGGGTGTGTAACTTCAGCTCTGCAGGATTGCATGGGGTAAAACTTGTTACCAACACATGTGAACCATTGCTACATTGTAGGTTGTGATCATTTTGCCCCACTGAAGCCCATGTATCTGACCTTACGTGCCTTTTGAACTAGGAGAATCGGGCTAATTTATTAATGATGATAATTATAATGTATCTGTAC**AGCACTTTTTACATTTGCGAAGTGCT**TTCCAATCCATGTTAGTTACTAGTTATTACAGCTGTAAGGATAAAACACGTCATGTGGATTCATTTTGAATTGGTGCTATTGGTATTTCCTCTGTTATTGCTAATAAATGAAAATGGTGGTATGaa**agaaatggtggtcatttct**aagaataggaggaaatagaacactgata**agcaattaatactgaagaagattgct**gcagtatgaaactcactgtaacactctcttatcacgtta**gcttttatgtccattactaaggcaaaagc**cccagtggaatcaagattcaaacttcatatcattagaatgggcagccaggaggttgctgcgtcactttgtagttacactatttcagttttctctaataagaattgcaacaaacccatttttatcacaagagcccttctaggcatctttttcattacttgtttagcccttgcacctctctacaattggaaagccaagtgacaccccaaagaaatgaacttttaggtcagggatagtggctcttgcctgtaatctcagaactttggaaggccaaggccagaggatggcttgaggccaggagttcaagatcagcctgggcagcatagcgagatcccat**ctgtacaaaaagaaataacttttattgagtacag**ggtgtatgccagccacactactgcataggttcagctgt**gacaaatagattgttttgtc**cttttgtagttaaatttgtataaatataagctttagatgctgtaggttctcagttaggagcaattttgccctcctggggacatttggcaacgtctgaagtcattgttgcttttcattacttgaagtggagggtgctattggcatctactgggtagggggcaacgatgctgtgaaacacctggcaatgcacaggacagccccctataacaaaggatcttacagcccaacatgtcaatg**tctcaaatcaagattgaga**aaccttgagatagagtgaccatataatttgtgtagtccacttttgaaatggaacaggagacaccattcgttattacttcaggattcaagatgtaaactgggactg**tcccaggctaactgggatttatagggactctataaat**aaactggtatagctgtgccactaactagatgg

Uppercase: PCDHACT-2

Lowercase: Flanking sequence[1000bp]

Red & Bold & Underline: Stem-loop [222]

Blue: Heptamer[368]

Green: Nonamer [49]

id-TRDC[C_gene_segment]

atgatcattgttatcctctgattaaagactcaaaggaaactatacagtagtcagtcccctacagaagaatgagtattcacctaaaggcagaaacatgtaccacctaccttcagcagatccctttccaccctttccagcagatct**ggaagggatggcacccttcc**gggaccccaacactgccatc**ctcagaacctctgag**ccagttatgcagtccttctcagaccctcgattctttaagtgacagaatgcatagcatgagcacctgccctgatctaccatccaagggcccatgcatttacaaaaaaccaactggtgcagagagatattcagcctgacctagattgtccagaggattgattaataaaacacagataatcg**gctggatgcagtggttcacacctgtaattccagc**actttgggaggccaaggtgggcggatcacctgaggtcgggagttcgagaccagcctgaccaacatggagaaaccccatctctactaaaaataaaaaaaat**tagctgggcatggtgatgcatgcctgtaatcccagcta**ctcgggaggctgaggcaggagaatcacttgaacccgggaggcggaggttgcagtgagccgagattgcaccattgcactccagcctgggcaacaagagtgaaactctatctcaaaaaaaaagaaaaagaaaaaaaagaaagaaaaagaaaaaagccacagataatcgaaacccagatgatccagaaacttctttgtagttgtcattttccatctgtg**cttccagccctatctcctagcagctcctaactggaag**tcaaacttactgcctttcatttctgctccccttgcctgtcctttcacagaggtggtaataagcccaggtcactaacaggatgcatggaggtctggattgtagtgtttggctccagggtaatcgaggtaatcaccactgtttaacccccacaaagttgtga**ataatcatctcacctaataagttgattat**atttgcagGAAGTCAGCCTCATACCAAACCATCCGTTTTTGTCATGAAAAATGGAACAAATGTCGCTTGTCTGGTGAAGGAATTCTACCCCAAGGATATAAGAATAAATCTCGTGTCATCCAAGAAGATAACAGAGTTTGATCCTGCTATTGTCATCTCTCCCAGTGGGAAGTACAATGCTGTCAAGCTTGGTAAATATGAAGATTCAAATTCAGTGACATGTTCAGTTCAACACGACAATAAAACTGTGCACTCCACTGACTTTGAAGTGAAGACAGATTCTACAGGTAGGCCATTTCTAGCTTCAAGGAGCTGGAGATTATGGGGAACAAGAATTGGGTGAAAGGGAAGTTAGAGATGTAACTGTGGACAAATCATTCTCAGTATAGCATCATGCTGGAAATAAGACTTAGGCCCAACTATAG**CCTGCCATTGGCAGG**GGAGGGAAATGCTTGTCATCCC**TAAGATGGAATCTAAAATAAAGCCCATCTTA**TTTCTTCCTCATCTCTCCTCTTTACCTACCACTGTCCCCTTCATACTAGACTCTGGGATTGAAAGTCCTCGTGCATTCTAATCCAGTGCTAAATTCCAACAAAGGGCAATGCGGCCTATTGTGGGGCTAAGAATCCTAGTTCTTCCCGCATATTGGTGTGCCTGGATGTTGACCTCATGGTACAATGAAAAAGGCTAGAATGGGAAGCATGTTGTATAATTAGCT**ATGTGACCTTCACCAAGTCACAT**CCTCTGGTCCTCAGTTCACTCACCCAAAGTCTCCTCCAGTTTTAAAAGCCTACAATCCTGTGAGCCTCTTCATTCCCAATGTAACCCTGACCACTGCTGTTTGTTCCAGATCACGTAAAACCAAAGGAAACTGAAAACACAAAGC**AACCTTCAAAGAGCTGCCATAAACCCAAAGGTT**AGTTCAAATCAAAGGGCCAACTTCAGAATCAAGGGTTAAAGCAAACTCTGTAATTGTCCACTGGGGCCAAAATGTATCAGATTTCAAAAGAAAACACAAACCATGCTGAGCCCAGTTGGTTCTGCAGAGCCTCTGACTCTCCCAGCAAGGCTTACCTGGCTGTCGGCATCGGTTTCCCAAGCTTGCCTGGCTGGGTGTCAGCAGCAGCCTAAGGTCTCTGACGGCTA**TCTGGCACCATGCTGGACTTGCAGGCCAGA**AAGAACTAGGCAGACTTGGGGCAGGCTAATATCAGTGTTGCAGGCCCAATTTCTGGAAGGGAATTTAATCTCTCCCTGAAGCCACCCTCTTCACTCTTTTTCAGCCATAGTTCATACCGAGAAGGTGAACATGATGTCCCTCACAGTGCTTGGGCTACGAATGCTGTTTGCAAAGACTGTTGCCGTCAATTTTCTCTTGACTGCCAAGTTATTTTTCTTGTAAGG**TAAGAATTAGCCGCTTCTTA**TTCCTATCTCTACCCACACCATGCTGCACATGGGGAAAGGGGATTTAGAAATGGCTAAGAAACCAAGAAGCATTGGAAAGGGAAAATAGAGACATACTTAGAATTAATAAGGTAGGCTATCTCAGCTACTCAGCAAAACAGCACCCCATTTCCTGTTATCCTTTCATTTTGAGTCAGACTGAAACCCATTAGGTAGTATAATGTACTATTTTTCTAATATACCAGAGAATAACCCTCACTTTCACTCCAGGGAGAAAACT**GAAGATAAATAAGGAGAATTCATCTTC**TTTACTGCTATTTTGTTCATCGACCAAGAGACCATGGGTTATTTGCAAGGCCATAAGTAAAGCTTTGCCTGAAGACTATCCAGAGGAAGCAGAACCTATCTTTGTCCTGATAAAGGGAATGCCAGCTCACACTCCTTAATCATAGGACCAGCCTGTCTA**TGTGTGGATCAGCCTGTCTTTGGGATGAACCTACACACA**GCATTAG**TAGTGAACAGCTCACTA**TTAGCTACTCAGGATGCATACACAGTTAGGTTTGAGATGTCAATTATTTGCCTGTTACTTGCATTCCGTTCTCAGGTAATCTCATACACGAAATTGAAACCAGGGCCTGGGTTTGCGT**GTGAGGCTTCTCCCCCTCAC**TCCTTCTCCAATACTCACAAATGCTCATCAACCCCAGCTTCTGT**TCTAGGAACAACCCTAGA**CTCAGAGTACT**TGACTGATCTGACCCAGACAGTAAACATACCCAGTCA**TCTCAAAAAGGGGGTCCAGCTCAC**CTGTTATTTTACTAACAG**TCCCAGGGACACAGGCAGTGTGGCTGGGTAAAAAGAGAGCTATACTAAGAGTCAGAAGATCTGGGTTTTATCCTGATTTTGACACTAACTTACATGTTACCTTAGCCAGGTTTCTTCATCTCTCTGGGCCTGGGT**TTTTGCATCTGCAAAA**CAAATGAGTTAAAAAAAATAATAACCAAAGGAGAACAAAAGAGTCTCATGACTTGTGCCAACACTG**ATAGCTGAACTAGCTGAGCTATTGCCATGTGTACATGGAAAGTGAGCCACAATAG**ATTAGTGATGTCTGCAATGGTACAGTATAGCAAGTACCGGCCAACCCCAGACTACATGACCCCTACGATTCCTTCTAGCACTGGCAGTCTATTCTATAACAACTTTCTTCACTGCAGGCTGACTGGCATGAGGAAGCTACACTCCTGAAGAAACCAAAGGCTTACAAAAATGCATCTCCTTGGCTTCTGACTTCTTTGTGATTCAAGTTGACCTGTCATAGCCTTGTTAAAATGGCTGCTAGCCAAACCACTTTTTCTTCAAAGACAACAAACCCAGCTCATCCTCCAGCTTGATGGGAAGACAAAAGTCCTGGGGAAGGGGGGTTTATGTCCTAACTGCTTTGTATGCTGTTTTATAAAGGGATAGAAGGATAT**AAAAAGATATAGGACTCTTTTT**TTACTCCTACAAGTGATACACTTTGAAAATGATGTTTTGTTCCTTTTG**ACTTTCTTTACCTTTTGAAGTAGAAAGT**GGGAACCAACAGGTTCACAGCTTCATTCCTCATGAGGCAAATAGGCCTTGGGAGAAGAAGAGCGGGTGCCCTTTTATCTAAACATGGAAGGCTCTGCTCAACTGAGCACTAGATTTGCTACAAACCAGCATCATCTTCTTCCTCCTGTCCTCACGGCTTGTCCCACCCTCTATGTTCACTTCAGGAGCCACACTAGAGATTCTGCATGGCGTGGAGGAGGACAAAGTTTCAGCACTTTCTGCCTCTCCTAATACTTTA**CAAATGAGATTACATTTG**AATTTGCTAATACTTTATGAGCAGGCAATGAGGTTTCCAAAATCTCATCTAAA**TACTCTCCAATCTATTAGCAAAAATCAGAGTA**AAATACAGAGGAAAGGCACTGCTTTC**TGTTAATTGATTTAACA**TGCATGAATTAGCTCCCTCTGAGTTCCAGGCACTATGCTGAGAGTACAAAGAAGACACAAGTCTG**CTTTCAAGCAACTCACTGTGAAAG**TGTTTTTGAAGGGAGGAACAGAAATGAGACCCCTATCTTTCCCTA**TAAAAACAACATTTTTA**CTGTCTTTTGCCTGCCAATCTGTATTTGAAACCATTGGACACTGATTCTCTGGCCTGGGACTTTGGCATTGATGGTTTTCTGCCTTTCTTCTCAGCCTCTGCCTCTATTGCATTTATTAAACTGCATTGTGTGCacctcgcctctggctttactctttgcagatcaccacagggggaaactcagctctgtgagctcactattagtcagccaaaagccaaattgaacttatgggtcacactgcactttcctgagccccaagcctacagaggccctgccttggagacccagcctcgggtttcctgctgcct**ctgctgatgccgctgttacagcaagagcaacagcag**cacctgcagc**gtggccagtgacctctgctactgccacccaggaaggccac**agccttttccaggagcactgctc**aggtgcagagccctagcaacccattattgtggttctctcagcacct**atgtggagcatcaagtcgggggttatgaggttacaagtgaaagagtga**ctgtcgccaacaactccacttgccatccccgacag**caaactcccccatcatgtctactacaccttgtatgtgtaaggaatgctgatttccaaaacattcacgagttaccttgtgtaatattagggtgtttcctctgaagttgttaggaacaaaaagatcatattct**catttttcagataaaacaaaaatg**ccctctcctcaaaaaagtatgtatcatgtccagagtcattcagtgaggcagggatattgctgggagcagaaccaaggtctcaagattttcagcccaaagttcctgctataagaacatactgcatccccaaatgaggacaagatgtcaaagtgagtatcagctagaatgaacttcaggattgtcccatatttagcatataaatgagccaccctctagcccac**tatgccagaaacaaaaaaaatcctggcata**tcacaggaccatgaatttcccacaccaaaaggagcctccagggaagattt**aaggcaagagccaggagggattatggctgcctt**ctgatctggggccatgatatgtgaactgtgagatggacatgtgtcacactcagtaggcattagaacagaagccccagaaagccaagccacatgcccaa

Uppercase: TRDC

Lowercase: Flanking sequence[1000bp]

Red & Bold & Underline: Stem-loop [36]

Blue: Heptamer[52]

Green: Nonamer [12]

id-IGHG4[C_gene_segment]

gaaatggggcctccctgtggcctgggggtcctggcaccatgcagggtggggagggccaagggcaggtgcaaggctcctacctg**tgctggggggcctgggttgagcccagca**gggaccttgccgggggaagctctggagagagggaggaggtgggctggtggccgagaaggccaggccagggctgggagggtgaggttgtggtgactga**gcctccagaagtaatgcaggacactgggaggc**agggggcatccaggcactcagggccctgacctgggctgctgcacactggggctaaggggaaaggaggggagaggctgaggaggaggctccaggaggctattccaaggcagggggttccggggccctggggctgaagggcgccgaccctatgcagtgtctggc**ccctctgctgcacagaagaaaagggccttggagggcagaggg**caggctatgaccag**ggccctgggcaagtcaggcccactcactagcggagggcc**acgctggggcggcagggtcaggagcttcaggggactcgggggacccacgagaagccatctgagaacagtgtccactggtcaagccaggcacccataaaaggctggagtggggccaatgggcatgagccgtccctgaggtggcaccgatggccagagctgaggccaagctagagacactggactgtgctgactcc**cggcaggcacagagcgctgacctggctgccg**agccccgccccctagg**ctgcaggggtgcctgcag**aagggcaccacagggccaccggtcctgcaagctttctggggcaggccgggcctgactttggctgggggcagggagggggctaaggtgacgcaggtggcgccagccaggcgcacacccaatgcccgtgagcccagacactggaccctgcatggaccat**cgcagatagacaagaaccgaggggcctctgcg**ccctgggcccagctctgtcccacaccgcggtcacatggcaccacctctcttgcagCTTCCACCAAGGGCCCATCGGTCTTCCCCCTGGCGCCCTGCTCCAGGAGCACCTCCGAGAGCACAGCCGCCCTGGGCTGCCTGGTCAAGGACTACTTCCCCGAACCGGTGACGGTGTCGTGGAACTCAGGCGCCCTGACCAGCGGCGTGCACACCTTCCCGGCTGTCCTACAGTCCTCAGGACTCTACTCCCTCAGCAGCGTGGTGAC**CGTGCCCTCCAGCAGCTTGGGCACG**AAGACCTACACCTGCAACGTAGATCACAAGCCCAGCAA**CACCAAGGTGGACAAGAGAGTTGGTG**AGAGG**CCAGCACAGGGAGGGAGGGTGTCTGCTGG**AAG**CCAGGCTCAGCCCTCCTGCCTGG**ACGCACCCCGGCTGTGCAGCCCCAGCCCAGGGCAGCAAGGCAGGCCCCATCTGTCTCCTCACCCGGAGGCCTCTGACCACCCCACTCATGCTCAGGGAGAGGGTCTTCTGGATTTTTCCACCAGGCTCCGGGCAGCCACAGGCTGGA**TGCCCCTACCCCAGGCCCTGCGCATACAGGGGCAGGTGCTGCGCTCAGACCTGCC**AAGAGCCATATCCGGGAGGACCCTGCCCCTGACCTAAGCCCACCCCAAAGGCCAAACTCTCCACTCCCTCAG**CTCAGACACCTTCTCTCCTCCCAGATCTGAG**TAACTCCCAATCTTCTCTCTGCAGAGTCCAAATATGGTCCCCCATGCCCATCATGCCCAGGTAAGCCAACCCAGGCCTCGCCCTCCAGCTCAAGGCGGGACAGGTGCCCTAGAGTAGCCTGCATCCAGGGACAGGCCCCAGCCG**GGTGCTGACGCATCCACCTCCATCTCTTCCTCAGCACC**TGAGTTCC**TGGGGGGACCATCAGTCTTCCTGTTCCCCCCA**AAACCCAAGGACACTCTCATGATCTCCCGGACCCCTGAGG**TCACGTGCGTGGTGGTGGACGTGA**GCCAGGAAGACCCCGAGGTCCAGTTCAACTGGTACGTGGATGGCGTGGAGGTGCATAATGCCAAGACAAAGCCGCGGGAGGAGCAGTTCAACAGCACGTACCGTGTGGTCAGCGTCCTCACC**GTCCTGCACCAGGAC**TGGCTGAACGGCAAGGAGTACAAGTGCAAGGTCTCCAACAAAGGCCTCCCGTCCTCCATCGAGAAAACCATCTCCAAAGCCAAAGGTGGGACCCACGGGGTGCGAGGGCCACATGG**ACAGAGGTCAGCTCGGCCCACCCTCTGCCCTGGGAGTGACCGCTGT**GCCAACCTCTGTCCCTACA**GGGCAGCCCCGAGAGCCACAGGTGTACACCCTGCCC**CCATCCCAGGAGGAGATGACCAAGAAC**CAGGTCAGCCTGACCTG**CCTGGTCAAAGGCTTCTACCCCAGCGACATCGCCGTGGAGTGGGAGAGCAATGGGCAGCCGGAGAACAACTACAAGACCACGCCTCCCGTGCTGGACTCCGACGGCTCCTTCTTCCTCTACAGCAGGCTCACCGTGGACAAGAGCAGGTGGCAGGAGGGGAATGTCTT**CTCATGCTCCGTGATGCATGAGGCTCTGCACAACCACTACACACAGAAGAGCCTC**TCCCTGTCTCTGGGTAAATGAgtgccagggccggcaagcccccgctccccgggctct**cggggtcgcgcgaggatgcttggcacgtaccccg**tgtacatacttcccgggc**gcccagcatggaaataaagcacccagcgctgccctgggc**ccctgcgagactgtgatggttctttccacggg**tcaggccgagtctgaggcctga**gtggcatgagggaggcagagcgggtcccactgtccccacact**ggcccaggctgtgcaggtgtgcctgggcc**gcctagggtggggctcagccaggggctgccctcggcagggtgggggatttgccag**cgtggccctccctccagcagcacctgccctgggctgggccacg**agaagccctaggagcccctgg**ggacagacacacagcccctgcctctgtaggagactgtcc**tgttctgtgagcgccctgtcctccgaccc**gcatgcccactcgggggcatgc**ctagtccatgtgcgtagggacaggccctccctcacccatctacccccacggcactaacccctggcagccctgcccagcctcgaacccacatggggacacaaccgactccggggacatgcactctcgggccctgtggagggactggtccagatgcccacacacacactcagcccagacccgttcaacaaaccccgcactgaggt**tggccggccacacggcca**ccacacacacacgtgcacgcctcacacacggagcctcacccgggcgaaccgcacagcacccagaccagagcaaggtcctcgcacacgtgaacactcctcagacacaggcccccacgagccccacgcggcacctcaaggcccacgagccgctcggcagcttctccacatgctgaccagctcagacaaacccagccctcctctcacaaggtg**cccctgcagccgccacacacacagggg**aacacacgccacgtcgcgtccctggcactggcccacgtcccaatacagcccttccctgcagctggggtcacatgaggggtg

Uppercase: IGHG4

Lowercase: Flanking sequence[1000bp]

Red & Bold & Underline: Stem-loop [32]

Blue: Heptamer[46]

Green: Nonamer [1]

id-IGHG3[C_gene_segment]

aatggggcctccctgtggcctgggggtcctggcaccacgcagggtggggagggccaagggcaggtgcaaggctcctacctg**tgctggggggcctgggttgagcccagca**gggaccttgccgggggaagctctggagagagggaggaggtgggctggtggctgagaaggccaggccagggctgggagggtgacggtgtggtgactga**gcctccagaagtaatgcaggacactgggaggc**agggggcatccaggcactcagggccctgacctgggctgctgcacactggggctaaggggaaaggaggggagaggctgaggaggaggctcccggggcgatattccaaggcagggggttccggggccctggggctgaagggcgccgaccctatgcagtgtctggc**ccctctgctgcacagaagaaaagggccttggagggcagaggg**caggctatgaccag**ggccctgggcaagtcaggcccactcactagcggagggcc**acgctggggcggcagggtcaggagcttcaggggactcaggggacccacgagaagccatctgagaacagtgtccactggtcaagccaggcacccataaaaggctggagtggggccaatgggcatgagccgtccctgaggtggcaccgatggccagagctgaggccaagctagaggccctggactgtgctgactcc**cggcagacacagagcgctgacctggctgccg**agccccgcctcctagg**ctgcaggggtgcctgcag**aagggcaccacagggccaccggtcctgcaagctttctggggcgggccgg**gcctgaccttggctttggggcagggagggggctaaggtgaggc**aggtggcgccagccaggcgcacacccaatgcccgtgagcccagacactggaccctgcctgga**ccctcgtggatagacaagaaccgaggg**gcctctgcgccctgggcccagctctgtcccacaccgcagtcacatggcgccatctctcttgcagCTTCCACCAAGGGCCCATCGGTCTTCCCCCTGGCGCCCTGCTCCAGGAGCACCTCTGGGGGCACAGCGGCCCTGGGCTGCCTGGTCAAGGACTACTTCCCAGAACCGGTGACGGTGTCGTGGAACTCAGGCGCCCTGACCAGCGGCGTGCACACCTTCCCGGCTGTCCTACAGTCCTCAGGACTCTACTCCCTCAGCAGCGTGGTGACC**GTGCCCTCCAGCAGCTTGGGCAC**CCAGACCTACACCTGCAACGTGAATCACAAGCCCAGCAA**CACCAAGGTGGACAAGAGAGTTGGTG**AGAGGCCAGCGCAGGGAGGGAGGGTGTCTGCTGGAAG**CCAGGCTCAGCCCTCCTGCCTGG**ACGCATCCCGGCTGTGCAGTCCCAGCCCAGGGCACCAAGGCAGGCCCCGTCTGACTCCTCACCCGGAGGCCTCTGCCCGCCCCACTCATGCTCAGGGAGAGGGTCTTCTGGCTTTTTCCACCAGGCTCCGGGCAGGCACAGGCTGGA**TGCCCCTACCCCAGGCCCTTCACACACAGGGGCA**GGTGCTGCGCTCAGAGCTGCCAAGAGCCATATCCAGGAGGACCCTGCCCCTGACCTAAGCCCACCCCAAAGGCCAAACTCTCTACTCACTCAG**CTCAGATACCTTCTCTCTTCCCAGATCTGAG**TAACTCCCAATCTTCTCTCTGCAGAGCTCAAAACCCCACTTGGTGACACAACTCACACATGCCCACGGTGCCCAGGTAAGCCAGCCCAGGCCTCGCCCTCCAGCTCAAGGCGGGACAAGAGCCCTAGAGT**GGCCTGAGTCCAGGGACAGGCC**CCAGCAGGGTGCTGACGCATCCACCTCCATCCCAGATCCCCGTAACTCCCAATCTTCTCTCTGCAGAGCCCAAATCTTGTGACACACCTCCCCCGTGCCCACGGTGCCCAGGTAAGCCAGCCCAGGCCTCGCCCTCCAGCTCAAGGCAGGACAAGAGCCCTAGAGTGGCCTGAGTCCAGGGACAGGCCCCAGCAGGGTGCTGACGCGTCCACCTCCATCCCAGATCCCCGTAACTCCCAATCTTCTCTCTGCAGAGCCCAAATCTTGTGACACACCTCCCCCATGCCCACGGTGCCCAGGTAAGCCAGCCCAGGCCTCGCCCTCCAGCTCAAGGCGGGACAAGAGCCCTAGAGTGGCCTGAGTCCAGGGACAGGCCCCAGCAGGGTGCTGACGCATCCACCTCCATCCCAGATCCCCGTAACTCCCAATCTTCTCTCTGCAGAGCCCAAATCTTGTGACACACCTCCCCCGTGCCCAAGGTGCCCAGGTAAGCCAGCCCAGGCCTCGCCCTCCAGCTCAAGGCAGGACAGGTGCCCTAGAGTGGCCTGCATCCAGGGACAGGTCCCAGTCG**GGTGCTGACACATCTGCCTCCATCTCTTCCTCAGCACC**TGAACTCCTGGGAGGACCGTCAGTCTTCCTCTTCCCCCCAAAACCCAAGGATACCCTTATGATTTCCCGGACCCCTGAGG**TCACGTGCGTGGTGGTGGACGTGA**GCCACGAAGACCCCGAGGTCCAGTTCAAGTGGTACGTGGACGGCGTGGAGGTGCATAATGCCAAGACAAAGCCGCGGGAGGAGCAGTACAACAGCACGTTCCGTGTGGTCAGCGTCCTCACC**GTCCTGCACCAGGAC**TGGCTGAACGGCAAGGAGTACAAGTGCAAGGTCTCCAACAAAGCCCTCCCAGCCCCCATCGAGAAAACCATCTCCAAAACCAAAGGTGGGACCCGCGGGGTATGAGGGCCACATGGA**CAGAGGCCAGCTTGACCCACCCTCTG**CCCTGGGAGTGACCGCTGTGCCAACCT**CTGTCCCTACAGGACAG**CCCCGAGAACCACAGGTGTACACCCTGCCCCCATCCCGGGAGGAGATGACCAAGAAC**CAGGTCAGCCTGACCTG**CCTGGTCAAAGGCTTCTACCCCAGCGACATCGCCGTGGAGTGGGAGAGCAGCGGGCAGCCGGAGAACAACTACAACACCACGCCTCCCATGCTGGACTCCGACGGCTCCTTCTTCCTCTACAGCAAGCTCACCGTGGACAAGAGCAGGTGGCAGCAGGGGAACATCTT**CTCATGCTCCGTGATGCATGAGGCTCTGCACAACCGCTTCACGCAGAAGAGCCTC**TCCCTGTCTCCGGGTAAATGAGTGCGACGGCCGGCAAGCCCCCGCTCCCCGGGCTCT**CGGGGTCGCGCGAGGATGCTTGGCACGTACCCCG**TGTACATACTTCCCGGGCACCCAGCATGGAAATAAAGCACCCAGCGCTGCCCTGGGCCCCTGCGAGACTGTGATGGTTCTTTCCACGGG**TCAGGCCGAGTCTGAGGCCTGA**GTGGCATGAGGGAGGCAGAGCGGGTCCCACTGTCCCCACACT**GGCCCAGGCTGTGCAGGTGTGCCTGGGCC**GCCTAGGGTGGGGCTCAGCCAGGGGCTGCCCTCGGCAGGGTGGGGGATTTGCCAG**CGTGGCCCTCCCTCCAGCAGCAGCTGCCCTGGGCTGGGCCACG**GGAAGCCCTAGGAGCCCCTGG**GGACAGACACACAGCCCCTGCCTCTGTAGGAGACTGTCC**TGTCCTGTGAGCGCCCTGTCCTCCGACCC**GCATGCCCACTCGGGGGCATGC**CTAGTCCATGTGCGTAGGGACAGGCCCTCCCTCACCCATCTACCCCCACGGCACTAACCCCTGGCAGCCCTGCCCAGCCTCGCACCCGCATGGGGACACAACCGACTCCGGGGACATGCACTCTCGGGCCCTGTGGAGAGACTGGTCCAGATGCCCACACACACACTCAGCCCAGACCCGTTCAACAAACCCCGCACTGAGGT**TGGCCGGCCACACGGCCA**CCACACACACACGTGCACGCCTCACACACGGAGCCTCACCCGGGCGAACCGCACAGCACCCAGACCAGAGCAAGGTCCTCGCACACGTGAACACTCCTCGGACACAGGCCCCCACGAGCCCCACGCGGCACCTCAAGGCCCACGAGCCGCTCGGCAGCTTCTCCACATGCTGACCAGCTCAGACAAACCCAGCCCTCCTCTCACAAGGTGCCCCTGCAGCCGCCACACACACACAGGCCCCCACACACAGGGGAACACACGCCACGTCGCGTCCCTGGCACTGGCCCACTTCCCAATACAGCCCTTCCCTGCAGCTGGGGTCACATGAGGTGTGGGCTTCACCATCCTCCTGCCCTCTGGGCCTCAGGGAGGGACACGGGAGACGGGGAGTGGGTCCTGCTGAGGGCCAGGTCGCTATCTAGGGCCGGGTGTGTGGCTGAGTCCCGGGGCCAAAGCTGGTGCCCAGGGCGGGCAGCTGTGGGGAGCTGACCTCAGGACACTGTTGGCCCATCCCGGCCGGGCCCTACATCCTGGGTCCTGCCACAGAGGGAATCACCCCCAGAGGCCCGAGCCCAGCAGGACACAGCACTGACCACCCTCTTCCTGTCCAGAGCTGCAACTGGAGGAGAGCTGTGCGGAGGCGCAGGACGGGGAGCTGGACGGGCTGTGGACGACCATCACCATCTTCATCACACTCTTCCTGTTAAGCGTGTGCTACAGTGCCACCGTCACCTTCT**TCAAGGTCGGCCGCACGTTGTCCCCAGCTGTCCTTGA**CATTGTCCTCCATGCTGTCACACACTGTCCCTGACACTGTCCCCAGGCTGTCCCCACCTGTCCCTGACACTGTCCCCCACGCTCTCACAAACTGTCCCTCACACTGTCCCCCATGCTGTCACAAACTGTCACTGACACTGTCCCCCATGCTATCCCCACCTGTCCCTGACACTGTCCCTGACACTGTCTCTCATGCTGTCCCCACTCATCTGCGACACTGTACCCCACGCTGTCCCCACTTGTCCTCAACAATGTCCCCCATGCTGTCCCCACCTGTCCCTGATGCTGTCCCCCACACTGTCCCAATCTGTCCCCACCACTCTCCCCCACGCTGTCCCCACCTGTCCCTGACACTGTCCCCCATGCCATCCCCATCTGTCCCGACAATGTCCCCAGGGTGTCCCCAGCTGTCCCTGATGCTGTCCCCCACACTGTCCCCACCTCTCCCTGACGCTGTCCCCCACGTGGTCCCCACTTGTCCCTGATGCTGTCCCCCACACTGTCCCCACCTGTCCCTGACACTGTCCCCCATGCCATCCCCATCTGTCCCGACAATGTCCCTATGGTGTCCCCAGCTGTCCCTGATGCTGTCCCCCACACTGTCCCCACCTGTCCCTGACGCTGTCCCCCACACTGTCCCCACCTCCCCCTGACACTGTCCCCCACACTGTCCCCACCTCTCCCTAACACTGTCCCACACACTGTCCCCTCCTGTCCCCAACACTTTCCCCCATGCTGTCCCCACCAGTCCCCAACACTGTACACCATGCTTTTCCCACCTGTCCCCAACACTGTCCCCCATGCTGTCCCCTCCTGTCCCCAACAATGTCCCCCATGCTGTTTCCTCCTGTCCCCAACACTGTCCGCCACTCTGTTTCCTCCTTTCCCTGACACTGTCCCCCACTCTGTCCCCACCTGTAGCCAACACTATCCCCTACGCTGTCTCCACCTGTCCCTGATGCTGTCCCCCACACTGTCCCCACTCCTCCCTGACACTGTCCCCTATGCTGTCCCCACCGGTTCCTAACACTGTCCCCCACACTGTCCCTACCTGTCCCCGACACTTTCTCCCATGCTGTTCCCACGTGTCTCCAACACTGTCCCCCACACAGTCTCCACCTGTCCCTGACACTGTCCCCCATGCTGTCCTCACCCATCTCTGACACTGTACACATACTGTCCCCACCTGTCCCTGATGCTGTCCTCCATGATGTCCCCACCTCTCCCTGACACTGTCACCCATGCTGTCCCCACCTGCCCCTGACACTCTCCTCCACGCTGTTCTCACCTGTCCCCAACACTCTCCCCCACACTGTCTCCACCTGTCCCTGACACTGTCCTCCACGCTGTCCCCACCTATCCCTGACACTGTCCCCCATGCTGTCCTCACCTGTCCCCAACACTCTCCTCCACACTGTCCTCACCTGTCCCCAACACTCTCCCCCCACACTGTCTCAACCTGTCCCTGACACTGTCCCCCATGCTGTCCTCACCTGTCCCTGACACTGTCCCCCATGCTGTCCTCACCTGTCTCTGACACTGTCCCCCGTGCTGTCCCCACCTGACACTATCTTCTGTGCTGTCCACATGCTGTTGCTGCCCTGGCTCTGCTCTCCATGT**CCAGGCCTCAGAGCAGGCAGTGGTGAGGCCCTGG**CACATGGGTGGCATGAGGGGCCGGATAGGCCTCAGGGGCAGGGCTGTGGCCTGGGTGGCCTG**AGGGGTGAGCAGGCCTCGGGGGCAGGGCTGTGGCCTCGCTCACCCCT**GTGCTGTGCCTTGCCTACAGGTGAAGTGGATCTTCTCCTCGGTGGTGGACCTGAAGCAGACCATCATCCCCGACTATAGGAACATGATTGGGCAGGGGGCCTAGggccaccctctgcggggtgtccagggccacccagatcccacacacgagccgtgggccatgctcagccaccacccaggccacaactgcccccgacctcaccgccctcaaccccatggctctctgtctttgcagtcgccctctgagccctgacacgccccccttccagaccctgtgcatagc**aggtctaccccagacct**ccgctgcttggtgcatgcagggagct**ggggaccaggtgtcccc**tcagcaggatgtccctgccctccagaccgcca**gatgctcacacaaaaggaggcagtgaccagcatc**cgaggcccccacccagg**caggagctggccctggagccaaccccgtccacgccagcctcctg**aacacaggcgtggtttccagatggtgagtgggagcatcagccgccaaggtag**ggaagccacagcaccatcaggccctgttggggaggcttcc**gagagctgc**gaaggctcactcagacggccttc**ctcccagcccgcagccagccagcctc**cattccgggcactcccgtgaactcctgacatgaggaatg**aggttgttctgatttcaagcaaagaacgctgctctctggctcctgggaacagtctcggtg**ccagcaccaccccttggctgcctgcctacactgctgg**attctcgggtggaactggacccgcagggacagccag**ccccagagtccgcactgggg**agagaaggggccaggcccaggacactgccacctcccacccactccagtccaccgagatcactcagagaagagcctgggccatgtggccactgcaggagccccacagtgcaagagtgaggatagcccaaggaaggg**ctgggcatctgcccag**acaggcctcccagag**aaggctggtgaccaggtcccaggcgggcaagactcagcctt**ggtg**gggcctgaggacagaggaggccc**aggagcatcggggagagaggtggagggacaccgggagagccaggagcgtggacacag

Uppercase: IGHG3

Lowercase: Flanking sequence[1000bp]

Red & Bold & Underline: Stem-loop [44]

Blue: Heptamer[124]

Green: Nonamer [2]

id-IGLC3[C_gene_segment]

aagccttgtt**ctgttctggcctcctcagtctgggttcttgtcggaacag**ctttgcccttgggttacctgggttccatctcctggggaattgggaacaaggggtctgagggaggcacctcctgggagactttagaaggacccagtgccctcggggctgatgctcgggaatcacagagctgggacccagagccaggatccagacccagaatgaggtaggaggtggaggggctg**ccctgggcgtctgggggctgccaggg**actgagc**cctgagccagcctgagactcagg**aaaccccgtcaggagggagaagggagaagcagactctggacaccagaaagccaggggaagg**gtcacaaaaggagtggatgtgac**ggaagggcgggctcctgggtctcttcagaacatat**cccctgtgcccagggg**gatcagaggggcagagtccactgcgtgaaag**ccccactgctatgaccaggtagccgggacgtgggg**tggatgccagaaaagactccacggaataaga**gagagcccaggacagcaggcaggctctc**cgatccccccaggcccttgccccatacacgggctccagaacacacatttggctggaacagcctgagggaccaaaaggccccagtatcccacagagctgaggagccaggccagaaaagtaaccccagagttcgctgtgcaggagagacacagagctctctttatctgtcaggatggcaggaggggacagggtcagggcactgagggtcagatgtcggtgtggggggccaaggccccgagagatctcaggacaggtggtcaggtgtctaaggtaaaacagctccccgtgcagatcaggacatagtggaaaacaccctgacccctctgcctggcatagaccttcagacacagagcccctgaacaagggcaccccaac**acctcatcatatactgaggtcaggggctccccaggtggacaccaggactctgaccccctg**cccctcatccaccccgcagGTCAGCCCAAGGCTGCCCCCTCGGTCACTCTGTTCCCACCCTCCTCTGAGGAGCTTCAAGCCAACAAGGCCACACTGGTGTGTCTCATAAGTGACTTCTACCCGGGAGCCGTGACAGTGGCCTGGAAGGCAGATAGCAGCCCCGTCAAGGCGGGAGTGGAGACCACCACACCCTCCAAACAAAGCAACAACAAGTACGCGGCCAGCAGCTACCTGAGCCTGACGCCTGAGCAGTGGAAGTCCCACAAAAGCTACAGCTGCCAGGTCACGCATGAAGGGAGCACCGTGGAGAAGACAGTGGCCCCTACAGAATGTTCATAGgttctcatccctcaccccccaccacgggagactagagctgcaggatcccag**gggaggggtctctcctccc**accccaaggcatcaagcccttctccctgcactcaataaaccctcaataaatattctcattgtcaatcagaaatcttgttttatctcattttttcttttctcacatataattcctagcc**ttccctgggttctcaatttacggtggagggaa**ttctgcacccagtgggaaagtcacccaaggga**ggaggcttacagcctcc**ccgagtcatctct**ctggaaggtccttcctcttccag**tcaccccttccccaactctccaccataccc**ctgagcctccagcctggcctcagctcag**accagtcccacaccctcctcaa**ttttacttctcaataaagacctgatcatgtaaaa**cccagtttccaatgtgtcgtctgtgtctggtcatgtgcctgtgctgaagggtcactgctct**gggacaggaggcagtttcaggtgagatcccatgtccc**cgtcatcccacac**cccacccaacctgccaggaaaccgggtggg**ctccctgtgccagggggaaccatgttccagagcagaaagttgtccctgcagagtggtccctgaaatgcagttcttgcccacctgggaaggatgtggagcctagtgaggacagagtggtggccctgagcagggcatcggggagaaacgaggagtgttccaggaccccctgctt**tgggctagagacagaaaacccttgagccca**ggccaagatcagagcagaa**acagggttgaacttccctgt**cccatccatgatacccagttaggagaccatttactaggtgccatcaccttacgttacattacaacattacgtgattgtgccatcacccgggagacatgaaaaaggctggaaaatggaacccttc**agtgtagtttacactttcacaatgtacgttagctatgaaag**atgctgacaagtcctgcagttggaaaacagttca

Uppercase: IGLC3

Lowercase: Flanking sequence[1000bp]

Red & Bold & Underline: Stem-loop [21]

Blue: Heptamer[33]

Green: Nonamer [2]

id-IGHD-2[C_gene_segment]

atggggagaagaggagggtcatccagaatttgggaaagcagggcgacagtttct**gccccaagggagaagggaaggaggatggggc**caccgccacaccagatgaccttgcgtaccaggccaaagaacgggaacacctggc**cccacctgagcagcaatagtcagtgtggtggtggg**cagacatgggtggaggcagggggtgagtagaaggttagactaagacggagcacctggggcctcca**gggacccaggcaagaaccctgcacttgctcagctgccctgggtacc**caggtctccaggaaagtgaggctgagagccaagcc**cagcaggcagccacacattctgggacctgccaccctacagcctgctg**t**ccatgagtaacaccccttacaaggggccaggtcagctcatgg**gtttat**cccaggcagcagagcccctggg**gccaggaatcaggga**gaggagcatccaatcccaccagctcctc**cggagccactca**gagggccagacgcatggccctc**cacagggaccccatggcccctgcagggcagctgaggacccgtggctggg**agcctgggcaggagggtcatacagccctaggcccgttgctcccaggct**tgagt**gcccctcctcccctcaggggc**ccaagggaagtgggttccagagaggttgggggcagcagggaaggtggaggtcccaggaatgc**ccagaggggcaccaaagcctctggagggaagacccctcc**cttccaggagctctcggcaacaagagcccagggtccacaaagccacaggtcccactcggttattctgactcacaacacaggagcggcagcaggggcattcgtgttcacgggccacttggtcagccccgctcaccctgggcactcctcctgggccccttttccctgccttccctgtc**accctgctgccagggt**cctctgccctgccctgccccttgtcctcagagcctccagcctcagactcccactgtgtctgtcttccagCACCCACCAAGGCTCCGGATGTGTTCCCCATCATATCAGGGTGCAGACACCCAAAGGATAACAGCCCTGTGGTCCTGGCATGCTTGATAACTGG**GTACCACCCAACGTCCGTGACTGTCACCTGGTAC**ATGGGGACACAGAGCCAGCCCCAGAGAACCTTCCCTGAGATACAAAGACGGGACAGCTACTACATGACAAGCAGCCAGCTCTCCACCCCCCTCCAGCAGTGGCGCCAAGGCGAGTACAAATGCGTGGTCCAGCACACC**GCCAGCAAGAGTAAGAAGGAGATCTTCCGCTGGC**CAGGTAGGTCGCACCGGAGATCACCCAGAAGGGCCCCCCAGGACCCCCAGCACCTTCCACTCAGGGCCTGACCACAAAGACAGAAGCAAGGGCTGGGCTGTGAGGCAACCCCCACCTCCCCCTCAGAGCACGTTCCTCCCCCTTCACCCTGTATCCACCCCTCCGGACCCTCCCCATCTCAGTCCCTCCGCTCCCTCTCTCTGAGGCCCATCTCCCAATACCCAGATCACTTTCCTTCCAGACCCTTCCCTCAGTGTGCACGGAGGCAGCTTGCCCAGCAAAGGTGACTGTCTAGTGGGCTTCCCACAGCCAAGCTCCCACCCCATGCTGCGGCCCCTCCCTTCTTCCTGCTTGGCTGCCTGTGCCCCCCACCTGCCTGTCCACAACCCAGCCTCTGGTACATCCATGCCCTCTGCCCTCAGCCTCACCTGCACTTTTCCTTGGATTTCAGAGTCTCCAAAGGCACAGGCCTCCTCAG**TGCCCACTGCACAACCCCAAGCAGAGGGCA**GCCTCGCCAAGGCAACCACAGCCCCAGCCACCACCCGTAACACAGGTGAGAAGCCCCTTCCCTGCACACTCCACCCCCACCCACCTGCTCATTCCTCAGCCGCCTCCTCCAGGCAGCCCTTCATAA**CTCCTTGTCTGAGTCTCCAAGTCACACTTTGGTAAGGAG**AGGGACACTGAACGGACCTCTAACAAACACCTACTGCCAGCCAGCCCC**AGTCTGGGGGCCAGCAGATGCCAAACAACCAGCAGACT**CCCAGA**GCAGACCTGGGCCGGCTCCCTGGCCCATGGACCCAGCTCTGC**CTC**GCTGAGCTGAGGCATGGGCTCTCAGCGCAGC**CTCACATAGAGCCACCCTGCCGAGGCAGTCCGGCTTGCAGACTCACA**GGTCACTTGGGCCGCAGCAGCCCCTCCCCGTGACC**CTCGCCTCCCGCCCGCCCCAGCCTGGCTCTCTCCAAGTGTTGGATCTTGGTGGCCAGCCTGCTTCTCACCCTCA**CCCTGCCTGCCACCTCAGAATGGCAGGG**GAAAGAGGGCCCTCACCAAGAACTTTATCTGAGAAGTCTGAGGCTTGTGACTCTGACCTGCCTGAGATGTCCATGTGGCCGGGGGGACGGGTTCAGTGTTCGGGAGAACTCGGGTACGTGCCTGACTTTCTCTGAGTAGGGCAGGAAGCTGTTAGGAGAAGCAGCAGTGAGGTGGGCTGGACCAACAGGCAGAATGACTGTCCCTCAGCCACCCTCTGGGATGTGGGTCAAGCTCTGACAAAGG**CATGGCACAGCCATGGTGGCCCCTGCTTGGATGAGTGGCCAC**GGTGCCCTCACCCTGGGCCAGAATCTGCCTCCACTCTGCAGGTGCAGAAACACGACATTCCCGTCTCTAAACACACCTAGCTCCTAGGCTTGGGGTGGGCCTATCAAATGCAGGGAGATGGACACAGCACAAGGGCCAGAGCTTCCCATGAGAAAGGTGAGGGCAGCTGCTCCCTGACCCGGGCATCTGCACTTGTCCC**TCTCCACCCTCCTCATGGGCAGTGGAGA**CTCAGCAACAAAACAAGTTGAGTGCATTAGCAGCCAGCTCTGG**AGCCAAGTCACTCACCCCACGGCCTTGGCTGCTGGTGGAGGGGCCTTCCCCTGGGCAGCC**TCCAAGAAGACAGCCAAGTGCTCTTACTCAGACCACGGCGCTGCTTCCTGGCACCTCGATTTCCCACAACAACATGGGGTGCAGACAGGCTAG**GGCCCCCTGCCCTGGGGCC**TGGACGGCATCCAGTTAAAGATGACCCTTCACGGGCGGTGC**CTGAGGTGTGCTGACCTCAG**CAGCTAAGCCCTCAGGTCTGGTCTGCACTGCCCCACCTGGAGGACCCAACTGACCCAGACACAGCCAGGGTTATGGCATGACCCCGTGGACGGTGACCCAC**AGGCCAGATGCAGCCGGGGGCTGTTTTGTGTGGCCT**AGAAATGTCTTTACAGTTGTAGTGGGATGGAGGAGGAAGAGGAAGAGAGGAGGGGAGAGGAAAGCAGGGAAGGGGAAAAAGAGGAGTTCAATGCAACCCCAAAAGCCAGAACAGTTTTGAGCTGAAAGAACAAGGCAGGAAACATCCCAGTACCTGACTTCAAAACATACTATAAAGCAGTTGTAATCAAAACAGGATCATAAAAACAGACACACAGACCCATGGAACAGAAAAGCGAGCCCAGAAATAAATCTACATGCTTGCAGT**CCATTGATTTTCAACAAAGGCACCAGGAAAACACAATGG**GGAGAGGA**CAGTTTCCTCAATAAATAGTGCTGGGGAAACTG**GATATCCATGTGCAGACTAATGAAACTACACAAAAATCAATTGAAAACAGTCTAGGCCAGGCGCGGTGGCTCATGCCGGTAATCCCAGCACTTTGGGAGGCCGAGACAGGCGGATCACCTGAGGTCAGGAGTTC**GAGACCAGCTTGGCCAACATGGCGAAACCCGGTCTC**CACTAAAAATACAAAAATTAGCACATGGTGGCCTACGTCTGTTATCCCAGCTTTTCAGGAGGCTGAGGCAGGAGAATCGCTTGAATCCGGGAGGTGAAGGTTGCAGGGAGCCAAGATTGCGCCACTGCATTCCAGCCTGGGCAATGGAGCGAGACTGTCTCAAAAAAAAAAAAAAAAAAAAGAAAAGAAAACAGTCTAAAGGTTTAACTGAACAGATAAAGCTACTAGAAGAAAACATAGGGGGAAAACTCCATGACATTAGTCTGAGCAACGAT**TTTTGGATATGATCCCAAAA**GCTCAGGCAGCACTAGTCACAAAAGCCAAGATACAGAACCAACCTAAGCACCCCTCAGCAGATGCACAGGTAAAGAAAATGTGGTACGTATGGGGCACAATGGAATACGATTCAGCCTTTAAAAACAGTGAAATTCTGTCATTGGCAACAATGTAGATGAACCTGAAGGA**CACTTATGCTAAGTG**AAATAAGCCAGGCACAGAAGGAGCAATACTGCATGATTGCACTTACATCTGGCAGGTTAAAAAGGCAAACTCTTAGAGGCAGACAGTAGAGAGGTGGTGCCAGGGAGCGGGCACTGGTGGCTGGGGAGATGTTGGTCAAAGGGCACAAAACTGCAGTTGGGAGGAATTAGTTCAGGACATCCCTTGTACATGGGGACAGTGGTTAGTAACAACGGATTGTATCCTTGAAAACCGCTAAGAAAATAGTTTTTAAGTGTTCTTGACACAAAAAGTGACACGTATGTGAGATACTGCATGGTCATTAGCTGGATTTAGCCATTC**CACAATGTACACATATTTCAAACATTGTG**TTGTATATGATAAACATGTAT**AATTTTTGTCAATTAAAAATT**TTTAGGAAGAGGAGGAGAAGAGAAGAAGAAGGAGAAGGAGAAAGAGGAACAAGAAGAGAGAGAGACAAAGACACCAGGTTTTTTCTGACCCCTGGGCTATCAAAACACCTATTGCCCAATAACTAGTTGGCCGTTGGTGCCCTAAACTATTGAAGCGATTGCTGTTATGTGGATGGGCCCCGGACACTTAGAAACTCGTGACCCCTGA**GGACCCCCACGAGGACAGTCAGGGTCC**CCCCGAACTCAGGGAGCACTGAGGAAGGAGCTCTTAGAGGCGT**GGGGCCCCTCAGGCCCCTCAGAGGGCTCTGCCACATGGGTCAGGGGCAGGCTGAGGGG**GAGTCCCAGGCTCCATGCCCAGCCTCTGTGCCTCTGACCAGGGTGTCCCCCACACCGCCTCCTCC**CCAGTGCCCTCCACTGGCCACACCTGGCCAG**AAGCTGGGGAGAGGAGAGCACAGTGGTTAAGTCAGTCCCTGCAGGGAGACGGCACCAGAAAAACCTGGC**CTGTGGATGAGTCCCGGCCTGGCAGCCACAGAGCAGAGAGCTCTG**GAAGCAACGAAGGCCCGAGTCTGCTCAGGGAAGAGCGGGCAGCAGCCCCAGGGCCGGACAGTGACCAAGAGTGGCACCGCCCATGGCTCAACGGGTCTTTGCC**CACAGATCCCCCAGCCCCTGGAGACAGGGTCTGTGTGCCTGGCCGTGCAGGCAGGCACCACA**CTCAGGGGGAGGCCACTGTGGAGCTCTGTGCAGAGCCCCGGG**CGGGAGCCTACTGCTCCCG**AAGGTCCGGCCACAGCTGCTCTCGTTTGCTCTCCCCTGCAGAGTGTCCGAGCCACACCCAGCCTCTTGGCGTCTACCTGCTAACCCCTGCAGTGCAGGACCT**GTGGCTCCGGGACAAAGCCACCTTCACCTGCTTCGTGGTGGGCAGTGACCTGAAGG**ATGCTCACCTGACCTGGGAGGTGGCTGGGAAGGTCCCCACAGGGGGCGTGGAGGAAGGGCTGCTGGAGCGGCACAGCA**ACGGCTCCCAGAGCCAGCACAGCCGT**CTGACCCTGCCC**AGGTCCTTGTGGAACGCGGGGACCT**CCGTCACCTGCACACTGAACCATCCCAGCCTCCCACCCCAGAGGTTGATGGCGCTGAGAGAACCCGGTGAGCCTGGCTCCCAGGTGGGGAGACGAGGGTGCCCACAGCCTGCTGACCCCTACGCCTGCCCCAGGGCC**ATGACCCCAGCTGGGCCCCAGCAGCACCGGTCAT**CCTCCACAGGAAAGGAGAAGGGAGGCACCAGCACC**CTGGCCGGCCCCACTTCTCTCCCAGTGCCCCCGTGGCCAG**AGCCTGACAGCCTCCCCCACCTCCCCGCAGCTGCGCAGGCACCCGTCAAGCTTTCCCTGAACCTGCTG**GCCTCGTCTGACCCTCCCGAGGC**GGCCTCGTGGCTCCTGTGTGAGGTGTCTGGCTTCTCGCCCCCCAACA**TCCTCCTGATGTGGCTGGAGGA**CCAGCGTGAGGTGAACACTTCTGGGTTTGCCCCCGCACGCCCCCCTCCACA**GCCCAGGAGCACCACGTTCTGGGC**CTGGAGTGTGCTGCGTGTCCCAGCCCCGCCCAGCCCTCAGCCAGCCACCTACACGTGTGTGGTCAGCCACGAGGACTCCCGGACTCTGCTCAACGCCAGCCGGAGCCTAGAAGTCAGCTGTGAGTCACCCCCAGGCCCAGGGTTGGGACGGGGACTCTGAGGGGGGCCATAAGGAGCTGGAATCCATACTAGGCAGGGGTGGGCACTGGGC**AGGGGCGGGGCTAGGCTGTCCTGGGCACACAGGCCCCT**TCTCGGTGTCCGG**CAGGAGCACAGACTTCCCAGTACTCCTG**GGCCATGGATGTCCCAGCGTCCATCCTTGCTGTCCACACCACGTGCTGGCCCAGGCTGGCTGGCACAGTGTAAGAGGTGGATACAACCCCTCGCCGTGCCCTGAGGAGTGGCGGTTTCCTCCCAAGACATTCCCCACGGCTGGGTGCTGGGCACAGGCCTTCCCTGGT**GTGACCGTGAATGTGGTCAC**CCTGAACAGCTGCCCTCTCTGGGGACAT**CTGACTGTCCAAGACCACAGTCAG**CA**CCTCTGGGAGCCAGAGGGGTCTCCAGAGACCCC**CAGATGTCAGGCTTGGGCTCAGTGCCCAGCGAAAGGTCAGCCCCACACATGCCCATAATGGGCGCCCACCCAGAGTGACAGCCCCCAGCCTCCTGCCAGGCCCACCCTTTTCCGCCCCCTTGAGGCATGGCACACAGACCAGTGCG**CCCACTGCCCGAGCATGGCCCCAGTGGG**ATGTGGTGGCCACGAGGGGCTGTACACA**CAGCAGGAGGCTGTCCGCCCTGCTCAGGGCCTGCTG**CCTATGCCCCAGCTGTCCAGCCAAGGGAGGCATGGAAGGGCCCCTGGTGTAAGCTGGAGCCAGGCACCCAGGCCCCCGGCCACCCTGCAGAGCCAAGGAAAGGAAGACACCCAAGTCAACAAGGGGCAGGGCTGAGGGCTGTCCCAGGCTCTTTTGGCCCGAG**GGGCTGCCAGCAGCCC**TGACCCGGCATGGGCCTTCCCCAGAAGCGACC**CTGTGAGGTGGCCTCACAG**AGAACCCCCTCTGAGGACAGTGTCTGACCCTGCCTGCCTCACACAGATGGGCCCCACAGCAGTGGGCAACCTGGGGGGCAGCAGCCCAACCTGACC**CTGCAGGGACTGCCCCCTGCAG**CAGCAGCTGCTTCTCAGTCCCCCAACCTCCCTGTCCCCGCCAGAGGGTCTTCCCCGAAGCT**GCAGCCCCAACCCATGGCTGC**CCACCTGGAACCGGGACTCCCTGTCCACTGCCCCCTCCCCTTCG**GGGCCCCATCTGTGCTGGGGCCC**AGGTTCGGCCTACAGATTCCCATCATTGCCATGGCCTCCTGACCTTGCCTATCCACCCCCAACCACCGGCTCCATGCTGACCCTCCCCCAGGCTCCCACGCCCAGCTGGCCGGCCATCCCCAGGCACAGACAGTCTGGGA**TCTCACAGGTTAGCCTGGACCATCCACCTGGCCAGACCTGGGAGA**GGCTGGAAGCT**GCCCTGCCACCATGCTCCAGGGC**CCCAGGTTGCAGTACTATGGGGTGA**GGGTGTGTGTGCACACCC**GTGTGTACCTAGGATATCCGAGTGTACC**CTTGTGCCCCCAAGCACAAG**TCTCCCTCCCAGGCAGTGAGGCCCAGATGGTGCAGTGGTTAGAGCTGAGGCTTATCCCACAGAGAACCCTGGCGCCTTGGTCAAGGAAGCCCCTATGCCTTTCTTGCCTCGATTTCCCCTCTTGT**CTGCTGAGCCAGCAG**GGGCCACGTCCTGGGCTGCTGTGAGGAGGAAGTGAGTTGGTGCTAGGAGGGGCTCCTGTGTGTGCATGGGCGGGA**GGGGTGCAGGTATCTGAGCACCCC**GGTCTCCACTTGAGAGAGCAGGGCAGGAGCTCCCTGACCCACCCAGACTACACACGCTGTGTCCACGTGTCTCACATTATCTGTGGCAGAGGATCCGGCTTCTTTCTCAATTTCCAGTTCTTC**ACAAAGCAATGCCTTTGT**AAAATGCAATAAGAAATACTAGAAAAATGATATGAACAGAAAGACACGCCGATTTTTTGTTATTAGATGTAACAGACCATGGCCCCATGAAATGAtcccggaccagatccgtccacacccgccactcagcagctctggccgagctcacagtacaaccacaataaactcttgttgaatgaactct**aggaagtctgtgacgtggctggttcttgtcaatgcttcct**gcctgcccacaggctcttcctcgtggatggggctgtgcttgccatggaagcgtttttcccggcctaggcttgccttgggccccactgccgtctccagctggagatgaccttctatacacacatttgctcatgacagacccttgcttagcccccttccatggctccctcctgctgct**gggataaaatcaccttgcctggatatccc**ctcctgggcccctttccaccctccttagtcagcacccccagttcagggcacctgctttccccgctgcggagaagccactctctccttgctgcccggctgtgtctt**gccttccacaccttgtcacagtggccacttcctaaggaaggc**ctccctgtgtgcaggtgtgcagaagtgccccagc**ctcccgtcacctttgtcacgggag**cccaatccatgagagtctatggttctgtctgt**ctgccccactcagggcag**cgacaagtccaggcggggaggacacagtaggcagagatttgtcgaggggacatatgagcaagagggtgaggctgggagctccctggagataaccacgcctcctgggaagactcgccgtcatttcag**ctccacgctgtgcgggggtgggtggag**gggtagcctggccctcatgaccagggagcttctcactcagcccccgttcctccccagacctggccatgacccccctgatccctcagagcaaggatgagaacagcgatgactacacgacctttgatgatgtgggcagcct**gtggaccaccctgtccac**gtttgtggccctcttcatcctcaccctcctctacagcggcattgtcactttcatcaaggtcaggggagcggccaggctctcagtgaccctcggggtgggtg

Uppercase: IGHD-2

Lowercase: Flanking sequence[1000bp]

Red & Bold & Underline: Stem-loop [83]

Blue: Heptamer[146]

Green: Nonamer [10]

id-PCDHACT[C_gene_segment]

gggccacctcaatctccgcccatgaaaacgcatctagaggag**tgtcacaagtttttcacagtgaca**tttttgcttactgatacaagacagtgatggtgactgatgatgtcccagtgatttctgagtagcttctaaccagcacataactcccccaacagtctttaagtctttaggtgcccatattttcctctttgttctcccccttcagactgagagtttgtagagagagggcaacagatcttttcaatacacaactaatgcaaaatgtatcaggtttttcttggatttcagctactccctgttaaacaatcagagcttagtgaacagtgattcagtgaggagggaaagcactcaggaaggagcaggaacaagtacaagtttatgaaaaggattagctagcaaaacaaggt**caaactctgcaatagtttg**ttttcctctccctagtatccctcttcaatcagaaaagagactgttatcagttgctggtgttatgactgggcacatccgccctgggtcaaatatg**ctgcagtctgcaaagccagcagcagattgcag**tcctctgcagtccagccaggccagcagaacttgtgtagccatgtgccctgttatagctgtaatactgaattgggaatgttcccttaatggggcacttgagggc**aaaagcagtgaaagctttt**cctttctcaaagcagac**tgttcttcccgtagtgttttaagaaca**cagacatgtattgggcaaggcaaagccaaaggtggcctttacaagattattaaatctggtcttc**cagggtatctaatctgtgtgaggaccctg**atgaactaattttcttctaaagtgctatatatgtagatatcatcatagagttacacatgaaatggctcattcaatacttttttagatgcctggaaatatttaagggagtaactaatcaattagcagcattcctgggagaacattgtcttgtcattttaacagaaaactctctttgtgattttgcagCCACGACAGCCCAACCCTGACTGGCGTTACTCTGCCTCCCTGAGAGCAGGCATGCACAGGTATGTATTTCCCTCCTCATTCACTCAGAAGTAACCTTAACTTGGTATGGCTCAGATAAACTGCATCTCCATAGGCCAGAAGCAGCTGTCAAAACTAAAAAGCTTTAGGTACTTTGCCAGGAAAATGCAATTATTTTGTCCCCATGTTTATTCCTTGAAAGATCGCAAATGGTCAG**TGCCAGATGCTTATCAAGTGCTGGCA**TATAAGAGTCCTCTGTAAAATCACAGAAACAGGCTGCTATGTATTTTCTCCCATCAAAATTTCTACAGGGAAGTAATTTCAACCTCCTTCATCCCTCTCTACCTATGCTTTCTTTTCCTCCTTTAAAAACTGTAATTAATACTCATGCTTTGAGACTTGGG**TACATTGTGCAATGTA**TACATACATGTTGTCTACCTTGTTTTTTTTTTAATCTCACATTGGCTATTACATCCTATTACCCTCAATAATTGATTGCTATTGTTGTTTGTGTTCACACCTATTAGAGCCTCCTCATCTTTCCCATCTGTTGCTATCTTATTGTCATCAATG**ACATGGTTCTTCAGAAGATGAGCCATGT**AAAGGGCTCCAAATCTAGCTTACTTTAAATTAACCTAGAGTAACGGTATTAGTCTAAGACTCAGATTAAATATAATTTTGCTTTCTCTACTTTGTCTCTCTGACAGTAATTGATTAACTACCATTATTTCTGGAGGTGATCCAGTATCCATGCCATGGGGC**CAAATAAAAGATTCATTATTTG**CGAATGTCTTTGGAAACCAAATGGGAAGGACCAAGAAACAAATGATCACAACTATCAAAAGGATTTAATTTTAAAGAAGAAATAATCTTCAAACTTAAGCCCCTCAAAATATCTGGGCAACTATCAAGTGAATATTCACCAAACTTAGATCAGTTCGTAAAGAGAAAGCCTACAAAGTATGTGTAGAG**TTAATGTGAAATTAGTTTTAGCCCATTAA**AATGCATTAGATTGAAATAAATTAACATACTCTCAAGCATTACAAATGAG**CAGAAATCATTACATTGGGTGCTATTTCTGATTCAGAAGCAATCAG**TGAAGGGCTGAAGATTAGTAGTTGGCTTGGTAAGATGTCACATTGGAACCTGGGTCATAATTTTAGGCCAGAAACATTCATGCATATACCAGAATATTAGGTATCAGAAGAAATTCTTTATATTTATTAGAG**ACCAACTTGTGCTTTTGCCTGCATCTGAGCTGTTGGT**GGAGACATGCAATGGGTAAAAGCATGGTTTACAGTAC**CAACTCTTGAAAAGTACCAAAGCTATGAGTTG**TGCCTTAAAAACTACATTTGAAATAAAACATTAAAACATAACTTCCTGGACTGGGCGCGGTGGCTCACACCTGTAATCCCAGCACTTTGGGAGGCCGAGGTGGGCAGATCATGAGGTCAAGAGATCGAGACCATCCTGGACAACACGGTGAAACCCTGTCTCTACTAAAAATACAAAAAT**TAGCTGGGCGTGATGGCATGTGCCTGTAGTTCCAGCTA**CTAGGGAGGCTGAGGCAGGAGAATCGCTTGAACCCGGGAGGCGGAAGT**TGCAGTGAGCCAAGATCGAGCCACTGCA**CTCCAGCCTGGCGACAGTGCGAGACTCTGTCTCAGAAGAATAAATAAATGAATAAAATAACATAACTTCCTTATCCCA**TTTTCAAATTGAAAA**AAAAAAGCCAAATGTGCTCCTATTCGGGTTTCAATTAAGATATTATGAGATTTGAGTAGGGTAAGAAATAAAATAAAAATTGAA**ATTAAAATGCCATTTCTTTTTTGCATTGTAAT**ACATTGAACATATTAAATGAGTTGTGAACCTAAATAATACTAATCTTTTTCGTATGTGTGCTTGGGTGTTCTCGGTCTTTCCAGTCTTGGACATCATGTAACTATTCTTTAAAAAATTCTGCTTTGAGCTGAGCTGGCTCCAGGATAGTTACACCTTCATGAATCTGACTGAGCCCACACAATTTGCTAGTAGGATCCAGGAACACTTGAAGGCTGTTAATATTTGGGGAAAAAAAACAGATAATTCTAGAGTGTAGACAAGGGGAAGAATAGTAAAAGGTCAGAGTTTAATGAGTGAATTTCTACTGGATATGTTGTTTGAAGTCAAAGAGTGAGA**AAACATTGAACTTATATGTTGCCTTCCCTCTAATAGTTCAAGTTT**GCCTGCTCTGTTGCCTCATATAACCCCTTTAGTCAGT**AGTCTAAATTTTATTTTAGAAATTTAACT**TTCAAGATAAGCAAATGTCTAGTTTAAAAGGGTCCTCTAGTCTAGGTGTAGTGGCTCATGCCTGTAATCCCAGCACTTTGGGAGGGTGAGGCAGGTGGATCACTTGAGGTCAGGAGTTCAAGACCAGCCTGGTCAATATGGTGAAAACCTGTCTCTACTAAAAATACAAAAATGAGC**CAGGCATGGTGGCGGGTGCCTG**TAGTCCCAGCTACTTGGGAGGCTGAGGCAGGAGAATTGCTTGAACCTGGGAGGCAGAGGTTGCAGTGAGCTGAGATCGTGCCACTGTACTCCAGCCTGAGTGACAGAGTGAGGCTTTGTCTCAAAAATAAATAAAATAAAA**CAAAATGTTCCTCTAATTTTG**ATGAGGGTTTTCTTGGACATTTTCTCTTAGGATCCCACTTATTTCTTCCTTCCTTTCTTCCTTCCTCCCTTCCATCATTCATTCATTCATTCATTCATTCATCCAACAAATATTTGAGAGATTAATATGAGTT**AGTATTAGACATACATAAATGAATACT**GCACCATGTTCTCTTTTCCCTTGAACAGTTTATGTTCTATCTCTGCTTGCCTCTAAAGGTCTCCCAGTTTGTATCTCACTCCCAGCAATGTTTTATGCTGAATTAATCTCTTCTGAGCGGGGATCTGTGAGTGGTGGTGATCAAGTTTCTCTAGTCTCAGGAAATATAGGGTGGGTCATCTATGCATAAAAGATATAGAAAGAGTAAAATAGAAAATAAGGTTAAGAATTAATTAGATAGCCA**AATTGGAACAATACTCCAATT**ATC**AGAAAATATTTTAGTGTGTTTTCT**TCTTTAGAGTAGAGAACCTAGGAACAAGAGAACCTGCAAGAGA**GGCTTGGAACTTTTGAGAACAAGCC**CTCCTCATCTGCTTCAGTATCGAGATGTTAAAATGGCTTAGTCCTCTGATGGGCTTCCTGTTAGATTTAGTGAGCGCCACATGGCGTTAATAAAAAACAGAATTGCCATAAAGATAGAACATGTGTGTTCCTGGAATAGTATAGCAGGCAATAAGTAAGTCAGCAATGCTTCTGCAGTTTATGCAGGGTGA**CTGCTCAGCAGTAATTGCTTCAGTTCAAGCATGAGCAG**AATGTGTTAGCTGCAGCCCTGGCTTCATAGTTGTAAGCAATTTCTGAGGGTGGAAGAAGAGATGGGAAAGAATTTATGATCTAACCGTTATCTGGGTCTGTGTGTTTATTCAGCTCTGTGCACCTAGAGGAGGCTGGCATTCTACGGGCTGGTCCAGGAGGGCCTGATCAGCAGTGGCCAACAGTATCCAGTGCAACACCAGGTAAAGAGCTGGGGTCTCTCCATTCTTTCTTGGTTTCTGGAAAGTGATCAGATGACCTACTTTTGTAAGATCAGGAATGTTGATGGCTCTTTTTCTTTTATATTTTTGTTATTCCCTTTTTTCCATACATACATGATTTCCTTACATATATGATTATTTTGATTTTATACCTAATGCTCTTCAGGAGTTGAAAAAGGATAACAAGGAAAGTGTGTGTGCACGCATGTGTGCATGTGTGTGTGTGTGTATGAAGTTTTTGGGGTTTGTTTGTTTGTTTAAATCAGGTACCTTTCAAATGCTTAGGTCATCCAAGCCATGCAGAGAAGATCTGGTGGCCTTATGCACAGAGATGACACTGTTAACAAAGATCTTTTGGTTGAAGACTATGGAAACCCACCCAAAGTAGTAAGGAAAGAAAGAGAAAAAGAAGGAAAGAAAGAAAAAGGAAGGAAGGAAGGAAGATGGTTTCTCATGGAAGTGGAAAATTATCGGAACCAAGGCATTGTTTTGAGTTCAAGTC**AAAGTCAATCTGCTTCTCTCTGCACATCAACAACATTCTGCAGACTGACTTT**TAGTGCC**TTGGCATGCATGTTGCCAA**ACATGACCGCCTCACAATTTCTTAGTTTAGAGGGATAATAGGGACTA**TTTCCTAATCCAAACTTTCAGGAAA**GAGAACCTGCTAAGTTGTGTAAAAAACCTAATGGCTGG**GTGAGTATAGGAAAATTGCTTAAACTCAC**ATTTGCTTGGTATCCTTATTCTTGCCCTATCACTAAAGCAGGGTCATGTAATAGAAATATGGCTTTGGAGGCCCATCGCTGTGGCAGTTTTCAGAAAGGGAAGATTAAGTGTTGGTAGAGACCACAAATTGTGTCTACTCTAATCCTCTATTAATAGAACATCATGATGATAATAGTAGTTACTAATTATTAAGCCATAATATGCCTAGACACTGTGCCAAGTACATTGTATGTGTGGTCTCATTTATTCCTTAAGTCAAGCTTGCAAGGGATTTATTACATCTATTCTACATATGAGAAACTGGAGAGGCAGAGAGATTAAGAAATGTACCCAAGTTCACGTAGCTTGTAAGTGGCAGAGGGTAGGATTCAAACCCAGATGTGTTTAGTTTCAAATCCAAGTATATCTAGCACTTATATTCATAACATGGCTGGCTTGCAATAATCCATTCAAATTCAAATACATATCTACATACATAACAGATGACAGAATGTGTGTGTAAAAGGTTTTTTCCCAAA**AATAATCAGATGCCTTTCAAATTATT**AAGTAACATGCAGCTAAGGGCCCATTTTTACTGGCAACTTTAAGGGCATTCGTTGATTCTAATCAGCCAGGATTTGCTATTTATGGATGTTGCACAA**TTCAACTAAAAGTCACATTTGTCCAAAAAATATACGAGTTGAA**GCAATTCATTAGAGAGCTAATATTGCCAGATTGCAAGGGGAAAAACATAAAATAGTTCATTGACAAATCTGTACCCTCAGTGCCAACGATGGAGTGAAGAAATGATGGAGGAGGAAGTGGTTTTAGACTGCCAAGTGTTGCAGGATGTGGAGGCATC**TGGGAAGGTGAGAACTTCCCA**AAGAAGCCACG**TGAAATCATGACTTTCTACCTTGCCTTTATTTCA**GAGTTTTTCTTGCTCTTATGGAGGCATTGTAGGTCGACCTGGTAAGCCACAAACTAACTTTGAATACATTCTCCCTCCCATTGGTGATGCTGGTTGGTGTG**TATTCCTAGGCAAATGTGGAATA**GGAACC**ATGTATTGATATTTCATACAT**CTGGCCAAGTCCCTCTTTCAGATTCAAAAAATGTTGAGAAC**CTATCTTTTTTACAGAGATAG**AGAAGGGGATCTCCCTTGTTCCCTTTCTTACTGTCCCAGCCCCTCTTGTATAACCCATTTTATCCAGAACTGTGCCTGGCTGCTGATGCATGAGT**CACAGTCTTCATGGACTGTG**CTGGATAGAGCTTACATCTTCCAACTACTCCATGGCAACCTAATCATACTTTTCAATACATACCTCTGCATCAGTGGTGTAAAGTTAAAGGGATTCTCTGCCTTCTCCCTGTCCTTCTGGTACTTTTAGGTTTTTAGGACTCAATATATGTTCTGCACTGCTTGGAGGGAATATGGCATAAAGATTAAGATTATGATTTAGAGTCAGATTTGAGTTGAATTCTAATCCCAAGCTTACTTGCTGGGTGAGCATAGACAAACTGCCTGAATTCATATTTTCTTAATTACCCTTTCTGTAAATTGGGTGTAGTAATAATAATAACACCTATTTTATTGAGTTACCATGAGAACTAAAGGAGAAAAAAAGAACTGAGCATAGTGCTTGACATATAGTTAATAAATGTCTAATCTTTTTTTTTTGA**GACAGAGTCTCGCTCTGTC**CCCCAGGCTGGAGTCCAGTGGCACGATATCGGCTCACAGCAACCTCTGCCTCCTGGGTTCAAGTGATTCTCCTGC**CTCAGCCTCCTGAGTAGCTGAG**ACTACAGGCGTGTGCCACCAAGCCTGGCTAATTTTTTGTGTTTTTAGTAAAGACGGGGTTTCACCGTGTTAGCCAGGATTGTCTCAATCTCCTGACCTCGTAATCCGCCTGCCTCGGTC**TCCCAAAGTGTTGGGA**TTA**CAGGCGTGAGCCACCGCGCCTG**GCCTAATCTTCTTACTCTTTTTTCTTTCTGGACTACTTTTCTGCAATCTATGATATAGTGTTGGCTGATAGCCTGGTGGCCAGAATTCAGTAGTTCTCATTTGCAGGCCCAG**ATATAGACCCTCTGAGGTTATCTGGGTCTATAT**AATCCAGTCACCCCAACTGTTCCCCTGGAAATGGAGTGAGGAGGATTTATTAGTTGCTGCCTGAAGAAAAGGGAAATGCTCCAAAAAATTTGGTTGTTTCCAGACTCAAATAGAGCCTGCCTTTCATTGATTCTGTTGCCCTTAAAGCTTCACGGTGAAGATGCAGTTGCTTCCAAA**AGGCTTCTTTCTGGTGCCTAAGCCT**CCTTATACTTGCTTCAGAGCCCTTTCCGTGAACCAGCTGTGTATTGCTCTTCTCATCCCAACACTTGCAATGGCTGAATAAAGGAAGTGGGGCCTGCCTTACGCTAATCCTCGTTCATATGTGTTTCTTAAAGTTATTTTTCCTTCACTGATGAATTCCTTTTTTTTTTTTTTTTTTTTGAGACAGTCTCGCTCTGTCGCCCAGGCTGGAGTGCAGTGGCACGATCTCAGCTCACAACAAGCTCTGCCTCCCGGGTTCATGCCACTCTCCTGCCTCAGCCTCCTGAGTAGCTGGGACTACAGGCGCCCGCCACCACTCCCGTCTAATTTTTTGTATTTTTAGTAGAGCCGGGGTTTCACTGTGTTAGCCAGGATGGTCTCAATCTCCTGACCTCGTGATCCGCCCACCTCAGCCTCCCAAAGTGCTGGGATTA**CAGGCGTGAGCCACTGCGCCTG**GCCTCACTGATGAA**TTCTTTTGCTTTTTAAAGAA**ACTGTTCATTTATTTTCACAGTCTGCAAAAGCCTAAGATAAA**GATGCCACACTCTGAAAGGATCAACAAGGGCATC**ACCAAGTAATGTTTTCTGCAGGATAAACAAG**TCAGGCATTAAATTGGTTAATCCTGA**TTACTGGCCCCTTTCTCTAGCCTCCCCTCTGTGTGAGCAGACCCGGACCACAGGCTTTCTTATTTCCTTTCAGCTTCCCTTGAGACTGAGCAGAGAGAGAAAATTAGCTAAATCAGGAATGCAGGGAATACAGTTGCAGCCTCTTCTTCAGATGGAGGAATGCGTTTTGGGGGGAGGGACATTAAAGGGCCAGTCG**CTCATGTTACAGCTCTTTTTAACTTCATGAG**TACTAATGCCCTGAAGAGGTTTTAATGAA**TGCCCTCTTGTGATCAGTTCCTAGGGCA**ATCCCAGGTTATAAAAGGACTGCCCCTGCCTGTGAGGGAACTG**GCCTGGCTTCAGTGGGCCAGGC**TGCTTTGTTATCTGTTATTGGTTTTTCCAGCTCCTCTTTCTACATTTCAAGGGATCTCAGGCCTTACCTAAGGCAACAGTACATTAGTTTTAGAGTGGGAGATGCTCACAGTTTTCAGAAGAGTTCAGAAAGTTTCAAAACACACAGCACTGCAGAAGATAACATTATAGCTTCTCAAGACCCCAGGGGATCTGGGACTAAACAGTGAAAGATTAATTAGGTAGCGGAAGCC**ACTAAGGCAGTGAGTCTTAGT**TA**GAGAACTTTGGTTTAGACAATGGTTCTC**AAAGGGGCAGCAACACCAACAATACCCGGAAACTTGTTAGAAATGCAAATTCTTAGGCCCTATCCTAGACTAATGAATCAGAAATTCTCAGGATGGAGGCTGGGTGTGGTCGCTCA**TGCCTGTAATTCCAGCACTTTGGGAGGCCAAGGCAGGCA**GATCACTTGAGGTCATGAGTTCGAGACCAGCCTGGTCAACATGGCGAAACCCCATCTCTACTAAAGTTACAAAAATG**AGCTGGGCGTGGTGGCAGGTGCCTGTAATCCCAGCT**ACTCGGGAGGCTGAGGCAGGAGAATTGCTTGAACTCGGGAGGTGGAGGT**TGCAGTGAGCTGAGATTGCACCACTGCA**CTCCAGCCTGGGTGACAGAGTGAGACTCCATCTCAAAAAAAAAAAAAATAATAAATAAAGAAAGAAAGACATTCTCAGGAATGGGACCCGGCAGTCTATGTTTTAACAAGCCTTCTATGTGATACCAATGTACTGTGAAGTTTTAAGAACTGGTCTAAGGTAAATATTCCTGAGGTTGTCTTATATCATTACAGGGTCAGAATGCATGCAAGGAAGCCATCTGTTTATGGTTCTTGTGAGAAGCAGGGGGCCTTTCCCCATGCCCGAGAGATAATTGTTAAGAGCTCAAGCTTGGGAGTCAGTGACCCTTTCTGAATTCTACCTCTGCCACTC**AGTAATTGTATGTTCCTGGGACCATTACT**TAACTTTCCTG**ATTCTCAGTTTCTTTCTCTATAAAATGGGGAGAAT**AGTGGTGTCTACCTTATAGGGTTCTTGGAAGAATTAGATGAGATAATGCACACATATTGCAGAATCTGAAGAACAATCAGGGTTTAGT**AAATAGTAGCTATTT**TTAAATGATTTTTCCAGGTATGAGTCTATCCTACAGCTTCAAAATTTAGACCCAGGTTGTTCTGAGTA**TTCTCTGTGGGAAGAATCTGCTATAGAGAA**GATTTTTTTAAAGTGCCTGTCTCTTTGTTTCCTTAGGGGATTGCTTTTGCCCTG**ATTTGCCACATCTCTTTACTCTGTGGAAAAT**GGACAGTTTATGTGCCCTAGTTTTATATGGGGATTTATATTCTTAATTGTCTCAAGGATTCTTACCTGTCTGACAAAACCAACTCC**CCATGGAAAGACTCCATGG**AGACTCCATCTCTGATCCTTCCCCAGAAAGAAAGCAT**GATTCTTAAGTTTTTTAGAATC**TGTTTAGGAGCACTGTCAA**CATGAATTTTTCTATTTCATG**AGTGAGTGCAGCCTCGGGCCTTGTTGGAGAATTTAGAAAGCATGCTGTTCCACTAACCTGTTCAACCTCAACTTCTGCCGT**TGTCATAGCAATGACA**GTCCTGGGAGGTGTGTGCATATCCTTATTAGGAAAAAAAAATGAGATCAGGGATCTATGTGAGTGGGGCAGCTCCCGCCTGTGAGTATCCTTC**GCTGTCACCTGCCATCTGACAGC**CCAGGAGTGCCAGCTTGGCTTGGCTTTCTCTACCCGAG**GAAAGTAAGTCCTTTTAAGATGCACTTTTACTTTC**TGGGGTTGTGAAACTCATTGTGTTTGCCAGAGTTCTCTTCGCAGCTTATGTAAAGAATTTGTTTGTTTTGGATTGACCTGAAGGGAGGAAGCAAGGGTGTGGGA**AGGGGAATTAGCATCCCCT**ACCTAGGAGAAGCCATGAAGCTTACTTAAGTCTCTGCTGGCTCCATCCATTCATGACTTTCTTCATCTTCTTCTTGGGAAAACACTCTGTACCTTCCA**CTTTTAATGGTCATGTAAATAAAAG**ACTAGAATGGAGATGTCCTGGTTTTCTGAAAT**TAAACATTTTTGTTTA**TGAATAGACTCTAAGATAATTCTTTCCCTAGACGCTCTGTATTTTCTTGGACCTCTCATTTGCCCCATAGTAAT**TATTCTAGGATTGGTGGCCTGGGCAGAATA**CAG**TCATGGTTAAGACCATGA**TACATAATAGAAGAATTTCTTCCGTAACAGTAGCCCCAAACAGAGATCACGTGTCTCCTGAAACC**TATCCAGTCTCATGTGCACTCAGATGTACGATCCTGGATA**TAACATTTTAGAGGGTGTAGTGAGGAGAAT**AGAACAACACTGTTCT**CTTAGGCTGCAGTTTCCTTCAAGCCCCACGATGGAGAGAAGCAGGCTAATGCAGGGTAAGGAAATAGAGACTTATTATAGTTTCTTCAGTTATCATTGATTCCTTTAATATGCCAGCTACATTGAGGCAATACAGTCTGGCCTCTGAAAACCTTGTCAGGAGAAACAACTTTTGAATATATCTTAGAAAAATAAGACACTTTATCCCTTCTTTTGTTTAAGAGTGTTGCATAAAACAGGAGAGTTTCTGAATTACCCTCCCTTCTAGCTTTCTTTATACACCATTCTTTGTAGCTGAAGTTTTAAGCCCCTTGTCAAAAGGGAGATTCAAGTTTCTGCTGGGGACATCTCAGTGTCACAAAGGGCCAAGGAAGTAGGGTCTCACCAACTTTGCCACCTGACTCAGCTCAA**AGGTGATAGGTCACCT**GTGTGACTGAGAGCTTTGTGGGAAAGATGATATGGAGGGTGGAGAGTCTGCACTTCACTCTGTGGGAAAAATTGCTGAGGTTGTTTAGAATTGTTAGGCTTGGAACTCCTCCTGGGAGGTGCTTGAACAAGAAAGACCTGCTTATAACCTGAGTTGAGGGCAGAGGAGGAAGATAGTTTGTAATTCCTTTACGTTTTGTGGCTCCGGCAAGCCCTGTCCTCAGCCTCACTGACACAAGTAAACTAAAA**ATGAAAGTCTGTCCTAGTGACAGCAAGGGTCTTTCAT**GGCAATATTTTAAAAGAAACTCTGCCCAGATTTCAAAGGAACGTGAAAATTTTATCTTCAGAGGCAGTCAGCTTTGCAGTTGAAAAGGCCATTGCCTAATCTGGAGAAACATATTCAGTTCAAGCACTGGCTAGAGACTAGAGGCCCCCAGGAAAGGGCCATAAGATTGGTCACTGCCAGGAAGCCTGTCAATGAGTGTGTGGACTGGAGAGGAATCTTTCTCTGCCTCAGCACTTGGAGTCTCCGTTATTCACAATCAGAAAAAGAGGGAGAGCAGAGATAGATGACATTCCACGTTTTTCTCGGTGAGACCAAGAGCTTCCTCAGCAACAGCCCTGCCTGAAATTGTCTATCTCCAAGGGCGTGGGCTAGGCAAAGGATGGGAGAAGTCAGTCCTGAAGATGGTAATACTTAAAGGAAATAAGGGGGACAAAGGATGCTTGTGCTACTATTGAAACAGGAAATTGAGAGCTCTGTAGAAGTCAGACTCAAGAGGCATAATAATAGAAAGTTAGGGTATGAAAAGGTGACTTTTAAGAACCAAATGTGGACCCGAAAGAGAACAAAGAGAAGTTTATTGTAACTAGCGTTGTATTCCTTGTCTGCTGGATACCAA**ACAATGTACCCCGGGTCTTGAGATTATCGATGCCATTGT**CTGTAAAAAACCAACCAAACAAACTTTAAAAATAAGTAAAGCCTGCCCTGTACAGAAAATCTAATAAAGCAAATTTGT**TGGTGATGTTCCAAGAAGGTTCACCA**GCTTGGGCTTTCAACCAGCATTGACCCAATCTTGTGTCCAGAGCTGTTGCTGTAGGTGGTATCATGTATCAAGCTGAATAGTGAGGTGACATTCCCCAAGTCTTCTCTCTTGTCTTTTTTGACTGTGCATCATAGATACTGTATTACCCAAGAACACACACCTGTTCGATCTCTTTTCCTATAATCACCTGGAGTCAGATGAATACCGTAAAGGTCTGTGGTAAGACTAGAAGCCTCAGCCAGTATGAATGATTTACATTAGATGCACACCAAGCTGCTTTCGGAGAGTCCAGATTTATGGGGATAAGAGCATCACT**AGGTATATCAACAGCCCTAGGGTGGATACCT**TTGAGCCTGTAAATTTGGCTCTGTGTTGGACGATGAACCATGGAATACAGAAGAAGCTCCCTTCTCTCAGCTAAAGCCTATTAGCAAAAATAGAGCCCTGAGGACATTGTATTTTAAGTGTATGTTCCAGTGAGTGGGTCGTTGCTGGGTCAAGTTTTATAAGTCTTTAAATCCCATACTGTTGCCTCCTGGACTGTAGAAATAG**TCTCTTAGAACAAGAGA**AAGGAAGACAATAGCTACCATTTATTGAGAACTGTGATAAGCACTTTACATATGTTGCATACTTAACTTTGAGAGAAGTACATTATTATTCCCATGTTTCAGATAAAAAAATTAAAGCTCA**TAAGAATTAAGTGGCTTTCTTA**CATTCACACAGCTAGTAAAAATGTTTTTTGCTCACCATTGTATTCC**CATTGTCAAGCAGGATGCCTAGCACACAATG**ATGCTCAATAAATTTTGTTAAATTCATTAATAAATGCAGTGGTAAAGGCAGTATTTGAACTCATGTCTGCTTGCTGCTGAACTCTATACACTTAACACATTACTATTATCTTGTCCCGCATGATACATGAAGGGAATAGCTTGGTAACTTGGAAAAGACTATTTACTGTCTGAGTTCTAGGCCAACTGTAGGGTGTCTGTATTTATTTCCCATTATGCTATAACAAATTAACCATAAACTTCATGGCTTAAAACAACACAATGTATTATCTTACAGTTCTGGAAGTCAGAAGTCCA**CAGTGAGTTTCACTG**GCTAAAATCAAGGTGTGTTCTTGCCACTATATAACAAGGATCCACTTTCCTTTAGTTTCCAATAACAGGCTCCTTATTTCTACCTGAGCCCTCACTGGCAGCACCTTTAATGCCATATTTGTAATAACAATCTGTTCATGACAATTTAGGTTTGCTCCTCACTTTTTTCTGAGTCCTCT**CTAGTAGAGCCATTAATATGCATATTGCTACTAG**CAGCCTGTTCAAGGTAATGTAGGCTCTTTCTATCATGCTCTTCAAAATTCTTCTAGACTCTGCCCATTTCCCAATCTCAAGGCCACTTCCGCATTTTTAGGCATTTATAACGGCAGAACCGCACTTCCAGATACCAAAATCTGTATTAGTTTCCTAGGGCTGCCATAACAAATTAACACAAAATCCCTTTTGCCATGTAACATAACATATTCACAAGTTCTAAAATTAGAATGCAGTCATTTTGGTGGGGCCATTAT**TCTGCCTACCACCCGGTCTTTCATGTTCAGGCAGA**GGTGGCTCTGTGTGTGTGTGTGTGTGTGTGTGTGTGTGTGTGTGTAAGTTAATTTCAGTAGAGAATGAGCTAGAGTAGAGAAAGAGAATTAAGGTGAGATGTGTCATGGAAGAACAGTGACTGATGATGCTAACTTTGCTCAATCAAGAAGTGTGATCCATTTAAATCATGCTTTCAGTGATCTATCCAATCAGATAAACTACTCTC**CCTTCCTGGGAAGAGTAAGGAAGG**AAGTAGAGCTAAAGATGAAAGTTCTTTTCGTAGCACATCCCTGCAAAGGATGGGAGTATTGTTTTGGTGGGCATTCCCTTTTCTAAGAGCAAAGATGGAAATGTGGAGAGAGGAGAAAAATGGTTCCCATTACATTATTGTGTTCTGGA**CTTAGAGATATTGTTCTGCCCCAGGTCTAAG**AATATTGTTCC**AAGAGTTGGGAGCAGGCAGCAGAGGACAGTACTCTT**TAGATCACCCAGAGGCCAATCTGTAAGGATTCAATCCTGGGGCATGGTAGCTAATGGAAAAGTCACTGTCACAAGTGATGCCAGGAGATGGGGCAACACATATTGCCTGCCTTGTATGCATCTAGCGTCTGTGTCTAAGTGAAAGCAGATTATATCCAGAACTTCCCTGGAAAATAGAAGAAAGAGCCTGATTGGTGTGTATGTGTGTGTTAGGGGTTGAGG**GGTGGGTAATGTTCCTGCCAGTATTCGTAAATCCCACC**TGCTTTTCCTGTGATGCTTCCTGTGTTGGGGATAGGAGGGTGGGATGGTTCAGGGTCAATTTTATGGATCCATATGTATCTAAGGATGTGTTTGTTTTTTGGGAACATTTTGTATAACAACCAGTTTCACTCTAGCTCCCTCCTTGATATTCTAAACAAGCTGGATATCTCCAAAGCTTAGGCCC**TCTCCATACTTACAGCAGCTCTGACTGGAGA**TTAAATCACCCCTTATACTGCTGACAATGATTAGGCCATGGGACCCATGAAAGAGCCTCCCCAGAGTCCAGCATCCCCTGTGGGCTCCGTGTGTCATCAAGATGTCAATCCAA**CTGCTGTTCCTGCAGTCTGCAATCAGCAG**GGCTATGTTTATTCAGCGGTCAGTGTCACATCAATTTCCTTCTGTTGCAACAAGTATAAATGGATTCTAAATATTTGCCTTTGGGAAATTT**CTTTAGAGGGAAACTCACTAAAGCTAATTTTTTTAGCTTT**TTGGTATGTTCTCTCTGAATCTGGGGATTTAGATATATAAATTAGCTTCTTTGGTCTTTTCTTGCCCAGGGTCACAACCTTGCCTCCAGGATAATACCTTCTA**AGTGTTTGTGATTTGAAGGGCACTACGAAGATCCTCACAAAAACT**ACCTCCCAGCCGGGTCCCTGAAACTCCATCATTAACCACCTTCATCAGCATTTCTCTTTTAAAATCCTTATTCATTCCTGTTCTCCTTCTTTCTTTCTCACACACACACACACACACACACACACACACACACACACACACACACGGGGAGAGAGAGACAGAGAGAGAGAGAGATAGAGAAGTGAAGTATATAGTATCCTTTCTAGGGATGCTTTTCTTGGCTTGGCTCCA**ATAGAGAATTTTGTTGGGACCCTCTAT**ATATAGTCGTGT**ACCAAATAATGACACTTTGGT**TAACAATGGACCACATATGTGACAGCTGTCCCATAAGATTGTAATACCATATTTTTACTGTACCTTTTCTATGTTTAGA**TACACAAATACTTGCCATTATATTACAATTGCGCAGTATTTTGTA**CAGTAACATGCTGTGCAGGTTTGTAGCCTAGGAGCAA**TAGGCTATACCATATAGCCTA**GGTATGTAGTAGGCTATGCCATCTAGGTTTGTGTAAGTACGCTCTATGATGTCCATACAACAA**AAATGCCTAGTGATGCATTT**CTCAGAACATATCCCCATTGTTAAGTGATACATGACTGTAGTCATCATCAAACATTTATTAAGCACTTAGGTCAGGCCAGGCTCTGTTCTAGGTGATGCAGATATAATATTAAATATAACACAGTCCTGCCCTTATAAATCTAATGGTGAAGGGAGA**AATGTAAAGAAATATGATGGAAATATTACATTGTCATGTGATAAGGGCGACAAT**AATG**CCCTACGTAGGTAGGG**TCAACGAAGGGGAAGGAAGCTGGGACAAACCACCAGGGCCTGTTGGTCCAGAATGGGACCCAGGGTCTGTCTATGTTATAATCAATTCAAACCCTAGGTAAATAAGGTAAGCTAGGCTGCCTTTCTTGAGACAGTCCCCAGATTGTTTTCACAGGGCCCAAACACTCTCAGCAACCATGAACAGTGGATGTTGTGGGACCTCAGAAGATGGAATTGGAAATTCAGGGGACTAAGTCAGAGATTATTTGACATTTAATACTGGATATTGAGGCCGGGCGCAGTGGCTCACACCTGTGATCCCAGCACTTTGGGAGGCTGAGGCGGGTGGATTGCCTGAA**CCCAGGAGTTTGAGAACAGCCTGGG**CAACATGGCAAAACCCCATCTCTACAAAAAAAATATAAAAATTAGCCGGGTGTGGTGGCATGCACTTGTAGT**CTCAGCTACTTGGGAGGCTGAG**GCACGAGAGTCACTTGAACCTGGAAGGTGGAGGT**TGCAGTGAGCTGAGATCACACTACTGCA**CTCCAGCCTGGGTGACAGAA**TGAGACCCTGTCTCA**AAAAAAAATTTAAAATACTGGGTATTGCAGGATGGCAAGTAGGATGTGGAGAGGAAGAGAATTCCAGGCAAAGGAAACAATGCGTGCATGCAAAAACAAAATACAAAAAACTGAATTGTGTAGTCTGGGTTAATAAGATGAGTGCATTTGATGGATGTGGTAGAATAAAGAGATGTTAGGAAATGAGATCAGAAATGTAGACTGGAATCAGATTTTGTAGGACCTTAAACATCCTGCTAAATAATTTGTAGTTTCATTTGTTGGCCATGGAGACCCACTAGAAGGTTAGTTGGTTTGTTTATATTT**AGGAAAGAAGATTCTTTTAAGTTTCCT**TAACTCTACTGAATTTTCAAAAATCAACCTACAGCAATCCAAACTTATCTGTGGAGATACATTCCAAGACCCTCAGTGGATGCCTGAAACCTCATTTAGTACTGACCTTTATATACATTATGTTCTT**TCATAGGTGCACCTATGA**TAAAGTTTAATTTATAAATTAG**GCACAATACTCTTGTGC**TTTGGGGCCA**GTATTAAGTAAAATAAGGGTACTTGAATAC**AAGCCCTTTGATACCAAAACAGCCAGTCGGATACCAAGACACTTTCTAAGTGAACTAACAGGTGAGTAGTGTAGACGGCATGGATAGGTCGGACAGAGGGATGATTCACGTACTGGGCAGGATAAAGCGGGATGGCTCGAGATTTCATCACATTACTCAGAACAGCTTGCAATTTAAAACTTATGAATTGTTTATTTTTGAAATTTTTCATTTAATATTTTCAGACTAAGGTTGACTGTGGGTAACTGAAGCCTCAGAAATCAAACCTCAAATAAGGGGCGATCACTGTACTGAATAATATGTCAAGATGAGCTATGAGCTTTTAACTTAGTGTTTTTCTTCTCCCAAACCAAATATTGGAAACTTTGGAGTGTTTAGAAAAGGAGAATCGAAAAGGGAAGTAAATGCAGCATTTTTTTTAATTGTTAAATAAAGGGCTGGGCTTGGAAATGTTCTTAGGTAATCCTGGTGGATTTGGTATATCATGTCTGAGAAAATGTGGAGAGTGAAGGGCAGGTTTAAAATTTGTCTCATGTTAGTTTCTGAAGGGAGATAGCCTGGCATTGCACAGTGCCTCACATTTCTATCGTATCACTTAATAATTGTGCAACCTGTGCTTCAGATTTATCAATGTATAAAAGAGGAGAGACCATAGGCTGTTGTAAGTATTAAATGACATAATGCACGCAAAATGCTTAGA**ACCATGCTTAGTTCATGGT**AGCCACTCAAAAATGTCGTCAGAACTATTACTACTCAGTTTAATGCTCCT**AGCACTTAACACAGTGCT**TGGCACACAGTAAAGAGGATTCAAGATGTTTTTGCTAGATTACATGGAATGGTATGGAAGTGTGCAGGGTGGTGTAGAGGGTTCTCCCATTATCTTGCCTTCCTTATACTGCAATGTGCTGACAGCACCTCCATTTTATACCCTCAATGCAAGGATATCACTACTTGTCTAGAAATGGTTGCTCAAGGCTGAGAAGTGA**CAGCAACACAGAGCCTGGGATTGCTG**CCAAATTTCAAGAAGCACAGATTGTAACAAAGTAAGGGGGGAGGGTAGAGAAGAAAAGTTTGAAAACCCAACCAAAGTCAGAAGCCATTTTGTTGTCGG**CTGAAATAATGCCTCATAGAGTTTCAG**GTGGGGCAGGCAGTGTGAAAACTTTGGGAATAGTCAGTGTCAAGTTGTGGTGCTAAGGGAGGAACCCAGTAGTGCTCAGGC**CTGGGCAAGTCTGGCTTCTTGGGGCCCAGCATTTAGAAAGGCCCTGTGTGTTTTGATATTATGCTG**TCACTATCTTGAAGTTCTTGATAATTTGTTAACAAGGGCCGCCTCCCCCCATTTTCATTTGGCAAATTTCACTGGTT**CCTGCAAATTATGTAGTTAATCCTGGGTGCAGG**CTAGTTTTTGAACCATGTTGGGCTAGTA**TTCTTCTGAACCTCTATCTCTTTGATTTAATTACAGAAGAA**ACAAAAGTCATAACAAATTATATCTGTGCAGGACTAAAATAAAATGAGTAGTGCCTCACTCCCTACATCCAATCATGCTTTCCAAAAGTAACCACTACCAATAATTTAGCGTGCATCTTCAGAATTCTTTATATGCCTACATTTTAATG**TATTTTTAAATAAAATA**GAATTACATTGTTGAAGTAAATATTA**AAAAATATAGCATTTTT**GCATAGAAATATTTATTTCCCTCAATCCCACTCCCCTTTGGTGAAGCAACCACTGTTAACAATTTTGTGTGTATCCTCCAATATTTTTTTGAAATTAATGAACTTTACTTTTTAGAGCAGTTTTAGGTTCACAGGAAAGTTGAGTAGAA**AGTGCAGAGTTCTCATGCACT**CCTGCTCTCCCACATACACAATGCCCCCACACCCCCGCCACAGTGACATCCTGCCCAGAGTGGTACATTCATGACAATTGATGAAACTATATTGACATCATTATTGAAACTATCATTACCACCCAACTTCCATGTTTACTTTAGGGTTCACTCTTCCAAATATTTCTTGCATCACTTAACAATGGGGATACACTGAGAAATGTGTCTTTAGGCAGTTTTTTCGTTGTACAA**ATATCATAGAATGTACTTACATAAACCTAGATGATAT**AGCATACTACACACTTAGGCTCTATCGTATGGCCTATTTCTCCTAGGCTACAAACCTGTATATCATGTTACTATACTGAATACTGTAGGCAATTTT**TACACAATGATAAGTATTTGTGTA**TCTCAACATACCTAAACGCAGAAAAAGTACAATAAAAATACATTATTATAATCTTACAGGACAACTGTCATATGTGTGGTCCGTTGTTGACTGAAACATGATTATGGGGCGCATGACTATATAGAGATTTTCAAAAAGCAAAATGGGATAATGCAATATGTATTATTATACAGCTTGTGTGTGTGTGTGTGTGTGTGTGTGTGTATGTTAACAAACACCTTTCTACGTCAGTACATATAGATTTGCCACAATCTTCTTGAGTAAGGATATAGCACTATGTTGTATACCTATATTATAATTTATCCAATTTGCTGTTGGTATCTATGTTTTCTAAACAATACAATATACATTCTTATACATATTTCTTATGCATGCTTGCTAGT**ATTTATACAGGATAAAT**TTCAAGAAGTAGAATTACAGAATCATAGGATATAAAAATATCAAATTGGCAAATTGGCATTCACGTACCTGTGGTTGGACTGCACTTGTTACATACTTCCCTCTGAGCCTTCCATCCCCACCTCGAGCTAGTGCTAT**ATGTCACTTAACTCAGTGACAT**CATCATCAACAGACTTAG**CCTCTGCAGTTGAAAATGCTAGAGCAAACAGAGG**AGAAAATTTACTTGTGAATCATAATAGCTAACCTGTACTGAACAGTTAAGCCATGTGCCAAGTATTATTCTAAGCACTTTACAAGTATTAACTCCTTTAATCTGTATAACCACACCCAGAAATTTGTGCATTATTATACTCATTTTACTGCTAAGAAAACTGACACCCATAGGATTAAA**TCAGATCACACATTTAGTAAGGGCACCAGGATCTGA**AGCAGGATTGTTTGACTTCTGAGTCTGTGCTCTTAACCACTGCACACACCGTCTCTAGAAAGTTTAATGCCATCTTTATGCCAAAGTTTATCTACCTGGTTTATCCTTTAACACTATTATTGTATTTATTCATTTACTTGTCTTTTCCTCCCACTGTGCTGTAACTCTTTGAGAACAGGGTACTTGTCCTAATATTATTCATATTTATATCCCTAATTCCTAATTTAATGTCTAATTTATTGTAAATAAGTTTTTTTTTTTTGAGACAGAGTTTTGCTCTTGTTGCCCAGGCTGGAGTGCAATGGCACAATCTCAGCTCACTGCAACCTCTGCCTCCTGGGTTCAAGTGATTCTCCTGCCTCAGCCTCCCAAGTAGCTGGGATTACAAGCTTGCACCACCACGCTAGGCTAATTTTGTATTTTTAGAAGAGACACAGTTTCACTATGTT**GGTCAGGCTGGTCTGGAACCCCTGACC**TCAGGTGATCTGCCTGCCTTGGCCTCCCAAAGTGCTGGATTACTGGTGTGAGCCACATGCCTGGCCTTGTAAATAATAAGTTTAGTTGAATAAATAACAATGCCTCCGGGAGGTAGCTATTATATCCATT**TTACAGATGAAGAAACTGTAA**GTCAATGTTAATCAAATAGTATCCTAGAAAATAGTAGAAAGCAGAATGCTGGAGCTGAGATTTGAACCCAAGACTTTTGATACTTCGTCCAGTGTGCTTTCCACCATGCCTAAGTAGTCTCCTTCACTTCCTCCTTCAGAGGGCTATGGAGAGTAACCTAGCAACCATTTCTAAGCTGGAAAATGTCACAGCCGGAAGTCTTCAGTCCCCTAGAAGGAAAAGAGCCTGATGGGGAGAGGGTCCTTGGAAAAGAGAATTTCTGGAAGTACTACATTTGAGAATAGGTGGTTAAAGTGGGAGTTGGATTAGTAAAGGAATACTGCATAGTAGTAATACCCATTCTTTATTTCAGAATTTCTTCATGACAGACATTGATCTGTGTGATTTATAAGCCTTGTCTCTTTTTAATGTTCACAATCCCATTAGATGGTTATTGTTATCTTTGTTTTGCATATGAAAGAATGGGAGCTAAGAGGCCAAGTACCTTG**CCTCTTATTCGTTCAACGAATAAGTGG**TGAAGCAGATTCCAACTCAAGTCTATGTTACCCAAGAACCTACATTTTAAGCATTTTGT**TACCCCCTGGATATGACAGCCAATGAAGAGGGGGTA**TTTTGAGAAGAGACTATAA**AGGGAAATTGCCTTCCCT**ACA**TCCTGGGGGACCTTATCAACCAGGA**AACAAGGTAGAGAAAACTGTGCAGCC**TGAGCCCTGCTGGGTTGCGGGGGGCTCA**CAGAAAGAAGAAATGTGATTTTTTTTTAGCTAACTACGGAGACCAGCCATTCCAATGTTTGAATCTGGGTTCGCAGCACATGATGTCTTTATACTCTTAACCTAGAAATGGCAGAGTTATTTTGGGCACAAAGCAAGAGCTGTGGCTTTAAAAATATGCCAAGTGTATTTATCTCTTCCGCTCCAAGATTACTGAAAATTAGCCCAGCTGTAGCTTGGGACACCAAACAGCCAAAAAATCTTCTTCCAGCTCAACTCATTCCACCCAAAGAGGGTGAAA**TACCCAGGAGTAGCAGCTCTAGCGGCCTCTGGGTA**GTGGTATTA**GATTGGCCTCCCCATTGCTAAGCCTGACATCCAATC**ACACACACACCACTCTCCCAGCTGCTCTGTAGATCACAATGCTAGGCCTGTGAATGGAGCTCAACTCCGTCTCTTCCCTCATCCCCAAGGCTTCACAAGAAGTAAACAAAACAAAC**AAAAACTATTTGATTATTGAGCCAAGGAGTCAATGTGAGAATAGTTTTT**CACCTTCATTATCAAATGCCTGTGTGGAGCTGAATGTGGTGGCTGACACCTGTGATCCTAGCACTTTGGGGAGGCAGAGGTGGGAATATTGCTTGAGGCCAGGAGTTCAAGACCAGCTTGGGCAACACAGCAAGGCCCTATCTCTCTCTCTGTCTCTCTCTGTCTCTCTCTCTCTCTCTCTCTCTCTCTCTCTCTC**TATATATATATATATATATATATATA**TTTTTTTTTTTTTTTTTTTTTTTGAGACAGAGTTTTGCTCTTGTTGCCCAAGCTGGAGTGCAATGGTAAGATCTCGGCTCACTGCAACCTCCTTCTCCAGGGTTCAAGTGATTCTCATGCCTCAAACTCCCGAG**TAGCTGGGATTACAGGTGCCTGCCACCATGCCCAGCTA**ATTTTTGTATTTTTAGTAGAGACAGGGTTTCACCATGTTGGGCAGGCTGGTCTCGAACTCCTGACCT**CAGGTGATCCACCTG**CCTCTGCCTCCCAAAGTGCTGGGATTA**CAGGCATGAGCCACTGTGCCTG**GCCCCTATCTCTGTATATTAAAAAAAAAAAATCTTAGCCAGGCATAGTGGCGCACACCTGTATTCCTAGCTACTCAGAAGGCTGAGGTGGGAGGATCACTTGAGTCCAGGAGATCCAGTCTGGCAGTCAGCCATGACCTCACCATTGTACTCCAACCTGGGCAACAGATATAGACGCTGTCTCTAAAAAAAAAAATCCTGTGTGATTTAGGAC**AAATTATCTTGCTGTAATTT**AAGCCTTCATATTCCTTTTTTATA**AAATAAATATGCTTTAAATATGTATTT**AGATATACACTTATAATAATAGC**TTTAATTAATTGTAATTAAA**TCATTTAAAATTAAATACATGCAAAATACCCCATAAACTGTCTAATCCATAATAAGCAATCAATAGTCCTTAAACAAATGAATCTTCTGTTGCTCTGATCTTAAT**AAGTAAAATTTAACGAGTTTTTACTT**TGCACCATGCACTTATGCTATGTGCTGTATAAGGATAGTTACATTTAATCTATAAATCAACCCTATGGGGCGGGCACTCTTATGAACTTTTTTTACGGATGAAAACTGAGGCCCAGAGATATGAAATAATTTGCCAA**ACATCACCATAATTTACATGATGT**AGCCAGGTTTAAGCCTACTATTCTGATTACAGAGCCTAAGATCCTACAGAGAACAGGGAGTATATTTTTACATTCCCTTGTTTCCA**TGGAAAAGTCTTTCCA**CTGTCAATTGAAGGACTAAGCAGCAGCGGGGAAGTGCTGGAAATGCTAGCCCAGGTGGGCTTAGCTTTCTCCCTCTCTCTCTGATCCTGG**ACAGGATTTGGGTTTCAGACCACGATTCTCCTGT**GTTTTGTGGGCTGTGATTACTCAGATTAGGTTTGCCCAATTCATAAAGGACTGGCGGGGGTTGGGGGTGAAGGTGTGGGAGAAGGCGGAGCTTGTCCAGTCGGTCCAACAAACCCCACAGATGGCGAAATAGGGGGCGGGGAAGGAGCTTGAGATATTTTACAACCTGGGCTGTTTCAGTGGTTGATGGCGGATGGTTTTTGCCTTTTGGTCAGCCTGAGTATATACTATCATTCTACAATCGGCCAAATTCTGACAGAGAGGGAGACAGAGAGAGAGGTTGATTAAATTGATGCCCAAAACCAAGAAGGAGCAAAGAAATCGGGGCTGTTTGAAAAAGACTGCAGTGGCTGACTTGGGTGGTGAGCGGAAATAAGGAGGAGGGAGAGGCGGGGTGTCTCTGCGCGGAAAGCCTGGAAGTTCACTTGCAAACACAGAATCTGCACAGCCTTCGGTGCCCTGACTCTTTCCTGGGCATCCAGAGGCAGCAGCAGCCGCCAGCGCCAAAGAACGAGCAGTCCAGGGGCTGGGCCGGAAAC**GGCTATAATCATTTAATAGCC**TTTGCCGGCTGCACTGACTTAGCAGAGTGGGCGGTAGGCAGGCTCCAGAGTGCTGTCTGGCAAGATAGTCCCCGGCTTTAATCAAAATGATGGGTTTTCTGGA**AGCTCTTTATTAACCTCAGCACTGAAATCCCAGAGCT**GGTAACAAAGGGATGAATGGGGAGCAAAGGGGCGGGGCCGAAACC**TGGAGGCCGGGCTGCATCCGCACCCCTTCCCCCACCTCCA**CTCCTTTCAACTCATTCTGGCTTAGGGCTCCTGCTGGCATCTCTGTGCCTCCCAAATAGTAGTAACAAAACAGGCAATAACCATAATAATTGGCACATT**TGTATAACGCTTTAGCATTTTCAAAGGATGACCTTGTTATACA**GCTCAGGATCT**GAGTCTCCTAGCTGGAGTCAGACTC**CCTAGATTTGAGTCCCAGGATCCACCAGTTAGTGACCATGTGACCATTAGTCCTTAGTCCCCTCATCTATAAAACAGACACATAACACAGACAGAGATACTAGAAATGTCTACTTCATAGGGCTCTTGTCAGGACCAAAACCTATACGACGTGCAGATTTGTTTAACTCTGTTGCACACACATAGGAAGGGCACAGTAAATGTTAA**CTACATCGACAACCCTGTGATGTAG**ACAGGAAGGGGATCATCTCCATTTTATGGATGAGGAAACTGAGGCTCAGAGATATTTATGTAAATGGCCC**AGGACTATGCACTAGTGAGAGAACCAGGGCAAGTCCT**CAAGAACAGAAAGAGAACAAAGCAAGATGAAGAGAAGATCAGAAAAGTGAACACCCTCCAACACCCTCCAGCACATGCCCCTCTGAGAGCTTTCCTGAAAATAGTATCCTTGGTCAAAATGAGTTTATGTTCTATTAGGGAAAGTCAGTGTAAATCAGAAAAGCCCTCCTGGCCTTAACAGAACAGTTC**CAAATGCAGATGAGGGTGAGTTTAACAGGCCTGGCATTTG**CTTCACAATCTTCTGGCCCTTTCCTGGCATTTGCCTTGGCAAAGACTCTGACCTGATCAATCCTAGTC**CCTGAGGCTCAACTCCATCAACTCAGG**CAGCCAGGGTTAGTTTAGCATGAAAGAGGAAAGCTGGAAATTTTGCCAAAAAGATTCC**TGGGCAGTGCCTAAGGGAAGTGCCCA**AATTGGATTATAGGATTACATGAAGTGGCCAGCTACTTCCAGAGGGCAGGGTTTTTTGTTTGTTTGCTCTGTCACCCAGGCTGGAGTGCAGTGGTGCAATCTCAGCTCACTGAAACCTCCGCCTCCCGGGTTCGAGTGATTCTTATGCCTCAGCCTCCCAAGTAGATGAAATTACAGGCGTGCACCACCACAGTCTCGCTAATTTTTATAGTTTTAGTAGAGATGGGGTTTCACCATGTTCCCTAGGCTGGT**CTTGAACTCCTGGCTTCAAG**TGATCCACCTGCCTCAGACTCCCAAAGTGCTGGGATTTTAGATGTGAGCCACCATTCCCGGCCCCAGAGGGCAGTTTTTAAAGTAGAAGTGAAGATCTGTATGCATTTA**TTAAAATATATGTTGTTCTGATTTTAA**AATATATCCCTACCAATTGTGAAGATATCGGCTAATCCAATAAAAAAGCAAA**ACCATATTTTGTATAATTATGGT**CGTATTCTGTTAAATAAACTTTTAAATCCTATTGGGTTGTAATCTGTAGTCTGGGAAGGGCTCTGCCTAACGA**TTCAGACCCCTACCAGATTTATATGTGTCTGAA**ATCCTCCTTCTAGCCTCTTAACAGGCCCATTCATTTGGGTCTTGACTGCAT**CCTCAGTCTTGTCTTTGCCTGAGG**GTGAGCTAGGACCCTGGGAGCATAAGGGAGGGGACTTGCCGGAGATCCTAGAAGGGAAAGGAGGCAGCACTGAAAGAAGAAACATTTCCTTGATTGATCATTTGCTGATACTGGCCCCTGGTTGGGGTTAGGGGTAGAAATGTGCTTCTTCCGTTTTCATCTTCTTCAAAAGGGAGTATCTCCATGATTCTGCCCCAAAGGCATGACATTTTATAGGCAAAGCCAGCCAAGTGTCTTGAGTGCTCTTAACCAAAAGGAATTCAGATTAGGAGGTCAGTCAGTGTTTGTCAGATCCTTAAGCTTTTGTTTTACTGGAATGAGTCACAGTCTACATGCTGCTGAGCTCTCAGAGATGTTTGTTTTATACAACAACCAGAACAGGTGAGGCACA**GTGGTCTGTGAGGGACTGGAGAGACCAC**ACCTTGTTCTGCTCTGCGGAAGCTGGACATGTGG**AGGAGGCACCTGACTAGATCTCTGCCTCCTGGAGTTTAGGCTGAGTCATATAGAACCAGGA**AGCTCAGTGACATGTTAAAGTGGATTAACTC**TTGGCAGTCCTGCTGTGAGGGGCTCCCTCTGCCAA**TACACACAGCCATTAATGTCCTTTAACTGTGCAAGATGAACATATCTCTGTGT**TCTGCATCTCCAGATGACAGTGCTTAGGCCTCATGCAGA**GTCCTGGTTATGGTTGAAGAAAAATTTCCATTTTGGGCTCTGAG**GGCAACTAAAGGACTGTGGAGTGGTGCTGAAACCCAGTTTTAGGTGCC**GAATCAGAGGTTTTTAAATACATTTCTCTCTTT**TGTCTCAGTCTCTTAGAGACA**GGACCTGTATTTTAAAGGTAAACAGGCAGATTCTGGCTGAGCTCAATTGCAGA**TTTGATTAACTTAGATAGATCAAA**GTCATTAGTCTCA**GAGAAAAAATTTGTTTCTC**ATCCCTAAAGTGCTATTGTGTCAGCTCTGCCAGGGTGTAAGGAAAAGGGTTTGGA**AGAGAGAAGAAAATTTCTTACCCTCTCT**CGTCACTGCCTGCTGTAAATTTGCAGTATCATTATCTTGGTCTCCTAAGCCCTGAGAGCTGCATTATATATAATTGCCCATATGTGATACCATTTACTGAATTCTTGCATTGTGCTAAGGATCAAGCTTAGTACTTTACATGCATTACATCATTGTCTTTAGGGACCCCAAAAGTGCCTCATTGGAGAGTTGTTTGGTCTGCTAAAGAGTGGGTACCACTTTCACTTCTCACATCAGTCCTTTCATTCATTCATCCAGCAAGTATTTACTAAACACCTGTTATGGGCCAGGAATTGTGCTAGGCACTGGTGATACATTGGTGAATAAACAGATACATTTTTTGCCTTTGTGAATCTTACAGTGGTAGAGAATAATAG**GCCAAGGGGGTGCTGCTTGGC**TAGGAATGTCATAGAATGCCTTTCCAAGGAGGTTACATTTAATACAGACTTGAAGAGTGAGGAGTCATGCTAAGAATGGATGAGAGGCTCACGCCTGTAATCCCAGCACTTTGGGAGGCCGAGACGGGCGGATCATGAGGTCAGGAGATCGAGACCATCCTGGCTAACACGGTGAAACCCCGTCTCTACTAAAAATACAAAAATTAGCCGGGCATGGTGGCGCGTGCCTGTAGTCCCAGCTACACAGGAGGCTGAGGCAGGAGAATGGCGTGAACCCGGGAGGCGGAGCT**TGCAGTGAGTCGAGATCGCGCCACTGCA**CTCCAGCCTGGGCGACAGAGCGAAACTCCGTCTCAAAAAAAAAAAAAAAAAAAAAAAAAAAAAAGAATGGATGAGAAATCATTCAAGACAAAAGCAAACCAGATGTGTAAAGATCCTGAGGCAAAAACAACTCCAAGGAGCCAGGTATGGTGGCCTGTAGTCCCAGCCACTTGGGAGGCTGAGACAGGAGGGTCGATTGAGTCCAGGAGTTTGAGGTTACATCAAGCAATGATTGCACCACTGCACTTCAGCCTGGTTGACAGAGTGAGAACCTATCTCTAA**CAAACAACAATAAAAAAACAATTCCAAAGAGTGTTTG**AGGATCTGAAAGAAGGCCTGTATGGTTGGAGTATAATAGTAAGAGGGAGATTTGTAGATGAGAGTGGAGAAGAAATAAAAATCAGATTATTGAAGGGACTTATGTATCATGGTAAGGAGTTTTTTTTTTTTTTCTCAAGGCAGTAGAAAGCTTAAGCAGAGGAGTG**ATATAATCTGATTTATAT**TTTTAAAAGAGTTTGCTATATGTATGTTATGCCTCATTTTTTTTTAAATTTTTTATTTTTAGATGGAGTCTTGCTCTGTTGCCCAGGCTGGAC**TGCAGTGGCACGATCTCAGCTCACTGCA**AGCTCCACTTCCCGGGTTCATGCCATTCTCCTGCCTCAGCCTCCTGAGTAGCTGGGACTATAGGCGCCCACCACCACGCCCGGCTAAGTTTTTTCTATTTTTTAGTAGAGACGCGGTTTCACTGTGTTAGCCAGGATGGTCTCAATCTCCTGACCTCGAGATCTGCCTGTCTCGGCCTCCCAAAGTGCTGGGATTACATGTGTGAGCCACCGCGCCTGGCTGTTATACATCAGTTTTTAAAGAGAGACATTAAGAAATAAAGATGACTCTGGCTACTGTGTGGAGGGTTGGAGAGAAGCAAAAGTGGAAGCAGACTGAATAAGGAGACTATTGCTGCAATTCATATAAGAGATGATGGTGTCCTGAAAGAGTGGTGGCAGTGAAAATTGGGAGAAGAGGACAGAG**TCAAGATATATTTAGGAGGCAGAAATGACAGGTCTTGA**TGATGTATTATAAATGGAGAATGAAGAATAGAATAGAGAAAAATGAATAATTAGCTTTGAACTTTCTGGCTTGAGAAATGGG**GTAAATGGGGTGTAATTTAC**TGAAACTGGAAAGATTTAGAAAGGAATAGATTCGAGG**AATCAAGAGTTTGATT**TCAGATTTCTTCAGTTTGGGATGCCCATGAGATTTCCAACTGGGGATACCAGATAGGCAGTTATACATGAGATTGGAGCACAGA**GAGATGTGGGCTTAAAATATAAGTCTGCATCTC**ATCAGCTTATTCATATGGTATTTATATCTATGGATATGGCTGAGATCCCAAAAGGAGAGAGTGAAGAGAAAATAGAGAAGAGAGTCTAGGGCCAAACCCAAGGAAGCTTCAACACACAGATGAGGAGACTGACAAAGGAAACTGTCAAAGAACAGTCAGAGAGAAAGGTCAGGAGAATGTTTTGATGGTGGGGGCCAGAATATGCTGTCCCAAAATAAGAAGGATTGTTGAGCTGAAGGCAAGTTAAAAGAAGC**AGATACAGGCTGGGTGCAGTGGGCTCATGCCTGTAATCT**TAGCATTTTGGGAGGCTAAAGTGGACAGATTGCCTGAGCTCAGGAGTTCGAGACCAGCCTGGGCAACATGATGGAACACCATCTCTACTAAAATACAAAAAAAAAAAAAAAAAAAAAAATTAGC**CAGGCATGGTGGCATGTGCCTG**TAGTCCCAGCTACTCAGGAGGCTGAGGCACGAGAATTACTTGGACCTAGGAGGCAGAGAC**TGCAGTGAGCTGATATCTCGCCACTGCA**CTCCAGCTTGGGCAACAGAGCAAGGCTCTATCTCAAATTTAAAAAATAATAATCATGATAATAATAAAAGAAGCAGATATAGGAGAGGTCTTCTGCCCTCCCTGTATTTGCCTAAAAAACCATAAATTTACAAAGACAAAAGTTATCCTACTTCCAC**CTCCCTCCTCTGCCTCCCACCACCAGGGAG**AACAAAGGTTAACCACTGAAGATAACTTTGGACTCTTATTGGCCTGGAAATGGTACTGCTTTACAAATTAGCCTTTATCTGCCATTCATTTGCCTTCCCCCAAGTAGCTACCCATTAGAGACTCAAAGTCCTTAAAGGTCTTTTCCTTTGTCTTACACTTCTTTAAAAATTTATTGTTCTTTGTTGAAGATGCTATATAAGCTGGAATTCTAAGCCACCTTTTTGAGAACTGCTCATTCTCTGGGTGTCTGT**CATGTATATATGAAATGTACATG**TTAATAAACTTCTGTTTCCTTTTTTTTCTTGTTAATCTGCCTTTTGTAACAGGGGTCCAGTCCATCTAAGAACTTATTGGGG**TTATTCTACATGACAAAGAACTCATGAGAATAA**GTGTTTCAAGAAGGGGATGACAAATGCTGCTGAGAGGTTTGATAAGATGAGGACTAAAATGTGACCATTGATTGTGGAAGTAATATAATGAACTTGACATGAGTTATGTTAGAAA**ATTACTGCTCAGGGACACCAAATTAGAGGAGACTGAGAAGTAAT**AGGAAATTGAAATAGCAGTTGTACCCAACCC**TAAACTGTAATTGAACATAAACAGTTTA**TTTGATGGAGCTTTTCACGTGTCAACCAAAGGAGCAGTGTTAGATACATAA**ACATTGCTATGATGTCAGAGTTCCAATGT**CATGGCCACTGGGATCACTTTGCCCAGACAGACCATTACCCTTCCTCTCCAGCTC**TGACTTCTACTCTTCTAGAAGTCA**CTGGTATACTTTATGGTGTGTCTTCCAATCAGACTCTTGGGAATGTCTTTTATTGAAAACTCCTGTGGATGCATAATCATTTTCCCAAGACTCAGGGCAGATTTCACCTCCTCTGGAACCCCATTAACTTTCTCAAGTATAATTAACAATTTCTTCTTCTGGAGTCCCACAATACTTTACATATACTTTAGTTATTGCATGTATTAAGTTGGTT**TCTAGTTGCTTGAGTGTATGTCCAACTAGA**CTGTGAGCACACTGAGGGAAGGAATTTGGCTTGGAAAGTAAAATTGGCTCCCAGTGTTTTATCTAAACAAATAGGCTCAATTTACAACAGGATTCCATCCTCTTACGCTGTGTATTCCCATCTCTTCCATGCTGCATCCCACCACCCTTCAATGTTATTACAATA**AAATTAAGATCAAATAATTT**ATTCAGCTAATTTTTCTTGTTTTGGACAAAATAGACATCCTAATACATTTATAGCCAAAGTTTAATCTAGACACTAAAAGGAGCATATTTTGCCTTTAGGCTCTGTATTTCCCATCCCGTTCCCAGTCAAGATAATCACAACTGTA**TTCTTTAGGAAATTCTTCACAAAGAA**CCAAACATATGTTACTATGAAACTAGATTCTTGGTATCCTGGTGAAAAAACCTGCCCTCTTGCTTGTCTGTAAATACTGGCCTTGGCTGGGTGTGGTAGCTCATGTCTATGATTCCAACACTTTGGGAGGCCAAGGTGGGAGGATCTCTTGAGCTTCAGTGTTTGAGACCAGCCTGGGCAACATAGTGAGATCCCATTTCTATAAAATTTTTTTTAAAAAGCTAGCCTGGCATGGGAGCTTGTGCCTGTAGTTCCAGCTACTTGGGAGGCTAAGATGGGAGGATTGATTGAGCACAGGAGGTCG**AGGCTGCAGTGACTGCATTTCAGCCT**GGGTGACAGAGGGAAATCCTGTCTCAAAAAAATTAAACAAATAAATAAATAAGTAAACACTTGCCTTGCCCTCAGACTTGAACAAACAACTCGTGATTTTTCTGGGGAGGTTCAGCCTGCCTATGCAGTACTCCTGTACTCTACCAGCAGTGTGGCATCAAGAGCATGTGTTGACCCTGTTAATGATTTGTAATGTTTTGTCTTTCAGAACCAGAGGCAGGAGAAGTGTCCCCTCCAGTCGGTGCGGGTGTCAACAGCAACAGCTGGACCTTTAAATACGGACCA**GGCAACCCCAAACAATCCGGTCCCGGTGAGTTGCC**CGACAAATTCATTATCCCAGGAT**CTCCTGCAATCATCTCCATCCGGCAGGAG**CCTACTAACAGCCAAATTGACAAAAGTGACTTCATAACCTTCGGCAAAAAGGAGGAGACCAAGAAAAAGAAGAAAAAGAAGAAGGGTAACAAGACCCAGGAGAAAAAAGAGAAAGGGAACAGCACGACTGACAACAGTGACCAGTGAGGTCCTCAAATGGAAACAAGCCACTTAGCCAGTTTTTGTAATAATGGCAAATCTCTCC**CATGTAGCAATTCCCTGCTCCTTTTTCCTATCTACATG**AGCCCTCTTAGAGACCTCAGAAATCTGCAGAAAGTTCCCTGTGTCTGTCTAGAACGCATTTAACAGGTTTTGTCGTAAAAGCTTTACTAAGTCTGGTGTTAACTCTTTCTCTCCACTCTGGCTTGTTTTCAGAACCTAAAAAGCAGACCCAAGTTTCCTT**TCTCCTCCGCCGCAAAGGAGA**GGCTTCCCAGCCCCGCCAGTGAGAGGTTGGACTCTCTGCCCTGTGCTCCGGGGATCCTGTCTTGATGACACTTGCAGGGCAGGCTGAAAAGTTTTGAGATTGAGCAGCTTGGGAGTTTGTGGCCACTGGGTATGTGTGGCTACCGCGGGTATGCGAGTGCCAGATATTGGCTGAGACGAGCCAGCTTAGACTAATTGGTACAAGGAAGGCAAGAAAACAAAGACAAATAAACAGCGGAAGTTATCAGTATGGAGGGGAAGTGTAAACTTAAAGGGACCAGACTTTCTAAATCTTACAACTCAAGAGGTGGCAGCCACCCTCTAGGAGACAAAACTACCCCCACTGACAAGG**CTTTAGGAGACCCTAAAG**TCTGTTGGCTGTGACGTCATTATACCTAAAATCTGCATCATACCTGCAAGCCAACAGTTCAGTGTTTTAACAGAGAACCACCCTG**GGAAACAGAAGCAGATCTGATGTGTTTCC**TATACATGTCCTGTGCTCACTTTATTAAAAATTCTTTTGCACACAATGTTTAT**GAAAAGGCCAGATCCTTTTC**CAATACTTATGCAAAAGCAAAAGAAAACCCCGACACCTCACCTTTCGCTGTTTGTTGTTTCAT**AGATTTATTTAAAAAAAGAGAAAGTCTATAGCTATAAATCT**TTAAAGAGAAATATGAATACAATTCCCCTAAACTCTCCTCAAAAGAGAATTCAGTCTACAGCCATTTAAATGATCATTGCTGC**TACAGAAGTGCTTTAAGAGAATTGCCTGAAACATCTGTA**TTATATCGGCCACCTGCCAATCACAGCTTTACTCTTTCAGGTCACTCTGGGGCTGCCTCTTGCATGTATTACTAAATAAAATGATCTCTCTTTCTCTCTCTCTCTCTCTTTTCTAAGAAACAATTATGTGCACTTTGATACACAACCTTCTCTAACCAACTATATATCAAGACCCAAAAATTGA**AGAAAAATATTGTTTTCT**CATACAGTGAGCAGATTTTTCAATCTACTAATTCTGTGACTTGTCTTGGTGTGCTAGCCTACACCTTCTCTTTGGTTTAGTTTTCCTTTTCTATAACACTCTGAATTGCTAATCTTACTAACACCTATGATGTTACCTGAAATCAATCTCCCATATGTATGCTGTATGCTATGCTAAGACTCCTGAAATATACTTACTCTGTGCTTGTGTATGTGAATGTTAATGCAACTATTACCTAGAGTGAACTTTAAGCTTTATTGTTGAATGTAATTCCATTATATTTCCTTTTGTACACCTGTGAAAAAGTGGAGTAGTGTTTTTTTAACCATTGTTAATCAGCTTTTGTGTAT**GAAAGACACAGTAAAATTTCTTTC**TTAAATCAAGATACTGGTGATTCAAGGAATTTTATTTATGGTCCAGCCAAGAGCCATCTCGTGCCAAGACTTCTGCTGGCAAGGGAATGGATAAAGCTGTTTTGTTCTAGTAACAATTT**TGGAATGAATACTGACAATATTCCA**TGAGGGTGTGCAAGCACAAATTTTACCAATCTGACCTCTTTGAAGTTGCAGAATGCTTTGAAATTCTAATGGTATCTGAAATATCAGCTCATAGAAAG**TAACAAAATTTGCTGTCACCTTAAATAAGACATTTTAATTTTGTTA**TAATGTACAATTTAGAAGTTTGATTAA**TTATATTATCTATTTAGGCATTAATATAA**AAGAGGTAGGAGTCTGTT**ATTTAAAAAAAGCATTAAAT**TTAAAAAAAAACTGTCTTGTCTACTTTTAGCTTCATTCTCCCATATTTTGAAGGGTGTGTAACTTCAGCTCTGCAGGATTGCATGGGGTAAAACTTGTTACCAACACATGTGAACCATTGCTACATTGTAGGTTGTGATCATTTTGCCCCACTGAAGCCCATGTATCTGACCTTACGTGCCTTTTGAACTAGGAGAATCGGGCTAATTTATTAATGATGATAATTATAATGTATCTGTAC**AGCACTTTTTACATTTGCGAAGTGCT**TTCCAATCCATGTTAGTTACTAGTTATTACAGCTGTAAGGATAAAACACGTCATGTGGATTCATTTTGAATTGGTGCTATTGGTATTTCCTCTGTTATTGCTAATAAATGAAAATGGTGGTATGaa**agaaatggtggtcatttct**aagaataggaggaaatagaacactgata**agcaattaatactgaagaagattgct**gcagtatgaaactcactgtaacactctcttatcacgtta**gcttttatgtccattactaaggcaaaagc**cccagtggaatcaagattcaaacttcatatcattagaatgggcagccaggaggttgctgcgtcactttgtagttacactatttcagttttctctaataagaattgcaacaaacccatttttatcacaagagcccttctaggcatctttttcattacttgtttagcccttgcacctctctacaattggaaagccaagtgacaccccaaagaaatgaacttttaggtcagggatagtggctcttgcctgtaatctcagaactttggaaggccaaggccagaggatggcttgaggccaggagttcaagatcagcctgggcagcatagcgagatcccat**ctgtacaaaaagaaataacttttattgagtacag**ggtgtatgccagccacactactgcataggttcagctgt**gacaaatagattgttttgtc**cttttgtagttaaatttgtataaatataagctttagatgctgtaggttctcagttaggagcaattttgccctcctggggacatttggcaacgtctgaagtcattgttgcttttcattacttgaagtggagggtgctattggcatctactgggtagggggcaacgatgctgtgaaacacctggcaatgcacaggacagccccctataacaaaggatcttacagcccaacatgtcaatg**tctcaaatcaagattgaga**aaccttgagatagagtgaccatataatttgtgtagtccacttttgaaatggaacaggagacaccattcgttattacttcaggattcaagatgtaaactgggactg**tcccaggctaactgggatttatagggactctataaat**aaactggtatagctgtgccactaactagatgg

Uppercase: PCDHACT

Lowercase: Flanking sequence[1000bp]

Red & Bold & Underline: Stem-loop [222]

Blue: Heptamer[368]

Green: Nonamer [49]

id-IGHM-2[C_gene_segment]

tgggctatactgggcttagctgggctgggctatactgggcttagctgggctgggctatactgggcttagctgggctgg**gctgagctgagatggtcttaggtggtctgagctcagc**taggctgggctgagctggtctg**agctcatctgagttgggctgagct**gagcttggctttgctgagctggggtggggtgggctgggctggattgagctggcctgggctgggatgaactggattgagctggcctgggctgggatgaactggaggacatggcactgggccaatcttcatgatcttgttggacatagatggatagcctcagctgagtctacactgcgttccccatcacactcaccctccctatactcact**cccaggcctgggttgtctgcctggg**gagacttcagggtagctggagtgtgactgagctggg**ggcagcagaagctgggctggagggactctattggctgcc**tgcggggtgtgtggctccaggcttcacattcaggtatgcaacctgggccctccagctgcatgtgctgggagctgagtgtgtg**cagcacctacgtgctg**atgcctcgggggaaagcaggcctggtccacccaaacctgagccctcagccat**tctgagcagggagccaggggcagtcaggcctcaga**gtgcagcagggcagccagctgaatggtggcagggatggctcagcctgctccaggagaccccaggtctgtccaggtgttcagtgctgggccctgcagcaggatgggc**tgaggcctgcagccccagcagccttggacaaagacctgaggcctca**ccacggccccgccacccctgatagccatgacagtctgggctttggaggcctgcaggtgggctcggccttggtggggcagccacagcgggacgcaagtagtgagggcactcagaacgccactcagccccgacaggcagggcac**gaggaggcagctcctc**acc**ctccctttctcttttgtcctgcgggtcctcagGGAG**TGCATCCGCCCCAACCCTTTTCCCCCTCGTCTCCTGTGAGAATTCCCCGTCGGATACGAGCAGCGTGGCCGTTGGCTGCCTCGCACAGGACTTCCTTCCCGACTCCATCACTTTCTCCTGGAAATACAAGAACAACTCTGACATCAGCAGCACCCGGGGCTTCCCATCAGTCCTGAGAGGGGGCAAGTACGCAGCCACCTCACAGGTGCTGCTGCCTTCCAAGGACGTCATGCAGGGCACAGACGAACACGTGGTGTGCAAAGTCCAGCACCCCAACGGCAACAAAGAAAAGAACGTGCCTCTTCCAGGTGAGGGCCGGGCCCAGCCACCGGGACAGAGAGGGAGCCGAAGGGGGCGGGAGTGGCGGGCACCGGGCTGACACGTGTCC**CTCACTGCAGTGATTGCTGAGCTGCCTCCCAAAGTGAG**CGTCTTCGTCCCACCCCGCGACGGCTTCTTCGGCAACCCCCGCAAGTCCAAGCTC**ATCTGCCAGGCCACGGGTTTCAGTCCCCGGCAGAT**TCAGGTGTCCTGGCTGCGCGAGGGGAAGCAGGTGGGGTCTGGCGTCACCACGGACCAGGTGCAGGCTGAGGCCAAAGAGTCTGGGCCCACGACCTACAAGGTGACCAGCACACTGACCATCAAAGAGAGCGACTGGCTCGGCCAGAGCATGTTCACCTGCCGCGTGGATCACAGGGGCCTGACCTTCCAGCAGAATGCGTCCTCCATGTGTGTCCCCGGTGAGTGACCTGTCCCCAGGGGCAGCACCCACCGACACACAGGGGTCCACTCGGGTCTGGCATTCGCCACCCCGGATGCAGCCATCTACTCCCTGAG**CCTTGGCTTCCCAGAGCGGCCAAGG**G**CAGGGGCTCGGGCGGCAGGACCCCTG**GGCTC**GGCAGAGGCAGTTGCTACTCTTTGGGTGGGAACCATGCCTCCGCC**CACATCCACACCTGCCCCACCTCTGACTCCCTTC**TCTTGACTCCAGATCAAGA**CACAGCCATCCGGGTCTTCGCCATCCCCCCATCCTTTGCCAGCATCTTCCTCACCAAGTCCACCAAGTTGACCTGCC**TGGTCACAGACCTGACCACCTATGACAGCGTGACCA**TCTCCTGGACCCGCCAGAATGGCGAAGCTGTGAAAACCCACACCAACATCTCCGAGAGCCACCCCAATGCCACTTTCAGCGCCGTGGGTGAGGCCAGCATCTGCGAGGATGACTGGAATTCCGGGGAGAGGTTCACGTGCACCGTGACCCACACAGACCTGCCCTCGCCACTGAAGCAGACCATCTCCCGGCCCAAGGGTAGGCCCCACTCTTGCCCCTCTTCCTGCACTCCCTGGGACCTCCCTTGGCCTCTGGGGCATGGTGGAAAGCACCCCTCAC**TCCCCCGTTGTCTGGGCAACTGGGGA**AAAGGGGACTCAACC**CCAGCCCACAGGCTGG**TCCCCCCACTGCCCCGCCCTCACCACCATCTCTGTTCACAGG**GGTGGCCCTGCACAGGCCCGATGTCTACTTGCTGCCACC**AGCCCGGGAGCAGCTGAACCTGCGGGAGTCGGCCACCATCACGTGCCTGGTGACGGGCTTCTCTCCCGCGGACGTCTTCGTGCAGTGGATGCAGAGGGGGCAGCCCTTGTCCCCGGAGAAGTATGTGACCAGCGCCCCAATGCCTGAGCCCCAGGCCCCAGGCCGGTACTTCGCCCACAGCATCCTGA**CCGTGTCCGAAGAGGAATGGAACACGG**GGGAGACCTACACCTGCGTGGTGGCCCATGAGGCCCTGCCCAACAGGGTCACCGAGAGGACCGTGGACAAGTCCACCGGTAAACCCACCCTGTACAACGTGTCCCTGGTCATGTCCGACACAGCTGGCACCTGCTACTGAccctgctggcctgcccacaggctcggggcggctggccgctctgtgtgtgc**atgcaaactaaccgtgtcaacggggtgagatgttgcat**cttataaaattagaaataaaaagatccattcaaaagatactggtcctgagtgcacgatgctctggcctactggggcggcggctgtgctgcacccaccctgcgcctcccctgcagaacaccttcctccacagcccccacccctgcctcacccacctgcgtgcctcagtggcttctagaaacccctgaattccctgcagctgctcacagcaggctgacctcagacttgccattcctcctactgcttccagaaagaaagctgaaagcaaggccacacgtatacaggcagcacacagg**catgtgtggatacacatg**gacagacacggacacacacaaacacatggacacacagagacgtgctaacccatgggcaca**cacatacacagacatggacccacacacaaacatatgtg**gacacacatgtacaaacatgcacaggcacacaaagagaacactgactacaggcacacacacacacgggcacacacatggatatgtgcacacatggacacataca**tgtgcaggacatgcaca**cacacagacacactagcacagaggcatacacacacagacacacacattcacaaacac**acatgtgcatgcaaacacacacacatgt**acagacacaagtacatggacacatgcacacccagagacacactgacacagacacacaggagcatgtgatacactaacacgtggacacacacgtctacccacaggcacacaacagatggacacgcgtacacagacatgcacacacccacaggcacaacacgtgcgcatgccggccggcccccgccaacattctcccagggccctgccggatactctgtccctgcagcagtttgctccctgcgctgtgctggcaccggggctttgggcccaggctctgcttgtccttctgtctctgct

Uppercase: IGHM-2

Lowercase: Flanking sequence[1000bp]

Red & Bold & Underline: Stem-loop [25]

Blue: Heptamer[63]

Green: Nonamer [1]

id-IGHA2-2[C_gene_segment]

gagctagactgggctgagctgggtgagcttaggtggactgagctgggctgggctgggctgagctgagctgagctgggctgggctgggctgggatgagctgtactgagctgccctggggtgggctgggctgagctgggctgagctgggctgggctggactgagctggactgagctggactggcctgggctgggctgggctggatgagctgaggtggctgctaatgtgggaaggaggccgtgggttgagtgtgactccacctgcagagccctgagcccagctgtgttcttaggggttctgagggccacgcagctctgttgcaccatgattctgtcttctctctt**gcccactgcctgaaggaaatttggagtgggc**tgggcccagagctcccctgggtaa**caggccctgtcctggagggcctg**gcagggacatggcttag**cctgttggcctctagtcccgagacctcataggccacagg**ggtccactgtggcttgtttgggcctggggtggggctcatggagtggtgggtgttggactgagactctgaccagggacaggggg**atggggtcacagccaagccactccacccctaccccat**gcacacagcactcagagcccaggccccctcctcagagcccccaccaaaatcctctctaggggcaggggaaagagcaagatatgtcccccacccagagcaggaactggggtcagggagctcaggggactcagccactccatggcagagccctgtttaatataacttgtgtctgggatggcctgggtcagaggccctatctaaggagcatgttcagaaactgtgtcgctgggatgag**acagctgggtccaaccgcaggcccatggtgcaggagctgt**gtaaccttggggctgtcaccaggcctctctgtgctgggttcctccagtgta**gaggagaggcaggtacagcctgtcctc**ctggggacatggcatgagggccgcgtcctcacagcgcattctgtgttccagCATCCCCGACCAGCCCCAAGGTCTTCCCGCTGAGCCTCGACAGCACCCCCCAAGATGGGAACGTGGTCGTCGCATGCCTGGTCCAGGGCTTCTTCCCCCAGGAGCCACTCAGTGTGACCTGGAGCGAAAGCGGACAGAACGTGACCGCCAGAAACTTCCCACCTAGCCAGGATGCCTCCGGGGACCTGTACACCACGAGCAGCCAGCTGACCCTGCCGGCCACACAGTGCCCAGACGGCAAGTCCGTGACATGCCACGTGAAGCACTACACGAATCCCAGCCAGGATGTGACTGTGC**CCTGCCCAGGTCAGAGGGCAGG**CTGGGGAGTGGGGCGGGGCCACCCCGTCCTGCCCTGACACTGCGCCTGCACCCGTGTTCCCCACAGGGAGCCGCCCCTTCACTCACACCAGAGTGGACCGCGGGCCGAGCCCCAGGAGGTGGTGGTGGACAGGCCAGGAGGGGCGAGGCGGGGGCACGGGGAAGGGCGTTCTGACCAGCTCAGGCCATCTCTCCACTCCAGTTCCCCCACCTCCCCCATGCTGCCACCCCCGACTGTCGCTGCACCGACCGGCCCTCGAGGACCTGCTCTT**AGGTTCAGAAGCGAACCT**CACGTGCACACTGACCGGCCTGAGAGATGCCTCTGGTGCCACCTTCACCTGGACGCCCTCAAGTGGGAAGAGCGCTGTTCAAGGACCACCTGAGCGTGACCTCTGTGGCTGCTACAGCGTGTCCAGTGTCCTGCC**TGGCTGTGCCCAGCCA**TGGAACCATGGGGAGACCTTCACCTGCACTGCTGCCCACCCCGAGTTGAAGACCCCACTAACCGCCAACATCACAAAATCCGGTGGGTCCAGACCCTGCTCGGGGCCCTGCTCAGTGCTCTGGTTTGCAAAGCATATTCCCGGCCTGCCTCCTCCCTCCCAATCCTGGGCTCCAGTGCTCATGCCAAGTACAGAGGGAAACTGAGGCAGGCTGAGGGGCCAGGACACAGCCCAGGGTGCCCACCAGAGCAGA**GGGGCTCTCTCATCCCCTGCCCAGCCCC**CTGACCTGGCTCTCTACCCTCCAGGAAACACATTCCGGCCCGAGGTCCACCTGCTGCCGCCGCCGTCGGAGGAGCTGGCCCTGAACGAGCTGGTGACGCTG**ACGTGCCTGGCACGT**GGCTTCAGCCCCAAGGATGTGCTGGTTCGCTGGCTGCAGGGGTCACAGGAGCTGCCCCGCGAGAAGTACCTGACTTGGGCATCCCGGCAGGAGCCCAGCCAGGGCACCACCACCTTCGCTGTGACCAGCATACTGCGCGTGGCAGCCGAGGACTGGA**AGAAGGGGGACACCTTCT**CCTGCATGGTGGGCCACGAGGCCCTGCCGCTGGCCTTCACACAGAAGACCATCGACCGCTTGGCGGGTAAACCCACCCATGTCAATGTGTCTGTTGTCATGGCGGAGGTGGACGGCACCTGCTACTGAgccgcccgcctgtccccacccctgaataaactccatgctcccccaagcagccccacgcttccatccggcgcctgtctgtccatcctcagggtctcagcacttgggaaag**ggccagggcatggacagggaagaataccccctgccctgagcc**tcggggggcccctggcacccccatgagactttccaccctggtgtgagtgtgagttgtgagtgtgagagtgtgtggtgcaggaggcctcgctggtgtgagatcttaggtctgccaaggcaggcacagcccaggatgggttctgagagacgca**catgccccggacagttctgagtgagcagtggcatg**gccgtttgtccctgagagagccgcctctggctgtagctgggagggaatagggagggtaaaaggagcaggctagccaagaaaggcgcaggtagtggcaggagcggcgagggagtgaggggctggactcca**gggccccactgggaggacaagctccaggagggccc**ca**ccaccctagtgggtgg**gcctcaggacgtcccactgacgcatgcagg**aaggggcacctcccctt**aaccacactgctctgtacggggcacgtgggcacacatgcacactcacactcacatatacgcctgagccctgcaggag**tggaacgttcacagcccagacccagttcca**gaaaagcca**ggggagtcccctcccaagcccccaagctcagcctgctcccc**caggcccctctggcttccctgtgtttccactgtgcacagatcaggcaccaactccacagacccctc**ccaggcagcccctgctccctgcctgg**ccaagtctcccatcccttcctaagcccaactaggacccaaagcatagacagggaggggccgcgtggggtggcatcagaa**gcaggccagtgagacagggcctgc**ccagggccctctgcatgcctctggcttctg**cctggggctcccagg**agtgaaagaacagtcccacaaccactgtggggacacc

Uppercase: IGHA2-2

Lowercase: Flanking sequence[1000bp]

Red & Bold & Underline: Stem-loop [22]

Blue: Heptamer[64]

Green: Nonamer [3]

id-TRBC1[C_gene_segment]

gcacagtaaattgtggtttcttccactcctcatgtgtcttcagatgaaatcattctttccaataatcccagaaattctttctctcctctctcaagcatgtgaaaaggtccagagctctgcagtgtgagctttctactgaaatggcccttggactttgtggttcattcatactcagtggtctagcttgtactacttttgagaatgcaaagcttaactgtggacggattccaatcctggccaggcagggttgctggacactctgagagaagaaagggtt**aatcccatgaccatcaacttccatgggatt**tcagccatcctggacaagctaccacac**cctcctgccccaggggaggagg**aaatgtggaccatcccatcagatattgaccaggtttggctctttaaagagggttacatgcaagaaaataaaattttttaaaaaggtgctgggcaggtgggggactcagatgtaatggaaaagtgt**cttttctagaaaagaaaag**ctaattctaatatgtgtcactaccccacgagacaaatatatacatcttgatttaaaaaaggaaaat**tataattagaaaaagtcaatttagttattgtaattata**ccactaatgagagtttcctacctcgagtttcaggattacatagccatgcaccaagcaaggctttgaa**aaataaagatacacagataaattattt**ggatagatgatcagacaagcctcagtaaaaacagccaagacaatcaggatataatgtgaccataggaagctggggagacagtaggcaatgtgcatccatgggacagcatagaaaggaggggcaaagtggagagagagcaacagacactgggatggtgaccccaaaacaatgagggcctagaatgacatagttgtgcttcattacggcccattccca**gggctctctctcacacacacagagccc**ctaccagaaccagacagctctcagagcaaccctggctccaacccctcttccctttccagAGGACCTGAACAAGGTGTTCCCACCCGAGGTCGCTGTGTTTGAGCCATCAGAAGCAGAGATCTCC**CACACCCAAAAGGCCACACTGGTGTG**CCTGGCCACAGGCTTCTTCCCTGACCACGTGGAGCTGAGCTGGTGGGTGAATGGGAAGGAGGTGCACAGTGGGGTCAGCACGGACCCGCAGCCCCTCAAGGAGCAGCCCGCCCTCAATGACTCCAGATACTGCCTGAGCAGCCGCCTGAGGGTCTCGGCCACCTTCTGGCAGAACCCCCGCAACCACTTCCGCTGTCAAGTCCAGTTCTACGGGCTCTCGGAGAATGACGAGTGGACCCAGGATAGGGCCAAACCCGTCACCCAGATCGTCAGCGCCGAGGCCTGGGGTAGAGCAGGTGAGTGGGGCCTGGGGAGATGCCTGGAGGAGATTAGGTGAGACCA**GCTACCAGGGAAAATGGAAAGATCCAGGTAGC**AGACAAGACTAGATCCAAAAAGAAAGGAACCAGCGCACACCATGAAG**GAGAATTGGGCACCTGTGGTTCATTCTTCTCCCAGATTCTC**AGCCCAACAGAGCCAAGCAGCTGGGTCCCCTTTCTATGTGGCCTGTGTAACTCTCATCTGGGTGGTGCCCCCCAGCCCCCTCAGTGCT**GCCACATGCCATGGATTGCAAGGACAATGTGGC**TGACATCTGCATGGCAGAAGAAAGGAGGTGCTGGGCTGTCAGAGGAAGCTGGTCTGGGCCTGGGAGTCTGTGCCAACTGCAAATCTGACTTTACTTTTAATTGCCTAT**GAAAATAAGGTCTCTCATTTATTTTC**CTCTCCCTGCTTTCTTTCAGACTGTGGCTTTACCTCGGGTAAGTAAGCCCTTCCTTTTCCTCTCCCTCTCTCA**TGGTTCTTGACCTAGAACCA**AGGCATGAAGAACTCACAGACACTGGAGGGTGGAGGGTGGGAGAGACCAGAGCTACCTGTGC**ACAGGTACCCACCTGT**CCTTCCTCCGTGCCAACAGTGTCCTACCAGCAAGGGGTCCTGTCTGCCACCATCCTCTATGAGATCCTGCTAGGGAAGGCCACCCTGTATGCTGTGCTGGTCAGCGCCCTTGTGTTGATGGCCATGGTAAGCAGGAGGGCAGGATGGGGCCAGCAGGCTGGAGGTGACACACTGACACCAAGCACCCAGAAGTATAGAGTCCCTGCCAGGATTGGAGCTGGGCAGTAGGGAGGGAAGAGATTTCATTC**AGGTGCCTCAGAAGATAACTTGCACCT**CTGTAGGATCACAGTGGAAGGGTCATGCTGGGAAGGAGAAGCTGGAGTCACCAGAAAACCCAATGGATGTTGTGATGAGCCTTACTATTTGTGTGGTCAATGGGCCCTACTACTTTCTCTCAATCCTCACAACTCCTGGCTCTTAATAACCCCCAAAA**CTTTCTCTTCTGCAGGTCAAGAGAAAG**GATTTCTGAaggcagccctggaagtggagttaggagcttctaacccgtcatggtttcaatacacattcttcttttgccagcgcttctgaagagctgctctcacctctctgcatcccaatagatatcccccta**tgtgcatgcacacctgcaca**ctcacggctgaaatctccctaacccagggggaccttagcatgcctaagtgactaaaccaataaaaatgttctggtctggcctgactctgacttgtgaatgtctggatagctccttggctgtctctgaactccctgtgactctccccattcagtcaggatagaaacaagaggtattcaaggaaaatgcagactcttcacgtaagagggatgaggggcccaccttgagatcaatagcagaggttaa**ttcagcgtgaaaggcagtgatgggagctgaa**gaggttacttctagaacagtctaggaagacacagatgttg**agtataggaattttctatatccaactatact**gttctgcccaggaaagacgtgctcagaggaagagccaacctatacaggtgtgttcaccc**tccagctggccatgtccccgtgactaacaaagctgga**ttccataagcatcacccaccttccttgcagctttcttattgagcactccattca**tcttcattggttcaccaagttgatttcccagctccaaagaaga**gaggctctgacttgcaaacttattttcaat**ggaagatgtgtcttcc**ggtttaagttacccatctgtttataaatctctctctagtgaattaaacca**gaatgaaaatgtcccctaatcattc**ctggaaggttagaaaataaaggtatctaaaactgagaatcagccccattcctacttctagaattccttcaaaagctccttctgttgtctcactgtcaccatggtgatggagtcccaaatcccaaaggtggcacagaagaccggatgattatccttgtctccttccacactctcctcacttctctcatccctgaagcc

Uppercase: TRBC1

Lowercase: Flanking sequence[1000bp]

Red & Bold & Underline: Stem-loop [22]

Blue: Heptamer[44]

Green: Nonamer [2]

id-IGHGP[C_gene_segment]

aaatatata**tttttcaaggtgaaaaa**aaaaaaagaaaaacctgccataatgaagagcagaccaatattccagaaaactgtcact**ttaacagagaagaccaaattctagtttcacatgaactgttaa**t**attaaagctaattttaat**taaaccttataaataattccatccag**gctgggcaaggtggcttacacctgtagtcccagc**attccaggagggtggatcacttaa**gcccaggagtttgagatcagcctgggc**aacatggggaagccctgtctctacaaaaaatacaaaaaaaaaattggccaggcatggtggcacgtgcccgtagtcccagctactt**gggaggctgaggtgggaggattgcttgagccctcccacctcaacctccc**tcagcct**cagtgctgagtgccactgcactg**ctgcctaggcaacacagtgggaccctgtctcaaaaataaataaataaataaatccggccaggtgcagtggctcatgcctgtaa**tcccaacactttggga**ggcctaggcgggtggatcatgaggtcaggagttcgagaccagcctggccaacgtggtgaaacccctgtctctactaaaactataaaaat**tagctgggcgtggtggtgggcacctgtaatcccagcta**ctctggaggctgaggcaggagaatcgcttgaacccaagaggcagaggttgcagtgaaccgagatcacaccattgcactccagcttaggcaacaagagtgaaactctgtctcaaaaaaataaataaataaatccaatcacagccgcctttgaccacataagatcccttttccacaatccttttacaactttttatttgtttgttttttgttttgttttgttttttgtttttgtttttgttttttttgagacaaagcactgacctggctgccgagccccgccccctagg**ctgcaggggtgcctgcag**aagggcaccacagggccaccagtcctgcaagctttctggggcaggccggGCCTGACTTTGGCTTTGGGGCAGGGAGGGGGCTAAGGTGAGGCAGGTGGCACCAGCCAGGTGCACACTCAATGCCCGTGAGCCCAGACACTGGACCCTGCCTGGA**CCCTCGCGGATAGACAAGAACCGAGGG**GCCTCTGCACCCTGGGCCCAGCTCTGTCCCACACCGCGGTCACATGGCACCACCTCTCTTGCAGCCTCCACCAAGGGCCCATCGGTCTTCCCCCTG**GTGCCCTCCTCCAGGAGCGTCTCTGAGGGCAC**AGCGGCCCTGGGCTGCCTGGTCAAGGACTACTTCCCCGAACCGGTGACGGTGTCGTGGAACTCAGGGGCCCTGACCAGAAGCGTGCACACCTTCCCGGCTGTCCTACAGTCCTCAGGACTCTACTCCCTCAGCAGCGTGGTGACC**GTGCCCTCCAGCAGCTTGGGCAC**CCAGACCTACACCTGCAACGTAGATCACAAGCCCAGCAA**CACCAAGGTGGACAAGACAGTTGGTG**AGAGG**CCAGCACAGGGAGGGAGGGTGTCTGCTGG**AAG**CCAGGCTCAGCCCTCTTGCCTGG**ACGTACCCCGGCTGTGCAGCCCCAGTCCAGGGCAGCAAGGCAGGCCCCATCTGTCTCCTCACCCGGAGGCCTCTGCCCGCCCCACTCATGCTCAGGGAGAGGGTCTTCTGGCTTTTTCCACCAGGCTCCAGGCAGCCACAGGCTGGAA**GCCCCTACCCCAGGCCCTGCGCACAAAGGGGC**AGGTGCTGCACTTAGACTGGCCAAGAGCCATATCCGGGAAGACCCTGCCCCTGACCTAAGCCCACCCCAAAGGCCAAGATCTCCACTCCCTCAG**CTCAGACACCTCTCCTCCCAGATCTGAG**TAACTCCCAATCTTCTCTCTGCAGAGCCCAAAACCCCATGTTGTGACACAACTCA**CACATGCCCACCATGTG**CAAGTAAGCCAGCCCAGGCCTCGCCCTCCAGCTCAAGGCGGGACAGGTGCCCTAGAGTAGCCTGCGTCCAGGGACAGGCCCCAACCGGG**TGCTGACACGTCCGCCTCCATCTCTTCCTCAGCA**ACTGAACCCC**TGGGGGGACCGTCAGTCTTCCTCTTCCCCCCA**AAACCCAAGGATACCCTCATGATCTCCCGGACCCCTGAGG**TCACGTGCGTGGTGGTGGACGTGA**GCCACGAAGACCCTGAGGTCAAGTTCAACTGGTACGTGGACGGCGTGGAGGTGCATAATGCCAAGACAAAGCCGTGGGAGGAGCAGTACAACAGCACGTACCATGTGGTCAGCGTCCTCACCGTCGTGCACCAGAACTGGCTGAACGGCAAGGAGTACAAGTGCAAGGTCTCCAACAAAGGCCTCCCAGCCCCCATCGAGAAAACCATCTCCAAAACCAAAGGTGGGACCCACGGAGCGCGAAGGCCACGTGGA**CAGAGGCCGGCTTGGCCCACCCTCTG**CCCTGGGAGTGACCGCTGTACCAACCTCTGTCCCTACA**GGGCAGCCCCGAGAACCACAGGTGTACACCCTGCCC**CCATCCCAGAAGATGACCAAGAAC**CAGGTCACCCTGACCTG**CCTGGTCAAAGGCTTCTACCCCAGCGACATCGCCGTGGAGTGGGAGAGCAATGGGCAGCCGGAGAACAACTACAAGACCACGCCTCCCATGCTGGACTCCAACGGCTCCTTCTTCCTCTATAGCAAGCTCACCGTGGACAAGAGCAGGTGGCAGCAGGGGAACGTCTT**CTCATGCTCCGTGATGCATGAG**GGTCTGCAGAACCACTACACGCAGAAGAGCCTCTCCCTGTCCCCGGGGTAAatga**gtgcgacggccggcaagcccccgctccccgggctctcgcggtcgcac**gaggatgcttggcacgtaccccgtctacatacttcccaggcacccagcatggaaataaagcacccaccactgccctgggcccctgcgagactgtgatggttctttccacggg**tcaggccgagtctgaggcctga**gtggcatgagggaggcagagcgggtcccactgtccccacact**ggcccaggctgtgcaggtgtgcctgggcc**gcctagggtggggctcagccaggggctgccctcggcagggtgggggatttgccagcgtggccctccctccagcagcacctgccctgggctgagccacgagaagccctaggagcccctgg**ggacagacacacagcccctgcctctgtaggagactgtcc**tgttctgtgagcgccctgtcctccgaccccc**catgcccactcgggggcatg**cctagtccatgtgcgtagggacaggccctccctcacccatctacccccacggcactaacccctggcagccctgcccagcctcgcacccgcatggggacacaaccgactccggggacatgcactctcgggccctatggagggactggtccagatgcccacacacacactcagcccagacccgttcaacaaaccccgcactgaggt**tggccggccacacggcca**ccacacacacacgtgcacgcctcacacacggagcctcacccgggcgaactgcacagcacccagaccagagcaaggtcctcgcacacgtgaacactcctcggacacaggcccctacaagccccatgcggcacctcaaggcccacgagcctctcggcagcttctccacttgctgaccagctcagacaaacccagtcctcctctcacaaagtg**cccctgcagccgccacacacacacagggg**atcacacaccacgtcacgtccctggccctggcccacttcccaatacagcccttccctgctcctggggtcacatg

Uppercase: IGHGP

Lowercase: Flanking sequence[1000bp]

Red & Bold & Underline: Stem-loop [33]

Blue: Heptamer[50]

Green: Nonamer [2]

id-IGLC5[C_gene_segment]

ctaacgaactttgtgcaagggaaactgaggccccatctcatgagggagagggaacaaggggctcgaaggagtgaccacctggtggactttagaaggacctgaaaccctcagagccaagataggggaatgaaaactcagagtctcagggccc**agtcccctggactgtgggact**ctggatccaggctgggaacaaggtaggaggtgcaggggcctctccaggtttctg**tgggctcccagggagagagccctgagctggcctgggaccca**tgaagccctgtcaggagggacgggaaggctctggacatgaaggagccaggtgaagtgtcacgaaaggccatggcattcagggaggtggctgatgggtctctgtgggaggcacccctagaagcaggaacccctgagttcaccgacaggcatatcccaaggcagaaaaactgtagattggccctaaacacagagagactctaacacagactccacagacaaagagcccaggacagacaga**cagtgggacttgggtgagcaaaggccctgactccactg**cagaagatccaggagagacggat**gtgggtacaaacaagagctcttacgtgagagacccac**tctcccccaacccagagcagctgtgtcaggtgagaaaagtttccagagtgaatctaacaagaggctcacagagctcagagaacacagccagagccggatcacgtcaggatgacagggtgaggcagcgccaggcaattgaaggtccactgtctatacaaaagcccaagtcc**ttggaagatttccaa**acaggcgatgggcaggggccaagctcagaggttttctgactgacagagttcttttaggaacatgatgtcacacccttctgagacagtgtggaccacatctgagaggtcctgcactggta**ctggggggcctgaacactcctcatgcactgaggtcaggggctccccag**gtggacaccaggactctgacctcctgcccctcatccattctgcagGTCCGCTCAAGGCCACCCCCTTGGTCACTCTGTTCCCGCCCTCCTCTGAGGAGCTCCAAGCCAACAAGGCCATGCTGGTGTGTCTCATAAATGA**CTTCTACCCAGGAGCCATAGAAG**GAAAATGGCACCCTAGTCACCAAGGGCATAGAGACAACCACACCCTCCACACAGAGCAACAACAACTATGCGGCCAGCAGCTACCTGAGCCTGACGCCCGAGCAGTGGAAGTCCCACAGAAGCTACAGCTGCCAGGTCACGCACAAAGAAAGTACCATGGAGAAGACAATGGCCCATGCAGAATGTTCTT**AGgcccccgaccctcaccccacccacaggggcct**ggagctgcaggatcccagggcagaggtgttccctcccaccccaagtcatccagtccttctgccgacacccaataaaccctcaataaatgtcctccttgtcaatcagaaaacatgctgtctgctcatttttgtttatccactttccatcctaaatttttttatc**tccaaagatcaacagaatcctttgga**cttgtgacaatgcagcctgggaccatcttcgtgtttctctgagccatactttttccatcaccttatccccatagaatgttccagaacaaggaaatagtcttcagaaggaaccagacatcttcattcttacccacgtctctgttgtgcttcctgctgccctgtgagcacgtccacctctggctg**cccggggacacaccctccttacatccctatggccccggg**agaacatctcagctcccttctttgttgtcctcggccctcactgtccagatgccatccctctctcaaggtgtttgtccatccccctccaaggccatgtggtcaccccacctgtatttcccctcatcaaagcctagagcatctccccttctccatgggcacactctgttcccgtcatcccagggaccctgccctcagcccccacaggccctccctataataatcactgatgggatgcttaccccctggaccctggtcactcaccccacatccccctctgggtgcctctcatcccacatgtgtcaccggatgccccagctgtattcaacattcctaggggacaagggtcggctctgctcctcactgacaccattatgaatgcctaatgtcctgctgtgcacaagttgtcaattgactggtgagttcaaagtcagattctggggttatgccaggagcttgggtgac**gctgattcttcttctgaagaactcagc**atgggatctgagctataatacacacacacgtgtacacacacacaatgcacataaacacacatgtccat

Uppercase: IGLC5

Lowercase: Flanking sequence[1000bp]

Red & Bold & Underline: Stem-loop [11]

Blue: Heptamer[36]

Green: Nonamer [4]

id-IGLC2[C_gene_segment]

aagccttgtt**ctgttctggcctcctcagtctgggttcttgtcggaacag**ctttgtccttgggttacctgggttccatctcctggggaattgggaacaaggggtctgagggaggcacctcctgggagactttagaaggacccagtgccctcggggctgatgctcgggaatcacagagctgggacccagagccaggatccagacccagaatgaggtaggaggtggaggggctg**ccctgggcgtctgggggctgccaggg**actgagc**cctgagccagcctgagactcagg**aaaccccgtcaggagggagaagggagaagcagactctggacaccagaaagccaggggaagg**gtcacaaaaggagtggatgtgac**ggaagggcgggctcctgggtctcttcagaacatat**cccctgtgcccagggg**gatcagaggggcagagtccactgcgtgaaag**ccccactgctatgaccaggtagccgggacgtgggg**tggatgccagaaaagactccacggaataaga**gagagcccaggacagcaggcaggctctc**cgatccccccaggcccttgccccatacacgggctccagaacacacatttggctggaacagcctgagggaccaaaaggccccagtatcccacagagctgaggagccaggccagaaaagtaaccccagagttcgctgtgcaggggagacacagagctctctttatctgtcaggatggcaggaggggacagggtcagggcgctgagggtcagatgtcggtgttgggggccaaggccccgagagatctcaggacaggtggtcaggtgtctaaggtaaaacagctccccgtgcaga**tcagggcatagtggaaaacaccctga**cccctctgcctggcatagaccttcagacacagagcccctgaacaagggcaccccaac**acctcatcatatactgaggtcaggggctccccaggtggacaccaggactctgaccccctg**cccctcatccaccccgcagGTCAGCCCAAGGCTGCCCCCTCGGTCACTCTGTTCCCGCCCTCCTCTGAGGAGCTTCAAGCCAACAAGGCCACACTGGTGTGTCTCATAAGTGACTTCTACCCGGGAGCCGTGACAGTGGCCTGGAAGGCAGATAGCAGCCCCGTCAAGGCGGGAGTGGAGACCACCACACCCTCCAAACAAAGCAACAACAAGTACGCGGCCAGCAGCTATCTGAGCCTGACGCCTGAGCAGTGGAAGTCCCACAGAAGCTACAGCTGCCAGGTCACGCATGAAGGGAGCACCGTGGAGAAGACAGTGGCCCCTACAGAATGTTCATAGgttctcaaccctcaccccccaccacgggagactagagctgcaggatcccag**gggaggggtctctcctccc**accccaaggcatcaagcccttctccctgcactcaataaaccctcaataaatattctcattgtcaatcagaaatcttgttttatctcattttttcttttctcacatataattcctagcctttc**ctgggttctcaatttgtggtggaaagaaccctgaacccag**tgggaaagttgcctatgtgaaggggttctcagttccctgggcatctctgcaggtaaggccttcctcacccagacaccccttcctcagctctccactgtacccctgagcca**ccagcctcgcctggctgg**gaccaggggggtgtcacactctcctagattctgcctttcaacagaaacctaaccacgcatcacacggcacttctcgcatgccttctgtgtctgctccagtctctgggctaaagagttgctggtccgggacaggggataggtccgctcttggtca**gatgccaggtccctgccatggcatc**cctgaccctatgcaacaagccagtgactctggtgagctctctgtgtcaggagaatccatgatccagagtttcatattgtcctgcaagcatctggtgggctgtagctcttgccaaactgggaaataccatggcccagcatcaggatgcaggacagtccggagagggaaatcaggagaagtgaaggggtctctgggg**agcccagatgtgggct**agaggcagaagtaagggtgaagagcacctatgagtcaatgtcatggtctcagcaggaacacagttgaaaatccccattcca**cacaagaccgtttagcaggaaaggagtccatacttgtg**ctgccaccaggatgtcctgagaagccttggagaatgaaacatacag**gtgcatttcctagacttgacaatgcac**gttagccaagtaaaggcaatgaaaagttctctactagggaaataatttcctgtggtaa

Uppercase: IGLC2

Lowercase: Flanking sequence[1000bp]

Red & Bold & Underline: Stem-loop [17]

Blue: Heptamer[25]

Green: Nonamer [5]

id-IGHG2[C_gene_segment]

aaatggggcctccctgtggcctgggggtcctggcaccacgcagggtggggagggccaagggcaggtgcaaggctcctacctg**tgctggggggcctgggttgagcccagca**gggaccttgccgggggaagctctggagagagggaggaggtgggctggtggccgagaaggccaggccagggctgggagggtgaggttgtggtgactga**gcctccagaagtaatgcaggacactgggaggc**agggggcatccaggcactcagggccctgacctgggctgctgcacactggggctaaggggaaaggaggggagaggctgaggaggaggctccaggaggctattccaaggcagggggttccggggccctggggctgaagggcgccgaccctatgcagtgtctggc**ccctctgctgcacagaagaaaagggccttggagggcagaggg**caggctatgaccag**ggccctgggcaagtcaggcccactcactagcggagggcc**acgctggggcggcagggtcaggagcttcaggggactcgggggacccacgagaagccatctgagaacagtgtccactggtcaagccaggcacccataaaaggctggagtggggccaatgggcatgagccgtccctgaggtggcaccgatggccagagctgaggccaagctagaggccctggactgtgctgactcc**cggcaggcacagagcgctgacctggctgccg**agccccgcctcctagg**ctgcaggggtgcctgcag**aagggcaccacagggccaccggtcctgcaagctttctggggcaggccgggcctgactttggctttggggcagggagggggctaaggtgacgcaggtggcgccagccaggtgcacacccaatgcccgtgagcccagacactggaccctgcctgga**ccctcgcagatagacaagaaccgaggg**gcctctgcgccctgggcccagctctgtcccacaccgcggtcacatggcaccacctctcttgcagCCTCCACCAAGGGCCCATCGGTCTTCCCCCTGGCGCCCTGCTCCAGGAGCACCTCCGAGAGCACAGCGGCCCTGGGCTGCCTGGTCAAGGACTACTTCCCCGAACCGGTGACGGTGTCGTGGAACTCAGGCGCTCTGACCAGCGGCGTGCACACCTTCCCGGCTGTCCTACAGTCCTCAGGACTCTACTCCCTCAGCAGCGTGGTGACCGTGCCCTCCAGCAACTTCGGCACCCAGACCTACACCTGCAACGTAGATCACAAGCCCAGCAA**CACCAAGGTGGACAAGACAGTTGGTG**AGAGGCCAGCTCAGGGAGGGAGGGTGTCTGCTGGAAG**CCAGGCTCAGCCCTCCTGCCTGG**ACGCACCCCGGCTGTGCAGCCCCAGCCCAGGGCAGCAAGGCAGGCCCCATCTGTCTCCTCACCCGGAGGCCTCTGCCCGCCCCACTCATGCTCAGGGAGAGGGTCTTCTGGCTTTTTCCACCAGGCTCCAGGCAGGCACAGGCTGGG**TGCCCCTACCCCAGGCCCTTCACACACAGGGGCAGGTGCTTGGCTCAGACCTGCC**AAAAGCCATATCCGGGAGGACCCTGCCCCTGACCTAAGCCGACCCCAAAGGCCAAACTGTCCACTCCCTCAG**CTCGGACACCTTCTCTCCTCCCAGATCCGAG**TAACTCCCAATCTTCTCTCTGCAGAGCGCAAATGTTGTGTCGAGTGCCCACCGTGCCCAGGTAAGCCAGCCCAGGCCTCGCCCTCCAGCTCAAGGCGGGACAGGTGCCCTAGAGTAGCCTGCATCCAGGGACAGACCCCAGCTG**GGTGCTGACACGTCCACCTCCATCTCTTCCTCAGCACC**ACCTGTGGCAGGACCGTCAGTCTTCCTCTTCCCCCCAAAACCCAAGGACACCCTCATGATCTCCCGGACCCCTGAGG**TCACGTGCGTGGTGGTGGACGTGA**GCCACGAAGACCCCGAGGTCCAGTTCAACTGGTACGTGGACGGCGTGGAGGTGCATAATGCCAAGACAAAGCCACGGGAGGAGCAGTTCAACAGCACGTTCCGTGTGGTCAGCGTCCTCACCGTCGTGCACCAGGACTGGCTGAACGGCAAGGAGTACAAGTGCAAGGTCTCCAACAAAGGCCTCCCAGCCCCCATCGAGAAAACCATCTCCAAAACCAAAGGTGGGACCCGCGGGGTATGAGGGCCACATGGA**CAGAGGCCGGCTCGGCCCACCCTCTG**CCCTGGGAGTGACCGCTGTGCCAACCTCTGTCCCTACA**GGGCAGCCCCGAGAACCACAGGTGTACACCCTGCCC**CCATCCCGGGAGGAGATGACCAAGAAC**CAGGTCAGCCTGACCTG**CCTGGTCAAAGGCTTCTACCCCAGCGACA**TCTCCGTGGAGTGGGAGAGCAATGGGCAGCCGGAGA**ACAACTACAAGACCACACCTCCCATGCTGGACTCCGACGGCTCCTTCTTCCTCTACAGCAAGCTCACCGTGGACAAGAGCAGGTGGCAGCAGGGGAACGTCTT**CTCATGCTCCGTGATGCATGAGGCTCTGCACAACCACTACACACAGAAGAGCCTC**TCCCTGTCTCCGGGTAAATGAgtgccacggccggcaagcccccgctccccaggctct**cggggtcgcgcgaggatgcttggcacgtaccccg**tctacatacttcccgggcacccagcatggaaataaagcacccagcgctgccctgggcccctgcgagactgtgatggttctttccgtggg**tcaggccgagtctgaggcctga**gtggcatgagggaggcagagcgggttccactgtccccacact**ggcccaggctgtgcaggtgtgcctgggcc**gcctagggtggggctcagccaggggctgccctcggcagggtgggggatttgccag**cgtggccctccctccagcagcagctgccctgggctgggccacg**ggaagccctaggagcccctgg**ggacagacacacagcccctgcctctgtaggagactgtcc**tgtcctgtgagcgccctgtcctccgacctc**catgcccactcgggggcatg**cctagtccatgtgcgtagggacaggccctccctcacccatctacccccacggcactaacccctggctgccctgcccagcctcgcacccgcatggggacacaaccgactccggggacatgcactctcgggccctgtggagggactggtccagatgcccacacacaca**ctcagcccagacccgttcaacaaaccccgcgctgag**gt**tggccggccacacggcca**ccacacacacacgtgcacgcctcacacacggagcctcacccgggcgaaccgcacagcacccagaccagagcaaggtcctcgcacacgtgaacactcctcagacacaggcccccacgagccccacgcggcacctcaaggcccacgagccgctcggcagcttctccacatgctgacctgctcagacaaacccagccctcctctcacaaggtgcccctgcagccgccacacacacacaggcccccacacacaggggaacacacgccacgtcgcgtccctggcactggcccacttcccaatgccgcccttccctgcagctga

Uppercase: IGHG2

Lowercase: Flanking sequence[1000bp]

Red & Bold & Underline: Stem-loop [28]

Blue: Heptamer[53]

Green: Nonamer [2]

id-TRGC2[C_gene_segment]

agtatgtacatgcggaagtagattctctttaaaacaagtgactgtgtatgttaaaaataaaagtgaacaaaatggctaggcatggtggctcacacctataatcccaggatttgggaggctgaggcaggcagatcacttgagcgcaggagttttaaaccagcctggtcaatatggtgaaatatgtctctagaaaaaacaaatattagccaggcatggtgatgcatacctgtagtcccagctacttgggaggctgaggtgggaggatcacttgagcctgggaagtcgagactgcagtgagctatgatcttgccgctgcattccaacctggacgacagagcaagccccagtctcaacaacaacaacaaaaattttgacattgattcatatgggaaataagataaataacaagataaatatggtgcatcgcagaatcagttaaatgaagagtggaagtacacgaaattgcagaacatgagaatgtgtcatatttggccaaagaacataagttacaaaggatggaaaagggaagtgagaaaaggaccaaagagtctcatctgtagagagatcattaaggttttctgaccttccactttcccctgccatccaccttgaaaacctgcttcactgtgatgaaacaaaagagatttaaaaataaaataaatatgctcatgaactttggaagccctggtaggaggcagttaaaaatcacactcatcacagcatgtgcagaataaacaaaggccaggttttcgtccagcatctgacacttgagagctgtgacttttggcaagttatttaatgtcttgattctcttttcaacatctgtaaaataagcacaataataagtactgtgcagcctatcctggatgaaaggccgcggtggacaccagctcaatggcatcttctctttttatggttatgtactaggccactccaaaccgtgcaatgtgtgtgtttctctaatgattcttttaaactcatatttcatttctccccatagATAAACAACTTGATGCAGATGTTTCCCCCAAGCCCACTATTTTTCTTCCTTCGATTGCTGAAACAAAACTCCAGAAGGCTGGAACATACCTTTGTCTTCTTGAGAAATTTTTCCCAGATATTATTAAGATACATTGGCAAGAAAAGAAGAGCAACACGATTCTGGGATCCCAGGAGGGGAACACCATGAAGACTAACGACACATACATGAAATTTAGCTGGTTAACGGTGCCAGAAGAGTCACTGGACAAAGAACACAGATGTATCGTCAGACATGAGAATAATAAAAACGGAATTGATCAAGAAATTATCTTTCCTCCAATAAAGACAGG**TATGTGTTTACACATA**TCATCTGTCAGAACACTTCTT**TGAAAGTGAATGCTGCATTTTTTCCTTTCA**GTATTAATGAAAAACATAAATCTTTCTTAAAAATTGTTA**CATTTAATGGTAGCGTAAATG**CCCTGCTACTTTTCTATAGAATTAAAATGGTATAGGTTTTGGAGAAAACAAAATTGAAAAAGTTGCTGAAGGTTTGTCAGCCTCAGCTCCATTATCCAAAATAAGAAAGTCACGTGCTGGTTTTTAGG**GTTGTTAGATGGATTAAAGAAACAACATACACAGAAGCATCTAGCAAC**GTGACACGTGGTAAACGC**TCAAAAAGTGTTCTCCCTTCTTTTGA**TGACTTTACTTGATCAGGAAATAACATATATATGTCTTTCAGGAATGTTCTGCCCAAGCAGGAGAGTCACTCACCTCAATCTTGCTACCCACAAAGTTTAACCTAAAAACAACGGGTTCATTGTTGACAAAATAATGTTTATCTGA**AGATAACTGTAGATCATATTTATCT**GT**AGATAATGTTTATCT**GTGGAGTGTGG**CTCTACAAAACATAGAATAGTCTTGGTCACTGCAGTTTTATAGAG**GCCTTGGGTTTTTCAGAGTTTCATTTTATATATCACCATAAAGTAACATTTCATAATTACAGGTTGGTAAGGCTTACATGTACAAACATTCTTCCATTTTCCATAATAAA**TGCATTTCCTGCCATTGGTGAATGCA**GCT**CAATAAACATTTATTG**TACAATTATGACACGCCAGGCTTAGTGGAAATGTGGATGAACAGACAAGGATGAGTT**ACTGTCCTAAGGATGATGCATGACAGT**GCAGAGAATATACTCTCTTCCTGATCACTCAGGGTCACTCATGATTCATGCGCGAGGTCCCAAAACAGTGCCTTTGATGCAGATTCTGTACATCTCTAGACGATTGGTCCAAGGGCTGAATGTGCTCTGGCCCAGTGGTCCAGTCTGTCACTATATGTCAACATCCTGAATATGAACATAACAGTCCAACATCTCAAGAGTGGGCATG**AAAAGGACTCATTTTGTGCTTTTTCCTGTGGTTAACAAGTCCTTTT**TAGCCTGGGGGAACAAGCATTAACAAAATGTTTGAAGATCTTTGCCACGTACCATTCCAAATTTCTAGGGTAAGTCTTTAGCTTTTCAGATCCTGAGTTTCTGCAATGATCAAATGTGATTTGGACAGTTGCGTTGACTTTCTCCTGGGGCTATAATGGAGTGCAAAGGAAACAATGGCAGGGAAAATGCTTGCTTTCAAAATGGTAGCATGGATGTGTTCATTCGTGTAGTTACTGTATTAGGTATAGCCTTTCCTGAAACTAACTGAAGTGGGGTTATAAAAACAGTCCCAATTTTCTATTTC**CTTTGCTGAGACACAAAGAGGAGACAAAAGAGCAAAG**CTTGAGGGTAGTTTTACCACTGTGCTTAAGTGTTCTGATTTTTCCAGTGATCAGGGTGAAATAAAAAGCATAGTAAGTTCCAGGGCAGTGAATACCATACAGGAGACAAGTTACAGTTTTATAATGTG**TTTTACTTTACACTAAATTCTAAAAGTAAAA**TGTCTTTTTTTTTTTCCGAGACAGAGTTTCACTCTTGTAGCCCAGGCAGGAGTGCTATGGTGTGATCTCGGCTCACAGCAACCTCCACCTCCCAGTTTCAAGCGATTCTTCTGCCTCAGCCTCCCGAGAAGTTGAAATTACAGGTGCCTGGCACCATATCTCGCTAATTATTCTATTTTTAGTAGAGATCGGGTTTTACCATGTTGGCCAGGCTGGTCTCGAACTCCTGACTTCAAGTGATCCACCCGCCTCAGCCTCCCAAAGTGCTGGGATTACAGGTGTGAGTCACTGTGCCGGACCTAACAGTAAAATGTCTTTCATGTGCTTCTCAAGGCAACT**ACATTAAGGAGGACACATCTCTTAATGT**CATTCTACAGTAGATTTCTAAT**GCTCTTTCTTGGAAGTTTGTTTTTCTGAGAAGAGC**TAAAA**ATATAATAACATGGAAGTGATCATATTATAT**AATCAATGAAGTGCTTTCAAAGGAGATAAAACTAACCTGGTCTGCATTTGCAACCAGCCTTGATTGAGAGAGAGAGAACTCAGGATACACTTAGAGATTTTATTATGGGGAATAGTTACTTTATTCATTTTACCTCAATCAATGCATGGAAATAAGTGACAGTCA**TTTTCATTTATCTTTTAATAAATAAAGTCACCATGAGGAAAATGAAAA**CCCATTAAAGTCAGTCCTTAAAGATATTTGGACATGCAGACATGATAACTAACATTTCCATTCGTGAGACTTACCCAAAACCTATACCTCAAGTCCATTTCTTAGAATACATGAAATAAAGATCTCAGTGAGTGTATAAAACTGCACACCAGAATCATATCCGTATAGACAAGAATACATCTACTAGAAAAATATAAACC**AAAACACCAAGGTGACTCTGTTTT**TTTCTGTTTTAAAATATGTTGTCTTTGTATGC**ATGTTTGCTTCTTCCTTTTTTTTTTTAAACAT**CGCAGATAAATTCAACTCTCACCTCAGTT**GAGAGAGAACTGTCAATGTGACTTGGCCTCTCTCTTTCTAGTCCCAGAAAGA**ATTGCACTGAAATGCTGAGCTCCTGTAATAAAAATGACCATTTGCTGAGAGTAATTAACATACTGAAAGAGATTTTCTTAGAATAGTGCACAATGGCCCAATGGTGACATTATATTGTCTCTTTATAAATTATTTTCTATCTATTTCTGTGGATTATTTCTACAAAGCACTTTTCATATGTCCAATTCCTTTTATTCCCCTACAAGTACTGACTGACTACTGGCTCTGCTGTTCACTGATATGACTTTCGGCAAGTTGCCTGCACTTTTTAAACGTTATTTCCTCATTCAGAACATGGGGCCATACAAAATACAACTCACTTCAGTGTTATTGGGGAATTAAACAAA**TAAATGCATGGGAAGCATTTA**ACATAGTGCCTGACACAATAATGAGCACTCAGTAGATGTTAGCTTTTATTAATATTGTTGTTGCTATGTCCAGAAACACTATACCTCCAG**AAAATCATGGGTACTTGCTGGGGACGTTGGGGATATGCATGATTTT**GAAAGGAGTGACTGCTCTTTACTGCTCAGATGAGAAATTTTTCTAAGCCAGACTCCTTCAAACATGTAAGATTCTGTTGTGGATTCTAGGACTGAAAGAATTCTTGGCCGAGTGTGGTGGCTTATCCTGGTAATCTCATCATTTGGGAGGACAAGGCAGGAAGATTGCTTGAGCCCAGGAGTTGGAAACAAGCCTGGACAACATGGCGAAACCCTGTCTCTACAAAAAATACAAACATTAGCTGGTCA**TGGGAGTGAGTGCCTGTACTCCCA**GCTA**CTCAGGAGGCTAAGATAGGAGGATCACCTGAG**CCTGGGCAGTTTGAGGTTTCAGTGAGCCGTGATGACACCATACTATACTCCACTCCAGCCTGGGTGACAGTGACATCCTGCCT**CAAAAAAACCCCCAAAATTATTCTTTTTG**CTGATTTCATG**TCAGCAGTGTGTGCTGA**AGGCTGTAAAGTAGCC**ACTTGTTCTGTTTATTTTTCCATTGAACAAGT**ATTTATCAAAAACGTACTTTGTGGAAGGCACTGTGCTAGGAACTATGCATACAGAAGGAAAACCAAATGTTCTTGGATACTACACTCCAG**TTGTGATAAAAAAGAAAAAAGTATTCTTCACAA**ACTTCAACATTTTGATGTGCAAAAACATAATATATGAATTAGATCTACCTAACTACACAGAATTAGACCAATTATTTCTGGGATTATGGGCTCATATTTTTAATAACTGTCCTCCTACCTCTCTGTTGACAGGTTTTATAAATATTCATTTAATTACACACAGTCACAGACACACTCAGACACACACACATACACACACACACACACCTTGACAAATAATGGGCATGAACAATTGACTGGTACTTGCTCTCATTCTTCTAGATGTCACCACAGTGGATCCCAAATACAATTATTCAAAGGATGCAAATGGTAAGTTTTTGTGTTTTTTATTTCCTCCTGATCATTTTAAGTTTTGAACTTCTCTGGCTTGA**AAAATCAGGGAATGGATTTT**GCTAGGTTGGATGCTGCAGAATGGACCTAATCATATTTTAAATTAGTCCCTCTTTTTCTAGGAGTTGTATTAACAAACCTAACTACTGCTTCATGTAAGAGATG**ACTGTAAATTGAAGGGTACAGT**GATATGCTTTCAGTTATTTC**AAAAAACAGACTTTACTCATCCATGTGTCTTTTTTCTTTTCTTTTTTTT**CTTTTTTGAGACGGAGTCTCGCTCTGTTGAACAGGCTGGATTGCAGTGACGCGATCTCACCTCACTACAACCTCCGCCT**CTGGAGTTCAAGCGATTCTCCAG**CCTCAGCTTCTCAAGTAGCTGGGACTACAGGCACATGCCACCATGTCCGGGTCATCTTTGTATTTTTAGCAGAGACCGGGTTTCACTATGTTGGCCAGGCTGGTCTAGAATTCCTG**ACTTCGTGATCTGCCCCCTCAGCCCTCCGAAGT**GCTGGGATTACAGACGTGAGTCACTGTGCCCGGCCTAACAGTAAAATGTCTTTCATGCGCTTCTCAAGGCAACTACGTTAAGGAGGACACTTCTCTTAATGTCATTCTACAGTAGATTTCTAATGCTCTTTCTTGGAAGTTTGTTTTTCTGAGAAAAGCTAAAAATATAACATGGAAGTGATCATATTGTATAATCAATGAAGTGCTTTTCAAGGAGATAAAACTAATCTGGTCCACGTTTGCAACCAACCTTGATTGAGAGAGAGAGAGAACTCAGGATACACTTGGAGATTTTATTATGGGGAATAGTTACTTTATTCTTTTTTCCTCAATCAATTCATGGAAATAAGTGATAGTCATA**TTCATTTATCTTTTAATAAATGAA**GTCACCATGAGGAAAATAAAAAGACATTGAAAACCCATTAAAGTTAGCCCTTAAAGATATTTGGACATGCAGACTTGATAACTAACGTTTGCATT**CTTGAGACTTACCCAAAACCCATACCTCAAG**TCCATGTTTTTAGAATTCATGAAATAAAGATCTCAGTGAGTGCATAAAATTGCGCACCAGAATCATATCCGTATAGACAAGAACACATCTACTAGAAAAATAATAAACC**AACACACCAATGCAACTGTGTT**TTCTTCTGTTTTAAAATATGTTGTCTTTGTATGCATGTTTGCTTCTTCCTTTTTTTTTTTTAACATCACAGATAAATTCAA**CTCTCACCTCAGGTTTTATTGAGAG**AACTGTCAATGTGACTTGGCCTCTG**TCTTTCTAGTCCCAGAAAGA**ATCGCACTGAAATGCTGAGCTCCTGTAATAAAAATGACCATTTGCTGAGAGTAATTAACATACTGAAAGAGATTTTCTTAGAGTACACAATGGTGACATTATATTGTCTCTTTATAAATAACTTTCTATCTATTTCTGTGGATTATTCCTACAAAGTACTTTTCATATGTCCAGTTTCTTTTCTTCCCCTACAACTACCGTCTGAATACTGGCTCTGCTATTTGCTGATATGATTCTCGGCAAGTTGCCTGCACTTTTTAAACTTTATTTCCTCATTCAGAACATGGGGCCAT**GTAATACTCATGTACGTGAGTATTAC**GTAATAATGCTCACTTAAGTGTTACTGGGGAATTAAACAAAA**AAATGCATGGCAAGCATTT**AACATAGTGCCTGACACAATAATGAGCACTCAGTAGATGTTAGATTTTATTAATATTGTTGTTGTTATGTCCGGAAACACTATACCTCCAGA**AAATCATGGGTACTTGCTTGGGATGTTGGGGATATGCATGATTT**GGAAAGGTATGACTGCTTTTT**TCTGCTTAGATGAGAAATTTTTCTAAGCCAGA**CTCCTTCAAATATGTAAGATTCTGTTGTGGATTCTAGGACGGAAAGAATTCTTGGTCAGGTGTGGTTTCTTATCCCTGTAATCCCAGAATTTTGGGAGGACAAGGCAGGAAGATTGCTTGA**GCCCAGGAGTTTGAAACCAGCCTGGGC**AACAAGACGAAACCCTGTCTCTACAAAAGTACATAAATTAGCTTGGCTTGGTGGTGTGTGCCTGTATTACCAGCTATTCGGGAGACTGAGATGGGAGGATCTCCTGAACCTGTGAAGTTTGAGGCTTCAGTGAGCCGTGATGACACCATACTATACTCGACTCCAGCCTGTGCGACAGTGAGACTCTGCGTCAAAAA**AAAAACCCCAAAATTATTGTTTTT**GCTGATTTCAGG**TCAGCAGTGTGTGCTGA**AGGGTGTAAAGTAGCC**ACTTGATCAGTTTATTTTTCCACTGAACAAGTATTTATCAAAAACATACTTTGTGGTCTGTTTTTGATAAATA**AAAAGGCACTGTGCTAGGAGCCATGAATACAGAAGGAAAACCAAATGTTCTTGGATACTACACTCCAGTTGTGATAAAAAAGA**AAAATGTATTCTTCACGAACTTCAACATTTT**GATATGCAAAAACATAGTATATAAATTAGATCTAC**CTGATTACGTAGAATCAG**ACCAATTATTTCTGGAATTGAGGGCTCATATTTTTAATAACTGTCCTCCTGCCTCTCTGTTGACAGGTTTTATAAATATTCATTTAATTACACACACACACACACACACCTTGACAAATAATGGACATGAACAATTGACTAGTACTTGCTCTCATTCTTCTAGATGTCATCACAATGGATCCCAAAGACAATTGGT**CAAAAGATGCAAATGGTAAGCTTTTG**TGTTTTTCCTTTCCTCCTGATCATTTTAAGTTTTGAACTTCTCTGGCTTGAAAAATCAGGGAATGGGCCGGGTGCGGTGGCTCACGCCTGTAATCCCAGCACTTTGGGAGGCCGAGGCGGGCGGATCACGAGGTCAGGAGATCGAGACCATCCCGGCTAAAACGGTGAAACCCCGTCTCTACTAAAAATACAAAAAATTAGCCGGGCTTAGTGGCGGGCGCCTGTAGTCCCAGCTACTTGGGAGGCTGAGGCAGGAGAATGGCGTGAACCCGGGAGGCGGAGCT**TGCAGTGAGCCGAGATTGCGCCACTGCA**CTCCACTCCAGCCTGGGCGACAGAGCGAGACTCCGTCTCAAAAAAAAAAAAAAAAAAAAAAAAAGA**AAAATCAGGGAATGGATTTT**G**CTAGGTTGGATGCTGCAGAATGGACCTAG**TGATATTTTAAATTAGTCCCTCTTTTTCTAGGAGTTGTATTAACAAACCTAACTACTGCTTCGGGTATGAGATG**ACTGTAAATTAGAGGGTACAGT**GATATGCTTTCAGTTATTTC**AAAAAACAGACTTTATTCATCCGTCTGTCTTTT**TTTTTTTTTTTTTTTTTTTTTTT**TGAGACGGAGGAGTCTCA**CTCTATCACCCAGGCTGGAGTGCAGTGGCGCGATCTCGGCTCACCATAACCTCCGCCTTACTGGTTCAAGCGATTCTCCAGCCTCAGCTTCTCAAGTAGCTGGGACTACAGGTGCACACCACCATACCTGGCTAATTTTTGTATTTTTAATAGAGATGGGGTTTCACCACGCTGGCCAGGATGGTCTTGAATTCTTGACCTCGTGATCTGCCCCCTCGGGCTCCCAAACTTCTGGGATTATAGGCGTGAGCCACTGTGCCCGGCCTTCTGTCTTTTGTTATAATGACTGGGGAA**AACATGATACCATGTT**GCTTCTTGAGTTGTTTTGTTTTAGTCTTTGGTCTTTGCTAGTAGCTAATAACACGAACTAGTGTTTATCAAGTGCTTTTTACACAGAAGGGCTTGTTCTGCATTTTCTAGTTTAATCATCTTAATACTCCTATAAAGTAGTACAATATATTTTCTCCCATTTTACAGTCCCTTTAAAGTAAATAACTATAAAAATCCCTTATACATGTCACACAGCTAGGTCTGGCATTTCAAATCAGGACATCAAACAAAGAATTCGTGCAGTTACTAAGTCCT**CTATTTTTTCTACAATAGAAAAAATAG**CAAGAATTACAGATAGCAAGACATTACAAGGCAGGAATCTGAAACGAAAGGGACATAATGTGGGGCTGGGTGGGTGCATGAGCTTTGCAGACTAGACTTTCATTCCAGCTCTTTTAATGATTAGGTGTAAGTGACCTACATTTTGTGAGTAACAGTTTTCTCATCAGCCAACTAAGAATAATTACACCAGATTCACAGTTATTGAAGAGATAAGGGCATGAATGTGAGA**TGTCTGGCGTAGGGTATCTCATTTAGCAGACA**CAGAATGAATACTTGTTTCTGGCTTTTTCTCTCTACATATGCACAAAGAATGTGACTAGAAGCATTGGCTCTAGCCCTGCTCAACTTTCCTCTATTTCCAATACCAAGGGGCTCTGACTTAGGCTGCCACACCAGGCAAGGAGGGGCAGTACCACCTCACTTGACCAAGGGCAGGGAGTCACGGACACATCACTTCCTGAGATCCTTTTCCACACCAAGGACTG**ATGTTTCTGGAATTCTCACTTTATGAAGACAAAACAT**ATAAATGGAAATTTCTGCAGGAAGAGACTCACTCTTGTAGCTCATTGAGTAGGCACTAGTGGTCCACCCCCACTGTCTTTACTTATTCCTTGACATCACATATCTCTTGTAAAACCTCAAATAATGTTAAATGCAATCACCCAATAATAGCATAGCCATAATTAGAGGCATTTAGGAAAGACAGGTGAGTGTGCCACAACTACCTAACACATCAGCAAATCTGGATTAACCACTTTCTTTGATTTTCCACAATGCAACCTTACTTTTTAATAGTTGGGAATGTTCTAAGTGAATTTAGCAGAGGTTGTTAATCAACTTGAAAGCTGAATTCTGACTTGTCTGACTCTTGGTGGTGCTGGTAGCAGTAGATGTTTACTTTTAGGTTTTGGTGGTGGTGGAATATCACTTCAACGTAAATCATCAGAAATAAGTATTTGTGAACCCCTCTCGCATTAATATATCTTATTCTGTAAAAAGAACATGTGCAATTTCTCTTAGATACACTACTGCTGCAGCTCACAAACACCTCTGCATATTACACGTACCTCCTCCTGCTCCTCAAGAGTGTGGTCTATTTTGCCATCATCACCTGCTGTCTGCTTAGAAGAACGGCTTTCTGCTGCAATGGAGAGAAATCATAAcagacggtggcacaaggaggccatcttt**tcctcatcggttattgtccctagaagcgtcttctgagga**tctagttgggctttctttctgggtttgggccatttcagttctcatgtgtgtactattctat**cattattgtataatg**gttttcaaaccagtgggcac**acagagaacctcactctgt**aataacaatgaggaatagccatggcgatctccagcaccaatctctccatgttttccacagctcctccagccaacccaaatagcgcctgctatagtgtagacagcctgcggcttctagccttgtccctctcttagtgttctttaatcagataactgcctggaagcctttcattttacacgccctgaagcagtcttctttgctagttgaattatgtggtgtgtttttccgtaataagcaaaataaatttaaaaaaatgaaaagttgacttttgtccatggtattttaa**ttggatgacatcaaattgaacatccaa**ggtaagaaacaacatggcaattgggctgtggaattctgtattggttgt**aagaatggtccaacaccccatttctaattctt**tccctgagatcgtggttatcacaccttctaagaggaactacaaccaaacgaaggagcccatgtggcttctgtttgaaaggtc**accagagtcagattcatctggt**ttaggacattccagtggctataggacactatctactgtgacgcgtaccgtgtgagctcagctgtagagtgtttcgcagacactgtgtttcctgatc**ctcacgatacccccgtgag**a**gctgccctaaaagcagagaggcagc**gtgatggagaggttcagcacatgctctctgat**cccaggaatcctggg**tatggtgtttcgtatctgtgtgacctcaggtgagttccaggaaatctatgtgccataatctcctcatgtaaaatgaagttataatgccccatttcctgaagttatgtggattagatgagttaatgacacctgg

Uppercase: TRGC2

Lowercase: Flanking sequence[1000bp]

Red & Bold & Underline: Stem-loop [71]

Blue: Heptamer[135]

Green: Nonamer [15]

id-IGHE[C_gene_segment]

tccagctttgctgagctaaactggaccgggctaaattgatctggactgaccattctcacctggctaagaggagctgagtcagaagcaagctggttgagctggctggactgaaataagagtttgctgcctgcaaggggaggtcctgggctgacctgggccaggctgaaccaggctggcttagagtgaacttcagagggcgactcccccggtaggccagtctcagctgaacttggctgtcccggtgggcagagcggggctggatactgtgattttgggggtacctagagcagacttcaagaccaagctaaactgggctccaggggcaggatgggctggggacttgggactccaggccaggggcgaagggccacgctgtacagaccgcactatctgggccagggttctgtggtgggagggactgactgcctggggcatcagggcaagtcttccc**gccctcccctagaggtcaggggtgggc**agagcaccatgggggtctggcaggtcaggtgagggctgctgtgatggggagatccaggcttggcactcaagagcccgaggagctgagaccacagccttggggggttggggtcagggttggagggcaggcagaccatccaccat**gagcccagagagagtttgaagggggagggctctggggtcccaggccccatggggtccctggg**tttcagcctaggggcatggcccagtgtctctgctcctgagtgcccaccgtgcagcacttgcaggggg**aggctggggtcatcctggaggcaccccccttcctgagcccagcct**gatgatagtggctgagcaacagcttctggtgggggaatgggggccctgggagccgcc**ctgggcctggggattgtggggaaaaaggcccag**aatgagcctggccatctggatccctgccacggggtccccagctcc**cccatccaggccccccaggcctgatggg**cgctggcct**gaggctggcactgactaggttctgtcctcacagCCTC**CACACAGAGCCCATCCGTCTTCCCCTTGACCCGCTGCTGCAAAAACATTCCCTCCAATGCCACCTCCGTGACTCTGGGCTGCCTGGCCACGGGCTACTTCCC**GGAGCCGGTGATGGTGACCTGGGACACAGGCTCC**CTCAACGGGACAACTATGACCTTACCAGCCACCACCCTCACGCTCTCTGGTCACTATGCCAC**CATCAGCTTGCTGACCGTCTCGGGTGCGTGGGCCAAGCAGATG**TTCACCTGCCGTGTGGCACACACTCCATCGTCCACAGACTGGGTCGACAACAAAACCTTCAGCGGTAAGAGAGGGCCAAGCTCAGAGACCACAGTTCCCAGGAG**TGCCAGGCTGAGGGCTGGCA**GAGTGGGCAGGGGTTGAGGGGG**TGGGTGGGCTCAAACGTGGGAACACCCA**GCATGCCTGGGGACCCGGGCCAGGACGCGGGGGCAAGAGGAGGGCACAC**AGAGCTCAGAGAGGCCAACAACCCTCATGACCACCAGCTCT**CCCCCAGTCTGCTCCAGGGA**CTTCACCCCGCCCACCGTGAAG**ATCTTACAGTCGTCCTGCGACGGCGGCGGGCACTTCCCCCCGACCATCCAGCTCCTGTGCCTCGTCTCTGGGTACACCCCAGGGACTATCAACATCACCTGGCTGGAGGACGGGCAGGTCATGGACGTGGACTTGTCCACCGCCTCTACCACGCAG**GAGGGTGAGCTGGCCTCCACACAAAGCGAGCTCACCCTCAGCCAGAAGCACTGGCTG**TCAGACCGCACCTACACCTGCCAGGTCACCTATCAAGGTCACACCTTTGAGGACAGCACCAAGAAGTGTGCAGGTACGTT**CCCACCTGCCCTGGTGGCCGCCACGGAGGCCAGAGAAGAGGGGCGGGTGGG**CCTCACACAGCCCTCCGGTGTACCACAGATTCCAACCCGAGAGGGGTGAGCGCCTACCTAAGCCGGCCCAGCCCGTTCGACCTGTTCATCCGCAAGTCGCCCACGATCACCTGTCTGGTGGTGGACCTGGCACCCAGCAAG**GGGACCGTGAACCTGACCTGGTCCC**GGGCCAGTGGGAAGCCTGTGAACCACTCCACCAGAAAGGAGGAGAAGCAGCGCAATGGCACGTTAACCGTCACGTCCACCCTGCCGGTGGGCACCCGAGACTGGATCGAGGGGGAGACCTACCAGTGCAGGGTGACCCACCCCCACCTGCCCAGGGCCCTCATGCGGTCCACGACCAAGACCAGCGGTGAGCCATGGGCAGGCCGGGGTCGTGGGGGAAGGGAGGGAGCGAGTGAGCGGGGCCCGGGCTGACCCCACGTCTGGCCACAGGCCCGCGTGCTG**CCCCGGAAGTCTATGCGTTTGCGACGCCGGAGTGGCCGGGG**AGCCGGGACAAGCGCACCCTCGCCTGCCTGATCCAGAACTTCATGCCTGAGGACATCTCGGTGCAGTG**GCTGCACAACGAGGTGCAGC**TCCCGGACGCCCGGCACAGCACGACGCAGCCCCGCAAGACCAAGGGCTCCGGCTTCTTCGTCTTCAGCCGCCTGGAGGTGACCAGGGCCGAATGGGAGCAGAAAGATGAGTTCATCTGCCGTGCAGTCCATGAGGCAGCAAGCCCCTC**ACAGACCGTCCAGCGAGCGGTGTCTGT**AAATCCCGGTAAATGAcgtactcctgcctccctc**cctcccagggctccatccagctgtgcagtggggagg**actggccagaccttctgtccactgttgcaatgaccccaggaagctacccccaataaactgtgcctgctcagagccccaggtacacccattcttgggagcgggcagggctgtgggcaggtgcatcttggcacagaggaatgggccccccaggaggggcagtgggaggaggtgggcagggctgagtccccccaggagaggcggtgggaggaggtgggcagggctgaggtgccactcatccatctgccttcgtgtcagggttatttgtcaaacagcatatctgcagggactcatcacagctaccccgggccctctctgcccccact**ctgggtctaccccctccaaggagtccaaagacccag**gggaggtcctcagggaaggggcaaggg**agcccccacagccctctctcttgggggct**tggcttctacccccctggacaggagcccctgcacccccaggtatagatgggcacacaggcccctccaggtggaaaaacagccctaagtgaaaccccca**cacagacacacacgacccgacagccctcgcccaagtctgtg**ccactggcgttcgcctctctgccctgtcccgccttgccgagtcct**ggccccagcaccggggcc**ggtggagccgagcccactcacaccccgcagcctccgccaccctgccctgtgggcacaccaggcccaggtcagcagccaggccccctctcctactgccccccaccgccccttggtccatcctgaatcggcccccaggggatcgccagcctcacacacccagtctcgcccactcacgcctcactcaaggcaca**gctgtgcacacactaggccccatagcaactccacagc**accctgtaccaccaccagggcgccatagacacccca**cacgtggtcacacgtg**gcccacactccgcctctcacgctgcctccagcgaggctactgccaagcc

Uppercase: IGHE

Lowercase: Flanking sequence[1000bp]

Red & Bold & Underline: Stem-loop [27]

Blue: Heptamer[58]

Green: Nonamer [1]

id-IGLC7[C_gene_segment]

tctgtgggtcccagttacggggctgcattaaacacagtgacaggaggcctttgactgaggacttggagagatgggggaggaaatggcaggaggacaaagatagaggaagaatattccgtgagaaggtggccccacagcgctgggtcacacgccatcccccaagacaggcagga**caccacagacagggtggtg**ggtctcagaaaactcaggccctaaacgtggatgcttaccaa**ttcctccactggaggaa**gacctcagagcagatgcccaggacagggacttctggtagggacggtgactgggacgggtgcctgtttgtcagggaaaacccactggagagtcagatcccccagataacttctcacgacatggagactctttcgaacagacaaagctccacgttcagctcagggagtaaaaaaaaaatgcctcaaatggaggcctttgatctactggaatccagcccccaggactgacaccctgtctcaccag**gcagcccagaggggtctctgcagggaggtggggtgggggctgc**aatgatggcaccagggagatgt**gtgggtaagaaacccac**tccctgtgagagagaagagcctgaacccaggaccaacagctgccctgcatgaagagatgagaacaaggggaactggtaggaggtgttcagacagacacccccaagatagacaaatacccagggtgagatgtggtcctggactccatcccatccagtgtggagccagcaccggtgggggtctataggtgatggaaaatatgaaaaagagacagatccaaga**gggggtctgtgaccccc**aa**gagtgggggcaactcccatctgacagcaagtgtctccactc**accgctgacct**gacctcagtccagcaagggtccggcctgaggtc**cctgccctgggccttagtcccatacccacttcaagactgaggt**caggggctccccaggtggacaccaggactctgaccccctg**cccctcatccaccccgcagGTCAGCCCAAGGCTGCCCCCTCGGTCACTCTGTTCCCACCCTCCTCTGAGGAGCTTCAAGCCAACAAGGCCACACTGGTGTGTCTCGTAAGTGACTTCAACCCGGGAGCCGTGACAGTGGCCTGGAAGGCAGATGGCAGCCCCGTCAAGGTGGGAGTGGAGACCACCAAACCCTCCAAACAAAGCAACAACAAGTATGCGGCCAGCAGCTACCTGAGCCTGACGCCCGAGCAGTGGAAGTCCCACAGAAGCTACAGCTGCCGGGTCACGCATGAAGGGAGCACCGTGGAGAAGACAGTGGCCCCTGCAGAATGCTCTT**AGgcccccgaccctcaccccacccacaggggcct**ggagctgcaggttcccaggggaggggtctctgcccccatcccaagtcatccagcccttctcaataaatatcctcatcgtcaacgagaaatcctgctccctctcttcttttcttatctcacacataatttgaggcctcccctgggttctcagtgt**tgggtgggggaatcctggcaccca**gtgagaaagtggccctgaggga**gaggctcatagcctc**ccggggtg**tctcctggtgaaagaggcccggataggaga**agtctgatcactgaacaccagtccctctgccctttcattcctcccccttctcctcaaagcatagccccccacctccctgcctcctgcctgga**tggagttgtctctggctgggactcca**gttacaccctcta**tctctgccctaacagaga**cacccttcacccaccatctcttttcccagcctgaaaaccct**gctccaagcctggagc**ctctttgcccctggcccttgcccccggaatgccctcctccctctgtgcccagctcagctcccacccaccctcaccccctccctgtcatccctgagagccagactgtccaggacactccagcaccactgacttctcaagctgttcaatgagggacccacctcg**attcctcacctgacagcaggttcagacacttaagaggaat**gggaggaagccaggaaggagactcacacgaaaggcctacaggaggctcgggacacttggaaaagataggctctgagctccctaggaaccagtgcaaggagggtaataagcacttcaccacttctgtaaatttaaacaaaccctttgctgaagggaaggtggtgttctcaatgctggggccagtcatctcgtcacaactaaaagtgggaacatgttctatccaaggcctcagtagtaatgggatgtgaaccttgacacacgcacaggtaactatctggcagcccttagacatggtgattctctcaagt

Uppercase: IGLC7

Lowercase: Flanking sequence[1000bp]

Red & Bold & Underline: Stem-loop [16]

Blue: Heptamer[23]

Green: Nonamer [2]

id-IGHEP1-2[C_gene_segment]

tgtcacccccaggacagggacagccaacccagagccgggagggagggtggggaggcggca**gcagggagctgtcctgagctccactgc**gcaactggctgatcttggcaagtccgagctgggtggactgaggggggcttggctgagtggactagactgagacgggcctaacagactgagctgaggcgagctgggtgggctgagagggctac**cctgtcccttagaggacagg**tggccaagctgggctgtcctgagccagggcgatcggggctggcccgggccaggcgggtttagctgagttgagtgagtggactgggtagagggaaatgagctaggctcagctgagctaggcttgagctgggttatcctaagccctaaggtggactgagctgggctgagctggacttatctggggag**cagggcaaagtcaggctgagctgaggtggcctgccctg**ggtg**gtccaggattgagttaagctgaattaggctgacctggac**ttgactggacttggttgaaataagctgggccgacacaggagtagggacaagctacagttctctacttaggataaaatgggtgctcgtggactatccgggctgaaggagaccaagctggggtatta**cctgctgagcttacctgacctggcctgagttcagcagg**gctgcgctgagctggacagacctg**agccaagcttagctggttgggctgagtaagctgggct**gagctaaatgggattgagctgaggagggctaggctgggggagagacctgacgacggacagggttaaaagctggagtgagcaggccttaaattattgaa**ctaaattgggctggggtgatctgaatttag**ctgggatgagctgggctgggctgaactgtgcccacgtgaactgggctaaactaggctcgcctgagtgga**ctcagctgggttggtctcaactgggttcagctgag**ctgggctcggctagactacactgggttcagctgacactacactgggttcAACCCGAGAGGGGTGAGCGCCTACCTAAGCCGGCCCAGCCCGTTCGACCTGTTCATCCGCAAGTCGCCCACGATCACCTGTCTGGTGGTGGACCTGGCACCCAGCAAGTGGACCGTGAACCTGACCTGGTCCCGGGCCAGTGGGAAGCCTGTGAACCACTCCACCAGAAAGGAGGAGAAGCAGCGCAATGGCACGTTAACCGTCACGTCCACCCTGCCGGTGGGCACCCGAGACTGGATCGAAGGGGAGACCTACCAGTGCAGGGTGACCCACCCCCACCTGCCCAGGGCCCTCGTGCGGTCCACGACCAAGACCAGCGGTGAGCCACGGGCAGGCCGGGGTCGTGGGGGGAGGGAGGGAGCGAGTGAGCG**GGGCCTGGGCTGACCCCACGTCTGGCCACAGGCCC**GCGTGCTGCCCCGGAAGTCTATGCGTTTGCGACGCCGGAGTGGCTGGGGAGCCGGGACAAGCGCACCCTCACCTGCCTGATCCAGAACTTCATGCCTGAGGACATCTCGGTGCAGTG**GCTGCACAACGAGGTGCAGC**TCCCGGACGCCCGGCACAGCACGACGCAGCCCCGCAAGACCAAGGGCTCCGGCTTCTTCATCTTCAGCCGCCTGGAGGTGACCAGGGCCGAGTGGGAGCAGAAAGATGAGTTCATCTGCCGTGCAGTCCATGAGGCAGCGAGCCCCTC**ACAGACCGTCCAGCGAGCGGTGTCTGT**AAATCCCGGTAAATGAcgtactcctgcctccct**ccctcccagggctccgtccagctgtgcagtggggaggg**ctggccagaccttctgtc**cactgttgcaacgaccccaggaagctacccccaataaacagtg**cctgctcagagcccagggtacacccgttcttgggagcgggcagggctgtgggcaggtgcatcttggcacagaggaatgggccccccaggaggggcagtgggaggaggtgggcagggctgagtccccccaggagaggtggtgggaggaggtgggcagggctgaggtgccactcatccatctgccttcgtgtcagggttatttgtcaaacagcgtatctgcagggactcatcacagctaccccgggccctctctgcccccactctcggtctaccccctccaaggagtccaaagacccaggggaggtcctcagggaaggggcaaggg**agccccgacagccctctctcttgggggct**tggcttctacccccctggacaggagcccctgcacccccaggtatagatgggcacacaggcccctccaggtagaaaaacagccctaagtgaaaccccca**cacagacacacacgacccgacagccctcgcccaagtctgtg**ccactggcgttcgcctctctgccctgtcccaccttgccgagtc**ctggccccagcaccggggccag**tggagccgagcccactcacaccccgcagcctccgccaccccgccctgtgggcacaccaggcccaggtcagcagccaggccccctctcctactgccccccaccgccccttggtccatcctgaatcggcctccaggggatcgccagcctcacacacccagtctcgcccactcacgcctcactcaaggcaca**gctgtgcacacactaggccccatagcaactccacagc**accctgtaccaccaccagggcgccatagacacccca**cacgtggtcacacgtg**gcccacactccgcctcccaccctgcctccagcgaggctactgccaagcc

Uppercase: IGHEP1-2

Lowercase: Flanking sequence[1000bp]

Red & Bold & Underline: Stem-loop [18]

Blue: Heptamer[27]

Green: Nonamer [0]

id-TRBC1-2[C_gene_segment]

gcacagtaaattgtggtttcttccactcctcatgtgtcttcagatgaaatcattctttccaataatcccagaaattctttctctcctctctcaagcatgtgaaaaggtccagagctctgcagtgtgagctttctactgaaatggcccttggactttgtggttcattcatactcagtggtctagcttgtactacttttgagaatgcaaagcttaactgtggacggattccaatcctggccaggcagggttgctggacactctgagagaagaaagggtt**aatcccatgaccatcaacttccatgggatt**tcagccatcctggacaagctaccacac**cctcctgccccaggggaggagg**aaatgtggaccatcccatcagatattgaccaggtttggctctttaaagagggttacatgcaagaaaataaaattttttaaaaaggtgctgggcaggtgggggactcagatgtaatggaaaagtgt**cttttctagaaaagaaaag**ctaattctaatatgtgtcactaccccacgagacaaatatatacatcttgatttaaaaaaggaaaat**tataattagaaaaagtcaatttagttattgtaattata**ccactaatgagagtttcctacctcgagtttcaggattacatagccatgcaccaagcaaggctttgaa**aaataaagatacacagataaattattt**ggatagatgatcagacaagcctcagtaaaaacagccaagacaatcaggatataatgtgaccataggaagctggggagacagtaggcaatgtgcatccatgggacagcatagaaaggaggggcaaagtggagagagagcaacagacactgggatggtgaccccaaaacaatgagggcctagaatgacatagttgtgcttcattacggcccattccca**gggctctctctcacacacacagagccc**ctaccagaaccagacagctctcagagcaaccctggctccaacccctcttccctttccagAGGACCTGAACAAGGTGTTCCCACCCGAGGTCGCTGTGTTTGAGCCATCAGAAGCAGAGATCTCC**CACACCCAAAAGGCCACACTGGTGTG**CCTGGCCACAGGCTTCTTCCCCGACCACGTGGAGCTGAGCTGGTGGGTGAATGGGAAGGAGGTGCACAGTGGGGTCAGCACGGACCCGCAGCCCCTCAAGGAGCAGCCCGCCCTCAATGACTCCAGATACTGCCTGAGCAGCCGCCTGAGGGTCTCGGCCACCTTCTGGCAGAACCCCCGCAACCACTTCCGCTGTCAAGTCCAGTTCTACGGGCTCTCGGAGAATGACGAGTGGACCCAGGATAGGGCCAAACCCGTCACCCAGATCGTCAGCGCCGAGGCCTGGGGTAGAGCAGGTGAGTGGGGCCTGGGGAGATGCCTGGAGGAGATTAGGTGAGACCA**GCTACCAGGGAAAATGGAAAGATCCAGGTAGC**AGACAAGACTAGATCCAAAAAGAAAGGAACCAGCGCACACCATGAAG**GAGAATTGGGCACCTGTGGTTCATTCTTCTCCCAGATTCTC**AGCCCAACAGAGCCAAGCAGCTGGGTCCCCTTTCTATGTGGCCTGTGTAACTCTCATCTGGGTGGTGCCCCCCATCCCCCTCAGTGCT**GCCACATGCCATGGATTGCAAGGACAATGTGGC**TGACATCTGCATGGCAGAAGAAAGGAGGTGCTGGGCTGTCAGAGGAAGCTGGTCTGGGCCTGGGAGTCTGTGCCAACTGCAAATCTGACTTTACTTTTAATTGCCTAT**GAAAATAAGGTCTCTCATTTATTTTC**CTCTCCCTGCTTTCTTTCAGACTGTGGCTTTACCTCGGGTAAGTAAGCCCTTCCTTTTCCTCTCCCTCTCTCA**TGGTTCTTGACCTAGAACCA**AGGCATGAAGAACTCACAGACACTGGAGGGTGGAGGGTGGGAGAGACCAGAGCTACCTGTGC**ACAGGTACCCACCTGT**CCTTCCTCCGTGCCAACAGTGTCCTACCAGCAAGGGGTCCTGTCTGCCACCATCCTCTATGAGATCCTGCTAGGGAAGGCCACCCTGTATGCTGTGCTGGTCAGCGCCCTTGTGTTGATGGCCATGGTAAGCAGGAGGGCAGGATGGGGCCAGCAGGCTGGAGGTGACACACTGACACCAAGCACCCAGAAGTATAGAGTCCCTGCCAGGATTGGAGCTGGGCAGTAGGGAGGGAAGAGATTTCATTC**AGGTGCCTCAGAAGATAACTTGCACCT**CTGTAGGATCACAGTGGAAGGGTCATGCTGGGAAGGAGAAGCTGGAGTCACCAGAAAACCCAATGGATGTTGTGATGAGCCTTACTATTTGTGTGGTCAATGGGCCCTACTACTTTCTCTCAATCCTCACAACTCCTGGCTCTTAATAACCCCCAAAA**CTTTCTCTTCTGCAGGTCAAGAGAAAG**GATTTCTGAaggcagccctggaagtggagttaggagcttctaacccgtcatggtttcaatacacattcttcttttgccagcgcttctgaagagctgctctcacctctctgcatcccaatagatatcccccta**tgtgcatgcacacctgcaca**ctcacggctgaaatctccctaacccagggggaccttagcatgcctaagtgactaaaccaataaaaatgttctggtctggcctgactctgacttgtgaatgtctggatagctccttggctgtctctgaactccctgtgactctccccattcagtcaggatagaaacaagaggtattcaaggaaaatgcagactcttcacgtaagagggatgaggggcccaccttgagatcaatagcagaagttaa**ttcagcgtgaaaggcagtgatgggagctgaa**gaggttacttctagaacagtctaggaagacacagatgttg**agtataggaattttctatatccaactatact**gttctgcccaggaaagacgtgctcagaggaagagccaacctatacaggtgtgttcaccc**tccagctggccatgtccccgtgactaacaaagctgga**ttccataagcatcacccaccttccttgcagctttcttattgagcactccattca**tcttcattggttcaccaagttgatttcccagctccaaagaaga**gaggctctgacttgcaaacttattttcaat**ggaagatgtgtcttcc**ggtttaagttacccatctgtttataaatctctctctagtgaattaaacca**gaatgaaaatgtcccctaatcattc**ctggaaggttagaaaataaaggtatctaaaactgagaatcagccccattcctacttctagaattccttcaaaagctccttctgttgtctcactgtcaccatggtgatggagtcccaaatcccaaaggtggcacagaagaccggatgattatccttgtctccttccacactctcctcacttctctcatccctgaagcc

Uppercase: TRBC1-2

Lowercase: Flanking sequence[1000bp]

Red & Bold & Underline: Stem-loop [22]

Blue: Heptamer[44]

Green: Nonamer [2]

id-IGHA1-2[C_gene_segment]

gggctgagctgtactaagctggcctgggctgggctgagctgtactgagctggcctgggctgggttgacctgggctgagctggactaagctgggctgacctgggctgggatgggatgggctaggatgacttgggctggactgggcgggactgagctggactggcctgggctgagctgggctgggctggactgagctggactggcctgggctgggctgggcgggatgggctgaggtggctgctaatgtgggaaagaggccgtgggttgagtgtgattccacctgcagagccctgagcccagctgtgttcttaggggttctgagggccacgcagctctgttgcaccatgattctgtcttctctctt**gcccactgcctgaaggaaatttggagtgggc**tgggcccagagctcccctgtatag**caggccctgtcctggagggcctg**gcagggacatggcttag**cctgttggcctctagtcccgagacctcataggccacagg**ggtccactgtggcttgtttgggcctggggtggggctcatggagtggtgggtgttggactgagactctgaccagggacaggggg**atggggtcacagccaagccactccacccctaccccat**gcacacagcactcagagcccagaccctctcctaagagcccccaccaaaatcctctctaggggcaggggatagagcaagacatgtcccccacccagagcaggggctgcggtcagggagctcaggggactcagccactccatggcagagccctgtttaatacaacttgtgtctgggatggcctgaatcagagaccctatctaaggagcatgttcagaaaccatgttgctgggatcagacagcagggtccaactgcaggcctgtggtgcaggagctgtgtgaccatggggctgtcaccag**gcctctctgtgctgggttcctccagtatagaggagaggc**agtatagaggagagggccgcgtcctcacagtgcattctgtgttccagCATCCCCGACCAGCCCCAAGGTCTTCCCGCTGAGCCTCTGCAGCACCCAGCCAGATGGGAACGTGGTCATCGCCTGCCTGGTCCAGGGCTTCTTCCCCCAGGAGCCACTCAGTGTGACCTGGAGCGAAAGCGGACAGGGCGTGACCGCCAGAAACTTCCCACCCAGCCAGGATGCCTCCGGGGACCTGTACACCACGAGCAGCCAGCTGACCC**TGCCGGCCACACAGTGCCTAGCCGGCA**AGTCCGTGACATGCCACGTGAAGCACTACACGAATCCCAGCCAGGATGTGACTGTGC**CCTGCCCAGGTCAGAGGGCAGG**CTGGGGAGTGGGGCGGGGCCACCCCGTCGTGCCCTGACACTGCGCCTGCACCCGTGTTCCCCACAGGGAGCCGCCCCTTCACTCACACCAGAGTGGACCGCGGGCCGAGCCCCAGGAGGTGGTGGTGGACAGGCCAGGAGGGGCGAGGCGGGGGCATGGGGAAGTATGTGCTGACCAGCTCAGGCCATCTCTCCACTCCAGTTCCCTCAACTCCACCTACCCCATCTCCCTCAACTCCACCTACCCCATCTCCCTCATGCTGCCACCCCCGACTGTCACTGCACCGACCGGCCCTCGAGGACCTGCTCTT**AGGTTCAGAAGCGAACCT**CACGTGCACACTGACCGG**CCTGAGAGATGCCTCAGGTGTCACCTTCACCTG**GACGCCCTCAAGTGGGAAGAGCGCTGTTCAAGGACCACCTGAGCGTGACCTCTGTGGCTGCTACAGCGTGTCCAGTGTCCTGCCGGGCTGTGCCGAGCCATGGAACCATGGGAAGACCTTCACTTGCACTGCTGCCTACCCCGAGTCCAAGACCCCGCTAACCGCCACCCTCTCAAAATCCGGTGGGTCCAGACCCTGCTCGGGGCCCTGCTCAGTGCTCTGGTTTGCAAAGCATATTCCTGGCCTGCCTCCTCCCTCCCAATCCTGGGCTCCAGTGCTCATGCCAAGTACAGAGGGAAACTGAGGCAGGCTGAGGGGCCAGGACACAGCCCAGGGTGCCCACCAGAGCAGA**GGGGCTCTCTCATCCCCTGCCCAGCCCC**CTGACCTGGCTCTCTACCCTCCAGGAAACACATTCCGGCCCGAGGTCCACCTGCTGCCGCCGCCGTCGGAGGAGCTGGCCCTGAACGAGCTGGTGACGCTGACGTGCCTGGCACGCGGCTTCAGCCCCAAGGATGTGCTGGTTCGCTGGCTGCAGGGGTCACAGGAGCTGCCCCGCGAGAAGTACCTGACTTGGGCATCCCGGCAGGAGCCCAGCCAGGGCACCACCACCTTCGCTGTGACCAGCATACTGCGCGTGGCAGCCGAGGACTGGA**AGAAGGGGGACACCTTCT**CCTGCATGGTGGGCCACGAGGCCCTGCCGCTGGCCTTCACACAGAAGACCATCGACCGCTTGGCGGGTAAACCCACCCATGTCAATGTGTCTGTTGTCATGGCGGAGGTGGACGGCACCTGCTACTGAgccgcccgcctgtccccacccctgaataaactccatgctcccccaagcagccccacgcttccatccggcgcctgtctgtccatcctcagggtctcagcacttgggaaagggcc**agggcatggacagggaagaataccccctgccct**cagcctc**ggggggcccctggcacccccc**tgagcctttccaccctggtgtgagtgtgagttgtgagtgtgagagtgtgtggtgcaggaggcctcgctggtgtgagatcttaggtctgccaaggcaggcacagcccaggatgggttctgagagatgca**catgccccggacagttctgagtgagcagtggcatg**gccgtttgtccctgagagagccgcctctggctgtagctgggagggaatagggagggtaaaaggagcaggctagccaagaaaggcgcaggtagtggcaggagcggcgagggagtgaggggctggactcca**gggccccactgggaggacaagctccaggagggccc**ca**ccaccctagtgggtgg**gcctcaggacgtcccactgacgcatgcagg**aaggggcacctcccctt**aaccacactgctctgtacggggcacgtgggcacaggtgcacactcacactcacatatacgcctgagccctgcaggagcggaacgttcacagcccagacccagttccagaaaa**gccaggggagtcccctcccaagcccccaagctcagcctgctcccctaggc**ccctctggcttccctgtgtttccactgtgcacagatcaggcaccaactccacagacccctc**ccaggcagcccctgctccctgcctgg**ccaagtctcccatcccttcctaagcccaactaggacccaaagcatagacagggaggggccacgtggggtggcatcagaa**gcaggccagtgagacagggcctgc**ccagggccctctgcatgcctctggcttctg**cctggggctcccagg**agtgtaagaacagtcccacaaccactgtggggacacc

Uppercase: IGHA1-2

Lowercase: Flanking sequence[1000bp]

Red & Bold & Underline: Stem-loop [22]

Blue: Heptamer[63]

Green: Nonamer [3]

id-IGHG4-2[C_gene_segment]

gaaatggggcctccctgtggcctgggggtcctggcaccatgcagggtggggagggccaagggcaggtgcaaggctcctacctg**tgctggggggcctgggttgagcccagca**gggaccttgccgggggaagctctggagagagggaggaggtgggctggtggccgagaaggccaggccagggctgggagggtgaggttgtggtgactga**gcctccagaagtaatgcaggacactgggaggc**agggggcatccaggcactcagggccctgacctgggctgctgcacactggggctaaggggaaaggaggggagaggctgaggaggaggctccaggaggctattccaaggcagggggttccggggccctggggctgaagggcgccgaccctatgcagtgtctggc**ccctctgctgcacagaagaaaagggccttggagggcagaggg**caggctatgaccag**ggccctgggcaagtcaggcccactcactagcggagggcc**acgctggggcggcagggtcaggagcttcaggggactcgggggacccacgagaagccatctgagaacagtgtccactggtcaagccaggcacccataaaaggctggagtggggccaatgggcatgagccgtccctgaggtggcaccgatggccagagctgaggccaagctagagacactggactgtgctgactcc**cggcaggcacagagcgctgacctggctgccg**agccccgccccctagg**ctgcaggggtgcctgcag**aagggcaccacagggccaccggtcctgcaagctttctggggcaggccgggcctgactttggctgggggcagggagggggctaaggtgacgcaggtggcgccagccaggcgcacacccaatgcccgtgagcccagacactggaccctgcatggaccat**cgcagatagacaagaaccgaggggcctctgcg**ccctgggcccagctctgtcccacaccgcggtcacatggcaccacctctcttgcagCTTCCACCAAGGGCCCATCGGTCTTCCCCCTGGCGCCCTGCTCCAGGAGCACCTCCGAGAGCACAGCCGCCCTGGGCTGCCTGGTCAAGGACTACTTCCCCGAACCGGTGACGGTGTCGTGGAACTCAGGCGCCCTGACCAGCGGCGTGCACACCTTCCCGGCTGTCCTACAGTCCTCAGGACTCTACTCCCTCAGCAGCGTGGTGAC**CGTGCCCTCCAGCAGCTTGGGCACG**AAGACCTACACCTGCAACGTAGATCACAAGCCCAGCAA**CACCAAGGTGGACAAGAGAGTTGGTG**AGAGG**CCAGCACAGGGAGGGAGGGTGTCTGCTGG**AAG**CCAGGCTCAGCCCTCCTGCCTGG**ACGCACCCCGGCTGTGCAGCCCCAGCCCAGGGCAGCAAGGCAGGCCCCATCTGTCTCCTCACCCGGAGGCCTCTGACCACCCCACTCATGCTCAGGGAGAGGGTCTTCTGGATTTTTCCACCAGGCTCCGGGCAGCCACAGGCTGGA**TGCCCCTACCCCAGGCCCTGCGCATACAGGGGCAGGTGCTGCGCTCAGACCTGCC**AAGAGCCATATCCGGGAGGACCCTGCCCCTGACCTAAGCCCACCCCAAAGGCCAAACTCTCCACTCCCTCAG**CTCAGACACCTTCTCTCCTCCCAGATCTGAG**TAACTCCCAATCTTCTCTCTGCAGAGTCCAAATATGGTCCCCCATGCCCATCATGCCCAGGTAAGCCAACCCAGGCCTCGCCCTCCAGCTCAAGGCGGGACAGGTGCCCTAGAGTAGCCTGCATCCAGGGACAGGCCCCAGCCG**GGTGCTGACGCATCCACCTCCATCTCTTCCTCAGCACC**TGAGTTCC**TGGGGGGACCATCAGTCTTCCTGTTCCCCCCA**AAACCCAAGGACACTCTCATGATCTCCCGGACCCCTGAGG**TCACGTGCGTGGTGGTGGACGTGA**GCCAGGAAGACCCCGAGGTCCAGTTCAACTGGTACGTGGATGGCGTGGAGGTGCATAATGCCAAGACAAAGCCGCGGGAGGAGCAGTTCAACAGCACGTACCGTGTGGTCAGCGTCCTCACC**GTCCTGCACCAGGAC**TGGCTGAACGGCAAGGAGTACAAGTGCAAGGTCTCCAACAAAGGCCTCCCGTCCTCCATCGAGAAAACCATCTCCAAAGCCAAAGGTGGGACCCACGGGGTGCGAGGGCCACATGG**ACAGAGGTCAGCTCGGCCCACCCTCTGCCCTGGGAGTGACCGCTGT**GCCAACCTCTGTCCCTACA**GGGCAGCCCCGAGAGCCACAGGTGTACACCCTGCCC**CCATCCCAGGAGGAGATGACCAAGAAC**CAGGTCAGCCTGACCTG**CCTGGTCAAAGGCTTCTACCCCAGCGACATCGCCGTGGAGTGGGAGAGCAATGGGCAGCCGGAGAACAACTACAAGACCACGCCTCCCGTGCTGGACTCCGACGGCTCCTTCTTCCTCTACAGCAGGCTCACCGTGGACAAGAGCAGGTGGCAGGAGGGGAATGTCTT**CTCATGCTCCGTGATGCATGAGGCTCTGCACAACCACTACACACAGAAGAGCCTC**TCCCTGTCTCTGGGTAAATGAgtgccagggccggcaagcccccgctccccgggctct**cggggtcgcgcgaggatgcttggcacgtaccccg**tgtacatacttcccgggc**gcccagcatggaaataaagcacccagcgctgccctgggc**ccctgcgagactgtgatggttctttccacggg**tcaggccgagtctgaggcctga**gtggcatgagggaggcagagcgggtcccactgtccccacact**ggcccaggctgtgcaggtgtgcctgggcc**gcctagggtggggctcagccaggggctgccctcggcagggtgggggatttgccag**cgtggccctccctccagcagcacctgccctgggctgggccacg**agaagccctaggagcccctgg**ggacagacacacagcccctgcctctgtaggagactgtcc**tgttctgtgagcgccctgtcctccgaccc**gcatgcccactcgggggcatgc**ctagtccatgtgcgtagggacaggccctccctcacccatctacccccacggcactaacccctggcagccctgcccagcctcgaacccacatggggacacaaccgactccggggacatgcactctcgggccctgtggagggactggtccagatgcccacacacacactcagcccagacccgttcaacaaaccccgcactgaggt**tggccggccacacggcca**ccacacacacacgtgcacgcctcacacacggagcctcacccgggcgaaccgcacagcacccagaccagagcaaggtcctcgcacacgtgaacactcctcagacacaggcccccacgagccccacgcggcacctcaaggcccacgagccgctcggcagcttctccacatgctgaccagctcagacaaacccagccctcctctcacaaggtg**cccctgcagccgccacacacacagggg**aacacacgccacgtcgcgtccctggcactggcccacgtcccaatacagcccttccctgcagctggggtcacatgaggggtg

Uppercase: IGHG4-2

Lowercase: Flanking sequence[1000bp]

Red & Bold & Underline: Stem-loop [32]

Blue: Heptamer[46]

Green: Nonamer [1]

id-IGHA2[C_gene_segment]

gagctagactgggctgagctgggtgagcttaggtggactgagctgggctgggctgggctgagctgagctgagctgggctgggctgggctgggatgagctgtactgagctgccctggggtgggctgggctgagctgggctgagctgggctgggctggactgagctggactgagctggactggcctgggctgggctgggctggatgagctgaggtggctgctaatgtgggaaggaggccgtgggttgagtgtgactccacctgcagagccctgagcccagctgtgttcttaggggttctgagggccacgcagctctgttgcaccatgattctgtcttctctctt**gcccactgcctgaaggaaatttggagtgggc**tgggcccagagctcccctgggtaa**caggccctgtcctggagggcctg**gcagggacatggcttag**cctgttggcctctagtcccgagacctcataggccacagg**ggtccactgtggcttgtttgggcctggggtggggctcatggagtggtgggtgttggactgagactctgaccagggacaggggg**atggggtcacagccaagccactccacccctaccccat**gcacacagcactcagagcccaggccccctcctcagagcccccaccaaaatcctctctaggggcaggggaaagagcaagacatgtcccccacccagagcaggaactggggtcagggagctcaggggactcagccactccatggcagagccctgtttaatataacttgtgtctgggatggcctgggtcagaggccctatctaaggagcatgttcagaaactgtgtcgctgggatgag**acagctgggtccaaccgcaggcccatggtgcaggagctgt**gtaaccttggggctgtcaccaggcctctctgtgctgggttcctccagtgta**gaggagaggcaggtacagcctgtcctc**ctggggacatggcatgagggccgcgtcctcacagcgcattctgtgttccagCATCCCCGACCAGCCCCAAGGTCTTCCCGCTGAGCCTCGACAGCACCCCCCAAGATGGGAACGTGGTCGTCGCATGCCTGGTCCAGGGCTTCTTCCCCCAGGAGCCACTCAGTGTGACCTGGAGCGAAAGCGGACAGAACGTGACCGCCAGAAACTTCCCACCTAGCCAGGATGCCTCCGGGGACCTGTACACCACGAGCAGCCAGCTGACCCTGCCGGCCACACAGTGCCCAGACGGCAAGTCCGTGACATGCCACGTGAAGCACTACACGAATTCCAGCCAGGATGTGACTGTGCCCTGCCGAGGTCAG**AGGGCAGGCTGGGGAGTGGGGCGGGGCCACCCCGTCCTGCCCT**GACACTGCGCCTGCACCCGTGTTCCCCACAGGGAGCCGCCCCTTCACTCACACCAGAGTGGACCGCGGGCCGAGCCCCAGGAGGTGGTGGTGGACAGGCCAGGAGGGGCGAGGCGGGGGCACGGGGAAGGGCGTTCTGACCAGCTCAGGCCATCTCTCCACTCCAGTTCCCCCACCTCCCCCATGCTGCCACCCCCGACTGTCGCTGCACCGACCGGCCCTCGAGGACCTGCTCTT**AGGTTCAGAAGCGAACCT**CACGTGCACACTGACCGGCCTGAGAGATGCCTCTGGTGCCACCTTCACCTGGACGCCCTCAAGTGGGAAGAGCGCTGTTCAAGGACCACCTGAGCGTGACCTCTGTGGCTGCTACAGCGTGTCCAGTGTCCTGCC**TGGCTGTGCCCAGCCA**TGGAACCATGGGGAGACCTTCACCTGCACTGCTGCCCACCCCGAGTTGAAGACCCCACTAACCGCCAACATCACAAAATCCGGTGGGTCCAGACCCTGCTCGGGGCCCTGCTCAGTGCTCTGGTTTGCAAAGCATATTCCCGGCCTGCCTCCTCCCTCCCAATCCTGGGCTCCAGTGCTCATGCCAAGTACAGAGGGAAACTGAGGCAGGCTGAGGGGCCAGGACACAGCCCAGGGTGCCCACCAGAGCAGA**GGGGCTCTCTCATCCCCTGCCCAGCCCC**CTGACCTGGCTCTCTACCCTCCAGGAAACACATTCCGGCCCGAGGTCCACCTGCTGCCGCCGCCGTCGGAGGAGCTGGCCCTGAACGAGCTGGTGACGCTG**ACGTGCCTGGCACGT**GGCTTCAGCCCCAAGGATGTGCTGGTTCGCTGGCTGCAGGGGTCACAGGAGCTGCCCCGCGAGAAGTACCTGACTTGGGCATCCCGGCAGGAGCCCAGCCAGGGCACCACCACCTACGCTGTAACCAGCATACTGCGCGTGGCAGCTGAGGACTGGA**AGAAGGGGGAGACCTTCT**CCTGCATGGTGGGCCACGAGGCCCTGCCGCTGGCCTTCACACAGAAGACCATCGACCGCATGGCGGGTAAACCCACCCACATCAATGTGTCTGTTGTCATGGCGG**AGGCGGATGGCACCTGCTACTGAgccgcccgcct**gtccccacccctgaataaactccatgctcccccaagcagccccacgcttccatccggcgcctgtctgtccatcctcagggtctcagcacttgggaaag**ggccagggcatggacagggaagaataccccctgccctgagcc**tcggggggcccctggcacccccatgagactttccaccctggtgtgagtgtgagttgtgagtgtgagagtgtgtggtgcaggaggcctcgctggtgtgagatcttaggtctgccaaggcaggcacagcccaggatgggttctgagagacgca**catgccccggacagttctgagtgagcagtggcatg**gccgtttgtccctgagagagccgcctctggctgtagctgggagggaatagggagggtaaaaggagcaggctagccaagaaaggcgcaggtagtggcaggagtggcgagggagtgaggggctggactcca**gggccccactgggaggacaagctccaggagggccc**ca**ccaccctagtgggtgg**gcctcaggacgtcccactgacgcatgcagg**aaggggcacctcccctt**aaccacactgctctgtacggggcacgtgggcacacatgcacactcacactcacatatacgcctgagccctgcaggag**tggaacgttcacagcccagacccagttcca**gaaaagcca**ggggagtcccctcccaagcccccaagctcagcctgctcccc**caggcccctctggct**tccctgtgtttccactgtgcacagctcaggga**ccaactccacagacccctc**ccaggcagcccctgctccctgcctgg**ccaagtctcccatcccttcctaagcccaactaggacccaaagcatagacagggaggggccgcgtggggtggcatcagaa**gcaggccagtgagacagggcctgc**ccagggccctctgcatgcctctggcttctg**cctggggctcccagg**agtgaaagaacagtcccacaaccactgtggggacacc

Uppercase: IGHA2

Lowercase: Flanking sequence[1000bp]

Red & Bold & Underline: Stem-loop [24]

Blue: Heptamer[64]

Green: Nonamer [3]

id-IGHG3-2[C_gene_segment]

aatggggcctccctgtggcctgggggtcctggcaccacgcagggtggggagggccaagggcaggtgcaaggctcctacctg**tgctggggggcctgggttgagcccagca**gggaccttgccgggggaagctctggagagagggaggaggtgggctggtggctgagaaggccaggccagggctgggagggtgacggtgtggtgactga**gcctccagaagtaatgcaggacactgggaggc**agggggcatccaggcactcagggccctgacctgggctgctgcacactggggctaaggggaaaggaggggagaggctgaggaggaggctcccggggcgatattccaaggcagggggttccggggccctggggctgaagggcgccgaccctatgcagtgtctggc**ccctctgctgcacagaagaaaagggccttggagggcagaggg**caggctatgaccag**ggccctgggcaagtcaggcccactcactagcggagggcc**acgctggggcggcagggtcaggagcttcaggggactcaggggacccacgagaagccatctgagaacagtgtccactggtcaagccaggcacccataaaaggctggagtggggccaatgggcatgagccgtccctgaggtggcaccgatggccagagctgaggccaagctagaggccctggactgtgctgactcc**cggcagacacagagcgctgacctggctgccg**agccccgcctcctagg**ctgcaggggtgcctgcag**aagggcaccacagggccaccggtcctgcaagctttctggggcgggccgg**gcctgaccttggctttggggcagggagggggctaaggtgaggc**aggtggcgccagccaggcgcacacccaatgcccgtgagcccagacactggaccctgcctgga**ccctcgtggatagacaagaaccgaggg**gcctctgcgccctgggcccagctctgtcccacaccgcagtcacatggcgccatctctcttgcagCTTCCACCAAGGGCCCATCGGTCTTCCCCCTGGCGCCCTGCTCCAGGAGCACCTCTGGGGGCACAGCGGCCCTGGGCTGCCTGGTCAAGGACTACTTCCCAGAACCGGTGACGGTGTCGTGGAACTCAGGCGCCCTGACCAGCGGCGTGCACACCTTCCCGGCTGTCCTACAGTCCTCAGGACTCTACTCCCTCAGCAGCGTGGTGACC**GTGCCCTCCAGCAGCTTGGGCAC**CCAGACCTACACCTGCAACGTGAATCACAAGCCCAGCAA**CACCAAGGTGGACAAGAGAGTTGGTG**AGAGGCCAGCGCAGGGAGGGAGGGTGTCTGCTGGAAG**CCAGGCTCAGCCCTCCTGCCTGG**ACGCATCCCGGCTGTGCAGTCCCAGCCCAGGGCACCAAGGCAGGCCCCGTCTGACTCCTCACCCGGAGGCCTCTGCCCGCCCCACTCATGCTCAGGGAGAGGGTCTTCTGGCTTTTTCCACCAGGCTCCGGGCAGGCACAGGCTGGA**TGCCCCTACCCCAGGCCCTTCACACACAGGGGCA**GGTGCTGCGCTCAGAGCTGCCAAGAGCCATATCCAGGAGGACCCTGCCCCTGACCTAAGCCCACCCCAAAGGCCAAACTCTCTACTCACTCAG**CTCAGATACCTTCTCTCTTCCCAGATCTGAG**TAACTCCCAATCTTCTCTCTGCAGAGCTCAAAACCCCACTTGGTGACACAACTCACACATGCCCACGGTGCCCAGGTAAGCCAGCCCAGGCCTCGCCCTCCAGCTCAAGGCGGGACAAGAGCCCTAGAGT**GGCCTGAGTCCAGGGACAGGCC**CCAGCAGGGTGCTGACGCATCCACCTCCATCCCAGATCCCCGTAACTCCCAATCTTCTCTCTGCAGAGCCCAAATCTTGTGACACACCTCCCCCGTGCCCACGGTGCCCAGGTAAGCCAGCCCAGGCCTCGCCCTCCAGCTCAAGGCAGGACAAGAGCCCTAGAGTGGCCTGAGTCCAGGGACAGGCCCCAGCAGGGTGCTGACGCGTCCACCTCCATCCCAGATCCCCGTAACTCCCAATCTTCTCTCTGCAGAGCCCAAATCTTGTGACACACCTCCCCCATGCCCACGGTGCCCAGGTAAGCCAGCCCAGGCCTCGCCCTCCAGCTCAAGGCGGGACAAGAGCCCTAGAGTGGCCTGAGTCCAGGGACAGGCCCCAGCAGGGTGCTGACGCATCCACCTCCATCCCAGATCCCCGTAACTCCCAATCTTCTCTCTGCAGAGCCCAAATCTTGTGACACACCTCCCCCGTGCCCAAGGTGCCCAGGTAAGCCAGCCCAGGCCTCGCCCTCCAGCTCAAGGCAGGACAGGTGCCCTAGAGTGGCCTGCATCCAGGGACAGGTCCCAGTCG**GGTGCTGACACATCTGCCTCCATCTCTTCCTCAGCACC**TGAACTCCTGGGAGGACCGTCAGTCTTCCTCTTCCCCCCAAAACCCAAGGATACCCTTATGATTTCCCGGACCCCTGAGG**TCACGTGCGTGGTGGTGGACGTGA**GCCACGAAGACCCCGAGGTCCAGTTCAAGTGGTACGTGGACGGCGTGGAGGTGCATAATGCCAAGACAAAGCCGCGGGAGGAGCAGTACAACAGCACGTTCCGTGTGGTCAGCGTCCTCACC**GTCCTGCACCAGGAC**TGGCTGAACGGCAAGGAGTACAAGTGCAAGGTCTCCAACAAAGCCCTCCCAGCCCCCATCGAGAAAACCATCTCCAAAACCAAAGGTGGGACCCGCGGGGTATGAGGGCCACATGGA**CAGAGGCCAGCTTGACCCACCCTCTG**CCCTGGGAGTGACCGCTGTGCCAACCT**CTGTCCCTACAGGACAG**CCCCGAGAACCACAGGTGTACACCCTGCCCCCATCCCGGGAGGAGATGACCAAGAAC**CAGGTCAGCCTGACCTG**CCTGGTCAAAGGCTTCTACCCCAGCGACATCGCCGTGGAGTGGGAGAGCAGCGGGCAGCCGGAGAACAACTACAACACCACGCCTCCCATGCTGGACTCCGACGGCTCCTTCTTCCTCTACAGCAAGCTCACCGTGGACAAGAGCAGGTGGCAGCAGGGGAACATCTT**CTCATGCTCCGTGATGCATGAGGCTCTGCACAACCGCTTCACGCAGAAGAGCCTC**TCCCTGTCTCCGGGTAAATGAgtgcgacggccggcaagcccccgctccccgggctct**cggggtcgcgcgaggatgcttggcacgtaccccg**tgtacatacttcccgggcacccagcatggaaataaagcacccagcgctgccctgggcccctgcgagactgtgatggttctttccacggg**tcaggccgagtctgaggcctga**gtggcatgagggaggcagagcgggtcccactgtccccacact**ggcccaggctgtgcaggtgtgcctgggcc**gcctagggtggggctcagccaggggctgccctcggcagggtgggggatttgccag**cgtggccctccctccagcagcagctgccctgggctgggccacg**ggaagccctaggagcccctgg**ggacagacacacagcccctgcctctgtaggagactgtcc**tgtcctgtgagcgccctgtcctccgaccc**gcatgcccactcgggggcatgc**ctagtccatgtgcgtagggacaggccctccctcacccatctacccccacggcactaacccctggcagccctgcccagcctcgcacccgcatggggacacaaccgactccggggacatgcactctcgggccctgtggagagactggtccagatgcccacacacacactcagcccagacccgttcaacaaaccccgcactgaggt**tggccggccacacggcca**ccacacacacacgtgcacgcctcacacacggagcctcacccgggcgaaccgcacagcacccagaccagagcaaggtcctcgcacacgtgaacactcctcggacacaggcccccacgagccccacgcggcacctcaaggcccacgagccgctcggcagcttctccacatgctgaccagctcagacaaacccagccctcctctcacaaggtgcccctgcagccgccacacacacacaggcccccacacacaggggaacacacgccacgtcgcgtccctggcactggcccacttcccaatacagcccttccctgcagctgg

Uppercase: IGHG3-2

Lowercase: Flanking sequence[1000bp]

Red & Bold & Underline: Stem-loop [29]

Blue: Heptamer[56]

Green: Nonamer [1]

id-IGHM-4[C_gene_segment]

tgggctatactgggcttagctgggctgggctatactgggcttagctgggctgggctatactgggcttagctgggctgg**gctgagctgagatggtcttaggtggtctgagctcagc**taggctgggctgagctggtctg**agctcatctgagttgggctgagct**gagcttggctttgctgagctggggtggggtgggctgggctggattgagctggcctgggctgggatgaactggattgagctggcctgggctgggatgaactggaggacatggcactgggccaatcttcatgatcttgttggacatagatggatagcctcagctgagtctacactgcgttccccatcacactcaccctccctatactcact**cccaggcctgggttgtctgcctggg**gagacttcagggtagctggagtgtgactgagctggg**ggcagcagaagctgggctggagggactctattggctgcc**tgcggggtgtgtggctccaggcttcacattcaggtatgcaacctgggccctccagctgcatgtgctgggagctgagtgtgtg**cagcacctacgtgctg**atgcctcgggggaaagcaggcctggtccacccaaacctgagccctcagccat**tctgagcagggagccaggggcagtcaggcctcaga**gtgcagcagggcagccagctgaatggtggcagggatggctcagcctgctccaggagaccccaggtctgtccaggtgttcagtgctgggccctgcagcaggatgggc**tgaggcctgcagccccagcagccttggacaaagacctgaggcctca**ccacggccccgccacccctgatagccatgacagtctgggctttggaggcctgcaggtgggctcggccttggtggggcagccacagcgggacgcaagtagtgagggcactcagaacgccactcagccccgacaggcagggcac**gaggaggcagctcctc**acc**ctccctttctcttttgtcctgcgggtcctcagGGAG**TGCATCCGCCCCAACCCTTTTCCCCCTCGTCTCCTGTGAGAATTCCCCGTCGGATACGAGCAGCGTGGCCGTTGGCTGCCTCGCACAGGACTTCCTTCCCGACTCCATCACTTTCTCCTGGAAATACAAGAACAACTCTGACATCAGCAGCACCCGGGGCTTCCCATCAGTCCTGAGAGGGGGCAAGTACGCAGCCACCTCACAGGTGCTGCTGCCTTCCAAGGACGTCATGCAGGGCACAGACGAACACGTGGTGTGCAAAGTCCAGCACCCCAACGGCAACAAAGAAAAGAACGTGCCTCTTCCAGGTGAGGGCCGGGCCCAGCCACCGGGACAGAGAGGGAGCCGAAGGGGGCGGGAGTGGCGGGCACCGGGCTGACACGTGTCC**CTCACTGCAGTGATTGCTGAGCTGCCTCCCAAAGTGAG**CGTCTTCGTCCCACCCCGCGACGGCTTCTTCGGCAACCCCCGCAAGTCCAAGCTC**ATCTGCCAGGCCACGGGTTTCAGTCCCCGGCAGAT**TCAGGTGTCCTGGCTGCGCGAGGGGAAGCAGGTGGGGTCTGGCGTCACCACGGACCAGGTGCAGGCTGAGGCCAAAGAGTCTGGGCCCACGACCTACAAGGTGACCAGCACACTGACCATCAAAGAGAGCGACTGGCTCGGCCAGAGCATGTTCACCTGCCGCGTGGATCACAGGGGCCTGACCTTCCAGCAGAATGCGTCCTCCATGTGTGTCCCCGGTGAGTGACCTGTCCCCAGGGGCAGCACCCACCGACACACAGGGGTCCACTCGGGTCTGGCATTCGCCACCCCGGATGCAGCCATCTACTCCCTGAG**CCTTGGCTTCCCAGAGCGGCCAAGG**G**CAGGGGCTCGGGCGGCAGGACCCCTG**GGCTC**GGCAGAGGCAGTTGCTACTCTTTGGGTGGGAACCATGCCTCCGCC**CACATCCACACCTGCCCCACCTCTGACTCCCTTC**TCTTGACTCCAGATCAAGA**CACAGCCATCCGGGTCTTCGCCATCCCCCCATCCTTTGCCAGCATCTTCCTCACCAAGTCCACCAAGTTGACCTGCC**TGGTCACAGACCTGACCACCTATGACAGCGTGACCA**TCTCCTGGACCCGCCAGAATGGCGAAGCTGTGAAAACCCACACCAACATCTCCGAGAGCCACCCCAATGCCACTTTCAGCGCCGTGGGTGAGGCCAGCATCTGCGAGGATGACTGGAATTCCGGGGAGAGGTTCACGTGCACCGTGACCCACACAGACCTGCCCTCGCCACTGAAGCAGACCATCTCCCGGCCCAAGGGTAGGCCCCACTCTTGCCCCTCTTCCTGCACTCCCTGGGACCTCCCTTGGCCTCTGGGGCATGGTGGAAAGCACCCCTCAC**TCCCCCGTTGTCTGGGCAACTGGGGA**AAAGGGGACTCAACC**CCAGCCCACAGGCTGG**TCCCCCCACTGCCCCGCCCTCACCACCATCTCTGTTCACAGG**GGTGGCCCTGCACAGGCCCGATGTCTACTTGCTGCCACC**AGCCCGGGAGCAGCTGAACCTGCGGGAGTCGGCCACCATCACGTGCCTGGTGACGGGCTTCTCTCCCGCGGACGTCTTCGTGCAGTGGATGCAGAGGGGGCAGCCCTTGTCCCCGGAGAAGTATGTGACCAGCGCCCCAATGCCTGAGCCCCAGGCCCCAGGCCGGTACTTCGCCCACAGCATCCTGA**CCGTGTCCGAAGAGGAATGGAACACGG**GGGAGACCTACACCTGCGTGGTGGCCCATGAGGCCCTGCCCAACAGGGTCACCGAGAGGACCGTGGACAAGTCCACCGGTAAACCCACCCTGTACAACGTGTCCCTGGTCATGTCCGACACAGCTGGCACCTGCTACTGAccctgctggcctgcccacaggctcggggcggctggccgctctgtgtgtgc**atgcaaactaaccgtgtcaacggggtgagatgttgcat**cttataaaattagaaataaaaagatccattcaaaagatactggtcctgagtgcacgatgctctggcctactggggcggcggctgtgctgcacccaccctgcgcctcccctgcagaacaccttcctccacagcccccacccctgcctcacccacctgcgtgcctcagtggcttctagaaacccctgaattccctgcagctgctcacagcaggctgacctcagacttgccattcctcctactgcttccagaaagaaagctgaaagcaaggccacacgtatacaggcagcacacagg**catgtgtggatacacatg**gacagacacggacacacacaaacacatggacacacagagacgtgctaacccatgggcaca**cacatacacagacatggacccacacacaaacatatgtg**gacacacatgtacaaacatgcacaggcacacaaagagaacactgactacaggcacacacacacacgggcacacacatggatatgtgcacacatggacacataca**tgtgcaggacatgcaca**cacacagacacactagcacagaggcatacacacacagacacacacattcacaaacac**acatgtgcatgcaaacacacacacatgt**acagacacaagtacatggacacatgcacacccagagacacactgacacagacacacaggagcatgtgatacactaacacgtggacacacacgtctacccacaggcacacaacagatggacacgcgtacacagacatgcacacacccacaggcacaacacgtgcgcatgccggccggcccccgccaacattctcccagggccctgccggatactctgtccctgcagcagtttgctccctgcgctgtgctggcaccggggctttgggcccaggctctgcttgtccttctgtctctgct

Uppercase: IGHM-4

Lowercase: Flanking sequence[1000bp]

Red & Bold & Underline: Stem-loop [25]

Blue: Heptamer[63]

Green: Nonamer [1]

id-IGLC1[C_gene_segment]

gcctgtgctgggtcatgaggacatggggacacagagggacgggtgagactgggtgaggtgccagaatccaaccctcccaggacagtcaccagaaaggagacagtctcttagggcagagatgtgtctgtccctggagccccgtcacctctggggcccagtgtctctctgttcacggatcggcctcctgccttcctcaaagggcatgttagactcaggaaatgaccagaggggagtgaatgaggggtgcagagaactccatggctaccaggtgaagtttggggtcatcacaggctgctggggtggg**cctgggggctgctgagtctcatagtctgtgggagcagccccagg**aacagctgaggtgaagggttctgtggtcgggcttgtggagacaggaaacatctcag**agcctcagaggagccctgaggct**tgtctaggtgg**agcccactccttgccaggagagccaagtgggct**gggctggggcagagcccggtgcctgtgagggataggaagctccagttcaaa**gcaggcttgggtctccccacacactgcctgc**caggacagtcctacaggatgagcaggggacccacagttcacggaggaggctctaggtcctggaagaataaagtgggtgatggaggggggtatagggatggaaatgagggatccaggggtcaaggccagattctaaactcagactccagagatcagagaagaaggaac**acagcctgccctgggtatatggagaaattgaggctgt**agaggagaggggctgggccaggacacctgtgaaaggtgacttgggagggctcctaggaaggcacagagctgtctgctctccacagggcatgagtggaaaggatggggaaagaagaggagagaaccccgggtggaccggatggccacactgtgaaccctcccagagactttagaca**gagagaggggctccacaacaccccggtattctgtctgccctctctc**acccccttccctgtccacacagGTCAGCCCAAGGCCAACCCCACTGTCACTCTGTTCCCGCCCTCCTCTGAGGAGCTCCAAGCCAACAAGGCCACACTAGTGTGTCTGATCAGTGACTTCTACCCGGGAGCTGTGACAGTGGCCTGGAAGGCAGATGGCAGCCCCGTCAAGGCGGGAGTGGAGACCACCAAACCCTCCAAACAGAGCAACAACAAGTACGCGGCCAGCAGCTACCTGAGCCTGACGCCCGAGCAGTGGAAGTCCCACAGAAGCTACAGCTGCCAGGTCACGCATGAAGGGAGCACCGTGGAGAAGACAGTGGCCCCTACAGAATGTTCATAGgttcccaactctaaccccacccacgggagcctggagctgcaggatccca**ggggaggggtctctctcccc**atcccaagtcatccagcccttctccctgcactcatgaaaccccaataaatatcctcattgacaaccagaaatcttgttttatctcattttttttctcacataaattgctagcctccccggggttctcagtgtggggtacagggaattctgcacccagtgtgaaaatcacccaaggga**ggaggctcacagcctcc**ctgagtcatctccccagagggtccttcctctcccagtcaccccttctccaactctccactgtacccctgagctaccagtctggcatcagttcagaccagtcccacaccctcctaaa**ttttacttctcaataaatacctgatcatgtaaaa**cgcagcatttctaatgtgcagtctctgtctggtcatgtgtctgggctgaagggtcactgctcagggacagggggcagttccaggtgagatcccatgtctccgtcatcccacaccccacccaacctgc**cagggaaccgggtgagctccctgtgccagtgggaactgcaatccaaggcaca**aaattgtcctgcagtccttgcccacctgggaaggg**acaggggcccagtgagaggtttgctggcgccctgt**ggggagattcaggagaaatgaagggggtccccggagaccagatgagggctagaggcagaaataatggaaaaaggacacccttgactcaaggccacggtctcagcaggaacagaaggtgaaattccccattgcatacgaggaaccagtcaggagagtgtttactgggtgagggataaataactgtgctgccactgggaacttgtaaaaacattgggaaaggaaacatgcaagtgtctttctaagacttgtacaatggacattggctaagtaaacatactgacaagtcctgcactagggaaccagtttaatatgatgagccacagcatatccaaaagcat

Uppercase: IGLC1

Lowercase: Flanking sequence[1000bp]

Red & Bold & Underline: Stem-loop [12]

Blue: Heptamer[40]

Green: Nonamer [4]
