## Supplementary Data 5 for "Adaptive immunity: from CRISPR to CRIHSP?"

In the following tables we show the IDs of the VDJ fragments, the presence or absence of stem-loop structures (StemL) in the vicinity of the heptamer and the nonamer, the presence is denoted by "+" and the absence is denoted by "-", and the distances of the StemL from the heptamer and the nonamer are shown (bp), as well as the sequences of the heptamer and the nonamer. For D fragments, we used yes or no to indicate whether the D fragment is located within the stem-loop structure.

| IGHV | ID | StemL | bp | Heptamer | H-N bp | Nonamer | bp | StemL |
|---|---|---|---|---|---|---|---|---|
| | IGHV3-9 | - | | CAGTGAG | 7 | acaaaaacc | 3 | + |
| | IGHV2-70 | + | 0 | cacagag | 23 | acaagaacc | | - |
| | IGHVIII-67-3-2 | + | 0 | cacagcg | 22 | acagaaacc | | - |
| | IGHV5-10-1 | + | 0 | cacagtg | 23 | ctaaaaccc | | - |
| | IGHVIII-67-2-2 | - | | cacatga | 22 | aataaacc | | - |
| | IGHV3-16-2 | - | | tcctgtg | 23 | acacaaacc | 0 | + |
| | IGHV7-4-1 | | +0 | cacagtg | 23 | tcaaaaacc | 0 | + |
| | IGHVIII-13-1 | + | 0 | cacagtg | 23 | acaccaacc | | - |
| | IGHVIII-16-1-2 | - | | cacagga | 24 | acagaaaaa | | - |
| | IGHV3-75-2 | - | | cgcagtg | 23 | acacaaacc | | - |
| | IGHV3-73-2 | | | cacagtg | 23 | acacaaacc | 0- | + |
| | IGHV3-23-2 | + | 0 | cacagtg | 23 | acacaaacc | | - |
| | IGHV6-1 | + | 0 | cacagtg | 23 | acacaaacc | | |
| | IGHVIII-11-1 | - | | ctcactgag | | | | - |
| | IGHVIV-44-1 | - | | cactgtg | 23 | acacaaacc | | - |
| | IGHV3-21-2 | + | 0 | cacagtg | 23 | | | - |
| | IGHV3-13-2 | + | 0 | cacagtg | 23 | acacaaacc | 1 | + |
| | IGHV5-10-1-2 | + | 0 | cacagtg | 23 | ctaaaaccc | | - |
| | IGHV7-56 | - | | caccgtg | 23 | ttagaaacc | | - |
| | IGHVIII-13-1-2 | + | 0 | cacagtg | 23 | acaccaacc | | - |
| | IGHVII-43-1-2 | + | 0 | CACAGCC | 38 | tgtttttga | | - |
| | IGHV3-35-2 | | | cactgtg | 23 | acacaaacc | 1 | + |
| | IGHV5-78-2 | - | | cagagtg | | | | - |
| | IGHV3-35 | - | | cactgtg | 23 | acacaaacc | 1 | + |
| | IGHV3-37-2 | - | | catggtg | 23 | ccagaaacc | 1 | + |
| | IGHVIII-5-2-2 | + | 0 | cacagtg | 22 | acgcaaact | | - |
| | IGHV3-53 | + | 0 | cacagtg | 23 | acacaaacc | 0 | + |
| | IGHV4-30-2 | - | | cacaatg | 23 | acacaaacc | 0 | + |
| | IGHV1-68 | + | 0 | CACGGTG | 23 | TCAGGAACC | 0 | + |
| | IGHV4-61-2 | + | 0 | cacagtg | 23 | acacaaacc | 0 | + |
| | IGHV3-15 | - | | cacagtg | 23 | acacaaacc | 0 | + |
| | IGHVII-74-1-2 | - | | | | | | - |
| | IGHVII-60- | - | | cagagtg | 23 | acccaaacc | | - |

| | | | | | | | | |
|---|---|---|---|---|---|---|---|---|
| | 1-2 | | | | | | | |
| | IGHV1-24-2 | + | 0 | cacagtg | 23 | tcagaaacc | | - |
| | IGHVIII-82 | - | | catagga | 24 | acacaaaat | | - |
| | IGHV7-27-2 | - | | cacagtg | | | | - |
| | IGHV2-5 | - | | cacaaag | 23 | acaaaaacc | | - |
| | IGHV3-66-2 | + | 0 | cacagtg | 23 | | 0 | + |
| | IGHVIII-22-2 | - | | cactgtc | | | | - |
| | IGHVII-30-1-2 | + | 0 | cacagtg | 23 | acccaagcc | 0 | + |
| | IGHV2-70 | + | 0 | cacagag | 23 | acaagaacc | | - |
| | IGHVIV-44-1-2 | - | | cactgtg | 23 | | | - |
| | IGHV3-43-2 | + | 0 | cacagtg | 23 | | 3 | + |
| | IGHV4-61 | + | 0 | cacagtg | 23 | acacaaacc | 0 | + |
| | IGHVII-43-1 | + | 0 | CACACAGCC | 38 | tgtttttga | | - |
| | IGHV3OR16-17 | + | 0 | CACAGCG | 24 | cacaaacc | | - |
| | IGHV7-40-2 | - | | cacagtg | 23 | tcagaaacc | | - |
| | IGHV3-54 | - | | ccaggta | 23 | acacagaat | | - |
| | IGHV3-63 | - | | CCAAGTG | 23 | acacaaaat | 0 | + |
| | IGHV3-29-2 | - | | ccaagtg | 23 | | 0 | + |
| | IGHV4-4 | + | 0 | cacagtg | 23 | acacaaacc | 0 | + |
| | IGHVII-74-1 | | | | | | | |
| | IGHV3-65-2 | - | | CACAGTG | 23 | ACACAAACC | 0 | + |
| | IGHV6-1-2 | + | 0 | cacagtg | 23 | acacaaacc | | - |
| | IGHV3-22-2 | - | | cacagtg | 23 | acacaaacc | 0 | + |
| | IGHVIII-38-1 | + | 0 | cacagtg | 23 | acacaaaag | 0 | + |
| | IGHV3-11-2 | + | 0 | cacagtg | 23 | acacaaacc | 0 | + |
| | IGHVII-40-1 | - | | cacagga | | | | - |
| | IGHV3-11 | + | 0 | Cacagtg | 23 | acacaaacc | 0 | + |
| | IGHV3-60-2 | 1 | | cgcagtg | 20 | acacaaacc | | - |
| | IGHV3-66 | 0 | + | cacagtg | 23 | acacaaacc | 0 | + |
| | IGHV1-2-2 | 0 | + | cacagtg | 23 | tcagaaacc | 0 | + |
| | IGHV4-80 | 0 | + | cacagga | 48 | gctttctga | | |
| | IGHV3-41-2 | - | | CAcagtg | 23 | acacaaacc | | - |
| | IGHV3-36-2 | - | | cattgtg | | | | - |
| | IGHV3-50 | - | | ccaaatg | 23 | acacaaaat | 3 | + |
| | IGHVIII-26-1-2 | - | | cacaggg | 24 | acacaaaaa | 0 | + |
| | IGHV2-70D | + | 0 | cacagag | 23 | acaagaacc | | - |
| | IGHV7-27 | - | | cacagtg | | | | - |
| | IGHV4-39 | + | 0 | cacagtg | 23 | acaaaaacc | 0 | + |
| | IGHV4-39-2 | + | 0 | cacagtg | 23 | acaaaaacc | 0 | + |
| | IGHV7-56-2 | - | | caccgtg | 23 | ttagaaacc | | - |
| | IGHVII-1-1 | + | 5 | tgctgtg | | | | - |
| | IGHVIII-44 | - | | | | | | - |
| | IGHV3-30-2-2 | - | | ccaggta | 23 | acacagttt | | - |
| | IGHVII-33-1 | + | 0- | cacagtg | 23 | acccaagcc | 0 | + |
| | IGHV3-38-2 | + | 0 | tacacag | 23 | acacaaacc | 1 | + |

| | | | | | | | | |
|---|---|---|---|---|---|---|---|---|
| | IGHV1-45-2 | + | 0 | cacagtg | 23 | tcagaaacc | | - |
| | IGHV8-51-1-2 | - | | catcgtg | 23 | agacagact | | - |
| | IGHV3-64 | + | 0 | cacagtg | 23 | gcagaaacc | 1 | + |
| | IGHV4-30-2-2 | - | | cacaatg | 23 | acacaaacc | 0 | + |
| | IGHV2-26 | + | 0 | cacagag | 23 | acaagaacc | | - |
| | IGHVIII-25-1-2 | - | | ctcactg | | | | - |
| | IGHV1-58 | + | 0 | cacagtg | 23 | tcagaaacg | | - |
| | IGHV7-40 | - | | cacagtg | 23 | tcagaaacc | | - |
| | IGHVII-65-1 | - | | cacaacg | 23 | atacaaacc | | - |
| | IGHV1-17-2 | + | 0 | cacagtg | 23 | tcagaaacc | 0 | + |
| | IGHV5-78 | - | | cagagtg | | | | - |
| | IGHV3-47-2 | + | 0 | cacagtg | 23 | atacaaact | 0 | + |
| | IGHVIII-2-1-2 | + | 0 | cacagtg | 23 | acacaaagc | 0 | + |
| | IGHVII-62-1 | + | 0 | cacagtg | 23 | accaaaacc | 0 | + |
| | IGHV3-13 | + | 0 | cacagtg | 23 | acacaaacc | 1 | + |
| | IGHV3-64-2 | + | 0 | cacagtg | 23 | gcagaaacc | 1 | + |
| | IGHVIII-67-3 | + | 0 | cacagcg | 22 | acagaaacc | 0 | + |
| | IGHV3-69-1 | + | 0 | cacagtg | 23 | acacaaacc | | - |
| | IGHVII-30-21-2 | + | 0 | cattgtg | 7 | acacaaacc | 1 | + |
| | IGHV3-74 | + | 0 | cacagtg | 23 | acacaaacc | | - |
| | IGHV4-4-2 | + | 0 | cacagtg | 23 | acacaaacc | 0 | + |
| | IGHV3-21 | + | 0 | cacagtg | 23 | acacaaacc | | - |
| | IGHV3-76 | - | | cacagtg | 23 | tcacaaacc | | - |
| | IGHV1-46[ | + | 0 | cacagtg | 23 | tcagaaacc | 0 | + |
| | IGHV3-54-2 | - | | ccaggta | 23 | acacagaat | | - |
| | IGHV3-15-2 | | | cacagtg | 23 | acacaaacc | 0 | + |
| | IGHV3-57 | + | 0 | cacagga | 24 | acacaaaaatc | | - |
| | IGHV3-69-1-2 | + | 0 | cacagtg | 23 | acacaaacc | | - |
| | IGHV3-33-3 | + | 0 | cacagtg | 23 | acacaaacc | 1 | + |
| | IGHV1-12-2 | + | 0 | CAGTGAA | | | | - |
| | IGHV3-33-2 | - | | ccaggta | 23 | acacagttt | | - |
| | IGHVII-40-1-2 | - | | cacagga | | | | - |
| | IGHV3-42 | + | 5 | cagtgag | 22 | | 1 | + |
| | IGHV3-53-2 | + | 0 | cacagtg | 23 | acacaaacc | 0 | + |
| | IGHVII-28-1-2 | + | 0 | cgcaatg | 23 | acacaaccc | 1 | + |
| | IGHV3-79 | - | | CCAGGTA | 23 | acacagagg | | - |
| | IGHV3-29 | - | | ccaagtg | 23 | acacaaaat | 0 | + |
| | IGHVII-67-1 | + | 0 | tactgtg | | | | - |
| | IGHV3-41 | - | | CAcagtg | 23 | acacaaacc | | - |
| | IGHV1-14 | + | 0 | cacagtg | 23 | tcagaaatc | | - |
| | IGHVII-60-1 | - | | cagagtg | 23 | acccaaacc | | - |
| | IGHVII-1-1-2 | + | 5 | tgctgtg | | | | - |

| | | | | | | | | |
|---|---|---|---|---|---|---|---|---|
| | IGHV3-62 | + | 0 | CTGTGTA | 42 | acacaaacc | 0 | + |
| | IGHVII-46-1 | | | | | | | |
| | IGHV1-69D | + | 0 | cacagtg | 23 | tcagaaacc | | - |
| | IGHVIII-38-1-2 | | 0 | cacagtg | 23 | acacaaaag | 0 | + |
| | IGHV1-18 | + | 0 | cacagtg | 23 | tcagaaacc | 0 | + |
| | IGHV4-59 | + | 0 | cacagtg | 23 | acaaaaacc | 0 | + |
| | IGHV2-70-2 | + | 0 | cacagag | 23 | acaagaacc | | - |
| | IGHVIII-16-1 | - | | cacagga | 24 | acagaaaaa | | - |
| | IGHV3-57-2 | + | 0 | cacagga | 24 | acacaaaaatc | | - |
| | IGHV3-75 | - | | cgcagtg | | acacaaacc | | - |
| | IGHV3-19-2 | - | | cactgtg | 23 | acacaaacc | 1 | + |
| | IGHVII-33-1-2 | + | 0 | cacagtg | 23 | acccaagcc | 0 | + |
| | IGHV3-47 | + | 0 | cacagtg | 23 | atacaaact | 0 | + |
| | IGHVII-44-2 | + | 0 | aacagtg | 23 | acataaacc | 0 | + |
| | IGHV3-72-2 | - | | cacagcg | 23 | acacaaacc | 0 | + |
| | IGHVII-15-1 | + | 0 | cattgtg | 7 | acacaaacc | 0 | + |
| | IGHV3-72 | - | | cacagcg | 23 | acacaaacc | 0 | + |
| | IGHV3-33-2-2 | - | | ccaggta | 23 | acacagttt | | - |
| | IGHV3-22 | - | | cacagtg | 23 | acacaaacc | 0 | + |
| | IGHV3-19 | - | | cactgtg | 23 | acacaaacc | 1 | + |
| | IGHVIII-44-2 | - | | | | acacaaacc | | - |
| | IGHVIII-5-1 | - | | cacatga | 17 | | | - |
| | IGHV3-38 | + | 0 | tacacag | 23 | acacaaacc | 1 | + |
| | IGHV3-30 | + | 0 | cacagtg | 23 | acacaaacc | 1 | + |
| | IGHVIII-47-1 | - | | cacggtg | 23 | acacaaacc | 4 | + |
| | IGHV3-32 | + | 0 | ccaagtg | 23 | acacaacat | 0 | + |
| | IGHV3-37 | - | | catggtg | 23 | ccagaaacc | 1 | + |
| | IGHV3OR16-8 | + | 0 | CACAGCG | 24 | cacaaacc | | - |
| | IGHV3-6-2 | - | | tacggta | 23 | acacaaacc | | - |
| | IGHVII-51-2-2 | - | | aacagaa | 21 | acacaaact | | - |
| | IGHV1-18-2 | + | 0 | cacagtg | 13 | tcagaaacc | 0 | + |
| | IGHV1-58-2 | + | 0 | cacagtg | 23 | tcagaaacg | | - |
| | IGHV3-7-2 | + | 0 | cacagtg | 23 | acacaaacc | | - |
| | IGHV3-7 | + | 0 | cacagtg | 23 | acacaaacc | | - |
| | IGHVII-51-2 | - | | aacagaa | 21 | acacaaact | | - |
| | IGHV3-48 | + | 0 | cacagtg | 23 | acacaaacc | | - |
| | IGHV3-36 | - | | cattgtg | | | | - |
| | IGHVII-78-1 | + | 0 | cacagtg | 23 | acccaaacc | 0 | + |
| | IGHV3-32-2 | + | 0 | cacaaca | | | | - |
| | IGHV3-25-2 | - | | cacagtg | 23 | acacaaacc | 0 | + |
| | IGHV4-55 | + | 0 | cacagtg | 23 | acacaaacc | 0 | + |
| | IGHV3-74-2 | + | 0 | cacagtg | 23 | acacaaacc | | - |
| | IGHV1-68-2 | + | 0 | CACGGTG | 23 | TCAGGAACC | 0 | + |
| | IGHV3-52 | + | 0 | cacagtg | 23 | acacaaacc | 0 | + |

| | | | | | | | | |
|---|---|---|---|---|---|---|---|---|
| | IGHV4-28-2 | + | 0 | cacagtg | 23 | acacaaacc | 0 | + |
| | IGHV5-51 | + | 0 | cacagtg | 23 | ctaaaaccc | | - |
| | IGHVIII-47-1-2 | - | | cacggtg | 23 | acacaaacc | 4 | + |
| | IGHVIII-2-1 | + | 0 | cacagtg | 23 | acacaaagc | 0 | + |
| | IGHV3-64D-2 | + | 0 | cacagtg | 23 | acacaaacc | 3 | + |
| | IGHVIII-67-4 | + | 0 | cacagga | 24 | acacaaaaa | | - |
| | IGHVIII-22-2-2 | + | 4 | cactatg | | | | - |
| | IGHV4-34-2 | + | 0 | cacagtg | 23 | acaaaaacc | 0 | + |
| | IGHV1-67 | + | 0 | cacagtg | 23 | tcagtaacc | | - |
| | IGHVIII-5-1-2 | - | | cacatga | 17 | acacaaacc | | - |
| | IGHV3-20 | + | 0 | cacagtg | 23 | acacaaacg | 3 | + |
| | IGHVIII-67-2 | - | | cacatga | 22 | acataaacc | | - |
| | IGHV1-67-2 | + | 0 | cacagtg | 23 | tcagtaacc | | - |
| | IGHVII-53-1-2 | - | | cacagta | 23 | acccaaacc | 1 | + |
| | IGHVII-22-1-2 | - | | cacagcg | 24 | acactctac | 0 | + |
| | IGHV1-45 | + | 0 | cacagtg | 23 | tcagaaacc | | - |
| | IGHV3-6 | - | | tacggta | 23 | acacaaacc | | - |
| | IGHV3-16 | - | | tcctgtg | 23 | acacaaacc | 0 | + |
| | IGHVIII-25-1 | - | | ctcactg | | | | - |
| | IGHV1-14-2 | + | 0 | cacagtg | 23 | tcagaaatc | | - |
| | IGHV3-60 | + | 0 | CTGTGTG | 32 | acacaaacc | | - |
| | IGHVII-26-2 | - | | ctcagtg | 23 | acacaaacc | 0 | + |
| | IGHV1-3-2 | + | 0 | cacagtg | 23 | tcagaaacc | | - |
| | IGHV7-34-1-2 | + | 0 | cacagtg | 23 | tcagaaagc | | - |
| | IGHVIII-76-1-2 | - | | cacaggg | 27 | acacaaacc | 0 | + |
| | IGHVIII-76-1 | - | | cacaggg | 27 | acacaaacc | 0 | + |
| | IGHV5-51-2 | + | 0 | cacagtg | 23 | ctaaaaccc | | - |
| | IGHVII-15-1-2 | + | 0 | cattgtg | 7 | acacaaacc | 0 | + |
| | IGHV1-69D-2 | + | 0 | cacagtg | 23 | tcagaaacc | | - |
| | IGHV7-34-1 | + | 0 | cacagtg | 23 | tcagaaagc | | - |
| | IGHV1-24 | + | 0 | cacagtg | 23 | tcagaaacc | | - |
| | IGHV1-3 | + | 0 | cacagtg | 23 | tcagaaacc | - | |
| | IGHVII-20-1 | - | | CACAGTG | 23 | ACACAAACC | | - |
| | IGHV1-69-2-2 | + | 0 | cacagtg | 23 | tcagaaacc | | - |
| | IGHVIII-5-2 | + | 0 | cacagtg | 22 | acgcaaact | | - |
| | IGHV4-34 | + | 0 | cacagtg | 23 | acaaaaacc | 0 | + |
| | IGHV3-49 | - | | cacagtg | 23 | acacaaacc | 0 | + |
| | IGHV4-59-2 | + | 0 | cacagtg | 23 | acaaaaacc | 0 | + |
| | IGHVIII-26- | - | | cacaggg | 11 | acacaaaaa | 0 | + |

| | | | | | | | | |
|---|---|---|---|---|---|---|---|---|
| | 1 | | | | | tc | | |
| | IGHVII-65-1-2 | - | | cacaacg | 23 | atacaaacc | | - |
| | IGHVII-30-21 | + | 0 | cattgtg | 7 | acacaaacc | 1 | + |
| | IGHV1-69-2 | + | 0 | cacagtg | 23 | tcagaaacc | | - |
| | IGHVII-49-1-2 | - | | tacagca | 23 | aaacaaacc | 0 | + |
| | IGHV1-69 | + | 0 | cacagtg | 23 | tcagaaacc | | - |
| | IGHV7-4-1-2 | + | 0 | cacagtg | 23 | tcaaaaacc | 0 | + |
| | IGHVII-30-1 | + | 0 | cacagtg | 23 | acccaagcc | 0 | + |
| | IGHVII-62-1-2 | + | 0 | cacagtg | 23 | accaaaacc | 0 | + |
| | IGHV3-50-2 | - | | ccaaatg | 23 | acacaaaat | 3 | + |
| | IGHVII-44-2-2 | + | 0 | aacagtg | 23 | acataaacc | 0 | + |
| | IGHV8-51-1 | - | | catcgtg | 23 | agacagact | | - |
| | IGHVII-20-1-2 | - | | CACAGTG | 23 | ACACAAACC | | - |
| | IGHV2-5-2 | - | | cacaaag | 23 | acaaaaacc | | - |
| | IGHV2-26-2 | + | 0 | cacagag | 23 | acaagaacc | | - |
| | IGHV3-25 | - | | cacagtg | 23 | acacaaacc | - | + |
| | IGHV3-48-2 | + | 0 | cacagtg | 23 | acacaaacc | | - |
| | IGHV3-20-2 | + | 0 | cacagtg | 23 | acacaaacg | 3 | + |
| | IGHV3-30-2[ | - | | ccaggta | 23 | acacagttt | | - |
| | IGHV3-71-2 | + | 0 | cacagtg | 23 | acacaaacc | 0 | - |
| | IGHV3-76-2 | - | | cacagtg | 23 | tcacaaacc | | - |
| | IGHVIII-67-4-2 | + | 0 | cacagga | 24 | acacaaaaatc | | - |
| | IGHV1-46-2 | + | 0 | cacagtg | 23 | tcagaaacc | 0 | + |
| | IGHV3-43 | + | 0 | cacagtg | 23 | acaaaaacc | 3 | + |
| | IGHV1-2 | + | 0 | cacagtg | 23 | tcagaaacc | 0 | + |
| | IGHVII-28-1 | + | 0 | cgcaatg | 23 | acacaaccc | 1 | + |
| | IGHVII-22-1 | - | | cacagcg | 24 | acactctac | 0 | + |
| | IGHVII-46-1-2 | - | | | | | | - |
| | IGHV1-17 | + | 0 | cacagtg | 23 | tcagaaacc | 0 | + |
| | IGHV7-81 | + | 0 | caccatg | 23 | tcagaaatc | 0 | + |
| | IGHV3-73 | - | | cacagtg | 23 | acacaaacc | 0 | + |
| | IGHVII-26-2-2 | - | | ctcagtg | 23 | acacaaacc | 0 | + |
| | IGHVII-67-1-2 | + | 0 | tactgtg | | | | - |
| | IGHVII-49-1 | - | | tacagca | 23 | aaacaaacc | 0 | + |
| | IGHV3-63-2 | - | | CCAAGTG | 23 | acacaaaat | 0 | + |
| | IGHV3-42-2 | + | 3 | GAcagtg | 24 | acacaaatc | 1 | + |
| | IGHV3-33 | + | 0 | cacagtg | 23 | | 1 | + |
| | IGHV3-71 | + | 0 | cacagtg | 23 | acacaaacc | 0 | + |
| | IGHV1-69-3 | + | 0 | cacagtg | 23 | tcagaaacc | | - |
| | IGHV3-65 | - | | CACAGTG | 23 | ACACAAACC | 0 | + |
| | IGHV3-64D | + | 0 | cacagtg | 23 | acacaaacc | 3 | + |
| | IGHVII-53-1 | | - | cacagta | 23 | acccaaacc | 1 | + |
| | IGHV3-49-2 | - | | cacagtg | 23 | acacaaacc | 0 | + |

| | | | | | | | | |
|---|---|---|---|---|---|---|---|---|
| | IGHV3-52-2 | + | 0 | cacagtg | 23 | acacaaacc | 0 | + |
| | IGHV4-55-2 | + | 0 | cacagtg | 23 | acacaaacc | 0 | + |
| | IGHVIII-11-1-2 | - | | ctcactg | | | | - |
| | IGHV3-62-2 | + | 0 | CTGTGTA | 42 | acacaaacc | 0 | + |
| | IGHV3-23 | + | 0 | cacagtg | 23 | acacaaacc | | - |
| | IGHV4-28 | | | cacagtg | 23 | acacaaacc | 0 | + |
| | IGHV1-12 | + | 0 | CAGTGAA | | | | |
| | IGHV3-30-3 | + | 0 | cacagtg | 23 | acacaaacc | 1 | + |
| IGHJ | ID | StemL | bp | Nonamer | H-N bp | Heptamer | bp | StemL |
| | IGHJ1 | + | 0 | ggtttctgt | 22 | caccgtg | - | - |
| | IGHJ1-2 | + | 0 | ggtttctgt | 22 | caccgtg | - | - |
| | IGHJ1P | - | - | - | - | - | - | - |
| | IGHJ1P-2 | - | - | - | - | - | - | - |
| | IGHJ2 | - | - | tgtttttgt | 22 | ggctgtg | - | - |
| | IGHJ2-2 | - | - | tgtttttgt | 22 | ggctgtg | - | - |
| | IGHJ2P | - | - | - | - | - | - | - |
| | IGHJ2P-2 | - | - | - | - | - | - | - |
| | IGHJ3 | + | 0 | ggtttatgt | 23 | ccctgtg | 0 | + |
| | IGHJ3-2 | + | 0 | ggtttatgt | 23 | ccctgtg | 0 | + |
| | IGHJ3P | - | - | - | - | - | - | - |
| | IGHJ3P-2 | - | - | - | - | - | - | - |
| | IGHJ4 | - | - | ggtttttgt | 23 | caatgtg | - | - |
| | IGHJ4-2 | - | - | ggtttttgt | 23 | caatgtg | - | - |
| | IGHJ5 | - | - | - | - | caatgtg | - | - |
| | IGHJ5-2 | - | - | - | - | caatgtg | - | - |
| | IGHJ6 | - | - | ggtttttgt | 22 | cattgtg | - | - |
| | IGHJ6-2 | - | - | ggtttttgt | 22 | cattgtg | - | - |
| | IGHJ1 | + | 0 | ggtttctgt | 22 | caccgtg | - | - |
| | IGHJ1-2 | + | 0 | ggtttctgt | 22 | caccgtg | - | - |
| | IGHJ1P | - | - | - | - | - | - | - |
| | IGHJ1P-2 | - | - | - | - | - | - | - |
| IGKV | ID | StemL | bp | Heptamer | H-N bp | Nonamer | bp | StemL |
| | IGKV1-5-3 | - | | cacagtg | 12 | acataaacc | | - |
| | IGKV1D-27 | + | 0 | cactgtg | 12 | acataaacc | | - |
| | IGKV1-27 | + | 0 | cactgtg | 12 | acataaacc | | - |
| | IGKV6-21-3 | - | | cactgtg | 12 | acaaaaact | 1 | + |
| | IGKV1D-42 | - | | cacaggg | | | | |
| | IGKV3-31-3 | + | 0 | cacagtg | | | | - |
| | IGKV2-19-2 | - | | cacagtg | 12 | atacaaacc | | - |
| | IGKV1-16 | - | | cacagtg | 12 | acataaacc | | - |
| | IGKV2-18-3 | - | | cacagtg | 12 | acagaaacc | 0 | + |
| | IGKV2-10 | - | | cacaatg | 12 | acacaaacc | | - |
| | IGKV1-32 | - | | cacagtg | | | | - |
| | IGKV1D-22 | | | | | | | |
| | IGKV2D-19 | - | | cacagtg | 12 | atacaaacc | | - |
| | IGKV2D-24 | - | | cacagtg | 12 | acaaaaacc | | - |
| | IGKV1D-43 | - | | cacagtg | 12 | acaaaaacc | | - |
| | IGKV1-39 | - | | cacagtg | 12 | acataaacc | | - |
| | IGKV2D-23 | + | 1 | cacaatg | 12 | acaaaagcc | | - |
| | IGKV1-35-3 | - | | cacagtg | | | | - |

| | | | | | | | | |
|---|---|---|---|---|---|---|---|---|
| | IGKV1-37 | - | | cacagtg | 12 | acataaacc | | - |
| | IGKV2D-28 | - | | cacagtg | 12 | acagaaacc | 0 | + |
| | IGKV2-38-2 | + | 0 | cacagtg | 12 | acacaaacc | | - |
| | IGKV3D-34 | - | | cacagta | 12 | acaaaaact | 0 | + |
| | IGKV1-12-3 | - | | cacagtg | 12 | acataaacc | | - |
| | IGKV1D-37 | - | | cacagtg | 12 | acataaacc | | - |
| | IGKV1-8-3 | - | | cacagtg | 12 | acaaaaacc | | - |
| | IGKV2-40 | + | 0 | cacagtg | 12 | acagaaacc | 0 | + |
| | IGKV3-34 | - | | cacagta | 12 | acaaaaact | 0 | + |
| | IGKV3-20 | | | cacagtg | 12 | acaaaaacc | 0 | + |
| | IGKV6D-21 | | | cactgtg | 12 | acaaaaact | 1 | + |
| | IGKV2D-38 | + | 0 | cacagtg | 12 | acacaaacc | | - |
| | IGKV1D-16 | | | cacagtg | 12 | acataaacc | 2 | + |
| | IGKV1D-32 | - | | cacagtg | | | | - |
| | IGKV1D-13 | - | | cacagtg | 12 | acataaacc | | - |
| | IGKV1-6 | - | | cacagtg | 12 | acagaaacc | | - |
| | IGKV1-22-3 | + | 0 | TACAGCA | | | | - |
| | IGKV1-32-3 | - | | cacagtg | | | | - |
| | IGKV2-40-2 | + | 0 | cacagtg | 12 | acagaaacc | 0 | + |
| | IGKV3-11-3 | - | | cacagtg | 12 | acaaaaacc | | - |
| | IGKV1-35 | - | | cacagtg | | | | - |
| | IGKV2D-40 | + | 0 | cacagtg | 12 | acagaaacc | 0 | +- |
| | IGKV2D-30 | - | | cacagtg | 12 | acaaaaacc | 4 | + |
| | IGKV1-33-3 | - | | cacagtg | | | | - |
| | IGKV2-14 | - | | cacaatg | 12 | acacaaacc | | - |
| | IGKV1D-8 | + | 0 | cacagtg | 12 | acaaaaacc | 0 | + |
| | IGKV1D-17 | - | | cacagtg | 12 | acataaacc | | - |
| | IGKV2D-36 | - | | | | | | - |
| | IGKV2-23 | + | 0 | CACACAG | | | | - |
| | IGKV1-17 | - | | cacagtg | 12 | acataaacc | | - |
| | IGKV1D-39 | - | | cacagtg | 12 | acataaacc | | - |
| | IGKV2D-29 | - | | cacagtg | 12 | acagaaacc | 5 | + |
| | IGKV1-13 | - | | catagtg | 12 | acataaacc | 2 | + |
| | IGKV1-5 | - | | cacagtg | 12 | acataaacc- | | - |
| | IGKV3-31 | + | 0 | cacagtg | | | | - |
| | IGKV2D-10 | - | | cacaatg | 12 | acacaaacc | | - |
| | IGKV3-7-3 | - | | cacagtg | 12 | acaaaaacc | | - |
| | IGKV2-19 | - | | cacagtg | 12 | atacaaacc | | - |
| | IGKV1-27-3 | + | 0 | cactgtg | 12 | acataaacc | | - |
| | IGKV3-25-3 | - | | CACTGTT | 41 | acaaaaacc | | - |
| | IGKV2-38 | + | 0 | cacagtg | 12 | acacaaacc | | - |
| | IGKV2D-14 | - | | cacaatg | 12 | acacaaacc | | - |
| | IGKV3-20-3 | - | | cacagtg | 12 | acaaaaacc | 0 | + |
| | IGKV2-18 | | | cacagtg | 12 | acagaaacc | 0 | + |
| | IGKV3-15 | - | | cacagtg | 12 | acaaaaacc | | - |
| | IGKV2-36 | - | | | | | | - |
| | IGKV1-37-3 | - | | cacagtg | 12 | acataaacc | | - |
| | IGKV2-26 | + | 1 | cacagtg | 13 | cacaaacc | | - |
| | IGKV3D-11 | - | | cacagtg | 12 | acaaaaacc | | - |
| | IGKV2-4-3 | + | 0 | cacacagtg | 12 | acacaaacc | 5 | + |
| | IGKV1-9 | - | | cacagtg | 12 | acataaacc | | - |
| | IGKV3-15-3 | - | | cacagtg | 12 | acaaaaacc | | - |

| | | | | | | | | |
|---|---|---|---|---|---|---|---|---|
| | IGKV2-14-3 | - | | cacaatg | 12 | acacaaacc | | - |
| | IGKV3-7 | - | | cacagtg | 12 | acaaaaacc | | - |
| | IGKV1-33 | - | | cacagtg | | | | - |
| | IGKV2-26-3 | + | 1 | cacagtg | 13 | cacaaacc | | - |
| | IGKV7-3-3 | = | | cacagtg | 12 | acaaaaacc | | - |
| | IGKV2-4 | + | 0 | cacagtg | 12 | acacaaacc | 5 | + |
| | IGKV2-24 | - | | cacagtg | 12 | acaaaaacc | | - |
| | IGKV3D-25 | - | | CACTGTT | 41 | acaaaaacc | | - |
| | IGKV1-39-3 | - | | cacagtg | 12 | acataaacc | | - |
| | IGKV7-3 | - | | cacagtg | 12 | acaaaaacc | | - |
| | IGKV6-21 | - | | cactgtg | 12 | acaaaaact | 1 | + |
| | IGKV2-28-3 | - | | cacagtg | 12 | acagaaacc | 0 | + |
| | IGKV1-9-3 | - | | cacagtg | 13 | acataaacc | | - |
| | IGKV2-10-3 | - | | cacaatg | 12 | acacaaacc | | - |
| | IGKV1D-12 | - | | cacagtg | 12 | acataaacc | | - |
| | IGKV1-13-3 | - | | catagtg | 12 | acataaacc | 2 | + |
| | IGKV2-30-3 | + | 0 | cacagtg | 12 | acaaaaacc | 0 | + |
| | IGKV2-30 | + | 0 | cacagtg | 12 | acaaaaacc | 0 | + |
| | IGKV2-36-2 | -- | | | | | | - |
| | IGKV5-2-3 | - | | cacagtg | 12 | acaaaaacc | | - |
| | IGKV1D-35 | + | 0 | ctttgtg | | | | - |
| | IGKV5-2 | - | | cacagtg | 12 | acaaaaacc | | - |
| | IGKV1-17-3 | - | | cacagtg | 12 | acataaacc | | - |
| | IGKV2-28 | - | | cacagtg | 12 | acagaaacc | 0 | + |
| | IGKV1-8 | - | | cacagtg | 12 | acaaaaacc | | - |
| | IGKV1-6-3 | - | | cacagtg | 12 | acagaaacc | | - |
| | IGKV3D-20 | - | | cacagtg | | acaaaaacc | 0 | + |
| | IGKV2-24-3 | - | | cacagtg | 12 | acaaaaacc | | - |
| | IGKV3D-15 | - | | cacagtg | 12 | acaaaaacc | | - |
| | IGKV3-34-3 | - | | cacagta | 12 | acaaaaact | 0 | + |
| | IGKV2-29-3 | - | | cacagtg | 12 | acagaaacc | 5 | + |
| | IGKV1-22 | + | 0 | TACAGCA | | | | - |
| | IGKV4-1 | - | | cacagtg | 12 | acacaaacc | | - |
| | IGKV2-29 | - | | cacagtg | 12 | | 5 | + |
| | IGKV2-23-3 | - | | CACAGCT | 24 | GTTTTCTGC | 2 | + |
| | IGKV3-11 | - | | cacagtg | 12 | acaaaaacc | | - |
| | IGKV3D-31 | + | 0 | cacagtg | | | | - |
| | IGKV1-12 | - | | cacagtg | 12 | acataaacc | | - |
| | IGKV1-16-3 | - | | cacagtg | 12 | acataaacc | | - |
| | IGKV1D-33 | - | | cacagtg | | | | - |
| | IGKV2D-18 | - | | cacagtg | 12 | acagaaacc | 0 | + |
| | IGKV3D-7 | - | | cacagtg | 12 | acaaaaacc | | - |
| | IGKV6D-41 | - | | cactgtg | | | | - |
| | IGKV2D-26 | + | 1 | cacagtg | 13 | cacaaacc | | - |
| | IGKV3-25 | - | | CACTGTT | 41 | acaaaaacc | | - |
| IGKJ | ID | StemL | bp | Nonamer | H-N bp | Heptamer | bp | StemL |
| | IGKJ1 | - | - | ggtttttgt | 23 | cactgtg | - | - |
| | IGKJ2 | - | - | agtttttgt | 23 | cattgtg | - | - |
| | IGKJ3 | - | - | ggtttttgt | 22 | cactgtg | 0 | + |
| | IGKJ4 | - | - | ggtttttgt | 23 | cactgtg | - | - |

| | | | | | | | | |
|---|---|---|---|---|---|---|---|---|
| | IGKJ5 | + | 0 | gatttttgt | 23 | cactgtg | - | - |
| IGLV | ID | StemL | bp | Heptamer | H-N bp | Nonamer | bp | StemL |
| | IGLV2-11 | - | | cacagtg | 23 | accaaaacc | 0 | + |
| | IGLV3-7 | + | 0 | cacagtg | 23 | acacaaacc | | - |
| | IGLV2-28 | - | | cacagtg | | | | - |
| | IGLV3-10 | - | | cacagtg | 23 | acacaaacc | | - |
| | IGLV4-69 | - | | cacagtg | 23 | acagaaacc | | - |
| | IGLV8OR8-1 | | | | | | | |
| | IGLVV-66 | | | | | acaaaaact | 1 | + |
| | IGLVV-58 | - | | cagtgaa | 12 | acaaaaact | | - |
| | IGLVIV-66-1 | + | 0 | cactgtg | | | | - |
| | IGLV3-12 | - | | cacggtg | 23 | acaaaaaca | 1 | + |
| | IGLV7-43 | + | 0 | cacagtg | 23 | acataaacc | | - |
| | IGLV3-1 | | | cacagtg | 23 | acagaaacc | | |
| | IGLV10-67 | - | | cacagcg | | | | - |
| | IGLV2-23 | - | | cacagtg | 23 | accaaaacc | | - |
| | IGLV3-27 | - | | cacagtg | 23 | acacaaacc | 0 | + |
| | IGLV5-45 | - | | cacagtg | 23 | acaaaaacc | | - |
| | IGLV7-46 | - | | cacagtg | 23 | acataaacc | | - |
| | IGLV2-8 | - | | cacagtg | | | | - |
| | IGLV3-29 | - | | | | | | - |
| | IGLV1-50 | - | | CAcacag | | | | - |
| | IGLVIV-53 | - | | cactgtg | | | | - |
| | IGLV1-62 | - | | | | acaagaacc | 5 | + |
| | IGLV3-21 | + | 0 | cacggtg | 23 | acaaaaaca | | - |
| | IGLV3-9 | + | 0 | cacagta | 23 | acacaaacc | | - |
| | IGLV1-51 | - | | cacagtg | 23 | acaagaacc | | - |
| | IGLV1-36 | - | | cacagtg | 23 | acaagaacc | 0 | + |
| | IGLV3-24 | + | 0 | cacagtg | 23 | acacaaacc | 0 | + |
| | IGLVI-42 | - | | aacagtg | | | | - |
| | IGLV3-32 | + | 0 | cacagtg | 23 | acacaaacc | 0 | + |
| | IGLV5-37-2 | + | 2 | cacagcc | | | | - |
| | IGLVI-56 | - | | ctctgtg | | | | - |
| | IGLV5-39 | - | | tttcactg | 19 | acaaaaaca | | - |
| | IGLV5-48 | - | | cacagtg | 45 | ggttcttgt | | - |
| | IGLV7-35 | - | | cacagtg | 23 | acataaacc | | - |
| | IGLVI-63 | - | | cacagtg | | | | - |
| | IGLVI-38 | + | 0 | cacggtg | | | | - |
| | IGLV2-18 | - | | cacagag | 23 | accaaaacc | 0 | + |
| | IGLVIV-65 | - | | CACAGCA | 34 | | 0 | + |
| | IGLV2-5 | - | | cacagtg | | | | - |
| | IGLV3-2 | + | 0 | cacagtg | 22 | atgcaaacc | 0 | + |
| | IGLV3-17 | + | 0 | cacagtg | | | | - |
| | IGLVI-20 | - | | caccgtg | 23 | acccaaacc | | - |
| | IGLVVI-25-1 | + | 0 | CACTGTT | 35 | accaaaacc | | - |
| | IGLVVI-22-1 | + | 0 | CCAAGTG | 13 | accaaaacc | | - |
| | IGLVIV-64 | -- | | CAcacag | | | | - |
| | IGLV6-57 | - | | cacagtg | 23 | acagaaact | | 0+ |
| | IGLV10-54 | - | | cacagtg | | | | - |
| | IGLV3-25 | - | | cacagtg | 23 | acataaacc | | - |
| | IGLV5-37 | - | | cacagtg | 23 | acaaaaacc | | - |

| | | | | | | | | |
|---|---|---|---|---|---|---|---|---|
| | IGLV3-19 | + | 0 | cacattg | 23 | acagaaacc | 0 | + |
| | IGLV7-43-2 | + | 0 | CACAGTG | 23 | ACATAAACC | | - |
| | IGLV3-30 | - | | cacagtg | | | | - |
| | IGLV3-15 | + | 0 | cacagtg | 23 | acacaaaaacc | | - |
| | IGLV5-45-2 | - | | CACAGTG | 23 | ACAAAAACC | | - |
| | IGLV11-55 | - | | cacagtg | 19 | acaaaaacc | | - |
| | IGLV1-47 | - | | cacagtg | 23 | acaagaacc | 0 | + |
| | IGLV1-44 | - | | cacagtg | 23 | acaagaacc | 5 | + |
| | IGLV4-60 | - | | cacagtg | 23 | acaaaatcc | 0 | + |
| | IGLV8-61 | + | 0 | cacagtg | | | | - |
| | IGLV3-13 | - | | cacagtg | 23 | acacaaatc | 0 | + |
| | IGLV3-26 | + | 0 | cacaatg | | | | - |
| | IGLV1-40 | - | | cacagtg | 23 | acaagaacc | 5 | + |
| | IGLVI-68 | - | | ctctgtg | | | | - |
| | IGLV2-33 | - | | catagtg | 23 | accaaaacc | 0 | + |
| | IGLV3-16 | - | | cacagtg | 23 | acataaacc | | - |
| | IGLVIV-59 | - | | TAcacag | | | | - |
| | IGLV1-41 | - | | cacagtg | 23 | acaagaacc | 5 | + |
| | IGLV3-31 | - | | | | acaaaaacc | | - |
| | IGLV4-3 | - | | cacagtg | | | | - |
| | IGLVVII-41-1 | - | | ttcactg | | | | - |
| | IGLV2-34 | - | | cacagtg | 22 | accaaaacc | 0 | + |
| | IGLV2-14 | - | | cacagtg | 23 | accaaaacc | 0 | + |
| | IGLV3-4 | + | 0 | catagtg | 24 | cacaaacc | 4 | + |
| | IGLV9-49 | - | | cacagtg | 22 | acaaaaacc | | - |
| | IGLV5-52 | + | 0 | cacagtg | | | | - |
| | IGLV3-6 | - | | cacagtg | 23 | acacaaacc | 0 | + |
| | IGLVI-70 | - | | cacagtg | 23 | acaaaaacc | | - |
| | IGLV3-22 | | | ctcagtg | 23 | acacaaact | 0 | + |
| IGLJ | ID | StemL | bp | Nonamer | H-N bp | Heptamer | bp | StemL |
| | IGLJ1 | + | 0 | ggttttggt | 12 | cactgtg | - | - |
| | IGLJ2 | - | - | ggttttttgt | 12 | cacagtg | - | - |
| | IGLJ3 | - | - | ggttttttgt | 12 | cacagtg | - | - |
| | IGLJ4 | - | - | - | - | - | - | - |
| | IGLJ5 | - | - | ggttttttgt | 12 | cacagca | - | - |
| | IGLJ6 | - | - | ggtttgtgt | 12 | cacagtg | - | - |
| | IGLJ7 | - | - | ggtttgtgt | 12 | cactgtg | - | - |
| TRAV | ID | StemL | bp | Heptamer | H-N bp | Nonamer | bp | StemL |
| | TRAV32 | + | - | - | - | - | - | - |
| | TRAV8-5 | + | 0 | cacagtg | 22 | acacaaact | - | - |
| | TRAV40 | + | 0 | cactgtg | 22 | acaaaaacc | - | - |
| | TRAV26-1 | - | - | - | - | - | - | - |
| | TRAV12-1 | + | 0 | cacagtg | 23 | acccaaacc | - | - |
| | TRAV13-1 | - | - | cacattg | 23 | acacaaacc | - | - |
| | TRAV37 | - | - | - | - | - | - | - |
| | TRAV17 | + | 0 | cacagtg | - | - | - | - |
| | TRAV20 | + | 0 | cacagcg | - | - | - | - |

| | | | | | | | | |
|---|---|---|---|---|---|---|---|---|
| | TRAV38-1 | + | 0 | cacaatg | 23 | acagaaacc | - | - |
| | TRAV26-2 | - | - | cacagtg | - | - | - | - |
| | TRAV34 | - | - | cacagcg | - | - | - | - |
| | TRAV28 | - | - | cactgtg | - | - | 2 | + |
| | TRAV11 | - | - | - | - | - | - | - |
| | TRAV9-1 | + | 0 | cacagtg | - | - | - | - |
| | TRAV24 | + | 0 | cacagtg | - | - | - | - |
| | TRAV8-7 | - | - | gactgtg | 23 | acacaaact | 0 | + |
| | TRAV41 | - | - | cacagtg | - | - | - | - |
| | TRAV3 | - | - | cacactg(mus) | 23 | acacaaact | - | - |
| | TRAV31 | - | - | cactgtg | - | - | - | - |
| | TRAV19 | + | 0 | cacagtg | 23 | acaaaaacc | 0 | + |
| | TRAV15 | + | 0 | cacaggg | - | - | - | - |
| | TRAV36DV7 | + | 0 | cacagtg | - | - | - | - |
| | TRAV14DV4 | + | 0 | cacagtg | 23 | acaaaagcc | 0 | + |
| | TRAV8-3 | + | 0 | cacagtg | 23 | acacaaact | - | - |
| | TRAV8-2 | + | 0 | cacagtg | - | - | - | - |
| | TRAV25 | + | 0 | cacagtg | - | - | - | - |
| | TRAV27 | + | 0 | cacagtg | 23 | acccaaacc | - | - |
| | TRAV18 | + | 0 | cagagtg | 24 | cacaaacc | - | - |
| | TRAV2 | - | - | - | - | acacagagg | - | - |
| | TRAV4 | + | 0 | cacagtg | - | - | - | - |
| | TRAV23DV6 | - | - | cacagtg | 23 | acccaaacc | - | - |
| | TRAV21 | + | 0 | cacagtg | - | - | - | - |
| | TRAV8-6 | - | - | cacagtg | 22 | acacaaacc | 0 | + |
| | TRAV7 | + | 0 | cacagta | - | - | - | - |
| | TRAV12-3 | + | 0 | cacagtg | 23 | acccaaacc | - | - |
| | TRAV22 | + | 3 | cacagtg | 23 | acacaaacc | - | - |
| | TRAV1-1 | + | 0 | cacagtg | - | - | - | - |
| | TRAV13-2 | - | - | cacattg | 23 | acccaaacc | - | - |
| | TRAV35 | + | 0 | cacagtg | - | - | - | - |
| | TRAV12-2 | - | - | cacagtg | 23 | acccaaacc | - | - |
| | TRAV6 | - | - | cacagta | - | - | - | - |
| | TRAV16 | + | 0 | cacagta | 22 | acacaaacc | - | - |
| | TRAV29DV5 | + | 0 | cacagtg | - | - | - | - |
| | TRAV8-1 | - | - | cacagtg | 23 | acacaaact | - | - |
| | TRAV5 | - | - | cacattg | 23 | acccaaacc | - | - |
| | TRAV33 | + | - | - | - | - | - | - |
| | TRAV8-4 | - | - | cacagtg | 22 | acataaacc | 0 | + |
| | TRAV10 | - | - | cactgtg | 23 | atgcaaacc | - | - |
| | TRAV1-2 | - | - | cacggtg | - | - | - | - |
| | TRAV38-2DV8 | + | 0 | cacagtg | 23 | acagaaacc | - | - |
| | TRAV9-2 | + | 2 | cacagtg | - | - | - | - |
| | TRAV30 | - | - | cacagtg | - | - | - | - |
| | TRAV39 | - | - | cacagtg | 23 | acccaaacc | 0 | + |
| TRAJ | ID | StemL | bp | Nonamer | H-N bp | Heptamer | bp | StemL |
| | TRAJ10 | - | - | - | - | cactgtg | - | - |
| | TRAJ43 | - | - | ggtttttgt | 12 | tactgtg | 0 | + |
| | TRAJ60 | - | - | - | - | cactatg | - | - |
| | TRAJ41 | - | - | - | - | cactgtg | - | - |
| | TRAJ24 | + | 1 | ccattttgt | 12 | cacagtg | 0 | + |
| | TRAJ32 | - | - | - | - | gactgtg | - | - |

| | | | | | | | | |
|---|---|---|---|---|---|---|---|---|
| | TRAJ55 | - | - | - | - | - | - | - |
| | TRAJ29 | + | 0 | ggtttttgt | 12 | cactgtg | - | - |
| | TRAJ46 | - | - | - | - | - | - | + |
| | TRAJ40 | - | - | ggtttatgt | 12 | cactgtg | 0 | + |
| | TRAJ42 | - | - | - | - | gactgtg | - | - |
| | TRAJ51 | - | - | - | - | - | - | + |
| | TRAJ20 | - | - | ggtttgtgt | 12 | cactgtg | 0 | + |
| | TRAJ25 | - | - | ggtttttga | 12 | cactatg | - | - |
| | TRAJ7 | + | 0 | ggtttttgt | 10 | cacagtg | - | - |
| | TRAJ21 | - | - | - | - | catggtg | - | - |
| | TRAJ38 | - | - | ggttttggt | 12 | gactgtg | - | - |
| | TRAJ57 | - | - | - | - | - | - | - |
| | TRAJ18 | - | - | - | - | cattgtg | - | - |
| | TRAJ49 | - | - | ggtttttgt | 12 | cacagtg | - | - |
| | TRAJ30 | + | 0 | agtttttgt | 12 | cacagtg | - | - |
| | TRAJ23 | - | - | tgtttttga | 12 | cacagtg | 0 | + |
| | TRAJ35 | - | - | ggtttttgt | 12 | cattgtg | - | - |
| | TRAJ61 | - | - | ggtttttgt | 12 | tcctgtg | - | - |
| | TRAJ26 | - | - | ggtttttgc | 12 | cactgtg | - | - |
| | TRAJ39 | - | - | ggtttttgc | 12 | cactgtg | - | - |
| | TRAJ5 | - | - | ggattttgt | 12 | - | - | - |
| | TRAJ53 | - | - | - | - | ggctgtg | - | - |
| | TRAJ31 | - | - | - | - | tgctgtg | - | - |
| | TRAJ4 | - | - | - | - | - | - | - |
| | TRAJ19 | - | - | - | - | - | - | - |
| | TRAJ36 | - | - | - | - | cactgtg | - | - |
| | TRAJ45 | - | - | - | - | cagagtg | - | - |
| | TRAJ27 | - | - | - | - | gactgtg | - | - |
| | TRAJ50 | - | - | - | - | ggctgtg | 0 | + |
| | TRAJ47 | - | - | tgtttttgt | 12 | cgctgtg | 0 | + |
| | TRAJ54 | - | - | agtttctgt | 12 | - | - | - |
| | TRAJ44 | - | - | ggtttctgt | 12 | cacagtg | - | - |
| | TRAJ17 | + | 5 | ggtttttgc | 12 | cattgtg | - | - |
| | TRAJ58 | - | - | ggtttttgc | 12 | cacagtg | 0 | + |
| | TRAJ37 | + | 2 | agtttttgt | 12 | - | - | - |
| | TRAJ12 | - | - | tgtttttga | 12 | cactgtg | 0 | + |
| TRGV | ID | StemL | bp | Heptamer | H-N bp | Nonamer | bp | StemL |
| | TRGVB | + | 0 | cacagca | - | - | - | - |
| | TRGV9 | + | 0 | cacagca | - | - | - | - |
| | TRGV3 | + | 0 | cacagtg | 24 | tgaaaatc | 3 | + |
| | TRGV2 | + | 0 | cacagtg | 24 | tgaaaatc | 3 | + |
| | TRGV8 | + | 0 | cacagtg | 24 | tgaaaatc | 3 | + |
| | TRGVA | - | - | - | - | - | - | - |
| | TRGV10 | - | - | - | - | - | - | - |
| | TRGV4 | + | 0 | cacagtg | - | - | - | - |
| | TRGV6 | + | 0 | cagagtg | - | - | - | - |
| | TRGV5 | + | 0 | cacagtg | 24 | tgaaaatc | - | - |
| | TRGV11 | + | 0 | cacagtg | 23 | acagaaact | 0 | + |
| | TRGV7 | + | 0 | cacagtg | 24 | tgaaaatc | 3 | + |
| | TRGV1 | + | 0 | cacagtg | 10 | tgaaaatc | - | - |
| | TRGV5P | + | 0 | cacagtg | - | - | - | - |
| TRGJ | ID | StemL | bp | Nonamer | H-N bp | Heptamer | bp | StemL |
| | TRGJP1 | - | - | gatttttgt | 12 | - | - | - |

| | | | | | | | | |
|---|---|---|---|---|---|---|---|---|
| | TRGJP | - | - | - | - | - | - | - |
| | TRGJ2 | - | - | agtttttga | 12 | cactgtg | - | - |
| | TRGJ1 | - | - | agtttttga | 12 | cactgtg | - | - |
| | TRGJP2 | - | - | gatttttgt | 12 | - | - | - |
| TRBV | ID | StemL | bp | Heptamer | H-N bp | Nonamer | bp | StemL |
| | TRBV26OR9-2 | - | - | cacagca | - | - | - | - |
| | TRBV23OR9-2 | - | - | cacagca | 23 | acacaaact | - | - |
| | TRBV10-3-2 | - | - | cacagtg | - | - | - | - |
| | TRBV11-2-2 | + | 0 | cacagtg | 23 | gcagaaaac | - | - |
| | TRBV18-2 | - | - | cacattg | - | - | - | - |
| | TRBV5-4 | + | 0 | cacagcc | - | - | - | - |
| | TRBV6-7 | - | - | cacagcg | - | - | - | - |
| | TRBV9 | + | 0 | cacagcc | - | - | - | - |
| | TRBV11-3 | + | 0 | cacagtg | 23 | gcagaaaac | - | - |
| | TRBV13 | + | 0 | cacagcc | 23 | acccaaacc | 0 | + |
| | TRBV6-7-2 | - | - | cacagcg | - | - | - | - |
| | TRBVA-2 | + | 0 | cacagca | 23 | acaaaatgg | - | - |
| | TRBV20-1 | - | - | cacagcg | 23 | gcaagaacc | - | - |
| | TRBV25-1 | - | - | cacagtg | - | - | - | - |
| | TRBV25OR9-2 | - | - | cacagtg | - | - | - | - |
| | TRBV19 | - | - | cacagtg | - | - | - | |
| | TRBV7-1-2 | - | - | cacagca | - | - | - | - |
| | TRBV2 | - | - | cacagcc | - | - | - | - |
| | TRBV7-9 | + | 0 | cacagca | 23 | tcacaaacc | - | - |
| | TRBV22OR9-2 | - | - | cacagtg | - | - | 3 | + |
| | TRBV19-2 | - | - | cacagtg | - | - | - | - |
| | TRBV5-8 | + | 0 | cacagcc | - | - | 3 | + |
| | TRBV15 | + | 0 | cacagag | - | - | 0 | + |
| | TRBV18 | - | - | cacattg | - | - | - | - |
| | TRBV7-6 | - | - | cacagtg | 23 | tcacaaacc | - | - |
| | TRBV29OR9-2 | - | - | cacagtg | 23 | gcaagaacc | - | - |
| | TRBV20OR9-2 | - | - | cacagcg | 23 | gcaagaacc | - | - |
| | TRBV12-3-2 | + | 0 | cacagcg | 23 | gcagaaaac | - | - |
| | TRBV7-8 | + | 0 | cacagca | 23 | tcacaaacc | - | - |
| | TRBV25-1-2 | - | - | cacagtg | - | - | - | - |
| | TRBV6-1 | - | - | cacagcg | - | - | - | - |
| | TRBV10-2-2 | - | - | cacagtg | - | - | - | - |
| | TRBV6-5 | - | - | cacagcg | - | - | - | - |
| | TRBV21-1 | + | 0 | cacagtg | 22 | acacaaact | - | - |
| | TRBV17-2 | + | - | - | - | - | - | - |
| | TRBV10-1 | - | - | cacagtg | - | - | - | - |
| | TRBV5-5-2 | + | 0 | cacagcc | - | - | 0 | + |
| | TRBV3-1-2 | + | 0 | cacagcc | 24 | cacaaacc | - | - |
| | TRBV21-1-2 | + | 0 | cacagtg | 22 | acacaaact | - | - |
| | TRBV4-3 | + | 0 | cacagcc | 23 | gcagaaacc | 0 | + |
| | TRBV8-2-2 | - | - | - | - | - | - | - |
| | TRBV23-1-2 | - | - | cacagca | 23 | acacaaact | - | - |
| | TRBV7-3-2 | - | - | cacagca | - | - | - | - |

| | | | | | | | | |
|---|---|---|---|---|---|---|---|---|
| | TRBV11-3-2 | + | 0 | cacagtg | 23 | gcagaaaac | - | - |
| | TRBV12-2-2 | - | - | cacagcg | - | - | - | - |
| | TRBV5-1-2 | + | 0 | cacagcc | 24 | cacaaacc | - | - |
| | TRBV8-1 | + | - | - | - | - | - | - |
| | TRBVAOR9-2 | + | 0 | cacagca | 23 | acaaaatgg | - | - |
| | TRBV6-4 | + | 0 | cacagtg | - | - | - | - |
| | TRBV6-3 | - | - | cacagtg | - | - | - | - |
| | TRBV4-2-2 | + | 0 | cacagcc | - | - | - | - |
| | TRBV17 | + | - | - | - | - | - | - |
| | TRBV10-2 | - | - | cacagtg | - | - | - | - |
| | TRBV9-2 | + | 0 | cacagcc | - | - | - | - |
| | TRBV6-3-2 | - | - | cacagtg | - | - | - | - |
| | TRBV7-2-2 | + | 0 | cacagca | - | - | - | - |
| | TRBV12-2 | - | - | cacagcg | - | - | - | - |
| | TRBVA | + | 0 | cacagca | 23 | acaaaatgg | - | - |
| | TRBV6-1-2 | - | - | cacagcg | - | - | - | - |
| | TRBV23-1 | - | - | cacagca | 23 | acacaaact | - | - |
| | TRBV2-2 | - | - | cacagcc | - | - | - | - |
| | TRBV27-2 | + | 0 | cacagtg | 23 | acaaaaaca | - | - |
| | TRBV12-3 | + | 0 | cacagcg | 23 | gcagaaaac | - | - |
| | TRBV10-3 | - | - | cacagtg | - | - | - | - |
| | TRBV11-1-2 | + | 0 | cacagcg | - | - | - | - |
| | TRBV11-1 | + | 0 | cacagcg | - | - | - | - |
| | TRBV16 | - | - | cacagtg | - | - | - | - |
| | TRBV21OR9-2 | - | - | cacagtg | 22 | acacaaact | - | - |
| | TRBV3-2 | + | 0 | cacagcc | 24 | cacaaacc | 0 | + |
| | TRBVB | - | - | - | - | - | - | - |
| | TRBV14 | - | - | cacagtg | 23 | gcaaaacca | - | - |
| | TRBV6-9 | - | - | cacagcg | - | - | - | - |
| | TRBV29-1-2 | - | - | cacagtg | 23 | gcaagaacc | 1 | + |
| | TRBV26-2 | + | - | - | - | - | - | - |
| | TRBV22-1-2 | + | 0 | cacaatg | - | - | - | - |
| | TRBV16-3 | - | - | cacagtg | - | - | - | - |
| | TRBV6-6-2 | - | - | cacagcg | - | - | - | - |
| | TRBV11-2 | + | 0 | cacagtg | 23 | gcagaaaac | - | - |
| | TRBV5-1 | + | 0 | cacagcc | 24 | cacaaacc | - | - |
| | TRBV6-5-2 | - | - | cacagcg | - | - | - | - |
| | TRBV8-2 | - | - | - | - | - | - | - |
| | TRBV29-1 | - | - | cacagtg | 23 | gcaagaacc | 1 | + |
| | TRBV7-1 | - | - | cacagca | - | - | - | - |
| | TRBV7-4 | + | 0 | cacagcg | 23 | tcacaaacc | - | - |
| | TRBV26 | + | - | - | - | - | - | - |
| | TRBV4-1-2 | + | 0 | cacagcc | 23 | gcagaaacc | 0 | + |
| | TRBV5-6-2 | + | 0 | cacagcc | - | - | - | - |
| | TRBV27 | + | 0 | cacagtg | 23 | acaaaaaca | - | - |
| | TRBV1-2 | + | 0 | cacagcc | 24 | cacaaacc | 2 | + |
| | TRBV5-7-2 | + | 0 | cacagcc | - | - | - | - |
| | TRBV7-5-2 | + | 0 | cacagtg | 23 | tcacaaacc | - | - |
| | TRBV10-1-2 | - | - | cacagtg | - | - | - | - |
| | TRBV12-1 | - | - | cacagca | 23 | gcagaaacc | - | - |
| | TRBV6-6 | - | - | cacagcg | - | - | - | - |
| | TRBV8-1-2 | + | - | - | - | - | - | - |
| | TRBV5-2-2 | - | - | - | - | - | - | - |

| | | | | | | | | |
|---|---|---|---|---|---|---|---|---|
| | TRBV28 | + | 0 | cacagcg | - | - | - | - |
| | TRBV12-1-2 | - | - | cacagca | 23 | gcagaaacc | - | - |
| | TRBV30 | - | - | cactgag | 21 | gcaaaaacc | 0 | + |
| | TRBV5-6 | + | 0 | cacagcc | - | - | - | - |
| | TRBV7-9-2 | + | 0 | cacagca | 23 | tcacaaacc | - | - |
| | TRBV28-2 | + | 0 | cacagcg | - | - | - | - |
| | TRBV15-2 | + | 0 | cacagag | - | - | 0 | + |
| | TRBV6-2 | - | - | cacagtg | - | - | - | - |
| | TRBV5-3-2 | + | 0 | cacagcc | - | - | 3 | + |
| | TRBV7-3 | - | - | cacagca | - | - | - | - |
| | TRBV7-6-2 | + | 0 | cacagca | 23 | tcacaaacc | - | - |
| | TRBV14-2 | - | - | cacagtg | 23 | gcaaaacca | - | - |
| | TRBV7-7 | + | 0 | cacagca | 23 | tcacaaacc | - | - |
| | TRBV24-1 | + | 0 | cacagtg | 23 | acagaaaga | - | - |
| | TRBV24OR9-2 | - | - | cacagtg | 23 | acagaaaga | - | - |
| | TRBV4-1 | + | 0 | cacagcc | 23 | gcagaaacc | 0 | + |
| | TRBV6-8 | - | - | cacagcg | - | - | - | - |
| | TRBV7-4-2 | + | 0 | cacagcg | 23 | tcacaaacc | - | - |
| | TRBV4-2 | - | - | cacagcc | 23 | gcagaaacc | 0 | + |
| | TRBV12-5-2 | + | 0 | cacagcg | 23 | gcagaaacc | - | - |
| | TRBV3-1 | + | 0 | cacagcc | 24 | cacaaacc | - | - |
| | TRBV7-7-2 | + | 0 | cacagca | 23 | tcacaaacc | - | - |
| | TRBV5-4-2 | + | 0 | cacagcc | - | - | - | - |
| | TRBV5-5 | + | 0 | cacagcc | - | - | 0 | + |
| | TRBV24-1-2 | + | 0 | cacagtg | 23 | acagaaaga | - | - |
| | TRBV12-4 | + | 0 | cacagcg | 23 | gcagaaacc | 0 | + |
| | TRBV12-4-2 | + | 0 | cacagcg | 23 | gcagaaaac | - | - |
| | TRBV13-2 | + | 0 | cacagcc | 23 | acccaaacc | 0 | + |
| | TRBV7-2 | + | 0 | cacagca | - | - | - | - |
| | TRBVB-2 | - | - | - | - | - | - | - |
| | TRBV5-2 | - | - | - | - | - | - | - |
| | TRBV6-8-2 | - | - | cacagcg | - | - | - | - |
| | TRBV7-5 | - | - | cacagtg | 23 | tcacaaacc | - | - |
| | TRBV12-5 | + | 0 | cacagcg | 23 | gcagaaacc | - | - |
| | TRBV30-2 | - | - | cactgag | 21 | gcaaaaacc | 0 | + |
| | TRBV6-4-2 | + | 0 | cacagtg | - | - | - | - |
| | TRBV5-3 | + | 0 | cacagcc | - | - | 3 | + |
| | TRBV20-1-2 | - | - | cacagcg | 23 | gcaagaacc | - | - |
| | TRBV5-7 | + | 0 | cacagcc | - | - | - | - |
| | TRBV1 | + | 0 | cacagcc | 24 | cacaaacc | 2 | + |
| | TRBV22-1 | + | 0 | cacaatg | - | - | - | - |
| TRBJ | ID | StemL | bp | Nonamer | H-N bp | Heptamer | bp | StemL |
| | TRBJ1-4-2 | + | 0 | ggttttcct | 12 | tgttgtg | - | - |
| | TRBJ1-3-2 | - | - | ggttttgaa | 12 | ggctgtg | - | - |
| | TRBJ2-6 | - | - | ggtttttgc | 12 | ggctgtg | - | - |
| | TRBJ1-5 | - | - | gggtttgcc | 12 | cactgtg | - | - |
| | TRBJ2-7-2 | - | - | ggtttgcat | 12 | ctccgtg | - | - |
| | TRBJ2-2 | - | - | ggtttgcgc | 12 | ggctgtg | - | - |
| | TRBJ2-3-2 | - | - | ggtttttgt | 12 | ggctgtg | - | - |
| | TRBJ2-5 | - | - | ggtttttgt | 12 | ggccgtg | - | - |
| | TRBJ1-1 | - | - | gattttcac | 12 | cactgtg | - | - |
| | TRBJ1-3 | - | - | ggttttgaa | 12 | ggctgtg | - | - |
| | TRBJ2-2-2 | - | - | ggtttgcgc | 12 | ggctgtg | - | - |

| | | | | | | | | |
|---|---|---|---|---|---|---|---|---|
| | TRBJ2-1-2 | - | - | gaattctgg | 12 | cactgtg | 0 | + |
| | TRBJ2-6-2 | - | - | ggtttttgc | 12 | ggctgtg | - | - |
| | TRBJ2-3 | - | - | ggtttttgt | 12 | ggctgtg | - | - |
| | TRBJ2-2P-2 | - | - | - | - | ggctgtg | - | - |
| | TRBJ1-5-2 | - | - | gggtttgcc | 12 | cactgtg | - | - |
| | TRBJ2-7 | - | - | ggtttgcat | 12 | ctccgtg | - | - |
| | TRBJ2-2P | - | - | - | - | ggctgtg | - | - |
| | TRBJ1-4 | + | 0 | ggttttcct | 12 | tgttgtg | - | - |
| | TRBJ1-1-2 | - | - | gattttcac | 12 | cactgtg | - | - |
| | TRBJ2-5-2 | - | - | ggtttttgt | 12 | ggccgtg | - | - |
| | TRBJ2-4 | - | - | agtttctgt | 12 | ggctgtg | - | - |
| | TRBJ1-2-2 | - | - | ccttttaga | 12 | ttatgtg | - | - |
| | TRBJ1-6 | - | - | gggttttat | 12 | agctgtg | - | - |
| | TRBJ1-2 | - | - | ccttttaga | 12 | ttatgtg | - | - |
| | TRBJ1-6-2 | - | - | gggttttat | 12 | agctgtg | - | - |
| | TRBJ2-1 | - | - | gaattctgg | 12 | cactgtg | 0 | + |
| | TRBJ2-4-2 | - | - | agtttctgt | 12 | ggctgtg | - | - |
| TRDV | ID | StemL | bp | Heptamer | H-N bp | Nonamer | bp | StemL |
| | TRDV3 | - | - | cactatg | 23 | acacaaact | 0 | + |
| | TRDV2 | - | - | - | - | - | - | - |
| | TRDV1 | - | | cacagtg | 23 | acaaaaacc | - | - |
| TRDJ | ID | StemL | bp | Nonamer | H-N bp | Heptamer | bp | StemL |
| | TRDJ3 | - | - | gttacctgt | 12 | taatgtg | - | - |
| | TRDJ1 | - | - | ggtttttgg | 12 | tgctgtg | - | - |
| | TRDJ4 | - | - | - | - | agctgtg | - | - |
| | TRDJ2 | - | - | - | - | - | - | - |

| IGHD | ID | Nonamer | bp | Heptamer | D-StemL | Heptamer | bp | Nonamer |
|---|---|---|---|---|---|---|---|---|
| | IGHD3-3-2 | ggtttgggg | 12 | cactgtg | yes | cacagtg | 12 | tcaaaaacc |
| | IGHD4-4 | gctttttgt | 12 | tactgtg | yes | cacagtg | 12 | gcaaaaact |
| | IGHD2-15 | ggattttgt | 12 | cactgtg | yes | cacagtg | 12 | tcccaaagc |
| | IGHD5-12 | ggttattgt | 12 | gactgtg | yes | cacagtg | 12 | gcagcaacc |
| | IGHD6-19-2 | ggtttctga | 12 | cacagtg | no | cacagtg | 12 | ccagaaacc |
| | IGHD1-1-2 | / | / | cacggtg | yes | caccgtg | 4 | aaactgtgt |
| | IGHD3-22 | / | / | cactgtg | yes | cacagtg | 12 | tcaaaaact |
| | IGHD5-24-2 | ggttattgt | 12 | ggccgtg | yes | cacagtg | 12 | gcagcaacc |
| | IGHD2-15-2 | ggattttgt | 12 | cactgtg | yes | cacagtg | 12 | tcccaaagc |
| | IGHD5-24 | ggttattgt | 12 | ggccgtg | yes | cacagtg | 12 | gcagcaacc |
| | IGHD3-16-2 | / | / | cactgtg | no | cacagca | 12 | tcagaaacc |
| | IGHD6-6 | aaacaaacc | 25 | cacagtg | yes | cacagtg | 12 | ccagaaacc |
| | IGHD4-11 | gctttttgt | 12 | tgctgtg | no | catagtg | 12 | gcaaaaact |

| | | | | | | | | |
|---|---|---|---|---|---|---|---|---|
| | IGHD2-2-2 | ggattttgt | 12 | cactgtg | yes | cacagtg | 12 | tcccaaagc |
| | IGHD3-9-2 | / | / | cactgtg | yes | cacagtg | 12 | tcaaaaacc |
| | IGHD6-13-2 | ggtttctga | 12 | cacagtg | yes | cacagtg | 12 | ccagaaacc |
| | IGHD3-10-2 | ggtttgggg | 12 | cactgtg | yes | cacagtg | 12 | tcaaaaacc |
| | IGHD2-8-2 | ggattttgt | 12 | cactgtg | yes | cacagtg | 12 | tcccaaagc |
| | IGHD3-22-2 | / | / | cactgtg | yes | cacagtg | 12 | tcaaaaact |
| | IGHD2-8 | ggattttgt | 12 | cactgtg | yes | cacagtg | 12 | tcccaaagc |
| | IGHD1-14 | ggattccga | 12 | cacagcg | no | cactgtc | 12 | tcaaaaact |
| | IGHD6-19 | ggtttctga | 12 | cacagtg | no | cacagtg | 12 | ccagaaacc |
| | IGHD3-16 | / | / | cactgtg | no | cacagca | 12 | tcagaaacc |
| | IGHD5-12-2 | ggttattgt | 12 | gactgtg | yes | cacagtg | 12 | gcagcaacc |
| | IGHD1-14-2 | ggattccga | 12 | cacagcg | no | cactgtc | 12 | tcaaaaact |
| | IGHD4-17 | gctttttgt | 12 | tactgtg | yes | cacagtg | 12 | gcaaaaact |
| | IGHD1-26-2 | ggattctga | 12 | cacggtg | no | cactgtg | - | - |
| | IGHD2-21-2 | ggattttgt | 12 | cactgtg | yes | cacagtg | 12 | tcctaaagc |
| | IGHD1-1 | / | / | cacggtg | yes | caccgtg | 4 | aaactgtgt |
| | IGHD2-2 | ggattttgt | 12 | cactgtg | yes | cacagtg | 12 | tcccaaagc |
| | IGHD3-9 | / | / | cactgtg | yes | cacagtg | 12 | tcaaaaacc |
| | IGHD4-23-2 | gctttttgt | 12 | tgctgtg | no | cacagtg | 12 | gcaaaaact |
| | IGHD5-18 | ggttattgt | 12 | gactgtg | yes | cacagtg | 12 | gcagcaacc |
| | IGHD5-5-2 | ggttattgt | 12 | gactgtg | yes | cacagtg | 12 | gcagcaacc |
| | IGHD4-17-2 | gctttttgt | 12 | tactgtg | yes | cacagtg | 12 | gcaaaaact |
| | IGHD2-21 | ggattttgt | 12 | cactgtg | yes | cacagtg | 12 | tcctaaagc |
| | IGHD7-27-2 | / | / | cactgtg | yes | cacagtg | 12 | acaaaaacc |
| | IGHD1-7-2 | ggattctga | 12 | cacagtg | yes | cactgtg | 12 | tccaaaacg |
| | IGHD6-6-2 | aaacaaacc | 25 | cacagtg | yes | cacagtg | 12 | ccagaaacc |
| | IGHD6-13 | ggtttctga | 12 | cacagtg | yes | cacagtg | 12 | ccagaaacc |
| | IGHD1-20-2 | ggattctga | 12 | cacagtg | no | caccgtg | 4 | aaactgtgt |
| | IGHD5-18-2 | ggttattgt | 12 | gactgtg | yes | cacagtg | 12 | gcagcaacc |
| | IGHD3-10 | ggtttgggg | 12 | cactgtg | yes | cacagtg | 12 | tcaaaaacc |

| | | | | | | | | |
|---|---|---|---|---|---|---|---|---|
| | IGHD6-25-2 | ggtttctga | 12 | cacagtc | yes | cacaatg | 12 | acagaaacc |
| | IGHD7-27 | / | / | cactgtg | yes | cacagtg | 12 | acaaaaacc |
| | IGHD4-23 | gctttttgt | 12 | tgctgtg | no | cacagtg | 12 | gcaaaaact |
| | IGHD4-4-2 | gctttttgt | 12 | tactgtg | yes | cacagtg | 12 | gcaaaaact |
| | IGHD3-3 | ggtttgggg | 12 | cactgtg | yes | cacagtg | 12 | tcaaaaacc |
| | IGHD6-25 | ggtttctga | 12 | cacagtc | yes | cacaatg | 12 | acagaaacc |
| | IGHD5-5 | ggttattgt | 12 | gactgtg | yes | cacagtg | 12 | gcagcaacc |
| | IGHD1-20 | ggattctga | 12 | cacagtg | no | caccgtg | 4 | aaactgtgt |
| | IGHD4-11-2 | gctttttgt | 12 | tgctgtg | no | catagtg | 12 | gcaaaaact |
| | IGHD1-26 | ggattctga | 12 | cacggtg | no | cactgtg | - | - |
| | IGHD1-7 | ggattctga | 12 | cacagtg | yes | cactgtg | 12 | tccaaaacg |
| TRBD | TRBD1 | tgtttttgt | 12 | cattgtg | yes | cacaatg | 23 | acaaaaacc |
| | TRBD2-2 | catttttgt | 12 | cattgtg | no | cacgatg | - | - |
| | TRBD2 | catttttgt | 12 | cattgtg | no | cacgatg | - | - |
| | TRBD1-2 | tgtttttgt | 12 | cattgtg | yes | cacaatg | 23 | acaaaaacc |
| TRDD | TRDD1 | - | - | - | no | - | - | - |
| | TRDD3 | agtttttgt | 12 | cactgtg | yes | cacagtg | 23 | acaaaaact |
| | TRDD2 | ggtttttat | 12 | cattgtg | no | cacacag | 23 | ccaaaaaca |
