## Supplementary Data 6 for "Adaptive immunity: from CRISPR to CRIHSP?"

**Heptamer:**

'CACGGTG', 'CGCAGTG', 'TGTTGTG', 'CAGAGTG', 'CAGTGAG', 'CATAGTG', 'CACACAG', 'CTCAGTG', 'CACATTA', 'CAATGTG', 'CACGATG', 'CACAGTA', 'CACAAAG', 'GACTGTG', 'GGCTGTG', 'CCAGGTA', 'CACAATG', 'CATCGTG', 'CACAACG', 'TACGGTA', 'CACAGCT', 'CACATAC', 'CACAACA', 'CACAGTG', 'CATTGTG', 'CACCGTG', 'AACAGAA', 'CACAGCA', 'TACAGCA', 'CACAGCC', 'CCAAATG', 'CCAAGTG', 'CACATTG', 'CAGTGAC', 'CACGGCC', 'CACAGGG', 'TACACAG', 'TTATGTG', 'CATGGTG', 'CGCAATG', 'CACAGGA', 'CTCCGTG', 'CACGGAG', 'CACAGTC', 'CACCATG', 'CACAGAG', 'CACTGTC', 'GGCCGTG', 'CACATAA', 'CATAGGA', 'CAGTGAA', 'AACAGTG', 'CCCTGTG', 'CACTGTG', 'TCCTGTG', 'CACATGA', 'GACAGAA', 'CGCTGTG', 'CACTGAG', 'CACTATG', 'CACAGCG', 'AGCTGTG'

**Nonamer:**

'ACAAAAACT', 'ACGCAAACT', 'ACACAAATC', 'ATTGAAACC', 'GTGAAAATC', 'ATAAAACCC', 'TCAAAAACC', 'ACACAGTTT', 'ACAAAAATG', 'GCAGCAACC', 'CCAAAAACC', 'ACCCAAACC', 'TCAGAAACC', 'TCAGGAACC', 'TCAGTAACC', 'ACACCAACC', 'ACACAAACG', 'AGACAGACT', 'ACACAACAT', 'ATACAAACC', 'CAGAAACC, 'ACACAGAGG', 'ACAAAAAGC', 'ACAGAAACC', 'TCCCAAAGC', 'TAGATAACC', 'CCCCAAACC', 'ACAATAACC', 'ACAAAAGCC', 'GCAAAAACC', 'CAAAGAACC', 'ACATAAACC', 'CTAAAACCC', 'TCAAAAACA', 'ACACAAACT', 'GCCAAAAAC', 'AAACAAACC', 'GCAAAACCA', 'ACAGAAAGA', 'ACAAAAACA', 'TCTAAAAGG', 'CCAAAAACA', 'ACACTCTAC', 'CCAGAAACC', 'TCCAAAACG', 'TCGGAATCC', 'ACAAAATCC', 'GGCAAACCC', 'CACAAACC', 'ACACAACCC', 'TTAGAAACC', 'ATGCAAACC', 'ACAGGTAAC', 'ACCCAAGCC', 'ACACAGATT', 'TGAAAATC ', 'TCACAAACC', 'GCAAGAACC', 'ACACAGAAT', 'ACACAAAAA', 'GCAGAAAAC', 'ACACAAAAG', 'ACAAAATGG', 'TCAGAATCC', 'ACAAGAACC', 'ACAAAAACC', 'ATACAAACT', 'AGGAAAACC', 'TCAGAAATC', 'GCAAAAACT', 'ACAAAACCA', 'ATAAAAACC', 'TTCAAAACC', 'ACACAAACC', 'TCAGAAAGC', 'ACACAAAGC', 'ACCAAAACC', 'ACAGAAAAA', 'CCAGAATTC', 'TCAATAAAA', 'GCAGAAACC', 'AAAGAACCG', 'TCAGAAACG', 'TCAAAAACT', 'TCCTAAAGC', 'ACAGAAACT', 'ACACAAAAT', 'GCGCAAACC', ' TCAAGAACA ', ' ACAAAAATC '
