## Supplementary Data 7 for "Adaptive immunity: from CRISPR to CRIHSP?"

**Supplementary Table 1**

The PCR primers we used to amplify the CRIHSP, non-CRIHSP and Ctrl sites in Figures 4F, 4G and 4H were provided as following:

| AR-CRIHSP-1F | 5′-TCCATAGAAGATTCTAGAAAAATGGCAGCAGGGGAGAGACAG-3′ |
| --- | --- |
| AR-CRIHSP-1R | 5′-GATTTAAATTCGAATTCAAACCGCCAAGGCCGAGCCTGAAG-3′ |
| AR-CRIHSP-2F | 5′-CCTCCATAGAAGATTCTAGAGGTGCCCTTCCTCCCACGTC-3′ |
| AR-CRIHSP-2R | 5′-CGATTTAAATTCGAATTCAGAGGGCCACCACACCCTGCGGATC-3′ |
| AR-CRIHSP-4F | 5′-CCTCCATAGAAGATTCTAGAGGCCTTGAAGGCATTTTGGAAATC-3′ |
| AR-CRIHSP-4R | 5′-TCCGATTTAAATTCGAATTCGAGCTGACTGGGGCAGAATTCACTC-3′ |
| AR-CRIHSP-6F | 5′-GACCTCCATAGAAGATTCTAGACCAGCCTGCCCCAGTCTAACCT-3′ |
| AR-CRIHSP-6R | 5′- ATCCGATTTAAATTCGAATTCCCACACCATCCACAGGCACCAAAT-3′ |
| AR-CRIHSP-7F | 5′-CCTCCATAGAAGATTCTAGAAAAAGCTGGGCCTGGTGGAGCG-3′ |
| AR-CRIHSP-7R | 5′-TCCGATTTAAATTCGAATTCTGCACACACTAGGCCCCATAGCAAC-3′ |
| AR-CRIHSP-8F | 5′-CCTCCATAGAAGATTCTAGAAAAGCAAGGCATCGTCTTGTACACT-3′ |
| AR-CRIHSP-8R | 5′-GATCCGATTTAAATTCGAATTCTCACAATGGTCAAGGTCGCACCC-3′ |
| AR-nonCRIHSP-3F | 5′-GACCTCCATAGAAGATTCTAGAGTCCCTGTTGATCTTCCGTTTTT-3′ |
| AR-nonCRIHSP-3R | 5′-ATCCGATTTAAATTCGAATTCCCAGGCATCCAGAATATGTCCTTG-3′ |
| AR-nonCRIHSP-4F | 5′-GACCTCCATAGAAGATTCTAGATTCTTCATCACAGTAGGGTGCAT-3′ |
| AR-nonCRIHSP-4R | 5′-ATCCGATTTAAATTCGAATTCCCTGCTTTGCTGTCTCACTTGG-3′ |
| AR-nonCRIHSP-IGHTY2F | 5′-AGAAGATTCTAGAGCTAGCGGAGAGCTATGATGTCACCAC-3′ |
| AR-nonCRIHSP-IGHTY2R | 5′-CGCGGATCCGATTTAAATTCGAATTCTACAGGGAAGAGCTTTTTG-3′ |
| AR-nonCRIHSP-IGHTY3F | 5′-ATTCTAGAGCTAGCGATCATTAAGTTTCATTCATTAGGAC-3′ |
| AR-nonCRIHSP-IGHTY3R | 5′-TTTAAATTCGAATTAATACCCTGAAAACCGAGTACACGG-3′ |
| AR-nonCRIHSP-IGHTY4F | 5′-ATTCTAGAGCTAGCGACCCAGATAAGACGACGGTG-3′ |
| AR-nonCRIHSP-IGHTY4R | 5′-CCGATTTAAATTCGAATTACTCCTGTCTTCTGGTCACT-3′ |
| AR-nonCRIHSP-IGHTY5F | 5′-ATTCTAGAGCTAGCGCTCTTTTGTACTTTGTCCCT-3′ |
| AR-nonCRIHSP-IGHTY5R | 5′-ATCCGATTTAAATTCGAATTGCCTTTGAAAATAAATCCTT-3′ |
| Ctrl-nonCRIHSP-UNJH-1F | 5′-TCCATAGAAGATTCTAGATGGACCAGGCATGGTGGTTTACGC-3′ |
| Ctrl-nonCRIHSP-UNJH-1R | 5′-TCCGATTTAAATTCGAATTCCTCACTATGTGCCAGGCATTGTTC-3′ |
| Ctrl-nonCRIHSP-UNJH-2F | 5′-CCTCCATAGAAGATTCTAGACCCTGATTGTTTTCCTCACCTTCTT-3′ |
| Ctrl-nonCRIHSP-UNJH-2R | 5′-TCCGATTTAAATTCGAATTCCAGACCACACATATAACAGTGGTCC-3′ |
| Ctrl-nonCRIHSP-UNJH-3F | 5′-CTCCATAGAAGATTCTAGACGGGGCTGACATTTTCCTGGCA-3′ |
| Ctrl-nonCRIHSP-UNJH-3R | 5′-TCCGATTTAAATTCGAATTCCATTTATTCAATAGCTCTGGGCCTG-3′ |
| Ctrl-nonCRIHSP-UNJH-4F | 5′-GACCTCCATAGAAGATTCTAGACTTACGAGCTCCCCTTGTGTGG-3′ |
| Ctrl-nonCRIHSP-UNJH-4R | 5′-ATCCGATTTAAATTCGAATTCCCCACTTCCTGCAAGGGTTATAAT-3′ |
| Ctrl-nonCRIHSP-UNJH-5F | 5′-GACCTCCATAGAAGATTCTAGAACCTGCGTCTCCTTCTAAGCTC-3′ |
| Ctrl-nonCRIHSP-UNJH-5R | 5′-CCGATTTAAATTCGAATTCCCTGTTTGCCTGTCCGGGTTGC-3′ |
| Ctrl-nonCRIHSP-NCR2F | 5′-ATTCTAGAGCTAGCGTGGCAAACACTAAGAACCGAGCAGA-3′ |
| Ctrl-nonCRIHSP-NCR2R | 5′-CCGATTTAAATTCGAATTCCAAGCATAGGAATCCACAACACCG-3′ |
