## Supplementary Data 8 for "Adaptive immunity: from CRISPR to CRIHSP?"

```
from Bio import SeqIO
import svgwrite
import plotly as py
import plotly.express as px
import plotly.graph_objs as go
from plotly.graph_objs import Scatter

Target_file_name=input("input target file name:")
DNA_Seq_dict = SeqIO.to_dict(SeqIO.parse(open(Target_file_name),'fasta')) # 直接转为字典格
式
for gene_name in DNA_Seq_dict:
    DNA_Seq= DNA_Seq_dict[gene_name].seq

#把所有 stem-loop 序列的(5',3')位置依次放在列表中
list_final_location=[]
list_final_location_5=[]
list_final_location_3=[]
for i in range(0, len(DNA_Seq)-60): #len(DNA_Seq)
    Fragment=DNA_Seq[i:i+7]
    Pre_DNA_Seq_R=DNA_Seq[i+7:i+60]
    DNA_Seq_R= Pre_DNA_Seq_R.reverse_complement()

    list_a=[]
    for j in range(0,43):
        list_a.append(DNA_Seq_R[-8-j:-1-j])

    for seq in range(0,len(list_a)):
        flag=0
        for base in range(0,len(Fragment)):
            if Fragment[base]==list_a[seq][base]:
                flag += 1
        if flag/len(Fragment) > 0.89:
            location_5_find = i #DNA_Seq.find(Fragment)
            location_3_ = list_a[seq].reverse_complement()
            location_3_find = Pre_DNA_Seq_R.find(location_3_) + 7 + i + 7
            list_final_location_5.append(location_5_find)
            list_final_location_3.append(location_3_find)
            if len(list_final_location_3)>=2:
                if list_final_location_3[-1]-list_final_location_3[-2] < 10:
                    list_final_location_3[-2]=max(list_final_location_3[-1],
list_final_location_3[-2])
                    list_final_location_3.remove(list_final_location_3[-1])
                    list_final_location_5.remove(list_final_location_5[-1])
```

```
for k in range(0,len(list_final_location_5)):
    list_final_location.append((list_final_location_5[k],list_final_location_3[k]))

#记录 stemloop 5'端 3'端的位置及相应序列：
Stemloop_sequences = []
for u in range(0,len(list_final_location)):
    Stemloop_sequences.append(list_final_location[u])
    Stemloop_sequences.append(DNA_Seq[list_final_location[u][0]:list_final_location[u][1]])
list_final_location_num = len(list_final_location)
with open('Stemloop_location_sequences.txt', 'w') as OUT:

OUT.write("{}\n{}\n{}\n".format(list_final_location_num,list_final_location,Stemloop_sequences))

#plotly 作图在线性基因上标注 stemloop 5'端的位置
data_5=[]
for t in list_final_location_5:
    list_plotly_x = [t,t]
    list_plotly_y = [0,10]
    trace_5=Scatter(x=list_plotly_x, y=list_plotly_y,mode="lines",\
                    line={'width':0.01,'color':'red'})
    data_5.append(trace_5)
layout = go.Layout(plot_bgcolor='rgba(0,0,0,0)')
fig = go.Figure(data=data_5, layout=layout)
py.offline.plot(fig,filename='stemloop5location.html')

# 定义一个函数，名为 check_element，接受两个参数：lst 和 x
def check_element(lst, x):
    # 创建一个空列表，用于存储每个子列表的检查结果
    result = []
    # 遍历 lst 中的每个子列表
    for sublist in lst:
    # 使用 in 运算符检查 x 是否在子列表中
        if x in sublist:
            # 如果在子列表中，返回 1，并添加到 result 列表中
            result.append(1)
        else:
            # 如果不在子列表中，返回 0，并添加到 result 列表中
            result.append(0)
    # 使用 sum 函数对 result 列表求和，并返回求和结果
    return sum(result)

#依次查找回文序列相似（90%以上）的 stemloop 并将相似序列的 5'端位置依次放在列表中
##plotly 可视化作图准备，将序列相同的回文序列 5'端放在一个列表中
```

```
list_plotly_whole = []
for tuple in range(0,len(list_final_location)):
    square_x=[]
    for tuple_behind in range(tuple+1,len(list_final_location)):
        flag=0
        for base in range(0,10):
            if
DNA_Seq[list_final_location[tuple][0]:list_final_location[tuple][1]][base]==DNA_Seq[list_final_loc
ation[tuple_behind][0]:list_final_location[tuple_behind][1]][base]:
                flag += 1
        if flag/10 > 0.89:
            if check_element(list_plotly_whole, list_final_location[tuple][0])<1 and
list_final_location[tuple][0] not in square_x:
                square_x.append(list_final_location[tuple][0])
            if check_element(list_plotly_whole, list_final_location[tuple_behind][0])<1 and
list_final_location[tuple_behind][0] not in square_x:
                square_x.append(list_final_location[tuple_behind][0])

    if len(square_x)!=0:
        list_plotly_whole.append(square_x)
    square_x=[]

# 创建一个空列表，用于存储每个子列表的元素个数
count = []
# 遍历 lst 中的每个子列表
for sublist in list_plotly_whole:
# 使用 len 函数计算子列表的长度，并添加到 count 列表中
    count.append(len(sublist))
total = sum(count)
HSP = total/2

for r in range(len(list_plotly_whole)):
    if len(list_plotly_whole[r])%2!= 0:
        list_plotly_whole[r].append(list_plotly_whole[r][-1])
with open('list_plotly_whole.txt', 'w') as OUT:
        OUT.write(">Seq{}\n{}\n{}\n{}\n{}\n".format('list_plotly_whole', count, total, HSP,
list_plotly_whole))

##plotly 可视化作图，列表中的元素是两个赋予同一种颜色，两个或更多，则颜色随机分配，
相同的序列颜色相同
data=[]
for s in range(len(list_plotly_whole)):
    list_x=[]
    list_y=[]
```

```
        for t in range(0,len(list_plotly_whole[s]),2):
            list_x.append(list_plotly_whole[s][t])
            list_y.append(list_plotly_whole[s][t+1])
        if len(list_x)<2:
            trace=Scatter(x=list_x, y=list_y,mode="markers",\
                          marker={'color':'green'})
            data.append(trace)
        if len(list_x)>=2:
            trace=Scatter(x=list_x, y=list_y,mode="markers")
            data.append(trace)
py.offline.plot(data,filename='similarstemlooplocation.html')
```
